# Rhpn2 promotes zebrafish melanoma development and aggressiveness *in vivo*

**DOI:** 10.64898/2026.07.03.736252

**Authors:** Mana Alavi, Alexia Gybels, Laura Gulizia, Katarzyna Konobrocka, Garnik Hovhannisyan, Sena Bekar, Camille Perazzolo, Sumeet Pal Singh, Isabelle Pirson

## Abstract

Melanoma, one of the most metastatic and multidrug resistant cancer, is the first leading cause of death from skin cancer. This complex disease requires identification of additional cooperating events that contribute to progression, invasion and metastasis to reinforce therapeutics. RhoGTPases play key roles in cancer development and metastasis. Rhophilin-2 (RHPN2), a Rho effector, is amplified in various human cancers and its role in melanoma remains unexplored. Here, we combined knock-down experiments in human melanoma cells, with knock-out and overexpression experiments in zebrafish to uncover the roles of RHPN2 in melanoma development. We show that in human melanoma cells RHPN2 contributes to growth, and to clonogenic, migratory and invasive properties of the cells. Using NRAS^Q61L^ and BRAF^V600E^ zebrafish models, we provide the first *in vivo* evidence that Rhpn2 promotes melanoma onset and development. Histological analysis of the Rhpn2 deficient tumors showed decreased cellular density and absence of primary cilia structures at the invasive tumor/stroma borders. Transcriptomic profiling of the Rhpn2-KO melanoma revealed increased expression of the IFN1-responsive genes and modulation of genes involved in lipid metabolism and cilia function. Together these findings position RHPN2 as a modulator of melanoma, offering new perspectives in considering it as a target to impair the development of the tumor.

## Introduction

Cutaneous melanoma, through its high metastatic potential and drug resistance, is the first leading cause of death related to skin cancer (Tasdogan et al., 2025). Context-dependent genetic mutations in melanocytes, that activate growth-promoting MAPK pathways (BRAF, NRAS) or attenuate tumor-suppressive mechanisms (P53 or PTEN), are driving factors of melanoma development (Centeno et al., 2023). Over the last decade, treatment has improved considerably with therapies targeting BRAF and MEK and the development of immune checkpoint inhibitors. However, due to cell plasticity and very dynamic tumor microenvironment (TME), most patients develop resistance to therapies. Melanoma development is a complex and heterogeneous process that also requires additional cooperating events that contribute to progression, invasion and metastasis and could represent new therapeutical targets (Wouters et al., 2020).

As main regulators of cytoskeleton dynamics, vesicular trafficking, and cell cycle progression regulation (Etienne-Manneville and Hall, 2002), RHO GTPases play key roles in cancer development, epithelial-to-mesenchymal transition (EMT) and cellular invasion (Clayton and Ridley, 2020; Crosas-Molist et al., 2022; Lee et al., 2025). RHOA/C signaling promotes bleb-based amoeboid migration which enables the cell to squeeze into pre-existing spaces and deform the ECM without the need for significant pericellular proteolysis (Wyckoff et al., 2006). RHOB specifically controls membrane blebbing in melanoma cells as well as their amoeboid migration in 3D-collagen (Gong et al., 2018). All the best characterized RHOA/B/C GTPases effectors ― ROCK, DIAPH1, and PKN/PRK — were also described as actors in melanoma migration and invasion (Arang et al., 2023; Carreira et al., 2006; Kümper et al., 2016). We and others identified Rhophilin-2 (RHPN2), as a RHOB effector and RHOA interactor (Mircescu et al., 2002; Peck et al., 2002).

RHPN2 is a highly conserved scaffold protein expressed in mouse and human secretory epithelia (Behrends et al., 2005; Mircescu et al., 2002). It is composed of different protein interaction domains (HR-1, Bro1 and PDZ) and was first shown to regulate actin-cytoskeleton, and vesicular trafficking (Peck et al., 2002; Steuve et al., 2006). Knock-out mice were previously generated, presenting with no macroscopic or tissues histological abnormalities, but these were not further analyzed in pathological conditions (Behrends et al., 2005). More recently, using different human cell lines, evidence has begun to emerge that RHPN2 could play a sustaining role in cancer cells growth and invasion processes. RHPN2 sustains glioblastoma EMT gene expression through RhoA activation (Danussi et al., 2013). Stable knock-down or overexpression of RHPN2 in lung and ovarian cancer cell lines, respectively, decreased or increased the growth of cell lines and the tumor volume in mice xenografts experiments (Xiao et al., 2020; Yu et al., 2020; Zhang et al., 2025). An intronic variant in the *rhpn2* gene was recently suggested as a cancer-associated germline risk (Kellman et al., 2025) and, in an *in silico* study, RHPN2 was proposed, but not biologically tested, as a candidate gene essential for melanoma cell survival (Pyatnitskiy et al., 2015). However, the impact of RHPN2 on *in vivo* tumor development, and particularly its role in melanoma remains unexplored.

In the present work, we show that RHPN2 KD in human melanoma cells significantly reduces growth capacities and viability. We pursue with *in vivo* experiments using zebrafish, now a well-established model for studying cancer initiation/progression and drug discovery (Letrado et al., 2018) especially in melanoma studies (Wojciechowska et al., 2016). Using constitutively active NRAS and BRAF melanoma models, we revealed, for the first time *in vivo,* that RHPN2 promotes melanoma onset and progression, that its deficiency decreases tubulin acetylation at the tumor-stroma interface and causes an decrease of some ciliary genes expression and regulation in IFN-response gene and lipid metabolism expression in melanoma cells. Taken together, our data suggest that RHPN2 is a novel regulator capable of controlling this highly aggressive cancer.

## Results

### RHPN2 is required for human melanoma cell proliferation and invasion

To address whether RHPN2 could influence melanoma cell behavior, we first measured RHPN2 expression levels in different melanoma cell lines A375 (originating from primary tumour), and A2058, SKMEL2 and SKMEL28 (originating from metastatic sites) by western blotting. We observed higher level of expression of RHPN2 in A2058 and A375 (Fig.1A). These cell lines were further used for knock-down experiments by targeting RHPN2 expression levels through lentivirus-mediated short hairpin RNA (shRNA). The reduction of RHPN2 protein expression was confirmed by immunoblotting in three different clones stably expressing RHPN2-shRNA (Fig.1B). Then, through MTT assays at different time-points, we evaluated the impact of RHPN2 silencing on cell growth, and showed a significant decrease in cell viability upon RHPN2 knock-down (KD) after 72h and 96h in all the clones tested compared to control cells (Fig.1C). Moreover, the colony formation experiments also demonstrated that downregulating RHPN2 decreased the number and the size of colonies in both A375 and A2058 (Fig.1D). Notably, the decrease observed in both experiments was proportional to RHPN2 knock-down in respective clones. This decrease in growth correlated with a significant decrease of pERK levels in both cells lines upon RHPN2 KD (Fig.1E) but no modulation in pAkt was noticed. Further, we showed, through transwell assay that both motility and invasive properties of RHPN2-KD cells were impaired compared to control cells (Fig.1F). These data indicate that RHPN2, like in other cancers, supports the growth and invasion of human melanoma cells *in vitro*.

**Figure 1.**
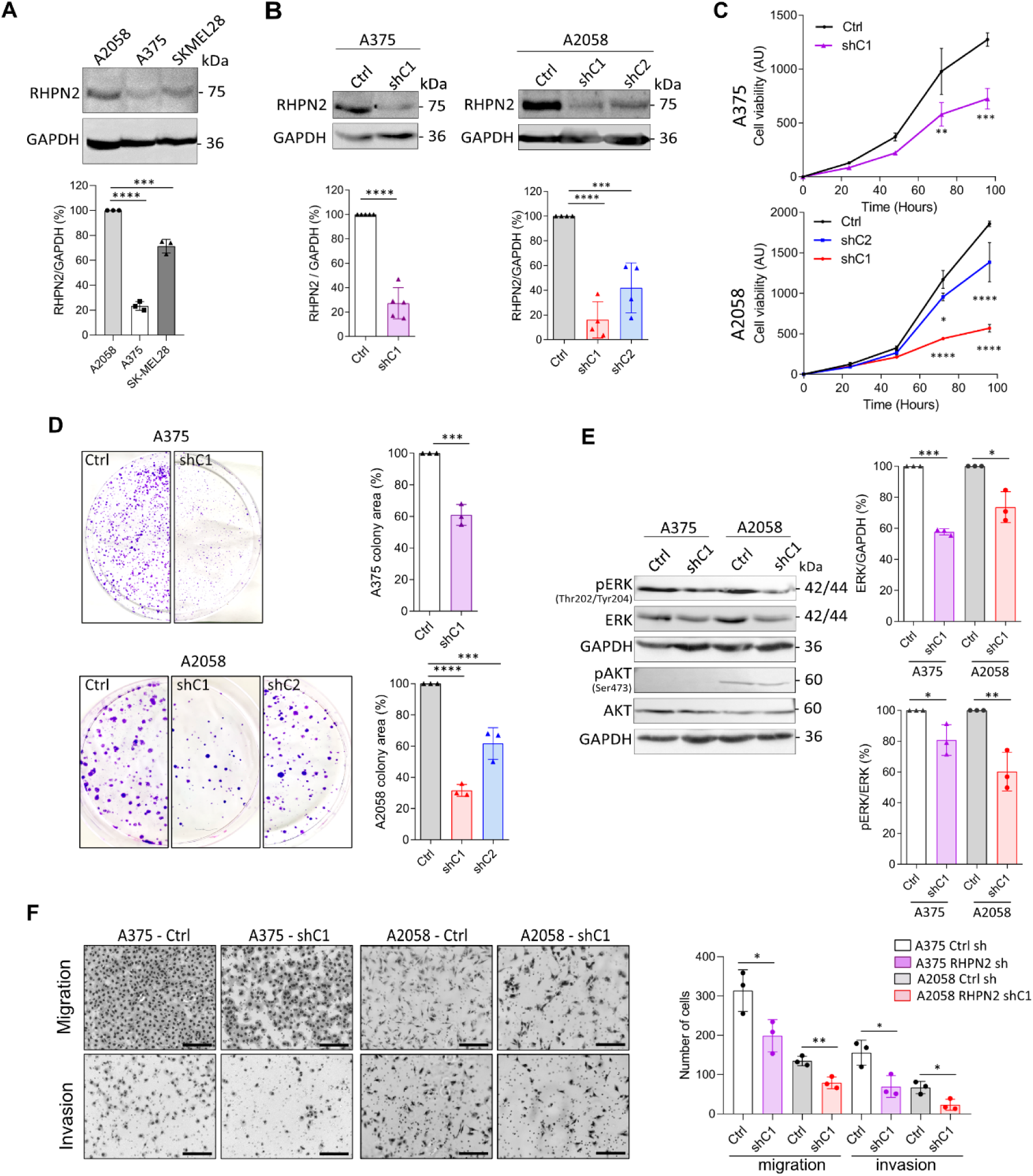
Rhpn2 knock-down in human melanoma cells decreases their clonogenic, viability and invasion properties. **A-** RHPN2 expression in melanoma cell by Western blotting (N=3; A375 vs A2058, ****p < 0.0001; SKMEL28 vs A2058, ***p = 0.0002, One-way Anova). **B-** Down regulation of RHPN2 expression in stable shRNA-RHPN2 clones (C) compared to shRNA-control (Ctrl) by Western blotting. Graphs represent the RHPN2 levels normalized to GAPDH. (N=4; A375 C1 vs Ctrl, ****p < 0.0001, unpaired T-test; A2058 C1 vs Ctrl, ****p < 0.0001; C2 vs Ctrl, ***p = 0.0006, One-way Anova). **C-** Viability tests in the presence of WST-8 for 4 days (N=2; A375, C1 vs Ctrl, **p = 0.0036 (72h **p=0.0044; 96h ***p=0004); A2058, **, p = 0.0013 (72h C1 *p=0.039; 96h **** p < 0.0001; C2 72h and 96h p < 0.0001); Two-way Anova Interaction). **D-** Clonogenic tendency of the clones after two weeks (N=3; A375 C1 vs Ctrl, ***p = 0.0005, unpaired T-test; A2058, C1 vs Ctrl, ****p < 0.0001; C2 vs Ctrl, ***p = 0.0005; One-way Anova). **E-** Erk and pErk quantification by Western blotting (N=3; Erk/GAPDH, A375 C1 vs Ctrl, **** p < 0.0001, A2058, C1 vs Ctrl, *p < 0.01; pErk/Erk, A375 C1 vs Ctrl, * p = 0.0288, A2058, C1 vs Ctrl, **p = 0.0054; unpaired T-test). **F-** Migration and invasion properties of the clones in transwell assays for 22h (N=3; Migration : A375 C1 vs Ctrl, *p = 0.04, A2058 C1 vs Ctrl, ** p = 0.0073, unpaired T-test; Invasion : A375 C1 vs Ctrl, *p = 0.0237, A2058 C1 vs Ctrl, *p = 0.0223, unpaired T-test). Data are represented as mean ± SD.

### Loss of Rhpn2 in zebrafish impairs NRAS^Q61L^-driven melanoma development

To gain insight into the biological role of Rhpn2 on tumor development *in vivo*, we used the zebrafish model system. Zebrafish is a well-established model for studying tumor development through potent genetic models and the striking conservation of cancer-related pathways between human and zebrafish allows extrapolation of results obtained in fish back to humans (Hason and Bartůněk, 2019). In zebrafish, *rhpn2* gene is unique and well conserved in sequence and in synteny between human and zebrafish (Fig.S1A). We first generated constitutive Rhpn2 knock-out (KO) zebrafish models using CRISPR/cas9 mediated mutagenesis. Two single guide RNA targeting exon 1 were used and two mutant alleles were obtained in F1 and validated by Sanger sequencing (Fig.S1B) respectively inducing a start loss (*rhpn2*^SL4/SL4^) and a premature stop codon (*rhpn2*^Δ2/Δ2^) in *rhpn2* gene (Fig.S1B; Fig.2A).

**Figure 2.**
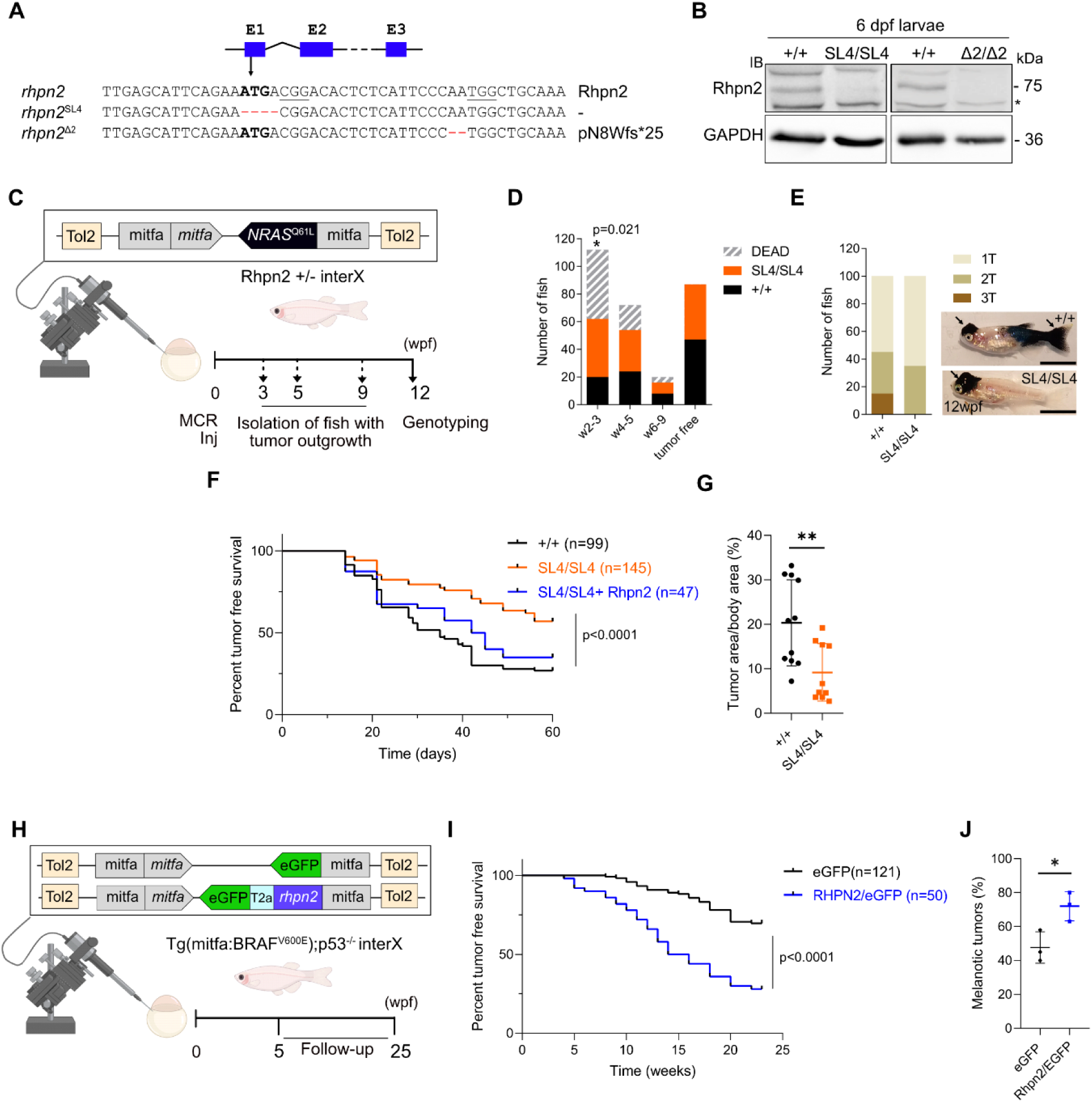
*In vivo*, Rhpn2 promotes the development of NRAS- and BRAF-dependent melanoma in zebrafish. **A-** Genomic sequence of *rhpn2* start codon (bold) region with sgRNA target PAM sequences underlined. The *rhpn2* mutant alleles present respectively a 4 base pairs deletion (start codon loss, SL4), and a 2 base pairs deletion (frameshift and premature stop codon 25 aa downstream, Δ2). **B-** Immunoblotting of 35 µg total protein extracts from 6 dpf larvae of the indicated genotype with ZF-Rhpn2 specific antibody. 75 kDa Rhpn2 protein was not detected. **C-** Schematic representation of the experiment workflow. The MCR-NRAS vector, injected in one-cell embryos allows tumor development. Fish presenting tumor are isolated at week 3, week 5 and week 9 and all genotyped at 12 wpf. **D-**The proportions of each genotype were significantly different than expected for the fish having developed tumor at 3 wpf compared to their tumor free siblings (N=3). Chi-square analysis : w2-3 (n = 112, +/+ vs SL4/SL4, *p = 0.021), w4-5 (n = 72, +/+ vs SL4/SL4, ns, p = 0,3), w6-9 (n = 20, +/+ vs SL4/SL4, ns, p = 0,52). **E-** The SL4/SL4 fish developed less tumors than +/+. **F-** A smaller proportion of *SL4/SL4* fish (orange) develop melanoma than +/+ (black) and this phenotype is rescued by coinjection MCR-Rhpn2 (blue). (N = 6; at 60 dpf WT (n=99), 73.12%; Rhpn2-KO (n=145), 43.1%; ****p<0,0001 Log-rank (Mantel-Cox) test). **G-** Tumor area measurements of SL4/SL4 (n = 10) compared with +/+ (n = 11) zebrafish. Statistical analysis, SL4/SL4 vs +/+, **p = 0,0062; T-test. **H-** Experiment workflow. The MCR-eGFP or MCR-Rhpn2/eGFP vectors, injected in one-cell embryos allow melanocyte rescue and tumor development in Tg(*mitfa*:BRAF^V600E^); p53-/- fish. Tumor appearance is followed-up till 6 mpf. I**-**A higher proportion of *Rhpn2-OE* fish (blue) develop melanoma than eGFP-OE (black). Statistical analysis: ****p<0,0001 Log-rank (Mantel-Cox) test. (N=4)**. J-** The proportion of melanotic tumors is higher in *Rhpn2-OE* fish. (N=3; *p = 0,01; T-test).

Adult heterozygous offsprings followed the expected Mendelian inheritance ratios (Fig.S1C) and homozygous mutants were viable, fertile and displayed no noticeable growth or developmental defects (Fig.S1E-G). Immunoblotting was performed on total protein extracts from 6 days post fertilization (dpf) larvae of both KO-lines and we confirmed protein deficiency (Fig.2B). This was done using an antibody designed against zebrafish Rhpn2, previously tested by western blot on recombinant proteins (Fig.S1G). The following experiments were done using *rhpn2*^SL4/SL4^ mutants.

To determine whether Rhpn2 may have an impact on melanoma growth *in vivo*, we first used the MiniCoopR system (Iyengar et al., 2012), which makes it possible to generate melanomas in fish within a few weeks. The MiniCoopR-NRAS^Q61L^ vector (MCR-NRAS) allows chimeric rescue of melanocytes in the melanocytes-deficient *mitfa*^-/-^ (nacre) background by reexpression of the mitfa under the control of its own promoter. Through simultaneous coexpression of the oncogene NRAS^Q61L^, tumors appear within the first weeks post fertilization (wpf). We microinjected the MCR-NRAS vector into one-cell stage embryos obtained from *rhpn2*^SL4/+^ nacre fish intercrosses (Fig.2C).

Tumor outgrowth was monitored weekly and fish presenting with nodular tumors were isolated at 3, 5, and 7 wpf. At 12 wpf, through genotyping, we observed a significant smaller proportion of WT fish compared to *rhpn2*^SL4/SL4^ in the population having developed tumors in early time-points (w2-3) (Fig.2D). This disequilibrium was not observed in tumor-free sibling animals. We also noticed that the tumor nodules were less numerous in *rhpn2*^SL4/SL4^ fish compared to their *rhpn2*^+/+^ siblings. However, when the genotyping of the fish was performed earlier, at 6 wpf instead of 12 wpf, we do not observe the significant decrease in live WT fish presenting nodular tumors (Fig.S2A). On the contrary, we observed an inverse increase, even if not significative. At 9wpf, the live fish that presented a rescue without tumor development enriched for Rhpn2-KO genotype, confirming that the tumor development was less frequent in *rhpn2*^SL4/SL4^ fish compared to their *rhpn2*^+/+^ siblings. This was also confirmed by the number of tumors per fish at 6 wpf (Fig.S2B). These results suggested that WT fish developed more aggressive tumors causing higher death rate compared to Rhpn2-KO fish.

To confirm this hypothesis, and analyze the tumor development earlier, we subsequently microinjected MCR-*NRAS* into embryos obtained seperately from intercrosses of *rhpn2*^SL4/SL4^ and intercrosses of *rhpn2*^+/+^ siblings. We first ensured that there was no difference in 1 dpf survival rate of the embryos of both genotypes (Fig.S2C) nor in the number of fish exhibiting melanocyte rescue at 7 dpf between Rhpn2-KO and WT fish (Fig.S2D). After injection, the tumor appearance was monitored daily and quantified using Kaplan-Meyer curves for both genotypes (Fig.2F). We observed that Rhpn2 deficiency significantly affected the number of fish developing tumors (at 60 dpf WT: 73.12%; Rhpn2-KO: 43.1%). Tumor size normalized to the fish body area was also significantly reduced in 6 wpf *rhpn2*^SL4/SL4^ compared to *rhpn2*^+/+^ siblings (Fig.2G). In these experiments, the entire organism is deficient in Rhpn2, making it difficult to delineate if Rhpn2 affects melanocytes or its microenvironment. To address this, we performed a melanocyte-specific rescue of Rhpn2 expression through coinjection of a MCR vector expressing Rhpn2 under the control of the mitfa promoter (MCR-Rhpn2) alongside with the MCR-NRAS in *rhpn2*^SL4/SL4^ embryos. Under these conditions, the tumor-free survival curve revealed a reversal of the phenotype (Fig. 2F, blue curve; rescued: 65%), suggesting, at least in part, a melanocyte-intrinsic effect of Rhpn2 expression during the early stages of tumor development. These results confirmed faster development of the tumor in Rhpn2^+/+^ fish as it was observed in Rhpn2^+/-^ intercrosses when WT fish presenting tumors died sooner than KO (Fig. 2D).

### Melanocytic specific overexpression of Rhpn2 promotes BRAFV600E-dependent melanoma development

To assess whether the observed effect of Rhpn2 extends to another genetic context of melanoma development, the Tg(*mitfa:BRAF^V600E^*);*p53*^-/-^ model was used (hereafter referred as BRAF) to overexpress Rhpn2 (Rhpn2-OE) in melanocytes using MCR- *rhpn2-T2a-eGFP* (MCR-Rhpn2) vector and MCR-*eGFP* as control (Fig.2H). We followed weekly the development of tumors in Rhpn2 expressing fish compared to MCR-eGFP injected control condition (Fig.2I). We confirmed the sustaining role of Rhpn2 expression on tumor development (at 22 wpf WT (n=121), 30.3 %; Rhpn2-OE (n=50), 72%; p<0.0001; N=4). The percentage of melanotic tumors observed in Rhpn2-OE fish was also higher than with MCR-eGFP individuals (Fig. 2J). When Rhpn2 was knocked-out in this model, the development of the tumor was also impaired in Tg(*mitfa:BRAF^V600E^*); *p53*^-/-^; *rhpn2*^-/-^ fish compared to *rhpn2*^+/+^ with 15% (9/59) in *rhpn2*^-/-^ and 23% (4/17) in *rhpn2*^+/+^ (data not shown).

### Lower cellular density and absence of acetylated-tubulin structures at the tumor boundaries in *rhpn2^SL4/SL4^* NRAS^Q61L^-driven melanoma

Histological analyses through hematoxylin/eosin staining were then performed on 6 µm sections of 6 wpf zebrafish bearing NRAS^Q61L^-driven tumors in both *rhpn2*^SL4/SL4^ and *rhpn2*^+/+^ fish. In *rhpn2*^SL4/SL4^ tumors compared to WT tumors, we observed a significant decrease in cellular density, a feature of tumor aggressiveness (Fig. 3A-B). This trend appeared already in tumors from 4 wpf fish, even if not statistically significant (Fig. 3B). Increased aggressiveness often correlates with higher mechanical forces at the tumor boundaries (Linke et al., 2024) and cytoskeletal reorganization mediated by RHO GTPases. Interestingly, reappearance of cilia at the junction of melanoma and stroma in zebrafish was identified as a characteristic of the tumor boundaries in zebrafish melanoma (Hunter et al., 2021). In addition, Rhpn2 was recently shown to play a role in stereocilia genesis and function in the inner ear of zebrafish (Dai et al., 2026) suggesting its involvement in environment sensing. We thus addressed the presence of ciliary structures by immunofluorescence analysis of the tumors using acetylated tubulin as a cilia marker. Our data revealed complete absence of acetylated tubulin signal at the tumor–stroma interface in *rhpn2*^SL4/SL4^ tumors compared to WT.

**Figure 3.**
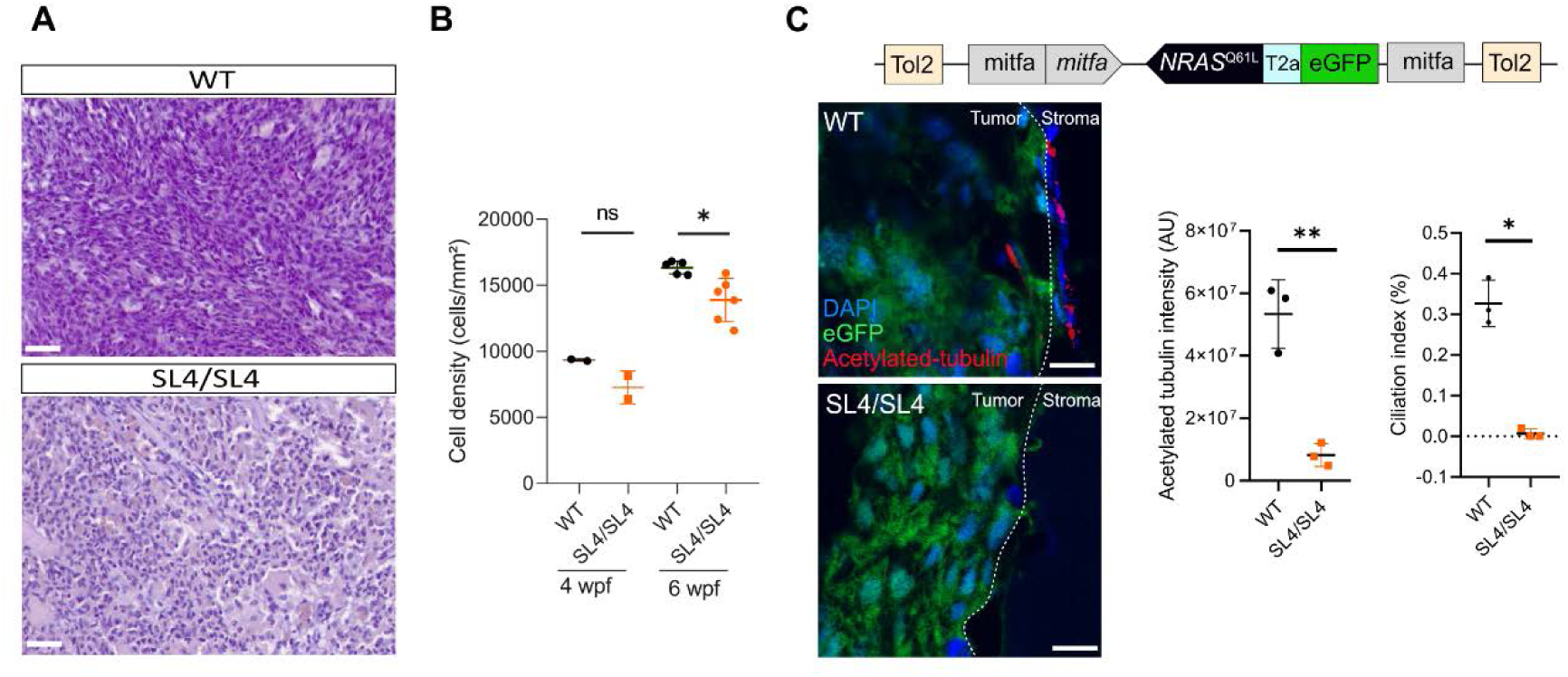
**A-** Hematoxylin/eosin staining of WT and SL4/SL4 tumors at 6 wpf. Scale bar 25µm. **B-** Quantification of tumor cellular density (15 fields/tumor) (at 4 wpf, ns, p = 0.333; at 6 wpf in WT (n = 5) and KO (n = 6) fish (*p = 0.013) (Welch test). **C-** Confocal images of sections of eGFP-expressing NRAS-dependent tumors from 6 wpf WT and *rhpn2*^SL4/SL4^ fish centered on tumor (green)/stroma boundaries and stained with anti-acetylated tubulin (red), and DAPI (blue). Scale bar 20µm. Quantification of acetylated-tubulin signal following background subtraction (n=3) (**p = 0.002; Student’s t-test). Quantification of Ciliation index (n=3) (*p = 0.048; Student’s t-test).

### Rhpn2 loss of expression generates an IFN response in the melanoma cells and decreases the expression of some ciliary genes

To gain a better understanding in the mechanisms of action and to assess how Rhpn2 influences tumor development, we subsequently performed a transcriptomic analysis of the *NRAS*-melanoma cells from *rhpn2*^SL4/SL4^ and *rhpn2*^+/+^ fish. We took advantage of the coinjection of a mitfa:mCherry Tol2 vector alongside with MCR-NRAS vector in the embryos to express fluorescence in melanoma cells. This allowed cell sorting and bulk RNA sequencing of the cells from *rhpn2*^SL4/SL4^ and *rhpn2*^+/+^ tumors. The melanoma nature of the sorted cells was confirmed by the expression of the hallmark genes in melanoma as mitfa, Sox10 or tyr (Fig. 4A). Despite high heterogeneity in the transcriptional states among the samples, we could identify specific differentially expressed genes (DEGs) (Fig. 4B-C). We observed that the relevant upgulated genes, in *rhpn2*^SL4/SL4^ melanoma cells, are interferon-stimulated genes (ISGs) immunity related e.g. Interferon-Induced protein with Tetratricopeptide Repeats 8 (*ifit8*), Interferon Alpha Inducible Protein 27 2/3 (*ifi27.2, ifi27.3*), Myxovirus resistance protein A (*mxa*), DExH-box helicase 58 (*dhx58*) and Radical S-Adenosyl Methionine Domain Containing 2 (*rsad2*) consistent with activation of defense pathways (Fig. 4D).

**Figure 4.**
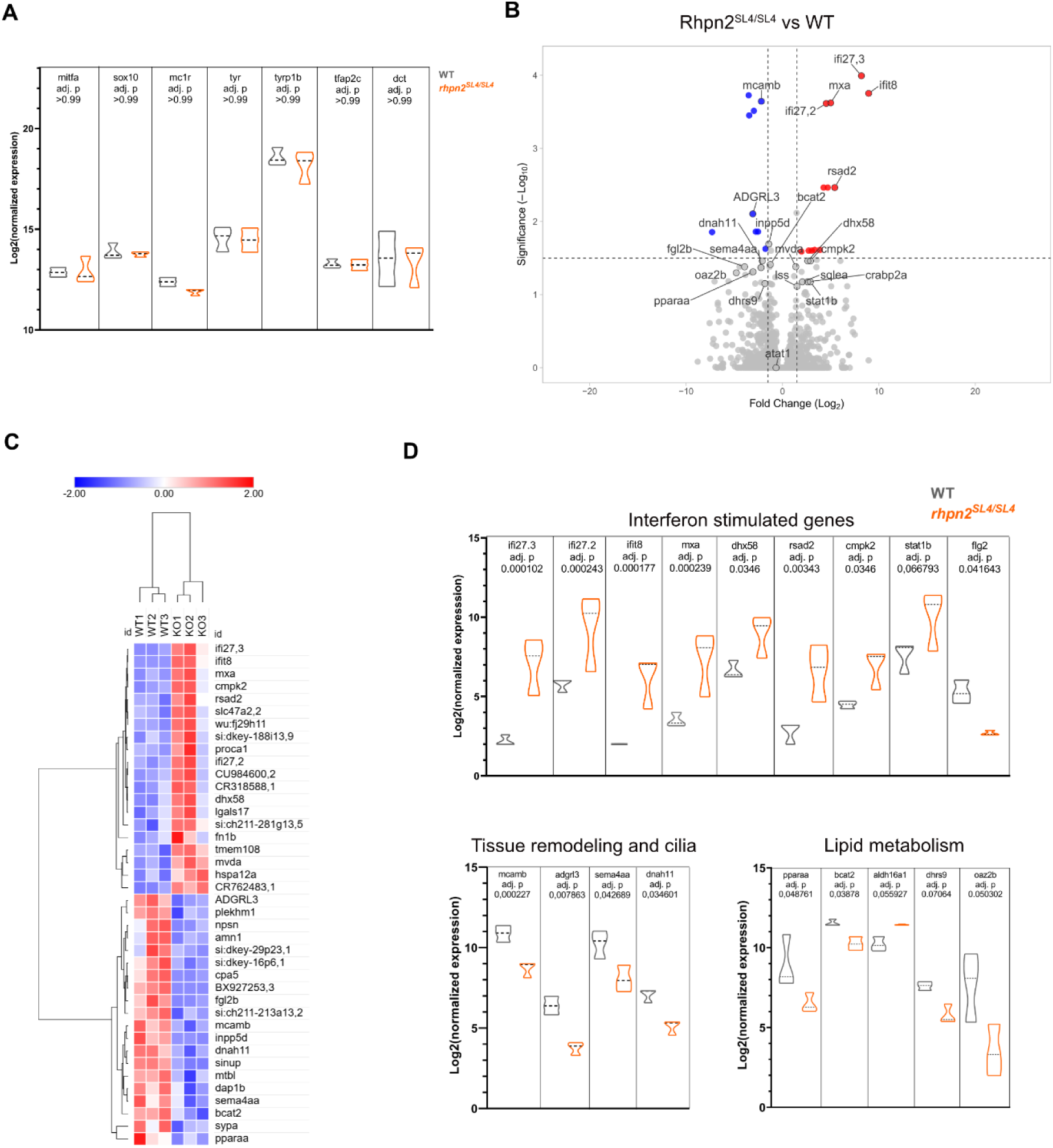
Rhpn2 loss decreases expression of ciliary genes and tubulin acetylation at the tumor/stroma interface. **A-** Violin plot depicting the relative expression of representative melanoma marker genes in WT and *rhpn2^SL4/SL4^* tumors. **B-** Distribution of differentially expressed genes (DEGs) shown as log2 fold change of expression levels and -log10 adjusted p-values. (Adj p-value<0.05, FC>2) **C-** Heat map of the significant DEGs (Adj p-value<0.05). **D-** Graphs representing normalized expression of selected DEGs. Upper panel: genes documented as interferon stimulated genes. Intermediate panel: selection of genes encoding channels related to tissue remodeling and cilium. Lower panel: selection of genes related to lipid metabolism.

Among the top downregulated genes in *rhpn2*^SL4/SL4^ melanoma we found proteins related to tissue remodeling and cilia e.g. Dynein axonemal heavy chain 11 (*dnah11*), Melanoma cell adhesion molecule b (*mcamb*), Semaphorine 4A (*sema4aa*) and Adhesion G Protein-Coupled Receptor L3 (*adgrl3*) and proteins involved in lipid metabolism e.g. Peroxisome Proliferator Activated Receptor Alpha (*pparaa*), Branched Chain Amino Acid Transaminase 2 (bcat2) (Fig. 4D).

## Discussion

Although Rhophilin-2 (RHPN2) was identified twenty-five years ago as a RHO GTPases associated protein its biological functions remain elusive (Mircescu et al., 2002; Peck et al., 2002). Here, we provide the first functional evidence demonstrating that RHPN2 acts as an *in vivo* pro-tumorigenic factor in melanoma, using complementary experiments in human cells and in zebrafish as models.

First identified as an actin cytoskeleton and vesicular trafficking regulator (Steuve et al., 2006), several studies have further proposed RHPN2 as a potential tumor associated-gene. However, to date, the role of RHPN2 in melanoma has not been characterized, although an *in-silico* study suggested that it is important for melanoma survival (Pyatnitskiy et al., 2015). In the present study, *in vitro*, using A375 and A2058 human melanoma cell lines, we showed that RHPN2 knock-down reduces cell viability and pERK levels. We also observed that RHPN2 knock-down reduced migration/invasion capacities and the clonogenic potential of the cells. These reinforced the hypothesis that RHPN2 could act as a regulator of melanoma aggressiveness, limiting both metastatic dissemination and tumor growth properties. RHPN2 knock-down experiments previously showed that RHPN2 is required for growth and invasion in ovarian cancer and lung adenocarcinoma (Xiao et al., 2020; Yan et al., 2021; Yu et al., 2020; Zhang et al., 2025) even if the molecular mechanisms impacted seem to be different. In lung adenocarcinoma, RHPN2 expression is essential for the transcription factor Kruppel-like factor 5 (KLF5) effects on tumor progression (Zhang et al., 2025) and confers resistance to glutamine depletion of the tumor (Xiao et al., 2020). In ovarian tumor, RHPN2 depletion blocks the cells in G1 phase and decreases integrin β1 signaling and metalloproteinases secretion (Yan et al., 2021). In glioblastoma, a cancer affecting glial cells, which like melanocytes, are derived from neural crest, RHPN2 was also previously shown to promote EMT (Danussi et al., 2013). Our findings in human melanoma cell lines thus support the idea that RHPN2 influences tumor cell growth and invasive properties, showing however for the first time, a decrease in Erk signaling when RHPN2 is knocked down. As BRAF and NRAS, both MAPKs activators, are the most frequently activated genes in melanoma, the direct molecular role of RHPN2 in this cascade would be interesting to address in future experiments.

Reported human tumor xenografts of both lung and ovarian cancer experiments have shown that RHPN2 overexpression increases tumor volume in mice (Xiao et al., 2020; Yu et al., 2020). However, here, we evidence, for the first time, the role of Rhpn2 in a primary tumor model of induced tumorigenesis, taking advantage of zebrafish, a well-established model for studying cancer development, especially in melanoma studies (Letrado et al., 2018; Wojciechowska et al., 2016). Using first NRAS^Q61L^ zebrafish melanoma models (Iyengar et al., 2012), to rapidly develop tumors (within weeks) in a mosaic fashion, we demonstrate that loss of Rhpn2 significantly reduces tumor development delaying tumor onset and reducing tumor burden. This effect is at least in part melanocyte intrinsic as shown by the rescue experiment performed by early re-re-expression of Rhpn2 in the fish embryo melanocytes. Wild-type animals displayed a higher frequency of tumors at early stages than Rhpn2-deficient animals indicating that Rhpn2 may facilitate the establishment of a tumor-permissive state rather than simply accelerating tumor growth. In this model, we also observed decreased cellular density in Rhpn2-KO fish correlating with lower melanoma aggressiveness. In parallel, we confirmed the positive impact of Rhpn2 on tumor development of *mitfa:BRAF^V600E^; p53^-/-^* stable transgenic fish model (Patton et al., 2005) in which tumors develop slower (within months). A decrease in tumors appeareance was also observed in *mitfa:BRAF^V600E^;p53^-/-^* ;*Rhpn2^-/-^* fish compared to *Rhpn2^+/+^.* The effects of Rhpn2 loss or overexpression were reproduced in both NRAS and BRAF/p53 melanoma models highlighting its potential relevance across genetically diverse melanoma subtypes during early tumor initiation and expansion.

Elevated cellular density is a hallmark of malignancy and tumor aggressiveness, reinforcing compressive stress, tissue invasion, and metastasis (Linke et al., 2024). We observed that Rhpn2 deficiency decreases cell density of the tumor. This is in accordance with impaired proliferative and/or migratory signaling observed in cell lines which could be linked to alterations in MAPK and Rho pathways. Consistently, zebrafish melanoma studies have shown that MEK inhibition suppresses melanocyte hyperplasia, linking MAPK signaling directly to tumor cell density (Fernandez Del Ama et al., 2016). Increased RhoA activity in melanoma has also been associated with enhanced proliferation, metabolic activity, and invasive phenotypes, particularly under therapeutic stress (Murali et al., 2024). Rhpn2 was proposed as a positive (Danussi et al., 2013) or negative (Xiao et al., 2020) regulator of RhoA in a way depending on the system. In melanoma, Rhpn2 deficiency could thus impair Rho-dependent actomyosin dynamics, leading to reduced cell cohesion within the tumor mass. Fibrous matrix and cells within a tumour experience tensile forces as adjacent cells proliferate or when additional matrix is deposited between cells (Johnson et al., 2025). This effect is most prominent at the tumour periphery (Mpekris et al., 2015). Interestingly, in zebrafish higher tumor cell density may be associated with the emergence of a specialized “interface” cell state where cancer cells interact with the surrounding microenvironment (Hunter et al., 2021). Primary cilia at dense, mechanically constrained tumor borders may function as mechanosensory hubs that integrate microenvironmental signals with RHO-dependent cytoskeletal remodeling (Collinson and Tanos, 2025; Johnson et al., 2025; Wang and Dynlacht, 2018). Our data showed that Rhpn2 is necessary to observe these ciliary structures at the interface. It has been reported that the loss of Rhpn2 in zebrafish leads shorter stereocilia, and vestibular abnormalities of inner ear stereocilia (Dai, Yubei et al., 2026) through a direct interaction with RhoA. Although stereocilia and primary cilia are fundamentally different structures—the latter being microtubule-based organelles and the former being actin-rich cellular protrusions—both contribute to the ability of cells to sense and respond to their environment (Nekooki-Machida and Hagiwara, 2020).

Despite the high heterogeneity observed between tumors, transcriptomic analysis of Rhpn2-deficient NRAS-driven melanomas identified a modest decreased expression of DNAH11, a ciliary axoneme protein and of mcamb which coordinates apical-basal and planar cell polarity and ciliogenesis during morphogenesis (Gao et al., 2017). Nevertheless, this experiment was performed on all the tumor melanocytes, representing only in minority the interface cells expressing cilia. In future work, it would be interesting to perform spatial transcriptomic analysis to elucidate Rhpn2 impact at the tumor interface.

Interestingly, we showed in Rhpn2-deficient tumors, the marked upregulation of *dhx58* (*lgp2*) together with multiple interferon-stimulated genes (*stat1, ifit8, mxa/mx1, rsad2, and ifi27*) suggesting an activation of type I interferon response. Type I interferon signaling can be triggered not only by viral infection but also by endogenous nucleic acids released during cellular stress, genomic instability, or organelle dysfunction in cancer (Schoggins, 2019). Several danger-sensing mechanisms could be involved, including the cGAS/STING pathway and the RIG-I-like receptors (RLRs) (Ishikawa and Barber, 2008; Liu et al., 2025) both known to be important in melanoma (Motwani et al., 2019; Oscherwitz et al., 2023). Dhrx58 plays a critical role as a regulator of RLRs in nucleic acids cellular responses (Bek et al., 2019).

Accordingly, several of the induced genes point toward a potential link between RHPN2 loss, mitochondrial stress, and lipid metabolism reprogramming. Mitochondrial dysfunction is a recognized source of endogenous danger signals capable of activating interferon pathways through the release of mitochondrial nucleic acids into the cytosol. Cmpk2 regulates mitochondrial function and inflammasome activation under inflammatory stress (Zhao et al., 2022), Rsad2 can disrupt mitochondrial lipid metabolism and antiviral stress responses (Chen et al., 2025; Majhi et al., 2025), and PPARα controls mitochondrial fatty acid oxidation and energy metabolism in cancer (Van den bossche et al., 2025). Together with the induction of ifi27, which has been associated with cellular stress, apoptosis, and inflammatory responses, and the elevated expression of mx1 and ifit-family genes, these findings support the hypothesis that RHPN2 deficiency causes broad cellular perturbations affecting the cytoskeleton, cilia, mitochondrial homeostasis, and metabolic pathways, ultimately leading to chronic activation of an interferon-mediated stress response that may contribute to the reduced growth observed in Rhpn2-KO tumors.

Altogether, our study identifies for the first time Rhpn2 as a regulator of melanoma progression that integrates cytoskeletal remodeling, MAPK signaling, and tumor–microenvironment interactions. By controlling tumor initiation, aggressiveness, ciliogenesis, and immune-related pathways, Rhpn2 emerges as a promising target for therapeutic in melanoma.

## Material and methods

### Cell culture and transfection

The A375 (ATCC-CRL-1619), A2058 (ATCC-CRL-3601) and SKML28 (ATCC-HTB- 72) cell lines were obtained from the American Type Culture Collection (ATCC, Manassas, VA, USA). They were maintained in Dulbecco’s Modified Eagle Medium (DMEM) supplemented with 10% fetal bovine serum (FBS), 1 mM sodium pyruvate, 100 U/mL penicillin, 100 μg/mL streptomycin, and 5 μg/mL amphotericin B. All components were obtained from ThermoFisher Scientific (Waltham, MA, US). All cell lines were maintained at 37°C in a humidified incubator with 5% CO₂. When necessary, cells were counted using hemocytometer. For recombinant protein expression, 2,5 × 10^6^ HEK293T cells were transfected with 3 µg of each plasmid using X-tremeGENE 9 (Roche Life Science, Basel, Switzerland) according to the manufacturer’s instructions. Cells were harvested and analyzed 48 h after transfection.

### Viral particles production and transduction

Lenti-X HEK293T cells were seeded at a density of 18 × 10⁶ cells per T225 flasks in complete medium 20-24 hours prior to transfection with 195 μL of PEI (1μg/μL) (company, country) and the following plasmids: pMD2.G (3.4 μg), PsPAX2 (6.8 μg), and target (13.6μg) in serum-free DMEM. Forty-eight hours post-transfection, the culture medium was collected, 0.45 μm filtered and concentrated by PEGx800 (25% in NaCl 2.5M) (1:3) during 24h. Centrifugation is performed at 4000 g for 1h at 4 °C. The viral pellet was resuspended in a volume corresponding to 1/100 of the initial supernatant volume. The resuspended viral particles were incubated on ice for 20 minutes before being used immediately or stored at −80°C until further use. Virus concentrations were titrated by serial dilutions. Selection of stably transduced melanoma cells was carried out in culture medium supplemented with 2 mg/ml G418 (ant-gn-5, InvivoGen, San Diego, CA, USA) or 1μg/μLPURO (#108321-42-2, InvivoGen, San Diego, CA, US). After 10 to 15 days of selection, cells were used for further experiments. For individual colonies, after 10 to 15 days of selection, clones were transferred to 24-well plates, grown to confluency, trypsinized, and further expanded.

### Viability assays

Cell viability was assessed using the MTT assay. Cells were seeded in 96-well plates at a density of 1.5 x 10^3^ per well and incubated at 37°C for 1 week. 10 μL of WST-8/CCK-8 tetrazolium salt (Abcam) (final concentration 0.5 mg/mL) was added to the cells daily, followed by a 2-hours incubation. The amount of produced formazan was measured by absorbance at 450 nm using a spectrophotometer (Bio-Rad Model 680 Microplate Reader). All experiments were performed in triplicate wells and repeated in at least three independent experiments.

### Colony formation assay

Control cells and cells expressing the desired shRNA were seeded in 6-well plates at a density of 1 x 10^3^ cells in 3-cm dishes and incubated at 37°C for 2 weeks to allow colony formation. The cells were washed with PBS and fixed with 4% paraformaldehyde for 5 minutes. Then the cells were washed three times with PBS, stained with 0.1% crystal violet for 2 minutes, thoroughly washed with distilled water and dried. The number and size of the colonies were evaluated using ImageJ software. Experiments were performed in triplicates.

### Cell migration and invasion assays

Cell migration and invasion assays were performed using 24-well Transwell chamber assays (8 µm pore size) (VWR 734-0038 and VWR 734-1047, Corning, Corning, NY, Leuven, Belgium), as described in the manufacturer’s instructions. 4 × 10^4^ cells were seeded into the upper compartment containing 500 µL of medium without serum whereas the lower chamber contained 750 µL of complete medium with FBS as attractant. After 20 h of incubation, the cells were fixed, stained (Differential Quik III Stain Kit, Polysciences, Inc., Warrington, PA, USA) and counted.

### Western blot

For Western blotting analysis, culture medium was removed, and the cells were washed in cold phosphate-buffered saline (PBS) and lyzed. For fish, 30 larvae were sampled after the experiments, euthanized by immersion in 0.25 mg/mL of buffered MS-222 (Sigma) and lyzed with thorough pipetting. Lysis buffer was composed of 50 mM Tris-HCl, pH 6.8, containing 12% sucrose, 2% SDS, 0.004% bromophenol blue, 20 mM dithiothreitol (DTT), and 1 x cOmplete™ Protease Inhibitor Cocktail (Roche, Basel, Switzerland). The lysates were cleared by centrifugation at 15,000 g at 4°C and the total protein concentration was measured using the PierceR 660 nm Protein assay reagent (22660, Thermo Scientific, Waltham, MA, US). The samples of total cell extracts (30 µg of proteins) were denaturated at 95°C in 1x Laemmli Buffer, resolved on a 8% SDS-PAGE gel and transferred to a nitrocellulose membrane (Amersham, GE Healthcare). Proteins were revealed by enhanced chemiluminescence (ECL) using SuperSignal West Pico Atto or Pico (Thermo Fisher Scientific, Waltham, MA, USA) and visualized using Fusion Solo S software (Vilber Lourmat, Marne-la-Vallée, France).

### Zebrafish husbandry

Zebrafish were maintained under standard laboratory conditions, accordancing to the FELASA guidelines and regulations (Aleström et al., 2020). All experimental procedures were approved by the ULB ethical committee for Animal Welfare (CEBEA) (Protocols 866N and 867N). The lines used were: Tg(mitfa:BRAF^V600E^); mitfa^-/-^ ; p53^M214K^ (lf) (Christian Lawrence, Boston Children’s Hospital, Boston, USA) (here referred to as: BRAF/p53). Stable Tg(mitfa:BRAF^V600E^), mitfa^-/-^, p53^-/-^, rhpn2^-/-^ fish were generated by successive outcrosses followed by several intercrosses. The term “adult” fish refers to animals aged between 6 months and 8 months old. *rhpn2^Δ2/+^ or rhpn2^SL4/+^* heterozygous fish were respectively intercrossed and their corresponding adult offspring were analyzed for morphometric measurements. The body length was measured using a ruler. All measurements were performed before genotyping and mendelian ratio inheritance confirmation.

### Generation of *rhpn2^-/-^* mutant zebrafish

The *rhpn2* (ENSDARG00000014577) knock-out mutant lines were generated using the CRISPR/Cas9 system. Two single guides RNA (sgRNAs) were identified (sgRNA1 5’ GGTTGAGCATTCAGAAATGA 3’ and sgRNA2 5’ CGGACACTCTCATTCCCAA 3’) and selected for their highest on-target efficiency and lowest predicted off-target score using Sequence Scan for CRISPR software (http://crispr.dfci.harvard.edu/SSC/) and CRISPR scan (http://www.cirsprscan.org/) targeting the first exon of *rhpn2*(Xu et al., 2015). The DNA template for the sgRNA synthesis was produced using the PCR-based short-oligo method as described previously (Talbot and Amacher, 2014). The resulting PCR product was purified by phenol-chloroform extraction and used for in vitro transcription using SP6 RNA polymerase (NEB, M0207). The shRNA was purified using the High Pure PCR Cleanup Microkit (Roche, 498395500). 60 pg sgRNA and 100 pg Cas9 protein (PNA Bio) were co-injected into one-cell stage wild-type embryos. The genotyping of both embryos and adults was performed using the following primers : *rhpn2* fw: 5’-ATCAACAGACATCCGAGAACC-3’, and *rhpn2* rv: 5’-TCTCAAAACTACAGTGACGGTAACA -3’. Mutations were validated by PCR and Sanger sequencing (Eurofins, Luxembourg City, Luxembourg). For both lines, individuals from at least the F3 generation were used in experiments.

### Generation and analysis of melanoma models

The *mitfa*:*NRAS^Q61L^* MiniCoopR (here referred to as MCR:NRAS), received from Hurstone lab (Badrock and Hurlstone, 2021), together with TOL2 transposase mRNA (20 pg), was microinjected in the yolk of single-cell embryos at a final concentration of 50 pg per embryo. Embryos were subsequently screened for pigmentation rescue at 5 dpf and closely monitored for apparent tumor onset and selection for further analysis.

For FACS purification of tumor cells MCR:NRAS was co-injected with mitfa:mCherry TOL2 vector for concomitant expression of fluorescence.

Evaluation of the impact of rhpn2 loss-of-function on tumor onset was achieved by intercrossing Tg(mitfa:BRAF^V600E^), p53^-/-^, rhpn2^+/-^ nacre fish. For Rhpn2 overexpression experiments, Tg(BRAF), p53^-/-^ nacre embryos were injected by MCR vector expressing eGFP or eGFP-T2a-Rhpn2 (Genescript customed) under the control of the mitfa promoter. Embryos were subsequently screened for pigmentation rescue at 5 dpf and monitored for tumor appearance from week 4 to week 25 and timing of tumor onset reported.

### Tumor histological analyses

Tumor-bearing fish were collected 3 weeks after tumor onset (6 wpf) and embedded in paraffin using Leica EG1150 H machine or in OCT for cryopreservation. Briefly, fish were euthanized and fixed with PFA4%. The next day, fish were decalcified by o/n incubation in 0.5M EDTA pH8 at 4°C. Samples were quickly rinsed and refixed in neutral buffered formalin solution (formaldehyde 10%, NaH_2_PO_4_.2H_2_O 4.9 g/L, Na_2_HPO_4_ 6.5g/L) at 4°C. Samples are then dehydrated by successive ethanol baths (70% 1h, 95% 1h, 100% 1h, 100% 2h), cleared in toluene o/n at room temperature before being embedded in paraffin at 60°C (Paraffin Paraplast, Leica Biosystems). Paraffin sections (6µm) were obtained using Leica RM2245 manual rotating microtome following manufacturer’s instructions. Paraffin blocks were cooled in an ice–water mixture prior to sectioning. Tissue sections were floated on a 45 °C water bath for a few seconds to allow proper flattening and were then mounted onto coated slides (VWR). The slides were dried horizontally at 50 °C and subsequently incubated overnight at 37 °C prior to further processing.

### Fluorescent immunostaining

Tumor-bearing fish were fixed in 4% PFA 48h. After PBS wash, they were treated o/n in 0.5M EDTA pH8 at 4°C. The samples were then refixed in 10% NBF pH7.2 (Formaldéhyde 4%, monobasic sodium phosphate 0.4%, dibasic sodium phosphate 0.65%, and methanol 1.5%) at room temperature o/n. The dissected tumors were then incubated o/n at 4°c in 30% sucrose and further embedded in OCT, snap-frozen, cryosectioned (8 µm, CM3050 S Leica, Wetzlar, Germany) and heated at 65°c for 30 minutes. Sections were air-dried for 1 hour at room temperature prior to staining. Tissue sections were outlined using a hydrophobic barrier pen (DAKO pen) and washed twice for 5 min in 1× PBS. Sections were then permeabilized for 15 min in PBS-0.5% Triton X-100 solution, followed by washes in PBS-0.1% Tween-20. Sections were blocked for 1 h at room temperature in blocking buffer (10 mg/mL BSA, 1% goat serum in 1× PBS) and incubated overnight at 4°C with the primary antibody. After washing, sections are incubated for 1 h with the appropriate secondary antibody in the dark. Nuclei were counterstained with DAPI (1:10,000) and sections were mounted using Fluo G medium. A minimum of 3 individuals were imaged for each condition. The primary and secondary antibodies used are the following : Anti-acetyl tubuline (mouse) (T6793 Sigma Aldrich;1/2000), Alexa Fluor 647-conjugated anti-mouse IgG (A-21239, Invitrogen; 1/500), Alexa Fluor 488-conjugated anti-rabbit IgG (A-21206, Invitrogen; 1/500). Immunostained cryosections were imaged using a Zeiss LSM 780 inverted confocal microscope, with Plan Apochromat 18 20x/0.8 M27 and LD C Apochromat 40x/1.1 W korr M27 objective. Quantification of acetylated tubulin+ cells was performed using ImageJ software.

### Hematoxylin and Eosin Staining of Paraffin Sections

Paraffin-embedded tissue sections were first deparaffinized and rehydrated through sequential 5-min washes in toluene, 100% ethanol, 90% ethanol, 70% ethanol, running tap water, and distilled water. Sections were depigmented by incubation in a solution containing 3% H₂O₂ and 1% KOH (Merck) for 15–45 min. Sections were stained with Hematoxylin (Mayer’s hematoxylin) for 4 min, followed by rinsing in running tap water until clear. Slides were then blued by immersion in 70% ethanol and chloridric acid solution. After rinsing in running water for 2 min, sections were counterstained with alcoholic Eosin. Slides were subsequently rinsed in running water for 5–10 min until the desired pink coloration was obtained. Sections were dehydrated through graded ethanol solutions (70% and 90%), followed by two washes in 100% isopropanol (2 min each) and two washes in toluene (5 min each). Slides were air-dried and coverslipped using dibutylphthalate polystyrene xylene mounting medium (DPX) (Merck). Whole slides were imaged using a Hamamatsu NanoZoomer digital slide scanner and visualized with NanoZoomer Slide software to evaluate tumor histological features. Qpath software was used for cell density quantification.

### Tumor cell sorting and RNA extraction

Fish were euthanized in 500 mg/l solution of tricaine (MS-222) (Sigma-Aldrich, E10521). Tumors (control and *rhpn2*^-/-^) were dissected and digested with 0.075 mg/ml Liberase (Roche-05401119001) at 33°C for 30 minutes in 0.9X Dulbeccoʹs Phosphate Buffered Saline (DPBS). Mechanical dissociation was performed during this process using a syringe with a 26 G needle (B.Braun, Omnifix 100 Duo). Cell suspensions were centrifuged (300 g) in 2% fetal bovine serum diluted in 1X DPBS (FACS buffer). Then, cells were resuspended in FACS buffer and filtered using a sterile 40 µm nylon mesh (VWR, Radnor, PA, USA). Cell sorting of mCherry^+^ cells was performed on a FACS ARIA (Becton Dickinson, San Jose, CA, USA) using DRQ7 (3 µM, Thermofisher scientific, D15105) to exclude non-viable cells. Sorting time did not exceed 15 min. Total RNA was extracted from approximately 150 000 cells using Relia Prep RNA tissue Miniprep system (Promega, Madison, WI, USA) according to the manufacturer’s instructions and sent for mRNA sequencing (Novogene-Europe, Munich, Germany).

### Bulk RNA sequencing and data analysis

FastQ data files were processed with the same approach for all datasets. Trimmed and filtered reads are mapped against the reference zebrafish genome GRCz11.95 using Hisat2. Differential gene expression analysis and downstream functional enrichment were performed using iDEP (version 1.1) (Ge et al., 2018).

### Statistical Analysis

Normality was assessed Statistical analysis was performed using GraphPad Prism software (version 8 GraphPad Software, San Diego, CA). When comparing multiple groups, we used a two-way ANOVA test followed by a multiple comparison test. No data was excluded from the analysis. The results are considered significant when p < 0.05 and are displayed with the mean and standard deviation (SD). Data display and sample sizes used for the statistical analysis are specified in the caption of the figures.

## DATA availability

## Acknowledgements

We thank Michiel Martens from the ULB Limif Platform, as well as Christine Dubois from the FACS facility. Finally, we thank Mustapha Chaouni and Kamel Gharbi for their daily maintenance of the fishroom facility. This project was funded by the Fonds National de la Recherche Scientifique (CDR: J0158.22F, J0170.24, Télévie grant :7-4592-24F). We are grateful to Adeline Augereau, Pierre Gillotay for their precious advices and fruitful discussions.

## Author contributions

Mana Alavi and Alexia Gybels: Conceptualization; Data curation; Formal analysis; Investigation; Visualization; Methodology; Writing—original draft; Writing—review and editing.

Sena Bekar, Katarzina Konobrocka, Laura Gulizia: Data curation; Formal analysis; Investigation; Visualization.

Garnik Hovhannisyan : Resources; Methodology; Imaging.

Camille Perazzolo : Technical assistance; Methodology.

Sumeet Pal Singh: Resources; Software; Formal analysis; Methodology; Writing-review and editing.

Isabelle Pirson: Conceptualization; Resources; Data curation; Software; Formal analysis; Supervision; Funding acquisition; Validation; Investigation; Visualization; Methodology; Writing—original draft; Project administration; Writing—review and editing.

## References

Aleström, P., D’Angelo, L., Midtlyng, P.J., Schorderet, D.F., Schulte-Merker, S., Sohm, F., Warner, S., 2020. Zebrafish: Housing and husbandry recommendations. Lab. Anim. 54, 213–224. 10.1177/0023677219869037

Arang, N., Lubrano, S., Ceribelli, M., Rigiracciolo, D.C., Saddawi-Konefka, R., Faraji, F., Ramirez, S.I., Kim, D., Tosto, F.A., Stevenson, E., Zhou, Y., Wang, Z., Bogomolovas, J., Molinolo, A.A., Swaney, D.L., Krogan, N.J., Yang, J., Coma, S., Pachter, J.A., Aplin, A.E., Alessi, D.R., Thomas, C.J., Gutkind, J.S., 2023. High-throughput chemogenetic drug screening reveals PKC-RhoA/PKN as a targetable signaling vulnerability in GNAQ-driven uveal melanoma. Cell Rep. Med. 4, 101244. 10.1016/j.xcrm.2023.101244

Badrock, A.P., Hurlstone, A., 2021. Dissecting Oncogenic RAS Signaling in Melanoma Development in Genetically Engineered Zebrafish Models, in: Rubio, I., Prior, I. (Eds.), Ras Activity and Signaling: Methods and Protocols. Springer US, New York, NY, pp. 411–422. 10.1007/978-1-0716-1190-6_25

Behrends, J., Clément, S., Pajak, B., Pohl, V., Maenhaut, C., Dumont, J.E., Schurmans, S., 2005. Normal thyroid structure and function in rhophilin 2-deficient mice. Mol. Cell. Biol. 25, 2846–2852. 10.1128/MCB.25.7.2846-2852.2005

Bek, S., Stritzke, F., Wintges, A., Nedelko, T., Böhmer, D.F.R., Fischer, J.C., Haas, T., Poeck, H., Heidegger, S., 2019. Targeting intrinsic RIG-I signaling turns melanoma cells into type I interferon-releasing cellular antitumor vaccines. Oncoimmunology 8, e1570779. 10.1080/2162402X.2019.1570779

Carreira, S., Goodall, J., Denat, L., Rodriguez, M., Nuciforo, P., Hoek, K.S., Testori, A., Larue, L., Goding, C.R., 2006. Mitf regulation of Dia1 controls melanoma proliferation and invasiveness. Genes Dev. 20, 3426–3439. 10.1101/gad.406406

Centeno, P.P., Pavet, V., Marais, R., 2023. The journey from melanocytes to melanoma. Nat. Rev. Cancer 23, 372–390. 10.1038/s41568-023-00565-7

Chen, S., Ye, J., Lin, Y., Chen, W., Huang, S., Yang, Q., Qian, H., Gao, S., Hua, C., 2025. Crucial Roles of RSAD2/viperin in Immunomodulation, Mitochondrial Metabolism and Autoimmune Diseases. Inflammation 48, 520–540. 10.1007/s10753-024-02076-5

Clayton, N.S., Ridley, A.J., 2020. Targeting Rho GTPase Signaling Networks in Cancer. Front. Cell Dev. Biol. 8. 10.3389/fcell.2020.00222

Collinson, R., Tanos, B., 2025. Primary cilia and cancer: a tale of many faces. Oncogene 44, 1551–1566. 10.1038/s41388-025-03416-x

Crosas-Molist, E., Samain, R., Kohlhammer, L., Orgaz, J.L., George, S.L., Maiques, O., Barcelo, J., Sanz-Moreno, V., 2022. Rho GTPase signaling in cancer progression and dissemination. Physiol. Rev. 102, 455–510. 10.1152/physrev.00045.2020

Dai, Y., Li, Q., Deng, J., Wu, S., Zhang, G., Hu, Y., Shen, Y., Liu, D., Wu, H., Gong, J., 2026. Rhpn2 regulates the development and function of vestibular sensory hair cells through the RhoA signaling in zebrafish. J. Genet. Genomics Yi Chuan Xue Bao 53, 131–142. 10.1016/j.jgg.2025.04.006

Danussi, C., Akavia, U.D., Niola, F., Jovic, A., Lasorella, A., Pe’er, D., Iavarone, A., 2013. RHPN2 drives mesenchymal transformation in malignant glioma by triggering RhoA activation. Cancer Res. 73, 5140–5150. 10.1158/0008-5472.CAN-13-1168-T

Etienne-Manneville, S., Hall, A., 2002. Rho GTPases in cell biology. Nature 420, 629–635. 10.1038/nature01148

Fernandez Del Ama, L., Jones, M., Walker, P., Chapman, A., Braun, J.A., Mohr, J., Hurlstone, A.F.L., 2016. Reprofiling using a zebrafish melanoma model reveals drugs cooperating with targeted therapeutics. Oncotarget 7, 40348–40361. 10.18632/oncotarget.9613

Gao, Q., Zhang, J., Wang, X., Liu, Y., He, R., Liu, X., Wang, F., Feng, J., Yang, D., Wang, Z., Meng, A., Yan, X., 2017. The signalling receptor MCAM coordinates apical-basal polarity and planar cell polarity during morphogenesis. Nat. Commun. 8, 15279. 10.1038/ncomms15279

Ge, S.X., Son, E.W., Yao, R., 2018. iDEP: an integrated web application for differential expression and pathway analysis of RNA-Seq data. BMC Bioinformatics 19, 534. 10.1186/s12859-018-2486-6

Gong, X., Didan, Y., Lock, J.G., Strömblad, S., 2018. KIF13A-regulated RhoB plasma membrane localization governs membrane blebbing and blebby amoeboid cell migration. EMBO J. 37, e98994. 10.15252/embj.201898994

Hason, M., Bartůněk, P., 2019. Zebrafish Models of Cancer-New Insights on Modeling Human Cancer in a Non-Mammalian Vertebrate. Genes 10, 935. 10.3390/genes10110935

Hunter, M.V., Moncada, R., Weiss, J.M., Yanai, I., White, R.M., 2021. Spatially resolved transcriptomics reveals the architecture of the tumor-microenvironment interface. Nat. Commun. 12, 6278. 10.1038/s41467-021-26614-z

Ishikawa, H., Barber, G.N., 2008. STING is an endoplasmic reticulum adaptor that facilitates innate immune signalling. Nature 455, 674–678. 10.1038/nature07317

Iyengar, S., Houvras, Y., Ceol, C.J., 2012. Screening for melanoma modifiers using a zebrafish autochthonous tumor model. J. Vis. Exp. JoVE e50086. 10.3791/50086

Johnson, A.M., Froman-Glover, C., Mistry, A., Yaddanapudi, K., Chen, J., 2025. The impact of compression and confinement in tumor growth and progression: emerging concepts in cancer mechanobiology. Front. Mater. 12. 10.3389/fmats.2025.1492438

Kellman, L.N., Neela, P.H., Srinivasan, S., Siprashvili, Z., Shanderson, R.L., Hong, A.W., Rao, D., Porter, D.F., Reynolds, D.L., Meyers, R.M., Guo, M.G., Yang, X., Zhao, Y., Wozniak, G.G., Donohue, L.K.H., Shenoy, R., Ko, L.A., Nguyen, D.T., Mondal, S., Garcia, O.S., Elcavage, L.E., Elfaki, I., Abell, N.S., Tao, S., Lopez, C.M., Montgomery, S.B., Khavari, P.A., 2025. Functional analysis of cancer-associated germline risk variants. Nat. Genet. 57, 718–728. 10.1038/s41588-024-02070-5

Kümper, S., Mardakheh, F.K., McCarthy, A., Yeo, M., Stamp, G.W., Paul, A., Worboys, J., Sadok, A., Jørgensen, C., Guichard, S., Marshall, C.J., 2016. Rho-associated kinase (ROCK) function is essential for cell cycle progression, senescence and tumorigenesis. eLife 5, e12994. 10.7554/eLife.12203

Lee, H., Lee, S., Seo, Y., Kim, D., Oh, Y., Jin, J., Hyeon, B., Han, Y., Kim, H., Lee, Y.J., Kim, H.M., Lee, G., Cho, K.-H., Do Heo, W., 2025. A Rho GTPase-effector ensemble governs cell migration behavior. Nat. Commun. 16, 9637. 10.1038/s41467-025-64635-0

Letrado, P., de Miguel, I., Lamberto, I., Díez-Martínez, R., Oyarzabal, J., 2018. Zebrafish: Speeding Up the Cancer Drug Discovery Process. Cancer Res. 78, 6048–6058. 10.1158/0008-5472.CAN-18-1029

Linke, J.A., Munn, L.L., Jain, R.K., 2024. Compressive stresses in cancer: characterization and implications for tumour progression and treatment. Nat. Rev. Cancer 24, 768–791. 10.1038/s41568-024-00745-z

Liu, Y., Tong, X., Wang, R., Li, Z.G., Xie, Z., Wang, D., Gu, W., Li, K., 2025. Cytokine-independent induction of LGP2/DHX58 in viral infection. J. Gen. Virol. 106, 002173. 10.1099/jgv.0.002173

Majhi, S., Roy, P., Jo, M., Liu, J., Hurto, R., Freddolino, L., Marsh, E.N.G., 2025. Viperin expression leads to downregulation of mitochondrial genes through misincorporation of ddhCTP by mitochondrial RNA polymerase. J. Biol. Chem. 301, 108359. 10.1016/j.jbc.2025.108359

Mircescu, H., Steuve, S., Savonet, V., Degraef, C., Mellor, H., Dumont, J.E., Maenhaut, C., Pirson, I., 2002. Identification and characterization of a novel activated RhoB binding protein containing a PDZ domain whose expression is specifically modulated in thyroid cells by cAMP. Eur. J. Biochem. 269, 6241–6249.

Motwani, M., Pesiridis, S., Fitzgerald, K.A., 2019. DNA sensing by the cGAS-STING pathway in health and disease. Nat. Rev. Genet. 20, 657–674. 10.1038/s41576-019-0151-1

Mpekris, F., Angeli, S., Pirentis, A.P., Stylianopoulos, T., 2015. Stress-mediated progression of solid tumors: effect of mechanical stress on tissue oxygenation, cancer cell proliferation, and drug delivery. Biomech. Model. Mechanobiol. 14, 1391–1402. 10.1007/s10237-015-0682-0

Murali, V.S., Rajendran, D., Isogai, T., DeBerardinis, R.J., Danuser, G., 2024. RhoA activation promotes glucose uptake to elevate proliferation in MAPK inhibitor resistant melanoma cells. 10.1101/2024.01.09.574940

Oscherwitz, M., Jiminez, V., Terhaar, H., Yusuf, N., 2023. Modulation of Skin Cancer by the Stimulator of Interferon Genes. Genes 14, 1794. 10.3390/genes14091794

Patton, E.E., Widlund, H.R., Kutok, J.L., Kopani, K.R., Amatruda, J.F., Murphey, R.D., Berghmans, S., Mayhall, E.A., Traver, D., Fletcher, C.D.M., Aster, J.C., Granter, S.R., Look, A.T., Lee, C., Fisher, D.E., Zon, L.I., 2005. BRAF mutations are sufficient to promote nevi formation and cooperate with p53 in the genesis of melanoma. Curr. Biol. CB 15, 249–254. 10.1016/j.cub.2005.01.031

Peck, J.W., Oberst, M., Bouker, K.B., Bowden, E., Burbelo, P.D., 2002. The RhoA-binding protein, rhophilin-2, regulates actin cytoskeleton organization. J. Biol. Chem. 277, 43924–43932. 10.1074/jbc.M203569200

Pyatnitskiy, M., Karpov, D., Poverennaya, E., Lisitsa, A., Moshkovskii, S., 2015. Bringing Down Cancer Aircraft: Searching for Essential Hypomutated Proteins in Skin Melanoma. PloS One 10, e0142819. 10.1371/journal.pone.0142819

Schoggins, J.W., 2019. Interferon-Stimulated Genes: What Do They All Do? Annu. Rev. Virol. 6, 567–584. 10.1146/annurev-virology-092818-015756

Steuve, S., Devosse, T., Lauwers, E., Vanderwinden, J.-M., André, B., Courtoy, P.J., Pirson, I., 2006. Rhophilin-2 is targeted to late-endosomal structures of the vesicular machinery in the presence of activated RhoB. Exp. Cell Res. 312, 3981–3989. 10.1016/j.yexcr.2006.08.028

Talbot, J.C., Amacher, S.L., 2014. A streamlined CRISPR pipeline to reliably generate zebrafish frameshifting alleles. Zebrafish 11, 583–585. 10.1089/zeb.2014.1047

Tasdogan, A., Sullivan, R.J., Katalinic, A., Lebbe, C., Whitaker, D., Puig, S., van de Poll-Franse, L.V., Massi, D., Schadendorf, D., 2025. Cutaneous melanoma. Nat. Rev. Dis. Primer 11, 23. 10.1038/s41572-025-00603-8

Van den bossche, V., Vignau, J., Vigneron, E., Rizzi, I., Zaryouh, H., Wouters, A., Ambroise, J., Van Laere, S., Beyaert, S., Helaers, R., van Marcke, C., Mignion, L., Lepicard, E.Y., Jordan, B.F., Guilbaud, C., Lowyck, O., Dahou, H., Mendola, A., Desgres, M., Aubert, L., Gerin, I., Bommer, G.T., Boidot, R., Vermonden, P., Warnant, A., Larondelle, Y., Machiels, J.-P., Feron, O., Schmitz, S., Corbet, C., 2025. PPARα-mediated lipid metabolism reprogramming supports anti-EGFR therapy resistance in head and neck squamous cell carcinoma. Nat. Commun. 16, 1237. 10.1038/s41467-025-56675-3

Wang, L., Dynlacht, B.D., 2018. The regulation of cilium assembly and disassembly in development and disease. Development 145, dev151407. 10.1242/dev.151407

Wojciechowska, S., van Rooijen, E., Ceol, C., Patton, E.E., White, R.M., 2016. Generation and analysis of zebrafish melanoma models. Methods Cell Biol. 134, 531–549. 10.1016/bs.mcb.2016.03.008

Wouters, J., Kalender-Atak, Z., Minnoye, L., Spanier, K.I., De Waegeneer, M., Bravo González-Blas, C., Mauduit, D., Davie, K., Hulselmans, G., Najem, A., Dewaele, M., Pedri, D., Rambow, F., Makhzami, S., Christiaens, V., Ceyssens, F., Ghanem, G., Marine, J.-C., Poovathingal, S., Aerts, S., 2020. Robust gene expression programs underlie recurrent cell states and phenotype switching in melanoma. Nat. Cell Biol. 22, 986–998. 10.1038/s41556-020-0547-3

Wyckoff, J.B., Pinner, S.E., Gschmeissner, S., Condeelis, J.S., Sahai, E., 2006. ROCK- and myosin-dependent matrix deformation enables protease-independent tumor-cell invasion in vivo. Curr. Biol. CB 16, 1515–1523. 10.1016/j.cub.2006.05.065

Xiao, D., He, Jiaxi, Guo, Z., He, H., Yang, S., Huang, L., Pan, H., He, Jianxing, 2020. Rhophilin-2 Upregulates Glutamine Synthetase by Stabilizing c-Myc Protein and Confers Resistance to Glutamine Deprivation in Lung Cancer. Front. Oncol. 10, 571384. 10.3389/fonc.2020.571384

Xu, H., Xiao, T., Chen, C.-H., Li, W., Meyer, C.A., Wu, Q., Wu, D., Cong, L., Zhang, F., Liu, J.S., Brown, M., Liu, X.S., 2015. Sequence determinants of improved CRISPR sgRNA design. Genome Res. 25, 1147–1157. 10.1101/gr.191452.115

Yan, L., He, Z., Li, W., Liu, N., Gao, S., 2021. P76RBE silencing inhibits ovarian cancer cell proliferation, migration, and invasion via suppressing the integrin β1/NF-κB pathway. Cell Cycle Georget. Tex 20, 1875–1889. 10.1080/15384101.2021.1963910

Yu, F., Qiao, P., Yin, G., Sun, Y., Yu, X., Sun, X., Chu, Y., Wang, Y., 2020. RHPN2 Promotes Malignant Cell Behaviours in Ovarian Cancer by Activating STAT3 Signalling. OncoTargets Ther. 13, 11517–11527. 10.2147/OTT.S272752

Zhang, T., Wang, R.-Q., Yang, Y.-B., Zhao, W.-J., Zhang, Q.-D., Ma, X.-L., Guo, J.-J., Su, Q., Shen, X.-Y., Lu, Y.-B., Su, F., Hou, X.-M., 2025. KLF5 facilitates lung adenocarcinoma metastasis by regulating the epithelial-mesenchymal transition pathway through RHPN2. J. Transl. Med. 23, 1078. 10.1186/s12967-025-07150-6

Zhao, M., Su, H.-Z., Zeng, Y.-H., Sun, Y., Guo, X.-X., Li, Y.-L., Wang, C., Zhao, Z.-Y., Huang, X.-J., Lin, K.-J., Ye, Z.-L., Lin, B.-W., Hong, S., Zheng, J., Liu, Y.-B., Yao, X.-P., Yang, D., Lu, Y.-Q., Chen, H.-Z., Zuo, E., Yang, G., Wang, H.-T., Huang, C.-W., Lin, X.-H., Cen, Z., Lai, L.-L., Zhang, Y.-K., Li, X., Lai, T., Lin, J., Zuo, D.-D., Lin, M.-T., Liou, C.-W., Kong, Q.-X., Yan, C.-Z., Xiong, Z.-Q., Wang, N., Luo, W., Zhao, C.-P., Cheng, X., Chen, W.-J., 2022. Loss of function of CMPK2 causes mitochondria deficiency and brain calcification. Cell Discov. 8, 128. 10.1038/s41421-022-00475-2

