## Supplementary figures for "Rhpn2 promotes zebrafish melanoma development and aggressiveness *in vivo*"

### Table of content

Supplementary Figures 1 and 2 and legends

#### Figure S1

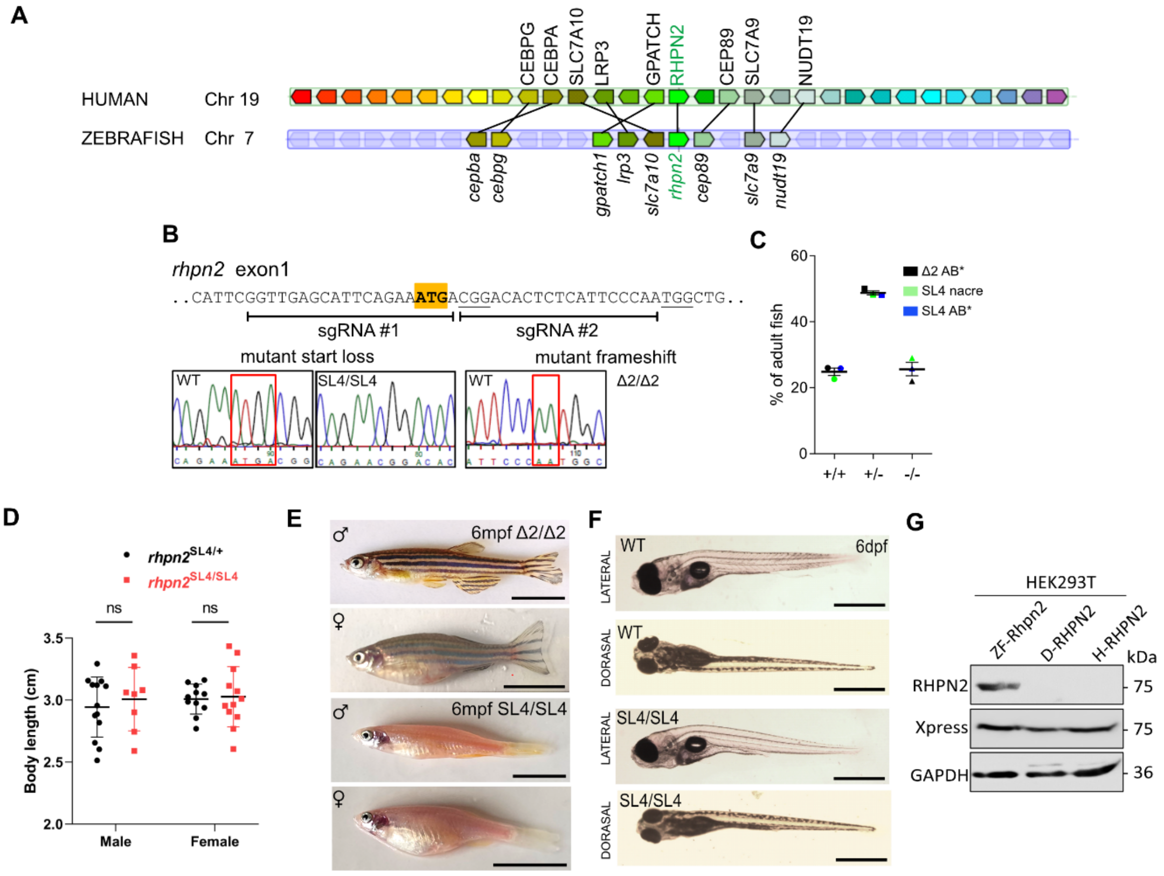


**Supplementary Figure 1. A-** Conservation of rhpn2 genes synteny between human and zebrafish using genomicus software (https://www.genomicus.biologie.ens.fr). Related orthologues are connected between chromosomes. **B-** Genomic sequence of *rhpn2* exon1 with sgRNA target sequences adjacent to the underlined PAM detailed. Sanger sequences respectively show a 4 bp and a 2 pd deletion (boxed in red). **C-** Genotypic analysis of adult fish from heterozygous intercrosses. A normal mendelian inheritance ratio is observed. Each dot color represents an independent cross (N = 3, at least 25 fish genotyped per cross, means: +/+, 24.8%; +/SL4, 48.8%; SL4/SL4, 25.6%). **D-** Body length measurements of adult males and females with *rhpn2*$^{SL4/SL4}$ loss-of-function mutations, compared to their respective heterozygous siblings (males, n = 18 for SL4/+ and n= 8 for SL4/SL4; females, n = 11 for SL4/+ and n= 13 for SL4/SL4). Statistical analysis: males : SL4/SL4 vs SL4/+: ns, p=0,42 ; females SL4/SL4 vs SL4/+: ns, p=0.59, unpaired T-test). **E-** Adult *rhpn2*$^{Δ2/Δ2}$ or *rhpn2*$^{SL4/SL4}$; *mitfa*$^{-/-}$ zebrafish show no macroscopic defects. Scale bar: 1 cm. **F-** No phenotypic differences were observed between 6 dpf WT or *rhpn2*$^{Δ2/Δ2}$ or *rhpn2*$^{SL4/SL4}$ larvae. Scale bar: 500 µm. **G-** Western blotting of 35 µg of total protein extract from HEK293T expressing zebrafish (ZF), dog (D) and human (H) RHPN2 with the antibody directed to a zebrafish C-terminal peptide.

**Figure S2**

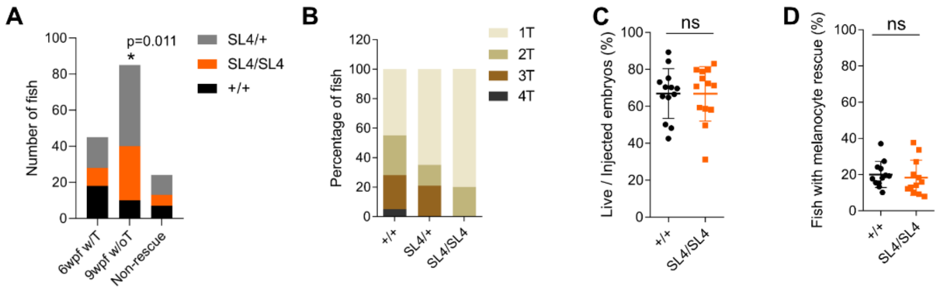


**Supplementary Figure 2. A-** The proportions of each genotype among fish having early developed tumors compared to their tumor free siblings were evaluated. Chi-square analysis : w6 with tumor (n = 45, ns p = 0.054), w9 w/o tumor (n = 85 , *p = 0,011), no rescue (n = 24, ns p = 0.417). **B-** The SL4/SL4 fish developed a lower number of tumor than +/+. **C-** No differences were observed in the percentage of live fish 1 day post-injection between +/+ and SL4/SL4 larvae (Unpaired t-Test, ns p = 0.98). **D-** No differences were observed between +/+ and SL4/SL4 larvae in the percentage of 5 dpf fish presenting rescued pigmentation (Unpaired t-Test, ns p = 0.616).
