## Supplementary table for "Rhpn2 promotes zebrafish melanoma development and aggressiveness *in vivo*"

### Program:featureCounts v2.1.1; Command:"featureCounts" "-p" "-T" "10" "--extraAttributes" "gene_name" "-a" "/home/singhlab/Documents/Genome/Danio_rerio/Ensembl/GRCz11/Annotation/Danio_rerio.GRCz11.110.gtf" "-o" "RAWdata_All_features_GRCz11.tsv" "RAWdata_Hisat2_Mapping/K0_MKRN250022282-1A_22VV33LT4_L7.hisat.bam" "RAWdata_Hisat2_Mapping/K1_MKRN250024061-1A_22VTJNLT4_L7.hisat.bam" "RAWdata_Hisat2_Mapping/K1_MKRN250024061-1A_22VVKVLT4_L8.hisat.bam" "RAWdata_Hisat2_Mapping/K2_MKRN250022283-1A_22VV33LT4_L7.hisat.bam" "RAWdata_Hisat2_Mapping/W1_MKRN250022279-1A_22VV33LT4_L7.hisat.bam" "RAWdata_Hisat2_Mapping/W2_MKRN250022280-1A_22VV33LT4_L7.hisat.bam" "RAWdata_Hisat2_Mapping/W3_MKRN250022281-1A_22VV33LT4_L7.hisat.bam"

| Geneid | Chr | Start | End | Strand | Length | gene_name | KO RAWdata | K1 RAWdata | K2 RAWdata | W1 RAWdata | W2 RAWdata | W3_RAWdata_Hisat2_Mapping/W3_MKRN250022281-1A_22VV33LT4_L7.hisat.bam |
|---|---|---|---|---|---|---|---|---|---|---|---|---|
| ENSDARG000 | 4;4;4 | 30402837:304 | 30402893:304 | +;+;+ | 423 | NA | 0 | 0 | 0 | 0 | 0 | 0 |
| ENSDARG000 | 4;4;4;4;4;4;4; | 1722899:1722 | 1722927:1722 | +;+;+;+;+;+;+; | 2014 | fgfr1op2 | 548 | 487 | 474 | 740 | 450 | 665 |
| ENSDARG000 | 4;4 | 40758511:407 | 40758849:407 | +;+ | 660 | NA | 0 | 0 | 0 | 0 | 1 | 0 |
| ENSDARG000 | 4;4;4;4;4;4;4; | 11099041:110 | 11099241:110 | -;-;-;-;-;-;-; | 3091 | NA | 0 | 1 | 0 | 0 | 0 | 0 |
| ENSDARG000 | 4;4;4;4;4;4;4; | 40835393:408 | 40835453:408 | +;+;+;+;+;+;+; | 2179 | NA | 0 | 0 | 0 | 0 | 0 | 6 |
| ENSDARG000 | 4;4;4 | 30409750:304 | 30410604:304 | -;-;- | 1096 | NA | 20 | 12 | 22 | 0 | 10 | 16 |
| ENSDARG000 | 4;4;4;4;4;4;4; | 11112580:111 | 11112626:111 | -;-;-;-;-;-;-; | 3646 | NA | 0 | 2 | 0 | 0 | 0 | 0 |
| ENSDARG000 | 4;4;4;4;4;4;4; | 8213063:8213 | 8213357:8213 | +;+;+;+;+;+;+; | 4126 | ERC1 | 352 | 223 | 336 | 281 | 310 | 285 |
| ENSDARG000 | 4;4 | 30428468:304 | 30430293:304 | -;- | 1976 | NA | 9 | 10 | 16 | 0 | 8 | 18 |
| ENSDARG000 | 4;4;4;4;4;4;4; | 8376362:8376 | 8376526:8376 | +;+;+;+;+;+;+; | 763 | NA | 0 | 0 | 0 | 0 | 0 | 1 |
| ENSDARG000 | 4;4 | 40839314:408 | 40839531:408 | +;+ | 461 | NA | 0 | 4 | 0 | 0 | 0 | 6 |
| ENSDARG000 | 4;4;4 | 30454496:304 | 30454538:304 | +;+;+ | 1774 | zgc:173716 | 5 | 28 | 38 | 0 | 12 | 14 |
| ENSDARG000 | 4;4;4 | 40846483:408 | 40847768:408 | -;-;- | 1466 | NA | 12 | 14 | 20 | 12 | 8 | 0 |
| ENSDARG000 | 4;4 | 40858450:408 | 40858599:408 | +;+ | 204 | NA | 0 | 0 | 0 | 0 | 1 | 0 |
| ENSDARG000 | 4;4;4;4;4;4;4; | 1732825:1736 | 1735043:1736 | -;-;-;-;-;-;-; | 3807 | TM7SF3 | 4181 | 3447 | 3392 | 3788 | 2240 | 4520 |
| ENSDARG000 | 4;4;4;4;4;4;4; | 5215950:5218 | 5216294:5218 | +;+;+;+;+;+;+; | 1973 | NA | 0 | 0 | 2 | 0 | 0 | 0 |
| ENSDARG000 | 4;4;4;4;4;4 | 40880433:408 | 40881466:408 | -;-;-;-;-;- | 1237 | zgc:174650 | 2 | 0 | 0 | 0 | 0 | 6 |
| ENSDARG000 | 4;4;4;4;4;4;4 | 19700308:197 | 19700433:197 | +;+;+;+;+;+;+ | 1231 | pax4 | 0 | 0 | 0 | 0 | 0 | 0 |
| ENSDARG000 | 4;4;4;4;4;4;4; | 13449775:134 | 13450109:134 | +;+;+;+;+;+;+; | 6622 | dyrk2 | 821 | 581 | 807 | 664 | 752 | 941 |
| ENSDARG000 | 4;4;4;4 | 22365061:224 | 22366063:224 | +;+;+;+ | 2519 | lhfpl3 | 2 | 0 | 4 | 12 | 8 | 0 |
| ENSDARG000 | 4;4;4;4;4;4;4 | 11124613:111 | 11124859:111 | -;-;-;-;-;-;- | 714 | NA | 0 | 0 | 0 | 0 | 0 | 0 |
| ENSDARG000 | 4;4;4;4;4;4;4; | 11124614:111 | 11124859:111 | -;-;-;-;-;-;-; | 2898 | NA | 8 | 8 | 8 | 46 | 0 | 51 |
| ENSDARG000 | 4;4;4;4;4;4;4; | 49765093:497 | 49765752:497 | -;-;-;-;-;-;-; | 4453 | NA | 103 | 70 | 113 | 22 | 37 | 58 |
| ENSDARG000 | 4;4;4;4;4;4;4; | 40971399:409 | 40971469:409 | -;-;-;-;-;-;-;- | 526 | NA | 0 | 0 | 0 | 0 | 0 | 0 |
| ENSDARG000 | 4;4;4;4;4;4;4; | 41057692:410 | 41058695:410 | -;-;-;-;-;-;-;- | 2249 | NA | 103 | 59 | 27 | 10 | 29 | 33 |
| ENSDARG000 | 4;4;4;4;4;4;4 | 40953320:409 | 40953368:409 | +;+;+;+;+;+;+ | 2715 | NA | 17 | 61 | 107 | 8 | 42 | 43 |
| ENSDARG000 | 4;4;4;4;4;4;4; | 8532580:8569 | 8532910:8569 | +;+;+;+;+;+;+; | 2065 | wnt5b | 18 | 12 | 22 | 10 | 32 | 8 |
| ENSDARG000 | 4;4;4;4;4;4;4; | 41157276:411 | 41157425:411 | +;+;+;+;+;+;+; | 1738 | NA | 0 | 2 | 0 | 0 | 0 | 2 |
| ENSDARG000 | 4;4;4;4;4;4 | 13499264:134 | 13499717:134 | -;-;-;-;-;- | 1243 | si:ch211-266a | 6 | 0 | 0 | 0 | 6 | 0 |
| ENSDARG000 | 4;4;4;4;4 | 41082518:410 | 41082573:410 | +;+;+;+;+ | 1277 | zgc:173721 | 12 | 2 | 6 | 6 | 15 | 6 |
| ENSDARG000 | 4;4 | 40970545:409 | 40970606:409 | +;+ | 717 | NA | 0 | 0 | 0 | 0 | 0 | 0 |
| ENSDARG000 | 4 | 8609517 | 8611841 | - | 2325 | fbxl14b | 292 | 442 | 565 | 514 | 667 | 360 |
| ENSDARG000 | 4;4 | 8634704:8637 | 8634841:8638 | -;- | 1411 | NA | 2 | 0 | 0 | 0 | 0 | 0 |
| ENSDARG000 | 4;4;4 | 30521650:305 | 30523033:305 | -;-;- | 1614 | NA | 0 | 0 | 0 | 0 | 0 | 0 |
| ENSDARG000 | 4;4 | 16001238:160 | 16001398:160 | +;+ | 662 | NA | 0 | 0 | 0 | 0 | 0 | 0 |
| ENSDARG000 | 4;4 | 16003900:160 | 16004097:160 | -;- | 369 | NA | 0 | 0 | 0 | 0 | 0 | 0 |
| ENSDARG000 | 4;4;4;4;4;4;4; | 5225552:5228 | 5226137:5228 | -;-;-;-;-;-;-; | 2348 | ADA2 | 136 | 99 | 134 | 58 | 117 | 175 |
| ENSDARG000 | 4 | 30645966 | 30646791 | + | 826 | NA | 0 | 0 | 0 | 0 | 0 | 0 |
| ENSDARG000 | 4;4;4;4;4 | 30707282:307 | 30707866:307 | -;-;-;-;- | 791 | NA | 2 | 2 | 0 | 0 | 4 | 0 |
| ENSDARG000 | 4;4;4;4;4;4;4; | 5240126:5242 | 5240578:5242 | -;-;-;-;-;-;-;- | 1330 | atp6v1e1b | 4041 | 4536 | 3991 | 4273 | 3894 | 3380 |
| ENSDARG000 | 4;4 | 16013301:160 | 16013500:160 | -;- | 371 | NA | 0 | 1 | 0 | 0 | 0 | 1 |
| ENSDARG000 | 4;4;4;4;4 | 16019215:160 | 16019394:160 | +;+;+;+;+ | 1101 | NA | 1 | 2 | 0 | 11 | 20 | 20 |
| ENSDARG000 | 4;4;4;4;4;4;4; | 3358383:3367 | 3358492:3367 | +;+;+;+;+;+;+; | 1722 | NAMPT | 422 | 816 | 484 | 294 | 456 | 571 |
| ENSDARG000 | 4;4 | 16040114:160 | 16040233:160 | -;- | 303 | NA | 0 | 0 | 0 | 0 | 0 | 0 |
| ENSDARG000 | 4;4 | 16042239:160 | 16042406:160 | +;+ | 368 | NA | 0 | 0 | 0 | 0 | 0 | 4 |
| ENSDARG000 | 4;4;4;4;4 | 49819352:498 | 49819464:498 | -;-;-;-;- | 327 | NA | 0 | 0 | 0 | 0 | 0 | 0 |
| ENSDARG000 | 4;4 | 16051600:160 | 16051767:160 | +;+ | 368 | NA | 0 | 0 | 0 | 0 | 0 | 4 |
| ENSDARG000 | 4;4 | 16060556:160 | 16060723:160 | +;+ | 368 | NA | 0 | 0 | 0 | 0 | 0 | 0 |
| ENSDARG000 | 4;4;4;4;4;4;4 | 30775376:307 | 30775448:307 | +;+;+;+;+;+;+ | 1133 | zgc:172014 | 2 | 22 | 22 | 16 | 4 | 16 |
| ENSDARG000 | 4;4 | 16069577:160 | 16069744:160 | +;+ | 368 | NA | 0 | 0 | 0 | 0 | 0 | 0 |
| ENSDARG000 | 4;4;4;4;4;4;4; | 49821234:498 | 49826308:498 | -;-;-;-;-;-;-; | 11189 | zgc:173709 | 123 | 98 | 108 | 35 | 57 | 82 |
| ENSDARG000 | 4;4;4;4;4;4;4; | 11135048:111 | 11135251:111 | +;+;+;+;+;+;+; | 5044 | CRACR2A | 2 | 2 | 5 | 16 | 4 | 19 |
| ENSDARG000 | 4;4;4;4;4;4;4; | 41202536:412 | 41202701:412 | -;-;-;-;-;-;-;- | 1050 | NA | 0 | 10 | 6 | 3 | 4 | 21 |
| ENSDARG000 | 4;4 | 49942510:499 | 49942772:499 | +;+ | 482 | NA | 0 | 0 | 0 | 0 | 0 | 0 |
| ENSDARG000 | 4;4;4;4;4;4;4; | 11144600:111 | 11148725:111 | -;-;-;-;-;-;-; | 6147 | prmt8b | 0 | 0 | 2 | 0 | 0 | 0 |
| ENSDARG000 | 4;4;4 | 41216522:412 | 41216580:412 | +;+;+ | 811 | si:ch211-181 | 0 | 0 | 0 | 6 | 3 | 0 |
| ENSDARG000 | 4;4;4;4;4;4;4; | 25497104:254 | 25497601:254 | -;-;-;-;-;-;-; | 1844 | rbm17 | 318 | 372 | 458 | 277 | 479 | 310 |
| ENSDARG000 | 4;4;4;4 | 13507584:135 | 13507742:135 | -;-;-;- | 570 | ifng1 | 0 | 18 | 0 | 2 | 8 | 109 |
| ENSDARG000 | 4;4;4;4;4;4;4; | 41212032:412 | 41212231:412 | -;-;-;-;-;-;-;- | 951 | NA | 0 | 0 | 0 | 3 | 0 | 2 |
| ENSDARG000 | 4;4;4;4;4;4;4; | 76291003:762 | 76291966:762 | -;-;-;-;-;-;-; | 4043 | NA | 66 | 38 | 90 | 17 | 33 | 27 |
| ENSDARG000 | 4;4;4;4;4;4;4; | 12074776:120 | 12075313:120 | -;-;-;-;-;-;-; | 3074 | braf | 412 | 307 | 431 | 362 | 425 | 560 |
| ENSDARG000 | 4;4;4;4;4;4;4; | 22424451:224 | 22425779:224 | -;-;-;-;-;-;-; | 5497 | KMT2E | 1927 | 1424 | 2588 | 2154 | 2374 | 1786 |
| ENSDARG000 | 4;4;4;4 | 13517446:135 | 13517541:135 | -;-;-;- | 513 | ifng1r | 5 | 11 | 0 | 0 | 4 | 0 |
| ENSDARG000 | 4;4;4;4 | 13535054:135 | 13535210:135 | -;-;-;- | 639 | il26 | 0 | 0 | 0 | 0 | 0 | 0 |
| ENSDARG000 | 4;4;4;4;4;4;4; | 30785713:307 | 30785820:307 | +;+;+;+;+;+;+; | 3353 | NA | 10 | 10 | 7 | 4 | 0 | 8 |
| ENSDARG000 | 4;4;4;4;4;4;4; | 49969199:499 | 49969798:499 | -;-;-;-;-;-;-; | 3573 | NA | 13 | 21 | 72 | 13 | 8 | 248 |
| ENSDARG000 | 4;4;4;4 | 1757462:1758 | 1757573:1758 | +;+;+;+ | 823 | med21 | 186 | 239 | 260 | 222 | 224 | 156 |
| ENSDARG000 | 4;4 | 50153271:501 | 50154156:501 | -;- | 1041 | NA | 0 | 0 | 3 | 0 | 14 | 0 |
| ENSDARG000 | 4;4;4;4;4;4;4; | 50260361:502 | 50260496:502 | +;+;+;+;+;+;+; | 3960 | NA | 276 | 208 | 433 | 198 | 425 | 768 |
| ENSDARG000 | 4;4;4 | 50293905:503 | 50295051:503 | -;-;- | 1391 | NA | 3 | 7 | 21 | 0 | 23 | 34 |
| ENSDARG000 | 4;4;4;4;4;4;4; | 30884012:308 | 30885243:308 | -;-;-;-;-;-;-; | 5310 | NA | 26 | 10 | 8 | 12 | 25 | 18 |
| ENSDARG000 | 4;4;4;4;4;4;4; | 50201633:502 | 50202912:502 | -;-;-;-;-;-;-;- | 2326 | NA | 76 | 40 | 48 | 4 | 50 | 60 |
| ENSDARG000 | 4;4;4;4;4;4;4; | 19710657:197 | 19712140:197 | -;-;-;-;-;-;-; | 3589 | hgfa | 2 | 2 | 2 | 4 | 2 | 2 |
| ENSDARG000 | 4;4;4 | 30925543:309 | 30926174:309 | -;-;- | 941 | NA | 3 | 0 | 0 | 0 | 2 | 0 |
| ENSDARG000 | 4;4;4;4;4;4;4 | 3389866:3394 | 3390401:3394 | +;+;+;+;+;+;+ | 1592 | SYPL1 | 129 | 311 | 52 | 168 | 31 | 398 |
| ENSDARG000 | 4;4;4;4;4 | 13544595:135 | 13544678:135 | -;-;-;-;- | 642 | il22 | 0 | 0 | 0 | 0 | 0 | 8 |
| ENSDARG000 | 4;4;4 | 3396322:3397 | 3396745:3398 | -;-;- | 773 | NA | 9 | 4 | 1 | 0 | 0 | 1 |
| ENSDARG000 | 4;4;4;4;4;4;4; | 16076719:160 | 16079026:160 | -;-;-;-;-;-;-; | 5951 | atp2b1a | 560 | 583 | 701 | 216 | 774 | 398 |
| ENSDARG000 | 4;4;4;4;4;4 | 8709082:8710 | 8709133:8710 | -;-;-;-;-;- | 348 | NA | 0 | 0 | 0 | 4 | 0 | 0 |
| ENSDARG000 | 4;4;4 | 41227127:412 | 41227152:412 | +;+;+ | 781 | NA | 0 | 6 | 2 | 0 | 4 | 2 |
| ENSDARG000 | 4;4;4;4;4;4;4; | 8670662:8680 | 8670748:8681 | +;+;+;+;+;+;+; | 6070 | adipor2 | 818 | 890 | 890 | 1113 | 955 | 880 |
| ENSDARG000 | 4;4;4;4;4;4 | 50305577:503 | 50305720:503 | +;+;+;+;+;+ | 2236 | NA | 12 | 8 | 2 | 0 | 0 | 0 |
| ENSDARG000 | 4;4;4 | 31085014:310 | 31085135:310 | +;+;+ | 710 | NA | 1 | 12 | 6 | 6 | 2 | 4 |
| ENSDARG000 | 4;4;4;4;4;4;4; | 13551400:135 | 13551569:135 | -;-;-;-;-;-;-; | 2374 | mdm1 | 78 | 62 | 67 | 34 | 58 | 84 |
| ENSDARG000 | 4;4 | 50401133:504 | 50401177:504 | +;+ | 900 | NA | 1 | 0 | 0 | 0 | 0 | 0 |
| ENSDARG000 | 4;4 | 31095829:310 | 31097390:310 | -;- | 1668 | NA | 28 | 20 | 48 | 14 | 30 | 47 |
| ENSDARG000 | 4;4 | 41311345:413 | 41311523:413 | +;+ | 692 | NA | 0 | 0 | 2 | 0 | 0 | 0 |
| ENSDARG000 | 4;4;4;4;4;4;4; | 11172964:111 | 11173156:111 | +;+;+;+;+;+;+; | 3170 | tspan11 | 7 | 5 | 9 | 0 | 8 | 13 |
| ENSDARG000 | 4;4;4 | 41315144:413 | 41317063:413 | -;-;- | 2122 | NA | 89 | 48 | 55 | 7 | 31 | 31 |
| ENSDARG000 | 4;4;4 | 41385580:413 | 41385626:413 | +;+;+ | 829 | NA | 0 | 0 | 0 | 0 | 0 | 2 |
| ENSDARG000 | 4;4;4;4;4;4;4; | 1761029:1762 | 1761134:1762 | +;+;+;+;+;+;+; | 1869 | pmpcb | 270 | 240 | 494 | 176 | 222 | 168 |
| ENSDARG000 | 4;4 | 11179509:111 | 11180043:111 | -;- | 785 | NA | 0 | 0 | 0 | 0 | 0 | 0 |
| ENSDARG000 | 4;4 | 41449611:414 | 41449805:414 | -;- | 497 | NA | 0 | 3 | 1 | 0 | 0 | 1 |
| ENSDARG000 | 4;4 | 41471349:414 | 41471487:414 | +;+ | 701 | NA | 0 | 0 | 0 | 0 | 0 | 0 |
| ENSDARG000 | 4;4;4;4;4;4;4; | 31247968:312 | 31248279:312 | -;-;-;-;-;-;-; | 2463 | zgc:173720 | 81 | 110 | 132 | 20 | 49 | 69 |
| ENSDARG000 | 4;4;4;4;4;4;4; | 76304911:763 | 76304944:763 | +;+;+;+;+;+;+; | 3718 | NA | 114 | 89 | 170 | 32 | 275 | 198 |
| ENSDARG000 | 4;4 | 31293362:312 | 31293452:312 | +;+ | 1136 | NA | 0 | 0 | 0 | 4 | 0 | 2 |
| ENSDARG000 | 4;4;4;4;4;4;4; | 1766333:1771 | 1771308:1771 | -;-;-;-;-;-;-; | 6766 | dnajc2 | 360 | 617 | 851 | 373 | 612 | 376 |
| ENSDARG000 | 4;4;4;4 | 50410452:504 | 50410627:504 | +;+;+;+ | 690 | NA | 4 | 9 | 4 | 0 | 0 | 6 |
| ENSDARG000 | 4;4;4;4;4;4 | 50426233:504 | 50426691:504 | -;-;-;-;-;- | 1997 | NA | 0 | 0 | 0 | 0 | 0 | 0 |
| ENSDARG000 | 4;4;4 | 50463619:504 | 50463648:504 | +;+;+ | 1887 | NA | 58 | 40 | 20 | 6 | 54 | 55 |
| ENSDARG000 | 4;4;4;4 | 50572302:505 | 50572389:505 | +;+;+;+ | 308 | NA | 8 | 2 | 3 | 0 | 5 | 0 |
| ENSDARG000 | 4;4;4 | 50701550:507 | 50702513:507 | -;-;- | 1160 | NA | 0 | 4 | 0 | 0 | 0 | 4 |
| ENSDARG000 | 4;4;4;4;4;4;4 | 22480169:224 | 22480353:224 | +;+;+;+;+;+;+ | 1095 | ndufb2 | 591 | 580 | 828 | 612 | 702 | 579 |
| ENSDARG000 | 4;4;4;4;4;4 | 3438510:3438 | 3438596:3438 | +;+;+;+;+;+ | 1627 | NA | 57 | 29 | 91 | 70 | 81 | 66 |
| ENSDARG000 | 4;4;4 | 41516714:415 | 41518002:415 | -;-;- | 1503 | NA | 6 | 0 | 7 | 0 | 2 | 3 |
| ENSDARG000 | 4;4 | 3446011:3455 | 3446459:3456 | -;- | 903 | NA | 0 | 1 | 1 | 0 | 0 | 0 |
| ENSDARG000 | 4;4;4;4;4;4;4; | 3455146:3455 | 3455358:3455 | +;+;+;+;+;+;+; | 2926 | znf800b | 232 | 197 | 254 | 169 | 192 | 186 |
| ENSDARG000 | 4;4;4 | 50726871:507 | 50727021:507 | +;+;+ | 1569 | NA | 0 | 0 | 0 | 0 | 0 | 0 |
| ENSDARG000 | 4;4;4;4;4;4;4; | 25607101:256 | 25607179:256 | +;+;+;+;+;+;+; | 2261 | acot14 | 12 | 0 | 4 | 2 | 0 | 4 |
| ENSDARG000 | 4;4;4;4;4;4;4; | 31327980:313 | 31328647:313 | +;+;+;+;+;+;+; | 5219 | NA | 461 | 304 | 487 | 143 | 390 | 364 |
| ENSDARG000 | 4;4 | 5249494:5253 | 5249522:5253 | +;+ | 859 | si:ch211-214i | 6 | 8 | 9 | 0 | 6 | 4 |
| ENSDARG000 | 4;4 | 50755309:507 | 50756882:507 | -;- | 1730 | NA | 14 | 34 | 70 | 10 | 29 | 60 |
| ENSDARG000 | 4;4;4;4 | 50776144:507 | 50776675:507 | -;-;-;- | 731 | NA | 0 | 0 | 0 | 0 | 2 | 0 |
| ENSDARG000 | 4;4;4;4;4;4;4; | 19768679:197 | 19768798:197 | -;-;-;-;-;-;-; | 3928 | cacna2d1a | 254 | 44 | 76 | 7 | 6 | 89 |
| ENSDARG000 | 4;4;4;4;4;4;4; | 22484830:224 | 22485489:224 | -;-;-;-;-;-;-; | 3466 | kdm7aa | 166 | 275 | 1094 | 718 | 372 | 108 |
| ENSDARG000 | 4;4 | 11202941:112 | 11203035:112 | +;+ | 194 | NA | 2 | 4 | 0 | 0 | 4 | 0 |
| ENSDARG000 | 4;4;4 | 41528590:415 | 41528681:415 | +;+;+ | 2253 | NA | 7 | 6 | 23 | 10 | 0 | 0 |
| ENSDARG000 | 4;4;4;4;4;4;4; | 41555642:415 | 41556694:415 | -;-;-;-;-;-;-; | 3866 | NA | 6 | 7 | 6 | 0 | 0 | 6 |
| ENSDARG000 | 4;4;4 | 11245332:112 | 11245450:112 | -;-;- | 539 | NA | 0 | 0 | 0 | 0 | 0 | 0 |
| ENSDARG000 | 4;4;4;4;4;4;4; | 8849816:8852 | 8852209:8852 | -;-;-;-;-;-;-; | 4214 | MPPED1 | 0 | 0 | 2 | 0 | 0 | 0 |
| ENSDARG000 | 4;4;4;4;4;4;4; | 5260941:5263 | 5261435:5263 | -;-;-;-;-;-;-; | 3267 | NA | 0 | 0 | 6 | 4 | 6 | 4 |
| ENSDARG000 | 4;4;4 | 50891961:508 | 50892109:508 | +;+;+ | 728 | NA | 3 | 0 | 4 | 0 | 0 | 2 |
| ENSDARG000 | 4;4;4;4;4;4;4 | 76336061:763 | 76336095:763 | +;+;+;+;+;+;+ | 4684 | NA | 104 | 31 | 58 | 24 | 117 | 172 |
| ENSDARG000 | 4;4;4;4;4;4;4; | 5255041:5255 | 5255211:5255 | +;+;+;+;+;+;+; | 978 | ccdc167 | 342 | 340 | 354 | 418 | 450 | 362 |
| ENSDARG000 | 4;4 | 31427055:314 | 31427330:314 | +;+ | 706 | NA | 0 | 0 | 0 | 0 | 0 | 2 |
| ENSDARG000 | 4;4;4;4;4 | 31450683:314 | 31450722:314 | +;+;+;+;+ | 2377 | NA | 10 | 23 | 30 | 0 | 11 | 22 |
| ENSDARG000 | 4;4 | 11285929:112 | 11286089:112 | +;+ | 916 | NA | 0 | 0 | 0 | 2 | 0 | 0 |
| ENSDARG000 | 4;4;4;4;4;4;4; | 31535449:316 | 31536131:316 | -;-;-;-;-;-;-; | 5914 | NA | 44 | 21 | 46 | 6 | 44 | 23 |
| ENSDARG000 | 4;4;4;4;4;4 | 76349919:763 | 76349925:763 | +;+;+;+;+;+ | 2714 | NA | 44 | 14 | 18 | 4 | 44 | 38 |
| ENSDARG000 | 4;4;4;4;4;4;4; | 50920712:509 | 50921304:509 | -;-;-;-;-;-;-; | 4923 | NA | 325 | 321 | 556 | 104 | 285 | 334 |
| ENSDARG000 | 4;4;4;4;4;4;4; | 31634604:316 | 31635129:316 | -;-;-;-;-;-;-; | 3198 | NA | 0 | 4 | 2 | 0 | 18 | 21 |
| ENSDARG000 | 4;4;4 | 50970601:509 | 50970677:509 | +;+;+ | 887 | NA | 3 | 6 | 20 | 0 | 4 | 8 |
| ENSDARG000 | 4;4;4;4;4 | 41675203:416 | 41675320:416 | -;-;-;-;- | 632 | NA | 4 | 2 | 4 | 0 | 0 | 6 |
| ENSDARG000 | 4;4;4;4 | 31555894:315 | 31555939:315 | -;-;-;- | 315 | NA | 4 | 11 | 10 | 2 | 9 | 12 |
| ENSDARG000 | 4;4;4;4;4;4;4; | 31535449:316 | 31536131:316 | -;-;-;-;-;-;-; | 6027 | NA | 25 | 67 | 41 | 32 | 115 | 102 |
| ENSDARG000 | 4;4 | 41602678:416 | 41603863:416 | -;- | 1335 | NA | 1 | 0 | 4 | 0 | 0 | 4 |
| ENSDARG000 | 4;4;4 | 41770710:417 | 41772698:417 | -;-;- | 2180 | NA | 0 | 12 | 6 | 0 | 3 | 0 |
| ENSDARG000 | 4;4;4;4;4;4;4; | 41598006:416 | 41598024:416 | -;-;-;-;-;-;-;- | 2033 | NA | 8 | 13 | 41 | 4 | 13 | 13 |
| ENSDARG000 | 4;4;4;4;4;4;4; | 31653917:316 | 31654749:316 | -;-;-;-;-;-;-; | 6312 | NA | 27 | 160 | 41 | 52 | 284 | 231 |
| ENSDARG000 | 4;4;4;4;4;4;4; | 76362874:763 | 76364433:763 | -;-;-;-;-;-;-; | 2769 | NA | 16 | 24 | 4 | 0 | 63 | 30 |
| ENSDARG000 | 4;4;4 | 41789497:417 | 41789545:417 | +;+;+ | 1143 | NA | 6 | 17 | 23 | 0 | 4 | 7 |
| ENSDARG000 | 4;4;4;4;4;4;4; | 3478049:3478 | 3478184:3478 | +;+;+;+;+;+;+; | 3498 | GRM8 | 2 | 6 | 17 | 2 | 0 | 0 |
| ENSDARG000 | 4;4;4;4;4;4;4; | 41994717:419 | 41994802:419 | +;+;+;+;+;+;+; | 2370 | NA | 43 | 2 | 45 | 16 | 2 | 14 |
| ENSDARG000 | 4;4;4;4;4;4;4; | 51026304:510 | 51026357:510 | -;-;-;-;-;-;-; | 2783 | NA | 21 | 29 | 44 | 6 | 43 | 31 |
| ENSDARG000 | 4;4;4 | 42009286:420 | 42009802:420 | -;-;- | 820 | NA | 0 | 0 | 0 | 0 | 0 | 0 |
| ENSDARG000 | 4;4;4;4;4;4;4; | 41834258:418 | 41834426:418 | +;+;+;+;+;+;+; | 3757 | NA | 92 | 44 | 145 | 79 | 275 | 339 |
| ENSDARG000 | 4;4;4;4;4;4;4; | 9769050:9769 | 9769212:9769 | -;-;-;-;-;-;-; | 2239 | svopl | 6 | 0 | 4 | 0 | 0 | 0 |
| ENSDARG000 | 4;4;4;4;4 | 41816778:418 | 41818383:418 | -;-;-;-;- | 1859 | NA | 2 | 6 | 6 | 0 | 20 | 22 |
| ENSDARG000 | 4 | 306036 | 308006 | + | 1971 | msgn1 | 0 | 4 | 8 | 6 | 20 | 0 |
| ENSDARG000 | 4;4;4;4;4 | 41915340:419 | 41915406:419 | +;+;+;+;+ | 941 | NA | 0 | 0 | 0 | 0 | 0 | 0 |
| ENSDARG000 | 4;4;4;4;4;4;4; | 13568469:135 | 13568610:135 | +;+;+;+;+;+;+; | 2042 | calua | 657 | 1050 | 1469 | 406 | 736 | 504 |
| ENSDARG000 | 4;4;4;4;4;4;4; | 22613625:226 | 22613785:226 | +;+;+;+;+;+;+; | 2141 | SEMA3C | 26 | 71 | 1621 | 893 | 9455 | 790 |
| ENSDARG000 | 4;4;4;4;4;4;4; | 25627147:256 | 25627287:256 | +;+;+;+;+;+;+; | 2060 | acot15 | 6 | 0 | 48 | 4 | 10 | 2 |
| ENSDARG000 | 4;4;4;4;4 | 12102267:121 | 12104106:121 | -;-;-;-;- | 2227 | mrps33 | 279 | 249 | 456 | 202 | 362 | 295 |
| ENSDARG000 | 4;4;4;4;4;4;4; | 76377227:763 | 76378556:763 | -;-;-;-;-;-;-; | 1985 | zgc:123107 | 28 | 64 | 22 | 16 | 156 | 46 |
| ENSDARG000 | 4;4;4;4;4;4;4; | 11311315:113 | 11311819:113 | +;+;+;+;+;+;+; | 4617 | sema3aa | 20395 | 9543 | 12275 | 15088 | 21071 | 8507 |
| ENSDARG000 | 4;4;4;4;4;4;4; | 8906893:8907 | 8907881:8907 | -;-;-;-;-;-;-; | 4580 | scube1 | 2 | 8 | 25 | 16 | 10 | 24 |
| ENSDARG000 | 4;4;4;4;4;4;4; | 308688:30869 | 308812:30881 | +;+;+;+;+;+;+; | 2102 | si:ch211-51e1 | 2027 | 2598 | 2818 | 1429 | 2255 | 1938 |
| ENSDARG000 | 4;4;4;4;4;4;4; | 51542447:515 | 51542512:515 | +;+;+;+;+;+;+; | 4316 | NA | 50 | 3 | 7 | 0 | 1 | 0 |

| | | | | | | | | | | | | |
|---|---|---|---|---|---|---|---|---|---|---|---|---|
| ENSDARG000 | 4;4 | 76387059:763 | 76387761:763 | -;- | 759 | NA | 0 | 0 | 0 | 0 | 0 | 0 |
| ENSDARG000 | 4;4 | 76390924:763 | 76391729:763 | -;- | 862 | NA | 0 | 0 | 0 | 0 | 0 | 0 |
| ENSDARG000 | 4;4;4 | 76426421:764 | 76426504:764 | +;+;+ | 1733 | zgc:172128 | 0 | 0 | 0 | 0 | 0 | 0 |
| ENSDARG000 | 4;4;4;4;4 | 76403698:764 | 76403819:764 | +;+;+;+;+ | 5673 | NA | 18 | 26 | 40 | 24 | 62 | 22 |
| ENSDARG000 | 4;4;4;4;4;4;4; | 76403698:764 | 76404727:764 | +;+;+;+;+;+;+; | 8589 | NA | 39 | 32 | 33 | 14 | 96 | 62 |
| ENSDARG000 | 4;4;4 | 76416268:764 | 76416337:764 | +;+;+ | 4900 | zgc:113209 | 77 | 90 | 90 | 14 | 125 | 100 |
| ENSDARG000 | 4;4;4;4;4 | 13577665:135 | 13578449:135 | -;-;-;-;- | 1777 | opn1sw1 | 0 | 0 | 0 | 2 | 6 | 0 |
| ENSDARG000 | 4;4;4;4;4;4;4; | 51669730:516 | 51669903:516 | -;-;-;-;-;-;-;-; | 3896 | NA | 44 | 10 | 33 | 3 | 25 | 35 |
| ENSDARG000 | 4;4;4;4;4;4;4; | 12111154:121 | 12111563:121 | +;+;+;+;+;+;+; | 10605 | tmem178bb | 22 | 5 | 10 | 8 | 11 | 4 |
| ENSDARG000 | 4;4;4;4;4;4;4; | 42019169:420 | 42019221:420 | +;+;+;+;+;+;+; | 1684 | NA | 0 | 0 | 0 | 4 | 28 | 26 |
| ENSDARG000 | 4;4;4;4;4;4;4; | 12122794:121 | 12123471:121 | -;-;-;-;-;-;-;- | 1289 | NA | 0 | 0 | 0 | 0 | 0 | 0 |
| ENSDARG000 | 4;4;4;4;4 | 16337156:163 | 16338896:163 | -;-;-;-;- | 2951 | kera | 12 | 2 | 22 | 0 | 0 | 6 |
| ENSDARG000 | 4 | 51876285 | 51876639 | + | 355 | NA | 0 | 11 | 0 | 0 | 6 | 18 |
| ENSDARG000 | 4;4;4;4;4;4;4; | 19946294:199 | 19950751:199 | -;-;-;-;-;-;-;-; | 6485 | drd4-rs | 0 | 0 | 0 | 0 | 0 | 0 |
| ENSDARG000 | 4;4;4;4;4;4;4; | 17707008:177 | 17708384:177 | -;-;-;-;-;-;-;- | 2563 | chpt1 | 506 | 809 | 746 | 296 | 825 | 520 |
| ENSDARG000 | 4;4;4;4;4;4;4; | 16317296:163 | 16321005:163 | -;-;-;-;-;-;-;-; | 6645 | epyc | 19 | 34 | 12 | 11 | 49 | 27 |
| ENSDARG000 | 4;4;4;4;4;4;4; | 31943831:319 | 31944534:319 | -;-;-;-;-;-;-;-; | 4304 | NA | 389 | 287 | 170 | 108 | 349 | 391 |
| ENSDARG000 | 4;4;4;4;4;4;4; | 22658472:226 | 22658932:226 | -;-;-;-;-;-;-;-; | 1880 | cd36 | 0 | 6 | 141 | 10 | 18 | 30 |
| ENSDARG000 | 4;4;4;4;4;4;4 | 31949672:319 | 31949720:319 | +;+;+;+;+;+;+ | 756 | NA | 3 | 2 | 3 | 0 | 5 | 0 |
| ENSDARG000 | 4;4;4 | 42031217:420 | 42032442:420 | -;-;- | 1444 | NA | 37 | 12 | 17 | 0 | 43 | 37 |
| ENSDARG000 | 4;4 | 3980247:3997 | 3981298:3998 | +;+ | 1883 | gpr37b | 0 | 20 | 0 | 1 | 6 | 26 |
| ENSDARG000 | 4;4;4;4;4;4 | 1788748:1791 | 1790063:1791 | -;-;-;-;-;- | 1981 | nudt4b | 38 | 98 | 116 | 56 | 48 | 54 |
| ENSDARG000 | 4;4;4;4;4 | 51971787:519 | 51971935:519 | +;+;+;+;+ | 1018 | NA | 0 | 0 | 0 | 0 | 0 | 0 |
| ENSDARG000 | 4;4;4;4;4;4;4; | 76440462:764 | 76440511:764 | +;+;+;+;+;+;+; | 5132 | NA | 40 | 16 | 19 | 0 | 32 | 42 |
| ENSDARG000 | 4;4;4;4 | 16315692:163 | 16315888:163 | +;+;+;+ | 1196 | si:dkeyp-38g8 | 4 | 16 | 13 | 6 | 2 | 7 |
| ENSDARG000 | 4;4;4;4;4;4;4 | 76455044:764 | 76455105:764 | +;+;+;+;+;+;+ | 3294 | zgc:113119 | 148 | 52 | 56 | 14 | 121 | 110 |
| ENSDARG000 | 4;4;4;4;4;4;4; | 52031199:520 | 52031242:520 | +;+;+;+;+;+;+; | 3134 | NA | 6 | 0 | 2 | 1 | 0 | 0 |
| ENSDARG000 | 4;4;4;4 | 1809562:1813 | 1810326:1813 | -;-;-;- | 1288 | ube2nb | 1034 | 1613 | 1397 | 1222 | 1366 | 1273 |
| ENSDARG000 | 4;4;4;4;4;4;4; | 52050410:520 | 52052598:520 | -;-;-;-;-;-;-;-; | 6503 | NA | 2 | 18 | 2 | 23 | 35 | 64 |
| ENSDARG000 | 4;4;4 | 76466751:764 | 76466785:764 | +;+;+ | 1475 | zgc:153116 | 35 | 28 | 10 | 2 | 57 | 44 |
| ENSDARG000 | 4;4;4;4;4;4;4; | 76468839:764 | 76468866:764 | -;-;-;-;-;-;-;-; | 3769 | NA | 158 | 74 | 84 | 20 | 211 | 160 |
| ENSDARG000 | 4;4;4;4;4;4;4; | 4079418:4088 | 4079776:4088 | +;+;+;+;+;+;+; | 2805 | waslb | 464 | 239 | 336 | 322 | 376 | 270 |
| ENSDARG000 | 4;4 | 6643421:6644 | 6643563:6647 | +;+ | 3035 | gpr85 | 56 | 38 | 292 | 236 | 124 | 156 |
| ENSDARG000 | 4;4;4 | 6678443:6687 | 6678489:6687 | +;+;+ | 479 | NA | 0 | 0 | 0 | 0 | 0 | 0 |
| ENSDARG000 | 4;4;4 | 42065748:420 | 42067434:420 | -;-;- | 1870 | zgc:174696 | 0 | 0 | 0 | 2 | 10 | 20 |
| ENSDARG000 | 4;4;4;4;4;4;4; | 25651720:256 | 25651786:256 | +;+;+;+;+;+;+; | 1950 | NA | 0 | 0 | 5 | 2 | 0 | 0 |
| ENSDARG000 | 4;4;4;4;4 | 42078825:420 | 42078974:420 | +;+;+;+;+ | 2439 | NA | 4 | 6 | 17 | 0 | 0 | 2 |
| ENSDARG000 | 4;4 | 32042094:320 | 32042174:320 | -;- | 311 | NA | 0 | 0 | 0 | 0 | 0 | 0 |
| ENSDARG000 | 4;4;4 | 20063279:200 | 20063382:200 | +;+;+ | 2870 | gcc1 | 530 | 417 | 515 | 305 | 597 | 457 |
| ENSDARG000 | 4;4;4;4 | 20051478:200 | 20051522:200 | +;+;+;+ | 979 | lamtor4 | 502 | 618 | 452 | 469 | 443 | 647 |
| ENSDARG000 | 4;4 | 42140245:421 | 42140325:421 | -;- | 659 | NA | 0 | 0 | 0 | 0 | 0 | 0 |
| ENSDARG000 | 4;4;4;4;4 | 20049260:200 | 20049794:200 | -;-;-;-;- | 926 | atp6v1f | 1356 | 1198 | 836 | 1155 | 1026 | 1117 |
| ENSDARG000 | 4;4;4;4;4;4 | 42152636:421 | 42152850:421 | -;-;-;-;-;- | 1918 | NA | 3 | 7 | 0 | 2 | 12 | 4 |
| ENSDARG000 | 4;4;4 | 42175261:421 | 42175326:421 | +;+;+ | 1491 | NA | 0 | 0 | 0 | 0 | 0 | 0 |
| ENSDARG000 | 4;4;4;4;4;4;4; | 6735053:6736 | 6735113:6737 | +;+;+;+;+;+;+; | 2251 | tmem168a | 708 | 483 | 1120 | 590 | 982 | 768 |
| ENSDARG000 | 4;4;4;4;4;4;4; | 13586455:135 | 13586971:135 | +;+;+;+;+;+;+; | 4193 | tnpo3 | 1208 | 1539 | 1703 | 835 | 1505 | 1360 |
| ENSDARG000 | 4;4;4;4;4 | 17807837:178 | 17808489:178 | -;-;-;-;- | 1095 | spic | 0 | 0 | 0 | 0 | 0 | 0 |
| ENSDARG000 | 4;4;4;4;4;4 | 17801328:178 | 17802105:178 | -;-;-;-;-;- | 1234 | NA | 0 | 0 | 0 | 0 | 0 | 0 |
| ENSDARG000 | 4;4;4 | 17793013:178 | 17793067:178 | +;+;+ | 2108 | NA | 2 | 0 | 0 | 0 | 2 | 0 |
| ENSDARG000 | 4;4 | 17736443:177 | 17737010:177 | -;- | 607 | NA | 0 | 0 | 0 | 0 | 0 | 0 |
| ENSDARG000 | 4;4;4;4;4 | 17821390:178 | 17822149:178 | -;-;-;-;- | 1033 | dnajb9b | 30 | 75 | 198 | 52 | 112 | 39 |
| ENSDARG000 | 4;4;4;4;4;4;4; | 32174137:321 | 32174308:321 | -;-;-;-;-;-;-;- | 2074 | zgc:113135 | 32 | 10 | 63 | 14 | 17 | 49 |
| ENSDARG000 | 4;4;4;4;4;4;4; | 17726841:177 | 17727476:177 | -;-;-;-;-;-;-;-; | 4273 | mybpc1 | 16 | 3 | 0 | 7 | 6 | 0 |
| ENSDARG000 | 4;4;4;4;4;4;4; | 32384950:323 | 32384998:323 | +;+;+;+;+;+;+; | 1156 | NA | 0 | 0 | 2 | 0 | 0 | 5 |
| ENSDARG000 | 4;4;4;4 | 52116776:521 | 52117899:521 | -;-;-;- | 1509 | NA | 0 | 2 | 0 | 0 | 4 | 9 |
| ENSDARG000 | 4;4;4;4;4;4;4; | 12243698:122 | 12245678:122 | -;-;-;-;-;-;-;-; | 4501 | cnot4b | 945 | 908 | 1139 | 780 | 900 | 1048 |
| ENSDARG000 | 4;4;4;4;4;4;4; | 9011448:9011 | 9012314:9011 | +;+;+;+;+;+;+; | 3671 | samm50l | 518 | 688 | 512 | 616 | 508 | 598 |
| ENSDARG000 | 4;4;4;4;4;4;4; | 20071128:200 | 20072570:200 | -;-;-;-;-;-;-;-; | 3123 | dennd6b | 92 | 100 | 160 | 98 | 97 | 96 |
| ENSDARG000 | 4;4;4;4;4 | 4115778:4117 | 4115804:4118 | -;-;-;-;- | 1839 | LMOD2 | 6 | 0 | 0 | 0 | 0 | 0 |
| ENSDARG000 | 4;4;4;4;4;4;4; | 42379152:423 | 42381861:423 | -;-;-;-;-;-;-;-; | 6735 | NA | 11 | 3 | 3 | 0 | 11 | 4 |
| ENSDARG000 | 4;4;4 | 42288973:422 | 42289813:422 | -;-;- | 1069 | NA | 0 | 0 | 1 | 0 | 0 | 0 |
| ENSDARG000 | 4;4;4;4;4;4;4; | 11375894:113 | 11376142:113 | +;+;+;+;+;+;+; | 19060 | NA | 196 | 200 | 71 | 56 | 71 | 38 |
| ENSDARG000 | 4;4;4;4;4;4;4; | 42195992:421 | 42200249:422 | -;-;-;-;-;-;-;-; | 9570 | NA | 62 | 11 | 35 | 4 | 15 | 29 |
| ENSDARG000 | 4;4;4;4;4 | 1821637:1821 | 1821834:1822 | -;-;-;-;- | 624 | mrpl42 | 210 | 218 | 327 | 250 | 226 | 299 |
| ENSDARG000 | 4;4;4;4;4;4;4; | 325028:32540 | 325259:32549 | -;-;-;-;-;-;-;-; | 1199 | mrps10 | 302 | 284 | 562 | 222 | 282 | 294 |
| ENSDARG000 | 4;4;4;4 | 52153452:521 | 52153819:521 | +;+;+;+ | 666 | NA | 0 | 0 | 0 | 0 | 0 | 0 |
| ENSDARG000 | 4;4;4;4;4 | 52158933:521 | 52159191:521 | -;-;-;-;- | 604 | NA | 0 | 0 | 0 | 0 | 0 | 0 |
| ENSDARG000 | 4;4;4 | 52148759:521 | 52149393:521 | -;-;- | 814 | NA | 0 | 0 | 1 | 0 | 4 | 0 |
| ENSDARG000 | 4;4;4;4;4;4;4; | 52162176:521 | 52162420:521 | +;+;+;+;+;+;+; | 1045 | NA | 0 | 2 | 10 | 0 | 0 | 1 |
| ENSDARG000 | 4;4;4;4;4;4;4; | 42338178:423 | 42338318:423 | +;+;+;+;+;+;+; | 496 | NA | 0 | 0 | 0 | 0 | 4 | 0 |
| ENSDARG000 | 4;4;4;4;4;4;4; | 11401018:114 | 11401290:114 | -;-;-;-;-;-;-;-; | 1387 | NA | 0 | 0 | 0 | 0 | 0 | 0 |
| ENSDARG000 | 4;4;4;4;4;4 | 42295479:422 | 42295532:422 | +;+;+;+;+;+ | 1673 | NA | 0 | 0 | 2 | 0 | 4 | 8 |
| ENSDARG000 | 4;4;4;4;4 | 42334850:423 | 42334986:423 | -;-;-;-;- | 674 | NA | 0 | 0 | 4 | 0 | 0 | 2 |
| ENSDARG000 | 4;4;4;4;4;4;4; | 32467854:324 | 32469757:324 | -;-;-;-;-;-;-;- | 2143 | NA | 8 | 14 | 14 | 4 | 18 | 4 |
| ENSDARG000 | 4;4;4;4;4;4;4; | 32429579:324 | 32432040:324 | -;-;-;-;-;-;-;-; | 6852 | NA | 0 | 27 | 2 | 0 | 11 | 2 |
| ENSDARG000 | 4;4;4;4;4;4;4; | 4120570:4120 | 4120739:4120 | -;-;-;-;-;-;-;- | 1755 | asb15b | 0 | 0 | 0 | 0 | 2 | 0 |
| ENSDARG000 | 4;4;4;4;4;4;4; | 32429579:324 | 32432040:324 | -;-;-;-;-;-;-;-; | 6614 | NA | 31 | 35 | 15 | 2 | 28 | 6 |
| ENSDARG000 | 4;4;4;4;4;4;4; | 22680442:226 | 22680869:226 | +;+;+;+;+;+;+; | 2431 | gnai1 | 1912 | 2341 | 2212 | 2482 | 1293 | 1384 |
| ENSDARG000 | 4;4;4;4;4;4;4; | 9813753:9814 | 9814251:9814 | -;-;-;-;-;-;-;-; | 2414 | parp12b | 366 | 293 | 360 | 248 | 396 | 504 |
| ENSDARG000 | 4;4 | 52177103:521 | 52177258:521 | +;+ | 1342 | NA | 4 | 8 | 4 | 0 | 16 | 6 |
| ENSDARG000 | 4;4 | 52212057:522 | 52212198:522 | +;+ | 1815 | NA | 8 | 0 | 2 | 0 | 0 | 18 |
| ENSDARG000 | 4;4;4;4;4 | 52234156:522 | 52234195:522 | +;+;+;+;+ | 1795 | NA | 6 | 4 | 23 | 2 | 12 | 2 |
| ENSDARG000 | 4;4 | 52339255:523 | 52339771:523 | +;+ | 632 | NA | 0 | 0 | 0 | 0 | 0 | 0 |
| ENSDARG000 | 4;4;4;4;4;4;4; | 9784301:9788 | 9788030:9788 | -;-;-;-;-;-;-;-; | 8078 | NA | 6 | 36 | 24 | 17 | 20 | 20 |
| ENSDARG000 | 4;4 | 4166344:4166 | 4166381:4167 | +;+ | 612 | NA | 0 | 0 | 0 | 0 | 0 | 0 |
| ENSDARG000 | 4;4;4;4;4;4;4; | 6799794:6800 | 6800218:6800 | -;-;-;-;-;-;-;-; | 2603 | ifrd1 | 674 | 920 | 998 | 647 | 1069 | 510 |
| ENSDARG000 | 4;4;4;4;4;4;4; | 1825839:1827 | 1827508:1827 | -;-;-;-;-;-;-;-; | 3440 | pwp1 | 649 | 628 | 899 | 457 | 531 | 600 |
| ENSDARG000 | 4;4;4;4;4;4;4; | 13605600:136 | 13606005:136 | -;-;-;-;-;-;-;-; | 2661 | irf5 | 78 | 4 | 22 | 1 | 74 | 55 |
| ENSDARG000 | 4;4;4;4 | 14826299:148 | 14826410:148 | +;+;+;+ | 776 | shisal1a | 0 | 0 | 0 | 0 | 0 | 0 |
| ENSDARG000 | 4;4;4;4;4;4;4; | 331351:33227 | 331520:33296 | +;+;+;+;+;+;+; | 6019 | tulp4a | 1993 | 1091 | 1916 | 1333 | 1885 | 1426 |
| ENSDARG000 | 4;4;4;4;4;4;4; | 20082475:200 | 20082852:200 | +;+;+;+;+;+;+; | 6653 | ppp6r2a | 1605 | 1373 | 1861 | 1214 | 2169 | 1760 |
| ENSDARG000 | 4;4 | 14895506:148 | 14896767:148 | -;- | 1296 | si:dkey-14k9.: | 104 | 83 | 106 | 58 | 111 | 127 |
| ENSDARG000 | 4;4;4;4;4;4;4; | 52347226:523 | 52347485:523 | +;+;+;+;+;+;+; | 1065 | NA | 0 | 2 | 0 | 0 | 0 | 10 |
| ENSDARG000 | 4;4;4;4 | 4224406:4224 | 4224654:4224 | -;-;-;- | 616 | NA | 10 | 3 | 2 | 2 | 7 | 6 |
| ENSDARG000 | 4;4;4;4;4;4;4; | 25678113:256 | 25678184:256 | +;+;+;+;+;+;+; | 3122 | acot17 | 2 | 0 | 2 | 0 | 6 | 2 |
| ENSDARG000 | 4;4;4;4;4 | 5304208:5307 | 5305107:5307 | -;-;-;-;- | 3165 | NA | 0 | 0 | 8 | 4 | 12 | 0 |
| ENSDARG000 | 4;4;4;4;4;4;4; | 5317483:5318 | 5317613:5319 | +;+;+;+;+;+;+; | 1819 | si:ch211-214i | 1783 | 2021 | 3695 | 3731 | 2271 | 2376 |
| ENSDARG000 | 4;4 | 52379250:523 | 52379604:523 | -;- | 384 | NA | 0 | 0 | 0 | 0 | 0 | 0 |
| ENSDARG000 | 4;4 | 52356485:523 | 52356634:523 | +;+ | 1633 | zgc:173705 | 18 | 5 | 9 | 12 | 8 | 11 |
| ENSDARG000 | 4;4 | 52504406:525 | 52504751:525 | -;- | 455 | NA | 0 | 0 | 0 | 0 | 4 | 0 |
| ENSDARG000 | 4;4 | 52511697:525 | 52512370:525 | +;+ | 853 | NA | 0 | 0 | 4 | 0 | 2 | 0 |
| ENSDARG000 | 4;4;4;4;4 | 52614984:526 | 52616165:526 | -;-;-;-;- | 1357 | NA | 10 | 13 | 10 | 0 | 2 | 6 |
| ENSDARG000 | 4 | 52671243 | 52672302 | - | 1060 | NA | 0 | 0 | 0 | 0 | 0 | 0 |
| ENSDARG000 | 4;4;4;4;4;4;4; | 32772213:327 | 32772295:327 | +;+;+;+;+;+;+; | 3738 | NA | 85 | 122 | 54 | 46 | 207 | 212 |
| ENSDARG000 | 4 | 52700533 | 52701516 | + | 984 | NA | 0 | 0 | 0 | 0 | 0 | 0 |
| ENSDARG000 | 4;4;4;4;4;4;4; | 32649955:326 | 32650104:326 | +;+;+;+;+;+;+; | 4967 | NA | 15 | 15 | 23 | 22 | 14 | 20 |
| ENSDARG000 | 4;4;4 | 32716854:327 | 32717202:327 | -;-;- | 592 | NA | 10 | 2 | 0 | 0 | 0 | 0 |
| ENSDARG000 | 4;4;4;4;4;4;4; | 9028819:9030 | 9029007:9030 | +;+;+;+;+;+;+; | 3075 | ALDH1L2 | 163 | 465 | 1013 | 351 | 283 | 339 |
| ENSDARG000 | 4;4;4;4;4 | 42403742:424 | 42404490:424 | -;-;-;-;- | 1354 | NA | 2 | 5 | 2 | 0 | 0 | 1 |
| ENSDARG000 | 4;4;4;4;4;4;4; | 14899720:149 | 14900177:149 | +;+;+;+;+;+;+; | 5581 | akr1b1.1 | 910 | 514 | 988 | 149 | 1110 | 2856 |
| ENSDARG000 | 4;4;4;4;4;4;4; | 4234994:4235 | 4235387:4235 | -;-;-;-;-;-;-;-; | 1944 | cd9b | 284 | 128 | 106 | 56 | 262 | 364 |
| ENSDARG000 | 4;4;4;4;4;4;4; | 12292274:122 | 12292371:122 | +;+;+;+;+;+;+; | 3473 | mkrn1 | 523 | 354 | 591 | 434 | 452 | 545 |
| ENSDARG000 | 4;4;4;4;4;4;4; | 42412212:424 | 42412398:424 | +;+;+;+;+;+;+; | 4006 | NA | 82 | 40 | 78 | 166 | 545 | 788 |
| ENSDARG000 | 4;4;4;4;4;4;4; | 14906608:149 | 14907207:149 | -;-;-;-;-;-;-;-; | 1592 | akr1b1.2 | 339 | 404 | 508 | 407 | 335 | 434 |
| ENSDARG000 | 4 | 12308976 | 12312976 | + | 4001 | TMEM121B | 4 | 2 | 2 | 4 | 6 | 2 |
| ENSDARG000 | 4;4;4 | 42556555:425 | 42556633:425 | +;+;+ | 1605 | NA | 3 | 10 | 8 | 0 | 10 | 6 |
| ENSDARG000 | 4;4;4 | 33012407:330 | 33012554:330 | +;+;+ | 664 | NA | 0 | 0 | 0 | 0 | 0 | 0 |
| ENSDARG000 | 4;4 | 33017081:330 | 33017209:330 | +;+ | 368 | NA | 0 | 0 | 0 | 0 | 0 | 0 |
| ENSDARG000 | 4;4;4;4;4;4;4; | 22730795:227 | 22731007:227 | -;-;-;-;-;-;-;-; | 2953 | nup107 | 668 | 1099 | 1412 | 642 | 1111 | 750 |
| ENSDARG000 | 4;4;4;4;4;4;4; | 357810:35781 | 358094:35809 | +;+;+;+;+;+;+; | 3399 | tmem181 | 1369 | 1201 | 1407 | 1422 | 1342 | 1424 |
| ENSDARG000 | 4;4;4;4;4;4;4 | 76705830:767 | 76705878:767 | +;+;+;+;+;+;+ | 998 | ms4a17a.7 | 31 | 2 | 2 | 2 | 15 | 10 |
| ENSDARG000 | 4;4;4;4;4;4;4 | 76722754:767 | 76722798:767 | +;+;+;+;+;+;+ | 1316 | ms4a17a.3 | 40 | 0 | 0 | 0 | 10 | 12 |
| ENSDARG000 | 4;4;4;4;4;4;4; | 33057844:330 | 33058112:330 | -;-;-;-;-;-;-;-; | 3925 | NA | 4 | 4 | 28 | 0 | 6 | 4 |
| ENSDARG000 | 4;4;4;4;4;4;4; | 13615731:136 | 13616305:136 | +;+;+;+;+;+;+; | 3626 | ccdc87 | 4 | 0 | 20 | 2 | 10 | 10 |
| ENSDARG000 | 4;4;4;4;4;4;4; | 76735113:767 | 76735153:767 | +;+;+;+;+;+;+; | 1035 | ms4a17a.6 | 58 | 12 | 22 | 14 | 82 | 130 |
| ENSDARG000 | 4;4;4;4;4;4;4; | 5327632:5327 | 5327763:5327 | -;-;-;-;-;-;-;-; | 1634 | osgep | 267 | 280 | 367 | 241 | 259 | 318 |
| ENSDARG000 | 4;4;4;4;4;4;4; | 6833583:6833 | 6833777:6833 | +;+;+;+;+;+;+; | 10697 | dock4b | 614 | 425 | 1095 | 428 | 1194 | 386 |
| ENSDARG000 | 4;4;4;4;4;4;4; | 76745015:767 | 76745054:767 | +;+;+;+;+;+;+; | 1054 | ms4a17a.10 | 50 | 35 | 38 | 12 | 252 | 100 |
| ENSDARG000 | 4;4;4;4;4;4;4; | 9824063:9824 | 9824481:9824 | -;-;-;-;-;-;-;-; | 3581 | NA | 0 | 34 | 36 | 18 | 4 | 13 |
| ENSDARG000 | 4;4;4;4;4;4;4; | 52739681:527 | 52739787:527 | -;-;-;-;-;-;-;-; | 1380 | NA | 12 | 0 | 0 | 20 | 2 | 10 |
| ENSDARG000 | 4;4;4;4;4;4;4; | 20101992:201 | 20103502:201 | -;-;-;-;-;-;-;-; | 2193 | fam3c | 786 | 722 | 797 | 632 | 969 | 853 |
| ENSDARG000 | 4;4;4;4;4;4;4; | 76755294:767 | 76755467:767 | +;+;+;+;+;+;+; | 2516 | ms4a17a.12 | 161 | 0 | 0 | 0 | 12 | 2 |
| ENSDARG000 | 4;4 | 6903549:6903 | 6903730:6904 | +;+ | 639 | NA | 0 | 0 | 0 | 0 | 0 | 0 |
| ENSDARG000 | 4;4;4;4;4;4;4 | 76752707:767 | 76752750:767 | +;+;+;+;+;+;+ | 1717 | ms4a17a.4 | 76 | 4 | 0 | 12 | 0 | 4 |
| ENSDARG000 | 4;4;4;4;4;4;4; | 52766525:527 | 52767094:527 | -;-;-;-;-;-;-;-; | 4082 | NA | 24 | 8 | 10 | 2 | 26 | 20 |
| ENSDARG000 | 4;4;4;4;4 | 16349042:163 | 16352450:163 | -;-;-;-;- | 4670 | lum | 218 | 16 | 27 | 2 | 239 | 78 |
| ENSDARG000 | 4;4 | 1844336:1844 | 1844380:1845 | +;+ | 419 | NA | 0 | 0 | 0 | 0 | 0 | 2 |
| ENSDARG000 | 4;4;4;4;4;4;4; | 12317461:123 | 12319278:123 | -;-;-;-;-;-;-;-; | 3069 | il17ra1a | 30 | 6 | 16 | 0 | 32 | 10 |
| ENSDARG000 | 4;4 | 33095657:331 | 33095806:331 | +;+ | 1620 | NA | 12 | 27 | 0 | 6 | 24 | 25 |
| ENSDARG000 | 4;4;4;4;4;4;4; | 76775837:767 | 76775881:767 | +;+;+;+;+;+;+; | 1699 | si:ch73-56d1: | 91 | 24 | 32 | 64 | 19 | 12 |
| ENSDARG000 | 4;4;4 | 33139862:331 | 33139955:331 | +;+;+ | 348 | NA | 14 | 2 | 6 | 0 | 4 | 0 |
| ENSDARG000 | 4;4;4;4;4;4;4; | 20110921:201 | 20111895:201 | -;-;-;-;-;-;-;-; | 2909 | NA | 0 | 0 | 0 | 8 | 4 | 0 |
| ENSDARG000 | 4;4;4;4;4;4;4; | 11458078:114 | 11458572:114 | +;+;+;+;+;+;+; | 1589 | ankrd16 | 62 | 48 | 67 | 80 | 47 | 117 |
| ENSDARG000 | 4;4;4;4;4;4;4; | 13625892:136 | 13625987:136 | +;+;+;+;+;+;+; | 3574 | PNPLA8 | 219 | 214 | 270 | 180 | 256 | 147 |
| ENSDARG000 | 4;4;4;4;4;4;4; | 386435:38714 | 387213:38721 | -;-;-;-;-;-;-;-; | 1065 | dynlt1b | 91 | 41 | 75 | 31 | 51 | 81 |
| ENSDARG000 | 4 | 13625982 | 13626831 | - | 850 | NA | 0 | 0 | 0 | 0 | 6 | 0 |
| ENSDARG000 | 4;4;4;4;4;4;4; | 76787274:767 | 76787341:767 | +;+;+;+;+;+;+; | 819 | NA | 50 | 0 | 0 | 7 | 0 | 6 |
| ENSDARG000 | 4;4;4;4;4;4 | 33179354:331 | 33179394:331 | -;-;-;-;-;- | 1488 | NA | 0 | 0 | 0 | 0 | 0 | 3 |
| ENSDARG000 | 4;4;4 | 33191390:331 | 33191536:331 | +;+;+ | 3147 | NA | 4 | 4 | 16 | 4 | 12 | 10 |
| ENSDARG000 | 4;4;4 | 43060711:430 | 43062016:430 | -;-;- | 1544 | NA | 0 | 0 | 1 | 0 | 0 | 10 |
| ENSDARG000 | 4;4 | 33225161:332 | 33225295:332 | +;+ | 1980 | NA | 30 | 17 | 22 | 28 | 18 | 16 |
| ENSDARG000 | 4;4;4;4;4 | 43048527:430 | 43050143:430 | -;-;-;-;- | 1947 | NA | 6 | 4 | 8 | 0 | 34 | 26 |
| ENSDARG000 | 4;4;4 | 33310760:333 | 33312119:333 | -;-;- | 1602 | NA | 0 | 0 | 0 | 0 | 0 | 0 |
| ENSDARG000 | 4;4;4;4;4;4;4; | 16368762:163 | 16370126:163 | -;-;-;-;-;-;-;-; | 2569 | dcn | 382 | 38 | 70 | 26 | 348 | 162 |
| ENSDARG000 | 4;4;4;4;4;4;4; | 9851810:9851 | 9854565:9854 | -;-;-;-;-;-;-;-; | 6000 | fbxo18 | 458 | 437 | 492 | 380 | 447 | 494 |
| ENSDARG000 | 4;4;4;4;4;4;4; | 76838989:768 | 76839632:768 | +;+;+;+;+;+;+; | 9042 | NA | 164 | 3 | 23 | 4 | 70 | 63 |
| ENSDARG000 | 4 | 76863480 | 76863920 | + | 441 | NA | 8 | 0 | 0 | 0 | 9 | 20 |
| ENSDARG000 | 4;4;4;4;4 | 76850983:768 | 76851171:768 | +;+;+;+;+ | 1288 | NA | 46 | 13 | 35 | 6 | 57 | 15 |
| ENSDARG000 | 4;4;4;4;4 | 76839169:768 | 76839361:768 | +;+;+;+;+ | 1114 | NA | 0 | 0 | 0 | 0 | 0 | 0 |
| ENSDARG000 | 4;4;4 | 76847462:768 | 76847713:768 | +;+;+ | 534 | NA | 6 | 6 | 5 | 1 | 7 | 25 |
| ENSDARG000 | 4;4;4;4;4;4;4; | 52814643:528 | 52815166:528 | -;-;-;-;-;-;-;- | 3403 | NA | 78 | 20 | 41 | 6 | 12 | 28 |
| ENSDARG000 | 4;4;4;4;4;4;4; | 9843923:9845 | 9844376:9845 | -;-;-;-;-;-;-;-; | 1585 | glt8d2 | 16 | 24 | 91 | 40 | 38 | 17 |

| | | | | | | | | | | | | |
|---|---|---|---|---|---|---|---|---|---|---|---|---|
| ENSDARG000 | 4:4:4:4:4:4:4: | 14917502:149 | 14918018:149 | -:-:-:-:-:-:-:-: | 2763 | prdm4 | 411 | 174 | 269 | 207 | 203 | 295 |
| ENSDARG000 | 4;4;4 | 33334774:333 | 33334924:333 | +;+;+ | 2177 | NA | 2 | 4 | 0 | 0 | 0 | 2 |
| ENSDARG000 | 4:4:4:4:4:4:4: | 391297:39137 | 391409:39140 | +:+:+:+:+:+:+: | 1281 | rpl18a | 14297 | 13207 | 23074 | 20604 | 13995 | 18551 |
| ENSDARG000 | 4:4:4:4:4:4:4: | 9836465:9837 | 9836640:9837 | +:+:+:+:+:+:+: | 2764 | hsp90b1 | 8167 | 18454 | 34465 | 10826 | 15463 | 6878 |
| ENSDARG000 | 4;4;4 | 33364466:333 | 33365080:333 | -;-;- | 799 | NA | 6 | 11 | 19 | 4 | 0 | 6 |
| ENSDARG000 | 4:4:4:4:4:4:4: | 76902513:769 | 76902649:769 | +:+:+:+:+:+:+: | 2587 | NA | 21 | 2 | 12 | 0 | 15 | 18 |
| ENSDARG000 | 4;4;4;4 | 9051588:9052 | 9052449:9052 | -;-;-;- | 1228 | nopchap1 | 103 | 169 | 165 | 83 | 152 | 112 |
| ENSDARG000 | 4;4 | 76918058:769 | 76918104:769 | +;+ | 544 | NA | 14 | 6 | 19 | 0 | 0 | 6 |
| ENSDARG000 | 4:4:4:4:4:4:4: | 25691642:256 | 25691686:256 | +:+:+:+:+:+:+: | 2468 | acot18 | 86 | 176 | 184 | 250 | 207 | 145 |
| ENSDARG000 | 4;4 | 52956204:529 | 52956349:529 | -;- | 193 | NA | 0 | 0 | 0 | 0 | 0 | 0 |
| ENSDARG000 | 4;4 | 52955173:529 | 52955404:529 | +;+ | 832 | NA | 0 | 0 | 0 | 0 | 0 | 0 |
| ENSDARG000 | 4;4;4 | 52959625:529 | 52959701:529 | +;+;+ | 1616 | NA | 2 | 16 | 6 | 4 | 2 | 19 |
| ENSDARG000 | 4:4:4:4:4:4:4: | 76926059:769 | 76926504:769 | +:+:+:+:+:+:+: | 7825 | NA | 964 | 18 | 10 | 3 | 742 | 865 |
| ENSDARG000 | 4;4;4 | 33373100:333 | 33373181:333 | +;+;+ | 1247 | NA | 0 | 0 | 0 | 0 | 0 | 0 |
| ENSDARG000 | 4:4:4:4:4:4:4: | 12358822:123 | 12358869:123 | +:+:+:+:+:+:+: | 916 | NA | 0 | 0 | 0 | 0 | 1 | 0 |
| ENSDARG000 | 4:4:4:4:4:4:4: | 20124863:201 | 20125948:201 | -:-:-:-:-:-:-:-: | 4116 | CEP83 | 282 | 205 | 266 | 168 | 200 | 190 |
| ENSDARG000 | 4:4:4:4:4:4:4: | 1878573:1880 | 1880380:1880 | -:-:-:-:-:-:-:-: | 4340 | ANO6 | 54 | 50 | 141 | 52 | 121 | 92 |
| ENSDARG000 | 4:4:4:4:4:4:4: | 12348568:123 | 12348608:123 | +:+:+:+:+:+:+: | 1062 | NA | 0 | 0 | 0 | 2 | 0 | 0 |
| ENSDARG000 | 4:4:4:4:4:4:4: | 12333168:123 | 12333285:123 | +:+:+:+:+:+:+: | 1502 | NA | 0 | 0 | 0 | 0 | 1 | 0 |
| ENSDARG000 | 4;4;4 | 33399196:334 | 33400311:334 | -;-;- | 1360 | NA | 4 | 8 | 10 | 0 | 0 | 14 |
| ENSDARG000 | 4;4;4;4;4 | 33420247:334 | 33420389:334 | +;+;+;+;+ | 2459 | NA | 40 | 9 | 31 | 0 | 4 | 0 |
| ENSDARG000 | 4;4;4;4;4;4 | 12333236:123 | 12333285:123 | +;+;+;+;+;+ | 911 | NA | 0 | 0 | 2 | 2 | 0 | 0 |
| ENSDARG000 | 4;4;4 | 43074375:430 | 43074521:430 | +;+;+ | 3238 | NA | 2 | 2 | 4 | 0 | 14 | 8 |
| ENSDARG000 | 4;4 | 17836098:178 | 17836162:178 | -;- | 927 | avpr2l | 0 | 0 | 0 | 0 | 0 | 0 |
| ENSDARG000 | 4;4;4;4;4;4 | 33461796:334 | 33461835:334 | +;+;+;+;+;+ | 1822 | NA | 0 | 6 | 0 | 2 | 0 | 0 |
| ENSDARG000 | 4:4:4:4:4:4:4: | 4267451:4270 | 4267528:4270 | +:+:+:+:+:+:+: | 6895 | ano2b | 4 | 9 | 6 | 0 | 2 | 14 |
| ENSDARG000 | 4;4;4 | 43156473:431 | 43157658:431 | -;-;- | 1428 | NA | 26 | 10 | 24 | 0 | 28 | 3 |
| ENSDARG000 | 4;4;4 | 4321168:4327 | 4321232:4328 | -;-;- | 331 | NA | 0 | 0 | 0 | 4 | 0 | 0 |
| ENSDARG000 | 4:4:4:4:4:4:4: | 52991210:529 | 52991270:529 | -:-:-:-:-:-:-:-: | 2366 | NA | 100 | 85 | 419 | 208 | 155 | 77 |
| ENSDARG000 | 4:4:4:4:4:4:4: | 9055488:9057 | 9055570:9058 | +:+:+:+:+:+:+: | 2275 | SLC41A2 | 436 | 340 | 504 | 370 | 504 | 543 |
| ENSDARG000 | 4 | 43192407 | 43193184 | - | 778 | NA | 0 | 0 | 0 | 0 | 0 | 0 |
| ENSDARG000 | 4:4:4:4:4:4:4: | 53034573:530 | 53035178:530 | -:-:-:-:-:-:-:-: | 5341 | NA | 13 | 65 | 19 | 10 | 46 | 54 |
| ENSDARG000 | 4:4:4:4:4:4:4: | 1921011:1921 | 1921042:1921 | -;-;-;-;-;-;-;- | 1728 | NA | 0 | 0 | 0 | 0 | 0 | 1 |
| ENSDARG000 | 4:4:4:4:4:4:4: | 43225728:432 | 43225795:432 | +:+:+:+:+:+:+: | 4997 | NA | 18 | 7 | 30 | 2 | 9 | 3 |
| ENSDARG000 | 4;4;4;4 | 20141191:201 | 20143181:201 | -;-;-;- | 3097 | lrrc17 | 334 | 18 | 28 | 12 | 108 | 48 |
| ENSDARG000 | 4;4;4;4;4 | 76988215:769 | 76989606:769 | -;-;-;-;- | 1827 | NA | 16 | 0 | 0 | 0 | 22 | 42 |
| ENSDARG000 | 4:4:4:4:4:4:4: | 76973771:769 | 76973960:769 | +:+:+:+:+:+:+: | 3465 | NA | 52 | 8 | 15 | 3 | 44 | 78 |
| ENSDARG000 | 4:4:4:4:4:4:4: | 43245188:432 | 43245267:432 | +:+:+:+:+:+:+: | 3840 | NA | 4 | 1 | 4 | 1 | 9 | 2 |
| ENSDARG000 | 4;4 | 33524237:335 | 33524386:335 | -;- | 201 | NA | 0 | 0 | 0 | 0 | 0 | 0 |
| ENSDARG000 | 4;4 | 33525559:335 | 33525600:335 | +;+ | 192 | NA | 0 | 0 | 0 | 0 | 0 | 0 |
| ENSDARG000 | 4:4:4:4:4:4:4: | 1937522:1937 | 1937767:1938 | +:+:+:+:+:+:+: | 1181 | NA | 0 | 0 | 0 | 0 | 0 | 0 |
| ENSDARG000 | 4:4:4:4:4:4:4: | 33537372:335 | 33537482:335 | +:+:+:+:+:+:+: | 5144 | NA | 22 | 9 | 16 | 12 | 6 | 10 |
| ENSDARG000 | 4;4 | 33537813:335 | 33537910:335 | +;+ | 377 | NA | 0 | 0 | 0 | 0 | 0 | 0 |
| ENSDARG000 | 4 | 7222080 | 7222367 | - | 288 | NA | 42 | 71 | 19 | 0 | 72 | 69 |
| ENSDARG000 | 4;4 | 16536173:165 | 16539431:165 | -;- | 3627 | btg1 | 7244 | 6125 | 6320 | 8614 | 5011 | 3571 |
| ENSDARG000 | 4;4;4;4 | 7229236:7230 | 7229623:7230 | -;-;-;- | 1011 | NA | 0 | 0 | 0 | 0 | 1 | 1 |
| ENSDARG000 | 4:4:4:4:4:4:4: | 13696537:136 | 13696701:136 | +:+:+:+:+:+:+: | 4505 | nrcama | 4 | 15 | 1 | 16 | 9 | 4 |
| ENSDARG000 | 4:4:4:4:4:4:4: | 11464255:114 | 11464402:114 | +:+:+:+:+:+:+: | 2542 | gdi2 | 13165 | 12687 | 12698 | 12228 | 11635 | 10447 |
| ENSDARG000 | 4;4;4;4 | 400376:40052 | 400452:40087 | +;+;+;+ | 871 | NA | 2 | 5 | 3 | 0 | 0 | 0 |
| ENSDARG000 | 4:4:4:4:4:4:4: | 53095146:530 | 53095228:530 | +:+:+:+:+:+:+: | 3184 | NA | 42 | 29 | 22 | 7 | 26 | 53 |
| ENSDARG000 | 4;4;4;4 | 402626:41668 | 402712:41684 | +;+;+;+ | 530 | NA | 0 | 2 | 0 | 0 | 0 | 0 |
| ENSDARG000 | 4;4;4 | 77122707:771 | 77125160:771 | -;-;- | 2613 | NA | 0 | 0 | 0 | 0 | 0 | 0 |
| ENSDARG000 | 4:4:4:4:4:4:4: | 77160027:771 | 77160072:771 | +:+:+:+:+:+:+: | 2607 | NA | 24 | 12 | 8 | 26 | 44 | 47 |
| ENSDARG000 | 4;4;4 | 9103182:9128 | 9105263:9128 | -;-;- | 2432 | chst11 | 377 | 1268 | 3274 | 1533 | 2954 | 2271 |
| ENSDARG000 | 4:4:4:4:4:4:4: | 11479705:114 | 11479865:114 | +:+:+:+:+:+:+: | 1852 | asb13a.1 | 50 | 30 | 68 | 72 | 60 | 108 |
| ENSDARG000 | 4;4;4;4;4 | 77141970:771 | 77142141:771 | +;+;+;+;+ | 4688 | NA | 94 | 106 | 242 | 68 | 179 | 200 |
| ENSDARG000 | 4;4 | 77151419:771 | 77151688:771 | +;+ | 4138 | NA | 16 | 20 | 42 | 22 | 16 | 64 |
| ENSDARG000 | 4;4;4;4;4 | 77053467:770 | 77053740:770 | +;+;+;+;+ | 5083 | NA | 40 | 40 | 49 | 18 | 93 | 62 |
| ENSDARG000 | 4;4;4 | 77136884:771 | 77139502:771 | -;-;- | 2778 | NA | 0 | 0 | 0 | 0 | 0 | 0 |
| ENSDARG000 | 4;4 | 77047067:770 | 77047164:770 | +;+ | 2960 | NA | 122 | 43 | 38 | 2 | 112 | 131 |
| ENSDARG000 | 4:4:4:4:4:4:4: | 77038392:770 | 77038552:770 | +:+:+:+:+:+:+: | 5775 | NA | 20 | 0 | 9 | 0 | 6 | 24 |
| ENSDARG000 | 4:4:4:4:4:4:4: | 77060861:770 | 77060904:770 | +:+:+:+:+:+:+: | 1389 | NA | 16 | 11 | 29 | 14 | 33 | 28 |
| ENSDARG000 | 4;4;4;4;4;4 | 4376498:4377 | 4376907:4377 | +;+;+;+;+;+ | 780 | NA | 0 | 0 | 0 | 0 | 0 | 0 |
| ENSDARG000 | 4:4:4:4:4:4:4: | 17843717:178 | 17843870:178 | +:+:+:+:+:+:+: | 4934 | apaf1 | 255 | 339 | 345 | 265 | 218 | 198 |
| ENSDARG000 | 4;4;4 | 77118052:771 | 77120492:771 | -;-;- | 2600 | NA | 0 | 0 | 0 | 0 | 0 | 0 |
| ENSDARG000 | 4:4:4:4:4:4:4: | 53115165:531 | 53115218:531 | +:+:+:+:+:+:+: | 4266 | NA | 51 | 91 | 34 | 15 | 89 | 68 |
| ENSDARG000 | 4;4 | 77092909:771 | 77093189:771 | +;+ | 8189 | NA | 24 | 24 | 70 | 38 | 46 | 72 |
| ENSDARG000 | 4;4;4 | 77076645:770 | 77076699:770 | +;+;+ | 3942 | NA | 20 | 3 | 6 | 4 | 26 | 14 |
| ENSDARG000 | 4:4:4:4:4:4:4: | 77126495:771 | 77129758:771 | -:-:-:-:-:-:-:-: | 4145 | NA | 0 | 0 | 0 | 0 | 0 | 0 |
| ENSDARG000 | 4:4:4:4:4:4:4: | 77021784:770 | 77021838:770 | +:+:+:+:+:+:+: | 6063 | NA | 8 | 0 | 0 | 4 | 0 | 10 |
| ENSDARG000 | 4:4:4:4:4:4:4: | 20198294:201 | 20198394:201 | -:-:-:-:-:-:-:-: | 2521 | GSAP | 189 | 96 | 264 | 300 | 210 | 136 |
| ENSDARG000 | 4:4:4:4:4:4:4: | 20177526:201 | 20177578:201 | +:+:+:+:+:+:+: | 3370 | ccdc146 | 8 | 3 | 8 | 5 | 42 | 28 |
| ENSDARG000 | 4:4:4:4:4:4:4: | 20224099:202 | 20224382:202 | -:-:-:-:-:-:-:-: | 1756 | stk38l | 16 | 10 | 33 | 8 | 31 | 50 |
| ENSDARG000 | 4;4;4 | 77132096:771 | 77134542:771 | -;-;- | 2606 | NA | 0 | 0 | 0 | 0 | 0 | 0 |
| ENSDARG000 | 4:4:4:4:4:4:4: | 7391110:7391 | 7391140:7391 | +:+:+:+:+:+:+: | 5524 | tnni4a | 457 | 421 | 833 | 1616 | 991 | 1046 |
| ENSDARG000 | 4:4:4:4:4:4:4: | 22755465:227 | 22758977:227 | -:-:-:-:-:-:-:-: | 7614 | magi2a | 606 | 491 | 678 | 428 | 820 | 226 |
| ENSDARG000 | 4;4;4;4;4;4 | 77113391:771 | 77115830:771 | -;-;-;-;-;- | 2599 | NA | 0 | 0 | 0 | 0 | 0 | 0 |
| ENSDARG000 | 4;4;4;4;4 | 43273634:432 | 43275232:432 | -;-;-;-;- | 1802 | NA | 0 | 8 | 3 | 0 | 0 | 6 |
| ENSDARG000 | 4:4:4:4:4:4:4: | 5333988:5333 | 5334089:5334 | +:+:+:+:+:+:+: | 3244 | apex1 | 904 | 956 | 2095 | 721 | 1100 | 719 |
| ENSDARG000 | 4;4;4 | 43295390:433 | 43297303:433 | -;-;- | 2143 | NA | 24 | 6 | 56 | 8 | 12 | 22 |
| ENSDARG000 | 4;4;4;4;4 | 43376293:433 | 43376391:433 | -;-;-;-;- | 645 | NA | 0 | 0 | 0 | 0 | 0 | 0 |
| ENSDARG000 | 4:4:4:4:4:4:4: | 14932537:149 | 14933778:149 | -:-:-:-:-:-:-:-: | 3803 | slc26a5 | 656 | 488 | 786 | 444 | 806 | 392 |
| ENSDARG000 | 4;4;4;4;4 | 43361785:433 | 43363004:433 | -;-;-;-;- | 1889 | NA | 2 | 0 | 4 | 12 | 18 | 74 |
| ENSDARG000 | 4;4;4;4;4 | 612363:61344 | 612571:61370 | +;+;+;+;+;+ | 2283 | pthlha | 4 | 0 | 0 | 0 | 0 | 0 |
| ENSDARG000 | 4;4 | 1950849:1951 | 1951091:1951 | +;+ | 564 | NA | 0 | 0 | 0 | 0 | 0 | 0 |
| ENSDARG000 | 4;4 | 20178694:201 | 20180541:201 | -;- | 2524 | fgl2a | 322 | 544 | 451 | 312 | 1121 | 709 |
| ENSDARG000 | 4:4:4:4:4:4:4: | 33574463:335 | 33574517:335 | +:+:+:+:+:+:+: | 6446 | NA | 5 | 111 | 2 | 40 | 140 | 41 |
| ENSDARG000 | 4:4:4:4:4:4:4: | 20176143:201 | 20176290:201 | -:-:-:-:-:-:-:-: | 609 | sinup | 6 | 3 | 4 | 32 | 49 | 31 |
| ENSDARG000 | 4;4 | 4397503:4431 | 4398438:4431 | -;- | 1070 | ntf3 | 106 | 66 | 44 | 47 | 74 | 101 |
| ENSDARG000 | 4;4;4 | 43392655:433 | 43392708:433 | +;+;+ | 1507 | NA | 0 | 0 | 0 | 0 | 12 | 6 |
| ENSDARG000 | 4;4;4 | 53159800:531 | 53161329:531 | -;-;- | 1721 | NA | 0 | 2 | 0 | 0 | 0 | 10 |
| ENSDARG000 | 4:4:4:4:4:4:4: | 9177997:9178 | 9178040:9178 | +:+:+:+:+:+:+: | 1122 | nfyba | 647 | 533 | 713 | 725 | 600 | 551 |
| ENSDARG000 | 4;4;4 | 53183373:531 | 53183438:531 | +;+;+ | 1817 | NA | 0 | 0 | 0 | 0 | 0 | 0 |
| ENSDARG000 | 4:4:4:4:4:4:4: | 14926948:149 | 14927072:149 | +:+:+:+:+:+:+: | 1456 | psmc2 | 2853 | 2607 | 2652 | 1765 | 2869 | 1850 |
| ENSDARG000 | 4:4:4:4:4:4:4: | 33720803:337 | 33723347:337 | -:-:-:-:-:-:-:-: | 5669 | NA | 3 | 0 | 17 | 0 | 6 | 6 |
| ENSDARG000 | 4:4:4:4:4:4:4: | 646154:64774 | 646234:64789 | +:+:+:+:+:+:+: | 3653 | xpot | 1361 | 2135 | 3343 | 1646 | 1855 | 1399 |
| ENSDARG000 | 4:4:4:4:4:4:4: | 1958240:1959 | 1958482:1960 | +:+:+:+:+:+:+: | 1286 | NA | 0 | 0 | 0 | 0 | 5 | 0 |
| ENSDARG000 | 4 | 33811110 | 33812019 | + | 910 | NA | 0 | 0 | 0 | 0 | 0 | 0 |
| ENSDARG000 | 4;4;4 | 43408004:434 | 43408095:434 | +;+;+ | 1655 | NA | 2 | 3 | 1 | 0 | 0 | 8 |
| ENSDARG000 | 4 | 43464313 | 43465197 | + | 885 | NA | 0 | 0 | 0 | 0 | 0 | 0 |
| ENSDARG000 | 4:4:4:4:4:4:4: | 16615026:166 | 16616271:166 | -:-:-:-:-:-:-:-: | 3626 | caprin2 | 2 | 0 | 0 | 0 | 0 | 4 |
| ENSDARG000 | 4:4:4:4:4:4:4: | 33862438:338 | 33862491:338 | +:+:+:+:+:+:+: | 3820 | NA | 40 | 134 | 71 | 10 | 126 | 123 |
| ENSDARG000 | 4:4:4:4:4:4:4: | 53246788:532 | 53246914:532 | +:+:+:+:+:+:+: | 6228 | NA | 23 | 7 | 23 | 2 | 5 | 23 |
| ENSDARG000 | 4:4:4:4:4:4:4: | 33818711:338 | 33818802:338 | +:+:+:+:+:+:+: | 3028 | NA | 6 | 19 | 8 | 14 | 26 | 26 |
| ENSDARG000 | 4;4;4 | 53257960:532 | 53258079:532 | -;-;- | 539 | NA | 0 | 0 | 0 | 0 | 0 | 0 |
| ENSDARG000 | 4;4;4 | 43632188:436 | 43633249:436 | -;-;- | 1266 | NA | 0 | 0 | 0 | 0 | 6 | 2 |
| ENSDARG000 | 4:4:4:4:4:4:4: | 43522680:435 | 43522709:435 | +:+:+:+:+:+:+: | 9127 | NA | 36 | 12 | 37 | 4 | 18 | 10 |
| ENSDARG000 | 4:4:4:4:4:4:4: | 1988460:1989 | 1989651:1989 | -:-:-:-:-:-:-:-: | 2657 | FAR2 | 172 | 82 | 118 | 238 | 157 | 133 |
| ENSDARG000 | 4:4:4:4:4:4:4: | 43695179:436 | 43695281:436 | +:+:+:+:+:+:+: | 3917 | NA | 6 | 1 | 6 | 8 | 91 | 85 |
| ENSDARG000 | 4:4:4:4:4:4:4: | 4484651:4488 | 4485968:4488 | -:-:-:-:-:-:-:-: | 3526 | TBC1D30 | 294 | 152 | 130 | 128 | 144 | 197 |
| ENSDARG000 | 4;4;4;4;4 | 43678985:436 | 43679050:436 | +;+;+;+;+ | 1905 | NA | 4 | 7 | 9 | 2 | 28 | 10 |
| ENSDARG000 | 4:4:4:4:4:4:4: | 43648421:436 | 43652282:436 | -:-:-:-:-:-:-:-: | 10841 | NA | 6 | 3 | 2 | 6 | 170 | 80 |
| ENSDARG000 | 4:4:4:4:4:4:4: | 43606820:436 | 43607387:436 | -:-:-:-:-:-:-:-: | 3330 | NA | 30 | 2 | 22 | 2 | 10 | 25 |
| ENSDARG000 | 4:4:4:4:4:4:4: | 34119824:341 | 34119921:341 | +:+:+:+:+:+:+: | 3868 | NA | 19 | 27 | 11 | 10 | 46 | 29 |
| ENSDARG000 | 4;4 | 34343524:343 | 34343596:343 | +;+ | 363 | NA | 0 | 0 | 0 | 0 | 0 | 0 |
| ENSDARG000 | 4;4;4;4;4 | 34359175:343 | 34359696:343 | -;-;-;-;- | 1472 | NA | 0 | 0 | 0 | 0 | 0 | 0 |
| ENSDARG000 | 4;4 | 23122744:231 | 23123165:231 | -;- | 484 | NA | 251 | 288 | 106 | 140 | 251 | 124 |
| ENSDARG000 | 4;4;4;4;4 | 23117557:231 | 23118159:231 | +;+;+;+;+ | 1354 | slc35e3 | 841 | 587 | 415 | 391 | 550 | 285 |
| ENSDARG000 | 4;4;4;4;4;4 | 53327149:533 | 53328683:533 | -;-;-;-;-;- | 3923 | NA | 6 | 6525 | 25 | 523 | 845 | 19949 |
| ENSDARG000 | 4:4:4:4:4:4:4: | 34369818:343 | 34369930:343 | +:+:+:+:+:+:+: | 5486 | NA | 25 | 18 | 15 | 4 | 12 | 8 |
| ENSDARG000 | 4;4;4 | 34389152:343 | 34389737:343 | -;-;- | 751 | NA | 0 | 0 | 0 | 0 | 0 | 0 |
| ENSDARG000 | 4:4:4:4:4:4:4: | 7415337:7415 | 7415437:7415 | +:+:+:+:+:+:+: | 1831 | cald1a | 165 | 1338 | 843 | 682 | 1438 | 801 |
| ENSDARG000 | 4:4:4:4:4:4:4: | 25704579:257 | 25704629:257 | +:+:+:+:+:+:+: | 5575 | lamb1b | 79213 | 41325 | 51325 | 53077 | 61438 | 55461 |
| ENSDARG000 | 4:4:4:4:4:4:4: | 12365775:123 | 12365815:123 | +:+:+:+:+:+:+: | 1206 | NA | 0 | 0 | 0 | 0 | 0 | 0 |
| ENSDARG000 | 4:4:4:4:4:4:4: | 53976731:539 | 53976815:539 | +:+:+:+:+:+:+: | 712 | NA | 2 | 0 | 10 | 0 | 0 | 0 |
| ENSDARG000 | 4:4:4:4:4:4:4: | 77207693:772 | 77207912:772 | -:-:-:-:-:-:-:-: | 2123 | psmb10 | 21 | 1419 | 1583 | 501 | 533 | 461 |
| ENSDARG000 | 4;4;4;4;4 | 77218965:772 | 77223015:772 | -;-;-;-;- | 4451 | NA | 44 | 38 | 47 | 6 | 65 | 64 |
| ENSDARG000 | 4;4;4;4;4;4 | 77229328:772 | 77231278:772 | -;-;-;-;-;- | 3236 | NA | 41 | 16 | 43 | 2 | 116 | 152 |
| ENSDARG000 | 4:4:4:4:4:4:4: | 9184481:9187 | 9184755:9187 | -:-:-:-:-:-:-:-: | 2630 | hcfc2 | 175 | 254 | 266 | 90 | 236 | 238 |
| ENSDARG000 | 4;4;4 | 43730608:437 | 43730777:437 | -;-;- | 518 | NA | 0 | 0 | 0 | 0 | 0 | 0 |
| ENSDARG000 | 4:4:4:4:4:4:4: | 43730606:437 | 43730777:437 | -:-:-:-:-:-:-:-: | 9001 | NA | 39 | 14 | 4 | 1 | 14 | 6 |
| ENSDARG000 | 4:4:4:4:4:4:4: | 13733838:137 | 13734038:137 | +:+:+:+:+:+:+: | 4786 | cntn1b | 0 | 0 | 2 | 0 | 0 | 0 |
| ENSDARG000 | 4;4;4 | 9880647:9891 | 9882162:9891 | -;-;- | 1824 | adm2a | 24 | 295 | 1099 | 704 | 90 | 42 |
| ENSDARG000 | 4:4:4:4:4:4:4: | 54268618:542 | 54269212:542 | -:-:-:-:-:-:-:-: | 1850 | NA | 0 | 0 | 0 | 0 | 2 | 0 |
| ENSDARG000 | 4:4:4:4:4:4:4: | 43904209:439 | 43904243:439 | -:-:-:-:-:-:-:-: | 2283 | NA | 48 | 16 | 19 | 20 | 46 | 71 |
| ENSDARG000 | 4;4;4;4;4;4 | 9901116:9904 | 9901914:9904 | -;-;-;-;-;- | 1419 | NA | 2 | 11 | 0 | 14 | 23 | 6 |
| ENSDARG000 | 4:4:4:4:4:4:4: | 11483335:114 | 11483529:114 | +:+:+:+:+:+:+: | 1598 | asb13a.2 | 20 | 8 | 12 | 4 | 32 | 24 |
| ENSDARG000 | 4:4:4:4:4:4:4: | 4509996:4510 | 4510343:4510 | +:+:+:+:+:+:+: | 2660 | gnsa | 1005 | 662 | 790 | 1246 | 911 | 927 |
| ENSDARG000 | 4:4:4:4:4:4:4: | 34443650:344 | 34444308:344 | -:-:-:-:-:-:-:-: | 5025 | NA | 6 | 3 | 14 | 2 | 9 | 12 |
| ENSDARG000 | 4;4 | 4514438:4516 | 4514456:4516 | -;- | 394 | NA | 0 | 0 | 0 | 0 | 0 | 0 |
| ENSDARG000 | 4:4:4:4:4:4:4: | 34553067:345 | 34553596:345 | -:-:-:-:-:-:-:-: | 3938 | NA | 135 | 398 | 136 | 48 | 256 | 308 |
| ENSDARG000 | 4:4:4:4:4:4:4: | 34599803:345 | 34599995:345 | +:+:+:+:+:+:+: | 1355 | NA | 0 | 0 | 0 | 0 | 0 | 0 |
| ENSDARG000 | 4:4:4:4:4:4:4: | 34468977:344 | 34469601:344 | -;-;-;-;-;-;-;- | 3227 | NA | 4 | 4 | 12 | 2 | 20 | 3 |
| ENSDARG000 | 4:4:4:4:4:4:4: | 16631483:166 | 16635800:166 | -:-:-:-:-:-:-:-: | 4852 | sinhcaf | 345 | 327 | 414 | 530 | 425 | 698 |
| ENSDARG000 | 4;4 | 34429101:344 | 34429694:344 | +;+ | 709 | NA | 0 | 0 | 1 | 0 | 0 | 0 |
| ENSDARG000 | 4;4;4;4;4 | 12381735:123 | 12383105:123 | -;-;-;-;- | 1875 | rergla | 9 | 4 | 0 | 0 | 2 | 0 |
| ENSDARG000 | 4:4:4:4:4:4:4: | 34753619:347 | 34754453:347 | -:-:-:-:-:-:-:-: | 3671 | NA | 10 | 3 | 14 | 0 | 12 | 9 |
| ENSDARG000 | 4:4:4:4:4:4:4: | 34417403:344 | 34417998:344 | +:+:+:+:+:+:+: | 10214 | NA | 38 | 12 | 14 | 4 | 17 | 17 |
| ENSDARG000 | 4:4:4:4:4:4:4: | 5341592:5341 | 5341788:5341 | +:+:+:+:+:+:+: | 8146 | zgc:113263 | 432 | 313 | 414 | 472 | 431 | 286 |
| ENSDARG000 | 4:4:4:4:4:4:4: | 5362909:5365 | 5367984:5365 | -:-:-:-:-:-:-:-: | 6373 | NA | 37 | 39 | 47 | 11 | 26 | 45 |
| ENSDARG000 | 4;4 | 681501:68297 | 681645:68346 | +;+ | 636 | NA | 0 | 0 | 0 | 0 | 0 | 0 |
| ENSDARG000 | 4:4:4:4:4:4:4: | 676710:67882 | 676947:67894 | -:-:-:-:-:-:-:-: | 1890 | rtcb | 766 | 792 | 1349 | 372 | 609 | 549 |
| ENSDARG000 | 4:4:4:4:4:4:4: | 2041067:2045 | 2045578:2045 | -:-:-:-:-:-:-:-: | 7197 | cpsf6 | 2374 | 2276 | 3551 | 2372 | 2637 | 2122 |
| ENSDARG000 | 4:4:4:4:4:4:4: | 17886838:178 | 17888207:178 | -:-:-:-:-:-:-:-: | 6628 | anks1b | 4514 | 3299 | 4829 | 3597 | 6237 | 5312 |
| ENSDARG000 | 4:4:4:4:4:4:4: | 23125689:231 | 23125787:231 | +:+:+:+:+:+:+: | 3894 | mdm2 | 4392 | 7637 | 1688 | 3009 | 6701 | 1584 |
| ENSDARG000 | 4:4:4:4:4:4:4: | 16649542:166 | 16651157:166 | -:-:-:-:-:-:-:-: | 5612 | dennd5b | 551 | 368 | 304 | 355 | 641 | 425 |
| ENSDARG000 | 4;4;4 | 44454270:444 | 44454447:444 | +;+;+ | 500 | NA | 0 | 1 | 0 | 0 | 0 | 0 |
| ENSDARG000 | 4:4:4:4:4:4:4: | 77349261:773 | 77349642:773 | -:-:-:-:-:-:-:-: | 2425 | NA | 0 | 0 | 0 | 0 | 0 | 0 |
| ENSDARG000 | 4:4:4:4:4:4:4: | 54316804:543 | 54321154:543 | -:-:-:-:-:-:-:-: | 8199 | NA | 19 | 11 | 9 | 28 | 10 | 43 |
| ENSDARG000 | 4:4:4:4:4:4:4: | 9911043:9911 | 9911725:9911 | +:+:+:+:+:+:+: | 11009 | sbf1 | 1933 | 2309 | 2816 | 2179 | 2783 | 2396 |
| ENSDARG000 | 4;4 | 44499731:445 | 44499904:445 | -;- | 348 | NA | 0 | 0 | 0 | 0 | 0 | 0 |
| ENSDARG000 | 4:4:4:4:4:4:4: | 44470531:444 | 44471239:444 | -:-:-:-:-:-:-:-: | 4732 | NA | 77 | 81 | 51 | 56 | 82 | 46 |

| | | | | | | | | | | | | |
|---|---|---|---|---|---|---|---|---|---|---|---|---|
| ENSDARG000 | 4:4:4:4:4:4:4: | 77389668:773 | 77390991:773 | -:-:-:-:-:-:-:-: | 3339 | slco1d1 | 0 | 2 | 6 | 0 | 6 | 10 |
| ENSDARG000 | 4:4:4:4:4:4:4: | 54355719:543 | 54356450:543 | -:-:-:-:-:-:-:-: | 3716 | NA | 28 | 21 | 14 | 0 | 28 | 37 |
| ENSDARG000 | 4;4;4;4 | 44514525:445 | 44514609:445 | -;-;-;- | 357 | NA | 0 | 0 | 0 | 0 | 0 | 0 |
| ENSDARG000 | 4;4 | 9207722:9212 | 9211129:9214 | -;- | 5174 | avpr1ab | 0 | 2 | 0 | 0 | 0 | 0 |
| ENSDARG000 | 4:4:4:4:4:4:4: | 77471853:774 | 77471929:774 | -;-;-;-;-;-;-;- | 4829 | NA | 33 | 6 | 29 | 13 | 12 | 9 |
| ENSDARG000 | 4 | 77709678 | 77712195 | + | 2518 | NA | 4 | 280 | 120 | 112 | 52 | 64 |
| ENSDARG000 | 4:4:4:4:4:4:4: | 11573568:115 | 11575267:115 | -:-:-:-:-:-:-:-: | 3759 | net1 | 2071 | 1367 | 2226 | 2767 | 2394 | 2279 |
| ENSDARG000 | 4;4;4;4;4;4;4 | 77681389:776 | 77681855:776 | +;+;+;+;+;+;+ | 3076 | gbp4 | 14 | 0 | 0 | 0 | 0 | 0 |
| ENSDARG000 | 4:4:4:4:4:4:4: | 9279515:9279 | 9280315:9280 | +:+:+:+:+:+:+: | 5199 | srgap1b | 12 | 6 | 2 | 0 | 4 | 14 |
| ENSDARG000 | 4:4:4:4:4:4:4: | 12401146:124 | 12401280:124 | -:-:-:-:-:-:-:-: | 2680 | arhgef39 | 36 | 172 | 165 | 66 | 146 | 86 |
| ENSDARG000 | 4:4:4:4:4:4:4: | 54480250:544 | 54480315:544 | +:+:+:+:+:+:+: | 3890 | si:ch211-227 | 3 | 9 | 15 | 16 | 7 | 11 |
| ENSDARG000 | 4:4:4:4:4:4:4: | 686157:68641 | 686183:68653 | +:+:+:+:+:+:+: | 1721 | fbxo7 | 1646 | 1311 | 787 | 677 | 761 | 904 |
| ENSDARG000 | 4:4:4:4:4:4:4: | 54434814:544 | 54434876:544 | +:+:+:+:+:+:+: | 4963 | NA | 19 | 9 | 25 | 4 | 1 | 6 |
| ENSDARG000 | 4;4;4;4;4;4;4 | 54461588:544 | 54461645:544 | +;+;+;+;+;+;+ | 1905 | NA | 1 | 0 | 3 | 0 | 0 | 0 |
| ENSDARG000 | 4;4;4;4 | 77620775:776 | 77622825:776 | -;-;-;- | 3018 | PTCHD3 | 0 | 0 | 12 | 0 | 36 | 0 |
| ENSDARG000 | 4;4;4;4 | 77675291:776 | 77676945:776 | -;-;-;- | 2359 | NA | 2 | 0 | 5 | 0 | 0 | 0 |
| ENSDARG000 | 4;4;4;4 | 77651736:776 | 77653571:776 | -;-;-;- | 2541 | hspa12a.3 | 0 | 0 | 5 | 0 | 1 | 0 |
| ENSDARG000 | 4:4:4:4:4:4:4: | 77657660:776 | 77657827:776 | +:+:+:+:+:+:+: | 2244 | si:dkey-61p9. | 4 | 4 | 0 | 0 | 2 | 0 |
| ENSDARG000 | 4;4;4;4;4;4;4 | 25754725:257 | 25756669:257 | -;-;-;-;-;-;- | 3507 | tmcc3 | 1840 | 1521 | 1324 | 1454 | 1990 | 1108 |
| ENSDARG000 | 4;4;4;4;4;4 | 20255160:202 | 20255445:202 | +;+;+;+;+;+ | 1501 | LRTM2 | 330 | 173 | 289 | 223 | 762 | 182 |
| ENSDARG000 | 4;4;4 | 34798625:347 | 34798944:347 | -;-;- | 585 | NA | 0 | 0 | 0 | 0 | 0 | 0 |
| ENSDARG000 | 4:4:4:4:4:4:4: | 77626530:776 | 77627184:776 | -:-:-:-:-:-:-:-: | 3410 | NA | 10 | 0 | 4 | 0 | 0 | 6 |
| ENSDARG000 | 4;4;4;4;4;4 | 77580532:775 | 77580704:775 | +;+;+;+;+;+ | 1582 | NA | 0 | 1 | 0 | 0 | 0 | 0 |
| ENSDARG000 | 4;4;4;4;4;4 | 77590387:775 | 77595945:775 | -;-;-;-;-;- | 6318 | fasn2 | 0 | 0 | 0 | 17 | 48 | 0 |
| ENSDARG000 | 4;4;4 | 44568015:445 | 44569169:445 | -;-;- | 1351 | NA | 10 | 6 | 48 | 10 | 39 | 24 |
| ENSDARG000 | 4;4 | 44577872:445 | 44578026:445 | +;+ | 1006 | NA | 4 | 8 | 31 | 18 | 7 | 19 |
| ENSDARG000 | 4;4;4;4;4;4 | 701612:70576 | 702140:70589 | -;-;-;-;-;- | 1178 | NA | 31 | 32 | 23 | 19 | 10 | 23 |
| ENSDARG000 | 4;4;4 | 44599987:446 | 44600853:446 | -;-;- | 1056 | NA | 67 | 53 | 97 | 8 | 32 | 39 |
| ENSDARG000 | 4;4;4 | 44666809:446 | 44667518:446 | -;-;- | 900 | NA | 0 | 6 | 2 | 0 | 1 | 2 |
| ENSDARG000 | 4;4;4 | 44701347:447 | 44701443:447 | +;+;+ | 1290 | NA | 0 | 0 | 0 | 0 | 0 | 0 |
| ENSDARG000 | 4:4:4:4:4:4:4: | 4526248:4528 | 4527891:4528 | -;-;-;-;-;-;-;- | 2763 | zgc:194209 | 40 | 12 | 66 | 46 | 38 | 93 |
| ENSDARG000 | 4:4:4:4:4:4:4: | 44713555:447 | 44713614:447 | +:+:+:+:+:+:+: | 4270 | NA | 99 | 118 | 193 | 16 | 117 | 239 |
| ENSDARG000 | 4;4;4 | 44778716:447 | 44779769:447 | -;-;- | 1239 | NA | 24 | 30 | 29 | 4 | 35 | 36 |
| ENSDARG000 | 4;4;4;4;4;4 | 44804898:448 | 44804982:448 | +;+;+;+;+;+ | 1375 | NA | 29 | 24 | 37 | 0 | 28 | 32 |
| ENSDARG000 | 4;4 | 44858444:448 | 44858586:448 | +;+ | 747 | NA | 31 | 49 | 101 | 74 | 212 | 255 |
| ENSDARG000 | 4:4:4:4:4:4:4: | 16710001:167 | 16710473:167 | +:+:+:+:+:+:+: | 2905 | pkp2 | 23 | 8 | 2 | 8 | 8 | 0 |
| ENSDARG000 | 4;4 | 44879202:448 | 44879335:448 | +;+ | 1762 | NA | 0 | 4 | 3 | 0 | 0 | 0 |
| ENSDARG000 | 4:4:4:4:4:4:4: | 7508316:7508 | 7508379:7508 | +:+:+:+:+:+:+: | 1765 | tnnt2e | 3 | 10 | 7 | 4 | 10 | 14 |
| ENSDARG000 | 4;4;4;4;4 | 44852270:448 | 44854265:448 | -;-;-;-;- | 2197 | znf1137 | 12 | 11 | 6 | 0 | 54 | 11 |
| ENSDARG000 | 4:4:4:4:4:4:4: | 13810811:138 | 13811101:138 | +:+:+:+:+:+:+: | 2589 | pdzrn4 | 0 | 0 | 0 | 0 | 0 | 2 |
| ENSDARG000 | 4;4;4;4;4 | 34853066:348 | 34853355:348 | -;-;-;-;- | 493 | NA | 0 | 0 | 0 | 0 | 0 | 0 |
| ENSDARG000 | 4:4:4:4:4:4:4: | 23223881:232 | 23224099:232 | +:+:+:+:+:+:+: | 2985 | TRHDE | 21 | 8 | 32 | 2 | 38 | 14 |
| ENSDARG000 | 4:4:4:4:4:4:4: | 737539:74037 | 737935:74045 | -:-:-:-:-:-:-:-: | 3128 | agbl5 | 731 | 387 | 469 | 610 | 348 | 579 |
| ENSDARG000 | 4;4;4;4 | 25829841:258 | 25830095:258 | -;-;-;- | 494 | ndufa12 | 463 | 570 | 507 | 246 | 434 | 280 |
| ENSDARG000 | 4:4:4:4:4:4:4: | 35167118:351 | 35167184:351 | +:+:+:+:+:+:+: | 5306 | NA | 9 | 2 | 24 | 8 | 10 | 10 |
| ENSDARG000 | 4;4;4 | 736395:73702 | 736496:73714 | +;+;+ | 331 | ost4 | 265 | 276 | 267 | 80 | 159 | 181 |
| ENSDARG000 | 4:4:4:4:4:4:4: | 35227135:352 | 35227316:352 | +:+:+:+:+:+:+: | 6738 | NA | 2 | 0 | 3 | 0 | 0 | 0 |
| ENSDARG000 | 4;4;4;4;4;4;4 | 54618332:546 | 54618423:546 | +;+;+;+;+;+;+ | 459 | NA | 0 | 0 | 0 | 0 | 0 | 0 |
| ENSDARG000 | 4:4:4:4:4:4:4: | 20271463:202 | 20271593:202 | -:-:-:-:-:-:-:-: | 2288 | CACNA2D4 | 0 | 0 | 0 | 0 | 1 | 2 |
| ENSDARG000 | 4:4:4:4:4:4:4: | 54583639:546 | 54583683:546 | +:+:+:+:+:+:+: | 8282 | NA | 22 | 8 | 21 | 5 | 13 | 9 |
| ENSDARG000 | 4:4:4:4:4:4:4: | 13905522:139 | 13906961:139 | -:-:-:-:-:-:-:-: | 2110 | yaf2 | 220 | 251 | 413 | 254 | 338 | 292 |
| ENSDARG000 | 4:4:4:4:4:4:4: | 35130120:351 | 35130710:351 | +:+:+:+:+:+:+: | 14908 | NA | 67 | 17 | 62 | 9 | 15 | 11 |
| ENSDARG000 | 4:4:4:4:4:4:4: | 5378338:5380 | 5378981:5380 | +:+:+:+:+:+:+: | 2673 | selenoo2 | 2 | 1 | 2 | 28 | 6 | 18 |
| ENSDARG000 | 4:4:4:4:4:4:4: | 54568329:545 | 54572372:545 | -:-:-:-:-:-:-:-: | 7063 | NA | 18 | 324 | 18 | 41 | 325 | 163 |
| ENSDARG000 | 4;4;4 | 9339067:9346 | 9340048:9346 | -;-;- | 1726 | kics2 | 220 | 201 | 221 | 173 | 257 | 201 |
| ENSDARG000 | 4:4:4:4:4:4:4: | 4540223:4540 | 4540670:4540 | -:-:-:-:-:-:-:-: | 1679 | RASSF3 | 343 | 329 | 338 | 216 | 355 | 294 |
| ENSDARG000 | 4:4:4:4:4:4:4: | 13923927:139 | 13925737:139 | -:-:-:-:-:-:-:-: | 2460 | zcrb1 | 86 | 106 | 194 | 90 | 116 | 162 |
| ENSDARG000 | 4;4 | 4585101:4585 | 4585204:4585 | +;+ | 684 | NA | 0 | 0 | 0 | 0 | 0 | 0 |
| ENSDARG000 | 4:4:4:4:4:4:4: | 13901458:139 | 13901557:139 | +:+:+:+:+:+:+: | 2494 | pphln1 | 482 | 587 | 771 | 385 | 568 | 500 |
| ENSDARG000 | 4;4;4;4;4 | 44954486:449 | 44954594:449 | +;+;+;+;+ | 563 | NA | 0 | 0 | 0 | 0 | 2 | 0 |
| ENSDARG000 | 4:4:4:4:4:4:4: | 44997263:449 | 44997962:449 | -:-:-:-:-:-:-:-: | 4113 | NA | 112 | 42 | 70 | 30 | 106 | 62 |
| ENSDARG000 | 4:4:4:4:4:4:4: | 14957360:149 | 14957698:149 | +:+:+:+:+:+:+: | 1762 | tspan33a | 62 | 2 | 8 | 12 | 55 | 14 |
| ENSDARG000 | 4:4:4:4:4:4:4: | 25838345:258 | 25840107:258 | -:-:-:-:-:-:-:-: | 4254 | nr2c1 | 214 | 348 | 462 | 206 | 333 | 332 |
| ENSDARG000 | 4;4;4;4;4;4;4 | 2059692:2060 | 2059918:2060 | +;+;+;+;+;+;+ | 1066 | yeats4 | 237 | 239 | 329 | 242 | 275 | 194 |
| ENSDARG000 | 4;4;4;4;4;4 | 2092683:2093 | 2092784:2093 | +;+;+;+;+;+ | 1542 | frs2a | 137 | 113 | 167 | 83 | 183 | 192 |
| ENSDARG000 | 4:4:4:4:4:4:4: | 2060524:2147 | 2061093:2147 | -:-:-:-:-:-:-:-: | 3163 | kcnc2 | 0 | 0 | 0 | 2 | 0 | 0 |
| ENSDARG000 | 4;4;4;4;4;4;4 | 13892773:138 | 13893429:138 | -;-;-;-;-;-;- | 1826 | gxylt1b | 18 | 12 | 8 | 6 | 0 | 12 |
| ENSDARG000 | 4:4:4:4:4:4:4: | 12612145:126 | 12612939:126 | +:+:+:+:+:+:+: | 3455 | lmo3 | 0 | 0 | 0 | 0 | 0 | 0 |
| ENSDARG000 | 4:4:4:4:4:4:4: | 23440404:234 | 23440547:234 | -:-:-:-:-:-:-:-: | 3051 | TRHDE | 1 | 0 | 0 | 0 | 0 | 0 |
| ENSDARG000 | 4;4 | 54718804:547 | 54719060:547 | +;+ | 687 | NA | 0 | 0 | 2 | 0 | 0 | 0 |
| ENSDARG000 | 4;4 | 54720270:547 | 54720460:547 | +;+ | 583 | NA | 0 | 0 | 0 | 0 | 0 | 0 |
| ENSDARG000 | 4;4 | 54741215:547 | 54741463:547 | -;- | 850 | NA | 0 | 2 | 2 | 0 | 0 | 0 |
| ENSDARG000 | 4:4:4:4:4:4:4: | 16725960:167 | 16726070:167 | +:+:+:+:+:+:+: | 2747 | tcp11l2 | 2635 | 708 | 932 | 1012 | 1716 | 2213 |
| ENSDARG000 | 4:4:4:4:4:4:4: | 77966055:779 | 77966368:779 | +:+:+:+:+:+:+: | 5819 | ikbip | 58 | 150 | 197 | 41 | 188 | 111 |
| ENSDARG000 | 4:4:4:4:4:4:4: | 4712410:4712 | 4715314:4712 | -:-:-:-:-:-:-:-: | 4632 | creb3l2 | 841 | 492 | 849 | 703 | 1161 | 677 |
| ENSDARG000 | 4:4:4:4:4:4:4: | 20302832:203 | 20303643:203 | -:-:-:-:-:-:-:-: | 3252 | dcp1b | 313 | 192 | 313 | 138 | 276 | 263 |
| ENSDARG000 | 4:4:4:4:4:4:4: | 77834796:778 | 77835027:778 | +:+:+:+:+:+:+: | 4274 | NA | 45 | 32 | 44 | 35 | 107 | 58 |
| ENSDARG000 | 4:4:4:4:4:4:4: | 10003523:100 | 10003633:100 | +:+:+:+:+:+:+: | 1902 | NA | 96 | 137 | 398 | 93 | 289 | 170 |
| ENSDARG000 | 4:4:4:4:4:4:4: | 77808579:778 | 77808782:778 | +:+:+:+:+:+:+: | 5512 | NA | 81 | 47 | 61 | 6 | 55 | 141 |
| ENSDARG000 | 4;4 | 77975221:779 | 77976788:779 | -;- | 1929 | ckap4 | 410 | 521 | 764 | 102 | 739 | 166 |
| ENSDARG000 | 4:4:4:4:4:4:4: | 77957611:779 | 77957775:779 | +:+:+:+:+:+:+: | 2035 | arfgap3 | 64 | 24 | 68 | 58 | 10 | 22 |
| ENSDARG000 | 4:4:4:4:4:4:4: | 18207655:182 | 18209464:182 | -:-:-:-:-:-:-:-: | 6300 | uhrf1bp1l | 202 | 156 | 281 | 81 | 226 | 161 |
| ENSDARG000 | 4:4:4:4:4:4:4: | 764100:76439 | 764557:76455 | -:-:-:-:-:-:-:-: | 3217 | tmem214 | 3941 | 3371 | 4216 | 2991 | 4170 | 2690 |
| ENSDARG000 | 4;4 | 16768961:167 | 16769123:167 | +;+ | 457 | NA | 0 | 2 | 0 | 0 | 0 | 13 |
| ENSDARG000 | 4;4 | 16777715:167 | 16777877:167 | +;+ | 365 | NA | 0 | 0 | 0 | 3 | 0 | 3 |
| ENSDARG000 | 4;4;4;4 | 78008298:780 | 78008372:780 | +;+;+;+ | 330 | NA | 254 | 495 | 607 | 317 | 353 | 392 |
| ENSDARG000 | 4:4:4:4:4:4:4: | 77973876:779 | 77973932:779 | +:+:+:+:+:+:+: | 2122 | terfa | 298 | 475 | 642 | 564 | 408 | 604 |
| ENSDARG000 | 4:4:4:4:4:4:4: | 23740655:237 | 23741303:237 | -:-:-:-:-:-:-:-: | 2093 | cpm | 24 | 30 | 34 | 68 | 32 | 23 |
| ENSDARG000 | 4;4;4;4 | 54754825:547 | 54755262:547 | -;-;-;- | 960 | NA | 0 | 0 | 0 | 0 | 0 | 0 |
| ENSDARG000 | 4:4:4:4:4:4:4: | 16787040:167 | 16787119:167 | +:+:+:+:+:+:+: | 1051 | golt1ba | 181 | 207 | 246 | 120 | 275 | 184 |
| ENSDARG000 | 4;4;4;4 | 54759510:547 | 54760032:547 | -;-;-;- | 818 | NA | 0 | 0 | 0 | 0 | 0 | 0 |
| ENSDARG000 | 4:4:4:4:4:4:4: | 16783972:167 | 16783976:167 | -:-:-:-:-:-:-:-: | 2593 | gys2 | 4 | 0 | 19 | 0 | 0 | 6 |
| ENSDARG000 | 4:4:4:4:4:4:4: | 45085360:450 | 45086793:450 | -:-:-:-:-:-:-:-: | 4801 | NA | 121 | 305 | 76 | 69 | 25 | 32 |
| ENSDARG000 | 4:4:4:4:4:4:4: | 35552485:355 | 35552588:355 | +:+:+:+:+:+:+: | 4165 | NA | 0 | 12 | 4 | 4 | 21 | 16 |
| ENSDARG000 | 4:4:4:4:4:4:4: | 4752622:4753 | 4753100:4753 | -:-:-:-:-:-:-:-: | 1037 | ssbp1 | 172 | 220 | 362 | 120 | 164 | 134 |
| ENSDARG000 | 4:4:4:4:4:4:4: | 45033922:450 | 45034022:450 | -:-:-:-:-:-:-:-: | 5107 | NA | 34 | 10 | 8 | 0 | 7 | 3 |
| ENSDARG000 | 4:4:4:4:4:4:4: | 54864061:550 | 54864101:550 | +:+:+:+:+:+:+: | 4530 | NA | 50 | 29 | 15 | 7 | 52 | 42 |
| ENSDARG000 | 4;4;4;4;4;4;4 | 11610718:116 | 11611332:116 | -;-;-;-;-;-;- | 1475 | NA | 6 | 28 | 79 | 26 | 5 | 6 |
| ENSDARG000 | 4:4:4:4:4:4:4: | 35576329:355 | 35576462:355 | +:+:+:+:+:+:+: | 7292 | NA | 118 | 104 | 92 | 40 | 127 | 96 |
| ENSDARG000 | 4;4;4 | 54955852:549 | 54956578:549 | -;-;- | 940 | NA | 0 | 144 | 0 | 0 | 122 | 327 |
| ENSDARG000 | 4;4;4 | 54973580:549 | 54973618:549 | +;+;+ | 1417 | NA | 34 | 19 | 18 | 22 | 52 | 28 |
| ENSDARG000 | 4:4:4:4:4:4:4: | 9360532:9360 | 9360885:9360 | +:+:+:+:+:+:+: | 3958 | ipo8 | 1785 | 1113 | 3047 | 3775 | 1437 | 1829 |
| ENSDARG000 | 4;4;4 | 5450499:5451 | 5450646:5451 | +;+;+ | 501 | NA | 2 | 4 | 0 | 1 | 0 | 2 |
| ENSDARG000 | 4;4;4 | 5456500:5456 | 5456814:5457 | -;-;- | 569 | NA | 16 | 7 | 43 | 36 | 13 | 20 |
| ENSDARG000 | 4;4 | 5420906:5421 | 5421015:5421 | +;+ | 393 | NA | 0 | 1 | 5 | 10 | 0 | 2 |
| ENSDARG000 | 4;4;4;4;4 | 55035164:550 | 55036352:550 | -;-;-;-;- | 1407 | NA | 0 | 0 | 2 | 2 | 0 | 0 |
| ENSDARG000 | 4;4;4 | 16784553:167 | 16784879:167 | +;+;+ | 1321 | stmp1 | 817 | 915 | 1115 | 868 | 1012 | 1029 |
| ENSDARG000 | 4;4;4;4 | 5421785:5421 | 5423035:5423 | -;-;-;- | 1479 | NA | 11 | 5 | 29 | 41 | 15 | 20 |
| ENSDARG000 | 4:4:4:4:4:4:4: | 54862025:548 | 54862115:548 | +:+:+:+:+:+:+: | 3106 | NA | 5 | 17 | 12 | 5 | 22 | 38 |
| ENSDARG000 | 4:4:4:4:4:4:4: | 5392094:5392 | 5392140:5392 | +:+:+:+:+:+:+: | 8657 | NA | 4 | 32 | 14 | 26 | 38 | 6 |
| ENSDARG000 | 4:4:4:4:4:4:4: | 77761901:777 | 77762034:777 | +:+:+:+:+:+:+: | 4710 | NA | 233 | 19 | 24 | 23 | 432 | 252 |
| ENSDARG000 | 4;4;4;4;4 | 45070770:450 | 45070899:450 | +;+;+;+;+ | 944 | NA | 13 | 14 | 12 | 4 | 17 | 8 |
| ENSDARG000 | 4:4:4:4:4:4:4: | 16794418:167 | 16794423:167 | +:+:+:+:+:+:+: | 671 | spx | 0 | 3 | 2 | 0 | 8 | 12 |
| ENSDARG000 | 4;4;4 | 55005641:550 | 55005912:550 | +;+;+ | 842 | NA | 4 | 3 | 4 | 0 | 0 | 2 |
| ENSDARG000 | 4;4;4;4 | 54948890:549 | 54948951:549 | +;+;+;+ | 464 | NA | 0 | 28 | 0 | 7 | 19 | 38 |
| ENSDARG000 | 4;4;4;4 | 45075758:450 | 45076340:450 | -;-;-;- | 814 | NA | 0 | 0 | 0 | 0 | 0 | 0 |
| ENSDARG000 | 4;4;4 | 54899568:549 | 54899597:549 | +;+;+ | 834 | NA | 0 | 103 | 46 | 16 | 104 | 28 |
| ENSDARG000 | 4;4;4;4 | 77735212:777 | 77735390:777 | +;+;+;+ | 1006 | NA | 776 | 0 | 0 | 0 | 684 | 789 |
| ENSDARG000 | 4;4;4 | 45038072:450 | 45038139:450 | +;+;+ | 751 | NA | 0 | 0 | 0 | 0 | 0 | 0 |
| ENSDARG000 | 4;4;4;4 | 5454006:5454 | 5454456:5454 | -;-;-;- | 667 | NA | 36 | 29 | 39 | 17 | 28 | 72 |
| ENSDARG000 | 4:4:4:4:4:4:4: | 77740228:777 | 77740691:777 | +:+:+:+:+:+:+: | 5186 | NA | 2 | 0 | 6 | 0 | 13 | 6 |
| ENSDARG000 | 4:4:4:4:4:4:4: | 77933084:779 | 77933193:779 | +:+:+:+:+:+:+: | 3441 | pacsin2 | 670 | 1011 | 819 | 614 | 896 | 776 |
| ENSDARG000 | 4:4:4:4:4:4:4: | 35948813:359 | 35949399:359 | -:-:-:-:-:-:-:-: | 3857 | NA | 6 | 3 | 0 | 0 | 0 | 0 |
| ENSDARG000 | 4:4:4:4:4:4:4: | 10017049:100 | 10017111:100 | +:+:+:+:+:+:+: | 2306 | NA | 10 | 22 | 12 | 62 | 19 | 46 |
| ENSDARG000 | 4;4;4;4;4;4;4 | 35851089:358 | 35851750:358 | -;-;-;-;-;-;- | 3445 | zgc:174653 | 2 | 0 | 0 | 6 | 2 | 0 |
| ENSDARG000 | 4 | 36016763 | 36017577 | - | 815 | NA | 2 | 2 | 0 | 0 | 0 | 14 |
| ENSDARG000 | 4;4;4 | 35876704:358 | 35876791:358 | +;+;+ | 1583 | NA | 1 | 1 | 2 | 0 | 3 | 0 |
| ENSDARG000 | 4;4;4 | 35982274:359 | 35984242:359 | -;-;- | 2259 | NA | 0 | 2 | 0 | 0 | 0 | 0 |
| ENSDARG000 | 4:4:4:4:4:4:4: | 35960020:359 | 35960523:359 | -:-:-:-:-:-:-:-: | 3782 | NA | 0 | 0 | 1 | 0 | 1 | 0 |
| ENSDARG000 | 4:4:4:4:4:4:4: | 20318127:203 | 20318184:203 | +:+:+:+:+:+:+: | 6852 | cacna1c | 23 | 6 | 2 | 0 | 8 | 68 |
| ENSDARG000 | 4;4;4;4;4 | 77880221:778 | 77880351:778 | +;+;+;+;+ | 2870 | NA | 22 | 17 | 34 | 21 | 35 | 42 |
| ENSDARG000 | 4;4;4 | 35993527:359 | 35993618:359 | +;+;+ | 1561 | NA | 4 | 4 | 0 | 0 | 11 | 5 |
| ENSDARG000 | 4;4;4;4;4 | 35900918:359 | 35900953:359 | +;+;+;+;+ | 1207 | NA | 0 | 0 | 0 | 0 | 0 | 0 |
| ENSDARG000 | 4:4:4:4:4:4:4: | 77919987:779 | 77920086:779 | +:+:+:+:+:+:+: | 2064 | ttll1 | 220 | 325 | 308 | 240 | 388 | 240 |
| ENSDARG000 | 4:4:4:4:4:4:4: | 14971239:149 | 14972141:149 | +:+:+:+:+:+:+: | 6212 | smo | 405 | 354 | 1102 | 518 | 780 | 390 |
| ENSDARG000 | 4:4:4:4:4:4:4: | 23771034:237 | 23776099:237 | -:-:-:-:-:-:-:-: | 6416 | usp6nl | 260 | 224 | 307 | 139 | 317 | 416 |
| ENSDARG000 | 4:4:4:4:4:4:4: | 4758001:4758 | 4758578:4758 | -:-:-:-:-:-:-:-: | 2404 | NA | 0 | 0 | 0 | 4 | 0 | 0 |
| ENSDARG000 | 4;4;4;4;4;4;4 | 12718965:127 | 12720639:127 | -;-;-;-;-;-;- | 2150 | mgst1.2 | 46 | 11 | 8 | 2 | 50 | 22 |
| ENSDARG000 | 4:4:4:4:4:4:4: | 12715695:127 | 12716340:127 | -;-;-;-;-;-;-;- | 1537 | mgst1.1 | 2 | 0 | 0 | 6 | 12 | 14 |
| ENSDARG000 | 4;4;4;4;4;4 | 2210317:2213 | 2210593:2214 | -;-;-;-;-;- | 1012 | NA | 47 | 49 | 76 | 126 | 50 | 68 |
| ENSDARG000 | 4:4:4:4:4:4:4: | 18281954:182 | 18283226:182 | -:-:-:-:-:-:-:-: | 5096 | PLXNC1 | 266 | 181 | 203 | 217 | 610 | 485 |
| ENSDARG000 | 4:4:4:4:4:4:4: | 783644:78530 | 785403:78540 | -:-:-:-:-:-:-:-: | 2619 | mapre3b | 677 | 740 | 755 | 758 | 595 | 546 |
| ENSDARG000 | 4:4:4:4:4:4:4: | 25859445:258 | 25862612:258 | -:-:-:-:-:-:-:-: | 7743 | fgd6 | 440 | 288 | 474 | 297 | 496 | 467 |
| ENSDARG000 | 4;4 | 36083288:360 | 36083437:360 | +;+ | 1006 | NA | 0 | 4 | 2 | 0 | 0 | 0 |
| ENSDARG000 | 4;4;4;4;4;4;4 | 55164069:551 | 55164130:551 | -;-;-;-;-;-;- | 639 | NA | 5 | 4 | 18 | 0 | 5 | 17 |
| ENSDARG000 | 4;4;4;4;4 | 4782721:4784 | 4783456:4784 | -;-;-;-;- | 1637 | zgc:162331 | 2 | 0 | 0 | 0 | 4 | 0 |
| ENSDARG000 | 4;4;4;4;4;4;4 | 55165697:551 | 55166130:551 | -;-;-;-;-;-;- | 1153 | NA | 6 | 0 | 1 | 0 | 0 | 0 |
| ENSDARG000 | 4;4;4 | 55172715:551 | 55173878:551 | -;-;- | 1406 | NA | 0 | 10 | 0 | 0 | 2 | 2 |
| ENSDARG000 | 4:4:4:4:4:4:4: | 36132112:361 | 36134961:361 | -:-:-:-:-:-:-:-: | 3114 | NA | 0 | 8 | 0 | 0 | 6 | 4 |
| ENSDARG000 | 4:4:4:4:4:4:4: | 55188228:552 | 55188815:552 | -;-;-;-;-;-;-;- | 3259 | NA | 42 | 19 | 31 | 11 | 25 | 5 |
| ENSDARG000 | 4;4;4 | 36150332:361 | 36150407:361 | +;+;+ | 1390 | NA | 8 | 6 | 30 | 0 | 9 | 2 |
| ENSDARG000 | 4;4;4 | 55421931:554 | 55421984:554 | +;+;+ | 221 | NA | 0 | 0 | 0 | 0 | 0 | 0 |
| ENSDARG000 | 4;4;4;4;4 | 55284199:552 | 55284372:552 | +;+;+;+;+ | 1394 | NA | 9 | 2 | 11 | 1 | 24 | 2 |
| ENSDARG000 | 4;4 | 55273988:552 | 55274036:552 | +;+ | 199 | NA | 1 | 0 | 0 | 0 | 0 | 0 |
| ENSDARG000 | 4;4;4 | 55258968:552 | 55259232:552 | -;-;- | 2465 | NA | 0 | 0 | 0 | 0 | 0 | 0 |
| ENSDARG000 | 4:4:4:4:4:4:4: | 55187370:551 | 55188815:551 | -:-:-:-:-:-:-:-: | 5959 | NA | 4 | 0 | 10 | 5 | 6 | 1 |
| ENSDARG000 | 4:4:4:4:4:4:4: | 2228754:2230 | 2228782:2230 | +:+:+:+:+:+:+: | 2169 | cmah | 9 | 6 | 20 | 24 | 16 | 46 |
| ENSDARG000 | 4;4;4;4;4;4;4 | 55313082:553 | 55313406:553 | -;-;-;-;-;-;- | 1876 | NA | 4 | 10 | 6 | 2 | 6 | 5 |
| ENSDARG000 | 4:4:4:4:4:4:4: | 36204590:362 | 36204868:362 | +:+:+:+:+:+:+: | 3247 | NA | 10 | 4 | 14 | 10 | 4 | 4 |
| ENSDARG000 | 4:4:4:4:4:4:4: | 23886726:238 | 23886820:238 | -:-:-:-:-:-:-:-: | 3783 | celf2 | 2649 | 2122 | 3988 | 1880 | 3600 | 2777 |
| ENSDARG000 | 4:4:4:4:4:4:4: | 11690923:116 | 11691036:116 | +:+:+:+:+:+:+: | 4480 | mkln1 | 669 | 641 | 830 | 532 | 639 | 602 |
| ENSDARG000 | 4:4:4:4:4:4:4: | 14981854:149 | 14981964:149 | +:+:+:+:+:+:+: | 2610 | cax1 | 4 | 2 | 469 | 84 | 162 | 124 |

| | | | | | | | | | | | | |
|---|---|---|---|---|---|---|---|---|---|---|---|---|
| ENSDARG000 | 4;4;4;4;4;4;4; | 9394426:9400 | 9394838:9400 | +;+;+;+;+;+;+; | 9056 | TMTC1 | 96 | 161 | 96 | 88 | 136 | 116 |
| ENSDARG000 | 4;4;4;4;4;4;4; | 36473781:364 | 36475792:364 | -;-;-;-;-;-;-;- | 2702 | NA | 1 | 2 | 3 | 0 | 5 | 24 |
| ENSDARG000 | 4;4 | 18383253:184 | 18383548:184 | -;- | 603 | cradd | 50 | 38 | 43 | 10 | 61 | 25 |
| ENSDARG000 | 4;4;4;4;4 | 45107809:451 | 45108658:451 | -;-;-;-;- | 1376 | NA | 2 | 2 | 7 | 0 | 14 | 0 |
| ENSDARG000 | 4;4;4;4;4;4;4; | 36616118:366 | 36616176:366 | +;+;+;+;+;+;+; | 3688 | si:dkev-151g2 | 8 | 4 | 7 | 0 | 8 | 8 |
| ENSDARG000 | 4;4;4;4;4;4;4; | 12755136:127 | 12755530:127 | -;-;-;-;-;-;-;- | 1444 | dera | 665 | 714 | 931 | 474 | 642 | 728 |
| ENSDARG000 | 4;4;4;4;4;4;4; | 5531583:5531 | 5531667:5531 | +;+;+;+;+;+;+; | 682 | si:dkev-14d8. | 0 | 0 | 0 | 0 | 0 | 0 |
| ENSDARG000 | 4;4;4;4;4;4;4; | 2252123:2252 | 2252313:2252 | +;+;+;+;+;+;+; | 938 | NA | 25 | 59 | 35 | 4 | 36 | 108 |
| ENSDARG000 | 4;4;4;4;4;4;4; | 798696:80287 | 800548:80305 | -;-;-;-;-;-;-;-; | 3615 | dpysl5b | 16 | 38 | 74 | 218 | 99 | 55 |
| ENSDARG000 | 4;4;4;4;4;4;4; | 14991789:149 | 14993554:149 | -;-;-;-;-;-;-;-; | 3018 | klhdc10 | 472 | 376 | 549 | 368 | 602 | 413 |
| ENSDARG000 | 4;4;4;4;4;4;4; | 5537101:5537 | 5537185:5537 | +;+;+;+;+;+;+; | 755 | si:dkev-14d8. | 0 | 0 | 0 | 0 | 0 | 0 |
| ENSDARG000 | 4;4;4;4;4 | 45128966:451 | 45129546:451 | -;-;-;-;- | 1570 | NA | 9 | 10 | 6 | 2 | 12 | 8 |
| ENSDARG000 | 4;4;4;4;4;4;4; | 5506946:5506 | 5507170:5507 | +;+;+;+;+;+;+; | 2530 | mapk11 | 65 | 87 | 51 | 46 | 86 | 233 |
| ENSDARG000 | 4;4;4;4;4;4;4; | 36789137:367 | 36789388:367 | -;-;-;-;-;-;-;-; | 2803 | NA | 0 | 0 | 1 | 0 | 5 | 0 |
| ENSDARG000 | 4;4 | 36875204:368 | 36875479:368 | +;+ | 704 | NA | 0 | 0 | 4 | 0 | 0 | 0 |
| ENSDARG000 | 4;4;4;4;4 | 4832144:4832 | 4832379:4832 | -;-;-;-;- | 644 | coa6 | 34 | 21 | 36 | 32 | 47 | 53 |
| ENSDARG000 | 4;4 | 36764181:371 | 36765304:371 | -;- | 1556 | NA | 11 | 12 | 38 | 22 | 6 | 4 |
| ENSDARG000 | 4;4;4;4;4;4;4; | 5523690:5525 | 5524109:5525 | +;+;+;+;+;+;+; | 1695 | mapk12b | 15 | 13 | 3 | 2 | 7 | 23 |
| ENSDARG000 | 4;4;4;4;4;4;4; | 4834085:4849 | 4834289:4849 | +;+;+;+;+;+;+; | 5047 | PTPRZ1 | 80 | 70 | 489 | 6 | 531 | 30 |
| ENSDARG000 | 4;4;4 | 55450711:554 | 55451857:554 | -;-;- | 1346 | NA | 10 | 10 | 6 | 0 | 6 | 4 |
| ENSDARG000 | 4;4;4;4;4;4 | 45147601:451 | 45147679:451 | +;+;+;+;+;+ | 550 | NA | 0 | 0 | 0 | 0 | 1 | 8 |
| ENSDARG000 | 4;4;4;4;4 | 45148652:451 | 45151713:451 | +;+;+;+;+ | 3786 | NA | 38 | 26 | 38 | 6 | 54 | 34 |
| ENSDARG000 | 4;4 | 55501017:555 | 55501598:555 | +;+ | 709 | NA | 0 | 0 | 0 | 0 | 0 | 0 |
| ENSDARG000 | 4;4;4 | 45181362:451 | 45182149:451 | -;-;- | 1028 | NA | 27 | 23 | 34 | 18 | 16 | 26 |
| ENSDARG000 | 4;4;4;4;4;4;4; | 4803698:4813 | 4804242:4813 | +;+;+;+;+;+;+; | 2862 | slc13a4 | 0 | 0 | 0 | 8 | 4 | 0 |
| ENSDARG000 | 4;4;4;4;4;4;4; | 2258935:2259 | 2259166:2259 | -;-;-;-;-;-;-;-; | 1267 | krr1 | 317 | 304 | 397 | 213 | 244 | 238 |
| ENSDARG000 | 4;4;4;4;4 | 55507920:555 | 55508268:555 | -;-;-;-;- | 1709 | NA | 2 | 12 | 22 | 2 | 24 | 24 |
| ENSDARG000 | 4;4 | 55509070:555 | 55509163:555 | -;- | 541 | NA | 0 | 0 | 0 | 0 | 0 | 0 |
| ENSDARG000 | 4;4;4;4;4;4;4; | 16828752:168 | 16829722:168 | -;-;-;-;-;-;-;-; | 1985 | ldhba | 12503 | 11927 | 16516 | 11664 | 14049 | 18341 |
| ENSDARG000 | 4;4;4;4;4;4 | 45195238:452 | 45195267:452 | +;+;+;+;+;+ | 1968 | NA | 18 | 56 | 75 | 5 | 91 | 83 |
| ENSDARG000 | 4;4;4;4;4 | 45195202:451 | 45195267:451 | +;+;+;+;+ | 1853 | NA | 25 | 4 | 25 | 6 | 8 | 18 |
| ENSDARG000 | 4;4;4;4;4 | 18432182:184 | 18433319:184 | -;-;-;-;- | 1791 | socs2 | 43 | 16 | 25 | 25 | 45 | 53 |
| ENSDARG000 | 4;4;4 | 45274792:452 | 45274938:452 | +;+;+ | 1414 | NA | 6 | 10 | 11 | 0 | 3 | 2 |
| ENSDARG000 | 4;4 | 18419542:184 | 18419556:184 | +;+ | 916 | NA | 6 | 3 | 18 | 0 | 0 | 30 |
| ENSDARG000 | 4;4;4;4;4 | 45292204:452 | 45293806:452 | -;-;-;-;- | 1806 | NA | 9 | 22 | 25 | 6 | 25 | 25 |
| ENSDARG000 | 4;4;4;4;4;4;4; | 20480399:204 | 20480758:204 | -;-;-;-;-;-;-;-; | 1772 | napepld | 216 | 174 | 130 | 297 | 176 | 136 |
| ENSDARG000 | 4;4;4 | 18453702:184 | 18454591:184 | -;-;- | 985 | sco2 | 0 | 0 | 0 | 0 | 0 | 0 |
| ENSDARG000 | 4;4;4;4;4 | 55593296:555 | 55593344:555 | +;+;+;+;+ | 1760 | NA | 8 | 9 | 24 | 10 | 7 | 8 |
| ENSDARG000 | 4;4;4;4;4;4;4; | 18441988:184 | 18442080:184 | +;+;+;+;+;+;+; | 5576 | ncaph2 | 198 | 536 | 563 | 186 | 386 | 250 |
| ENSDARG000 | 4;4;4;4;4;4;4; | 55604974:556 | 55605059:556 | +;+;+;+;+;+;+; | 2370 | NA | 11 | 5 | 27 | 2 | 27 | 26 |
| ENSDARG000 | 4;4;4;4;4;4;4; | 10365857:103 | 10366328:103 | +;+;+;+;+;+;+; | 6132 | kcnd2 | 0 | 0 | 0 | 0 | 0 | 6 |
| ENSDARG000 | 4;4;4;4;4;4;4; | 12769951:127 | 12770659:127 | -;-;-;-;-;-;-;-; | 4197 | HELB | 178 | 281 | 286 | 154 | 206 | 258 |
| ENSDARG000 | 4;4 | 15006216:150 | 15006404:150 | -;- | 230 | NA | 0 | 0 | 0 | 0 | 0 | 0 |
| ENSDARG000 | 4;4;4;4;4;4;4; | 15006217:150 | 15006392:150 | +;+;+;+;+;+;+; | 2411 | zc3hc1 | 243 | 520 | 525 | 258 | 586 | 309 |
| ENSDARG000 | 4;4;4 | 55631467:556 | 55634020:556 | -;-;- | 2756 | NA | 0 | 0 | 0 | 0 | 0 | 0 |
| ENSDARG000 | 4;4 | 842010:84429 | 842121:84503 | +;+ | 851 | NA | 38 | 2 | 2 | 25 | 20 | 16 |
| ENSDARG000 | 4;4;4;4;4;4;4; | 45321432:453 | 45321486:453 | -;-;-;-;-;-;-;-; | 2686 | NA | 6 | 1 | 9 | 0 | 9 | 12 |
| ENSDARG000 | 4;4;4;4;4;4;4; | 818928:82339 | 819615:82473 | -;-;-;-;-;-;-;-; | 2824 | sobpb | 99 | 10 | 28 | 135 | 39 | 54 |
| ENSDARG000 | 4;4;4;4;4;4;4; | 25912308:259 | 25912574:259 | +;+;+;+;+;+;+; | 2943 | vezt | 195 | 220 | 194 | 197 | 337 | 221 |
| ENSDARG000 | 4;4 | 45357558:453 | 45357982:453 | +;+ | 1424 | NA | 36 | 46 | 66 | 52 | 42 | 46 |
| ENSDARG000 | 4;4;4;4;4;4;4; | 16844973:168 | 16845394:168 | -;-;-;-;-;-;-;-; | 1421 | slc25a3a | 2 | 0 | 2 | 0 | 2 | 10 |
| ENSDARG000 | 4;4;4;4;4;4;4; | 45388995:453 | 45389086:453 | +;+;+;+;+;+;+; | 4078 | NA | 263 | 60 | 109 | 23 | 42 | 90 |
| ENSDARG000 | 4;4;4;4 | 18489207:184 | 18489334:184 | +;+;+;+ | 562 | NA | 0 | 0 | 0 | 2 | 0 | 0 |
| ENSDARG000 | 4;4;4;4;4;4;4; | 20486041:204 | 20486306:204 | +;+;+;+;+;+;+; | 2639 | ints13 | 505 | 463 | 575 | 447 | 523 | 454 |
| ENSDARG000 | 4;4;4;4;4;4;4; | 2267641:2269 | 2268006:2269 | +;+;+;+;+;+;+; | 1831 | NA | 246 | 123 | 918 | 98 | 271 | 523 |
| ENSDARG000 | 4;4 | 45423547:454 | 45424794:454 | -;- | 1432 | NA | 0 | 0 | 0 | 0 | 0 | 0 |
| ENSDARG000 | 4;4;4;4;4;4;4; | 55663384:556 | 55664249:556 | -;-;-;-;-;-;-;-; | 3876 | NA | 16 | 24 | 2 | 0 | 0 | 6 |
| ENSDARG000 | 4 | 55724043 | 55724313 | + | 271 | NA | 0 | 1 | 0 | 0 | 0 | 1 |
| ENSDARG000 | 4;4;4;4;4;4;4; | 9467049:9467 | 9467206:9467 | +;+;+;+;+;+;+; | 2874 | ovch1 | 0 | 4 | 3 | 6 | 1 | 0 |
| ENSDARG000 | 4 | 55727476 | 55727632 | + | 157 | NA | 1 | 0 | 0 | 0 | 0 | 0 |
| ENSDARG000 | 4;4;4;4;4;4 | 25054510:250 | 25055895:250 | -;-;-;-;-;- | 3047 | gata3 | 38 | 22 | 8 | 16 | 30 | 54 |
| ENSDARG000 | 4;4 | 45439770:454 | 45439881:454 | +;+ | 334 | NA | 0 | 0 | 0 | 0 | 0 | 0 |
| ENSDARG000 | 4;4;4 | 45441669:454 | 45441832:454 | +;+;+ | 1171 | NA | 29 | 0 | 2 | 0 | 0 | 0 |
| ENSDARG000 | 4;4;4;4;4;4 | 18494857:184 | 18494910:184 | +;+;+;+;+;+ | 726 | NA | 0 | 0 | 0 | 0 | 0 | 1 |
| ENSDARG000 | 4;4;4 | 45458619:454 | 45458668:454 | +;+;+ | 696 | NA | 6 | 0 | 0 | 0 | 0 | 0 |
| ENSDARG000 | 4;4;4;4;4;4;4; | 10574822:105 | 10575605:105 | -;-;-;-;-;-;-;- | 1851 | tspan12 | 580 | 18 | 16 | 6 | 45 | 61 |
| ENSDARG000 | 4;4;4 | 45484774:454 | 45484807:454 | +;+;+ | 482 | NA | 0 | 0 | 0 | 0 | 0 | 0 |
| ENSDARG000 | 4;4;4;4;4;4;4; | 13963499:139 | 13965031:139 | -;-;-;-;-;-;-;-; | 3531 | prickle1b | 711 | 696 | 1369 | 955 | 1258 | 1019 |
| ENSDARG000 | 4;4;4;4;4;4;4; | 2482046:2490 | 2482139:2490 | +;+;+;+;+;+;+; | 2186 | zdhhc17 | 488 | 320 | 369 | 317 | 407 | 403 |
| ENSDARG000 | 4;4;4;4;4 | 45504471:455 | 45504521:455 | +;+;+;+;+ | 1789 | NA | 6 | 10 | 0 | 2 | 3 | 6 |
| ENSDARG000 | 4 | 55728130 | 55730004 | + | 1875 | NA | 2 | 0 | 1 | 0 | 0 | 1 |
| ENSDARG000 | 4 | 55777234 | 55779108 | - | 1875 | NA | 0 | 1 | 0 | 0 | 0 | 4 |
| ENSDARG000 | 4 | 55779603 | 55779776 | - | 174 | NA | 0 | 0 | 0 | 0 | 0 | 0 |
| ENSDARG000 | 4 | 55782926 | 55783379 | - | 454 | NA | 1 | 0 | 0 | 0 | 0 | 2 |
| ENSDARG000 | 4 | 55793431 | 55795305 | - | 1875 | NA | 0 | 0 | 0 | 0 | 0 | 0 |
| ENSDARG000 | 4 | 55795803 | 55795959 | - | 157 | NA | 0 | 0 | 0 | 0 | 0 | 0 |
| ENSDARG000 | 4 | 55799122 | 55799572 | - | 451 | NA | 0 | 0 | 0 | 0 | 0 | 0 |
| ENSDARG000 | 4 | 55808991 | 55810865 | - | 1875 | NA | 0 | 0 | 0 | 0 | 0 | 0 |
| ENSDARG000 | 4 | 55811363 | 55811519 | - | 157 | NA | 0 | 0 | 0 | 0 | 0 | 0 |
| ENSDARG000 | 4 | 55827254 | 55827702 | - | 449 | NA | 0 | 0 | 0 | 0 | 0 | 1 |
| ENSDARG000 | 4;4;4;4;4;4 | 55864895:558 | 55864966:558 | +;+;+;+;+;+ | 433 | NA | 0 | 0 | 0 | 0 | 0 | 0 |
| ENSDARG000 | 4;4;4;4;4;4;4 | 15011341:150 | 15011512:150 | +;+;+;+;+;+;+ | 2758 | ube2h | 1781 | 1106 | 1902 | 1576 | 2209 | 1827 |
| ENSDARG000 | 4;4;4;4;4;4;4; | 55871735:558 | 55871784:558 | +;+;+;+;+;+;+; | 3423 | NA | 182 | 108 | 259 | 53 | 135 | 78 |
| ENSDARG000 | 4;4;4;4 | 55980166:559 | 55980599:559 | -;-;-;- | 959 | NA | 0 | 0 | 0 | 0 | 0 | 0 |
| ENSDARG000 | 4;4;4;4;4;4;4; | 11732156:117 | 11734909:117 | -;-;-;-;-;-;-;-; | 4160 | podxl | 210 | 18 | 34 | 10 | 56 | 112 |
| ENSDARG000 | 4;4 | 26006442:260 | 26006614:260 | -;- | 339 | NA | 0 | 0 | 0 | 0 | 0 | 0 |
| ENSDARG000 | 4;4;4;4;4;4;4; | 25950372:259 | 25950510:259 | +;+;+;+;+;+;+; | 1419 | metap2a | 15 | 29 | 33 | 40 | 26 | 23 |
| ENSDARG000 | 4;4;4 | 45556878:455 | 45557022:455 | +;+;+ | 528 | NA | 0 | 0 | 0 | 0 | 0 | 0 |
| ENSDARG000 | 4;4;4;4;4;4;4; | 861444:86204 | 862136:86213 | -;-;-;-;-;-;-;-; | 9736 | NA | 278 | 50 | 134 | 138 | 110 | 90 |
| ENSDARG000 | 4;4;4;4;4;4 | 25084228:250 | 25084804:250 | -;-;-;-;-;- | 3275 | taf3 | 972 | 941 | 1161 | 1030 | 1197 | 867 |
| ENSDARG000 | 4;4;4 | 45610254:456 | 45610408:456 | +;+;+ | 1353 | NA | 26 | 3 | 14 | 4 | 0 | 4 |
| ENSDARG000 | 4;4;4;4;4;4 | 46615995:466 | 46616274:466 | -;-;-;-;-;- | 642 | NA | 47 | 16 | 61 | 6 | 36 | 2 |
| ENSDARG000 | 4;4 | 46681803:466 | 46682019:466 | -;- | 473 | NA | 0 | 0 | 0 | 0 | 0 | 0 |
| ENSDARG000 | 4;4;4;4;4;4;4; | 16857722:168 | 16858556:168 | -;-;-;-;-;-;-;-; | 2587 | tmpoa | 379 | 534 | 638 | 301 | 438 | 372 |
| ENSDARG000 | 4;4;4;4;4;4;4; | 46842926:468 | 46845877:468 | -;-;-;-;-;-;-;-; | 9700 | NA | 57 | 27 | 51 | 10 | 34 | 32 |
| ENSDARG000 | 4;4;4;4;4;4;4; | 10604110:106 | 10604287:106 | +;+;+;+;+;+;+; | 2155 | ing3 | 931 | 620 | 783 | 721 | 582 | 1004 |
| ENSDARG000 | 4;4;4;4;4;4;4; | 46845345:468 | 46845877:468 | -;-;-;-;-;-;-;-; | 3513 | NA | 98 | 31 | 32 | 14 | 24 | 38 |
| ENSDARG000 | 4;4;4 | 47178124:471 | 47178174:471 | +;+;+ | 638 | NA | 0 | 0 | 0 | 0 | 0 | 0 |
| ENSDARG000 | 4;4;4;4;4;4;4; | 46771835:467 | 46771874:467 | +;+;+;+;+;+;+; | 1041 | NA | 56 | 22 | 50 | 16 | 57 | 54 |
| ENSDARG000 | 4;4;4;4;4;4;4; | 46890749:468 | 46891341:468 | -;-;-;-;-;-;-;-; | 3849 | NA | 581 | 182 | 518 | 280 | 373 | 502 |
| ENSDARG000 | 4;4;4;4;4;4;4; | 9478500:9486 | 9478675:9487 | +;+;+;+;+;+;+; | 2582 | lmf2b | 1148 | 2861 | 3005 | 2618 | 2419 | 1073 |
| ENSDARG000 | 4 | 55991165 | 55992044 | + | 880 | NA | 0 | 0 | 0 | 0 | 0 | 0 |
| ENSDARG000 | 4;4;4;4;4;4;4; | 46812891:468 | 46812987:468 | +;+;+;+;+;+;+; | 3398 | NA | 63 | 43 | 6 | 43 | 137 | 163 |
| ENSDARG000 | 4;4;4;4;4;4;4; | 46770787:468 | 46771310:468 | -;-;-;-;-;-;-;-; | 5473 | NA | 308 | 71 | 340 | 89 | 407 | 249 |
| ENSDARG000 | 4;4;4 | 56150518:561 | 56151550:561 | -;-;- | 1330 | NA | 0 | 11 | 0 | 0 | 1 | 0 |
| ENSDARG000 | 4;4;4;4 | 11765486:117 | 11765723:117 | -;-;-;- | 649 | NA | 24 | 7 | 13 | 16 | 39 | 23 |
| ENSDARG000 | 4;4;4;4;4;4;4; | 56110201:561 | 56110783:561 | -;-;-;-;-;-;-;-; | 3158 | NA | 15 | 7 | 5 | 2 | 2 | 10 |
| ENSDARG000 | 4;4;4;4;4;4;4; | 11762097:117 | 11762126:117 | +;+;+;+;+;+;+; | 895 | NA | 9 | 2 | 0 | 0 | 4 | 9 |
| ENSDARG000 | 4;4 | 56058846:560 | 56060855:560 | -;- | 2158 | NA | 0 | 9 | 0 | 0 | 0 | 0 |
| ENSDARG000 | 4;4;4;4;4;4;4; | 46981658:469 | 46982018:469 | +;+;+;+;+;+;+; | 4224 | NA | 1054 | 568 | 1754 | 515 | 1109 | 1298 |
| ENSDARG000 | 4;4;4;4;4;4;4; | 56090842:560 | 56091461:560 | -;-;-;-;-;-;-;-; | 4088 | NA | 18 | 19 | 18 | 16 | 12 | 18 |
| ENSDARG000 | 4;4;4;4;4;4;4; | 12785964:127 | 12785988:127 | -;-;-;-;-;-;-;-; | 2348 | b2m | 8133 | 12073 | 5388 | 6340 | 5859 | 6064 |
| ENSDARG000 | 4;4;4 | 46808017:468 | 46808836:468 | -;-;- | 942 | NA | 0 | 2 | 2 | 18 | 0 | 6 |
| ENSDARG000 | 4;4;4;4;4;4 | 56059807:560 | 56060704:560 | -;-;-;-;-;- | 2840 | NA | 12 | 4 | 4 | 0 | 5 | 10 |
| ENSDARG000 | 4;4 | 47174366:471 | 47175119:471 | +;+ | 805 | NA | 0 | 0 | 0 | 0 | 0 | 0 |
| ENSDARG000 | 4;4;4;4;4;4;4; | 25173299:251 | 25173434:251 | -;-;-;-;-;-;-;- | 1039 | atp5f1c | 3591 | 3891 | 4754 | 2896 | 3745 | 2966 |
| ENSDARG000 | 4;4;4;4;4;4;4; | 20507105:205 | 20507352:205 | -;-;-;-;-;-;-;- | 1681 | rassf8b | 140 | 98 | 139 | 100 | 103 | 79 |
| ENSDARG000 | 4;4;4;4;4;4 | 14107775:141 | 14107883:141 | -;-;-;-;-;- | 886 | NA | 0 | 1 | 10 | 2 | 2 | 2 |
| ENSDARG000 | 4;4;4;4 | 14106511:141 | 14107153:141 | +;+;+;+ | 1264 | NA | 2 | 8 | 10 | 79 | 2 | 10 |
| ENSDARG000 | 4;4;4;4;4;4;4; | 12782134:127 | 12782504:127 | -;-;-;-;-;-;-;-; | 2262 | IRAK3 | 234 | 73 | 83 | 76 | 58 | 98 |
| ENSDARG000 | 4;4;4;4;4;4;4; | 16876831:168 | 16877522:168 | -;-;-;-;-;-;-;-; | 1823 | strap | 980 | 1281 | 1560 | 1000 | 1244 | 1096 |
| ENSDARG000 | 4;4;4;4;4;4;4; | 18500012:185 | 18500454:185 | -;-;-;-;-;-;-;-; | 2611 | rint1 | 282 | 312 | 421 | 300 | 318 | 220 |
| ENSDARG000 | 4;4;4 | 7545930:7549 | 7546060:7549 | +;+;+ | 482 | NA | 47 | 32 | 31 | 26 | 58 | 101 |
| ENSDARG000 | 4;4;4;4 | 26127177:261 | 26127297:261 | -;-;-;- | 597 | NA | 0 | 0 | 0 | 0 | 0 | 0 |
| ENSDARG000 | 4;4;4;4;4;4 | 37230587:372 | 37231968:372 | -;-;-;-;-;- | 1593 | NA | 12 | 8 | 2 | 0 | 4 | 0 |
| ENSDARG000 | 4;4;4;4;4;4;4; | 7653607:7656 | 7654007:7656 | -;-;-;-;-;-;-;-; | 2183 | lta4h | 365 | 563 | 659 | 331 | 753 | 572 |
| ENSDARG000 | 4;4;4;4;4;4;4 | 14116366:141 | 14116888:141 | +;+;+;+;+;+;+ | 1645 | NA | 7 | 19 | 14 | 49 | 19 | 9 |
| ENSDARG000 | 4;4;4;4;4;4;4; | 14118978:141 | 14119128:141 | -;-;-;-;-;-;-;- | 1067 | NA | 0 | 1 | 4 | 0 | 1 | 5 |
| ENSDARG000 | 4;4;4;4;4 | 37239091:372 | 37239125:372 | +;+;+;+;+ | 1657 | znf973 | 93 | 44 | 82 | 11 | 33 | 45 |
| ENSDARG000 | 4;4;4 | 56193313:562 | 56194380:562 | -;-;- | 1293 | NA | 2 | 2 | 4 | 0 | 6 | 10 |
| ENSDARG000 | 4;4;4;4;4;4;4; | 37267210:372 | 37267273:372 | -;-;-;-;-;-;-;-; | 2369 | NA | 10 | 6 | 12 | 8 | 4 | 16 |
| ENSDARG000 | 4;4;4 | 912563:91292 | 912722:91298 | +;+;+ | 558 | ripply2 | 4 | 4 | 21 | 16 | 10 | 2 |
| ENSDARG000 | 4;4;4 | 7650003:7650 | 7650248:7650 | -;-;- | 639 | NA | 73 | 32 | 35 | 2 | 12 | 25 |
| ENSDARG000 | 4;4 | 18514457:185 | 18515332:185 | +;+ | 1753 | NA | 0 | 0 | 0 | 0 | 0 | 0 |
| ENSDARG000 | 4;4;4;4;4 | 18592743:185 | 18594230:185 | -;-;-;-;- | 2357 | cdkn1ba | 785 | 981 | 1987 | 921 | 1743 | 930 |
| ENSDARG000 | 4;4;4;4;4;4;4; | 9508505:9508 | 9508713:9509 | +;+;+;+;+;+;+; | 1845 | kitlgb | 30 | 22 | 32 | 14 | 22 | 25 |
| ENSDARG000 | 4;4;4;4;4;4;4; | 2511629:2513 | 2512192:2514 | -;-;-;-;-;-;-;-; | 2711 | csrp2 | 6387 | 6970 | 3620 | 11707 | 2572 | 4898 |
| ENSDARG000 | 4;4 | 11779997:117 | 11780095:117 | -;- | 302 | NA | 3 | 0 | 0 | 0 | 1 | 9 |
| ENSDARG000 | 4;4;4;4;4;4;4; | 15081385:150 | 15082891:150 | -;-;-;-;-;-;-;-; | 3013 | nrf1 | 578 | 576 | 749 | 524 | 764 | 438 |
| ENSDARG000 | 4;4;4;4 | 56244131:562 | 56244511:562 | -;-;-;- | 943 | NA | 0 | 0 | 0 | 0 | 0 | 0 |
| ENSDARG000 | 4;4;4;4;4;4;4; | 26303613:263 | 26306398:263 | -;-;-;-;-;-;-;- | 5575 | st8sia1 | 36 | 90 | 79 | 44 | 28 | 39 |
| ENSDARG000 | 4;4;4;4;4;4 | 11791197:117 | 11791247:117 | +;+;+;+;+;+ | 530 | NA | 0 | 0 | 0 | 0 | 1 | 0 |
| ENSDARG000 | 4;4;4;4;4;4;4; | 10616626:106 | 10617625:106 | +;+;+;+;+;+;+; | 4043 | CPED1 | 288 | 217 | 103 | 395 | 319 | 273 |
| ENSDARG000 | 4;4;4;4;4;4;4; | 9524805:9528 | 9525981:9528 | -;-;-;-;-;-;-;- | 2043 | NA | 2 | 10 | 4 | 4 | 12 | 10 |
| ENSDARG000 | 4;4;4 | 14159321:141 | 14159592:141 | -;-;- | 421 | NA | 0 | 0 | 0 | 0 | 1 | 0 |
| ENSDARG000 | 4;4;4;4;4;4 | 14167042:141 | 14167344:141 | -;-;-;-;-;- | 1180 | NA | 8 | 24 | 13 | 31 | 6 | 30 |
| ENSDARG000 | 4;4;4;4;4;4;4; | 56306039:563 | 56306632:563 | -;-;-;-;-;-;-;-; | 1938 | NA | 4 | 0 | 0 | 2 | 0 | 4 |
| ENSDARG000 | 4;4;4;4;4;4;4; | 26357221:263 | 26357400:263 | +;+;+;+;+;+;+; | 2481 | tnni1d | 6 | 0 | 0 | 0 | 0 | 0 |
| ENSDARG000 | 4;4;4;4;4;4;4; | 18597162:185 | 18597568:185 | -;-;-;-;-;-;-;-; | 2593 | cdpf1 | 472 | 386 | 506 | 372 | 873 | 423 |
| ENSDARG000 | 4;4;4 | 37406676:374 | 37407428:374 | +;+;+ | 958 | NA | 0 | 0 | 0 | 0 | 0 | 0 |
| ENSDARG000 | 4;4;4 | 56315530:563 | 56315558:563 | +;+;+ | 1874 | zgc:174275 | 40 | 24 | 82 | 8 | 28 | 52 |
| ENSDARG000 | 4;4;4 | 14170459:141 | 14170730:141 | -;-;- | 421 | NA | 0 | 0 | 0 | 0 | 0 | 0 |
| ENSDARG000 | 4;4;4;4;4;4;4; | 56379274:563 | 56380083:563 | +;+;+;+;+;+;+; | 5819 | NA | 0 | 0 | 0 | 0 | 0 | 2 |
| ENSDARG000 | 4;4;4 | 37503994:375 | 37504151:375 | +;+;+ | 1598 | NA | 55 | 23 | 46 | 20 | 32 | 20 |
| ENSDARG000 | 4;4;4;4;4;4;4; | 37455405:374 | 37455463:374 | +;+;+;+;+;+;+; | 3565 | NA | 14 | 8 | 8 | 2 | 4 | 0 |
| ENSDARG000 | 4;4;4;4;4;4;4; | 20812900:208 | 20813173:208 | +;+;+;+;+;+;+; | 8267 | nav3 | 750 | 240 | 701 | 332 | 612 | 835 |
| ENSDARG000 | 4;4;4;4;4 | 37476367:374 | 37476845:374 | -;-;-;-;- | 1782 | NA | 0 | 0 | 0 | 0 | 0 | 0 |
| ENSDARG000 | 4;4;4;4;4;4;4; | 56379907:563 | 56380083:563 | +;+;+;+;+;+;+; | 3593 | NA | 35 | 6 | 8 | 5 | 14 | 36 |
| ENSDARG000 | 4;4;4;4;4;4;4; | 26390851:263 | 26391516:263 | -;-;-;-;-;-;-;-; | 2847 | NA | 32 | 0 | 0 | 4 | 0 | 0 |
| ENSDARG000 | 4;4;4;4;4;4;4; | 929557:93172 | 929930:93195 | -;-;-;-;-;-;-;-; | 1827 | sim1b | 4 | 0 | 0 | 6 | 2 | 0 |
| ENSDARG000 | 4;4;4 | 56508620:565 | 56509205:565 | -;-;- | 751 | NA | 0 | 0 | 0 | 0 | 0 | 0 |
| ENSDARG000 | 4;4;4;4;4;4;4; | 11804675:118 | 11805705:118 | -;-;-;-;-;-;-;-; | 1907 | cyb5r3 | 1578 | 1646 | 1521 | 2193 | 1693 | 2457 |

| | | | | | | | | | | | | |
|---|---|---|---|---|---|---|---|---|---|---|---|---|
| ENSDARG000 | 4;4 | 14174239:141 | 14174299:141 | -;- | 393 | NA | 0 | 0 | 0 | 0 | 0 | 0 |
| ENSDARG000 | 4;4;4 | 14180997:141 | 14181269:141 | -;-;- | 422 | NA | 0 | 0 | 0 | 0 | 0 | 1 |
| ENSDARG000 | 4;4;4;4;4;4;4; | 14176521:141 | 14176823:141 | -;-;-;-;-;-;-;- | 1553 | NA | 2 | 4 | 7 | 6 | 7 | 2 |
| ENSDARG000 | 4;4 | 14174367:141 | 14174463:141 | +;+ | 245 | NA | 0 | 0 | 0 | 0 | 1 | 0 |
| ENSDARG000 | 4;4;4;4 | 9536860:9538 | 9537050:9538 | +;+;+;+ | 628 | lsm8 | 238 | 219 | 278 | 170 | 194 | 158 |
| ENSDARG000 | 4;4;4;4;4;4;4; | 15135239:151 | 15135716:151 | -;-;-;-;-;-;-;-; | 1124 | NA | 6 | 14 | 2 | 28 | 16 | 9 |
| ENSDARG000 | 4;4;4 | 37577527:375 | 37578307:375 | -;-;- | 1078 | NA | 0 | 0 | 0 | 0 | 6 | 2 |
| ENSDARG000 | 4;4;4;4;4 | 12819272:128 | 12819946:128 | -;-;-;-;- | 1415 | hmga2 | 10 | 58 | 113 | 58 | 92 | 79 |
| ENSDARG000 | 4;4;4;4;4;4;4; | 25187912:251 | 25188367:251 | -;-;-;-;-;-;-;-; | 3261 | itih2 | 0 | 0 | 27 | 0 | 0 | 2 |
| ENSDARG000 | 4;4;4;4;4;4;4; | 5568688:5587 | 5568858:5588 | -;-;-;-;-;-;-;-; | 3004 | vegfab | 321 | 2210 | 7709 | 5382 | 1350 | 249 |
| ENSDARG000 | 4;4;4;4;4;4;4; | 2530841:2532 | 2532184:2532 | -;-;-;-;-;-;-;-; | 3544 | e2f7 | 94 | 287 | 255 | 144 | 189 | 220 |
| ENSDARG000 | 4 | 47257854 | 47262290 | + | 4437 | crestin | 5943 | 1602 | 8430 | 1253 | 610 | 7694 |
| ENSDARG000 | 4;4;4;4;4;4;4; | 15207433:152 | 15207578:152 | +;+;+;+;+;+;+; | 697 | NA | 14 | 2 | 2 | 2 | 17 | 0 |
| ENSDARG000 | 4;4;4;4;4;4;4; | 15217929:152 | 15220988:152 | -;-;-;-;-;-;-;-; | 8676 | plxna4 | 136 | 249 | 578 | 230 | 296 | 131 |
| ENSDARG000 | 4;4;4;4;4 | 47303276:473 | 47303942:473 | -;-;-;-;- | 1007 | NA | 0 | 0 | 4 | 0 | 0 | 0 |
| ENSDARG000 | 4;4;4;4;4;4;4; | 25181572:251 | 25181872:251 | +;+;+;+;+;+;+; | 1678 | kin | 160 | 203 | 259 | 140 | 238 | 169 |
| ENSDARG000 | 4;4;4;4;4;4 | 56590242:565 | 56590681:565 | -;-;-;-;-;- | 1340 | NA | 0 | 0 | 0 | 0 | 0 | 0 |
| ENSDARG000 | 4;4;4;4;4;4;4 | 18609539:186 | 18610103:186 | -;-;-;-;-;-;- | 1953 | pparaa | 62 | 131 | 90 | 1458 | 332 | 213 |
| ENSDARG000 | 4;4;4;4;4;4;4; | 37676163:376 | 37676858:376 | -;-;-;-;-;-;-;-; | 3641 | NA | 156 | 385 | 162 | 68 | 274 | 250 |
| ENSDARG000 | 4;4;4 | 47385355:473 | 47385444:473 | +;+;+ | 1579 | NA | 28 | 8 | 12 | 0 | 12 | 20 |
| ENSDARG000 | 4;4;4;4;4;4;4; | 26496489:264 | 26497081:264 | +;+;+;+;+;+;+; | 8626 | NA | 317 | 104 | 408 | 216 | 38 | 427 |
| ENSDARG000 | 4;4;4;4;4;4;4; | 47436126:474 | 47436230:474 | +;+;+;+;+;+;+; | 5899 | NA | 33 | 10 | 29 | 0 | 8 | 24 |
| ENSDARG000 | 4;4;4 | 47656992:476 | 47657138:476 | +;+;+ | 2791 | NA | 42 | 31 | 30 | 20 | 14 | 32 |
| ENSDARG000 | 4;4;4;4;4;4;4; | 56651458:566 | 56651639:566 | -;-;-;-;-;-;-;-; | 3688 | NA | 47 | 17 | 53 | 4 | 22 | 22 |
| ENSDARG000 | 4;4;4;4;4;4;4; | 10706755:107 | 10707032:107 | +;+;+;+;+;+;+; | 2058 | wnt16 | 30 | 16 | 4 | 6 | 8 | 28 |
| ENSDARG000 | 4;4;4;4;4;4;4; | 21129752:211 | 21129812:211 | +;+;+;+;+;+;+; | 5301 | syt1a | 4 | 2 | 0 | 8 | 30 | 28 |
| ENSDARG000 | 4;4;4 | 56771551:567 | 56772194:567 | +;+;+ | 933 | NA | 0 | 0 | 0 | 0 | 0 | 16 |
| ENSDARG000 | 4;4;4;4;4;4 | 37701476:377 | 37702709:377 | -;-;-;-;-;- | 1447 | NA | 0 | 57 | 19 | 15 | 74 | 48 |
| ENSDARG000 | 4;4;4;4;4;4;4 | 56833071:568 | 56833177:568 | +;+;+;+;+;+;+ | 3193 | NA | 7 | 4 | 10 | 9 | 7 | 9 |
| ENSDARG000 | 4;4;4;4;4;4;4; | 9555280:9555 | 9556055:9556 | -;-;-;-;-;-;-;-; | 5558 | SHANK3 | 80 | 4 | 36 | 12 | 40 | 216 |
| ENSDARG000 | 4;4;4;4;4;4;4; | 11813778:118 | 11816863:118 | -;-;-;-;-;-;-;-; | 6223 | btbd11a | 172 | 90 | 198 | 282 | 216 | 204 |
| ENSDARG000 | 4;4;4;4;4;4;4; | 12890668:128 | 12894153:128 | -;-;-;-;-;-;-;-; | 5206 | msrb3 | 255 | 168 | 113 | 150 | 152 | 146 |
| ENSDARG000 | 4;4;4;4;4;4;4; | 11946875:119 | 11947209:119 | +;+;+;+;+;+;+; | 2389 | NA | 0 | 0 | 1 | 0 | 0 | 0 |
| ENSDARG000 | 4;4;4;4;4 | 1054708:1060 | 1056005:1060 | -;-;-;-;- | 1699 | mtpn | 3385 | 3718 | 2742 | 3059 | 3853 | 3425 |
| ENSDARG000 | 4;4;4;4;4;4;4; | 47736069:477 | 47736154:477 | +;+;+;+;+;+;+; | 7296 | NA | 40 | 25 | 55 | 11 | 73 | 48 |
| ENSDARG000 | 4;4;4;4;4 | 37710093:377 | 37710151:377 | +;+;+;+;+ | 906 | NA | 0 | 9 | 0 | 2 | 4 | 0 |
| ENSDARG000 | 4;4;4;4;4;4;4; | 5642696:5642 | 5642873:5642 | +;+;+;+;+;+;+; | 986 | mrps18a | 562 | 907 | 1422 | 1452 | 901 | 458 |
| ENSDARG000 | 4;4;4;4;4;4;4 | 56889552:568 | 56891700:568 | -;-;-;-;-;-;- | 2562 | NA | 0 | 0 | 0 | 0 | 0 | 0 |
| ENSDARG000 | 4;4;4;4;4;4;4 | 18702129:187 | 18702211:187 | +;+;+;+;+;+;+ | 1099 | NA | 2135 | 496 | 5791 | 3801 | 9980 | 2118 |
| ENSDARG000 | 4;4;4;4;4;4;4 | 37760928:377 | 37762161:377 | -;-;-;-;-;-;- | 2081 | NA | 0 | 52 | 0 | 6 | 134 | 107 |
| ENSDARG000 | 4;4;4;4;4;4;4; | 26972862:269 | 26973104:269 | +;+;+;+;+;+;+; | 6859 | NA | 0 | 0 | 2 | 4 | 0 | 0 |
| ENSDARG000 | 4;4;4;4;4;4;4; | 37783972:377 | 37784053:377 | +;+;+;+;+;+;+; | 5263 | NA | 27 | 4 | 7 | 0 | 12 | 5 |
| ENSDARG000 | 4;4;4 | 1146346:1283 | 1146475:1284 | +;+;+ | 1754 | chrm2a | 6 | 0 | 0 | 0 | 0 | 2 |
| ENSDARG000 | 4;4;4;4;4;4;4; | 47782411:477 | 47782590:477 | +;+;+;+;+;+;+; | 3551 | NA | 25 | 19 | 35 | 2 | 5 | 9 |
| ENSDARG000 | 4;4;4;4 | 1103158:1104 | 1103428:1104 | -;-;-;- | 842 | NA | 6 | 0 | 0 | 0 | 0 | 0 |
| ENSDARG000 | 4;4;4;4;4;4;4; | 18720785:187 | 18721059:187 | -;-;-;-;-;-;-;-; | 1717 | atxn10 | 189 | 218 | 368 | 401 | 1420 | 471 |
| ENSDARG000 | 4;4;4;4;4;4;4; | 14188327:141 | 14188896:141 | -;-;-;-;-;-;-;-; | 2242 | pus7l | 52 | 68 | 66 | 88 | 64 | 169 |
| ENSDARG000 | 4;4;4;4;4;4;4; | 47876171:478 | 47878214:478 | -;-;-;-;-;-;-;-; | 5167 | NA | 340 | 55 | 238 | 76 | 241 | 221 |
| ENSDARG000 | 4;4;4;4;4;4;4; | 48484490:484 | 48485323:484 | -;-;-;-;-;-;-;-; | 3504 | NA | 80 | 71 | 21 | 14 | 37 | 15 |
| ENSDARG000 | 4;4;4;4;4;4;4; | 2548783:2548 | 2549470:2549 | -;-;-;-;-;-;-;-; | 3836 | cog5 | 307 | 340 | 356 | 367 | 417 | 294 |
| ENSDARG000 | 4;4;4;4;4;4;4; | 37936249:379 | 37938666:379 | +;+;+;+;+;+;+; | 8583 | NA | 3 | 0 | 0 | 0 | 6 | 1 |
| ENSDARG000 | 4;4;4;4;4;4;4 | 48628562:486 | 48630708:486 | -;-;-;-;-;-;- | 2565 | NA | 6 | 4 | 3 | 14 | 2 | 8 |
| ENSDARG000 | 4;4;4;4;4;4;4; | 56915604:569 | 56915714:569 | +;+;+;+;+;+;+; | 914 | NA | 0 | 4 | 9 | 7 | 2 | 1 |
| ENSDARG000 | 4;4;4;4;4 | 2619132:2620 | 2619410:2621 | +;+;+;+;+ | 3156 | gpr22a | 0 | 0 | 0 | 4 | 0 | 0 |
| ENSDARG000 | 4;4;4;4;4;4;4; | 48693208:486 | 48693378:486 | -;-;-;-;-;-;-;- | 2817 | NA | 24 | 8 | 8 | 2 | 4 | 4 |
| ENSDARG000 | 4;4;4;4;4;4 | 38001640:380 | 38001734:380 | +;+;+;+;+;+ | 1446 | znf1051 | 1 | 0 | 0 | 0 | 8 | 0 |
| ENSDARG000 | 4;4;4;4 | 38032613:380 | 38032818:380 | -;-;-;- | 832 | NA | 0 | 0 | 0 | 0 | 0 | 0 |
| ENSDARG000 | 4;4;4;4;4 | 48769555:487 | 48769603:487 | +;+;+;+;+ | 2441 | NA | 24 | 30 | 21 | 0 | 16 | 25 |
| ENSDARG000 | 4;4;4;4;4;4;4 | 10749520:107 | 10750421:107 | -;-;-;-;-;-;- | 1376 | NA | 0 | 0 | 0 | 0 | 1 | 6 |
| ENSDARG000 | 4;4;4;4;4;4;4; | 56932843:569 | 56933465:569 | -;-;-;-;-;-;-;-; | 3516 | NA | 19 | 9 | 25 | 10 | 22 | 11 |
| ENSDARG000 | 4;4;4;4;4;4;4; | 10721795:107 | 10721974:107 | +;+;+;+;+;+;+; | 8087 | stab2 | 422 | 68 | 479 | 146 | 223 | 711 |
| ENSDARG000 | 4;4;4;4;4;4;4; | 1301436:1303 | 1303392:1303 | -;-;-;-;-;-;-;- | 2547 | ptn | 62 | 110 | 403 | 244 | 190 | 66 |
| ENSDARG000 | 4;4 | 56970503:569 | 56970815:569 | -;- | 371 | NA | 0 | 0 | 0 | 0 | 0 | 0 |
| ENSDARG000 | 4;4;4;4;4;4 | 56974252:569 | 56974378:569 | +;+;+;+;+;+ | 2961 | NA | 0 | 0 | 0 | 0 | 2 | 4 |
| ENSDARG000 | 4;4;4;4;4;4;4; | 21323006:213 | 21324167:213 | -;-;-;-;-;-;-;-; | 2265 | pawr | 46 | 32 | 53 | 10 | 40 | 6 |
| ENSDARG000 | 4;4;4;4;4;4;4; | 27081972:270 | 27082356:270 | -;-;-;-;-;-;-;-; | 1545 | creld2 | 196 | 396 | 552 | 100 | 346 | 172 |
| ENSDARG000 | 4;4;4;4;4 | 5645251:5647 | 5645584:5647 | -;-;-;-;- | 1138 | rsph9 | 4 | 6 | 3 | 0 | 4 | 2 |
| ENSDARG000 | 4;4;4;4;4 | 48802365:488 | 48802422:488 | +;+;+;+;+ | 1597 | znf1113 | 39 | 31 | 62 | 30 | 56 | 29 |
| ENSDARG000 | 4;4;4;4 | 9587968:9591 | 9589454:9591 | -;-;-;- | 1916 | cdnf | 6 | 2 | 14 | 8 | 4 | 26 |
| ENSDARG000 | 4;4;4 | 56995821:570 | 56996882:570 | -;-;- | 1242 | NA | 0 | 1 | 2 | 0 | 0 | 0 |
| ENSDARG000 | 4;4;4;4 | 38035942:380 | 38036153:380 | -;-;-;- | 514 | NA | 2 | 0 | 0 | 0 | 0 | 0 |
| ENSDARG000 | 4 | 38086141 | 38086998 | + | 858 | NA | 0 | 0 | 0 | 0 | 0 | 0 |
| ENSDARG000 | 4;4;4;4;4;4;4; | 48930297:489 | 48931537:489 | -;-;-;-;-;-;-;- | 2551 | NA | 6 | 10 | 23 | 0 | 16 | 19 |
| ENSDARG000 | 4;4;4 | 38103421:381 | 38103512:381 | +;+;+ | 2661 | NA | 7 | 0 | 0 | 0 | 0 | 0 |
| ENSDARG000 | 4;4;4;4;4;4;4; | 18759289:187 | 18759327:187 | -;-;-;-;-;-;-;-; | 1777 | NA | 65 | 57 | 66 | 56 | 109 | 76 |
| ENSDARG000 | 4;4;4;4;4;4;4; | 4930406:4930 | 4930576:4930 | -;-;-;-;-;-;-;-; | 658 | ndufa5 | 394 | 489 | 571 | 454 | 444 | 379 |
| ENSDARG000 | 4;4;4;4;4;4;4; | 57046954:570 | 57047232:570 | +;+;+;+;+;+;+; | 3634 | NA | 0 | 0 | 8 | 0 | 0 | 0 |
| ENSDARG000 | 4;4;4 | 49016956:490 | 49017250:490 | +;+;+ | 1386 | NA | 6 | 4 | 39 | 7 | 10 | 14 |
| ENSDARG000 | 4;4;4;4;4;4;4; | 49031364:490 | 49031781:490 | -;-;-;-;-;-;-;-; | 1854 | NA | 0 | 0 | 0 | 0 | 4 | 0 |
| ENSDARG000 | 4;4;4 | 49109355:491 | 49110775:491 | -;-;- | 1624 | NA | 10 | 2 | 9 | 0 | 7 | 1 |
| ENSDARG000 | 4;4 | 57093908:570 | 57094967:570 | +;+ | 1337 | NA | 10 | 10 | 14 | 8 | 2 | 0 |
| ENSDARG000 | 4;4;4;4;4;4;4; | 18782539:187 | 18782575:187 | +;+;+;+;+;+;+; | 2280 | slc26a4 | 8 | 10 | 2 | 4 | 18 | 10 |
| ENSDARG000 | 4;4;4;4;4 | 57099307:570 | 57099404:570 | +;+;+;+;+ | 521 | si:ch211-238 | 23 | 18 | 11 | 13 | 12 | 11 |
| ENSDARG000 | 4;4;4;4;4 | 57101749:571 | 57101812:571 | +;+;+;+;+ | 578 | NA | 13 | 6 | 2 | 4 | 8 | 2 |
| ENSDARG000 | 4;4;4 | 49129407:491 | 49129514:491 | -;-;- | 283 | NA | 30 | 12 | 30 | 1 | 11 | 9 |
| ENSDARG000 | 4;4 | 49151806:491 | 49152033:491 | +;+ | 552 | NA | 0 | 0 | 0 | 0 | 0 | 0 |
| ENSDARG000 | 4;4;4;4;4;4;4; | 57104534:571 | 57104585:571 | +;+;+;+;+;+;+; | 1629 | NA | 127 | 132 | 102 | 186 | 121 | 102 |
| ENSDARG000 | 4;4;4;4;4;4;4; | 5665182:5665 | 5665801:5665 | -;-;-;-;-;-;-;-; | 2958 | tmem63a | 218 | 215 | 372 | 227 | 329 | 225 |
| ENSDARG000 | 4;4;4 | 49196098:491 | 49196189:491 | +;+;+ | 1241 | NA | 2 | 2 | 2 | 0 | 0 | 0 |
| ENSDARG000 | 4;4;4;4;4;4;4; | 21488990:214 | 21490568:214 | -;-;-;-;-;-;-;-; | 4846 | ppp1r12a | 719 | 602 | 969 | 695 | 813 | 695 |
| ENSDARG000 | 4;4;4;4;4;4;4; | 16885854:168 | 16886000:168 | +;+;+;+;+;+;+; | 4131 | ETNK1 | 9126 | 3344 | 5378 | 7775 | 5549 | 10769 |
| ENSDARG000 | 4;4;4;4;4;4;4; | 57131456:571 | 57131556:571 | +;+;+;+;+;+;+; | 3303 | NA | 10 | 3 | 1 | 4 | 22 | 5 |
| ENSDARG000 | 4;4 | 38170708:381 | 38170747:381 | +;+ | 1345 | NA | 0 | 0 | 0 | 0 | 0 | 0 |
| ENSDARG000 | 4;4;4;4 | 49256109:492 | 49256773:492 | -;-;-;- | 900 | NA | 0 | 0 | 0 | 0 | 0 | 0 |
| ENSDARG000 | 4;4 | 38188797:381 | 38188943:381 | +;+ | 1485 | NA | 4 | 4 | 4 | 0 | 2 | 0 |
| ENSDARG000 | 4;4;4;4;4;4;4 | 9623461:9623 | 9623633:9623 | +;+;+;+;+;+;+ | 1239 | NA | 42 | 26 | 56 | 137 | 50 | 48 |
| ENSDARG000 | 4;4;4 | 38254699:382 | 38255860:382 | -;-;- | 1404 | NA | 18 | 16 | 16 | 14 | 0 | 2 |
| ENSDARG000 | 4;4;4;4;4;4;4; | 27100528:271 | 27100706:271 | +;+;+;+;+;+;+; | 1931 | alg12 | 241 | 392 | 381 | 211 | 378 | 377 |
| ENSDARG000 | 4;4;4;4;4;4;4; | 9592486:9592 | 9592604:9592 | +;+;+;+;+;+;+; | 2249 | hspa14 | 1219 | 1153 | 2002 | 1138 | 1643 | 1157 |
| ENSDARG000 | 4;4;4;4;4;4;4; | 49262131:492 | 49262259:492 | +;+;+;+;+;+;+; | 3520 | NA | 6 | 8 | 15 | 5 | 126 | 13 |
| ENSDARG000 | 4;4;4;4;4;4 | 57320332:573 | 57320747:573 | +;+;+;+;+;+ | 1866 | NA | 2 | 2 | 0 | 1 | 0 | 0 |
| ENSDARG000 | 4;4;4;4;4;4;4; | 7677318:7698 | 7677715:7699 | +;+;+;+;+;+;+; | 2335 | elk3 | 1112 | 1045 | 1159 | 903 | 1264 | 1094 |
| ENSDARG000 | 4;4;4;4;4 | 57348205:573 | 57348302:573 | +;+;+;+;+ | 522 | NA | 0 | 0 | 0 | 0 | 4 | 0 |
| ENSDARG000 | 4;4;4;4 | 49278098:492 | 49278997:492 | -;-;-;- | 1505 | NA | 10 | 4 | 4 | 2 | 8 | 6 |
| ENSDARG000 | 4;4;4;4;4;4;4; | 38265870:382 | 38267879:382 | -;-;-;-;-;-;-;-; | 5895 | NA | 11 | 5 | 14 | 2 | 12 | 33 |
| ENSDARG000 | 4;4;4;4;4;4;4; | 14193080:141 | 14194664:141 | -;-;-;-;-;-;-;-; | 2616 | twf1b | 226 | 243 | 224 | 132 | 258 | 162 |
| ENSDARG000 | 4;4;4;4;4;4 | 38267281:382 | 38267879:382 | -;-;-;-;-;- | 1729 | NA | 0 | 1 | 1 | 1 | 0 | 5 |
| ENSDARG000 | 4;4;4;4;4;4;4; | 9612574:9612 | 9612718:9612 | +;+;+;+;+;+;+; | 1956 | tmem243b | 1882 | 1231 | 1538 | 2512 | 1477 | 2820 |
| ENSDARG000 | 4;4 | 57351005:573 | 57353246:573 | -;- | 2424 | NA | 10 | 8 | 9 | 0 | 14 | 10 |
| ENSDARG000 | 4;4;4 | 57380165:574 | 57381232:574 | -;-;- | 1242 | NA | 2 | 3 | 0 | 0 | 0 | 1 |
| ENSDARG000 | 4;4;4;4;4 | 57419948:574 | 57420256:574 | +;+;+;+;+ | 1611 | NA | 2 | 26 | 0 | 3 | 20 | 28 |
| ENSDARG000 | 4;4;4 | 4938873:4939 | 4939062:4940 | +;+;+ | 776 | NA | 2 | 9 | 21 | 22 | 2 | 0 |
| ENSDARG000 | 4;4;4;4;4 | 57518790:575 | 57520374:575 | -;-;-;-;- | 1788 | zgc:174702 | 0 | 4 | 8 | 2 | 4 | 8 |
| ENSDARG000 | 4;4;4;4;4;4 | 9609905:9609 | 9609988:9609 | +;+;+;+;+;+ | 861 | meig1 | 48 | 50 | 45 | 58 | 26 | 36 |
| ENSDARG000 | 4;4 | 57516325:575 | 57516703:575 | +;+ | 597 | NA | 0 | 0 | 0 | 0 | 0 | 0 |
| ENSDARG000 | 4;4;4;4;4;4;4; | 9603306:9604 | 9604418:9604 | -;-;-;-;-;-;-;-; | 2903 | dclre1c | 278 | 166 | 272 | 209 | 376 | 381 |
| ENSDARG000 | 4;4;4;4;4;4;4; | 1516716:1530 | 1516792:1530 | +;+;+;+;+;+;+; | 5272 | slc38a4 | 40 | 6 | 36 | 8 | 36 | 12 |
| ENSDARG000 | 4;4;4;4;4;4;4 | 9630958:9630 | 9631130:9631 | +;+;+;+;+;+;+ | 1239 | NA | 4 | 0 | 0 | 10 | 4 | 6 |
| ENSDARG000 | 4;4;4;4 | 4963892:4963 | 4964031:4964 | +;+;+;+ | 1683 | NA | 9 | 24 | 84 | 137 | 19 | 85 |
| ENSDARG000 | 4;4;4;4;4;4;4; | 7718854:7720 | 7719003:7721 | -;-;-;-;-;-;-;-; | 2033 | cdk17 | 1322 | 962 | 1061 | 740 | 1438 | 890 |
| ENSDARG000 | 4;4;4 | 38410690:384 | 38411850:384 | -;-;- | 1458 | NA | 85 | 54 | 338 | 26 | 47 | 144 |
| ENSDARG000 | 4;4;4;4;4;4 | 38335552:383 | 38335722:383 | -;-;-;-;-;- | 1825 | NA | 0 | 2 | 8 | 0 | 0 | 4 |
| ENSDARG000 | 4;4;4;4;4;4;4; | 18804317:188 | 18804382:188 | +;+;+;+;+;+;+; | 2687 | slc26a3.2 | 4 | 8 | 6 | 4 | 10 | 12 |
| ENSDARG000 | 4;4 | 38475577:384 | 38476402:384 | -;- | 873 | NA | 8 | 0 | 0 | 0 | 0 | 1 |
| ENSDARG000 | 4;4;4;4 | 49457488:494 | 49457640:494 | +;+;+;+ | 870 | NA | 0 | 0 | 0 | 0 | 0 | 0 |
| ENSDARG000 | 4;4;4 | 38424204:384 | 38424280:384 | +;+;+ | 1362 | NA | 2 | 14 | 0 | 0 | 15 | 32 |
| ENSDARG000 | 4;4;4;4;4;4;4; | 38335535:385 | 38335729:385 | -;-;-;-;-;-;-;-; | 7759 | NA | 9 | 8 | 71 | 6 | 8 | 17 |
| ENSDARG000 | 4;4;4;4;4;4;4; | 12917164:129 | 12917715:129 | -;-;-;-;-;-;-;-; | 2988 | lemd3 | 1519 | 1342 | 1178 | 688 | 1213 | 848 |
| ENSDARG000 | 4;4;4;4;4;4;4; | 16982249:169 | 16985398:169 | -;-;-;-;-;-;-;-; | 5312 | sox5 | 8323 | 8203 | 12854 | 8273 | 13260 | 6270 |
| ENSDARG000 | 4;4;4;4;4;4 | 49501970:495 | 49502124:495 | +;+;+;+;+;+ | 1343 | NA | 10 | 27 | 50 | 6 | 14 | 26 |
| ENSDARG000 | 4;4;4;4;4;4;4; | 12013043:120 | 12013893:120 | +;+;+;+;+;+;+; | 3324 | cry1a | 955 | 482 | 445 | 608 | 943 | 873 |
| ENSDARG000 | 4;4;4;4;4;4 | 2637947:2638 | 2638245:2638 | +;+;+;+;+;+ | 1190 | dus4l | 152 | 186 | 200 | 90 | 172 | 82 |
| ENSDARG000 | 4;4;4 | 10769186:107 | 10769327:107 | +;+;+ | 330 | NA | 4 | 7 | 6 | 0 | 0 | 8 |
| ENSDARG000 | 4;4;4;4;4;4;4 | 10783815:107 | 10783853:107 | -;-;-;-;-;-;- | 474 | NA | 4 | 6 | 1 | 9 | 5 | 0 |
| ENSDARG000 | 4;4 | 10772097:107 | 10772246:107 | -;- | 385 | NA | 2 | 2 | 11 | 2 | 1 | 15 |
| ENSDARG000 | 4;4;4 | 10773580:107 | 10773669:107 | -;-;- | 330 | NA | 0 | 0 | 0 | 0 | 5 | 0 |
| ENSDARG000 | 4;4;4 | 27112399:271 | 27117228:271 | -;-;- | 4961 | zbed4 | 565 | 512 | 724 | 474 | 501 | 700 |
| ENSDARG000 | 4;4;4;4;4;4;4; | 27130412:271 | 27130672:271 | +;+;+;+;+;+;+; | 6725 | BRD1 | 662 | 339 | 663 | 351 | 691 | 542 |
| ENSDARG000 | 4;4;4;4;4;4;4; | 49550966:495 | 49556182:495 | -;-;-;-;-;-;-;-; | 8919 | NA | 1397 | 889 | 1043 | 824 | 1329 | 1620 |
| ENSDARG000 | 4;4;4;4;4;4;4; | 25220674:252 | 25220847:252 | +;+;+;+;+;+;+; | 3436 | ITIH5 | 40 | 4 | 14 | 16 | 12 | 79 |
| ENSDARG000 | 4;4;4;4;4;4;4; | 2655358:2657 | 2655478:2658 | +;+;+;+;+;+;+; | 1788 | bcap29 | 433 | 398 | 451 | 437 | 463 | 535 |
| ENSDARG000 | 4;4;4;4;4;4;4; | 27251865:272 | 27253248:272 | -;-;-;-;-;-;-;-; | 2403 | tafa5a | 0 | 2 | 0 | 0 | 0 | 6 |
| ENSDARG000 | 4;4;4;4;4;4;4; | 4971042:4971 | 4973234:4973 | -;-;-;-;-;-;-;-; | 4892 | NA | 22 | 16 | 26 | 24 | 12 | 18 |
| ENSDARG000 | 4;4;4;4;4;4;4; | 38547203:385 | 38547278:385 | +;+;+;+;+;+;+; | 2170 | NA | 325 | 135 | 76 | 22 | 47 | 139 |
| ENSDARG000 | 4;4;4;4;4;4;4; | 57564614:575 | 57564886:575 | -;-;-;-;-;-;-;-; | 1721 | NA | 0 | 0 | 0 | 0 | 0 | 0 |
| ENSDARG000 | 4;4;4;4;4;4;4; | 12931763:129 | 12931988:129 | +;+;+;+;+;+;+; | 1883 | wif1 | 345 | 311 | 100 | 178 | 50 | 308 |
| ENSDARG000 | 4;4 | 38564564:385 | 38564660:385 | -;- | 833 | NA | 60 | 31 | 19 | 2 | 17 | 32 |
| ENSDARG000 | 4;4;4 | 7817996:7820 | 7818079:7820 | +;+;+ | 913 | si:ch1073-67i | 272 | 72 | 34 | 10 | 106 | 56 |
| ENSDARG000 | 4 | 5741733 | 5744457 | + | 2725 | pou3f2a | 0 | 0 | 0 | 0 | 0 | 0 |
| ENSDARG000 | 4;4;4 | 38634554:386 | 38634579:386 | +;+;+ | 189 | NA | 0 | 0 | 0 | 0 | 0 | 0 |
| ENSDARG000 | 4;4;4 | 7822773:7824 | 7822879:7824 | +;+;+ | 1822 | si:ch1073-67i | 107 | 234 | 46 | 44 | 108 | 124 |
| ENSDARG000 | 4;4;4;4;4;4;4; | 7827261:7827 | 7827396:7827 | +;+;+;+;+;+;+; | 2995 | phyh | 782 | 626 | 569 | 360 | 1032 | 869 |
| ENSDARG000 | 4;4;4;4;4;4 | 5750854:5753 | 5751882:5753 | -;-;-;-;-;- | 2166 | FAXC | 15 | 4 | 6 | 4 | 10 | 6 |
| ENSDARG000 | 4;4;4;4;4;4;4; | 5764486:5768 | 5766526:5768 | -;-;-;-;-;-;-;-; | 2968 | ccnc | 686 | 643 | 838 | 798 | 793 | 744 |
| ENSDARG000 | 4;4 | 17244072:172 | 17244212:172 | -;- | 262 | NA | 0 | 0 | 0 | 0 | 0 | 1 |
| ENSDARG000 | 4;4;4;4;4;4;4; | 14247693:142 | 14250156:142 | -;-;-;-;-;-;-;-; | 5154 | NELL2 | 0 | 0 | 2 | 2 | 0 | 4 |
| ENSDARG000 | 4;4;4;4;4;4;4; | 18824959:188 | 18825086:188 | +;+;+;+;+;+;+; | 2244 | slc26a3.1 | 0 | 0 | 6 | 0 | 0 | 1 |
| ENSDARG000 | 4;4;4 | 1576889:1577 | 1576967:1577 | +;+;+ | 620 | NA | 0 | 0 | 0 | 0 | 0 | 0 |
| ENSDARG000 | 4;4;4;4;4;4;4; | 25251855:252 | 25252571:252 | -;-;-;-;-;-;-;-; | 1793 | tmem110l | 514 | 389 | 265 | 190 | 200 | 632 |
| ENSDARG000 | 4;4;4;4;4;4;4; | 17245001:172 | 17245043:172 | +;+;+;+;+;+;+; | 2325 | dnai7 | 0 | 0 | 6 | 0 | 14 | 6 |
| ENSDARG000 | 4;4 | 49587497:495 | 49588782:495 | -;- | 1439 | NA | 0 | 0 | 8 | 0 | 4 | 4 |
| ENSDARG000 | 4;4;4;4;4;4;4; | 15490215:154 | 15490818:154 | -;-;-;-;-;-;-;-; | 1827 | chchd3a | 450 | 410 | 590 | 402 | 543 | 508 |

| | | | | | | | | | | | | |
|---|---|---|---|---|---|---|---|---|---|---|---|---|
| ENSDARG000 | 4;4;4;4;4;4;4; | 38818194:388 | 38818957:388 | -;-;-;-;-;-;-;-; | 3750 | NA | 47 | 18 | 20 | 4 | 5 | 5 |
| ENSDARG000 | 4;4;4 | 49600697:496 | 49600775:496 | +;+;+ | 732 | NA | 0 | 0 | 0 | 0 | 0 | 0 |
| ENSDARG000 | 4;4;4;4;4;4;4; | 57580376:575 | 57580611:575 | +;+;+;+;+;+;+; | 996 | NA | 0 | 0 | 0 | 0 | 0 | 0 |
| ENSDARG000 | 4;4;4;4;4;4;4; | 57580303:575 | 57580611:575 | +;+;+;+;+;+;+; | 2685 | NA | 2 | 2 | 6 | 8 | 0 | 8 |
| ENSDARG000 | 4;4;4;4;4 | 7841627:7842 | 7841725:7842 | +;+;+;+;+ | 580 | ucmaa | 67 | 5 | 8 | 0 | 0 | 10 |
| ENSDARG000 | 4;4;4;4 | 49605855:496 | 49606087:496 | -;-;-;- | 776 | NA | 0 | 0 | 0 | 0 | 0 | 0 |
| ENSDARG000 | 4;4;4;4;4;4;4; | 21648010:216 | 21649201:216 | -;-;-;-;-;-;-;-; | 2581 | rps16 | 9556 | 9200 | 14457 | 12196 | 8683 | 13175 |
| ENSDARG000 | 4;4;4;4;4;4;4; | 7851129:7852 | 7851438:7852 | -;-;-;-;-;-;-;-; | 2837 | mcm10 | 32 | 151 | 200 | 48 | 178 | 56 |
| ENSDARG000 | 4;4 | 10793677:107 | 10793839:107 | -;- | 321 | NA | 0 | 0 | 0 | 0 | 0 | 53 |
| ENSDARG000 | 4;4;4;4;4;4;4; | 10811490:108 | 10813824:108 | -;-;-;-;-;-;-;-; | 5190 | NA | 1286 | 1177 | 2955 | 2740 | 1674 | 1942 |
| ENSDARG000 | 4;4;4;4;4;4;4; | 27626869:276 | 27628954:276 | -;-;-;-;-;-;-;-; | 4149 | tbc1d22a | 277 | 258 | 344 | 321 | 370 | 313 |
| ENSDARG000 | 4;4;4;4;4;4;4; | 1600034:1601 | 1600325:1601 | +;+;+;+;+;+;+; | 5401 | slc38a2 | 1064 | 3331 | 3782 | 2167 | 1096 | 1380 |
| ENSDARG000 | 4;4;4;4;4;4;4; | 15605844:156 | 15605923:156 | +;+;+;+;+;+;+; | 3102 | exoc4 | 571 | 520 | 694 | 481 | 670 | 442 |
| ENSDARG000 | 4;4;4;4;4;4 | 10790960:107 | 10791005:107 | -;-;-;-;-;- | 443 | NA | 0 | 0 | 0 | 0 | 0 | 9 |
| ENSDARG000 | 4;4;4;4;4 | 10816240:108 | 10816330:108 | +;+;+;+;+ | 927 | NA | 6 | 2 | 8 | 2 | 1 | 0 |
| ENSDARG000 | 4;4;4 | 49664256:496 | 49664345:496 | +;+;+ | 1272 | NA | 14 | 14 | 18 | 2 | 32 | 2 |
| ENSDARG000 | 4;4;4;4;4 | 49656481:497 | 49656829:497 | -;-;-;-;- | 2646 | NA | 9 | 33 | 18 | 6 | 15 | 32 |
| ENSDARG000 | 4;4 | 57709567:577 | 57709680:577 | +;+ | 548 | NA | 0 | 0 | 0 | 0 | 0 | 4 |
| ENSDARG000 | 4;4;4;4;4;4;4; | 15757435:157 | 15758142:157 | -;-;-;-;-;-;-;-; | 1156 | si:ch211-92l1 | 8 | 60 | 50 | 26 | 34 | 20 |
| ENSDARG000 | 4;4;4;4 | 57711173:577 | 57711644:577 | -;-;-;- | 1326 | NA | 24 | 0 | 24 | 20 | 8 | 10 |
| ENSDARG000 | 4;4;4 | 38937575:389 | 38938109:389 | +;+;+ | 1317 | NA | 1 | 2 | 2 | 0 | 12 | 0 |
| ENSDARG000 | 4;4;4;4 | 38981587:389 | 38981685:389 | +;+;+;+ | 432 | NA | 0 | 0 | 0 | 0 | 0 | 0 |
| ENSDARG000 | 4;4 | 21656245:216 | 21656582:216 | -;- | 722 | NA | 0 | 0 | 0 | 0 | 0 | 0 |
| ENSDARG000 | 4;4;4;4 | 21653609:216 | 21653654:216 | +;+;+;+ | 750 | NA | 4 | 1 | 2 | 2 | 0 | 0 |
| ENSDARG000 | 4;4 | 49727181:497 | 49727345:497 | -;- | 360 | NA | 0 | 0 | 0 | 0 | 0 | 0 |
| ENSDARG000 | 4;4 | 49725892:497 | 49725938:497 | +;+ | 401 | NA | 0 | 0 | 0 | 0 | 0 | 0 |
| ENSDARG000 | 4;4;4;4;4;4;4; | 27898833:279 | 27898983:279 | +;+;+;+;+;+;+; | 1950 | cerk | 193 | 88 | 193 | 50 | 203 | 210 |
| ENSDARG000 | 4;4;4;4 | 49748740:497 | 49748810:497 | +;+;+;+ | 860 | NA | 0 | 0 | 0 | 0 | 0 | 0 |
| ENSDARG000 | 4;4;4;4;4;4;4; | 2674898:2675 | 2675359:2675 | -;-;-;-;-;-;-;-; | 2804 | slco1c1 | 1147 | 1197 | 1618 | 1297 | 1423 | 1318 |
| ENSDARG000 | 4;4;4 | 21717793:217 | 21718302:217 | +;+;+ | 720 | myf6 | 0 | 0 | 1 | 0 | 0 | 0 |
| ENSDARG000 | 4;4 | 57855379:578 | 57856984:578 | -;- | 1748 | zgc:174314 | 6 | 14 | 30 | 4 | 12 | 52 |
| ENSDARG000 | 4;4;4;4;4 | 38990661:389 | 38990724:389 | +;+;+;+;+ | 578 | si:dkev-122c1 | 0 | 0 | 0 | 0 | 0 | 0 |
| ENSDARG000 | 4;4;4 | 12966640:129 | 12966857:129 | +;+;+ | 540 | vhll | 56 | 87 | 65 | 79 | 60 | 92 |
| ENSDARG000 | 4;4;4;4 | 57887079:578 | 57887278:578 | -;-;-;- | 666 | NA | 0 | 0 | 0 | 0 | 0 | 4 |
| ENSDARG000 | 4;4;4;4;4 | 57881965:578 | 57882040:578 | +;+;+;+;+ | 604 | NA | 0 | 0 | 0 | 0 | 0 | 0 |
| ENSDARG000 | 4;4;4;4;4;4 | 57880688:578 | 57880734:578 | -;-;-;-;-;- | 523 | NA | 0 | 0 | 0 | 0 | 0 | 0 |
| ENSDARG000 | 4;4;4 | 39055027:390 | 39055061:390 | +;+;+ | 1606 | NA | 0 | 33 | 0 | 6 | 46 | 83 |
| ENSDARG000 | 4;4;4;4;4;4;4 | 21741228:217 | 21741714:217 | +;+;+;+;+;+;+ | 1415 | myf5 | 4 | 0 | 8 | 0 | 0 | 4 |
| ENSDARG000 | 4;4;4;4;4;4;4; | 18836791:188 | 18838093:188 | -;-;-;-;-;-;-;-; | 1953 | cbll1 | 149 | 238 | 266 | 145 | 298 | 184 |
| ENSDARG000 | 4;4;4;4;4 | 39105046:391 | 39107472:391 | -;-;-;-;- | 2601 | NA | 0 | 0 | 0 | 0 | 0 | 0 |
| ENSDARG000 | 4;4;4;4;4;4;4; | 12971255:129 | 12972436:129 | -;-;-;-;-;-;-;-; | 1897 | tmbim4 | 566 | 419 | 601 | 570 | 443 | 682 |
| ENSDARG000 | 4;4;4 | 59748607:597 | 59748623:597 | +;+;+ | 2094 | znf1068 | 64 | 55 | 76 | 77 | 66 | 110 |
| ENSDARG000 | 4;4;4;4;4;4;4; | 39158005:391 | 39163228:391 | -;-;-;-;-;-;-;-; | 9546 | si:ch211-22k7 | 75 | 17 | 66 | 24 | 15 | 7 |
| ENSDARG000 | 4;4;4;4;4;4;4; | 59766049:597 | 59766775:597 | -;-;-;-;-;-;-;- | 3534 | NA | 302 | 221 | 301 | 56 | 247 | 213 |
| ENSDARG000 | 4;4;4;4;4 | 18839856:188 | 18840006:188 | -;-;-;-;- | 813 | MCAT | 0 | 0 | 0 | 0 | 0 | 0 |
| ENSDARG000 | 4;4;4;4 | 18846624:188 | 18847087:188 | -;-;-;- | 1380 | mcat | 32 | 37 | 20 | 28 | 37 | 9 |
| ENSDARG000 | 4;4;4;4;4;4;4; | 5025625:5026 | 5026371:5026 | -;-;-;-;-;-;-;-; | 2851 | ahcyl2b | 912 | 1066 | 1537 | 1008 | 1319 | 727 |
| ENSDARG000 | 4;4;4;4;4;4;4; | 59841099:598 | 59841158:598 | +;+;+;+;+;+;+; | 3867 | NA | 553 | 524 | 501 | 76 | 452 | 350 |
| ENSDARG000 | 4;4;4;4;4;4;4; | 18843015:188 | 18843125:188 | +;+;+;+;+;+;+; | 1262 | bik | 52 | 8 | 17 | 17 | 58 | 34 |
| ENSDARG000 | 4;4;4 | 57983841:579 | 57985359:579 | -;-;- | 1763 | NA | 0 | 2 | 4 | 0 | 4 | 0 |
| ENSDARG000 | 4;4;4;4;4;4 | 21751762:217 | 21753729:217 | -;-;-;-;-;- | 2723 | lin7a | 912 | 319 | 914 | 629 | 2599 | 592 |
| ENSDARG000 | 4;4;4;4;4;4;4; | 12033873:120 | 12034883:120 | -;-;-;-;-;-;-;-; | 2153 | si:dkev-222f8. | 2666 | 2135 | 1748 | 2950 | 1659 | 2826 |
| ENSDARG000 | 4;4;4 | 58057168:580 | 58057197:580 | +;+;+ | 2822 | si:dkev-57k17 | 24 | 25 | 93 | 2 | 26 | 12 |
| ENSDARG000 | 4;4;4;4;4;4;4; | 58240778:582 | 58241202:582 | +;+;+;+;+;+;+; | 4519 | NA | 24 | 28 | 17 | 6 | 60 | 40 |
| ENSDARG000 | 4;4;4;4;4;4;4; | 25300910:253 | 25301074:253 | +;+;+;+;+;+;+; | 3240 | SFMBT2 | 30 | 64 | 135 | 57 | 51 | 128 |
| ENSDARG000 | 4 | 12031958 | 12033866 | + | 1909 | tnnt2c | 24 | 37 | 9 | 41 | 25 | 52 |
| ENSDARG000 | 4;4;4;4;4;4;4; | 58272337:582 | 58272466:582 | +;+;+;+;+;+;+; | 4147 | NA | 6 | 1 | 10 | 8 | 2 | 15 |
| ENSDARG000 | 4;4;4;4;4;4 | 59889820:598 | 59889863:598 | +;+;+;+;+;+ | 2320 | NA | 12 | 8 | 21 | 14 | 6 | 6 |
| ENSDARG000 | 4;4;4;4;4;4;4; | 28062473:280 | 28064390:280 | -;-;-;-;-;-;-;-; | 3696 | GRAMD4 | 0 | 2 | 0 | 0 | 4 | 2 |
| ENSDARG000 | 4;4;4;4;4;4;4; | 7870384:7870 | 7871091:7871 | -;-;-;-;-;-;-;-; | 1628 | nudt5 | 609 | 639 | 1866 | 1231 | 654 | 699 |
| ENSDARG000 | 4;4;4 | 60001731:600 | 60001807:600 | +;+;+ | 1013 | NA | 4 | 2 | 0 | 0 | 0 | 0 |
| ENSDARG000 | 4;4;4 | 60062722:600 | 60063065:600 | -;-;- | 528 | NA | 0 | 2 | 6 | 1 | 35 | 0 |
| ENSDARG000 | 4;4;4;4;4;4;4; | 60048421:600 | 60049030:600 | -;-;-;-;-;-;-;- | 1569 | NA | 24 | 21 | 11 | 2 | 31 | 22 |
| ENSDARG000 | 4;4;4 | 60072177:600 | 60072358:600 | +;+;+ | 656 | NA | 12 | 3 | 8 | 2 | 9 | 11 |
| ENSDARG000 | 4;4;4;4;4 | 14343706:143 | 14343758:143 | +;+;+;+;+ | 971 | prl2 | 0 | 2 | 0 | 0 | 0 | 0 |
| ENSDARG000 | 4;4;4;4;4;4 | 60121087:601 | 60121204:601 | -;-;-;-;-;- | 912 | NA | 0 | 0 | 0 | 0 | 0 | 1 |
| ENSDARG000 | 4;4;4;4;4;4;4; | 21866851:218 | 21867651:218 | +;+;+;+;+;+;+; | 3759 | ACSS3 | 209 | 117 | 163 | 94 | 139 | 207 |
| ENSDARG000 | 4;4;4;4;4;4;4; | 39368978:393 | 39369062:393 | +;+;+;+;+;+;+; | 713 | NA | 6 | 0 | 8 | 2 | 0 | 18 |
| ENSDARG000 | 4;4;4;4;4;4;4 | 14360372:143 | 14360843:143 | +;+;+;+;+;+;+ | 2124 | NUAK1 | 38 | 60 | 261 | 33 | 45 | 84 |
| ENSDARG000 | 4;4;4 | 39428602:394 | 39430371:394 | -;-;- | 2063 | NA | 4 | 0 | 2 | 0 | 0 | 0 |
| ENSDARG000 | 4;4;4;4;4;4 | 12993913:129 | 12994182:129 | -;-;-;-;-;- | 658 | si:dkey-6a5.3 | 6 | 0 | 3 | 21 | 10 | 0 |
| ENSDARG000 | 4;4;4;4;4;4;4; | 39451635:394 | 39451762:394 | -;-;-;-;-;-;-;- | 798 | NA | 0 | 0 | 0 | 0 | 0 | 0 |
| ENSDARG000 | 4;4;4;4;4;4;4; | 12997407:129 | 12997585:129 | +;+;+;+;+;+;+; | 988 | NA | 43 | 7 | 14 | 27 | 27 | 9 |
| ENSDARG000 | 4;4;4;4;4;4;4; | 39372775:394 | 39372860:394 | -;-;-;-;-;-;-;-; | 4451 | NA | 15 | 7 | 29 | 3 | 14 | 21 |
| ENSDARG000 | 4;4;4 | 39482550:394 | 39482611:394 | +;+;+ | 751 | NA | 0 | 0 | 0 | 0 | 0 | 0 |
| ENSDARG000 | 4;4;4;4;4;4;4; | 17252596:172 | 17254283:172 | -;-;-;-;-;-;-;-; | 6003 | lrmp | 6 | 6 | 20 | 8 | 5 | 27 |
| ENSDARG000 | 4;4;4 | 60215803:602 | 60216450:602 | -;-;- | 870 | NA | 0 | 5 | 6 | 0 | 0 | 0 |
| ENSDARG000 | 4;4 | 14399472:144 | 14399679:144 | +;+ | 767 | NA | 0 | 0 | 0 | 0 | 0 | 4 |
| ENSDARG000 | 4;4;4;4;4;4;4; | 1619584:1620 | 1619751:1620 | +;+;+;+;+;+;+; | 7491 | NA | 2031 | 1754 | 2663 | 1748 | 2095 | 1797 |
| ENSDARG000 | 4;4;4;4;4;4;4; | 14433890:144 | 14433985:144 | -;-;-;-;-;-;-;-; | 454 | si:dkev-261e2 | 2 | 2 | 4 | 4 | 2 | 2 |
| ENSDARG000 | 4;4 | 60241780:602 | 60241914:602 | -;- | 288 | NA | 0 | 0 | 0 | 0 | 0 | 0 |
| ENSDARG000 | 4;4;4 | 60296027:602 | 60296153:602 | +;+;+ | 1144 | NA | 0 | 0 | 0 | 0 | 0 | 0 |
| ENSDARG000 | 4;4;4;4;4;4;4; | 60310050:603 | 60310710:603 | -;-;-;-;-;-;-;- | 1403 | NA | 12 | 7 | 44 | 0 | 0 | 0 |
| ENSDARG000 | 4;4;4;4;4;4;4; | 58298838:582 | 58298879:583 | +;+;+;+;+;+;+; | 956 | NA | 4 | 23 | 4 | 8 | 59 | 27 |
| ENSDARG000 | 4;4;4;4;4 | 60339527:603 | 60339783:603 | -;-;-;-;- | 1310 | NA | 6 | 0 | 32 | 0 | 2 | 8 |
| ENSDARG000 | 4;4;4 | 58363998:583 | 58364087:583 | +;+;+ | 1870 | NA | 4 | 36 | 11 | 1 | 66 | 52 |
| ENSDARG000 | 4;4;4;4 | 58400159:584 | 58400576:584 | +;+;+;+ | 1517 | NA | 0 | 0 | 0 | 0 | 0 | 0 |
| ENSDARG000 | 4;4;4;4;4;4;4; | 2752852:2752 | 2753092:2753 | -;-;-;-;-;-;-;-; | 4092 | PDE3A | 34 | 31 | 74 | 71 | 90 | 9 |
| ENSDARG000 | 4;4;4;4;4;4;4; | 60358706:603 | 60359628:603 | -;-;-;-;-;-;-;-; | 2234 | NA | 3 | 0 | 0 | 2 | 0 | 0 |
| ENSDARG000 | 4;4 | 39576200:395 | 39576427:395 | +;+ | 552 | NA | 0 | 0 | 0 | 0 | 0 | 0 |
| ENSDARG000 | 4;4 | 39643903:396 | 39644475:396 | -;- | 717 | NA | 0 | 0 | 0 | 0 | 0 | 0 |
| ENSDARG000 | 4;4;4 | 60381608:603 | 60381642:603 | +;+;+ | 1785 | NA | 8 | 21 | 36 | 2 | 16 | 12 |
| ENSDARG000 | 4;4;4;4;4;4;4; | 70317135:703 | 70317182:703 | +;+;+;+;+;+;+; | 3595 | NA | 205 | 11 | 16 | 15 | 168 | 75 |
| ENSDARG000 | 4;4;4;4;4;4;4 | 10888762:108 | 10889083:108 | +;+;+;+;+;+;+ | 4974 | syt10 | 0 | 0 | 0 | 0 | 0 | 0 |
| ENSDARG000 | 4;4;4;4;4;4 | 58446500:584 | 58446741:584 | -;-;-;-;-;- | 1442 | NA | 0 | 0 | 0 | 0 | 2 | 0 |
| ENSDARG000 | 4;4;4;4;4;4;4; | 28190013:281 | 28192662:281 | -;-;-;-;-;-;-;-; | 11760 | CELSR1 | 136 | 32 | 120 | 153 | 182 | 72 |
| ENSDARG000 | 4;4;4;4;4;4;4; | 15819728:158 | 15819978:158 | +;+;+;+;+;+;+; | 2549 | LRGUK | 8 | 0 | 0 | 0 | 0 | 4 |
| ENSDARG000 | 4;4;4 | 58576146:585 | 58576170:585 | +;+;+ | 1278 | NA | 0 | 0 | 0 | 0 | 3 | 0 |
| ENSDARG000 | 4;4 | 39669246:396 | 39669473:396 | +;+ | 549 | NA | 0 | 0 | 0 | 0 | 0 | 0 |
| ENSDARG000 | 4;4;4;4;4;4;4; | 39749453:397 | 39749530:397 | +;+;+;+;+;+;+; | 3332 | si:dkeyp-4c4. | 3 | 0 | 0 | 0 | 4 | 0 |
| ENSDARG000 | 4;4;4;4 | 13008680:130 | 13008785:130 | +;+;+;+ | 700 | NA | 22 | 6 | 24 | 18 | 39 | 16 |
| ENSDARG000 | 4;4;4 | 70420080:704 | 70420232:704 | +;+;+ | 805 | NA | 6 | 0 | 13 | 5 | 13 | 7 |
| ENSDARG000 | 4;4;4;4;4;4;4; | 70409414:704 | 70409606:704 | +;+;+;+;+;+;+; | 5364 | NA | 10 | 24 | 7 | 2 | 45 | 20 |
| ENSDARG000 | 4;4;4;4;4;4;4; | 25410293:254 | 25410410:254 | +;+;+;+;+;+;+; | 2394 | prkcq | 6 | 68 | 30 | 40 | 77 | 75 |
| ENSDARG000 | 4;4 | 70414733:704 | 70414947:704 | +;+ | 436 | NA | 0 | 0 | 0 | 0 | 0 | 0 |
| ENSDARG000 | 4;4;4 | 70351760:703 | 70351912:703 | +;+;+ | 1393 | NA | 98 | 2 | 1 | 0 | 50 | 76 |
| ENSDARG000 | 4;4;4;4;4;4;4; | 70341246:703 | 70341342:703 | +;+;+;+;+;+;+; | 3205 | NA | 63 | 11 | 11 | 11 | 33 | 41 |
| ENSDARG000 | 4;4 | 58964468:589 | 58964930:589 | +;+ | 702 | NA | 0 | 0 | 0 | 0 | 0 | 0 |
| ENSDARG000 | 4;4;4 | 58768257:587 | 58768550:587 | -;-;- | 536 | NA | 0 | 2 | 0 | 0 | 0 | 0 |
| ENSDARG000 | 4;4;4;4;4;4;4; | 21962316:219 | 21962374:219 | -;-;-;-;-;-;-;-; | 3713 | ppfia2 | 3021 | 2308 | 3102 | 3193 | 3543 | 3241 |
| ENSDARG000 | 4;4;4 | 58849254:588 | 58849738:588 | -;-;- | 770 | NA | 0 | 0 | 0 | 0 | 0 | 0 |
| ENSDARG000 | 4;4;4;4;4 | 58942335:589 | 58943643:589 | -;-;-;-;- | 2438 | NA | 8 | 0 | 1 | 0 | 0 | 0 |
| ENSDARG000 | 4;4;4;4 | 70380429:703 | 70380533:703 | +;+;+;+ | 758 | NA | 8 | 5 | 6 | 0 | 6 | 0 |
| ENSDARG000 | 4;4 | 58918869:589 | 58919094:589 | +;+ | 549 | NA | 0 | 0 | 0 | 0 | 0 | 0 |
| ENSDARG000 | 4;4;4 | 59041720:590 | 59042494:590 | -;-;- | 1036 | NA | 0 | 2 | 0 | 0 | 0 | 0 |
| ENSDARG000 | 4;4;4;4;4;4;4; | 60428812:604 | 60428864:604 | +;+;+;+;+;+;+; | 3535 | NA | 32 | 47 | 32 | 4 | 14 | 10 |
| ENSDARG000 | 4;4;4;4 | 59061652:590 | 59061688:590 | +;+;+;+ | 1247 | NA | 0 | 0 | 0 | 0 | 4 | 0 |
| ENSDARG000 | 4;4;4;4;4;4;4; | 1647098:1648 | 1648645:1648 | -;-;-;-;-;-;-;-; | 6475 | arid2 | 1128 | 687 | 1474 | 797 | 1351 | 818 |
| ENSDARG000 | 4;4;4;4;4;4;4; | 58673287:588 | 58673351:588 | -;-;-;-;-;-;-;- | 2004 | NA | 0 | 0 | 0 | 0 | 2 | 0 |
| ENSDARG000 | 4;4 | 39829175:398 | 39829496:398 | -;- | 506 | NA | 0 | 0 | 0 | 0 | 0 | 0 |
| ENSDARG000 | 4;4;4;4 | 58976357:589 | 58976572:589 | -;-;-;- | 489 | NA | 0 | 0 | 0 | 0 | 0 | 0 |
| ENSDARG000 | 4;4;4;4;4;4;4; | 58864222:588 | 58865530:588 | -;-;-;-;-;-;-;- | 2635 | NA | 1 | 6 | 3 | 2 | 4 | 4 |
| ENSDARG000 | 4;4;4;4;4;4;4; | 5776347:5777 | 5776643:5778 | +;+;+;+;+;+;+; | 2388 | ngs | 0 | 0 | 0 | 0 | 0 | 0 |
| ENSDARG000 | 4;4;4 | 58695514:586 | 58695664:586 | +;+;+ | 1725 | NA | 0 | 0 | 0 | 0 | 0 | 0 |
| ENSDARG000 | 4;4;4;4;4;4;4; | 7876197:7876 | 7876336:7876 | +;+;+;+;+;+;+; | 3317 | cdc123 | 270 | 224 | 538 | 374 | 262 | 276 |
| ENSDARG000 | 4;4 | 21966094:219 | 21966135:219 | +;+ | 1137 | NA | 6 | 4 | 0 | 0 | 0 | 0 |
| ENSDARG000 | 4;4;4;4;4;4;4; | 18852821:188 | 18855037:188 | -;-;-;-;-;-;-;-; | 2889 | rap1b | 4054 | 4341 | 3464 | 3393 | 3732 | 3458 |
| ENSDARG000 | 4;4;4;4 | 60515709:605 | 60516040:605 | -;-;-;- | 632 | NA | 0 | 0 | 0 | 0 | 0 | 0 |
| ENSDARG000 | 4;4;4;4;4;4;4; | 10926171:109 | 10927222:109 | -;-;-;-;-;-;-;-; | 3710 | NA | 0 | 0 | 0 | 0 | 0 | 2 |
| ENSDARG000 | 4;4;4;4;4;4;4; | 60482963:604 | 60483042:604 | +;+;+;+;+;+;+; | 5443 | NA | 43 | 9 | 30 | 4 | 27 | 16 |
| ENSDARG000 | 4;4;4;4;4;4;4 | 60547709:605 | 60547805:605 | +;+;+;+;+;+;+ | 2442 | NA | 8 | 1 | 20 | 0 | 8 | 5 |
| ENSDARG000 | 4;4;4 | 39919216:399 | 39920694:399 | -;-;- | 1737 | NA | 4 | 6 | 2 | 3 | 2 | 20 |
| ENSDARG000 | 4;4;4;4;4;4 | 14481835:144 | 14481922:144 | -;-;-;-;-;- | 1792 | NA | 8 | 12 | 28 | 16 | 8 | 10 |
| ENSDARG000 | 4;4;4;4;4;4;4; | 14449233:144 | 14451056:144 | -;-;-;-;-;-;-;-; | 7524 | plxnb2a.1 | 2020 | 1812 | 2262 | 1662 | 2100 | 1736 |
| ENSDARG000 | 4;4;4;4;4;4;4; | 10949389:109 | 10949480:109 | -;-;-;-;-;-;-;-; | 4832 | NA | 0 | 0 | 2 | 0 | 0 | 3 |
| ENSDARG000 | 4;4;4;4;4;4;4; | 60760283:607 | 60761075:607 | -;-;-;-;-;-;-;-; | 4142 | NA | 6 | 4 | 3 | 0 | 3 | 5 |
| ENSDARG000 | 4;4;4;4;4;4;4; | 10972585:109 | 10974944:109 | -;-;-;-;-;-;-;-; | 5146 | tmtc2a | 44 | 78 | 156 | 60 | 62 | 74 |
| ENSDARG000 | 4;4;4;4;4;4;4; | 39992253:399 | 39992912:399 | -;-;-;-;-;-;-;-; | 3997 | NA | 22 | 339 | 18 | 108 | 363 | 330 |
| ENSDARG000 | 4;4;4 | 60777691:607 | 60777752:607 | +;+;+ | 751 | NA | 0 | 0 | 0 | 0 | 0 | 0 |
| ENSDARG000 | 4;4;4;4;4;4;4; | 10959442:109 | 10960748:109 | -;-;-;-;-;-;-;-; | 6167 | NA | 0 | 0 | 0 | 0 | 0 | 14 |
| ENSDARG000 | 4 | 1685021 | 1685345 | + | 325 | NA | 4 | 2 | 10 | 0 | 4 | 18 |
| ENSDARG000 | 4;4;4;4;4;4;4; | 2923199:2926 | 2923359:2928 | -;-;-;-;-;-;-;-; | 2814 | aebp2 | 210 | 235 | 277 | 198 | 273 | 198 |
| ENSDARG000 | 4;4;4;4;4;4;4; | 1689312:1692 | 1690812:1692 | -;-;-;-;-;-;-;- | 2432 | gas2l3 | 138 | 112 | 206 | 153 | 90 | 204 |
| ENSDARG000 | 4;4;4 | 70434872:704 | 70435024:704 | +;+;+ | 790 | NA | 4 | 1 | 1 | 4 | 5 | 4 |
| ENSDARG000 | 4;4;4;4;4;4;4; | 28338642:283 | 28339267:283 | -;-;-;-;-;-;-;-; | 1833 | trmu | 105 | 136 | 178 | 41 | 140 | 117 |
| ENSDARG000 | 4;4;4;4;4;4;4; | 7888047:7943 | 7888504:7944 | +;+;+;+;+;+;+; | 2842 | camk1da | 45 | 0 | 60 | 0 | 24 | 34 |
| ENSDARG000 | 4;4;4;4;4;4;4; | 13035870:130 | 13036198:130 | -;-;-;-;-;-;-;-; | 4726 | grip1 | 5 | 6 | 10 | 0 | 56 | 0 |
| ENSDARG000 | 4;4 | 60811087:608 | 60811129:608 | -;- | 206 | NA | 0 | 0 | 0 | 0 | 0 | 0 |
| ENSDARG000 | 4;4;4 | 60811946:608 | 60812048:608 | -;-;- | 388 | NA | 0 | 2 | 0 | 0 | 0 | 0 |
| ENSDARG000 | 4;4;4 | 60859063:608 | 60859631:608 | -;-;- | 846 | NA | 6 | 0 | 3 | 0 | 0 | 5 |
| ENSDARG000 | 4;4 | 60894595:608 | 60894807:608 | +;+ | 369 | NA | 0 | 0 | 0 | 0 | 0 | 0 |
| ENSDARG000 | 4;4;4;4;4;4;4; | 60889489:608 | 60890065:608 | -;-;-;-;-;-;-;-; | 3682 | NA | 70 | 16 | 41 | 10 | 20 | 10 |
| ENSDARG000 | 4;4;4 | 9649283:9651 | 9649683:9651 | -;-;- | 700 | NA | 0 | 0 | 0 | 0 | 0 | 0 |
| ENSDARG000 | 4;4 | 18952482:189 | 18953705:189 | -;- | 1293 | NA | 110 | 70 | 112 | 34 | 40 | 105 |
| ENSDARG000 | 4;4;4;4 | 70570937:705 | 70571019:705 | +;+;+;+ | 1467 | NA | 0 | 0 | 4 | 8 | 2 | 0 |
| ENSDARG000 | 4;4;4;4;4;4;4; | 70573439:705 | 70574218:705 | -;-;-;-;-;-;-;-; | 2716 | NA | 0 | 0 | 0 | 0 | 2 | 0 |
| ENSDARG000 | 4;4;4;4;4;4;4; | 70566694:705 | 70569078:705 | -;-;-;-;-;-;-;- | 2964 | NA | 0 | 0 | 0 | 0 | 0 | 0 |
| ENSDARG000 | 4;4;4;4;4;4 | 70556298:705 | 70556369:705 | +;+;+;+;+;+ | 1698 | NA | 0 | 0 | 0 | 0 | 0 | 0 |
| ENSDARG000 | 4;4;4;4 | 15942075:159 | 15942209:159 | +;+;+;+ | 503 | NA | 2 | 4 | 0 | 0 | 0 | 0 |
| ENSDARG000 | 4;4 | 15944245:159 | 15944274:159 | +;+ | 89 | NA | 0 | 0 | 0 | 0 | 0 | 0 |
| ENSDARG000 | 4;4;4;4;4;4;4; | 59112311:591 | 59112547:591 | -;-;-;-;-;-;-;- | 2802 | NA | 1 | 17 | 4 | 6 | 8 | 7 |
| ENSDARG000 | 4;4;4;4;4;4;4; | 15889437:158 | 15889617:158 | +;+;+;+;+;+;+; | 7679 | NA | 2 | 4 | 4 | 0 | 2 | 1 |

| | | | | | | | | | | | |
|---|---|---|---|---|---|---|---|---|---|---|---|
| ENSDARG000 | 4;4 | 59146023:591 | 59146264:591 | -;- | 515 | NA | 0 | 0 | 0 | 0 | 0 | 0 |
| ENSDARG000 | 4;4 | 60934218:609 | 60934369:609 | +;+ | 1739 | NA | 4 | 12 | 5 | 8 | 1 | 0 |
| ENSDARG000 | 4;4 | 60939506:609 | 60940782:609 | -;- | 1648 | NA | 0 | 0 | 0 | 0 | 0 | 0 |
| ENSDARG000 | 4;4;4;4 | 59240496:592 | 59240903:592 | +;+;+;+ | 1461 | NA | 0 | 0 | 0 | 0 | 0 | 0 |
| ENSDARG000 | 4:4:4:4:4:4:4: | 60956214:609 | 60956316:609 | +:+:+:+:+:+:+: | 1545 | NA | 0 | 0 | 0 | 4 | 0 | 0 |
| ENSDARG000 | 4:4:4:4:4:4:4: | 59234719:592 | 59234776:592 | +:+:+:+:+:+:+: | 2098 | NA | 0 | 10 | 8 | 0 | 2 | 0 |
| ENSDARG000 | 4:4:4:4:4:4:4: | 9652216:9653 | 9654174:9654 | -:-:-:-:-:-:-:-: | 4219 | dmtf1 | 283 | 194 | 376 | 242 | 294 | 214 |
| ENSDARG000 | 4;4;4 | 59320243:593 | 59320334:593 | +;+;+ | 2163 | NA | 17 | 29 | 24 | 0 | 10 | 18 |
| ENSDARG000 | 4;4;4;4 | 60966693:609 | 60967138:609 | +;+;+;+ | 1143 | NA | 0 | 0 | 0 | 0 | 0 | 0 |
| ENSDARG000 | 4;4 | 22236358:222 | 22236802:222 | -;- | 661 | NA | 0 | 0 | 0 | 0 | 0 | 0 |
| ENSDARG000 | 4:4:4:4:4:4:4: | 59245846:592 | 59246192:592 | +:+:+:+:+:+:+: | 3750 | NA | 52 | 12 | 13 | 4 | 12 | 15 |
| ENSDARG000 | 4;4;4 | 22297839:222 | 22297918:222 | +;+;+ | 898 | llph | 650 | 779 | 930 | 390 | 546 | 517 |
| ENSDARG000 | 4;4;4;4;4;4 | 14636309:146 | 14636908:146 | -;-;-;-;-;- | 2274 | plxnb2a.2 | 4 | 2 | 3 | 1 | 10 | 5 |
| ENSDARG000 | 4:4:4:4:4:4:4: | 15954293:159 | 15954730:159 | +:+:+:+:+:+:+: | 1593 | NA | 0 | 0 | 0 | 0 | 0 | 0 |
| ENSDARG000 | 4;4 | 61044749:610 | 61044879:610 | -;- | 179 | NA | 0 | 0 | 0 | 0 | 0 | 0 |
| ENSDARG000 | 4;4 | 61047336:610 | 61047466:610 | -;- | 179 | NA | 0 | 0 | 0 | 0 | 0 | 0 |
| ENSDARG000 | 4;4;4 | 8001874:8012 | 8002692:8012 | -;-;- | 1444 | ccdc3a | 12 | 0 | 0 | 2 | 0 | 4 |
| ENSDARG000 | 4;4;4 | 59362813:593 | 59362963:593 | +;+;+ | 2840 | NA | 18 | 33 | 17 | 0 | 2 | 6 |
| ENSDARG000 | 4:4:4:4:4:4:4: | 40018440:400 | 40018500:400 | +:+:+:+:+:+:+: | 1726 | NA | 0 | 7 | 4 | 0 | 0 | 2 |
| ENSDARG000 | 4;4;4 | 28374628:283 | 28374950:283 | +;+;+ | 2019 | alg10 | 83 | 84 | 77 | 38 | 97 | 111 |
| ENSDARG000 | 4:4:4:4:4:4:4: | 28356606:283 | 28356837:283 | +:+:+:+:+:+:+: | 4913 | NA | 58 | 185 | 194 | 92 | 235 | 128 |
| ENSDARG000 | 4:4:4:4:4:4:4: | 28371732:283 | 28372252:283 | -;-;-;-;-;-;-;- | 1458 | tprkb | 41 | 35 | 53 | 22 | 40 | 30 |
| ENSDARG000 | 4:4:4:4:4:4:4: | 40022273:400 | 40022391:400 | +:+:+:+:+:+:+: | 551 | NA | 0 | 5 | 0 | 0 | 2 | 0 |
| ENSDARG000 | 4;4;4;4 | 28355058:283 | 28355144:283 | +;+;+;+ | 586 | NA | 0 | 0 | 0 | 0 | 0 | 0 |
| ENSDARG000 | 4:4:4:4:4:4:4: | 25472505:254 | 25474499:254 | -:-:-:-:-:-:-:-: | 3827 | pfkfb3 | 957 | 2396 | 7952 | 7744 | 1590 | 922 |
| ENSDARG000 | 4;4;4;4;4;4 | 40035116:400 | 40035179:400 | +;+;+;+;+;+ | 1129 | NA | 8 | 25 | 12 | 0 | 47 | 15 |
| ENSDARG000 | 4;4 | 59425539:594 | 59425690:594 | +;+ | 1658 | NA | 2 | 19 | 12 | 2 | 40 | 8 |
| ENSDARG000 | 4;4 | 59541396:595 | 59542073:595 | -;- | 828 | NA | 0 | 0 | 0 | 0 | 0 | 0 |
| ENSDARG000 | 4:4:4:4:4:4:4: | 40035564:400 | 40035734:400 | +:+:+:+:+:+:+: | 5211 | NA | 3 | 13 | 12 | 0 | 6 | 10 |
| ENSDARG000 | 4:4:4:4:4:4:4: | 5784970:5786 | 5786065:5786 | -:-:-:-:-:-:-:-: | 2731 | pgm3 | 282 | 259 | 400 | 220 | 330 | 260 |
| ENSDARG000 | 4;4;4 | 70633232:706 | 70633305:706 | +;+;+ | 995 | NA | 0 | 0 | 0 | 0 | 0 | 0 |
| ENSDARG000 | 4;4 | 61049133:610 | 61049263:610 | -;- | 179 | NA | 0 | 0 | 0 | 0 | 0 | 0 |
| ENSDARG000 | 4;4 | 61050930:610 | 61051060:610 | -;- | 179 | NA | 0 | 0 | 0 | 0 | 0 | 0 |
| ENSDARG000 | 4:4:4:4:4:4:4: | 18960914:189 | 18961265:189 | +:+:+:+:+:+:+: | 3576 | impdh1b | 2452 | 3078 | 3746 | 1257 | 2807 | 2775 |
| ENSDARG000 | 4:4:4:4:4:4:4: | 5094007:5097 | 5094482:5097 | -:-:-:-:-:-:-:-: | 2488 | tmem209 | 546 | 680 | 869 | 483 | 724 | 478 |
| ENSDARG000 | 4;4 | 61052727:610 | 61052857:610 | -;- | 179 | NA | 0 | 0 | 0 | 0 | 0 | 0 |
| ENSDARG000 | 4;4 | 61054524:610 | 61054654:610 | -;- | 179 | NA | 0 | 0 | 0 | 0 | 0 | 0 |
| ENSDARG000 | 4;4;4 | 59589201:595 | 59589242:595 | +;+;+ | 991 | NA | 0 | 4 | 0 | 0 | 4 | 0 |
| ENSDARG000 | 4:4:4:4:4:4:4: | 17279966:172 | 17280029:172 | +:+:+:+:+:+:+: | 2950 | bcat1 | 187 | 326 | 1045 | 637 | 123 | 393 |
| ENSDARG000 | 4;4;4;4 | 59635278:596 | 59635731:596 | -;-;-;- | 1060 | NA | 0 | 14 | 0 | 0 | 0 | 6 |
| ENSDARG000 | 4;4;4;4;4 | 40218600:402 | 40218622:402 | +;+;+;+;+ | 2598 | NA | 0 | 0 | 0 | 0 | 0 | 0 |
| ENSDARG000 | 4;4;4;4;4;4 | 14644076:146 | 14644678:146 | -;-;-;-;-;- | 2264 | plxnb2a.3 | 20 | 11 | 40 | 20 | 10 | 54 |
| ENSDARG000 | 4:4:4:4:4:4:4: | 70887089:708 | 70887139:708 | +:+:+:+:+:+:+: | 2922 | NA | 154 | 65 | 91 | 38 | 65 | 46 |
| ENSDARG000 | 4:4:4:4:4:4:4: | 70967639:709 | 70967745:709 | +:+:+:+:+:+:+: | 3636 | NA | 21 | 7 | 10 | 18 | 9 | 3 |
| ENSDARG000 | 4:4:4:4:4:4:4: | 70850583:708 | 70851379:708 | -:-:-:-:-:-:-:-: | 3434 | NA | 178 | 169 | 313 | 105 | 431 | 291 |
| ENSDARG000 | 4;4 | 61060200:610 | 61060475:610 | +;+ | 704 | NA | 8 | 0 | 0 | 0 | 4 | 8 |
| ENSDARG000 | 4;4 | 61083750:610 | 61083839:610 | -;- | 233 | NA | 0 | 0 | 0 | 0 | 1 | 0 |
| ENSDARG000 | 4:4:4:4:4:4:4: | 2948063:2948 | 2948846:2948 | -:-:-:-:-:-:-:-: | 5024 | plekha5 | 4482 | 2681 | 2813 | 3944 | 4118 | 2336 |
| ENSDARG000 | 4;4;4;4 | 61165722:611 | 61165829:611 | +;+;+;+ | 533 | NA | 0 | 0 | 0 | 0 | 0 | 0 |
| ENSDARG000 | 4;4;4;4 | 61171310:611 | 61171461:611 | +;+;+;+ | 837 | NA | 0 | 0 | 0 | 0 | 0 | 0 |
| ENSDARG000 | 4;4;4;4;4;4;4 | 40265583:402 | 40265641:402 | +;+;+;+;+;+;+ | 642 | NA | 5 | 0 | 11 | 4 | 7 | 0 |
| ENSDARG000 | 4;4 | 70957417:709 | 70957543:709 | -;- | 201 | NA | 0 | 0 | 0 | 0 | 0 | 0 |
| ENSDARG000 | 4;4 | 70950578:709 | 70950707:709 | -;- | 194 | NA | 2 | 0 | 0 | 0 | 0 | 0 |
| ENSDARG000 | 4;4;4;4;4;4 | 71105738:711 | 71105799:711 | -;-;-;-;-;- | 1817 | NA | 0 | 4 | 0 | 0 | 0 | 0 |
| ENSDARG000 | 4:4:4:4:4:4:4: | 71014655:710 | 71014720:710 | +:+:+:+:+:+:+: | 3962 | NA | 38 | 29 | 38 | 13 | 33 | 23 |
| ENSDARG000 | 4;4;4;4;4;4 | 61644499:616 | 61646165:616 | -;-;-;-;-;- | 1867 | NA | 2 | 8 | 6 | 2 | 3 | 2 |
| ENSDARG000 | 4:4:4:4:4:4:4: | 70712807:707 | 70712920:707 | +:+:+:+:+:+:+: | 4904 | NA | 36 | 17 | 25 | 7 | 32 | 12 |
| ENSDARG000 | 4:4:4:4:4:4:4: | 8016457:8018 | 8016644:8019 | +:+:+:+:+:+:+: | 2455 | optn | 1845 | 1408 | 1254 | 1409 | 1151 | 1014 |
| ENSDARG000 | 4;4;4 | 61685313:616 | 61686017:616 | -;-;- | 950 | NA | 2 | 29 | 17 | 0 | 13 | 11 |
| ENSDARG000 | 4:4:4:4:4:4:4: | 11054309:110 | 11055866:110 | -:-:-:-:-:-:-:-: | 5193 | NA | 39 | 48 | 48 | 32 | 72 | 98 |
| ENSDARG000 | 4:4:4:4:4:4:4: | 71018579:710 | 71019424:710 | +:+:+:+:+:+:+: | 6040 | NA | 122 | 73 | 160 | 38 | 70 | 43 |
| ENSDARG000 | 4:4:4:4:4:4:4: | 11035103:110 | 11035950:110 | -:-:-:-:-:-:-:-: | 2591 | mettl25 | 74 | 78 | 130 | 64 | 93 | 48 |
| ENSDARG000 | 4:4:4:4:4:4:4: | 70926424:709 | 70926467:709 | +:+:+:+:+:+:+: | 3371 | NA | 12 | 9 | 8 | 3 | 18 | 6 |
| ENSDARG000 | 4:4:4:4:4:4:4: | 70791887:707 | 70792058:707 | +:+:+:+:+:+:+: | 5416 | NA | 458 | 10 | 25 | 5 | 324 | 393 |
| ENSDARG000 | 4;4;4 | 59701066:597 | 59702435:597 | -;-;- | 1555 | NA | 22 | 24 | 38 | 6 | 16 | 4 |
| ENSDARG000 | 4;4;4 | 59711338:597 | 59711414:597 | +;+;+ | 1216 | NA | 0 | 0 | 0 | 0 | 2 | 0 |
| ENSDARG000 | 4;4;4;4 | 11053301:110 | 11053931:110 | +;+;+;+ | 1752 | ccdc59 | 3 | 10 | 8 | 14 | 6 | 6 |
| ENSDARG000 | 4:4:4:4:4:4:4: | 70782916:708 | 70782974:708 | +:+:+:+:+:+:+: | 6328 | NA | 18 | 5 | 14 | 2 | 19 | 33 |
| ENSDARG000 | 4:4:4:4:4:4:4: | 71058396:710 | 71058439:710 | +:+:+:+:+:+:+: | 3621 | NA | 63 | 57 | 45 | 11 | 29 | 40 |
| ENSDARG000 | 4;4;4 | 61693986:616 | 61694075:616 | +;+;+ | 1625 | NA | 2 | 0 | 17 | 0 | 4 | 4 |
| ENSDARG000 | 4:4:4:4:4:4:4: | 70842085:708 | 70843356:708 | -:-:-:-:-:-:-:-: | 1725 | NA | 20 | 2 | 11 | 0 | 20 | 24 |
| ENSDARG000 | 4:4:4:4:4:4:4: | 22300330:223 | 22301257:223 | -:-:-:-:-:-:-:-: | 2501 | hcls1 | 132 | 153 | 80 | 60 | 220 | 278 |
| ENSDARG000 | 4;4;4;4;4;4 | 71005025:710 | 71005395:710 | -;-;-;-;-;- | 865 | NA | 0 | 0 | 0 | 2 | 0 | 0 |
| ENSDARG000 | 4;4;4 | 61715860:617 | 61715953:617 | +;+;+ | 1541 | NA | 3 | 0 | 2 | 0 | 2 | 2 |
| ENSDARG000 | 4;4;4 | 61904277:619 | 61905188:619 | -;-;- | 1128 | NA | 0 | 0 | 6 | 0 | 0 | 2 |
| ENSDARG000 | 4:4:4:4:4:4:4: | 61865790:618 | 61865882:618 | +:+:+:+:+:+:+: | 1926 | NA | 7 | 1 | 2 | 0 | 14 | 8 |
| ENSDARG000 | 4;4 | 17370686:173 | 17370883:173 | +;+ | 487 | NA | 4 | 0 | 0 | 8 | 0 | 0 |
| ENSDARG000 | 4:4:4:4:4:4:4: | 17337763:173 | 17339427:173 | -:-:-:-:-:-:-:-: | 4071 | parpbp | 122 | 232 | 328 | 132 | 197 | 76 |
| ENSDARG000 | 4;4;4;4;4 | 40357513:403 | 40359113:403 | -;-;-;-;- | 1804 | NA | 3 | 6 | 0 | 2 | 37 | 12 |
| ENSDARG000 | 4;4;4;4 | 61979756:619 | 61980147:619 | -;-;-;- | 871 | NA | 0 | 3 | 3 | 0 | 4 | 0 |
| ENSDARG000 | 4:4:4:4:4:4:4: | 17327704:173 | 17327961:173 | +:+:+:+:+:+:+: | 3599 | nup37 | 126 | 342 | 320 | 182 | 292 | 212 |
| ENSDARG000 | 4:4:4:4:4:4:4: | 5796761:5798 | 5796783:5798 | +:+:+:+:+:+:+: | 3434 | NA | 0 | 0 | 14 | 4 | 10 | 0 |
| ENSDARG000 | 4;4;4 | 28418829:284 | 28418869:284 | +;+;+ | 2285 | znf1041 | 60 | 63 | 64 | 70 | 66 | 60 |
| ENSDARG000 | 4;4;4;4;4 | 28442262:284 | 28442302:284 | +;+;+;+;+ | 645 | NA | 0 | 0 | 0 | 0 | 2 | 1 |
| ENSDARG000 | 4:4:4:4:4:4:4: | 28473223:284 | 28473255:284 | +:+:+:+:+:+:+: | 3047 | NA | 19 | 2 | 0 | 0 | 2 | 10 |
| ENSDARG000 | 4;4;4 | 28442223:284 | 28442302:284 | +;+;+ | 1708 | NA | 88 | 74 | 65 | 29 | 84 | 52 |
| ENSDARG000 | 4:4:4:4:4:4:4: | 28384437:283 | 28384480:283 | +:+:+:+:+:+:+: | 3143 | NA | 12 | 38 | 30 | 24 | 44 | 22 |
| ENSDARG000 | 4;4;4 | 17336557:173 | 17336809:173 | +;+;+ | 580 | pmch | 0 | 0 | 0 | 0 | 0 | 0 |
| ENSDARG000 | 4;4;4;4;4 | 17310143:173 | 17310437:173 | +;+;+;+;+ | 1237 | igf1 | 4 | 4 | 3 | 0 | 2 | 2 |
| ENSDARG000 | 4;4;4;4;4;4 | 40427249:404 | 40427382:404 | +;+;+;+;+;+ | 1353 | NA | 4 | 3 | 1 | 0 | 2 | 2 |
| ENSDARG000 | 4:4:4:4:4:4:4: | 40373029:403 | 40373108:403 | +:+:+:+:+:+:+: | 5784 | NA | 171 | 229 | 167 | 6 | 201 | 74 |
| ENSDARG000 | 4;4 | 40402017:404 | 40403294:404 | -;- | 1435 | NA | 10 | 13 | 17 | 0 | 48 | 34 |
| ENSDARG000 | 4:4:4:4:4:4:4: | 61994037:619 | 61994122:619 | +:+:+:+:+:+:+: | 2366 | NA | 125 | 127 | 296 | 112 | 167 | 12 |
| ENSDARG000 | 4:4:4:4:4:4:4: | 22312856:223 | 22314183:223 | -:-:-:-:-:-:-:-: | 11949 | NA | 2084 | 1534 | 2084 | 1268 | 2943 | 1964 |
| ENSDARG000 | 4;4 | 8029968:8030 | 8030457:8030 | -;- | 513 | si:ch211-240l | 0 | 0 | 0 | 0 | 0 | 0 |
| ENSDARG000 | 4:4:4:4:4:4:4: | 40490896:404 | 40491036:404 | +:+:+:+:+:+:+: | 902 | NA | 0 | 2 | 0 | 2 | 1 | 0 |
| ENSDARG000 | 4:4:4:4:4:4:4: | 62028152:620 | 62030033:620 | -:-:-:-:-:-:-:-: | 2262 | NA | 0 | 2 | 10 | 0 | 5 | 1 |
| ENSDARG000 | 4;4;4 | 40502515:405 | 40502573:405 | +;+;+ | 659 | NA | 0 | 0 | 1 | 0 | 32 | 0 |
| ENSDARG000 | 4:4:4:4:4:4:4: | 40488564:404 | 40488729:404 | -;-;-;-;-;-;-;- | 976 | NA | 0 | 0 | 0 | 0 | 2 | 0 |
| ENSDARG000 | 4;4;4;4 | 40498020:405 | 40498219:405 | -;-;-;- | 663 | NA | 0 | 0 | 0 | 0 | 2 | 8 |
| ENSDARG000 | 4;4 | 8034381:8035 | 8034946:8035 | -;- | 589 | si:ch211-240l | 0 | 0 | 0 | 0 | 0 | 0 |
| ENSDARG000 | 4;4 | 8039791:8040 | 8040314:8040 | -;- | 569 | si:ch211-240l | 0 | 0 | 0 | 2 | 2 | 3 |
| ENSDARG000 | 4;4;4 | 8042805:8043 | 8043835:8043 | -;-;- | 1088 | si:ch211-240l | 0 | 0 | 2 | 2 | 0 | 6 |
| ENSDARG000 | 4:4:4:4:4:4:4: | 9696965:9698 | 9697946:9698 | -;-;-;-;-;-;-;- | 1883 | tspan9b | 1321 | 1004 | 1400 | 1018 | 1422 | 982 |
| ENSDARG000 | 4;4;4 | 28814850:288 | 28814878:288 | +;+;+ | 2644 | znf1059 | 82 | 76 | 72 | 37 | 72 | 99 |
| ENSDARG000 | 4:4:4:4:4:4:4: | 9724192:9724 | 9725085:9725 | -:-:-:-:-:-:-:-: | 6079 | mical3b | 711 | 1813 | 1084 | 1051 | 1256 | 1688 |
| ENSDARG000 | 4;4 | 5825368:5825 | 5825588:5826 | +;+ | 837 | NA | 0 | 0 | 0 | 0 | 0 | 0 |
| ENSDARG000 | 4:4:4:4:4:4:4: | 5807964:5809 | 5810077:5810 | -:-:-:-:-:-:-:-: | 3831 | foxm1 | 70 | 428 | 494 | 206 | 327 | 184 |
| ENSDARG000 | 4;4;4;4 | 22343093:223 | 22343362:223 | +;+;+;+ | 2310 | guca1a | 14 | 4 | 21 | 0 | 14 | 4 |
| ENSDARG000 | 4:4:4:4:4:4:4: | 28829894:288 | 28830475:288 | -;-;-;-;-;-;-;- | 3761 | NA | 211 | 65 | 217 | 173 | 189 | 270 |
| ENSDARG000 | 4:4:4:4:4:4:4: | 9669717:9674 | 9669911:9674 | +:+:+:+:+:+:+: | 3824 | elapor2b | 6 | 4 | 0 | 10 | 4 | 2 |
| ENSDARG000 | 4:4:4:4:4:4:4: | 8045564:8047 | 8049260:8047 | -:-:-:-:-:-:-:-: | 11790 | WNK1 | 1554 | 1102 | 1286 | 1155 | 957 | 1374 |
| ENSDARG000 | 4;4;4 | 28951558:289 | 28952795:289 | -;-;- | 1428 | zgc:174315 | 56 | 36 | 37 | 44 | 40 | 69 |
| ENSDARG000 | 4;4;4 | 62184754:621 | 62184844:621 | +;+;+ | 1948 | NA | 2 | 0 | 0 | 0 | 2 | 0 |
| ENSDARG000 | 4;4;4 | 28962774:289 | 28962824:289 | +;+;+ | 2463 | NA | 92 | 56 | 86 | 28 | 52 | 53 |
| ENSDARG000 | 4:4:4:4:4:4:4: | 13412030:134 | 13412317:134 | +:+:+:+:+:+:+: | 4824 | cand1 | 3198 | 2598 | 3839 | 2503 | 3519 | 2870 |
| ENSDARG000 | 4:4:4:4:4:4:4: | 28997595:289 | 28997703:289 | +:+:+:+:+:+:+: | 1213 | si:dkev-13e3. | 0 | 0 | 0 | 0 | 0 | 0 |
| ENSDARG000 | 4:4:4:4:4:4:4: | 14727018:147 | 14727429:147 | +:+:+:+:+:+:+: | 2230 | cmasa | 673 | 781 | 885 | 794 | 898 | 848 |
| ENSDARG000 | 4:4:4:4:4:4:4: | 40525075:405 | 40525123:405 | -:-:-:-:-:-:-:-: | 1187 | NA | 8 | 6 | 0 | 2 | 2 | 6 |
| ENSDARG000 | 4:4:4:4:4:4:4: | 62198397:621 | 62198983:621 | -:-:-:-:-:-:-:-: | 4879 | NA | 34 | 9 | 20 | 10 | 6 | 12 |
| ENSDARG000 | 4:4:4:4:4:4:4: | 13428993:134 | 13429060:134 | +:+:+:+:+:+:+: | 2145 | si:dkev-39a18 | 0 | 0 | 0 | 0 | 10 | 0 |
| ENSDARG000 | 4:4:4:4:4:4:4: | 40554704:405 | 40555396:405 | -:-:-:-:-:-:-:-: | 5295 | NA | 34 | 14 | 38 | 19 | 32 | 30 |
| ENSDARG000 | 4;4 | 14717502:147 | 14717851:147 | +;+ | 1251 | kcnj8 | 18 | 0 | 2 | 0 | 2 | 0 |
| ENSDARG000 | 4:4:4:4:4:4:4: | 14660769:146 | 14660940:146 | +:+:+:+:+:+:+: | 4879 | abcc9 | 141 | 28 | 12 | 4 | 38 | 59 |
| ENSDARG000 | 4:4:4:4:4:4:4: | 3343171:3346 | 3344747:3346 | -:-:-:-:-:-:-:-: | 4762 | pik3cg | 865 | 765 | 921 | 460 | 918 | 898 |
| ENSDARG000 | 4;4;4;4;4 | 62262253:622 | 62262305:622 | +;+;+;+;+ | 1633 | zgc:174704 | 1 | 0 | 0 | 0 | 0 | 0 |
| ENSDARG000 | 4;4;4;4;4;4 | 62337473:623 | 62337619:623 | +;+;+;+;+;+ | 1933 | NA | 11 | 10 | 13 | 0 | 7 | 8 |
| ENSDARG000 | 4;4 | 62346014:623 | 62346273:623 | -;- | 456 | NA | 0 | 0 | 0 | 0 | 0 | 0 |
| ENSDARG000 | 4;4;4 | 62431132:624 | 62431224:624 | +;+;+ | 958 | NA | 6 | 12 | 10 | 0 | 8 | 4 |
| ENSDARG000 | 4;4;4 | 62452535:624 | 62453788:624 | -;-;- | 1480 | NA | 0 | 0 | 0 | 0 | 0 | 0 |
| ENSDARG000 | 4:4:4:4:4:4:4: | 18985529:189 | 18989402:189 | -:-:-:-:-:-:-:-: | 5545 | NA | 26 | 14 | 142 | 53 | 36 | 56 |
| ENSDARG000 | 4;4;4 | 11066785:110 | 11066926:110 | -;-;- | 268 | NA | 0 | 0 | 0 | 0 | 0 | 0 |
| ENSDARG000 | 4:4:4:4:4:4:4: | 29032643:290 | 29033945:290 | -:-:-:-:-:-:-:-: | 6151 | NA | 48 | 4 | 51 | 24 | 2 | 6 |
| ENSDARG000 | 4:4:4:4:4:4:4: | 62491221:624 | 62491785:624 | -:-:-:-:-:-:-:-: | 4234 | NA | 28 | 13 | 10 | 4 | 14 | 7 |
| ENSDARG000 | 4;4;4;4 | 8168514:8206 | 8168773:8207 | +;+;+;+ | 1002 | ninj2 | 4 | 9 | 2 | 0 | 0 | 4 |
| ENSDARG000 | 4:4:4:4:4:4:4: | 62702193:627 | 62702787:627 | -:-:-:-:-:-:-:-: | 3388 | NA | 3 | 1 | 0 | 0 | 7 | 0 |
| ENSDARG000 | 4:4:4:4:4:4:4: | 11070864:110 | 11070913:110 | -:-:-:-:-:-:-:-: | 3176 | NA | 0 | 0 | 0 | 0 | 0 | 0 |
| ENSDARG000 | 4:4:4:4:4:4:4: | 62576422:625 | 62576666:625 | +:+:+:+:+:+:+: | 3953 | NA | 29 | 7 | 23 | 0 | 9 | 12 |
| ENSDARG000 | 4;4;4;4;4 | 5132951:5134 | 5133208:5134 | +;+;+;+;+ | 2538 | ccnd2b | 120 | 142 | 401 | 58 | 515 | 136 |
| ENSDARG000 | 4;4 | 62679430:626 | 62679604:626 | +;+ | 1653 | NA | 0 | 2 | 8 | 0 | 8 | 2 |
| ENSDARG000 | 4:4:4:4:4:4:4: | 62532285:625 | 62536494:625 | -:-:-:-:-:-:-:-: | 7807 | NA | 26 | 12 | 41 | 6 | 15 | 33 |
| ENSDARG000 | 4;4 | 29139757:291 | 29140629:291 | -;- | 1093 | NA | 9 | 10 | 1 | 1 | 2 | 6 |
| ENSDARG000 | 4;4;4 | 29262191:292 | 29262467:292 | -;-;- | 485 | NA | 0 | 28 | 0 | 0 | 0 | 8 |
| ENSDARG000 | 4;4;4;4;4 | 29206813:292 | 29207086:292 | +;+;+;+;+ | 1218 | NA | 0 | 4 | 0 | 0 | 4 | 8 |
| ENSDARG000 | 4;4;4;4 | 15968483:159 | 15968819:159 | +;+;+;+ | 1017 | si:dkev-117n7 | 10 | 0 | 0 | 0 | 2 | 0 |
| ENSDARG000 | 4;4 | 19031768:190 | 19032245:190 | -;- | 589 | lepb | 74 | 10 | 6 | 0 | 2 | 8 |
| ENSDARG000 | 4;4;4 | 19127973:191 | 19128086:191 | +;+;+ | 736 | NA | 14 | 2 | 10 | 0 | 0 | 1 |
| ENSDARG000 | 4 | 15981096 | 15981539 | + | 444 | ucn3l | 0 | 0 | 0 | 0 | 0 | 0 |
| ENSDARG000 | 4:4:4:4:4:4:4: | 22353465:223 | 22353729:223 | -:-:-:-:-:-:-:-: | 1671 | orc5 | 46 | 168 | 293 | 170 | 244 | 150 |
| ENSDARG000 | 4;4 | 29532731:295 | 29532872:295 | +;+ | 1528 | znf1140 | 12 | 12 | 14 | 6 | 16 | 18 |
| ENSDARG000 | 4;4;4;4;4;4 | 5156117:5158 | 5156272:5158 | +;+;+;+;+;+ | 1177 | tigarb | 141 | 202 | 199 | 134 | 224 | 126 |
| ENSDARG000 | 4;4;4;4;4;4 | 29620959:296 | 29621237:296 | +;+;+;+;+;+ | 3035 | NA | 3 | 0 | 4 | 0 | 2 | 1 |
| ENSDARG000 | 4:4:4:4:4:4:4: | 11079058:110 | 11079258:110 | -:-:-:-:-:-:-:-: | 1236 | NA | 0 | 1 | 0 | 0 | 0 | 0 |
| ENSDARG000 | 4;4;4;4 | 71235680:712 | 71235759:712 | -;-;-;- | 358 | NA | 10 | 0 | 2 | 0 | 0 | 9 |
| ENSDARG000 | 4;4 | 71154171:711 | 71154268:711 | +;+ | 971 | NA | 87 | 19 | 18 | 14 | 96 | 159 |
| ENSDARG000 | 4:4:4:4:4:4:4: | 71274311:712 | 71275001:712 | -:-:-:-:-:-:-:-: | 3510 | NA | 92 | 11 | 19 | 2 | 106 | 114 |
| ENSDARG000 | 4:4:4:4:4:4:4: | 71132873:711 | 71133564:711 | -;-;-;-;-;-;-;- | 1263 | NA | 0 | 0 | 0 | 0 | 0 | 0 |
| ENSDARG000 | 4:4:4:4:4:4:4: | 71132844:711 | 71133564:711 | -:-:-:-:-:-:-:-: | 6068 | NA | 31 | 26 | 26 | 4 | 6 | 11 |
| ENSDARG000 | 4:4:4:4:4:4:4: | 29702703:297 | 29703260:297 | +:+:+:+:+:+:+: | 5859 | NA | 313 | 76 | 143 | 92 | 150 | 106 |
| ENSDARG000 | 4;4;4 | 71229095:712 | 71229364:712 | -;-;- | 400 | NA | 10 | 0 | 0 | 0 | 14 | 0 |
| ENSDARG000 | 4:4:4:4:4:4:4: | 71194643:711 | 71195306:711 | -:-:-:-:-:-:-:-: | 3545 | NA | 33 | 5 | 18 | 6 | 12 | 8 |
| ENSDARG000 | 4;4;4 | 71281920:712 | 71281981:712 | +;+;+ | 751 | NA | 0 | 0 | 0 | 0 | 0 | 0 |

| | | | | | | | | | | | | |
|---|---|---|---|---|---|---|---|---|---|---|---|---|
| ENSDARG000 | 4;4;4 | 62801285:628 | 62801435:628 | +;+;+ | 1761 | NA | 0 | 0 | 0 | 0 | 0 | 0 |
| ENSDARG000 | 4;4;4;4;4;4;4; | 71221655:712 | 71221783:712 | -;-;-;-;-;-;-;-; | 3663 | NA | 32 | 23 | 24 | 10 | 16 | 7 |
| ENSDARG000 | 4;4;4;4;4;4;4; | 71164915:711 | 71165602:711 | -;-;-;-;-;-;-;-; | 4638 | NA | 227 | 35 | 57 | 15 | 225 | 126 |
| ENSDARG000 | 4;4;4;4;4;4;4; | 5832311:5832 | 5832500:5832 | +;+;+;+;+;+;+;+; | 1108 | rhno1 | 76 | 164 | 140 | 164 | 106 | 126 |
| ENSDARG000 | 4;4;4;4;4;4;4; | 62855162:628 | 62855273:628 | +;+;+;+;+;+;+;+; | 3349 | NA | 7 | 0 | 5 | 0 | 0 | 2 |
| ENSDARG000 | 4;4;4;4;4;4;4; | 15983541:159 | 15983680:159 | +;+;+;+;+;+;+;+; | 2978 | copg2 | 3128 | 3095 | 4500 | 2436 | 3710 | 2819 |
| ENSDARG000 | 4;4 | 29753347:297 | 29753652:297 | -;- | 420 | NA | 0 | 4 | 2 | 0 | 5 | 3 |
| ENSDARG000 | 4;4;4;4;4 | 5835393:5837 | 5835747:5837 | +;+;+;+;+ | 1705 | pex26 | 269 | 136 | 220 | 150 | 196 | 176 |
| ENSDARG000 | 4;4;4;4;4;4;4; | 11084209:110 | 11084741:110 | -;-;-;-;-;-;-;-; | 5587 | NA | 0 | 0 | 0 | 0 | 0 | 0 |
| ENSDARG000 | 4;4;4 | 29758289:297 | 29759522:297 | -;-;- | 1439 | znf1119 | 32 | 43 | 68 | 16 | 85 | 45 |
| ENSDARG000 | 4;4;4;4;4 | 62984730:629 | 62984966:629 | +;+;+;+;+ | 938 | NA | 0 | 0 | 0 | 0 | 0 | 0 |
| ENSDARG000 | 4;4;4 | 62986341:629 | 62986881:629 | -;-;- | 855 | NA | 0 | 0 | 0 | 0 | 0 | 0 |
| ENSDARG000 | 4;4;4;4;4;4;4; | 5842433:5842 | 5842632:5842 | +;+;+;+;+;+;+;+; | 1707 | NA | 206 | 254 | 144 | 124 | 122 | 130 |
| ENSDARG000 | 4;4;4;4;4;4;4; | 63253425:632 | 63254114:632 | +;+;+;+;+;+;+;+; | 4888 | NA | 2 | 0 | 0 | 0 | 6 | 2 |
| ENSDARG000 | 4;4;4 | 29820388:298 | 29820587:298 | -;-;- | 586 | NA | 0 | 0 | 0 | 0 | 0 | 0 |
| ENSDARG000 | 4;4;4;4;4;4;4; | 63544595:635 | 63548042:635 | -;-;-;-;-;-;-;-; | 6674 | NA | 41 | 17 | 23 | 8 | 13 | 11 |
| ENSDARG000 | 4;4;4;4;4 | 63466876:637 | 63467867:637 | -;-;-;-;- | 2535 | NA | 18 | 14 | 21 | 2 | 18 | 16 |
| ENSDARG000 | 4;4;4;4;4;4;4; | 17380578:173 | 17380894:173 | -;-;-;-;-;-;-;-; | 1635 | th2 | 16 | 4 | 4 | 3 | 4 | 86 |
| ENSDARG000 | 4;4 | 29828668:298 | 29829468:298 | -;- | 951 | NA | 0 | 0 | 0 | 0 | 0 | 0 |
| ENSDARG000 | 4;4;4;4;4;4;4 | 63484571:634 | 63484647:634 | +;+;+;+;+;+;+ | 463 | NA | 0 | 0 | 0 | 0 | 0 | 0 |
| ENSDARG000 | 4;4;4;4;4;4;4; | 19534833:195 | 19535175:195 | +;+;+;+;+;+;+;+; | 4421 | LRRC4 | 13 | 2 | 20 | 21 | 46 | 6 |
| ENSDARG000 | 4;4;4 | 29848173:298 | 29848266:298 | +;+;+ | 1461 | NA | 6 | 0 | 8 | 0 | 0 | 2 |
| ENSDARG000 | 4;4;4;4;4;4;4; | 19396945:194 | 19397865:194 | -;-;-;-;-;-;-;-; | 4220 | snd1 | 5314 | 6486 | 11104 | 6341 | 7242 | 5197 |
| ENSDARG000 | 4;4;4 | 5180650:5181 | 5181079:5181 | +;+;+ | 721 | fgf6b | 0 | 0 | 0 | 0 | 0 | 0 |
| ENSDARG000 | 4;4;4 | 29909870:299 | 29909964:299 | +;+;+ | 2321 | NA | 7 | 11 | 26 | 4 | 12 | 8 |
| ENSDARG000 | 4;4;4 | 29967255:299 | 29967337:299 | +;+;+ | 1427 | NA | 31 | 18 | 28 | 10 | 18 | 35 |
| ENSDARG000 | 4;4;4;4;4;4;4; | 30045493:300 | 30049247:300 | -;-;-;-;-;-;-;-; | 6328 | NA | 12 | 0 | 0 | 0 | 0 | 0 |
| ENSDARG000 | 4;4;4;4;4 | 63769987:637 | 63770065:637 | +;+;+;+;+ | 2840 | zgc:173713 | 12 | 25 | 8 | 7 | 20 | 84 |
| ENSDARG000 | 4;4;4 | 63790028:637 | 63790117:637 | +;+;+ | 1213 | NA | 7 | 14 | 0 | 0 | 14 | 14 |
| ENSDARG000 | 4;4;4 | 71332814:713 | 71333071:713 | -;-;- | 389 | NA | 0 | 0 | 0 | 0 | 0 | 0 |
| ENSDARG000 | 4;4;4;4;4;4;4; | 71304561:713 | 71304647:713 | -;-;-;-;-;-;-;-; | 3663 | NA | 90 | 45 | 37 | 18 | 28 | 68 |
| ENSDARG000 | 4;4;4 | 71322824:713 | 71322980:713 | +;+;+ | 562 | NA | 0 | 0 | 4 | 0 | 1 | 5 |
| ENSDARG000 | 4;4;4;4;4 | 63818212:638 | 63818299:638 | +;+;+;+;+ | 1667 | NA | 2 | 4 | 6 | 0 | 4 | 1 |
| ENSDARG000 | 4 | 63906609 | 63907480 | - | 872 | NA | 0 | 0 | 0 | 0 | 0 | 0 |
| ENSDARG000 | 4;4;4;4 | 63921795:639 | 63922114:639 | -;-;-;- | 749 | NA | 0 | 4 | 2 | 2 | 0 | 0 |
| ENSDARG000 | 4;4;4;4;4;4;4; | 30145984:301 | 30146576:301 | -;-;-;-;-;-;-;-; | 4772 | NA | 28 | 18 | 34 | 18 | 23 | 16 |
| ENSDARG000 | 4;4 | 63930285:639 | 63930641:639 | +;+ | 536 | NA | 0 | 0 | 0 | 0 | 0 | 0 |
| ENSDARG000 | 4;4 | 63928858:639 | 63928968:639 | +;+ | 1087 | NA | 0 | 3 | 0 | 0 | 0 | 4 |
| ENSDARG000 | 4;4;4;4;4;4;4; | 17391680:173 | 17392483:173 | -;-;-;-;-;-;-;-; | 2314 | pah | 35827 | 24520 | 21651 | 33747 | 32307 | 36276 |
| ENSDARG000 | 4;4;4;4 | 71353792:713 | 71354698:713 | -;-;-;- | 1297 | NA | 0 | 0 | 0 | 0 | 0 | 0 |
| ENSDARG000 | 4;4;4;4;4;4 | 5848229:5848 | 5848306:5848 | +;+;+;+;+;+ | 1925 | lyrm5a | 105 | 140 | 262 | 235 | 187 | 284 |
| ENSDARG000 | 4;4;4;4;4;4;4; | 5850225:5850 | 5850412:5850 | -;-;-;-;-;-;-;-; | 1979 | SLC5A8 | 4 | 9 | 35 | 46 | 25 | 29 |
| ENSDARG000 | 4;4;4 | 63973666:639 | 63975056:639 | -;-;- | 1592 | NA | 8 | 2 | 0 | 0 | 0 | 0 |
| ENSDARG000 | 4;4;4 | 63992019:639 | 63992070:639 | +;+;+ | 1390 | zgc:173607 | 0 | 0 | 2 | 4 | 0 | 0 |
| ENSDARG000 | 4;4;4 | 5193198:5193 | 5193516:5193 | +;+;+ | 919 | fgf23 | 0 | 0 | 0 | 0 | 0 | 0 |
| ENSDARG000 | 4;4;4;4;4;4;4; | 5205265:5207 | 5205568:5207 | +;+;+;+;+;+;+;+; | 2218 | NA | 0 | 2 | 2 | 3 | 0 | 0 |
| ENSDARG000 | 4;4;4;4;4;4;4; | 64025160:640 | 64025257:640 | +;+;+;+;+;+;+;+; | 4045 | NA | 2 | 0 | 0 | 0 | 8 | 2 |
| ENSDARG000 | 4;4;4;4;4;4;4; | 30209190:302 | 30209875:302 | -;-;-;-;-;-;-;-; | 3757 | NA | 14 | 10 | 6 | 2 | 35 | 34 |
| ENSDARG000 | 4;4;4;4;4;4;4; | 5196469:5197 | 5196741:5197 | +;+;+;+;+;+;+;+; | 1816 | rad51ap1 | 28 | 163 | 153 | 64 | 90 | 76 |
| ENSDARG000 | 4;4;4;4 | 64070952:640 | 64071466:640 | -;-;-;- | 1492 | NA | 0 | 7 | 5 | 0 | 14 | 7 |
| ENSDARG000 | 4;4;4 | 64102312:641 | 64102406:641 | +;+;+ | 1262 | NA | 0 | 2 | 20 | 0 | 0 | 0 |
| ENSDARG000 | 4;4;4;4;4 | 64133570:641 | 64134931:641 | -;-;-;-;- | 1578 | NA | 0 | 0 | 0 | 0 | 0 | 0 |
| ENSDARG000 | 4;4 | 30256991:302 | 30257523:302 | +;+ | 854 | NA | 6 | 6 | 7 | 0 | 18 | 25 |
| ENSDARG000 | 4;4;4 | 71383270:713 | 71383856:713 | -;-;- | 752 | NA | 0 | 0 | 0 | 0 | 0 | 0 |
| ENSDARG000 | 4;4;4;4;4;4;4; | 71366326:713 | 71366420:713 | +;+;+;+;+;+;+;+; | 3505 | NA | 126 | 9 | 8 | 4 | 30 | 53 |
| ENSDARG000 | 4;4;4 | 64147241:641 | 64147274:641 | +;+;+ | 1568 | NA | 0 | 1 | 3 | 0 | 0 | 6 |
| ENSDARG000 | 4;4 | 64178733:641 | 64179687:641 | -;- | 1040 | NA | 0 | 0 | 0 | 0 | 0 | 0 |
| ENSDARG000 | 4;4;4 | 64200907:642 | 64201060:642 | +;+;+ | 877 | NA | 0 | 0 | 0 | 0 | 0 | 0 |
| ENSDARG000 | 4;4;4 | 30262399:302 | 30263453:302 | -;-;- | 1298 | NA | 2 | 10 | 8 | 3 | 16 | 24 |
| ENSDARG000 | 4;4 | 17417111:174 | 17417855:174 | +;+ | 1977 | ascl1a | 0 | 7 | 8 | 14 | 2 | 0 |
| ENSDARG000 | 4;4;4;4;4;4 | 71401321:714 | 71401376:714 | +;+;+;+;+;+ | 521 | NA | 0 | 0 | 0 | 0 | 0 | 0 |
| ENSDARG000 | 4;4;4 | 64295728:642 | 64295881:642 | +;+;+ | 574 | NA | 0 | 0 | 0 | 0 | 0 | 0 |
| ENSDARG000 | 4;4 | 71414829:714 | 71415590:714 | -;- | 822 | NA | 0 | 0 | 0 | 0 | 0 | 0 |
| ENSDARG000 | 4;4;4;4;4;4;4; | 64318868:643 | 64319065:643 | +;+;+;+;+;+;+;+; | 2225 | NA | 37 | 0 | 83 | 12 | 7 | 26 |
| ENSDARG000 | 4;4;4;4;4 | 64328155:643 | 64329506:643 | -;-;-;-;- | 1579 | znf1147 | 0 | 4 | 0 | 0 | 2 | 8 |
| ENSDARG000 | 4;4 | 30348676:303 | 30348825:303 | -;- | 242 | NA | 2 | 1 | 5 | 0 | 11 | 0 |
| ENSDARG000 | 4;4;4 | 30354053:303 | 30355265:303 | -;-;- | 1455 | NA | 12 | 33 | 35 | 4 | 9 | 40 |
| ENSDARG000 | 4;4 | 30363762:303 | 30364051:303 | -;- | 453 | NA | 0 | 2 | 0 | 0 | 0 | 0 |
| ENSDARG000 | 4;4;4 | 30363474:303 | 30363553:303 | +;+;+ | 1269 | NA | 30 | 25 | 22 | 16 | 17 | 18 |
| ENSDARG000 | 4;4;4;4;4;4;4; | 71423591:714 | 71424158:714 | -;-;-;-;-;-;-;-; | 3376 | NA | 293 | 23 | 6 | 10 | 253 | 306 |
| ENSDARG000 | 4;4;4;4;4;4;4; | 6423041:6423 | 6423566:6423 | -;-;-;-;-;-;-;-; | 3951 | foxp2 | 24 | 6 | 31 | 44 | 23 | 16 |
| ENSDARG000 | 4;4;4;4;4;4;4; | 64338860:643 | 64338947:643 | +;+;+;+;+;+;+;+; | 1839 | NA | 47 | 37 | 25 | 6 | 26 | 20 |
| ENSDARG000 | 4;4 | 71443269:714 | 71443405:714 | -;- | 205 | NA | 9 | 2 | 0 | 0 | 22 | 6 |
| ENSDARG000 | 4;4;4;4;4;4;4; | 71460543:714 | 71461144:714 | -;-;-;-;-;-;-;-; | 5157 | NA | 93 | 23 | 46 | 13 | 46 | 83 |
| ENSDARG000 | 4;4;4 | 64386780:643 | 64386839:643 | +;+;+ | 333 | NA | 2 | 1 | 0 | 0 | 6 | 0 |
| ENSDARG000 | 4;4;4;4;4;4;4; | 15996234:159 | 15996618:159 | -;-;-;-;-;-;-;-; | 1468 | mest | 170 | 217 | 232 | 256 | 179 | 111 |
| ENSDARG000 | 4;4;4;4 | 6572364:6575 | 6573309:6575 | +;+;+;+ | 4865 | NA | 0 | 0 | 0 | 0 | 0 | 0 |
| ENSDARG000 | 4;4;4;4;4;4;4; | 6416278:6419 | 6416458:6419 | +;+;+;+;+;+;+;+; | 1434 | NA | 2 | 0 | 8 | 0 | 0 | 0 |
| ENSDARG000 | 4 | 6639292 | 6640415 | + | 1124 | NA | 1176 | 934 | 1086 | 1509 | 1314 | 1814 |
| ENSDARG000 | 4;4;4 | 6411080:6413 | 6411270:6413 | -;-;- | 392 | NA | 0 | 0 | 0 | 0 | 0 | 0 |
| ENSDARG000 | 4;4;4;4;4;4;4; | 17424855:174 | 17425388:174 | +;+;+;+;+;+;+;+; | 3578 | recql | 761 | 482 | 865 | 489 | 980 | 754 |
| ENSDARG000 | 4;4;4;4;4;4;4; | 6339572:6344 | 6344702:6344 | -;-;-;-;-;-;-;-; | 6431 | mdfic | 2510 | 923 | 1400 | 1952 | 1568 | 2742 |
| ENSDARG000 | 4;4;4 | 64395855:643 | 64396036:643 | +;+;+ | 656 | NA | 11 | 0 | 8 | 0 | 0 | 0 |
| ENSDARG000 | 4;4;4 | 64399553:644 | 64399890:644 | -;-;- | 584 | NA | 0 | 0 | 4 | 0 | 0 | 0 |
| ENSDARG000 | 4;4;4 | 64471942:644 | 64473333:644 | -;-;- | 1634 | NA | 0 | 3 | 9 | 0 | 8 | 0 |
| ENSDARG000 | 4;4;4 | 64530121:645 | 64531627:645 | -;-;- | 1743 | NA | 20 | 19 | 2 | 6 | 8 | 7 |
| ENSDARG000 | 4;4;4;4;4;4;4; | 71489538:714 | 71490111:714 | -;-;-;-;-;-;-;-; | 6685 | NA | 517 | 205 | 206 | 49 | 327 | 424 |
| ENSDARG000 | 4;4;4;4 | 64549970:645 | 64550116:645 | +;+;+;+ | 2281 | NA | 0 | 2 | 3 | 0 | 0 | 8 |
| ENSDARG000 | 4;4;4;4;4 | 64577406:645 | 64577493:645 | +;+;+;+;+ | 1844 | NA | 6 | 4 | 0 | 2 | 12 | 6 |
| ENSDARG000 | 4;4;4;4;4;4;4 | 64586627:645 | 64586843:645 | -;-;-;-;-;-;- | 1826 | NA | 15 | 8 | 6 | 5 | 2 | 16 |
| ENSDARG000 | 4;4;4;4;4;4;4; | 17539134:175 | 17539336:175 | +;+;+;+;+;+;+;+; | 1482 | itfg2 | 302 | 226 | 203 | 344 | 290 | 287 |
| ENSDARG000 | 4;4 | 64608808:646 | 64608937:646 | +;+ | 1246 | NA | 4 | 8 | 4 | 12 | 40 | 6 |
| ENSDARG000 | 4;4;4;4;4;4;4; | 71536326:715 | 71537076:715 | -;-;-;-;-;-;-;-; | 3406 | NA | 20 | 13 | 11 | 16 | 12 | 17 |
| ENSDARG000 | 4;4 | 64688888:646 | 64689779:646 | -;- | 1217 | NA | 14 | 83 | 48 | 7 | 74 | 67 |
| ENSDARG000 | 4;4 | 71561409:715 | 71561527:715 | -;- | 419 | NA | 0 | 0 | 0 | 0 | 0 | 0 |
| ENSDARG000 | 4;4;4;4;4;4;4; | 71578099:715 | 71578251:715 | +;+;+;+;+;+;+;+; | 6839 | NA | 827 | 9 | 18 | 4 | 816 | 1040 |
| ENSDARG000 | 4;4;4;4;4 | 71617362:716 | 71617405:716 | +;+;+;+;+ | 1332 | NA | 6 | 0 | 1 | 0 | 0 | 1 |
| ENSDARG000 | 4;4;4;4 | 64708721:647 | 64708926:647 | -;-;-;- | 832 | NA | 0 | 0 | 0 | 0 | 0 | 0 |
| ENSDARG000 | 4;4;4;4;4;4 | 17572495:175 | 17575065:175 | -;-;-;-;-;- | 3685 | nrip2 | 19 | 11 | 24 | 26 | 71 | 14 |
| ENSDARG000 | 4;4;4;4;4;4;4; | 71641813:716 | 71641856:716 | +;+;+;+;+;+;+;+; | 3090 | NA | 123 | 118 | 113 | 40 | 86 | 19 |
| ENSDARG000 | 4;4;4 | 64712035:647 | 64712266:647 | -;-;- | 384 | NA | 0 | 0 | 0 | 0 | 0 | 0 |
| ENSDARG000 | 4;4;4;4;4;4;4; | 64755464:647 | 64755555:647 | +;+;+;+;+;+;+;+; | 2679 | NA | 6 | 7 | 4 | 0 | 4 | 11 |
| ENSDARG000 | 4;4;4;4;4;4;4 | 64857878:649 | 64858358:649 | -;-;-;-;-;-;- | 1751 | NA | 8 | 2 | 8 | 0 | 12 | 2 |
| ENSDARG000 | 4;4;4 | 64857820:648 | 64858830:648 | -;-;- | 1256 | NA | 0 | 0 | 0 | 0 | 0 | 0 |
| ENSDARG000 | 4;4;4 | 17634311:176 | 17636657:176 | -;-;- | 3649 | klhl42 | 135 | 57 | 138 | 86 | 142 | 125 |
| ENSDARG000 | 4;4 | 64921810:649 | 64921956:649 | +;+ | 297 | NA | 0 | 0 | 0 | 0 | 0 | 0 |
| ENSDARG000 | 4;4;4;4;4;4;4; | 71689470:716 | 71689767:716 | -;-;-;-;-;-;-;-; | 1935 | NA | 167 | 72 | 174 | 25 | 107 | 42 |
| ENSDARG000 | 4;4;4 | 71855046:718 | 71855128:718 | +;+;+ | 2571 | NA | 0 | 0 | 0 | 0 | 0 | 0 |
| ENSDARG000 | 4;4;4;4 | 71871289:718 | 71872633:718 | -;-;-;- | 2033 | NA | 0 | 0 | 0 | 0 | 0 | 0 |
| ENSDARG000 | 4;4;4;4;4;4;4; | 71691806:716 | 71695491:716 | -;-;-;-;-;-;-;-; | 11659 | NA | 18 | 7 | 42 | 5 | 12 | 26 |
| ENSDARG000 | 4;4;4;4;4;4;4; | 71896596:718 | 71897178:718 | -;-;-;-;-;-;-;-; | 3543 | NA | 32 | 7 | 168 | 1 | 7 | 43 |
| ENSDARG000 | 4;4;4;4;4 | 65031177:650 | 65032823:650 | -;-;-;-;- | 1889 | NA | 4 | 0 | 0 | 0 | 0 | 4 |
| ENSDARG000 | 4;4;4;4;4;4;4; | 71819777:718 | 71819811:718 | +;+;+;+;+;+;+;+; | 2317 | NA | 0 | 0 | 0 | 0 | 0 | 0 |
| ENSDARG000 | 4;4;4;4;4;4;4; | 17671164:176 | 17671927:176 | +;+;+;+;+;+;+;+; | 5619 | gnptab | 605 | 477 | 530 | 360 | 690 | 562 |
| ENSDARG000 | 4;4;4;4;4;4;4; | 65043312:650 | 65043497:650 | -;-;-;-;-;-;-;-; | 1547 | NA | 0 | 0 | 0 | 0 | 0 | 0 |
| ENSDARG000 | 4;4;4;4;4;4;4 | 17655872:176 | 17656053:176 | +;+;+;+;+;+;+ | 2256 | washc3 | 628 | 598 | 644 | 532 | 557 | 733 |
| ENSDARG000 | 4;4;4;4;4;4;4 | 17662413:176 | 17662624:176 | -;-;-;-;-;-;- | 1016 | dram1 | 151 | 207 | 223 | 87 | 292 | 199 |
| ENSDARG000 | 4;4;4;4;4;4;4; | 17642731:176 | 17642862:176 | +;+;+;+;+;+;+;+; | 1993 | cwf19l1 | 314 | 257 | 398 | 228 | 460 | 321 |
| ENSDARG000 | 4;4;4;4;4;4;4; | 65057234:650 | 65057293:650 | +;+;+;+;+;+;+;+; | 4160 | NA | 42 | 22 | 95 | 4 | 33 | 47 |
| ENSDARG000 | 4;4;4 | 65081703:650 | 65084008:650 | -;-;- | 2536 | NA | 0 | 2 | 0 | 0 | 0 | 0 |
| ENSDARG000 | 4;4 | 65127317:651 | 65127492:651 | +;+ | 1939 | NA | 0 | 0 | 0 | 0 | 0 | 0 |
| ENSDARG000 | 4;4 | 65139267:651 | 65139444:651 | +;+ | 1402 | NA | 10 | 2 | 7 | 0 | 0 | 0 |
| ENSDARG000 | 4;4;4 | 65296666:653 | 65298143:653 | -;-;- | 1719 | NA | 20 | 16 | 25 | 8 | 30 | 1 |
| ENSDARG000 | 4;4;4;4;4;4;4; | 65167250:651 | 65167847:651 | -;-;-;-;-;-;-;-; | 5606 | NA | 42 | 149 | 22 | 15 | 259 | 244 |
| ENSDARG000 | 4;4;4 | 65306260:653 | 65306336:653 | +;+;+ | 1495 | NA | 0 | 3 | 0 | 0 | 10 | 34 |
| ENSDARG000 | 4;4;4;4;4;4 | 65271779:652 | 65271870:652 | +;+;+;+;+;+ | 1792 | NA | 1 | 4 | 2 | 0 | 10 | 0 |
| ENSDARG000 | 4;4;4 | 65332578:653 | 65332671:653 | +;+;+ | 1289 | NA | 0 | 0 | 0 | 0 | 0 | 2 |
| ENSDARG000 | 4 | 72430431 | 72431296 | + | 866 | NA | 0 | 4 | 4 | 0 | 0 | 0 |
| ENSDARG000 | 4;4;4 | 72436227:724 | 72439870:724 | -;-;- | 4221 | si:ch211-161r | 22 | 55 | 72 | 28 | 78 | 100 |
| ENSDARG000 | 4;4;4;4;4;4;4; | 65537216:656 | 65537539:656 | +;+;+;+;+;+;+;+; | 8607 | NA | 25 | 35 | 11 | 2 | 63 | 26 |
| ENSDARG000 | 4;4;4;4;4;4;4; | 65487457:654 | 65487553:654 | +;+;+;+;+;+;+;+; | 5085 | NA | 23 | 6 | 18 | 12 | 14 | 16 |
| ENSDARG000 | 4;4;4;4;4;4;4; | 65607540:656 | 65608000:656 | +;+;+;+;+;+;+;+; | 2830 | NA | 0 | 0 | 0 | 0 | 0 | 0 |
| ENSDARG000 | 4;4;4;4;4;4;4; | 72463842:724 | 72465186:724 | -;-;-;-;-;-;-;-; | 4226 | NA | 0 | 0 | 4 | 0 | 0 | 0 |
| ENSDARG000 | 4 | 72486529 | 72488135 | - | 1607 | NA | 0 | 0 | 0 | 0 | 0 | 0 |
| ENSDARG000 | 4;4;4;4 | 72500840:725 | 72501005:725 | -;-;-;- | 2104 | NA | 0 | 2 | 2 | 0 | 14 | 0 |
| ENSDARG000 | 4;4;4 | 72534789:725 | 72535991:725 | -;-;- | 1507 | zgc:175107 | 42 | 41 | 78 | 49 | 56 | 146 |
| ENSDARG000 | 4;4;4 | 72546478:725 | 72550390:725 | -;-;- | 4220 | NA | 34 | 66 | 127 | 24 | 126 | 114 |
| ENSDARG000 | 4;4;4;4;4 | 72603266:726 | 72604850:726 | -;-;-;-;- | 2366 | NA | 50 | 38 | 75 | 16 | 47 | 45 |
| ENSDARG000 | 4;4;4;4;4 | 72559042:725 | 72560669:725 | -;-;-;-;- | 1865 | NA | 28 | 82 | 82 | 20 | 115 | 50 |
| ENSDARG000 | 4;4;4;4;4;4;4; | 72578191:725 | 72578567:725 | +;+;+;+;+;+;+;+; | 4877 | NA | 56 | 48 | 91 | 22 | 102 | 34 |
| ENSDARG000 | 4;4;4;4;4;4;4; | 66017000:660 | 66017096:660 | +;+;+;+;+;+;+;+; | 2733 | NA | 2 | 0 | 2 | 1 | 2 | 0 |
| ENSDARG000 | 4;4;4 | 72612792:726 | 72612863:726 | +;+;+ | 1484 | NA | 15 | 25 | 38 | 12 | 16 | 23 |
| ENSDARG000 | 4;4;4;4;4;4;4; | 66447839:664 | 66447898:664 | +;+;+;+;+;+;+;+; | 3258 | NA | 0 | 0 | 0 | 0 | 14 | 0 |
| ENSDARG000 | 4;4;4;4 | 66618956:666 | 66620174:666 | -;-;-;- | 1438 | NA | 0 | 0 | 0 | 0 | 4 | 0 |
| ENSDARG000 | 4;4 | 66668652:666 | 66668710:666 | +;+ | 263 | NA | 0 | 0 | 0 | 0 | 0 | 0 |
| ENSDARG000 | 4;4 | 66800865:668 | 66801072:668 | +;+ | 816 | NA | 0 | 0 | 0 | 0 | 0 | 18 |
| ENSDARG000 | 4;4;4;4;4;4;4; | 66853199:668 | 66853322:668 | +;+;+;+;+;+;+;+; | 3668 | NA | 119 | 39 | 448 | 42 | 66 | 119 |
| ENSDARG000 | 4;4;4;4;4;4;4; | 73085993:730 | 73086060:730 | +;+;+;+;+;+;+;+; | 3577 | NA | 0 | 2 | 1 | 0 | 2 | 0 |
| ENSDARG000 | 4;4;4;4;4 | 66972965:669 | 66973615:669 | -;-;-;-;- | 2813 | NA | 14 | 4 | 24 | 10 | 0 | 4 |
| ENSDARG000 | 4;4;4;4;4;4;4 | 73115553:731 | 73118507:731 | -;-;-;-;-;-;- | 3690 | NA | 4 | 2 | 0 | 6 | 22 | 3 |
| ENSDARG000 | 4;4;4;4;4;4;4; | 67786391:677 | 67786980:677 | -;-;-;-;-;-;-;-; | 4425 | NA | 4 | 4 | 4 | 0 | 4 | 1 |
| ENSDARG000 | 4;4;4;4;4 | 73155796:731 | 73155859:731 | -;-;-;-;- | 1083 | NA | 4 | 0 | 0 | 0 | 0 | 0 |
| ENSDARG000 | 4;4;4;4;4;4;4; | 67784163:679 | 67787277:679 | -;-;-;-;-;-;-;-; | 7581 | NA | 10 | 7 | 1 | 0 | 3 | 4 |
| ENSDARG000 | 4;4;4;4;4;4;4; | 67994238:679 | 67994333:679 | +;+;+;+;+;+;+;+; | 8250 | NA | 13 | 5 | 2 | 17 | 26 | 27 |
| ENSDARG000 | 4;4;4;4;4;4;4; | 68475654:684 | 68475789:684 | +;+;+;+;+;+;+;+; | 4187 | NA | 45 | 26 | 88 | 47 | 63 | 72 |
| ENSDARG000 | 4;4;4;4;4 | 73241193:732 | 73245240:732 | -;-;-;-;- | 4980 | NA | 36 | 42 | 77 | 16 | 141 | 50 |
| ENSDARG000 | 4;4;4 | 73256872:732 | 73259179:732 | -;-;- | 2526 | wu:fc30c06 | 4 | 14 | 65 | 8 | 50 | 24 |
| ENSDARG000 | 4;4;4;4;4;4;4 | 73335421:733 | 73338383:733 | -;-;-;-;-;-;- | 4155 | NA | 38 | 12 | 96 | 28 | 80 | 35 |
| ENSDARG000 | 4;4;4;4;4 | 73306451:733 | 73310091:733 | -;-;-;-;- | 4681 | NA | 4 | 2 | 4 | 4 | 26 | 7 |
| ENSDARG000 | 4;4;4;4;4;4;4; | 73273857:732 | 73274195:732 | -;-;-;-;-;-;-;-; | 4614 | NA | 35 | 25 | 58 | 34 | 60 | 75 |
| ENSDARG000 | 4;4;4;4;4;4;4; | 68519945:685 | 68520013:685 | +;+;+;+;+;+;+;+; | 4000 | NA | 82 | 48 | 103 | 17 | 36 | 38 |
| ENSDARG000 | 4;4;4;4;4;4;4; | 68591587:685 | 68592227:685 | +;+;+;+;+;+;+;+; | 6175 | NA | 60 | 27 | 48 | 19 | 28 | 24 |

| | | | | | | | | | | | | |
|---|---|---|---|---|---|---|---|---|---|---|---|---|
| ENSDARG000 | 4;4;4;4;4;4;4;4; | 73382464:733 | 73383974:733 | -;-;-;-;-;-;-;-;- | 3556 | zgc:171727 | 29 | 83 | 74 | 37 | 62 | 57 |
| ENSDARG000 | 4;4;4;4;4 | 73398788:734 | 73400139:734 | -;-;-;-;- | 3374 | NA | 42 | 42 | 47 | 24 | 96 | 67 |
| ENSDARG000 | 4;4 | 73454634:734 | 73456231:734 | -;- | 1837 | NA | 60 | 80 | 69 | 50 | 105 | 108 |
| ENSDARG000 | 4;4;4 | 73472272:734 | 73474100:734 | -;-;- | 2209 | NA | 48 | 79 | 214 | 59 | 45 | 64 |
| ENSDARG000 | 4;4;4 | 73500848:735 | 73501039:735 | +;+;+ | 1585 | NA | 0 | 0 | 0 | 0 | 0 | 0 |
| ENSDARG000 | 4;4;4;4;4 | 73548320:735 | 73551862:735 | -;-;-;-;- | 3949 | NA | 26 | 46 | 40 | 44 | 48 | 50 |
| ENSDARG000 | 4;4;4;4;4;4;4;4; | 73491106:734 | 73491543:735 | -;-;-;-;-;-;-;-;- | 14040 | znf1015 | 87 | 182 | 235 | 127 | 314 | 253 |
| ENSDARG000 | 4;4;4;4;4;4;4;4; | 68614373:686 | 68614656:686 | +;+;+;+;+;+;+;+; | 3780 | zgc:174180 | 10 | 1 | 35 | 0 | 13 | 6 |
| ENSDARG000 | 4;4 | 68710181:687 | 68710286:687 | +;+ | 437 | NA | 0 | 0 | 0 | 0 | 0 | 0 |
| ENSDARG000 | 4;4 | 68717067:687 | 68717170:687 | +;+ | 435 | NA | 0 | 0 | 0 | 0 | 0 | 0 |
| ENSDARG000 | 4;4 | 68733298:687 | 68733721:687 | -;- | 955 | NA | 0 | 0 | 1 | 0 | 3 | 0 |
| ENSDARG000 | 4;4;4 | 73562952:735 | 73566098:735 | -;-;- | 3433 | NA | 10 | 46 | 42 | 22 | 34 | 30 |
| ENSDARG000 | 4;4;4 | 73573395:735 | 73576132:735 | -;-;- | 2946 | NA | 28 | 35 | 76 | 8 | 62 | 70 |
| ENSDARG000 | 4;4;4;4 | 68742914:687 | 68743623:687 | -;-;-;- | 974 | NA | 0 | 0 | 1 | 0 | 0 | 1 |
| ENSDARG000 | 4;4;4;4;4 | 68776476:687 | 68776517:687 | +;+;+;+;+ | 455 | NA | 0 | 0 | 0 | 0 | 0 | 0 |
| ENSDARG000 | 4;4;4;4;4;4;4;4; | 68821997:688 | 68822550:688 | -;-;-;-;-;-;-;-;- | 4961 | NA | 12 | 0 | 9 | 0 | 0 | 2 |
| ENSDARG000 | 4;4 | 73606482:736 | 73606913:736 | +;+ | 2232 | si:ch211-165i | 18 | 72 | 172 | 98 | 280 | 118 |
| ENSDARG000 | 4;4;4;4;4;4;4;4; | 73635284:736 | 73639746:736 | -;-;-;-;-;-;-;-;- | 4807 | si:ch73-266f2 | 64 | 153 | 166 | 60 | 148 | 238 |
| ENSDARG000 | 4;4;4;4;4;4;4;4; | 73651053:736 | 73651113:736 | +;+;+;+;+;+;+;+; | 1485 | znf989 | 14 | 22 | 46 | 8 | 23 | 12 |
| ENSDARG000 | 4;4;4;4;4;4;4;4; | 73672430:736 | 73672485:736 | +;+;+;+;+;+;+;+; | 1549 | NA | 18 | 40 | 34 | 22 | 8 | 28 |
| ENSDARG000 | 4;4;4 | 73683468:736 | 73683512:736 | +;+;+ | 1336 | zgc:173624 | 8 | 14 | 20 | 26 | 18 | 10 |
| ENSDARG000 | 4;4;4 | 73701801:737 | 73702630:737 | -;-;- | 1042 | NA | 6 | 10 | 29 | 18 | 14 | 17 |
| ENSDARG000 | 4;4;4;4;4;4;4 | 73727692:737 | 73731503:737 | -;-;-;-;-;-;- | 4530 | zgc:171551 | 15 | 2 | 4 | 2 | 3 | 0 |
| ENSDARG000 | 4;4;4;4 | 73740012:737 | 73742476:737 | -;-;-;- | 2760 | NA | 7 | 26 | 45 | 26 | 42 | 48 |
| ENSDARG000 | 4;4 | 73725609:737 | 73726395:737 | -;- | 1088 | NA | 0 | 0 | 0 | 0 | 0 | 4 |
| ENSDARG000 | 4;4;4;4;4 | 73711227:737 | 73715768:737 | -;-;-;-;- | 9373 | NA | 75 | 135 | 169 | 101 | 122 | 158 |
| ENSDARG000 | 4;4;4 | 73757963:737 | 73760923:737 | -;-;- | 3163 | NA | 53 | 40 | 93 | 24 | 52 | 75 |
| ENSDARG000 | 4;4;4;4;4;4;4;4; | 68856289:688 | 68860348:688 | -;-;-;-;-;-;-;-;- | 8701 | NA | 61 | 32 | 33 | 8 | 17 | 26 |
| ENSDARG000 | 4;4;4;4;4;4 | 73780345:737 | 73781936:737 | -;-;-;-;-;- | 1890 | NA | 4 | 10 | 24 | 14 | 15 | 35 |
| ENSDARG000 | 4;4;4;4;4;4 | 73796172:737 | 73797232:737 | -;-;-;-;-;- | 1369 | NA | 22 | 38 | 54 | 32 | 48 | 28 |
| ENSDARG000 | 4;4;4;4;4;4;4;4; | 68922112:689 | 68922198:689 | +;+;+;+;+;+;+;+; | 10716 | NA | 32 | 6 | 17 | 0 | 5 | 8 |
| ENSDARG000 | 4;4;4;4;4;4;4;4; | 69251882:692 | 69251971:692 | +;+;+;+;+;+;+;+; | 3794 | NA | 3 | 0 | 2 | 0 | 3 | 0 |
| ENSDARG000 | 4;4 | 69245481:692 | 69245619:692 | -;- | 206 | NA | 0 | 0 | 0 | 0 | 2 | 4 |
| ENSDARG000 | 4;4;4;4;4 | 73814116:738 | 73818481:738 | -;-;-;-;- | 4881 | NA | 22 | 34 | 35 | 32 | 68 | 34 |
| ENSDARG000 | 4;4 | 73847789:738 | 73849900:738 | -;- | 2316 | NA | 6 | 0 | 8 | 2 | 12 | 4 |
| ENSDARG000 | 4;4 | 73869433:738 | 73869500:738 | -;- | 419 | NA | 1 | 0 | 0 | 0 | 0 | 0 |
| ENSDARG000 | 4;4;4;4;4 | 73872345:738 | 73872406:738 | +;+;+;+;+ | 2793 | NA | 10 | 44 | 68 | 13 | 38 | 50 |
| ENSDARG000 | 4;4 | 73941991:739 | 73942190:739 | +;+ | 2275 | NA | 22 | 33 | 48 | 2 | 28 | 33 |
| ENSDARG000 | 4;4;4 | 69122156:691 | 69122902:691 | -;-;- | 921 | NA | 0 | 0 | 0 | 0 | 0 | 0 |
| ENSDARG000 | 4;4;4;4;4;4;4;4; | 69150355:691 | 69150393:691 | +;+;+;+;+;+;+;+; | 612 | NA | 2 | 0 | 2 | 0 | 0 | 9 |
| ENSDARG000 | 4;4;4;4;4 | 69162228:691 | 69162539:691 | -;-;-;-;- | 798 | NA | 26 | 3 | 37 | 12 | 16 | 21 |
| ENSDARG000 | 4;4;4;4;4;4 | 69078543:692 | 69078693:692 | +;+;+;+;+;+ | 2124 | NA | 23 | 14 | 34 | 2 | 29 | 31 |
| ENSDARG000 | 4;4;4;4;4 | 69190524:691 | 69190566:691 | +;+;+;+;+ | 1796 | NA | 18 | 0 | 2 | 0 | 0 | 2 |
| ENSDARG000 | 4;4;4 | 69375564:693 | 69375655:693 | +;+;+ | 1433 | NA | 0 | 2 | 0 | 0 | 0 | 0 |
| ENSDARG000 | 4;4;4 | 69409331:694 | 69409422:694 | +;+;+ | 1591 | NA | 2 | 14 | 4 | 0 | 10 | 0 |
| ENSDARG000 | 4;4 | 69427328:694 | 69427757:694 | -;- | 706 | NA | 0 | 0 | 0 | 0 | 0 | 0 |
| ENSDARG000 | 4;4;4 | 69510337:695 | 69510428:695 | +;+;+ | 1938 | NA | 3 | 4 | 2 | 8 | 0 | 0 |
| ENSDARG000 | 4;4;4 | 69545067:695 | 69546099:695 | -;-;- | 1276 | NA | 2 | 2 | 2 | 0 | 2 | 1 |
| ENSDARG000 | 4;4;4 | 69559692:695 | 69559783:695 | +;+;+ | 1892 | NA | 0 | 0 | 2 | 0 | 0 | 0 |
| ENSDARG000 | 4;4;4;4;4;4;4 | 69577362:695 | 69577458:695 | +;+;+;+;+;+;+ | 2576 | NA | 9 | 3 | 6 | 0 | 1 | 0 |
| ENSDARG000 | 4;4;4 | 69593339:696 | 69595366:696 | -;-;- | 2267 | NA | 0 | 0 | 0 | 0 | 0 | 3 |
| ENSDARG000 | 4;4;4;4;4;4;4;4; | 69712232:697 | 69712290:697 | +;+;+;+;+;+;+;+; | 3447 | NA | 139 | 52 | 79 | 18 | 38 | 51 |
| ENSDARG000 | 4;4;4;4;4;4;4;4; | 69801372:698 | 69801433:698 | +;+;+;+;+;+;+;+; | 2435 | NA | 0 | 0 | 0 | 0 | 0 | 0 |
| ENSDARG000 | 4;4;4 | 69823638:698 | 69823714:698 | +;+;+ | 2177 | NA | 0 | 2 | 0 | 0 | 0 | 0 |
| ENSDARG000 | 4;4 | 69818495:698 | 69819126:698 | -;- | 755 | NA | 0 | 0 | 0 | 0 | 0 | 0 |
| ENSDARG000 | 4;4;4 | 69782658:697 | 69784038:697 | -;-;- | 1626 | NA | 16 | 12 | 17 | 0 | 16 | 0 |
| ENSDARG000 | 4;4;4;4;4;4;4;4; | 69712223:697 | 69712290:697 | +;+;+;+;+;+;+;+; | 3841 | NA | 18 | 7 | 7 | 0 | 3 | 11 |
| ENSDARG000 | 4;4;4;4;4;4;4 | 74087575:740 | 74088113:740 | -;-;-;-;-;-;- | 1247 | rab21 | 774 | 1170 | 1064 | 526 | 1110 | 678 |
| ENSDARG000 | 4;4 | 74131530:741 | 74131762:741 | +;+ | 733 | NA | 0 | 2 | 0 | 4 | 0 | 0 |
| ENSDARG000 | 4;4;4;4;4;4;4;4; | 74226447:742 | 74227057:742 | +;+;+;+;+;+;+;+; | 6632 | PTPRR | 58 | 60 | 48 | 46 | 96 | 88 |
| ENSDARG000 | 4;4;4;4;4 | 69863019:698 | 69863045:698 | +;+;+;+;+ | 1486 | znf1117 | 4 | 2 | 4 | 4 | 2 | 2 |
| ENSDARG000 | 4;4;4 | 69888328:698 | 69888367:698 | +;+;+ | 946 | NA | 0 | 0 | 0 | 0 | 0 | 0 |
| ENSDARG000 | 4;4;4;4;4 | 69911256:699 | 69913005:699 | -;-;-;-;- | 1994 | NA | 18 | 10 | 0 | 2 | 0 | 10 |
| ENSDARG000 | 4;4;4 | 74551606:745 | 74551824:745 | +;+;+ | 2075 | KCNA1 | 0 | 0 | 0 | 0 | 0 | 2 |
| ENSDARG000 | 4 | 74564192 | 74565834 | + | 1643 | NA | 0 | 0 | 0 | 0 | 0 | 2 |
| ENSDARG000 | 4;4 | 70038196:700 | 70038369:700 | -;- | 303 | NA | 0 | 0 | 0 | 0 | 0 | 0 |
| ENSDARG000 | 4;4;4;4;4;4;4;4; | 70269825:702 | 70270405:702 | -;-;-;-;-;-;-;-;- | 4877 | NA | 20 | 5 | 18 | 2 | 15 | 23 |
| ENSDARG000 | 4;4;4;4;4;4;4;4; | 70015788:701 | 70016211:701 | +;+;+;+;+;+;+;+; | 1419 | NA | 3 | 20 | 22 | 10 | 9 | 7 |
| ENSDARG000 | 4;4;4;4;4;4;4;4; | 70171984:701 | 70172138:701 | +;+;+;+;+;+;+;+; | 3745 | NA | 122 | 5 | 28 | 7 | 84 | 106 |
| ENSDARG000 | 4;4;4;4;4;4;4;4; | 70010099:700 | 70010173:700 | +;+;+;+;+;+;+;+; | 5974 | NA | 58 | 19 | 31 | 5 | 37 | 38 |
| ENSDARG000 | 4;4;4 | 70127566:701 | 70128497:701 | -;-;- | 1225 | NA | 0 | 0 | 0 | 0 | 0 | 0 |
| ENSDARG000 | 4;4;4;4;4;4;4;4; | 74928675:749 | 74929487:749 | +;+;+;+;+;+;+;+; | 5575 | nup50 | 510 | 1376 | 1314 | 414 | 1300 | 760 |
| ENSDARG000 | 4;4;4;4;4;4;4;4; | 70086974:700 | 70087506:700 | -;-;-;-;-;-;-;-;- | 3250 | NA | 13 | 2 | 0 | 0 | 23 | 16 |
| ENSDARG000 | 4;4;4;4;4;4;4;4; | 70045731:700 | 70046440:700 | -;-;-;-;-;-;-;-;- | 4957 | NA | 37 | 5 | 13 | 5 | 15 | 17 |
| ENSDARG000 | 4;4;4;4;4;4;4;4; | 74970357:749 | 74970626:749 | -;-;-;-;-;-;-;-;- | 1568 | kiaa0930 | 26 | 86 | 96 | 26 | 56 | 21 |
| ENSDARG000 | 4;4;4;4;4;4;4;4; | 70245533:702 | 70246231:702 | -;-;-;-;-;-;-;-;- | 4948 | NA | 26 | 3 | 22 | 17 | 6 | 8 |
| ENSDARG000 | 4;4 | 70042955:700 | 70043128:700 | -;- | 200 | NA | 0 | 0 | 0 | 0 | 0 | 0 |
| ENSDARG000 | 4;4;4 | 70303503:703 | 70304665:703 | -;-;- | 1456 | NA | 0 | 0 | 4 | 16 | 2 | 0 |
| ENSDARG000 | 4;4;4;4;4;4;4;4; | 75138640:751 | 75139324:751 | -;-;-;-;-;-;-;-;- | 2993 | NA | 34 | 60 | 80 | 55 | 115 | 97 |
| ENSDARG000 | 4;4;4;4;4;4;4;4; | 75310097:753 | 75310483:753 | +;+;+;+;+;+;+;+; | 6307 | CPNE8 | 1694 | 522 | 1060 | 773 | 1518 | 1176 |
| ENSDARG000 | 4;4;4;4;4;4;4;4; | 75458537:754 | 75458743:754 | +;+;+;+;+;+;+;+; | 6538 | NA | 109 | 49 | 76 | 11 | 43 | 74 |
| ENSDARG000 | 4;4;4;4;4;4;4 | 76142882:761 | 76144643:761 | -;-;-;-;-;-;- | 2399 | NA | 3 | 0 | 0 | 0 | 6 | 0 |
| ENSDARG000 | 4;4;4;4;4;4;4;4; | 75601353:756 | 75604591:756 | -;-;-;-;-;-;-;-;- | 6892 | NA | 9 | 3 | 10 | 2 | 11 | 0 |
| ENSDARG000 | 4;4;4;4;4;4;4;4; | 75481536:754 | 75482220:754 | -;-;-;-;-;-;-;-;- | 4882 | NA | 21 | 10 | 12 | 1 | 15 | 23 |
| ENSDARG000 | 4;4;4;4;4;4;4;4; | 75631236:756 | 75631887:756 | -;-;-;-;-;-;-;-;- | 4072 | NA | 27 | 23 | 19 | 6 | 34 | 39 |
| ENSDARG000 | 4;4;4;4;4;4;4;4; | 76117415:761 | 76118036:761 | -;-;-;-;-;-;-;-;- | 3225 | NA | 10 | 4 | 11 | 5 | 7 | 6 |
| ENSDARG000 | 4;4;4;4;4;4;4;4; | 75526044:755 | 75527205:755 | -;-;-;-;-;-;-;-;- | 4138 | si:dkey-71l4.4 | 40 | 6 | 22 | 1 | 8 | 7 |
| ENSDARG000 | 4;4;4;4;4;4;4;4; | 75603917:756 | 75604591:756 | -;-;-;-;-;-;-;-;- | 4639 | NA | 63 | 10 | 5 | 2 | 44 | 26 |
| ENSDARG000 | 4;4;4;4;4;4;4;4; | 76001682:760 | 76002333:760 | -;-;-;-;-;-;-;-;- | 3621 | NA | 11 | 15 | 8 | 15 | 3 | 9 |
| ENSDARG000 | 4;4;4;4;4;4;4;4; | 76173376:761 | 76173651:761 | -;-;-;-;-;-;-;-;- | 5648 | NA | 26 | 13 | 26 | 4 | 13 | 20 |
| ENSDARG000 | 4;4;4;4;4;4;4;4; | 75537772:755 | 75537816:755 | +;+;+;+;+;+;+;+; | 4000 | NA | 30 | 12 | 33 | 9 | 20 | 16 |
| ENSDARG000 | 4;4;4;4;4;4;4;4; | 75781680:757 | 75782224:757 | -;-;-;-;-;-;-;-;- | 3446 | NA | 22 | 6 | 12 | 2 | 3 | 4 |
| ENSDARG000 | 4;4;4;4;4;4;4;4; | 75883021:758 | 75883653:758 | -;-;-;-;-;-;-;-;- | 3627 | NA | 20 | 9 | 7 | 5 | 18 | 13 |
| ENSDARG000 | 4;4;4;4;4;4;4 | 75666631:756 | 75667154:756 | -;-;-;-;-;-;- | 2486 | NA | 4 | 2 | 4 | 3 | 6 | 2 |
| ENSDARG000 | 4;4;4;4;4;4;4;4; | 76060322:760 | 76061363:760 | -;-;-;-;-;-;-;-;- | 5104 | NA | 58 | 19 | 25 | 11 | 16 | 5 |
| ENSDARG000 | 4;4;4;4;4;4;4 | 76255295:762 | 76255788:762 | -;-;-;-;-;-;- | 2949 | NA | 14 | 13 | 17 | 2 | 12 | 11 |
| ENSDARG000 | 4;4;4;4;4;4 | 75816428:758 | 75816601:758 | +;+;+;+;+;+ | 1218 | NA | 17 | 16 | 20 | 1 | 3 | 16 |
| ENSDARG000 | 4;4;4;4;4;4;4;4; | 75804584:758 | 75806956:758 | -;-;-;-;-;-;-;-;- | 7256 | NA | 14 | 15 | 55 | 6 | 10 | 11 |
| ENSDARG000 | 4;4;4;4;4;4;4;4; | 75690971:756 | 75691622:756 | -;-;-;-;-;-;-;-;- | 3874 | NA | 12 | 12 | 14 | 4 | 16 | 4 |
| ENSDARG000 | 4;4;4;4 | 76150837:761 | 76152243:761 | -;-;-;- | 2808 | NA | 0 | 2 | 0 | 0 | 0 | 0 |
| ENSDARG000 | 4;4;4;4;4;4;4;4; | 76048706:760 | 76049379:760 | -;-;-;-;-;-;-;-;- | 3800 | NA | 5 | 1 | 7 | 2 | 0 | 3 |
| ENSDARG000 | 4;4;4;4;4;4;4;4; | 75840975:759 | 75841501:759 | -;-;-;-;-;-;-;-;- | 5637 | NA | 35 | 5 | 59 | 4 | 13 | 25 |
| ENSDARG000 | 4;4;4;4;4;4;4;4; | 75840584:758 | 75841501:758 | -;-;-;-;-;-;-;-;- | 5437 | NA | 54 | 38 | 37 | 7 | 14 | 10 |
| ENSDARG000 | 4;4;4;4;4;4;4;4; | 76221239:762 | 76221927:762 | -;-;-;-;-;-;-;-;- | 3751 | NA | 23 | 39 | 28 | 11 | 30 | 25 |
| ENSDARG000 | 4;4;4;4;4;4;4;4; | 13698985:136 | 13698992:136 | -;-;-;-;-;-;-;-;- | 708 | NA | 0 | 0 | 0 | 0 | 0 | 0 |
| ENSDARG000 | 4;4;4;4;4;4;4;4; | 12979653:129 | 12979676:129 | +;+;+;+;+;+;+;+; | 228 | NA | 22 | 4 | 2 | 24 | 8 | 6 |
| ENSDARG000 | 4;4;4;4;4;4;4;4; | 76492691:764 | 76493264:764 | +;+;+;+;+;+;+;+; | 5156 | ms4a17a.17 | 74 | 2 | 4 | 4 | 41 | 76 |
| ENSDARG000 | 4;4;4 | 74002176:740 | 74002331:740 | +;+;+ | 1230 | si:cabz01069( | 2 | 16 | 20 | 24 | 46 | 39 |
| ENSDARG000 | 4;4 | 52188449:521 | 52189216:521 | -;- | 972 | NA | 0 | 2 | 1 | 0 | 10 | 0 |
| ENSDARG000 | 4;4;4 | 58016732:580 | 58016816:580 | +;+;+ | 963 | NA | 2 | 0 | 0 | 0 | 0 | 0 |
| ENSDARG000 | 4;4;4;4;4;4;4;4; | 90048:904441 | 90283:906281 | +;+;+;+;+;+;+;+; | 2808 | lrp6 | 961 | 491 | 800 | 539 | 824 | 653 |
| ENSDARG000 | 4;4;4;4;4;4 | 68087932:680 | 68087947:680 | +;+;+;+;+;+ | 3628 | NA | 5 | 0 | 0 | 0 | 14 | 15 |
| ENSDARG000 | 4;4;4;4;4;4;4;4; | 72723304:727 | 72723935:727 | +;+;+;+;+;+;+;+; | 6728 | rab3ip | 770 | 1205 | 2201 | 1014 | 3120 | 863 |
| ENSDARG000 | 4;4;4;4;4 | 36489448:365 | 36490247:365 | +;+;+;+;+ | 846 | NA | 0 | 0 | 0 | 0 | 0 | 0 |
| ENSDARG000 | 4;4;4;4;4;4;4;4; | 19903784:199 | 19907532:199 | -;-;-;-;-;-;-;-;- | 6135 | NA | 0 | 0 | 0 | 0 | 0 | 0 |
| ENSDARG000 | 4;4;4;4;4;4;4;4; | 63159757:631 | 63162046:631 | -;-;-;-;-;-;-;- | 3372 | NA | 41 | 21 | 30 | 11 | 4 | 2 |
| ENSDARG000 | 4;4;4;4;4;4;4;4; | 76692834:766 | 76692886:766 | +;+;+;+;+;+;+;+; | 1084 | ms4a17a.1 | 98 | 3 | 0 | 1 | 16 | 1 |
| ENSDARG000 | 4;4;4;4;4;4;4;4; | 56422336:564 | 56422534:564 | +;+;+;+;+;+;+;+; | 1338 | NA | 0 | 0 | 0 | 0 | 0 | 0 |
| ENSDARG000 | 4;4;4;4;4;4;4 | 8797197:8802 | 8797451:8802 | +;+;+;+;+;+;+ | 2801 | sult4a1 | 0 | 0 | 0 | 0 | 4 | 2 |
| ENSDARG000 | 4 | 77557112 | 77557279 | - | 168 | NA | 0 | 1 | 1 | 0 | 0 | 0 |
| ENSDARG000 | 4;4 | 36906436:369 | 36906520:369 | +;+ | 966 | NA | 4 | 6 | 9 | 2 | 0 | 2 |
| ENSDARG000 | 4;4;4;4;4;4;4;4; | 72134837:721 | 72135185:721 | -;-;-;-;-;-;-;-;- | 2709 | slco1f4 | 0 | 0 | 4 | 16 | 24 | 0 |
| ENSDARG000 | 4 | 37992753 | 37997390 | + | 4638 | NA | 0 | 2 | 0 | 0 | 0 | 0 |
| ENSDARG000 | 4;4;4 | 42450386:424 | 42450475:424 | +;+;+ | 345 | NA | 1 | 10 | 0 | 0 | 8 | 14 |
| ENSDARG000 | 4;4;4;4;4;4;4;4; | 78058635:780 | 78058778:780 | +;+;+;+;+;+;+;+; | 3178 | NA | 0 | 43 | 37 | 19 | 43 | 36 |
| ENSDARG000 | 4;4 | 73015260:730 | 73016101:730 | -;- | 927 | NA | 24 | 44 | 51 | 14 | 44 | 44 |
| ENSDARG000 | 4;4 | 50539777:505 | 50540687:505 | -;- | 996 | NA | 16 | 7 | 7 | 0 | 4 | 12 |
| ENSDARG000 | 4;4 | 1014548:1015 | 1014883:1015 | -;- | 675 | NA | 0 | 2 | 0 | 0 | 0 | 0 |
| ENSDARG000 | 4;4;4 | 34657691:346 | 34657775:346 | +;+;+ | 993 | NA | 4 | 0 | 2 | 0 | 0 | 1 |
| ENSDARG000 | 4 | 77319718 | 77319838 | + | 121 | 5S_rRNA | 0 | 0 | 0 | 0 | 0 | 0 |
| ENSDARG000 | 4 | 42645881 | 42646044 | + | 164 | U1 | 0 | 0 | 0 | 0 | 0 | 0 |
| ENSDARG000 | 4 | 35233297 | 35233413 | - | 117 | 5S_rRNA | 0 | 0 | 0 | 0 | 0 | 0 |
| ENSDARG000 | 4 | 61180974 | 61181137 | + | 164 | U1 | 0 | 0 | 0 | 0 | 0 | 0 |
| ENSDARG000 | 4 | 419659 | 419774 | + | 116 | 5S_rRNA | 0 | 0 | 0 | 0 | 0 | 0 |
| ENSDARG000 | 4 | 58818242 | 58818348 | - | 107 | U6 | 0 | 0 | 0 | 0 | 0 | 0 |
| ENSDARG000 | 4 | 45206709 | 45206827 | - | 119 | 5S_rRNA | 0 | 0 | 0 | 0 | 0 | 2 |
| ENSDARG000 | 4 | 38234160 | 38234266 | - | 107 | U6 | 0 | 0 | 0 | 0 | 0 | 0 |
| ENSDARG000 | 4 | 32263223 | 32263329 | + | 107 | U6 | 0 | 0 | 0 | 0 | 0 | 0 |
| ENSDARG000 | 4 | 77560655 | 77560810 | - | 156 | 5_8S_rRNA | 0 | 1 | 0 | 0 | 0 | 0 |
| ENSDARG000 | 4 | 40915566 | 40915672 | - | 107 | U6 | 0 | 0 | 0 | 0 | 0 | 0 |
| ENSDARG000 | 4 | 61444218 | 61444324 | - | 107 | U6 | 0 | 0 | 0 | 0 | 0 | 0 |
| ENSDARG000 | 4;4;4;4;4;4;4;4; | 74271067:742 | 74271461:742 | -;-;-;-;-;-;-;-;- | 6436 | ptprb | 1126 | 151 | 137 | 78 | 1517 | 1433 |
| ENSDARG000 | 4 | 69956050 | 69956213 | - | 164 | U1 | 0 | 0 | 0 | 0 | 0 | 0 |
| ENSDARG000 | 4 | 30993186 | 30993349 | - | 164 | U1 | 0 | 0 | 0 | 0 | 0 | 0 |
| ENSDARG000 | 4 | 50119341 | 50119489 | + | 149 | U1 | 1 | 0 | 0 | 0 | 0 | 1 |
| ENSDARG000 | 4 | 57971674 | 57971822 | + | 149 | U1 | 0 | 0 | 0 | 0 | 0 | 0 |
| ENSDARG000 | 4 | 23379262 | 23379378 | + | 117 | 5S_rRNA | 0 | 0 | 0 | 0 | 0 | 0 |
| ENSDARG000 | 4 | 38021474 | 38021637 | - | 164 | U1 | 2 | 0 | 0 | 0 | 0 | 0 |
| ENSDARG000 | 4 | 51396232 | 51396380 | - | 149 | U1 | 0 | 0 | 0 | 0 | 0 | 0 |
| ENSDARG000 | 4 | 58794970 | 58795076 | - | 107 | U6 | 0 | 0 | 0 | 0 | 0 | 0 |
| ENSDARG000 | 4 | 64799863 | 64799969 | + | 107 | U6 | 0 | 0 | 0 | 0 | 0 | 0 |
| ENSDARG000 | 4 | 55779607 | 55779760 | + | 154 | 5_8S_rRNA | 0 | 0 | 0 | 0 | 0 | 0 |
| ENSDARG000 | 4 | 51401779 | 51401885 | + | 107 | U6 | 0 | 0 | 0 | 0 | 0 | 0 |
| ENSDARG000 | 4 | 61459997 | 61460103 | - | 107 | U6 | 0 | 0 | 0 | 0 | 0 | 0 |
| ENSDARG000 | 4 | 31194936 | 31195042 | - | 107 | U6 | 0 | 0 | 0 | 0 | 0 | 0 |
| ENSDARG000 | 4 | 18654155 | 18654261 | - | 107 | dre-let-7a-3 | 0 | 2 | 1 | 7 | 4 | 5 |
| ENSDARG000 | 4 | 68564061 | 68564158 | - | 98 | dre-mir-738 | 0 | 0 | 0 | 0 | 0 | 0 |
| ENSDARG000 | 4 | 45591994 | 45592100 | + | 107 | U6 | 0 | 0 | 0 | 0 | 0 | 0 |
| ENSDARG000 | 4 | 64800383 | 64800489 | + | 107 | U6 | 0 | 0 | 0 | 0 | 0 | 0 |
| ENSDARG000 | 4 | 69152234 | 69152397 | + | 164 | U1 | 0 | 0 | 0 | 0 | 0 | 0 |
| ENSDARG000 | 4 | 48177977 | 48178083 | - | 107 | U6 | 0 | 0 | 0 | 0 | 0 | 0 |
| ENSDARG000 | 4 | 69488581 | 69488682 | + | 102 | U6 | 0 | 0 | 0 | 0 | 0 | 0 |

| | | | | | | | | | | | | |
|---|---|---|---|---|---|---|---|---|---|---|---|---|
| ENSDARG000 | 4 | 40898384 | 40898490 | - | 107 | U6 | 0 | 0 | 0 | 0 | 0 | 0 |
| ENSDARG000 | 4 | 35545432 | 35545546 | + | 115 | 5S_rRNA | 0 | 0 | 0 | 0 | 0 | 0 |
| ENSDARG000 | 4 | 28703350 | 28703422 | + | 73 | dre-mir-430c- | 0 | 0 | 0 | 0 | 0 | 0 |
| ENSDARG000 | 4 | 32279072 | 32279178 | + | 107 | U6 | 0 | 0 | 0 | 0 | 0 | 0 |
| ENSDARG000 | 4;4;4;4;4 | 4601815:4603 | 4602298:4603 | -;-;-;-;- | 1217 | NA | 365 | 349 | 415 | 642 | 276 | 206 |
| ENSDARG000 | 4 | 46467143 | 46467249 | - | 107 | U6 | 0 | 0 | 0 | 0 | 0 | 0 |
| ENSDARG000 | 4 | 52727108 | 52727284 | + | 177 | U1 | 15 | 0 | 17 | 11 | 3 | 16 |
| ENSDARG000 | 4 | 42643299 | 42643462 | + | 164 | U1 | 0 | 0 | 0 | 0 | 0 | 0 |
| ENSDARG000 | 4 | 64726277 | 64726383 | + | 107 | U6 | 0 | 0 | 0 | 0 | 0 | 0 |
| ENSDARG000 | 4 | 7185195 | 7185311 | - | 117 | 5S_rRNA | 0 | 0 | 0 | 0 | 0 | 0 |
| ENSDARG000 | 4 | 61954121 | 61954239 | + | 119 | 5S_rRNA | 0 | 0 | 0 | 0 | 0 | 0 |
| ENSDARG000 | 4 | 67547938 | 67548044 | - | 107 | U6 | 0 | 0 | 0 | 0 | 0 | 0 |
| ENSDARG000 | 4 | 47187316 | 47187422 | - | 107 | U6 | 0 | 0 | 0 | 0 | 0 | 0 |
| ENSDARG000 | 4 | 52587943 | 52588049 | + | 107 | U6 | 0 | 0 | 0 | 0 | 0 | 0 |
| ENSDARG000 | 4 | 45577960 | 45578066 | + | 107 | U6 | 0 | 0 | 0 | 0 | 0 | 0 |
| ENSDARG000 | 4 | 45570271 | 45570376 | + | 106 | U6 | 0 | 0 | 0 | 0 | 0 | 0 |
| ENSDARG000 | 4;4;4;4;4;4;4; | 72272050:722 | 72273119:722 | -;-;-;-;-;-;-;-; | 2383 | NA | 92 | 85 | 168 | 38 | 112 | 70 |
| ENSDARG000 | 4 | 32261642 | 32261743 | + | 102 | U6 | 0 | 0 | 0 | 0 | 0 | 0 |
| ENSDARG000 | 4 | 40911863 | 40911969 | - | 107 | U6 | 0 | 0 | 0 | 0 | 0 | 0 |
| ENSDARG000 | 4;4 | 52360358:523 | 52360372:523 | -;- | 762 | NA | 6 | 10 | 41 | 1 | 5 | 23 |
| ENSDARG000 | 4 | 61481649 | 61481812 | + | 164 | U1 | 0 | 0 | 0 | 0 | 0 | 0 |
| ENSDARG000 | 4 | 41338330 | 41338482 | - | 153 | U1 | 0 | 0 | 0 | 0 | 2 | 0 |
| ENSDARG000 | 4 | 31227559 | 31227627 | - | 69 | U6 | 0 | 0 | 0 | 0 | 0 | 0 |
| ENSDARG000 | 4;4;4 | 68353501:683 | 68355954:683 | -;-;- | 2731 | NA | 7 | 0 | 12 | 0 | 1 | 8 |
| ENSDARG000 | 4;4;4;4 | 20042807:200 | 20042953:200 | -;-;-;- | 365 | zgc:193726 | 0 | 0 | 0 | 0 | 0 | 4 |
| ENSDARG000 | 4 | 68046522 | 68046595 | - | 74 | U6 | 0 | 0 | 0 | 0 | 2 | 0 |
| ENSDARG000 | 4 | 28695576 | 28695657 | + | 82 | dre-mir-430a- | 0 | 0 | 0 | 0 | 0 | 0 |
| ENSDARG000 | 4;4;4;4 | 54225677:542 | 54225703:542 | +;+;+;+ | 2580 | NA | 6 | 1 | 6 | 1 | 0 | 0 |
| ENSDARG000 | 4 | 64785471 | 64785634 | - | 164 | U1 | 0 | 0 | 0 | 0 | 0 | 0 |
| ENSDARG000 | 4;4;4;4;4;4;4; | 73215536:732 | 73215654:732 | +;+;+;+;+;+;+; | 1595 | NA | 6 | 8 | 8 | 13 | 16 | 8 |
| ENSDARG000 | 4;4;4;4;4;4;4; | 269443:27303 | 272212:27318 | -;-;-;-;-;-;-;- | 3695 | si:ch73-252i1 | 2194 | 1131 | 1275 | 1318 | 1111 | 1010 |
| ENSDARG000 | 4;4 | 72235562:722 | 72236334:722 | +;+ | 789 | NA | 34 | 30 | 27 | 9 | 34 | 37 |
| ENSDARG000 | 4 | 31219259 | 31219365 | - | 107 | U6 | 0 | 0 | 0 | 0 | 0 | 0 |
| ENSDARG000 | 4 | 15074715 | 15074824 | + | 110 | dre-mir-96 | 0 | 0 | 0 | 0 | 0 | 0 |
| ENSDARG000 | 4 | 68047016 | 68047112 | - | 97 | U6 | 0 | 0 | 0 | 0 | 0 | 0 |
| ENSDARG000 | 4;4;4;4;4;4;4; | 74445535:744 | 74445678:744 | -;-;-;-;-;-;-;- | 1424 | NA | 0 | 0 | 5 | 2 | 29 | 0 |
| ENSDARG000 | 4 | 46481388 | 46481494 | - | 107 | U6 | 0 | 0 | 0 | 0 | 0 | 0 |
| ENSDARG000 | 4 | 57964669 | 57964775 | - | 107 | U6 | 0 | 0 | 0 | 0 | 0 | 0 |
| ENSDARG000 | 4 | 69492559 | 69492665 | + | 107 | U6 | 0 | 0 | 0 | 0 | 0 | 0 |
| ENSDARG000 | 4 | 61457918 | 61457991 | - | 74 | U6 | 0 | 0 | 0 | 0 | 0 | 0 |
| ENSDARG000 | 4 | 39364038 | 39364215 | - | 178 | U1 | 0 | 0 | 0 | 0 | 0 | 0 |
| ENSDARG000 | 4 | 67548983 | 67549089 | - | 107 | U6 | 0 | 0 | 0 | 0 | 0 | 0 |
| ENSDARG000 | 4 | 28708430 | 28708511 | + | 82 | dre-mir-430a- | 0 | 0 | 0 | 0 | 0 | 0 |
| ENSDARG000 | 4;4 | 61802304:618 | 61803953:618 | -;- | 1814 | NA | 57 | 33 | 68 | 22 | 39 | 36 |
| ENSDARG000 | 4 | 28707110 | 28707190 | + | 81 | dre-mir-430b- | 0 | 0 | 0 | 0 | 0 | 0 |
| ENSDARG000 | 4 | 28701587 | 28701659 | + | 73 | dre-mir-430c- | 0 | 0 | 0 | 0 | 0 | 0 |
| ENSDARG000 | 4 | 393766 | 393902 | + | 137 | NA | 0 | 0 | 5 | 4 | 2 | 0 |
| ENSDARG000 | 4 | 52586888 | 52586994 | + | 107 | U6 | 0 | 0 | 0 | 0 | 0 | 0 |
| ENSDARG000 | 4 | 10661561 | 10661677 | - | 117 | 5S_rRNA | 0 | 0 | 0 | 0 | 0 | 0 |
| ENSDARG000 | 4 | 31223488 | 31223594 | - | 107 | U6 | 0 | 0 | 0 | 0 | 0 | 0 |
| ENSDARG000 | 4 | 28695370 | 28695450 | + | 81 | dre-mir-430b- | 0 | 0 | 0 | 0 | 0 | 0 |
| ENSDARG000 | 4 | 32271151 | 32271257 | + | 107 | U6 | 0 | 0 | 0 | 0 | 0 | 0 |
| ENSDARG000 | 4 | 28706853 | 28706925 | + | 73 | dre-mir-430c- | 0 | 0 | 0 | 0 | 0 | 0 |
| ENSDARG000 | 4 | 28703813 | 28703894 | + | 82 | dre-mir-430a- | 0 | 0 | 0 | 0 | 0 | 0 |
| ENSDARG000 | 4 | 28432331 | 28432445 | - | 115 | 5S_rRNA | 0 | 0 | 0 | 0 | 0 | 0 |
| ENSDARG000 | 4 | 28700961 | 28701033 | + | 73 | dre-mir-430c- | 0 | 0 | 0 | 0 | 0 | 0 |
| ENSDARG000 | 4;4;4;4 | 72063970:720 | 72064461:720 | -;-;-;- | 940 | NA | 0 | 0 | 0 | 0 | 0 | 0 |
| ENSDARG000 | 4 | 42626690 | 42626853 | + | 164 | U1 | 0 | 0 | 0 | 0 | 0 | 0 |
| ENSDARG000 | 4;4;4 | 32824850:328 | 32826538:328 | -;-;- | 1879 | NA | 3 | 2 | 1 | 0 | 0 | 2 |
| ENSDARG000 | 4;4;4;4;4;4;4; | 2396024:2396 | 2396105:2396 | -;-;-;-;-;-;-;-; | 2046 | OSBPL8 | 51 | 22 | 61 | 16 | 51 | 52 |
| ENSDARG000 | 4 | 14725074 | 14725214 | + | 141 | U4 | 0 | 0 | 0 | 0 | 0 | 0 |
| ENSDARG000 | 4 | 65245933 | 65246038 | + | 106 | U6 | 0 | 0 | 0 | 0 | 0 | 0 |
| ENSDARG000 | 4 | 52737998 | 52738143 | + | 146 | U1 | 0 | 0 | 0 | 0 | 0 | 0 |
| ENSDARG000 | 4 | 28705347 | 28705427 | + | 81 | dre-mir-430b- | 0 | 0 | 0 | 0 | 0 | 0 |
| ENSDARG000 | 4;4 | 68181039:681 | 68185893:681 | -;- | 4889 | NA | 29 | 6 | 8 | 1 | 2 | 0 |
| ENSDARG000 | 4;4;4;4;4;4;4; | 78022547:780 | 78022710:780 | -;-;-;-;-;-;-;- | 994 | cct2 | 2252 | 5948 | 8243 | 4200 | 5384 | 5573 |
| ENSDARG000 | 4 | 57963615 | 57963721 | - | 107 | U6 | 0 | 0 | 0 | 0 | 0 | 0 |
| ENSDARG000 | 4 | 45558046 | 45558209 | - | 164 | U1 | 0 | 0 | 0 | 0 | 0 | 0 |
| ENSDARG000 | 4 | 28697921 | 28698002 | + | 82 | dre-mir-430a- | 0 | 0 | 0 | 0 | 0 | 0 |
| ENSDARG000 | 4 | 28695960 | 28696040 | + | 81 | dre-mir-430b- | 0 | 0 | 0 | 0 | 0 | 0 |
| ENSDARG000 | 4 | 18189406 | 18189520 | - | 115 | 5S_rRNA | 0 | 0 | 0 | 0 | 0 | 0 |
| ENSDARG000 | 4 | 64698800 | 64698906 | + | 107 | U6 | 0 | 0 | 0 | 0 | 0 | 0 |
| ENSDARG000 | 4;4 | 48550308:485 | 48550392:485 | +;+ | 1077 | NA | 0 | 0 | 0 | 0 | 0 | 0 |
| ENSDARG000 | 4 | 61482954 | 61483117 | + | 164 | U1 | 0 | 0 | 0 | 0 | 0 | 0 |
| ENSDARG000 | 4 | 48171637 | 48171742 | - | 106 | U6 | 0 | 0 | 0 | 0 | 0 | 0 |
| ENSDARG000 | 4 | 48174808 | 48174914 | - | 107 | U6 | 0 | 0 | 0 | 0 | 0 | 0 |
| ENSDARG000 | 4 | 28709454 | 28709534 | + | 81 | dre-mir-430b- | 0 | 0 | 0 | 0 | 0 | 0 |
| ENSDARG000 | 4;4;4;4;4 | 3806258:3807 | 3807031:3807 | -;-;-;-;- | 2369 | NA | 4 | 0 | 2 | 6 | 14 | 1 |
| ENSDARG000 | 4;4;4;4 | 75292781:752 | 75294426:752 | +;+;+;+ | 4335 | ABCD2 | 34 | 18 | 32 | 4 | 62 | 40 |
| ENSDARG000 | 4;4;4;4;4;4 | 958051:96092 | 959308:96100 | -;-;-;-;-;- | 1819 | atp23 | 142 | 160 | 173 | 98 | 180 | 138 |
| ENSDARG000 | 4;4 | 67050047:670 | 67050230:670 | +;+ | 2421 | znf1070 | 17 | 10 | 24 | 0 | 10 | 22 |
| ENSDARG000 | 4;4;4;4;4;4;4 | 76637548:766 | 76637598:766 | +;+;+;+;+;+;+ | 1034 | ms4a17a.8 | 32 | 14 | 2 | 0 | 33 | 38 |
| ENSDARG000 | 4;4;4 | 51458762:514 | 51459529:514 | -;-;- | 972 | NA | 0 | 4 | 4 | 4 | 2 | 8 |
| ENSDARG000 | 4;4;4 | 65745953:657 | 65746491:657 | -;-;- | 876 | NA | 14 | 10 | 2 | 4 | 27 | 35 |
| ENSDARG000 | 4;4;4 | 68562464:685 | 68562496:685 | +;+;+ | 888 | NA | 0 | 0 | 0 | 0 | 0 | 0 |
| ENSDARG000 | 4;4 | 48575332:485 | 48576626:485 | -;- | 1733 | NA | 0 | 0 | 1 | 0 | 0 | 4 |
| ENSDARG000 | 4;4 | 63075252:630 | 63076181:630 | -;- | 1044 | NA | 27 | 12 | 36 | 8 | 11 | 4 |
| ENSDARG000 | 4;4;4;4;4;4 | 1461079:1462 | 1461949:1462 | +;+;+;+;+;+ | 2256 | NA | 20 | 51 | 0 | 0 | 2 | 4 |
| ENSDARG000 | 4 | 70003003 | 70003166 | - | 164 | U1 | 0 | 0 | 0 | 0 | 0 | 0 |
| ENSDARG000 | 4 | 43367796 | 43374443 | - | 6648 | NA | 0 | 0 | 0 | 14 | 32 | 38 |
| ENSDARG000 | 4;4 | 29165840:292 | 29166653:292 | +;+ | 6561 | NA | 76 | 84 | 168 | 20 | 148 | 125 |
| ENSDARG000 | 4;4;4 | 77253851:772 | 77255093:772 | -;-;- | 1493 | zgc:162948 | 38 | 83 | 110 | 57 | 115 | 122 |
| ENSDARG000 | 4 | 51961329 | 51961447 | - | 119 | 5S_rRNA | 0 | 0 | 0 | 0 | 0 | 0 |
| ENSDARG000 | 4;4;4;4;4;4;4 | 71989418:719 | 71989888:719 | +;+;+;+;+;+;+ | 2154 | PARP11 | 10 | 16 | 12 | 0 | 39 | 16 |
| ENSDARG000 | 4;4;4 | 73973242:739 | 73973311:739 | +;+;+ | 3254 | NA | 0 | 0 | 0 | 2 | 25 | 0 |
| ENSDARG000 | 4;4;4;4;4;4;4; | 72092423:720 | 72093127:720 | -;-;-;-;-;-;-;-; | 2423 | slco1f1 | 0 | 0 | 1 | 0 | 12 | 0 |
| ENSDARG000 | 4;4;4;4;4;4;4; | 5906958:5911 | 5910727:5911 | -;-;-;-;-;-;-;-; | 4287 | arl1 | 998 | 1337 | 1281 | 911 | 1129 | 1144 |
| ENSDARG000 | 4;4 | 67136110:671 | 67137104:671 | -;- | 1080 | NA | 2 | 2 | 4 | 0 | 0 | 9 |
| ENSDARG000 | 4 | 1191527 | 1191644 | + | 118 | 5S_rRNA | 0 | 0 | 0 | 0 | 0 | 0 |
| ENSDARG000 | 4;4;4;4;4 | 20037521:200 | 20038711:200 | -;-;-;-;- | 1461 | NA | 2 | 4 | 21 | 12 | 5 | 7 |
| ENSDARG000 | 4;4;4;4 | 9543459:9547 | 9543467:9547 | -;-;-;- | 441 | NA | 0 | 0 | 3 | 0 | 0 | 0 |
| ENSDARG000 | 4 | 45563414 | 45563515 | + | 102 | U6 | 0 | 0 | 0 | 0 | 0 | 0 |
| ENSDARG000 | 4 | 20396049 | 20396170 | - | 122 | 5S_rRNA | 0 | 0 | 0 | 0 | 0 | 0 |
| ENSDARG000 | 4 | 45586412 | 45586518 | + | 107 | U6 | 0 | 0 | 0 | 0 | 0 | 0 |
| ENSDARG000 | 4 | 61182284 | 61182447 | + | 164 | U1 | 0 | 0 | 0 | 0 | 0 | 0 |
| ENSDARG000 | 4 | 28704927 | 28705008 | + | 82 | dre-mir-430a- | 0 | 0 | 0 | 0 | 0 | 0 |
| ENSDARG000 | 4 | 40922745 | 40922893 | + | 149 | U1 | 0 | 0 | 0 | 0 | 0 | 0 |
| ENSDARG000 | 4 | 46482444 | 46482550 | - | 107 | U6 | 0 | 0 | 0 | 0 | 0 | 0 |
| ENSDARG000 | 4 | 28703607 | 28703687 | + | 81 | dre-mir-430b- | 0 | 0 | 0 | 0 | 0 | 0 |
| ENSDARG000 | 4;4;4;4;4;4;4; | 61810270:618 | 61810354:618 | +;+;+;+;+;+;+; | 1089 | NA | 4 | 0 | 1 | 0 | 0 | 4 |
| ENSDARG000 | 4 | 42624108 | 42624271 | + | 164 | U1 | 0 | 0 | 0 | 0 | 0 | 0 |
| ENSDARG000 | 4 | 67581201 | 67581365 | + | 165 | U1 | 0 | 0 | 0 | 0 | 0 | 0 |
| ENSDARG000 | 4 | 69959938 | 69960101 | - | 164 | U1 | 0 | 0 | 0 | 0 | 0 | 0 |
| ENSDARG000 | 4 | 64786781 | 64786929 | - | 149 | U1 | 0 | 0 | 0 | 0 | 0 | 0 |
| ENSDARG000 | 4 | 58817185 | 58817291 | - | 107 | U6 | 0 | 0 | 0 | 0 | 0 | 0 |
| ENSDARG000 | 4 | 16213424 | 16213540 | - | 117 | 5S_rRNA | 0 | 0 | 0 | 0 | 0 | 0 |
| ENSDARG000 | 4 | 52591639 | 52591745 | + | 107 | U6 | 0 | 0 | 0 | 0 | 0 | 0 |
| ENSDARG000 | 4;4;4;4;4;4;4; | 75013307:750 | 75013722:750 | -;-;-;-;-;-;-;-; | 3149 | large1 | 156 | 135 | 293 | 130 | 259 | 197 |
| ENSDARG000 | 4 | 21841853 | 21841969 | + | 117 | 5S_rRNA | 0 | 0 | 0 | 0 | 0 | 2 |
| ENSDARG000 | 4 | 64800910 | 64801016 | + | 107 | U6 | 0 | 0 | 0 | 0 | 0 | 0 |
| ENSDARG000 | 4 | 28706690 | 28706771 | + | 82 | dre-mir-430a- | 0 | 0 | 0 | 0 | 0 | 0 |
| ENSDARG000 | 4;4;4;4;4;4;4; | 54472921:544 | 54472947:544 | -;-;-;-;-;-;-;- | 165 | NA | 0 | 1 | 3 | 0 | 0 | 0 |
| ENSDARG000 | 4 | 28705090 | 28705162 | + | 73 | dre-mir-430c- | 0 | 0 | 0 | 0 | 0 | 0 |
| ENSDARG000 | 4;4 | 51151834:511 | 51152375:511 | -;- | 627 | NA | 12 | 14 | 9 | 2 | 5 | 6 |
| ENSDARG000 | 4;4 | 32191378:321 | 32192159:321 | -;- | 987 | NA | 35 | 4 | 42 | 9 | 13 | 11 |
| ENSDARG000 | 4 | 58818772 | 58818878 | - | 107 | U6 | 0 | 0 | 0 | 0 | 0 | 0 |
| ENSDARG000 | 4;4;4;4;4 | 57505842:575 | 57506527:575 | +;+;+;+;+ | 849 | NA | 4 | 13 | 8 | 2 | 4 | 0 |
| ENSDARG000 | 4 | 60177193 | 60177356 | - | 164 | U1 | 0 | 0 | 0 | 0 | 0 | 0 |
| ENSDARG000 | 4 | 57782177 | 57782296 | - | 120 | U1 | 0 | 0 | 0 | 0 | 0 | 0 |
| ENSDARG000 | 4 | 31216612 | 31216718 | - | 107 | U6 | 0 | 0 | 0 | 0 | 0 | 0 |
| ENSDARG000 | 4 | 46485439 | 46485512 | - | 74 | U6 | 0 | 0 | 0 | 0 | 0 | 0 |
| ENSDARG000 | 4 | 50113179 | 50113285 | - | 107 | U6 | 0 | 0 | 0 | 0 | 0 | 0 |
| ENSDARG000 | 4 | 48184797 | 48184945 | + | 149 | U1 | 1 | 0 | 0 | 0 | 0 | 0 |
| ENSDARG000 | 4 | 58925847 | 58925965 | - | 119 | 5S_rRNA | 0 | 0 | 0 | 0 | 0 | 0 |
| ENSDARG000 | 4 | 30905272 | 30905435 | + | 164 | U1 | 0 | 0 | 0 | 0 | 1 | 0 |
| ENSDARG000 | 4 | 31199160 | 31199266 | - | 107 | U6 | 0 | 0 | 0 | 0 | 0 | 0 |
| ENSDARG000 | 4 | 61175662 | 61175730 | - | 69 | U6 | 0 | 0 | 0 | 0 | 0 | 0 |
| ENSDARG000 | 4 | 28702456 | 28702536 | + | 81 | dre-mir-430b- | 0 | 0 | 0 | 0 | 0 | 0 |
| ENSDARG000 | 4 | 6914716 | 6914851 | + | 136 | 5S_rRNA | 0 | 0 | 0 | 0 | 0 | 0 |
| ENSDARG000 | 4 | 38232079 | 38232152 | - | 74 | U6 | 0 | 0 | 0 | 0 | 0 | 0 |
| ENSDARG000 | 4 | 58794453 | 58794559 | - | 107 | U6 | 0 | 0 | 0 | 0 | 0 | 0 |
| ENSDARG000 | 4 | 64704994 | 64705100 | + | 107 | U6 | 0 | 0 | 0 | 0 | 0 | 0 |
| ENSDARG000 | 4;4;4;4;4;4;4; | 9679840:9695 | 9679860:9695 | +;+;+;+;+;+;+; | 300 | NA | 228 | 216 | 342 | 389 | 409 | 324 |
| ENSDARG000 | 4 | 28695113 | 28695185 | + | 73 | dre-mir-430c- | 0 | 0 | 0 | 0 | 0 | 0 |
| ENSDARG000 | 4;4;4 | 68393422:684 | 68394388:684 | +;+;+ | 1458 | NA | 8 | 6 | 18 | 1 | 7 | 9 |
| ENSDARG000 | 4 | 69489105 | 69489178 | + | 74 | U6 | 0 | 0 | 0 | 0 | 0 | 0 |
| ENSDARG000 | 4 | 55021058 | 55021176 | + | 119 | 5S_rRNA | 0 | 0 | 0 | 0 | 0 | 0 |
| ENSDARG000 | 4;4;4;4;4;4;4; | 184863:18486 | 184967:18496 | -;-;-;-;-;-;-;-; | 1202 | NA | 3153 | 1779 | 2634 | 2711 | 2198 | 2125 |
| ENSDARG000 | 4 | 26407146 | 26407263 | - | 118 | 5S_rRNA | 0 | 0 | 0 | 0 | 0 | 0 |
| ENSDARG000 | 4 | 68626746 | 68626862 | - | 117 | 5S_rRNA | 0 | 0 | 0 | 0 | 0 | 0 |
| ENSDARG000 | 4 | 52588998 | 52589104 | + | 107 | U6 | 0 | 0 | 0 | 0 | 0 | 0 |
| ENSDARG000 | 4 | 48174279 | 48174385 | - | 107 | U6 | 0 | 0 | 0 | 0 | 0 | 0 |
| ENSDARG000 | 4 | 31224544 | 31224650 | - | 107 | U6 | 0 | 0 | 0 | 0 | 0 | 0 |
| ENSDARG000 | 4 | 52756813 | 52756968 | - | 156 | U1 | 0 | 0 | 0 | 0 | 0 | 0 |
| ENSDARG000 | 4 | 45585882 | 45585988 | + | 107 | U6 | 0 | 0 | 0 | 0 | 0 | 0 |
| ENSDARG000 | 4 | 48176391 | 48176496 | - | 106 | U6 | 0 | 0 | 0 | 0 | 0 | 0 |
| ENSDARG000 | 4 | 61460522 | 61460628 | - | 107 | U6 | 0 | 0 | 0 | 0 | 0 | 0 |
| ENSDARG000 | 4 | 61447389 | 61447495 | - | 107 | U6 | 0 | 0 | 0 | 0 | 0 | 0 |
| ENSDARG000 | 4 | 28696854 | 28696926 | + | 73 | dre-mir-430c- | 0 | 0 | 0 | 0 | 0 | 0 |
| ENSDARG000 | 4 | 28703187 | 28703268 | + | 82 | dre-mir-430a- | 0 | 0 | 0 | 0 | 0 | 0 |
| ENSDARG000 | 4 | 67976543 | 67976659 | + | 117 | 5S_rRNA | 0 | 0 | 0 | 0 | 0 | 0 |
| ENSDARG000 | 4 | 55827086 | 55827183 | + | 98 | NA | 0 | 0 | 0 | 0 | 0 | 0 |

| | | | | | | | | | | | | |
|---|---|---|---|---|---|---|---|---|---|---|---|---|
| ENSDARG000 | 4;4;4;4;4;4 | 12702440:127 | 12702472:127 | +;+;+;+;+;+ | 144 | NA | 0 | 0 | 0 | 0 | 0 | 0 |
| ENSDARG000 | 4:4:4:4:4:4:4: | 35316044:354 | 35316128:354 | +:+:+:+:+:+:+: | 1212 | zgc:174698 | 0 | 0 | 6 | 0 | 0 | 2 |
| ENSDARG000 | 4 | 68039083 | 68039189 | - | 107 | U6 | 0 | 0 | 0 | 0 | 0 | 0 |
| ENSDARG000 | 4 | 52587414 | 52587520 | + | 107 | U6 | 0 | 0 | 0 | 0 | 0 | 0 |
| ENSDARG000 | 4 | 40905768 | 40905874 | - | 107 | U6 | 0 | 0 | 0 | 0 | 0 | 0 |
| ENSDARG000 | 4 | 18653148 | 18653241 | - | 94 | dre-let-7b | 13 | 30 | 36 | 64 | 68 | 38 |
| ENSDARG000 | 4;4;4;4;4;4 | 73051901:730 | 73051942:730 | +;+;+;+;+;+ | 2038 | zgc:152938 | 34 | 82 | 63 | 22 | 86 | 69 |
| ENSDARG000 | 4;4;4 | 4232562:4232 | 4232585:4232 | +;+;+ | 535 | smkr1 | 0 | 2 | 0 | 0 | 4 | 0 |
| ENSDARG000 | 4 | 60193434 | 60193597 | - | 164 | U1 | 0 | 0 | 0 | 0 | 0 | 0 |
| ENSDARG000 | 4;4;4;4;4;4 | 20545448:205 | 20545535:205 | +;+;+;+;+;+ | 3290 | NA | 0 | 0 | 0 | 0 | 6 | 24 |
| ENSDARG000 | 4;4 | 56711650:567 | 56712827:567 | -;- | 1263 | NA | 2 | 4 | 6 | 0 | 6 | 14 |
| ENSDARG000 | 4 | 64419246 | 64419395 | - | 150 | U1 | 12 | 0 | 14 | 0 | 7 | 7 |
| ENSDARG000 | 4 | 32276432 | 32276538 | + | 107 | U6 | 0 | 0 | 0 | 0 | 0 | 0 |
| ENSDARG000 | 4 | 66672025 | 66672131 | + | 107 | U6 | 0 | 0 | 0 | 0 | 0 | 0 |
| ENSDARG000 | 4 | 47191002 | 47191108 | - | 107 | U6 | 0 | 0 | 0 | 0 | 0 | 0 |
| ENSDARG000 | 4 | 51399667 | 51399768 | + | 102 | U6 | 0 | 0 | 0 | 0 | 0 | 0 |
| ENSDARG000 | 4 | 28700798 | 28700879 | + | 82 | dre-mir-430a- | 0 | 0 | 0 | 0 | 0 | 0 |
| ENSDARG000 | 4:4:4:4:4:4:4: | 2354892:2360 | 2355040:2360 | -:-:-:-:-:-:-:-: | 1398 | nap1l1 | 6036 | 7663 | 10413 | 7766 | 7662 | 8130 |
| ENSDARG000 | 4:4:4:4:4:4:4: | 892959:89909 | 892988:89910 | +:+:+:+:+:+:+: | 189 | NA | 0 | 0 | 0 | 3 | 0 | 0 |
| ENSDARG000 | 4 | 64121772 | 64123545 | - | 1774 | NA | 0 | 0 | 0 | 0 | 0 | 0 |
| ENSDARG000 | 4:4:4:4:4:4:4: | 31259:32438: | 31303:32494: | +:+:+:+:+:+:+: | 834 | phtf2 | 149 | 113 | 136 | 146 | 113 | 103 |
| ENSDARG000 | 4 | 42443560 | 42443627 | - | 68 | U6 | 0 | 0 | 0 | 0 | 1 | 0 |
| ENSDARG000 | 4:4:4:4:4:4:4: | 67318419:673 | 67318503:673 | +:+:+:+:+:+:+: | 1761 | NA | 11 | 2 | 16 | 0 | 9 | 6 |
| ENSDARG000 | 4:4:4:4:4:4:4: | 121709:12178 | 121836:12183 | +:+:+:+:+:+:+: | 1298 | crebl2 | 891 | 788 | 1060 | 1437 | 1012 | 1208 |
| ENSDARG000 | 4 | 57965196 | 57965302 | - | 107 | U6 | 0 | 0 | 0 | 0 | 0 | 0 |
| ENSDARG000 | 4 | 46463459 | 46463565 | - | 107 | U6 | 0 | 0 | 0 | 0 | 0 | 0 |
| ENSDARG000 | 4:4:4:4:4:4:4: | 77286165:772 | 77286174:772 | -:-:-:-:-:-:-:-: | 2139 | slco1f2 | 0 | 0 | 0 | 0 | 0 | 0 |
| ENSDARG000 | 4:4:4:4:4:4:4: | 26082530:260 | 26083187:260 | -:-:-:-:-:-:-:-: | 1945 | si:ch211-244 | 238 | 224 | 103 | 106 | 89 | 176 |
| ENSDARG000 | 4 | 61480344 | 61480507 | + | 164 | U1 | 0 | 0 | 0 | 0 | 0 | 0 |
| ENSDARG000 | 4;4 | 47768097:477 | 47768132:477 | +;+ | 345 | NA | 0 | 0 | 0 | 0 | 0 | 1 |
| ENSDARG000 | 4;4 | 53891317:539 | 53891498:539 | +;+ | 1589 | NA | 8 | 0 | 2 | 0 | 0 | 0 |
| ENSDARG000 | 4;4;4;4;4;4;4 | 107287:11205 | 110846:11217 | -;-;-;-;-;-;- | 4927 | dusp16 | 674 | 458 | 639 | 404 | 552 | 392 |
| ENSDARG000 | 4;4;4 | 41265205:412 | 41265350:412 | -;-;- | 942 | NA | 5 | 2 | 0 | 0 | 3 | 0 |
| ENSDARG000 | 4;4;4;4 | 56621120:566 | 56621395:566 | -;-;-;- | 1242 | NA | 4 | 11 | 0 | 2 | 7 | 26 |
| ENSDARG000 | 4:4:4:4:4:4:4: | 74141400:741 | 74141891:741 | +:+:+:+:+:+:+: | 4009 | trim24 | 389 | 461 | 697 | 414 | 604 | 562 |
| ENSDARG000 | 4 | 31233116 | 31233279 | + | 164 | U1 | 0 | 0 | 0 | 0 | 0 | 0 |
| ENSDARG000 | 4;4;4;4 | 30718997:307 | 30719081:307 | +;+;+;+ | 648 | NA | 16 | 10 | 14 | 1 | 12 | 20 |
| ENSDARG000 | 4;4 | 68564654:685 | 68564794:685 | -;- | 318 | NA | 0 | 0 | 0 | 0 | 0 | 0 |
| ENSDARG000 | 4 | 43143873 | 43144021 | - | 149 | U1 | 0 | 0 | 0 | 0 | 0 | 0 |
| ENSDARG000 | 4 | 63088692 | 63090306 | + | 1615 | NA | 2 | 0 | 0 | 0 | 0 | 0 |
| ENSDARG000 | 4 | 32257582 | 32257730 | - | 149 | U1 | 0 | 0 | 0 | 0 | 0 | 1 |
| ENSDARG000 | 4 | 28707457 | 28707529 | + | 73 | dre-mir-430c- | 0 | 0 | 0 | 0 | 0 | 0 |
| ENSDARG000 | 4 | 67568809 | 67568972 | + | 164 | U1 | 0 | 0 | 0 | 0 | 0 | 0 |
| ENSDARG000 | 4 | 58815603 | 58815709 | - | 107 | U6 | 0 | 0 | 0 | 0 | 0 | 0 |
| ENSDARG000 | 4:4:4:4:4:4:4: | 26053044:260 | 26053542:260 | +:+:+:+:+:+:+: | 7401 | scyl2 | 591 | 463 | 633 | 476 | 593 | 493 |
| ENSDARG000 | 4 | 51407598 | 51407704 | + | 107 | U6 | 0 | 0 | 0 | 0 | 0 | 0 |
| ENSDARG000 | 4 | 47188893 | 47188999 | - | 107 | U6 | 0 | 0 | 0 | 0 | 0 | 0 |
| ENSDARG000 | 4 | 50114764 | 50114870 | - | 107 | U6 | 0 | 0 | 0 | 0 | 0 | 0 |
| ENSDARG000 | 4:4:4:4:4:4:4: | 78010696:780 | 78010849:780 | -:-:-:-:-:-:-:-: | 1084 | nots | 0 | 6 | 16 | 10 | 8 | 12 |
| ENSDARG000 | 4 | 61202591 | 61202754 | + | 164 | U1 | 0 | 0 | 0 | 0 | 0 | 0 |
| ENSDARG000 | 4:4:4:4:4:4:4: | 51339670:513 | 51339754:513 | +:+:+:+:+:+:+: | 2163 | NA | 2 | 3 | 1 | 0 | 6 | 16 |
| ENSDARG000 | 4 | 32262166 | 32262272 | + | 107 | U6 | 0 | 0 | 0 | 0 | 0 | 0 |
| ENSDARG000 | 4 | 32275377 | 32275483 | + | 107 | U6 | 0 | 0 | 0 | 0 | 0 | 0 |
| ENSDARG000 | 4;4 | 286425:28730 | 286545:28742 | -;- | 240 | echdc3 | 17 | 6 | 17 | 4 | 22 | 8 |
| ENSDARG000 | 4:4:4:4:4:4:4: | 47636303:476 | 47636580:476 | +:+:+:+:+:+:+: | 5305 | NA | 18 | 16 | 31 | 8 | 46 | 25 |
| ENSDARG000 | 4 | 60153876 | 60154024 | - | 149 | U1 | 0 | 0 | 0 | 0 | 0 | 0 |
| ENSDARG000 | 4 | 48455954 | 48456117 | - | 164 | U1 | 0 | 0 | 0 | 0 | 0 | 0 |
| ENSDARG000 | 4 | 51394923 | 51395086 | - | 164 | U1 | 0 | 0 | 0 | 0 | 0 | 0 |
| ENSDARG000 | 4 | 57809976 | 57810049 | + | 74 | U6 | 0 | 0 | 0 | 0 | 0 | 0 |
| ENSDARG000 | 4 | 61507749 | 61507912 | + | 164 | U1 | 0 | 0 | 0 | 0 | 0 | 1 |
| ENSDARG000 | 4 | 14732983 | 14733123 | + | 141 | U4 | 5 | 2 | 6 | 10 | 0 | 3 |
| ENSDARG000 | 4 | 28697317 | 28697398 | + | 82 | dre-mir-430a- | 0 | 0 | 0 | 0 | 0 | 0 |
| ENSDARG000 | 4 | 57962556 | 57962662 | - | 107 | U6 | 0 | 0 | 0 | 0 | 0 | 0 |
| ENSDARG000 | 4 | 58810841 | 58810947 | - | 107 | U6 | 0 | 0 | 0 | 0 | 0 | 0 |
| ENSDARG000 | 4 | 51414467 | 51414573 | + | 107 | U6 | 0 | 0 | 0 | 0 | 0 | 0 |
| ENSDARG000 | 4 | 55724384 | 55724481 | - | 98 | NA | 0 | 0 | 0 | 0 | 0 | 0 |
| ENSDARG000 | 4 | 23226733 | 23226849 | - | 117 | 5S_rRNA | 0 | 0 | 0 | 0 | 0 | 0 |
| ENSDARG000 | 4 | 52593214 | 52593320 | + | 107 | U6 | 0 | 0 | 0 | 0 | 0 | 0 |
| ENSDARG000 | 4 | 48169539 | 48169645 | - | 107 | U6 | 1 | 0 | 0 | 0 | 0 | 0 |
| ENSDARG000 | 4 | 28709056 | 28709137 | + | 82 | dre-mir-430a- | 0 | 0 | 0 | 0 | 0 | 0 |
| ENSDARG000 | 4 | 52582450 | 52582598 | - | 149 | U1 | 0 | 0 | 0 | 0 | 0 | 0 |
| ENSDARG000 | 4:4:4:4:4:4:4: | 4626794:4628 | 4626951:4628 | -:-:-:-:-:-:-:-: | 2199 | DGKI | 46 | 61 | 161 | 139 | 92 | 109 |
| ENSDARG000 | 4;4;4;4;4 | 68456852:684 | 68456904:684 | +;+;+;+;+ | 1627 | NA | 0 | 0 | 4 | 0 | 0 | 0 |
| ENSDARG000 | 4:4:4:4:4:4:4: | 1469106:1471 | 1471326:1471 | -:-:-:-:-:-:-:-: | 3771 | NA | 376 | 295 | 319 | 431 | 368 | 433 |
| ENSDARG000 | 4 | 31221376 | 31221482 | - | 107 | U6 | 0 | 0 | 0 | 0 | 0 | 0 |
| ENSDARG000 | 4 | 52595846 | 52595952 | + | 107 | U6 | 0 | 0 | 0 | 0 | 0 | 0 |
| ENSDARG000 | 4:4:4:4:4:4:4: | 73203746:732 | 73203751:732 | -;-;-;-;-;-;- | 777 | NA | 44 | 60 | 52 | 3 | 47 | 64 |
| ENSDARG000 | 4 | 46700036 | 46700154 | + | 119 | 5S_rRNA | 0 | 0 | 0 | 0 | 0 | 0 |
| ENSDARG000 | 4 | 45559355 | 45559503 | - | 149 | U1 | 0 | 0 | 0 | 0 | 0 | 0 |
| ENSDARG000 | 4;4;4 | 48717758:487 | 48718483:487 | -;-;- | 1038 | NA | 8 | 2 | 18 | 0 | 7 | 4 |
| ENSDARG000 | 4 | 61731254 | 61731402 | + | 149 | U1 | 0 | 0 | 0 | 0 | 0 | 0 |
| ENSDARG000 | 4 | 65244885 | 65244991 | + | 107 | U6 | 0 | 0 | 0 | 0 | 0 | 0 |
| ENSDARG000 | 4 | 61204137 | 61204285 | + | 149 | U1 | 0 | 0 | 0 | 0 | 0 | 0 |
| ENSDARG000 | 4;4;4;4;4;4;4 | 16447485:164 | 16448574:164 | -;-;-;-;-;-;- | 3021 | NA | 0 | 0 | 0 | 0 | 0 | 0 |
| ENSDARG000 | 4 | 38237604 | 38237704 | - | 101 | U6 | 0 | 0 | 0 | 0 | 0 | 0 |
| ENSDARG000 | 4 | 48178506 | 48178612 | - | 107 | U6 | 0 | 0 | 0 | 0 | 0 | 0 |
| ENSDARG000 | 4;4;4;4;4;4 | 57271603:572 | 57271790:572 | -;-;-;-;-;- | 2600 | NA | 58 | 13 | 51 | 4 | 9 | 15 |
| ENSDARG000 | 4;4;4;4 | 47134868:471 | 47134977:471 | -;-;-;- | 1224 | NA | 0 | 9 | 9 | 0 | 4 | 3 |
| ENSDARG000 | 4 | 48171108 | 48171214 | - | 107 | U6 | 0 | 0 | 0 | 0 | 0 | 0 |
| ENSDARG000 | 4 | 28704197 | 28704277 | + | 81 | dre-mir-430b- | 0 | 0 | 0 | 0 | 0 | 0 |
| ENSDARG000 | 4 | 64704467 | 64704573 | + | 107 | U6 | 0 | 0 | 0 | 0 | 0 | 0 |
| ENSDARG000 | 4 | 41989850 | 41990027 | - | 178 | U1 | 0 | 0 | 1 | 0 | 0 | 0 |
| ENSDARG000 | 4 | 45564986 | 45565092 | + | 107 | U6 | 0 | 0 | 0 | 0 | 0 | 0 |
| ENSDARG000 | 4 | 28542021 | 28542135 | - | 115 | 5S_rRNA | 0 | 0 | 0 | 0 | 0 | 0 |
| ENSDARG000 | 4 | 48173219 | 48173325 | - | 107 | U6 | 0 | 0 | 0 | 0 | 0 | 0 |
| ENSDARG000 | 4 | 28697458 | 28697530 | + | 73 | dre-mir-430c- | 0 | 0 | 0 | 0 | 0 | 0 |
| ENSDARG000 | 4;4;4;4 | 74023396:740 | 74023669:740 | -;-;-;- | 2696 | NA | 2 | 1 | 3 | 0 | 1 | 3 |
| ENSDARG000 | 4;4;4;4 | 46524666:465 | 46524882:465 | -;-;-;- | 1587 | NA | 28 | 3 | 37 | 75 | 20 | 1 |
| ENSDARG000 | 4;4;4;4;4;4;4 | 74396786:743 | 74397254:743 | +;+;+;+;+;+;+ | 2435 | NA | 135 | 119 | 197 | 32 | 141 | 216 |
| ENSDARG000 | 4;4;4 | 68569204:685 | 68569406:685 | -;-;- | 246 | NA | 0 | 0 | 0 | 0 | 0 | 0 |
| ENSDARG000 | 4;4;4;4 | 43999348:440 | 43999548:440 | +;+;+;+ | 952 | NA | 0 | 2 | 0 | 0 | 0 | 0 |
| ENSDARG000 | 4;4 | 53781557:537 | 53781808:537 | +;+ | 1334 | NA | 0 | 0 | 0 | 0 | 0 | 0 |
| ENSDARG000 | 4;4;4 | 74044796:740 | 74045322:740 | -;-;- | 771 | NA | 11 | 5 | 7 | 0 | 0 | 3 |
| ENSDARG000 | 4 | 41339872 | 41340036 | - | 165 | U1 | 0 | 0 | 0 | 0 | 0 | 0 |
| ENSDARG000 | 4 | 64725232 | 64725338 | + | 107 | U6 | 0 | 0 | 0 | 0 | 0 | 0 |
| ENSDARG000 | 4 | 52595319 | 52595425 | + | 107 | U6 | 0 | 0 | 0 | 0 | 0 | 0 |
| ENSDARG000 | 4 | 68834616 | 68834731 | + | 116 | 5S_rRNA | 0 | 0 | 0 | 0 | 0 | 0 |
| ENSDARG000 | 4 | 32270096 | 32270202 | + | 107 | U6 | 0 | 0 | 0 | 0 | 0 | 0 |
| ENSDARG000 | 4 | 69492029 | 69492135 | + | 107 | U6 | 0 | 0 | 0 | 0 | 0 | 0 |
| ENSDARG000 | 4 | 69494142 | 69494248 | + | 107 | U6 | 0 | 0 | 0 | 0 | 0 | 0 |
| ENSDARG000 | 4 | 55494751 | 55498830 | + | 4080 | NA | 0 | 0 | 0 | 0 | 2 | 0 |
| ENSDARG000 | 4 | 42625399 | 42625562 | + | 164 | U1 | 0 | 0 | 0 | 0 | 0 | 0 |
| ENSDARG000 | 4:4:4:4:4:4:4: | 20576857:205 | 20576902:205 | +:+:+:+:+:+:+: | 1501 | NA | 0 | 0 | 0 | 0 | 0 | 0 |
| ENSDARG000 | 4 | 42658612 | 42658775 | + | 164 | U1 | 0 | 0 | 0 | 0 | 0 | 0 |
| ENSDARG000 | 4 | 57812851 | 57812957 | + | 107 | U6 | 0 | 0 | 0 | 0 | 0 | 0 |
| ENSDARG000 | 4 | 41985234 | 41985397 | + | 164 | U1 | 0 | 0 | 0 | 0 | 0 | 0 |
| ENSDARG000 | 4 | 45593038 | 45593144 | + | 107 | U6 | 0 | 0 | 0 | 0 | 0 | 0 |
| ENSDARG000 | 4 | 11566040 | 11566156 | + | 117 | 5S_rRNA | 0 | 0 | 0 | 0 | 0 | 0 |
| ENSDARG000 | 4 | 72422722 | 72422836 | - | 115 | 5S_rRNA | 3 | 17 | 9 | 11 | 12 | 17 |
| ENSDARG000 | 4 | 32264279 | 32264385 | + | 107 | U6 | 0 | 0 | 0 | 0 | 0 | 0 |
| ENSDARG000 | 4 | 57965723 | 57965829 | - | 107 | U6 | 0 | 0 | 0 | 0 | 0 | 0 |
| ENSDARG000 | 4 | 40918733 | 40918833 | - | 101 | U6 | 0 | 0 | 0 | 0 | 0 | 0 |
| ENSDARG000 | 4:4:4:4:4:4:4: | 49322310:493 | 49322414:493 | +:+:+:+:+:+:+: | 2339 | NA | 4 | 4 | 0 | 4 | 0 | 0 |
| ENSDARG000 | 4 | 28701844 | 28701924 | + | 81 | dre-mir-430b- | 0 | 0 | 0 | 0 | 0 | 0 |
| ENSDARG000 | 4 | 28701218 | 28701298 | + | 81 | dre-mir-430b- | 0 | 0 | 0 | 0 | 0 | 0 |
| ENSDARG000 | 4 | 13584483 | 13584581 | + | 99 | dre-mir-729 | 0 | 0 | 0 | 0 | 0 | 0 |
| ENSDARG000 | 4 | 34612088 | 34612206 | + | 119 | 5S_rRNA | 0 | 0 | 0 | 0 | 0 | 0 |
| ENSDARG000 | 4;4 | 3287819:3289 | 3288385:3290 | +;+ | 1740 | NA | 2 | 4 | 16 | 8 | 0 | 8 |
| ENSDARG000 | 4:4:4:4:4:4:4: | 76608282:766 | 76608862:766 | +:+:+:+:+:+:+: | 3052 | ms4a17a.9 | 162 | 26 | 22 | 8 | 177 | 210 |
| ENSDARG000 | 4 | 68043563 | 68043669 | - | 107 | U6 | 0 | 0 | 0 | 0 | 0 | 0 |
| ENSDARG000 | 4;4;4;4;4 | 75165958:751 | 75167611:751 | -;-;-;-;- | 2589 | NAA40 | 0 | 9 | 10 | 0 | 2 | 4 |
| ENSDARG000 | 4 | 46479806 | 46479912 | - | 107 | U6 | 0 | 0 | 0 | 0 | 0 | 0 |
| ENSDARG000 | 4 | 40913453 | 40913559 | - | 107 | U6 | 0 | 0 | 0 | 0 | 0 | 0 |
| ENSDARG000 | 4 | 38876767 | 38877924 | - | 1158 | NA | 16 | 16 | 3 | 0 | 6 | 8 |
| ENSDARG000 | 4;4;4;4;4;4;4 | 20566371:205 | 20566424:205 | +;+;+;+;+;+;+ | 2831 | NA | 0 | 0 | 0 | 4 | 0 | 19 |
| ENSDARG000 | 4:4:4:4:4:4:4: | 76659013:766 | 76659057:766 | +:+:+:+:+:+:+: | 1039 | ms4a17a.5 | 175 | 2 | 0 | 0 | 18 | 13 |
| ENSDARG000 | 4 | 57801344 | 57801507 | - | 164 | U1 | 0 | 0 | 0 | 0 | 0 | 0 |
| ENSDARG000 | 4 | 77100338 | 77100455 | - | 118 | 5S_rRNA | 0 | 0 | 0 | 0 | 0 | 0 |
| ENSDARG000 | 4;4 | 77267930:772 | 77273812:772 | -;- | 6151 | NA | 122 | 125 | 248 | 94 | 216 | 247 |
| ENSDARG000 | 4 | 31250316 | 31250464 | + | 149 | U1 | 2 | 2 | 3 | 0 | 2 | 0 |
| ENSDARG000 | 4 | 30974916 | 30975079 | - | 164 | U1 | 0 | 0 | 0 | 0 | 0 | 0 |
| ENSDARG000 | 4 | 51418162 | 51418268 | + | 107 | U6 | 0 | 0 | 0 | 0 | 0 | 0 |
| ENSDARG000 | 4 | 42437497 | 42437603 | - | 107 | U6 | 0 | 0 | 0 | 0 | 9 | 30 |
| ENSDARG000 | 4 | 44503489 | 44503605 | + | 117 | 5S_rRNA | 0 | 0 | 0 | 0 | 0 | 0 |
| ENSDARG000 | 4 | 40907355 | 40907461 | - | 107 | U6 | 0 | 0 | 0 | 0 | 0 | 0 |
| ENSDARG000 | 4 | 61445274 | 61445380 | - | 107 | U6 | 0 | 0 | 0 | 0 | 0 | 0 |
| ENSDARG000 | 4:4:4:4:4:4:4: | 17647547:176 | 17647570:176 | -;-;-;-;-;-;-;- | 189 | NA | 2 | 2 | 3 | 3 | 4 | 1 |
| ENSDARG000 | 4:4:4:4:4:4:4: | 51201875:512 | 51202111:512 | +:+:+:+:+:+:+: | 4984 | NA | 18 | 3 | 42 | 2 | 5 | 20 |
| ENSDARG000 | 4 | 46476112 | 46476218 | - | 107 | U6 | 0 | 0 | 0 | 0 | 0 | 0 |
| ENSDARG000 | 4 | 69957355 | 69957518 | - | 164 | U1 | 0 | 0 | 0 | 0 | 0 | 0 |
| ENSDARG000 | 4 | 28698305 | 28698385 | + | 81 | dre-mir-430b- | 0 | 0 | 0 | 0 | 0 | 0 |
| ENSDARG000 | 4 | 19047466 | 19047555 | - | 90 | dre-mir-129-1 | 0 | 0 | 0 | 0 | 0 | 0 |
| ENSDARG000 | 4:4:4:4:4:4:4: | 77322543:773 | 77322940:773 | -:-:-:-:-:-:-:-: | 2059 | slco1f3 | 8 | 0 | 0 | 0 | 0 | 0 |
| ENSDARG000 | 4:4:4:4:4:4:4: | 57262332:572 | 57265340:572 | -:-:-:-:-:-:-:-: | 5778 | NA | 7 | 25 | 16 | 1 | 51 | 14 |
| ENSDARG000 | 4:4:4:4:4:4:4: | 48519:49992: | 48560:50038: | -;-;-;-;-;-;- | 411 | NA | 8 | 22 | 17 | 0 | 23 | 14 |
| ENSDARG000 | 4 | 31193892 | 31193998 | - | 107 | U6 | 0 | 0 | 0 | 0 | 0 | 0 |
| ENSDARG000 | 4:4:4:4:4:4:4: | 14713008:147 | 14713017:147 | -:-:-:-:-:-:-:-: | 501 | NA | 21 | 3 | 11 | 2 | 3 | 18 |
| ENSDARG000 | 4 | 68567154 | 68567307 | - | 154 | 5_8S_rRNA | 0 | 0 | 0 | 0 | 0 | 0 |
| ENSDARG000 | 4 | 48458568 | 48458716 | - | 149 | U1 | 0 | 0 | 0 | 0 | 0 | 0 |
| ENSDARG000 | 4 | 32256272 | 32256435 | - | 164 | U1 | 0 | 0 | 0 | 0 | 0 | 0 |
| ENSDARG000 | 4 | 41352921 | 41353067 | - | 147 | U1 | 0 | 0 | 0 | 0 | 0 | 0 |
| ENSDARG000 | 4;4;4;4;4 | 75156398:751 | 75156413:751 | -;-;-;-;- | 1065 | NA | 1016 | 491 | 1029 | 783 | 875 | 567 |

| | | | | | | | | | | | | |
|---|---|---|---|---|---|---|---|---|---|---|---|---|
| ENSDARG000 | 4 | 32293872 | 32293945 | + | 74 | U6 | 0 | 0 | 0 | 0 | 0 | 0 |
| ENSDARG000 | 4 | 41360684 | 41360847 | - | 164 | U1 | 0 | 0 | 0 | 0 | 0 | 0 |
| ENSDARG000 | 4 | 30903981 | 30904129 | + | 149 | U1 | 0 | 0 | 0 | 0 | 0 | 0 |
| ENSDARG000 | 4 | 30906578 | 30906741 | + | 164 | U1 | 0 | 2 | 0 | 0 | 0 | 0 |
| ENSDARG000 | 4 | 42661194 | 42661339 | + | 146 | U1 | 2 | 0 | 0 | 0 | 0 | 0 |
| ENSDARG000 | 4 | 50115291 | 50115391 | - | 101 | U6 | 0 | 0 | 0 | 0 | 0 | 0 |
| ENSDARG000 | 4 | 53545298 | 53545475 | - | 178 | U1 | 0 | 0 | 0 | 0 | 0 | 0 |
| ENSDARG000 | 4 | 17061980 | 17062097 | + | 118 | 5S_rRNA | 0 | 0 | 0 | 0 | 0 | 0 |
| ENSDARG000 | 4 | 41366009 | 41366108 | + | 100 | U6 | 0 | 0 | 0 | 0 | 0 | 0 |
| ENSDARG000 | 4;4;4 | 39141908:391 | 39141992:391 | +;+;+ | 1287 | NA | 0 | 0 | 0 | 0 | 0 | 0 |
| ENSDARG000 | 4 | 33981766 | 33981839 | + | 74 | U6 | 0 | 0 | 0 | 0 | 0 | 0 |
| ENSDARG000 | 4 | 35735006 | 35735169 | - | 164 | U1 | 0 | 0 | 2 | 0 | 0 | 0 |
| ENSDARG000 | 4 | 33968210 | 33968352 | - | 143 | U1 | 0 | 0 | 0 | 0 | 0 | 0 |
| ENSDARG000 | 4 | 15075665 | 15075732 | + | 68 | dre-mir-182 | 0 | 0 | 0 | 0 | 0 | 0 |
| ENSDARG000 | 4 | 18916968 | 18917082 | + | 115 | 5S_rRNA | 0 | 0 | 0 | 0 | 0 | 0 |
| ENSDARG000 | 4 | 56332290 | 56332408 | + | 119 | 5S_rRNA | 0 | 0 | 0 | 0 | 0 | 0 |
| ENSDARG000 | 4;4;4 | 39244401:392 | 39244787:392 | -;-;- | 2540 | NA | 0 | 15 | 0 | 4 | 31 | 4 |
| ENSDARG000 | 4 | 42471811 | 42471959 | + | 149 | U1 | 0 | 0 | 0 | 0 | 0 | 0 |
| ENSDARG000 | 4 | 41361919 | 41362060 | - | 142 | U1 | 0 | 0 | 0 | 0 | 0 | 0 |
| ENSDARG000 | 4 | 53149224 | 53149340 | - | 117 | 5S_rRNA | 0 | 0 | 0 | 0 | 0 | 0 |
| ENSDARG000 | 4;4 | 77561226:775 | 77561629:775 | -;- | 432 | NA | 0 | 0 | 0 | 0 | 0 | 0 |
| ENSDARG000 | 4 | 41344483 | 41344646 | - | 164 | U1 | 0 | 0 | 0 | 0 | 0 | 0 |
| ENSDARG000 | 4 | 45566042 | 45566148 | + | 107 | U6 | 0 | 0 | 0 | 0 | 0 | 0 |
| ENSDARG000 | 4 | 47193629 | 47193735 | - | 107 | U6 | 0 | 0 | 0 | 0 | 0 | 0 |
| ENSDARG000 | 4 | 61446860 | 61446966 | - | 107 | U6 | 0 | 0 | 0 | 0 | 0 | 0 |
| ENSDARG000 | 4 | 52588468 | 52588574 | + | 107 | U6 | 0 | 0 | 0 | 0 | 0 | 0 |
| ENSDARG000 | 4;4;4;4;4;4 | 4376498:4377 | 4376907:4377 | -;-;-;-;-;- | 784 | NA | 0 | 0 | 0 | 0 | 0 | 0 |
| ENSDARG000 | 4 | 38233633 | 38233739 | - | 107 | U6 | 0 | 0 | 0 | 0 | 0 | 0 |
| ENSDARG000 | 4 | 45592511 | 45592617 | + | 107 | U6 | 0 | 0 | 0 | 0 | 0 | 0 |
| ENSDARG000 | 4 | 32269039 | 32269145 | + | 107 | U6 | 0 | 0 | 0 | 0 | 0 | 0 |
| ENSDARG000 | 4 | 61445800 | 61445906 | - | 107 | U6 | 0 | 0 | 0 | 0 | 0 | 0 |
| ENSDARG000 | 4 | 30995796 | 30995959 | - | 164 | U1 | 0 | 0 | 0 | 0 | 0 | 0 |
| ENSDARG000 | 4;4;4;4;4;4;4; | 5868034:5868 | 5868084:5868 | +;+;+;+;+;+;+; | 8385 | UTP20 | 718 | 653 | 1285 | 304 | 881 | 584 |
| ENSDARG000 | 4 | 28707316 | 28707397 | + | 82 | dre-mir-430a- | 0 | 0 | 0 | 0 | 0 | 0 |
| ENSDARG000 | 4;4;4;4;4 | 38589553:385 | 38589587:385 | -;-;-;-;- | 993 | NA | 6 | 1 | 2 | 0 | 0 | 3 |
| ENSDARG000 | 4 | 15074566 | 15074663 | + | 98 | dre-mir-183 | 0 | 0 | 0 | 0 | 0 | 0 |
| ENSDARG000 | 4;4;4;4;4;4;4; | 26036682:260 | 26036863:260 | +;+;+;+;+;+;+; | 4130 | DEPDC4 | 4 | 60 | 18 | 44 | 26 | 28 |
| ENSDARG000 | 4;4;4;4;4;4;4; | 26021749:260 | 26022728:260 | -;-;-;-;-;-;-;-; | 3055 | usp44 | 160 | 210 | 323 | 214 | 222 | 208 |
| ENSDARG000 | 4 | 48170583 | 48170689 | - | 107 | U6 | 0 | 0 | 0 | 0 | 0 | 0 |
| ENSDARG000 | 4 | 69494662 | 69494768 | + | 107 | U6 | 0 | 0 | 0 | 0 | 0 | 0 |
| ENSDARG000 | 4 | 64733363 | 64733469 | + | 107 | U6 | 0 | 0 | 0 | 0 | 0 | 0 |
| ENSDARG000 | 4 | 51409182 | 51409288 | + | 107 | U6 | 0 | 0 | 0 | 0 | 0 | 0 |
| ENSDARG000 | 4;4;4;4;4;4;4; | 66839620:668 | 66840233:668 | -;-;-;-;-;-;-;-; | 3136 | NA | 0 | 0 | 4 | 4 | 0 | 0 |
| ENSDARG000 | 4 | 28693834 | 28693915 | + | 82 | dre-mir-430a- | 0 | 0 | 0 | 0 | 0 | 0 |
| ENSDARG000 | 4 | 28705959 | 28706039 | + | 81 | dre-mir-430b- | 0 | 0 | 0 | 0 | 0 | 0 |
| ENSDARG000 | 4;4;4 | 69619:70135: | 69882:70272: | +;+;+ | 1400 | mansc1 | 0 | 0 | 2 | 0 | 0 | 0 |
| ENSDARG000 | 4 | 70004282 | 70004322 | - | 41 | U1 | 0 | 0 | 0 | 0 | 0 | 0 |
| ENSDARG000 | 4 | 48457264 | 48457427 | - | 164 | U1 | 0 | 0 | 0 | 0 | 0 | 0 |
| ENSDARG000 | 4 | 68064687 | 68064850 | + | 164 | U1 | 0 | 0 | 0 | 0 | 0 | 0 |
| ENSDARG000 | 4;4;4;4;4;4;4 | 76671012:766 | 76671056:766 | +;+;+;+;+;+;+ | 914 | ms4a17a.2 | 126 | 2 | 0 | 1 | 11 | 1 |
| ENSDARG000 | 4;4;4;4;4;4;4; | 50188524:501 | 50188742:501 | +;+;+;+;+;+;+; | 2186 | NA | 0 | 0 | 1 | 0 | 0 | 0 |
| ENSDARG000 | 4;4;4;4;4;4;4; | 5498230:5499 | 5498247:5499 | -;-;-;-;-;-;-;-; | 597 | NA | 0 | 0 | 0 | 6 | 0 | 0 |
| ENSDARG000 | 4;4;4;4;4;4;4; | 134412:13533 | 135241:13540 | -;-;-;-;-;-;-;-; | 3025 | tbk1 | 2240 | 1649 | 1783 | 1501 | 1574 | 1775 |
| ENSDARG000 | 4;4;4;4;4;4;4 | 42878556:428 | 42878672:428 | +;+;+;+;+;+;+ | 3000 | NA | 21 | 4 | 3 | 0 | 2 | 7 |
| ENSDARG000 | 4;4;4;4;4 | 288633:29025 | 288793:29039 | +;+;+;+;+ | 672 | NA | 6 | 37 | 40 | 10 | 17 | 12 |
| ENSDARG000 | 4;4;4;4 | 42604252:426 | 42604254:426 | +;+;+;+ | 330 | NA | 13 | 22 | 1 | 1 | 8 | 8 |
| ENSDARG000 | 4;4 | 49073583:490 | 49073667:490 | +;+ | 1080 | NA | 2 | 0 | 6 | 0 | 0 | 0 |
| ENSDARG000 | 4;4;4 | 32096811:320 | 32096818:320 | -;-;- | 177 | NA | 0 | 0 | 0 | 0 | 0 | 0 |
| ENSDARG000 | 4;4 | 55810436:558 | 55810463:558 | +;+ | 429 | NA | 0 | 0 | 0 | 0 | 0 | 0 |
| ENSDARG000 | 4;4;4;4;4 | 61840440:618 | 61842797:618 | +;+;+;+;+ | 5727 | NA | 48 | 21 | 19 | 21 | 15 | 8 |
| ENSDARG000 | 4;4;4;4;4;4 | 68678959:686 | 68678968:686 | -;-;-;-;-;- | 964 | NA | 0 | 0 | 2 | 0 | 0 | 0 |
| ENSDARG000 | 4;4;4;4 | 54519511:545 | 54519562:545 | +;+;+;+ | 615 | NA | 0 | 0 | 0 | 0 | 0 | 0 |
| ENSDARG000 | 4 | 46491191 | 46491354 | + | 164 | U1 | 0 | 0 | 0 | 0 | 0 | 0 |
| ENSDARG000 | 4 | 8503687 | 8503801 | + | 115 | 5S_rRNA | 0 | 0 | 0 | 0 | 0 | 0 |
| ENSDARG000 | 4 | 50113704 | 50113810 | - | 107 | U6 | 0 | 0 | 0 | 0 | 0 | 0 |
| ENSDARG000 | 4 | 50112649 | 50112755 | - | 107 | U6 | 0 | 0 | 0 | 0 | 0 | 0 |
| ENSDARG000 | 4;4;4;4;4 | 29773917:297 | 29774001:297 | +;+;+;+;+ | 822 | NA | 59 | 70 | 81 | 23 | 42 | 64 |
| ENSDARG000 | 4 | 62377750 | 62377806 | + | 57 | NA | 0 | 0 | 0 | 0 | 0 | 0 |
| ENSDARG000 | 4 | 40916096 | 40916202 | - | 107 | U6 | 0 | 0 | 0 | 0 | 0 | 0 |
| ENSDARG000 | 4 | 55798954 | 55799051 | + | 98 | NA | 0 | 0 | 0 | 0 | 0 | 0 |
| ENSDARG000 | 4;4;4;4;4;4;4 | 16323970:163 | 16324009:163 | +;+;+;+;+;+;+ | 549 | NA | 0 | 0 | 0 | 0 | 0 | 0 |
| ENSDARG000 | 4 | 28694219 | 28694299 | + | 81 | dre-mir-430b- | 0 | 0 | 0 | 0 | 0 | 0 |
| ENSDARG000 | 4 | 28702213 | 28702285 | + | 73 | dre-mir-430c- | 0 | 0 | 0 | 0 | 0 | 0 |
| ENSDARG000 | 4 | 31195993 | 31196099 | - | 107 | U6 | 0 | 0 | 0 | 0 | 0 | 0 |
| ENSDARG000 | 4 | 32289117 | 32289223 | + | 107 | U6 | 0 | 0 | 0 | 0 | 0 | 0 |
| ENSDARG000 | 4 | 52728428 | 52728572 | + | 145 | U1 | 1 | 0 | 0 | 0 | 0 | 0 |
| ENSDARG000 | 4 | 67556083 | 67556189 | - | 107 | U6 | 0 | 0 | 0 | 0 | 0 | 0 |
| ENSDARG000 | 4;4;4;4;4;4;4; | 20536120:205 | 20536335:205 | +;+;+;+;+;+;+; | 2012 | NA | 0 | 0 | 0 | 0 | 12 | 4 |
| ENSDARG000 | 4;4;4;4;4;4;4; | 74828055:748 | 74828312:748 | -;-;-;-;-;-;-;-; | 1401 | PHF21B | 0 | 2 | 0 | 0 | 0 | 2 |
| ENSDARG000 | 4;4;4;4;4 | 67588046:675 | 67588306:675 | -;-;-;-;- | 573 | NA | 6 | 4 | 6 | 4 | 16 | 3 |
| ENSDARG000 | 4 | 42657321 | 42657484 | + | 164 | U1 | 0 | 0 | 0 | 0 | 0 | 0 |
| ENSDARG000 | 4 | 38023028 | 38023176 | - | 149 | U1 | 0 | 0 | 1 | 5 | 1 | 0 |
| ENSDARG000 | 4 | 31239305 | 31239453 | + | 149 | U1 | 0 | 0 | 0 | 0 | 0 | 0 |
| ENSDARG000 | 4 | 51311287 | 51311450 | - | 164 | U1 | 0 | 0 | 0 | 0 | 0 | 0 |
| ENSDARG000 | 4 | 42642008 | 42642171 | + | 164 | U1 | 0 | 0 | 0 | 0 | 0 | 0 |
| ENSDARG000 | 4 | 64703408 | 64703514 | + | 107 | U6 | 0 | 0 | 0 | 0 | 0 | 0 |
| ENSDARG000 | 4 | 32282774 | 32282880 | + | 107 | U6 | 0 | 0 | 0 | 0 | 0 | 0 |
| ENSDARG000 | 4 | 68043053 | 68043159 | - | 107 | U6 | 0 | 0 | 0 | 0 | 0 | 0 |
| ENSDARG000 | 4 | 50102614 | 50102720 | - | 107 | U6 | 0 | 0 | 0 | 0 | 0 | 0 |
| ENSDARG000 | 4 | 31860679 | 31869702 | + | 9024 | NA | 3 | 0 | 0 | 0 | 0 | 0 |
| ENSDARG000 | 4;4;4 | 51160367:511 | 51160396:511 | +;+;+ | 1886 | NA | 151 | 69 | 98 | 16 | 83 | 64 |
| ENSDARG000 | 4 | 51404956 | 51405062 | + | 107 | U6 | 0 | 0 | 0 | 0 | 0 | 0 |
| ENSDARG000 | 4 | 42440107 | 42440212 | - | 106 | U6 | 0 | 0 | 0 | 0 | 8 | 6 |
| ENSDARG000 | 4 | 2349335 | 2350371 | - | 1037 | phlda1 | 1302 | 624 | 1028 | 771 | 1248 | 1000 |
| ENSDARG000 | 4 | 23451817 | 23451931 | - | 115 | 5S_rRNA | 0 | 0 | 0 | 0 | 0 | 0 |
| ENSDARG000 | 4 | 28705716 | 28705788 | + | 73 | dre-mir-430c- | 0 | 0 | 0 | 0 | 0 | 0 |
| ENSDARG000 | 4;4;4;4;4 | 67590576:675 | 67590747:675 | -;-;-;-;- | 521 | NA | 23 | 28 | 18 | 27 | 20 | 46 |
| ENSDARG000 | 4 | 61479039 | 61479202 | + | 164 | U1 | 0 | 0 | 0 | 0 | 0 | 0 |
| ENSDARG000 | 4 | 64312206 | 64312355 | - | 150 | U1 | 0 | 0 | 0 | 0 | 0 | 0 |
| ENSDARG000 | 4 | 52586361 | 52586467 | + | 107 | U6 | 0 | 0 | 0 | 0 | 0 | 0 |
| ENSDARG000 | 4 | 64725762 | 64725868 | + | 107 | U6 | 0 | 0 | 0 | 0 | 0 | 0 |
| ENSDARG000 | 4 | 42440636 | 42440742 | - | 107 | U6 | 0 | 0 | 0 | 6 | 14 | 31 |
| ENSDARG000 | 4 | 68035422 | 68035528 | - | 107 | U6 | 0 | 0 | 0 | 0 | 0 | 0 |
| ENSDARG000 | 4 | 28694950 | 28695031 | + | 82 | dre-mir-430a- | 0 | 0 | 0 | 0 | 0 | 0 |
| ENSDARG000 | 4;4 | 128697:13337 | 130435:13349 | -;- | 1857 | gpr19 | 114 | 79 | 178 | 380 | 225 | 192 |
| ENSDARG000 | 4;4;4;4;4;4;4; | 61330443:613 | 61330576:613 | -;-;-;-;-;-;-;-; | 2283 | NA | 4 | 6 | 10 | 7 | 0 | 3 |
| ENSDARG000 | 4;4;4;4;4;4;4 | 75200467:752 | 75201245:752 | +;+;+;+;+;+;+ | 1689 | SLC2A13 | 10 | 0 | 9 | 0 | 0 | 29 |
| ENSDARG000 | 4;4;4;4 | 17308:17681: | 17548:17772: | -;-;-;- | 510 | ptpn12 | 208 | 220 | 314 | 218 | 189 | 196 |
| ENSDARG000 | 4 | 64718416 | 64718489 | + | 74 | U6 | 0 | 0 | 0 | 0 | 0 | 0 |
| ENSDARG000 | 4;4;4 | 233812:23646 | 234302:23654 | -;-;- | 626 | RERG | 47 | 45 | 101 | 167 | 64 | 89 |
| ENSDARG000 | 4 | 7210755 | 7212875 | - | 2121 | LRRN3 | 531 | 518 | 479 | 153 | 626 | 542 |
| ENSDARG000 | 4;4;4;4;4;4;4; | 190547:19195 | 190641:19209 | -;-;-;-;-;-;-;-; | 2598 | NA | 5462 | 2562 | 4224 | 3896 | 3645 | 3117 |
| ENSDARG000 | 4 | 60164210 | 60164361 | - | 152 | U1 | 1 | 0 | 1 | 0 | 6 | 0 |
| ENSDARG000 | 4 | 33390222 | 33390384 | - | 163 | U1 | 0 | 0 | 0 | 0 | 0 | 0 |
| ENSDARG000 | 4 | 38237117 | 38237190 | - | 74 | U6 | 0 | 0 | 0 | 0 | 0 | 0 |
| ENSDARG000 | 4 | 58823314 | 58823462 | + | 149 | U1 | 0 | 0 | 0 | 0 | 0 | 0 |
| ENSDARG000 | 4 | 60194853 | 60194994 | - | 142 | U1 | 0 | 0 | 0 | 0 | 0 | 0 |
| ENSDARG000 | 4 | 42622832 | 42622980 | + | 149 | U1 | 0 | 0 | 0 | 0 | 0 | 0 |
| ENSDARG000 | 4 | 42651603 | 42651766 | + | 164 | U1 | 0 | 0 | 0 | 0 | 0 | 0 |
| ENSDARG000 | 4 | 554267 | 554384 | + | 118 | 5S_rRNA | 0 | 0 | 0 | 0 | 0 | 0 |
| ENSDARG000 | 4 | 7163758 | 7163874 | + | 117 | 5S_rRNA | 0 | 0 | 0 | 0 | 0 | 0 |
| ENSDARG000 | 4 | 7679979 | 7680095 | - | 117 | 5S_rRNA | 0 | 0 | 0 | 0 | 0 | 0 |
| ENSDARG000 | 4 | 39357666 | 39357785 | - | 120 | 5S_rRNA | 0 | 0 | 0 | 0 | 0 | 0 |
| ENSDARG000 | 4 | 68042524 | 68042630 | - | 107 | U6 | 0 | 0 | 0 | 0 | 0 | 0 |
| ENSDARG000 | 4 | 32291756 | 32291862 | + | 107 | U6 | 0 | 0 | 0 | 0 | 0 | 0 |
| ENSDARG000 | 4 | 32272738 | 32272844 | + | 107 | U6 | 0 | 0 | 0 | 0 | 0 | 0 |
| ENSDARG000 | 4 | 32288057 | 32288163 | + | 107 | U6 | 0 | 0 | 0 | 0 | 0 | 0 |
| ENSDARG000 | 4 | 48177447 | 48177553 | - | 107 | U6 | 0 | 0 | 0 | 0 | 0 | 0 |
| ENSDARG000 | 4 | 52581140 | 52581303 | - | 164 | U1 | 0 | 0 | 0 | 0 | 0 | 0 |
| ENSDARG000 | 4 | 33391775 | 33391921 | - | 147 | U1 | 0 | 0 | 0 | 0 | 0 | 0 |
| ENSDARG000 | 4;4;4;4;4;4;4; | 26095462:260 | 26097297:260 | -;-;-;-;-;-;-;-; | 3084 | si:ch211-244t | 480 | 673 | 158 | 312 | 376 | 450 |
| ENSDARG000 | 4 | 51408128 | 51408234 | + | 107 | U6 | 0 | 0 | 0 | 0 | 0 | 0 |
| ENSDARG000 | 4 | 52594802 | 52594908 | + | 107 | U6 | 0 | 0 | 0 | 0 | 0 | 0 |
| ENSDARG000 | 4 | 51413937 | 51414043 | + | 107 | U6 | 0 | 0 | 0 | 0 | 0 | 0 |
| ENSDARG000 | 4 | 30998406 | 30998555 | - | 150 | U1 | 0 | 0 | 0 | 0 | 2 | 0 |
| ENSDARG000 | 4 | 40916621 | 40916727 | - | 107 | U6 | 0 | 0 | 0 | 0 | 0 | 0 |
| ENSDARG000 | 4 | 57798235 | 57798398 | - | 164 | U1 | 0 | 0 | 0 | 0 | 0 | 0 |
| ENSDARG000 | 4 | 48173749 | 48173855 | - | 107 | U6 | 0 | 0 | 0 | 0 | 0 | 0 |
| ENSDARG000 | 4 | 61477742 | 61477897 | + | 156 | U1 | 0 | 0 | 0 | 0 | 0 | 0 |
| ENSDARG000 | 4 | 69153546 | 69153694 | + | 149 | U1 | 4 | 0 | 12 | 0 | 2 | 1 |
| ENSDARG000 | 4 | 31200217 | 31200323 | - | 107 | U6 | 0 | 0 | 0 | 0 | 0 | 0 |
| ENSDARG000 | 4 | 67557663 | 67557769 | - | 107 | U6 | 0 | 0 | 0 | 0 | 0 | 0 |
| ENSDARG000 | 4 | 50995331 | 50995399 | + | 69 | U6 | 0 | 0 | 0 | 0 | 0 | 0 |
| ENSDARG000 | 4 | 32262693 | 32262799 | + | 107 | U6 | 0 | 0 | 0 | 0 | 0 | 0 |
| ENSDARG000 | 4 | 68042047 | 68042120 | - | 74 | U6 | 0 | 0 | 0 | 0 | 0 | 0 |
| ENSDARG000 | 4 | 43570799 | 43570866 | - | 68 | U6 | 0 | 0 | 0 | 0 | 0 | 0 |
| ENSDARG000 | 4 | 50103659 | 50103765 | - | 107 | U6 | 0 | 0 | 0 | 0 | 0 | 0 |
| ENSDARG000 | 4 | 28707700 | 28707780 | + | 81 | dre-mir-430b- | 0 | 0 | 0 | 0 | 0 | 0 |
| ENSDARG000 | 4;4;4;4;4;4;4; | 1388965:1390 | 1389158:1390 | -;-;-;-;-;-;-;-; | 1413 | NA | 526 | 348 | 551 | 302 | 533 | 410 |
| ENSDARG000 | 4 | 60166049 | 60166212 | - | 164 | U1 | 0 | 0 | 0 | 0 | 0 | 0 |
| ENSDARG000 | 4 | 57802898 | 57803053 | - | 156 | U1 | 0 | 0 | 0 | 0 | 0 | 0 |
| ENSDARG000 | 4;4;4;4;4;4;4 | 76575585:765 | 76575624:765 | +;+;+;+;+;+;+ | 927 | ms4a17a.11 | 503 | 81 | 66 | 18 | 299 | 149 |
| ENSDARG000 | 4 | 68037008 | 68037114 | - | 107 | U6 | 0 | 0 | 0 | 0 | 0 | 0 |
| ENSDARG000 | 4 | 47194674 | 47194774 | - | 101 | U6 | 0 | 0 | 0 | 0 | 0 | 0 |
| ENSDARG000 | 4 | 47194154 | 47194260 | - | 107 | U6 | 0 | 0 | 0 | 0 | 0 | 0 |
| ENSDARG000 | 4 | 42438026 | 42438132 | - | 107 | U6 | 0 | 0 | 0 | 0 | 13 | 8 |
| ENSDARG000 | 4;4 | 68567783:685 | 68568183:685 | -;- | 429 | NA | 0 | 0 | 0 | 0 | 0 | 0 |
| ENSDARG000 | 4;4;4;4;4 | 72628893:726 | 72630424:726 | -;-;-;-;- | 2420 | NA | 32 | 60 | 97 | 8 | 75 | 52 |
| ENSDARG000 | 4 | 59835872 | 59836867 | - | 996 | NA | 16 | 0 | 2 | 8 | 2 | 0 |
| ENSDARG000 | 4;4;4;4;4;4;4; | 57034:57071: | 57046:58388: | -;-;-;-;-;-;-;-; | 2562 | NA | 392 | 175 | 569 | 379 | 266 | 287 |
| ENSDARG000 | 4 | 67567275 | 67567423 | + | 149 | U1 | 0 | 0 | 0 | 0 | 0 | 0 |
| ENSDARG000 | 4;4;4;4;4;4;4; | 3312852:3317 | 3312975:3317 | +;+;+;+;+;+;+; | 3576 | NA | 10 | 16 | 12 | 18 | 82 | 90 |

| | | | | | | | | | | | | |
|---|---|---|---|---|---|---|---|---|---|---|---|---|
| ENSDARG000 | 4 | 68055981 | 68056083 | - | 103 | U6 | 0 | 0 | 0 | 0 | 0 | 0 |
| ENSDARG000 | 4 | 28693975 | 28694047 | + | 73 | dre-mir-430c- | 0 | 0 | 0 | 0 | 0 | 0 |
| ENSDARG000 | 4:4:4:4:4:4:4: | 71987709:720 | 71989909:720 | -:-:-:-:-:-:-:-: | 5017 | pyroxd1 | 108 | 237 | 173 | 127 | 294 | 205 |
| ENSDARG000 | 4:4:4:4:4:4:4: | 8780179:878 | 8780780:878 | -:-:-:-:-:-:-:-: | 1891 | pnpla3 | 14 | 28 | 16 | 66 | 0 | 41 |
| ENSDARG000 | 4 | 60192143 | 60192306 | - | 164 | U1 | 0 | 0 | 0 | 0 | 0 | 0 |
| ENSDARG000 | 4 | 68062658 | 68062821 | + | 164 | U1 | 0 | 0 | 0 | 0 | 0 | 0 |
| ENSDARG000 | 4 | 76483254 | 76483368 | + | 115 | 5S_rRNA | 0 | 0 | 0 | 0 | 2 | 3 |
| ENSDARG000 | 4 | 61494699 | 61494862 | + | 164 | U1 | 0 | 0 | 0 | 0 | 0 | 0 |
| ENSDARG000 | 4;4;4;4 | 55813426:558 | 55814102:558 | -;-;-;- | 894 | NA | 0 | 2 | 17 | 0 | 0 | 10 |
| ENSDARG000 | 4 | 58654592 | 58654665 | - | 74 | U6 | 0 | 0 | 0 | 0 | 13 | 2 |
| ENSDARG000 | 4 | 51406015 | 51406121 | + | 107 | U6 | 0 | 0 | 0 | 0 | 0 | 0 |
| ENSDARG000 | 4 | 31198100 | 31198206 | - | 107 | U6 | 0 | 0 | 0 | 0 | 0 | 0 |
| ENSDARG000 | 4 | 42439064 | 42439170 | - | 107 | U6 | 0 | 0 | 0 | 4 | 5 | 12 |
| ENSDARG000 | 4;4 | 73185344:731 | 73188546:731 | -;- | 3249 | NA | 116 | 163 | 197 | 65 | 288 | 185 |
| ENSDARG000 | 4 | 31251606 | 31251769 | + | 164 | U1 | 0 | 0 | 0 | 0 | 0 | 0 |
| ENSDARG000 | 4 | 30985356 | 30985519 | - | 164 | U1 | 0 | 0 | 0 | 0 | 0 | 0 |
| ENSDARG000 | 4 | 7025167 | 7025283 | + | 117 | 5S_rRNA | 0 | 0 | 0 | 0 | 0 | 0 |
| ENSDARG000 | 4:4:4:4:4:4:4: | 57192681:571 | 57192724:571 | +:+:+:+:+:+:+: | 3129 | NA | 32 | 20 | 36 | 3 | 9 | 11 |
| ENSDARG000 | 4 | 51407069 | 51407175 | + | 107 | U6 | 0 | 0 | 0 | 0 | 0 | 0 |
| ENSDARG000 | 4:4:4:4:4:4:4: | 73224617:732 | 73226559:732 | +:+:+:+:+:+:+: | 2877 | NA | 99 | 73 | 97 | 52 | 92 | 116 |
| ENSDARG000 | 4 | 46462932 | 46463038 | - | 107 | U6 | 0 | 0 | 0 | 0 | 0 | 0 |
| ENSDARG000 | 4 | 40915039 | 40915145 | - | 107 | U6 | 0 | 0 | 0 | 0 | 0 | 0 |
| ENSDARG000 | 4 | 60181213 | 60181376 | - | 164 | U1 | 0 | 0 | 0 | 0 | 0 | 0 |
| ENSDARG000 | 4;4;4 | 36506439:365 | 36508228:365 | -;-;- | 1980 | NA | 0 | 10 | 0 | 0 | 3 | 12 |
| ENSDARG000 | 4 | 8134594 | 8134710 | + | 117 | 5S_rRNA | 0 | 0 | 0 | 0 | 0 | 0 |
| ENSDARG000 | 4 | 4690726 | 4690841 | + | 116 | 5S_rRNA | 0 | 0 | 0 | 0 | 0 | 0 |
| ENSDARG000 | 4 | 23419320 | 23419455 | - | 136 | 5S_rRNA | 0 | 0 | 0 | 0 | 0 | 0 |
| ENSDARG000 | 4 | 45570797 | 45570903 | + | 107 | U6 | 0 | 0 | 0 | 0 | 0 | 0 |
| ENSDARG000 | 4:4:4:4:4:4:4: | 40295896:402 | 40295980:402 | +:+:+:+:+:+:+: | 3298 | NA | 16 | 33 | 25 | 0 | 31 | 54 |
| ENSDARG000 | 4 | 42466419 | 42466521 | - | 103 | U6 | 0 | 0 | 0 | 0 | 0 | 0 |
| ENSDARG000 | 4;4;4 | 905483:90743 | 905873:90753 | -;-;- | 600 | mrap2b | 0 | 0 | 0 | 0 | 2 | 0 |
| ENSDARG000 | 4;4;4;4;4 | 43999348:440 | 43999548:440 | +;+;+;+;+ | 2663 | NA | 4 | 0 | 0 | 0 | 0 | 0 |
| ENSDARG000 | 4 | 61163721 | 61163827 | - | 107 | U6 | 0 | 0 | 0 | 0 | 0 | 0 |
| ENSDARG000 | 4 | 28708850 | 28708930 | + | 81 | dre-mir-430b- | 0 | 0 | 0 | 0 | 0 | 0 |
| ENSDARG000 | 4 | 53538926 | 53539045 | - | 120 | 5S_rRNA | 0 | 0 | 0 | 0 | 0 | 0 |
| ENSDARG000 | 4 | 47199396 | 47199555 | + | 160 | U1 | 0 | 0 | 0 | 0 | 0 | 0 |
| ENSDARG000 | 4:4:4:4:4:4:4: | 8739093:874 | 8742834:874 | -:-:-:-:-:-:-:-: | 5739 | ttll12 | 124 | 178 | 203 | 206 | 182 | 159 |
| ENSDARG000 | 4 | 57796681 | 57796844 | - | 164 | U1 | 0 | 0 | 0 | 0 | 0 | 0 |
| ENSDARG000 | 4:4:4:4:4:4:4: | 982028:98596 | 986478:98647 | -:-:-:-:-:-:-:-: | 5696 | naga | 1468 | 919 | 1103 | 2014 | 1951 | 972 |
| ENSDARG000 | 4 | 48186092 | 48186254 | + | 163 | U1 | 1 | 0 | 0 | 0 | 0 | 1 |
| ENSDARG000 | 4 | 67557133 | 67557239 | - | 107 | U6 | 0 | 0 | 0 | 0 | 0 | 0 |
| ENSDARG000 | 4;4;4;4 | 65508134:655 | 65508173:655 | -;-;-;- | 954 | NA | 0 | 0 | 0 | 0 | 0 | 0 |
| ENSDARG000 | 4 | 51424256 | 51424362 | + | 107 | U6 | 0 | 0 | 0 | 0 | 0 | 0 |
| ENSDARG000 | 4;4 | 64981411:650 | 64981570:650 | +;+ | 4401 | NA | 4 | 2 | 0 | 0 | 4 | 10 |
| ENSDARG000 | 4 | 47190477 | 47190583 | - | 107 | U6 | 0 | 0 | 0 | 0 | 0 | 0 |
| ENSDARG000 | 4 | 61441050 | 61441156 | - | 107 | U6 | 0 | 0 | 0 | 0 | 0 | 0 |
| ENSDARG000 | 4;4 | 16827390:168 | 16827690:168 | +;+ | 1036 | kiss2 | 0 | 0 | 2 | 0 | 0 | 0 |
| ENSDARG000 | 4 | 42630563 | 42630726 | + | 164 | U1 | 0 | 0 | 0 | 0 | 0 | 0 |
| ENSDARG000 | 4 | 64699330 | 64699436 | + | 107 | U6 | 0 | 0 | 0 | 0 | 0 | 0 |
| ENSDARG000 | 4 | 64794290 | 64794396 | + | 107 | U6 | 0 | 0 | 0 | 0 | 0 | 0 |
| ENSDARG000 | 4:4:4:4:4:4:4: | 6032640:603 | 6032861:603 | +:+:+:+:+:+:+: | 1410 | tfec | 256 | 155 | 934 | 66 | 328 | 399 |
| ENSDARG000 | 4 | 33976808 | 33976909 | + | 102 | U6 | 0 | 0 | 0 | 0 | 0 | 0 |
| ENSDARG000 | 4 | 51402839 | 51402945 | + | 107 | U6 | 0 | 0 | 0 | 0 | 0 | 0 |
| ENSDARG000 | 4;4;4 | 26449406:264 | 26449429:264 | +;+;+ | 399 | NA | 0 | 4 | 0 | 0 | 2 | 0 |
| ENSDARG000 | 4 | 68563695 | 68563862 | - | 168 | NA | 0 | 0 | 0 | 0 | 0 | 0 |
| ENSDARG000 | 4 | 58824609 | 58824772 | + | 164 | U1 | 0 | 0 | 0 | 0 | 0 | 0 |
| ENSDARG000 | 4 | 60170162 | 60170325 | - | 164 | U1 | 0 | 0 | 0 | 0 | 0 | 0 |
| ENSDARG000 | 4 | 50120632 | 50120795 | + | 164 | U1 | 0 | 0 | 0 | 0 | 0 | 0 |
| ENSDARG000 | 4 | 33966917 | 33967068 | - | 152 | U1 | 0 | 0 | 0 | 0 | 0 | 0 |
| ENSDARG000 | 4 | 58819301 | 58819402 | - | 102 | U6 | 0 | 0 | 0 | 0 | 0 | 0 |
| ENSDARG000 | 4 | 57781567 | 57781730 | - | 164 | U1 | 0 | 0 | 0 | 0 | 0 | 0 |
| ENSDARG000 | 4 | 58793926 | 58794032 | - | 107 | U6 | 0 | 0 | 0 | 0 | 0 | 0 |
| ENSDARG000 | 4 | 42638867 | 42639030 | + | 164 | U1 | 0 | 0 | 0 | 0 | 0 | 0 |
| ENSDARG000 | 4 | 42629272 | 42629435 | + | 164 | U1 | 0 | 0 | 0 | 0 | 0 | 0 |
| ENSDARG000 | 4 | 31212385 | 31212491 | - | 107 | U6 | 0 | 0 | 0 | 0 | 0 | 0 |
| ENSDARG000 | 4 | 25840786 | 25840902 | - | 117 | 5S_rRNA | 0 | 3 | 0 | 0 | 0 | 0 |
| ENSDARG000 | 4 | 67554499 | 67554605 | - | 107 | U6 | 0 | 0 | 0 | 0 | 0 | 0 |
| ENSDARG000 | 4 | 28700067 | 28700147 | + | 81 | dre-mir-430b- | 0 | 0 | 0 | 0 | 0 | 0 |
| ENSDARG000 | 4;4 | 55728109:557 | 55728509:557 | -;- | 429 | NA | 0 | 0 | 0 | 0 | 0 | 0 |
| ENSDARG000 | 4 | 2581916 | 2582031 | + | 116 | 5S_rRNA | 0 | 0 | 0 | 0 | 0 | 0 |
| ENSDARG000 | 4 | 65235102 | 65235265 | - | 164 | U1 | 0 | 0 | 0 | 0 | 0 | 0 |
| ENSDARG000 | 4 | 51403896 | 51404002 | + | 107 | U6 | 0 | 0 | 0 | 0 | 0 | 0 |
| ENSDARG000 | 4 | 45567630 | 45567736 | + | 107 | U6 | 0 | 0 | 0 | 0 | 0 | 0 |
| ENSDARG000 | 4 | 32267984 | 32268090 | + | 107 | U6 | 0 | 0 | 0 | 0 | 0 | 0 |
| ENSDARG000 | 4 | 51401252 | 51401358 | + | 107 | U6 | 0 | 0 | 0 | 0 | 0 | 0 |
| ENSDARG000 | 4 | 28697111 | 28697191 | + | 81 | dre-mir-430b- | 0 | 0 | 0 | 0 | 0 | 0 |
| ENSDARG000 | 4 | 28693371 | 28693443 | + | 73 | dre-mir-430c- | 0 | 0 | 0 | 0 | 0 | 0 |
| ENSDARG000 | 4 | 19097324 | 19097444 | + | 121 | 5S_rRNA | 0 | 0 | 0 | 0 | 0 | 0 |
| ENSDARG000 | 4 | 60175901 | 60176064 | - | 164 | U1 | 0 | 0 | 0 | 0 | 0 | 0 |
| ENSDARG000 | 4 | 67570358 | 67570521 | + | 164 | U1 | 0 | 0 | 0 | 0 | 0 | 0 |
| ENSDARG000 | 4 | 57238239 | 57238306 | + | 68 | U6 | 0 | 0 | 0 | 0 | 0 | 0 |
| ENSDARG000 | 4 | 32290172 | 32290278 | + | 107 | U6 | 0 | 0 | 0 | 0 | 0 | 0 |
| ENSDARG000 | 4 | 46478222 | 46478328 | - | 107 | U6 | 0 | 0 | 0 | 0 | 0 | 0 |
| ENSDARG000 | 4 | 61458415 | 61458521 | - | 107 | U6 | 0 | 0 | 0 | 0 | 0 | 0 |
| ENSDARG000 | 4:4:4:4:4:4:4: | 20237461:202 | 20238428:202 | -;-;-;-;-;-;-;- | 1469 | NA | 8 | 2 | 1 | 0 | 1 | 6 |
| ENSDARG000 | 4 | 72837670 | 72837786 | - | 117 | 5S_rRNA | 0 | 0 | 0 | 0 | 0 | 0 |
| ENSDARG000 | 4 | 67553973 | 67554076 | - | 104 | U6 | 0 | 0 | 0 | 0 | 0 | 0 |
| ENSDARG000 | 4;4;4;4;4;4 | 27018663:270 | 27018779:270 | +;+;+;+;+;+ | 1874 | NA | 0 | 0 | 0 | 1 | 0 | 0 |
| ENSDARG000 | 4;4;4 | 55781670:557 | 55782366:557 | -;-;- | 882 | NA | 0 | 0 | 0 | 2 | 0 | 0 |
| ENSDARG000 | 4 | 64279882 | 64284924 | - | 5043 | NA | 5 | 4 | 18 | 19 | 19 | 35 |
| ENSDARG000 | 4;4;4;4;4;4 | 74047033:740 | 74047526:740 | -;-;-;-;-;- | 2930 | NA | 18 | 30 | 41 | 32 | 61 | 49 |
| ENSDARG000 | 4;4 | 72348618:723 | 72349477:723 | -;- | 1049 | NA | 26 | 8 | 30 | 8 | 16 | 26 |
| ENSDARG000 | 4;4;4 | 29344957:293 | 29345293:293 | +;+;+ | 1527 | NA | 8 | 8 | 2 | 2 | 4 | 10 |
| ENSDARG000 | 4:4:4:4:4:4:4: | 966061:96794 | 966150:96813 | +:+:+:+:+:+:+: | 2220 | rpap3 | 347 | 349 | 467 | 325 | 500 | 270 |
| ENSDARG000 | 4:4:4:4:4:4:4: | 25558849:255 | 25558963:255 | +:+:+:+:+:+:+: | 1082 | il15ra | 85 | 108 | 117 | 55 | 127 | 113 |
| ENSDARG000 | 4;4;4 | 32566617:325 | 32566709:325 | +;+;+ | 1559 | NA | 0 | 12 | 5 | 0 | 12 | 46 |
| ENSDARG000 | 4:4:4:4:4:4:4: | 40541015:406 | 40541099:406 | +:+:+:+:+:+:+: | 1116 | NA | 0 | 2 | 0 | 0 | 0 | 0 |
| ENSDARG000 | 4 | 8141349 | 8141466 | - | 118 | 5S_rRNA | 0 | 0 | 0 | 0 | 0 | 0 |
| ENSDARG000 | 4 | 64714866 | 64714972 | + | 107 | U6 | 0 | 0 | 0 | 0 | 0 | 0 |
| ENSDARG000 | 4 | 47188366 | 47188472 | - | 107 | U6 | 0 | 0 | 0 | 0 | 0 | 0 |
| ENSDARG000 | 4 | 48170061 | 48170167 | - | 107 | U6 | 1 | 0 | 0 | 0 | 0 | 0 |
| ENSDARG000 | 4 | 42439593 | 42439699 | - | 107 | U6 | 0 | 0 | 0 | 2 | 7 | 2 |
| ENSDARG000 | 4 | 21743863 | 21743984 | + | 122 | NA | 0 | 0 | 0 | 0 | 0 | 0 |
| ENSDARG000 | 4;4 | 72205923:722 | 72205989:722 | +;+ | 5254 | zgc:165515 | 28 | 118 | 170 | 8 | 112 | 108 |
| ENSDARG000 | 4 | 31231570 | 31231725 | + | 156 | U1 | 0 | 0 | 0 | 0 | 0 | 0 |
| ENSDARG000 | 4 | 61729981 | 61730126 | + | 146 | U1 | 0 | 0 | 0 | 0 | 0 | 0 |
| ENSDARG000 | 4 | 45564459 | 45564565 | + | 107 | U6 | 0 | 0 | 0 | 0 | 0 | 0 |
| ENSDARG000 | 4 | 58814013 | 58814119 | - | 107 | U6 | 0 | 0 | 0 | 0 | 0 | 0 |
| ENSDARG000 | 4 | 53948449 | 53948612 | + | 164 | U1 | 5 | 4 | 13 | 1 | 3 | 2 |
| ENSDARG000 | 4 | 65244358 | 65244464 | + | 107 | U6 | 0 | 0 | 0 | 0 | 0 | 0 |
| ENSDARG000 | 4 | 28696691 | 28696772 | + | 82 | dre-mir-430a- | 0 | 0 | 0 | 0 | 0 | 0 |
| ENSDARG000 | 4 | 28697715 | 28697795 | + | 81 | dre-mir-430b- | 0 | 0 | 0 | 0 | 0 | 0 |
| ENSDARG000 | 4 | 28699198 | 28699270 | + | 73 | dre-mir-430c- | 0 | 0 | 0 | 0 | 0 | 0 |
| ENSDARG000 | 4 | 57795128 | 57795291 | - | 164 | U1 | 0 | 0 | 0 | 0 | 0 | 0 |
| ENSDARG000 | 4 | 64732316 | 64732422 | + | 107 | U6 | 0 | 0 | 0 | 0 | 0 | 0 |
| ENSDARG000 | 4 | 41980187 | 41980351 | + | 165 | U1 | 0 | 0 | 0 | 0 | 0 | 0 |
| ENSDARG000 | 4 | 47193104 | 47193210 | - | 107 | U6 | 0 | 0 | 0 | 0 | 0 | 0 |
| ENSDARG000 | 4 | 47192054 | 47192160 | - | 107 | U6 | 0 | 0 | 0 | 0 | 0 | 0 |
| ENSDARG000 | 4:4:4:4:4:4:4: | 151506:15205 | 152173:15217 | -:-:-:-:-:-:-:-: | 3170 | eps8a | 221 | 130 | 106 | 183 | 155 | 109 |
| ENSDARG000 | 4 | 58798141 | 58798247 | - | 107 | U6 | 0 | 0 | 0 | 0 | 0 | 0 |
| ENSDARG000 | 4 | 28709197 | 28709269 | + | 73 | dre-mir-430c- | 0 | 0 | 0 | 0 | 0 | 0 |
| ENSDARG000 | 4;4 | 47047575:470 | 47048507:470 | -;- | 987 | NA | 53 | 17 | 49 | 3 | 45 | 55 |
| ENSDARG000 | 4;4 | 29653496:296 | 29654195:296 | -;- | 1542 | NA | 118 | 65 | 118 | 36 | 83 | 97 |
| ENSDARG000 | 4;4;4 | 30299435:303 | 30299519:303 | +;+;+ | 882 | NA | 8 | 7 | 28 | 2 | 16 | 14 |
| ENSDARG000 | 4 | 61179684 | 61179832 | + | 149 | U1 | 0 | 0 | 0 | 0 | 0 | 0 |
| ENSDARG000 | 4 | 65236412 | 65236560 | - | 149 | U1 | 0 | 0 | 0 | 0 | 0 | 0 |
| ENSDARG000 | 4;4 | 37092809:370 | 37092893:370 | +;+ | 1323 | NA | 0 | 0 | 0 | 0 | 0 | 0 |
| ENSDARG000 | 4;4;4;4;4;4 | 71973211:719 | 71975495:719 | -;-;-;-;-;- | 3414 | ftr64 | 90 | 94 | 130 | 29 | 67 | 94 |
| ENSDARG000 | 4;4;4 | 67675471:677 | 67676032:677 | -;-;- | 1542 | NA | 2 | 0 | 0 | 12 | 2 | 0 |
| ENSDARG000 | 4;4;4;4;4 | 54798291:547 | 54798375:547 | +;+;+;+;+ | 1159 | NA | 2 | 0 | 16 | 0 | 4 | 5 |
| ENSDARG000 | 4;4 | 39742119:397 | 39742971:397 | +;+ | 4552 | NA | 0 | 2 | 0 | 0 | 2 | 0 |
| ENSDARG000 | 4:4:4:4:4:4:4: | 38344:38344: | 38512:38512: | +:+:+:+:+:+:+: | 2827 | PHTF2 | 493 | 415 | 612 | 690 | 601 | 366 |
| ENSDARG000 | 4;4 | 55758103:557 | 55758124:557 | +;+ | 306 | NA | 0 | 0 | 0 | 0 | 0 | 0 |
| ENSDARG000 | 4;4;4 | 55797866:557 | 55798562:557 | -;-;- | 882 | NA | 0 | 0 | 0 | 0 | 0 | 0 |
| ENSDARG000 | 4;4;4 | 77551100:775 | 77551304:775 | -;-;- | 291 | NA | 0 | 0 | 0 | 0 | 0 | 0 |
| ENSDARG000 | 4;4;4 | 70480701:704 | 70481905:704 | -;-;- | 1385 | NA | 2 | 10 | 4 | 0 | 3 | 0 |
| ENSDARG000 | 4:4:4:4:4:4:4: | 3792950:3792 | 3794176:3794 | -;-;-;-;-;-;-;- | 2683 | NA | 0 | 0 | 0 | 0 | 0 | 0 |
| ENSDARG000 | 4;4 | 77262694:772 | 77263418:772 | -;- | 801 | NA | 4 | 39 | 37 | 4 | 10 | 7 |
| ENSDARG000 | 4 | 12595680 | 12595804 | - | 125 | 5S_rRNA | 0 | 0 | 0 | 0 | 0 | 0 |
| ENSDARG000 | 4 | 27141179 | 27141303 | - | 125 | 5S_rRNA | 0 | 0 | 0 | 0 | 0 | 0 |
| ENSDARG000 | 4 | 45563929 | 45564035 | + | 107 | U6 | 0 | 0 | 0 | 0 | 0 | 0 |
| ENSDARG000 | 4 | 52585848 | 52585947 | + | 100 | U6 | 0 | 0 | 0 | 0 | 0 | 0 |
| ENSDARG000 | 4 | 37730119 | 37730282 | - | 164 | U1 | 0 | 0 | 0 | 0 | 0 | 0 |
| ENSDARG000 | 4 | 51299147 | 51299293 | + | 147 | U1 | 0 | 0 | 0 | 0 | 0 | 0 |
| ENSDARG000 | 4 | 28693628 | 28693708 | + | 81 | dre-mir-430b- | 0 | 0 | 0 | 0 | 0 | 0 |
| ENSDARG000 | 4 | 28699035 | 28699116 | + | 82 | dre-mir-430a- | 0 | 0 | 0 | 0 | 0 | 0 |
| ENSDARG000 | 4 | 60185112 | 60185275 | - | 164 | U1 | 0 | 0 | 0 | 0 | 0 | 0 |
| ENSDARG000 | 4 | 65245413 | 65245519 | + | 107 | U6 | 0 | 0 | 0 | 0 | 0 | 0 |
| ENSDARG000 | 4 | 28701424 | 28701505 | + | 82 | dre-mir-430a- | 0 | 0 | 0 | 0 | 0 | 0 |
| ENSDARG000 | 4 | 64702881 | 64702987 | + | 107 | U6 | 0 | 0 | 0 | 0 | 0 | 0 |
| ENSDARG000 | 4 | 41342927 | 41343090 | - | 164 | U1 | 0 | 0 | 0 | 0 | 0 | 0 |
| ENSDARG000 | 4 | 32264809 | 32264915 | + | 107 | U6 | 0 | 0 | 0 | 0 | 0 | 0 |
| ENSDARG000 | 4;4;4;4;4;4 | 77250504:772 | 77251784:772 | -;-;-;-;-;- | 1899 | zgc:174310 | 69 | 72 | 115 | 82 | 80 | 92 |
| ENSDARG000 | 4 | 30991881 | 30992044 | - | 164 | U1 | 0 | 0 | 0 | 0 | 0 | 0 |
| ENSDARG000 | 4 | 40924040 | 40924203 | + | 164 | U1 | 0 | 0 | 0 | 0 | 0 | 0 |
| ENSDARG000 | 4 | 45567102 | 45567208 | + | 107 | U6 | 0 | 0 | 0 | 0 | 0 | 0 |
| ENSDARG000 | 4 | 40897857 | 40897963 | - | 107 | U6 | 0 | 0 | 0 | 0 | 0 | 0 |
| ENSDARG000 | 4 | 61995745 | 61997189 | + | 1445 | NA | 16 | 6 | 2 | 0 | 4 | 0 |
| ENSDARG000 | 4;4;4 | 279669:28206 | 279949:28220 | +;+;+ | 866 | NA | 251 | 203 | 251 | 115 | 179 | 130 |
| ENSDARG000 | 4 | 11606758 | 11606855 | - | 98 | mir29a-1 | 2 | 2 | 3 | 0 | 0 | 0 |
| ENSDARG000 | 4:4:4:4:4:4:4: | 72668095:726 | 72668198:726 | +:+:+:+:+:+:+: | 3544 | NA | 22 | 6 | 21 | 194 | 301 | 8 |
| ENSDARG000 | 4;4 | 55778679:557 | 55778706:557 | +;+ | 429 | NA | 0 | 0 | 0 | 0 | 0 | 0 |

| | | | | | | | | | | | | |
|---|---|---|---|---|---|---|---|---|---|---|---|---|
| ENSDARG000 | 4 | 31252911 | 31253074 | + | 164 | U1 | 0 | 0 | 0 | 0 | 0 | 0 |
| ENSDARG000 | 4 | 38242945 | 38243108 | + | 164 | U1 | 0 | 0 | 0 | 0 | 0 | 0 |
| ENSDARG000 | 4 | 52591113 | 52591219 | + | 107 | U6 | 0 | 0 | 0 | 0 | 0 | 0 |
| ENSDARG000 | 4 | 52592164 | 52592270 | + | 107 | U6 | 0 | 0 | 0 | 0 | 0 | 0 |
| ENSDARG000 | 4 | 46481917 | 46482023 | - | 107 | U6 | 0 | 0 | 0 | 0 | 0 | 0 |
| ENSDARG000 | 4 | 40918203 | 40918309 | - | 107 | U6 | 0 | 0 | 0 | 0 | 0 | 0 |
| ENSDARG000 | 4:4:4:4:4:4:4: | 10066840:10 | 10067218:10 | +:+:+:+:+:+:+: | 8322 | FLNC | 9632 | 3943 | 5918 | 5567 | 6117 | 4080 |
| ENSDARG000 | 4:4:4:4:4:4:4: | 54095523:54 | 54095577:54 | -:-:-:-:-:-:-:-: | 2256 | NA | 1 | 10 | 1 | 0 | 29 | 21 |
| ENSDARG000 | 4 | 43140778 | 43140926 | - | 149 | U1 | 0 | 0 | 0 | 0 | 0 | 0 |
| ENSDARG000 | 4 | 45584825 | 45584931 | + | 107 | U6 | 0 | 0 | 0 | 0 | 0 | 0 |
| ENSDARG000 | 4;4;4 | 69223136:692 | 69223852:692 | -;-;- | 1491 | NA | 0 | 0 | 0 | 0 | 0 | 0 |
| ENSDARG000 | 4 | 63846615 | 63848305 | - | 1691 | NA | 0 | 6 | 0 | 0 | 2 | 1 |
| ENSDARG000 | 4 | 61512969 | 61513117 | + | 149 | U1 | 0 | 0 | 0 | 0 | 0 | 0 |
| ENSDARG000 | 4 | 50114234 | 50114340 | - | 107 | U6 | 0 | 0 | 0 | 0 | 0 | 0 |
| ENSDARG000 | 4 | 53958451 | 53958628 | - | 178 | U1 | 0 | 0 | 0 | 0 | 0 | 0 |
| ENSDARG000 | 4 | 11608196 | 11608283 | - | 88 | dre-mir-29b-2 | 0 | 0 | 0 | 3 | 0 | 0 |
| ENSDARG000 | 4 | 28699455 | 28699535 | + | 81 | dre-mir-430b- | 0 | 0 | 0 | 0 | 0 | 0 |
| ENSDARG000 | 4:4:4:4:4:4:4: | 58661331:586 | 58661414:586 | +:+:+:+:+:+:+: | 1103 | NA | 5 | 22 | 0 | 10 | 47 | 49 |
| ENSDARG000 | 4;4;4;4 | 59339234:594 | 59339732:594 | -;-;-;- | 1539 | NA | 1 | 1 | 0 | 0 | 0 | 0 |
| ENSDARG000 | 4:4:4:4:4:4:4: | 65160450:651 | 65160658:651 | -:-:-:-:-:-:-:-: | 2135 | NA | 0 | 19 | 1 | 6 | 32 | 27 |
| ENSDARG000 | 4 | 26570698 | 26570813 | + | 116 | 5S_rRNA | 0 | 0 | 0 | 0 | 0 | 0 |
| ENSDARG000 | 4 | 61183594 | 61183742 | + | 149 | U1 | 0 | 0 | 0 | 0 | 0 | 0 |
| ENSDARG000 | 4 | 58812958 | 58813064 | - | 107 | U6 | 0 | 0 | 0 | 0 | 0 | 0 |
| ENSDARG000 | 4 | 47192579 | 47192685 | - | 107 | U6 | 0 | 0 | 0 | 0 | 0 | 0 |
| ENSDARG000 | 4 | 57972964 | 57973127 | + | 164 | U1 | 0 | 0 | 0 | 0 | 0 | 0 |
| ENSDARG000 | 4 | 31213971 | 31214077 | - | 107 | U6 | 0 | 0 | 0 | 0 | 0 | 0 |
| ENSDARG000 | 4 | 42438554 | 42438660 | - | 107 | U6 | 0 | 0 | 0 | 2 | 22 | 29 |
| ENSDARG000 | 4 | 28695717 | 28695789 | + | 73 | dre-mir-430c- | 0 | 0 | 0 | 0 | 0 | 0 |
| ENSDARG000 | 4 | 61509054 | 61509217 | + | 164 | U1 | 0 | 0 | 0 | 0 | 0 | 0 |
| ENSDARG000 | 4 | 45565515 | 45565621 | + | 107 | U6 | 0 | 0 | 0 | 0 | 0 | 0 |
| ENSDARG000 | 4 | 51402311 | 51402417 | + | 107 | U6 | 0 | 0 | 0 | 0 | 0 | 0 |
| ENSDARG000 | 4:4:4:4:4:4:4: | 32156734:321 | 32156818:321 | +:+:+:+:+:+:+: | 660 | NA | 12 | 10 | 6 | 0 | 4 | 5 |
| ENSDARG000 | 4 | 51418692 | 51418798 | + | 107 | U6 | 0 | 0 | 0 | 0 | 0 | 0 |
| ENSDARG000 | 4;4;4;4;4;4;4 | 49308991:493 | 49309182:493 | -;-;-;-;-;-;- | 1231 | NA | 0 | 0 | 0 | 0 | 0 | 0 |
| ENSDARG000 | 4;4;4 | 71719248:717 | 71719652:717 | -;-;- | 1038 | NA | 0 | 8 | 4 | 2 | 2 | 0 |
| ENSDARG000 | 4;4;4 | 71735828:717 | 71735902:717 | +;+;+ | 1314 | NA | 4 | 0 | 1 | 0 | 0 | 0 |
| ENSDARG000 | 4 | 28699824 | 28699896 | + | 73 | dre-mir-430c- | 0 | 0 | 0 | 0 | 0 | 0 |
| ENSDARG000 | 4 | 30955341 | 30955504 | - | 164 | U1 | 0 | 0 | 0 | 0 | 0 | 0 |
| ENSDARG000 | 4 | 50989958 | 50990121 | - | 164 | U1 | 0 | 0 | 0 | 0 | 0 | 0 |
| ENSDARG000 | 4;4 | 60664235:606 | 60666196:606 | +;+ | 2224 | NA | 1 | 3 | 14 | 1 | 6 | 1 |
| ENSDARG000 | 4 | 64721299 | 64721405 | + | 107 | U6 | 0 | 0 | 0 | 0 | 0 | 0 |
| ENSDARG000 | 4 | 50102087 | 50102193 | - | 107 | U6 | 0 | 0 | 0 | 0 | 0 | 0 |
| ENSDARG000 | 4 | 37731429 | 37731577 | - | 149 | U1 | 0 | 0 | 0 | 0 | 0 | 0 |
| ENSDARG000 | 4 | 52012366 | 52012480 | + | 115 | 5S_rRNA | 0 | 0 | 0 | 0 | 0 | 0 |
| ENSDARG000 | 4 | 8587787 | 8587908 | - | 122 | 5S_rRNA | 0 | 0 | 0 | 0 | 0 | 0 |
| ENSDARG000 | 4 | 41346037 | 41346182 | - | 146 | U1 | 0 | 0 | 0 | 0 | 0 | 4 |
| ENSDARG000 | 4 | 51400192 | 51400298 | + | 107 | U6 | 0 | 0 | 0 | 0 | 0 | 0 |
| ENSDARG000 | 4 | 51400722 | 51400828 | + | 107 | U6 | 0 | 0 | 0 | 0 | 0 | 0 |
| ENSDARG000 | 4:4:4:4:4:4:4: | 76509294:765 | 76509341:765 | +:+:+:+:+:+:+: | 998 | ms4a17a.16 | 6 | 0 | 0 | 0 | 0 | 0 |
| ENSDARG000 | 4:4:4:4:4:4:4: | 72797711:727 | 72797946:727 | +:+:+:+:+:+:+: | 3301 | NA | 89 | 4 | 6 | 27 | 124 | 20 |
| ENSDARG000 | 4;4 | 46661522:466 | 46661606:466 | +;+ | 1152 | NA | 5 | 5 | 4 | 4 | 0 | 18 |
| ENSDARG000 | 4 | 42627981 | 42628144 | + | 164 | U1 | 0 | 0 | 0 | 0 | 0 | 0 |
| ENSDARG000 | 4 | 67582756 | 67582919 | + | 164 | U1 | 0 | 0 | 0 | 0 | 0 | 0 |
| ENSDARG000 | 4 | 69485075 | 69485239 | - | 165 | U1 | 0 | 0 | 0 | 0 | 0 | 0 |
| ENSDARG000 | 4 | 64698271 | 64698377 | + | 107 | U6 | 0 | 0 | 0 | 0 | 0 | 0 |
| ENSDARG000 | 4 | 57806997 | 57807066 | + | 70 | U6 | 0 | 0 | 0 | 0 | 0 | 0 |
| ENSDARG000 | 4;4 | 55794876:557 | 55794903:557 | +;+ | 429 | NA | 0 | 0 | 0 | 0 | 0 | 0 |
| ENSDARG000 | 4;4;4;4;4 | 62057454:620 | 62057550:620 | +;+;+;+;+ | 2338 | NA | 5 | 4 | 7 | 2 | 2 | 6 |
| ENSDARG000 | 4 | 57779867 | 57780030 | - | 164 | U1 | 0 | 0 | 0 | 0 | 0 | 0 |
| ENSDARG000 | 4 | 28115607 | 28115721 | + | 115 | 5S_rRNA | 0 | 0 | 0 | 0 | 0 | 0 |
| ENSDARG000 | 4 | 40917678 | 40917784 | - | 107 | U6 | 0 | 0 | 0 | 0 | 0 | 0 |
| ENSDARG000 | 4;4;4 | 77562651:775 | 77562855:775 | -;-;- | 291 | NA | 0 | 0 | 0 | 0 | 0 | 0 |
| ENSDARG000 | 4 | 64697744 | 64697847 | + | 104 | U6 | 0 | 0 | 0 | 0 | 0 | 0 |
| ENSDARG000 | 4 | 69493085 | 69493191 | + | 107 | U6 | 0 | 0 | 0 | 0 | 0 | 0 |
| ENSDARG000 | 4:4:4:4:4:4:4: | 34860183:348 | 34860267:348 | +:+:+:+:+:+:+: | 2407 | NA | 2 | 11 | 0 | 0 | 14 | 16 |
| ENSDARG000 | 4 | 42631854 | 42632017 | + | 164 | U1 | 0 | 0 | 0 | 0 | 0 | 0 |
| ENSDARG000 | 4 | 42640158 | 42640321 | + | 164 | U1 | 0 | 0 | 0 | 0 | 0 | 0 |
| ENSDARG000 | 4 | 14744576 | 14744691 | - | 116 | 5S_rRNA | 0 | 0 | 0 | 0 | 0 | 0 |
| ENSDARG000 | 4 | 31218729 | 31218835 | - | 107 | U6 | 0 | 0 | 0 | 0 | 0 | 0 |
| ENSDARG000 | 4 | 32279602 | 32279708 | + | 107 | U6 | 0 | 0 | 0 | 0 | 0 | 0 |
| ENSDARG000 | 4 | 55782758 | 55782855 | + | 98 | NA | 0 | 0 | 0 | 0 | 0 | 0 |
| ENSDARG000 | 4 | 28708593 | 28708665 | + | 73 | dre-mir-430c- | 0 | 0 | 0 | 0 | 0 | 0 |
| ENSDARG000 | 4:4:4:4:4:4:4: | 11787351:117 | 11787718:117 | -;-;-;-;-;-;-;- | 1522 | NA | 5 | 2 | 13 | 2 | 1 | 24 |
| ENSDARG000 | 4;4;4;4;4 | 12873982:128 | 12874632:128 | -;-;-;-;- | 930 | NA | 0 | 2 | 0 | 0 | 0 | 0 |
| ENSDARG000 | 4;4;4;4;4 | 13270792:132 | 13271884:132 | +;+;+;+;+ | 3127 | NA | 24 | 12 | 16 | 36 | 26 | 16 |
| ENSDARG000 | 4;4;4;4 | 17165644:171 | 17165916:171 | -;-;-;- | 1265 | NA | 45 | 23 | 52 | 42 | 65 | 28 |
| ENSDARG000 | 4:4:4:4:4:4:4: | 18469651:184 | 18469779:184 | +:+:+:+:+:+:+: | 2109 | NA | 0 | 0 | 1 | 2 | 0 | 0 |
| ENSDARG000 | 4;4;4 | 20531757:205 | 20531899:205 | +;+;+ | 421 | NA | 0 | 0 | 0 | 0 | 0 | 0 |
| ENSDARG000 | 4;4;4;4;4;4;4 | 21637724:216 | 21637789:216 | -;-;-;-;-;-;- | 843 | NA | 0 | 0 | 2 | 0 | 0 | 0 |
| ENSDARG000 | 4;4;4 | 22547576:225 | 22547593:225 | +;+;+ | 1184 | NA | 0 | 0 | 0 | 0 | 0 | 0 |
| ENSDARG000 | 4;4;4;4;4;4 | 30104535:301 | 30104685:301 | -;-;-;-;-;- | 4319 | NA | 4 | 2 | 6 | 0 | 4 | 16 |
| ENSDARG000 | 4;4;4 | 31148179:311 | 31148409:311 | -;-;- | 535 | NA | 0 | 0 | 0 | 0 | 0 | 0 |
| ENSDARG000 | 4;4;4;4 | 33796692:338 | 33796865:338 | -;-;-;- | 586 | NA | 0 | 0 | 0 | 0 | 0 | 0 |
| ENSDARG000 | 4;4;4 | 34200661:342 | 34200806:342 | +;+;+ | 526 | NA | 4 | 0 | 0 | 0 | 0 | 0 |
| ENSDARG000 | 4:4:4:4:4:4:4: | 35083725:350 | 35084254:350 | +:+:+:+:+:+:+: | 2814 | NA | 2 | 0 | 10 | 0 | 2 | 0 |
| ENSDARG000 | 4;4;4;4 | 37311797:373 | 37311867:373 | +;+;+;+ | 242 | NA | 0 | 0 | 0 | 0 | 0 | 0 |
| ENSDARG000 | 4;4;4;4;4 | 41540236:415 | 41540971:415 | -;-;-;-;- | 1572 | NA | 0 | 0 | 0 | 0 | 0 | 0 |
| ENSDARG000 | 4;4;4;4;4 | 42830392:428 | 42831144:428 | +;+;+;+;+ | 3520 | NA | 0 | 0 | 0 | 0 | 0 | 0 |
| ENSDARG000 | 4;4;4 | 45535334:455 | 45535606:455 | +;+;+ | 543 | NA | 14 | 2 | 0 | 1 | 0 | 0 |
| ENSDARG000 | 4;4;4 | 47275789:472 | 47275900:472 | -;-;- | 431 | NA | 0 | 0 | 0 | 0 | 0 | 0 |
| ENSDARG000 | 4;4;4;4 | 50513865:505 | 50513942:505 | -;-;-;- | 768 | NA | 0 | 0 | 0 | 0 | 0 | 0 |
| ENSDARG000 | 4;4;4 | 50622706:506 | 50622793:506 | +;+;+ | 1269 | NA | 0 | 0 | 0 | 0 | 0 | 0 |
| ENSDARG000 | 4;4;4 | 52488291:524 | 52488450:524 | +;+;+ | 389 | NA | 0 | 0 | 0 | 0 | 0 | 0 |
| ENSDARG000 | 4:4:4:4:4:4:4: | 53672375:536 | 53672619:536 | -;-;-;-;-;-;-;- | 864 | NA | 0 | 0 | 0 | 0 | 0 | 0 |
| ENSDARG000 | 4;4;4;4 | 53675056:536 | 53675557:536 | -;-;-;- | 1048 | NA | 0 | 0 | 0 | 0 | 0 | 0 |
| ENSDARG000 | 4;4;4;4;4 | 54181734:541 | 54182224:541 | +;+;+;+;+ | 2554 | NA | 0 | 0 | 0 | 0 | 0 | 8 |
| ENSDARG000 | 4;4;4 | 56001936:560 | 56002080:560 | -;-;- | 331 | NA | 4 | 0 | 4 | 0 | 0 | 2 |
| ENSDARG000 | 4;4;4;4 | 56577460:565 | 56577743:565 | -;-;-;- | 790 | NA | 0 | 0 | 0 | 0 | 0 | 0 |
| ENSDARG000 | 4;4;4;4;4 | 57478707:574 | 57478883:574 | +;+;+;+;+ | 622 | NA | 0 | 0 | 0 | 0 | 0 | 1 |
| ENSDARG000 | 4;4;4;4;4 | 57642367:576 | 57642403:576 | +;+;+;+;+ | 975 | NA | 9 | 0 | 0 | 0 | 0 | 0 |
| ENSDARG000 | 4;4;4 | 58139396:581 | 58139587:581 | -;-;- | 625 | NA | 11 | 2 | 2 | 0 | 4 | 2 |
| ENSDARG000 | 4;4;4;4 | 59731272:597 | 59731512:597 | -;-;-;- | 599 | NA | 0 | 8 | 2 | 9 | 0 | 0 |
| ENSDARG000 | 4;4;4;4;4 | 60098668:600 | 60098958:600 | +;+;+;+;+ | 886 | NA | 0 | 0 | 0 | 0 | 0 | 0 |
| ENSDARG000 | 4:4:4:4:4:4:4: | 61071556:610 | 61071843:610 | -:-:-:-:-:-:-:-: | 1012 | NA | 0 | 3 | 0 | 2 | 0 | 1 |
| ENSDARG000 | 4;4;4;4;4 | 62302545:623 | 62302743:623 | +;+;+;+;+ | 1048 | NA | 20 | 1 | 12 | 1 | 0 | 0 |
| ENSDARG000 | 4:4:4:4:4:4:4: | 63177298:631 | 63177675:631 | +:+:+:+:+:+:+: | 1801 | NA | 17 | 0 | 10 | 22 | 4 | 0 |
| ENSDARG000 | 4;4;4;4 | 63829645:638 | 63829785:638 | +;+;+;+ | 589 | NA | 0 | 0 | 0 | 2 | 0 | 0 |
| ENSDARG000 | 4;4;4;4 | 63853097:638 | 63853158:638 | -;-;-;- | 441 | NA | 0 | 0 | 0 | 0 | 0 | 0 |
| ENSDARG000 | 4;4;4;4;4 | 66922672:669 | 66922756:669 | -;-;-;-;- | 380 | NA | 0 | 0 | 0 | 0 | 0 | 0 |
| ENSDARG000 | 4;4;4;4;4;4;4 | 70472455:704 | 70473477:704 | -;-;-;-;-;-;- | 1930 | NA | 0 | 0 | 0 | 0 | 0 | 0 |
| ENSDARG000 | 4;4;4 | 71953615:719 | 71954041:719 | -;-;- | 908 | NA | 59 | 202 | 507 | 90 | 279 | 182 |
| ENSDARG000 | 4;4;4 | 72304336:723 | 72305554:723 | -;-;- | 1682 | NA | 3 | 16 | 15 | 0 | 10 | 34 |
| ENSDARG000 | 4;4;4;4;4 | 74028579:740 | 74028727:740 | -;-;-;-;- | 777 | NA | 0 | 0 | 0 | 0 | 0 | 0 |
| ENSDARG000 | 4;4;4 | 75285319:752 | 75285473:752 | -;-;- | 444 | NA | 0 | 0 | 0 | 0 | 0 | 0 |
| ENSDARG000 | 7:7:7:7:7:7:7: | 528593:5295 | 529852:5298 | +:+:+:+:+:+:+: | 5075 | NA | 54 | 546 | 294 | 1005 | 70 | 552 |
| ENSDARG000 | 7;7;7;7;7;7 | 67817256:678 | 67817416:678 | -;-;-;-;-;- | 1124 | NA | 2 | 1 | 0 | 0 | 0 | 0 |
| ENSDARG000 | 7:7:7:7:7:7:7: | 29502874:295 | 29502956:295 | +:+:+:+:+:+:+: | 1278 | NA | 0 | 0 | 0 | 0 | 10 | 0 |
| ENSDARG000 | 7:7:7:7:7:7:7: | 37742299:377 | 37742425:377 | +:+:+:+:+:+:+: | 2077 | brd7 | 891 | 724 | 1330 | 527 | 793 | 624 |
| ENSDARG000 | 7:7:7:7:7:7:7: | 742517:7433 | 743647:7436 | -:-:-:-:-:-:-:-: | 2006 | drap1 | 1654 | 2162 | 2396 | 1700 | 2293 | 1597 |
| ENSDARG000 | 7:7:7:7:7:7:7: | 67845942:678 | 67846198:678 | -:-:-:-:-:-:-:-: | 2001 | NA | 2 | 0 | 0 | 0 | 2 | 0 |
| ENSDARG000 | 7:7:7:7:7:7:7: | 72882446:728 | 72882489:728 | +:+:+:+:+:+:+: | 8222 | ADGRL3 | 6 | 10 | 16 | 64 | 142 | 52 |
| ENSDARG000 | 7;7;7 | 756942:7612 | 757034:7616 | +;+;+ | 2042 | C7H11orf68 | 177 | 170 | 240 | 188 | 194 | 94 |
| ENSDARG000 | 7:7:7:7:7:7:7: | 57108823:571 | 57109372:571 | +:+:+:+:+:+:+: | 6277 | enosf1 | 570 | 303 | 429 | 271 | 557 | 591 |
| ENSDARG000 | 7;7;7 | 5676779:5678 | 5677017:5678 | -;-;- | 737 | NA | 1 | 0 | 0 | 0 | 0 | 0 |
| ENSDARG000 | 7:7:7:7:7:7:7: | 808916:8089 | 810006:8100 | -:-:-:-:-:-:-:-: | 2037 | CABP4 | 0 | 0 | 0 | 0 | 0 | 0 |
| ENSDARG000 | 7 | 18247037 | 18247691 | - | 655 | NA | 0 | 0 | 0 | 0 | 0 | 0 |
| ENSDARG000 | 7 | 57341805 | 57342998 | + | 1194 | NA | 4 | 5 | 31 | 4 | 39 | 0 |
| ENSDARG000 | 7;7 | 68365760:683 | 68366080:683 | +;+ | 908 | NA | 0 | 0 | 0 | 0 | 0 | 1 |
| ENSDARG000 | 7;7 | 18364176:183 | 18367267:183 | +;+ | 3092 | cd248a | 208 | 44 | 32 | 6 | 22 | 78 |
| ENSDARG000 | 7:7:7:7:7:7:7: | 68214935:682 | 68221374:682 | -;-;-;-;-;-;-;-;- | 16288 | zfhx3b | 1540 | 594 | 1914 | 1291 | 1784 | 1740 |
| ENSDARG000 | 7:7:7:7:7:7:7: | 49619375:496 | 49621472:496 | -:-:-:-:-:-:-:-: | 3297 | NA | 316 | 212 | 224 | 172 | 252 | 211 |
| ENSDARG000 | 7;7;7 | 51752239:517 | 51755041:517 | -;-;- | 3082 | leap2 | 0 | 2 | 0 | 0 | 0 | 0 |
| ENSDARG000 | 7:7:7:7:7:7:7: | 828289:8378 | 829225:8382 | +:+:+:+:+:+:+: | 2485 | NA | 78 | 37 | 52 | 17 | 51 | 66 |
| ENSDARG000 | 7;7;7;7;7;7 | 73295890:732 | 73295985:732 | +;+;+;+;+;+ | 2572 | NA | 2 | 0 | 2 | 4 | 14 | 8 |
| ENSDARG000 | 7:7:7:7:7:7:7: | 29509255:295 | 29509297:295 | +:+:+:+:+:+:+: | 1459 | NA | 0 | 0 | 0 | 0 | 0 | 0 |
| ENSDARG000 | 7;7;7;7;7;7;7 | 854237:8547 | 854979:8548 | +;+;+;+;+;+;+ | 3391 | NA | 84 | 30 | 42 | 16 | 20 | 28 |
| ENSDARG000 | 7;7;7;7;7;7 | 30371893:303 | 30372059:303 | +;+;+;+;+;+ | 2554 | aqp9b | 0 | 0 | 6 | 0 | 0 | 16 |
| ENSDARG000 | 7 | 41459653 | 41460435 | + | 783 | NA | 0 | 0 | 0 | 0 | 0 | 0 |
| ENSDARG000 | 7;7;7;7;7;7 | 864492:8683 | 864656:8686 | -;-;-;-;-;- | 2292 | NA | 44 | 22 | 18 | 12 | 32 | 6 |
| ENSDARG000 | 7:7:7:7:7:7:7: | 41436488:414 | 41439189:414 | -:-:-:-:-:-:-:-: | 4611 | smarcd3b | 1908 | 845 | 1042 | 2044 | 1284 | 933 |
| ENSDARG000 | 7;7;7;7 | 18412621:184 | 18416863:184 | -;-;-;- | 4507 | NA | 0 | 0 | 0 | 0 | 0 | 0 |
| ENSDARG000 | 7;7;7;7;7;7;7 | 18462242:184 | 18462730:184 | -;-;-;-;-;-;- | 4210 | znf16l | 286 | 272 | 460 | 182 | 436 | 250 |
| ENSDARG000 | 7;7;7 | 7662195:7674 | 7666956:7674 | -;-;- | 5293 | mfap3l | 2220 | 1963 | 2239 | 1960 | 1603 | 2167 |
| ENSDARG000 | 7:7:7:7:7:7:7: | 73308566:733 | 73308797:733 | +:+:+:+:+:+:+: | 2980 | NA | 14 | 16 | 35 | 66 | 42 | 28 |
| ENSDARG000 | 7:7:7:7:7:7:7: | 7675685:7675 | 7675714:7675 | -:-:-:-:-:-:-:-: | 1800 | AADAT | 40 | 46 | 157 | 55 | 78 | 44 |
| ENSDARG000 | 7:7:7:7:7:7:7: | 5699624:5700 | 5700057:5700 | -:-:-:-:-:-:-:-: | 3077 | NA | 0 | 29 | 17 | 88 | 75 | 40 |
| ENSDARG000 | 7:7:7:7:7:7:7: | 73332486:733 | 73332748:733 | +:+:+:+:+:+:+: | 2133 | SLC24A2 | 0 | 0 | 4 | 0 | 0 | 2 |
| ENSDARG000 | 7:7:7:7:7:7:7: | 24979833:249 | 24981088:249 | -:-:-:-:-:-:-:-: | 3411 | rcor2 | 6 | 30 | 56 | 16 | 28 | 12 |
| ENSDARG000 | 7:7:7:7:7:7:7: | 31924983:319 | 31926565:319 | -:-:-:-:-:-:-:-: | 3011 | bdnf | 293 | 335 | 248 | 202 | 136 | 95 |
| ENSDARG000 | 7:7:7:7:7:7:7: | 34241319:342 | 34241892:342 | -:-:-:-:-:-:-:-: | 3525 | dis3l | 161 | 96 | 132 | 60 | 194 | 135 |
| ENSDARG000 | 7;7 | 5737873:5784 | 5737981:5784 | +;+ | 217 | NA | 0 | 0 | 0 | 0 | 0 | 0 |
| ENSDARG000 | 7:7:7:7:7:7:7: | 34236238:342 | 34236368:342 | +:+:+:+:+:+:+: | 5700 | tipin | 197 | 230 | 280 | 116 | 288 | 233 |
| ENSDARG000 | 7:7:7:7:7:7:7: | 5776656:5777 | 5776749:5777 | -;-;-;-;-;-;-;- | 2392 | NA | 0 | 4 | 0 | 1 | 4 | 13 |
| ENSDARG000 | 7;7;7;7;7;7;7 | 5739921:5744 | 5740166:5745 | -;-;-;-;-;-;- | 1938 | NA | 0 | 8 | 2 | 27 | 15 | 7 |
| ENSDARG000 | 7:7:7:7:7:7:7: | 62320134:624 | 62321649:624 | +:+:+:+:+:+:+: | 4444 | STIM2 | 98 | 52 | 84 | 28 | 50 | 36 |
| ENSDARG000 | 7;7 | 73423306:734 | 73424055:734 | -;- | 772 | NA | 0 | 0 | 0 | 0 | 0 | 0 |
| ENSDARG000 | 7;7;7;7;7 | 7696665:7700 | 7697015:7701 | +;+;+;+;+ | 1751 | ino80b | 294 | 228 | 335 | 323 | 290 | 212 |
| ENSDARG000 | 7;7;7;7;7;7;7 | 57432318:574 | 57432569:574 | +;+;+;+;+;+;+ | 1180 | drd4b | 309 | 44 | 142 | 64 | 134 | 65 |
| ENSDARG000 | 7:7:7:7:7:7:7: | 51774502:517 | 51774706:517 | +:+:+:+:+:+:+: | 1981 | got2b | 1538 | 2354 | 2814 | 1761 | 1763 | 1978 |
| ENSDARG000 | 7:7:7:7:7:7:7: | 57465814:574 | 57470361:574 | -:-:-:-:-:-:-:-: | 6641 | sirt3 | 201 | 108 | 155 | 84 | 177 | 327 |
| ENSDARG000 | 7;7;7;7 | 68540585:685 | 68540669:685 | +;+;+;+ | 2106 | NA | 4 | 0 | 2 | 0 | 0 | 4 |
| ENSDARG000 | 7:7:7:7:7:7:7: | 13039426:130 | 13040672:130 | +:+:+:+:+:+:+: | 14847 | syt7b | 20 | 18 | 34 | 2 | 278 | 0 |

| | | | | | | | | | | | | |
|---|---|---|---|---|---|---|---|---|---|---|---|---|
| ENSDARG000 | 7;7;7;7;7 | 51768343:517 | 51769835:517 | -;-;-;-;- | 2348 | bmp15 | 2 | 0 | 0 | 0 | 0 | 6 |
| ENSDARG000 | 7;7;7 | 21787507:217 | 21787723:217 | +;+;+ | 1309 | tmem88b | 3 | 0 | 8 | 0 | 0 | 0 |
| ENSDARG000 | 7;7;7 | 73444325:734 | 73444605:734 | +;+;+ | 810 | NA | 0 | 0 | 0 | 0 | 0 | 0 |
| ENSDARG000 | 7;7;7;7;7;7;7; | 40536928:405 | 40538690:405 | -;-;-;-;-;-;-;-; | 4494 | dnajb6b | 84 | 64 | 76 | 46 | 66 | 40 |
| ENSDARG000 | 7;7;7;7;7;7 | 21785595:217 | 21786226:217 | -;-;-;-;-;- | 1911 | NA | 0 | 0 | 0 | 0 | 0 | 0 |
| ENSDARG000 | 7;7;7;7;7;7;7; | 30406150:304 | 30411251:304 | -;-;-;-;-;-;-;-; | 7623 | adam10a | 10223 | 6312 | 7991 | 7210 | 9713 | 13091 |
| ENSDARG000 | 7;7;7 | 73447724:734 | 73447978:734 | +;+;+ | 1158 | cbln6 | 0 | 0 | 0 | 0 | 0 | 0 |
| ENSDARG000 | 7 | 30475836 | 30476382 | + | 547 | NA | 0 | 0 | 0 | 0 | 0 | 0 |
| ENSDARG000 | 7;7;7;7;7;7;7; | 30392613:303 | 30392848:303 | +;+;+;+;+;+;+; | 2140 | lipca | 0 | 0 | 4 | 0 | 0 | 0 |
| ENSDARG000 | 7;7;7;7;7;7;7; | 28253644:282 | 28253723:282 | +;+;+;+;+;+;+; | 1259 | NA | 18 | 0 | 2 | 0 | 8 | 6 |
| ENSDARG000 | 7;7;7;7;7;7;7; | 18474563:184 | 18475202:184 | -;-;-;-;-;-;-;-; | 4882 | NA | 182 | 196 | 368 | 314 | 266 | 244 |
| ENSDARG000 | 7;7;7 | 73453486:734 | 73453795:734 | +;+;+ | 825 | cbln8 | 0 | 0 | 8 | 0 | 2 | 0 |
| ENSDARG000 | 7;7;7;7;7;7;7; | 28374947:283 | 28375803:283 | -;-;-;-;-;-;-;-; | 2386 | stk33 | 2 | 0 | 0 | 0 | 0 | 0 |
| ENSDARG000 | 7;7;7;7;7;7;7; | 20017211:200 | 20017342:200 | +;+;+;+;+;+;+; | 3492 | BCL6B | 176 | 14 | 106 | 32 | 142 | 98 |
| ENSDARG000 | 7;7;7;7;7;7;7; | 26297169:262 | 26300568:263 | -;-;-;-;-;-;-;-; | 5309 | zgc:77439 | 453 | 218 | 433 | 194 | 742 | 488 |
| ENSDARG000 | 7;7;7;7;7;7;7; | 19978434:199 | 19980106:199 | -;-;-;-;-;-;-;-; | 4535 | NA | 446 | 330 | 479 | 300 | 499 | 467 |
| ENSDARG000 | 7;7;7;7;7;7;7; | 52949847:529 | 52950027:529 | +;+;+;+;+;+;+; | 3931 | cdkn2aip | 124 | 148 | 175 | 123 | 186 | 244 |
| ENSDARG000 | 7;7;7;7;7;7;7; | 49638413:496 | 49640482:496 | -;-;-;-;-;-;-;-; | 3051 | hrasb | 194 | 242 | 230 | 233 | 143 | 247 |
| ENSDARG000 | 7;7;7 | 62735810:627 | 62736106:627 | -;-;- | 517 | NA | 0 | 0 | 0 | 0 | 0 | 0 |
| ENSDARG000 | 7;7;7;7;7;7;7; | 52929964:529 | 52933374:529 | -;-;-;-;-;-;-;-; | 5910 | tubgcp4 | 352 | 506 | 777 | 420 | 646 | 511 |
| ENSDARG000 | 7;7;7;7;7;7;7 | 63324888:633 | 63328905:633 | +;+;+;+;+;+;+ | 5325 | PCDH7 | 12 | 6 | 34 | 0 | 38 | 0 |
| ENSDARG000 | 7;7 | 32022061:320 | 32022666:320 | -;- | 697 | NA | 0 | 0 | 0 | 0 | 0 | 0 |
| ENSDARG000 | 7;7;7;7;7;7;7; | 32021669:320 | 32021756:320 | +;+;+;+;+;+;+; | 2207 | mettl15 | 54 | 77 | 67 | 42 | 61 | 26 |
| ENSDARG000 | 7;7;7;7;7 | 21841037:218 | 21841297:218 | +;+;+;+;+ | 1654 | tm4sf5 | 0 | 0 | 0 | 0 | 0 | 0 |
| ENSDARG000 | 7;7;7;7;7;7;7; | 38667066:386 | 38667246:386 | +;+;+;+;+;+;+; | 1846 | mtch2 | 2588 | 2413 | 2745 | 1898 | 3025 | 2619 |
| ENSDARG000 | 7;7;7;7;7;7;7 | 34256520:342 | 34257866:342 | -;-;-;-;-;-;- | 2431 | NA | 52 | 6 | 2 | 14 | 20 | 6 |
| ENSDARG000 | 7;7;7;7;7;7;7; | 73488035:735 | 73488278:735 | +;+;+;+;+;+;+; | 6047 | NA | 32 | 28 | 88 | 37 | 55 | 55 |
| ENSDARG000 | 7;7 | 63637143:636 | 63637295:636 | -;- | 384 | NA | 0 | 0 | 0 | 0 | 0 | 0 |
| ENSDARG000 | 7 | 34290051 | 34293073 | + | 3023 | fibinb | 72 | 106 | 318 | 142 | 291 | 36 |
| ENSDARG000 | 7;7;7;7;7;7;7; | 31976764:319 | 31977099:319 | -;-;-;-;-;-;-;-; | 3197 | kif18a | 52 | 276 | 328 | 161 | 224 | 106 |
| ENSDARG000 | 7;7;7;7;7;7;7; | 16033273:160 | 16034009:160 | +;+;+;+;+;+;+; | 2404 | immp1l | 39 | 41 | 65 | 44 | 47 | 26 |
| ENSDARG000 | 7;7;7 | 21859337:218 | 21859509:218 | +;+;+ | 2440 | si:dkev-85k7.1 | 128 | 280 | 1564 | 1615 | 402 | 131 |
| ENSDARG000 | 7;7;7;7;7;7;7; | 37761524:377 | 37767903:377 | -;-;-;-;-;-;-;-; | 10297 | adcy7 | 62 | 53 | 196 | 32 | 142 | 85 |
| ENSDARG000 | 7;7;7;7;7;7;7; | 5833060:5834 | 5833164:5834 | -;-;-;-;-;-;-;- | 2415 | NA | 1 | 2 | 0 | 0 | 0 | 6 |
| ENSDARG000 | 7;7;7 | 37851111:378 | 37851196:378 | +;+;+ | 778 | NA | 0 | 0 | 0 | 0 | 0 | 0 |
| ENSDARG000 | 7;7;7;7;7;7;7; | 37861247:378 | 37863069:378 | -;-;-;-;-;-;-;-; | 4297 | tent4b | 481 | 385 | 678 | 433 | 628 | 411 |
| ENSDARG000 | 7;7;7;7;7;7;7; | 29518396:295 | 29518670:295 | -;-;-;-;-;-;-;-; | 1561 | anxa2b | 6 | 0 | 0 | 0 | 0 | 0 |
| ENSDARG000 | 7;7;7;7;7;7;7; | 7708296:7710 | 7711856:7711 | -;-;-;-;-;-;-;-; | 6806 | intu | 224 | 171 | 294 | 212 | 143 | 272 |
| ENSDARG000 | 7;7;7;7 | 13353080:133 | 13354990:133 | -;-;-;- | 2373 | sdhaf2 | 462 | 391 | 603 | 362 | 538 | 432 |
| ENSDARG000 | 7;7;7;7;7 | 28406134:284 | 28408405:284 | -;-;-;-;- | 2662 | rerglb | 18 | 0 | 2 | 4 | 24 | 23 |
| ENSDARG000 | 7;7;7;7;7;7;7; | 65103874:651 | 65105335:651 | -;-;-;-;-;-;-;-; | 3188 | tmem41b | 329 | 394 | 272 | 129 | 392 | 170 |
| ENSDARG000 | 7;7;7;7;7;7;7; | 24069421:240 | 24069604:240 | +;+;+;+;+;+;+; | 6155 | acin1b | 3090 | 2827 | 3826 | 2344 | 4047 | 3496 |
| ENSDARG000 | 7;7;7;7;7;7;7; | 34296789:342 | 34296932:342 | +;+;+;+;+;+;+; | 3880 | bbox1 | 114 | 296 | 1148 | 1254 | 236 | 264 |
| ENSDARG000 | 7;7;7;7;7;7;7; | 57539146:575 | 57539306:575 | +;+;+;+;+;+;+; | 2850 | pde5ab | 111 | 36 | 155 | 92 | 468 | 184 |
| ENSDARG000 | 7;7;7;7;7;7;7; | 39410180:394 | 39410451:394 | +;+;+;+;+;+;+; | 4213 | NA | 319 | 58 | 106 | 61 | 139 | 148 |
| ENSDARG000 | 7;7;7;7;7;7;7; | 21876548:218 | 21876766:218 | -;-;-;-;-;-;-;-; | 2132 | mettl3 | 154 | 222 | 258 | 136 | 216 | 90 |
| ENSDARG000 | 7;7;7;7;7;7;7; | 24049776:240 | 24049958:240 | +;+;+;+;+;+;+; | 5497 | efs | 475 | 533 | 539 | 370 | 464 | 366 |
| ENSDARG000 | 7;7;7;7;7 | 57629370:576 | 57629686:576 | -;-;-;-;- | 872 | mad2l1 | 284 | 771 | 712 | 258 | 485 | 303 |
| ENSDARG000 | 7;7;7 | 18549669:185 | 18550685:185 | -;-;- | 1726 | msantd1 | 70 | 94 | 188 | 80 | 126 | 98 |
| ENSDARG000 | 7;7 | 24058768:240 | 24060183:240 | -;- | 1562 | NA | 4 | 6 | 2 | 2 | 4 | 0 |
| ENSDARG000 | 7;7;7;7;7;7;7; | 28423996:284 | 28425207:284 | -;-;-;-;-;-;-;-; | 4605 | NA | 8 | 16 | 19 | 20 | 3 | 25 |
| ENSDARG000 | 7;7;7 | 44705349:447 | 44705538:447 | +;+;+ | 853 | NA | 8 | 6 | 8 | 2 | 16 | 2 |
| ENSDARG000 | 7;7;7;7;7;7;7; | 69142757:691 | 69148480:691 | -;-;-;-;-;-;-;-; | 9433 | USP10 | 3777 | 4774 | 6219 | 3592 | 5945 | 4537 |
| ENSDARG000 | 7;7;7 | 44713135:447 | 44713270:447 | +;+;+ | 2417 | NA | 758 | 735 | 2145 | 490 | 1874 | 566 |
| ENSDARG000 | 7;7;7;7;7;7;7; | 4384863:4384 | 4385232:4385 | +;+;+;+;+;+;+; | 4203 | slc12a10.3 | 0 | 4 | 8 | 12 | 4 | 3 |
| ENSDARG000 | 7;7;7 | 39437384:394 | 39437841:394 | -;-;- | 539 | NA | 1 | 0 | 0 | 3 | 10 | 0 |
| ENSDARG000 | 7;7 | 44730625:447 | 44730730:447 | +;+ | 872 | NA | 8 | 5 | 16 | 14 | 10 | 28 |
| ENSDARG000 | 7;7;7;7;7;7;7; | 18557939:185 | 18560999:185 | -;-;-;-;-;-;-;-; | 5506 | dtx4a | 190 | 102 | 107 | 96 | 183 | 158 |
| ENSDARG000 | 7;7;7;7;7;7;7; | 49654588:496 | 49654707:496 | +;+;+;+;+;+;+; | 4406 | rassf7b | 38 | 31 | 18 | 16 | 40 | 8 |
| ENSDARG000 | 7;7;7;7;7;7;7; | 5949085:5949 | 5950775:5950 | -;-;-;-;-;-;-;-; | 2813 | NA | 2 | 0 | 0 | 0 | 0 | 0 |
| ENSDARG000 | 7;7;7;7;7 | 73593814:736 | 73594042:736 | +;+;+;+;+ | 7953 | znf219 | 18 | 12 | 6 | 0 | 6 | 6 |
| ENSDARG000 | 7;7;7;7;7;7;7; | 20030888:200 | 20031335:200 | +;+;+;+;+;+;+; | 5777 | SLC16A13 | 112 | 61 | 188 | 88 | 214 | 120 |
| ENSDARG000 | 7;7;7;7;7;7;7; | 6317866:6320 | 6318023:6320 | +;+;+;+;+;+;+; | 2249 | NA | 0 | 0 | 0 | 0 | 0 | 0 |
| ENSDARG000 | 7;7;7 | 13363898:133 | 13364243:133 | -;-;- | 532 | polr2l | 358 | 749 | 750 | 275 | 498 | 378 |
| ENSDARG000 | 7 | 6457759 | 6458222 | - | 464 | zgc:154164 | 68 | 14 | 47 | 19 | 39 | 2 |
| ENSDARG000 | 7 | 6467050 | 6467510 | - | 461 | NA | 6 | 6 | 5 | 2 | 0 | 0 |
| ENSDARG000 | 7 | 6363702 | 6364168 | - | 467 | zgc:112234 | 12 | 0 | 4 | 1 | 4 | 6 |
| ENSDARG000 | 7;7;7;7;7;7;7; | 30532636:305 | 30539049:305 | -;-;-;-;-;-;-;-; | 9994 | sltm | 2640 | 1939 | 3359 | 2845 | 2589 | 2337 |
| ENSDARG000 | 7;7;7;7;7;7;7; | 5885800:6070 | 5886100:6070 | -;-;-;-;-;-;-;-; | 3622 | dusp19b | 111 | 104 | 110 | 112 | 73 | 86 |
| ENSDARG000 | 7;7;7 | 16166558:161 | 16166714:161 | +;+;+ | 876 | NA | 0 | 2 | 2 | 2 | 2 | 0 |
| ENSDARG000 | 7;7 | 51802374:518 | 51802523:518 | -;- | 657 | NA | 0 | 0 | 0 | 0 | 0 | 0 |
| ENSDARG000 | 7;7;7;7;7;7;7; | 51816321:518 | 51816589:518 | +;+;+;+;+;+;+; | 2497 | si:ch211-122f | 944 | 854 | 619 | 509 | 900 | 829 |
| ENSDARG000 | 7;7 | 51825557:518 | 51825708:518 | +;+ | 359 | NA | 0 | 0 | 0 | 0 | 3 | 0 |
| ENSDARG000 | 7;7;7;7;7;7;7; | 51795667:517 | 51795838:517 | +;+;+;+;+;+;+; | 4781 | slc38a7 | 174 | 100 | 105 | 84 | 166 | 199 |
| ENSDARG000 | 7;7;7;7;7;7;7; | 6480212:6486 | 6480254:6486 | +;+;+;+;+;+;+; | 3426 | NA | 2 | 46 | 26 | 30 | 116 | 104 |
| ENSDARG000 | 7;7;7;7;7;7;7 | 73630751:736 | 73631002:736 | +;+;+;+;+;+;+ | 4963 | NA | 112 | 132 | 189 | 76 | 114 | 80 |
| ENSDARG000 | 7;7;7;7;7;7;7; | 65121743:651 | 65122079:651 | +;+;+;+;+;+;+; | 5559 | ipo7 | 6983 | 4941 | 7587 | 4527 | 5073 | 5849 |
| ENSDARG000 | 7 | 5906327 | 5906995 | + | 669 | zgc:112234 | 25 | 6 | 6 | 1 | 9 | 4 |
| ENSDARG000 | 7;7;7;7;7;7;7; | 6066772:6067 | 6067179:6067 | -;-;-;-;-;-;-;-; | 4220 | NA | 14 | 308 | 43 | 773 | 363 | 88 |
| ENSDARG000 | 7;7;7;7;7 | 6029084:6030 | 6030231:6031 | -;-;-;-;- | 2653 | si:dkev-273g1 | 22 | 4 | 62 | 12 | 4 | 18 |
| ENSDARG000 | 7;7;7 | 21887307:218 | 21887427:218 | +;+;+ | 856 | pop7 | 270 | 250 | 232 | 248 | 246 | 198 |
| ENSDARG000 | 7 | 5904020 | 5904598 | + | 579 | hist1h2a11 | 0 | 0 | 2 | 0 | 0 | 0 |
| ENSDARG000 | 7 | 6469133 | 6470431 | - | 1299 | zgc:110425 | 0 | 0 | 0 | 0 | 0 | 4 |
| ENSDARG000 | 7 | 6443547 | 6444011 | - | 465 | zgc:112234 | 0 | 0 | 2 | 0 | 0 | 0 |
| ENSDARG000 | 7 | 5965611 | 5966108 | + | 498 | NA | 8 | 14 | 14 | 0 | 2 | 0 |
| ENSDARG000 | 7 | 6349530 | 6351021 | - | 1492 | si:ch73-368i2 | 50 | 34 | 68 | 24 | 62 | 14 |
| ENSDARG000 | 7;7;7;7;7;7;7; | 6085485:6090 | 6085826:6090 | -;-;-;-;-;-;-;-; | 7640 | NA | 0 | 1 | 10 | 0 | 0 | 7 |
| ENSDARG000 | 7;7;7;7;7;7;7; | 73670137:736 | 73670493:736 | +;+;+;+;+;+;+; | 2454 | btr12 | 16 | 0 | 0 | 6 | 4 | 0 |
| ENSDARG000 | 7 | 6340617 | 6340998 | - | 382 | NA | 2 | 5 | 0 | 0 | 0 | 1 |
| ENSDARG000 | 7 | 6465468 | 6466119 | - | 652 | zgc:112234 | 150 | 33 | 62 | 4 | 0 | 47 |
| ENSDARG000 | 7;7;7;7;7;7;7; | 6385664:6387 | 6385935:6387 | +;+;+;+;+;+;+; | 2768 | NA | 2 | 34 | 14 | 48 | 26 | 0 |
| ENSDARG000 | 7;7 | 13375565:133 | 13377784:133 | -;- | 2490 | NA | 14 | 18 | 24 | 142 | 44 | 98 |
| ENSDARG000 | 7;7;7;7;7;7;7; | 30493684:304 | 30494502:304 | +;+;+;+;+;+;+; | 6363 | mindy2 | 206 | 87 | 240 | 62 | 221 | 184 |
| ENSDARG000 | 7 | 21917857 | 21918877 | + | 1021 | ufsp1 | 22 | 25 | 36 | 12 | 40 | 48 |
| ENSDARG000 | 7 | 6356388 | 6356875 | - | 488 | zgc:112234 | 22 | 6 | 22 | 2 | 12 | 2 |
| ENSDARG000 | 7;7;7;7;7;7;7; | 21890473:218 | 21891864:218 | -;-;-;-;-;-;-;-; | 2407 | epoa | 12 | 51 | 224 | 14 | 16 | 46 |
| ENSDARG000 | 7 | 6416636 | 6417339 | - | 704 | si:ch73-368i2 | 0 | 2 | 0 | 13 | 2 | 0 |
| ENSDARG000 | 7;7;7;7;7;7;7; | 73690366:736 | 73691015:736 | +;+;+;+;+;+;+; | 5958 | NA | 0 | 0 | 0 | 0 | 0 | 0 |
| ENSDARG000 | 7 | 5910467 | 5910989 | + | 523 | NA | 0 | 0 | 0 | 6 | 0 | 0 |
| ENSDARG000 | 7;7;7;7;7;7;7; | 20109968:201 | 20110502:201 | +;+;+;+;+;+;+; | 3470 | agfg2 | 43 | 89 | 71 | 159 | 92 | 94 |
| ENSDARG000 | 7 | 6426736 | 6427198 | - | 463 | hist1h2a6 | 0 | 2 | 0 | 0 | 0 | 0 |
| ENSDARG000 | 7 | 5909271 | 5909676 | + | 406 | NA | 4 | 2 | 8 | 6 | 8 | 0 |
| ENSDARG000 | 7 | 5908077 | 5908583 | + | 507 | NA | 0 | 0 | 16 | 0 | 0 | 0 |
| ENSDARG000 | 7 | 6468174 | 6468611 | - | 438 | NA | 6 | 0 | 4 | 0 | 0 | 0 |
| ENSDARG000 | 7 | 6425130 | 6425549 | - | 420 | NA | 6 | 2 | 4 | 0 | 0 | 4 |
| ENSDARG000 | 7 | 6460425 | 6460821 | - | 397 | NA | 0 | 4 | 0 | 4 | 0 | 2 |
| ENSDARG000 | 7 | 5977707 | 5978723 | + | 1017 | NA | 37 | 8 | 26 | 10 | 0 | 14 |
| ENSDARG000 | 7 | 5948383 | 5948685 | + | 303 | NA | 0 | 0 | 0 | 0 | 0 | 0 |
| ENSDARG000 | 7;7;7;7;7;7;7; | 6564653:6564 | 6565047:6565 | -;-;-;-;-;-;-;-; | 1843 | KCNJ10 | 0 | 0 | 0 | 0 | 0 | 0 |
| ENSDARG000 | 7;7;7 | 20147202:201 | 20147859:201 | +;+;+ | 7027 | NA | 62 | 38 | 200 | 44 | 300 | 108 |
| ENSDARG000 | 7;7 | 6400350:6401 | 6400575:6401 | -;- | 291 | NA | 0 | 0 | 0 | 0 | 0 | 0 |
| ENSDARG000 | 7;7 | 20132336:201 | 20132915:201 | -;- | 774 | NA | 0 | 0 | 0 | 0 | 0 | 2 |
| ENSDARG000 | 7 | 6430573 | 6431158 | - | 586 | NA | 0 | 2 | 2 | 0 | 0 | 0 |
| ENSDARG000 | 7;7;7;7;7;7;7; | 34317401:343 | 34319002:343 | -;-;-;-;-;-;-;-; | 7797 | MADD | 477 | 268 | 436 | 405 | 446 | 380 |
| ENSDARG000 | 7 | 5976613 | 5977242 | + | 630 | hist2h3ca1 | 2 | 0 | 4 | 0 | 2 | 0 |
| ENSDARG000 | 7;7;7;7;7;7;7; | 20058857:200 | 20059579:200 | -;-;-;-;-;-;-;-; | 3603 | acap1 | 628 | 792 | 845 | 922 | 1074 | 1185 |
| ENSDARG000 | 7 | 5905091 | 5905740 | + | 650 | NA | 0 | 0 | 4 | 0 | 6 | 0 |
| ENSDARG000 | 7 | 6366829 | 6367347 | - | 519 | NA | 0 | 2 | 0 | 0 | 2 | 2 |
| ENSDARG000 | 7 | 6463626 | 6464225 | - | 600 | hist1h2a2 | 1 | 4 | 0 | 0 | 0 | 2 |
| ENSDARG000 | 7 | 6438122 | 6438544 | - | 423 | NA | 0 | 1 | 0 | 0 | 0 | 2 |
| ENSDARG000 | 7 | 6409215 | 6409499 | - | 285 | NA | 0 | 0 | 0 | 0 | 0 | 0 |
| ENSDARG000 | 7;7;7;7;7;7;7; | 5920665:5922 | 5920784:5922 | +;+;+;+;+;+;+; | 4201 | NA | 0 | 12 | 6 | 12 | 9 | 2 |
| ENSDARG000 | 7;7;7;7;7;7;7; | 25000060:250 | 25000433:250 | +;+;+;+;+;+;+; | 2171 | naa40 | 419 | 624 | 816 | 426 | 647 | 555 |
| ENSDARG000 | 7;7;7;7;7;7;7; | 38680034:386 | 38680096:386 | +;+;+;+;+;+;+; | 1860 | psmc3 | 2379 | 2698 | 2689 | 1198 | 2374 | 1403 |
| ENSDARG000 | 7;7;7;7;7;7;7; | 5878622:5881 | 5879227:5881 | -;-;-;-;-;-;-;-; | 1683 | NA | 4 | 2 | 0 | 0 | 2 | 16 |
| ENSDARG000 | 7 | 6357559 | 6357952 | - | 394 | NA | 2 | 0 | 2 | 0 | 0 | 0 |
| ENSDARG000 | 7 | 6428579 | 6429031 | - | 453 | zgc:112234 | 2 | 5 | 6 | 4 | 0 | 0 |
| ENSDARG000 | 7 | 6432132 | 6432854 | - | 723 | NA | 0 | 10 | 10 | 8 | 14 | 4 |
| ENSDARG000 | 7 | 5975194 | 5975991 | + | 798 | si:dkev-23a13 | 0 | 0 | 0 | 0 | 0 | 0 |
| ENSDARG000 | 7 | 6444608 | 6445129 | - | 522 | NA | 0 | 0 | 0 | 0 | 0 | 0 |
| ENSDARG000 | 7 | 5968225 | 5968991 | + | 767 | zgc:163040 | 8 | 0 | 2 | 0 | 0 | 13 |
| ENSDARG000 | 7 | 6344680 | 6345507 | - | 828 | H3C15 | 2 | 6 | 0 | 0 | 2 | 6 |
| ENSDARG000 | 7 | 5959203 | 5959818 | + | 616 | NA | 0 | 0 | 0 | 0 | 2 | 1 |
| ENSDARG000 | 7;7;7;7;7;7 | 16176847:161 | 16177394:161 | -;-;-;-;-;- | 1603 | NA | 4 | 0 | 26 | 10 | 28 | 10 |
| ENSDARG000 | 7 | 6447299 | 6448182 | - | 884 | NA | 0 | 1 | 8 | 8 | 8 | 0 |
| ENSDARG000 | 7 | 6355000 | 6355459 | - | 460 | NA | 0 | 0 | 2 | 0 | 0 | 0 |
| ENSDARG000 | 7 | 5964296 | 5965015 | + | 720 | si:dkev-23a13 | 0 | 2 | 0 | 0 | 0 | 0 |
| ENSDARG000 | 7 | 6445595 | 6446172 | - | 578 | NA | 0 | 0 | 0 | 0 | 0 | 0 |
| ENSDARG000 | 7 | 6361991 | 6362460 | - | 470 | NA | 24 | 12 | 1 | 7 | 3 | 6 |
| ENSDARG000 | 7 | 5911670 | 5912374 | + | 705 | NA | 0 | 4 | 14 | 0 | 0 | 2 |
| ENSDARG000 | 7;7;7;7;7;7;7; | 40583939:405 | 40585997:405 | -;-;-;-;-;-;-;-; | 5648 | ube3c | 1066 | 756 | 1056 | 448 | 1081 | 676 |
| ENSDARG000 | 7;7;7;7;7;7 | 6652967:6658 | 6653217:6658 | +;+;+;+;+;+ | 3076 | pnp5a | 3351 | 1667 | 2389 | 2058 | 4247 | 3142 |
| ENSDARG000 | 7;7;7;7;7;7;7; | 44773912:447 | 44774176:447 | -;-;-;-;-;-;-;-; | 1829 | nae1 | 822 | 911 | 996 | 524 | 1034 | 760 |
| ENSDARG000 | 7 | 6429632 | 6430101 | - | 470 | zgc:173552 | 0 | 1 | 0 | 3 | 0 | 4 |
| ENSDARG000 | 7 | 5913030 | 5913529 | + | 500 | NA | 0 | 0 | 0 | 0 | 1 | 0 |
| ENSDARG000 | 7 | 6346119 | 6346859 | - | 741 | NA | 0 | 17 | 20 | 0 | 0 | 0 |
| ENSDARG000 | 7;7;7 | 18606798:186 | 18609700:186 | -;-;- | 3041 | NA | 22 | 50 | 32 | 8 | 138 | 153 |
| ENSDARG000 | 7;7;7;7;7;7;7; | 16185376:161 | 16185953:161 | -;-;-;-;-;-;-;-; | 1735 | NA | 0 | 0 | 107 | 48 | 208 | 52 |
| ENSDARG000 | 7 | 5955160 | 5956028 | + | 869 | NA | 2 | 7 | 7 | 7 | 3 | 6 |
| ENSDARG000 | 7 | 6415501 | 6415991 | - | 491 | NA | 4 | 12 | 8 | 0 | 6 | 5 |
| ENSDARG000 | 7;7;7;7;7;7;7; | 6127741:6128 | 6128122:6128 | -;-;-;-;-;-;-;-; | 3869 | NA | 0 | 15 | 13 | 11 | 4 | 20 |
| ENSDARG000 | 7;7 | 32168360:321 | 32169157:321 | -;- | 897 | NA | 0 | 2 | 2 | 0 | 0 | 2 |
| ENSDARG000 | 7 | 6367904 | 6368406 | - | 503 | NA | 2 | 2 | 0 | 0 | 0 | 2 |
| ENSDARG000 | 7 | 6352164 | 6352569 | - | 406 | NA | 17 | 6 | 20 | 2 | 16 | 1 |
| ENSDARG000 | 7 | 6459019 | 6459481 | - | 463 | zgc:112234 | 158 | 22 | 60 | 20 | 63 | 31 |
| ENSDARG000 | 7 | 6441160 | 6441865 | - | 706 | NA | 0 | 0 | 1 | 0 | 0 | 0 |
| ENSDARG000 | 7;7;7;7;7;7;7; | 6524041:6526 | 6525776:6526 | -;-;-;-;-;-;-;- | 2897 | NA | 12 | 12 | 4 | 0 | 18 | 8 |
| ENSDARG000 | 7 | 5956937 | 5957597 | + | 661 | hist1h2a7 | 14 | 3 | 2 | 2 | 2 | 0 |
| ENSDARG000 | 7 | 5960491 | 5961503 | + | 1013 | si:dkev-23a13 | 9 | 0 | 2 | 0 | 0 | 8 |
| ENSDARG000 | 7;7;7;7;7;7 | 73719444:737 | 73719535:737 | +;+;+;+;+;+ | 578 | NA | 35 | 98 | 5 | 46 | 69 | 72 |
| ENSDARG000 | 7;7;7;7;7;7;7; | 7783113:7783 | 7783662:7783 | -;-;-;-;-;-;-;-; | 2087 | trmt10b | 56 | 43 | 59 | 89 | 103 | 108 |
| ENSDARG000 | 7;7;7;7;7;7;7; | 73717581:737 | 73718695:737 | -;-;-;-;-;-;-;-; | 2513 | casq1b | 0 | 0 | 2 | 0 | 6 | 0 |

| | | | | | | | | | | | | |
|---|---|---|---|---|---|---|---|---|---|---|---|---|
| ENSDARG000 | 7;7;7;7;7;7;7; | 13382852:134 | 13383601:134 | +;+;+;+;+;+;+; | 6488 | DAGLA | 836 | 550 | 723 | 3296 | 1046 | 2007 |
| ENSDARG000 | 7;7;7 | 52962632:529 | 52962978:529 | -;-;- | 627 | cart3 | 66 | 989 | 3282 | 2226 | 1101 | 4619 |
| ENSDARG000 | 7;7;7;7;7;7;7; | 32369026:323 | 32369259:323 | +;+;+;+;+;+;+; | 6857 | lgr4 | 28 | 0 | 52 | 2 | 40 | 40 |
| ENSDARG000 | 7;7;7;7;7;7;7; | 41491493:414 | 41491996:414 | -;-;-;-;-;-;-; | 4233 | NFX1 | 928 | 864 | 1107 | 814 | 1020 | 844 |
| ENSDARG000 | 7;7;7;7;7;7;7; | 28464794:284 | 28465476:284 | -;-;-;-;-;-;-; | 4483 | dennd2b | 2953 | 3118 | 3857 | 3167 | 4564 | 2205 |
| ENSDARG000 | 7;7;7;7;7;7;7; | 49695904:496 | 49696069:496 | +;+;+;+;+;+;+; | 4674 | NA | 778 | 798 | 415 | 266 | 674 | 472 |
| ENSDARG000 | 7;7;7;7;7;7;7 | 21920662:219 | 21921342:219 | -;-;-;-;-;-;- | 999 | NA | 0 | 0 | 0 | 0 | 14 | 6 |
| ENSDARG000 | 7;7;7 | 7806682:7808 | 7807119:7808 | -;-;- | 697 | cxcl19 | 206 | 19 | 84 | 162 | 153 | 182 |
| ENSDARG000 | 7;7;7;7;7;7;7; | 16196903:161 | 16199174:161 | -;-;-;-;-;-;-; | 3374 | btr05 | 0 | 0 | 0 | 0 | 0 | 0 |
| ENSDARG000 | 7;7;7;7;7;7;7; | 53007450:530 | 53007628:530 | +;+;+;+;+;+;+; | 3106 | NA | 22 | 22 | 14 | 12 | 22 | 1 |
| ENSDARG000 | 7;7;7 | 7819156:7822 | 7820926:7823 | -;-;- | 2130 | cxcl8b.3 | 0 | 4 | 6 | 1 | 4 | 4 |
| ENSDARG000 | 7;7;7;7 | 7839612:7842 | 7839998:7842 | -;-;-;- | 714 | cxcl8b.1 | 32 | 34 | 25 | 53 | 129 | 86 |
| ENSDARG000 | 7;7;7;7;7;7;7; | 18617982:186 | 18620684:186 | -;-;-;-;-;-;-; | 4572 | coro1b | 48 | 93 | 95 | 52 | 144 | 162 |
| ENSDARG000 | 7;7;7;7;7;7;7; | 38685654:386 | 38685674:386 | +;+;+;+;+;+;+; | 5649 | syt13 | 12 | 56 | 714 | 116 | 646 | 92 |
| ENSDARG000 | 7;7;7;7 | 1442059:1442 | 1442202:1442 | +;+;+;+ | 659 | NA | 340 | 177 | 110 | 120 | 315 | 140 |
| ENSDARG000 | 7;7 | 21927586:219 | 21928218:219 | -;- | 975 | NA | 6 | 0 | 4 | 2 | 0 | 0 |
| ENSDARG000 | 7;7;7;7;7;7;7; | 38687268:386 | 38687623:386 | +;+;+;+;+;+;+; | 1929 | NA | 0 | 8 | 0 | 0 | 4 | 0 |
| ENSDARG000 | 7;7;7;7;7;7;7; | 38695719:386 | 38696570:386 | -;-;-;-;-;-;- | 1144 | cd59 | 20 | 2 | 0 | 2 | 62 | 16 |
| ENSDARG000 | 7;7;7;7;7 | 1449999:1451 | 1450191:1451 | +;+;+;+;+ | 948 | NA | 246 | 128 | 168 | 160 | 110 | 108 |
| ENSDARG000 | 7;7;7;7;7;7;7; | 26326462:263 | 26326641:263 | +;+;+;+;+;+;+; | 13728 | zanl | 15 | 2 | 6 | 0 | 10 | 2 |
| ENSDARG000 | 7;7;7 | 38685545:386 | 38687616:386 | -;-;- | 2250 | aplnr2 | 5 | 1 | 4 | 0 | 17 | 28 |
| ENSDARG000 | 7;7;7;7;7;7;7; | 57725708:577 | 57726022:577 | +;+;+;+;+;+;+; | 4019 | camk2d1 | 60 | 36 | 14 | 30 | 252 | 12 |
| ENSDARG000 | 7;7;7;7;7;7;7; | 38703338:387 | 38705150:387 | -;-;-;-;-;-;-; | 3078 | fbxo3 | 315 | 202 | 324 | 187 | 308 | 203 |
| ENSDARG000 | 7;7;7;7;7;7;7; | 8040720:8041 | 8040929:8041 | -;-;-;-;-;-;-; | 1901 | NA | 28 | 25 | 23 | 16 | 81 | 64 |
| ENSDARG000 | 7;7;7;7;7;7;7; | 7906806:7906 | 7909006:7909 | -;-;-;-;-;-;-; | 6997 | NA | 17 | 0 | 0 | 0 | 13 | 4 |
| ENSDARG000 | 7;7;7 | 1457379:1467 | 1457554:1468 | +;+;+ | 522 | emc4 | 10 | 1 | 8 | 2 | 5 | 8 |
| ENSDARG000 | 7;7;7;7;7;7;7; | 7988986:7993 | 7990821:7993 | -;-;-;-;-;-;-; | 5502 | NA | 18 | 2 | 4 | 8 | 4 | 0 |
| ENSDARG000 | 7;7;7;7;7;7;7; | 73801377:738 | 73804851:738 | +;+;+;+;+;+;+; | 8078 | NA | 1387 | 1125 | 1260 | 1120 | 1238 | 1355 |
| ENSDARG000 | 7;7;7;7;7;7;7; | 7906806:7906 | 7909006:7909 | -;-;-;-;-;-;-; | 8915 | NA | 104 | 34 | 51 | 10 | 80 | 50 |
| ENSDARG000 | 7;7;7;7;7;7;7 | 69187585:691 | 69187701:691 | +;+;+;+;+;+;+ | 1800 | marveld3 | 18 | 4 | 0 | 4 | 16 | 6 |
| ENSDARG000 | 7;7;7;7;7;7;7 | 34453185:344 | 34453354:344 | +;+;+;+;+;+;+ | 548 | NA | 0 | 0 | 0 | 0 | 0 | 0 |
| ENSDARG000 | 7;7;7;7;7;7;7; | 13491452:134 | 13491867:134 | +;+;+;+;+;+;+; | 5006 | arih1l | 650 | 688 | 707 | 496 | 906 | 734 |
| ENSDARG000 | 7;7;7;7;7;7;7;7 | 21929461:219 | 21929851:219 | +;+;+;+;+;+;+;+ | 6933 | NA | 20 | 32 | 106 | 0 | 72 | 0 |
| ENSDARG000 | 7;7;7;7;7;7;7; | 24100155:241 | 24100772:241 | -;-;-;-;-;-;-;- | 3788 | AJUBA | 725 | 344 | 483 | 376 | 504 | 445 |
| ENSDARG000 | 7;7;7;7;7 | 73924257:739 | 73924293:739 | +;+;+;+;+ | 1671 | NA | 499 | 516 | 543 | 480 | 408 | 593 |
| ENSDARG000 | 7;7;7;7;7;7;7; | 16205609:162 | 16207389:162 | -;-;-;-;-;-;-; | 3045 | btr06 | 0 | 0 | 0 | 0 | 0 | 0 |
| ENSDARG000 | 7;7;7 | 32475343:324 | 32475471:324 | +;+;+ | 253 | NA | 0 | 0 | 0 | 0 | 0 | 0 |
| ENSDARG000 | 7;7 | 32538063:325 | 32538372:325 | -;- | 570 | NA | 0 | 0 | 0 | 0 | 8 | 0 |
| ENSDARG000 | 7;7;7;7;7;7;7; | 29538628:295 | 29541414:295 | -;-;-;-;-;-;-; | 4487 | rorab | 1643 | 1574 | 1632 | 881 | 1707 | 2378 |
| ENSDARG000 | 7;7 | 49715750:497 | 49716407:497 | +;+ | 1273 | ascl1b | 0 | 0 | 0 | 0 | 0 | 0 |
| ENSDARG000 | 7;7;7;7;7;7;7; | 18657008:186 | 18659042:186 | -;-;-;-;-;-;-; | 4049 | NA | 108 | 4 | 0 | 14 | 12 | 2 |
| ENSDARG000 | 7;7;7 | 49797528:497 | 49797570:498 | -;-;- | 2064 | FJX1 | 1666 | 935 | 864 | 460 | 1140 | 712 |
| ENSDARG000 | 7;7;7;7;7;7;7; | 4425348:4425 | 4425486:4425 | -;-;-;-;-;-;-; | 3102 | slc12a10.1 | 0 | 0 | 18 | 4 | 26 | 2 |
| ENSDARG000 | 7;7;7;7;7;7;7; | 49800478:498 | 49800681:498 | +;+;+;+;+;+;+; | 4224 | PAMR1 | 116 | 59 | 158 | 136 | 164 | 138 |
| ENSDARG000 | 7;7;7 | 32595981:325 | 32598856:325 | -;-;- | 3353 | KCNA4 | 10 | 2 | 4 | 0 | 12 | 0 |
| ENSDARG000 | 7;7;7;7;7;7;7; | 21940857:219 | 21945593:219 | -;-;-;-;-;-;-; | 6392 | nlgn2a | 34 | 6 | 98 | 31 | 91 | 20 |
| ENSDARG000 | 7;7;7;7;7;7;7; | 25015151:250 | 25015585:250 | +;+;+;+;+;+;+; | 4051 | b3gat3 | 296 | 322 | 418 | 124 | 357 | 252 |
| ENSDARG000 | 7;7;7;7;7;7;7; | 65177132:651 | 65178312:651 | -;-;-;-;-;-;-; | 8619 | NA | 14 | 20 | 26 | 6 | 34 | 28 |
| ENSDARG000 | 7;7;7;7;7 | 32605705:326 | 32605730:326 | +;+;+;+;+ | 1907 | fshb | 0 | 0 | 0 | 0 | 0 | 0 |
| ENSDARG000 | 7;7 | 73961361:739 | 73961674:739 | +;+ | 667 | NA | 9 | 12 | 10 | 2 | 2 | 2 |
| ENSDARG000 | 7;7;7 | 40638210:406 | 40638840:406 | +;+;+ | 1469 | mnx1 | 0 | 0 | 0 | 0 | 0 | 0 |
| ENSDARG000 | 7;7;7;7;7;7;7; | 53059299:530 | 53060738:530 | -;-;-;-;-;-;-; | 5085 | cdh1 | 12379 | 10089 | 8643 | 13920 | 8068 | 10700 |
| ENSDARG000 | 7;7;7;7;7;7;7; | 34472363:344 | 34472765:344 | -;-;-;-;-;-;-;- | 1232 | NA | 0 | 2 | 0 | 0 | 2 | 0 |
| ENSDARG000 | 7;7;7;7;7;7;7; | 16231363:162 | 16232579:162 | -;-;-;-;-;-;- | 2422 | NA | 0 | 0 | 0 | 0 | 4 | 0 |
| ENSDARG000 | 7 | 51904909 | 51905343 | - | 435 | NA | 0 | 0 | 0 | 0 | 2 | 0 |
| ENSDARG000 | 7;7 | 51922694:519 | 51923227:519 | -;- | 1125 | NA | 0 | 0 | 0 | 0 | 1 | 0 |
| ENSDARG000 | 7;7;7;7;7;7;7; | 1473929:1481 | 1474227:1481 | +;+;+;+;+;+;+; | 2124 | lpcat4 | 27 | 34 | 24 | 50 | 28 | 30 |
| ENSDARG000 | 7;7;7;7;7;7;7; | 49862837:498 | 49863114:498 | +;+;+;+;+;+;+; | 3763 | slc1a2a | 21 | 0 | 0 | 0 | 4 | 0 |
| ENSDARG000 | 7;7;7;7;7;7;7; | 13527396:135 | 13528359:135 | +;+;+;+;+;+;+; | 12173 | PLEKHO2 | 1134 | 915 | 1030 | 934 | 1332 | 1308 |
| ENSDARG000 | 7;7;7;7;7;7;7 | 44802353:448 | 44802586:448 | +;+;+;+;+;+;+ | 1873 | ca7 | 0 | 0 | 0 | 0 | 0 | 0 |
| ENSDARG000 | 7;7;7;7;7;7;7 | 34487833:344 | 34488167:344 | +;+;+;+;+;+;+ | 1748 | cln6a | 960 | 745 | 937 | 742 | 1008 | 888 |
| ENSDARG000 | 7;7;7;7;7;7;7; | 37898618:378 | 37898895:378 | -;-;-;-;-;-;-; | 2607 | heatr3 | 312 | 288 | 628 | 355 | 592 | 383 |
| ENSDARG000 | 7;7;7;7;7;7;7; | 18702534:187 | 18705536:187 | -;-;-;-;-;-;-; | 4517 | trmt10a | 132 | 147 | 140 | 104 | 130 | 100 |
| ENSDARG000 | 7;7;7;7;7 | 32624684:326 | 32625069:326 | -;-;-;-;- | 949 | arl14ep | 384 | 529 | 395 | 258 | 372 | 419 |
| ENSDARG000 | 7;7;7;7;7;7;7; | 8267985:8270 | 8271154:8271 | -;-;-;-;-;-;-; | 6476 | f13a1b | 257 | 502 | 532 | 679 | 1638 | 1052 |
| ENSDARG000 | 7;7;7;7;7;7;7; | 40644769:406 | 40645220:406 | -;-;-;-;-;-;-; | 3279 | nom1 | 476 | 439 | 673 | 95 | 466 | 287 |
| ENSDARG000 | 7;7 | 16348739:163 | 16349149:163 | -;- | 601 | NA | 6 | 1 | 10 | 10 | 4 | 2 |
| ENSDARG000 | 7;7;7;7;7;7;7; | 16348835:163 | 16349089:163 | +;+;+;+;+;+;+; | 3126 | mpped2a | 287 | 317 | 423 | 217 | 551 | 316 |
| ENSDARG000 | 7;7;7 | 8324506:8328 | 8324675:8329 | +;+;+ | 7434 | NA | 74 | 72 | 179 | 213 | 165 | 171 |
| ENSDARG000 | 7;7;7;7;7;7;7; | 24114694:241 | 24114919:241 | +;+;+;+;+;+;+; | 695 | mrpl52 | 64 | 88 | 119 | 44 | 89 | 68 |
| ENSDARG000 | 7;7;7;7;7;7;7; | 29628398:296 | 29631307:296 | -;-;-;-;-;-;-; | 14988 | VPS13C | 22088 | 9735 | 15985 | 16385 | 28019 | 14726 |
| ENSDARG000 | 7;7;7;7;7;7;7; | 41517173:415 | 41520033:415 | -;-;-;-;-;-;-; | 4614 | plxdc2b | 1692 | 785 | 613 | 3107 | 1788 | 4407 |
| ENSDARG000 | 7;7;7;7;7;7;7; | 25033924:250 | 25034009:250 | +;+;+;+;+;+;+; | 1594 | sb:cb1058 | 4 | 12 | 22 | 8 | 48 | 36 |
| ENSDARG000 | 7;7;7;7;7;7;7; | 32629674:326 | 32629851:326 | +;+;+;+;+;+;+; | 1958 | si:ch73-314g1 | 134 | 80 | 135 | 69 | 140 | 125 |
| ENSDARG000 | 7;7;7 | 22313533:223 | 22313648:223 | +;+;+ | 4071 | tmem102 | 345 | 338 | 388 | 227 | 365 | 313 |
| ENSDARG000 | 7;7;7;7;7;7;7; | 32644673:326 | 32646133:326 | -;-;-;-;-;-;-; | 2475 | spi1b | 150 | 144 | 299 | 149 | 405 | 413 |
| ENSDARG000 | 7;7;7;7 | 22293894:222 | 22294120:222 | +;+;+;+ | 847 | tmem256 | 147 | 169 | 140 | 115 | 132 | 149 |
| ENSDARG000 | 7;7;7;7;7;7;7; | 53199763:532 | 53199865:532 | +;+;+;+;+;+;+; | 2488 | NA | 7 | 8 | 0 | 0 | 4 | 2 |
| ENSDARG000 | 7;7;7;7;7;7;7; | 34496113:344 | 34496467:344 | -;-;-;-;-;-;-; | 1540 | tex9 | 10 | 16 | 5 | 2 | 26 | 24 |
| ENSDARG000 | 7;7;7;7;7;7;7; | 53152108:531 | 53152239:531 | +;+;+;+;+;+;+; | 8675 | NA | 82 | 13 | 29 | 22 | 72 | 80 |
| ENSDARG000 | 7;7;7;7;7;7;7; | 34492744:344 | 34492809:344 | +;+;+;+;+;+;+; | 663 | calml4a | 2 | 0 | 4 | 0 | 0 | 0 |
| ENSDARG000 | 7;7;7;7;7;7;7; | 53170128:531 | 53170256:531 | +;+;+;+;+;+;+; | 2542 | NA | 51 | 2 | 10 | 3 | 10 | 12 |
| ENSDARG000 | 7;7;7;7;7;7;7; | 65240227:652 | 65240400:652 | +;+;+;+;+;+;+; | 2563 | bco1l | 21 | 19 | 10 | 42 | 20 | 32 |
| ENSDARG000 | 7;7;7;7;7;7;7; | 24124289:241 | 24126057:241 | -;-;-;-;-;-;-; | 3509 | mmp14a | 388 | 64 | 206 | 88 | 230 | 140 |
| ENSDARG000 | 7;7;7;7;7;7;7; | 26366451:263 | 26369646:263 | -;-;-;-;-;-;-; | 5070 | trappc14 | 439 | 348 | 456 | 262 | 326 | 418 |
| ENSDARG000 | 7;7;7;7;7;7;7; | 13582256:135 | 13582434:135 | +;+;+;+;+;+;+; | 3380 | ankdd1a | 100 | 133 | 70 | 50 | 103 | 64 |
| ENSDARG000 | 7;7;7 | 1498482:1501 | 1498742:1502 | -;-;- | 1126 | NA | 0 | 0 | 0 | 0 | 0 | 0 |
| ENSDARG000 | 7;7;7 | 4474880:4477 | 4474965:4478 | +;+;+ | 638 | NA | 0 | 0 | 2 | 0 | 4 | 0 |
| ENSDARG000 | 7;7;7;7;7;7;7; | 16509201:165 | 16509347:165 | +;+;+;+;+;+;+; | 3093 | zdhhc13 | 530 | 600 | 720 | 448 | 729 | 480 |
| ENSDARG000 | 7;7;7;7;7;7;7; | 51950614:519 | 51950954:519 | -;-;-;-;-;-;-; | 1741 | cyp2x10.2 | 0 | 0 | 0 | 0 | 0 | 0 |
| ENSDARG000 | 7;7;7;7;7;7;7; | 16548219:165 | 16549076:165 | -;-;-;-;-;-;-; | 1456 | csrp3 | 5 | 9 | 7 | 13 | 3 | 17 |
| ENSDARG000 | 7;7;7;7;7;7;7; | 22330466:223 | 22330662:223 | +;+;+;+;+;+;+; | 1521 | fgf11a | 0 | 0 | 0 | 0 | 0 | 0 |
| ENSDARG000 | 7;7 | 1505507:1508 | 1505677:1508 | +;+ | 591 | nop10 | 325 | 386 | 491 | 273 | 478 | 364 |
| ENSDARG000 | 7;7;7 | 1521834:1522 | 1522064:1522 | +;+;+ | 2208 | NA | 1453 | 717 | 605 | 727 | 1206 | 684 |
| ENSDARG000 | 7;7;7 | 44895614:448 | 44895802:448 | +;+;+ | 983 | NA | 0 | 2 | 0 | 0 | 2 | 0 |
| ENSDARG000 | 7;7;7;7;7;7;7; | 44857271:448 | 44860837:448 | -;-;-;-;-;-;-; | 5814 | NA | 0 | 0 | 6 | 0 | 10 | 2 |
| ENSDARG000 | 7;7;7 | 4483378:4483 | 4483405:4484 | +;+;+ | 616 | NA | 0 | 0 | 5 | 0 | 0 | 0 |
| ENSDARG000 | 7;7;7;7 | 8358547:8359 | 8358578:8359 | +;+;+;+ | 1710 | jac7 | 4 | 6 | 40 | 80 | 85 | 208 |
| ENSDARG000 | 7;7;7;7;7;7;7; | 34506795:345 | 34507232:345 | +;+;+;+;+;+;+; | 8031 | RFX7 | 114 | 82 | 240 | 170 | 274 | 74 |
| ENSDARG000 | 7;7;7;7;7;7;7; | 1530024:1534 | 1530133:1534 | +;+;+;+;+;+;+; | 1652 | tox4b | 232 | 132 | 293 | 110 | 190 | 270 |
| ENSDARG000 | 7;7;7;7 | 4493556:4493 | 4493592:4494 | +;+;+;+ | 769 | NA | 0 | 0 | 0 | 0 | 0 | 0 |
| ENSDARG000 | 7;7;7;7;7;7 | 38036841:380 | 38037048:380 | -;-;-;-;-;- | 900 | NA | 48 | 18 | 8 | 8 | 12 | 12 |
| ENSDARG000 | 7;7;7;7;7;7 | 38089650:380 | 38089793:380 | +;+;+;+;+;+ | 2621 | cebpg | 391 | 579 | 1506 | 1450 | 581 | 572 |
| ENSDARG000 | 7;7;7 | 18721547:187 | 18721910:187 | -;-;- | 5030 | C7H4orf54 | 0 | 0 | 0 | 0 | 0 | 0 |
| ENSDARG000 | 7;7;7;7;7 | 8367627:8367 | 8367746:8368 | +;+;+;+;+ | 1648 | zgc:172143 | 4 | 0 | 4 | 0 | 10 | 0 |
| ENSDARG000 | 7 | 38084986 | 38087865 | - | 2880 | cebpa | 172 | 216 | 112 | 89 | 404 | 178 |
| ENSDARG000 | 7;7;7;7;7;7;7; | 18878973:188 | 18879044:188 | +;+;+;+;+;+;+; | 5974 | NA | 334 | 418 | 384 | 166 | 352 | 268 |
| ENSDARG000 | 7;7;7;7;7;7;7; | 24153070:241 | 24153127:241 | +;+;+;+;+;+;+; | 3542 | NA | 3414 | 2623 | 2475 | 2236 | 3069 | 2626 |
| ENSDARG000 | 7;7;7;7;7;7;7; | 49882750:498 | 49883414:498 | -;-;-;-;-;-;-; | 1970 | NA | 248 | 14 | 2 | 16 | 16 | 6 |
| ENSDARG000 | 7;7;7;7;7;7;7; | 28556166:285 | 28560595:285 | -;-;-;-;-;-;-; | 5208 | tmem9b | 2369 | 2194 | 2321 | 2784 | 1959 | 2555 |
| ENSDARG000 | 7;7;7;7;7;7;7; | 18779555:187 | 18780380:187 | -;-;-;-;-;-;-; | 3239 | mllt3 | 12 | 68 | 74 | 42 | 63 | 52 |
| ENSDARG000 | 7;7;7;7 | 26393751:264 | 26398908:264 | -;-;-;- | 5823 | gal3st4 | 3434 | 1993 | 3096 | 2149 | 2625 | 3849 |
| ENSDARG000 | 7;7 | 8373025:8374 | 8374570:8374 | -;- | 1651 | aep1 | 6 | 2 | 0 | 6 | 10 | 0 |
| ENSDARG000 | 7;7;7 | 8379833:8380 | 8381092:8381 | -;-;- | 1322 | jac10 | 10 | 4 | 0 | 0 | 20 | 0 |
| ENSDARG000 | 7;7;7;7;7;7;7; | 34549198:345 | 34549541:345 | +;+;+;+;+;+;+; | 6027 | FHOD1 | 66 | 154 | 42 | 282 | 646 | 50 |
| ENSDARG000 | 7;7 | 8406686:8407 | 8407670:8408 | -;- | 1108 | jac11 | 4 | 2 | 6 | 6 | 16 | 8 |
| ENSDARG000 | 7;7;7;7;7;7;7; | 30561588:305 | 30563296:305 | -;-;-;-;-;-;-; | 5144 | rnf111 | 1037 | 802 | 1263 | 829 | 1306 | 926 |
| ENSDARG000 | 7;7;7 | 8415494:8415 | 8416767:8416 | -;-;- | 1337 | NA | 96 | 76 | 123 | 62 | 203 | 76 |
| ENSDARG000 | 7;7;7;7;7;7;7; | 13625929:136 | 13626495:136 | -;-;-;-;-;-;-; | 1407 | spg21 | 1089 | 944 | 1035 | 851 | 1203 | 1116 |
| ENSDARG000 | 7;7;7;7 | 29731254:297 | 29731400:297 | +;+;+;+ | 726 | NA | 0 | 0 | 0 | 0 | 12 | 0 |
| ENSDARG000 | 7 | 8428954 | 8429919 | + | 966 | NA | 0 | 2 | 2 | 0 | 0 | 0 |
| ENSDARG000 | 7;7;7;7;7;7;7; | 57886148:578 | 57887534:578 | -;-;-;-;-;-;-; | 14058 | ank2b | 4411 | 2137 | 3536 | 2855 | 5688 | 7330 |
| ENSDARG000 | 7;7 | 8437206:8438 | 8438420:8438 | -;- | 1347 | si:dkeyp-32g1 | 254 | 20 | 4 | 54 | 42 | 52 |
| ENSDARG000 | 7;7;7;7;7;7;7; | 1550966:1550 | 1551709:1551 | +;+;+;+;+;+;+; | 9516 | supt16h | 1571 | 2760 | 3971 | 1756 | 2622 | 2181 |
| ENSDARG000 | 7;7;7;7;7;7;7; | 22543963:225 | 22544117:225 | +;+;+;+;+;+;+; | 5002 | chrnb1 | 0 | 0 | 0 | 0 | 0 | 0 |
| ENSDARG000 | 7;7;7 | 20239915:202 | 20241346:202 | -;-;- | 1499 | si:ch73-335l2 | 1689 | 557 | 1314 | 1750 | 538 | 1046 |
| ENSDARG000 | 7;7;7;7;7 | 4499264:4499 | 4499324:4499 | +;+;+;+;+ | 1166 | si:dkev-83f18. | 0 | 0 | 0 | 0 | 0 | 0 |
| ENSDARG000 | 7;7;7;7;7;7;7; | 41632928:416 | 41636012:416 | -;-;-;-;-;-;-; | 8462 | malrd1 | 10 | 6 | 7 | 10 | 8 | 14 |
| ENSDARG000 | 7;7;7;7;7;7;7; | 26429941:264 | 26432740:264 | -;-;-;-;-;-;-;- | 3132 | her8a | 10 | 6 | 160 | 18 | 8 | 108 |
| ENSDARG000 | 7;7;7;7;7 | 44960188:449 | 44961202:449 | -;-;-;-;- | 1747 | rrad | 56 | 60 | 70 | 87 | 45 | 76 |
| ENSDARG000 | 7;7;7;7;7;7;7; | 39444843:394 | 39444908:394 | +;+;+;+;+;+;+; | 2791 | tnnt3b | 28 | 8 | 8 | 14 | 26 | 32 |
| ENSDARG000 | 7;7;7;7;7;7;7; | 65261354:652 | 65261669:652 | +;+;+;+;+;+;+; | 2183 | bco1 | 2381 | 638 | 1207 | 570 | 1058 | 104 |
| ENSDARG000 | 7;7;7;7;7;7;7; | 49901483:499 | 49901655:499 | +;+;+;+;+;+;+; | 873 | apip | 655 | 557 | 698 | 624 | 588 | 664 |
| ENSDARG000 | 7;7 | 38104703:381 | 38104781:381 | +;+ | 556 | NA | 0 | 0 | 0 | 0 | 0 | 0 |
| ENSDARG000 | 7;7;7;7;7;7;7; | 26451869:264 | 26452972:264 | -;-;-;-;-;-;- | 1997 | tm4sf21b | 0 | 0 | 2 | 0 | 0 | 3 |
| ENSDARG000 | 7;7;7;7;7;7;7; | 30626378:306 | 30626587:306 | +;+;+;+;+;+;+; | 1688 | ccnb2 | 16 | 144 | 131 | 36 | 84 | 80 |
| ENSDARG000 | 7 | 26445742 | 26446726 | - | 985 | NA | 1 | 2 | 8 | 4 | 0 | 4 |
| ENSDARG000 | 7;7;7;7;7;7;7; | 22585447:225 | 22585727:225 | +;+;+;+;+;+;+; | 3174 | chrnb1l | 0 | 2 | 0 | 0 | 0 | 0 |
| ENSDARG000 | 7;7;7;7;7;7;7; | 26446020:264 | 26446777:264 | -;-;-;-;-;-;- | 2311 | NA | 0 | 3 | 19 | 9 | 23 | 25 |
| ENSDARG000 | 7;7;7;7;7;7;7; | 4524059:4582 | 4524430:4583 | +;+;+;+;+;+;+; | 2378 | NA | 0 | 0 | 0 | 0 | 0 | 0 |
| ENSDARG000 | 7;7;7;7 | 29745906:297 | 29746030:297 | +;+;+;+ | 752 | NA | 0 | 0 | 0 | 0 | 0 | 3 |
| ENSDARG000 | 7;7;7;7 | 29757774:297 | 29757900:297 | +;+;+;+ | 758 | NA | 0 | 0 | 0 | 0 | 0 | 0 |
| ENSDARG000 | 7;7;7;7 | 4552513:4552 | 4552549:4552 | +;+;+;+ | 1091 | NA | 8 | 0 | 0 | 0 | 0 | 0 |
| ENSDARG000 | 7;7;7;7;7;7;7; | 4538334:4538 | 4538401:4538 | +;+;+;+;+;+;+; | 1204 | si:dkev-83f18. | 0 | 0 | 0 | 0 | 0 | 0 |
| ENSDARG000 | 7;7;7;7;7;7;7; | 24165371:241 | 24165598:241 | +;+;+;+;+;+;+; | 2324 | NA | 0 | 0 | 0 | 0 | 0 | 2 |
| ENSDARG000 | 7;7;7;7;7;7;7; | 4523051:4523 | 4523107:4523 | +;+;+;+;+;+;+; | 1230 | NA | 0 | 0 | 0 | 0 | 6 | 0 |
| ENSDARG000 | 7;7;7;7;7;7;7; | 24172987:241 | 24173795:241 | -;-;-;-;-;-;-; | 2514 | zgc:153151 | 22 | 14 | 30 | 2 | 33 | 10 |
| ENSDARG000 | 7;7;7;7;7 | 29772789:297 | 29772866:297 | +;+;+;+;+ | 762 | NA | 0 | 3 | 0 | 0 | 0 | 0 |
| ENSDARG000 | 7;7 | 20247871:202 | 20247983:202 | +;+ | 767 | si:ch73-335l2 | 0 | 1 | 0 | 2 | 1 | 2 |
| ENSDARG000 | 7;7;7;7;7 | 4572727:4572 | 4572796:4572 | +;+;+;+;+ | 1210 | si:dkev-83f18. | 0 | 0 | 0 | 0 | 0 | 0 |
| ENSDARG000 | 7;7;7 | 20247871:202 | 20247983:202 | +;+;+ | 1182 | NA | 0 | 0 | 0 | 0 | 0 | 0 |
| ENSDARG000 | 7;7 | 38144619:381 | 38144648:381 | +;+ | 660 | NA | 0 | 0 | 0 | 0 | 0 | 0 |
| ENSDARG000 | 7;7;7;7 | 29792766:297 | 29792911:297 | +;+;+;+ | 722 | NA | 2 | 1 | 2 | 0 | 0 | 3 |
| ENSDARG000 | 7;7;7;7 | 29806350:298 | 29806401:298 | +;+;+;+ | 695 | NA | 0 | 0 | 0 | 0 | 1 | 0 |
| ENSDARG000 | 7;7;7;7;7;7;7; | 34620418:346 | 34620876:346 | +;+;+;+;+;+;+; | 4528 | slc9a5 | 0 | 2 | 0 | 0 | 2 | 0 |
| ENSDARG000 | 7;7 | 8456999:8457 | 8457130:8458 | +;+ | 1375 | jac4 | 258 | 10 | 2 | 40 | 48 | 40 |
| ENSDARG000 | 7;7;7;7;7;7;7; | 41713920:417 | 41716144:417 | -;-;-;-;-;-;-; | 3305 | arl8 | 279 | 362 | 518 | 451 | 601 | 446 |
| ENSDARG000 | 7;7;7 | 50053339:501 | 50053392:501 | +;+;+ | 4447 | LRRC4C | 0 | 0 | 0 | 4 | 2 | 0 |
| ENSDARG000 | 7;7;7;7;7;7;7; | 20260172:202 | 20260688:202 | +;+;+;+;+;+;+; | 4848 | dvl2 | 1076 | 773 | 1646 | 1292 | 1023 | 1028 |
| ENSDARG000 | 7;7;7;7;7;7;7; | 44965773:449 | 44967390:449 | -;-;-;-;-;-;-; | 2241 | ciao2b | 1043 | 949 | 1130 | 709 | 849 | 1051 |
| ENSDARG000 | 7;7;7;7 | 29822214:298 | 29822291:298 | +;+;+;+ | 821 | NA | 0 | 0 | 0 | 0 | 1 | 0 |
| ENSDARG000 | 7;7;7;7 | 29836386:298 | 29836463:298 | +;+;+;+ | 720 | NA | 5 | 0 | 0 | 0 | 4 | 0 |

| | | | | | | | | | | | | |
|---|---|---|---|---|---|---|---|---|---|---|---|---|
| ENSDARG000 | 7;7;7;7;7;7;7; | 38158604:381 | 38159120:381 | -;-;-;-;-;-;-;-; | 4779 | ABCC12 | 64 | 72 | 164 | 150 | 50 | 38 |
| ENSDARG000 | 7;7;7;7;7;7;7; | 50228015:502 | 50229332:502 | -;-;-;-;-;-;-;-; | 2297 | hsd17b12b | 1219 | 1363 | 1564 | 744 | 2167 | 1393 |
| ENSDARG000 | 7 | 8469459 | 8470860 | - | 1402 | NA | 0 | 0 | 26 | 0 | 0 | 0 |
| ENSDARG000 | 7;7;7;7;7;7;7; | 53215678:532 | 53215948:532 | +;+;+;+;+;+;+; | 2492 | NA | 4 | 3 | 4 | 0 | 10 | 4 |
| ENSDARG000 | 7;7;7;7;7;7;7; | 32693890:326 | 32694044:326 | +;+;+;+;+;+;+; | 3061 | slc39a13 | 858 | 1334 | 2519 | 960 | 2531 | 946 |
| ENSDARG000 | 7;7;7;7;7 | 38744924:387 | 38745010:387 | +;+;+;+;+ | 2474 | znf408 | 46 | 33 | 94 | 32 | 14 | 66 |
| ENSDARG000 | 7;7;7;7;7;7;7; | 52093291:520 | 52093631:520 | -;-;-;-;-;-;-;-; | 1715 | NA | 0 | 0 | 0 | 0 | 0 | 0 |
| ENSDARG000 | 7;7;7;7;7;7;7; | 16576246:165 | 16576994:165 | -;-;-;-;-;-;-;-; | 3798 | e2f8 | 186 | 591 | 545 | 318 | 346 | 224 |
| ENSDARG000 | 7 | 18989241 | 18990075 | + | 835 | zgc:194252 | 10 | 17 | 10 | 35 | 27 | 28 |
| ENSDARG000 | 7;7 | 8489333:849( | 8490566:849( | -;- | 1300 | jac6 | 0 | 1 | 147 | 0 | 11 | 4 |
| ENSDARG000 | 7;7;7;7;7;7;7; | 25042078:250 | 25042105:250 | +;+;+;+;+;+;+; | 1962 | si:dkev-23i12. | 206 | 257 | 714 | 332 | 565 | 438 |
| ENSDARG000 | 7;7;7;7;7;7;7 | 1630275:163 | 1630387:163 | +;+;+;+;+;+;+ | 883 | NA | 0 | 2 | 0 | 0 | 0 | 0 |
| ENSDARG000 | 7;7;7;7;7;7;7; | 32746914:327 | 32750934:327 | -;-;-;-;-;-;- | 5053 | GAS2 | 0 | 0 | 0 | 0 | 0 | 0 |
| ENSDARG000 | 7;7 | 8502801:850 | 8504035:850 | -;- | 1301 | loc564660 | 6 | 2 | 161 | 7 | 11 | 2 |
| ENSDARG000 | 7;7;7;7 | 32722227:327 | 32722861:327 | +;+;+;+ | 8311 | NA | 2 | 0 | 4 | 0 | 0 | 0 |
| ENSDARG000 | 7;7;7;7;7;7;7 | 18993828:189 | 18994091:189 | -;-;-;-;-;-;- | 947 | hacd4 | 10 | 82 | 74 | 82 | 29 | 157 |
| ENSDARG000 | 7;7;7;7 | 1812480:1817 | 1812759:1817 | +;+;+;+ | 757 | NA | 0 | 0 | 0 | 0 | 0 | 0 |
| ENSDARG000 | 7;7 | 8513993:8515 | 8515230:8515 | -;- | 1304 | jac9 | 2 | 0 | 155 | 0 | 15 | 6 |
| ENSDARG000 | 7;7;7;7;7;7;7; | 53254234:532 | 53254378:532 | +;+;+;+;+;+;+; | 1946 | trip4 | 229 | 247 | 262 | 146 | 333 | 130 |
| ENSDARG000 | 7;7;7;7;7;7;7; | 28569266:285 | 28570214:285 | -;-;-;-;-;-;-;-; | 3969 | scube2 | 14 | 16 | 26 | 32 | 12 | 49 |
| ENSDARG000 | 7;7;7;7;7;7;7; | 1938278:193 | 1938334:193 | -;-;-;-;-;-;-;-; | 1287 | NA | 0 | 0 | 0 | 0 | 0 | 0 |
| ENSDARG000 | 7;7 | 50277537:502 | 50278263:502 | -;- | 802 | NA | 0 | 0 | 0 | 0 | 0 | 0 |
| ENSDARG000 | 7;7;7;7 | 26459360:264 | 26461072:264 | -;-;-;- | 2288 | MBLAC1 | 18 | 38 | 59 | 30 | 44 | 44 |
| ENSDARG000 | 7;7;7;7;7;7;7; | 26466826:264 | 26466979:264 | +;+;+;+;+;+;+; | 2433 | mpdu1b | 594 | 798 | 1044 | 527 | 1239 | 706 |
| ENSDARG000 | 7;7;7 | 8532327:853 | 8532413:853 | -;-;- | 359 | NA | 0 | 0 | 0 | 0 | 0 | 0 |
| ENSDARG000 | 7;7;7;7;7;7 | 22616212:226 | 22616684:226 | +;+;+;+;+;+ | 2015 | cldn7a | 36 | 4 | 76 | 0 | 50 | 2 |
| ENSDARG000 | 7;7;7 | 1996444:200 | 1996702:200 | -;-;- | 468 | NA | 0 | 0 | 0 | 0 | 0 | 0 |
| ENSDARG000 | 7;7;7;7;7;7;7; | 1980829:198( | 1980992:198( | -;-;-;-;-;-;-;-; | 1375 | zgc:171424 | 0 | 0 | 0 | 0 | 0 | 0 |
| ENSDARG000 | 7;7;7;7;7;7;7; | 50295715:502 | 50297055:502 | -;-;-;-;-;-;-;-; | 5211 | ttc17 | 592 | 501 | 656 | 328 | 518 | 554 |
| ENSDARG000 | 7;7;7;7;7;7;7; | 24187581:241 | 24188147:241 | -;-;-;-;-;-;-;-; | 1853 | si:ch211-216 | 22 | 16 | 8 | 10 | 9 | 2 |
| ENSDARG000 | 7;7 | 8543317:854 | 8543363:854 | +;+ | 1284 | jac8 | 0 | 6 | 0 | 0 | 0 | 0 |
| ENSDARG000 | 7;7 | 8570332:857 | 8571574:857 | -;- | 1289 | NA | 6 | 105 | 173 | 58 | 337 | 112 |
| ENSDARG000 | 7;7;7 | 40866815:408 | 40866926:408 | -;-;- | 927 | NA | 0 | 0 | 0 | 0 | 0 | 6 |
| ENSDARG000 | 7;7;7 | 40884012:408 | 40884630:408 | +;+;+ | 4551 | shha | 0 | 0 | 0 | 0 | 0 | 0 |
| ENSDARG000 | 7;7 | 8598995:860 | 8599944:860 | -;- | 1016 | NA | 27 | 763 | 832 | 398 | 2009 | 939 |
| ENSDARG000 | 7;7 | 8623945:862 | 8624040:862 | +;+ | 631 | NA | 0 | 0 | 0 | 0 | 0 | 0 |
| ENSDARG000 | 7;7;7;7;7 | 45040526:450 | 45040759:450 | -;-;-;-;- | 1603 | vstm2b | 0 | 0 | 0 | 0 | 0 | 0 |
| ENSDARG000 | 7;7;7 | 38756326:387 | 38756827:387 | -;-;- | 718 | NA | 0 | 0 | 0 | 0 | 0 | 0 |
| ENSDARG000 | 7;7;7;7;7;7;7; | 22629902:226 | 22630710:226 | -;-;-;-;-;-;-;-; | 1751 | NA | 116 | 140 | 275 | 170 | 209 | 187 |
| ENSDARG000 | 7;7;7;7;7;7;7; | 38762043:387 | 38762138:387 | +;+;+;+;+;+;+; | 3972 | arhgap1 | 2238 | 1383 | 2085 | 1962 | 2394 | 1791 |
| ENSDARG000 | 7;7;7;7;7;7;7; | 65309738:653 | 65309975:653 | +;+;+;+;+;+;+; | 1584 | vat1l | 0 | 0 | 2 | 10 | 0 | 0 |
| ENSDARG000 | 7;7;7;7;7;7;7; | 8670398:867( | 8670496:867( | +;+;+;+;+;+;+; | 966 | NA | 0 | 2 | 0 | 0 | 7 | 0 |
| ENSDARG000 | 7;7 | 22632049:226 | 22632231:226 | +;+ | 237 | NA | 0 | 0 | 0 | 0 | 0 | 0 |
| ENSDARG000 | 7;7;7;7 | 22637515:226 | 22637712:226 | +;+;+;+ | 598 | si:dkev-112a7 | 0 | 0 | 0 | 0 | 0 | 0 |
| ENSDARG000 | 7;7;7;7;7;7;7; | 65296901:652 | 65297096:652 | +;+;+;+;+;+;+; | 1121 | bola3 | 58 | 84 | 107 | 62 | 47 | 84 |
| ENSDARG000 | 7;7;7;7;7;7;7; | 38750871:387 | 38751021:387 | +;+;+;+;+;+;+; | 3136 | f2 | 4 | 2 | 39 | 18 | 9 | 4 |
| ENSDARG000 | 7;7;7;7;7 | 41748693:417 | 41749490:417 | +;+;+;+;+ | 2925 | hrh3 | 4 | 8 | 2 | 6 | 0 | 2 |
| ENSDARG000 | 7;7;7;7;7;7;7; | 53641257:536 | 53641875:536 | -;-;-;-;-;-;-;-; | 1774 | chrna7a | 6 | 23 | 35 | 0 | 18 | 0 |
| ENSDARG000 | 7;7 | 45400716:454 | 45401008:454 | +;+ | 3965 | uqcrfs1 | 2505 | 3152 | 2922 | 1655 | 2705 | 1868 |
| ENSDARG000 | 7;7;7;7;7;7;7; | 34687969:346 | 34688695:346 | +;+;+;+;+;+;+; | 9634 | NA | 501 | 805 | 1117 | 495 | 1668 | 296 |
| ENSDARG000 | 7;7;7;7;7;7;7; | 24196575:241 | 24198635:241 | -;-;-;-;-;-;-;-; | 3017 | gmpr2 | 1247 | 1486 | 1447 | 997 | 1496 | 1132 |
| ENSDARG000 | 7;7 | 24196542:242 | 24196667:242 | +;+ | 376 | NA | 0 | 0 | 0 | 0 | 0 | 0 |
| ENSDARG000 | 7;7;7;7;7;7;7; | 30632229:306 | 30633263:306 | -;-;-;-;-;-;-;-; | 5154 | MYO1E | 62 | 40 | 113 | 89 | 155 | 38 |
| ENSDARG000 | 7;7;7;7;7;7;7; | 69449814:694 | 69450447:694 | +;+;+;+;+;+;+; | 2546 | ctdnep1b | 164 | 119 | 206 | 132 | 216 | 140 |
| ENSDARG000 | 7 | 19007840 | 19008579 | + | 740 | NA | 0 | 0 | 0 | 0 | 0 | 0 |
| ENSDARG000 | 7;7;7;7 | 29850789:298 | 29850914:298 | +;+;+;+ | 694 | NA | 0 | 0 | 0 | 0 | 0 | 2 |
| ENSDARG000 | 7;7;7;7;7;7;7; | 20298333:202 | 20298603:202 | +;+;+;+;+;+;+; | 3604 | acadvl | 2110 | 1728 | 2179 | 1561 | 1998 | 1550 |
| ENSDARG000 | 7;7;7;7;7;7;7; | 53754653:537 | 53755126:537 | +;+;+;+;+;+;+; | 6254 | neo1a | 1331 | 555 | 1250 | 458 | 1122 | 752 |
| ENSDARG000 | 7;7;7;7;7 | 45651479:456 | 45651519:456 | -;-;-;-;- | 547 | NA | 0 | 0 | 0 | 0 | 0 | 0 |
| ENSDARG000 | 7;7;7;7;7;7;7; | 13684012:136 | 13685145:136 | +;+;+;+;+;+;+; | 4773 | pdcd7 | 191 | 148 | 256 | 180 | 183 | 198 |
| ENSDARG000 | 7;7;7;7;7;7;7; | 40900212:409 | 40907213:409 | -;-;-;-;-;-;-;-; | 10699 | RBM33 | 1600 | 1111 | 1956 | 1854 | 1781 | 1630 |
| ENSDARG000 | 7 | 19014767 | 19015511 | + | 745 | NA | 0 | 0 | 0 | 0 | 0 | 0 |
| ENSDARG000 | 7 | 40948875 | 40949190 | - | 316 | NA | 0 | 0 | 0 | 5 | 0 | 0 |
| ENSDARG000 | 7;7;7;7;7 | 19033883:190 | 19034097:190 | -;-;-;-;- | 491 | fcer1g | 45 | 28 | 19 | 8 | 113 | 64 |
| ENSDARG000 | 7;7;7;7;7;7;7; | 19043750:190 | 19043915:190 | -;-;-;-;-;-;-;-; | 1593 | ndufs2 | 1820 | 1955 | 2136 | 1234 | 1678 | 1542 |
| ENSDARG000 | 7;7 | 19028827:190 | 19028885:190 | -;- | 603 | NA | 30 | 21 | 44 | 24 | 121 | 114 |
| ENSDARG000 | 7 | 19026314 | 19027070 | + | 757 | NA | 0 | 0 | 0 | 0 | 12 | 0 |
| ENSDARG000 | 7;7 | 26494583:264 | 26496250:264 | -;- | 2561 | sox19b | 2 | 0 | 0 | 0 | 0 | 0 |
| ENSDARG000 | 7;7;7 | 38193343:381 | 38193387:381 | +;+;+ | 910 | NA | 0 | 0 | 4 | 7 | 2 | 0 |
| ENSDARG000 | 7 | 13748687 | 13750108 | - | 1422 | KBTBD13 | 0 | 0 | 0 | 0 | 0 | 0 |
| ENSDARG000 | 7 | 34786591 | 34788033 | + | 1443 | si:dkev-148a1 | 4 | 4 | 0 | 0 | 8 | 0 |
| ENSDARG000 | 7;7;7 | 34789353:347 | 34789387:347 | +;+;+ | 1358 | si:dkev-148a1 | 6 | 6 | 2 | 14 | 32 | 20 |
| ENSDARG000 | 7;7;7;7;7;7;7; | 52122224:521 | 52122405:521 | +;+;+;+;+;+;+; | 2305 | NA | 0 | 0 | 0 | 0 | 0 | 0 |
| ENSDARG000 | 7;7;7;7;7;7;7; | 45828129:458 | 45829464:458 | -;-;-;-;-;-;-;-; | 3340 | shcbp1 | 658 | 1440 | 1255 | 626 | 1332 | 672 |
| ENSDARG000 | 7 | 38202538 | 38203393 | + | 856 | NA | 0 | 2 | 0 | 0 | 0 | 6 |
| ENSDARG000 | 7;7;7;7;7;7;7; | 69470142:694 | 69470531:694 | +;+;+;+;+;+;+; | 1336 | gabarapb | 2754 | 1800 | 2753 | 2642 | 2722 | 2487 |
| ENSDARG000 | 7;7;7 | 38229070:382 | 38229624:382 | +;+;+ | 725 | NA | 0 | 0 | 0 | 0 | 0 | 0 |
| ENSDARG000 | 7;7;7;7;7;7;7; | 50353297:503 | 50353776:503 | -;-;-;-;-;-;-;-; | 2391 | prc1b | 350 | 983 | 690 | 337 | 655 | 374 |
| ENSDARG000 | 7;7 | 38245927:382 | 38245956:382 | +;+ | 280 | NA | 0 | 0 | 0 | 0 | 0 | 0 |
| ENSDARG000 | 7;7;7;7;7;7;7; | 32811907:328 | 32816071:328 | -;-;-;-;-;-;-;-; | 5680 | slc17a6b | 0 | 20 | 68 | 4 | 0 | 0 |
| ENSDARG000 | 7;7;7;7;7;7;7; | 8694557:870( | 8695204:870( | -;-;-;-;-;-;-;-; | 1422 | tex261 | 304 | 398 | 430 | 206 | 391 | 297 |
| ENSDARG000 | 7;7;7 | 38249858:382 | 38250100:382 | +;+;+ | 2677 | NA | 95 | 136 | 189 | 62 | 130 | 94 |
| ENSDARG000 | 7;7 | 45862631:458 | 45862800:458 | +;+ | 447 | NA | 2 | 0 | 0 | 0 | 0 | 0 |
| ENSDARG000 | 7;7;7;7;7;7;7; | 45963767:459 | 45963857:459 | +;+;+;+;+;+;+; | 1366 | pop4 | 113 | 240 | 237 | 84 | 183 | 156 |
| ENSDARG000 | 7;7;7;7;7 | 54092291:540 | 54092643:540 | -;-;-;-;- | 751 | rps17 | 6794 | 6750 | 8261 | 7397 | 5516 | 9221 |
| ENSDARG000 | 7;7;7;7;7;7;7; | 2021025:202 | 2021459:202 | -;-;-;-;-;-;-;-; | 3056 | NA | 2 | 2 | 8 | 28 | 0 | 4 |
| ENSDARG000 | 7;7;7;7;7;7;7; | 52135791:521 | 52135964:521 | +;+;+;+;+;+;+; | 2172 | si:dkev-286h2 | 0 | 6 | 2 | 0 | 2 | 0 |
| ENSDARG000 | 7;7;7;7;7;7;7; | 41812190:418 | 41812619:418 | +;+;+;+;+;+;+; | 3190 | orc6 | 54 | 238 | 188 | 146 | 234 | 156 |
| ENSDARG000 | 7;7;7;7 | 24205034:242 | 24205196:242 | +;+;+;+ | 654 | nedd8 | 742 | 918 | 866 | 670 | 798 | 665 |
| ENSDARG000 | 7;7;7;7;7;7 | 4592676:459 | 4592718:459 | +;+;+;+;+;+ | 1543 | NA | 0 | 0 | 0 | 0 | 0 | 0 |
| ENSDARG000 | 7;7;7;7;7;7;7; | 38258749:382 | 38258789:382 | +;+;+;+;+;+;+; | 1215 | cnep1r1 | 195 | 226 | 205 | 146 | 257 | 212 |
| ENSDARG000 | 7;7;7;7;7;7;7; | 58179513:581 | 58179641:581 | +;+;+;+;+;+;+; | 1335 | ggh | 443 | 342 | 416 | 340 | 495 | 598 |
| ENSDARG000 | 7;7;7;7;7;7;7; | 41007773:410 | 41010171:410 | -;-;-;-;-;-;-;-; | 3939 | insig1 | 1075 | 1250 | 959 | 761 | 460 | 689 |
| ENSDARG000 | 7;7;7;7;7 | 4620586:462( | 4620608:462( | +;+;+;+;+ | 1520 | zgc:162060 | 0 | 0 | 0 | 0 | 0 | 0 |
| ENSDARG000 | 7;7;7;7;7;7;7; | 8722566:872 | 8722921:872 | -;-;-;-;-;-;-;-; | 2166 | NA | 18 | 20 | 26 | 8 | 78 | 40 |
| ENSDARG000 | 7;7;7;7;7;7;7; | 41757462:417 | 41758508:417 | -;-;-;-;-;-;-;-; | 4625 | vps35 | 4684 | 4759 | 4724 | 4636 | 5715 | 4222 |
| ENSDARG000 | 7;7;7;7;7;7;7; | 26510827:265 | 26512081:265 | -;-;-;-;-;-;-;-; | 3544 | eif4a1a | 4492 | 7248 | 10754 | 2557 | 9249 | 4031 |
| ENSDARG000 | 7;7;7;7;7;7;7; | 20344032:203 | 20344255:203 | +;+;+;+;+;+;+; | 1435 | ponzr1 | 156 | 181 | 284 | 78 | 236 | 290 |
| ENSDARG000 | 7;7;7;7;7;7;7; | 40963311:409 | 40963412:409 | +;+;+;+;+;+;+; | 1119 | si:dkev-151p1 | 4 | 0 | 2 | 0 | 2 | 0 |
| ENSDARG000 | 7;7;7;7;7;7;7; | 50395002:503 | 50395211:503 | +;+;+;+;+;+;+; | 1315 | hddc3 | 194 | 351 | 311 | 256 | 220 | 266 |
| ENSDARG000 | 7;7 | 40988901:409 | 40991012:409 | -;- | 2910 | en2a | 0 | 0 | 0 | 0 | 0 | 0 |
| ENSDARG000 | 7;7;7;7;7;7;7; | 50376097:503 | 50376894:503 | -;-;-;-;-;-;-;-; | 2634 | vps33b | 376 | 301 | 423 | 318 | 389 | 388 |
| ENSDARG000 | 7;7;7;7;7;7;7; | 2084587:208 | 2090014:209( | -;-;-;-;-;-;-;-; | 6477 | NA | 4675 | 1716 | 1378 | 1640 | 2495 | 3236 |
| ENSDARG000 | 7;7;7;7;7;7 | 52150698:521 | 52150996:521 | -;-;-;-;-;- | 1133 | NA | 74 | 66 | 96 | 31 | 81 | 36 |
| ENSDARG000 | 7;7;7;7;7;7;7; | 41821311:418 | 41822231:418 | -;-;-;-;-;-;-;-; | 4010 | mylk3 | 0 | 4 | 0 | 0 | 0 | 0 |
| ENSDARG000 | 7;7;7;7;7;7;7; | 2120630:212( | 2121043:212 | -;-;-;-;-;-;-;-; | 1861 | NA | 12 | 0 | 3 | 8 | 0 | 0 |
| ENSDARG000 | 7;7;7;7;7;7;7; | 2145304:214 | 2145423:214 | -;-;-;-;-;-;-;-; | 1917 | NA | 4 | 0 | 0 | 0 | 0 | 10 |
| ENSDARG000 | 7;7;7;7;7;7 | 52154215:521 | 52154418:521 | +;+;+;+;+;+ | 1492 | tmem208 | 361 | 492 | 426 | 290 | 310 | 284 |
| ENSDARG000 | 7;7;7;7;7 | 4865367:486 | 4865410:486 | +;+;+;+;+ | 1148 | NA | 0 | 0 | 0 | 0 | 0 | 0 |
| ENSDARG000 | 7;7;7;7;7;7;7; | 29863548:298 | 29863634:298 | +;+;+;+;+;+;+; | 9837 | tln2a | 56 | 82 | 90 | 80 | 80 | 128 |
| ENSDARG000 | 7;7;7;7;7;7;7; | 24208611:242 | 24212510:242 | -;-;-;-;-;-;-;-; | 6116 | slc7a8a | 548 | 315 | 988 | 622 | 1634 | 878 |
| ENSDARG000 | 7;7;7;7;7 | 4718382:471 | 4718420:471 | +;+;+;+;+ | 1140 | si:ch211-225 | 0 | 0 | 0 | 0 | 0 | 0 |
| ENSDARG000 | 7;7;7;7;7;7;7; | 4642009:464 | 4642069:464 | +;+;+;+;+;+;+; | 1261 | si:dkev-83f18. | 0 | 0 | 0 | 0 | 0 | 0 |
| ENSDARG000 | 7;7;7;7;7;7;7; | 20334345:203 | 20335428:203 | -;-;-;-;-;-;-;-; | 2208 | NA | 51 | 59 | 101 | 97 | 74 | 89 |
| ENSDARG000 | 7;7;7;7 | 4839451:484 | 4839808:484 | -;-;-;- | 1087 | NA | 0 | 0 | 0 | 0 | 0 | 0 |
| ENSDARG000 | 7;7;7;7;7 | 24210071:242 | 24210229:242 | +;+;+;+;+ | 762 | NA | 0 | 0 | 0 | 0 | 0 | 0 |
| ENSDARG000 | 7;7;7;7;7 | 4897673:489 | 4897711:489 | +;+;+;+;+ | 1143 | NA | 0 | 0 | 0 | 0 | 0 | 0 |
| ENSDARG000 | 7;7 | 45975537:459 | 45975647:459 | +;+ | 1225 | plekhf1 | 237 | 44 | 29 | 34 | 96 | 77 |
| ENSDARG000 | 7;7;7;7;7;7;7; | 54100664:541 | 54105544:541 | -;-;-;-;-;-;-;-; | 7221 | csnk1g1 | 668 | 712 | 800 | 626 | 877 | 620 |
| ENSDARG000 | 7;7;7;7;7 | 4732597:473 | 4732696:473 | +;+;+;+;+ | 1207 | NA | 0 | 0 | 0 | 0 | 0 | 0 |
| ENSDARG000 | 7;7;7;7;7;7;7 | 4682458:468 | 4682517:468 | +;+;+;+;+;+;+ | 1237 | NA | 0 | 0 | 0 | 0 | 0 | 0 |
| ENSDARG000 | 7;7;7;7;7;7;7; | 4657987:465 | 4658044:465 | +;+;+;+;+;+;+; | 1677 | si:dkev-83f18. | 4 | 0 | 0 | 0 | 0 | 0 |
| ENSDARG000 | 7;7;7;7;7;7;7; | 28612671:286 | 28613001:286 | +;+;+;+;+;+;+; | 2155 | slc7a6os | 706 | 677 | 836 | 535 | 980 | 503 |
| ENSDARG000 | 7;7;7;7;7;7;7; | 30725473:307 | 30725815:307 | +;+;+;+;+;+;+; | 5340 | mtmr10 | 266 | 159 | 253 | 236 | 294 | 226 |
| ENSDARG000 | 7;7;7;7;7 | 4795707:479 | 4795763:479 | +;+;+;+;+ | 1161 | NA | 0 | 0 | 0 | 0 | 0 | 0 |
| ENSDARG000 | 7;7;7 | 4880423:488( | 4880459:488( | +;+;+ | 867 | NA | 0 | 0 | 0 | 0 | 2 | 0 |
| ENSDARG000 | 7;7;7;7;7 | 2180238:218 | 2181945:218 | -;-;-;-;- | 2586 | NA | 0 | 0 | 2 | 0 | 0 | 0 |
| ENSDARG000 | 7;7;7;7;7;7;7; | 34794829:347 | 34794945:347 | +;+;+;+;+;+;+; | 3608 | esrp2 | 168 | 0 | 4 | 10 | 6 | 30 |
| ENSDARG000 | 7;7;7;7;7 | 4624926:462 | 4624986:462 | +;+;+;+;+ | 1140 | NA | 0 | 0 | 0 | 0 | 4 | 0 |
| ENSDARG000 | 7;7;7;7;7;7;7; | 58259455:582 | 58259672:582 | -;-;-;-;-;-;-;- | 1269 | sdcbp | 4 | 24 | 17 | 34 | 4 | 18 |
| ENSDARG000 | 7;7;7;7;7 | 4776790:477 | 4776849:477 | +;+;+;+;+ | 1144 | NA | 0 | 0 | 0 | 0 | 0 | 0 |
| ENSDARG000 | 7;7;7;7;7;7;7; | 16806473:168 | 16806688:168 | +;+;+;+;+;+;+; | 11638 | NAV2 | 20 | 12 | 28 | 30 | 36 | 8 |
| ENSDARG000 | 7;7;7;7;7 | 4824871:482 | 4824924:482 | +;+;+;+;+ | 1139 | NA | 0 | 0 | 0 | 0 | 0 | 0 |
| ENSDARG000 | 7;7;7;7;7 | 45988091:459 | 45989066:459 | -;-;-;-;- | 1165 | C7H19orf12 | 8 | 2 | 10 | 0 | 17 | 40 |
| ENSDARG000 | 7;7;7;7;7;7;7; | 4718378:482 | 4718420:482 | +;+;+;+;+;+;+; | 1833 | NA | 0 | 0 | 0 | 0 | 0 | 0 |
| ENSDARG000 | 7;7;7;7;7;7;7 | 45992048:459 | 45992897:459 | -;-;-;-;-;-;- | 1451 | C7H19orf12 | 6 | 4 | 0 | 4 | 24 | 22 |
| ENSDARG000 | 7;7;7 | 45993716:459 | 45994208:459 | -;-;- | 754 | C7H19orf12 | 0 | 0 | 0 | 0 | 0 | 12 |
| ENSDARG000 | 7;7;7;7;7 | 4648398:464 | 4648468:464 | +;+;+;+;+ | 1156 | si:dkev-83f18. | 0 | 0 | 0 | 0 | 0 | 0 |
| ENSDARG000 | 7;7;7;7;7;7;7; | 4671241:467 | 4671301:467 | +;+;+;+;+;+;+; | 1188 | si:ch211-225 | 0 | 0 | 0 | 0 | 4 | 0 |
| ENSDARG000 | 7;7 | 2259413:225 | 2259590:226 | +;+ | 4293 | NA | 0 | 4 | 2 | 2 | 1 | 2 |
| ENSDARG000 | 7;7;7;7;7;7;7; | 2214348:221 | 2214683:221 | -;-;-;-;-;-;-;-; | 2642 | NA | 0 | 6 | 2 | 6 | 0 | 0 |
| ENSDARG000 | 7;7;7;7;7;7;7 | 4911590:492 | 4911626:492 | +;+;+;+;+;+;+ | 1249 | NA | 0 | 0 | 0 | 0 | 0 | 0 |
| ENSDARG000 | 7;7;7;7;7;7;7; | 32834142:328 | 32836513:328 | -;-;-;-;-;-;-;-; | 5569 | ano5b | 727 | 728 | 857 | 745 | 948 | 1030 |
| ENSDARG000 | 7;7 | 2236317:223 | 2236531:224 | +;+ | 3482 | zgc:172065 | 0 | 0 | 0 | 0 | 1 | 0 |
| ENSDARG000 | 7;7;7;7;7;7;7; | 19070464:190 | 19077523:190 | -;-;-;-;-;-;-;-; | 9034 | kirrel1a | 1090 | 1344 | 2062 | 1189 | 2623 | 986 |
| ENSDARG000 | 7;7;7;7;7;7;7 | 2244859:224 | 2245875:224 | +;+;+;+;+;+;+ | 4842 | zgc:172075 | 0 | 0 | 0 | 1 | 0 | 0 |
| ENSDARG000 | 7;7;7 | 58686860:586 | 58686915:586 | +;+;+ | 1323 | penkb | 8 | 6 | 8 | 30 | 2 | 0 |
| ENSDARG000 | 7;7;7;7;7;7;7; | 4625915:469 | 4626230:469 | +;+;+;+;+;+;+; | 2344 | NA | 0 | 0 | 0 | 0 | 0 | 0 |
| ENSDARG000 | 7;7 | 2228276:223 | 2228493:223 | +;+ | 2353 | NA | 6 | 0 | 0 | 0 | 0 | 0 |
| ENSDARG000 | 7;7;7;7;7 | 4911404:491 | 4911452:491 | +;+;+;+;+ | 1170 | NA | 0 | 0 | 0 | 0 | 0 | 0 |
| ENSDARG000 | 7;7;7;7;7;7;7; | 69478914:694 | 69479183:694 | +;+;+;+;+;+;+; | 2543 | spata6l | 51 | 64 | 6 | 102 | 70 | 196 |
| ENSDARG000 | 7;7;7;7;7 | 4702269:470 | 4702312:470 | +;+;+;+;+ | 1163 | si:ch211-225 | 0 | 0 | 0 | 0 | 0 | 0 |
| ENSDARG000 | 7;7;7;7;7;7;7; | 26518843:265 | 26519364:265 | -;-;-;-;-;-;-;-; | 2273 | senp3b | 304 | 549 | 514 | 276 | 382 | 360 |
| ENSDARG000 | 7;7;7;7;7;7 | 22657566:226 | 22657755:226 | +;+;+;+;+;+ | 1537 | ponzr5 | 0 | 0 | 0 | 0 | 4 | 0 |
| ENSDARG000 | 7;7;7;7;7;7;7; | 65673885:656 | 65674140:656 | +;+;+;+;+;+;+; | 2024 | parvab | 460 | 1072 | 1016 | 977 | 1228 | 605 |
| ENSDARG000 | 7;7;7;7;7;7 | 22664936:226 | 22665317:226 | -;-;-;-;-;- | 796 | NA | 126 | 14 | 78 | 68 | 48 | 40 |
| ENSDARG000 | 7;7;7;7;7;7;7; | 58699718:586 | 58699870:587 | +;+;+;+;+;+;+; | 1656 | sdr16c5b | 1408 | 1469 | 1296 | 1224 | 821 | 1353 |
| ENSDARG000 | 7;7;7;7;7;7;7; | 19153320:191 | 19154096:191 | -;-;-;-;-;-;-;-; | 2126 | il13ra1 | 271 | 238 | 339 | 377 | 152 | 224 |
| ENSDARG000 | 7;7;7;7;7;7 | 22675475:226 | 22675533:226 | +;+;+;+;+;+ | 866 | ponzr3 | 0 | 0 | 0 | 0 | 0 | 0 |
| ENSDARG000 | 7;7;7;7;7;7;7; | 28623294:286 | 28626109:286 | -;-;-;-;-;-;-;-; | 4369 | slc7a6 | 48 | 6 | 19 | 30 | 8 | 16 |
| ENSDARG000 | 7;7;7 | 41080649:410 | 41083023:410 | -;-;- | 3510 | htr5ab | 0 | 0 | 2 | 0 | 2 | 0 |
| ENSDARG000 | 7;7;7;7;7;7;7; | 52158818:521 | 52159028:521 | +;+;+;+;+;+;+; | 3378 | katnb1 | 674 | 643 | 1059 | 788 | 913 | 707 |

| | | | | | | | | | | | | |
|---|---|---|---|---|---|---|---|---|---|---|---|---|
| ENSDARG000 | 7;7;7;7 | 22680560:226 | 22680591:226 | +;+;+;+ | 1109 | ponzr4 | 0 | 0 | 0 | 0 | 0 | 0 |
| ENSDARG000 | 7:7:7:7:7:7:7: | 22688781:226 | 22688880:226 | +:+:+:+:+:+:+: | 2519 | ugt5g1 | 28 | 6 | 6 | 0 | 20 | 28 |
| ENSDARG000 | 7 | 41112901 | 41113295 | + | 395 | NA | 0 | 0 | 0 | 0 | 0 | 0 |
| ENSDARG000 | 7;7;7;7 | 50407575:504 | 50408023:504 | -;-;-;- | 757 | hypk | 919 | 1554 | 1690 | 900 | 1175 | 990 |
| ENSDARG000 | 7:7:7:7:7:7:7: | 50399627:504 | 50399852:504 | +:+:+:+:+:+:+: | 1813 | mfap1 | 847 | 626 | 894 | 540 | 795 | 757 |
| ENSDARG000 | 7:7:7:7:7:7:7: | 38265363:382 | 38265445:382 | +:+:+:+:+:+:+: | 3292 | gpatch1 | 604 | 430 | 694 | 445 | 672 | 521 |
| ENSDARG000 | 7;7;7;7;7;7 | 26534131:265 | 26534232:265 | +;+;+;+;+;+ | 1251 | NA | 25 | 12 | 15 | 12 | 10 | 10 |
| ENSDARG000 | 7;7 | 65756041:657 | 65756232:657 | +;+ | 1335 | NA | 29 | 6 | 16 | 1 | 14 | 8 |
| ENSDARG000 | 7 | 52175958 | 52177419 | + | 1462 | NA | 2 | 2 | 14 | 6 | 2 | 4 |
| ENSDARG000 | 7;7 | 2289715:2292 | 2291314:2292 | -;- | 1707 | si:dkev-187i1 | 8 | 0 | 0 | 0 | 2 | 0 |
| ENSDARG000 | 7:7:7:7:7:7:7: | 30772991:307 | 30773242:307 | -:-:-:-:-:-:-: | 2248 | mphosph10 | 424 | 598 | 803 | 270 | 520 | 289 |
| ENSDARG000 | 7:7:7:7:7:7:7: | 65813301:658 | 65813448:658 | +:+:+:+:+:+:+: | 9546 | TEAD1 | 1269 | 717 | 851 | 757 | 1225 | 728 |
| ENSDARG000 | 7:7:7:7:7:7:7: | 25047219:250 | 25047684:250 | -:-:-:-:-:-:-: | 1322 | arl2 | 461 | 530 | 446 | 398 | 485 | 360 |
| ENSDARG000 | 7;7;7;7 | 58720601:587 | 58720948:587 | -;-;-;- | 650 | chchd7 | 154 | 162 | 182 | 64 | 163 | 223 |
| ENSDARG000 | 7;7 | 2304846:2309 | 2308934:2309 | -;- | 4201 | NA | 2 | 0 | 0 | 0 | 0 | 0 |
| ENSDARG000 | 7;7;7 | 8844285:8876 | 8844853:8877 | -;-;- | 1099 | vax2 | 0 | 0 | 0 | 0 | 0 | 0 |
| ENSDARG000 | 7;7;7 | 2407857:2410 | 2408260:2410 | -;-;- | 597 | NA | 0 | 0 | 0 | 0 | 0 | 0 |
| ENSDARG000 | 7;7;7;7;7 | 30779761:307 | 30779816:307 | +;+;+;+;+ | 1696 | mcee | 178 | 250 | 349 | 117 | 278 | 272 |
| ENSDARG000 | 7:7:7:7:7:7:7: | 14211322:142 | 14211363:142 | +:+:+:+:+:+:+: | 4553 | fes | 70 | 26 | 16 | 14 | 42 | 36 |
| ENSDARG000 | 7:7:7:7:7:7:7: | 24251367:242 | 24252427:242 | +:+:+:+:+:+:+: | 11731 | SVEP1 | 78 | 0 | 24 | 3 | 64 | 16 |
| ENSDARG000 | 7;7;7;7;7;7;7 | 38787831:387 | 38788250:387 | -;-;-;-;-;-;- | 1245 | NA | 0 | 8 | 4 | 14 | 3 | 4 |
| ENSDARG000 | 7:7:7:7:7:7:7: | 34823533:348 | 34828431:348 | -:-:-:-:-:-:-: | 8571 | NFATC3 | 633 | 361 | 608 | 429 | 521 | 495 |
| ENSDARG000 | 7;7;7 | 17023289:170 | 17023355:170 | +;+;+ | 875 | NA | 0 | 0 | 0 | 0 | 0 | 0 |
| ENSDARG000 | 7;7;7;7 | 17024602:170 | 17025279:170 | -;-;-;- | 1433 | dbx1a | 4 | 0 | 0 | 0 | 0 | 0 |
| ENSDARG000 | 7;7 | 34827523:348 | 34828042:348 | +;+ | 780 | NA | 0 | 0 | 0 | 0 | 2 | 0 |
| ENSDARG000 | 7;7;7;7;7 | 58730201:587 | 58730333:587 | +;+;+;+;+ | 5036 | plag1 | 106 | 16 | 33 | 10 | 36 | 84 |
| ENSDARG000 | 7;7;7;7;7;7 | 8890366:8898 | 8894989:8898 | -;-;-;-;-;- | 5301 | ADISSP | 118 | 433 | 452 | 274 | 308 | 266 |
| ENSDARG000 | 7:7:7:7:7:7:7: | 50475703:504 | 50477691:504 | -:-:-:-:-:-:-: | 3900 | NA | 22 | 0 | 11 | 0 | 20 | 16 |
| ENSDARG000 | 7:7:7:7:7:7:7: | 46003449:460 | 46003569:460 | +:+:+:+:+:+:+: | 3062 | si:ch211-260 | 448 | 304 | 257 | 425 | 298 | 279 |
| ENSDARG000 | 7:7:7:7:7:7:7: | 50448478:504 | 50448546:504 | +:+:+:+:+:+:+: | 4171 | SERINC4 | 2 | 0 | 6 | 0 | 10 | 0 |
| ENSDARG000 | 7;7;7;7;7 | 58745987:587 | 58746088:587 | +;+;+;+;+ | 523 | rps20 | 13940 | 17668 | 21357 | 18212 | 13319 | 16434 |
| ENSDARG000 | 7:7:7:7:7:7:7: | 46009862:460 | 46010099:460 | -;-;-;-;-;-;- | 1649 | zgc:162297 | 0 | 0 | 4 | 0 | 5 | 12 |
| ENSDARG000 | 7:7:7:7:7:7:7: | 54222156:542 | 54222522:542 | +:+:+:+:+:+:+: | 2778 | pacsin3 | 612 | 551 | 617 | 493 | 911 | 1482 |
| ENSDARG000 | 7;7 | 2427600:2430 | 2429206:2430 | -;- | 1713 | zgc:66473 | 12 | 2 | 0 | 0 | 0 | 0 |
| ENSDARG000 | 7:7:7:7:7:7:7: | 28648706:286 | 28650653:286 | -:-:-:-:-:-:-: | 3221 | NA | 164 | 85 | 110 | 122 | 116 | 206 |
| ENSDARG000 | 7:7:7:7:7:7:7: | 22702224:227 | 22702273:227 | +:+:+:+:+:+:+: | 5106 | NA | 14 | 0 | 2 | 0 | 0 | 0 |
| ENSDARG000 | 7 | 2455344 | 2456229 | + | 886 | NA | 0 | 0 | 0 | 0 | 0 | 0 |
| ENSDARG000 | 7;7;7;7;7 | 8967974:8974 | 8968588:8974 | -;-;-;-;- | 1057 | NA | 453 | 366 | 337 | 102 | 86 | 196 |
| ENSDARG000 | 7;7;7 | 2467057:2467 | 2467160:2467 | +;+;+ | 1046 | NA | 0 | 0 | 0 | 0 | 0 | 0 |
| ENSDARG000 | 7:7:7:7:7:7:7: | 19179819:191 | 19183540:191 | -:-:-:-:-:-:-: | 9973 | dock11 | 3752 | 2195 | 2972 | 1820 | 4399 | 3692 |
| ENSDARG000 | 7:7:7:7:7:7:7: | 32897723:328 | 32897761:328 | +:+:+:+:+:+:+: | 1871 | tssc4 | 521 | 558 | 752 | 694 | 672 | 575 |
| ENSDARG000 | 7:7:7:7:7:7:7: | 28668979:286 | 28671275:286 | -:-:-:-:-:-:-: | 4306 | adgrg1 | 36 | 36 | 59 | 190 | 36 | 152 |
| ENSDARG000 | 7;7;7;7;7;7;7 | 25049545:250 | 25049616:250 | +;+;+;+;+;+;+ | 1843 | batf2 | 238 | 165 | 378 | 108 | 106 | 58 |
| ENSDARG000 | 7:7:7:7:7:7:7: | 17063761:170 | 17063846:170 | +:+:+:+:+:+:+: | 4857 | htatip2 | 268 | 211 | 249 | 160 | 398 | 268 |
| ENSDARG000 | 7:7:7:7:7:7:7: | 25049577:250 | 25049616:250 | +:+:+:+:+:+:+: | 2978 | PPP2R5B | 12 | 6 | 12 | 0 | 2 | 6 |
| ENSDARG000 | 7 | 34963748 | 34965654 | + | 1907 | ddx28 | 145 | 155 | 236 | 171 | 208 | 146 |
| ENSDARG000 | 7:7:7:7:7:7:7: | 41146560:411 | 41146699:411 | +:+:+:+:+:+:+: | 2450 | puf60b | 2105 | 2578 | 3514 | 1965 | 3081 | 2004 |
| ENSDARG000 | 7;7;7 | 2489969:2490 | 2490103:2490 | +;+;+ | 1820 | NA | 0 | 0 | 0 | 0 | 0 | 0 |
| ENSDARG000 | 7:7:7:7:7:7:7: | 2538071:2541 | 2538265:2541 | +:+:+:+:+:+:+: | 1076 | NA | 0 | 0 | 0 | 0 | 0 | 0 |
| ENSDARG000 | 7:7:7:7:7:7:7: | 9008372:9011 | 9011324:9011 | +:+:+:+:+:+:+: | 5414 | helz2c | 6 | 4 | 0 | 0 | 4 | 0 |
| ENSDARG000 | 7;7;7 | 34979173:350 | 34980037:350 | -;-;- | 2034 | galr1b | 0 | 2 | 0 | 4 | 4 | 0 |
| ENSDARG000 | 7:7:7:7:7:7:7: | 22718251:227 | 22718845:227 | +:+:+:+:+:+:+: | 4106 | fxr2 | 1203 | 1288 | 1392 | 893 | 1435 | 1152 |
| ENSDARG000 | 7:7:7:7:7:7:7: | 38278860:382 | 38278997:382 | +:+:+:+:+:+:+: | 4607 | lrp3 | 77 | 26 | 91 | 28 | 114 | 96 |
| ENSDARG000 | 7:7:7:7:7:7:7: | 30787903:308 | 30788034:308 | +:+:+:+:+:+:+: | 4425 | APBA2 | 28 | 6 | 14 | 0 | 8 | 12 |
| ENSDARG000 | 7:7:7:7:7:7:7: | 26545502:265 | 26545753:265 | +:+:+:+:+:+:+: | 3545 | tnk1 | 40 | 8 | 8 | 34 | 10 | 0 |
| ENSDARG000 | 7:7:7:7:7:7:7: | 38793832:387 | 38794656:387 | -:-:-:-:-:-:-: | 2580 | atg13 | 1177 | 962 | 1329 | 1210 | 1105 | 1089 |
| ENSDARG000 | 7:7:7:7:7:7:7: | 24357231:243 | 24357545:243 | -:-:-:-:-:-:-: | 1568 | txn | 276 | 425 | 451 | 89 | 303 | 446 |
| ENSDARG000 | 7:7:7:7:7:7:7: | 69496293:695 | 69497658:695 | -:-:-:-:-:-:-: | 2978 | slc1a1 | 4 | 6 | 4 | 2 | 2 | 10 |
| ENSDARG000 | 7:7:7:7:7:7:7: | 46019780:460 | 46020223:460 | +:+:+:+:+:+:+: | 2615 | ccne1 | 102 | 225 | 238 | 60 | 228 | 115 |
| ENSDARG000 | 7;7;7;7 | 65965300:659 | 65965390:659 | +;+;+;+ | 769 | NA | 0 | 0 | 0 | 2 | 0 | 0 |
| ENSDARG000 | 7;7;7 | 2980248:3009 | 2981426:3010 | -;-;- | 1361 | zgc:171695 | 0 | 0 | 0 | 0 | 0 | 0 |
| ENSDARG000 | 7:7:7:7:7:7:7: | 3125746:3125 | 3125861:3126 | +:+:+:+:+:+:+: | 1415 | NA | 0 | 0 | 0 | 0 | 0 | 0 |
| ENSDARG000 | 7;7;7 | 66027991:660 | 66028084:660 | +;+;+ | 334 | NA | 0 | 0 | 0 | 0 | 0 | 0 |
| ENSDARG000 | 7:7:7:7:7:7:7: | 58843700:588 | 58843845:588 | +:+:+:+:+:+:+: | 939 | lypla1 | 95 | 91 | 63 | 71 | 65 | 120 |
| ENSDARG000 | 7;7;7;7 | 25103965:251 | 25104114:251 | +;+;+;+ | 678 | NA | 0 | 0 | 0 | 0 | 0 | 0 |
| ENSDARG000 | 7;7;7;7;7;7;7 | 35062132:350 | 35063068:350 | -;-;-;-;-;-;- | 1588 | ctrb.3 | 0 | 0 | 4 | 2 | 10 | 0 |
| ENSDARG000 | 7;7;7 | 25109700:251 | 25110674:251 | -;-;- | 1263 | gpha2 | 0 | 0 | 0 | 0 | 2 | 0 |
| ENSDARG000 | 7;7;7 | 54605640:546 | 54606518:546 | +;+;+ | 1887 | fgf3 | 4 | 0 | 0 | 0 | 0 | 4 |
| ENSDARG000 | 7:7:7:7:7:7:7: | 24366962:243 | 24367252:243 | -;-;-;-;-;-;- | 1209 | NA | 4 | 14 | 22 | 32 | 12 | 13 |
| ENSDARG000 | 7:7:7:7:7:7:7: | 28709931:287 | 28710206:287 | -:-:-:-:-:-:-: | 1695 | NA | 2 | 0 | 1 | 0 | 0 | 2 |
| ENSDARG000 | 7;7;7 | 54617060:546 | 54617690:546 | +;+;+ | 3170 | fgf4 | 0 | 0 | 2 | 0 | 4 | 4 |
| ENSDARG000 | 7;7;7;7;7;7;7 | 54651109:546 | 54651274:546 | +;+;+;+;+;+;+ | 1660 | lto1 | 66 | 108 | 95 | 55 | 48 | 80 |
| ENSDARG000 | 7:7:7:7:7:7:7: | 19334314:193 | 19334850:193 | -:-:-:-:-:-:-: | 2527 | ntn4 | 135 | 42 | 26 | 78 | 48 | 91 |
| ENSDARG000 | 7;7;7;7;7;7 | 54642005:546 | 54642289:546 | +;+;+;+;+;+ | 1032 | fgf19 | 0 | 0 | 0 | 0 | 0 | 0 |
| ENSDARG000 | 7;7;7 | 50766094:507 | 50766287:507 | +;+;+ | 608 | cartl | 84 | 8 | 4 | 8 | 16 | 14 |
| ENSDARG000 | 7;7;7;7;7 | 54609479:546 | 54609755:546 | -;-;-;-;- | 1316 | NA | 0 | 0 | 0 | 0 | 0 | 0 |
| ENSDARG000 | 7:7:7:7:7:7:7: | 17106160:171 | 17106681:171 | +:+:+:+:+:+:+: | 2974 | prmt3 | 287 | 232 | 400 | 128 | 302 | 187 |
| ENSDARG000 | 7:7:7:7:7:7:7: | 66048102:660 | 66048205:660 | +:+:+:+:+:+:+: | 3645 | bmal1b | 128 | 321 | 248 | 114 | 84 | 243 |
| ENSDARG000 | 7;7;7;7;7;7;7 | 35068036:350 | 35068098:350 | +;+;+;+;+;+;+ | 869 | ctrb.2 | 0 | 0 | 0 | 0 | 2 | 4 |
| ENSDARG000 | 7:7:7:7:7:7:7: | 9290929:9290 | 9291168:9291 | +:+:+:+:+:+:+: | 1803 | snrpa1 | 401 | 515 | 945 | 429 | 811 | 478 |
| ENSDARG000 | 7;7;7 | 3171585:3178 | 3171665:3178 | +;+;+ | 1291 | NA | 0 | 0 | 0 | 0 | 0 | 0 |
| ENSDARG000 | 7;7;7;7;7;7;7 | 25120226:251 | 25120294:251 | -;-;-;-;-;-;- | 599 | MAJIN | 0 | 0 | 2 | 0 | 0 | 0 |
| ENSDARG000 | 7:7:7:7:7:7:7: | 41873080:418 | 41873794:418 | -:-:-:-:-:-:-: | 1324 | C7H16orf87 | 428 | 496 | 701 | 592 | 789 | 743 |
| ENSDARG000 | 7:7:7:7:7:7:7: | 38315647:383 | 38317164:383 | -:-:-:-:-:-:-: | 3025 | slc7a10a | 2348 | 1107 | 1818 | 1059 | 1729 | 1150 |
| ENSDARG000 | 7:7:7:7:7:7:7: | 14291323:142 | 14291595:142 | +:+:+:+:+:+:+: | 2183 | rhcga | 0 | 0 | 0 | 0 | 1 | 2 |
| ENSDARG000 | 7;7;7;7 | 22767678:227 | 22767724:227 | +;+;+;+ | 781 | ponzr6 | 8 | 14 | 8 | 26 | 48 | 56 |
| ENSDARG000 | 7;7;7 | 38315664:383 | 38316277:383 | +;+;+ | 1803 | NA | 0 | 0 | 1 | 0 | 2 | 0 |
| ENSDARG000 | 7:7:7:7:7:7:7: | 52290812:522 | 52297107:522 | -:-:-:-:-:-:-: | 8505 | SLCO3A1 | 0 | 0 | 1 | 0 | 0 | 6 |
| ENSDARG000 | 7;7;7;7 | 19374683:193 | 19374872:193 | +;+;+;+ | 623 | snrpf | 474 | 913 | 962 | 458 | 558 | 626 |
| ENSDARG000 | 7:7:7:7:7:7:7: | 19397129:194 | 19401424:194 | -:-:-:-:-:-:-: | 6673 | NA | 156 | 49 | 92 | 71 | 60 | 60 |
| ENSDARG000 | 7:7:7:7:7:7:7: | 3229828:3230 | 3229955:3230 | +:+:+:+:+:+:+: | 796 | NA | 0 | 0 | 0 | 0 | 0 | 0 |
| ENSDARG000 | 7;7 | 22776174:227 | 22776566:227 | -;- | 568 | NA | 6 | 2 | 2 | 0 | 0 | 4 |
| ENSDARG000 | 7:7:7:7:7:7:7: | 19389304:193 | 19389510:193 | +:+:+:+:+:+:+: | 2516 | amdhd1 | 24 | 0 | 63 | 0 | 25 | 26 |
| ENSDARG000 | 7:7:7:7:7:7:7: | 28724910:287 | 28725043:287 | +:+:+:+:+:+:+: | 3844 | ccdc102a | 106 | 85 | 173 | 156 | 206 | 306 |
| ENSDARG000 | 7;7 | 50810249:508 | 50810724:508 | -;- | 648 | NA | 0 | 0 | 0 | 0 | 0 | 0 |
| ENSDARG000 | 7:7:7:7:7:7:7: | 50796422:508 | 50796886:508 | +:+:+:+:+:+:+: | 2965 | NA | 15 | 20 | 0 | 8 | 14 | 9 |
| ENSDARG000 | 7;7 | 22788718:227 | 22789970:227 | -;- | 1553 | NA | 836 | 801 | 1324 | 1629 | 582 | 668 |
| ENSDARG000 | 7:7:7:7:7:7:7: | 30923283:309 | 30923665:309 | -:-:-:-:-:-:-: | 1432 | sord | 123 | 207 | 194 | 110 | 156 | 109 |
| ENSDARG000 | 7;7;7;7 | 14314386:143 | 14314714:143 | +;+;+;+ | 734 | NA | 0 | 0 | 0 | 0 | 0 | 0 |
| ENSDARG000 | 7;7;7 | 52364748:523 | 52365278:523 | +;+;+ | 1614 | gnrhr2 | 0 | 0 | 0 | 0 | 0 | 0 |
| ENSDARG000 | 7:7:7:7:7:7:7: | 3292468:3292 | 3292572:3292 | +:+:+:+:+:+:+: | 756 | NA | 0 | 0 | 0 | 0 | 0 | 0 |
| ENSDARG000 | 7;7 | 25126629:251 | 25126874:251 | +;+ | 1604 | zgc:101765 | 171 | 162 | 210 | 100 | 240 | 172 |
| ENSDARG000 | 7;7;7;7;7 | 38808027:388 | 38808092:388 | +;+;+;+;+ | 1438 | harbi1 | 144 | 95 | 124 | 40 | 112 | 146 |
| ENSDARG000 | 7:7:7:7:7:7:7: | 25129520:251 | 25130338:251 | -:-:-:-:-:-:-: | 1359 | badb | 155 | 133 | 197 | 140 | 196 | 119 |
| ENSDARG000 | 7;7 | 3332091:3333 | 3332214:3333 | +;+ | 179 | NA | 0 | 0 | 0 | 0 | 0 | 0 |
| ENSDARG000 | 7;7;7;7 | 3357973:3358 | 3358207:3358 | +;+;+;+ | 1430 | NA | 0 | 14 | 10 | 21 | 4 | 83 |
| ENSDARG000 | 7;7 | 3375546:3380 | 3377133:3380 | -;- | 1632 | NA | 0 | 0 | 0 | 0 | 0 | 2 |
| ENSDARG000 | 7:7:7:7:7:7:7: | 17229282:172 | 17229553:172 | +:+:+:+:+:+:+: | 7760 | slc6a5 | 32 | 18 | 32 | 0 | 27 | 16 |
| ENSDARG000 | 7:7:7:7:7:7:7: | 39463050:394 | 39464119:394 | -:-:-:-:-:-:-: | 10444 | OTOG | 26 | 8 | 36 | 10 | 6 | 0 |
| ENSDARG000 | 7:7:7:7:7:7:7: | 35075847:350 | 35076086:350 | +:+:+:+:+:+:+: | 1918 | ctrb.1 | 0 | 0 | 0 | 0 | 1 | 0 |
| ENSDARG000 | 7:7:7:7:7:7:7: | 24375070:243 | 24376309:243 | -:-:-:-:-:-:-: | 2706 | ptgr1.1 | 3764 | 7405 | 2012 | 4291 | 2600 | 4184 |
| ENSDARG000 | 7:7:7:7:7:7:7: | 26567864:265 | 26568048:265 | -:-:-:-:-:-:-: | 1780 | NA | 0 | 0 | 0 | 0 | 0 | 0 |
| ENSDARG000 | 7:7:7:7:7:7:7: | 38821438:388 | 38824774:388 | -:-:-:-:-:-:-: | 5780 | phf21aa | 22 | 26 | 119 | 28 | 31 | 14 |
| ENSDARG000 | 7:7:7:7:7:7:7: | 29951997:299 | 29952418:299 | +:+:+:+:+:+:+: | 3308 | tpma | 16 | 16 | 12 | 38 | 52 | 18 |
| ENSDARG000 | 7;7;7;7;7 | 14353922:143 | 14354469:143 | -;-;-;-;- | 1485 | NA | 0 | 2 | 0 | 0 | 0 | 0 |
| ENSDARG000 | 7;7;7;7;7;7;7 | 38811800:388 | 38811903:388 | +;+;+;+;+;+;+ | 2817 | zgc:110699 | 0 | 0 | 30 | 12 | 0 | 0 |
| ENSDARG000 | 7;7;7;7;7;7;7 | 46035854:460 | 46035963:460 | +;+;+;+;+;+;+ | 1697 | NA | 0 | 0 | 0 | 16 | 0 | 0 |
| ENSDARG000 | 7:7:7:7:7:7:7: | 20431633:204 | 20433894:204 | -:-:-:-:-:-:-: | 4510 | NA | 2 | 1 | 0 | 10 | 0 | 4 |
| ENSDARG000 | 7:7:7:7:7:7:7: | 20383841:203 | 20384481:203 | +:+:+:+:+:+:+: | 6943 | NA | 37 | 30 | 24 | 23 | 32 | 33 |
| ENSDARG000 | 7:7:7:7:7:7:7: | 35080490:350 | 35081599:350 | -:-:-:-:-:-:-: | 1676 | agrp | 0 | 0 | 0 | 0 | 0 | 0 |
| ENSDARG000 | 7:7:7:7:7:7:7: | 54668303:546 | 54673237:546 | -:-:-:-:-:-:-: | 6621 | ccnd1 | 4241 | 6452 | 9250 | 7155 | 21360 | 10963 |
| ENSDARG000 | 7 | 52377189 | 52379021 | + | 1833 | NA | 0 | 0 | 0 | 0 | 0 | 0 |
| ENSDARG000 | 7:7:7:7:7:7:7: | 52371112:523 | 52373039:523 | -:-:-:-:-:-:-: | 3915 | NA | 0 | 0 | 0 | 0 | 2 | 0 |
| ENSDARG000 | 7:7:7:7:7:7:7: | 30930653:309 | 30930727:309 | +:+:+:+:+:+:+: | 9182 | TJP1 | 4045 | 2319 | 3380 | 2466 | 2280 | 3416 |
| ENSDARG000 | 7:7:7:7:7:7:7: | 32901658:329 | 32901843:329 | +:+:+:+:+:+:+: | 4117 | NA | 50 | 4 | 10 | 9 | 20 | 10 |
| ENSDARG000 | 7;7;7;7;7;7;7 | 29962559:299 | 29962708:299 | +;+;+;+;+;+;+ | 3263 | FBXO22 | 497 | 604 | 1050 | 521 | 735 | 460 |
| ENSDARG000 | 7:7:7:7:7:7:7: | 26580501:265 | 26582788:265 | -:-:-:-:-:-:-: | 3667 | plscr3b | 1696 | 1472 | 3173 | 1212 | 1980 | 2552 |
| ENSDARG000 | 7;7;7;7 | 30926289:309 | 30926529:309 | +;+;+;+ | 1523 | NA | 0 | 2 | 4 | 0 | 0 | 0 |
| ENSDARG000 | 7;7;7;7;7;7;7 | 30929459:309 | 30930012:309 | -;-;-;-;-;-;- | 1364 | terb2 | 8 | 5 | 8 | 19 | 0 | 2 |
| ENSDARG000 | 7:7:7:7:7:7:7: | 28901786:289 | 28902107:289 | +:+:+:+:+:+:+: | 7012 | DOK4 | 12 | 14 | 12 | 48 | 14 | 12 |
| ENSDARG000 | 7;7;7;7;7 | 38357915:383 | 38358241:383 | -;-;-;-;- | 922 | NA | 1 | 0 | 0 | 0 | 0 | 0 |
| ENSDARG000 | 7:7:7:7:7:7:7: | 38349665:383 | 38349864:383 | +:+:+:+:+:+:+: | 4190 | rhpn2 | 195 | 19 | 77 | 193 | 53 | 68 |
| ENSDARG000 | 7:7:7:7:7:7:7: | 22792132:227 | 22792154:227 | +:+:+:+:+:+:+: | 2569 | rbm4.3 | 6626 | 6364 | 9161 | 4997 | 7280 | 4811 |
| ENSDARG000 | 7:7:7:7:7:7:7: | 41887429:418 | 41887910:418 | +:+:+:+:+:+:+: | 4612 | gpt2 | 1095 | 1881 | 1997 | 962 | 2090 | 1249 |
| ENSDARG000 | 7:7:7:7:7:7:7: | 71278625:712 | 71279143:712 | -:-:-:-:-:-:-: | 2974 | dhx15 | 2392 | 2354 | 3617 | 1771 | 3011 | 2054 |
| ENSDARG000 | 7:7:7:7:7:7:7: | 50822465:508 | 50822621:508 | +:+:+:+:+:+:+: | 7782 | per1b | 4758 | 2598 | 3506 | 2979 | 5272 | 3650 |
| ENSDARG000 | 7:7:7:7:7:7:7: | 35086369:350 | 35086588:350 | +:+:+:+:+:+:+: | 2145 | atp6v0d1 | 2574 | 2533 | 2776 | 2836 | 2903 | 2650 |
| ENSDARG000 | 7:7:7:7:7:7:7: | 59046524:590 | 59046995:590 | -;-;-;-;-;-;- | 1451 | chmp5b | 1926 | 1917 | 1725 | 1756 | 1857 | 1877 |
| ENSDARG000 | 7;7 | 3427806:3429 | 3429396:3429 | -;- | 1696 | NA | 0 | 0 | 0 | 0 | 14 | 10 |
| ENSDARG000 | 7:7:7:7:7:7:7: | 46252993:462 | 46253519:462 | +:+:+:+:+:+:+: | 7671 | ZNF536 | 16 | 19 | 18 | 18 | 28 | 163 |
| ENSDARG000 | 7:7:7:7:7:7:7: | 22796083:227 | 22796314:227 | +:+:+:+:+:+:+: | 3054 | rbm4.2 | 2263 | 1909 | 2532 | 1344 | 2334 | 1577 |
| ENSDARG000 | 7:7:7:7:7:7:7: | 66112194:661 | 66113167:661 | -:-:-:-:-:-:-: | 2193 | BTBD10 | 1399 | 1677 | 1813 | 1366 | 1480 | 1069 |
| ENSDARG000 | 7:7:7:7:7:7:7: | 3442823:3442 | 3442915:3442 | +:+:+:+:+:+:+: | 1912 | NA | 0 | 0 | 0 | 0 | 0 | 2 |
| ENSDARG000 | 7;7;7;7;7 | 26613183:266 | 26613309:266 | +;+;+;+;+ | 775 | NA | 0 | 0 | 0 | 0 | 0 | 0 |
| ENSDARG000 | 7:7:7:7:7:7:7: | 14393059:143 | 14395053:143 | -:-:-:-:-:-:-: | 5851 | kif7 | 420 | 416 | 512 | 498 | 436 | 425 |
| ENSDARG000 | 7:7:7:7:7:7:7: | 39537790:395 | 39538444:395 | -:-:-:-:-:-:-: | 2048 | slc22a18 | 142 | 59 | 100 | 104 | 142 | 203 |
| ENSDARG000 | 7:7:7:7:7:7:7: | 26616241:266 | 26617620:266 | -:-:-:-:-:-:-: | 2593 | phf23b | 452 | 520 | 526 | 376 | 627 | 376 |
| ENSDARG000 | 7:7:7:7:7:7:7: | 17332123:173 | 17332275:173 | -:-:-:-:-:-:-: | 1324 | nitr8 | 8 | 0 | 0 | 0 | 0 | 0 |
| ENSDARG000 | 7;7;7;7;7;7;7 | 17266904:172 | 17266945:172 | -;-;-;-;-;-;- | 782 | NA | 0 | 0 | 0 | 0 | 0 | 0 |
| ENSDARG000 | 7:7:7:7:7:7:7: | 17358084:173 | 17358319:173 | -;-;-;-;-;-;- | 943 | NA | 68 | 0 | 2 | 0 | 0 | 0 |
| ENSDARG000 | 7:7:7:7:7:7:7: | 32926760:329 | 32928692:329 | -:-:-:-:-:-:-: | 4649 | PKP3 | 32 | 46 | 17 | 18 | 68 | 8 |
| ENSDARG000 | 7:7:7:7:7:7:7: | 19424857:194 | 19425110:194 | +:+:+:+:+:+:+: | 7784 | NA | 8 | 6 | 8 | 4 | 8 | 6 |
| ENSDARG000 | 7:7:7:7:7:7:7: | 17255705:172 | 17255876:172 | +:+:+:+:+:+:+: | 2079 | nitr9 | 20 | 0 | 0 | 0 | 0 | 0 |
| ENSDARG000 | 7:7:7:7:7:7:7: | 46727891:467 | 46732822:467 | -;-;-;-;-;-;- | 5785 | tshz3b | 796 | 1006 | 1485 | 744 | 1461 | 1059 |
| ENSDARG000 | 7;7;7 | 46782320:468 | 46782606:468 | +;+;+ | 761 | NA | 2 | 14 | 22 | 10 | 40 | 19 |
| ENSDARG000 | 7:7:7:7:7:7:7: | 24391129:243 | 24391181:243 | +:+:+:+:+:+:+: | 3927 | poln | 0 | 4 | 8 | 4 | 2 | 20 |
| ENSDARG000 | 7 | 17346399 | 17347139 | + | 741 | NA | 0 | 0 | 0 | 0 | 0 | 0 |
| ENSDARG000 | 7 | 17353030 | 17353772 | + | 743 | NA | 2 | 0 | 0 | 0 | 0 | 0 |
| ENSDARG000 | 7;7;7;7;7 | 17261616:172 | 17261775:172 | -;-;-;-;- | 998 | NA | 0 | 0 | 0 | 0 | 0 | 0 |
| ENSDARG000 | 7:7:7:7:7:7:7: | 32983440:329 | 32986572:329 | -:-:-:-:-:-:-: | 4599 | cd81a | 27891 | 25705 | 28904 | 26770 | 31167 | 35447 |
| ENSDARG000 | 7:7:7:7:7:7:7: | 17255707:172 | 17255876:172 | +:+:+:+:+:+:+: | 940 | NA | 43 | 0 | 0 | 0 | 0 | 0 |
| ENSDARG000 | 7:7:7:7:7:7:7: | 29012033:290 | 29012236:290 | +:+:+:+:+:+:+: | 2719 | dnaaf1 | 2 | 6 | 0 | 0 | 0 | 0 |

| | | | | | | | | | | | |
|---|---|---|---|---|---|---|---|---|---|---|---|
| ENSDARG000 | 7;7;7;7;7;7 | 17374560:173 | 17374602:173 | +;+;+;+;+;+ | 585 | nitr7a | 4 | 0 | 0 | 0 | 0 | 0 |
| ENSDARG000 | 7 | 17356482 | 17357237 | + | 756 | NA | 12 | 0 | 0 | 0 | 0 | 0 |
| ENSDARG000 | 7;7;7;7;7;7;7 | 17253149:172 | 17253219:172 | -;-;-;-;-;-;-;- | 1280 | nitr12 | 2 | 0 | 0 | 0 | 0 | 2 |
| ENSDARG000 | 7;7;7;7;7;7 | 9308625:9312 | 9308836:9312 | +;+;+;+;+;+ | 3587 | selenos | 882 | 1522 | 1905 | 1636 | 1880 | 2271 |
| ENSDARG000 | 7;7;7;7;7;7;7 | 24390939:243 | 24391181:243 | +;+;+;+;+;+;+ | 2529 | haus3 | 151 | 504 | 332 | 244 | 300 | 227 |
| ENSDARG000 | 7;7 | 47155993:471 | 47157316:471 | -;- | 1448 | NA | 0 | 0 | 0 | 0 | 0 | 0 |
| ENSDARG000 | 7;7;7;7;7 | 35112052:351 | 35113567:351 | -;-;-;-;- | 2592 | hsd11b2 | 10 | 14 | 6 | 14 | 62 | 0 |
| ENSDARG000 | 7;7;7 | 47162928:472 | 47164681:472 | -;-;- | 2005 | NA | 0 | 0 | 0 | 0 | 0 | 0 |
| ENSDARG000 | 7;7;7;7;7;7;7 | 47243564:472 | 47243737:472 | +;+;+;+;+;+;+ | 4473 | ZNF507 | 154 | 92 | 197 | 116 | 212 | 148 |
| ENSDARG000 | 7;7;7 | 29015846:290 | 29017808:290 | -;-;- | 2412 | nrn1lb | 0 | 0 | 4 | 0 | 0 | 0 |
| ENSDARG000 | 7;7;7 | 47226092:472 | 47226381:472 | -;-;- | 637 | NA | 0 | 0 | 0 | 0 | 0 | 0 |
| ENSDARG000 | 7;7;7;7;7;7;7 | 59156413:591 | 59157280:591 | -;-;-;-;-;-;-;- | 3355 | haus6 | 170 | 546 | 517 | 217 | 442 | 337 |
| ENSDARG000 | 7;7;7;7;7;7;7 | 35191220:351 | 35191829:351 | +;+;+;+;+;+;+ | 4136 | zdhhc1 | 218 | 237 | 386 | 365 | 438 | 396 |
| ENSDARG000 | 7;7;7;7 | 54926474:549 | 54927013:549 | -;-;-;- | 821 | NA | 0 | 0 | 0 | 0 | 1 | 0 |
| ENSDARG000 | 7;7;7;7;7;7;7 | 22801465:228 | 22801554:228 | +;+;+;+;+;+;+ | 1705 | rbm4.1 | 746 | 570 | 987 | 440 | 1270 | 650 |
| ENSDARG000 | 7;7;7;7 | 14536083:145 | 14536377:145 | +;+;+;+ | 1828 | mrpl46 | 210 | 276 | 362 | 94 | 368 | 144 |
| ENSDARG000 | 7;7;7;7;7;7;7 | 52389958:523 | 52391120:523 | -;-;-;-;-;-;-;- | 3257 | MYZAP | 4 | 4 | 4 | 6 | 22 | 2 |
| ENSDARG000 | 7;7;7;7;7;7;7 | 14512000:145 | 14512307:145 | -;-;-;-;-;-;-;- | 1029 | mrps11 | 359 | 524 | 617 | 230 | 405 | 376 |
| ENSDARG000 | 7;7;7;7;7;7;7 | 26629084:266 | 26629291:266 | +;+;+;+;+;+;+ | 856 | hsbp1a | 244 | 535 | 259 | 852 | 155 | 593 |
| ENSDARG000 | 7;7;7 | 55112922:551 | 55113138:551 | +;+;+ | 1393 | snai3 | 2 | 0 | 0 | 0 | 0 | 0 |
| ENSDARG000 | 7;7;7 | 9326234:9343 | 9326832:9344 | +;+;+ | 3921 | chsy1 | 1690 | 1384 | 903 | 1661 | 2146 | 1514 |
| ENSDARG000 | 7;7;7;7 | 3461500:3465 | 3463952:3465 | -;-;-;- | 4454 | NA | 22 | 0 | 0 | 0 | 6 | 0 |
| ENSDARG000 | 7;7;7;7;7;7;7 | 55149001:551 | 55149081:551 | +;+;+;+;+;+;+ | 2844 | NA | 20 | 0 | 0 | 4 | 0 | 0 |
| ENSDARG000 | 7;7;7;7;7;7;7 | 38897836:388 | 38898309:388 | +;+;+;+;+;+;+ | 3907 | creb3l1 | 44 | 76 | 14 | 6 | 24 | 0 |
| ENSDARG000 | 7;7;7 | 66136409:661 | 66136721:661 | -;-;- | 488 | pth1b | 0 | 0 | 0 | 0 | 0 | 0 |
| ENSDARG000 | 7;7;7;7;7;7;7 | 20459671:204 | 20460244:204 | -;-;-;-;-;-;-;- | 1374 | cnpy4 | 104 | 92 | 98 | 92 | 143 | 92 |
| ENSDARG000 | 7;7;7;7;7;7;7 | 3483334:3483 | 3483337:3483 | +;+;+;+;+;+;+ | 5644 | NA | 0 | 0 | 0 | 6 | 0 | 0 |
| ENSDARG000 | 7;7;7;7;7;7;7 | 9526268:9533 | 9526537:9533 | -;-;-;-;-;-;-;- | 6534 | LRRK1 | 120 | 42 | 333 | 104 | 187 | 195 |
| ENSDARG000 | 7;7;7;7;7;7;7 | 55292959:552 | 55292998:552 | +;+;+;+;+;+;+ | 1799 | ctu2 | 176 | 174 | 304 | 190 | 179 | 191 |
| ENSDARG000 | 7;7;7;7;7;7;7 | 19482084:194 | 19482122:194 | +;+;+;+;+;+;+ | 2203 | NA | 680 | 653 | 855 | 334 | 1842 | 624 |
| ENSDARG000 | 7;7;7;7;7;7;7 | 50849142:508 | 50849418:508 | +;+;+;+;+;+;+ | 2600 | pcolceb | 122 | 82 | 174 | 196 | 398 | 24 |
| ENSDARG000 | 7;7;7;7;7;7;7 | 25135739:251 | 25135895:251 | +;+;+;+;+;+;+ | 4170 | exoc6b | 945 | 644 | 749 | 777 | 749 | 1032 |
| ENSDARG000 | 7;7;7;7;7 | 47273352:472 | 47273607:472 | -;-;-;-;- | 768 | NA | 0 | 0 | 0 | 0 | 2 | 0 |
| ENSDARG000 | 7;7;7;7;7;7;7 | 26709251:267 | 26709346:267 | +;+;+;+;+;+;+ | 4123 | cd82a | 762 | 339 | 956 | 716 | 218 | 1088 |
| ENSDARG000 | 7;7;7;7;7;7;7 | 41903540:419 | 41904549:419 | -;-;-;-;-;-;-;- | 2343 | dnaja2a | 4760 | 4580 | 5815 | 3590 | 3989 | 3600 |
| ENSDARG000 | 7;7;7;7;7;7;7 | 26649319:266 | 26649397:266 | +;+;+;+;+;+;+ | 1999 | tp53i11a | 132 | 93 | 187 | 74 | 164 | 188 |
| ENSDARG000 | 7;7;7;7;7;7;7 | 35229645:352 | 35229851:352 | +;+;+;+;+;+;+ | 3184 | tppp3 | 14 | 22 | 12 | 38 | 54 | 18 |
| ENSDARG000 | 7 | 50876581 | 50883433 | - | 6853 | PAQR9 | 38 | 54 | 64 | 44 | 70 | 28 |
| ENSDARG000 | 7;7;7;7 | 59169081:591 | 59169289:591 | +;+;+;+ | 834 | ostc | 1177 | 1795 | 2004 | 1163 | 1673 | 1260 |
| ENSDARG000 | 7;7;7;7;7;7;7 | 35245607:352 | 35246023:352 | +;+;+;+;+;+;+ | 5453 | AMFR | 23 | 15 | 4 | 24 | 50 | 56 |
| ENSDARG000 | 7;7;7;7;7;7;7 | 39553046:395 | 39553866:395 | -;-;-;-;-;-;-;- | 5657 | ptprja | 408 | 145 | 437 | 730 | 1128 | 283 |
| ENSDARG000 | 7;7;7;7;7;7;7 | 35268054:352 | 35268175:352 | +;+;+;+;+;+;+ | 1573 | NA | 335 | 228 | 697 | 375 | 615 | 1586 |
| ENSDARG000 | 7;7;7;7;7;7;7 | 9721385:9724 | 9722119:9724 | -;-;-;-;-;-;-;- | 2256 | aldh1a3 | 73 | 2 | 3 | 0 | 32 | 0 |
| ENSDARG000 | 7;7;7;7;7 | 59212666:592 | 59212969:592 | +;+;+;+;+ | 2467 | dok1b | 686 | 540 | 701 | 593 | 771 | 905 |
| ENSDARG000 | 7;7;7;7;7;7;7 | 22809905:228 | 22810296:228 | +;+;+;+;+;+;+ | 3009 | sf1 | 3691 | 3489 | 5056 | 3468 | 4663 | 3328 |
| ENSDARG000 | 7;7;7;7;7;7;7 | 66259046:662 | 66259837:662 | -;-;-;-;-;-;-;- | 2833 | GALNT18 | 8 | 0 | 3 | 0 | 6 | 10 |
| ENSDARG000 | 7;7 | 71335815:713 | 71335918:713 | +;+ | 1125 | sod3a | 0 | 0 | 0 | 0 | 0 | 0 |
| ENSDARG000 | 7;7;7 | 50890078:508 | 50890163:508 | -;-;- | 282 | NA | 0 | 0 | 0 | 0 | 0 | 0 |
| ENSDARG000 | 7;7;7;7;7 | 9840994:9858 | 9845276:9859 | -;-;-;-;- | 5278 | asb7 | 207 | 150 | 200 | 120 | 214 | 242 |
| ENSDARG000 | 7;7;7;7 | 3532393:3532 | 3532438:3532 | +;+;+;+ | 1181 | NA | 0 | 0 | 0 | 0 | 0 | 0 |
| ENSDARG000 | 7;7 | 38456426:384 | 38457782:384 | -;- | 1743 | zgc:165481 | 0 | 4 | 0 | 9 | 0 | 0 |
| ENSDARG000 | 7;7;7;7;7;7;7 | 38395197:383 | 38395481:383 | +;+;+;+;+;+;+ | 3799 | cep89 | 99 | 42 | 73 | 38 | 68 | 14 |
| ENSDARG000 | 7;7;7;7;7;7 | 20467549:204 | 20467798:204 | +;+;+;+;+;+ | 480 | NA | 2 | 0 | 0 | 0 | 0 | 0 |
| ENSDARG000 | 7;7;7;7;7;7;7 | 20471315:204 | 20471440:204 | +;+;+;+;+;+;+ | 3170 | NA | 0 | 0 | 0 | 0 | 0 | 0 |
| ENSDARG000 | 7;7;7 | 24457191:244 | 24462425:244 | -;-;- | 5836 | nat8l | 14 | 207 | 158 | 14 | 70 | 54 |
| ENSDARG000 | 7;7;7;7;7;7 | 14995849:149 | 14996784:149 | -;-;-;-;-;- | 1350 | NA | 0 | 0 | 0 | 0 | 0 | 0 |
| ENSDARG000 | 7;7;7;7;7;7 | 24485048:244 | 24486996:244 | -;-;-;-;-;- | 2413 | C7H4orf48 | 402 | 476 | 507 | 617 | 408 | 507 |
| ENSDARG000 | 7;7;7;7;7;7 | 50910959:509 | 50911820:509 | -;-;-;-;-;- | 1452 | ankrd46b | 173 | 198 | 160 | 100 | 178 | 154 |
| ENSDARG000 | 7;7;7;7;7;7;7 | 3558030:3558 | 3558081:3558 | +;+;+;+;+;+;+ | 2717 | si:dkev-192d1 | 0 | 0 | 0 | 8 | 0 | 10 |
| ENSDARG000 | 7;7;7;7;7;7;7 | 29024282:290 | 29024435:290 | +;+;+;+;+;+;+ | 3936 | cog4 | 620 | 548 | 710 | 276 | 735 | 296 |
| ENSDARG000 | 7;7;7;7;7;7 | 71535045:715 | 71535436:715 | +;+;+;+;+;+ | 1243 | tyms | 233 | 1546 | 1168 | 466 | 941 | 538 |
| ENSDARG000 | 7;7;7;7;7;7;7 | 9873750:9873 | 9873884:9873 | +;+;+;+;+;+;+ | 2126 | NA | 56 | 36 | 58 | 56 | 80 | 41 |
| ENSDARG000 | 7;7;7;7;7;7;7 | 41160225:411 | 41160383:411 | +;+;+;+;+;+;+ | 9356 | scrib | 8029 | 4948 | 8511 | 9558 | 9072 | 7783 |
| ENSDARG000 | 7;7;7;7;7;7;7 | 9904627:9922 | 9904693:9922 | +;+;+;+;+;+;+ | 5173 | CERS3 | 0 | 0 | 0 | 0 | 0 | 0 |
| ENSDARG000 | 7;7;7;7;7;7;7 | 52431747:524 | 52432859:524 | -;-;-;-;-;-;-;- | 5012 | CGNL1 | 202 | 32 | 62 | 76 | 48 | 94 |
| ENSDARG000 | 7;7;7;7;7;7;7 | 20503344:205 | 20503408:205 | +;+;+;+;+;+;+ | 866 | NA | 8 | 0 | 0 | 0 | 0 | 0 |
| ENSDARG000 | 7;7;7;7;7;7;7 | 47287307:472 | 47287353:472 | +;+;+;+;+;+;+ | 8702 | DPY19L3 | 666 | 388 | 689 | 496 | 744 | 899 |
| ENSDARG000 | 7;7;7;7;7;7;7 | 19495379:194 | 19496270:194 | +;+;+;+;+;+;+ | 4437 | si:ch211-212k | 38 | 61 | 83 | 47 | 46 | 100 |
| ENSDARG000 | 7;7;7;7;7 | 47385216:473 | 47387628:473 | -;-;-;-;- | 3440 | NA | 0 | 0 | 0 | 0 | 0 | 0 |
| ENSDARG000 | 7;7;7;7;7;7;7 | 31145386:311 | 31146073:311 | +;+;+;+;+;+;+ | 3719 | ENTREP2 | 2 | 0 | 2 | 0 | 2 | 0 |
| ENSDARG000 | 7;7;7;7;7;7;7 | 50920437:509 | 50923751:509 | -;-;-;-;-;-;-;- | 8456 | NA | 18 | 4 | 10 | 14 | 16 | 0 |
| ENSDARG000 | 7;7;7;7;7;7;7 | 25323742:253 | 25324698:253 | +;+;+;+;+;+;+ | 3876 | cyp26b1 | 137 | 256 | 466 | 312 | 1455 | 28 |
| ENSDARG000 | 7;7;7;7;7;7;7 | 15087193:150 | 15091549:150 | -;-;-;-;-;-;-;- | 12888 | si:dkev-172h2 | 3427 | 1777 | 1965 | 2159 | 2192 | 1821 |
| ENSDARG000 | 7;7;7;7;7 | 20512419:205 | 20512475:205 | +;+;+;+;+ | 873 | NA | 52 | 32 | 4 | 6 | 10 | 34 |
| ENSDARG000 | 7;7;7;7;7;7;7 | 25339756:254 | 25339974:254 | -;-;-;-;-;-;-;- | 1194 | NA | 0 | 0 | 0 | 0 | 0 | 0 |
| ENSDARG000 | 7;7;7;7;7;7;7 | 59247447:592 | 59249189:592 | -;-;-;-;-;-;-;- | 3315 | m1ap | 8 | 0 | 2 | 0 | 36 | 14 |
| ENSDARG000 | 7 | 41254621 | 41255247 | + | 627 | NA | 4 | 0 | 0 | 0 | 0 | 0 |
| ENSDARG000 | 7;7;7;7;7;7;7 | 41275201:412 | 41275536:412 | -;-;-;-;-;-;-;- | 2582 | NA | 2 | 17 | 66 | 9 | 9 | 4 |
| ENSDARG000 | 7;7;7;7;7;7;7 | 35282885:352 | 35284195:352 | -;-;-;-;-;-;-;- | 4692 | SLC12A4 | 2627 | 1483 | 2107 | 1617 | 2566 | 2923 |
| ENSDARG000 | 7;7 | 66536017:665 | 66536447:665 | +;+ | 572 | NA | 0 | 0 | 0 | 0 | 0 | 0 |
| ENSDARG000 | 7;7;7;7;7;7;7 | 39624728:396 | 39624847:396 | +;+;+;+;+;+;+ | 8327 | PTPN5 | 66 | 10 | 112 | 8 | 34 | 0 |
| ENSDARG000 | 7;7;7;7;7;7;7 | 3597413:3601 | 3597679:3601 | +;+;+;+;+;+;+ | 6934 | si:dkev-192d1 | 0 | 0 | 0 | 0 | 0 | 0 |
| ENSDARG000 | 7;7;7;7;7;7;7 | 66565572:665 | 66565971:665 | +;+;+;+;+;+;+ | 2679 | NA | 46 | 0 | 0 | 20 | 14 | 12 |
| ENSDARG000 | 7;7;7;7;7;7;7 | 38962207:389 | 38962649:389 | +;+;+;+;+;+;+ | 9008 | DGKZ | 1146 | 962 | 1089 | 962 | 1192 | 749 |
| ENSDARG000 | 7;7;7;7;7;7;7 | 55518519:555 | 55519129:555 | +;+;+;+;+;+;+ | 2432 | CDT1 | 66 | 136 | 202 | 40 | 118 | 87 |
| ENSDARG000 | 7;7;7;7;7 | 20518218:205 | 20518360:205 | +;+;+;+;+ | 2137 | zgc:103586 | 8 | 2 | 4 | 16 | 81 | 0 |
| ENSDARG000 | 7;7;7;7;7 | 3678848:3679 | 3679235:3679 | -;-;-;-;- | 1185 | NA | 0 | 0 | 0 | 0 | 0 | 0 |
| ENSDARG000 | 7;7;7;7;7 | 3654800:3655 | 3655210:3655 | -;-;-;-;- | 1208 | si:ch211-282i | 0 | 0 | 0 | 0 | 4 | 0 |
| ENSDARG000 | 7;7;7 | 3664574:3669 | 3666692:3669 | -;-;- | 2214 | NA | 0 | 0 | 0 | 0 | 0 | 0 |
| ENSDARG000 | 7;7;7;7;7;7;7 | 55530532:555 | 55531485:555 | -;-;-;-;-;-;-;- | 2742 | aprt | 312 | 298 | 459 | 296 | 368 | 336 |
| ENSDARG000 | 7;7;7;7;7;7;7 | 3622558:3716 | 3623361:3717 | -;-;-;-;-;-;- | 2335 | NA | 2 | 4 | 4 | 0 | 0 | 0 |
| ENSDARG000 | 7;7;7;7;7 | 3643979:3644 | 3644408:3644 | -;-;-;-;- | 1271 | si:ch211-282i | 0 | 0 | 0 | 0 | 0 | 2 |
| ENSDARG000 | 7;7;7;7;7;7 | 3623003:3627 | 3623361:3627 | -;-;-;-;-;- | 1213 | NA | 0 | 0 | 0 | 0 | 0 | 0 |
| ENSDARG000 | 7;7;7;7;7;7;7 | 6684123:6687 | 6684872:6687 | -;-;-;-;-;-;-;- | 3182 | capn1 | 851 | 567 | 823 | 720 | 587 | 600 |
| ENSDARG000 | 7;7;7;7;7 | 3684657:3687 | 3689074:3689 | -;-;-;-;- | 4965 | NA | 6 | 0 | 0 | 0 | 0 | 0 |
| ENSDARG000 | 7;7;7;7;7 | 3632351:3637 | 3632718:3637 | -;-;-;-;- | 1142 | NA | 0 | 0 | 0 | 0 | 0 | 0 |
| ENSDARG000 | 7;7;7 | 3698790:3699 | 3699151:3699 | -;-;- | 945 | NA | 0 | 0 | 0 | 0 | 0 | 0 |
| ENSDARG000 | 7;7;7;7;7;7 | 10502631:105 | 10503076:105 | -;-;-;-;-;- | 1424 | mthfs | 138 | 102 | 242 | 80 | 78 | 120 |
| ENSDARG000 | 7;7;7;7;7;7;7 | 71547747:715 | 71548218:715 | +;+;+;+;+;+;+ | 1902 | adcyap1a | 0 | 0 | 0 | 0 | 0 | 0 |
| ENSDARG000 | 7;7;7;7;7;7;7 | 38978002:389 | 38978071:389 | -;-;-;-;-;-;- | 393 | NA | 0 | 0 | 4 | 0 | 2 | 2 |
| ENSDARG000 | 7;7;7;7;7;7;7 | 33028452:330 | 33029188:330 | +;+;+;+;+;+;+ | 6573 | kcnc1a | 6 | 0 | 0 | 2 | 0 | 0 |
| ENSDARG000 | 7;7;7;7;7;7;7 | 59505081:595 | 59505387:595 | -;-;-;-;-;-;-;- | 1274 | slx1b | 204 | 192 | 281 | 215 | 275 | 238 |
| ENSDARG000 | 7;7;7;7;7;7;7 | 20524064:205 | 20524117:205 | +;+;+;+;+;+;+ | 2303 | slc3a2a | 585 | 223 | 484 | 152 | 1022 | 1370 |
| ENSDARG000 | 7;7;7;7;7;7;7 | 24508123:245 | 24508388:245 | -;-;-;-;-;-;-;- | 1752 | faah2b | 0 | 0 | 0 | 0 | 0 | 0 |
| ENSDARG000 | 7;7;7;7;7;7;7 | 66739166:667 | 66739449:667 | +;+;+;+;+;+;+ | 4012 | IRAG1 | 16 | 6 | 12 | 2 | 24 | 18 |
| ENSDARG000 | 7;7;7;7;7;7;7 | 42206543:422 | 42206908:422 | +;+;+;+;+;+;+ | 4646 | PHKB | 538 | 446 | 515 | 340 | 793 | 749 |
| ENSDARG000 | 7;7;7;7;7;7;7 | 24491966:244 | 24492110:244 | +;+;+;+;+;+;+ | 5632 | nelfa | 390 | 371 | 581 | 472 | 428 | 382 |
| ENSDARG000 | 7;7;7;7;7;7;7 | 4931366:4931 | 4931440:4931 | +;+;+;+;+;+;+ | 1892 | si:dkey-28d5.1 | 8 | 0 | 0 | 2 | 2 | 0 |
| ENSDARG000 | 7;7;7;7;7;7 | 66822229:668 | 66822566:668 | +;+;+;+;+;+ | 1472 | lyve1a | 157 | 24 | 74 | 96 | 71 | 126 |
| ENSDARG000 | 7;7;7;7;7;7;7 | 24520518:245 | 24520811:245 | +;+;+;+;+;+;+ | 2511 | btr09 | 146 | 254 | 149 | 159 | 49 | 106 |
| ENSDARG000 | 7;7;7;7;7;7;7 | 47417529:474 | 47420210:474 | -;-;-;-;-;-;-;- | 10842 | NA | 44 | 6 | 0 | 10 | 0 | 108 |
| ENSDARG000 | 7;7;7;7;7;7;7 | 31339129:313 | 31339484:313 | +;+;+;+;+;+;+ | 2815 | GOLM2 | 3560 | 2807 | 3347 | 2633 | 5224 | 2520 |
| ENSDARG000 | 7;7;7;7;7;7;7 | 38507006:385 | 38507119:385 | +;+;+;+;+;+;+ | 2327 | SLC7A9 | 30 | 6 | 20 | 44 | 59 | 0 |
| ENSDARG000 | 7;7;7;7;7;7;7 | 6756206:6756 | 6756372:6756 | +;+;+;+;+;+;+ | 2677 | sf3b2 | 2449 | 2361 | 3277 | 1789 | 2855 | 1600 |
| ENSDARG000 | 7;7;7 | 26750471:267 | 26750608:267 | +;+;+ | 379 | NA | 0 | 2 | 0 | 2 | 0 | 7 |
| ENSDARG000 | 7;7;7;7;7;7;7 | 39664055:396 | 39664307:396 | +;+;+;+;+;+;+ | 8443 | kiaa0232 | 4022 | 2413 | 3179 | 1864 | 2966 | 2392 |
| ENSDARG000 | 7;7;7;7;7;7;7 | 33106332:331 | 33107160:331 | -;-;-;-;-;-;-;- | 2173 | tph1b | 4 | 0 | 4 | 0 | 0 | 0 |
| ENSDARG000 | 7;7;7;7;7;7;7 | 25558275:255 | 25561213:255 | -;-;-;-;-;-;-;- | 9317 | dysf | 208 | 171 | 1268 | 324 | 226 | 294 |
| ENSDARG000 | 7;7;7;7;7;7;7 | 26762717:267 | 26763181:267 | +;+;+;+;+;+;+ | 1994 | tspan18a | 1621 | 1172 | 1233 | 1341 | 390 | 3474 |
| ENSDARG000 | 7;7;7;7;7;7;7 | 59649399:596 | 59649648:596 | +;+;+;+;+;+;+ | 1490 | rpl34 | 12428 | 12124 | 19150 | 19235 | 11618 | 19018 |
| ENSDARG000 | 7;7;7;7;7;7;7 | 20577694:205 | 20577745:205 | -;-;-;-;-;-;-;- | 3266 | zpax4 | 0 | 2 | 8 | 0 | 2 | 4 |
| ENSDARG000 | 7;7;7;7;7;7;7 | 66836476:668 | 66839699:668 | -;-;-;-;-;-;-;- | 5615 | AMPD3 | 2 | 2 | 4 | 0 | 17 | 12 |
| ENSDARG000 | 7;7;7;7;7;7;7 | 20536258:205 | 20536728:205 | -;-;-;-;-;-;-;- | 1379 | NA | 0 | 0 | 0 | 0 | 0 | 0 |
| ENSDARG000 | 7;7;7;7;7;7;7 | 10562118:105 | 10562297:105 | +;+;+;+;+;+;+ | 1709 | zfand6 | 354 | 294 | 423 | 227 | 326 | 328 |
| ENSDARG000 | 7;7;7;7;7;7;7 | 20563305:205 | 20564069:205 | +;+;+;+;+;+;+ | 3409 | NA | 8 | 2 | 0 | 4 | 2 | 14 |
| ENSDARG000 | 7;7;7;7;7;7;7 | 22823889:228 | 22824290:228 | +;+;+;+;+;+;+ | 5071 | pygmb | 220 | 669 | 3151 | 798 | 1288 | 442 |
| ENSDARG000 | 7;7;7;7;7;7;7 | 20535869:205 | 20535891:205 | +;+;+;+;+;+;+ | 1701 | zgc:158423 | 0 | 0 | 6 | 0 | 2 | 4 |
| ENSDARG000 | 7;7;7;7;7 | 4938463:4938 | 4938511:4938 | +;+;+;+;+ | 1162 | NA | 0 | 0 | 0 | 0 | 0 | 0 |
| ENSDARG000 | 7;7;7;7;7;7 | 29992889:299 | 29993039:299 | +;+;+;+;+;+ | 2181 | isl2b | 0 | 0 | 0 | 2 | 4 | 0 |
| ENSDARG000 | 7;7;7;7;7;7 | 17393794:173 | 17393856:173 | +;+;+;+;+;+ | 833 | nitr7b | 50 | 0 | 0 | 0 | 0 | 0 |
| ENSDARG000 | 7;7;7;7;7;7;7 | 35348471:353 | 35351777:353 | -;-;-;-;-;-;-;- | 5398 | SLC6A2 | 62 | 3 | 0 | 84 | 40 | 72 |
| ENSDARG000 | 7;7;7;7;7;7;7 | 51103232:511 | 51103493:511 | +;+;+;+;+;+;+ | 6455 | col4a5 | 36 | 0 | 24 | 2 | 38 | 0 |
| ENSDARG000 | 7 | 29041465 | 29043075 | - | 1611 | thap11 | 124 | 100 | 178 | 98 | 229 | 106 |
| ENSDARG000 | 7;7;7;7 | 29044888:290 | 29045217:290 | +;+;+;+ | 2347 | gfod2 | 130 | 118 | 85 | 142 | 143 | 185 |
| ENSDARG000 | 7;7;7;7;7;7 | 25702443:257 | 25703003:257 | +;+;+;+;+;+ | 1967 | NA | 20 | 4 | 0 | 0 | 0 | 0 |
| ENSDARG000 | 7;7;7;7 | 66874770:668 | 66876020:668 | -;-;-;- | 1687 | adma | 90 | 158 | 208 | 612 | 85 | 74 |
| ENSDARG000 | 7;7;7;7;7;7;7 | 33120167:331 | 33120573:331 | -;-;-;-;-;-;-;- | 2149 | mns1 | 28 | 20 | 58 | 80 | 36 | 38 |
| ENSDARG000 | 7;7;7;7;7;7;7 | 55543351:555 | 55543757:555 | -;-;-;-;-;-;-;- | 2033 | galns | 1079 | 722 | 1025 | 1004 | 864 | 843 |
| ENSDARG000 | 7;7;7;7;7;7;7 | 19502744:195 | 19502777:195 | +;+;+;+;+;+;+ | 2290 | NA | 54 | 43 | 93 | 50 | 64 | 76 |
| ENSDARG000 | 7;7;7;7;7;7;7 | 17400445:174 | 17400776:174 | -;-;-;-;-;-;-;- | 1802 | nitr3a | 14 | 18 | 0 | 0 | 10 | 0 |
| ENSDARG000 | 7;7;7;7;7 | 26841136:268 | 26841735:268 | -;-;-;-;- | 1392 | si:ch211-107r | 106 | 96 | 130 | 110 | 122 | 84 |
| ENSDARG000 | 7;7;7;7;7;7;7 | 17412992:174 | 17413114:174 | -;-;-;-;-;-;-;- | 1842 | nitr3b | 4 | 14 | 3 | 0 | 1 | 2 |
| ENSDARG000 | 7;7;7;7;7;7;7 | 35385472:353 | 35386393:353 | -;-;-;-;-;-;-;- | 3792 | lpcat2 | 62 | 26 | 18 | 16 | 69 | 42 |
| ENSDARG000 | 7 | 4966092 | 4966337 | + | 246 | NA | 0 | 0 | 0 | 0 | 2 | 0 |
| ENSDARG000 | 7;7;7;7;7;7;7 | 66884291:668 | 66884624:668 | +;+;+;+;+;+;+ | 7112 | SBF2 | 1838 | 1502 | 1534 | 1575 | 1211 | 1089 |
| ENSDARG000 | 7;7;7;7;7;7;7 | 30051488:300 | 30051607:300 | +;+;+;+;+;+;+ | 3530 | pstpip1b | 14 | 40 | 66 | 44 | 97 | 40 |
| ENSDARG000 | 7;7;7;7;7;7 | 29053217:290 | 29053560:290 | +;+;+;+;+;+ | 2215 | NA | 459 | 457 | 560 | 339 | 588 | 405 |
| ENSDARG000 | 7;7;7;7;7;7 | 17624185:176 | 17624860:176 | -;-;-;-;-;- | 1439 | nitr6a | 0 | 0 | 0 | 0 | 0 | 0 |
| ENSDARG000 | 7;7;7;7;7;7;7 | 31374383:313 | 31374515:313 | +;+;+;+;+;+;+ | 6519 | ctdspl2b | 298 | 499 | 668 | 383 | 449 | 404 |
| ENSDARG000 | 7;7;7;7;7;7;7 | 4971938:4971 | 4971986:4971 | +;+;+;+;+;+;+ | 1249 | si:dkev-81n2.1 | 0 | 0 | 0 | 0 | 0 | 0 |
| ENSDARG000 | 7;7;7;7;7;7;7 | 6969909:7048 | 6970046:7048 | +;+;+;+;+;+;+ | 6533 | actn3b | 12 | 12 | 6 | 2 | 28 | 16 |
| ENSDARG000 | 7;7 | 52551760:525 | 52552194:525 | -;- | 528 | NA | 0 | 6 | 0 | 4 | 0 | 0 |
| ENSDARG000 | 7;7;7;7;7;7;7 | 52516706:525 | 52519236:525 | -;-;-;-;-;-;-;- | 6495 | tcf12 | 1596 | 1279 | 1820 | 1053 | 2687 | 1143 |
| ENSDARG000 | 7;7;7;7;7;7;7 | 59677273:596 | 59677730:596 | +;+;+;+;+;+;+ | 2746 | trmt44 | 202 | 207 | 157 | 64 | 233 | 150 |
| ENSDARG000 | 7;7;7;7;7;7 | 17633113:176 | 17633275:176 | -;-;-;-;-;- | 840 | NA | 0 | 0 | 0 | 0 | 0 | 0 |
| ENSDARG000 | 7;7;7 | 42386533:423 | 42387059:423 | -;-;- | 1060 | NA | 18 | 4 | 6 | 0 | 0 | 0 |
| ENSDARG000 | 7;7;7;7;7;7;7 | 42367467:423 | 42367585:423 | +;+;+;+;+;+;+ | 862 | NA | 30 | 12 | 14 | 0 | 0 | 0 |
| ENSDARG000 | 7;7;7;7;7;7;7 | 6990510:6991 | 6990555:6991 | +;+;+;+;+;+;+ | 2118 | ccs | 184 | 110 | 141 | 80 | 295 | 204 |
| ENSDARG000 | 7;7;7;7;7;7;7 | 39679944:396 | 39680355:396 | +;+;+;+;+;+;+ | 7070 | TBC1D14 | 1103 | 926 | 1061 | 474 | 798 | 882 |
| ENSDARG000 | 7;7;7;7;7;7;7 | 6941583:6941 | 6942164:6942 | +;+;+;+;+;+;+ | 11166 | rbm14b | 1067 | 629 | 845 | 614 | 1296 | 1104 |
| ENSDARG000 | 7;7;7;7;7;7;7 | 19509888:195 | 19510129:195 | -;-;-;-;-;-;-;- | 3231 | MAN2B2 | 2220 | 1559 | 1308 | 1127 | 1816 | 1288 |
| ENSDARG000 | 7;7;7;7;7;7;7 | 33130639:331 | 33130755:331 | +;+;+;+;+;+;+ | 2210 | zgc:153219 | 264 | 222 | 378 | 422 | 391 | 362 |

| | | | | | | | | | | | | |
|---|---|---|---|---|---|---|---|---|---|---|---|---|
| ENSDARG000 | 7;7;7 | 48219331:482 | 48219367:482 | -;-;- | 301 | NA | 0 | 0 | 0 | 0 | 0 | 0 |
| ENSDARG000 | 7;7;7;7;7;7;7; | 4987530:4987 | 4987572:4987 | +;+;+;+;+;+;+; | 1153 | si:dkev-81n2. | 0 | 0 | 0 | 0 | 0 | 0 |
| ENSDARG000 | 7;7;7;7;7;7;7; | 33136533:331 | 33136548:331 | +;+;+;+;+;+;+; | 1315 | NA | 0 | 0 | 0 | 0 | 0 | 0 |
| ENSDARG000 | 7;7;7;7;7;7;7; | 15266093:152 | 15266214:152 | +;+;+;+;+;+;+; | 4734 | sv2ba | 4 | 2 | 39 | 0 | 14 | 6 |
| ENSDARG000 | 7;7;7;7;7;7 | 38529263:385 | 38530049:385 | +;+;+;+;+;+ | 2386 | nudt19 | 341 | 299 | 582 | 253 | 255 | 51 |
| ENSDARG000 | 7;7;7;7;7;7;7; | 33130661:331 | 33130794:331 | +;+;+;+;+;+;+; | 1227 | NA | 0 | 0 | 0 | 0 | 0 | 0 |
| ENSDARG000 | 7;7;7;7;7;7;7; | 25858380:258 | 25858648:258 | +;+;+;+;+;+;+; | 4081 | mtmr1a | 110 | 116 | 137 | 113 | 156 | 169 |
| ENSDARG000 | 7;7;7;7;7;7;7; | 31395525:313 | 31396043:313 | +;+;+;+;+;+;+; | 7241 | NA | 2 | 0 | 1 | 0 | 0 | 2 |
| ENSDARG000 | 7;7;7;7;7;7;7; | 26844261:268 | 26844425:268 | +;+;+;+;+;+;+; | 3480 | ext2 | 2522 | 2066 | 2248 | 1634 | 2311 | 2157 |
| ENSDARG000 | 7;7;7;7;7;7;7; | 29065915:290 | 29066302:290 | +;+;+;+;+;+;+; | 2221 | vrk3 | 354 | 338 | 418 | 183 | 310 | 288 |
| ENSDARG000 | 7;7;7;7;7;7;7; | 25878553:258 | 25881952:258 | -;-;-;-;-;-;-; | 4647 | cd99l2 | 1024 | 1738 | 1551 | 744 | 1446 | 1085 |
| ENSDARG000 | 7;7;7;7;7;7;7; | 71664624:716 | 71664921:716 | +;+;+;+;+;+;+; | 3011 | NA | 4 | 2 | 10 | 0 | 40 | 2 |
| ENSDARG000 | 7;7;7;7;7;7;7; | 35410050:354 | 35411000:354 | -;-;-;-;-;-;-; | 3033 | mmp2 | 1104 | 500 | 1026 | 532 | 3472 | 206 |
| ENSDARG000 | 7;7;7;7;7;7 | 17641188:176 | 17641292:176 | -;-;-;-;-;- | 713 | NA | 0 | 0 | 0 | 0 | 0 | 0 |
| ENSDARG000 | 7;7;7;7;7;7;7; | 48240707:482 | 48242015:482 | -;-;-;-;-;-;-; | 3572 | cpeb1b | 0 | 0 | 0 | 0 | 0 | 0 |
| ENSDARG000 | 7;7;7;7;7;7;7; | 42460302:424 | 42460602:424 | +;+;+;+;+;+;+; | 7458 | ADAMTS18 | 617 | 98 | 6 | 6 | 208 | 16 |
| ENSDARG000 | 7;7 | 4998793:4998 | 4998829:4999 | +;+ | 387 | NA | 0 | 0 | 0 | 0 | 0 | 0 |
| ENSDARG000 | 7;7;7;7;7;7;7; | 55637379:556 | 55637714:556 | -;-;-;-;-;-;-; | 1085 | pabpn1l | 0 | 3 | 0 | 1 | 0 | 0 |
| ENSDARG000 | 7;7;7;7;7;7;7; | 60111458:601 | 60111812:601 | +;+;+;+;+;+;+; | 3245 | hspa12b | 49 | 16 | 33 | 3 | 77 | 124 |
| ENSDARG000 | 7;7;7;7;7;7;7 | 30075245:300 | 30077832:300 | -;-;-;-;-;-;- | 3381 | tspan3b | 0 | 4 | 6 | 0 | 15 | 18 |
| ENSDARG000 | 7;7;7;7;7;7;7; | 39166460:391 | 39166588:391 | +;+;+;+;+;+;+; | 2750 | mdka | 170 | 28 | 86 | 14 | 66 | 96 |
| ENSDARG000 | 7;7;7;7;7 | 3744945:3745 | 3745319:3745 | -;-;-;-;- | 1166 | si:ch211-282i | 0 | 0 | 0 | 0 | 0 | 0 |
| ENSDARG000 | 7;7;7;7;7 | 5007310:5007 | 5007350:5007 | +;+;+;+;+ | 1168 | si:ch211-272h | 0 | 0 | 0 | 0 | 0 | 0 |
| ENSDARG000 | 7;7;7;7;7;7;7; | 3755739:3755 | 3756108:3756 | -;-;-;-;-;-;-;- | 1204 | si:ch211-282i | 0 | 0 | 0 | 0 | 0 | 0 |
| ENSDARG000 | 7;7;7;7;7;7;7; | 71688710:716 | 71690161:716 | -;-;-;-;-;-;-; | 6959 | MYOM1 | 0 | 2 | 1 | 0 | 2 | 0 |
| ENSDARG000 | 7;7;7;7;7;7;7; | 10701732:107 | 10701976:107 | +;+;+;+;+;+;+; | 8155 | arnt2 | 480 | 239 | 502 | 713 | 458 | 415 |
| ENSDARG000 | 7;7;7;7;7;7;7; | 22853788:228 | 22854082:228 | +;+;+;+;+;+;+; | 6823 | CLCN5 | 370 | 342 | 663 | 354 | 524 | 362 |
| ENSDARG000 | 7;7;7;7 | 26893688:268 | 26897487:268 | -;-;-;- | 4928 | ALX4 | 2127 | 789 | 9708 | 675 | 3048 | 6462 |
| ENSDARG000 | 7;7;7;7;7;7;7; | 19528265:195 | 19528392:195 | +;+;+;+;+;+;+; | 1650 | ttc5 | 182 | 192 | 264 | 190 | 212 | 200 |
| ENSDARG000 | 7;7;7;7;7;7;7; | 19538425:195 | 19538793:195 | -;-;-;-;-;-;-;- | 999 | CCNB1IP1 | 0 | 0 | 0 | 0 | 0 | 0 |
| ENSDARG000 | 7;7;7;7;7;7;7; | 38535934:385 | 38537884:385 | -;-;-;-;-;-;-; | 4055 | celf1 | 1891 | 1978 | 2666 | 1957 | 2818 | 2491 |
| ENSDARG000 | 7;7;7;7;7;7;7; | 43131218:431 | 43131711:431 | -;-;-;-;-;-;-; | 4306 | NA | 6 | 12 | 29 | 58 | 0 | 2 |
| ENSDARG000 | 7;7 | 35514655:355 | 35515199:355 | +;+ | 2938 | NA | 0 | 0 | 0 | 0 | 0 | 0 |
| ENSDARG000 | 7;7;7;7;7;7;7; | 35435258:354 | 35435447:354 | +;+;+;+;+;+;+; | 2421 | NA | 0 | 0 | 0 | 0 | 2 | 0 |
| ENSDARG000 | 7;7;7;7;7;7;7; | 17648931:176 | 17649351:176 | -;-;-;-;-;-;-; | 3149 | map4k2 | 32 | 6 | 15 | 31 | 35 | 44 |
| ENSDARG000 | 7;7;7;7;7;7;7; | 20587506:205 | 20587848:205 | +;+;+;+;+;+;+; | 1693 | NA | 0 | 0 | 0 | 0 | 0 | 0 |
| ENSDARG000 | 7;7;7;7;7;7;7; | 35511732:355 | 35512305:355 | -;-;-;-;-;-;-; | 2340 | irx6a | 0 | 0 | 0 | 0 | 0 | 0 |
| ENSDARG000 | 7 | 38539306 | 38540433 | - | 1128 | NA | 431 | 369 | 625 | 499 | 868 | 980 |
| ENSDARG000 | 7;7;7;7;7;7;7; | 60148452:601 | 60149086:601 | -;-;-;-;-;-;-; | 2226 | cct7 | 9514 | 9469 | 12726 | 7778 | 9138 | 9228 |
| ENSDARG000 | 7;7 | 43466635:434 | 43466671:434 | +;+ | 409 | NA | 0 | 0 | 0 | 4 | 0 | 0 |
| ENSDARG000 | 7;7 | 43468336:434 | 43468377:434 | -;- | 279 | NA | 0 | 0 | 0 | 0 | 0 | 0 |
| ENSDARG000 | 7;7;7 | 30085149:300 | 30085441:300 | -;-;- | 679 | nmbb | 4 | 0 | 0 | 0 | 0 | 0 |
| ENSDARG000 | 7;7;7 | 39706004:397 | 39706128:397 | +;+;+ | 778 | ccl36.1 | 7 | 22 | 0 | 15 | 52 | 205 |
| ENSDARG000 | 7;7;7;7;7;7;7; | 31425976:314 | 31426087:314 | +;+;+;+;+;+;+; | 2023 | eif3jb | 766 | 1050 | 1259 | 555 | 1003 | 798 |
| ENSDARG000 | 7;7;7;7;7 | 15308219:153 | 15308313:153 | +;+;+;+;+ | 1120 | mespba | 6 | 0 | 0 | 2 | 0 | 0 |
| ENSDARG000 | 7;7;7;7;7;7;7; | 15308042:153 | 15308126:153 | +;+;+;+;+;+;+; | 2521 | mespaa | 0 | 7 | 6 | 0 | 14 | 0 |
| ENSDARG000 | 7;7 | 43489894:434 | 43490022:434 | +;+ | 651 | NA | 14 | 0 | 4 | 0 | 0 | 2 |
| ENSDARG000 | 7;7;7;7;7;7;7; | 56098590:561 | 56098637:561 | +;+;+;+;+;+;+; | 2643 | cdh15 | 6 | 0 | 2 | 0 | 0 | 0 |
| ENSDARG000 | 7;7;7;7;7;7;7; | 24528414:245 | 24528585:245 | +;+;+;+;+;+;+; | 3664 | TESK1 | 20 | 16 | 50 | 26 | 28 | 28 |
| ENSDARG000 | 7;7;7;7;7;7;7; | 71764665:717 | 71764956:717 | +;+;+;+;+;+;+; | 1459 | myl12.1 | 6641 | 9973 | 9500 | 7745 | 7700 | 7980 |
| ENSDARG000 | 7;7;7;7;7;7;7; | 3831626:3831 | 3832002:3832 | -;-;-;-;-;-;-;- | 1227 | si:dkev-88n24 | 0 | 0 | 0 | 0 | 0 | 0 |
| ENSDARG000 | 7;7;7;7;7;7;7; | 3848298:3848 | 3848667:3848 | -;-;-;-;-;-;-; | 1200 | si:dkev-88n24 | 1 | 0 | 0 | 0 | 0 | 0 |
| ENSDARG000 | 7;7;7;7;7 | 3802249:3802 | 3803886:3803 | -;-;-;-;- | 1979 | zgc:195077 | 0 | 0 | 0 | 0 | 0 | 0 |
| ENSDARG000 | 7;7;7;7;7;7;7; | 30109990:301 | 30110679:301 | -;-;-;-;-;-;-; | 6122 | NA | 0 | 0 | 0 | 0 | 0 | 0 |
| ENSDARG000 | 7;7;7;7;7;7;7; | 3814931:3815 | 3815474:3815 | -;-;-;-;-;-;-; | 1356 | si:dkev-88n24 | 0 | 0 | 0 | 0 | 0 | 0 |
| ENSDARG000 | 7;7;7;7;7;7;7; | 67183718:671 | 67183833:671 | -;-;-;-;-;-;-; | 1831 | SWAP70 | 24 | 12 | 0 | 0 | 41 | 32 |
| ENSDARG000 | 7;7;7 | 19552381:195 | 19552682:195 | +;+;+ | 4443 | NA | 0 | 0 | 10 | 12 | 0 | 0 |
| ENSDARG000 | 7;7;7;7;7;7;7; | 3783875:3783 | 3784289:3784 | -;-;-;-;-;-;-; | 1278 | si:ch73-335d1 | 0 | 0 | 0 | 0 | 0 | 0 |
| ENSDARG000 | 7;7;7;7 | 39713215:397 | 39713374:397 | +;+;+;+ | 384 | NA | 0 | 0 | 0 | 0 | 0 | 0 |
| ENSDARG000 | 7;7;7;7;7;7;7; | 5016717:5018 | 5019399:5019 | -;-;-;-;-;-;-;- | 3881 | ltb4r | 136 | 192 | 41 | 21 | 89 | 77 |
| ENSDARG000 | 7;7;7 | 5020958:5023 | 5020989:5023 | +;+;+ | 680 | NA | 0 | 0 | 0 | 0 | 0 | 0 |
| ENSDARG000 | 7;7;7 | 20600674:206 | 20602307:206 | -;-;- | 2884 | NA | 54 | 40 | 128 | 26 | 41 | 10 |
| ENSDARG000 | 7;7;7;7;7;7;7; | 26998169:269 | 26998361:269 | +;+;+;+;+;+;+; | 3396 | caprin1a | 1337 | 1179 | 1785 | 780 | 1704 | 1220 |
| ENSDARG000 | 7;7;7;7;7;7;7; | 33145529:331 | 33145575:331 | +;+;+;+;+;+;+; | 1322 | itln2 | 0 | 0 | 0 | 0 | 0 | 0 |
| ENSDARG000 | 7;7;7;7;7;7 | 20629411:206 | 20629872:206 | +;+;+;+;+;+ | 3633 | NA | 492 | 525 | 765 | 384 | 583 | 368 |
| ENSDARG000 | 7;7;7;7;7;7;7; | 22911610:229 | 22917042:229 | -;-;-;-;-;-;-; | 6425 | tnfsf10l | 52 | 16 | 96 | 2 | 82 | 40 |
| ENSDARG000 | 7;7;7;7;7;7;7; | 5049909:5052 | 5052832:5052 | -;-;-;-;-;-;-;- | 3986 | ltb4r2a | 34 | 0 | 0 | 1 | 0 | 4 |
| ENSDARG000 | 7;7;7 | 5052336:5052 | 5052382:5052 | +;+;+ | 903 | NA | 0 | 0 | 0 | 0 | 0 | 0 |
| ENSDARG000 | 7 | 39715712 | 39716073 | - | 362 | NA | 4 | 0 | 0 | 1 | 5 | 1 |
| ENSDARG000 | 7;7;7 | 7151832:7238 | 7151955:7238 | +;+;+ | 9074 | GAL3ST3 | 0 | 0 | 0 | 0 | 0 | 0 |
| ENSDARG000 | 7;7;7;7;7;7;7; | 29080684:290 | 29080871:290 | +;+;+;+;+;+;+; | 2082 | acd | 32 | 102 | 164 | 82 | 155 | 105 |
| ENSDARG000 | 7;7;7;7;7;7;7; | 39720317:397 | 39720485:397 | +;+;+;+;+;+;+; | 1345 | NA | 0 | 0 | 0 | 2 | 6 | 2 |
| ENSDARG000 | 7;7;7;7;7;7;7; | 19562045:195 | 19562246:195 | +;+;+;+;+;+;+; | 2395 | slc7a7 | 148 | 85 | 148 | 89 | 532 | 204 |
| ENSDARG000 | 7;7;7;7;7;7;7; | 24593209:245 | 24593469:245 | -;-;-;-;-;-;-; | 3188 | pik3c3 | 2110 | 2313 | 2107 | 1469 | 2181 | 2395 |
| ENSDARG000 | 7;7 | 39730429:397 | 39731689:397 | -;- | 2504 | ccdc96 | 31 | 26 | 35 | 14 | 53 | 25 |
| ENSDARG000 | 7;7;7;7;7;7;7; | 43603957:436 | 43604397:436 | -;-;-;-;-;-;-;- | 1129 | NA | 0 | 0 | 0 | 24 | 0 | 0 |
| ENSDARG000 | 7;7;7;7;7;7;7; | 7279903:7312 | 7279958:7312 | +;+;+;+;+;+;+; | 10581 | NA | 2 | 10 | 0 | 0 | 0 | 0 |
| ENSDARG000 | 7;7;7;7;7;7 | 5134670:5140 | 5137701:5140 | -;-;-;-;-;- | 3270 | NA | 11 | 0 | 2 | 18 | 27 | 11 |
| ENSDARG000 | 7;7;7;7;7;7;7 | 10911396:109 | 10912173:109 | +;+;+;+;+;+;+ | 3082 | abhd17c | 206 | 358 | 352 | 196 | 438 | 264 |
| ENSDARG000 | 7;7;7;7;7;7;7; | 56152267:561 | 56152913:561 | -;-;-;-;-;-;-; | 3109 | SPG7 | 559 | 418 | 620 | 290 | 682 | 474 |
| ENSDARG000 | 7;7;7;7;7;7;7; | 60255852:602 | 60259751:602 | -;-;-;-;-;-;-; | 8147 | plcb3 | 727 | 333 | 855 | 417 | 895 | 921 |
| ENSDARG000 | 7;7 | 39180876:392 | 39183346:392 | -;- | 2864 | CHRM4 | 0 | 0 | 0 | 0 | 0 | 0 |
| ENSDARG000 | 7;7;7;7;7;7;7; | 31432140:314 | 31434917:314 | -;-;-;-;-;-;-;- | 4130 | cilp | 85 | 40 | 59 | 9 | 207 | 69 |
| ENSDARG000 | 7;7;7 | 51219008:512 | 51222311:512 | -;-;- | 7078 | NA | 808 | 516 | 788 | 646 | 666 | 850 |
| ENSDARG000 | 7;7;7;7;7;7;7; | 33151731:331 | 33151834:331 | +;+;+;+;+;+;+; | 2215 | NA | 19 | 2 | 36 | 26 | 34 | 37 |
| ENSDARG000 | 7;7;7;7;7;7;7; | 20645121:206 | 20645316:206 | +;+;+;+;+;+;+; | 1718 | sat2a.2 | 5 | 7 | 11 | 2 | 8 | 5 |
| ENSDARG000 | 7;7;7 | 5155968:5159 | 5157252:5160 | -;-;- | 1451 | zgc:195075 | 98 | 40 | 37 | 34 | 62 | 33 |
| ENSDARG000 | 7;7;7;7;7;7;7; | 71776905:717 | 71777049:717 | +;+;+;+;+;+;+; | 1869 | NSUN6 | 67 | 46 | 128 | 14 | 78 | 14 |
| ENSDARG000 | 7;7;7;7;7;7;7; | 24573721:245 | 24573984:245 | +;+;+;+;+;+;+; | 2867 | NA | 11 | 2 | 8 | 13 | 4 | 12 |
| ENSDARG000 | 7;7 | 39738505:397 | 39738840:397 | +;+ | 3503 | tada2b | 198 | 150 | 247 | 177 | 286 | 216 |
| ENSDARG000 | 7;7;7;7;7;7;7; | 48252638:482 | 48253607:482 | -;-;-;-;-;-;-; | 2609 | rbpms2b | 54 | 4 | 6 | 4 | 11 | 13 |
| ENSDARG000 | 7;7;7;7;7 | 35706765:357 | 35707759:357 | -;-;-;-;- | 1927 | irx5a | 14 | 0 | 12 | 2 | 8 | 4 |
| ENSDARG000 | 7;7;7;7;7;7;7; | 19605830:196 | 19606667:196 | -;-;-;-;-;-;-; | 1833 | NA | 24 | 12 | 6 | 0 | 4 | 8 |
| ENSDARG000 | 7;7;7;7;7;7;7; | 19600262:196 | 19600463:196 | +;+;+;+;+;+;+; | 2031 | casp23 | 62 | 55 | 12 | 10 | 18 | 50 |
| ENSDARG000 | 7;7;7;7;7;7;7; | 35679424:356 | 35679577:356 | +;+;+;+;+;+;+; | 1479 | NA | 0 | 0 | 0 | 0 | 1 | 0 |
| ENSDARG000 | 7;7;7;7;7;7;7; | 19588608:195 | 19588855:195 | -;-;-;-;-;-;-; | 1551 | oxa1l | 534 | 532 | 685 | 555 | 417 | 410 |
| ENSDARG000 | 7;7;7 | 5180261:5187 | 5185980:5187 | -;-;- | 5952 | NA | 168 | 124 | 116 | 46 | 48 | 96 |
| ENSDARG000 | 7;7;7;7;7;7;7; | 5199667:5200 | 5200004:5200 | -;-;-;-;-;-;-; | 2114 | NA | 2 | 0 | 11 | 6 | 0 | 0 |
| ENSDARG000 | 7;7;7;7;7;7;7; | 7378920:7380 | 7380010:7380 | -;-;-;-;-;-;- | 2135 | zgc:101810 | 95 | 11 | 0 | 0 | 14 | 2 |
| ENSDARG000 | 7;7;7;7;7;7 | 39744244:397 | 39745479:397 | -;-;-;-;-;- | 1955 | grpel1 | 709 | 917 | 1047 | 619 | 660 | 575 |
| ENSDARG000 | 7;7;7;7 | 22968756:229 | 22969344:229 | -;-;-;- | 2451 | NA | 29 | 57 | 188 | 47 | 139 | 81 |
| ENSDARG000 | 7;7;7;7;7;7;7; | 22981441:229 | 22982412:229 | +;+;+;+;+;+;+; | 4833 | ccnb3 | 132 | 422 | 748 | 148 | 549 | 118 |
| ENSDARG000 | 7;7;7;7;7;7;7; | 11197022:111 | 11197129:111 | +;+;+;+;+;+;+; | 6351 | CEMIP | 960 | 571 | 289 | 1534 | 512 | 1278 |
| ENSDARG000 | 7;7;7;7;7;7;7; | 51267478:512 | 51268197:512 | -;-;-;-;-;-;-; | 4345 | gc2 | 0 | 0 | 2 | 0 | 0 | 4 |
| ENSDARG000 | 7;7;7;7;7;7;7; | 15341333:153 | 15341714:153 | -;-;-;-;-;-;-; | 3117 | NA | 1465 | 3530 | 368 | 2963 | 526 | 880 |
| ENSDARG000 | 7;7;7 | 31454001:314 | 31454335:314 | -;-;- | 541 | NA | 0 | 0 | 0 | 0 | 0 | 0 |
| ENSDARG000 | 7;7;7;7;7;7;7; | 30135768:301 | 30140073:301 | -;-;-;-;-;-;-; | 6107 | FRMD5 | 6 | 0 | 6 | 6 | 0 | 0 |
| ENSDARG000 | 7;7;7 | 5209616:5211 | 5210120:5211 | -;-;- | 929 | NA | 0 | 0 | 0 | 2 | 0 | 0 |
| ENSDARG000 | 7;7;7;7;7;7;7; | 67220374:672 | 67220667:672 | -;-;-;-;-;-;-; | 2208 | znf143a | 308 | 317 | 315 | 175 | 321 | 351 |
| ENSDARG000 | 7;7;7;7;7;7;7; | 39360797:393 | 39360975:393 | +;+;+;+;+;+;+; | 1209 | acp2 | 1078 | 577 | 708 | 790 | 915 | 1173 |
| ENSDARG000 | 7;7;7;7;7;7;7; | 3856998:3857 | 3857375:3857 | -;-;-;-;-;-;-; | 1518 | si:dkev-88n24 | 0 | 0 | 0 | 0 | 0 | 0 |
| ENSDARG000 | 7;7;7;7;7 | 33158383:331 | 33158903:331 | -;-;-;-;- | 1437 | NA | 44 | 30 | 42 | 36 | 86 | 76 |
| ENSDARG000 | 7;7;7;7 | 3878593:3883 | 3879013:3884 | -;-;-;- | 845 | NA | 0 | 0 | 0 | 0 | 0 | 0 |
| ENSDARG000 | 7;7;7;7;7;7;7; | 31494741:315 | 31501954:315 | -;-;-;-;-;-;-; | 9849 | IGDCC3 | 2962 | 1667 | 2256 | 2893 | 2051 | 2600 |
| ENSDARG000 | 7;7;7;7;7;7;7; | 29096194:290 | 29096414:290 | +;+;+;+;+;+;+; | 2523 | ogfod1 | 399 | 431 | 468 | 202 | 341 | 290 |
| ENSDARG000 | 7;7 | 43857191:438 | 43857535:438 | +;+ | 881 | NA | 0 | 0 | 0 | 0 | 0 | 0 |
| ENSDARG000 | 7;7;7 | 33168919:331 | 33169067:331 | +;+;+ | 925 | NA | 14 | 38 | 15 | 9 | 24 | 67 |
| ENSDARG000 | 7;7;7;7;7;7;7; | 43760149:437 | 43761967:437 | -;-;-;-;-;-;-; | 4718 | cdh11 | 644 | 284 | 1775 | 4009 | 2757 | 1800 |
| ENSDARG000 | 7;7;7;7;7;7;7; | 39757608:397 | 39757723:397 | +;+;+;+;+;+;+; | 8045 | SORCS2 | 1993 | 706 | 1529 | 579 | 824 | 633 |
| ENSDARG000 | 7;7;7;7;7;7;7; | 41295974:412 | 41296199:412 | +;+;+;+;+;+;+; | 926 | si:dkev-86l18. | 0 | 0 | 5 | 2 | 4 | 0 |
| ENSDARG000 | 7;7 | 29108956:291 | 29112173:291 | -;- | 3534 | fbxl8 | 94 | 54 | 66 | 72 | 101 | 85 |
| ENSDARG000 | 7;7;7;7;7;7;7 | 51314734:513 | 51316104:513 | -;-;-;-;-;-;- | 2040 | rasl11a | 10 | 8 | 12 | 0 | 10 | 2 |
| ENSDARG000 | 7;7;7 | 7401692:7406 | 7405039:7407 | -;-;- | 4083 | six7 | 0 | 0 | 0 | 0 | 0 | 0 |
| ENSDARG000 | 7;7;7;7;7 | 3889073:3892 | 3893862:3893 | -;-;-;-;- | 5212 | NA | 0 | 0 | 0 | 0 | 0 | 0 |
| ENSDARG000 | 7;7;7;7;7;7;7; | 39373234:393 | 39373347:393 | -;-;-;-;-;-;-;- | 1284 | NA | 129 | 80 | 42 | 65 | 64 | 100 |
| ENSDARG000 | 7;7;7;7;7 | 3910249:3910 | 3910612:3911 | -;-;-;-;- | 1176 | NA | 2 | 0 | 0 | 0 | 0 | 0 |
| ENSDARG000 | 7;7;7;7 | 7477398:7482 | 7478939:7482 | -;-;-;- | 1847 | pfdn2 | 1008 | 2061 | 2119 | 1131 | 1250 | 1003 |
| ENSDARG000 | 7;7;7;7;7;7;7; | 36035432:360 | 36035434:360 | +;+;+;+;+;+;+; | 5923 | irx3a | 40 | 4 | 2 | 2 | 8 | 24 |
| ENSDARG000 | 7;7 | 7466049:7473 | 7469753:7474 | -;- | 5080 | b4gat1 | 273 | 353 | 511 | 292 | 343 | 406 |
| ENSDARG000 | 7;7;7;7;7;7;7; | 7495034:7495 | 7495394:7495 | +;+;+;+;+;+;+; | 1497 | nit1 | 92 | 42 | 46 | 32 | 78 | 101 |
| ENSDARG000 | 7;7;7;7;7;7;7; | 33176517:331 | 33177671:331 | -;-;-;-;-;-;-; | 4762 | itga11a | 29 | 4 | 14 | 2 | 2 | 0 |
| ENSDARG000 | 7;7;7;7;7;7;7; | 51324834:513 | 51324974:513 | +;+;+;+;+;+;+; | 5359 | NA | 804 | 785 | 844 | 875 | 728 | 933 |
| ENSDARG000 | 7;7;7;7;7;7;7; | 33172066:331 | 33172109:331 | +;+;+;+;+;+;+; | 1860 | ITLN1 | 0 | 0 | 0 | 0 | 0 | 0 |
| ENSDARG000 | 7;7;7;7;7;7;7; | 31625662:316 | 31629243:316 | -;-;-;-;-;-;-; | 7058 | IGDCC4 | 23 | 50 | 226 | 68 | 194 | 16 |
| ENSDARG000 | 7;7;7;7;7;7;7; | 67253561:672 | 67253848:672 | +;+;+;+;+;+;+; | 2597 | NA | 0 | 0 | 0 | 0 | 0 | 0 |
| ENSDARG000 | 7;7;7;7;7;7;7; | 67307273:673 | 67310873:673 | -;-;-;-;-;-;-; | 4624 | psmd7 | 707 | 745 | 853 | 359 | 836 | 638 |
| ENSDARG000 | 7;7;7;7 | 23013946:230 | 23013993:230 | +;+;+;+ | 383 | NA | 0 | 0 | 0 | 0 | 0 | 0 |
| ENSDARG000 | 7;7;7;7;7;7;7; | 23292133:233 | 23292385:233 | +;+;+;+;+;+;+; | 3916 | htr2cl1 | 10 | 2 | 19 | 12 | 38 | 12 |
| ENSDARG000 | 7;7;7 | 30175656:301 | 30176652:301 | -;-;- | 1506 | tmed3 | 406 | 276 | 623 | 254 | 460 | 296 |
| ENSDARG000 | 7;7;7;7;7;7;7; | 67252880:672 | 67253848:672 | +;+;+;+;+;+;+; | 8068 | NA | 13 | 2 | 16 | 12 | 4 | 7 |
| ENSDARG000 | 7;7;7;7;7;7;7; | 38573917:385 | 38574037:385 | -;-;-;-;-;-;-; | 1277 | NA | 80 | 52 | 47 | 126 | 12 | 36 |
| ENSDARG000 | 7;7;7;7;7;7;7; | 56472568:564 | 56472666:564 | +;+;+;+;+;+;+; | 3032 | ist1 | 2251 | 1925 | 2247 | 1613 | 2458 | 1925 |
| ENSDARG000 | 7;7 | 44002012:440 | 44002425:440 | -;- | 492 | NA | 0 | 0 | 0 | 0 | 0 | 0 |
| ENSDARG000 | 7;7;7;7;7;7;7; | 27021368:270 | 27025708:270 | -;-;-;-;-;-;-; | 6280 | nucb2a | 2977 | 1805 | 2936 | 2247 | 2765 | 3136 |
| ENSDARG000 | 7;7;7;7;7;7;7; | 39385540:393 | 39385541:393 | +;+;+;+;+;+;+; | 2688 | tnni2b.2 | 0 | 3 | 0 | 0 | 0 | 0 |
| ENSDARG000 | 7;7;7;7;7;7;7; | 5305381:5309 | 5305518:5309 | -;-;-;-;-;-;-;- | 1055 | NA | 17 | 60 | 222 | 49 | 353 | 148 |
| ENSDARG000 | 7;7;7;7;7;7 | 20656942:206 | 20657519:206 | +;+;+;+;+;+ | 2172 | tnfsf12 | 81 | 26 | 28 | 30 | 123 | 50 |
| ENSDARG000 | 7;7 | 44396899:443 | 44397379:443 | +;+ | 955 | NA | 0 | 0 | 0 | 0 | 0 | 0 |
| ENSDARG000 | 7;7;7;7;7;7;7; | 17691847:176 | 17692937:176 | -;-;-;-;-;-;-; | 2659 | men1 | 146 | 230 | 348 | 140 | 268 | 158 |
| ENSDARG000 | 7;7;7;7;7;7 | 3922236:3922 | 3922400:3923 | -;-;-;-;-;- | 1150 | NA | 0 | 0 | 0 | 0 | 0 | 0 |
| ENSDARG000 | 7;7;7;7;7;7;7; | 36110972:361 | 36113098:361 | -;-;-;-;-;-;-; | 4295 | FTO | 184 | 145 | 218 | 147 | 142 | 124 |
| ENSDARG000 | 7;7;7;7;7;7;7; | 15373412:153 | 15376614:153 | -;-;-;-;-;-;-; | 7158 | ANPEP | 2279 | 711 | 880 | 10512 | 1817 | 4768 |
| ENSDARG000 | 7;7;7;7;7;7;7; | 60396957:604 | 60397018:604 | +;+;+;+;+;+;+; | 2902 | brms1 | 635 | 529 | 855 | 593 | 1037 | 707 |
| ENSDARG000 | 7;7;7;7;7;7;7; | 5324061:5329 | 5328262:5329 | -;-;-;-;-;-;-; | 5862 | vangl2 | 3023 | 3580 | 5065 | 3642 | 4117 | 4436 |
| ENSDARG000 | 7;7;7;7;7;7;7; | 11376325:113 | 11376750:113 | -;-;-;-;-;-;-; | 1370 | mesd | 787 | 906 | 980 | 513 | 827 | 718 |
| ENSDARG000 | 7;7;7;7;7;7;7; | 60359347:603 | 60360107:603 | +;+;+;+;+;+;+; | 2782 | ppp1r14bb | 1048 | 914 | 1766 | 636 | 2251 | 920 |
| ENSDARG000 | 7;7;7;7;7;7;7; | 30178253:301 | 30178347:301 | +;+;+;+;+;+;+; | 4418 | si:ch211-107c | 44 | 82 | 75 | 56 | 63 | 74 |
| ENSDARG000 | 7;7;7;7;7;7;7; | 40081630:400 | 40081778:400 | +;+;+;+;+;+;+; | 2897 | maf1b | 620 | 770 | 871 | 1156 | 708 | 677 |
| ENSDARG000 | 7;7;7;7;7;7;7; | 44445595:444 | 44445719:444 | +;+;+;+;+;+;+; | 6772 | cdh5 | 74 | 38 | 154 | 24 | 55 | 30 |
| ENSDARG000 | 7;7 | 51387644:513 | 51387741:513 | +;+ | 322 | NA | 0 | 0 | 0 | 0 | 0 | 0 |
| ENSDARG000 | 7;7;7;7;7;7;7; | 51364290:513 | 51365170:513 | -;-;-;-;-;-;-; | 3890 | arhgap36 | 69 | 95 | 48 | 54 | 86 | 39 |
| ENSDARG000 | 7;7 | 30186636:301 | 30189646:301 | -;- | 3696 | NA | 0 | 0 | 0 | 0 | 0 | 0 |

| | | | | | | | | | | | | |
|---|---|---|---|---|---|---|---|---|---|---|---|---|
| ENSDARG000 | 7 | 11390462 | 11393193 | + | 2732 | TLNRD1 | 306 | 414 | 424 | 316 | 386 | 286 |
| ENSDARG000 | 7:7:7:7:7:7:7: | 40058757:40( | 40060873:40( | -:-:-:-:-:-:-:-: | 6697 | shrprbck1r | 1223 | 871 | 1290 | 906 | 690 | 1016 |
| ENSDARG000 | 7:7:7:7:7:7:7: | 56517402:565 | 56517502:565 | +:+:+:+:+:+:+: | 1562 | dhodh | 86 | 185 | 242 | 114 | 248 | 189 |
| ENSDARG000 | 7;7 | 19613811:196 | 19614176:196 | -;- | 891 | zgc:194655 | 43 | 39 | 10 | 18 | 0 | 21 |
| ENSDARG000 | 7:7:7:7:7:7:7: | 11595264:115 | 11595574:115 | -:-:-:-:-:-:-:-: | 935 | STARD5 | 27 | 22 | 37 | 2 | 11 | 16 |
| ENSDARG000 | 7:7:7:7:7:7:7: | 44484853:444 | 44484880:444 | +:+:+:+:+:+:+: | 9671 | NA | 8 | 14 | 28 | 0 | 8 | 10 |
| ENSDARG000 | 7;7 | 11604834:116 | 11605012:116 | -;- | 683 | NA | 12 | 7 | 2 | 6 | 17 | 2 |
| ENSDARG000 | 7:7:7:7:7:7:7: | 11459235:114 | 11459410:114 | +:+:+:+:+:+:+: | 8504 | IL16 | 3428 | 1765 | 2633 | 1542 | 2935 | 2499 |
| ENSDARG000 | 7:7:7:7:7:7:7: | 3930742:3936 | 3936980:3936 | -:-:-:-:-:-:-:-: | 8040 | ndrg2 | 1540 | 1114 | 1225 | 1799 | 2170 | 1492 |
| ENSDARG000 | 7:7:7:7:7:7:7: | 5380320:5383 | 5383575:5383 | -:-:-:-:-:-:-:-: | 8007 | NA | 1571 | 824 | 1613 | 1125 | 1744 | 2010 |
| ENSDARG000 | 7:7:7:7:7:7:7: | 48288762:482 | 48288870:482 | +:+:+:+:+:+:+: | 1682 | oaz2b | 6 | 0 | 40 | 216 | 903 | 36 |
| ENSDARG000 | 7;7;7;7;7;7 | 7555456:7557 | 7555579:7557 | +;+;+;+;+;+ | 927 | NA | 0 | 4 | 0 | 0 | 0 | 0 |
| ENSDARG000 | 7;7;7 | 48266129:482 | 48267110:482 | -;-;- | 1316 | NA | 0 | 2 | 0 | 0 | 0 | 2 |
| ENSDARG000 | 7:7:7:7:7:7:7: | 71792290:717 | 71794797:717 | -:-:-:-:-:-:-:-: | 4511 | cacnb2a | 391 | 320 | 597 | 380 | 497 | 443 |
| ENSDARG000 | 7;7;7;7;7;7;7 | 7570701:757( | 7570777:757( | +;+;+;+;+;+;+ | 834 | NA | 0 | 0 | 0 | 0 | 0 | 0 |
| ENSDARG000 | 7:7:7:7:7:7:7: | 31776055:317 | 31778232:317 | -:-:-:-:-:-:-:-: | 3502 | nap1l4b | 1955 | 1857 | 2350 | 2172 | 1997 | 2445 |
| ENSDARG000 | 7;7;7;7;7;7 | 29115890:291 | 29116017:291 | +;+;+;+;+;+ | 1302 | tradd | 241 | 276 | 198 | 153 | 229 | 250 |
| ENSDARG000 | 7;7;7 | 41303572:413 | 41304041:413 | +;+;+ | 2096 | rgs9bp | 0 | 0 | 0 | 0 | 0 | 0 |
| ENSDARG000 | 7:7:7:7:7:7:7: | 19615056:196 | 19615146:196 | +:+:+:+:+:+:+: | 1329 | NA | 152 | 165 | 34 | 26 | 34 | 64 |
| ENSDARG000 | 7:7:7:7:7:7:7: | 7511914:7543 | 7512028:7543 | +:+:+:+:+:+:+: | 6141 | CLCN3 | 2980 | 2572 | 2996 | 2779 | 3312 | 2803 |
| ENSDARG000 | 7;7 | 44593756:445 | 44594613:445 | +;+ | 3410 | NA | 26 | 33 | 18 | 15 | 25 | 76 |
| ENSDARG000 | 7:7:7:7:7:7:7: | 41313568:413 | 41313971:413 | +:+:+:+:+:+:+: | 1417 | MCUR1 | 247 | 288 | 272 | 250 | 232 | 358 |
| ENSDARG000 | 7:7:7:7:7:7:7: | 27041315:27( | 27041398:27( | +:+:+:+:+:+:+: | 812 | rps13 | 8908 | 8580 | 14203 | 11822 | 9039 | 12521 |
| ENSDARG000 | 7:7:7:7:7:7:7: | 17716601:177 | 17716862:177 | +:+:+:+:+:+:+: | 5443 | rtn3 | 3129 | 1814 | 3120 | 3091 | 5096 | 3344 |
| ENSDARG000 | 7:7:7:7:7:7:7: | 29133321:291 | 29133699:291 | +:+:+:+:+:+:+: | 2396 | gnao1b | 52 | 125 | 110 | 81 | 124 | 175 |
| ENSDARG000 | 7;7;7 | 60551133:605 | 60551662:605 | +;+;+ | 3298 | NA | 0 | 0 | 0 | 0 | 0 | 0 |
| ENSDARG000 | 7:7:7:7:7:7:7: | 39399747:394 | 39399772:394 | +:+:+:+:+:+:+: | 1460 | tnni2b.1 | 4 | 0 | 0 | 0 | 12 | 0 |
| ENSDARG000 | 7:7:7:7:7:7:7: | 23495986:234 | 23496211:234 | +:+:+:+:+:+:+: | 5996 | zgc:109889 | 1294 | 1319 | 1309 | 1699 | 1723 | 1087 |
| ENSDARG000 | 7:7:7:7:7:7:7: | 60457138:604 | 60459231:604 | -:-:-:-:-:-:-:-: | 5842 | pcxb | 0 | 2 | 0 | 0 | 12 | 2 |
| ENSDARG000 | 7:7:7:7:7:7:7: | 33279108:332 | 33279876:332 | +:+:+:+:+:+:+: | 4160 | CORO2B | 3692 | 4429 | 4230 | 3108 | 6062 | 2495 |
| ENSDARG000 | 7;7 | 60590632:605 | 60590964:605 | +;+ | 517 | NA | 0 | 0 | 0 | 0 | 0 | 0 |
| ENSDARG000 | 7:7:7:7:7:7:7: | 51467822:514 | 51473027:514 | -:-:-:-:-:-:-:-: | 9561 | NHSL2 | 1064 | 738 | 473 | 590 | 529 | 684 |
| ENSDARG000 | 7:7:7:7:7:7:7: | 67325933:673 | 67326005:673 | +:+:+:+:+:+:+: | 3783 | NA | 32 | 26 | 74 | 81 | 31 | 104 |
| ENSDARG000 | 7:7:7:7:7:7:7: | 56577522:565 | 56577586:565 | +:+:+:+:+:+:+: | 3296 | NA | 0 | 0 | 4 | 0 | 0 | 0 |
| ENSDARG000 | 7;7;7 | 29148714:291 | 29148837:291 | +;+;+ | 357 | mtbl | 454 | 187 | 606 | 928 | 1301 | 1634 |
| ENSDARG000 | 7:7:7:7:7:7:7: | 15446229:154 | 15447258:154 | +:+:+:+:+:+:+: | 12373 | igf1rb | 560 | 249 | 608 | 1062 | 633 | 488 |
| ENSDARG000 | 7;7;7;7 | 48297842:483 | 48297976:483 | +;+;+;+ | 2834 | slc25a44b | 208 | 258 | 228 | 244 | 370 | 266 |
| ENSDARG000 | 7:7:7:7:7:7:7: | 24627076:246 | 24632084:246 | -:-:-:-:-:-:-:-: | 6720 | rgp1 | 344 | 248 | 178 | 134 | 315 | 266 |
| ENSDARG000 | 7:7:7:7:7:7:7: | 38577869:385 | 38578158:385 | +:+:+:+:+:+:+: | 1341 | ptpmt1 | 137 | 146 | 182 | 90 | 88 | 129 |
| ENSDARG000 | 7:7:7:7:7:7:7: | 24645186:246 | 24645215:246 | +:+:+:+:+:+:+: | 3969 | GBA2 | 916 | 1057 | 971 | 810 | 1633 | 1101 |
| ENSDARG000 | 7:7:7:7:7:7:7: | 11689691:116 | 11689911:116 | +:+:+:+:+:+:+: | 2302 | NA | 0 | 0 | 0 | 0 | 0 | 0 |
| ENSDARG000 | 7;7;7;7;7 | 29153018:291 | 29154094:291 | +;+;+;+;+ | 2856 | tmppe | 565 | 440 | 298 | 288 | 524 | 447 |
| ENSDARG000 | 7;7;7 | 24628546:246 | 24628673:246 | +;+;+ | 796 | MSMP | 0 | 0 | 0 | 0 | 0 | 0 |
| ENSDARG000 | 7;7;7;7;7;7 | 56602521:566 | 56603696:566 | -;-;-;-;-;- | 1980 | sult5a1 | 48 | 12 | 12 | 16 | 87 | 40 |
| ENSDARG000 | 7:7:7:7:7:7:7: | 29163762:291 | 29164096:291 | +:+:+:+:+:+:+: | 2228 | SLC38A8 | 46 | 14 | 10 | 20 | 12 | 20 |
| ENSDARG000 | 7 | 29163592 | 29164818 | - | 1227 | exosc6 | 114 | 271 | 272 | 63 | 262 | 117 |
| ENSDARG000 | 7:7:7:7:7:7:7: | 36467315:364 | 36467390:364 | +:+:+:+:+:+:+: | 2797 | aktip | 733 | 669 | 747 | 567 | 769 | 726 |
| ENSDARG000 | 7:7:7:7:7:7:7: | 41322407:413 | 41322470:413 | +:+:+:+:+:+:+: | 3184 | dph2 | 205 | 283 | 365 | 179 | 233 | 192 |
| ENSDARG000 | 7;7 | 5494565:5514 | 5494599:552( | +;+ | 6843 | MYADM | 1250 | 1783 | 1022 | 1824 | 890 | 1880 |
| ENSDARG000 | 7:7:7:7:7:7:7: | 41329748:413 | 41331700:413 | -:-:-:-:-:-:-:-: | 3407 | ncf2 | 22 | 40 | 56 | 71 | 207 | 107 |
| ENSDARG000 | 7;7;7;7;7;7;7 | 4162994:4167 | 4163107:4167 | +;+;+;+;+;+;+ | 12562 | NA | 22 | 1 | 4 | 0 | 0 | 6 |
| ENSDARG000 | 7:7:7:7:7:7:7: | 19625622:196 | 19625850:196 | -:-:-:-:-:-:-:-: | 7105 | NA | 5568 | 3952 | 6488 | 4892 | 5352 | 4936 |
| ENSDARG000 | 7;7 | 5529967:5534 | 5530160:5536 | +;+ | 1456 | NA | 6 | 0 | 0 | 6 | 8 | 0 |
| ENSDARG000 | 7;7;7;7 | 5530154:5541 | 5530160:5541 | +;+;+;+ | 3263 | NA | 0 | 6 | 33 | 26 | 0 | 0 |
| ENSDARG000 | 7:7:7:7:7:7:7: | 33345624:333 | 33346542:333 | -:-:-:-:-:-:-:-: | 2522 | anp32a | 2384 | 3106 | 3581 | 2452 | 2892 | 3055 |
| ENSDARG000 | 7:7:7:7:7:7:7: | 25913225:259 | 25913287:259 | +:+:+:+:+:+:+: | 1917 | hmgb3a | 3680 | 2922 | 2992 | 2532 | 3415 | 2896 |
| ENSDARG000 | 7:7:7:7:7:7:7: | 25920792:259 | 25920999:259 | +:+:+:+:+:+:+: | 5245 | arrb2b | 7019 | 6143 | 6590 | 8704 | 7173 | 5320 |
| ENSDARG000 | 7;7 | 12061787:12( | 12064501:12( | -;- | 3216 | mex3b | 390 | 142 | 274 | 98 | 208 | 136 |
| ENSDARG000 | 7;7;7;7 | 48319916:483 | 48320116:483 | +;+;+;+ | 921 | crabp1b | 0 | 2 | 0 | 0 | 0 | 0 |
| ENSDARG000 | 7:7:7:7:7:7:7: | 29176924:291 | 29177159:291 | +:+:+:+:+:+:+: | 2406 | aph1b | 2409 | 3028 | 2258 | 1889 | 2405 | 2219 |
| ENSDARG000 | 7:7:7:7:7:7:7: | 52712807:527 | 52712930:527 | +:+:+:+:+:+:+: | 3468 | NA | 505 | 489 | 549 | 295 | 541 | 549 |
| ENSDARG000 | 7:7:7:7:7:7:7: | 17760713:177 | 17767030:177 | -:-:-:-:-:-:-:-: | 8270 | NA | 110 | 70 | 136 | 80 | 146 | 125 |
| ENSDARG000 | 7:7:7:7:7:7:7: | 4193869:4193 | 4193910:4193 | +:+:+:+:+:+:+: | 5670 | NA | 11 | 0 | 0 | 0 | 0 | 0 |
| ENSDARG000 | 7:7:7:7:7:7:7: | 44604835:446 | 44604944:446 | +:+:+:+:+:+:+: | 841 | NA | 16 | 19 | 29 | 0 | 33 | 56 |
| ENSDARG000 | 7;7;7;7;7;7 | 40093662:40( | 40094001:40( | +;+;+;+;+;+ | 3779 | NA | 138 | 88 | 151 | 70 | 112 | 60 |
| ENSDARG000 | 7:7:7:7:7:7:7: | 4194055:4243 | 4194091:4243 | +:+:+:+:+:+:+: | 1413 | NA | 0 | 0 | 0 | 0 | 0 | 0 |
| ENSDARG000 | 7:7:7:7:7:7:7: | 4219234:4219 | 4219297:4219 | +:+:+:+:+:+:+: | 4628 | NA | 0 | 0 | 4 | 0 | 0 | 0 |
| ENSDARG000 | 7:7:7:7:7:7:7: | 56703254:567 | 56703372:567 | +:+:+:+:+:+:+: | 1746 | nqo1 | 1828 | 914 | 586 | 1102 | 977 | 960 |
| ENSDARG000 | 7:7:7:7:7:7:7: | 61281844:612 | 61282494:612 | -:-:-:-:-:-:-:-: | 1397 | rwdd | 510 | 334 | 574 | 350 | 435 | 533 |
| ENSDARG000 | 7:7:7:7:7:7:7: | 44599313:445 | 44600007:446 | -:-:-:-:-:-:-:-: | 2271 | tk2 | 156 | 162 | 148 | 155 | 144 | 137 |
| ENSDARG000 | 7:7:7:7:7:7:7: | 30201611:302 | 30201746:302 | +:+:+:+:+:+:+: | 2077 | wdr76 | 8 | 138 | 123 | 110 | 134 | 48 |
| ENSDARG000 | 7;7;7 | 5556575:5559 | 5557059:5559 | -;-;- | 966 | NA | 0 | 0 | 0 | 0 | 1 | 1 |
| ENSDARG000 | 7;7;7;7;7 | 5556407:5556 | 5556582:5557 | +;+;+;+;+ | 829 | NA | 0 | 0 | 1 | 6 | 1 | 0 |
| ENSDARG000 | 7:7:7:7:7:7:7: | 48345747:483 | 48348846:483 | -:-:-:-:-:-:-:-: | 8163 | NA | 796 | 894 | 1080 | 512 | 1148 | 905 |
| ENSDARG000 | 7;7;7;7;7;7 | 41340520:413 | 41340885:413 | +;+;+;+;+;+ | 5026 | garem | 448 | 383 | 366 | 313 | 478 | 374 |
| ENSDARG000 | 7:7:7:7:7:7:7: | 19680252:196 | 19682515:196 | -:-:-:-:-:-:-:-: | 5092 | NA | 332 | 211 | 336 | 114 | 322 | 282 |
| ENSDARG000 | 7:7:7:7:7:7:7: | 41348543:413 | 41349190:413 | -:-:-:-:-:-:-:-: | 12981 | PRKDC | 613 | 559 | 863 | 564 | 1221 | 757 |
| ENSDARG000 | 7:7:7:7:7:7:7: | 31796237:317 | 31796998:317 | -:-:-:-:-:-:-:-: | 3237 | cars1 | 519 | 782 | 1170 | 502 | 832 | 510 |
| ENSDARG000 | 7:7:7:7:7:7:7: | 51633815:516 | 51633969:516 | -:-:-:-:-:-:-:-: | 1669 | rps4x | 21878 | 19821 | 35894 | 30400 | 22240 | 32568 |
| ENSDARG000 | 7;7;7;7;7;7;7 | 38588866:385 | 38588949:385 | +;+;+;+;+;+;+ | 916 | ndufs3 | 608 | 869 | 948 | 666 | 756 | 735 |
| ENSDARG000 | 7:7:7:7:7:7:7: | 40102391:401 | 40102677:401 | -:-:-:-:-:-:-:-: | 3795 | NA | 0 | 0 | 1 | 0 | 2 | 14 |
| ENSDARG000 | 7:7:7:7:7:7:7: | 17782436:177 | 17782718:177 | +:+:+:+:+:+:+: | 6588 | NA | 412 | 252 | 379 | 411 | 442 | 429 |
| ENSDARG000 | 7:7:7:7:7:7:7: | 12140955:121 | 12146480:121 | -:-:-:-:-:-:-:-: | 8934 | EFL1 | 536 | 422 | 652 | 430 | 410 | 450 |
| ENSDARG000 | 7;7;7;7 | 12371061:123 | 12371188:123 | +;+;+;+ | 1526 | saxo2 | 151 | 113 | 159 | 98 | 83 | 79 |
| ENSDARG000 | 7:7:7:7:7:7:7: | 19738665:197 | 19738745:197 | +:+:+:+:+:+:+: | 3286 | tkfc | 656 | 589 | 508 | 560 | 724 | 558 |
| ENSDARG000 | 7;7 | 23561035:235 | 23562035:235 | -;- | 1231 | NA | 0 | 0 | 4 | 5 | 2 | 0 |
| ENSDARG000 | 7;7;7;7;7 | 23569736:235 | 23570853:235 | -;-;-;-;- | 1745 | NA | 0 | 0 | 0 | 0 | 0 | 0 |
| ENSDARG000 | 7:7:7:7:7:7:7: | 17805329:178 | 17805849:178 | -:-:-:-:-:-:-:-: | 2353 | ecsit | 181 | 196 | 289 | 111 | 273 | 182 |
| ENSDARG000 | 7:7:7:7:7:7:7: | 5566582:5567 | 5566951:5567 | +:+:+:+:+:+:+: | 2122 | NA | 0 | 0 | 6 | 4 | 0 | 2 |
| ENSDARG000 | 7;7;7;7;7;7;7 | 4278839:4283 | 4280444:4284 | +;+;+;+;+;+;+ | 5674 | NA | 0 | 0 | 0 | 0 | 0 | 0 |
| ENSDARG000 | 7:7:7:7:7:7:7: | 56751746:567 | 56752262:567 | -:-:-:-:-:-:-:-: | 2453 | csnk2a2a | 486 | 494 | 797 | 522 | 608 | 531 |
| ENSDARG000 | 7;7;7 | 4291707:4294 | 4292166:4295 | -;-;- | 936 | cbln11 | 8 | 0 | 2 | 2 | 0 | 0 |
| ENSDARG000 | 7:7:7:7:7:7:7: | 17816006:178 | 17816711:178 | +:+:+:+:+:+:+: | 7058 | EML3 | 4866 | 1903 | 4036 | 1642 | 3211 | 1494 |
| ENSDARG000 | 7;7;7;7;7 | 4299907:4303 | 4302508:4303 | -;-;-;-;- | 7057 | NA | 0 | 0 | 2 | 0 | 0 | 0 |
| ENSDARG000 | 7:7:7:7:7:7:7: | 24681278:246 | 24681811:246 | -:-:-:-:-:-:-:-: | 1341 | calb2b | 2 | 2 | 0 | 2 | 0 | 0 |
| ENSDARG000 | 7:7:7:7:7:7:7: | 61598572:615 | 61598682:615 | +:+:+:+:+:+:+: | 1560 | NA | 24 | 6 | 28 | 62 | 12 | 14 |
| ENSDARG000 | 7:7:7:7:7:7:7: | 61578660:615 | 61578818:615 | -:-:-:-:-:-:-:-: | 4404 | NA | 1009 | 881 | 1022 | 1123 | 953 | 678 |
| ENSDARG000 | 7;7;7;7;7 | 67451108:674 | 67451386:674 | +;+;+;+;+ | 1028 | gcshb | 21 | 0 | 4 | 6 | 8 | 0 |
| ENSDARG000 | 7:7:7:7:7:7:7: | 23577723:235 | 23577872:235 | +:+:+:+:+:+:+: | 5922 | NA | 66 | 20 | 12 | 14 | 30 | 68 |
| ENSDARG000 | 7:7:7:7:7:7:7: | 67429185:674 | 67429591:674 | +:+:+:+:+:+:+: | 5343 | kars1 | 6042 | 6483 | 9289 | 4751 | 6110 | 5227 |
| ENSDARG000 | 7:7:7:7:7:7:7: | 33372680:333 | 33372914:333 | +:+:+:+:+:+:+: | 4925 | glceb | 598 | 989 | 406 | 920 | 679 | 813 |
| ENSDARG000 | 7:7:7:7:7:7:7: | 27059330:27( | 27059412:27( | +:+:+:+:+:+:+: | 5688 | PLEKHA7 | 6 | 0 | 0 | 2 | 26 | 24 |
| ENSDARG000 | 7;7;7;7;7 | 56777668:567 | 56779260:567 | -;-;-;-;- | 4176 | NA | 0 | 0 | 0 | 0 | 0 | 0 |
| ENSDARG000 | 7;7;7 | 56830881:568 | 56831642:568 | -;-;- | 1100 | senp8 | 47 | 48 | 79 | 33 | 50 | 57 |
| ENSDARG000 | 7:7:7:7:7:7:7: | 29189100:291 | 29189726:291 | -:-:-:-:-:-:-:-: | 15074 | HERC1 | 5807 | 3060 | 3048 | 1653 | 2822 | 2200 |
| ENSDARG000 | 7;7 | 36549782:365 | 36552152:365 | -;- | 2701 | NA | 161 | 80 | 187 | 64 | 93 | 36 |
| ENSDARG000 | 7:7:7:7:7:7:7: | 36484598:364 | 36484715:365 | +:+:+:+:+:+:+: | 11835 | CHD9 | 3473 | 2040 | 2877 | 2022 | 2824 | 2564 |
| ENSDARG000 | 7:7:7:7:7:7:7: | 33457148:334 | 33457331:334 | +:+:+:+:+:+:+: | 3411 | paqr5b | 7 | 0 | 0 | 0 | 0 | 0 |
| ENSDARG000 | 7;7 | 51642671:516 | 51644181:516 | -;- | 1752 | cited1 | 4424 | 4845 | 5722 | 6039 | 6616 | 5727 |
| ENSDARG000 | 7:7:7:7:7:7:7: | 40122918:401 | 40123688:401 | -:-:-:-:-:-:-:-: | 4031 | DYNC2I1 | 126 | 98 | 113 | 50 | 102 | 86 |
| ENSDARG000 | 7;7;7;7;7 | 5573759:5582 | 5573815:5583 | +;+;+;+;+ | 570 | NA | 0 | 0 | 0 | 0 | 0 | 0 |
| ENSDARG000 | 7:7:7:7:7:7:7: | 5582771:5582 | 5582857:5583 | -:-:-:-:-:-:-:-: | 2958 | NA | 0 | 2 | 5 | 3 | 2 | 32 |
| ENSDARG000 | 7:7:7:7:7:7:7: | 15659280:156 | 15659387:156 | +:+:+:+:+:+:+: | 2009 | NA | 0 | 2 | 2 | 2 | 2 | 0 |
| ENSDARG000 | 7;7;7;7;7;7 | 33463902:335 | 33464462:335 | -;-;-;-;-;- | 1179 | NA | 0 | 0 | 0 | 0 | 0 | 0 |
| ENSDARG000 | 7:7:7:7:7:7:7: | 4332430:4337 | 4332675:4337 | +:+:+:+:+:+:+: | 6367 | slc12a10.2 | 0 | 0 | 1 | 0 | 2 | 2 |
| ENSDARG000 | 7:7:7:7:7:7:7: | 19762595:197 | 19762702:197 | +:+:+:+:+:+:+: | 4505 | NA | 6 | 0 | 33 | 6 | 40 | 10 |
| ENSDARG000 | 7:7:7:7:7:7:7: | 38600150:386 | 38603314:386 | -;-;-;-;-;-;-;- | 3907 | c1qtnf4 | 30 | 34 | 28 | 52 | 8 | 22 |
| ENSDARG000 | 7;7 | 24714840:247 | 24717496:247 | -;- | 2740 | cmtr2 | 234 | 131 | 244 | 135 | 240 | 260 |
| ENSDARG000 | 7;7 | 48460239:484 | 48460543:484 | +;+ | 3128 | lingo1b | 6 | 2 | 0 | 0 | 0 | 0 |
| ENSDARG000 | 7;7 | 4339514:4356 | 4339603:4357 | -;- | 559 | NA | 0 | 0 | 0 | 0 | 0 | 0 |
| ENSDARG000 | 7:7:7:7:7:7:7: | 12389866:123 | 12390653:123 | -:-:-:-:-:-:-:-: | 6385 | ADAMTSL3 | 30 | 28 | 90 | 24 | 48 | 26 |
| ENSDARG000 | 7:7:7:7:7:7:7: | 26006819:26( | 26006970:26( | +:+:+:+:+:+:+: | 4095 | NA | 1788 | 1070 | 1935 | 1222 | 1600 | 1286 |
| ENSDARG000 | 7:7:7:7:7:7:7: | 57088920:57( | 57089046:57( | +:+:+:+:+:+:+: | 2361 | scamp2l | 12 | 0 | 0 | 4 | 0 | 10 |
| ENSDARG000 | 7:7:7:7:7:7:7: | 5629313:563( | 5629596:563( | -;-;-;-;-;-;-;- | 2355 | NA | 0 | 216 | 24 | 302 | 267 | 60 |
| ENSDARG000 | 7:7:7:7:7:7:7: | 38635538:386 | 38635648:386 | -:-:-:-:-:-:-:-: | 1403 | c6ast4 | 0 | 0 | 0 | 0 | 0 | 0 |
| ENSDARG000 | 7:7:7:7:7:7:7: | 40148213:401 | 40148397:401 | +:+:+:+:+:+:+: | 4117 | ESYT2 | 14 | 12 | 6 | 16 | 12 | 10 |
| ENSDARG000 | 7:7:7:7:7:7:7: | 51656099:516 | 51656790:516 | -:-:-:-:-:-:-:-: | 2393 | hdac8 | 258 | 292 | 332 | 253 | 284 | 194 |
| ENSDARG000 | 7:7:7:7:7:7:7: | 61977050:619 | 61977776:619 | -:-:-:-:-:-:-:-: | 2621 | plaa | 760 | 783 | 940 | 404 | 941 | 508 |
| ENSDARG000 | 7:7:7:7:7:7:7: | 30209911:302 | 30211038:302 | -:-:-:-:-:-:-:-: | 3256 | ZNF710 | 906 | 899 | 1161 | 868 | 1018 | 842 |
| ENSDARG000 | 7:7:7:7:7:7:7: | 24729558:247 | 24729834:247 | +:+:+:+:+:+:+: | 6497 | NA | 923 | 338 | 658 | 583 | 372 | 680 |
| ENSDARG000 | 7:7:7:7:7:7:7: | 48555400:485 | 48556131:485 | +:+:+:+:+:+:+: | 4033 | kcnq1.1 | 0 | 2 | 0 | 0 | 2 | 4 |
| ENSDARG000 | 7:7:7:7:7:7:7: | 38642114:386 | 38642236:386 | -:-:-:-:-:-:-:-: | 920 | c6ast3 | 0 | 0 | 0 | 0 | 0 | 0 |
| ENSDARG000 | 7;7;7;7 | 19835569:198 | 19836128:198 | +;+;+;+ | 2536 | ovol1a | 20 | 0 | 0 | 0 | 6 | 0 |
| ENSDARG000 | 7:7:7:7:7:7:7: | 27250012:272 | 27250582:272 | +:+:+:+:+:+:+: | 9577 | sox6 | 6817 | 3893 | 4906 | 2921 | 8549 | 4849 |
| ENSDARG000 | 7;7;7 | 36625275:366 | 36625646:366 | -;-;- | 641 | NA | 0 | 0 | 0 | 0 | 0 | 0 |
| ENSDARG000 | 7;7;7 | 36642633:366 | 36643007:366 | -;-;- | 644 | NA | 0 | 0 | 0 | 0 | 0 | 0 |
| ENSDARG000 | 7:7:7:7:7:7:7: | 62080456:621 | 62080593:621 | +:+:+:+:+:+:+: | 3196 | rbpjb | 122 | 78 | 138 | 82 | 79 | 78 |
| ENSDARG000 | 7:7:7:7:7:7:7: | 23727768:237 | 23728530:237 | -:-:-:-:-:-:-:-: | 2689 | zgc:92429 | 0 | 4 | 16 | 8 | 0 | 6 |
| ENSDARG000 | 7:7:7:7:7:7:7: | 30240791:302 | 30241255:302 | +:+:+:+:+:+:+: | 6393 | SEMA4B | 16 | 26 | 135 | 14 | 20 | 12 |
| ENSDARG000 | 7;7;7;7;7 | 41401488:414 | 41401704:414 | -;-;-;-;- | 827 | NA | 0 | 4 | 2 | 4 | 26 | 4 |
| ENSDARG000 | 7;7;7;7;7 | 12799576:128 | 12803056:128 | -;-;-;-;- | 4142 | zgc:158785 | 0 | 0 | 0 | 0 | 0 | 4 |
| ENSDARG000 | 7 | 12835048 | 12836225 | + | 1178 | gjd6 | 0 | 0 | 0 | 0 | 0 | 0 |
| ENSDARG000 | 7:7:7:7:7:7:7: | 40205394:402 | 40205485:402 | +:+:+:+:+:+:+: | 4199 | ncapg2 | 116 | 498 | 637 | 180 | 380 | 253 |
| ENSDARG000 | 7:7:7:7:7:7:7: | 17907883:179 | 17908262:179 | +:+:+:+:+:+:+: | 5162 | mta2 | 2345 | 2213 | 3145 | 1546 | 2701 | 1286 |
| ENSDARG000 | 7 | 38649136 | 38649529 | + | 394 | NA | 0 | 0 | 0 | 0 | 0 | 0 |
| ENSDARG000 | 7;7 | 38649588:386 | 38650017:386 | -;- | 568 | NA | 0 | 0 | 0 | 0 | 2 | 0 |
| ENSDARG000 | 7:7:7:7:7:7:7: | 15818778:158 | 15819703:158 | -:-:-:-:-:-:-:-: | 2099 | RCN1 | 579 | 876 | 751 | 412 | 898 | 569 |
| ENSDARG000 | 7;7 | 36671512:366 | 36671784:366 | +;+ | 948 | NA | 0 | 0 | 0 | 0 | 0 | 0 |
| ENSDARG000 | 7;7;7;7;7;7 | 36671337:366 | 36671678:366 | -;-;-;-;-;- | 730 | NA | 2 | 0 | 0 | 0 | 0 | 0 |
| ENSDARG000 | 7:7:7:7:7:7:7: | 33644434:336 | 33647214:336 | -:-:-:-:-:-:-:-: | 6000 | tle3b | 2042 | 2195 | 1701 | 1313 | 2305 | 1271 |
| ENSDARG000 | 7;7 | 36698679:366 | 36699113:366 | +;+ | 952 | zgc:195356 | 0 | 0 | 0 | 0 | 12 | 0 |
| ENSDARG000 | 7;7 | 24814866:248 | 24815034:248 | +;+ | 527 | NA | 0 | 0 | 0 | 0 | 0 | 0 |
| ENSDARG000 | 7:7:7:7:7:7:7: | 24830576:248 | 24830788:248 | -:-:-:-:-:-:-:-: | 2039 | fam113 | 0 | 0 | 0 | 0 | 2 | 0 |
| ENSDARG000 | 7:7:7:7:7:7:7: | 51710354:517 | 51714032:517 | -:-:-:-:-:-:-:-: | 8109 | PHKA1 | 463 | 371 | 427 | 431 | 612 | 319 |
| ENSDARG000 | 7:7:7:7:7:7:7: | 12840556:128 | 12846702:128 | -:-:-:-:-:-:-:-: | 7245 | sh3gl3a | 100 | 68 | 75 | 62 | 278 | 62 |
| ENSDARG000 | 7;7;7;7 | 30270410:302 | 30270813:302 | -;-;-;- | 1081 | harbi2 | 176 | 77 | 110 | 118 | 118 | 201 |
| ENSDARG000 | 7:7:7:7:7:7:7: | 36898622:368 | 36899710:368 | +:+:+:+:+:+:+: | 4346 | TOX3 | 28 | 0 | 4 | 0 | 6 | 6 |
| ENSDARG000 | 7;7 | 24828870:248 | 24829052:248 | +;+ | 2113 | cskmt | 37 | 54 | 83 | 49 | 107 | 56 |
| ENSDARG000 | 7:7:7:7:7:7:7: | 24815809:248 | 24816353:248 | -:-:-:-:-:-:-:-: | 1282 | otub1b | 560 | 507 | 635 | 497 | 446 | 542 |
| ENSDARG000 | 7;7 | 19882066:198 | 19882625:199 | +;+ | 9392 | TMEM151A | 0 | 12 | 5 | 0 | 14 | 4 |
| ENSDARG000 | 7;7;7;7 | 19903924:199 | 19904227:199 | +;+;+;+ | 703 | si:ch211-285i | 0 | 0 | 0 | 0 | 0 | 1 |
| ENSDARG000 | 7:7:7:7:7:7:7: | 38653481:386 | 38654003:386 | -:-:-:-:-:-:-:-: | 1636 | npsn | 70 | 69 | 154 | 176 | 1411 | 1522 |

| | | | | | | | | | | | | |
|---|---|---|---|---|---|---|---|---|---|---|---|---|
| ENSDARG000 | 7;7 | 19882225:19 | 19882625:19 | +;+ | 572 | NA | 0 | 0 | 0 | 0 | 0 | 0 |
| ENSDARG000 | 7;7;7;7;7;7;7 | 19850889:198 | 19850967:198 | +;+;+;+;+;+;+ | 6110 | mus81 | 55 | 207 | 250 | 108 | 161 | 192 |
| ENSDARG000 | 7;7;7;7;7;7;7 | 41403690:414 | 41404804:414 | -;-;-;-;-;-;- | 3396 | abcf2b | 38 | 53 | 88 | 12 | 32 | 20 |
| ENSDARG000 | 7;7;7;7;7;7;7 | 48663031:486 | 48664141:486 | -;-;-;-;-;-;- | 2199 | cdkn1ca | 4 | 82 | 30 | 12 | 62 | 14 |
| ENSDARG000 | 7;7;7;7;7;7;7 | 62248514:622 | 62248738:622 | +;+;+;+;+;+;+ | 2122 | tbc1d19 | 89 | 60 | 121 | 61 | 134 | 140 |
| ENSDARG000 | 7;7;7;7;7;7;7 | 30275513:302 | 30276621:302 | -;-;-;-;-;-;- | 1325 | serf2b | 1852 | 1790 | 2101 | 1860 | 1633 | 2270 |
| ENSDARG000 | 7;7;7;7;7;7;7 | 26029672:260 | 26029900:260 | +;+;+;+;+;+;+ | 3595 | alox12 | 92 | 42 | 35 | 52 | 154 | 90 |
| ENSDARG000 | 7;7;7;7;7;7;7 | 48667081:486 | 48667266:486 | +;+;+;+;+;+;+ | 4675 | trpm5 | 0 | 2 | 3 | 0 | 8 | 0 |
| ENSDARG000 | 7;7 | 17947217:179 | 17947435:179 | +;+ | 1553 | cth1 | 0 | 0 | 0 | 7 | 0 | 2 |
| ENSDARG000 | 7;7;7;7;7;7;7 | 12927944:129 | 12928186:129 | +;+;+;+;+;+;+ | 1094 | fth1a | 12431 | 10768 | 15815 | 18614 | 14105 | 15819 |
| ENSDARG000 | 7;7;7;7;7;7;7 | 67467702:674 | 67467747:674 | +;+;+;+;+;+;+ | 1997 | rpl13 | 16403 | 15578 | 26166 | 23434 | 17240 | 22136 |
| ENSDARG000 | 7;7;7;7;7 | 26048033:260 | 26048559:260 | -;-;-;-;- | 1000 | rnaseka | 1976 | 2007 | 1644 | 1761 | 1733 | 1959 |
| ENSDARG000 | 7;7;7;7;7;7;7 | 44608224:446 | 44608597:446 | +;+;+;+;+;+;+ | 1705 | cmtm3 | 476 | 650 | 558 | 1118 | 1557 | 670 |
| ENSDARG000 | 7;7;7;7;7;7;7 | 12950507:129 | 12950527:129 | +;+;+;+;+;+;+ | 640 | saa | 1040 | 56 | 6 | 20 | 2 | 14 |
| ENSDARG000 | 7;7;7;7;7;7;7 | 44611623:446 | 44612248:446 | -;-;-;-;-;-;- | 884 | NA | 7 | 0 | 1 | 8 | 4 | 10 |
| ENSDARG000 | 7;7;7;7;7;7 | 72012397:720 | 72012605:720 | +;+;+;+;+;+ | 6915 | ywhae2 | 5895 | 6228 | 7767 | 1333 | 6608 | 5125 |
| ENSDARG000 | 7;7;7;7;7;7;7 | 12959846:129 | 12959963:129 | -;-;-;-;-;-;- | 505 | rplp2l | 7031 | 8220 | 10723 | 8470 | 6711 | 9179 |
| ENSDARG000 | 7;7;7;7;7;7;7 | 48705227:487 | 48705326:487 | +;+;+;+;+;+;+ | 993 | NA | 0 | 0 | 0 | 0 | 4 | 0 |
| ENSDARG000 | 7;7;7;7;7;7;7 | 29278580:292 | 29282120:292 | -;-;-;-;-;-;- | 4432 | dapk2a | 207 | 108 | 258 | 156 | 188 | 124 |
| ENSDARG000 | 7;7;7;7;7;7;7 | 17953589:179 | 17953785:179 | +;+;+;+;+;+;+ | 3778 | taf6l | 420 | 397 | 426 | 364 | 433 | 433 |
| ENSDARG000 | 7;7;7;7 | 41421998:414 | 41422606:414 | +;+;+;+ | 3401 | CHPF2 | 839 | 638 | 964 | 668 | 959 | 847 |
| ENSDARG000 | 7;7;7;7;7;7 | 17973097:179 | 17973158:179 | +;+;+;+;+;+ | 1343 | NA | 0 | 2 | 2 | 0 | 0 | 0 |
| ENSDARG000 | 7;7;7;7 | 30282389:302 | 30282615:302 | +;+;+;+ | 2685 | polr2m | 112 | 173 | 282 | 132 | 206 | 144 |
| ENSDARG000 | 7;7;7;7;7;7;7 | 48719924:487 | 48723792:487 | -;-;-;-;-;-;- | 4938 | traf6 | 762 | 865 | 1029 | 554 | 840 | 818 |
| ENSDARG000 | 7;7;7;7;7;7;7 | 26049818:260 | 26050151:260 | +;+;+;+;+;+;+ | 1636 | NA | 207 | 183 | 153 | 124 | 130 | 156 |
| ENSDARG000 | 7;7;7;7;7;7;7 | 23748971:237 | 23757023:237 | -;-;-;-;-;-;- | 10098 | NA | 2987 | 1738 | 2690 | 1407 | 2918 | 2324 |
| ENSDARG000 | 7;7;7;7;7;7;7 | 12976173:129 | 12976321:129 | +;+;+;+;+;+;+ | 1700 | NA | 10 | 0 | 0 | 0 | 18 | 0 |
| ENSDARG000 | 7;7;7;7;7;7;7 | 67486807:674 | 67486916:674 | +;+;+;+;+;+;+ | 2916 | CPNE7 | 2 | 0 | 6 | 0 | 6 | 4 |
| ENSDARG000 | 7;7;7 | 26058772:260 | 26061610:260 | +;+;+ | 2839 | NA | 11 | 21 | 37 | 12 | 67 | 6 |
| ENSDARG000 | 7;7;7 | 37060405:370 | 37060659:370 | +;+;+ | 962 | NA | 0 | 0 | 0 | 0 | 0 | 0 |
| ENSDARG000 | 7;7;7;7;7;7;7 | 37359004:373 | 37359108:373 | +;+;+;+;+;+;+ | 6858 | sall1a | 188 | 32 | 50 | 6 | 12 | 22 |
| ENSDARG000 | 7;7;7;7;7;7;7 | 27455321:274 | 27455400:274 | +;+;+;+;+;+;+ | 2822 | CYP2R1 | 4 | 0 | 2 | 0 | 44 | 0 |
| ENSDARG000 | 7;7;7;7;7;7;7 | 52753067:527 | 52753592:527 | +;+;+;+;+;+;+ | 1655 | mtfmt | 186 | 173 | 142 | 160 | 174 | 170 |
| ENSDARG000 | 7;7;7;7;7;7;7 | 27680556:276 | 27681114:276 | -;-;-;-;-;-;- | 1673 | calca | 0 | 0 | 0 | 0 | 6 | 0 |
| ENSDARG000 | 7;7;7;7;7 | 26077153:260 | 26079906:260 | -;-;-;-;- | 4795 | ache | 24 | 16 | 60 | 39 | 53 | 10 |
| ENSDARG000 | 7;7;7;7;7 | 26065218:260 | 26065381:260 | +;+;+;+;+ | 2618 | NA | 2 | 2 | 2 | 0 | 13 | 0 |
| ENSDARG000 | 7;7;7;7;7;7;7 | 52761841:527 | 52761889:527 | +;+;+;+;+;+;+ | 7683 | PPIP5K1 | 382 | 200 | 427 | 240 | 576 | 415 |
| ENSDARG000 | 7;7;7;7;7 | 26073114:260 | 26073618:260 | -;-;-;-;- | 1574 | C10H17orf49 | 282 | 296 | 565 | 425 | 597 | 480 |
| ENSDARG000 | 7;7;7 | 37420980:374 | 37423079:374 | -;-;- | 2570 | NA | 0 | 0 | 0 | 0 | 0 | 0 |
| ENSDARG000 | 7;7;7;7;7;7;7 | 33798131:338 | 33802296:338 | -;-;-;-;-;-;- | 8699 | UACA | 124 | 6 | 10 | 12 | 22 | 18 |
| ENSDARG000 | 7;7;7;7;7;7 | 29293452:292 | 29293853:292 | +;+;+;+;+;+ | 2180 | NA | 75 | 18 | 51 | 40 | 66 | 68 |
| ENSDARG000 | 7;7 | 29293759:292 | 29293860:292 | -;- | 675 | NA | 0 | 5 | 14 | 49 | 35 | 16 |
| ENSDARG000 | 7;7 | 72032484:720 | 72033514:720 | -;- | 2773 | NA | 6 | 8 | 27 | 0 | 10 | 4 |
| ENSDARG000 | 7;7 | 67636769:676 | 67638735:676 | -;- | 2466 | NA | 0 | 0 | 0 | 0 | 0 | 0 |
| ENSDARG000 | 7;7;7;7;7;7;7 | 48761646:487 | 48762017:487 | +;+;+;+;+;+;+ | 6229 | NA | 0 | 14 | 110 | 32 | 264 | 158 |
| ENSDARG000 | 7;7;7;7;7;7;7 | 31838320:318 | 31838388:318 | +;+;+;+;+;+;+ | 5870 | mybpc3 | 0 | 0 | 0 | 0 | 0 | 0 |
| ENSDARG000 | 7;7;7;7;7;7;7 | 29301497:293 | 29301764:293 | +;+;+;+;+;+;+ | 3162 | rab8b | 2104 | 2061 | 2229 | 1116 | 1513 | 1551 |
| ENSDARG000 | 7;7;7;7;7;7;7 | 37514780:375 | 37516290:375 | -;-;-;-;-;-;- | 4639 | CYLD | 636 | 564 | 792 | 997 | 1601 | 545 |
| ENSDARG000 | 7;7;7;7;7 | 67699009:676 | 67700104:677 | +;+;+;+;+ | 2432 | zgc:162592 | 0 | 0 | 0 | 0 | 0 | 0 |
| ENSDARG000 | 7;7;7;7 | 44628681:446 | 44629344:446 | -;-;-;- | 1610 | cmtm4 | 145 | 102 | 140 | 290 | 238 | 90 |
| ENSDARG000 | 7;7;7;7;7;7;7 | 38661107:386 | 38662002:386 | -;-;-;-;-;-;- | 1613 | c6ast1 | 42 | 16 | 6 | 4 | 26 | 2 |
| ENSDARG000 | 7;7 | 44671925:446 | 44672095:446 | +;+ | 416 | NA | 5 | 3 | 4 | 0 | 3 | 0 |
| ENSDARG000 | 7;7;7;7;7;7;7 | 30326184:303 | 30327753:303 | -;-;-;-;-;-;- | 3734 | aldh1a2 | 275 | 124 | 142 | 80 | 216 | 102 |
| ENSDARG000 | 7;7;7;7;7;7;7 | 44674956:446 | 44678009:446 | -;-;-;-;-;-;- | 4489 | dync1li2 | 15790 | 9676 | 6036 | 8452 | 8282 | 9061 |
| ENSDARG000 | 7;7 | 67730148:677 | 67730900:677 | -;- | 894 | NA | 0 | 0 | 0 | 0 | 10 | 3 |
| ENSDARG000 | 7;7;7;7;7 | 67733759:677 | 67734108:677 | +;+;+;+;+ | 1798 | cyb5b | 857 | 938 | 1191 | 323 | 958 | 620 |
| ENSDARG000 | 7 | 72101890 | 72103391 | + | 1502 | NA | 15 | 15 | 0 | 2 | 23 | 130 |
| ENSDARG000 | 7 | 67747252 | 67748015 | + | 764 | NA | 195 | 82 | 456 | 140 | 312 | 234 |
| ENSDARG000 | 7;7;7;7;7;7;7 | 48788147:487 | 48789038:487 | -;-;-;-;-;-;- | 2589 | mfge8a | 142 | 53 | 294 | 188 | 375 | 204 |
| ENSDARG000 | 7;7;7;7;7;7;7 | 19909774:199 | 19911141:199 | -;-;-;-;-;-;- | 2333 | tmem179ba | 3110 | 3641 | 5536 | 3902 | 4707 | 5169 |
| ENSDARG000 | 7;7;7;7;7;7;7 | 72276511:722 | 72276624:722 | +;+;+;+;+;+;+ | 1419 | tollip | 622 | 588 | 693 | 640 | 466 | 570 |
| ENSDARG000 | 7;7;7;7;7;7;7 | 72291181:722 | 72291525:722 | -;-;-;-;-;-;- | 17245 | NA | 872 | 5 | 240 | 18 | 72 | 44 |
| ENSDARG000 | 7;7;7;7;7;7;7 | 40228422:402 | 40228630:402 | +;+;+;+;+;+;+ | 4722 | ptprn2 | 0 | 0 | 6 | 14 | 6 | 0 |
| ENSDARG000 | 7 | 33879753 | 33881275 | - | 1523 | NA | 0 | 0 | 2 | 4 | 11 | 0 |
| ENSDARG000 | 7;7;7;7;7;7;7 | 26100024:261 | 26100173:261 | +;+;+;+;+;+;+ | 5533 | NA | 4 | 44 | 214 | 192 | 94 | 94 |
| ENSDARG000 | 7;7;7;7;7;7;7 | 33882597:338 | 33886894:338 | -;-;-;-;-;-;- | 6002 | PIAS1 | 122 | 54 | 148 | 105 | 76 | 151 |
| ENSDARG000 | 7;7;7;7;7;7;7 | 17992455:179 | 17993002:179 | +;+;+;+;+;+;+ | 17462 | NA | 47 | 14 | 126 | 40 | 74 | 98 |
| ENSDARG000 | 7;7;7;7;7 | 19930564:199 | 19932025:199 | -;-;-;-;- | 3561 | prox3 | 143 | 157 | 422 | 292 | 257 | 167 |
| ENSDARG000 | 7;7;7;7;7;7;7 | 27700274:277 | 27703118:277 | -;-;-;-;-;-;- | 5924 | PDE3B | 560 | 235 | 411 | 303 | 348 | 425 |
| ENSDARG000 | 7;7;7;7;7;7;7 | 19953756:199 | 19955118:199 | -;-;-;-;-;-;- | 2566 | snx15 | 624 | 408 | 729 | 523 | 662 | 542 |
| ENSDARG000 | 7 | 27797821 | 27799035 | + | 1215 | NA | 1 | 2 | 2 | 0 | 3 | 1 |
| ENSDARG000 | 7;7;7;7;7;7;7 | 37555568:375 | 37556230:375 | -;-;-;-;-;-;- | 3862 | nod2 | 232 | 170 | 197 | 470 | 291 | 240 |
| ENSDARG000 | 7;7;7;7;7;7;7 | 29312376:293 | 29314580:293 | -;-;-;-;-;-;- | 6196 | trpm1a | 14 | 6 | 14 | 2 | 11 | 12 |
| ENSDARG000 | 7;7;7;7;7;7;7 | 48805534:488 | 48805697:488 | +;+;+;+;+;+;+ | 3706 | cpt1aa | 2048 | 1701 | 2019 | 715 | 1794 | 2137 |
| ENSDARG000 | 7;7;7 | 23854361:238 | 23854951:238 | -;-;- | 1225 | NA | 0 | 0 | 0 | 0 | 0 | 0 |
| ENSDARG000 | 7;7;7;7;7;7;7 | 21321317:213 | 21321465:213 | +;+;+;+;+;+;+ | 14042 | DNAH2 | 4 | 2 | 8 | 0 | 16 | 0 |
| ENSDARG000 | 7;7;7 | 23865336:238 | 23865742:238 | -;-;- | 1247 | zgc:162171 | 0 | 6 | 12 | 4 | 4 | 0 |
| ENSDARG000 | 7;7;7;7;7;7;7 | 24840701:248 | 24841354:248 | -;-;-;-;-;-;- | 4166 | NA | 142 | 43 | 56 | 223 | 134 | 200 |
| ENSDARG000 | 7;7;7 | 23875269:238 | 23875688:238 | +;+;+ | 2434 | rab39bb | 10 | 18 | 10 | 18 | 41 | 30 |
| ENSDARG000 | 7;7;7;7;7;7;7 | 24862471:248 | 24863147:248 | -;-;-;-;-;-;-;- | 2154 | NA | 0 | 0 | 0 | 0 | 0 | 0 |
| ENSDARG000 | 7;7;7;7;7;7;7 | 67748939:677 | 67749405:677 | +;+;+;+;+;+;+ | 9226 | dhx38 | 568 | 407 | 761 | 234 | 669 | 494 |
| ENSDARG000 | 7;7;7;7;7;7 | 23886756:238 | 23888239:238 | -;-;-;-;-;- | 2213 | vbp1 | 990 | 1051 | 1454 | 1180 | 926 | 1216 |
| ENSDARG000 | 7;7;7;7;7 | 26116103:261 | 26116955:261 | -;-;-;-;- | 1833 | NA | 114 | 92 | 173 | 136 | 142 | 104 |
| ENSDARG000 | 7;7;7;7;7;7;7 | 26132665:261 | 26132752:261 | +;+;+;+;+;+;+ | 10055 | NAT16 | 11 | 6 | 53 | 6 | 5 | 7 |
| ENSDARG000 | 7;7;7;7;7;7;7 | 33954099:339 | 33955527:339 | -;-;-;-;-;-;- | 3875 | skor1a | 4 | 4 | 0 | 2 | 8 | 0 |
| ENSDARG000 | 7;7;7;7;7;7;7 | 24881680:248 | 24881856:248 | +;+;+;+;+;+;+ | 2646 | krcp | 40 | 393 | 422 | 222 | 288 | 244 |
| ENSDARG000 | 7;7;7;7;7;7;7 | 27802480:278 | 27802569:278 | +;+;+;+;+;+;+ | 4040 | copb1 | 2891 | 3247 | 5108 | 3242 | 4169 | 2635 |
| ENSDARG000 | 7;7;7;7;7;7;7 | 37582281:375 | 37586369:375 | -;-;-;-;-;-;- | 5120 | nkd1 | 60 | 8 | 16 | 18 | 18 | 8 |
| ENSDARG000 | 7;7;7;7;7;7;7 | 23902484:239 | 23903647:239 | -;-;-;-;-;-;- | 1901 | NA | 4 | 0 | 0 | 2 | 0 | 0 |
| ENSDARG000 | 7;7;7;7;7 | 26167420:261 | 26167641:261 | +;+;+;+;+ | 1384 | si:ch211-196f | 2 | 10 | 0 | 4 | 10 | 26 |
| ENSDARG000 | 7;7;7;7;7;7 | 48870496:488 | 48870680:488 | +;+;+;+;+;+ | 2090 | igf2a | 24 | 12 | 10 | 2 | 27 | 57 |
| ENSDARG000 | 7;7;7;7 | 29351499:293 | 29351829:293 | -;-;-;- | 760 | gtf2a2 | 234 | 332 | 394 | 250 | 264 | 280 |
| ENSDARG000 | 7;7;7;7;7;7;7 | 26172396:261 | 26172695:261 | +;+;+;+;+;+;+ | 2745 | nat16l | 214 | 167 | 214 | 185 | 103 | 182 |
| ENSDARG000 | 7;7;7;7;7;7;7 | 48999723:489 | 48999918:489 | +;+;+;+;+;+;+ | 6002 | NA | 337 | 298 | 364 | 161 | 355 | 516 |
| ENSDARG000 | 7;7;7;7;7;7;7 | 27834130:278 | 27834470:278 | +;+;+;+;+;+;+ | 4832 | rras2 | 1627 | 1633 | 1484 | 1728 | 1345 | 1224 |
| ENSDARG000 | 7;7;7;7 | 29359988:293 | 29360073:293 | -;-;-;- | 756 | NA | 2 | 0 | 1 | 0 | 5 | 0 |
| ENSDARG000 | 7;7;7;7 | 23907692:239 | 23907970:239 | +;+;+;+ | 2327 | syt4 | 2 | 8 | 4 | 2 | 8 | 0 |
| ENSDARG000 | 7;7;7;7;7;7;7 | 72345925:723 | 72346199:723 | -;-;-;-;-;-;-;- | 1530 | NA | 0 | 12 | 2 | 0 | 0 | 2 |
| ENSDARG000 | 7;7;7;7;7;7;7 | 24888910:248 | 24889833:248 | +;+;+;+;+;+;+ | 5488 | MARK2 | 1677 | 1022 | 1530 | 1285 | 2001 | 1468 |
| ENSDARG000 | 7;7;7;7;7;7;7 | 33970698:339 | 33973678:339 | -;-;-;-;-;-;- | 4512 | map2k5 | 502 | 404 | 546 | 264 | 656 | 445 |
| ENSDARG000 | 7;7;7;7;7;7;7 | 23940933:239 | 23940976:239 | +;+;+;+;+;+;+ | 1592 | NA | 0 | 0 | 0 | 0 | 6 | 4 |
| ENSDARG000 | 7;7;7;7 | 29413895:294 | 29413980:294 | -;-;-;- | 756 | NA | 0 | 0 | 6 | 2 | 4 | 1 |
| ENSDARG000 | 7;7;7;7 | 29390067:293 | 29390151:293 | -;-;-;- | 755 | NA | 2 | 0 | 0 | 0 | 7 | 0 |
| ENSDARG000 | 7;7;7;7 | 29378430:293 | 29378515:293 | -;-;-;- | 756 | NA | 0 | 0 | 0 | 0 | 1 | 0 |
| ENSDARG000 | 7;7;7;7 | 29431710:294 | 29431796:294 | -;-;-;- | 695 | NA | 0 | 0 | 2 | 0 | 0 | 0 |
| ENSDARG000 | 7;7;7;7;7 | 29402489:294 | 29402646:294 | -;-;-;-;- | 846 | NA | 4 | 0 | 1 | 0 | 2 | 0 |
| ENSDARG000 | 7;7;7;7;7;7;7 | 23967332:239 | 23967486:239 | -;-;-;-;-;-;- | 6405 | NA | 34 | 30 | 50 | 60 | 61 | 36 |
| ENSDARG000 | 7;7 | 26224211:262 | 26224427:262 | +;+ | 3953 | NA | 92 | 128 | 2020 | 366 | 470 | 2258 |
| ENSDARG000 | 7;7;7;7;7;7;7 | 27898091:278 | 27898364:278 | +;+;+;+;+;+;+ | 10918 | TUB | 567 | 20 | 48 | 1406 | 28 | 1394 |
| ENSDARG000 | 7;7;7;7;7;7;7 | 49078890:490 | 49079598:490 | +;+;+;+;+;+;+ | 4784 | B4GALNT4 | 1 | 0 | 0 | 6 | 0 | 0 |
| ENSDARG000 | 7;7;7 | 28058202:280 | 28058520:280 | +;+;+ | 522 | NA | 0 | 0 | 0 | 0 | 0 | 0 |
| ENSDARG000 | 7;7;7;7;7;7 | 28047279:280 | 28050215:280 | -;-;-;-;-;- | 3616 | RIC3 | 0 | 0 | 0 | 0 | 0 | 20 |
| ENSDARG000 | 7;7;7;7;7;7;7 | 49272702:492 | 49273711:492 | +;+;+;+;+;+;+ | 5462 | ABTB2 | 156 | 386 | 867 | 666 | 772 | 1232 |
| ENSDARG000 | 7;7;7;7;7;7;7 | 28109364:281 | 28110198:281 | -;-;-;-;-;-;-;- | 1821 | lmo1 | 4 | 0 | 4 | 0 | 4 | 0 |
| ENSDARG000 | 7 | 29461060 | 29463253 | + | 2194 | foxb1b | 0 | 0 | 2 | 0 | 0 | 0 |
| ENSDARG000 | 7;7;7;7;7;7 | 29461816:294 | 29462250:294 | -;-;-;-;-;- | 1104 | NA | 0 | 2 | 0 | 0 | 0 | 0 |
| ENSDARG000 | 7;7;7;7;7;7;7 | 34102661:341 | 34107450:341 | -;-;-;-;-;-;-;- | 6173 | smad3a | 162 | 188 | 279 | 108 | 209 | 126 |
| ENSDARG000 | 7;7;7;7;7;7;7 | 23997518:239 | 23997918:239 | +;+;+;+;+;+;+ | 1320 | NA | 0 | 0 | 0 | 0 | 0 | 0 |
| ENSDARG000 | 7 | 24002096 | 24005268 | - | 3173 | NA | 2 | 0 | 6 | 0 | 0 | 0 |
| ENSDARG000 | 7;7;7;7;7;7;7 | 26245061:262 | 26246195:262 | -;-;-;-;-;-;- | 1853 | ap1s1 | 422 | 534 | 495 | 233 | 576 | 425 |
| ENSDARG000 | 7;7;7;7;7;7;7 | 31902333:319 | 31906032:319 | -;-;-;-;-;-;- | 5055 | lin7c | 2048 | 1422 | 2286 | 1346 | 2038 | 2393 |
| ENSDARG000 | 7;7;7;7;7;7;7 | 49324103:493 | 49325765:493 | -;-;-;-;-;-;- | 4368 | BRSK2 | 4 | 4 | 24 | 22 | 18 | 18 |
| ENSDARG000 | 7;7;7;7;7;7;7 | 34227075:342 | 34227105:342 | +;+;+;+;+;+;+ | 2680 | lctla | 22 | 44 | 124 | 232 | 16 | 272 |
| ENSDARG000 | 7;7;7;7;7 | 34182963:341 | 34184191:341 | -;-;-;-;- | 2781 | smad6a | 50 | 6 | 40 | 10 | 55 | 8 |
| ENSDARG000 | 7;7;7;7;7;7 | 34222433:342 | 34222672:342 | -;-;-;-;-;- | 734 | crybgx | 4 | 0 | 0 | 0 | 0 | 0 |
| ENSDARG000 | 7;7 | 24006875:240 | 24007670:240 | +;+ | 3420 | homezb | 542 | 554 | 660 | 444 | 619 | 510 |
| ENSDARG000 | 7;7;7;7;7;7;7 | 72644051:726 | 72644197:726 | -;-;-;-;-;-;- | 5712 | NA | 980 | 929 | 1995 | 839 | 1105 | 705 |
| ENSDARG000 | 7;7;7;7;7;7;7 | 21632100:216 | 21632513:216 | +;+;+;+;+;+;+ | 10097 | KDM6B | 3102 | 2754 | 4120 | 2372 | 4316 | 3278 |
| ENSDARG000 | 7;7;7;7;7;7;7 | 26265584:262 | 26266130:262 | -;-;-;-;-;-;-;- | 1789 | serpine1 | 713 | 123 | 96 | 132 | 646 | 218 |
| ENSDARG000 | 7;7;7;7;7 | 24015507:240 | 24016046:240 | -;-;-;-;- | 1175 | cideb | 2 | 0 | 0 | 0 | 0 | 0 |
| ENSDARG000 | 7;7 | 24018851:240 | 24019576:240 | +;+ | 1593 | NA | 0 | 0 | 0 | 0 | 0 | 0 |
| ENSDARG000 | 7;7;7;7;7;7;7 | 24039464:240 | 24039780:240 | -;-;-;-;-;-;- | 2344 | dhrs4 | 116 | 224 | 229 | 207 | 182 | 235 |
| ENSDARG000 | 7;7;7;7;7;7;7 | 24023653:240 | 24023971:240 | +;+;+;+;+;+;+ | 2329 | tinf2 | 74 | 161 | 152 | 70 | 144 | 157 |
| ENSDARG000 | 7;7;7;7;7;7;7 | 52830254:528 | 52834794:528 | -;-;-;-;-;-;- | 12455 | MAP1A | 30 | 36 | 74 | 44 | 222 | 40 |
| ENSDARG000 | 7;7;7;7;7;7;7 | 71463106:714 | 71463222:714 | -;-;-;-;-;-;- | 2884 | ACO1 | 1116 | 436 | 1064 | 991 | 849 | 836 |
| ENSDARG000 | 7;7;7;7;7;7;7 | 71955486:719 | 71955700:719 | +;+;+;+;+;+;+ | 3431 | lrp4 | 1128 | 490 | 634 | 417 | 662 | 783 |
| ENSDARG000 | 7;7;7;7;7;7;7 | 19817306:198 | 19817325:198 | +;+;+;+;+;+;+ | 1476 | NA | 2 | 0 | 7 | 0 | 0 | 0 |
| ENSDARG000 | 7;7;7 | 65387374:653 | 65387479:653 | +;+;+ | 3364 | CLEC3A | 596 | 2340 | 602 | 1026 | 1087 | 171 |
| ENSDARG000 | 7 | 47372795 | 47381146 | - | 8352 | NA | 0 | 0 | 0 | 0 | 0 | 0 |
| ENSDARG000 | 7;7;7;7;7;7;7 | 64378068:643 | 64378397:643 | -;-;-;-;-;-;- | 5419 | NA | 28 | 38 | 8 | 6 | 54 | 18 |
| ENSDARG000 | 7;7;7;7;7;7;7 | 71575995:715 | 71578151:715 | -;-;-;-;-;-;-;- | 3498 | METTL4 | 175 | 115 | 122 | 48 | 129 | 100 |
| ENSDARG000 | 7;7;7;7;7;7;7 | 53413018:534 | 53413821:534 | +;+;+;+;+;+;+ | 10961 | znf609b | 1889 | 1528 | 1899 | 1480 | 1976 | 2049 |
| ENSDARG000 | 7;7;7;7;7;7;7 | 52887701:528 | 52887761:528 | +;+;+;+;+;+;+ | 5291 | tp53bp1 | 963 | 1145 | 1863 | 1259 | 1718 | 1409 |
| ENSDARG000 | 7;7;7;7;7;7;7 | 61843532:618 | 61846802:618 | -;-;-;-;-;-;- | 7310 | NA | 10 | 0 | 20 | 10 | 8 | 28 |
| ENSDARG000 | 7;7 | 57677120:576 | 57677914:576 | +;+ | 4917 | ARSJ | 54 | 16 | 24 | 0 | 234 | 16 |
| ENSDARG000 | 7;7;7;7;7;7 | 54475134:544 | 54475480:544 | +;+;+;+;+;+ | 1533 | zgc:153993 | 398 | 439 | 483 | 350 | 452 | 359 |
| ENSDARG000 | 7;7;7;7;7;7;7 | 59200481:592 | 59203305:592 | -;-;-;-;-;-;- | 4068 | nagk | 326 | 383 | 308 | 366 | 350 | 304 |
| ENSDARG000 | 7;7;7;7;7;7;7 | 71437142:714 | 71437970:714 | -;-;-;-;-;-;- | 2458 | sepsecs | 82 | 175 | 137 | 72 | 176 | 172 |
| ENSDARG000 | 7;7;7;7;7;7;7 | 31154282:311 | 31154364:311 | -;-;-;-;-;-;- | 600 | NA | 0 | 0 | 0 | 0 | 0 | 0 |
| ENSDARG000 | 7;7;7;7;7;7;7 | 10592152:105 | 10592255:105 | +;+;+;+;+;+;+ | 1058 | fah | 82 | 124 | 71 | 20 | 271 | 176 |
| ENSDARG000 | 7;7;7;7;7;7;7 | 65533307:655 | 65533567:655 | +;+;+;+;+;+;+ | 3526 | MICAL2 | 20 | 0 | 0 | 4 | 16 | 16 |
| ENSDARG000 | 7;7;7;7;7 | 20826240:208 | 20831254:208 | -;-;-;-;- | 5587 | cldn15a | 0 | 0 | 8 | 4 | 6 | 0 |
| ENSDARG000 | 7;7;7;7;7;7;7 | 34293635:342 | 34293778:342 | -;-;-;-;-;-;- | 804 | NA | 0 | 0 | 0 | 0 | 0 | 0 |
| ENSDARG000 | 7 | 21830686 | 21830850 | - | 165 | U1 | 0 | 0 | 0 | 0 | 0 | 0 |
| ENSDARG000 | 7 | 14094794 | 14094911 | - | 118 | 5S_rRNA | 0 | 0 | 0 | 0 | 0 | 0 |
| ENSDARG000 | 7 | 42812055 | 42812169 | - | 115 | 5S_rRNA | 0 | 0 | 0 | 0 | 0 | 0 |
| ENSDARG000 | 7;7;7;7;7;7 | 17467551:174 | 17467590:174 | +;+;+;+;+;+ | 999 | nitr1b | 0 | 0 | 0 | 0 | 0 | 1 |
| ENSDARG000 | 7;7;7;7;7;7;7 | 50641919:506 | 50644537:506 | -;-;-;-;-;-;- | 7680 | iqgap1 | 8007 | 6502 | 7427 | 4757 | 11513 | 6997 |
| ENSDARG000 | 7 | 1813535 | 1813653 | + | 119 | 5S_rRNA | 0 | 0 | 0 | 0 | 0 | 0 |
| ENSDARG000 | 7 | 7778963 | 7779175 | + | 213 | U3 | 0 | 0 | 0 | 0 | 0 | 0 |
| ENSDARG000 | 7 | 25045410 | 25045479 | + | 70 | SNORD30 | 0 | 0 | 4 | 1 | 2 | 0 |
| ENSDARG000 | 7;7;7;7;7;7;7 | 20876303:208 | 20876849:208 | +;+;+;+;+;+;+ | 3723 | GIGYF1 | 441 | 224 | 429 | 278 | 581 | 665 |

| | | | | | | | | | | | | |
|---|---|---|---|---|---|---|---|---|---|---|---|---|
| ENSDARG000 | 7:7:7:7:7:7:7: | 21180747:212 | 21181040:212 | +:+:+:+:+:+:+: | 5808 | serpinh2 | 176 | 62 | 182 | 122 | 171 | 148 |
| ENSDARG000 | 7 | 68089813 | 68089928 | + | 116 | 5S_rRNA | 0 | 0 | 0 | 0 | 0 | 0 |
| ENSDARG000 | 7;7;7;7 | 74134010:741 | 74134301:741 | +;+;+;+ | 3032 | cldnd1a | 174 | 188 | 248 | 115 | 160 | 140 |
| ENSDARG000 | 7;7;7 | 57404701:574 | 57404883:574 | +;+;+ | 611 | NA | 110 | 8 | 2 | 90 | 0 | 0 |
| ENSDARG000 | 7 | 50947555 | 50947674 | + | 120 | 5S_rRNA | 0 | 0 | 0 | 0 | 0 | 0 |
| ENSDARG000 | 7 | 65151737 | 65151917 | + | 181 | SNORA23 | 0 | 4 | 0 | 0 | 0 | 0 |
| ENSDARG000 | 7 | 59153778 | 59153894 | - | 117 | 5S_rRNA | 0 | 0 | 0 | 0 | 0 | 0 |
| ENSDARG000 | 7:7:7:7:7:7:7: | 6915609:6916 | 6915654:6917 | +:+:+:+:+:+:+: | 879 | CCS | 3 | 14 | 13 | 23 | 0 | 4 |
| ENSDARG000 | 7 | 66640915 | 66641031 | + | 117 | 5S_rRNA | 0 | 0 | 0 | 0 | 0 | 0 |
| ENSDARG000 | 7 | 19956078 | 19956192 | - | 115 | 5S_rRNA | 0 | 0 | 6 | 0 | 0 | 0 |
| ENSDARG000 | 7 | 14281093 | 14281225 | + | 133 | 5S_rRNA | 0 | 0 | 0 | 0 | 0 | 0 |
| ENSDARG000 | 7:7:7:7:7:7:7: | 65398161:653 | 65398404:653 | +:+:+:+:+:+:+: | 6029 | usp47 | 1138 | 1130 | 1368 | 1061 | 1670 | 827 |
| ENSDARG000 | 7 | 27825945 | 27826061 | + | 117 | 5S_rRNA | 0 | 0 | 0 | 0 | 0 | 0 |
| ENSDARG000 | 7 | 11603780 | 11603894 | - | 115 | 5S_rRNA | 0 | 0 | 4 | 0 | 0 | 0 |
| ENSDARG000 | 7 | 25045848 | 25045974 | + | 127 | SNORD22 | 0 | 0 | 0 | 0 | 0 | 0 |
| ENSDARG000 | 7 | 61015820 | 61015938 | - | 119 | 5S_rRNA | 0 | 0 | 0 | 0 | 0 | 0 |
| ENSDARG000 | 7 | 10941100 | 10941214 | - | 115 | 5S_rRNA | 0 | 0 | 0 | 0 | 0 | 0 |
| ENSDARG000 | 7 | 5296 | 5449 | + | 154 | 5_8S_rRNA | 41 | 47 | 230 | 57 | 12 | 146 |
| ENSDARG000 | 7;7;7;7;7;7;7 | 18121722:181 | 18123149:181 | -;-;-;-;-;-;- | 2609 | PELI3 | 166 | 109 | 244 | 539 | 120 | 240 |
| ENSDARG000 | 7 | 73263751 | 73263822 | - | 72 | dre-mir-31 | 0 | 0 | 0 | 0 | 0 | 0 |
| ENSDARG000 | 7 | 53641042 | 53641178 | - | 137 | 5S_rRNA | 5 | 0 | 3 | 0 | 2 | 0 |
| ENSDARG000 | 7 | 40870631 | 40870748 | + | 118 | 5S_rRNA | 0 | 0 | 0 | 0 | 0 | 0 |
| ENSDARG000 | 7:7:7:7:7:7:7: | 69732435:697 | 69732506:697 | -:-:-:-:-:-:-:-: | 2805 | USP53 | 162 | 48 | 33 | 56 | 36 | 40 |
| ENSDARG000 | 7 | 41493017 | 41493131 | - | 115 | 5S_rRNA | 0 | 0 | 1 | 0 | 0 | 0 |
| ENSDARG000 | 7:7:7:7:7:7:7: | 41947739:419 | 41947785:419 | -:-:-:-:-:-:-:-: | 1777 | NETO2 | 68 | 12 | 4 | 30 | 57 | 58 |
| ENSDARG000 | 7 | 53113556 | 53113672 | - | 117 | 5S_rRNA | 3 | 1 | 1 | 8 | 28 | 11 |
| ENSDARG000 | 7:7:7:7:7:7:7: | 59335082:594 | 59335169:594 | +:+:+:+:+:+:+: | 2301 | SEMA4F | 342 | 12 | 107 | 62 | 72 | 67 |
| ENSDARG000 | 7:7:7:7:7:7:7: | 17566111:175 | 17568229:175 | -:-:-:-:-:-:-:-: | 3071 | nitr5 | 0 | 0 | 1 | 13 | 2 | 42 |
| ENSDARG000 | 7;7;7;7;7;7;7 | 47143303:471 | 47146344:471 | -;-;-;-;-;-;- | 4041 | NA | 0 | 0 | 0 | 0 | 0 | 0 |
| ENSDARG000 | 7;7;7 | 59553040:595 | 59554954:595 | -;-;- | 2489 | BOLA2 | 250 | 248 | 392 | 242 | 374 | 441 |
| ENSDARG000 | 7:7:7:7:7:7:7: | 54323944:543 | 54324141:543 | -:-:-:-:-:-:-:-: | 2778 | ano1a | 140 | 38 | 30 | 14 | 97 | 59 |
| ENSDARG000 | 7;7;7;7;7;7 | 17534485:175 | 17534524:175 | +;+;+;+;+;+ | 987 | nitr1k | 0 | 0 | 0 | 0 | 0 | 0 |
| ENSDARG000 | 7:7:7:7:7:7:7: | 1062972:1062 | 1063113:1063 | -:-:-:-:-:-:-:-: | 4661 | dctn1a | 2978 | 1409 | 1521 | 1217 | 1975 | 2004 |
| ENSDARG000 | 7;7;7;7;7;7;7 | 69242104:692 | 69242459:692 | -;-;-;-;-;-;- | 1557 | phlpp2 | 46 | 31 | 40 | 39 | 46 | 18 |
| ENSDARG000 | 7 | 40485510 | 40485590 | + | 81 | dre-mir-153a | 0 | 0 | 0 | 0 | 0 | 0 |
| ENSDARG000 | 7:7:7:7:7:7:7: | 56889879:569 | 56890721:569 | +:+:+:+:+:+:+: | 7569 | myo9aa | 3311 | 1901 | 2971 | 2161 | 2536 | 2100 |
| ENSDARG000 | 7:7:7:7:7:7:7: | 56253914:563 | 56253994:563 | +:+:+:+:+:+:+: | 8603 | ANKRD11 | 1286 | 909 | 1861 | 1098 | 1811 | 1263 |
| ENSDARG000 | 7;7 | 56651759:566 | 56652505:566 | +;+ | 1608 | NA | 0 | 0 | 0 | 0 | 0 | 0 |
| ENSDARG000 | 7;7;7;7;7;7;7 | 72630369:726 | 72630719:726 | +;+;+;+;+;+;+ | 1107 | NA | 11 | 13 | 22 | 0 | 4 | 25 |
| ENSDARG000 | 7;7 | 56676838:566 | 56677153:566 | +;+ | 5712 | zgc:194679 | 0 | 0 | 0 | 0 | 0 | 0 |
| ENSDARG000 | 7:7:7:7:7:7:7: | 71586485:715 | 71586652:715 | +:+:+:+:+:+:+: | 5949 | SMCHD1 | 393 | 226 | 618 | 321 | 530 | 369 |
| ENSDARG000 | 7;7;7;7;7;7;7 | 7469762:7471 | 7469787:7471 | +;+;+;+;+;+;+ | 147 | NA | 0 | 1 | 2 | 0 | 0 | 0 |
| ENSDARG000 | 7:7:7:7:7:7:7: | 57612905:576 | 57612937:576 | +:+:+:+:+:+:+: | 183 | NA | 5 | 0 | 9 | 3 | 56 | 9 |
| ENSDARG000 | 7;7;7;7;7;7;7 | 59020972:590 | 59021204:590 | +;+;+;+;+;+;+ | 3702 | hacd1 | 14 | 70 | 133 | 36 | 86 | 54 |
| ENSDARG000 | 7:7:7:7:7:7:7: | 61480296:614 | 61480349:614 | +:+:+:+:+:+:+: | 4481 | NA | 1259 | 676 | 1962 | 542 | 3713 | 2466 |
| ENSDARG000 | 7:7:7:7:7:7:7: | 15736230:157 | 15736465:157 | +:+:+:+:+:+:+: | 3661 | NA | 10 | 14 | 37 | 16 | 60 | 39 |
| ENSDARG000 | 7;7;7;7;7;7 | 6908131:6919 | 6908156:6919 | -;-;-;-;-;- | 123 | NA | 0 | 2 | 4 | 8 | 0 | 0 |
| ENSDARG000 | 7;7;7;7;7;7;7 | 65472578:654 | 65474249:654 | -;-;-;-;-;-;- | 2644 | dkk3a | 0 | 2 | 2 | 0 | 0 | 0 |
| ENSDARG000 | 7 | 25291033 | 25291152 | - | 120 | 5S_rRNA | 0 | 0 | 0 | 0 | 0 | 0 |
| ENSDARG000 | 7:7:7:7:7:7:7: | 18075504:180 | 18075581:180 | +:+:+:+:+:+:+: | 2172 | NA | 40 | 83 | 69 | 91 | 232 | 114 |
| ENSDARG000 | 7 | 29336996 | 29337119 | - | 124 | dre-mir-204-2 | 0 | 0 | 0 | 0 | 0 | 0 |
| ENSDARG000 | 7 | 50287933 | 50288019 | - | 87 | dre-mir-129-2 | 0 | 0 | 0 | 0 | 0 | 0 |
| ENSDARG000 | 7 | 73827805 | 73829243 | + | 1439 | zgc:173587 | 9 | 8 | 12 | 9 | 18 | 7 |
| ENSDARG000 | 7:7:7:7:7:7:7: | 15905691:159 | 15906643:159 | -:-:-:-:-:-:-:-: | 7304 | elp4 | 187 | 152 | 232 | 150 | 194 | 170 |
| ENSDARG000 | 7;7 | 3158104:3159 | 3158135:3160 | +;+ | 1074 | NA | 0 | 0 | 0 | 4 | 0 | 0 |
| ENSDARG000 | 7 | 43563124 | 43563240 | - | 117 | 5S_rRNA | 0 | 0 | 0 | 0 | 0 | 0 |
| ENSDARG000 | 7 | 47058656 | 47058770 | + | 115 | 5S_rRNA | 0 | 0 | 0 | 0 | 0 | 0 |
| ENSDARG000 | 7 | 68310824 | 68310941 | + | 118 | 5S_rRNA | 0 | 0 | 0 | 0 | 0 | 0 |
| ENSDARG000 | 7 | 26513822 | 26513960 | - | 139 | NA | 0 | 0 | 0 | 0 | 0 | 0 |
| ENSDARG000 | 7:7:7:7:7:7:7: | 69528850:695 | 69529279:695 | +:+:+:+:+:+:+: | 1976 | RAP1GDS1 | 258 | 99 | 304 | 100 | 320 | 272 |
| ENSDARG000 | 7;7;7;7;7;7 | 73713799:737 | 73713847:737 | -;-;-;-;-;- | 856 | NA | 0 | 6 | 0 | 0 | 0 | 0 |
| ENSDARG000 | 7:7:7:7:7:7:7: | 13824150:138 | 13824318:138 | +:+:+:+:+:+:+: | 1645 | abhd2a | 36 | 22 | 66 | 28 | 10 | 40 |
| ENSDARG000 | 7 | 59696033 | 59696149 | + | 117 | 5S_rRNA | 0 | 0 | 0 | 0 | 0 | 2 |
| ENSDARG000 | 7:7:7:7:7:7:7: | 61050329:610 | 61050684:610 | +:+:+:+:+:+:+: | 10417 | NWD2 | 34 | 30 | 14 | 10 | 36 | 54 |
| ENSDARG000 | 7;7;7;7;7;7 | 55223955:552 | 55224020:552 | -;-;-;-;-;- | 729 | RNF166 | 47 | 12 | 35 | 26 | 55 | 40 |
| ENSDARG000 | 7:7:7:7:7:7:7: | 7841951:7842 | 7841987:7842 | +:+:+:+:+:+:+: | 180 | NA | 10 | 0 | 0 | 0 | 6 | 1 |
| ENSDARG000 | 7:7:7:7:7:7:7: | 20965880:209 | 20966549:209 | +:+:+:+:+:+:+: | 1959 | efnb3b | 135 | 85 | 159 | 34 | 150 | 80 |
| ENSDARG000 | 7 | 28157554 | 28157668 | - | 115 | 5S_rRNA | 0 | 0 | 0 | 0 | 0 | 0 |
| ENSDARG000 | 7 | 30438905 | 30439021 | + | 117 | 5S_rRNA | 0 | 0 | 0 | 0 | 0 | 0 |
| ENSDARG000 | 7 | 51597522 | 51597636 | - | 115 | 5S_rRNA | 0 | 0 | 0 | 0 | 0 | 2 |
| ENSDARG000 | 7 | 65336322 | 65336440 | + | 119 | 5S_rRNA | 0 | 0 | 0 | 0 | 0 | 0 |
| ENSDARG000 | 7 | 41761261 | 41761379 | + | 119 | 5S_rRNA | 0 | 0 | 2 | 0 | 0 | 0 |
| ENSDARG000 | 7 | 10248072 | 10248187 | + | 116 | 5S_rRNA | 0 | 0 | 0 | 0 | 0 | 0 |
| ENSDARG000 | 7 | 27912938 | 27913055 | + | 118 | 5S_rRNA | 0 | 0 | 0 | 0 | 0 | 0 |
| ENSDARG000 | 7:7:7:7:7:7:7: | 40658003:406 | 40658062:406 | +:+:+:+:+:+:+: | 1596 | lmbr1 | 67 | 70 | 59 | 63 | 52 | 105 |
| ENSDARG000 | 7 | 69490235 | 69490425 | + | 191 | U2 | 0 | 0 | 0 | 0 | 0 | 0 |
| ENSDARG000 | 7 | 26071426 | 26071508 | + | 83 | dre-mir-457a | 0 | 0 | 0 | 0 | 0 | 0 |
| ENSDARG000 | 7 | 14450524 | 14450620 | - | 97 | dre-mir-7a-2 | 0 | 0 | 0 | 0 | 0 | 0 |
| ENSDARG000 | 7 | 59157559 | 59157689 | - | 131 | SCARNA8 | 3 | 6 | 13 | 3 | 5 | 1 |
| ENSDARG000 | 7:7:7:7:7:7:7: | 69088015:690 | 69088066:690 | -:-:-:-:-:-:-:-: | 1539 | CRISPLD2 | 26 | 8 | 8 | 2 | 14 | 6 |
| ENSDARG000 | 7 | 12172129 | 12172246 | + | 118 | 5S_rRNA | 0 | 0 | 0 | 0 | 0 | 0 |
| ENSDARG000 | 7 | 46226244 | 46226360 | - | 117 | 5S_rRNA | 0 | 0 | 0 | 0 | 0 | 0 |
| ENSDARG000 | 7 | 32681640 | 32681755 | + | 116 | 5S_rRNA | 0 | 0 | 3 | 0 | 0 | 0 |
| ENSDARG000 | 7 | 38678418 | 38678532 | + | 115 | 5S_rRNA | 0 | 0 | 0 | 0 | 0 | 0 |
| ENSDARG000 | 7 | 61632769 | 61632883 | + | 115 | 5S_rRNA | 0 | 0 | 0 | 0 | 0 | 0 |
| ENSDARG000 | 7;7;7;7;7;7 | 17588689:175 | 17588817:175 | -;-;-;-;-;- | 894 | NA | 0 | 0 | 0 | 0 | 0 | 0 |
| ENSDARG000 | 7;7;7;7;7;7 | 64961892:649 | 64963630:649 | -;-;-;-;-;- | 2813 | sinhcafl | 64 | 208 | 286 | 277 | 77 | 174 |
| ENSDARG000 | 7 | 52940360 | 52940482 | - | 123 | 5S_rRNA | 6 | 2 | 2 | 0 | 2 | 2 |
| ENSDARG000 | 7:7:7:7:7:7:7: | 55005229:550 | 55005307:550 | -:-:-:-:-:-:-:-: | 2325 | tpcn2 | 121 | 38 | 138 | 47 | 139 | 144 |
| ENSDARG000 | 7 | 10683487 | 10683607 | + | 121 | 5S_rRNA | 0 | 0 | 0 | 0 | 0 | 0 |
| ENSDARG000 | 7 | 48498587 | 48498702 | - | 116 | 5S_rRNA | 0 | 0 | 0 | 0 | 0 | 0 |
| ENSDARG000 | 7 | 68678939 | 68679055 | + | 117 | 5S_rRNA | 0 | 0 | 0 | 0 | 0 | 0 |
| ENSDARG000 | 7:7:7:7:7:7:7: | 69308270:693 | 69308632:693 | -:-:-:-:-:-:-:-: | 2798 | ap1g1 | 1025 | 1006 | 1403 | 830 | 1345 | 930 |
| ENSDARG000 | 7:7:7:7:7:7:7: | 58139372:581 | 58144370:581 | -:-:-:-:-:-:-:-: | 7825 | NSMAF | 482 | 331 | 511 | 316 | 512 | 590 |
| ENSDARG000 | 7;7;7 | 65939098:659 | 65939958:659 | +;+;+ | 1278 | RASSF10 | 38 | 24 | 8 | 24 | 10 | 28 |
| ENSDARG000 | 7:7:7:7:7:7:7: | 48041790:480 | 48042568:480 | -:-:-:-:-:-:-:-: | 2937 | NA | 10 | 16 | 153 | 55 | 57 | 89 |
| ENSDARG000 | 7;7;7;7;7;7 | 1045637:1047 | 1045756:1047 | +;+;+;+;+;+ | 1983 | epdl1 | 160 | 83 | 96 | 80 | 204 | 291 |
| ENSDARG000 | 7:7:7:7:7:7:7: | 39255781:392 | 39256513:392 | -:-:-:-:-:-:-:-: | 5656 | ambra1a | 354 | 227 | 297 | 193 | 375 | 276 |
| ENSDARG000 | 7:7:7:7:7:7:7: | 40715756:407 | 40716151:407 | -;-;-;-;-;-;-;- | 1267 | rnf32 | 0 | 0 | 0 | 0 | 0 | 0 |
| ENSDARG000 | 7:7:7:7:7:7:7: | 6879534:6879 | 6879676:6879 | +:+:+:+:+:+:+: | 2319 | prkd4 | 0 | 30 | 0 | 0 | 18 | 22 |
| ENSDARG000 | 7;7;7;7;7 | 20859437:208 | 20860018:208 | -;-;-;-;- | 985 | fis1 | 421 | 586 | 666 | 579 | 428 | 462 |
| ENSDARG000 | 7;7;7;7;7;7 | 71385513:713 | 71385704:713 | -;-;-;-;-;- | 1316 | NA | 7 | 20 | 0 | 0 | 13 | 25 |
| ENSDARG000 | 7:7:7:7:7:7:7: | 2797795:2798 | 2797948:2798 | +:+:+:+:+:+:+: | 1752 | NA | 0 | 0 | 0 | 0 | 0 | 0 |
| ENSDARG000 | 7;7;7;7 | 13896769:139 | 13897902:139 | -;-;-;- | 1885 | slc39a1 | 2447 | 1874 | 1380 | 1561 | 1772 | 1904 |
| ENSDARG000 | 7 | 13340143 | 13340268 | + | 126 | 5S_rRNA | 0 | 0 | 0 | 0 | 0 | 0 |
| ENSDARG000 | 7:7:7:7:7:7:7: | 64936903:649 | 64938002:649 | +:+:+:+:+:+:+: | 2928 | NA | 36 | 37 | 28 | 30 | 53 | 36 |
| ENSDARG000 | 7:7:7:7:7:7:7: | 17466011:174 | 17466020:174 | -:-:-:-:-:-:-:-: | 282 | NA | 0 | 0 | 0 | 0 | 0 | 2 |
| ENSDARG000 | 7 | 45740492 | 45740608 | - | 117 | 5S_rRNA | 0 | 0 | 0 | 0 | 0 | 0 |
| ENSDARG000 | 7 | 27043715 | 27043842 | + | 128 | NA | 0 | 1 | 1 | 1 | 0 | 0 |
| ENSDARG000 | 7 | 16732674 | 16732790 | + | 117 | 5S_rRNA | 0 | 0 | 0 | 0 | 0 | 0 |
| ENSDARG000 | 7;7;7;7;7;7 | 7658:7658:93 | 8093:8041:94 | -;-;-;-;-;- | 635 | NA | 390 | 381 | 6655 | 520 | 130 | 2103 |
| ENSDARG000 | 7;7;7;7;7;7 | 71517094:715 | 71517280:715 | -;-;-;-;-;- | 1101 | ACER2 | 46 | 36 | 50 | 67 | 60 | 34 |
| ENSDARG000 | 7 | 50686781 | 50686897 | - | 117 | 5S_rRNA | 0 | 0 | 0 | 0 | 0 | 0 |
| ENSDARG000 | 7 | 33412304 | 33412420 | - | 117 | 5S_rRNA | 0 | 0 | 0 | 0 | 0 | 0 |
| ENSDARG000 | 7 | 14379544 | 14379631 | - | 88 | dre-mir-9-6 | 0 | 0 | 0 | 0 | 0 | 0 |
| ENSDARG000 | 7 | 42290978 | 42291094 | - | 117 | 5S_rRNA | 0 | 0 | 0 | 0 | 0 | 0 |
| ENSDARG000 | 7:7:7:7:7:7:7: | 17559929:175 | 17559987:175 | +:+:+:+:+:+:+: | 2130 | nitr1o | 0 | 0 | 0 | 0 | 0 | 0 |
| ENSDARG000 | 7 | 11213408 | 11213524 | + | 117 | 5S_rRNA | 0 | 0 | 0 | 0 | 0 | 0 |
| ENSDARG000 | 7 | 604530 | 604645 | + | 116 | 5S_rRNA | 0 | 0 | 0 | 0 | 0 | 0 |
| ENSDARG000 | 7 | 51108524 | 51108638 | + | 115 | 5S_rRNA | 0 | 0 | 0 | 0 | 0 | 0 |
| ENSDARG000 | 7;7;7;7;7 | 1139805:1141 | 1140509:1141 | -;-;-;-;- | 1602 | NA | 1 | 0 | 0 | 0 | 0 | 0 |
| ENSDARG000 | 7:7:7:7:7:7:7: | 69653981:696 | 69654051:696 | +:+:+:+:+:+:+: | 7217 | frem1a | 26 | 0 | 322 | 20 | 104 | 4 |
| ENSDARG000 | 7 | 66198624 | 66198740 | - | 117 | 5S_rRNA | 0 | 0 | 0 | 0 | 0 | 0 |
| ENSDARG000 | 7 | 48754170 | 48754286 | - | 117 | 5S_rRNA | 0 | 0 | 0 | 0 | 0 | 0 |
| ENSDARG000 | 7 | 30449415 | 30449530 | - | 116 | 5S_rRNA | 0 | 0 | 0 | 0 | 0 | 4 |
| ENSDARG000 | 7:7:7:7:7:7:7: | 53541173:535 | 53541357:535 | +:+:+:+:+:+:+: | 1756 | NA | 1095 | 620 | 843 | 1148 | 278 | 1111 |
| ENSDARG000 | 7 | 12719781 | 12719911 | - | 131 | 5S_rRNA | 0 | 0 | 0 | 0 | 0 | 0 |
| ENSDARG000 | 7;7 | 73819078:738 | 73819894:738 | +;+ | 1241 | NA | 312 | 62 | 228 | 64 | 154 | 90 |
| ENSDARG000 | 7 | 70949727 | 70949843 | + | 117 | 5S_rRNA | 0 | 0 | 0 | 0 | 0 | 0 |
| ENSDARG000 | 7;7 | 7653;7847 | 7823;7987 | +;+ | 312 | NA | 1 | 2 | 31 | 2 | 1 | 26 |
| ENSDARG000 | 7 | 16203995 | 16204113 | + | 119 | 5S_rRNA | 0 | 0 | 0 | 0 | 0 | 0 |
| ENSDARG000 | 7 | 68180335 | 68180449 | - | 115 | 5S_rRNA | 0 | 0 | 0 | 0 | 0 | 0 |
| ENSDARG000 | 7;7;7;7;7;7 | 73838557:738 | 73839073:738 | -;-;-;-;-;- | 1065 | caap1 | 343 | 314 | 405 | 288 | 423 | 296 |
| ENSDARG000 | 7 | 73845326 | 73845736 | - | 411 | zgc:173552 | 2 | 8 | 10 | 0 | 4 | 10 |
| ENSDARG000 | 7;7 | 7900626:7904 | 7900730:7904 | -;- | 546 | NA | 0 | 0 | 0 | 0 | 0 | 1 |
| ENSDARG000 | 7;7;7;7;7;7;7 | 61230013:612 | 61231938:612 | -;-;-;-;-;-;- | 2670 | rell1 | 739 | 421 | 486 | 425 | 683 | 1024 |
| ENSDARG000 | 7:7:7:7:7:7:7: | 58751504:587 | 58751723:587 | +:+:+:+:+:+:+: | 4678 | zgc:56231 | 2 | 4 | 0 | 0 | 2 | 18 |
| ENSDARG000 | 7;7;7;7;7;7 | 17597884:175 | 17598030:175 | -;-;-;-;-;- | 743 | NA | 0 | 0 | 0 | 12 | 0 | 0 |
| ENSDARG000 | 7 | 42935126 | 42939263 | + | 4138 | NA | 2 | 2 | 0 | 0 | 0 | 4 |
| ENSDARG000 | 7:7:7:7:7:7:7: | 15871156:158 | 15871529:158 | +:+:+:+:+:+:+: | 7064 | pax6b | 6 | 4 | 4 | 0 | 2 | 0 |
| ENSDARG000 | 7:7:7:7:7:7:7: | 20673287:206 | 20673390:206 | -:-:-:-:-:-:-:-: | 6024 | CHD3 | 2081 | 1090 | 1852 | 1291 | 1898 | 1609 |
| ENSDARG000 | 7;7;7 | 62004048:620 | 62004213:620 | +;+;+ | 560 | smim20 | 142 | 150 | 147 | 105 | 130 | 190 |
| ENSDARG000 | 7;7;7;7;7;7;7 | 72577898:725 | 72578034:725 | +;+;+;+;+;+;+ | 789 | mrpl21 | 161 | 255 | 269 | 170 | 240 | 190 |
| ENSDARG000 | 7;7;7 | 58856044:588 | 58856229:588 | -;-;- | 696 | NA | 4 | 0 | 0 | 2 | 8 | 0 |
| ENSDARG000 | 7 | 61099850 | 61099966 | + | 117 | 5S_rRNA | 0 | 0 | 0 | 0 | 0 | 0 |
| ENSDARG000 | 7 | 21823363 | 21823527 | - | 165 | U1 | 0 | 0 | 0 | 0 | 0 | 0 |
| ENSDARG000 | 7 | 1203121 | 1203239 | - | 119 | 5S_rRNA | 0 | 0 | 0 | 0 | 0 | 0 |
| ENSDARG000 | 7:7:7:7:7:7:7: | 59068601:590 | 59070286:590 | -:-:-:-:-:-:-:-: | 7390 | DENND4C | 2213 | 1411 | 1590 | 1943 | 1572 | 1796 |
| ENSDARG000 | 7:7:7:7:7:7:7: | 56615554:566 | 56615764:566 | +:+:+:+:+:+:+: | 1749 | dpep1 | 0 | 0 | 0 | 0 | 0 | 0 |
| ENSDARG000 | 7 | 54485565 | 54485685 | - | 121 | 5S_rRNA | 0 | 0 | 0 | 0 | 0 | 0 |
| ENSDARG000 | 7 | 67473392 | 67473474 | + | 83 | NA | 10 | 8 | 6 | 0 | 18 | 3 |
| ENSDARG000 | 7 | 67471357 | 67471438 | + | 82 | SNORD68 | 1 | 7 | 5 | 8 | 13 | 4 |
| ENSDARG000 | 7 | 57171618 | 57171739 | - | 122 | 5S_rRNA | 0 | 0 | 1 | 0 | 0 | 0 |
| ENSDARG000 | 7:7:7:7:7:7:7: | 15618127:156 | 15618230:156 | +:+:+:+:+:+:+: | 1433 | LRRC28 | 84 | 91 | 88 | 135 | 58 | 86 |
| ENSDARG000 | 7:7:7:7:7:7:7: | 13918349:139 | 13918638:139 | +:+:+:+:+:+:+: | 6302 | NA | 88 | 677 | 633 | 425 | 1721 | 276 |
| ENSDARG000 | 7 | 16114300 | 16114428 | - | 129 | 5S_rRNA | 0 | 0 | 0 | 0 | 0 | 0 |
| ENSDARG000 | 7;7;7 | 11666686:116 | 11666824:116 | -;-;- | 636 | NA | 0 | 0 | 0 | 0 | 0 | 0 |
| ENSDARG000 | 7:7:7:7:7:7:7: | 1330501:1331 | 1330607:1332 | -;-;-;-;-;-;-;- | 1881 | NA | 22 | 35 | 11 | 7 | 48 | 25 |
| ENSDARG000 | 7:7:7:7:7:7:7: | 72402279:724 | 72402409:724 | -:-:-:-:-:-:-:-: | 2319 | NA | 6 | 0 | 0 | 4 | 0 | 6 |
| ENSDARG000 | 7:7:7:7:7:7:7: | 69905307:699 | 69905887:699 | +:+:+:+:+:+:+: | 4532 | sec24d | 453 | 569 | 720 | 304 | 925 | 322 |
| ENSDARG000 | 7:7:7:7:7:7:7: | 637597:64348 | 638273:64366 | -:-:-:-:-:-:-:-: | 2661 | MAP3K11 | 314 | 275 | 335 | 234 | 305 | 254 |
| ENSDARG000 | 7 | 62556472 | 62556592 | - | 121 | 5S_rRNA | 0 | 0 | 0 | 0 | 0 | 0 |

| | | | | | | | | | | | | |
|---|---|---|---|---|---|---|---|---|---|---|---|---|
| ENSDARG000 | 7 | 29931525 | 29931639 | + | 115 | 5S_rRNA | 0 | 0 | 0 | 0 | 0 | 0 |
| ENSDARG000 | 7 | 55183300 | 55183416 | - | 117 | 5S_rRNA | 0 | 0 | 0 | 0 | 0 | 0 |
| ENSDARG000 | 7 | 71908214 | 71908328 | + | 115 | 5S_rRNA | 0 | 0 | 0 | 0 | 0 | 0 |
| ENSDARG000 | 7 | 7354319 | 7354445 | - | 127 | 5S_rRNA | 0 | 0 | 0 | 0 | 0 | 0 |
| ENSDARG000 | 7:7:7:7:7:7:7: | 59572379:595 | 59576082:595 | -:-:-:-:-:-:-:-:-: | 5440 | larp7 | 495 | 476 | 662 | 325 | 1088 | 733 |
| ENSDARG000 | 7 | 73846609 | 73846995 | - | 387 | NA | 0 | 14 | 1 | 0 | 8 | 1 |
| ENSDARG000 | 7;7;7;7;7 | 6420435:6422 | 6420494:6422 | +;+;+;+;+ | 132 | NA | 0 | 0 | 0 | 0 | 0 | 0 |
| ENSDARG000 | 7;7;7;7;7;7 | 17456686:174 | 17456767:174 | +;+;+;+;+;+ | 1032 | nitr1a | 0 | 0 | 0 | 0 | 0 | 1 |
| ENSDARG000 | 7 | 25045182 | 25045246 | + | 65 | SNORD29 | 0 | 0 | 0 | 0 | 0 | 0 |
| ENSDARG000 | 7:7:7:7:7:7:7: | 61764040:617 | 61764227:617 | +:+:+:+:+:+:+: | 3731 | acox3 | 249 | 239 | 318 | 234 | 412 | 325 |
| ENSDARG000 | 7 | 52526304 | 52526430 | - | 127 | 5S_rRNA | 0 | 0 | 0 | 0 | 0 | 4 |
| ENSDARG000 | 7 | 42313774 | 42313888 | + | 115 | 5S_rRNA | 0 | 0 | 0 | 0 | 0 | 0 |
| ENSDARG000 | 7;7;7;7;7;7 | 72594954:725 | 72595140:725 | -;-;-;-;-;- | 1398 | MAPK8IP1 | 0 | 2 | 0 | 0 | 0 | 0 |
| ENSDARG000 | 7 | 73855687 | 73856073 | - | 387 | NA | 4 | 24 | 2 | 0 | 8 | 14 |
| ENSDARG000 | 7 | 26070874 | 26071007 | + | 134 | dre-mir-15b | 0 | 0 | 0 | 0 | 0 | 0 |
| ENSDARG000 | 7;7;7;7 | 69841017:698 | 69841098:698 | +;+;+;+ | 369 | NA | 0 | 0 | 0 | 0 | 0 | 0 |
| ENSDARG000 | 7 | 31485485 | 31485601 | - | 117 | 5S_rRNA | 0 | 0 | 0 | 0 | 0 | 0 |
| ENSDARG000 | 7;7;7;7;7;7 | 17443567:174 | 17443609:174 | +;+;+;+;+;+ | 942 | nitr2b | 0 | 0 | 0 | 0 | 0 | 0 |
| ENSDARG000 | 7 | 39234995 | 39235112 | - | 118 | 5S_rRNA | 0 | 0 | 0 | 0 | 0 | 0 |
| ENSDARG000 | 7;7 | 54313489:543 | 54313760:543 | -;- | 1471 | FADD | 247 | 161 | 145 | 112 | 221 | 134 |
| ENSDARG000 | 7;7 | 2969661:2970 | 2969852:2970 | +;+ | 348 | NA | 0 | 0 | 0 | 0 | 2 | 0 |
| ENSDARG000 | 7;7;7;7;7;7 | 17520427:175 | 17520493:175 | +;+;+;+;+;+ | 987 | nitr1i | 0 | 0 | 0 | 0 | 0 | 0 |
| ENSDARG000 | 7;7;7;7 | 74006756:740 | 74010351:740 | -;-;-;- | 4697 | NA | 0 | 0 | 4 | 0 | 12 | 4 |
| ENSDARG000 | 7 | 55101440 | 55101556 | - | 117 | 5S_rRNA | 0 | 0 | 0 | 0 | 0 | 0 |
| ENSDARG000 | 7 | 4208061 | 4208177 | + | 117 | 5S_rRNA | 0 | 0 | 0 | 0 | 0 | 0 |
| ENSDARG000 | 7 | 69893435 | 69893563 | + | 129 | 5S_rRNA | 0 | 0 | 0 | 0 | 0 | 0 |
| ENSDARG000 | 7 | 44906299 | 44906413 | - | 115 | 5S_rRNA | 0 | 0 | 0 | 0 | 0 | 0 |
| ENSDARG000 | 7 | 72267099 | 72269049 | - | 1951 | NA | 0 | 5 | 0 | 0 | 4 | 8 |
| ENSDARG000 | 7 | 7772165 | 7772378 | + | 214 | NA | 0 | 0 | 0 | 0 | 0 | 0 |
| ENSDARG000 | 7 | 4077734 | 4077871 | - | 138 | 5S_rRNA | 0 | 0 | 0 | 0 | 0 | 0 |
| ENSDARG000 | 7 | 66937450 | 66937564 | + | 115 | 5S_rRNA | 0 | 0 | 0 | 0 | 0 | 0 |
| ENSDARG000 | 7 | 21836676 | 21836866 | - | 191 | U2 | 0 | 0 | 0 | 0 | 0 | 0 |
| ENSDARG000 | 7 | 29009161 | 29010135 | + | 975 | NA | 0 | 0 | 0 | 0 | 0 | 0 |
| ENSDARG000 | 7:7:7:7:7:7:7: | 25825667:258 | 25825820:258 | +:+:+:+:+:+:+: | 2136 | mtm1 | 244 | 177 | 230 | 109 | 216 | 214 |
| ENSDARG000 | 7 | 50258060 | 50258175 | - | 116 | 5S_rRNA | 0 | 0 | 0 | 0 | 0 | 0 |
| ENSDARG000 | 7 | 49227366 | 49227486 | - | 121 | 5S_rRNA | 0 | 0 | 0 | 0 | 0 | 0 |
| ENSDARG000 | 7 | 41582544 | 41582659 | + | 116 | 5S_rRNA | 0 | 0 | 0 | 0 | 0 | 0 |
| ENSDARG000 | 7:7:7:7:7:7:7: | 64978738:649 | 64978921:649 | -:-:-:-:-:-:-:-:-: | 4094 | DENND5A | 2386 | 1908 | 2774 | 2003 | 3631 | 1818 |
| ENSDARG000 | 7;7 | 58823406:588 | 58825405:588 | -;- | 2313 | sox32 | 0 | 0 | 0 | 0 | 2 | 6 |
| ENSDARG000 | 7 | 24076711 | 24076821 | + | 111 | NA | 0 | 0 | 1 | 2 | 0 | 0 |
| ENSDARG000 | 7 | 36095269 | 36096582 | - | 1314 | NA | 0 | 0 | 0 | 0 | 0 | 0 |
| ENSDARG000 | 7;7;7 | 52811291:528 | 52811300:528 | +;+;+ | 234 | NA | 0 | 0 | 0 | 0 | 0 | 0 |
| ENSDARG000 | 7:7:7:7:7:7:7: | 70578985:705 | 70579088:705 | -:-:-:-:-:-:-:-:-: | 2580 | PPARGC1A | 2 | 0 | 14 | 0 | 18 | 4 |
| ENSDARG000 | 7;7;7;7;7;7;7 | 61876641:618 | 61877441:618 | +;+;+;+;+;+;+ | 3179 | NA | 0 | 0 | 0 | 0 | 0 | 0 |
| ENSDARG000 | 7;7;7;7;7;7;7 | 59033902:590 | 59034171:590 | +;+;+;+;+;+;+ | 2241 | FASTKD3 | 221 | 218 | 165 | 170 | 187 | 220 |
| ENSDARG000 | 7;7;7 | 1306723:1310 | 1307848:1310 | +;+;+ | 1444 | NA | 0 | 0 | 0 | 0 | 0 | 0 |
| ENSDARG000 | 7:7:7:7:7:7:7: | 59004234:590 | 59004247:590 | -:-:-:-:-:-:-:-:-: | 228 | NA | 20 | 7 | 18 | 8 | 22 | 13 |
| ENSDARG000 | 7:7:7:7:7:7:7: | 14632157:146 | 14632887:146 | +:+:+:+:+:+:+: | 4313 | NTRK3 | 0 | 1 | 2 | 4 | 0 | 0 |
| ENSDARG000 | 7:7:7:7:7:7:7: | 1383027:1388 | 1383512:1388 | +:+:+:+:+:+:+: | 7088 | NA | 825 | 704 | 1408 | 1429 | 1070 | 925 |
| ENSDARG000 | 7:7:7:7:7:7:7: | 20779026:207 | 20782111:207 | -:-:-:-:-:-:-:-:-: | 4449 | dnajc3b | 1415 | 1490 | 1546 | 786 | 1416 | 1109 |
| ENSDARG000 | 7 | 29155395 | 29155511 | - | 117 | 5S_rRNA | 0 | 0 | 0 | 0 | 0 | 0 |
| ENSDARG000 | 7:7:7:7:7:7:7: | 18176162:181 | 18176597:181 | +:+:+:+:+:+:+: | 1232 | RCE1 | 114 | 127 | 170 | 76 | 158 | 118 |
| ENSDARG000 | 7:7:7:7:7:7:7: | 57146542:571 | 57146690:571 | -:-:-:-:-:-:-:-:-: | 4983 | ADAMTS7 | 696 | 985 | 2493 | 1096 | 2860 | 1082 |
| ENSDARG000 | 7 | 7772165 | 7772377 | + | 213 | U3 | 0 | 0 | 0 | 0 | 0 | 0 |
| ENSDARG000 | 7 | 25791229 | 25791355 | - | 127 | 5S_rRNA | 0 | 0 | 0 | 0 | 0 | 0 |
| ENSDARG000 | 7:7:7:7:7:7:7: | 13639473:136 | 13639885:136 | +:+:+:+:+:+:+: | 4888 | clpxa | 1082 | 791 | 818 | 911 | 970 | 905 |
| ENSDARG000 | 7 | 7778963 | 7779176 | + | 214 | NA | 0 | 0 | 0 | 0 | 0 | 0 |
| ENSDARG000 | 7;7 | 69020928:690 | 69022011:690 | -;- | 1373 | NA | 0 | 0 | 0 | 0 | 0 | 0 |
| ENSDARG000 | 7;7;7;7;7 | 2849020:2861 | 2849181:2861 | +;+;+;+;+ | 2577 | NA | 0 | 0 | 0 | 0 | 0 | 0 |
| ENSDARG000 | 7:7:7:7:7:7:7: | 74150839:741 | 74150952:741 | +:+:+:+:+:+:+: | 693 | ppp1cbl | 2491 | 2008 | 2927 | 4292 | 2625 | 2693 |
| ENSDARG000 | 7;7;7;7;7;7 | 72272918:722 | 72273912:722 | -;-;-;-;-;- | 1628 | slc35c1 | 92 | 63 | 69 | 68 | 61 | 98 |
| ENSDARG000 | 7:7:7:7:7:7:7: | 72040939:720 | 72041049:720 | -:-:-:-:-:-:-:-:-: | 1563 | NA | 0 | 4 | 0 | 0 | 0 | 4 |
| ENSDARG000 | 7:7:7:7:7:7:7: | 55950229:559 | 55950333:559 | +:+:+:+:+:+:+: | 2337 | ACSF3 | 78 | 84 | 52 | 34 | 102 | 56 |
| ENSDARG000 | 7:7:7:7:7:7:7: | 58892856:588 | 58893094:588 | -:-:-:-:-:-:-:-:-: | 4323 | MRC1 | 111 | 106 | 650 | 152 | 1370 | 688 |
| ENSDARG000 | 7:7:7:7:7:7:7: | 71350469:713 | 71351179:713 | -:-:-:-:-:-:-:-:-: | 2030 | ccdc149a | 10 | 2 | 14 | 96 | 18 | 0 |
| ENSDARG000 | 7 | 49185938 | 49186052 | - | 115 | 5S_rRNA | 0 | 0 | 0 | 0 | 0 | 0 |
| ENSDARG000 | 7 | 72215450 | 72215568 | - | 119 | 5S_rRNA | 7 | 1 | 1 | 0 | 0 | 2 |
| ENSDARG000 | 7 | 34080184 | 34080298 | + | 115 | 5S_rRNA | 0 | 0 | 0 | 0 | 0 | 0 |
| ENSDARG000 | 7:7:7:7:7:7:7: | 55660263:556 | 55660601:556 | -:-:-:-:-:-:-:-:-: | 1910 | CBFA2T3 | 7 | 2 | 4 | 0 | 34 | 14 |
| ENSDARG000 | 7;7;7;7;7;7;7 | 11614144:116 | 11615002:116 | -;-;-;-;-;-;- | 1611 | NA | 7 | 4 | 2 | 0 | 2 | 0 |
| ENSDARG000 | 7:7:7:7:7:7:7: | 69959141:699 | 69959188:699 | -:-:-:-:-:-:-:-:-: | 1327 | KCNIP4 | 4 | 0 | 18 | 0 | 4 | 0 |
| ENSDARG000 | 7;7;7 | 71503356:715 | 71504286:715 | +;+;+ | 1904 | ppp1r15a | 9177 | 2549 | 9037 | 7469 | 2261 | 2607 |
| ENSDARG000 | 7;7;7;7;7 | 51788669:517 | 51788782:517 | -;-;-;-;- | 297 | NA | 0 | 0 | 0 | 0 | 0 | 0 |
| ENSDARG000 | 7;7;7 | 1872184:1875 | 1872445:1875 | -;-;- | 668 | NA | 0 | 0 | 0 | 0 | 0 | 0 |
| ENSDARG000 | 7 | 9343638 | 9343754 | - | 117 | 5S_rRNA | 0 | 0 | 0 | 0 | 0 | 0 |
| ENSDARG000 | 7 | 4970757 | 4970885 | - | 129 | 5S_rRNA | 0 | 0 | 0 | 0 | 0 | 0 |
| ENSDARG000 | 7 | 11527252 | 11527371 | - | 120 | 5S_rRNA | 0 | 0 | 0 | 0 | 0 | 0 |
| ENSDARG000 | 7 | 50030675 | 50030791 | - | 117 | 5S_rRNA | 0 | 0 | 0 | 0 | 0 | 0 |
| ENSDARG000 | 7 | 37693838 | 37693952 | - | 115 | 5S_rRNA | 0 | 0 | 0 | 0 | 0 | 0 |
| ENSDARG000 | 7 | 48306510 | 48306645 | - | 136 | 5S_rRNA | 0 | 0 | 0 | 0 | 0 | 0 |
| ENSDARG000 | 7 | 58748945 | 58749013 | + | 69 | SNORD54 | 0 | 0 | 0 | 4 | 0 | 0 |
| ENSDARG000 | 7 | 25763184 | 25763298 | - | 115 | 5S_rRNA | 0 | 0 | 0 | 0 | 0 | 0 |
| ENSDARG000 | 7;7;7;7;7;7 | 17494169:174 | 17494250:174 | +;+;+;+;+;+ | 1031 | nitr1f | 0 | 0 | 0 | 0 | 0 | 0 |
| ENSDARG000 | 7;7 | 73701410:737 | 73701951:737 | -;- | 570 | NA | 0 | 0 | 0 | 0 | 0 | 0 |
| ENSDARG000 | 7;7 | 73822054:738 | 73822363:738 | +;+ | 348 | NA | 4 | 25 | 8 | 4 | 2 | 22 |
| ENSDARG000 | 7 | 30189592 | 30189670 | - | 79 | dre-mir-184-2 | 0 | 0 | 0 | 0 | 0 | 0 |
| ENSDARG000 | 7;7;7 | 1157610:1161 | 1157768:1162 | -;-;- | 568 | NA | 0 | 0 | 0 | 0 | 0 | 0 |
| ENSDARG000 | 7:7:7:7:7:7:7: | 26680868:266 | 26680877:266 | -;-;-;-;-;-;-;- | 201 | NA | 0 | 0 | 0 | 0 | 5 | 0 |
| ENSDARG000 | 7 | 5189778 | 5189896 | + | 119 | 5S_rRNA | 0 | 0 | 1 | 0 | 0 | 0 |
| ENSDARG000 | 7 | 24263623 | 24263739 | + | 117 | 5S_rRNA | 0 | 0 | 0 | 0 | 0 | 0 |
| ENSDARG000 | 7 | 22937129 | 22937248 | - | 120 | 5S_rRNA | 0 | 0 | 2 | 0 | 0 | 0 |
| ENSDARG000 | 7 | 25046139 | 25046208 | + | 70 | SNORD31 | 0 | 0 | 7 | 2 | 1 | 1 |
| ENSDARG000 | 7 | 62836290 | 62836406 | + | 117 | 5S_rRNA | 0 | 0 | 0 | 0 | 0 | 0 |
| ENSDARG000 | 7 | 64150332 | 64152087 | - | 1756 | NA | 0 | 0 | 0 | 0 | 0 | 0 |
| ENSDARG000 | 7 | 16841999 | 16842116 | - | 118 | 5S_rRNA | 0 | 0 | 0 | 0 | 0 | 0 |
| ENSDARG000 | 7;7;7;7;7;7;7 | 70267090:702 | 70268473:702 | -;-;-;-;-;-;- | 2184 | ADGRA3 | 1129 | 712 | 1005 | 931 | 1174 | 770 |
| ENSDARG000 | 7 | 43332628 | 43332742 | - | 115 | 5S_rRNA | 0 | 0 | 0 | 0 | 0 | 0 |
| ENSDARG000 | 7;7;7;7;7;7 | 58270474:582 | 58273389:582 | -;-;-;-;-;- | 4305 | NA | 10 | 6 | 23 | 0 | 46 | 32 |
| ENSDARG000 | 7:7:7:7:7:7:7: | 4085115:4088 | 4089639:4089 | -:-:-:-:-:-:-:-:-: | 5493 | zgc:55733 | 1153 | 820 | 994 | 891 | 1255 | 880 |
| ENSDARG000 | 7;7 | 58772209:587 | 58774376:587 | -;- | 2848 | sox17 | 8 | 0 | 0 | 0 | 8 | 0 |
| ENSDARG000 | 7:7:7:7:7:7:7: | 72132047:721 | 72132431:721 | -:-:-:-:-:-:-:-:-: | 4180 | SIPA1 | 10 | 12 | 7 | 2 | 43 | 55 |
| ENSDARG000 | 7 | 60095683 | 60096318 | - | 636 | NA | 0 | 0 | 0 | 0 | 0 | 0 |
| ENSDARG000 | 7;7;7;7 | 69641197:696 | 69641326:696 | -;-;-;- | 1098 | NA | 2 | 4 | 16 | 0 | 4 | 0 |
| ENSDARG000 | 7:7:7:7:7:7:7: | 18100996:181 | 18101281:181 | +:+:+:+:+:+:+: | 2970 | rab1ba | 2553 | 1928 | 2092 | 1549 | 2222 | 1794 |
| ENSDARG000 | 7:7:7:7:7:7:7: | 60079302:600 | 60079360:600 | +:+:+:+:+:+:+: | 3587 | etnppl | 6 | 2 | 4 | 8 | 10 | 0 |
| ENSDARG000 | 7 | 4427247 | 4427365 | - | 119 | 5S_rRNA | 0 | 0 | 0 | 0 | 0 | 0 |
| ENSDARG000 | 7:7:7:7:7:7:7: | 66634167:666 | 66634398:666 | +:+:+:+:+:+:+: | 4032 | eif4g2a | 4202 | 5273 | 5576 | 2051 | 5448 | 2880 |
| ENSDARG000 | 7 | 23285035 | 23285151 | + | 117 | 5S_rRNA | 0 | 0 | 0 | 0 | 0 | 0 |
| ENSDARG000 | 7 | 44369563 | 44369679 | + | 117 | 5S_rRNA | 0 | 0 | 0 | 0 | 0 | 0 |
| ENSDARG000 | 7:7:7:7:7:7:7: | 72003301:720 | 72003616:720 | +:+:+:+:+:+:+: | 950 | psmd9 | 351 | 320 | 409 | 137 | 279 | 252 |
| ENSDARG000 | 7 | 73852283 | 73852594 | - | 312 | NA | 6 | 8 | 0 | 0 | 6 | 0 |
| ENSDARG000 | 7;7 | 73847702:738 | 73848046:738 | -;- | 692 | zgc:163061 | 40 | 36 | 47 | 28 | 20 | 16 |
| ENSDARG000 | 7:7:7:7:7:7:7: | 13988075:139 | 13988413:139 | +:+:+:+:+:+:+: | 7295 | furina | 1433 | 403 | 478 | 578 | 562 | 692 |
| ENSDARG000 | 7;7;7;7;7 | 64739078:647 | 64743165:647 | -;-;-;-;- | 4841 | zdhhc21 | 214 | 208 | 130 | 170 | 294 | 306 |
| ENSDARG000 | 7 | 68329989 | 68330107 | - | 119 | 5S_rRNA | 0 | 0 | 0 | 0 | 0 | 0 |
| ENSDARG000 | 7 | 17862425 | 17862540 | - | 116 | 5S_rRNA | 0 | 0 | 7 | 0 | 0 | 0 |
| ENSDARG000 | 7:7:7:7:7:7:7: | 55633483:556 | 55633541:556 | +:+:+:+:+:+:+: | 2142 | trappc2l | 240 | 272 | 211 | 234 | 226 | 204 |
| ENSDARG000 | 7;7;7;7;7;7 | 17433836:174 | 17433912:174 | +;+;+;+;+;+ | 1085 | nitr2a | 0 | 0 | 0 | 0 | 0 | 0 |
| ENSDARG000 | 7;7;7;7;7;7 | 19016023:190 | 19016063:190 | +;+;+;+;+;+ | 171 | NA | 0 | 0 | 0 | 0 | 0 | 0 |
| ENSDARG000 | 7;7;7;7;7;7 | 56453646:564 | 56453782:564 | +;+;+;+;+;+ | 2922 | SLC22A31 | 153 | 228 | 435 | 188 | 593 | 264 |
| ENSDARG000 | 7;7;7;7;7;7;7 | 3872189:3872 | 3872261:3872 | +;+;+;+;+;+;+ | 804 | NA | 0 | 81 | 17 | 32 | 2 | 37 |
| ENSDARG000 | 7 | 44668003 | 44668117 | - | 115 | 5S_rRNA | 0 | 0 | 0 | 0 | 0 | 0 |
| ENSDARG000 | 7 | 47860077 | 47860193 | + | 117 | 5S_rRNA | 0 | 0 | 0 | 0 | 0 | 0 |
| ENSDARG000 | 7;7;7;7;7;7 | 20771754:207 | 20772417:207 | -;-;-;-;-;- | 2001 | cyb5d1 | 16 | 0 | 6 | 4 | 8 | 0 |
| ENSDARG000 | 7 | 42778000 | 42778114 | + | 115 | 5S_rRNA | 0 | 0 | 0 | 0 | 0 | 0 |
| ENSDARG000 | 7 | 33853188 | 33853307 | + | 120 | 5S_rRNA | 0 | 0 | 0 | 0 | 0 | 0 |
| ENSDARG000 | 7;7;7;7;7;7 | 17543487:175 | 17543562:175 | +;+;+;+;+;+ | 987 | NA | 0 | 0 | 0 | 0 | 6 | 0 |
| ENSDARG000 | 7:7:7:7:7:7:7: | 41981366:419 | 41981425:419 | -:-:-:-:-:-:-:-:-: | 1800 | ITFG1 | 443 | 362 | 432 | 366 | 571 | 670 |
| ENSDARG000 | 7;7;7;7;7;7 | 17485782:174 | 17485821:174 | +;+;+;+;+;+ | 975 | NA | 0 | 0 | 0 | 0 | 0 | 0 |
| ENSDARG000 | 7:7:7:7:7:7:7: | 69829442:698 | 69830670:698 | -:-:-:-:-:-:-:-:-: | 1801 | myoz2a | 2 | 0 | 1 | 0 | 0 | 0 |
| ENSDARG000 | 7:7:7:7:7:7:7: | 62229075:622 | 62229196:622 | -:-:-:-:-:-:-:-:-: | 1475 | CCKAR | 0 | 4 | 3 | 6 | 16 | 0 |
| ENSDARG000 | 7:7:7:7:7:7:7: | 69601812:696 | 69602064:696 | -:-:-:-:-:-:-:-:-: | 1389 | tspan5a | 811 | 459 | 1155 | 345 | 1191 | 622 |
| ENSDARG000 | 7;7;7;7;7;7 | 73829477:738 | 73829676:738 | -;-;-;-;-;- | 1314 | zgc:92594 | 87 | 40 | 113 | 137 | 44 | 65 |
| ENSDARG000 | 7;7 | 10351038:103 | 10353341:103 | +;+ | 2608 | MINAR1 | 0 | 0 | 0 | 0 | 0 | 0 |
| ENSDARG000 | 7;7;7;7;7;7;7 | 25896619:259 | 25896641:259 | -;-;-;-;-;-;- | 153 | NA | 1 | 0 | 2 | 2 | 2 | 6 |
| ENSDARG000 | 7;7;7 | 68384854:684 | 68385042:684 | -;-;- | 240 | NA | 0 | 0 | 0 | 0 | 0 | 0 |
| ENSDARG000 | 7:7:7:7:7:7:7: | 778297:77943 | 778959:77953 | -:-:-:-:-:-:-:-:-: | 1775 | rela | 283 | 213 | 317 | 163 | 298 | 200 |
| ENSDARG000 | 7;7;7;7;7 | 71494246:714 | 71494390:714 | -;-;-;-;- | 564 | NA | 4107 | 1288 | 5742 | 10920 | 2426 | 2524 |
| ENSDARG000 | 7:7:7:7:7:7:7: | 69019851:690 | 69019965:690 | +:+:+:+:+:+:+: | 1392 | NA | 2 | 0 | 8 | 0 | 0 | 0 |
| ENSDARG000 | 7 | 28280171 | 28280285 | - | 115 | 5S_rRNA | 0 | 0 | 0 | 0 | 0 | 0 |
| ENSDARG000 | 7 | 26120631 | 26120736 | + | 106 | NA | 0 | 0 | 0 | 0 | 0 | 0 |
| ENSDARG000 | 7;7;7;7;7;7 | 56735195:567 | 56735948:567 | +;+;+;+;+;+ | 1597 | MATCAP1 | 183 | 201 | 133 | 106 | 224 | 165 |
| ENSDARG000 | 7 | 28578449 | 28578565 | - | 117 | 5S_rRNA | 0 | 0 | 0 | 0 | 0 | 0 |
| ENSDARG000 | 7:7:7:7:7:7:7: | 13756215:137 | 13756727:137 | +:+:+:+:+:+:+: | 4625 | rasl12 | 72 | 23 | 41 | 1 | 16 | 25 |
| ENSDARG000 | 7 | 15132977 | 15133091 | + | 115 | 5S_rRNA | 0 | 0 | 0 | 0 | 0 | 0 |
| ENSDARG000 | 7;7;7;7;7;7 | 74141297:741 | 74141362:741 | +;+;+;+;+;+ | 483 | NA | 33 | 23 | 34 | 22 | 42 | 10 |
| ENSDARG000 | 7:7:7:7:7:7:7: | 61817629:618 | 61817797:618 | -:-:-:-:-:-:-:-:-: | 1422 | HTRA3 | 123 | 125 | 12 | 43 | 6 | 41 |
| ENSDARG000 | 7:7:7:7:7:7:7: | 9189547:9192 | 9189685:9192 | +:+:+:+:+:+:+: | 1590 | PCSK6 | 53 | 6 | 6 | 0 | 12 | 6 |
| ENSDARG000 | 7 | 42153410 | 42153524 | - | 115 | 5S_rRNA | 0 | 0 | 0 | 0 | 0 | 0 |
| ENSDARG000 | 7 | 24074861 | 24074975 | + | 115 | NA | 0 | 0 | 0 | 0 | 0 | 0 |
| ENSDARG000 | 7 | 25043010 | 25043080 | + | 71 | SNORD31 | 0 | 0 | 0 | 0 | 0 | 0 |
| ENSDARG000 | 7 | 42720405 | 42720519 | + | 115 | 5S_rRNA | 0 | 0 | 0 | 0 | 0 | 0 |
| ENSDARG000 | 7:7:7:7:7:7:7: | 61906903:619 | 61907045:619 | +:+:+:+:+:+:+: | 4588 | tdrd7b | 3206 | 3310 | 2300 | 2112 | 1524 | 1336 |
| ENSDARG000 | 7;7;7 | 73780936:737 | 73781107:737 | +;+;+ | 393 | PEA15 | 981 | 432 | 818 | 382 | 958 | 660 |
| ENSDARG000 | 7 | 25046348 | 25046421 | + | 74 | SNORD31 | 0 | 0 | 7 | 0 | 4 | 0 |
| ENSDARG000 | 7;7;7;7;7;7 | 69356661:693 | 69357549:693 | +;+;+;+;+;+ | 2163 | ZNF821 | 21 | 11 | 22 | 2 | 30 | 10 |
| ENSDARG000 | 7:7:7:7:7:7:7: | 13867722:138 | 13868344:138 | -:-:-:-:-:-:-:-:-: | 1562 | rlbp1a | 0 | 0 | 0 | 0 | 0 | 2 |
| ENSDARG000 | 7:7:7:7:7:7:7: | 69428831:694 | 69429453:694 | +:+:+:+:+:+:+: | 3980 | elp5 | 172 | 168 | 341 | 245 | 220 | 136 |
| ENSDARG000 | 7;7;7 | 69415393:694 | 69416688:694 | -;-;- | 2667 | ATXN1L | 115 | 38 | 207 | 84 | 149 | 60 |
| ENSDARG000 | 7;7;7;7;7;7 | 17295053:172 | 17295154:172 | -;-;-;-;-;- | 585 | nitr11a | 0 | 0 | 0 | 0 | 0 | 0 |
| ENSDARG000 | 7;7;7 | 25903399:259 | 25903413:259 | -;-;- | 72 | NA | 0 | 2 | 1 | 0 | 2 | 0 |
| ENSDARG000 | 7:7:7:7:7:7:7: | 74084203:740 | 74084374:740 | -;-;-;-;-;-;-;- | 1928 | TYRP1 | 54463 | 53217 | 14605 | 47287 | 33593 | 36581 |

| | | | | | | | | | | | |
|---|---|---|---|---|---|---|---|---|---|---|---|
| ENSDARG000 | 7;7;7;7;7;7 | 61184104:611 | 61184794:611 | +;+;+;+;+;+ | 1517 | zgc:194930 | 20 | 6 | 4 | 0 | 8 | 8 |
| ENSDARG000 | 7:7:7:7:7:7:7: | 9981757:9981 | 9981822:9981 | +:+:+:+:+:+:+: | 3150 | ADAMTS17 | 0 | 6 | 6 | 0 | 152 | 0 |
| ENSDARG000 | 7:7:7:7:7:7:7: | 20917966:209 | 20918172:209 | +:+:+:+:+:+:+: | 2034 | wrap53 | 162 | 183 | 372 | 167 | 231 | 144 |
| ENSDARG000 | 7 | 73814951 | 73815262 | - | 312 | NA | 0 | 0 | 0 | 0 | 0 | 0 |
| ENSDARG000 | 7 | 2012801 | 2012916 | - | 116 | 5S_rRNA | 0 | 0 | 0 | 0 | 0 | 0 |
| ENSDARG000 | 7 | 28461463 | 28461578 | + | 116 | 5S_rRNA | 0 | 0 | 0 | 0 | 0 | 0 |
| ENSDARG000 | 7:7:7:7:7:7:7: | 71423612:714 | 71424429:714 | -;-;-;-;-;-;-;- | 2009 | lgi2a | 0 | 6 | 12 | 0 | 8 | 8 |
| ENSDARG000 | 7 | 47961177 | 47961293 | - | 117 | 5S_rRNA | 0 | 0 | 0 | 0 | 0 | 0 |
| ENSDARG000 | 7 | 34978351 | 34978469 | - | 119 | 5S_rRNA | 0 | 0 | 0 | 0 | 0 | 0 |
| ENSDARG000 | 7 | 62578520 | 62578635 | + | 116 | 5S_rRNA | 0 | 0 | 0 | 0 | 0 | 0 |
| ENSDARG000 | 7 | 54990518 | 54990633 | + | 116 | 5S_rRNA | 0 | 0 | 0 | 2 | 0 | 0 |
| ENSDARG000 | 7 | 67476155 | 67476231 | + | 77 | NA | 7 | 12 | 23 | 1 | 14 | 3 |
| ENSDARG000 | 7;7 | 56554001:565 | 56554875:565 | -;- | 1152 | NA | 0 | 0 | 0 | 0 | 0 | 0 |
| ENSDARG000 | 7;7;7;7 | 72381111:723 | 72381267:723 | -;-;-;- | 660 | NA | 0 | 0 | 0 | 0 | 0 | 0 |
| ENSDARG000 | 7:7:7:7:7:7:7: | 17574435:175 | 17579272:175 | -:-:-:-:-:-:-:-: | 5858 | nitr4a | 0 | 0 | 0 | 0 | 0 | 0 |
| ENSDARG000 | 7;7;7;7;7;7 | 17551908:175 | 17551967:175 | +;+;+;+;+;+ | 1007 | nitr1m | 0 | 0 | 0 | 0 | 0 | 0 |
| ENSDARG000 | 7 | 73850906 | 73851280 | - | 375 | NA | 13 | 2 | 8 | 4 | 0 | 7 |
| ENSDARG000 | 7 | 73854066 | 73854476 | - | 411 | zgc:173552 | 18 | 93 | 41 | 25 | 65 | 16 |
| ENSDARG000 | 7:7:7:7:7:7:7: | 66655729:666 | 66656403:666 | -:-:-:-:-:-:-:-: | 3873 | ctr9 | 1332 | 1123 | 1771 | 902 | 1595 | 940 |
| ENSDARG000 | 7 | 12853103 | 12853217 | + | 115 | 5S_rRNA | 0 | 0 | 0 | 0 | 0 | 0 |
| ENSDARG000 | 7 | 24072381 | 24072484 | + | 104 | NA | 0 | 0 | 0 | 0 | 0 | 0 |
| ENSDARG000 | 7:7:7:7:7:7:7: | 34414194:344 | 34419113:344 | -:-:-:-:-:-:-:-: | 7339 | nr1h3 | 892 | 878 | 1109 | 660 | 900 | 773 |
| ENSDARG000 | 7:7:7:7:7:7:7: | 427503:45933 | 429141:45958 | +:+:+:+:+:+:+: | 3522 | NRXN2 | 20 | 296 | 124 | 298 | 18 | 219 |
| ENSDARG000 | 7:7:7:7:7:7:7: | 50548074:505 | 50548279:505 | -:-:-:-:-:-:-:-: | 2048 | NA | 878 | 351 | 779 | 163 | 1422 | 313 |
| ENSDARG000 | 7:7:7:7:7:7:7: | 55328420:553 | 55328675:553 | -:-:-:-:-:-:-:-: | 7311 | PIEZO1 | 576 | 655 | 459 | 176 | 663 | 602 |
| ENSDARG000 | 7:7:7:7:7:7:7: | 60161002:601 | 60161507:601 | +:+:+:+:+:+:+: | 8335 | slc8a4b | 2 | 0 | 0 | 2 | 36 | 62 |
| ENSDARG000 | 7 | 64383483 | 64383599 | - | 117 | 5S_rRNA | 0 | 0 | 0 | 0 | 0 | 0 |
| ENSDARG000 | 7 | 39136650 | 39136774 | - | 125 | U5 | 0 | 1 | 0 | 0 | 0 | 0 |
| ENSDARG000 | 7:7:7:7:7:7:7: | 10895883:109 | 10895913:109 | -:-:-:-:-:-:-:-: | 315 | NA | 0 | 0 | 0 | 0 | 0 | 0 |
| ENSDARG000 | 7;7;7;7 | 70338270:703 | 70338418:703 | +;+;+;+ | 1288 | gba3 | 0 | 56 | 6 | 18 | 45 | 11 |
| ENSDARG000 | 7;7;7;7;7 | 58837286:588 | 58837711:588 | -;-;-;-;- | 999 | mrpl15 | 485 | 600 | 772 | 406 | 498 | 422 |
| ENSDARG000 | 7:7:7:7:7:7:7: | 71809723:718 | 71809750:718 | +:+:+:+:+:+:+: | 282 | NA | 4 | 0 | 4 | 1 | 4 | 0 |
| ENSDARG000 | 7:7:7:7:7:7:7: | 58188174:581 | 58188324:581 | -:-:-:-:-:-:-:-: | 2340 | ASPH | 470 | 402 | 485 | 297 | 631 | 569 |
| ENSDARG000 | 7 | 13452269 | 13452389 | - | 121 | 5S_rRNA | 0 | 0 | 0 | 0 | 0 | 0 |
| ENSDARG000 | 7 | 56068166 | 56068280 | - | 115 | 5S_rRNA | 0 | 0 | 0 | 0 | 0 | 0 |
| ENSDARG000 | 7 | 26028659 | 26028773 | + | 115 | 5S_rRNA | 0 | 0 | 0 | 0 | 0 | 0 |
| ENSDARG000 | 7;7;7;7 | 22678346:226 | 22678370:226 | -;-;-;- | 111 | NA | 0 | 0 | 0 | 0 | 0 | 0 |
| ENSDARG000 | 7;7;7;7;7 | 22281666:223 | 22281679:223 | -;-;-;-;- | 105 | NA | 0 | 0 | 2 | 0 | 0 | 0 |
| ENSDARG000 | 7;7;7;7 | 1912805:1914 | 1913209:1914 | -;-;-;- | 1082 | NA | 0 | 0 | 0 | 0 | 0 | 0 |
| ENSDARG000 | 7;7;7 | 59739164:597 | 59739831:597 | +;+;+ | 1020 | NA | 0 | 0 | 0 | 0 | 0 | 0 |
| ENSDARG000 | 7:7:7:7:7:7:7: | 72222542:722 | 72222728:722 | -:-:-:-:-:-:-:-: | 2515 | RASGRP2 | 19 | 6 | 7 | 7 | 16 | 28 |
| ENSDARG000 | 7;7;7;7 | 72441515:724 | 72442873:724 | +;+;+;+ | 3606 | NA | 0 | 12 | 4 | 0 | 2 | 8 |
| ENSDARG000 | 7:7:7:7:7:7:7: | 72460911:724 | 72460943:724 | +:+:+:+:+:+:+: | 13226 | HECTD4 | 658 | 287 | 777 | 326 | 787 | 418 |
| ENSDARG000 | 7 | 6697973 | 6698091 | + | 119 | 5S_rRNA | 0 | 3 | 0 | 0 | 0 | 0 |
| ENSDARG000 | 7 | 23362162 | 23362245 | + | 84 | dre-mir-730 | 0 | 0 | 0 | 0 | 0 | 0 |
| ENSDARG000 | 7 | 29005622 | 29006398 | + | 777 | NA | 0 | 0 | 0 | 0 | 0 | 0 |
| ENSDARG000 | 7;7;7;7;7;7 | 68886106:688 | 68886364:688 | -;-;-;-;-;- | 1360 | NA | 0 | 0 | 0 | 0 | 0 | 0 |
| ENSDARG000 | 7;7;7 | 886685:88891 | 886833:88919 | +;+;+ | 1258 | NA | 0 | 0 | 0 | 0 | 0 | 0 |
| ENSDARG000 | 7:7:7:7:7:7:7: | 936128:94238 | 936250:94245 | +:+:+:+:+:+:+: | 10786 | NA | 20 | 2 | 0 | 14 | 16 | 0 |
| ENSDARG000 | 7:7:7:7:7:7:7: | 980292:98029 | 980462:98046 | +:+:+:+:+:+:+: | 5517 | NA | 26 | 0 | 0 | 20 | 0 | 4 |
| ENSDARG000 | 7;7;7;7;7 | 1123824:1127 | 1125633:1127 | -;-;-;-;- | 2567 | NA | 255 | 98 | 105 | 95 | 149 | 156 |
| ENSDARG000 | 7;7;7 | 1772769:1774 | 1772821:1774 | +;+;+ | 378 | NA | 0 | 0 | 0 | 0 | 0 | 0 |
| ENSDARG000 | 7;7;7 | 2352951:2353 | 2353013:2353 | +;+;+ | 277 | NA | 0 | 0 | 0 | 0 | 0 | 0 |
| ENSDARG000 | 7;7;7 | 2591979:2606 | 2592134:2606 | +;+;+ | 2369 | NA | 0 | 0 | 0 | 0 | 0 | 0 |
| ENSDARG000 | 7;7;7 | 2623061:2632 | 2623192:2632 | +;+;+ | 2933 | NA | 0 | 0 | 0 | 0 | 0 | 0 |
| ENSDARG000 | 7;7;7;7;7 | 11656567:116 | 11656767:116 | -;-;-;-;- | 916 | NA | 0 | 0 | 0 | 0 | 0 | 0 |
| ENSDARG000 | 7;7;7;7;7;7 | 16075216:160 | 16075233:160 | -;-;-;-;-;- | 1520 | NA | 0 | 0 | 0 | 0 | 0 | 0 |
| ENSDARG000 | 7;7;7;7;7 | 20234414:202 | 20238684:202 | -;-;-;-;- | 4562 | NA | 0 | 0 | 2 | 8 | 0 | 0 |
| ENSDARG000 | 7;7;7;7;7 | 35585636:355 | 35585934:355 | +;+;+;+;+ | 1928 | NA | 0 | 0 | 0 | 0 | 0 | 0 |
| ENSDARG000 | 7:7:7:7:7:7:7: | 42703718:427 | 42703799:427 | +:+:+:+:+:+:+: | 3596 | NA | 0 | 0 | 0 | 0 | 0 | 0 |
| ENSDARG000 | 7:7:7:7:7:7:7: | 45914757:459 | 45915035:459 | +:+:+:+:+:+:+: | 1150 | NA | 0 | 0 | 0 | 0 | 0 | 0 |
| ENSDARG000 | 7;7;7;7 | 48979664:489 | 48979812:489 | -;-;-;- | 685 | NA | 0 | 0 | 0 | 0 | 0 | 0 |
| ENSDARG000 | 7;7;7;7;7;7;7 | 55053248:550 | 55055031:550 | -;-;-;-;-;-;- | 2883 | nadsyn1 | 93 | 74 | 92 | 70 | 84 | 70 |
| ENSDARG000 | 7;7;7;7;7 | 58309898:583 | 58310374:583 | +;+;+;+;+ | 2244 | NA | 10 | 28 | 9 | 4 | 0 | 0 |
| ENSDARG000 | 7:7:7:7:7:7:7: | 59750111:597 | 59750808:597 | +:+:+:+:+:+:+: | 6066 | NA | 0 | 0 | 0 | 0 | 0 | 0 |
| ENSDARG000 | 7;7;7;7;7;7 | 67916111:679 | 67916459:679 | -;-;-;-;-;- | 854 | NA | 0 | 0 | 0 | 0 | 0 | 0 |
| ENSDARG000 | 7;7;7 | 69868210:698 | 69868767:698 | -;-;- | 4129 | NA | 0 | 6 | 6 | 0 | 4 | 0 |
| ENSDARG000 | 7;7;7;7;7 | 72606759:726 | 72606782:726 | +;+;+;+;+ | 821 | NA | 0 | 0 | 6 | 2 | 1 | 0 |
| ENSDARG000 | 7;7;7 | 72612470:726 | 72612698:726 | +;+;+ | 592 | NA | 6 | 0 | 0 | 0 | 0 | 0 |
| ENSDARG000 | 7;7;7;7;7 | 73167923:731 | 73168372:731 | +;+;+;+;+ | 2048 | NA | 0 | 0 | 0 | 0 | 0 | 0 |
| ENSDARG000 | 5;5;5;5;5 | 67463027:674 | 67463298:674 | -;-;-;-;- | 822 | si:dkev-251i1 | 217 | 130 | 329 | 80 | 194 | 438 |
| ENSDARG000 | 5;5 | 28282449:282 | 28282629:282 | +;+ | 563 | NA | 20 | 15 | 16 | 35 | 24 | 18 |
| ENSDARG000 | 5;5;5 | 28271412:282 | 28271521:282 | +;+;+ | 1823 | vamp8 | 70 | 26 | 24 | 42 | 108 | 152 |
| ENSDARG000 | 5:5:5:5:5:5:5: | 41774366:417 | 41774593:417 | +:+:+:+:+:+:+: | 2964 | NA | 2 | 13 | 0 | 0 | 4 | 0 |
| ENSDARG000 | 5:5:5:5:5:5:5: | 37087583:370 | 37087772:370 | +:+:+:+:+:+:+: | 1359 | tagln2 | 848 | 500 | 124 | 304 | 308 | 303 |
| ENSDARG000 | 5;5;5;5;5;5;5 | 69856153:698 | 69857152:698 | +;+;+;+;+;+;+ | 1899 | ugt2a6 | 351 | 4 | 212 | 7 | 271 | 5 |
| ENSDARG000 | 5:5:5:5:5:5:5: | 33275150:332 | 33275997:332 | -:-:-:-:-:-:-:-: | 1502 | surf6 | 5 | 14 | 17 | 2 | 9 | 3 |
| ENSDARG000 | 5:5:5:5:5:5:5: | 33264842:332 | 33267116:332 | -:-:-:-:-:-:-:-: | 6953 | kyat1 | 100 | 176 | 182 | 130 | 106 | 160 |
| ENSDARG000 | 5:5:5:5:5:5:5: | 15667874:157 | 15668473:157 | +:+:+:+:+:+:+: | 12640 | NA | 14 | 12 | 127 | 34 | 24 | 108 |
| ENSDARG000 | 5:5:5:5:5:5:5: | 22969635:229 | 22969763:229 | +:+:+:+:+:+:+: | 1647 | hmgn7 | 4243 | 5572 | 7328 | 3565 | 6564 | 4963 |
| ENSDARG000 | 5:5:5:5:5:5:5: | 29714786:297 | 29715141:297 | +:+:+:+:+:+:+: | 2886 | DDX31 | 164 | 186 | 350 | 130 | 183 | 195 |
| ENSDARG000 | 5:5:5:5:5:5:5: | 67474119:674 | 67474508:674 | -:-:-:-:-:-:-:-: | 2637 | UTP14C | 378 | 656 | 898 | 352 | 693 | 413 |
| ENSDARG000 | 5;5;5;5;5;5 | 54545432:545 | 54545765:545 | -;-;-;-;-;- | 847 | ssna1 | 46 | 88 | 156 | 84 | 111 | 76 |
| ENSDARG000 | 5:5:5:5:5:5:5: | 22974019:229 | 22974148:229 | +:+:+:+:+:+:+: | 5141 | BRWD3 | 458 | 437 | 534 | 264 | 592 | 431 |
| ENSDARG000 | 5:5:5:5:5:5:5: | 9179578:9179 | 9179881:9179 | -:-:-:-:-:-:-:-: | 4450 | NA | 10 | 4 | 8 | 2 | 0 | 0 |
| ENSDARG000 | 5;5;5;5;5 | 25025053:250 | 25025743:250 | -;-;-;-;- | 1238 | gbgt1l5 | 10 | 12 | 6 | 0 | 26 | 60 |
| ENSDARG000 | 5;5;5;5 | 54501105:545 | 54503012:545 | +;+;+;+ | 2445 | NA | 14 | 2 | 14 | 2 | 54 | 18 |
| ENSDARG000 | 5:5:5:5:5:5:5: | 22956894:229 | 22961572:229 | -:-:-:-:-:-:-:-: | 6063 | NA | 34 | 28 | 60 | 20 | 48 | 104 |
| ENSDARG000 | 5:5:5:5:5:5:5: | 33281105:332 | 33281300:332 | +:+:+:+:+:+:+: | 1293 | surf1 | 295 | 248 | 365 | 295 | 476 | 346 |
| ENSDARG000 | 5:5:5:5:5:5:5: | 69868911:698 | 69869794:698 | +:+:+:+:+:+:+: | 2756 | ugt2a5 | 183 | 118 | 117 | 253 | 133 | 119 |
| ENSDARG000 | 5;5;5;5;5;5 | 4366431:4366 | 4366662:4366 | +;+;+;+;+;+ | 1418 | sat1a.2 | 2422 | 2633 | 3196 | 2717 | 1330 | 1386 |
| ENSDARG000 | 5:5:5:5:5:5:5: | 57210237:572 | 57210304:572 | +:+:+:+:+:+:+: | 2413 | pja2 | 530 | 356 | 556 | 474 | 496 | 470 |
| ENSDARG000 | 5:5:5:5:5:5:5: | 51615025:516 | 51617279:516 | -;-;-;-;-;-;- | 3576 | otpb | 0 | 0 | 0 | 12 | 7 | 8 |
| ENSDARG000 | 5:5:5:5:5:5:5: | 32206378:322 | 32206418:322 | +:+:+:+:+:+:+: | 5990 | myhz2 | 6 | 0 | 4 | 0 | 5 | 11 |
| ENSDARG000 | 5;5;5;5;5 | 30986512:309 | 30986675:309 | +;+;+;+;+ | 2172 | ube2g1a | 440 | 466 | 456 | 228 | 654 | 362 |
| ENSDARG000 | 5;5;5;5 | 4921092:4921 | 4922648:4922 | -;-;-;- | 1603 | zbtb43 | 20 | 0 | 62 | 14 | 50 | 28 |
| ENSDARG000 | 5:5:5:5:5:5:5: | 5032833:5035 | 5033374:5035 | -:-:-:-:-:-:-:-: | 2063 | lmx1ba | 0 | 0 | 0 | 22 | 0 | 2 |
| ENSDARG000 | 5:5:5:5:5:5:5: | 35875085:358 | 35875224:358 | -:-:-:-:-:-:-:-: | 2593 | NA | 0 | 0 | 0 | 0 | 0 | 0 |
| ENSDARG000 | 5;5;5;5;5 | 51739181:517 | 51740026:517 | -;-;-;-;- | 1764 | lhfpl2b | 949 | 1152 | 1821 | 1233 | 2248 | 1196 |
| ENSDARG000 | 5:5:5:5:5:5:5: | 15819651:158 | 15819801:158 | +:+:+:+:+:+:+: | 4545 | hspb8 | 20 | 6 | 61 | 12 | 36 | 2 |
| ENSDARG000 | 5:5:5:5:5:5:5: | 43798345:437 | 43798436:437 | -:-:-:-:-:-:-:-: | 1019 | smn1 | 130 | 164 | 270 | 195 | 127 | 159 |
| ENSDARG000 | 5:5:5:5:5:5:5: | 69878693:698 | 69878879:698 | +:+:+:+:+:+:+: | 3112 | ythdc1 | 1054 | 602 | 1210 | 868 | 1307 | 1098 |
| ENSDARG000 | 5:5:5:5:5:5:5: | 28297250:282 | 28297393:282 | +:+:+:+:+:+:+: | 1976 | vamp5 | 2373 | 1370 | 668 | 537 | 592 | 773 |
| ENSDARG000 | 5:5:5:5:5:5:5: | 5171254:5175 | 5172276:5175 | -:-:-:-:-:-:-:-: | 2310 | mvb12ba | 154 | 170 | 219 | 85 | 149 | 166 |
| ENSDARG000 | 5:5:5:5:5:5:5: | 39706345:397 | 39708726:397 | -:-:-:-:-:-:-:-: | 4691 | rasgef1ba | 674 | 1165 | 607 | 381 | 1738 | 907 |
| ENSDARG000 | 5:5:5:5:5:5:5: | 28308774:283 | 28308999:283 | +:+:+:+:+:+:+: | 13969 | DNAH10 | 116 | 53 | 66 | 28 | 62 | 24 |
| ENSDARG000 | 5:5:5:5:5:5:5: | 57243568:572 | 57244431:572 | -:-:-:-:-:-:-:-: | 3483 | FER | 478 | 289 | 434 | 230 | 493 | 394 |
| ENSDARG000 | 5:5:5:5:5:5:5: | 26234425:262 | 26234823:262 | -:-:-:-:-:-:-:-: | 3923 | erap1b | 2084 | 1403 | 1044 | 464 | 1692 | 779 |
| ENSDARG000 | 5:5:5:5:5:5:5: | 21181047:211 | 21181178:211 | +:+:+:+:+:+:+: | 1803 | arhgap25 | 34 | 25 | 18 | 16 | 112 | 92 |
| ENSDARG000 | 5:5:5:5:5:5:5: | 59494079:595 | 59494314:595 | +:+:+:+:+:+:+: | 5557 | GTF2IRD1 | 2098 | 2424 | 3530 | 3665 | 3364 | 2270 |
| ENSDARG000 | 5:5:5:5:5:5:5: | 37101393:371 | 37102238:371 | -:-:-:-:-:-:-:-: | 2155 | il13ra2 | 152 | 10 | 64 | 30 | 48 | 56 |
| ENSDARG000 | 5:5:5:5:5:5:5: | 65491390:654 | 65491457:654 | +:+:+:+:+:+:+: | 3656 | KCNT1 | 0 | 4 | 0 | 0 | 0 | 7 |
| ENSDARG000 | 5:5:5:5:5:5:5: | 38440099:384 | 38441201:384 | -:-:-:-:-:-:-:-: | 2557 | chrne | 3 | 0 | 0 | 0 | 41 | 4 |
| ENSDARG000 | 5:5:5:5:5:5:5: | 46207017:462 | 46207046:462 | -:-:-:-:-:-:-:-: | 1089 | ssbp2a | 15 | 6 | 0 | 0 | 0 | 0 |
| ENSDARG000 | 5:5:5:5:5:5:5: | 38388443:383 | 38391132:383 | -:-:-:-:-:-:-:-: | 5520 | PLD2 | 727 | 715 | 857 | 455 | 723 | 637 |
| ENSDARG000 | 5;5;5 | 29746355:297 | 29747326:297 | -;-;- | 1960 | barhl1a | 5 | 11 | 18 | 11 | 27 | 20 |
| ENSDARG000 | 5:5:5:5:5:5:5: | 25044311:250 | 25046532:250 | -:-:-:-:-:-:-:-: | 6341 | PNPLA7 | 1087 | 399 | 715 | 1065 | 873 | 891 |
| ENSDARG000 | 5:5:5:5:5:5:5: | 43808838:438 | 43809067:438 | -:-:-:-:-:-:-:-: | 2132 | mccc2 | 1129 | 1165 | 1170 | 766 | 1252 | 1103 |
| ENSDARG000 | 5:5:5:5:5:5:5: | 12908781:129 | 12908920:129 | +:+:+:+:+:+:+: | 1921 | ppil2 | 445 | 440 | 464 | 414 | 544 | 450 |
| ENSDARG000 | 5:5:5:5:5:5:5: | 51774983:517 | 51776197:517 | -:-:-:-:-:-:-:-: | 3358 | homer1b | 0 | 10 | 10 | 14 | 0 | 4 |
| ENSDARG000 | 5;5;5;5;5;5;5 | 67533180:675 | 67534826:675 | -;-;-;-;-;-;- | 4184 | zbtb20 | 255 | 263 | 467 | 215 | 587 | 158 |
| ENSDARG000 | 5;5 | 21211135:212 | 21211534:212 | +;+ | 1458 | bmp10 | 0 | 2 | 0 | 0 | 4 | 2 |
| ENSDARG000 | 5;5;5;5;5 | 43807003:438 | 43807164:438 | +;+;+;+;+ | 684 | hspb11 | 24 | 42 | 31 | 24 | 36 | 52 |
| ENSDARG000 | 5:5:5:5:5:5:5: | 54555567:545 | 54555754:545 | +:+:+:+:+:+:+: | 2647 | anapc2 | 214 | 310 | 296 | 206 | 316 | 184 |
| ENSDARG000 | 5;5;5;5;5 | 36271016:362 | 36271401:362 | -;-;-;-;- | 1026 | efnb1 | 700 | 555 | 859 | 562 | 503 | 700 |
| ENSDARG000 | 5;5;5 | 36415978:364 | 36416703:364 | +;+;+ | 1206 | nalf2 | 8 | 5 | 20 | 7 | 2 | 0 |
| ENSDARG000 | 5:5:5:5:5:5:5: | 41793001:417 | 41793226:417 | +:+:+:+:+:+:+: | 2053 | bcl7a | 582 | 468 | 547 | 357 | 674 | 677 |
| ENSDARG000 | 5:5:5:5:5:5:5: | 21225017:212 | 21225100:212 | +:+:+:+:+:+:+: | 4005 | NA | 192 | 204 | 248 | 119 | 171 | 361 |
| ENSDARG000 | 5;5;5;5 | 41800082:418 | 41800506:418 | -;-;-;- | 999 | alkbh2 | 150 | 202 | 168 | 115 | 146 | 116 |
| ENSDARG000 | 5;5;5;5 | 15841523:158 | 15844015:158 | -;-;-;- | 3223 | NA | 0 | 0 | 0 | 0 | 0 | 0 |
| ENSDARG000 | 5;5;5 | 21258218:212 | 21258317:212 | +;+;+ | 485 | NA | 0 | 3 | 0 | 0 | 0 | 0 |
| ENSDARG000 | 5:5:5:5:5:5:5: | 19385503:193 | 19386449:193 | -:-:-:-:-:-:-:-: | 2945 | foxn4 | 3 | 4 | 0 | 0 | 0 | 0 |
| ENSDARG000 | 5;5;5 | 21270450:212 | 21270597:212 | +;+;+ | 666 | NA | 0 | 4 | 2 | 2 | 0 | 8 |
| ENSDARG000 | 5:5:5:5:5:5:5: | 36439405:364 | 36439836:364 | +:+:+:+:+:+:+: | 1216 | eda | 2 | 2 | 2 | 0 | 0 | 0 |
| ENSDARG000 | 5;5;5 | 57320113:573 | 57320338:573 | +;+;+ | 1069 | atp6v1g1 | 5496 | 6278 | 4725 | 6208 | 4891 | 5696 |
| ENSDARG000 | 5;5;5;5;5 | 21274165:212 | 21274268:212 | +;+;+;+;+ | 649 | NA | 0 | 0 | 0 | 0 | 0 | 0 |
| ENSDARG000 | 5:5:5:5:5:5:5: | 63322093:633 | 63322702:633 | +:+:+:+:+:+:+: | 3633 | NA | 176 | 226 | 221 | 218 | 330 | 345 |
| ENSDARG000 | 5:5:5:5:5:5:5: | 30996322:309 | 30996331:309 | +:+:+:+:+:+:+: | 4918 | ankfy1 | 1371 | 2326 | 2395 | 1827 | 2529 | 2025 |
| ENSDARG000 | 5;5 | 36505473:365 | 36505662:365 | +;+ | 495 | NA | 0 | 0 | 0 | 0 | 0 | 0 |
| ENSDARG000 | 5:5:5:5:5:5:5: | 5302932:5307 | 5303024:5307 | -:-:-:-:-:-:-:-: | 1538 | NA | 0 | 0 | 0 | 0 | 0 | 0 |
| ENSDARG000 | 5:5:5:5:5:5:5: | 15944411:159 | 15945356:159 | -:-:-:-:-:-:-:-: | 2373 | xbp1 | 5130 | 5737 | 12068 | 7762 | 17994 | 13443 |
| ENSDARG000 | 5:5:5:5:5:5:5: | 69907222:699 | 69908256:699 | -:-:-:-:-:-:-:-: | 2619 | fktn | 289 | 275 | 282 | 253 | 448 | 298 |
| ENSDARG000 | 5;5;5 | 21288227:212 | 21288363:212 | +;+;+ | 321 | sh2d1aa | 0 | 0 | 6 | 2 | 5 | 3 |
| ENSDARG000 | 5:5:5:5:5:5:5: | 29752604:297 | 29752884:297 | -:-:-:-:-:-:-:-: | 1064 | cfap77 | 0 | 0 | 0 | 6 | 0 | 2 |
| ENSDARG000 | 5:5:5:5:5:5:5: | 37132016:371 | 37132135:371 | -:-:-:-:-:-:-:-: | 2152 | LRCH2 | 347 | 308 | 447 | 383 | 181 | 438 |
| ENSDARG000 | 5;5;5;5;5;5;5 | 46277593:462 | 46277937:462 | +;+;+;+;+;+;+ | 1844 | zgc:110626 | 172 | 108 | 230 | 132 | 106 | 263 |
| ENSDARG000 | 5;5;5;5;5 | 36513605:365 | 36513757:365 | +;+;+;+;+ | 2132 | wnt11f2 | 2 | 0 | 0 | 0 | 0 | 0 |
| ENSDARG000 | 5;5;5;5;5 | 67662430:676 | 67662798:676 | +;+;+;+;+ | 1158 | NA | 12 | 36 | 7 | 8 | 6 | 6 |
| ENSDARG000 | 5:5:5:5:5:5:5: | 63340637:633 | 63340679:633 | +:+:+:+:+:+:+: | 1811 | NA | 0 | 6 | 0 | 4 | 0 | 36 |
| ENSDARG000 | 5:5:5:5:5:5:5: | 51848756:518 | 51848966:518 | +:+:+:+:+:+:+: | 8745 | cmya5 | 4 | 0 | 4 | 0 | 16 | 0 |
| ENSDARG000 | 5:5:5:5:5:5:5: | 23107804:231 | 23108169:231 | -;-;-;-;-;-;-;- | 1510 | uprt | 432 | 482 | 1131 | 753 | 532 | 604 |
| ENSDARG000 | 5:5:5:5:5:5:5: | 57328535:573 | 57328708:573 | +:+:+:+:+:+:+: | 4740 | slc31a1 | 985 | 1209 | 1899 | 1129 | 1532 | 1268 |
| ENSDARG000 | 5;5;5 | 23118470:231 | 23118531:231 | +;+;+ | 6759 | NEXMIF | 2 | 0 | 20 | 6 | 20 | 0 |
| ENSDARG000 | 5:5:5:5:5:5:5: | 51833132:518 | 51833407:518 | +:+:+:+:+:+:+: | 2475 | tent2 | 443 | 340 | 427 | 191 | 360 | 384 |
| ENSDARG000 | 5:5:5:5:5:5:5: | 59553636:595 | 59554053:595 | +:+:+:+:+:+:+: | 4181 | dtx2 | 472 | 835 | 2301 | 1109 | 3667 | 833 |
| ENSDARG000 | 5:5:5:5:5:5:5: | 21295747:212 | 21297328:212 | -:-:-:-:-:-:-:-: | 9037 | TENM1 | 1080 | 163 | 733 | 288 | 72 | 1382 |
| ENSDARG000 | 5:5:5:5:5:5:5: | 26248380:262 | 26248660:262 | +:+:+:+:+:+:+: | 2853 | naaladl1 | 0 | 3 | 0 | 0 | 0 | 0 |
| ENSDARG000 | 5:5:5:5:5:5:5: | 67671207:676 | 67673583:676 | -:-:-:-:-:-:-:-: | 4378 | NA | 1687 | 1294 | 2547 | 1725 | 1875 | 1774 |
| ENSDARG000 | 5;5;5 | 37267247:372 | 37267354:372 | -;-;- | 258 | NA | 0 | 0 | 6 | 0 | 0 | 0 |
| ENSDARG000 | 5:5:5:5:5:5:5: | 69926766:699 | 69927537:699 | -:-:-:-:-:-:-:-: | 5042 | ugt2a1 | 76 | 34 | 26 | 12 | 57 | 37 |
| ENSDARG000 | 5:5:5:5:5:5:5: | 15992655:160 | 15993128:160 | +:+:+:+:+:+:+: | 3196 | znrf3 | 180 | 247 | 314 | 148 | 718 | 390 |
| ENSDARG000 | 5:5:5:5:5:5:5: | 67736822:677 | 67737586:677 | -:-:-:-:-:-:-:-: | 2049 | b4galt4 | 305 | 201 | 219 | 98 | 352 | 226 |

| | | | | | | | | | | | | |
|---|---|---|---|---|---|---|---|---|---|---|---|---|
| ENSDARG000 | 5;5;5;5;5;5;5; | 37277242:372 | 37277597:372 | +;+;+;+;+;+;+; | 1902 | NA | 466 | 365 | 243 | 225 | 227 | 276 |
| ENSDARG000 | 5;5;5;5;5;5;5; | 36530836:365 | 36531117:365 | +;+;+;+;+;+;+; | 2482 | igbp1 | 527 | 447 | 542 | 230 | 691 | 410 |
| ENSDARG000 | 5;5;5;5;5;5;5; | 63359520:633 | 63359685:633 | +;+;+;+;+;+;+; | 2997 | ttf1.1 | 60 | 94 | 156 | 80 | 86 | 79 |
| ENSDARG000 | 5;5;5 | 25072894:250 | 25072973:250 | +;+;+ | 640 | mrpl41 | 224 | 236 | 334 | 163 | 199 | 228 |
| ENSDARG000 | 5;5;5;5;5;5;5 | 46293850:462 | 46294183:462 | +;+;+;+;+;+;+ | 1628 | si:ch211-130r | 2 | 0 | 0 | 2 | 1 | 0 |
| ENSDARG000 | 5;5;5;5;5;5;5; | 63375386:633 | 63376247:633 | +;+;+;+;+;+;+; | 2329 | ttf1.2 | 41 | 65 | 102 | 47 | 94 | 41 |
| ENSDARG000 | 5;5;5;5;5;5;5; | 65613292:656 | 65614914:656 | -;-;-;-;-;-;-;-; | 6106 | camsap1b | 436 | 430 | 574 | 362 | 599 | 452 |
| ENSDARG000 | 5;5;5;5;5;5;5; | 5398966:5406 | 5399113:5406 | +;+;+;+;+;+;+; | 1900 | MAPKAP1 | 151 | 141 | 215 | 171 | 207 | 176 |
| ENSDARG000 | 5;5;5;5;5;5;5; | 25077212:250 | 25077818:250 | -;-;-;-;-;-;-;-; | 1864 | dph7 | 288 | 309 | 384 | 221 | 351 | 221 |
| ENSDARG000 | 5;5;5;5;5;5;5; | 19413838:194 | 19414068:194 | +;+;+;+;+;+;+; | 3349 | MYO1H | 0 | 0 | 0 | 0 | 0 | 3 |
| ENSDARG000 | 5;5;5;5;5;5;5; | 32221755:322 | 32221783:322 | +;+;+;+;+;+;+; | 5983 | myhc4 | 2 | 2 | 0 | 3 | 6 | 4 |
| ENSDARG000 | 5;5;5;5 | 31032149:310 | 31032338:310 | -;-;-;- | 965 | cyb5d2 | 82 | 62 | 60 | 56 | 74 | 90 |
| ENSDARG000 | 5;5;5;5;5;5 | 39875646:398 | 39875908:398 | +;+;+;+;+;+ | 1970 | NA | 0 | 0 | 0 | 0 | 0 | 0 |
| ENSDARG000 | 5;5;5;5;5 | 60157256:601 | 60157904:601 | -;-;-;-;- | 1104 | NA | 0 | 2 | 4 | 1 | 0 | 0 |
| ENSDARG000 | 5;5;5;5;5;5;5; | 23136544:231 | 23136839:231 | +;+;+;+;+;+;+; | 2572 | prps1a | 2973 | 2940 | 4099 | 1592 | 3486 | 2405 |
| ENSDARG000 | 5;5;5;5 | 38462121:384 | 38462360:384 | +;+;+;+ | 1067 | gltpd2b | 0 | 0 | 2 | 0 | 0 | 0 |
| ENSDARG000 | 5;5;5;5;5;5;5; | 33287611:332 | 33287704:332 | +;+;+;+;+;+;+; | 2337 | med22 | 383 | 343 | 408 | 314 | 526 | 496 |
| ENSDARG000 | 5;5;5;5;5;5;5; | 33284144:332 | 33284281:332 | -;-;-;-;-;-;-;-; | 1478 | rpl7a | 29983 | 28606 | 48398 | 42032 | 31724 | 40945 |
| ENSDARG000 | 5;5;5;5;5;5;5; | 57363187:573 | 57363381:573 | -;-;-;-;-;-;-;- | 1140 | NA | 17 | 59 | 70 | 51 | 14 | 46 |
| ENSDARG000 | 5;5;5;5;5;5;5; | 39981479:399 | 39981632:399 | -;-;-;-;-;-;-;-; | 1317 | NA | 0 | 0 | 0 | 0 | 0 | 0 |
| ENSDARG000 | 5;5;5;5;5;5;5; | 60518611:605 | 60519253:605 | +;+;+;+;+;+;+; | 6443 | tmem132e | 258 | 100 | 136 | 214 | 122 | 191 |
| ENSDARG000 | 5;5;5;5;5;5;5 | 16128214:161 | 16128845:161 | -;-;-;-;-;-;- | 1546 | coq5 | 64 | 126 | 128 | 67 | 98 | 74 |
| ENSDARG000 | 5;5;5;5;5;5;5; | 38475531:384 | 38476938:384 | -;-;-;-;-;-;-;-; | 2559 | atp1b2b | 0 | 0 | 0 | 0 | 0 | 0 |
| ENSDARG000 | 5;5;5;5;5;5;5; | 63390315:633 | 63390415:633 | +;+;+;+;+;+;+; | 2474 | rab14 | 7046 | 5882 | 5436 | 7172 | 5557 | 5252 |
| ENSDARG000 | 5;5;5;5;5;5;5; | 39994283:399 | 39994575:399 | +;+;+;+;+;+;+; | 1655 | tmem175 | 153 | 139 | 121 | 127 | 83 | 174 |
| ENSDARG000 | 5;5;5;5;5;5;5; | 25084385:250 | 25084598:250 | +;+;+;+;+;+;+; | 893 | paxx | 154 | 138 | 105 | 74 | 114 | 150 |
| ENSDARG000 | 5;5;5;5;5;5;5; | 67835595:678 | 67835690:678 | +;+;+;+;+;+;+; | 3205 | si:ch211-271 | 1488 | 1224 | 1731 | 870 | 1723 | 1096 |
| ENSDARG000 | 5;5;5;5;5;5;5; | 6670945:6672 | 6670954:6673 | +;+;+;+;+;+;+; | 2992 | pxna | 671 | 685 | 906 | 662 | 573 | 727 |
| ENSDARG000 | 5;5;5;5;5;5;5; | 60934345:609 | 60934614:609 | +;+;+;+;+;+;+; | 1138 | NA | 0 | 0 | 0 | 0 | 0 | 0 |
| ENSDARG000 | 5;5;5;5;5;5;5; | 60919378:609 | 60919747:609 | +;+;+;+;+;+;+; | 2622 | DOC2B | 2 | 2 | 0 | 0 | 17 | 4 |
| ENSDARG000 | 5;5;5;5;5;5;5; | 57407515:574 | 57408067:574 | -;-;-;-;-;-;-;-; | 1982 | NA | 0 | 0 | 0 | 0 | 0 | 0 |
| ENSDARG000 | 5;5;5;5;5;5;5; | 31036911:310 | 31037507:310 | +;+;+;+;+;+;+; | 11299 | ZZEF1 | 1065 | 808 | 1255 | 549 | 1234 | 918 |
| ENSDARG000 | 5;5;5;5;5;5;5; | 54711027:547 | 54711701:547 | -;-;-;-;-;-;-;-; | 2363 | ccnb1 | 466 | 1743 | 1871 | 726 | 1153 | 524 |
| ENSDARG000 | 5;5;5;5;5;5;5; | 998296:10135 | 998751:10136 | -;-;-;-;-;-;-;- | 1562 | mbd2 | 126 | 161 | 247 | 244 | 143 | 158 |
| ENSDARG000 | 5 | 46327205 | 46329880 | - | 2676 | NA | 234 | 273 | 262 | 102 | 306 | 193 |
| ENSDARG000 | 5;5;5;5;5;5;5; | 41826958:418 | 41827281:418 | -;-;-;-;-;-;-;-; | 13502 | NA | 14 | 4 | 0 | 12 | 6 | 16 |
| ENSDARG000 | 5;5;5;5;5;5;5; | 26263128:262 | 26269124:262 | -;-;-;-;-;-;-;-; | 10116 | arvcfb | 256 | 172 | 273 | 44 | 177 | 171 |
| ENSDARG000 | 5;5;5;5 | 65670605:656 | 65672422:656 | -;-;-;- | 2431 | NA | 0 | 0 | 6 | 0 | 0 | 0 |
| ENSDARG000 | 5;5;5;5 | 9218318:9219 | 9218677:9219 | +;+;+;+ | 868 | NA | 2 | 4 | 0 | 8 | 4 | 9 |
| ENSDARG000 | 5;5;5;5;5;5;5; | 1055042:1055 | 1055115:1055 | +;+;+;+;+;+;+; | 1136 | rnf185 | 586 | 474 | 663 | 579 | 524 | 506 |
| ENSDARG000 | 5;5;5;5;5;5;5; | 19436483:194 | 19439572:194 | -;-;-;-;-;-;-;-; | 4213 | kctd10 | 1250 | 1436 | 1499 | 1230 | 1112 | 1294 |
| ENSDARG000 | 5;5;5;5;5;5;5; | 16151944:161 | 16152942:161 | -;-;-;-;-;-;-;-; | 2783 | kremen1 | 60 | 32 | 78 | 66 | 48 | 12 |
| ENSDARG000 | 5;5;5 | 9224051:9231 | 9224108:9231 | +;+;+ | 952 | NA | 6 | 6 | 8 | 14 | 0 | 1 |
| ENSDARG000 | 5;5;5;5;5;5;5; | 46424437:464 | 46424808:464 | +;+;+;+;+;+;+; | 2013 | NA | 0 | 2 | 6 | 2 | 0 | 0 |
| ENSDARG000 | 5;5;5;5 | 69950882:699 | 69951197:699 | +;+;+;+ | 1393 | dnajc25 | 228 | 249 | 275 | 72 | 450 | 100 |
| ENSDARG000 | 5;5;5;5;5;5;5; | 37290260:372 | 37290398:372 | +;+;+;+;+;+;+; | 3321 | zgc:163098 | 919 | 703 | 1179 | 907 | 1063 | 1450 |
| ENSDARG000 | 5;5 | 6722310:6728 | 6722375:6728 | +;+ | 372 | NA | 0 | 0 | 0 | 0 | 0 | 0 |
| ENSDARG000 | 5;5;5;5;5;5;5; | 6726155:6726 | 6728702:6726 | -;-;-;-;-;-;-;-; | 3755 | ostf1 | 1505 | 1558 | 1226 | 1360 | 1539 | 1657 |
| ENSDARG000 | 5;5 | 21736359:217 | 21736686:217 | -;- | 621 | NA | 0 | 0 | 0 | 0 | 0 | 0 |
| ENSDARG000 | 5;5;5;5;5;5;5; | 24047292:240 | 24047909:240 | +;+;+;+;+;+;+; | 2120 | ctdnep1a | 232 | 284 | 250 | 230 | 218 | 229 |
| ENSDARG000 | 5;5;5;5;5;5;5; | 43849529:438 | 43849651:438 | -;-;-;-;-;-;-;-; | 2089 | NA | 10 | 12 | 36 | 2 | 20 | 0 |
| ENSDARG000 | 5;5;5;5;5;5;5; | 46490622:464 | 46494664:464 | -;-;-;-;-;-;-;-; | 4928 | hapln1a | 324 | 98 | 3532 | 1728 | 352 | 105 |
| ENSDARG000 | 5;5;5;5;5;5;5; | 9246018:9246 | 9246561:9246 | +;+;+;+;+;+;+; | 4366 | NA | 22 | 36 | 42 | 81 | 88 | 120 |
| ENSDARG000 | 5;5;5;5;5;5;5; | 36551470:365 | 36551724:365 | -;-;-;-;-;-;-;-; | 6394 | taf1 | 1128 | 849 | 1156 | 590 | 1161 | 1079 |
| ENSDARG000 | 5;5;5;5;5;5;5; | 28398449:283 | 28398502:283 | +;+;+;+;+;+;+; | 7047 | nsmfb | 8 | 10 | 8 | 4 | 26 | 12 |
| ENSDARG000 | 5;5;5;5;5;5;5; | 40009958:400 | 40010093:400 | -;-;-;-;-;-;-;-; | 1750 | CDS1 | 66 | 97 | 176 | 63 | 205 | 123 |
| ENSDARG000 | 5;5 | 33332548:333 | 33332896:333 | +;+ | 2028 | NA | 0 | 0 | 2 | 0 | 0 | 13 |
| ENSDARG000 | 5;5;5 | 33339762:333 | 33339976:333 | +;+;+ | 996 | ptges | 18 | 0 | 0 | 0 | 4 | 3 |
| ENSDARG000 | 5;5;5;5;5;5;5; | 69981453:699 | 69981582:699 | +;+;+;+;+;+;+; | 1526 | NA | 0 | 0 | 2 | 0 | 0 | 0 |
| ENSDARG000 | 5;5;5 | 29784172:297 | 29784260:297 | +;+;+ | 853 | NA | 4 | 12 | 95 | 16 | 52 | 96 |
| ENSDARG000 | 5;5;5;5;5;5 | 29794058:297 | 29794143:297 | +;+;+;+;+;+ | 1621 | thy1 | 1966 | 852 | 1121 | 2262 | 558 | 2904 |
| ENSDARG000 | 5;5;5;5;5;5;5; | 29803380:298 | 29803466:298 | +;+;+;+;+;+;+; | 2074 | usf1l | 782 | 697 | 929 | 788 | 559 | 601 |
| ENSDARG000 | 5;5;5;5;5 | 67869785:678 | 67870223:678 | -;-;-;-;- | 1052 | tagln3a | 2 | 8 | 0 | 4 | 3 | 0 |
| ENSDARG000 | 5;5;5;5;5 | 41853438:418 | 41853474:418 | -;-;-;-;- | 616 | NA | 0 | 2 | 0 | 0 | 0 | 10 |
| ENSDARG000 | 5;5;5;5;5;5;5; | 16428411:164 | 16432206:164 | -;-;-;-;-;-;-;-; | 7514 | piwil2 | 0 | 4 | 19 | 0 | 0 | 4 |
| ENSDARG000 | 5;5;5;5;5;5;5; | 25088622:250 | 25089270:250 | -;-;-;-;-;-;-;-; | 8892 | ABCA2 | 403 | 275 | 252 | 214 | 236 | 311 |
| ENSDARG000 | 5;5;5 | 1072930:1074 | 1073996:1074 | -;-;- | 1532 | NA | 48 | 36 | 71 | 72 | 87 | 30 |
| ENSDARG000 | 5;5;5;5;5;5;5; | 63400366:634 | 63400747:634 | -;-;-;-;-;-;-;-; | 5342 | NA | 888 | 815 | 1032 | 1893 | 1188 | 879 |
| ENSDARG000 | 5;5;5;5;5;5;5; | 6755466:6755 | 6755677:6755 | +;+;+;+;+;+;+; | 7740 | NA | 16 | 6 | 12 | 0 | 12 | 0 |
| ENSDARG000 | 5;5;5;5;5;5;5 | 24059598:240 | 24059921:240 | +;+;+;+;+;+;+ | 1434 | gabarapa | 2020 | 1557 | 1616 | 2017 | 1779 | 2133 |
| ENSDARG000 | 5;5;5;5;5;5;5; | 28526569:285 | 28526885:285 | -;-;-;-;-;-;-;-; | 1507 | zgc:162952 | 15 | 21 | 10 | 10 | 18 | 19 |
| ENSDARG000 | 5;5;5;5;5;5;5; | 67884087:678 | 67887448:678 | -;-;-;-;-;-;-;-; | 4869 | abhd10a | 174 | 178 | 228 | 210 | 106 | 274 |
| ENSDARG000 | 5;5;5;5;5;5;5; | 57417873:574 | 57418425:574 | -;-;-;-;-;-;-;-; | 1993 | NA | 26 | 22 | 14 | 6 | 16 | 34 |
| ENSDARG000 | 5;5;5;5;5;5;5; | 19448078:194 | 19448245:194 | +;+;+;+;+;+;+; | 3217 | ube3b | 237 | 218 | 378 | 214 | 183 | 298 |
| ENSDARG000 | 5;5;5;5;5;5;5; | 9297585:9300 | 9297933:9300 | +;+;+;+;+;+;+; | 2201 | NA | 0 | 0 | 0 | 0 | 0 | 0 |
| ENSDARG000 | 5 | 28555905 | 28559870 | - | 3966 | NA | 258 | 224 | 242 | 222 | 224 | 360 |
| ENSDARG000 | 5;5;5;5;5;5;5; | 72106837:721 | 72111421:721 | -;-;-;-;-;-;-;-; | 6822 | smyd1a | 96 | 174 | 143 | 84 | 134 | 91 |
| ENSDARG000 | 5;5;5;5;5;5;5; | 40052930:400 | 40053050:400 | -;-;-;-;-;-;-;-; | 10573 | WDFY3 | 700 | 487 | 664 | 458 | 1257 | 754 |
| ENSDARG000 | 5;5;5;5;5;5;5; | 21852714:218 | 21853376:218 | -;-;-;-;-;-;-;-; | 4216 | smarca1 | 449 | 366 | 328 | 769 | 632 | 809 |
| ENSDARG000 | 5;5;5;5;5 | 33488860:334 | 33489046:334 | +;+;+;+;+ | 665 | NA | 0 | 0 | 0 | 0 | 0 | 0 |
| ENSDARG000 | 5;5;5;5;5;5;5 | 61336781:613 | 61337093:613 | -;-;-;-;-;-;- | 1057 | NA | 0 | 0 | 0 | 0 | 0 | 0 |
| ENSDARG000 | 5;5;5;5;5 | 33498253:334 | 33498442:334 | +;+;+;+;+ | 702 | NA | 0 | 0 | 0 | 0 | 0 | 0 |
| ENSDARG000 | 5;5;5;5;5;5 | 41857450:418 | 41857508:418 | -;-;-;-;-;- | 832 | NA | 0 | 0 | 0 | 0 | 0 | 8 |
| ENSDARG000 | 5;5;5;5;5;5;5; | 65679595:656 | 65681127:656 | -;-;-;-;-;-;-;-; | 7908 | notch1b | 54 | 22 | 36 | 27 | 58 | 62 |
| ENSDARG000 | 5;5 | 1079423:1082 | 1079580:1084 | +;+ | 1733 | NA | 99 | 155 | 173 | 55 | 117 | 160 |
| ENSDARG000 | 5;5;5;5;5 | 38604877:386 | 38605401:386 | -;-;-;-;- | 1009 | znhit1 | 212 | 214 | 228 | 130 | 142 | 110 |
| ENSDARG000 | 5;5 | 9348284:9348 | 9348555:9349 | +;+ | 655 | tal2 | 4 | 0 | 2 | 10 | 0 | 0 |
| ENSDARG000 | 5;5;5;5;5;5;5; | 38596630:385 | 38597068:385 | +;+;+;+;+;+;+; | 1434 | mogat3b | 136 | 182 | 115 | 129 | 58 | 87 |
| ENSDARG000 | 5;5;5;5;5;5;5; | 23151169:231 | 23151414:231 | +;+;+;+;+;+;+; | 1636 | tbx5b | 0 | 0 | 0 | 0 | 0 | 0 |
| ENSDARG000 | 5;5;5;5;5;5;5; | 37343643:373 | 37343695:373 | +;+;+;+;+;+;+; | 3260 | klhl13 | 5 | 12 | 0 | 9 | 2 | 1 |
| ENSDARG000 | 5;5;5;5;5;5;5; | 6796291:6796 | 6796401:6796 | +;+;+;+;+;+;+; | 2128 | me2 | 644 | 637 | 594 | 346 | 872 | 547 |
| ENSDARG000 | 5;5;5;5;5;5;5; | 37379825:374 | 37380844:374 | +;+;+;+;+;+;+; | 3973 | NA | 0 | 0 | 0 | 0 | 2 | 0 |
| ENSDARG000 | 5;5;5;5;5 | 33503108:335 | 33503300:335 | +;+;+;+;+ | 651 | NA | 0 | 0 | 0 | 0 | 0 | 0 |
| ENSDARG000 | 5;5;5 | 41874958:418 | 41875074:418 | -;-;- | 623 | NA | 0 | 2 | 0 | 0 | 0 | 2 |
| ENSDARG000 | 5;5;5;5;5;5;5; | 1239657:1243 | 1240621:1243 | -;-;-;-;-;-;-;-; | 2946 | NA | 116 | 111 | 294 | 34 | 180 | 132 |
| ENSDARG000 | 5;5;5;5;5;5;5; | 23176493:231 | 23177629:231 | -;-;-;-;-;-;-;- | 1392 | NA | 1 | 1 | 2 | 0 | 1 | 0 |
| ENSDARG000 | 5;5;5;5;5;5;5; | 24063046:240 | 24063578:240 | +;+;+;+;+;+;+; | 2476 | gps2 | 480 | 484 | 605 | 429 | 580 | 511 |
| ENSDARG000 | 5;5;5;5;5;5 | 33519943:335 | 33519973:335 | +;+;+;+;+;+ | 961 | ms4a17c.1 | 6 | 4 | 0 | 0 | 11 | 0 |
| ENSDARG000 | 5;5;5;5;5;5;5; | 61349983:613 | 61350025:613 | +;+;+;+;+;+;+; | 1499 | NA | 256 | 295 | 403 | 147 | 438 | 279 |
| ENSDARG000 | 5;5;5 | 72129708:721 | 72131676:721 | -;-;- | 2579 | spra | 1516 | 2995 | 1534 | 1399 | 2201 | 1419 |
| ENSDARG000 | 5;5;5;5;5;5;5; | 52000163:520 | 52000337:520 | -;-;-;-;-;-;-;-; | 1311 | cdk7 | 321 | 264 | 530 | 332 | 422 | 286 |
| ENSDARG000 | 5;5;5;5;5;5;5; | 40200818:402 | 40201340:402 | -;-;-;-;-;-;-;- | 1578 | si:dkey-193c2 | 237 | 214 | 312 | 170 | 271 | 160 |
| ENSDARG000 | 5;5;5;5;5;5;5; | 9428876:9431 | 9429608:9431 | +;+;+;+;+;+;+; | 1925 | ugt2b6 | 421 | 130 | 5 | 163 | 413 | 168 |
| ENSDARG000 | 5;5;5;5;5;5;5; | 28560847:285 | 28562305:285 | -;-;-;-;-;-;-;-; | 7899 | tncb | 72 | 21 | 38 | 2 | 37 | 10 |
| ENSDARG000 | 5;5;5;5;5;5;5; | 19477555:194 | 19477610:194 | +;+;+;+;+;+;+; | 1509 | NA | 48 | 9 | 0 | 45 | 26 | 0 |
| ENSDARG000 | 5;5;5;5;5;5;5; | 32246872:322 | 32246909:322 | +;+;+;+;+;+;+; | 5915 | NA | 0 | 0 | 2 | 3 | 4 | 3 |
| ENSDARG000 | 5;5;5;5;5;5;5; | 40219964:402 | 40220200:402 | +;+;+;+;+;+;+; | 9993 | NA | 310 | 152 | 325 | 122 | 276 | 264 |
| ENSDARG000 | 5;5;5;5;5;5;5; | 26686279:266 | 26686676:266 | +;+;+;+;+;+;+; | 6238 | tango2 | 518 | 467 | 484 | 335 | 502 | 566 |
| ENSDARG000 | 5;5;5;5;5;5;5; | 19494198:194 | 19494486:194 | +;+;+;+;+;+;+; | 1775 | mvk | 34 | 106 | 84 | 34 | 108 | 40 |
| ENSDARG000 | 5;5;5;5;5;5;5; | 46658045:466 | 46662432:466 | -;-;-;-;-;-;-;-; | 5549 | EDIL3 | 378 | 272 | 1869 | 727 | 1311 | 94 |
| ENSDARG000 | 5;5;5;5;5;5;5; | 19489581:194 | 19489845:194 | -;-;-;-;-;-;-;-; | 945 | mmab | 110 | 208 | 149 | 85 | 148 | 151 |
| ENSDARG000 | 5;5;5;5;5;5;5; | 51947150:519 | 51950471:519 | -;-;-;-;-;-;-;-; | 4891 | serinc5 | 4162 | 2304 | 1839 | 2708 | 2438 | 2726 |
| ENSDARG000 | 5;5;5;5;5;5;5; | 9441198:9445 | 9445240:9445 | -;-;-;-;-;-;-;-; | 8237 | GAK | 2565 | 1558 | 2073 | 1517 | 3480 | 1923 |
| ENSDARG000 | 5;5 | 37504309:375 | 37505390:375 | +;+ | 4162 | NA | 350 | 306 | 403 | 298 | 356 | 374 |
| ENSDARG000 | 5;5;5;5;5;5;5; | 1278092:1278 | 1278332:1278 | +;+;+;+;+;+;+; | 5369 | dnm1a | 0 | 4 | 2 | 0 | 2 | 10 |
| ENSDARG000 | 5;5;5;5;5;5;5; | 43860564:438 | 43864585:438 | -;-;-;-;-;-;-;-; | 7023 | DAB2 | 997 | 334 | 329 | 122 | 368 | 432 |
| ENSDARG000 | 5;5;5;5;5;5;5; | 13037106:130 | 13038142:130 | -;-;-;-;-;-;-;- | 1731 | ypel1 | 703 | 131 | 192 | 177 | 364 | 251 |
| ENSDARG000 | 5;5;5;5;5;5;5; | 63629000:636 | 63629760:636 | -;-;-;-;-;-;-;- | 1543 | surf4l | 560 | 776 | 903 | 488 | 663 | 413 |
| ENSDARG000 | 5;5;5;5;5;5;5; | 57430675:574 | 57431227:574 | -;-;-;-;-;-;-;-; | 1926 | NA | 8 | 4 | 6 | 10 | 20 | 6 |
| ENSDARG000 | 5;5;5;5;5;5;5; | 43870389:438 | 43870572:438 | +;+;+;+;+;+;+; | 1700 | zgc:112966 | 0 | 1 | 0 | 0 | 0 | 18 |
| ENSDARG000 | 5;5;5;5 | 16496273:164 | 16496727:164 | +;+;+;+ | 1204 | NA | 0 | 0 | 2 | 8 | 6 | 0 |
| ENSDARG000 | 5;5;5;5;5;5;5; | 54716670:547 | 54716941:547 | -;-;-;-;-;-;-;-; | 2815 | slc30a5 | 715 | 891 | 1049 | 644 | 999 | 1029 |
| ENSDARG000 | 5;5 | 67909644:679 | 67910558:679 | -;- | 1379 | gsx1 | 0 | 0 | 0 | 0 | 0 | 0 |
| ENSDARG000 | 5;5 | 40272848:402 | 40273869:402 | -;- | 2472 | snx18a | 161 | 301 | 475 | 106 | 435 | 408 |
| ENSDARG000 | 5;5;5;5;5;5;5; | 31168096:311 | 31168213:311 | +;+;+;+;+;+;+; | 5899 | ATP2A3 | 237 | 84 | 61 | 54 | 208 | 255 |
| ENSDARG000 | 5;5;5;5;5;5;5; | 16677264:166 | 16677985:166 | -;-;-;-;-;-;-;-; | 3595 | atad1a | 296 | 199 | 381 | 242 | 287 | 227 |
| ENSDARG000 | 5;5;5;5;5;5;5; | 63648164:636 | 63648864:636 | -;-;-;-;-;-;-;-; | 1976 | ciz1b | 78 | 66 | 94 | 116 | 52 | 64 |
| ENSDARG000 | 5;5;5;5;5;5;5; | 33537789:335 | 33538314:335 | -;-;-;-;-;-;-;-; | 2383 | NA | 17 | 8 | 8 | 2 | 4 | 12 |
| ENSDARG000 | 5;5;5;5 | 67971627:679 | 67971794:679 | +;+;+;+ | 1027 | mtif3 | 113 | 157 | 302 | 174 | 197 | 200 |
| ENSDARG000 | 5;5;5;5;5;5;5; | 40294595:402 | 40294637:402 | +;+;+;+;+;+;+; | 1090 | arl15a | 0 | 0 | 0 | 6 | 0 | 0 |
| ENSDARG000 | 5;5;5;5;5;5;5; | 61361815:613 | 61362254:613 | +;+;+;+;+;+;+; | 3197 | castor2 | 1886 | 1557 | 2318 | 3323 | 2708 | 2450 |
| ENSDARG000 | 5 | 40296551 | 40297003 | - | 453 | hspb3 | 0 | 0 | 0 | 0 | 0 | 0 |
| ENSDARG000 | 5;5;5;5;5;5;5; | 6854345:6854 | 6854781:6854 | +;+;+;+;+;+;+; | 1693 | elac1 | 298 | 317 | 541 | 219 | 158 | 306 |
| ENSDARG000 | 5;5;5;5;5;5;5; | 9556943:9556 | 9557289:9557 | -;-;-;-;-;-;-;-; | 2043 | NA | 143 | 48 | 143 | 40 | 115 | 132 |
| ENSDARG000 | 5;5;5;5;5;5;5; | 72137331:721 | 72137524:721 | +;+;+;+;+;+;+; | 1390 | exosc2 | 174 | 309 | 370 | 110 | 300 | 128 |
| ENSDARG000 | 5;5;5;5;5;5;5; | 13104851:131 | 13105133:131 | -;-;-;-;-;-;-;-; | 2171 | mapk1 | 383 | 334 | 587 | 121 | 635 | 807 |
| ENSDARG000 | 5;5;5;5;5;5;5; | 67980187:679 | 67980801:679 | -;-;-;-;-;-;-;-; | 1575 | gtf3aa | 166 | 341 | 423 | 129 | 216 | 219 |
| ENSDARG000 | 5;5;5 | 47623564:476 | 47624032:476 | -;-;- | 693 | NA | 0 | 0 | 0 | 0 | 0 | 0 |
| ENSDARG000 | 5;5;5;5;5;5;5; | 23180505:231 | 23180621:231 | +;+;+;+;+;+;+; | 941 | NA | 0 | 0 | 0 | 0 | 0 | 0 |
| ENSDARG000 | 5;5;5;5;5;5;5; | 37649206:376 | 37649391:376 | +;+;+;+;+;+;+; | 7544 | SPTBN2 | 153 | 56 | 306 | 58 | 284 | 229 |
| ENSDARG000 | 5;5 | 21891305:219 | 21891376:219 | +;+ | 376 | NA | 87 | 38 | 72 | 108 | 73 | 138 |
| ENSDARG000 | 5;5;5;5;5;5;5; | 9699922:9699 | 9700129:9700 | -;-;-;-;-;-;-;-; | 2953 | NA | 1680 | 512 | 600 | 429 | 1198 | 569 |
| ENSDARG000 | 5;5;5;5;5;5;5; | 33611951:336 | 33612554:336 | +;+;+;+;+;+;+; | 2327 | ttll11 | 248 | 205 | 137 | 117 | 116 | 258 |
| ENSDARG000 | 5;5;5;5;5;5;5; | 38612134:386 | 38612180:386 | +;+;+;+;+;+;+; | 712 | si:ch211-271 | 348 | 328 | 234 | 124 | 228 | 304 |
| ENSDARG000 | 5;5;5 | 47726197:477 | 47726284:477 | -;-;- | 387 | cox7c | 1391 | 1793 | 1252 | 795 | 996 | 1156 |
| ENSDARG000 | 5;5;5;5;5;5;5; | 38619813:386 | 38619880:386 | +;+;+;+;+;+;+; | 932 | si:ch211-271 | 132 | 222 | 132 | 31 | 19 | 132 |
| ENSDARG000 | 5;5;5;5;5;5;5; | 33679559:336 | 33680312:336 | -;-;-;-;-;-;-;-; | 4648 | dab2ipb | 548 | 319 | 367 | 382 | 525 | 377 |
| ENSDARG000 | 5;5;5;5;5;5;5; | 57442271:574 | 57442397:574 | +;+;+;+;+;+;+; | 1812 | prpf4 | 318 | 441 | 513 | 255 | 315 | 325 |
| ENSDARG000 | 5;5;5;5;5;5;5; | 41878252:418 | 41880403:418 | -;-;-;-;-;-;-;-; | 9569 | ncor1 | 5479 | 4812 | 5754 | 5421 | 6164 | 5134 |
| ENSDARG000 | 5;5;5;5;5;5;5; | 38631119:386 | 38631193:386 | +;+;+;+;+;+;+; | 1148 | NA | 128 | 145 | 149 | 40 | 23 | 92 |
| ENSDARG000 | 5;5 | 57480014:574 | 57480202:574 | +;+ | 3165 | NA | 60 | 0 | 6 | 2 | 50 | 2 |
| ENSDARG000 | 5;5;5;5;5 | 33879807:338 | 33880404:338 | -;-;-;-;- | 1277 | si:dkeyp-72a4 | 0 | 0 | 0 | 0 | 0 | 10 |
| ENSDARG000 | 5;5;5;5;5;5;5; | 68059924:680 | 68066996:680 | -;-;-;-;-;-;-;-; | 7663 | spag7 | 1413 | 1407 | 1329 | 1180 | 1770 | 1524 |
| ENSDARG000 | 5;5;5;5;5;5;5; | 57453830:574 | 57454282:574 | -;-;-;-;-;-;-;-; | 1147 | rnf224 | 1 | 2 | 4 | 0 | 10 | 2 |
| ENSDARG000 | 5;5;5;5;5 | 33784860:337 | 33784978:337 | -;-;-;-;- | 880 | NA | 1 | 1 | 0 | 0 | 1 | 0 |
| ENSDARG000 | 5;5;5;5;5;5 | 21922534:219 | 21922586:219 | +;+;+;+;+;+ | 1254 | NA | 289 | 160 | 248 | 251 | 158 | 420 |
| ENSDARG000 | 5;5;5;5;5;5;5; | 6875272:6878 | 6875617:6878 | +;+;+;+;+;+;+; | 1624 | SMAD4 | 216 | 136 | 257 | 171 | 252 | 205 |
| ENSDARG000 | 5;5;5;5;5;5;5; | 16797209:168 | 16801017:168 | -;-;-;-;-;-;-;-; | 6456 | GALNT9 | 0 | 0 | 5 | 0 | 0 | 0 |
| ENSDARG000 | 5;5;5;5;5;5;5; | 1432074:1433 | 1432745:1433 | -;-;-;-;-;-;-;-; | 4015 | golga2 | 1102 | 762 | 1074 | 613 | 939 | 733 |
| ENSDARG000 | 5;5;5;5 | 36595149:365 | 36596992:365 | -;-;-;- | 2174 | rhogc | 728 | 749 | 815 | 687 | 824 | 628 |
| ENSDARG000 | 5;5;5;5;5;5;5; | 24086227:240 | 24086395:240 | +;+;+;+;+;+;+; | 2737 | tp53 | 1437 | 3261 | 3055 | 2641 | 3353 | 2186 |
| ENSDARG000 | 5;5;5;5;5;5;5; | 25366677:253 | 25366835:253 | -;-;-;-;-;-;-;-; | 1594 | rorb | 6 | 1 | 12 | 0 | 4 | 2 |

| | | | | | | | | | | | | |
|---|---|---|---|---|---|---|---|---|---|---|---|---|
| ENSDARG000 | 5;5;5;5 | 32260502:322 | 32260546:322 | +;+;+;+ | 651 | NA | 0 | 0 | 0 | 0 | 0 | 0 |
| ENSDARG000 | 5;5;5;5;5;5;5; | 32262499:322 | 32262574:322 | -;-;-;-;-;-;-;-; | 5961 | NA | 0 | 0 | 0 | 6 | 13 | 2 |
| ENSDARG000 | 5;5;5;5;5;5;5; | 23190642:231 | 23190747:231 | +;+;+;+;+;+;+; | 1185 | CFAP73 | 8 | 2 | 1 | 16 | 5 | 21 |
| ENSDARG000 | 5;5;5 | 17040621:170 | 17040889:170 | -;-;- | 445 | NA | 0 | 0 | 0 | 0 | 0 | 0 |
| ENSDARG000 | 5;5;5;5;5;5;5; | 43965078:439 | 43965787:439 | +;+;+;+;+;+;+; | 6025 | TMEM8B | 1 | 0 | 0 | 2 | 4 | 3 |
| ENSDARG000 | 5;5;5;5;5;5;5; | 9894143:9896 | 9897328:9897 | -;-;-;-;-;-;-;- | 3988 | tor4ab | 0 | 2 | 21 | 0 | 12 | 0 |
| ENSDARG000 | 5;5;5;5;5;5;5; | 17624463:176 | 17625131:176 | +;+;+;+;+;+;+; | 5248 | NA | 1147 | 1182 | 3287 | 1382 | 3081 | 2818 |
| ENSDARG000 | 5;5;5;5;5 | 44064764:440 | 44064953:440 | +;+;+;+;+ | 1181 | NA | 0 | 0 | 0 | 0 | 0 | 0 |
| ENSDARG000 | 5;5;5;5;5 | 17599870:176 | 17600518:176 | -;-;-;-;- | 1163 | NA | 2 | 2 | 0 | 4 | 2 | 3 |
| ENSDARG000 | 5;5;5;5;5;5;5; | 28755475:287 | 28755693:287 | -;-;-;-;-;-;-;-; | 1791 | TRAF2 | 157 | 49 | 233 | 89 | 53 | 140 |
| ENSDARG000 | 5;5;5;5;5;5;5; | 21939733:219 | 21941089:219 | -;-;-;-;-;-;-;-; | 3109 | NA | 36 | 13 | 10 | 22 | 48 | 32 |
| ENSDARG000 | 5;5;5;5;5;5;5; | 26728115:267 | 26728173:267 | +;+;+;+;+;+;+; | 1230 | tmem150aa | 1374 | 810 | 718 | 909 | 727 | 736 |
| ENSDARG000 | 5;5;5;5;5;5;5; | 38641933:386 | 38642112:386 | -;-;-;-;-;-;-;-; | 1583 | NA | 0 | 8 | 12 | 4 | 0 | 6 |
| ENSDARG000 | 5;5;5;5;5;5;5; | 57489253:574 | 57490553:574 | -;-;-;-;-;-;-;-; | 2683 | pisd | 545 | 577 | 980 | 724 | 627 | 614 |
| ENSDARG000 | 5;5;5;5;5;5;5; | 68076913:680 | 68077689:680 | -;-;-;-;-;-;-;-; | 1667 | slc25a11 | 1319 | 1447 | 1772 | 1369 | 1696 | 1169 |
| ENSDARG000 | 5;5;5;5;5;5;5; | 7279104:7281 | 7279111:7281 | +;+;+;+;+;+;+; | 3113 | UNC5C | 0 | 14 | 6 | 6 | 0 | 1 |
| ENSDARG000 | 5;5;5;5;5 | 40485503:404 | 40485620:404 | +;+;+;+;+ | 629 | ndufs4 | 378 | 438 | 608 | 327 | 503 | 425 |
| ENSDARG000 | 5;5;5;5;5;5;5; | 63668735:636 | 63669036:636 | +;+;+;+;+;+;+; | 3068 | dnm1b | 14 | 4 | 22 | 20 | 1 | 57 |
| ENSDARG000 | 5;5;5;5;5;5;5; | 25536616:255 | 25537565:255 | -;-;-;-;-;-;-;-; | 2282 | MFSD3 | 229 | 101 | 154 | 53 | 214 | 87 |
| ENSDARG000 | 5;5;5;5;5;5;5; | 61432541:614 | 61433531:614 | -;-;-;-;-;-;-;-; | 2133 | ncf1 | 215 | 238 | 265 | 234 | 296 | 349 |
| ENSDARG000 | 5;5;5;5;5;5;5 | 44272964:442 | 44274162:442 | -;-;-;-;-;-;- | 2016 | NA | 16 | 0 | 0 | 0 | 0 | 0 |
| ENSDARG000 | 5;5 | 19712011:197 | 19712122:197 | +;+ | 1584 | fam222aa | 44 | 18 | 57 | 3 | 25 | 0 |
| ENSDARG000 | 5;5;5;5;5;5;5; | 61404497:614 | 61404719:614 | -;-;-;-;-;-;-;-; | 1725 | rcc1l | 394 | 385 | 326 | 172 | 449 | 272 |
| ENSDARG000 | 5;5;5;5;5;5;5; | 9983350:9986 | 9983545:9986 | -;-;-;-;-;-;-;-; | 1363 | wdr31 | 3 | 14 | 24 | 4 | 24 | 38 |
| ENSDARG000 | 5;5;5;5;5 | 23196067:231 | 23196544:231 | -;-;-;-;- | 1689 | IQCD | 64 | 30 | 84 | 71 | 10 | 117 |
| ENSDARG000 | 5;5;5;5;5;5;5; | 47862636:478 | 47863529:478 | +;+;+;+;+;+;+; | 4915 | RASA1 | 1210 | 1174 | 1432 | 787 | 1456 | 1103 |
| ENSDARG000 | 5;5;5;5;5;5;5; | 10014604:100 | 10014708:100 | +;+;+;+;+;+;+; | 2868 | slc2a11b | 4760 | 4066 | 3804 | 4175 | 11571 | 5113 |
| ENSDARG000 | 5;5;5;5;5;5;5; | 40504910:405 | 40507633:405 | -;-;-;-;-;-;-;-; | 3529 | fsta | 20 | 4 | 0 | 8 | 4 | 0 |
| ENSDARG000 | 5;5;5;5;5;5;5; | 54765702:547 | 54769678:547 | -;-;-;-;-;-;-;-; | 7005 | pik3r1 | 2851 | 1946 | 3519 | 2591 | 3398 | 3449 |
| ENSDARG000 | 5;5;5;5;5;5;5; | 47963940:479 | 47964137:479 | -;-;-;-;-;-;-;-; | 1298 | ccnh | 144 | 146 | 206 | 130 | 168 | 169 |
| ENSDARG000 | 5;5;5;5;5;5;5; | 57641554:576 | 57641898:576 | +;+;+;+;+;+;+; | 2644 | cryabb | 0 | 0 | 0 | 0 | 21 | 0 |
| ENSDARG000 | 5;5;5;5;5;5;5; | 65880890:658 | 65885083:658 | -;-;-;-;-;-;-;-; | 6772 | ADAMTSL2 | 0 | 1 | 0 | 0 | 0 | 0 |
| ENSDARG000 | 5;5;5;5;5 | 57629207:576 | 57629676:576 | -;-;-;-;- | 793 | hspb2 | 0 | 0 | 0 | 0 | 0 | 0 |
| ENSDARG000 | 5;5;5;5;5;5;5; | 72145468:721 | 72145603:721 | +;+;+;+;+;+;+; | 3374 | ABL1 | 1725 | 1322 | 1994 | 1734 | 1592 | 1191 |
| ENSDARG000 | 5;5;5;5;5;5;5; | 13181029:131 | 13181687:131 | -;-;-;-;-;-;-;- | 1924 | ppm1f | 76 | 52 | 95 | 48 | 142 | 74 |
| ENSDARG000 | 5;5;5;5;5 | 38654215:386 | 38654277:386 | +;+;+;+;+ | 814 | NA | 88 | 249 | 157 | 175 | 0 | 92 |
| ENSDARG000 | 5;5;5;5;5;5;5; | 19795977:197 | 19796259:197 | -;-;-;-;-;-;-;-; | 2741 | trpv4 | 365 | 664 | 535 | 792 | 347 | 697 |
| ENSDARG000 | 5;5;5;5;5;5;5; | 28770273:287 | 28770693:287 | +;+;+;+;+;+;+; | 3295 | trafd1 | 603 | 440 | 590 | 394 | 562 | 340 |
| ENSDARG000 | 5;5;5;5;5;5;5; | 36599107:365 | 36599343:365 | +;+;+;+;+;+;+; | 2612 | micu2 | 488 | 437 | 507 | 604 | 378 | 504 |
| ENSDARG000 | 5;5;5;5;5;5;5; | 68408346:684 | 68408512:684 | -;-;-;-;-;-;-;-; | 4421 | ephb4a | 493 | 273 | 289 | 152 | 270 | 299 |
| ENSDARG000 | 5;5;5;5;5;5;5; | 40557232:405 | 40558330:405 | -;-;-;-;-;-;-;-; | 2729 | NA | 0 | 0 | 0 | 0 | 0 | 0 |
| ENSDARG000 | 5;5;5;5;5;5;5; | 23201370:232 | 23201458:232 | +;+;+;+;+;+;+; | 4039 | TPCN1 | 4666 | 5350 | 11033 | 8871 | 9871 | 10121 |
| ENSDARG000 | 5;5 | 40713349:407 | 40713898:407 | +;+ | 660 | NA | 0 | 0 | 0 | 0 | 0 | 0 |
| ENSDARG000 | 5;5;5 | 40699830:407 | 40699887:407 | +;+;+ | 925 | NA | 0 | 2 | 2 | 10 | 0 | 0 |
| ENSDARG000 | 5;5;5 | 37720777:377 | 37720978:377 | +;+;+ | 1481 | cdc42ep2 | 16 | 10 | 4 | 0 | 38 | 0 |
| ENSDARG000 | 5 | 63739863 | 63741137 | + | 1275 | NA | 0 | 10 | 2 | 8 | 0 | 16 |
| ENSDARG000 | 5;5;5;5;5;5;5; | 44346691:443 | 44346887:443 | +;+;+;+;+;+;+; | 2676 | tars1 | 2689 | 3108 | 5747 | 3576 | 2287 | 2342 |
| ENSDARG000 | 5;5;5;5;5;5;5; | 13211452:132 | 13214959:132 | -;-;-;-;-;-;-;-; | 5806 | TOP3B | 233 | 300 | 311 | 69 | 341 | 205 |
| ENSDARG000 | 5;5;5;5;5 | 38662125:386 | 38662167:386 | +;+;+;+;+ | 764 | NA | 325 | 469 | 217 | 137 | 120 | 191 |
| ENSDARG000 | 5;5;5;5;5;5;5; | 65946222:659 | 65946474:659 | +;+;+;+;+;+;+; | 2029 | mymk | 0 | 0 | 2 | 0 | 0 | 0 |
| ENSDARG000 | 5;5;5 | 44372168:443 | 44372479:443 | +;+;+ | 848 | NA | 41 | 66 | 23 | 21 | 23 | 72 |
| ENSDARG000 | 5;5;5;5;5;5 | 33940215:339 | 33941209:339 | -;-;-;-;-;- | 1562 | zgc:63972 | 87 | 16 | 64 | 20 | 56 | 97 |
| ENSDARG000 | 5;5;5;5;5;5;5; | 10055980:100 | 10056195:100 | -;-;-;-;-;-;-;-; | 1816 | chek2 | 75 | 179 | 221 | 75 | 184 | 120 |
| ENSDARG000 | 5;5;5;5;5;5;5; | 17780475:177 | 17780600:177 | +;+;+;+;+;+;+; | 2741 | chfr | 478 | 327 | 615 | 340 | 483 | 427 |
| ENSDARG000 | 5;5;5;5;5;5;5; | 10046643:100 | 10046849:100 | +;+;+;+;+;+;+; | 3379 | gstt2 | 352 | 317 | 218 | 259 | 442 | 354 |
| ENSDARG000 | 5;5;5;5;5 | 19850828:198 | 19851768:198 | -;-;-;-;- | 1463 | gltpa | 131 | 252 | 212 | 170 | 142 | 316 |
| ENSDARG000 | 5;5;5;5;5;5;5; | 7488064:7488 | 7488575:7488 | -;-;-;-;-;-;-;-; | 2277 | bmpr1ba | 8 | 17 | 6 | 2 | 22 | 0 |
| ENSDARG000 | 5;5;5;5;5;5;5; | 25578181:255 | 25578366:255 | -;-;-;-;-;-;-;-; | 1642 | anxa1a | 18998 | 12319 | 3900 | 20249 | 10544 | 21477 |
| ENSDARG000 | 5;5;5;5 | 63749368:637 | 63749498:637 | +;+;+;+ | 526 | NA | 21 | 23 | 22 | 4 | 38 | 87 |
| ENSDARG000 | 5;5;5;5;5;5;5; | 65956918:659 | 65957081:659 | -;-;-;-;-;-;-;-; | 2212 | NA | 2 | 0 | 2 | 0 | 0 | 0 |
| ENSDARG000 | 5;5;5;5;5;5;5; | 28797771:287 | 28798661:287 | +;+;+;+;+;+;+; | 3148 | NA | 0 | 0 | 6 | 0 | 0 | 0 |
| ENSDARG000 | 5;5;5;5 | 38673168:386 | 38673488:386 | +;+;+;+ | 1060 | NA | 33 | 38 | 19 | 10 | 28 | 34 |
| ENSDARG000 | 5;5;5 | 23242370:232 | 23242600:232 | +;+;+ | 1875 | agtr2 | 116 | 12 | 8 | 7 | 22 | 20 |
| ENSDARG000 | 5;5;5;5 | 22021142:220 | 22021904:220 | -;-;-;- | 1583 | asb12a | 609 | 735 | 638 | 789 | 693 | 652 |
| ENSDARG000 | 5;5;5;5;5;5;5; | 37744625:377 | 37744686:377 | +;+;+;+;+;+;+; | 2375 | dpf2 | 221 | 291 | 359 | 294 | 339 | 193 |
| ENSDARG000 | 5;5;5;5;5;5;5; | 57714902:577 | 57715686:577 | +;+;+;+;+;+;+; | 1924 | ufc1 | 417 | 600 | 518 | 248 | 396 | 352 |
| ENSDARG000 | 5;5;5;5;5;5 | 40728613:407 | 40729756:407 | -;-;-;-;-;- | 2303 | isl1a | 0 | 0 | 0 | 0 | 0 | 4 |
| ENSDARG000 | 5;5;5;5;5;5;5; | 63767376:637 | 63767448:637 | +;+;+;+;+;+;+; | 6905 | NA | 495 | 400 | 356 | 157 | 175 | 1748 |
| ENSDARG000 | 5;5;5;5;5;5;5; | 57693472:576 | 57694038:576 | -;-;-;-;-;-;-;-; | 1584 | si:dkev-71l1.1 | 6 | 10 | 11 | 5 | 16 | 0 |
| ENSDARG000 | 5;5;5;5;5;5;5; | 26756021:267 | 26756950:267 | -;-;-;-;-;-;-;-; | 1749 | rnf181 | 401 | 533 | 569 | 727 | 430 | 541 |
| ENSDARG000 | 5;5;5;5;5;5;5; | 23256187:232 | 23256241:232 | +;+;+;+;+;+;+; | 2957 | NA | 22 | 2 | 0 | 0 | 12 | 4 |
| ENSDARG000 | 5;5;5;5;5;5;5 | 40835601:408 | 40835839:408 | +;+;+;+;+;+;+ | 2269 | si:dkey-3h3.3 | 142 | 226 | 254 | 122 | 207 | 194 |
| ENSDARG000 | 5;5;5;5;5;5;5; | 38679623:386 | 38679671:386 | +;+;+;+;+;+;+; | 1589 | si:dkev-58f10. | 19 | 84 | 45 | 19 | 7 | 55 |
| ENSDARG000 | 5;5;5;5;5;5;5; | 72264192:722 | 72264863:722 | -;-;-;-;-;-;-;-; | 2167 | tbx5a | 0 | 0 | 0 | 0 | 0 | 0 |
| ENSDARG000 | 5;5;5;5;5;5 | 10084100:100 | 10084453:100 | +;+;+;+;+;+ | 963 | hscb | 48 | 60 | 116 | 40 | 48 | 34 |
| ENSDARG000 | 5;5;5;5;5;5;5; | 7722017:7722 | 7722711:7723 | -;-;-;-;-;-;-;-; | 6231 | pdlim5a | 4 | 4 | 0 | 0 | 0 | 0 |
| ENSDARG000 | 5;5;5;5;5;5;5; | 38679623:386 | 38679671:386 | +;+;+;+;+;+;+; | 1465 | si:dkev-58f10. | 353 | 619 | 503 | 211 | 132 | 359 |
| ENSDARG000 | 5;5 | 38697496:387 | 38698108:387 | -;- | 841 | NA | 0 | 0 | 0 | 0 | 0 | 0 |
| ENSDARG000 | 5;5 | 17848665:178 | 17849536:178 | -;- | 949 | NA | 2 | 0 | 0 | 0 | 0 | 0 |
| ENSDARG000 | 5;5;5;5;5;5;5; | 68765883:687 | 68767818:687 | -;-;-;-;-;-;-;-; | 5654 | mepcea | 634 | 416 | 771 | 390 | 612 | 573 |
| ENSDARG000 | 5;5;5;5;5;5;5; | 1911814:1911 | 1911858:1911 | +;+;+;+;+;+;+; | 1972 | si:ch73-55i23 | 54 | 9 | 46 | 10 | 98 | 97 |
| ENSDARG000 | 5;5;5;5;5;5;5; | 10232193:102 | 10233331:102 | -;-;-;-;-;-;-;-; | 2578 | si:ch73-42k18 | 2 | 0 | 0 | 0 | 2 | 0 |
| ENSDARG000 | 5;5;5;5;5;5 | 28812453:288 | 28812497:288 | +;+;+;+;+;+ | 1477 | zgc:174260 | 0 | 0 | 10 | 0 | 0 | 0 |
| ENSDARG000 | 5;5;5;5;5;5;5; | 31283576:313 | 31284182:313 | +;+;+;+;+;+;+; | 3208 | camkk1a | 32 | 63 | 91 | 39 | 46 | 65 |
| ENSDARG000 | 5;5 | 10265648:102 | 10265861:102 | +;+ | 486 | NA | 0 | 0 | 0 | 0 | 0 | 0 |
| ENSDARG000 | 5;5;5;5;5;5;5; | 29820266:298 | 29820489:298 | +;+;+;+;+;+;+; | 1428 | f11r.2 | 12 | 0 | 0 | 0 | 0 | 2 |
| ENSDARG000 | 5;5;5;5;5;5;5; | 40844968:408 | 40846244:408 | -;-;-;-;-;-;-;-; | 4122 | parp8 | 60 | 140 | 261 | 116 | 427 | 175 |
| ENSDARG000 | 5;5;5;5;5;5;5 | 72316803:723 | 72318564:723 | -;-;-;-;-;-;- | 3872 | tbx3a | 38 | 0 | 0 | 0 | 14 | 0 |
| ENSDARG000 | 5;5;5;5;5;5;5; | 7853463:7861 | 7853675:7861 | -;-;-;-;-;-;-;-; | 2221 | opn4xb | 531 | 55 | 298 | 123 | 16 | 815 |
| ENSDARG000 | 5;5;5;5;5;5;5; | 19867794:198 | 19868243:198 | +;+;+;+;+;+;+; | 2043 | tchp | 174 | 140 | 110 | 130 | 116 | 100 |
| ENSDARG000 | 5;5;5;5;5;5;5; | 17881074:178 | 17882583:178 | -;-;-;-;-;-;-;-; | 3914 | zdhhc8b | 118 | 176 | 228 | 74 | 143 | 218 |
| ENSDARG000 | 5;5;5;5 | 28824612:288 | 28824962:288 | +;+;+;+ | 831 | NA | 0 | 0 | 0 | 0 | 0 | 0 |
| ENSDARG000 | 5;5;5;5;5;5;5; | 34066591:340 | 34066859:340 | -;-;-;-;-;-;-;-; | 1864 | fibcd1b | 2 | 61 | 52 | 73 | 10 | 18 |
| ENSDARG000 | 5;5 | 10336135:103 | 10336412:103 | +;+ | 336 | NA | 0 | 0 | 0 | 0 | 0 | 0 |
| ENSDARG000 | 5;5;5;5;5;5;5; | 65970235:659 | 65970468:659 | +;+;+;+;+;+;+; | 1917 | slc2a8 | 322 | 234 | 319 | 173 | 237 | 360 |
| ENSDARG000 | 5;5;5;5;5;5;5; | 26765241:267 | 26765310:267 | +;+;+;+;+;+;+; | 2776 | NA | 217 | 115 | 157 | 374 | 120 | 280 |
| ENSDARG000 | 5;5 | 7989210:7995 | 7989282:7995 | +;+ | 495 | GDNF | 0 | 0 | 0 | 0 | 0 | 0 |
| ENSDARG000 | 5;5;5;5;5;5;5; | 37785152:377 | 37785409:377 | +;+;+;+;+;+;+; | 3823 | myo1ca | 1992 | 2210 | 1977 | 4817 | 1574 | 3650 |
| ENSDARG000 | 5;5;5;5 | 61459422:614 | 61459528:614 | +;+;+;+ | 631 | polr2j | 233 | 301 | 362 | 282 | 299 | 244 |
| ENSDARG000 | 5;5;5;5;5 | 54938634:549 | 54938996:549 | +;+;+;+;+ | 2279 | cd180 | 26 | 0 | 14 | 0 | 50 | 8 |
| ENSDARG000 | 5;5 | 47977617:480 | 47977673:480 | +;+ | 690 | NA | 0 | 0 | 0 | 0 | 0 | 0 |
| ENSDARG000 | 5;5;5;5;5;5;5; | 13353866:133 | 13353994:133 | +;+;+;+;+;+;+; | 1733 | ccl19a.1 | 63 | 10 | 10 | 0 | 52 | 24 |
| ENSDARG000 | 5;5;5;5;5;5;5; | 13308139:133 | 13308243:133 | +;+;+;+;+;+;+; | 1062 | ydjc | 130 | 135 | 177 | 55 | 74 | 111 |
| ENSDARG000 | 5;5;5;5;5;5;5; | 63913310:639 | 63913481:639 | -;-;-;-;-;-;-;-; | 5118 | pappab | 12 | 0 | 0 | 0 | 0 | 8 |
| ENSDARG000 | 5;5;5 | 13308531:133 | 13309575:133 | -;-;- | 1513 | sdf2l1 | 278 | 757 | 893 | 206 | 620 | 310 |
| ENSDARG000 | 5;5 | 57722263:577 | 57723149:577 | -;- | 1161 | gig2p | 106 | 89 | 45 | 61 | 55 | 70 |
| ENSDARG000 | 5;5;5;5;5;5;5; | 54945021:549 | 54945087:549 | -;-;-;-;-;-;-;-; | 3954 | NA | 126 | 113 | 124 | 75 | 175 | 104 |
| ENSDARG000 | 5;5;5;5;5;5;5; | 13337507:133 | 13339259:133 | -;-;-;-;-;-;-;-; | 2455 | ube2l3b | 3691 | 3960 | 3753 | 2207 | 3923 | 2866 |
| ENSDARG000 | 5;5;5 | 38720140:387 | 38720235:387 | +;+;+ | 491 | NA | 0 | 27 | 10 | 16 | 0 | 10 |
| ENSDARG000 | 5;5;5;5;5;5;5; | 28830388:288 | 28830448:288 | +;+;+;+;+;+;+; | 1502 | serpina7 | 0 | 0 | 32 | 0 | 0 | 0 |
| ENSDARG000 | 5;5;5;5;5;5;5; | 40931658:409 | 40931824:409 | -;-;-;-;-;-;-;-; | 7360 | NA | 75 | 118 | 563 | 71 | 251 | 24 |
| ENSDARG000 | 5;5;5;5;5;5 | 61467618:614 | 61467757:614 | +;+;+;+;+;+ | 957 | alkbh4 | 116 | 99 | 82 | 78 | 76 | 102 |
| ENSDARG000 | 5;5;5;5;5;5;5; | 22027588:220 | 22028959:220 | -;-;-;-;-;-;-;-; | 3569 | mtmr8 | 362 | 272 | 476 | 429 | 442 | 563 |
| ENSDARG000 | 5;5;5;5;5;5;5; | 34407763:344 | 34408132:344 | +;+;+;+;+;+;+; | 5075 | NA | 57 | 14 | 44 | 4 | 30 | 14 |
| ENSDARG000 | 5;5;5;5;5;5;5 | 10488937:104 | 10489795:104 | -;-;-;-;-;-;- | 1492 | NA | 11 | 1 | 0 | 0 | 4 | 0 |
| ENSDARG000 | 5;5;5;5;5;5;5; | 1950660:1952 | 1950872:1952 | -;-;-;-;-;-;-;-; | 1173 | rplp0 | 44691 | 40380 | 71595 | 59555 | 56537 | 62325 |
| ENSDARG000 | 5;5;5;5;5 | 57726425:577 | 57726755:577 | +;+;+;+;+ | 1905 | fdxacb1 | 110 | 24 | 239 | 83 | 253 | 98 |
| ENSDARG000 | 5;5;5;5 | 68817034:688 | 68817156:688 | -;-;-;- | 495 | C5H2orf68 | 50 | 55 | 35 | 4 | 39 | 20 |
| ENSDARG000 | 5;5;5;5 | 68788658:687 | 68790137:687 | -;-;-;- | 1780 | her1 | 14 | 2 | 4 | 0 | 10 | 0 |
| ENSDARG000 | 5;5;5;5;5;5;5 | 65991152:659 | 65991293:659 | +;+;+;+;+;+;+ | 1142 | lcn15 | 39 | 10 | 6 | 0 | 4 | 6 |
| ENSDARG000 | 5;5;5;5;5 | 68807170:688 | 68807341:688 | +;+;+;+;+ | 1221 | her7 | 14 | 2 | 8 | 18 | 0 | 2 |
| ENSDARG000 | 5;5;5;5;5;5 | 37837245:378 | 37837367:378 | +;+;+;+;+;+ | 890 | epd | 0 | 0 | 0 | 0 | 0 | 146 |
| ENSDARG000 | 5;5;5;5;5;5;5; | 24102499:241 | 24103040:241 | -;-;-;-;-;-;-;-; | 1376 | NA | 0 | 0 | 0 | 0 | 0 | 0 |
| ENSDARG000 | 5;5;5;5;5;5;5 | 41996889:419 | 41997246:419 | +;+;+;+;+;+;+ | 1550 | pigl | 203 | 156 | 121 | 100 | 183 | 183 |
| ENSDARG000 | 5;5;5;5;5;5;5; | 28848870:288 | 28848949:288 | +;+;+;+;+;+;+; | 1507 | zgc:174259 | 0 | 0 | 8 | 0 | 0 | 0 |
| ENSDARG000 | 5;5;5;5 | 38752287:387 | 38752460:387 | +;+;+;+ | 780 | cxcl11.8 | 2 | 0 | 0 | 0 | 0 | 0 |
| ENSDARG000 | 5;5;5;5;5;5;5; | 42070695:420 | 42070912:420 | -;-;-;-;-;-;-;- | 699 | cxcl11.6 | 0 | 0 | 0 | 0 | 7 | 1 |
| ENSDARG000 | 5;5;5;5;5;5;5; | 38743644:387 | 38743684:387 | +;+;+;+;+;+;+; | 1666 | NA | 154 | 134 | 85 | 45 | 140 | 123 |
| ENSDARG000 | 5;5;5;5 | 42076672:420 | 42077004:420 | -;-;-;- | 608 | cxcl11.6 | 4 | 0 | 0 | 0 | 7 | 0 |
| ENSDARG000 | 5;5;5;5;5;5;5; | 42022826:420 | 42022853:420 | -;-;-;-;-;-;-;-; | 1179 | CENPV | 48 | 31 | 23 | 27 | 32 | 11 |
| ENSDARG000 | 5;5;5;5 | 55119510:551 | 55120182:551 | +;+;+;+ | 1116 | NA | 0 | 0 | 0 | 0 | 2 | 14 |
| ENSDARG000 | 5;5;5;5;5;5;5; | 38726854:387 | 38727038:387 | +;+;+;+;+;+;+; | 1286 | si:dkev-58f10. | 94 | 235 | 157 | 131 | 6 | 90 |
| ENSDARG000 | 5;5 | 42064144:420 | 42064249:420 | +;+ | 1650 | UBB | 4724 | 5518 | 4103 | 4650 | 2766 | 2911 |
| ENSDARG000 | 5;5;5;5;5;5;5; | 52012883:520 | 52013216:520 | +;+;+;+;+;+;+; | 990 | NA | 0 | 0 | 0 | 6 | 0 | 7 |
| ENSDARG000 | 5;5;5;5;5;5;5; | 19886055:198 | 19886639:198 | -;-;-;-;-;-;-;-; | 2938 | git2a | 370 | 372 | 797 | 554 | 394 | 489 |
| ENSDARG000 | 5;5;5;5;5;5;5; | 29831195:298 | 29831226:298 | +;+;+;+;+;+;+; | 1886 | f11r.1 | 254 | 34 | 20 | 36 | 93 | 96 |
| ENSDARG000 | 5;5;5;5;5;5;5; | 28857969:288 | 28858013:288 | +;+;+;+;+;+;+; | 1322 | NA | 0 | 1 | 3 | 0 | 0 | 0 |
| ENSDARG000 | 5;5;5;5;5;5;5; | 55160348:551 | 55160717:551 | -;-;-;-;-;-;-;-; | 1879 | srek1 | 215 | 113 | 221 | 262 | 249 | 348 |
| ENSDARG000 | 5;5;5;5;5;5;5; | 61475346:614 | 61475463:614 | +;+;+;+;+;+;+; | 2778 | lrwd1 | 74 | 172 | 268 | 96 | 138 | 118 |
| ENSDARG000 | 5;5;5;5;5;5;5; | 36611128:366 | 36611363:366 | +;+;+;+;+;+;+; | 2056 | nova1 | 394 | 220 | 259 | 190 | 505 | 141 |
| ENSDARG000 | 5;5;5;5;5;5 | 26786969:267 | 26788015:267 | -;-;-;-;-;- | 1861 | ccdc117 | 480 | 416 | 372 | 447 | 328 | 452 |
| ENSDARG000 | 5;5;5;5;5;5;5; | 31439390:314 | 31442189:314 | -;-;-;-;-;-;-;-; | 4617 | p2rx1 | 0 | 0 | 0 | 0 | 20 | 16 |
| ENSDARG000 | 5;5;5 | 1997765:1997 | 1998117:1999 | -;-;- | 1724 | nupr1a | 1142 | 975 | 890 | 1632 | 1024 | 1880 |
| ENSDARG000 | 5;5;5;5;5;5;5; | 48027981:480 | 48030100:480 | -;-;-;-;-;-;-;-; | 3502 | tmem161b | 264 | 265 | 380 | 159 | 368 | 307 |
| ENSDARG000 | 5;5 | 10507826:105 | 10507949:105 | -;- | 300 | NA | 0 | 0 | 0 | 0 | 0 | 0 |
| ENSDARG000 | 5;5 | 10511958:105 | 10512015:105 | +;+ | 309 | NA | 0 | 0 | 0 | 0 | 0 | 0 |
| ENSDARG000 | 5;5;5;5;5;5 | 2002804:2002 | 2003044:2003 | +;+;+;+;+;+ | 1508 | vkorc1l1 | 440 | 380 | 501 | 560 | 491 | 364 |
| ENSDARG000 | 5 | 64167620 | 64168415 | - | 796 | cmlc1 | 0 | 0 | 0 | 0 | 0 | 0 |
| ENSDARG000 | 5;5;5;5;5;5 | 66025336:660 | 66028560:660 | -;-;-;-;-;- | 3311 | nrarpb | 98 | 18 | 40 | 30 | 28 | 55 |
| ENSDARG000 | 5;5;5;5;5;5;5; | 1965296:1968 | 1965625:1968 | +;+;+;+;+;+;+; | 4243 | dhx33 | 44 | 70 | 109 | 38 | 48 | 42 |
| ENSDARG000 | 5;5;5;5;5;5;5; | 24116428:241 | 24116962:241 | -;-;-;-;-;-;-;-; | 1706 | capga | 0 | 44 | 82 | 24 | 68 | 14 |
| ENSDARG000 | 5;5;5;5;5;5;5 | 37840914:378 | 37841117:378 | +;+;+;+;+;+;+ | 2881 | panx1b | 2 | 0 | 4 | 0 | 14 | 18 |
| ENSDARG000 | 5;5 | 1983570:1986 | 1983643:1986 | +;+ | 338 | NA | 0 | 0 | 0 | 0 | 0 | 0 |
| ENSDARG000 | 5;5;5;5;5;5;5 | 41082534:410 | 41086842:410 | -;-;-;-;-;-;- | 5047 | golph3a | 1464 | 1007 | 1266 | 877 | 1937 | 1473 |
| ENSDARG000 | 5;5;5;5;5;5 | 10719302:107 | 10719519:107 | +;+;+;+;+;+ | 784 | NA | 0 | 0 | 1 | 0 | 0 | 0 |
| ENSDARG000 | 5;5;5;5;5;5;5; | 57743815:577 | 57743985:577 | +;+;+;+;+;+;+; | 2548 | alg9 | 724 | 659 | 662 | 342 | 684 | 578 |
| ENSDARG000 | 5;5;5 | 10720894:107 | 10721446:107 | -;-;- | 869 | NA | 0 | 0 | 3 | 0 | 0 | 0 |
| ENSDARG000 | 5;5;5;5;5;5;5 | 52039067:520 | 52039546:520 | +;+;+;+;+;+;+ | 5089 | SETBP1 | 124 | 106 | 217 | 84 | 187 | 210 |
| ENSDARG000 | 5;5;5;5;5;5;5; | 36654817:366 | 36654958:366 | +;+;+;+;+;+;+; | 1809 | capns1a | 1582 | 1600 | 1823 | 1924 | 1484 | 2032 |
| ENSDARG000 | 5;5;5;5;5;5;5; | 64177484:641 | 64177867:641 | -;-;-;-;-;-;-;- | 1638 | ak5l | 10 | 0 | 14 | 8 | 6 | 7 |
| ENSDARG000 | 5;5;5;5;5;5;5; | 52055227:520 | 52055343:520 | +;+;+;+;+;+;+; | 913 | NA | 0 | 0 | 5 | 2 | 2 | 4 |
| ENSDARG000 | 5;5 | 31548312:315 | 31548459:315 | +;+ | 185 | NA | 0 | 0 | 0 | 0 | 0 | 0 |

| | | | | | | | | | | | | |
|---|---|---|---|---|---|---|---|---|---|---|---|---|
| ENSDARG000 | 5;5;5;5;5;5;5; | 29851433:298 | 29851560:298 | +;+;+;+;+;+;+; | 1932 | UBASH3B | 561 | 431 | 712 | 413 | 626 | 470 |
| ENSDARG000 | 5;5;5;5;5;5 | 31480342:315 | 31480997:315 | +;+;+;+;+;+ | 2112 | NA | 173 | 40 | 41 | 20 | 58 | 50 |
| ENSDARG000 | 5;5;5;5;5 | 10747294:107 | 10747450:107 | +;+;+;+;+ | 1368 | NA | 0 | 0 | 2 | 0 | 0 | 0 |
| ENSDARG000 | 5;5;5;5;5;5;5; | 44654535:446 | 44655270:446 | +;+;+;+;+;+;+; | 7453 | dapk1 | 364 | 439 | 294 | 202 | 538 | 428 |
| ENSDARG000 | 5;5;5;5;5;5;5; | 23268121:232 | 23269011:232 | -;-;-;-;-;-;-;-; | 2150 | plp1b | 14 | 0 | 14 | 0 | 112 | 70 |
| ENSDARG000 | 5;5;5;5;5;5;5; | 18012154:180 | 18012227:180 | +;+;+;+;+;+;+; | 2907 | ascc2 | 425 | 401 | 491 | 221 | 450 | 432 |
| ENSDARG000 | 5;5;5;5;5;5;5; | 28870372:288 | 28873179:288 | -;-;-;-;-;-;-;-;- | 3891 | NA | 90 | 108 | 510 | 188 | 426 | 83 |
| ENSDARG000 | 5;5;5;5;5;5;5; | 26795465:267 | 26795765:267 | +;+;+;+;+;+;+; | 2114 | tcn2 | 914 | 1092 | 1566 | 3617 | 1579 | 3933 |
| ENSDARG000 | 5;5;5;5 | 36666715:366 | 36666798:366 | +;+;+;+ | 2587 | zgc:153990 | 403 | 445 | 461 | 268 | 541 | 338 |
| ENSDARG000 | 5;5 | 22067570:220 | 22067935:220 | +;+ | 1108 | shisa2a | 0 | 2 | 0 | 0 | 42 | 4 |
| ENSDARG000 | 5;5;5;5;5;5;5; | 36650064:366 | 36650331:366 | +;+;+;+;+;+;+; | 1561 | alkbh6 | 90 | 122 | 144 | 81 | 172 | 87 |
| ENSDARG000 | 5;5;5;5;5;5;5; | 26824303:268 | 26824308:268 | -;-;-;-;-;-;-;-; | 978 | si:ch211-102 | 207 | 123 | 233 | 187 | 253 | 320 |
| ENSDARG000 | 5;5;5;5;5;5;5; | 26805608:268 | 26806270:268 | +;+;+;+;+;+;+; | 3283 | RABL6 | 16200 | 17627 | 13699 | 21896 | 13371 | 23320 |
| ENSDARG000 | 5;5;5;5;5;5;5; | 8005911:8009 | 8007036:8009 | -;-;-;-;-;-;-;-; | 8317 | nipbla | 1120 | 803 | 1286 | 945 | 1213 | 1064 |
| ENSDARG000 | 5;5;5;5 | 10765611:107 | 10768258:107 | -;-;-;- | 3275 | rtn4r | 0 | 4 | 0 | 0 | 0 | 0 |
| ENSDARG000 | 5;5;5;5;5;5 | 42081884:420 | 42082337:420 | -;-;-;-;-;- | 1284 | NA | 0 | 14 | 0 | 0 | 6 | 2 |
| ENSDARG000 | 5;5;5;5;5 | 42092227:420 | 42092297:420 | +;+;+;+;+ | 2079 | ubb | 3858 | 7318 | 2298 | 2720 | 2213 | 3910 |
| ENSDARG000 | 5;5;5;5;5;5;5; | 68826514:688 | 68826657:688 | +;+;+;+;+;+;+; | 1856 | usp39 | 299 | 454 | 464 | 327 | 379 | 378 |
| ENSDARG000 | 5;5;5;5;5 | 64203526:642 | 64208106:642 | -;-;-;-;- | 5571 | NA | 38 | 42 | 445 | 24 | 338 | 135 |
| ENSDARG000 | 5;5;5;5;5;5;5; | 32281815:322 | 32281914:322 | -;-;-;-;-;-;-;-; | 6018 | myhz1.2 | 4 | 0 | 2 | 0 | 12 | 0 |
| ENSDARG000 | 5;5;5;5;5;5;5; | 55221593:552 | 55221703:552 | +;+;+;+;+;+;+; | 3386 | tmc2a | 0 | 0 | 12 | 0 | 4 | 0 |
| ENSDARG000 | 5;5;5;5 | 31605322:316 | 31605526:316 | +;+;+;+ | 1100 | si:dkev-220k2 | 11 | 21 | 0 | 8 | 26 | 22 |
| ENSDARG000 | 5;5;5;5;5;5;5; | 2066262:2074 | 2069856:2074 | -;-;-;-;-;-;-;-; | 5381 | GUSB | 5902 | 1467 | 1890 | 2257 | 6807 | 2056 |
| ENSDARG000 | 5;5 | 64349580:643 | 64350108:643 | -;- | 789 | fam78aa | 2 | 0 | 16 | 22 | 12 | 0 |
| ENSDARG000 | 5;5;5;5;5 | 64356637:643 | 64356716:643 | +;+;+;+;+ | 2201 | NA | 0 | 0 | 0 | 0 | 0 | 0 |
| ENSDARG000 | 5;5;5;5;5 | 37854685:378 | 37854731:378 | +;+;+;+;+ | 485 | ins | 0 | 0 | 0 | 0 | 0 | 0 |
| ENSDARG000 | 5;5;5;5;5;5;5; | 66043820:660 | 66044460:660 | +;+;+;+;+;+;+; | 7957 | jak2b | 882 | 702 | 1050 | 831 | 906 | 768 |
| ENSDARG000 | 5;5;5;5;5;5;5; | 19933356:199 | 19933657:199 | +;+;+;+;+;+;+; | 2066 | ankrd13a | 137 | 169 | 134 | 84 | 152 | 189 |
| ENSDARG000 | 5;5;5;5;5;5;5; | 42097608:420 | 42097773:420 | +;+;+;+;+;+;+; | 2337 | p2rx5 | 3 | 0 | 2 | 0 | 6 | 1 |
| ENSDARG000 | 5;5;5;5;5;5;5; | 52167986:521 | 52168101:521 | +;+;+;+;+;+;+; | 2702 | slc14a2 | 16 | 0 | 0 | 0 | 4 | 38 |
| ENSDARG000 | 5;5;5;5;5;5;5; | 41105602:411 | 41106756:411 | -;-;-;-;-;-;-;-; | 2868 | mtmr12 | 962 | 844 | 534 | 978 | 577 | 2531 |
| ENSDARG000 | 5;5;5;5;5;5;5; | 37864705:378 | 37867500:378 | -;-;-;-;-;-;-;-; | 8331 | NA | 1005 | 938 | 1124 | 622 | 1780 | 1265 |
| ENSDARG000 | 5;5;5;5;5;5;5; | 24128908:241 | 24129132:241 | +;+;+;+;+;+;+; | 4123 | si:ch211-114 | 2730 | 2357 | 3899 | 3292 | 3663 | 4090 |
| ENSDARG000 | 5;5;5;5;5;5;5; | 34549365:345 | 34549555:345 | +;+;+;+;+;+;+; | 2513 | aif1l | 184 | 232 | 272 | 470 | 220 | 302 |
| ENSDARG000 | 5;5;5;5;5;5;5; | 41124098:411 | 41126343:411 | -;-;-;-;-;-;-;-; | 5354 | zfr | 3534 | 2996 | 3076 | 2072 | 3205 | 3015 |
| ENSDARG000 | 5;5;5;5;5;5;5 | 22073710:220 | 22074687:220 | -;-;-;-;-;-;- | 2052 | zc4h2 | 287 | 281 | 424 | 302 | 370 | 358 |
| ENSDARG000 | 5;5;5;5;5;5;5; | 11724428:117 | 11727932:117 | -;-;-;-;-;-;-;-; | 6173 | nf2a | 30 | 42 | 99 | 79 | 106 | 56 |
| ENSDARG000 | 5;5;5;5;5;5;5; | 61508418:615 | 61508778:615 | +;+;+;+;+;+;+; | 2831 | bckdk | 1074 | 2917 | 2770 | 2551 | 1979 | 2081 |
| ENSDARG000 | 5;5;5;5;5;5;5; | 38771779:387 | 38772533:387 | -;-;-;-;-;-;-;-; | 1443 | NA | 1 | 0 | 0 | 0 | 0 | 4 |
| ENSDARG000 | 5;5;5;5;5;5;5; | 2815021:2815 | 2815067:2815 | +;+;+;+;+;+;+; | 1962 | hpda | 0 | 0 | 112 | 2 | 0 | 0 |
| ENSDARG000 | 5;5;5 | 61556172:615 | 61556240:615 | +;+;+ | 878 | orai2 | 733 | 982 | 1511 | 1481 | 339 | 711 |
| ENSDARG000 | 5;5;5;5;5;5;5 | 30081241:300 | 30081750:300 | -;-;-;-;-;-;- | 933 | sdhda | 44 | 38 | 86 | 64 | 88 | 42 |
| ENSDARG000 | 5;5;5;5 | 52472744:524 | 52472889:524 | -;-;-;- | 372 | NA | 0 | 0 | 1 | 0 | 0 | 0 |
| ENSDARG000 | 5;5;5;5;5;5;5; | 30052714:300 | 30057201:300 | -;-;-;-;-;-;-;-; | 6785 | BCO2 | 886 | 705 | 1313 | 868 | 816 | 1065 |
| ENSDARG000 | 5;5;5;5;5;5;5; | 31640881:316 | 31645250:316 | -;-;-;-;-;-;-;-; | 6174 | sh3glb2b | 2549 | 1989 | 2119 | 3111 | 1707 | 2628 |
| ENSDARG000 | 5;5 | 42122810:421 | 42123532:421 | -;- | 904 | emc6 | 643 | 954 | 977 | 480 | 866 | 629 |
| ENSDARG000 | 5;5;5;5;5;5;5; | 48110155:481 | 48110462:481 | -;-;-;-;-;-;-;-; | 1533 | NA | 0 | 10 | 0 | 0 | 6 | 0 |
| ENSDARG000 | 5;5 | 64368770:643 | 64369318:643 | +;+ | 2709 | plpp7a | 15 | 12 | 27 | 18 | 12 | 15 |
| ENSDARG000 | 5;5 | 19961988:199 | 19963181:199 | -;- | 1351 | NA | 20 | 8 | 30 | 6 | 8 | 6 |
| ENSDARG000 | 5;5;5;5;5;5;5; | 42124706:421 | 42124923:421 | +;+;+;+;+;+;+; | 1505 | SHPK | 54 | 57 | 104 | 57 | 92 | 30 |
| ENSDARG000 | 5;5;5;5;5;5;5; | 68845048:688 | 68845896:688 | -;-;-;-;-;-;-;-; | 1822 | sftpbb | 208 | 205 | 184 | 163 | 331 | 355 |
| ENSDARG000 | 5;5 | 52502966:525 | 52504739:525 | -;- | 1990 | MYORG | 0 | 0 | 0 | 0 | 0 | 0 |
| ENSDARG000 | 5;5;5 | 22087656:220 | 22087916:220 | -;-;- | 645 | NA | 2 | 0 | 0 | 0 | 0 | 0 |
| ENSDARG000 | 5;5;5;5;5;5;5; | 57773222:577 | 57773377:577 | +;+;+;+;+;+;+; | 2883 | ppp2r1ba | 5605 | 5906 | 7180 | 5223 | 6761 | 5072 |
| ENSDARG000 | 5;5;5;5;5;5;5; | 52509108:525 | 52509169:525 | -;-;-;-;-;-;-;-; | 2077 | mamdc2a | 32 | 12 | 41 | 12 | 54 | 0 |
| ENSDARG000 | 5;5;5 | 25585869:255 | 25585961:255 | +;+;+ | 364 | NA | 12 | 12 | 0 | 7 | 8 | 5 |
| ENSDARG000 | 5;5;5;5;5;5;5; | 11812080:118 | 11812319:118 | +;+;+;+;+;+;+; | 1989 | FBXO21 | 132 | 102 | 155 | 112 | 124 | 151 |
| ENSDARG000 | 5;5;5;5 | 25586321:255 | 25586344:255 | -;-;-;- | 364 | NA | 1 | 0 | 4 | 1 | 0 | 0 |
| ENSDARG000 | 5;5;5;5;5;5;5; | 25603761:256 | 25607545:256 | -;-;-;-;-;-;-;-; | 6018 | cyp1d1 | 81 | 23 | 21 | 30 | 20 | 30 |
| ENSDARG000 | 5;5;5;5;5 | 22098591:221 | 22099155:221 | +;+;+;+;+ | 2516 | ZC3H12B | 127 | 104 | 166 | 108 | 99 | 141 |
| ENSDARG000 | 5;5;5;5;5;5;5; | 38786481:387 | 38787289:387 | -;-;-;-;-;-;-;-; | 2613 | cnot6l | 730 | 441 | 742 | 638 | 787 | 884 |
| ENSDARG000 | 5;5 | 30089715:300 | 30089950:300 | +;+ | 687 | timm8b | 428 | 291 | 339 | 150 | 406 | 220 |
| ENSDARG000 | 5;5;5;5;5;5;5; | 55265274:552 | 55265648:552 | +;+;+;+;+;+;+; | 5759 | NA | 4 | 2 | 0 | 0 | 10 | 2 |
| ENSDARG000 | 5;5;5;5;5;5;5; | 25596234:255 | 25596282:255 | +;+;+;+;+;+;+; | 1192 | NA | 4 | 24 | 5 | 14 | 1 | 7 |
| ENSDARG000 | 5;5;5;5;5;5;5; | 13394543:133 | 13394777:133 | +;+;+;+;+;+;+; | 2900 | NA | 60 | 39 | 94 | 84 | 78 | 56 |
| ENSDARG000 | 5;5;5;5;5;5;5; | 66250856:662 | 66250977:662 | +;+;+;+;+;+;+; | 1809 | NA | 423 | 333 | 816 | 453 | 790 | 842 |
| ENSDARG000 | 5;5;5 | 68878124:688 | 68878411:688 | +;+;+ | 1263 | spaw | 2 | 8 | 12 | 29 | 3 | 15 |
| ENSDARG000 | 5;5 | 55293099:552 | 55293232:552 | -;- | 486 | NA | 0 | 0 | 0 | 0 | 0 | 0 |
| ENSDARG000 | 5;5 | 38786068:387 | 38786140:387 | +;+ | 324 | NA | 9 | 19 | 2 | 0 | 8 | 4 |
| ENSDARG000 | 5;5 | 3501859:3544 | 3502094:3545 | +;+ | 1374 | ywhag1 | 2310 | 723 | 741 | 884 | 550 | 753 |
| ENSDARG000 | 5;5 | 8246603:8250 | 8246671:8252 | +;+ | 1958 | NA | 30 | 132 | 18 | 32 | 236 | 69 |
| ENSDARG000 | 5;5;5;5;5;5;5; | 19969794:199 | 19969919:199 | -;-;-;-;-;-;-;-; | 1852 | slc6a4b | 0 | 0 | 0 | 0 | 0 | 0 |
| ENSDARG000 | 5;5;5;5;5;5;5; | 18040054:180 | 18040067:180 | -;-;-;-;-;-;-;-; | 1614 | NA | 90 | 40 | 49 | 78 | 201 | 63 |
| ENSDARG000 | 5;5;5;5;5 | 64397454:643 | 64397811:643 | +;+;+;+;+ | 1041 | edf1 | 2148 | 2364 | 2711 | 2327 | 2073 | 2626 |
| ENSDARG000 | 5;5;5;5;5;5;5; | 34578479:345 | 34578753:345 | +;+;+;+;+;+;+; | 2815 | gfm2 | 502 | 239 | 342 | 168 | 908 | 476 |
| ENSDARG000 | 5;5;5;5;5;5;5; | 37890521:378 | 37890646:378 | +;+;+;+;+;+;+; | 3146 | tmprss4b | 0 | 0 | 6 | 0 | 0 | 0 |
| ENSDARG000 | 5;5;5;5;5;5;5 | 52596850:525 | 52596954:525 | +;+;+;+;+;+;+ | 1300 | ptar1 | 192 | 117 | 177 | 83 | 190 | 184 |
| ENSDARG000 | 5;5;5;5;5;5;5; | 24138446:241 | 24138581:241 | -;-;-;-;-;-;-;-; | 1972 | mrps31 | 372 | 530 | 698 | 411 | 472 | 472 |
| ENSDARG000 | 5;5;5;5;5;5;5; | 34600714:346 | 34601109:346 | -;-;-;-;-;-;-;-; | 2498 | hexb | 5686 | 3476 | 3755 | 4256 | 5641 | 4938 |
| ENSDARG000 | 5;5 | 68891515:688 | 68891843:688 | -;- | 830 | NA | 0 | 0 | 0 | 8 | 0 | 0 |
| ENSDARG000 | 5;5;5;5;5;5;5; | 28999474:290 | 28999476:290 | -;-;-;-;-;-;-;-; | 8373 | WHRN | 8 | 0 | 20 | 8 | 90 | 0 |
| ENSDARG000 | 5;5;5;5;5;5;5; | 37888095:378 | 37888358:378 | -;-;-;-;-;-;-;-; | 1836 | tmprss13b | 3 | 0 | 0 | 1 | 5 | 0 |
| ENSDARG000 | 5;5;5;5;5;5;5; | 8284358:8284 | 8284553:8284 | +;+;+;+;+;+;+; | 5981 | NA | 32 | 19 | 21 | 10 | 0 | 10 |
| ENSDARG000 | 5;5;5;5;5;5;5; | 28972935:289 | 28973336:289 | +;+;+;+;+;+;+; | 5245 | stxbp1b | 0 | 0 | 2 | 0 | 10 | 4 |
| ENSDARG000 | 5;5;5;5;5;5;5; | 11840905:118 | 11841088:118 | +;+;+;+;+;+;+; | 2298 | tesca | 26 | 12 | 10 | 12 | 6 | 14 |
| ENSDARG000 | 5;5;5;5;5;5;5; | 42136359:421 | 42136478:421 | +;+;+;+;+;+;+; | 3046 | trpv1 | 3240 | 6830 | 2902 | 7871 | 5468 | 12076 |
| ENSDARG000 | 5;5;5;5;5;5;5; | 37884022:378 | 37884178:378 | -;-;-;-;-;-;-;-; | 508 | fxyd2 | 0 | 0 | 47 | 20 | 82 | 0 |
| ENSDARG000 | 5;5;5;5;5;5 | 24156170:241 | 24156265:241 | +;+;+;+;+;+ | 2301 | slc25a15b | 238 | 267 | 349 | 183 | 300 | 111 |
| ENSDARG000 | 5;5;5;5;5;5;5; | 22115250:221 | 22116077:221 | -;-;-;-;-;-;-;-; | 2642 | las1l | 294 | 340 | 471 | 290 | 384 | 365 |
| ENSDARG000 | 5;5;5;5;5;5;5; | 3568016:3568 | 3568554:3568 | -;-;-;-;-;-;-;-;- | 1663 | hspb1 | 208 | 414 | 1138 | 1266 | 177 | 206 |
| ENSDARG000 | 5 | 64407896 | 64408695 | + | 800 | NA | 49 | 72 | 75 | 56 | 83 | 45 |
| ENSDARG000 | 5;5;5;5;5;5;5; | 26847190:268 | 26847253:268 | +;+;+;+;+;+;+; | 1141 | imp4 | 320 | 524 | 624 | 172 | 520 | 450 |
| ENSDARG000 | 5;5;5;5;5;5;5; | 52625975:526 | 52626211:526 | +;+;+;+;+;+;+; | 7521 | APBA1 | 59 | 114 | 94 | 20 | 106 | 62 |
| ENSDARG000 | 5;5;5;5;5;5;5; | 66282256:662 | 66282735:662 | -;-;-;-;-;-;-;-; | 1359 | prlrb | 0 | 0 | 7 | 4 | 10 | 0 |
| ENSDARG000 | 5;5;5;5;5;5;5; | 48179340:481 | 48183748:481 | -;-;-;-;-;-;-;-; | 7103 | mef2cb | 1362 | 713 | 896 | 982 | 1977 | 2568 |
| ENSDARG000 | 5;5;5 | 61583655:615 | 61584209:615 | -;-;- | 1406 | pex12 | 102 | 92 | 154 | 90 | 192 | 79 |
| ENSDARG000 | 5;5;5;5;5;5;5; | 11916928:119 | 11917709:119 | -;-;-;-;-;-;-;-; | 2238 | rnft2 | 46 | 59 | 88 | 46 | 57 | 44 |
| ENSDARG000 | 5;5;5;5;5;5;5; | 11943792:119 | 11944426:119 | +;+;+;+;+;+;+; | 1598 | spring1 | 219 | 195 | 193 | 94 | 204 | 240 |
| ENSDARG000 | 5;5;5;5;5;5;5; | 8355647:8356 | 8356213:8357 | -;-;-;-;-;-;-;-; | 5684 | spef2 | 14 | 12 | 12 | 4 | 8 | 8 |
| ENSDARG000 | 5;5;5;5;5;5;5; | 30131275:301 | 30131311:301 | -;-;-;-;-;-;-;-; | 2332 | zbtb44 | 110 | 27 | 41 | 20 | 59 | 61 |
| ENSDARG000 | 5;5;5;5;5;5;5; | 30155834:301 | 30156399:301 | -;-;-;-;-;-;-;-; | 2676 | ADAMTS8 | 56 | 0 | 10 | 0 | 8 | 8 |
| ENSDARG000 | 5;5;5 | 11965113:119 | 11965782:119 | -;-;- | 851 | NA | 2 | 0 | 4 | 0 | 0 | 4 |
| ENSDARG000 | 5;5;5;5;5;5;5; | 25624819:256 | 25625057:256 | -;-;-;-;-;-;-;-; | 2679 | tmc2b | 0 | 0 | 0 | 4 | 0 | 0 |
| ENSDARG000 | 5;5;5 | 61601290:616 | 61604763:616 | -;-;- | 3739 | drll.3 | 2 | 2 | 33 | 8 | 0 | 0 |
| ENSDARG000 | 5;5;5;5;5;5;5; | 68899473:689 | 68900896:689 | -;-;-;-;-;-;-;-; | 7540 | ank1a | 12 | 23 | 28 | 7 | 110 | 28 |
| ENSDARG000 | 5;5;5;5;5;5;5; | 66310495:663 | 66310520:663 | +;+;+;+;+;+;+; | 3078 | NA | 58 | 12 | 48 | 14 | 35 | 54 |
| ENSDARG000 | 5;5;5;5;5;5;5; | 32298624:322 | 32298716:322 | -;-;-;-;-;-;-;-; | 6010 | myhz1.1 | 4 | 0 | 0 | 4 | 20 | 4 |
| ENSDARG000 | 5;5;5;5;5;5;5; | 13427826:134 | 13427902:134 | +;+;+;+;+;+;+; | 2658 | sec14l8 | 2156 | 285 | 425 | 351 | 787 | 594 |
| ENSDARG000 | 5;5;5;5;5 | 24165141:241 | 24165501:241 | -;-;-;-;- | 968 | tm4sf21a | 11 | 36 | 16 | 6 | 35 | 12 |
| ENSDARG000 | 5 | 12030872 | 12031174 | - | 303 | NA | 0 | 0 | 0 | 0 | 0 | 0 |
| ENSDARG000 | 5;5;5;5;5;5;5; | 11982364:119 | 11986739:119 | -;-;-;-;-;-;-;-; | 6609 | castor1 | 122 | 98 | 203 | 206 | 95 | 316 |
| ENSDARG000 | 5;5;5;5;5;5;5; | 68900506:689 | 68900896:689 | -;-;-;-;-;-;-;-; | 3587 | ank1a | 0 | 4 | 0 | 2 | 8 | 0 |
| ENSDARG000 | 5;5;5;5 | 3611498:3617 | 3612665:3617 | -;-;-;- | 3002 | NA | 0 | 0 | 0 | 0 | 2 | 8 |
| ENSDARG000 | 5;5;5;5;5;5;5; | 52745401:527 | 52745664:527 | -;-;-;-;-;-;-;-; | 3021 | ENTREP1 | 215 | 176 | 219 | 122 | 217 | 144 |
| ENSDARG000 | 5;5;5;5;5;5;5; | 64411516:644 | 64414161:644 | -;-;-;-;-;-;-;-; | 5015 | brd3b | 1466 | 1286 | 2087 | 1137 | 1939 | 1764 |
| ENSDARG000 | 5;5;5 | 61615663:616 | 61617272:616 | -;-;- | 1875 | drll.2 | 0 | 3 | 14 | 25 | 6 | 0 |
| ENSDARG000 | 5;5;5;5;5;5;5 | 24179307:241 | 24179511:241 | +;+;+;+;+;+;+ | 2689 | mpdu1a | 14 | 42 | 42 | 141 | 37 | 48 |
| ENSDARG000 | 5;5;5;5;5;5;5; | 3862586:3864 | 3862638:3864 | -;-;-;-;-;-;-;-; | 849 | NA | 0 | 0 | 0 | 0 | 0 | 0 |
| ENSDARG000 | 5;5;5;5;5;5;5; | 20115774:201 | 20116511:201 | -;-;-;-;-;-;-;-;- | 1395 | pxmp2 | 173 | 186 | 170 | 154 | 260 | 163 |
| ENSDARG000 | 5;5 | 42164079:421 | 42165784:421 | -;- | 1829 | fam222ba | 705 | 589 | 698 | 518 | 677 | 753 |
| ENSDARG000 | 5;5;5;5;5;5;5; | 30179010:301 | 30179948:301 | +;+;+;+;+;+;+; | 2877 | adamts15a | 81 | 8 | 2 | 20 | 26 | 58 |
| ENSDARG000 | 5;5;5 | 61630696:616 | 61632308:616 | -;-;- | 1878 | drll.1 | 0 | 8 | 65 | 51 | 55 | 0 |
| ENSDARG000 | 5;5;5;5;5;5;5; | 55371248:553 | 55375282:553 | -;-;-;-;-;-;-;-; | 5301 | prune2 | 408 | 0 | 2 | 0 | 2 | 0 |
| ENSDARG000 | 5;5;5;5;5;5;5; | 31693316:316 | 31693943:316 | -;-;-;-;-;-;-;-; | 2119 | PIP5KL1 | 95 | 95 | 148 | 135 | 179 | 175 |
| ENSDARG000 | 5;5 | 42252209:422 | 42252293:422 | -;- | 255 | NA | 0 | 0 | 0 | 0 | 0 | 0 |
| ENSDARG000 | 5;5;5;5;5;5;5; | 12032802:120 | 12033203:120 | -;-;-;-;-;-;-;-; | 1472 | nipsnap1 | 0 | 0 | 8 | 0 | 4 | 0 |
| ENSDARG000 | 5;5;5;5;5;5;5 | 44833313:448 | 44833917:448 | -;-;-;-;-;-;- | 1592 | fbp1a | 277 | 194 | 170 | 243 | 197 | 572 |
| ENSDARG000 | 5;5;5 | 44812921:448 | 44812970:448 | +;+;+ | 576 | NA | 0 | 0 | 0 | 1 | 0 | 0 |
| ENSDARG000 | 5;5;5;5;5;5;5; | 44804791:448 | 44804911:448 | +;+;+;+;+;+;+; | 2083 | ctsla | 32870 | 29487 | 25013 | 30012 | 33411 | 28284 |
| ENSDARG000 | 5;5;5;5;5;5;5; | 30351233:303 | 30351271:303 | -;-;-;-;-;-;-;-; | 2651 | snx19a | 2 | 7 | 17 | 14 | 8 | 16 |
| ENSDARG000 | 5;5;5;5;5;5;5; | 57814165:578 | 57814861:578 | -;-;-;-;-;-;-;-; | 2961 | sik2a | 2 | 2 | 0 | 0 | 0 | 0 |
| ENSDARG000 | 5;5;5;5;5;5;5 | 44814064:448 | 44814383:448 | -;-;-;-;-;-;- | 1298 | fbp2 | 0 | 0 | 0 | 0 | 0 | 0 |
| ENSDARG000 | 5;5;5 | 61645156:616 | 61646766:616 | -;-;- | 1862 | drl | 0 | 2 | 38 | 87 | 6 | 0 |
| ENSDARG000 | 5;5;5;5;5;5;5; | 22133153:221 | 22133535:221 | +;+;+;+;+;+;+; | 3780 | msna | 8170 | 10689 | 9672 | 6888 | 10448 | 6770 |
| ENSDARG000 | 5;5;5;5;5;5;5; | 20125934:201 | 20126515:201 | -;-;-;-;-;-;-;-; | 1811 | usp30 | 33 | 64 | 53 | 31 | 64 | 101 |
| ENSDARG000 | 5;5 | 48659747:486 | 48660132:486 | -;- | 828 | mblac2 | 27 | 33 | 36 | 30 | 10 | 46 |
| ENSDARG000 | 5;5;5;5;5;5;5 | 52801620:528 | 52801932:528 | -;-;-;-;-;-;- | 1474 | zgc:158260 | 3 | 0 | 4 | 0 | 4 | 1 |
| ENSDARG000 | 5;5;5 | 66347006:663 | 66347233:663 | -;-;- | 369 | cplx4b | 0 | 0 | 0 | 0 | 1 | 7 |
| ENSDARG000 | 5;5;5;5;5;5;5; | 18047111:180 | 18047411:180 | +;+;+;+;+;+;+; | 4394 | HIRA | 651 | 678 | 743 | 300 | 694 | 474 |
| ENSDARG000 | 5;5;5;5;5;5;5; | 42258994:422 | 42259248:422 | +;+;+;+;+;+;+; | 2071 | eral1 | 214 | 258 | 277 | 132 | 273 | 302 |
| ENSDARG000 | 5;5;5;5;5;5;5; | 38825225:388 | 38825278:388 | +;+;+;+;+;+;+; | 1042 | mrpl1 | 192 | 165 | 231 | 140 | 190 | 152 |
| ENSDARG000 | 5;5;5;5;5;5;5; | 25652727:256 | 25653160:256 | -;-;-;-;-;-;-;-; | 3315 | TMC1 | 4 | 10 | 2 | 0 | 12 | 4 |
| ENSDARG000 | 5;5;5;5;5;5;5; | 26856253:268 | 26856456:268 | +;+;+;+;+;+;+; | 7665 | araf | 1772 | 1065 | 2007 | 1181 | 1876 | 1583 |
| ENSDARG000 | 5;5;5 | 48672553:486 | 48673812:486 | -;-;- | 1700 | lysmd3 | 54 | 27 | 17 | 47 | 25 | 11 |
| ENSDARG000 | 5;5;5;5;5;5;5; | 48666485:486 | 48666642:486 | +;+;+;+;+;+;+; | 3887 | polr3g | 4 | 0 | 10 | 0 | 0 | 6 |
| ENSDARG000 | 5;5;5;5;5;5;5; | 41143563:411 | 41143640:411 | +;+;+;+;+;+;+; | 3010 | sub1b | 507 | 736 | 1162 | 567 | 933 | 637 |
| ENSDARG000 | 5;5;5 | 13521081:135 | 13521125:135 | +;+;+ | 791 | NA | 22 | 8 | 12 | 18 | 21 | 8 |
| ENSDARG000 | 5;5;5;5;5;5;5; | 13472234:134 | 13473599:134 | +;+;+;+;+;+;+; | 2337 | CNNM4 | 177 | 104 | 147 | 81 | 171 | 70 |
| ENSDARG000 | 5;5;5;5 | 31714358:317 | 31714951:317 | -;-;-;- | 845 | dpm2 | 402 | 450 | 519 | 493 | 374 | 421 |
| ENSDARG000 | 5;5;5;5;5;5;5; | 3891485:3891 | 3891669:3891 | +;+;+;+;+;+;+; | 1004 | rpain | 120 | 88 | 77 | 98 | 158 | 102 |
| ENSDARG000 | 5;5;5;5;5;5;5; | 64476118:644 | 64476794:644 | -;-;-;-;-;-;-;-;- | 1721 | rxrab | 329 | 144 | 291 | 233 | 389 | 334 |
| ENSDARG000 | 5;5;5;5 | 24199180:241 | 24200442:242 | -;-;-;- | 2170 | sox19a | 12 | 4 | 0 | 22 | 0 | 4 |
| ENSDARG000 | 5;5;5;5;5;5;5; | 55402072:554 | 55402301:554 | -;-;-;-;-;-;-;-; | 889 | NA | 0 | 0 | 0 | 0 | 0 | 0 |
| ENSDARG000 | 5;5;5;5;5;5;5; | 20147830:201 | 20147861:201 | +;+;+;+;+;+;+; | 1647 | SVOP | 4 | 3 | 2 | 0 | 0 | 6 |
| ENSDARG000 | 5;5;5;5;5;5;5; | 66353589:663 | 66353656:663 | +;+;+;+;+;+;+; | 1938 | emb | 7 | 1 | 0 | 7 | 1 | 4 |
| ENSDARG000 | 5;5;5;5;5 | 30384554:303 | 30384711:303 | +;+;+;+;+ | 1421 | zgc:158412 | 4 | 2 | 0 | 0 | 24 | 2 |
| ENSDARG000 | 5;5;5;5;5;5;5; | 30392148:303 | 30392657:303 | +;+;+;+;+;+;+; | 4474 | stk36 | 75 | 61 | 163 | 92 | 155 | 131 |
| ENSDARG000 | 5;5;5;5;5;5;5; | 29142797:291 | 29143239:291 | -;-;-;-;-;-;-;-; | 1354 | noxa1 | 15 | 0 | 2 | 0 | 0 | 0 |
| ENSDARG000 | 5;5;5;5;5;5;5; | 24220157:242 | 24220841:242 | -;-;-;-;-;-;-;-; | 2387 | senp3a | 106 | 149 | 200 | 98 | 192 | 206 |
| ENSDARG000 | 5;5 | 30390893:303 | 30391389:303 | -;- | 551 | NA | 0 | 0 | 0 | 1 | 0 | 0 |
| ENSDARG000 | 5;5;5;5;5;5 | 24202623:242 | 24202685:242 | +;+;+;+;+;+ | 1552 | NA | 0 | 0 | 2 | 0 | 0 | 1 |
| ENSDARG000 | 5;5;5;5;5;5;5; | 24208413:242 | 24209544:242 | -;-;-;-;-;-;-;-; | 2276 | eif4a1b | 7762 | 7489 | 12999 | 10981 | 9966 | 11213 |

| | | | | | | | | | | | | |
|---|---|---|---|---|---|---|---|---|---|---|---|---|
| ENSDARG000 | 5;5;5;5;5;5;5; | 61657010:616 | 61657832:616 | +;+;+;+;+;+;+; | 4810 | ap2b1 | 4927 | 4221 | 6102 | 4755 | 5907 | 5440 |
| ENSDARG000 | 5;5;5;5;5;5;5; | 8417184:841 | 8418812:841 | -;-;-;-;-;-;-;-; | 4081 | lifra | 318 | 226 | 520 | 264 | 430 | 302 |
| ENSDARG000 | 5;5;5;5;5;5;5; | 36693859:366 | 36694124:366 | +;+;+;+;+;+;+; | 3094 | dlb | 0 | 0 | 0 | 0 | 12 | 8 |
| ENSDARG000 | 5;5;5;5;5;5;5; | 32323775:323 | 32324128:323 | -;-;-;-;-;-;-;-; | 3242 | dab2 | 60 | 6 | 164 | 46 | 210 | 128 |
| ENSDARG000 | 5;5;5;5;5 | 52971425:529 | 52971748:529 | -;-;-;-;- | 1364 | mrps30 | 369 | 502 | 580 | 337 | 505 | 400 |
| ENSDARG000 | 5;5;5;5;5;5;5; | 29159777:291 | 29160248:291 | +;+;+;+;+;+;+; | 2228 | dpp7 | 2760 | 1929 | 2804 | 1858 | 2645 | 2701 |
| ENSDARG000 | 5;5;5 | 42271810:422 | 42272550:422 | -;-;- | 741 | NA | 8 | 7 | 2 | 2 | 17 | 13 |
| ENSDARG000 | 5;5;5;5;5;5;5; | 64732036:647 | 64732335:647 | +;+;+;+;+;+;+; | 2300 | olfm1a | 0 | 0 | 0 | 0 | 0 | 0 |
| ENSDARG000 | 5;5;5;5;5;5;5; | 31723404:317 | 31724447:317 | -;-;-;-;-;-;-;-; | 2502 | eeig1b | 354 | 384 | 386 | 387 | 666 | 292 |
| ENSDARG000 | 5;5;5;5;5;5;5 | 23283890:232 | 23286454:232 | -;-;-;-;-;-;- | 4438 | nlgn3b | 0 | 0 | 0 | 0 | 0 | 0 |
| ENSDARG000 | 5;5;5;5;5;5;5; | 42280372:422 | 42280685:422 | +;+;+;+;+;+;+; | 3080 | tbx16l | 0 | 2 | 3 | 0 | 0 | 0 |
| ENSDARG000 | 5;5;5;5;5;5;5; | 25680842:256 | 25680977:256 | +;+;+;+;+;+;+; | 2682 | zfand5a | 2417 | 1486 | 2320 | 2960 | 2554 | 1561 |
| ENSDARG000 | 5;5;5;5;5;5 | 57886049:578 | 57886400:578 | -;-;-;-;-;- | 1037 | LAYN | 0 | 0 | 0 | 0 | 0 | 0 |
| ENSDARG000 | 5;5;5;5;5 | 23331153:233 | 23331265:233 | -;-;-;-;- | 567 | NA | 0 | 0 | 0 | 0 | 0 | 0 |
| ENSDARG000 | 5;5;5;5;5;5;5; | 41199458:411 | 41199584:411 | +;+;+;+;+;+;+; | 1753 | NA | 0 | 0 | 0 | 0 | 0 | 0 |
| ENSDARG000 | 5;5;5;5;5;5;5; | 13549603:135 | 13549785:135 | +;+;+;+;+;+;+; | 3164 | ckap2l | 132 | 372 | 494 | 262 | 342 | 181 |
| ENSDARG000 | 5;5;5;5;5;5;5; | 12065957:120 | 12066001:120 | -;-;-;-;-;-;-;-; | 1437 | NA | 0 | 0 | 2 | 4 | 2 | 0 |
| ENSDARG000 | 5;5 | 13563298:135 | 13564444:135 | -;- | 1504 | NA | 10 | 13 | 8 | 13 | 48 | 8 |
| ENSDARG000 | 5;5;5;5;5;5;5; | 20170960:201 | 20171298:201 | -;-;-;-;-;-;-;-; | 1295 | dao.2 | 0 | 4 | 13 | 1 | 4 | 6 |
| ENSDARG000 | 5;5;5;5;5;5;5; | 8492640:849 | 8492778:849 | +;+;+;+;+;+;+; | 2346 | NA | 0 | 2 | 4 | 10 | 14 | 12 |
| ENSDARG000 | 5;5 | 3935917:393 | 3935984:393 | -;- | 165 | NA | 0 | 4 | 0 | 0 | 0 | 0 |
| ENSDARG000 | 5;5;5;5;5 | 3927989:392 | 3928132:392 | +;+;+;+;+ | 663 | znhit3 | 40 | 46 | 91 | 18 | 32 | 22 |
| ENSDARG000 | 5;5;5;5;5;5;5; | 3914403:391 | 3914745:391 | -;-;-;-;-;-;-;-; | 1172 | c1qbp | 1626 | 1462 | 1646 | 1019 | 1794 | 1306 |
| ENSDARG000 | 5;5 | 13643167:136 | 13646054:136 | -;- | 2888 | purba | 1400 | 1585 | 1962 | 1217 | 2058 | 1460 |
| ENSDARG000 | 5;5;5;5;5;5;5; | 48683447:487 | 48683767:487 | +;+;+;+;+;+;+; | 19174 | adgrv1 | 42 | 16 | 8 | 6 | 16 | 14 |
| ENSDARG000 | 5;5;5;5 | 30414250:304 | 30414757:304 | -;-;-;- | 2336 | atf5a | 70 | 214 | 605 | 952 | 86 | 223 |
| ENSDARG000 | 5;5;5;5;5 | 53009083:530 | 53009247:530 | +;+;+;+;+ | 692 | hint2 | 114 | 168 | 156 | 137 | 156 | 156 |
| ENSDARG000 | 5;5 | 42301124:423 | 42301316:423 | -;- | 375 | NA | 0 | 0 | 0 | 0 | 0 | 0 |
| ENSDARG000 | 5;5;5;5;5;5;5; | 57901168:579 | 57902103:579 | -;-;-;-;-;-;-;-; | 1784 | hoatz | 2 | 2 | 5 | 2 | 0 | 0 |
| ENSDARG000 | 5;5;5;5;5;5;5 | 55588389:555 | 55588582:555 | -;-;-;-;-;-;- | 1745 | gnaq | 147 | 174 | 206 | 164 | 213 | 190 |
| ENSDARG000 | 5;5;5;5;5;5;5; | 23353570:233 | 23353739:233 | -;-;-;-;-;-;-;-; | 3377 | gria3a | 0 | 0 | 17 | 0 | 0 | 2 |
| ENSDARG000 | 5;5;5;5;5;5;5; | 31779911:317 | 31780234:317 | +;+;+;+;+;+;+; | 1755 | slc25a25b | 75 | 236 | 310 | 60 | 170 | 103 |
| ENSDARG000 | 5;5;5;5;5;5;5; | 42308955:423 | 42309132:423 | +;+;+;+;+;+;+; | 1589 | NA | 0 | 0 | 0 | 0 | 0 | 0 |
| ENSDARG000 | 5;5;5;5;5;5;5; | 66366865:663 | 66366959:663 | -;-;-;-;-;-;-;-; | 3948 | hip1rb | 14 | 2 | 21 | 18 | 10 | 4 |
| ENSDARG000 | 5;5;5;5;5;5;5; | 13647288:136 | 13647368:136 | +;+;+;+;+;+;+; | 1359 | h2az2a | 678 | 992 | 1411 | 651 | 934 | 737 |
| ENSDARG000 | 5;5;5;5;5;5;5; | 41260478:412 | 41262047:412 | -;-;-;-;-;-;-;- | 3297 | npr3 | 56 | 0 | 2 | 0 | 0 | 4 |
| ENSDARG000 | 5;5;5;5;5;5;5; | 38837429:388 | 38837590:388 | +;+;+;+;+;+;+; | 13772 | fras1 | 140 | 10 | 10 | 0 | 40 | 10 |
| ENSDARG000 | 5;5;5;5;5;5;5; | 12100126:121 | 12100684:121 | -;-;-;-;-;-;-;-; | 5681 | nos1 | 58 | 42 | 110 | 180 | 161 | 226 |
| ENSDARG000 | 5;5;5;5;5;5;5; | 53580846:535 | 53581676:535 | +;+;+;+;+;+;+; | 7094 | taf6 | 453 | 475 | 743 | 412 | 663 | 532 |
| ENSDARG000 | 5;5;5;5;5;5;5; | 69140656:691 | 69141528:691 | -;-;-;-;-;-;-;-; | 3618 | ganaba | 717 | 632 | 976 | 640 | 1101 | 680 |
| ENSDARG000 | 5;5;5 | 53714794:537 | 53715785:537 | -;-;- | 1241 | NA | 0 | 0 | 0 | 0 | 0 | 2 |
| ENSDARG000 | 5;5 | 53581489:536 | 53581676:536 | +;+ | 853 | NA | 0 | 0 | 0 | 0 | 0 | 2 |
| ENSDARG000 | 5;5;5;5;5;5;5; | 12272632:122 | 12272638:122 | -;-;-;-;-;-;-;-; | 6402 | KSR2 | 6 | 6 | 16 | 2 | 2 | 2 |
| ENSDARG000 | 5;5;5;5 | 36730726:367 | 36730946:367 | +;+;+;+ | 938 | ppp1r14aa | 8 | 0 | 0 | 8 | 0 | 4 |
| ENSDARG000 | 5;5 | 69183255:691 | 69183293:691 | +;+ | 712 | NA | 6 | 16 | 59 | 12 | 24 | 36 |
| ENSDARG000 | 5;5 | 12272448:122 | 12272638:122 | -;- | 418 | NA | 0 | 0 | 0 | 0 | 0 | 0 |
| ENSDARG000 | 5;5;5;5;5;5;5; | 42322847:423 | 42323342:423 | +;+;+;+;+;+;+; | 1753 | NA | 0 | 73 | 3 | 0 | 13 | 145 |
| ENSDARG000 | 5;5 | 12515402:125 | 12515490:125 | +;+ | 1141 | zgc:171242 | 6 | 0 | 0 | 0 | 0 | 0 |
| ENSDARG000 | 5;5 | 12503494:125 | 12503634:125 | -;- | 469 | NA | 0 | 0 | 0 | 0 | 0 | 0 |
| ENSDARG000 | 5;5 | 42333171:423 | 42333782:423 | -;- | 692 | NA | 0 | 0 | 0 | 0 | 0 | 0 |
| ENSDARG000 | 5;5;5;5;5;5;5; | 8533929:853 | 8538795:853 | -;-;-;-;-;-;-;-; | 10034 | RICTOR | 532 | 382 | 636 | 380 | 627 | 591 |
| ENSDARG000 | 5;5;5;5 | 13671614:136 | 13672538:136 | -;-;-;- | 2436 | zgc:65851 | 0 | 0 | 2 | 0 | 0 | 0 |
| ENSDARG000 | 5;5;5;5;5 | 22177033:222 | 22177182:222 | +;+;+;+;+ | 973 | fdx1b | 2 | 0 | 2 | 0 | 0 | 0 |
| ENSDARG000 | 5;5;5;5;5;5;5 | 36752943:367 | 36753044:367 | +;+;+;+;+;+;+ | 2043 | NA | 71 | 13 | 24 | 26 | 30 | 80 |
| ENSDARG000 | 5;5;5;5;5;5;5; | 69199198:692 | 69200217:692 | -;-;-;-;-;-;-;- | 2035 | mat2ab | 130 | 332 | 483 | 130 | 265 | 304 |
| ENSDARG000 | 5;5;5;5;5;5;5; | 29181725:291 | 29181989:291 | -;-;-;-;-;-;-;-; | 2723 | man1b1b | 325 | 375 | 581 | 278 | 571 | 411 |
| ENSDARG000 | 5;5;5;5;5;5;5; | 41322783:413 | 41323051:413 | +;+;+;+;+;+;+; | 2479 | arid3c | 10 | 0 | 2 | 0 | 6 | 2 |
| ENSDARG000 | 5;5;5;5;5;5;5; | 30427198:304 | 30427338:304 | -;-;-;-;-;-;-;-; | 6108 | PHLDB1 | 61 | 6 | 14 | 18 | 95 | 34 |
| ENSDARG000 | 5;5;5;5;5;5 | 13706959:137 | 13707260:137 | -;-;-;-;-;- | 1282 | NA | 102 | 34 | 68 | 50 | 95 | 54 |
| ENSDARG000 | 5;5;5;5;5;5 | 13767743:137 | 13768145:137 | -;-;-;-;-;- | 931 | snrnp27 | 620 | 527 | 571 | 563 | 500 | 548 |
| ENSDARG000 | 5;5;5;5;5;5;5; | 18431401:184 | 18431612:184 | -;-;-;-;-;-;-;-; | 2156 | TMEM132C | 4 | 40 | 76 | 21 | 118 | 38 |
| ENSDARG000 | 5;5;5;5;5;5;5; | 55602720:556 | 55604761:556 | -;-;-;-;-;-;-;-; | 4012 | hnrpkl | 1421 | 2511 | 2530 | 1543 | 2409 | 1865 |
| ENSDARG000 | 5;5;5;5;5;5;5; | 53824959:538 | 53825143:538 | +;+;+;+;+;+;+; | 2915 | sat2b | 220 | 142 | 192 | 148 | 213 | 135 |
| ENSDARG000 | 5;5;5;5;5;5;5; | 18827314:188 | 18828176:188 | -;-;-;-;-;-;-;-; | 2817 | aacs | 244 | 251 | 173 | 162 | 116 | 256 |
| ENSDARG000 | 5;5;5;5;5;5;5; | 69232230:692 | 69232331:692 | -;-;-;-;-;-;-;-; | 2203 | INPP5J | 0 | 1 | 5 | 3 | 8 | 0 |
| ENSDARG000 | 5;5;5;5 | 23479562:234 | 23479637:234 | -;-;-;- | 669 | NA | 0 | 0 | 0 | 0 | 0 | 0 |
| ENSDARG000 | 5;5;5;5;5;5 | 23468045:234 | 23468113:234 | -;-;-;-;-;- | 564 | NA | 0 | 0 | 0 | 0 | 0 | 0 |
| ENSDARG000 | 5;5;5;5;5;5;5; | 23457112:234 | 23457348:234 | -;-;-;-;-;-;-;-; | 2398 | NA | 15 | 41 | 66 | 40 | 26 | 19 |
| ENSDARG000 | 5;5;5;5;5;5;5; | 61862199:618 | 61862330:618 | +;+;+;+;+;+;+; | 6493 | SYMPK | 1152 | 1237 | 2070 | 901 | 1172 | 1055 |
| ENSDARG000 | 5;5;5;5;5;5;5; | 22264051:222 | 22264319:222 | +;+;+;+;+;+;+; | 2212 | NA | 10 | 4 | 37 | 0 | 15 | 7 |
| ENSDARG000 | 5;5;5;5;5;5;5 | 24234466:242 | 24234525:242 | -;-;-;-;-;-;- | 802 | plscr3a | 0 | 2 | 5 | 10 | 22 | 0 |
| ENSDARG000 | 5;5;5;5;5;5;5; | 64814482:648 | 64814832:648 | -;-;-;-;-;-;-;- | 1021 | lix1 | 0 | 0 | 0 | 0 | 0 | 0 |
| ENSDARG000 | 5;5;5;5;5;5;5; | 25708369:257 | 25708985:257 | -;-;-;-;-;-;-;-; | 1935 | gda | 593 | 385 | 245 | 219 | 894 | 182 |
| ENSDARG000 | 5;5;5;5;5;5 | 34622320:346 | 34622657:346 | +;+;+;+;+;+ | 13077 | ENC1 | 3260 | 1733 | 1469 | 2132 | 1351 | 7227 |
| ENSDARG000 | 5;5;5 | 18904968:189 | 18905664:189 | -;-;- | 1088 | bri3bp | 503 | 634 | 625 | 223 | 821 | 619 |
| ENSDARG000 | 5;5;5;5;5;5;5; | 36768674:367 | 36768822:367 | +;+;+;+;+;+;+; | 2505 | NA | 8 | 4 | 6 | 2 | 64 | 10 |
| ENSDARG000 | 5;5;5;5;5;5;5; | 41476443:414 | 41476529:414 | +;+;+;+;+;+;+; | 2807 | pias2 | 711 | 515 | 806 | 647 | 674 | 721 |
| ENSDARG000 | 5;5;5;5 | 22363015:223 | 22363492:223 | -;-;-;- | 1056 | GPR83 | 0 | 0 | 0 | 0 | 0 | 0 |
| ENSDARG000 | 5;5;5;5;5;5 | 22370167:223 | 22370461:223 | +;+;+;+;+;+ | 1410 | NA | 0 | 2 | 0 | 0 | 7 | 7 |
| ENSDARG000 | 5;5;5;5;5;5;5; | 8616667:861 | 8616689:861 | -;-;-;-;-;-;-;-; | 894 | NA | 0 | 2 | 0 | 0 | 0 | 3 |
| ENSDARG000 | 5;5;5;5;5;5;5; | 29199292:292 | 29199476:292 | -;-;-;-;-;-;-;-; | 3614 | NA | 0 | 5 | 8 | 0 | 0 | 0 |
| ENSDARG000 | 5;5;5;5;5;5;5; | 32345187:323 | 32345288:323 | +;+;+;+;+;+;+; | 2345 | c9 | 0 | 0 | 67 | 2 | 0 | 0 |
| ENSDARG000 | 5;5;5;5;5;5;5; | 31811662:318 | 31812166:318 | +;+;+;+;+;+;+; | 1911 | uap1l1 | 3052 | 1670 | 1641 | 1864 | 2697 | 2963 |
| ENSDARG000 | 5;5;5;5;5;5;5; | 8661414:866 | 8661756:866 | -;-;-;-;-;-;-;-; | 2302 | zgc:153352 | 64 | 104 | 74 | 17 | 108 | 90 |
| ENSDARG000 | 5;5;5 | 32342243:323 | 32342730:323 | -;-;- | 1767 | NA | 0 | 0 | 0 | 0 | 0 | 0 |
| ENSDARG000 | 5;5;5;5;5;5;5; | 31819436:318 | 31819489:318 | +;+;+;+;+;+;+; | 1544 | zdhhc12b | 171 | 69 | 67 | 294 | 128 | 189 |
| ENSDARG000 | 5;5;5;5;5;5 | 64840656:648 | 64840806:648 | +;+;+;+;+;+ | 7829 | LRRC8A | 472 | 391 | 828 | 634 | 774 | 366 |
| ENSDARG000 | 5;5;5;5 | 57924611:579 | 57925029:579 | +;+;+;+ | 1236 | btg4 | 0 | 0 | 0 | 0 | 0 | 0 |
| ENSDARG000 | 5;5;5;5;5;5;5; | 3943157:394 | 3943487:394 | -;-;-;-;-;-;-;-; | 2883 | MYO19 | 247 | 139 | 194 | 117 | 268 | 35 |
| ENSDARG000 | 5;5;5;5;5;5;5; | 44846280:448 | 44846491:448 | +;+;+;+;+;+;+; | 9244 | NA | 3776 | 1652 | 1712 | 2497 | 1594 | 2788 |
| ENSDARG000 | 5;5;5;5;5;5;5; | 13781968:137 | 13786065:137 | -;-;-;-;-;-;-;-; | 6839 | add2 | 12 | 8 | 70 | 7 | 64 | 107 |
| ENSDARG000 | 5;5;5;5;5;5;5; | 26869580:268 | 26870322:268 | -;-;-;-;-;-;-;-; | 1878 | zgc:64051 | 120 | 295 | 327 | 169 | 710 | 469 |
| ENSDARG000 | 5;5;5;5;5;5;5; | 24240318:242 | 24240641:242 | -;-;-;-;-;-;-;- | 2020 | phf23a | 367 | 312 | 478 | 416 | 611 | 305 |
| ENSDARG000 | 5;5 | 66676356:666 | 66676793:667 | -;- | 3865 | MN1 | 14 | 14 | 24 | 8 | 81 | 24 |
| ENSDARG000 | 5;5;5 | 53878433:538 | 53878940:538 | -;-;- | 887 | NA | 0 | 0 | 0 | 0 | 4 | 0 |
| ENSDARG000 | 5;5;5;5;5;5 | 64851986:648 | 64852068:648 | +;+;+;+;+;+ | 635 | ppil3 | 34 | 96 | 106 | 124 | 88 | 75 |
| ENSDARG000 | 5;5 | 53878846:538 | 53879074:538 | -;- | 537 | NA | 0 | 4 | 0 | 0 | 0 | 0 |
| ENSDARG000 | 5;5;5;5;5;5;5; | 53908609:539 | 53910027:539 | -;-;-;-;-;-;-;- | 2210 | cldn15b | 131 | 141 | 164 | 164 | 171 | 180 |
| ENSDARG000 | 5;5;5;5;5;5;5; | 37918810:379 | 37919525:379 | -;-;-;-;-;-;-;-; | 1878 | scn4bb | 2 | 0 | 0 | 0 | 0 | 6 |
| ENSDARG000 | 5;5 | 53929512:539 | 53929690:539 | -;- | 519 | NA | 0 | 0 | 0 | 0 | 0 | 0 |
| ENSDARG000 | 5;5;5 | 53876169:538 | 53876268:538 | +;+;+ | 743 | naa38 | 243 | 216 | 215 | 169 | 199 | 199 |
| ENSDARG000 | 5;5 | 4006837:400 | 4007103:401 | +;+ | 1862 | pigw | 106 | 142 | 162 | 100 | 136 | 140 |
| ENSDARG000 | 5;5;5;5;5;5;5; | 23488055:234 | 23489467:234 | -;-;-;-;-;-;-;-; | 5371 | stag2a | 1882 | 1551 | 1738 | 1584 | 1755 | 1716 |
| ENSDARG000 | 5;5;5;5;5;5;5; | 8684652:869 | 8685538:869 | -;-;-;-;-;-;-;-; | 2293 | paip1 | 591 | 545 | 555 | 683 | 711 | 636 |
| ENSDARG000 | 5;5;5 | 42361013:423 | 42361114:423 | +;+;+ | 1816 | NA | 0 | 34 | 10 | 6 | 10 | 91 |
| ENSDARG000 | 5;5;5;5;5;5;5; | 30502249:305 | 30503677:305 | -;-;-;-;-;-;-;-; | 3038 | arcn1a | 1140 | 1199 | 1636 | 777 | 1326 | 998 |
| ENSDARG000 | 5;5;5;5;5;5 | 26896103:268 | 26896215:268 | +;+;+;+;+;+ | 976 | NA | 0 | 14 | 0 | 0 | 0 | 0 |
| ENSDARG000 | 5;5;5;5;5 | 69278089:692 | 69278267:692 | +;+;+;+;+ | 862 | selenom | 156 | 591 | 389 | 226 | 252 | 219 |
| ENSDARG000 | 5;5;5;5;5;5;5; | 42402536:424 | 42402662:424 | +;+;+;+;+;+;+; | 1462 | NA | 0 | 0 | 0 | 0 | 0 | 0 |
| ENSDARG000 | 5;5;5;5;5;5;5; | 58075301:580 | 58075501:580 | -;-;-;-;-;-;-;-; | 1692 | drd2b | 40 | 16 | 197 | 68 | 357 | 88 |
| ENSDARG000 | 5;5;5;5;5;5;5; | 66733289:667 | 66734230:667 | -;-;-;-;-;-;-;-; | 2825 | kat5b | 237 | 154 | 209 | 54 | 202 | 177 |
| ENSDARG000 | 5;5;5;5;5;5;5; | 34827614:348 | 34830367:348 | -;-;-;-;-;-;-;-; | 8667 | ARHGEF28 | 64 | 48 | 212 | 98 | 102 | 46 |
| ENSDARG000 | 5;5;5;5;5;5;5; | 42388464:423 | 42388590:423 | +;+;+;+;+;+;+; | 1462 | NA | 0 | 0 | 0 | 0 | 0 | 0 |
| ENSDARG000 | 5;5;5;5;5;5;5; | 42395655:423 | 42395781:423 | +;+;+;+;+;+;+; | 1462 | NA | 0 | 0 | 0 | 0 | 0 | 0 |
| ENSDARG000 | 5;5;5;5;5;5;5; | 42381273:423 | 42381399:423 | +;+;+;+;+;+;+; | 1462 | NA | 0 | 0 | 0 | 0 | 0 | 0 |
| ENSDARG000 | 5;5;5 | 34920395:349 | 34920443:349 | +;+;+ | 386 | NA | 0 | 0 | 0 | 0 | 0 | 0 |
| ENSDARG000 | 5;5;5 | 34918991:349 | 34919142:349 | -;-;- | 367 | NA | 0 | 0 | 0 | 0 | 0 | 0 |
| ENSDARG000 | 5;5 | 55646312:556 | 55647069:556 | -;- | 901 | NA | 12 | 16 | 46 | 45 | 40 | 68 |
| ENSDARG000 | 5;5;5;5;5;5;5; | 55626693:556 | 55627049:556 | +;+;+;+;+;+;+; | 2616 | ntrk2b | 0 | 0 | 11 | 2 | 15 | 6 |
| ENSDARG000 | 5;5;5;5;5;5;5; | 64856666:648 | 64856921:648 | +;+;+;+;+;+;+; | 1071 | zgc:101858 | 614 | 508 | 766 | 627 | 638 | 536 |
| ENSDARG000 | 5;5;5;5;5;5;5; | 37948283:379 | 37949161:379 | -;-;-;-;-;-;-;-; | 2368 | mpzl2b | 131 | 143 | 83 | 47 | 81 | 61 |
| ENSDARG000 | 5;5;5;5;5;5;5 | 42379517:423 | 42379789:423 | +;+;+;+;+;+;+ | 1247 | NA | 0 | 0 | 0 | 0 | 0 | 0 |
| ENSDARG000 | 5;5;5;5;5;5;5; | 22406672:224 | 22406758:224 | +;+;+;+;+;+;+; | 1212 | NA | 267 | 0 | 232 | 115 | 0 | 0 |
| ENSDARG000 | 5;5;5;5;5;5;5; | 69289990:692 | 69293324:692 | -;-;-;-;-;-;-;-; | 4778 | NA | 41 | 159 | 88 | 44 | 68 | 36 |
| ENSDARG000 | 5;5;5;5 | 26949739:269 | 26949832:269 | +;+;+;+ | 576 | NA | 0 | 0 | 0 | 0 | 0 | 0 |
| ENSDARG000 | 5;5;5;5;5;5;5; | 26930062:269 | 26930833:269 | -;-;-;-;-;-;-;-; | 2299 | NA | 9 | 0 | 0 | 0 | 0 | 0 |
| ENSDARG000 | 5;5;5;5;5;5;5; | 31830476:318 | 31830678:318 | -;-;-;-;-;-;-;-; | 3901 | NA | 226 | 328 | 207 | 148 | 190 | 151 |
| ENSDARG000 | 5;5;5;5;5;5;5; | 24245682:242 | 24245730:242 | +;+;+;+;+;+;+; | 2609 | atp6v1aa | 1038 | 1162 | 1180 | 644 | 1349 | 812 |
| ENSDARG000 | 5;5;5;5;5;5;5; | 8832299:884 | 8836501:884 | -;-;-;-;-;-;-;-; | 8835 | NA | 12 | 8 | 32 | 10 | 32 | 18 |
| ENSDARG000 | 5;5;5;5;5 | 53998978:539 | 53999064:539 | -;-;-;-;- | 636 | NA | 0 | 0 | 0 | 0 | 0 | 0 |
| ENSDARG000 | 5;5;5;5;5;5;5; | 12528693:125 | 12528811:125 | +;+;+;+;+;+;+; | 1798 | rfc5 | 350 | 655 | 1028 | 583 | 604 | 651 |
| ENSDARG000 | 5;5;5;5;5;5;5 | 58372164:583 | 58372316:583 | +;+;+;+;+;+;+ | 969 | nrgna | 4 | 20 | 185 | 102 | 33 | 114 |
| ENSDARG000 | 5;5;5;5;5;5;5; | 18912950:189 | 18913822:189 | -;-;-;-;-;-;-;-; | 2352 | poli | 123 | 78 | 123 | 98 | 160 | 110 |
| ENSDARG000 | 5;5 | 24252459:242 | 24252849:242 | -;- | 488 | NA | 0 | 0 | 0 | 0 | 0 | 0 |
| ENSDARG000 | 5;5 | 54024166:540 | 54024418:540 | -;- | 520 | NA | 0 | 0 | 0 | 0 | 0 | 0 |
| ENSDARG000 | 5;5;5;5;5;5;5; | 12549551:125 | 12550023:125 | -;-;-;-;-;-;-;- | 1523 | wsb2 | 170 | 96 | 63 | 112 | 73 | 142 |
| ENSDARG000 | 5 | 58416335 | 58417513 | + | 1179 | NA | 0 | 66 | 68 | 29 | 0 | 50 |
| ENSDARG000 | 5;5;5;5 | 25726479:257 | 25728545:257 | -;-;-;- | 2743 | C5H9orf85 | 376 | 331 | 432 | 234 | 368 | 443 |
| ENSDARG000 | 5;5;5;5;5;5;5; | 58421536:584 | 58421944:584 | +;+;+;+;+;+;+; | 1842 | NA | 74 | 45 | 100 | 50 | 62 | 38 |
| ENSDARG000 | 5;5;5;5;5 | 27039024:270 | 27039169:270 | -;-;-;-;- | 977 | NA | 0 | 4 | 6 | 0 | 8 | 0 |
| ENSDARG000 | 5;5;5;5;5;5;5; | 4016271:401 | 4016379:401 | +;+;+;+;+;+;+; | 4011 | ggnbp2 | 634 | 784 | 1091 | 684 | 750 | 692 |
| ENSDARG000 | 5;5;5;5;5;5;5 | 55719757:557 | 55720543:557 | -;-;-;-;-;-;- | 1866 | kcmf1 | 393 | 491 | 709 | 350 | 792 | 323 |
| ENSDARG000 | 5;5;5;5;5;5;5; | 48985861:489 | 48988086:489 | -;-;-;-;-;-;-;-; | 3806 | arrdc3a | 897 | 674 | 1256 | 1422 | 1382 | 943 |
| ENSDARG000 | 5;5;5;5;5;5;5; | 8919698:892 | 8919827:892 | +;+;+;+;+;+;+; | 2453 | agpat9l | 56 | 8 | 32 | 4 | 32 | 10 |
| ENSDARG000 | 5;5;5;5;5;5;5; | 55763276:557 | 55764621:557 | -;-;-;-;-;-;-;-; | 2611 | camk4 | 42 | 34 | 81 | 14 | 119 | 60 |
| ENSDARG000 | 5;5;5;5;5 | 37966505:379 | 37966583:379 | +;+;+;+;+ | 743 | pafah1b2 | 152 | 171 | 152 | 79 | 150 | 123 |
| ENSDARG000 | 5;5;5;5;5;5 | 22435786:224 | 22436002:224 | -;-;-;-;-;- | 601 | NA | 0 | 0 | 0 | 0 | 0 | 6 |
| ENSDARG000 | 5;5;5;5 | 13839316:138 | 13839817:138 | -;-;-;- | 1122 | figla | 0 | 0 | 0 | 0 | 0 | 0 |
| ENSDARG000 | 5;5;5;5;5;5;5; | 39087364:390 | 39087448:390 | +;+;+;+;+;+;+; | 1322 | anxa3a | 2 | 2 | 0 | 0 | 0 | 8 |
| ENSDARG000 | 5;5;5;5;5;5;5; | 20188438:201 | 20189151:201 | -;-;-;-;-;-;-;-; | 1684 | dao.1 | 0 | 0 | 2 | 0 | 0 | 0 |
| ENSDARG000 | 5;5;5;5;5;5;5; | 36832079:368 | 36832578:368 | -;-;-;-;-;-;-;- | 1584 | ckma | 18 | 26 | 12 | 78 | 46 | 22 |
| ENSDARG000 | 5;5;5;5;5;5;5; | 23518399:235 | 23519784:235 | -;-;-;-;-;-;-;-; | 7393 | acsl4b | 1337 | 1113 | 1056 | 1611 | 1748 | 3926 |
| ENSDARG000 | 5;5;5;5;5;5;5; | 66807179:668 | 66808047:668 | -;-;-;-;-;-;-;-; | 3145 | stip1 | 2867 | 3207 | 4082 | 2978 | 3018 | 2744 |
| ENSDARG000 | 5;5;5;5;5;5;5; | 44944778:449 | 44945135:449 | +;+;+;+;+;+;+; | 3500 | dmrt1 | 48 | 12 | 54 | 76 | 22 | 28 |
| ENSDARG000 | 5;5;5;5;5 | 64861368:648 | 64861593:648 | -;-;-;-;- | 3684 | zbtb26 | 106 | 90 | 152 | 92 | 162 | 76 |
| ENSDARG000 | 5;5;5;5;5;5;5 | 23576328:235 | 23576612:235 | -;-;-;-;-;-;- | 1114 | cwc15 | 10 | 15 | 0 | 4 | 5 | 2 |
| ENSDARG000 | 5;5;5;5;5;5;5; | 22465086:224 | 22465451:224 | -;-;-;-;-;-;-;-; | 4239 | NA | 10 | 0 | 8 | 22 | 9 | 14 |
| ENSDARG000 | 5;5;5;5;5;5;5 | 14165567:141 | 14166572:141 | -;-;-;-;-;-;- | 1944 | NA | 44 | 2 | 0 | 26 | 20 | 0 |
| ENSDARG000 | 5;5;5;5;5;5;5; | 66755748:667 | 66755927:667 | -;-;-;-;-;-;-;-; | 1926 | FERMT3 | 42 | 53 | 62 | 37 | 117 | 89 |
| ENSDARG000 | 5;5;5;5;5;5;5; | 54035570:540 | 54037952:540 | -;-;-;-;-;-;-;-; | 3424 | NA | 0 | 0 | 0 | 0 | 0 | 0 |
| ENSDARG000 | 5 | 32360336 | 32363372 | - | 3037 | gas1b | 118 | 32 | 220 | 40 | 272 | 116 |
| ENSDARG000 | 5;5 | 58676282:586 | 58676968:586 | -;- | 782 | NA | 0 | 0 | 0 | 0 | 0 | 0 |

| | | | | | | | | | | | | |
|---|---|---|---|---|---|---|---|---|---|---|---|---|
| ENSDARG000 | 5;5;5;5;5;5;5; | 31860043:318 | 31860163:318 | +;+;+;+;+;+;+; | 1424 | iscub | 2242 | 852 | 680 | 1368 | 978 | 1128 |
| ENSDARG000 | 5;5;5;5;5;5;5; | 14306475:143 | 14311630:143 | -;-;-;-;-;-;-;- | 11443 | tet3 | 2264 | 2575 | 6287 | 3673 | 3423 | 2650 |
| ENSDARG000 | 5;5;5;5;5;5;5; | 41485277:414 | 41486119:414 | -;-;-;-;-;-;-;- | 3669 | EEF2 | 96 | 239 | 1203 | 1478 | 95 | 527 |
| ENSDARG000 | 5;5;5;5;5;5;5; | 20197138:201 | 20197564:201 | -;-;-;-;-;-;-;- | 1384 | dao.3 | 4 | 2 | 2 | 0 | 10 | 0 |
| ENSDARG000 | 5;5;5;5;5;5;5; | 29296396:292 | 29296926:292 | -;-;-;-;-;-;-;- | 7205 | cacna1ba | 0 | 4 | 4 | 0 | 0 | 0 |
| ENSDARG000 | 5;5 | 14368100:143 | 14368467:143 | -;- | 539 | NA | 0 | 0 | 0 | 0 | 0 | 0 |
| ENSDARG000 | 5;5;5 | 24262230:242 | 24264878:242 | -;-;- | 3208 | NA | 282 | 180 | 277 | 151 | 490 | 0 |
| ENSDARG000 | 5;5;5;5;5;5;5; | 32380026:323 | 32381674:323 | -;-;-;-;-;-;-;- | 3453 | fbxw2 | 635 | 828 | 973 | 534 | 856 | 758 |
| ENSDARG000 | 5;5;5;5;5;5;5; | 25733774:257 | 25734136:257 | +;+;+;+;+;+;+; | 6233 | abhd17b | 2191 | 2489 | 1765 | 1861 | 2453 | 2880 |
| ENSDARG000 | 5;5;5;5;5;5 | 18925367:189 | 18925823:189 | +;+;+;+;+;+ | 2384 | pgam5 | 1002 | 882 | 1204 | 707 | 1146 | 1130 |
| ENSDARG000 | 5;5;5;5;5;5;5; | 64869176:648 | 64869387:648 | -;-;-;-;-;-;-;- | 1522 | krt1-c5 | 0 | 0 | 0 | 0 | 0 | 0 |
| ENSDARG000 | 5;5;5;5 | 8959728:8960 | 8960312:8960 | -;-;-;- | 873 | si:ch211-155r | 0 | 0 | 0 | 0 | 0 | 0 |
| ENSDARG000 | 5;5;5;5;5;5;5 | 61919217:619 | 61919476:619 | +;+;+;+;+;+;+ | 1775 | RSPH6A | 0 | 12 | 48 | 24 | 12 | 0 |
| ENSDARG000 | 5;5;5;5;5;5;5; | 27137473:272 | 27137936:272 | +;+;+;+;+;+;+; | 3239 | UNC5D | 0 | 0 | 2 | 0 | 0 | 1 |
| ENSDARG000 | 5;5;5;5 | 37978501:379 | 37978528:379 | +;+;+;+ | 974 | apoa1a | 1 | 2 | 32 | 16 | 5 | 12 |
| ENSDARG000 | 5;5 | 54091223:540 | 54091395:540 | -;- | 302 | NA | 0 | 0 | 0 | 0 | 0 | 0 |
| ENSDARG000 | 5;5;5;5;5;5;5; | 12590403:125 | 12590570:125 | +;+;+;+;+;+;+; | 1373 | si:dkev-98f17. | 26 | 10 | 12 | 7 | 16 | 27 |
| ENSDARG000 | 5;5;5;5 | 24290643:242 | 24291530:242 | -;-;-;- | 1041 | NA | 0 | 0 | 0 | 0 | 0 | 0 |
| ENSDARG000 | 5;5;5;5;5;5;5; | 58695838:586 | 58697169:586 | -;-;-;-;-;-;-;- | 7004 | NA | 4549 | 2707 | 3672 | 2830 | 3814 | 2631 |
| ENSDARG000 | 5;5;5;5;5;5;5 | 24282570:242 | 24282691:242 | +;+;+;+;+;+;+ | 1308 | zdhhc23a | 0 | 0 | 0 | 0 | 0 | 2 |
| ENSDARG000 | 5;5;5 | 54106010:541 | 54106052:541 | +;+;+ | 857 | NA | 0 | 0 | 0 | 0 | 0 | 0 |
| ENSDARG000 | 5;5;5;5;5;5;5; | 34981584:349 | 34981663:349 | +;+;+;+;+;+;+; | 1656 | ankra2 | 212 | 336 | 368 | 361 | 343 | 364 |
| ENSDARG000 | 5;5;5;5;5;5;5; | 25760112:257 | 25760374:257 | +;+;+;+;+;+;+; | 7173 | cemip2 | 1407 | 1180 | 1332 | 1131 | 1396 | 1334 |
| ENSDARG000 | 5;5;5;5;5;5;5; | 32402026:324 | 32403964:324 | -;-;-;-;-;-;-;- | 3127 | ncs1a | 16 | 20 | 16 | 41 | 30 | 54 |
| ENSDARG000 | 5;5;5;5;5;5;5; | 49002612:490 | 49004252:490 | -;-;-;-;-;-;-;- | 3241 | NA | 0 | 2 | 0 | 0 | 0 | 0 |
| ENSDARG000 | 5;5;5;5;5;5 | 36850650:368 | 36851216:368 | +;+;+;+;+;+ | 1718 | nccrp1 | 992 | 737 | 633 | 618 | 788 | 914 |
| ENSDARG000 | 5;5;5;5;5;5;5; | 37982117:379 | 37982179:379 | -;-;-;-;-;-;-;- | 2212 | NA | 177 | 172 | 256 | 94 | 207 | 162 |
| ENSDARG000 | 5;5;5;5;5 | 37981800:379 | 37982179:379 | -;-;-;-;- | 1019 | NA | 39 | 32 | 27 | 29 | 66 | 12 |
| ENSDARG000 | 5;5;5;5;5;5;5; | 22515347:225 | 22515947:225 | -;-;-;-;-;-;-;- | 1926 | glra4b | 0 | 0 | 0 | 0 | 0 | 0 |
| ENSDARG000 | 5 | 12713258 | 12713920 | - | 663 | cldn26 | 0 | 0 | 0 | 0 | 0 | 0 |
| ENSDARG000 | 5;5 | 31872049:318 | 31873160:318 | -;- | 1533 | tmem119b | 4659 | 798 | 202 | 1055 | 430 | 1337 |
| ENSDARG000 | 5;5;5;5;5 | 22510639:225 | 22510800:225 | +;+;+;+;+ | 951 | rpl36a | 9724 | 9033 | 13383 | 11793 | 7286 | 13552 |
| ENSDARG000 | 5;5;5;5;5;5;5; | 30518036:305 | 30518645:305 | +;+;+;+;+;+;+; | 2600 | hmbsa | 462 | 645 | 659 | 386 | 552 | 625 |
| ENSDARG000 | 5;5;5;5;5;5;5; | 54161362:541 | 54162236:541 | -;-;-;-;-;-;-;- | 2442 | grk1b | 0 | 0 | 0 | 0 | 0 | 0 |
| ENSDARG000 | 5;5;5;5;5;5;5; | 14375108:143 | 14376183:143 | -;-;-;-;-;-;-;- | 2299 | ap3m2 | 3696 | 4675 | 4476 | 3698 | 3400 | 5607 |
| ENSDARG000 | 5;5;5;5;5;5;5; | 39099172:390 | 39099912:390 | +;+;+;+;+;+;+; | 5624 | bmp2k | 575 | 658 | 1009 | 601 | 794 | 976 |
| ENSDARG000 | 5;5;5 | 54159265:541 | 54159772:541 | -;-;- | 936 | NA | 0 | 0 | 0 | 0 | 0 | 0 |
| ENSDARG000 | 5;5;5;5;5;5;5; | 54110853:541 | 54115899:541 | -;-;-;-;-;-;-;- | 7678 | NA | 6 | 22 | 14 | 10 | 72 | 40 |
| ENSDARG000 | 5;5 | 39142856:391 | 39142940:391 | +;+ | 1690 | NA | 0 | 0 | 0 | 0 | 0 | 0 |
| ENSDARG000 | 5;5;5;5;5;5;5; | 55866769:558 | 55867020:558 | -;-;-;-;-;-;-;- | 2917 | wdr36 | 683 | 1083 | 1464 | 592 | 1113 | 740 |
| ENSDARG000 | 5;5 | 45000597:450 | 45001182:450 | +;+ | 1920 | dmrt3a | 0 | 0 | 2 | 8 | 4 | 4 |
| ENSDARG000 | 5;5;5;5;5;5;5 | 45007962:450 | 45008137:450 | +;+;+;+;+;+;+ | 4333 | dmrt2a | 10 | 4 | 0 | 0 | 5 | 2 |
| ENSDARG000 | 5;5;5;5;5;5;5 | 4054704:4131 | 4055184:4132 | +;+;+;+;+;+;+ | 1791 | dhrs11a | 57 | 8 | 40 | 44 | 60 | 50 |
| ENSDARG000 | 5;5;5;5;5;5;5; | 61931337:619 | 61933000:619 | -;-;-;-;-;-;-;- | 2511 | napaa | 574 | 472 | 786 | 744 | 675 | 535 |
| ENSDARG000 | 5;5;5;5;5;5;5; | 12724046:127 | 12724634:127 | -;-;-;-;-;-;-;- | 3558 | lztr1 | 2985 | 965 | 1001 | 1524 | 1529 | 1266 |
| ENSDARG000 | 5;5;5;5;5;5;5; | 31880586:318 | 31884644:318 | -;-;-;-;-;-;-;- | 6074 | CORO1C | 970 | 1561 | 1380 | 895 | 941 | 1060 |
| ENSDARG000 | 5 | 32791245 | 32793481 | + | 2237 | ier5l | 5241 | 4951 | 5393 | 4425 | 4793 | 5966 |
| ENSDARG000 | 5;5;5;5;5;5;5; | 49019182:490 | 49019603:490 | -;-;-;-;-;-;-;- | 2140 | si:dkev-172m | 510 | 609 | 798 | 488 | 516 | 570 |
| ENSDARG000 | 5;5;5;5;5;5;5; | 20255568:202 | 20255623:202 | +;+;+;+;+;+;+; | 2016 | SSH1 | 348 | 364 | 218 | 173 | 387 | 311 |
| ENSDARG000 | 5;5;5;5;5;5;5; | 24305877:243 | 24305904:243 | +;+;+;+;+;+;+; | 1424 | ctsll | 0 | 0 | 0 | 0 | 6 | 0 |
| ENSDARG000 | 5;5;5;5 | 61941610:619 | 61941876:619 | +;+;+;+ | 4243 | NA | 586 | 351 | 448 | 300 | 518 | 430 |
| ENSDARG000 | 5;5;5;5;5;5;5; | 36870737:368 | 36871042:368 | +;+;+;+;+;+;+; | 2685 | slc8a2a | 0 | 10 | 0 | 0 | 19 | 4 |
| ENSDARG000 | 5;5 | 38009624:380 | 38009767:380 | +;+ | 499 | NA | 0 | 0 | 0 | 0 | 0 | 0 |
| ENSDARG000 | 5;5;5;5;5;5;5; | 4150814:4151 | 4151742:4151 | -;-;-;-;-;-;-;- | 2570 | NA | 0 | 0 | 4 | 0 | 36 | 0 |
| ENSDARG000 | 5;5;5;5;5;5;5; | 18943043:189 | 18943811:189 | -;-;-;-;-;-;-;- | 3812 | ankle2 | 192 | 181 | 282 | 138 | 234 | 178 |
| ENSDARG000 | 5;5 | 38022577:380 | 38022709:380 | +;+ | 486 | NA | 0 | 0 | 0 | 0 | 0 | 0 |
| ENSDARG000 | 5;5;5;5;5;5;5; | 14394323:143 | 14398521:143 | -;-;-;-;-;-;-;- | 7796 | kat6a | 1937 | 1430 | 2428 | 1805 | 2228 | 1731 |
| ENSDARG000 | 5;5;5;5;5;5;5; | 67079540:670 | 67079749:670 | -;-;-;-;-;-;-;- | 2270 | rps6ka4 | 256 | 297 | 276 | 135 | 307 | 197 |
| ENSDARG000 | 5;5;5;5;5;5;5; | 41496490:414 | 41496728:414 | +;+;+;+;+;+;+; | 2925 | fancg | 62 | 153 | 203 | 98 | 108 | 75 |
| ENSDARG000 | 5;5 | 24329983:243 | 24332272:243 | -;- | 2948 | znf703 | 1030 | 425 | 861 | 1038 | 1311 | 1145 |
| ENSDARG000 | 5;5;5;5;5;5;5; | 32793886:327 | 32798957:327 | -;-;-;-;-;-;-;- | 6294 | ptpa | 1011 | 1012 | 1129 | 900 | 966 | 840 |
| ENSDARG000 | 5;5;5;5;5;5;5; | 45138934:451 | 45139029:451 | +;+;+;+;+;+;+; | 6267 | smarca2 | 2917 | 2114 | 3420 | 3092 | 3465 | 2584 |
| ENSDARG000 | 5;5;5;5;5;5;5; | 64900223:649 | 64900624:649 | +;+;+;+;+;+;+; | 2480 | ptgs1 | 8 | 2 | 10 | 0 | 20 | 0 |
| ENSDARG000 | 5;5;5;5;5;5;5; | 23585215:235 | 23586015:235 | -;-;-;-;-;-;-;- | 2069 | fam76b | 635 | 757 | 844 | 556 | 713 | 441 |
| ENSDARG000 | 5;5;5;5;5;5;5; | 8964926:8964 | 8965485:8965 | +;+;+;+;+;+;+; | 4237 | tnksb | 1768 | 1478 | 1717 | 1447 | 1634 | 1230 |
| ENSDARG000 | 5;5;5;5;5;5;5; | 20319519:203 | 20319668:203 | +;+;+;+;+;+;+; | 5204 | coro1ca | 8458 | 5819 | 5978 | 5628 | 5538 | 5845 |
| ENSDARG000 | 5;5;5;5;5;5;5; | 25945246:259 | 25945248:259 | +;+;+;+;+;+;+; | 5208 | TRPM3 | 0 | 2 | 0 | 0 | 2 | 0 |
| ENSDARG000 | 5;5;5;5;5;5 | 64893630:648 | 64893815:648 | -;-;-;-;-;- | 768 | LHX6 | 0 | 0 | 0 | 0 | 0 | 0 |
| ENSDARG000 | 5;5;5;5;5;5;5; | 27404946:274 | 27405039:274 | +;+;+;+;+;+;+; | 1647 | hdr | 133 | 63 | 418 | 52 | 79 | 461 |
| ENSDARG000 | 5;5;5 | 49734238:497 | 49736006:497 | -;-;- | 1946 | NA | 8 | 0 | 0 | 4 | 0 | 2 |
| ENSDARG000 | 5;5;5;5;5 | 25964182:259 | 25964414:259 | -;-;-;-;- | 569 | NA | 2 | 0 | 0 | 0 | 0 | 0 |
| ENSDARG000 | 5;5;5 | 58836354:588 | 58837094:588 | -;-;- | 1140 | tlcd5b | 6 | 4 | 4 | 0 | 2 | 12 |
| ENSDARG000 | 5 | 27432958 | 27434351 | + | 1394 | histh1l1 | 7538 | 7106 | 7790 | 11161 | 4396 | 7902 |
| ENSDARG000 | 5;5;5;5;5;5 | 38035337:380 | 38035526:380 | -;-;-;-;-;- | 958 | NA | 0 | 0 | 2 | 0 | 0 | 0 |
| ENSDARG000 | 5 | 27429872 | 27430247 | + | 376 | zgc:153409 | 26 | 2 | 2 | 0 | 0 | 12 |
| ENSDARG000 | 5;5;5;5;5;5;5; | 27421025:274 | 27421116:274 | +;+;+;+;+;+;+; | 1316 | cyb561a3a | 296 | 261 | 202 | 221 | 334 | 224 |
| ENSDARG000 | 5;5;5;5;5;5;5; | 42835082:428 | 42835651:428 | -;-;-;-;-;-;-;- | 2472 | flot2a | 3351 | 2570 | 3975 | 2943 | 2959 | 4857 |
| ENSDARG000 | 5;5;5;5;5;5;5; | 58850626:588 | 58853855:588 | -;-;-;-;-;-;-;- | 5105 | NA | 72 | 241 | 526 | 90 | 235 | 48 |
| ENSDARG000 | 5;5;5;5;5;5;5; | 34987448:349 | 34987831:349 | -;-;-;-;-;-;-;- | 1112 | btf3 | 8541 | 8955 | 13861 | 15025 | 8640 | 13049 |
| ENSDARG000 | 5;5;5;5;5;5;5; | 55920892:559 | 55922092:559 | -;-;-;-;-;-;-;- | 2053 | slc25a46 | 248 | 369 | 306 | 200 | 408 | 226 |
| ENSDARG000 | 5;5;5;5;5;5;5; | 4209163:4209 | 4209859:4209 | -;-;-;-;-;-;-;- | 1219 | NA | 8 | 85 | 150 | 78 | 318 | 172 |
| ENSDARG000 | 5;5;5;5;5;5;5; | 61953086:619 | 61953117:619 | -;-;-;-;-;-;-;- | 2575 | NA | 0 | 0 | 0 | 0 | 0 | 29 |
| ENSDARG000 | 5;5;5;5;5;5;5; | 18965494:189 | 18966806:189 | -;-;-;-;-;-;-;- | 5826 | golga3 | 724 | 717 | 1317 | 782 | 839 | 662 |
| ENSDARG000 | 5;5;5;5;5;5;5; | 24510619:245 | 24512319:245 | -;-;-;-;-;-;-;- | 2765 | tia1l | 3646 | 4193 | 4935 | 2835 | 4494 | 3582 |
| ENSDARG000 | 5;5;5 | 38040407:380 | 38040485:380 | +;+;+ | 574 | si:dkev-111e8 | 0 | 0 | 0 | 0 | 0 | 0 |
| ENSDARG000 | 5;5;5 | 49744713:497 | 49745916:497 | +;+;+ | 2994 | nr2f1a | 76 | 8 | 6 | 32 | 26 | 36 |
| ENSDARG000 | 5;5;5;5 | 67126774:671 | 67130454:671 | -;-;-;- | 4796 | rom1a | 0 | 0 | 0 | 0 | 0 | 0 |
| ENSDARG000 | 5;5;5;5;5;5;5; | 38060959:380 | 38061177:380 | -;-;-;-;-;-;-;- | 1060 | si:dkev-111e8 | 4 | 1 | 0 | 2 | 4 | 4 |
| ENSDARG000 | 5;5;5;5;5;5;5 | 39154617:391 | 39155837:391 | -;-;-;-;-;-;- | 2566 | paqr3a | 468 | 467 | 299 | 182 | 486 | 580 |
| ENSDARG000 | 5;5;5;5;5;5;5; | 12743640:127 | 12743846:127 | +;+;+;+;+;+;+; | 7154 | pole | 193 | 718 | 864 | 204 | 733 | 567 |
| ENSDARG000 | 5;5;5;5;5;5;5; | 69622005:696 | 69622132:696 | +;+;+;+;+;+;+; | 2525 | vps33a | 912 | 798 | 840 | 750 | 972 | 852 |
| ENSDARG000 | 5;5;5;5 | 20428838:204 | 20428917:204 | +;+;+;+ | 755 | NA | 2 | 2 | 0 | 0 | 2 | 0 |
| ENSDARG000 | 5;5;5;5;5;5;5; | 69641166:696 | 69641367:696 | -;-;-;-;-;-;-;- | 1269 | eif2b1 | 373 | 483 | 865 | 555 | 479 | 380 |
| ENSDARG000 | 5 | 20441653 | 20442314 | + | 662 | NA | 0 | 0 | 0 | 0 | 0 | 0 |
| ENSDARG000 | 5;5;5 | 67161442:671 | 67161675:671 | -;-;- | 523 | NA | 0 | 0 | 0 | 0 | 0 | 0 |
| ENSDARG000 | 5;5;5;5;5;5;5; | 4223515:4223 | 4223581:4223 | -;-;-;-;-;-;-;- | 818 | NA | 17 | 84 | 106 | 73 | 302 | 110 |
| ENSDARG000 | 5;5;5;5;5;5;5; | 54280135:542 | 54280690:542 | +;+;+;+;+;+;+; | 1244 | rangrf | 99 | 120 | 198 | 128 | 162 | 138 |
| ENSDARG000 | 5;5 | 26075230:260 | 26075939:260 | +;+ | 1460 | klf9 | 164 | 139 | 64 | 136 | 141 | 245 |
| ENSDARG000 | 5;5;5;5;5;5;5; | 14477282:144 | 14478874:144 | -;-;-;-;-;-;-;- | 3723 | si:ch211-244 | 205 | 536 | 637 | 308 | 630 | 290 |
| ENSDARG000 | 5;5 | 49755257:497 | 49755467:497 | +;+ | 668 | NA | 6 | 2 | 0 | 0 | 0 | 0 |
| ENSDARG000 | 5;5;5;5;5;5;5; | 49768223:497 | 49770079:497 | -;-;-;-;-;-;-;- | 3032 | fam172a | 1017 | 622 | 928 | 848 | 1232 | 1187 |
| ENSDARG000 | 5 | 30534702 | 30535327 | - | 626 | h2ax | 62 | 210 | 247 | 84 | 184 | 54 |
| ENSDARG000 | 5;5;5;5;5;5;5; | 23597140:235 | 23597177:235 | +;+;+;+;+;+;+; | 2264 | kat5a | 496 | 226 | 582 | 560 | 399 | 507 |
| ENSDARG000 | 5;5;5;5;5 | 67175283:671 | 67175941:671 | -;-;-;-;- | 1655 | cdca5 | 46 | 308 | 230 | 140 | 211 | 100 |
| ENSDARG000 | 5;5;5;5;5;5;5; | 22550614:225 | 22551226:225 | -;-;-;-;-;-;-;- | 2865 | aifm1 | 813 | 1023 | 1403 | 578 | 1251 | 853 |
| ENSDARG000 | 5;5;5;5;5;5;5; | 32815745:328 | 32816258:328 | +;+;+;+;+;+;+; | 5364 | crata | 478 | 426 | 496 | 546 | 382 | 474 |
| ENSDARG000 | 5;5;5;5;5;5;5; | 50084936:502 | 50086821:502 | -;-;-;-;-;-;-;- | 6111 | NA | 122 | 77 | 71 | 70 | 88 | 116 |
| ENSDARG000 | 5;5;5;5;5;5;5; | 59030096:590 | 59030292:590 | +;+;+;+;+;+;+; | 6315 | rabep1 | 1039 | 811 | 1028 | 668 | 1057 | 840 |
| ENSDARG000 | 5;5;5;5;5;5;5; | 32831561:328 | 32831625:328 | +;+;+;+;+;+;+; | 3298 | NA | 32 | 32 | 333 | 390 | 7 | 10 |
| ENSDARG000 | 5;5;5;5;5;5;5; | 20443888:204 | 20444445:204 | -;-;-;-;-;-;-;- | 1100 | ISCU | 494 | 382 | 521 | 765 | 488 | 693 |
| ENSDARG000 | 5;5;5;5;5;5;5; | 31914906:319 | 31918884:319 | -;-;-;-;-;-;-;- | 6361 | SSH1 | 96 | 52 | 84 | 10 | 56 | 18 |
| ENSDARG000 | 5;5 | 26079178:260 | 26079354:260 | +;+ | 594 | NA | 36 | 25 | 5 | 44 | 36 | 36 |
| ENSDARG000 | 5;5;5;5;5;5;5; | 50371509:503 | 50371736:503 | +;+;+;+;+;+;+; | 3821 | NA | 82 | 40 | 86 | 30 | 24 | 94 |
| ENSDARG000 | 5;5;5;5;5;5;5; | 26082434:260 | 26082888:260 | -;-;-;-;-;-;-;- | 1066 | fam219ab | 66 | 89 | 46 | 42 | 9 | 104 |
| ENSDARG000 | 5;5;5;5;5;5;5; | 4232196:4235 | 4235671:4235 | -;-;-;-;-;-;-;- | 4321 | NA | 0 | 0 | 0 | 0 | 0 | 0 |
| ENSDARG000 | 5;5;5;5;5;5;5; | 30537640:305 | 30538129:305 | -;-;-;-;-;-;-;- | 2732 | c2cd2l | 93 | 63 | 89 | 57 | 109 | 69 |
| ENSDARG000 | 5;5;5;5;5;5;5; | 69650148:696 | 69650292:696 | +;+;+;+;+;+;+; | 1271 | gtf2h3 | 167 | 295 | 447 | 237 | 354 | 306 |
| ENSDARG000 | 5;5;5;5;5;5 | 61976511:619 | 61976734:619 | +;+;+;+;+;+ | 1641 | EHD2 | 4 | 0 | 0 | 0 | 20 | 0 |
| ENSDARG000 | 5;5;5;5;5;5;5; | 55934129:559 | 55934187:559 | +;+;+;+;+;+;+; | 2188 | tmem150ab | 154 | 257 | 252 | 85 | 307 | 153 |
| ENSDARG000 | 5;5;5;5 | 39504136:395 | 39504320:395 | +;+;+;+ | 2502 | prdm8b | 49 | 2 | 34 | 30 | 34 | 10 |
| ENSDARG000 | 5;5;5;5;5 | 54297319:542 | 54297547:542 | -;-;-;-;- | 955 | slc25a35 | 38 | 49 | 55 | 35 | 87 | 90 |
| ENSDARG000 | 5;5;5;5;5;5;5; | 67188245:671 | 67188530:671 | -;-;-;-;-;-;-;- | 4767 | CLIP1 | 966 | 500 | 802 | 414 | 869 | 580 |
| ENSDARG000 | 5;5;5;5;5;5;5; | 42943972:429 | 42944133:429 | -;-;-;-;-;-;-;- | 1482 | grsf1 | 262 | 584 | 518 | 316 | 477 | 294 |
| ENSDARG000 | 5;5 | 38088594:380 | 38091779:380 | -;- | 3538 | NA | 0 | 0 | 0 | 0 | 3 | 0 |
| ENSDARG000 | 5 | 34997763 | 34999553 | + | 1791 | foxd1 | 0 | 2 | 13 | 0 | 4 | 6 |
| ENSDARG000 | 5;5;5 | 38096729:381 | 38097701:381 | +;+;+ | 2367 | si:ch211-284 | 84 | 86 | 132 | 51 | 73 | 75 |
| ENSDARG000 | 5;5;5;5;5;5;5; | 24520437:245 | 24520548:245 | +;+;+;+;+;+;+; | 4703 | dgcr8 | 1594 | 1071 | 1935 | 1426 | 2202 | 2075 |
| ENSDARG000 | 5;5;5;5;5;5;5; | 19006701:190 | 19008564:190 | -;-;-;-;-;-;-;- | 3010 | ranbp1 | 2234 | 3464 | 5246 | 2054 | 3466 | 2847 |
| ENSDARG000 | 5;5;5;5;5;5;5; | 35123662:351 | 35126075:351 | -;-;-;-;-;-;-;- | 5879 | fcho2 | 1118 | 728 | 892 | 455 | 1113 | 951 |
| ENSDARG000 | 5;5;5;5;5 | 54304494:543 | 54304744:543 | -;-;-;-;- | 651 | tmem141 | 102 | 102 | 68 | 90 | 128 | 207 |
| ENSDARG000 | 5;5;5;5 | 38104901:381 | 38105606:381 | -;-;-;- | 1328 | NA | 5 | 24 | 28 | 10 | 7 | 8 |
| ENSDARG000 | 5;5 | 35121048:351 | 35121094:351 | +;+ | 1262 | NA | 30 | 10 | 0 | 2 | 9 | 7 |
| ENSDARG000 | 5;5;5;5;5;5;5 | 54313824:543 | 54314056:543 | -;-;-;-;-;-;- | 696 | NA | 0 | 0 | 0 | 0 | 18 | 0 |
| ENSDARG000 | 5;5;5 | 69662132:696 | 69662259:696 | +;+;+ | 774 | NA | 0 | 0 | 0 | 0 | 4 | 0 |
| ENSDARG000 | 5;5;5;5;5;5;5; | 38110384:381 | 38111134:381 | -;-;-;-;-;-;-;- | 2330 | NA | 29 | 38 | 48 | 26 | 32 | 46 |
| ENSDARG000 | 5;5;5 | 69668643:696 | 69668808:696 | +;+;+ | 935 | NA | 38 | 16 | 46 | 26 | 36 | 43 |
| ENSDARG000 | 5;5;5;5;5;5;5; | 31931834:319 | 31931978:319 | -;-;-;-;-;-;-;- | 1379 | NA | 0 | 0 | 0 | 0 | 0 | 0 |
| ENSDARG000 | 5;5;5;5;5;5;5; | 65009764:650 | 65009950:650 | -;-;-;-;-;-;-;- | 1650 | anxa1c | 58 | 6 | 0 | 0 | 20 | 0 |
| ENSDARG000 | 5;5;5 | 39537195:395 | 39537411:395 | +;+;+ | 678 | fgf5 | 0 | 0 | 0 | 0 | 0 | 0 |
| ENSDARG000 | 5;5;5;5;5;5;5; | 41521043:415 | 41521186:415 | -;-;-;-;-;-;-;- | 1098 | akr1a1a | 0 | 2 | 0 | 0 | 4 | 0 |
| ENSDARG000 | 5;5;5;5;5;5;5; | 50912729:509 | 50913443:509 | +;+;+;+;+;+;+; | 7016 | cert1a | 803 | 426 | 377 | 496 | 671 | 345 |
| ENSDARG000 | 5;5;5;5;5;5;5; | 61998641:619 | 61998766:619 | +;+;+;+;+;+;+; | 857 | selenow1 | 619 | 364 | 359 | 561 | 589 | 627 |
| ENSDARG000 | 5;5;5;5;5;5;5; | 36895545:368 | 36895674:368 | +;+;+;+;+;+;+; | 2250 | srsf7a | 1897 | 2118 | 2781 | 1583 | 2248 | 1531 |
| ENSDARG000 | 5;5;5;5;5;5;5; | 41510387:415 | 41510529:415 | +;+;+;+;+;+;+; | 3097 | vcp | 11027 | 11546 | 11716 | 4958 | 12417 | 6800 |
| ENSDARG000 | 5;5;5;5;5;5;5; | 30588856:305 | 30589010:305 | +;+;+;+;+;+;+; | 1295 | dpagt1 | 383 | 417 | 594 | 326 | 588 | 431 |
| ENSDARG000 | 5;5;5 | 55976013:559 | 55976682:559 | -;-;- | 843 | NA | 0 | 0 | 0 | 1 | 0 | 0 |
| ENSDARG000 | 5;5;5;5;5;5;5; | 55984270:559 | 55984403:559 | +;+;+;+;+;+;+; | 1533 | fkbp6 | 6 | 0 | 0 | 0 | 2 | 0 |
| ENSDARG000 | 5;5;5;5;5;5;5; | 30596477:305 | 30596601:305 | +;+;+;+;+;+;+; | 5192 | hinfp | 130 | 144 | 243 | 120 | 210 | 128 |
| ENSDARG000 | 5;5;5;5;5;5;5; | 45746457:457 | 45748851:457 | -;-;-;-;-;-;-;- | 6822 | slc4a4a | 4 | 8 | 4 | 24 | 50 | 20 |
| ENSDARG000 | 5;5;5;5;5;5;5; | 4253929:4257 | 4254233:4257 | -;-;-;-;-;-;-;- | 3030 | srrt | 1341 | 1785 | 2530 | 1154 | 2400 | 1264 |
| ENSDARG000 | 5;5;5;5;5;5;5; | 20453874:204 | 20454168:204 | +;+;+;+;+;+;+; | 4754 | sart3 | 802 | 1143 | 1307 | 645 | 1020 | 814 |
| ENSDARG000 | 5;5;5;5;5;5;5; | 69702602:697 | 69702885:697 | -;-;-;-;-;-;-;- | 1189 | dguok | 80 | 108 | 117 | 86 | 95 | 115 |
| ENSDARG000 | 5;5;5;5;5;5;5; | 67250959:672 | 67255069:672 | -;-;-;-;-;-;-;- | 5552 | zcchc8 | 372 | 428 | 462 | 291 | 496 | 426 |
| ENSDARG000 | 5;5;5;5;5;5;5; | 24532401:245 | 24532700:245 | -;-;-;-;-;-;-;- | 2466 | trmt2a | 157 | 251 | 448 | 152 | 310 | 199 |
| ENSDARG000 | 5;5;5;5;5;5;5 | 30607838:306 | 30607992:306 | +;+;+;+;+;+;+ | 965 | NA | 0 | 0 | 0 | 0 | 0 | 0 |
| ENSDARG000 | 5;5;5;5;5;5;5; | 31944270:319 | 31944354:319 | +;+;+;+;+;+;+; | 2701 | UNG | 1 | 6 | 0 | 0 | 0 | 0 |
| ENSDARG000 | 5;5;5 | 39667272:396 | 39667781:396 | +;+;+ | 1601 | bmp3 | 3 | 2 | 0 | 0 | 10 | 0 |
| ENSDARG000 | 5;5;5;5;5;5;5; | 39563301:395 | 39563430:395 | +;+;+;+;+;+;+; | 1127 | cfap299 | 0 | 0 | 0 | 0 | 6 | 0 |
| ENSDARG000 | 5;5;5;5;5;5;5; | 54336898:543 | 54338400:543 | -;-;-;-;-;-;-;- | 2290 | zmynd19 | 310 | 608 | 527 | 293 | 559 | 344 |
| ENSDARG000 | 5;5;5;5;5;5;5; | 19021786:190 | 19022855:190 | -;-;-;-;-;-;-;- | 3914 | NA | 317 | 255 | 468 | 684 | 316 | 604 |
| ENSDARG000 | 5;5;5;5;5;5;5; | 59089699:590 | 59089996:590 | -;-;-;-;-;-;-;- | 2410 | nup88 | 474 | 594 | 760 | 396 | 632 | 479 |
| ENSDARG000 | 5;5;5;5;5;5;5; | 12807902:128 | 12809539:128 | -;-;-;-;-;-;-;- | 3557 | p2rx2 | 13 | 5 | 0 | 6 | 26 | 0 |

| | | | | | | | | | | | | |
|---|---|---|---|---|---|---|---|---|---|---|---|---|
| ENSDARG000 | 5;5;5 | 38153360:381 | 38153412:381 | +;+;+ | 716 | NA | 0 | 0 | 0 | 0 | 0 | 0 |
| ENSDARG000 | 5;5;5;5;5;5;5; | 38123633:381 | 38124065:381 | -;-;-;-;-;-;-;-; | 4522 | gucy2d | 2 | 2 | 2 | 2 | 2 | 0 |
| ENSDARG000 | 5;5 | 26100788:261 | 26103644:261 | -;- | 3142 | NA | 12 | 9 | 20 | 34 | 10 | 32 |
| ENSDARG000 | 5;5;5;5;5;5;5; | 62009765:620 | 62009951:620 | +;+;+;+;+;+;+; | 2109 | AKAP10 | 568 | 491 | 549 | 423 | 461 | 446 |
| ENSDARG000 | 5;5;5;5;5;5;5; | 69708528:697 | 69709183:697 | -;-;-;-;-;-;-;-; | 1722 | mob1a | 2075 | 1930 | 1800 | 3011 | 1817 | 2750 |
| ENSDARG000 | 5;5;5;5 | 9037650:9040 | 9037960:9040 | +;+;+;+ | 1745 | si:ch211-155r | 82 | 40 | 6 | 14 | 76 | 22 |
| ENSDARG000 | 5;5;5;5;5;5;5; | 31959954:319 | 31960136:319 | +;+;+;+;+;+;+; | 3064 | NA | 0 | 0 | 6 | 2 | 4 | 7 |
| ENSDARG000 | 5;5;5;5;5;5 | 20485624:204 | 20487545:204 | -;-;-;-;-;- | 2431 | ficd | 208 | 211 | 484 | 244 | 366 | 292 |
| ENSDARG000 | 5;5;5;5;5;5;5; | 69716458:697 | 69716750:697 | +;+;+;+;+;+;+; | 2386 | mthfd2 | 4226 | 3069 | 3486 | 1854 | 3039 | 2359 |
| ENSDARG000 | 5;5;5;5;5;5;5; | 45889932:458 | 45891445:458 | -;-;-;-;-;-;-;-; | 2650 | ifngr2 | 1349 | 1301 | 791 | 505 | 1046 | 1657 |
| ENSDARG000 | 5;5;5;5;5;5;5; | 14529101:145 | 14530631:145 | -;-;-;-;-;-;-;-; | 3940 | sema4c | 4162 | 2458 | 2516 | 3571 | 1455 | 2553 |
| ENSDARG000 | 5;5;5;5;5;5;5; | 29491104:294 | 29491845:294 | -;-;-;-;-;-;-;-; | 3867 | ehmt1a | 684 | 503 | 800 | 367 | 689 | 654 |
| ENSDARG000 | 5;5;5 | 56023186:560 | 56023206:560 | +;+;+ | 1767 | FZD9 | 8 | 1 | 5 | 4 | 6 | 0 |
| ENSDARG000 | 5;5;5;5;5;5;5 | 27488975:274 | 27489109:274 | +;+;+;+;+;+;+ | 1792 | sfrp1a | 161 | 4 | 34 | 44 | 45 | 132 |
| ENSDARG000 | 5;5;5;5;5;5;5; | 27434601:274 | 27434785:274 | +;+;+;+;+;+;+; | 6460 | loxl2b | 18181 | 7195 | 9056 | 46036 | 19116 | 11333 |
| ENSDARG000 | 5 | 27437481 | 27438812 | - | 1332 | drd7 | 6 | 43 | 161 | 369 | 14 | 17 |
| ENSDARG000 | 5;5;5;5;5;5;5; | 56090851:560 | 56091126:560 | -;-;-;-;-;-;-;-; | 2066 | mks1 | 81 | 82 | 74 | 46 | 60 | 58 |
| ENSDARG000 | 5;5;5;5;5;5;5; | 26111596:261 | 26111696:261 | -;-;-;-;-;-;-;- | 973 | ela3l | 0 | 0 | 0 | 0 | 6 | 0 |
| ENSDARG000 | 5;5;5;5;5;5;5; | 42951852:429 | 42952028:429 | +;+;+;+;+;+;+; | 2313 | mob1ba | 1310 | 1172 | 1307 | 1251 | 1673 | 1219 |
| ENSDARG000 | 5;5 | 20542992:205 | 20543054:205 | +;+ | 845 | NA | 35 | 21 | 36 | 24 | 47 | 131 |
| ENSDARG000 | 5;5;5;5;5;5;5; | 65022287:650 | 65022966:650 | -;-;-;-;-;-;-;-; | 1704 | anxa1b | 58 | 2 | 0 | 12 | 24 | 2 |
| ENSDARG000 | 5;5;5 | 20570573:205 | 20570647:205 | +;+;+ | 1399 | cmklr1 | 75 | 33 | 100 | 44 | 273 | 148 |
| ENSDARG000 | 5 | 20599658 | 20600537 | + | 880 | CMKLR1 | 4 | 0 | 0 | 0 | 4 | 5 |
| ENSDARG000 | 5;5;5;5;5;5;5; | 23608026:236 | 23608335:236 | -;-;-;-;-;-;-;-; | 2132 | mtmr2 | 557 | 689 | 710 | 633 | 669 | 525 |
| ENSDARG000 | 5;5;5;5;5;5 | 54400971:544 | 54401438:544 | +;+;+;+;+;+ | 1485 | bspry | 14 | 3 | 8 | 12 | 16 | 8 |
| ENSDARG000 | 5;5;5;5;5 | 22579975:225 | 22580295:225 | +;+;+;+;+ | 1130 | tnfsf10l4 | 38 | 0 | 13 | 1 | 24 | 20 |
| ENSDARG000 | 5;5;5;5;5;5;5; | 19094153:190 | 19094186:190 | +;+;+;+;+;+;+; | 5347 | NA | 169 | 149 | 410 | 309 | 220 | 183 |
| ENSDARG000 | 5;5;5;5;5;5;5; | 9046813:9047 | 9047762:9047 | +;+;+;+;+;+;+; | 5052 | IDUA | 456 | 321 | 354 | 221 | 424 | 342 |
| ENSDARG000 | 5;5;5;5;5;5;5; | 39679767:396 | 39679862:396 | -;-;-;-;-;-;-;-; | 2526 | prkg2 | 2 | 0 | 0 | 0 | 1 | 0 |
| ENSDARG000 | 5;5;5;5;5;5;5; | 12836913:128 | 12837136:128 | +;+;+;+;+;+;+; | 2546 | pes | 1401 | 1296 | 2079 | 934 | 1641 | 1266 |
| ENSDARG000 | 5;5 | 20613466:206 | 20614431:206 | +;+ | 966 | CMKLR1 | 2 | 6 | 2 | 0 | 20 | 2 |
| ENSDARG000 | 5;5;5;5;5;5;5; | 31995683:320 | 31995912:320 | +;+;+;+;+;+;+; | 2352 | SCAI | 66 | 26 | 26 | 24 | 121 | 54 |
| ENSDARG000 | 5;5;5;5;5;5 | 43006422:430 | 43006600:430 | +;+;+;+;+;+ | 1806 | aqp3a | 28076 | 1092 | 1158 | 1127 | 28078 | 11559 |
| ENSDARG000 | 5;5;5;5;5;5;5; | 32865364:328 | 32865486:328 | -;-;-;-;-;-;-;-; | 2166 | lrsam1 | 384 | 191 | 291 | 157 | 264 | 320 |
| ENSDARG000 | 5;5;5;5;5;5;5; | 26121393:261 | 26121475:261 | +;+;+;+;+;+;+; | 2140 | bco2l | 4 | 0 | 6 | 0 | 6 | 0 |
| ENSDARG000 | 5;5;5;5;5;5;5; | 65040228:650 | 65040298:650 | +;+;+;+;+;+;+; | 2226 | pmpca | 704 | 751 | 1207 | 532 | 937 | 714 |
| ENSDARG000 | 5;5;5;5;5;5;5; | 15044307:150 | 15044902:150 | -;-;-;-;-;-;-;-; | 1977 | NA | 4 | 0 | 0 | 0 | 0 | 0 |
| ENSDARG000 | 5;5;5;5;5;5;5; | 20627200:206 | 20628182:206 | -;-;-;-;-;-;-;- | 2349 | wscd2 | 10 | 2 | 4 | 0 | 4 | 13 |
| ENSDARG000 | 5;5;5;5;5;5;5; | 31983098:319 | 31983257:319 | +;+;+;+;+;+;+; | 1773 | ppp6c | 1523 | 1777 | 1972 | 1623 | 1493 | 1464 |
| ENSDARG000 | 5;5;5;5;5;5;5; | 62052538:620 | 62053064:620 | +;+;+;+;+;+;+; | 1015 | NA | 0 | 0 | 0 | 0 | 0 | 0 |
| ENSDARG000 | 5;5;5;5;5;5;5; | 29516844:295 | 29517485:295 | -;-;-;-;-;-;-;- | 2028 | arrdc1a | 71 | 0 | 0 | 4 | 14 | 22 |
| ENSDARG000 | 5;5;5;5;5;5;5; | 62340799:623 | 62341202:623 | +;+;+;+;+;+;+; | 1269 | aspa | 148 | 24 | 103 | 90 | 161 | 92 |
| ENSDARG000 | 5;5;5;5 | 59116068:591 | 59116442:591 | +;+;+;+ | 4335 | NA | 82 | 80 | 174 | 234 | 139 | 148 |
| ENSDARG000 | 5;5;5 | 20693724:206 | 20693792:206 | +;+;+ | 760 | NA | 8 | 0 | 0 | 0 | 8 | 8 |
| ENSDARG000 | 5;5;5;5;5;5;5; | 15202495:152 | 15202621:152 | +;+;+;+;+;+;+; | 2470 | tbx1 | 188 | 4 | 6 | 10 | 15 | 85 |
| ENSDARG000 | 5;5;5;5;5;5;5; | 22590326:225 | 22594253:225 | -;-;-;-;-;-;-;-; | 5342 | nono | 1445 | 2101 | 2799 | 1293 | 2908 | 1376 |
| ENSDARG000 | 5;5;5;5;5;5;5; | 54422941:544 | 54422979:544 | +;+;+;+;+;+;+; | 2370 | NA | 502 | 298 | 717 | 281 | 395 | 412 |
| ENSDARG000 | 5;5;5;5;5;5;5; | 45895613:458 | 45895925:458 | +;+;+;+;+;+;+; | 2985 | POLK | 105 | 148 | 174 | 70 | 152 | 125 |
| ENSDARG000 | 5;5 | 15186823:152 | 15186952:152 | -;- | 1006 | NA | 0 | 0 | 0 | 0 | 0 | 0 |
| ENSDARG000 | 5;5;5;5;5;5;5; | 20786772:207 | 20788797:207 | -;-;-;-;-;-;-;-; | 4143 | si:ch211-225t | 56 | 54 | 168 | 18 | 225 | 80 |
| ENSDARG000 | 5;5;5;5;5;5;5; | 27583117:275 | 27583576:275 | +;+;+;+;+;+;+; | 1274 | ZMAT4 | 0 | 0 | 0 | 0 | 0 | 0 |
| ENSDARG000 | 5;5;5;5 | 9070382:9070 | 9070738:9070 | -;-;-;- | 629 | atp5meb | 1061 | 1421 | 1343 | 995 | 930 | 1213 |
| ENSDARG000 | 5;5;5;5;5;5 | 59234421:592 | 59234585:592 | +;+;+;+;+;+ | 884 | prkrip1 | 158 | 196 | 250 | 90 | 145 | 184 |
| ENSDARG000 | 5;5;5;5;5;5;5; | 54446262:544 | 54446293:544 | -;-;-;-;-;-;-;-; | 5090 | fbxw5 | 817 | 653 | 978 | 646 | 837 | 790 |
| ENSDARG000 | 5;5;5 | 38167810:381 | 38168723:381 | -;-;- | 1056 | NA | 0 | 25 | 24 | 0 | 0 | 19 |
| ENSDARG000 | 5;5;5;5;5;5;5; | 22585269:225 | 22585289:225 | -;-;-;-;-;-;-;-; | 1203 | casp22 | 61 | 46 | 29 | 20 | 31 | 34 |
| ENSDARG000 | 5;5;5;5;5;5;5; | 54416258:544 | 54416970:544 | -;-;-;-;-;-;-;-; | 1736 | coq4 | 190 | 230 | 324 | 258 | 225 | 227 |
| ENSDARG000 | 5;5;5;5;5 | 43064941:430 | 43065331:430 | -;-;-;-;- | 1692 | NA | 10 | 2 | 0 | 0 | 6 | 2 |
| ENSDARG000 | 5;5;5;5;5;5 | 59252701:592 | 59252824:592 | +;+;+;+;+;+ | 1582 | upk3b | 55 | 6 | 16 | 0 | 13 | 20 |
| ENSDARG000 | 5;5;5;5;5;5;5; | 24543862:245 | 24544485:245 | +;+;+;+;+;+;+; | 4283 | atp6v0a2b | 7211 | 6042 | 4535 | 6382 | 6053 | 7201 |
| ENSDARG000 | 5;5 | 9078198:9088 | 9078768:9090 | -;- | 1916 | kcnv2b | 0 | 4 | 0 | 0 | 0 | 10 |
| ENSDARG000 | 5 | 27866649 | 27867657 | - | 1009 | tcima | 1933 | 310 | 812 | 368 | 551 | 528 |
| ENSDARG000 | 5;5;5;5;5;5;5; | 62611400:626 | 62611555:626 | +;+;+;+;+;+;+; | 975 | abr | 357 | 192 | 334 | 194 | 513 | 120 |
| ENSDARG000 | 5;5 | 35398745:354 | 35399635:354 | +;+ | 1494 | ptger4b | 6 | 12 | 26 | 38 | 41 | 12 |
| ENSDARG000 | 5;5;5;5;5;5;5; | 69733096:697 | 69733225:697 | +;+;+;+;+;+;+; | 1160 | arl6ip4 | 387 | 470 | 734 | 421 | 750 | 402 |
| ENSDARG000 | 5;5;5;5;5 | 35421766:354 | 35422965:354 | -;-;-;-;- | 1735 | ttc33 | 171 | 193 | 164 | 146 | 134 | 180 |
| ENSDARG000 | 5;5;5;5;5;5;5; | 50972485:509 | 50973825:509 | -;-;-;-;-;-;-;-; | 4304 | hmgcra | 656 | 1255 | 1808 | 734 | 336 | 342 |
| ENSDARG000 | 5;5;5;5;5;5;5 | 4298636:4299 | 4299008:4299 | +;+;+;+;+;+;+ | 1039 | prdx4 | 829 | 1447 | 2932 | 1250 | 1651 | 1217 |
| ENSDARG000 | 5;5;5;5;5;5;5; | 32882688:328 | 32882859:328 | +;+;+;+;+;+;+; | 1570 | rpl12 | 13300 | 15092 | 23894 | 21220 | 14583 | 21731 |
| ENSDARG000 | 5;5;5;5;5;5;5; | 26131690:261 | 26132561:261 | +;+;+;+;+;+;+; | 2805 | si:ch211-214i | 168 | 132 | 124 | 67 | 127 | 116 |
| ENSDARG000 | 5;5;5 | 56246736:562 | 56247108:562 | +;+;+ | 1072 | NA | 0 | 0 | 0 | 0 | 0 | 0 |
| ENSDARG000 | 5;5;5;5 | 56256188:562 | 56256258:562 | +;+;+;+ | 302 | NA | 0 | 0 | 0 | 0 | 0 | 0 |
| ENSDARG000 | 5;5;5;5 | 9112291:9112 | 9112434:9113 | +;+;+;+ | 786 | NA | 0 | 0 | 0 | 0 | 0 | 0 |
| ENSDARG000 | 5;5;5;5;5;5;5; | 43427841:434 | 43428700:434 | +;+;+;+;+;+;+; | 3172 | rhobtb4 | 92 | 89 | 30 | 116 | 63 | 94 |
| ENSDARG000 | 5;5;5;5 | 30614532:306 | 30614783:306 | -;-;-;- | 812 | tcnbb | 126 | 68 | 22 | 44 | 51 | 100 |
| ENSDARG000 | 5;5;5 | 62642924:626 | 62643035:626 | +;+;+ | 796 | NA | 42 | 33 | 101 | 50 | 46 | 22 |
| ENSDARG000 | 5;5;5;5;5;5;5 | 12888260:128 | 12888903:128 | +;+;+;+;+;+;+ | 5744 | gal3st1a | 3115 | 667 | 996 | 1377 | 650 | 2077 |
| ENSDARG000 | 5;5;5 | 23630384:236 | 23630543:236 | +;+;+ | 1404 | gja2 | 0 | 0 | 0 | 0 | 0 | 0 |
| ENSDARG000 | 5;5;5 | 62699748:627 | 62699859:627 | +;+;+ | 791 | NA | 20 | 8 | 3 | 2 | 10 | 7 |
| ENSDARG000 | 5;5;5;5 | 23622177:236 | 23622348:236 | +;+;+;+ | 1516 | gjb1a | 6 | 10 | 10 | 2 | 4 | 0 |
| ENSDARG000 | 5;5;5;5;5;5;5; | 27897504:278 | 27898393:278 | +;+;+;+;+;+;+; | 3839 | adam28 | 172 | 38 | 109 | 60 | 149 | 72 |
| ENSDARG000 | 5;5;5;5;5;5;5; | 23641833:236 | 23644260:236 | -;-;-;-;-;-;-;-; | 5193 | tbc1d8b | 390 | 353 | 601 | 274 | 591 | 390 |
| ENSDARG000 | 5;5;5 | 38176040:381 | 38176742:381 | -;-;- | 851 | si:ch211-284e | 0 | 6 | 7 | 2 | 0 | 0 |
| ENSDARG000 | 5;5;5;5;5;5;5; | 15222011:152 | 15222500:152 | -;-;-;-;-;-;-;-; | 1438 | gnb1l | 14 | 44 | 45 | 18 | 42 | 27 |
| ENSDARG000 | 5;5;5;5 | 67268110:672 | 67268371:672 | +;+;+;+ | 2725 | si:ch211-110r | 31 | 61 | 46 | 30 | 28 | 69 |
| ENSDARG000 | 5;5;5;5;5;5 | 69747417:697 | 69747447:697 | +;+;+;+;+;+ | 1148 | mtrfr | 214 | 156 | 298 | 128 | 247 | 190 |
| ENSDARG000 | 5;5;5;5 | 9134187:9134 | 9134312:9134 | +;+;+;+ | 768 | NA | 0 | 0 | 0 | 0 | 0 | 0 |
| ENSDARG000 | 5;5;5 | 62707833:627 | 62707944:627 | +;+;+ | 796 | NA | 3 | 0 | 2 | 4 | 5 | 0 |
| ENSDARG000 | 5;5;5;5 | 30617915:306 | 30618164:306 | -;-;-;- | 655 | tcnba | 1 | 0 | 0 | 0 | 0 | 0 |
| ENSDARG000 | 5;5;5;5;5;5;5; | 59278193:592 | 59278657:592 | +;+;+;+;+;+;+; | 7508 | rasa4 | 40 | 164 | 193 | 388 | 340 | 50 |
| ENSDARG000 | 5;5;5;5;5;5;5; | 30624183:306 | 30624671:306 | +;+;+;+;+;+;+; | 2546 | ABCG4 | 0 | 0 | 2 | 0 | 0 | 0 |
| ENSDARG000 | 5;5 | 56258431:562 | 56258542:562 | +;+ | 351 | NA | 0 | 0 | 0 | 0 | 0 | 0 |
| ENSDARG000 | 5;5;5;5;5 | 56268436:562 | 56269051:562 | +;+;+;+;+ | 2074 | lhx1b | 0 | 2 | 0 | 0 | 0 | 0 |
| ENSDARG000 | 5;5;5;5;5;5 | 62726349:627 | 62726390:627 | +;+;+;+;+;+ | 898 | zswim7 | 40 | 78 | 64 | 28 | 34 | 56 |
| ENSDARG000 | 5;5;5 | 51026563:510 | 51026626:510 | +;+;+ | 1796 | gcnt4a | 48 | 4 | 76 | 18 | 56 | 12 |
| ENSDARG000 | 5;5;5;5;5;5;5; | 35458346:354 | 35458461:354 | +;+;+;+;+;+;+; | 774 | ERLIN2 | 0 | 0 | 0 | 0 | 0 | 0 |
| ENSDARG000 | 5;5;5;5;5;5;5; | 35458190:354 | 35458461:354 | +;+;+;+;+;+;+; | 1622 | fbp1b | 0 | 0 | 43 | 20 | 30 | 0 |
| ENSDARG000 | 5;5;5;5;5;5 | 26138313:261 | 26138487:261 | +;+;+;+;+;+ | 1734 | dhfr | 40 | 203 | 187 | 84 | 140 | 155 |
| ENSDARG000 | 5;5;5;5;5;5;5; | 35463688:354 | 35463838:354 | +;+;+;+;+;+;+; | 5752 | erlin2 | 238 | 293 | 368 | 291 | 249 | 323 |
| ENSDARG000 | 5;5;5;5;5;5;5; | 20823409:208 | 20823602:208 | +;+;+;+;+;+;+; | 3796 | limk2 | 132 | 8 | 40 | 10 | 76 | 28 |
| ENSDARG000 | 5;5;5;5;5 | 62781626:627 | 62782371:627 | -;-;-;-;- | 1602 | adora2b | 29 | 2 | 12 | 2 | 8 | 2 |
| ENSDARG000 | 5;5;5;5;5;5;5; | 43530388:435 | 43530739:435 | +;+;+;+;+;+;+; | 2759 | NA | 113 | 3 | 15 | 0 | 20 | 10 |
| ENSDARG000 | 5;5;5;5;5;5 | 15311591:153 | 15311714:153 | -;-;-;-;-;- | 620 | TXNRD2 | 0 | 0 | 0 | 0 | 0 | 0 |
| ENSDARG000 | 5;5;5;5;5;5;5; | 15311591:153 | 15311714:153 | -;-;-;-;-;-;-;-; | 1800 | TXNRD2 | 25 | 48 | 51 | 22 | 56 | 57 |
| ENSDARG000 | 5;5;5;5;5;5 | 38182601:381 | 38182647:381 | +;+;+;+;+;+ | 528 | NA | 0 | 0 | 0 | 0 | 0 | 0 |
| ENSDARG000 | 5;5;5;5;5;5;5; | 65065968:650 | 65066234:650 | -;-;-;-;-;-;-;-; | 2750 | inpp5e | 195 | 215 | 282 | 140 | 273 | 228 |
| ENSDARG000 | 5;5;5 | 38193453:381 | 38194092:381 | -;-;- | 783 | si:ch211-284e | 58 | 81 | 33 | 22 | 30 | 17 |
| ENSDARG000 | 5;5;5;5;5;5;5; | 9147497:9149 | 9147557:9149 | +;+;+;+;+;+;+; | 909 | rchy1 | 366 | 437 | 474 | 319 | 385 | 304 |
| ENSDARG000 | 5;5;5;5;5;5;5; | 4332220:4332 | 4332450:4332 | +;+;+;+;+;+;+; | 1758 | sat1a.1 | 410 | 82 | 425 | 169 | 90 | 79 |
| ENSDARG000 | 5;5;5;5;5;5;5; | 62804252:628 | 62807563:628 | -;-;-;-;-;-;-;-; | 7814 | SPECC1 | 104 | 336 | 511 | 520 | 284 | 220 |
| ENSDARG000 | 5;5;5;5 | 41542436:415 | 41542669:415 | -;-;-;- | 1184 | dnajb5 | 6 | 16 | 3 | 0 | 0 | 4 |
| ENSDARG000 | 5;5;5;5;5;5;5 | 23677420:236 | 23678148:236 | -;-;-;-;-;-;- | 1981 | rnf128a | 306 | 86 | 208 | 168 | 152 | 186 |
| ENSDARG000 | 5;5;5;5;5;5;5; | 56277866:562 | 56277985:562 | +;+;+;+;+;+;+; | 2330 | aatf | 281 | 389 | 511 | 187 | 345 | 345 |
| ENSDARG000 | 5;5;5;5;5;5;5; | 45918680:459 | 45918724:459 | +;+;+;+;+;+;+; | 1490 | ANKDD1B | 1 | 3 | 7 | 0 | 2 | 0 |
| ENSDARG000 | 5;5;5;5;5;5;5; | 30680804:306 | 30680885:306 | +;+;+;+;+;+;+; | 1994 | nherf4b | 0 | 2 | 0 | 0 | 0 | 0 |
| ENSDARG000 | 5;5;5;5;5;5;5; | 45938316:459 | 45938907:459 | -;-;-;-;-;-;-;-; | 2252 | NA | 16 | 105 | 108 | 66 | 78 | 26 |
| ENSDARG000 | 5;5;5;5;5;5;5; | 29548441:295 | 29549507:295 | -;-;-;-;-;-;-;-; | 2907 | nelfb | 482 | 502 | 476 | 258 | 534 | 308 |
| ENSDARG000 | 5;5;5;5;5;5;5; | 43617262:436 | 43617869:436 | -;-;-;-;-;-;-;-; | 2016 | rtkna | 544 | 264 | 461 | 372 | 631 | 491 |
| ENSDARG000 | 5;5 | 26156079:261 | 26156281:261 | +;+ | 1647 | ankrd34bb | 0 | 0 | 4 | 10 | 2 | 10 |
| ENSDARG000 | 5;5;5;5;5;5;5; | 69785990:697 | 69786108:697 | +;+;+;+;+;+;+; | 1906 | kmt5ab | 216 | 474 | 664 | 286 | 370 | 320 |
| ENSDARG000 | 5;5;5;5;5;5;5; | 26188264:261 | 26189616:261 | -;-;-;-;-;-;-;-; | 3244 | rad17 | 180 | 170 | 203 | 125 | 161 | 233 |
| ENSDARG000 | 5;5;5;5 | 32888119:328 | 32888738:328 | -;-;-;- | 1277 | pole3 | 369 | 794 | 903 | 452 | 741 | 500 |
| ENSDARG000 | 5;5;5;5;5;5;5; | 32060881:320 | 32061407:320 | -;-;-;-;-;-;-;- | 1101 | arpc5lb | 261 | 168 | 232 | 84 | 176 | 140 |
| ENSDARG000 | 5;5;5;5;5;5;5; | 26162401:261 | 26162749:261 | -;-;-;-;-;-;-;-; | 1804 | fam151b | 56 | 60 | 106 | 22 | 87 | 106 |
| ENSDARG000 | 5;5;5;5;5;5;5; | 24616591:246 | 24616802:246 | -;-;-;-;-;-;-;-; | 863 | NA | 4 | 0 | 4 | 26 | 61 | 20 |
| ENSDARG000 | 5;5;5;5;5;5;5; | 29563819:295 | 29563896:295 | -;-;-;-;-;-;-;- | 1362 | NA | 2 | 18 | 13 | 13 | 5 | 21 |
| ENSDARG000 | 5;5;5;5;5;5;5; | 15350954:153 | 15351135:153 | +;+;+;+;+;+;+; | 2453 | pebp1 | 270 | 467 | 465 | 487 | 731 | 195 |
| ENSDARG000 | 5;5;5;5;5;5;5; | 59392183:593 | 59393005:593 | +;+;+;+;+;+;+; | 5182 | CLIP2 | 476 | 706 | 833 | 352 | 908 | 258 |
| ENSDARG000 | 5;5;5;5;5;5;5; | 24571750:245 | 24571892:245 | -;-;-;-;-;-;-;-; | 3438 | osbp2b | 578 | 529 | 875 | 702 | 802 | 951 |
| ENSDARG000 | 5;5;5;5;5;5;5; | 22605530:226 | 22606601:226 | -;-;-;-;-;-;-;-; | 2032 | zgc:113208 | 24 | 66 | 347 | 362 | 520 | 523 |
| ENSDARG000 | 5;5;5;5;5;5;5; | 26171564:261 | 26171874:261 | -;-;-;-;-;-;-;-; | 1494 | ccdc125 | 26 | 2 | 4 | 22 | 0 | 42 |
| ENSDARG000 | 5;5;5;5;5;5;5; | 63088740:630 | 63088824:630 | -;-;-;-;-;-;-;-; | 1197 | NA | 409 | 586 | 209 | 288 | 386 | 193 |
| ENSDARG000 | 5;5;5 | 9166395:9170 | 9166846:9170 | -;-;- | 586 | NA | 8 | 11 | 0 | 2 | 11 | 23 |
| ENSDARG000 | 5;5;5;5;5;5;5; | 41582643:415 | 41582825:415 | -;-;-;-;-;-;-;-; | 7599 | epg5 | 801 | 347 | 606 | 342 | 564 | 538 |
| ENSDARG000 | 5;5;5 | 9173538:9179 | 9174199:9179 | -;-;- | 797 | NA | 30 | 24 | 0 | 11 | 4 | 2 |
| ENSDARG000 | 5;5;5;5;5;5;5; | 27924347:279 | 27925190:279 | -;-;-;-;-;-;-;-; | 2914 | ppp3cca | 275 | 226 | 338 | 196 | 278 | 252 |
| ENSDARG000 | 5;5;5;5;5;5;5; | 51111343:511 | 51111621:511 | +;+;+;+;+;+;+; | 3201 | pomt1 | 391 | 382 | 506 | 293 | 669 | 287 |
| ENSDARG000 | 5;5;5;5;5;5;5; | 19261391:192 | 19261433:192 | +;+;+;+;+;+;+; | 3735 | NA | 703 | 521 | 821 | 1235 | 373 | 1674 |
| ENSDARG000 | 5;5;5;5;5;5;5; | 65086668:650 | 65087291:650 | +;+;+;+;+;+;+; | 2149 | ptrh1 | 80 | 193 | 295 | 117 | 158 | 130 |
| ENSDARG000 | 5;5;5;5;5;5;5; | 38200807:382 | 38201056:382 | +;+;+;+;+;+;+; | 2936 | NA | 196 | 74 | 106 | 132 | 64 | 709 |
| ENSDARG000 | 5;5;5 | 30699260:306 | 30699410:306 | +;+;+ | 515 | atp5l | 2341 | 2374 | 2657 | 2036 | 2166 | 2030 |
| ENSDARG000 | 5;5;5;5;5;5;5; | 29571803:295 | 29574391:295 | -;-;-;-;-;-;-;-; | 3981 | entpd2a.1 | 220 | 26 | 28 | 16 | 50 | 86 |
| ENSDARG000 | 5;5;5;5;5;5;5; | 23702295:237 | 23703605:237 | -;-;-;-;-;-;-;-; | 2068 | gbgt1l1 | 0 | 0 | 2 | 0 | 0 | 0 |
| ENSDARG000 | 5;5;5;5;5;5;5; | 67275888:672 | 67276612:672 | -;-;-;-;-;-;-;-; | 2823 | rsrc2 | 703 | 671 | 901 | 491 | 959 | 905 |
| ENSDARG000 | 5;5;5;5;5;5;5; | 15359995:153 | 15361475:153 | -;-;-;-;-;-;-;-; | 4702 | TAOK3 | 1234 | 767 | 1500 | 1855 | 1214 | 1228 |
| ENSDARG000 | 5;5 | 65100409:651 | 65100946:651 | +;+ | 1488 | NA | 0 | 0 | 0 | 2 | 0 | 0 |
| ENSDARG000 | 5;5;5 | 63197955:632 | 63198122:632 | +;+;+ | 854 | NA | 0 | 0 | 0 | 0 | 0 | 0 |
| ENSDARG000 | 5;5;5;5;5;5;5; | 63177355:631 | 63177458:631 | -;-;-;-;-;-;-;-; | 6969 | TECTA | 19 | 4 | 22 | 9 | 28 | 10 |
| ENSDARG000 | 5;5;5;5;5;5;5; | 32076109:320 | 32076350:320 | +;+;+;+;+;+;+; | 884 | zmat5 | 72 | 140 | 98 | 155 | 67 | 154 |
| ENSDARG000 | 5;5;5;5;5 | 65112481:651 | 65113527:651 | -;-;-;-;- | 1782 | tor2a | 192 | 122 | 191 | 190 | 193 | 268 |
| ENSDARG000 | 5;5;5;5;5;5;5; | 45976130:459 | 45976431:459 | +;+;+;+;+;+;+; | 2620 | sv2ca | 0 | 0 | 0 | 0 | 0 | 0 |
| ENSDARG000 | 5;5;5;5;5;5;5; | 32086448:320 | 32086575:320 | -;-;-;-;-;-;-;-; | 646 | CABP7 | 0 | 0 | 0 | 0 | 0 | 0 |
| ENSDARG000 | 5;5;5;5;5;5;5; | 43782267:437 | 43782306:437 | +;+;+;+;+;+;+; | 4149 | nos2a | 138 | 6 | 6 | 9 | 2 | 6 |
| ENSDARG000 | 5;5;5;5;5;5;5; | 41639693:416 | 41639957:416 | -;-;-;-;-;-;-;-; | 2161 | riok2 | 588 | 540 | 825 | 448 | 708 | 523 |
| ENSDARG000 | 5;5;5;5 | 27999886:280 | 28000733:280 | -;-;-;- | 1208 | vps37ba | 834 | 465 | 501 | 417 | 519 | 376 |
| ENSDARG000 | 5;5;5;5;5 | 30702084:307 | 30702386:307 | -;-;-;-;- | 944 | ift22 | 22 | 43 | 48 | 35 | 38 | 42 |
| ENSDARG000 | 5;5;5;5;5 | 32914321:329 | 32914591:329 | +;+;+;+;+ | 1855 | ccn1l1 | 0 | 0 | 0 | 0 | 0 | 0 |
| ENSDARG000 | 5;5;5;5;5;5 | 20866572:208 | 20867956:208 | -;-;-;-;-;- | 2094 | pik3ip1 | 314 | 265 | 567 | 1553 | 152 | 448 |
| ENSDARG000 | 5;5;5;5;5 | 23721515:237 | 23722205:237 | -;-;-;-;- | 999 | NA | 0 | 0 | 0 | 0 | 0 | 4 |
| ENSDARG000 | 5;5;5;5;5;5 | 23738167:237 | 23739275:237 | -;-;-;-;-;- | 1488 | gbgt1l2 | 4 | 0 | 2 | 0 | 0 | 6 |
| ENSDARG000 | 5;5;5;5;5;5;5; | 38215924:382 | 38218203:382 | -;-;-;-;-;-;-;-; | 4466 | slc12a9 | 2343 | 2311 | 2390 | 2511 | 2956 | 3746 |
| ENSDARG000 | 5;5;5;5;5;5;5; | 32924669:329 | 32925036:329 | +;+;+;+;+;+;+; | 2444 | lmo4a | 109 | 112 | 445 | 47 | 148 | 193 |

| | | | | | | | | | | | | |
|---|---|---|---|---|---|---|---|---|---|---|---|---|
| ENSDARG000 | 5:5:5:5:5:5:5: | 32129482:321 | 32129931:321 | -:-:-:-:-:-:-:-: | 1267 | NA | 1 | 0 | 2 | 0 | 0 | 0 |
| ENSDARG000 | 5;5 | 46196147:461 | 46196283:461 | +;+ | 1309 | f2r | 43 | 4 | 30 | 9 | 11 | 26 |
| ENSDARG000 | 5:5:5:5:5:5:5: | 46077710:460 | 46078021:460 | +:+:+:+:+:+:+: | 9208 | iqgap2 | 496 | 146 | 171 | 333 | 156 | 808 |
| ENSDARG000 | 5:5:5:5:5:5:5: | 56308622:563 | 56309919:563 | -:-:-:-:-:-:-:-: | 9447 | acaca | 3612 | 2864 | 2914 | 1197 | 2673 | 1981 |
| ENSDARG000 | 5:5:5:5:5:5:5: | 19309877:193 | 19311950:193 | +:+:+:+:+:+:+: | 12362 | NA | 924 | 806 | 1303 | 1098 | 1029 | 1369 |
| ENSDARG000 | 5:5:5:5:5:5:5: | 35526757:355 | 35527425:355 | -:-:-:-:-:-:-:-: | 1399 | si:dkev-204l1 | 8 | 45 | 86 | 58 | 104 | 14 |
| ENSDARG000 | 5:5:5:5:5:5:5: | 54482325:544 | 54483378:544 | -:-:-:-:-:-:-:-: | 2461 | alad | 276 | 278 | 388 | 286 | 458 | 232 |
| ENSDARG000 | 5:5:5:5:5:5:5: | 63248266:632 | 63248774:632 | -:-:-:-:-:-:-:-: | 1752 | TBCEL | 131 | 140 | 211 | 115 | 100 | 92 |
| ENSDARG000 | 5:5:5:5:5:5:5: | 65124168:651 | 65124749:651 | -:-:-:-:-:-:-:-: | 2896 | sh2d3cb | 45 | 58 | 36 | 45 | 37 | 36 |
| ENSDARG000 | 5;5 | 22633188:226 | 22633631:226 | +;+ | 1333 | mtnr1c | 0 | 2 | 14 | 14 | 59 | 6 |
| ENSDARG000 | 5:5:5:5:5:5:5: | 29597645:295 | 29599983:295 | -:-:-:-:-:-:-:-: | 5788 | grin1b | 4 | 0 | 8 | 2 | 28 | 2 |
| ENSDARG000 | 5;5;5 | 22677786:226 | 22677956:226 | +;+;+ | 1055 | vma21 | 930 | 1398 | 1166 | 964 | 1835 | 1154 |
| ENSDARG000 | 5 | 23757933 | 23758587 | - | 655 | NA | 0 | 0 | 0 | 0 | 0 | 0 |
| ENSDARG000 | 5;5;5 | 23768377:237 | 23769009:237 | -;-;- | 991 | NA | 0 | 0 | 0 | 0 | 0 | 0 |
| ENSDARG000 | 5:5:5:5:5:5:5: | 65157576:651 | 65157774:651 | +:+:+:+:+:+:+: | 1576 | cfap157 | 0 | 2 | 2 | 0 | 2 | 0 |
| ENSDARG000 | 5:5:5:5:5:5:5: | 51164440:511 | 51169233:511 | -:-:-:-:-:-:-:-: | 9947 | snapc4 | 305 | 219 | 462 | 276 | 438 | 362 |
| ENSDARG000 | 5;5 | 38250988:382 | 38252146:382 | -;- | 2253 | NA | 174 | 261 | 230 | 348 | 189 | 227 |
| ENSDARG000 | 5;5;5;5;5;5;5 | 23776680:237 | 23777406:237 | -;-;-;-;-;-;- | 1218 | gbgt1l8 | 4 | 0 | 0 | 0 | 0 | 0 |
| ENSDARG000 | 5;5;5;5;5;5;5 | 30707133:307 | 30708048:307 | -;-;-;-;-;-;- | 2237 | ftr82 | 110 | 5 | 19 | 16 | 30 | 14 |
| ENSDARG000 | 5:5:5:5:5:5:5: | 20886282:208 | 20887008:208 | -:-:-:-:-:-:-:-: | 6035 | NA | 314 | 175 | 366 | 144 | 372 | 197 |
| ENSDARG000 | 5;5;5;5 | 33018764:330 | 33019980:330 | -;-;-;- | 1586 | zgc:55461 | 42 | 116 | 193 | 142 | 252 | 84 |
| ENSDARG000 | 5:5:5:5:5:5:5: | 35561607:356 | 35563174:356 | +:+:+:+:+:+:+: | 3177 | AR | 432 | 500 | 1330 | 417 | 2192 | 489 |
| ENSDARG000 | 5;5;5;5 | 32135202:321 | 32135836:321 | -;-;-;- | 869 | NA | 0 | 0 | 0 | 2 | 0 | 0 |
| ENSDARG000 | 5:5:5:5:5:5:5: | 28018246:280 | 28018926:280 | -:-:-:-:-:-:-:-: | 2742 | abcb9 | 100 | 14 | 58 | 236 | 210 | 190 |
| ENSDARG000 | 5:5:5:5:5:5:5: | 38263240:382 | 38263875:382 | +:+:+:+:+:+:+: | 3053 | gnb2 | 1618 | 2187 | 2086 | 2699 | 1462 | 1795 |
| ENSDARG000 | 5:5:5:5:5:5:5: | 54497475:544 | 54497501:544 | +:+:+:+:+:+:+: | 1186 | tmem203 | 98 | 102 | 150 | 76 | 122 | 119 |
| ENSDARG000 | 5:5:5:5:5:5:5: | 63302660:633 | 63302939:633 | +:+:+:+:+:+:+: | 2261 | NA | 108 | 180 | 179 | 250 | 214 | 181 |
| ENSDARG000 | 5;5;5 | 24723405:247 | 24723675:247 | -;-;- | 612 | rasl10a | 6 | 2 | 2 | 0 | 11 | 0 |
| ENSDARG000 | 5:5:5:5:5:5:5: | 63294865:632 | 63295238:632 | -:-:-:-:-:-:-:-: | 2678 | gsnb | 236 | 23 | 38 | 85 | 54 | 91 |
| ENSDARG000 | 5;5;5;5 | 19334206:193 | 19334261:193 | -;-;-;- | 720 | NA | 0 | 0 | 4 | 0 | 0 | 0 |
| ENSDARG000 | 5:5:5:5:5:5:5: | 15495351:154 | 15495558:154 | +:+:+:+:+:+:+: | 1568 | suds3 | 651 | 719 | 942 | 762 | 655 | 782 |
| ENSDARG000 | 5:5:5:5:5:5:5: | 63288599:632 | 63289559:632 | +:+:+:+:+:+:+: | 2135 | NA | 96 | 169 | 181 | 147 | 176 | 159 |
| ENSDARG000 | 5:5:5:5:5:5:5: | 28041967:280 | 28042213:280 | +:+:+:+:+:+:+: | 1257 | ogfod2 | 26 | 15 | 11 | 14 | 0 | 8 |
| ENSDARG000 | 5;5;5;5 | 20922100:209 | 20922163:209 | +;+;+;+ | 1037 | si:dkev-174n2 | 150 | 138 | 174 | 133 | 149 | 127 |
| ENSDARG000 | 5:5:5:5:5:5:5: | 28045764:280 | 28048752:280 | -:-:-:-:-:-:-:-: | 6692 | SLC4A5 | 8 | 0 | 0 | 0 | 0 | 0 |
| ENSDARG000 | 5:5:5:5:5:5:5: | 28030954:280 | 28033413:280 | -:-:-:-:-:-:-:-: | 3936 | zgc:113436 | 122 | 115 | 140 | 80 | 111 | 178 |
| ENSDARG000 | 5:5:5:5:5:5:5: | 51199943:512 | 51200510:512 | -:-:-:-:-:-:-:-: | 1782 | entr1 | 373 | 270 | 270 | 232 | 277 | 277 |
| ENSDARG000 | 5:5:5:5:5:5:5: | 32141790:321 | 32141979:321 | +:+:+:+:+:+:+: | 1801 | tescb | 14 | 16 | 32 | 2 | 26 | 42 |
| ENSDARG000 | 5:5:5:5:5:5:5: | 67315816:673 | 67319658:673 | -:-:-:-:-:-:-:-: | 6755 | mlxip | 747 | 644 | 1237 | 831 | 519 | 1082 |
| ENSDARG000 | 5:5:5:5:5:5:5: | 51226846:512 | 51227284:512 | +:+:+:+:+:+:+: | 5275 | ubac1 | 696 | 785 | 596 | 622 | 654 | 1218 |
| ENSDARG000 | 5:5:5:5:5:5:5: | 22791686:227 | 22791804:227 | +:+:+:+:+:+:+: | 2683 | npas2 | 146 | 202 | 152 | 29 | 274 | 191 |
| ENSDARG000 | 5:5:5:5:5:5:5: | 36899691:369 | 36899739:369 | +:+:+:+:+:+:+: | 2390 | hnrnpl | 2966 | 2794 | 4129 | 1994 | 3399 | 2171 |
| ENSDARG000 | 5;5;5;5 | 69795204:698 | 69796549:698 | -;-;-;- | 2009 | rilpl2 | 802 | 714 | 404 | 480 | 564 | 696 |
| ENSDARG000 | 5:5:5:5:5:5:5: | 23788592:237 | 23789796:237 | -:-:-:-:-:-:-:-: | 1936 | gbgt1l4 | 2 | 0 | 0 | 0 | 0 | 0 |
| ENSDARG000 | 5:5:5:5:5:5:5: | 20931252:209 | 20935016:209 | -:-:-:-:-:-:-:-: | 5919 | camk2b1 | 1725 | 1360 | 1358 | 1380 | 1576 | 1992 |
| ENSDARG000 | 5:5:5:5:5:5:5: | 30723129:307 | 30723925:307 | +:+:+:+:+:+:+: | 2549 | NA | 0 | 2 | 0 | 4 | 0 | 0 |
| ENSDARG000 | 5:5:5:5:5:5:5: | 33028083:330 | 33029768:330 | -:-:-:-:-:-:-:-: | 3439 | NA | 50 | 50 | 62 | 30 | 64 | 156 |
| ENSDARG000 | 5:5:5:5:5:5:5: | 22836364:228 | 22836485:228 | +:+:+:+:+:+:+: | 1586 | NA | 17 | 53 | 55 | 34 | 66 | 19 |
| ENSDARG000 | 5:5:5:5:5:5:5: | 69808763:698 | 69808821:698 | +:+:+:+:+:+:+: | 1909 | FSD1L | 223 | 118 | 151 | 155 | 177 | 210 |
| ENSDARG000 | 5:5:5:5:5:5:5: | 24756496:247 | 24758033:247 | -:-:-:-:-:-:-:-: | 4878 | ap1b1 | 1931 | 1762 | 2068 | 1316 | 2225 | 1556 |
| ENSDARG000 | 5;5;5;5;5;5 | 32155141:321 | 32155769:321 | -;-;-;-;-;- | 870 | m17 | 52 | 38 | 24 | 24 | 79 | 79 |
| ENSDARG000 | 5:5:5:5:5:5:5: | 22851857:228 | 22852303:228 | -;-;-;-;-;-;-;- | 1761 | apool | 307 | 381 | 439 | 337 | 443 | 419 |
| ENSDARG000 | 5:5:5:5:5:5:5: | 51248784:512 | 51249082:512 | +:+:+:+:+:+:+: | 5205 | msh3 | 48 | 208 | 178 | 100 | 168 | 142 |
| ENSDARG000 | 5:5:5:5:5:5:5: | 19337108:193 | 19337472:193 | +:+:+:+:+:+:+: | 8331 | ACACB | 1162 | 656 | 623 | 767 | 830 | 818 |
| ENSDARG000 | 5:5:5:5:5:5:5: | 38301906:383 | 38306044:383 | -:-:-:-:-:-:-:-: | 8663 | mink1 | 4084 | 2468 | 2420 | 2526 | 1830 | 3203 |
| ENSDARG000 | 5;5 | 65174672:651 | 65174750:651 | +;+ | 495 | NA | 7 | 0 | 0 | 0 | 0 | 2 |
| ENSDARG000 | 5:5:5:5:5:5:5: | 32162684:321 | 32162833:321 | +:+:+:+:+:+:+: | 2149 | NA | 0 | 0 | 0 | 0 | 0 | 0 |
| ENSDARG000 | 5:5:5:5:5:5:5: | 41645479:416 | 41646021:416 | -:-:-:-:-:-:-:-: | 4204 | atxn2 | 948 | 567 | 712 | 304 | 1032 | 691 |
| ENSDARG000 | 5:5:5:5:5:5:5: | 30741730:307 | 30742363:307 | +:+:+:+:+:+:+: | 5985 | ftr83 | 44 | 2 | 0 | 6 | 12 | 8 |
| ENSDARG000 | 5:5:5:5:5:5:5: | 67351219:673 | 67351635:673 | -:-:-:-:-:-:-:-: | 1527 | unga | 102 | 261 | 216 | 92 | 184 | 170 |
| ENSDARG000 | 5:5:5:5:5:5:5: | 22871718:228 | 22874038:228 | -:-:-:-:-:-:-:-: | 5450 | NA | 36 | 28 | 34 | 30 | 60 | 158 |
| ENSDARG000 | 5:5:5:5:5:5:5: | 33067283:330 | 33067285:330 | +:+:+:+:+:+:+: | 1386 | NPFFR2 | 4 | 0 | 0 | 0 | 0 | 0 |
| ENSDARG000 | 5:5:5:5:5:5:5: | 35682681:356 | 35683256:356 | -:-:-:-:-:-:-:-: | 4207 | ophn1 | 960 | 733 | 1014 | 595 | 2072 | 699 |
| ENSDARG000 | 5;5;5;5;5 | 26199537:262 | 26199746:262 | +;+;+;+;+ | 1087 | AK6 | 350 | 346 | 332 | 188 | 392 | 308 |
| ENSDARG000 | 5:5:5:5:5:5:5: | 28134688:281 | 28135625:281 | -;-;-;-;-;-;-;- | 1929 | zgc:110329 | 52 | 35 | 100 | 21 | 84 | 77 |
| ENSDARG000 | 5;5;5;5 | 56474733:564 | 56474873:564 | +;+;+;+ | 2572 | NA | 0 | 0 | 0 | 0 | 0 | 0 |
| ENSDARG000 | 5:5:5:5:5:5:5: | 51443009:515 | 51443668:515 | +:+:+:+:+:+:+: | 4119 | rasgrf2b | 2 | 2 | 0 | 0 | 0 | 0 |
| ENSDARG000 | 5;5;5;5;5;5;5 | 56504717:565 | 56508192:565 | -;-;-;-;-;-;- | 5470 | tbx2a | 2237 | 294 | 1636 | 700 | 768 | 1900 |
| ENSDARG000 | 5:5:5:5:5:5:5: | 26204561:262 | 26204677:262 | +:+:+:+:+:+:+: | 3276 | marveld2b | 38 | 2 | 5 | 0 | 0 | 6 |
| ENSDARG000 | 5;5 | 51482777:514 | 51483175:514 | -;- | 783 | NA | 2 | 0 | 0 | 0 | 0 | 0 |
| ENSDARG000 | 5:5:5:5:5:5:5: | 33199998:332 | 33200149:332 | -:-:-:-:-:-:-:-: | 935 | NA | 19 | 4 | 0 | 0 | 4 | 0 |
| ENSDARG000 | 5:5:5:5:5:5:5: | 36913423:369 | 36914134:369 | -:-:-:-:-:-:-:-: | 1955 | kptn | 387 | 279 | 374 | 336 | 278 | 394 |
| ENSDARG000 | 5:5:5:5:5:5:5: | 65180792:651 | 65180938:651 | -:-:-:-:-:-:-:-: | 4972 | col27a1b | 31 | 4 | 4 | 6 | 4 | 0 |
| ENSDARG000 | 5;5;5;5;5;5;5 | 21033771:210 | 21034191:210 | -;-;-;-;-;-;- | 1259 | NA | 107 | 65 | 41 | 47 | 29 | 39 |
| ENSDARG000 | 5;5 | 23818446:238 | 23819027:238 | +;+ | 709 | NA | 0 | 0 | 0 | 0 | 0 | 0 |
| ENSDARG000 | 5;5;5;5;5 | 21059459:210 | 21059577:210 | -;-;-;-;- | 583 | NA | 46 | 20 | 36 | 16 | 29 | 43 |
| ENSDARG000 | 5;5 | 36934568:369 | 36935328:369 | -;- | 825 | NA | 0 | 0 | 0 | 0 | 0 | 0 |
| ENSDARG000 | 5;5;5 | 36932718:369 | 36933240:369 | +;+;+ | 2381 | crx | 0 | 0 | 1 | 0 | 0 | 0 |
| ENSDARG000 | 5:5:5:5:5:5:5: | 33222124:332 | 33222650:332 | -:-:-:-:-:-:-:-: | 4354 | NA | 2 | 0 | 0 | 0 | 0 | 0 |
| ENSDARG000 | 5;5;5;5 | 67371650:673 | 67371836:673 | +;+;+;+ | 4528 | sebox | 0 | 0 | 0 | 0 | 0 | 2 |
| ENSDARG000 | 5:5:5:5:5:5:5: | 30759232:307 | 30759291:307 | +:+:+:+:+:+:+: | 1712 | ruvbl2 | 480 | 633 | 938 | 450 | 592 | 544 |
| ENSDARG000 | 5:5:5:5:5:5:5: | 23839770:238 | 23840485:238 | -;-;-;-;-;-;-;- | 974 | GBGT1 | 0 | 0 | 0 | 0 | 0 | 0 |
| ENSDARG000 | 5:5:5:5:5:5:5: | 28160503:281 | 28161465:281 | +:+:+:+:+:+:+: | 1837 | tacr1a | 0 | 0 | 0 | 0 | 0 | 4 |
| ENSDARG000 | 5:5:5:5:5:5:5: | 24816248:248 | 24821393:248 | -;-;-;-;-;-;-;- | 6357 | NA | 3737 | 3480 | 2115 | 2427 | 3668 | 1828 |
| ENSDARG000 | 5:5:5:5:5:5:5: | 26212621:262 | 26212778:262 | +:+:+:+:+:+:+: | 2389 | oclnb | 70 | 0 | 0 | 2 | 8 | 6 |
| ENSDARG000 | 5;5;5;5;5;5;5 | 23849647:238 | 23850471:238 | -;-;-;-;-;-;- | 1316 | gbgt1l3 | 0 | 0 | 5 | 0 | 0 | 0 |
| ENSDARG000 | 5;5 | 21100356:211 | 21100712:211 | +;+ | 2082 | hs3st1l2 | 12 | 12 | 4 | 6 | 8 | 0 |
| ENSDARG000 | 5:5:5:5:5:5:5: | 37032038:370 | 37032844:370 | +:+:+:+:+:+:+: | 3243 | nfkbib | 780 | 706 | 936 | 564 | 642 | 666 |
| ENSDARG000 | 5:5:5:5:5:5:5: | 29652198:296 | 29652574:296 | +:+:+:+:+:+:+: | 8752 | tsc1a | 918 | 610 | 778 | 489 | 848 | 614 |
| ENSDARG000 | 5;5;5;5;5;5;5 | 29669938:296 | 29670288:296 | -;-;-;-;-;-;- | 992 | spaca9 | 0 | 0 | 0 | 0 | 0 | 0 |
| ENSDARG000 | 5:5:5:5:5:5:5: | 29671681:296 | 29671755:296 | +:+:+:+:+:+:+: | 2611 | ak8 | 0 | 0 | 0 | 0 | 4 | 2 |
| ENSDARG000 | 5:5:5:5:5:5:5: | 30802798:308 | 30807014:308 | -:-:-:-:-:-:-:-: | 7014 | spns2 | 112 | 8 | 60 | 4 | 128 | 110 |
| ENSDARG000 | 5;5;5 | 37043549:370 | 37043705:370 | -;-;- | 386 | NA | 0 | 0 | 0 | 0 | 0 | 0 |
| ENSDARG000 | 5:5:5:5:5:5:5: | 37040853:370 | 37041397:370 | -:-:-:-:-:-:-:-: | 1717 | NA | 0 | 0 | 0 | 0 | 0 | 0 |
| ENSDARG000 | 5;5;5;5 | 30780600:307 | 30780662:307 | +;+;+;+ | 412 | NA | 0 | 0 | 0 | 0 | 0 | 0 |
| ENSDARG000 | 5:5:5:5:5:5:5: | 67390115:673 | 67391189:673 | +:+:+:+:+:+:+: | 4994 | ebf2 | 8 | 7 | 11 | 0 | 15 | 6 |
| ENSDARG000 | 5:5:5:5:5:5:5: | 33238545:332 | 33241045:332 | -:-:-:-:-:-:-:-: | 3939 | prkaa1 | 634 | 514 | 601 | 398 | 754 | 493 |
| ENSDARG000 | 5;5;5;5;5 | 23874796:238 | 23875526:238 | -;-;-;-;- | 985 | NA | 16 | 10 | 1 | 8 | 10 | 18 |
| ENSDARG000 | 5:5:5:5:5:5:5: | 21144269:211 | 21144444:211 | +:+:+:+:+:+:+: | 2822 | cds2 | 6482 | 7021 | 6767 | 6658 | 8469 | 7240 |
| ENSDARG000 | 5;5;5;5;5;5 | 56914891:569 | 56915468:569 | -;-;-;-;-;- | 1894 | ppm1db | 215 | 144 | 156 | 80 | 268 | 215 |
| ENSDARG000 | 5:5:5:5:5:5:5: | 23888314:238 | 23888615:238 | -:-:-:-:-:-:-:-: | 3445 | NA | 65 | 31 | 26 | 26 | 54 | 48 |
| ENSDARG000 | 5:5:5:5:5:5:5: | 28259655:282 | 28259755:282 | +:+:+:+:+:+:+: | 2684 | ncaph | 152 | 603 | 730 | 227 | 386 | 303 |
| ENSDARG000 | 5:5:5:5:5:5:5: | 26221011:262 | 26221328:262 | +:+:+:+:+:+:+: | 2381 | gtf2h2 | 214 | 193 | 288 | 56 | 217 | 202 |
| ENSDARG000 | 5:5:5:5:5:5:5: | 24873196:248 | 24873898:248 | -:-:-:-:-:-:-:-: | 3172 | ewsr1b | 3751 | 4052 | 5307 | 3833 | 4876 | 3678 |
| ENSDARG000 | 5:5:5:5:5:5:5: | 35744623:357 | 35744713:357 | +:+:+:+:+:+:+: | 1616 | yipf6 | 631 | 707 | 771 | 719 | 833 | 767 |
| ENSDARG000 | 5:5:5:5:5:5:5: | 51594209:515 | 51594325:515 | +:+:+:+:+:+:+: | 1828 | ckmt2b | 0 | 0 | 10 | 8 | 8 | 0 |
| ENSDARG000 | 5;5;5;5 | 37056818:370 | 37056955:370 | +;+;+;+ | 991 | tppp2 | 2 | 0 | 0 | 0 | 0 | 20 |
| ENSDARG000 | 5;5;5;5 | 23913585:239 | 23913703:239 | +;+;+;+ | 4697 | ercc6l | 190 | 978 | 864 | 284 | 705 | 372 |
| ENSDARG000 | 5;5 | 33258289:332 | 33258773:332 | -;- | 700 | ifitm1 | 230 | 2 | 10 | 34 | 10 | 116 |
| ENSDARG000 | 5;5 | 30929047:309 | 30929247:309 | -;- | 684 | NA | 0 | 0 | 0 | 0 | 0 | 0 |
| ENSDARG000 | 5;5;5;5;5;5 | 37068223:370 | 37068333:370 | +;+;+;+;+;+ | 1783 | NA | 0 | 0 | 0 | 0 | 0 | 0 |
| ENSDARG000 | 5:5:5:5:5:5:5: | 35786141:357 | 35786593:357 | +:+:+:+:+:+:+: | 5965 | stard8 | 523 | 395 | 734 | 280 | 635 | 336 |
| ENSDARG000 | 5:5:5:5:5:5:5: | 30956981:309 | 30960210:309 | -:-:-:-:-:-:-:-: | 5103 | spns3 | 1696 | 1080 | 1377 | 1171 | 1900 | 1559 |
| ENSDARG000 | 5:5:5:5:5:5:5: | 24031314:240 | 24032522:240 | -:-:-:-:-:-:-:-: | 2523 | pdha1a | 1809 | 2380 | 2225 | 1307 | 1890 | 1960 |
| ENSDARG000 | 5:5:5:5:5:5:5: | 23957572:239 | 23958752:239 | -:-:-:-:-:-:-:-: | 4974 | map7d2a | 0 | 9 | 52 | 20 | 76 | 47 |
| ENSDARG000 | 5:5:5:5:5:5:5: | 24009668:240 | 24011020:240 | -:-:-:-:-:-:-:-: | 3693 | rps6ka3a | 619 | 525 | 694 | 624 | 917 | 714 |
| ENSDARG000 | 5;5;5;5 | 67448664:674 | 67448774:674 | +;+;+;+ | 900 | si:dkev-251i1 | 2 | 0 | 0 | 0 | 11 | 20 |
| ENSDARG000 | 5;5;5;5;5;5;5 | 24002598:240 | 24003058:240 | -;-;-;-;-;-;- | 950 | eif1axa | 936 | 1154 | 1394 | 824 | 1615 | 729 |
| ENSDARG000 | 5;5;5;5;5 | 24882633:248 | 24883053:248 | +;+;+;+;+ | 1707 | rhbdd3 | 78 | 40 | 56 | 49 | 128 | 84 |
| ENSDARG000 | 5:5:5:5:5:5:5: | 24887219:248 | 24888123:248 | -:-:-:-:-:-:-:-: | 3112 | NA | 57 | 15 | 12 | 4 | 30 | 58 |
| ENSDARG000 | 5:5:5:5:5:5:5: | 56927703:569 | 56927905:569 | -:-:-:-:-:-:-:-: | 2304 | C5H2orf42 | 22 | 51 | 52 | 18 | 37 | 41 |
| ENSDARG000 | 5:5:5:5:5:5:5: | 56950861:569 | 56952343:569 | -:-:-:-:-:-:-:-: | 3416 | tia1 | 1850 | 2444 | 1804 | 952 | 2257 | 1390 |
| ENSDARG000 | 5:5:5:5:5:5:5: | 56965177:569 | 56965358:569 | +:+:+:+:+:+:+: | 1978 | tmem232 | 10 | 21 | 4 | 10 | 6 | 22 |
| ENSDARG000 | 5;5;5 | 57167932:571 | 57168449:571 | -;-;- | 635 | NA | 0 | 0 | 0 | 0 | 0 | 0 |
| ENSDARG000 | 5:5:5:5:5:5:5: | 57005838:570 | 57006339:570 | -:-:-:-:-:-:-:-: | 4233 | man2a1 | 999 | 703 | 930 | 448 | 932 | 776 |
| ENSDARG000 | 5;5 | 37487687:374 | 37488496:374 | +;+ | 1541 | NA | 283 | 287 | 273 | 190 | 124 | 381 |
| ENSDARG000 | 5:5:5:5:5:5:5: | 52937583:529 | 52938253:529 | -;-;-;-;-;-;-;- | 2091 | zfpl1 | 384 | 394 | 552 | 462 | 458 | 486 |
| ENSDARG000 | 5;5;5;5;5;5;5 | 250078:25561 | 250200:25617 | +;+;+;+;+;+;+ | 1485 | trabd2a | 1 | 0 | 10 | 0 | 17 | 0 |
| ENSDARG000 | 5:5:5:5:5:5:5: | 50817382:508 | 50817459:508 | -:-:-:-:-:-:-:-: | 4758 | SKIC3 | 270 | 227 | 282 | 118 | 316 | 242 |
| ENSDARG000 | 5:5:5:5:5:5:5: | 6515741:6516 | 6516359:6516 | -:-:-:-:-:-:-:-: | 4814 | tnks1bp1 | 166 | 12 | 14 | 24 | 12 | 6 |
| ENSDARG000 | 5;5;5 | 828663:83215 | 828740:83231 | +;+;+ | 672 | NA | 21 | 7 | 108 | 4 | 1 | 150 |
| ENSDARG000 | 5;5 | 71983005:719 | 71984155:719 | -;- | 1521 | fam78ab | 6 | 4 | 14 | 7 | 33 | 37 |
| ENSDARG000 | 5:5:5:5:5:5:5: | 23083555:230 | 23083703:230 | +:+:+:+:+:+:+: | 1374 | zdhhc15a | 10 | 20 | 44 | 73 | 15 | 50 |
| ENSDARG000 | 5;5;5;5 | 72383890:723 | 72385233:723 | -;-;-;- | 1798 | wbp1 | 359 | 276 | 400 | 319 | 263 | 229 |
| ENSDARG000 | 5:5:5:5:5:5:5: | 10003458:100 | 10003637:100 | -;-;-;-;-;-;-;- | 1759 | NA | 0 | 0 | 1 | 0 | 0 | 0 |
| ENSDARG000 | 5;5 | 30381936:303 | 30382780:303 | -;- | 902 | gig2o | 351 | 456 | 355 | 186 | 288 | 94 |
| ENSDARG000 | 5 | 12366209 | 12366325 | - | 117 | 5S_rRNA | 0 | 0 | 0 | 0 | 0 | 0 |
| ENSDARG000 | 5:5:5:5:5:5:5: | 45322734:453 | 45323082:453 | +:+:+:+:+:+:+: | 3778 | adamts3 | 17 | 6 | 8 | 4 | 0 | 2 |
| ENSDARG000 | 5;5;5;5;5;5;5 | 1624359:1642 | 1624699:1642 | +;+;+;+;+;+;+ | 1258 | PPP1CA | 1504 | 1106 | 1073 | 933 | 1181 | 908 |
| ENSDARG000 | 5;5;5;5;5;5 | 6350328:6355 | 6352057:6356 | -;-;-;-;-;- | 2300 | zgc:73226 | 0 | 14 | 4 | 4 | 2 | 14 |
| ENSDARG000 | 5;5;5;5 | 6617401:6617 | 6617490:6618 | +;+;+;+ | 1458 | msi1a | 14 | 38 | 37 | 2 | 7 | 6 |
| ENSDARG000 | 5 | 61353794 | 61353917 | + | 124 | NA | 0 | 9 | 19 | 4 | 25 | 6 |
| ENSDARG000 | 5;5 | 61784262:617 | 61784554:617 | -;- | 946 | NA | 0 | 0 | 0 | 0 | 0 | 0 |
| ENSDARG000 | 5;5;5;5;5;5 | 8769069:8769 | 8770052:8770 | -;-;-;-;-;- | 1739 | fgf10b | 4 | 0 | 0 | 0 | 0 | 0 |
| ENSDARG000 | 5 | 62723233 | 62723673 | + | 441 | NA | 0 | 4 | 0 | 0 | 0 | 0 |
| ENSDARG000 | 5:5:5:5:5:5:5: | 11864429:118 | 11864847:118 | -:-:-:-:-:-:-:-: | 2231 | fbxw8 | 246 | 273 | 251 | 236 | 269 | 299 |
| ENSDARG000 | 5:5:5:5:5:5:5: | 50898849:508 | 50898872:508 | +:+:+:+:+:+:+: | 2058 | arsk | 96 | 73 | 210 | 100 | 247 | 49 |
| ENSDARG000 | 5:5:5:5:5:5:5: | 8767290:8767 | 8767302:8767 | +:+:+:+:+:+:+: | 288 | NA | 0 | 0 | 0 | 0 | 0 | 0 |
| ENSDARG000 | 5 | 32885205 | 32885331 | + | 127 | NA | 9 | 0 | 5 | 2 | 0 | 20 |
| ENSDARG000 | 5:5:5:5:5:5:5: | 61788819:617 | 61790291:617 | -;-;-;-;-;-;-;- | 2355 | im:7138535 | 147 | 148 | 187 | 104 | 113 | 218 |
| ENSDARG000 | 5 | 43123176 | 43123291 | - | 116 | U5 | 0 | 0 | 0 | 0 | 0 | 0 |
| ENSDARG000 | 5:5:5:5:5:5:5: | 72066835:720 | 72067029:720 | -;-;-;-;-;-;-;- | 1626 | DQX1 | 260 | 205 | 202 | 135 | 184 | 123 |
| ENSDARG000 | 5 | 19008646 | 19008774 | - | 129 | SNORA77B | 0 | 0 | 2 | 0 | 3 | 0 |
| ENSDARG000 | 5 | 27094224 | 27094338 | - | 115 | 5S_rRNA | 0 | 0 | 0 | 0 | 0 | 0 |
| ENSDARG000 | 5 | 55607744 | 55607847 | - | 104 | dre-mir-7b | 1 | 0 | 8 | 14 | 16 | 3 |
| ENSDARG000 | 5:5:5:5:5:5:5: | 43072951:430 | 43073187:430 | +:+:+:+:+:+:+: | 1963 | wdr54 | 113 | 121 | 140 | 94 | 176 | 157 |
| ENSDARG000 | 5:5:5:5:5:5:5: | 52844681:528 | 52845656:528 | +:+:+:+:+:+:+: | 6302 | scarb2a | 6184 | 5644 | 10009 | 4972 | 6860 | 8387 |
| ENSDARG000 | 5:5:5:5:5:5:5: | 33301005:333 | 33301103:333 | +:+:+:+:+:+:+: | 4996 | usp20 | 675 | 514 | 710 | 569 | 748 | 546 |
| ENSDARG000 | 5:5:5:5:5:5:5: | 33356425:333 | 33356575:333 | -:-:-:-:-:-:-:-: | 3048 | garnl3 | 456 | 65 | 129 | 32 | 1209 | 271 |
| ENSDARG000 | 5;5 | 58990583:589 | 58993625:589 | -;- | 3203 | mis12 | 54 | 261 | 297 | 170 | 128 | 100 |
| ENSDARG000 | 5 | 48170098 | 48170184 | - | 87 | dre-mir-9-5 | 0 | 0 | 0 | 0 | 0 | 0 |
| ENSDARG000 | 5;5;5;5;5;5;5 | 2693601:2696 | 2693949:2696 | +;+;+;+;+;+;+ | 1396 | rabepk | 79 | 141 | 162 | 109 | 147 | 160 |

| | | | | | | | | | | | | |
|---|---|---|---|---|---|---|---|---|---|---|---|---|
| ENSDARG000 | 5 | 61352331 | 61352465 | + | 135 | NA | 0 | 0 | 0 | 0 | 0 | 0 |
| ENSDARG000 | 5;5;5 | 64319590:643 | 64319703:643 | +;+;+ | 258 | NA | 0 | 17 | 4 | 0 | 19 | 7 |
| ENSDARG000 | 5 | 2212672 | 2212788 | - | 117 | 5S_rRNA | 0 | 0 | 0 | 0 | 0 | 0 |
| ENSDARG000 | 5 | 60463393 | 60463507 | - | 115 | 5S_rRNA | 0 | 0 | 0 | 0 | 0 | 0 |
| ENSDARG000 | 5;5;5;5;5;5;5; | 1493767:149 | 1494034:149 | +;+;+;+;+;+;+; | 1961 | haus4 | 40 | 189 | 124 | 74 | 116 | 60 |
| ENSDARG000 | 5;5;5;5;5;5;5; | 8196264:819 | 8196409:819 | +;+;+;+;+;+;+; | 3259 | lmbrd2a | 169 | 142 | 257 | 243 | 170 | 234 |
| ENSDARG000 | 5 | 13296307 | 13296422 | + | 116 | dre-mir-454b | 0 | 0 | 0 | 0 | 0 | 0 |
| ENSDARG000 | 5;5;5 | 1515938:1519 | 1516266:1519 | +;+;+ | 1906 | ddrgk1 | 160 | 149 | 326 | 178 | 147 | 145 |
| ENSDARG000 | 5 | 7972759 | 7972879 | - | 121 | 5S_rRNA | 0 | 0 | 0 | 0 | 0 | 0 |
| ENSDARG000 | 5 | 60146271 | 60146389 | - | 119 | 5S_rRNA | 0 | 0 | 0 | 0 | 0 | 0 |
| ENSDARG000 | 5 | 29990147 | 29990221 | + | 75 | mir100-2 | 0 | 0 | 0 | 0 | 0 | 0 |
| ENSDARG000 | 5 | 38798204 | 38798318 | - | 115 | 5S_rRNA | 0 | 0 | 0 | 0 | 0 | 0 |
| ENSDARG000 | 5;5;5;5;5;5;5; | 69458694:694 | 69459280:694 | -;-;-;-;-;-;-;-; | 3004 | NA | 299 | 359 | 511 | 171 | 603 | 270 |
| ENSDARG000 | 5 | 69656394 | 69656508 | - | 115 | 5S_rRNA | 8 | 0 | 3 | 0 | 6 | 8 |
| ENSDARG000 | 5 | 66496949 | 66497066 | + | 118 | 5S_rRNA | 0 | 0 | 0 | 0 | 1 | 1 |
| ENSDARG000 | 5;5;5;5;5 | 13243446:132 | 13243460:132 | +;+;+;+;+ | 1264 | NA | 0 | 0 | 0 | 0 | 0 | 0 |
| ENSDARG000 | 5;5;5;5;5 | 69124146:691 | 69124256:691 | +;+;+;+;+ | 885 | NA | 0 | 0 | 0 | 2 | 0 | 0 |
| ENSDARG000 | 5 | 32886472 | 32886598 | + | 127 | NA | 0 | 6 | 3 | 6 | 4 | 5 |
| ENSDARG000 | 5 | 63985444 | 63985560 | - | 117 | 5S_rRNA | 0 | 0 | 0 | 0 | 0 | 0 |
| ENSDARG000 | 5 | 41004148 | 41004262 | - | 115 | 5S_rRNA | 0 | 0 | 0 | 0 | 0 | 0 |
| ENSDARG000 | 5;5;5;5;5 | 72375523:723 | 72375589:723 | -;-;-;-;- | 498 | MTFP1 | 102 | 141 | 120 | 46 | 83 | 70 |
| ENSDARG000 | 5 | 43124399 | 43124514 | - | 116 | U5 | 0 | 0 | 0 | 0 | 0 | 0 |
| ENSDARG000 | 5 | 29533717 | 29534748 | - | 1032 | NA | 0 | 4 | 0 | 0 | 6 | 78 |
| ENSDARG000 | 5 | 32637648 | 32637765 | + | 118 | 5S_rRNA | 0 | 0 | 0 | 0 | 0 | 0 |
| ENSDARG000 | 5 | 7632695 | 7632819 | - | 125 | 5S_rRNA | 0 | 0 | 0 | 0 | 0 | 0 |
| ENSDARG000 | 5 | 7431425 | 7431523 | - | 99 | U6 | 0 | 0 | 0 | 0 | 0 | 0 |
| ENSDARG000 | 5 | 42489838 | 42489932 | - | 95 | NA | 0 | 0 | 0 | 0 | 0 | 0 |
| ENSDARG000 | 5;5;5;5;5;5;5; | 3819190:3821 | 3819402:3821 | -;-;-;-;-;-;-;-; | 1753 | NA | 0 | 0 | 12 | 0 | 12 | 10 |
| ENSDARG000 | 5;5;5 | 32321313:323 | 32322399:323 | -;-;- | 1392 | hspb15 | 0 | 0 | 8 | 0 | 0 | 0 |
| ENSDARG000 | 5;5;5;5;5;5;5; | 52190513:521 | 52193688:521 | -;-;-;-;-;-;-;-; | 6977 | lnpep | 2298 | 1768 | 2449 | 1586 | 2493 | 2288 |
| ENSDARG000 | 5;5;5;5;5;5;5; | 51127953:511 | 51128361:511 | -;-;-;-;-;-;-;-; | 3541 | NA | 1 | 16 | 7 | 2 | 12 | 20 |
| ENSDARG000 | 5;5;5;5;5;5;5; | 11290851:113 | 11290960:113 | +;+;+;+;+;+;+; | 2523 | SCARF2 | 1698 | 895 | 1538 | 576 | 2472 | 1436 |
| ENSDARG000 | 5 | 54685175 | 54685710 | + | 536 | pmchl | 0 | 0 | 0 | 0 | 0 | 0 |
| ENSDARG000 | 5;5;5;5;5;5;5; | 2409554:2413 | 2410631:2413 | -;-;-;-;-;-;-;-; | 7035 | CIT | 476 | 923 | 1018 | 344 | 746 | 290 |
| ENSDARG000 | 5 | 42272074 | 42272162 | - | 89 | dre-mir-144 | 0 | 0 | 0 | 0 | 0 | 0 |
| ENSDARG000 | 5;5 | 20112032:201 | 20112182:201 | +;+ | 1141 | isg15 | 1212 | 3634 | 2476 | 1814 | 116 | 1095 |
| ENSDARG000 | 5 | 29998762 | 29998847 | + | 86 | dre-mir-125b- | 0 | 0 | 0 | 0 | 0 | 0 |
| ENSDARG000 | 5;5;5;5;5;5;5; | 2241723:2247 | 2243242:2247 | -;-;-;-;-;-;-;- | 2294 | ca4a | 0 | 0 | 0 | 0 | 0 | 0 |
| ENSDARG000 | 5;5;5;5;5;5;5; | 1833741:1839 | 1833894:1839 | -;-;-;-;-;-;-;-;- | 1539 | rcl1 | 380 | 318 | 580 | 312 | 455 | 455 |
| ENSDARG000 | 5;5;5;5 | 42892196:428 | 42892742:428 | -;-;-;- | 1558 | cxcl11.1 | 16 | 16 | 2 | 14 | 20 | 14 |
| ENSDARG000 | 5;5;5;5;5;5;5; | 9377005:9382 | 9377737:9383 | +;+;+;+;+;+;+; | 2393 | ugt2b1 | 31 | 8 | 1 | 20 | 11 | 7 |
| ENSDARG000 | 5;5;5;5;5;5;5; | 1109098:1113 | 1109254:1113 | +;+;+;+;+;+;+; | 4245 | YTHDC2 | 173 | 113 | 128 | 54 | 118 | 98 |
| ENSDARG000 | 5;5;5;5;5;5;5; | 68529728:685 | 68529834:685 | -;-;-;-;-;-;-;-; | 2124 | DLG4 | 0 | 13 | 7 | 0 | 0 | 5 |
| ENSDARG000 | 5;5;5;5;5 | 70155935:701 | 70156035:701 | +;+;+;+;+ | 1640 | rgs3a | 3696 | 3432 | 6689 | 6100 | 5531 | 5299 |
| ENSDARG000 | 5;5 | 15690639:156 | 15691648:156 | -;- | 1245 | NA | 0 | 2 | 0 | 0 | 0 | 0 |
| ENSDARG000 | 5;5;5;5;5;5;5; | 66203154:662 | 66203315:662 | -;-;-;-;-;-;- | 1696 | coq2 | 163 | 147 | 143 | 122 | 126 | 196 |
| ENSDARG000 | 5 | 41183136 | 41183252 | - | 117 | 5S_rRNA | 0 | 0 | 0 | 0 | 0 | 0 |
| ENSDARG000 | 5;5;5;5;5;5 | 51001985:510 | 51002307:510 | +;+;+;+;+;+ | 1771 | ankrd31 | 0 | 0 | 0 | 0 | 0 | 0 |
| ENSDARG000 | 5;5;5;5;5;5;5; | 23045096:230 | 23045115:230 | +;+;+;+;+;+;+; | 5283 | NA | 668 | 698 | 1106 | 431 | 933 | 719 |
| ENSDARG000 | 5 | 62887128 | 62887243 | + | 116 | 5S_rRNA | 0 | 0 | 0 | 0 | 0 | 0 |
| ENSDARG000 | 5;5;5;5;5;5;5; | 58550291:585 | 58550399:585 | +;+;+;+;+;+;+; | 2214 | pou2f3 | 373 | 14 | 2 | 6 | 20 | 2 |
| ENSDARG000 | 5 | 66537217 | 66537334 | - | 118 | 5S_rRNA | 0 | 0 | 0 | 0 | 0 | 0 |
| ENSDARG000 | 5 | 29214530 | 29214646 | + | 117 | 5S_rRNA | 0 | 0 | 0 | 0 | 0 | 0 |
| ENSDARG000 | 5 | 11849377 | 11849491 | + | 115 | 5S_rRNA | 0 | 0 | 0 | 0 | 0 | 0 |
| ENSDARG000 | 5 | 42402176 | 42402270 | - | 95 | NA | 0 | 0 | 0 | 0 | 0 | 0 |
| ENSDARG000 | 5 | 60606983 | 60607099 | - | 117 | 5S_rRNA | 0 | 0 | 0 | 0 | 0 | 0 |
| ENSDARG000 | 5;5;5;5;5 | 1983495:1992 | 1983505:1992 | +;+;+;+;+ | 105 | NA | 0 | 0 | 0 | 0 | 0 | 0 |
| ENSDARG000 | 5 | 69183390 | 69183522 | + | 133 | NA | 1 | 4 | 4 | 0 | 3 | 6 |
| ENSDARG000 | 5 | 7198757 | 7199998 | - | 1242 | NA | 0 | 0 | 0 | 0 | 0 | 0 |
| ENSDARG000 | 5 | 42468244 | 42468338 | - | 95 | NA | 0 | 0 | 0 | 0 | 0 | 0 |
| ENSDARG000 | 5;5;5;5;5 | 68326858:683 | 68327904:683 | -;-;-;-;- | 2047 | h1m | 0 | 0 | 0 | 0 | 0 | 0 |
| ENSDARG000 | 5;5;5;5;5;5;5; | 8111350:8111 | 8111886:8111 | -;-;-;-;-;-;-;-; | 2053 | slc1a3a | 74 | 126 | 337 | 77 | 158 | 198 |
| ENSDARG000 | 5;5;5;5;5;5;5; | 16360209:163 | 16360984:163 | -;-;-;-;-;-;-;-; | 2593 | slc39a14 | 96 | 44 | 90 | 68 | 118 | 29 |
| ENSDARG000 | 5;5;5;5;5;5;5; | 51079504:510 | 51079597:510 | +;+;+;+;+;+;+; | 1849 | fam169aa | 8 | 0 | 0 | 0 | 8 | 0 |
| ENSDARG000 | 5;5;5;5;5;5;5 | 25304499:253 | 25304694:253 | +;+;+;+;+;+;+ | 1546 | carnmt1 | 95 | 51 | 118 | 70 | 116 | 168 |
| ENSDARG000 | 5 | 43121393 | 43121508 | - | 116 | U5 | 0 | 0 | 0 | 0 | 0 | 0 |
| ENSDARG000 | 5;5;5;5;5 | 66132394:661 | 66132499:661 | +;+;+;+;+ | 1532 | zgc:114041 | 14 | 8 | 30 | 38 | 8 | 16 |
| ENSDARG000 | 5;5;5;5;5;5;5; | 51885121:518 | 51887022:518 | -;-;-;-;-;-;-;-; | 3065 | mtx3 | 130 | 56 | 110 | 55 | 59 | 138 |
| ENSDARG000 | 5;5;5 | 52256885:522 | 52259976:522 | -;-;- | 4052 | rgmb | 16133 | 6430 | 5832 | 7109 | 6830 | 3218 |
| ENSDARG000 | 5;5;5;5;5;5;5; | 69597166:695 | 69597221:695 | -;-;-;-;-;-;-;-; | 2409 | aldh2.2 | 1152 | 1371 | 1308 | 1148 | 1343 | 2278 |
| ENSDARG000 | 5 | 15734525 | 15734580 | - | 56 | SNORD63 | 0 | 0 | 0 | 0 | 0 | 0 |
| ENSDARG000 | 5 | 51555333 | 51555464 | + | 132 | 5S_rRNA | 0 | 0 | 0 | 0 | 0 | 0 |
| ENSDARG000 | 5;5;5;5;5;5;5; | 32499070:324 | 32500609:325 | -;-;-;-;-;-;-;-; | 2083 | ntmt1 | 249 | 414 | 540 | 252 | 286 | 243 |
| ENSDARG000 | 5;5;5;5 | 38517486:385 | 38517869:385 | -;-;-;- | 2516 | NA | 2 | 0 | 0 | 0 | 0 | 2 |
| ENSDARG000 | 5;5;5 | 4523633:4528 | 4523761:4528 | -;-;- | 306 | cst14b.1 | 2776 | 1051 | 1330 | 1952 | 992 | 993 |
| ENSDARG000 | 5 | 42395295 | 42395389 | - | 95 | NA | 0 | 0 | 0 | 0 | 0 | 0 |
| ENSDARG000 | 5 | 14245149 | 14245280 | - | 132 | 5S_rRNA | 0 | 0 | 0 | 0 | 0 | 0 |
| ENSDARG000 | 5 | 6929034 | 6929148 | - | 115 | 5S_rRNA | 0 | 0 | 0 | 0 | 0 | 0 |
| ENSDARG000 | 5;5;5;5;5;5;5 | 42657474:426 | 42657556:426 | -;-;-;-;-;-;- | 1044 | NA | 0 | 0 | 0 | 0 | 10 | 4 |
| ENSDARG000 | 5 | 33284908 | 33284983 | - | 76 | SNORD36 | 0 | 0 | 0 | 0 | 0 | 0 |
| ENSDARG000 | 5;5;5;5;5;5;5; | 60847823:608 | 60847981:608 | +;+;+;+;+;+;+; | 4152 | lig3 | 1138 | 954 | 1143 | 681 | 1035 | 979 |
| ENSDARG000 | 5 | 69021859 | 69021962 | - | 104 | dre-mir-2188 | 0 | 0 | 0 | 0 | 0 | 0 |
| ENSDARG000 | 5 | 13299331 | 13299414 | + | 84 | dre-mir-130b | 0 | 0 | 0 | 0 | 0 | 0 |
| ENSDARG000 | 5;5;5;5;5;5;5; | 872299:8895 | 872397:8896 | +;+;+;+;+;+;+; | 1713 | fubp3 | 897 | 883 | 1138 | 568 | 792 | 546 |
| ENSDARG000 | 5 | 23222701 | 23222819 | - | 119 | 5S_rRNA | 3 | 7 | 43 | 33 | 0 | 30 |
| ENSDARG000 | 5;5;5;5;5;5;5; | 71924175:719 | 71925101:719 | +;+;+;+;+;+;+; | 5963 | NA | 624 | 390 | 826 | 525 | 734 | 490 |
| ENSDARG000 | 5 | 61259952 | 61260067 | + | 116 | 5S_rRNA | 0 | 0 | 0 | 0 | 0 | 0 |
| ENSDARG000 | 5 | 42504242 | 42504336 | - | 95 | NA | 0 | 0 | 0 | 0 | 0 | 0 |
| ENSDARG000 | 5;5;5;5;5;5;5 | 67768162:677 | 67769620:677 | -;-;-;-;-;-;- | 3716 | CASR | 0 | 0 | 0 | 0 | 0 | 0 |
| ENSDARG000 | 5;5;5;5;5;5;5; | 70345518:703 | 70346326:703 | +;+;+;+;+;+;+; | 5582 | pappaa | 6 | 2 | 4 | 4 | 15 | 4 |
| ENSDARG000 | 5 | 23910333 | 23910449 | + | 117 | 5S_rRNA | 0 | 0 | 4 | 0 | 0 | 0 |
| ENSDARG000 | 5;5;5;5;5;5;5 | 35270859:352 | 35274473:352 | -;-;-;-;-;-;- | 9643 | map1b | 2 | 36 | 70 | 68 | 18 | 30 |
| ENSDARG000 | 5;5;5;5;5;5;5; | 12562188:125 | 12563319:125 | -;-;-;-;-;-;-;-;- | 3036 | vsig10 | 48 | 0 | 0 | 6 | 4 | 4 |
| ENSDARG000 | 5 | 46738081 | 46738197 | + | 117 | 5S_rRNA | 0 | 0 | 0 | 0 | 0 | 0 |
| ENSDARG000 | 5;5;5;5;5;5;5; | 53479576:534 | 53479616:534 | +;+;+;+;+;+;+; | 1606 | NA | 0 | 0 | 0 | 0 | 0 | 0 |
| ENSDARG000 | 5;5;5;5 | 57646919:576 | 57647687:576 | -;-;-;- | 1147 | cfap68 | 0 | 0 | 0 | 0 | 12 | 0 |
| ENSDARG000 | 5;5;5;5;5;5;5; | 13373593:133 | 13373709:133 | +;+;+;+;+;+;+; | 610 | ccl19a.2 | 146 | 2 | 0 | 0 | 10 | 0 |
| ENSDARG000 | 5 | 64857844 | 64857958 | + | 115 | 5S_rRNA | 2 | 3 | 0 | 0 | 0 | 1 |
| ENSDARG000 | 5;5;5;5 | 42882679:428 | 42883071:428 | -;-;-;- | 744 | NA | 12 | 16 | 2 | 2 | 24 | 8 |
| ENSDARG000 | 5 | 33285526 | 33285597 | - | 72 | SNORD24 | 2 | 1 | 12 | 5 | 10 | 8 |
| ENSDARG000 | 5;5;5;5;5;5;5; | 67785991:677 | 67788313:677 | -;-;-;-;-;-;-;-; | 3174 | eif4e1b | 0 | 0 | 0 | 0 | 0 | 0 |
| ENSDARG000 | 5;5;5 | 42542252:425 | 42543498:425 | -;-;- | 1856 | NA | 0 | 8 | 5 | 10 | 0 | 0 |
| ENSDARG000 | 5;5;5;5;5;5;5; | 42912966:429 | 42913496:429 | +;+;+;+;+;+;+; | 1809 | rufy3 | 276 | 235 | 267 | 276 | 205 | 329 |
| ENSDARG000 | 5;5;5;5;5;5;5; | 26884676:268 | 26885984:268 | -;-;-;-;-;-;- | 2252 | lman2lb | 708 | 1278 | 1008 | 594 | 896 | 591 |
| ENSDARG000 | 5;5;5;5;5;5;5; | 68228268:682 | 68228350:682 | -;-;-;-;-;-;-;-; | 1683 | KIF1C | 26 | 15 | 35 | 14 | 19 | 42 |
| ENSDARG000 | 5;5;5;5;5;5;5; | 67812062:678 | 67812442:678 | +;+;+;+;+;+;+; | 2741 | zgc:175280 | 0 | 2 | 18 | 34 | 50 | 0 |
| ENSDARG000 | 5;5;5;5;5 | 58665648:586 | 58665676:586 | +;+;+;+;+ | 486 | zgc:194948 | 0 | 0 | 0 | 0 | 0 | 0 |
| ENSDARG000 | 5;5;5;5;5;5;5; | 61799629:618 | 61799848:618 | +;+;+;+;+;+;+; | 2541 | NA | 2918 | 2370 | 3698 | 2551 | 3597 | 2847 |
| ENSDARG000 | 5 | 16898923 | 16899047 | - | 125 | 5S_rRNA | 0 | 0 | 0 | 0 | 0 | 0 |
| ENSDARG000 | 5;5;5;5 | 4533244:4545 | 4533574:4545 | +;+;+;+ | 867 | NA | 204 | 62 | 54 | 134 | 185 | 94 |
| ENSDARG000 | 5 | 47748248 | 47748370 | - | 123 | 5S_rRNA | 0 | 0 | 0 | 0 | 0 | 0 |
| ENSDARG000 | 5;5;5;5;5;5;5 | 58996765:589 | 58997310:589 | +;+;+;+;+;+;+ | 4971 | derl2 | 533 | 656 | 786 | 418 | 938 | 596 |
| ENSDARG000 | 5 | 462121 | 462235 | - | 115 | 5S_rRNA | 0 | 0 | 1 | 3 | 0 | 0 |
| ENSDARG000 | 5;5;5 | 817016:82051 | 817027:82066 | +;+;+ | 606 | NA | 13 | 15 | 529 | 20 | 6 | 318 |
| ENSDARG000 | 5;5;5 | 4564233:4565 | 4564853:4565 | +;+;+ | 2937 | NA | 2 | 9 | 6 | 2 | 41 | 94 |
| ENSDARG000 | 5;5;5;5;5 | 20103085:201 | 20103443:201 | +;+;+;+;+ | 804 | gatc | 282 | 226 | 349 | 441 | 356 | 380 |
| ENSDARG000 | 5;5;5;5;5;5;5; | 71252856:712 | 71254954:712 | -;-;-;-;-;-;- | 3516 | brinp1 | 0 | 0 | 0 | 0 | 0 | 2 |
| ENSDARG000 | 5 | 33284556 | 33284626 | - | 71 | SNORD36 | 0 | 0 | 0 | 0 | 0 | 0 |
| ENSDARG000 | 5 | 67011063 | 67011179 | + | 117 | 5S_rRNA | 0 | 0 | 0 | 0 | 0 | 0 |
| ENSDARG000 | 5;5;5;5;5;5;5; | 72377851:723 | 72377968:723 | +;+;+;+;+;+;+; | 1011 | lman2la | 131 | 259 | 241 | 202 | 179 | 144 |
| ENSDARG000 | 5;5;5;5;5;5;5; | 45613024:456 | 45613861:456 | -;-;-;-;-;-;- | 2066 | NPFFR2 | 0 | 4 | 0 | 0 | 2 | 0 |
| ENSDARG000 | 5;5;5;5;5;5;5 | 60886103:608 | 60886338:608 | +;+;+;+;+;+;+ | 1397 | tmem248 | 512 | 408 | 572 | 240 | 561 | 575 |
| ENSDARG000 | 5 | 23568347 | 23568462 | + | 116 | 5S_rRNA | 6 | 2 | 11 | 16 | 0 | 72 |
| ENSDARG000 | 5;5;5;5;5;5;5; | 66433287:664 | 66433459:664 | +;+;+;+;+;+;+; | 6873 | NA | 168 | 931 | 1346 | 459 | 1067 | 485 |
| ENSDARG000 | 5 | 4942598 | 4942712 | - | 115 | 5S_rRNA | 0 | 0 | 0 | 0 | 0 | 0 |
| ENSDARG000 | 5;5;5;5;5;5;5; | 31748:32037: | 31863:32260: | -;-;-;-;-;-;-;-; | 3600 | dcc | 0 | 0 | 6 | 0 | 6 | 0 |
| ENSDARG000 | 5 | 43217368 | 43217483 | - | 116 | U5 | 0 | 0 | 0 | 0 | 0 | 0 |
| ENSDARG000 | 5 | 42409361 | 42409455 | - | 95 | NA | 0 | 0 | 0 | 0 | 0 | 0 |
| ENSDARG000 | 5;5;5;5;5;5;5; | 71715703:717 | 71715709:717 | -;-;-;-;-;-;-;-; | 1902 | gpsm1b | 208 | 343 | 421 | 239 | 243 | 281 |
| ENSDARG000 | 5;5;5;5;5;5;5; | 69504910:695 | 69506334:695 | -;-;-;-;-;-;-;-; | 2475 | tmem116 | 43 | 4 | 61 | 4 | 43 | 29 |
| ENSDARG000 | 5;5 | 20421539:204 | 20421711:204 | +;+ | 1554 | selplg | 145 | 161 | 164 | 211 | 363 | 325 |
| ENSDARG000 | 5;5;5;5;5;5;5; | 337215:33815 | 337339:33831 | +;+;+;+;+;+;+; | 3023 | rnf170 | 1041 | 609 | 908 | 775 | 1078 | 530 |
| ENSDARG000 | 5;5;5;5;5 | 62319217:623 | 62319583:623 | +;+;+;+;+ | 8579 | pho | 252 | 404 | 320 | 114 | 309 | 389 |
| ENSDARG000 | 5 | 34237907 | 34238023 | - | 117 | 5S_rRNA | 0 | 0 | 0 | 0 | 0 | 0 |
| ENSDARG000 | 5;5;5;5;5;5;5; | 61839707:618 | 61839866:618 | +;+;+;+;+;+;+; | 3445 | NA | 42 | 30 | 95 | 74 | 0 | 70 |
| ENSDARG000 | 5 | 5352071 | 5352187 | + | 117 | 5S_rRNA | 0 | 0 | 0 | 0 | 0 | 0 |
| ENSDARG000 | 5 | 11900557 | 11900674 | + | 118 | 5S_rRNA | 0 | 0 | 0 | 0 | 0 | 0 |
| ENSDARG000 | 5 | 37053878 | 37053998 | + | 121 | 5S_rRNA | 0 | 0 | 0 | 0 | 0 | 0 |
| ENSDARG000 | 5;5;5;5;5 | 5689476:5689 | 5689594:5690 | +;+;+;+;+ | 846 | TRAF2 | 2 | 6 | 10 | 14 | 0 | 0 |
| ENSDARG000 | 5;5;5 | 66157117:661 | 66158014:661 | -;-;- | 1248 | mboat4 | 0 | 4 | 2 | 2 | 3 | 10 |
| ENSDARG000 | 5 | 52225684 | 52225805 | + | 122 | 5S_rRNA | 4 | 4 | 4 | 32 | 0 | 9 |
| ENSDARG000 | 5;5;5;5;5;5;5; | 68007172:680 | 68007329:680 | -;-;-;-;-;-;- | 1302 | NA | 0 | 0 | 0 | 14 | 2 | 0 |
| ENSDARG000 | 5 | 59755144 | 59755261 | + | 118 | 5S_rRNA | 0 | 0 | 0 | 0 | 0 | 0 |
| ENSDARG000 | 5 | 1960073 | 1960200 | - | 128 | SNORA63 | 4 | 2 | 0 | 0 | 1 | 0 |
| ENSDARG000 | 5;5;5;5;5;5;5; | 51909740:519 | 51909818:519 | +;+;+;+;+;+;+; | 6244 | thbs4a | 24 | 12 | 0 | 2 | 10 | 1 |
| ENSDARG000 | 5 | 65250379 | 65250474 | - | 96 | dre-mir-455-1 | 0 | 0 | 0 | 0 | 0 | 0 |
| ENSDARG000 | 5;5;5;5;5;5;5; | 35202063:352 | 35205171:352 | -;-;-;-;-;-;-;-; | 5804 | TNPO1 | 636 | 698 | 710 | 642 | 707 | 722 |
| ENSDARG000 | 5;5;5;5;5;5 | 58679071:586 | 58679237:586 | +;+;+;+;+;+ | 1219 | foxr1 | 2 | 0 | 0 | 0 | 0 | 0 |
| ENSDARG000 | 5;5;5;5;5;5;5; | 2651412:2651 | 2652249:2652 | -;-;-;-;-;-;- | 2822 | RAB35 | 115 | 140 | 115 | 89 | 135 | 83 |
| ENSDARG000 | 5;5;5;5;5;5;5; | 69541667:695 | 69542017:695 | -;-;-;-;-;-;-;-; | 1179 | kiss1rb | 13 | 12 | 5 | 0 | 13 | 6 |
| ENSDARG000 | 5;5;5;5;5;5 | 6489260:6497 | 6489711:6498 | -;-;-;-;-;- | 1028 | crybb3 | 4 | 2 | 0 | 0 | 0 | 0 |
| ENSDARG000 | 5;5;5;5 | 26921033:269 | 26921526:269 | +;+;+;+ | 1080 | tm2d2 | 312 | 267 | 254 | 165 | 351 | 357 |
| ENSDARG000 | 5;5;5;5;5;5;5; | 37316683:373 | 37316771:373 | -;-;-;-;-;-;-;-; | 2715 | wdr44 | 1349 | 868 | 1203 | 953 | 1047 | 885 |
| ENSDARG000 | 5;5;5;5;5;5;5; | 4382378:4384 | 4382592:4384 | -;-;-;-;-;-;-;-;- | 880 | apooa | 253 | 319 | 447 | 320 | 448 | 190 |
| ENSDARG000 | 5;5;5;5;5;5;5 | 69686368:696 | 69686538:696 | +;+;+;+;+;+;+ | 2149 | znf1005 | 18 | 63 | 34 | 60 | 35 | 69 |
| ENSDARG000 | 5;5;5;5;5;5;5; | 23622717:236 | 23622730:236 | -;-;-;-;-;-;-;-;- | 204 | NA | 0 | 0 | 3 | 0 | 0 | 0 |
| ENSDARG000 | 5 | 43728137 | 43728269 | - | 133 | 5S_rRNA | 0 | 0 | 0 | 0 | 0 | 0 |
| ENSDARG000 | 5;5;5 | 15523008:155 | 15523053:155 | -;-;- | 318 | NA | 0 | 0 | 0 | 7 | 0 | 0 |
| ENSDARG000 | 5;5;5;5;5;5;5; | 64277604:642 | 64277838:642 | +;+;+;+;+;+;+; | 1971 | QSOX2 | 312 | 379 | 371 | 211 | 284 | 175 |
| ENSDARG000 | 5 | 42497045 | 42497139 | - | 95 | NA | 0 | 0 | 0 | 0 | 0 | 0 |
| ENSDARG000 | 5;5;5 | 2680455:2686 | 2681163:2686 | -;-;- | 1343 | gng10 | 106 | 119 | 126 | 159 | 108 | 171 |
| ENSDARG000 | 5 | 51810809 | 51810925 | + | 117 | 5S_rRNA | 0 | 0 | 0 | 0 | 0 | 0 |

| | | | | | | | | | | | | |
|---|---|---|---|---|---|---|---|---|---|---|---|---|
| ENSDARG000 | 5:5:5:5:5:5:5: | 4482437:4482 | 4482793:4482 | -:-:-:-:-:-:-:-: | 929 | zgc:153704 | 374 | 8 | 0 | 85 | 46 | 2 |
| ENSDARG000 | 5 | 997812 | 997928 | - | 117 | 5S_rRNA | 0 | 0 | 0 | 0 | 0 | 0 |
| ENSDARG000 | 5:5:5:5:5:5:5: | 57673141:576 | 57674737:576 | -;-;-;-;-;-;-;- | 2649 | crfb12 | 12 | 0 | 0 | 16 | 0 | 0 |
| ENSDARG000 | 5:5:5:5:5:5:5: | 36538535:365 | 36538921:365 | -:-:-:-:-:-:-:-: | 1264 | zgc:158432 | 0 | 2 | 6 | 4 | 4 | 0 |
| ENSDARG000 | 5;5;5;5;5;5;5 | 36944533:369 | 36946610:369 | -;-;-;-;-;-;- | 2437 | h3f3c | 6272 | 4356 | 6377 | 4952 | 7960 | 4757 |
| ENSDARG000 | 5 | 33286587 | 33286658 | - | 72 | SNORD24 | 0 | 0 | 0 | 1 | 0 | 0 |
| ENSDARG000 | 5;5;5;5;5 | 11502578:115 | 11502708:115 | -;-;-;-;- | 648 | CABP7 | 0 | 0 | 0 | 0 | 0 | 0 |
| ENSDARG000 | 5:5:5:5:5:5:5: | 8712798:8718 | 8713177:8718 | -:-:-:-:-:-:-:-: | 3894 | mybbp1a | 1298 | 1277 | 2408 | 820 | 1349 | 1057 |
| ENSDARG000 | 5:5:5:5:5:5:5: | 71995056:719 | 71995078:720 | +:+:+:+:+:+:+: | 897 | PLPP7 | 0 | 2 | 0 | 0 | 0 | 0 |
| ENSDARG000 | 5:5:5:5:5:5:5: | 50869091:508 | 50869956:508 | +:+:+:+:+:+:+: | 6580 | nol6 | 680 | 929 | 1411 | 476 | 1117 | 8290 |
| ENSDARG000 | 5 | 12192898 | 12193014 | - | 117 | 5S_rRNA | 0 | 0 | 0 | 0 | 0 | 0 |
| ENSDARG000 | 5 | 68617958 | 68618089 | - | 132 | 5S_rRNA | 0 | 0 | 0 | 0 | 0 | 0 |
| ENSDARG000 | 5 | 17029298 | 17029414 | + | 117 | 5S_rRNA | 0 | 0 | 0 | 0 | 0 | 0 |
| ENSDARG000 | 5 | 43284630 | 43284745 | - | 116 | U5 | 0 | 0 | 0 | 0 | 0 | 0 |
| ENSDARG000 | 5 | 58037772 | 58037886 | + | 115 | 5S_rRNA | 0 | 0 | 0 | 0 | 0 | 0 |
| ENSDARG000 | 5 | 43224699 | 43224814 | - | 116 | U5 | 0 | 0 | 0 | 0 | 0 | 0 |
| ENSDARG000 | 5 | 29990452 | 29990544 | + | 93 | dre-let-7a-4 | 0 | 0 | 0 | 0 | 0 | 0 |
| ENSDARG000 | 5 | 42271971 | 42272030 | - | 60 | dre-mir-451 | 0 | 0 | 0 | 0 | 0 | 0 |
| ENSDARG000 | 5:5:5:5:5:5:5: | 66220601:662 | 66220663:662 | +:+:+:+:+:+:+: | 1923 | mfsd10 | 936 | 1882 | 3380 | 2875 | 3671 | 5459 |
| ENSDARG000 | 5:5:5:5:5:5:5: | 66170479:661 | 66170754:661 | +:+:+:+:+:+:+: | 2946 | gldc | 214 | 234 | 462 | 236 | 278 | 726 |
| ENSDARG000 | 5:5:5:5:5:5:5: | 480119:48396 | 480173:48428 | +:+:+:+:+:+:+: | 3539 | tek | 308 | 86 | 330 | 310 | 468 | 175 |
| ENSDARG000 | 5:5:5:5:5:5:5: | 58687541:586 | 58687745:586 | +:+:+:+:+:+:+: | 1894 | cenatac | 105 | 85 | 215 | 121 | 203 | 159 |
| ENSDARG000 | 5;5;5 | 44455754:444 | 44456502:444 | -;-;- | 3337 | gas1a | 214 | 191 | 429 | 233 | 373 | 391 |
| ENSDARG000 | 5 | 43480856 | 43480972 | - | 117 | 5S_rRNA | 0 | 0 | 0 | 0 | 0 | 0 |
| ENSDARG000 | 5 | 46609029 | 46609143 | + | 115 | 5S_rRNA | 0 | 0 | 0 | 0 | 0 | 0 |
| ENSDARG000 | 5:5:5:5:5:5:5: | 33295678:332 | 33295920:332 | -;-;-;-;-;-;-;- | 1103 | C5H9orf78 | 343 | 226 | 403 | 266 | 370 | 317 |
| ENSDARG000 | 5:5:5:5:5:5:5: | 9360394:9360 | 9360473:9360 | +:+:+:+:+:+:+: | 963 | NA | 6 | 0 | 4 | 3 | 5 | 2 |
| ENSDARG000 | 5 | 43120171 | 43120286 | - | 116 | U5 | 0 | 0 | 0 | 0 | 0 | 0 |
| ENSDARG000 | 5;5 | 6955900:6958 | 6956575:6962 | +;+ | 4593 | NA | 236 | 205 | 293 | 243 | 216 | 272 |
| ENSDARG000 | 5;5;5 | 4575800:4576 | 4575956:4576 | +;+;+ | 1685 | NA | 8 | 4 | 0 | 4 | 50 | 38 |
| ENSDARG000 | 5:5:5:5:5:5:5: | 23024033:230 | 23024184:230 | +:+:+:+:+:+:+: | 1453 | chmp1b | 2070 | 1160 | 1364 | 1303 | 1148 | 1204 |
| ENSDARG000 | 5;5 | 9677305:9677 | 9677456:9678 | -;- | 732 | NA | 30 | 38 | 43 | 30 | 87 | 39 |
| ENSDARG000 | 5;5 | 821952:82211 | 822092:82228 | -;- | 312 | NA | 0 | 0 | 16 | 1 | 0 | 6 |
| ENSDARG000 | 5:5:5:5:5:5:5: | 72194444:721 | 72194712:721 | +:+:+:+:+:+:+: | 4309 | ddx54 | 407 | 603 | 968 | 344 | 629 | 384 |
| ENSDARG000 | 5:5:5:5:5:5:5: | 68112194:681 | 68112295:681 | +:+:+:+:+:+:+: | 1170 | KCNAB3 | 0 | 0 | 14 | 0 | 0 | 0 |
| ENSDARG000 | 5;5;5;5;5;5;5 | 35955209:359 | 35955667:359 | +;+;+;+;+;+;+ | 4712 | steep1 | 432 | 424 | 470 | 430 | 394 | 356 |
| ENSDARG000 | 5 | 41163655 | 41163771 | + | 117 | 5S_rRNA | 0 | 0 | 0 | 0 | 0 | 0 |
| ENSDARG000 | 5;5 | 65871688:658 | 65872503:658 | +;+ | 1275 | FAM163B | 0 | 3 | 4 | 16 | 0 | 9 |
| ENSDARG000 | 5 | 61580599 | 61580697 | + | 99 | NA | 0 | 1 | 0 | 0 | 0 | 4 |
| ENSDARG000 | 5;5;5;5 | 69426654:694 | 69429856:694 | -;-;-;- | 3628 | isca1 | 445 | 270 | 360 | 407 | 373 | 388 |
| ENSDARG000 | 5 | 42475439 | 42475533 | - | 95 | NA | 0 | 0 | 0 | 0 | 0 | 0 |
| ENSDARG000 | 5 | 8722928 | 8723043 | - | 116 | 5S_rRNA | 0 | 0 | 0 | 0 | 0 | 0 |
| ENSDARG000 | 5;5;5;5 | 72087619:720 | 72087670:720 | +;+;+;+ | 697 | oxt | 2 | 0 | 2 | 0 | 5 | 0 |
| ENSDARG000 | 5:5:5:5:5:5:5: | 6954162:6970 | 6954267:6970 | +:+:+:+:+:+:+: | 1686 | NA | 10 | 2 | 9 | 13 | 6 | 2 |
| ENSDARG000 | 5 | 63709618 | 63709683 | - | 66 | NA | 0 | 0 | 0 | 0 | 0 | 0 |
| ENSDARG000 | 5:5:5:5:5:5:5: | 58397646:583 | 58397697:583 | +:+:+:+:+:+:+: | 1272 | HEPACAM | 0 | 0 | 0 | 0 | 0 | 6 |
| ENSDARG000 | 5 | 1837149 | 1837260 | - | 112 | dre-mir-101b | 1 | 0 | 0 | 0 | 0 | 0 |
| ENSDARG000 | 5;5;5;5;5;5 | 29789123:297 | 29789143:297 | -;-;-;-;-;- | 147 | NA | 6 | 4 | 39 | 27 | 28 | 33 |
| ENSDARG000 | 5;5;5 | 57359542:573 | 57359933:573 | -;-;- | 558 | cdc26 | 185 | 248 | 257 | 209 | 145 | 218 |
| ENSDARG000 | 5;5;5;5 | 42895592:428 | 42897061:428 | -;-;-;- | 2083 | cxcl20 | 14 | 6 | 14 | 0 | 56 | 2 |
| ENSDARG000 | 5;5;5;5;5 | 9408901:9409 | 9409023:9409 | +;+;+;+;+ | 825 | NA | 0 | 0 | 0 | 0 | 0 | 0 |
| ENSDARG000 | 5;5;5;5;5;5;5 | 55543580:555 | 55544600:555 | -;-;-;-;-;-;- | 1937 | gna14 | 44 | 4 | 0 | 0 | 8 | 0 |
| ENSDARG000 | 5:5:5:5:5:5:5: | 40336828:403 | 40336860:403 | -:-:-:-:-:-:-:-: | 231 | NA | 1 | 5 | 1 | 0 | 0 | 0 |
| ENSDARG000 | 5;5 | 63078198:630 | 63081130:630 | -;- | 2991 | NA | 8 | 14 | 20 | 14 | 12 | 12 |
| ENSDARG000 | 5 | 58212161 | 58212284 | + | 124 | 5S_rRNA | 0 | 0 | 0 | 0 | 0 | 0 |
| ENSDARG000 | 5;5 | 46146084:461 | 46148218:461 | -;- | 2975 | F2RL2 | 26 | 1 | 0 | 2 | 0 | 8 |
| ENSDARG000 | 5:5:5:5:5:5:5: | 58487368:584 | 58487436:584 | +:+:+:+:+:+:+: | 2664 | NA | 0 | 2 | 4 | 0 | 6 | 0 |
| ENSDARG000 | 5;5 | 1219187:1223 | 1219423:1223 | +;+ | 816 | bbln | 165 | 202 | 86 | 256 | 172 | 150 |
| ENSDARG000 | 5 | 2564145 | 2564261 | + | 117 | 5S_rRNA | 0 | 0 | 0 | 0 | 0 | 0 |
| ENSDARG000 | 5 | 31944607 | 31944722 | + | 116 | 5S_rRNA | 0 | 0 | 0 | 0 | 0 | 0 |
| ENSDARG000 | 5 | 6531067 | 6531182 | - | 116 | 5S_rRNA | 0 | 0 | 0 | 0 | 0 | 0 |
| ENSDARG000 | 5;5;5;5;5 | 8256574:8264 | 8256677:8264 | +;+;+;+;+ | 2274 | NA | 84 | 161 | 167 | 24 | 32 | 108 |
| ENSDARG000 | 5 | 57922284 | 57922367 | - | 84 | dre-mir-34c | 0 | 0 | 0 | 0 | 0 | 0 |
| ENSDARG000 | 5;5;5;5;5;5 | 68267908:682 | 68267994:682 | +;+;+;+;+;+ | 600 | tnfsf13 | 0 | 3 | 0 | 0 | 2 | 0 |
| ENSDARG000 | 5 | 51102010 | 51104671 | + | 2662 | zgc:194398 | 10 | 4 | 8 | 0 | 6 | 2 |
| ENSDARG000 | 5;5 | 25311309:253 | 25311816:253 | +;+ | 1215 | wu:fa19b12 | 85 | 86 | 143 | 110 | 157 | 117 |
| ENSDARG000 | 5 | 35823418 | 35823548 | - | 131 | 5S_rRNA | 0 | 0 | 0 | 0 | 0 | 0 |
| ENSDARG000 | 5;5;5;5 | 4806851:4818 | 4807782:4818 | +;+;+;+ | 3513 | angptl2a | 154 | 4 | 1221 | 38 | 10 | 19 |
| ENSDARG000 | 5;5;5 | 42467867:424 | 42468426:424 | +;+;+ | 1014 | NA | 0 | 0 | 0 | 0 | 0 | 0 |
| ENSDARG000 | 5 | 22234407 | 22234505 | - | 99 | mir223 | 0 | 0 | 1 | 2 | 2 | 8 |
| ENSDARG000 | 5:5:5:5:5:5:5: | 7919878:7919 | 7920029:7920 | -:-:-:-:-:-:-:-: | 2961 | EGFLAM | 0 | 0 | 0 | 0 | 6 | 8 |
| ENSDARG000 | 5:5:5:5:5:5:5: | 2742232:2742 | 2742320:2742 | -:-:-:-:-:-:-:-: | 1974 | LZTS3 | 19 | 25 | 40 | 14 | 29 | 35 |
| ENSDARG000 | 5 | 13288828 | 13288906 | + | 79 | dre-mir-130c- | 0 | 0 | 0 | 0 | 0 | 0 |
| ENSDARG000 | 5 | 65600554 | 65600664 | + | 111 | 5S_rRNA | 0 | 0 | 0 | 0 | 0 | 0 |
| ENSDARG000 | 5 | 57922520 | 57922604 | - | 85 | dre-mir-34b | 0 | 0 | 0 | 0 | 0 | 0 |
| ENSDARG000 | 5 | 1181409 | 1181524 | - | 116 | 5S_rRNA | 0 | 0 | 0 | 2 | 0 | 0 |
| ENSDARG000 | 5;5;5 | 61738276:617 | 61738719:617 | +;+;+ | 1042 | RASL10B | 0 | 0 | 0 | 0 | 0 | 0 |
| ENSDARG000 | 5;5;5;5;5;5;5 | 62287716:622 | 62292605:622 | -;-;-;-;-;-;- | 5692 | NA | 10 | 12 | 4 | 8 | 4 | 16 |
| ENSDARG000 | 5:5:5:5:5:5:5: | 23994038:239 | 23994053:239 | +:+:+:+:+:+:+: | 162 | NA | 1 | 0 | 0 | 0 | 0 | 0 |
| ENSDARG000 | 5:5:5:5:5:5:5: | 51156808:511 | 51157165:511 | -:-:-:-:-:-:-:-: | 1923 | CARD9 | 135 | 122 | 81 | 48 | 79 | 174 |
| ENSDARG000 | 5;5 | 22459087:224 | 22459261:224 | +;+ | 370 | NA | 133 | 78 | 88 | 50 | 134 | 49 |
| ENSDARG000 | 5 | 16445349 | 16445465 | - | 117 | 5S_rRNA | 0 | 0 | 0 | 0 | 0 | 0 |
| ENSDARG000 | 5:5:5:5:5:5:5: | 39386225:394 | 39389444:394 | -:-:-:-:-:-:-:-: | 5216 | antxr2a | 1249 | 283 | 1287 | 351 | 3231 | 421 |
| ENSDARG000 | 5;5;5;5;5 | 56119975:561 | 56120564:561 | +;+;+;+;+ | 1391 | mrm1 | 64 | 63 | 51 | 10 | 50 | 46 |
| ENSDARG000 | 5:5:5:5:5:5:5: | 457729:45821 | 457812:45834 | +:+:+:+:+:+:+: | 1752 | ift74 | 1714 | 1818 | 2059 | 1273 | 2182 | 1120 |
| ENSDARG000 | 5;5;5 | 69417462:694 | 69417989:694 | -;-;- | 903 | NA | 0 | 0 | 2 | 0 | 0 | 0 |
| ENSDARG000 | 5:5:5:5:5:5:5: | 69486184:694 | 69486853:694 | -:-:-:-:-:-:-:-: | 1783 | psat1 | 1850 | 2115 | 3234 | 2428 | 937 | 1527 |
| ENSDARG000 | 5:5:5:5:5:5:5: | 64994085:649 | 64994271:649 | -:-:-:-:-:-:-:-: | 1194 | anxa1d | 7 | 4 | 0 | 0 | 8 | 2 |
| ENSDARG000 | 5:5:5:5:5:5:5: | 16315637:163 | 16315737:163 | -:-:-:-:-:-:-:-: | 2169 | SUSD2 | 52 | 59 | 142 | 46 | 56 | 53 |
| ENSDARG000 | 5;5 | 36974931:369 | 36975039:369 | +;+ | 968 | gjd1a | 14 | 0 | 0 | 0 | 0 | 0 |
| ENSDARG000 | 5;5;5;5;5;5;5 | 9417409:9417 | 9418141:9418 | +;+;+;+;+;+;+ | 1732 | ugt2b5 | 22 | 4 | 0 | 6 | 7 | 3 |
| ENSDARG000 | 5:5:5:5:5:5:5: | 68037629:680 | 68038062:680 | -:-:-:-:-:-:-:-: | 1798 | NA | 552 | 336 | 470 | 416 | 558 | 486 |
| ENSDARG000 | 5:5:5:5:5:5:5: | 27414770:274 | 27414800:274 | -:-:-:-:-:-:-:-: | 204 | NA | 0 | 0 | 0 | 3 | 3 | 4 |
| ENSDARG000 | 5 | 42192566 | 42192680 | + | 115 | 5S_rRNA | 0 | 0 | 0 | 0 | 0 | 1 |
| ENSDARG000 | 5 | 12927438 | 12927554 | + | 117 | 5S_rRNA | 0 | 0 | 0 | 0 | 0 | 0 |
| ENSDARG000 | 5 | 55671466 | 55671582 | + | 117 | 5S_rRNA | 0 | 0 | 0 | 0 | 0 | 0 |
| ENSDARG000 | 5 | 12610802 | 12610883 | + | 82 | dre-mir-22b | 0 | 0 | 0 | 0 | 0 | 0 |
| ENSDARG000 | 5 | 3683053 | 3684402 | - | 1350 | NA | 0 | 0 | 0 | 0 | 0 | 0 |
| ENSDARG000 | 5;5 | 69536091:695 | 69536719:695 | -;- | 674 | zgc:153044 | 38 | 44 | 24 | 55 | 42 | 38 |
| ENSDARG000 | 5;5;5;5;5;5;5 | 431994:43199 | 432064:43206 | +;+;+;+;+;+;+ | 719 | thap1 | 131 | 64 | 85 | 118 | 142 | 120 |
| ENSDARG000 | 5 | 39324768 | 39324884 | - | 117 | 5S_rRNA | 0 | 0 | 0 | 0 | 0 | 0 |
| ENSDARG000 | 5;5;5;5;5;5;5 | 1589458:1597 | 1589535:1597 | +;+;+;+;+;+;+ | 1982 | MYL2 | 2 | 0 | 0 | 0 | 0 | 0 |
| ENSDARG000 | 5 | 824709 | 825920 | - | 1212 | NA | 299 | 284 | 3362 | 659 | 148 | 1420 |
| ENSDARG000 | 5 | 46046172 | 46046286 | - | 115 | 5S_rRNA | 0 | 0 | 0 | 0 | 0 | 0 |
| ENSDARG000 | 5 | 6875920 | 6876034 | + | 115 | 5S_rRNA | 0 | 0 | 0 | 0 | 0 | 0 |
| ENSDARG000 | 5 | 13289353 | 13289433 | + | 81 | dre-mir-301b | 0 | 0 | 0 | 0 | 0 | 0 |
| ENSDARG000 | 5:5:5:5:5:5:5: | 71699893:716 | 71699961:716 | -:-:-:-:-:-:-:-: | 904 | ak1 | 1283 | 1580 | 2525 | 3361 | 1446 | 1914 |
| ENSDARG000 | 5;5;5 | 1089600:1091 | 1089735:1091 | +;+;+ | 744 | NA | 33 | 26 | 15 | 18 | 28 | 30 |
| ENSDARG000 | 5:5:5:5:5:5:5: | 1877464:1877 | 1877505:1877 | +:+:+:+:+:+:+: | 2044 | zgc:101699 | 59 | 103 | 68 | 6 | 142 | 107 |
| ENSDARG000 | 5 | 1952652 | 1952778 | - | 127 | SNORA63 | 2 | 12 | 15 | 6 | 5 | 2 |
| ENSDARG000 | 5 | 39029811 | 39029925 | - | 115 | 5S_rRNA | 0 | 0 | 0 | 0 | 0 | 0 |
| ENSDARG000 | 5;5;5;5 | 57653459:576 | 57653504:576 | -;-;-;- | 485 | mia | 621 | 706 | 277 | 91 | 2905 | 200 |
| ENSDARG000 | 5;5;5;5;5 | 7564644:7564 | 7564812:7565 | +;+;+;+;+ | 918 | NA | 0 | 0 | 26 | 1 | 0 | 9 |
| ENSDARG000 | 5 | 24521682 | 24521750 | + | 69 | dre-mir-1306 | 0 | 0 | 0 | 0 | 0 | 0 |
| ENSDARG000 | 5 | 15743137 | 15743253 | + | 117 | 5S_rRNA | 0 | 0 | 0 | 0 | 0 | 0 |
| ENSDARG000 | 5 | 42380913 | 42381007 | - | 95 | NA | 0 | 0 | 0 | 0 | 0 | 0 |
| ENSDARG000 | 5:5:5:5:5:5:5: | 5708576:5711 | 5708686:5712 | -:-:-:-:-:-:-:-: | 5025 | NA | 395 | 66 | 110 | 105 | 92 | 277 |
| ENSDARG000 | 5;5;5 | 50788329:507 | 50788589:507 | +;+;+ | 525 | NA | 0 | 0 | 0 | 0 | 0 | 0 |
| ENSDARG000 | 5:5:5:5:5:5:5: | 63458263:634 | 63458416:634 | -:-:-:-:-:-:-:-: | 5115 | NA | 0 | 0 | 20 | 0 | 0 | 2 |
| ENSDARG000 | 5 | 24489875 | 24489974 | - | 100 | 5S_rRNA | 0 | 0 | 0 | 0 | 0 | 0 |
| ENSDARG000 | 5:5:5:5:5:5:5: | 45677781:456 | 45677934:456 | +:+:+:+:+:+:+: | 1807 | gc | 2 | 0 | 12 | 0 | 4 | 0 |
| ENSDARG000 | 5:5:5:5:5:5:5: | 69574677:695 | 69575045:695 | -:-:-:-:-:-:-:-: | 1918 | aldh2.1 | 1170 | 1175 | 1088 | 1255 | 1253 | 1498 |
| ENSDARG000 | 5 | 62085817 | 62085934 | + | 118 | 5S_rRNA | 0 | 0 | 0 | 0 | 0 | 0 |
| ENSDARG000 | 5 | 61351368 | 61351501 | + | 134 | NA | 14 | 7 | 2 | 0 | 4 | 9 |
| ENSDARG000 | 5 | 487859 | 487975 | - | 117 | 5S_rRNA | 8 | 0 | 0 | 8 | 2 | 0 |
| ENSDARG000 | 5 | 42674156 | 42674250 | + | 95 | NA | 0 | 0 | 0 | 0 | 0 | 0 |
| ENSDARG000 | 5:5:5:5:5:5:5: | 70262153:702 | 70262601:702 | +:+:+:+:+:+:+: | 4705 | znf618 | 451 | 430 | 642 | 230 | 691 | 421 |
| ENSDARG000 | 5 | 9823754 | 9824908 | - | 1155 | zgc:158343 | 580 | 293 | 492 | 546 | 340 | 673 |
| ENSDARG000 | 5:5:5:5:5:5:5: | 62304529:623 | 62309221:623 | -;-;-;-;-;-;-;- | 5696 | C5H11orf54 | 614 | 409 | 376 | 292 | 444 | 689 |
| ENSDARG000 | 5:5:5:5:5:5:5: | 52226807:522 | 52227304:522 | -:-:-:-:-:-:-:-: | 3225 | erap2 | 469 | 714 | 520 | 157 | 638 | 801 |
| ENSDARG000 | 5;5 | 393738:39535 | 394924:39601 | +;+ | 1850 | june | 291 | 124 | 236 | 164 | 108 | 98 |
| ENSDARG000 | 5:5:5:5:5:5:5: | 70539703:705 | 70540007:705 | -:-:-:-:-:-:-:-: | 1110 | NA | 0 | 0 | 0 | 4 | 0 | 0 |
| ENSDARG000 | 5:5:5:5:5:5:5: | 72094502:720 | 72094880:720 | -:-:-:-:-:-:-:-: | 2051 | thnsl2 | 24 | 54 | 73 | 36 | 54 | 20 |
| ENSDARG000 | 5;5;5;5;5 | 63509761:635 | 63511604:635 | -;-;-;-;- | 3027 | prdm12b | 0 | 0 | 6 | 0 | 0 | 2 |
| ENSDARG000 | 5 | 60775881 | 60775998 | + | 118 | 5S_rRNA | 0 | 0 | 0 | 0 | 0 | 0 |
| ENSDARG000 | 5 | 39174600 | 39174716 | + | 117 | 5S_rRNA | 0 | 0 | 0 | 0 | 0 | 0 |
| ENSDARG000 | 5 | 21293130 | 21293245 | - | 116 | 5S_rRNA | 0 | 0 | 0 | 0 | 0 | 0 |
| ENSDARG000 | 5 | 43216145 | 43216260 | - | 116 | U5 | 0 | 0 | 0 | 0 | 0 | 0 |
| ENSDARG000 | 5 | 32993786 | 32993891 | + | 106 | 5S_rRNA | 0 | 0 | 0 | 0 | 0 | 0 |
| ENSDARG000 | 5 | 42482642 | 42482736 | - | 95 | NA | 0 | 0 | 0 | 0 | 0 | 0 |
| ENSDARG000 | 5 | 25972068 | 25972160 | + | 93 | dre-mir-204-1 | 0 | 0 | 0 | 0 | 0 | 0 |
| ENSDARG000 | 5:5:5:5:5:5:5: | 3094981:3095 | 3095424:3095 | -:-:-:-:-:-:-:-: | 1697 | hsf5 | 0 | 0 | 2 | 0 | 10 | 6 |
| ENSDARG000 | 5:5:5:5:5:5:5: | 6401427:6408 | 6401468:6408 | -;-;-;-;-;-;-;- | 714 | NA | 15 | 42 | 71 | 42 | 42 | 29 |
| ENSDARG000 | 5:5:5:5:5:5:5: | 11407504:114 | 11407664:114 | +:+:+:+:+:+:+: | 1836 | zgc:112294 | 0 | 0 | 0 | 0 | 0 | 0 |
| ENSDARG000 | 5;5;5;5;5;5 | 50761236:507 | 50761298:507 | +;+;+;+;+;+ | 171 | NA | 0 | 0 | 0 | 0 | 0 | 0 |
| ENSDARG000 | 5 | 43285851 | 43285966 | - | 116 | U5 | 0 | 0 | 0 | 0 | 0 | 0 |
| ENSDARG000 | 5 | 9405283 | 9406062 | + | 780 | NA | 0 | 0 | 0 | 0 | 0 | 0 |
| ENSDARG000 | 5;5 | 22011575:220 | 22017322:220 | -;- | 5826 | amer1 | 800 | 510 | 764 | 698 | 838 | 710 |
| ENSDARG000 | 5:5:5:5:5:5:5: | 50442112:504 | 50446191:504 | -:-:-:-:-:-:-:-: | 7180 | MCTP1 | 60 | 5 | 37 | 131 | 82 | 257 |
| ENSDARG000 | 5 | 20408836 | 20408952 | - | 117 | 5S_rRNA | 3 | 0 | 0 | 0 | 0 | 1 |
| ENSDARG000 | 5;5;5 | 66761506:667 | 66761518:667 | +;+;+ | 300 | NA | 0 | 0 | 0 | 0 | 0 | 0 |
| ENSDARG000 | 5 | 5615331 | 5615446 | + | 116 | 5S_rRNA | 0 | 0 | 0 | 0 | 0 | 0 |
| ENSDARG000 | 5:5:5:5:5:5:5: | 38686046:386 | 38688509:386 | -;-;-;-;-;-;-;- | 3208 | NA | 419 | 279 | 206 | 100 | 346 | 314 |
| ENSDARG000 | 5;5;5;5;5 | 63002872:630 | 63003180:630 | -;-;-;-;- | 1231 | NA | 156 | 261 | 439 | 192 | 368 | 279 |
| ENSDARG000 | 5;5;5;5 | 58520266:585 | 58520490:585 | +;+;+;+ | 804 | OAF | 79 | 30 | 20 | 44 | 30 | 43 |
| ENSDARG000 | 5;5 | 833599:83376 | 833739:83393 | -;- | 312 | NA | 0 | 0 | 3 | 0 | 0 | 0 |
| ENSDARG000 | 5:5:5:5:5:5:5: | 4436405:4438 | 4436471:4438 | +:+:+:+:+:+:+: | 2151 | NA | 414 | 144 | 327 | 215 | 438 | 240 |
| ENSDARG000 | 5;5;5;5;5;5;5 | 3184755:3191 | 3185242:3192 | -;-;-;-;-;-;- | 1080 | NA | 0 | 4 | 2 | 1 | 2 | 0 |
| ENSDARG000 | 5;5;5;5 | 57619943:576 | 57622003:576 | -;-;-;- | 2782 | zgc:193711 | 6 | 0 | 0 | 0 | 2 | 0 |
| ENSDARG000 | 5:5:5:5:5:5:5: | 5614620:5619 | 5617383:5619 | -:-:-:-:-:-:-:-: | 5027 | RABL6 | 180 | 126 | 189 | 102 | 122 | 131 |
| ENSDARG000 | 5:5:5:5:5:5:5: | 62374092:624 | 62375270:624 | +:+:+:+:+:+:+: | 2690 | ABR | 881 | 279 | 725 | 422 | 942 | 214 |
| ENSDARG000 | 5:5:5:5:5:5:5: | 25317061:253 | 25317119:253 | +:+:+:+:+:+:+: | 6336 | trpm6 | 281 | 103 | 126 | 116 | 122 | 301 |

| | | | | | | | | | | | | |
|---|---|---|---|---|---|---|---|---|---|---|---|---|
| ENSDARG000 | 5:5:5:5:5:5:5: | 32490238:324 | 32490337:324 | +:+:+:+:+:+:+: | 2096 | ndor1 | 174 | 251 | 268 | 76 | 293 | 215 |
| ENSDARG000 | 5 | 45201332 | 45201448 | + | 117 | 5S_rRNA | 0 | 0 | 0 | 0 | 0 | 0 |
| ENSDARG000 | 5 | 48553086 | 48553203 | + | 118 | 5S_rRNA | 0 | 0 | 0 | 0 | 0 | 0 |
| ENSDARG000 | 5 | 34833978 | 34834093 | - | 116 | 5S_rRNA | 0 | 0 | 0 | 0 | 0 | 0 |
| ENSDARG000 | 5 | 42659776 | 42659870 | + | 95 | NA | 0 | 0 | 0 | 0 | 0 | 0 |
| ENSDARG000 | 5 | 42388104 | 42388198 | - | 95 | NA | 0 | 0 | 0 | 0 | 0 | 0 |
| ENSDARG000 | 5:5:5:5:5:5:5: | 71662162:716 | 71662716:716 | -:-:-:-:-:-:-: | 2047 | NA | 52 | 26 | 130 | 69 | 40 | 66 |
| ENSDARG000 | 5;5;5;5;5;5 | 38732861:387 | 38732886:387 | -;-;-;-;-;- | 129 | NA | 0 | 0 | 2 | 0 | 0 | 0 |
| ENSDARG000 | 5;5;5;5 | 62988916:629 | 62989463:629 | +;+;+;+ | 895 | TIMM22 | 157 | 142 | 163 | 69 | 155 | 114 |
| ENSDARG000 | 5 | 8635268 | 8635392 | - | 125 | 5S_rRNA | 0 | 0 | 0 | 0 | 0 | 0 |
| ENSDARG000 | 5 | 836137 | 836290 | - | 154 | 5_8S_rRNA | 2 | 0 | 3 | 0 | 0 | 1 |
| ENSDARG000 | 5:5:5:5:5:5:5: | 36641200:366 | 36641225:366 | -:-:-:-:-:-:-: | 513 | NA | 2 | 4 | 0 | 0 | 0 | 6 |
| ENSDARG000 | 5:5:5:5:5:5:5: | 35253457:352 | 35253888:352 | -:-:-:-:-:-:-: | 1389 | ptcd2 | 46 | 91 | 129 | 72 | 85 | 88 |
| ENSDARG000 | 5;5;5;5;5;5;5 | 26913120:269 | 26913684:269 | +;+;+;+;+;+;+ | 2242 | NA | 2 | 1172 | 12 | 6 | 22 | 6 |
| ENSDARG000 | 5 | 47975758 | 47977361 | + | 1604 | NA | 130 | 70 | 88 | 54 | 106 | 176 |
| ENSDARG000 | 5;5;5;5 | 10011922:100 | 10012038:100 | -;-;-;- | 558 | NA | 0 | 0 | 0 | 0 | 0 | 0 |
| ENSDARG000 | 5:5:5:5:5:5:5: | 52881744:528 | 52883247:528 | -:-:-:-:-:-:-: | 3362 | PPEF2 | 2 | 0 | 0 | 0 | 0 | 3 |
| ENSDARG000 | 5;5;5;5;5;5 | 54666099:546 | 54666230:546 | -;-;-;-;-;- | 1087 | NA | 0 | 0 | 0 | 0 | 0 | 0 |
| ENSDARG000 | 5 | 61351920 | 61351995 | + | 76 | NA | 0 | 0 | 0 | 0 | 0 | 0 |
| ENSDARG000 | 5 | 1376773 | 1376868 | - | 96 | dre-mir-199-2 | 0 | 0 | 0 | 0 | 0 | 0 |
| ENSDARG000 | 5;5;5;5;5;5 | 1203767:1207 | 1205919:1207 | -;-;-;-;-;- | 2816 | surf2 | 175 | 263 | 266 | 207 | 253 | 170 |
| ENSDARG000 | 5;5;5;5 | 42875850:428 | 42877293:428 | -;-;-;- | 1756 | NA | 4 | 20 | 8 | 2 | 0 | 6 |
| ENSDARG000 | 5 | 30166456 | 30166572 | + | 117 | 5S_rRNA | 0 | 0 | 0 | 0 | 0 | 0 |
| ENSDARG000 | 5;5;5;5;5;5 | 72108241:721 | 72108317:721 | +;+;+;+;+;+ | 584 | fabp1a | 0 | 0 | 0 | 0 | 0 | 0 |
| ENSDARG000 | 5 | 42666965 | 42667059 | + | 95 | NA | 0 | 0 | 0 | 0 | 0 | 0 |
| ENSDARG000 | 5:5:5:5:5:5:5: | 72172299:721 | 72172634:721 | -;-;-;-;-;-;-;- | 2415 | rab14l | 918 | 938 | 1141 | 1200 | 907 | 1068 |
| ENSDARG000 | 5 | 14642152 | 14642269 | + | 118 | 5S_rRNA | 0 | 0 | 2 | 0 | 0 | 0 |
| ENSDARG000 | 5 | 59746283 | 59746397 | - | 115 | 5S_rRNA | 0 | 0 | 0 | 0 | 0 | 0 |
| ENSDARG000 | 5:5:5:5:5:5:5: | 20030414:200 | 20030497:200 | +:+:+:+:+:+:+: | 3579 | SGSM1 | 0 | 0 | 2 | 0 | 0 | 15 |
| ENSDARG000 | 5:5:5:5:5:5:5: | 54580962:545 | 54581016:545 | +:+:+:+:+:+:+: | 3277 | NA | 276 | 86 | 453 | 142 | 151 | 55 |
| ENSDARG000 | 5;5;5;5;5;5 | 1180703:1185 | 1180969:1185 | -;-;-;-;-;- | 1384 | surf4 | 511 | 688 | 712 | 490 | 677 | 467 |
| ENSDARG000 | 5;5;5;5 | 28497956:285 | 28498015:285 | +;+;+;+ | 588 | nfr | 0 | 0 | 0 | 8 | 0 | 0 |
| ENSDARG000 | 5;5 | 35502290:355 | 35502717:355 | +;+ | 804 | NA | 0 | 0 | 0 | 0 | 0 | 0 |
| ENSDARG000 | 5;5;5;5;5 | 3118231:3120 | 3118958:3120 | +;+;+;+;+ | 2288 | supt4h1 | 377 | 424 | 522 | 276 | 341 | 330 |
| ENSDARG000 | 5:5:5:5:5:5:5: | 315183:31518 | 315311:31531 | -:-:-:-:-:-:-: | 1687 | HOOK3 | 746 | 710 | 793 | 423 | 744 | 618 |
| ENSDARG000 | 5 | 1138670 | 1138785 | + | 116 | 5S_rRNA | 0 | 0 | 0 | 0 | 0 | 0 |
| ENSDARG000 | 5:5:5:5:5:5:5: | 72023719:720 | 72023955:720 | -:-:-:-:-:-:-: | 6361 | PRRC2B | 4899 | 3118 | 6875 | 2973 | 6265 | 3488 |
| ENSDARG000 | 5;5;5;5;5 | 37817870:378 | 37818382:378 | -;-;-;-;- | 1051 | NA | 10 | 6 | 3 | 1 | 0 | 0 |
| ENSDARG000 | 5:5:5:5:5:5:5: | 38425279:384 | 38425317:384 | +:+:+:+:+:+:+: | 273 | NA | 0 | 0 | 1 | 0 | 3 | 0 |
| ENSDARG000 | 5 | 32885760 | 32885886 | + | 127 | NA | 0 | 0 | 0 | 0 | 0 | 0 |
| ENSDARG000 | 5 | 1962282 | 1962409 | - | 128 | SNORA63 | 0 | 0 | 0 | 0 | 0 | 0 |
| ENSDARG000 | 5:5:5:5:5:5:5: | 60873055:608 | 60873474:608 | -:-:-:-:-:-:-: | 1431 | rad51d | 12 | 92 | 70 | 31 | 58 | 29 |
| ENSDARG000 | 5:5:5:5:5:5:5: | 71802014:718 | 71802083:718 | +:+:+:+:+:+:+: | 1331 | LHX3 | 0 | 0 | 1 | 0 | 0 | 0 |
| ENSDARG000 | 5:5:5:5:5:5:5: | 72407526:724 | 72407593:724 | -:-:-:-:-:-:-: | 1347 | PAX8 | 0 | 0 | 6 | 0 | 4 | 0 |
| ENSDARG000 | 5;5 | 5666864:5669 | 5667033:5669 | -;- | 300 | NA | 16 | 19 | 8 | 3 | 25 | 24 |
| ENSDARG000 | 5:5:5:5:5:5:5: | 1668478:1672 | 1668658:1672 | -:-:-:-:-:-:-: | 2985 | MCC | 466 | 186 | 534 | 336 | 404 | 485 |
| ENSDARG000 | 5:5:5:5:5:5:5: | 32460453:324 | 32461452:324 | -:-:-:-:-:-:-: | 2559 | gpr107 | 1132 | 900 | 934 | 669 | 1216 | 1131 |
| ENSDARG000 | 5 | 72089395 | 72089511 | - | 117 | 5S_rRNA | 0 | 0 | 0 | 0 | 4 | 0 |
| ENSDARG000 | 5:5:5:5:5:5:5: | 2713779:2715 | 2714583:2715 | -;-;-;-;-;-;-;- | 2339 | hspa5 | 13663 | 35597 | 81477 | 37286 | 31399 | 10952 |
| ENSDARG000 | 5:5:5:5:5:5:5: | 13870340:138 | 13870500:138 | +:+:+:+:+:+:+: | 3832 | hk2 | 625 | 940 | 806 | 220 | 706 | 336 |
| ENSDARG000 | 5 | 38892622 | 38892759 | - | 138 | 5S_rRNA | 0 | 0 | 0 | 0 | 0 | 0 |
| ENSDARG000 | 5 | 61094808 | 61094926 | - | 119 | 5S_rRNA | 0 | 0 | 0 | 0 | 0 | 0 |
| ENSDARG000 | 5 | 20077694 | 20077810 | + | 117 | 5S_rRNA | 0 | 0 | 0 | 0 | 0 | 0 |
| ENSDARG000 | 5 | 68144905 | 68145021 | + | 117 | 5S_rRNA | 0 | 0 | 0 | 0 | 0 | 0 |
| ENSDARG000 | 5 | 25426275 | 25426404 | - | 130 | 5S_rRNA | 0 | 0 | 0 | 0 | 0 | 0 |
| ENSDARG000 | 5:5:5:5:5:5:5: | 57658898:576 | 57658982:576 | +:+:+:+:+:+:+: | 1925 | bicdl2 | 32 | 4 | 0 | 0 | 2 | 14 |
| ENSDARG000 | 5:5:5:5:5:5:5: | 69553686:695 | 69553979:695 | -:-:-:-:-:-:-: | 1772 | mapkapk5 | 263 | 292 | 416 | 183 | 275 | 321 |
| ENSDARG000 | 5;5;5;5;5;5;5 | 23035645:230 | 23035741:230 | +;+;+;+;+;+;+ | 1875 | CNGA2 | 111 | 47 | 313 | 73 | 262 | 163 |
| ENSDARG000 | 5:5:5:5:5:5:5: | 58455488:584 | 58455666:584 | +:+:+:+:+:+:+: | 3098 | slc37a2 | 2600 | 2264 | 1892 | 1936 | 2524 | 2355 |
| ENSDARG000 | 5;5;5 | 35105085:351 | 35105184:351 | -;-;- | 717 | TMEM174 | 0 | 2 | 12 | 6 | 23 | 0 |
| ENSDARG000 | 5 | 50575790 | 50575906 | - | 117 | 5S_rRNA | 0 | 0 | 0 | 0 | 0 | 0 |
| ENSDARG000 | 5 | 46652479 | 46652595 | - | 117 | 5S_rRNA | 0 | 0 | 0 | 0 | 0 | 0 |
| ENSDARG000 | 5 | 32170170 | 32170286 | - | 117 | 5S_rRNA | 0 | 0 | 0 | 0 | 0 | 0 |
| ENSDARG000 | 5 | 13532874 | 13532988 | - | 115 | 5S_rRNA | 0 | 0 | 0 | 0 | 0 | 0 |
| ENSDARG000 | 5 | 43283405 | 43283520 | - | 116 | U5 | 0 | 0 | 0 | 0 | 0 | 0 |
| ENSDARG000 | 5:5:5:5:5:5:5: | 12637823:126 | 12637865:126 | -:-:-:-:-:-:-: | 2947 | AIFM3 | 0 | 0 | 0 | 0 | 0 | 0 |
| ENSDARG000 | 5:5:5:5:5:5:5: | 1933131:1933 | 1933589:1934 | +:+:+:+:+:+:+: | 2974 | NA | 0 | 2 | 1 | 0 | 8 | 2 |
| ENSDARG000 | 5 | 23673259 | 23673375 | + | 117 | 5S_rRNA | 0 | 0 | 0 | 0 | 0 | 0 |
| ENSDARG000 | 5 | 20674933 | 20675050 | + | 118 | 5S_rRNA | 0 | 0 | 0 | 0 | 0 | 0 |
| ENSDARG000 | 5 | 56066682 | 56066796 | - | 115 | 5S_rRNA | 0 | 0 | 0 | 0 | 0 | 0 |
| ENSDARG000 | 5 | 7148291 | 7148407 | - | 117 | 5S_rRNA | 0 | 0 | 0 | 0 | 0 | 0 |
| ENSDARG000 | 5:5:5:5:5:5:5: | 11242291:112 | 11244748:112 | -:-:-:-:-:-:-: | 3511 | NA | 74 | 19 | 6 | 28 | 44 | 22 |
| ENSDARG000 | 5:5:5:5:5:5:5: | 37517800:375 | 37518455:375 | +:+:+:+:+:+:+: | 5483 | NA | 841 | 228 | 116 | 184 | 1067 | 663 |
| ENSDARG000 | 5:5:5:5:5:5:5: | 28915389:289 | 28918843:289 | -:-:-:-:-:-:-: | 5542 | niban2b | 2300 | 918 | 1182 | 746 | 1088 | 1650 |
| ENSDARG000 | 5 | 65101371 | 65101450 | + | 80 | dre-mir-219-2 | 0 | 0 | 0 | 0 | 0 | 0 |
| ENSDARG000 | 5 | 50780985 | 50781623 | - | 639 | zgc:194908 | 2 | 0 | 4 | 0 | 0 | 6 |
| ENSDARG000 | 5 | 2473511 | 2473627 | + | 117 | 5S_rRNA | 0 | 0 | 0 | 0 | 0 | 0 |
| ENSDARG000 | 5 | 3170736 | 3170826 | + | 91 | dre-mir-142a | 0 | 0 | 0 | 0 | 0 | 0 |
| ENSDARG000 | 5;5;5 | 2772289:2774 | 2772641:2775 | -;-;- | 587 | NA | 133 | 50 | 45 | 35 | 88 | 111 |
| ENSDARG000 | 5;5;5;5;5 | 3258123:3258 | 3258671:3258 | -;-;-;-;- | 686 | NA | 0 | 0 | 0 | 0 | 0 | 0 |
| ENSDARG000 | 5:5:5:5:5:5:5: | 3311008:3311 | 3311274:3311 | -:-:-:-:-:-:-: | 3281 | NA | 0 | 0 | 0 | 0 | 2 | 0 |
| ENSDARG000 | 5;5;5;5 | 3448497:3457 | 3448774:3457 | +;+;+;+ | 502 | NA | 0 | 0 | 0 | 0 | 0 | 0 |
| ENSDARG000 | 5;5;5 | 3660658:3664 | 3662245:3664 | -;-;- | 1778 | NA | 0 | 0 | 0 | 0 | 2 | 0 |
| ENSDARG000 | 5;5;5;5;5 | 4640459:4640 | 4641057:4642 | -;-;-;-;- | 2573 | NA | 0 | 0 | 0 | 0 | 0 | 0 |
| ENSDARG000 | 5;5;5;5;5 | 4685675:4689 | 4686632:4690 | -;-;-;-;- | 3319 | NA | 0 | 0 | 1 | 0 | 0 | 5 |
| ENSDARG000 | 5;5;5 | 4714732:4729 | 4714948:4729 | -;-;- | 618 | NA | 0 | 0 | 4 | 0 | 0 | 0 |
| ENSDARG000 | 5;5;5 | 8637804:8639 | 8637911:8639 | -;-;- | 359 | NA | 0 | 0 | 2 | 0 | 0 | 1 |
| ENSDARG000 | 5;5;5 | 9287035:9287 | 9287196:9287 | +;+;+ | 464 | NA | 0 | 0 | 0 | 0 | 0 | 0 |
| ENSDARG000 | 5;5;5 | 9292071:9294 | 9292330:9294 | +;+;+ | 639 | NA | 0 | 0 | 0 | 0 | 0 | 0 |
| ENSDARG000 | 5;5;5;5 | 10379293:104 | 10379548:104 | +;+;+;+ | 612 | NA | 0 | 0 | 0 | 0 | 0 | 0 |
| ENSDARG000 | 5;5;5;5 | 11278426:112 | 11280031:112 | -;-;-;- | 2059 | NA | 30 | 6 | 8 | 9 | 36 | 26 |
| ENSDARG000 | 5;5;5;5;5;5 | 13603536:136 | 13603593:136 | +;+;+;+;+;+ | 4865 | NA | 165 | 39 | 0 | 110 | 34 | 10 |
| ENSDARG000 | 5;5;5;5;5;5;5 | 29258016:292 | 29258764:292 | -;-;-;-;-;-;- | 1619 | NA | 0 | 0 | 0 | 0 | 0 | 0 |
| ENSDARG000 | 5;5;5 | 32626010:327 | 32627607:327 | -;-;- | 1770 | NA | 0 | 0 | 4 | 0 | 0 | 2 |
| ENSDARG000 | 5;5;5;5;5 | 42357256:423 | 42357474:423 | +;+;+;+;+ | 708 | NA | 1 | 20 | 21 | 0 | 24 | 37 |
| ENSDARG000 | 5:5:5:5:5:5:5: | 43822113:438 | 43824638:438 | -:-:-:-:-:-:-: | 7548 | NA | 39 | 60 | 158 | 6 | 65 | 48 |
| ENSDARG000 | 5;5;5;5;5 | 47066818:471 | 47067352:471 | -;-;-;-;- | 945 | NA | 0 | 0 | 0 | 0 | 0 | 0 |
| ENSDARG000 | 5;5;5 | 52438392:524 | 52438678:524 | +;+;+ | 426 | NA | 0 | 0 | 0 | 0 | 0 | 0 |
| ENSDARG000 | 5;5;5;5 | 52923261:529 | 52925732:529 | -;-;-;- | 2836 | NA | 0 | 2 | 0 | 0 | 0 | 0 |
| ENSDARG000 | 5;5;5;5;5;5;5 | 59605750:596 | 59606351:596 | +;+;+;+;+;+;+ | 3675 | NA | 8 | 0 | 0 | 0 | 0 | 0 |
| ENSDARG000 | 5;5;5 | 64656728:646 | 64658975:646 | -;-;- | 2897 | NA | 0 | 0 | 0 | 0 | 0 | 2 |
| ENSDARG000 | 5;5;5 | 68205952:682 | 68209651:682 | -;-;- | 4249 | NA | 81 | 62 | 71 | 68 | 50 | 88 |
| ENSDARG000 | 5;5;5;5;5 | 68259919:682 | 68260522:682 | +;+;+;+;+ | 1947 | NA | 0 | 0 | 6 | 0 | 0 | 0 |
| ENSDARG000 | 5;5;5 | 69319978:693 | 69320565:693 | -;-;- | 1231 | NA | 5 | 14 | 12 | 13 | 23 | 8 |
| ENSDARG000 | 3;3 | 20913310:209 | 20913467:209 | +;+ | 1704 | NA | 0 | 0 | 0 | 0 | 0 | 0 |
| ENSDARG000 | 3:3:3:3:3:3:3: | 14571813:145 | 14572116:145 | +:+:+:+:+:+:+: | 2267 | ZNF653 | 82 | 38 | 75 | 32 | 60 | 79 |
| ENSDARG000 | 3;3;3 | 29664357:296 | 29664500:296 | +;+;+ | 1673 | NA | 0 | 1 | 0 | 0 | 0 | 0 |
| ENSDARG000 | 3;3;3;3;3;3;3 | 29655677:296 | 29656426:296 | -;-;-;-;-;-;- | 2627 | NA | 13 | 9 | 23 | 8 | 7 | 8 |
| ENSDARG000 | 3:3:3:3:3:3:3: | 18437397:184 | 18437569:184 | +:+:+:+:+:+:+: | 6248 | tbc1d16 | 247 | 226 | 302 | 148 | 440 | 222 |
| ENSDARG000 | 3:3:3:3:3:3:3: | 29654961:296 | 29656426:296 | -:-:-:-:-:-:-: | 2389 | NA | 4 | 23 | 17 | 11 | 22 | 22 |
| ENSDARG000 | 3:3:3:3:3:3:3: | 20947560:209 | 20950585:209 | -;-;-;-;-;-;-;- | 3332 | ndufa4a | 3200 | 2996 | 3467 | 2840 | 3203 | 3012 |
| ENSDARG000 | 3;3;3;3 | 36424055:364 | 36424177:364 | +;+;+;+ | 1063 | SMIM22 | 8 | 3 | 14 | 2 | 18 | 20 |
| ENSDARG000 | 3:3:3:3:3:3:3: | 26734162:267 | 26734455:267 | +:+:+:+:+:+:+: | 3004 | NA | 156 | 126 | 119 | 74 | 113 | 242 |
| ENSDARG000 | 3:3:3:3:3:3:3: | 31118026:311 | 31119276:311 | -:-:-:-:-:-:-: | 12276 | smg1 | 3730 | 3361 | 4822 | 2935 | 4094 | 3288 |
| ENSDARG000 | 3:3:3:3:3:3:3: | 16240984:162 | 16241426:162 | -:-:-:-:-:-:-: | 2042 | NA | 1707 | 2113 | 522 | 2866 | 608 | 1556 |
| ENSDARG000 | 3:3:3:3:3:3:3: | 20963162:209 | 20963953:209 | -:-:-:-:-:-:-: | 2578 | slc35b1 | 570 | 1043 | 1868 | 670 | 1373 | 711 |
| ENSDARG000 | 3;3;3;3;3;3 | 823084:82689 | 823600:82698 | +;+;+;+;+;+ | 1264 | NA | 103 | 29 | 24 | 13 | 68 | 68 |
| ENSDARG000 | 3:3:3:3:3:3:3: | 25059502:250 | 25060088:250 | -:-:-:-:-:-:-: | 2130 | xpnpep3 | 113 | 176 | 160 | 100 | 136 | 144 |
| ENSDARG000 | 3:3:3:3:3:3:3: | 36427988:364 | 36429049:364 | -:-:-:-:-:-:-: | 2542 | rogdi | 414 | 358 | 450 | 336 | 506 | 528 |
| ENSDARG000 | 3;3;3;3;3;3;3 | 16265924:162 | 16266098:162 | +;+;+;+;+;+;+ | 4582 | NA | 23030 | 26488 | 12416 | 26723 | 16844 | 21567 |
| ENSDARG000 | 3;3 | 25055976:250 | 25056154:250 | +;+ | 380 | NA | 0 | 1 | 2 | 0 | 0 | 2 |
| ENSDARG000 | 3:3:3:3:3:3:3: | 7839880:7840 | 7840395:7840 | -:-:-:-:-:-:-: | 4217 | ubn2b | 516 | 680 | 745 | 276 | 668 | 468 |
| ENSDARG000 | 3;3 | 29687567:296 | 29687631:296 | +;+ | 439 | NA | 12 | 0 | 0 | 30 | 5 | 0 |
| ENSDARG000 | 3:3:3:3:3:3:3: | 29674308:296 | 29675570:296 | -:-:-:-:-:-:-: | 3071 | NA | 571 | 811 | 919 | 385 | 48 | 31 |
| ENSDARG000 | 3:3:3:3:3:3:3: | 14611299:146 | 14611477:146 | +:+:+:+:+:+:+: | 1747 | tspan35 | 16 | 6 | 22 | 12 | 19 | 6 |
| ENSDARG000 | 3;3;3 | 842733:84749 | 843266:84808 | +;+;+ | 1427 | NA | 0 | 10 | 0 | 0 | 0 | 4 |
| ENSDARG000 | 3:3:3:3:3:3:3: | 938438:94005 | 938948:94078 | +:+:+:+:+:+:+: | 3105 | NA | 0 | 0 | 0 | 1 | 2 | 0 |
| ENSDARG000 | 3:3:3:3:3:3:3: | 18555899:185 | 18556166:185 | +:+:+:+:+:+:+: | 3191 | cbx4 | 186 | 67 | 119 | 90 | 230 | 82 |
| ENSDARG000 | 3;3;3;3 | 29714775:298 | 29715477:298 | +;+;+;+ | 2994 | cacng2a | 8 | 0 | 0 | 0 | 2 | 2 |
| ENSDARG000 | 3;3 | 29713211:297 | 29714238:297 | -;- | 1236 | NA | 0 | 0 | 0 | 0 | 6 | 0 |
| ENSDARG000 | 3;3;3;3;3;3;3 | 29835271:298 | 29835964:298 | -;-;-;-;-;-;- | 1133 | NA | 0 | 0 | 0 | 0 | 0 | 3 |
| ENSDARG000 | 3;3;3;3;3 | 1005059:1005 | 1005103:1005 | +;+;+;+;+ | 2648 | zgc:172051 | 6 | 0 | 50 | 0 | 106 | 296 |
| ENSDARG000 | 3;3;3;3;3 | 26751861:267 | 26754369:267 | -;-;-;-;- | 3407 | rab40c | 797 | 755 | 1000 | 771 | 1030 | 1074 |
| ENSDARG000 | 3:3:3:3:3:3:3: | 29867213:298 | 29867330:298 | -:-:-:-:-:-:-: | 1698 | rpl3 | 48042 | 48000 | 78517 | 70661 | 53350 | 74464 |
| ENSDARG000 | 3;3;3 | 3971436:3975 | 3976133:3976 | -;-;- | 4895 | NA | 18 | 0 | 2 | 0 | 2 | 4 |
| ENSDARG000 | 3:3:3:3:3:3:3: | 20983395:209 | 20986128:209 | -:-:-:-:-:-:-: | 4329 | FAM117A | 188 | 104 | 195 | 106 | 176 | 166 |
| ENSDARG000 | 3:3:3:3:3:3:3: | 14641962:146 | 14642250:146 | +:+:+:+:+:+:+: | 3009 | zgc:158403 | 11 | 5 | 13 | 15 | 24 | 39 |
| ENSDARG000 | 3;3;3;3;3 | 4047789:4048 | 4047861:4049 | +;+;+;+;+ | 10403 | NA | 27 | 16 | 2 | 0 | 1 | 0 |
| ENSDARG000 | 3;3;3;3;3 | 18567560:185 | 18567679:185 | +;+;+;+;+ | 2763 | cbx8a | 375 | 293 | 471 | 428 | 427 | 527 |
| ENSDARG000 | 3;3;3 | 4225128:4225 | 4225291:4226 | +;+;+ | 365 | NA | 0 | 0 | 0 | 0 | 0 | 0 |
| ENSDARG000 | 3;3;3;3;3 | 31396149:313 | 31396865:313 | +;+;+;+;+ | 897 | NA | 0 | 0 | 0 | 0 | 0 | 0 |
| ENSDARG000 | 3:3:3:3:3:3:3: | 36441306:364 | 36443059:364 | -:-:-:-:-:-:-: | 3433 | glyr1 | 1442 | 1574 | 1833 | 1485 | 1655 | 1750 |
| ENSDARG000 | 3:3:3:3:3:3:3: | 1015867:1016 | 1015956:1016 | +:+:+:+:+:+:+: | 5566 | NA | 0 | 1 | 6 | 0 | 14 | 31 |
| ENSDARG000 | 3:3:3:3:3:3:3: | 31600593:316 | 31600809:316 | +:+:+:+:+:+:+: | 2796 | ccdc43 | 400 | 552 | 592 | 278 | 572 | 468 |
| ENSDARG000 | 3:3:3:3:3:3:3: | 26792546:267 | 26793481:267 | -:-:-:-:-:-:-: | 2845 | pigq | 301 | 229 | 316 | 190 | 271 | 262 |
| ENSDARG000 | 3;3;3;3;3 | 4239106:4250 | 4241111:4250 | -;-;-;-;- | 2831 | NA | 0 | 0 | 0 | 0 | 0 | 0 |
| ENSDARG000 | 3:3:3:3:3:3:3: | 25087605:250 | 25087720:250 | +:+:+:+:+:+:+: | 1984 | st13 | 3278 | 2866 | 3228 | 3652 | 2682 | 3661 |
| ENSDARG000 | 3;3;3;3 | 21025153:210 | 21025789:210 | +;+;+;+ | 4101 | zwi | 8 | 6 | 8 | 20 | 34 | 14 |
| ENSDARG000 | 3:3:3:3:3:3:3: | 21018053:210 | 21018568:210 | -:-:-:-:-:-:-: | 1776 | RUNDC3A | 20 | 78 | 55 | 26 | 95 | 30 |
| ENSDARG000 | 3;3;3 | 26813058:268 | 26813249:268 | +;+;+ | 1154 | socs1a | 557 | 434 | 473 | 265 | 234 | 475 |
| ENSDARG000 | 3:3:3:3:3:3:3: | 29893966:298 | 29894053:298 | -:-:-:-:-:-:-: | 717 | rpl27 | 8551 | 10647 | 14008 | 11209 | 8450 | 11809 |
| ENSDARG000 | 3;3;3;3;3;3 | 7904390:7907 | 7904961:7908 | -;-;-;-;-;- | 1619 | trim35-21 | 0 | 0 | 1 | 7 | 2 | 0 |
| ENSDARG000 | 3;3;3;3 | 12440099:124 | 12440281:124 | +;+;+;+ | 3808 | vasnb | 76 | 40 | 60 | 0 | 120 | 9 |
| ENSDARG000 | 3:3:3:3:3:3:3: | 29872819:298 | 29873769:298 | -:-:-:-:-:-:-: | 2223 | slc25a39 | 278 | 254 | 328 | 152 | 299 | 362 |
| ENSDARG000 | 3;3;3 | 4266289:4267 | 4266330:4267 | +;+;+ | 1003 | NA | 0 | 0 | 0 | 0 | 0 | 0 |
| ENSDARG000 | 3;3 | 4269354:4270 | 4270138:4270 | +;+ | 1053 | NA | 0 | 0 | 0 | 0 | 2 | 0 |
| ENSDARG000 | 3:3:3:3:3:3:3: | 31606781:316 | 31606809:316 | -:-:-:-:-:-:-: | 3899 | NA | 18 | 36 | 6 | 15 | 49 | 2 |
| ENSDARG000 | 3;3;3;3;3 | 18572279:185 | 18572491:185 | -;-;-;-;- | 998 | aqp8b | 0 | 0 | 0 | 2 | 0 | 0 |
| ENSDARG000 | 3:3:3:3:3:3:3: | 26823788:268 | 26825675:268 | -:-:-:-:-:-:-: | 6539 | CLEC16A | 941 | 694 | 1455 | 1448 | 1470 | 1165 |
| ENSDARG000 | 3:3:3:3:3:3:3: | 7920211:7938 | 7920685:7938 | -:-:-:-:-:-:-: | 1464 | NA | 1 | 0 | 0 | 0 | 0 | 0 |
| ENSDARG000 | 3:3:3:3:3:3:3: | 25101960:251 | 25102158:251 | -:-:-:-:-:-:-: | 591 | rbx1 | 966 | 1376 | 1012 | 752 | 941 | 670 |
| ENSDARG000 | 3 | 31621774 | 31624213 | + | 2440 | fzd2 | 1495 | 460 | 881 | 450 | 822 | 1067 |

| | | | | | | | | | | | | |
|---|---|---|---|---|---|---|---|---|---|---|---|---|
| ENSDARG000 | 3;3 | 14691312:146 | 14692333:146 | -;- | 1448 | GADD45GIP1 | 320 | 357 | 346 | 182 | 416 | 186 |
| ENSDARG000 | 3;3 | 26905024:269 | 26905833:269 | +;+ | 1252 | dexi | 0 | 0 | 0 | 0 | 0 | 0 |
| ENSDARG000 | 3;3;3;3 | 4294936:4299 | 4296983:4300 | -;-;-;- | 2901 | NA | 0 | 0 | 0 | 0 | 0 | 0 |
| ENSDARG000 | 3;3;3;3;3;3;3; | 14768364:147 | 14768487:147 | +;+;+;+;+;+;+; | 2883 | nfixb | 26 | 16 | 41 | 12 | 71 | 10 |
| ENSDARG000 | 3;3;3;3;3;3;3; | 18579806:185 | 18580111:185 | +;+;+;+;+;+;+; | 3170 | arhgap17b | 605 | 320 | 452 | 490 | 382 | 531 |
| ENSDARG000 | 3;3;3;3;3;3;3; | 31662126:316 | 31662447:316 | +;+;+;+;+;+;+; | 3210 | NA | 60 | 22 | 22 | 22 | 12 | 46 |
| ENSDARG000 | 3;3 | 4346854:4347 | 4346932:4347 | +;+ | 782 | NA | 0 | 0 | 0 | 0 | 0 | 0 |
| ENSDARG000 | 3;3;3;3;3 | 18634685:186 | 18634846:186 | +;+;+;+;+ | 1370 | NA | 23 | 4 | 17 | 17 | 14 | 23 |
| ENSDARG000 | 3;3;3;3;3;3 | 7943202:7946 | 7943958:7946 | -;-;-;-;-;- | 1828 | trim35-24 | 2 | 0 | 0 | 0 | 0 | 0 |
| ENSDARG000 | 3;3;3;3;3;3;3; | 4403980:4405 | 4404072:4406 | +;+;+;+;+;+;+; | 6363 | NA | 4 | 0 | 4 | 0 | 0 | 0 |
| ENSDARG000 | 3;3 | 4420902:4421 | 4421094:4421 | -;- | 279 | NA | 0 | 0 | 0 | 0 | 0 | 0 |
| ENSDARG000 | 3;3;3;3;3;3;3; | 1030479:1031 | 1030521:1031 | +;+;+;+;+;+;+; | 1947 | zgc:172253 | 0 | 1 | 90 | 0 | 140 | 281 |
| ENSDARG000 | 3;3;3;3;3;3;3; | 1037946:1039 | 1038014:1039 | +;+;+;+;+;+;+; | 2201 | NA | 0 | 0 | 0 | 0 | 0 | 0 |
| ENSDARG000 | 3;3;3;3;3 | 1037959:1039 | 1038014:1039 | +;+;+;+;+ | 1929 | NA | 0 | 0 | 0 | 0 | 4 | 10 |
| ENSDARG000 | 3;3;3;3;3;3;3; | 12593558:125 | 12593611:125 | +;+;+;+;+;+;+; | 6878 | ABCA3 | 2952 | 1217 | 3091 | 3713 | 4861 | 3855 |
| ENSDARG000 | 3;3;3;3;3;3;3; | 31705454:317 | 31707219:317 | -;-;-;-;-;-;-; | 3814 | ccdc47 | 2706 | 3001 | 3556 | 1876 | 4054 | 3327 |
| ENSDARG000 | 3;3;3;3;3;3;3; | 21044154:210 | 21044533:210 | -;-;-;-;-;-;-; | 1762 | tlcd3ba | 73 | 82 | 98 | 89 | 86 | 109 |
| ENSDARG000 | 3;3;3;3;3;3;3; | 36459790:364 | 36462904:364 | -;-;-;-;-;-;-; | 5905 | ppl | 269 | 28 | 12 | 26 | 124 | 36 |
| ENSDARG000 | 3;3;3 | 1096831:1101 | 1096923:1102 | +;+;+ | 2079 | NA | 844 | 577 | 641 | 682 | 564 | 364 |
| ENSDARG000 | 3;3;3;3;3;3 | 7952790:7956 | 7953289:7956 | -;-;-;-;-;- | 1589 | trim35-23 | 3 | 0 | 0 | 4 | 3 | 0 |
| ENSDARG000 | 3;3;3;3;3;3;3; | 25108968:251 | 25110222:251 | -;-;-;-;-;-;-; | 3781 | NA | 4810 | 54 | 67 | 535 | 46 | 254 |
| ENSDARG000 | 3;3;3;3;3;3;3; | 16389020:163 | 16389407:163 | -;-;-;-;-;-;-; | 3366 | eftud2 | 953 | 1185 | 1885 | 467 | 1275 | 862 |
| ENSDARG000 | 3;3;3;3;3;3;3; | 18655673:186 | 18656146:186 | -;-;-;-;-;-;-; | 2405 | slc5a11 | 114 | 194 | 183 | 166 | 182 | 266 |
| ENSDARG000 | 3;3;3;3;3;3;3; | 26908978:269 | 26909355:269 | -;-;-;-;-;-;-; | 3810 | NA | 42 | 38 | 62 | 15 | 62 | 49 |
| ENSDARG000 | 3;3;3;3;3;3 | 7964606:7966 | 7965095:7966 | -;-;-;-;-;- | 1516 | trim35-22 | 12 | 1 | 2 | 2 | 0 | 7 |
| ENSDARG000 | 3;3;3;3;3;3;3; | 15068321:150 | 15068888:150 | -;-;-;-;-;-;-; | 1878 | nme4 | 199 | 339 | 633 | 420 | 272 | 656 |
| ENSDARG000 | 3;3;3;3;3;3;3; | 4444233:4444 | 4444738:4444 | -;-;-;-;-;-;-; | 2521 | NA | 22 | 16 | 17 | 4 | 8 | 20 |
| ENSDARG000 | 3;3;3;3;3;3;3; | 26931710:269 | 26932490:269 | -;-;-;-;-;-;-; | 1708 | nubp1 | 247 | 279 | 320 | 178 | 257 | 265 |
| ENSDARG000 | 3;3;3;3;3;3;3; | 31717261:317 | 31717790:317 | +;+;+;+;+;+;+; | 2877 | dcaf7 | 1663 | 1547 | 1965 | 1480 | 1916 | 1581 |
| ENSDARG000 | 3;3;3;3;3;3;3; | 15085294:150 | 15085936:150 | -;-;-;-;-;-;-; | 4303 | xpo6 | 1779 | 1431 | 2185 | 2266 | 2283 | 1969 |
| ENSDARG000 | 3;3;3;3;3 | 29902221:299 | 29903941:299 | -;-;-;-;- | 2717 | RUNDC1 | 0 | 2 | 6 | 0 | 12 | 6 |
| ENSDARG000 | 3;3;3;3;3;3;3; | 1107102:1113 | 1107331:1113 | +;+;+;+;+;+;+; | 4383 | srebf2 | 3493 | 1649 | 2769 | 1574 | 1822 | 1526 |
| ENSDARG000 | 3;3;3;3;3;3;3; | 23334967:233 | 23335091:233 | -;-;-;-;-;-;-; | 2622 | RAPGEFL1 | 25 | 8 | 25 | 24 | 14 | 4 |
| ENSDARG000 | 3;3;3;3;3;3;3; | 25121156:251 | 25121556:251 | +;+;+;+;+;+;+; | 2601 | napsa | 14682 | 6228 | 5897 | 17762 | 7372 | 13544 |
| ENSDARG000 | 3;3;3;3;3;3;3; | 23247325:232 | 23247540:232 | +;+;+;+;+;+;+; | 9157 | PPP1R9B | 1625 | 1256 | 1451 | 940 | 1695 | 1748 |
| ENSDARG000 | 3;3;3;3;3;3;3; | 36515376:365 | 36515538:365 | +;+;+;+;+;+;+; | 6307 | NA | 2 | 0 | 8 | 0 | 2 | 8 |
| ENSDARG000 | 3;3;3;3;3;3;3; | 7987111:7989 | 7987584:7989 | -;-;-;-;-;-;-; | 1498 | NA | 3 | 6 | 8 | 0 | 0 | 1 |
| ENSDARG000 | 3;3 | 25132020:251 | 25132295:251 | -;- | 319 | NA | 0 | 0 | 0 | 0 | 0 | 0 |
| ENSDARG000 | 3;3;3 | 27045647:270 | 27046010:270 | -;-;- | 669 | NA | 2 | 1 | 2 | 1 | 7 | 0 |
| ENSDARG000 | 3;3;3;3;3;3;3 | 27027781:270 | 27027900:270 | +;+;+;+;+;+;+ | 2817 | emp2 | 816 | 677 | 1230 | 3349 | 1457 | 1509 |
| ENSDARG000 | 3;3;3;3;3;3;3; | 4458390:4460 | 4459571:4460 | -;-;-;-;-;-;-; | 4054 | zc3h7ba | 633 | 742 | 336 | 384 | 343 | 390 |
| ENSDARG000 | 3;3;3;3;3 | 8006777:8012 | 8007322:8012 | -;-;-;-;- | 1015 | NA | 4 | 0 | 0 | 0 | 0 | 0 |
| ENSDARG000 | 3;3;3;3;3;3;3; | 31728871:317 | 31730067:317 | -;-;-;-;-;-;-; | 4933 | kcnh6a | 14 | 0 | 10 | 4 | 8 | 6 |
| ENSDARG000 | 3;3;3;3;3;3 | 8030592:8033 | 8031667:8033 | -;-;-;-;-;- | 2153 | trim35-19 | 4 | 6 | 22 | 0 | 10 | 12 |
| ENSDARG000 | 3;3;3;3;3;3;3; | 29914427:299 | 29915409:299 | -;-;-;-;-;-;-; | 4227 | grna | 7755 | 2169 | 708 | 1794 | 6441 | 1614 |
| ENSDARG000 | 3;3;3;3;3;3;3; | 18675716:186 | 18677411:186 | -;-;-;-;-;-;-; | 7951 | GRID2IP | 16 | 2 | 10 | 6 | 2 | 0 |
| ENSDARG000 | 3;3 | 29929582:299 | 29929645:299 | +;+ | 216 | NA | 0 | 0 | 0 | 0 | 0 | 0 |
| ENSDARG000 | 3;3;3;3;3;3;3; | 15132629:151 | 15134755:151 | -;-;-;-;-;-;-; | 3201 | antkmt | 127 | 128 | 175 | 140 | 244 | 138 |
| ENSDARG000 | 3;3;3;3 | 1132908:1141 | 1135743:1141 | -;-;-;- | 3459 | NA | 166 | 87 | 204 | 87 | 119 | 92 |
| ENSDARG000 | 3;3;3;3;3;3;3; | 36589427:365 | 36590074:365 | -;-;-;-;-;-;-; | 2577 | RRN3 | 505 | 755 | 874 | 421 | 852 | 574 |
| ENSDARG000 | 3;3;3;3;3;3;3; | 12710350:127 | 12710561:127 | +;+;+;+;+;+;+; | 2335 | cyp2k18 | 8 | 0 | 24 | 20 | 4 | 16 |
| ENSDARG000 | 3;3 | 8046984:8048 | 8047064:8048 | -;- | 122 | NA | 0 | 0 | 0 | 0 | 0 | 0 |
| ENSDARG000 | 3;3;3;3 | 8049263:8049 | 8049380:8049 | -;-;-;- | 465 | NA | 0 | 0 | 0 | 0 | 0 | 0 |
| ENSDARG000 | 3;3;3;3;3;3;3; | 27058461:270 | 27059044:270 | -;-;-;-;-;-;-; | 1876 | NA | 291 | 266 | 399 | 228 | 466 | 363 |
| ENSDARG000 | 3;3;3;3;3;3;3; | 1150348:1150 | 1150526:1150 | +;+;+;+;+;+;+; | 1599 | nol12 | 188 | 292 | 304 | 169 | 214 | 136 |
| ENSDARG000 | 3;3;3;3;3;3;3; | 31792441:317 | 31793782:317 | -;-;-;-;-;-;-;- | 2651 | gfap | 14 | 0 | 32 | 14 | 4 | 0 |
| ENSDARG000 | 3;3;3;3;3;3;3; | 18714143:187 | 18714645:187 | -;-;-;-;-;-;-; | 2361 | e4f1 | 165 | 129 | 141 | 96 | 185 | 119 |
| ENSDARG000 | 3;3;3;3;3;3 | 25136153:251 | 25138869:251 | -;-;-;-;-;- | 3708 | mief1 | 724 | 620 | 716 | 980 | 699 | 702 |
| ENSDARG000 | 3;3;3;3;3;3;3; | 31815184:318 | 31817042:318 | -;-;-;-;-;-;-; | 5100 | MAP3K3 | 695 | 254 | 529 | 336 | 583 | 507 |
| ENSDARG000 | 3;3;3;3 | 8139316:8143 | 8139376:8144 | -;-;-;- | 826 | NA | 0 | 0 | 0 | 0 | 0 | 0 |
| ENSDARG000 | 3;3;3;3;3;3;3; | 27071749:270 | 27071831:270 | +;+;+;+;+;+;+; | 9558 | GRIN2A | 0 | 6 | 22 | 8 | 20 | 0 |
| ENSDARG000 | 3;3;3;3;3;3;3; | 8118244:8118 | 8122429:8122 | -;-;-;-;-;-;-; | 5639 | NA | 0 | 0 | 0 | 0 | 0 | 0 |
| ENSDARG000 | 3;3;3;3;3;3;3; | 23428953:234 | 23430383:234 | -;-;-;-;-;-;-; | 3767 | casc3 | 1210 | 1190 | 1456 | 808 | 1097 | 1038 |
| ENSDARG000 | 3;3;3;3 | 25154078:251 | 25154159:251 | +;+;+;+ | 5367 | NA | 3337 | 1528 | 8456 | 494 | 2984 | 5938 |
| ENSDARG000 | 3;3;3;3;3;3;3; | 29941777:299 | 29941836:299 | +;+;+;+;+;+;+; | 2044 | ifi35 | 703 | 426 | 241 | 106 | 398 | 238 |
| ENSDARG000 | 3;3 | 25166805:251 | 25167481:251 | +;+ | 1663 | mpst | 1845 | 673 | 575 | 1015 | 719 | 1117 |
| ENSDARG000 | 3;3;3;3;3;3;3; | 4502066:4504 | 4502172:4504 | +;+;+;+;+;+;+; | 1899 | rangap1a | 805 | 1152 | 1955 | 1324 | 1079 | 1328 |
| ENSDARG000 | 3;3 | 36605149:366 | 36605370:366 | -;- | 832 | si:dkeyp-72e1 | 0 | 0 | 1 | 0 | 0 | 2 |
| ENSDARG000 | 3;3;3;3 | 36610085:366 | 36610854:366 | -;-;-;- | 1666 | NA | 2 | 0 | 4 | 0 | 8 | 18 |
| ENSDARG000 | 3;3;3;3 | 15149540:151 | 15150114:151 | -;-;-;- | 881 | si:dkey-282h2 | 34 | 16 | 34 | 8 | 67 | 37 |
| ENSDARG000 | 3;3;3;3;3;3;3; | 36616713:366 | 36617044:366 | +;+;+;+;+;+;+; | 4258 | pdxdc1 | 714 | 839 | 691 | 513 | 1041 | 590 |
| ENSDARG000 | 3;3;3;3;3;3;3; | 15193521:151 | 15193973:151 | -;-;-;-;-;-;- | 1948 | hirip3 | 170 | 435 | 452 | 285 | 503 | 318 |
| ENSDARG000 | 3;3;3;3;3;3;3; | 21066315:210 | 21068396:210 | -;-;-;-;-;-;-; | 4973 | nlk1 | 1076 | 731 | 844 | 570 | 671 | 688 |
| ENSDARG000 | 3;3;3;3;3;3;3; | 12718100:127 | 12718374:127 | +;+;+;+;+;+;+; | 1731 | NA | 0 | 0 | 0 | 6 | 0 | 0 |
| ENSDARG000 | 3;3;3;3;3;3;3; | 25191366:251 | 25191509:251 | +;+;+;+;+;+;+; | 4385 | il2rb | 20 | 16 | 8 | 6 | 32 | 28 |
| ENSDARG000 | 3;3;3 | 8224622:8224 | 8224681:8224 | +;+;+ | 906 | NA | 0 | 0 | 0 | 0 | 0 | 0 |
| ENSDARG000 | 3;3;3;3;3;3;3 | 15213980:152 | 15215124:152 | -;-;-;-;-;-;- | 2087 | kctd13 | 298 | 274 | 268 | 296 | 256 | 299 |
| ENSDARG000 | 3;3;3;3;3;3;3; | 18740950:187 | 18741630:187 | -;-;-;-;-;-;-; | 2910 | zgc:113333 | 70 | 18 | 54 | 44 | 67 | 34 |
| ENSDARG000 | 3;3;3;3;3;3;3; | 1167026:1167 | 1167219:1167 | +;+;+;+;+;+;+; | 2348 | triobpb | 784 | 630 | 705 | 897 | 585 | 562 |
| ENSDARG000 | 3;3;3;3 | 31850851:318 | 31851317:318 | -;-;-;- | 866 | limd2 | 199 | 210 | 179 | 78 | 173 | 81 |
| ENSDARG000 | 3;3;3;3;3;3;3; | 23462819:234 | 23465313:234 | -;-;-;-;-;-;-; | 4706 | msl1a | 368 | 309 | 481 | 350 | 484 | 624 |
| ENSDARG000 | 3;3;3;3;3;3;3; | 36646054:366 | 36646513:366 | +;+;+;+;+;+;+; | 2601 | gspt1l | 6558 | 7350 | 10295 | 5431 | 6902 | 6919 |
| ENSDARG000 | 3;3;3;3;3 | 25216790:252 | 25216823:252 | +;+;+;+;+ | 1411 | C1QTNF6 | 150 | 2 | 8 | 69 | 24 | 31 |
| ENSDARG000 | 3;3;3;3;3;3;3 | 23471180:234 | 23471662:234 | -;-;-;-;-;-;- | 1663 | NA | 272 | 201 | 206 | 188 | 239 | 208 |
| ENSDARG000 | 3;3;3;3;3;3;3; | 29949984:299 | 29951931:299 | -;-;-;-;-;-;-; | 3487 | bcat2 | 1669 | 1103 | 1108 | 2210 | 3202 | 3477 |
| ENSDARG000 | 3;3;3;3;3;3;3; | 31879442:318 | 31880877:318 | -;-;-;-;-;-;-; | 4032 | ddx42 | 1064 | 957 | 1130 | 653 | 1284 | 1308 |
| ENSDARG000 | 3;3;3;3;3;3;3; | 36667802:366 | 36667863:366 | -;-;-;-;-;-;-; | 6619 | MYH11 | 339 | 330 | 438 | 267 | 242 | 241 |
| ENSDARG000 | 3;3;3;3;3;3;3; | 4534566:4535 | 4534840:4536 | -;-;-;-;-;-;-; | 2037 | ftr43 | 82 | 148 | 26 | 40 | 46 | 28 |
| ENSDARG000 | 3;3;3;3;3;3;3; | 36671585:366 | 36671801:366 | +;+;+;+;+;+;+; | 2811 | nde1 | 1052 | 1267 | 1199 | 1282 | 1170 | 1140 |
| ENSDARG000 | 3;3;3;3;3;3;3; | 8229466:8234 | 8229864:8234 | -;-;-;-;-;-;-; | 2113 | cyp2k22 | 0 | 0 | 6 | 0 | 80 | 0 |
| ENSDARG000 | 3;3;3;3;3;3;3; | 23488652:234 | 23488959:234 | +;+;+;+;+;+;+; | 2926 | nr1d1 | 5190 | 2313 | 5199 | 2348 | 6528 | 3923 |
| ENSDARG000 | 3;3;3;3;3;3;3; | 15232181:152 | 15232668:152 | -;-;-;-;-;-;-; | 3262 | NA | 8 | 0 | 4 | 0 | 6 | 2 |
| ENSDARG000 | 3;3;3;3;3;3;3; | 23498437:235 | 23499709:235 | -;-;-;-;-;-;-; | 2431 | NA | 84 | 208 | 272 | 100 | 128 | 112 |
| ENSDARG000 | 3;3;3;3;3;3;3; | 25269861:252 | 25270810:252 | -;-;-;-;-;-;-; | 3463 | ddx5 | 18986 | 13092 | 19375 | 8187 | 22371 | 9428 |
| ENSDARG000 | 3;3;3;3;3;3;3; | 21096027:211 | 21096902:211 | -;-;-;-;-;-;-; | 2466 | NA | 373 | 330 | 381 | 227 | 395 | 263 |
| ENSDARG000 | 3;3;3;3;3;3;3; | 18756231:187 | 18756322:187 | +;+;+;+;+;+;+; | 2271 | narfl | 220 | 181 | 254 | 142 | 234 | 212 |
| ENSDARG000 | 3;3;3 | 16449490:164 | 16449888:164 | +;+;+ | 2743 | NA | 0 | 0 | 0 | 0 | 0 | 0 |
| ENSDARG000 | 3;3;3;3;3;3;3; | 29969143:299 | 29970504:299 | -;-;-;-;-;-;-; | 2632 | hsd17b14 | 380 | 231 | 182 | 190 | 274 | 276 |
| ENSDARG000 | 3;3;3;3;3;3;3; | 8256231:8280 | 8256790:8281 | -;-;-;-;-;-;-; | 2178 | trim35-9 | 0 | 0 | 0 | 0 | 0 | 0 |
| ENSDARG000 | 3;3 | 18771528:187 | 18771682:187 | +;+ | 815 | NA | 20 | 10 | 9 | 12 | 2 | 5 |
| ENSDARG000 | 3;3;3 | 1182315:1184 | 1182464:1185 | +;+;+ | 827 | ndufa6 | 627 | 619 | 534 | 430 | 482 | 312 |
| ENSDARG000 | 3;3;3;3;3;3;3; | 18774511:187 | 18775418:187 | -;-;-;-;-;-;-; | 1767 | hagh | 3059 | 1645 | 1421 | 3369 | 1491 | 2549 |
| ENSDARG000 | 3;3;3;3;3;3;3; | 8261132:8261 | 8261319:8261 | -;-;-;-;-;-;-; | 1562 | NA | 0 | 0 | 4 | 0 | 0 | 0 |
| ENSDARG000 | 3;3;3;3;3;3;3; | 5019502:5019 | 5019981:5019 | -;-;-;-;-;-;-;- | 1308 | polr3h | 678 | 613 | 423 | 328 | 496 | 333 |
| ENSDARG000 | 3;3 | 27289608:272 | 27290400:272 | -;- | 944 | NA | 0 | 0 | 0 | 0 | 0 | 0 |
| ENSDARG000 | 3;3;3 | 25237399:252 | 25239116:252 | -;-;- | 2063 | mgat3a | 10 | 0 | 0 | 0 | 12 | 20 |
| ENSDARG000 | 3;3;3;3;3;3;3; | 12732382:127 | 12732648:127 | +;+;+;+;+;+;+; | 3437 | cyp2k19 | 0 | 1 | 15 | 2 | 12 | 0 |
| ENSDARG000 | 3;3;3;3;3;3;3; | 36713477:367 | 36713647:367 | +;+;+;+;+;+;+; | 6719 | ABCC1 | 882 | 1155 | 1347 | 530 | 817 | 800 |
| ENSDARG000 | 3;3;3 | 1187244:1189 | 1187285:1190 | +;+;+ | 238 | NA | 0 | 1 | 0 | 0 | 0 | 0 |
| ENSDARG000 | 3;3;3;3;3 | 1186540:1187 | 1186639:1187 | -;-;-;-;- | 534 | NA | 4 | 1 | 8 | 3 | 3 | 7 |
| ENSDARG000 | 3;3;3;3;3;3 | 21125293:211 | 21126218:211 | -;-;-;-;-;- | 1563 | cdipt | 1285 | 1220 | 1395 | 1400 | 1205 | 1497 |
| ENSDARG000 | 3;3;3;3;3;3;3; | 16473006:164 | 16473133:164 | -;-;-;-;-;-;-; | 1866 | NA | 2 | 0 | 2 | 4 | 0 | 14 |
| ENSDARG000 | 3;3;3;3;3;3;3; | 36744518:367 | 36745186:367 | -;-;-;-;-;-;-; | 3758 | NA | 428 | 18 | 107 | 156 | 28 | 255 |
| ENSDARG000 | 3;3;3;3;3;3;3; | 36807642:368 | 36807998:368 | -;-;-;-;-;-;-; | 5175 | NA | 48 | 2 | 10 | 19 | 1 | 35 |
| ENSDARG000 | 3;3 | 29980603:299 | 29980748:299 | +;+ | 871 | NA | 0 | 0 | 0 | 2 | 6 | 0 |
| ENSDARG000 | 3;3 | 5035028:5040 | 5035229:5040 | +;+ | 513 | NA | 82 | 42 | 24 | 106 | 38 | 22 |
| ENSDARG000 | 3;3 | 30022078:300 | 30022184:300 | -;- | 881 | NA | 2 | 0 | 2 | 0 | 0 | 0 |
| ENSDARG000 | 3;3;3;3;3;3;3; | 27595037:275 | 27596617:275 | -;-;-;-;-;-;-;- | 2637 | hapstr1a | 529 | 525 | 441 | 244 | 631 | 382 |
| ENSDARG000 | 3;3;3;3;3;3;3; | 16513219:165 | 16513535:165 | -;-;-;-;-;-;-; | 2061 | eps8l1a | 52 | 0 | 24 | 22 | 46 | 10 |
| ENSDARG000 | 3;3;3;3;3;3;3; | 12744083:127 | 12744295:127 | +;+;+;+;+;+;+; | 1558 | NA | 0 | 1 | 0 | 0 | 4 | 0 |
| ENSDARG000 | 3;3;3;3;3;3;3; | 36836711:368 | 36838183:368 | -;-;-;-;-;-;-; | 2104 | ramp2 | 199 | 131 | 156 | 216 | 74 | 328 |
| ENSDARG000 | 3;3;3;3;3;3;3; | 1191370:1195 | 1193195:1195 | -;-;-;-;-;-;-; | 3525 | fam234b | 418 | 457 | 524 | 543 | 474 | 310 |
| ENSDARG000 | 3;3;3;3;3;3;3; | 21138540:211 | 21143343:211 | -;-;-;-;-;-;-; | 7864 | NA | 2428 | 2152 | 3099 | 2751 | 3138 | 3374 |
| ENSDARG000 | 3;3;3;3;3;3;3; | 8290595:8291 | 8291007:8291 | +;+;+;+;+;+;+; | 1387 | NA | 0 | 10 | 6 | 16 | 62 | 12 |
| ENSDARG000 | 3;3;3;3 | 5052822:5056 | 5054053:5056 | -;-;-;- | 2014 | tefb | 624 | 565 | 516 | 782 | 760 | 446 |
| ENSDARG000 | 3;3 | 8297757:8298 | 8297897:8298 | -;- | 353 | NA | 0 | 0 | 8 | 0 | 31 | 0 |
| ENSDARG000 | 3;3;3;3;3;3;3; | 15271943:152 | 15272125:152 | +;+;+;+;+;+;+; | 1944 | NA | 0 | 0 | 0 | 0 | 0 | 0 |
| ENSDARG000 | 3;3;3;3;3;3;3; | 36859701:368 | 36859905:368 | +;+;+;+;+;+;+; | 4594 | ezh1 | 645 | 526 | 551 | 614 | 503 | 1321 |
| ENSDARG000 | 3;3;3;3;3;3;3; | 31919042:319 | 31919204:319 | -;-;-;-;-;-;-; | 1454 | rnf113a | 162 | 165 | 270 | 78 | 180 | 126 |
| ENSDARG000 | 3;3;3;3;3;3;3; | 30044011:300 | 30044169:300 | -;-;-;-;-;-;-; | 5036 | PPFIA3 | 4 | 4 | 4 | 2 | 36 | 1 |
| ENSDARG000 | 3;3;3;3;3;3;3; | 31897029:318 | 31897154:318 | +;+;+;+;+;+;+; | 2305 | strada | 138 | 136 | 161 | 62 | 169 | 128 |
| ENSDARG000 | 3;3;3;3;3;3;3; | 5102656:5102 | 5106312:5106 | -;-;-;-;-;-;-; | 9635 | MYH9 | 5980 | 5362 | 6435 | 4030 | 6632 | 3770 |
| ENSDARG000 | 3;3;3;3;3;3;3; | 12755535:127 | 12755874:127 | +;+;+;+;+;+;+; | 1806 | cyp2k17 | 0 | 0 | 0 | 0 | 0 | 0 |
| ENSDARG000 | 3;3;3;3;3;3;3; | 5083407:5087 | 5083469:5087 | +;+;+;+;+;+;+; | 8966 | NA | 47 | 2 | 0 | 4 | 2 | 0 |
| ENSDARG000 | 3;3;3;3;3;3;3; | 23518137:235 | 23524605:235 | -;-;-;-;-;-;-; | 9028 | igf2bp1 | 3701 | 2001 | 4243 | 3254 | 3152 | 5169 |
| ENSDARG000 | 3;3;3;3;3;3;3; | 27642078:276 | 27642128:276 | +;+;+;+;+;+;+; | 2604 | NA | 0 | 0 | 2 | 0 | 10 | 0 |
| ENSDARG000 | 3;3;3;3;3;3;3; | 8323586:8323 | 8323972:8323 | -;-;-;-;-;-;-; | 2352 | RBFOX3 | 0 | 2 | 8 | 24 | 10 | 42 |
| ENSDARG000 | 3;3;3;3;3;3;3; | 27606505:276 | 27607612:276 | +;+;+;+;+;+;+; | 7047 | USP7 | 2120 | 2346 | 3037 | 1473 | 2899 | 2538 |
| ENSDARG000 | 3;3;3;3 | 36895465:368 | 36895611:368 | +;+;+;+ | 895 | NA | 0 | 0 | 0 | 0 | 0 | 0 |
| ENSDARG000 | 3;3;3;3;3;3;3; | 36904039:369 | 36904238:369 | -;-;-;-;-;-;-; | 4158 | NA | 4 | 5 | 0 | 14 | 0 | 4 |
| ENSDARG000 | 3;3;3;3;3;3;3; | 27642386:276 | 27643759:276 | -;-;-;-;-;-;-; | 2048 | si:ch211-157c | 45 | 0 | 0 | 14 | 14 | 12 |
| ENSDARG000 | 3;3;3;3;3;3;3; | 12764894:127 | 12765157:127 | +;+;+;+;+;+;+; | 1872 | cyp2k16 | 6 | 0 | 0 | 0 | 0 | 0 |
| ENSDARG000 | 3;3;3;3;3;3;3; | 21189766:211 | 21189892:211 | +;+;+;+;+;+;+; | 1710 | zgc:123295 | 8 | 6 | 2 | 0 | 0 | 0 |
| ENSDARG000 | 3;3;3;3;3;3;3; | 15296824:152 | 15296875:152 | +;+;+;+;+;+;+; | 1163 | cabp5b | 2 | 12 | 4 | 4 | 108 | 4 |
| ENSDARG000 | 3;3;3;3;3;3;3; | 1209006:1211 | 1209093:1211 | +;+;+;+;+;+;+; | 1533 | POLDIP3 | 687 | 590 | 818 | 473 | 602 | 393 |
| ENSDARG000 | 3;3;3;3;3;3;3; | 30140580:301 | 30141412:301 | -;-;-;-;-;-;-; | 2198 | nucb1 | 1351 | 988 | 1144 | 870 | 1217 | 1102 |
| ENSDARG000 | 3;3;3;3;3;3;3; | 16566361:165 | 16566600:165 | +;+;+;+;+;+;+; | 2332 | NA | 53 | 52 | 56 | 3 | 40 | 109 |
| ENSDARG000 | 3;3;3;3;3;3;3; | 12784460:127 | 12784765:127 | +;+;+;+;+;+;+; | 1954 | cyp2k8 | 15 | 2 | 8 | 0 | 44 | 4 |
| ENSDARG000 | 3;3;3;3;3;3;3; | 36972298:369 | 36972720:369 | +;+;+;+;+;+;+; | 6375 | PLEKHH3 | 2578 | 2465 | 3371 | 2046 | 2438 | 1530 |
| ENSDARG000 | 3;3;3;3 | 30153966:301 | 30154509:301 | -;-;-;- | 1373 | ccndx | 0 | 0 | 0 | 0 | 4 | 0 |
| ENSDARG000 | 3;3;3;3;3;3;3 | 1219344:1219 | 1219450:1219 | +;+;+;+;+;+;+ | 1005 | rrp7a | 278 | 238 | 522 | 251 | 385 | 307 |
| ENSDARG000 | 3;3;3;3;3;3;3; | 37059803:370 | 37061008:370 | -;-;-;-;-;-;-; | 3226 | tubg1 | 359 | 611 | 583 | 466 | 735 | 580 |
| ENSDARG000 | 3;3;3;3;3;3 | 21200763:212 | 21200922:212 | +;+;+;+;+;+ | 1148 | zgc:112038 | 0 | 0 | 1 | 0 | 12 | 0 |
| ENSDARG000 | 3;3;3;3;3;3 | 21201428:212 | 21202122:212 | -;-;-;-;-;- | 1199 | NA | 0 | 4 | 2 | 4 | 2 | 0 |
| ENSDARG000 | 3;3;3;3;3;3;3; | 8825382:8826 | 8825406:8827 | +;+;+;+;+;+;+; | 1568 | NA | 7 | 6 | 7 | 0 | 3 | 4 |
| ENSDARG000 | 3;3;3;3;3;3;3; | 15332268:153 | 15333131:153 | -;-;-;-;-;-;-; | 1831 | pitpnbl | 875 | 994 | 966 | 472 | 717 | 799 |
| ENSDARG000 | 3 | 18795570 | 18796559 | + | 990 | fahd1 | 133 | 135 | 120 | 175 | 173 | 116 |
| ENSDARG000 | 3;3;3;3;3;3;3; | 30166946:301 | 30167622:301 | -;-;-;-;-;-;-; | 2438 | tbc1d17 | 636 | 469 | 522 | 619 | 474 | 530 |
| ENSDARG000 | 3;3;3;3;3;3;3; | 23581614:235 | 23584607:235 | -;-;-;-;-;-;-; | 5489 | ube2z | 1141 | 1056 | 994 | 882 | 1030 | 903 |
| ENSDARG000 | 3;3;3;3;3;3;3; | 31925067:319 | 31925252:319 | +;+;+;+;+;+;+; | 2705 | snrnp70 | 2468 | 2204 | 2986 | 1744 | 2868 | 2066 |
| ENSDARG000 | 3;3;3;3;3;3;3; | 18798431:188 | 18801278:188 | -;-;-;-;-;-;-; | 3290 | msrb1a | 245 | 108 | 302 | 296 | 354 | 282 |
| ENSDARG000 | 3;3;3;3;3;3;3; | 37083765:370 | 37084170:370 | +;+;+;+;+;+;+; | 5641 | NA | 594 | 550 | 749 | 576 | 420 | 649 |

| | | | | | | | | | | | | |
|---|---|---|---|---|---|---|---|---|---|---|---|---|
| ENSDARG000 | 3;3;3;3;3;3 | 21225750:212 | 21225817:212 | +;+;+;+;+;+ | 1240 | zgc:153968 | 0 | 0 | 0 | 0 | 0 | 0 |
| ENSDARG000 | 3;3;3;3;3;3;3; | 37112693:371 | 37112824:371 | +;+;+;+;+;+;+; | 961 | psmc3ip | 12 | 17 | 24 | 18 | 26 | 23 |
| ENSDARG000 | 3;3 | 8873204:8873 | 8873347:8873 | -;- | 210 | NA | 0 | 0 | 0 | 0 | 0 | 0 |
| ENSDARG000 | 3;3;3;3;3;3;3; | 25281947:252 | 25283009:252 | -;-;-;-;-;-;-; | 3399 | smurf2 | 748 | 599 | 720 | 532 | 754 | 482 |
| ENSDARG000 | 3;3;3;3;3;3;3; | 37135516:371 | 37136925:371 | -;-;-;-;-;-;-; | 2222 | mlx | 313 | 287 | 320 | 260 | 244 | 309 |
| ENSDARG000 | 3;3;3;3;3 | 23599290:235 | 23599613:235 | -;-;-;-;- | 862 | atp5mc1 | 2822 | 3571 | 3568 | 2372 | 3417 | 2224 |
| ENSDARG000 | 3;3;3;3;3;3;3; | 31933893:319 | 31934129:319 | +;+;+;+;+;+;+; | 1589 | lin7b | 148 | 76 | 95 | 88 | 382 | 66 |
| ENSDARG000 | 3;3 | 8933639:8934 | 8934147:8934 | -;- | 554 | NA | 0 | 0 | 0 | 0 | 0 | 0 |
| ENSDARG000 | 3;3;3;3;3;3;3; | 12816362:128 | 12816630:128 | +;+;+;+;+;+;+; | 2125 | cyp2k6 | 0 | 2 | 8 | 2 | 2 | 0 |
| ENSDARG000 | 3;3;3;3;3;3;3; | 9071806:9072 | 9071840:9072 | +;+;+;+;+;+;+; | 2354 | NA | 1 | 0 | 0 | 0 | 1 | 2 |
| ENSDARG000 | 3;3;3;3;3;3;3; | 1223824:1224 | 1223990:1224 | +;+;+;+;+;+;+; | 1176 | wbp2nl | 3297 | 1935 | 2479 | 2918 | 2528 | 1814 |
| ENSDARG000 | 3;3;3;3;3;3;3; | 12828824:128 | 12828878:128 | +;+;+;+;+;+;+; | 1660 | NA | 6 | 16 | 0 | 0 | 1 | 29 |
| ENSDARG000 | 3;3;3;3;3 | 31945604:319 | 31945856:319 | +;+;+;+;+ | 1118 | zgc:193811 | 2 | 0 | 2 | 0 | 2 | 0 |
| ENSDARG000 | 3;3;3;3;3;3 | 30190419:301 | 30190613:301 | +;+;+;+;+;+ | 2379 | NA | 2073 | 1554 | 3245 | 3318 | 2067 | 2808 |
| ENSDARG000 | 3;3;3;3;3;3;3; | 31953145:319 | 31953185:319 | +;+;+;+;+;+;+; | 2638 | kcnc3a | 0 | 0 | 4 | 0 | 0 | 0 |
| ENSDARG000 | 3;3;3;3;3 | 12835370:128 | 12835423:128 | +;+;+;+;+ | 668 | NA | 2 | 0 | 0 | 0 | 1 | 0 |
| ENSDARG000 | 3;3 | 21240594:212 | 21241530:212 | -;- | 1190 | tcap | 11 | 1 | 2 | 2 | 14 | 1 |
| ENSDARG000 | 3;3 | 1228132:1232 | 1230246:1232 | -;- | 2393 | NA | 160 | 115 | 124 | 192 | 113 | 112 |
| ENSDARG000 | 3;3;3;3;3 | 30202863:302 | 30204284:302 | -;-;-;-;- | 1746 | zgc:195001 | 2 | 2 | 0 | 0 | 2 | 0 |
| ENSDARG000 | 3;3;3;3;3;3;3; | 15358459:153 | 15358816:153 | +;+;+;+;+;+;+; | 4210 | sh2b1 | 1199 | 934 | 1344 | 793 | 978 | 909 |
| ENSDARG000 | 3;3;3;3;3;3;3; | 1260350:1261 | 1261112:1261 | -;-;-;-;-;-;-; | 8828 | NA | 2500 | 1766 | 2711 | 2017 | 2228 | 2254 |
| ENSDARG000 | 3;3;3;3;3;3;3; | 16612574:166 | 16612663:166 | +;+;+;+;+;+;+; | 1820 | slc17a7a | 0 | 0 | 0 | 0 | 3 | 0 |
| ENSDARG000 | 3;3 | 1260360:1268 | 1260407:1268 | +;+ | 616 | NA | 0 | 0 | 0 | 0 | 0 | 0 |
| ENSDARG000 | 3;3 | 23654233:236 | 23654990:236 | +;+ | 1170 | hoxb13a | 0 | 0 | 0 | 0 | 0 | 0 |
| ENSDARG000 | 3;3;3;3;3;3 | 23641935:236 | 23642781:236 | -;-;-;-;-;- | 1320 | eve1 | 0 | 0 | 0 | 0 | 0 | 0 |
| ENSDARG000 | 3;3;3;3;3;3 | 32074806:320 | 32075305:320 | -;-;-;-;-;- | 1271 | baxb | 244 | 360 | 246 | 278 | 157 | 215 |
| ENSDARG000 | 3;3;3;3 | 5265101:5267 | 5265875:5267 | -;-;-;- | 1236 | txn2 | 816 | 928 | 1243 | 691 | 1245 | 789 |
| ENSDARG000 | 3;3;3;3;3 | 5297493:5300 | 5297676:5301 | +;+;+;+;+ | 740 | NA | 0 | 0 | 0 | 0 | 0 | 0 |
| ENSDARG000 | 3;3;3;3;3;3;3; | 18807006:188 | 18807571:188 | +;+;+;+;+;+;+; | 7225 | tnpo2a | 5299 | 6268 | 9047 | 5445 | 7132 | 6204 |
| ENSDARG000 | 3;3;3;3;3;3;3; | 30241237:302 | 30241541:302 | -;-;-;-;-;-;-; | 1286 | emc10 | 1189 | 1223 | 1281 | 779 | 1211 | 1036 |
| ENSDARG000 | 3;3;3;3;3;3 | 12848123:128 | 12848176:128 | +;+;+;+;+;+ | 902 | NA | 0 | 4 | 0 | 0 | 0 | 10 |
| ENSDARG000 | 3;3;3;3;3 | 16643253:166 | 16643375:166 | -;-;-;-;- | 648 | NA | 0 | 0 | 0 | 0 | 0 | 0 |
| ENSDARG000 | 3;3;3;3;3;3;3; | 37196258:371 | 37196532:371 | +;+;+;+;+;+;+; | 6855 | FMNL1 | 144 | 147 | 116 | 108 | 292 | 322 |
| ENSDARG000 | 3;3;3;3;3;3;3; | 25369965:253 | 25370210:253 | -;-;-;-;-;-;-; | 1933 | kpna2 | 1723 | 4010 | 3632 | 1981 | 2790 | 1497 |
| ENSDARG000 | 3;3;3;3;3;3;3; | 37235924:372 | 37239969:372 | -;-;-;-;-;-;-; | 9262 | SRCIN1 | 4 | 8 | 0 | 16 | 24 | 14 |
| ENSDARG000 | 3;3;3;3 | 21251627:212 | 21252259:212 | -;-;-;- | 1366 | syngr1a | 9775 | 8925 | 9991 | 7744 | 11564 | 9470 |
| ENSDARG000 | 3;3;3;3;3;3;3; | 1378098:1378 | 1378328:1378 | -;-;-;-;-;-;-; | 1383 | ddx47 | 288 | 320 | 607 | 190 | 327 | 345 |
| ENSDARG000 | 3;3;3;3;3 | 5331428:5339 | 5331612:5339 | +;+;+;+;+ | 741 | NA | 0 | 0 | 0 | 0 | 0 | 0 |
| ENSDARG000 | 3 | 12861158 | 12862480 | + | 1323 | KCNJ2 | 0 | 0 | 0 | 0 | 0 | 0 |
| ENSDARG000 | 3;3;3;3;3;3;3; | 27664864:276 | 27665357:276 | +;+;+;+;+;+;+; | 6186 | clcn7 | 2758 | 2864 | 3312 | 2604 | 3409 | 3066 |
| ENSDARG000 | 3;3 | 23669267:236 | 23669689:236 | +;+ | 1417 | hoxb10a | 16 | 95 | 2 | 10 | 6 | 0 |
| ENSDARG000 | 3;3;3;3;3;3;3; | 32080403:320 | 32080699:320 | +;+;+;+;+;+;+; | 2850 | aldh16a1 | 2810 | 2533 | 3480 | 1330 | 1308 | 870 |
| ENSDARG000 | 3;3;3;3;3;3;3; | 15394428:153 | 15394711:153 | +;+;+;+;+;+;+; | 3681 | atxn2l | 649 | 889 | 1279 | 376 | 1003 | 500 |
| ENSDARG000 | 3;3;3;3 | 37574885:375 | 37575039:375 | +;+;+;+ | 1953 | wnt9b | 0 | 2 | 0 | 0 | 0 | 14 |
| ENSDARG000 | 3;3;3;3;3;3;3; | 37558491:375 | 37560636:375 | -;-;-;-;-;-;-; | 2847 | arf2a | 9133 | 11032 | 9640 | 7429 | 10552 | 8180 |
| ENSDARG000 | 3;3;3;3;3;3;3; | 21280658:212 | 21283962:212 | -;-;-;-;-;-;-; | 4809 | FAM171A2 | 4279 | 3655 | 2705 | 4211 | 4609 | 3762 |
| ENSDARG000 | 3;3;3;3;3;3;3; | 32099507:320 | 32099949:321 | +;+;+;+;+;+;+; | 1493 | fthl27 | 5976 | 7369 | 8638 | 6030 | 6832 | 5022 |
| ENSDARG000 | 3;3;3;3;3;3;3 | 1395079:1397 | 1396129:1397 | -;-;-;-;-;-;- | 2056 | NA | 531 | 426 | 880 | 557 | 780 | 370 |
| ENSDARG000 | 3;3 | 23677351:236 | 23678014:236 | +;+ | 2577 | hoxb9a | 26 | 927 | 76 | 72 | 36 | 128 |
| ENSDARG000 | 3;3;3;3;3;3 | 12884117:128 | 12885191:128 | -;-;-;-;-;- | 1548 | BTBD17 | 2 | 6 | 4 | 0 | 2 | 4 |
| ENSDARG000 | 3;3;3;3;3;3 | 12893141:128 | 12894307:128 | -;-;-;-;-;- | 1833 | ccdc137 | 268 | 311 | 460 | 137 | 334 | 236 |
| ENSDARG000 | 3 | 23686198 | 23686700 | - | 503 | NA | 0 | 0 | 0 | 0 | 0 | 0 |
| ENSDARG000 | 3;3 | 23687909:236 | 23688711:236 | +;+ | 1817 | hoxb8a | 6 | 8 | 8 | 0 | 18 | 194 |
| ENSDARG000 | 3;3;3;3;3;3;3; | 23691847:236 | 23692270:236 | +;+;+;+;+;+;+; | 1573 | hoxb7a | 150 | 739 | 800 | 786 | 417 | 1924 |
| ENSDARG000 | 3;3;3;3;3;3;3; | 37634574:376 | 37635859:376 | -;-;-;-;-;-;-; | 1509 | NA | 0 | 2 | 0 | 0 | 2 | 0 |
| ENSDARG000 | 3;3;3;3;3;3;3; | 5344999:5349 | 5350035:5350 | -;-;-;-;-;-;-; | 6744 | NA | 80 | 48 | 49 | 10 | 68 | 70 |
| ENSDARG000 | 3;3;3;3;3;3;3; | 12924930:129 | 12925983:129 | -;-;-;-;-;-;-; | 2363 | pdgfab | 93 | 218 | 137 | 60 | 186 | 102 |
| ENSDARG000 | 3;3;3;3;3;3 | 37649436:376 | 37649621:376 | +;+;+;+;+;+ | 1483 | gosr2 | 742 | 1275 | 1446 | 1095 | 858 | 657 |
| ENSDARG000 | 3;3;3;3;3;3;3; | 32118670:321 | 32118911:321 | +;+;+;+;+;+;+; | 999 | fthl31 | 23 | 14 | 2 | 8 | 23 | 46 |
| ENSDARG000 | 3;3;3;3;3;3;3; | 5366371:5377 | 5366642:5378 | -;-;-;-;-;-;-; | 2149 | NA | 59 | 81 | 0 | 16 | 24 | 8 |
| ENSDARG000 | 3;3;3;3;3;3;3; | 25378801:253 | 25380610:253 | -;-;-;-;-;-;-; | 11050 | BPTF | 2421 | 1609 | 3201 | 1448 | 2971 | 2236 |
| ENSDARG000 | 3;3;3;3;3;3;3 | 21379057:213 | 21379105:213 | +;+;+;+;+;+;+ | 778 | NA | 0 | 0 | 0 | 0 | 0 | 0 |
| ENSDARG000 | 3;3;3;3;3;3;3; | 21350509:213 | 21351144:213 | -;-;-;-;-;-;-; | 3755 | itga2b | 0 | 2 | 2 | 0 | 2 | 6 |
| ENSDARG000 | 3;3;3;3;3;3;3; | 16662209:166 | 16662855:166 | +;+;+;+;+;+;+; | 6532 | zgc:55558 | 700 | 740 | 769 | 544 | 706 | 665 |
| ENSDARG000 | 3;3;3;3;3;3;3; | 23697997:236 | 23698465:236 | +;+;+;+;+;+;+; | 6575 | hoxb3a | 224 | 609 | 643 | 498 | 345 | 1006 |
| ENSDARG000 | 3;3;3;3;3;3;3; | 37662444:376 | 37668801:376 | -;-;-;-;-;-;-; | 7886 | gpatch8 | 3522 | 1814 | 3545 | 2591 | 4524 | 3884 |
| ENSDARG000 | 3;3 | 23707691:237 | 23708366:237 | +;+ | 1790 | hoxb5a | 77 | 130 | 120 | 154 | 75 | 257 |
| ENSDARG000 | 3;3 | 23752150:237 | 23752711:237 | +;+ | 1832 | hoxb2a | 32 | 57 | 39 | 60 | 36 | 67 |
| ENSDARG000 | 3;3 | 23703704:237 | 23704133:237 | +;+ | 989 | hoxb6a | 58 | 34 | 82 | 100 | 77 | 144 |
| ENSDARG000 | 3;3;3;3 | 23710839:237 | 23710939:237 | +;+;+;+ | 1336 | hoxb4a | 64 | 61 | 76 | 51 | 44 | 139 |
| ENSDARG000 | 3;3;3;3;3;3;3; | 23701915:237 | 23702231:237 | -;-;-;-;-;-;-; | 1412 | NA | 12 | 20 | 29 | 11 | 7 | 37 |
| ENSDARG000 | 3 | 23740362 | 23740665 | - | 304 | NA | 0 | 0 | 6 | 0 | 0 | 0 |
| ENSDARG000 | 3;3 | 23768898:237 | 23769570:237 | +;+ | 1507 | hoxb1a | 0 | 0 | 2 | 6 | 0 | 13 |
| ENSDARG000 | 3;3;3;3;3;3;3; | 27713610:277 | 27713680:277 | +;+;+;+;+;+;+; | 3355 | arhgdig | 3577 | 2562 | 2815 | 2789 | 3958 | 3093 |
| ENSDARG000 | 3;3;3;3;3 | 5403848:5404 | 5404525:5404 | -;-;-;-;- | 1414 | vmo1a | 378 | 471 | 4 | 88 | 65 | 42 |
| ENSDARG000 | 3;3;3;3;3;3;3; | 15455319:154 | 15456277:154 | -;-;-;-;-;-;-; | 2609 | spns1 | 3178 | 2995 | 4611 | 2922 | 3646 | 5707 |
| ENSDARG000 | 3;3;3;3;3;3;3; | 37707432:377 | 37707500:377 | +;+;+;+;+;+;+; | 2846 | NA | 415 | 261 | 614 | 489 | 458 | 423 |
| ENSDARG000 | 3;3;3;3;3;3;3; | 18840810:188 | 18841149:188 | +;+;+;+;+;+;+; | 5896 | TMEM104 | 276 | 145 | 152 | 122 | 157 | 178 |
| ENSDARG000 | 3;3;3;3;3;3;3; | 25432419:254 | 25437579:254 | -;-;-;-;-;-;-; | 6241 | grb2b | 1338 | 1298 | 1160 | 1082 | 1364 | 1104 |
| ENSDARG000 | 3;3;3;3;3;3;3; | 13004657:130 | 13005603:130 | -;-;-;-;-;-;-; | 1993 | prkar1b | 1779 | 676 | 304 | 1343 | 453 | 643 |
| ENSDARG000 | 3;3;3 | 30317853:303 | 30317976:303 | +;+;+ | 493 | NA | 0 | 0 | 0 | 0 | 0 | 0 |
| ENSDARG000 | 3;3;3;3;3;3;3; | 30257582:302 | 30257775:302 | +;+;+;+;+;+;+; | 4869 | mybpc2a | 32 | 66 | 68 | 74 | 129 | 88 |
| ENSDARG000 | 3;3;3;3;3;3;3; | 32142382:321 | 32142626:321 | +;+;+;+;+;+;+; | 1834 | syt5a | 0 | 2 | 0 | 0 | 20 | 101 |
| ENSDARG000 | 3;3;3;3 | 1405504:1405 | 1405635:1405 | -;-;-;- | 372 | bglap | 0 | 0 | 0 | 0 | 0 | 0 |
| ENSDARG000 | 3;3;3;3;3;3;3 | 27770110:277 | 27770232:277 | +;+;+;+;+;+;+ | 1131 | eci1 | 247 | 328 | 438 | 288 | 426 | 384 |
| ENSDARG000 | 3;3;3;3 | 1405504:1405 | 1405635:1405 | -;-;-;- | 324 | NA | 0 | 0 | 0 | 0 | 0 | 0 |
| ENSDARG000 | 3;3;3;3 | 1414870:1433 | 1415319:1433 | -;-;-;- | 637 | mgp | 0 | 6 | 0 | 0 | 0 | 2 |
| ENSDARG000 | 3;3;3 | 25503442:255 | 25503588:255 | +;+;+ | 4741 | MCHR1 | 19 | 14 | 62 | 28 | 8 | 0 |
| ENSDARG000 | 3;3;3;3;3;3;3; | 32157140:321 | 32157382:321 | -;-;-;-;-;-;-; | 1282 | tnnt1 | 80 | 34 | 79 | 49 | 1 | 5 |
| ENSDARG000 | 3;3;3;3;3;3 | 5420406:5428 | 5420969:5428 | -;-;-;-;-;- | 1584 | trim35-7 | 0 | 0 | 5 | 0 | 4 | 2 |
| ENSDARG000 | 3;3;3;3;3;3;3; | 25546672:255 | 25548265:255 | -;-;-;-;-;-;-; | 4404 | NA | 0 | 0 | 6 | 0 | 0 | 0 |
| ENSDARG000 | 3;3;3;3;3;3;3; | 37757385:377 | 37757690:377 | -;-;-;-;-;-;-; | 1475 | NA | 4 | 0 | 0 | 0 | 0 | 0 |
| ENSDARG000 | 3;3 | 25572778:255 | 25572824:255 | +;+ | 565 | NA | 0 | 0 | 0 | 0 | 0 | 0 |
| ENSDARG000 | 3;3;3;3;3;3 | 5474559:5474 | 5474643:5474 | +;+;+;+;+;+ | 833 | NA | 0 | 0 | 0 | 0 | 0 | 0 |
| ENSDARG000 | 3;3;3;3;3;3;3; | 15476536:154 | 15477006:154 | -;-;-;-;-;-;-; | 1441 | nfatc2ip | 225 | 327 | 448 | 388 | 365 | 420 |
| ENSDARG000 | 3;3;3;3;3;3;3; | 21669545:217 | 21670497:217 | +;+;+;+;+;+;+; | 9998 | CRHR1 | 0 | 64 | 46 | 32 | 36 | 59 |
| ENSDARG000 | 3;3;3;3;3;3;3; | 27778975:277 | 27779327:277 | -;-;-;-;-;-;-; | 1309 | dnase1 | 6 | 0 | 0 | 0 | 0 | 0 |
| ENSDARG000 | 3;3;3;3 | 5504990:5505 | 5505360:5505 | +;+;+;+ | 954 | NA | 0 | 0 | 8 | 0 | 0 | 0 |
| ENSDARG000 | 3;3;3;3;3;3;3; | 32172534:321 | 32172958:321 | -;-;-;-;-;-;-; | 2210 | pih1d1 | 219 | 156 | 271 | 162 | 194 | 222 |
| ENSDARG000 | 3;3;3 | 5514677:5516 | 5516395:5516 | -;-;- | 1909 | NA | 0 | 0 | 0 | 0 | 0 | 0 |
| ENSDARG000 | 3;3;3;3;3;3;3; | 5538485:5538 | 5538614:5539 | -;-;-;-;-;-;-; | 2022 | trim35-31 | 42 | 38 | 2 | 4 | 14 | 3 |
| ENSDARG000 | 3;3;3;3;3;3;3; | 16687830:166 | 16688051:166 | -;-;-;-;-;-;-; | 3561 | pold1 | 268 | 648 | 894 | 203 | 527 | 433 |
| ENSDARG000 | 3;3;3;3;3;3;3; | 13190012:131 | 13190272:131 | +;+;+;+;+;+;+; | 4223 | sun1b | 1254 | 850 | 1243 | 772 | 1234 | 893 |
| ENSDARG000 | 3;3;3;3;3;3;3; | 5575313:5575 | 5575405:5575 | +;+;+;+;+;+;+; | 1182 | NA | 78 | 90 | 87 | 23 | 85 | 88 |
| ENSDARG000 | 3;3;3 | 23935073:239 | 23935236:239 | +;+;+ | 753 | NA | 0 | 0 | 0 | 0 | 0 | 0 |
| ENSDARG000 | 3;3;3;3;3;3 | 24072720:240 | 24074908:240 | -;-;-;-;-;- | 3303 | nfe2l1a | 192 | 173 | 208 | 171 | 241 | 208 |
| ENSDARG000 | 3;3;3 | 23934882:240 | 23935143:240 | -;-;- | 485 | NA | 0 | 0 | 0 | 0 | 0 | 0 |
| ENSDARG000 | 3;3;3;3;3;3;3; | 24050043:240 | 24050103:240 | +;+;+;+;+;+;+; | 2509 | cbx1a | 781 | 1452 | 1646 | 1278 | 1448 | 1553 |
| ENSDARG000 | 3;3;3;3;3;3;3; | 18936006:189 | 18936628:189 | +;+;+;+;+;+;+; | 2104 | si:ch211-198r | 0 | 0 | 0 | 0 | 0 | 0 |
| ENSDARG000 | 3;3;3;3;3;3;3; | 25682884:256 | 25683361:256 | -;-;-;-;-;-;-; | 1212 | NA | 0 | 0 | 0 | 0 | 0 | 0 |
| ENSDARG000 | 3;3;3;3;3;3;3 | 27786601:277 | 27786840:277 | +;+;+;+;+;+;+ | 1652 | nat15 | 337 | 348 | 430 | 253 | 387 | 334 |
| ENSDARG000 | 3;3;3;3;3;3;3; | 5619058:5623 | 5619277:5623 | -;-;-;-;-;-;-; | 1587 | ddx39ab | 1470 | 1809 | 3582 | 1130 | 1931 | 1326 |
| ENSDARG000 | 3;3;3;3;3;3;3; | 25708549:257 | 25711960:257 | -;-;-;-;-;-;-; | 5447 | ntn1b | 8 | 0 | 2 | 0 | 18 | 28 |
| ENSDARG000 | 3;3;3 | 25745636:257 | 25745685:257 | +;+;+ | 138 | NA | 0 | 0 | 0 | 0 | 0 | 0 |
| ENSDARG000 | 3;3 | 25752221:257 | 25752288:257 | +;+ | 588 | NA | 0 | 0 | 0 | 0 | 0 | 0 |
| ENSDARG000 | 3;3;3;3;3;3 | 5647772:5652 | 5648520:5652 | -;-;-;-;-;- | 2574 | NA | 0 | 0 | 0 | 0 | 0 | 2 |
| ENSDARG000 | 3;3;3;3;3;3;3; | 27798094:277 | 27798246:277 | +;+;+;+;+;+;+; | 1556 | carhsp1 | 52 | 18 | 58 | 25 | 27 | 102 |
| ENSDARG000 | 3;3;3;3;3;3;3; | 37766473:377 | 37767604:377 | -;-;-;-;-;-;-; | 2046 | NA | 4 | 0 | 4 | 0 | 49 | 12 |
| ENSDARG000 | 3;3;3;3;3 | 32186134:321 | 32186768:321 | -;-;-;-;- | 834 | NA | 0 | 0 | 19 | 0 | 20 | 6 |
| ENSDARG000 | 3;3;3;3;3;3;3; | 37776489:377 | 37777933:377 | -;-;-;-;-;-;-; | 2061 | baxa | 1180 | 1584 | 1309 | 573 | 993 | 674 |
| ENSDARG000 | 3;3;3 | 21929942:219 | 21929992:219 | +;+;+ | 888 | NA | 0 | 0 | 0 | 0 | 0 | 0 |
| ENSDARG000 | 3;3;3;3;3;3;3; | 19005187:190 | 19006777:190 | -;-;-;-;-;-;-; | 4287 | GRIN2C | 0 | 0 | 0 | 0 | 0 | 0 |
| ENSDARG000 | 3;3;3;3;3;3;3; | 15489725:154 | 15490076:154 | -;-;-;-;-;-;-; | 1637 | sgf29 | 204 | 244 | 354 | 264 | 304 | 351 |
| ENSDARG000 | 3;3;3;3;3;3;3; | 32228901:322 | 32229091:322 | -;-;-;-;-;-;-; | 2934 | NA | 43 | 20 | 52 | 20 | 14 | 28 |
| ENSDARG000 | 3;3;3;3;3;3;3; | 37787229:377 | 37787313:377 | +;+;+;+;+;+;+; | 927 | rps11 | 9429 | 9029 | 13759 | 12014 | 8750 | 14037 |
| ENSDARG000 | 3;3;3;3;3;3;3; | 27832500:278 | 27836720:278 | -;-;-;-;-;-;-; | 6209 | abat | 2796 | 1528 | 3001 | 2232 | 2418 | 2121 |
| ENSDARG000 | 3;3;3;3;3;3;3; | 37790351:377 | 37790610:377 | +;+;+;+;+;+;+; | 2243 | NA | 18 | 66 | 80 | 46 | 50 | 22 |
| ENSDARG000 | 3;3;3;3;3;3;3; | 19152222:191 | 19152454:191 | -;-;-;-;-;-;-; | 3598 | rfx1a | 141 | 76 | 84 | 34 | 112 | 74 |
| ENSDARG000 | 3;3;3;3;3;3 | 38540380:385 | 38540481:385 | +;+;+;+;+;+ | 974 | NA | 0 | 0 | 0 | 0 | 0 | 0 |
| ENSDARG000 | 3;3;3;3 | 38525979:385 | 38526894:385 | -;-;-;- | 1827 | NA | 4 | 0 | 4 | 0 | 0 | 0 |
| ENSDARG000 | 3;3;3;3;3;3;3; | 37824268:378 | 37824689:378 | +;+;+;+;+;+;+; | 3155 | asic2 | 2 | 8 | 4 | 4 | 0 | 0 |
| ENSDARG000 | 3;3;3;3 | 1492174:1494 | 1492305:1495 | +;+;+;+ | 3205 | sox10 | 15379 | 11526 | 17161 | 16625 | 15490 | 12090 |
| ENSDARG000 | 3;3;3;3;3;3;3; | 24094581:240 | 24094645:240 | +;+;+;+;+;+;+; | 2255 | copz2 | 580 | 488 | 768 | 599 | 793 | 640 |
| ENSDARG000 | 3;3;3;3;3;3;3; | 27884685:278 | 27884851:278 | -;-;-;-;-;-;-; | 1315 | mettl22 | 62 | 102 | 102 | 80 | 143 | 84 |
| ENSDARG000 | 3;3;3;3;3;3;3; | 13423935:134 | 13425003:134 | -;-;-;-;-;-;-; | 2267 | dhps | 334 | 300 | 453 | 242 | 391 | 283 |
| ENSDARG000 | 3;3;3;3;3;3;3; | 16747317:167 | 16747444:167 | -;-;-;-;-;-;-; | 992 | aspdh | 0 | 0 | 2 | 0 | 0 | 0 |
| ENSDARG000 | 3;3;3;3;3;3;3; | 16722014:167 | 16722288:167 | +;+;+;+;+;+;+; | 6271 | gys1 | 1398 | 1933 | 4209 | 2695 | 2058 | 1931 |
| ENSDARG000 | 3;3 | 21999006:220 | 22000073:220 | -;- | 1154 | NA | 0 | 0 | 0 | 0 | 0 | 0 |
| ENSDARG000 | 3;3;3;3;3;3;3; | 38647987:386 | 38651082:386 | -;-;-;-;-;-;-; | 4910 | MPP3 | 0 | 0 | 0 | 0 | 6 | 0 |
| ENSDARG000 | 3;3;3;3;3;3;3; | 1511199:1512 | 1511527:1512 | -;-;-;-;-;-;-; | 1328 | polr2f | 442 | 506 | 436 | 312 | 496 | 352 |
| ENSDARG000 | 3;3;3;3;3;3;3; | 38705425:387 | 38707807:387 | -;-;-;-;-;-;-; | 5851 | NA | 10 | 38 | 100 | 8 | 40 | 14 |
| ENSDARG000 | 3;3 | 30354768:303 | 30355287:303 | +;+ | 642 | NA | 0 | 0 | 0 | 0 | 0 | 0 |
| ENSDARG000 | 3;3;3;3;3;3;3; | 30369765:303 | 30369814:303 | -;-;-;-;-;-;-; | 4695 | LRRC4B | 4 | 0 | 10 | 0 | 26 | 0 |
| ENSDARG000 | 3;3;3;3;3;3;3; | 38762891:387 | 38767385:387 | -;-;-;-;-;-;-; | 5617 | ZNF281 | 356 | 312 | 477 | 218 | 395 | 308 |
| ENSDARG000 | 3;3;3;3;3;3;3; | 22035863:220 | 22036260:220 | +;+;+;+;+;+;+; | 6871 | cdc27 | 1655 | 1751 | 3480 | 3257 | 2122 | 2181 |
| ENSDARG000 | 3;3;3;3;3;3 | 1554302:1554 | 1554384:1555 | +;+;+;+;+;+ | 2095 | NA | 4 | 1 | 3 | 4 | 1 | 0 |
| ENSDARG000 | 3;3;3;3;3;3;3; | 30454353:304 | 30454853:304 | -;-;-;-;-;-;-; | 2759 | med25 | 611 | 505 | 620 | 482 | 674 | 562 |
| ENSDARG000 | 3;3 | 1572850:1575 | 1573214:1576 | +;+ | 680 | NA | 0 | 0 | 0 | 0 | 0 | 0 |
| ENSDARG000 | 3;3;3;3 | 1576753:1586 | 1576949:1586 | +;+;+;+ | 2641 | NA | 0 | 0 | 0 | 0 | 0 | 0 |
| ENSDARG000 | 3;3;3;3;3;3;3; | 22057109:220 | 22057255:220 | +;+;+;+;+;+;+; | 7019 | KANSL1 | 1318 | 1181 | 1623 | 1623 | 1467 | 1361 |
| ENSDARG000 | 3;3;3 | 38793499:388 | 38794482:388 | +;+;+ | 2024 | NA | 2 | 2 | 0 | 0 | 1 | 0 |
| ENSDARG000 | 3;3;3;3;3;3;3; | 27982269:279 | 27986247:279 | -;-;-;-;-;-;-; | 6442 | rbfox1 | 132 | 68 | 196 | 66 | 88 | 240 |
| ENSDARG000 | 3;3;3;3;3;3;3; | 32294745:322 | 32296973:322 | -;-;-;-;-;-;-; | 5469 | NA | 66 | 10 | 261 | 308 | 50 | 92 |
| ENSDARG000 | 3;3;3;3;3;3;3; | 38912557:389 | 38913449:389 | -;-;-;-;-;-;-; | 2267 | si:dkev-106c1 | 0 | 1 | 0 | 0 | 0 | 0 |
| ENSDARG000 | 3;3;3;3;3;3;3 | 39099716:391 | 39099820:391 | +;+;+;+;+;+;+ | 1695 | tmem98 | 0 | 0 | 0 | 0 | 0 | 0 |
| ENSDARG000 | 3;3;3 | 9984718:9991 | 9985427:9991 | -;-;- | 1257 | NA | 0 | 0 | 0 | 0 | 0 | 0 |
| ENSDARG000 | 3;3;3;3;3;3;3; | 6178347:6180 | 6181547:6181 | -;-;-;-;-;-;-; | 4260 | NA | 0 | 0 | 0 | 0 | 0 | 0 |
| ENSDARG000 | 3;3;3;3;3 | 15505275:155 | 15505497:155 | +;+;+;+;+ | 794 | nupr1b | 32 | 18 | 58 | 168 | 96 | 46 |
| ENSDARG000 | 3;3;3;3;3;3;3; | 39112898:391 | 39113295:391 | -;-;-;-;-;-;-; | 5167 | NA | 32 | 4 | 84 | 24 | 6 | 0 |
| ENSDARG000 | 3;3;3;3;3 | 30500968:305 | 30501326:305 | +;+;+;+;+ | 2987 | NA | 6 | 18 | 12 | 4 | 56 | 18 |
| ENSDARG000 | 3;3;3;3;3;3;3; | 24134418:241 | 24134628:241 | +;+;+;+;+;+;+; | 3292 | NA | 24 | 42 | 99 | 96 | 49 | 64 |
| ENSDARG000 | 3;3;3;3;3;3;3; | 1782815:1783 | 1782896:1783 | +;+;+;+;+;+;+; | 3757 | NA | 105 | 63 | 4 | 22 | 13 | 0 |

| | | | | | | | | | | | | |
|---|---|---|---|---|---|---|---|---|---|---|---|---|
| ENSDARG000 | 3;3;3 | 13440900:134 | 13440940:134 | +;+;+ | 542 | gng14 | 20 | 9 | 19 | 14 | 29 | 23 |
| ENSDARG000 | 3;3;3 | 16762483:167 | 16762630:167 | +;+;+ | 2782 | tmem86b | 0 | 0 | 12 | 0 | 2 | 0 |
| ENSDARG000 | 3;3;3;3;3;3;3; | 13445772:134 | 13445797:134 | -;-;-;-;-;-;-;-; | 1971 | NA | 162 | 152 | 243 | 122 | 166 | 138 |
| ENSDARG000 | 3;3;3;3;3;3;3 | 39175860:391 | 39175973:391 | -;-;-;-;-;-;- | 939 | NA | 6 | 0 | 0 | 0 | 0 | 0 |
| ENSDARG000 | 3;3;3;3;3;3;3; | 1842616:1842 | 1842783:1842 | -;-;-;-;-;-;-;-; | 2527 | MICALL1 | 87 | 49 | 80 | 25 | 73 | 8 |
| ENSDARG000 | 3;3;3;3;3;3;3; | 19216567:192 | 19216889:192 | +;+;+;+;+;+;+; | 2284 | NA | 229 | 237 | 458 | 353 | 281 | 260 |
| ENSDARG000 | 3;3;3;3;3;3;3; | 15550522:155 | 15550859:155 | +;+;+;+;+;+;+; | 1424 | DOC2A | 0 | 2 | 0 | 0 | 6 | 0 |
| ENSDARG000 | 3;3 | 19207176:192 | 19207408:192 | +;+ | 721 | rln3a | 0 | 0 | 0 | 14 | 2 | 0 |
| ENSDARG000 | 3;3;3 | 24161869:241 | 24161982:241 | +;+;+ | 635 | NA | 6 | 0 | 0 | 0 | 0 | 6 |
| ENSDARG000 | 3;3 | 19207234:192 | 19207367:192 | -;- | 197 | NA | 0 | 0 | 0 | 0 | 0 | 0 |
| ENSDARG000 | 3;3 | 16771797:167 | 16772066:167 | +;+ | 920 | NA | 0 | 0 | 0 | 0 | 0 | 0 |
| ENSDARG000 | 3;3;3;3;3;3;3; | 32325143:323 | 32325418:323 | -;-;-;-;-;-;-;-; | 815 | si:dkev-16p21 | 28 | 22 | 62 | 26 | 82 | 78 |
| ENSDARG000 | 3;3;3;3;3 | 16780455:167 | 16782467:167 | -;-;-;-;- | 2115 | NA | 194 | 80 | 66 | 34 | 260 | 4 |
| ENSDARG000 | 3;3;3;3 | 24180633:241 | 24180697:241 | +;+;+;+ | 555 | NA | 6 | 3 | 4 | 0 | 1 | 4 |
| ENSDARG000 | 3;3 | 15622547:156 | 15623141:156 | -;- | 816 | NA | 77 | 54 | 75 | 63 | 121 | 84 |
| ENSDARG000 | 3;3;3 | 15627268:156 | 15629581:156 | -;-;- | 3319 | NA | 537 | 443 | 403 | 346 | 501 | 661 |
| ENSDARG000 | 3;3;3;3;3;3;3; | 30515365:305 | 30515764:305 | -;-;-;-;-;-;-;-; | 3140 | SYT3 | 8 | 15 | 30 | 10 | 32 | 2 |
| ENSDARG000 | 3;3;3;3;3 | 32341917:323 | 32342000:323 | +;+;+;+;+ | 1399 | prl | 0 | 0 | 0 | 0 | 4 | 0 |
| ENSDARG000 | 3;3;3;3;3;3;3; | 39181387:391 | 39182136:391 | -;-;-;-;-;-;-;-; | 2523 | retsat.2 | 474 | 209 | 320 | 214 | 640 | 861 |
| ENSDARG000 | 3;3;3;3;3 | 24189804:241 | 24189887:241 | +;+;+;+;+ | 1598 | prr15la | 28 | 26 | 12 | 42 | 24 | 16 |
| ENSDARG000 | 3;3;3;3;3;3;3; | 39191815:391 | 39192532:391 | -;-;-;-;-;-;-;-; | 2416 | retsat.1 | 0 | 3 | 8 | 0 | 6 | 0 |
| ENSDARG000 | 3;3;3;3;3;3;3 | 1862871:1862 | 1868525:1868 | -;-;-;-;-;-;- | 6602 | NA | 0 | 0 | 10 | 0 | 2 | 2 |
| ENSDARG000 | 3;3;3;3;3;3;3; | 25825043:258 | 25825126:258 | +;+;+;+;+;+;+; | 2669 | NA | 4 | 10 | 4 | 119 | 14 | 61 |
| ENSDARG000 | 3;3;3 | 24197934:241 | 24198093:241 | +;+;+ | 1981 | atf4b | 7283 | 8640 | 10056 | 13884 | 6417 | 6794 |
| ENSDARG000 | 3;3 | 25844151:258 | 25844358:258 | +;+ | 553 | NA | 0 | 0 | 0 | 0 | 0 | 1 |
| ENSDARG000 | 3;3;3;3;3;3;3; | 16841942:168 | 16842015:168 | +;+;+;+;+;+;+; | 2858 | stk17al | 58 | 69 | 10 | 28 | 84 | 98 |
| ENSDARG000 | 3;3;3;3;3;3;3 | 13559199:135 | 13559266:135 | +;+;+;+;+;+;+ | 507 | NA | 0 | 2 | 0 | 0 | 0 | 0 |
| ENSDARG000 | 3;3;3;3 | 24201956:242 | 24202279:242 | -;-;-;- | 1748 | NA | 176 | 126 | 164 | 270 | 93 | 205 |
| ENSDARG000 | 3;3;3;3 | 13569806:135 | 13569863:135 | +;+;+;+ | 361 | NA | 0 | 2 | 0 | 2 | 20 | 2 |
| ENSDARG000 | 3;3;3;3;3;3;3; | 15635946:156 | 15636168:156 | -;-;-;-;-;-;-;-; | 1765 | NA | 476 | 218 | 160 | 329 | 182 | 163 |
| ENSDARG000 | 3;3;3 | 1880882:1881 | 1881496:1881 | -;-;- | 730 | NA | 0 | 0 | 126 | 0 | 48 | 133 |
| ENSDARG000 | 3;3;3;3;3;3;3; | 1930066:1933 | 1930098:1933 | -;-;-;-;-;-;-;-; | 876 | NA | 33 | 9 | 16 | 0 | 5 | 9 |
| ENSDARG000 | 3;3;3;3;3;3;3 | 39200082:392 | 39200242:392 | -;-;-;-;-;-;- | 744 | NA | 0 | 0 | 2 | 0 | 0 | 0 |
| ENSDARG000 | 3;3;3;3;3;3;3 | 1931815:1931 | 1935549:1935 | -;-;-;-;-;-;- | 4528 | zgc:152753 | 0 | 6 | 20 | 0 | 68 | 100 |
| ENSDARG000 | 3;3;3;3;3;3;3; | 30665195:306 | 30666106:306 | -;-;-;-;-;-;-;-; | 2429 | NA | 2096 | 2094 | 1808 | 1164 | 1558 | 2122 |
| ENSDARG000 | 3;3;3;3;3;3;3; | 39238096:392 | 39239401:392 | -;-;-;-;-;-;-;-; | 3012 | nmt1a | 3324 | 3557 | 4495 | 2274 | 4089 | 3090 |
| ENSDARG000 | 3;3 | 6204184:6211 | 6205004:6211 | -;- | 1057 | NA | 0 | 6 | 2 | 12 | 12 | 24 |
| ENSDARG000 | 3;3;3 | 25849560:258 | 25849729:258 | +;+;+ | 1938 | mfsd6l | 9 | 0 | 0 | 0 | 0 | 0 |
| ENSDARG000 | 3;3;3;3 | 16874401:168 | 16874493:168 | +;+;+;+ | 978 | NA | 5 | 12 | 16 | 0 | 10 | 0 |
| ENSDARG000 | 3;3;3;3;3;3;3; | 6221205:6222 | 6221516:6223 | -;-;-;-;-;-;-;- | 1918 | NA | 2 | 1 | 2 | 0 | 0 | 0 |
| ENSDARG000 | 3;3;3;3;3;3;3; | 39260082:392 | 39264179:392 | -;-;-;-;-;-;-;-; | 6706 | plcd3a | 30 | 18 | 101 | 42 | 24 | 12 |
| ENSDARG000 | 3;3;3;3;3;3;3; | 32356833:323 | 32358091:323 | -;-;-;-;-;-;-;-; | 2442 | prmt1 | 3628 | 4732 | 6083 | 2181 | 5163 | 3648 |
| ENSDARG000 | 3;3;3;3;3;3;3; | 6291635:6291 | 6292297:6292 | +;+;+;+;+;+;+; | 3307 | NA | 0 | 0 | 0 | 0 | 0 | 0 |
| ENSDARG000 | 3;3;3;3;3;3;3; | 39259430:392 | 39259491:392 | +;+;+;+;+;+;+; | 690 | NA | 0 | 0 | 0 | 0 | 0 | 0 |
| ENSDARG000 | 3;3;3 | 16900243:169 | 16900874:169 | -;-;- | 840 | NA | 0 | 0 | 0 | 0 | 0 | 0 |
| ENSDARG000 | 3;3;3;3;3;3;3; | 10152092:101 | 10152226:101 | +;+;+;+;+;+;+; | 2141 | cbx2 | 56 | 98 | 218 | 145 | 110 | 112 |
| ENSDARG000 | 3;3;3;3;3;3;3; | 25857008:258 | 25857162:258 | +;+;+;+;+;+;+; | 2115 | zgc:171844 | 0 | 0 | 10 | 0 | 0 | 0 |
| ENSDARG000 | 3;3;3;3;3;3;3; | 6325895:6325 | 6326395:6326 | +;+;+;+;+;+;+; | 1784 | zgc:136767 | 0 | 0 | 0 | 0 | 0 | 0 |
| ENSDARG000 | 3;3;3;3;3;3;3 | 24220354:242 | 24220460:242 | +;+;+;+;+;+;+ | 1210 | tmem150b | 0 | 2 | 2 | 0 | 2 | 0 |
| ENSDARG000 | 3;3;3;3;3;3;3; | 24207243:242 | 24207560:242 | +;+;+;+;+;+;+; | 1832 | adsl | 780 | 1276 | 1336 | 1073 | 1295 | 773 |
| ENSDARG000 | 3;3;3;3;3 | 30694401:306 | 30694426:306 | +;+;+;+;+ | 375 | NA | 0 | 1 | 15 | 14 | 6 | 7 |
| ENSDARG000 | 3;3;3;3;3;3;3; | 16922226:169 | 16922309:169 | +;+;+;+;+;+;+; | 3556 | atp6v0a1a | 1050 | 746 | 1005 | 582 | 787 | 1135 |
| ENSDARG000 | 3;3;3;3 | 2013324:2016 | 2014489:2016 | -;-;-;- | 2420 | NA | 0 | 0 | 0 | 0 | 18 | 22 |
| ENSDARG000 | 3;3;3;3;3;3;3; | 15646192:156 | 15647822:156 | -;-;-;-;-;-;-;-; | 6087 | zgc:66474 | 723 | 604 | 554 | 508 | 654 | 551 |
| ENSDARG000 | 3;3 | 39371873:394 | 39372200:394 | -;- | 596 | NA | 39 | 45 | 59 | 52 | 36 | 36 |
| ENSDARG000 | 3;3;3;3;3;3;3; | 39313094:393 | 39317643:393 | -;-;-;-;-;-;-;-; | 8042 | NA | 2195 | 1541 | 1713 | 1638 | 1707 | 2324 |
| ENSDARG000 | 3;3 | 16976095:170 | 16977088:170 | +;+ | 2156 | CAVIN1 | 6 | 0 | 2 | 0 | 0 | 0 |
| ENSDARG000 | 3;3;3;3;3;3;3; | 13582019:135 | 13582600:135 | -;-;-;-;-;-;-;-; | 1784 | tufm | 1630 | 1879 | 1908 | 1025 | 1749 | 1156 |
| ENSDARG000 | 3;3;3;3;3;3;3; | 39472342:394 | 39475430:394 | -;-;-;-;-;-;-;-; | 4748 | zgc:100868 | 99 | 8 | 0 | 20 | 24 | 6 |
| ENSDARG000 | 3;3;3 | 2083860:2087 | 2085040:2087 | -;-;- | 1841 | NA | 0 | 0 | 0 | 0 | 0 | 0 |
| ENSDARG000 | 3;3;3;3;3;3;3; | 30714111:307 | 30714151:307 | -;-;-;-;-;-;-;-; | 2865 | NA | 0 | 0 | 9 | 0 | 1 | 6 |
| ENSDARG000 | 3;3 | 15669952:156 | 15671725:156 | -;- | 2097 | zgc:66443 | 202 | 165 | 273 | 167 | 266 | 170 |
| ENSDARG000 | 3;3;3;3;3;3;3; | 13600714:136 | 13600916:136 | +;+;+;+;+;+;+; | 1877 | hspbp1 | 170 | 258 | 321 | 335 | 202 | 86 |
| ENSDARG000 | 3;3;3;3;3;3 | 2329708:2329 | 2330261:2330 | -;-;-;-;-;- | 1939 | NA | 0 | 0 | 0 | 0 | 0 | 6 |
| ENSDARG000 | 3;3;3;3;3;3;3; | 39540014:395 | 39540074:395 | +;+;+;+;+;+;+; | 2123 | zgc:165423 | 0 | 0 | 0 | 2 | 0 | 0 |
| ENSDARG000 | 3;3;3;3;3;3;3; | 6360774:6360 | 6361201:6361 | +;+;+;+;+;+;+; | 1288 | NA | 0 | 0 | 0 | 0 | 0 | 0 |
| ENSDARG000 | 3;3;3;3 | 39555762:395 | 39555968:395 | +;+;+;+ | 1935 | NA | 112 | 104 | 118 | 100 | 89 | 50 |
| ENSDARG000 | 3;3;3;3;3;3;3; | 2572267:2572 | 2572463:2572 | -;-;-;-;-;-;-;-; | 1510 | NA | 0 | 4 | 118 | 0 | 0 | 0 |
| ENSDARG000 | 3;3;3;3;3;3;3; | 19245804:192 | 19245910:192 | +;+;+;+;+;+;+; | 6327 | smarca4a | 4432 | 3497 | 5262 | 1938 | 4459 | 2787 |
| ENSDARG000 | 3;3;3;3;3 | 15694215:156 | 15694370:156 | -;-;-;-;- | 1150 | NA | 0 | 0 | 6 | 0 | 0 | 0 |
| ENSDARG000 | 3;3;3;3;3 | 15691303:157 | 15691334:157 | +;+;+;+;+ | 659 | NA | 29 | 4 | 4 | 11 | 0 | 2 |
| ENSDARG000 | 3;3;3;3;3;3;3; | 15708419:157 | 15708860:157 | -;-;-;-;-;-;-;-; | 3150 | mvp | 11199 | 6300 | 8284 | 7096 | 6061 | 5020 |
| ENSDARG000 | 3;3;3;3;3;3;3; | 39566999:395 | 39567076:395 | +;+;+;+;+;+;+; | 1871 | aldoaa | 13948 | 31536 | 51719 | 65558 | 18483 | 21200 |
| ENSDARG000 | 3;3;3;3;3;3;3; | 6380160:6388 | 6380627:6388 | -;-;-;-;-;-;-;- | 1302 | jpt1b | 2500 | 1840 | 3079 | 2884 | 2304 | 2813 |
| ENSDARG000 | 3;3;3;3;3 | 13624815:136 | 13625056:136 | +;+;+;+;+ | 1611 | pglyrp6 | 1 | 6 | 0 | 3 | 2 | 3 |
| ENSDARG000 | 3;3;3;3;3;3;3; | 39579393:395 | 39579686:395 | +;+;+;+;+;+;+; | 1873 | cln3 | 356 | 387 | 417 | 448 | 310 | 451 |
| ENSDARG000 | 3;3;3;3;3;3;3; | 32365811:323 | 32366545:323 | +;+;+;+;+;+;+; | 5797 | AP2A1 | 5360 | 4853 | 6072 | 4071 | 5588 | 4082 |
| ENSDARG000 | 3;3;3 | 10179191:101 | 10179695:101 | -;-;- | 818 | NA | 0 | 4 | 4 | 0 | 12 | 0 |
| ENSDARG000 | 3;3;3;3;3;3;3; | 25907266:259 | 25907419:259 | +;+;+;+;+;+;+; | 2385 | tom1 | 69 | 56 | 162 | 34 | 163 | 94 |
| ENSDARG000 | 3;3;3;3;3;3;3; | 2602341:2602 | 2602743:2602 | -;-;-;-;-;-;-;-; | 2757 | NA | 37 | 14 | 68 | 17 | 21 | 20 |
| ENSDARG000 | 3;3;3;3;3;3;3; | 24275597:242 | 24276638:242 | +;+;+;+;+;+;+; | 7212 | NA | 80 | 18 | 12 | 4 | 54 | 52 |
| ENSDARG000 | 3;3;3;3;3;3;3; | 24437363:244 | 24439004:244 | -;-;-;-;-;-;-;-; | 3101 | NA | 0 | 6 | 7 | 6 | 38 | 12 |
| ENSDARG000 | 3 | 10310090 | 10312232 | - | 2143 | nog1 | 0 | 0 | 0 | 0 | 10 | 0 |
| ENSDARG000 | 3;3;3;3 | 28444706:284 | 28445062:284 | -;-;-;- | 493 | NA | 0 | 6 | 0 | 0 | 0 | 0 |
| ENSDARG000 | 3;3;3;3;3;3;3; | 28502419:285 | 28502867:285 | +;+;+;+;+;+;+; | 4427 | NA | 1786 | 913 | 1403 | 2612 | 1272 | 834 |
| ENSDARG000 | 3;3;3;3;3;3;3 | 3181205:3181 | 3181387:3182 | -;-;-;-;-;-;- | 1120 | NA | 0 | 0 | 0 | 0 | 0 | 0 |
| ENSDARG000 | 3;3;3;3;3;3;3; | 24314697:243 | 24318626:243 | -;-;-;-;-;-;-;-; | 6120 | csf2rb | 92 | 284 | 270 | 140 | 738 | 457 |
| ENSDARG000 | 3;3;3;3;3;3;3; | 13637383:136 | 13637627:136 | +;+;+;+;+;+;+; | 8359 | NA | 596 | 310 | 612 | 156 | 774 | 610 |
| ENSDARG000 | 3;3;3;3;3;3;3; | 39600562:396 | 39600678:396 | +;+;+;+;+;+;+; | 2555 | prpsap2 | 623 | 799 | 1325 | 776 | 764 | 872 |
| ENSDARG000 | 3;3;3 | 24438616:244 | 24438749:244 | +;+;+ | 803 | NA | 0 | 0 | 0 | 0 | 0 | 0 |
| ENSDARG000 | 3;3 | 24297748:243 | 24297841:243 | +;+ | 300 | NA | 0 | 0 | 0 | 0 | 0 | 0 |
| ENSDARG000 | 3;3;3;3;3;3;3; | 24458204:244 | 24458357:244 | +;+;+;+;+;+;+; | 1732 | CBX6 | 102 | 104 | 143 | 37 | 159 | 82 |
| ENSDARG000 | 3;3;3;3;3;3;3; | 24297482:243 | 24297841:243 | +;+;+;+;+;+;+; | 1590 | pvalb6 | 50 | 316 | 98 | 44 | 100 | 72 |
| ENSDARG000 | 3;3;3;3;3;3;3 | 30888126:308 | 30891331:308 | -;-;-;-;-;-;- | 4544 | ppp1caa | 3255 | 3345 | 4546 | 2561 | 3384 | 2636 |
| ENSDARG000 | 3;3;3;3;3;3;3; | 30868326:308 | 30873770:308 | -;-;-;-;-;-;-;-; | 6627 | kmt5c | 957 | 741 | 1103 | 712 | 1036 | 917 |
| ENSDARG000 | 3;3;3;3;3;3;3; | 6423274:6423 | 6423878:6423 | -;-;-;-;-;-;-;- | 869 | sumo2b | 1902 | 2264 | 3450 | 2336 | 2000 | 2098 |
| ENSDARG000 | 3;3 | 13830599:138 | 13831525:138 | +;+ | 1358 | ptger1b | 3 | 0 | 0 | 0 | 4 | 8 |
| ENSDARG000 | 3;3;3;3;3;3;3; | 10535957:105 | 10536699:105 | -;-;-;-;-;-;-;-; | 3209 | elac2 | 531 | 438 | 815 | 313 | 599 | 592 |
| ENSDARG000 | 3;3;3;3;3;3;3; | 22158024:221 | 22162010:221 | -;-;-;-;-;-;-;-; | 8021 | NA | 10 | 16 | 10 | 2 | 52 | 14 |
| ENSDARG000 | 3;3;3;3;3 | 39619571:396 | 39620113:396 | -;-;-;-;- | 5242 | NA | 10 | 2 | 2 | 11 | 10 | 34 |
| ENSDARG000 | 3;3;3;3;3 | 39630930:396 | 39631728:396 | -;-;-;-;- | 1419 | grapa | 31 | 20 | 39 | 18 | 56 | 27 |
| ENSDARG000 | 3;3 | 39627461:396 | 39627581:396 | +;+ | 673 | NA | 4 | 0 | 6 | 2 | 14 | 4 |
| ENSDARG000 | 3;3;3;3;3;3;3; | 17030665:170 | 17030852:170 | +;+;+;+;+;+;+; | 3442 | stat3 | 2639 | 2659 | 3853 | 2584 | 2182 | 1990 |
| ENSDARG000 | 3;3;3;3;3 | 3200912:3205 | 3202056:3206 | -;-;-;-;- | 2796 | NA | 0 | 0 | 0 | 0 | 0 | 0 |
| ENSDARG000 | 3;3;3;3 | 25951407:259 | 25956383:259 | -;-;-;- | 5384 | NA | 18 | 242 | 2 | 52 | 178 | 100 |
| ENSDARG000 | 3;3;3;3 | 25984754:259 | 25986711:259 | -;-;-;- | 2278 | NA | 60 | 205 | 276 | 133 | 361 | 204 |
| ENSDARG000 | 3;3;3;3;3;3;3; | 6443992:6443 | 6444151:6444 | +;+;+;+;+;+;+; | 1363 | NUP85 | 166 | 263 | 428 | 204 | 296 | 224 |
| ENSDARG000 | 3;3;3;3;3;3;3; | 25990052:259 | 25990379:259 | +;+;+;+;+;+;+; | 1722 | gcat | 406 | 526 | 488 | 389 | 425 | 552 |
| ENSDARG000 | 3;3;3;3;3;3;3; | 39656188:396 | 39656705:396 | +;+;+;+;+;+;+; | 8415 | EPN2 | 1390 | 1029 | 1826 | 1554 | 1830 | 1400 |
| ENSDARG000 | 3;3;3;3;3;3;3; | 3313950:3313 | 3314146:3314 | -;-;-;-;-;-;-;-; | 1330 | NA | 2 | 0 | 15 | 0 | 0 | 0 |
| ENSDARG000 | 3;3;3;3;3 | 30921246:309 | 30921507:309 | +;+;+;+;+ | 2474 | cldni | 1083 | 60 | 2 | 50 | 98 | 232 |
| ENSDARG000 | 3;3;3;3;3;3;3; | 30926364:309 | 30926709:309 | -;-;-;-;-;-;-;-; | 1813 | coro1a | 568 | 804 | 814 | 781 | 873 | 1213 |
| ENSDARG000 | 3;3;3;3;3;3 | 28634467:286 | 28635197:286 | -;-;-;-;-;- | 2369 | fbxl16 | 28 | 12 | 13 | 28 | 4 | 32 |
| ENSDARG000 | 3;3;3;3;3 | 28718817:287 | 28720163:287 | -;-;-;-;- | 2170 | gsg1l | 4 | 0 | 4 | 0 | 1 | 2 |
| ENSDARG000 | 3;3;3 | 30947964:309 | 30948459:309 | -;-;- | 1114 | paqr4a | 102 | 78 | 49 | 20 | 142 | 52 |
| ENSDARG000 | 3;3;3;3;3;3;3; | 19299309:192 | 19299366:192 | +;+;+;+;+;+;+; | 5052 | ldlra | 48 | 4 | 4 | 18 | 6 | 16 |
| ENSDARG000 | 3;3;3;3;3;3;3; | 28695762:286 | 28696536:286 | -;-;-;-;-;-;-;-; | 1487 | NA | 8 | 1 | 0 | 0 | 0 | 6 |
| ENSDARG000 | 3;3;3;3;3;3;3; | 22222244:222 | 22223997:222 | -;-;-;-;-;-;-;-; | 3130 | myl4 | 8 | 2 | 0 | 24 | 0 | 6 |
| ENSDARG000 | 3;3;3;3;3 | 32403758:324 | 32404021:324 | +;+;+;+;+ | 2039 | NA | 96 | 102 | 70 | 175 | 88 | 39 |
| ENSDARG000 | 3;3;3;3;3;3;3; | 28649967:286 | 28650020:286 | +;+;+;+;+;+;+; | 1244 | NA | 6 | 8 | 3 | 12 | 13 | 13 |
| ENSDARG000 | 3;3;3;3;3;3;3; | 22242269:222 | 22242630:222 | +;+;+;+;+;+;+; | 8024 | scn4ab | 220 | 327 | 218 | 46 | 196 | 924 |
| ENSDARG000 | 3;3 | 6508565:6510 | 6508778:6510 | +;+ | 282 | NA | 0 | 0 | 0 | 0 | 0 | 0 |
| ENSDARG000 | 3;3;3;3;3;3;3; | 6490520:6490 | 6492502:6492 | -;-;-;-;-;-;-;-; | 4404 | GGA3 | 226 | 183 | 295 | 88 | 240 | 180 |
| ENSDARG000 | 3;3;3;3;3;3;3; | 30968176:309 | 30968610:309 | +;+;+;+;+;+;+; | 3701 | prf1.9 | 26 | 28 | 4 | 6 | 47 | 42 |
| ENSDARG000 | 3;3;3;3;3;3;3; | 3335167:3340 | 3340963:3340 | -;-;-;-;-;-;-;-; | 9994 | NA | 380 | 323 | 340 | 252 | 357 | 338 |
| ENSDARG000 | 3;3;3;3 | 30980852:309 | 30981583:309 | +;+;+;+ | 2175 | prf1.1 | 0 | 4 | 0 | 0 | 12 | 6 |
| ENSDARG000 | 3;3;3 | 6553362:6554 | 6554029:6554 | -;-;- | 777 | NA | 0 | 0 | 0 | 0 | 0 | 0 |
| ENSDARG000 | 3;3;3;3;3;3;3; | 6556960:6556 | 6563356:6563 | -;-;-;-;-;-;-;-; | 6804 | NA | 0 | 0 | 0 | 12 | 10 | 46 |
| ENSDARG000 | 3;3;3;3;3;3;3; | 39691525:396 | 39692397:396 | -;-;-;-;-;-;-;-; | 1581 | b9d1 | 112 | 112 | 133 | 139 | 88 | 95 |
| ENSDARG000 | 3;3;3;3;3;3;3; | 25999477:260 | 25999747:260 | +;+;+;+;+;+;+; | 2617 | mcm5 | 764 | 2879 | 2875 | 1123 | 2552 | 1668 |
| ENSDARG000 | 3;3;3;3;3;3;3; | 13879446:138 | 13879635:138 | +;+;+;+;+;+;+; | 1884 | farsa | 1337 | 1755 | 3076 | 1812 | 1226 | 1182 |
| ENSDARG000 | 3;3;3;3;3;3;3; | 32410746:324 | 32411068:324 | +;+;+;+;+;+;+; | 2802 | rras | 1969 | 2214 | 1778 | 1385 | 2041 | 1818 |
| ENSDARG000 | 3;3;3;3;3;3;3; | 13842554:138 | 13842632:138 | +;+;+;+;+;+;+; | 6000 | ilf3b | 4665 | 4628 | 5926 | 2365 | 4806 | 3723 |
| ENSDARG000 | 3;3;3;3;3;3;3; | 30998362:309 | 30998374:309 | -;-;-;-;-;-;-;-; | 933 | NA | 13 | 3 | 9 | 24 | 4 | 17 |
| ENSDARG000 | 3;3 | 6600294:6607 | 6600777:6609 | -;- | 916 | si:ch73-59f11 | 30 | 14 | 0 | 9 | 6 | 6 |
| ENSDARG000 | 3;3;3;3;3;3;3; | 39759130:397 | 39759140:397 | +;+;+;+;+;+;+; | 8345 | NA | 8 | 2 | 4 | 0 | 4 | 0 |
| ENSDARG000 | 3;3 | 39945865:399 | 39946081:399 | -;- | 606 | NA | 0 | 0 | 0 | 0 | 0 | 0 |
| ENSDARG000 | 3;3;3;3;3 | 22327738:223 | 22327800:223 | +;+;+;+;+ | 1197 | gh1 | 0 | 4 | 0 | 0 | 0 | 0 |
| ENSDARG000 | 3;3;3;3;3;3 | 39958407:399 | 39958838:399 | +;+;+;+;+;+ | 1666 | CCDC97 | 168 | 236 | 252 | 158 | 208 | 152 |
| ENSDARG000 | 3;3;3;3;3;3;3; | 39970355:399 | 39972686:399 | -;-;-;-;-;-;-;-; | 5963 | llgl1 | 72 | 22 | 50 | 34 | 146 | 54 |
| ENSDARG000 | 3;3;3;3;3;3;3 | 15736811:157 | 15736909:157 | +;+;+;+;+;+;+ | 1763 | phb | 2218 | 3032 | 3268 | 1373 | 2190 | 1675 |
| ENSDARG000 | 3;3;3;3;3;3;3; | 22362687:223 | 22362969:223 | -;-;-;-;-;-;-;-; | 823 | ifnphi1 | 23 | 27 | 65 | 84 | 19 | 5 |
| ENSDARG000 | 3;3;3;3;3;3;3; | 26011482:260 | 26011959:260 | -;-;-;-;-;-;-;-; | 1737 | hmox1a | 308 | 470 | 136 | 303 | 515 | 234 |
| ENSDARG000 | 3;3;3;3;3;3;3; | 22375596:223 | 22376480:223 | +;+;+;+;+;+;+; | 6050 | NA | 110 | 32 | 27 | 29 | 65 | 63 |
| ENSDARG000 | 3;3;3;3;3;3;3; | 17251542:172 | 17251608:172 | +;+;+;+;+;+;+; | 2634 | stat5a | 284 | 192 | 393 | 235 | 381 | 414 |
| ENSDARG000 | 3;3;3;3;3;3;3 | 31039923:310 | 31040016:310 | +;+;+;+;+;+;+ | 714 | cox6a2 | 2 | 2 | 22 | 10 | 55 | 18 |
| ENSDARG000 | 3;3;3;3;3;3;3; | 22341043:223 | 22341055:223 | -;-;-;-;-;-;-;-; | 598 | ifnphi2 | 0 | 0 | 0 | 0 | 0 | 0 |
| ENSDARG000 | 3;3;3 | 6639796:6659 | 6640046:6659 | +;+;+ | 6673 | cygb1 | 182 | 100 | 438 | 112 | 94 | 178 |
| ENSDARG000 | 3;3;3;3;3;3;3 | 22335030:223 | 22335282:223 | +;+;+;+;+;+;+ | 1400 | rdm1 | 32 | 96 | 69 | 42 | 28 | 37 |
| ENSDARG000 | 3;3;3;3;3;3;3 | 31042585:310 | 31045511:310 | -;-;-;-;-;-;- | 5335 | NA | 344 | 218 | 327 | 208 | 340 | 354 |
| ENSDARG000 | 3;3;3;3;3;3;3; | 40056485:400 | 40057095:400 | -;-;-;-;-;-;-;-; | 13044 | NA | 0 | 0 | 0 | 0 | 4 | 2 |
| ENSDARG000 | 3;3 | 31058464:310 | 31058698:310 | +;+ | 1041 | NA | 52 | 67 | 63 | 41 | 116 | 42 |
| ENSDARG000 | 3;3;3;3;3;3;3; | 6692369:6692 | 6692586:6693 | -;-;-;-;-;-;-;-; | 1389 | NA | 55 | 56 | 101 | 78 | 72 | 36 |
| ENSDARG000 | 3;3;3;3;3 | 24482999:244 | 24483712:244 | +;+;+;+;+ | 1614 | NA | 0 | 0 | 4 | 0 | 10 | 0 |
| ENSDARG000 | 3;3;3;3;3;3;3; | 32416948:324 | 32417370:324 | +;+;+;+;+;+;+; | 1674 | prrg2 | 20 | 6 | 2 | 0 | 10 | 4 |
| ENSDARG000 | 3;3;3;3;3;3;3; | 40153819:401 | 40154534:401 | -;-;-;-;-;-;-;-; | 2163 | drg2 | 713 | 619 | 971 | 365 | 745 | 538 |
| ENSDARG000 | 3;3;3;3;3;3;3; | 15773991:157 | 15774269:157 | +;+;+;+;+;+;+; | 2408 | ZNF652 | 435 | 308 | 406 | 213 | 381 | 276 |
| ENSDARG000 | 3;3;3;3;3;3;3; | 26019426:260 | 26019537:260 | +;+;+;+;+;+;+; | 2623 | foxred2 | 707 | 454 | 897 | 426 | 801 | 608 |
| ENSDARG000 | 3;3 | 15755723:157 | 15755981:157 | +;+ | 447 | NA | 5 | 4 | 14 | 6 | 7 | 2 |
| ENSDARG000 | 3;3;3;3;3;3;3; | 31063206:310 | 31063995:310 | -;-;-;-;-;-;-;-; | 1488 | NA | 163 | 106 | 170 | 53 | 102 | 126 |
| ENSDARG000 | 3;3;3;3;3;3;3; | 3386221:3386 | 3386309:3386 | -;-;-;-;-;-;-;-; | 4517 | NA | 0 | 0 | 2 | 0 | 0 | 0 |
| ENSDARG000 | 3;3;3;3;3;3;3; | 19336286:193 | 19336674:193 | +;+;+;+;+;+;+; | 2971 | kri1 | 307 | 378 | 601 | 200 | 349 | 370 |

| | | | | | | | | | | | | |
|---|---|---|---|---|---|---|---|---|---|---|---|---|
| ENSDARG000 | 3;3;3;3 | 24511959:245 | 24512026:245 | +;+;+;+ | 2076 | dnal4a | 166 | 79 | 211 | 126 | 154 | 205 |
| ENSDARG000 | 3;3;3;3;3;3;3; | 40162399:401 | 40162625:401 | +;+;+;+;+;+;+; | 1564 | gfer | 240 | 489 | 443 | 870 | 411 | 232 |
| ENSDARG000 | 3;3;3 | 31066787:310 | 31066950:310 | -;-;- | 321 | NA | 0 | 0 | 0 | 0 | 0 | 0 |
| ENSDARG000 | 3;3;3;3;3;3;3; | 31069276:310 | 31070705:310 | -;-;-;-;-;-;-;- | 1852 | elob | 2173 | 2355 | 1925 | 1706 | 2210 | 2090 |
| ENSDARG000 | 3;3;3;3 | 40170216:401 | 40170382:401 | +;+;+;+ | 2817 | syngr3a | 236 | 323 | 394 | 560 | 46 | 442 |
| ENSDARG000 | 3;3;3;3;3;3;3; | 28869610:288 | 28869750:288 | -;-;-;-;-;-;-;-; | 1884 | eef2kmt | 87 | 64 | 169 | 80 | 146 | 167 |
| ENSDARG000 | 3;3;3;3;3;3;3; | 28851175:288 | 28851528:288 | -;-;-;-;-;-;-;- | 1319 | C3H16orf89 | 0 | 0 | 0 | 0 | 0 | 0 |
| ENSDARG000 | 3;3;3;3;3;3;3; | 28801366:289 | 28801625:289 | -;-;-;-;-;-;-;-; | 1453 | NA | 0 | 1 | 0 | 0 | 2 | 7 |
| ENSDARG000 | 3;3;3;3;3;3;3; | 6720858:6722 | 6723338:6722 | -;-;-;-;-;-;-;-; | 7671 | MAST1 | 35 | 41 | 92 | 148 | 38 | 138 |
| ENSDARG000 | 3;3;3;3;3;3;3 | 28822065:288 | 28824608:288 | -;-;-;-;-;-;- | 3664 | NA | 38 | 0 | 2 | 0 | 10 | 0 |
| ENSDARG000 | 3;3 | 17395239:174 | 17395522:174 | +;+ | 387 | NA | 0 | 0 | 0 | 0 | 0 | 0 |
| ENSDARG000 | 3;3;3;3;3;3;3; | 17346502:173 | 17346617:173 | +;+;+;+;+;+;+; | 3518 | NA | 32 | 49 | 165 | 158 | 92 | 216 |
| ENSDARG000 | 3;3;3;3;3;3;3; | 28831450:288 | 28831675:288 | +;+;+;+;+;+;+; | 5993 | flr | 220 | 148 | 184 | 181 | 173 | 220 |
| ENSDARG000 | 3;3;3;3;3;3;3; | 24537023:245 | 24537127:245 | +;+;+;+;+;+;+; | 2879 | NA | 72 | 250 | 224 | 94 | 214 | 162 |
| ENSDARG000 | 3;3;3;3 | 15805917:158 | 15805977:158 | +;+;+;+ | 2098 | phospho1 | 26 | 17 | 14 | 14 | 24 | 4 |
| ENSDARG000 | 3;3;3 | 15817644:158 | 15817838:158 | +;+;+ | 3246 | natd1 | 802 | 583 | 585 | 706 | 416 | 661 |
| ENSDARG000 | 3;3 | 28899595:289 | 28899800:289 | -;- | 1865 | NA | 0 | 0 | 0 | 0 | 0 | 0 |
| ENSDARG000 | 3;3;3;3;3;3;3; | 28860283:288 | 28860582:288 | +;+;+;+;+;+;+; | 1797 | alg1 | 293 | 274 | 387 | 271 | 355 | 388 |
| ENSDARG000 | 3;3;3;3;3;3;3; | 26035550:260 | 26036194:260 | +;+;+;+;+;+;+; | 1792 | ankrd54 | 489 | 612 | 771 | 508 | 598 | 416 |
| ENSDARG000 | 3;3;3;3;3;3;3; | 32425202:324 | 32425430:324 | +;+;+;+;+;+;+; | 9242 | PRR12 | 766 | 495 | 1143 | 727 | 1060 | 573 |
| ENSDARG000 | 3;3;3;3;3;3;3; | 31080680:310 | 31081197:310 | -;-;-;-;-;-;-;-; | 1747 | zgc:153292 | 129 | 103 | 111 | 180 | 169 | 226 |
| ENSDARG000 | 3;3;3;3;3;3;3; | 10592597:105 | 10592824:105 | -;-;-;-;-;-;-;-; | 1615 | map2k4a | 416 | 282 | 504 | 357 | 380 | 448 |
| ENSDARG000 | 3;3;3;3;3;3;3; | 32458720:324 | 32459579:324 | -;-;-;-;-;-;-;-; | 3091 | NA | 0 | 0 | 0 | 0 | 0 | 0 |
| ENSDARG000 | 3;3;3;3;3;3;3; | 26044969:260 | 26045012:260 | +;+;+;+;+;+;+; | 1963 | hmgxb4a | 8 | 0 | 50 | 0 | 6 | 4 |
| ENSDARG000 | 3;3;3;3;3;3;3; | 19355869:193 | 19355947:193 | +;+;+;+;+;+;+; | 1602 | spc24 | 70 | 226 | 366 | 144 | 220 | 148 |
| ENSDARG000 | 3;3;3;3;3;3;3; | 24556640:245 | 24556907:245 | +;+;+;+;+;+;+; | 1928 | NA | 69 | 85 | 118 | 36 | 55 | 82 |
| ENSDARG000 | 3;3;3;3;3;3;3; | 40185253:401 | 40186205:401 | -;-;-;-;-;-;-;-; | 3446 | znf598 | 850 | 873 | 1035 | 341 | 1026 | 612 |
| ENSDARG000 | 3;3;3;3;3;3;3; | 24571163:245 | 24571307:245 | +;+;+;+;+;+;+; | 5053 | NA | 22 | 8 | 0 | 0 | 4 | 15 |
| ENSDARG000 | 3;3 | 17537352:175 | 17537427:175 | +;+ | 483 | hcrt | 0 | 0 | 4 | 14 | 0 | 0 |
| ENSDARG000 | 3;3;3;3;3;3;3; | 17547532:175 | 17547607:175 | +;+;+;+;+;+;+; | 6683 | kcnh4a | 2 | 12 | 31 | 42 | 49 | 49 |
| ENSDARG000 | 3;3;3;3;3;3;3; | 40206031:402 | 40207701:402 | -;-;-;-;-;-;-;-; | 5403 | flii | 1292 | 1380 | 1323 | 962 | 1034 | 948 |
| ENSDARG000 | 3;3 | 31087509:310 | 31087789:310 | +;+ | 1595 | NA | 46 | 28 | 18 | 24 | 65 | 36 |
| ENSDARG000 | 3;3;3;3;3;3;3; | 24595922:245 | 24596504:245 | +;+;+;+;+;+;+; | 2291 | NA | 0 | 0 | 0 | 0 | 0 | 0 |
| ENSDARG000 | 3;3;3;3;3;3;3; | 40235288:402 | 40235539:402 | -;-;-;-;-;-;-;-; | 3526 | TOP3A | 262 | 230 | 254 | 211 | 223 | 327 |
| ENSDARG000 | 3;3;3;3;3;3;3; | 22432069:224 | 22432521:224 | +;+;+;+;+;+;+; | 10388 | NA | 891 | 439 | 503 | 649 | 992 | 783 |
| ENSDARG000 | 3;3 | 7263209:7271 | 7264413:7271 | -;- | 1406 | NA | 0 | 0 | 43 | 66 | 30 | 22 |
| ENSDARG000 | 3;3 | 22426714:224 | 22426986:224 | +;+ | 633 | coa3b | 13 | 0 | 2 | 8 | 2 | 0 |
| ENSDARG000 | 3;3 | 7276432:7283 | 7277422:7283 | -;- | 1178 | zgc:173577 | 2 | 8 | 20 | 20 | 4 | 2 |
| ENSDARG000 | 3;3;3;3 | 15828426:158 | 15828557:158 | +;+;+;+ | 1327 | tmem11 | 643 | 1042 | 2036 | 1160 | 838 | 674 |
| ENSDARG000 | 3 | 7295464 | 7296189 | - | 726 | NA | 18 | 13 | 32 | 10 | 16 | 17 |
| ENSDARG000 | 3;3;3;3 | 31098063:311 | 31098302:311 | +;+;+;+ | 3588 | NA | 66 | 5 | 18 | 73 | 190 | 236 |
| ENSDARG000 | 3;3;3;3;3;3;3; | 31104891:311 | 31105814:311 | -;-;-;-;-;-;-;-; | 1880 | arl6ip1 | 8000 | 7420 | 6092 | 8822 | 10141 | 9221 |
| ENSDARG000 | 3;3;3;3;3;3 | 7299141:7303 | 7305003:7305 | -;-;-;-;-;- | 6448 | zgc:113443 | 22 | 23 | 15 | 33 | 25 | 35 |
| ENSDARG000 | 3;3 | 40255408:402 | 40257521:402 | +;+ | 2605 | smcr8a | 420 | 356 | 384 | 247 | 281 | 178 |
| ENSDARG000 | 3;3 | 31093455:310 | 31093764:310 | +;+ | 2455 | NA | 61 | 105 | 152 | 40 | 118 | 112 |
| ENSDARG000 | 3;3 | 7557916:7570 | 7558566:7570 | -;- | 765 | NA | 66 | 31 | 56 | 16 | 23 | 31 |
| ENSDARG000 | 3;3;3;3;3;3;3; | 40260929:402 | 40261409:402 | -;-;-;-;-;-;-;-; | 1898 | shmt1 | 2115 | 2689 | 4019 | 1975 | 1180 | 1153 |
| ENSDARG000 | 3;3;3 | 19364948:193 | 19367285:193 | -;-;- | 2443 | s1pr5a | 64 | 22 | 35 | 22 | 24 | 24 |
| ENSDARG000 | 3;3;3;3 | 26064091:260 | 26064168:260 | +;+;+;+ | 1506 | si:dkeyp-69e1 | 1780 | 1285 | 673 | 817 | 1024 | 1097 |
| ENSDARG000 | 3;3;3;3;3;3;3; | 19557604:195 | 19558202:195 | -;-;-;-;-;-;-;- | 1733 | zgc:163079 | 4 | 0 | 0 | 0 | 0 | 0 |
| ENSDARG000 | 3;3;3;3;3;3 | 40277704:402 | 40278202:402 | -;-;-;-;-;- | 1421 | pdap1b | 670 | 866 | 1375 | 1076 | 585 | 694 |
| ENSDARG000 | 3;3;3;3 | 40284598:402 | 40285007:402 | +;+;+;+ | 1185 | bud31 | 813 | 850 | 1229 | 820 | 741 | 696 |
| ENSDARG000 | 3;3;3;3;3;3;3; | 3402223:3402 | 3402307:3402 | -;-;-;-;-;-;-;-; | 4530 | zgc:171446 | 0 | 0 | 0 | 0 | 0 | 0 |
| ENSDARG000 | 3;3;3;3;3 | 7617353:7617 | 7617535:7617 | +;+;+;+;+ | 1567 | rtbdn | 0 | 0 | 0 | 0 | 0 | 0 |
| ENSDARG000 | 3 | 19609638 | 19610990 | + | 1353 | rprml | 0 | 0 | 0 | 0 | 0 | 0 |
| ENSDARG000 | 3 | 7654655 | 7656059 | - | 1405 | junbb | 45781 | 18785 | 39231 | 34513 | 13373 | 18180 |
| ENSDARG000 | 3;3;3;3;3;3;3; | 24603923:246 | 24604026:246 | +;+;+;+;+;+;+; | 1192 | NA | 84 | 101 | 12 | 6 | 7 | 10 |
| ENSDARG000 | 3;3;3;3;3;3;3; | 17615599:176 | 17615728:176 | +;+;+;+;+;+;+; | 2593 | rab5c | 6628 | 9039 | 8191 | 7724 | 9414 | 6933 |
| ENSDARG000 | 3;3 | 32474661:324 | 32474713:324 | +;+ | 576 | NA | 0 | 0 | 0 | 0 | 0 | 0 |
| ENSDARG000 | 3;3;3;3;3;3;3; | 19665319:196 | 19665521:196 | +;+;+;+;+;+;+; | 1475 | mettl2a | 182 | 221 | 296 | 74 | 248 | 134 |
| ENSDARG000 | 3;3 | 26123736:261 | 26123843:261 | +;+ | 2358 | hsp70.3 | 1364 | 1113 | 4434 | 13514 | 101 | 314 |
| ENSDARG000 | 3;3;3;3;3;3;3; | 26081343:260 | 26081379:260 | +;+;+;+;+;+;+; | 4331 | atp2a1 | 14 | 7 | 12 | 4 | 26 | 16 |
| ENSDARG000 | 3;3;3;3;3;3;3; | 32477782:324 | 32477969:324 | +;+;+;+;+;+;+; | 5783 | NA | 1941 | 1642 | 2214 | 1270 | 2108 | 1620 |
| ENSDARG000 | 3;3;3;3;3;3;3; | 40289418:402 | 40289817:402 | +;+;+;+;+;+;+; | 1874 | cpsf4 | 337 | 397 | 442 | 286 | 488 | 416 |
| ENSDARG000 | 3;3;3;3 | 40296785:402 | 40297273:402 | -;-;-;- | 997 | atp5mf | 1199 | 1731 | 1497 | 907 | 1185 | 1123 |
| ENSDARG000 | 3;3;3;3;3;3 | 26119603:261 | 26121798:261 | -;-;-;-;-;- | 2302 | hsp70.2 | 815 | 649 | 3174 | 8789 | 103 | 87 |
| ENSDARG000 | 3;3;3;3;3;3 | 26112780:261 | 26112989:261 | -;-;-;-;-;- | 2297 | hsp70.1 | 547 | 496 | 1442 | 7248 | 36 | 39 |
| ENSDARG000 | 3 | 26108033 | 26109322 | - | 1290 | foxd7 | 0 | 6 | 0 | 2 | 0 | 0 |
| ENSDARG000 | 3;3;3;3;3;3;3 | 15849391:158 | 15849852:158 | -;-;-;-;-;-;- | 1232 | dhrs7b | 88 | 8 | 46 | 8 | 17 | 10 |
| ENSDARG000 | 3;3;3;3;3;3;3; | 17653784:176 | 17654064:176 | +;+;+;+;+;+;+; | 2915 | kat2a | 192 | 178 | 225 | 35 | 208 | 183 |
| ENSDARG000 | 3;3;3;3;3;3;3; | 32492467:324 | 32492745:324 | +;+;+;+;+;+;+; | 4659 | trpm4a | 26 | 6 | 16 | 0 | 2 | 2 |
| ENSDARG000 | 3;3;3;3;3;3;3; | 40315410:403 | 40315628:403 | +;+;+;+;+;+;+; | 2554 | slc29a4a | 0 | 0 | 4 | 6 | 0 | 2 |
| ENSDARG000 | 3;3;3;3;3;3;3; | 7771420:7771 | 7771517:7771 | +;+;+;+;+;+;+; | 1164 | NA | 1229 | 467 | 337 | 641 | 655 | 404 |
| ENSDARG000 | 3;3 | 40397862:404 | 40398286:404 | -;- | 1074 | NA | 4 | 0 | 10 | 6 | 0 | 10 |
| ENSDARG000 | 3;3;3;3;3;3;3; | 7763114:7771 | 7763158:7771 | +;+;+;+;+;+;+; | 2453 | hook2 | 4722 | 1798 | 1369 | 2394 | 2045 | 1644 |
| ENSDARG000 | 3;3;3;3;3;3;3; | 40407352:404 | 40407602:404 | +;+;+;+;+;+;+; | 13023 | TNRC18 | 2322 | 883 | 1845 | 810 | 890 | 1434 |
| ENSDARG000 | 3;3;3;3;3;3;3 | 32526263:325 | 32527417:325 | +;+;+;+;+;+;+ | 3453 | NA | 16 | 4 | 0 | 6 | 1 | 0 |
| ENSDARG000 | 3;3;3;3;3;3;3; | 24612191:246 | 24613324:246 | +;+;+;+;+;+;+; | 3078 | zgc:171506 | 40 | 28 | 19 | 38 | 18 | 12 |
| ENSDARG000 | 3;3;3;3;3;3;3; | 15859612:158 | 15863405:158 | -;-;-;-;-;-;-;-; | 7333 | CRAMP1 | 486 | 342 | 754 | 705 | 512 | 534 |
| ENSDARG000 | 3;3;3;3;3;3 | 22817155:228 | 22817465:228 | -;-;-;-;-;- | 1091 | cyb561 | 836 | 317 | 428 | 140 | 322 | 227 |
| ENSDARG000 | 3;3;3;3 | 28939759:289 | 28939813:289 | +;+;+;+ | 868 | lgals1l1 | 22 | 0 | 9 | 0 | 6 | 14 |
| ENSDARG000 | 3;3;3;3;3;3;3; | 22578369:225 | 22578722:225 | +;+;+;+;+;+;+; | 11775 | TANC2 | 1593 | 620 | 1467 | 564 | 919 | 882 |
| ENSDARG000 | 3;3;3;3;3;3;3 | 24634481:246 | 24634662:246 | +;+;+;+;+;+;+ | 869 | NA | 4 | 0 | 0 | 0 | 0 | 8 |
| ENSDARG000 | 3;3;3;3;3;3;3; | 28953274:289 | 28953362:289 | +;+;+;+;+;+;+; | 2149 | lgals2a | 4084 | 8055 | 7421 | 8705 | 9525 | 7112 |
| ENSDARG000 | 3;3;3;3;3;3;3; | 32535986:325 | 32536545:325 | -;-;-;-;-;-;-;-; | 2147 | rcn3 | 94 | 329 | 494 | 154 | 713 | 105 |
| ENSDARG000 | 3;3;3;3;3 | 15893039:158 | 15893218:158 | +;+;+;+;+ | 2146 | jpt2 | 3380 | 3832 | 4871 | 3402 | 4162 | 3888 |
| ENSDARG000 | 3;3;3;3;3;3;3; | 40526107:405 | 40526792:405 | -;-;-;-;-;-;-;-; | 2169 | actb2 | 147013 | 173301 | 246679 | 364811 | 199982 | 270221 |
| ENSDARG000 | 3;3;3;3;3;3;3; | 32532000:325 | 32532632:325 | +;+;+;+;+;+;+; | 2251 | nosip | 140 | 161 | 236 | 195 | 255 | 221 |
| ENSDARG000 | 3;3;3;3;3;3;3; | 24640946:246 | 24641235:246 | +;+;+;+;+;+;+; | 2249 | sp100.4 | 511 | 575 | 551 | 331 | 667 | 541 |
| ENSDARG000 | 3;3;3;3;3;3 | 10637330:106 | 10637520:106 | +;+;+;+;+;+ | 839 | tmem220 | 88 | 114 | 68 | 96 | 132 | 153 |
| ENSDARG000 | 3;3;3;3;3 | 13911950:139 | 13916073:139 | -;-;-;-;- | 4785 | NA | 0 | 0 | 0 | 0 | 2 | 0 |
| ENSDARG000 | 3;3;3;3;3;3;3; | 29082267:291 | 29082496:291 | +;+;+;+;+;+;+; | 8058 | NA | 24 | 27 | 116 | 180 | 130 | 46 |
| ENSDARG000 | 3;3;3;3;3;3;3; | 19685873:196 | 19686456:196 | +;+;+;+;+;+;+; | 4080 | tlk2 | 945 | 783 | 1114 | 550 | 1165 | 988 |
| ENSDARG000 | 3;3;3;3;3;3;3; | 32661765:326 | 32662122:326 | +;+;+;+;+;+;+; | 2277 | NA | 596 | 454 | 465 | 610 | 406 | 542 |
| ENSDARG000 | 3;3;3;3;3 | 40576447:405 | 40577381:405 | +;+;+;+;+ | 3077 | fscn1a | 3053 | 4701 | 3968 | 3186 | 5462 | 3707 |
| ENSDARG000 | 3;3;3;3;3 | 13929860:139 | 13930013:139 | +;+;+;+;+ | 1226 | syce2 | 11 | 5 | 8 | 44 | 14 | 22 |
| ENSDARG000 | 3;3;3;3;3;3;3; | 40600770:406 | 40605677:406 | -;-;-;-;-;-;-;-; | 7531 | RNF216 | 638 | 587 | 625 | 554 | 619 | 564 |
| ENSDARG000 | 3;3 | 22854673:228 | 22854857:228 | +;+ | 862 | NA | 16 | 4 | 9 | 14 | 5 | 4 |
| ENSDARG000 | 3;3;3 | 7822948:7828 | 7823670:7829 | +;+;+ | 1527 | trir | 500 | 660 | 458 | 326 | 613 | 486 |
| ENSDARG000 | 3;3;3;3;3;3;3; | 10652075:106 | 10652173:106 | -;-;-;-;-;-;-;-; | 1862 | NA | 0 | 0 | 0 | 0 | 0 | 0 |
| ENSDARG000 | 3;3;3;3;3;3;3; | 15907297:159 | 15907754:159 | +;+;+;+;+;+;+; | 5496 | mapk8ip3 | 2474 | 4516 | 3447 | 4438 | 7449 | 4418 |
| ENSDARG000 | 3;3;3 | 34140507:341 | 34140639:341 | +;+;+ | 3053 | si:dkey-204f1 | 13223 | 13589 | 15374 | 18716 | 14684 | 18161 |
| ENSDARG000 | 3;3;3;3;3;3;3; | 32671146:326 | 32671204:326 | +;+;+;+;+;+;+; | 1817 | rnf25 | 841 | 518 | 708 | 827 | 530 | 881 |
| ENSDARG000 | 3;3;3;3;3;3;3; | 34149337:341 | 34149482:341 | +;+;+;+;+;+;+; | 3334 | carm1 | 2041 | 2647 | 3762 | 1746 | 2931 | 1972 |
| ENSDARG000 | 3 | 40679931 | 40681001 | - | 1071 | NA | 0 | 0 | 0 | 0 | 0 | 0 |
| ENSDARG000 | 3;3;3;3 | 41726360:417 | 41726428:417 | +;+;+;+ | 1490 | chst12a | 198 | 106 | 177 | 130 | 171 | 169 |
| ENSDARG000 | 3;3;3;3;3;3;3; | 41784542:417 | 41786233:417 | -;-;-;-;-;-;-;- | 2151 | grifin | 0 | 4 | 0 | 16 | 10 | 10 |
| ENSDARG000 | 3;3;3;3;3;3;3; | 13937426:139 | 13938232:139 | -;-;-;-;-;-;-;-; | 2190 | gcdhb | 190 | 111 | 165 | 222 | 191 | 112 |
| ENSDARG000 | 3;3;3;3;3;3;3; | 32546499:325 | 32546602:325 | +;+;+;+;+;+;+; | 1829 | pax10 | 8 | 5 | 19 | 5 | 23 | 4 |
| ENSDARG000 | 3;3;3;3;3;3;3; | 40690212:406 | 40692333:406 | -;-;-;-;-;-;-;-; | 4409 | smurf1 | 600 | 896 | 1282 | 1232 | 935 | 750 |
| ENSDARG000 | 3;3;3;3;3;3;3; | 22905341:229 | 22905491:229 | +;+;+;+;+;+;+; | 3740 | HDAC5 | 306 | 135 | 230 | 174 | 180 | 107 |
| ENSDARG000 | 3;3;3;3;3;3;3 | 17867126:178 | 17869083:178 | -;-;-;-;-;-;- | 2591 | nkiras2 | 462 | 281 | 392 | 343 | 352 | 476 |
| ENSDARG000 | 3;3;3;3;3;3;3; | 17744339:178 | 17744579:178 | +;+;+;+;+;+;+; | 2220 | ZNF385C | 2 | 7 | 14 | 18 | 0 | 6 |
| ENSDARG000 | 3;3;3;3;3;3;3; | 32681274:326 | 32681648:326 | +;+;+;+;+;+;+; | 2952 | kat8 | 399 | 386 | 432 | 258 | 363 | 379 |
| ENSDARG000 | 3;3;3;3;3;3;3; | 17878124:178 | 17878528:178 | +;+;+;+;+;+;+; | 4712 | dnajc7 | 2021 | 2293 | 2488 | 1540 | 2398 | 1953 |
| ENSDARG000 | 3;3;3;3;3;3;3; | 24681855:246 | 24682027:246 | +;+;+;+;+;+;+; | 3459 | slc25a17 | 169 | 218 | 208 | 164 | 306 | 325 |
| ENSDARG000 | 3;3;3;3;3;3;3; | 32689707:326 | 32690167:326 | +;+;+;+;+;+;+; | 2694 | NA | 0 | 0 | 4 | 0 | 6 | 0 |
| ENSDARG000 | 3;3;3 | 10702884:107 | 10702943:107 | -;-;- | 790 | NA | 0 | 0 | 0 | 0 | 6 | 0 |
| ENSDARG000 | 3;3;3;3;3;3;3; | 29445473:294 | 29445595:294 | +;+;+;+;+;+;+; | 1829 | grap2a | 290 | 183 | 972 | 134 | 944 | 1048 |
| ENSDARG000 | 3;3;3;3;3;3;3; | 10728970:107 | 10729219:107 | -;-;-;-;-;-;-;-; | 1436 | zgc:112965 | 0 | 0 | 0 | 0 | 0 | 0 |
| ENSDARG000 | 3;3;3;3;3;3;3; | 40791720:407 | 40791801:407 | +;+;+;+;+;+;+; | 1312 | NA | 2 | 0 | 24 | 2 | 10 | 0 |
| ENSDARG000 | 3;3;3 | 32570759:325 | 32571415:325 | -;-;- | 1009 | NA | 0 | 2 | 4 | 0 | 2 | 8 |
| ENSDARG000 | 3;3;3;3;3;3;3; | 32571929:325 | 32572288:325 | +;+;+;+;+;+;+; | 1753 | NA | 60 | 14 | 31 | 16 | 47 | 29 |
| ENSDARG000 | 3;3;3;3;3;3;3; | 3415921:3415 | 3416005:3416 | -;-;-;-;-;-;-;-; | 4543 | zgc:171445 | 0 | 0 | 0 | 0 | 0 | 0 |
| ENSDARG000 | 3;3;3;3;3;3;3; | 40796110:407 | 40796279:407 | +;+;+;+;+;+;+; | 1687 | arpc1a | 4301 | 4840 | 4533 | 3244 | 5285 | 3316 |
| ENSDARG000 | 3;3;3;3;3;3;3; | 29469283:294 | 29469744:294 | +;+;+;+;+;+;+; | 4656 | NA | 32 | 18 | 128 | 10 | 49 | 36 |
| ENSDARG000 | 3;3;3;3;3;3;3; | 41917499:419 | 41918025:419 | +;+;+;+;+;+;+; | 3099 | lfng | 18665 | 10532 | 3998 | 11081 | 5476 | 5339 |
| ENSDARG000 | 3;3;3;3;3;3;3; | 26157073:261 | 26158594:261 | -;-;-;-;-;-;-;-; | 3064 | rabep2 | 226 | 321 | 250 | 170 | 372 | 315 |
| ENSDARG000 | 3;3;3 | 10762721:107 | 10763022:107 | -;-;- | 560 | NA | 0 | 0 | 0 | 0 | 0 | 0 |
| ENSDARG000 | 3;3;3;3;3;3;3; | 40809011:408 | 40809152:408 | +;+;+;+;+;+;+; | 1708 | arpc1b | 904 | 1459 | 1614 | 1204 | 2377 | 2475 |
| ENSDARG000 | 3;3;3;3;3;3;3; | 24708685:247 | 24708844:247 | +;+;+;+;+;+;+; | 4689 | NA | 89 | 99 | 181 | 78 | 211 | 176 |
| ENSDARG000 | 3;3;3;3;3;3;3; | 41955358:419 | 41962731:419 | -;-;-;-;-;-;-;-; | 9797 | TTYH3 | 1772 | 2699 | 3198 | 2455 | 4111 | 4248 |
| ENSDARG000 | 3;3;3;3;3 | 11072958:110 | 11073147:110 | +;+;+;+;+ | 1030 | mrps7 | 338 | 319 | 440 | 220 | 393 | 232 |
| ENSDARG000 | 3;3;3;3;3;3;3; | 40821657:408 | 40822197:408 | -;-;-;-;-;-;-;-; | 1728 | wipi2 | 779 | 598 | 1141 | 680 | 702 | 788 |
| ENSDARG000 | 3;3;3;3;3;3;3; | 42090418:420 | 42090532:420 | +;+;+;+;+;+;+; | 2753 | snx8a | 1711 | 2423 | 2413 | 2959 | 2528 | 3117 |
| ENSDARG000 | 3;3;3;3 | 42121166:421 | 42121972:421 | -;-;-;- | 1264 | nudt1 | 71 | 142 | 165 | 114 | 136 | 101 |
| ENSDARG000 | 3;3;3 | 240636:24148 | 240791:24163 | +;+;+ | 1684 | zgc:174288 | 243 | 203 | 12 | 248 | 155 | 74 |
| ENSDARG000 | 3;3 | 42126062:421 | 42126834:421 | +;+ | 1255 | MRM2 | 4 | 8 | 14 | 4 | 2 | 14 |
| ENSDARG000 | 3;3;3;3;3;3;3; | 346530:34934 | 346699:34961 | -;-;-;-;-;-;-;-; | 2785 | NA | 1436 | 713 | 480 | 855 | 848 | 329 |
| ENSDARG000 | 3;3;3 | 42184493:421 | 42185273:421 | -;-;- | 906 | zgc:194207 | 0 | 2 | 6 | 12 | 4 | 0 |
| ENSDARG000 | 3;3;3 | 42194210:421 | 42194708:421 | +;+;+ | 769 | NA | 0 | 0 | 0 | 0 | 0 | 0 |
| ENSDARG000 | 3;3;3;3;3;3;3; | 32587010:325 | 32587440:325 | -;-;-;-;-;-;-;-; | 1642 | tspan4b | 109 | 28 | 68 | 24 | 104 | 82 |
| ENSDARG000 | 3;3;3;3;3;3;3 | 42194210:421 | 42194671:421 | -;-;-;-;-;-;- | 1037 | NA | 0 | 0 | 0 | 0 | 0 | 0 |
| ENSDARG000 | 3;3 | 42203476:422 | 42203677:422 | +;+ | 242 | NA | 0 | 0 | 4 | 0 | 6 | 0 |
| ENSDARG000 | 3;3;3;3;3;3 | 24973025:249 | 24974469:249 | -;-;-;-;-;- | 2138 | desi1a | 910 | 821 | 744 | 705 | 954 | 826 |
| ENSDARG000 | 3;3 | 42212710:422 | 42212772:422 | -;- | 749 | NA | 4 | 1 | 6 | 26 | 32 | 25 |
| ENSDARG000 | 3;3;3;3;3;3;3; | 32631372:326 | 32631538:326 | +;+;+;+;+;+;+; | 4519 | srrm2 | 1200 | 1221 | 1647 | 1139 | 1557 | 1025 |
| ENSDARG000 | 3;3;3;3;3;3;3; | 40841897:408 | 40842021:408 | +;+;+;+;+;+;+; | 1664 | mmd2a | 152 | 76 | 140 | 74 | 93 | 115 |
| ENSDARG000 | 3;3;3;3;3;3;3; | 26173069:261 | 26175715:261 | -;-;-;-;-;-;-;-; | 4400 | NA | 430 | 834 | 742 | 201 | 508 | 276 |
| ENSDARG000 | 3;3;3;3;3;3;3; | 42923275:429 | 42923688:429 | +;+;+;+;+;+;+; | 2965 | tmem184a | 42 | 1 | 2 | 0 | 15 | 43 |
| ENSDARG000 | 3;3;3;3;3;3;3; | 40876553:408 | 40876823:408 | -;-;-;-;-;-;-;-; | 2439 | ap5z1 | 338 | 299 | 348 | 339 | 361 | 395 |
| ENSDARG000 | 3;3;3;3;3;3;3; | 34174004:341 | 34174511:341 | -;-;-;-;-;-;-;-; | 1724 | yipf2 | 480 | 319 | 491 | 364 | 380 | 338 |
| ENSDARG000 | 3;3;3;3;3 | 42964045:429 | 42965193:429 | -;-;-;-;- | 1345 | mafk | 424 | 253 | 458 | 463 | 251 | 309 |
| ENSDARG000 | 3;3;3;3;3;3;3; | 15990547:159 | 15991182:159 | -;-;-;-;-;-;-;-; | 1190 | nme3 | 909 | 955 | 1050 | 822 | 827 | 773 |
| ENSDARG000 | 3;3;3;3;3;3;3; | 32698424:327 | 32698566:327 | +;+;+;+;+;+;+; | 2823 | fus | 799 | 874 | 638 | 459 | 710 | 572 |
| ENSDARG000 | 3;3;3;3;3;3 | 24986145:249 | 24986385:249 | +;+;+;+;+;+ | 2064 | cbx7a | 518 | 871 | 1446 | 2390 | 323 | 608 |
| ENSDARG000 | 3;3 | 32599819:326 | 32601842:326 | -;- | 2336 | NA | 102 | 41 | 16 | 23 | 72 | 151 |
| ENSDARG000 | 3;3 | 32607780:326 | 32610468:326 | -;- | 2779 | NA | 80 | 103 | 80 | 84 | 246 | 332 |
| ENSDARG000 | 3;3;3;3;3;3;3; | 40901743:409 | 40902712:409 | -;-;-;-;-;-;-;- | 3200 | foxk1 | 1034 | 845 | 1482 | 1080 | 1246 | 1084 |
| ENSDARG000 | 3;3;3 | 42918846:429 | 42919015:429 | +;+;+ | 815 | psmg3 | 210 | 358 | 420 | 424 | 234 | 372 |
| ENSDARG000 | 3;3;3;3;3;3;3; | 32620575:326 | 32620682:326 | +;+;+;+;+;+;+; | 1737 | NA | 0 | 0 | 0 | 0 | 0 | 0 |
| ENSDARG000 | 3;3;3;3;3;3;3; | 32594357:326 | 32594420:326 | +;+;+;+;+;+;+; | 1161 | NA | 0 | 0 | 0 | 0 | 0 | 0 |

| | | | | | | | | | | | | |
|---|---|---|---|---|---|---|---|---|---|---|---|---|
| ENSDARG000 | 3;3;3;3;3;3 | 11079404:11 | 11080223:11 | -;-;-;-;-;- | 1387 | mif4gdb | 62 | 44 | 96 | 32 | 102 | 66 |
| ENSDARG000 | 3;3;3;3;3 | 22984098:22 | 22984260:22 | +;+;+;+;+ | 1938 | lsm12a | 519 | 721 | 810 | 499 | 694 | 424 |
| ENSDARG000 | 3:3:3:3:3:3:3: | 42999277:42 | 42999356:42 | +:+:+:+:+:+:+: | 6866 | INTS1 | 1353 | 927 | 1996 | 1395 | 1484 | 1341 |
| ENSDARG000 | 3:3:3:3:3:3:3: | 40936094:40 | 40936907:40 | -:-:-:-:-:-:-: | 2316 | cyp3c4 | 0 | 0 | 24 | 0 | 0 | 8 |
| ENSDARG000 | 3:3:3:3:3:3:3: | 14157090:14 | 14157436:14 | +:+:+:+:+:+:+: | 7057 | NA | 26 | 18 | 28 | 14 | 6 | 24 |
| ENSDARG000 | 3:3:3:3:3:3:3: | 29487300:29 | 29490032:29 | -:-:-:-:-:-:-: | 4063 | NA | 12 | 10 | 6 | 5 | 50 | 54 |
| ENSDARG000 | 3;3 | 22999849:23 | 23000485:23 | +;+ | 1492 | NA | 10 | 0 | 0 | 0 | 0 | 8 |
| ENSDARG000 | 3;3;3;3 | 16003122:16 | 16003692:16 | -;-;-;- | 1018 | mrps34 | 388 | 502 | 457 | 292 | 506 | 286 |
| ENSDARG000 | 3:3:3:3:3:3:3: | 40947961:40 | 40948614:40 | -:-:-:-:-:-:-: | 2225 | cyp3c3 | 18 | 0 | 1 | 0 | 0 | 0 |
| ENSDARG000 | 3:3:3:3:3:3:3: | 23029484:23 | 23030594:23 | +:+:+:+:+:+:+: | 6297 | nags | 2 | 1 | 0 | 0 | 0 | 0 |
| ENSDARG000 | 3:3:3:3:3:3:3: | 23045214:23 | 23045245:23 | +:+:+:+:+:+:+: | 3112 | NA | 0 | 2 | 6 | 16 | 4 | 12 |
| ENSDARG000 | 3:3:3:3:3:3:3: | 383848:3838 | 384116:3841 | -:-:-:-:-:-:-: | 1388 | mhc1zja | 460 | 513 | 200 | 431 | 285 | 95 |
| ENSDARG000 | 3;3 | 14259115:14 | 14259294:14 | +;+ | 571 | NA | 0 | 0 | 0 | 0 | 0 | 0 |
| ENSDARG000 | 3:3:3:3:3:3:3: | 40958870:40 | 40958974:40 | -:-:-:-:-:-:-: | 1699 | NA | 0 | 0 | 0 | 0 | 0 | 0 |
| ENSDARG000 | 3:3:3:3:3:3:3: | 32714157:32 | 32714408:32 | +:+:+:+:+:+:+: | 8708 | NA | 1164 | 924 | 1463 | 813 | 1269 | 1035 |
| ENSDARG000 | 3:3:3:3:3:3:3: | 40969242:40 | 40969579:40 | -:-:-:-:-:-:-: | 2217 | cyp3c1 | 296 | 41 | 124 | 92 | 50 | 164 |
| ENSDARG000 | 3;3;3 | 43086548:43 | 43086816:43 | +;+;+ | 1107 | NA | 0 | 0 | 0 | 0 | 0 | 0 |
| ENSDARG000 | 3;3;3;3;3;3;3 | 16018202:16 | 16022350:16 | -;-;-;-;-;-;- | 5071 | spsb3a | 650 | 510 | 630 | 560 | 561 | 779 |
| ENSDARG000 | 3;3 | 402886:4053 | 402947:4056 | -;- | 282 | NA | 0 | 0 | 0 | 0 | 0 | 0 |
| ENSDARG000 | 3;3;3 | 19813248:19 | 19813464:19 | -;-;- | 361 | NA | 2 | 3 | 1 | 0 | 5 | 0 |
| ENSDARG000 | 3:3:3:3:3:3:3: | 24996930:24 | 25000974:24 | -:-:-:-:-:-:-: | 10800 | EP300 | 2007 | 928 | 1700 | 1110 | 2025 | 1621 |
| ENSDARG000 | 3:3:3:3:3:3:3: | 14388010:14 | 14388304:14 | +:+:+:+:+:+:+: | 4321 | tlcd4b | 76 | 14 | 73 | 78 | 130 | 84 |
| ENSDARG000 | 3:3:3:3:3:3:3: | 43102010:43 | 43102390:43 | +:+:+:+:+:+:+: | 3773 | micall2a | 62 | 0 | 2 | 6 | 0 | 20 |
| ENSDARG000 | 3:3:3:3:3:3:3: | 17910569:17 | 17910803:17 | +:+:+:+:+:+:+: | 1815 | odad4 | 12 | 4 | 6 | 0 | 0 | 2 |
| ENSDARG000 | 3;3;3 | 11101585:11 | 11101940:11 | +;+;+ | 2128 | sstr5 | 0 | 0 | 6 | 0 | 8 | 2 |
| ENSDARG000 | 3;3;3;3;3 | 19850937:19 | 19851435:19 | -;-;-;-;- | 1214 | rnd2 | 16 | 18 | 0 | 0 | 0 | 0 |
| ENSDARG000 | 3:3:3:3:3:3:3: | 40978997:40 | 40980536:40 | -:-:-:-:-:-:-: | 7747 | SDK1 | 10 | 2 | 12 | 6 | 16 | 4 |
| ENSDARG000 | 3 | 14399225 | 14399652 | + | 428 | NA | 0 | 0 | 0 | 0 | 0 | 0 |
| ENSDARG000 | 3;3;3 | 11358518:11 | 11358608:11 | -;-;- | 495 | NA | 1 | 6 | 0 | 0 | 0 | 0 |
| ENSDARG000 | 3:3:3:3:3:3:3: | 3432960:343 | 3433284:343 | -:-:-:-:-:-:-: | 4813 | si:dkev-46g23 | 0 | 0 | 0 | 0 | 0 | 0 |
| ENSDARG000 | 3;3;3 | 420342:4210 | 420522:4211 | +;+;+ | 805 | NA | 2 | 7 | 6 | 20 | 7 | 0 |
| ENSDARG000 | 3:3:3:3:3:3:3: | 11397269:11 | 11397481:11 | -:-:-:-:-:-:-: | 2146 | NA | 223 | 15 | 93 | 84 | 12 | 10 |
| ENSDARG000 | 3;3 | 34180835:34 | 34181082:34 | +;+ | 1120 | timm29 | 205 | 235 | 302 | 156 | 235 | 224 |
| ENSDARG000 | 3:3:3:3:3:3:3: | 32743946:32 | 32744021:32 | +:+:+:+:+:+:+: | 2378 | hsd3b7 | 108 | 68 | 80 | 72 | 108 | 98 |
| ENSDARG000 | 3:3:3:3:3:3:3: | 34220194:34 | 34220394:34 | +:+:+:+:+:+:+: | 1446 | slc25a23b | 14 | 5 | 24 | 26 | 8 | 7 |
| ENSDARG000 | 3:3:3:3:3:3:3: | 34234029:34 | 34234148:34 | +:+:+:+:+:+:+: | 1946 | znf207a | 1399 | 1026 | 1631 | 812 | 1056 | 806 |
| ENSDARG000 | 3;3;3;3;3 | 41531279:41 | 41531599:41 | -;-;-;-;- | 527 | si:ch211-222 | 0 | 0 | 0 | 0 | 0 | 0 |
| ENSDARG000 | 3:3:3:3:3:3:3: | 32789605:32 | 32790047:32 | +:+:+:+:+:+:+: | 5739 | tbc1d10b | 916 | 839 | 908 | 568 | 1023 | 938 |
| ENSDARG000 | 3:3:3:3:3:3:3: | 11779689:11 | 11781412:11 | -;-;-;-;-;-;-;- | 3489 | FOXK2 | 747 | 1049 | 1643 | 916 | 1068 | 731 |
| ENSDARG000 | 3:3:3:3:3:3:3: | 26186153:26 | 26187198:26 | -:-:-:-:-:-:-: | 1847 | ypel3 | 149 | 212 | 115 | 207 | 116 | 324 |
| ENSDARG000 | 3;3;3;3;3;3 | 19914332:19 | 19915063:19 | +;+;+;+;+;+ | 4318 | vat1 | 450 | 165 | 603 | 172 | 1003 | 532 |
| ENSDARG000 | 3;3;3;3;3;3;3 | 431250:4366 | 431400:4367 | +;+;+;+;+;+;+ | 838 | CCDC134 | 0 | 0 | 0 | 0 | 0 | 0 |
| ENSDARG000 | 3:3:3:3:3:3:3: | 41558682:41 | 41558827:41 | +:+:+:+:+:+:+: | 4667 | CARD11 | 52 | 32 | 38 | 12 | 66 | 72 |
| ENSDARG000 | 3:3:3:3:3:3:3: | 431208:4366 | 431400:4367 | +:+:+:+:+:+:+: | 6001 | si:ch73-308m | 435 | 405 | 199 | 315 | 213 | 25 |
| ENSDARG000 | 3:3:3:3:3:3:3: | 41714966:41 | 41715198:41 | +:+:+:+:+:+:+: | 2663 | eif3ba | 8541 | 8455 | 10473 | 6161 | 9451 | 9443 |
| ENSDARG000 | 3 | 43650041 | 43653791 | + | 3751 | NA | 12 | 8 | 25 | 15 | 54 | 16 |
| ENSDARG000 | 3:3:3:3:3:3:3: | 43373867:43 | 43374002:43 | +:+:+:+:+:+:+: | 5397 | zfand2a | 426 | 350 | 444 | 655 | 350 | 214 |
| ENSDARG000 | 3;3;3;3 | 41647637:41 | 41648363:41 | +;+;+;+ | 4379 | gna12a | 3502 | 3837 | 4809 | 3162 | 4443 | 3652 |
| ENSDARG000 | 3:3:3:3:3:3:3: | 43578975:43 | 43581605:43 | -:-:-:-:-:-:-: | 5840 | axin1 | 1058 | 1180 | 1683 | 1350 | 1373 | 1108 |
| ENSDARG000 | 3:3:3:3:3:3:3: | 29510818:29 | 29511095:29 | +:+:+:+:+:+:+: | 4898 | rac2 | 376 | 917 | 639 | 744 | 2005 | 1039 |
| ENSDARG000 | 3:3:3:3:3:3:3: | 17933132:17 | 17933340:17 | +:+:+:+:+:+:+: | 6136 | NA | 3644 | 6565 | 6140 | 4705 | 4811 | 5906 |
| ENSDARG000 | 3:3:3:3:3:3:3: | 43347517:43 | 43347603:43 | -;-;-;-;-;-;- | 1759 | snn | 203 | 121 | 118 | 127 | 138 | 115 |
| ENSDARG000 | 3;3;3;3 | 43757181:43 | 43758007:43 | -;-;-;- | 1325 | pla2g10 | 37 | 24 | 41 | 51 | 56 | 39 |
| ENSDARG000 | 3;3;3 | 43352556:43 | 43353865:43 | -;-;- | 1923 | UNCX | 6 | 4 | 0 | 0 | 11 | 4 |
| ENSDARG000 | 3:3:3:3:3:3:3: | 32813048:32 | 32813984:32 | -:-:-:-:-:-:-: | 1529 | mylpfa | 20 | 36 | 6 | 20 | 58 | 14 |
| ENSDARG000 | 3;3 | 16055009:16 | 16055061:16 | +;+ | 648 | NA | 0 | 1 | 2 | 1 | 1 | 0 |
| ENSDARG000 | 3;3;3;3;3;3;3 | 16044680:16 | 16045821:16 | -;-;-;-;-;-;- | 1629 | atp6v0ca | 15374 | 17835 | 13970 | 13980 | 13504 | 13532 |
| ENSDARG000 | 3;3;3;3;3;3 | 43460696:43 | 43460854:43 | +;+;+;+;+;+ | 5000 | galr2b | 20 | 8 | 8 | 4 | 2 | 18 |
| ENSDARG000 | 3:3:3:3:3:3:3: | 43774369:43 | 43774464:43 | +:+:+:+:+:+:+: | 3998 | zc3h7a | 1196 | 1522 | 1787 | 2770 | 1496 | 1447 |
| ENSDARG000 | 3;3 | 43711165:43 | 43711486:43 | +;+ | 1140 | NA | 1706 | 733 | 1608 | 1645 | 1428 | 1592 |
| ENSDARG000 | 3;3;3;3;3;3 | 20001608:20 | 20001965:20 | +;+;+;+;+;+ | 1705 | asb16 | 0 | 0 | 0 | 0 | 0 | 0 |
| ENSDARG000 | 3 | 29600917 | 29602479 | + | 1563 | NA | 0 | 0 | 0 | 0 | 0 | 0 |
| ENSDARG000 | 3;3;3;3;3;3;3 | 32821205:32 | 32825994:32 | -;-;-;-;-;-;- | 5373 | pheta2 | 52 | 34 | 52 | 8 | 120 | 58 |
| ENSDARG000 | 3;3;3;3;3;3;3 | 14463941:14 | 14464078:14 | +;+;+;+;+;+;+ | 1080 | CNN1 | 0 | 0 | 0 | 0 | 3 | 0 |
| ENSDARG000 | 3:3:3:3:3:3:3: | 26193592:26 | 26193937:26 | -:-:-:-:-:-:-: | 1577 | gdpd3a | 140 | 6 | 20 | 12 | 34 | 10 |
| ENSDARG000 | 3;3;3;3 | 14496319:14 | 14497278:14 | -;-;-;- | 1232 | elof1 | 1078 | 1162 | 1371 | 1138 | 1430 | 1322 |
| ENSDARG000 | 3:3:3:3:3:3:3: | 26211974:26 | 26212672:26 | -:-:-:-:-:-:-: | 2217 | mapk3 | 1610 | 1501 | 1301 | 1100 | 1152 | 1162 |
| ENSDARG000 | 3:3:3:3:3:3:3: | 32651697:32 | 32652098:32 | +:+:+:+:+:+:+: | 1525 | thoc6 | 91 | 170 | 194 | 52 | 120 | 162 |
| ENSDARG000 | 3:3:3:3:3:3:3: | 32832042:32 | 32832602:32 | +:+:+:+:+:+:+: | 1888 | cd2bp2 | 278 | 625 | 589 | 260 | 362 | 259 |
| ENSDARG000 | 3;3 | 16064878:16 | 16065253:16 | -;- | 656 | NA | 2 | 32 | 14 | 60 | 14 | 96 |
| ENSDARG000 | 3:3:3:3:3:3:3: | 34258755:34 | 34265424:34 | -:-:-:-:-:-:-: | 12765 | TNRC6C | 2356 | 1283 | 2387 | 1456 | 3194 | 2340 |
| ENSDARG000 | 3:3:3:3:3:3:3: | 14499075:14 | 14499188:14 | +:+:+:+:+:+:+: | 2247 | tmem205 | 539 | 543 | 421 | 349 | 474 | 466 |
| ENSDARG000 | 3:3:3:3:3:3:3: | 11926030:11 | 11926311:11 | +:+:+:+:+:+:+: | 2902 | dnaja3a | 406 | 389 | 606 | 207 | 455 | 361 |
| ENSDARG000 | 3;3;3;3;3 | 12345280:12 | 12345443:12 | -;-;-;-;- | 598 | pam16 | 138 | 258 | 341 | 216 | 260 | 162 |
| ENSDARG000 | 3;3 | 43828545:43 | 43830348:43 | -;- | 2164 | NA | 11 | 24 | 7 | 12 | 5 | 16 |
| ENSDARG000 | 3;3;3;3;3;3;3 | 43866785:43 | 43866841:43 | -;-;-;-;-;-;- | 833 | NA | 0 | 0 | 0 | 0 | 0 | 0 |
| ENSDARG000 | 3:3:3:3:3:3:3: | 32842794:32 | 32842892:32 | +:+:+:+:+:+:+: | 4462 | NA | 460 | 230 | 270 | 157 | 295 | 303 |
| ENSDARG000 | 3:3:3:3:3:3:3: | 12322170:12 | 12322359:12 | +:+:+:+:+:+:+: | 1515 | glis2b | 105 | 148 | 315 | 41 | 191 | 86 |
| ENSDARG000 | 3:3:3:3:3:3:3: | 43835558:43 | 43836426:43 | -:-:-:-:-:-:-: | 5715 | NA | 0 | 0 | 0 | 0 | 12 | 0 |
| ENSDARG000 | 3:3:3:3:3:3:3: | 11867701:11 | 11867886:11 | -:-:-:-:-:-:-: | 2915 | coro7 | 272 | 244 | 334 | 157 | 406 | 287 |
| ENSDARG000 | 3;3;3 | 32855157:32 | 32855913:32 | -;-;- | 1239 | NA | 83 | 22 | 10 | 18 | 22 | 24 |
| ENSDARG000 | 3:3:3:3:3:3:3: | 43825785:43 | 43826040:43 | +:+:+:+:+:+:+: | 2374 | litaf | 396 | 206 | 169 | 362 | 223 | 236 |
| ENSDARG000 | 3:3:3:3:3:3:3: | 26223376:26 | 26223526:26 | +:+:+:+:+:+:+: | 1443 | nudt9 | 365 | 596 | 348 | 489 | 340 | 476 |
| ENSDARG000 | 3:3:3:3:3:3:3: | 29640837:29 | 29640879:29 | +:+:+:+:+:+:+: | 3082 | eif3d | 6710 | 5307 | 9561 | 7148 | 6655 | 7861 |
| ENSDARG000 | 3:3:3:3:3:3:3: | 43878456:43 | 43878627:43 | -:-:-:-:-:-:-: | 901 | NA | 0 | 0 | 0 | 0 | 0 | 0 |
| ENSDARG000 | 3:3:3:3:3:3:3: | 11958592:11 | 11958719:11 | -:-:-:-:-:-:-: | 1231 | NA | 330 | 189 | 201 | 108 | 346 | 213 |
| ENSDARG000 | 3:3:3:3:3:3:3: | 16072966:16 | 16074390:16 | -:-:-:-:-:-:-: | 2468 | lasp1 | 2155 | 2442 | 3575 | 2684 | 2929 | 2933 |
| ENSDARG000 | 3:3:3:3:3:3:3: | 43942032:43 | 43942398:43 | -:-:-:-:-:-:-: | 2086 | ubfd1 | 459 | 548 | 579 | 392 | 496 | 515 |
| ENSDARG000 | 3;3;3;3;3;3 | 23092762:23 | 23092903:23 | +;+;+;+;+;+ | 658 | gngt2a | 364 | 351 | 195 | 390 | 346 | 270 |
| ENSDARG000 | 3:3:3:3:3:3:3: | 44048907:44 | 44049828:44 | -:-:-:-:-:-:-: | 2508 | il4r.1 | 413 | 381 | 349 | 275 | 217 | 254 |
| ENSDARG000 | 3:3:3:3:3:3:3: | 32862730:32 | 32862770:32 | +:+:+:+:+:+:+: | 2082 | rusf1 | 252 | 381 | 392 | 391 | 424 | 343 |
| ENSDARG000 | 3;3;3;3;3 | 14512670:14 | 14512876:14 | +;+;+;+;+ | 1647 | rab3db | 0 | 0 | 4 | 8 | 18 | 6 |
| ENSDARG000 | 3:3:3:3:3:3:3: | 14543953:14 | 14544117:14 | +:+:+:+:+:+:+: | 1655 | epor | 0 | 0 | 2 | 0 | 14 | 6 |
| ENSDARG000 | 3:3:3:3:3:3:3: | 26236279:26 | 26236690:26 | -;-;-;-;-;-;-;- | 1822 | ppp4ca | 213 | 307 | 382 | 196 | 310 | 264 |
| ENSDARG000 | 3:3:3:3:3:3:3: | 44062576:44 | 44062685:44 | +:+:+:+:+:+:+: | 1584 | nsmce1 | 34 | 45 | 63 | 32 | 22 | 34 |
| ENSDARG000 | 3:3:3:3:3:3:3: | 17951790:17 | 17951927:17 | +:+:+:+:+:+:+: | 5219 | aclya | 1618 | 1638 | 1966 | 931 | 2297 | 1369 |
| ENSDARG000 | 3;3;3 | 14565761:14 | 14566301:14 | -;-;- | 1227 | swsap1 | 4 | 30 | 70 | 2 | 36 | 34 |
| ENSDARG000 | 3:3:3:3:3:3:3: | 34452336:34 | 34456599:34 | -:-:-:-:-:-:-: | 7569 | septin9a | 13401 | 13153 | 14302 | 19479 | 18841 | 17410 |
| ENSDARG000 | 3:3:3:3:3:3:3: | 44081620:44 | 44081814:44 | -:-:-:-:-:-:-: | 1918 | kdm8 | 72 | 100 | 137 | 54 | 116 | 106 |
| ENSDARG000 | 3 | 20063509 | 20064130 | + | 622 | NA | 18 | 15 | 42 | 18 | 25 | 16 |
| ENSDARG000 | 3:3:3:3:3:3:3: | 20041907:20 | 20044036:20 | -:-:-:-:-:-:-: | 4527 | ubtf | 2522 | 2095 | 2332 | 1500 | 2357 | 2316 |
| ENSDARG000 | 3;3;3;3;3 | 629707:6303 | 630252:6304 | -;-;-;-;- | 1318 | NA | 0 | 1 | 0 | 0 | 5 | 2 |
| ENSDARG000 | 3;3;3 | 495748:4984 | 495961:4985 | -;-;- | 436 | NA | 0 | 0 | 0 | 0 | 0 | 0 |
| ENSDARG000 | 3;3;3;3;3;3 | 650429:6505 | 650458:6506 | -;-;-;-;-;- | 752 | NA | 0 | 0 | 4 | 0 | 0 | 0 |
| ENSDARG000 | 3;3;3 | 472158:4731 | 472249:4733 | +;+;+ | 1109 | zgc:194659 | 272 | 27 | 45 | 180 | 25 | 3 |
| ENSDARG000 | 3:3:3:3:3:3:3: | 464127:4641 | 464185:4641 | +:+:+:+:+:+:+: | 1909 | dicp1.9 | 14 | 16 | 5 | 4 | 54 | 0 |
| ENSDARG000 | 3:3:3:3:3:3:3: | 23096728:23 | 23096845:23 | +:+:+:+:+:+:+: | 1359 | NA | 4 | 2 | 0 | 0 | 1 | 0 |
| ENSDARG000 | 3;3;3;3;3 | 720222:7205 | 720311:7210 | -;-;-;-;- | 1367 | NA | 0 | 0 | 0 | 0 | 2 | 6 |
| ENSDARG000 | 3:3:3:3:3:3:3: | 26244353:26 | 26244666:26 | +:+:+:+:+:+:+: | 6063 | NA | 240 | 514 | 716 | 363 | 539 | 384 |
| ENSDARG000 | 3;3;3 | 32870185:32 | 32871644:32 | -;-;- | 1813 | zgc:113090 | 130 | 91 | 113 | 136 | 50 | 66 |
| ENSDARG000 | 3:3:3:3:3:3:3: | 16134376:16 | 16134966:16 | -:-:-:-:-:-:-: | 1572 | arl5c | 331 | 107 | 240 | 275 | 151 | 250 |
| ENSDARG000 | 3;3;3;3;3;3;3 | 18023975:18 | 18024512:18 | -;-;-;-;-;-;- | 1474 | si:ch73-141c | 380 | 242 | 290 | 172 | 528 | 199 |
| ENSDARG000 | 3:3:3:3:3:3:3: | 23221047:23 | 23221255:23 | +:+:+:+:+:+:+: | 6021 | col1a1a | 2921 | 633 | 1589 | 370 | 7433 | 480 |
| ENSDARG000 | 3;3;3;3;3;3 | 34587988:34 | 34589525:34 | -;-;-;-;-;- | 2987 | si:ch211-165 | 210 | 208 | 229 | 260 | 291 | 335 |
| ENSDARG000 | 3:3:3:3:3:3:3: | 34597741:34 | 34597865:34 | -;-;-;-;-;-;-;- | 901 | nmrk1 | 81 | 18 | 94 | 42 | 54 | 50 |
| ENSDARG000 | 3:3:3:3:3:3:3: | 23220270:23 | 23221553:23 | -;-;-;-;-;-;-;- | 2615 | NA | 0 | 0 | 0 | 0 | 0 | 0 |
| ENSDARG000 | 3:3:3:3:3:3:3: | 32889796:32 | 32890227:32 | -:-:-:-:-:-:-: | 3310 | kat7a | 634 | 359 | 487 | 444 | 546 | 571 |
| ENSDARG000 | 3;3;3;3;3;3 | 18038972:18 | 18040755:18 | -;-;-;-;-;- | 2558 | hsd17b1 | 7 | 8 | 0 | 0 | 0 | 0 |
| ENSDARG000 | 3:3:3:3:3:3:3: | 18046776:18 | 18047138:18 | +:+:+:+:+:+:+: | 2569 | mettl26 | 254 | 187 | 302 | 191 | 289 | 220 |
| ENSDARG000 | 3:3:3:3:3:3:3: | 34601852:34 | 34602006:34 | -:-:-:-:-:-:-: | 1539 | zgc:163143 | 40 | 22 | 74 | 24 | 38 | 42 |
| ENSDARG000 | 3:3:3:3:3:3:3: | 26267725:26 | 26267966:26 | +:+:+:+:+:+:+: | 1792 | ADAP2 | 18 | 24 | 18 | 10 | 86 | 48 |
| ENSDARG000 | 3:3:3:3:3:3:3: | 20065861:20 | 20066549:20 | -:-:-:-:-:-:-: | 3556 | slc4a1a | 94 | 40 | 620 | 8 | 1516 | 104 |
| ENSDARG000 | 3;3;3;3;3 | 34623560:34 | 34623661:34 | -;-;-;-;- | 503 | tac4 | 0 | 0 | 0 | 0 | 0 | 0 |
| ENSDARG000 | 3;3;3;3;3;3 | 34652144:34 | 34655124:34 | -;-;-;-;-;- | 3359 | znfl1 | 0 | 0 | 9 | 27 | 0 | 28 |
| ENSDARG000 | 3:3:3:3:3:3:3: | 16157120:16 | 16157821:16 | -:-:-:-:-:-:-: | 2889 | cacnb1 | 32 | 24 | 12 | 89 | 13 | 69 |
| ENSDARG000 | 3;3;3 | 34670076:34 | 34670554:34 | +;+;+ | 1195 | dlx4a | 0 | 0 | 2 | 0 | 12 | 0 |
| ENSDARG000 | 3;3 | 44541322:44 | 44541976:44 | -;- | 1269 | hs3st4 | 0 | 0 | 0 | 0 | 0 | 2 |
| ENSDARG000 | 3;3 | 44947355:45 | 44947884:45 | +;+ | 1149 | hs3st2 | 0 | 0 | 0 | 0 | 0 | 0 |
| ENSDARG000 | 3:3:3:3:3:3:3: | 34688022:34 | 34688227:34 | -:-:-:-:-:-:-: | 2720 | thraa | 172 | 249 | 166 | 226 | 327 | 114 |
| ENSDARG000 | 3;3;3;3;3;3 | 45261387:45 | 45262523:45 | -;-;-;-;-;- | 2035 | kctd5a | 1397 | 1670 | 1136 | 944 | 1635 | 983 |
| ENSDARG000 | 3;3;3 | 34683096:34 | 34683268:34 | +;+;+ | 648 | dusp3b | 0 | 0 | 0 | 0 | 0 | 0 |
| ENSDARG000 | 3:3:3:3:3:3:3: | 45283391:45 | 45286894:45 | -:-:-:-:-:-:-: | 5149 | pdpk1a | 238 | 180 | 213 | 312 | 219 | 278 |
| ENSDARG000 | 3:3:3:3:3:3:3: | 26288981:26 | 26289322:26 | +:+:+:+:+:+:+: | 3325 | rhot1a | 598 | 873 | 832 | 402 | 1006 | 753 |
| ENSDARG000 | 3;3;3;3;3 | 45315477:45 | 45315692:45 | -;-;-;-;- | 762 | il21r.2 | 0 | 2 | 0 | 6 | 2 | 21 |
| ENSDARG000 | 3:3:3:3:3:3:3: | 45343782:45 | 45345003:45 | -;-;-;-;-;-;-;- | 2398 | il21r.1 | 106 | 532 | 266 | 3076 | 272 | 489 |
| ENSDARG000 | 3;3;3;3;3 | 32906082:32 | 32906599:32 | -;-;-;-;- | 1169 | G6PC1 | 0 | 0 | 2 | 2 | 8 | 0 |
| ENSDARG000 | 3;3;3;3;3 | 18062871:18 | 18063311:18 | +;+;+;+;+ | 3018 | ANKRD40 | 518 | 609 | 552 | 439 | 656 | 338 |
| ENSDARG000 | 3:3:3:3:3:3:3: | 32924185:32 | 32924892:32 | -:-:-:-:-:-:-: | 3152 | aoc2 | 334 | 142 | 293 | 100 | 430 | 498 |
| ENSDARG000 | 3:3:3:3:3:3:3: | 45364849:45 | 45365031:45 | +:+:+:+:+:+:+: | 2228 | ube2ia | 1252 | 1423 | 1094 | 1182 | 1352 | 1369 |
| ENSDARG000 | 3:3:3:3:3:3:3: | 32953075:32 | 32954738:32 | -:-:-:-:-:-:-: | 3001 | casp6b.1 | 20 | 4 | 2 | 0 | 2 | 0 |
| ENSDARG000 | 3:3:3:3:3:3:3: | 45401472:45 | 45401986:45 | +:+:+:+:+:+:+: | 4180 | BAIAP3 | 6 | 20 | 41 | 85 | 0 | 455 |
| ENSDARG000 | 3;3;3;3;3;3 | 34767541:34 | 34767894:34 | -;-;-;-;-;- | 819 | NA | 0 | 0 | 0 | 0 | 0 | 0 |
| ENSDARG000 | 3;3;3;3;3;3 | 26322596:26 | 26322722:26 | +;+;+;+;+;+ | 767 | NA | 9 | 14 | 0 | 52 | 5 | 0 |
| ENSDARG000 | 3:3:3:3:3:3:3: | 34766989:34 | 34767894:34 | -:-:-:-:-:-:-: | 3373 | nsfa | 0 | 0 | 0 | 2 | 4 | 0 |
| ENSDARG000 | 3;3 | 18097700:18 | 18098146:18 | +;+ | 1944 | NA | 2 | 0 | 6 | 0 | 0 | 0 |
| ENSDARG000 | 3:3:3:3:3:3:3: | 45602345:45 | 45602875:45 | +:+:+:+:+:+:+: | 3164 | NA | 0 | 0 | 0 | 0 | 0 | 3 |
| ENSDARG000 | 3:3:3:3:3:3:3: | 20103746:20 | 20104525:20 | -;-;-;-;-;-;-;- | 1147 | selenow2b | 82 | 6 | 4 | 10 | 8 | 2 |
| ENSDARG000 | 3;3;3 | 18152487:18 | 18153055:18 | -;-;- | 912 | NA | 0 | 0 | 0 | 0 | 1 | 0 |
| ENSDARG000 | 3:3:3:3:3:3:3: | 18187330:18 | 18187696:18 | -;-;-;-;-;-;-;- | 1263 | NA | 0 | 0 | 0 | 0 | 0 | 0 |
| ENSDARG000 | 3;3 | 18189685:18 | 18189877:18 | +;+ | 966 | NA | 0 | 0 | 0 | 0 | 4 | 0 |
| ENSDARG000 | 3;3 | 18185616:18 | 18187696:18 | -;- | 2279 | tob1a | 639 | 310 | 431 | 490 | 621 | 450 |
| ENSDARG000 | 3:3:3:3:3:3:3: | 32961488:32 | 32962320:32 | -;-;-;-;-;-;-;- | 1578 | casp6a | 8 | 8 | 12 | 6 | 4 | 0 |
| ENSDARG000 | 3:3:3:3:3:3:3: | 26331243:26 | 26331615:26 | -:-:-:-:-:-:-: | 1543 | C3H17orf75 | 289 | 200 | 192 | 234 | 175 | 233 |
| ENSDARG000 | 3;3 | 45687266:45 | 45687682:45 | +;+ | 3695 | gpr146 | 292 | 196 | 357 | 184 | 153 | 516 |
| ENSDARG000 | 3;3;3;3;3 | 45756605:45 | 45756788:45 | +;+;+;+;+ | 2820 | gper1 | 0 | 0 | 0 | 0 | 0 | 2 |
| ENSDARG000 | 3:3:3:3:3:3:3: | 45776370:45 | 45776750:45 | -:-:-:-:-:-:-: | 809 | h3f3a | 1375 | 2866 | 3983 | 1556 | 2629 | 1517 |
| ENSDARG000 | 3:3:3:3:3:3:3: | 45778810:45 | 45778874:45 | +:+:+:+:+:+:+: | 7406 | UNKL | 855 | 535 | 871 | 787 | 883 | 1243 |
| ENSDARG000 | 3;3;3;3;3;3 | 45818626:45 | 45818653:45 | -;-;-;-;-;- | 695 | NA | 0 | 0 | 0 | 0 | 0 | 0 |
| ENSDARG000 | 3;3;3 | 45845007:45 | 45848048:45 | -;-;- | 3081 | IGFALS | 6 | 0 | 2 | 0 | 0 | 0 |
| ENSDARG000 | 3 | 26342768 | 26343648 | + | 881 | NA | 188 | 682 | 694 | 288 | 553 | 332 |
| ENSDARG000 | 3:3:3:3:3:3:3: | 32969204:32 | 32969906:32 | -:-:-:-:-:-:-: | 1806 | casp6b.2 | 0 | 0 | 2 | 0 | 4 | 0 |
| ENSDARG000 | 3;3;3;3 | 46043406:46 | 46044362:46 | -;-;-;- | 2075 | shisa9b | 0 | 0 | 0 | 0 | 0 | 0 |

| | | | | | | | | | | | |
|---|---|---|---|---|---|---|---|---|---|---|---|
| ENSDARG000 | 3;3;3;3;3;3;3; | 26345732:263 | 26345821:263 | +;+;+;+;+;+;+; | 1315 | rps15a | 9671 | 10190 | 14355 | 11009 | 7780 | 12582 |
| ENSDARG000 | 3;3;3;3;3;3;3; | 46224589:462 | 46224842:462 | -;-;-;-;-;-;-;- | 1230 | NA | 0 | 0 | 0 | 0 | 0 | 0 |
| ENSDARG000 | 3;3;3;3;3;3;3; | 18303407:183 | 18303537:183 | -;-;-;-;-;-;-;- | 4576 | spag9a | 162 | 65 | 76 | 21 | 112 | 71 |
| ENSDARG000 | 3;3;3;3 | 20108221:201 | 20110652:201 | -;-;-;- | 2694 | selenow2a | 1946 | 1789 | 2104 | 1344 | 2026 | 1990 |
| ENSDARG000 | 3;3;3;3 | 33045883:330 | 33045989:330 | +;+;+;+ | 1080 | NA | 0 | 0 | 8 | 5 | 0 | 0 |
| ENSDARG000 | 3;3;3;3;3;3;3; | 46271911:462 | 46272321:462 | +;+;+;+;+;+;+; | 6014 | NA | 148 | 91 | 184 | 88 | 123 | 68 |
| ENSDARG000 | 3;3;3;3;3 | 46377305:463 | 46377611:463 | -;-;-;-;- | 980 | snrnp25 | 243 | 258 | 296 | 212 | 317 | 270 |
| ENSDARG000 | 3;3;3 | 46380086:463 | 46380346:463 | +;+;+ | 690 | polr3k | 206 | 311 | 342 | 154 | 167 | 170 |
| ENSDARG000 | 3;3;3;3;3;3;3; | 46387860:463 | 46389154:463 | -;-;-;-;-;-;-;- | 1996 | cdip1 | 117 | 107 | 159 | 151 | 203 | 203 |
| ENSDARG000 | 3;3;3;3;3;3;3; | 16229911:162 | 16229971:162 | +;+;+;+;+;+;+; | 975 | rpl19 | 16866 | 16229 | 25855 | 27952 | 19612 | 25724 |
| ENSDARG000 | 3;3;3;3;3;3;3; | 33065613:330 | 33067213:330 | -;-;-;-;-;-;-;- | 4212 | rarab | 1246 | 1309 | 1258 | 1023 | 1513 | 1576 |
| ENSDARG000 | 3;3;3;3;3 | 46425262:464 | 46425381:464 | +;+;+;+;+ | 1387 | sc:d0202 | 0 | 0 | 0 | 0 | 0 | 0 |
| ENSDARG000 | 3;3;3;3;3;3;3; | 26418451:264 | 26418773:264 | -;-;-;-;-;-;-;- | 2868 | xylt1 | 243 | 264 | 328 | 130 | 254 | 1810 |
| ENSDARG000 | 3;3;3;3;3;3;3; | 46444625:464 | 46444736:464 | +;+;+;+;+;+;+; | 3379 | si:ch211-66e2 | 58 | 6 | 22 | 16 | 69 | 76 |
| ENSDARG000 | 3;3;3;3;3;3;3; | 46457450:464 | 46457583:464 | +;+;+;+;+;+;+; | 1691 | NA | 2 | 0 | 14 | 0 | 46 | 2 |
| ENSDARG000 | 3;3;3;3;3;3;3; | 46479705:464 | 46480029:464 | +;+;+;+;+;+;+; | 8411 | TYK2 | 1312 | 1279 | 2309 | 1693 | 1238 | 1026 |
| ENSDARG000 | 3;3;3;3;3;3 | 20156956:201 | 20157269:201 | +;+;+;+;+;+ | 9139 | NGFR | 14 | 0 | 4 | 6 | 52 | 10 |
| ENSDARG000 | 3 | 33300522 | 33301321 | + | 800 | hspb9 | 0 | 0 | 0 | 0 | 0 | 0 |
| ENSDARG000 | 3;3;3;3;3;3;3; | 34801146:348 | 34804668:348 | -;-;-;-;-;-;-;- | 5127 | psmd11a | 924 | 1159 | 1380 | 426 | 1064 | 566 |
| ENSDARG000 | 3;3;3 | 33300418:333 | 33300709:333 | -;-;- | 754 | NA | 0 | 0 | 0 | 0 | 0 | 0 |
| ENSDARG000 | 3;3 | 20404912:204 | 20405292:204 | +;+ | 884 | NXPH3 | 0 | 0 | 0 | 0 | 0 | 0 |
| ENSDARG000 | 3;3;3;3;3;3;3 | 33340939:333 | 33341085:333 | +;+;+;+;+;+;+ | 846 | pyya | 0 | 2 | 0 | 0 | 0 | 0 |
| ENSDARG000 | 3;3 | 33344076:333 | 33344158:333 | -;- | 186 | NA | 0 | 0 | 0 | 0 | 0 | 0 |
| ENSDARG000 | 3;3;3;3;3;3;3; | 33345347:333 | 33345488:333 | +;+;+;+;+;+;+; | 2521 | mpp2a | 2 | 0 | 0 | 0 | 0 | 4 |
| ENSDARG000 | 3 | 34821327 | 34822320 | + | 994 | cdk5r1a | 0 | 0 | 0 | 0 | 0 | 0 |
| ENSDARG000 | 3;3;3;3;3;3;3; | 18376269:183 | 18377779:183 | -;-;-;-;-;-;-;- | 2422 | kdelr2a | 2632 | 3538 | 4722 | 2068 | 4687 | 2996 |
| ENSDARG000 | 3;3;3;3;3;3;3; | 20658203:206 | 20659205:206 | -;-;-;-;-;-;-;- | 2550 | spop | 607 | 646 | 742 | 408 | 786 | 538 |
| ENSDARG000 | 3;3;3;3;3;3;3; | 46559639:465 | 46559995:465 | +;+;+;+;+;+;+; | 4335 | raver1 | 77 | 244 | 409 | 95 | 271 | 200 |
| ENSDARG000 | 3;3 | 33384282:333 | 33384490:333 | +;+ | 311 | NA | 0 | 0 | 0 | 0 | 0 | 0 |
| ENSDARG000 | 3;3;3;3 | 33389023:333 | 33390386:333 | -;-;-;- | 1801 | NA | 0 | 0 | 0 | 0 | 0 | 0 |
| ENSDARG000 | 3;3;3;3;3;3;3; | 46628885:467 | 46630105:467 | +;+;+;+;+;+;+; | 7511 | PDE4A | 28 | 6 | 43 | 84 | 46 | 82 |
| ENSDARG000 | 3;3;3;3;3;3 | 18389363:183 | 18389527:183 | +;+;+;+;+;+ | 1099 | NA | 6 | 0 | 10 | 8 | 30 | 26 |
| ENSDARG000 | 3;3;3;3;3;3;3; | 33395942:333 | 33396880:333 | -;-;-;-;-;-;-;- | 2447 | abi3a | 8 | 20 | 6 | 4 | 72 | 62 |
| ENSDARG000 | 3;3 | 46635527:466 | 46635614:466 | +;+ | 1127 | NA | 0 | 0 | 2 | 0 | 0 | 0 |
| ENSDARG000 | 3;3;3;3;3;3;3; | 34846201:348 | 34846517:348 | +;+;+;+;+;+;+; | 3914 | NA | 70 | 31 | 28 | 189 | 16 | 0 |
| ENSDARG000 | 3;3 | 46757611:467 | 46757707:467 | -;- | 514 | NA | 0 | 0 | 0 | 0 | 0 | 0 |
| ENSDARG000 | 3;3;3;3;3;3;3; | 46762703:467 | 46762776:467 | +;+;+;+;+;+;+; | 5968 | prkcsh | 3047 | 3109 | 4490 | 2236 | 3944 | 2709 |
| ENSDARG000 | 3;3;3;3;3;3;3; | 18398876:183 | 18399454:183 | +;+;+;+;+;+;+; | 1453 | rps2 | 45740 | 45115 | 78024 | 69718 | 56806 | 64549 |
| ENSDARG000 | 3 | 34947638 | 34948034 | - | 397 | NA | 0 | 0 | 0 | 0 | 0 | 0 |
| ENSDARG000 | 3;3;3;3;3;3;3; | 34919810:349 | 34920225:349 | +;+;+;+;+;+;+; | 2003 | CA10 | 0 | 10 | 6 | 14 | 0 | 0 |
| ENSDARG000 | 3;3;3;3;3;3;3 | 33420214:334 | 33422071:334 | -;-;-;-;-;-;- | 2291 | ccdc103 | 18 | 38 | 16 | 6 | 10 | 12 |
| ENSDARG000 | 3;3;3;3;3;3;3; | 46800190:468 | 46803378:468 | -;-;-;-;-;-;-;- | 4629 | elavl3 | 10 | 0 | 0 | 0 | 1 | 6 |
| ENSDARG000 | 3;3;3;3;3;3;3; | 34985895:349 | 34985957:349 | +;+;+;+;+;+;+; | 2158 | smarce1 | 3400 | 3978 | 5762 | 3407 | 4214 | 3164 |
| ENSDARG000 | 3;3;3;3;3;3;3; | 50032394:500 | 50032511:500 | -;-;-;-;-;-;-;- | 1253 | SLC25A10 | 0 | 1 | 10 | 0 | 0 | 0 |
| ENSDARG000 | 3;3;3;3;3;3;3; | 33423986:334 | 33424112:334 | -;-;-;-;-;-;-;- | 1622 | rpl23 | 8718 | 10772 | 13992 | 11835 | 8206 | 12517 |
| ENSDARG000 | 3;3;3;3;3 | 33432191:334 | 33433937:334 | -;-;-;-;- | 3026 | NA | 0 | 0 | 0 | 0 | 0 | 0 |
| ENSDARG000 | 3;3;3;3 | 18405136:184 | 18405350:184 | -;-;-;- | 733 | ndufb10 | 839 | 979 | 965 | 799 | 908 | 1046 |
| ENSDARG000 | 3;3;3;3;3;3;3; | 50054187:500 | 50054822:500 | -;-;-;-;-;-;-;- | 1287 | mrpl12 | 532 | 799 | 958 | 321 | 721 | 456 |
| ENSDARG000 | 3;3;3;3;3;3;3; | 50076793:500 | 50077432:500 | -;-;-;-;-;-;-;- | 2968 | hgs | 1630 | 1355 | 1634 | 1179 | 1627 | 1206 |
| ENSDARG000 | 3;3;3;3;3;3;3; | 18424244:184 | 18424325:184 | +;+;+;+;+;+;+; | 1439 | NA | 96 | 60 | 172 | 153 | 202 | 108 |
| ENSDARG000 | 3;3;3;3;3;3;3; | 50069204:500 | 50069939:500 | +;+;+;+;+;+;+; | 1656 | zgc:103625 | 23 | 25 | 16 | 12 | 20 | 15 |
| ENSDARG000 | 3;3;3;3;3;3;3; | 33440615:334 | 33441059:334 | +;+;+;+;+;+;+; | 3219 | gtpbp1 | 953 | 1204 | 3390 | 3275 | 1028 | 782 |
| ENSDARG000 | 3;3;3 | 50123507:501 | 50124537:501 | -;-;- | 1729 | cldnk | 6 | 0 | 0 | 0 | 20 | 4 |
| ENSDARG000 | 3;3;3;3;3;3;3; | 50127726:501 | 50129113:501 | -;-;-;-;-;-;-;- | 3121 | btr02 | 0 | 0 | 0 | 6 | 4 | 4 |
| ENSDARG000 | 3;3;3;3;3;3;3; | 33461075:334 | 33462245:334 | -;-;-;-;-;-;-;- | 3576 | sgsm3 | 2460 | 1458 | 2063 | 4165 | 1667 | 1542 |
| ENSDARG000 | 3;3;3;3 | 50141691:501 | 50143657:501 | -;-;-;- | 2415 | NA | 26 | 12 | 4 | 9 | 2 | 12 |
| ENSDARG000 | 3;3;3;3;3;3;3; | 50155245:501 | 50155459:501 | +;+;+;+;+;+;+; | 2452 | rsad1 | 25 | 15 | 37 | 12 | 38 | 30 |
| ENSDARG000 | 3;3;3;3;3;3;3; | 50155245:501 | 50155674:501 | +;+;+;+;+;+;+; | 6493 | NA | 469 | 424 | 508 | 317 | 495 | 438 |
| ENSDARG000 | 3;3 | 50175166:501 | 50176666:501 | -;- | 2075 | rasd2a | 0 | 0 | 0 | 0 | 0 | 0 |
| ENSDARG000 | 3;3;3 | 33681518:336 | 33681620:336 | -;-;- | 352 | NA | 0 | 0 | 0 | 0 | 0 | 0 |
| ENSDARG000 | 3;3;3;3;3;3;3; | 50271962:502 | 50272440:502 | -;-;-;-;-;-;-;- | 1818 | arl16 | 98 | 54 | 62 | 26 | 66 | 56 |
| ENSDARG000 | 3;3;3;3;3;3;3; | 35005062:350 | 35006043:350 | +;+;+;+;+;+;+; | 7074 | prkcbb | 14 | 23 | 14 | 24 | 78 | 70 |
| ENSDARG000 | 3;3;3;3;3;3;3; | 50310684:503 | 50310918:503 | +;+;+;+;+;+;+; | 2669 | gas7a | 54 | 37 | 123 | 6 | 29 | 73 |
| ENSDARG000 | 3;3;3;3;3 | 50441444:504 | 50441620:504 | -;-;-;-;- | 1131 | rcvrna | 0 | 0 | 0 | 0 | 1 | 0 |
| ENSDARG000 | 3;3;3;3;3;3;3 | 50537127:505 | 50537277:505 | +;+;+;+;+;+;+ | 803 | NA | 0 | 0 | 0 | 0 | 0 | 0 |
| ENSDARG000 | 3;3;3;3 | 35298078:353 | 35299422:353 | +;+;+;+ | 4350 | CACNG3 | 4 | 6 | 60 | 2 | 6 | 8 |
| ENSDARG000 | 3;3 | 50728859:507 | 50729483:507 | -;- | 650 | NA | 12 | 8 | 0 | 30 | 10 | 7 |
| ENSDARG000 | 3;3;3;3;3;3;3; | 50602866:506 | 50603383:506 | +;+;+;+;+;+;+; | 2055 | NA | 32 | 4 | 2 | 14 | 6 | 6 |
| ENSDARG000 | 3;3;3;3;3;3;3; | 50861176:508 | 50861712:508 | -;-;-;-;-;-;-;- | 1227 | pmp22a | 1959 | 1026 | 1496 | 1139 | 769 | 795 |
| ENSDARG000 | 3;3;3;3;3;3;3; | 35406395:354 | 35407163:354 | +;+;+;+;+;+;+; | 6965 | rbbp6 | 2605 | 1990 | 3694 | 3214 | 3515 | 2978 |
| ENSDARG000 | 3;3;3;3 | 50951203:509 | 50952853:509 | -;-;-;- | 2123 | cdc42ep4a | 393 | 500 | 650 | 376 | 499 | 484 |
| ENSDARG000 | 3;3;3;3;3;3;3; | 35495570:354 | 35495823:354 | +;+;+;+;+;+;+; | 10457 | TNRC6A | 4739 | 3929 | 5407 | 3790 | 5365 | 4143 |
| ENSDARG000 | 3;3;3;3;3;3;3; | 51030667:510 | 51030905:510 | -;-;-;-;-;-;-;- | 6561 | SDK2 | 10 | 12 | 35 | 12 | 72 | 14 |
| ENSDARG000 | 3;3;3;3;3 | 51417422:514 | 51417641:514 | +;+;+;+;+ | 1711 | NA | 381 | 448 | 1288 | 728 | 1596 | 822 |
| ENSDARG000 | 3;3;3;3;3;3;3; | 51563695:515 | 51564016:515 | +;+;+;+;+;+;+; | 2791 | ttyh2l | 5625 | 4873 | 3857 | 4527 | 3593 | 5607 |
| ENSDARG000 | 3;3;3;3;3;3;3; | 51660158:516 | 51660223:516 | +;+;+;+;+;+;+; | 623 | CHMP6 | 0 | 4 | 14 | 0 | 0 | 18 |
| ENSDARG000 | 3;3;3;3;3;3;3; | 35535121:355 | 35536159:355 | -;-;-;-;-;-;-;- | 1537 | ndufab1b | 635 | 922 | 838 | 592 | 612 | 586 |
| ENSDARG000 | 3;3;3;3;3;3;3; | 33773156:337 | 33773776:337 | -;-;-;-;-;-;-;- | 7101 | CACNA1A | 789 | 392 | 828 | 652 | 698 | 457 |
| ENSDARG000 | 3;3;3;3;3;3;3; | 51899578:519 | 51900906:519 | -;-;-;-;-;-;-;- | 6021 | AATK | 62 | 30 | 282 | 16 | 154 | 44 |
| ENSDARG000 | 3;3;3;3;3;3;3; | 51684963:516 | 51685132:516 | +;+;+;+;+;+;+; | 7762 | baiap2a | 1865 | 468 | 174 | 505 | 537 | 798 |
| ENSDARG000 | 3;3 | 52078798:520 | 52078864:520 | +;+ | 763 | FUT9 | 1 | 2 | 2 | 0 | 2 | 0 |
| ENSDARG000 | 3;3;3 | 52064001:520 | 52064487:520 | -;-;- | 670 | NA | 4 | 2 | 0 | 4 | 6 | 4 |
| ENSDARG000 | 3;3;3;3;3;3;3; | 35542067:355 | 35542347:355 | +;+;+;+;+;+;+; | 1714 | rpusd1 | 105 | 122 | 153 | 62 | 110 | 168 |
| ENSDARG000 | 3;3;3;3;3 | 52092599:520 | 52092620:520 | +;+;+;+;+ | 2116 | FUT9 | 169 | 136 | 99 | 92 | 135 | 120 |
| ENSDARG000 | 3;3;3;3;3;3;3; | 52145511:521 | 52145665:521 | +;+;+;+;+;+;+; | 2009 | NA | 44 | 11 | 16 | 26 | 32 | 19 |
| ENSDARG000 | 3;3;3;3;3;3;3; | 52215045:522 | 52215746:522 | -;-;-;-;-;-;-;- | 1399 | NA | 1 | 75 | 12 | 19 | 20 | 14 |
| ENSDARG000 | 3;3;3;3;3 | 52367302:523 | 52367365:523 | +;+;+;+;+ | 704 | NA | 0 | 12 | 2 | 0 | 4 | 4 |
| ENSDARG000 | 3;3;3;3 | 52442102:524 | 52442343:524 | +;+;+;+ | 778 | NA | 31 | 43 | 7 | 34 | 102 | 51 |
| ENSDARG000 | 3;3;3;3 | 52467879:524 | 52468041:524 | +;+;+;+ | 617 | NA | 0 | 2 | 7 | 7 | 17 | 4 |
| ENSDARG000 | 3;3;3;3 | 52473670:524 | 52473823:524 | +;+;+;+ | 353 | NA | 0 | 1 | 6 | 0 | 7 | 7 |
| ENSDARG000 | 3;3;3;3;3;3;3; | 52475058:524 | 52475124:524 | +;+;+;+;+;+;+; | 1783 | NA | 158 | 186 | 16 | 376 | 738 | 281 |
| ENSDARG000 | 3;3 | 52528606:525 | 52529553:525 | +;+ | 1335 | ptger1c | 0 | 0 | 0 | 0 | 4 | 0 |
| ENSDARG000 | 3;3;3;3;3;3;3; | 52545014:525 | 52545253:525 | +;+;+;+;+;+;+; | 2906 | slc27a1a | 1004 | 852 | 1150 | 564 | 1069 | 904 |
| ENSDARG000 | 3;3;3;3;3;3 | 35550870:355 | 35551367:355 | -;-;-;-;-;- | 1028 | dctn5 | 830 | 1187 | 870 | 959 | 860 | 1067 |
| ENSDARG000 | 3;3;3;3;3;3 | 52607039:526 | 52607716:526 | -;-;-;-;-;- | 1551 | NA | 0 | 0 | 0 | 0 | 2 | 0 |
| ENSDARG000 | 3;3;3;3 | 52641737:526 | 52642294:526 | -;-;-;- | 1249 | NA | 26 | 16 | 22 | 10 | 28 | 28 |
| ENSDARG000 | 3;3;3;3;3;3;3; | 52647144:526 | 52647347:526 | -;-;-;-;-;-;-;- | 2503 | zgc:113210 | 230 | 147 | 190 | 108 | 202 | 153 |
| ENSDARG000 | 3;3;3;3;3;3;3; | 52667699:526 | 52668686:526 | -;-;-;-;-;-;-;- | 1664 | NA | 123 | 101 | 194 | 28 | 104 | 148 |
| ENSDARG000 | 3;3;3 | 35589212:355 | 35590428:355 | -;-;- | 1663 | gng13b | 8 | 0 | 22 | 24 | 20 | 14 |
| ENSDARG000 | 3;3;3;3 | 52676577:526 | 52677538:526 | -;-;-;- | 1585 | NA | 90 | 88 | 113 | 75 | 114 | 94 |
| ENSDARG000 | 3;3;3;3;3;3 | 52684556:526 | 52684628:526 | +;+;+;+;+;+ | 1948 | pgls | 249 | 230 | 436 | 44 | 310 | 292 |
| ENSDARG000 | 3;3;3;3;3;3;3; | 52702700:527 | 52703140:527 | +;+;+;+;+;+;+; | 2422 | COLGALT1 | 391 | 197 | 409 | 84 | 600 | 382 |
| ENSDARG000 | 3;3;3;3;3;3;3; | 52737565:527 | 52737816:527 | +;+;+;+;+;+;+; | 3576 | NA | 58 | 78 | 68 | 32 | 147 | 77 |
| ENSDARG000 | 3;3;3;3;3;3;3; | 52806347:528 | 52806484:528 | +;+;+;+;+;+;+; | 1340 | NA | 0 | 0 | 0 | 0 | 2 | 0 |
| ENSDARG000 | 3;3;3;3;3;3;3; | 35608385:356 | 35608551:356 | +;+;+;+;+;+;+; | 3914 | traf7 | 399 | 416 | 480 | 465 | 554 | 373 |
| ENSDARG000 | 3;3;3;3;3;3;3; | 52899801:529 | 52900232:529 | +;+;+;+;+;+;+; | 4083 | atp13a1 | 1460 | 1370 | 1777 | 817 | 2128 | 1106 |
| ENSDARG000 | 3;3;3;3;3;3;3; | 35632773:356 | 35643582:356 | -;-;-;-;-;-;-;- | 15937 | CASKIN1 | 1124 | 454 | 1042 | 1788 | 1238 | 1173 |
| ENSDARG000 | 3;3;3;3;3;3;3; | 52999962:530 | 53001074:530 | +;+;+;+;+;+;+; | 4379 | pbx4 | 2003 | 1805 | 3969 | 1493 | 2211 | 1581 |
| ENSDARG000 | 3;3;3;3;3 | 53081027:530 | 53083610:530 | -;-;-;-;- | 3638 | lpar2a | 24 | 28 | 16 | 50 | 16 | 12 |
| ENSDARG000 | 3;3;3;3;3;3;3; | 53116172:531 | 53116237:531 | +;+;+;+;+;+;+; | 6124 | brd4 | 1655 | 1861 | 2286 | 1866 | 2178 | 1425 |
| ENSDARG000 | 3;3;3;3 | 53228684:532 | 53228783:532 | +;+;+;+ | 614 | pet100 | 156 | 161 | 239 | 158 | 105 | 186 |
| ENSDARG000 | 3;3;3;3;3;3;3; | 53240562:532 | 53240616:532 | +;+;+;+;+;+;+; | 2529 | stxbp2 | 1553 | 965 | 1321 | 950 | 1322 | 1110 |
| ENSDARG000 | 3;3;3;3;3;3;3; | 53317040:533 | 53317179:533 | +;+;+;+;+;+;+; | 2903 | xab2 | 659 | 617 | 1025 | 534 | 814 | 481 |
| ENSDARG000 | 3;3;3;3;3;3;3; | 33921971:339 | 33922682:339 | -;-;-;-;-;-;-;- | 2796 | gtf2f1 | 697 | 697 | 661 | 314 | 636 | 503 |
| ENSDARG000 | 3;3;3;3;3;3;3; | 53457192:534 | 53457781:534 | -;-;-;-;-;-;-;- | 2163 | nr5a5 | 0 | 0 | 0 | 0 | 2 | 0 |
| ENSDARG000 | 3;3;3;3 | 33918510:339 | 33918894:339 | +;+;+;+ | 1286 | alkbh7 | 67 | 164 | 142 | 90 | 116 | 128 |
| ENSDARG000 | 3;3;3;3;3;3;3; | 53526025:535 | 53531476:535 | -;-;-;-;-;-;-;- | 11783 | notch3 | 864 | 81 | 56 | 68 | 222 | 331 |
| ENSDARG000 | 3;3;3;3;3;3;3; | 35812040:358 | 35812126:358 | +;+;+;+;+;+;+; | 3292 | crlf3 | 758 | 776 | 1040 | 900 | 674 | 702 |
| ENSDARG000 | 3;3;3;3;3;3;3; | 53687645:536 | 53688425:536 | -;-;-;-;-;-;-;- | 1789 | rdh8a | 4118 | 1874 | 1230 | 1295 | 1021 | 1309 |
| ENSDARG000 | 3;3;3;3;3;3;3; | 53773256:538 | 53773403:538 | +;+;+;+;+;+;+; | 6759 | col5a3a | 38 | 18 | 4 | 0 | 52 | 42 |
| ENSDARG000 | 3;3;3;3;3;3;3; | 54012708:540 | 54013048:540 | +;+;+;+;+;+;+; | 3759 | olfm2a | 0 | 0 | 0 | 0 | 4 | 0 |
| ENSDARG000 | 3;3;3;3;3;3 | 54524227:545 | 54524924:545 | -;-;-;-;-;- | 1983 | angptl6 | 0 | 0 | 10 | 0 | 0 | 0 |
| ENSDARG000 | 3;3;3;3;3;3;3; | 54502350:545 | 54505214:545 | -;-;-;-;-;-;-;- | 5532 | si:ch73-208g1 | 47 | 34 | 10 | 18 | 75 | 63 |
| ENSDARG000 | 3;3;3;3;3;3;3; | 54553931:545 | 54554083:545 | +;+;+;+;+;+;+; | 1872 | ppan | 358 | 572 | 824 | 260 | 538 | 372 |
| ENSDARG000 | 3;3 | 54572725:545 | 54572765:545 | +;+ | 1178 | p2ry11 | 6 | 4 | 6 | 0 | 6 | 0 |
| ENSDARG000 | 3;3;3;3;3;3;3; | 54581987:545 | 54582114:545 | +;+;+;+;+;+;+; | 1076 | eif3g | 2700 | 2744 | 4482 | 3878 | 3059 | 3825 |
| ENSDARG000 | 3;3;3;3;3;3;3; | 35849392:358 | 35851398:358 | -;-;-;-;-;-;-;- | 4728 | suz12a | 393 | 662 | 687 | 553 | 1160 | 656 |
| ENSDARG000 | 3;3;3;3;3;3;3; | 54591218:545 | 54591540:545 | -;-;-;-;-;-;-;- | 4896 | dnmt1 | 492 | 1891 | 2238 | 680 | 2062 | 1271 |
| ENSDARG000 | 3;3;3;3;3;3;3; | 36023268:360 | 36023661:360 | -;-;-;-;-;-;-;- | 2495 | rab11fip4a | 0 | 10 | 11 | 32 | 8 | 16 |
| ENSDARG000 | 3;3;3;3;3;3;3; | 54744069:547 | 54744432:547 | +;+;+;+;+;+;+; | 1720 | NA | 288 | 521 | 855 | 283 | 854 | 502 |
| ENSDARG000 | 3;3;3;3;3;3;3 | 54761569:547 | 54761758:547 | +;+;+;+;+;+;+ | 1253 | NA | 204 | 226 | 398 | 129 | 264 | 77 |
| ENSDARG000 | 3;3;3;3;3;3;3; | 33949246:339 | 33950074:339 | -;-;-;-;-;-;-;- | 5297 | NA | 0 | 0 | 0 | 0 | 2 | 0 |
| ENSDARG000 | 3;3;3;3;3;3;3; | 54776117:547 | 54776569:547 | +;+;+;+;+;+;+; | 2450 | NA | 29 | 51 | 10 | 28 | 27 | 25 |
| ENSDARG000 | 3;3;3 | 54887782:548 | 54888086:548 | +;+;+ | 1446 | aanat2 | 236 | 180 | 142 | 66 | 127 | 94 |
| ENSDARG000 | 3;3;3;3;3;3;3; | 54896331:548 | 54897386:548 | -;-;-;-;-;-;-;- | 3753 | rhbdf1a | 4053 | 3818 | 6350 | 3749 | 7524 | 4976 |
| ENSDARG000 | 3;3;3 | 55031685:550 | 55032208:550 | +;+;+ | 1234 | mpg | 23 | 24 | 45 | 42 | 32 | 8 |
| ENSDARG000 | 3;3;3;3;3;3;3; | 55045346:550 | 55046261:550 | -;-;-;-;-;-;-;- | 2958 | nprl3 | 271 | 211 | 313 | 225 | 289 | 220 |
| ENSDARG000 | 3;3;3 | 55091230:550 | 55091433:550 | -;-;- | 650 | hbba2 | 14 | 5 | 22 | 16 | 64 | 0 |
| ENSDARG000 | 3;3;3 | 55093581:550 | 55093716:550 | +;+;+ | 564 | NA | 6 | 12 | 10 | 28 | 44 | 0 |
| ENSDARG000 | 3;3;3 | 55098457:550 | 55098701:550 | -;-;- | 617 | hbba1 | 16 | 19 | 110 | 25 | 213 | 48 |
| ENSDARG000 | 3;3;3 | 55100045:551 | 55100254:551 | +;+;+ | 651 | hbaa1 | 2230 | 1374 | 6055 | 832 | 17793 | 4324 |
| ENSDARG000 | 3;3;3;3;3;3;3 | 36119782:361 | 36120240:361 | -;-;-;-;-;-;- | 2109 | coil | 110 | 187 | 169 | 110 | 288 | 136 |
| ENSDARG000 | 3;3;3;3;3;3;3 | 55103445:551 | 55104011:551 | -;-;-;-;-;-;- | 843 | hbba1 | 377 | 121 | 647 | 78 | 2031 | 407 |
| ENSDARG000 | 3;3;3 | 55105066:551 | 55105280:551 | +;+;+ | 657 | si:ch211-5k11 | 38 | 13 | 109 | 6 | 297 | 78 |
| ENSDARG000 | 3;3;3;3;3 | 55114097:551 | 55114569:551 | +;+;+;+;+ | 684 | hbbe1.1 | 0 | 0 | 2 | 0 | 3 | 0 |
| ENSDARG000 | 3;3;3 | 55120330:551 | 55120568:551 | -;-;- | 633 | hbae1.1 | 0 | 0 | 0 | 0 | 2 | 0 |
| ENSDARG000 | 3;3;3;3;3 | 55122662:551 | 55123134:551 | +;+;+;+;+ | 684 | hbbe1.2 | 0 | 0 | 0 | 0 | 4 | 0 |
| ENSDARG000 | 3;3;3 | 55127549:551 | 55127750:551 | -;-;- | 547 | NA | 0 | 0 | 6 | 0 | 2 | 0 |
| ENSDARG000 | 3;3;3 | 55131289:551 | 55131437:551 | +;+;+ | 582 | hbbe1.3 | 2 | 0 | 16 | 0 | 21 | 0 |
| ENSDARG000 | 3;3;3 | 55138413:551 | 55138629:551 | -;-;- | 555 | NA | 0 | 0 | 23 | 0 | 4 | 3 |
| ENSDARG000 | 3;3;3;3;3;3;3; | 36136530:361 | 36136974:361 | -;-;-;-;-;-;-;- | 3013 | engase | 217 | 180 | 183 | 106 | 200 | 136 |
| ENSDARG000 | 3;3;3;3;3;3;3; | 55146418:551 | 55146814:551 | -;-;-;-;-;-;-;- | 787 | hbae3 | 0 | 0 | 0 | 0 | 8 | 0 |
| ENSDARG000 | 3;3;3;3;3;3;3; | 36127287:361 | 36127469:361 | +;+;+;+;+;+;+; | 4668 | scpep1 | 2760 | 2820 | 2458 | 1512 | 3249 | 2259 |
| ENSDARG000 | 3;3;3;3;3;3;3; | 55161627:551 | 55162382:551 | -;-;-;-;-;-;-;- | 4312 | kank2 | 3388 | 2556 | 2608 | 4313 | 4056 | 3117 |
| ENSDARG000 | 3 | 34017838 | 34017876 | - | 39 | NA | 0 | 0 | 0 | 0 | 0 | 0 |
| ENSDARG000 | 3 | 33975926 | 33975939 | - | 14 | NA | 0 | 0 | 0 | 0 | 0 | 0 |
| ENSDARG000 | 3 | 34058679 | 34058717 | - | 39 | NA | 0 | 0 | 0 | 0 | 0 | 0 |
| ENSDARG000 | 3;3 | 34074289:340 | 34074594:340 | -;- | 349 | NA | 0 | 0 | 2 | 0 | 0 | 0 |
| ENSDARG000 | 3;3;3;3 | 34050922:340 | 34051232:340 | -;-;-;- | 379 | NA | 0 | 0 | 0 | 0 | 0 | 0 |
| ENSDARG000 | 3;3;3;3;3;3;3; | 55218597:552 | 55219220:552 | -;-;-;-;-;-;-;- | 7448 | DOCK6 | 922 | 802 | 1318 | 882 | 1980 | 1120 |
| ENSDARG000 | 3;3 | 34078200:340 | 34078421:340 | +;+ | 534 | NA | 0 | 0 | 0 | 0 | 0 | 0 |
| ENSDARG000 | 3;3 | 34088852:340 | 34089160:340 | -;- | 367 | NA | 0 | 0 | 0 | 0 | 0 | 0 |

| | | | | | | | | | | | | |
|---|---|---|---|---|---|---|---|---|---|---|---|---|
| ENSDARG000 | 3;3;3;3;3 | 34074278:34 | 34074289:34 | -;-;-;-;- | 419 | NA | 0 | 0 | 0 | 0 | 0 | 0 |
| ENSDARG000 | 3 | 33980241 | 33980251 | - | 11 | NA | 0 | 0 | 0 | 0 | 0 | 0 |
| ENSDARG000 | 3;3 | 34023165:34 | 34023476:34 | -;- | 359 | NA | 0 | 0 | 1 | 0 | 0 | 0 |
| ENSDARG000 | 3;3;3;3;3;3;3 | 34036619:34 | 34036900:34 | -;-;-;-;-;-;- | 417 | NA | 0 | 0 | 0 | 0 | 0 | 0 |
| ENSDARG000 | 3;3 | 34034048:34 | 34034356:34 | -;- | 361 | NA | 0 | 0 | 0 | 0 | 0 | 0 |
| ENSDARG000 | 3;3 | 34048205:34 | 34048516:34 | -;- | 355 | NA | 0 | 0 | 0 | 0 | 0 | 0 |
| ENSDARG000 | 3;3 | 34037481:34 | 34037753:34 | -;- | 319 | NA | 0 | 0 | 0 | 0 | 0 | 0 |
| ENSDARG000 | 3;3 | 34092458:34 | 34092751:34 | -;- | 380 | NA | 0 | 0 | 0 | 0 | 0 | 4 |
| ENSDARG000 | 3;3 | 34080570:34 | 34080875:34 | -;- | 369 | NA | 0 | 0 | 0 | 0 | 0 | 0 |
| ENSDARG000 | 3;3 | 34072012:34 | 34072320:34 | -;- | 352 | NA | 0 | 0 | 0 | 0 | 0 | 0 |
| ENSDARG000 | 3;3 | 34038753:34 | 34039059:34 | -;- | 350 | NA | 0 | 0 | 0 | 0 | 0 | 0 |
| ENSDARG000 | 3;3 | 34031577:34 | 34031880:34 | -;- | 344 | NA | 0 | 0 | 0 | 0 | 0 | 0 |
| ENSDARG000 | 3;3 | 34084861:34 | 34085166:34 | -;- | 395 | NA | 0 | 0 | 0 | 0 | 0 | 0 |
| ENSDARG000 | 3;3 | 34076378:34 | 34076674:34 | -;- | 343 | NA | 0 | 0 | 0 | 0 | 0 | 0 |
| ENSDARG000 | 3 | 33970681 | 33970734 | - | 54 | NA | 0 | 0 | 0 | 0 | 0 | 0 |
| ENSDARG000 | 3;3 | 34047179:34 | 34047478:34 | -;- | 343 | NA | 0 | 0 | 0 | 0 | 0 | 0 |
| ENSDARG000 | 3;3;3;3;3;3;3; | 55288200:55 | 55288551:55 | +;+;+;+;+;+;+; | 1007 | angptl8 | 0 | 2 | 2 | 0 | 10 | 6 |
| ENSDARG000 | 3;3 | 34094760:34 | 34095068:34 | -;- | 372 | NA | 0 | 0 | 0 | 0 | 0 | 0 |
| ENSDARG000 | 3;3 | 34070022:34 | 34070327:34 | -;- | 349 | NA | 0 | 0 | 0 | 0 | 0 | 0 |
| ENSDARG000 | 3;3 | 34086041:34 | 34086337:34 | -;- | 340 | NA | 0 | 0 | 0 | 0 | 0 | 0 |
| ENSDARG000 | 3;3 | 34028457:34 | 34028756:34 | -;- | 343 | NA | 0 | 0 | 0 | 0 | 0 | 0 |
| ENSDARG000 | 3;3 | 34039630:34 | 34039935:34 | -;- | 349 | NA | 0 | 0 | 0 | 0 | 0 | 0 |
| ENSDARG000 | 3 | 33977959 | 33977969 | - | 11 | NA | 0 | 0 | 0 | 0 | 0 | 0 |
| ENSDARG000 | 3;3 | 55401951:55 | 55402448:55 | -;- | 592 | mafga | 16 | 23 | 48 | 32 | 42 | 37 |
| ENSDARG000 | 3;3 | 34059697:34 | 34059998:34 | -;- | 382 | NA | 0 | 0 | 0 | 0 | 0 | 0 |
| ENSDARG000 | 3 | 33979568 | 33979579 | - | 12 | NA | 0 | 0 | 0 | 0 | 0 | 0 |
| ENSDARG000 | 3 | 33970381 | 33970431 | - | 51 | NA | 0 | 0 | 0 | 0 | 0 | 0 |
| ENSDARG000 | 3 | 34000364 | 34000409 | - | 46 | NA | 0 | 0 | 0 | 0 | 0 | 0 |
| ENSDARG000 | 3 | 34091300 | 34091356 | - | 57 | NA | 0 | 0 | 0 | 0 | 0 | 0 |
| ENSDARG000 | 3;3 | 34089983:34 | 34090294:34 | -;- | 358 | NA | 0 | 0 | 0 | 0 | 0 | 0 |
| ENSDARG000 | 3;3 | 34066113:34 | 34066415:34 | -;- | 346 | NA | 0 | 0 | 0 | 0 | 11 | 0 |
| ENSDARG000 | 3;3 | 34055269:34 | 34055580:34 | -;- | 393 | NA | 0 | 0 | 0 | 0 | 0 | 0 |
| ENSDARG000 | 3;3;3;3;3;3;3; | 33994684:33 | 33994820:33 | -;-;-;-;-;-;-;-; | 1676 | NA | 0 | 6 | 0 | 4 | 0 | 0 |
| ENSDARG000 | 3;3;3;3;3;3;3; | 55420176:55 | 55424795:55 | -;-;-;-;-;-;-;-; | 8186 | NA | 1629 | 888 | 744 | 2211 | 1674 | 1708 |
| ENSDARG000 | 3;3 | 34069166:34 | 34069468:34 | -;- | 346 | NA | 0 | 2 | 0 | 0 | 22 | 0 |
| ENSDARG000 | 3;3 | 34052395:34 | 34052704:34 | -;- | 362 | NA | 0 | 0 | 0 | 0 | 0 | 0 |
| ENSDARG000 | 3 | 33970138 | 33970190 | - | 53 | NA | 0 | 0 | 0 | 0 | 0 | 0 |
| ENSDARG000 | 3 | 34017270 | 34017279 | - | 10 | NA | 0 | 0 | 0 | 0 | 0 | 0 |
| ENSDARG000 | 3;3 | 34030460:34 | 34030785:34 | -;- | 390 | NA | 0 | 0 | 0 | 0 | 0 | 0 |
| ENSDARG000 | 3 | 34015518 | 34015533 | - | 16 | NA | 0 | 0 | 0 | 0 | 0 | 0 |
| ENSDARG000 | 3;3 | 55482160:55 | 55482415:55 | +;+ | 777 | si:dkev-188h1 | 0 | 13 | 12 | 0 | 2 | 14 |
| ENSDARG000 | 3;3 | 34035192:34 | 34035500:34 | -;- | 358 | NA | 0 | 0 | 0 | 0 | 0 | 0 |
| ENSDARG000 | 3;3 | 34063955:34 | 34064805:34 | -;- | 900 | NA | 0 | 0 | 0 | 0 | 0 | 0 |
| ENSDARG000 | 3;3 | 34061400:34 | 34061728:34 | -;- | 410 | NA | 0 | 0 | 0 | 0 | 0 | 0 |
| ENSDARG000 | 3 | 33970948 | 33970998 | - | 51 | NA | 0 | 0 | 0 | 0 | 0 | 0 |
| ENSDARG000 | 3 | 33979335 | 33979345 | - | 11 | NA | 0 | 0 | 0 | 0 | 0 | 0 |
| ENSDARG000 | 3;3 | 34086948:34 | 34087256:34 | -;- | 371 | NA | 0 | 0 | 0 | 0 | 0 | 0 |
| ENSDARG000 | 3;3 | 34053623:34 | 34053925:34 | -;- | 384 | NA | 0 | 0 | 0 | 0 | 0 | 0 |
| ENSDARG000 | 3;3 | 34026706:34 | 34027009:34 | -;- | 374 | NA | 0 | 0 | 0 | 0 | 0 | 0 |
| ENSDARG000 | 3;3;3;3 | 34081851:34 | 34082132:34 | -;-;-;- | 414 | NA | 0 | 0 | 0 | 0 | 0 | 0 |
| ENSDARG000 | 3 | 34000919 | 34000963 | - | 45 | NA | 0 | 0 | 0 | 0 | 0 | 0 |
| ENSDARG000 | 3;3;3;3;3;3;3; | 55487261:55 | 55488365:55 | -;-;-;-;-;-;-;-; | 4970 | tex2 | 1484 | 755 | 745 | 1476 | 1235 | 1555 |
| ENSDARG000 | 3;3 | 34021654:34 | 34021950:34 | -;- | 337 | NA | 0 | 0 | 0 | 0 | 0 | 0 |
| ENSDARG000 | 3;3;3;3 | 34066951:34 | 34067232:34 | -;-;-;- | 369 | NA | 0 | 0 | 0 | 0 | 0 | 0 |
| ENSDARG000 | 3 | 33971247 | 33971297 | - | 51 | NA | 0 | 0 | 0 | 0 | 0 | 0 |
| ENSDARG000 | 3;3;3;3;3;3;3; | 33963247:33 | 33963389:33 | -;-;-;-;-;-;-;-; | 2982 | NA | 14 | 7 | 18 | 8 | 164 | 34 |
| ENSDARG000 | 3;3;3;3 | 36160979:36 | 36161201:36 | +;+;+;+ | 1506 | slc16a3a | 2 | 2 | 16 | 0 | 18 | 4 |
| ENSDARG000 | 3;3;3;3;3;3;3; | 55627948:55 | 55631988:55 | -;-;-;-;-;-;-;-; | 7075 | axin2 | 513 | 281 | 249 | 262 | 469 | 253 |
| ENSDARG000 | 3;3 | 56099256:56 | 56099594:56 | +;+ | 782 | NA | 0 | 0 | 0 | 0 | 0 | 0 |
| ENSDARG000 | 3;3;3;3;3;3;3; | 56163276:56 | 56163867:56 | +;+;+;+;+;+;+; | 2363 | NA | 2 | 0 | 2 | 8 | 12 | 6 |
| ENSDARG000 | 3;3;3;3;3;3;3; | 56366395:56 | 56366559:56 | +;+;+;+;+;+;+; | 1950 | CACNG5 | 0 | 0 | 0 | 0 | 0 | 0 |
| ENSDARG000 | 3;3;3;3;3;3;3; | 56574623:56 | 56574837:56 | +;+;+;+;+;+;+; | 1977 | rac1b | 556 | 608 | 592 | 479 | 547 | 569 |
| ENSDARG000 | 3;3 | 56411141:56 | 56411701:56 | -;- | 706 | NA | 0 | 0 | 0 | 0 | 0 | 0 |
| ENSDARG000 | 3;3;3;3;3;3;3; | 56475776:56 | 56475857:56 | -;-;-;-;-;-;-;- | 979 | CYTH3 | 0 | 0 | 0 | 0 | 0 | 4 |
| ENSDARG000 | 3;3;3;3;3;3;3; | 56408840:56 | 56409689:56 | -;-;-;-;-;-;-;-; | 2531 | CYTH3 | 20 | 0 | 10 | 0 | 5 | 15 |
| ENSDARG000 | 3;3;3;3;3;3;3; | 36173203:36 | 36175208:36 | -;-;-;-;-;-;-;-; | 3647 | csnk1da | 3052 | 2266 | 2733 | 2434 | 2853 | 2688 |
| ENSDARG000 | 3;3;3;3;3;3;3; | 56601829:56 | 56601998:56 | -;-;-;-;-;-;-;-; | 1179 | stk19 | 12 | 3 | 17 | 22 | 10 | 34 |
| ENSDARG000 | 3;3;3;3;3;3;3; | 56861059:56 | 56861575:56 | -;-;-;-;-;-;-;-; | 1818 | zgc:112148 | 56 | 41 | 40 | 209 | 98 | 70 |
| ENSDARG000 | 3 | 56876280 | 56877918 | + | 1639 | NA | 4 | 0 | 8 | 0 | 68 | 0 |
| ENSDARG000 | 3;3;3;3;3 | 56882832:56 | 56888152:56 | -;-;-;-;- | 6103 | USH1G | 0 | 307 | 0 | 0 | 2 | 2 |
| ENSDARG000 | 3;3;3;3;3;3;3; | 56905103:56 | 56905451:56 | +;+;+;+;+;+;+; | 2060 | OTOP2 | 2 | 10 | 2 | 22 | 4 | 2 |
| ENSDARG000 | 3;3;3;3;3;3;3; | 56921990:56 | 56922591:56 | -;-;-;-;-;-;-;- | 1160 | NA | 14 | 24 | 18 | 26 | 33 | 15 |
| ENSDARG000 | 3;3 | 57423137:57 | 57424971:57 | -;- | 1935 | socs3a | 44094 | 13923 | 28925 | 25901 | 14199 | 34633 |
| ENSDARG000 | 3;3;3;3;3;3;3; | 57462228:57 | 57465978:57 | -;-;-;-;-;-;-;-; | 5061 | cyth1a | 1543 | 1465 | 1550 | 1417 | 1663 | 1447 |
| ENSDARG000 | 3;3;3;3;3;3;3; | 57583656:57 | 57585179:57 | -;-;-;-;-;-;-;-; | 4956 | usp36 | 1280 | 1253 | 2363 | 1243 | 1654 | 1726 |
| ENSDARG000 | 3;3;3 | 57636374:57 | 57636456:57 | +;+;+ | 402 | NA | 0 | 0 | 0 | 0 | 0 | 0 |
| ENSDARG000 | 3;3;3;3;3;3;3; | 57632965:57 | 57634505:57 | -;-;-;-;-;-;-;-; | 3530 | timp2b | 1424 | 1384 | 934 | 573 | 912 | 518 |
| ENSDARG000 | 3;3;3;3;3 | 57699246:57 | 57700530:57 | -;-;-;-;- | 1977 | lgals3bp.3 | 2 | 30 | 12 | 2 | 6 | 6 |
| ENSDARG000 | 3;3;3;3;3 | 57718367:57 | 57719630:57 | -;-;-;-;- | 1977 | lgals3bp.2 | 0 | 1 | 2 | 0 | 0 | 0 |
| ENSDARG000 | 3;3;3;3;3 | 57741933:57 | 57743207:57 | -;-;-;-;- | 1994 | lgals3bp.1 | 272 | 445 | 46 | 67 | 127 | 102 |
| ENSDARG000 | 3;3;3;3;3;3;3 | 57747716:57 | 57748642:57 | -;-;-;-;-;-;- | 1942 | cant1a | 893 | 830 | 546 | 966 | 785 | 864 |
| ENSDARG000 | 3;3;3;3 | 57766627:57 | 57769545:57 | -;-;-;- | 3500 | syngr2a | 1594 | 1156 | 1184 | 1119 | 1086 | 1569 |
| ENSDARG000 | 3;3;3;3;3;3;3; | 57779673:57 | 57779861:57 | +;+;+;+;+;+;+; | 2583 | NA | 14 | 8 | 14 | 0 | 10 | 31 |
| ENSDARG000 | 3;3;3;3;3;3;3 | 57788761:57 | 57788867:57 | +;+;+;+;+;+;+ | 744 | NA | 0 | 0 | 0 | 0 | 0 | 0 |
| ENSDARG000 | 3;3;3;3;3;3;3; | 57825938:57 | 57826066:57 | +;+;+;+;+;+;+; | 573 | cenpx | 120 | 222 | 166 | 108 | 126 | 100 |
| ENSDARG000 | 3;3;3;3;3;3;3; | 57997980:57 | 57998087:57 | +;+;+;+;+;+;+; | 2726 | pycr1a | 119 | 79 | 56 | 208 | 112 | 106 |
| ENSDARG000 | 3;3;3;3;3;3;3; | 36222799:36 | 36222875:36 | +;+;+;+;+;+;+; | 2730 | dus1l | 87 | 273 | 225 | 104 | 202 | 169 |
| ENSDARG000 | 3;3;3;3 | 58039261:58 | 58039389:58 | -;-;-;- | 672 | NA | 0 | 0 | 0 | 0 | 0 | 0 |
| ENSDARG000 | 3;3 | 58057614:58 | 58058172:58 | -;- | 712 | NA | 0 | 0 | 0 | 0 | 0 | 0 |
| ENSDARG000 | 3;3;3;3 | 58067750:58 | 58067925:58 | -;-;-;- | 483 | NA | 0 | 0 | 0 | 0 | 0 | 0 |
| ENSDARG000 | 3;3;3;3;3;3 | 58092212:58 | 58092321:58 | +;+;+;+;+;+ | 879 | NA | 2 | 2 | 0 | 0 | 2 | 0 |
| ENSDARG000 | 3;3;3;3;3;3;3; | 58104899:58 | 58105029:58 | -;-;-;-;-;-;-;- | 1098 | NA | 4 | 8 | 6 | 0 | 0 | 2 |
| ENSDARG000 | 3;3 | 58108431:58 | 58108825:58 | -;- | 641 | si:dkeyp-79f1 | 7 | 0 | 14 | 0 | 0 | 6 |
| ENSDARG000 | 3;3;3;3;3;3;3; | 58119571:58 | 58119643:58 | +;+;+;+;+;+;+; | 1538 | NA | 8 | 2 | 0 | 0 | 0 | 4 |
| ENSDARG000 | 3;3;3;3;3;3;3; | 58133227:58 | 58133357:58 | +;+;+;+;+;+;+; | 2715 | NA | 215 | 194 | 238 | 153 | 243 | 249 |
| ENSDARG000 | 3;3;3 | 58158595:58 | 58160030:58 | -;-;- | 1706 | snu13a | 146 | 267 | 370 | 46 | 234 | 184 |
| ENSDARG000 | 3;3;3;3;3;3;3; | 58167288:58 | 58167378:58 | +;+;+;+;+;+;+; | 2059 | uqcrc2a | 951 | 1020 | 1074 | 1049 | 1006 | 988 |
| ENSDARG000 | 3 | 58188903 | 58189429 | - | 527 | NA | 255 | 137 | 462 | 1 | 207 | 258 |
| ENSDARG000 | 3;3;3;3;3;3;3; | 58354852:58 | 58355103:58 | +;+;+;+;+;+;+; | 1877 | NA | 6 | 2 | 0 | 8 | 0 | 2 |
| ENSDARG000 | 3;3;3;3;3;3;3; | 58379450:58 | 58379688:58 | +;+;+;+;+;+;+; | 1705 | SDR42E2 | 0 | 0 | 0 | 0 | 0 | 0 |
| ENSDARG000 | 3;3;3;3;3;3 | 58430678:58 | 58431290:58 | -;-;-;-;-;- | 1549 | CDR2 | 8 | 4 | 2 | 22 | 0 | 0 |
| ENSDARG000 | 3;3;3;3;3;3 | 58455428:58 | 58455802:58 | +;+;+;+;+;+ | 1735 | mfsd13al | 48 | 49 | 53 | 57 | 102 | 68 |
| ENSDARG000 | 3 | 58469935 | 58470519 | + | 585 | NA | 14 | 10 | 14 | 16 | 16 | 18 |
| ENSDARG000 | 3;3;3;3;3;3 | 58472305:58 | 58472486:58 | +;+;+;+;+;+ | 5235 | NA | 176 | 128 | 253 | 150 | 206 | 160 |
| ENSDARG000 | 3;3 | 58623923:58 | 58624379:58 | -;- | 655 | NA | 94 | 75 | 96 | 33 | 55 | 49 |
| ENSDARG000 | 3;3;3;3;3;3;3; | 58633692:58 | 58634598:58 | -;-;-;-;-;-;-;-; | 1808 | dhrs7ca | 0 | 0 | 0 | 0 | 0 | 0 |
| ENSDARG000 | 3;3 | 58654063:58 | 58654199:58 | -;- | 906 | NA | 0 | 0 | 0 | 0 | 0 | 0 |
| ENSDARG000 | 3;3;3;3;3;3;3; | 58654598:58 | 58654709:58 | +;+;+;+;+;+;+; | 2844 | NA | 2 | 0 | 10 | 2 | 12 | 19 |
| ENSDARG000 | 3;3 | 58669650:58 | 58669770:58 | -;- | 1231 | NA | 0 | 0 | 0 | 0 | 0 | 0 |
| ENSDARG000 | 3;3 | 58659756:58 | 58659876:58 | -;- | 1303 | NA | 0 | 0 | 0 | 0 | 0 | 0 |
| ENSDARG000 | 3;3 | 34098259:34 | 34098572:34 | -;- | 376 | NA | 0 | 0 | 0 | 0 | 0 | 0 |
| ENSDARG000 | 3;3 | 34101631:34 | 34101936:34 | -;- | 369 | NA | 0 | 0 | 0 | 0 | 0 | 0 |
| ENSDARG000 | 3;3;3;3;3 | 34095514:34 | 34095564:34 | -;-;-;-;- | 419 | NA | 0 | 0 | 0 | 0 | 0 | 0 |
| ENSDARG000 | 3;3 | 34102957:34 | 34103246:34 | -;- | 336 | NA | 0 | 0 | 0 | 0 | 0 | 0 |
| ENSDARG000 | 3;3 | 34096513:34 | 34096833:34 | -;- | 367 | NA | 0 | 0 | 0 | 0 | 0 | 0 |
| ENSDARG000 | 3;3 | 58694176:58 | 58694304:58 | +;+ | 545 | NA | 14 | 0 | 14 | 0 | 13 | 5 |
| ENSDARG000 | 3;3;3;3;3;3 | 58702320:58 | 58702441:58 | +;+;+;+;+;+ | 2913 | NA | 126 | 40 | 173 | 29 | 147 | 112 |
| ENSDARG000 | 3;3 | 34106046:34 | 34106345:34 | -;- | 343 | NA | 0 | 0 | 0 | 0 | 0 | 0 |
| ENSDARG000 | 3;3 | 34107197:34 | 34107505:34 | -;- | 398 | NA | 0 | 0 | 0 | 0 | 0 | 0 |
| ENSDARG000 | 3;3;3;3;3;3;3; | 58714741:58 | 58715312:58 | -;-;-;-;-;-;-;-; | 4347 | radil2a | 38 | 4 | 28 | 26 | 50 | 12 |
| ENSDARG000 | 3;3 | 34111957:34 | 34112274:34 | +;+ | 495 | si:dkev-204f1 | 0 | 0 | 0 | 0 | 0 | 0 |
| ENSDARG000 | 3;3;3;3;3;3;3; | 58805790:58 | 58806308:58 | -;-;-;-;-;-;-;-; | 2039 | luc7l3 | 989 | 934 | 1418 | 755 | 1261 | 1041 |
| ENSDARG000 | 3;3 | 34113410:34 | 34113715:34 | -;- | 349 | NA | 0 | 0 | 0 | 0 | 0 | 0 |
| ENSDARG000 | 3;3 | 58833306:58 | 58833339:58 | +;+ | 334 | NA | 0 | 1 | 1 | 0 | 3 | 1 |
| ENSDARG000 | 3;3 | 34115438:34 | 34115749:34 | -;- | 358 | NA | 0 | 0 | 0 | 0 | 0 | 0 |
| ENSDARG000 | 3;3;3;3 | 60034194:60 | 60034315:60 | +;+;+;+ | 6295 | NA | 78 | 107 | 154 | 41 | 101 | 105 |
| ENSDARG000 | 3;3;3;3;3 | 60146309:60 | 60149225:60 | -;-;-;-;- | 3579 | NA | 2 | 6 | 16 | 4 | 4 | 0 |
| ENSDARG000 | 3;3;3;3;3;3;3; | 60007703:60 | 60007850:60 | +;+;+;+;+;+;+; | 1082 | si:ch211-110 | 7 | 2 | 2 | 0 | 16 | 12 |
| ENSDARG000 | 3;3;3;3;3;3;3; | 59411956:59 | 59412105:59 | +;+;+;+;+;+;+; | 2160 | NA | 242 | 174 | 169 | 182 | 268 | 176 |
| ENSDARG000 | 3;3;3;3;3;3;3; | 60129848:60 | 60130407:60 | -;-;-;-;-;-;-;-; | 1338 | NA | 5 | 2 | 8 | 10 | 33 | 9 |
| ENSDARG000 | 3;3;3;3;3;3;3; | 59986120:59 | 59986450:59 | -;-;-;-;-;-;-;-; | 3167 | NA | 513 | 324 | 585 | 446 | 675 | 497 |
| ENSDARG000 | 3;3;3;3;3;3;3; | 36234060:36 | 36234476:36 | -;-;-;-;-;-;-;-; | 2383 | gps1 | 924 | 907 | 1249 | 696 | 989 | 904 |
| ENSDARG000 | 3;3;3;3;3;3;3; | 60285115:60 | 60286202:60 | -;-;-;-;-;-;-;-; | 4893 | NA | 42 | 83 | 37 | 61 | 61 | 32 |
| ENSDARG000 | 3;3;3;3 | 60314274:60 | 60314826:60 | -;-;-;- | 853 | NA | 115 | 345 | 261 | 295 | 137 | 97 |
| ENSDARG000 | 3;3;3;3;3;3;3; | 60338988:60 | 60339591:60 | -;-;-;-;-;-;-;-; | 5018 | NA | 1418 | 1919 | 1585 | 907 | 1542 | 1078 |
| ENSDARG000 | 3;3;3;3;3;3;3; | 60957512:60 | 60957535:60 | +;+;+;+;+;+;+; | 6661 | helz | 2074 | 1639 | 2986 | 1472 | 2717 | 1509 |
| ENSDARG000 | 3;3;3;3 | 61043774:61 | 61044082:61 | -;-;-;- | 763 | CACNG1 | 0 | 0 | 0 | 0 | 0 | 0 |
| ENSDARG000 | 3;3;3;3 | 61078057:61 | 61079616:61 | -;-;-;- | 2249 | cacng4b | 0 | 2 | 7 | 8 | 0 | 2 |
| ENSDARG000 | 3;3;3;3;3;3;3; | 61108356:61 | 61109102:61 | -;-;-;-;-;-;-;- | 1465 | aimp2 | 1494 | 1797 | 3557 | 2267 | 1392 | 1471 |
| ENSDARG000 | 3;3;3;3;3 | 61154219:61 | 61154486:61 | -;-;-;-;- | 840 | pvalb8 | 8 | 0 | 0 | 0 | 6 | 0 |
| ENSDARG000 | 3;3;3;3;3;3;3; | 61174750:61 | 61176357:61 | -;-;-;-;-;-;-;- | 1976 | pvalb4 | 2 | 23 | 3 | 0 | 6 | 13 |
| ENSDARG000 | 3;3;3;3;3;3;3; | 34120191:34 | 34120331:34 | +;+;+;+;+;+;+; | 1985 | aldh3b1 | 116 | 98 | 84 | 44 | 74 | 84 |
| ENSDARG000 | 3;3;3;3;3;3;3; | 61193890:61 | 61194182:61 | -;-;-;-;-;-;-;- | 669 | pvalb1 | 10 | 26 | 10 | 18 | 52 | 22 |
| ENSDARG000 | 3;3;3;3;3;3;3; | 61208544:61 | 61209477:61 | -;-;-;-;-;-;-;-; | 1855 | pvalb5 | 0 | 0 | 0 | 0 | 0 | 0 |
| ENSDARG000 | 3;3;3;3;3;3;3; | 61221194:61 | 61221664:61 | +;+;+;+;+;+;+; | 8767 | lmtk2 | 1183 | 1174 | 1399 | 605 | 1644 | 1370 |
| ENSDARG000 | 3;3;3;3;3;3;3; | 61292935:61 | 61293171:61 | -;-;-;-;-;-;-;-; | 3584 | TECPR1 | 76 | 21 | 153 | 90 | 176 | 130 |
| ENSDARG000 | 3;3 | 61342019:61 | 61343702:61 | -;- | 2041 | NA | 230 | 120 | 179 | 75 | 185 | 288 |
| ENSDARG000 | 3;3 | 61349155:61 | 61352084:61 | -;- | 3120 | NA | 136 | 64 | 116 | 78 | 159 | 196 |
| ENSDARG000 | 3;3 | 61357949:61 | 61360314:61 | -;- | 3529 | NA | 425 | 389 | 376 | 263 | 500 | 349 |
| ENSDARG000 | 3;3;3 | 61382190:61 | 61384106:61 | -;-;- | 2088 | NA | 293 | 319 | 310 | 169 | 291 | 215 |
| ENSDARG000 | 3;3;3;3;3 | 61371434:61 | 61373433:61 | -;-;-;-;- | 3686 | NA | 100 | 136 | 171 | 114 | 173 | 186 |
| ENSDARG000 | 3;3;3;3 | 61364830:61 | 61366515:61 | -;-;-;- | 2014 | NA | 150 | 138 | 146 | 45 | 173 | 243 |
| ENSDARG000 | 3;3;3 | 61391636:61 | 61392067:61 | +;+;+ | 2990 | bri3 | 7645 | 6213 | 5476 | 8228 | 6108 | 8499 |
| ENSDARG000 | 3;3;3;3;3;3;3 | 61531215:61 | 61531381:61 | +;+;+;+;+;+;+ | 2442 | NA | 78 | 0 | 100 | 59 | 53 | 87 |
| ENSDARG000 | 3;3;3;3;3;3 | 61522759:61 | 61523089:61 | +;+;+;+;+;+ | 1939 | NA | 0 | 0 | 0 | 8 | 0 | 0 |
| ENSDARG000 | 3;3;3;3;3 | 61540527:61 | 61540864:61 | +;+;+;+;+ | 1636 | zgc:113295 | 157 | 50 | 132 | 101 | 120 | 88 |
| ENSDARG000 | 3;3;3;3;3;3 | 36248851:36 | 36249958:36 | -;-;-;-;-;- | 1849 | rac3a | 4 | 42 | 36 | 50 | 8 | 14 |
| ENSDARG000 | 3;3;3;3;3;3;3; | 61566374:61 | 61566828:61 | -;-;-;-;-;-;-;-; | 2346 | nptx2a | 4 | 0 | 8 | 19 | 0 | 0 |
| ENSDARG000 | 3;3;3;3;3;3;3; | 62353650:62 | 62353788:62 | +;+;+;+;+;+;+; | 1075 | iqck | 15 | 4 | 38 | 20 | 49 | 21 |
| ENSDARG000 | 3;3;3;3;3;3;3; | 34129505:34 | 34130080:34 | -;-;-;-;-;-;-;-; | 1584 | clpp | 495 | 598 | 665 | 396 | 598 | 364 |
| ENSDARG000 | 3;3;3;3 | 62367307:62 | 62367785:62 | -;-;-;- | 2087 | gprc5ba | 2 | 4 | 20 | 6 | 17 | 10 |
| ENSDARG000 | 3;3;3;3;3;3;3; | 62382661:62 | 62384623:62 | -;-;-;-;-;-;-;-; | 3411 | proza | 2407 | 860 | 1841 | 1577 | 2088 | 2318 |

| | | | | | | | | | | | | |
|---|---|---|---|---|---|---|---|---|---|---|---|---|
| ENSDARG000 | 3;3;3 | 62397718:623 | 62398893:623 | -;-;- | 1902 | sox8b | 4 | 4 | 12 | 0 | 12 | 12 |
| ENSDARG000 | 3;3 | 62432901:624 | 62433144:624 | +;+ | 509 | NA | 85 | 69 | 86 | 50 | 48 | 67 |
| ENSDARG000 | 3;3;3;3;3 | 62521041:625 | 62523641:625 | -;-;-;-;- | 3511 | sox9b | 70 | 4 | 38 | 4 | 54 | 52 |
| ENSDARG000 | 3;3;3;3;3;3;3; | 36262593:362 | 36265545:362 | -;-;-;-;-;-;-;-; | 3964 | prkar1aa | 7738 | 5558 | 6806 | 5666 | 6222 | 7277 |
| ENSDARG000 | 3;3;3;3;3;3;3; | 36284986:362 | 36285354:362 | +;+;+;+;+;+;+; | 2458 | wipi1 | 822 | 594 | 762 | 818 | 506 | 455 |
| ENSDARG000 | 3;3;3;3;3;3;3; | 36275633:362 | 36275824:362 | +;+;+;+;+;+;+; | 4595 | zgc:86896 | 74 | 20 | 20 | 7 | 32 | 28 |
| ENSDARG000 | 3;3;3;3;3;3 | 36313532:363 | 36313889:363 | +;+;+;+;+;+ | 4909 | slc16a6b | 932 | 1043 | 1001 | 707 | 1712 | 779 |
| ENSDARG000 | 3;3;3;3 | 36332579:363 | 36333569:363 | -;-;-;- | 1574 | gna13b | 765 | 720 | 970 | 711 | 878 | 736 |
| ENSDARG000 | 3;3;3;3;3;3;3; | 36366053:363 | 36366777:363 | +;+;+;+;+;+;+; | 4317 | rgs9a | 2 | 8 | 23 | 73 | 14 | 12 |
| ENSDARG000 | 3;3;3;3;3;3;3; | 36401105:364 | 36402984:364 | -;-;-;-;-;-;-;-; | 5351 | COG1 | 258 | 335 | 370 | 225 | 411 | 272 |
| ENSDARG000 | 3;3;3;3;3;3;3; | 10914113:109 | 10914256:109 | -;-;-;-;-;-;-;-; | 1694 | NA | 0 | 0 | 12 | 3 | 0 | 2 |
| ENSDARG000 | 3;3;3;3 | 11585298:115 | 11587175:115 | -;-;-;- | 2945 | hlfa | 2301 | 1369 | 1169 | 1616 | 2513 | 2072 |
| ENSDARG000 | 3;3;3 | 6278752:628( | 6280065:628( | -;-;- | 1484 | NA | 0 | 0 | 0 | 0 | 0 | 0 |
| ENSDARG000 | 3;3 | 7001608:700( | 7001719:701 | +;+ | 5192 | NA | 49 | 62 | 68 | 33 | 39 | 14 |
| ENSDARG000 | 3;3;3;3;3;3;3; | 13511984:135 | 13511988:135 | +;+;+;+;+;+;+; | 624 | JMJD8 | 40 | 71 | 54 | 49 | 63 | 39 |
| ENSDARG000 | 3;3;3;3;3;3;3; | 22334012:223 | 22334026:223 | +;+;+;+;+;+;+; | 1275 | ifnphi3 | 0 | 1 | 4 | 16 | 0 | 0 |
| ENSDARG000 | 3;3;3;3;3;3;3; | 21385251:213 | 21385421:213 | -;-;-;-;-;-;-;-; | 1035 | NA | 0 | 0 | 0 | 0 | 0 | 0 |
| ENSDARG000 | 3;3;3 | 15176737:151 | 15177648:151 | +;+;+ | 1866 | TSPAN10 | 4234 | 2760 | 2183 | 4066 | 4719 | 5456 |
| ENSDARG000 | 3;3;3;3;3;3;3; | 32979969:329 | 32980342:329 | -;-;-;-;-;-;-;-; | 3654 | NA | 16 | 31 | 24 | 17 | 127 | 85 |
| ENSDARG000 | 3;3 | 13482542:134 | 13482550:134 | +;+ | 5042 | NA | 0 | 0 | 0 | 0 | 0 | 0 |
| ENSDARG000 | 3;3;3;3;3;3 | 5882130:588( | 5883022:589( | -;-;-;-;-;- | 5851 | NA | 1257 | 1395 | 410 | 482 | 353 | 728 |
| ENSDARG000 | 3;3;3;3;3;3;3; | 19732173:197 | 19732284:197 | +;+;+;+;+;+;+; | 4422 | MRC2 | 29 | 6 | 22 | 8 | 70 | 8 |
| ENSDARG000 | 3 | 30694783 | 30694872 | + | 90 | NA | 0 | 0 | 0 | 0 | 0 | 0 |
| ENSDARG000 | 3;3;3;3;3;3;3; | 17689419:176 | 17689592:176 | +;+;+;+;+;+;+; | 2052 | dhx58 | 716 | 923 | 204 | 64 | 84 | 150 |
| ENSDARG000 | 3;3;3 | 7133388:713 | 7133539:713 | -;-;- | 627 | NA | 16 | 10 | 109 | 37 | 39 | 97 |
| ENSDARG000 | 3;3;3;3;3;3;3; | 12128117:121 | 12132906:121 | -;-;-;-;-;-;-;-; | 5766 | srl | 0 | 0 | 0 | 0 | 6 | 4 |
| ENSDARG000 | 3;3;3;3 | 27929505:279 | 27930115:279 | +;+;+;+ | 1945 | TMEM114 | 2 | 6 | 0 | 0 | 0 | 0 |
| ENSDARG000 | 3;3;3;3;3;3 | 29881035:298 | 29881056:298 | +;+;+;+;+;+ | 141 | NA | 0 | 0 | 0 | 0 | 4 | 7 |
| ENSDARG000 | 3;3;3 | 1637358:163 | 1637379:163 | +;+;+ | 1236 | NA | 3 | 0 | 0 | 0 | 0 | 0 |
| ENSDARG000 | 3;3;3;3;3;3 | 3810919:381 | 3811002:381 | +;+;+;+;+;+ | 609 | PMM1 | 39 | 40 | 112 | 217 | 52 | 74 |
| ENSDARG000 | 3;3;3;3;3;3 | 19487707:194 | 19488953:194 | -;-;-;-;-;- | 1910 | NA | 8 | 1 | 0 | 0 | 0 | 0 |
| ENSDARG000 | 3;3;3;3;3 | 31851004:318 | 31851022:318 | +;+;+;+;+ | 120 | NA | 0 | 0 | 0 | 0 | 0 | 0 |
| ENSDARG000 | 3;3;3;3;3;3;3; | 40492:40492: | 41195:41195: | -;-;-;-;-;-;-;-; | 1177 | NA | 40 | 29 | 1 | 20 | 17 | 26 |
| ENSDARG000 | 3;3;3;3;3;3;3; | 3641429:365( | 3642235:365( | +;+;+;+;+;+;+; | 2351 | plbd1a | 738 | 351 | 368 | 315 | 550 | 572 |
| ENSDARG000 | 3;3;3;3;3;3;3; | 1323732:133 | 1325245:133 | -;-;-;-;-;-;-;- | 2315 | tmem106c | 241 | 187 | 177 | 142 | 223 | 243 |
| ENSDARG000 | 3 | 39649445 | 39649562 | - | 118 | 5S_rRNA | 0 | 0 | 0 | 0 | 0 | 0 |
| ENSDARG000 | 3;3 | 9379877:941( | 9380820:942 | +;+ | 6815 | GPR139 | 2 | 0 | 0 | 0 | 0 | 2 |
| ENSDARG000 | 3;3;3 | 58218661:582 | 58222046:582 | -;-;- | 4170 | NA | 60 | 71 | 26 | 180 | 52 | 46 |
| ENSDARG000 | 3 | 40088685 | 40088801 | - | 117 | 5S_rRNA | 0 | 0 | 0 | 0 | 0 | 0 |
| ENSDARG000 | 3 | 9041989 | 9042095 | + | 107 | U6 | 0 | 0 | 0 | 0 | 0 | 0 |
| ENSDARG000 | 3;3;3;3 | 11371786:113 | 11372003:113 | -;-;-;- | 1056 | NA | 0 | 1 | 2 | 0 | 0 | 0 |
| ENSDARG000 | 3;3 | 7111704:711 | 7114108:711 | -;- | 2511 | NA | 152 | 80 | 181 | 186 | 84 | 156 |
| ENSDARG000 | 3;3;3;3;3;3;3; | 19446997:194 | 19447109:194 | +;+;+;+;+;+;+; | 1496 | zgc:123297 | 0 | 2 | 0 | 0 | 0 | 0 |
| ENSDARG000 | 3;3 | 1979271:198 | 1981317:198 | -;- | 2185 | NA | 0 | 0 | 0 | 0 | 0 | 13 |
| ENSDARG000 | 3;3;3;3;3;3;3; | 53491105:534 | 53491499:534 | -;-;-;-;-;-;-;-; | 1263 | shfl | 12 | 28 | 8 | 10 | 21 | 16 |
| ENSDARG000 | 3;3;3 | 3016408:301 | 3019310:301 | -;-;- | 3071 | zgc:172090 | 8 | 20 | 2 | 0 | 32 | 0 |
| ENSDARG000 | 3;3 | 1735214:173 | 1735540:173 | +;+ | 2272 | NA | 0 | 0 | 139 | 0 | 25 | 18 |
| ENSDARG000 | 3;3;3;3;3 | 31647540:31( | 31648372:31( | -;-;-;-;- | 1619 | NA | 0 | 0 | 0 | 0 | 0 | 0 |
| ENSDARG000 | 3 | 35960989 | 35961086 | + | 98 | dre-mir-365-2 | 6 | 4 | 0 | 0 | 13 | 0 |
| ENSDARG000 | 3 | 24095957 | 24096036 | + | 80 | dre-mir-152 | 0 | 0 | 0 | 0 | 0 | 0 |
| ENSDARG000 | 3;3;3;3;3;3;3; | 49576662:495 | 49576735:495 | +;+;+;+;+;+;+; | 2266 | NA | 3 | 0 | 0 | 0 | 0 | 0 |
| ENSDARG000 | 3;3;3;3;3;3;3; | 1303862:130 | 1304000:130 | -;-;-;-;-;-;-;-; | 4365 | GRIN2B | 0 | 2 | 12 | 10 | 4 | 18 |
| ENSDARG000 | 3 | 34158875 | 34159072 | + | 198 | SNORA74 | 0 | 0 | 2 | 0 | 0 | 0 |
| ENSDARG000 | 3 | 22568987 | 22569106 | + | 120 | 5S_rRNA | 0 | 0 | 0 | 0 | 0 | 0 |
| ENSDARG000 | 3;3;3 | 709393:7995 | 709605:7998 | +;+;+ | 866 | NA | 7 | 6 | 0 | 4 | 16 | 9 |
| ENSDARG000 | 3 | 18917461 | 18917583 | + | 123 | 5S_rRNA | 0 | 0 | 0 | 0 | 0 | 0 |
| ENSDARG000 | 3 | 30800541 | 30800658 | - | 118 | 5S_rRNA | 0 | 0 | 0 | 0 | 0 | 0 |
| ENSDARG000 | 3 | 43716827 | 43716915 | + | 89 | dre-mir-365-1 | 0 | 0 | 0 | 0 | 0 | 0 |
| ENSDARG000 | 3;3;3;3 | 10743236:107 | 10743287:107 | -;-;-;- | 1077 | NA | 0 | 0 | 0 | 0 | 0 | 0 |
| ENSDARG000 | 3;3;3;3;3;3;3; | 62173396:621 | 62173687:621 | -;-;-;-;-;-;-;-; | 2694 | tbl3 | 776 | 880 | 1307 | 452 | 863 | 666 |
| ENSDARG000 | 3 | 42497814 | 42504515 | + | 6702 | NA | 3 | 2 | 6 | 0 | 0 | 0 |
| ENSDARG000 | 3 | 23720651 | 23720745 | + | 95 | dre-mir-10c | 0 | 0 | 0 | 0 | 0 | 0 |
| ENSDARG000 | 3 | 9037246 | 9037352 | + | 107 | U6 | 0 | 0 | 0 | 0 | 0 | 0 |
| ENSDARG000 | 3 | 49077762 | 49077879 | - | 118 | 5S_rRNA | 0 | 0 | 0 | 0 | 0 | 0 |
| ENSDARG000 | 3 | 19349719 | 19349817 | + | 99 | NA | 1 | 0 | 3 | 0 | 0 | 0 |
| ENSDARG000 | 3 | 21616920 | 21617040 | - | 121 | 5S_rRNA | 0 | 0 | 0 | 0 | 0 | 0 |
| ENSDARG000 | 3 | 3598555 | 3603348 | + | 4794 | NA | 24 | 30 | 0 | 2 | 10 | 2 |
| ENSDARG000 | 3;3;3 | 60851049:608 | 60851197:608 | -;-;- | 1026 | DXO | 64 | 28 | 65 | 48 | 83 | 52 |
| ENSDARG000 | 3;3;3;3;3 | 60716904:607 | 60717112:607 | +;+;+;+;+ | 4773 | foxj1a | 0 | 0 | 8 | 10 | 28 | 10 |
| ENSDARG000 | 3 | 19312991 | 19313107 | + | 117 | 5S_rRNA | 0 | 0 | 0 | 0 | 0 | 0 |
| ENSDARG000 | 3;3;3;3;3;3;3; | 56699187:567 | 56699354:567 | +;+;+;+;+;+;+; | 16192 | rnf213a | 8013 | 3488 | 3042 | 1614 | 3218 | 3357 |
| ENSDARG000 | 3 | 62416556 | 62417677 | - | 1122 | SSTR2 | 0 | 0 | 0 | 0 | 0 | 0 |
| ENSDARG000 | 3 | 9914272 | 9914389 | - | 118 | 5S_rRNA | 0 | 0 | 0 | 0 | 0 | 0 |
| ENSDARG000 | 3;3;3;3;3;3;3; | 48584684:485 | 48587251:485 | -;-;-;-;-;-;-;-; | 3596 | ndel1b | 263 | 223 | 371 | 355 | 363 | 190 |
| ENSDARG000 | 3;3;3;3;3;3;3; | 49757578:497 | 49760694:497 | -;-;-;-;-;-;-;-; | 4712 | NA | 16 | 2 | 12 | 8 | 4 | 0 |
| ENSDARG000 | 3;3;3;3;3;3 | 48221413:482 | 48221610:482 | +;+;+;+;+;+ | 1335 | fn3krp | 41 | 45 | 82 | 43 | 122 | 74 |
| ENSDARG000 | 3;3;3;3;3;3 | 2105598:210 | 2105704:210 | -;-;-;-;-;- | 1970 | NA | 0 | 0 | 0 | 0 | 0 | 0 |
| ENSDARG000 | 3 | 5697609 | 5697732 | - | 124 | 5S_rRNA | 0 | 0 | 0 | 0 | 3 | 0 |
| ENSDARG000 | 3 | 37787818 | 37787902 | + | 85 | NA | 0 | 8 | 16 | 2 | 16 | 2 |
| ENSDARG000 | 3 | 43714699 | 43714783 | + | 85 | dre-mir-193b | 0 | 0 | 0 | 0 | 0 | 0 |
| ENSDARG000 | 3 | 35743184 | 35743256 | + | 73 | NA | 0 | 0 | 0 | 0 | 0 | 0 |
| ENSDARG000 | 3;3;3;3;3 | 60589157:605 | 60589935:605 | +;+;+;+;+ | 1523 | mettl23 | 66 | 98 | 221 | 143 | 86 | 41 |
| ENSDARG000 | 3;3;3;3;3;3;3; | 31875772:318 | 31877437:318 | -;-;-;-;-;-;-;- | 2909 | zp3e | 0 | 2 | 12 | 0 | 0 | 0 |
| ENSDARG000 | 3 | 9036190 | 9036296 | + | 107 | U6 | 0 | 0 | 0 | 0 | 0 | 0 |
| ENSDARG000 | 3;3;3;3;3;3;3; | 9634777:964 | 9642035:964 | -;-;-;-;-;-;-;-; | 12266 | CREBBP | 2554 | 1673 | 3393 | 2121 | 2980 | 2239 |
| ENSDARG000 | 3;3 | 3693417:369 | 3693779:369 | +;+ | 1764 | NA | 0 | 0 | 0 | 0 | 0 | 2 |
| ENSDARG000 | 3;3 | 32139666:321 | 32139770:321 | +;+ | 279 | NA | 0 | 0 | 0 | 0 | 0 | 10 |
| ENSDARG000 | 3 | 3935115 | 3939785 | - | 4671 | NA | 0 | 0 | 2 | 0 | 0 | 0 |
| ENSDARG000 | 3;3;3 | 866044:8692 | 866461:8698 | +;+;+ | 3050 | NA | 0 | 0 | 1 | 0 | 0 | 2 |
| ENSDARG000 | 3;3;3;3 | 3046909:304 | 3047060:304 | +;+;+;+ | 5782 | NA | 16 | 44 | 0 | 2 | 36 | 0 |
| ENSDARG000 | 3;3;3;3;3 | 24793921:248 | 24794381:248 | -;-;-;-;- | 2098 | NA | 0 | 0 | 0 | 0 | 0 | 0 |
| ENSDARG000 | 3;3;3;3;3;3;3; | 726000:7264 | 726054:7267 | +;+;+;+;+;+;+; | 918 | dicp1.17 | 0 | 7 | 0 | 2 | 37 | 9 |
| ENSDARG000 | 3;3;3;3;3;3;3; | 55684449:556 | 55685021:556 | -;-;-;-;-;-;-;-; | 3618 | cep112 | 58 | 134 | 188 | 48 | 132 | 93 |
| ENSDARG000 | 3;3 | 17715685:177 | 17716084:177 | -;- | 405 | NA | 0 | 0 | 0 | 0 | 0 | 0 |
| ENSDARG000 | 3 | 41312022 | 41312143 | - | 122 | 5S_rRNA | 0 | 0 | 0 | 0 | 0 | 0 |
| ENSDARG000 | 3 | 27407096 | 27407211 | + | 116 | 5S_rRNA | 0 | 0 | 0 | 0 | 0 | 0 |
| ENSDARG000 | 3 | 33424192 | 33424328 | - | 137 | NA | 2 | 3 | 2 | 0 | 4 | 0 |
| ENSDARG000 | 3 | 31788521 | 31788637 | + | 117 | 5S_rRNA | 0 | 0 | 0 | 0 | 1 | 0 |
| ENSDARG000 | 3 | 33761549 | 33762726 | + | 1178 | ier2a | 14924 | 7818 | 12310 | 17517 | 7117 | 12588 |
| ENSDARG000 | 3 | 36170943 | 36171968 | - | 1026 | NA | 0 | 0 | 0 | 0 | 0 | 0 |
| ENSDARG000 | 3;3;3;3;3;3;3; | 56645710:566 | 56645772:566 | +;+;+;+;+;+;+; | 5691 | NA | 0 | 0 | 0 | 0 | 0 | 0 |
| ENSDARG000 | 3;3;3;3;3;3;3; | 48638594:486 | 48639069:486 | -;-;-;-;-;-;-;-; | 2877 | ARHGAP44 | 27 | 24 | 38 | 16 | 32 | 26 |
| ENSDARG000 | 3;3;3;3;3 | 58608169:586 | 58608385:586 | -;-;-;-;- | 3246 | NA | 267 | 298 | 251 | 200 | 198 | 123 |
| ENSDARG000 | 3;3 | 2669813:267 | 2670510:267 | +;+ | 1642 | NA | 0 | 0 | 0 | 0 | 0 | 0 |
| ENSDARG000 | 3;3;3;3;3;3 | 47322494:473 | 47322934:473 | +;+;+;+;+;+ | 1613 | ppap2d | 204 | 259 | 486 | 212 | 556 | 244 |
| ENSDARG000 | 3;3;3 | 62317990:623 | 62318100:623 | +;+;+ | 2027 | znf1124 | 95 | 68 | 78 | 75 | 62 | 83 |
| ENSDARG000 | 3;3 | 532862:5396 | 532966:5456 | +;+ | 6166 | NA | 6 | 0 | 2 | 0 | 0 | 0 |
| ENSDARG000 | 3;3;3;3 | 5008971:501( | 5009133:501( | -;-;-;- | 1220 | CSDC2 | 1227 | 470 | 481 | 975 | 526 | 192 |
| ENSDARG000 | 3;3;3;3;3;3;3; | 66842:66845: | 66944:66944: | -;-;-;-;-;-;-;-; | 4588 | zgc:165518 | 0 | 0 | 0 | 0 | 2 | 6 |
| ENSDARG000 | 3;3;3;3;3 | 48561112:485 | 48561739:485 | +;+;+;+;+ | 2293 | SLC16A5 | 0 | 2 | 4 | 0 | 0 | 0 |
| ENSDARG000 | 3;3;3;3;3;3;3 | 1689220:169 | 1689269:169 | +;+;+;+;+;+;+ | 3035 | NA | 6 | 8 | 0 | 4 | 4 | 0 |
| ENSDARG000 | 3 | 54067523 | 54067639 | + | 117 | 5S_rRNA | 0 | 0 | 0 | 0 | 0 | 0 |
| ENSDARG000 | 3;3;3 | 47281236:472 | 47284303:472 | -;-;- | 3475 | tnfsf18 | 10 | 7 | 4 | 8 | 20 | 16 |
| ENSDARG000 | 3 | 3383090 | 3383207 | - | 118 | 5S_rRNA | 0 | 0 | 0 | 0 | 0 | 0 |
| ENSDARG000 | 3;3;3;3;3;3 | 6469754:647 | 6469913:647 | +;+;+;+;+;+ | 717 | NUP85 | 134 | 198 | 301 | 154 | 240 | 137 |
| ENSDARG000 | 3;3;3;3;3;3;3 | 60862441:608 | 60862754:608 | -;-;-;-;-;-;- | 786 | PITPNC1 | 0 | 8 | 10 | 0 | 12 | 2 |
| ENSDARG000 | 3;3;3;3;3;3;3; | 47498433:474 | 47499328:474 | -;-;-;-;-;-;-;-; | 5574 | ADGRL1 | 0 | 0 | 0 | 0 | 0 | 2 |
| ENSDARG000 | 3;3 | 7245170:725 | 7246842:725 | -;- | 1683 | NA | 14 | 0 | 16 | 5 | 2 | 10 |
| ENSDARG000 | 3;3;3;3;3;3;3; | 53476155:534 | 53476345:534 | -;-;-;-;-;-;-;- | 1842 | NA | 20 | 10 | 58 | 4 | 56 | 8 |
| ENSDARG000 | 3;3 | 2070039:207 | 2072311:207 | -;- | 2356 | NA | 0 | 0 | 0 | 0 | 0 | 0 |
| ENSDARG000 | 3;3 | 7123912:712 | 7124783:712 | -;- | 1015 | NA | 42 | 41 | 162 | 102 | 42 | 137 |
| ENSDARG000 | 3;3;3 | 62205858:622 | 62205912:622 | +;+;+ | 759 | znf1024 | 68 | 53 | 70 | 73 | 95 | 148 |
| ENSDARG000 | 3 | 4755681 | 4760384 | - | 4704 | NA | 0 | 0 | 13 | 0 | 4 | 6 |
| ENSDARG000 | 3 | 27083298 | 27083414 | - | 117 | 5S_rRNA | 0 | 0 | 0 | 0 | 0 | 0 |
| ENSDARG000 | 3 | 9039887 | 9039993 | + | 107 | U6 | 0 | 0 | 0 | 0 | 0 | 0 |
| ENSDARG000 | 3 | 9038830 | 9038936 | + | 107 | U6 | 0 | 0 | 0 | 0 | 0 | 0 |
| ENSDARG000 | 3;3;3 | 1749793:175( | 1750201:175( | +;+;+ | 1783 | NA | 0 | 0 | 4 | 0 | 14 | 3 |
| ENSDARG000 | 3;3;3 | 49150822:491 | 49151350:491 | -;-;- | 1593 | dnajb1a | 274 | 120 | 208 | 140 | 269 | 226 |
| ENSDARG000 | 3 | 30248739 | 30249036 | + | 298 | Metazoa_SRP | 0 | 8 | 103 | 10 | 0 | 60 |
| ENSDARG000 | 3;3 | 62339264:623 | 62339318:623 | +;+ | 945 | NA | 77 | 38 | 30 | 33 | 44 | 82 |
| ENSDARG000 | 3;3;3;3 | 49521106:495 | 49521345:495 | +;+;+;+ | 2458 | crb3a | 28 | 8 | 34 | 10 | 31 | 2 |
| ENSDARG000 | 3;3;3;3;3;3;3; | 59880317:598 | 59880577:598 | +;+;+;+;+;+;+; | 2937 | alyref | 3124 | 5278 | 4508 | 2780 | 3459 | 2041 |
| ENSDARG000 | 3 | 23673377 | 23673455 | + | 79 | mir196b | 0 | 0 | 0 | 0 | 0 | 0 |
| ENSDARG000 | 3;3;3;3 | 8758286:876 | 8759887:876 | -;-;-;- | 2093 | NA | 376 | 213 | 248 | 18 | 151 | 94 |
| ENSDARG000 | 3;3;3;3;3;3;3; | 61401311:614 | 61402288:614 | -;-;-;-;-;-;-;-; | 2592 | baiap2l1b | 20 | 14 | 100 | 138 | 10 | 24 |
| ENSDARG000 | 3;3;3;3;3;3;3; | 59899452:599 | 59899577:599 | +;+;+;+;+;+;+; | 2202 | arhgdia | 7564 | 8836 | 8640 | 7398 | 8830 | 7996 |
| ENSDARG000 | 3;3;3;3;3;3;3; | 62064805:62( | 62064938:62( | -;-;-;-;-;-;-;-; | 2826 | srebf1 | 1724 | 1085 | 1052 | 380 | 1027 | 806 |
| ENSDARG000 | 3;3;3;3;3;3;3; | 31201542:312 | 31201738:312 | -;-;-;-;-;-;-;-; | 1537 | SYT17 | 3 | 0 | 4 | 2 | 0 | 0 |
| ENSDARG000 | 3 | 45635077 | 45635186 | + | 110 | dre-mir-2195 | 0 | 0 | 0 | 0 | 0 | 0 |
| ENSDARG000 | 3;3 | 56565300:565 | 56565945:565 | -;- | 915 | SMIM10 | 108 | 133 | 144 | 196 | 114 | 168 |
| ENSDARG000 | 3;3;3;3;3;3;3 | 49425793:494 | 49426351:494 | -;-;-;-;-;-;- | 1681 | samd1a | 168 | 304 | 256 | 126 | 258 | 170 |
| ENSDARG000 | 3;3 | 7152770:717( | 7156159:717( | -;- | 3533 | NA | 75 | 90 | 89 | 108 | 88 | 120 |
| ENSDARG000 | 3;3;3;3;3;3;3; | 49459828:494 | 49460521:494 | -;-;-;-;-;-;-;-; | 2199 | prkacaa | 3349 | 1808 | 1818 | 1494 | 2074 | 1744 |
| ENSDARG000 | 3;3;3;3;3;3;3; | 32933663:329 | 32933930:329 | +;+;+;+;+;+;+; | 4525 | NA | 1123 | 762 | 1154 | 898 | 1063 | 1043 |
| ENSDARG000 | 3 | 29868635 | 29868732 | - | 98 | NA | 6 | 7 | 17 | 3 | 11 | 4 |
| ENSDARG000 | 3 | 57991074 | 57991985 | + | 912 | MYADML2 | 0 | 0 | 0 | 0 | 8 | 0 |
| ENSDARG000 | 3;3;3;3;3;3;3; | 57038033:57( | 57038329:57( | +;+;+;+;+;+;+; | 8941 | NA | 247 | 115 | 523 | 91 | 416 | 214 |
| ENSDARG000 | 3;3;3;3;3;3 | 49043917:49( | 49044157:49( | +;+;+;+;+;+ | 1874 | zgc:92161 | 31 | 26 | 2 | 6 | 36 | 26 |
| ENSDARG000 | 3 | 18867835 | 18867952 | + | 118 | 5S_rRNA | 0 | 0 | 0 | 0 | 0 | 0 |
| ENSDARG000 | 3;3 | 60044780:60( | 60044882:60( | +;+ | 1659 | NA | 37 | 21 | 31 | 17 | 36 | 10 |
| ENSDARG000 | 3 | 33948669 | 33948788 | - | 120 | 5S_rRNA | 0 | 0 | 0 | 0 | 0 | 0 |
| ENSDARG000 | 3 | 10213098 | 10213220 | - | 123 | 5S_rRNA | 0 | 0 | 0 | 0 | 0 | 0 |
| ENSDARG000 | 3;3;3 | 196316:2055 | 198966:2056 | -;-;- | 3479 | NA | 6 | 0 | 10 | 2 | 0 | 8 |
| ENSDARG000 | 3 | 9040417 | 9040523 | + | 107 | U6 | 0 | 0 | 0 | 0 | 0 | 0 |
| ENSDARG000 | 3 | 29167020 | 29167147 | + | 128 | 5S_rRNA | 0 | 0 | 0 | 0 | 0 | 0 |
| ENSDARG000 | 3;3;3;3 | 59070401:59( | 59070559:59( | -;-;-;- | 516 | prpsap1 | 1354 | 757 | 1137 | 549 | 751 | 777 |
| ENSDARG000 | 3;3 | 31257172:312 | 31257858:312 | -;- | 821 | NA | 6 | 0 | 0 | 0 | 6 | 0 |
| ENSDARG000 | 3;3;3;3;3;3 | 58197025:581 | 58197190:581 | -;-;-;-;-;- | 2625 | NA | 234 | 161 | 55 | 399 | 248 | 256 |
| ENSDARG000 | 3;3;3;3;3;3;3; | 12977:12977: | 14075:14075: | -;-;-;-;-;-;-;- | 1626 | zgc:110249 | 77 | 60 | 0 | 112 | 32 | 25 |
| ENSDARG000 | 3 | 55299043 | 55299158 | + | 116 | 5S_rRNA | 0 | 0 | 0 | 0 | 0 | 0 |
| ENSDARG000 | 3;3 | 832788:8340 | 832815:8361 | +;+ | 2073 | NA | 0 | 0 | 1 | 0 | 0 | 4 |
| ENSDARG000 | 3;3;3;3;3;3;3; | 15425785:154 | 15426133:154 | -;-;-;-;-;-;-;-; | 1420 | lat | 2 | 14 | 4 | 14 | 16 | 44 |

| | | | | | | | | | | | |
|---|---|---|---|---|---|---|---|---|---|---|---|
| ENSDARG000 | 3;3;3 | 3681116:3685 | 3681155:3685 | +;+;+ | 1162 | art4 | 2 | 0 | 0 | 2 | 0 | 0 |
| ENSDARG000 | 3;3;3;3 | 32066983:320 | 32067551:320 | -;-;-;- | 953 | NA | 0 | 14 | 0 | 0 | 0 | 10 |
| ENSDARG000 | 3;3;3 | 7435802:7436 | 7435980:7436 | -;-;- | 987 | NA | 12 | 9 | 0 | 17 | 6 | 21 |
| ENSDARG000 | 3 | 37788476 | 37788560 | + | 85 | NA | 0 | 2 | 1 | 0 | 0 | 0 |
| ENSDARG000 | 3;3;3;3;3;3;3; | 59951010:599 | 59952400:599 | -;-;-;-;-;-;-;-; | 2358 | cdr2l | 336 | 281 | 273 | 270 | 456 | 278 |
| ENSDARG000 | 3 | 33407588 | 33407710 | + | 123 | 5S_rRNA | 0 | 0 | 0 | 0 | 0 | 0 |
| ENSDARG000 | 3;3;3;3;3;3 | 60575113:605 | 60579742:605 | -;-;-;-;-;- | 5710 | jmjd6 | 420 | 708 | 1057 | 724 | 639 | 422 |
| ENSDARG000 | 3;3;3;3;3;3;3; | 53352018:533 | 53352889:533 | +;+;+;+;+;+;+; | 5002 | NA | 48 | 18 | 54 | 14 | 74 | 14 |
| ENSDARG000 | 3 | 39182746 | 39182862 | + | 117 | 5S_rRNA | 0 | 0 | 0 | 0 | 0 | 0 |
| ENSDARG000 | 3;3;3;3 | 49559261:495 | 49560649:495 | -;-;-;- | 1750 | tubb4bl | 2 | 2 | 28 | 0 | 6 | 0 |
| ENSDARG000 | 3;3 | 6895555:6902 | 6896606:6902 | -;- | 1157 | NA | 41 | 45 | 40 | 15 | 49 | 50 |
| ENSDARG000 | 3 | 9041462 | 9041568 | + | 107 | U6 | 0 | 0 | 0 | 0 | 0 | 0 |
| ENSDARG000 | 3 | 32571309 | 32571371 | + | 63 | mir150 | 0 | 0 | 0 | 0 | 0 | 0 |
| ENSDARG000 | 3;3;3;3;3;3;3; | 9588705:9588 | 9588890:9588 | +;+;+;+;+;+;+; | 3039 | trap1 | 929 | 978 | 1308 | 564 | 1169 | 853 |
| ENSDARG000 | 3 | 7545733 | 7546740 | - | 1008 | NA | 18 | 16 | 18 | 36 | 19 | 66 |
| ENSDARG000 | 3;3;3;3;3;3;3; | 3454610:3454 | 3454792:3455 | +;+;+;+;+;+;+; | 4233 | NA | 0 | 0 | 0 | 2 | 0 | 0 |
| ENSDARG000 | 3;3 | 146153:14731 | 146940:14736 | -;- | 843 | NA | 14 | 24 | 7 | 10 | 30 | 4 |
| ENSDARG000 | 3;3;3 | 13332870:133 | 13332999:133 | +;+;+ | 393 | NA | 0 | 0 | 1 | 0 | 0 | 0 |
| ENSDARG000 | 3;3;3 | 4606718:4608 | 4607076:4608 | -;-;- | 1386 | NA | 87 | 37 | 38 | 16 | 12 | 53 |
| ENSDARG000 | 3;3 | 54616015:546 | 54618019:546 | -;- | 2370 | s1pr2 | 2132 | 2428 | 2520 | 1381 | 2803 | 1821 |
| ENSDARG000 | 3;3;3;3;3;3 | 49103210:491 | 49103742:491 | -;-;-;-;-;- | 2070 | NA | 2 | 0 | 0 | 0 | 3 | 5 |
| ENSDARG000 | 3 | 20679923 | 20680038 | + | 116 | 5S_rRNA | 0 | 0 | 0 | 0 | 0 | 0 |
| ENSDARG000 | 3 | 55483513 | 55483647 | + | 135 | NA | 0 | 6 | 3 | 12 | 8 | 0 |
| ENSDARG000 | 3;3;3 | 59287569:592 | 59287607:592 | -;-;- | 243 | NA | 0 | 0 | 0 | 0 | 0 | 0 |
| ENSDARG000 | 3;3;3;3 | 9485208:9487 | 9485927:9487 | -;-;-;- | 807 | NA | 0 | 0 | 0 | 0 | 4 | 0 |
| ENSDARG000 | 3;3;3;3;3;3;3; | 40672242:406 | 40672248:406 | +;+;+;+;+;+;+; | 1551 | NA | 0 | 0 | 0 | 0 | 0 | 1 |
| ENSDARG000 | 3 | 39884819 | 39884956 | + | 138 | 5S_rRNA | 0 | 0 | 0 | 0 | 0 | 0 |
| ENSDARG000 | 3;3;3;3;3;3;3; | 9542168:9542 | 9542272:9542 | -;-;-;-;-;-;-;-; | 3250 | NA | 0 | 0 | 0 | 0 | 0 | 0 |
| ENSDARG000 | 3;3;3 | 11202622:112 | 11202725:112 | -;-;- | 795 | NA | 0 | 0 | 0 | 0 | 0 | 0 |
| ENSDARG000 | 3 | 26663048 | 26663165 | - | 118 | 5S_rRNA | 0 | 0 | 0 | 0 | 0 | 0 |
| ENSDARG000 | 3;3 | 8158039:8199 | 8159549:8199 | -;- | 1841 | NA | 0 | 0 | 11 | 1 | 15 | 21 |
| ENSDARG000 | 3;3;3;3;3;3 | 49021079:490 | 49021246:490 | +;+;+;+;+;+ | 801 | zgc:163083 | 0 | 0 | 0 | 0 | 0 | 0 |
| ENSDARG000 | 3;3 | 301479:30188 | 301751:30222 | +;+ | 612 | KDELR3 | 0 | 0 | 0 | 0 | 0 | 0 |
| ENSDARG000 | 3;3;3;3;3;3;3; | 43955864:439 | 43956135:439 | +;+;+;+;+;+;+; | 3829 | NA | 4135 | 2874 | 3671 | 4267 | 3248 | 4935 |
| ENSDARG000 | 3;3;3 | 48253202:482 | 48255925:482 | -;-;- | 4932 | znf750 | 60 | 0 | 0 | 0 | 7 | 0 |
| ENSDARG000 | 3;3;3;3;3;3;3 | 49126103:491 | 49126254:491 | -;-;-;-;-;-;- | 1348 | gipc1 | 244 | 131 | 199 | 79 | 157 | 213 |
| ENSDARG000 | 3;3;3;3 | 32977861:329 | 32977919:329 | -;-;-;- | 405 | NA | 28 | 28 | 50 | 15 | 151 | 51 |
| ENSDARG000 | 3;3 | 7395174:7405 | 7396341:7405 | -;- | 1369 | zgc:173517 | 2 | 0 | 0 | 0 | 2 | 0 |
| ENSDARG000 | 3;3;3;3 | 52892671:528 | 52895519:528 | -;-;-;- | 3641 | NA | 67 | 104 | 120 | 54 | 148 | 58 |
| ENSDARG000 | 3;3;3;3 | 49504919:495 | 49505616:495 | -;-;-;- | 1356 | asf1ba | 42 | 290 | 406 | 136 | 234 | 214 |
| ENSDARG000 | 3 | 53689310 | 53689428 | - | 119 | 5S_rRNA | 3 | 0 | 4 | 0 | 0 | 1 |
| ENSDARG000 | 3;3;3;3;3;3 | 62140077:621 | 62140265:621 | +;+;+;+;+;+ | 654 | GID4 | 141 | 118 | 237 | 124 | 157 | 189 |
| ENSDARG000 | 3;3;3;3;3 | 52275691:522 | 52275813:522 | +;+;+;+;+ | 609 | NA | 0 | 1 | 13 | 2 | 4 | 4 |
| ENSDARG000 | 3;3;3;3;3 | 59453467:594 | 59456233:594 | -;-;-;-;- | 4237 | nptx1l | 604 | 735 | 1200 | 913 | 497 | 911 |
| ENSDARG000 | 3 | 9031619 | 9031767 | - | 149 | U1 | 0 | 0 | 0 | 0 | 0 | 0 |
| ENSDARG000 | 3;3;3;3;3;3 | 49097775:490 | 49097943:490 | +;+;+;+;+;+ | 1220 | zgc:123284 | 3 | 3 | 0 | 0 | 0 | 0 |
| ENSDARG000 | 3 | 7463168 | 7464250 | - | 1083 | NA | 0 | 2 | 14 | 0 | 10 | 7 |
| ENSDARG000 | 3;3 | 4911207:4911 | 4911817:4911 | -;- | 711 | NA | 0 | 0 | 0 | 0 | 0 | 0 |
| ENSDARG000 | 3 | 47195137 | 47195234 | - | 98 | dre-mir-199-3 | 0 | 2 | 0 | 0 | 0 | 0 |
| ENSDARG000 | 3;3;3;3;3;3;3; | 23063718:230 | 23063939:230 | +;+;+;+;+;+;+; | 1651 | NA | 2082 | 1670 | 582 | 1031 | 1613 | 1272 |
| ENSDARG000 | 3;3;3;3;3;3;3; | 8506061:8506 | 8506428:8507 | -;-;-;-;-;-;-;-; | 1413 | NA | 0 | 0 | 1 | 0 | 0 | 10 |
| ENSDARG000 | 3;3;3;3;3;3;3; | 59117136:591 | 59117434:591 | +;+;+;+;+;+;+; | 1530 | MGAT5B | 0 | 0 | 0 | 0 | 0 | 0 |
| ENSDARG000 | 3;3;3;3;3;3;3; | 3765831:3765 | 3765885:3766 | -;-;-;-;-;-;-;-; | 3075 | NA | 2121 | 1091 | 1865 | 402 | 1637 | 701 |
| ENSDARG000 | 3;3;3;3 | 57820913:578 | 57820999:578 | +;+;+;+ | 856 | NA | 0 | 4 | 0 | 0 | 0 | 0 |
| ENSDARG000 | 3;3 | 62271320:622 | 62271374:622 | +;+ | 1185 | zgc:174624 | 71 | 45 | 80 | 93 | 48 | 56 |
| ENSDARG000 | 3 | 39957358 | 39957474 | - | 117 | 5S_rRNA | 0 | 0 | 0 | 0 | 0 | 0 |
| ENSDARG000 | 3 | 37788203 | 37788289 | + | 87 | NA | 0 | 2 | 0 | 0 | 1 | 2 |
| ENSDARG000 | 3;3;3;3;3;3;3; | 49397115:493 | 49397698:493 | +;+;+;+;+;+;+; | 4673 | TECR | 0 | 0 | 4 | 0 | 0 | 0 |
| ENSDARG000 | 3 | 34910280 | 34910399 | + | 120 | 5S_rRNA | 0 | 0 | 0 | 0 | 0 | 0 |
| ENSDARG000 | 3 | 28859284 | 28859400 | - | 117 | 5S_rRNA | 0 | 0 | 0 | 0 | 0 | 0 |
| ENSDARG000 | 3;3;3;3;3;3;3; | 3610140:3610 | 3610307:3610 | +;+;+;+;+;+;+; | 1831 | pick1 | 190 | 167 | 170 | 106 | 189 | 165 |
| ENSDARG000 | 3 | 37788861 | 37788945 | + | 85 | NA | 2 | 5 | 14 | 4 | 14 | 3 |
| ENSDARG000 | 3;3 | 49010987:490 | 49012151:490 | -;- | 1738 | NA | 4 | 0 | 0 | 0 | 0 | 2 |
| ENSDARG000 | 3;3;3 | 59851537:598 | 59852625:598 | +;+;+ | 5258 | NA | 0 | 0 | 0 | 0 | 0 | 0 |
| ENSDARG000 | 3;3;3;3;3;3;3; | 62161184:621 | 62161323:621 | +;+;+;+;+;+;+; | 3049 | noxo1a | 36 | 25 | 31 | 66 | 22 | 40 |
| ENSDARG000 | 3;3;3;3;3;3 | 10407458:104 | 10407535:104 | -;-;-;-;-;- | 1160 | pctp | 115 | 102 | 93 | 53 | 116 | 116 |
| ENSDARG000 | 3;3;3;3;3 | 40293506:402 | 40293514:402 | +;+;+;+;+ | 111 | NA | 0 | 0 | 4 | 0 | 0 | 3 |
| ENSDARG000 | 3;3;3;3 | 49379774:493 | 49379841:493 | -;-;-;- | 345 | NA | 0 | 0 | 0 | 0 | 0 | 0 |
| ENSDARG000 | 3 | 1207098 | 1207250 | - | 153 | U12 | 0 | 0 | 0 | 0 | 0 | 0 |
| ENSDARG000 | 3;3;3;3 | 57268363:572 | 57268864:572 | +;+;+;+ | 4543 | NA | 0 | 0 | 5 | 0 | 2 | 0 |
| ENSDARG000 | 3;3 | 14099693:141 | 14101933:141 | -;- | 2478 | nfil3-6 | 90 | 476 | 62 | 98 | 421 | 32 |
| ENSDARG000 | 3 | 47460488 | 47460603 | + | 116 | 5S_rRNA | 0 | 0 | 0 | 0 | 0 | 0 |
| ENSDARG000 | 3;3;3;3;3;3;3; | 33570583:335 | 33570989:335 | -;-;-;-;-;-;-;-; | 2531 | ELFN2 | 6 | 8 | 2 | 0 | 6 | 0 |
| ENSDARG000 | 3;3 | 4113551:4114 | 4113915:4118 | +;+ | 4740 | NA | 0 | 0 | 0 | 0 | 0 | 0 |
| ENSDARG000 | 3;3;3;3 | 32125452:321 | 32125556:321 | +;+;+;+ | 1682 | fthl29 | 5 | 0 | 0 | 0 | 4 | 46 |
| ENSDARG000 | 3;3;3;3;3;3;3; | 11977271:119 | 11977351:119 | -;-;-;-;-;-;-;-; | 3153 | NA | 619 | 329 | 668 | 675 | 736 | 2 |
| ENSDARG000 | 3;3;3;3;3;3;3; | 41629605:416 | 41629619:416 | -;-;-;-;-;-;-;-; | 387 | NA | 6 | 13 | 6 | 5 | 8 | 20 |
| ENSDARG000 | 3;3;3;3;3;3;3; | 17855981:178 | 17855993:178 | +;+;+;+;+;+;+; | 165 | NA | 3 | 0 | 0 | 0 | 0 | 1 |
| ENSDARG000 | 3;3 | 7141360:7147 | 7142486:7147 | -;- | 1233 | NA | 107 | 59 | 90 | 69 | 101 | 88 |
| ENSDARG000 | 3 | 3573696 | 3578297 | + | 4602 | NA | 13 | 50 | 0 | 78 | 39 | 14 |
| ENSDARG000 | 3;3 | 1942219:1942 | 1942320:1943 | +;+ | 1184 | zgc:165583 | 0 | 0 | 346 | 0 | 190 | 214 |
| ENSDARG000 | 3;3 | 7509323:7524 | 7511441:7524 | -;- | 2306 | znf1001 | 54 | 38 | 99 | 73 | 97 | 120 |
| ENSDARG000 | 3;3;3;3;3;3;3; | 54431093:544 | 54435460:544 | -;-;-;-;-;-;-;-; | 6150 | trip10a | 1741 | 1571 | 1915 | 1297 | 2056 | 1930 |
| ENSDARG000 | 3;3;3;3;3;3;3; | 60607241:606 | 60607336:606 | +;+;+;+;+;+;+; | 1062 | mfsd11 | 108 | 66 | 108 | 56 | 144 | 55 |
| ENSDARG000 | 3 | 23817922 | 23818046 | - | 125 | 5S_rRNA | 0 | 0 | 0 | 0 | 0 | 0 |
| ENSDARG000 | 3;3 | 10969712:109 | 10970067:109 | -;- | 705 | NA | 7 | 1 | 8 | 1 | 6 | 1 |
| ENSDARG000 | 3 | 9496077 | 9496853 | - | 777 | NA | 0 | 0 | 0 | 0 | 0 | 0 |
| ENSDARG000 | 3 | 36967871 | 36967993 | - | 123 | 5S_rRNA | 0 | 0 | 0 | 0 | 0 | 0 |
| ENSDARG000 | 3;3;3;3;3;3;3; | 9332984:9333 | 9333306:9333 | +;+;+;+;+;+;+; | 3883 | NA | 0 | 0 | 0 | 0 | 0 | 0 |
| ENSDARG000 | 3;3 | 15442412:154 | 15443472:154 | -;- | 1395 | si:dkey-56d12 | 124 | 30 | 143 | 0 | 66 | 49 |
| ENSDARG000 | 3 | 29867404 | 29867540 | - | 137 | NA | 5 | 11 | 42 | 8 | 41 | 11 |
| ENSDARG000 | 3;3 | 9737490:9738 | 9737558:9738 | -;- | 1023 | NA | 1 | 2 | 0 | 5 | 0 | 0 |
| ENSDARG000 | 3 | 52828912 | 52829006 | + | 95 | NA | 0 | 0 | 0 | 0 | 0 | 0 |
| ENSDARG000 | 3;3 | 3519607:3527 | 3520161:3528 | +;+ | 1148 | NA | 0 | 4 | 6 | 20 | 6 | 10 |
| ENSDARG000 | 3;3;3;3;3;3;3 | 562333:56518 | 562965:56525 | -;-;-;-;-;-;- | 1450 | dicp1.3-4 | 0 | 4 | 20 | 1 | 21 | 46 |
| ENSDARG000 | 3;3;3;3;3;3;3; | 47114328:471 | 47114608:471 | -;-;-;-;-;-;-;-; | 1358 | qtrt1 | 106 | 114 | 222 | 96 | 162 | 119 |
| ENSDARG000 | 3;3;3;3;3 | 31249885:312 | 31251471:312 | -;-;-;-;- | 1820 | apnl | 12 | 0 | 0 | 0 | 24 | 8 |
| ENSDARG000 | 3;3 | 58984463:589 | 58986465:589 | +;+ | 2501 | NA | 98 | 103 | 89 | 59 | 100 | 121 |
| ENSDARG000 | 3 | 19333504 | 19333631 | - | 128 | 5S_rRNA | 0 | 0 | 0 | 0 | 0 | 0 |
| ENSDARG000 | 3;3;3;3;3;3;3; | 62216727:622 | 62217576:622 | +;+;+;+;+;+;+; | 10745 | NA | 322 | 309 | 255 | 354 | 265 | 331 |
| ENSDARG000 | 3 | 18270043 | 18274691 | - | 4649 | NA | 0 | 0 | 0 | 0 | 0 | 0 |
| ENSDARG000 | 3;3 | 58655375:586 | 58655441:586 | +;+ | 384 | NA | 0 | 0 | 0 | 0 | 0 | 0 |
| ENSDARG000 | 3 | 9031370 | 9031520 | + | 151 | U1 | 0 | 0 | 0 | 0 | 0 | 0 |
| ENSDARG000 | 3 | 9024485 | 9024552 | - | 68 | U6 | 0 | 0 | 0 | 0 | 0 | 0 |
| ENSDARG000 | 3 | 56354644 | 56355966 | - | 1323 | NA | 0 | 0 | 0 | 0 | 0 | 0 |
| ENSDARG000 | 3;3;3 | 60684909:606 | 60685732:606 | -;-;- | 1076 | ubald2 | 750 | 262 | 543 | 603 | 365 | 532 |
| ENSDARG000 | 3 | 13406451 | 13406538 | + | 88 | dre-mir-27c | 3 | 1 | 3 | 15 | 0 | 5 |
| ENSDARG000 | 3;3 | 2754885:2756 | 2755198:2757 | +;+ | 968 | NA | 0 | 0 | 0 | 0 | 0 | 0 |
| ENSDARG000 | 3 | 49132341 | 49132477 | + | 137 | 5S_rRNA | 0 | 0 | 0 | 0 | 0 | 0 |
| ENSDARG000 | 3;3;3;3;3;3;3; | 11032192:110 | 11032560:110 | -;-;-;-;-;-;-;-; | 2094 | NA | 0 | 5 | 0 | 0 | 2 | 0 |
| ENSDARG000 | 3;3;3;3;3;3;3; | 45195219:451 | 45196705:451 | -;-;-;-;-;-;-;-; | 4310 | USP31 | 319 | 217 | 262 | 208 | 311 | 275 |
| ENSDARG000 | 3;3;3;3;3;3;3; | 612432:61331 | 612575:61364 | +;+;+;+;+;+;+; | 1392 | NA | 2 | 0 | 0 | 0 | 0 | 0 |
| ENSDARG000 | 3;3;3;3;3;3 | 6993107:7081 | 6994155:7081 | -;-;-;-;-;- | 2423 | NA | 83 | 44 | 70 | 58 | 59 | 93 |
| ENSDARG000 | 3;3 | 60812851:608 | 60818825:608 | -;- | 6630 | anks4b | 59 | 70 | 158 | 77 | 187 | 78 |
| ENSDARG000 | 3;3 | 58828341:588 | 58828708:588 | -;- | 463 | NA | 0 | 0 | 0 | 0 | 0 | 0 |
| ENSDARG000 | 3 | 52620058 | 52620175 | + | 118 | 5S_rRNA | 0 | 0 | 0 | 0 | 0 | 0 |
| ENSDARG000 | 3;3;3 | 20012891:200 | 20013059:200 | +;+;+ | 1144 | tmub2 | 288 | 181 | 418 | 152 | 193 | 236 |
| ENSDARG000 | 3 | 7040363 | 7041760 | + | 1398 | NA | 46 | 34 | 85 | 31 | 26 | 87 |
| ENSDARG000 | 3;3;3;3;3;3 | 33745014:337 | 33745758:337 | +;+;+;+;+;+ | 1650 | NACC1 | 802 | 974 | 1123 | 1013 | 1109 | 362 |
| ENSDARG000 | 3;3 | 32129632:321 | 32129736:321 | +;+ | 279 | NA | 0 | 0 | 0 | 0 | 0 | 5 |
| ENSDARG000 | 3;3;3;3 | 47222096:472 | 47225053:472 | -;-;-;- | 3938 | tmed1a | 32 | 2 | 12 | 6 | 6 | 10 |
| ENSDARG000 | 3;3;3;3;3;3;3; | 58565263:585 | 58565428:585 | -;-;-;-;-;-;-;-; | 3308 | NA | 76 | 101 | 78 | 82 | 99 | 70 |
| ENSDARG000 | 3;3;3;3;3;3;3; | 4571886:4572 | 4572016:4573 | -;-;-;-;-;-;-;-; | 1283 | NA | 10 | 6 | 3 | 6 | 5 | 1 |
| ENSDARG000 | 3;3;3;3;3;3 | 48206469:482 | 48206674:482 | +;+;+;+;+;+ | 2476 | NA | 0 | 2 | 10 | 4 | 54 | 11 |
| ENSDARG000 | 3;3 | 3740729:3741 | 3740785:3741 | +;+ | 603 | NA | 0 | 0 | 0 | 0 | 2 | 4 |
| ENSDARG000 | 3;3 | 58663942:586 | 58664091:586 | +;+ | 1294 | NA | 0 | 0 | 0 | 0 | 0 | 0 |
| ENSDARG000 | 3;3;3;3 | 32135037:321 | 32135147:321 | +;+;+;+ | 534 | NA | 0 | 1 | 0 | 1 | 3 | 9 |
| ENSDARG000 | 3 | 13406226 | 13406323 | + | 98 | dre-mir-23a-2 | 0 | 0 | 0 | 17 | 0 | 5 |
| ENSDARG000 | 3;3;3;3 | 27706113:277 | 27706189:277 | -;-;-;- | 252 | NA | 0 | 0 | 0 | 0 | 0 | 0 |
| ENSDARG000 | 3;3;3;3;3;3;3; | 57948377:579 | 57949457:579 | +;+;+;+;+;+;+; | 2246 | notum1a | 2 | 2 | 4 | 0 | 0 | 2 |
| ENSDARG000 | 3;3;3;3;3;3;3; | 60761811:607 | 60761871:607 | +;+;+;+;+;+;+; | 1514 | tsen54 | 246 | 173 | 278 | 323 | 214 | 226 |
| ENSDARG000 | 3 | 6038203 | 6038338 | + | 136 | SNORA62 | 0 | 0 | 0 | 0 | 0 | 2 |
| ENSDARG000 | 3 | 61185660 | 61187105 | + | 1446 | NA | 0 | 0 | 0 | 0 | 0 | 0 |
| ENSDARG000 | 3 | 14244666 | 14244784 | + | 119 | 5S_rRNA | 0 | 0 | 0 | 0 | 0 | 0 |
| ENSDARG000 | 3 | 26194344 | 26194459 | + | 116 | 5S_rRNA | 0 | 0 | 0 | 2 | 0 | 0 |
| ENSDARG000 | 3 | 344817 | 344937 | - | 121 | 5S_rRNA | 0 | 0 | 0 | 0 | 0 | 0 |
| ENSDARG000 | 3;3;3;3;3;3;3; | 49551580:495 | 49551900:495 | -;-;-;-;-;-;-;-; | 1572 | NA | 2 | 1 | 52 | 0 | 2 | 0 |
| ENSDARG000 | 3;3;3 | 3281734:3295 | 3284962:3296 | -;-;- | 4717 | NA | 0 | 0 | 0 | 0 | 0 | 0 |
| ENSDARG000 | 3 | 60451500 | 60451628 | - | 129 | 5S_rRNA | 0 | 0 | 0 | 0 | 0 | 0 |
| ENSDARG000 | 3;3 | 13469704:134 | 13469745:134 | +;+ | 1557 | zgc:77748 | 0 | 0 | 4 | 0 | 0 | 0 |
| ENSDARG000 | 3;3;3;3 | 48473346:484 | 48473852:484 | +;+;+;+ | 2035 | metrnla | 1997 | 1663 | 2547 | 2332 | 1987 | 2153 |
| ENSDARG000 | 3;3 | 3729821:3731 | 3731404:3731 | -;- | 1585 | NA | 7 | 0 | 7 | 0 | 6 | 2 |
| ENSDARG000 | 3;3;3;3 | 11568523:115 | 11568558:115 | +;+;+;+ | 276 | NA | 5 | 3 | 0 | 1 | 3 | 0 |
| ENSDARG000 | 3 | 34197694 | 34197771 | + | 78 | dre-mir-181c | 0 | 0 | 0 | 0 | 0 | 0 |
| ENSDARG000 | 3;3;3;3;3;3;3 | 62196672:621 | 62196981:621 | +;+;+;+;+;+;+ | 1178 | sco1 | 270 | 330 | 365 | 204 | 308 | 303 |
| ENSDARG000 | 3 | 36756033 | 36756149 | - | 117 | 5S_rRNA | 0 | 0 | 0 | 0 | 0 | 0 |
| ENSDARG000 | 3 | 3139240 | 3139974 | + | 735 | NA | 56 | 56 | 0 | 25 | 12 | 0 |
| ENSDARG000 | 3;3 | 62290024:622 | 62290078:622 | +;+ | 852 | zgc:173575 | 23 | 17 | 12 | 25 | 13 | 42 |
| ENSDARG000 | 3 | 33450100 | 33450219 | + | 120 | 5S_rRNA | 0 | 0 | 0 | 0 | 0 | 0 |
| ENSDARG000 | 3 | 10187033 | 10187151 | - | 119 | 5S_rRNA | 0 | 0 | 0 | 0 | 0 | 0 |
| ENSDARG000 | 3;3;3;3;3;3;3; | 11306799:113 | 11306927:113 | -;-;-;-;-;-;-;-; | 1864 | NA | 19 | 7 | 2 | 28 | 2 | 8 |
| ENSDARG000 | 3 | 9035676 | 9035776 | + | 101 | U6 | 0 | 0 | 0 | 0 | 0 | 0 |
| ENSDARG000 | 3;3;3;3;3;3;3; | 4047789:4048 | 4047861:4048 | +;+;+;+;+;+;+; | 1803 | NA | 4 | 0 | 0 | 0 | 0 | 0 |
| ENSDARG000 | 3;3;3;3;3 | 155659:15817 | 155711:15869 | +;+;+;+;+ | 1930 | NA | 0 | 0 | 0 | 0 | 0 | 4 |
| ENSDARG000 | 3;3;3;3;3;3;3; | 14317802:143 | 14317939:143 | +;+;+;+;+;+;+; | 1763 | plppr2a | 2 | 0 | 6 | 0 | 0 | 0 |
| ENSDARG000 | 3 | 3545825 | 3550612 | + | 4788 | NA | 16 | 94 | 8 | 40 | 27 | 14 |
| ENSDARG000 | 3;3;3;3;3;3;3; | 162315:16231 | 163804:16380 | -;-;-;-;-;-;-;-; | 2511 | kdelr3 | 132 | 118 | 204 | 179 | 240 | 161 |
| ENSDARG000 | 3;3;3 | 58231560:582 | 58231665:582 | +;+;+ | 504 | NA | 43 | 71 | 62 | 70 | 106 | 61 |
| ENSDARG000 | 3 | 9036717 | 9036823 | + | 107 | U6 | 0 | 0 | 0 | 0 | 0 | 0 |
| ENSDARG000 | 3;3;3;3;3;3;3; | 4576089:4577 | 4576775:4577 | -;-;-;-;-;-;-;-; | 2860 | NA | 24 | 25 | 12 | 11 | 6 | 14 |
| ENSDARG000 | 3;3;3;3;3;3;3; | 18413461:184 | 18413946:184 | +;+;+;+;+;+;+; | 1785 | GAA | 61 | 79 | 118 | 31 | 219 | 78 |
| ENSDARG000 | 3;3;3;3;3;3;3; | 44002300:440 | 44003151:440 | -;-;-;-;-;-;-;-; | 2355 | NA | 309 | 267 | 302 | 228 | 135 | 199 |

| | | | | | | | | | | | |
|---|---|---|---|---|---|---|---|---|---|---|---|
| ENSDARG000 | 3;3;3 | 5490104:549 | 5490150:549 | +;+;+ | 3230 | NA | 0 | 0 | 14 | 0 | 11 | 0 |
| ENSDARG000 | 3;3;3;3;3;3;3; | 48234445:482 | 48234613:482 | +;+;+;+;+;+;+; | 3576 | TBCD | 339 | 339 | 380 | 207 | 516 | 337 |
| ENSDARG000 | 3;3;3 | 2200895:220 | 2202695:220 | -;-;- | 1953 | NA | 0 | 0 | 0 | 0 | 0 | 0 |
| ENSDARG000 | 3;3 | 3113543:311 | 3114234:311 | +;+ | 732 | NA | 0 | 0 | 0 | 4 | 0 | 0 |
| ENSDARG000 | 3;3;3;3;3;3;3; | 60177774:602 | 60177930:602 | +;+;+;+;+;+;+; | 2726 | xylt2 | 350 | 261 | 598 | 276 | 602 | 169 |
| ENSDARG000 | 3 | 14888052 | 14888168 | - | 117 | 5S_rRNA | 0 | 0 | 0 | 0 | 0 | 0 |
| ENSDARG000 | 3 | 37075698 | 37075813 | - | 116 | 5S_rRNA | 0 | 0 | 0 | 0 | 0 | 0 |
| ENSDARG000 | 3 | 8389555 | 8389671 | + | 117 | 5S_rRNA | 0 | 0 | 0 | 0 | 0 | 0 |
| ENSDARG000 | 3;3;3;3;3;3;3; | 3852614:385 | 3852694:385 | +;+;+;+;+;+;+; | 4633 | NA | 8 | 0 | 0 | 0 | 0 | 0 |
| ENSDARG000 | 3 | 9028494 | 9028642 | + | 149 | U1 | 0 | 0 | 0 | 0 | 0 | 0 |
| ENSDARG000 | 3;3;3;3;3;3 | 325757:3308 | 327588:3309 | -;-;-;-;-;- | 2865 | mhc1zfa | 14 | 9 | 11 | 0 | 13 | 7 |
| ENSDARG000 | 3;3;3;3;3 | 3992590:399 | 3993035:399 | +;+;+;+;+ | 8613 | NA | 32 | 0 | 0 | 2 | 15 | 0 |
| ENSDARG000 | 3;3 | 62327332:62 | 62327386:62 | +;+ | 930 | zgc:113407 | 81 | 62 | 104 | 127 | 57 | 58 |
| ENSDARG000 | 3;3;3;3;3;3;3 | 12202982:12 | 12203131:12 | -;-;-;-;-;-;- | 1602 | TFAP4 | 228 | 340 | 425 | 231 | 399 | 382 |
| ENSDARG000 | 3;3;3;3 | 47245764:47 | 47246294:47 | +;+;+;+ | 3537 | tnfsf14 | 0 | 10 | 0 | 0 | 30 | 4 |
| ENSDARG000 | 3 | 2113367 | 2113483 | - | 117 | 5S_rRNA | 0 | 0 | 0 | 0 | 0 | 0 |
| ENSDARG000 | 3;3;3;3;3;3;3 | 11548516:11 | 11548793:11 | +;+;+;+;+;+;+ | 1781 | mmd | 366 | 342 | 320 | 190 | 479 | 402 |
| ENSDARG000 | 3;3;3;3 | 2719243:288 | 2719243:288 | +;+;+;+ | 905 | NA | 0 | 0 | 0 | 0 | 0 | 0 |
| ENSDARG000 | 3;3;3;3 | 32112004:32 | 32112158:32 | +;+;+;+ | 888 | fthl30 | 5 | 0 | 0 | 0 | 3 | 16 |
| ENSDARG000 | 3;3;3;3 | 19460991:19 | 19461066:19 | +;+;+;+ | 1227 | pde6ga | 0 | 0 | 0 | 0 | 0 | 0 |
| ENSDARG000 | 3 | 24655643 | 24660328 | + | 4686 | NA | 505 | 284 | 138 | 82 | 99 | 171 |
| ENSDARG000 | 3;3;3;3;3;3;3; | 54849083:54 | 54849684:54 | +;+;+;+;+;+;+; | 6604 | mgrn1b | 1142 | 736 | 925 | 582 | 985 | 903 |
| ENSDARG000 | 3;3 | 4997545:499 | 4997661:499 | +;+ | 240 | NA | 16 | 23 | 62 | 148 | 29 | 23 |
| ENSDARG000 | 3;3;3;3;3;3;3; | 3469600:347 | 3469832:347 | +;+;+;+;+;+;+; | 2534 | NA | 0 | 0 | 0 | 0 | 0 | 0 |
| ENSDARG000 | 3;3;3;3;3;3;3; | 12554801:12 | 12554934:12 | +;+;+;+;+;+;+; | 3414 | ccnf | 365 | 578 | 925 | 459 | 860 | 539 |
| ENSDARG000 | 3;3;3 | 1724941:172 | 1725003:172 | +;+;+ | 1663 | NA | 0 | 5 | 82 | 0 | 42 | 17 |
| ENSDARG000 | 3;3;3;3;3;3 | 48965923:48 | 48966446:48 | -;-;-;-;-;- | 1620 | NA | 4 | 0 | 0 | 0 | 0 | 0 |
| ENSDARG000 | 3;3;3;3;3 | 31177972:31 | 31178140:31 | +;+;+;+;+ | 1249 | clec19a | 17 | 0 | 0 | 18 | 14 | 10 |
| ENSDARG000 | 3;3;3;3 | 24675029:24 | 24675936:24 | -;-;-;- | 1476 | NA | 4 | 0 | 0 | 0 | 0 | 0 |
| ENSDARG000 | 3 | 24403149 | 24403269 | - | 121 | 5S_rRNA | 0 | 0 | 0 | 0 | 0 | 0 |
| ENSDARG000 | 3;3;3;3;3;3 | 3489456:348 | 3489657:349 | -;-;-;-;-;- | 780 | NA | 0 | 4 | 0 | 0 | 0 | 0 |
| ENSDARG000 | 3;3;3 | 580351:5834 | 580385:5838 | -;-;- | 414 | NA | 8 | 0 | 0 | 0 | 4 | 8 |
| ENSDARG000 | 3;3;3;3;3;3;3; | 62126981:62 | 62127044:62 | +;+;+;+;+;+;+; | 1728 | drc3 | 2 | 2 | 12 | 22 | 20 | 10 |
| ENSDARG000 | 3;3;3;3;3;3;3; | 47130912:47 | 47131272:47 | +;+;+;+;+;+;+; | 3398 | dnm2a | 1307 | 1015 | 1344 | 746 | 1507 | 976 |
| ENSDARG000 | 3;3;3;3;3;3;3; | 3666138:366 | 3666141:366 | -;-;-;-;-;-;-;- | 843 | NA | 0 | 0 | 0 | 0 | 0 | 0 |
| ENSDARG000 | 3 | 18892650 | 18892766 | + | 117 | 5S_rRNA | 0 | 0 | 0 | 0 | 0 | 0 |
| ENSDARG000 | 3;3;3;3;3;3 | 57234093:57 | 57234110:57 | +;+;+;+;+;+ | 1467 | fscn2a | 0 | 0 | 0 | 0 | 0 | 0 |
| ENSDARG000 | 3;3;3;3;3;3;3; | 61896514:61 | 61896613:61 | +;+;+;+;+;+;+; | 1662 | NA | 19 | 7 | 12 | 55 | 27 | 13 |
| ENSDARG000 | 3;3 | 48986671:48 | 48990139:48 | -;- | 4087 | NA | 8 | 33 | 0 | 0 | 6 | 6 |
| ENSDARG000 | 3;3;3;3;3;3;3; | 5778954:578 | 5779157:578 | -;-;-;-;-;-;-;-; | 2730 | pkn1b | 90 | 32 | 45 | 38 | 109 | 119 |
| ENSDARG000 | 3;3;3;3;3;3 | 4555758:455 | 4555861:455 | -;-;-;-;-;- | 1239 | NA | 33 | 4 | 6 | 3 | 3 | 3 |
| ENSDARG000 | 3;3;3;3;3;3;3; | 4644031:464 | 4644576:464 | -;-;-;-;-;-;-;- | 1500 | slc25a38a | 10 | 22 | 106 | 14 | 8 | 44 |
| ENSDARG000 | 3;3 | 2971852:297 | 2972001:297 | +;+ | 2473 | NA | 4 | 4 | 1 | 0 | 0 | 0 |
| ENSDARG000 | 3 | 59051503 | 59052994 | + | 1492 | rasd4 | 0 | 0 | 0 | 0 | 8 | 6 |
| ENSDARG000 | 3;3;3;3 | 58506242:58 | 58507096:58 | -;-;-;- | 1654 | NA | 4 | 2 | 0 | 0 | 0 | 8 |
| ENSDARG000 | 3;3;3 | 60594138:60 | 60596850:60 | -;-;- | 3553 | srsf2b | 321 | 442 | 476 | 284 | 516 | 218 |
| ENSDARG000 | 3;3;3;3;3;3;3; | 32603256:32 | 32603298:32 | -;-;-;-;-;-;-;-; | 222 | NA | 0 | 0 | 0 | 0 | 0 | 0 |
| ENSDARG000 | 3;3 | 62219158:62 | 62219212:62 | +;+ | 933 | NA | 42 | 36 | 41 | 14 | 22 | 52 |
| ENSDARG000 | 3;3;3;3;3;3;3; | 46919522:46 | 46919639:46 | -;-;-;-;-;-;-;-; | 4210 | NA | 0 | 0 | 0 | 0 | 16 | 2 |
| ENSDARG000 | 3;3;3;3;3 | 59864872:59 | 59864969:59 | +;+;+;+;+ | 1146 | mcrip1 | 340 | 435 | 467 | 470 | 439 | 450 |
| ENSDARG000 | 3;3;3;3;3;3;3; | 60828813:60 | 60829579:60 | +;+;+;+;+;+;+; | 1634 | NA | 57 | 106 | 97 | 18 | 117 | 62 |
| ENSDARG000 | 3 | 5326140 | 5326261 | - | 122 | 5S_rRNA | 0 | 0 | 0 | 0 | 0 | 0 |
| ENSDARG000 | 3;3 | 58656312:58 | 58656472:58 | +;+ | 594 | NA | 0 | 0 | 0 | 0 | 0 | 0 |
| ENSDARG000 | 3 | 35959912 | 35960010 | + | 99 | dre-mir-193a- | 0 | 1 | 0 | 0 | 4 | 0 |
| ENSDARG000 | 3 | 44888037 | 44888153 | + | 117 | 5S_rRNA | 0 | 0 | 0 | 0 | 0 | 0 |
| ENSDARG000 | 3;3;3;3;3;3;3; | 13523773:13 | 13526064:13 | -;-;-;-;-;-;-;-; | 3775 | amdhd2 | 1418 | 1438 | 1249 | 1486 | 1403 | 1460 |
| ENSDARG000 | 3;3;3;3;3 | 1352555:136 | 1352726:136 | -;-;-;-;- | 2442 | NA | 129 | 79 | 190 | 33 | 209 | 266 |
| ENSDARG000 | 3;3;3;3;3;3;3; | 52953489:52 | 52954473:52 | +;+;+;+;+;+;+; | 1943 | NA | 58 | 267 | 291 | 98 | 188 | 113 |
| ENSDARG000 | 3;3;3;3;3;3;3; | 62091298:62 | 62097582:62 | -;-;-;-;-;-;-;-; | 7664 | TOM1L2 | 1620 | 865 | 1368 | 1637 | 1566 | 1616 |
| ENSDARG000 | 3 | 61420079 | 61421085 | + | 1007 | NA | 0 | 0 | 0 | 0 | 0 | 0 |
| ENSDARG000 | 3 | 51548133 | 51548250 | - | 118 | 5S_rRNA | 0 | 0 | 0 | 0 | 0 | 0 |
| ENSDARG000 | 3;3;3;3;3;3 | 49074008:49 | 49074176:49 | +;+;+;+;+;+ | 945 | zgc:112146 | 0 | 0 | 0 | 0 | 0 | 0 |
| ENSDARG000 | 3;3;3 | 7056679:709 | 7057712:709 | -;-;- | 1716 | NA | 21 | 21 | 95 | 24 | 26 | 75 |
| ENSDARG000 | 3;3 | 47552353:47 | 47552362:47 | +;+ | 213 | NA | 0 | 0 | 0 | 0 | 0 | 0 |
| ENSDARG000 | 3;3;3;3;3;3;3; | 42918143:42 | 42918151:42 | -;-;-;-;-;-;-;-; | 345 | NA | 180 | 139 | 194 | 234 | 135 | 151 |
| ENSDARG000 | 3;3;3;3 | 1895523:190 | 1896298:190 | -;-;-;- | 922 | zgc:101806 | 0 | 0 | 10 | 0 | 32 | 3 |
| ENSDARG000 | 3;3;3;3;3 | 62000822:62 | 62001857:62 | +;+;+;+;+ | 9170 | NA | 0 | 0 | 0 | 14 | 0 | 6 |
| ENSDARG000 | 3 | 29770142 | 29770258 | - | 117 | 5S_rRNA | 0 | 0 | 0 | 0 | 0 | 0 |
| ENSDARG000 | 3;3;3;3 | 32105175:32 | 32105402:32 | +;+;+;+ | 889 | fthl28 | 49 | 40 | 31 | 24 | 74 | 29 |
| ENSDARG000 | 3 | 23382492 | 23382608 | - | 117 | 5S_rRNA | 0 | 0 | 0 | 0 | 0 | 0 |
| ENSDARG000 | 3;3 | 2959343:296 | 2959538:296 | +;+ | 967 | NA | 0 | 0 | 0 | 0 | 0 | 0 |
| ENSDARG000 | 3;3;3;3;3;3;3; | 634682:6371 | 634740:6373 | +;+;+;+;+;+;+; | 965 | NA | 2 | 0 | 2 | 5 | 11 | 1 |
| ENSDARG000 | 3 | 9457762 | 9458445 | - | 684 | NA | 0 | 0 | 0 | 0 | 0 | 0 |
| ENSDARG000 | 3 | 34359215 | 34359332 | + | 118 | 5S_rRNA | 0 | 0 | 0 | 0 | 0 | 0 |
| ENSDARG000 | 3 | 3718608 | 3723494 | - | 4887 | NA | 6 | 0 | 2 | 0 | 3 | 4 |
| ENSDARG000 | 3;3;3;3;3;3 | 19468777:19 | 19469196:19 | +;+;+;+;+;+ | 4327 | NA | 6 | 4 | 12 | 0 | 22 | 76 |
| ENSDARG000 | 3 | 29552641 | 29552762 | - | 122 | 5S_rRNA | 0 | 0 | 0 | 0 | 1 | 0 |
| ENSDARG000 | 3;3;3;3;3;3;3; | 19621034:19 | 19621154:19 | +;+;+;+;+;+;+; | 2535 | itgb3a | 64 | 6 | 2 | 0 | 6 | 8 |
| ENSDARG000 | 3;3;3;3;3;3;3 | 9431501:943 | 9432939:943 | -;-;-;-;-;-;- | 2915 | rnps1 | 48 | 46 | 22 | 34 | 52 | 33 |
| ENSDARG000 | 3;3 | 8173070:817 | 8174152:817 | -;- | 1125 | NA | 0 | 0 | 0 | 0 | 6 | 0 |
| ENSDARG000 | 3;3;3 | 19507415:19 | 19508176:19 | -;-;- | 1495 | NA | 0 | 0 | 0 | 0 | 0 | 0 |
| ENSDARG000 | 3;3;3;3;3;3;3 | 53097587:53 | 53097824:53 | -;-;-;-;-;-;- | 1107 | NA | 0 | 0 | 2 | 0 | 0 | 0 |
| ENSDARG000 | 3 | 29251949 | 29252065 | - | 117 | 5S_rRNA | 0 | 0 | 0 | 2 | 0 | 8 |
| ENSDARG000 | 3;3;3 | 54833658:54 | 54837888:54 | -;-;- | 4551 | ubald1b | 903 | 685 | 896 | 815 | 524 | 516 |
| ENSDARG000 | 3 | 4726621 | 4731396 | + | 4776 | NA | 2 | 0 | 0 | 0 | 4 | 0 |
| ENSDARG000 | 3;3;3;3;3;3;3; | 59784632:59 | 59784724:59 | +;+;+;+;+;+;+; | 5360 | pecam1a | 284 | 24 | 44 | 2 | 33 | 20 |
| ENSDARG000 | 3 | 16373012 | 16373127 | + | 116 | 5S_rRNA | 0 | 0 | 0 | 0 | 0 | 0 |
| ENSDARG000 | 3 | 9040947 | 9041053 | + | 107 | U6 | 0 | 0 | 0 | 0 | 0 | 0 |
| ENSDARG000 | 3 | 51907486 | 51907562 | - | 77 | dre-mir-338-[illegible] | 0 | 0 | 0 | 0 | 0 | 0 |
| ENSDARG000 | 3 | 24745766 | 24745881 | - | 116 | 5S_rRNA | 0 | 0 | 0 | 0 | 0 | 0 |
| ENSDARG000 | 3;3;3;3;3;3;3; | 20025709:20 | 20026774:20 | -;-;-;-;-;-;-;-; | 2245 | atxn7l3a | 283 | 190 | 220 | 152 | 230 | 145 |
| ENSDARG000 | 3;3;3;3;3;3;3 | 13487760:13 | 13488678:13 | -;-;-;-;-;-;- | 1820 | slc25a19 | 84 | 85 | 80 | 103 | 93 | 71 |
| ENSDARG000 | 3;3 | 7098948:716 | 7100140:716 | -;- | 1316 | NA | 92 | 19 | 72 | 34 | 42 | 80 |
| ENSDARG000 | 3;3;3;3;3;3 | 53511936:53 | 53512032:53 | +;+;+;+;+;+ | 933 | NA | 0 | 0 | 0 | 0 | 0 | 0 |
| ENSDARG000 | 3 | 42866083 | 42866200 | - | 118 | 5S_rRNA | 0 | 0 | 0 | 0 | 0 | 0 |
| ENSDARG000 | 3 | 49815197 | 49815287 | - | 91 | NA | 0 | 0 | 0 | 0 | 0 | 0 |
| ENSDARG000 | 3;3;3;3;3 | 60525982:60 | 60526677:60 | -;-;-;-;- | 2083 | si:ch73-366l1 | 26 | 16 | 58 | 14 | 28 | 18 |
| ENSDARG000 | 3;3;3;3;3;3 | 48549294:48 | 48550878:48 | -;-;-;-;-;- | 2084 | atp5pd | 2422 | 2212 | 3048 | 1801 | 2321 | 1912 |
| ENSDARG000 | 3 | 41500409 | 41500525 | + | 117 | 5S_rRNA | 0 | 0 | 0 | 0 | 0 | 0 |
| ENSDARG000 | 3 | 3099492 | 3100238 | + | 747 | NA | 2 | 0 | 0 | 5 | 10 | 0 |
| ENSDARG000 | 3;3;3;3;3;3;3; | 5991765:599 | 5991896:599 | -;-;-;-;-;-;-;-; | 2016 | MCOLN1 | 36 | 2 | 36 | 4 | 52 | 18 |
| ENSDARG000 | 3;3 | 48842918:48 | 48843939:48 | +;+ | 1641 | hs3st3b1a | 32 | 0 | 4 | 0 | 39 | 0 |
| ENSDARG000 | 3;3;3;3 | 60047032:60 | 60047105:60 | +;+;+;+ | 654 | NA | 23 | 15 | 28 | 18 | 36 | 14 |
| ENSDARG000 | 3;3;3;3 | 314376:3151 | 315008:3154 | -;-;-;- | 1204 | NA | 22 | 22 | 17 | 18 | 16 | 2 |
| ENSDARG000 | 3;3;3;3;3;3;3 | 906519:9065 | 906729:9067 | +;+;+;+;+;+;+ | 1810 | NA | 26 | 24 | 32 | 6 | 2 | 5 |
| ENSDARG000 | 3;3;3;3;3 | 1589564:158 | 1589659:158 | +;+;+;+;+ | 512 | NA | 0 | 0 | 0 | 0 | 0 | 0 |
| ENSDARG000 | 3;3;3;3;3;3 | 4626684:463 | 4630626:463 | -;-;-;-;-;- | 4925 | NA | 0 | 0 | 0 | 0 | 0 | 0 |
| ENSDARG000 | 3;3;3 | 5718913:572 | 5719198:572 | -;-;- | 797 | NA | 0 | 6 | 0 | 0 | 0 | 0 |
| ENSDARG000 | 3;3;3 | 5744408:574 | 5744601:574 | -;-;- | 424 | NA | 0 | 0 | 0 | 0 | 0 | 0 |
| ENSDARG000 | 3;3;3 | 8094011:809 | 8094100:809 | +;+;+ | 4669 | NA | 6 | 6 | 0 | 0 | 0 | 0 |
| ENSDARG000 | 3;3;3;3;3 | 9475575:947 | 9475881:947 | -;-;-;-;- | 829 | NA | 0 | 0 | 2 | 0 | 0 | 0 |
| ENSDARG000 | 3;3;3 | 9744455:974 | 9744642:974 | -;-;- | 1318 | NA | 0 | 0 | 0 | 0 | 0 | 0 |
| ENSDARG000 | 3;3;3 | 9754014:975 | 9754437:975 | -;-;- | 836 | NA | 0 | 0 | 0 | 0 | 0 | 0 |
| ENSDARG000 | 3;3;3;3;3 | 13149550:13 | 13150513:13 | +;+;+;+;+ | 3482 | NA | 36 | 32 | 35 | 28 | 46 | 28 |
| ENSDARG000 | 3;3;3 | 14242119:14 | 14243308:14 | +;+;+ | 2040 | NA | 0 | 0 | 0 | 0 | 0 | 0 |
| ENSDARG000 | 3;3;3 | 18972408:18 | 18973072:18 | -;-;- | 1602 | NA | 0 | 0 | 0 | 0 | 0 | 0 |
| ENSDARG000 | 3;3;3;3;3 | 19986341:19 | 19989595:19 | -;-;-;-;- | 6024 | NA | 394 | 104 | 337 | 70 | 305 | 84 |
| ENSDARG000 | 3;3;3 | 27294344:27 | 27294824:27 | +;+;+ | 640 | NA | 0 | 0 | 0 | 0 | 0 | 0 |
| ENSDARG000 | 3;3;3;3;3;3;3 | 27319816:27 | 27319938:27 | +;+;+;+;+;+;+ | 1005 | NA | 0 | 0 | 0 | 0 | 0 | 0 |
| ENSDARG000 | 3;3;3 | 29406013:29 | 29406167:29 | +;+;+ | 584 | NA | 0 | 0 | 0 | 0 | 0 | 0 |
| ENSDARG000 | 3;3;3 | 32194752:32 | 32194943:32 | -;-;- | 438 | NA | 1 | 1 | 4 | 0 | 2 | 12 |
| ENSDARG000 | 3;3;3;3;3 | 32196192:32 | 32196611:32 | -;-;-;-;- | 672 | NA | 0 | 0 | 4 | 0 | 7 | 0 |
| ENSDARG000 | 3;3;3;3 | 41396123:41 | 41396506:41 | -;-;-;- | 1679 | NA | 0 | 0 | 0 | 0 | 0 | 0 |
| ENSDARG000 | 3;3;3;3 | 45833840:45 | 45834326:45 | -;-;-;- | 1700 | NA | 0 | 0 | 0 | 0 | 0 | 0 |
| ENSDARG000 | 3;3;3;3;3 | 48461685:48 | 48462614:48 | -;-;-;-;- | 1363 | NA | 0 | 0 | 0 | 0 | 0 | 0 |
| ENSDARG000 | 3;3;3;3 | 56619195:56 | 56619323:56 | +;+;+;+ | 424 | NA | 0 | 0 | 0 | 0 | 0 | 0 |
| ENSDARG000 | 3;3;3;3;3 | 58210305:58 | 58210630:58 | -;-;-;-;- | 653 | NA | 21 | 21 | 0 | 45 | 50 | 29 |
| ENSDARG000 | 3;3;3;3 | 58344862:58 | 58344864:58 | +;+;+;+ | 1070 | NA | 0 | 0 | 0 | 0 | 0 | 0 |
| ENSDARG000 | 3;3;3;3;3 | 58961904:58 | 58962168:58 | +;+;+;+;+ | 966 | NA | 210 | 66 | 53 | 30 | 123 | 26 |
| ENSDARG000 | 3;3;3;3 | 60646736:60 | 60647298:60 | +;+;+;+ | 1133 | NA | 0 | 1 | 1 | 2 | 10 | 0 |
| ENSDARG000 | 6;6;6;6;6;6;6; | 40482330:40 | 40482453:40 | +;+;+;+;+;+;+; | 1978 | NA | 6 | 0 | 0 | 0 | 6 | 20 |
| ENSDARG000 | 6;6;6;6;6 | 34028532:34 | 34028650:34 | +;+;+;+;+ | 1161 | si:ch73-185c | 0 | 0 | 16 | 4 | 23 | 0 |
| ENSDARG000 | 6;6;6;6;6;6;6; | 9204099:920 | 9204118:920 | +;+;+;+;+;+;+; | 1311 | TBX19 | 0 | 0 | 0 | 0 | 0 | 0 |
| ENSDARG000 | 6;6;6;6;6 | 52242139:52 | 52242271:52 | +;+;+;+;+ | 5413 | zc3h10 | 328 | 257 | 364 | 204 | 284 | 182 |
| ENSDARG000 | 6;6;6;6;6;6;6; | 39279104:39 | 39279385:39 | +;+;+;+;+;+;+; | 2257 | NA | 2 | 0 | 0 | 0 | 0 | 0 |
| ENSDARG000 | 6;6;6;6;6;6;6; | 16673793:16 | 16674683:16 | -;-;-;-;-;-;-;-; | 4434 | NOMO1 | 2442 | 2339 | 3611 | 1444 | 3123 | 1978 |
| ENSDARG000 | 6;6;6 | 34213971:34 | 34214758:34 | -;-;- | 1435 | dmrt2b | 0 | 0 | 0 | 0 | 0 | 0 |
| ENSDARG000 | 6;6 | 37308716:37 | 37309567:37 | +;+ | 1206 | PTGER3 | 38 | 5 | 18 | 37 | 12 | 2 |
| ENSDARG000 | 6;6;6;6;6;6;6; | 34511886:34 | 34512491:34 | +;+;+;+;+;+;+; | 9207 | lrp8 | 92 | 84 | 186 | 10 | 150 | 165 |
| ENSDARG000 | 6 | 37334559 | 37334963 | + | 405 | NA | 0 | 0 | 0 | 0 | 0 | 0 |
| ENSDARG000 | 6;6;6;6;6;6;6; | 10700996:10 | 10702327:10 | -;-;-;-;-;-;-;-; | 2892 | NA | 304 | 314 | 254 | 227 | 157 | 212 |
| ENSDARG000 | 6;6 | 19137841:19 | 19138060:19 | -;- | 553 | NA | 0 | 0 | 0 | 0 | 0 | 0 |
| ENSDARG000 | 6;6 | 19146424:19 | 19146644:19 | -;- | 554 | NA | 0 | 0 | 1 | 0 | 0 | 0 |
| ENSDARG000 | 6;6;6;6 | 19163923:19 | 19164097:19 | -;-;-;- | 716 | NA | 0 | 0 | 0 | 0 | 4 | 21 |
| ENSDARG000 | 6;6;6;6;6;6;6; | 40523370:40 | 40523442:40 | +;+;+;+;+;+;+; | 2908 | prkcda | 62 | 10 | 34 | 16 | 120 | 88 |
| ENSDARG000 | 6;6;6;6;6;6;6; | 52263236:52 | 52263583:52 | +;+;+;+;+;+;+; | 5100 | NA | 1124 | 596 | 835 | 846 | 1923 | 1402 |
| ENSDARG000 | 6;6;6;6;6;6;6; | 41255485:41 | 41255938:41 | +;+;+;+;+;+;+; | 5754 | NA | 0 | 4 | 103 | 201 | 2 | 0 |
| ENSDARG000 | 6;6;6;6;6;6;6; | 9225594:922 | 9226418:922 | -;-;-;-;-;-;-;-; | 2747 | iqcb1 | 4 | 8 | 12 | 14 | 14 | 8 |
| ENSDARG000 | 6;6;6;6;6;6;6; | 6275401:627 | 6275783:627 | -;-;-;-;-;-;-;-; | 2646 | mtif2 | 266 | 229 | 356 | 222 | 326 | 282 |
| ENSDARG000 | 6;6;6;6;6;6;6; | 12462079:12 | 12462211:12 | +;+;+;+;+;+;+; | 2863 | nr4a2b | 1422 | 101 | 291 | 336 | 184 | 88 |
| ENSDARG000 | 6;6;6;6;6;6;6; | 39290777:39 | 39290900:39 | +;+;+;+;+;+;+; | 2141 | NA | 109 | 81 | 23 | 44 | 32 | 44 |
| ENSDARG000 | 6;6;6;6;6;6;6; | 28205296:28 | 28205678:28 | +;+;+;+;+;+;+; | 4736 | LSM2 | 6 | 4 | 8 | 0 | 3 | 8 |
| ENSDARG000 | 6 | 30221834 | 30221970 | - | 137 | NA | 0 | 0 | 0 | 0 | 0 | 0 |
| ENSDARG000 | 6;6;6;6;6;6;6; | 37389683:37 | 37390311:37 | -;-;-;-;-;-;-;-; | 1861 | cthl | 4 | 4 | 14 | 0 | 31 | 12 |
| ENSDARG000 | 6;6;6;6;6;6;6; | 19263433:19 | 19264392:19 | -;-;-;-;-;-;-;-; | 2255 | zgc:174863 | 2916 | 2571 | 2732 | 2329 | 2443 | 3013 |
| ENSDARG000 | 6;6;6;6;6;6 | 28205284:28 | 28205370:28 | +;+;+;+;+;+ | 752 | smx5 | 0 | 0 | 0 | 0 | 0 | 0 |
| ENSDARG000 | 6;6;6;6;6;6;6; | 41446541:41 | 41446748:41 | +;+;+;+;+;+;+; | 1342 | cplane2 | 34 | 88 | 50 | 87 | 18 | 50 |
| ENSDARG000 | 6;6;6;6;6;6;6; | 34823721:34 | 34826971:34 | -;-;-;-;-;-;-;-; | 4707 | mier1a | 240 | 280 | 372 | 240 | 307 | 302 |
| ENSDARG000 | 6;6;6;6;6 | 9241121:924 | 9241198:924 | +;+;+;+;+ | 777 | NA | 1 | 2 | 3 | 3 | 7 | 4 |
| ENSDARG000 | 6;6;6;6;6;6;6; | 9245087:924 | 9245231:924 | -;-;-;-;-;-;-;-; | 1264 | iqcb1 | 0 | 0 | 0 | 0 | 0 | 0 |
| ENSDARG000 | 6;6 | 40545865:40 | 40545887:40 | +;+ | 362 | NA | 0 | 0 | 0 | 0 | 0 | 0 |
| ENSDARG000 | 6;6;6;6;6;6;6; | 41452979:41 | 41453426:41 | +;+;+;+;+;+;+; | 2200 | wdr82 | 434 | 763 | 786 | 465 | 734 | 572 |

| | | | | | | | | | | | | |
|---|---|---|---|---|---|---|---|---|---|---|---|---|
| ENSDARG000 | 6;6;6;6;6;6; | 30491129:305 | 30491497:305 | +;+;+;+;+;+;+; | 4168 | wwc3 | 1657 | 1009 | 1064 | 734 | 1235 | 1122 |
| ENSDARG000 | 6;6;6;6;6;6; | 39308498:393 | 39309533:393 | -;-;-;-;-;-;-;- | 2469 | krt4 | 1498 | 569 | 140 | 508 | 376 | 506 |
| ENSDARG000 | 6;6;6;6;6;6; | 37409558:374 | 37410132:374 | -;-;-;-;-;-;-; | 1828 | cth | 516 | 462 | 687 | 292 | 553 | 566 |
| ENSDARG000 | 6;6;6;6;6;6; | 52339548:523 | 52339860:523 | -;-;-;-;-;-;-; | 1536 | eif6 | 1239 | 1419 | 2039 | 904 | 1326 | 1060 |
| ENSDARG000 | 6;6;6;6;6;6; | 9268857:9271 | 9269813:9271 | -;-;-;-;-;-;-; | 2897 | NA | 76 | 68 | 106 | 36 | 153 | 122 |
| ENSDARG000 | 6;6;6;6;6;6; | 13871641:138 | 13872047:138 | -;-;-;-;-;-;-; | 2252 | desmb | 2 | 0 | 0 | 20 | 10 | 13 |
| ENSDARG000 | 6;6;6;6;6;6; | 12482599:125 | 12482712:125 | +;+;+;+;+;+;+; | 2778 | stk24b | 999 | 664 | 1022 | 818 | 914 | 793 |
| ENSDARG000 | 6;6;6;6;6;6; | 10721765:107 | 10722440:107 | -;-;-;-;-;-;-; | 4681 | SP3 | 803 | 555 | 809 | 523 | 764 | 733 |
| ENSDARG000 | 6;6;6;6;6;6; | 41463785:414 | 41464098:414 | +;+;+;+;+;+;+; | 2677 | twf2a | 6 | 0 | 2 | 16 | 4 | 649 |
| ENSDARG000 | 6;6 | 10727344:107 | 10727894:107 | +;+ | 1017 | NA | 9 | 8 | 6 | 1 | 9 | 8 |
| ENSDARG000 | 6;6;6;6;6;6; | 6301205:6303 | 6301617:6303 | -;-;-;-;-;-;-; | 4730 | CCDC88A | 351 | 32 | 163 | 94 | 107 | 86 |
| ENSDARG000 | 6;6;6;6;6;6; | 28216487:282 | 28216727:282 | -;-;-;-;-;-;-;- | 2314 | bcl6aa | 404 | 162 | 150 | 303 | 108 | 68 |
| ENSDARG000 | 6;6;6;6;6;6; | 40554551:405 | 40554859:405 | +;+;+;+;+;+;+; | 4199 | ddi2 | 1019 | 1027 | 1284 | 486 | 1285 | 879 |
| ENSDARG000 | 6;6;6;6;6;6; | 41503854:415 | 41504380:415 | +;+;+;+;+;+;+; | 1731 | cish | 216 | 70 | 210 | 34 | 99 | 205 |
| ENSDARG000 | 6;6;6;6 | 6376395:6384 | 6376818:6384 | -;-;-;- | 1083 | NA | 15 | 0 | 0 | 4 | 0 | 0 |
| ENSDARG000 | 6;6;6;6;6;6; | 39336731:393 | 39337295:393 | -;-;-;-;-;-;-;- | 1917 | zgc:158846 | 0 | 4 | 28 | 9 | 18 | 20 |
| ENSDARG000 | 6;6;6;6;6;6; | 6390471:6390 | 6390522:6390 | -;-;-;-;-;-;-; | 2524 | NA | 2 | 0 | 0 | 0 | 6 | 0 |
| ENSDARG000 | 6;6;6;6;6;6; | 6445065:6459 | 6445846:6459 | -;-;-;-;-;-;-; | 5176 | NA | 217 | 180 | 102 | 248 | 76 | 199 |
| ENSDARG000 | 6;6;6;6 | 52350443:523 | 52350554:523 | +;+;+;+ | 5771 | NA | 83 | 32 | 35 | 38 | 104 | 75 |
| ENSDARG000 | 6;6;6;6;6;6; | 41510135:415 | 41510817:415 | -;-;-;-;-;-;-; | 1811 | hemk1 | 380 | 342 | 506 | 131 | 334 | 536 |
| ENSDARG000 | 6;6;6;6;6;6; | 52362395:523 | 52364816:523 | -;-;-;-;-;-;-; | 4126 | MMP24 | 8 | 0 | 6 | 2 | 0 | 2 |
| ENSDARG000 | 6;6;6;6;6;6; | 40563848:405 | 40564153:405 | +;+;+;+;+;+;+; | 2071 | NA | 0 | 0 | 2 | 0 | 6 | 2 |
| ENSDARG000 | 6 | 28222989 | 28224964 | + | 1976 | NA | 134 | 20 | 148 | 36 | 52 | 46 |
| ENSDARG000 | 6;6;6;6;6;6; | 40572545:405 | 40572710:405 | -;-;-;-;-;-;-; | 633 | tomm6 | 415 | 724 | 688 | 272 | 413 | 346 |
| ENSDARG000 | 6;6;6;6;6;6 | 40578219:405 | 40578695:405 | -;-;-;-;-;- | 934 | tspo | 262 | 257 | 369 | 400 | 426 | 578 |
| ENSDARG000 | 6;6;6;6;6;6; | 39360377:393 | 39360956:393 | +;+;+;+;+;+;+; | 1889 | krt18b | 78181 | 65979 | 76891 | 128797 | 52743 | 121753 |
| ENSDARG000 | 6;6;6;6 | 40573365:405 | 40573533:405 | +;+;+;+ | 711 | uqcc2 | 225 | 369 | 375 | 291 | 281 | 262 |
| ENSDARG000 | 6;6;6;6;6 | 19294852:193 | 19295774:193 | -;-;-;-;- | 1403 | jpt1a | 7 | 38 | 6 | 86 | 12 | 48 |
| ENSDARG000 | 6;6;6;6;6;6; | 7721292:7725 | 7724020:7726 | -;-;-;-;-;-;-; | 4266 | slc25a38b | 540 | 369 | 427 | 224 | 666 | 360 |
| ENSDARG000 | 6;6;6;6;6;6; | 37431169:374 | 37431677:374 | -;-;-;-;-;-;-; | 1299 | nit2 | 219 | 256 | 234 | 245 | 278 | 202 |
| ENSDARG000 | 6;6;6;6;6;6 | 19306192:193 | 19307868:193 | -;-;-;-;-;- | 2311 | NA | 0 | 2 | 0 | 0 | 0 | 0 |
| ENSDARG000 | 6;6;6;6 | 52468139:524 | 52469093:524 | -;-;-;- | 1874 | fam83c | 22 | 6 | 4 | 2 | 2 | 5 |
| ENSDARG000 | 6;6;6;6;6;6; | 28294113:283 | 28294337:283 | +;+;+;+;+;+;+; | 6399 | lpp | 1596 | 1176 | 1898 | 1661 | 1380 | 2076 |
| ENSDARG000 | 6;6;6;6;6 | 30625945:306 | 30631104:306 | -;-;-;-;- | 5987 | lurap1 | 42 | 16 | 18 | 36 | 32 | 12 |
| ENSDARG000 | 6;6;6;6;6;6; | 13896264:138 | 13896407:138 | +;+;+;+;+;+;+; | 1931 | dnpep | 799 | 1029 | 1192 | 1166 | 1005 | 896 |
| ENSDARG000 | 6;6;6;6 | 52637308:526 | 52639009:526 | -;-;-;- | 2754 | GDF5 | 4 | 2 | 0 | 0 | 0 | 6 |
| ENSDARG000 | 6;6;6;6 | 52663097:526 | 52664759:526 | -;-;-;- | 2306 | sdc4 | 25870 | 16737 | 19674 | 28718 | 13088 | 21697 |
| ENSDARG000 | 6;6 | 25808140:258 | 25808350:258 | +;+ | 445 | NA | 0 | 0 | 1 | 2 | 8 | 1 |
| ENSDARG000 | 6;6;6;6;6;6; | 12556427:125 | 12557006:125 | -;-;-;-;-;-;-; | 2631 | slc15a1b | 4 | 0 | 0 | 2 | 0 | 0 |
| ENSDARG000 | 6;6;6;6;6;6; | 34847276:348 | 34848708:348 | -;-;-;-;-;-;-; | 2351 | slc35d1a | 445 | 356 | 288 | 203 | 396 | 282 |
| ENSDARG000 | 6;6;6;6;6;6; | 10741266:107 | 10742156:107 | -;-;-;-;-;-;-; | 2238 | ola1 | 3547 | 3804 | 5541 | 4432 | 3819 | 5409 |
| ENSDARG000 | 6;6;6;6;6;6 | 30668098:306 | 30668683:306 | +;+;+;+;+;+;+ | 2090 | TTC22 | 34 | 14 | 23 | 14 | 19 | 19 |
| ENSDARG000 | 6;6;6;6;6;6; | 25933736:259 | 25934650:259 | -;-;-;-;-;-;-; | 1776 | lmo4b | 296 | 41 | 96 | 36 | 51 | 90 |
| ENSDARG000 | 6;6 | 52718973:527 | 52719082:527 | +;+ | 277 | NA | 0 | 0 | 0 | 0 | 0 | 0 |
| ENSDARG000 | 6;6;6;6 | 52715708:527 | 52716837:527 | -;-;-;- | 1704 | oser1 | 577 | 529 | 585 | 726 | 605 | 578 |
| ENSDARG000 | 6;6;6;6;6;6; | 19315715:193 | 19315981:193 | -;-;-;-;-;-;-; | 2264 | gga3a | 831 | 881 | 1251 | 962 | 809 | 708 |
| ENSDARG000 | 6;6;6 | 6491013:6498 | 6491341:6498 | +;+;+ | 2605 | bcl11ab | 2144 | 2266 | 1531 | 1322 | 1801 | 2704 |
| ENSDARG000 | 6;6;6;6;6;6; | 39370587:393 | 39370730:393 | +;+;+;+;+;+;+; | 4153 | NA | 406 | 232 | 323 | 430 | 213 | 542 |
| ENSDARG000 | 6;6;6;6;6;6; | 41554794:415 | 41555772:415 | +;+;+;+;+;+;+; | 6570 | SRGAP3 | 404 | 249 | 358 | 296 | 578 | 428 |
| ENSDARG000 | 6;6 | 6662479:6665 | 6662667:6666 | +;+ | 1610 | zgc:113227 | 85 | 67 | 95 | 19 | 93 | 104 |
| ENSDARG000 | 6;6;6;6;6;6; | 52790049:528 | 52790064:528 | +;+;+;+;+;+;+; | 3760 | matn4 | 18 | 4 | 89 | 0 | 14 | 8 |
| ENSDARG000 | 6;6;6;6;6;6; | 40587551:405 | 40587981:405 | +;+;+;+;+;+;+; | 4859 | FRS3 | 4 | 0 | 6 | 0 | 0 | 0 |
| ENSDARG000 | 6;6;6;6;6;6; | 30676995:306 | 30677755:306 | -;-;-;-;-;-;-; | 2024 | ttc4 | 228 | 493 | 578 | 196 | 444 | 361 |
| ENSDARG000 | 6;6;6;6;6;6; | 37441789:374 | 37442354:374 | +;+;+;+;+;+;+; | 1991 | GTPBP6 | 18 | 44 | 88 | 60 | 38 | 84 |
| ENSDARG000 | 6;6;6;6;6;6; | 52757433:527 | 52757756:527 | -;-;-;-;-;-;-; | 1690 | RBPJL | 0 | 0 | 0 | 0 | 0 | 0 |
| ENSDARG000 | 6;6;6;6;6;6; | 1553442:1556 | 1556895:1556 | -;-;-;-;-;-;-; | 5021 | NA | 134 | 108 | 140 | 33 | 128 | 92 |
| ENSDARG000 | 6;6;6;6;6 | 16722014:167 | 16722199:167 | -;-;-;-;- | 842 | NA | 0 | 0 | 0 | 0 | 0 | 0 |
| ENSDARG000 | 6;6;6;6;6;6; | 9289802:9318 | 9290457:9318 | +;+;+;+;+;+;+; | 8912 | kalrnb | 112 | 2 | 95 | 50 | 148 | 36 |
| ENSDARG000 | 6;6;6;6;6;6; | 37451341:374 | 37451905:374 | -;-;-;-;-;-;-; | 1504 | NA | 13 | 12 | 16 | 16 | 28 | 23 |
| ENSDARG000 | 6;6;6;6;6;6; | 28752943:287 | 28753069:287 | +;+;+;+;+;+;+; | 1587 | tprg1 | 50 | 14 | 24 | 3 | 52 | 36 |
| ENSDARG000 | 6;6;6 | 6669369:6671 | 6669933:6671 | -;-;- | 1537 | si:dkev-261i1 | 125 | 139 | 151 | 102 | 101 | 132 |
| ENSDARG000 | 6;6;6;6;6;6; | 39392442:393 | 39393598:393 | -;-;-;-;-;-;-; | 6036 | scn8ab | 10 | 2 | 0 | 0 | 6 | 4 |
| ENSDARG000 | 6;6;6;6;6;6; | 40629066:406 | 40629313:406 | +;+;+;+;+;+;+; | 2533 | slc6a11a | 2 | 0 | 0 | 0 | 12 | 2 |
| ENSDARG000 | 6;6;6;6;6;6 | 16734086:167 | 16734110:167 | -;-;-;-;-;- | 750 | NA | 0 | 0 | 0 | 0 | 0 | 0 |
| ENSDARG000 | 6;6;6;6;6;6; | 34868156:348 | 34868751:348 | +;+;+;+;+;+;+; | 4023 | il23r | 0 | 5 | 4 | 2 | 9 | 10 |
| ENSDARG000 | 6;6;6;6;6;6; | 7735806:7735 | 7736862:7736 | -;-;-;-;-;-;-; | 7144 | myh9a | 804 | 215 | 226 | 184 | 578 | 454 |
| ENSDARG000 | 6;6;6;6;6;6; | 37460269:374 | 37461732:374 | -;-;-;-;-;-;-; | 2680 | NA | 16 | 40 | 28 | 4 | 42 | 14 |
| ENSDARG000 | 6;6;6;6 | 16736871:167 | 16736934:167 | +;+;+;+ | 374 | NA | 0 | 0 | 0 | 0 | 0 | 0 |
| ENSDARG000 | 6;6;6;6;6;6; | 19334365:193 | 19335264:193 | -;-;-;-;-;-;-; | 1520 | mif4gda | 156 | 133 | 248 | 192 | 222 | 147 |
| ENSDARG000 | 6;6;6;6;6;6; | 34908597:349 | 34908858:349 | +;+;+;+;+;+;+; | 2379 | il12rb2 | 29 | 68 | 239 | 158 | 205 | 142 |
| ENSDARG000 | 6;6;6;6;6;6; | 6736925:6736 | 6738097:6738 | -;-;-;-;-;-;-; | 2875 | ccdc85al | 32 | 32 | 88 | 16 | 34 | 36 |
| ENSDARG000 | 6;6;6;6;6;6; | 10758181:107 | 10758465:107 | -;-;-;-;-;-;-; | 2640 | GPR155 | 179 | 133 | 153 | 164 | 175 | 143 |
| ENSDARG000 | 6;6;6;6;6;6; | 30689239:306 | 30689371:306 | +;+;+;+;+;+;+; | 2932 | dnai4 | 3 | 4 | 8 | 24 | 4 | 2 |
| ENSDARG000 | 6;6;6;6;6;6; | 1567446:1567 | 1569332:1569 | -;-;-;-;-;-;-; | 4796 | NA | 395 | 193 | 472 | 443 | 322 | 329 |
| ENSDARG000 | 6;6;6 | 52853384:528 | 52854596:528 | +;+;+ | 1223 | NA | 0 | 0 | 0 | 0 | 0 | 0 |
| ENSDARG000 | 6 | 52859495 | 52861521 | + | 2027 | or90a1 | 0 | 0 | 0 | 0 | 0 | 0 |
| ENSDARG000 | 6;6;6;6;6;6; | 41808673:418 | 41808856:418 | +;+;+;+;+;+;+; | 2213 | rad18 | 104 | 252 | 226 | 76 | 156 | 150 |
| ENSDARG000 | 6;6;6;6;6;6; | 26063962:260 | 26064191:260 | -;-;-;-;-;-;-; | 1566 | HS2ST1 | 174 | 185 | 276 | 161 | 250 | 284 |
| ENSDARG000 | 6;6;6 | 30684696:306 | 30685694:306 | -;-;- | 1711 | pars2 | 22 | 76 | 83 | 30 | 76 | 56 |
| ENSDARG000 | 6;6;6 | 39499623:395 | 39500150:395 | +;+;+ | 938 | NA | 10 | 6 | 3 | 24 | 21 | 2 |
| ENSDARG000 | 6;6;6 | 39506043:395 | 39506617:395 | +;+;+ | 1012 | mettl7a.3 | 0 | 0 | 0 | 0 | 0 | 0 |
| ENSDARG000 | 6;6;6;6;6;6 | 16801893:168 | 16801961:168 | -;-;-;-;-;- | 902 | NA | 0 | 0 | 0 | 0 | 0 | 0 |
| ENSDARG000 | 6;6;6;6;6;6; | 39505565:395 | 39510548:395 | -;-;-;-;-;-;-; | 6009 | atf1 | 1217 | 1042 | 1541 | 1145 | 1341 | 845 |
| ENSDARG000 | 6;6 | 52864455:528 | 52864724:528 | +;+ | 1079 | NA | 0 | 0 | 0 | 0 | 0 | 0 |
| ENSDARG000 | 6;6;6 | 39493864:394 | 39494404:394 | +;+;+ | 1838 | mettl7a.1 | 182 | 183 | 289 | 238 | 242 | 270 |
| ENSDARG000 | 6 | 52869892 | 52871963 | + | 2072 | NA | 0 | 0 | 0 | 0 | 0 | 0 |
| ENSDARG000 | 6;6;6;6;6;6; | 39491846:394 | 39492318:394 | -;-;-;-;-;-;-; | 956 | NA | 0 | 0 | 0 | 0 | 0 | 0 |
| ENSDARG000 | 6;6 | 52873822:528 | 52873884:528 | +;+ | 2342 | or90b2 | 0 | 0 | 4 | 0 | 0 | 0 |
| ENSDARG000 | 6 | 52877147 | 52879068 | + | 1922 | NA | 0 | 0 | 0 | 0 | 0 | 0 |
| ENSDARG000 | 6;6;6;6;6;6; | 40647788:406 | 40648091:406 | -;-;-;-;-;-;-; | 1383 | ppil1 | 156 | 196 | 239 | 70 | 184 | 110 |
| ENSDARG000 | 6 | 52880292 | 52882674 | + | 2383 | or90e2 | 0 | 0 | 0 | 0 | 0 | 0 |
| ENSDARG000 | 6;6 | 37474977:374 | 37475158:374 | +;+ | 514 | NA | 0 | 2 | 4 | 2 | 6 | 7 |
| ENSDARG000 | 6 | 32699359 | 32703317 | - | 3959 | mafaa | 0 | 4 | 4 | 0 | 2 | 2 |
| ENSDARG000 | 6;6;6;6;6;6; | 40661703:406 | 40661861:406 | +;+;+;+;+;+;+; | 1850 | eno1b | 32 | 310 | 981 | 356 | 188 | 102 |
| ENSDARG000 | 6;6;6;6;6;6; | 34931597:349 | 34932680:349 | -;-;-;-;-;-;-; | 2344 | serbp1a | 25232 | 25403 | 35473 | 26281 | 27388 | 34207 |
| ENSDARG000 | 6 | 52889353 | 52891181 | + | 1829 | or137-9 | 0 | 0 | 0 | 0 | 0 | 0 |
| ENSDARG000 | 6 | 52890832 | 52891194 | + | 363 | NA | 0 | 0 | 0 | 0 | 0 | 0 |
| ENSDARG000 | 6;6;6;6;6;6; | 10781146:107 | 10783718:107 | -;-;-;-;-;-;-; | 4280 | NA | 2040 | 2397 | 2336 | 2311 | 2429 | 2564 |
| ENSDARG000 | 6 | 52891947 | 52893701 | + | 1755 | NA | 0 | 0 | 0 | 0 | 0 | 0 |
| ENSDARG000 | 6 | 52895008 | 52897484 | + | 2477 | NA | 0 | 0 | 0 | 0 | 0 | 0 |
| ENSDARG000 | 6 | 52903302 | 52905209 | + | 1908 | NA | 0 | 0 | 0 | 0 | 0 | 0 |
| ENSDARG000 | 6;6;6;6;6;6; | 13920479:139 | 13920771:139 | +;+;+;+;+;+;+; | 3550 | PTPRN | 651 | 311 | 363 | 240 | 241 | 456 |
| ENSDARG000 | 6 | 52911338 | 52913806 | + | 2469 | NA | 0 | 0 | 0 | 0 | 0 | 0 |
| ENSDARG000 | 6 | 52918537 | 52919757 | + | 1221 | or90d3 | 0 | 0 | 0 | 0 | 0 | 0 |
| ENSDARG000 | 6;6;6;6;6;6; | 37478765:374 | 37480119:374 | -;-;-;-;-;-;-; | 3612 | jade3 | 344 | 487 | 548 | 358 | 461 | 263 |
| ENSDARG000 | 6;6;6;6;6;6; | 32717438:327 | 32717919:327 | -;-;-;-;-;-;-; | 4086 | NA | 101 | 116 | 187 | 112 | 112 | 130 |
| ENSDARG000 | 6 | 52924103 | 52925772 | + | 1670 | or90b1 | 0 | 0 | 0 | 0 | 0 | 0 |
| ENSDARG000 | 6 | 52927651 | 52929994 | + | 2344 | NA | 0 | 0 | 0 | 0 | 0 | 0 |
| ENSDARG000 | 6 | 52931841 | 52933122 | + | 1282 | NA | 0 | 0 | 0 | 0 | 0 | 0 |
| ENSDARG000 | 6 | 52938534 | 52940148 | + | 1615 | NA | 0 | 0 | 0 | 0 | 0 | 0 |
| ENSDARG000 | 6;6;6;6;6 | 41956355:419 | 41956908:419 | +;+;+;+;+ | 1886 | oxtra | 0 | 0 | 0 | 0 | 0 | 0 |
| ENSDARG000 | 6;6;6;6;6;6; | 32834007:328 | 32834384:328 | +;+;+;+;+;+;+; | 2923 | NA | 297 | 202 | 193 | 78 | 184 | 172 |
| ENSDARG000 | 6;6 | 41996066:420 | 41996726:420 | -;- | 1017 | cav3 | 0 | 0 | 0 | 0 | 0 | 0 |
| ENSDARG000 | 6 | 52942895 | 52945294 | + | 2400 | or90h4 | 0 | 0 | 0 | 0 | 0 | 0 |
| ENSDARG000 | 6;6;6;6;6;6; | 12593986:125 | 12595797:125 | -;-;-;-;-;-;-; | 8469 | NA | 80 | 55 | 82 | 16 | 89 | 85 |
| ENSDARG000 | 6;6;6;6;6;6 | 16804761:168 | 16804807:168 | +;+;+;+;+;+ | 750 | NA | 0 | 0 | 0 | 0 | 0 | 0 |
| ENSDARG000 | 6;6;6;6 | 26305223:263 | 26305659:263 | +;+;+;+ | 2362 | NA | 10 | 2 | 8 | 4 | 2 | 0 |
| ENSDARG000 | 6;6;6;6;6;6; | 19344749:193 | 19345223:193 | -;-;-;-;-;-;-; | 1497 | grb2a | 354 | 365 | 282 | 294 | 319 | 271 |
| ENSDARG000 | 6;6;6;6;6;6; | 26309968:263 | 26310002:263 | +;+;+;+;+;+;+; | 3466 | dzip1l | 0 | 0 | 6 | 0 | 0 | 0 |
| ENSDARG000 | 6;6;6;6;6;6; | 52947186:529 | 52947444:529 | +;+;+;+;+;+;+; | 3438 | NA | 552 | 335 | 255 | 96 | 289 | 226 |
| ENSDARG000 | 6;6;6;6;6;6; | 10812337:108 | 10813134:108 | -;-;-;-;-;-;-; | 2280 | chrna1 | 2 | 10 | 12 | 6 | 12 | 0 |
| ENSDARG000 | 6;6;6;6;6;6; | 28795780:287 | 28795811:287 | +;+;+;+;+;+;+; | 3536 | tp63 | 244 | 9 | 8 | 21 | 61 | 26 |
| ENSDARG000 | 6;6;6;6;6;6; | 6779114:6779 | 6779247:6779 | +;+;+;+;+;+;+; | 4357 | sec23b | 1114 | 1061 | 1558 | 916 | 1222 | 763 |
| ENSDARG000 | 6;6 | 42031836:420 | 42033937:420 | -;- | 2597 | NA | 50 | 32 | 38 | 113 | 62 | 19 |
| ENSDARG000 | 6;6;6;6;6;6; | 40671336:406 | 40671452:406 | +;+;+;+;+;+;+; | 4359 | NA | 775 | 496 | 584 | 256 | 758 | 746 |
| ENSDARG000 | 6;6;6 | 53041180:530 | 53042130:530 | -;-;- | 1103 | inka1b | 1126 | 563 | 976 | 1434 | 930 | 612 |
| ENSDARG000 | 6;6;6;6;6;6; | 37513911:375 | 37515623:375 | -;-;-;-;-;-;-; | 2956 | rp2 | 349 | 294 | 313 | 344 | 447 | 319 |
| ENSDARG000 | 6;6 | 30703828:307 | 30704185:307 | +;+ | 753 | insl5a | 0 | 1 | 1 | 7 | 0 | 4 |
| ENSDARG000 | 6 | 26557979 | 26559921 | - | 1943 | sox14 | 0 | 0 | 0 | 0 | 0 | 0 |
| ENSDARG000 | 6;6;6;6;6;6; | 13993500:139 | 13993913:139 | -;-;-;-;-;-;-; | 1792 | gca | 813 | 1092 | 788 | 814 | 770 | 767 |
| ENSDARG000 | 6 | 39524924 | 39525505 | - | 582 | NA | 222 | 132 | 114 | 141 | 112 | 165 |
| ENSDARG000 | 6;6;6;6;6;6; | 42038358:420 | 42039322:420 | -;-;-;-;-;-;-; | 2456 | parp3 | 1174 | 2024 | 1602 | 1108 | 2277 | 1649 |
| ENSDARG000 | 6;6;6;6;6 | 26572001:265 | 26572107:265 | +;+;+;+;+ | 817 | NA | 0 | 0 | 0 | 0 | 0 | 0 |
| ENSDARG000 | 6;6;6;6;6;6; | 39531135:395 | 39531953:395 | -;-;-;-;-;-;-; | 5454 | NA | 792 | 338 | 587 | 386 | 444 | 395 |
| ENSDARG000 | 6;6;6;6;6;6; | 53050166:530 | 53051020:530 | -;-;-;-;-;-;-; | 2618 | gnai2b | 64 | 44 | 36 | 38 | 180 | 85 |
| ENSDARG000 | 6;6;6;6;6;6; | 7977611:8041 | 7977995:8041 | +;+;+;+;+;+;+; | 3452 | palm3 | 28 | 8 | 0 | 6 | 12 | 0 |
| ENSDARG000 | 6;6;6;6;6;6; | 32882821:328 | 32883151:328 | +;+;+;+;+;+;+; | 2172 | NA | 741 | 715 | 213 | 298 | 984 | 649 |
| ENSDARG000 | 6;6;6;6;6;6; | 18074521:180 | 18074703:180 | -;-;-;-;-;-;-; | 1998 | SEC14L1 | 367 | 236 | 369 | 228 | 332 | 394 |
| ENSDARG000 | 6 | 19383267 | 19384304 | + | 1038 | MCHR1 | 0 | 0 | 0 | 0 | 0 | 0 |
| ENSDARG000 | 6;6;6;6 | 40689357:406 | 40690630:406 | -;-;-;- | 3962 | perm1 | 285 | 166 | 266 | 137 | 307 | 238 |
| ENSDARG000 | 6;6;6;6;6;6 | 16854742:168 | 16855038:168 | -;-;-;-;-;- | 1100 | NA | 0 | 0 | 0 | 0 | 0 | 0 |
| ENSDARG000 | 6;6;6;6;6;6; | 1831111:1831 | 1831392:1831 | -;-;-;-;-;-;-; | 3064 | DLGAP4 | 51 | 58 | 115 | 57 | 48 | 75 |
| ENSDARG000 | 6;6;6;6;6;6; | 30709926:307 | 30715896:307 | -;-;-;-;-;-;-; | 9258 | SGIP1 | 2616 | 1504 | 2626 | 5795 | 2178 | 3733 |
| ENSDARG000 | 6;6;6;6;6;6; | 10830844:108 | 10831150:108 | -;-;-;-;-;-;-; | 1040 | atp5mc3b | 5433 | 6878 | 8010 | 6278 | 6302 | 4837 |
| ENSDARG000 | 6;6;6;6;6;6; | 42051480:420 | 42051991:420 | -;-;-;-;-;-;-; | 3363 | GRM2 | 0 | 0 | 0 | 0 | 0 | 0 |
| ENSDARG000 | 6;6;6;6;6;6; | 19481394:194 | 19482412:194 | -;-;-;-;-;-;-; | 3635 | usp43a | 6 | 12 | 0 | 0 | 6 | 6 |
| ENSDARG000 | 6;6;6;6;6;6; | 34943351:349 | 34944133:349 | -;-;-;-;-;-;-; | 2018 | hsd17b7 | 66 | 36 | 66 | 50 | 56 | 27 |
| ENSDARG000 | 6;6;6;6;6;6; | 40707162:407 | 40708793:407 | -;-;-;-;-;-;-; | 2830 | si:ch211-157 | 144 | 161 | 271 | 268 | 150 | 250 |
| ENSDARG000 | 6;6;6;6;6;6; | 40714811:407 | 40714866:407 | +;+;+;+;+;+;+; | 1236 | NA | 0 | 0 | 0 | 0 | 0 | 0 |
| ENSDARG000 | 6;6;6;6;6;6; | 40715637:407 | 40718341:407 | -;-;-;-;-;-;-; | 5122 | kbtbd12 | 2 | 6 | 0 | 0 | 22 | 13 |
| ENSDARG000 | 6;6;6;6;6;6; | 53147446:531 | 53152980:531 | -;-;-;-;-;-;-; | 7022 | SLC38A3 | 8 | 8 | 42 | 32 | 22 | 20 |
| ENSDARG000 | 6;6 | 26653913:267 | 26653981:267 | +;+ | 217 | NA | 0 | 0 | 0 | 0 | 0 | 0 |
| ENSDARG000 | 6;6;6;6;6;6; | 53263854:532 | 53263979:532 | -;-;-;-;-;-;-; | 3942 | rbm5 | 3082 | 1961 | 5161 | 5233 | 4347 | 4312 |
| ENSDARG000 | 6;6;6;6;6;6; | 18142623:181 | 18142838:181 | +;+;+;+;+;+;+; | 2298 | NA | 15 | 2 | 24 | 0 | 15 | 37 |
| ENSDARG000 | 6;6;6;6;6;6; | 34960025:349 | 34963934:349 | -;-;-;-;-;-;-; | 6564 | DDR2 | 3188 | 1564 | 3372 | 3040 | 2426 | 1804 |
| ENSDARG000 | 6;6;6;6;6;6; | 39613357:396 | 39615334:396 | -;-;-;-;-;-;-; | 3487 | atf7b | 897 | 721 | 1361 | 1397 | 1129 | 1132 |
| ENSDARG000 | 6;6;6;6;6;6; | 37529383:375 | 37529746:375 | +;+;+;+;+;+;+; | 5182 | slc9a7 | 714 | 463 | 637 | 532 | 934 | 665 |
| ENSDARG000 | 6;6;6;6;6;6 | 16867518:168 | 16867542:168 | -;-;-;-;-;- | 750 | NA | 0 | 0 | 0 | 0 | 0 | 0 |
| ENSDARG000 | 6;6;6;6;6 | 16870004:168 | 16870371:168 | +;+;+;+;+ | 877 | NA | 0 | 0 | 0 | 0 | 0 | 0 |
| ENSDARG000 | 6;6;6;6;6;6; | 2030703:2030 | 2030923:2030 | +;+;+;+;+;+;+; | 7524 | quo | 111 | 48 | 106 | 18 | 10 | 42 |
| ENSDARG000 | 6;6;6 | 18196961:181 | 18198188:181 | -;-;- | 1774 | ppp1r27b | 8 | 11 | 2 | 2 | 0 | 4 |
| ENSDARG000 | 6;6;6;6;6;6; | 32966750:329 | 32967758:329 | -;-;-;-;-;-;-; | 2804 | ADCYAP1R1 | 0 | 0 | 0 | 0 | 2 | 0 |

| | | | | | | | | | | | |
|---|---|---|---|---|---|---|---|---|---|---|---|
| ENSDARG000 | 6:6:6:6:6:6:6 | 19667619:196 | 19668051:196 | -:-:-:-:-:-:-:-:-: | 2512 | cfap52 | 2 | 0 | 0 | 0 | 0 | 0 |
| ENSDARG000 | 6;6 | 53288018:532 | 53288250:532 | +;+ | 1383 | NA | 2 | 0 | 0 | 0 | 0 | 4 |
| ENSDARG000 | 6:6:6:6:6:6:6 | 26874839:268 | 26876051:268 | -:-:-:-:-:-:-:-:-: | 5644 | hdlbpa | 24747 | 27066 | 46626 | 39501 | 30830 | 33790 |
| ENSDARG000 | 6:6:6:6:6:6:6 | 53304977:533 | 53307329:533 | -:-:-:-:-:-:-:-:-: | 3973 | gnb1b | 7678 | 9571 | 8278 | 7653 | 9277 | 6717 |
| ENSDARG000 | 6;6;6;6;6 | 16893171:168 | 16893195:168 | -;-;-;-;- | 705 | NA | 0 | 0 | 0 | 0 | 0 | 0 |
| ENSDARG000 | 6;6;6;6;6;6 | 33070766:330 | 33071385:330 | -;-;-;-;-;- | 1205 | NA | 0 | 0 | 0 | 0 | 0 | 0 |
| ENSDARG000 | 6:6:6:6:6:6:6 | 14030903:140 | 14033334:140 | -:-:-:-:-:-:-:-:-: | 4118 | etv5b | 2921 | 2002 | 2842 | 1625 | 2825 | 1856 |
| ENSDARG000 | 6:6:6:6:6:6:6 | 53349966:533 | 53350262:533 | +:+:+:+:+:+:+: | 933 | C6H3orf18 | 250 | 47 | 135 | 98 | 235 | 122 |
| ENSDARG000 | 6:6:6:6:6:6:6 | 8079974:808 | 8080010:808 | +:+:+:+:+:+:+: | 3366 | NA | 9 | 9 | 28 | 4 | 20 | 10 |
| ENSDARG000 | 6:6:6:6:6:6:6 | 35033457:350 | 35036916:350 | -:-:-:-:-:-:-:-:-: | 6082 | uap1 | 651 | 648 | 726 | 517 | 940 | 569 |
| ENSDARG000 | 6;6;6;6;6;6 | 53373498:533 | 53378377:533 | -;-;-;-;-;- | 5217 | NA | 14 | 18 | 49 | 34 | 90 | 110 |
| ENSDARG000 | 6;6;6 | 8129543:813 | 8129793:813 | +;+;+ | 2525 | klf1 | 6 | 0 | 27 | 4 | 102 | 6 |
| ENSDARG000 | 6:6:6:6:6:6:6 | 53394125:533 | 53394537:533 | -:-:-:-:-:-:-:-:-: | 2430 | mst1 | 27 | 16 | 21 | 14 | 62 | 26 |
| ENSDARG000 | 6:6:6:6:6:6:6 | 40745437:407 | 40745788:407 | -:-:-:-:-:-:-:-:-: | 6228 | NA | 10 | 16 | 6 | 16 | 0 | 16 |
| ENSDARG000 | 6;6;6;6 | 42145626:421 | 42146596:421 | -;-;-;- | 1928 | tex264a | 1482 | 933 | 965 | 754 | 1070 | 1176 |
| ENSDARG000 | 6;6;6;6;6 | 16895685:168 | 16896043:168 | +;+;+;+;+ | 868 | NA | 0 | 0 | 0 | 0 | 0 | 0 |
| ENSDARG000 | 6:6:6:6:6:6:6 | 18203464:182 | 18204694:182 | -:-:-:-:-:-:-:-:-: | 2852 | p4hb | 5778 | 7247 | 12719 | 5411 | 9423 | 6731 |
| ENSDARG000 | 6;6;6 | 8172227:817 | 8172479:818 | +;+;+ | 4331 | nfil3-5 | 744 | 1831 | 1219 | 1092 | 318 | 540 |
| ENSDARG000 | 6:6:6:6:6:6:6 | 40723554:407 | 40723761:407 | +:+:+:+:+:+:+: | 2978 | slc26a6l | 2 | 2 | 34 | 8 | 30 | 2 |
| ENSDARG000 | 6;6;6 | 53429228:534 | 53429594:534 | +;+;+ | 2288 | abhd14b | 160 | 172 | 254 | 168 | 300 | 216 |
| ENSDARG000 | 6;6 | 37603968:376 | 37605012:376 | -;- | 2825 | CHST7 | 2748 | 1276 | 2733 | 1306 | 2461 | 1856 |
| ENSDARG000 | 6;6;6;6;6;6 | 40775800:407 | 40776455:407 | +;+;+;+;+;+ | 2133 | NA | 0 | 8 | 0 | 0 | 0 | 0 |
| ENSDARG000 | 6 | 40769517 | 40771813 | - | 2297 | h1-10 | 3123 | 3992 | 4139 | 8860 | 2027 | 3367 |
| ENSDARG000 | 6:6:6:6:6:6:6 | 40743367:407 | 40743369:407 | -:-:-:-:-:-:-:-:-: | 2154 | arpc4 | 1818 | 2635 | 2130 | 1670 | 2599 | 1956 |
| ENSDARG000 | 6:6:6:6:6:6:6 | 40730869:407 | 40735550:407 | -:-:-:-:-:-:-:-:-: | 6560 | P4HTM | 8 | 8 | 44 | 26 | 28 | 64 |
| ENSDARG000 | 6:6:6:6:6:6:6 | 30851545:308 | 30854230:308 | -:-:-:-:-:-:-:-:-: | 5574 | PDE4B | 3936 | 4299 | 4480 | 2739 | 2453 | 2029 |
| ENSDARG000 | 6;6;6;6;6 | 54063032:540 | 54063652:540 | -;-;-;-;- | 1700 | hyal1 | 314 | 214 | 212 | 165 | 498 | 335 |
| ENSDARG000 | 6;6;6;6;6 | 16946724:169 | 16946748:169 | -;-;-;-;- | 579 | NA | 0 | 0 | 0 | 0 | 0 | 0 |
| ENSDARG000 | 6:6:6:6:6:6:6 | 42291308:422 | 42291526:422 | -:-:-:-:-:-:-:-:-: | 4916 | fancd2 | 70 | 475 | 621 | 259 | 530 | 316 |
| ENSDARG000 | 6 | 54078703 | 54080052 | + | 1350 | fhit | 233 | 153 | 332 | 200 | 351 | 319 |
| ENSDARG000 | 6:6:6:6:6:6:6 | 19689464:196 | 19689565:196 | +:+:+:+:+:+:+: | 1505 | stx8 | 323 | 268 | 222 | 162 | 290 | 349 |
| ENSDARG000 | 6:6:6:6:6:6:6 | 8210115:821 | 8211317:821 | -:-:-:-:-:-:-:-:-: | 3874 | NA | 44 | 29 | 79 | 32 | 16 | 54 |
| ENSDARG000 | 6:6:6:6:6:6:6 | 10836147:108 | 10839940:108 | -:-:-:-:-:-:-:-:-: | 5218 | lnpk | 302 | 274 | 341 | 238 | 432 | 356 |
| ENSDARG000 | 6:6:6:6:6:6:6 | 39633065:396 | 39637574:396 | -:-:-:-:-:-:-:-:-: | 6940 | NA | 1053 | 1325 | 1841 | 687 | 2020 | 1400 |
| ENSDARG000 | 6;6;6;6;6;6;6 | 54093057:540 | 54093894:540 | -;-;-;-;-;-;- | 2063 | HYAL2 | 435 | 136 | 130 | 316 | 537 | 569 |
| ENSDARG000 | 6;6;6;6;6;6;6 | 54117004:541 | 54117423:541 | -;-;-;-;-;-;- | 829 | tusc2a | 91 | 98 | 56 | 111 | 54 | 68 |
| ENSDARG000 | 6:6:6:6:6:6:6 | 33076839:330 | 33076961:330 | +:+:+:+:+:+:+: | 3428 | pomgnt1 | 358 | 301 | 478 | 221 | 494 | 448 |
| ENSDARG000 | 6:6:6:6:6:6:6 | 54142311:541 | 54142500:541 | +:+:+:+:+:+:+: | 4079 | hmga1b | 58 | 82 | 103 | 42 | 258 | 26 |
| ENSDARG000 | 6:6:6:6:6:6:6 | 35052721:350 | 35053087:350 | +:+:+:+:+:+:+: | 2259 | uhmk1 | 344 | 182 | 314 | 169 | 359 | 375 |
| ENSDARG000 | 6;6;6;6 | 6797520:679 | 6797758:679 | +;+;+;+ | 894 | NA | 113 | 89 | 111 | 124 | 74 | 176 |
| ENSDARG000 | 6;6;6;6;6 | 54154807:541 | 54155064:541 | +;+;+;+;+ | 931 | nudt3b | 82 | 66 | 109 | 107 | 153 | 92 |
| ENSDARG000 | 6:6:6:6:6:6:6 | 12853655:128 | 12854095:128 | +:+:+:+:+:+:+: | 1942 | FAM117B | 0 | 0 | 0 | 0 | 0 | 0 |
| ENSDARG000 | 6:6:6:6:6:6:6 | 39661588:396 | 39662255:396 | -:-:-:-:-:-:-:-:-: | 1444 | pfdn5 | 1224 | 1642 | 1641 | 1520 | 1292 | 1546 |
| ENSDARG000 | 6:6:6:6:6:6:6 | 12797648:128 | 12798636:128 | -:-:-:-:-:-:-:-:-: | 5060 | bmpr2a | 650 | 566 | 667 | 451 | 746 | 640 |
| ENSDARG000 | 6:6:6:6:6:6:6 | 26948093:269 | 26948201:269 | +:+:+:+:+:+:+: | 6225 | FARP2 | 127 | 24 | 74 | 8 | 32 | 52 |
| ENSDARG000 | 6;6 | 39666347:396 | 39666553:396 | +;+ | 434 | NA | 0 | 0 | 0 | 0 | 2 | 0 |
| ENSDARG000 | 6:6:6:6:6:6:6 | 2093206:209 | 2093427:209 | +:+:+:+:+:+:+: | 3330 | tgm2b | 84 | 0 | 48 | 10 | 22 | 10 |
| ENSDARG000 | 6:6:6:6:6:6:6 | 54174696:541 | 54174859:541 | -:-:-:-:-:-:-:-:-: | 1083 | rps10 | 14594 | 15910 | 23629 | 23933 | 17330 | 21386 |
| ENSDARG000 | 6:6:6:6:6:6:6 | 19804048:198 | 19809000:198 | -;-;-;-;-;-;-;- | 7005 | ntn1a | 91 | 12 | 18 | 29 | 29 | 49 |
| ENSDARG000 | 6;6;6;6;6;6 | 16949735:169 | 16949861:169 | +;+;+;+;+;+ | 899 | NA | 0 | 0 | 0 | 0 | 0 | 0 |
| ENSDARG000 | 6:6:6:6:6:6:6 | 35106115:351 | 35106659:351 | -:-:-:-:-:-:-:-:-: | 3626 | NOS1AP | 12 | 0 | 20 | 8 | 6 | 0 |
| ENSDARG000 | 6:6:6:6:6:6:6 | 6802142:680 | 6802624:680 | +:+:+:+:+:+:+: | 5165 | dtd1 | 368 | 451 | 796 | 480 | 493 | 416 |
| ENSDARG000 | 6;6 | 35099897:351 | 35102930:351 | -;- | 3345 | NA | 56 | 2 | 10 | 2 | 10 | 12 |
| ENSDARG000 | 6:6:6:6:6:6:6 | 54187643:542 | 54187694:542 | +:+:+:+:+:+:+: | 2945 | pacsin1b | 12 | 24 | 24 | 16 | 74 | 85 |
| ENSDARG000 | 6:6:6:6:6:6:6 | 37623693:376 | 37623950:376 | +:+:+:+:+:+:+: | 4406 | tubgcp5 | 220 | 458 | 474 | 320 | 356 | 256 |
| ENSDARG000 | 6;6;6;6;6 | 16980025:169 | 16980349:169 | -;-;-;-;- | 855 | NA | 0 | 0 | 0 | 0 | 0 | 0 |
| ENSDARG000 | 6;6;6;6;6;6;6 | 18244220:182 | 18245197:182 | -;-;-;-;-;-;- | 1312 | anapc11 | 256 | 299 | 266 | 155 | 193 | 152 |
| ENSDARG000 | 6:6:6:6:6:6:6 | 28914233:289 | 28915493:289 | -:-:-:-:-:-:-:-:-: | 3273 | tbc1d23 | 876 | 853 | 1083 | 720 | 1011 | 981 |
| ENSDARG000 | 6:6:6:6:6:6:6 | 10871659:108 | 10872112:108 | +:+:+:+:+:+:+: | 1488 | mtx2 | 521 | 602 | 492 | 298 | 488 | 298 |
| ENSDARG000 | 6;6;6;6;6 | 54325008:543 | 54326863:543 | -;-;-;-;- | 2759 | ilrun | 967 | 725 | 959 | 864 | 918 | 1039 |
| ENSDARG000 | 6:6:6:6:6:6:6 | 39703232:397 | 39703476:397 | -:-:-:-:-:-:-:-:-: | 2440 | NA | 42 | 61 | 60 | 12 | 197 | 61 |
| ENSDARG000 | 6:6:6:6:6:6:6 | 54358529:543 | 54358923:543 | +:+:+:+:+:+:+: | 3489 | NA | 26 | 0 | 12 | 10 | 6 | 10 |
| ENSDARG000 | 6;6;6;6;6 | 9457966:945 | 9458104:945 | -;-;-;-;- | 891 | NA | 0 | 0 | 4 | 0 | 4 | 2 |
| ENSDARG000 | 6;6;6;6;6;6 | 16982613:169 | 16982659:169 | +;+;+;+;+;+ | 750 | NA | 0 | 0 | 0 | 0 | 0 | 0 |
| ENSDARG000 | 6;6 | 39700200:397 | 39700387:397 | +;+ | 543 | NA | 0 | 0 | 0 | 0 | 0 | 0 |
| ENSDARG000 | 6:6:6:6:6:6:6 | 39668850:396 | 39669427:396 | -:-:-:-:-:-:-:-:-: | 7188 | NA | 145 | 642 | 686 | 335 | 492 | 287 |
| ENSDARG000 | 6:6:6:6:6:6:6 | 2145564:214 | 2146058:214 | -:-:-:-:-:-:-:-:-: | 2402 | tgm5l | 22 | 8 | 8 | 22 | 18 | 8 |
| ENSDARG000 | 6:6:6:6:6:6:6 | 35362225:353 | 35362346:353 | +:+:+:+:+:+:+: | 2633 | rgs4 | 62 | 14 | 158 | 64 | 116 | 32 |
| ENSDARG000 | 6:6:6:6:6:6:6 | 9471367:947 | 9471498:947 | -;-;-;-;-;-;-;- | 1945 | NA | 307 | 434 | 226 | 102 | 325 | 104 |
| ENSDARG000 | 6;6 | 42338309:423 | 42338573:423 | +;+ | 866 | gpx1b | 0 | 4 | 4 | 10 | 14 | 0 |
| ENSDARG000 | 6:6:6:6:6:6:6 | 8263548:826 | 8265385:826 | -:-:-:-:-:-:-:-:-: | 5672 | SLC44A2 | 572 | 704 | 784 | 776 | 744 | 900 |
| ENSDARG000 | 6:6:6:6:6:6:6 | 8250382:825 | 8250623:825 | -:-:-:-:-:-:-:-:-: | 1846 | odad3 | 4 | 3 | 5 | 2 | 4 | 4 |
| ENSDARG000 | 6:6:6:6:6:6:6 | 19933763:199 | 19933976:199 | +:+:+:+:+:+:+: | 4883 | PIK3R5 | 72 | 88 | 40 | 22 | 180 | 100 |
| ENSDARG000 | 6;6;6;6;6 | 17032975:170 | 17033205:170 | -;-;-;-;- | 836 | NA | 0 | 0 | 0 | 0 | 0 | 0 |
| ENSDARG000 | 6:6:6:6:6:6:6 | 9542448:954 | 9542962:954 | -:-:-:-:-:-:-:-:-: | 2745 | MAP3K2 | 337 | 119 | 238 | 121 | 253 | 209 |
| ENSDARG000 | 6;6;6;6;6;6 | 54430413:544 | 54430905:544 | -;-;-;-;-;- | 917 | snrpc | 421 | 718 | 695 | 350 | 615 | 463 |
| ENSDARG000 | 6:6:6:6:6:6:6 | 6822592:682 | 6822696:682 | +:+:+:+:+:+:+: | 2677 | RIN2 | 94 | 58 | 147 | 189 | 294 | 97 |
| ENSDARG000 | 6;6;6;6;6 | 12908959:129 | 12909107:129 | -;-;-;-;- | 575 | NA | 0 | 0 | 0 | 0 | 0 | 0 |
| ENSDARG000 | 6;6;6;6 | 54438355:544 | 54438766:544 | -;-;-;- | 926 | sys1 | 316 | 359 | 363 | 409 | 356 | 377 |
| ENSDARG000 | 6:6:6:6:6:6:6 | 12870744:128 | 12871370:128 | -:-:-:-:-:-:-:-:-: | 3141 | ical1 | 521 | 359 | 487 | 477 | 287 | 383 |
| ENSDARG000 | 6;6;6;6;6;6 | 17050800:170 | 17050824:170 | -;-;-;-;-;- | 750 | NA | 0 | 0 | 0 | 0 | 0 | 0 |
| ENSDARG000 | 6;6;6;6;6 | 54496808:544 | 54496924:544 | +;+;+;+;+ | 1176 | NA | 4 | 2 | 3 | 6 | 20 | 4 |
| ENSDARG000 | 6;6;6 | 12922002:129 | 12922186:129 | +;+;+ | 1726 | cxcr4a | 118 | 40 | 2 | 4 | 48 | 73 |
| ENSDARG000 | 6;6;6 | 35384191:353 | 35384337:353 | +;+;+ | 419 | NA | 0 | 0 | 0 | 6 | 0 | 0 |
| ENSDARG000 | 6:6:6:6:6:6:6 | 35387367:353 | 35389942:353 | -;-;-;-;-;-;-;- | 3255 | rgs5a | 107 | 2 | 8 | 24 | 60 | 18 |
| ENSDARG000 | 6:6:6:6:6:6:6 | 2174082:219 | 2174437:219 | +:+:+:+:+:+:+: | 2071 | ACVR1B | 383 | 209 | 440 | 308 | 221 | 284 |
| ENSDARG000 | 6:6:6:6:6:6:6 | 18251140:182 | 18251304:182 | +:+:+:+:+:+:+: | 3548 | ccdc40 | 19 | 172 | 335 | 412 | 185 | 37 |
| ENSDARG000 | 6:6:6:6:6:6:6 | 14086688:140 | 14088373:140 | -:-:-:-:-:-:-:-:-: | 3500 | cacnb4b | 673 | 412 | 717 | 1259 | 665 | 897 |
| ENSDARG000 | 6;6 | 18266596:182 | 18266974:182 | -;- | 638 | NA | 0 | 2 | 3 | 2 | 2 | 0 |
| ENSDARG000 | 6;6 | 18290802:182 | 18291020:182 | -;- | 338 | NA | 0 | 0 | 0 | 0 | 0 | 0 |
| ENSDARG000 | 6;6;6;6;6;6;6 | 18254400:182 | 18254447:182 | +;+;+;+;+;+;+ | 680 | NA | 0 | 10 | 49 | 19 | 9 | 0 |
| ENSDARG000 | 6:6:6:6:6:6:6 | 54538948:545 | 54539061:545 | +:+:+:+:+:+:+: | 2166 | NA | 0 | 0 | 0 | 0 | 0 | 0 |
| ENSDARG000 | 6:6:6:6:6:6:6 | 42345005:423 | 42346764:423 | -:-:-:-:-:-:-:-:-: | 4871 | NA | 1179 | 1503 | 1828 | 608 | 1668 | 1023 |
| ENSDARG000 | 6:6:6:6:6:6:6 | 33104544:331 | 33105689:331 | -:-:-:-:-:-:-:-:-: | 1935 | TSPAN1 | 0 | 0 | 0 | 0 | 0 | 4 |
| ENSDARG000 | 6:6:6:6:6:6:6 | 9568149:956 | 9569173:956 | -;-;-;-;-;-;-;- | 2623 | cyp27c1 | 35 | 16 | 4 | 2 | 4 | 0 |
| ENSDARG000 | 6:6:6:6:6:6:6 | 6850821:685 | 6851230:685 | +:+:+:+:+:+:+: | 6017 | rapgef4b | 4 | 2 | 8 | 0 | 48 | 28 |
| ENSDARG000 | 6;6;6;6;6 | 35443430:354 | 35444307:354 | -;-;-;-;- | 1306 | rgs16 | 2 | 0 | 4 | 0 | 2 | 4 |
| ENSDARG000 | 6:6:6:6:6:6:6 | 28946041:289 | 28946400:289 | -:-:-:-:-:-:-:-:-: | 1981 | tomm70a | 1316 | 1440 | 1993 | 918 | 1511 | 1463 |
| ENSDARG000 | 6:6:6:6:6:6:6 | 9595414:959 | 9595834:959 | -:-:-:-:-:-:-:-:-: | 2759 | ercc3 | 321 | 411 | 455 | 278 | 430 | 377 |
| ENSDARG000 | 6:6:6:6:6:6:6 | 54576520:545 | 54576686:545 | +:+:+:+:+:+:+: | 1938 | tead3b | 454 | 322 | 612 | 521 | 270 | 258 |
| ENSDARG000 | 6;6;6 | 31069530:310 | 31069768:310 | -;-;- | 901 | NA | 40 | 22 | 2 | 7 | 4 | 2 |
| ENSDARG000 | 6:6:6:6:6:6:6 | 35459369:354 | 35462541:354 | -:-:-:-:-:-:-:-:-: | 4501 | rgs8 | 0 | 0 | 0 | 0 | 8 | 6 |
| ENSDARG000 | 6:6:6:6:6:6:6 | 9630523:963 | 9631260:963 | -:-:-:-:-:-:-:-:-: | 2075 | wdr12 | 410 | 485 | 722 | 324 | 529 | 440 |
| ENSDARG000 | 6;6;6;6;6;6 | 17053650:170 | 17053696:170 | +;+;+;+;+;+ | 750 | NA | 0 | 0 | 0 | 0 | 0 | 0 |
| ENSDARG000 | 6;6;6;6 | 10888798:108 | 10891210:108 | -;-;-;- | 3002 | nfe2l2b | 12 | 24 | 177 | 245 | 8 | 26 |
| ENSDARG000 | 6;6 | 10903494:109 | 10903594:109 | +;+ | 487 | NA | 0 | 0 | 0 | 0 | 0 | 0 |
| ENSDARG000 | 6;6;6;6;6 | 17099672:171 | 17099958:171 | -;-;-;-;- | 966 | NA | 0 | 0 | 0 | 0 | 0 | 0 |
| ENSDARG000 | 6:6:6:6:6:6:6 | 27041594:270 | 27042015:270 | -:-:-:-:-:-:-:-:-: | 2118 | stk25a | 28 | 4 | 18 | 8 | 12 | 18 |
| ENSDARG000 | 6:6:6:6:6:6:6 | 23712911:237 | 23712953:237 | +:+:+:+:+:+:+: | 5534 | pik3r6a | 221 | 434 | 582 | 463 | 967 | 364 |
| ENSDARG000 | 6;6;6;6;6;6;6 | 2222764:222 | 2222796:222 | +;+;+;+;+;+;+ | 586 | NA | 0 | 0 | 0 | 0 | 0 | 0 |
| ENSDARG000 | 6:6:6:6:6:6:6 | 2206495:221 | 2208789:221 | -:-:-:-:-:-:-:-:-: | 3405 | prkag1 | 799 | 840 | 1135 | 826 | 736 | 650 |
| ENSDARG000 | 6;6;6 | 10910589:109 | 10911232:109 | -;-;- | 963 | cycsb | 2371 | 4022 | 3704 | 1534 | 2679 | 1593 |
| ENSDARG000 | 6:6:6:6:6:6:6 | 54680730:546 | 54681164:546 | +:+:+:+:+:+:+: | 4524 | NA | 638 | 621 | 717 | 616 | 1261 | 610 |
| ENSDARG000 | 6:6:6:6:6:6:6 | 39737408:397 | 39740172:397 | -:-:-:-:-:-:-:-:-: | 5690 | pfkmb | 0 | 2 | 16 | 10 | 4 | 4 |
| ENSDARG000 | 6;6;6;6;6;6 | 17110835:171 | 17110859:171 | -;-;-;-;-;- | 750 | NA | 0 | 0 | 0 | 0 | 0 | 0 |
| ENSDARG000 | 6;6;6;6;6;6 | 17113677:171 | 17113723:171 | +;+;+;+;+;+ | 750 | NA | 0 | 0 | 0 | 0 | 0 | 0 |
| ENSDARG000 | 6:6:6:6:6:6:6 | 35675784:356 | 35677723:356 | -:-:-:-:-:-:-:-:-: | 3739 | brinp3a.1 | 10 | 0 | 0 | 2 | 10 | 0 |
| ENSDARG000 | 6:6:6:6:6:6:6 | 31184058:311 | 31187129:311 | -:-:-:-:-:-:-:-:-: | 6038 | lepr | 728 | 484 | 568 | 1261 | 248 | 485 |
| ENSDARG000 | 6:6:6:6:6:6:6 | 54711306:547 | 54711613:547 | +:+:+:+:+:+:+: | 2833 | NA | 128 | 36 | 4 | 36 | 100 | 142 |
| ENSDARG000 | 6;6 | 35794799:357 | 35795014:357 | +;+ | 277 | NA | 0 | 0 | 0 | 0 | 0 | 0 |
| ENSDARG000 | 6;6;6;6;6;6 | 17181157:171 | 17181203:171 | +;+;+;+;+;+ | 750 | NA | 0 | 0 | 0 | 0 | 0 | 0 |
| ENSDARG000 | 6:6:6:6:6:6:6 | 54761141:547 | 54761488:547 | -:-:-:-:-:-:-:-:-: | 1162 | NA | 0 | 0 | 0 | 0 | 0 | 0 |
| ENSDARG000 | 6;6;6;6;6 | 17208022:172 | 17208127:172 | +;+;+;+;+ | 807 | NA | 0 | 0 | 0 | 0 | 0 | 0 |
| ENSDARG000 | 6;6;6;6;6 | 17204785:172 | 17204978:172 | -;-;-;-;- | 807 | NA | 0 | 0 | 0 | 0 | 0 | 0 |
| ENSDARG000 | 6;6 | 17119279:172 | 17119576:172 | -;- | 468 | NA | 0 | 0 | 0 | 0 | 0 | 0 |
| ENSDARG000 | 6;6;6;6;6;6 | 17178321:171 | 17178345:171 | -;-;-;-;-;- | 750 | NA | 0 | 0 | 0 | 0 | 0 | 0 |
| ENSDARG000 | 6:6:6:6:6:6:6 | 23809163:238 | 23809601:238 | +:+:+:+:+:+:+: | 2344 | glulb | 23985 | 21296 | 32216 | 34468 | 38394 | 36231 |
| ENSDARG000 | 6:6:6:6:6:6:6 | 28970513:289 | 28970622:289 | -:-:-:-:-:-:-:-:-: | 2147 | glmnb | 0 | 0 | 0 | 0 | 0 | 0 |
| ENSDARG000 | 6;6;6 | 9651703:965 | 9651831:965 | +;+;+ | 527 | NA | 0 | 0 | 0 | 0 | 0 | 0 |
| ENSDARG000 | 6;6;6 | 10923105:109 | 10924192:109 | -;-;- | 1184 | ccr7 | 0 | 2 | 0 | 0 | 6 | 2 |
| ENSDARG000 | 6:6:6:6:6:6:6 | 33537267:335 | 33537685:335 | +:+:+:+:+:+:+: | 7998 | pik3r3b | 1625 | 1120 | 1456 | 508 | 2033 | 1406 |
| ENSDARG000 | 6:6:6:6:6:6:6 | 42369999:423 | 42371772:423 | -:-:-:-:-:-:-:-:-: | 2644 | emc3 | 1400 | 1816 | 2319 | 1347 | 2162 | 1811 |
| ENSDARG000 | 6:6:6:6:6:6:6 | 54888493:549 | 54889450:549 | +:+:+:+:+:+:+: | 8179 | NAV1 | 1738 | 335 | 1513 | 599 | 1751 | 958 |
| ENSDARG000 | 6:6:6:6:6:6:6 | 37655078:376 | 37655298:376 | +:+:+:+:+:+:+: | 5137 | cyfip1 | 3364 | 2959 | 3223 | 2536 | 3205 | 2789 |
| ENSDARG000 | 6:6:6:6:6:6:6 | 18298444:183 | 18298899:183 | +:+:+:+:+:+:+: | 8203 | NA | 60 | 4 | 0 | 12 | 0 | 2 |
| ENSDARG000 | 6:6:6:6:6:6:6 | 27062512:270 | 27063183:270 | +:+:+:+:+:+:+: | 2939 | boka | 298 | 18 | 16 | 24 | 21 | 112 |
| ENSDARG000 | 6:6:6:6:6:6:6 | 6924637:692 | 6924735:692 | +:+:+:+:+:+:+: | 4745 | map3k20a | 42 | 6 | 22 | 10 | 44 | 12 |
| ENSDARG000 | 6:6:6:6:6:6:6 | 55032439:550 | 55032819:550 | +:+:+:+:+:+:+: | 6175 | mybphb | 8 | 28 | 8 | 0 | 7 | 8 |
| ENSDARG000 | 6;6;6;6;6;6 | 40794015:407 | 40794119:407 | +;+;+;+;+;+ | 1955 | gata2b | 2 | 0 | 0 | 0 | 2 | 0 |
| ENSDARG000 | 6;6;6;6 | 40795898:407 | 40796238:407 | -;-;-;- | 539 | NA | 0 | 0 | 0 | 0 | 0 | 0 |
| ENSDARG000 | 6;6;6;6;6;6;6 | 40822587:408 | 40822932:408 | -;-;-;-;-;-;- | 2093 | eefsec | 259 | 328 | 368 | 174 | 331 | 196 |
| ENSDARG000 | 6:6:6:6:6:6:6 | 12986853:129 | 12986916:129 | +:+:+:+:+:+:+: | 1476 | si:ch73-71c2( | 222 | 161 | 235 | 202 | 136 | 199 |
| ENSDARG000 | 6:6:6:6:6:6:6 | 12968101:129 | 12968287:129 | +:+:+:+:+:+:+: | 3218 | mcm6 | 967 | 3130 | 3751 | 1079 | 3204 | 1734 |
| ENSDARG000 | 6:6:6:6:6:6:6 | 35910498:359 | 35911819:359 | -:-:-:-:-:-:-:-:-: | 2339 | NA | 0 | 0 | 0 | 0 | 2 | 0 |
| ENSDARG000 | 6:6:6:6:6:6:6 | 10943188:109 | 10945444:109 | -:-:-:-:-:-:-:-:-: | 3848 | notum2 | 0 | 0 | 4 | 8 | 2 | 0 |
| ENSDARG000 | 6:6:6:6:6:6:6 | 55327909:553 | 55328176:553 | -:-:-:-:-:-:-:-:-: | 3718 | pcif1 | 116 | 114 | 102 | 59 | 133 | 145 |
| ENSDARG000 | 6;6 | 23887314:238 | 23887732:238 | +;+ | 1938 | ZNF648 | 0 | 0 | 0 | 2 | 20 | 0 |
| ENSDARG000 | 6;6;6 | 2227632:222 | 2228169:222 | +;+;+ | 672 | NA | 0 | 0 | 0 | 4 | 0 | 0 |
| ENSDARG000 | 6:6:6:6:6:6:6 | 23931236:239 | 23931296:239 | +:+:+:+:+:+:+: | 1254 | gadd45ab | 1865 | 692 | 473 | 1110 | 887 | 1472 |
| ENSDARG000 | 6:6:6:6:6:6:6 | 31279542:312 | 31283842:312 | -:-:-:-:-:-:-:-:-: | 7977 | DNAJC6 | 331 | 112 | 50 | 34 | 36 | 76 |
| ENSDARG000 | 6:6:6:6:6:6:6 | 23891230:238 | 23891238:238 | -:-:-:-:-:-:-:-:-: | 3654 | NA | 20 | 56 | 68 | 31 | 78 | 48 |
| ENSDARG000 | 6;6;6;6;6;6;6 | 23955374:239 | 23955952:239 | -;-;-;-;-;-;- | 942 | GNG12 | 84 | 28 | 19 | 80 | 16 | 38 |
| ENSDARG000 | 6;6 | 39812475:398 | 39814009:398 | +;+ | 2711 | C1QL4 | 0 | 0 | 0 | 0 | 0 | 0 |
| ENSDARG000 | 6;6;6;6;6;6 | 29004062:290 | 29004240:290 | -;-;-;-;-;- | 1228 | gfi1ab | 6 | 10 | 14 | 10 | 21 | 52 |
| ENSDARG000 | 6:6:6:6:6:6:6 | 55357188:553 | 55357278:553 | +:+:+:+:+:+:+: | 1923 | pltp | 3159 | 315 | 1719 | 892 | 1532 | 477 |
| ENSDARG000 | 6:6:6:6:6:6:6 | 27112884:271 | 27113108:271 | -:-:-:-:-:-:-:-:-: | 1837 | agxta | 0 | 0 | 6 | 2 | 8 | 0 |
| ENSDARG000 | 6:6:6:6:6:6:6 | 9664206:966 | 9664638:966 | -:-:-:-:-:-:-:-:-: | 2245 | nop58 | 1176 | 2547 | 3584 | 982 | 2099 | 1560 |
| ENSDARG000 | 6:6:6:6:6:6:6 | 40843786:408 | 40844123:408 | -:-:-:-:-:-:-:-:-: | 2092 | RFT1 | 368 | 376 | 481 | 238 | 397 | 427 |
| ENSDARG000 | 6:6:6:6:6:6:6 | 39836474:398 | 39836604:398 | +:+:+:+:+:+:+: | 5184 | SMARCC2 | 660 | 580 | 758 | 342 | 872 | 620 |
| ENSDARG000 | 6;6;6 | 40841156:408 | 40841343:408 | -;-;- | 598 | mustn1a | 14 | 0 | 19 | 14 | 0 | 6 |
| ENSDARG000 | 6;6;6 | 27108048:271 | 27108116:271 | +;+;+ | 353 | NA | 0 | 0 | 0 | 0 | 0 | 0 |
| ENSDARG000 | 6:6:6:6:6:6:6 | 27126620:271 | 27126850:271 | -:-:-:-:-:-:-:-:-: | 1528 | zgc:103559 | 0 | 0 | 0 | 0 | 0 | 0 |

| | | | | | | | | | | | | |
|---|---|---|---|---|---|---|---|---|---|---|---|---|
| ENSDARG000 | 6:6:6:6:6:6: | 40832635:408 | 40832922:408 | +:+:+:+:+:+:+: | 1898 | ruvbl1 | 763 | 1051 | 1655 | 844 | 1294 | 904 |
| ENSDARG000 | 6;6;6;6;6 | 9677420:9678 | 9677563:9678 | +;+;+;+;+ | 1436 | sumo1 | 1216 | 1158 | 1367 | 1453 | 1289 | 1117 |
| ENSDARG000 | 6:6:6:6:6:6: | 55396807:553 | 55397436:553 | -:-:-:-:-:-:-:-: | 2045 | ctsa | 1724 | 1937 | 1764 | 1150 | 2867 | 1859 |
| ENSDARG000 | 6;6;6;6;6;6 | 10982786:109 | 10983578:109 | -;-;-;-;-;- | 1469 | coq7 | 184 | 155 | 206 | 214 | 130 | 170 |
| ENSDARG000 | 6:6:6:6:6:6: | 2235893:2236 | 2235919:2236 | +:+:+:+:+:+:+: | 1119 | NA | 2 | 0 | 0 | 0 | 0 | 0 |
| ENSDARG000 | 6;6;6;6 | 10988764:110 | 10988879:110 | +;+;+;+ | 443 | NA | 0 | 2 | 12 | 0 | 6 | 2 |
| ENSDARG000 | 6;6;6 | 35916500:359 | 35916688:359 | +;+;+ | 860 | NA | 0 | 0 | 0 | 0 | 0 | 0 |
| ENSDARG000 | 6:6:6:6:6:6: | 27090800:270 | 27090882:270 | +:+:+:+:+:+:+: | 1827 | atg4b | 367 | 386 | 742 | 664 | 429 | 229 |
| ENSDARG000 | 6:6:6:6:6:6: | 42377911:423 | 42379122:423 | -:-:-:-:-:-:-:-: | 2903 | sec61a1l | 610 | 949 | 1410 | 504 | 1132 | 768 |
| ENSDARG000 | 6:6:6:6:6:6: | 14172884:141 | 14173701:141 | -:-:-:-:-:-:-:-: | 3648 | inpp4ab | 104 | 31 | 121 | 67 | 119 | 75 |
| ENSDARG000 | 6;6;6;6;6;6 | 24055421:240 | 24056860:240 | -;-;-;-;-;- | 2322 | dr1 | 508 | 576 | 705 | 354 | 580 | 435 |
| ENSDARG000 | 6;6;6;6;6 | 27104962:271 | 27105187:271 | -;-;-;-;- | 908 | dtymk | 53 | 220 | 203 | 78 | 114 | 95 |
| ENSDARG000 | 6;6;6;6;6;6 | 17264905:172 | 17264929:172 | -;-;-;-;-;- | 750 | NA | 2 | 0 | 0 | 0 | 0 | 0 |
| ENSDARG000 | 6:6:6:6:6:6: | 36116704:361 | 36116780:361 | +:+:+:+:+:+:+: | 1078 | NA | 4 | 1 | 2 | 0 | 0 | 1 |
| ENSDARG000 | 6:6:6:6:6:6: | 8314451:8314 | 8314563:8314 | +:+:+:+:+:+:+: | 4170 | gcdha | 460 | 396 | 377 | 259 | 508 | 520 |
| ENSDARG000 | 6:6:6:6:6:6: | 55428924:554 | 55429247:554 | +:+:+:+:+:+:+: | 1945 | ncoa5 | 1608 | 1180 | 2118 | 1386 | 1837 | 1695 |
| ENSDARG000 | 6;6 | 42383194:423 | 42383330:423 | +;+ | 167 | NA | 0 | 0 | 0 | 0 | 0 | 0 |
| ENSDARG000 | 6:6:6:6:6:6: | 2619893:2619 | 2620244:2620 | -:-:-:-:-:-:-:-: | 1239 | hyi | 300 | 179 | 207 | 261 | 125 | 246 |
| ENSDARG000 | 6;6;6;6;6;6 | 18364048:183 | 18364270:183 | -;-;-;-;-;- | 849 | si:dkev-31g6. | 50 | 62 | 164 | 42 | 31 | 8 |
| ENSDARG000 | 6;6 | 33813288:338 | 33813560:338 | +;+ | 912 | NA | 0 | 0 | 0 | 0 | 0 | 0 |
| ENSDARG000 | 6:6:6:6:6:6: | 33663666:336 | 33667607:336 | -:-:-:-:-:-:-:-: | 9451 | MAST2 | 98 | 22 | 122 | 50 | 171 | 349 |
| ENSDARG000 | 6;6;6;6;6 | 18359306:183 | 18359467:183 | +;+;+;+;+ | 1660 | cbx8b | 259 | 213 | 212 | 110 | 215 | 207 |
| ENSDARG000 | 6:6:6:6:6:6: | 6972953:6972 | 6974084:6974 | -:-:-:-:-:-:-:-: | 1709 | tuba8l4 | 5441 | 14573 | 14365 | 11474 | 12176 | 7017 |
| ENSDARG000 | 6:6:6:6:6:6: | 11026354:110 | 11026907:110 | +:+:+:+:+:+:+: | 6446 | NA | 74 | 14 | 156 | 24 | 114 | 134 |
| ENSDARG000 | 6;6;6;6;6;6 | 24076954:240 | 24077371:240 | -;-;-;-;-;- | 1237 | RDH8 | 0 | 2 | 4 | 0 | 0 | 0 |
| ENSDARG000 | 6:6:6:6:6:6: | 29011911:290 | 29012267:290 | -:-:-:-:-:-:-:-: | 4855 | EVI5 | 231 | 226 | 257 | 311 | 331 | 241 |
| ENSDARG000 | 6;6;6 | 33817691:338 | 33818137:338 | +;+;+ | 682 | NA | 0 | 0 | 0 | 0 | 0 | 0 |
| ENSDARG000 | 6;6;6;6 | 6989454:6990 | 6990434:6990 | -;-;-;- | 1409 | si:ch211-114r | 907 | 1937 | 2041 | 1372 | 1858 | 1106 |
| ENSDARG000 | 6:6:6:6:6:6: | 2951645:2951 | 2951869:2951 | +:+:+:+:+:+:+: | 9026 | ptprfa | 106 | 25 | 39 | 0 | 82 | 64 |
| ENSDARG000 | 6:6:6:6:6:6: | 36127505:361 | 36132147:361 | -:-:-:-:-:-:-:-: | 6737 | brinp3a.2 | 420 | 406 | 695 | 353 | 666 | 386 |
| ENSDARG000 | 6 | 6995294 | 6997346 | - | 2053 | NA | 42 | 104 | 164 | 154 | 171 | 55 |
| ENSDARG000 | 6:6:6:6:6:6: | 55449300:554 | 55449926:554 | -:-:-:-:-:-:-:-: | 3863 | NA | 4 | 0 | 0 | 0 | 0 | 0 |
| ENSDARG000 | 6:6:6:6:6:6: | 42409082:424 | 42410421:424 | -:-:-:-:-:-:-:-: | 2793 | ip6k2a | 712 | 358 | 700 | 564 | 1598 | 1015 |
| ENSDARG000 | 6;6;6;6 | 55563650:555 | 55563720:555 | +;+;+;+ | 750 | NA | 0 | 0 | 0 | 0 | 1 | 0 |
| ENSDARG000 | 6 | 40874206 | 40875253 | + | 1048 | zgc:112163 | 14 | 42 | 20 | 56 | 26 | 39 |
| ENSDARG000 | 6;6;6;6;6;6 | 17267681:172 | 17267727:172 | +;+;+;+;+;+ | 750 | NA | 0 | 0 | 0 | 0 | 0 | 0 |
| ENSDARG000 | 6:6:6:6:6:6: | 40922572:409 | 40922962:409 | +:+:+:+:+:+:+: | 4293 | eif4enif1 | 566 | 365 | 623 | 226 | 562 | 442 |
| ENSDARG000 | 6:6:6:6:6:6: | 18367388:183 | 18368252:183 | +:+:+:+:+:+:+: | 2594 | dgke | 414 | 210 | 297 | 117 | 257 | 265 |
| ENSDARG000 | 6:6:6:6:6:6: | 24090950:240 | 24091571:240 | -:-:-:-:-:-:-:-: | 2837 | scinla | 540 | 572 | 596 | 1795 | 519 | 1266 |
| ENSDARG000 | 6;6;6;6;6 | 17281907:172 | 17283008:172 | -;-;-;-;- | 1582 | NA | 54 | 18 | 30 | 83 | 28 | 48 |
| ENSDARG000 | 6:6:6:6:6:6: | 40877394:408 | 40878043:408 | -:-:-:-:-:-:-:-: | 2388 | sirt4 | 128 | 121 | 66 | 58 | 90 | 122 |
| ENSDARG000 | 6:6:6:6:6:6: | 6999487:6999 | 7000043:7000 | -:-:-:-:-:-:-:-: | 2526 | bin1b | 123 | 178 | 220 | 172 | 222 | 130 |
| ENSDARG000 | 6:6:6:6:6:6: | 40884615:408 | 40884853:408 | -:-:-:-:-:-:-:-: | 2415 | sxph | 77 | 6 | 0 | 0 | 26 | 0 |
| ENSDARG000 | 6:6:6:6:6:6: | 42475730:424 | 42475976:424 | +:+:+:+:+:+:+: | 4722 | NA | 46 | 2 | 26 | 0 | 22 | 8 |
| ENSDARG000 | 6;6;6 | 17612044:176 | 17612305:176 | +;+;+ | 950 | NA | 0 | 0 | 0 | 0 | 0 | 2 |
| ENSDARG000 | 6:6:6:6:6:6: | 40898877:408 | 40900380:409 | -:-:-:-:-:-:-:-: | 6540 | NA | 157 | 164 | 187 | 107 | 173 | 189 |
| ENSDARG000 | 6:6:6:6:6:6: | 39874177:398 | 39874591:398 | -:-:-:-:-:-:-:-: | 2836 | myl6 | 1152 | 1109 | 871 | 1151 | 774 | 1069 |
| ENSDARG000 | 6;6 | 8339298:8342 | 8339742:8346 | +;+ | 4652 | NA | 0 | 0 | 0 | 0 | 6 | 0 |
| ENSDARG000 | 6:6:6:6:6:6: | 11070261:110 | 11071371:110 | -:-:-:-:-:-:-:-: | 2618 | sumo3a | 1532 | 2775 | 3180 | 1479 | 2534 | 2190 |
| ENSDARG000 | 6:6:6:6:6:6: | 12996038:129 | 12996416:129 | -:-:-:-:-:-:-:-: | 1689 | tmbim1a | 1110 | 865 | 1003 | 1315 | 643 | 670 |
| ENSDARG000 | 6:6:6:6:6:6: | 31350101:313 | 31353724:313 | -:-:-:-:-:-:-:-: | 4493 | ak4 | 854 | 1402 | 1552 | 2661 | 1415 | 1487 |
| ENSDARG000 | 6;6;6;6 | 33875137:338 | 33876057:338 | -;-;-;- | 1074 | tmem69 | 111 | 124 | 120 | 105 | 128 | 102 |
| ENSDARG000 | 6:6:6:6:6:6: | 42545797:425 | 42545992:425 | +:+:+:+:+:+:+: | 2915 | CAMKV | 0 | 0 | 0 | 0 | 0 | 0 |
| ENSDARG000 | 6;6 | 9703721:9707 | 9706226:9707 | -;- | 2846 | fzd7b | 1478 | 1258 | 1628 | 1739 | 1510 | 1596 |
| ENSDARG000 | 6;6;6;6;6;6 | 17972241:179 | 17972571:179 | -;-;-;-;-;- | 1277 | NA | 0 | 0 | 0 | 0 | 0 | 0 |
| ENSDARG000 | 6:6:6:6:6:6: | 17725788:177 | 17727598:177 | -:-:-:-:-:-:-:-: | 6673 | RPTOR | 590 | 522 | 651 | 417 | 733 | 530 |
| ENSDARG000 | 6;6 | 55819038:558 | 55819353:558 | +;+ | 1082 | NA | 54 | 65 | 54 | 28 | 75 | 55 |
| ENSDARG000 | 6;6 | 55814757:558 | 55815138:558 | -;- | 679 | NA | 0 | 0 | 2 | 0 | 0 | 0 |
| ENSDARG000 | 6:6:6:6:6:6: | 39897543:398 | 39898599:398 | -:-:-:-:-:-:-:-: | 1869 | tsfm | 86 | 194 | 165 | 42 | 185 | 66 |
| ENSDARG000 | 6;6;6;6;6;6 | 42604241:426 | 42604493:426 | +;+;+;+;+;+ | 816 | NA | 16 | 24 | 38 | 24 | 44 | 30 |
| ENSDARG000 | 6;6;6;6;6 | 14301214:143 | 14301666:143 | +;+;+;+;+ | 1359 | TMEM182 | 0 | 0 | 0 | 0 | 0 | 0 |
| ENSDARG000 | 6:6:6:6:6:6: | 55909795:559 | 55911530:559 | -:-:-:-:-:-:-:-: | 3734 | eya2 | 22 | 14 | 34 | 3 | 2 | 10 |
| ENSDARG000 | 6:6:6:6:6:6: | 36281005:362 | 36281336:362 | +:+:+:+:+:+:+: | 2390 | NA | 18 | 14 | 12 | 0 | 13 | 4 |
| ENSDARG000 | 6;6;6 | 14316794:143 | 14317063:143 | -;-;- | 495 | NA | 0 | 0 | 0 | 0 | 0 | 0 |
| ENSDARG000 | 6:6:6:6:6:6: | 9747108:9748 | 9747856:9748 | -:-:-:-:-:-:-:-: | 2115 | cdk15 | 2457 | 2135 | 2226 | 2014 | 2306 | 2340 |
| ENSDARG000 | 6;6;6;6;6 | 42618902:426 | 42619009:426 | +;+;+;+;+ | 1012 | NA | 6 | 2 | 1 | 4 | 4 | 0 |
| ENSDARG000 | 6:6:6:6:6:6: | 18388327:183 | 18389920:183 | -:-:-:-:-:-:-:-: | 3213 | trim25 | 1390 | 1561 | 1240 | 905 | 1162 | 889 |
| ENSDARG000 | 6;6;6;6;6;6 | 11078693:110 | 11079157:110 | -;-;-;-;-;- | 1227 | pttg1ipa | 14 | 3 | 6 | 2 | 6 | 34 |
| ENSDARG000 | 6:6:6:6:6:6: | 29086594:290 | 29086693:290 | +:+:+:+:+:+:+: | 1060 | rpl5b | 16513 | 15314 | 26797 | 24194 | 16649 | 24765 |
| ENSDARG000 | 6 | 11077114 | 11077609 | - | 496 | NA | 16 | 1 | 2 | 0 | 10 | 23 |
| ENSDARG000 | 6:6:6:6:6:6: | 8355122:8357 | 8356109:8357 | -:-:-:-:-:-:-:-: | 1974 | acp5a | 58 | 28 | 28 | 36 | 64 | 40 |
| ENSDARG000 | 6;6;6;6;6 | 7059999:7059 | 7060908:7060 | -;-;-;-;- | 1786 | ihhb | 0 | 6 | 4 | 14 | 0 | 0 |
| ENSDARG000 | 6:6:6:6:6:6: | 56076595:560 | 56077009:560 | +:+:+:+:+:+:+: | 4324 | rtf2 | 628 | 407 | 696 | 614 | 458 | 510 |
| ENSDARG000 | 6:6:6:6:6:6: | 39905021:399 | 39905448:399 | +:+:+:+:+:+:+: | 1394 | endou | 0 | 0 | 0 | 0 | 0 | 0 |
| ENSDARG000 | 6;6;6;6;6;6 | 42635960:426 | 42636246:426 | +;+;+;+;+;+ | 1296 | NA | 0 | 0 | 16 | 12 | 6 | 18 |
| ENSDARG000 | 6:6:6:6:6:6: | 102506:10324 | 102680:10348 | +:+:+:+:+:+:+: | 3078 | NA | 894 | 1036 | 613 | 238 | 502 | 212 |
| ENSDARG000 | 6;6;6;6;6;6 | 56097971:561 | 56099801:561 | -;-;-;-;-;- | 3290 | GCNT7 | 10 | 8 | 10 | 0 | 18 | 28 |
| ENSDARG000 | 6;6;6;6;6;6 | 42646411:426 | 42646663:426 | +;+;+;+;+;+ | 1250 | NA | 0 | 1 | 6 | 0 | 0 | 12 |
| ENSDARG000 | 6:6:6:6:6:6: | 39913853:399 | 39914033:399 | -:-:-:-:-:-:-:-: | 2387 | sumf1 | 338 | 228 | 333 | 225 | 307 | 266 |
| ENSDARG000 | 6;6 | 24140757:241 | 24141155:241 | -;- | 808 | NA | 14750 | 15391 | 14019 | 28111 | 17820 | 7735 |
| ENSDARG000 | 6:6:6:6:6:6: | 33881372:338 | 33881612:338 | +:+:+:+:+:+:+: | 7379 | GPBP1L1 | 1984 | 1740 | 2355 | 1360 | 2260 | 1696 |
| ENSDARG000 | 6:6:6:6:6:6: | 33897624:338 | 33898589:338 | +:+:+:+:+:+:+: | 2558 | NA | 2 | 0 | 0 | 0 | 0 | 0 |
| ENSDARG000 | 6:6:6:6:6:6: | 29102942:291 | 29103826:291 | -;-;-;-;-;-;- | 1511 | DIPK1A | 1495 | 391 | 518 | 829 | 1204 | 926 |
| ENSDARG000 | 6:6:6:6:6:6: | 3280939:3282 | 3281011:3282 | +:+:+:+:+:+:+: | 6709 | NA | 684 | 331 | 817 | 450 | 684 | 697 |
| ENSDARG000 | 6:6:6:6:6:6: | 18418651:184 | 18418920:184 | +:+:+:+:+:+:+: | 3319 | rab11fip4b | 105 | 90 | 70 | 78 | 99 | 217 |
| ENSDARG000 | 6:6:6:6:6:6: | 42656722:426 | 42657185:426 | +:+:+:+:+:+:+: | 923 | NA | 12 | 28 | 22 | 32 | 20 | 40 |
| ENSDARG000 | 6:6:6:6:6:6: | 7085889:7085 | 7086177:7086 | -:-:-:-:-:-:-:-: | 1818 | nhej1 | 80 | 95 | 180 | 52 | 128 | 82 |
| ENSDARG000 | 6:6:6:6:6:6: | 24152816:241 | 24153349:241 | -:-:-:-:-:-:-:-: | 4948 | NA | 223 | 254 | 250 | 260 | 289 | 194 |
| ENSDARG000 | 6:6:6:6:6:6: | 29144776:291 | 29145776:291 | -:-:-:-:-:-:-:-: | 4583 | zbtb11 | 534 | 794 | 1155 | 619 | 886 | 545 |
| ENSDARG000 | 6:6:6:6:6:6: | 11101751:111 | 11101961:111 | -:-:-:-:-:-:-:-: | 7237 | NA | 4 | 2 | 5 | 2 | 0 | 4 |
| ENSDARG000 | 6:6:6:6:6:6: | 13024448:130 | 13024636:130 | +:+:+:+:+:+:+: | 1577 | pnkd | 155 | 189 | 129 | 207 | 168 | 165 |
| ENSDARG000 | 6:6:6:6:6:6: | 56141852:561 | 56141909:561 | +:+:+:+:+:+:+: | 5091 | tfap2c | 11835 | 8838 | 9563 | 6994 | 13798 | 9394 |
| ENSDARG000 | 6;6;6;6 | 29184137:291 | 29184610:291 | -;-;-;- | 1040 | dpt | 4 | 4 | 6 | 2 | 14 | 2 |
| ENSDARG000 | 6:6:6:6:6:6: | 42666694:426 | 42667004:426 | +:+:+:+:+:+:+: | 4750 | NA | 1030 | 1023 | 1646 | 937 | 1176 | 1037 |
| ENSDARG000 | 6;6;6;6;6;6 | 24318031:243 | 24318425:243 | -;-;-;-;-;- | 1336 | NA | 4 | 0 | 0 | 0 | 0 | 0 |
| ENSDARG000 | 6:6:6:6:6:6: | 57401178:574 | 57401707:574 | -:-:-:-:-:-:-:-: | 5709 | ZNFX1 | 1910 | 1657 | 413 | 397 | 498 | 416 |
| ENSDARG000 | 6:6:6:6:6:6: | 7109954:7112 | 7110632:7112 | -:-:-:-:-:-:-:-: | 1706 | atg3 | 1461 | 1411 | 1513 | 1154 | 1720 | 1320 |
| ENSDARG000 | 6:6:6:6:6:6: | 29217392:292 | 29217632:292 | +:+:+:+:+:+:+: | 2354 | atp1b1a | 5185 | 4019 | 5886 | 2567 | 6892 | 3740 |
| ENSDARG000 | 6;6;6;6;6;6 | 24329275:243 | 24330292:243 | -;-;-;-;-;- | 2207 | EPHX4 | 0 | 0 | 0 | 0 | 0 | 11 |
| ENSDARG000 | 6;6 | 57431319:574 | 57431407:574 | +;+ | 247 | NA | 0 | 0 | 0 | 0 | 0 | 0 |
| ENSDARG000 | 6;6;6;6;6;6 | 57478337:574 | 57478521:574 | -;-;-;-;-;- | 963 | snrpb | 2136 | 3031 | 4653 | 2396 | 2980 | 2115 |
| ENSDARG000 | 6:6:6:6:6:6: | 7123980:7124 | 7124498:7124 | +:+:+:+:+:+:+: | 2242 | GPR161 | 66 | 137 | 157 | 32 | 91 | 66 |
| ENSDARG000 | 6:6:6:6:6:6: | 57436314:574 | 57436434:574 | -:-:-:-:-:-:-:-: | 2626 | ddx27 | 850 | 834 | 1562 | 525 | 1008 | 663 |
| ENSDARG000 | 6;6;6;6;6 | 7157468:7157 | 7158203:7157 | -;-;-;-;- | 1263 | HS6ST3 | 0 | 2 | 0 | 0 | 0 | 0 |
| ENSDARG000 | 6;6 | 57479251:574 | 57479749:574 | +;+ | 929 | NA | 7 | 6 | 27 | 18 | 18 | 14 |
| ENSDARG000 | 6:6:6:6:6:6: | 31368071:313 | 31368274:313 | +:+:+:+:+:+:+: | 6712 | jak1 | 3433 | 2649 | 3500 | 2503 | 2240 | 1823 |
| ENSDARG000 | 6;6;6;6;6 | 42693114:426 | 42693191:426 | +;+;+;+;+ | 1213 | NA | 0 | 0 | 0 | 0 | 0 | 0 |
| ENSDARG000 | 6;6;6 | 18441331:184 | 18441439:184 | +;+;+ | 847 | NA | 0 | 2 | 0 | 4 | 2 | 2 |
| ENSDARG000 | 6:6:6:6:6:6: | 15036828:150 | 15037272:150 | -:-:-:-:-:-:-:-: | 3070 | tgfbrap1 | 169 | 150 | 194 | 124 | 278 | 184 |
| ENSDARG000 | 6;6;6;6;6 | 7240403:7242 | 7240634:7242 | +;+;+;+;+ | 816 | NA | 0 | 0 | 0 | 0 | 0 | 0 |
| ENSDARG000 | 6:6:6:6:6:6: | 9793495:9793 | 9793537:9793 | +:+:+:+:+:+:+: | 7299 | ALS2 | 1435 | 576 | 930 | 726 | 783 | 664 |
| ENSDARG000 | 6:6:6:6:6:6: | 13039951:130 | 13040164:130 | +:+:+:+:+:+:+: | 1377 | catip | 18 | 6 | 16 | 50 | 23 | 2 |
| ENSDARG000 | 6:6:6:6:6:6: | 112579:11359 | 112842:11375 | +:+:+:+:+:+:+: | 1776 | ap1m2 | 48 | 10 | 6 | 22 | 24 | 2 |
| ENSDARG000 | 6:6:6:6:6:6: | 29288006:292 | 29288210:292 | +:+:+:+:+:+:+: | 1783 | zgc:172121 | 38 | 60 | 63 | 42 | 77 | 44 |
| ENSDARG000 | 6:6:6:6:6:6: | 18524444:185 | 18524736:185 | -:-:-:-:-:-:-:-: | 2280 | utp6 | 626 | 652 | 927 | 502 | 824 | 720 |
| ENSDARG000 | 6:6:6:6:6:6: | 42818963:428 | 42819513:428 | +:+:+:+:+:+:+: | 4882 | sema3fa | 6 | 2 | 2 | 0 | 12 | 6 |
| ENSDARG000 | 6:6:6:6:6:6: | 57491794:574 | 57492411:574 | -:-:-:-:-:-:-:-: | 3513 | itcha | 334 | 345 | 324 | 132 | 447 | 278 |
| ENSDARG000 | 6:6:6:6:6:6: | 8363629:8365 | 8366464:8366 | -:-:-:-:-:-:-:-: | 6527 | ilf3a | 1516 | 1425 | 2407 | 1299 | 1885 | 1829 |
| ENSDARG000 | 6:6:6:6:6:6: | 29227489:292 | 29227584:292 | -:-:-:-:-:-:-:-: | 1536 | nme7 | 114 | 162 | 179 | 108 | 181 | 114 |
| ENSDARG000 | 6:6:6:6:6:6: | 39923052:399 | 39923176:399 | +:+:+:+:+:+:+: | 9101 | ITPR1 | 162 | 149 | 248 | 281 | 222 | 212 |
| ENSDARG000 | 6:6:6:6:6:6: | 37733608:377 | 37734626:377 | -:-:-:-:-:-:-:-: | 2000 | nipa2 | 332 | 351 | 290 | 168 | 349 | 277 |
| ENSDARG000 | 6:6:6:6:6:6: | 13045885:130 | 13046028:130 | +:+:+:+:+:+:+: | 4280 | casp8 | 868 | 1067 | 860 | 1176 | 1124 | 556 |
| ENSDARG000 | 6:6:6:6:6:6: | 24364678:243 | 24365897:243 | -:-:-:-:-:-:-:-: | 5669 | brdt | 897 | 549 | 1050 | 1027 | 967 | 806 |
| ENSDARG000 | 6;6 | 18520859:185 | 18521014:185 | +;+ | 898 | NA | 188 | 155 | 176 | 180 | 79 | 153 |
| ENSDARG000 | 6;6;6;6;6;6 | 36287538:362 | 36288148:362 | -;-;-;-;-;- | 930 | NA | 8 | 0 | 10 | 0 | 0 | 8 |
| ENSDARG000 | 6:6:6:6:6:6: | 7249531:7249 | 7249709:7249 | +:+:+:+:+:+:+: | 3355 | dzip1 | 18 | 24 | 72 | 86 | 63 | 6 |
| ENSDARG000 | 6:6:6:6:6:6: | 15071399:150 | 15071754:150 | -:-:-:-:-:-:-:-: | 2280 | fhl2b | 12 | 0 | 3 | 2 | 0 | 0 |
| ENSDARG000 | 6:6:6:6:6:6: | 42918933:429 | 42919201:429 | +:+:+:+:+:+:+: | 1967 | gnat1 | 0 | 6 | 0 | 0 | 8 | 0 |
| ENSDARG000 | 6:6:6:6:6:6: | 40951227:409 | 40953208:409 | +:+:+:+:+:+:+: | 7175 | PATZ1 | 544 | 242 | 489 | 162 | 418 | 464 |
| ENSDARG000 | 6;6;6;6;6;6 | 120181:12030 | 120233:12045 | +;+;+;+;+;+ | 631 | CDKN2D | 0 | 38 | 20 | 2 | 20 | 12 |
| ENSDARG000 | 6;6;6;6;6 | 124172:12690 | 129137:12913 | -;-;-;-;- | 5519 | NA | 66 | 46 | 18 | 48 | 44 | 48 |
| ENSDARG000 | 6;6;6 | 15127507:151 | 15127693:151 | +;+;+ | 555 | dph3 | 26 | 46 | 33 | 16 | 29 | 12 |
| ENSDARG000 | 6:6:6:6:6:6: | 57541776:575 | 57541976:575 | +:+:+:+:+:+:+: | 1533 | NECAB3 | 0 | 2 | 0 | 0 | 2 | 0 |
| ENSDARG000 | 6:6:6:6:6:6: | 36381709:363 | 36381991:363 | +:+:+:+:+:+:+: | 1983 | rhcgl1 | 54 | 0 | 2 | 12 | 10 | 10 |
| ENSDARG000 | 6:6:6:6:6:6: | 11249706:112 | 11250503:112 | +:+:+:+:+:+:+: | 6033 | atg9a | 646 | 476 | 435 | 372 | 415 | 604 |
| ENSDARG000 | 6:6:6:6:6:6: | 27146671:271 | 27147042:271 | +:+:+:+:+:+:+: | 9258 | KIF1A | 262 | 116 | 80 | 10 | 222 | 110 |
| ENSDARG000 | 6:6:6:6:6:6: | 40992409:409 | 40992925:409 | +:+:+:+:+:+:+: | 3072 | tgfa | 26 | 20 | 74 | 11 | 52 | 62 |
| ENSDARG000 | 6:6:6:6:6:6: | 40975655:409 | 40978186:409 | -:-:-:-:-:-:-:-: | 4296 | NA | 8 | 0 | 2 | 12 | 70 | 0 |
| ENSDARG000 | 6 | 57630414 | 57631164 | - | 751 | NA | 397 | 335 | 662 | 424 | 1047 | 1105 |
| ENSDARG000 | 6:6:6:6:6:6: | 29295310:292 | 29295874:292 | -:-:-:-:-:-:-:-: | 1946 | BIVM | 59 | 48 | 69 | 67 | 110 | 65 |
| ENSDARG000 | 6:6:6:6:6:6: | 57634360:576 | 57634999:576 | -:-:-:-:-:-:-:-: | 2699 | cbfa2t2 | 1600 | 746 | 2434 | 1512 | 1973 | 1496 |
| ENSDARG000 | 6:6:6:6:6:6: | 33911154:339 | 33911359:339 | -:-:-:-:-:-:-:-: | 2496 | nasp | 307 | 1175 | 1574 | 476 | 1310 | 730 |
| ENSDARG000 | 6;6;6;6;6;6 | 15145123:152 | 15145300:152 | +;+;+;+;+;+;+ | 5404 | nck2b | 190 | 186 | 186 | 152 | 207 | 88 |
| ENSDARG000 | 6;6;6;6 | 15250672:152 | 15250697:152 | +;+;+;+ | 533 | NA | 0 | 0 | 0 | 0 | 0 | 2 |
| ENSDARG000 | 6:6:6:6:6:6: | 3334392:3334 | 3334562:3334 | +:+:+:+:+:+:+: | 2330 | st3gal3a | 200 | 270 | 378 | 282 | 479 | 348 |
| ENSDARG000 | 6:6:6:6:6:6: | 57900237:579 | 57901241:579 | -:-:-:-:-:-:-:-: | 2567 | tox2 | 20 | 10 | 50 | 20 | 40 | 44 |
| ENSDARG000 | 6:6:6:6:6:6: | 42932757:429 | 42934240:429 | -:-:-:-:-:-:-:-: | 3454 | edem1 | 1374 | 1462 | 1425 | 916 | 1524 | 1388 |
| ENSDARG000 | 6:6:6:6:6:6: | 24398907:244 | 24399174:244 | +:+:+:+:+:+:+: | 3008 | tgfbr3 | 58 | 4 | 12 | 8 | 16 | 26 |
| ENSDARG000 | 6;6;6;6;6 | 36459897:364 | 36460116:364 | +;+;+;+;+ | 2277 | lrrc15 | 208 | 96 | 148 | 122 | 172 | 202 |
| ENSDARG000 | 6:6:6:6:6:6: | 15268685:152 | 15268818:152 | +:+:+:+:+:+:+: | 714 | ecrg4b | 0 | 0 | 0 | 0 | 0 | 0 |
| ENSDARG000 | 6:6:6:6:6:6: | 13096912:130 | 13097014:130 | +:+:+:+:+:+:+: | 3331 | dytn | 21 | 2 | 14 | 14 | 27 | 50 |
| ENSDARG000 | 6:6:6:6:6:6: | 131998:13199 | 133039:13303 | -:-:-:-:-:-:-:-: | 3200 | keap1b | 738 | 561 | 662 | 393 | 713 | 412 |
| ENSDARG000 | 6:6:6:6:6:6: | 58280936:582 | 58281111:582 | +:+:+:+:+:+:+: | 4673 | RALGAPB | 1162 | 898 | 1187 | 803 | 1295 | 960 |
| ENSDARG000 | 6;6 | 9873911:9881 | 9874473:9881 | -;- | 702 | NA | 0 | 0 | 0 | 0 | 0 | 0 |
| ENSDARG000 | 6:6:6:6:6:6: | 9867426:9870 | 9867558:9870 | +:+:+:+:+:+:+: | 2630 | MPP4 | 13 | 0 | 0 | 7 | 2 | 0 |
| ENSDARG000 | 6;6;6;6;6;6 | 36465347:364 | 36465420:364 | +;+;+;+;+;+ | 6058 | NA | 2 | 0 | 6 | 4 | 50 | 32 |
| ENSDARG000 | 6;6 | 36479193:365 | 36479510:365 | -;- | 538 | NA | 0 | 0 | 0 | 0 | 0 | 0 |
| ENSDARG000 | 6;6;6 | 140121:14012 | 141564:14033 | -;-;- | 1577 | s1pr5b | 20 | 4 | 0 | 20 | 4 | 0 |
| ENSDARG000 | 6:6:6:6:6:6: | 24660817:246 | 24661854:246 | +:+:+:+:+:+:+: | 11490 | ZNF644 | 318 | 90 | 154 | 152 | 232 | 190 |
| ENSDARG000 | 6:6:6:6:6:6: | 7287102:7288 | 7287462:7288 | -;-;-;-;-;-;-;- | 951 | NA | 0 | 0 | 0 | 0 | 0 | 0 |
| ENSDARG000 | 6;6;6;6;6 | 7299194:7299 | 7299435:7299 | -;-;-;-;- | 881 | cldn10a | 0 | 4 | 2 | 0 | 0 | 0 |
| ENSDARG000 | 6;6;6;6;6;6 | 36551083:365 | 36551915:365 | -;-;-;-;-;- | 1538 | her6 | 203 | 100 | 176 | 136 | 64 | 262 |

| | | | | | | | | | | | | |
|---|---|---|---|---|---|---|---|---|---|---|---|---|
| ENSDARG000 | 6;6;6 | 7313548:7319 | 7314194:7319 | -;-;- | 804 | NA | 3 | 0 | 0 | 0 | 0 | 0 |
| ENSDARG000 | 6;6;6;6;6;6 | 8393378:8395 | 8396015:8396 | -;-;-;-;-;- | 3846 | ptger1a | 0 | 0 | 4 | 0 | 0 | 2 |
| ENSDARG000 | 6;6;6;6;6;6 | 29305190:293 | 29305440:293 | +;+;+;+;+;+ | 1355 | si:ch211-201l | 12 | 2 | 6 | 0 | 26 | 2 |
| ENSDARG000 | 6;6;6;6 | 15373153:154 | 15374095:154 | +;+;+;+ | 2560 | tmtops2a | 2 | 0 | 2 | 0 | 0 | 0 |
| ENSDARG000 | 6;6;6;6;6;6;6 | 15437132:154 | 15437801:154 | -;-;-;-;-;-;- | 2633 | st6gal2b | 6 | 1 | 8 | 0 | 0 | 12 |
| ENSDARG000 | 6;6;6;6;6;6;6 | 58406014:584 | 58406309:584 | +;+;+;+;+;+;+ | 6501 | KCNQ2 | 0 | 0 | 0 | 2 | 0 | 2 |
| ENSDARG000 | 6;6;6;6;6;6;6 | 13136602:131 | 13137218:131 | -;-;-;-;-;-;- | 3707 | ADAM23 | 10 | 16 | 16 | 8 | 9 | 9 |
| ENSDARG000 | 6;6;6;6;6;6;6 | 58508420:585 | 58508483:585 | +;+;+;+;+;+;+ | 1653 | NA | 0 | 2 | 0 | 0 | 0 | 0 |
| ENSDARG000 | 6;6;6 | 369991:37483 | 370065:37497 | +;+;+ | 346 | NA | 123 | 154 | 147 | 101 | 79 | 144 |
| ENSDARG000 | 6;6;6;6;6;6;6 | 8453398:8454 | 8454212:8454 | -;-;-;-;-;-;- | 3741 | NA | 388 | 271 | 805 | 603 | 367 | 522 |
| ENSDARG000 | 6;6;6;6;6;6;6 | 31430791:314 | 31435085:314 | -;-;-;-;-;-;- | 6831 | RAVER2 | 34 | 8 | 38 | 14 | 51 | 24 |
| ENSDARG000 | 6;6;6;6;6;6;6 | 7322587:7329 | 7322777:7329 | +;+;+;+;+;+;+ | 4305 | abcc4 | 839 | 638 | 555 | 329 | 1024 | 537 |
| ENSDARG000 | 6;6;6;6;6 | 9893554:9893 | 9893645:9893 | +;+;+;+;+ | 788 | NA | 0 | 0 | 0 | 0 | 0 | 0 |
| ENSDARG000 | 6;6;6;6;6 | 41014791:410 | 41019232:410 | -;-;-;-;- | 5478 | hdhd3 | 34 | 32 | 46 | 32 | 46 | 78 |
| ENSDARG000 | 6;6;6;6;6;6;6 | 24742871:247 | 24742900:247 | +;+;+;+;+;+;+ | 966 | NA | 0 | 0 | 0 | 0 | 0 | 0 |
| ENSDARG000 | 6;6;6;6;6;6;6 | 58522557:585 | 58522933:585 | +;+;+;+;+;+;+ | 1694 | arfrp1 | 183 | 215 | 290 | 164 | 255 | 217 |
| ENSDARG000 | 6 | 377959 | 379248 | - | 1290 | NA | 0 | 0 | 0 | 0 | 4 | 0 |
| ENSDARG000 | 6;6;6;6;6;6;6 | 42951960:429 | 42952902:429 | -;-;-;-;-;-;- | 1772 | arl8ba | 1381 | 1450 | 1603 | 1334 | 1377 | 1850 |
| ENSDARG000 | 6;6;6;6;6;6;6 | 27275716:272 | 27275785:272 | +;+;+;+;+;+;+ | 2111 | scly | 155 | 134 | 205 | 54 | 248 | 165 |
| ENSDARG000 | 6;6;6;6;6 | 24817852:248 | 24818803:248 | +;+;+;+;+ | 1864 | barhl2 | 0 | 0 | 0 | 0 | 0 | 0 |
| ENSDARG000 | 6;6;6;6;6;6;6 | 11272286:112 | 11272671:112 | -;-;-;-;-;-;- | 13210 | pcnt | 1878 | 1041 | 1477 | 1248 | 1638 | 1136 |
| ENSDARG000 | 6;6;6;6;6;6;6 | 58543336:585 | 58543539:585 | +;+;+;+;+;+;+ | 3658 | STMN3 | 987 | 264 | 6 | 50 | 264 | 26 |
| ENSDARG000 | 6;6;6;6;6;6;6 | 41023580:410 | 41024063:410 | -;-;-;-;-;-;- | 1413 | trub2 | 192 | 213 | 202 | 174 | 238 | 329 |
| ENSDARG000 | 6;6;6;6;6;6;6 | 37745719:377 | 37746477:377 | -;-;-;-;-;-;- | 1559 | nipa1 | 34 | 60 | 148 | 96 | 82 | 140 |
| ENSDARG000 | 6;6;6;6;6 | 42975278:429 | 42976074:429 | -;-;-;-;- | 1540 | tnfrsf18 | 120 | 52 | 12 | 36 | 138 | 56 |
| ENSDARG000 | 6;6;6;6;6;6;6 | 58569808:585 | 58569974:585 | +;+;+;+;+;+;+ | 9119 | NA | 5881 | 2314 | 624 | 606 | 1397 | 1655 |
| ENSDARG000 | 6;6;6 | 25145265:251 | 25145339:251 | -;-;- | 296 | NA | 0 | 0 | 0 | 0 | 0 | 0 |
| ENSDARG000 | 6;6;6;6;6;6;6 | 9898683:9898 | 9898834:9899 | +;+;+;+;+;+;+ | 1645 | tmem237b | 69 | 33 | 104 | 48 | 60 | 100 |
| ENSDARG000 | 6;6;6;6;6;6;6 | 392815:39531 | 392875:39538 | +;+;+;+;+;+;+ | 1752 | cyth4b | 44 | 28 | 68 | 7 | 105 | 85 |
| ENSDARG000 | 6;6;6;6;6;6;6 | 15526769:155 | 15527713:155 | -;-;-;-;-;-;- | 3244 | NA | 231 | 108 | 189 | 174 | 306 | 429 |
| ENSDARG000 | 6;6;6;6;6;6;6 | 41057943:410 | 41058541:410 | -;-;-;-;-;-;- | 6889 | NA | 51 | 45 | 34 | 34 | 133 | 145 |
| ENSDARG000 | 6;6;6;6;6;6;6 | 41038757:410 | 41039252:410 | +;+;+;+;+;+;+ | 6278 | entpd8 | 32 | 2 | 15 | 6 | 6 | 14 |
| ENSDARG000 | 6;6;6;6;6;6;6 | 39993605:399 | 39994835:399 | -;-;-;-;-;-;- | 3066 | pfkfb4b | 508 | 845 | 391 | 309 | 602 | 431 |
| ENSDARG000 | 6;6;6;6;6;6;6 | 43001964:430 | 43002689:430 | -;-;-;-;-;-;- | 1741 | NA | 16 | 25 | 4 | 10 | 26 | 42 |
| ENSDARG000 | 6 | 7414215 | 7415343 | + | 1129 | sox21a | 4 | 22 | 42 | 2 | 74 | 20 |
| ENSDARG000 | 6;6;6;6 | 58622831:586 | 58622948:586 | +;+;+;+ | 5629 | sp7 | 6 | 8 | 4 | 12 | 16 | 0 |
| ENSDARG000 | 6;6;6;6;6;6;6 | 25157331:251 | 25158313:251 | -;-;-;-;-;-;- | 2558 | znf326 | 2 | 4 | 0 | 4 | 2 | 0 |
| ENSDARG000 | 6;6;6 | 43015916:430 | 43016340:430 | +;+;+ | 1046 | tcta | 276 | 266 | 304 | 204 | 360 | 295 |
| ENSDARG000 | 6;6;6;6;6;6;6 | 27304503:273 | 27304694:273 | +;+;+;+;+;+;+ | 3078 | ESPNL | 6 | 0 | 0 | 1 | 2 | 2 |
| ENSDARG000 | 6;6;6;6;6;6;6 | 58698475:587 | 58699025:587 | +;+;+;+;+;+;+ | 4577 | igsf8 | 420 | 488 | 580 | 317 | 498 | 348 |
| ENSDARG000 | 6;6;6;6;6;6;6 | 58772209:587 | 58772285:587 | -;-;-;-;-;-;- | 3266 | NA | 71 | 71 | 126 | 76 | 154 | 105 |
| ENSDARG000 | 6;6;6;6;6;6;6 | 37752781:377 | 37752861:377 | +;+;+;+;+;+;+ | 14609 | HERC2 | 2164 | 921 | 2194 | 1018 | 2549 | 1386 |
| ENSDARG000 | 6;6;6;6;6;6;6 | 29313220:293 | 29314264:293 | -;-;-;-;-;-;- | 6496 | TMEM131 | 1913 | 1316 | 2210 | 1521 | 2619 | 2453 |
| ENSDARG000 | 6;6;6;6;6;6;6 | 43019642:430 | 43020727:430 | -;-;-;-;-;-;- | 2357 | glyctk | 340 | 204 | 200 | 227 | 250 | 216 |
| ENSDARG000 | 6;6;6;6;6 | 25167250:251 | 25174224:251 | -;-;-;-;- | 7127 | lrrc8db | 232 | 149 | 241 | 318 | 268 | 229 |
| ENSDARG000 | 6;6;6 | 25189837:251 | 25192077:251 | -;-;- | 2494 | lrrc8c | 1977 | 876 | 1165 | 1138 | 1135 | 1609 |
| ENSDARG000 | 6 | 25188023 | 25188783 | - | 761 | NA | 749 | 332 | 457 | 743 | 389 | 595 |
| ENSDARG000 | 6;6;6;6;6;6;6 | 58737168:587 | 58737688:587 | -;-;-;-;-;-;- | 2810 | soat2 | 22 | 7 | 25 | 10 | 42 | 25 |
| ENSDARG000 | 6;6;6;6;6;6;6 | 8470598:8470 | 8470996:8470 | -;-;-;-;-;-;- | 3928 | NA | 62 | 97 | 36 | 64 | 148 | 233 |
| ENSDARG000 | 6;6;6;6;6;6;6 | 7421898:7422 | 7422072:7422 | +;+;+;+;+;+;+ | 1531 | ccdc65 | 4 | 4 | 2 | 0 | 6 | 0 |
| ENSDARG000 | 6;6;6;6;6;6;6 | 31590751:315 | 31592055:315 | -;-;-;-;-;-;- | 4930 | CACHD1 | 1002 | 473 | 837 | 445 | 758 | 425 |
| ENSDARG000 | 6;6;6;6;6;6;6 | 11369135:113 | 11369291:113 | +;+;+;+;+;+;+ | 1438 | zgc:112416 | 0 | 0 | 0 | 0 | 0 | 0 |
| ENSDARG000 | 6;6;6;6;6;6;6 | 27338512:273 | 27339239:273 | +;+;+;+;+;+;+ | 2691 | KLHL30 | 0 | 0 | 0 | 0 | 0 | 0 |
| ENSDARG000 | 6;6;6;6;6;6;6 | 36717985:367 | 36721412:367 | -;-;-;-;-;-;- | 6833 | opa1 | 1390 | 1413 | 1851 | 1075 | 1419 | 1372 |
| ENSDARG000 | 6;6 | 41086608:410 | 41086969:410 | -;- | 522 | NA | 0 | 0 | 0 | 0 | 0 | 0 |
| ENSDARG000 | 6;6;6;6;6;6;6 | 41080207:410 | 41080371:410 | -;-;-;-;-;-;- | 1222 | srsf3a | 0 | 0 | 0 | 0 | 0 | 0 |
| ENSDARG000 | 6 | 3459026 | 3459340 | + | 315 | NA | 0 | 0 | 0 | 0 | 0 | 0 |
| ENSDARG000 | 6;6;6;6;6;6;6 | 41080003:410 | 41080371:410 | -;-;-;-;-;-;- | 1643 | srsf3a | 40 | 36 | 52 | 42 | 30 | 32 |
| ENSDARG000 | 6;6;6;6;6;6;6 | 40031007:400 | 40031133:400 | -;-;-;-;-;-;- | 968 | NA | 4 | 0 | 0 | 0 | 0 | 4 |
| ENSDARG000 | 6;6;6;6;6 | 9920919:9921 | 9921007:9921 | -;-;-;-;- | 800 | NA | 0 | 5 | 0 | 6 | 0 | 2 |
| ENSDARG000 | 6;6;6;6;6;6;6 | 414653:41552 | 416458:41645 | -;-;-;-;-;-;- | 2579 | fam83fb | 20 | 0 | 0 | 6 | 2 | 4 |
| ENSDARG000 | 6;6;6;6;6;6 | 40039534:400 | 40039814:400 | -;-;-;-;-;- | 1719 | NA | 0 | 0 | 0 | 0 | 0 | 0 |
| ENSDARG000 | 6;6;6 | 40043615:400 | 40043744:400 | +;+;+ | 532 | NA | 0 | 0 | 0 | 0 | 0 | 0 |
| ENSDARG000 | 6;6;6;6;6;6;6 | 40045035:400 | 40045788:400 | -;-;-;-;-;-;- | 2677 | uroc1 | 0 | 0 | 5 | 3 | 3 | 0 |
| ENSDARG000 | 6;6;6;6;6;6;6 | 3473657:3474 | 3473911:3474 | +;+;+;+;+;+;+ | 2589 | rad54l | 88 | 463 | 513 | 182 | 325 | 288 |
| ENSDARG000 | 6;6;6;6;6;6;6 | 3492972:3493 | 3493304:3493 | -;-;-;-;-;-;- | 2772 | NA | 154 | 88 | 195 | 62 | 261 | 229 |
| ENSDARG000 | 6;6 | 43076516:430 | 43079709:430 | -;- | 3497 | lrrn1 | 282 | 16 | 72 | 54 | 54 | 26 |
| ENSDARG000 | 6;6;6;6;6;6;6 | 7428613:7429 | 7429461:7429 | -;-;-;-;-;-;- | 2331 | fkbp11 | 78 | 99 | 152 | 49 | 100 | 76 |
| ENSDARG000 | 6;6;6;6;6;6;6 | 27361978:273 | 27362449:273 | +;+;+;+;+;+;+ | 7151 | NA | 8 | 2 | 2 | 0 | 0 | 0 |
| ENSDARG000 | 6;6 | 7429311:7435 | 7429541:7435 | +;+ | 451 | NA | 0 | 0 | 0 | 0 | 0 | 0 |
| ENSDARG000 | 6;6;6;6;6;6;6 | 9952678:9953 | 9952861:9953 | +;+;+;+;+;+;+ | 1751 | cyp20a1 | 905 | 921 | 885 | 904 | 919 | 852 |
| ENSDARG000 | 6;6;6;6;6;6;6 | 43222524:432 | 43226562:432 | -;-;-;-;-;-;- | 5192 | trnt1 | 461 | 621 | 800 | 394 | 661 | 558 |
| ENSDARG000 | 6;6;6;6;6;6;6 | 15604759:156 | 15606448:156 | -;-;-;-;-;-;- | 3340 | mlpha | 25505 | 15027 | 17552 | 25168 | 15553 | 19077 |
| ENSDARG000 | 6;6;6;6;6;6;6 | 43220191:432 | 43220212:432 | -;-;-;-;-;-;- | 8123 | FRMD4B | 87 | 52 | 111 | 94 | 62 | 72 |
| ENSDARG000 | 6;6;6;6;6;6;6 | 11397269:113 | 11397837:113 | +;+;+;+;+;+;+ | 3850 | NA | 433 | 474 | 592 | 278 | 386 | 478 |
| ENSDARG000 | 6;6;6;6;6;6;6 | 18531932:185 | 18532531:185 | +;+;+;+;+;+;+ | 4057 | suz12b | 1110 | 1338 | 1369 | 1372 | 1555 | 1282 |
| ENSDARG000 | 6;6 | 43346929:433 | 43347362:433 | -;- | 548 | NA | 20 | 1 | 13 | 2 | 5 | 6 |
| ENSDARG000 | 6;6 | 43218851:432 | 43219415:432 | -;- | 1591 | NA | 0 | 1 | 0 | 4 | 0 | 6 |
| ENSDARG000 | 6;6;6 | 43234213:432 | 43234646:432 | +;+;+ | 1466 | arl6ip5a | 253 | 248 | 169 | 222 | 337 | 288 |
| ENSDARG000 | 6;6;6;6 | 43342179:433 | 43342355:433 | +;+;+;+ | 1372 | NA | 14 | 4 | 12 | 2 | 12 | 20 |
| ENSDARG000 | 6;6;6;6;6;6;6 | 58832155:588 | 58832358:588 | +;+;+;+;+;+;+ | 2050 | dctn2 | 4881 | 4236 | 3695 | 4685 | 4364 | 3627 |
| ENSDARG000 | 6;6;6;6;6;6;6 | 38132511:381 | 38132789:381 | -;-;-;-;-;-;- | 1773 | NA | 2 | 0 | 0 | 0 | 0 | 0 |
| ENSDARG000 | 6;6;6;6;6;6;6 | 41096058:410 | 41096248:410 | +;+;+;+;+;+;+ | 4388 | fkbp5 | 1267 | 3343 | 307 | 1249 | 2785 | 315 |
| ENSDARG000 | 6;6;6;6;6;6;6 | 40060409:400 | 40061406:400 | -;-;-;-;-;-;- | 1324 | chchd4b | 32 | 42 | 44 | 29 | 74 | 81 |
| ENSDARG000 | 6;6;6;6;6;6;6 | 8490322:8490 | 8490570:8491 | -;-;-;-;-;-;- | 2769 | pglyrp2 | 0 | 0 | 8 | 0 | 0 | 8 |
| ENSDARG000 | 6;6;6;6 | 7444899:7449 | 7445091:7449 | +;+;+;+ | 645 | arf3b | 46 | 36 | 23 | 20 | 18 | 20 |
| ENSDARG000 | 6;6;6;6;6;6;6 | 38381957:384 | 38382461:384 | +;+;+;+;+;+;+ | 3940 | GABRB3 | 2 | 4 | 4 | 0 | 4 | 2 |
| ENSDARG000 | 6;6;6;6;6;6;6 | 427317:42731 | 428979:42803 | -;-;-;-;-;-;- | 2741 | grap2b | 52 | 64 | 133 | 78 | 278 | 366 |
| ENSDARG000 | 6;6;6;6;6;6;6 | 38366936:383 | 38367208:383 | -;-;-;-;-;-;- | 2016 | GABRA5 | 0 | 2 | 0 | 0 | 10 | 0 |
| ENSDARG000 | 6;6;6;6;6;6;6 | 58871600:588 | 58871910:588 | -;-;-;-;-;-;- | 4975 | NA | 473 | 335 | 607 | 479 | 543 | 368 |
| ENSDARG000 | 6;6;6;6;6;6;6 | 25213934:252 | 25213975:252 | +;+;+;+;+;+;+ | 1551 | KYAT3 | 1 | 0 | 0 | 0 | 1 | 2 |
| ENSDARG000 | 6;6;6;6 | 58915889:589 | 58916005:589 | +;+;+;+ | 1388 | ddit3 | 1393 | 919 | 2169 | 2291 | 658 | 884 |
| ENSDARG000 | 6 | 7453355 | 7453831 | + | 477 | NA | 22 | 32 | 5 | 15 | 16 | 38 |
| ENSDARG000 | 6;6;6;6;6;6;6 | 3509036:3509 | 3509334:3509 | -;-;-;-;-;-;- | 818 | uqcrh | 1564 | 1865 | 2196 | 1435 | 1688 | 1779 |
| ENSDARG000 | 6;6;6;6;6;6;6 | 40065201:400 | 40066437:400 | -;-;-;-;-;-;- | 2586 | sec61a1 | 6301 | 7733 | 17979 | 13078 | 9727 | 7855 |
| ENSDARG000 | 6 | 15651977 | 15653494 | - | 1518 | trim63a | 16 | 0 | 26 | 4 | 12 | 4 |
| ENSDARG000 | 6;6;6;6;6;6;6 | 15762647:157 | 15762880:157 | +;+;+;+;+;+;+ | 3396 | iqca1 | 2 | 0 | 0 | 0 | 0 | 0 |
| ENSDARG000 | 6;6;6;6;6;6;6 | 7466223:7466 | 7466707:7466 | +;+;+;+;+;+;+ | 6761 | ERBB3 | 111 | 4 | 132 | 26 | 82 | 91 |
| ENSDARG000 | 6;6 | 15751982:157 | 15753814:157 | -;- | 1953 | ackr3b | 198 | 30 | 168 | 30 | 178 | 258 |
| ENSDARG000 | 6;6;6;6;6;6;6 | 3516695:3516 | 3517138:3517 | +;+;+;+;+;+;+ | 2004 | faah | 91 | 86 | 88 | 21 | 186 | 34 |
| ENSDARG000 | 6;6;6;6;6;6;6 | 9970733:9970 | 9970935:9970 | +;+;+;+;+;+;+ | 2300 | abi2b | 1323 | 1054 | 1476 | 1217 | 1228 | 1255 |
| ENSDARG000 | 6;6;6;6;6;6;6 | 27451879:274 | 27452498:274 | +;+;+;+;+;+;+ | 5472 | NA | 4 | 0 | 0 | 8 | 6 | 0 |
| ENSDARG000 | 6;6;6;6;6;6;6 | 41118910:411 | 41119755:411 | -;-;-;-;-;-;- | 2752 | slc6a22.1 | 178 | 152 | 82 | 56 | 160 | 36 |
| ENSDARG000 | 6;6;6;6;6;6;6 | 11422044:114 | 11422338:114 | -;-;-;-;-;-;- | 1113 | NA | 0 | 2 | 0 | 0 | 0 | 2 |
| ENSDARG000 | 6;6;6;6;6;6;6 | 29386728:293 | 29386947:293 | +;+;+;+;+;+;+ | 2072 | actl6a | 511 | 904 | 1140 | 362 | 864 | 670 |
| ENSDARG000 | 6;6;6;6;6;6;6 | 8499865:8507 | 8501059:8507 | -;-;-;-;-;-;- | 7107 | myh11a | 268 | 228 | 92 | 181 | 346 | 386 |
| ENSDARG000 | 6;6;6;6;6;6;6 | 58921240:589 | 58921613:589 | -;-;-;-;-;-;- | 3174 | mars1 | 2382 | 2181 | 3160 | 1402 | 2094 | 1205 |
| ENSDARG000 | 6;6;6;6;6;6;6 | 40089743:400 | 40091095:400 | -;-;-;-;-;-;- | 2765 | ip6k2b | 1555 | 1323 | 2059 | 1542 | 3726 | 2578 |
| ENSDARG000 | 6;6 | 40174216:401 | 40174775:401 | -;- | 903 | NA | 0 | 0 | 0 | 0 | 0 | 0 |
| ENSDARG000 | 6;6;6;6;6;6;6 | 36795225:367 | 36795261:367 | +;+;+;+;+;+;+ | 2782 | NA | 2 | 0 | 0 | 0 | 0 | 0 |
| ENSDARG000 | 6;6;6;6;6;6;6 | 18544791:185 | 18545556:185 | +;+;+;+;+;+;+ | 7921 | atad5b | 151 | 439 | 358 | 255 | 314 | 385 |
| ENSDARG000 | 6;6;6;6;6;6;6 | 25257728:252 | 25257769:252 | +;+;+;+;+;+;+ | 6571 | kyat3.2 | 2869 | 318 | 455 | 323 | 3998 | 5580 |
| ENSDARG000 | 6;6;6;6;6;6;6 | 41144155:411 | 41144357:411 | +;+;+;+;+;+;+ | 2408 | slc6a22.2 | 8 | 40 | 8 | 0 | 38 | 34 |
| ENSDARG000 | 6;6;6;6;6;6;6 | 18554819:185 | 18555677:185 | -;-;-;-;-;-;- | 1803 | tefm | 128 | 317 | 220 | 172 | 351 | 212 |
| ENSDARG000 | 6;6;6;6;6;6;6 | 515181:51897 | 515249:51915 | +;+;+;+;+;+;+ | 3039 | NA | 0 | 20 | 18 | 2 | 20 | 12 |
| ENSDARG000 | 6;6 | 31804665:318 | 31805057:318 | -;- | 572 | NA | 7 | 0 | 2 | 0 | 0 | 0 |
| ENSDARG000 | 6;6;6;6;6;6;6 | 11438972:114 | 11439218:114 | +;+;+;+;+;+;+ | 4896 | NA | 24 | 0 | 16 | 0 | 60 | 0 |
| ENSDARG000 | 6;6;6;6;6;6;6 | 31772182:317 | 31775186:317 | -;-;-;-;-;-;- | 4716 | ROR1 | 260 | 2 | 118 | 22 | 299 | 484 |
| ENSDARG000 | 6;6 | 31770121:317 | 31770147:317 | +;+ | 283 | NA | 0 | 0 | 0 | 0 | 0 | 0 |
| ENSDARG000 | 6;6;6 | 59350457:593 | 59351422:593 | -;-;- | 1728 | fam210b | 124 | 68 | 90 | 116 | 106 | 70 |
| ENSDARG000 | 6;6;6;6;6;6;6 | 59370821:593 | 59373234:593 | -;-;-;-;-;-;- | 3302 | PPP1R16B | 90 | 56 | 58 | 20 | 77 | 106 |
| ENSDARG000 | 6;6;6;6;6;6;6 | 27514465:275 | 27515177:275 | +;+;+;+;+;+;+ | 3984 | ryk | 1667 | 1392 | 2537 | 1607 | 2519 | 2517 |
| ENSDARG000 | 6;6;6;6;6;6;6 | 59808677:598 | 59808868:598 | +;+;+;+;+;+;+ | 2354 | caskb | 995 | 1221 | 1132 | 1111 | 965 | 1028 |
| ENSDARG000 | 6;6;6;6;6;6;6 | 13189563:131 | 13192234:131 | -;-;-;-;-;-;- | 5679 | eef1b2 | 2529 | 2316 | 4195 | 2977 | 2822 | 3666 |
| ENSDARG000 | 6;6;6;6;6;6;6 | 38626684:386 | 38627353:386 | +;+;+;+;+;+;+ | 5696 | ATP10A | 250 | 125 | 234 | 68 | 304 | 202 |
| ENSDARG000 | 6;6;6;6;6;6;6 | 3827751:3827 | 3828042:3828 | +;+;+;+;+;+;+ | 5063 | gad1b | 4 | 0 | 3 | 2 | 4 | 4 |
| ENSDARG000 | 6;6;6;6;6;6;6 | 7533601:7536 | 7533854:7536 | +;+;+;+;+;+;+ | 2476 | pa2g4a | 3266 | 3802 | 6060 | 2420 | 4159 | 3689 |
| ENSDARG000 | 6;6;6;6;6;6;6 | 40184235:401 | 40185686:401 | -;-;-;-;-;-;- | 10973 | NA | 58 | 16 | 18 | 6 | 14 | 68 |
| ENSDARG000 | 6;6;6;6;6 | 41181869:411 | 41182309:411 | +;+;+;+;+ | 1574 | opn1mw1 | 6 | 0 | 4 | 0 | 6 | 0 |
| ENSDARG000 | 6;6;6;6;6;6;6 | 38759546:387 | 38759705:387 | -;-;-;-;-;-;- | 1771 | NA | 0 | 0 | 0 | 0 | 0 | 0 |
| ENSDARG000 | 6;6;6;6;6;6 | 36821621:368 | 36821912:368 | +;+;+;+;+;+ | 1972 | tmem45a | 20 | 70 | 170 | 89 | 54 | 16 |
| ENSDARG000 | 6;6;6;6;6 | 8591585:8592 | 8591949:8592 | -;-;-;-;- | 848 | cep20 | 80 | 45 | 55 | 54 | 54 | 18 |
| ENSDARG000 | 6;6;6;6;6;6;6 | 33918813:339 | 33918973:339 | -;-;-;-;-;-;- | 1756 | akr1a1b | 784 | 565 | 676 | 704 | 768 | 935 |
| ENSDARG000 | 6;6;6;6;6;6;6 | 59832786:598 | 59833084:598 | +;+;+;+;+;+;+ | 5959 | ddx3xb | 6673 | 7382 | 9530 | 4031 | 8466 | 4834 |
| ENSDARG000 | 6;6;6;6;6;6 | 8598428:8598 | 8598585:8598 | +;+;+;+;+;+ | 1410 | ube2g2 | 510 | 393 | 632 | 337 | 488 | 518 |
| ENSDARG000 | 6;6;6 | 15895634:158 | 15895722:158 | +;+;+ | 706 | NA | 0 | 0 | 0 | 0 | 0 | 0 |
| ENSDARG000 | 6;6;6;6;6;6;6 | 33926829:339 | 33927137:339 | -;-;-;-;-;-;- | 1308 | prpf38a | 372 | 557 | 521 | 368 | 486 | 402 |
| ENSDARG000 | 6;6;6;6;6;6;6 | 33949280:339 | 33949962:339 | -;-;-;-;-;-;- | 2355 | mpl | 0 | 0 | 0 | 0 | 10 | 3 |
| ENSDARG000 | 6;6;6;6;6;6;6 | 36839509:368 | 36839825:368 | +;+;+;+;+;+;+ | 1395 | sft2d2a | 0 | 0 | 0 | 0 | 0 | 0 |
| ENSDARG000 | 6;6;6;6;6;6;6 | 43390387:433 | 43390429:433 | +;+;+;+;+;+;+ | 2282 | mitfa | 13234 | 4954 | 7773 | 7078 | 7342 | 7307 |
| ENSDARG000 | 6;6;6;6;6;6;6 | 33979523:339 | 33984471:339 | -;-;-;-;-;-;- | 9153 | tie1 | 230 | 20 | 164 | 18 | 126 | 124 |
| ENSDARG000 | 6;6;6 | 15938621:159 | 15938777:159 | +;+;+ | 581 | NA | 0 | 0 | 0 | 0 | 0 | 0 |
| ENSDARG000 | 6;6;6;6;6;6 | 29397554:293 | 29397769:293 | -;-;-;-;-;- | 906 | mrpl47 | 304 | 285 | 398 | 287 | 357 | 264 |
| ENSDARG000 | 6;6;6;6;6;6;6 | 3864040:3864 | 3864223:3864 | +;+;+;+;+;+;+ | 2013 | gorasp2 | 1691 | 1809 | 2332 | 1378 | 2294 | 1626 |
| ENSDARG000 | 6;6;6;6;6;6;6 | 33931095:339 | 33931921:339 | +;+;+;+;+;+;+ | 7125 | orc1 | 154 | 436 | 417 | 172 | 453 | 244 |
| ENSDARG000 | 6;6;6 | 15993146:159 | 15993224:159 | +;+;+ | 555 | NA | 0 | 0 | 0 | 0 | 0 | 0 |
| ENSDARG000 | 6;6;6;6;6;6;6 | 7553085:7553 | 7553264:7553 | +;+;+;+;+;+;+ | 7721 | MYH10 | 2512 | 3647 | 6692 | 2678 | 3742 | 2413 |
| ENSDARG000 | 6;6;6;6;6;6;6 | 584632:58846 | 584847:58859 | +;+;+;+;+;+;+ | 2899 | zgc:92360 | 84 | 21 | 14 | 26 | 50 | 90 |
| ENSDARG000 | 6;6;6;6;6;6;6 | 32010183:320 | 32010481:320 | -;-;-;-;-;-;- | 2008 | pgm1 | 1958 | 3898 | 7584 | 9561 | 3377 | 2965 |
| ENSDARG000 | 6;6;6;6;6;6;6 | 11497646:114 | 11498053:114 | -;-;-;-;-;-;- | 1548 | GULP1 | 1 | 0 | 6 | 0 | 18 | 0 |
| ENSDARG000 | 6;6;6;6;6 | 41186320:411 | 41186746:411 | +;+;+;+;+ | 1661 | opn1mw2 | 0 | 0 | 0 | 0 | 0 | 0 |
| ENSDARG000 | 6;6 | 16031189:160 | 16032108:160 | +;+ | 2095 | gbx2 | 4 | 0 | 0 | 0 | 0 | 0 |
| ENSDARG000 | 6;6;6;6;6;6;6 | 38773376:387 | 38773519:387 | +;+;+;+;+;+;+ | 4104 | ube3a | 1113 | 1316 | 1546 | 706 | 1464 | 1200 |
| ENSDARG000 | 6;6;6;6 | 18594977:185 | 18595637:185 | -;-;-;- | 1226 | cd7al | 22 | 15 | 11 | 29 | 154 | 65 |
| ENSDARG000 | 6;6;6;6;6;6 | 29402997:294 | 29403140:294 | +;+;+;+;+;+ | 858 | ndufb5 | 702 | 955 | 1034 | 629 | 894 | 586 |
| ENSDARG000 | 6;6;6;6;6;6;6 | 18569453:185 | 18569676:185 | +;+;+;+;+;+;+ | 2707 | rhot1b | 79 | 370 | 312 | 452 | 189 | 306 |
| ENSDARG000 | 6;6;6;6;6 | 41191482:411 | 41191893:411 | +;+;+;+;+ | 1379 | opn1mw3 | 0 | 0 | 0 | 0 | 0 | 0 |
| ENSDARG000 | 6;6;6;6;6;6;6 | 8622228:8626 | 8622339:8626 | +;+;+;+;+;+;+ | 2200 | TSPEAR | 6 | 0 | 0 | 0 | 0 | 0 |
| ENSDARG000 | 6;6;6;6;6 | 43438074:434 | 43438529:434 | -;-;-;-;- | 1887 | NA | 8 | 0 | 11 | 12 | 18 | 0 |
| ENSDARG000 | 6 | 598669 | 599688 | + | 1020 | NA | 0 | 0 | 0 | 0 | 0 | 0 |
| ENSDARG000 | 6;6;6;6;6;6;6 | 38791900:387 | 38791963:387 | -;-;-;-;-;-;- | 3020 | CNGA3 | 2 | 4 | 0 | 0 | 0 | 0 |
| ENSDARG000 | 6;6;6;6;6 | 41200500:412 | 41200934:412 | +;+;+;+;+ | 1379 | opn1mw4 | 0 | 0 | 0 | 0 | 0 | 4 |

| | | | | | | | | | | | | |
|---|---|---|---|---|---|---|---|---|---|---|---|---|
| ENSDARG000 | 6;6;6;6;6;6;6 | 29410986:294 | 29411396:294 | +;+;+;+;+;+;+;+ | 5644 | usp13 | 8 | 26 | 16 | 6 | 12 | 38 |
| ENSDARG000 | 6;6;6;6;6;6;6 | 27620687:276 | 27621291:276 | -;-;-;-;-;-;-;- | 947 | NA | 0 | 42 | 44 | 37 | 8 | 50 |
| ENSDARG000 | 6;6;6;6;6;6;6 | 59854224:598 | 59854630:598 | +;+;+;+;+;+;+;+ | 5935 | kdm6al | 643 | 458 | 769 | 363 | 648 | 578 |
| ENSDARG000 | 6;6;6;6;6;6;6 | 38845676:388 | 38846043:388 | +;+;+;+;+;+;+;+ | 4737 | stk35l | 1350 | 2128 | 2204 | 2421 | 2621 | 1169 |
| ENSDARG000 | 6;6;6;6;6;6 | 10015525:100 | 10016561:100 | -;-;-;-;-;- | 1859 | itm2ba | 10476 | 7700 | 9308 | 9057 | 11881 | 9087 |
| ENSDARG000 | 6;6;6;6;6 | 41223204:412 | 41223992:412 | -;-;-;-;- | 1410 | synprb | 0 | 0 | 0 | 0 | 0 | 0 |
| ENSDARG000 | 6;6;6;6;6;6;6 | 13206516:132 | 13206559:132 | +;+;+;+;+;+;+;+ | 2515 | ndufs1 | 0 | 0 | 2 | 0 | 0 | 0 |
| ENSDARG000 | 6;6;6;6;6;6;6 | 40329308:403 | 40329958:403 | -;-;-;-;-;-;-;- | 2518 | ttll3 | 104 | 16 | 13 | 60 | 5 | 54 |
| ENSDARG000 | 6;6;6;6;6;6;6 | 13207139:132 | 13207215:132 | +;+;+;+;+;+;+;+ | 9842 | ino80db | 388 | 304 | 343 | 328 | 415 | 424 |
| ENSDARG000 | 6;6;6;6;6;6;6 | 38879961:388 | 38880275:388 | +;+;+;+;+;+;+;+ | 3861 | bin2b | 600 | 700 | 447 | 687 | 825 | 857 |
| ENSDARG000 | 6;6;6;6;6;6;6 | 43450221:434 | 43450503:434 | +;+;+;+;+;+;+;+ | 2334 | zgc:113054 | 182 | 92 | 210 | 68 | 168 | 159 |
| ENSDARG000 | 6;6;6;6;6;6;6 | 36877968:368 | 36878113:368 | +;+;+;+;+;+;+;+ | 5902 | NA | 32 | 6 | 6 | 7 | 31 | 16 |
| ENSDARG000 | 6;6;6;6;6;6;6 | 3878492:3879 | 3879385:3879 | -;-;-;-;-;-;-;- | 3219 | tlk1b | 999 | 932 | 1085 | 835 | 1029 | 1031 |
| ENSDARG000 | 6;6;6;6;6;6;6 | 59913147:599 | 59913227:599 | +;+;+;+;+;+;+;+ | 1963 | aamp | 2066 | 1974 | 3347 | 2625 | 1774 | 2126 |
| ENSDARG000 | 6;6;6;6;6;6;6 | 36844127:368 | 36844351:368 | +;+;+;+;+;+;+;+ | 4437 | chd1l | 408 | 576 | 534 | 319 | 459 | 452 |
| ENSDARG000 | 6;6;6 | 44400757:444 | 44400807:444 | +;+;+ | 248 | NA | 0 | 0 | 0 | 0 | 0 | 0 |
| ENSDARG000 | 6;6;6;6;6;6;6 | 44243070:442 | 44246895:442 | -;-;-;-;-;-;-;- | 6191 | pdzrn3b | 1022 | 66 | 589 | 236 | 498 | 400 |
| ENSDARG000 | 6;6;6;6;6;6;6 | 27624023:276 | 27624146:276 | +;+;+;+;+;+;+;+ | 2350 | SLCO2A1 | 74 | 10 | 11 | 2 | 21 | 46 |
| ENSDARG000 | 6;6;6;6;6;6;6 | 16043318:160 | 16047255:160 | -;-;-;-;-;-;-;- | 7360 | AGAP1 | 405 | 465 | 609 | 413 | 565 | 362 |
| ENSDARG000 | 6;6;6;6;6;6;6 | 8652310:8654 | 8653046:8654 | +;+;+;+;+;+;+;+ | 4651 | NA | 470 | 331 | 614 | 410 | 486 | 404 |
| ENSDARG000 | 6;6;6;6 | 43491635:434 | 43492138:434 | -;-;-;- | 936 | mdfic2 | 0 | 0 | 0 | 0 | 0 | 0 |
| ENSDARG000 | 6;6;6 | 43499071:434 | 43499104:434 | +;+;+ | 241 | NA | 0 | 0 | 0 | 0 | 0 | 0 |
| ENSDARG000 | 6;6;6;6;6;6;6 | 44812444:448 | 44812869:448 | +;+;+;+;+;+;+;+ | 4414 | NA | 4 | 51 | 375 | 6 | 12 | 8 |
| ENSDARG000 | 6;6;6;6;6;6;6 | 60036767:600 | 60036792:600 | +;+;+;+;+;+;+ | 1032 | NA | 34 | 4 | 2 | 6 | 2 | 1 |
| ENSDARG000 | 6;6;6;6;6;6;6 | 45114613:451 | 45114856:451 | +;+;+;+;+;+;+;+ | 2773 | NA | 0 | 10 | 85 | 12 | 14 | 20 |
| ENSDARG000 | 6;6;6;6;6;6;6 | 60043053:600 | 60044494:600 | -;-;-;-;-;-;-;- | 2434 | ccndbp1 | 363 | 399 | 445 | 372 | 416 | 454 |
| ENSDARG000 | 6;6;6;6;6;6;6 | 3961319:3961 | 3962225:3962 | -;-;-;-;-;-;-;- | 2880 | slc25a12 | 3862 | 2167 | 2525 | 2629 | 2730 | 2524 |
| ENSDARG000 | 6;6;6;6;6;6 | 40383424:403 | 40385079:403 | -;-;-;-;-;- | 2156 | si:dkev-28n18 | 20 | 8 | 3 | 1 | 0 | 13 |
| ENSDARG000 | 6;6;6;6;6;6;6 | 40354424:403 | 40354998:403 | +;+;+;+;+;+;+;+ | 5289 | SLC4A8 | 3325 | 447 | 188 | 1418 | 259 | 1804 |
| ENSDARG000 | 6;6;6;6;6;6 | 38896158:388 | 38896395:388 | +;+;+;+;+;+ | 1625 | slc48a1b | 481 | 694 | 368 | 932 | 510 | 633 |
| ENSDARG000 | 6;6;6;6;6;6;6 | 45494227:455 | 45494352:455 | +;+;+;+;+;+;+;+ | 3479 | cntn4 | 0 | 0 | 0 | 0 | 2 | 2 |
| ENSDARG000 | 6;6;6;6;6;6;6 | 43602143:436 | 43602446:436 | -;-;-;-;-;-;-;- | 8357 | foxp1b | 55 | 12 | 56 | 8 | 49 | 30 |
| ENSDARG000 | 6;6;6;6;6;6;6 | 603278:60328 | 603532:60353 | -;-;-;-;-;-;-;- | 675 | lgals2b | 10 | 0 | 14 | 0 | 8 | 0 |
| ENSDARG000 | 6;6 | 48380245:483 | 48380431:483 | -;- | 812 | NA | 264 | 195 | 290 | 288 | 218 | 333 |
| ENSDARG000 | 6;6;6;6;6;6;6 | 32029231:320 | 32029702:320 | -;-;-;-;-;-;-;- | 2988 | efcab7 | 201 | 155 | 246 | 245 | 225 | 244 |
| ENSDARG000 | 6;6;6;6;6;6;6 | 60055168:600 | 60055292:600 | +;+;+;+;+;+;+;+ | 2895 | pck1 | 2 | 8 | 12 | 4 | 176 | 0 |
| ENSDARG000 | 6;6;6;6;6;6;6 | 11680726:116 | 11681157:116 | +;+;+;+;+;+;+;+ | 1926 | CALCRL | 19 | 0 | 0 | 0 | 0 | 66 |
| ENSDARG000 | 6;6;6;6;6;6;6 | 18605644:186 | 18606303:186 | -;-;-;-;-;-;-;- | 1934 | znf207b | 734 | 621 | 1170 | 594 | 959 | 845 |
| ENSDARG000 | 6;6;6;6;6 | 48382697:483 | 48384878:483 | -;-;-;-;- | 2745 | rhoca | 1556 | 1399 | 1463 | 939 | 1223 | 1819 |
| ENSDARG000 | 6;6;6 | 13276022:133 | 13278563:133 | -;-;- | 4375 | KCNJ3 | 4 | 4 | 8 | 0 | 16 | 20 |
| ENSDARG000 | 6;6;6;6;6;6;6 | 13347848:133 | 13348244:133 | -;-;-;-;-;-;-;- | 1893 | arl6ip6 | 125 | 139 | 134 | 73 | 182 | 92 |
| ENSDARG000 | 6;6;6;6;6;6;6 | 60076456:600 | 60076719:600 | -;-;-;-;-;-;- | 811 | pmepa1 | 595 | 345 | 421 | 295 | 184 | 252 |
| ENSDARG000 | 6;6;6;6;6;6;6 | 27896657:278 | 27896743:278 | +;+;+;+;+;+;+;+ | 2635 | cep63 | 370 | 274 | 323 | 174 | 694 | 411 |
| ENSDARG000 | 6 | 8691408 | 8691993 | + | 586 | NA | 11 | 2 | 10 | 8 | 14 | 7 |
| ENSDARG000 | 6;6;6;6;6;6 | 60112200:601 | 60112311:601 | +;+;+;+;+;+ | 1635 | prelid3b | 2216 | 1760 | 2380 | 1150 | 3108 | 1872 |
| ENSDARG000 | 6;6;6;6;6;6;6 | 38905047:389 | 38905695:389 | -;-;-;-;-;-;-;- | 3212 | NA | 14 | 28 | 109 | 130 | 76 | 66 |
| ENSDARG000 | 6;6;6;6;6;6 | 8698677:8702 | 8699009:8702 | -;-;-;-;-;- | 1181 | nabp1b | 0 | 0 | 0 | 0 | 0 | 0 |
| ENSDARG000 | 6;6;6;6;6;6;6 | 60125033:601 | 60125259:601 | +;+;+;+;+;+;+;+ | 1754 | aurka | 122 | 466 | 376 | 188 | 224 | 171 |
| ENSDARG000 | 6;6;6;6;6;6;6 | 27667359:276 | 27667535:276 | +;+;+;+;+;+;+;+ | 3511 | rab6ba | 13 | 14 | 0 | 2 | 2 | 16 |
| ENSDARG000 | 6;6;6;6 | 27884372:278 | 27885215:278 | -;-;-;- | 1465 | im:7152348 | 64 | 12 | 21 | 22 | 124 | 52 |
| ENSDARG000 | 6;6;6;6;6;6;6 | 48425366:484 | 48426796:484 | -;-;-;-;-;-;-;- | 3082 | NA | 110 | 46 | 111 | 64 | 139 | 106 |
| ENSDARG000 | 6;6;6;6;6;6;6 | 32046202:320 | 32046272:320 | +;+;+;+;+;+;+;+ | 851 | itgb3bp | 28 | 148 | 80 | 55 | 51 | 35 |
| ENSDARG000 | 6;6;6;6 | 8705149:8708 | 8705245:8708 | +;+;+;+ | 789 | NA | 0 | 0 | 0 | 0 | 6 | 0 |
| ENSDARG000 | 6;6;6;6;6;6;6 | 8756177:8759 | 8759985:8760 | -;-;-;-;-;-;-;- | 5514 | TMEFF2 | 1280 | 1556 | 2206 | 1828 | 1584 | 649 |
| ENSDARG000 | 6;6 | 8707812:8735 | 8710416:8736 | -;- | 3403 | cavin2b | 3107 | 3042 | 970 | 3889 | 1761 | 1338 |
| ENSDARG000 | 6;6;6 | 11725451:117 | 11725684:117 | -;-;- | 412 | NA | 0 | 0 | 0 | 0 | 0 | 0 |
| ENSDARG000 | 6;6;6;6;6;6;6 | 45843193:458 | 45845132:458 | -;-;-;-;-;-;-;- | 4779 | rbm19 | 548 | 533 | 790 | 414 | 653 | 414 |
| ENSDARG000 | 6;6;6;6;6;6;6 | 11760749:117 | 11761080:117 | +;+;+;+;+;+;+;+ | 2315 | zswim2 | 0 | 0 | 0 | 0 | 0 | 0 |
| ENSDARG000 | 6;6 | 16394629:163 | 16394711:163 | +;+ | 453 | NA | 0 | 0 | 0 | 0 | 0 | 0 |
| ENSDARG000 | 6;6;6;6;6;6;6 | 7659576:7660 | 7660934:7660 | -;-;-;-;-;-;-;- | 5085 | ubn2a | 547 | 337 | 662 | 433 | 484 | 541 |
| ENSDARG000 | 6;6;6;6 | 16400049:164 | 16400318:164 | -;-;-;- | 840 | faimb | 0 | 0 | 0 | 0 | 2 | 0 |
| ENSDARG000 | 6;6;6;6;6;6;6 | 29476933:294 | 29478013:294 | -;-;-;-;-;-;-;- | 3789 | PEX5L | 0 | 38 | 63 | 135 | 6 | 154 |
| ENSDARG000 | 6;6;6;6;6;6;6 | 3984841:3985 | 3985289:3985 | -;-;-;-;-;-;-;- | 1237 | unc50 | 440 | 333 | 533 | 536 | 541 | 493 |
| ENSDARG000 | 6;6;6;6;6;6;6 | 13390289:133 | 13391426:133 | -;-;-;-;-;-;-;- | 5851 | NA | 202 | 66 | 343 | 98 | 180 | 405 |
| ENSDARG000 | 6;6;6;6;6;6;6 | 16406723:164 | 16408813:164 | +;+;+;+;+;+;+ | 3904 | NA | 12 | 2 | 4 | 2 | 4 | 0 |
| ENSDARG000 | 6 | 43903209 | 43907321 | + | 4113 | gpr27 | 58 | 36 | 60 | 122 | 54 | 72 |
| ENSDARG000 | 6;6;6 | 43918529:439 | 43919088:439 | -;-;- | 972 | prok2 | 0 | 0 | 0 | 0 | 0 | 0 |
| ENSDARG000 | 6;6;6;6;6;6;6 | 40429773:404 | 40430206:404 | -;-;-;-;-;-;-;- | 1890 | ccdc174 | 272 | 222 | 331 | 278 | 238 | 273 |
| ENSDARG000 | 6;6;6;6;6;6;6 | 38954733:389 | 38955315:389 | -;-;-;-;-;-;-;- | 2128 | vdrb | 99 | 37 | 63 | 148 | 75 | 161 |
| ENSDARG000 | 6;6;6;6;6;6;6 | 48618512:486 | 48618624:486 | +;+;+;+;+;+;+ | 1334 | TAFA3 | 0 | 4 | 0 | 0 | 0 | 14 |
| ENSDARG000 | 6;6;6;6;6;6;6 | 40422224:404 | 40422455:404 | -;-;-;-;-;-;-;- | 1510 | NA | 100 | 58 | 52 | 86 | 22 | 149 |
| ENSDARG000 | 6;6;6;6;6;6;6 | 10037252:100 | 10037369:100 | +;+;+;+;+;+;+;+ | 2111 | sucla2 | 1426 | 1315 | 1413 | 914 | 1755 | 1167 |
| ENSDARG000 | 6;6;6;6;6;6 | 48720617:487 | 48720672:487 | +;+;+;+;+;+ | 1375 | NA | 0 | 0 | 0 | 0 | 0 | 0 |
| ENSDARG000 | 6;6;6;6;6;6;6 | 11765341:117 | 11766771:117 | -;-;-;-;-;-;-;- | 4497 | NA | 0 | 1 | 1 | 0 | 0 | 6 |
| ENSDARG000 | 6;6;6;6;6;6;6 | 11765541:117 | 11766771:117 | -;-;-;-;-;-;-;- | 2515 | FAM171B | 0 | 0 | 2 | 0 | 0 | 0 |
| ENSDARG000 | 6;6;6;6;6;6;6 | 18623492:186 | 18625226:186 | -;-;-;-;-;-;-;- | 3321 | rhbdl3 | 3 | 8 | 0 | 20 | 0 | 6 |
| ENSDARG000 | 6;6;6;6;6;6;6 | 11781231:117 | 11782081:117 | -;-;-;-;-;-;-;- | 3230 | marchf7 | 320 | 172 | 307 | 216 | 185 | 242 |
| ENSDARG000 | 6;6;6;6;6;6;6 | 16436572:164 | 16436706:164 | -;-;-;-;-;-;-;- | 1672 | SLC19A2 | 92 | 103 | 148 | 37 | 172 | 75 |
| ENSDARG000 | 6;6;6 | 43979711:439 | 43980597:439 | -;-;- | 1219 | NA | 0 | 0 | 0 | 0 | 0 | 0 |
| ENSDARG000 | 6;6;6;6;6;6;6 | 48978202:489 | 48978333:489 | +;+;+;+;+;+;+;+ | 2634 | slc16a1a | 0 | 2 | 0 | 0 | 2 | 6 |
| ENSDARG000 | 6;6;6;6;6;6;6 | 36942966:369 | 36943324:369 | +;+;+;+;+;+;+ | 2472 | negr1 | 4 | 0 | 14 | 0 | 0 | 0 |
| ENSDARG000 | 6;6;6;6;6 | 44006709:440 | 44009847:440 | -;-;-;-;- | 3809 | rybpb | 892 | 541 | 587 | 548 | 757 | 760 |
| ENSDARG000 | 6;6;6;6;6;6;6 | 44089189:441 | 44092654:441 | -;-;-;-;-;-;-;- | 4704 | shq1 | 289 | 190 | 265 | 166 | 344 | 258 |
| ENSDARG000 | 6;6;6;6;6;6;6 | 45874595:458 | 45875277:458 | -;-;-;-;-;-;-;- | 3709 | epha2a | 160 | 42 | 44 | 12 | 20 | 38 |
| ENSDARG000 | 6;6;6;6;6;6;6 | 32065094:320 | 32065611:320 | -;-;-;-;-;-;-;- | 3172 | alg6 | 1268 | 1095 | 1338 | 854 | 1109 | 1081 |
| ENSDARG000 | 6;6;6;6;6;6;6 | 29693492:296 | 29693735:296 | +;+;+;+;+;+;+;+ | 1148 | pde6d | 87 | 109 | 100 | 95 | 135 | 114 |
| ENSDARG000 | 6;6;6;6;6;6;6 | 9107063:9108 | 9107241:9108 | +;+;+;+;+;+;+;+ | 3463 | vps16 | 2326 | 1870 | 2139 | 1837 | 2249 | 1944 |
| ENSDARG000 | 6;6;6;6;6;6;6 | 39011703:390 | 39014703:390 | -;-;-;-;-;-;-;- | 7794 | NA | 2598 | 1231 | 1697 | 2623 | 2440 | 2457 |
| ENSDARG000 | 6;6;6;6 | 4001317:4002 | 4001730:4002 | +;+;+;+ | 1529 | NA | 0 | 2 | 1 | 0 | 0 | 2 |
| ENSDARG000 | 6;6;6;6;6 | 16468776:164 | 16469151:164 | +;+;+;+;+ | 2762 | zgc:161969 | 613 | 616 | 859 | 458 | 734 | 694 |
| ENSDARG000 | 6;6;6;6;6;6;6 | 45918981:459 | 45919034:459 | +;+;+;+;+;+;+ | 6214 | H6PD | 814 | 248 | 558 | 280 | 753 | 694 |
| ENSDARG000 | 6;6;6;6;6;6;6 | 13458257:134 | 13458843:134 | -;-;-;-;-;-;-;- | 3594 | MYLK | 0 | 2 | 0 | 0 | 10 | 4 |
| ENSDARG000 | 6;6;6;6;6;6;6 | 7688491:7688 | 7688897:7688 | -;-;-;-;-;-;-;- | 2767 | ttc26 | 102 | 72 | 120 | 77 | 117 | 96 |
| ENSDARG000 | 6;6;6 | 37193908:371 | 37194239:371 | +;+;+ | 524 | NA | 0 | 0 | 4 | 0 | 0 | 0 |
| ENSDARG000 | 6;6;6;6;6;6;6 | 4033832:4061 | 4034076:4061 | +;+;+;+;+;+;+;+ | 3653 | SEMA5B | 6 | 4 | 23 | 0 | 11 | 2 |
| ENSDARG000 | 6;6 | 27938489:279 | 27939822:279 | -;- | 1707 | NA | 0 | 1 | 2 | 0 | 0 | 0 |
| ENSDARG000 | 6;6;6;6;6;6;6 | 45932241:459 | 45932378:459 | +;+;+;+;+;+;+ | 731 | rbp7b | 4 | 0 | 6 | 1 | 0 | 4 |
| ENSDARG000 | 6;6;6;6;6;6;6 | 7709662:7712 | 7713491:7713 | -;-;-;-;-;-;-;- | 4848 | rpsa | 34644 | 33663 | 57233 | 52948 | 39929 | 46076 |
| ENSDARG000 | 6;6;6;6;6;6;6 | 29707129:297 | 29707187:297 | +;+;+;+;+;+;+;+ | 1511 | phb2b | 409 | 824 | 958 | 267 | 740 | 326 |
| ENSDARG000 | 6;6;6 | 10077756:100 | 10078303:100 | +;+;+ | 1552 | HTR2A | 8 | 0 | 0 | 0 | 0 | 16 |
| ENSDARG000 | 6;6 | 10087230:100 | 10088408:100 | -;- | 1207 | NA | 4 | 2 | 0 | 1 | 0 | 0 |
| ENSDARG000 | 6;6;6;6;6;6;6 | 9130989:9133 | 9131121:9133 | +;+;+;+;+;+;+ | 1490 | rgn | 156 | 279 | 332 | 200 | 219 | 296 |
| ENSDARG000 | 6;6;6;6;6;6;6 | 45934682:459 | 45934792:459 | +;+;+;+;+;+;+;+ | 1041 | cenps | 72 | 149 | 149 | 59 | 116 | 67 |
| ENSDARG000 | 6;6;6;6;6;6;6 | 44163727:441 | 44164103:441 | +;+;+;+;+;+;+ | 2168 | gxylt2 | 660 | 322 | 303 | 332 | 541 | 322 |
| ENSDARG000 | 6;6;6;6 | 49061077:490 | 49061324:490 | -;-;-;- | 1005 | si:ch211-105i | 22 | 17 | 2 | 0 | 14 | 20 |
| ENSDARG000 | 6;6;6;6;6;6;6 | 49052741:490 | 49052817:490 | +;+;+;+;+;+;+;+ | 3482 | sycp1 | 9 | 1 | 7 | 3 | 19 | 4 |
| ENSDARG000 | 6;6;6;6;6;6;6 | 44197099:441 | 44197472:441 | +;+;+;+;+;+;+;+ | 4461 | ppp4r2b | 761 | 802 | 1028 | 600 | 789 | 741 |
| ENSDARG000 | 6;6;6;6;6;6;6 | 37301341:373 | 37301641:373 | +;+;+;+;+;+;+;+ | 2408 | zranb2 | 1163 | 948 | 1796 | 1006 | 1258 | 1296 |
| ENSDARG000 | 6;6;6;6;6;6;6 | 27991943:279 | 27992159:279 | +;+;+;+;+;+;+;+ | 4246 | amotl2a | 839 | 423 | 1090 | 1421 | 1019 | 960 |
| ENSDARG000 | 6;6;6;6;6;6;6 | 10094061:100 | 10094167:100 | +;+;+;+;+;+;+;+ | 4129 | ATP7B | 18 | 4 | 19 | 6 | 8 | 30 |
| ENSDARG000 | 6;6;6;6;6;6;6 | 612330:61233 | 612763:61276 | +;+;+;+;+;+;+;+ | 6609 | NA | 4 | 0 | 20 | 0 | 40 | 0 |
| ENSDARG000 | 6;6;6;6;6;6 | 16478099:164 | 16478258:164 | -;-;-;-;-;- | 903 | psmb3 | 1866 | 1760 | 2025 | 1097 | 1810 | 1313 |
| ENSDARG000 | 6;6;6;6 | 16493755:164 | 16493822:164 | +;+;+;+ | 805 | NA | 134 | 96 | 114 | 95 | 115 | 111 |
| ENSDARG000 | 6;6;6;6;6;6;6 | 11829867:118 | 11830006:118 | +;+;+;+;+;+;+;+ | 7337 | BAZ2B | 1264 | 615 | 1132 | 847 | 945 | 921 |
| ENSDARG000 | 6;6;6;6;6;6 | 45937777:459 | 45939567:459 | -;-;-;-;-;- | 2798 | tardbpb | 2560 | 2186 | 3161 | 1574 | 2734 | 2091 |
| ENSDARG000 | 6;6 | 10140554:101 | 10142871:101 | -;- | 2415 | B3GALT1 | 92 | 2 | 6 | 2 | 2 | 10 |
| ENSDARG000 | 6;6;6;6;6;6;6 | 10170241:101 | 10173258:101 | -;-;-;-;-;-;-;- | 13117 | XIRP2 | 171 | 62 | 66 | 52 | 82 | 56 |
| ENSDARG000 | 6;6;6;6;6;6;6 | 18714213:187 | 18719045:187 | -;-;-;-;-;-;-;- | 9950 | NA | 2101 | 1638 | 3007 | 2623 | 3488 | 1998 |
| ENSDARG000 | 6;6;6;6;6;6;6 | 13506841:135 | 13507129:135 | +;+;+;+;+;+;+;+ | 2010 | gmppab | 378 | 568 | 496 | 451 | 462 | 302 |
| ENSDARG000 | 6;6 | 11815638:118 | 11820746:118 | -;- | 5402 | NA | 20 | 33 | 17 | 30 | 52 | 16 |
| ENSDARG000 | 6;6;6;6;6;6;6 | 4255319:4274 | 4255369:4274 | +;+;+;+;+;+;+;+ | 8423 | NBEAL1 | 519 | 258 | 885 | 385 | 767 | 448 |
| ENSDARG000 | 6;6;6;6 | 40437987:404 | 40438134:404 | +;+;+;+ | 599 | ghrl | 0 | 0 | 0 | 0 | 0 | 0 |
| ENSDARG000 | 6;6;6;6;6;6;6 | 9142731:9142 | 9142832:9143 | +;+;+;+;+;+;+ | 1497 | NA | 0 | 20 | 30 | 26 | 108 | 30 |
| ENSDARG000 | 6;6;6;6;6;6;6 | 39064824:390 | 39065190:390 | -;-;-;-;-;-;-;- | 2456 | eif4bb | 3161 | 3453 | 5627 | 3900 | 3406 | 3378 |
| ENSDARG000 | 6 | 32092032 | 32093830 | - | 1799 | foxd3 | 1709 | 2558 | 1579 | 1152 | 2842 | 1334 |
| ENSDARG000 | 6;6;6;6;6;6;6 | 45951893:459 | 45952683:459 | -;-;-;-;-;-;-;- | 6036 | PTPRG | 50 | 18 | 0 | 6 | 14 | 22 |
| ENSDARG000 | 6;6;6;6;6;6;6 | 28018390:280 | 28018452:280 | +;+;+;+;+;+;+;+ | 4417 | sap130a | 658 | 530 | 650 | 458 | 776 | 482 |
| ENSDARG000 | 6;6;6 | 40451270:404 | 40451840:404 | -;-;- | 1184 | vhl | 110 | 135 | 271 | 325 | 113 | 108 |
| ENSDARG000 | 6;6;6;6;6;6;6 | 40439858:404 | 40440241:404 | -;-;-;-;-;-;-;- | 2716 | tatdn2 | 13 | 2 | 23 | 0 | 6 | 0 |
| ENSDARG000 | 6;6 | 40448634:404 | 40449574:404 | +;+ | 1120 | NA | 0 | 0 | 1 | 2 | 0 | 10 |
| ENSDARG000 | 6;6;6;6;6;6;6 | 9155252:9155 | 9155353:9155 | +;+;+;+;+;+;+;+ | 3007 | zgc:64065 | 2 | 17 | 46 | 22 | 222 | 28 |
| ENSDARG000 | 6;6;6;6;6;6;6 | 16509818:165 | 16510162:165 | -;-;-;-;-;-;-;- | 9928 | UNC80 | 4 | 9 | 25 | 22 | 23 | 4 |
| ENSDARG000 | 6;6;6;6;6;6;6 | 13606410:136 | 13607320:136 | +;+;+;+;+;+;+;+ | 2316 | asic4b | 0 | 0 | 1 | 4 | 0 | 0 |
| ENSDARG000 | 6;6;6;6;6;6;6 | 10248332:102 | 10249527:102 | -;-;-;-;-;-;-;- | 6477 | scn1lab | 32 | 6 | 86 | 102 | 4 | 8 |
| ENSDARG000 | 6;6;6;6;6;6;6 | 13698813:136 | 13700583:137 | -;-;-;-;-;-;- | 3317 | CHPF | 200 | 522 | 847 | 634 | 1187 | 782 |
| ENSDARG000 | 6;6;6;6;6 | 13726844:137 | 13727180:137 | +;+;+;+;+ | 1346 | NA | 12 | 20 | 46 | 96 | 47 | 45 |
| ENSDARG000 | 6 | 13742899 | 13744681 | + | 1783 | cdk5r2a | 22 | 0 | 0 | 4 | 0 | 0 |
| ENSDARG000 | 6;6;6;6;6;6;6 | 29754141:297 | 29754205:297 | +;+;+;+;+;+;+;+ | 1544 | ptmaa | 750 | 344 | 416 | 272 | 647 | 571 |
| ENSDARG000 | 6;6 | 13740479:137 | 13740641:137 | -;- | 280 | NA | 0 | 0 | 0 | 0 | 0 | 0 |
| ENSDARG000 | 6;6;6;6;6;6;6 | 40456798:404 | 40458129:404 | -;-;-;-;-;-;-;- | 2526 | crbn | 318 | 243 | 278 | 264 | 270 | 270 |
| ENSDARG000 | 6;6;6;6;6;6;6 | 4370935:4371 | 4371486:4371 | +;+;+;+;+;+;+;+ | 6277 | RBM26 | 562 | 609 | 910 | 399 | 777 | 473 |
| ENSDARG000 | 6;6;6;6;6;6;6 | 39085969:390 | 39086054:390 | +;+;+;+;+;+;+;+ | 1573 | prss60.1 | 0 | 2 | 0 | 0 | 0 | 0 |
| ENSDARG000 | 6;6;6;6;6;6;6 | 49063289:490 | 49063707:490 | -;-;-;-;-;-;-;- | 2367 | slc5a8l | 0 | 2 | 1 | 8 | 6 | 10 |
| ENSDARG000 | 6;6;6;6;6;6;6 | 10333920:103 | 10334047:103 | +;+;+;+;+;+;+;+ | 3003 | NA | 687 | 200 | 52 | 499 | 26 | 201 |
| ENSDARG000 | 6;6;6;6;6;6;6 | 39098397:390 | 39098545:390 | +;+;+;+;+;+;+;+ | 1595 | prss60.2 | 2 | 0 | 0 | 0 | 0 | 0 |
| ENSDARG000 | 6;6;6;6;6;6;6 | 9163007:9163 | 9163357:9163 | +;+;+;+;+;+;+;+ | 3241 | zgc:112023 | 334 | 213 | 260 | 302 | 313 | 363 |
| ENSDARG000 | 6;6;6 | 29826493:298 | 29828770:298 | -;-;- | 2428 | NA | 1 | 2 | 6 | 0 | 21 | 0 |
| ENSDARG000 | 6;6;6;6 | 29821617:298 | 29821722:298 | +;+;+;+ | 792 | NA | 0 | 0 | 0 | 0 | 0 | 0 |
| ENSDARG000 | 6;6;6 | 46168017:461 | 46168745:461 | -;-;- | 924 | NA | 0 | 0 | 0 | 0 | 0 | 0 |
| ENSDARG000 | 6;6;6 | 40467562:404 | 40467694:404 | +;+;+ | 806 | zgc:152986 | 0 | 0 | 0 | 0 | 0 | 0 |
| ENSDARG000 | 6;6;6;6 | 9175886:9181 | 9176304:9181 | +;+;+;+ | 1559 | NA | 2 | 2 | 6 | 0 | 6 | 10 |
| ENSDARG000 | 6;6;6;6;6;6;6 | 29842213:298 | 29842284:298 | +;+;+;+;+;+;+ | 556 | NA | 0 | 0 | 8 | 15 | 19 | 11 |
| ENSDARG000 | 6;6;6;6;6;6;6 | 49081141:490 | 49081197:490 | +;+;+;+;+;+;+;+ | 833 | tshba | 0 | 0 | 0 | 0 | 0 | 0 |
| ENSDARG000 | 6;6;6;6;6;6 | 32093936:320 | 32094143:320 | +;+;+;+;+;+ | 2331 | NA | 159 | 120 | 173 | 188 | 271 | 172 |
| ENSDARG000 | 6;6;6;6 | 19040507:190 | 19040827:190 | -;-;-;- | 892 | si:rp71-81e14 | 2 | 10 | 0 | 3 | 8 | 8 |
| ENSDARG000 | 6;6;6;6 | 46309795:463 | 46309979:463 | +;+;+;+ | 1488 | NA | 256 | 188 | 72 | 164 | 91 | 238 |
| ENSDARG000 | 6;6;6;6;6;6;6 | 18935815:189 | 18937974:189 | -;-;-;-;-;-;-;- | 4486 | septin9b | 2342 | 2485 | 2286 | 2327 | 1929 | 3125 |
| ENSDARG000 | 6;6;6;6;6;6;6 | 39108858:391 | 39108996:391 | +;+;+;+;+;+;+;+ | 1276 | NA | 2 | 2 | 1 | 0 | 0 | 0 |
| ENSDARG000 | 6;6 | 32195891:321 | 32196381:321 | -;- | 528 | NA | 0 | 0 | 0 | 0 | 0 | 0 |
| ENSDARG000 | 6;6;6;6;6;6;6 | 46258820:462 | 46259163:462 | +;+;+;+;+;+;+;+ | 2887 | cfap20dc | 0 | 0 | 0 | 0 | 0 | 4 |
| ENSDARG000 | 6;6;6;6;6;6;6 | 9191981:9192 | 9192179:9192 | +;+;+;+;+;+;+;+ | 1766 | sft2d2b | 4 | 0 | 0 | 8 | 0 | 6 |
| ENSDARG000 | 6;6 | 29846519:298 | 29846931:298 | +;+ | 846 | NA | 0 | 0 | 0 | 0 | 2 | 0 |

| | | | | | | | | | | | | |
|---|---|---|---|---|---|---|---|---|---|---|---|---|
| ENSDARG000 | 6;6;6;6;6;6;6; | 13780205:137 | 13780747:137 | -;-;-;-;-;-;-;-;-; | 1702 | cryba2a | 0 | 0 | 0 | 0 | 0 | 0 |
| ENSDARG000 | 6;6;6 | 46294361:463 | 46295106:463 | -;-;- | 931 | NA | 0 | 0 | 0 | 0 | 0 | 0 |
| ENSDARG000 | 6;6;6;6;6;6;6; | 13779857:137 | 13779940:137 | +;+;+;+;+;+;+;+; | 3427 | tmem198b | 1259 | 963 | 1120 | 1244 | 1949 | 1308 |
| ENSDARG000 | 6;6;6 | 10519295:105 | 10519587:105 | +;+;+ | 830 | NA | 0 | 0 | 0 | 0 | 0 | 0 |
| ENSDARG000 | 6;6;6;6;6;6;6; | 10450000:104 | 10450078:104 | +;+;+;+;+;+;+;+; | 4653 | NA | 0 | 4 | 0 | 0 | 0 | 0 |
| ENSDARG000 | 6;6 | 19046987:190 | 19047312:190 | +;+ | 641 | NA | 0 | 0 | 0 | 8 | 0 | 0 |
| ENSDARG000 | 6;6;6;6;6;6;6; | 49095636:490 | 49095721:490 | +;+;+;+;+;+;+;+; | 3164 | slc25a55a | 2316 | 3343 | 3314 | 1662 | 3901 | 2420 |
| ENSDARG000 | 6;6 | 39114345:391 | 39114471:391 | +;+ | 1822 | gpr84 | 110 | 140 | 105 | 80 | 198 | 178 |
| ENSDARG000 | 6;6;6;6;6;6;6; | 46320693:463 | 46320857:463 | +;+;+;+;+;+;+;+; | 1823 | NA | 69 | 26 | 25 | 16 | 16 | 46 |
| ENSDARG000 | 6;6;6;6;6;6;6; | 46353910:463 | 46354038:463 | -;-;-;-;-;-;-;-;-; | 1749 | CAMK1 | 0 | 0 | 7 | 2 | 15 | 9 |
| ENSDARG000 | 6;6;6;6;6;6;6; | 39123544:391 | 39127656:391 | -;-;-;-;-;-;-;-;-; | 6618 | stat2 | 4625 | 3720 | 1050 | 721 | 1153 | 1632 |
| ENSDARG000 | 6;6 | 4416849:4433 | 4417135:4433 | -;- | 396 | NA | 0 | 2 | 0 | 2 | 0 | 0 |
| ENSDARG000 | 6;6;6;6;6;6;6; | 32300018:323 | 32301986:323 | -;-;-;-;-;-;-;-;-; | 4166 | atg4c | 793 | 755 | 884 | 723 | 773 | 788 |
| ENSDARG000 | 6;6;6;6;6;6;6; | 4872883:4872 | 4873484:4873 | +;+;+;+;+;+;+;+; | 3942 | PCDH9 | 1888 | 731 | 1139 | 768 | 1284 | 926 |
| ENSDARG000 | 6;6 | 39162294:391 | 39164668:391 | -;- | 2612 | NA | 2994 | 1213 | 1215 | 2429 | 2327 | 1103 |
| ENSDARG000 | 6;6;6;6;6;6;6; | 29860776:298 | 29861537:298 | +;+;+;+;+;+;+;+; | 6056 | dlg1a | 1321 | 1063 | 1159 | 1566 | 1237 | 1369 |
| ENSDARG000 | 6;6;6;6;6;6;6; | 28051978:280 | 28052130:280 | +;+;+;+;+;+;+;+; | 7559 | MYO7B | 2 | 35 | 20 | 24 | 10 | 22 |
| ENSDARG000 | 6;6;6;6;6;6;6 | 39184236:391 | 39184282:391 | +;+;+;+;+;+;+ | 830 | tac3b | 4 | 3 | 6 | 5 | 9 | 4 |
| ENSDARG000 | 6;6;6 | 5700659:5709 | 5701047:5709 | -;-;- | 579 | NA | 0 | 0 | 0 | 0 | 0 | 0 |
| ENSDARG000 | 6;6;6 | 6181909:6182 | 6182233:6182 | -;-;- | 539 | NA | 0 | 0 | 0 | 0 | 0 | 0 |
| ENSDARG000 | 6;6 | 32320838:323 | 32320883:323 | +;+ | 1452 | NA | 0 | 0 | 0 | 0 | 2 | 1 |
| ENSDARG000 | 6;6;6;6;6;6;6; | 46406565:464 | 46406681:464 | +;+;+;+;+;+;+;+; | 6348 | pbrm1l | 1324 | 1143 | 1339 | 892 | 1582 | 1109 |
| ENSDARG000 | 6;6;6;6;6;6;6; | 49107048:491 | 49109660:491 | -;-;-;-;-;-;-;-;-; | 3506 | tspan2a | 84 | 291 | 515 | 300 | 297 | 409 |
| ENSDARG000 | 6;6;6;6;6;6;6; | 12197186:121 | 12201441:122 | -;-;-;-;-;-;-;-;-; | 8029 | NA | 1096 | 734 | 467 | 700 | 594 | 1167 |
| ENSDARG000 | 6;6;6;6;6;6;6; | 49169420:491 | 49170402:491 | -;-;-;-;-;-;-;-;-; | 1715 | ngfb | 0 | 0 | 0 | 2 | 0 | 6 |
| ENSDARG000 | 6;6;6;6;6;6;6 | 39190402:391 | 39191195:391 | -;-;-;-;-;-;- | 2092 | c1galt1lb | 390 | 322 | 399 | 251 | 686 | 468 |
| ENSDARG000 | 6;6;6;6;6;6;6; | 6231012:6231 | 6232390:6232 | -;-;-;-;-;-;-;-;-; | 2852 | rtn4a | 2487 | 2589 | 4305 | 1768 | 3103 | 2678 |
| ENSDARG000 | 6;6;6;6 | 49255706:492 | 49255830:492 | +;+;+;+ | 971 | NA | 0 | 0 | 0 | 0 | 0 | 0 |
| ENSDARG000 | 6;6;6;6 | 49310908:493 | 49311400:493 | -;-;-;- | 1181 | NA | 0 | 0 | 0 | 0 | 0 | 0 |
| ENSDARG000 | 6;6;6;6;6;6 | 49541278:495 | 49541424:495 | -;-;-;-;-;- | 773 | NA | 74 | 98 | 110 | 46 | 126 | 74 |
| ENSDARG000 | 6;6;6;6;6;6;6; | 10586019:105 | 10586266:105 | -;-;-;-;-;-;-;-;-; | 14597 | NA | 28 | 30 | 45 | 88 | 30 | 84 |
| ENSDARG000 | 6;6;6;6;6;6;6; | 49551614:495 | 49551928:495 | +;+;+;+;+;+;+;+; | 5676 | rab22a | 645 | 626 | 779 | 702 | 856 | 631 |
| ENSDARG000 | 6;6;6;6;6;6;6; | 39202685:392 | 39203080:392 | -;-;-;-;-;-;-;-;-; | 2519 | os9 | 1597 | 1225 | 1775 | 923 | 1811 | 1541 |
| ENSDARG000 | 6;6;6 | 10691940:106 | 10691965:106 | +;+;+ | 639 | NA | 0 | 0 | 0 | 0 | 0 | 0 |
| ENSDARG000 | 6;6;6;6;6;6;6; | 12299188:123 | 12300071:123 | -;-;-;-;-;-;-;-;-; | 1598 | si:dkev-276i7. | 845 | 771 | 865 | 1176 | 760 | 901 |
| ENSDARG000 | 6;6;6;6;6;6;6; | 46431848:464 | 46432029:464 | +;+;+;+;+;+;+;+; | 3231 | stau1 | 3928 | 2728 | 3153 | 2339 | 3508 | 3344 |
| ENSDARG000 | 6;6;6;6;6;6 | 49650170:496 | 49652911:496 | -;-;-;-;-;- | 3884 | APCDD1L | 1532 | 908 | 1864 | 3563 | 1977 | 926 |
| ENSDARG000 | 6;6;6;6;6;6 | 49583153:496 | 49583465:496 | +;+;+;+;+;+ | 1965 | vapb | 1137 | 988 | 1154 | 915 | 1054 | 1071 |
| ENSDARG000 | 6;6;6;6;6;6;6; | 39222598:392 | 39222850:392 | +;+;+;+;+;+;+;+; | 3685 | B4GALNT1 | 447 | 108 | 90 | 212 | 165 | 511 |
| ENSDARG000 | 6;6;6;6;6;6;6; | 12316438:123 | 12316616:123 | +;+;+;+;+;+;+;+; | 786 | NA | 40 | 18 | 21 | 70 | 2 | 20 |
| ENSDARG000 | 6;6;6;6;6;6;6; | 39256170:392 | 39256406:392 | -;-;-;-;-;-;-;-;-; | 2342 | NA | 2 | 0 | 0 | 0 | 0 | 48 |
| ENSDARG000 | 6;6;6;6;6;6;6; | 49722970:497 | 49723420:497 | +;+;+;+;+;+;+;+; | 4816 | STX16 | 876 | 787 | 829 | 536 | 1086 | 1070 |
| ENSDARG000 | 6;6 | 28115383:281 | 28115965:281 | -;- | 900 | NA | 0 | 0 | 0 | 0 | 0 | 0 |
| ENSDARG000 | 6;6;6;6;6;6;6; | 32410848:324 | 32410885:324 | +;+;+;+;+;+;+;+; | 5731 | KANK4 | 1675 | 625 | 966 | 1167 | 1991 | 1196 |
| ENSDARG000 | 6 | 28124393 | 28125412 | + | 1020 | GPR17 | 53 | 116 | 0 | 209 | 4 | 16 |
| ENSDARG000 | 6;6;6;6;6;6;6; | 32326074:323 | 32326247:323 | +;+;+;+;+;+;+;+; | 7497 | DOCK7 | 6007 | 4848 | 8212 | 7289 | 8325 | 4332 |
| ENSDARG000 | 6;6;6;6;6;6;6; | 49742164:497 | 49742423:497 | +;+;+;+;+;+;+;+; | 2193 | npepl1 | 2496 | 2525 | 2614 | 2769 | 2626 | 2155 |
| ENSDARG000 | 6;6;6;6;6;6;6; | 30091811:300 | 30092073:300 | +;+;+;+;+;+;+;+; | 2840 | MELTF | 6 | 0 | 0 | 0 | 16 | 0 |
| ENSDARG000 | 6;6;6;6;6 | 46458416:464 | 46458639:464 | -;-;-;-;- | 1106 | NA | 0 | 2 | 0 | 10 | 0 | 0 |
| ENSDARG000 | 6;6;6;6;6;6;6 | 32344197:323 | 32345407:323 | -;-;-;-;-;-;- | 2539 | angptl3 | 10 | 0 | 0 | 2 | 2 | 0 |
| ENSDARG000 | 6;6;6;6;6;6;6; | 32394502:324 | 32394906:324 | -;-;-;-;-;-;-;-;-; | 3752 | usp1 | 301 | 560 | 636 | 455 | 584 | 538 |
| ENSDARG000 | 6;6;6 | 49761229:497 | 49764009:497 | -;-;- | 3207 | NA | 379 | 279 | 431 | 462 | 401 | 579 |
| ENSDARG000 | 6;6 | 32415749:324 | 32415949:324 | -;- | 524 | NA | 4 | 0 | 4 | 0 | 14 | 18 |
| ENSDARG000 | 6;6;6;6;6;6;6; | 49771372:497 | 49771758:497 | +;+;+;+;+;+;+;+; | 2274 | ctsz | 3984 | 4073 | 2551 | 2264 | 4562 | 3129 |
| ENSDARG000 | 6;6;6;6;6 | 46481024:464 | 46481438:464 | +;+;+;+;+ | 1188 | NA | 0 | 0 | 0 | 0 | 4 | 0 |
| ENSDARG000 | 6;6 | 12326267:123 | 12326380:123 | +;+ | 740 | NA | 0 | 0 | 0 | 0 | 5 | 0 |
| ENSDARG000 | 6;6;6;6;6 | 46507609:465 | 46509537:465 | -;-;-;-;- | 3099 | KCNB1 | 0 | 0 | 0 | 0 | 2 | 0 |
| ENSDARG000 | 6;6;6;6;6;6;6; | 28140816:281 | 28141274:281 | +;+;+;+;+;+;+;+; | 3899 | INPP5D | 160 | 144 | 182 | 431 | 384 | 371 |
| ENSDARG000 | 6;6;6;6;6;6;6; | 46568119:465 | 46568433:465 | -;-;-;-;-;-;-;-;-; | 2019 | ptgis | 53 | 34 | 20 | 30 | 56 | 96 |
| ENSDARG000 | 6;6;6;6;6;6;6; | 30113750:301 | 30113960:301 | +;+;+;+;+;+;+;+; | 2865 | lrrc40 | 390 | 321 | 381 | 271 | 385 | 396 |
| ENSDARG000 | 6;6 | 46668717:466 | 46669755:466 | +;+ | 1532 | oxtrb | 0 | 0 | 4 | 0 | 0 | 0 |
| ENSDARG000 | 6;6;6;6;6;6;6; | 49781884:497 | 49782632:497 | -;-;-;-;-;-;-;-;-; | 2553 | nelfcd | 429 | 414 | 447 | 252 | 574 | 379 |
| ENSDARG000 | 6;6;6;6;6;6;6; | 46697710:466 | 46698008:466 | +;+;+;+;+;+;+;+; | 2823 | NA | 8 | 8 | 8 | 10 | 38 | 28 |
| ENSDARG000 | 6;6;6;6;6;6;6; | 12379666:123 | 12380169:123 | -;-;-;-;-;-;-;-;-; | 4720 | gpd2 | 742 | 445 | 573 | 321 | 409 | 612 |
| ENSDARG000 | 6;6;6;6;6;6;6 | 46725540:467 | 46726442:467 | -;-;-;-;-;-;- | 1964 | tarbp2 | 371 | 339 | 310 | 212 | 333 | 294 |
| ENSDARG000 | 6;6 | 49859980:498 | 49861176:498 | +;+ | 1492 | NA | 0 | 0 | 0 | 0 | 0 | 0 |
| ENSDARG000 | 6;6;6;6;6;6;6; | 49811831:498 | 49812520:498 | -;-;-;-;-;-;-;-;-; | 2841 | GNAS | 1680 | 1208 | 1466 | 1430 | 1835 | 1451 |
| ENSDARG000 | 6;6;6;6 | 46736251:467 | 46737545:467 | -;-;-;- | 2372 | zgc:66479 | 30 | 12 | 22 | 14 | 12 | 8 |
| ENSDARG000 | 6;6;6;6;6;6;6; | 46743246:467 | 46743424:467 | -;-;-;-;-;-;-;-;-; | 4637 | NA | 0 | 12 | 34 | 36 | 0 | 0 |
| ENSDARG000 | 6;6;6;6;6;6;6; | 30124197:301 | 30124839:301 | -;-;-;-;-;-;-;-;-; | 5086 | LRRC7 | 4 | 10 | 33 | 14 | 14 | 20 |
| ENSDARG000 | 6;6;6;6 | 49881864:498 | 49881992:498 | +;+;+;+ | 1473 | tubb1 | 6 | 0 | 37 | 0 | 31 | 8 |
| ENSDARG000 | 6;6 | 49919936:499 | 49920633:499 | -;- | 807 | NA | 3 | 4 | 6 | 4 | 3 | 2 |
| ENSDARG000 | 6;6;6;6;6;6 | 49893488:498 | 49893622:498 | -;-;-;-;-;- | 1099 | atp5f1e | 734 | 997 | 724 | 479 | 386 | 516 |
| ENSDARG000 | 6;6 | 49918254:499 | 49918619:499 | -;- | 539 | NA | 2 | 0 | 0 | 0 | 0 | 0 |
| ENSDARG000 | 6;6;6;6;6;6;6; | 49926115:499 | 49926251:499 | +;+;+;+;+;+;+;+; | 1922 | ahcy | 5019 | 4403 | 6884 | 4835 | 4938 | 4757 |
| ENSDARG000 | 6;6;6;6;6 | 49901465:499 | 49901799:499 | +;+;+;+;+ | 2011 | chmp4ba | 647 | 495 | 738 | 821 | 500 | 501 |
| ENSDARG000 | 6;6;6;6;6;6;6; | 46771654:467 | 46772149:467 | -;-;-;-;-;-;-;-;-; | 4311 | NA | 0 | 0 | 0 | 0 | 0 | 0 |
| ENSDARG000 | 6;6;6;6;6;6;6; | 46843592:468 | 46844405:468 | -;-;-;-;-;-;-;-;-; | 7605 | NA | 0 | 0 | 0 | 0 | 0 | 0 |
| ENSDARG000 | 6;6;6;6 | 49975910:499 | 49976347:499 | -;-;-;- | 728 | asip1 | 2 | 0 | 0 | 12 | 4 | 0 |
| ENSDARG000 | 6;6;6;6;6;6;6; | 49995781:499 | 49995858:499 | +;+;+;+;+;+;+;+; | 1290 | eif2s2 | 3957 | 4438 | 6633 | 3735 | 4509 | 3728 |
| ENSDARG000 | 6;6;6;6;6;6;6; | 50021410:500 | 50022667:500 | -;-;-;-;-;-;-;-;-; | 2891 | raly | 2 | 10 | 18 | 2 | 4 | 0 |
| ENSDARG000 | 6;6;6;6;6;6;6; | 47424266:474 | 47424714:474 | +;+;+;+;+;+;+;+; | 6230 | NA | 170 | 268 | 1891 | 2624 | 469 | 1711 |
| ENSDARG000 | 6;6;6;6;6;6;6; | 50381347:503 | 50381719:503 | +;+;+;+;+;+;+;+; | 2051 | cyc1 | 2178 | 3177 | 3528 | 2090 | 2527 | 2040 |
| ENSDARG000 | 6;6;6;6;6;6;6; | 47529881:475 | 47531196:475 | -;-;-;-;-;-;-;-;-; | 5565 | MAGI3 | 10 | 0 | 0 | 2 | 4 | 0 |
| ENSDARG000 | 6;6;6 | 50401620:504 | 50402324:504 | -;-;- | 954 | romo1 | 1352 | 1349 | 1695 | 2522 | 1269 | 2077 |
| ENSDARG000 | 6;6;6;6;6;6;6; | 50393047:503 | 50393228:503 | +;+;+;+;+;+;+;+; | 2033 | ergic3 | 509 | 625 | 874 | 387 | 657 | 514 |
| ENSDARG000 | 6;6;6;6;6;6;6; | 47809755:478 | 47813917:478 | -;-;-;-;-;-;-;-;-; | 7352 | LRIG2 | 2380 | 1598 | 1973 | 1789 | 2031 | 2461 |
| ENSDARG000 | 6;6;6;6;6 | 50451337:504 | 50451931:504 | +;+;+;+;+ | 2496 | mych | 7148 | 6608 | 10216 | 8553 | 5338 | 5645 |
| ENSDARG000 | 6;6;6;6;6;6;6; | 6270686:6270 | 6270763:6270 | +;+;+;+;+;+;+;+; | 1067 | rps27a | 13527 | 16861 | 21026 | 17835 | 13574 | 19687 |
| ENSDARG000 | 6;6;6;6;6;6;6; | 50580940:505 | 50582611:505 | -;-;-;-;-;-;-;-;-; | 3848 | mtss1 | 1359 | 762 | 828 | 1273 | 825 | 996 |
| ENSDARG000 | 6;6;6;6;6;6 | 50695381:506 | 50696559:506 | -;-;-;-;-;- | 1820 | osgn1 | 915 | 786 | 589 | 687 | 596 | 862 |
| ENSDARG000 | 6;6;6;6;6;6;6; | 50713800:507 | 50714496:507 | -;-;-;-;-;-;-;-;-; | 1891 | pigu | 251 | 295 | 360 | 149 | 268 | 423 |
| ENSDARG000 | 6;6;6;6;6;6;6; | 50742315:507 | 50743045:507 | -;-;-;-;-;-;-;-;-; | 1502 | NA | 12 | 0 | 16 | 5 | 6 | 5 |
| ENSDARG000 | 6;6;6;6;6;6;6; | 47843760:478 | 47843998:478 | +;+;+;+;+;+;+;+; | 2801 | padi2 | 58 | 24 | 18 | 46 | 39 | 30 |
| ENSDARG000 | 6;6 | 50754501:507 | 50755303:507 | -;- | 1232 | NA | 0 | 0 | 0 | 0 | 0 | 2 |
| ENSDARG000 | 6;6;6;6;6;6;6; | 47869397:478 | 47870506:478 | -;-;-;-;-;-;-;-;-; | 3477 | RAP1GAP | 44 | 22 | 106 | 383 | 47 | 144 |
| ENSDARG000 | 6;6;6;6;6;6;6; | 51020918:510 | 51021131:510 | -;-;-;-;-;-;-;-;-; | 4495 | PTPRT | 0 | 0 | 0 | 0 | 0 | 0 |
| ENSDARG000 | 6;6;6;6;6;6;6; | 51009981:510 | 51010236:510 | -;-;-;-;-;-;-;-;-; | 2101 | PTPRT | 0 | 0 | 0 | 0 | 0 | 0 |
| ENSDARG000 | 6;6;6;6 | 47979305:480 | 47979481:480 | -;-;-;- | 449 | NA | 0 | 0 | 0 | 0 | 0 | 0 |
| ENSDARG000 | 6;6;6;6;6;6;6; | 48041759:480 | 48042223:480 | +;+;+;+;+;+;+;+; | 3078 | NA | 351 | 280 | 252 | 298 | 315 | 193 |
| ENSDARG000 | 6;6;6;6;6;6;6; | 48068695:480 | 48068964:480 | -;-;-;-;-;-;-;-;-; | 1903 | SLC2A1 | 769 | 2111 | 5859 | 4668 | 1453 | 565 |
| ENSDARG000 | 6;6;6;6;6;6;6; | 51529208:515 | 51532337:515 | -;-;-;-;-;-;-;- | 4549 | NA | 490 | 330 | 487 | 273 | 458 | 572 |
| ENSDARG000 | 6;6;6;6;6;6;6; | 48154954:481 | 48155094:481 | +;+;+;+;+;+;+;+; | 3644 | phc2b | 650 | 625 | 588 | 643 | 356 | 793 |
| ENSDARG000 | 6;6;6;6;6 | 48253449:482 | 48253794:482 | +;+;+;+;+ | 1771 | wnt2ba | 0 | 0 | 0 | 0 | 4 | 0 |
| ENSDARG000 | 6;6;6;6;6;6;6; | 48290125:482 | 48290331:482 | -;-;-;-;-;-;-;-;-; | 1739 | st7l | 194 | 214 | 246 | 140 | 230 | 244 |
| ENSDARG000 | 6;6;6;6;6;6;6; | 51549203:515 | 51549539:515 | -;-;-;-;-;-;-;-;-; | 3438 | rbl1 | 491 | 525 | 640 | 287 | 588 | 679 |
| ENSDARG000 | 6;6;6;6;6;6;6; | 51575924:515 | 51581278:515 | -;-;-;-;-;-;-;-;-; | 14200 | NA | 1008 | 710 | 1225 | 528 | 1204 | 1004 |
| ENSDARG000 | 6;6;6;6;6;6;6; | 48322250:483 | 48324152:483 | -;-;-;-;-;-;-;-;-; | 2885 | capza1a | 2623 | 2693 | 2417 | 2373 | 1975 | 2689 |
| ENSDARG000 | 6;6;6;6;6;6;6; | 48348415:483 | 48348603:483 | +;+;+;+;+;+;+;+; | 3444 | MOV10 | 3859 | 1968 | 1800 | 1962 | 1970 | 3244 |
| ENSDARG000 | 6;6;6;6;6;6;6; | 51676227:517 | 51676456:517 | +;+;+;+;+;+;+;+; | 4122 | ripor3 | 6 | 6 | 26 | 0 | 37 | 18 |
| ENSDARG000 | 6;6 | 51769760:517 | 51770614:517 | -;- | 955 | blcap | 613 | 693 | 917 | 923 | 643 | 683 |
| ENSDARG000 | 6;6;6;6 | 51773873:517 | 51774129:517 | +;+;+;+ | 1201 | TMEM74B | 2 | 0 | 0 | 0 | 0 | 0 |
| ENSDARG000 | 6;6;6;6;6;6;6 | 51796296:517 | 51796561:517 | -;-;-;-;-;-;- | 1286 | psmf1 | 670 | 575 | 503 | 485 | 498 | 378 |
| ENSDARG000 | 6;6;6;6;6 | 51824596:518 | 51824688:518 | +;+;+;+;+ | 1129 | rspo4 | 0 | 0 | 0 | 0 | 0 | 0 |
| ENSDARG000 | 6;6;6;6;6;6;6; | 51932267:519 | 51932874:519 | +;+;+;+;+;+;+;+; | 2271 | ANGPT4 | 2 | 6 | 2 | 0 | 10 | 4 |
| ENSDARG000 | 6;6 | 52064304:520 | 52064907:520 | +;+ | 4800 | NA | 0 | 0 | 0 | 0 | 2 | 0 |
| ENSDARG000 | 6;6;6;6;6;6 | 52081752:520 | 52082930:520 | -;-;-;-;-;- | 2392 | RIMS4 | 0 | 0 | 0 | 0 | 0 | 0 |
| ENSDARG000 | 6;6;6;6;6;6;6; | 52212574:522 | 52212710:522 | +;+;+;+;+;+;+;+; | 3851 | ywhaba | 9192 | 12026 | 12517 | 9433 | 11099 | 9800 |
| ENSDARG000 | 6;6;6;6;6;6;6; | 52226682:522 | 52227406:522 | -;-;-;-;-;-;-;-;-; | 2990 | tomm34 | 374 | 519 | 388 | 152 | 398 | 232 |
| ENSDARG000 | 6;6;6;6 | 52235441:522 | 52235486:522 | +;+;+;+ | 415 | cox6c | 1231 | 1664 | 1589 | 1171 | 1241 | 1130 |
| ENSDARG000 | 6;6;6 | 14949950:149 | 14953352:149 | +;+;+ | 3403 | pou3f3b | 2 | 0 | 12 | 0 | 2 | 0 |
| ENSDARG000 | 6;6;6;6;6;6;6; | 22119189:221 | 22121055:221 | -;-;-;-;-;-;-;-;-; | 4004 | gga1 | 1488 | 1691 | 1901 | 1465 | 2197 | 1475 |
| ENSDARG000 | 6;6;6;6;6;6;6; | 21874932:218 | 21878548:218 | -;-;-;-;-;-;-;-;-; | 10935 | NA | 311 | 246 | 471 | 346 | 704 | 82 |
| ENSDARG000 | 6;6;6;6;6;6;6; | 21470261:214 | 21472469:214 | -;-;-;-;-;-;-;-;- | 3779 | cacng1a | 2 | 2 | 4 | 0 | 12 | 0 |
| ENSDARG000 | 6;6;6;6;6;6;6; | 55299511:552 | 55299628:552 | -;-;-;-;-;-;-;-;- | 489 | DNTTIP1 | 153 | 112 | 158 | 115 | 111 | 98 |
| ENSDARG000 | 6;6;6;6;6;6;6; | 55221490:552 | 55222737:552 | -;-;-;-;-;-;-;-;-; | 3746 | NA | 12 | 10 | 2 | 7 | 18 | 4 |
| ENSDARG000 | 6;6 | 168962:16983 | 169059:17299 | +;+ | 3253 | NA | 14 | 6 | 19 | 0 | 14 | 28 |
| ENSDARG000 | 6;6;6;6;6;6;6; | 37894220:378 | 37894258:378 | +;+;+;+;+;+;+;+; | 2551 | OCA2 | 899 | 352 | 42 | 326 | 49 | 1561 |
| ENSDARG000 | 6;6;6;6;6;6;6; | 22068589:220 | 22069157:220 | +;+;+;+;+;+;+;+; | 5863 | aldh1l1 | 80 | 380 | 1511 | 1708 | 498 | 160 |
| ENSDARG000 | 6;6;6;6 | 60024117:600 | 60024192:600 | -;-;-;- | 735 | slain1b | 0 | 0 | 0 | 0 | 0 | 0 |
| ENSDARG000 | 6;6;6;6;6;6;6 | 55287823:552 | 55288188:552 | -;-;-;-;-;-;- | 926 | ube2c | 575 | 1032 | 890 | 466 | 670 | 507 |
| ENSDARG000 | 6;6;6;6;6;6;6 | 21001264:210 | 21001507:210 | +;+;+;+;+;+;+ | 4026 | gjc4a.1 | 26 | 6 | 2 | 24 | 4 | 6 |
| ENSDARG000 | 6;6;6;6;6;6;6; | 56638688:566 | 56638970:566 | -;-;-;-;-;-;-;-;-; | 2367 | CDH22 | 42 | 4 | 12 | 63 | 4 | 42 |
| ENSDARG000 | 6;6;6;6;6;6;6 | 9945303:9945 | 9945395:9945 | -;-;-;-;-;-;- | 1259 | zp2l2 | 0 | 0 | 2 | 0 | 0 | 0 |
| ENSDARG000 | 6;6;6 | 269204:27098 | 269230:27130 | +;+;+ | 1406 | atf4a | 9656 | 12023 | 15804 | 12879 | 10526 | 8372 |
| ENSDARG000 | 6 | 6124783 | 6124897 | - | 115 | 5S_rRNA | 0 | 0 | 0 | 0 | 0 | 0 |
| ENSDARG000 | 6;6;6;6;6 | 55282629:552 | 55282727:552 | +;+;+;+;+ | 1219 | zgc:109913 | 0 | 0 | 0 | 0 | 0 | 0 |
| ENSDARG000 | 6;6;6;6;6;6 | 3680651:3685 | 3681757:3685 | +;+;+;+;+;+ | 4255 | klhl41b | 0 | 0 | 2 | 0 | 12 | 8 |
| ENSDARG000 | 6 | 20132532 | 20132629 | + | 98 | NA | 0 | 0 | 0 | 0 | 0 | 0 |
| ENSDARG000 | 6;6;6;6;6;6;6; | 20954400:209 | 20954562:209 | +;+;+;+;+;+;+;+; | 6664 | stk11ip | 784 | 488 | 668 | 1116 | 778 | 1165 |
| ENSDARG000 | 6;6;6;6;6;6;6; | 3716666:3717 | 3717114:3717 | +;+;+;+;+;+;+;+; | 2595 | ssb | 1629 | 2094 | 3209 | 1211 | 2281 | 1728 |
| ENSDARG000 | 6 | 8484901 | 8485015 | - | 115 | 5S_rRNA | 0 | 0 | 0 | 0 | 0 | 0 |
| ENSDARG000 | 6 | 40946357 | 40946475 | - | 119 | 5S_rRNA | 0 | 0 | 0 | 0 | 0 | 0 |
| ENSDARG000 | 6 | 48612034 | 48612150 | - | 117 | 5S_rRNA | 0 | 0 | 0 | 0 | 0 | 0 |
| ENSDARG000 | 6 | 15544055 | 15544170 | + | 116 | 5S_rRNA | 0 | 0 | 0 | 0 | 0 | 0 |
| ENSDARG000 | 6 | 33726047 | 33726148 | - | 102 | 5S_rRNA | 0 | 0 | 0 | 0 | 0 | 0 |
| ENSDARG000 | 6 | 55555262 | 55555396 | - | 135 | 5S_rRNA | 0 | 0 | 0 | 0 | 0 | 0 |
| ENSDARG000 | 6 | 20299551 | 20299648 | - | 98 | NA | 0 | 0 | 0 | 0 | 0 | 0 |
| ENSDARG000 | 6 | 20647227 | 20647333 | - | 107 | U6 | 0 | 0 | 0 | 0 | 0 | 0 |
| ENSDARG000 | 6 | 30787928 | 30788044 | + | 117 | 5S_rRNA | 1 | 0 | 0 | 0 | 0 | 0 |
| ENSDARG000 | 6 | 49122130 | 49122255 | + | 126 | 5S_rRNA | 0 | 0 | 0 | 0 | 0 | 0 |
| ENSDARG000 | 6 | 17182138 | 17182235 | - | 98 | NA | 0 | 0 | 0 | 0 | 0 | 0 |
| ENSDARG000 | 6 | 34644253 | 34644370 | - | 118 | 5S_rRNA | 0 | 0 | 0 | 0 | 0 | 0 |
| ENSDARG000 | 6 | 49412754 | 49414235 | + | 1482 | KCNA2 | 0 | 0 | 4 | 0 | 0 | 0 |
| ENSDARG000 | 6 | 20337144 | 20337241 | - | 98 | NA | 0 | 0 | 0 | 0 | 0 | 0 |
| ENSDARG000 | 6 | 17268662 | 17268759 | - | 98 | NA | 0 | 0 | 0 | 0 | 0 | 0 |
| ENSDARG000 | 6;6;6;6;6;6 | 49529014:495 | 49529103:495 | -;-;-;-;-;- | 756 | NA | 132 | 83 | 148 | 70 | 115 | 106 |
| ENSDARG000 | 6;6;6;6;6;6;6; | 21809633:218 | 21809946:218 | -;-;-;-;-;-;-;-;-; | 3663 | SETD5 | 351 | 262 | 400 | 169 | 449 | 388 |
| ENSDARG000 | 6 | 54483686 | 54483802 | + | 117 | 5S_rRNA | 0 | 0 | 0 | 0 | 0 | 0 |
| ENSDARG000 | 6;6;6 | 21633286:216 | 21637947:216 | -;-;- | 5891 | NA | 0 | 0 | 0 | 0 | 0 | 12 |
| ENSDARG000 | 6;6;6;6;6;6;6 | 21202639:212 | 21202734:212 | +;+;+;+;+;+;+ | 1761 | cidec | 0 | 2 | 4 | 0 | 0 | 6 |
| ENSDARG000 | 6;6;6;6 | 22679610:226 | 22680350:226 | +;+;+;+ | 4407 | NA | 72 | 98 | 42 | 48 | 0 | 30 |

| | | | | | | | | | | | | |
|---|---|---|---|---|---|---|---|---|---|---|---|---|
| ENSDARG000 | 6 | 19716423 | 19716541 | - | 119 | 5S_rRNA | 0 | 0 | 0 | 0 | 0 | 0 |
| ENSDARG000 | 6;6;6;6;6 | 15899609:158 | 15899686:158 | +;+;+;+;+ | 891 | NA | 0 | 0 | 0 | 0 | 0 | 0 |
| ENSDARG000 | 6;6;6;6;6;6;6; | 21588205:215 | 21589672:215 | -;-;-;-;-;-;-;-; | 3542 | ppp1r12c | 797 | 762 | 752 | 540 | 766 | 1092 |
| ENSDARG000 | 6;6;6;6;6;6;6; | 54838269:548 | 54838589:548 | -;-;-;-;-;-;-;-; | 877 | csrp1b | 32 | 4 | 56 | 2 | 24 | 0 |
| ENSDARG000 | 6;6;6;6;6;6;6; | 55852274:558 | 55852831:558 | -;-;-;-;-;-;-;-; | 2118 | cyp24a1 | 4 | 0 | 2 | 0 | 0 | 0 |
| ENSDARG000 | 6;6;6;6;6;6;6; | 21684296:216 | 21684361:216 | +;+;+;+;+;+;+; | 3635 | rhebl1 | 1103 | 1413 | 2285 | 1367 | 1607 | 1494 |
| ENSDARG000 | 6;6;6;6;6;6;6; | 1764469:1764 | 1765244:1765 | -;-;-;-;-;-;-;- | 1581 | zgc:158417 | 0 | 0 | 8 | 0 | 0 | 0 |
| ENSDARG000 | 6;6;6;6 | 20921542:209 | 20922497:209 | -;-;-;- | 1748 | igfbp2a | 2 | 17 | 46 | 310 | 30 | 6 |
| ENSDARG000 | 6;6;6;6;6;6;6 | 21053548:210 | 21053828:210 | +;+;+;+;+;+;+ | 4505 | gjc4a.2 | 22 | 4 | 0 | 0 | 0 | 0 |
| ENSDARG000 | 6;6;6;6;6 | 54280365:542 | 54280543:542 | -;-;-;-;- | 978 | SPDEF | 2 | 0 | 0 | 0 | 0 | 0 |
| ENSDARG000 | 6;6;6;6;6;6;6; | 20647155:206 | 20647396:206 | +;+;+;+;+;+;+; | 3862 | SLC19A1 | 765 | 1009 | 1024 | 648 | 812 | 673 |
| ENSDARG000 | 6;6;6;6 | 49508805:495 | 49508933:495 | -;-;-;- | 412 | rplp2 | 5116 | 6250 | 7592 | 7801 | 5397 | 7505 |
| ENSDARG000 | 6 | 20136059 | 20136156 | - | 98 | NA | 0 | 0 | 0 | 0 | 0 | 0 |
| ENSDARG000 | 6 | 16983589 | 16983686 | - | 98 | NA | 0 | 0 | 0 | 0 | 0 | 0 |
| ENSDARG000 | 6 | 47467910 | 47467992 | + | 83 | dre-mir-29b-1 | 0 | 0 | 2 | 19 | 0 | 1 |
| ENSDARG000 | 6;6;6;6;6;6;6; | 28339662:283 | 28340758:283 | -;-;-;-;-;-;-;- | 2413 | NA | 0 | 0 | 35 | 32 | 0 | 0 |
| ENSDARG000 | 6;6;6;6;6;6;6; | 46981486:470 | 46981887:470 | -;-;-;-;-;-;-;-; | 3092 | GRM4 | 2 | 12 | 0 | 10 | 8 | 0 |
| ENSDARG000 | 6;6;6;6;6;6;6; | 20764848:207 | 20768446:207 | -;-;-;-;-;-;-;-; | 9190 | NA | 2 | 74 | 38 | 24 | 0 | 16 |
| ENSDARG000 | 6 | 9523373 | 9523489 | - | 117 | 5S_rRNA | 0 | 0 | 0 | 0 | 4 | 0 |
| ENSDARG000 | 6;6;6;6;6;6;6 | 21395051:214 | 21395271:214 | +;+;+;+;+;+;+ | 1849 | cacng5a | 0 | 0 | 0 | 0 | 0 | 0 |
| ENSDARG000 | 6;6;6;6;6;6;6 | 59984772:599 | 59984850:599 | +;+;+;+;+;+;+ | 708 | NA | 49 | 35 | 62 | 48 | 40 | 18 |
| ENSDARG000 | 6 | 17355068 | 17355165 | - | 98 | NA | 0 | 0 | 0 | 0 | 0 | 0 |
| ENSDARG000 | 6 | 13333133 | 13334222 | + | 1090 | rprmb | 0 | 0 | 0 | 0 | 0 | 0 |
| ENSDARG000 | 6;6;6 | 28203:28586: | 28393:28754: | +;+;+ | 1232 | NA | 537 | 409 | 677 | 360 | 658 | 381 |
| ENSDARG000 | 6 | 53669511 | 53669630 | + | 120 | 5S_rRNA | 0 | 0 | 0 | 0 | 0 | 0 |
| ENSDARG000 | 6 | 57263510 | 57263645 | - | 136 | 5S_rRNA | 0 | 0 | 0 | 0 | 0 | 0 |
| ENSDARG000 | 6 | 42162650 | 42162774 | + | 125 | 5S_rRNA | 0 | 0 | 0 | 0 | 0 | 0 |
| ENSDARG000 | 6 | 38087495 | 38087611 | + | 117 | 5S_rRNA | 0 | 0 | 0 | 0 | 0 | 0 |
| ENSDARG000 | 6 | 17114658 | 17114755 | - | 98 | NA | 0 | 0 | 0 | 0 | 0 | 0 |
| ENSDARG000 | 6 | 17205192 | 17205289 | + | 98 | NA | 0 | 0 | 0 | 0 | 0 | 0 |
| ENSDARG000 | 6 | 38597669 | 38597784 | - | 116 | 5S_rRNA | 0 | 0 | 0 | 0 | 0 | 0 |
| ENSDARG000 | 6 | 38377091 | 38377211 | + | 121 | 5S_rRNA | 0 | 0 | 0 | 0 | 0 | 0 |
| ENSDARG000 | 6;6;6;6;6;6;6; | 22104565:221 | 22105297:221 | +;+;+;+;+;+;+; | 3378 | ddx17 | 1649 | 1504 | 3091 | 2995 | 2518 | 4553 |
| ENSDARG000 | 6;6;6;6;6;6;6; | 60018233:600 | 60018336:600 | -;-;-;-;-;-;-;-; | 387 | NA | 34 | 22 | 15 | 26 | 27 | 26 |
| ENSDARG000 | 6;6;6;6;6;6;6 | 21576164:215 | 21576679:215 | -;-;-;-;-;-;- | 1084 | si:dkev-43k4.: | 54 | 47 | 67 | 118 | 66 | 80 |
| ENSDARG000 | 6 | 41194687 | 41194801 | + | 115 | 5S_rRNA | 0 | 0 | 0 | 0 | 0 | 0 |
| ENSDARG000 | 6;6;6;6;6;6;6; | 153146:15455 | 153350:15529 | +;+;+;+;+;+;+; | 2720 | zglp1 | 6 | 0 | 8 | 0 | 0 | 0 |
| ENSDARG000 | 6 | 17459301 | 17459398 | - | 98 | NA | 0 | 0 | 0 | 0 | 0 | 0 |
| ENSDARG000 | 6 | 17381485 | 17381582 | + | 98 | NA | 0 | 0 | 0 | 0 | 0 | 0 |
| ENSDARG000 | 6;6;6;6;6;6;6; | 1724889:1725 | 1724935:1725 | +;+;+;+;+;+;+; | 1320 | NA | 114 | 69 | 83 | 35 | 65 | 79 |
| ENSDARG000 | 6 | 16950495 | 16950592 | - | 98 | NA | 0 | 0 | 0 | 0 | 0 | 0 |
| ENSDARG000 | 6;6;6;6;6;6;6; | 23122789:231 | 23122897:231 | +;+;+;+;+;+;+; | 2931 | acox1 | 1097 | 781 | 743 | 550 | 1187 | 1270 |
| ENSDARG000 | 6;6;6 | 13201358:132 | 13201473:132 | +;+;+ | 252 | NA | 0 | 0 | 0 | 0 | 0 | 0 |
| ENSDARG000 | 6;6;6;6;6;6;6; | 34038963:340 | 34039139:340 | +;+;+;+;+;+;+; | 3606 | ap1m3 | 35 | 4 | 4 | 8 | 18 | 10 |
| ENSDARG000 | 6 | 54082522 | 54082656 | + | 135 | 5S_rRNA | 2 | 0 | 0 | 3 | 0 | 2 |
| ENSDARG000 | 6;6;6 | 46341306:463 | 46341463:463 | +;+;+ | 590 | FAM107A | 30 | 25 | 4 | 0 | 4 | 4 |
| ENSDARG000 | 6 | 3710865 | 3713097 | + | 2233 | phospho2 | 63 | 69 | 68 | 30 | 80 | 26 |
| ENSDARG000 | 6 | 13800093 | 13800172 | - | 80 | dre-mir-375-1 | 0 | 0 | 0 | 0 | 0 | 0 |
| ENSDARG000 | 6 | 46025846 | 46025963 | + | 118 | 5S_rRNA | 0 | 0 | 0 | 0 | 0 | 0 |
| ENSDARG000 | 6 | 54507105 | 54507178 | - | 74 | NA | 0 | 0 | 0 | 0 | 0 | 0 |
| ENSDARG000 | 6;6;6;6 | 49520124:495 | 49520807:495 | -;-;-;- | 1445 | rps26l | 6846 | 8322 | 11516 | 8638 | 6684 | 8976 |
| ENSDARG000 | 6 | 24296039 | 24301324 | - | 5286 | NA | 6 | 6 | 0 | 2 | 5 | 8 |
| ENSDARG000 | 6 | 48604823 | 48604937 | + | 115 | 5S_rRNA | 0 | 0 | 0 | 0 | 0 | 0 |
| ENSDARG000 | 6;6;6;6;6;6;6; | 1494817:1498 | 1494969:1498 | -;-;-;-;-;-;-;- | 920 | chchd6b | 723 | 825 | 765 | 653 | 740 | 624 |
| ENSDARG000 | 6 | 11202009 | 11202126 | - | 118 | 5S_rRNA | 0 | 0 | 0 | 0 | 0 | 0 |
| ENSDARG000 | 6 | 21824400 | 21824516 | + | 117 | 5S_rRNA | 0 | 0 | 0 | 0 | 0 | 0 |
| ENSDARG000 | 6;6;6;6;6;6;6; | 21861087:218 | 21862705:218 | -;-;-;-;-;-;-;-; | 2319 | si:dkev-19e4.! | 780 | 832 | 1159 | 724 | 904 | 932 |
| ENSDARG000 | 6;6;6;6;6;6;6 | 1808039:1812 | 1808887:1812 | -;-;-;-;-;-;- | 1630 | NA | 10 | 23 | 35 | 28 | 18 | 4 |
| ENSDARG000 | 6 | 18517970 | 18518067 | - | 98 | dre-mir-193a- | 0 | 0 | 0 | 0 | 0 | 0 |
| ENSDARG000 | 6;6;6;6;6;6;6; | 14980761:149 | 14981208:149 | +;+;+;+;+;+;+; | 3134 | mrps9 | 392 | 470 | 547 | 358 | 485 | 321 |
| ENSDARG000 | 6;6 | 54459413:544 | 54460040:544 | -;- | 1017 | NA | 0 | 0 | 0 | 0 | 0 | 1 |
| ENSDARG000 | 6 | 46749506 | 46749624 | + | 119 | 5S_rRNA | 0 | 0 | 0 | 0 | 0 | 0 |
| ENSDARG000 | 6;6;6;6;6;6;6; | 1285832:1288 | 1285927:1288 | -;-;-;-;-;-;-;-; | 5658 | PLXNA1 | 1175 | 484 | 714 | 491 | 419 | 803 |
| ENSDARG000 | 6 | 47278139 | 47278257 | + | 119 | 5S_rRNA | 0 | 0 | 0 | 0 | 0 | 0 |
| ENSDARG000 | 6 | 53032996 | 53033111 | - | 116 | 5S_rRNA | 0 | 0 | 0 | 0 | 0 | 0 |
| ENSDARG000 | 6 | 17208761 | 17208858 | - | 98 | NA | 0 | 0 | 0 | 0 | 0 | 0 |
| ENSDARG000 | 6 | 16805744 | 16805838 | - | 95 | NA | 0 | 0 | 0 | 0 | 0 | 0 |
| ENSDARG000 | 6;6;6 | 23616893:236 | 23618099:236 | -;-;- | 1609 | aanat1 | 0 | 4 | 18 | 0 | 0 | 14 |
| ENSDARG000 | 6 | 58793883 | 58793997 | + | 115 | 5S_rRNA | 0 | 0 | 0 | 0 | 0 | 2 |
| ENSDARG000 | 6 | 30202921 | 30203037 | - | 117 | 5S_rRNA | 0 | 0 | 0 | 0 | 0 | 0 |
| ENSDARG000 | 6 | 11774617 | 11774732 | + | 116 | 5S_rRNA | 0 | 0 | 0 | 2 | 0 | 0 |
| ENSDARG000 | 6;6;6;6;6;6;6; | 13083665:130 | 13083690:130 | -;-;-;-;-;-;-;-; | 4824 | casp20 | 215 | 37 | 31 | 9 | 44 | 145 |
| ENSDARG000 | 6 | 24842170 | 24842286 | - | 117 | 5S_rRNA | 0 | 0 | 0 | 0 | 0 | 0 |
| ENSDARG000 | 6 | 40625304 | 40625420 | - | 117 | 5S_rRNA | 0 | 0 | 0 | 0 | 0 | 0 |
| ENSDARG000 | 6;6;6;6;6;6;6; | 15355244:153 | 15355259:153 | -;-;-;-;-;-;-;-; | 399 | NA | 0 | 0 | 0 | 0 | 2 | 0 |
| ENSDARG000 | 6 | 50135319 | 50135437 | - | 119 | 5S_rRNA | 0 | 0 | 0 | 0 | 0 | 0 |
| ENSDARG000 | 6 | 24785827 | 24785943 | - | 117 | 5S_rRNA | 0 | 0 | 0 | 0 | 0 | 0 |
| ENSDARG000 | 6;6;6;6;6;6 | 48862:55415: | 49113:55809: | +;+;+;+;+;+ | 3286 | NA | 517 | 304 | 582 | 253 | 514 | 242 |
| ENSDARG000 | 6 | 11040128 | 11040242 | - | 115 | 5S_rRNA | 0 | 0 | 0 | 0 | 0 | 0 |
| ENSDARG000 | 6;6;6;6;6 | 7795602:7798 | 7795784:7798 | -;-;-;-;- | 936 | PLPPR2 | 0 | 0 | 0 | 0 | 0 | 0 |
| ENSDARG000 | 6;6;6 | 21740672:217 | 21741077:217 | +;+;+ | 648 | LHFPL4 | 0 | 1 | 0 | 0 | 0 | 0 |
| ENSDARG000 | 6;6;6;6;6;6;6; | 20678890:206 | 20679953:206 | -;-;-;-;-;-;-;-; | 3571 | NA | 96 | 50 | 40 | 36 | 130 | 28 |
| ENSDARG000 | 6;6;6;6;6;6;6; | 3693441:3696 | 3693546:3696 | +;+;+;+;+;+;+; | 2595 | ppig | 522 | 915 | 1236 | 539 | 723 | 642 |
| ENSDARG000 | 6;6;6;6 | 1787160:1794 | 1787311:1794 | +;+;+;+ | 2957 | myl9b | 230 | 827 | 935 | 1980 | 632 | 430 |
| ENSDARG000 | 6 | 11018416 | 11018532 | + | 117 | 5S_rRNA | 0 | 0 | 0 | 0 | 0 | 0 |
| ENSDARG000 | 6 | 3165608 | 3165722 | + | 115 | 5S_rRNA | 0 | 0 | 0 | 0 | 0 | 0 |
| ENSDARG000 | 6;6;6 | 55212928:552 | 55213086:552 | +;+;+ | 1565 | manbal | 428 | 348 | 465 | 334 | 512 | 449 |
| ENSDARG000 | 6 | 36744134 | 36744251 | - | 118 | 5S_rRNA | 0 | 0 | 0 | 0 | 0 | 0 |
| ENSDARG000 | 6 | 22540982 | 22541076 | - | 95 | NA | 0 | 0 | 0 | 0 | 0 | 0 |
| ENSDARG000 | 6;6;6;6;6;6;6; | 21992820:219 | 21993625:219 | +;+;+;+;+;+;+; | 2412 | thumpd3 | 334 | 517 | 675 | 485 | 582 | 230 |
| ENSDARG000 | 6;6;6 | 72040:72264: | 72048:72503: | +;+;+ | 525 | crygm4 | 0 | 0 | 0 | 0 | 0 | 0 |
| ENSDARG000 | 6;6;6;6;6;6 | 60004871:600 | 60004882:600 | +;+;+;+;+;+ | 132 | NA | 3 | 0 | 0 | 0 | 0 | 0 |
| ENSDARG000 | 6 | 19392860 | 19392977 | - | 118 | 5S_rRNA | 0 | 0 | 0 | 0 | 0 | 0 |
| ENSDARG000 | 6;6;6;6;6;6;6; | 21095918:210 | 21096232:210 | +;+;+;+;+;+;+; | 9444 | spega | 0 | 0 | 0 | 0 | 0 | 0 |
| ENSDARG000 | 6 | 57769542 | 57769658 | - | 117 | 5S_rRNA | 0 | 0 | 0 | 0 | 0 | 0 |
| ENSDARG000 | 6;6;6;6;6;6;6; | 1161224:1162 | 1162415:1162 | -;-;-;-;-;-;-;-; | 3116 | txnrd3 | 3914 | 6973 | 5185 | 2018 | 4503 | 2274 |
| ENSDARG000 | 6 | 47602599 | 47602715 | - | 117 | 5S_rRNA | 0 | 0 | 0 | 0 | 0 | 0 |
| ENSDARG000 | 6 | 17317557 | 17317653 | + | 97 | NA | 0 | 0 | 0 | 0 | 0 | 0 |
| ENSDARG000 | 6 | 29096849 | 29096945 | + | 97 | SNORD21 | 15 | 5 | 5 | 7 | 6 | 37 |
| ENSDARG000 | 6;6;6;6;6;6;6; | 25312387:253 | 25315627:253 | -;-;-;-;-;-;-;-; | 8138 | pkn2b | 1091 | 627 | 910 | 444 | 850 | 668 |
| ENSDARG000 | 6;6;6;6;6;6;6; | 12006557:120 | 12006954:120 | +;+;+;+;+;+;+; | 1464 | wdsub1 | 479 | 191 | 296 | 222 | 505 | 320 |
| ENSDARG000 | 6;6;6;6;6;6;6 | 181142:18172 | 181259:18180 | +;+;+;+;+;+;+ | 1038 | TAB1 | 280 | 188 | 431 | 365 | 246 | 262 |
| ENSDARG000 | 6 | 6071053 | 6071169 | - | 117 | 5S_rRNA | 0 | 0 | 0 | 0 | 0 | 0 |
| ENSDARG000 | 6;6;6;6;6;6;6; | 475264:48721 | 475525:48733 | +;+;+;+;+;+;+; | 1254 | NA | 0 | 5 | 0 | 0 | 4 | 10 |
| ENSDARG000 | 6;6;6;6;6;6;6; | 46878889:468 | 46879464:468 | -;-;-;-;-;-;-;-; | 2290 | igfn1.1 | 0 | 0 | 0 | 0 | 0 | 0 |
| ENSDARG000 | 6 | 36733709 | 36733826 | - | 118 | 5S_rRNA | 0 | 0 | 0 | 0 | 0 | 0 |
| ENSDARG000 | 6;6;6;6;6;6;6; | 23010242:230 | 23011069:230 | +;+;+;+;+;+;+; | 3522 | POLG2 | 72 | 94 | 142 | 56 | 160 | 142 |
| ENSDARG000 | 6 | 36990570 | 36990684 | + | 115 | 5S_rRNA | 0 | 0 | 0 | 0 | 0 | 0 |
| ENSDARG000 | 6 | 40653129 | 40653238 | - | 110 | dre-mir-429b | 0 | 0 | 0 | 0 | 0 | 0 |
| ENSDARG000 | 6 | 16896978 | 16897074 | - | 97 | NA | 0 | 0 | 0 | 0 | 0 | 0 |
| ENSDARG000 | 6 | 36769939 | 36770069 | - | 131 | 5S_rRNA | 0 | 0 | 0 | 0 | 0 | 0 |
| ENSDARG000 | 6;6;6;6;6;6;6; | 49585461:495 | 49585476:495 | -;-;-;-;-;-;-;-; | 321 | NA | 0 | 0 | 2 | 7 | 0 | 0 |
| ENSDARG000 | 6;6;6;6;6;6;6; | 45346808:453 | 45346925:453 | +;+;+;+;+;+;+; | 6577 | NA | 12 | 44 | 222 | 64 | 14 | 50 |
| ENSDARG000 | 6 | 16737867 | 16737964 | - | 98 | NA | 0 | 0 | 0 | 0 | 0 | 0 |
| ENSDARG000 | 6 | 9668277 | 9668361 | - | 85 | NA | 0 | 2 | 1 | 0 | 0 | 0 |
| ENSDARG000 | 6;6;6;6;6;6;6; | 27696323:276 | 27696349:276 | -;-;-;-;-;-;-;- | 210 | NA | 0 | 0 | 0 | 0 | 0 | 0 |
| ENSDARG000 | 6 | 10867024 | 10867141 | + | 118 | dre-mir-10d | 0 | 0 | 0 | 0 | 0 | 0 |
| ENSDARG000 | 6 | 13980945 | 13981092 | + | 148 | dre-mir-153b | 2 | 0 | 4 | 0 | 2 | 2 |
| ENSDARG000 | 6 | 50064180 | 50064296 | + | 117 | 5S_rRNA | 0 | 0 | 0 | 0 | 0 | 0 |
| ENSDARG000 | 6 | 68220 | 69904 | + | 1685 | sowahcb | 18 | 12 | 0 | 4 | 10 | 2 |
| ENSDARG000 | 6 | 6215794 | 6215910 | + | 117 | 5S_rRNA | 0 | 0 | 0 | 0 | 0 | 0 |
| ENSDARG000 | 6 | 5684181 | 5684295 | + | 115 | 5S_rRNA | 0 | 0 | 0 | 0 | 0 | 0 |
| ENSDARG000 | 6 | 29329810 | 29329927 | + | 118 | 5S_rRNA | 0 | 0 | 0 | 1 | 0 | 5 |
| ENSDARG000 | 6 | 45605280 | 45605396 | + | 117 | 5S_rRNA | 0 | 0 | 0 | 0 | 0 | 0 |
| ENSDARG000 | 6 | 41454791 | 41454873 | + | 83 | dre-let-7j | 0 | 0 | 0 | 0 | 2 | 4 |
| ENSDARG000 | 6 | 41997963 | 41998096 | + | 134 | 5S_rRNA | 0 | 0 | 0 | 0 | 0 | 0 |
| ENSDARG000 | 6;6;6;6;6;6;6; | 22337081:223 | 22337218:223 | +;+;+;+;+;+;+; | 1760 | uqcrc1 | 3086 | 3648 | 3827 | 2272 | 3351 | 2932 |
| ENSDARG000 | 6 | 47974879 | 47975014 | + | 136 | 5S_rRNA | 0 | 0 | 0 | 0 | 0 | 0 |
| ENSDARG000 | 6 | 20174884 | 20174981 | + | 98 | NA | 0 | 0 | 0 | 0 | 0 | 0 |
| ENSDARG000 | 6;6;6;6;6;6;6; | 53503273:535 | 53503698:535 | -;-;-;-;-;-;-;-; | 1668 | cacna2d2a | 7 | 0 | 10 | 8 | 0 | 0 |
| ENSDARG000 | 6 | 55109962 | 55110076 | + | 115 | 5S_rRNA | 0 | 0 | 0 | 0 | 0 | 0 |
| ENSDARG000 | 6 | 31365446 | 31365526 | + | 81 | dre-mir-101a | 0 | 0 | 0 | 0 | 0 | 0 |
| ENSDARG000 | 6;6;6;6;6;6;6; | 59426148:594 | 59431334:594 | -;-;-;-;-;-;-;-; | 7333 | gli1 | 10 | 2 | 6 | 0 | 0 | 6 |
| ENSDARG000 | 6;6;6;6;6;6;6 | 10028886:100 | 10029364:100 | -;-;-;-;-;-;- | 1126 | nudt15 | 46 | 32 | 29 | 34 | 86 | 40 |
| ENSDARG000 | 6 | 58009067 | 58009195 | + | 129 | 5S_rRNA | 0 | 0 | 0 | 0 | 0 | 0 |
| ENSDARG000 | 6;6;6;6;6;6 | 344423:34459 | 344825:34482 | -;-;-;-;-;- | 724 | pde6ha | 19 | 8 | 10 | 12 | 20 | 2 |
| ENSDARG000 | 6 | 22141746 | 22141862 | + | 117 | 5S_rRNA | 0 | 0 | 0 | 0 | 0 | 0 |
| ENSDARG000 | 6;6;6;6;6;6;6; | 2124469:2125 | 2126661:2125 | -;-;-;-;-;-;-;- | 3371 | VSTM2L | 0 | 1 | 4 | 0 | 0 | 0 |
| ENSDARG000 | 6;6;6;6;6;6;6; | 21525543:215 | 21526637:215 | -;-;-;-;-;-;-;-; | 2413 | psmd12 | 2233 | 2404 | 2413 | 1189 | 2552 | 1634 |
| ENSDARG000 | 6;6;6;6;6;6;6 | 2421028:2424 | 2423107:2424 | -;-;-;-;-;-;- | 3445 | med8 | 258 | 242 | 309 | 367 | 245 | 288 |
| ENSDARG000 | 6;6;6;6;6;6;6; | 44584775:445 | 44587606:445 | -;-;-;-;-;-;-;-; | 6398 | NA | 4 | 0 | 0 | 6 | 0 | 0 |
| ENSDARG000 | 6;6;6;6;6;6;6; | 2271559:2279 | 2271779:2279 | +;+;+;+;+;+;+; | 1938 | pbx1b | 421 | 174 | 595 | 158 | 468 | 83 |
| ENSDARG000 | 6;6;6;6 | 25282671:252 | 25282795:252 | +;+;+;+ | 528 | NA | 7 | 0 | 0 | 0 | 7 | 2 |
| ENSDARG000 | 6;6;6;6;6;6 | 6215042:6217 | 6215051:6217 | +;+;+;+;+;+ | 162 | NA | 0 | 2 | 0 | 0 | 0 | 0 |
| ENSDARG000 | 6;6;6;6;6;6 | 149405:14971 | 149613:14977 | +;+;+;+;+;+ | 1137 | fdx2 | 312 | 394 | 369 | 225 | 260 | 297 |
| ENSDARG000 | 6;6;6;6 | 1769905:1772 | 1770410:1773 | -;-;-;- | 1371 | zgc:113442 | 8 | 2 | 8 | 0 | 2 | 0 |
| ENSDARG000 | 6;6;6;6;6;6;6; | 22025806:220 | 22027645:220 | +;+;+;+;+;+;+; | 3696 | NA | 1029 | 674 | 784 | 662 | 1162 | 731 |
| ENSDARG000 | 6 | 56539009 | 56539125 | - | 117 | 5S_rRNA | 0 | 0 | 0 | 0 | 0 | 0 |
| ENSDARG000 | 6 | 1779397 | 1780633 | - | 1237 | NA | 191 | 245 | 257 | 260 | 192 | 224 |
| ENSDARG000 | 6 | 20409557 | 20409654 | - | 98 | NA | 0 | 0 | 0 | 0 | 0 | 0 |
| ENSDARG000 | 6;6;6;6;6;6;6; | 59642695:596 | 59642826:596 | +;+;+;+;+;+;+; | 2806 | R3HDM2 | 112 | 53 | 99 | 60 | 63 | 83 |
| ENSDARG000 | 6;6;6 | 39536596:395 | 39536928:395 | +;+;+ | 384 | NA | 0 | 2 | 0 | 0 | 8 | 1 |
| ENSDARG000 | 6;6;6;6;6;6;6; | 3640381:3648 | 3640935:3648 | +;+;+;+;+;+;+; | 3426 | NA | 0 | 0 | 0 | 0 | 1 | 0 |
| ENSDARG000 | 6;6;6;6;6;6 | 13083146:130 | 13083354:130 | +;+;+;+;+;+ | 1255 | L3HYPDH | 250 | 228 | 238 | 183 | 220 | 119 |
| ENSDARG000 | 6;6 | 59557501:595 | 59558252:595 | -;- | 1140 | NA | 0 | 0 | 0 | 0 | 0 | 0 |
| ENSDARG000 | 6;6;6;6;6 | 3730843:3809 | 3730985:3810 | +;+;+;+;+ | 2092 | NA | 0 | 0 | 0 | 0 | 0 | 2 |
| ENSDARG000 | 6;6;6;6;6;6;6; | 54805136:548 | 54807417:548 | -;-;-;-;-;-;-;-; | 2888 | tnni1b | 6 | 0 | 0 | 0 | 0 | 0 |
| ENSDARG000 | 6;6;6;6;6;6;6; | 59176470:591 | 59176502:591 | +;+;+;+;+;+;+; | 1479 | shmt2 | 716 | 601 | 1824 | 681 | 792 | 980 |
| ENSDARG000 | 6 | 37615366 | 37615483 | - | 118 | 5S_rRNA | 0 | 0 | 0 | 0 | 0 | 0 |
| ENSDARG000 | 6 | 29469991 | 29470105 | + | 115 | 5S_rRNA | 0 | 0 | 0 | 0 | 0 | 0 |
| ENSDARG000 | 6;6;6 | 214269:21616 | 215991:21641 | -;-;- | 2173 | NA | 18 | 12 | 2 | 2 | 16 | 21 |
| ENSDARG000 | 6;6;6;6;6;6;6; | 59991076:599 | 59991079:599 | +;+;+;+;+;+;+; | 999 | GPR180 | 80 | 73 | 120 | 24 | 85 | 39 |

| | | | | | | | | | | | | |
|---|---|---|---|---|---|---|---|---|---|---|---|---|
| ENSDARG000 | 6:6:6:6:6:6:6: | 23057311:230 | 23057363:230 | +:+:+:+:+:+:+: | 8722 | evpla | 154 | 6 | 12 | 12 | 69 | 24 |
| ENSDARG000 | 6:6:6:6:6:6:6: | 30430591:304 | 30430688:304 | +:+:+:+:+:+:+: | 8247 | NA | 1233 | 1634 | 1941 | 1058 | 1540 | 1133 |
| ENSDARG000 | 6;6;6;6 | 34138386:341 | 34138625:341 | -;-;-;- | 614 | cfap144 | 0 | 0 | 0 | 0 | 0 | 0 |
| ENSDARG000 | 6;6 | 55277419:552 | 55277463:552 | +;+ | 726 | NA | 0 | 0 | 4 | 0 | 0 | 0 |
| ENSDARG000 | 6;6;6;6 | 21513363:215 | 21516690:215 | -;-;-;- | 4014 | NA | 8 | 2 | 6 | 0 | 60 | 22 |
| ENSDARG000 | 6;6;6;6;6 | 48206535:482 | 48206620:482 | +;+;+;+;+ | 2301 | CTTNBP2NL | 432 | 388 | 488 | 326 | 503 | 252 |
| ENSDARG000 | 6:6:6:6:6:6:6: | 52521467:525 | 52522066:525 | -;-;-;-;-;-;-;-;- | 1409 | uqcc1 | 370 | 410 | 481 | 318 | 303 | 273 |
| ENSDARG000 | 6 | 57080758 | 57080872 | + | 115 | 5S_rRNA | 0 | 0 | 0 | 0 | 0 | 0 |
| ENSDARG000 | 6:6:6:6:6:6:6: | 296130:31908 | 296255:31922 | +:+:+:+:+:+:+: | 3274 | rbfox2 | 5147 | 4051 | 4224 | 4429 | 4138 | 4005 |
| ENSDARG000 | 6 | 59685145 | 59685261 | - | 117 | 5S_rRNA | 0 | 0 | 0 | 0 | 0 | 0 |
| ENSDARG000 | 6 | 52923373 | 52923489 | - | 117 | 5S_rRNA | 0 | 0 | 0 | 0 | 0 | 0 |
| ENSDARG000 | 6 | 17385032 | 17385129 | - | 98 | NA | 0 | 0 | 0 | 0 | 0 | 0 |
| ENSDARG000 | 6 | 52756334 | 52757276 | - | 943 | NA | 0 | 0 | 0 | 0 | 0 | 0 |
| ENSDARG000 | 6 | 51075056 | 51075171 | + | 116 | 5S_rRNA | 0 | 0 | 0 | 0 | 0 | 0 |
| ENSDARG000 | 6 | 40667347 | 40667463 | - | 117 | 5S_rRNA | 0 | 0 | 2 | 0 | 0 | 0 |
| ENSDARG000 | 6 | 18516311 | 18516407 | - | 97 | dre-mir-365-3 | 0 | 0 | 0 | 6 | 0 | 0 |
| ENSDARG000 | 6 | 33443334 | 33443455 | - | 122 | 5S_rRNA | 0 | 0 | 0 | 0 | 3 | 0 |
| ENSDARG000 | 6 | 17051039 | 17051135 | + | 97 | NA | 0 | 0 | 0 | 0 | 0 | 0 |
| ENSDARG000 | 6 | 9062276 | 9062391 | - | 116 | 5S_rRNA | 0 | 0 | 0 | 0 | 0 | 0 |
| ENSDARG000 | 6;6 | 60142619:601 | 60145308:601 | -;- | 3495 | slc32a1 | 0 | 0 | 0 | 0 | 0 | 0 |
| ENSDARG000 | 6:6:6:6:6:6:6: | 3816241:3816 | 3816441:3816 | +:+:+:+:+:+:+: | 1284 | erich2 | 0 | 6 | 10 | 6 | 6 | 0 |
| ENSDARG000 | 6;6 | 21068783:210 | 21069516:210 | -;- | 1044 | inha | 0 | 0 | 0 | 0 | 0 | 0 |
| ENSDARG000 | 6 | 36511812 | 36511930 | + | 119 | 5S_rRNA | 0 | 0 | 0 | 0 | 0 | 0 |
| ENSDARG000 | 6 | 20333347 | 20333444 | + | 98 | NA | 0 | 0 | 0 | 0 | 0 | 0 |
| ENSDARG000 | 6 | 8771476 | 8771596 | + | 121 | 5S_rRNA | 0 | 0 | 0 | 0 | 0 | 0 |
| ENSDARG000 | 6;6 | 54481941:544 | 54482625:544 | -;- | 1017 | NA | 0 | 2 | 0 | 0 | 0 | 0 |
| ENSDARG000 | 6 | 4658945 | 4659059 | - | 115 | 5S_rRNA | 0 | 0 | 0 | 0 | 0 | 0 |
| ENSDARG000 | 6;6;6 | 59967994:599 | 59968194:599 | +;+;+ | 1647 | zgc:65895 | 314 | 189 | 280 | 149 | 488 | 130 |
| ENSDARG000 | 6 | 7717788 | 7717929 | - | 142 | SNORA62 | 6 | 2 | 8 | 5 | 4 | 2 |
| ENSDARG000 | 6:6:6:6:6:6:6: | 22326624:223 | 22326694:223 | +:+:+:+:+:+:+: | 1560 | rae1 | 618 | 915 | 955 | 469 | 949 | 630 |
| ENSDARG000 | 6:6:6:6:6:6:6: | 19025384:190 | 19025435:190 | +:+:+:+:+:+:+: | 225 | NA | 0 | 0 | 0 | 0 | 0 | 0 |
| ENSDARG000 | 6 | 20464402 | 20464499 | - | 98 | NA | 0 | 0 | 0 | 0 | 0 | 0 |
| ENSDARG000 | 6:6:6:6:6:6:6: | 23026170:230 | 23026833:230 | +:+:+:+:+:+:+: | 6568 | srp68 | 1170 | 1334 | 2192 | 1140 | 1873 | 1248 |
| ENSDARG000 | 6 | 25435093 | 25435221 | + | 129 | 5S_rRNA | 0 | 0 | 0 | 0 | 0 | 0 |
| ENSDARG000 | 6:6:6:6:6:6:6: | 47484084:474 | 47484147:474 | -:-:-:-:-:-:-:-: | 1080 | NA | 28 | 27 | 30 | 30 | 46 | 39 |
| ENSDARG000 | 6 | 12414215 | 12414329 | - | 115 | 5S_rRNA | 0 | 0 | 0 | 0 | 0 | 0 |
| ENSDARG000 | 6 | 1270561 | 1270675 | - | 115 | 5S_rRNA | 0 | 0 | 0 | 0 | 0 | 0 |
| ENSDARG000 | 6;6;6;6;6;6 | 57741039:577 | 57741321:577 | +;+;+;+;+;+ | 1164 | SNTA1 | 14 | 10 | 44 | 2 | 76 | 28 |
| ENSDARG000 | 6;6;6;6 | 59935950:599 | 59936590:599 | -;-;-;- | 957 | NA | 464 | 315 | 484 | 442 | 525 | 227 |
| ENSDARG000 | 6;6 | 202367:20385 | 202702:20529 | +;+ | 1770 | MGAT3 | 173 | 85 | 201 | 56 | 178 | 126 |
| ENSDARG000 | 6;6;6;6;6;6;6 | 17789481:177 | 17789605:177 | +;+;+;+;+;+;+ | 819 | NA | 2 | 1 | 0 | 2 | 0 | 1 |
| ENSDARG000 | 6:6:6:6:6:6:6: | 3934738:3935 | 3934845:3935 | +:+:+:+:+:+:+: | 1887 | DYNC1I2 | 1268 | 1261 | 1300 | 1267 | 972 | 758 |
| ENSDARG000 | 6:6:6:6:6:6:6: | 1748463:1748 | 1748714:1748 | -:-:-:-:-:-:-:-: | 1503 | orc4 | 112 | 440 | 560 | 187 | 376 | 248 |
| ENSDARG000 | 6:6:6:6:6:6:6: | 4229360:4229 | 4229909:4229 | +:+:+:+:+:+:+: | 5655 | STRADB | 268 | 220 | 400 | 219 | 358 | 275 |
| ENSDARG000 | 6;6;6;6;6;6 | 4528631:4531 | 4529110:4531 | +;+;+;+;+;+ | 2043 | NA | 8 | 30 | 12 | 22 | 28 | 8 |
| ENSDARG000 | 6 | 8153813 | 8154866 | + | 1054 | nfil3-3 | 18 | 4 | 5 | 0 | 0 | 2 |
| ENSDARG000 | 6 | 7174967 | 7175083 | - | 117 | 5S_rRNA | 0 | 0 | 0 | 0 | 0 | 0 |
| ENSDARG000 | 6 | 7717164 | 7717302 | - | 139 | SNORA62 | 0 | 0 | 3 | 0 | 0 | 0 |
| ENSDARG000 | 6 | 55650119 | 55650251 | - | 133 | 5S_rRNA | 0 | 0 | 0 | 0 | 0 | 0 |
| ENSDARG000 | 6 | 58441344 | 58441460 | + | 117 | 5S_rRNA | 0 | 0 | 0 | 0 | 0 | 0 |
| ENSDARG000 | 6:6:6:6:6:6:6: | 2163826:2163 | 2168371:2167 | -;-;-;-;-;-;-;-;- | 5330 | pxmp4 | 338 | 426 | 502 | 360 | 438 | 370 |
| ENSDARG000 | 6:6:6:6:6:6:6: | 31684:31714: | 31968:32040: | +:+:+:+:+:+:+: | 6646 | MARF1 | 1305 | 744 | 978 | 770 | 1174 | 905 |
| ENSDARG000 | 6:6:6:6:6:6:6: | 12036951:120 | 12038127:120 | -:-:-:-:-:-:-:-: | 6044 | NA | 38 | 8 | 2 | 22 | 26 | 12 |
| ENSDARG000 | 6:6:6:6:6:6:6: | 21536131:215 | 21537023:215 | +:+:+:+:+:+:+: | 3398 | NA | 405 | 271 | 407 | 224 | 578 | 315 |
| ENSDARG000 | 6;6;6;6 | 264927:26573 | 265264:26586 | +;+;+;+ | 2110 | NA | 501 | 358 | 550 | 350 | 523 | 420 |
| ENSDARG000 | 6;6;6;6 | 1219279:1219 | 1219500:1219 | -;-;-;- | 2472 | NA | 0 | 0 | 0 | 0 | 0 | 0 |
| ENSDARG000 | 6;6;6;6;6 | 55174744:551 | 55174870:551 | +;+;+;+;+ | 774 | SYT2 | 0 | 0 | 0 | 0 | 0 | 6 |
| ENSDARG000 | 6 | 30484005 | 30485009 | - | 1005 | cldn33b | 22 | 6 | 8 | 6 | 0 | 0 |
| ENSDARG000 | 6;6 | 2457455:2463 | 2457745:2464 | +;+ | 520 | NA | 3 | 3 | 0 | 0 | 0 | 1 |
| ENSDARG000 | 6:6:6:6:6:6:6: | 4185107:4185 | 4185869:4185 | -:-:-:-:-:-:-:-: | 3390 | TRAK2 | 1748 | 1442 | 1985 | 3253 | 2134 | 1645 |
| ENSDARG000 | 6;6;6;6;6;6 | 42500:45997: | 42612:46058: | -;-;-;-;-;- | 858 | bmerb1 | 27 | 12 | 25 | 37 | 24 | 4 |
| ENSDARG000 | 6 | 46870184 | 46870298 | + | 115 | 5S_rRNA | 0 | 0 | 0 | 0 | 0 | 0 |
| ENSDARG000 | 6 | 57133975 | 57134089 | + | 115 | 5S_rRNA | 0 | 0 | 0 | 0 | 0 | 0 |
| ENSDARG000 | 6 | 17515775 | 17515872 | - | 98 | NA | 0 | 0 | 0 | 0 | 0 | 0 |
| ENSDARG000 | 6 | 59136671 | 59136787 | - | 117 | 5S_rRNA | 0 | 0 | 0 | 0 | 0 | 0 |
| ENSDARG000 | 6:6:6:6:6:6:6: | 4160579:4160 | 4161871:4161 | +:+:+:+:+:+:+: | 3222 | NA | 10 | 8 | 4 | 2 | 6 | 1 |
| ENSDARG000 | 6 | 54506458 | 54506586 | - | 129 | dre-let-7a-6 | 6 | 12 | 9 | 17 | 9 | 3 |
| ENSDARG000 | 6 | 6879927 | 6880049 | + | 123 | 5S_rRNA | 0 | 0 | 0 | 0 | 0 | 0 |
| ENSDARG000 | 6;6;6 | 22597362:226 | 22597888:226 | +;+;+ | 2335 | cygb2 | 0 | 0 | 0 | 0 | 0 | 0 |
| ENSDARG000 | 6:6:6:6:6:6:6: | 22957709:229 | 22958625:229 | -:-:-:-:-:-:-:-: | 3530 | nol11 | 764 | 838 | 1207 | 543 | 924 | 758 |
| ENSDARG000 | 6:6:6:6:6:6:6: | 21697128:217 | 21700380:217 | -:-:-:-:-:-:-:-: | 5230 | MTMR14 | 669 | 495 | 468 | 329 | 569 | 551 |
| ENSDARG000 | 6;6;6;6 | 20910551:209 | 20911186:209 | +;+;+;+ | 2072 | igfbp5a | 0 | 24 | 27 | 3 | 0 | 6 |
| ENSDARG000 | 6;6;6;6;6;6;6 | 59255136:592 | 59255207:592 | -;-;-;-;-;-;- | 694 | NA | 0 | 0 | 0 | 0 | 0 | 0 |
| ENSDARG000 | 6:6:6:6:6:6:6: | 1535303:1538 | 1535936:1538 | -:-:-:-:-:-:-:-: | 1686 | tpra1 | 453 | 233 | 348 | 492 | 315 | 318 |
| ENSDARG000 | 6 | 43353271 | 43361226 | + | 7956 | NA | 61 | 0 | 39 | 0 | 6 | 30 |
| ENSDARG000 | 6 | 9672680 | 9672763 | - | 84 | NA | 0 | 0 | 0 | 0 | 0 | 0 |
| ENSDARG000 | 6 | 37439255 | 37439370 | - | 116 | 5S_rRNA | 0 | 0 | 0 | 0 | 0 | 0 |
| ENSDARG000 | 6 | 23394843 | 23394899 | - | 57 | NA | 0 | 0 | 0 | 0 | 0 | 0 |
| ENSDARG000 | 6 | 45355409 | 45355525 | + | 117 | 5S_rRNA | 0 | 0 | 0 | 0 | 0 | 0 |
| ENSDARG000 | 6 | 46228117 | 46228232 | - | 116 | 5S_rRNA | 0 | 0 | 0 | 0 | 0 | 0 |
| ENSDARG000 | 6 | 37447028 | 37447143 | - | 116 | 5S_rRNA | 3 | 0 | 4 | 0 | 4 | 0 |
| ENSDARG000 | 6;6 | 4539953:4540 | 4540434:4543 | +;+ | 2732 | pou4f1 | 6 | 0 | 0 | 6 | 2 | 0 |
| ENSDARG000 | 6;6;6;6;6 | 3563890:3566 | 3564435:3566 | -;-;-;-;- | 1430 | dmbx1b | 0 | 0 | 0 | 0 | 0 | 0 |
| ENSDARG000 | 6 | 55547135 | 55547270 | - | 136 | 5S_rRNA | 0 | 0 | 0 | 0 | 0 | 0 |
| ENSDARG000 | 6:6:6:6:6:6:6: | 21157195:211 | 21157784:211 | -:-:-:-:-:-:-:-: | 6339 | obsl1a | 0 | 10 | 30 | 12 | 9 | 0 |
| ENSDARG000 | 6;6;6;6;6;6 | 23107602:231 | 23107913:231 | -;-;-;-;-;- | 647 | ten1 | 48 | 82 | 78 | 110 | 116 | 26 |
| ENSDARG000 | 6 | 17054631 | 17054727 | - | 97 | NA | 0 | 0 | 0 | 0 | 0 | 0 |
| ENSDARG000 | 6:6:6:6:6:6:6: | 22043127:220 | 22043503:220 | -:-:-:-:-:-:-:-: | 2051 | cfap100 | 7 | 4 | 5 | 5 | 7 | 15 |
| ENSDARG000 | 6;6 | 30456788:304 | 30456874:304 | +;+ | 902 | NA | 0 | 0 | 10 | 0 | 0 | 0 |
| ENSDARG000 | 6:6:6:6:6:6:6: | 21227621:212 | 21228203:212 | +:+:+:+:+:+:+: | 6143 | prkcaa | 1138 | 762 | 682 | 872 | 990 | 1051 |
| ENSDARG000 | 6:6:6:6:6:6:6: | 59944488:599 | 59944568:599 | +:+:+:+:+:+:+: | 1547 | nufip1 | 124 | 253 | 431 | 184 | 254 | 252 |
| ENSDARG000 | 6:6:6:6:6:6:6: | 840434:84043 | 844869:84486 | -:-:-:-:-:-:-:-: | 7622 | zeb2b | 607 | 32 | 114 | 68 | 70 | 146 |
| ENSDARG000 | 6 | 52304431 | 52304621 | + | 191 | U2 | 0 | 0 | 0 | 0 | 0 | 0 |
| ENSDARG000 | 6:6:6:6:6:6:6: | 22352223:223 | 22352626:223 | -:-:-:-:-:-:-:-: | 1537 | nprl2 | 301 | 269 | 312 | 196 | 290 | 403 |
| ENSDARG000 | 6 | 55744383 | 55744498 | + | 116 | 5S_rRNA | 0 | 0 | 0 | 0 | 0 | 0 |
| ENSDARG000 | 6;6;6 | 49628431:496 | 49628527:496 | -;-;- | 174 | NA | 0 | 0 | 0 | 0 | 3 | 0 |
| ENSDARG000 | 6 | 13765310 | 13765427 | - | 118 | 5S_rRNA | 0 | 0 | 0 | 0 | 0 | 0 |
| ENSDARG000 | 6 | 20178436 | 20178533 | - | 98 | NA | 0 | 0 | 0 | 0 | 0 | 0 |
| ENSDARG000 | 6 | 29089559 | 29089648 | + | 90 | SNORD21 | 2 | 0 | 0 | 0 | 1 | 0 |
| ENSDARG000 | 6:6:6:6:6:6:6: | 83873:84073: | 83971:84178: | +:+:+:+:+:+:+: | 4695 | SMARCA4 | 64 | 32 | 42 | 22 | 42 | 28 |
| ENSDARG000 | 6 | 44132968 | 44133082 | + | 115 | 5S_rRNA | 0 | 0 | 0 | 0 | 0 | 0 |
| ENSDARG000 | 6 | 40656003 | 40656085 | - | 83 | dre-mir-141 | 0 | 0 | 0 | 0 | 0 | 0 |
| ENSDARG000 | 6;6;6;6;6;6;6 | 59029485:590 | 59029514:590 | +;+;+;+;+;+;+ | 735 | NA | 20 | 13 | 11 | 35 | 30 | 10 |
| ENSDARG000 | 6 | 22537132 | 22537229 | + | 98 | NA | 0 | 0 | 0 | 0 | 0 | 0 |
| ENSDARG000 | 6 | 47156302 | 47156416 | - | 115 | 5S_rRNA | 0 | 0 | 0 | 0 | 0 | 0 |
| ENSDARG000 | 6 | 9673032 | 9673122 | - | 91 | SNORD70B | 0 | 0 | 0 | 0 | 0 | 0 |
| ENSDARG000 | 6 | 16871306 | 16871403 | - | 98 | NA | 0 | 0 | 0 | 0 | 0 | 0 |
| ENSDARG000 | 6 | 40656204 | 40656288 | - | 85 | dre-mir-200c | 0 | 0 | 0 | 0 | 0 | 0 |
| ENSDARG000 | 6 | 7718273 | 7718414 | - | 142 | SNORA62 | 0 | 2 | 0 | 0 | 0 | 0 |
| ENSDARG000 | 6 | 54814921 | 54815037 | - | 117 | 5S_rRNA | 0 | 0 | 0 | 0 | 0 | 0 |
| ENSDARG000 | 6;6;6;6;6 | 237924:24201 | 241170:24231 | -;-;-;-;- | 4393 | NA | 46 | 125 | 78 | 64 | 172 | 199 |
| ENSDARG000 | 6;6;6;6 | 808649:82042 | 808844:82092 | -;-;-;- | 972 | NA | 0 | 0 | 0 | 1 | 0 | 0 |
| ENSDARG000 | 6;6;6;6 | 1262612:1270 | 1262647:1271 | -;-;-;- | 1725 | NA | 0 | 0 | 0 | 0 | 0 | 0 |
| ENSDARG000 | 6:6:6:6:6:6:6: | 2491733:2492 | 2492021:2492 | +:+:+:+:+:+:+: | 3225 | NA | 44 | 21 | 38 | 59 | 41 | 48 |
| ENSDARG000 | 6:6:6:6:6:6:6: | 2519973:2519 | 2520216:2520 | +:+:+:+:+:+:+: | 4983 | NA | 328 | 116 | 191 | 180 | 214 | 215 |
| ENSDARG000 | 6;6;6 | 5938477:5960 | 5938759:5961 | +;+;+ | 802 | NA | 0 | 0 | 0 | 0 | 0 | 0 |
| ENSDARG000 | 6;6;6;6;6;6 | 20202009:202 | 20202085:202 | -;-;-;-;-;- | 418 | NA | 0 | 0 | 4 | 0 | 0 | 0 |
| ENSDARG000 | 6;6;6 | 20284289:202 | 20284466:202 | -;-;- | 1088 | NA | 0 | 0 | 0 | 0 | 0 | 0 |
| ENSDARG000 | 6;6;6;6 | 22462474:224 | 22462561:224 | +;+;+;+ | 2514 | NA | 18 | 18 | 38 | 30 | 42 | 22 |
| ENSDARG000 | 6;6;6;6;6;6 | 23620640:236 | 23620675:236 | +;+;+;+;+;+ | 1966 | NA | 23 | 8 | 4 | 8 | 10 | 49 |
| ENSDARG000 | 6:6:6:6:6:6:6: | 24310761:243 | 24311748:243 | +:+:+:+:+:+:+: | 2686 | NA | 8 | 25 | 55 | 44 | 61 | 30 |
| ENSDARG000 | 6;6;6;6;6 | 32521642:325 | 32521874:325 | +;+;+;+;+ | 654 | NA | 0 | 0 | 0 | 0 | 0 | 0 |
| ENSDARG000 | 6;6;6 | 32573996:325 | 32574024:325 | -;-;- | 241 | NA | 0 | 0 | 0 | 0 | 0 | 0 |
| ENSDARG000 | 6;6;6 | 34442409:344 | 34442491:344 | +;+;+ | 498 | NA | 0 | 0 | 0 | 0 | 0 | 6 |
| ENSDARG000 | 6;6;6;6 | 34487776:344 | 34487947:344 | +;+;+;+ | 1128 | NA | 0 | 0 | 0 | 0 | 0 | 6 |
| ENSDARG000 | 6;6;6 | 38518219:385 | 38518282:385 | +;+;+ | 333 | NA | 0 | 0 | 0 | 0 | 0 | 0 |
| ENSDARG000 | 6;6;6;6;6;6 | 40475711:404 | 40475943:404 | +;+;+;+;+;+ | 2702 | NA | 4 | 4 | 0 | 0 | 0 | 2 |
| ENSDARG000 | 6;6;6;6 | 41909373:419 | 41909732:419 | +;+;+;+ | 684 | NA | 0 | 0 | 0 | 0 | 0 | 0 |
| ENSDARG000 | 2:2:2:2:2:2:2: | 6042039:6042 | 6042272:6042 | -:-:-:-:-:-:-:-: | 1146 | czib | 395 | 531 | 592 | 384 | 488 | 450 |
| ENSDARG000 | 2;2 | 11976261:119 | 11978455:119 | -;- | 2795 | NA | 0 | 0 | 0 | 0 | 2 | 0 |
| ENSDARG000 | 2;2;2;2 | 49799470:497 | 49799687:497 | +;+;+;+ | 1157 | NA | 0 | 0 | 0 | 0 | 0 | 0 |
| ENSDARG000 | 2:2:2:2:2:2:2: | 11946554:119 | 11946685:119 | +:+:+:+:+:+:+: | 2749 | cdc73 | 940 | 950 | 1251 | 869 | 1072 | 1112 |
| ENSDARG000 | 2:2:2:2:2:2:2: | 37775655:377 | 37775991:377 | -:-:-:-:-:-:-:-: | 4201 | NFATC4 | 126 | 22 | 164 | 185 | 164 | 10 |
| ENSDARG000 | 2:2:2:2:2:2:2: | 44049081:440 | 44049305:440 | -:-:-:-:-:-:-:-: | 1796 | cadm3 | 14 | 8 | 6 | 2 | 14 | 12 |
| ENSDARG000 | 2:2:2:2:2:2:2: | 38554260:385 | 38554305:385 | +:+:+:+:+:+:+: | 2600 | CDH24 | 6 | 6 | 6 | 0 | 4 | 4 |
| ENSDARG000 | 2:2:2:2:2:2:2: | 23295415:232 | 23297078:232 | -:-:-:-:-:-:-:-: | 3364 | NIBAN1 | 62 | 32 | 100 | 82 | 173 | 93 |
| ENSDARG000 | 2;2;2;2 | 27386617:273 | 27387052:273 | +;+;+;+ | 1732 | si:ch73-382f3 | 40 | 61 | 76 | 42 | 90 | 44 |
| ENSDARG000 | 2;2;2;2;2;2 | 45643122:456 | 45643244:456 | +;+;+;+;+;+ | 1336 | NA | 0 | 2 | 0 | 0 | 4 | 0 |
| ENSDARG000 | 2;2 | 26286465:262 | 26286743:262 | +;+ | 760 | NA | 0 | 2 | 0 | 8 | 0 | 1 |
| ENSDARG000 | 2:2:2:2:2:2:2: | 27394798:273 | 27394883:273 | +:+:+:+:+:+:+: | 1603 | selenop2 | 0 | 2 | 4 | 0 | 0 | 0 |
| ENSDARG000 | 2;2;2;2;2;2;2 | 55833425:558 | 55834022:558 | -;-;-;-;-;-;- | 1489 | rx2 | 0 | 0 | 0 | 0 | 0 | 0 |
| ENSDARG000 | 2:2:2:2:2:2:2: | 31830183:318 | 31830631:318 | -;-;-;-;-;-;-;- | 1136 | NA | 491 | 607 | 798 | 322 | 484 | 618 |
| ENSDARG000 | 2:2:2:2:2:2:2: | 49805892:498 | 49805955:498 | +:+:+:+:+:+:+: | 2686 | wdr48b | 995 | 470 | 486 | 299 | 489 | 454 |
| ENSDARG000 | 2:2:2:2:2:2:2: | 2181709:2181 | 2181846:2181 | +:+:+:+:+:+:+: | 2314 | NA | 6 | 4 | 6 | 0 | 8 | 0 |
| ENSDARG000 | 2:2:2:2:2:2:2: | 16885285:168 | 16886168:168 | -:-:-:-:-:-:-:-: | 3709 | CLCN2 | 0 | 0 | 2 | 2 | 0 | 0 |
| ENSDARG000 | 2:2:2:2:2:2:2: | 55914699:559 | 55915173:559 | +:+:+:+:+:+:+: | 10170 | ATCAY | 2 | 0 | 2 | 0 | 0 | 0 |
| ENSDARG000 | 2:2:2:2:2:2:2: | 36664173:366 | 36664902:366 | -:-:-:-:-:-:-:-: | 2879 | ccnl1b | 1017 | 810 | 1153 | 558 | 1289 | 928 |
| ENSDARG000 | 2:2:2:2:2:2:2: | 4207209:4207 | 4207239:4207 | +:+:+:+:+:+:+: | 2579 | gata6 | 0 | 0 | 2 | 0 | 0 | 0 |
| ENSDARG000 | 2:2:2:2:2:2:2: | 45655294:456 | 45656489:456 | -:-:-:-:-:-:-:-: | 2466 | prpf38b | 826 | 952 | 1375 | 666 | 1179 | 667 |
| ENSDARG000 | 2:2:2:2:2:2:2: | 10183640:101 | 10187119:101 | -:-:-:-:-:-:-:-: | 4421 | dmbx1a | 0 | 0 | 0 | 6 | 4 | 0 |
| ENSDARG000 | 2;2;2;2;2;2 | 24352497:243 | 24352890:243 | +;+;+;+;+;+ | 1348 | NA | 0 | 2 | 0 | 0 | 0 | 0 |
| ENSDARG000 | 2:2:2:2:2:2:2: | 55982300:559 | 55983100:559 | +:+:+:+:+:+:+: | 2644 | nmrk2 | 0 | 0 | 0 | 0 | 0 | 0 |
| ENSDARG000 | 2:2:2:2:2:2:2: | 35240485:352 | 35241090:352 | +:+:+:+:+:+:+: | 5335 | tnr | 2 | 0 | 0 | 0 | 6 | 6 |
| ENSDARG000 | 2;2;2;2;2;2;2 | 37804235:378 | 37804433:378 | +;+;+;+;+;+;+ | 1867 | ltb4r2b | 9 | 2 | 0 | 19 | 5 | 0 |
| ENSDARG000 | 2:2:2:2:2:2:2: | 55890673:558 | 55893849:558 | -:-:-:-:-:-:-:-: | 4451 | cplx4c | 0 | 0 | 0 | 0 | 0 | 0 |
| ENSDARG000 | 2:2:2:2:2:2:2: | 26288301:262 | 26288383:262 | +:+:+:+:+:+:+: | 4314 | ptbp1a | 3626 | 3060 | 4283 | 2347 | 5032 | 3386 |
| ENSDARG000 | 2:2:2:2:2:2:2: | 2223837:2223 | 2224012:2224 | +:+:+:+:+:+:+: | 1108 | tmie | 0 | 0 | 0 | 0 | 0 | 0 |
| ENSDARG000 | 2;2;2;2;2 | 24365452:243 | 24366135:243 | -;-;-;-;- | 2395 | plvapa | 36 | 34 | 118 | 36 | 78 | 42 |
| ENSDARG000 | 2:2:2:2:2:2:2: | 32743807:327 | 32744030:327 | +:+:+:+:+:+:+: | 2122 | klhl18 | 143 | 280 | 178 | 138 | 221 | 155 |
| ENSDARG000 | 2:2:2:2:2:2:2: | 27403300:274 | 27403497:274 | +:+:+:+:+:+:+: | 1330 | elovl8a | 0 | 0 | 0 | 0 | 0 | 0 |
| ENSDARG000 | 2:2:2:2:2:2:2: | 38742338:387 | 38742495:387 | +:+:+:+:+:+:+: | 7921 | carmil3 | 107 | 4 | 20 | 16 | 28 | 10 |
| ENSDARG000 | 2;2;2;2;2;2 | 37815687:378 | 37815789:378 | +;+;+;+;+;+ | 1983 | sdr39u1 | 261 | 330 | 299 | 173 | 341 | 251 |

| | | | | | | | | | | | | |
|---|---|---|---|---|---|---|---|---|---|---|---|---|
| ENSDARG000 | 2;2;2;2;2;2;2; | 44192652:441 | 44193031:441 | -;-;-;-;-;-;-;-; | 1032 | sdhc | 658 | 804 | 1031 | 1088 | 768 | 866 |
| ENSDARG000 | 2;2;2;2;2;2;2; | 49860722:498 | 49861004:498 | +;+;+;+;+;+;+; | 6136 | COA1 | 204 | 99 | 188 | 110 | 142 | 98 |
| ENSDARG000 | 2;2;2;2;2;2;2; | 24374305:243 | 24374415:243 | +;+;+;+;+;+;+; | 3106 | nr2f6a | 393 | 435 | 613 | 405 | 872 | 381 |
| ENSDARG000 | 2;2;2 | 49855746:498 | 49855808:498 | +;+;+ | 820 | NA | 2 | 0 | 0 | 8 | 0 | 2 |
| ENSDARG000 | 2;2;2;2;2;2;2; | 49844630:498 | 49844998:498 | -;-;-;-;-;-;-;-; | 1227 | blvra | 315 | 321 | 126 | 214 | 450 | 277 |
| ENSDARG000 | 2;2;2;2;2;2;2 | 6051916:6054 | 6052987:6054 | -;-;-;-;-;-;- | 1822 | uck2b | 160 | 404 | 454 | 162 | 303 | 238 |
| ENSDARG000 | 2;2;2;2;2;2;2; | 12019341:120 | 12019507:120 | -;-;-;-;-;-;-;-; | 3671 | NA | 0 | 0 | 2 | 0 | 2 | 0 |
| ENSDARG000 | 2;2 | 27484559:274 | 27484812:274 | -;- | 442 | NA | 0 | 4 | 10 | 6 | 6 | 7 |
| ENSDARG000 | 2;2;2 | 32758663:327 | 32758771:327 | +;+;+ | 791 | NA | 0 | 0 | 0 | 0 | 2 | 0 |
| ENSDARG000 | 2;2;2;2;2;2;2; | 27491066:275 | 27491599:275 | +;+;+;+;+;+;+; | 6228 | zswim5 | 874 | 950 | 1217 | 817 | 2016 | 772 |
| ENSDARG000 | 2;2;2 | 31833997:318 | 31834358:318 | +;+;+ | 1437 | epdr1 | 287 | 1093 | 715 | 722 | 1046 | 1025 |
| ENSDARG000 | 2;2;2;2;2;2;2; | 20604775:206 | 20605323:206 | +;+;+;+;+;+;+; | 3963 | OLFML2B | 38 | 180 | 217 | 110 | 99 | 123 |
| ENSDARG000 | 2;2;2;2;2;2;2; | 2290148:2290 | 2290855:2290 | -;-;-;-;-;-;-;-; | 2114 | pth1r | 3 | 4 | 50 | 2 | 98 | 4 |
| ENSDARG000 | 2;2;2;2 | 10280645:102 | 10280815:102 | +;+;+;+ | 1253 | gadd45aa | 382 | 286 | 315 | 377 | 374 | 422 |
| ENSDARG000 | 2;2;2;2;2;2;2; | 8586415:8586 | 8587613:8587 | -;-;-;-;-;-;-;-; | 2465 | si:ch211-71m | 265 | 388 | 134 | 180 | 78 | 230 |
| ENSDARG000 | 2;2;2;2;2;2;2; | 23351454:233 | 23351495:233 | +;+;+;+;+;+;+; | 2097 | rnf2 | 1060 | 970 | 1204 | 990 | 938 | 1010 |
| ENSDARG000 | 2;2;2;2;2;2;2; | 31838442:318 | 31838538:318 | +;+;+;+;+;+;+; | 2665 | stard3nl | 691 | 521 | 708 | 675 | 682 | 760 |
| ENSDARG000 | 2;2;2;2;2;2 | 30531726:305 | 30531981:305 | +;+;+;+;+;+ | 1457 | ANKRD33B | 0 | 1 | 3 | 0 | 4 | 0 |
| ENSDARG000 | 2;2 | 20632221:206 | 20632258:206 | +;+ | 333 | NA | 0 | 0 | 0 | 0 | 0 | 0 |
| ENSDARG000 | 2;2;2;2;2;2;2; | 45665493:456 | 45666254:456 | -;-;-;-;-;-;-;-; | 2099 | eeig2a | 54 | 60 | 114 | 70 | 81 | 47 |
| ENSDARG000 | 2;2;2;2;2;2;2; | 24400725:244 | 24403011:244 | -;-;-;-;-;-;-;-; | 5600 | NA | 0 | 0 | 2 | 0 | 10 | 0 |
| ENSDARG000 | 2;2;2;2;2;2;2; | 24386453:243 | 24386687:243 | -;-;-;-;-;-;-;-; | 2761 | zgc:154006 | 2 | 3 | 2 | 0 | 8 | 6 |
| ENSDARG000 | 2;2;2;2;2 | 10287133:102 | 10288560:102 | -;-;-;-;- | 2061 | gng12a | 420 | 443 | 544 | 502 | 302 | 399 |
| ENSDARG000 | 2;2;2;2;2;2;2; | 32761130:327 | 32761503:327 | -;-;-;-;-;-;-;-; | 2541 | bfsp2 | 2 | 0 | 6 | 26 | 36 | 18 |
| ENSDARG000 | 2;2;2 | 38892631:389 | 38892812:389 | +;+;+ | 1346 | NA | 0 | 0 | 0 | 0 | 0 | 0 |
| ENSDARG000 | 2;2;2;2 | 26303627:264 | 26304478:264 | +;+;+;+ | 2654 | efna2a | 4 | 0 | 0 | 0 | 18 | 0 |
| ENSDARG000 | 2;2;2 | 10386479:103 | 10386862:103 | +;+;+ | 701 | NA | 0 | 0 | 0 | 0 | 0 | 0 |
| ENSDARG000 | 2;2;2;2;2;2;2; | 10396579:104 | 10397171:104 | -;-;-;-;-;-;-;-; | 2735 | scinlb | 760 | 1482 | 1882 | 642 | 2247 | 1648 |
| ENSDARG000 | 2;2;2;2;2;2 | 26434936:264 | 26435756:264 | -;-;-;-;-;- | 949 | NA | 0 | 26 | 12 | 0 | 0 | 14 |
| ENSDARG000 | 2;2;2;2;2 | 17055069:170 | 17055123:170 | +;+;+;+;+ | 576 | thpo | 0 | 0 | 0 | 0 | 8 | 0 |
| ENSDARG000 | 2;2;2;2;2 | 38924975:389 | 38925284:389 | +;+;+;+;+ | 1569 | rem2 | 0 | 0 | 2 | 0 | 2 | 0 |
| ENSDARG000 | 2;2 | 4281682:4288 | 4281967:4288 | +;+ | 758 | NA | 6 | 0 | 0 | 0 | 0 | 0 |
| ENSDARG000 | 2;2;2;2;2;2;2; | 10342177:103 | 10343075:103 | -;-;-;-;-;-;-;-; | 2545 | wls | 508 | 244 | 207 | 307 | 242 | 261 |
| ENSDARG000 | 2;2;2;2 | 35482156:354 | 35482245:354 | +;+;+;+ | 1031 | NA | 25 | 17 | 27 | 26 | 38 | 18 |
| ENSDARG000 | 2;2;2 | 31856242:318 | 31856582:318 | -;-;- | 734 | NA | 0 | 2 | 10 | 0 | 11 | 5 |
| ENSDARG000 | 2;2;2;2;2;2;2; | 49884696:498 | 49884807:498 | -;-;-;-;-;-;-;-; | 4716 | hecw1b | 2 | 6 | 20 | 0 | 22 | 2 |
| ENSDARG000 | 2;2;2;2;2;2;2; | 17096161:170 | 17097796:170 | -;-;-;-;-;-;-;-; | 4123 | pif1 | 14 | 70 | 122 | 26 | 32 | 30 |
| ENSDARG000 | 2;2;2;2;2;2;2; | 23379401:233 | 23380379:233 | -;-;-;-;-;-;-;-; | 4000 | ivns1abpb | 3607 | 3246 | 4249 | 5040 | 5693 | 3498 |
| ENSDARG000 | 2;2;2;2;2;2;2; | 37836821:378 | 37837280:378 | +;+;+;+;+;+;+; | 2629 | parp2 | 561 | 754 | 568 | 612 | 658 | 410 |
| ENSDARG000 | 2;2;2;2;2;2;2; | 12104389:121 | 12110276:121 | -;-;-;-;-;-;-;-; | 9150 | GPR158 | 0 | 0 | 2 | 0 | 0 | 0 |
| ENSDARG000 | 2;2 | 8634240:8634 | 8634552:8635 | -;- | 700 | NA | 0 | 0 | 0 | 0 | 0 | 0 |
| ENSDARG000 | 2;2;2;2;2;2;2; | 8617816:8617 | 8617854:8617 | -;-;-;-;-;-;-;-; | 887 | NA | 0 | 0 | 0 | 8 | 0 | 0 |
| ENSDARG000 | 2;2;2;2;2;2;2; | 15300088:153 | 15300385:153 | -;-;-;-;-;-;-;-; | 1618 | tecrl2b | 0 | 0 | 0 | 0 | 0 | 0 |
| ENSDARG000 | 2;2;2;2;2;2;2; | 37827567:378 | 37828106:378 | -;-;-;-;-;-;-;-; | 1844 | NA | 331 | 217 | 293 | 244 | 289 | 249 |
| ENSDARG000 | 2;2 | 36691771:366 | 36692316:366 | +;+ | 1163 | PTX3 | 2 | 0 | 9 | 0 | 15 | 4 |
| ENSDARG000 | 2;2 | 35512002:355 | 35512076:355 | +;+ | 461 | NA | 1 | 0 | 0 | 0 | 0 | 0 |
| ENSDARG000 | 2;2;2;2;2;2;2; | 35509418:355 | 35510173:355 | -;-;-;-;-;-;-;-; | 4270 | tnn | 58 | 0 | 7 | 2 | 10 | 6 |
| ENSDARG000 | 2;2;2;2;2;2;2; | 20699481:207 | 20700015:207 | -;-;-;-;-;-;-;-; | 1838 | NA | 0 | 2 | 0 | 0 | 4 | 4 |
| ENSDARG000 | 2;2;2;2;2;2;2; | 32780138:327 | 32780807:327 | +;+;+;+;+;+;+; | 1662 | zgc:136930 | 18 | 0 | 18 | 102 | 60 | 24 |
| ENSDARG000 | 2;2;2;2;2;2;2; | 20658259:206 | 20661407:206 | -;-;-;-;-;-;-;-; | 4439 | dusp12 | 72 | 100 | 108 | 94 | 110 | 126 |
| ENSDARG000 | 2;2;2;2;2;2;2; | 30554256:305 | 30556532:305 | -;-;-;-;-;-;-;-; | 6215 | ctnnd2b | 14 | 6 | 84 | 38 | 54 | 120 |
| ENSDARG000 | 2;2;2;2;2;2;2; | 38985489:389 | 38987223:389 | -;-;-;-;-;-;-;- | 2362 | si:ch211-119 | 0 | 12 | 0 | 0 | 8 | 8 |
| ENSDARG000 | 2;2 | 17181777:171 | 17182488:171 | +;+ | 1602 | ptger4c | 0 | 0 | 0 | 0 | 0 | 0 |
| ENSDARG000 | 2;2;2;2;2;2;2 | 8644422:8644 | 8644601:8644 | -;-;-;-;-;-;- | 1077 | si:ch211-71m | 8 | 0 | 0 | 0 | 0 | 0 |
| ENSDARG000 | 2;2;2;2;2;2;2; | 42382472:423 | 42385634:423 | -;-;-;-;-;-;-;-; | 5788 | slco5a1a | 590 | 300 | 446 | 424 | 461 | 402 |
| ENSDARG000 | 2;2;2;2;2;2;2; | 27556168:275 | 27556367:275 | -;-;-;-;-;-;-;-; | 1292 | urod | 350 | 407 | 448 | 186 | 436 | 345 |
| ENSDARG000 | 2;2;2 | 32796873:327 | 32796987:327 | +;+;+ | 1856 | ccr9a | 170 | 266 | 190 | 128 | 993 | 468 |
| ENSDARG000 | 2;2;2;2;2;2;2; | 36721357:367 | 36721511:367 | +;+;+;+;+;+;+; | 1251 | pdcd10b | 578 | 729 | 592 | 635 | 576 | 559 |
| ENSDARG000 | 2;2;2;2;2;2;2; | 26463114:264 | 26463916:264 | -;-;-;-;-;-;-;-; | 1922 | rabggtb | 641 | 577 | 649 | 324 | 516 | 462 |
| ENSDARG000 | 2;2;2;2;2;2;2; | 36701322:367 | 36701754:367 | +;+;+;+;+;+;+; | 2488 | golim4b | 42 | 36 | 73 | 108 | 58 | 101 |
| ENSDARG000 | 2;2;2 | 23406784:234 | 23407030:234 | -;-;- | 786 | NA | 1 | 0 | 2 | 4 | 1 | 0 |
| ENSDARG000 | 2 | 31861353 | 31861398 | - | 46 | NA | 4 | 0 | 0 | 10 | 0 | 2 |
| ENSDARG000 | 2 | 31861919 | 31861973 | - | 55 | NA | 0 | 0 | 0 | 0 | 0 | 0 |
| ENSDARG000 | 2 | 31862569 | 31862617 | - | 49 | NA | 0 | 0 | 0 | 0 | 0 | 0 |
| ENSDARG000 | 2;2;2 | 23438411:234 | 23438660:234 | -;-;- | 789 | NA | 5 | 0 | 4 | 1 | 7 | 0 |
| ENSDARG000 | 2 | 31863717 | 31863765 | - | 49 | NA | 0 | 1 | 0 | 0 | 0 | 0 |
| ENSDARG000 | 2;2;2;2 | 17200320:172 | 17200427:172 | -;-;-;- | 566 | NA | 2 | 0 | 0 | 0 | 0 | 0 |
| ENSDARG000 | 2;2;2;2;2;2;2; | 24410494:244 | 24414238:244 | -;-;-;-;-;-;-;-; | 6314 | kcnn1a | 0 | 0 | 0 | 0 | 0 | 4 |
| ENSDARG000 | 2;2;2;2;2;2;2; | 56012016:560 | 56012784:560 | +;+;+;+;+;+;+; | 5255 | loxl5b | 0 | 0 | 0 | 4 | 2 | 0 |
| ENSDARG000 | 2;2;2;2;2;2;2; | 12255568:122 | 12255934:122 | +;+;+;+;+;+;+; | 2301 | prtfdc1b | 110 | 301 | 490 | 217 | 331 | 471 |
| ENSDARG000 | 2 | 31864066 | 31864111 | - | 46 | NA | 0 | 0 | 0 | 4 | 0 | 0 |
| ENSDARG000 | 2 | 31864399 | 31864450 | - | 52 | NA | 0 | 0 | 2 | 0 | 0 | 0 |
| ENSDARG000 | 2 | 31865457 | 31865501 | - | 45 | NA | 0 | 0 | 0 | 0 | 0 | 4 |
| ENSDARG000 | 2;2 | 31866722:318 | 31867022:318 | -;- | 344 | NA | 0 | 1 | 2 | 8 | 14 | 1 |
| ENSDARG000 | 2;2 | 31869121:318 | 31869418:318 | -;- | 350 | NA | 0 | 10 | 0 | 13 | 1 | 2 |
| ENSDARG000 | 2;2 | 31873021:318 | 31873316:318 | -;- | 339 | NA | 0 | 0 | 0 | 0 | 0 | 0 |
| ENSDARG000 | 2;2 | 31874398:318 | 31874701:318 | -;- | 347 | NA | 0 | 0 | 0 | 0 | 0 | 0 |
| ENSDARG000 | 2 | 31877471 | 31877825 | - | 355 | NA | 0 | 0 | 0 | 0 | 0 | 0 |
| ENSDARG000 | 2;2;2;2;2;2 | 56060400:560 | 56066445:560 | -;-;-;-;-;- | 7925 | NA | 544 | 243 | 489 | 728 | 176 | 250 |
| ENSDARG000 | 2;2;2 | 35591423:355 | 35591523:355 | +;+;+ | 728 | mrps14 | 302 | 416 | 357 | 230 | 305 | 344 |
| ENSDARG000 | 2;2;2;2;2;2;2; | 32812402:328 | 32812606:328 | -;-;-;-;-;-;-;-; | 3153 | PRPF4B | 549 | 406 | 494 | 223 | 556 | 540 |
| ENSDARG000 | 2;2;2;2;2;2;2; | 4303125:4303 | 4303761:4303 | +;+;+;+;+;+;+; | 1911 | cables1 | 122 | 62 | 116 | 37 | 132 | 74 |
| ENSDARG000 | 2;2;2 | 23475723:234 | 23475969:234 | -;-;- | 780 | NA | 60 | 0 | 9 | 52 | 34 | 0 |
| ENSDARG000 | 2;2;2;2;2;2;2 | 22480074:224 | 22480826:224 | -;-;-;-;-;-;- | 1609 | gtf2b | 679 | 789 | 720 | 632 | 759 | 640 |
| ENSDARG000 | 2;2 | 49984159:499 | 49985246:499 | -;- | 1113 | ccr12a | 22 | 66 | 92 | 70 | 280 | 222 |
| ENSDARG000 | 2;2;2 | 23511594:235 | 23511840:235 | -;-;- | 780 | NA | 0 | 0 | 1 | 0 | 0 | 0 |
| ENSDARG000 | 2;2;2;2;2;2;2; | 15341689:153 | 15344311:153 | -;-;-;-;-;-;-;-; | 3457 | OLFM3 | 8 | 0 | 0 | 0 | 2 | 0 |
| ENSDARG000 | 2;2;2;2;2;2 | 35595454:355 | 35595596:355 | +;+;+;+;+;+ | 1072 | cacybp | 572 | 775 | 844 | 662 | 680 | 812 |
| ENSDARG000 | 2;2;2;2;2;2;2; | 8663108:8666 | 8663873:8666 | -;-;-;-;-;-;-;-; | 2574 | miga1 | 181 | 191 | 298 | 120 | 268 | 135 |
| ENSDARG000 | 2;2;2;2;2;2;2; | 17266759:172 | 17267749:172 | -;-;-;-;-;-;-;-; | 3256 | st3gal3b | 1636 | 1040 | 1281 | 892 | 1679 | 1443 |
| ENSDARG000 | 2;2;2;2;2;2;2; | 111919:11214 | 112007:11226 | +;+;+;+;+;+;+; | 1757 | fggy | 115 | 104 | 199 | 177 | 168 | 115 |
| ENSDARG000 | 2;2;2;2;2;2;2; | 8649293:8649 | 8649448:8649 | +;+;+;+;+;+;+; | 4976 | fubp1 | 6074 | 5221 | 9141 | 4850 | 6783 | 6032 |
| ENSDARG000 | 2;2;2;2;2;2;2; | 38997357:389 | 38997731:389 | -;-;-;-;-;-;-;-; | 4194 | copb2 | 4517 | 4042 | 6544 | 3377 | 5285 | 3039 |
| ENSDARG000 | 2 | 56125174 | 56131312 | - | 6139 | jund | 1886 | 470 | 938 | 1093 | 573 | 1066 |
| ENSDARG000 | 2;2 | 31880194:318 | 31880504:318 | -;- | 354 | NA | 0 | 0 | 0 | 0 | 0 | 0 |
| ENSDARG000 | 2;2;2;2 | 118225:11854 | 119215:11921 | -;-;-;- | 1151 | znfl2a | 12 | 20 | 38 | 12 | 74 | 6 |
| ENSDARG000 | 2;2;2 | 36815484:368 | 36818132:368 | -;-;- | 3043 | SLITRK3 | 0 | 0 | 0 | 0 | 0 | 0 |
| ENSDARG000 | 2;2 | 31882411:318 | 31882711:318 | -;- | 338 | NA | 0 | 0 | 0 | 0 | 0 | 0 |
| ENSDARG000 | 2;2;2 | 23537868:235 | 23537974:235 | -;-;- | 646 | NA | 0 | 0 | 0 | 0 | 0 | 0 |
| ENSDARG000 | 2;2 | 31885581:318 | 31885903:318 | -;- | 366 | NA | 0 | 0 | 0 | 0 | 0 | 0 |
| ENSDARG000 | 2;2 | 31888975:318 | 31889276:318 | -;- | 345 | NA | 0 | 0 | 0 | 0 | 0 | 0 |
| ENSDARG000 | 2;2;2 | 22495274:224 | 22495381:225 | +;+;+ | 5389 | NA | 76 | 100 | 77 | 77 | 112 | 136 |
| ENSDARG000 | 2;2;2;2 | 124433:12524 | 125059:12537 | -;-;-;- | 1299 | igfbp1b | 0 | 0 | 2 | 0 | 0 | 0 |
| ENSDARG000 | 2;2;2;2;2;2;2; | 27577470:275 | 27579294:275 | -;-;-;-;-;-;-;-; | 3300 | lyn | 74 | 139 | 98 | 85 | 194 | 238 |
| ENSDARG000 | 2;2;2;2;2;2;2; | 35603637:356 | 35603948:356 | +;+;+;+;+;+;+; | 3466 | plk3 | 4400 | 751 | 2516 | 2511 | 1774 | 1841 |
| ENSDARG000 | 2;2;2 | 23540185:235 | 23540273:235 | -;-;- | 639 | NA | 0 | 0 | 0 | 0 | 0 | 0 |
| ENSDARG000 | 2;2;2;2 | 45918939:459 | 45919309:459 | +;+;+;+ | 1031 | NA | 0 | 0 | 0 | 0 | 0 | 0 |
| ENSDARG000 | 2;2;2;2;2;2;2; | 56139941:561 | 56140119:561 | +;+;+;+;+;+;+; | 5806 | pgpep1 | 248 | 206 | 283 | 114 | 330 | 174 |
| ENSDARG000 | 2;2;2;2;2;2;2; | 20725684:207 | 20725811:207 | +;+;+;+;+;+;+; | 1867 | rxrga | 251 | 626 | 336 | 252 | 537 | 216 |
| ENSDARG000 | 2;2;2;2;2 | 4490604:4490 | 4491356:4491 | -;-;-;-;- | 1120 | adipoqb | 0 | 0 | 0 | 0 | 0 | 0 |
| ENSDARG000 | 2;2;2 | 23542636:235 | 23542882:235 | -;-;- | 780 | NA | 2 | 0 | 3 | 6 | 1 | 0 |
| ENSDARG000 | 2;2;2;2 | 39016066:390 | 39016755:390 | -;-;-;- | 1086 | rbp2b | 0 | 0 | 8 | 0 | 0 | 0 |
| ENSDARG000 | 2;2;2;2;2;2;2; | 12302527:123 | 12302861:123 | +;+;+;+;+;+;+; | 7781 | ARHGAP21 | 1458 | 1094 | 1274 | 1431 | 1069 | 1860 |
| ENSDARG000 | 2;2;2;2;2;2;2; | 42419841:424 | 42422432:424 | -;-;-;-;-;-;-;-; | 5460 | ESYT2 | 1006 | 947 | 1329 | 1478 | 943 | 1128 |
| ENSDARG000 | 2;2;2;2;2;2;2; | 6092058:6092 | 6092406:6092 | -;-;-;-;-;-;-;-; | 1411 | zte38 | 0 | 2 | 0 | 14 | 2 | 0 |
| ENSDARG000 | 2;2;2;2;2;2;2; | 44203075:442 | 44203857:442 | -;-;-;-;-;-;-;-; | 3907 | atp1a2a | 4 | 7 | 3 | 16 | 8 | 4 |
| ENSDARG000 | 2;2;2;2;2;2;2; | 136346:13634 | 136720:13672 | -;-;-;-;-;-;-;-; | 2539 | adcy1b | 4 | 16 | 0 | 4 | 4 | 0 |
| ENSDARG000 | 2;2;2;2;2;2;2; | 10467799:104 | 10468643:104 | -;-;-;-;-;-;-;-; | 5145 | ccdc18 | 1570 | 2415 | 2254 | 5068 | 4410 | 2892 |
| ENSDARG000 | 2;2;2;2;2;2;2; | 6067371:6067 | 6067398:6067 | +;+;+;+;+;+;+; | 2330 | aldh9a1b | 353 | 421 | 439 | 496 | 496 | 390 |
| ENSDARG000 | 2;2 | 24495804:245 | 24497777:245 | -;- | 3581 | NA | 2 | 3 | 4 | 42 | 2 | 9 |
| ENSDARG000 | 2;2 | 24487320:244 | 24487971:244 | -;- | 932 | insl3 | 0 | 1 | 0 | 4 | 2 | 0 |
| ENSDARG000 | 2;2;2;2;2;2;2; | 4729155:4733 | 4730486:4733 | -;-;-;-;-;-;-;-; | 5514 | NA | 107 | 58 | 196 | 18 | 74 | 68 |
| ENSDARG000 | 2;2;2;2;2;2 | 24484275:244 | 24484394:244 | +;+;+;+;+;+ | 1204 | NA | 0 | 0 | 0 | 0 | 0 | 0 |
| ENSDARG000 | 2;2;2;2;2;2;2; | 49987088:499 | 49988785:499 | -;-;-;-;-;-;-;-; | 4184 | si:ch211-106 | 4 | 14 | 8 | 0 | 33 | 22 |
| ENSDARG000 | 2;2;2;2;2;2;2; | 56168619:561 | 56169773:561 | +;+;+;+;+;+;+; | 8615 | upf1 | 2685 | 2270 | 3885 | 2368 | 3070 | 2734 |
| ENSDARG000 | 2;2;2;2;2;2;2; | 27611956:276 | 27614093:276 | -;-;-;-;-;-;-;-; | 4511 | NA | 206 | 204 | 302 | 136 | 318 | 245 |
| ENSDARG000 | 2;2;2;2;2;2 | 22506364:225 | 22507321:225 | -;-;-;-;-;- | 4411 | znf644a | 388 | 321 | 625 | 312 | 486 | 520 |
| ENSDARG000 | 2;2;2 | 23568907:235 | 23569013:235 | -;-;- | 646 | NA | 0 | 0 | 1 | 0 | 0 | 0 |
| ENSDARG000 | 2;2;2 | 23571230:235 | 23571318:235 | -;-;- | 639 | NA | 0 | 0 | 0 | 0 | 0 | 0 |
| ENSDARG000 | 2;2;2;2;2;2;2; | 30721070:307 | 30721207:307 | +;+;+;+;+;+;+; | 2820 | si:dkev-94e7. | 0 | 2 | 0 | 2 | 0 | 2 |
| ENSDARG000 | 2;2;2 | 30713868:307 | 30714779:307 | -;-;- | 1199 | GPR141 | 0 | 2 | 0 | 0 | 0 | 13 |
| ENSDARG000 | 2;2;2;2 | 39021282:390 | 39021388:390 | +;+;+;+ | 448 | rbp1.2 | 0 | 0 | 0 | 0 | 0 | 0 |
| ENSDARG000 | 2;2;2 | 23573682:235 | 23573928:235 | -;-;- | 780 | NA | 0 | 0 | 2 | 3 | 0 | 0 |
| ENSDARG000 | 2;2;2;2;2 | 56339058:563 | 56339221:563 | -;-;-;-;- | 496 | uba52 | 10846 | 11522 | 17301 | 14061 | 11106 | 14010 |
| ENSDARG000 | 2;2;2;2;2;2;2; | 32826235:328 | 32826499:328 | +;+;+;+;+;+;+; | 2414 | NA | 154 | 89 | 104 | 71 | 102 | 108 |
| ENSDARG000 | 2;2;2;2;2;2;2; | 20793982:207 | 20794022:207 | +;+;+;+;+;+;+; | 1999 | prg4a | 30 | 26 | 19 | 67 | 39 | 34 |
| ENSDARG000 | 2;2 | 4812694:4813 | 4812753:4814 | -;- | 402 | NA | 0 | 0 | 0 | 0 | 0 | 0 |
| ENSDARG000 | 2;2;2;2;2;2;2; | 46589798:465 | 46590155:465 | +;+;+;+;+;+;+; | 2085 | EPHB1 | 0 | 0 | 0 | 0 | 4 | 0 |
| ENSDARG000 | 2;2;2;2;2;2;2; | 26470070:264 | 26470430:264 | -;-;-;-;-;-;-;-; | 2085 | acadm | 1769 | 1255 | 1862 | 1315 | 1945 | 2165 |
| ENSDARG000 | 2;2;2;2;2;2;2; | 20811677:208 | 20811935:208 | -;-;-;-;-;-;-;-; | 7333 | TPR | 1528 | 1721 | 2668 | 1057 | 2211 | 1570 |
| ENSDARG000 | 2;2;2;2 | 39028337:390 | 39028864:390 | -;-;-;- | 968 | rbp1.1 | 0 | 0 | 0 | 0 | 0 | 0 |
| ENSDARG000 | 2;2 | 39036182:390 | 39036762:390 | +;+ | 1646 | NA | 0 | 0 | 0 | 0 | 0 | 0 |
| ENSDARG000 | 2;2;2;2;2;2;2; | 4849269:4849 | 4850005:4850 | -;-;-;-;-;-;-;-; | 2925 | tfr1a | 64 | 55 | 199 | 36 | 258 | 92 |
| ENSDARG000 | 2;2;2;2;2;2;2; | 44259482:442 | 44261109:442 | -;-;-;-;-;-;-;-; | 3838 | mpz | 32 | 2 | 4 | 0 | 132 | 52 |
| ENSDARG000 | 2;2;2;2;2;2;2; | 31891335:318 | 31892469:318 | -;-;-;-;-;-;-;-; | 3651 | amph | 2 | 0 | 4 | 0 | 4 | 0 |
| ENSDARG000 | 2;2;2 | 23599896:236 | 23600002:236 | -;-;- | 645 | NA | 0 | 4 | 0 | 0 | 3 | 2 |
| ENSDARG000 | 2;2;2;2;2;2;2; | 22531185:225 | 22531377:225 | +;+;+;+;+;+;+; | 3817 | HFM1 | 6 | 10 | 36 | 24 | 10 | 24 |
| ENSDARG000 | 2;2;2;2;2;2;2; | 42496498:424 | 42496775:424 | -;-;-;-;-;-;-;-; | 4753 | NA | 1030 | 488 | 776 | 2466 | 554 | 647 |
| ENSDARG000 | 2;2;2;2;2;2;2; | 12406713:124 | 12408108:124 | -;-;-;-;-;-;-;-; | 3893 | dpp6b | 4 | 0 | 4 | 4 | 0 | 2 |
| ENSDARG000 | 2;2 | 23602463:236 | 23602560:236 | -;- | 456 | NA | 18 | 2 | 2 | 1 | 12 | 4 |
| ENSDARG000 | 2;2;2;2;2;2;2; | 258698:25869 | 259697:25969 | +;+;+;+;+;+;+; | 7154 | PHLPP1 | 1551 | 1153 | 2167 | 1844 | 2012 | 1812 |
| ENSDARG000 | 2;2 | 30732886:307 | 30733027:307 | -;- | 778 | NA | 0 | 0 | 0 | 0 | 0 | 0 |
| ENSDARG000 | 2;2;2 | 23604654:236 | 23604900:236 | -;-;- | 786 | NA | 21 | 0 | 15 | 10 | 22 | 2 |
| ENSDARG000 | 2;2;2;2;2;2;2; | 50062165:500 | 50062728:500 | +;+;+;+;+;+;+; | 3690 | NA | 509 | 566 | 684 | 431 | 718 | 591 |
| ENSDARG000 | 2;2;2;2;2;2;2; | 36828243:368 | 36828285:368 | +;+;+;+;+;+;+; | 4110 | NRDC | 1907 | 1797 | 2255 | 1088 | 2080 | 1505 |
| ENSDARG000 | 2;2;2;2;2;2;2; | 23613040:236 | 23613330:236 | +;+;+;+;+;+;+; | 1443 | rpp40 | 148 | 151 | 202 | 58 | 151 | 176 |
| ENSDARG000 | 2;2;2;2;2;2;2; | 10603550:106 | 10608390:106 | -;-;-;-;-;-;-;-; | 6664 | mtf2 | 89 | 300 | 210 | 114 | 198 | 234 |
| ENSDARG000 | 2;2;2;2;2;2;2; | 56356652:563 | 56356858:563 | -;-;-;-;-;-;-;-; | 1221 | NA | 4 | 30 | 16 | 22 | 6 | 56 |
| ENSDARG000 | 2;2 | 39059635:390 | 39059887:390 | +;+ | 2094 | NA | 4 | 0 | 0 | 0 | 0 | 0 |
| ENSDARG000 | 2;2;2;2;2;2;2; | 39062754:390 | 39063368:390 | -;-;-;-;-;-;-;-; | 1960 | NMNAT3 | 21 | 49 | 44 | 26 | 102 | 48 |
| ENSDARG000 | 2;2 | 35612621:356 | 35612799:356 | +;+ | 572 | NA | 1 | 0 | 0 | 5 | 0 | 0 |
| ENSDARG000 | 2;2;2;2;2;2;2; | 24507597:245 | 24507816:245 | +;+;+;+;+;+;+; | 1612 | rps28 | 5188 | 4940 | 6076 | 5039 | 3476 | 5076 |

| | | | | | | | | | | | | |
|---|---|---|---|---|---|---|---|---|---|---|---|---|
| ENSDARG000 | 2;2;2;2;2;2;2; | 56463167:564 | 56463410:564 | +;+;+;+;+;+;+; | 1176 | rab11bb | 80 | 88 | 96 | 74 | 26 | 58 |
| ENSDARG000 | 2;2;2;2;2;2;2; | 30739680:307 | 30740172:307 | +;+;+;+;+;+;+; | 3381 | tcea1 | 936 | 1298 | 1607 | 1024 | 1440 | 976 |
| ENSDARG000 | 2;2;2;2;2;2;2; | 27855102:278 | 27855230:278 | +;+;+;+;+;+;+; | 3137 | buc | 10 | 12 | 0 | 0 | 10 | 16 |
| ENSDARG000 | 2;2;2;2;2;2;2; | 30752501:307 | 30756974:307 | -;-;-;-;-;-;-; | 5569 | rgs20 | 6 | 0 | 4 | 2 | 4 | 0 |
| ENSDARG000 | 2;2 | 47038403:470 | 47039063:470 | -;- | 1020 | NA | 0 | 0 | 0 | 0 | 0 | 0 |
| ENSDARG000 | 2;2;2;2 | 56487668:564 | 56488258:564 | +;+;+;+ | 1086 | NA | 48 | 45 | 18 | 3 | 16 | 4 |
| ENSDARG000 | 2;2;2;2;2 | 32835679:328 | 32835880:328 | +;+;+;+;+ | 742 | NA | 0 | 0 | 0 | 0 | 2 | 0 |
| ENSDARG000 | 2;2 | 4894873:489 | 4894936:489 | +;+ | 536 | NA | 210 | 196 | 179 | 191 | 144 | 200 |
| ENSDARG000 | 2;2;2;2;2;2;2; | 8717155:871 | 8717388:871 | +;+;+;+;+;+;+; | 3907 | usp33 | 826 | 1264 | 1544 | 1146 | 1212 | 896 |
| ENSDARG000 | 2;2;2;2 | 56497894:564 | 56498091:564 | +;+;+;+ | 586 | NA | 67 | 59 | 35 | 6 | 35 | 27 |
| ENSDARG000 | 2;2;2;2;2;2;2; | 35615547:356 | 35616116:356 | +;+;+;+;+;+;+; | 1648 | ankrd45 | 1 | 4 | 7 | 1 | 2 | 0 |
| ENSDARG000 | 2;2;2;2;2;2;2; | 37851124:378 | 37851658:378 | -;-;-;-;-;-;-; | 1501 | NA | 240 | 92 | 29 | 16 | 30 | 92 |
| ENSDARG000 | 2;2;2;2;2;2;2; | 17450228:174 | 17451434:174 | -;-;-;-;-;-;-; | 4745 | NA | 684 | 411 | 615 | 509 | 596 | 474 |
| ENSDARG000 | 2;2;2;2;2;2;2; | 35732652:357 | 35733751:357 | +;+;+;+;+;+;+; | 5991 | RASAL2 | 4042 | 3657 | 4246 | 4165 | 5857 | 4201 |
| ENSDARG000 | 2 | 32843133 | 32846324 | + | 3192 | opn4.1 | 4 | 0 | 6 | 2 | 18 | 0 |
| ENSDARG000 | 2;2;2;2;2;2;2; | 283764:2841 | 284402:2844 | -;-;-;-;-;-;-; | 2867 | MEP1B | 28 | 2 | 8 | 7 | 11 | 13 |
| ENSDARG000 | 2;2;2 | 35728589:357 | 35729290:357 | -;-;- | 1018 | NA | 0 | 0 | 6 | 0 | 0 | 0 |
| ENSDARG000 | 2;2 | 35615832:356 | 35616098:356 | -;- | 333 | NA | 0 | 0 | 0 | 0 | 0 | 0 |
| ENSDARG000 | 2;2;2;2;2;2;2; | 6104378:610 | 6107020:610 | -;-;-;-;-;-;-; | 3479 | prdx1 | 4888 | 7977 | 6387 | 6534 | 7346 | 5147 |
| ENSDARG000 | 2;2;2 | 32846602:328 | 32846841:328 | +;+;+ | 1331 | tmem53 | 66 | 97 | 116 | 132 | 138 | 68 |
| ENSDARG000 | 2;2 | 47112696:471 | 47113110:471 | +;+ | 504 | NA | 0 | 0 | 0 | 0 | 0 | 0 |
| ENSDARG000 | 2;2;2;2;2;2;2; | 56508451:565 | 56513961:565 | -;-;-;-;-;-;-; | 8051 | PIP5K1C | 95 | 70 | 62 | 46 | 63 | 50 |
| ENSDARG000 | 2;2 | 47148933:471 | 47149301:471 | +;+ | 467 | NA | 0 | 0 | 0 | 0 | 0 | 0 |
| ENSDARG000 | 2;2 | 47249179:472 | 47249302:472 | +;+ | 168 | NA | 0 | 0 | 0 | 0 | 0 | 0 |
| ENSDARG000 | 2;2;2;2;2;2;2; | 50104811:501 | 50105703:501 | -;-;-;-;-;-;-; | 3628 | exoc3 | 1724 | 1400 | 1396 | 1291 | 1414 | 1155 |
| ENSDARG000 | 2;2;2;2;2 | 37866880:378 | 37868730:378 | -;-;-;-;- | 2873 | zgc:66427 | 767 | 900 | 908 | 515 | 1073 | 796 |
| ENSDARG000 | 2;2 | 32854216:328 | 32854888:328 | +;+ | 2061 | NA | 0 | 0 | 0 | 0 | 0 | 0 |
| ENSDARG000 | 2;2;2;2;2;2;2 | 23622788:236 | 23624596:236 | -;-;-;-;-;-;- | 3970 | CDY2B | 284 | 356 | 422 | 292 | 385 | 388 |
| ENSDARG000 | 2;2 | 295899:2986 | 296351:2999 | +;+ | 1729 | NA | 55 | 33 | 62 | 39 | 36 | 70 |
| ENSDARG000 | 2;2;2;2;2;2;2; | 23677179:237 | 23677326:237 | +;+;+;+;+;+;+; | 7185 | oxsr1a | 1860 | 1904 | 2668 | 1437 | 2357 | 2876 |
| ENSDARG000 | 2;2;2;2;2;2;2; | 27877908:278 | 27878598:278 | -;-;-;-;-;-;-; | 2323 | gad3 | 0 | 0 | 0 | 0 | 0 | 0 |
| ENSDARG000 | 2;2;2;2;2;2;2; | 15389193:153 | 15391831:153 | -;-;-;-;-;-;-; | 8491 | col11a1b | 328 | 52 | 226 | 128 | 298 | 24 |
| ENSDARG000 | 2;2;2;2;2;2;2; | 50133594:501 | 50134484:501 | -;-;-;-;-;-;-; | 1780 | ahrrb | 204 | 40 | 49 | 6 | 43 | 172 |
| ENSDARG000 | 2;2;2;2;2;2;2; | 8778490:877 | 8779203:877 | +;+;+;+;+;+;+; | 6570 | zzz3 | 1042 | 1022 | 1107 | 719 | 1244 | 800 |
| ENSDARG000 | 2;2;2;2;2;2;2; | 31948352:319 | 31948621:319 | +;+;+;+;+;+;+; | 6331 | ankhb | 0 | 12 | 18 | 14 | 26 | 8 |
| ENSDARG000 | 2;2;2;2;2 | 10642047:106 | 10642431:106 | +;+;+;+;+ | 3113 | dipk1aa | 48 | 66 | 75 | 54 | 117 | 42 |
| ENSDARG000 | 2;2;2;2;2;2;2; | 31942390:319 | 31942419:319 | +;+;+;+;+;+;+; | 1938 | otulinb | 234 | 169 | 248 | 210 | 263 | 209 |
| ENSDARG000 | 2;2;2;2;2;2;2; | 32866587:328 | 32866783:328 | +;+;+;+;+;+;+; | 2083 | NA | 0 | 0 | 0 | 0 | 0 | 0 |
| ENSDARG000 | 2;2;2;2;2;2;2; | 47345041:473 | 47345436:473 | -;-;-;-;-;-;-; | 2125 | NA | 0 | 0 | 0 | 0 | 0 | 0 |
| ENSDARG000 | 2;2;2;2;2;2;2; | 24536762:245 | 24537359:245 | +;+;+;+;+;+;+; | 2704 | angptl4 | 4161 | 4937 | 5782 | 12245 | 11938 | 2053 |
| ENSDARG000 | 2;2;2;2;2;2;2; | 39549376:395 | 39549532:395 | -;-;-;-;-;-;-;- | 1361 | NA | 0 | 0 | 0 | 0 | 0 | 0 |
| ENSDARG000 | 2;2 | 301898:3045 | 302403:3067 | +;+ | 2690 | NA | 65 | 57 | 91 | 94 | 90 | 76 |
| ENSDARG000 | 2;2;2;2;2;2;2; | 39108339:392 | 39108516:392 | +;+;+;+;+;+;+; | 9327 | clstn2a | 0 | 0 | 0 | 0 | 0 | 2 |
| ENSDARG000 | 2;2;2;2 | 36862473:368 | 36862837:368 | +;+;+;+ | 1028 | NA | 0 | 10 | 2 | 0 | 10 | 0 |
| ENSDARG000 | 2;2 | 12798366:128 | 12798783:128 | +;+ | 2091 | NA | 1 | 42 | 22 | 16 | 27 | 12 |
| ENSDARG000 | 2;2;2;2;2;2;2; | 47370323:473 | 47370528:473 | -;-;-;-;-;-;-; | 2919 | pax3a | 2023 | 1986 | 2379 | 1415 | 2499 | 1679 |
| ENSDARG000 | 2;2;2;2;2;2;2; | 42592572:425 | 42592742:425 | +;+;+;+;+;+;+; | 1008 | NA | 0 | 0 | 0 | 0 | 0 | 0 |
| ENSDARG000 | 2;2;2;2;2;2;2 | 36895025:368 | 36895231:368 | -;-;-;-;-;-;- | 922 | NA | 3 | 0 | 0 | 0 | 0 | 0 |
| ENSDARG000 | 2;2;2;2;2;2;2; | 36898982:369 | 36899154:369 | +;+;+;+;+;+;+; | 1595 | NA | 0 | 0 | 0 | 0 | 3 | 8 |
| ENSDARG000 | 2;2;2;2;2;2;2; | 26479676:264 | 26480289:264 | +;+;+;+;+;+;+; | 3825 | hectd3 | 1279 | 1813 | 1350 | 1097 | 1674 | 1927 |
| ENSDARG000 | 2;2;2;2;2;2;2; | 17508323:175 | 17508527:175 | +;+;+;+;+;+;+; | 1539 | NA | 0 | 0 | 0 | 0 | 0 | 0 |
| ENSDARG000 | 2;2 | 33038085:330 | 33038682:330 | -;- | 1303 | NA | 0 | 2 | 0 | 0 | 0 | 0 |
| ENSDARG000 | 2;2 | 57058078:570 | 57058174:570 | -;- | 301 | NA | 0 | 0 | 0 | 0 | 0 | 0 |
| ENSDARG000 | 2;2;2;2;2;2;2; | 6126086:612 | 6126905:612 | +;+;+;+;+;+;+; | 2811 | FZR1 | 395 | 329 | 335 | 298 | 319 | 420 |
| ENSDARG000 | 2;2;2;2;2 | 307665:3141 | 308212:3146 | +;+;+;+;+ | 2350 | zgc:113452 | 102 | 86 | 125 | 98 | 78 | 97 |
| ENSDARG000 | 2;2;2;2;2;2;2; | 10687276:106 | 10687441:106 | -;-;-;-;-;-;-;- | 1039 | rpl5a | 11911 | 12445 | 20231 | 15304 | 13282 | 18607 |
| ENSDARG000 | 2;2;2;2;2;2;2; | 4900213:490 | 4903764:490 | -;-;-;-;-;-;-; | 8675 | dlg1b | 1083 | 545 | 1280 | 890 | 1663 | 1174 |
| ENSDARG000 | 2;2;2;2;2;2;2; | 37874771:378 | 37874919:378 | +;+;+;+;+;+;+; | 1146 | cbln13 | 2 | 0 | 2 | 0 | 8 | 2 |
| ENSDARG000 | 2;2;2 | 47432087:474 | 47432787:474 | -;-;- | 882 | NA | 0 | 0 | 2 | 2 | 0 | 2 |
| ENSDARG000 | 2;2;2;2;2 | 17540720:175 | 17540801:175 | +;+;+;+;+ | 887 | NA | 0 | 0 | 0 | 0 | 0 | 0 |
| ENSDARG000 | 2;2;2;2;2;2;2; | 44301276:443 | 44301549:443 | -;-;-;-;-;-;-; | 3561 | lig1 | 251 | 1237 | 1419 | 418 | 1115 | 638 |
| ENSDARG000 | 2;2;2;2;2;2;2; | 17524278:175 | 17524374:175 | +;+;+;+;+;+;+; | 1425 | NA | 0 | 4 | 0 | 0 | 0 | 0 |
| ENSDARG000 | 2;2;2;2;2;2;2; | 17569644:175 | 17569735:175 | +;+;+;+;+;+;+; | 2683 | NA | 0 | 0 | 0 | 0 | 0 | 0 |
| ENSDARG000 | 2 | 27932781 | 27933864 | + | 1084 | mc4r | 0 | 0 | 0 | 0 | 0 | 0 |
| ENSDARG000 | 2;2;2;2;2;2;2; | 17556177:175 | 17556381:175 | +;+;+;+;+;+;+; | 1608 | NA | 2 | 0 | 0 | 0 | 0 | 0 |
| ENSDARG000 | 2;2;2;2;2;2;2; | 10709557:107 | 10709888:107 | +;+;+;+;+;+;+; | 4448 | NA | 2 | 8 | 10 | 8 | 16 | 14 |
| ENSDARG000 | 2;2;2;2;2;2;2; | 57076079:570 | 57076687:570 | -;-;-;-;-;-;-;- | 1698 | slc25a42 | 113 | 77 | 110 | 96 | 143 | 66 |
| ENSDARG000 | 2;2 | 39618951:396 | 39619067:396 | +;+ | 1449 | zgc:136870 | 46 | 36 | 42 | 7 | 94 | 30 |
| ENSDARG000 | 2;2;2 | 39584066:396 | 39585153:396 | -;-;- | 2346 | spsb4a | 218 | 281 | 317 | 217 | 438 | 210 |
| ENSDARG000 | 2;2;2;2;2;2;2; | 47471647:474 | 47472127:474 | +;+;+;+;+;+;+; | 6680 | acsl3b | 4929 | 9609 | 7093 | 3208 | 6192 | 7064 |
| ENSDARG000 | 2;2;2;2;2;2;2; | 8845840:885 | 8846524:885 | -;-;-;-;-;-;-; | 2559 | ak5 | 2 | 2 | 42 | 18 | 48 | 8 |
| ENSDARG000 | 2;2;2;2;2;2;2 | 317338:3173 | 317561:3175 | +;+;+;+;+;+;+ | 1597 | NA | 108 | 102 | 94 | 55 | 116 | 116 |
| ENSDARG000 | 2;2;2;2;2;2;2; | 50199738:502 | 50199935:502 | -;-;-;-;-;-;-; | 2585 | mcm6l | 2 | 2 | 0 | 4 | 2 | 4 |
| ENSDARG000 | 2;2;2;2;2;2;2; | 20866898:208 | 20866958:208 | +;+;+;+;+;+;+; | 1669 | odr4 | 210 | 209 | 274 | 142 | 179 | 226 |
| ENSDARG000 | 2;2;2;2;2 | 37879812:378 | 37880408:378 | -;-;-;-;- | 1112 | hbl4 | 4 | 0 | 0 | 0 | 8 | 6 |
| ENSDARG000 | 2;2;2;2;2;2;2; | 57149002:571 | 57149089:571 | +;+;+;+;+;+;+; | 2845 | eef2b | 100654 | 85083 | 167230 | 131234 | 117062 | 135790 |
| ENSDARG000 | 2;2;2;2;2;2;2; | 24548249:245 | 24549756:245 | -;-;-;-;-;-;-; | 2452 | cnn2 | 3449 | 4975 | 2604 | 4323 | 3079 | 3746 |
| ENSDARG000 | 2;2 | 327081:3305 | 327592:3344 | +;+ | 4382 | zgc:174263 | 343 | 288 | 394 | 242 | 315 | 429 |
| ENSDARG000 | 2;2;2;2;2;2;2; | 37887277:378 | 37887933:378 | -;-;-;-;-;-;-; | 1055 | mbl2 | 0 | 0 | 0 | 0 | 4 | 0 |
| ENSDARG000 | 2;2 | 50202638:502 | 50202663:502 | +;+ | 360 | NA | 11 | 0 | 6 | 0 | 2 | 0 |
| ENSDARG000 | 2;2;2;2;2;2;2; | 33051730:330 | 33053008:330 | +;+;+;+;+;+;+; | 9433 | rnf220a | 644 | 672 | 758 | 434 | 725 | 784 |
| ENSDARG000 | 2;2;2 | 37892606:378 | 37893201:378 | -;-;- | 885 | NA | 10 | 0 | 4 | 0 | 0 | 0 |
| ENSDARG000 | 2;2;2;2 | 27975554:279 | 27975730:279 | +;+;+;+ | 3260 | NA | 18 | 0 | 14 | 5 | 8 | 0 |
| ENSDARG000 | 2;2;2;2;2;2;2; | 27957642:279 | 27958115:279 | -;-;-;-;-;-;-; | 2077 | C2H5orf22 | 96 | 100 | 117 | 54 | 93 | 70 |
| ENSDARG000 | 2;2;2;2;2;2;2; | 39731284:397 | 39734836:397 | -;-;-;-;-;-;-; | 5032 | slc25a36a | 962 | 629 | 1474 | 1191 | 911 | 1383 |
| ENSDARG000 | 2;2;2;2;2;2;2; | 6150855:615 | 6152218:615 | -;-;-;-;-;-;-; | 4655 | NA | 0 | 4 | 2 | 0 | 5 | 8 |
| ENSDARG000 | 2;2;2;2;2;2;2; | 42605252:426 | 42605493:426 | -;-;-;-;-;-;-; | 7061 | MYO10 | 573 | 765 | 1008 | 711 | 1422 | 1010 |
| ENSDARG000 | 2;2;2;2;2;2;2; | 35854242:358 | 35854350:358 | +;+;+;+;+;+;+; | 4118 | dhx9 | 2023 | 1625 | 2522 | 1053 | 2399 | 1454 |
| ENSDARG000 | 2;2;2;2;2 | 6181383:618 | 6182051:618 | +;+;+;+;+ | 2156 | NA | 8 | 4 | 11 | 2 | 9 | 29 |
| ENSDARG000 | 2;2;2;2 | 20882954:208 | 20885661:208 | -;-;-;- | 2971 | pdca | 4 | 15 | 22 | 3 | 18 | 8 |
| ENSDARG000 | 2;2;2;2;2;2;2; | 10766744:107 | 10766817:107 | +;+;+;+;+;+;+; | 1681 | gfi1aa | 0 | 10 | 26 | 6 | 6 | 6 |
| ENSDARG000 | 2;2;2;2;2;2;2; | 12847602:128 | 12848323:128 | +;+;+;+;+;+;+; | 17128 | KMT2C | 1176 | 685 | 1376 | 883 | 1315 | 1022 |
| ENSDARG000 | 2;2;2 | 335949:3376 | 336197:3377 | -;-;- | 804 | NA | 0 | 0 | 0 | 0 | 0 | 0 |
| ENSDARG000 | 2;2;2 | 342478:3440 | 343216:3441 | -;-;- | 1056 | si:dkey-33c14 | 32 | 9 | 0 | 0 | 6 | 0 |
| ENSDARG000 | 2;2;2 | 352097:3522 | 352206:3523 | -;-;- | 332 | NA | 0 | 0 | 0 | 0 | 9 | 0 |
| ENSDARG000 | 2;2;2;2;2;2;2; | 20916497:209 | 20917170:209 | -;-;-;-;-;-;-; | 2373 | ptgs2a | 405 | 44 | 6 | 100 | 50 | 58 |
| ENSDARG000 | 2;2;2;2;2 | 32016256:320 | 32016443:320 | +;+;+;+;+ | 1822 | mycb | 5395 | 2839 | 5238 | 4577 | 2645 | 3352 |
| ENSDARG000 | 2;2;2;2;2;2;2; | 354424:3573 | 357185:3574 | -;-;-;-;-;-;-;- | 4149 | NA | 443 | 374 | 425 | 324 | 406 | 508 |
| ENSDARG000 | 2;2;2;2;2;2;2; | 32019707:320 | 32019831:320 | +;+;+;+;+;+;+; | 1709 | NA | 52 | 55 | 159 | 110 | 113 | 78 |
| ENSDARG000 | 2;2;2;2;2;2;2; | 22592139:225 | 22592253:225 | +;+;+;+;+;+;+; | 2299 | septin2 | 3093 | 4535 | 4693 | 3183 | 5323 | 4530 |
| ENSDARG000 | 2;2;2;2;2;2 | 37894849:378 | 37895501:378 | -;-;-;-;-;- | 1015 | NA | 0 | 0 | 4 | 4 | 6 | 0 |
| ENSDARG000 | 2;2;2;2;2;2;2; | 36907136:369 | 36907327:369 | -;-;-;-;-;-;-;- | 1402 | ifi44c1 | 27 | 7 | 2 | 13 | 20 | 12 |
| ENSDARG000 | 2;2 | 47541737:475 | 47541778:475 | -;- | 144 | NA | 0 | 0 | 0 | 0 | 0 | 0 |
| ENSDARG000 | 2;2;2;2;2;2;2; | 28049046:280 | 28051237:280 | -;-;-;-;-;-;-; | 4258 | cdh6 | 616 | 36 | 24 | 58 | 74 | 242 |
| ENSDARG000 | 2;2;2;2;2;2;2; | 30786773:307 | 30786825:307 | +;+;+;+;+;+;+; | 2151 | atp6v1h | 3960 | 2831 | 3238 | 2817 | 2797 | 2334 |
| ENSDARG000 | 2;2;2;2;2;2;2; | 9061885:906 | 9062024:906 | +;+;+;+;+;+;+; | 1482 | pigk | 190 | 176 | 233 | 66 | 343 | 170 |
| ENSDARG000 | 2;2;2;2;2;2;2; | 20925899:209 | 20925943:209 | +;+;+;+;+;+;+; | 2591 | pla2g4aa | 104 | 71 | 129 | 104 | 219 | 213 |
| ENSDARG000 | 2;2;2;2;2;2;2; | 36914408:369 | 36915371:369 | -;-;-;-;-;-;-;- | 2161 | ifi44c2 | 76 | 128 | 15 | 35 | 42 | 36 |
| ENSDARG000 | 2;2;2;2;2;2;2; | 6212205:621 | 6212355:621 | -;-;-;-;-;-;-; | 1581 | NA | 0 | 0 | 0 | 0 | 0 | 0 |
| ENSDARG000 | 2;2;2;2;2;2 | 39926961:399 | 39927317:399 | -;-;-;-;-;- | 931 | NA | 6 | 0 | 0 | 0 | 0 | 0 |
| ENSDARG000 | 2;2;2;2;2;2;2; | 36917613:369 | 36921679:369 | -;-;-;-;-;-;-;- | 7678 | NA | 718 | 746 | 1086 | 1181 | 1305 | 872 |
| ENSDARG000 | 2;2 | 366429:3678 | 366761:3696 | +;+ | 2127 | NA | 60 | 56 | 94 | 28 | 102 | 32 |
| ENSDARG000 | 2;2;2 | 6243144:624 | 6243388:624 | +;+;+ | 855 | gng5 | 889 | 1200 | 1290 | 1343 | 830 | 1094 |
| ENSDARG000 | 2;2;2;2;2 | 47581997:475 | 47582105:475 | +;+;+;+;+ | 2701 | scg2b | 12 | 4 | 0 | 0 | 0 | 0 |
| ENSDARG000 | 2;2;2;2;2 | 47587238:475 | 47587787:475 | -;-;-;-;- | 1079 | ap1s3b | 64 | 0 | 10 | 2 | 14 | 20 |
| ENSDARG000 | 2;2;2;2;2;2;2; | 57239978:572 | 57240835:572 | -;-;-;-;-;-;-; | 1856 | mbd3a | 925 | 1026 | 1358 | 956 | 1005 | 849 |
| ENSDARG000 | 2;2;2;2;2;2;2; | 10790875:108 | 10791736:108 | -;-;-;-;-;-;-; | 2539 | rpap2 | 114 | 171 | 214 | 45 | 168 | 110 |
| ENSDARG000 | 2;2;2;2;2;2;2; | 37897079:378 | 37897133:379 | +;+;+;+;+;+;+; | 1747 | NA | 85 | 32 | 64 | 58 | 84 | 40 |
| ENSDARG000 | 2;2;2;2 | 24558127:245 | 24560752:245 | -;-;-;- | 3745 | crsp7 | 468 | 370 | 511 | 282 | 655 | 392 |
| ENSDARG000 | 2;2;2;2 | 20967673:209 | 20967930:209 | +;+;+;+ | 3433 | arpc5a | 2639 | 3078 | 2840 | 1792 | 3015 | 2541 |
| ENSDARG000 | 2;2 | 371759:3720 | 371836:3724 | +;+ | 408 | NA | 2 | 0 | 0 | 1 | 2 | 4 |
| ENSDARG000 | 2;2;2;2;2;2;2; | 44348473:443 | 44348625:443 | +;+;+;+;+;+;+; | 3406 | zgc:152670 | 19 | 12 | 2 | 0 | 6 | 20 |
| ENSDARG000 | 2;2;2;2;2;2;2; | 6253246:625 | 6253807:625 | +;+;+;+;+;+;+; | 1855 | zp3b | 22 | 34 | 22 | 59 | 36 | 50 |
| ENSDARG000 | 2;2;2;2 | 57268781:572 | 57269158:572 | -;-;-;- | 609 | si:ch1073-325 | 852 | 1254 | 1211 | 826 | 1022 | 1119 |
| ENSDARG000 | 2;2;2;2;2;2;2; | 23731194:237 | 23731568:237 | +;+;+;+;+;+;+; | 1781 | NA | 2 | 0 | 0 | 0 | 10 | 0 |
| ENSDARG000 | 2;2;2;2;2;2;2 | 32115851:321 | 32116458:321 | -;-;-;-;-;-;- | 1352 | NA | 0 | 0 | 6 | 0 | 1 | 0 |
| ENSDARG000 | 2;2;2;2;2;2;2; | 47623202:476 | 47623317:476 | +;+;+;+;+;+;+; | 1131 | NA | 0 | 0 | 0 | 0 | 8 | 0 |
| ENSDARG000 | 2;2;2;2;2;2;2; | 23749614:237 | 23752300:237 | -;-;-;-;-;-;-;- | 8291 | xirp1 | 0 | 2 | 6 | 2 | 18 | 0 |
| ENSDARG000 | 2;2;2;2;2;2;2 | 374365:3753 | 375167:3753 | -;-;-;-;-;-;- | 2302 | wrnip1 | 185 | 259 | 238 | 120 | 205 | 136 |
| ENSDARG000 | 2;2;2;2;2;2;2; | 9160947:916 | 9161193:916 | -;-;-;-;-;-;-;- | 1173 | ST6GALNAC5 | 0 | 0 | 1 | 0 | 12 | 0 |
| ENSDARG000 | 2;2;2;2;2;2;2; | 40067808:400 | 40068861:400 | -;-;-;-;-;-;-; | 4952 | epha4a | 42 | 34 | 20 | 0 | 52 | 80 |
| ENSDARG000 | 2;2;2;2;2;2;2 | 36939749:369 | 36940154:369 | +;+;+;+;+;+;+ | 1485 | gadd45bb | 1530 | 567 | 1163 | 678 | 1358 | 999 |
| ENSDARG000 | 2;2 | 28599070:285 | 28599522:285 | -;- | 715 | NA | 24 | 0 | 2 | 4 | 0 | 2 |
| ENSDARG000 | 2;2;2;2;2;2;2; | 57310523:573 | 57310838:573 | -;-;-;-;-;-;-; | 3102 | tcf3a | 349 | 144 | 330 | 134 | 241 | 148 |
| ENSDARG000 | 2;2;2;2;2;2;2; | 36990024:369 | 36990553:369 | -;-;-;-;-;-;-;- | 1151 | gng7 | 133 | 115 | 152 | 113 | 132 | 171 |
| ENSDARG000 | 2;2;2 | 20980162:209 | 20980400:209 | -;-;- | 536 | lyrm4 | 90 | 132 | 128 | 144 | 78 | 91 |
| ENSDARG000 | 2;2;2;2;2 | 9330146:933 | 9330936:933 | -;-;-;-;- | 1707 | st6galnac3 | 300 | 292 | 315 | 197 | 346 | 341 |
| ENSDARG000 | 2;2;2;2;2;2;2; | 35880600:359 | 35881338:359 | +;+;+;+;+;+;+; | 6133 | lamc1 | 12593 | 8453 | 13326 | 8942 | 16899 | 6418 |
| ENSDARG000 | 2;2;2;2;2;2;2; | 50229183:502 | 50229915:502 | -;-;-;-;-;-;-; | 3711 | cul1a | 1398 | 1470 | 1710 | 1089 | 1693 | 1221 |
| ENSDARG000 | 2;2;2;2;2;2;2; | 24700922:247 | 24701342:247 | +;+;+;+;+;+;+; | 9546 | pik3r2 | 3497 | 1691 | 3236 | 2575 | 4733 | 3621 |
| ENSDARG000 | 2;2;2;2;2;2;2; | 28672152:286 | 28672344:286 | +;+;+;+;+;+;+; | 1800 | nadsyn1 | 125 | 168 | 221 | 121 | 201 | 129 |
| ENSDARG000 | 2;2;2;2;2;2;2; | 28648888:286 | 28651688:286 | -;-;-;-;-;-;-; | 3978 | dhcr7 | 346 | 230 | 404 | 287 | 286 | 286 |
| ENSDARG000 | 2;2;2;2;2;2;2; | 5371492:538 | 5371606:538 | +;+;+;+;+;+;+; | 971 | NA | 32 | 34 | 149 | 202 | 159 | 42 |
| ENSDARG000 | 2;2;2;2;2;2;2; | 42708674:427 | 42709270:427 | +;+;+;+;+;+;+; | 2574 | NA | 74 | 99 | 6 | 4 | 10 | 25 |
| ENSDARG000 | 2;2;2;2;2;2;2; | 394166:4109 | 394359:4110 | +;+;+;+;+;+;+; | 2021 | MYLK4 | 0 | 2 | 0 | 0 | 0 | 1 |
| ENSDARG000 | 2 | 6261413 | 6262441 | - | 1029 | arl14 | 10 | 1 | 16 | 20 | 15 | 10 |
| ENSDARG000 | 2;2 | 33272564:332 | 33273038:332 | -;- | 528 | NA | 0 | 0 | 0 | 0 | 0 | 0 |
| ENSDARG000 | 2;2;2 | 20991985:209 | 20993444:209 | -;-;- | 1790 | nrn1b | 0 | 6 | 2 | 0 | 0 | 2 |
| ENSDARG000 | 2;2;2;2;2;2;2; | 9513160:951 | 9513378:951 | -;-;-;-;-;-;-; | 1202 | cope | 1783 | 2149 | 2417 | 1454 | 2142 | 1366 |
| ENSDARG000 | 2 | 37059304 | 37059966 | - | 663 | diras1b | 0 | 0 | 0 | 0 | 0 | 0 |
| ENSDARG000 | 2;2;2;2 | 6265768:627 | 6267295:627 | -;-;-;- | 2709 | ppm1la | 1802 | 1293 | 1593 | 2132 | 1312 | 2327 |
| ENSDARG000 | 2;2;2;2 | 23773078:237 | 23773208:237 | -;-;-;- | 318 | NA | 0 | 0 | 0 | 0 | 2 | 0 |
| ENSDARG000 | 2;2;2 | 33326522:333 | 33326680:333 | +;+;+ | 1625 | klf17 | 4 | 0 | 24 | 0 | 18 | 6 |
| ENSDARG000 | 2;2;2;2;2;2;2; | 47644041:476 | 47644681:476 | -;-;-;-;-;-;-; | 2958 | cul3a | 1246 | 1391 | 2001 | 1380 | 1711 | 1251 |
| ENSDARG000 | 2;2;2;2;2;2;2; | 42715995:427 | 42716222:427 | +;+;+;+;+;+;+; | 1747 | NA | 4 | 4 | 9 | 0 | 0 | 2 |
| ENSDARG000 | 2;2;2;2;2 | 6303639:630 | 6303674:630 | +;+;+;+;+ | 1520 | otol1a | 11 | 0 | 12 | 35 | 10 | 24 |
| ENSDARG000 | 2 | 686873 | 689047 | - | 2175 | foxc1a | 28 | 0 | 2 | 0 | 0 | 10 |
| ENSDARG000 | 2;2;2;2;2;2;2; | 21000334:210 | 21000656:210 | +;+;+;+;+;+;+; | 8211 | RREB1 | 800 | 771 | 1357 | 1110 | 1166 | 987 |
| ENSDARG000 | 2;2;2;2;2;2;2; | 57416844:574 | 57417860:574 | -;-;-;-;-;-;-; | 2608 | map2k2b | 433 | 267 | 488 | 212 | 454 | 382 |
| ENSDARG000 | 2;2;2;2;2;2;2; | 32215272:322 | 32215913:322 | -;-;-;-;-;-;-; | 2463 | fam49ba | 84 | 78 | 76 | 32 | 195 | 100 |
| ENSDARG000 | 2;2;2;2;2;2;2; | 35991137:359 | 35991210:359 | +;+;+;+;+;+;+; | 3934 | NA | 348 | 303 | 48 | 46 | 32 | 32 |
| ENSDARG000 | 2;2;2;2;2;2;2; | 10821127:108 | 10821160:108 | +;+;+;+;+;+;+; | 2701 | glmna | 243 | 324 | 276 | 260 | 274 | 258 |

| | | | | | | | | | | | | |
|---|---|---|---|---|---|---|---|---|---|---|---|---|
| ENSDARG000 | 2;2;2;2;2;2;2;2; | 37089362:370 | 37089707:370 | -;-;-;-;-;-;-;- | 1159 | zgc:101744 | 8 | 10 | 26 | 12 | 26 | 6 |
| ENSDARG000 | 2;2;2 | 5397064:5397 | 5398993:5399 | -;-;- | 2439 | NA | 4 | 12 | 2 | 5 | 12 | 12 |
| ENSDARG000 | 2;2;2;2;2 | 23790748:237 | 23790988:237 | +;+;+;+;+ | 3138 | NA | 360 | 466 | 349 | 573 | 283 | 274 |
| ENSDARG000 | 2;2;2;2;2;2;2;2; | 37110504:371 | 37110758:371 | +;+;+;+;+;+;+;+; | 1770 | slc1a8b | 4 | 2 | 0 | 2 | 4 | 0 |
| ENSDARG000 | 2;2 | 42724404:427 | 42724551:427 | +;+ | 2450 | basp1 | 0 | 2 | 6 | 0 | 22 | 8 |
| ENSDARG000 | 2 | 5406236 | 5407872 | + | 1637 | sft2d3 | 130 | 252 | 195 | 177 | 428 | 242 |
| ENSDARG000 | 2;2;2;2;2;2;2;2; | 696244:6990 | 698115:6990 | -;-;-;-;-;-;-;- | 3063 | NA | 157 | 191 | 186 | 102 | 199 | 162 |
| ENSDARG000 | 2;2;2;2;2;2;2;2; | 23808640:238 | 23808830:238 | +;+;+;+;+;+;+;+; | 5133 | gorasp1a | 1192 | 1140 | 1599 | 716 | 1193 | 941 |
| ENSDARG000 | 2;2;2;2;2;2;2;2; | 33335911:333 | 33336095:333 | +;+;+;+;+;+;+;+; | 10324 | slc6a9 | 114 | 81 | 26 | 26 | 50 | 81 |
| ENSDARG000 | 2;2;2;2;2;2;2;2; | 28889936:289 | 28890191:289 | +;+;+;+;+;+;+;+; | 2756 | cdh10a | 28 | 44 | 95 | 11 | 63 | 24 |
| ENSDARG000 | 2;2;2;2;2;2;2;2; | 26498446:264 | 26498557:264 | +;+;+;+;+;+;+;+; | 1711 | rps8a | 17220 | 13824 | 24369 | 21254 | 15825 | 23630 |
| ENSDARG000 | 2;2;2;2;2;2;2;2; | 15612755:156 | 15612959:156 | +;+;+;+;+;+;+;+; | 1698 | amy2a | 2 | 0 | 0 | 0 | 4 | 0 |
| ENSDARG000 | 2;2;2;2;2;2;2;2; | 50276755:502 | 50279236:502 | -;-;-;-;-;-;-;-; | 6638 | cntnap2b | 5 | 6 | 4 | 22 | 8 | 0 |
| ENSDARG000 | 2;2;2;2;2;2;2;2; | 33422411:334 | 33423311:334 | -;-;-;-;-;-;-;-; | 2435 | ccdc24 | 2 | 1 | 1 | 12 | 0 | 31 |
| ENSDARG000 | 2;2;2 | 9552456:9553 | 9552846:9553 | +;+;+ | 1948 | dnajb4 | 360 | 249 | 197 | 128 | 318 | 260 |
| ENSDARG000 | 2;2;2;2;2;2;2;2; | 24762567:247 | 24762798:247 | +;+;+;+;+;+;+;+; | 1166 | ifi30 | 310 | 432 | 568 | 466 | 840 | 513 |
| ENSDARG000 | 2;2;2;2;2;2;2;2; | 9529488:9529 | 9529745:9529 | -;-;-;-;-;-;-;-; | 1771 | slc25a24l | 2 | 0 | 2 | 0 | 0 | 0 |
| ENSDARG000 | 2;2 | 44366972:443 | 44367041:443 | -;- | 295 | NA | 0 | 0 | 0 | 0 | 0 | 0 |
| ENSDARG000 | 2;2 | 44373221:443 | 44373250:443 | +;+ | 486 | NA | 0 | 0 | 0 | 0 | 0 | 0 |
| ENSDARG000 | 2;2;2;2;2;2;2 | 44383175:443 | 44386372:443 | -;-;-;-;-;-;- | 3929 | pip4p1a | 282 | 261 | 189 | 109 | 239 | 205 |
| ENSDARG000 | 2;2;2;2;2;2;2;2; | 6321623:6323 | 6322706:6323 | -;-;-;-;-;-;-;-; | 4267 | si:dkev-119f1. | 276 | 309 | 350 | 336 | 333 | 331 |
| ENSDARG000 | 2;2;2;2;2;2;2;2; | 57447105:574 | 57447480:574 | -;-;-;-;-;-;-;-; | 1856 | pias4b | 0 | 0 | 0 | 0 | 0 | 0 |
| ENSDARG000 | 2;2;2;2;2;2;2;2; | 22617675:226 | 22617983:226 | -;-;-;-;-;-;-;-; | 1924 | stk25b | 984 | 914 | 1128 | 1005 | 1170 | 911 |
| ENSDARG000 | 2;2;2;2;2;2;2;2; | 5410432:5410 | 5411433:5411 | -;-;-;-;-;-;-;-; | 5060 | wdr33 | 1485 | 1530 | 1858 | 1236 | 1589 | 1311 |
| ENSDARG000 | 2;2;2;2;2;2;2;2; | 42776224:427 | 42776708:427 | -;-;-;-;-;-;-;-; | 2744 | adcy8 | 2 | 6 | 10 | 0 | 17 | 9 |
| ENSDARG000 | 2;2;2;2;2 | 24770435:247 | 24770998:247 | +;+;+;+;+ | 1563 | mpv17l2 | 178 | 231 | 286 | 70 | 160 | 196 |
| ENSDARG000 | 2;2;2;2;2;2;2;2; | 37123338:371 | 37125883:371 | -;-;-;-;-;-;-;-; | 3520 | elavl1a | 3732 | 4686 | 5175 | 3012 | 5337 | 3449 |
| ENSDARG000 | 2;2;2;2;2;2 | 42773349:427 | 42773375:427 | +;+;+;+;+;+ | 1564 | NA | 0 | 0 | 0 | 0 | 0 | 0 |
| ENSDARG000 | 2;2 | 713080:7151 | 713939:7164 | -;- | 2173 | foxf2a | 0 | 0 | 0 | 4 | 0 | 0 |
| ENSDARG000 | 2;2;2;2 | 30878864:308 | 30878993:308 | +;+;+;+ | 1486 | oprk1 | 0 | 0 | 2 | 0 | 0 | 0 |
| ENSDARG000 | 2;2;2;2;2;2 | 9560740:9580 | 9561196:9580 | +;+;+;+;+;+ | 1775 | gipc2 | 54 | 26 | 54 | 42 | 42 | 12 |
| ENSDARG000 | 2;2;2;2;2 | 23823622:238 | 23823794:238 | +;+;+;+;+ | 1072 | vopp1a | 1134 | 339 | 565 | 273 | 638 | 714 |
| ENSDARG000 | 2;2;2;2;2;2;2;2; | 28995776:290 | 28996152:290 | +;+;+;+;+;+;+;+; | 2944 | cdh12a | 0 | 0 | 0 | 0 | 0 | 0 |
| ENSDARG000 | 2;2;2;2;2;2;2;2; | 13056802:130 | 13057023:130 | +;+;+;+;+;+;+;+; | 2241 | NA | 130 | 35 | 84 | 34 | 45 | 133 |
| ENSDARG000 | 2;2;2;2;2;2;2;2; | 36026183:360 | 36027005:360 | -;-;-;-;-;-;-;-; | 1761 | nmnat2 | 0 | 15 | 0 | 0 | 4 | 10 |
| ENSDARG000 | 2;2;2;2;2 | 6475567:6478 | 6476447:6478 | -;-;-;-;- | 1279 | rgs13 | 691 | 140 | 88 | 368 | 410 | 441 |
| ENSDARG000 | 2;2 | 720779:7207 | 722156:7221 | -;- | 1392 | foxq1a | 172 | 2 | 0 | 10 | 16 | 42 |
| ENSDARG000 | 2;2;2;2;2;2;2;2; | 26504969:265 | 26505105:265 | -;-;-;-;-;-;-;-; | 1133 | BEST4 | 0 | 2 | 2 | 0 | 8 | 0 |
| ENSDARG000 | 2;2;2;2;2;2;2 | 9592319:9598 | 9594094:9598 | -;-;-;-;-;-;- | 2308 | txndc12 | 1067 | 951 | 1275 | 477 | 1334 | 787 |
| ENSDARG000 | 2;2;2;2;2;2;2;2; | 6511499:6511 | 6512410:6512 | -;-;-;-;-;-;-;- | 1465 | NA | 4 | 0 | 0 | 0 | 0 | 0 |
| ENSDARG000 | 2;2 | 17707492:177 | 17707673:177 | +;+ | 248 | NA | 9 | 10 | 25 | 20 | 11 | 16 |
| ENSDARG000 | 2;2;2;2;2;2;2;2; | 17578320:175 | 17580060:175 | -;-;-;-;-;-;-;-; | 8404 | ptprfb | 1691 | 1117 | 4350 | 1334 | 5621 | 1396 |
| ENSDARG000 | 2;2 | 26518813:265 | 26519428:265 | -;- | 711 | NA | 0 | 2 | 0 | 0 | 0 | 0 |
| ENSDARG000 | 2;2;2;2;2;2;2;2; | 36046126:360 | 36046292:360 | +;+;+;+;+;+;+;+; | 7632 | smg7 | 1812 | 1772 | 2623 | 2180 | 2003 | 1910 |
| ENSDARG000 | 2;2;2;2;2;2;2;2; | 23856045:238 | 23856344:238 | -;-;-;-;-;-;-;- | 1703 | tgfbr1a | 117 | 53 | 95 | 80 | 112 | 70 |
| ENSDARG000 | 2;2;2;2;2;2;2;2; | 40623015:406 | 40623576:406 | -;-;-;-;-;-;-;-; | 1194 | NA | 0 | 0 | 0 | 0 | 0 | 0 |
| ENSDARG000 | 2;2;2;2;2;2;2;2; | 44400770:444 | 44401185:444 | -;-;-;-;-;-;-;-; | 1337 | dnaaf6 | 2 | 0 | 0 | 0 | 0 | 0 |
| ENSDARG000 | 2;2;2 | 40901997:411 | 40902717:411 | -;-;- | 1458 | hs6st1a | 0 | 4 | 12 | 0 | 0 | 5 |
| ENSDARG000 | 2;2;2;2;2 | 5446087:5446 | 5446196:5447 | +;+;+;+;+ | 1530 | DUSP28 | 0 | 41 | 24 | 126 | 53 | 51 |
| ENSDARG000 | 2 | 36087769 | 36087779 | + | 11 | NA | 0 | 0 | 0 | 0 | 0 | 0 |
| ENSDARG000 | 2;2;2;2;2;2;2 | 26529447:265 | 26529874:265 | -;-;-;-;-;-;- | 1689 | NA | 37 | 0 | 8 | 2 | 6 | 0 |
| ENSDARG000 | 2;2;2;2;2;2;2;2; | 29249204:292 | 29249633:292 | +;+;+;+;+;+;+;+; | 4613 | cdh18a | 2 | 2 | 2 | 0 | 4 | 0 |
| ENSDARG000 | 2;2;2;2;2;2;2;2; | 37134281:371 | 37134376:371 | +;+;+;+;+;+;+;+; | 2212 | pex19 | 1692 | 1380 | 1722 | 1125 | 1617 | 1440 |
| ENSDARG000 | 2;2;2;2;2;2;2;2; | 21063660:210 | 21063865:210 | +;+;+;+;+;+;+;+; | 1955 | riok1 | 469 | 498 | 681 | 304 | 552 | 544 |
| ENSDARG000 | 2;2 | 32355428:323 | 32355465:323 | -;- | 200 | NA | 0 | 0 | 0 | 0 | 0 | 0 |
| ENSDARG000 | 2;2;2;2;2;2;2;2; | 32263215:322 | 32268190:322 | -;-;-;-;-;-;-;-; | 9140 | asap1a | 843 | 992 | 1009 | 417 | 1031 | 971 |
| ENSDARG000 | 2 | 36088047 | 36088056 | + | 10 | NA | 4 | 0 | 1 | 2 | 0 | 3 |
| ENSDARG000 | 2 | 36088431 | 36088479 | + | 49 | NA | 0 | 0 | 3 | 0 | 2 | 0 |
| ENSDARG000 | 2;2;2;2;2;2 | 6535865:6540 | 6536803:6540 | -;-;-;-;-;- | 1694 | rgs18 | 6 | 18 | 2 | 4 | 56 | 50 |
| ENSDARG000 | 2 | 36088658 | 36088717 | + | 60 | NA | 0 | 0 | 0 | 0 | 0 | 0 |
| ENSDARG000 | 2;2;2;2;2;2;2;2; | 854444:8553 | 854834:8554 | -;-;-;-;-;-;-;- | 1633 | irf4a | 12 | 12 | 16 | 4 | 20 | 6 |
| ENSDARG000 | 2;2;2 | 36089660:360 | 36089950:360 | +;+;+ | 437 | NA | 0 | 2 | 8 | 10 | 0 | 16 |
| ENSDARG000 | 2;2;2;2 | 6885852:6886 | 6885975:6886 | +;+;+;+ | 1030 | rgs5b | 478 | 76 | 80 | 69 | 101 | 265 |
| ENSDARG000 | 2;2;2 | 6875751:6917 | 6876215:6917 | -;-;- | 828 | NA | 0 | 0 | 0 | 0 | 0 | 0 |
| ENSDARG000 | 2;2;2;2;2;2;2;2; | 50862527:508 | 50862655:508 | +;+;+;+;+;+;+;+; | 9277 | adcyap1r1a | 4 | 2 | 2 | 2 | 0 | 2 |
| ENSDARG000 | 2;2;2;2;2;2;2 | 37136662:371 | 37136955:371 | -;-;-;-;-;-;- | 1370 | tspan37 | 258 | 232 | 212 | 309 | 196 | 286 |
| ENSDARG000 | 2;2;2;2;2 | 36092953:360 | 36093532:360 | +;+;+;+;+ | 853 | NA | 0 | 0 | 0 | 0 | 0 | 2 |
| ENSDARG000 | 2;2;2;2;2;2;2;2; | 30894595:308 | 30894759:308 | +;+;+;+;+;+;+;+; | 1107 | CHMP5 | 16 | 35 | 43 | 43 | 48 | 45 |
| ENSDARG000 | 2;2;2;2;2;2;2;2; | 29477383:294 | 29478197:294 | -;-;-;-;-;-;-;-; | 1919 | cahz | 90 | 77 | 412 | 74 | 1099 | 226 |
| ENSDARG000 | 2;2;2;2;2;2;2 | 57801960:578 | 57801984:578 | +;+;+;+;+;+;+ | 991 | NA | 0 | 2 | 0 | 0 | 0 | 0 |
| ENSDARG000 | 2;2;2;2;2;2;2;2; | 15776156:157 | 15776256:157 | +;+;+;+;+;+;+;+; | 4945 | NTNG1 | 0 | 0 | 0 | 0 | 0 | 0 |
| ENSDARG000 | 2;2;2;2;2 | 15833553:158 | 15833641:158 | +;+;+;+;+ | 564 | NA | 0 | 0 | 0 | 2 | 0 | 0 |
| ENSDARG000 | 2;2;2;2;2;2;2;2; | 24781026:247 | 24781079:247 | +;+;+;+;+;+;+;+; | 6992 | NA | 214 | 112 | 192 | 522 | 283 | 320 |
| ENSDARG000 | 2;2;2;2;2;2 | 9684305:9689 | 9684963:9689 | -;-;-;-;-;- | 1301 | selenot1a | 1156 | 1790 | 2004 | 1531 | 1792 | 1771 |
| ENSDARG000 | 2;2 | 13185302:132 | 13188226:132 | -;- | 4017 | BCL2 | 4 | 12 | 80 | 88 | 12 | 207 |
| ENSDARG000 | 2;2;2;2;2;2;2 | 47900506:479 | 47900565:479 | -;-;-;-;-;-;- | 1212 | NA | 24 | 84 | 54 | 40 | 86 | 35 |
| ENSDARG000 | 2;2;2;2;2;2;2;2; | 6926100:6927 | 6926725:6927 | +;+;+;+;+;+;+;+; | 2015 | NA | 0 | 0 | 0 | 0 | 4 | 0 |
| ENSDARG000 | 2 | 36094994 | 36095056 | + | 63 | NA | 0 | 0 | 0 | 0 | 0 | 0 |
| ENSDARG000 | 2 | 36095406 | 36095465 | + | 60 | NA | 0 | 0 | 0 | 0 | 0 | 0 |
| ENSDARG000 | 2;2;2;2;2;2;2;2; | 5456541:5456 | 5458099:5458 | -;-;-;-;-;-;-;-; | 2589 | PROC | 0 | 0 | 0 | 0 | 0 | 0 |
| ENSDARG000 | 2;2;2;2;2;2;2;2; | 57825432:578 | 57825788:578 | -;-;-;-;-;-;-;-; | 1756 | sf3a2 | 986 | 1242 | 1341 | 998 | 1163 | 834 |
| ENSDARG000 | 2;2;2;2;2;2;2;2; | 5457388:5460 | 5458099:5460 | -;-;-;-;-;-;-;-; | 3115 | proca | 0 | 0 | 0 | 0 | 2 | 0 |
| ENSDARG000 | 2;2;2;2 | 22640215:226 | 22640797:226 | +;+;+;+ | 1509 | bokb | 4 | 0 | 2 | 0 | 0 | 0 |
| ENSDARG000 | 2 | 36095905 | 36095967 | + | 63 | NA | 0 | 0 | 0 | 0 | 0 | 0 |
| ENSDARG000 | 2;2;2;2;2;2;2;2; | 44426609:444 | 44426861:444 | +;+;+;+;+;+;+;+; | 2161 | NA | 0 | 0 | 8 | 0 | 2 | 0 |
| ENSDARG000 | 2 | 6958513 | 6959134 | + | 622 | NA | 0 | 0 | 0 | 0 | 0 | 0 |
| ENSDARG000 | 2;2;2;2;2;2;2;2; | 26576853:265 | 26577480:265 | -;-;-;-;-;-;-;-; | 3270 | ndc1 | 475 | 928 | 1289 | 579 | 1249 | 876 |
| ENSDARG000 | 2;2;2;2;2;2;2;2; | 24867534:248 | 24868088:248 | +;+;+;+;+;+;+;+; | 3038 | rab3aa | 8 | 4 | 6 | 22 | 4 | 0 |
| ENSDARG000 | 2;2;2;2;2;2;2;2; | 2927463:2927 | 2927657:2927 | -;-;-;-;-;-;-;-; | 3132 | NA | 2 | 2 | 4 | 6 | 0 | 6 |
| ENSDARG000 | 2;2;2;2;2;2;2;2; | 2957515:2967 | 2957624:2967 | +;+;+;+;+;+;+;+; | 3464 | pik3r3a | 694 | 794 | 593 | 456 | 1098 | 371 |
| ENSDARG000 | 2;2;2;2;2;2;2;2; | 881954:8823 | 882679:8826 | -;-;-;-;-;-;-;-; | 1435 | dusp22b | 258 | 230 | 248 | 165 | 124 | 234 |
| ENSDARG000 | 2;2;2;2;2;2;2;2; | 13223755:132 | 13225338:132 | -;-;-;-;-;-;-;-; | 2797 | kdsr | 595 | 841 | 801 | 424 | 727 | 814 |
| ENSDARG000 | 2;2;2;2;2;2;2;2; | 6963296:6965 | 6963375:6965 | +;+;+;+;+;+;+;+; | 2845 | DDR2 | 28 | 0 | 6 | 0 | 0 | 0 |
| ENSDARG000 | 2;2;2;2;2;2;2 | 30904069:309 | 30904161:309 | +;+;+;+;+;+;+ | 1298 | bag1 | 555 | 717 | 650 | 654 | 629 | 749 |
| ENSDARG000 | 2 | 36096367 | 36096429 | + | 63 | NA | 0 | 0 | 0 | 0 | 0 | 0 |
| ENSDARG000 | 2;2 | 24885987:248 | 24886245:248 | +;+ | 423 | NA | 0 | 0 | 0 | 0 | 0 | 0 |
| ENSDARG000 | 2;2;2;2;2;2;2;2; | 9702053:9704 | 9702993:9705 | -;-;-;-;-;-;-;-; | 3009 | dvl3a | 526 | 654 | 824 | 460 | 698 | 408 |
| ENSDARG000 | 2;2;2;2;2;2;2;2; | 22659787:226 | 22659976:226 | +;+;+;+;+;+;+;+; | 1910 | zgc:161973 | 2 | 0 | 0 | 0 | 1 | 0 |
| ENSDARG000 | 2;2;2;2;2;2;2 | 30910134:309 | 30910575:309 | -;-;-;-;-;-;- | 932 | myl12.2 | 360 | 419 | 602 | 694 | 420 | 501 |
| ENSDARG000 | 2 | 36096728 | 36096787 | + | 60 | NA | 0 | 0 | 0 | 0 | 0 | 0 |
| ENSDARG000 | 2 | 36097414 | 36097469 | + | 56 | NA | 0 | 0 | 0 | 0 | 0 | 0 |
| ENSDARG000 | 2 | 36097984 | 36098047 | + | 64 | NA | 0 | 0 | 0 | 0 | 0 | 0 |
| ENSDARG000 | 2;2;2;2;2;2;2;2; | 22652981:226 | 22653545:226 | -;-;-;-;-;-;-;-; | 2065 | thap4 | 212 | 161 | 225 | 267 | 132 | 218 |
| ENSDARG000 | 2;2;2 | 33457310:334 | 33457844:334 | +;+;+ | 1117 | zgc:113531 | 321 | 320 | 211 | 197 | 392 | 409 |
| ENSDARG000 | 2;2;2;2;2;2;2;2; | 47905735:479 | 47906285:479 | +;+;+;+;+;+;+;+; | 3423 | NA | 115 | 1 | 61 | 23 | 39 | 35 |
| ENSDARG000 | 2;2;2;2 | 47927026:479 | 47927648:479 | +;+;+;+ | 1112 | NA | 3 | 0 | 32 | 0 | 0 | 28 |
| ENSDARG000 | 2;2;2;2;2;2;2;2; | 41524238:415 | 41524362:415 | +;+;+;+;+;+;+;+; | 3706 | acvr1l | 1601 | 1389 | 1733 | 1154 | 1629 | 1610 |
| ENSDARG000 | 2;2;2;2;2;2;2;2; | 22676743:226 | 22677408:226 | -;-;-;-;-;-;-;-; | 2211 | agxtb | 0 | 0 | 45 | 36 | 178 | 0 |
| ENSDARG000 | 2;2;2;2;2;2;2 | 47916204:479 | 47916873:479 | -;-;-;-;-;-;- | 1912 | NA | 121 | 142 | 166 | 93 | 188 | 128 |
| ENSDARG000 | 2;2;2;2;2;2;2;2; | 29491314:295 | 29491544:295 | +;+;+;+;+;+;+;+; | 3474 | dlgap1a | 0 | 0 | 0 | 0 | 2 | 0 |
| ENSDARG000 | 2;2;2;2;2;2;2;2; | 37176814:371 | 37177011:371 | +;+;+;+;+;+;+;+; | 4337 | copa | 5782 | 5272 | 7453 | 4242 | 6359 | 3831 |
| ENSDARG000 | 2 | 36098341 | 36098397 | + | 57 | NA | 0 | 0 | 0 | 0 | 0 | 0 |
| ENSDARG000 | 2;2;2;2 | 26591968:265 | 26592717:265 | -;-;-;- | 998 | zgc:113691 | 2 | 18 | 9 | 8 | 4 | 24 |
| ENSDARG000 | 2;2;2;2;2;2;2 | 21076585:210 | 21078029:210 | -;-;-;-;-;-;- | 2381 | nebl | 8 | 6 | 22 | 14 | 4 | 19 |
| ENSDARG000 | 2;2;2;2;2;2;2;2; | 15993389:159 | 15994049:159 | -;-;-;-;-;-;-;-; | 3766 | vav3b | 27 | 1350 | 422 | 1026 | 1836 | 14 |
| ENSDARG000 | 2;2;2;2;2;2;2;2; | 10863942:108 | 10864521:108 | -;-;-;-;-;-;-;-; | 2241 | cdc7 | 32 | 171 | 145 | 94 | 118 | 79 |
| ENSDARG000 | 2 | 36098750 | 36098812 | + | 63 | NA | 0 | 0 | 0 | 0 | 0 | 0 |
| ENSDARG000 | 2;2;2;2;2;2;2;2; | 37207461:372 | 37207543:372 | +;+;+;+;+;+;+;+; | 1098 | apoda.2 | 0 | 0 | 26 | 1 | 28 | 0 |
| ENSDARG000 | 2 | 36098969 | 36099028 | + | 60 | NA | 0 | 0 | 0 | 0 | 0 | 0 |
| ENSDARG000 | 2;2;2;2;2;2;2;2; | 6999369:7014 | 6999730:7014 | +;+;+;+;+;+;+;+; | 3889 | xpr1b | 659 | 546 | 671 | 332 | 534 | 540 |
| ENSDARG000 | 2 | 36099210 | 36099272 | + | 63 | NA | 0 | 0 | 0 | 0 | 0 | 0 |
| ENSDARG000 | 2 | 36099905 | 36099964 | + | 60 | NA | 0 | 0 | 0 | 0 | 0 | 0 |
| ENSDARG000 | 2;2;2;2;2;2;2;2; | 44512324:445 | 44512909:445 | +;+;+;+;+;+;+;+; | 4818 | NA | 34 | 170 | 206 | 66 | 146 | 62 |
| ENSDARG000 | 2 | 36100207 | 36100269 | + | 63 | NA | 0 | 0 | 0 | 0 | 0 | 0 |
| ENSDARG000 | 2;2;2;2;2 | 37207799:372 | 37208293:372 | -;-;-;-;- | 853 | apoda.1 | 876 | 736 | 4266 | 274 | 913 | 2315 |
| ENSDARG000 | 2 | 36100465 | 36100527 | + | 63 | NA | 0 | 0 | 0 | 0 | 0 | 0 |
| ENSDARG000 | 2 | 36101377 | 36101442 | + | 66 | NA | 0 | 0 | 0 | 0 | 0 | 0 |
| ENSDARG000 | 2;2;2;2;2;2;2;2; | 13255986:132 | 13257225:132 | -;-;-;-;-;-;-;-; | 3006 | vps4b | 2600 | 1953 | 2035 | 1906 | 2401 | 2390 |
| ENSDARG000 | 2;2;2;2;2;2;2;2; | 50906097:509 | 50906515:509 | -;-;-;-;-;-;-;-; | 1742 | ghrhra | 0 | 0 | 2 | 0 | 0 | 0 |
| ENSDARG000 | 2;2;2 | 57879347:578 | 57879647:578 | -;-;- | 1108 | tpgs1 | 481 | 432 | 398 | 427 | 378 | 392 |
| ENSDARG000 | 2;2;2;2;2;2;2;2; | 5480125:5482 | 5482256:5482 | -;-;-;-;-;-;-;-; | 3385 | saga | 0 | 0 | 0 | 2 | 0 | 2 |
| ENSDARG000 | 2;2;2;2;2;2;2;2; | 26598801:266 | 26599767:266 | -;-;-;-;-;-;-;-; | 2249 | yipf1 | 261 | 212 | 319 | 237 | 294 | 330 |
| ENSDARG000 | 2;2;2 | 44522856:445 | 44523113:445 | -;-;- | 389 | NA | 0 | 0 | 0 | 0 | 0 | 0 |
| ENSDARG000 | 2;2;2 | 57913490:579 | 57914010:579 | -;-;- | 1378 | NA | 30 | 16 | 17 | 17 | 53 | 36 |
| ENSDARG000 | 2;2;2;2;2;2;2;2; | 57892666:578 | 57894203:578 | -;-;-;-;-;-;-;- | 2931 | NA | 172 | 105 | 87 | 73 | 143 | 113 |
| ENSDARG000 | 2;2;2;2;2;2;2;2; | 2994052:2994 | 2994158:2994 | +;+;+;+;+;+;+;+; | 2138 | ipp | 372 | 222 | 342 | 73 | 227 | 247 |
| ENSDARG000 | 2;2;2;2;2;2;2;2; | 57848844:578 | 57849055:578 | +;+;+;+;+;+;+;+; | 4948 | plekhj1 | 662 | 768 | 754 | 595 | 623 | 733 |
| ENSDARG000 | 2;2;2 | 13298051:132 | 13298483:132 | -;-;- | 623 | NA | 2 | 1 | 0 | 0 | 1 | 7 |
| ENSDARG000 | 2;2;2 | 24896992:248 | 24897072:248 | -;-;- | 637 | NA | 0 | 0 | 0 | 0 | 0 | 0 |
| ENSDARG000 | 2 | 36101762 | 36101824 | + | 63 | NA | 0 | 0 | 0 | 0 | 0 | 0 |
| ENSDARG000 | 2 | 36102061 | 36102123 | + | 63 | NA | 0 | 0 | 0 | 0 | 0 | 0 |
| ENSDARG000 | 2 | 36102608 | 36102670 | + | 63 | NA | 0 | 0 | 0 | 0 | 0 | 0 |
| ENSDARG000 | 2 | 36102854 | 36102916 | + | 63 | NA | 0 | 0 | 0 | 0 | 0 | 0 |
| ENSDARG000 | 2;2;2;2 | 47935476:479 | 47936331:479 | +;+;+;+ | 1348 | NA | 46 | 44 | 16 | 48 | 4 | 27 |
| ENSDARG000 | 2;2;2 | 33697340:336 | 33697718:336 | +;+;+ | 639 | NA | 0 | 0 | 0 | 0 | 0 | 0 |
| ENSDARG000 | 2;2;2;2;2;2;2;2; | 33681199:336 | 33682369:336 | -;-;-;-;-;-;-;- | 1955 | atp6v0b | 2903 | 3795 | 2641 | 1846 | 3154 | 3412 |
| ENSDARG000 | 2;2;2;2;2;2;2;2; | 33476766:334 | 33477238:334 | -;-;-;-;-;-;-;-; | 1986 | b4galt2 | 780 | 669 | 718 | 215 | 1323 | 1070 |
| ENSDARG000 | 2;2;2;2 | 24900571:249 | 24900747:249 | -;-;-;- | 699 | NA | 0 | 0 | 0 | 0 | 0 | 0 |
| ENSDARG000 | 2;2 | 29710857:297 | 29711451:297 | +;+ | 798 | ptf1a | 0 | 0 | 0 | 0 | 0 | 0 |
| ENSDARG000 | 2;2;2;2;2 | 33692165:336 | 33692356:336 | +;+;+;+;+ | 1337 | NA | 0 | 0 | 0 | 0 | 0 | 0 |
| ENSDARG000 | 2;2;2;2;2 | 5508542:5510 | 5508799:5510 | -;-;-;-;- | 933 | cldn15la | 4 | 0 | 0 | 2 | 2 | 0 |
| ENSDARG000 | 2;2;2;2;2;2;2;2; | 42942552:429 | 42943243:429 | -;-;-;-;-;-;-;-; | 3451 | oc90 | 0 | 0 | 0 | 0 | 0 | 0 |
| ENSDARG000 | 2;2 | 29712697:297 | 29712772:297 | +;+ | 525 | NA | 0 | 0 | 0 | 0 | 0 | 0 |
| ENSDARG000 | 2;2;2 | 24902808:249 | 24903061:249 | -;-;- | 801 | NA | 0 | 0 | 0 | 0 | 0 | 0 |
| ENSDARG000 | 2;2;2;2;2;2;2;2; | 29717786:297 | 29717884:297 | -;-;-;-;-;-;-;-; | 1682 | NA | 0 | 0 | 0 | 0 | 0 | 2 |
| ENSDARG000 | 2;2;2;2;2;2 | 36114192:361 | 36114257:361 | +;+;+;+;+;+ | 905 | NA | 0 | 0 | 0 | 0 | 0 | 0 |
| ENSDARG000 | 2;2;2;2 | 50967947:509 | 50968056:509 | +;+;+;+ | 471 | NA | 0 | 0 | 0 | 0 | 0 | 0 |
| ENSDARG000 | 2 | 36104161 | 36104223 | + | 63 | NA | 0 | 0 | 0 | 0 | 0 | 0 |
| ENSDARG000 | 2 | 36113846 | 36113908 | + | 63 | NA | 0 | 0 | 0 | 0 | 0 | 0 |
| ENSDARG000 | 2 | 36105789 | 36105854 | + | 66 | NA | 0 | 0 | 0 | 0 | 0 | 0 |
| ENSDARG000 | 2 | 36119542 | 36119604 | + | 63 | NA | 0 | 0 | 0 | 0 | 0 | 0 |
| ENSDARG000 | 2 | 36120881 | 36120946 | + | 66 | NA | 0 | 0 | 0 | 0 | 0 | 0 |

| | | | | | | | | | | | | |
|---|---|---|---|---|---|---|---|---|---|---|---|---|
| ENSDARG000 | 2 | 36108374 | 36108436 | + | 63 | NA | 0 | 0 | 0 | 0 | 0 | 0 |
| ENSDARG000 | 2 | 36116237 | 36116302 | + | 66 | NA | 0 | 0 | 0 | 0 | 0 | 0 |
| ENSDARG000 | 2 | 36108185 | 36108247 | + | 63 | NA | 0 | 0 | 0 | 0 | 0 | 0 |
| ENSDARG000 | 2 | 36117004 | 36117063 | + | 60 | NA | 0 | 0 | 0 | 0 | 0 | 0 |
| ENSDARG000 | 2 | 36105476 | 36105538 | + | 63 | NA | 0 | 0 | 0 | 0 | 0 | 0 |
| ENSDARG000 | 2 | 36121720 | 36121776 | + | 57 | NA | 0 | 0 | 0 | 0 | 0 | 0 |
| ENSDARG000 | 2 | 36120620 | 36120676 | + | 57 | NA | 0 | 0 | 0 | 0 | 0 | 0 |
| ENSDARG000 | 2 | 36120263 | 36120325 | + | 63 | NA | 0 | 0 | 0 | 0 | 0 | 0 |
| ENSDARG000 | 2 | 36112872 | 36112934 | + | 63 | NA | 0 | 0 | 0 | 0 | 0 | 0 |
| ENSDARG000 | 2 | 36114754 | 36114819 | + | 66 | NA | 0 | 0 | 0 | 0 | 0 | 0 |
| ENSDARG000 | 2 | 36109592 | 36109657 | + | 66 | NA | 0 | 0 | 0 | 0 | 0 | 0 |
| ENSDARG000 | 2 | 36118795 | 36118857 | + | 63 | NA | 0 | 0 | 0 | 0 | 0 | 0 |
| ENSDARG000 | 2 | 36122310 | 36122372 | + | 63 | NA | 0 | 0 | 0 | 0 | 0 | 0 |
| ENSDARG000 | 2 | 36123056 | 36123115 | + | 60 | NA | 0 | 0 | 0 | 0 | 0 | 0 |
| ENSDARG000 | 2 | 36119123 | 36119188 | + | 66 | NA | 0 | 0 | 0 | 0 | 0 | 0 |
| ENSDARG000 | 2;2;2;2;2 | 36103309:361 | 36103378:361 | +;+;+;+;+ | 399 | NA | 0 | 0 | 0 | 0 | 0 | 0 |
| ENSDARG000 | 2 | 36103931 | 36103993 | + | 63 | NA | 0 | 0 | 0 | 0 | 0 | 0 |
| ENSDARG000 | 2 | 36122605 | 36122664 | + | 60 | NA | 0 | 0 | 0 | 0 | 0 | 0 |
| ENSDARG000 | 2:2:2:2:2:2:2: | 9746437:9746 | 9746770:9746 | -:-:-:-:-:-:-:-: | 989 | NA | 0 | 0 | 1 | 0 | 3 | 0 |
| ENSDARG000 | 2 | 36117657 | 36117719 | + | 63 | NA | 0 | 0 | 0 | 0 | 0 | 0 |
| ENSDARG000 | 2;2 | 47942191:479 | 47942253:479 | +;+ | 668 | NA | 27 | 23 | 16 | 0 | 5 | 7 |
| ENSDARG000 | 2 | 36115571 | 36115627 | + | 57 | NA | 0 | 0 | 0 | 0 | 0 | 0 |
| ENSDARG000 | 2 | 36105192 | 36105254 | + | 63 | NA | 0 | 0 | 0 | 0 | 0 | 0 |
| ENSDARG000 | 2 | 36118397 | 36118462 | + | 66 | NA | 0 | 0 | 0 | 0 | 0 | 0 |
| ENSDARG000 | 2;2;2;2;2;2 | 36109002:361 | 36109071:361 | +;+;+;+;+;+ | 907 | NA | 0 | 0 | 0 | 0 | 0 | 0 |
| ENSDARG000 | 2 | 36124223 | 36124282 | + | 60 | NA | 0 | 0 | 0 | 0 | 0 | 0 |
| ENSDARG000 | 2 | 36123871 | 36123931 | + | 61 | NA | 0 | 0 | 0 | 0 | 0 | 0 |
| ENSDARG000 | 2 | 36107575 | 36107637 | + | 63 | NA | 0 | 0 | 0 | 0 | 0 | 0 |
| ENSDARG000 | 2 | 36110886 | 36110942 | + | 57 | NA | 0 | 0 | 0 | 0 | 0 | 0 |
| ENSDARG000 | 2 | 36107285 | 36107344 | + | 60 | NA | 0 | 0 | 0 | 0 | 0 | 0 |
| ENSDARG000 | 2 | 36113177 | 36113236 | + | 60 | NA | 0 | 0 | 0 | 0 | 0 | 0 |
| ENSDARG000 | 2 | 36115205 | 36115267 | + | 63 | NA | 0 | 0 | 0 | 0 | 0 | 0 |
| ENSDARG000 | 2;2 | 24013250:240 | 24013423:240 | +;+ | 440 | NA | 0 | 0 | 0 | 0 | 0 | 0 |
| ENSDARG000 | 2 | 36124755 | 36124817 | + | 63 | NA | 0 | 0 | 0 | 0 | 0 | 0 |
| ENSDARG000 | 2 | 36104569 | 36104634 | + | 66 | NA | 0 | 0 | 0 | 0 | 0 | 0 |
| ENSDARG000 | 2:2:2:2:2:2:2: | 23936954:239 | 23938040:239 | -:-:-:-:-:-:-:-: | 4796 | col15a1b | 644 | 876 | 1276 | 309 | 4937 | 1210 |
| ENSDARG000 | 2 | 36107846 | 36107911 | + | 66 | NA | 0 | 0 | 0 | 0 | 0 | 0 |
| ENSDARG000 | 2 | 36106071 | 36106133 | + | 63 | NA | 0 | 0 | 0 | 0 | 0 | 0 |
| ENSDARG000 | 2;2;2;2;2;2 | 36121373:361 | 36121436:361 | +;+;+;+;+;+ | 896 | NA | 0 | 0 | 0 | 0 | 0 | 0 |
| ENSDARG000 | 2 | 36110158 | 36110223 | + | 66 | NA | 0 | 0 | 0 | 0 | 0 | 0 |
| ENSDARG000 | 2 | 36111650 | 36111715 | + | 66 | NA | 0 | 0 | 0 | 0 | 0 | 0 |
| ENSDARG000 | 2 | 36117394 | 36117456 | + | 63 | NA | 0 | 0 | 0 | 0 | 2 | 0 |
| ENSDARG000 | 2;2;2;2;2 | 36112268:361 | 36112327:361 | +;+;+;+;+ | 981 | NA | 0 | 0 | 0 | 0 | 0 | 0 |
| ENSDARG000 | 2 | 36116544 | 36116606 | + | 63 | NA | 0 | 0 | 0 | 0 | 0 | 0 |
| ENSDARG000 | 2 | 36110489 | 36110551 | + | 63 | NA | 0 | 0 | 0 | 0 | 0 | 0 |
| ENSDARG000 | 2 | 36117949 | 36118011 | + | 63 | NA | 0 | 0 | 0 | 0 | 0 | 0 |
| ENSDARG000 | 2 | 36106830 | 36106892 | + | 63 | NA | 0 | 0 | 0 | 0 | 0 | 0 |
| ENSDARG000 | 2 | 36111275 | 36111337 | + | 63 | NA | 0 | 0 | 0 | 0 | 0 | 0 |
| ENSDARG000 | 2 | 24931677 | 24932753 | + | 1077 | AGTR1 | 4 | 0 | 3 | 0 | 2 | 2 |
| ENSDARG000 | 2;2;2 | 3416683:3417 | 3417279:3417 | -;-;- | 1350 | iba57 | 108 | 100 | 150 | 212 | 100 | 150 |
| ENSDARG000 | 2;2;2;2;2;2;2 | 41563073:415 | 41563860:415 | -;-;-;-;-;-;- | 2098 | ZNF622 | 613 | 694 | 951 | 517 | 749 | 605 |
| ENSDARG000 | 2:2:2:2:2:2:2: | 904362:90573 | 904437:90591 | +:+:+:+:+:+:+: | 2194 | ripk1l | 391 | 256 | 342 | 106 | 346 | 169 |
| ENSDARG000 | 2;2 | 3412076:3413 | 3412497:3416 | -;- | 3387 | NA | 14 | 11 | 42 | 8 | 23 | 24 |
| ENSDARG000 | 2:2:2:2:2:2:2: | 41546271:415 | 41546663:415 | -:-:-:-:-:-:-:-: | 1833 | d2hgdh | 41 | 76 | 117 | 42 | 126 | 99 |
| ENSDARG000 | 2;2;2;2;2;2 | 3403264:3405 | 3403861:3405 | +;+;+;+;+;+ | 1776 | jmjd4 | 47 | 27 | 27 | 51 | 50 | 42 |
| ENSDARG000 | 2;2;2;2;2;2;2 | 47944967:479 | 47945263:479 | +;+;+;+;+;+;+ | 2032 | NA | 6 | 0 | 0 | 10 | 4 | 1 |
| ENSDARG000 | 2:2:2:2:2:2:2: | 3387390:3387 | 3387962:3387 | -:-:-:-:-:-:-:-: | 2355 | snap47 | 231 | 110 | 204 | 113 | 186 | 170 |
| ENSDARG000 | 2:2:2:2:2:2:2: | 5521671:5521 | 5521904:5521 | +:+:+:+:+:+:+: | 2175 | NA | 17 | 0 | 2 | 0 | 13 | 3 |
| ENSDARG000 | 2:2:2:2:2:2:2: | 21090317:210 | 21090472:210 | +:+:+:+:+:+:+: | 2783 | NA | 56 | 97 | 148 | 32 | 67 | 71 |
| ENSDARG000 | 2:2:2:2:2:2:2: | 24944040:249 | 24944243:249 | -:-:-:-:-:-:-:-: | 4371 | HLTF | 228 | 401 | 360 | 282 | 389 | 456 |
| ENSDARG000 | 2;2 | 7080302:7080 | 7080457:7080 | -;- | 346 | NA | 0 | 0 | 0 | 0 | 0 | 0 |
| ENSDARG000 | 2;2;2;2;2;2 | 7106837:7107 | 7106954:7107 | +;+;+;+;+;+ | 1703 | NA | 8 | 0 | 11 | 2 | 6 | 2 |
| ENSDARG000 | 2:2:2:2:2:2:2: | 7075220:7076 | 7075582:7076 | +:+:+:+:+:+:+: | 2562 | cdc14aa | 2 | 4 | 4 | 0 | 0 | 0 |
| ENSDARG000 | 2:2:2:2:2:2:2: | 30916188:309 | 30916301:309 | +:+:+:+:+:+:+: | 5969 | myom1a | 0 | 2 | 4 | 9 | 20 | 0 |
| ENSDARG000 | 2 | 47955160 | 47957673 | - | 2514 | fzd8b | 2 | 0 | 0 | 10 | 0 | 0 |
| ENSDARG000 | 2;2;2;2;2;2;2 | 24936766:249 | 24936887:249 | +;+;+;+;+;+;+ | 2624 | gyg1a | 1069 | 1417 | 2540 | 1968 | 1614 | 2025 |
| ENSDARG000 | 2;2;2;2;2 | 47986662:479 | 47987374:479 | -;-;-;-;- | 1141 | si:ch211-77g1 | 0 | 0 | 2 | 0 | 0 | 0 |
| ENSDARG000 | 2:2:2:2:2:2:2: | 16160906:161 | 16161045:161 | +:+:+:+:+:+:+: | 1737 | selenoj | 1156 | 54 | 536 | 784 | 823 | 648 |
| ENSDARG000 | 2:2:2:2:2:2:2: | 32357226:323 | 32358283:323 | -:-:-:-:-:-:-:-: | 4084 | ubtfl | 1477 | 989 | 1271 | 907 | 1551 | 1745 |
| ENSDARG000 | 2:2:2:2:2:2:2: | 50999477:509 | 50999773:509 | +:+:+:+:+:+:+: | 5750 | eef1da | 3519 | 2607 | 3869 | 4093 | 2794 | 4688 |
| ENSDARG000 | 2;2;2;2;2;2;2 | 48073972:480 | 48074464:480 | +;+;+;+;+;+;+ | 1553 | NA | 4 | 4 | 0 | 2 | 2 | 8 |
| ENSDARG000 | 2;2;2;2 | 44545912:445 | 44545988:445 | +;+;+;+ | 1060 | mterf4 | 60 | 30 | 60 | 48 | 44 | 87 |
| ENSDARG000 | 2 | 3435888 | 3437862 | - | 1975 | cyp8b1.1 | 0 | 0 | 6 | 0 | 0 | 6 |
| ENSDARG000 | 2;2;2;2 | 935952:93674 | 935984:93692 | +;+;+;+ | 1828 | si:ch211-241e | 97 | 69 | 97 | 68 | 49 | 177 |
| ENSDARG000 | 2;2 | 13376099:133 | 13376446:133 | -;- | 1211 | NMUR1 | 0 | 0 | 0 | 0 | 0 | 0 |
| ENSDARG000 | 2;2;2;2 | 33722547:337 | 33722738:337 | +;+;+;+ | 1067 | NA | 0 | 0 | 0 | 0 | 0 | 0 |
| ENSDARG000 | 2;2 | 18405569:184 | 18405726:184 | +;+ | 469 | NA | 0 | 0 | 0 | 0 | 0 | 0 |
| ENSDARG000 | 2;2;2;2 | 29764395:297 | 29765073:297 | -;-;-;- | 1295 | NA | 10 | 0 | 0 | 0 | 0 | 0 |
| ENSDARG000 | 2;2;2;2 | 10878406:108 | 10878626:108 | +;+;+;+ | 961 | tceanc2 | 35 | 22 | 22 | 46 | 26 | 4 |
| ENSDARG000 | 2:2:2:2:2:2:2: | 21164930:211 | 21165941:211 | -:-:-:-:-:-:-:-: | 2303 | bmi1b | 146 | 195 | 298 | 206 | 245 | 115 |
| ENSDARG000 | 2;2;2 | 29842588:298 | 29843044:298 | -;-;- | 540 | NA | 0 | 0 | 0 | 0 | 0 | 0 |
| ENSDARG000 | 2:2:2:2:2:2:2: | 29795833:298 | 29796010:298 | +:+:+:+:+:+:+: | 8501 | NA | 124 | 6 | 4 | 4 | 36 | 32 |
| ENSDARG000 | 2:2:2:2:2:2:2: | 9755422:9757 | 9755587:9757 | +:+:+:+:+:+:+: | 1823 | pcyt1aa | 324 | 179 | 333 | 205 | 277 | 260 |
| ENSDARG000 | 2:2:2:2:2:2:2: | 13462305:134 | 13462548:134 | +:+:+:+:+:+:+: | 3165 | CFAP57 | 0 | 0 | 4 | 4 | 4 | 6 |
| ENSDARG000 | 2;2;2;2;2;2 | 57925518:579 | 57925555:579 | -;-;-;-;-;- | 650 | NA | 44 | 53 | 49 | 8 | 29 | 20 |
| ENSDARG000 | 2:2:2:2:2:2:2: | 44571200:445 | 44571391:445 | +:+:+:+:+:+:+: | 2264 | klhl24a | 768 | 283 | 416 | 340 | 660 | 847 |
| ENSDARG000 | 2 | 3452590 | 3454556 | + | 1967 | cyp8b1.2 | 2 | 4 | 0 | 0 | 6 | 0 |
| ENSDARG000 | 2;2 | 3468069:3468 | 3470019:3470 | +;+ | 1951 | cyp8b1.3 | 4 | 0 | 0 | 0 | 10 | 2 |
| ENSDARG000 | 2:2:2:2:2:2:2: | 48089057:480 | 48089253:480 | +:+:+:+:+:+:+: | 3846 | pitrm1 | 838 | 940 | 1161 | 536 | 960 | 586 |
| ENSDARG000 | 2:2:2:2:2:2:2: | 43001512:430 | 43002042:430 | -:-:-:-:-:-:-:-: | 3711 | KCNQ3 | 0 | 12 | 10 | 0 | 2 | 14 |
| ENSDARG000 | 2:2:2:2:2:2:2: | 18698388:187 | 18703514:187 | -:-:-:-:-:-:-:-: | 7713 | pbx1a | 18 | 4 | 18 | 10 | 54 | 2 |
| ENSDARG000 | 2;2;2 | 33726862:337 | 33727241:337 | +;+;+ | 640 | NA | 0 | 0 | 0 | 0 | 0 | 0 |
| ENSDARG000 | 2;2;2;2 | 33747509:337 | 33747700:337 | +;+;+;+ | 1067 | NA | 0 | 0 | 0 | 0 | 0 | 0 |
| ENSDARG000 | 2;2 | 3472832:3496 | 3472971:3498 | +;+ | 2074 | gjc2 | 0 | 0 | 4 | 0 | 2 | 0 |
| ENSDARG000 | 2:2:2:2:2:2:2: | 57953239:579 | 57957178:579 | -:-:-:-:-:-:-:-: | 6915 | NA | 56 | 66 | 62 | 54 | 166 | 112 |
| ENSDARG000 | 2;2;2 | 3502430:3502 | 3502565:3502 | +;+;+ | 917 | SERP1 | 1987 | 1836 | 2485 | 2586 | 1704 | 2750 |
| ENSDARG000 | 2:2:2:2:2:2:2: | 13694450:136 | 13695044:136 | +:+:+:+:+:+:+: | 5174 | ebna1bp2 | 852 | 892 | 1163 | 575 | 933 | 754 |
| ENSDARG000 | 2;2;2;2;2;2;2 | 26604773:266 | 26605409:266 | +;+;+;+;+;+;+ | 1537 | lrrc42 | 27 | 54 | 96 | 38 | 45 | 75 |
| ENSDARG000 | 2:2:2:2:2:2:2: | 10881168:108 | 10881650:108 | -:-:-:-:-:-:-:-: | 1853 | CDCP2 | 15 | 13 | 2 | 0 | 11 | 12 |
| ENSDARG000 | 2:2:2:2:2:2:2: | 7120582:7123 | 7123334:7124 | -:-:-:-:-:-:-:-: | 4150 | extl2 | 160 | 205 | 365 | 244 | 368 | 263 |
| ENSDARG000 | 2;2;2 | 33751490:337 | 33751868:337 | +;+;+ | 639 | NA | 0 | 0 | 0 | 0 | 0 | 0 |
| ENSDARG000 | 2:2:2:2:2:2:2: | 5543901:5546 | 5544272:5546 | -:-:-:-:-:-:-:-: | 2696 | parla | 69 | 84 | 66 | 47 | 95 | 75 |
| ENSDARG000 | 2;2;2 | 33756378:337 | 33756756:337 | +;+;+ | 645 | NA | 0 | 0 | 2 | 0 | 0 | 0 |
| ENSDARG000 | 2;2;2;2;2;2;2 | 10914023:109 | 10914528:109 | +;+;+;+;+;+;+ | 1506 | mrpl37 | 659 | 666 | 907 | 490 | 708 | 497 |
| ENSDARG000 | 2:2:2:2:2:2:2: | 26612574:266 | 26613183:266 | -:-:-:-:-:-:-:-: | 1594 | tmem59 | 2236 | 1728 | 1612 | 2097 | 2358 | 2118 |
| ENSDARG000 | 2:2:2:2:2:2:2: | 944341:94483 | 944743:94493 | -:-:-:-:-:-:-:-: | 4539 | NA | 248 | 46 | 14 | 24 | 16 | 62 |
| ENSDARG000 | 2:2:2:2:2:2:2: | 13721888:137 | 13722356:137 | +:+:+:+:+:+:+: | 3178 | ZBTB41 | 165 | 109 | 157 | 184 | 164 | 252 |
| ENSDARG000 | 2:2:2:2:2:2:2: | 13783393:137 | 13783507:137 | -:-:-:-:-:-:-:-: | 1467 | NA | 0 | 0 | 0 | 5 | 7 | 4 |
| ENSDARG000 | 2;2;2 | 948068:94965 | 948321:95009 | +;+;+ | 745 | NA | 0 | 0 | 0 | 0 | 0 | 0 |
| ENSDARG000 | 2:2:2:2:2:2:2: | 16173999:161 | 16174397:161 | +:+:+:+:+:+:+: | 3051 | sh3glb1b | 2835 | 3223 | 6306 | 4068 | 6649 | 4695 |
| ENSDARG000 | 2:2:2:2:2:2:2: | 29902294:299 | 29903310:299 | -:-:-:-:-:-:-:-: | 4637 | paxip1 | 539 | 523 | 911 | 356 | 574 | 634 |
| ENSDARG000 | 2:2:2:2:2:2:2: | 9789811:9789 | 9790606:9790 | -:-:-:-:-:-:-:-: | 2300 | ap2m1a | 10166 | 9197 | 9879 | 8200 | 9750 | 7653 |
| ENSDARG000 | 2;2 | 5563077:5577 | 5563613:5581 | +;+ | 4922 | NA | 24 | 32 | 18 | 16 | 12 | 36 |
| ENSDARG000 | 2;2;2;2;2 | 33796207:337 | 33796432:337 | +;+;+;+;+ | 1101 | kiss1ra | 0 | 0 | 0 | 0 | 0 | 0 |
| ENSDARG000 | 2:2:2:2:2:2:2: | 16198735:161 | 16199741:161 | -:-:-:-:-:-:-:-: | 11937 | NA | 3012 | 3850 | 3033 | 1790 | 3082 | 4220 |
| ENSDARG000 | 2:2:2:2:2:2:2: | 44615000:446 | 44615796:446 | +:+:+:+:+:+:+: | 4957 | YEATS2 | 310 | 222 | 439 | 223 | 399 | 378 |
| ENSDARG000 | 2:2:2:2:2:2:2: | 43098237:430 | 43098505:430 | -:-:-:-:-:-:-:-: | 2659 | dnaaf11 | 42 | 18 | 28 | 38 | 22 | 36 |
| ENSDARG000 | 2:2:2:2:2:2:2: | 44666705:446 | 44666869:446 | +:+:+:+:+:+:+: | 2475 | NA | 91 | 98 | 145 | 127 | 136 | 116 |
| ENSDARG000 | 2:2:2:2:2:2:2: | 18988407:189 | 18988506:189 | +:+:+:+:+:+:+: | 2176 | glula | 254 | 164 | 225 | 184 | 308 | 83 |
| ENSDARG000 | 2:2:2:2:2:2:2: | 24027744:240 | 24031286:240 | -:-:-:-:-:-:-:-: | 8353 | NA | 1187 | 1447 | 1076 | 1088 | 1683 | 2007 |
| ENSDARG000 | 2:2:2:2:2:2:2: | 3533117:3533 | 3533286:3533 | +:+:+:+:+:+:+: | 5942 | gpt | 20 | 110 | 136 | 136 | 338 | 98 |
| ENSDARG000 | 2:2:2:2:2:2:2: | 41577507:415 | 41583205:415 | -:-:-:-:-:-:-:-: | 7405 | SSBP4 | 527 | 409 | 961 | 321 | 964 | 616 |
| ENSDARG000 | 2:2:2:2:2:2:2: | 43128362:431 | 43128837:431 | -:-:-:-:-:-:-:-: | 1738 | NA | 160 | 95 | 132 | 73 | 115 | 221 |
| ENSDARG000 | 2:2:2:2:2:2:2: | 21174998:211 | 21175346:211 | +:+:+:+:+:+:+: | 2265 | dnajc1 | 1363 | 2048 | 2858 | 1853 | 1916 | 1510 |
| ENSDARG000 | 2:2:2:2:2:2:2: | 7132281:7132 | 7132420:7132 | +:+:+:+:+:+:+: | 1604 | zgc:110366 | 158 | 122 | 141 | 130 | 156 | 126 |
| ENSDARG000 | 2;2;2;2;2;2 | 43131147:431 | 43132377:431 | -;-;-;-;-;- | 2490 | ftr14 | 377 | 231 | 268 | 206 | 263 | 275 |
| ENSDARG000 | 2:2:2:2:2:2:2: | 5587495:5601 | 5587573:5601 | +:+:+:+:+:+:+: | 3311 | fgf12a | 4 | 8 | 0 | 0 | 4 | 0 |
| ENSDARG000 | 2:2:2:2:2:2:2: | 37227011:372 | 37227228:372 | +:+:+:+:+:+:+: | 2180 | samd7 | 4 | 1 | 0 | 2 | 6 | 0 |
| ENSDARG000 | 2;2 | 990043:99068 | 990589:99103 | +;+ | 903 | NA | 11 | 0 | 0 | 0 | 0 | 0 |
| ENSDARG000 | 2;2;2 | 44697568:447 | 44697812:447 | -;-;- | 938 | MAP6D1 | 0 | 1 | 3 | 1 | 2 | 0 |
| ENSDARG000 | 2:2:2:2:2:2:2: | 33926606:339 | 33927123:339 | +:+:+:+:+:+:+: | 5018 | kif2c | 378 | 953 | 1142 | 910 | 773 | 438 |
| ENSDARG000 | 2;2;2;2 | 19119847:191 | 19120274:191 | -;-;-;- | 746 | kncn | 0 | 0 | 0 | 0 | 0 | 0 |
| ENSDARG000 | 2;2 | 19100422:191 | 19105440:191 | -;- | 5058 | TMEM275 | 0 | 0 | 0 | 0 | 0 | 4 |
| ENSDARG000 | 2:2:2:2:2:2:2: | 43144938:431 | 43145233:431 | -:-:-:-:-:-:-:-: | 1430 | NA | 11 | 8 | 12 | 0 | 14 | 18 |
| ENSDARG000 | 2:2:2:2:2:2:2: | 30960351:309 | 30960564:309 | +:+:+:+:+:+:+: | 4923 | LPIN2 | 1029 | 1367 | 3042 | 5027 | 1700 | 1055 |
| ENSDARG000 | 2:2:2:2:2:2:2: | 22694382:227 | 22694490:227 | +:+:+:+:+:+:+: | 5584 | KIF1A | 8 | 6 | 12 | 0 | 10 | 47 |
| ENSDARG000 | 2:2:2:2:2:2:2: | 37245382:372 | 37245487:372 | +:+:+:+:+:+:+: | 3083 | sec62 | 1584 | 1739 | 2038 | 1426 | 2184 | 1846 |
| ENSDARG000 | 2;2;2 | 19152372:191 | 19153283:191 | -;-;- | 1037 | si:dkev-225f2 | 0 | 2 | 16 | 0 | 0 | 0 |
| ENSDARG000 | 2:2:2:2:2:2:2: | 19163965:191 | 19164236:191 | +:+:+:+:+:+:+: | 2111 | elovl1a | 44 | 24 | 98 | 10 | 52 | 2 |
| ENSDARG000 | 2:2:2:2:2:2:2: | 44725550:447 | 44727470:447 | -:-:-:-:-:-:-:-: | 3790 | acsm3 | 4 | 2 | 22 | 0 | 0 | 10 |
| ENSDARG000 | 2:2:2:2:2:2:2: | 29925561:299 | 29925828:299 | +:+:+:+:+:+:+: | 3058 | htr5aa | 4 | 0 | 0 | 0 | 0 | 0 |
| ENSDARG000 | 2:2:2:2:2:2:2: | 5709424:5711 | 5710000:5711 | -;-;-;-;-;-;-;- | 1937 | cwc25 | 587 | 492 | 748 | 554 | 287 | 426 |
| ENSDARG000 | 2:2:2:2:2:2:2: | 58887599:588 | 58888034:588 | -:-:-:-:-:-:-:-: | 3026 | MAU2 | 945 | 815 | 1020 | 786 | 1047 | 729 |
| ENSDARG000 | 2;2 | 29976419:299 | 29977235:299 | +;+ | 1825 | en2b | 3 | 4 | 34 | 4 | 3 | 2 |
| ENSDARG000 | 2:2:2:2:2:2:2: | 10934479:109 | 10935302:109 | -:-:-:-:-:-:-:-: | 2834 | ssbp3a | 3073 | 2353 | 2715 | 3922 | 2041 | 2013 |
| ENSDARG000 | 2;2 | 3595333:3597 | 3596180:3597 | +;+ | 1430 | c1ql3b | 4 | 6 | 22 | 0 | 8 | 2 |
| ENSDARG000 | 2;2;2 | 24963861:249 | 24964793:249 | -;-;- | 1736 | NA | 1008 | 1167 | 577 | 930 | 911 | 1020 |
| ENSDARG000 | 2;2 | 36187180:361 | 36187517:361 | -;- | 378 | NA | 8 | 4 | 0 | 2 | 2 | 0 |
| ENSDARG000 | 2 | 36189586 | 36189919 | - | 334 | NA | 0 | 0 | 0 | 0 | 0 | 0 |
| ENSDARG000 | 2:2:2:2:2:2:2: | 7149588:7155 | 7155290:7155 | -:-:-:-:-:-:-:-: | 9300 | RC3H1 | 1516 | 1329 | 1892 | 949 | 1939 | 1423 |
| ENSDARG000 | 2;2;2;2;2;2;2 | 41835025:418 | 41835576:418 | -;-;-;-;-;-;- | 2868 | keap1a | 915 | 885 | 855 | 599 | 1127 | 624 |
| ENSDARG000 | 2;2;2;2;2;2;2 | 51037175:510 | 51037841:510 | -;-;-;-;-;-;- | 1934 | NA | 0 | 0 | 0 | 0 | 0 | 0 |
| ENSDARG000 | 2;2 | 36190807:361 | 36191129:361 | -;- | 366 | NA | 7 | 4 | 0 | 7 | 0 | 0 |
| ENSDARG000 | 2;2 | 36193346:361 | 36193689:361 | -;- | 445 | NA | 21 | 22 | 5 | 12 | 2 | 8 |
| ENSDARG000 | 2:2:2:2:2:2:2: | 43159171:431 | 43159895:431 | -:-:-:-:-:-:-:-: | 1760 | NA | 1255 | 335 | 803 | 616 | 410 | 389 |
| ENSDARG000 | 2:2:2:2:2:2:2: | 29985664:299 | 29986660:299 | -:-:-:-:-:-:-:-: | 1861 | cnpy1 | 316 | 510 | 1046 | 342 | 694 | 330 |
| ENSDARG000 | 2;2;2;2 | 14007938:140 | 14008451:140 | -;-;-;- | 1338 | NA | 0 | 6 | 2 | 0 | 24 | 0 |
| ENSDARG000 | 2:2:2:2:2:2:2: | 9821757:9821 | 9821836:9821 | +:+:+:+:+:+:+: | 2160 | anxa13l | 1869 | 1113 | 985 | 1630 | 1149 | 2105 |
| ENSDARG000 | 2:2:2:2:2:2:2: | 41876595:418 | 41876930:418 | +:+:+:+:+:+:+: | 2700 | crlf1a | 462 | 601 | 303 | 358 | 226 | 336 |
| ENSDARG000 | 2:2:2:2:2:2:2: | 3603589:3603 | 3604005:3604 | -:-:-:-:-:-:-:-: | 1784 | pter | 128 | 132 | 232 | 265 | 182 | 126 |
| ENSDARG000 | 2:2:2:2:2:2:2: | 29995590:299 | 29995629:299 | +:+:+:+:+:+:+: | 2444 | NA | 0 | 0 | 1 | 0 | 0 | 2 |
| ENSDARG000 | 2:2:2:2:2:2:2: | 48119494:481 | 48121833:481 | -:-:-:-:-:-:-:-: | 5259 | PFKP | 1843 | 3243 | 6016 | 4599 | 3146 | 2212 |

| | | | | | | | | | | | | |
|---|---|---|---|---|---|---|---|---|---|---|---|---|
| ENSDARG000 | 2:2:2:2:2:2:2:2: | 26623377:266 | 26623871:266 | -:-:-:-:-:-:-:-: | 4081 | u2surp | 2670 | 2094 | 4271 | 3354 | 3300 | 3697 |
| ENSDARG000 | 2;2;2;2 | 41926707:419 | 41927060:419 | +;+;+;+ | 1163 | odf3l2a | 0 | 2 | 0 | 0 | 0 | 0 |
| ENSDARG000 | 2;2 | 36195585:361 | 36195961:361 | -;- | 426 | NA | 10 | 20 | 3 | 0 | 2 | 10 |
| ENSDARG000 | 2;2 | 24990398:249 | 24992592:249 | -;- | 2411 | slc35g2a | 20 | 3 | 4 | 2 | 10 | 8 |
| ENSDARG000 | 2;2 | 51071699:510 | 51071772:510 | +;+ | 385 | NA | 0 | 1 | 0 | 0 | 2 | 0 |
| ENSDARG000 | 2;2;2;2;2;2;2 | 51065777:510 | 51066370:510 | -;-;-;-;-;-;- | 1863 | ftr67 | 484 | 220 | 329 | 355 | 302 | 371 |
| ENSDARG000 | 2 | 36196420 | 36196737 | - | 318 | NA | 1 | 0 | 1 | 0 | 0 | 1 |
| ENSDARG000 | 2;2 | 36198657:361 | 36198997:361 | -;- | 387 | NA | 2 | 10 | 7 | 2 | 4 | 0 |
| ENSDARG000 | 2;2 | 36200154:362 | 36200476:362 | -;- | 396 | NA | 2 | 1 | 7 | 2 | 0 | 0 |
| ENSDARG000 | 2:2:2:2:2:2:2:2: | 44749201:447 | 44750127:447 | -:-:-:-:-:-:-:-: | 5618 | ncapd2 | 560 | 1371 | 2019 | 781 | 1357 | 616 |
| ENSDARG000 | 2;2 | 5726353:5728 | 5726715:5728 | -;- | 539 | sst2 | 0 | 0 | 0 | 0 | 0 | 0 |
| ENSDARG000 | 2:2:2:2:2:2:2:2: | 59015878:590 | 59016016:590 | +:+:+:+:+:+:+:+: | 940 | NA | 10 | 4 | 14 | 4 | 2 | 0 |
| ENSDARG000 | 2:2:2:2:2:2:2:2: | 37267241:372 | 37268153:372 | -:-:-:-:-:-:-:-: | 2517 | nadkb | 621 | 573 | 666 | 426 | 494 | 571 |
| ENSDARG000 | 2;2 | 5726629:5728 | 5726741:5729 | +;+ | 608 | NA | 0 | 0 | 0 | 0 | 0 | 0 |
| ENSDARG000 | 2:2:2:2:2:2:2:2: | 7192966:7196 | 7193225:7197 | +:+:+:+:+:+:+:+: | 4073 | NA | 4 | 0 | 0 | 0 | 0 | 0 |
| ENSDARG000 | 2;2 | 36201297:362 | 36201619:362 | -;- | 411 | NA | 0 | 0 | 0 | 0 | 0 | 0 |
| ENSDARG000 | 2;2 | 14009630:140 | 14010008:140 | -;- | 951 | NA | 2 | 8 | 2 | 0 | 4 | 0 |
| ENSDARG000 | 2;2;2 | 21206878:212 | 21207298:212 | -;-;- | 629 | NA | 6 | 2 | 0 | 0 | 10 | 0 |
| ENSDARG000 | 2;2;2;2;2;2 | 41936603:419 | 41936820:419 | -;-;-;-;-;- | 1036 | ebi3 | 50 | 4 | 22 | 16 | 16 | 41 |
| ENSDARG000 | 2:2:2:2:2:2:2:2: | 21185903:211 | 21186291:211 | +:+:+:+:+:+:+:+: | 1765 | si:dkev-29d8.: | 46 | 12 | 16 | 15 | 17 | 18 |
| ENSDARG000 | 2:2:2:2:2:2:2:2: | 30986996:309 | 30987324:309 | -:-:-:-:-:-:-:-: | 3304 | EMILIN2 | 802 | 296 | 408 | 1422 | 1117 | 1140 |
| ENSDARG000 | 2;2;2 | 14021956:140 | 14022058:140 | -;-;- | 412 | NA | 0 | 0 | 0 | 0 | 0 | 0 |
| ENSDARG000 | 2;2;2;2;2;2;2 | 5734870:5740 | 5736428:5740 | -;-;-;-;-;-;- | 4818 | NYAP2 | 0 | 4 | 0 | 0 | 4 | 0 |
| ENSDARG000 | 2:2:2:2:2:2:2:2: | 21229909:212 | 21230319:212 | +:+:+:+:+:+:+:+: | 2812 | NA | 2 | 0 | 0 | 0 | 6 | 2 |
| ENSDARG000 | 2 | 36205716 | 36206038 | - | 323 | NA | 8 | 4 | 51 | 17 | 25 | 0 |
| ENSDARG000 | 2:2:2:2:2:2:2:2: | 32391897:323 | 32394663:323 | -:-:-:-:-:-:-:-: | 6745 | slc4a2a | 1668 | 922 | 1552 | 2197 | 1896 | 1282 |
| ENSDARG000 | 2:2:2:2:2:2:2:2: | 59041081:590 | 59041218:590 | +:+:+:+:+:+:+:+: | 2118 | stk11 | 469 | 450 | 606 | 259 | 602 | 328 |
| ENSDARG000 | 2:2:2:2:2:2:2:2: | 43204919:432 | 43205314:432 | +:+:+:+:+:+:+:+: | 5376 | pard3ab | 781 | 331 | 640 | 382 | 870 | 445 |
| ENSDARG000 | 2;2;2;2;2 | 51088448:510 | 51091328:510 | -;-;-;-;- | 3435 | THEM6 | 54 | 98 | 105 | 18 | 118 | 48 |
| ENSDARG000 | 2:2:2:2:2:2:2:2: | 7213146:7213 | 7213445:7213 | +:+:+:+:+:+:+:+: | 3004 | NA | 0 | 0 | 0 | 2 | 0 | 0 |
| ENSDARG000 | 2;2 | 36206362:362 | 36206695:362 | -;- | 377 | NA | 16 | 10 | 70 | 30 | 10 | 2 |
| ENSDARG000 | 2;2;2 | 51098549:511 | 51099057:511 | -;-;- | 1125 | them6 | 6 | 29 | 12 | 28 | 9 | 48 |
| ENSDARG000 | 2 | 36209068 | 36209415 | - | 348 | NA | 24 | 3 | 60 | 14 | 42 | 13 |
| ENSDARG000 | 2:2:2:2:2:2:2:2: | 1001560:1002 | 1002222:1002 | +:+:+:+:+:+:+:+: | 7378 | NA | 10 | 0 | 46 | 28 | 10 | 24 |
| ENSDARG000 | 2:2:2:2:2:2:2:2: | 3630307:3631 | 3631645:3631 | -:-:-:-:-:-:-:-: | 3177 | MMP16 | 0 | 2 | 0 | 2 | 2 | 0 |
| ENSDARG000 | 2:2:2:2:2:2:2:2: | 14207559:142 | 14208522:142 | -:-:-:-:-:-:-:-: | 1415 | dynlt5 | 0 | 14 | 8 | 4 | 0 | 4 |
| ENSDARG000 | 2;2;2;2;2;2 | 51183320:511 | 51183416:511 | +;+;+;+;+;+ | 2554 | LRRC24 | 0 | 0 | 0 | 0 | 0 | 0 |
| ENSDARG000 | 2;2;2 | 41945975:419 | 41946325:419 | -;-;- | 471 | NA | 0 | 0 | 0 | 0 | 0 | 0 |
| ENSDARG000 | 2;2;2 | 59163700:591 | 59163844:591 | +;+;+ | 341 | NA | 0 | 0 | 0 | 0 | 0 | 0 |
| ENSDARG000 | 2:2:2:2:2:2:2:2: | 25108569:251 | 25109061:251 | -:-:-:-:-:-:-:-: | 2228 | pccb | 1212 | 1139 | 1319 | 871 | 1253 | 1055 |
| ENSDARG000 | 2;2;2;2;2 | 51220283:512 | 51220846:512 | -;-;-;-;- | 947 | pigrl4.2 | 10 | 2 | 0 | 3 | 4 | 18 |
| ENSDARG000 | 2;2 | 59171268:591 | 59171473:591 | +;+ | 449 | NA | 0 | 0 | 0 | 0 | 0 | 0 |
| ENSDARG000 | 2;2 | 31101088:311 | 31101249:311 | +;+ | 281 | NA | 0 | 0 | 0 | 0 | 0 | 0 |
| ENSDARG000 | 2;2 | 31107929:311 | 31108090:311 | +;+ | 281 | NA | 0 | 0 | 0 | 0 | 0 | 0 |
| ENSDARG000 | 2:2:2:2:2:2:2:2: | 25019387:250 | 25019536:250 | +:+:+:+:+:+:+:+: | 7677 | STAG1 | 486 | 356 | 572 | 235 | 630 | 528 |
| ENSDARG000 | 2;2 | 36212897:362 | 36213289:362 | -;- | 483 | NA | 19 | 8 | 39 | 23 | 26 | 5 |
| ENSDARG000 | 2;2 | 36213715:362 | 36214031:362 | -;- | 357 | NA | 2 | 6 | 14 | 12 | 8 | 0 |
| ENSDARG000 | 2;2 | 36214831:362 | 36215186:362 | -;- | 462 | NA | 2 | 8 | 0 | 0 | 0 | 0 |
| ENSDARG000 | 2;2 | 36216397:362 | 36216776:362 | -;- | 470 | NA | 3 | 0 | 7 | 1 | 0 | 11 |
| ENSDARG000 | 2:2:2:2:2:2:2:2: | 41949186:419 | 41949541:419 | -:-:-:-:-:-:-:-: | 6814 | NA | 7439 | 3443 | 4530 | 3735 | 4380 | 8320 |
| ENSDARG000 | 2:2:2:2:2:2:2:2: | 22765492:227 | 22765785:227 | +:+:+:+:+:+:+:+: | 3013 | ESPNL | 0 | 0 | 0 | 0 | 2 | 0 |
| ENSDARG000 | 2;2 | 36217875:362 | 36218215:362 | -;- | 431 | NA | 5 | 3 | 8 | 0 | 3 | 8 |
| ENSDARG000 | 2:2:2:2:2:2:2:2: | 38087413:380 | 38088597:380 | -:-:-:-:-:-:-:-: | 8826 | chd8 | 1493 | 1180 | 2071 | 830 | 1778 | 1192 |
| ENSDARG000 | 2;2;2;2;2 | 7238549:7241 | 7242092:7242 | -;-;-;-;- | 4280 | zgc:153115 | 736 | 1405 | 1260 | 549 | 1418 | 718 |
| ENSDARG000 | 2;2;2 | 24076013:240 | 24076240:240 | -;-;- | 505 | NA | 0 | 0 | 0 | 0 | 0 | 0 |
| ENSDARG000 | 2:2:2:2:2:2:2:2: | 24080694:240 | 24081125:240 | +:+:+:+:+:+:+:+: | 5168 | kcnh2a | 0 | 0 | 0 | 0 | 4 | 2 |
| ENSDARG000 | 2;2 | 36218876:362 | 36219204:362 | -;- | 372 | NA | 2 | 4 | 3 | 4 | 12 | 4 |
| ENSDARG000 | 2:2:2:2:2:2:2:2: | 59174872:591 | 59175249:591 | -;-;-;-;-;-;-;- | 1707 | NA | 8 | 14 | 7 | 2 | 2 | 8 |
| ENSDARG000 | 2:2:2:2:2:2:2:2: | 51251589:512 | 51251799:512 | -:-:-:-:-:-:-:-: | 1223 | NA | 0 | 0 | 0 | 0 | 0 | 0 |
| ENSDARG000 | 2:2:2:2:2:2:2:2: | 33969212:339 | 33969398:339 | -:-:-:-:-:-:-:-: | 4853 | ptch2 | 118 | 30 | 24 | 92 | 43 | 2 |
| ENSDARG000 | 2:2:2:2:2:2:2:2: | 19217586:192 | 19218266:192 | -:-:-:-:-:-:-:-: | 2601 | cdc20 | 116 | 1049 | 533 | 244 | 400 | 556 |
| ENSDARG000 | 2:2:2:2:2:2:2:2: | 14276639:142 | 14278863:142 | -:-:-:-:-:-:-:-: | 5324 | NA | 48 | 132 | 106 | 76 | 67 | 83 |
| ENSDARG000 | 2:2:2:2:2:2:2:2: | 5793908:5795 | 5794160:5795 | +:+:+:+:+:+:+:+: | 1574 | slc35d1b | 210 | 161 | 203 | 56 | 127 | 254 |
| ENSDARG000 | 2:2:2:2:2:2:2:2: | 7279646:7279 | 7279719:7279 | +:+:+:+:+:+:+:+: | 1193 | NA | 0 | 0 | 0 | 0 | 0 | 0 |
| ENSDARG000 | 2;2 | 36219587:362 | 36219924:362 | -;- | 378 | NA | 10 | 5 | 15 | 5 | 6 | 5 |
| ENSDARG000 | 2;2 | 36221823:362 | 36222195:362 | -;- | 419 | NA | 0 | 0 | 0 | 2 | 18 | 4 |
| ENSDARG000 | 2;2 | 36223759:362 | 36224106:362 | -;- | 378 | NA | 2 | 0 | 0 | 0 | 2 | 0 |
| ENSDARG000 | 2;2;2 | 51273795:512 | 51274106:512 | -;-;- | 683 | NA | 0 | 0 | 0 | 0 | 0 | 0 |
| ENSDARG000 | 2;2;2;2;2 | 14394120:143 | 14394174:143 | -;-;-;-;- | 409 | NA | 7 | 15 | 9 | 5 | 11 | 15 |
| ENSDARG000 | 2:2:2:2:2:2:2:2: | 31296644:312 | 31299733:312 | -:-:-:-:-:-:-:-: | 3791 | adcyap1b | 8 | 125 | 316 | 192 | 18 | 542 |
| ENSDARG000 | 2;2 | 36226781:362 | 36227070:362 | -;- | 330 | NA | 8 | 0 | 0 | 2 | 4 | 0 |
| ENSDARG000 | 2;2 | 7288097:7289 | 7288745:7289 | -;- | 1009 | si:dkevp-106c | 0 | 0 | 2 | 0 | 0 | 0 |
| ENSDARG000 | 2;2 | 36228816:362 | 36229189:362 | -;- | 417 | NA | 7 | 10 | 2 | 20 | 1 | 1 |
| ENSDARG000 | 2;2 | 36229529:362 | 36229851:362 | -;- | 366 | NA | 0 | 2 | 5 | 1 | 0 | 5 |
| ENSDARG000 | 2:2:2:2:2:2:2:2: | 31363286:313 | 31363923:313 | -:-:-:-:-:-:-:-: | 2085 | clul1 | 10 | 9 | 10 | 5 | 1 | 22 |
| ENSDARG000 | 2:2:2:2:2:2:2:2: | 31271082:312 | 31271280:312 | +:+:+:+:+:+:+:+: | 6578 | yes1 | 9620 | 6903 | 8340 | 6695 | 10231 | 8169 |
| ENSDARG000 | 2:2:2:2:2:2:2:2: | 44781185:447 | 44781686:447 | +:+:+:+:+:+:+:+: | 4735 | EEF1AKMT4 | 76 | 42 | 83 | 50 | 44 | 24 |
| ENSDARG000 | 2;2 | 44923799:449 | 44924182:449 | +;+ | 675 | NA | 0 | 0 | 0 | 0 | 0 | 0 |
| ENSDARG000 | 2;2;2 | 48182604:481 | 48183236:481 | -;-;- | 919 | NA | 0 | 2 | 4 | 0 | 0 | 0 |
| ENSDARG000 | 2;2 | 36231242:362 | 36231540:362 | -;- | 348 | NA | 0 | 0 | 0 | 0 | 8 | 0 |
| ENSDARG000 | 2;2 | 36234304:362 | 36234680:362 | -;- | 420 | NA | 25 | 11 | 1 | 12 | 20 | 7 |
| ENSDARG000 | 2;2 | 36235043:362 | 36235347:362 | -;- | 348 | NA | 0 | 0 | 0 | 0 | 4 | 1 |
| ENSDARG000 | 2:2:2:2:2:2:2:2: | 7295515:7296 | 7295603:7296 | +:+:+:+:+:+:+:+: | 6581 | NA | 6 | 0 | 4 | 20 | 14 | 10 |
| ENSDARG000 | 2;2;2 | 16365768:163 | 16366209:163 | -;-;- | 3284 | NA | 779 | 674 | 626 | 277 | 384 | 770 |
| ENSDARG000 | 2:2:2:2:2:2:2:2: | 59190666:592 | 59191040:592 | -:-:-:-:-:-:-:-: | 1774 | NA | 1 | 0 | 2 | 0 | 2 | 22 |
| ENSDARG000 | 2:2:2:2:2:2:2:2: | 51281856:512 | 51281897:512 | -:-:-:-:-:-:-:-: | 1769 | pigrl3.5 | 0 | 0 | 0 | 0 | 0 | 0 |
| ENSDARG000 | 2:2:2:2:2:2:2:2: | 22795494:227 | 22795715:227 | +:+:+:+:+:+:+:+: | 3003 | rab6bb | 99 | 132 | 56 | 228 | 50 | 186 |
| ENSDARG000 | 2;2;2;2;2;2 | 51281779:512 | 51281897:512 | -;-;-;-;-;- | 877 | NA | 0 | 0 | 0 | 0 | 0 | 0 |
| ENSDARG000 | 2;2 | 51300617:513 | 51300926:513 | -;- | 631 | NA | 0 | 0 | 0 | 0 | 0 | 0 |
| ENSDARG000 | 2;2 | 44943396:449 | 44945065:449 | -;- | 2125 | camk2n1a | 12 | 108 | 117 | 274 | 70 | 64 |
| ENSDARG000 | 2;2 | 36238336:362 | 36238712:362 | -;- | 420 | NA | 3 | 6 | 7 | 4 | 5 | 2 |
| ENSDARG000 | 2;2 | 36241408:362 | 36241750:362 | -;- | 383 | NA | 9 | 0 | 0 | 4 | 0 | 0 |
| ENSDARG000 | 2;2 | 36242122:362 | 36242444:362 | -;- | 366 | NA | 2 | 0 | 3 | 9 | 4 | 12 |
| ENSDARG000 | 2;2;2;2;2;2;2 | 51321773:513 | 51321891:513 | -;-;-;-;-;-;- | 1690 | NA | 0 | 1 | 0 | 0 | 6 | 0 |
| ENSDARG000 | 2:2:2:2:2:2:2:2: | 14444276:144 | 14444303:144 | -:-:-:-:-:-:-:-: | 2336 | PDE4B | 9 | 5 | 74 | 184 | 167 | 29 |
| ENSDARG000 | 2;2 | 36244910:362 | 36245253:362 | -;- | 390 | NA | 0 | 0 | 0 | 0 | 0 | 0 |
| ENSDARG000 | 2;2 | 36247073:362 | 36247401:362 | -;- | 369 | NA | 20 | 6 | 6 | 1 | 8 | 0 |
| ENSDARG000 | 2 | 36249400 | 36249731 | - | 332 | NA | 0 | 0 | 0 | 0 | 0 | 2 |
| ENSDARG000 | 2;2;2 | 14489942:144 | 14490454:144 | -;-;- | 745 | NA | 0 | 0 | 15 | 2 | 4 | 0 |
| ENSDARG000 | 2:2:2:2:2:2:2:2: | 3696038:3696 | 3696273:3696 | +:+:+:+:+:+:+:+: | 8594 | egfra | 170 | 70 | 169 | 18 | 66 | 30 |
| ENSDARG000 | 2:2:2:2:2:2:2:2: | 32475794:324 | 32476376:324 | -:-:-:-:-:-:-:-: | 1901 | ttc19 | 125 | 182 | 128 | 70 | 140 | 166 |
| ENSDARG000 | 2:2:2:2:2:2:2:2: | 37307326:373 | 37307568:373 | -:-:-:-:-:-:-:-: | 2202 | SKIL | 72 | 55 | 60 | 37 | 86 | 87 |
| ENSDARG000 | 2;2 | 36251713:362 | 36252065:362 | -;- | 393 | NA | 0 | 0 | 0 | 0 | 0 | 0 |
| ENSDARG000 | 2:2:2:2:2:2:2:2: | 59221948:592 | 59222325:592 | -;-;-;-;-;-;-;- | 3579 | NA | 2 | 0 | 0 | 0 | 1 | 3 |
| ENSDARG000 | 2:2:2:2:2:2:2:2: | 5811482:5817 | 5814049:5817 | -:-:-:-:-:-:-:-: | 3766 | mier1b | 214 | 207 | 246 | 168 | 288 | 238 |
| ENSDARG000 | 2;2 | 36253604:362 | 36253923:362 | -;- | 357 | NA | 4 | 0 | 0 | 0 | 0 | 2 |
| ENSDARG000 | 2;2 | 36254217:362 | 36254572:362 | -;- | 396 | NA | 0 | 0 | 0 | 0 | 0 | 8 |
| ENSDARG000 | 2;2;2;2;2 | 44960876:449 | 44966236:449 | -;-;-;-;- | 5818 | mul1a | 168 | 293 | 313 | 247 | 180 | 344 |
| ENSDARG000 | 2:2:2:2:2:2:2:2: | 42005217:420 | 42005324:420 | +:+:+:+:+:+:+:+: | 2410 | NA | 1275 | 594 | 522 | 370 | 435 | 531 |
| ENSDARG000 | 2;2;2;2;2 | 51383977:513 | 51384308:513 | -;-;-;-;- | 711 | NA | 0 | 0 | 0 | 0 | 0 | 0 |
| ENSDARG000 | 2:2:2:2:2:2:2:2: | 16384285:163 | 16384689:163 | -:-:-:-:-:-:-:-: | 8355 | ATR | 469 | 43393 | 39427 | 23690 | 22314 | 46728 |
| ENSDARG000 | 2 | 51415503 | 51415811 | - | 309 | NA | 0 | 0 | 0 | 0 | 2 | 0 |
| ENSDARG000 | 2;2 | 36256312:362 | 36256643:362 | -;- | 396 | NA | 0 | 0 | 0 | 0 | 6 | 2 |
| ENSDARG000 | 2:2:2:2:2:2:2:2: | 11028923:110 | 11029292:110 | +:+:+:+:+:+:+:+: | 2575 | acot11a | 26 | 79 | 107 | 128 | 21 | 26 |
| ENSDARG000 | 2:2:2:2:2:2:2:2: | 21309272:213 | 21309862:213 | +:+:+:+:+:+:+:+: | 3714 | NA | 1066 | 328 | 498 | 233 | 413 | 52 |
| ENSDARG000 | 2;2 | 21312742:213 | 21313000:213 | -;- | 529 | NA | 0 | 0 | 0 | 0 | 0 | 0 |
| ENSDARG000 | 2;2;2;2;2 | 59245240:592 | 59245617:592 | -;-;-;-;- | 928 | NA | 9 | 5 | 4 | 0 | 0 | 14 |
| ENSDARG000 | 2 | 51429173 | 51429490 | - | 318 | NA | 0 | 0 | 0 | 0 | 0 | 0 |
| ENSDARG000 | 2;2 | 36258891:362 | 36259247:362 | -;- | 397 | NA | 0 | 0 | 0 | 6 | 0 | 0 |
| ENSDARG000 | 2;2 | 36259718:362 | 36260049:362 | -;- | 399 | NA | 4 | 0 | 0 | 0 | 0 | 0 |
| ENSDARG000 | 2;2 | 36261170:362 | 36261489:362 | -;- | 381 | NA | 0 | 0 | 0 | 0 | 4 | 0 |
| ENSDARG000 | 2;2;2;2;2;2;2 | 7698446:7702 | 7699291:7702 | -;-;-;-;-;-;- | 1618 | dcun1d1 | 313 | 408 | 354 | 182 | 357 | 246 |
| ENSDARG000 | 2;2 | 51451103:514 | 51451413:514 | -;- | 639 | NA | 0 | 0 | 0 | 0 | 0 | 0 |
| ENSDARG000 | 2;2;2;2 | 51461380:514 | 51461580:514 | -;-;-;- | 668 | NA | 0 | 0 | 0 | 0 | 0 | 0 |
| ENSDARG000 | 2;2 | 36263197:362 | 36263506:362 | -;- | 371 | NA | 0 | 0 | 0 | 0 | 0 | 0 |
| ENSDARG000 | 2:2:2:2:2:2:2:2: | 22808946:228 | 22809092:228 | +:+:+:+:+:+:+:+: | 2306 | aire | 8 | 57 | 10 | 9 | 14 | 6 |
| ENSDARG000 | 2;2 | 14779104:147 | 14779374:147 | +;+ | 690 | NA | 28 | 10 | 14 | 0 | 0 | 0 |
| ENSDARG000 | 2;2;2;2 | 38121415:381 | 38121758:381 | -;-;-;- | 1107 | cbln12 | 2 | 0 | 4 | 0 | 2 | 0 |
| ENSDARG000 | 2;2 | 5838938:5840 | 5839582:5840 | -;- | 795 | NA | 0 | 12 | 28 | 20 | 0 | 0 |
| ENSDARG000 | 2:2:2:2:2:2:2:2: | 48203278:482 | 48203972:482 | -:-:-:-:-:-:-:-: | 4696 | gigyf2 | 1337 | 1152 | 2052 | 832 | 1376 | 1025 |
| ENSDARG000 | 2:2:2:2:2:2:2:2: | 32494212:324 | 32494587:324 | -:-:-:-:-:-:-:-: | 1250 | faim2a | 270 | 160 | 298 | 213 | 299 | 278 |
| ENSDARG000 | 2:2:2:2:2:2:2:2: | 42020915:420 | 42021610:420 | -:-:-:-:-:-:-:-: | 2687 | gbp1 | 1563 | 1692 | 1204 | 362 | 1232 | 997 |
| ENSDARG000 | 2;2;2;2 | 51592392:515 | 51592516:515 | -;-;-;- | 818 | NA | 0 | 0 | 0 | 0 | 2 | 0 |
| ENSDARG000 | 2;2 | 25124498:251 | 25125496:251 | +;+ | 4040 | msl2a | 306 | 419 | 544 | 260 | 440 | 490 |
| ENSDARG000 | 2:2:2:2:2:2:2:2: | 44972720:449 | 44972953:449 | +:+:+:+:+:+:+:+: | 2277 | alg3 | 230 | 284 | 349 | 184 | 337 | 278 |
| ENSDARG000 | 2:2:2:2:2:2:2:2: | 32505554:325 | 32506092:325 | -:-:-:-:-:-:-:-: | 2414 | abcf2a | 1464 | 3178 | 2713 | 555 | 4020 | 1384 |
| ENSDARG000 | 2:2:2:2:2:2:2:2: | 37365551:373 | 37367146:373 | -:-:-:-:-:-:-:-: | 3778 | prkci | 1711 | 2170 | 2157 | 1427 | 1874 | 2072 |
| ENSDARG000 | 2:2:2:2:2:2:2:2: | 34008798:340 | 34009120:340 | -:-:-:-:-:-:-:-: | 2342 | eif2b3 | 474 | 487 | 665 | 440 | 478 | 333 |
| ENSDARG000 | 2:2:2:2:2:2:2:2: | 38127986:381 | 38128501:381 | -;-;-;-;-;-;-;- | 1964 | NA | 6 | 22 | 17 | 47 | 32 | 20 |
| ENSDARG000 | 2;2 | 19252270:192 | 19252834:192 | -;- | 638 | NA | 0 | 0 | 1 | 2 | 0 | 0 |
| ENSDARG000 | 2;2 | 51610584:516 | 51610912:516 | -;- | 673 | NA | 0 | 0 | 0 | 0 | 0 | 0 |
| ENSDARG000 | 2;2;2;2;2;2 | 21326957:213 | 21327717:213 | -;-;-;-;-;- | 2528 | klhl40a | 84 | 4 | 30 | 65 | 67 | 0 |
| ENSDARG000 | 2:2:2:2:2:2:2:2: | 19236677:192 | 19236795:192 | +:+:+:+:+:+:+:+: | 6483 | CC2D1B | 810 | 773 | 758 | 722 | 646 | 939 |
| ENSDARG000 | 2:2:2:2:2:2:2:2: | 21333529:213 | 21338295:213 | -:-:-:-:-:-:-:-: | 6344 | hhatla | 7200 | 322 | 3630 | 2609 | 4091 | 224 |
| ENSDARG000 | 2:2:2:2:2:2:2:2: | 14790294:147 | 14791773:147 | -:-:-:-:-:-:-:-: | 2746 | NA | 0 | 32 | 25 | 161 | 10 | 23 |
| ENSDARG000 | 2:2:2:2:2:2:2:2: | 25131349:251 | 25132836:251 | -:-:-:-:-:-:-:-: | 2393 | nceh1a | 6 | 1 | 4 | 7 | 0 | 11 |
| ENSDARG000 | 2:2:2:2:2:2:2:2: | 22851832:228 | 22851960:228 | +:+:+:+:+:+:+:+: | 2985 | amotl2b | 3914 | 3255 | 3038 | 3106 | 5792 | 2561 |
| ENSDARG000 | 2;2;2 | 25130523:251 | 25130831:251 | +;+;+ | 1661 | NA | 0 | 0 | 0 | 0 | 0 | 0 |
| ENSDARG000 | 2:2:2:2:2:2:2:2: | 59255193:592 | 59255570:592 | -;-;-;-;-;-;-;- | 1628 | NA | 3 | 16 | 3 | 1 | 0 | 3 |
| ENSDARG000 | 2:2:2:2:2:2:2:2: | 51613453:516 | 51614354:516 | -:-:-:-:-:-:-:-: | 1836 | NA | 16 | 0 | 16 | 0 | 4 | 0 |
| ENSDARG000 | 2:2:2:2:2:2:2:2: | 7715810:7715 | 7715872:7715 | +:+:+:+:+:+:+:+: | 1910 | eif4a2 | 819 | 1430 | 2416 | 2600 | 1906 | 2091 |
| ENSDARG000 | 2:2:2:2:2:2:2:2: | 5841108:5843 | 5841207:5844 | +:+:+:+:+:+:+:+: | 5183 | DIS3L2 | 214 | 316 | 427 | 235 | 371 | 214 |
| ENSDARG000 | 2;2;2;2 | 9848636:9850 | 9850476:9850 | -;-;-;- | 2961 | NA | 18 | 12 | 0 | 8 | 8 | 0 |
| ENSDARG000 | 2;2 | 25158342:251 | 25158465:251 | -;- | 574 | NA | 0 | 0 | 0 | 0 | 0 | 0 |
| ENSDARG000 | 2;2;2;2;2 | 38147761:381 | 38147862:381 | +;+;+;+;+ | 6324 | NA | 136 | 73 | 172 | 65 | 86 | 83 |
| ENSDARG000 | 2:2:2:2:2:2:2:2: | 26647472:266 | 26647740:266 | +:+:+:+:+:+:+:+: | 1923 | ttpa | 58 | 0 | 72 | 32 | 34 | 12 |
| ENSDARG000 | 2;2 | 36264323:362 | 36264624:362 | -;- | 342 | NA | 0 | 0 | 0 | 0 | 0 | 0 |
| ENSDARG000 | 2:2:2:2:2:2:2:2: | 43469241:434 | 43469707:434 | +:+:+:+:+:+:+:+: | 2026 | nrp1b | 36 | 22 | 26 | 24 | 28 | 64 |
| ENSDARG000 | 2;2 | 14938586:149 | 14938879:149 | -;- | 723 | NA | 0 | 0 | 0 | 0 | 0 | 2 |
| ENSDARG000 | 2 | 9862833 | 9863388 | - | 556 | NA | 44 | 64 | 92 | 148 | 20 | 54 |
| ENSDARG000 | 2:2:2:2:2:2:2:2: | 24177006:241 | 24177242:241 | +:+:+:+:+:+:+:+: | 7610 | NA | 10 | 14 | 18 | 18 | 16 | 24 |
| ENSDARG000 | 2:2:2:2:2:2:2:2: | 38161318:381 | 38161783:381 | +:+:+:+:+:+:+:+: | 4473 | mmp14b | 3177 | 4338 | 4735 | 2044 | 3614 | 2469 |

| | | | | | | | | | | | | |
|---|---|---|---|---|---|---|---|---|---|---|---|---|
| ENSDARG000 | 2;2;2 | 51640591:516 | 51641038:516 | -;-;- | 890 | dad1 | 1260 | 1413 | 2137 | 1190 | 1530 | 1268 |
| ENSDARG000 | 2;2;2;2;2;2;2; | 11069055:110 | 11069899:110 | -;-;-;-;-;-;-;- | 2250 | fam151a | 36 | 0 | 13 | 2 | 16 | 6 |
| ENSDARG000 | 2;2 | 36266386:362 | 36266691:362 | -;- | 346 | NA | 0 | 0 | 0 | 0 | 0 | 0 |
| ENSDARG000 | 2;2;2;2;2;2;2; | 25198648:252 | 25198715:252 | +;+;+;+;+;+;+; | 6371 | PPP2R3A | 51 | 38 | 44 | 62 | 19 | 45 |
| ENSDARG000 | 2;2 | 36269807:362 | 36270170:362 | -;- | 404 | NA | 0 | 0 | 0 | 0 | 0 | 0 |
| ENSDARG000 | 2;2;2 | 14970007:149 | 14970325:149 | +;+;+ | 832 | NA | 846 | 0 | 348 | 157 | 191 | 641 |
| ENSDARG000 | 2;2 | 36273162:362 | 36273471:362 | -;- | 350 | NA | 0 | 0 | 0 | 0 | 0 | 0 |
| ENSDARG000 | 2;2 | 36276985:362 | 36277360:362 | -;- | 413 | NA | 0 | 0 | 0 | 0 | 0 | 0 |
| ENSDARG000 | 2;2;2;2;2;2;2; | 59278841:592 | 59279218:592 | -;-;-;-;-;-;-;- | 2073 | NA | 1 | 0 | 0 | 4 | 0 | 0 |
| ENSDARG000 | 2;2;2;2;2;2;2; | 51645164:516 | 51645315:516 | +;+;+;+;+;+;+; | 1551 | abhd4 | 403 | 207 | 189 | 240 | 230 | 214 |
| ENSDARG000 | 2;2;2;2;2;2;2; | 5887626:5887 | 5887936:5887 | +;+;+;+;+;+;+; | 3600 | ACAP2 | 28 | 23 | 8 | 3 | 14 | 19 |
| ENSDARG000 | 2;2;2;2;2;2;2; | 21356242:213 | 21356896:213 | +;+;+;+;+;+;+; | 1642 | ctdsplb | 3013 | 1160 | 1648 | 1677 | 1803 | 746 |
| ENSDARG000 | 2;2;2;2;2;2;2; | 9868601:9869 | 9869233:9869 | -;-;-;-;-;-;-; | 2087 | abi1b | 60 | 72 | 185 | 206 | 42 | 87 |
| ENSDARG000 | 2;2;2;2;2;2;2; | 11100613:111 | 11104806:111 | -;-;-;-;-;-;-; | 6002 | cryz | 219 | 852 | 461 | 479 | 509 | 648 |
| ENSDARG000 | 2;2;2;2;2;2;2; | 31382590:313 | 31383038:313 | +;+;+;+;+;+;+; | 4850 | colec12 | 212 | 93 | 86 | 299 | 61 | 177 |
| ENSDARG000 | 2;2;2 | 14982383:149 | 14982502:149 | -;-;- | 435 | HCCS | 114 | 12 | 55 | 14 | 32 | 59 |
| ENSDARG000 | 2;2 | 36278980:362 | 36279326:362 | -;- | 387 | NA | 0 | 0 | 0 | 0 | 0 | 0 |
| ENSDARG000 | 2;2;2;2;2;2 | 11119315:111 | 11119671:111 | +;+;+;+;+;+ | 1488 | tyw3 | 42 | 67 | 74 | 40 | 49 | 93 |
| ENSDARG000 | 2;2 | 37405351:374 | 37405499:374 | -;- | 741 | NA | 0 | 0 | 2 | 0 | 0 | 0 |
| ENSDARG000 | 2;2;2;2;2;2;2; | 11083311:110 | 11084588:110 | -;-;-;-;-;-;-;- | 2761 | NA | 16 | 4 | 16 | 0 | 6 | 2 |
| ENSDARG000 | 2;2 | 36292896:362 | 36293205:362 | -;- | 380 | NA | 0 | 3 | 9 | 0 | 35 | 2 |
| ENSDARG000 | 2;2 | 36293820:362 | 36294140:362 | -;- | 358 | NA | 0 | 0 | 0 | 0 | 0 | 0 |
| ENSDARG000 | 2;2 | 36297301:362 | 36297652:362 | -;- | 389 | NA | 0 | 0 | 0 | 0 | 0 | 0 |
| ENSDARG000 | 2;2;2;2;2;2;2; | 59295260:592 | 59295637:592 | -;-;-;-;-;-;-;- | 1629 | NA | 3 | 9 | 7 | 1 | 2 | 12 |
| ENSDARG000 | 2;2;2;2;2;2;2; | 44984509:449 | 44985118:449 | -;-;-;-;-;-;-; | 4043 | NA | 0 | 2 | 6 | 2 | 16 | 0 |
| ENSDARG000 | 2;2;2;2;2;2;2; | 42039036:420 | 42039361:420 | -;-;-;-;-;-;-; | 2950 | NA | 207 | 213 | 297 | 120 | 178 | 174 |
| ENSDARG000 | 2;2 | 7749394:7757 | 7750989:7757 | -;- | 1810 | b3gnt5b | 8 | 20 | 10 | 4 | 10 | 44 |
| ENSDARG000 | 2 | 48263750 | 48264602 | + | 853 | NA | 6 | 0 | 4 | 2 | 6 | 14 |
| ENSDARG000 | 2;2;2;2;2;2;2; | 34112100:341 | 34112215:341 | +;+;+;+;+;+;+; | 3029 | klhl20 | 660 | 685 | 591 | 361 | 721 | 530 |
| ENSDARG000 | 2;2 | 36302279:363 | 36302580:363 | -;- | 336 | NA | 0 | 0 | 0 | 0 | 0 | 0 |
| ENSDARG000 | 2;2 | 36303393:363 | 36303727:363 | -;- | 375 | NA | 0 | 0 | 0 | 0 | 0 | 0 |
| ENSDARG000 | 2;2;2;2;2;2 | 48284464:482 | 48285503:482 | -;-;-;-;-;- | 1857 | itm2cb | 407 | 206 | 441 | 296 | 248 | 474 |
| ENSDARG000 | 2;2;2;2;2;2;2 | 14992879:149 | 14993275:149 | +;+;+;+;+;+;+ | 1383 | NA | 88 | 43 | 61 | 94 | 20 | 268 |
| ENSDARG000 | 2 | 48282590 | 48284160 | + | 1571 | lpar5a | 2 | 14 | 24 | 2 | 6 | 28 |
| ENSDARG000 | 2;2;2;2;2;2;2; | 48288461:483 | 48288517:483 | +;+;+;+;+;+;+; | 1497 | hes6 | 174 | 139 | 212 | 160 | 276 | 120 |
| ENSDARG000 | 2;2;2;2;2;2;2 | 37424261:374 | 37424649:374 | +;+;+;+;+;+;+ | 1218 | NA | 23 | 16 | 35 | 40 | 60 | 71 |
| ENSDARG000 | 2;2 | 36304603:363 | 36304946:363 | -;- | 381 | NA | 0 | 0 | 0 | 0 | 0 | 2 |
| ENSDARG000 | 2;2 | 36306186:363 | 36306510:363 | -;- | 397 | NA | 0 | 0 | 0 | 0 | 0 | 0 |
| ENSDARG000 | 2;2;2;2 | 1124512:1126 | 1124670:1126 | +;+;+;+ | 613 | polr2d | 418 | 430 | 579 | 348 | 522 | 292 |
| ENSDARG000 | 2;2;2;2;2;2;2; | 7764671:7772 | 7768806:7772 | -;-;-;-;-;-;-; | 5999 | tbl1xr1b | 492 | 549 | 879 | 432 | 684 | 628 |
| ENSDARG000 | 2;2;2 | 5927255:5927 | 5927341:5927 | +;+;+ | 3969 | NA | 0 | 0 | 0 | 10 | 8 | 0 |
| ENSDARG000 | 2;2 | 36308826:363 | 36309118:363 | -;- | 339 | NA | 0 | 0 | 0 | 0 | 0 | 0 |
| ENSDARG000 | 2;2;2;2;2;2;2; | 37449294:374 | 37449831:374 | -;-;-;-;-;-;-; | 2596 | NA | 475 | 227 | 583 | 814 | 442 | 697 |
| ENSDARG000 | 2;2;2;2;2;2;2; | 25315591:253 | 25315744:253 | +;+;+;+;+;+;+; | 8860 | fndc3ba | 768 | 448 | 689 | 460 | 778 | 342 |
| ENSDARG000 | 2;2;2;2;2 | 5933902:5936 | 5937105:5937 | -;-;-;-;- | 3650 | tmem125b | 16 | 0 | 0 | 0 | 0 | 0 |
| ENSDARG000 | 2;2;2;2;2;2;2; | 32515846:325 | 32516582:325 | -;-;-;-;-;-;-; | 2520 | SMARCD3 | 22 | 12 | 25 | 14 | 28 | 34 |
| ENSDARG000 | 2;2;2;2;2;2;2; | 3823813:3823 | 3823919:3823 | +;+;+;+;+;+;+; | 5092 | npc1 | 2526 | 1966 | 4242 | 1651 | 2990 | 2715 |
| ENSDARG000 | 2;2;2;2;2;2;2; | 1714734:1750 | 1714809:1750 | +;+;+;+;+;+;+; | 5288 | ADGRL2 | 59 | 57 | 110 | 6 | 131 | 39 |
| ENSDARG000 | 2;2 | 36312612:363 | 36312963:363 | -;- | 413 | NA | 0 | 0 | 0 | 0 | 0 | 0 |
| ENSDARG000 | 2;2 | 36316513:363 | 36316868:363 | -;- | 414 | NA | 0 | 0 | 0 | 0 | 0 | 0 |
| ENSDARG000 | 2;2 | 36318102:363 | 36318421:363 | -;- | 375 | NA | 0 | 0 | 0 | 0 | 0 | 0 |
| ENSDARG000 | 2;2;2;2;2;2 | 1714640:1750 | 1714809:1750 | +;+;+;+;+;+ | 1751 | NA | 0 | 2 | 3 | 4 | 6 | 0 |
| ENSDARG000 | 2;2 | 36320760:363 | 36321095:363 | -;- | 385 | NA | 0 | 0 | 0 | 0 | 0 | 0 |
| ENSDARG000 | 2;2;2;2;2;2;2; | 19280829:192 | 19281954:192 | -;-;-;-;-;-;-; | 5138 | ZFYVE9 | 804 | 389 | 688 | 588 | 629 | 538 |
| ENSDARG000 | 2;2 | 36324496:363 | 36324865:363 | -;- | 416 | NA | 0 | 0 | 0 | 0 | 0 | 0 |
| ENSDARG000 | 2;2 | 36327607:363 | 36327977:363 | -;- | 423 | NA | 0 | 0 | 0 | 0 | 0 | 0 |
| ENSDARG000 | 2;2;2;2;2;2;2; | 9918678:9918 | 9918739:9919 | +;+;+;+;+;+;+; | 1347 | NA | 2 | 0 | 12 | 16 | 0 | 2 |
| ENSDARG000 | 2 | 36332018 | 36332331 | - | 314 | NA | 0 | 0 | 0 | 0 | 0 | 0 |
| ENSDARG000 | 2;2 | 36597322:365 | 36597619:365 | -;- | 392 | NA | 0 | 0 | 1 | 0 | 4 | 2 |
| ENSDARG000 | 2;2;2;2;2;2;2; | 31437547:314 | 31437741:314 | +;+;+;+;+;+;+; | 3717 | stam | 1516 | 1636 | 1935 | 1615 | 1434 | 1556 |
| ENSDARG000 | 2;2;2;2;2;2;2; | 51665527:516 | 51669611:516 | -;-;-;-;-;-;-; | 6386 | NA | 510 | 2010 | 1777 | 464 | 2054 | 1087 |
| ENSDARG000 | 2;2;2;2;2;2 | 45049290:450 | 45049448:450 | +;+;+;+;+;+ | 897 | NA | 0 | 0 | 0 | 0 | 6 | 0 |
| ENSDARG000 | 2;2 | 36598554:365 | 36598864:365 | -;- | 351 | NA | 4 | 0 | 0 | 0 | 0 | 2 |
| ENSDARG000 | 2;2 | 36600510:366 | 36600813:366 | -;- | 344 | NA | 8 | 0 | 2 | 8 | 3 | 0 |
| ENSDARG000 | 2;2 | 36602489:366 | 36602793:366 | -;- | 345 | NA | 0 | 0 | 0 | 4 | 0 | 0 |
| ENSDARG000 | 2;2;2 | 45068366:450 | 45068428:450 | +;+;+ | 712 | NA | 0 | 0 | 8 | 14 | 2 | 0 |
| ENSDARG000 | 2;2 | 36604152:366 | 36604455:366 | -;- | 353 | NA | 0 | 0 | 0 | 0 | 0 | 0 |
| ENSDARG000 | 2;2;2;2;2;2;2; | 14998612:149 | 14999021:149 | -;-;-;-;-;-;-; | 1929 | hccsa.1 | 366 | 266 | 374 | 226 | 226 | 367 |
| ENSDARG000 | 2;2;2;2;2;2;2; | 22880605:228 | 22881094:228 | -;-;-;-;-;-;-; | 7330 | NA | 10 | 59 | 49 | 60 | 122 | 22 |
| ENSDARG000 | 2;2 | 45099174:450 | 45099185:450 | -;- | 236 | NA | 0 | 0 | 0 | 0 | 0 | 0 |
| ENSDARG000 | 2;2;2;2;2;2 | 59312776:593 | 59313153:593 | -;-;-;-;-;- | 1584 | NA | 25 | 30 | 23 | 12 | 19 | 19 |
| ENSDARG000 | 2;2;2;2;2;2;2; | 59322130:593 | 59322280:593 | +;+;+;+;+;+;+; | 1195 | NA | 0 | 4 | 7 | 8 | 0 | 2 |
| ENSDARG000 | 2;2;2;2 | 9946121:9946 | 9946346:9946 | +;+;+;+ | 6943 | tsen15 | 127 | 102 | 233 | 245 | 186 | 184 |
| ENSDARG000 | 2;2;2;2;2;2;2; | 26655744:266 | 26656403:266 | +;+;+;+;+;+;+; | 5275 | asph | 1797 | 3061 | 4066 | 1904 | 4803 | 2482 |
| ENSDARG000 | 2;2;2;2;2;2;2; | 45139470:451 | 45139966:451 | -;-;-;-;-;-;-; | 2804 | CAPN10 | 194 | 298 | 385 | 445 | 304 | 467 |
| ENSDARG000 | 2;2;2;2;2;2;2; | 38225427:382 | 38225789:382 | +;+;+;+;+;+;+; | 1555 | C2H14orf119 | 286 | 270 | 260 | 232 | 271 | 349 |
| ENSDARG000 | 2;2 | 26667680:266 | 26667716:266 | +;+ | 776 | NA | 4 | 6 | 19 | 5 | 12 | 8 |
| ENSDARG000 | 2;2;2;2;2;2;2 | 42072231:420 | 42075251:420 | +;+;+;+;+;+;+ | 4112 | VCPIP1 | 290 | 266 | 320 | 114 | 377 | 323 |
| ENSDARG000 | 2;2;2 | 30050541:300 | 30051775:300 | -;-;- | 2019 | shhb | 0 | 0 | 0 | 0 | 0 | 4 |
| ENSDARG000 | 2;2;2;2;2;2;2; | 16449627:164 | 16449813:164 | +;+;+;+;+;+;+; | 1454 | zgc:110269 | 463 | 374 | 487 | 482 | 362 | 313 |
| ENSDARG000 | 2;2;2;2;2 | 42088456:420 | 42088893:420 | -;-;-;-;- | 876 | NA | 0 | 0 | 0 | 0 | 0 | 0 |
| ENSDARG000 | 2;2;2;2;2;2;2; | 38197820:381 | 38198788:381 | -;-;-;-;-;-;-; | 7500 | NA | 4383 | 4084 | 4576 | 3824 | 4489 | 3831 |
| ENSDARG000 | 2;2;2;2;2;2;2; | 36608387:366 | 36609306:366 | +;+;+;+;+;+;+; | 3604 | pak2a | 1865 | 2151 | 2087 | 1376 | 2164 | 1743 |
| ENSDARG000 | 2 | 19362798 | 19363161 | - | 364 | NA | 0 | 0 | 0 | 4 | 0 | 0 |
| ENSDARG000 | 2;2;2;2;2;2;2; | 19505263:195 | 19505482:195 | +;+;+;+;+;+;+; | 1512 | pimr141 | 0 | 0 | 0 | 0 | 0 | 0 |
| ENSDARG000 | 2;2;2;2;2;2;2; | 19506311:195 | 19506364:195 | -;-;-;-;-;-;-;- | 1008 | NA | 0 | 0 | 0 | 0 | 0 | 0 |
| ENSDARG000 | 2;2;2;2;2;2;2; | 36646451:366 | 36646804:366 | +;+;+;+;+;+;+; | 2488 | klhl6 | 6 | 14 | 28 | 17 | 74 | 100 |
| ENSDARG000 | 2 | 24261217 | 24261296 | + | 80 | NA | 0 | 0 | 0 | 0 | 0 | 0 |
| ENSDARG000 | 2;2;2;2;2;2;2; | 24252987:242 | 24256646:242 | -;-;-;-;-;-;-; | 9980 | myh7 | 16 | 0 | 0 | 4 | 0 | 0 |
| ENSDARG000 | 2;2;2;2;2;2;2; | 36606019:366 | 36606180:366 | +;+;+;+;+;+;+; | 1181 | ing5b | 66 | 202 | 224 | 94 | 133 | 90 |
| ENSDARG000 | 2;2;2 | 1939391:1940 | 1939619:1941 | +;+;+ | 782 | exosc4 | 294 | 377 | 497 | 161 | 419 | 248 |
| ENSDARG000 | 2;2;2;2;2;2;2; | 30103131:301 | 30103367:301 | +;+;+;+;+;+;+; | 1533 | dnajb6a | 809 | 712 | 706 | 677 | 646 | 748 |
| ENSDARG000 | 2;2;2 | 1945211:1948 | 1945557:1948 | +;+;+ | 3859 | NA | 102 | 46 | 43 | 68 | 60 | 46 |
| ENSDARG000 | 2;2;2;2;2;2;2; | 42095487:420 | 42096133:420 | -;-;-;-;-;-;-; | 1995 | adhfe1 | 94 | 128 | 203 | 275 | 168 | 135 |
| ENSDARG000 | 2;2;2;2;2;2;2; | 5948534:5948 | 5948854:5948 | +;+;+;+;+;+;+; | 2006 | slc1a7a | 6 | 0 | 4 | 8 | 0 | 10 |
| ENSDARG000 | 2;2;2;2;2;2;2; | 1958604:1958 | 1958886:1958 | +;+;+;+;+;+;+; | 3469 | NA | 24 | 30 | 24 | 6 | 12 | 36 |
| ENSDARG000 | 2;2;2;2;2;2;2; | 51737097:517 | 51737226:517 | -;-;-;-;-;-;-; | 4113 | NA | 565 | 186 | 41 | 193 | 295 | 182 |
| ENSDARG000 | 2;2;2;2;2;2;2; | 9970942:9973 | 9971051:9973 | +;+;+;+;+;+;+; | 1777 | sec22ba | 273 | 427 | 641 | 358 | 552 | 296 |
| ENSDARG000 | 2;2;2 | 37460640:374 | 37460888:374 | -;-;- | 1126 | NA | 144 | 93 | 156 | 206 | 118 | 227 |
| ENSDARG000 | 2;2;2;2 | 31454562:314 | 31454840:314 | +;+;+;+ | 1084 | tmem236 | 0 | 0 | 14 | 0 | 2 | 0 |
| ENSDARG000 | 2;2;2;2;2;2;2; | 48309600:483 | 48310740:483 | -;-;-;-;-;-;-; | 5418 | per2 | 886 | 1200 | 1098 | 841 | 1008 | 746 |
| ENSDARG000 | 2;2;2;2;2 | 25551457:255 | 25551761:255 | -;-;-;-;- | 1108 | ghsra | 0 | 0 | 0 | 0 | 0 | 0 |
| ENSDARG000 | 2;2;2;2;2;2;2; | 9990491:1000 | 9990562:1000 | +;+;+;+;+;+;+; | 1968 | slc35a3b | 4 | 6 | 2 | 0 | 2 | 18 |
| ENSDARG000 | 2;2;2;2;2;2;2; | 19518520:195 | 19518611:195 | -;-;-;-;-;-;-; | 1393 | pimr52 | 0 | 0 | 0 | 0 | 0 | 0 |
| ENSDARG000 | 2;2;2;2;2;2;2; | 9958081:9958 | 9959263:9959 | -;-;-;-;-;-;-; | 1867 | zgc:55943 | 585 | 620 | 851 | 630 | 1030 | 608 |
| ENSDARG000 | 2;2;2;2;2;2;2; | 31515853:315 | 31517866:315 | -;-;-;-;-;-;-; | 6190 | mrc1b | 130 | 601 | 569 | 445 | 2147 | 3086 |
| ENSDARG000 | 2;2;2;2;2;2;2; | 31475772:314 | 31476082:314 | +;+;+;+;+;+;+; | 2079 | cacnb2b | 2 | 2 | 2 | 0 | 13 | 0 |
| ENSDARG000 | 2;2;2 | 37463416:374 | 37463459:374 | -;-;- | 526 | NA | 1 | 6 | 0 | 16 | 0 | 0 |
| ENSDARG000 | 2;2;2;2;2;2;2; | 15035090:150 | 15036359:150 | -;-;-;-;-;-;-; | 2195 | si:dkey-10f21 | 139 | 100 | 122 | 130 | 279 | 90 |
| ENSDARG000 | 2;2;2;2;2 | 7818368:7822 | 7818481:7822 | +;+;+;+;+ | 1182 | kcnmb2 | 0 | 0 | 0 | 0 | 0 | 0 |
| ENSDARG000 | 2;2;2;2 | 45191049:451 | 45191281:451 | +;+;+;+ | 632 | ccl20a.3 | 14 | 2 | 0 | 8 | 0 | 6 |
| ENSDARG000 | 2;2;2;2;2;2;2; | 42114310:421 | 42114578:421 | -;-;-;-;-;-;-; | 1704 | trim55a | 2 | 78 | 62 | 20 | 32 | 40 |
| ENSDARG000 | 2;2;2;2;2;2;2 | 3881000:3882 | 3881080:3882 | +;+;+;+;+;+;+ | 714 | NA | 0 | 0 | 0 | 0 | 0 | 0 |
| ENSDARG000 | 2;2;2;2;2;2;2; | 11205795:112 | 11205981:112 | +;+;+;+;+;+;+; | 2138 | lhx8a | 0 | 0 | 0 | 0 | 0 | 0 |
| ENSDARG000 | 2;2;2;2;2;2;2; | 7831914:7832 | 7832486:7832 | -;-;-;-;-;-;-;- | 1760 | NA | 2 | 0 | 0 | 0 | 0 | 0 |
| ENSDARG000 | 2;2 | 42112503:421 | 42112551:421 | +;+ | 690 | NA | 0 | 0 | 0 | 0 | 0 | 0 |
| ENSDARG000 | 2;2;2;2;2;2;2; | 34128738:341 | 34129524:341 | -;-;-;-;-;-;-;- | 2114 | cenpl | 34 | 267 | 181 | 66 | 101 | 38 |
| ENSDARG000 | 2;2;2;2;2;2;2; | 34138376:341 | 34138864:341 | +;+;+;+;+;+;+; | 2784 | DARS2 | 230 | 289 | 397 | 204 | 444 | 227 |
| ENSDARG000 | 2;2;2;2;2;2;2; | 51888033:518 | 51889258:518 | -;-;-;-;-;-;-; | 5799 | scn12aa | 0 | 0 | 10 | 0 | 0 | 0 |
| ENSDARG000 | 2;2;2;2;2;2;2; | 37469576:374 | 37470147:374 | -;-;-;-;-;-;-; | 1459 | DAPK3 | 612 | 303 | 507 | 339 | 563 | 297 |
| ENSDARG000 | 2;2;2;2;2;2;2; | 16460321:164 | 16460634:164 | +;+;+;+;+;+;+; | 2430 | agfg1b | 745 | 652 | 939 | 794 | 687 | 732 |
| ENSDARG000 | 2;2;2;2;2;2;2; | 26682913:266 | 26683169:266 | +;+;+;+;+;+;+; | 1314 | impa1 | 738 | 654 | 561 | 396 | 652 | 398 |
| ENSDARG000 | 2;2;2;2;2;2;2; | 1988036:1989 | 1988239:1990 | +;+;+;+;+;+;+; | 2758 | ssx2ipa | 82 | 122 | 158 | 92 | 118 | 80 |
| ENSDARG000 | 2;2;2;2;2;2;2; | 6005199:6011 | 6005897:6011 | -;-;-;-;-;-;-; | 2394 | scp2a | 220 | 190 | 397 | 162 | 186 | 214 |
| ENSDARG000 | 2;2;2;2;2;2;2 | 52065884:520 | 52065997:520 | +;+;+;+;+;+;+ | 1616 | shda | 68 | 68 | 67 | 59 | 88 | 104 |
| ENSDARG000 | 2;2;2;2;2;2;2; | 45300512:453 | 45300650:453 | +;+;+;+;+;+;+; | 6850 | CAMSAP2 | 1516 | 607 | 1016 | 800 | 1076 | 842 |
| ENSDARG000 | 2;2;2;2;2;2;2; | 52049239:520 | 52049412:520 | +;+;+;+;+;+;+; | 2089 | yju2 | 376 | 379 | 485 | 192 | 353 | 310 |
| ENSDARG000 | 2;2;2;2;2;2;2; | 7847480:7847 | 7847768:7848 | +;+;+;+;+;+;+; | 3763 | NA | 36 | 22 | 272 | 21 | 86 | 87 |
| ENSDARG000 | 2;2 | 48395488:483 | 48395785:483 | -;- | 384 | NA | 0 | 0 | 0 | 0 | 0 | 1 |
| ENSDARG000 | 2;2;2;2;2;2;2; | 30116486:301 | 30117565:301 | -;-;-;-;-;-;-; | 4245 | trpa1a | 229 | 346 | 306 | 379 | 266 | 466 |
| ENSDARG000 | 2;2 | 30126414:301 | 30126466:301 | +;+ | 200 | NA | 0 | 0 | 0 | 0 | 0 | 0 |
| ENSDARG000 | 2;2;2;2;2;2;2; | 45382433:453 | 45382615:453 | +;+;+;+;+;+;+; | 3121 | WDR47 | 219 | 116 | 172 | 116 | 219 | 84 |
| ENSDARG000 | 2;2;2;2;2;2;2 | 4019538:4023 | 4020795:4023 | -;-;-;-;-;-;- | 1805 | rab18b | 560 | 601 | 488 | 348 | 457 | 377 |
| ENSDARG000 | 2;2;2;2;2;2;2; | 15048410:150 | 15048719:150 | +;+;+;+;+;+;+; | 2213 | cnn3b | 4395 | 3506 | 3192 | 4149 | 3793 | 2184 |
| ENSDARG000 | 2;2;2;2;2;2;2 | 19522082:195 | 19522474:195 | +;+;+;+;+;+;+ | 1374 | pimr49 | 0 | 0 | 0 | 0 | 0 | 0 |
| ENSDARG000 | 2;2;2;2 | 16482338:164 | 16482447:164 | +;+;+;+ | 968 | fbxo36b | 8 | 2 | 26 | 4 | 13 | 4 |
| ENSDARG000 | 2;2 | 21605547:216 | 21606036:216 | +;+ | 1420 | mplkip | 123 | 183 | 158 | 94 | 112 | 166 |
| ENSDARG000 | 2;2;2;2;2;2 | 42135719:421 | 42135987:421 | +;+;+;+;+;+ | 1707 | CYP7B1 | 91 | 57 | 90 | 56 | 117 | 90 |
| ENSDARG000 | 2;2;2 | 32561183:325 | 32562113:325 | -;-;- | 1625 | tmub1 | 358 | 420 | 481 | 354 | 537 | 418 |
| ENSDARG000 | 2;2;2 | 48405272:484 | 48405616:484 | -;-;- | 766 | NA | 0 | 0 | 0 | 0 | 0 | 0 |
| ENSDARG000 | 2;2;2;2;2;2;2; | 21608352:216 | 21610247:216 | -;-;-;-;-;-;-;- | 2939 | ralab | 799 | 901 | 881 | 648 | 669 | 749 |
| ENSDARG000 | 2;2;2;2;2;2;2; | 25560556:255 | 25560680:255 | +;+;+;+;+;+;+; | 4461 | pld1a | 4292 | 3672 | 4775 | 6458 | 7548 | 5371 |
| ENSDARG000 | 2;2;2;2;2;2;2; | 32575685:325 | 32579359:325 | -;-;-;-;-;-;-; | 7016 | fastk | 988 | 1013 | 997 | 351 | 1052 | 1003 |
| ENSDARG000 | 2;2;2;2;2;2;2; | 42156286:421 | 42156811:421 | -;-;-;-;-;-;-; | 2645 | slc39a6 | 344 | 253 | 339 | 287 | 345 | 190 |
| ENSDARG000 | 2;2;2 | 48413946:484 | 48414297:484 | -;-;- | 773 | NA | 0 | 0 | 0 | 0 | 0 | 0 |
| ENSDARG000 | 2;2;2;2 | 2010277:2013 | 2011585:2013 | -;-;-;- | 1644 | tubb2 | 0 | 0 | 0 | 0 | 0 | 0 |
| ENSDARG000 | 2;2;2;2;2;2;2; | 16502780:165 | 16504585:165 | -;-;-;-;-;-;-; | 2415 | NA | 4 | 7 | 3 | 2 | 8 | 4 |
| ENSDARG000 | 2;2;2;2;2;2;2; | 37480669:374 | 37480783:374 | +;+;+;+;+;+;+; | 2851 | sppl2 | 1214 | 842 | 1332 | 1000 | 1726 | 1156 |
| ENSDARG000 | 2;2;2;2;2;2;2; | 48545804:485 | 48546139:485 | -;-;-;-;-;-;-;- | 1915 | si:dkeyp-94g1 | 6 | 2 | 4 | 0 | 0 | 0 |
| ENSDARG000 | 2;2;2;2;2;2;2; | 52122663:521 | 52122873:521 | +;+;+;+;+;+;+; | 1103 | theg | 0 | 0 | 0 | 0 | 0 | 0 |
| ENSDARG000 | 2;2;2;2;2;2;2; | 30177476:301 | 30177532:301 | -;-;-;-;-;-;-;- | 1039 | rpl7 | 19909 | 18886 | 32009 | 29407 | 21062 | 30635 |
| ENSDARG000 | 2;2;2;2;2;2;2; | 19559047:195 | 19559266:195 | +;+;+;+;+;+;+; | 1497 | pimr140 | 0 | 0 | 0 | 0 | 0 | 0 |
| ENSDARG000 | 2;2;2 | 21620177:216 | 21621878:216 | -;-;- | 1779 | C2H7orf25 | 172 | 353 | 270 | 100 | 324 | 200 |
| ENSDARG000 | 2;2;2 | 19560092:195 | 19560147:195 | -;-;- | 333 | NA | 0 | 0 | 0 | 0 | 0 | 0 |
| ENSDARG000 | 2;2;2;2;2 | 19539680:195 | 19539901:195 | -;-;-;-;- | 920 | NA | 0 | 0 | 0 | 0 | 0 | 0 |
| ENSDARG000 | 2;2;2;2;2;2;2; | 30144979:301 | 30145307:301 | +;+;+;+;+;+;+; | 4749 | NA | 58 | 66 | 49 | 113 | 102 | 140 |
| ENSDARG000 | 2;2;2;2;2;2;2; | 2024610:2029 | 2026820:2029 | -;-;-;-;-;-;-; | 4150 | slc22a23 | 20 | 14 | 60 | 18 | 59 | 14 |
| ENSDARG000 | 2;2;2;2 | 21634128:216 | 21634486:216 | +;+;+;+ | 2371 | fbxl7 | 38 | 4 | 12 | 16 | 4 | 4 |
| ENSDARG000 | 2;2;2;2;2;2;2; | 43705593:437 | 43705596:437 | -;-;-;-;-;-;-; | 3283 | KIF5B | 5970 | 3841 | 4243 | 3168 | 3921 | 3281 |
| ENSDARG000 | 2;2 | 11337040:113 | 11337566:113 | -;- | 620 | NA | 150 | 33 | 196 | 12 | 93 | 83 |
| ENSDARG000 | 2;2 | 11330724:113 | 11330899:113 | -;- | 269 | NA | 0 | 0 | 7 | 1 | 0 | 2 |
| ENSDARG000 | 2;2;2;2;2;2;2; | 11216814:112 | 11217070:112 | -;-;-;-;-;-;-; | 2664 | SLC44A5 | 0 | 0 | 0 | 0 | 0 | 2 |

| | | | | | | | | | | | | |
|---|---|---|---|---|---|---|---|---|---|---|---|---|
| ENSDARG000 | 2;2;2;2;2;2 | 15100742:151 | 15100942:151 | +;+;+;+;+;+ | 1353 | f3b | 2247 | 1371 | 1726 | 3841 | 1233 | 1824 |
| ENSDARG000 | 2;2 | 15102103:151 | 15102269:151 | -;- | 247 | NA | 85 | 60 | 58 | 170 | 62 | 81 |
| ENSDARG000 | 2;2;2;2;2;2;2; | 7923964:793 | 7925195:793 | -;-;-;-;-;-;-;-; | 4322 | ephb3a | 276 | 159 | 1305 | 112 | 443 | 684 |
| ENSDARG000 | 2;2;2;2;2;2;2; | 37504244:375 | 37505463:375 | -;-;-;-;-;-;-;-; | 4244 | arhgef18a | 36 | 0 | 14 | 2 | 34 | 16 |
| ENSDARG000 | 2;2;2;2;2;2;2; | 53357953:533 | 53358016:533 | +;+;+;+;+;+;+; | 1233 | NA | 0 | 0 | 0 | 0 | 0 | 0 |
| ENSDARG000 | 2;2;2;2;2;2;2; | 24274293:242 | 24274444:242 | -;-;-;-;-;-;-;-; | 6941 | myh7l | 0 | 2 | 10 | 0 | 0 | 0 |
| ENSDARG000 | 2;2 | 59335781:593 | 59336385:593 | -;- | 729 | NA | 0 | 0 | 2 | 0 | 0 | 0 |
| ENSDARG000 | 2;2 | 16508824:165 | 16509796:165 | -;- | 1243 | NA | 0 | 6 | 3 | 2 | 4 | 17 |
| ENSDARG000 | 2;2;2;2;2;2;2; | 4074412:4075 | 4075400:4075 | +;+;+;+;+;+;+; | 5551 | mib1 | 2037 | 1996 | 2431 | 2107 | 2672 | 2046 |
| ENSDARG000 | 2;2;2;2;2;2;2; | 31599963:316 | 31600993:316 | -;-;-;-;-;-;-;-; | 9405 | NA | 36 | 66 | 48 | 42 | 48 | 159 |
| ENSDARG000 | 2;2;2;2;2;2;2; | 31597841:315 | 31598319:315 | +;+;+;+;+;+;+; | 4345 | NA | 6 | 17 | 24 | 11 | 51 | 28 |
| ENSDARG000 | 2;2;2;2 | 19601696:196 | 19601917:196 | -;-;-;- | 770 | NA | 0 | 0 | 0 | 0 | 0 | 0 |
| ENSDARG000 | 2;2;2;2;2;2;2; | 19574496:195 | 19574587:195 | -;-;-;-;-;-;-;-; | 1546 | pimr51 | 0 | 0 | 0 | 0 | 0 | 0 |
| ENSDARG000 | 2;2;2;2;2;2;2; | 31611034:316 | 31611153:316 | -;-;-;-;-;-;-;-; | 1456 | NA | 5 | 6 | 7 | 0 | 10 | 14 |
| ENSDARG000 | 2;2;2;2 | 32604847:326 | 32604889:326 | +;+;+;+ | 821 | NA | 0 | 2 | 0 | 2 | 4 | 0 |
| ENSDARG000 | 2;2;2;2;2;2 | 59339650:593 | 59340027:593 | -;-;-;-;-;- | 1566 | NA | 3 | 4 | 0 | 4 | 0 | 4 |
| ENSDARG000 | 2;2;2;2;2;2;2; | 19578079:195 | 19578470:195 | +;+;+;+;+;+;+; | 1388 | pimr50 | 0 | 0 | 0 | 0 | 0 | 0 |
| ENSDARG000 | 2;2;2;2;2;2;2 | 19628134:196 | 19628225:196 | -;-;-;-;-;-;- | 1372 | NA | 0 | 0 | 0 | 0 | 0 | 0 |
| ENSDARG000 | 2;2;2;2;2;2 | 16512861:165 | 16512896:165 | -;-;-;-;-;- | 914 | NA | 0 | 1 | 2 | 2 | 0 | 0 |
| ENSDARG000 | 2;2;2;2;2;2;2; | 54696042:547 | 54696191:547 | +;+;+;+;+;+;+; | 6536 | ankrd12 | 1418 | 979 | 1514 | 1224 | 1147 | 1739 |
| ENSDARG000 | 2;2;2;2 | 16536453:165 | 16537222:165 | +;+;+;+ | 3379 | grk7a | 6 | 0 | 2 | 0 | 4 | 0 |
| ENSDARG000 | 2;2;2;2;2;2;2 | 45441007:454 | 45441036:454 | +;+;+;+;+;+;+ | 1149 | NA | 14 | 0 | 30 | 8 | 6 | 0 |
| ENSDARG000 | 2;2;2;2 | 32610878:326 | 32611040:326 | -;-;-;- | 513 | NA | 62 | 28 | 86 | 6 | 32 | 34 |
| ENSDARG000 | 2;2;2;2;2;2;2; | 31614533:316 | 31614758:316 | +;+;+;+;+;+;+; | 3496 | NA | 19 | 135 | 37 | 79 | 17 | 109 |
| ENSDARG000 | 2;2;2;2;2;2;2; | 2110271:2116 | 2110733:2116 | +;+;+;+;+;+;+; | 7439 | DSP | 631 | 123 | 215 | 206 | 258 | 185 |
| ENSDARG000 | 2;2;2;2;2;2;2 | 32621677:326 | 32622128:326 | -;-;-;-;-;-;- | 1710 | si:dkeyp-73d8 | 0 | 0 | 20 | 0 | 0 | 0 |
| ENSDARG000 | 2;2;2 | 34217643:342 | 34217726:342 | -;-;- | 611 | NA | 0 | 0 | 0 | 0 | 0 | 0 |
| ENSDARG000 | 2;2;2;2;2;2 | 30182431:301 | 30183066:301 | +;+;+;+;+;+ | 1785 | rdh10b | 236 | 268 | 204 | 185 | 394 | 225 |
| ENSDARG000 | 2;2;2;2 | 32636508:326 | 32636771:326 | -;-;-;- | 706 | NA | 0 | 0 | 0 | 0 | 0 | 2 |
| ENSDARG000 | 2;2 | 45454415:454 | 45454876:454 | +;+ | 2253 | NA | 8 | 0 | 0 | 0 | 0 | 0 |
| ENSDARG000 | 2;2;2;2;2;2;2; | 16547706:165 | 16549016:165 | -;-;-;-;-;-;-;-; | 2805 | atp1b3a | 428 | 763 | 1151 | 1009 | 1105 | 842 |
| ENSDARG000 | 2;2;2;2;2;2;2; | 43742574:437 | 43743123:437 | -;-;-;-;-;-;-;-; | 1747 | ftr16 | 220 | 123 | 60 | 86 | 68 | 132 |
| ENSDARG000 | 2;2;2;2;2;2;2; | 26692358:266 | 26692528:266 | -;-;-;-;-;-;-;-; | 2353 | NA | 225 | 132 | 118 | 129 | 219 | 124 |
| ENSDARG000 | 2;2;2;2;2;2;2; | 45479841:454 | 45481569:454 | +;+;+;+;+;+;+; | 4143 | NA | 109 | 45 | 90 | 57 | 81 | 88 |
| ENSDARG000 | 2;2;2;2;2;2;2; | 45511459:455 | 45511662:455 | +;+;+;+;+;+;+; | 1570 | NA | 0 | 0 | 0 | 0 | 0 | 0 |
| ENSDARG000 | 2;2;2;2;2;2;2; | 45548890:455 | 45549147:455 | +;+;+;+;+;+;+; | 2684 | FNDC7 | 2 | 0 | 0 | 0 | 0 | 0 |
| ENSDARG000 | 2;2;2;2;2;2;2; | 22931159:229 | 22933640:229 | -;-;-;-;-;-;-;-; | 6424 | sap130b | 469 | 593 | 776 | 223 | 830 | 420 |
| ENSDARG000 | 2;2;2;2;2;2;2; | 45491032:454 | 45491889:454 | -;-;-;-;-;-;-;-; | 3741 | gpsm2 | 444 | 921 | 861 | 522 | 744 | 646 |
| ENSDARG000 | 2;2;2;2;2;2;2; | 25816843:258 | 25816936:258 | +;+;+;+;+;+;+; | 3172 | slc2a2 | 9 | 2 | 37 | 20 | 40 | 4 |
| ENSDARG000 | 2;2;2;2;2;2;2 | 32642338:326 | 32642922:326 | -;-;-;-;-;-;- | 1084 | si:dkeyp-73d8 | 0 | 0 | 0 | 0 | 0 | 0 |
| ENSDARG000 | 2;2;2;2;2;2;2; | 59367429:593 | 59367655:593 | -;-;-;-;-;-;-;- | 3455 | NA | 0 | 2 | 8 | 0 | 0 | 4 |
| ENSDARG000 | 2;2;2;2;2;2;2; | 25657958:256 | 25658449:256 | +;+;+;+;+;+;+; | 6996 | tnika | 4245 | 3009 | 3478 | 3642 | 3418 | 1218 |
| ENSDARG000 | 2;2;2;2;2;2;2; | 11483072:114 | 11483272:114 | -;-;-;-;-;-;-;-; | 1962 | sdr16c5a | 150 | 132 | 186 | 98 | 134 | 176 |
| ENSDARG000 | 2;2;2;2;2;2;2; | 34380982:343 | 34382101:343 | -;-;-;-;-;-;-;-; | 3117 | brinp2 | 0 | 0 | 2 | 0 | 0 | 2 |
| ENSDARG000 | 2;2;2;2;2;2;2; | 42176922:421 | 42177223:421 | +;+;+;+;+;+;+; | 1436 | NA | 2 | 0 | 2 | 0 | 0 | 1 |
| ENSDARG000 | 2;2;2;2;2;2;2; | 21765670:217 | 21767774:217 | -;-;-;-;-;-;-;-; | 11233 | CHD7 | 2717 | 1180 | 3042 | 1644 | 2974 | 1856 |
| ENSDARG000 | 2;2;2;2;2;2;2; | 15109528:151 | 15109901:151 | -;-;-;-;-;-;-;-; | 2389 | abcd3b | 96 | 189 | 130 | 88 | 203 | 110 |
| ENSDARG000 | 2;2;2;2;2;2;2; | 8031234:8031 | 8031504:8031 | +;+;+;+;+;+;+; | 5078 | PCOLCE2 | 2 | 10 | 30 | 7 | 26 | 42 |
| ENSDARG000 | 2;2;2;2;2 | 25814552:258 | 25814747:258 | -;-;-;-;- | 841 | NA | 2 | 1 | 0 | 0 | 0 | 0 |
| ENSDARG000 | 2;2;2;2;2;2 | 30193747:301 | 30195078:301 | -;-;-;-;-;- | 2005 | ube2wa | 157 | 112 | 165 | 164 | 129 | 174 |
| ENSDARG000 | 2;2;2 | 11508691:115 | 11509789:115 | -;-;- | 1436 | penka | 0 | 0 | 8 | 2 | 32 | 20 |
| ENSDARG000 | 2;2;2;2;2;2;2; | 54798689:548 | 54798783:548 | +;+;+;+;+;+;+; | 5549 | twsg1a | 204 | 182 | 375 | 214 | 386 | 156 |
| ENSDARG000 | 2;2;2;2;2;2;2; | 38231695:382 | 38232512:382 | -;-;-;-;-;-;-;-; | 4415 | NA | 8 | 44 | 96 | 88 | 348 | 204 |
| ENSDARG000 | 2;2;2;2;2;2;2; | 27010439:270 | 27010639:270 | +;+;+;+;+;+;+; | 2772 | cdh7a | 36 | 6 | 0 | 0 | 2 | 2 |
| ENSDARG000 | 2;2;2;2;2;2;2; | 8059957:8060 | 8062058:8062 | -;-;-;-;-;-;-;-; | 4090 | pls1 | 38 | 53 | 88 | 46 | 142 | 58 |
| ENSDARG000 | 2;2;2;2;2;2 | 16585549:165 | 16585981:165 | +;+;+;+;+;+ | 2143 | NA | 28 | 50 | 50 | 0 | 8 | 12 |
| ENSDARG000 | 2;2;2;2;2 | 59382428:593 | 59382487:593 | -;-;-;-;- | 1139 | ftr39p | 0 | 4 | 0 | 4 | 1 | 0 |
| ENSDARG000 | 2;2 | 2171371:2172 | 2171811:2172 | -;- | 742 | NA | 30 | 16 | 19 | 29 | 13 | 37 |
| ENSDARG000 | 2;2;2;2;2;2;2; | 2168547:2168 | 2168648:2168 | +;+;+;+;+;+;+; | 3739 | higd1a | 2300 | 3653 | 2297 | 3055 | 3103 | 1153 |
| ENSDARG000 | 2;2;2;2;2;2;2; | 55270566:552 | 55270888:552 | -;-;-;-;-;-;-;-; | 2563 | rab8a | 694 | 646 | 1045 | 459 | 910 | 857 |
| ENSDARG000 | 2;2 | 27000542:270 | 27000747:270 | +;+ | 373 | NA | 0 | 0 | 0 | 0 | 0 | 0 |
| ENSDARG000 | 2;2 | 42191592:421 | 42192077:421 | +;+ | 1293 | cavin4a | 2 | 0 | 0 | 0 | 0 | 0 |
| ENSDARG000 | 2;2;2;2;2 | 11633253:116 | 11633800:116 | -;-;-;-;- | 1644 | bpnt2 | 462 | 528 | 468 | 220 | 601 | 369 |
| ENSDARG000 | 2;2 | 10056100:100 | 10057097:100 | -;- | 2338 | FAM78B | 0 | 0 | 0 | 0 | 0 | 0 |
| ENSDARG000 | 2;2 | 11648163:116 | 11648314:116 | +;+ | 246 | NA | 0 | 0 | 0 | 0 | 0 | 0 |
| ENSDARG000 | 2;2;2;2;2 | 30249977:302 | 30250227:302 | +;+;+;+;+ | 1070 | TMEM70 | 88 | 82 | 98 | 61 | 170 | 124 |
| ENSDARG000 | 2;2;2;2;2;2;2; | 30244376:302 | 30246474:302 | -;-;-;-;-;-;-;-; | 3750 | elocb | 984 | 1230 | 1471 | 1023 | 1375 | 1083 |
| ENSDARG000 | 2;2;2;2;2;2;2; | 42200390:422 | 42200687:422 | +;+;+;+;+;+;+; | 1513 | ift57 | 186 | 145 | 218 | 172 | 120 | 87 |
| ENSDARG000 | 2;2;2;2;2;2;2 | 19633493:196 | 19633882:196 | +;+;+;+;+;+;+ | 1370 | NA | 0 | 0 | 0 | 0 | 0 | 0 |
| ENSDARG000 | 2;2;2;2;2;2;2; | 37544951:375 | 37546435:375 | -;-;-;-;-;-;-;-; | 6541 | insra | 302 | 275 | 761 | 304 | 631 | 490 |
| ENSDARG000 | 2;2;2;2;2;2;2; | 11651176:116 | 11651714:116 | -;-;-;-;-;-;-;-; | 1699 | mffb | 66 | 106 | 121 | 46 | 125 | 92 |
| ENSDARG000 | 2;2;2;2;2;2;2; | 55300425:553 | 55302554:553 | -;-;-;-;-;-;-;-; | 3388 | tpm4b | 93 | 72 | 126 | 90 | 186 | 112 |
| ENSDARG000 | 2;2;2;2;2;2;2; | 38261748:382 | 38261857:382 | +;+;+;+;+;+;+; | 1339 | dhrs1 | 490 | 685 | 718 | 594 | 746 | 869 |
| ENSDARG000 | 2;2;2;2;2;2;2; | 34572690:346 | 34573450:346 | +;+;+;+;+;+;+; | 7127 | ASTN1 | 175 | 102 | 68 | 66 | 66 | 68 |
| ENSDARG000 | 2 | 8112449 | 8114608 | + | 2160 | chst2a | 1386 | 1193 | 1294 | 995 | 1040 | 1827 |
| ENSDARG000 | 2;2;2;2;2;2;2; | 43749221:437 | 43749354:437 | +;+;+;+;+;+;+; | 3954 | arhgap12a | 42 | 2 | 19 | 27 | 92 | 45 |
| ENSDARG000 | 2;2 | 38271392:382 | 38271526:382 | +;+ | 2117 | homeza | 492 | 127 | 343 | 263 | 151 | 109 |
| ENSDARG000 | 2;2;2;2 | 8261109:8274 | 8261372:8275 | +;+;+;+ | 1990 | dipk2aa | 371 | 290 | 413 | 859 | 221 | 657 |
| ENSDARG000 | 2;2;2;2;2;2 | 11670270:116 | 11670865:116 | +;+;+;+;+;+ | 1968 | NA | 0 | 0 | 2 | 0 | 0 | 0 |
| ENSDARG000 | 2;2;2;2;2 | 19640952:196 | 19640997:196 | +;+;+;+;+ | 1448 | rwdd3 | 86 | 53 | 73 | 34 | 57 | 117 |
| ENSDARG000 | 2;2;2;2;2;2;2; | 38277275:382 | 38277413:382 | -;-;-;-;-;-;-;-; | 846 | NA | 0 | 0 | 0 | 0 | 0 | 0 |
| ENSDARG000 | 2;2;2;2;2;2;2; | 42212948:422 | 42213028:422 | +;+;+;+;+;+;+; | 1649 | tmem71 | 720 | 492 | 654 | 449 | 521 | 904 |
| ENSDARG000 | 2;2;2;2;2;2 | 48817958:488 | 48818053:488 | -;-;-;-;-;- | 538 | NA | 0 | 0 | 0 | 0 | 0 | 3 |
| ENSDARG000 | 2;2;2;2;2;2;2; | 10063747:100 | 10063773:100 | +;+;+;+;+;+;+; | 1614 | cmpk | 1327 | 2233 | 1887 | 1179 | 1665 | 1293 |
| ENSDARG000 | 2;2;2;2;2;2;2; | 16597011:165 | 16597188:165 | +;+;+;+;+;+;+; | 6418 | xrn1 | 868 | 777 | 1601 | 661 | 972 | 863 |
| ENSDARG000 | 2;2;2;2;2;2;2 | 48950787:489 | 48950880:489 | -;-;-;-;-;-;- | 1282 | KCNJ9 | 0 | 0 | 0 | 0 | 0 | 0 |
| ENSDARG000 | 2;2;2;2;2;2;2; | 32650729:326 | 32654653:326 | -;-;-;-;-;-;-;-; | 6883 | puf60a | 1247 | 1779 | 2511 | 1178 | 1868 | 1548 |
| ENSDARG000 | 2;2;2 | 27321014:273 | 27321086:273 | +;+;+ | 905 | NA | 0 | 0 | 0 | 0 | 0 | 0 |
| ENSDARG000 | 2;2;2;2;2;2;2; | 27318174:273 | 27318778:273 | -;-;-;-;-;-;-;-; | 2760 | tmx3a | 0 | 0 | 0 | 0 | 0 | 0 |
| ENSDARG000 | 2;2;2;2;2;2;2; | 42228741:422 | 42228857:422 | -;-;-;-;-;-;-;-; | 870 | apom | 0 | 0 | 56 | 0 | 0 | 0 |
| ENSDARG000 | 2;2;2;2;2;2 | 27311863:273 | 27315438:273 | -;-;-;-;-;- | 3735 | DSEL | 2 | 18 | 20 | 32 | 44 | 14 |
| ENSDARG000 | 2;2;2 | 30269306:302 | 30269571:302 | +;+;+ | 527 | NA | 0 | 0 | 0 | 0 | 0 | 0 |
| ENSDARG000 | 2;2;2 | 30276217:302 | 30276470:302 | +;+;+ | 515 | NA | 0 | 0 | 0 | 0 | 0 | 0 |
| ENSDARG000 | 2;2;2;2;2;2;2; | 19777146:197 | 19777253:197 | +;+;+;+;+;+;+; | 3077 | ptbp2b | 355 | 340 | 613 | 311 | 471 | 396 |
| ENSDARG000 | 2;2;2;2;2 | 30281444:302 | 30281710:302 | +;+;+;+;+ | 1050 | NA | 0 | 0 | 0 | 0 | 0 | 0 |
| ENSDARG000 | 2;2;2;2;2;2;2; | 11685742:117 | 11685922:117 | +;+;+;+;+;+;+; | 9510 | greb1l | 18 | 13 | 105 | 34 | 44 | 40 |
| ENSDARG000 | 2;2;2;2;2;2;2; | 34986201:349 | 34986807:349 | -;-;-;-;-;-;-;-; | 6727 | PAPPA2 | 17000 | 6118 | 1927 | 10166 | 1527 | 3058 |
| ENSDARG000 | 2;2;2;2;2;2;2; | 15128027:151 | 15128518:151 | +;+;+;+;+;+;+; | 7081 | arhgap29b | 462 | 260 | 349 | 408 | 561 | 434 |
| ENSDARG000 | 2;2;2;2;2;2;2; | 30290719:302 | 30291208:302 | -;-;-;-;-;-;-;-; | 3921 | jph1b | 0 | 0 | 0 | 0 | 2 | 0 |
| ENSDARG000 | 2;2;2;2;2;2;2; | 42236118:422 | 42236748:422 | +;+;+;+;+;+;+; | 2584 | NA | 20 | 78 | 34 | 0 | 45 | 45 |
| ENSDARG000 | 2;2 | 11760826:117 | 11761562:117 | -;- | 861 | NA | 0 | 0 | 0 | 0 | 4 | 0 |
| ENSDARG000 | 2;2 | 24304854:243 | 24305502:243 | +;+ | 1469 | fitm1l | 2 | 0 | 0 | 0 | 0 | 0 |
| ENSDARG000 | 2;2;2;2;2;2;2; | 22970140:229 | 22971553:229 | -;-;-;-;-;-;-;-; | 8980 | mllt1b | 269 | 389 | 424 | 213 | 463 | 404 |
| ENSDARG000 | 2;2;2;2;2 | 49007592:490 | 49009058:490 | -;-;-;-;- | 1930 | cdc34b | 825 | 961 | 733 | 801 | 579 | 827 |
| ENSDARG000 | 2;2;2;2;2;2;2; | 30368800:303 | 30368879:303 | +;+;+;+;+;+;+; | 4240 | PI15 | 8 | 0 | 0 | 0 | 0 | 0 |
| ENSDARG000 | 2;2;2;2;2 | 42247560:422 | 42248132:422 | +;+;+;+;+ | 1062 | NA | 5 | 0 | 0 | 0 | 9 | 1 |
| ENSDARG000 | 2;2;2;2;2;2;2; | 8490588:8490 | 8490885:8490 | -;-;-;-;-;-;-;-; | 2536 | NA | 0 | 0 | 0 | 0 | 1 | 0 |
| ENSDARG000 | 2;2;2;2;2;2;2; | 22993958:229 | 22994515:229 | -;-;-;-;-;-;-;-; | 1720 | znf414 | 13 | 7 | 7 | 16 | 7 | 0 |
| ENSDARG000 | 2;2;2;2;2;2;2; | 21821683:218 | 21823201:218 | -;-;-;-;-;-;-;-; | 2587 | rab2a | 4241 | 4210 | 4397 | 3656 | 4255 | 4007 |
| ENSDARG000 | 2;2;2;2;2;2;2; | 38282647:382 | 38282747:382 | -;-;-;-;-;-;-;-; | 1166 | zp3f.1 | 0 | 2 | 0 | 0 | 0 | 0 |
| ENSDARG000 | 2;2 | 22986282:229 | 22986455:229 | +;+ | 786 | NA | 0 | 0 | 0 | 0 | 0 | 0 |
| ENSDARG000 | 2;2;2 | 45593783:455 | 45594012:455 | +;+;+ | 539 | NA | 0 | 0 | 0 | 0 | 0 | 0 |
| ENSDARG000 | 2;2;2 | 45619949:456 | 45620159:456 | +;+;+ | 493 | NA | 0 | 0 | 0 | 0 | 0 | 0 |
| ENSDARG000 | 2;2;2;2;2 | 49039159:490 | 49039328:490 | +;+;+;+;+ | 1552 | abhd17aa | 124 | 146 | 136 | 48 | 256 | 88 |
| ENSDARG000 | 2;2 | 45636034:456 | 45636241:456 | +;+ | 284 | NA | 0 | 0 | 0 | 0 | 0 | 0 |
| ENSDARG000 | 2;2;2;2;2;2;2; | 10087334:100 | 10088000:100 | -;-;-;-;-;-;-;- | 2312 | bcl6ab | 1211 | 1058 | 1474 | 1680 | 1876 | 783 |
| ENSDARG000 | 2;2;2;2;2;2;2; | 25839193:258 | 25839457:258 | +;+;+;+;+;+;+; | 3128 | eif5a2 | 11568 | 15118 | 19602 | 11652 | 12883 | 16464 |
| ENSDARG000 | 2;2;2;2;2;2;2; | 35116627:351 | 35117184:351 | +;+;+;+;+;+;+; | 2822 | cop1 | 801 | 572 | 748 | 536 | 971 | 745 |
| ENSDARG000 | 2;2 | 10127762:101 | 10127975:101 | +;+ | 374 | insl5b | 0 | 10 | 0 | 0 | 1 | 1 |
| ENSDARG000 | 2;2;2;2;2;2;2; | 16652238:166 | 16652858:166 | +;+;+;+;+;+;+; | 3132 | gk5 | 504 | 495 | 421 | 118 | 312 | 252 |
| ENSDARG000 | 2;2;2;2;2;2;2; | 49232559:492 | 49232682:492 | +;+;+;+;+;+;+; | 2812 | NA | 0 | 0 | 0 | 0 | 8 | 0 |
| ENSDARG000 | 2;2;2 | 42318012:423 | 42318602:423 | +;+;+ | 864 | NA | 4 | 4 | 0 | 0 | 0 | 2 |
| ENSDARG000 | 2;2;2;2;2;2;2; | 42294944:422 | 42295537:422 | +;+;+;+;+;+;+; | 4413 | NA | 0 | 2 | 0 | 0 | 4 | 5 |
| ENSDARG000 | 2;2;2;2;2;2;2; | 42260021:422 | 42260611:422 | +;+;+;+;+;+;+; | 4013 | NA | 0 | 9 | 10 | 3 | 5 | 0 |
| ENSDARG000 | 2;2;2;2;2 | 42312889:423 | 42313075:423 | -;-;-;-;- | 863 | NA | 25 | 10 | 8 | 6 | 0 | 0 |
| ENSDARG000 | 2;2;2;2;2 | 42330828:423 | 42331058:423 | +;+;+;+;+ | 877 | NA | 0 | 7 | 4 | 1 | 12 | 0 |
| ENSDARG000 | 2;2;2;2;2 | 42271830:422 | 42272423:422 | +;+;+;+;+ | 883 | NA | 3 | 0 | 0 | 2 | 0 | 0 |
| ENSDARG000 | 2;2;2;2;2;2;2; | 30379650:303 | 30379836:303 | +;+;+;+;+;+;+; | 1979 | crispld1b | 0 | 0 | 0 | 0 | 2 | 4 |
| ENSDARG000 | 2;2;2;2;2;2;2; | 37656078:376 | 37658034:376 | -;-;-;-;-;-;-;-; | 3342 | mfsd14a2 | 956 | 826 | 917 | 783 | 952 | 808 |
| ENSDARG000 | 2;2;2 | 38373272:383 | 38373499:383 | +;+;+ | 1308 | psmb5 | 2243 | 3611 | 3646 | 1271 | 3552 | 2330 |
| ENSDARG000 | 2;2;2;2;2;2;2; | 21855036:218 | 21855401:218 | +;+;+;+;+;+;+; | 2174 | ca8 | 31 | 124 | 72 | 50 | 112 | 80 |
| ENSDARG000 | 2;2;2;2;2;2;2; | 43851983:438 | 43852060:438 | +;+;+;+;+;+;+; | 2318 | NA | 6 | 8 | 0 | 8 | 2 | 24 |
| ENSDARG000 | 2;2;2;2;2;2;2; | 43807230:438 | 43807774:438 | -;-;-;-;-;-;-;-; | 4420 | NA | 205 | 140 | 226 | 131 | 189 | 167 |
| ENSDARG000 | 2;2;2;2;2;2;2; | 24310635:243 | 24311432:243 | -;-;-;-;-;-;-;-; | 2797 | trnau1apb | 1440 | 1237 | 2407 | 1898 | 1903 | 2040 |
| ENSDARG000 | 2;2;2 | 42344304:423 | 42344858:423 | +;+;+ | 829 | NA | 0 | 0 | 2 | 0 | 0 | 0 |
| ENSDARG000 | 2;2;2;2;2;2;2 | 10134345:101 | 10134757:101 | +;+;+;+;+;+;+ | 1488 | ahsg2 | 0 | 0 | 36 | 0 | 0 | 2 |
| ENSDARG000 | 2;2;2;2;2;2;2; | 27379861:273 | 27380527:273 | -;-;-;-;-;-;-;-; | 1901 | toe1 | 221 | 197 | 285 | 157 | 274 | 252 |
| ENSDARG000 | 2;2;2;2;2;2;2; | 49417900:494 | 49418227:494 | +;+;+;+;+;+;+; | 9352 | rorcb | 212 | 822 | 140 | 193 | 80 | 262 |
| ENSDARG000 | 2;2;2;2;2;2;2; | 27330461:273 | 27330909:273 | +;+;+;+;+;+;+; | 4324 | tesk2 | 170 | 134 | 110 | 156 | 187 | 226 |
| ENSDARG000 | 2;2;2;2;2;2;2 | 42354643:423 | 42356149:423 | -;-;-;-;-;-;- | 2092 | NA | 267 | 126 | 183 | 129 | 184 | 147 |
| ENSDARG000 | 2;2;2;2;2;2;2; | 16696052:166 | 16696115:166 | +;+;+;+;+;+;+; | 2424 | ppp1r7 | 453 | 716 | 564 | 916 | 540 | 684 |
| ENSDARG000 | 2;2;2;2;2;2 | 25852679:258 | 25852806:258 | -;-;-;-;-;- | 984 | NA | 0 | 0 | 0 | 0 | 0 | 0 |
| ENSDARG000 | 2;2;2;2;2;2;2; | 42361373:423 | 42361819:423 | -;-;-;-;-;-;-;-; | 2354 | gtpbp4 | 5297 | 4298 | 9660 | 7070 | 4673 | 8578 |
| ENSDARG000 | 2;2;2;2;2;2;2; | 19824190:198 | 19825900:198 | -;-;-;-;-;-;-;-; | 5383 | dpydb | 0 | 2 | 6 | 14 | 22 | 0 |
| ENSDARG000 | 2;2;2;2;2;2;2; | 31665836:316 | 31665998:316 | +;+;+;+;+;+;+; | 804 | NA | 16 | 2 | 0 | 16 | 42 | 34 |
| ENSDARG000 | 2;2;2;2;2;2;2; | 15203322:152 | 15203440:152 | +;+;+;+;+;+;+; | 7620 | ABCA4 | 0 | 6 | 0 | 2 | 0 | 0 |
| ENSDARG000 | 2;2;2;2;2;2;2; | 31686365:316 | 31686397:316 | -;-;-;-;-;-;-;-; | 4697 | LRRCC1 | 86 | 162 | 240 | 45 | 196 | 70 |
| ENSDARG000 | 2;2;2;2;2;2;2; | 31676790:316 | 31677975:316 | -;-;-;-;-;-;-;-; | 3275 | e2f5 | 2 | 6 | 0 | 6 | 2 | 0 |
| ENSDARG000 | 2;2;2;2;2;2;2; | 49457449:494 | 49457853:494 | +;+;+;+;+;+;+; | 2053 | sh3gl1a | 52 | 112 | 10 | 90 | 50 | 24 |
| ENSDARG000 | 2;2 | 31671545:316 | 31671593:316 | +;+ | 1150 | ACKR4 | 12 | 0 | 0 | 0 | 4 | 4 |
| ENSDARG000 | 2;2;2;2;2;2;2; | 25860344:258 | 25860464:258 | +;+;+;+;+;+;+; | 5803 | slc7a14a | 0 | 0 | 0 | 0 | 0 | 0 |
| ENSDARG000 | 2;2;2;2;2;2;2; | 37684922:376 | 37686419:376 | -;-;-;-;-;-;-;-; | 8000 | NA | 124 | 108 | 176 | 58 | 199 | 100 |
| ENSDARG000 | 2;2;2;2;2;2;2; | 21982911:219 | 21983394:219 | +;+;+;+;+;+;+; | 6470 | tox | 54 | 40 | 11 | 0 | 60 | 28 |
| ENSDARG000 | 2;2;2;2;2;2;2; | 11848510:118 | 11855665:118 | -;-;-;-;-;-;-;-; | 8371 | abhd3 | 282 | 316 | 288 | 110 | 514 | 263 |
| ENSDARG000 | 2;2 | 30423428:304 | 30423809:304 | +;+ | 3056 | socs6b | 394 | 466 | 464 | 389 | 581 | 429 |
| ENSDARG000 | 2;2;2;2;2;2;2; | 10147029:101 | 10147397:101 | +;+;+;+;+;+;+; | 2823 | pfn2a | 9253 | 11852 | 14145 | 8126 | 11887 | 9915 |
| ENSDARG000 | 2;2;2;2;2;2;2; | 16710883:167 | 16710976:167 | +;+;+;+;+;+;+; | 2576 | ubxn7 | 371 | 379 | 442 | 411 | 349 | 302 |
| ENSDARG000 | 2;2 | 20146139:201 | 20146183:201 | -;- | 344 | NA | 0 | 0 | 0 | 0 | 0 | 0 |
| ENSDARG000 | 2;2;2;2;2;2;2; | 23006792:230 | 23006880:230 | +;+;+;+;+;+;+; | 4008 | mknk2a | 93 | 559 | 392 | 82 | 424 | 590 |
| ENSDARG000 | 2;2;2;2;2;2;2; | 49522178:495 | 49522304:495 | +;+;+;+;+;+;+; | 2299 | stap2a | 0 | 2 | 0 | 2 | 2 | 0 |
| ENSDARG000 | 2;2;2 | 25997426:259 | 25998157:259 | -;-;- | 1316 | cldn11a | 37 | 0 | 0 | 20 | 6 | 20 |
| ENSDARG000 | 2;2;2 | 30458706:304 | 30458903:304 | -;-;- | 657 | cbln2a | 2 | 6 | 2 | 2 | 4 | 0 |

| | | | | | | | | | | | | |
|---|---|---|---|---|---|---|---|---|---|---|---|---|
| ENSDARG000 | 2;2;2;2;2;2;2 | 49555768:495 | 49556022:495 | -;-;-;-;-;-;- | 1024 | NA | 0 | 0 | 0 | 0 | 0 | 0 |
| ENSDARG000 | 2;2;2;2;2;2;2; | 49569017:495 | 49569095:495 | +;+;+;+;+;+;+; | 2983 | sema4e | 30 | 16 | 4 | 12 | 26 | 32 |
| ENSDARG000 | 2;2;2;2;2;2 | 20262479:202 | 20264373:202 | -;-;-;-;-;- | 2961 | plppr5a | 1149 | 2711 | 1618 | 1290 | 3486 | 1051 |
| ENSDARG000 | 2;2;2;2;2;2;2; | 30463825:304 | 30464103:304 | +;+;+;+;+;+;+; | 5586 | NETO1 | 4 | 4 | 0 | 0 | 0 | 4 |
| ENSDARG000 | 2;2;2;2;2;2 | 10160484:101 | 10160616:101 | +;+;+;+;+;+ | 1394 | nsun4 | 84 | 80 | 119 | 71 | 126 | 87 |
| ENSDARG000 | 2;2;2;2;2;2 | 22150134:221 | 22150812:221 | -;-;-;-;-;- | 1987 | cyp7a1 | 2 | 0 | 2 | 4 | 18 | 0 |
| ENSDARG000 | 2;2;2 | 26034175:260 | 26034280:260 | +;+;+ | 753 | NA | 0 | 0 | 0 | 0 | 0 | 0 |
| ENSDARG000 | 2;2;2;2;2;2;2; | 20331445:203 | 20332201:203 | +;+;+;+;+;+;+; | 5650 | PLPPR4 | 2 | 0 | 4 | 6 | 0 | 8 |
| ENSDARG000 | 2;2 | 37750016:377 | 37750091:377 | +;+ | 1072 | MYADM | 95 | 121 | 308 | 96 | 219 | 146 |
| ENSDARG000 | 2;2;2;2;2;2 | 37762924:377 | 37763535:377 | +;+;+;+;+;+ | 1847 | NA | 0 | 10 | 2 | 0 | 0 | 0 |
| ENSDARG000 | 2;2;2;2;2;2;2; | 43894368:438 | 43894470:438 | +;+;+;+;+;+;+; | 3271 | NA | 981 | 771 | 1166 | 741 | 986 | 1235 |
| ENSDARG000 | 2;2;2;2;2;2;2; | 30480907:304 | 30481406:304 | +;+;+;+;+;+;+; | 1047 | atpsckmt | 72 | 85 | 135 | 48 | 95 | 93 |
| ENSDARG000 | 2;2;2;2;2;2;2; | 16780643:167 | 16780707:167 | +;+;+;+;+;+;+; | 5009 | tfa | 74 | 16 | 770 | 16 | 24 | 0 |
| ENSDARG000 | 2;2;2;2;2;2;2; | 20406399:204 | 20406480:204 | +;+;+;+;+;+;+; | 7806 | PALMD | 0 | 0 | 0 | 0 | 0 | 0 |
| ENSDARG000 | 2;2;2;2;2;2;2; | 23039041:230 | 23039218:230 | +;+;+;+;+;+;+; | 3324 | csnk1g2a | 1110 | 1029 | 1194 | 997 | 1223 | 1176 |
| ENSDARG000 | 2;2;2;2;2;2;2; | 26054113:260 | 26054509:260 | -;-;-;-;-;-;-;- | 2049 | NA | 6 | 4 | 0 | 0 | 0 | 0 |
| ENSDARG000 | 2;2;2;2;2 | 22208404:222 | 22210924:222 | -;-;-;-;- | 2711 | fam110b | 722 | 897 | 545 | 854 | 704 | 720 |
| ENSDARG000 | 2;2;2;2;2;2;2; | 11923615:119 | 11923938:119 | +;+;+;+;+;+;+; | 3007 | ro60 | 468 | 211 | 387 | 356 | 296 | 576 |
| ENSDARG000 | 2;2;2;2;2;2;2; | 26060528:261 | 26061092:261 | +;+;+;+;+;+;+; | 4090 | NA | 0 | 0 | 8 | 0 | 0 | 0 |
| ENSDARG000 | 2;2;2;2 | 20430366:204 | 20430682:204 | +;+;+;+ | 3018 | NA | 0 | 0 | 0 | 0 | 0 | 0 |
| ENSDARG000 | 2;2;2;2;2;2;2; | 31781843:317 | 31782300:317 | -;-;-;-;-;-;-;- | 1953 | NA | 163 | 82 | 89 | 97 | 124 | 103 |
| ENSDARG000 | 2;2 | 31784172:317 | 31784332:317 | +;+ | 803 | NA | 4 | 0 | 3 | 2 | 5 | 2 |
| ENSDARG000 | 2;2;2;2;2;2;2; | 20444477:204 | 20444555:204 | -;-;-;-;-;-;-;- | 1758 | FRRS1 | 0 | 0 | 0 | 0 | 0 | 0 |
| ENSDARG000 | 2;2;2;2;2;2;2; | 23081247:230 | 23081672:230 | +;+;+;+;+;+;+; | 6476 | mfsd12a | 73 | 59 | 50 | 32 | 158 | 109 |
| ENSDARG000 | 2;2 | 23084639:230 | 23085212:230 | -;- | 975 | NA | 0 | 0 | 0 | 0 | 0 | 0 |
| ENSDARG000 | 2;2;2;2;2 | 32701820:327 | 32701921:327 | +;+;+;+;+ | 456 | NA | 1 | 0 | 4 | 0 | 0 | 0 |
| ENSDARG000 | 2;2;2;2;2;2;2; | 32676002:326 | 32676503:326 | -;-;-;-;-;-;-;- | 3370 | NRBP2 | 320 | 192 | 248 | 232 | 282 | 182 |
| ENSDARG000 | 2;2;2;2 | 24317465:243 | 24317607:243 | +;+;+;+ | 937 | mrpl34 | 266 | 307 | 405 | 306 | 276 | 330 |
| ENSDARG000 | 2;2;2;2;2;2;2; | 31806602:318 | 31807220:318 | +;+;+;+;+;+;+; | 3337 | ranbp9 | 1189 | 905 | 1096 | 1044 | 921 | 1043 |
| ENSDARG000 | 2;2;2;2;2;2;2; | 26179096:261 | 26179464:261 | +;+;+;+;+;+;+; | 3820 | plppr3a | 206 | 198 | 33 | 78 | 99 | 131 |
| ENSDARG000 | 2;2;2;2;2;2;2; | 43920461:439 | 43920576:439 | +;+;+;+;+;+;+; | 1830 | si:ch211-195 | 307 | 214 | 306 | 140 | 212 | 372 |
| ENSDARG000 | 2;2 | 24348589:243 | 24349061:243 | +;+ | 533 | NA | 0 | 0 | 0 | 0 | 0 | 0 |
| ENSDARG000 | 2;2;2;2;2;2;2; | 24321751:243 | 24322244:243 | -;-;-;-;-;-;-;- | 3462 | ANO8 | 12 | 15 | 29 | 28 | 59 | 45 |
| ENSDARG000 | 2;2;2;2;2 | 43940925:439 | 43941075:439 | -;-;-;-;- | 993 | NA | 11 | 14 | 8 | 0 | 10 | 10 |
| ENSDARG000 | 2;2;2;2;2;2;2; | 30489846:304 | 30490094:304 | +;+;+;+;+;+;+; | 4748 | marchf6 | 2053 | 1668 | 1995 | 1260 | 1997 | 1832 |
| ENSDARG000 | 2;2;2;2;2;2;2; | 26237322:262 | 26237438:262 | +;+;+;+;+;+;+; | 2759 | palm1b | 6168 | 3842 | 2481 | 3678 | 3295 | 5109 |
| ENSDARG000 | 2;2 | 26238769:262 | 26239074:262 | -;- | 428 | NA | 0 | 0 | 0 | 0 | 0 | 1 |
| ENSDARG000 | 2;2;2;2;2;2;2; | 20472150:204 | 20472208:204 | +;+;+;+;+;+;+; | 5224 | agla | 965 | 1395 | 2889 | 1847 | 1588 | 1237 |
| ENSDARG000 | 2 | 23130389 | 23132134 | + | 1746 | NA | 0 | 0 | 0 | 0 | 0 | 0 |
| ENSDARG000 | 2;2;2;2;2;2;2; | 16798652:167 | 16798808:167 | +;+;+;+;+;+;+; | 10190 | eif4g1a | 6315 | 7594 | 11460 | 3478 | 9258 | 5274 |
| ENSDARG000 | 2;2;2;2 | 23156047:231 | 23156102:231 | +;+;+;+ | 1058 | NA | 10 | 0 | 2 | 0 | 12 | 8 |
| ENSDARG000 | 2;2;2;2 | 23165250:231 | 23166121:231 | -;-;-;- | 1908 | prrx1a | 189 | 70 | 2 | 16 | 14 | 439 |
| ENSDARG000 | 2;2;2;2;2;2;2; | 43961626:439 | 43963549:439 | -;-;-;-;-;-;-;- | 3955 | kirrel1b | 6 | 0 | 4 | 0 | 6 | 0 |
| ENSDARG000 | 2;2;2;2;2;2;2; | 23222939:232 | 23223077:232 | +;+;+;+;+;+;+; | 3963 | kifap3b | 170 | 1134 | 392 | 540 | 286 | 528 |
| ENSDARG000 | 2;2;2;2;2;2;2; | 20555394:205 | 20556306:205 | -;-;-;-;-;-;-;- | 3731 | NA | 16 | 16 | 72 | 16 | 52 | 0 |
| ENSDARG000 | 2;2;2;2;2 | 20539402:205 | 20539482:205 | +;+;+;+;+ | 2899 | NA | 4 | 8 | 44 | 48 | 50 | 14 |
| ENSDARG000 | 2;2;2;2;2;2;2; | 49608796:496 | 49609917:496 | -;-;-;-;-;-;-;- | 3143 | cactin | 544 | 480 | 593 | 290 | 644 | 354 |
| ENSDARG000 | 2;2;2;2;2;2;2; | 49631043:496 | 49631097:496 | +;+;+;+;+;+;+; | 925 | NA | 12 | 0 | 6 | 2 | 35 | 14 |
| ENSDARG000 | 2;2;2 | 49644803:496 | 49644943:496 | +;+;+ | 939 | NA | 0 | 2 | 0 | 0 | 0 | 0 |
| ENSDARG000 | 2;2 | 49646958:496 | 49647048:496 | +;+ | 402 | NA | 0 | 0 | 0 | 0 | 0 | 0 |
| ENSDARG000 | 2;2 | 49666756:496 | 49666922:496 | -;- | 190 | NA | 0 | 0 | 0 | 0 | 0 | 0 |
| ENSDARG000 | 2;2 | 49767578:497 | 49767770:497 | -;- | 536 | NA | 0 | 0 | 0 | 0 | 0 | 0 |
| ENSDARG000 | 2;2;2;2;2;2 | 49792508:497 | 49792572:497 | -;-;-;-;-;- | 286 | NA | 0 | 0 | 16 | 2 | 16 | 0 |
| ENSDARG000 | 2;2;2;2;2;2;2; | 57206248:572 | 57206409:572 | -;-;-;-;-;-;-;- | 1770 | BTBD2 | 382 | 238 | 334 | 242 | 310 | 281 |
| ENSDARG000 | 2;2;2;2;2;2;2; | 38318000:383 | 38318094:383 | -;-;-;-;-;-;-;- | 1533 | NA | 2 | 0 | 11 | 0 | 20 | 5 |
| ENSDARG000 | 2;2;2;2 | 54919257:549 | 54919323:549 | -;-;-;- | 1008 | NA | 0 | 1 | 0 | 0 | 0 | 0 |
| ENSDARG000 | 2;2;2;2;2;2;2; | 176711:17671 | 178342:1783 | -;-;-;-;-;-;-;- | 1969 | zgc:113518 | 112 | 98 | 94 | 116 | 112 | 136 |
| ENSDARG000 | 2;2;2;2 | 19104231:191 | 19104255:191 | +;+;+;+ | 114 | NA | 0 | 0 | 0 | 0 | 0 | 0 |
| ENSDARG000 | 2 | 31804582 | 31805604 | + | 1023 | rnf182 | 0 | 2 | 0 | 0 | 16 | 0 |
| ENSDARG000 | 2;2;2;2;2;2;2; | 15586632:155 | 15586713:155 | +;+;+;+;+;+;+; | 1572 | NA | 2 | 2 | 0 | 2 | 2 | 0 |
| ENSDARG000 | 2;2;2;2;2;2;2; | 37975026:379 | 37975130:379 | +;+;+;+;+;+;+; | 2325 | NA | 124 | 53 | 47 | 36 | 67 | 104 |
| ENSDARG000 | 2;2;2 | 23482798:234 | 23482845:234 | +;+;+ | 423 | NA | 2 | 0 | 2 | 4 | 2 | 2 |
| ENSDARG000 | 2;2;2;2;2;2;2; | 28453338:284 | 28454192:284 | +;+;+;+;+;+;+; | 2703 | MMP15 | 1782 | 959 | 1253 | 639 | 1987 | 690 |
| ENSDARG000 | 2 | 33541928 | 33543046 | + | 1119 | NA | 0 | 0 | 6 | 0 | 0 | 0 |
| ENSDARG000 | 2;2 | 57504073:575 | 57505105:575 | +;+ | 1326 | ONECUT3 | 0 | 0 | 0 | 0 | 0 | 0 |
| ENSDARG000 | 2 | 731491 | 732864 | - | 1374 | NA | 0 | 0 | 0 | 0 | 0 | 0 |
| ENSDARG000 | 2 | 23257777 | 23257897 | + | 121 | 5S_rRNA | 0 | 0 | 0 | 0 | 0 | 0 |
| ENSDARG000 | 2;2;2;2 | 2737422:2742 | 2737960:2743 | +;+;+;+ | 1435 | aqp1a.1 | 96 | 30 | 1209 | 731 | 139 | 279 |
| ENSDARG000 | 2;2;2;2;2;2;2; | 38025260:380 | 38025337:380 | +;+;+;+;+;+;+; | 1726 | hnrnpc | 868 | 987 | 1449 | 961 | 1369 | 920 |
| ENSDARG000 | 2;2;2;2;2 | 58077662:580 | 58078109:580 | -;-;-;-;- | 667 | fcer1gl | 147 | 182 | 238 | 138 | 349 | 408 |
| ENSDARG000 | 2;2;2;2;2;2;2; | 56679214:566 | 56683698:566 | -;-;-;-;-;-;-;- | 7859 | ARHGAP45 | 522 | 217 | 593 | 342 | 1035 | 350 |
| ENSDARG000 | 2;2;2;2;2;2;2; | 24638367:246 | 24638449:246 | +;+;+;+;+;+;+; | 3948 | MAST3 | 1897 | 1121 | 1557 | 1368 | 2873 | 1218 |
| ENSDARG000 | 2;2;2;2;2;2;2; | 56891858:568 | 56892126:568 | +;+;+;+;+;+;+; | 1898 | tm6sf2a | 0 | 0 | 0 | 0 | 2 | 0 |
| ENSDARG000 | 2 | 33900435 | 33900512 | + | 78 | dre-mir-23a-3 | 0 | 0 | 0 | 0 | 2 | 0 |
| ENSDARG000 | 2;2;2;2;2;2;2; | 1522721:1523 | 1522802:1523 | -;-;-;-;-;-;-;- | 2177 | dab1b | 0 | 0 | 0 | 0 | 1 | 0 |
| ENSDARG000 | 2 | 56838681 | 56838799 | + | 119 | 5S_rRNA | 0 | 0 | 0 | 0 | 0 | 0 |
| ENSDARG000 | 2;2 | 7601996:7604 | 7602378:7605 | -;- | 606 | NA | 47 | 24 | 75 | 56 | 45 | 35 |
| ENSDARG000 | 2 | 14655134 | 14656669 | - | 1536 | NA | 661 | 364 | 381 | 486 | 619 | 693 |
| ENSDARG000 | 2;2 | 36346930:363 | 36349825:363 | -;- | 3058 | NA | 0 | 0 | 0 | 0 | 4 | 0 |
| ENSDARG000 | 2;2;2;2;2;2;2; | 54641644:546 | 54641861:546 | +;+;+;+;+;+;+; | 911 | ndufv2 | 691 | 682 | 776 | 493 | 636 | 644 |
| ENSDARG000 | 2;2;2;2;2 | 5131964:5132 | 5132614:5132 | -;-;-;-;- | 1034 | ptmab | 4918 | 2915 | 6159 | 3045 | 4733 | 3552 |
| ENSDARG000 | 2 | 46288860 | 46288996 | + | 137 | 5S_rRNA | 0 | 0 | 0 | 0 | 0 | 0 |
| ENSDARG000 | 2;2;2;2;2;2 | 55980227:559 | 55980260:559 | -;-;-;-;-;- | 132 | NA | 0 | 0 | 0 | 0 | 0 | 0 |
| ENSDARG000 | 2;2;2;2;2;2;2; | 5178172:5180 | 5178560:5180 | -;-;-;-;-;-;-;- | 1332 | phb2a | 1294 | 1393 | 1846 | 753 | 1125 | 977 |
| ENSDARG000 | 2 | 7721212 | 7721390 | + | 179 | SNORA81 | 0 | 4 | 1 | 0 | 2 | 1 |
| ENSDARG000 | 2;2;2;2;2;2;2 | 56937548:569 | 56937599:569 | +;+;+;+;+;+;+ | 735 | NA | 6 | 2 | 28 | 2 | 4 | 2 |
| ENSDARG000 | 2;2 | 927204:93099 | 927428:93163 | +;+ | 872 | NA | 48 | 57 | 76 | 37 | 51 | 85 |
| ENSDARG000 | 2;2;2;2;2;2;2; | 59527149:595 | 59527215:595 | +;+;+;+;+;+;+; | 1950 | NA | 0 | 0 | 0 | 2 | 0 | 0 |
| ENSDARG000 | 2;2;2;2;2;2;2; | 51509663:515 | 51510305:515 | -;-;-;-;-;-;-;- | 2123 | pigrl2.3 | 4 | 32 | 26 | 0 | 52 | 42 |
| ENSDARG000 | 2;2;2 | 48620155:486 | 48620200:486 | +;+;+ | 639 | NA | 0 | 0 | 0 | 0 | 0 | 2 |
| ENSDARG000 | 2 | 33902325 | 33902455 | + | 131 | dre-mir-27a | 2 | 4 | 1 | 0 | 0 | 0 |
| ENSDARG000 | 2 | 19628697 | 19628794 | + | 98 | NA | 0 | 0 | 0 | 0 | 0 | 0 |
| ENSDARG000 | 2;2;2;2;2;2;2; | 58513535:585 | 58513642:585 | -;-;-;-;-;-;-;- | 2514 | epb41l3b | 3543 | 2521 | 3236 | 3978 | 2027 | 1818 |
| ENSDARG000 | 2;2 | 16487443:164 | 16487491:164 | +;+ | 1089 | NA | 27 | 2 | 46 | 0 | 31 | 7 |
| ENSDARG000 | 2 | 41421606 | 41421722 | - | 117 | 5S_rRNA | 0 | 0 | 0 | 0 | 0 | 0 |
| ENSDARG000 | 2;2;2;2;2 | 58444873:584 | 58445267:584 | -;-;-;-;- | 1455 | APCDD1 | 4 | 4 | 0 | 0 | 3 | 0 |
| ENSDARG000 | 2 | 32666324 | 32666459 | - | 136 | NA | 0 | 0 | 0 | 0 | 0 | 0 |
| ENSDARG000 | 2;2 | 27867830:278 | 27867982:278 | +;+ | 1492 | RNF152 | 0 | 6 | 16 | 2 | 16 | 31 |
| ENSDARG000 | 2 | 45629707 | 45630823 | - | 1117 | NA | 0 | 0 | 0 | 0 | 0 | 0 |
| ENSDARG000 | 2;2;2;2;2;2;2; | 9629558:9631 | 9629747:9632 | -;-;-;-;-;-;-;- | 1816 | SCHIP1 | 0 | 0 | 0 | 0 | 0 | 0 |
| ENSDARG000 | 2;2;2;2;2;2;2; | 27469669:274 | 27469838:274 | -;-;-;-;-;-;-;- | 2138 | NA | 2 | 0 | 0 | 3 | 1 | 6 |
| ENSDARG000 | 2;2;2;2;2;2;2 | 53501750:535 | 53502017:535 | -;-;-;-;-;-;- | 1274 | pnhd | 0 | 0 | 0 | 0 | 0 | 0 |
| ENSDARG000 | 2;2;2;2;2;2;2; | 212059:21230 | 212230:21242 | +;+;+;+;+;+;+; | 3522 | DHX30 | 255 | 159 | 252 | 56 | 210 | 88 |
| ENSDARG000 | 2;2;2;2;2;2;2; | 27651984:276 | 27652032:276 | +;+;+;+;+;+;+; | 4417 | tmem68 | 341 | 310 | 266 | 148 | 402 | 270 |
| ENSDARG000 | 2 | 7720939 | 7721009 | + | 71 | NA | 0 | 0 | 0 | 0 | 0 | 0 |
| ENSDARG000 | 2;2 | 36490105:364 | 36490397:364 | -;- | 342 | NA | 0 | 0 | 0 | 0 | 0 | 0 |
| ENSDARG000 | 2 | 10905062 | 10905179 | - | 118 | 5S_rRNA | 0 | 0 | 0 | 0 | 0 | 0 |
| ENSDARG000 | 2;2;2 | 51790494:517 | 51790603:517 | -;-;- | 285 | NA | 0 | 0 | 15 | 0 | 14 | 23 |
| ENSDARG000 | 2 | 6367702 | 6367816 | + | 115 | 5S_rRNA | 0 | 0 | 0 | 0 | 0 | 0 |
| ENSDARG000 | 2;2;2;2;2;2;2; | 56642138:566 | 56642446:566 | -;-;-;-;-;-;-;- | 882 | gpx4b | 1694 | 1989 | 2458 | 1544 | 2416 | 1644 |
| ENSDARG000 | 2 | 44564378 | 44564494 | + | 117 | 5S_rRNA | 0 | 0 | 0 | 0 | 0 | 5 |
| ENSDARG000 | 2 | 17657958 | 17658076 | + | 119 | 5S_rRNA | 0 | 0 | 0 | 0 | 0 | 0 |
| ENSDARG000 | 2;2;2;2;2;2 | 68789:70137 | 69011:70322 | +;+;+;+;+;+ | 633 | NA | 38 | 6 | 1 | 15 | 2 | 6 |
| ENSDARG000 | 2;2;2;2;2;2;2; | 33664387:336 | 33664400:336 | +;+;+;+;+;+;+; | 368 | NA | 1 | 0 | 1 | 2 | 14 | 2 |
| ENSDARG000 | 2 | 50706439 | 50707494 | - | 1056 | NA | 0 | 0 | 4 | 0 | 0 | 4 |
| ENSDARG000 | 2;2;2 | 51774062:517 | 51774174:517 | -;-;- | 288 | NA | 0 | 0 | 0 | 2 | 0 | 0 |
| ENSDARG000 | 2 | 7723394 | 7723583 | + | 190 | NA | 2 | 2 | 1 | 0 | 4 | 3 |
| ENSDARG000 | 2;2 | 51520262:515 | 51520578:515 | -;- | 634 | NA | 0 | 0 | 13 | 0 | 0 | 34 |
| ENSDARG000 | 2 | 41211978 | 41212094 | - | 117 | 5S_rRNA | 0 | 0 | 0 | 0 | 0 | 0 |
| ENSDARG000 | 2;2;2;2;2;2;2; | 5338759:5340 | 5338975:5340 | -;-;-;-;-;-;-;- | 1317 | MFN1 | 0 | 2 | 6 | 0 | 19 | 6 |
| ENSDARG000 | 2;2;2;2 | 37971353:379 | 37971417:379 | +;+;+;+ | 1741 | apol1 | 404 | 222 | 100 | 343 | 207 | 426 |
| ENSDARG000 | 2;2;2;2;2;2;2; | 58345157:583 | 58347596:583 | -;-;-;-;-;-;-;- | 4265 | NA | 246 | 247 | 535 | 148 | 389 | 261 |
| ENSDARG000 | 2;2;2 | 7663942:7664 | 7664405:7664 | -;-;- | 672 | NA | 8 | 298 | 2002 | 1428 | 1145 | 8486 |
| ENSDARG000 | 2;2;2;2;2 | 31762852:317 | 31764887:317 | -;-;-;-;- | 2530 | and2 | 2 | 2 | 0 | 4 | 10 | 14 |
| ENSDARG000 | 2 | 19579582 | 19579679 | - | 98 | NA | 0 | 0 | 0 | 0 | 0 | 0 |
| ENSDARG000 | 2 | 18017000 | 18022765 | + | 5766 | NA | 12 | 0 | 6 | 0 | 0 | 2 |
| ENSDARG000 | 2;2;2;2;2;2;2; | 31719439:317 | 31719605:317 | -;-;-;-;-;-;-;- | 1378 | NA | 193 | 40 | 26 | 4 | 25 | 14 |
| ENSDARG000 | 2;2;2;2;2;2;2; | 22416275:224 | 22417166:224 | +;+;+;+;+;+;+; | 3889 | PKN2 | 318 | 383 | 380 | 349 | 310 | 424 |
| ENSDARG000 | 2;2;2;2 | 51765179:517 | 51765529:517 | -;-;-;- | 553 | NA | 1084 | 162 | 356 | 107 | 19 | 84 |
| ENSDARG000 | 2 | 11006657 | 11006790 | + | 134 | 5S_rRNA | 0 | 2 | 0 | 0 | 2 | 0 |
| ENSDARG000 | 2;2;2;2;2;2;2; | 40830768:408 | 40833235:408 | -;-;-;-;-;-;-;- | 7702 | uggt1 | 758 | 1256 | 1256 | 282 | 1472 | 926 |
| ENSDARG000 | 2 | 8189285 | 8189400 | + | 116 | 5S_rRNA | 0 | 0 | 0 | 0 | 0 | 0 |
| ENSDARG000 | 2;2;2;2;2;2;2; | 54844755:548 | 54845238:548 | +;+;+;+;+;+;+; | 4071 | rnf126 | 245 | 204 | 362 | 136 | 247 | 180 |
| ENSDARG000 | 2 | 26501033 | 26501103 | + | 71 | SNORD38A | 0 | 0 | 0 | 0 | 0 | 0 |
| ENSDARG000 | 2;2;2;2;2;2;2; | 2818645:2849 | 2819518:2849 | +;+;+;+;+;+;+; | 5176 | rock1 | 1857 | 1455 | 1785 | 825 | 1473 | 1319 |
| ENSDARG000 | 2;2 | 3516913:3519 | 3517769:3519 | +;+ | 1393 | NA | 29 | 22 | 24 | 50 | 55 | 28 |
| ENSDARG000 | 2 | 42962228 | 42962344 | - | 117 | 5S_rRNA | 0 | 0 | 0 | 0 | 0 | 0 |
| ENSDARG000 | 2 | 32293401 | 32293516 | - | 116 | 5S_rRNA | 0 | 0 | 0 | 0 | 0 | 0 |
| ENSDARG000 | 2 | 30181130 | 30181219 | - | 90 | NA | 0 | 0 | 8 | 0 | 14 | 2 |
| ENSDARG000 | 2;2;2;2;2;2;2; | 54586647:545 | 54586774:545 | -;-;-;-;-;-;-;- | 2043 | acss2l | 25 | 14 | 16 | 12 | 16 | 104 |
| ENSDARG000 | 2 | 37836619 | 37836920 | + | 302 | RNaseP_nuc | 0 | 2 | 0 | 0 | 0 | 0 |
| ENSDARG000 | 2 | 14660356 | 14660546 | + | 191 | U2 | 0 | 0 | 0 | 0 | 0 | 0 |
| ENSDARG000 | 2;2;2;2;2 | 37959040:379 | 37959198:379 | -;-;-;-;- | 942 | NA | 0 | 0 | 2 | 0 | 0 | 0 |
| ENSDARG000 | 2;2;2;2;2 | 40181633:401 | 40184626:401 | -;-;-;-;- | 3582 | si:ch211-122 | 0 | 0 | 0 | 0 | 0 | 0 |
| ENSDARG000 | 2;2;2;2;2;2;2; | 52332022:523 | 52332141:523 | -;-;-;-;-;-;-;- | 1023 | tle2c | 156 | 90 | 132 | 176 | 185 | 137 |
| ENSDARG000 | 2;2;2;2;2;2 | 58377395:584 | 58377790:584 | +;+;+;+;+;+ | 4853 | vapal | 683 | 577 | 792 | 598 | 892 | 545 |
| ENSDARG000 | 2;2;2;2;2;2;2; | 1470070:1471 | 1470395:1471 | -;-;-;-;-;-;-;- | 2699 | si:ch211-188 | 4 | 2 | 1 | 0 | 0 | 3 |
| ENSDARG000 | 2;2;2;2;2;2;2; | 2904080:2905 | 2904231:2905 | +;+;+;+;+;+;+; | 1832 | eif2b5 | 769 | 710 | 975 | 631 | 746 | 574 |
| ENSDARG000 | 2;2;2;2;2;2;2; | 21452822:214 | 21452928:214 | +;+;+;+;+;+;+; | 3457 | afg3l2 | 754 | 745 | 1009 | 537 | 801 | 730 |
| ENSDARG000 | 2;2;2;2 | 38288308:382 | 38288439:382 | +;+;+;+ | 972 | nedd8l | 0 | 0 | 0 | 0 | 0 | 0 |
| ENSDARG000 | 2 | 7394790 | 7394908 | + | 119 | 5S_rRNA | 0 | 0 | 0 | 0 | 0 | 0 |
| ENSDARG000 | 2 | 59038195 | 59038312 | - | 118 | 5S_rRNA | 0 | 0 | 0 | 0 | 0 | 0 |
| ENSDARG000 | 2 | 44774275 | 44774595 | - | 321 | NA | 0 | 0 | 2 | 0 | 0 | 8 |
| ENSDARG000 | 2;2;2;2;2;2 | 45100963:451 | 45101212:451 | +;+;+;+;+;+ | 849 | NA | 0 | 0 | 0 | 0 | 0 | 0 |
| ENSDARG000 | 2;2;2;2;2;2;2; | 52847049:528 | 52847157:528 | +;+;+;+;+;+;+; | 1868 | NA | 0 | 0 | 0 | 0 | 0 | 0 |
| ENSDARG000 | 2;2;2;2;2;2;2; | 1453034:1456 | 1453277:1456 | -;-;-;-;-;-;-;- | 1848 | GPAA1 | 1071 | 504 | 700 | 515 | 591 | 403 |
| ENSDARG000 | 2;2;2 | 198565:20416 | 202264:20427 | -;-;- | 4334 | NA | 130 | 100 | 146 | 82 | 108 | 162 |
| ENSDARG000 | 2 | 56042512 | 56042630 | - | 119 | 5S_rRNA | 0 | 0 | 0 | 0 | 0 | 0 |
| ENSDARG000 | 2 | 51850157 | 51850271 | - | 115 | 5S_rRNA | 0 | 0 | 0 | 0 | 0 | 0 |
| ENSDARG000 | 2;2 | 48496207:484 | 48496646:484 | -;- | 864 | NA | 0 | 0 | 0 | 2 | 0 | 0 |
| ENSDARG000 | 2;2;2;2;2;2;2; | 45116819:451 | 45117590:451 | -;-;-;-;-;-;-;- | 1630 | eif4e2 | 110 | 438 | 264 | 226 | 364 | 462 |
| ENSDARG000 | 2;2;2;2;2;2;2; | 1594344:1599 | 1598676:1599 | -;-;-;-;-;-;-;- | 5263 | prkacbb | 1961 | 1487 | 1759 | 1192 | 1910 | 1648 |

| | | | | | | | | | | | | |
|---|---|---|---|---|---|---|---|---|---|---|---|---|
| ENSDARG000 | 2;2;2;2;2 | 243778:2469 | 243838:2470 | +;+;+;+;+ | 357 | NA | 2 | 2 | 2 | 0 | 0 | 0 |
| ENSDARG000 | 2;2;2;2;2;2 | 59135878:591 | 59137390:591 | -;-;-;-;-;- | 3155 | NA | 135 | 56 | 124 | 92 | 74 | 98 |
| ENSDARG000 | 2;2;2;2 | 27670284:276 | 27673534:276 | -;-;-;- | 4674 | xkr4 | 2 | 2 | 0 | 2 | 12 | 0 |
| ENSDARG000 | 2 | 57103401 | 57103485 | - | 85 | dre-mir-7148 | 0 | 0 | 0 | 0 | 0 | 0 |
| ENSDARG000 | 2;2;2;2 | 55199721:552 | 55199740:552 | +;+;+;+ | 663 | NA | 20 | 17 | 16 | 18 | 22 | 21 |
| ENSDARG000 | 2 | 59102169 | 59102252 | - | 84 | dre-mir-9-7 | 0 | 0 | 0 | 0 | 0 | 0 |
| ENSDARG000 | 2;2;2;2;2;2;2; | 27811038:278 | 27812594:278 | -;-;-;-;-;-;-;- | 5264 | NA | 555 | 351 | 451 | 266 | 392 | 319 |
| ENSDARG000 | 2;2;2;2;2;2;2; | 55365727:553 | 55365951:553 | +;+;+;+;+;+;+; | 1727 | LPL | 0 | 0 | 0 | 0 | 0 | 0 |
| ENSDARG000 | 2;2;2;2;2;2;2; | 13907452:139 | 13907552:139 | +;+;+;+;+;+;+; | 1977 | cpdp | 695 | 270 | 400 | 188 | 457 | 303 |
| ENSDARG000 | 2 | 39370811 | 39370927 | - | 117 | 5S_rRNA | 0 | 0 | 0 | 0 | 0 | 0 |
| ENSDARG000 | 2;2;2;2;2;2;2; | 38342298:383 | 38343009:383 | -;-;-;-;-;-;-;- | 2459 | prmt5 | 711 | 1107 | 1737 | 461 | 1247 | 695 |
| ENSDARG000 | 2 | 9406558 | 9406675 | - | 118 | 5S_rRNA | 2 | 0 | 0 | 0 | 0 | 0 |
| ENSDARG000 | 2 | 20757582 | 20757699 | - | 118 | 5S_rRNA | 0 | 1 | 0 | 0 | 2 | 0 |
| ENSDARG000 | 2 | 9746858 | 9746993 | - | 136 | NA | 0 | 0 | 0 | 0 | 0 | 0 |
| ENSDARG000 | 2 | 3174870 | 3174995 | + | 126 | NA | 0 | 0 | 0 | 0 | 0 | 0 |
| ENSDARG000 | 2;2;2;2;2;2;2 | 9621289:9621 | 9621408:9621 | -;-;-;-;-;-;- | 564 | NA | 2 | 0 | 0 | 2 | 6 | 6 |
| ENSDARG000 | 2;2;2;2;2;2;2; | 47708853:477 | 47709112:477 | +;+;+;+;+;+;+; | 2838 | mbnl1 | 628 | 791 | 1064 | 729 | 1173 | 773 |
| ENSDARG000 | 2;2;2;2;2;2;2; | 55783896:557 | 55785659:557 | -;-;-;-;-;-;-;- | 2917 | calr3b | 2476 | 5253 | 6421 | 1260 | 4306 | 2729 |
| ENSDARG000 | 2;2;2;2;2;2 | 54043678:540 | 54043791:540 | -;-;-;-;-;- | 912 | NA | 184 | 109 | 131 | 190 | 151 | 177 |
| ENSDARG000 | 2 | 16944272 | 16944391 | - | 120 | 5S_rRNA | 0 | 0 | 0 | 0 | 0 | 0 |
| ENSDARG000 | 2;2;2 | 63298:64038: | 65533:65529: | -;-;- | 2259 | zgc:153913 | 2 | 4 | 28 | 26 | 25 | 28 |
| ENSDARG000 | 2 | 27039688 | 27039804 | - | 117 | 5S_rRNA | 0 | 0 | 0 | 0 | 0 | 0 |
| ENSDARG000 | 2 | 19519085 | 19519182 | + | 98 | NA | 0 | 0 | 0 | 0 | 0 | 0 |
| ENSDARG000 | 2;2;2;2;2;2;2 | 22363824:223 | 22364394:223 | +;+;+;+;+;+;+ | 5095 | hs2st1a | 534 | 1440 | 1240 | 765 | 1601 | 1717 |
| ENSDARG000 | 2;2;2 | 28414830:284 | 28415685:284 | -;-;- | 1288 | NA | 3 | 4 | 0 | 0 | 4 | 0 |
| ENSDARG000 | 2 | 7723870 | 7723941 | + | 72 | NA | 0 | 4 | 2 | 0 | 0 | 3 |
| ENSDARG000 | 2 | 42586738 | 42587035 | + | 298 | RNaseP_nuc | 0 | 0 | 0 | 0 | 0 | 0 |
| ENSDARG000 | 2;2;2;2;2;2;2; | 42871831:428 | 42872741:428 | +;+;+;+;+;+;+; | 4414 | efr3a | 159 | 128 | 82 | 71 | 172 | 103 |
| ENSDARG000 | 2;2 | 58631830:586 | 58632481:586 | -;- | 2254 | NA | 0 | 10 | 8 | 0 | 0 | 0 |
| ENSDARG000 | 2 | 24261217 | 24261296 | - | 80 | dre-mir-736 | 0 | 0 | 0 | 0 | 0 | 0 |
| ENSDARG000 | 2 | 50473426 | 50473542 | - | 117 | 5S_rRNA | 0 | 0 | 0 | 0 | 0 | 0 |
| ENSDARG000 | 2;2;2;2;2;2;2; | 55665095:556 | 55665391:556 | +;+;+;+;+;+;+; | 6004 | klf2b | 481 | 40 | 74 | 138 | 156 | 382 |
| ENSDARG000 | 2 | 25155842 | 25155958 | - | 117 | 5S_rRNA | 0 | 0 | 0 | 0 | 0 | 0 |
| ENSDARG000 | 2 | 40544946 | 40545061 | - | 116 | 5S_rRNA | 0 | 0 | 0 | 0 | 0 | 0 |
| ENSDARG000 | 2;2 | 36496636:364 | 36496928:364 | -;- | 342 | NA | 0 | 0 | 0 | 2 | 0 | 0 |
| ENSDARG000 | 2;2;2;2;2;2;2; | 38287358:382 | 38290242:382 | -;-;-;-;-;-;-;- | 5652 | ap1g2 | 458 | 249 | 434 | 133 | 281 | 277 |
| ENSDARG000 | 2;2;2;2;2;2;2; | 36632676:366 | 36632725:366 | -;-;-;-;-;-;-;- | 186 | NA | 6 | 0 | 2 | 0 | 5 | 0 |
| ENSDARG000 | 2;2;2;2;2;2;2; | 2772447:2774 | 2772623:2774 | +;+;+;+;+;+;+; | 2094 | thoc1 | 654 | 657 | 1121 | 458 | 830 | 678 |
| ENSDARG000 | 2;2;2;2;2;2;2; | 2630188:2632 | 2630247:2632 | -;-;-;-;-;-;-;- | 1734 | NA | 3759 | 3620 | 3992 | 4175 | 3986 | 3513 |
| ENSDARG000 | 2;2;2;2 | 40194452:401 | 40194522:401 | -;-;-;- | 371 | ccl34a.3 | 10 | 54 | 84 | 65 | 245 | 50 |
| ENSDARG000 | 2;2;2 | 4402765:4405 | 4403125:4405 | +;+;+ | 1407 | bambib | 269 | 220 | 502 | 732 | 896 | 362 |
| ENSDARG000 | 2;2;2;2;2;2;2; | 1218758:1220 | 1218916:1220 | -;-;-;-;-;-;-;- | 1713 | ABCF3 | 433 | 450 | 553 | 193 | 516 | 308 |
| ENSDARG000 | 2;2 | 53204750:532 | 53205906:532 | +;+ | 2012 | zgc:165603 | 0 | 0 | 0 | 0 | 0 | 0 |
| ENSDARG000 | 2;2;2;2;2;2;2; | 51783120:517 | 51783272:517 | +;+;+;+;+;+;+; | 1081 | crygn1 | 2 | 0 | 0 | 0 | 0 | 0 |
| ENSDARG000 | 2;2;2;2;2;2;2; | 53880669:538 | 53880678:538 | +;+;+;+;+;+;+; | 189 | NA | 0 | 0 | 0 | 0 | 0 | 0 |
| ENSDARG000 | 2 | 13885155 | 13886842 | - | 1688 | NA | 0 | 0 | 0 | 0 | 0 | 0 |
| ENSDARG000 | 2;2;2;2;2;2;2; | 56811549:568 | 56811914:568 | -;-;-;-;-;-;-;- | 4440 | sbno2a | 137 | 61 | 206 | 103 | 199 | 161 |
| ENSDARG000 | 2;2;2 | 36932796:369 | 36932903:369 | -;-;- | 321 | CARTPT | 0 | 4 | 2 | 0 | 0 | 0 |
| ENSDARG000 | 2;2;2;2;2;2;2 | 52469854:524 | 52470838:524 | -;-;-;-;-;-;- | 2381 | gna11b | 761 | 895 | 1119 | 882 | 1110 | 935 |
| ENSDARG000 | 2 | 14767477 | 14767626 | + | 150 | U2 | 0 | 0 | 0 | 0 | 0 | 0 |
| ENSDARG000 | 2;2 | 53909462:539 | 53909916:539 | -;- | 785 | NA | 0 | 0 | 2 | 0 | 0 | 0 |
| ENSDARG000 | 2;2;2;2;2;2;2; | 52406565:524 | 52407159:524 | -;-;-;-;-;-;-;- | 1512 | chico | 519 | 231 | 241 | 333 | 598 | 212 |
| ENSDARG000 | 2;2;2;2;2;2 | 97692:97793: | 97757:98319: | -;-;-;-;-;- | 1869 | angptl1b | 0 | 4 | 0 | 0 | 0 | 0 |
| ENSDARG000 | 2;2;2 | 22409249:224 | 22409958:224 | +;+;+ | 3812 | zgc:56628 | 0 | 28 | 12 | 4 | 2 | 20 |
| ENSDARG000 | 2 | 40056899 | 40057016 | + | 118 | 5S_rRNA | 0 | 0 | 0 | 0 | 0 | 0 |
| ENSDARG000 | 2 | 18502308 | 18502424 | + | 117 | 5S_rRNA | 0 | 0 | 0 | 0 | 0 | 0 |
| ENSDARG000 | 2;2;2;2;2 | 57609291:576 | 57609854:576 | -;-;-;-;- | 2241 | NA | 0 | 0 | 0 | 0 | 0 | 0 |
| ENSDARG000 | 2;2 | 58256276:582 | 58257344:582 | -;- | 1265 | foxl2b | 0 | 0 | 0 | 0 | 0 | 0 |
| ENSDARG000 | 2 | 41036076 | 41036191 | + | 116 | 5S_rRNA | 0 | 0 | 0 | 0 | 0 | 0 |
| ENSDARG000 | 2 | 50709366 | 50709482 | - | 117 | 5S_rRNA | 0 | 0 | 0 | 0 | 0 | 0 |
| ENSDARG000 | 2;2;2 | 51477200:514 | 51477537:514 | -;-;- | 723 | NA | 20 | 3 | 11 | 10 | 30 | 39 |
| ENSDARG000 | 2;2;2;2 | 2761882:2765 | 2762256:2765 | +;+;+;+ | 833 | aqp1a.2 | 0 | 0 | 5 | 0 | 0 | 8 |
| ENSDARG000 | 2;2;2;2 | 46411079:464 | 46411094:464 | +;+;+;+ | 768 | NA | 0 | 0 | 2 | 0 | 0 | 0 |
| ENSDARG000 | 2;2;2;2 | 38034325:380 | 38034602:380 | -;-;-;- | 642 | NA | 0 | 0 | 0 | 0 | 0 | 0 |
| ENSDARG000 | 2;2;2;2;2 | 53483322:534 | 53485505:534 | -;-;-;-;- | 2941 | admp | 0 | 0 | 0 | 0 | 0 | 0 |
| ENSDARG000 | 2;2;2;2;2;2;2; | 58115123:581 | 58115158:581 | -;-;-;-;-;-;-;- | 1797 | NA | 790 | 598 | 555 | 310 | 561 | 423 |
| ENSDARG000 | 2;2;2;2;2;2;2; | 1478225:1480 | 1478235:1480 | -;-;-;-;-;-;-;- | 2329 | c8b | 2 | 0 | 8 | 0 | 0 | 0 |
| ENSDARG000 | 2;2;2;2;2;2 | 38731696:387 | 38732364:387 | +;+;+;+;+;+ | 6251 | NA | 8 | 32 | 2 | 0 | 2 | 0 |
| ENSDARG000 | 2;2;2;2;2;2;2; | 2470687:2470 | 2470940:2470 | +;+;+;+;+;+;+; | 1150 | myl13 | 2 | 4 | 0 | 0 | 10 | 2 |
| ENSDARG000 | 2 | 3125958 | 3126074 | - | 117 | 5S_rRNA | 0 | 0 | 0 | 0 | 0 | 0 |
| ENSDARG000 | 2;2;2;2;2;2;2; | 50722439:507 | 50722496:507 | +;+;+;+;+;+;+; | 4618 | FYCO1 | 118 | 84 | 369 | 95 | 281 | 176 |
| ENSDARG000 | 2;2;2;2;2 | 3106237:3107 | 3106716:3107 | -;-;-;-;- | 1098 | wnt3a | 0 | 0 | 2 | 0 | 0 | 2 |
| ENSDARG000 | 2;2;2;2;2 | 22353422:223 | 22354001:223 | -;-;-;-;- | 964 | selenof | 1326 | 2333 | 1903 | 1075 | 2380 | 1327 |
| ENSDARG000 | 2;2;2;2 | 9961868:9961 | 9961919:9962 | -;-;-;- | 393 | NA | 2 | 2 | 2 | 0 | 0 | 0 |
| ENSDARG000 | 2 | 30751977 | 30752109 | - | 133 | 5S_rRNA | 0 | 0 | 0 | 0 | 0 | 0 |
| ENSDARG000 | 2 | 47852969 | 47853084 | - | 116 | 5S_rRNA | 0 | 0 | 0 | 0 | 0 | 0 |
| ENSDARG000 | 2 | 19634994 | 19635091 | - | 98 | NA | 0 | 0 | 0 | 0 | 0 | 0 |
| ENSDARG000 | 2;2;2;2;2;2;2; | 24578053:245 | 24578342:245 | -;-;-;-;-;-;-;- | 1774 | crtc1a | 107 | 58 | 206 | 73 | 167 | 153 |
| ENSDARG000 | 2 | 17451656 | 17452724 | + | 1069 | NA | 0 | 0 | 0 | 0 | 0 | 0 |
| ENSDARG000 | 2;2;2;2 | 40181633:401 | 40182128:401 | +;+;+;+ | 1328 | NA | 0 | 0 | 0 | 0 | 0 | 0 |
| ENSDARG000 | 2;2;2;2;2;2;2; | 4044636:4047 | 4046536:4047 | -;-;-;-;-;-;-;- | 4004 | yme1l1b | 484 | 490 | 457 | 340 | 569 | 373 |
| ENSDARG000 | 2;2;2;2;2 | 21485492:214 | 21485707:214 | +;+;+;+;+ | 4118 | inhbab | 26 | 4 | 6 | 0 | 0 | 0 |
| ENSDARG000 | 2 | 9875529 | 9875645 | + | 117 | 5S_rRNA | 1 | 0 | 0 | 0 | 0 | 0 |
| ENSDARG000 | 2;2;2;2;2;2;2 | 5300405:5305 | 5300461:5305 | +;+;+;+;+;+;+ | 750 | GNB4 | 317 | 317 | 321 | 220 | 252 | 225 |
| ENSDARG000 | 2;2;2 | 1174778:1184 | 1175518:1185 | -;-;- | 1191 | NA | 0 | 0 | 0 | 0 | 0 | 0 |
| ENSDARG000 | 2;2;2;2;2;2;2; | 52810862:528 | 52811179:528 | -;-;-;-;-;-;-;- | 2313 | ralbp1 | 524 | 263 | 576 | 292 | 430 | 373 |
| ENSDARG000 | 2;2;2;2 | 53582357:535 | 53587203:535 | -;-;-;- | 5160 | NA | 12 | 2 | 0 | 0 | 22 | 4 |
| ENSDARG000 | 2;2;2;2;2;2;2; | 58777364:587 | 58777432:587 | -;-;-;-;-;-;-;- | 2343 | PTPRS | 64 | 56 | 141 | 51 | 127 | 92 |
| ENSDARG000 | 2 | 11243886 | 11244007 | + | 122 | 5S_rRNA | 0 | 0 | 0 | 0 | 0 | 0 |
| ENSDARG000 | 2 | 15868425 | 15868540 | + | 116 | 5S_rRNA | 0 | 0 | 0 | 0 | 0 | 0 |
| ENSDARG000 | 2 | 14675795 | 14675985 | + | 191 | U2 | 0 | 0 | 0 | 0 | 0 | 0 |
| ENSDARG000 | 2;2;2;2;2;2;2 | 52782056:527 | 52782153:527 | -;-;-;-;-;-;- | 591 | NA | 354 | 186 | 337 | 276 | 300 | 178 |
| ENSDARG000 | 2;2;2;2;2;2;2; | 46032678:460 | 46032989:460 | +;+;+;+;+;+;+; | 2426 | gpc1b | 725 | 642 | 739 | 401 | 1092 | 319 |
| ENSDARG000 | 2 | 45159855 | 45162467 | + | 2613 | NA | 6 | 0 | 239 | 84 | 244 | 0 |
| ENSDARG000 | 2;2;2;2;2;2;2; | 51493515:514 | 51493765:514 | -;-;-;-;-;-;-;- | 1626 | pigrl2.4 | 6 | 9 | 10 | 0 | 37 | 20 |
| ENSDARG000 | 2;2;2 | 37963266:379 | 37963734:379 | -;-;- | 816 | cbln9 | 0 | 4 | 0 | 0 | 0 | 0 |
| ENSDARG000 | 2 | 3428357 | 3429454 | + | 1098 | ACKR2 | 10 | 0 | 2 | 0 | 11 | 0 |
| ENSDARG000 | 2;2;2;2;2;2;2; | 50391331:503 | 50391881:503 | +;+;+;+;+;+;+; | 1658 | viml | 577 | 1121 | 186 | 994 | 1052 | 292 |
| ENSDARG000 | 2 | 53686071 | 53686188 | + | 118 | 5S_rRNA | 0 | 0 | 0 | 0 | 0 | 0 |
| ENSDARG000 | 2 | 15173267 | 15173383 | + | 117 | 5S_rRNA | 0 | 0 | 0 | 0 | 0 | 0 |
| ENSDARG000 | 2 | 12224423 | 12224539 | - | 117 | 5S_rRNA | 0 | 0 | 0 | 0 | 0 | 0 |
| ENSDARG000 | 2 | 37826736 | 37827038 | + | 303 | RNaseP_nuc | 0 | 2 | 9 | 0 | 0 | 9 |
| ENSDARG000 | 2 | 38370535 | 38371942 | + | 1408 | psmb11a | 0 | 0 | 0 | 0 | 2 | 1 |
| ENSDARG000 | 2;2;2;2 | 2621400:2623 | 2621886:2623 | -;-;-;- | 813 | NA | 101 | 108 | 128 | 58 | 120 | 119 |
| ENSDARG000 | 2;2;2;2;2;2;2; | 21387902:213 | 21388616:213 | -;-;-;-;-;-;-;- | 3042 | plcd1b | 76 | 6 | 8 | 11 | 36 | 6 |
| ENSDARG000 | 2;2;2;2;2;2;2 | 45172838:451 | 45172940:451 | -;-;-;-;-;-;- | 180 | NA | 0 | 0 | 0 | 0 | 0 | 0 |
| ENSDARG000 | 2;2;2;2;2;2;2; | 2790573:2792 | 2791285:2792 | -;-;-;-;-;-;-;- | 2200 | usp14 | 1622 | 1636 | 2051 | 732 | 1910 | 1203 |
| ENSDARG000 | 2;2;2;2;2 | 59154588:591 | 59154653:591 | -;-;-;-;- | 864 | NA | 5 | 12 | 6 | 1 | 0 | 3 |
| ENSDARG000 | 2;2;2;2;2;2;2; | 54387710:543 | 54387785:543 | +;+;+;+;+;+;+; | 6086 | rab12 | 394 | 461 | 529 | 211 | 524 | 368 |
| ENSDARG000 | 2;2;2;2;2;2;2; | 18866:19165: | 18934:19280: | -;-;-;-;-;-;-;- | 1230 | NA | 0 | 0 | 0 | 5 | 2 | 0 |
| ENSDARG000 | 2 | 40595375 | 40595494 | - | 120 | 5S_rRNA | 0 | 0 | 0 | 0 | 0 | 0 |
| ENSDARG000 | 2 | 16615940 | 16616055 | - | 116 | 5S_rRNA | 0 | 0 | 0 | 0 | 0 | 0 |
| ENSDARG000 | 2;2;2;2;2 | 1264971:1269 | 1265117:1269 | -;-;-;-;- | 3720 | ZNF521 | 6 | 0 | 2 | 4 | 10 | 14 |
| ENSDARG000 | 2 | 19523586 | 19523683 | - | 98 | NA | 0 | 0 | 0 | 0 | 0 | 0 |
| ENSDARG000 | 2;2;2;2 | 43537846:435 | 43538154:435 | -;-;-;- | 832 | ITGB1 | 1435 | 1113 | 1401 | 982 | 1119 | 897 |
| ENSDARG000 | 2;2;2;2;2 | 37953571:379 | 37954570:379 | -;-;-;-;- | 1462 | cbln10 | 0 | 0 | 2 | 0 | 0 | 0 |
| ENSDARG000 | 2 | 4571013 | 4571103 | - | 91 | NA | 0 | 0 | 0 | 0 | 0 | 0 |
| ENSDARG000 | 2;2;2;2;2;2;2; | 43678039:436 | 43678176:436 | -;-;-;-;-;-;-;- | 2875 | NA | 5 | 6 | 4 | 2 | 5 | 18 |
| ENSDARG000 | 2;2;2;2;2;2;2; | 43577294:435 | 43577460:435 | -;-;-;-;-;-;-;- | 1563 | itgb1b | 3361 | 2926 | 3222 | 2381 | 2504 | 2264 |
| ENSDARG000 | 2;2;2;2;2;2;2; | 43638983:436 | 43639584:436 | -;-;-;-;-;-;-;- | 2383 | itgb1b.2 | 4 | 6 | 1 | 6 | 6 | 0 |
| ENSDARG000 | 2;2;2;2 | 38055529:380 | 38055924:380 | +;+;+;+ | 2370 | si:rp71-1g18. | 254 | 134 | 217 | 138 | 210 | 244 |
| ENSDARG000 | 2;2;2;2;2 | 3062737:3070 | 3067338:3070 | -;-;-;-;- | 5093 | arf1 | 2624 | 2944 | 3281 | 2446 | 2631 | 2680 |
| ENSDARG000 | 2;2;2;2;2;2 | 55249838:552 | 55249855:552 | +;+;+;+;+;+ | 543 | CIB3 | 0 | 0 | 0 | 0 | 2 | 0 |
| ENSDARG000 | 2;2;2 | 7419986:7427 | 7424503:7427 | -;-;- | 5107 | asip2b | 6 | 18 | 235 | 19 | 108 | 90 |
| ENSDARG000 | 2;2;2;2;2;2;2; | 58739111:587 | 58739254:587 | +;+;+;+;+;+;+; | 810 | NA | 210 | 149 | 144 | 134 | 134 | 70 |
| ENSDARG000 | 2 | 22434430 | 22434546 | - | 117 | 5S_rRNA | 0 | 0 | 0 | 0 | 0 | 0 |
| ENSDARG000 | 2;2;2;2;2;2;2; | 55429988:554 | 55430189:554 | +;+;+;+;+;+;+; | 4889 | ap1m1 | 3050 | 2858 | 3594 | 2221 | 3494 | 2828 |
| ENSDARG000 | 2;2;2;2;2;2;2; | 53065039:530 | 53065625:530 | -;-;-;-;-;-;-;- | 1601 | sirt6 | 190 | 174 | 136 | 98 | 153 | 102 |
| ENSDARG000 | 2;2;2;2 | 54015958:540 | 54016119:540 | -;-;-;- | 1352 | ABHD8 | 0 | 0 | 0 | 0 | 8 | 0 |
| ENSDARG000 | 2;2;2;2;2;2;2; | 58026496:580 | 58026765:580 | -;-;-;-;-;-;-;- | 2134 | NA | 10 | 0 | 0 | 0 | 0 | 0 |
| ENSDARG000 | 2;2 | 6908662:6908 | 6908676:6908 | -;- | 18 | NA | 0 | 0 | 0 | 0 | 0 | 0 |
| ENSDARG000 | 2;2;2 | 58877162:588 | 58877326:588 | +;+;+ | 1248 | NA | 285 | 144 | 318 | 164 | 306 | 90 |
| ENSDARG000 | 2;2 | 48478399:485 | 48478867:485 | -;- | 934 | NA | 0 | 0 | 0 | 0 | 0 | 0 |
| ENSDARG000 | 2 | 24505975 | 24506090 | - | 116 | 5S_rRNA | 0 | 0 | 0 | 2 | 0 | 0 |
| ENSDARG000 | 2;2;2;2 | 58008980:580 | 58009444:580 | +;+;+;+ | 3069 | si:ch211-155 | 257 | 112 | 179 | 150 | 208 | 247 |
| ENSDARG000 | 2;2;2;2;2;2;2; | 1428696:1430 | 1428841:1430 | -;-;-;-;-;-;-;- | 1910 | rpe65a | 3588 | 30 | 4 | 33 | 0 | 4 |
| ENSDARG000 | 2 | 42423684 | 42423799 | + | 116 | 5S_rRNA | 1 | 0 | 2 | 0 | 4 | 0 |
| ENSDARG000 | 2 | 19575059 | 19575156 | + | 98 | NA | 0 | 0 | 0 | 0 | 0 | 0 |
| ENSDARG000 | 2 | 36753649 | 36753758 | + | 110 | dre-mir-2198 | 2 | 0 | 0 | 0 | 0 | 0 |
| ENSDARG000 | 2;2;2;2 | 56265791:562 | 56265792:562 | +;+;+;+ | 336 | NA | 5 | 4 | 11 | 2 | 3 | 0 |
| ENSDARG000 | 2;2;2;2;2;2;2; | 53720028:537 | 53720159:537 | +;+;+;+;+;+;+; | 2018 | ctnnbl1 | 684 | 709 | 852 | 428 | 703 | 470 |
| ENSDARG000 | 2;2;2;2;2;2;2; | 43619555:436 | 43619660:436 | -;-;-;-;-;-;-;- | 2451 | itgb1b.1 | 41 | 7 | 6 | 0 | 40 | 26 |
| ENSDARG000 | 2;2 | 26499502:264 | 26499613:264 | -;- | 122 | NA | 0 | 0 | 0 | 0 | 0 | 0 |
| ENSDARG000 | 2;2;2;2;2 | 58221163:582 | 58221502:582 | +;+;+;+;+ | 1242 | KCNJ10 | 26 | 32 | 0 | 4 | 8 | 2 |
| ENSDARG000 | 2;2;2;2;2;2;2 | 58841181:588 | 58841247:588 | +;+;+;+;+;+;+ | 1305 | cirbpa | 10097 | 17144 | 18460 | 12527 | 14279 | 12680 |
| ENSDARG000 | 2 | 50590332 | 50590448 | + | 117 | 5S_rRNA | 0 | 0 | 0 | 7 | 5 | 0 |
| ENSDARG000 | 2;2;2;2;2;2;2; | 9536021:9536 | 9536039:9536 | +;+;+;+;+;+;+; | 186 | NA | 0 | 2 | 1 | 0 | 0 | 4 |
| ENSDARG000 | 2;2;2;2 | 58173047:581 | 58174786:581 | -;-;-;- | 5550 | NA | 12 | 17 | 12 | 12 | 14 | 28 |
| ENSDARG000 | 2;2;2 | 48638827:486 | 48638947:486 | +;+;+ | 747 | NA | 0 | 0 | 4 | 0 | 3 | 33 |
| ENSDARG000 | 2;2;2;2;2;2;2 | 56657804:566 | 56658362:566 | +;+;+;+;+;+;+ | 1135 | POLR2E | 0 | 0 | 0 | 0 | 0 | 0 |
| ENSDARG000 | 2;2;2;2;2;2;2; | 1348281:1348 | 1349159:1349 | -;-;-;-;-;-;-;- | 2085 | ss18 | 730 | 844 | 1292 | 890 | 1017 | 742 |
| ENSDARG000 | 2 | 14670997 | 14671173 | + | 177 | U2 | 0 | 0 | 0 | 0 | 0 | 0 |
| ENSDARG000 | 2 | 9747302 | 9747431 | - | 130 | NA | 0 | 0 | 0 | 0 | 0 | 0 |
| ENSDARG000 | 2 | 21375467 | 21375594 | + | 128 | dre-mir-26a-2 | 2 | 2 | 0 | 0 | 0 | 0 |
| ENSDARG000 | 2 | 16824759 | 16824819 | + | 61 | NA | 0 | 0 | 0 | 0 | 0 | 0 |
| ENSDARG000 | 2;2;2;2;2;2;2; | 1486822:1487 | 1487207:1487 | +;+;+;+;+;+;+; | 2320 | c8a | 4 | 2 | 10 | 46 | 0 | 10 |
| ENSDARG000 | 2 | 45332137 | 45336975 | - | 4839 | SAMD9 | 26 | 0 | 2 | 0 | 2 | 0 |
| ENSDARG000 | 2;2;2 | 2503396:2503 | 2503516:2503 | +;+;+ | 294 | crhr2 | 0 | 0 | 0 | 0 | 0 | 0 |
| ENSDARG000 | 2;2;2;2;2 | 28385115:283 | 28385463:283 | -;-;-;-;- | 930 | NA | 0 | 2 | 0 | 0 | 0 | 0 |
| ENSDARG000 | 2;2;2 | 4383061:4383 | 4383176:4383 | +;+;+ | 1042 | NA | 162 | 119 | 120 | 47 | 154 | 120 |
| ENSDARG000 | 2;2;2;2;2;2;2; | 1202347:1203 | 1202456:1203 | +;+;+;+;+;+;+; | 1359 | NA | 0 | 0 | 0 | 0 | 0 | 0 |
| ENSDARG000 | 2;2 | 22564928:225 | 22569635:225 | -;- | 5621 | SOX14 | 0 | 0 | 0 | 4 | 0 | 7 |

| | | | | | | | | | | | | |
|---|---|---|---|---|---|---|---|---|---|---|---|---|
| ENSDARG000 | 2 | 33569922 | 33570039 | - | 118 | 5S_rRNA | 0 | 0 | 0 | 0 | 0 | 0 |
| ENSDARG000 | 2 | 1575184 | 1575290 | + | 107 | 5S_rRNA | 0 | 0 | 0 | 0 | 0 | 0 |
| ENSDARG000 | 2;2;2;2;2;2 | 52232630:522 | 52233014:522 | +;+;+;+;+;+ | 1185 | PLPP2 | 20 | 1 | 8 | 0 | 23 | 12 |
| ENSDARG000 | 2;2 | 48447353:484 | 48447730:484 | -;- | 782 | NA | 0 | 0 | 0 | 0 | 0 | 0 |
| ENSDARG000 | 2;2 | 36499886:365 | 36500169:365 | -;- | 333 | NA | 0 | 2 | 4 | 0 | 8 | 4 |
| ENSDARG000 | 2;2;2;2;2;2;2; | 40294313:403 | 40294444:403 | +;+;+;+;+;+;+; | 3311 | epha4b | 38 | 6 | 8 | 10 | 12 | 52 |
| ENSDARG000 | 2 | 48746683 | 48746812 | - | 130 | 5S_rRNA | 0 | 0 | 0 | 0 | 0 | 0 |
| ENSDARG000 | 2;2;2;2;2;2;2; | 49667767:496 | 49668052:496 | +;+;+;+;+;+;+; | 7120 | NA | 14 | 0 | 0 | 6 | 20 | 6 |
| ENSDARG000 | 2 | 32667132 | 32667265 | - | 134 | NA | 0 | 0 | 0 | 0 | 0 | 0 |
| ENSDARG000 | 2;2;2;2;2;2;2; | 57121596:571 | 57121808:571 | -;-;-;-;-;-;-;- | 1443 | ARMC6 | 206 | 184 | 264 | 175 | 299 | 158 |
| ENSDARG000 | 2 | 4114118 | 4114216 | - | 99 | dre-mir-133a- | 0 | 0 | 0 | 0 | 0 | 0 |
| ENSDARG000 | 2;2;2;2;2;2;2; | 3044992:305 | 3045373:305 | +;+;+;+;+;+;+; | 2859 | C2H1orf35 | 188 | 286 | 435 | 155 | 268 | 287 |
| ENSDARG000 | 2;2;2;2;2;2;2; | 2563192:257 | 2563278:257 | +;+;+;+;+;+;+; | 1005 | CRHR2 | 0 | 0 | 0 | 0 | 4 | 0 |
| ENSDARG000 | 2;2;2 | 48748254:487 | 48748340:487 | -;-;- | 249 | NA | 0 | 0 | 1 | 6 | 10 | 0 |
| ENSDARG000 | 2;2;2;2;2;2;2; | 38039857:38 | 38040246:38 | +;+;+;+;+;+;+; | 2979 | casq1a | 0 | 0 | 0 | 0 | 4 | 0 |
| ENSDARG000 | 2;2 | 47869807:478 | 47869920:478 | -;- | 705 | NA | 88 | 50 | 55 | 45 | 71 | 105 |
| ENSDARG000 | 2 | 14665168 | 14665353 | + | 186 | U2 | 0 | 0 | 0 | 0 | 0 | 0 |
| ENSDARG000 | 2 | 37295088 | 37296159 | + | 1072 | gpr160 | 2 | 0 | 0 | 0 | 8 | 0 |
| ENSDARG000 | 2;2;2 | 37995176:379 | 37995716:379 | -;-;- | 765 | pcp4l1 | 4 | 17 | 7 | 0 | 4 | 10 |
| ENSDARG000 | 2;2;2;2;2;2;2; | 3026802:302 | 3029513:302 | -;-;-;-;-;-;-; | 3613 | guk1a | 1565 | 1770 | 3410 | 2074 | 1970 | 2517 |
| ENSDARG000 | 2;2;2;2;2;2;2; | 7557912:756 | 7558341:756 | +;+;+;+;+;+;+; | 1598 | ripk2 | 62 | 102 | 123 | 78 | 96 | 96 |
| ENSDARG000 | 2;2;2;2;2;2;2 | 3986083:398 | 3986207:398 | +;+;+;+;+;+;+ | 1224 | mkxb | 5 | 0 | 19 | 0 | 9 | 6 |
| ENSDARG000 | 2;2;2;2;2;2;2; | 58186846:581 | 58187269:581 | -;-;-;-;-;-;-; | 1138 | pnp5b | 498 | 1323 | 2701 | 2126 | 730 | 461 |
| ENSDARG000 | 2 | 22897285 | 22897401 | - | 117 | 5S_rRNA | 0 | 0 | 0 | 0 | 0 | 0 |
| ENSDARG000 | 2 | 10206001 | 10206118 | + | 118 | 5S_rRNA | 0 | 0 | 0 | 0 | 0 | 0 |
| ENSDARG000 | 2 | 48564830 | 48564946 | - | 117 | 5S_rRNA | 0 | 0 | 0 | 0 | 0 | 0 |
| ENSDARG000 | 2 | 9688055 | 9688185 | - | 131 | 5S_rRNA | 2 | 0 | 0 | 0 | 1 | 4 |
| ENSDARG000 | 2 | 45202223 | 45202331 | - | 109 | dre-mir-181b- | 0 | 0 | 6 | 0 | 2 | 0 |
| ENSDARG000 | 2;2;2;2 | 54482603:544 | 54483940:544 | +;+;+;+ | 5120 | NA | 2 | 0 | 0 | 0 | 0 | 0 |
| ENSDARG000 | 2;2 | 13688624:136 | 13691673:136 | -;- | 3127 | NA | 85 | 48 | 53 | 25 | 60 | 85 |
| ENSDARG000 | 2 | 20145741 | 20145825 | - | 85 | dre-mir-137-1 | 0 | 0 | 0 | 0 | 0 | 0 |
| ENSDARG000 | 2;2;2;2 | 40199002:401 | 40199237:401 | -;-;-;- | 530 | ccl34a.4 | 451 | 607 | 875 | 599 | 1926 | 1171 |
| ENSDARG000 | 2;2;2;2;2;2;2; | 38002717:38 | 38002753:38 | +;+;+;+;+;+;+; | 1824 | DCAF8 | 623 | 469 | 488 | 347 | 451 | 432 |
| ENSDARG000 | 2;2;2;2;2;2;2; | 41504752:415 | 41504795:415 | -;-;-;-;-;-;-;- | 1149 | otomp | 0 | 4 | 13 | 6 | 12 | 33 |
| ENSDARG000 | 2;2 | 36509085:365 | 36509377:365 | -;- | 342 | NA | 0 | 0 | 0 | 0 | 0 | 0 |
| ENSDARG000 | 2;2;2;2;2;2;2; | 205763:2058 | 206091:2060 | +;+;+;+;+;+;+; | 1537 | zgc:113293 | 0 | 0 | 4 | 0 | 0 | 0 |
| ENSDARG000 | 2;2;2;2;2;2;2; | 36715510:367 | 36715530:367 | -;-;-;-;-;-;-; | 234 | NA | 1 | 0 | 0 | 4 | 6 | 3 |
| ENSDARG000 | 2;2;2;2;2;2;2; | 16846772:168 | 16847053:168 | +;+;+;+;+;+;+; | 1865 | fam131aa | 4 | 23 | 26 | 26 | 12 | 44 |
| ENSDARG000 | 2;2;2;2;2;2;2; | 53114476:531 | 53114523:531 | +;+;+;+;+;+;+; | 2380 | NA | 0 | 0 | 0 | 0 | 0 | 0 |
| ENSDARG000 | 2;2;2;2;2;2;2 | 6734108:676 | 6734936:676 | +;+;+;+;+;+;+ | 4701 | NA | 94 | 0 | 0 | 0 | 4 | 2 |
| ENSDARG000 | 2 | 56430847 | 56431298 | - | 452 | ctxn1 | 0 | 0 | 0 | 0 | 0 | 0 |
| ENSDARG000 | 2;2;2;2;2;2;2; | 55751966:557 | 55752193:557 | -;-;-;-;-;-;-; | 1608 | NA | 74 | 66 | 110 | 100 | 86 | 62 |
| ENSDARG000 | 2 | 58636676 | 58636792 | - | 117 | 5S_rRNA | 0 | 0 | 0 | 0 | 0 | 0 |
| ENSDARG000 | 2 | 45073794 | 45073910 | - | 117 | 5S_rRNA | 0 | 0 | 0 | 0 | 0 | 0 |
| ENSDARG000 | 2;2;2;2;2 | 105748:1078 | 105960:1080 | +;+;+;+;+ | 904 | TADA1 | 164 | 155 | 251 | 120 | 253 | 116 |
| ENSDARG000 | 2 | 26645317 | 26645398 | - | 82 | dre-mir-124-1 | 0 | 0 | 0 | 0 | 0 | 0 |
| ENSDARG000 | 2;2;2;2;2;2;2 | 53459873:534 | 53460047:534 | -;-;-;-;-;-;- | 846 | NA | 6 | 0 | 0 | 0 | 4 | 0 |
| ENSDARG000 | 2;2 | 36493849:364 | 36494138:364 | -;- | 339 | NA | 0 | 0 | 0 | 0 | 0 | 0 |
| ENSDARG000 | 2 | 50718112 | 50719170 | - | 1059 | XCR1 | 0 | 0 | 0 | 0 | 0 | 2 |
| ENSDARG000 | 2;2 | 4631335:464 | 4631669:464 | -;- | 769 | NA | 0 | 0 | 0 | 0 | 0 | 0 |
| ENSDARG000 | 2 | 13050225 | 13051517 | + | 1293 | NA | 0 | 2 | 4 | 0 | 0 | 0 |
| ENSDARG000 | 2;2;2;2;2;2 | 58914674:589 | 58915007:589 | -;-;-;-;-;- | 1041 | NA | 256 | 192 | 290 | 129 | 257 | 199 |
| ENSDARG000 | 2 | 10597810 | 10600950 | - | 3141 | NA | 6 | 8 | 8 | 20 | 24 | 20 |
| ENSDARG000 | 2;2;2;2;2 | 22566082:225 | 22566652:225 | -;-;-;-;- | 1217 | NA | 0 | 0 | 0 | 0 | 0 | 0 |
| ENSDARG000 | 2;2;2 | 48687528:486 | 48687873:486 | +;+;+ | 681 | NA | 0 | 1 | 4 | 0 | 0 | 0 |
| ENSDARG000 | 2 | 26499649 | 26499754 | + | 106 | SNORD46 | 0 | 0 | 0 | 0 | 0 | 0 |
| ENSDARG000 | 2 | 47260637 | 47260753 | - | 117 | 5S_rRNA | 0 | 0 | 0 | 0 | 0 | 0 |
| ENSDARG000 | 2;2;2;2;2;2 | 184818:1878 | 189440:1894 | -;-;-;-;-;- | 4987 | znf1014 | 194 | 158 | 274 | 178 | 248 | 333 |
| ENSDARG000 | 2 | 46009507 | 46012744 | + | 3238 | NA | 0 | 2 | 2 | 0 | 0 | 0 |
| ENSDARG000 | 2;2 | 58414598:584 | 58414691:584 | -;- | 162 | NA | 0 | 0 | 0 | 0 | 0 | 0 |
| ENSDARG000 | 2;2 | 194742:1973 | 196291:1976 | -;- | 1868 | NA | 150 | 123 | 160 | 164 | 110 | 142 |
| ENSDARG000 | 2 | 42998802 | 42998919 | + | 118 | 5S_rRNA | 0 | 0 | 0 | 4 | 0 | 0 |
| ENSDARG000 | 2;2;2;2;2;2;2; | 45081489:45 | 45081624:45 | +;+;+;+;+;+;+; | 2520 | chrng | 8 | 0 | 0 | 0 | 8 | 4 |
| ENSDARG000 | 2;2;2;2;2;2;2 | 27830436:278 | 27830792:278 | +;+;+;+;+;+;+ | 1157 | NA | 0 | 2 | 0 | 0 | 0 | 0 |
| ENSDARG000 | 2;2;2;2;2;2;2; | 38285414:382 | 38285799:382 | -;-;-;-;-;-;-; | 1469 | zp3f.2 | 2 | 4 | 0 | 4 | 2 | 0 |
| ENSDARG000 | 2;2;2;2;2;2;2; | 54364273:543 | 54364413:543 | -;-;-;-;-;-;-; | 933 | napgb | 57 | 22 | 36 | 57 | 64 | 32 |
| ENSDARG000 | 2 | 2301358 | 2301434 | - | 77 | dre-mir-460 | 0 | 0 | 0 | 0 | 0 | 0 |
| ENSDARG000 | 2;2;2;2;2;2;2; | 20763289:207 | 20763300:207 | -;-;-;-;-;-;-; | 381 | NA | 2 | 21 | 1 | 24 | 21 | 13 |
| ENSDARG000 | 2;2 | 45330452:453 | 45330607:453 | -;- | 348 | NA | 4 | 0 | 0 | 0 | 0 | 0 |
| ENSDARG000 | 2 | 583451 | 583567 | - | 117 | 5S_rRNA | 0 | 0 | 0 | 0 | 0 | 0 |
| ENSDARG000 | 2 | 26501768 | 26501900 | + | 133 | NA | 3 | 17 | 13 | 5 | 12 | 19 |
| ENSDARG000 | 2 | 14713394 | 14713538 | + | 145 | U2 | 0 | 0 | 0 | 0 | 0 | 0 |
| ENSDARG000 | 2;2;2 | 51499900:515 | 51500321:515 | -;-;- | 795 | NA | 2 | 4 | 0 | 0 | 21 | 10 |
| ENSDARG000 | 2;2;2;2;2;2;2; | 53619268:536 | 53619420:536 | +;+;+;+;+;+;+; | 4422 | ranbp3b | 762 | 825 | 976 | 647 | 960 | 675 |
| ENSDARG000 | 2 | 45744863 | 45744979 | - | 117 | 5S_rRNA | 0 | 0 | 0 | 0 | 0 | 0 |
| ENSDARG000 | 2 | 23679873 | 23679989 | + | 117 | 5S_rRNA | 0 | 0 | 0 | 0 | 0 | 0 |
| ENSDARG000 | 2;2;2;2;2;2;2; | 43585571:435 | 43587734:435 | -;-;-;-;-;-;-; | 4485 | epc1b | 999 | 693 | 951 | 798 | 1018 | 965 |
| ENSDARG000 | 2;2;2;2;2;2;2; | 45696743:456 | 45696899:456 | +;+;+;+;+;+;+; | 1371 | NA | 2 | 0 | 19 | 0 | 41 | 0 |
| ENSDARG000 | 2;2;2 | 48458703:484 | 48459411:484 | -;-;- | 1464 | NA | 0 | 0 | 0 | 2 | 0 | 0 |
| ENSDARG000 | 2;2;2;2;2;2;2 | 50608099:506 | 50608229:506 | +;+;+;+;+;+;+ | 1835 | neurod6b | 0 | 0 | 0 | 0 | 2 | 0 |
| ENSDARG000 | 2;2;2;2;2;2;2; | 50477779:504 | 50477879:504 | +;+;+;+;+;+;+; | 1842 | PDE1C | 0 | 0 | 0 | 21 | 22 | 0 |
| ENSDARG000 | 2;2;2;2;2;2;2 | 48519594:485 | 48521371:485 | -;-;-;-;-;-;- | 3680 | NA | 0 | 0 | 1 | 0 | 0 | 0 |
| ENSDARG000 | 2;2;2;2;2;2;2; | 51818039:518 | 51818522:518 | +;+;+;+;+;+;+; | 1977 | ACVR2B | 37 | 52 | 67 | 16 | 43 | 44 |
| ENSDARG000 | 2 | 44763082 | 44763375 | - | 294 | NA | 0 | 4 | 7 | 5 | 11 | 19 |
| ENSDARG000 | 2 | 56847743 | 56847859 | + | 117 | 5S_rRNA | 0 | 0 | 0 | 0 | 0 | 0 |
| ENSDARG000 | 2;2;2;2 | 3201345:325 | 3201491:325 | +;+;+;+ | 1147 | wnt9a | 0 | 0 | 0 | 4 | 6 | 0 |
| ENSDARG000 | 2 | 16829689 | 16829764 | + | 76 | NA | 0 | 0 | 0 | 0 | 0 | 0 |
| ENSDARG000 | 2;2 | 49366684:493 | 49367565:493 | -;- | 891 | NA | 0 | 2 | 0 | 0 | 8 | 0 |
| ENSDARG000 | 2;2;2;2;2;2;2; | 9983079:998 | 9983360:998 | -;-;-;-;-;-;-; | 904 | imp3 | 201 | 226 | 258 | 171 | 245 | 204 |
| ENSDARG000 | 2 | 26085162 | 26085279 | - | 118 | 5S_rRNA | 0 | 0 | 0 | 0 | 0 | 0 |
| ENSDARG000 | 2 | 14724960 | 14725146 | + | 187 | U2 | 0 | 0 | 0 | 0 | 0 | 0 |
| ENSDARG000 | 2;2 | 2918759:292 | 2919655:292 | +;+ | 1431 | NA | 2 | 6 | 0 | 0 | 0 | 2 |
| ENSDARG000 | 2 | 4116314 | 4116400 | - | 87 | dre-mir-1-2 | 0 | 0 | 0 | 0 | 0 | 0 |
| ENSDARG000 | 2;2;2;2;2 | 53890006:538 | 53890463:538 | -;-;-;-;- | 962 | capsla | 0 | 0 | 0 | 0 | 8 | 0 |
| ENSDARG000 | 2 | 13123566 | 13123682 | + | 117 | 5S_rRNA | 0 | 0 | 1 | 0 | 0 | 0 |
| ENSDARG000 | 2 | 7720559 | 7720629 | + | 71 | NA | 4 | 5 | 0 | 5 | 5 | 7 |
| ENSDARG000 | 2;2;2;2;2;2;2; | 38068069:38 | 38068266:38 | -;-;-;-;-;-;-;- | 2198 | NA | 525 | 398 | 530 | 310 | 610 | 313 |
| ENSDARG000 | 2;2;2;2;2;2;2; | 5245692:524 | 5245794:524 | +;+;+;+;+;+;+; | 1551 | NA | 232 | 79 | 61 | 1 | 82 | 41 |
| ENSDARG000 | 2;2;2;2;2;2;2; | 54327160:543 | 54327212:543 | +;+;+;+;+;+;+; | 1764 | NA | 0 | 0 | 0 | 0 | 2 | 0 |
| ENSDARG000 | 2 | 35831329 | 35831445 | + | 117 | 5S_rRNA | 0 | 1 | 1 | 0 | 0 | 0 |
| ENSDARG000 | 2 | 27233666 | 27233781 | - | 116 | 5S_rRNA | 0 | 0 | 0 | 0 | 0 | 0 |
| ENSDARG000 | 2 | 51458113 | 51458228 | - | 116 | 5S_rRNA | 0 | 0 | 0 | 0 | 0 | 0 |
| ENSDARG000 | 2 | 14994393 | 14994490 | - | 98 | NA | 0 | 0 | 0 | 0 | 0 | 0 |
| ENSDARG000 | 2;2;2;2;2;2;2; | 54086436:541 | 54086499:541 | +;+;+;+;+;+;+; | 4062 | PIEZO2 | 2 | 0 | 0 | 0 | 0 | 0 |
| ENSDARG000 | 2;2 | 58868109:588 | 58869038:588 | -;- | 1344 | fem1a | 568 | 433 | 511 | 457 | 595 | 267 |
| ENSDARG000 | 2;2;2;2;2;2;2; | 7668573:766 | 7669061:766 | -;-;-;-;-;-;-; | 3262 | CCDC39 | 0 | 2 | 4 | 0 | 0 | 0 |
| ENSDARG000 | 2 | 54727306 | 54727426 | - | 121 | 5S_rRNA | 0 | 0 | 0 | 0 | 0 | 0 |
| ENSDARG000 | 2 | 6766293 | 6766407 | - | 115 | 5S_rRNA | 0 | 0 | 0 | 0 | 0 | 0 |
| ENSDARG000 | 2;2;2 | 7487592:749 | 7487672:749 | -;-;- | 289 | NA | 0 | 3 | 8 | 0 | 4 | 13 |
| ENSDARG000 | 2;2;2 | 12819999:128 | 12820611:128 | -;-;- | 1315 | NA | 0 | 0 | 0 | 2 | 0 | 0 |
| ENSDARG000 | 2;2;2 | 21882881:219 | 21883038:219 | +;+;+ | 532 | NA | 0 | 0 | 0 | 0 | 0 | 0 |
| ENSDARG000 | 2;2;2 | 27218149:272 | 27218276:272 | +;+;+ | 353 | NA | 0 | 0 | 0 | 0 | 0 | 0 |
| ENSDARG000 | 2;2;2 | 28165967:281 | 28167277:281 | -;-;- | 1692 | NA | 0 | 0 | 0 | 3 | 0 | 0 |
| ENSDARG000 | 2;2;2;2 | 28372943:283 | 28373653:283 | -;-;-;- | 1076 | NA | 10 | 28 | 20 | 9 | 31 | 16 |
| ENSDARG000 | 2;2;2;2;2 | 30277334:302 | 30278326:302 | +;+;+;+;+ | 1394 | NA | 2 | 0 | 0 | 0 | 0 | 0 |
| ENSDARG000 | 2;2;2;2;2;2;2 | 31100444:311 | 31100688:311 | +;+;+;+;+;+;+ | 1220 | NA | 0 | 0 | 0 | 0 | 0 | 0 |
| ENSDARG000 | 2;2;2 | 37926380:379 | 37926751:379 | +;+;+ | 797 | NA | 15 | 6 | 15 | 7 | 31 | 14 |
| ENSDARG000 | 2;2;2 | 40786665:407 | 40786751:407 | -;-;- | 431 | NA | 0 | 0 | 0 | 0 | 0 | 0 |
| ENSDARG000 | 2;2;2 | 41162103:412 | 41162274:412 | -;-;- | 569 | NA | 0 | 0 | 0 | 0 | 0 | 8 |
| ENSDARG000 | 2;2;2 | 41230920:412 | 41231125:412 | +;+;+ | 902 | NA | 0 | 0 | 0 | 0 | 0 | 0 |
| ENSDARG000 | 2;2;2 | 41972537:419 | 41973599:419 | -;-;- | 1966 | NA | 716 | 286 | 551 | 281 | 518 | 751 |
| ENSDARG000 | 2;2;2 | 43509502:435 | 43509946:435 | -;-;- | 1311 | NA | 6 | 4 | 4 | 0 | 8 | 0 |
| ENSDARG000 | 2;2;2 | 47312577:473 | 47312763:473 | +;+;+ | 1117 | NA | 0 | 2 | 2 | 0 | 0 | 0 |
| ENSDARG000 | 2;2;2;2;2;2;2; | 52154778:521 | 52155105:521 | +;+;+;+;+;+;+; | 1522 | NA | 6 | 0 | 0 | 0 | 0 | 6 |
| ENSDARG000 | 2;2;2 | 54305610:543 | 54305849:543 | +;+;+ | 3400 | NA | 0 | 0 | 0 | 2 | 0 | 0 |
| ENSDARG000 | 2;2;2 | 54430632:544 | 54431348:544 | +;+;+ | 3311 | NA | 0 | 0 | 0 | 0 | 0 | 0 |
| ENSDARG000 | 2;2;2;2;2;2;2; | 55725568:557 | 55726436:557 | -;-;-;-;-;-;-;- | 1503 | NA | 45 | 25 | 47 | 30 | 36 | 29 |
| ENSDARG000 | 2;2;2;2;2 | 58272078:582 | 58276711:582 | -;-;-;-;- | 5998 | NA | 2 | 0 | 10 | 0 | 2 | 8 |
| ENSDARG000 | 2;2;2;2;2;2;2; | 58995304:589 | 58996084:589 | +;+;+;+;+;+;+; | 5033 | NA | 0 | 0 | 0 | 0 | 0 | 6 |
| ENSDARG000 | 1;1;1;1;1;1;1; | 27977297:279 | 27977653:279 | +;+;+;+;+;+;+; | 3099 | sugt1 | 885 | 954 | 981 | 712 | 1149 | 900 |
| ENSDARG000 | 1;1;1;1;1;1;1 | 38398696:384 | 38399229:384 | -;-;-;-;-;-;- | 1232 | hpgd | 8 | 0 | 10 | 10 | 62 | 2 |
| ENSDARG000 | 1;1 | 56943492:569 | 56944389:569 | -;- | 1016 | NA | 3 | 1 | 0 | 0 | 0 | 2 |
| ENSDARG000 | 1;1;1;1;1;1;1; | 44707054:447 | 44707201:447 | -;-;-;-;-;-;-;- | 746 | pold4 | 216 | 160 | 253 | 172 | 189 | 294 |
| ENSDARG000 | 1;1;1;1;1;1;1; | 31809173:318 | 31809570:318 | -;-;-;-;-;-;-; | 2744 | nt5c2b | 592 | 446 | 360 | 643 | 509 | 884 |
| ENSDARG000 | 1;1;1;1;1;1;1; | 7574351:757 | 7574671:757 | -;-;-;-;-;-;-; | 2291 | mxb | 189 | 551 | 6 | 4 | 0 | 0 |
| ENSDARG000 | 1;1;1;1;1;1;1; | 20733102:207 | 20734331:207 | -;-;-;-;-;-;-; | 4633 | tll1 | 16 | 6 | 46 | 12 | 23 | 18 |
| ENSDARG000 | 1 | 56950797 | 56951369 | - | 573 | NA | 0 | 0 | 0 | 0 | 0 | 0 |
| ENSDARG000 | 1;1;1;1;1;1 | 56955349:569 | 56955379:569 | +;+;+;+;+;+ | 1186 | NA | 0 | 0 | 0 | 0 | 0 | 0 |
| ENSDARG000 | 1;1;1;1;1;1;1; | 17527342:175 | 17527453:175 | +;+;+;+;+;+;+; | 2992 | casp3a | 315 | 287 | 375 | 365 | 306 | 320 |
| ENSDARG000 | 1;1;1;1;1;1 | 56972616:569 | 56972648:569 | +;+;+;+;+;+ | 1060 | NA | 0 | 0 | 0 | 0 | 0 | 0 |
| ENSDARG000 | 1;1;1;1;1;1;1; | 25886437:258 | 25886605:258 | -;-;-;-;-;-;-; | 3319 | USP53 | 4 | 0 | 28 | 0 | 12 | 11 |
| ENSDARG000 | 1;1;1;1;1;1;1; | 53881844:538 | 53882377:538 | -;-;-;-;-;-;-; | 2668 | capn9 | 44 | 4 | 4 | 4 | 10 | 6 |
| ENSDARG000 | 1;1;1;1;1;1;1; | 40237276:402 | 40237520:402 | +;+;+;+;+;+;+; | 2352 | faah2a | 158 | 166 | 102 | 99 | 191 | 194 |
| ENSDARG000 | 1;1;1;1;1;1;1; | 22647756:226 | 22648211:226 | -;-;-;-;-;-;-; | 1720 | uchl1 | 2240 | 2885 | 1490 | 5321 | 1873 | 2498 |
| ENSDARG000 | 1;1;1;1;1;1;1; | 51191049:511 | 51191141:511 | +;+;+;+;+;+;+; | 2157 | btbd3a | 447 | 253 | 381 | 304 | 403 | 388 |
| ENSDARG000 | 1;1;1;1;1;1;1 | 58608978:586 | 58609064:586 | +;+;+;+;+;+;+ | 2138 | NA | 82 | 89 | 127 | 103 | 111 | 84 |
| ENSDARG000 | 1;1;1;1;1;1;1; | 58613972:586 | 58614060:586 | +;+;+;+;+;+;+; | 2179 | NA | 50 | 78 | 125 | 69 | 103 | 116 |
| ENSDARG000 | 1;1;1;1;1;1;1; | 45616067:456 | 45617938:456 | -;-;-;-;-;-;-; | 3172 | septin3 | 2 | 2 | 8 | 0 | 28 | 27 |
| ENSDARG000 | 1;1;1;1;1;1 | 56992262:569 | 56992316:569 | +;+;+;+;+;+ | 1218 | NA | 0 | 0 | 0 | 1 | 0 | 0 |
| ENSDARG000 | 1;1;1;1;1;1;1; | 45570059:455 | 45571034:455 | -;-;-;-;-;-;-; | 4360 | atf7ip | 2239 | 1546 | 1772 | 1557 | 2819 | 2047 |
| ENSDARG000 | 1;1;1;1;1;1;1; | 46888033:468 | 46888508:468 | -;-;-;-;-;-;-; | 1435 | pdxkb | 296 | 267 | 300 | 63 | 349 | 240 |
| ENSDARG000 | 1;1;1;1;1;1;1; | 20839482:208 | 20840257:208 | -;-;-;-;-;-;-; | 6373 | cpe | 912 | 34 | 140 | 4001 | 114 | 118 |
| ENSDARG000 | 1;1;1;1;1;1 | 57004178:570 | 57004238:570 | +;+;+;+;+;+ | 1157 | NA | 0 | 0 | 0 | 0 | 0 | 0 |
| ENSDARG000 | 1;1;1;1 | 12135129:121 | 12135190:121 | +;+;+;+ | 304 | spink4 | 0 | 0 | 0 | 0 | 0 | 6 |
| ENSDARG000 | 1;1;1;1;1 | 43686251:436 | 43686477:436 | +;+;+;+;+ | 1633 | cisd2 | 320 | 398 | 483 | 215 | 443 | 326 |
| ENSDARG000 | 1;1;1;1;1;1;1; | 49568335:495 | 49568497:495 | +;+;+;+;+;+;+; | 4745 | col17a1a | 36 | 0 | 0 | 0 | 6 | 10 |
| ENSDARG000 | 1;1 | 57013640:570 | 57013851:570 | +;+ | 699 | NA | 0 | 2 | 0 | 0 | 0 | 0 |
| ENSDARG000 | 1;1;1 | 58620953:586 | 58621034:586 | +;+;+ | 589 | zgc:174890 | 12 | 7 | 16 | 6 | 7 | 11 |
| ENSDARG000 | 1;1;1;1;1;1;1; | 59067978:590 | 59068002:590 | +;+;+;+;+;+;+; | 1470 | MFAP4 | 19 | 8 | 40 | 1 | 2 | 8 |
| ENSDARG000 | 1;1;1;1;1;1;1; | 35494837:354 | 35495277:354 | +;+;+;+;+;+;+; | 5166 | gab1 | 1347 | 1062 | 1127 | 929 | 1345 | 774 |
| ENSDARG000 | 1;1;1;1;1;1;1; | 29862074:298 | 29862162:298 | +;+;+;+;+;+;+; | 3119 | pcca | 861 | 855 | 1257 | 436 | 1188 | 725 |
| ENSDARG000 | 1;1;1;1;1;1;1 | 46948010:469 | 46948129:469 | +;+;+;+;+;+;+ | 980 | TMPRSS3 | 57 | 23 | 8 | 33 | 21 | 39 |

| | | | | | | | | | | | |
|---|---|---|---|---|---|---|---|---|---|---|---|
| ENSDARG000 | 1:1:1:1:1:1:1:1: | 25917591:259 | 25918504:259 | -:-:-:-:-:-:-:-: | 1606 | myoz2b | 0 | 0 | 0 | 0 | 0 | 0 |
| ENSDARG000 | 1;1;1;1;1;1;1 | 245076:25160 | 245952:25172 | +;+;+;+;+;+;+ | 4056 | grk1a | 0 | 10 | 4 | 0 | 6 | 4 |
| ENSDARG000 | 1:1:1:1:1:1:1:1: | 57040472:570 | 57040504:570 | +:+:+:+:+:+:+:+: | 1695 | NA | 0 | 0 | 0 | 0 | 0 | 0 |
| ENSDARG000 | 1;1;1;1;1;1;1 | 40263671:402 | 40263814:402 | +;+;+;+;+;+;+ | 1088 | TXN | 5548 | 7875 | 7880 | 4822 | 6297 | 5493 |
| ENSDARG000 | 1;1;1;1;1;1 | 20916551:209 | 20917353:209 | -;-;-;-;-;- | 1598 | msmo1 | 416 | 161 | 311 | 381 | 130 | 199 |
| ENSDARG000 | 1:1:1:1:1:1:1:1: | 1712140:1712 | 1712356:1712 | +:+:+:+:+:+:+:+: | 4285 | atp1a1a.1 | 20286 | 15979 | 21890 | 12597 | 21678 | 15639 |
| ENSDARG000 | 1:1:1:1:1:1:1:1: | 57050899:570 | 57050964:570 | +:+:+:+:+:+:+:+: | 1841 | NA | 0 | 0 | 0 | 0 | 0 | 0 |
| ENSDARG000 | 1;1;1;1;1;1;1 | 59101838:591 | 59102095:591 | -;-;-;-;-;-;- | 748 | NA | 0 | 0 | 2 | 0 | 0 | 0 |
| ENSDARG000 | 1:1:1:1:1:1:1:1: | 43695033:436 | 43695596:436 | -:-:-:-:-:-:-:-: | 6090 | SLC9B2 | 48 | 8 | 55 | 26 | 63 | 65 |
| ENSDARG000 | 1:1:1:1:1:1:1:1: | 59073203:590 | 59073343:590 | +:+:+:+:+:+:+:+: | 1212 | mfap4.2 | 124 | 10 | 87 | 83 | 167 | 31 |
| ENSDARG000 | 1:1:1:1:1:1:1:1: | 1789357:1789 | 1789429:1789 | +:+:+:+:+:+:+:+: | 3311 | atp1a1a.2 | 2 | 2 | 0 | 32 | 0 | 2 |
| ENSDARG000 | 1:1:1:1:1:1:1:1: | 43714149:437 | 43714477:437 | -:-:-:-:-:-:-:-: | 1719 | bdh2 | 0 | 0 | 0 | 0 | 0 | 0 |
| ENSDARG000 | 1;1;1;1;1;1;1 | 59094416:590 | 59094805:590 | -;-;-;-;-;-;- | 985 | MFAP4 | 0 | 0 | 0 | 0 | 8 | 7 |
| ENSDARG000 | 1:1:1:1:1:1:1:1: | 59090583:590 | 59090749:590 | +:+:+:+:+:+:+:+: | 1150 | mfap4.1 | 117 | 46 | 108 | 38 | 151 | 64 |
| ENSDARG000 | 1:1:1:1:1:1:1:1: | 20932052:209 | 20932602:209 | -:-:-:-:-:-:-:-: | 2612 | KLHL2 | 304 | 383 | 338 | 181 | 332 | 221 |
| ENSDARG000 | 1:1:1:1:1:1:1:1: | 59075795:590 | 59076159:590 | -:-:-:-:-:-:-:-: | 2171 | MFAP4 | 0 | 6 | 2 | 0 | 4 | 1 |
| ENSDARG000 | 1:1:1:1:1:1:1:1: | 59073203:590 | 59073343:590 | +:+:+:+:+:+:+:+: | 864 | MFAP4 | 0 | 0 | 2 | 0 | 0 | 4 |
| ENSDARG000 | 1;1 | 20929437:209 | 20929678:209 | -;- | 404 | NA | 0 | 0 | 0 | 0 | 0 | 0 |
| ENSDARG000 | 1;1 | 20936711:209 | 20936931:209 | +;+ | 688 | NA | 5 | 2 | 0 | 8 | 0 | 16 |
| ENSDARG000 | 1;1;1 | 10636064:106 | 10637820:106 | -;-;- | 2291 | zgc:174945 | 0 | 2 | 0 | 0 | 2 | 6 |
| ENSDARG000 | 1;1;1;1;1 | 52518176:525 | 52518543:525 | +;+;+;+;+ | 3875 | tacr3l | 4 | 0 | 0 | 0 | 0 | 0 |
| ENSDARG000 | 1;1;1 | 57118783:571 | 57119130:571 | +;+;+ | 741 | NA | 0 | 0 | 0 | 0 | 0 | 0 |
| ENSDARG000 | 1:1:1:1:1:1:1:1: | 44710955:447 | 44711023:447 | +:+:+:+:+:+:+:+: | 2679 | SSRP1 | 1030 | 806 | 948 | 629 | 1156 | 675 |
| ENSDARG000 | 1:1:1:1:1:1:1:1: | 12141876:121 | 12141962:121 | -;-;-;-;-;-;-;- | 1123 | NA | 4 | 6 | 4 | 2 | 16 | 40 |
| ENSDARG000 | 1;1 | 38665754:387 | 38665996:387 | +;+ | 552 | NA | 0 | 0 | 0 | 0 | 0 | 0 |
| ENSDARG000 | 1:1:1:1:1:1:1:1: | 38649141:386 | 38651830:386 | -:-:-:-:-:-:-:-: | 4527 | gpm6ab | 13056 | 8757 | 12703 | 9823 | 15138 | 9985 |
| ENSDARG000 | 1:1:1:1:1:1:1:1: | 7632119:7634 | 7632380:7634 | -:-:-:-:-:-:-:-: | 2039 | efnb2b | 41 | 19 | 13 | 20 | 12 | 9 |
| ENSDARG000 | 1;1;1;1;1;1 | 55264025:552 | 55264656:552 | -;-;-;-;-;- | 960 | si:ch211-286l | 900 | 843 | 1716 | 967 | 1391 | 1512 |
| ENSDARG000 | 1;1 | 10662312:106 | 10662372:106 | +;+ | 541 | NA | 0 | 0 | 0 | 0 | 0 | 0 |
| ENSDARG000 | 1;1;1 | 10667447:106 | 10667542:106 | -;-;- | 766 | NA | 29 | 16 | 27 | 2 | 2 | 14 |
| ENSDARG000 | 1;1;1;1;1;1 | 7662091:7662 | 7663038:7663 | -;-;-;-;-;- | 2290 | arglu1b | 927 | 756 | 1208 | 1019 | 1273 | 1371 |
| ENSDARG000 | 1:1:1:1:1:1:1:1: | 10642322:106 | 10645554:106 | -:-:-:-:-:-:-:-: | 3877 | NA | 8 | 2 | 6 | 2 | 0 | 8 |
| ENSDARG000 | 1;1;1;1;1;1;1 | 25959084:259 | 25960129:259 | -;-;-;-;-;-;- | 4530 | NA | 0 | 0 | 0 | 0 | 0 | 0 |
| ENSDARG000 | 1;1;1;1;1 | 10672843:106 | 10673164:106 | -;-;-;-;- | 1220 | NA | 1 | 1 | 11 | 0 | 0 | 2 |
| ENSDARG000 | 1;1 | 57128562:571 | 57129179:571 | -;- | 738 | NA | 4 | 0 | 0 | 0 | 0 | 0 |
| ENSDARG000 | 1;1;1;1;1;1;1 | 19303241:193 | 19303444:193 | +;+;+;+;+;+;+ | 3993 | NA | 81 | 66 | 72 | 219 | 48 | 156 |
| ENSDARG000 | 1:1:1:1:1:1:1:1: | 46963993:469 | 46964226:469 | -:-:-:-:-:-:-:-: | 1516 | PKNOX1 | 2600 | 1048 | 479 | 605 | 782 | 625 |
| ENSDARG000 | 1:1:1:1:1:1:1:1: | 22654875:226 | 22654950:226 | +:+:+:+:+:+:+:+: | 1913 | anxa5b | 7991 | 7484 | 4554 | 10412 | 4485 | 9354 |
| ENSDARG000 | 1:1:1:1:1:1:1:1: | 51202984:512 | 51203429:512 | -:-:-:-:-:-:-:-: | 2840 | esf1 | 464 | 542 | 844 | 303 | 679 | 385 |
| ENSDARG000 | 1;1;1;1;1;1;1 | 55279124:552 | 55279453:552 | +;+;+;+;+;+;+ | 2563 | NA | 2 | 4 | 38 | 2 | 34 | 81 |
| ENSDARG000 | 1;1 | 22663723:226 | 22664383:226 | -;- | 773 | NA | 7 | 0 | 2 | 6 | 14 | 3 |
| ENSDARG000 | 1 | 9230244 | 9231952 | - | 1709 | NA | 2 | 8 | 4 | 0 | 12 | 4 |
| ENSDARG000 | 1:1:1:1:1:1:1:1: | 53911834:539 | 53912729:539 | -:-:-:-:-:-:-:-: | 2220 | taf5l | 119 | 129 | 237 | 72 | 193 | 159 |
| ENSDARG000 | 1;1;1 | 31864404:318 | 31865522:318 | +;+;+ | 1972 | inab | 0 | 0 | 0 | 0 | 0 | 0 |
| ENSDARG000 | 1:1:1:1:1:1:1:1: | 53919110:539 | 53919381:539 | +:+:+:+:+:+:+:+: | 3659 | nup133 | 380 | 601 | 1138 | 206 | 768 | 529 |
| ENSDARG000 | 1:1:1:1:1:1:1:1: | 51229327:512 | 51232570:512 | -:-:-:-:-:-:-:-: | 6820 | KIF16B | 190 | 263 | 222 | 106 | 400 | 227 |
| ENSDARG000 | 1:1:1:1:1:1:1:1: | 12172322:121 | 12172419:121 | +:+:+:+:+:+:+:+: | 5458 | stra6l | 0 | 0 | 14 | 0 | 16 | 8 |
| ENSDARG000 | 1:1:1:1:1:1:1:1: | 23596731:235 | 23597486:235 | -;-;-;-;-;-;-;- | 913 | NA | 0 | 0 | 0 | 0 | 0 | 0 |
| ENSDARG000 | 1;1;1;1;1 | 57198387:571 | 57198447:571 | +;+;+;+;+ | 746 | NA | 0 | 0 | 0 | 0 | 0 | 0 |
| ENSDARG000 | 1:1:1:1:1:1:1:1: | 51221320:512 | 51221583:512 | +:+:+:+:+:+:+:+: | 1216 | ndufaf5 | 470 | 359 | 464 | 307 | 407 | 353 |
| ENSDARG000 | 1;1;1;1;1;1 | 57208005:572 | 57208065:572 | +;+;+;+;+;+ | 1217 | NA | 0 | 0 | 0 | 0 | 0 | 0 |
| ENSDARG000 | 1;1;1;1;1;1 | 57151926:571 | 57151986:571 | +;+;+;+;+;+ | 1143 | NA | 0 | 0 | 0 | 0 | 0 | 0 |
| ENSDARG000 | 1;1;1 | 57145072:571 | 57145140:571 | +;+;+ | 240 | NA | 0 | 0 | 0 | 0 | 0 | 0 |
| ENSDARG000 | 1:1:1:1:1:1:1:1: | 57132140:571 | 57132180:571 | +:+:+:+:+:+:+:+: | 1479 | NA | 0 | 0 | 0 | 0 | 0 | 0 |
| ENSDARG000 | 1;1;1;1;1 | 57176256:571 | 57176288:571 | +;+;+;+;+ | 949 | NA | 0 | 0 | 0 | 0 | 0 | 0 |
| ENSDARG000 | 1:1:1:1:1:1:1:1: | 17537451:175 | 17539789:175 | -:-:-:-:-:-:-:-: | 4809 | acsl1a | 1667 | 352 | 867 | 2104 | 1303 | 1549 |
| ENSDARG000 | 1:1:1:1:1:1:1:1: | 57218301:572 | 57218357:572 | +:+:+:+:+:+:+:+: | 1644 | NA | 0 | 0 | 0 | 0 | 0 | 0 |
| ENSDARG000 | 1 | 57170145 | 57172294 | - | 2150 | NA | 23 | 12 | 19 | 14 | 22 | 15 |
| ENSDARG000 | 1;1;1;1 | 57187794:571 | 57187867:571 | +;+;+;+ | 754 | NA | 1 | 0 | 0 | 0 | 0 | 0 |
| ENSDARG000 | 1:1:1:1:1:1:1:1: | 40291054:402 | 40291263:402 | +:+:+:+:+:+:+:+: | 2883 | NA | 10 | 0 | 0 | 8 | 6 | 0 |
| ENSDARG000 | 1;1;1;1;1 | 57235896:572 | 57236147:572 | +;+;+;+;+ | 888 | NA | 0 | 0 | 0 | 0 | 0 | 65 |
| ENSDARG000 | 1:1:1:1:1:1:1:1: | 55293424:552 | 55293857:552 | +:+:+:+:+:+:+:+: | 2302 | zgc:172106 | 2 | 0 | 541 | 180 | 372 | 491 |
| ENSDARG000 | 1:1:1:1:1:1:1:1: | 9234928:9235 | 9235103:9235 | -:-:-:-:-:-:-:-: | 1696 | zgc:136472 | 0 | 17 | 11 | 6 | 22 | 5 |
| ENSDARG000 | 1:1:1:1:1:1:1:1: | 35614089:356 | 35615188:356 | -:-:-:-:-:-:-:-: | 9823 | frem3 | 24 | 2 | 14 | 10 | 8 | 2 |
| ENSDARG000 | 1:1:1:1:1:1:1:1: | 28031697:280 | 28031918:280 | +:+:+:+:+:+:+:+: | 2417 | chst10 | 432 | 279 | 427 | 421 | 531 | 480 |
| ENSDARG000 | 1:1:1:1:1:1:1:1: | 10683843:106 | 10684189:106 | +:+:+:+:+:+:+:+: | 905 | mpc2a | 8 | 5 | 16 | 12 | 10 | 12 |
| ENSDARG000 | 1:1:1:1:1:1:1:1: | 26025292:260 | 26025531:260 | -:-:-:-:-:-:-:-: | 1088 | NA | 64 | 2 | 32 | 14 | 30 | 52 |
| ENSDARG000 | 1:1:1:1:1:1:1:1: | 26011479:260 | 26013270:260 | -;-;-;-;-;-;-;- | 2714 | NA | 1 | 0 | 0 | 0 | 0 | 0 |
| ENSDARG000 | 1:1:1:1:1:1:1:1: | 31942961:319 | 31943227:319 | +:+:+:+:+:+:+:+: | 2392 | anos1a | 24 | 2 | 12 | 8 | 25 | 4 |
| ENSDARG000 | 1:1:1:1:1:1:1:1: | 26036123:260 | 26036719:260 | -:-:-:-:-:-:-:-: | 2010 | gpank1 | 94 | 66 | 97 | 89 | 61 | 85 |
| ENSDARG000 | 1:1:1:1:1:1:1:1: | 19332837:193 | 19332914:193 | +:+:+:+:+:+:+:+: | 1969 | tyrp1b | 476667 | 318709 | 187222 | 441351 | 347415 | 349881 |
| ENSDARG000 | 1:1:1:1:1:1:1:1: | 26046148:260 | 26047406:260 | -:-:-:-:-:-:-:-: | 2945 | pdcd4a | 3826 | 2406 | 3611 | 3410 | 3526 | 4693 |
| ENSDARG000 | 1:1:1:1:1:1:1:1: | 260039:26003 | 260146:26014 | +:+:+:+:+:+:+:+: | 8469 | cenpe | 1212 | 2329 | 3115 | 1413 | 1983 | 1148 |
| ENSDARG000 | 1;1;1;1;1;1;1 | 46985884:469 | 46985915:469 | -;-;-;-;-;-;- | 1223 | NA | 0 | 4 | 2 | 10 | 14 | 12 |
| ENSDARG000 | 1;1;1;1;1;1 | 55304374:553 | 55304406:553 | +;+;+;+;+;+ | 997 | NA | 0 | 7 | 0 | 2 | 12 | 7 |
| ENSDARG000 | 1:1:1:1:1:1:1:1: | 52560549:525 | 52560706:525 | +:+:+:+:+:+:+:+: | 10133 | abca1a | 1725 | 990 | 1239 | 836 | 1836 | 1226 |
| ENSDARG000 | 1:1:1:1:1:1:1:1: | 40308077:403 | 40308255:403 | +:+:+:+:+:+:+:+: | 3741 | NA | 2 | 0 | 1 | 0 | 0 | 0 |
| ENSDARG000 | 1;1;1 | 7679328:7679 | 7680108:7680 | +;+;+ | 1576 | en1b | 12 | 6 | 4 | 0 | 10 | 10 |
| ENSDARG000 | 1:1:1:1:1:1:1:1: | 20980975:209 | 20981234:209 | -;-;-;-;-;-;-;- | 1419 | NA | 0 | 0 | 0 | 0 | 0 | 0 |
| ENSDARG000 | 1;1 | 7700023:7701 | 7700215:7701 | -;- | 329 | NA | 0 | 0 | 0 | 0 | 0 | 0 |
| ENSDARG000 | 1:1:1:1:1:1:1:1: | 5437390:5438 | 5437426:5438 | +:+:+:+:+:+:+:+: | 3286 | atic | 3519 | 1937 | 4000 | 1851 | 1630 | 2578 |
| ENSDARG000 | 1;1;1;1;1;1 | 22685961:226 | 22686343:226 | -;-;-;-;-;- | 1377 | fgfbp2b | 76 | 2 | 2 | 0 | 0 | 0 |
| ENSDARG000 | 1:1:1:1:1:1:1:1: | 22691256:226 | 22691429:226 | +:+:+:+:+:+:+:+: | 1534 | zmynd10 | 10 | 16 | 0 | 0 | 6 | 0 |
| ENSDARG000 | 1:1:1:1:1:1:1:1: | 7847787:7847 | 7848042:7848 | -:-:-:-:-:-:-:-: | 4032 | radil | 10 | 12 | 34 | 32 | 36 | 38 |
| ENSDARG000 | 1;1;1;1;1;1 | 5449777:5449 | 5450250:5450 | -;-;-;-;-;- | 1152 | mnx2b | 0 | 5 | 5 | 0 | 0 | 0 |
| ENSDARG000 | 1:1:1:1:1:1:1:1: | 55348537:553 | 55348580:553 | +:+:+:+:+:+:+:+: | 1720 | NA | 10 | 14 | 8 | 4 | 30 | 12 |
| ENSDARG000 | 1:1:1:1:1:1:1:1: | 5504128:5504 | 5505136:5505 | -:-:-:-:-:-:-:-: | 9157 | fn1b | 14169 | 1796 | 713 | 221 | 476 | 738 |
| ENSDARG000 | 1:1:1:1:1:1:1:1: | 29954690:299 | 29955256:299 | -;-;-;-;-;-;-;- | 1589 | march4l | 0 | 2 | 4 | 0 | 2 | 0 |
| ENSDARG000 | 1:1:1:1:1:1:1:1: | 44746154:447 | 44746467:447 | +:+:+:+:+:+:+:+: | 3179 | p2rx3b | 0 | 0 | 0 | 0 | 0 | 2 |
| ENSDARG000 | 1:1:1:1:1:1:1:1: | 47106268:471 | 47106751:471 | -:-:-:-:-:-:-:-: | 2027 | setd4 | 203 | 150 | 219 | 82 | 250 | 188 |
| ENSDARG000 | 1:1:1:1:1:1:1:1: | 19353048:193 | 19358344:193 | -:-:-:-:-:-:-:-: | 7235 | rbm47 | 874 | 584 | 825 | 422 | 1502 | 923 |
| ENSDARG000 | 1:1:1:1:1:1:1:1: | 46999441:469 | 46999893:469 | -:-:-:-:-:-:-:-: | 6364 | ITSN1 | 1318 | 802 | 1700 | 803 | 1436 | 1184 |
| ENSDARG000 | 1;1;1;1;1;1 | 10720294:107 | 10720430:107 | +;+;+;+;+;+ | 2279 | atp1b1b | 4 | 0 | 0 | 0 | 0 | 15 |
| ENSDARG000 | 1:1:1:1:1:1:1:1: | 40336160:403 | 40337552:403 | -:-:-:-:-:-:-:-: | 5755 | NA | 446 | 469 | 885 | 746 | 718 | 331 |
| ENSDARG000 | 1 | 32017437 | 32018115 | + | 679 | NA | 0 | 8 | 14 | 0 | 16 | 4 |
| ENSDARG000 | 1:1:1:1:1:1:1:1: | 47089501:470 | 47089751:470 | +:+:+:+:+:+:+:+: | 2396 | cryzl1 | 416 | 416 | 425 | 328 | 483 | 368 |
| ENSDARG000 | 1:1:1:1:1:1:1:1: | 7908282:7908 | 7908485:7908 | -:-:-:-:-:-:-:-: | 708 | MMD2 | 7 | 5 | 4 | 7 | 13 | 6 |
| ENSDARG000 | 1:1:1:1:1:1:1:1: | 20990877:209 | 20991136:209 | -;-;-;-;-;-;-;- | 1419 | NA | 0 | 0 | 0 | 0 | 0 | 0 |
| ENSDARG000 | 1;1 | 55476002:554 | 55476150:554 | +;+ | 602 | NA | 0 | 0 | 0 | 0 | 0 | 4 |
| ENSDARG000 | 1:1:1:1:1:1:1:1: | 22712931:227 | 22713997:227 | -:-:-:-:-:-:-:-: | 3936 | prom1b | 2 | 9 | 12 | 2 | 6 | 4 |
| ENSDARG000 | 1:1:1:1:1:1:1:1: | 49606518:496 | 49609250:496 | -:-:-:-:-:-:-:-: | 6150 | slkb | 440 | 220 | 393 | 178 | 498 | 440 |
| ENSDARG000 | 1:1:1:1:1:1:1:1: | 40562540:405 | 40563004:405 | +:+:+:+:+:+:+:+: | 2199 | scoca | 460 | 488 | 668 | 594 | 575 | 653 |
| ENSDARG000 | 1:1:1:1:1:1:1:1: | 23783349:237 | 23785062:237 | +:+:+:+:+:+:+:+: | 10246 | slit2 | 26 | 2 | 52 | 4 | 56 | 10 |
| ENSDARG000 | 1:1:1:1:1:1:1:1: | 45650285:456 | 45650901:456 | -:-:-:-:-:-:-:-: | 1713 | serhl | 294 | 495 | 428 | 415 | 505 | 692 |
| ENSDARG000 | 1:1:1:1:1:1:1:1: | 21000784:210 | 21001043:210 | -;-;-;-;-;-;-;- | 1419 | NA | 0 | 0 | 0 | 0 | 0 | 0 |
| ENSDARG000 | 1:1:1:1:1:1:1:1: | 32051581:320 | 32051700:320 | +:+:+:+:+:+:+:+: | 3038 | STS | 20 | 10 | 42 | 0 | 24 | 46 |
| ENSDARG000 | 1:1:1:1:1:1:1:1: | 55478653:554 | 55478835:554 | +:+:+:+:+:+:+:+: | 3178 | NA | 2 | 0 | 0 | 0 | 22 | 0 |
| ENSDARG000 | 1;1;1 | 9277846:9278 | 9277955:9278 | +;+;+ | 525 | NA | 0 | 0 | 0 | 0 | 0 | 1 |
| ENSDARG000 | 1:1:1:1:1:1:1:1: | 9258931:9261 | 9258962:9262 | -:-:-:-:-:-:-:-: | 2307 | ubn1 | 2 | 2 | 16 | 8 | 10 | 39 |
| ENSDARG000 | 1:1:1:1:1:1:1:1: | 35662474:356 | 35663804:356 | -:-:-:-:-:-:-:-: | 2449 | NA | 162 | 234 | 128 | 72 | 192 | 371 |
| ENSDARG000 | 1:1:1:1:1:1:1:1: | 44767657:447 | 44767693:447 | +:+:+:+:+:+:+:+: | 824 | si:dkey-9i23.4 | 2 | 8 | 2 | 0 | 6 | 14 |
| ENSDARG000 | 1:1:1:1:1:1:1:1: | 51312752:513 | 51312771:513 | +:+:+:+:+:+:+:+: | 5647 | MAST1 | 12 | 2 | 72 | 22 | 14 | 0 |
| ENSDARG000 | 1;1 | 57277945:572 | 57279141:572 | -;- | 1828 | NA | 10 | 0 | 0 | 0 | 0 | 2 |
| ENSDARG000 | 1;1;1;1;1;1;1 | 7924956:7925 | 7925380:7925 | -;-;-;-;-;-;- | 1215 | NA | 598 | 332 | 556 | 243 | 360 | 1107 |
| ENSDARG000 | 1:1:1:1:1:1:1:1: | 57269441:572 | 57269684:572 | +:+:+:+:+:+:+:+: | 3094 | NA | 0 | 2 | 2 | 0 | 0 | 0 |
| ENSDARG000 | 1:1:1:1:1:1:1:1: | 21010689:210 | 21010948:210 | -;-;-;-;-;-;-;- | 1419 | NA | 0 | 0 | 0 | 0 | 0 | 0 |
| ENSDARG000 | 1:1:1:1:1:1:1:1: | 19418380:194 | 19418982:194 | -:-:-:-:-:-:-:-: | 2087 | lap3 | 508 | 644 | 570 | 269 | 556 | 420 |
| ENSDARG000 | 1:1:1:1:1:1:1:1: | 12195700:121 | 12195772:121 | +:+:+:+:+:+:+:+: | 3912 | tdrd7a | 2 | 0 | 0 | 0 | 0 | 0 |
| ENSDARG000 | 1:1:1:1:1:1:1:1: | 38758261:387 | 38758510:387 | +:+:+:+:+:+:+:+: | 5454 | wdr17 | 6 | 26 | 72 | 43 | 34 | 17 |
| ENSDARG000 | 1:1:1:1:1:1:1:1: | 10771457:107 | 10771539:107 | +:+:+:+:+:+:+:+: | 1899 | CNGA3 | 0 | 0 | 0 | 0 | 0 | 0 |
| ENSDARG000 | 1;1;1;1 | 17593392:175 | 17593611:175 | +;+;+;+ | 951 | helt | 0 | 0 | 0 | 0 | 6 | 0 |
| ENSDARG000 | 1:1:1:1:1:1:1:1: | 59107633:591 | 59108081:591 | -;-;-;-;-;-;-;- | 921 | NA | 0 | 8 | 0 | 0 | 0 | 20 |
| ENSDARG000 | 1:1:1:1:1:1:1:1: | 53954230:539 | 53954417:539 | +:+:+:+:+:+:+:+: | 2606 | CCSAP | 0 | 0 | 0 | 0 | 0 | 2 |
| ENSDARG000 | 1:1:1:1:1:1:1:1: | 53945934:539 | 53946050:539 | +:+:+:+:+:+:+:+: | 1978 | acta1a | 8 | 4 | 0 | 2 | 2 | 4 |
| ENSDARG000 | 1:1:1:1:1:1:1:1: | 28067771:280 | 28068130:280 | -:-:-:-:-:-:-:-: | 1532 | snapc3 | 420 | 369 | 411 | 184 | 446 | 367 |
| ENSDARG000 | 1;1;1;1;1;1 | 44780849:447 | 44780876:447 | +;+;+;+;+;+ | 806 | si:dkey-9i23.5 | 2 | 0 | 1 | 0 | 6 | 0 |
| ENSDARG000 | 1:1:1:1:1:1:1:1: | 6408:6642:66 | 6760:6760:67 | -:-:-:-:-:-:-:-: | 1723 | rpl24 | 16944 | 17085 | 24413 | 21927 | 16513 | 17741 |
| ENSDARG000 | 1:1:1:1:1:1:1:1: | 10783670:107 | 10783825:107 | -:-:-:-:-:-:-:-: | 2633 | NA | 0 | 6 | 0 | 0 | 0 | 0 |
| ENSDARG000 | 1:1:1:1:1:1:1:1: | 55535827:555 | 55535901:555 | +:+:+:+:+:+:+:+: | 1483 | NA | 2 | 0 | 0 | 0 | 0 | 0 |
| ENSDARG000 | 1:1:1:1:1:1:1:1: | 11822:11992: | 12034:12034: | +:+:+:+:+:+:+:+: | 2656 | cep97 | 210 | 213 | 368 | 202 | 287 | 154 |
| ENSDARG000 | 1:1:1:1:1:1:1:1: | 40572757:405 | 40573724:405 | -:-:-:-:-:-:-:-: | 2956 | clgn | 752 | 1231 | 1463 | 824 | 1493 | 970 |
| ENSDARG000 | 1:1:1:1:1:1:1:1: | 59128019:591 | 59128441:591 | -:-:-:-:-:-:-:-: | 1112 | NA | 0 | 0 | 0 | 1 | 6 | 29 |
| ENSDARG000 | 1:1:1:1:1:1:1:1: | 30083578:300 | 30083750:300 | +:+:+:+:+:+:+:+: | 4593 | vps8 | 1002 | 1016 | 973 | 1369 | 1271 | 1246 |
| ENSDARG000 | 1;1;1;1;1;1 | 44784793:447 | 44785238:447 | -;-;-;-;-;- | 2339 | NA | 2 | 2 | 0 | 8 | 0 | 0 |
| ENSDARG000 | 1;1;1 | 44800298:448 | 44801179:448 | +;+;+ | 882 | TMEM187 | 84 | 50 | 97 | 84 | 100 | 143 |
| ENSDARG000 | 1;1;1;1;1;1 | 7936270:7936 | 7936439:7936 | -;-;-;-;-;- | 831 | NA | 10 | 6 | 19 | 8 | 10 | 8 |
| ENSDARG000 | 1;1;1;1;1 | 59136015:591 | 59136366:591 | -;-;-;-;- | 874 | mfap4.10 | 0 | 4 | 14 | 12 | 32 | 23 |
| ENSDARG000 | 1;1;1;1;1;1;1 | 17618015:176 | 17618514:176 | -;-;-;-;-;-;- | 1375 | NA | 2 | 0 | 0 | 0 | 0 | 2 |
| ENSDARG000 | 1:1:1:1:1:1:1:1: | 22770255:227 | 22772780:227 | -:-:-:-:-:-:-:-: | 4027 | tapt1b | 768 | 493 | 526 | 566 | 577 | 475 |
| ENSDARG000 | 1:1:1:1:1:1:1:1: | 59132757:591 | 59133115:591 | -:-:-:-:-:-:-:-: | 1786 | NA | 0 | 0 | 2 | 1 | 12 | 10 |
| ENSDARG000 | 1:1:1:1:1:1:1:1: | 53980120:539 | 53980586:539 | -:-:-:-:-:-:-:-: | 1554 | RHOU | 2 | 0 | 8 | 0 | 0 | 13 |
| ENSDARG000 | 1:1:1:1:1:1:1:1: | 21135012:211 | 21139482:211 | -:-:-:-:-:-:-:-: | 9252 | NA | 771 | 608 | 667 | 478 | 745 | 578 |
| ENSDARG000 | 1:1:1:1:1:1:1:1: | 45663727:456 | 45663779:456 | +:+:+:+:+:+:+:+: | 1479 | trappc5 | 670 | 737 | 835 | 663 | 707 | 698 |
| ENSDARG000 | 1:1:1:1:1:1:1:1: | 55563532:555 | 55563607:555 | +:+:+:+:+:+:+:+: | 1532 | NA | 0 | 0 | 0 | 0 | 0 | 0 |
| ENSDARG000 | 1;1;1;1;1;1;1 | 44802794:448 | 44803067:448 | -;-;-;-;-;-;- | 1050 | NA | 0 | 0 | 0 | 0 | 2 | 0 |
| ENSDARG000 | 1:1:1:1:1:1:1:1: | 32097179:320 | 32097634:320 | -:-:-:-:-:-:-:-: | 1364 | pnpla4 | 76 | 48 | 80 | 35 | 83 | 69 |
| ENSDARG000 | 1:1:1:1:1:1:1:1: | 57331813:573 | 57332056:573 | +:+:+:+:+:+:+:+: | 1694 | EPN3 | 24 | 12 | 60 | 38 | 8 | 24 |
| ENSDARG000 | 1:1:1:1:1:1:1:1: | 43801957:438 | 43802251:438 | -;-;-;-;-;-;-;- | 1261 | tacr3a | 0 | 0 | 6 | 0 | 0 | 0 |
| ENSDARG000 | 1:1:1:1:1:1:1:1: | 52632856:526 | 52633675:526 | +:+:+:+:+:+:+:+: | 2752 | NA | 470 | 402 | 387 | 313 | 451 | 418 |
| ENSDARG000 | 1:1:1:1:1:1:1:1: | 49651012:496 | 49651092:496 | +:+:+:+:+:+:+:+: | 2713 | tsga10 | 0 | 0 | 0 | 0 | 0 | 0 |
| ENSDARG000 | 1:1:1:1:1:1:1:1: | 22805398:228 | 22808921:228 | -;-;-;-;-;-;-;- | 4581 | ldb2b | 28 | 20 | 36 | 4 | 13 | 10 |
| ENSDARG000 | 1:1:1:1:1:1:1:1: | 55583116:555 | 55583158:555 | +:+:+:+:+:+:+:+: | 1464 | NA | 0 | 2 | 0 | 0 | 0 | 0 |
| ENSDARG000 | 1:1:1:1:1:1:1:1: | 35790082:357 | 35790761:357 | +:+:+:+:+:+:+:+: | 4784 | hhip | 42 | 21 | 16 | 4 | 12 | 126 |
| ENSDARG000 | 1:1:1:1:1:1:1:1: | 51386649:513 | 51386943:513 | +:+:+:+:+:+:+:+: | 4671 | atg4da | 613 | 538 | 890 | 343 | 765 | 722 |
| ENSDARG000 | 1:1:1:1:1:1:1:1: | 17652388:176 | 17652496:176 | -:-:-:-:-:-:-:-: | 1707 | NA | 0 | 0 | 0 | 0 | 0 | 0 |
| ENSDARG000 | 1:1:1:1:1:1:1:1: | 312784:31601 | 313378:31622 | -:-:-:-:-:-:-:-: | 2637 | gas6 | 340 | 523 | 221 | 128 | 230 | 182 |
| ENSDARG000 | 1;1;1;1;1;1 | 26071869:260 | 26072100:260 | +;+;+;+;+;+ | 3919 | MXD4 | 536 | 334 | 574 | 633 | 428 | 406 |
| ENSDARG000 | 1;1;1 | 9290103:9291 | 9290523:9291 | +;+;+ | 1902 | uncx4.1 | 0 | 0 | 0 | 0 | 2 | 2 |
| ENSDARG000 | 1;1;1;1;1;1;1 | 7946842:7947 | 7947271:7947 | -;-;-;-;-;-;- | 1221 | NA | 110 | 115 | 257 | 97 | 184 | 379 |
| ENSDARG000 | 1:1:1:1:1:1:1:1: | 59140583:591 | 59140946:591 | -;-;-;-;-;-;-;- | 855 | NA | 0 | 4 | 6 | 1 | 14 | 7 |
| ENSDARG000 | 1:1:1:1:1:1:1:1: | 55600504:556 | 55600882:556 | +:+:+:+:+:+:+:+: | 1257 | NA | 0 | 0 | 0 | 0 | 6 | 0 |
| ENSDARG000 | 1:1:1:1:1:1:1:1: | 43894238:438 | 43896032:438 | -:-:-:-:-:-:-:-: | 2594 | cnga1a | 0 | 0 | 0 | 0 | 0 | 0 |

| | | | | | | | | | | | |
|---|---|---|---|---|---|---|---|---|---|---|---|
| ENSDARG000 | 1:1:1:1:1:1:1:1: | 10809604:108 | 10810093:108 | -:-:-:-:-:-:-:-: | 1311 | crfb15 | 2 | 0 | 2 | 0 | 0 | 2 |
| ENSDARG000 | 1:1:1:1:1:1:1:1: | 7964276:796 | 7968003:796 | -:-:-:-:-:-:-:-: | 5472 | NA | 22 | 8 | 28 | 54 | 14 | 22 |
| ENSDARG000 | 1:1:1:1:1:1:1:1: | 54037077:540 | 54038223:540 | +:+:+:+:+:+:+:+: | 3520 | NA | 1238 | 741 | 1085 | 782 | 1092 | 833 |
| ENSDARG000 | 1;1;1;1;1 | 32316335:323 | 32316940:323 | -;-;-;-;- | 911 | NA | 0 | 2 | 0 | 0 | 0 | 0 |
| ENSDARG000 | 1:1:1:1:1:1:1:1: | 59142783:591 | 59143019:591 | +:+:+:+:+:+:+:+: | 2828 | GPR108 | 1279 | 1160 | 1443 | 1017 | 1397 | 1220 |
| ENSDARG000 | 1;1;1;1;1;1;1 | 32341484:323 | 32341816:323 | +;+;+;+;+;+;+ | 1634 | pudp | 164 | 130 | 191 | 165 | 243 | 182 |
| ENSDARG000 | 1:1:1:1:1:1:1:1: | 54017703:540 | 54017748:540 | -:-:-:-:-:-:-:-: | 962 | NA | 0 | 0 | 1 | 0 | 0 | 0 |
| ENSDARG000 | 1;1;1 | 54062510:540 | 54062763:540 | -;-;- | 612 | smdt1b | 376 | 312 | 389 | 166 | 367 | 344 |
| ENSDARG000 | 1:1:1:1:1:1:1:1: | 54013457:540 | 54013920:540 | +:+:+:+:+:+:+:+: | 4825 | dla | 0 | 0 | 0 | 0 | 6 | 2 |
| ENSDARG000 | 1:1:1:1:1:1:1:1: | 40613292:406 | 40613516:406 | +:+:+:+:+:+:+:+: | 3243 | naa15b | 1080 | 1197 | 1609 | 701 | 1356 | 900 |
| ENSDARG000 | 1;1;1;1;1;1;1 | 44814322:448 | 44814391:448 | +;+;+;+;+;+;+ | 1575 | ndufs8a | 1160 | 1585 | 1490 | 1274 | 1413 | 1355 |
| ENSDARG000 | 1:1:1:1:1:1:1:1: | 45671687:456 | 45671933:456 | +:+:+:+:+:+:+:+: | 5620 | mcoln1a | 2399 | 1509 | 1739 | 1977 | 2079 | 3169 |
| ENSDARG000 | 1;1;1;1;1;1;1 | 26099250:260 | 26099511:260 | +;+;+;+;+;+;+ | 708 | ndufb6 | 564 | 625 | 564 | 371 | 467 | 500 |
| ENSDARG000 | 1:1:1:1:1:1:1:1: | 19433004:194 | 19433352:194 | +:+:+:+:+:+:+:+: | 3682 | clockb | 402 | 383 | 582 | 309 | 451 | 531 |
| ENSDARG000 | 1:1:1:1:1:1:1:1: | 28089703:281 | 28089761:281 | +:+:+:+:+:+:+:+: | 1578 | psip1a | 224 | 257 | 325 | 294 | 350 | 334 |
| ENSDARG000 | 1:1:1:1:1:1:1:1: | 55608520:556 | 55608576:556 | +:+:+:+:+:+:+:+: | 1948 | NA | 0 | 0 | 0 | 0 | 2 | 0 |
| ENSDARG000 | 1:1:1:1:1:1:1:1: | 32521469:325 | 32521875:325 | +:+:+:+:+:+:+:+: | 5478 | nlgn4xa | 0 | 4 | 34 | 0 | 112 | 104 |
| ENSDARG000 | 1:1:1:1:1:1:1:1: | 52653626:526 | 52654081:526 | -:-:-:-:-:-:-:-: | 1451 | aptx | 92 | 132 | 177 | 88 | 230 | 154 |
| ENSDARG000 | 1:1:1:1:1:1:1:1: | 43901966:439 | 43902769:439 | -:-:-:-:-:-:-:-: | 1779 | enam | 166 | 74 | 56 | 8 | 168 | 2 |
| ENSDARG000 | 1:1:1:1:1:1:1:1: | 7993632:799 | 7994738:799 | -:-:-:-:-:-:-:-: | 1983 | NA | 0 | 1 | 0 | 0 | 0 | 0 |
| ENSDARG000 | 1:1:1:1:1:1:1:1: | 343292:3436 | 343872:3438 | -:-:-:-:-:-:-:-: | 2678 | pros1 | 337 | 197 | 141 | 128 | 96 | 260 |
| ENSDARG000 | 1:1:1:1:1:1:1:1: | 17677373:176 | 17677397:176 | -:-:-:-:-:-:-:-: | 1567 | NA | 3 | 7 | 7 | 0 | 13 | 5 |
| ENSDARG000 | 1:1:1:1:1:1:1:1: | 17681044:176 | 17681575:176 | -;-;-;-;-;-;-;- | 2033 | cfap97 | 234 | 200 | 306 | 272 | 255 | 226 |
| ENSDARG000 | 1:1:1:1:1:1:1:1: | 55643198:556 | 55643396:556 | +:+:+:+:+:+:+:+: | 1745 | NA | 0 | 0 | 0 | 0 | 0 | 0 |
| ENSDARG000 | 1;1;1;1 | 17676745:176 | 17676929:176 | +;+;+;+ | 1434 | slc25a4 | 6 | 0 | 0 | 0 | 2 | 0 |
| ENSDARG000 | 1;1;1;1 | 17695426:176 | 17695583:176 | +;+;+;+ | 1076 | ankrd37 | 73 | 456 | 1186 | 941 | 278 | 144 |
| ENSDARG000 | 1:1:1:1:1:1:1:1: | 17699471:176 | 17699882:176 | -:-:-:-:-:-:-:-: | 1964 | ufsp2 | 475 | 825 | 1035 | 672 | 612 | 566 |
| ENSDARG000 | 1:1:1:1:1:1:1:1: | 57347888:573 | 57348173:573 | +:+:+:+:+:+:+:+: | 2097 | b3gntl1 | 232 | 213 | 203 | 120 | 297 | 264 |
| ENSDARG000 | 1:1:1:1:1:1:1:1: | 1805294:180 | 1805378:180 | +:+:+:+:+:+:+:+: | 3267 | atp1a1a.3 | 131 | 117 | 102 | 55 | 124 | 43 |
| ENSDARG000 | 1 | 24076243 | 24078099 | + | 1857 | mab21l2 | 6 | 0 | 0 | 0 | 0 | 4 |
| ENSDARG000 | 1:1:1:1:1:1:1:1: | 24337062:243 | 24337098:243 | +:+:+:+:+:+:+:+: | 1030 | NA | 2 | 0 | 0 | 0 | 0 | 0 |
| ENSDARG000 | 1:1:1:1:1:1:1:1: | 44826316:448 | 44826622:448 | +:+:+:+:+:+:+:+: | 1938 | stx3a | 0 | 1 | 0 | 1 | 1 | 0 |
| ENSDARG000 | 1:1:1:1:1:1:1:1: | 23919418:239 | 23922115:239 | -:-:-:-:-:-:-:-: | 11203 | LRBA | 2006 | 1044 | 2380 | 868 | 2078 | 1150 |
| ENSDARG000 | 1;1;1;1;1;1;1 | 33215241:332 | 33215283:332 | +;+;+;+;+;+;+ | 901 | NA | 0 | 0 | 4 | 0 | 0 | 0 |
| ENSDARG000 | 1:1:1:1:1:1:1:1: | 22851261:228 | 22851886:228 | +:+:+:+:+:+:+:+: | 6386 | gtf2e2 | 671 | 769 | 813 | 600 | 884 | 632 |
| ENSDARG000 | 1:1:1:1:1:1:1:1: | 51407520:514 | 51407607:514 | +:+:+:+:+:+:+:+: | 4778 | actr2a | 3443 | 3932 | 4396 | 2925 | 3868 | 4101 |
| ENSDARG000 | 1:1:1:1:1:1:1:1: | 33217911:332 | 33218614:332 | +:+:+:+:+:+:+:+: | 4857 | PRKX | 124 | 141 | 136 | 42 | 156 | 104 |
| ENSDARG000 | 1:1:1:1:1:1:1:1: | 22845225:228 | 22845572:228 | -:-:-:-:-:-:-:-: | 1038 | qdprb.1 | 152 | 265 | 343 | 252 | 255 | 217 |
| ENSDARG000 | 1:1:1:1:1:1:1:1: | 38804961:388 | 38805595:388 | -:-:-:-:-:-:-:-: | 1972 | asb5b | 64 | 12 | 38 | 55 | 6 | 46 |
| ENSDARG000 | 1;1 | 22857753:228 | 22858571:228 | -;- | 918 | smim18 | 0 | 0 | 0 | 0 | 0 | 0 |
| ENSDARG000 | 1:1:1:1:1:1:1:1: | 12231478:122 | 12231594:122 | +:+:+:+:+:+:+:+: | 1266 | TMOD1 | 2 | 2 | 2 | 0 | 8 | 0 |
| ENSDARG000 | 1:1:1:1:1:1:1:1: | 9459760:946 | 9460994:946 | -:-:-:-:-:-:-:-: | 4018 | micall2b | 62 | 124 | 105 | 96 | 155 | 80 |
| ENSDARG000 | 1;1 | 28308616:283 | 28308663:283 | -;- | 206 | NA | 0 | 0 | 0 | 0 | 0 | 0 |
| ENSDARG000 | 1;1;1;1;1;1 | 26105141:261 | 26105333:261 | +;+;+;+;+;+ | 3648 | toporsa | 238 | 202 | 218 | 188 | 184 | 261 |
| ENSDARG000 | 1:1:1:1:1:1:1:1: | 8017069:801 | 8017596:801 | -;-;-;-;-;-;- | 1418 | NA | 1050 | 709 | 350 | 138 | 493 | 680 |
| ENSDARG000 | 1;1 | 28348958:283 | 28349569:283 | -;- | 709 | NA | 0 | 0 | 0 | 0 | 0 | 0 |
| ENSDARG000 | 1:1:1:1:1:1:1:1: | 45707219:457 | 45707508:457 | +:+:+:+:+:+:+:+: | 7762 | NA | 2102 | 1237 | 2473 | 1400 | 1502 | 1764 |
| ENSDARG000 | 1;1;1;1;1;1 | 35862550:358 | 35862769:358 | +;+;+;+;+;+ | 800 | NA | 0 | 0 | 0 | 0 | 0 | 0 |
| ENSDARG000 | 1:1:1:1:1:1:1:1: | 59154521:591 | 59154658:591 | +:+:+:+:+:+:+:+: | 818 | NA | 0 | 0 | 0 | 0 | 0 | 2 |
| ENSDARG000 | 1:1:1:1:1:1:1:1: | 45744118:457 | 45745074:457 | -:-:-:-:-:-:-:-: | 1939 | si:ch211-214 | 0 | 0 | 0 | 0 | 0 | 0 |
| ENSDARG000 | 1:1:1:1:1:1:1:1: | 35896488:358 | 35898560:358 | -:-:-:-:-:-:-:-: | 4125 | smad1 | 2123 | 1373 | 1894 | 1375 | 1577 | 1076 |
| ENSDARG000 | 1:1:1:1:1:1:1:1: | 33322174:333 | 33322466:333 | +:+:+:+:+:+:+:+: | 10545 | MXRA5 | 114 | 28 | 24 | 62 | 16 | 56 |
| ENSDARG000 | 1:1:1:1:1:1:1:1: | 8022320:802 | 8024123:802 | -:-:-:-:-:-:-:-: | 6455 | NA | 162 | 102 | 377 | 150 | 1275 | 505 |
| ENSDARG000 | 1:1:1:1:1:1:1:1: | 55662491:556 | 55662680:556 | +:+:+:+:+:+:+:+: | 1960 | NA | 0 | 0 | 3 | 0 | 0 | 0 |
| ENSDARG000 | 1:1:1:1:1:1:1:1: | 49682312:496 | 49682450:496 | +:+:+:+:+:+:+:+: | 2597 | NA | 0 | 0 | 0 | 8 | 6 | 4 |
| ENSDARG000 | 1:1:1:1:1:1:1:1: | 357518:3603 | 357900:3604 | -:-:-:-:-:-:-:-: | 2894 | RASA3 | 229 | 146 | 43 | 4 | 66 | 89 |
| ENSDARG000 | 1:1:1:1:1:1:1:1: | 28391631:283 | 28392229:283 | -:-:-:-:-:-:-:-: | 2044 | NA | 522 | 287 | 462 | 326 | 555 | 587 |
| ENSDARG000 | 1:1:1:1:1:1:1:1: | 8127675:813 | 8131569:813 | -;-;-;-;-;-;-;- | 6253 | FAM83G | 3685 | 2579 | 2760 | 2696 | 5225 | 4319 |
| ENSDARG000 | 1:1:1:1:1:1:1:1: | 8110562:811 | 8110672:811 | +:+:+:+:+:+:+:+: | 1914 | SLC5A10 | 11 | 7 | 8 | 30 | 39 | 8 |
| ENSDARG000 | 1:1:1:1:1:1:1:1: | 52662203:526 | 52662370:526 | +:+:+:+:+:+:+:+: | 2537 | osbp | 887 | 662 | 893 | 453 | 817 | 521 |
| ENSDARG000 | 1:1:1:1:1:1:1:1: | 33336923:333 | 33337850:333 | -:-:-:-:-:-:-:-: | 2694 | gyg2 | 18 | 51 | 4 | 2 | 16 | 30 |
| ENSDARG000 | 1:1:1:1:1:1:1:1: | 47116121:471 | 47117055:471 | -:-:-:-:-:-:-:-: | 2188 | mhc1zea | 953 | 1672 | 1673 | 578 | 1671 | 1112 |
| ENSDARG000 | 1:1:1:1:1:1:1:1: | 33353174:333 | 33356496:333 | -:-:-:-:-:-:-:-: | 5539 | cd99 | 6108 | 3482 | 2069 | 5630 | 4678 | 7392 |
| ENSDARG000 | 1:1:1:1:1:1:1:1: | 43909933:439 | 43910234:439 | -:-:-:-:-:-:-:-: | 950 | scpp5 | 0 | 18 | 0 | 0 | 2 | 0 |
| ENSDARG000 | 1:1:1:1:1:1:1:1: | 18716:18716: | 18912:18912: | +:+:+:+:+:+:+:+: | 1969 | NFKBIZ | 227 | 22 | 33 | 23 | 68 | 37 |
| ENSDARG000 | 1:1:1:1:1:1:1:1: | 57371447:573 | 57371658:573 | +:+:+:+:+:+:+:+: | 1144 | NA | 6 | 13 | 0 | 6 | 7 | 20 |
| ENSDARG000 | 1 | 28708241 | 28709951 | - | 1711 | NA | 0 | 0 | 0 | 0 | 0 | 0 |
| ENSDARG000 | 1:1:1:1:1:1:1:1: | 51444549:514 | 51450323:514 | -:-:-:-:-:-:-:-: | 7092 | SPRED2 | 1603 | 931 | 1363 | 1220 | 1739 | 1665 |
| ENSDARG000 | 1:1:1:1:1:1:1:1: | 12262980:122 | 12264120:122 | -:-:-:-:-:-:-:-: | 2132 | cplx2l | 0 | 40 | 8 | 82 | 10 | 98 |
| ENSDARG000 | 1:1:1:1:1:1:1:1: | 55703120:557 | 55703396:557 | +:+:+:+:+:+:+:+: | 1293 | NA | 0 | 0 | 0 | 0 | 0 | 0 |
| ENSDARG000 | 1:1:1:1:1:1:1:1: | 44838706:448 | 44838953:448 | +:+:+:+:+:+:+:+: | 5010 | kdm2aa | 1182 | 1242 | 1859 | 772 | 1652 | 1063 |
| ENSDARG000 | 1;1;1 | 49699903:497 | 49700053:497 | +;+;+ | 398 | NA | 0 | 0 | 0 | 0 | 0 | 0 |
| ENSDARG000 | 1;1;1;1;1 | 8165340:816 | 8167814:816 | -;-;-;-;- | 3206 | grapb | 306 | 8 | 2 | 161 | 94 | 162 |
| ENSDARG000 | 1:1:1:1:1:1:1:1: | 5576151:557 | 5576279:557 | +:+:+:+:+:+:+:+: | 1607 | cpo | 2 | 0 | 0 | 0 | 0 | 2 |
| ENSDARG000 | 1:1:1:1:1:1:1:1: | 57724157:577 | 57724511:577 | +:+:+:+:+:+:+:+: | 2929 | NA | 0 | 0 | 0 | 0 | 0 | 5 |
| ENSDARG000 | 1:1:1:1:1:1:1:1: | 10824351:108 | 10824848:108 | -:-:-:-:-:-:-:-: | 12133 | dmd | 34 | 62 | 190 | 34 | 102 | 42 |
| ENSDARG000 | 1:1:1:1:1:1:1:1: | 55723870:557 | 55724050:557 | +:+:+:+:+:+:+:+: | 1935 | NA | 0 | 0 | 0 | 0 | 0 | 0 |
| ENSDARG000 | 1:1:1:1:1:1:1:1: | 19473028:194 | 19473378:194 | -:-:-:-:-:-:-:-: | 3346 | kitb | 0 | 4 | 9 | 2 | 6 | 2 |
| ENSDARG000 | 1:1:1:1:1:1:1:1: | 24355515:243 | 24355607:243 | +:+:+:+:+:+:+:+: | 1874 | rps3a | 27142 | 25258 | 42279 | 35315 | 27200 | 37522 |
| ENSDARG000 | 1:1:1:1:1:1:1:1: | 47125610:471 | 47125715:471 | -:-:-:-:-:-:-:-: | 1835 | mhc1zda | 200 | 116 | 195 | 182 | 280 | 410 |
| ENSDARG000 | 1 | 49805384 | 49805818 | - | 435 | NA | 0 | 0 | 0 | 0 | 0 | 0 |
| ENSDARG000 | 1:1:1:1:1:1:1:1: | 8304904:830 | 8304996:830 | +:+:+:+:+:+:+:+: | 6170 | CACNA1H | 0 | 2 | 12 | 0 | 0 | 0 |
| ENSDARG000 | 1;1;1;1;1;1;1 | 24365582:243 | 24365671:243 | +;+;+;+;+;+;+ | 855 | NA | 0 | 1 | 7 | 0 | 0 | 0 |
| ENSDARG000 | 1:1:1:1:1:1:1:1: | 54065611:540 | 54066751:540 | -:-:-:-:-:-:-:-: | 1407 | NA | 2 | 0 | 6 | 10 | 4 | 6 |
| ENSDARG000 | 1:1:1:1:1:1:1:1: | 47140788:471 | 47141782:471 | -:-:-:-:-:-:-:-: | 2187 | mhc1zca | 9 | 0 | 6 | 12 | 8 | 22 |
| ENSDARG000 | 1:1:1:1:1:1:1:1: | 59218067:592 | 59218755:592 | -:-:-:-:-:-:-:-: | 2700 | akap8l | 3603 | 3012 | 3900 | 3111 | 3726 | 2513 |
| ENSDARG000 | 1;1;1 | 30333202:303 | 30333408:303 | -;-;- | 579 | NA | 0 | 0 | 0 | 0 | 0 | 0 |
| ENSDARG000 | 1;1;1;1;1 | 21242836:212 | 21243252:212 | -;-;-;-;- | 1620 | npy1r | 0 | 4 | 0 | 0 | 0 | 0 |
| ENSDARG000 | 1:1:1:1:1:1:1:1: | 47155424:471 | 47156453:471 | -;-;-;-;-;-;-;- | 2323 | mhc1zba | 16638 | 14235 | 13616 | 16050 | 14803 | 18324 |
| ENSDARG000 | 1:1:1:1:1:1:1:1: | 9499645:950 | 9501441:950 | -:-:-:-:-:-:-:-: | 4821 | NA | 609 | 564 | 1135 | 370 | 1016 | 466 |
| ENSDARG000 | 1:1:1:1:1:1:1:1: | 43917934:439 | 43918202:439 | -:-:-:-:-:-:-:-: | 1037 | scpp7 | 322 | 225 | 128 | 6 | 512 | 14 |
| ENSDARG000 | 1;1 | 30343714:303 | 30348028:303 | -;- | 4764 | C9H3orf70 | 6974 | 5635 | 6883 | 5777 | 9078 | 7787 |
| ENSDARG000 | 1;1;1;1;1;1;1 | 21309386:213 | 21309570:213 | +;+;+;+;+;+;+ | 909 | med28 | 282 | 246 | 426 | 106 | 254 | 266 |
| ENSDARG000 | 1:1:1:1:1:1:1:1: | 35956435:359 | 35956615:359 | +:+:+:+:+:+:+:+: | 1899 | mmaa | 212 | 214 | 274 | 108 | 247 | 152 |
| ENSDARG000 | 1:1:1:1:1:1:1:1: | 55730205:557 | 55730269:557 | +:+:+:+:+:+:+:+: | 3625 | NA | 12 | 14 | 61 | 133 | 32 | 12 |
| ENSDARG000 | 1:1:1:1:1:1:1:1: | 49814461:498 | 49815148:498 | +:+:+:+:+:+:+:+: | 4778 | lef1 | 86 | 107 | 252 | 52 | 391 | 194 |
| ENSDARG000 | 1;1;1;1;1 | 55745878:557 | 55746711:557 | -;-;-;-;- | 2118 | dnajb1b | 765 | 553 | 1378 | 2293 | 464 | 210 |
| ENSDARG000 | 1:1:1:1:1:1:1:1: | 21313165:213 | 21314097:213 | -:-:-:-:-:-:-:-: | 2350 | tmem144a | 225 | 146 | 218 | 204 | 317 | 305 |
| ENSDARG000 | 1;1;1;1;1 | 26127818:261 | 26127835:261 | -;-;-;-;- | 396 | NA | 71 | 33 | 61 | 43 | 98 | 317 |
| ENSDARG000 | 1:1:1:1:1:1:1:1: | 26110985:261 | 26111092:261 | +:+:+:+:+:+:+:+: | 1359 | mtap | 1054 | 1273 | 1543 | 1327 | 1354 | 1031 |
| ENSDARG000 | 1:1:1:1:1:1:1:1: | 12295142:122 | 12295307:122 | +:+:+:+:+:+:+:+: | 4146 | gne | 1459 | 1418 | 2164 | 1329 | 2201 | 1718 |
| ENSDARG000 | 1:1:1:1:1:1:1:1: | 52763979:527 | 52765184:527 | -:-:-:-:-:-:-:-: | 4214 | patl1 | 1679 | 1465 | 2141 | 1674 | 1726 | 1666 |
| ENSDARG000 | 1:1:1:1:1:1:1:1: | 28723817:287 | 28724445:287 | -:-:-:-:-:-:-:-: | 2514 | GK3 | 411 | 416 | 480 | 370 | 448 | 357 |
| ENSDARG000 | 1:1:1:1:1:1:1:1: | 51475094:514 | 51475460:514 | +:+:+:+:+:+:+:+: | 2937 | meis1a | 10 | 6 | 0 | 0 | 21 | 4 |
| ENSDARG000 | 1;1;1;1;1;1 | 57774454:577 | 57774577:577 | +;+;+;+;+;+ | 1155 | NA | 0 | 0 | 0 | 0 | 0 | 0 |
| ENSDARG000 | 1:1:1:1:1:1:1:1: | 54069450:540 | 54069561:540 | +:+:+:+:+:+:+:+: | 2318 | DCAF15 | 176 | 225 | 175 | 156 | 180 | 177 |
| ENSDARG000 | 1:1:1:1:1:1:1:1: | 24387659:243 | 24387992:243 | +:+:+:+:+:+:+:+: | 1126 | NA | 0 | 0 | 0 | 0 | 0 | 0 |
| ENSDARG000 | 1:1:1:1:1:1:1:1: | 38818268:388 | 38818547:388 | +:+:+:+:+:+:+:+: | 822 | spcs3 | 686 | 987 | 1257 | 584 | 1039 | 644 |
| ENSDARG000 | 1;1 | 40733921:407 | 40735227:407 | -;- | 1370 | zgc:153642 | 0 | 0 | 0 | 0 | 0 | 0 |
| ENSDARG000 | 1:1:1:1:1:1:1:1: | 40740951:407 | 40741760:407 | +:+:+:+:+:+:+:+: | 3444 | htra3a | 130 | 2 | 12 | 12 | 3 | 2 |
| ENSDARG000 | 1:1:1:1:1:1:1:1: | 1838164:183 | 1838243:183 | +:+:+:+:+:+:+:+: | 3820 | atp1a1a.5 | 0 | 7 | 0 | 0 | 4 | 8 |
| ENSDARG000 | 1:1:1:1:1:1:1:1: | 24376698:243 | 24378889:243 | -:-:-:-:-:-:-:-: | 5439 | sh3d19 | 137 | 4 | 2 | 16 | 14 | 12 |
| ENSDARG000 | 1:1:1:1:1:1:1:1: | 45754868:457 | 45755570:457 | +:+:+:+:+:+:+:+: | 7190 | PKN1 | 1849 | 2156 | 2608 | 1872 | 2316 | 2138 |
| ENSDARG000 | 1;1;1;1;1 | 57831922:578 | 57833108:578 | -;-;-;-;- | 2119 | NA | 7 | 2 | 0 | 6 | 0 | 8 |
| ENSDARG000 | 1:1:1:1:1:1:1:1: | 5606460:561 | 5608648:561 | -:-:-:-:-:-:-:-: | 5437 | nrp2a | 494 | 216 | 1523 | 294 | 801 | 709 |
| ENSDARG000 | 1:1:1:1:1:1:1:1: | 27690:27814: | 27955:27955: | +:+:+:+:+:+:+:+: | 1771 | eed | 263 | 375 | 609 | 333 | 440 | 405 |
| ENSDARG000 | 1;1;1;1;1;1;1 | 57927489:579 | 57927623:579 | +;+;+;+;+;+;+ | 2597 | NA | 4 | 2 | 12 | 0 | 4 | 0 |
| ENSDARG000 | 1;1;1;1;1;1;1 | 47165842:471 | 47165895:471 | +;+;+;+;+;+;+ | 4065 | cbr1 | 1267 | 923 | 1205 | 835 | 1710 | 1132 |
| ENSDARG000 | 1:1:1:1:1:1:1:1: | 43926153:439 | 43926779:439 | -;-;-;-;-;-;- | 1189 | ambn | 26 | 17 | 2 | 0 | 42 | 6 |
| ENSDARG000 | 1:1:1:1:1:1:1:1: | 40760448:407 | 40760634:407 | +:+:+:+:+:+:+:+: | 4474 | GPR78 | 4 | 0 | 0 | 0 | 2 | 2 |
| ENSDARG000 | 1;1;1 | 19515228:195 | 19515728:195 | +;+;+ | 950 | clrn2 | 0 | 6 | 0 | 0 | 7 | 0 |
| ENSDARG000 | 1;1;1;1;1 | 9626277:962 | 9627922:962 | -;-;-;-;- | 1820 | ugt5b5 | 8 | 3 | 0 | 9 | 1 | 20 |
| ENSDARG000 | 1;1;1;1;1 | 9557212:961 | 9557481:961 | +;+;+;+;+ | 4282 | ELFN1 | 18 | 32 | 6 | 6 | 6 | 54 |
| ENSDARG000 | 1:1:1:1:1:1:1:1: | 24382651:243 | 24382997:243 | +:+:+:+:+:+:+:+: | 1245 | QDPR | 0 | 0 | 0 | 0 | 0 | 0 |
| ENSDARG000 | 1;1;1;1;1 | 9645275:964 | 9646964:964 | -;-;-;-;- | 2279 | ugt5b1 | 87 | 71 | 11 | 67 | 37 | 73 |
| ENSDARG000 | 1;1;1;1;1 | 30422143:304 | 30422416:304 | +;+;+;+;+ | 1938 | tmem41ab | 0 | 2 | 0 | 4 | 2 | 0 |
| ENSDARG000 | 1:1:1:1:1:1:1:1: | 9630996:963 | 9633073:963 | -:-:-:-:-:-:-:-: | 2768 | ugt5b4 | 112 | 74 | 0 | 46 | 34 | 156 |
| ENSDARG000 | 1;1;1;1;1;1;1 | 8415983:842 | 8418943:842 | -;-;-;-;-;-;- | 3827 | syngr3b | 782 | 300 | 238 | 1068 | 650 | 987 |
| ENSDARG000 | 1:1:1:1:1:1:1:1: | 24376394:243 | 24376449:243 | +:+:+:+:+:+:+:+: | 849 | qdprb.4 | 0 | 0 | 0 | 0 | 0 | 0 |
| ENSDARG000 | 1;1;1;1 | 43945629:439 | 43946059:439 | -;-;-;- | 610 | NA | 0 | 0 | 0 | 0 | 0 | 0 |
| ENSDARG000 | 1;1;1 | 26183739:261 | 26183813:261 | -;-;- | 353 | NA | 0 | 0 | 0 | 0 | 0 | 0 |
| ENSDARG000 | 1;1;1;1;1 | 9629161:964 | 9629568:964 | -;-;-;-;- | 2272 | ugt5b2 | 253 | 453 | 93 | 201 | 111 | 470 |
| ENSDARG000 | 1:1:1:1:1:1:1:1: | 9635978:963 | 9637750:963 | -;-;-;-;-;-;-;- | 2105 | ugt5b3 | 48 | 23 | 11 | 19 | 39 | 68 |
| ENSDARG000 | 1:1:1:1:1:1:1:1: | 35985813:359 | 35986235:359 | +:+:+:+:+:+:+:+: | 8913 | zgc:152968 | 0 | 0 | 0 | 0 | 0 | 0 |
| ENSDARG000 | 1:1:1:1:1:1:1:1: | 40801448:408 | 40801824:408 | +:+:+:+:+:+:+:+: | 3852 | CPZ | 34 | 6 | 2 | 4 | 26 | 8 |
| ENSDARG000 | 1:1:1:1:1:1:1:1: | 26250729:262 | 26253236:262 | -:-:-:-:-:-:-:-: | 3971 | CXXC4 | 0 | 0 | 0 | 0 | 0 | 0 |
| ENSDARG000 | 1:1:1:1:1:1:1:1: | 22943551:229 | 22946180:229 | -:-:-:-:-:-:-:-: | 6812 | ADGRL3 | 92 | 32 | 96 | 56 | 44 | 98 |
| ENSDARG000 | 1:1:1:1:1:1:1:1: | 47178529:471 | 47178709:471 | +:+:+:+:+:+:+:+: | 3014 | morc3b | 88 | 24 | 40 | 0 | 6 | 38 |
| ENSDARG000 | 1;1;1;1;1;1 | 12335816:123 | 12336053:123 | +;+;+;+;+;+ | 1383 | nansa | 779 | 1122 | 1304 | 573 | 1055 | 740 |
| ENSDARG000 | 1:1:1:1:1:1:1:1: | 54088563:540 | 54089367:540 | -:-:-:-:-:-:-:-: | 3353 | RFX1 | 582 | 596 | 919 | 601 | 579 | 632 |
| ENSDARG000 | 1;1 | 6068436:608 | 6068537:608 | -;- | 542 | NA | 2 | 0 | 0 | 0 | 0 | 0 |
| ENSDARG000 | 1:1:1:1:1:1:1:1: | 26356205:263 | 26356750:263 | +:+:+:+:+:+:+:+: | 9834 | TET2 | 1054 | 563 | 1124 | 528 | 1158 | 1042 |
| ENSDARG000 | 1:1:1:1:1:1:1:1: | 26389153:263 | 26389577:263 | -:-:-:-:-:-:-:-: | 3535 | ppa2 | 208 | 234 | 264 | 144 | 216 | 299 |
| ENSDARG000 | 1:1:1:1:1:1:1:1: | 57966831:579 | 57967105:579 | +:+:+:+:+:+:+:+: | 6595 | NA | 59 | 116 | 64 | 19 | 113 | 50 |
| ENSDARG000 | 1:1:1:1:1:1:1:1: | 43958198:439 | 43958291:439 | -;-;-;-;-;-;-;- | 830 | odam | 0 | 0 | 0 | 0 | 0 | 0 |
| ENSDARG000 | 1 | 30439422 | 30441060 | - | 1639 | NA | 642 | 1459 | 2241 | 1360 | 2745 | 1517 |
| ENSDARG000 | 1:1:1:1:1:1:1:1: | 45839927:458 | 45840071:458 | +:+:+:+:+:+:+:+: | 1425 | map2k7 | 97 | 48 | 122 | 44 | 100 | 78 |
| ENSDARG000 | 1;1;1;1;1;1;1 | 36552:36727: | 36878:36878: | +;+;+;+;+;+;+ | 1272 | hikeshi | 308 | 384 | 437 | 322 | 370 | 358 |
| ENSDARG000 | 1:1:1:1:1:1:1:1: | 52792439:527 | 52792699:527 | +:+:+:+:+:+:+:+: | 3711 | smarca5 | 1605 | 2688 | 3311 | 978 | 3195 | 1192 |
| ENSDARG000 | 1;1 | 8507004:850 | 8507224:851 | -;- | 1401 | NA | 2 | 0 | 0 | 0 | 0 | 9 |
| ENSDARG000 | 1;1;1;1 | 8508753:851 | 8509789:851 | +;+;+;+ | 1556 | alkbh5 | 183 | 264 | 371 | 258 | 243 | 154 |
| ENSDARG000 | 1:1:1:1:1:1:1:1: | 30443193:304 | 30443392:304 | -:-:-:-:-:-:-:-: | 1932 | igf2bp2b | 607 | 1132 | 1689 | 694 | 2165 | 1123 |
| ENSDARG000 | 1;1;1 | 40861690:408 | 40861733:408 | +;+;+ | 668 | NA | 2 | 0 | 0 | 0 | 0 | 0 |
| ENSDARG000 | 1:1:1:1:1:1:1:1: | 8442071:844 | 8442153:844 | +:+:+:+:+:+:+:+: | 7334 | MYO15A | 12 | 2 | 11 | 2 | 12 | 3 |
| ENSDARG000 | 1;1 | 40909750:409 | 40910887:409 | -;- | 1475 | hmx4 | 0 | 0 | 0 | 0 | 0 | 0 |
| ENSDARG000 | 1:1:1:1:1:1:1:1: | 38850830:388 | 38851249:388 | -:-:-:-:-:-:-:-: | 2235 | vegfc | 31 | 2 | 22 | 2 | 24 | 10 |
| ENSDARG000 | 1:1:1:1:1:1:1:1: | 21333068:213 | 21333581:213 | -:-:-:-:-:-:-:-: | 3449 | gria2a | 29 | 28 | 14 | 45 | 62 | 26 |
| ENSDARG000 | 1;1 | 40912613:409 | 40913586:409 | -;- | 1555 | hmx1 | 0 | 0 | 0 | 0 | 0 | 0 |
| ENSDARG000 | 1:1:1:1:1:1:1:1: | 57981823:579 | 57981880:579 | +:+:+:+:+:+:+:+: | 1424 | NA | 0 | 0 | 0 | 0 | 0 | 6 |
| ENSDARG000 | 1;1;1;1;1 | 57997535:579 | 57998278:579 | -;-;-;-;- | 1131 | si:ch211-114 | 174 | 186 | 123 | 38 | 80 | 131 |
| ENSDARG000 | 1 | 40992682 | 40994259 | - | 1578 | adra2c | 0 | 0 | 0 | 4 | 0 | 0 |
| ENSDARG000 | 1:1:1:1:1:1:1:1: | 28832671:288 | 28835778:288 | -:-:-:-:-:-:-:-: | 3994 | timmdc1 | 533 | 415 | 438 | 375 | 537 | 536 |

| | | | | | | | | | | | |
|---|---|---|---|---|---|---|---|---|---|---|---|
| ENSDARG000 | 1;1;1 | 19535144:195 | 19535306:195 | +;+;+ | 887 | exoc1l | 0 | 2 | 2 | 0 | 1 | 2 |
| ENSDARG000 | 1;1;1;1;1;1;1 | 59237328:592 | 59238468:592 | -;-;-;-;-;-;- | 1623 | NA | 183 | 124 | 86 | 135 | 74 | 74 |
| ENSDARG000 | 1;1;1;1;1;1;1; | 19535120:195 | 19535306:195 | +;+;+;+;+;+;+; | 4260 | smc2 | 372 | 1648 | 2060 | 670 | 1594 | 742 |
| ENSDARG000 | 1;1;1;1;1;1;1; | 43969797:439 | 43970169:439 | -;-;-;-;-;-;-;-;- | 909 | scpp9 | 0 | 0 | 0 | 0 | 0 | 0 |
| ENSDARG000 | 1;1;1;1;1;1;1; | 418869:42017 | 419121:42030 | +;+;+;+;+;+;+; | 5496 | TPP2 | 960 | 596 | 716 | 308 | 641 | 641 |
| ENSDARG000 | 1;1;1 | 58005222:580 | 58006818:580 | -;-;- | 2062 | nxnl1 | 0 | 0 | 0 | 0 | 0 | 0 |
| ENSDARG000 | 1;1;1;1;1;1;1; | 58013076:580 | 58015098:580 | -;-;-;-;-;-;-;-;- | 3636 | colgalt1b | 877 | 432 | 2215 | 294 | 1308 | 695 |
| ENSDARG000 | 1;1;1;1;1;1;1; | 49909672:499 | 49910080:499 | -;-;-;-;-;-;-;-;- | 1498 | hadh | 453 | 572 | 579 | 558 | 635 | 644 |
| ENSDARG000 | 1;1 | 58003615:580 | 58003860:580 | +;+ | 540 | NA | 2 | 0 | 1 | 0 | 0 | 0 |
| ENSDARG000 | 1;1;1;1;1;1 | 43973880:439 | 43976715:439 | -;-;-;-;-;- | 4044 | NA | 6 | 10 | 18 | 16 | 14 | 4 |
| ENSDARG000 | 1;1;1;1;1;1 | 51537250:515 | 51537465:515 | +;+;+;+;+;+ | 2429 | etaa1a | 28 | 87 | 111 | 54 | 84 | 41 |
| ENSDARG000 | 1 | 54606354 | 54608032 | + | 1679 | NA | 51 | 14 | 18 | 22 | 22 | 30 |
| ENSDARG000 | 1;1;1;1;1;1;1; | 59244159:592 | 59245524:592 | -;-;-;-;-;-;-;-;- | 2537 | olfm2b | 187 | 56 | 96 | 138 | 86 | 90 |
| ENSDARG000 | 1;1;1;1;1;1;1; | 23161369:231 | 23161424:231 | +;+;+;+;+;+;+; | 6549 | NA | 0 | 0 | 0 | 0 | 0 | 0 |
| ENSDARG000 | 1;1;1;1;1;1;1; | 55752593:557 | 55752717:557 | +;+;+;+;+;+;+; | 1853 | tecrb | 1505 | 1599 | 1631 | 1508 | 1653 | 1255 |
| ENSDARG000 | 1;1;1;1;1;1;1; | 44870880:448 | 44870889:448 | -;-;-;-;-;-;-;-;- | 3349 | TCIRG1 | 3465 | 1343 | 3466 | 2473 | 1598 | 5246 |
| ENSDARG000 | 1;1;1;1;1;1;1; | 6135176:6135 | 6135334:6135 | +;+;+;+;+;+;+; | 2951 | abcb6a | 50 | 14 | 25 | 8 | 16 | 10 |
| ENSDARG000 | 1;1;1;1;1;1;1; | 43988715:439 | 43989581:439 | -;-;-;-;-;-;-;-;- | 2745 | zgc:66472 | 248 | 210 | 256 | 126 | 286 | 206 |
| ENSDARG000 | 1;1;1;1;1;1 | 54613757:546 | 54614775:546 | -;-;-;-;-;- | 2227 | tekt4 | 0 | 0 | 2 | 0 | 0 | 0 |
| ENSDARG000 | 1;1;1;1;1;1;1; | 53266504:532 | 53267398:532 | -;-;-;-;-;-;-;-;- | 2137 | znf330 | 234 | 453 | 947 | 354 | 644 | 484 |
| ENSDARG000 | 1;1;1;1;1;1;1; | 41131481:411 | 41131796:411 | +;+;+;+;+;+;+; | 2123 | lrpap1 | 1930 | 2249 | 2847 | 1626 | 2762 | 1895 |
| ENSDARG000 | 1;1;1;1;1;1;1; | 44839:45788: | 45550:46371: | -;-;-;-;-;-;-;-;- | 2775 | ildr1a | 108 | 5 | 8 | 16 | 6 | 13 |
| ENSDARG000 | 1;1;1;1;1;1;1; | 39313:39325: | 40061:40061: | -;-;-;-;-;-;-;-;- | 3111 | tmem39a | 659 | 758 | 1361 | 646 | 1214 | 649 |
| ENSDARG000 | 1;1;1;1;1;1;1; | 47331644:473 | 47331978:473 | +;+;+;+;+;+;+; | 7567 | bcl9 | 1033 | 780 | 1167 | 836 | 1203 | 806 |
| ENSDARG000 | 1;1;1;1;1;1 | 51557789:515 | 51559861:515 | -;-;-;-;-;- | 2795 | ppp3r1b | 1221 | 1517 | 2080 | 1406 | 1459 | 1636 |
| ENSDARG000 | 1;1;1;1;1;1;1; | 59252435:592 | 59252970:592 | -;-;-;-;-;-;-;-;- | 5692 | col5a3b | 50 | 6 | 16 | 0 | 8 | 5 |
| ENSDARG000 | 1;1;1;1;1;1;1; | 44003654:440 | 44003860:440 | +;+;+;+;+;+;+; | 1893 | NA | 0 | 0 | 0 | 0 | 0 | 0 |
| ENSDARG000 | 1;1;1;1;1;1;1; | 28874389:288 | 28874903:288 | -;-;-;-;-;-;-;-;- | 1690 | poglut1 | 262 | 363 | 510 | 326 | 465 | 348 |
| ENSDARG000 | 1;1;1;1;1;1;1; | 33383644:333 | 33384101:333 | +;+;+;+;+;+;+; | 5416 | dhrsx | 278 | 178 | 118 | 50 | 167 | 164 |
| ENSDARG000 | 1;1;1;1;1;1;1; | 53234693:532 | 53235119:532 | -;-;-;-;-;-;-;-;- | 1400 | il15 | 10 | 4 | 6 | 5 | 20 | 23 |
| ENSDARG000 | 1;1;1;1;1;1;1; | 52929185:529 | 52929297:529 | +;+;+;+;+;+;+; | 3895 | INPP4B | 40 | 42 | 239 | 6 | 125 | 80 |
| ENSDARG000 | 1;1;1;1;1;1;1; | 36017867:360 | 36022149:360 | -;-;-;-;-;-;-;-;- | 10828 | ZNF827 | 732 | 606 | 937 | 581 | 592 | 745 |
| ENSDARG000 | 1 | 59252435 | 59252973 | - | 539 | NA | 0 | 0 | 0 | 0 | 0 | 0 |
| ENSDARG000 | 1;1;1;1;1;1;1; | 12348213:123 | 12348506:123 | +;+;+;+;+;+;+; | 2396 | clta | 2993 | 3466 | 3259 | 2785 | 3014 | 2530 |
| ENSDARG000 | 1;1 | 47414734:474 | 47416856:474 | -;- | 2801 | gja5b | 3026 | 2116 | 2158 | 1461 | 4882 | 1725 |
| ENSDARG000 | 1;1;1;1;1;1;1; | 8521323:8521 | 8521426:8521 | +;+;+;+;+;+;+; | 3021 | mief2 | 230 | 195 | 391 | 431 | 166 | 177 |
| ENSDARG000 | 1;1;1;1 | 47446032:474 | 47446174:474 | +;+;+;+ | 2459 | gja8b | 4 | 0 | 2 | 0 | 4 | 0 |
| ENSDARG000 | 1;1 | 30551777:305 | 30552405:305 | +;+ | 1751 | gpr183b | 12 | 12 | 11 | 16 | 32 | 13 |
| ENSDARG000 | 1;1;1;1;1;1;1; | 30539971:305 | 30540656:305 | -;-;-;-;-;-;-;-;- | 1783 | ubac2 | 365 | 381 | 676 | 385 | 732 | 479 |
| ENSDARG000 | 1;1;1;1;1;1;1 | 1904419:1906 | 1904533:1906 | +;+;+;+;+;+;+ | 1137 | NA | 2 | 2 | 0 | 6 | 6 | 21 |
| ENSDARG000 | 1;1;1;1 | 1915967:1921 | 1916060:1922 | +;+;+;+ | 693 | NA | 2 | 4 | 34 | 8 | 25 | 4 |
| ENSDARG000 | 1;1;1;1;1;1;1 | 1896737:1897 | 1896848:1898 | +;+;+;+;+;+;+ | 1399 | NA | 0 | 2 | 0 | 0 | 1 | 10 |
| ENSDARG000 | 1;1;1;1;1;1 | 454315:45653 | 454696:45665 | -;-;-;-;-;- | 1008 | METTL21C | 0 | 0 | 0 | 0 | 0 | 0 |
| ENSDARG000 | 1;1;1;1;1;1;1; | 1870861:1874 | 1870899:1874 | -;-;-;-;-;-;-;-;- | 3159 | NA | 280 | 227 | 191 | 103 | 236 | 1004 |
| ENSDARG000 | 1;1;1;1;1;1;1; | 28898945:288 | 28899672:288 | -;-;-;-;-;-;-;-;- | 3501 | pwp2h | 718 | 699 | 1265 | 331 | 964 | 667 |
| ENSDARG000 | 1;1;1;1;1;1;1 | 44011472:440 | 44011587:440 | -;-;-;-;-;-;- | 1456 | NA | 99 | 70 | 109 | 77 | 80 | 133 |
| ENSDARG000 | 1;1;1;1;1;1 | 58039551:580 | 58040038:580 | +;+;+;+;+;+ | 3397 | trim35-1 | 138 | 146 | 200 | 114 | 170 | 104 |
| ENSDARG000 | 1;1;1;1;1;1;1; | 21423010:214 | 21423791:214 | -;-;-;-;-;-;-;-;- | 2421 | glrba | 16 | 6 | 2 | 56 | 4 | 26 |
| ENSDARG000 | 1;1 | 28903770:289 | 28903922:289 | +;+ | 330 | NA | 0 | 0 | 0 | 0 | 8 | 0 |
| ENSDARG000 | 1;1;1;1;1;1;1; | 23274710:232 | 23274860:232 | +;+;+;+;+;+;+; | 1746 | lias | 484 | 472 | 362 | 316 | 490 | 310 |
| ENSDARG000 | 1;1;1;1;1;1;1; | 466780:46678 | 467842:46784 | -;-;-;-;-;-;-;-;- | 1612 | mettl21e | 6 | 0 | 0 | 0 | 0 | 6 |
| ENSDARG000 | 1;1;1;1;1;1;1; | 6171585:6171 | 6171865:6171 | +;+;+;+;+;+;+; | 1802 | NA | 225 | 107 | 131 | 154 | 100 | 96 |
| ENSDARG000 | 1;1;1;1;1;1;1; | 23269503:232 | 23269887:232 | -;-;-;-;-;-;-;-;- | 1904 | rpl9 | 23 | 45 | 48 | 42 | 64 | 30 |
| ENSDARG000 | 1;1;1;1;1 | 54626491:546 | 54626543:546 | +;+;+;+;+ | 689 | si:ch211-202 | 0 | 2 | 0 | 0 | 0 | 0 |
| ENSDARG000 | 1;1;1;1;1;1;1; | 36194761:361 | 36194822:361 | +;+;+;+;+;+;+; | 747 | lsm6 | 523 | 883 | 709 | 372 | 622 | 482 |
| ENSDARG000 | 1;1;1;1;1;1;1; | 6161930:6162 | 6162152:6162 | +;+;+;+;+;+;+; | 5106 | cnppd1 | 807 | 691 | 988 | 766 | 753 | 890 |
| ENSDARG000 | 1;1;1 | 51593884:516 | 51594892:516 | -;-;- | 1420 | cnrip1a | 30 | 46 | 32 | 16 | 14 | 78 |
| ENSDARG000 | 1;1;1;1;1;1;1; | 41142881:411 | 41143733:411 | -;-;-;-;-;-;-;-;- | 2975 | DOK7 | 0 | 3 | 10 | 0 | 6 | 1 |
| ENSDARG000 | 1;1;1;1;1;1;1; | 58050905:580 | 58051565:580 | -;-;-;-;-;-;-;-;- | 1819 | caspb | 124 | 8 | 0 | 6 | 56 | 42 |
| ENSDARG000 | 1;1;1 | 44027611:440 | 44032854:440 | -;-;- | 6452 | NA | 84 | 52 | 94 | 118 | 94 | 128 |
| ENSDARG000 | 1;1;1;1;1;1;1; | 47459723:474 | 47459814:474 | +;+;+;+;+;+;+; | 1511 | fstl1a | 538 | 1222 | 2296 | 2713 | 1894 | 1879 |
| ENSDARG000 | 1;1;1;1;1;1;1; | 9739370:9740 | 9741326:9741 | -;-;-;-;-;-;-;-;- | 4116 | mad1l1 | 196 | 332 | 256 | 134 | 307 | 196 |
| ENSDARG000 | 1;1;1;1;1;1;1 | 17721853:177 | 17722562:177 | -;-;-;-;-;-;- | 1504 | pdlim3a | 0 | 0 | 0 | 0 | 0 | 0 |
| ENSDARG000 | 1;1;1 | 36436936:364 | 36437428:364 | +;+;+ | 3210 | pou4f2 | 0 | 0 | 0 | 5 | 0 | 0 |
| ENSDARG000 | 1;1;1;1 | 44040211:440 | 44041743:440 | -;-;-;- | 2400 | si:ch73-109d | 150 | 183 | 209 | 138 | 223 | 252 |
| ENSDARG000 | 1;1;1;1;1;1;1; | 39387876:393 | 39388411:393 | +;+;+;+;+;+;+; | 15558 | tenm3 | 686 | 624 | 2095 | 634 | 2632 | 430 |
| ENSDARG000 | 1;1;1;1;1;1;1; | 23243093:232 | 23243789:232 | -;-;-;-;-;-;-;-;- | 6860 | rfc1 | 326 | 698 | 778 | 347 | 730 | 485 |
| ENSDARG000 | 1;1;1;1;1;1;1; | 33547532:335 | 33548553:335 | -;-;-;-;-;-;-;-;- | 2056 | creb1a | 409 | 516 | 491 | 278 | 456 | 457 |
| ENSDARG000 | 1;1;1;1;1;1;1; | 44906517:449 | 44906673:449 | +;+;+;+;+;+;+; | 3733 | NA | 1511 | 1364 | 1213 | 1499 | 928 | 2574 |
| ENSDARG000 | 1;1 | 8534698:8539 | 8537534:8540 | +;+ | 3662 | smcr8b | 298 | 134 | 145 | 187 | 258 | 188 |
| ENSDARG000 | 1;1;1;1;1;1;1; | 8539671:8540 | 8540148:8540 | -;-;-;-;-;-;-;-;- | 3694 | zgc:112980 | 184 | 214 | 315 | 124 | 290 | 205 |
| ENSDARG000 | 1;1;1;1;1;1;1; | 45866692:458 | 45870285:458 | -;-;-;-;-;-;-;-;- | 7810 | PNPLA6 | 1065 | 643 | 652 | 596 | 800 | 689 |
| ENSDARG000 | 1;1;1 | 44925422:449 | 44926211:449 | -;-;- | 1429 | si:dkev-9i23.1 | 562 | 407 | 510 | 373 | 339 | 577 |
| ENSDARG000 | 1;1;1;1;1;1;1; | 41201586:412 | 41203193:412 | -;-;-;-;-;-;-;-;- | 7369 | rgs12b | 4302 | 3783 | 3112 | 4481 | 3311 | 2159 |
| ENSDARG000 | 1;1;1;1;1;1;1; | 23283532:232 | 23284265:232 | -;-;-;-;-;-;-;-;- | 2677 | ugdh | 973 | 712 | 861 | 556 | 1090 | 394 |
| ENSDARG000 | 1;1;1;1;1 | 49923402:499 | 49923594:499 | -;-;-;-;- | 1708 | cyp2u1 | 142 | 48 | 36 | 76 | 138 | 98 |
| ENSDARG000 | 1;1 | 39675706:396 | 39676132:396 | -;- | 482 | NA | 0 | 0 | 0 | 0 | 1 | 0 |
| ENSDARG000 | 1;1;1;1;1;1;1; | 54638126:546 | 54638195:546 | +;+;+;+;+;+;+; | 1718 | si:ch211-202 | 0 | 0 | 0 | 0 | 0 | 0 |
| ENSDARG000 | 1;1;1;1;1;1;1; | 51608742:516 | 51609178:516 | -;-;-;-;-;-;-;-;- | 891 | fbxo48 | 196 | 126 | 88 | 146 | 96 | 282 |
| ENSDARG000 | 1;1;1;1;1;1;1; | 58066109:580 | 58066209:580 | +;+;+;+;+;+;+; | 3059 | NA | 13 | 0 | 9 | 0 | 0 | 1 |
| ENSDARG000 | 1;1;1;1 | 44053641:440 | 44053974:440 | +;+;+;+ | 680 | NA | 65 | 27 | 55 | 28 | 46 | 36 |
| ENSDARG000 | 1;1;1;1;1;1;1; | 474085:47428 | 474685:47468 | -;-;-;-;-;-;-;-;- | 3414 | NA | 332 | 196 | 385 | 293 | 340 | 285 |
| ENSDARG000 | 1;1;1;1;1;1 | 47486104:474 | 47486747:474 | +;+;+;+;+;+ | 1328 | lrrc58a | 0 | 0 | 1 | 0 | 4 | 3 |
| ENSDARG000 | 1;1;1;1;1;1;1; | 26411496:264 | 26411747:264 | +;+;+;+;+;+;+; | 2764 | ARHGEF38 | 14 | 10 | 8 | 16 | 7 | 0 |
| ENSDARG000 | 1;1;1;1;1;1;1; | 49933652:499 | 49934325:499 | -;-;-;-;-;-;-;-;- | 2125 | sgms2a | 0 | 1 | 2 | 0 | 18 | 0 |
| ENSDARG000 | 1;1;1;1;1;1;1; | 54650048:546 | 54650075:546 | +;+;+;+;+;+;+; | 796 | NA | 4 | 0 | 4 | 0 | 2 | 0 |
| ENSDARG000 | 1;1;1 | 36447498:364 | 36447571:364 | -;-;- | 441 | NA | 0 | 0 | 0 | 0 | 0 | 0 |
| ENSDARG000 | 1;1;1;1;1;1;1; | 30585140:305 | 30585847:305 | -;-;-;-;-;-;-;-;- | 3060 | dachc | 148 | 206 | 257 | 238 | 170 | 158 |
| ENSDARG000 | 1;1;1;1;1;1;1; | 44074445:440 | 44074568:440 | -;-;-;-;-;-;-;-;- | 1517 | NA | 22 | 6 | 6 | 29 | 46 | 10 |
| ENSDARG000 | 1;1;1;1;1;1 | 33558555:335 | 33558730:335 | +;+;+;+;+;+ | 1986 | chmp2bb | 1659 | 1260 | 1272 | 930 | 1377 | 1345 |
| ENSDARG000 | 1;1;1;1;1;1;1; | 47499888:475 | 47500189:475 | +;+;+;+;+;+;+; | 1495 | stn1 | 93 | 182 | 208 | 102 | 118 | 181 |
| ENSDARG000 | 1;1;1;1;1;1;1; | 58079053:580 | 58079158:580 | +;+;+;+;+;+;+; | 3309 | si:ch211-114 | 15 | 0 | 0 | 0 | 0 | 0 |
| ENSDARG000 | 1;1;1;1;1;1;1 | 54673846:546 | 54674545:546 | +;+;+;+;+;+;+ | 2016 | gprc5bb | 0 | 4 | 8 | 0 | 0 | 0 |
| ENSDARG000 | 1;1;1;1;1;1;1; | 26438597:264 | 26439096:264 | -;-;-;-;-;-;- | 1994 | ints12 | 322 | 354 | 460 | 330 | 438 | 308 |
| ENSDARG000 | 1;1;1;1;1;1;1 | 28961653:289 | 28962983:289 | -;-;-;-;-;-;- | 2308 | gpm6ba | 0 | 2 | 40 | 4 | 108 | 0 |
| ENSDARG000 | 1;1;1;1;1;1;1; | 1941031:1949 | 1941339:1949 | +;+;+;+;+;+;+; | 3026 | NA | 22 | 6 | 2 | 0 | 10 | 0 |
| ENSDARG000 | 1 | 2025085 | 2027100 | + | 2016 | slc5a3a | 0 | 0 | 4 | 0 | 8 | 4 |
| ENSDARG000 | 1;1 | 2048047:2051 | 2049309:2051 | -;- | 1290 | oxgr1a.1 | 4 | 6 | 0 | 2 | 0 | 0 |
| ENSDARG000 | 1;1;1;1;1;1;1; | 17747656:177 | 17748010:177 | -;-;-;-;-;-;-;-;- | 4603 | NA | 47 | 18 | 32 | 66 | 38 | 48 |
| ENSDARG000 | 1;1;1;1;1;1;1; | 24469313:244 | 24469632:244 | +;+;+;+;+;+;+; | 6524 | FHIP1A | 79 | 12 | 346 | 23 | 27 | 105 |
| ENSDARG000 | 1 | 2060430 | 2061419 | - | 990 | oxgr1a.3 | 0 | 2 | 0 | 0 | 0 | 0 |
| ENSDARG000 | 1 | 2074513 | 2075529 | - | 1017 | oxgr1a.2 | 0 | 0 | 0 | 4 | 2 | 2 |
| ENSDARG000 | 1;1;1;1;1;1;1 | 51615506:516 | 51615825:516 | +;+;+;+;+;+;+ | 809 | NA | 92 | 36 | 118 | 92 | 156 | 37 |
| ENSDARG000 | 1;1;1;1;1;1;1; | 9860381:9863 | 9860748:9863 | +;+;+;+;+;+;+; | 1394 | pmm2 | 392 | 430 | 474 | 233 | 434 | 365 |
| ENSDARG000 | 1;1;1;1;1;1;1; | 11086957:110 | 11087165:110 | -;-;-;-;-;-;-;-;- | 5496 | NA | 226 | 808 | 1071 | 314 | 586 | 378 |
| ENSDARG000 | 1;1;1;1;1;1;1; | 21495850:215 | 21496079:215 | -;-;-;-;-;-;-;-;- | 4683 | WDR19 | 196 | 142 | 223 | 100 | 213 | 232 |
| ENSDARG000 | 1;1;1;1;1;1;1; | 47585700:475 | 47585776:475 | +;+;+;+;+;+;+; | 6887 | SH3PXD2A | 1490 | 460 | 1016 | 544 | 1670 | 1159 |
| ENSDARG000 | 1;1;1;1;1;1 | 44930457:449 | 44930614:449 | -;-;-;-;-;- | 795 | NA | 6 | 0 | 0 | 20 | 4 | 6 |
| ENSDARG000 | 1;1;1;1;1;1;1; | 54683655:546 | 54683752:546 | +;+;+;+;+;+;+; | 1970 | NA | 411 | 927 | 393 | 293 | 558 | 422 |
| ENSDARG000 | 1;1;1;1;1;1 | 60979:60987: | 61106:61106: | -;-;-;-;-;- | 737 | NA | 99 | 55 | 36 | 51 | 76 | 75 |
| ENSDARG000 | 1;1;1;1;1;1;1 | 55780128:557 | 55780380:557 | -;-;-;-;-;-;- | 697 | ndufb7 | 874 | 940 | 1164 | 894 | 937 | 931 |
| ENSDARG000 | 1;1;1;1;1;1;1; | 36612660:366 | 36612756:366 | +;+;+;+;+;+;+; | 3172 | ednraa | 374 | 353 | 950 | 1387 | 545 | 781 |
| ENSDARG000 | 1;1;1;1;1;1;1; | 19562752:195 | 19563646:195 | -;-;-;-;-;-;-;-;- | 6542 | ecpas | 1271 | 1536 | 1257 | 448 | 1349 | 1010 |
| ENSDARG000 | 1;1;1;1;1 | 30718246:307 | 30718584:307 | -;-;-;-;- | 1119 | mzt1 | 210 | 151 | 250 | 132 | 233 | 260 |
| ENSDARG000 | 1;1;1;1;1;1;1; | 58091659:580 | 58091692:580 | +;+;+;+;+;+;+; | 3417 | NA | 0 | 0 | 6 | 2 | 0 | 12 |
| ENSDARG000 | 1;1;1;1;1;1 | 47701924:477 | 47704217:477 | -;-;-;-;-;- | 3866 | neurl1aa | 74 | 16 | 246 | 10 | 66 | 96 |
| ENSDARG000 | 1;1;1;1;1;1;1; | 65705:65705: | 67548:67548: | -;-;-;-;-;-;-;-;- | 3915 | gpa33a | 510 | 22 | 16 | 48 | 78 | 36 |
| ENSDARG000 | 1;1;1;1;1;1;1; | 24520503:245 | 24521082:245 | -;-;-;-;-;-;-;-;- | 3855 | gatb | 172 | 160 | 250 | 132 | 194 | 198 |
| ENSDARG000 | 1;1 | 51673028:516 | 51673964:516 | -;- | 1639 | prokr1a | 0 | 0 | 0 | 0 | 6 | 0 |
| ENSDARG000 | 1;1;1;1;1;1;1; | 494297:49434 | 494382:49438 | +;+;+;+;+;+;+; | 3044 | blzf1 | 452 | 365 | 374 | 211 | 452 | 403 |
| ENSDARG000 | 1 | 24557414 | 24558749 | + | 1336 | dctpp1 | 0 | 0 | 0 | 0 | 0 | 0 |
| ENSDARG000 | 1;1;1;1;1;1;1; | 58105762:581 | 58106448:581 | -;-;-;-;-;-;-;-;- | 1993 | caspbl | 4 | 6 | 0 | 2 | 16 | 0 |
| ENSDARG000 | 1;1;1;1;1;1;1; | 2101541:2101 | 2101796:2101 | +;+;+;+;+;+;+; | 5784 | mbnl2 | 3716 | 3176 | 3983 | 3761 | 3518 | 4924 |
| ENSDARG000 | 1 | 33634186 | 33635304 | + | 1119 | NA | 0 | 0 | 0 | 0 | 0 | 0 |
| ENSDARG000 | 1;1;1;1;1 | 33645044:336 | 33645968:336 | -;-;-;-;- | 1113 | cldng | 19 | 4 | 8 | 0 | 10 | 2 |
| ENSDARG000 | 1;1;1;1;1;1;1; | 6189949:6189 | 6190246:6190 | +;+;+;+;+;+;+; | 2749 | retreg2 | 463 | 280 | 417 | 194 | 353 | 302 |
| ENSDARG000 | 1;1;1;1;1;1;1; | 9873295:9873 | 9873657:9873 | -;-;-;-;-;-;-;-;- | 1795 | ints15 | 258 | 220 | 347 | 154 | 340 | 240 |
| ENSDARG000 | 1;1;1;1;1;1;1; | 44141758:441 | 44142001:441 | -;-;-;-;-;-;-;-;- | 2190 | slc43a3a | 8 | 0 | 16 | 2 | 21 | 8 |
| ENSDARG000 | 1;1;1;1;1;1;1; | 51702087:517 | 51702500:517 | -;-;-;-;-;-;-;-;- | 1227 | snrpb2 | 402 | 408 | 521 | 238 | 378 | 313 |
| ENSDARG000 | 1;1;1;1;1;1 | 59293873:592 | 59294054:592 | +;+;+;+;+;+ | 2088 | rdh8b | 2 | 0 | 0 | 2 | 2 | 2 |
| ENSDARG000 | 1;1;1;1;1;1;1; | 49951707:499 | 49951799:499 | +;+;+;+;+;+;+; | 2652 | gstcd | 102 | 64 | 137 | 56 | 139 | 71 |
| ENSDARG000 | 1;1;1;1;1;1;1; | 33647682:336 | 33647818:336 | +;+;+;+;+;+;+; | 5977 | NA | 1226 | 1096 | 2138 | 1265 | 1931 | 1450 |
| ENSDARG000 | 1;1;1;1;1;1;1; | 36722122:367 | 36722589:367 | +;+;+;+;+;+;+; | 6549 | tmem184c | 565 | 456 | 537 | 364 | 490 | 572 |
| ENSDARG000 | 1;1;1;1;1;1;1; | 44127292:441 | 44127339:441 | +;+;+;+;+;+;+; | 1794 | cabp2a | 488 | 342 | 190 | 500 | 395 | 564 |
| ENSDARG000 | 1 | 45922699 | 45924004 | + | 1306 | lipt1 | 30 | 16 | 28 | 26 | 46 | 46 |
| ENSDARG000 | 1;1;1;1;1;1;1; | 8555323:8556 | 8555629:8557 | -;-;-;-;-;-;- | 2344 | PTCD1 | 264 | 247 | 369 | 140 | 252 | 246 |
| ENSDARG000 | 1;1;1;1;1;1;1; | 36746078:367 | 36746513:367 | -;-;-;-;-;-;-;-;- | 3102 | prmt9 | 130 | 115 | 155 | 115 | 123 | 73 |
| ENSDARG000 | 1;1;1;1;1;1;1; | 47831619:478 | 47831973:478 | +;+;+;+;+;+;+; | 3179 | cfap58 | 0 | 2 | 4 | 0 | 2 | 2 |
| ENSDARG000 | 1;1;1;1;1;1;1; | 55786986:557 | 55787339:557 | -;-;-;-;-;-;-;-;- | 2864 | zgc:136908 | 0 | 0 | 12 | 38 | 3 | 8 |
| ENSDARG000 | 1;1;1;1;1;1;1; | 54700657:547 | 54700777:547 | -;-;-;-;-;-;- | 612 | exosc1 | 530 | 450 | 609 | 582 | 452 | 731 |
| ENSDARG000 | 1;1;1;1;1;1;1; | 36771954:367 | 36772501:367 | +;+;+;+;+;+;+; | 4226 | arhgap10 | 565 | 399 | 531 | 268 | 579 | 549 |
| ENSDARG000 | 1;1;1;1;1;1;1; | 44935442:449 | 44935708:449 | -;-;-;-;-;-;-;-;- | 1144 | tmem176 | 202 | 52 | 188 | 36 | 26 | 52 |
| ENSDARG000 | 1;1;1;1;1;1;1; | 11107688:111 | 11107763:111 | +;+;+;+;+;+;+; | 1124 | knstrn | 142 | 236 | 358 | 170 | 168 | 84 |
| ENSDARG000 | 1;1;1;1;1;1;1; | 29058268:290 | 29059092:290 | -;-;-;-;-;-;-;-;- | 1503 | gemin8 | 359 | 363 | 440 | 278 | 394 | 316 |
| ENSDARG000 | 1;1;1;1;1;1;1; | 50085440:500 | 50085707:500 | +;+;+;+;+;+;+; | 2897 | npnta | 12 | 0 | 10 | 25 | 38 | 10 |
| ENSDARG000 | 1;1 | 17858380:178 | 17858878:178 | +;+ | 539 | NA | 0 | 0 | 0 | 0 | 0 | 0 |
| ENSDARG000 | 1;1;1;1;1;1;1; | 26444986:264 | 26445189:264 | +;+;+;+;+;+;+; | 4389 | g3bp2a | 968 | 808 | 1009 | 324 | 1276 | 698 |
| ENSDARG000 | 1;1;1;1;1;1;1; | 12364456:123 | 12364851:123 | -;-;-;-;-;-;-;-;- | 2964 | SCLT1 | 90 | 250 | 244 | 310 | 178 | 212 |
| ENSDARG000 | 1;1;1;1;1 | 17892944:178 | 17893145:178 | +;+;+;+;+ | 3161 | tlr3 | 26 | 6 | 18 | 0 | 20 | 18 |
| ENSDARG000 | 1;1;1;1;1;1;1; | 58116900:581 | 58117509:581 | +;+;+;+;+;+;+; | 4474 | NA | 22 | 0 | 6 | 0 | 4 | 2 |
| ENSDARG000 | 1;1;1;1;1;1;1; | 45925150:459 | 45925484:459 | +;+;+;+;+;+;+; | 8064 | NA | 1644 | 2790 | 4327 | 1882 | 2524 | 1682 |
| ENSDARG000 | 1;1;1;1;1;1;1; | 33668236:336 | 33668846:336 | +;+;+;+;+;+;+; | 3728 | arl13b | 358 | 302 | 248 | 103 | 350 | 234 |
| ENSDARG000 | 1;1;1;1;1;1;1; | 30723380:307 | 30723465:307 | +;+;+;+;+;+;+; | 3330 | bora | 24 | 98 | 170 | 54 | 58 | 52 |
| ENSDARG000 | 1;1;1;1;1;1;1; | 59304096:593 | 59305506:593 | -;-;-;-;-;-;-;-;- | 3478 | txndc11 | 212 | 240 | 348 | 179 | 345 | 154 |
| ENSDARG000 | 1;1;1;1;1;1;1; | 109798:11064 | 109928:11080 | +;+;+;+;+;+;+; | 2870 | f7l | 0 | 0 | 6 | 0 | 10 | 2 |
| ENSDARG000 | 1;1 | 108285:11057 | 108338:11080 | -;- | 282 | NA | 0 | 0 | 0 | 0 | 0 | 3 |
| ENSDARG000 | 1;1;1;1;1;1;1; | 12394205:123 | 12394304:123 | +;+;+;+;+;+;+; | 4273 | C1H4orf33 | 40 | 100 | 66 | 68 | 66 | 91 |
| ENSDARG000 | 1;1;1;1;1;1;1; | 46026457:460 | 46027059:460 | +;+;+;+;+;+;+; | 2092 | NA | 0 | 0 | 0 | 0 | 0 | 0 |
| ENSDARG000 | 1;1;1;1 | 8575307:8575 | 8575403:8575 | +;+;+;+ | 372 | NA | 1 | 0 | 3 | 1 | 0 | 0 |
| ENSDARG000 | 1 | 48921944 | 48922273 | - | 330 | NA | 2 | 0 | 0 | 0 | 0 | 0 |
| ENSDARG000 | 1;1;1;1;1;1;1 | 55817653:558 | 55818435:558 | -;-;-;-;-;-;- | 2570 | NA | 28 | 38 | 21 | 23 | 38 | 38 |
| ENSDARG000 | 1 | 48923328 | 48927049 | + | 3722 | NA | 0 | 0 | 0 | 0 | 0 | 0 |

| | | | | | | | | | | | | |
|---|---|---|---|---|---|---|---|---|---|---|---|---|
| ENSDARG000 | 1 | 46194333 | 46196107 | + | 1775 | sox1b | 0 | 0 | 0 | 0 | 0 | 0 |
| ENSDARG000 | 1;1;1;1;1;1 | 8587230:858 | 8587361:858 | +;+;+;+;+;+ | 1232 | NA | 48 | 33 | 62 | 26 | 47 | 46 |
| ENSDARG000 | 1;1;1;1;1 | 41318241:41 | 41318367:41 | +;+;+;+;+ | 902 | NA | 0 | 0 | 0 | 0 | 0 | 0 |
| ENSDARG000 | 1;1 | 8593560:859 | 8593613:859 | +;+ | 2526 | NA | 6 | 4 | 9 | 0 | 20 | 24 |
| ENSDARG000 | 1:1:1:1:1:1:1: | 41293964:412 | 41295591:412 | -:-:-:-:-:-:-:-: | 10780 | htt | 1699 | 1210 | 1876 | 1247 | 1754 | 1468 |
| ENSDARG000 | 1:1:1:1:1:1:1: | 508885:5090 | 509494:5094 | -:-:-:-:-:-:-:-: | 1820 | trmt10c | 102 | 138 | 222 | 108 | 152 | 106 |
| ENSDARG000 | 1:1:1:1:1:1:1: | 17900306:17 | 17900546:17 | +:+:+:+:+:+:+: | 2643 | cyp4v2a | 0 | 0 | 8 | 0 | 0 | 0 |
| ENSDARG000 | 1:1:1:1:1:1:1: | 9886991:988 | 9887089:988 | +:+:+:+:+:+:+: | 1840 | rgs11 | 6 | 12 | 0 | 0 | 0 | 0 |
| ENSDARG000 | 1;1;1;1;1;1 | 8615535:861 | 8616371:861 | -;-;-;-;-;- | 1813 | NA | 0 | 0 | 0 | 0 | 8 | 0 |
| ENSDARG000 | 1;1;1;1 | 54707667:54 | 54709514:54 | -;-;-;- | 2615 | pgam1b | 2192 | 2336 | 2012 | 2359 | 2334 | 1915 |
| ENSDARG000 | 1:1:1:1:1:1:1: | 8601935:860 | 8602071:860 | +:+:+:+:+:+:+: | 1791 | NA | 0 | 0 | 0 | 0 | 0 | 0 |
| ENSDARG000 | 1:1:1:1:1:1:1: | 46238774:46 | 46238939:46 | -:-:-:-:-:-:-:-: | 1878 | NA | 0 | 0 | 0 | 0 | 0 | 0 |
| ENSDARG000 | 1:1:1:1:1:1:1: | 6218632:621 | 6218829:621 | -:-:-:-:-:-:-:-: | 1674 | mdh1b | 14 | 32 | 16 | 14 | 16 | 4 |
| ENSDARG000 | 1:1:1:1:1:1:1: | 24870661:24 | 24871357:24 | -:-:-:-:-:-:-:-: | 3544 | FBXW7 | 255 | 146 | 195 | 146 | 244 | 129 |
| ENSDARG000 | 1:1:1:1:1:1:1: | 51712869:51 | 51713187:51 | -:-:-:-:-:-:-:-: | 1307 | rnaseh2a | 275 | 446 | 658 | 424 | 492 | 252 |
| ENSDARG000 | 1:1:1:1:1:1:1: | 58137020:58 | 58137192:58 | +:+:+:+:+:+:+: | 3259 | NA | 17 | 1 | 0 | 0 | 7 | 0 |
| ENSDARG000 | 1:1:1:1:1:1:1: | 55843552:55 | 55844863:55 | -:-:-:-:-:-:-:-: | 2666 | prkacab | 746 | 426 | 551 | 498 | 645 | 538 |
| ENSDARG000 | 1:1:1:1:1:1:1: | 25154076:25 | 25154660:25 | -:-:-:-:-:-:-:-: | 1423 | tmem154 | 36 | 68 | 54 | 50 | 43 | 56 |
| ENSDARG000 | 1:1:1:1:1:1:1: | 44173245:44 | 44173308:44 | +:+:+:+:+:+:+: | 5601 | CTNND1 | 2789 | 2374 | 2718 | 1727 | 2387 | 2177 |
| ENSDARG000 | 1:1:1:1:1:1:1: | 11124746:11 | 11125371:11 | -:-:-:-:-:-:-:-: | 7350 | SDK1 | 0 | 0 | 0 | 0 | 6 | 0 |
| ENSDARG000 | 1;1 | 46250113:46 | 46250490:46 | +;+ | 784 | NA | 2 | 0 | 0 | 0 | 0 | 0 |
| ENSDARG000 | 1:1:1:1:1:1:1: | 54721240:54 | 54721633:54 | -:-:-:-:-:-:-:-: | 1528 | cdc25d | 10 | 6 | 4 | 15 | 0 | 0 |
| ENSDARG000 | 1;1;1;1;1;1;1 | 58150000:58 | 58150218:58 | +;+;+;+;+;+;+ | 3096 | NA | 1 | 1 | 8 | 6 | 1 | 0 |
| ENSDARG000 | 1;1;1;1;1 | 8638162:863 | 8638610:864 | -;-;-;-;- | 2626 | fbxl18 | 166 | 226 | 218 | 92 | 194 | 146 |
| ENSDARG000 | 1:1:1:1:1:1:1: | 119960:1201 | 120036:1202 | +:+:+:+:+:+:+: | 1809 | f7 | 0 | 0 | 5 | 0 | 0 | 0 |
| ENSDARG000 | 1;1;1;1 | 48997724:49 | 48997953:49 | -;-;-;- | 650 | NA | 0 | 0 | 0 | 0 | 0 | 0 |
| ENSDARG000 | 1;1;1;1;1;1 | 39813229:39 | 39814049:39 | -;-;-;-;-;- | 1597 | dctd | 118 | 424 | 397 | 232 | 356 | 240 |
| ENSDARG000 | 1:1:1:1:1:1:1: | 39878634:39 | 39883387:39 | -:-:-:-:-:-:-:-: | 6293 | tmem134 | 1678 | 1949 | 1451 | 1308 | 2206 | 1915 |
| ENSDARG000 | 1:1:1:1:1:1:1: | 29068649:29 | 29068684:29 | +:+:+:+:+:+:+: | 3287 | cbsa | 50 | 102 | 187 | 20 | 98 | 62 |
| ENSDARG000 | 1;1;1;1;1 | 18251335:18 | 18252727:18 | -;-;-;-;- | 2095 | mtnr1aa | 2 | 0 | 0 | 0 | 2 | 0 |
| ENSDARG000 | 1;1;1;1;1;1 | 18187612:18 | 18187867:18 | -;-;-;-;-;- | 718 | NA | 0 | 0 | 0 | 0 | 0 | 0 |
| ENSDARG000 | 1:1:1:1:1:1:1: | 39859782:39 | 39859873:39 | +:+:+:+:+:+:+: | 2421 | irf2a | 507 | 677 | 675 | 270 | 570 | 481 |
| ENSDARG000 | 1;1;1;1 | 49222164:49 | 49222804:49 | -;-;-;- | 1101 | cxcl18b | 529 | 390 | 444 | 186 | 118 | 166 |
| ENSDARG000 | 1:1:1:1:1:1:1: | 44941031:44 | 44941085:44 | +:+:+:+:+:+:+: | 819 | si:dkev-9i23.1 | 52 | 32 | 54 | 35 | 14 | 38 |
| ENSDARG000 | 1:1:1:1:1:1:1: | 50198607:50 | 50198910:50 | -:-:-:-:-:-:-:-: | 3134 | tbck | 418 | 503 | 579 | 410 | 556 | 424 |
| ENSDARG000 | 1:1:1:1:1:1:1: | 6225493:622 | 6225575:622 | +:+:+:+:+:+:+: | 2149 | NA | 337 | 438 | 802 | 343 | 481 | 479 |
| ENSDARG000 | 1;1;1;1;1;1 | 51721851:51 | 51721942:51 | +;+;+;+;+;+ | 1018 | prdx2 | 8459 | 8059 | 9174 | 8603 | 10456 | 6537 |
| ENSDARG000 | 1;1;1;1;1;1;1 | 50293938:50 | 50294087:50 | +;+;+;+;+;+;+ | 1187 | aimp1a | 820 | 847 | 1137 | 470 | 839 | 764 |
| ENSDARG000 | 1:1:1:1:1:1:1: | 513986:5140 | 514219:5142 | +:+:+:+:+:+:+: | 1600 | txnl4b | 47 | 141 | 182 | 76 | 194 | 162 |
| ENSDARG000 | 1;1 | 49245229:49 | 49246190:49 | -;- | 1002 | NA | 0 | 0 | 0 | 0 | 0 | 0 |
| ENSDARG000 | 1:1:1:1:1:1:1: | 44949909:44 | 44950294:44 | +:+:+:+:+:+:+: | 5451 | vps37c | 1390 | 1503 | 1823 | 1343 | 1527 | 1273 |
| ENSDARG000 | 1;1;1;1;1 | 49266886:49 | 49266997:49 | +;+;+;+;+ | 1872 | caly | 2 | 18 | 2 | 2 | 10 | 0 |
| ENSDARG000 | 1;1;1;1;1;1 | 58160151:58 | 58160299:58 | +;+;+;+;+;+ | 3264 | NA | 9 | 0 | 8 | 0 | 1 | 2 |
| ENSDARG000 | 1;1;1;1;1 | 58176023:58 | 58176578:58 | +;+;+;+;+ | 1475 | NA | 13 | 9 | 11 | 8 | 5 | 3 |
| ENSDARG000 | 1:1:1:1:1:1:1: | 26467071:26 | 26467297:26 | +:+:+:+:+:+:+: | 5584 | uso1 | 1954 | 1948 | 2642 | 1615 | 2217 | 1854 |
| ENSDARG000 | 1;1;1;1;1;1 | 58182400:58 | 58182623:58 | +;+;+;+;+;+ | 3079 | NA | 0 | 0 | 0 | 0 | 0 | 1 |
| ENSDARG000 | 1;1;1 | 44992207:44 | 44992415:44 | +;+;+ | 385 | NA | 0 | 0 | 0 | 0 | 0 | 0 |
| ENSDARG000 | 1:1:1:1:1:1:1: | 44983825:44 | 44984053:44 | +:+:+:+:+:+:+: | 3814 | NA | 0 | 4 | 0 | 0 | 0 | 0 |
| ENSDARG000 | 1:1:1:1:1:1:1: | 46263473:46 | 46269765:46 | -:-:-:-:-:-:-:-: | 11102 | ATP11A | 19323 | 10076 | 14659 | 14002 | 17032 | 19803 |
| ENSDARG000 | 1:1:1:1:1:1:1: | 33697170:33 | 33697341:33 | +:+:+:+:+:+:+: | 2086 | nsun3 | 60 | 83 | 119 | 36 | 80 | 119 |
| ENSDARG000 | 1;1;1;1;1;1;1 | 53279861:53 | 53280476:53 | +;+;+;+;+;+;+ | 1810 | rnf150a | 16 | 32 | 64 | 42 | 96 | 46 |
| ENSDARG000 | 1;1;1;1;1 | 58194268:58 | 58196003:58 | +;+;+;+;+ | 2645 | NA | 0 | 1 | 0 | 0 | 0 | 1 |
| ENSDARG000 | 1;1 | 2222363:222 | 2222679:224 | +;+ | 555 | rap2ab | 130 | 35 | 43 | 72 | 85 | 125 |
| ENSDARG000 | 1:1:1:1:1:1:1: | 8649099:864 | 8649746:865 | -:-:-:-:-:-:-:-: | 2890 | actb1 | 69842 | 103793 | 101531 | 89589 | 99447 | 88190 |
| ENSDARG000 | 1:1:1:1:1:1:1: | 9919909:992 | 9920373:992 | -:-:-:-:-:-:-:-: | 2334 | pgap6 | 251 | 168 | 234 | 97 | 180 | 212 |
| ENSDARG000 | 1;1 | 46400612:46 | 46400719:46 | +;+ | 923 | NA | 0 | 0 | 0 | 0 | 1 | 1 |
| ENSDARG000 | 1:1:1:1:1:1:1: | 18363833:18 | 18364034:18 | -:-:-:-:-:-:-:-: | 2455 | klhl5 | 88 | 58 | 169 | 66 | 110 | 38 |
| ENSDARG000 | 1:1:1:1:1:1:1: | 127250:1272 | 127324:1273 | +:+:+:+:+:+:+: | 2467 | f7i | 0 | 0 | 2 | 0 | 0 | 6 |
| ENSDARG000 | 1;1 | 49352900:49 | 49353358:49 | +;+ | 1424 | msx1b | 36 | 0 | 2 | 0 | 4 | 0 |
| ENSDARG000 | 1:1:1:1:1:1:1: | 58205904:58 | 58205974:58 | +:+:+:+:+:+:+: | 1629 | NA | 0 | 0 | 0 | 0 | 0 | 0 |
| ENSDARG000 | 1;1;1;1 | 55878113:55 | 55878869:55 | -;-;-;- | 1610 | asf1bb | 232 | 462 | 786 | 250 | 490 | 444 |
| ENSDARG000 | 1:1:1:1:1:1:1: | 59314675:59 | 59314768:59 | +:+:+:+:+:+:+: | 2669 | parn | 1173 | 955 | 1182 | 939 | 1335 | 1106 |
| ENSDARG000 | 1;1;1;1;1;1 | 6357059:635 | 6357188:635 | +;+;+;+;+;+ | 1350 | NA | 31 | 42 | 45 | 25 | 53 | 69 |
| ENSDARG000 | 1;1;1;1 | 518853:5204 | 519127:5206 | -;-;-;- | 1167 | si:ch73-41e3. | 0 | 0 | 0 | 0 | 0 | 0 |
| ENSDARG000 | 1:1:1:1:1:1:1: | 524388:5246 | 524596:5248 | +:+:+:+:+:+:+: | 3118 | mrpl16 | 912 | 929 | 1061 | 586 | 1016 | 767 |
| ENSDARG000 | 1:1:1:1:1:1:1: | 25178106:25 | 25178163:25 | +:+:+:+:+:+:+: | 1906 | mnd1 | 57 | 128 | 257 | 81 | 170 | 124 |
| ENSDARG000 | 1;1;1 | 51732979:51 | 51734535:51 | -;-;- | 1557 | junba | 19761 | 8137 | 15708 | 17193 | 6173 | 10593 |
| ENSDARG000 | 1;1;1;1;1;1;1 | 49357695:49 | 49358049:49 | -;-;-;-;-;-;- | 582 | si:dkeyp-80c1 | 0 | 0 | 0 | 0 | 0 | 0 |
| ENSDARG000 | 1:1:1:1:1:1:1: | 9954489:995 | 9954773:995 | +:+:+:+:+:+:+: | 3406 | pdia2 | 8 | 2 | 4 | 0 | 28 | 22 |
| ENSDARG000 | 1;1;1;1;1 | 55891376:55 | 55891902:55 | -;-;-;-;- | 960 | NA | 18 | 22 | 6 | 24 | 36 | 28 |
| ENSDARG000 | 1:1:1:1:1:1:1: | 19602389:19 | 19602484:19 | +:+:+:+:+:+:+: | 3221 | fbxo10 | 32 | 41 | 48 | 34 | 36 | 52 |
| ENSDARG000 | 1:1:1:1:1:1:1: | 45012391:45 | 45016979:45 | -:-:-:-:-:-:-:-: | 6690 | NA | 289 | 296 | 341 | 179 | 405 | 304 |
| ENSDARG000 | 1:1:1:1:1:1:1: | 45043767:45 | 45043906:45 | -:-:-:-:-:-:-:-: | 1604 | rps6 | 22153 | 20338 | 36353 | 29692 | 23266 | 29929 |
| ENSDARG000 | 1:1:1:1:1:1:1: | 21540453:21 | 21540686:21 | -:-:-:-:-:-:-:-: | 4560 | ncapd3 | 168 | 753 | 935 | 340 | 668 | 417 |
| ENSDARG000 | 1;1;1;1;1;1 | 18550864:18 | 18550970:18 | +;+;+;+;+;+ | 749 | NA | 2 | 9 | 4 | 3 | 21 | 31 |
| ENSDARG000 | 1:1:1:1:1:1:1: | 45002971:45 | 45003089:45 | +:+:+:+:+:+:+: | 2467 | dnaja1 | 275 | 237 | 251 | 157 | 260 | 168 |
| ENSDARG000 | 1;1;1;1;1 | 8662530:866 | 8663370:866 | +;+;+;+;+ | 3781 | FSCN1 | 4 | 12 | 0 | 2 | 6 | 14 |
| ENSDARG000 | 1:1:1:1:1:1:1: | 51753125:51 | 51753362:51 | +:+:+:+:+:+:+: | 1069 | NA | 0 | 0 | 0 | 0 | 0 | 0 |
| ENSDARG000 | 1;1 | 51802535:51 | 51802708:51 | -;- | 314 | NA | 0 | 0 | 0 | 0 | 0 | 0 |
| ENSDARG000 | 1;1;1;1 | 6403551:641 | 6405472:641 | -;-;-;- | 2960 | klf7a | 286 | 123 | 144 | 291 | 160 | 237 |
| ENSDARG000 | 1:1:1:1:1:1:1: | 54737353:54 | 54737804:54 | +:+:+:+:+:+:+: | 3822 | pi4k2a | 2186 | 1685 | 1518 | 1392 | 1539 | 1288 |
| ENSDARG000 | 1:1:1:1:1:1:1: | 2260407:226 | 2260524:226 | +:+:+:+:+:+:+: | 4242 | kpnb3 | 8730 | 8586 | 9909 | 5508 | 9701 | 10707 |
| ENSDARG000 | 1:1:1:1:1:1:1: | 36935154:36 | 36935603:36 | -:-:-:-:-:-:-:-: | 3540 | nr3c2 | 146 | 34 | 14 | 27 | 32 | 76 |
| ENSDARG000 | 1:1:1:1:1:1:1: | 135903:1363 | 136147:1364 | +:+:+:+:+:+:+: | 2727 | f10 | 2459 | 1034 | 1108 | 2678 | 1787 | 944 |
| ENSDARG000 | 1;1;1 | 55903811:55 | 55903943:55 | -;-;- | 321 | NA | 0 | 2 | 0 | 0 | 0 | 0 |
| ENSDARG000 | 1:1:1:1:1:1:1: | 33969015:33 | 33969691:33 | +:+:+:+:+:+:+: | 5654 | EPHA6 | 0 | 2 | 0 | 0 | 14 | 0 |
| ENSDARG000 | 1;1;1 | 55908578:55 | 55908710:55 | -;-;- | 321 | NA | 0 | 0 | 0 | 0 | 0 | 0 |
| ENSDARG000 | 1:1:1:1:1:1:1: | 53321878:53 | 53321998:53 | +:+:+:+:+:+:+: | 4072 | TBC1D9 | 1716 | 1111 | 1994 | 1654 | 2705 | 1529 |
| ENSDARG000 | 1:1:1:1:1:1:1: | 18575747:18 | 18576261:18 | -:-:-:-:-:-:-:-: | 2273 | fam114a1 | 277 | 220 | 344 | 340 | 290 | 359 |
| ENSDARG000 | 1;1 | 51827046:51 | 51827613:51 | +;+ | 1294 | dand5 | 0 | 0 | 0 | 0 | 0 | 0 |
| ENSDARG000 | 1:1:1:1:1:1:1: | 49370549:49 | 49370616:49 | -:-:-:-:-:-:-:-: | 1405 | gsk3bb | 88 | 99 | 70 | 75 | 93 | 55 |
| ENSDARG000 | 1;1 | 51869067:51 | 51869506:51 | -;- | 772 | NA | 1 | 9 | 5 | 0 | 13 | 4 |
| ENSDARG000 | 1:1:1:1:1:1:1: | 6646529:664 | 6646741:664 | +:+:+:+:+:+:+: | 2365 | ube2f | 679 | 607 | 845 | 440 | 653 | 661 |
| ENSDARG000 | 1:1:1:1:1:1:1: | 51896147:51 | 51896215:51 | +:+:+:+:+:+:+: | 2531 | NFIX | 40 | 60 | 92 | 11 | 43 | 54 |
| ENSDARG000 | 1:1:1:1:1:1:1: | 144284:1449 | 144377:1450 | +:+:+:+:+:+:+: | 1519 | prozb | 0 | 8 | 24 | 24 | 0 | 0 |
| ENSDARG000 | 1;1;1;1;1;1 | 6640437:664 | 6640575:664 | +;+;+;+;+;+ | 950 | si:ch211-93g | 6 | 4 | 10 | 0 | 2 | 4 |
| ENSDARG000 | 1;1 | 44236341:44 | 44236488:44 | +;+ | 727 | NA | 0 | 0 | 0 | 0 | 0 | 0 |
| ENSDARG000 | 1;1;1 | 55920864:55 | 55921002:55 | -;-;- | 341 | NA | 0 | 0 | 0 | 0 | 0 | 0 |
| ENSDARG000 | 1:1:1:1:1:1:1: | 58221372:58 | 58221399:58 | +:+:+:+:+:+:+: | 3633 | NA | 310 | 171 | 271 | 187 | 176 | 98 |
| ENSDARG000 | 1;1;1 | 55925810:55 | 55925939:55 | -;-;- | 318 | NA | 0 | 2 | 4 | 0 | 4 | 8 |
| ENSDARG000 | 1;1;1 | 54758575:54 | 54759087:54 | -;-;- | 1012 | NA | 592 | 331 | 459 | 319 | 448 | 238 |
| ENSDARG000 | 1:1:1:1:1:1:1: | 54766943:54 | 54766992:54 | +:+:+:+:+:+:+: | 3748 | NA | 255 | 250 | 139 | 113 | 183 | 110 |
| ENSDARG000 | 1:1:1:1:1:1:1: | 52162642:52 | 52162793:52 | +:+:+:+:+:+:+: | 1438 | BEST2 | 6 | 0 | 14 | 0 | 0 | 0 |
| ENSDARG000 | 1;1;1;1;1;1;1 | 52130164:52 | 52130352:52 | +;+;+;+;+;+;+ | 2843 | rnf11a | 578 | 352 | 350 | 646 | 381 | 780 |
| ENSDARG000 | 1:1:1:1:1:1:1: | 52137528:52 | 52137812:52 | +:+:+:+:+:+:+: | 4613 | get3 | 2137 | 2050 | 2480 | 1929 | 2482 | 1854 |
| ENSDARG000 | 1;1;1;1;1;1 | 44239558:44 | 44239629:44 | +;+;+;+;+;+ | 910 | NA | 0 | 0 | 0 | 0 | 0 | 0 |
| ENSDARG000 | 1;1 | 52137392:52 | 52137731:52 | -;- | 578 | NA | 0 | 0 | 0 | 0 | 1 | 0 |
| ENSDARG000 | 1:1:1:1:1:1:1: | 532766:5328 | 532899:5328 | +:+:+:+:+:+:+: | 2648 | mrpl39 | 380 | 631 | 837 | 318 | 591 | 313 |
| ENSDARG000 | 1:1:1:1:1:1:1: | 52111273:52 | 52111532:52 | -;-;-;-;-;-;-;- | 1410 | rad23aa | 909 | 623 | 839 | 533 | 630 | 560 |
| ENSDARG000 | 1:1:1:1:1:1:1: | 58228542:58 | 58228564:58 | +:+:+:+:+:+:+: | 3987 | NA | 38 | 18 | 22 | 0 | 33 | 9 |
| ENSDARG000 | 1:1:1:1:1:1:1: | 30736449:30 | 30736851:30 | -:-:-:-:-:-:-:-: | 3247 | DIS3 | 402 | 499 | 736 | 258 | 696 | 422 |
| ENSDARG000 | 1:1:1:1:1:1:1: | 39898327:39 | 39902052:39 | -:-:-:-:-:-:-:-: | 10886 | STOX2 | 62 | 44 | 36 | 34 | 10 | 52 |
| ENSDARG000 | 1:1:1:1:1:1:1: | 29102011:29 | 29102428:29 | -:-:-:-:-:-:-:-: | 1515 | HSF2BP | 2 | 0 | 0 | 0 | 0 | 0 |
| ENSDARG000 | 1;1 | 44246467:44 | 44246600:44 | +;+ | 373 | NA | 0 | 0 | 0 | 0 | 0 | 0 |
| ENSDARG000 | 1;1;1;1;1 | 55938565:55 | 55939004:55 | -;-;-;-;- | 1131 | NA | 10 | 15 | 18 | 28 | 56 | 76 |
| ENSDARG000 | 1;1;1 | 29096881:29 | 29097203:29 | +;+;+ | 1417 | cryaa | 0 | 0 | 0 | 0 | 0 | 0 |
| ENSDARG000 | 1:1:1:1:1:1:1: | 8673061:867 | 8674221:867 | -:-:-:-:-:-:-:-: | 2918 | abcg2b | 14 | 4 | 4 | 0 | 16 | 2 |
| ENSDARG000 | 1:1:1:1:1:1:1: | 58242498:58 | 58242602:58 | +:+:+:+:+:+:+: | 1823 | NA | 0 | 14 | 5 | 0 | 0 | 4 |
| ENSDARG000 | 1;1;1;1;1;1 | 44249616:44 | 44249732:44 | +;+;+;+;+;+ | 955 | NA | 0 | 0 | 0 | 0 | 0 | 0 |
| ENSDARG000 | 1:1:1:1:1:1:1: | 49415281:49 | 49415748:49 | +:+:+:+:+:+:+: | 2701 | taf5 | 328 | 226 | 399 | 100 | 297 | 295 |
| ENSDARG000 | 1;1 | 44256883:44 | 44257065:44 | +;+ | 656 | NA | 0 | 0 | 0 | 0 | 0 | 0 |
| ENSDARG000 | 1;1;1;1;1;1 | 44260085:44 | 44260201:44 | +;+;+;+;+;+ | 955 | NA | 0 | 0 | 0 | 0 | 0 | 0 |
| ENSDARG000 | 1 | 11595798 | 11596829 | - | 1032 | NA | 0 | 0 | 0 | 0 | 0 | 0 |
| ENSDARG000 | 1;1 | 53364510:53 | 53365012:53 | +;+ | 688 | NA | 509 | 341 | 582 | 768 | 675 | 451 |
| ENSDARG000 | 1;1 | 55947389:55 | 55947495:55 | -;- | 211 | NA | 0 | 0 | 0 | 0 | 0 | 0 |
| ENSDARG000 | 1:1:1:1:1:1:1: | 49432137:49 | 49432236:49 | -;-;-;-;-;-;-;- | 898 | atp5md | 375 | 554 | 406 | 174 | 206 | 285 |
| ENSDARG000 | 1;1 | 11606010:11 | 11606903:11 | -;- | 1053 | NA | 1 | 0 | 0 | 0 | 0 | 0 |
| ENSDARG000 | 1;1 | 11615273:11 | 11616142:11 | -;- | 1023 | NA | 38 | 0 | 14 | 0 | 59 | 21 |
| ENSDARG000 | 1;1;1;1;1;1;1 | 58260886:58 | 58261106:58 | +;+;+;+;+;+;+ | 3154 | NA | 16 | 6 | 8 | 2 | 20 | 0 |
| ENSDARG000 | 1;1 | 44336897:44 | 44337104:44 | +;+ | 793 | NA | 0 | 0 | 0 | 0 | 0 | 0 |
| ENSDARG000 | 1;1;1;1 | 39981451:39 | 39982691:39 | -;-;-;- | 1783 | ing2 | 261 | 424 | 424 | 196 | 369 | 258 |
| ENSDARG000 | 1:1:1:1:1:1:1: | 45056371:45 | 45056529:45 | +:+:+:+:+:+:+: | 1915 | NA | 4 | 0 | 0 | 0 | 0 | 4 |
| ENSDARG000 | 1;1;1;1 | 50392716:50 | 50393263:50 | -;-;-;- | 1357 | dkk2 | 0 | 10 | 0 | 0 | 18 | 0 |
| ENSDARG000 | 1;1 | 39990547:39 | 39990656:39 | +;+ | 1218 | SOD3 | 8 | 4 | 4 | 0 | 12 | 6 |
| ENSDARG000 | 1;1;1;1;1 | 58282449:58 | 58284209:58 | +;+;+;+;+ | 2662 | NA | 1 | 0 | 8 | 0 | 2 | 0 |
| ENSDARG000 | 1:1:1:1:1:1:1: | 41454210:41 | 41454560:41 | +:+:+:+:+:+:+: | 2949 | si:ch211-89o | 0 | 0 | 0 | 0 | 0 | 0 |
| ENSDARG000 | 1:1:1:1:1:1:1: | 46404968:46 | 46405548:46 | +:+:+:+:+:+:+: | 6714 | tubgcp3 | 584 | 788 | 868 | 514 | 787 | 729 |
| ENSDARG000 | 1:1:1:1:1:1:1: | 34203817:34 | 34203945:34 | +:+:+:+:+:+:+: | 3299 | arl6 | 108 | 144 | 112 | 132 | 161 | 96 |
| ENSDARG000 | 1;1;1 | 11626252:11 | 11627373:11 | -;-;- | 1212 | NA | 93 | 0 | 29 | 0 | 69 | 45 |
| ENSDARG000 | 1:1:1:1:1:1:1: | 45080897:45 | 45080979:45 | +:+:+:+:+:+:+: | 3883 | NA | 0 | 0 | 0 | 0 | 0 | 0 |
| ENSDARG000 | 1:1:1:1:1:1:1: | 56107522:56 | 56107661:56 | -:-:-:-:-:-:-:-: | 2443 | NA | 0 | 0 | 10 | 4 | 70 | 16 |
| ENSDARG000 | 1;1 | 6665033:666 | 6665080:666 | +;+ | 920 | NA | 0 | 0 | 3 | 1 | 3 | 2 |
| ENSDARG000 | 1:1:1:1:1:1:1: | 50487586:50 | 50488089:50 | -:-:-:-:-:-:-:-: | 2377 | papss1 | 495 | 603 | 683 | 230 | 457 | 366 |
| ENSDARG000 | 1:1:1:1:1:1:1: | 54834119:54 | 54835894:54 | +:+:+:+:+:+:+: | 3176 | NA | 10 | 0 | 12 | 2 | 7 | 11 |
| ENSDARG000 | 1:1:1:1:1:1:1: | 39995008:39 | 39995175:39 | +:+:+:+:+:+:+: | 1920 | aip | 1469 | 1280 | 1593 | 1456 | 1653 | 1562 |
| ENSDARG000 | 1:1:1:1:1:1:1: | 54865552:54 | 54865729:54 | +:+:+:+:+:+:+: | 4279 | NA | 32 | 1 | 35 | 15 | 31 | 36 |
| ENSDARG000 | 1:1:1:1:1:1:1: | 18594146:18 | 18596388:18 | -;-;-;-;-;-;-;- | 3877 | klf3 | 98 | 20 | 20 | 16 | 30 | 62 |
| ENSDARG000 | 1;1 | 11641482:11 | 11642432:11 | -;- | 1032 | NA | 0 | 0 | 0 | 0 | 0 | 0 |
| ENSDARG000 | 1;1;1;1 | 59406300:59 | 59406435:59 | -;-;-;- | 612 | NA | 0 | 0 | 0 | 0 | 0 | 0 |
| ENSDARG000 | 1:1:1:1:1:1:1: | 59406243:59 | 59406435:59 | -:-:-:-:-:-:-:-: | 2266 | NA | 20 | 0 | 0 | 3 | 4 | 4 |
| ENSDARG000 | 1:1:1:1:1:1:1: | 37195465:37 | 37196657:37 | +:+:+:+:+:+:+: | 9529 | dclk2a | 944 | 1392 | 1244 | 1055 | 1004 | 1046 |
| ENSDARG000 | 1:1:1:1:1:1:1: | 53374454:53 | 53374556:53 | +:+:+:+:+:+:+: | 3098 | ucp1 | 2 | 4 | 323 | 311 | 333 | 15 |
| ENSDARG000 | 1:1:1:1:1:1:1: | 54889733:54 | 54889945:54 | +:+:+:+:+:+:+: | 2986 | zfyve27 | 1236 | 878 | 526 | 850 | 966 | 709 |
| ENSDARG000 | 1:1:1:1:1:1:1: | 26538446:26 | 26539436:26 | -:-:-:-:-:-:-:-: | 2781 | NA | 0 | 0 | 0 | 2 | 0 | 0 |
| ENSDARG000 | 1;1;1;1;1;1;1 | 8694196:869 | 8694546:869 | +;+;+;+;+;+;+ | 2146 | hapstr1b | 2668 | 2099 | 2251 | 2412 | 1810 | 2328 |
| ENSDARG000 | 1;1;1 | 12672979:12 | 12673219:12 | +;+;+ | 859 | NA | 2 | 0 | 0 | 0 | 0 | 0 |
| ENSDARG000 | 1:1:1:1:1:1:1: | 12675627:12 | 12676111:12 | -;-;-;-;-;-;-;- | 810 | NA | 0 | 3 | 0 | 0 | 2 | 1 |
| ENSDARG000 | 1;1;1;1;1;1;1 | 44340124:44 | 44340240:44 | +;+;+;+;+;+;+ | 1207 | NA | 0 | 0 | 0 | 0 | 0 | 0 |
| ENSDARG000 | 1;1;1;1;1;1;1 | 58290933:58 | 58291009:58 | +;+;+;+;+;+;+ | 2973 | NA | 10 | 4 | 7 | 6 | 30 | 9 |
| ENSDARG000 | 1:1:1:1:1:1:1: | 9972565:997 | 9973130:997 | -:-:-:-:-:-:-:-: | 1199 | NA | 0 | 0 | 4 | 0 | 6 | 0 |
| ENSDARG000 | 1;1;1;1;1 | 9966384:996 | 9966407:996 | +;+;+;+;+ | 679 | NA | 0 | 0 | 0 | 0 | 0 | 0 |
| ENSDARG000 | 1:1:1:1:1:1:1: | 40010103:40 | 40010170:40 | -:-:-:-:-:-:-:-: | 1585 | cdk2ap2 | 752 | 831 | 916 | 417 | 1076 | 565 |

| | | | | | | | | | | | | |
|---|---|---|---|---|---|---|---|---|---|---|---|---|
| ENSDARG000 | 1:1:1:1:1:1:1: | 21563311:215 | 21563444:215 | +:+:+:+:+:+:+: | 2022 | primpol | 52 | 281 | 364 | 110 | 289 | 162 |
| ENSDARG000 | 1;1 | 44347388:443 | 44347595:443 | +;+ | 793 | NA | 0 | 0 | 0 | 0 | 0 | 0 |
| ENSDARG000 | 1 | 30829997 | 30831196 | - | 1200 | NA | 2 | 0 | 1 | 0 | 0 | 1 |
| ENSDARG000 | 1 | 30840216 | 30841415 | - | 1200 | NA | 3 | 0 | 0 | 0 | 2 | 1 |
| ENSDARG000 | 1 | 30853969 | 30857925 | + | 3957 | NA | 0 | 0 | 38 | 0 | 130 | 0 |
| ENSDARG000 | 1:1:1:1:1:1:1: | 11670879:116 | 11671676:116 | -:-:-:-:-:-:-:-: | 2629 | brat1 | 96 | 232 | 244 | 230 | 198 | 142 |
| ENSDARG000 | 1:1:1:1:1:1:1: | 152007:15231 | 152405:15240 | -:-:-:-:-:-:-:-: | 2196 | pcid2 | 288 | 339 | 376 | 404 | 352 | 398 |
| ENSDARG000 | 1:1:1:1:1:1:1: | 26551034:265 | 26551248:265 | +:+:+:+:+:+:+: | 2291 | NA | 0 | 0 | 0 | 2 | 0 | 0 |
| ENSDARG000 | 1;1 | 30919336:309 | 30919670:309 | -;- | 831 | NA | 0 | 0 | 0 | 0 | 0 | 0 |
| ENSDARG000 | 1;1;1;1 | 9986675:9986 | 9986775:9987 | -;-;-;- | 625 | NA | 0 | 4 | 2 | 0 | 2 | 2 |
| ENSDARG000 | 1:1:1:1:1:1:1: | 19637312:196 | 19637599:196 | -:-:-:-:-:-:-:-: | 1908 | polr1e | 230 | 292 | 531 | 138 | 282 | 207 |
| ENSDARG000 | 1:1:1:1:1:1:1: | 45121393:451 | 45121637:451 | +:+:+:+:+:+:+: | 7790 | NA | 18 | 4 | 0 | 0 | 0 | 6 |
| ENSDARG000 | 1:1:1:1:1:1:1: | 8763429:8765 | 8767763:8767 | -:-:-:-:-:-:-:-: | 7880 | NA | 0 | 0 | 2 | 0 | 0 | 6 |
| ENSDARG000 | 1;1;1;1;1;1;1 | 58303892:583 | 58303976:583 | +;+;+;+;+;+;+ | 3112 | NA | 0 | 0 | 12 | 0 | 11 | 1 |
| ENSDARG000 | 1;1;1 | 59416186:594 | 59416232:594 | -;-;- | 363 | NA | 4 | 0 | 0 | 3 | 0 | 0 |
| ENSDARG000 | 1:1:1:1:1:1:1: | 25218177:252 | 25218415:252 | +:+:+:+:+:+:+: | 5410 | NA | 719 | 464 | 793 | 547 | 702 | 741 |
| ENSDARG000 | 1:1:1:1:1:1:1: | 40023640:400 | 40023856:400 | +:+:+:+:+:+:+: | 1679 | lgi2b | 214 | 274 | 184 | 223 | 93 | 424 |
| ENSDARG000 | 1;1;1;1;1;1;1 | 26573435:265 | 26573520:265 | -;-;-;-;-;-;- | 1050 | NA | 0 | 0 | 0 | 0 | 0 | 0 |
| ENSDARG000 | 1;1;1;1;1;1;1 | 44350628:443 | 44350744:443 | +;+;+;+;+;+;+ | 1207 | NA | 0 | 0 | 0 | 0 | 0 | 0 |
| ENSDARG000 | 1:1:1:1:1:1:1: | 29140884:291 | 29141340:291 | +:+:+:+:+:+:+: | 2478 | ankrd10a | 34 | 90 | 130 | 106 | 55 | 76 |
| ENSDARG000 | 1:1:1:1:1:1:1: | 6817292:6817 | 6817443:6817 | +:+:+:+:+:+:+: | 4613 | erbb4a | 0 | 0 | 12 | 0 | 4 | 0 |
| ENSDARG000 | 1:1:1:1:1:1:1: | 40034061:400 | 40034227:400 | +:+:+:+:+:+:+: | 2502 | ccdc149b | 136 | 127 | 284 | 413 | 143 | 144 |
| ENSDARG000 | 1;1;1;1;1;1;1 | 58461990:584 | 58462038:584 | +;+;+;+;+;+;+ | 3043 | NA | 7 | 0 | 4 | 0 | 0 | 0 |
| ENSDARG000 | 1;1;1;1;1 | 58393630:583 | 58395479:583 | +;+;+;+;+ | 2786 | NA | 12 | 3 | 1 | 4 | 28 | 12 |
| ENSDARG000 | 1:1:1:1:1:1:1: | 58442694:584 | 58442917:584 | +:+:+:+:+:+:+: | 3573 | zgc:194906 | 8 | 0 | 0 | 0 | 0 | 0 |
| ENSDARG000 | 1:1:1:1:1:1:1: | 12763461:127 | 12763970:127 | +:+:+:+:+:+:+: | 4553 | pcdh10a | 44422 | 22840 | 34050 | 26790 | 37055 | 32945 |
| ENSDARG000 | 1:1:1:1:1:1:1: | 11690360:116 | 11690420:116 | +:+:+:+:+:+:+: | 2007 | EIF3B | 0 | 0 | 10 | 4 | 30 | 2 |
| ENSDARG000 | 1:1:1:1:1:1:1: | 58375871:583 | 58375931:583 | +:+:+:+:+:+:+: | 1474 | NA | 0 | 0 | 0 | 0 | 0 | 0 |
| ENSDARG000 | 1:1:1:1:1:1:1: | 58353661:583 | 58353752:583 | +:+:+:+:+:+:+: | 3116 | NA | 8 | 5 | 10 | 13 | 3 | 1 |
| ENSDARG000 | 1:1:1:1:1:1:1: | 58470287:584 | 58470551:584 | +:+:+:+:+:+:+: | 3173 | NA | 3 | 2 | 0 | 5 | 31 | 0 |
| ENSDARG000 | 1:1:1:1:1:1:1: | 26577288:265 | 26577474:265 | +:+:+:+:+:+:+: | 2821 | NA | 0 | 0 | 0 | 4 | 0 | 0 |
| ENSDARG000 | 1:1:1:1:1:1:1: | 58400409:584 | 58400472:584 | +:+:+:+:+:+:+: | 1865 | NA | 0 | 0 | 0 | 0 | 0 | 0 |
| ENSDARG000 | 1:1:1:1:1:1:1: | 30946231:309 | 30946289:309 | +:+:+:+:+:+:+: | 1119 | metap1d | 91 | 105 | 226 | 42 | 123 | 194 |
| ENSDARG000 | 1;1;1;1;1 | 58456494:584 | 58458265:584 | +;+;+;+;+ | 2710 | NA | 0 | 4 | 0 | 0 | 1 | 0 |
| ENSDARG000 | 1;1;1;1;1 | 58323980:583 | 58325735:583 | +;+;+;+;+ | 2768 | NA | 0 | 0 | 0 | 0 | 0 | 0 |
| ENSDARG000 | 1;1;1 | 58370526:583 | 58370985:583 | +;+;+ | 808 | NA | 0 | 0 | 0 | 0 | 0 | 0 |
| ENSDARG000 | 1:1:1:1:1:1:1: | 58360935:583 | 58360998:583 | +:+:+:+:+:+:+: | 3363 | NA | 0 | 0 | 2 | 0 | 2 | 7 |
| ENSDARG000 | 1:1:1:1:1:1:1: | 52178916:521 | 52179178:521 | -;-;-;-;-;-;-;- | 1200 | rab3da | 14 | 22 | 42 | 6 | 18 | 22 |
| ENSDARG000 | 1:1:1:1:1:1:1: | 58312187:583 | 58312283:583 | +:+:+:+:+:+:+: | 4372 | NA | 0 | 0 | 0 | 6 | 16 | 4 |
| ENSDARG000 | 1:1:1:1:1:1:1: | 58424507:584 | 58424729:584 | +:+:+:+:+:+:+: | 2435 | NA | 0 | 0 | 0 | 0 | 0 | 0 |
| ENSDARG000 | 1:1:1:1:1:1:1: | 34243650:342 | 34243903:342 | +:+:+:+:+:+:+: | 5150 | SLC5A7 | 0 | 0 | 4 | 0 | 4 | 0 |
| ENSDARG000 | 1:1:1:1:1:1:1: | 45138905:451 | 45139254:451 | -:-:-:-:-:-:-:-: | 2064 | NA | 0 | 4 | 0 | 0 | 0 | 0 |
| ENSDARG000 | 1:1:1:1:1:1:1: | 9994811:9994 | 9994844:9994 | +:+:+:+:+:+:+: | 772 | NA | 2 | 20 | 18 | 12 | 155 | 80 |
| ENSDARG000 | 1:1:1:1:1:1:1: | 41465771:414 | 41465869:414 | +:+:+:+:+:+:+: | 1821 | paip2b | 2124 | 1929 | 2015 | 1474 | 1630 | 1734 |
| ENSDARG000 | 1:1:1:1:1:1:1: | 58332000:583 | 58332215:583 | +:+:+:+:+:+:+: | 3570 | NA | 0 | 0 | 0 | 0 | 0 | 2 |
| ENSDARG000 | 1;1 | 44357737:443 | 44357944:443 | +;+ | 793 | NA | 0 | 0 | 0 | 0 | 0 | 0 |
| ENSDARG000 | 1;1;1;1;1 | 59419086:594 | 59419124:594 | -;-;-;-;- | 618 | NA | 2 | 0 | 4 | 0 | 2 | 0 |
| ENSDARG000 | 1;1;1 | 8962624:8962 | 8963517:8963 | -;-;- | 964 | socs1b | 54 | 40 | 64 | 6 | 44 | 39 |
| ENSDARG000 | 1;1;1 | 8950874:8958 | 8950936:8958 | +;+;+ | 865 | NA | 0 | 0 | 0 | 0 | 0 | 2 |
| ENSDARG000 | 1:1:1:1:1:1:1: | 26605065:266 | 26605272:266 | +:+:+:+:+:+:+: | 3836 | coro2aa | 88 | 0 | 2 | 0 | 12 | 20 |
| ENSDARG000 | 1:1:1:1:1:1:1: | 40057412:400 | 40057860:400 | -;-;-;-;-;-;-;- | 2953 | NA | 487 | 304 | 561 | 404 | 623 | 326 |
| ENSDARG000 | 1;1;1;1 | 40095123:400 | 40095675:400 | -;-;-;- | 1137 | cntf | 28 | 36 | 43 | 13 | 36 | 10 |
| ENSDARG000 | 1:1:1:1:1:1:1: | 2301958:2301 | 2302137:2302 | +:+:+:+:+:+:+: | 5486 | FARP1 | 1484 | 902 | 1098 | 1990 | 2149 | 1827 |
| ENSDARG000 | 1:1:1:1:1:1:1: | 594584:59473 | 594772:59477 | +:+:+:+:+:+:+: | 1273 | jam2a | 62 | 10 | 20 | 30 | 24 | 27 |
| ENSDARG000 | 1;1;1 | 59520300:595 | 59520433:595 | +;+;+ | 874 | NA | 2 | 0 | 0 | 8 | 2 | 0 |
| ENSDARG000 | 1:1:1:1:1:1:1: | 56130222:561 | 56130482:561 | -:-:-:-:-:-:-:-: | 5069 | c3a.1 | 170 | 34 | 341 | 18 | 2 | 64 |
| ENSDARG000 | 1:1:1:1:1:1:1: | 53391345:533 | 53395913:533 | -:-:-:-:-:-:-:-: | 6018 | elmod2 | 678 | 650 | 1733 | 523 | 994 | 1500 |
| ENSDARG000 | 1 | 34295925 | 34297403 | + | 1479 | kctd12.2 | 300 | 455 | 128 | 50 | 114 | 74 |
| ENSDARG000 | 1;1;1;1 | 52203009:522 | 52203813:522 | -;-;-;- | 1711 | pld6 | 0 | 0 | 2 | 2 | 0 | 0 |
| ENSDARG000 | 1:1:1:1:1:1:1: | 50538839:505 | 50539657:505 | +:+:+:+:+:+:+: | 3336 | pkd2 | 398 | 279 | 404 | 230 | 512 | 257 |
| ENSDARG000 | 1;1;1 | 29167321:291 | 29167395:291 | -;-;- | 321 | NA | 0 | 0 | 0 | 0 | 0 | 0 |
| ENSDARG000 | 1;1;1;1;1;1;1 | 44360973:443 | 44361089:443 | +;+;+;+;+;+;+ | 1207 | NA | 0 | 0 | 0 | 0 | 0 | 0 |
| ENSDARG000 | 1:1:1:1:1:1:1: | 54908895:549 | 54909201:549 | +:+:+:+:+:+:+: | 3784 | golga7ba | 22 | 40 | 54 | 66 | 58 | 21 |
| ENSDARG000 | 1:1:1:1:1:1:1: | 45149844:451 | 45150307:451 | -:-:-:-:-:-:-:-: | 3626 | NA | 0 | 0 | 0 | 0 | 0 | 0 |
| ENSDARG000 | 1;1;1;1 | 2331637:2332 | 2332053:2332 | -;-;-;- | 703 | NA | 124 | 0 | 0 | 35 | 30 | 56 |
| ENSDARG000 | 1;1;1 | 34321574:343 | 34322149:343 | -;-;- | 833 | NA | 0 | 0 | 0 | 0 | 0 | 0 |
| ENSDARG000 | 1:1:1:1:1:1:1: | 12848061:128 | 12848109:128 | +:+:+:+:+:+:+: | 1038 | NA | 10 | 34 | 6 | 20 | 8 | 0 |
| ENSDARG000 | 1:1:1:1:1:1:1: | 10003193:100 | 10003553:100 | +:+:+:+:+:+:+: | 4090 | NA | 0 | 0 | 0 | 0 | 0 | 0 |
| ENSDARG000 | 1;1;1;1;1;1 | 19649545:196 | 19649652:196 | +;+;+;+;+;+ | 1635 | tmem192 | 251 | 375 | 518 | 334 | 398 | 362 |
| ENSDARG000 | 1;1 | 12931446:129 | 12931676:129 | +;+ | 1356 | NA | 120 | 28 | 32 | 4 | 54 | 6 |
| ENSDARG000 | 1 | 13002978 | 13003067 | + | 90 | NA | 0 | 0 | 0 | 0 | 0 | 0 |
| ENSDARG000 | 1;1;1;1;1;1;1 | 52214895:522 | 52215107:522 | -;-;-;-;-;-;- | 1010 | cnn1a | 0 | 0 | 0 | 0 | 0 | 0 |
| ENSDARG000 | 1;1;1;1;1 | 13077467:131 | 13077629:131 | -;-;-;-;- | 892 | NA | 0 | 0 | 0 | 0 | 0 | 0 |
| ENSDARG000 | 1;1 | 13087797:130 | 13088155:130 | +;+ | 1020 | NA | 0 | 41 | 2 | 6 | 4 | 2 |
| ENSDARG000 | 1;1;1;1 | 8974500:8975 | 8975196:8975 | -;-;-;- | 973 | NA | 80 | 93 | 68 | 78 | 77 | 80 |
| ENSDARG000 | 1:1:1:1:1:1:1: | 44395976:443 | 44396211:443 | +:+:+:+:+:+:+: | 2793 | unc93b1 | 56 | 57 | 85 | 118 | 166 | 149 |
| ENSDARG000 | 1;1 | 44390470:443 | 44390705:443 | +;+ | 834 | timm10 | 234 | 295 | 308 | 93 | 267 | 143 |
| ENSDARG000 | 1;1 | 29174195:291 | 29174307:291 | +;+ | 626 | NA | 0 | 0 | 0 | 0 | 0 | 0 |
| ENSDARG000 | 1:1:1:1:1:1:1: | 29178117:291 | 29178490:291 | +:+:+:+:+:+:+: | 1425 | zgc:173948 | 0 | 4 | 0 | 22 | 0 | 0 |
| ENSDARG000 | 1:1:1:1:1:1:1: | 59525648:595 | 59525873:595 | -:-:-:-:-:-:-:-: | 1721 | NA | 18 | 10 | 0 | 14 | 4 | 0 |
| ENSDARG000 | 1:1:1:1:1:1:1: | 11711226:117 | 11711531:117 | +:+:+:+:+:+:+: | 3142 | NA | 5 | 6 | 10 | 0 | 0 | 10 |
| ENSDARG000 | 1:1:1:1:1:1:1: | 29174504:291 | 29174765:291 | -:-:-:-:-:-:-:-: | 679 | NA | 0 | 0 | 0 | 0 | 0 | 0 |
| ENSDARG000 | 1;1 | 45170046:451 | 45170507:451 | +;+ | 515 | NA | 0 | 0 | 0 | 0 | 0 | 0 |
| ENSDARG000 | 1:1:1:1:1:1:1: | 54934522:549 | 54934878:549 | -:-:-:-:-:-:-:-: | 2178 | crtac1a | 12 | 2 | 70 | 23 | 92 | 26 |
| ENSDARG000 | 1:1:1:1:1:1:1: | 29183962:291 | 29184431:291 | +:+:+:+:+:+:+: | 2310 | NA | 198 | 241 | 317 | 140 | 282 | 266 |
| ENSDARG000 | 1:1:1:1:1:1:1: | 41478345:414 | 41478504:414 | +:+:+:+:+:+:+: | 1802 | NA | 20 | 12 | 0 | 0 | 2 | 10 |
| ENSDARG000 | 1:1:1:1:1:1:1: | 46493944:465 | 46494136:465 | +:+:+:+:+:+:+: | 4198 | MCF2L | 196 | 165 | 738 | 191 | 287 | 140 |
| ENSDARG000 | 1;1 | 8978908:8980 | 8979301:8980 | -;- | 647 | NA | 0 | 28 | 0 | 6 | 0 | 0 |
| ENSDARG000 | 1:1:1:1:1:1:1: | 21576674:215 | 21577016:215 | -:-:-:-:-:-:-:-: | 2226 | NA | 268 | 78 | 110 | 38 | 60 | 108 |
| ENSDARG000 | 1;1 | 52264131:522 | 52264749:522 | -;- | 1039 | NA | 16 | 2 | 0 | 10 | 0 | 0 |
| ENSDARG000 | 1;1;1;1;1 | 19667947:196 | 19668315:196 | -;-;-;-;- | 868 | apela | 0 | 0 | 12 | 2 | 2 | 17 |
| ENSDARG000 | 1:1:1:1:1:1:1: | 14497458:144 | 14502104:145 | -;-;-;-;-;-;-;- | 5150 | NA | 153 | 176 | 229 | 279 | 159 | 418 |
| ENSDARG000 | 1;1;1 | 30977936:309 | 30978808:309 | -;-;- | 1521 | dlx2b | 4 | 0 | 10 | 0 | 0 | 0 |
| ENSDARG000 | 1;1;1 | 52392511:523 | 52392568:523 | +;+;+ | 641 | NA | 12 | 0 | 0 | 0 | 0 | 0 |
| ENSDARG000 | 1:1:1:1:1:1:1: | 18642117:186 | 18642130:186 | -:-:-:-:-:-:-:-: | 7032 | TBC1D1 | 2433 | 2060 | 2477 | 2434 | 3050 | 2055 |
| ENSDARG000 | 1:1:1:1:1:1:1: | 16073887:161 | 16074007:161 | +:+:+:+:+:+:+: | 1137 | tusc3 | 327 | 546 | 693 | 578 | 526 | 478 |
| ENSDARG000 | 1;1;1 | 8984068:8988 | 8984156:8988 | +;+;+ | 2457 | tlr2 | 0 | 19 | 12 | 0 | 27 | 2 |
| ENSDARG000 | 1:1:1:1:1:1:1: | 34365845:343 | 34367324:343 | -:-:-:-:-:-:-:-: | 6560 | lmo7b | 12 | 28 | 36 | 4 | 32 | 14 |
| ENSDARG000 | 1:1:1:1:1:1:1: | 26660369:266 | 26660990:266 | -:-:-:-:-:-:-:-: | 2149 | anp32b | 2239 | 4194 | 4638 | 2145 | 3397 | 2899 |
| ENSDARG000 | 1;1 | 40107811:401 | 40107990:401 | +;+ | 3802 | NA | 218 | 206 | 231 | 278 | 260 | 316 |
| ENSDARG000 | 1:1:1:1:1:1:1: | 10037444:100 | 10038040:100 | -:-:-:-:-:-:-:-: | 1556 | rnf175 | 369 | 290 | 382 | 266 | 589 | 454 |
| ENSDARG000 | 1;1;1;1;1;1 | 52398205:523 | 52398779:523 | +;+;+;+;+;+ | 2036 | NA | 64 | 5 | 0 | 8 | 16 | 6 |
| ENSDARG000 | 1;1;1;1;1;1 | 19689749:196 | 19691618:196 | -;-;-;-;-;- | 2288 | NA | 0 | 0 | 0 | 0 | 0 | 0 |
| ENSDARG000 | 1:1:1:1:1:1:1: | 19708508:197 | 19708600:197 | +:+:+:+:+:+:+: | 1634 | NA | 0 | 0 | 0 | 0 | 0 | 0 |
| ENSDARG000 | 1:1:1:1:1:1:1: | 157793:15803 | 158136:15813 | +:+:+:+:+:+:+: | 2146 | cul4a | 1400 | 1096 | 1297 | 1467 | 1408 | 886 |
| ENSDARG000 | 1:1:1:1:1:1:1: | 610082:61032 | 610784:61131 | -:-:-:-:-:-:-:-: | 1542 | atp5pf | 1784 | 1679 | 1899 | 1300 | 1642 | 1328 |
| ENSDARG000 | 1:1:1:1:1:1:1: | 54948059:549 | 54948151:549 | +:+:+:+:+:+:+: | 1704 | NA | 540 | 532 | 291 | 352 | 492 | 247 |
| ENSDARG000 | 1;1;1 | 16393077:163 | 16393368:163 | -;-;- | 1107 | fgf20a | 0 | 0 | 0 | 0 | 0 | 2 |
| ENSDARG000 | 1;1;1 | 59533317:595 | 59533341:595 | +;+;+ | 1169 | sp6 | 8 | 0 | 0 | 0 | 12 | 0 |
| ENSDARG000 | 1:1:1:1:1:1:1: | 41487339:414 | 41487821:414 | +:+:+:+:+:+:+: | 3640 | NA | 267 | 267 | 278 | 219 | 344 | 248 |
| ENSDARG000 | 1;1;1 | 59542001:595 | 59542292:595 | -;-;- | 585 | NA | 9 | 0 | 0 | 0 | 8 | 2 |
| ENSDARG000 | 1:1:1:1:1:1:1: | 45173421:451 | 45174042:451 | -:-:-:-:-:-:-:-: | 2242 | zgc:111983 | 705 | 24 | 12 | 136 | 90 | 95 |
| ENSDARG000 | 1 | 11736420 | 11737595 | + | 1176 | NA | 0 | 0 | 2 | 0 | 0 | 0 |
| ENSDARG000 | 1;1;1;1;1;1 | 44421920:444 | 44423008:444 | -;-;-;-;-;- | 1717 | cryba1l2 | 0 | 0 | 0 | 0 | 0 | 0 |
| ENSDARG000 | 1;1 | 52429810:524 | 52430515:524 | -;- | 777 | zgc:194101 | 4 | 22 | 28 | 0 | 16 | 20 |
| ENSDARG000 | 1:1:1:1:1:1:1: | 50566256:505 | 50566529:505 | -:-:-:-:-:-:-:-: | 2116 | abcg2d | 0 | 2 | 0 | 6 | 0 | 0 |
| ENSDARG000 | 1:1:1:1:1:1:1: | 16396870:163 | 16397333:163 | +:+:+:+:+:+:+: | 5016 | MICU3 | 0 | 46 | 46 | 30 | 112 | 44 |
| ENSDARG000 | 1:1:1:1:1:1:1: | 54960189:549 | 54961109:549 | -:-:-:-:-:-:-:-: | 3162 | khsrp | 1383 | 2336 | 1508 | 741 | 1834 | 829 |
| ENSDARG000 | 1:1:1:1:1:1:1: | 31019165:310 | 31019414:310 | +:+:+:+:+:+:+: | 4759 | itga6b | 2654 | 766 | 2382 | 456 | 3119 | 1866 |
| ENSDARG000 | 1:1:1:1:1:1:1: | 40116572:401 | 40117792:401 | -:-:-:-:-:-:-:-: | 3051 | NA | 722 | 419 | 807 | 509 | 708 | 591 |
| ENSDARG000 | 1;1;1 | 6957096:6961 | 6957324:6961 | +;+;+ | 402 | NA | 0 | 0 | 0 | 0 | 0 | 0 |
| ENSDARG000 | 1:1:1:1:1:1:1: | 53454741:534 | 53455284:534 | -:-:-:-:-:-:-:-: | 2166 | zgc:66455 | 8 | 0 | 26 | 0 | 0 | 0 |
| ENSDARG000 | 1 | 7063837 | 7063943 | + | 107 | NA | 0 | 0 | 0 | 0 | 0 | 0 |
| ENSDARG000 | 1:1:1:1:1:1:1: | 25322445:253 | 25323240:253 | -:-:-:-:-:-:-:-: | 3089 | trim2a | 58 | 64 | 76 | 54 | 98 | 61 |
| ENSDARG000 | 1:1:1:1:1:1:1: | 11781544:117 | 11785937:117 | -:-:-:-:-:-:-:-: | 6346 | ttyh3b | 1024 | 1234 | 1423 | 426 | 1323 | 385 |
| ENSDARG000 | 1:1:1:1:1:1:1: | 7189856:7320 | 7190351:7321 | +:+:+:+:+:+:+: | 5307 | ERBB4 | 0 | 0 | 4 | 0 | 0 | 0 |
| ENSDARG000 | 1:1:1:1:1:1:1: | 37312580:373 | 37312888:373 | +:+:+:+:+:+:+: | 5536 | fam193a | 1046 | 953 | 1112 | 723 | 1179 | 876 |
| ENSDARG000 | 1 | 16460124 | 16461398 | - | 1275 | NA | 16 | 32 | 13 | 8 | 11 | 2 |
| ENSDARG000 | 1:1:1:1:1:1:1: | 19759370:197 | 19760938:197 | -:-:-:-:-:-:-:-: | 2173 | tma16 | 170 | 164 | 326 | 133 | 186 | 116 |
| ENSDARG000 | 1:1:1:1:1:1:1: | 41498188:414 | 41498485:414 | +:+:+:+:+:+:+: | 4404 | NA | 326 | 163 | 408 | 262 | 251 | 197 |
| ENSDARG000 | 1:1:1:1:1:1:1: | 9004719:9004 | 9005007:9005 | +:+:+:+:+:+:+: | 3024 | prkcba | 82 | 105 | 26 | 16 | 90 | 53 |
| ENSDARG000 | 1:1:1:1:1:1:1: | 26667872:266 | 26667984:266 | +:+:+:+:+:+:+: | 2440 | NA | 102 | 57 | 210 | 42 | 360 | 108 |
| ENSDARG000 | 1:1:1:1:1:1:1: | 21610471:216 | 21612542:216 | -;-;-;-;-;-;-;- | 3393 | zcchc7 | 268 | 282 | 444 | 207 | 395 | 279 |
| ENSDARG000 | 1:1:1:1:1:1:1: | 49435017:494 | 49435128:494 | +:+:+:+:+:+:+: | 5909 | pdcd11 | 3474 | 2637 | 4877 | 1533 | 3282 | 2268 |
| ENSDARG000 | 1;1;1;1;1 | 2446996:2447 | 2448591:2448 | -;-;-;-;- | 1810 | ggact.3 | 0 | 0 | 0 | 0 | 0 | 291 |
| ENSDARG000 | 1;1 | 2450140:2453 | 2452269:2453 | -;- | 2240 | ggact.2 | 0 | 0 | 0 | 0 | 0 | 200 |
| ENSDARG000 | 1:1:1:1:1:1:1: | 26676758:266 | 26676873:266 | +:+:+:+:+:+:+: | 5275 | NA | 395 | 1148 | 1188 | 556 | 815 | 518 |
| ENSDARG000 | 1:1:1:1:1:1:1: | 19764995:197 | 19765136:197 | +:+:+:+:+:+:+: | 2744 | fnip2 | 1865 | 1263 | 1626 | 1906 | 1692 | 1908 |
| ENSDARG000 | 1;1;1;1;1;1;1 | 41588071:415 | 41588285:415 | +;+;+;+;+;+;+ | 1052 | NA | 10 | 0 | 0 | 3 | 14 | 0 |
| ENSDARG000 | 1;1 | 19787281:197 | 19788005:197 | -;- | 827 | NA | 12 | 2 | 5 | 0 | 9 | 3 |
| ENSDARG000 | 1:1:1:1:1:1:1: | 26670828:266 | 26671206:266 | -:-:-:-:-:-:-:-: | 1736 | trmo | 54 | 69 | 123 | 39 | 70 | 60 |
| ENSDARG000 | 1;1 | 52433847:524 | 52434711:524 | -;- | 964 | NA | 10 | 13 | 34 | 4 | 192 | 54 |
| ENSDARG000 | 1;1 | 56180416:561 | 56180571:561 | +;+ | 1336 | crb3b | 0 | 0 | 6 | 0 | 14 | 14 |
| ENSDARG000 | 1:1:1:1:1:1:1: | 44439661:444 | 44439709:444 | +:+:+:+:+:+:+: | 2694 | crybb1l2 | 0 | 0 | 0 | 0 | 0 | 0 |
| ENSDARG000 | 1;1;1;1 | 40140498:401 | 40140554:401 | +;+;+;+ | 2803 | zgc:152658 | 172 | 313 | 34 | 0 | 16 | 42 |
| ENSDARG000 | 1;1;1;1;1;1;1 | 40148193:401 | 40148243:401 | +;+;+;+;+;+;+ | 1509 | zgc:194443 | 25 | 23 | 12 | 0 | 0 | 0 |
| ENSDARG000 | 1:1:1:1:1:1:1: | 23297623:232 | 23299086:232 | -:-:-:-:-:-:-:-: | 2287 | smim14 | 650 | 212 | 292 | 458 | 454 | 486 |
| ENSDARG000 | 1;1;1;1;1;1;1 | 19808408:198 | 19808614:198 | +;+;+;+;+;+;+ | 2709 | NA | 74 | 77 | 76 | 52 | 94 | 30 |
| ENSDARG000 | 1 | 53475074 | 53476758 | - | 1685 | b3gnt2a | 16 | 10 | 53 | 6 | 40 | 9 |
| ENSDARG000 | 1:1:1:1:1:1:1: | 53490669:534 | 53490724:534 | -:-:-:-:-:-:-:-: | 1793 | commd1 | 399 | 261 | 787 | 380 | 412 | 659 |
| ENSDARG000 | 1;1 | 52441080:524 | 52441921:524 | -;- | 962 | NA | 341 | 265 | 201 | 80 | 279 | 178 |
| ENSDARG000 | 1:1:1:1:1:1:1: | 54974191:549 | 54978741:549 | -:-:-:-:-:-:-:-: | 6252 | NA | 178 | 121 | 100 | 62 | 104 | 76 |
| ENSDARG000 | 1:1:1:1:1:1:1: | 16463079:164 | 16463523:164 | +:+:+:+:+:+:+: | 2077 | vps37a | 476 | 385 | 555 | 543 | 474 | 358 |
| ENSDARG000 | 1;1;1;1;1 | 55007272:550 | 55008468:550 | -;-;-;-;- | 1528 | zgc:136864 | 0 | 0 | 0 | 0 | 0 | 0 |
| ENSDARG000 | 1:1:1:1:1:1:1: | 56188094:561 | 56189076:561 | -:-:-:-:-:-:-:-: | 3656 | NA | 18 | 18 | 4 | 6 | 25 | 66 |
| ENSDARG000 | 1;1;1;1;1 | 55002583:550 | 55002660:550 | +;+;+;+;+ | 517 | NA | 0 | 0 | 0 | 0 | 0 | 0 |
| ENSDARG000 | 1;1 | 52446149:524 | 52447142:524 | -;- | 1071 | si:ch211-217 | 28 | 4 | 0 | 6 | 6 | 0 |
| ENSDARG000 | 1:1:1:1:1:1:1: | 55006945:550 | 55011802:550 | -:-:-:-:-:-:-:-: | 8053 | vps54 | 613 | 516 | 628 | 343 | 577 | 389 |
| ENSDARG000 | 1:1:1:1:1:1:1: | 25399514:254 | 25403507:254 | -:-:-:-:-:-:-:-: | 6086 | NA | 4 | 0 | 0 | 0 | 2 | 8 |
| ENSDARG000 | 1;1;1;1 | 52452010:524 | 52452367:524 | -;-;-;- | 1212 | ccdc86 | 505 | 702 | 1606 | 510 | 704 | 1089 |
| ENSDARG000 | 1:1:1:1:1:1:1: | 7517449:7517 | 7517496:7517 | +:+:+:+:+:+:+: | 3116 | lancl1 | 536 | 545 | 822 | 662 | 755 | 755 |
| ENSDARG000 | 1;1;1;1;1;1 | 37372919:373 | 37373502:373 | -;-;-;-;-;- | 1753 | NA | 48 | 111 | 198 | 40 | 114 | 137 |

| | | | | | | | | | | | | |
|---|---|---|---|---|---|---|---|---|---|---|---|---|
| ENSDARG000 | 1:1:1:1:1:1:1:1: | 11854710:118 | 11855897:118 | -:-:-:-:-:-:-:-: | 3497 | iqce | 108 | 152 | 226 | 56 | 190 | 157 |
| ENSDARG000 | 1:1:1:1:1:1:1:1: | 46598502:465 | 46599078:465 | +:+:+:+:+:+:+:+: | 3359 | cab39l | 1857 | 1821 | 2353 | 1833 | 2442 | 2398 |
| ENSDARG000 | 1:1:1:1:1:1:1:1: | 7556965:7556 | 7557049:7557 | +:+:+:+:+:+:+:+: | 1368 | pcnp | 498 | 171 | 706 | 518 | 637 | 606 |
| ENSDARG000 | 1:1:1:1:1:1:1:1: | 41596099:415 | 41596451:415 | +:+:+:+:+:+:+:+: | 2182 | NA | 196 | 138 | 176 | 148 | 176 | 186 |
| ENSDARG000 | 1:1:1:1:1:1:1:1: | 46617006:466 | 46617601:466 | -:-:-:-:-:-:-:-: | 2697 | cdadc1 | 242 | 142 | 170 | 158 | 133 | 175 |
| ENSDARG000 | 1:1:1:1:1:1:1:1: | 50594175:505 | 50595973:505 | -:-:-:-:-:-:-:-: | 3179 | ppm1kb | 607 | 537 | 817 | 1118 | 464 | 913 |
| ENSDARG000 | 1;1 | 2454094:2457 | 2455779:2457 | -;- | 1790 | ggact.1 | 0 | 0 | 0 | 0 | 0 | 120 |
| ENSDARG000 | 1:1:1:1:1:1:1:1: | 617620:62099 | 621475:62147 | -:-:-:-:-:-:-:-: | 6084 | appa | 10662 | 6627 | 9880 | 8499 | 10966 | 10157 |
| ENSDARG000 | 1:1:1:1:1:1:1:1: | 45205920:452 | 45206180:452 | -:-:-:-:-:-:-:-: | 1745 | ddx39aa | 1614 | 2032 | 2718 | 1322 | 2297 | 1676 |
| ENSDARG000 | 1;1;1;1;1;1 | 9082417:9086 | 9082627:9086 | +;+;+;+;+;+ | 2016 | NA | 0 | 0 | 0 | 0 | 0 | 0 |
| ENSDARG000 | 1:1:1:1:1:1:1:1: | 53515089:535 | 53515174:535 | +:+:+:+:+:+:+:+: | 2443 | NA | 0 | 2 | 0 | 0 | 0 | 0 |
| ENSDARG000 | 1;1 | 21721651:217 | 21722008:217 | -;- | 566 | NA | 0 | 0 | 0 | 0 | 0 | 0 |
| ENSDARG000 | 1:1:1:1:1:1:1:1: | 9100447:9100 | 9100607:9100 | -:-:-:-:-:-:-:-: | 3815 | NA | 57 | 12 | 53 | 0 | 3 | 32 |
| ENSDARG000 | 1:1:1:1:1:1:1:1: | 37379063:373 | 37381198:373 | -:-:-:-:-:-:-:-: | 3071 | scpp1 | 0 | 6 | 8 | 6 | 0 | 0 |
| ENSDARG000 | 1;1 | 29211540:292 | 29214788:292 | +;+ | 4190 | irs2a | 6 | 333 | 9 | 1234 | 1 | 294 |
| ENSDARG000 | 1;1 | 9113630:9114 | 9113748:9115 | -;- | 377 | NA | 0 | 0 | 0 | 0 | 0 | 0 |
| ENSDARG000 | 1;1;1;1;1 | 9105039:9106 | 9105081:9107 | -;-;-;-;- | 540 | NA | 0 | 0 | 0 | 0 | 0 | 0 |
| ENSDARG000 | 1:1:1:1:1:1:1:1: | 7538909:7540 | 7538937:7540 | +:+:+:+:+:+:+:+: | 1432 | mylz3 | 116 | 124 | 107 | 162 | 141 | 107 |
| ENSDARG000 | 1:1:1:1:1:1:1:1: | 40156907:401 | 40157135:401 | +:+:+:+:+:+:+:+: | 3624 | NA | 38 | 40 | 15 | 10 | 28 | 31 |
| ENSDARG000 | 1:1:1:1:1:1:1:1: | 21725821:217 | 21726350:217 | +:+:+:+:+:+:+:+: | 1718 | pax5 | 0 | 0 | 0 | 0 | 10 | 2 |
| ENSDARG000 | 1;1 | 2704604:2714 | 2705226:2714 | -;- | 662 | NA | 0 | 0 | 0 | 0 | 0 | 0 |
| ENSDARG000 | 1:1:1:1:1:1:1:1: | 19835803:198 | 19837114:198 | -:-:-:-:-:-:-:-: | 2728 | grhprb | 0 | 0 | 6 | 6 | 14 | 0 |
| ENSDARG000 | 1;1 | 4237208:4250 | 4237621:4250 | -;- | 504 | NA | 0 | 0 | 0 | 0 | 0 | 0 |
| ENSDARG000 | 1:1:1:1:1:1:1:1: | 52462068:524 | 52462200:524 | +:+:+:+:+:+:+:+: | 2321 | glb1 | 343 | 412 | 340 | 234 | 541 | 363 |
| ENSDARG000 | 1;1 | 29267841:292 | 29268052:292 | +;+ | 3262 | lig4 | 172 | 213 | 248 | 109 | 260 | 266 |
| ENSDARG000 | 1;1;1;1 | 56217388:562 | 56218601:562 | -;-;-;- | 1624 | zgc:65894 | 0 | 2 | 4 | 8 | 0 | 28 |
| ENSDARG000 | 1:1:1:1:1:1:1:1: | 11881559:118 | 11881641:118 | +:+:+:+:+:+:+:+: | 2753 | SNX8 | 4 | 0 | 10 | 0 | 0 | 0 |
| ENSDARG000 | 1;1;1 | 5273481:5275 | 5273957:5277 | +;+;+ | 2297 | spry2 | 3401 | 2736 | 4055 | 3476 | 3549 | 3568 |
| ENSDARG000 | 1:1:1:1:1:1:1:1: | 53657337:536 | 53657483:536 | -:-:-:-:-:-:-:-: | 3775 | XPO1 | 1387 | 1679 | 2882 | 1229 | 2852 | 1860 |
| ENSDARG000 | 1:1:1:1:1:1:1:1: | 41609254:416 | 41609316:416 | +:+:+:+:+:+:+:+: | 6728 | mogs | 1134 | 1174 | 1397 | 566 | 1353 | 861 |
| ENSDARG000 | 1:1:1:1:1:1:1:1: | 37391141:373 | 37391254:373 | +:+:+:+:+:+:+:+: | 4360 | sparcl1 | 0 | 2 | 2 | 0 | 2 | 10 |
| ENSDARG000 | 1:1:1:1:1:1:1:1: | 10051763:100 | 10051907:100 | +:+:+:+:+:+:+:+: | 2259 | fgb | 0 | 0 | 64 | 0 | 0 | 0 |
| ENSDARG000 | 1:1:1:1:1:1:1:1: | 23311617:233 | 23312134:233 | +:+:+:+:+:+:+:+: | 2673 | ube2kb | 1038 | 1293 | 1206 | 847 | 1398 | 1346 |
| ENSDARG000 | 1:1:1:1:1:1:1:1: | 16484211:164 | 16484740:164 | -:-:-:-:-:-:-:-: | 2593 | mtmr7b | 14 | 14 | 33 | 15 | 12 | 42 |
| ENSDARG000 | 1:1:1:1:1:1:1:1: | 55044658:550 | 55049554:550 | -:-:-:-:-:-:-:-: | 5919 | NA | 92 | 36 | 21 | 6 | 88 | 24 |
| ENSDARG000 | 1;1;1 | 26701610:267 | 26702992:267 | -;-;- | 1383 | NA | 0 | 0 | 0 | 0 | 0 | 0 |
| ENSDARG000 | 1:1:1:1:1:1:1:1: | 5354489:5363 | 5359003:5363 | -:-:-:-:-:-:-:-: | 5475 | ndfip2 | 3614 | 3059 | 3178 | 2749 | 3620 | 3459 |
| ENSDARG000 | 1;1;1;1;1 | 44491077:444 | 44491237:444 | +;+;+;+;+ | 3980 | rtn4rl2a | 12 | 20 | 4 | 0 | 22 | 24 |
| ENSDARG000 | 1:1:1:1:1:1:1:1: | 50613868:506 | 50614233:506 | +:+:+:+:+:+:+:+: | 3639 | NA | 455 | 560 | 106 | 87 | 173 | 134 |
| ENSDARG000 | 1;1 | 29311527:293 | 29312024:293 | +;+ | 777 | NA | 0 | 0 | 0 | 0 | 0 | 0 |
| ENSDARG000 | 1;1 | 29323374:293 | 29323561:293 | -;- | 655 | NA | 0 | 0 | 0 | 1 | 0 | 0 |
| ENSDARG000 | 1;1;1 | 29281764:293 | 29282980:293 | +;+;+ | 2844 | NALF1 | 0 | 0 | 0 | 0 | 0 | 0 |
| ENSDARG000 | 1;1 | 1313605:1315 | 1313691:1315 | -;- | 244 | NA | 0 | 0 | 0 | 0 | 0 | 0 |
| ENSDARG000 | 1;1;1;1;1 | 1599979:1606 | 1600208:1607 | +;+;+;+;+ | 1060 | urp2 | 0 | 0 | 0 | 0 | 12 | 0 |
| ENSDARG000 | 1:1:1:1:1:1:1:1: | 34496855:345 | 34497067:345 | +:+:+:+:+:+:+:+: | 5144 | klf12a | 296 | 344 | 282 | 236 | 172 | 147 |
| ENSDARG000 | 1;1 | 40186407:401 | 40186748:401 | -;- | 424 | NA | 0 | 0 | 0 | 0 | 0 | 0 |
| ENSDARG000 | 1;1;1;1;1 | 40199074:401 | 40199171:401 | +;+;+;+;+ | 1516 | si:ch211-113e | 4 | 0 | 4 | 0 | 0 | 0 |
| ENSDARG000 | 1:1:1:1:1:1:1:1: | 31071708:310 | 31072889:310 | -:-:-:-:-:-:-:-: | 5692 | NA | 190 | 86 | 106 | 58 | 34 | 163 |
| ENSDARG000 | 1:1:1:1:1:1:1:1: | 41666611:416 | 41666635:416 | +:+:+:+:+:+:+:+: | 3071 | FBXO41 | 32 | 112 | 53 | 34 | 34 | 108 |
| ENSDARG000 | 1;1 | 44534625:445 | 44534735:445 | -;- | 448 | NA | 0 | 0 | 0 | 0 | 0 | 0 |
| ENSDARG000 | 1:1:1:1:1:1:1:1: | 25442699:254 | 25443222:254 | -:-:-:-:-:-:-:-: | 2605 | arfip1 | 762 | 814 | 1190 | 835 | 1009 | 1052 |
| ENSDARG000 | 1:1:1:1:1:1:1:1: | 26740718:267 | 26741543:267 | -:-:-:-:-:-:-:-: | 1881 | sh3gl2a | 12 | 82 | 10 | 42 | 8 | 38 |
| ENSDARG000 | 1:1:1:1:1:1:1:1: | 44523516:445 | 44523899:445 | +:+:+:+:+:+:+:+: | 5682 | zdhhc5a | 1281 | 1097 | 1396 | 943 | 1198 | 1292 |
| ENSDARG000 | 1:1:1:1:1:1:1:1: | 40210840:402 | 40212459:402 | -:-:-:-:-:-:-:-: | 5499 | TESK1 | 153 | 82 | 111 | 52 | 137 | 109 |
| ENSDARG000 | 1:1:1:1:1:1:1:1: | 34593490:346 | 34597865:346 | -:-:-:-:-:-:-:-: | 11484 | KLF5 | 42 | 5 | 0 | 6 | 16 | 8 |
| ENSDARG000 | 1;1;1;1;1;1;1 | 9118100:9118 | 9118340:9118 | -;-;-;-;-;-;- | 918 | ndufab1a | 636 | 815 | 807 | 424 | 662 | 530 |
| ENSDARG000 | 1:1:1:1:1:1:1:1: | 45217425:452 | 45217807:452 | +:+:+:+:+:+:+:+: | 3027 | EVI5L | 151 | 57 | 92 | 95 | 184 | 98 |
| ENSDARG000 | 1:1:1:1:1:1:1:1: | 34616054:346 | 34616273:346 | -:-:-:-:-:-:-:-: | 2563 | pibf1 | 0 | 0 | 0 | 0 | 0 | 0 |
| ENSDARG000 | 1:1:1:1:1:1:1:1: | 55080797:550 | 55080926:550 | +:+:+:+:+:+:+:+: | 3020 | lgalsla | 495 | 501 | 104 | 168 | 272 | 60 |
| ENSDARG000 | 1:1:1:1:1:1:1:1: | 37697373:376 | 37697593:376 | -:-:-:-:-:-:-:-: | 781 | NA | 0 | 0 | 1 | 0 | 0 | 2 |
| ENSDARG000 | 1:1:1:1:1:1:1:1: | 1618994:1619 | 1619314:1619 | -:-:-:-:-:-:-:-: | 920 | NA | 6 | 2 | 18 | 8 | 36 | 32 |
| ENSDARG000 | 1:1:1:1:1:1:1:1: | 49481203:494 | 49484221:494 | -:-:-:-:-:-:-:-: | 4059 | zgc:175214 | 1411 | 574 | 755 | 792 | 844 | 1245 |
| ENSDARG000 | 1:1:1:1:1:1:1:1: | 45316063:453 | 45316611:453 | -;-;-;-;-;-;-;- | 1311 | NA | 2 | 2 | 0 | 6 | 0 | 28 |
| ENSDARG000 | 1;1;1 | 29444452:294 | 29444519:294 | +;+;+ | 517 | NA | 0 | 0 | 0 | 0 | 0 | 0 |
| ENSDARG000 | 1;1;1 | 29451465:294 | 29451534:294 | +;+;+ | 517 | NA | 0 | 0 | 0 | 0 | 0 | 0 |
| ENSDARG000 | 1:1:1:1:1:1:1:1: | 16548733:165 | 16548964:165 | +:+:+:+:+:+:+:+: | 5336 | NA | 162 | 272 | 335 | 179 | 502 | 186 |
| ENSDARG000 | 1;1;1;1;1 | 52481332:524 | 52481361:524 | +;+;+;+;+ | 1797 | cldnd1b | 723 | 776 | 801 | 887 | 667 | 671 |
| ENSDARG000 | 1;1;1;1;1;1;1 | 53686995:536 | 53687675:536 | -;-;-;-;-;-;- | 2775 | FAM161A | 2 | 2 | 16 | 16 | 8 | 40 |
| ENSDARG000 | 1;1;1;1;1 | 49501171:495 | 49501242:495 | -;-;-;-;- | 1011 | CALHM1 | 0 | 6 | 0 | 0 | 0 | 0 |
| ENSDARG000 | 1:1:1:1:1:1:1:1: | 21861387:218 | 21861891:218 | -:-:-:-:-:-:-:-: | 2730 | melk | 46 | 185 | 226 | 82 | 197 | 83 |
| ENSDARG000 | 1:1:1:1:1:1:1:1: | 41770842:417 | 41771796:417 | -:-:-:-:-:-:-:-: | 2046 | rnf24 | 151 | 261 | 452 | 446 | 323 | 92 |
| ENSDARG000 | 1:1:1:1:1:1:1:1: | 10063501:100 | 10063575:100 | -:-:-:-:-:-:-:-: | 3142 | fga | 0 | 0 | 83 | 2 | 0 | 0 |
| ENSDARG000 | 1;1 | 29507647:295 | 29508154:295 | +;+ | 615 | NA | 0 | 0 | 0 | 0 | 1 | 0 |
| ENSDARG000 | 1;1;1;1 | 29515160:295 | 29515312:295 | +;+;+;+ | 1251 | NA | 0 | 4 | 0 | 0 | 0 | 0 |
| ENSDARG000 | 1:1:1:1:1:1:1:1: | 11902579:119 | 11907812:119 | -:-:-:-:-:-:-:-: | 7765 | grk4 | 834 | 508 | 580 | 460 | 502 | 553 |
| ENSDARG000 | 1;1;1;1;1 | 29608128:296 | 29608823:296 | -;-;-;-;- | 1772 | cckbra | 20 | 10 | 20 | 6 | 8 | 5 |
| ENSDARG000 | 1:1:1:1:1:1:1:1: | 50639416:506 | 50639512:506 | +:+:+:+:+:+:+:+: | 3693 | herc3 | 301 | 281 | 287 | 223 | 355 | 250 |
| ENSDARG000 | 1;1;1;1;1 | 37701261:377 | 37701489:377 | +;+;+;+;+ | 1398 | NA | 2 | 2 | 0 | 0 | 0 | 0 |
| ENSDARG000 | 1:1:1:1:1:1:1:1: | 5402476:5402 | 5402683:5402 | +:+:+:+:+:+:+:+: | 1616 | tuba8l2 | 196 | 242 | 96 | 108 | 198 | 310 |
| ENSDARG000 | 1;1;1;1 | 29653947:296 | 29655034:296 | -;-;-;- | 1705 | NA | 23 | 18 | 40 | 20 | 46 | 28 |
| ENSDARG000 | 1:1:1:1:1:1:1:1: | 31110817:311 | 31110852:311 | +:+:+:+:+:+:+:+: | 2224 | eef1a1b | 21 | 0 | 88 | 0 | 46 | 20 |
| ENSDARG000 | 1:1:1:1:1:1:1:1: | 23323002:233 | 23323711:233 | -:-:-:-:-:-:-:-: | 5923 | pds5a | 2062 | 1925 | 2728 | 1568 | 2267 | 1712 |
| ENSDARG000 | 1:1:1:1:1:1:1:1: | 45332641:453 | 45332975:453 | -:-:-:-:-:-:-:-: | 1295 | f8a | 244 | 223 | 262 | 266 | 337 | 237 |
| ENSDARG000 | 1:1:1:1:1:1:1:1: | 9125023:9125 | 9125412:9125 | -:-:-:-:-:-:-:-: | 2013 | NA | 45 | 88 | 116 | 92 | 137 | 95 |
| ENSDARG000 | 1:1:1:1:1:1:1:1: | 52489962:524 | 52490721:524 | -:-:-:-:-:-:-:-: | 3536 | acy3.2 | 187 | 116 | 217 | 167 | 228 | 203 |
| ENSDARG000 | 1;1 | 49510209:495 | 49510312:495 | +;+ | 2064 | ITPRIP | 8 | 6 | 0 | 2 | 22 | 12 |
| ENSDARG000 | 1:1:1:1:1:1:1:1: | 1689775:1690 | 1689861:1690 | +:+:+:+:+:+:+:+: | 4471 | atp1a1a.4 | 0 | 14 | 603 | 68 | 1000 | 6 |
| ENSDARG000 | 1:1:1:1:1:1:1:1: | 58491504:584 | 58492145:584 | -:-:-:-:-:-:-:-: | 2196 | NA | 0 | 0 | 0 | 0 | 0 | 0 |
| ENSDARG000 | 1:1:1:1:1:1:1:1: | 53714734:537 | 53715125:537 | +:+:+:+:+:+:+:+: | 2167 | pus10 | 86 | 133 | 190 | 68 | 142 | 98 |
| ENSDARG000 | 1:1:1:1:1:1:1:1: | 29664336:296 | 29664558:296 | +:+:+:+:+:+:+:+: | 6135 | raph1b | 3007 | 1168 | 2040 | 1316 | 2463 | 1834 |
| ENSDARG000 | 1:1:1:1:1:1:1:1: | 29743442:297 | 29743944:297 | -:-:-:-:-:-:-:-: | 1021 | spp2 | 4 | 9 | 19 | 0 | 7 | 0 |
| ENSDARG000 | 1:1:1:1:1:1:1:1: | 52499412:524 | 52499929:524 | -:-:-:-:-:-:-:-: | 2053 | acy3.1 | 81 | 54 | 53 | 26 | 100 | 108 |
| ENSDARG000 | 1:1:1:1:1:1:1:1: | 44569282:445 | 44571906:445 | -;-;-;-;-;-;-;- | 3447 | tmx2b | 388 | 445 | 605 | 246 | 616 | 568 |
| ENSDARG000 | 1:1:1:1:1:1:1:1: | 53701801:537 | 53701929:537 | -:-:-:-:-:-:-:-: | 1914 | cct4 | 7108 | 7260 | 13160 | 8179 | 7580 | 7579 |
| ENSDARG000 | 1;1;1;1;1;1 | 45341934:453 | 45342132:453 | -;-;-;-;-;- | 1040 | NA | 36 | 20 | 17 | 2 | 36 | 26 |
| ENSDARG000 | 1:1:1:1:1:1:1:1: | 56229674:562 | 56229758:562 | +:+:+:+:+:+:+:+: | 5167 | c3a.2 | 0 | 0 | 44 | 10 | 0 | 0 |
| ENSDARG000 | 1:1:1:1:1:1:1:1: | 45323142:453 | 45323498:453 | +:+:+:+:+:+:+:+: | 1459 | emp1 | 13 | 9 | 4 | 7 | 0 | 4 |
| ENSDARG000 | 1:1:1:1:1:1:1:1: | 52495043:525 | 52495181:525 | +:+:+:+:+:+:+:+: | 1202 | NA | 0 | 0 | 0 | 0 | 0 | 0 |
| ENSDARG000 | 1:1:1:1:1:1:1:1: | 18779168:187 | 18780744:187 | -:-:-:-:-:-:-:-: | 3437 | pgm2 | 2548 | 1945 | 1594 | 1288 | 1785 | 2045 |
| ENSDARG000 | 1:1:1:1:1:1:1:1: | 5422439:5422 | 5422532:5422 | +:+:+:+:+:+:+:+: | 1940 | stk16 | 273 | 284 | 158 | 100 | 182 | 195 |
| ENSDARG000 | 1:1:1:1:1:1:1:1: | 19849168:198 | 19849208:198 | +:+:+:+:+:+:+:+: | 3502 | UBA6 | 498 | 491 | 709 | 213 | 587 | 545 |
| ENSDARG000 | 1;1 | 10118455:101 | 10119029:101 | +;+ | 1117 | lratb.2 | 277 | 0 | 0 | 16 | 0 | 0 |
| ENSDARG000 | 1;1;1;1;1 | 10110203:101 | 10110744:101 | +;+;+;+;+ | 1599 | LRAT | 122 | 6 | 17 | 61 | 16 | 8 |
| ENSDARG000 | 1;1;1 | 25491324:254 | 25491928:254 | -;-;- | 1213 | sfrp2 | 86 | 0 | 0 | 0 | 2 | 2 |
| ENSDARG000 | 1:1:1:1:1:1:1:1: | 10074343:100 | 10074943:100 | +:+:+:+:+:+:+:+: | 3972 | NA | 91 | 123 | 387 | 132 | 216 | 200 |
| ENSDARG000 | 1:1:1:1:1:1:1:1: | 26821963:268 | 26824453:268 | -:-:-:-:-:-:-:-: | 7477 | NA | 52 | 158 | 109 | 126 | 131 | 42 |
| ENSDARG000 | 1:1:1:1:1:1:1:1: | 41849152:418 | 41849355:418 | +:+:+:+:+:+:+:+: | 3811 | smox | 1216 | 1191 | 1215 | 796 | 2592 | 914 |
| ENSDARG000 | 1:1:1:1:1:1:1:1: | 49514224:495 | 49514464:495 | -:-:-:-:-:-:-:-: | 1603 | zp3c | 6 | 0 | 0 | 0 | 0 | 0 |
| ENSDARG000 | 1;1 | 31141317:311 | 31141380:311 | +;+ | 593 | NA | 20 | 3 | 12 | 9 | 0 | 8 |
| ENSDARG000 | 1;1 | 31288536:312 | 31288759:312 | -;- | 287 | NA | 0 | 0 | 0 | 0 | 0 | 0 |
| ENSDARG000 | 1:1:1:1:1:1:1:1: | 31134942:311 | 31136812:311 | -:-:-:-:-:-:-:-: | 4129 | NA | 1318 | 510 | 2270 | 395 | 1815 | 1058 |
| ENSDARG000 | 1;1;1 | 25555725:255 | 25556310:255 | -;-;- | 772 | NA | 0 | 0 | 0 | 0 | 0 | 0 |
| ENSDARG000 | 1:1:1:1:1:1:1:1: | 37752171:377 | 37752729:377 | +:+:+:+:+:+:+:+: | 5122 | galntl6 | 0 | 4 | 12 | 2 | 42 | 0 |
| ENSDARG000 | 1;1;1;1;1;1 | 25624758:256 | 25625008:256 | -;-;-;-;-;- | 3610 | NA | 0 | 0 | 0 | 0 | 0 | 0 |
| ENSDARG000 | 1:1:1:1:1:1:1:1: | 11977426:119 | 11977774:119 | +:+:+:+:+:+:+:+: | 2069 | tspan5b | 364 | 430 | 518 | 226 | 445 | 498 |
| ENSDARG000 | 1:1:1:1:1:1:1:1: | 46635236:466 | 46636406:466 | -:-:-:-:-:-:-:-: | 2750 | rcbtb1 | 420 | 303 | 314 | 244 | 419 | 392 |
| ENSDARG000 | 1:1:1:1:1:1:1:1: | 55095314:550 | 55095456:551 | +:+:+:+:+:+:+:+: | 2882 | aftpha | 618 | 620 | 654 | 498 | 669 | 446 |
| ENSDARG000 | 1:1:1:1:1:1:1:1: | 58508790:585 | 58508874:585 | +:+:+:+:+:+:+:+: | 2165 | NA | 0 | 0 | 0 | 0 | 0 | 0 |
| ENSDARG000 | 1:1:1:1:1:1:1:1: | 44582369:445 | 44582670:445 | +:+:+:+:+:+:+:+: | 1177 | med19b | 274 | 169 | 252 | 188 | 281 | 184 |
| ENSDARG000 | 1:1:1:1:1:1:1:1: | 21893289:218 | 21893381:218 | -:-:-:-:-:-:-:-: | 1169 | NA | 0 | 0 | 6 | 4 | 0 | 0 |
| ENSDARG000 | 1:1:1:1:1:1:1:1: | 50794669:507 | 50796292:507 | -:-:-:-:-:-:-:-: | 4802 | zgc:154142 | 0 | 0 | 2 | 2 | 2 | 0 |
| ENSDARG000 | 1:1:1:1:1:1:1:1: | 29759678:297 | 29759776:297 | +:+:+:+:+:+:+:+: | 1621 | cpb2 | 0 | 0 | 0 | 0 | 0 | 0 |
| ENSDARG000 | 1;1 | 58535306:585 | 58535489:585 | +;+ | 1231 | zgc:172091 | 0 | 0 | 0 | 0 | 0 | 0 |
| ENSDARG000 | 1:1:1:1:1:1:1:1: | 49525096:495 | 49525419:495 | -:-:-:-:-:-:-:-: | 3165 | herc56.4 | 324 | 297 | 75 | 9 | 3 | 34 |
| ENSDARG000 | 1:1:1:1:1:1:1:1: | 29750937:297 | 29752518:297 | -:-:-:-:-:-:-:-: | 2934 | alg11 | 595 | 530 | 567 | 430 | 630 | 618 |
| ENSDARG000 | 1:1:1:1:1:1:1:1: | 34695373:346 | 34695553:346 | +:+:+:+:+:+:+:+: | 1723 | gtf2f2a | 733 | 750 | 1035 | 643 | 1005 | 1011 |
| ENSDARG000 | 1:1:1:1:1:1:1:1: | 18810891:188 | 18811169:188 | -:-:-:-:-:-:-:-: | 584 | tmsb5 | 0 | 0 | 0 | 0 | 0 | 0 |
| ENSDARG000 | 1:1:1:1:1:1:1:1: | 25650917:256 | 25651066:256 | +:+:+:+:+:+:+:+: | 2192 | plrg1 | 678 | 745 | 1014 | 447 | 877 | 826 |
| ENSDARG000 | 1:1:1:1:1:1:1:1: | 38169512:381 | 38170085:381 | -:-:-:-:-:-:-:-: | 1470 | hmgb2a | 6 | 7 | 7 | 0 | 4 | 2 |
| ENSDARG000 | 1:1:1:1:1:1:1:1: | 34685405:346 | 34685624:346 | +:+:+:+:+:+:+:+: | 1714 | gpalpp1 | 418 | 364 | 580 | 290 | 450 | 430 |
| ENSDARG000 | 1:1:1:1:1:1:1:1: | 38142354:381 | 38143183:381 | +:+:+:+:+:+:+:+: | 8070 | galnt7 | 732 | 852 | 861 | 866 | 938 | 790 |
| ENSDARG000 | 1 | 9144958 | 9145896 | + | 939 | rasd3 | 0 | 0 | 0 | 0 | 2 | 0 |
| ENSDARG000 | 1:1:1:1:1:1:1:1: | 34707932:347 | 34708383:347 | -:-:-:-:-:-:-:-: | 1732 | tpt1 | 28950 | 28312 | 41752 | 43918 | 30024 | 49046 |
| ENSDARG000 | 1:1:1:1:1:1:1:1: | 27023687:270 | 27024415:270 | +:+:+:+:+:+:+:+: | 9574 | bnc2 | 1137 | 1320 | 1526 | 1352 | 1965 | 1823 |
| ENSDARG000 | 1;1;1;1 | 50847783:508 | 50850491:508 | -;-;-;- | 2939 | NA | 8 | 82 | 136 | 174 | 66 | 138 |
| ENSDARG000 | 1:1:1:1:1:1:1:1: | 10127163:101 | 10127308:101 | +:+:+:+:+:+:+:+: | 2403 | rbm46 | 2 | 0 | 0 | 0 | 0 | 0 |
| ENSDARG000 | 1:1:1:1:1:1:1:1: | 44591492:445 | 44593543:445 | -:-:-:-:-:-:-:-: | 3807 | slc43a1b | 18 | 22 | 20 | 22 | 26 | 22 |
| ENSDARG000 | 1:1:1:1:1:1:1:1: | 21937201:219 | 21937282:219 | +:+:+:+:+:+:+:+: | 5166 | kdm4c | 716 | 480 | 1210 | 1223 | 873 | 833 |
| ENSDARG000 | 1;1;1;1 | 55112801:551 | 55113613:551 | -;-;-;- | 1157 | NA | 168 | 248 | 322 | 284 | 191 | 213 |
| ENSDARG000 | 1:1:1:1:1:1:1:1: | 12009673:120 | 12009762:120 | +:+:+:+:+:+:+:+: | 5712 | slc24a2 | 288 | 40 | 34 | 112 | 192 | 56 |
| ENSDARG000 | 1:1:1:1:1:1:1:1: | 16621345:166 | 16621461:166 | +:+:+:+:+:+:+:+: | 6942 | pcm1 | 2156 | 1204 | 1913 | 1088 | 1835 | 1341 |
| ENSDARG000 | 1 | 46652615 | 46654528 | + | 1914 | arl11 | 0 | 0 | 0 | 0 | 4 | 0 |
| ENSDARG000 | 1;1;1;1;1;1 | 50921266:509 | 50921510:509 | +;+;+;+;+;+ | 1889 | otx2a | 6 | 0 | 0 | 0 | 0 | 0 |
| ENSDARG000 | 1:1:1:1:1:1:1:1: | 49532662:495 | 49533000:495 | -:-:-:-:-:-:-:-: | 3186 | herc56.3 | 311 | 389 | 26 | 12 | 13 | 3 |
| ENSDARG000 | 1;1;1;1 | 46660213:466 | 46660751:466 | -;-;-;- | 967 | ebpl | 18 | 106 | 108 | 74 | 59 | 63 |
| ENSDARG000 | 1;1;1;1;1 | 10221568:102 | 10221817:102 | +;+;+;+;+ | 1563 | npy2rl | 0 | 0 | 0 | 0 | 0 | 0 |
| ENSDARG000 | 1:1:1:1:1:1:1:1: | 41886830:418 | 41887089:418 | +:+:+:+:+:+:+:+: | 7622 | NA | 8 | 0 | 6 | 0 | 0 | 0 |
| ENSDARG000 | 1:1:1:1:1:1:1:1: | 58544659:585 | 58545801:585 | -:-:-:-:-:-:-:-: | 3131 | slc27a1b | 26 | 6 | 32 | 6 | 134 | 4 |
| ENSDARG000 | 1;1;1;1;1;1 | 55124296:551 | 55124567:551 | -;-;-;-;-;- | 886 | rab1aa | 620 | 613 | 835 | 401 | 714 | 307 |
| ENSDARG000 | 1;1 | 10270247:102 | 10270955:102 | -;- | 869 | NA | 0 | 0 | 0 | 0 | 0 | 0 |
| ENSDARG000 | 1;1 | 18811679:188 | 18812015:188 | +;+ | 1355 | slc25a51a | 0 | 0 | 2 | 1 | 2 | 0 |
| ENSDARG000 | 1:1:1:1:1:1:1:1: | 31508739:315 | 31509183:315 | -:-:-:-:-:-:-:-: | 1689 | cenpk | 52 | 244 | 194 | 82 | 224 | 118 |
| ENSDARG000 | 1:1:1:1:1:1:1:1: | 53735195:537 | 53735737:537 | -:-:-:-:-:-:-:-: | 2276 | NA | 417 | 426 | 565 | 426 | 284 | 304 |
| ENSDARG000 | 1;1;1;1;1;1 | 27530825:275 | 27531305:275 | -;-;-;-;-;- | 959 | NA | 0 | 0 | 0 | 0 | 0 | 0 |
| ENSDARG000 | 1:1:1:1:1:1:1:1: | 18841462:188 | 18842204:188 | -;-;-;-;-;-;-;- | 1034 | zgc:195282 | 4 | 18 | 12 | 8 | 25 | 18 |
| ENSDARG000 | 1:1:1:1:1:1:1:1: | 10297027:102 | 10297165:102 | +:+:+:+:+:+:+:+: | 893 | eif3f | 3205 | 3043 | 4876 | 3980 | 3162 | 4257 |
| ENSDARG000 | 1:1:1:1:1:1:1:1: | 27713349:277 | 27713393:277 | +:+:+:+:+:+:+:+: | 1277 | NA | 0 | 0 | 0 | 0 | 0 | 0 |
| ENSDARG000 | 1:1:1:1:1:1:1:1: | 9153141:9154 | 9153826:9154 | +:+:+:+:+:+:+:+: | 2618 | plk1 | 839 | 1986 | 1860 | 1063 | 1111 | 770 |
| ENSDARG000 | 1 | 41949303 | 41950304 | + | 1002 | NA | 0 | 0 | 0 | 0 | 0 | 0 |
| ENSDARG000 | 1:1:1:1:1:1:1:1: | 19930520:199 | 19930557:199 | +:+:+:+:+:+:+:+: | 3776 | NA | 511 | 208 | 348 | 256 | 292 | 289 |
| ENSDARG000 | 1;1 | 9157310:9162 | 9157453:9162 | -;- | 424 | NA | 0 | 0 | 0 | 0 | 0 | 0 |
| ENSDARG000 | 1:1:1:1:1:1:1:1: | 50968908:509 | 50969311:509 | +:+:+:+:+:+:+:+: | 2602 | mdh1aa | 33507 | 27260 | 31171 | 67557 | 50298 | 47124 |

| | | | | | | | | | | | | |
|---|---|---|---|---|---|---|---|---|---|---|---|---|
| ENSDARG000 | 1:1:1:1:1:1:1: | 44651727:446 | 44653763:446 | -:-:-:-:-:-:-:-: | 5496 | NA | 1032 | 976 | 894 | 664 | 1024 | 634 |
| ENSDARG000 | 1:1:1:1:1:1:1: | 46666696:466 | 46669177:466 | -:-:-:-:-:-:-:-: | 3998 | kpna3 | 1857 | 2413 | 2451 | 1260 | 2385 | 1740 |
| ENSDARG000 | 1;1;1;1;1;1 | 31519239:315 | 31519323:315 | +;+;+;+;+;+ | 846 | NA | 14 | 2 | 6 | 2 | 32 | 0 |
| ENSDARG000 | 1;1;1;1;1 | 12046544:120 | 12046637:120 | +;+;+;+;+ | 394 | NA | 0 | 12 | 0 | 0 | 12 | 2 |
| ENSDARG000 | 1:1:1:1:1:1:1: | 55137943:551 | 55138015:551 | +:+:+:+:+:+:+: | 1490 | mb | 66 | 4 | 55 | 2 | 126 | 20 |
| ENSDARG000 | 1;1 | 41957210:419 | 41961608:419 | -;- | 5643 | ADRA1D | 0 | 0 | 6 | 0 | 0 | 0 |
| ENSDARG000 | 1:1:1:1:1:1:1: | 7562316:7563 | 7563241:7563 | -:-:-:-:-:-:-:-: | 2646 | mxa | 270 | 418 | 34 | 4 | 14 | 6 |
| ENSDARG000 | 1;1 | 38193147:381 | 38193778:381 | -;- | 1641 | hand2 | 6 | 0 | 0 | 0 | 0 | 0 |
| ENSDARG000 | 1:1:1:1:1:1:1: | 58563957:585 | 58564446:585 | -:-:-:-:-:-:-:-: | 2298 | NA | 35 | 32 | 157 | 111 | 45 | 42 |
| ENSDARG000 | 1:1:1:1:1:1:1: | 18863060:188 | 18863185:188 | +:+:+:+:+:+:+: | 2843 | NA | 819 | 724 | 1153 | 1359 | 1366 | 805 |
| ENSDARG000 | 1:1:1:1:1:1:1: | 25669799:256 | 25670234:256 | -:-:-:-:-:-:-:-: | 2069 | fgg | 0 | 0 | 106 | 0 | 0 | 0 |
| ENSDARG000 | 1;1 | 31531990:315 | 31533044:315 | -;- | 1619 | lbx1b | 0 | 0 | 0 | 0 | 0 | 0 |
| ENSDARG000 | 1;1;1;1;1;1;1 | 53842344:538 | 53842576:538 | +;+;+;+;+;+;+ | 1850 | ifngr1 | 33 | 28 | 28 | 5 | 6 | 28 |
| ENSDARG000 | 1;1;1;1;1;1 | 38346043:383 | 38346851:383 | -;-;-;-;-;- | 1815 | fbxo8 | 271 | 239 | 235 | 247 | 230 | 263 |
| ENSDARG000 | 1:1:1:1:1:1:1: | 49547045:495 | 49547278:495 | -:-:-:-:-:-:-:-: | 3034 | herc56.2 | 142 | 170 | 14 | 8 | 8 | 20 |
| ENSDARG000 | 1;1;1;1 | 45351890:453 | 45352040:453 | +;+;+;+ | 981 | si:ch211-243a | 372 | 326 | 1336 | 132 | 437 | 728 |
| ENSDARG000 | 1:1:1:1:1:1:1: | 31573225:315 | 31573396:315 | +:+:+:+:+:+:+: | 1909 | slc2a15b | 25751 | 21663 | 19801 | 19988 | 27130 | 25175 |
| ENSDARG000 | 1:1:1:1:1:1:1: | 20063502:200 | 20064042:200 | -:-:-:-:-:-:-:-: | 1714 | mettl14 | 210 | 245 | 281 | 174 | 336 | 179 |
| ENSDARG000 | 1:1:1:1:1:1:1: | 56274613:562 | 56274753:562 | +:+:+:+:+:+:+: | 5209 | c3a.3 | 0 | 0 | 74 | 0 | 0 | 0 |
| ENSDARG000 | 1:1:1:1:1:1:1: | 27738647:277 | 27739090:277 | -:-:-:-:-:-:-:-: | 1660 | NA | 0 | 4 | 0 | 0 | 0 | 0 |
| ENSDARG000 | 1:1:1:1:1:1:1: | 12049229:120 | 12049489:120 | +:+:+:+:+:+:+: | 1752 | saraf | 741 | 664 | 1505 | 1599 | 835 | 665 |
| ENSDARG000 | 1:1:1:1:1:1:1: | 12061284:120 | 12062172:120 | -;-;-;-;-;-;-;- | 1599 | pla2g12a | 101 | 129 | 120 | 111 | 74 | 107 |
| ENSDARG000 | 1:1:1:1:1:1:1: | 58580412:585 | 58580515:585 | -:-:-:-:-:-:-:-: | 2301 | NA | 51 | 104 | 256 | 135 | 307 | 117 |
| ENSDARG000 | 1;1 | 42776553:427 | 42779075:427 | -;- | 2523 | LRRTM1 | 0 | 8 | 0 | 0 | 0 | 0 |
| ENSDARG000 | 1;1;1;1;1;1 | 42281308:422 | 42282121:422 | -;-;-;-;-;- | 1271 | NA | 0 | 0 | 3 | 0 | 0 | 0 |
| ENSDARG000 | 1:1:1:1:1:1:1: | 10318077:103 | 10318306:103 | +:+:+:+:+:+:+: | 1611 | pip4p1b | 244 | 150 | 118 | 135 | 119 | 197 |
| ENSDARG000 | 1:1:1:1:1:1:1: | 9169807:9169 | 9169977:9169 | -:-:-:-:-:-:-:-: | 2961 | NA | 234 | 108 | 141 | 105 | 118 | 215 |
| ENSDARG000 | 1:1:1:1:1:1:1: | 42224769:422 | 42225322:422 | +:+:+:+:+:+:+: | 16723 | ctnna2 | 149 | 168 | 118 | 260 | 276 | 72 |
| ENSDARG000 | 1:1:1:1:1:1:1: | 27808916:278 | 27809396:278 | +:+:+:+:+:+:+: | 2100 | pspc1 | 449 | 439 | 512 | 402 | 778 | 597 |
| ENSDARG000 | 1:1:1:1:1:1:1: | 23372470:233 | 23372892:233 | +:+:+:+:+:+:+: | 8782 | n4bp2 | 350 | 274 | 361 | 219 | 345 | 431 |
| ENSDARG000 | 1:1:1:1:1:1:1: | 10305611:103 | 10305782:103 | +:+:+:+:+:+:+: | 1713 | zgc:77880 | 34 | 36 | 46 | 28 | 50 | 28 |
| ENSDARG000 | 1;1 | 25696798:256 | 25697350:256 | +;+ | 1160 | lrata | 0 | 0 | 0 | 0 | 0 | 0 |
| ENSDARG000 | 1:1:1:1:1:1:1: | 16654254:166 | 16654785:166 | -:-:-:-:-:-:-:-: | 1946 | asah1b | 3219 | 2237 | 1506 | 2276 | 2156 | 2069 |
| ENSDARG000 | 1:1:1:1:1:1:1: | 55161083:551 | 55161475:551 | -:-:-:-:-:-:-:-: | 943 | pane1 | 196 | 286 | 194 | 80 | 218 | 116 |
| ENSDARG000 | 1 | 23398405 | 23398992 | + | 588 | rhoh | 6 | 12 | 2 | 0 | 0 | 6 |
| ENSDARG000 | 1:1:1:1:1:1:1: | 53865677:538 | 53866354:538 | -:-:-:-:-:-:-:-: | 2156 | ltv1 | 767 | 695 | 1171 | 584 | 672 | 551 |
| ENSDARG000 | 1:1:1:1:1:1:1: | 20069535:200 | 20069850:200 | +:+:+:+:+:+:+: | 5655 | PRSS12 | 333 | 106 | 148 | 64 | 164 | 131 |
| ENSDARG000 | 1:1:1:1:1:1:1: | 29766725:297 | 29766853:297 | +:+:+:+:+:+:+: | 5966 | zc3h13 | 1420 | 1147 | 1949 | 728 | 1880 | 1265 |
| ENSDARG000 | 1:1:1:1:1:1:1: | 45372562:453 | 45378947:453 | -:-:-:-:-:-:-:-: | 9345 | grin2bb | 2 | 4 | 21 | 2 | 7 | 13 |
| ENSDARG000 | 1:1:1:1:1:1:1: | 31654203:316 | 31654512:316 | -:-:-:-:-:-:-:-: | 1028 | dpcd | 90 | 109 | 82 | 72 | 91 | 51 |
| ENSDARG000 | 1:1:1:1:1:1:1: | 27867706:278 | 27868462:278 | +:+:+:+:+:+:+: | 3375 | dnajb14 | 220 | 278 | 292 | 202 | 249 | 283 |
| ENSDARG000 | 1:1:1:1:1:1:1: | 38362348:383 | 38362450:383 | +:+:+:+:+:+:+: | 2199 | cep44 | 26 | 107 | 96 | 40 | 88 | 53 |
| ENSDARG000 | 1:1:1:1:1:1:1: | 31638274:316 | 31638789:316 | +:+:+:+:+:+:+: | 2456 | fgf8b | 0 | 0 | 2 | 0 | 0 | 0 |
| ENSDARG000 | 1:1:1:1:1:1:1: | 31657842:316 | 31657902:316 | +:+:+:+:+:+:+: | 2102 | poll | 216 | 96 | 177 | 124 | 282 | 145 |
| ENSDARG000 | 1;1;1 | 25709813:257 | 25710025:257 | -;-;- | 572 | NA | 0 | 0 | 0 | 0 | 0 | 0 |
| ENSDARG000 | 1;1 | 25719891:257 | 25720161:257 | +;+ | 669 | si:dkev-30k22 | 0 | 0 | 0 | 0 | 0 | 0 |
| ENSDARG000 | 1 | 25732715 | 25733867 | + | 1153 | NPY2R | 0 | 0 | 0 | 0 | 0 | 0 |
| ENSDARG000 | 1:1:1:1:1:1:1: | 20147222:201 | 20147575:201 | -:-:-:-:-:-:-:-: | 3702 | NDST4 | 4 | 6 | 2 | 0 | 22 | 8 |
| ENSDARG000 | 1;1;1;1 | 58593247:585 | 58595774:585 | -;-;-;- | 3060 | NA | 132 | 187 | 200 | 136 | 202 | 121 |
| ENSDARG000 | 1:1:1:1:1:1:1: | 50997156:510 | 50997402:510 | +:+:+:+:+:+:+: | 1817 | ugp2a | 130 | 125 | 272 | 176 | 212 | 173 |
| ENSDARG000 | 1:1:1:1:1:1:1: | 25737767:257 | 25738303:257 | -:-:-:-:-:-:-:-: | 2574 | map9 | 38 | 40 | 46 | 24 | 34 | 22 |
| ENSDARG000 | 1:1:1:1:1:1:1: | 58601680:586 | 58601897:586 | +:+:+:+:+:+:+: | 2116 | NA | 50 | 44 | 73 | 36 | 100 | 21 |
| ENSDARG000 | 1:1:1:1:1:1:1: | 22002649:220 | 22002818:220 | +:+:+:+:+:+:+: | 12816 | DNAH6 | 12 | 37 | 62 | 48 | 24 | 61 |
| ENSDARG000 | 1:1:1:1:1:1:1: | 55169130:551 | 55171285:551 | -:-:-:-:-:-:-:-: | 3701 | zgc:158803 | 846 | 895 | 2207 | 1099 | 1314 | 908 |
| ENSDARG000 | 1:1:1:1:1:1:1: | 18965750:189 | 18966071:189 | +:+:+:+:+:+:+: | 3880 | limch1a | 46 | 50 | 19 | 8 | 67 | 49 |
| ENSDARG000 | 1:1:1:1:1:1:1: | 55191150:551 | 55191757:551 | -;-;-;-;-;-;-;- | 1642 | mri1 | 280 | 224 | 544 | 229 | 374 | 363 |
| ENSDARG000 | 1;1 | 12065156:120 | 12067196:120 | -;- | 2181 | NA | 0 | 0 | 4 | 0 | 0 | 4 |
| ENSDARG000 | 1:1:1:1:1:1:1: | 16673841:166 | 16675667:166 | -:-:-:-:-:-:-:-: | 3045 | frg1 | 330 | 232 | 364 | 172 | 343 | 238 |
| ENSDARG000 | 1:1:1:1:1:1:1: | 25783801:257 | 25784233:257 | +:+:+:+:+:+:+: | 2929 | gucy1a1 | 0 | 4 | 0 | 0 | 0 | 0 |
| ENSDARG000 | 1:1:1:1:1:1:1: | 46710647:467 | 46711185:467 | -;-;-;-;-;-;-;- | 1478 | spryd7a | 548 | 519 | 612 | 480 | 572 | 468 |
| ENSDARG000 | 1;1;1;1;1 | 20481109:204 | 20481231:204 | -;-;-;-;- | 1114 | NA | 0 | 0 | 0 | 0 | 0 | 0 |
| ENSDARG000 | 1;1;1;1;1;1;1 | 34743459:347 | 34743553:347 | -;-;-;-;-;-;- | 1075 | NA | 0 | 0 | 1 | 0 | 0 | 0 |
| ENSDARG000 | 1:1:1:1:1:1:1: | 12068800:120 | 12068877:120 | +:+:+:+:+:+:+: | 2166 | xpa | 81 | 71 | 102 | 29 | 61 | 32 |
| ENSDARG000 | 1:1:1:1:1:1:1: | 55239160:552 | 55239374:552 | +:+:+:+:+:+:+: | 2358 | NA | 8 | 9 | 20 | 6 | 2 | 5 |
| ENSDARG000 | 1;1;1;1;1 | 17330495:173 | 17331068:173 | -;-;-;-;- | 977 | NA | 0 | 0 | 0 | 0 | 0 | 0 |
| ENSDARG000 | 1;1 | 20593653:206 | 20593934:206 | +;+ | 459 | NA | 0 | 0 | 0 | 0 | 0 | 0 |
| ENSDARG000 | 1;1;1;1;1;1 | 20556067:205 | 20559944:205 | -;-;-;-;-;- | 5298 | ugt8 | 14 | 2 | 2 | 0 | 32 | 20 |
| ENSDARG000 | 1:1:1:1:1:1:1: | 23406403:234 | 23406947:234 | +:+:+:+:+:+:+: | 2618 | CHRNA9 | 0 | 0 | 0 | 0 | 0 | 2 |
| ENSDARG000 | 1;1;1;1 | 34748695:347 | 34749320:347 | -;-;-;- | 1089 | NA | 0 | 0 | 0 | 0 | 0 | 0 |
| ENSDARG000 | 1:1:1:1:1:1:1: | 56344651:563 | 56345325:563 | -:-:-:-:-:-:-:-: | 1898 | NA | 2 | 0 | 12 | 0 | 6 | 10 |
| ENSDARG000 | 1:1:1:1:1:1:1: | 10336681:103 | 10336824:103 | -:-:-:-:-:-:-:-: | 1091 | FRMD7 | 0 | 7 | 11 | 15 | 8 | 0 |
| ENSDARG000 | 1:1:1:1:1:1:1: | 9199031:9199 | 9199243:9199 | +:+:+:+:+:+:+: | 3982 | chtf18 | 86 | 477 | 400 | 156 | 238 | 188 |
| ENSDARG000 | 1:1:1:1:1:1:1: | 12105901:121 | 12108141:121 | -:-:-:-:-:-:-:-: | 4907 | mttp | 2 | 0 | 0 | 0 | 0 | 0 |
| ENSDARG000 | 1:1:1:1:1:1:1: | 9227081:9227 | 9227397:9227 | -;-;-;-;-;-;-;- | 573 | gng13a | 5 | 10 | 2 | 1 | 21 | 0 |
| ENSDARG000 | 1:1:1:1:1:1:1: | 51021279:510 | 51021938:510 | -:-:-:-:-:-:-:-: | 2413 | spast | 2839 | 2261 | 2694 | 3960 | 1829 | 1485 |
| ENSDARG000 | 1;1;1;1 | 31674297:316 | 31674563:316 | +;+;+;+ | 1539 | wbp1lb | 338 | 302 | 340 | 312 | 426 | 266 |
| ENSDARG000 | 1;1;1 | 19074326:190 | 19074747:190 | -;-;- | 872 | phox2ba | 0 | 10 | 2 | 20 | 8 | 10 |
| ENSDARG000 | 1;1 | 46725808:467 | 46726540:467 | -;- | 876 | NA | 36 | 7 | 29 | 46 | 36 | 18 |
| ENSDARG000 | 1:1:1:1:1:1:1: | 46837849:468 | 46838823:468 | -:-:-:-:-:-:-:-: | 2623 | agpat3 | 1316 | 1166 | 1695 | 746 | 2017 | 1864 |
| ENSDARG000 | 1;1;1;1;1;1;1 | 56801021:568 | 56801081:568 | +;+;+;+;+;+;+ | 1314 | NA | 0 | 2 | 0 | 0 | 0 | 0 |
| ENSDARG000 | 1:1:1:1:1:1:1: | 34763539:347 | 34763602:347 | +:+:+:+:+:+:+: | 1610 | zgc:172122 | 6 | 0 | 4 | 4 | 0 | 0 |
| ENSDARG000 | 1;1;1;1;1 | 46827891:468 | 46827915:468 | +;+;+;+;+ | 858 | NA | 0 | 1 | 0 | 2 | 0 | 0 |
| ENSDARG000 | 1;1 | 46830809:468 | 46831578:468 | -;- | 1280 | dleu7 | 0 | 0 | 3 | 0 | 0 | 0 |
| ENSDARG000 | 1:1:1:1:1:1:1: | 25801648:258 | 25801825:258 | +:+:+:+:+:+:+: | 2360 | gucy1b1 | 16 | 55 | 88 | 22 | 55 | 19 |
| ENSDARG000 | 1:1:1:1:1:1:1: | 29798140:298 | 29798434:298 | +:+:+:+:+:+:+: | 3232 | tm9sf2 | 5451 | 4474 | 5110 | 3495 | 5967 | 4636 |
| ENSDARG000 | 1:1:1:1:1:1:1: | 43652480:436 | 43652843:436 | -:-:-:-:-:-:-:-: | 1593 | suclg1 | 1432 | 1560 | 1657 | 1044 | 1678 | 1279 |
| ENSDARG000 | 1 | 55247370 | 55248496 | - | 1127 | nanos3 | 0 | 2 | 2 | 0 | 0 | 0 |
| ENSDARG000 | 1:1:1:1:1:1:1: | 23489269:234 | 23489360:234 | +:+:+:+:+:+:+: | 4337 | NA | 36 | 240 | 209 | 104 | 148 | 122 |
| ENSDARG000 | 1;1;1;1;1;1 | 10462692:104 | 10463665:104 | -;-;-;-;-;- | 1659 | cnpy3 | 270 | 360 | 315 | 125 | 394 | 288 |
| ENSDARG000 | 1:1:1:1:1:1:1: | 23506525:235 | 23507181:235 | -:-:-:-:-:-:-:-: | 4063 | fam184b | 1602 | 18 | 22 | 0 | 30 | 54 |
| ENSDARG000 | 1:1:1:1:1:1:1: | 10486093:104 | 10487757:104 | -:-:-:-:-:-:-:-: | 4956 | trpc5a | 0 | 0 | 0 | 0 | 0 | 0 |
| ENSDARG000 | 1;1;1;1;1 | 56813845:568 | 56813923:568 | +;+;+;+;+ | 1001 | NA | 0 | 0 | 0 | 0 | 0 | 0 |
| ENSDARG000 | 1:1:1:1:1:1:1: | 19083501:190 | 19083673:190 | +:+:+:+:+:+:+: | 1393 | exosc9 | 382 | 711 | 827 | 328 | 457 | 679 |
| ENSDARG000 | 1:1:1:1:1:1:1: | 10378706:103 | 10379282:103 | +:+:+:+:+:+:+: | 3021 | dachb | 14 | 12 | 10 | 0 | 14 | 2 |
| ENSDARG000 | 1;1;1;1;1 | 56823437:568 | 56823469:568 | +;+;+;+;+ | 954 | NA | 0 | 1 | 0 | 0 | 0 | 0 |
| ENSDARG000 | 1;1 | 17465956:174 | 17466097:174 | -;- | 201 | NA | 0 | 1 | 0 | 0 | 0 | 0 |
| ENSDARG000 | 1:1:1:1:1:1:1: | 17376922:173 | 17377021:173 | +:+:+:+:+:+:+: | 16411 | fat1a | 6004 | 2249 | 5443 | 3592 | 7684 | 5398 |
| ENSDARG000 | 1;1;1;1 | 56830798:568 | 56830882:568 | +;+;+;+ | 611 | NA | 0 | 0 | 0 | 0 | 0 | 0 |
| ENSDARG000 | 1;1;1 | 35317354:353 | 35317520:353 | -;-;- | 286 | NA | 0 | 0 | 0 | 0 | 0 | 0 |
| ENSDARG000 | 1:1:1:1:1:1:1: | 20635190:206 | 20635298:206 | +:+:+:+:+:+:+: | 2719 | spock3 | 14441 | 13145 | 4351 | 13123 | 4836 | 6556 |
| ENSDARG000 | 1:1:1:1:1:1:1: | 51039558:510 | 51039645:510 | +:+:+:+:+:+:+: | 630 | dpy30 | 415 | 579 | 564 | 566 | 400 | 488 |
| ENSDARG000 | 1;1;1;1;1;1 | 56843411:568 | 56843474:568 | +;+;+;+;+;+ | 1227 | NA | 0 | 0 | 0 | 0 | 0 | 0 |
| ENSDARG000 | 1:1:1:1:1:1:1: | 31706668:317 | 31706702:317 | +:+:+:+:+:+:+: | 1555 | as3mt | 145 | 193 | 300 | 230 | 285 | 183 |
| ENSDARG000 | 1;1;1;1;1 | 56852400:568 | 56852460:568 | +;+;+;+;+ | 1106 | NA | 0 | 2 | 0 | 0 | 0 | 0 |
| ENSDARG000 | 1;1;1;1;1 | 56859745:568 | 56859805:568 | +;+;+;+;+ | 974 | NA | 0 | 0 | 0 | 0 | 0 | 0 |
| ENSDARG000 | 1:1:1:1:1:1:1: | 55250107:552 | 55250232:552 | -:-:-:-:-:-:-:-: | 1835 | NA | 0 | 0 | 2 | 0 | 0 | 0 |
| ENSDARG000 | 1;1;1;1;1;1;1 | 56873359:568 | 56873428:568 | +;+;+;+;+;+;+ | 1512 | NA | 0 | 0 | 0 | 0 | 0 | 0 |
| ENSDARG000 | 1:1:1:1:1:1:1: | 35332924:353 | 35333044:353 | -:-:-:-:-:-:-:-: | 1110 | NA | 0 | 0 | 0 | 0 | 0 | 0 |
| ENSDARG000 | 1:1:1:1:1:1:1: | 56886214:568 | 56886284:568 | +:+:+:+:+:+:+: | 1558 | NA | 0 | 1 | 0 | 0 | 0 | 0 |
| ENSDARG000 | 1;1;1;1;1;1 | 25848231:258 | 25848714:258 | +;+;+;+;+;+ | 1337 | fabp2 | 6 | 0 | 4 | 0 | 2 | 0 |
| ENSDARG000 | 1:1:1:1:1:1:1: | 22236489:222 | 22237031:222 | -:-:-:-:-:-:-:-: | 1658 | NA | 0 | 0 | 0 | 0 | 0 | 0 |
| ENSDARG000 | 1:1:1:1:1:1:1: | 31725154:317 | 31726786:317 | +:+:+:+:+:+:+: | 5073 | CNNM2 | 14 | 20 | 34 | 22 | 60 | 6 |
| ENSDARG000 | 1;1;1;1;1;1 | 22388799:223 | 22389412:223 | -;-;-;-;-;- | 2128 | chrnb3a | 346 | 56 | 108 | 90 | 184 | 531 |
| ENSDARG000 | 1;1;1;1;1;1;1 | 29831759:298 | 29831864:298 | +;+;+;+;+;+;+ | 2425 | zic2b | 0 | 0 | 0 | 0 | 0 | 0 |
| ENSDARG000 | 1:1:1:1:1:1:1: | 35473219:354 | 35473525:354 | +:+:+:+:+:+:+: | 6283 | USP38 | 390 | 253 | 515 | 439 | 445 | 390 |
| ENSDARG000 | 1:1:1:1:1:1:1: | 23559478:235 | 23559529:235 | +:+:+:+:+:+:+: | 3721 | ncapg | 184 | 1113 | 1390 | 644 | 728 | 521 |
| ENSDARG000 | 1:1:1:1:1:1:1: | 23573224:235 | 23575006:235 | -:-:-:-:-:-:-:-: | 9183 | NA | 439 | 291 | 674 | 154 | 575 | 438 |
| ENSDARG000 | 1:1:1:1:1:1:1: | 19094548:190 | 19094608:190 | +:+:+:+:+:+:+: | 2562 | NA | 496 | 820 | 704 | 548 | 586 | 886 |
| ENSDARG000 | 1;1 | 25862966:258 | 25863005:258 | -;- | 594 | NA | 0 | 4 | 6 | 12 | 6 | 8 |
| ENSDARG000 | 1;1;1;1;1;1 | 56900580:569 | 56900620:569 | +;+;+;+;+;+ | 1208 | NA | 0 | 0 | 0 | 0 | 0 | 0 |
| ENSDARG000 | 1;1;1 | 22427135:224 | 22427492:224 | +;+;+ | 1173 | NA | 0 | 0 | 0 | 0 | 0 | 0 |
| ENSDARG000 | 1;1;1;1;1 | 56919742:569 | 56919790:569 | +;+;+;+;+ | 738 | NA | 0 | 0 | 0 | 0 | 0 | 0 |
| ENSDARG000 | 1:1:1:1:1:1:1: | 22481723:224 | 22484930:224 | -:-:-:-:-:-:-:-: | 4852 | chrna6 | 796 | 28 | 16 | 54 | 42 | 458 |
| ENSDARG000 | 1;1;1;1;1 | 56929873:569 | 56929929:569 | +;+;+;+;+ | 969 | NA | 0 | 0 | 0 | 0 | 0 | 0 |
| ENSDARG000 | 1:1:1:1:1:1:1: | 22542457:225 | 22546747:225 | -:-:-:-:-:-:-:-: | 6426 | NA | 1489 | 421 | 497 | 474 | 929 | 3142 |
| ENSDARG000 | 1:1:1:1:1:1:1: | 19122920:191 | 19123678:191 | -:-:-:-:-:-:-:-: | 7367 | ptprdb | 381 | 158 | 257 | 295 | 402 | 515 |
| ENSDARG000 | 1:1:1:1:1:1:1: | 51048975:510 | 51049500:510 | -:-:-:-:-:-:-:-: | 1840 | psen2 | 2072 | 1846 | 1407 | 1966 | 1702 | 1592 |
| ENSDARG000 | 1:1:1:1:1:1:1: | 51066671:510 | 51067105:510 | +:+:+:+:+:+:+: | 1775 | srd5a2a | 54 | 70 | 27 | 223 | 98 | 43 |
| ENSDARG000 | 1:1:1:1:1:1:1: | 51104984:511 | 51108220:511 | -:-:-:-:-:-:-:-: | 6678 | jag1a | 154 | 52 | 129 | 78 | 42 | 46 |
| ENSDARG000 | 1 | 7956030 | 7961696 | + | 5667 | NA | 0 | 0 | 0 | 0 | 0 | 0 |
| ENSDARG000 | 1;1;1;1 | 58967210:589 | 58967266:589 | -;-;-;- | 420 | NA | 8 | 4 | 0 | 17 | 14 | 20 |
| ENSDARG000 | 1;1 | 56463494:564 | 56463536:564 | +;+ | 941 | zgc:171452 | 2 | 0 | 0 | 0 | 0 | 0 |
| ENSDARG000 | 1;1;1;1;1;1;1 | 54178703:541 | 54178804:541 | -;-;-;-;-;-;- | 2235 | NTHL1 | 241 | 179 | 290 | 130 | 302 | 162 |
| ENSDARG000 | 1;1;1 | 14522913:145 | 14529137:145 | -;-;- | 6710 | NA | 107 | 117 | 126 | 150 | 59 | 131 |
| ENSDARG000 | 1;1;1;1 | 40724140:407 | 40724169:407 | +;+;+;+ | 615 | NA | 0 | 0 | 2 | 0 | 0 | 0 |
| ENSDARG000 | 1;1;1 | 57741577:577 | 57741947:577 | +;+;+ | 825 | NA | 0 | 0 | 0 | 0 | 0 | 0 |
| ENSDARG000 | 1;1;1;1;1 | 6307305:6317 | 6307635:6317 | +;+;+;+;+ | 2185 | NA | 17 | 10 | 17 | 4 | 15 | 35 |
| ENSDARG000 | 1;1;1 | 50159645:501 | 50159696:501 | -;-;- | 381 | NA | 0 | 0 | 0 | 0 | 0 | 0 |
| ENSDARG000 | 1;1;1 | 13930625:139 | 13930853:139 | +;+;+ | 1353 | NOCT | 18 | 10 | 6 | 28 | 20 | 4 |
| ENSDARG000 | 1 | 10363720 | 10363851 | + | 132 | 5S_rRNA | 0 | 0 | 0 | 0 | 0 | 0 |
| ENSDARG000 | 1 | 33967157 | 33967273 | + | 117 | 5S_rRNA | 0 | 0 | 0 | 0 | 0 | 0 |
| ENSDARG000 | 1:1:1:1:1:1:1: | 58981985:589 | 58982291:589 | +:+:+:+:+:+:+: | 1371 | thumpd1 | 200 | 241 | 435 | 148 | 236 | 160 |
| ENSDARG000 | 1 | 22191952 | 22192067 | - | 116 | 5S_rRNA | 0 | 0 | 0 | 0 | 0 | 0 |
| ENSDARG000 | 1:1:1:1:1:1:1: | 13957235:139 | 13958918:139 | -:-:-:-:-:-:-:-: | 3776 | elf2b | 1172 | 1361 | 2054 | 1487 | 1576 | 1222 |
| ENSDARG000 | 1;1;1;1;1 | 58762742:587 | 58763095:587 | -;-;-;-;- | 843 | NA | 0 | 0 | 8 | 0 | 0 | 10 |
| ENSDARG000 | 1:1:1:1:1:1:1: | 5798548:5817 | 5799085:5817 | -:-:-:-:-:-:-:-: | 3599 | pard3ba | 60 | 6 | 30 | 2 | 16 | 4 |
| ENSDARG000 | 1:1:1:1:1:1:1: | 55997173:559 | 55997212:559 | -:-:-:-:-:-:-:-: | 4929 | NA | 23 | 39 | 94 | 52 | 310 | 164 |
| ENSDARG000 | 1:1:1:1:1:1:1: | 58949548:589 | 58949865:589 | -:-:-:-:-:-:-:-: | 2006 | NA | 29 | 8 | 14 | 1 | 10 | 4 |
| ENSDARG000 | 1:1:1:1:1:1:1: | 40181404:401 | 40181412:401 | -:-:-:-:-:-:-:-: | 894 | NA | 0 | 0 | 0 | 0 | 0 | 0 |
| ENSDARG000 | 1;1;1;1;1 | 57744400:577 | 57744442:577 | +;+;+;+;+ | 861 | NA | 0 | 0 | 0 | 0 | 0 | 0 |
| ENSDARG000 | 1 | 42277523 | 42277650 | - | 128 | 5S_rRNA | 0 | 0 | 0 | 0 | 0 | 0 |
| ENSDARG000 | 1;1;1;1 | 11659861:116 | 11660272:116 | +;+;+;+ | 1246 | GNA12 | 0 | 1 | 25 | 0 | 0 | 3 |
| ENSDARG000 | 1:1:1:1:1:1:1: | 57521361:575 | 57521405:575 | +:+:+:+:+:+:+: | 1197 | tspan34 | 0 | 0 | 0 | 0 | 8 | 0 |
| ENSDARG000 | 1 | 45768416 | 45768530 | + | 115 | 5S_rRNA | 0 | 0 | 0 | 0 | 0 | 0 |
| ENSDARG000 | 1;1;1;1 | 658405:66146 | 660810:66162 | -;-;-;- | 2928 | cyyr1 | 237 | 358 | 691 | 217 | 563 | 254 |
| ENSDARG000 | 1 | 11380497 | 11380611 | + | 115 | 5S_rRNA | 0 | 0 | 0 | 0 | 0 | 0 |
| ENSDARG000 | 1;1 | 54115839:541 | 54116064:541 | +;+ | 894 | rln3b | 0 | 0 | 0 | 0 | 4 | 0 |
| ENSDARG000 | 1:1:1:1:1:1:1: | 58990121:589 | 58990347:589 | +:+:+:+:+:+:+: | 2925 | NA | 196 | 198 | 285 | 47 | 314 | 169 |
| ENSDARG000 | 1 | 46722379 | 46722518 | - | 140 | dre-mir-16b | 6 | 9 | 4 | 6 | 8 | 1 |
| ENSDARG000 | 1 | 30165953 | 30166065 | + | 113 | dre-mir-740 | 0 | 0 | 0 | 0 | 0 | 0 |
| ENSDARG000 | 1 | 2403103 | 2403224 | + | 122 | 5S_rRNA | 3 | 0 | 0 | 3 | 2 | 5 |
| ENSDARG000 | 1:1:1:1:1:1:1: | 35194454:351 | 35194608:351 | +:+:+:+:+:+:+: | 3651 | sc:d189 | 0 | 2 | 8 | 0 | 4 | 2 |

| | | | | | | | | | | | | |
|---|---|---|---|---|---|---|---|---|---|---|---|---|
| ENSDARG000 | 1 | 39837656 | 39837783 | + | 128 | 5S_rRNA | 0 | 0 | 0 | 0 | 0 | 0 |
| ENSDARG000 | 1 | 56363884 | 56363974 | - | 91 | dre-mir-722 | 0 | 0 | 0 | 0 | 0 | 0 |
| ENSDARG000 | 1 | 11469536 | 11469650 | + | 115 | 5S_rRNA | 0 | 0 | 0 | 0 | 0 | 0 |
| ENSDARG000 | 1;1;1 | 59569516:59! | 59571785:59! | -;-;- | 2270 | wfikkn1 | 0 | 0 | 0 | 0 | 0 | 0 |
| ENSDARG000 | 1 | 30227391 | 30227505 | + | 115 | 5S_rRNA | 0 | 0 | 0 | 0 | 0 | 0 |
| ENSDARG000 | 1 | 7801531 | 7801646 | - | 116 | 5S_rRNA | 0 | 0 | 0 | 0 | 0 | 0 |
| ENSDARG000 | 1 | 56595955 | 56596701 | - | 747 | NA | 2 | 0 | 0 | 0 | 0 | 0 |
| ENSDARG000 | 1 | 2275605 | 2275742 | + | 138 | NA | 1 | 0 | 0 | 1 | 0 | 2 |
| ENSDARG000 | 1:1:1:1:1:1:1:1: | 14417785:144 | 14418550:144 | -:-:-:-:-:-:-:-: | 1674 | NA | 54 | 68 | 88 | 31 | 99 | 50 |
| ENSDARG000 | 1 | 12735920 | 12736034 | + | 115 | 5S_rRNA | 0 | 0 | 0 | 0 | 0 | 0 |
| ENSDARG000 | 1 | 16458525 | 16458741 | + | 217 | NA | 0 | 0 | 0 | 0 | 0 | 0 |
| ENSDARG000 | 1:1:1:1:1:1:1:1: | 673163:6736( | 673284:6741! | -:-:-:-:-:-:-:-: | 2736 | ADAMTS1 | 127 | 34 | 23 | 10 | 103 | 25 |
| ENSDARG000 | 1 | 3298112 | 3298249 | - | 138 | dre-mir-17a-1 | 0 | 0 | 0 | 0 | 0 | 0 |
| ENSDARG000 | 1 | 43265123 | 43265241 | + | 119 | 5S_rRNA | 0 | 0 | 0 | 0 | 0 | 0 |
| ENSDARG000 | 1 | 14074307 | 14074375 | - | 69 | U6 | 0 | 0 | 0 | 0 | 0 | 0 |
| ENSDARG000 | 1 | 14723156 | 14723250 | + | 95 | NA | 0 | 0 | 0 | 0 | 0 | 0 |
| ENSDARG000 | 1 | 1575483 | 1575563 | + | 81 | dre-mir-459 | 0 | 0 | 0 | 0 | 0 | 0 |
| ENSDARG000 | 1;1;1 | 58628165:586 | 58628332:586 | +;+;+ | 1470 | NA | 89 | 79 | 142 | 86 | 119 | 120 |
| ENSDARG000 | 1;1 | 56533355:565 | 56533596:565 | -;- | 2037 | NA | 4 | 1 | 4 | 2 | 10 | 6 |
| ENSDARG000 | 1:1:1:1:1:1:1:1: | 15258641:152 | 15259224:152 | +:+:+:+:+:+:+: | 2647 | pi4k2b | 598 | 720 | 506 | 548 | 573 | 405 |
| ENSDARG000 | 1;1;1;1;1;1;1 | 28624186:286 | 28624419:286 | -;-;-;-;-;-;- | 1489 | ednrba | 15238 | 8772 | 13603 | 8079 | 16323 | 15145 |
| ENSDARG000 | 1;1 | 56413733:564 | 56415464:564 | -;- | 1987 | NA | 0 | 0 | 0 | 0 | 0 | 0 |
| ENSDARG000 | 1;1;1 | 56779495:567 | 56779552:567 | +;+;+ | 633 | NA | 0 | 0 | 0 | 0 | 0 | 0 |
| ENSDARG000 | 1:1:1:1:1:1:1:1: | 59163060:591 | 59163262:591 | -;-;-;-;-;-;-;- | 1641 | NA | 14 | 8 | 2 | 0 | 22 | 0 |
| ENSDARG000 | 1:1:1:1:1:1:1:1: | 200294:20094 | 200563:20126 | -;-;-;-;-;-;-;- | 1538 | adprhl1 | 11 | 3 | 1 | 11 | 14 | 17 |
| ENSDARG000 | 1:1:1:1:1:1:1:1: | 14163506:141 | 14163982:141 | -:-:-:-:-:-:-:-: | 2403 | camk2d2 | 103 | 33 | 240 | 32 | 601 | 62 |
| ENSDARG000 | 1;1;1 | 56447107:564 | 56447354:564 | +;+;+ | 6539 | NA | 0 | 0 | 0 | 0 | 2 | 6 |
| ENSDARG000 | 1:1:1:1:1:1:1:1: | 58857088:588 | 58857205:588 | -:-:-:-:-:-:-:-: | 1296 | NA | 1166 | 763 | 1146 | 547 | 888 | 432 |
| ENSDARG000 | 1 | 8579197 | 8586066 | + | 6870 | NA | 23 | 14 | 37 | 14 | 30 | 24 |
| ENSDARG000 | 1:1:1:1:1:1:1:1: | 48256950:483 | 48257248:483 | +:+:+:+:+:+:+: | 6993 | NA | 0 | 0 | 0 | 0 | 0 | 0 |
| ENSDARG000 | 1;1 | 14672440:146 | 14673564:146 | -;- | 1462 | NA | 30 | 9 | 35 | 12 | 0 | 58 |
| ENSDARG000 | 1 | 7918795 | 7918882 | - | 88 | NA | 0 | 0 | 0 | 0 | 0 | 0 |
| ENSDARG000 | 1:1:1:1:1:1:1:1: | 59555936:595 | 59556043:595 | -:-:-:-:-:-:-:-: | 3400 | NA | 120 | 109 | 125 | 106 | 407 | 428 |
| ENSDARG000 | 1;1 | 45644345:456 | 45644388:456 | -;- | 168 | NA | 0 | 0 | 1 | 0 | 2 | 3 |
| ENSDARG000 | 1;1;1;1;1;1 | 58912051:589 | 58912848:589 | -;-;-;-;-;- | 1396 | mfap4.4 | 0 | 0 | 0 | 0 | 1 | 4 |
| ENSDARG000 | 1;1;1 | 58735326:587 | 58735366:587 | +;+;+ | 393 | NA | 0 | 0 | 6 | 0 | 0 | 0 |
| ENSDARG000 | 1 | 14744645 | 14744739 | + | 95 | NA | 0 | 0 | 0 | 0 | 0 | 0 |
| ENSDARG000 | 1;1 | 56546605:565 | 56546634:565 | -;- | 732 | NA | 0 | 0 | 0 | 0 | 0 | 0 |
| ENSDARG000 | 1:1:1:1:1:1:1:1: | 14348399:143 | 14348512:143 | +:+:+:+:+:+:+: | 2226 | crmp1 | 14 | 0 | 2 | 7 | 1 | 0 |
| ENSDARG000 | 1 | 39513741 | 39513857 | + | 117 | 5S_rRNA | 0 | 0 | 0 | 0 | 0 | 0 |
| ENSDARG000 | 1 | 47914095 | 47914209 | + | 115 | 5S_rRNA | 0 | 0 | 0 | 0 | 0 | 0 |
| ENSDARG000 | 1 | 16768683 | 16768809 | + | 127 | 5S_rRNA | 0 | 0 | 0 | 0 | 0 | 0 |
| ENSDARG000 | 1 | 56253961 | 56254076 | - | 116 | 5S_rRNA | 0 | 0 | 0 | 0 | 0 | 0 |
| ENSDARG000 | 1 | 36154261 | 36154377 | + | 117 | 5S_rRNA | 0 | 0 | 0 | 0 | 0 | 0 |
| ENSDARG000 | 1 | 1218691 | 1218805 | + | 115 | 5S_rRNA | 0 | 0 | 0 | 0 | 0 | 0 |
| ENSDARG000 | 1 | 39609833 | 39609950 | + | 118 | 5S_rRNA | 0 | 0 | 0 | 0 | 0 | 0 |
| ENSDARG000 | 1 | 14593705 | 14593798 | - | 94 | NA | 0 | 0 | 0 | 0 | 0 | 0 |
| ENSDARG000 | 1;1;1;1;1;1;1 | 57787642:577 | 57787745:577 | +;+;+;+;+;+;+ | 1696 | NA | 0 | 0 | 0 | 0 | 0 | 0 |
| ENSDARG000 | 1 | 14701736 | 14701830 | + | 95 | NA | 0 | 0 | 0 | 0 | 0 | 0 |
| ENSDARG000 | 1:1:1:1:1:1:1:1: | 56036644:56( | 56036955:56( | -:-:-:-:-:-:-:-: | 5152 | c3a.6 | 0 | 0 | 44 | 2 | 0 | 17 |
| ENSDARG000 | 1 | 24825147 | 24825263 | - | 117 | 5S_rRNA | 0 | 0 | 0 | 0 | 0 | 0 |
| ENSDARG000 | 1;1 | 56553900:565 | 56554021:565 | -;- | 735 | NA | 8 | 0 | 0 | 1 | 4 | 0 |
| ENSDARG000 | 1 | 57147219 | 57147335 | - | 117 | 5S_rRNA | 0 | 0 | 0 | 0 | 0 | 0 |
| ENSDARG000 | 1;1;1;1;1;1;1 | 15623961:156 | 15624158:156 | -;-;-;-;-;-;- | 1373 | SGCZ | 0 | 2 | 0 | 0 | 0 | 0 |
| ENSDARG000 | 1 | 32706109 | 32706227 | + | 119 | 5S_rRNA | 0 | 0 | 0 | 0 | 0 | 0 |
| ENSDARG000 | 1:1:1:1:1:1:1:1: | 15293531:152 | 15293627:152 | -:-:-:-:-:-:-:-: | 3702 | DLC1 | 4122 | 3535 | 3521 | 3054 | 2730 | 3191 |
| ENSDARG000 | 1 | 51683519 | 51683618 | + | 100 | U6 | 0 | 0 | 0 | 0 | 0 | 0 |
| ENSDARG000 | 1:1:1:1:1:1:1:1: | 14020445:14( | 14020525:14( | +:+:+:+:+:+:+: | 1841 | hpf1 | 316 | 608 | 762 | 368 | 668 | 416 |
| ENSDARG000 | 1 | 41125286 | 41125402 | + | 117 | 5S_rRNA | 0 | 0 | 0 | 0 | 0 | 0 |
| ENSDARG000 | 1 | 42094624 | 42094739 | + | 116 | 5S_rRNA | 0 | 0 | 0 | 0 | 0 | 0 |
| ENSDARG000 | 1 | 14372680 | 14373891 | + | 1212 | DRD5 | 0 | 0 | 0 | 0 | 12 | 0 |
| ENSDARG000 | 1 | 4210498 | 4210652 | - | 155 | 5_8S_rRNA | 0 | 0 | 0 | 0 | 0 | 0 |
| ENSDARG000 | 1 | 14515445 | 14515539 | - | 95 | NA | 0 | 0 | 4 | 0 | 0 | 0 |
| ENSDARG000 | 1 | 9197859 | 9197966 | - | 108 | U6 | 0 | 0 | 0 | 0 | 0 | 0 |
| ENSDARG000 | 1:1:1:1:1:1:1:1: | 218524:21928 | 218910:21946 | +:+:+:+:+:+:+: | 1998 | TMCO3 | 136 | 141 | 131 | 171 | 134 | 129 |
| ENSDARG000 | 1:1:1:1:1:1:1:1: | 961607:96323 | 961665:96334 | +:+:+:+:+:+:+: | 1159 | n6amt1 | 192 | 207 | 222 | 128 | 200 | 246 |
| ENSDARG000 | 1 | 54186039 | 54186156 | + | 118 | 5S_rRNA | 0 | 0 | 0 | 0 | 0 | 0 |
| ENSDARG000 | 1:1:1:1:1:1:1:1: | 227241:23029 | 227378:23037 | +:+:+:+:+:+:+: | 3773 | tfdp1b | 193 | 318 | 447 | 142 | 312 | 197 |
| ENSDARG000 | 1;1 | 9071324:9074 | 9071347:9074 | +;+ | 177 | NA | 2 | 0 | 0 | 0 | 0 | 0 |
| ENSDARG000 | 1:1:1:1:1:1:1:1: | 26012668:26( | 26012707:26( | +:+:+:+:+:+:+: | 240 | NA | 0 | 0 | 0 | 0 | 0 | 0 |
| ENSDARG000 | 1:1:1:1:1:1:1:1: | 48581374:485 | 48583855:485 | -:-:-:-:-:-:-:-: | 3791 | NA | 0 | 0 | 6 | 0 | 2 | 0 |
| ENSDARG000 | 1;1 | 56755536:567 | 56756376:567 | +;+ | 2193 | NA | 0 | 0 | 0 | 0 | 0 | 0 |
| ENSDARG000 | 1:1:1:1:1:1:1:1: | 14466860:144 | 14466980:144 | +:+:+:+:+:+:+: | 7838 | NA | 572 | 357 | 722 | 741 | 452 | 1091 |
| ENSDARG000 | 1 | 24114525 | 24114558 | + | 34 | 5S_rRNA | 0 | 0 | 0 | 0 | 0 | 0 |
| ENSDARG000 | 1 | 11062524 | 11062647 | - | 124 | 5S_rRNA | 0 | 0 | 0 | 0 | 0 | 0 |
| ENSDARG000 | 1;1 | 52754085:527 | 52754111:527 | +;+ | 5371 | NA | 1282 | 2302 | 39 | 1565 | 44 | 10 |
| ENSDARG000 | 1 | 13782930 | 13783046 | - | 117 | 5S_rRNA | 0 | 0 | 0 | 0 | 0 | 0 |
| ENSDARG000 | 1 | 39985905 | 39986160 | - | 256 | RNase_MRP | 0 | 0 | 0 | 0 | 0 | 0 |
| ENSDARG000 | 1 | 41601035 | 41601152 | - | 118 | 5S_rRNA | 4 | 0 | 0 | 0 | 0 | 0 |
| ENSDARG000 | 1 | 3298006 | 3298088 | - | 83 | dre-mir-18a | 0 | 0 | 0 | 0 | 0 | 0 |
| ENSDARG000 | 1 | 14716016 | 14716110 | + | 95 | NA | 0 | 0 | 0 | 0 | 0 | 0 |
| ENSDARG000 | 1:1:1:1:1:1:1:1: | 15226268:152 | 15226520:152 | +:+:+:+:+:+:+: | 6059 | HGSNAT | 2776 | 3022 | 2574 | 1171 | 1842 | 2666 |
| ENSDARG000 | 1 | 26350526 | 26350641 | - | 116 | 5S_rRNA | 0 | 0 | 0 | 0 | 0 | 0 |
| ENSDARG000 | 1:1:1:1:1:1:1:1: | 691738:69466 | 692524:69483 | -;-;-;-;-;-;-;- | 2814 | adamts5 | 156 | 18 | 128 | 58 | 44 | 10 |
| ENSDARG000 | 1:1:1:1:1:1:1:1: | 208181:20897 | 208616:2093( | -;-;-;-;-;-;-;- | 1489 | dcun1d2b | 617 | 413 | 531 | 459 | 649 | 750 |
| ENSDARG000 | 1;1;1;1 | 15137901:151 | 15141412:151 | +;+;+;+ | 7522 | NA | 78 | 32 | 40 | 16 | 20 | 104 |
| ENSDARG000 | 1 | 53764107 | 53764223 | - | 117 | 5S_rRNA | 0 | 0 | 0 | 0 | 0 | 0 |
| ENSDARG000 | 1 | 22521212 | 22521326 | + | 115 | 5S_rRNA | 0 | 0 | 0 | 0 | 0 | 0 |
| ENSDARG000 | 1 | 6787690 | 6787804 | + | 115 | 5S_rRNA | 0 | 0 | 0 | 0 | 0 | 0 |
| ENSDARG000 | 1 | 46945027 | 46945143 | - | 117 | 5S_rRNA | 0 | 0 | 0 | 0 | 0 | 0 |
| ENSDARG000 | 1 | 14648999 | 14649069 | - | 71 | NA | 0 | 0 | 0 | 0 | 0 | 0 |
| ENSDARG000 | 1 | 43224477 | 43224591 | + | 115 | 5S_rRNA | 0 | 0 | 0 | 0 | 0 | 0 |
| ENSDARG000 | 1;1;1;1 | 58870054:588 | 58871577:588 | -;-;-;- | 2286 | NA | 589 | 282 | 377 | 150 | 371 | 162 |
| ENSDARG000 | 1;1;1;1;1;1 | 56502706:565 | 56502807:565 | +;+;+;+;+;+ | 5034 | NA | 0 | 2 | 0 | 9 | 4 | 4 |
| ENSDARG000 | 1:1:1:1:1:1:1:1: | 34954697:349 | 34956109:349 | -:-:-:-:-:-:-:-: | 3562 | NA | 0 | 6 | 2 | 10 | 20 | 12 |
| ENSDARG000 | 1:1:1:1:1:1:1:1: | 58922027:589 | 58922050:589 | +:+:+:+:+:+:+: | 1398 | TRIP10 | 10 | 12 | 18 | 6 | 6 | 0 |
| ENSDARG000 | 1:1:1:1:1:1:1:1: | 15204633:152 | 15204803:152 | +:+:+:+:+:+:+: | 2191 | itln1 | 0 | 13 | 0 | 8 | 38 | 12 |
| ENSDARG000 | 1:1:1:1:1:1:1:1: | 14454663:144 | 14454709:144 | +:+:+:+:+:+:+: | 5139 | rbpja | 531 | 486 | 666 | 526 | 554 | 505 |
| ENSDARG000 | 1:1:1:1:1:1:1:1: | 31374751:313 | 31374975:313 | -:-:-:-:-:-:-:-: | 4065 | NA | 0 | 10 | 13 | 8 | 2 | 4 |
| ENSDARG000 | 1:1:1:1:1:1:1:1: | 160302:16198 | 160382:16201 | -:-:-:-:-:-:-:-: | 297 | NA | 0 | 4 | 0 | 0 | 0 | 3 |
| ENSDARG000 | 1:1:1:1:1:1:1:1: | 23261857:232 | 23261875:232 | +:+:+:+:+:+:+: | 243 | NA | 3 | 0 | 1 | 0 | 1 | 0 |
| ENSDARG000 | 1;1;1 | 54124209:541 | 54124307:541 | +;+;+ | 372 | NA | 135 | 200 | 285 | 116 | 237 | 104 |
| ENSDARG000 | 1;1;1;1;1;1 | 36188184:361 | 36188203:361 | -;-;-;-;-;- | 117 | NA | 0 | 0 | 0 | 0 | 0 | 0 |
| ENSDARG000 | 1;1 | 56714284:567 | 56717701:567 | -;- | 3458 | NA | 0 | 0 | 0 | 0 | 0 | 0 |
| ENSDARG000 | 1 | 32513989 | 32514104 | + | 116 | 5S_rRNA | 0 | 0 | 0 | 0 | 0 | 0 |
| ENSDARG000 | 1;1;1;1;1;1 | 55264712:552 | 55264728:552 | +;+;+;+;+;+ | 135 | NA | 1 | 1 | 0 | 2 | 2 | 0 |
| ENSDARG000 | 1 | 16932586 | 16932801 | - | 216 | U3 | 0 | 0 | 0 | 0 | 0 | 0 |
| ENSDARG000 | 1;1;1;1 | 43985417:439 | 43986514:439 | -;-;-;- | 1767 | NA | 0 | 0 | 0 | 0 | 0 | 8 |
| ENSDARG000 | 1;1;1;1 | 14253118:142 | 14253334:142 | +;+;+;+ | 716 | NA | 230 | 150 | 178 | 266 | 314 | 178 |
| ENSDARG000 | 1 | 34122880 | 34122972 | + | 93 | dre-mir-734 | 0 | 0 | 0 | 0 | 0 | 0 |
| ENSDARG000 | 1 | 3297603 | 3297755 | - | 153 | dre-mir-20a | 0 | 0 | 0 | 0 | 0 | 0 |
| ENSDARG000 | 1:1:1:1:1:1:1:1: | 59353230:593 | 59355257:593 | -:-:-:-:-:-:-:-: | 3470 | NA | 16 | 14 | 42 | 30 | 49 | 22 |
| ENSDARG000 | 1:1:1:1:1:1:1:1: | 54282042:542 | 54282401:542 | -:-:-:-:-:-:-:-: | 13041 | NA | 1773 | 589 | 1422 | 874 | 1863 | 1093 |
| ENSDARG000 | 1:1:1:1:1:1:1:1: | 14382584:143 | 14383100:143 | +:+:+:+:+:+:+: | 3192 | fip1l1b | 1051 | 1142 | 1348 | 1117 | 1314 | 1116 |
| ENSDARG000 | 1;1;1;1;1;1 | 59192197:591 | 59192669:591 | -;-;-;-;-;- | 1786 | lpar2b | 43 | 2 | 10 | 4 | 8 | 9 |
| ENSDARG000 | 1;1;1;1;1;1;1 | 8001435:8004 | 8004202:8004 | -;-;-;-;-;-;- | 3813 | NA | 0 | 0 | 0 | 0 | 0 | 0 |
| ENSDARG000 | 1;1;1 | 59062603:59( | 59062807:59( | +;+;+ | 743 | pin1 | 935 | 1074 | 1274 | 1134 | 910 | 908 |
| ENSDARG000 | 1;1;1;1;1;1;1 | 53221858:532 | 53221882:532 | +;+;+;+;+;+;+ | 153 | NA | 0 | 1 | 1 | 0 | 1 | 3 |
| ENSDARG000 | 1;1;1;1;1;1;1 | 22842935:228 | 22842944:228 | +;+;+;+;+;+;+ | 147 | NA | 0 | 2 | 1 | 2 | 0 | 4 |
| ENSDARG000 | 1 | 33735018 | 33735134 | + | 117 | 5S_rRNA | 1 | 0 | 0 | 0 | 0 | 0 |
| ENSDARG000 | 1;1;1;1;1;1 | 57497959:574 | 57498736:574 | -;-;-;-;-;- | 1715 | zgc:171470 | 0 | 0 | 0 | 0 | 0 | 0 |
| ENSDARG000 | 1:1:1:1:1:1:1:1: | 59015237:59( | 59015346:59( | -;-;-;-;-;-;-;- | 1589 | ears2 | 146 | 158 | 248 | 168 | 199 | 156 |
| ENSDARG000 | 1 | 14337211 | 14337313 | + | 103 | NA | 0 | 0 | 0 | 0 | 0 | 0 |
| ENSDARG000 | 1 | 41311777 | 41311892 | - | 116 | 5S_rRNA | 0 | 0 | 0 | 0 | 0 | 0 |
| ENSDARG000 | 1 | 16941215 | 16941430 | - | 216 | U3 | 0 | 0 | 0 | 0 | 0 | 0 |
| ENSDARG000 | 1 | 34712513 | 34712583 | - | 71 | SNORD58 | 2 | 0 | 0 | 0 | 0 | 0 |
| ENSDARG000 | 1 | 14767576 | 14767670 | + | 95 | NA | 0 | 0 | 5 | 8 | 0 | 0 |
| ENSDARG000 | 1 | 3297367 | 3297451 | - | 85 | dre-mir-92a-1 | 0 | 0 | 0 | 0 | 0 | 0 |
| ENSDARG000 | 1 | 37123642 | 37123772 | + | 131 | 5S_rRNA | 0 | 0 | 0 | 0 | 0 | 0 |
| ENSDARG000 | 1 | 12431328 | 12431442 | + | 115 | 5S_rRNA | 0 | 0 | 0 | 0 | 0 | 0 |
| ENSDARG000 | 1 | 7142039 | 7142143 | - | 105 | U6 | 0 | 0 | 0 | 0 | 0 | 0 |
| ENSDARG000 | 1:1:1:1:1:1:1:1: | 14073891:14( | 14073998:14( | +:+:+:+:+:+:+: | 14544 | NA | 6 | 2 | 18 | 0 | 0 | 0 |
| ENSDARG000 | 1;1;1;1;1 | 58840889:588 | 58841179:588 | +;+;+;+;+ | 1043 | tmed1b | 211 | 235 | 330 | 118 | 270 | 203 |
| ENSDARG000 | 1:1:1:1:1:1:1:1: | 52121918:521 | 52121943:521 | -;-;-;-;-;-;-;- | 177 | NA | 0 | 0 | 0 | 0 | 2 | 0 |
| ENSDARG000 | 1:1:1:1:1:1:1:1: | 54198893:541 | 54199082:541 | +:+:+:+:+:+:+: | 6551 | TSC2 | 1989 | 1196 | 1806 | 1564 | 1889 | 1861 |
| ENSDARG000 | 1:1:1:1:1:1:1:1: | 46500436:465 | 46501503:465 | -;-;-;-;-;-;-;- | 1947 | si:busm1-105 | 50 | 19 | 168 | 136 | 18 | 28 |
| ENSDARG000 | 1:1:1:1:1:1:1:1: | 2712956:2789 | 2713361:2789 | +:+:+:+:+:+:+: | 2021 | gpc6a | 14 | 0 | 16 | 6 | 6 | 2 |
| ENSDARG000 | 1 | 10870117 | 10870241 | - | 125 | 5S_rRNA | 0 | 0 | 0 | 0 | 0 | 0 |
| ENSDARG000 | 1 | 7920119 | 7920206 | - | 88 | NA | 0 | 0 | 0 | 0 | 0 | 0 |
| ENSDARG000 | 1 | 57163027 | 57163141 | - | 115 | 5S_rRNA | 0 | 0 | 0 | 0 | 0 | 0 |
| ENSDARG000 | 1 | 25120089 | 25120205 | + | 117 | 5S_rRNA | 0 | 0 | 0 | 0 | 0 | 0 |
| ENSDARG000 | 1 | 44590267 | 44590381 | + | 115 | 5S_rRNA | 0 | 0 | 0 | 0 | 0 | 0 |
| ENSDARG000 | 1 | 576444 | 576521 | + | 78 | dre-mir-155 | 0 | 0 | 0 | 0 | 0 | 0 |
| ENSDARG000 | 1;1;1;1;1;1;1 | 38151262:381 | 38151289:381 | +;+;+;+;+;+;+ | 168 | NA | 0 | 1 | 1 | 0 | 0 | 3 |
| ENSDARG000 | 1;1 | 35949269:359 | 35950544:359 | +;+ | 1533 | NA | 21 | 26 | 9 | 14 | 11 | 34 |
| ENSDARG000 | 1;1;1;1;1 | 58977262:589 | 58977355:589 | +;+;+;+;+ | 1396 | eri2 | 28 | 37 | 40 | 23 | 40 | 16 |
| ENSDARG000 | 1 | 1050941 | 1051057 | - | 117 | 5S_rRNA | 0 | 0 | 0 | 0 | 0 | 0 |
| ENSDARG000 | 1;1 | 59064686:59( | 59064942:59( | -;- | 453 | NA | 0 | 0 | 0 | 0 | 0 | 4 |
| ENSDARG000 | 1;1;1;1 | 50773727:507 | 50774454:507 | -;-;-;- | 1355 | NA | 26 | 19 | 28 | 2 | 25 | 10 |
| ENSDARG000 | 1;1 | 4101741:415( | 4101889:4154 | +;+ | 3890 | slitrk6 | 44 | 70 | 110 | 228 | 58 | 96 |
| ENSDARG000 | 1 | 42974760 | 42974875 | - | 116 | 5S_rRNA | 0 | 0 | 0 | 0 | 0 | 0 |
| ENSDARG000 | 1;1;1 | 54154230:541 | 54154445:541 | -;-;- | 865 | NA | 10 | 2 | 4 | 0 | 1 | 16 |
| ENSDARG000 | 1:1:1:1:1:1:1:1: | 13260809:132 | 13262394:132 | -;-;-;-;-;-;-;- | 5254 | pcdh18a | 128 | 39 | 406 | 16 | 70 | 146 |
| ENSDARG000 | 1;1 | 56768263:567 | 56768500:567 | +;+ | 878 | NA | 0 | 0 | 0 | 0 | 0 | 0 |
| ENSDARG000 | 1;1 | 58677434:586 | 58678099:586 | +;+ | 1796 | NA | 143 | 113 | 193 | 157 | 181 | 155 |
| ENSDARG000 | 1 | 16162104 | 16162220 | + | 117 | 5S_rRNA | 0 | 0 | 0 | 0 | 0 | 0 |
| ENSDARG000 | 1;1;1;1;1 | 44374034:443 | 44374064:443 | -;-;-;-;- | 105 | NA | 0 | 0 | 0 | 0 | 0 | 0 |
| ENSDARG000 | 1:1:1:1:1:1:1:1: | 14432434:144 | 14432682:144 | +:+:+:+:+:+:+: | 2615 | slc34a2a | 4 | 2 | 13 | 4 | 10 | 12 |
| ENSDARG000 | 1:1:1:1:1:1:1:1: | 18234470:182 | 18234504:182 | +:+:+:+:+:+:+: | 399 | NA | 0 | 0 | 0 | 0 | 0 | 0 |
| ENSDARG000 | 1:1:1:1:1:1:1:1: | 38416106:384 | 38418398:384 | -;-;-;-;-;-;-;- | 3767 | glra3 | 2 | 6 | 327 | 54 | 38 | 8 |
| ENSDARG000 | 1 | 21254353 | 21254482 | + | 130 | 5S_rRNA | 0 | 0 | 0 | 0 | 0 | 0 |
| ENSDARG000 | 1 | 16937418 | 16937633 | - | 216 | U3 | 0 | 0 | 0 | 0 | 0 | 0 |

| | | | | | | | | | | | |
|---|---|---|---|---|---|---|---|---|---|---|---|
| ENSDARG000 | 1 | 34711932 | 34712003 | - | 72 | SNORD58 | 0 | 0 | 0 | 0 | 0 | 0 |
| ENSDARG000 | 1 | 19705699 | 19705832 | + | 134 | 5S_rRNA | 0 | 0 | 0 | 0 | 0 | 0 |
| ENSDARG000 | 1;1;1;1;1;1 | 14406098:14 | 14406376:14 | +;+;+;+;+;+ | 1909 | chic2 | 440 | 444 | 507 | 314 | 510 | 477 |
| ENSDARG000 | 1 | 47706127 | 47706244 | - | 118 | 5S_rRNA | 0 | 0 | 0 | 0 | 0 | 0 |
| ENSDARG000 | 1;1 | 22677429:22 | 22678243:22 | -;- | 866 | NA | 30 | 4 | 24 | 32 | 0 | 6 |
| ENSDARG000 | 1 | 6453404 | 6453519 | - | 116 | 5S_rRNA | 0 | 0 | 0 | 0 | 0 | 0 |
| ENSDARG000 | 1 | 39373802 | 39373911 | - | 110 | dre-mir-2197 | 0 | 0 | 0 | 0 | 0 | 0 |
| ENSDARG000 | 1;1;1;1 | 54194801:54 | 54194935:54 | +;+;+;+ | 489 | NA | 28 | 34 | 53 | 22 | 65 | 55 |
| ENSDARG000 | 1 | 12543359 | 12543475 | + | 117 | 5S_rRNA | 0 | 0 | 0 | 0 | 0 | 0 |
| ENSDARG000 | 1:1:1:1:1:1:1: | 34696503:34 | 34696505:34 | +:+:+:+:+:+:+: | 600 | NA | 97 | 61 | 100 | 55 | 136 | 168 |
| ENSDARG000 | 1;1;1;1 | 44056129:44 | 44056147:44 | +;+;+;+ | 438 | NA | 20 | 30 | 36 | 19 | 57 | 56 |
| ENSDARG000 | 1 | 57486529 | 57486644 | - | 116 | 5S_rRNA | 0 | 0 | 0 | 0 | 0 | 0 |
| ENSDARG000 | 1:1:1:1:1:1:1: | 58898124:58 | 58899916:58 | -:-:-:-:-:-:-:-: | 2583 | MFAP4 | 0 | 4 | 4 | 0 | 0 | 0 |
| ENSDARG000 | 1 | 59131292 | 59131406 | + | 115 | 5S_rRNA | 0 | 0 | 0 | 0 | 0 | 0 |
| ENSDARG000 | 1:1:1:1:1:1:1: | 57620964:57 | 57621035:57 | -:-:-:-:-:-:-:-: | 1461 | NA | 8 | 14 | 2 | 19 | 25 | 21 |
| ENSDARG000 | 1 | 7919480 | 7919567 | - | 88 | NA | 2 | 4 | 9 | 4 | 7 | 7 |
| ENSDARG000 | 1 | 14673777 | 14673871 | + | 95 | NA | 0 | 0 | 0 | 0 | 0 | 0 |
| ENSDARG000 | 1;1;1;1;1;1;1 | 14313850:14 | 14315029:14 | -;-;-;-;-;-;- | 2833 | WFS1 | 0 | 2 | 0 | 6 | 0 | 0 |
| ENSDARG000 | 1:1:1:1:1:1:1: | 176583:1765 | 176951:1768 | +:+:+:+:+:+:+: | 1194 | LAMP1 | 2911 | 1963 | 1692 | 2137 | 2641 | 1606 |
| ENSDARG000 | 1 | 23751099 | 23751213 | + | 115 | 5S_rRNA | 0 | 0 | 0 | 0 | 0 | 0 |
| ENSDARG000 | 1 | 14560391 | 14560485 | - | 95 | NA | 0 | 0 | 0 | 1 | 1 | 1 |
| ENSDARG000 | 1;1 | 14658801:14 | 14658835:14 | +;+ | 1269 | NA | 103 | 116 | 154 | 100 | 112 | 78 |
| ENSDARG000 | 1;1;1 | 52735484:52 | 52736007:52 | +;+;+ | 582 | NA | 0 | 0 | 12 | 0 | 3 | 0 |
| ENSDARG000 | 1 | 7988052 | 7989314 | + | 1263 | NA | 8 | 0 | 0 | 0 | 0 | 2 |
| ENSDARG000 | 1;1;1;1;1;1 | 14257158:14 | 14257280:14 | -;-;-;-;-;- | 528 | NA | 1 | 11 | 4 | 0 | 9 | 0 |
| ENSDARG000 | 1;1 | 59123750:59 | 59124257:59 | -;- | 677 | NA | 0 | 0 | 0 | 0 | 1 | 3 |
| ENSDARG000 | 1 | 47897269 | 47897385 | + | 117 | 5S_rRNA | 0 | 0 | 0 | 0 | 0 | 0 |
| ENSDARG000 | 1 | 34400240 | 34400354 | + | 115 | 5S_rRNA | 0 | 0 | 0 | 0 | 0 | 0 |
| ENSDARG000 | 1 | 16927724 | 16927939 | - | 216 | U3 | 0 | 0 | 0 | 0 | 0 | 0 |
| ENSDARG000 | 1 | 14694561 | 14694655 | + | 95 | NA | 0 | 0 | 0 | 0 | 0 | 0 |
| ENSDARG000 | 1;1 | 55452892:55 | 55453235:55 | +;+ | 646 | NA | 0 | 0 | 0 | 0 | 0 | 0 |
| ENSDARG000 | 1 | 34708497 | 34708627 | - | 131 | NA | 0 | 3 | 3 | 1 | 13 | 0 |
| ENSDARG000 | 1 | 32013688 | 32014487 | + | 800 | NA | 40 | 21 | 61 | 47 | 54 | 0 |
| ENSDARG000 | 1 | 23866878 | 23866965 | + | 88 | dre-mir-218a- | 0 | 0 | 0 | 0 | 0 | 0 |
| ENSDARG000 | 1 | 14737470 | 14737564 | + | 95 | NA | 0 | 0 | 0 | 0 | 0 | 0 |
| ENSDARG000 | 1 | 38858399 | 38859425 | + | 1027 | NA | 4 | 0 | 0 | 0 | 0 | 0 |
| ENSDARG000 | 1;1;1;1;1;1;1 | 32791920:32 | 32791986:32 | +;+;+;+;+;+;+ | 1050 | NA | 2 | 8 | 2 | 0 | 4 | 4 |
| ENSDARG000 | 1:1:1:1:1:1:1: | 54137089:54 | 54137333:54 | +:+:+:+:+:+:+: | 1413 | TNPO2 | 497 | 674 | 905 | 422 | 895 | 558 |
| ENSDARG000 | 1 | 37996459 | 37996577 | + | 119 | 5S_rRNA | 0 | 0 | 0 | 0 | 0 | 0 |
| ENSDARG000 | 1 | 58527632 | 58527756 | - | 125 | 5S_rRNA | 0 | 0 | 0 | 0 | 0 | 0 |
| ENSDARG000 | 1 | 16939166 | 16939381 | - | 216 | U3 | 0 | 0 | 0 | 0 | 0 | 0 |
| ENSDARG000 | 1 | 14639582 | 14639676 | - | 95 | NA | 0 | 0 | 0 | 0 | 0 | 0 |
| ENSDARG000 | 1 | 13875805 | 13875921 | - | 117 | 5S_rRNA | 0 | 0 | 0 | 0 | 0 | 0 |
| ENSDARG000 | 1;1;1;1;1;1;1 | 1362932:136 | 1364280:136 | -;-;-;-;-;-;- | 2220 | runx1 | 235 | 177 | 346 | 266 | 234 | 206 |
| ENSDARG000 | 1 | 50854364 | 50854478 | + | 115 | 5S_rRNA | 0 | 0 | 0 | 0 | 0 | 0 |
| ENSDARG000 | 1 | 16935395 | 16935611 | - | 217 | NA | 0 | 0 | 0 | 0 | 0 | 0 |
| ENSDARG000 | 1 | 21680364 | 21680478 | - | 115 | 5S_rRNA | 0 | 0 | 0 | 0 | 0 | 0 |
| ENSDARG000 | 1 | 16935396 | 16935611 | - | 216 | U3 | 0 | 0 | 0 | 0 | 0 | 0 |
| ENSDARG000 | 1 | 14708876 | 14708970 | + | 95 | NA | 0 | 0 | 0 | 0 | 0 | 0 |
| ENSDARG000 | 1 | 3759924 | 3760034 | + | 111 | 5S_rRNA | 0 | 0 | 0 | 0 | 0 | 0 |
| ENSDARG000 | 1 | 33001559 | 33001673 | - | 115 | 5S_rRNA | 0 | 0 | 0 | 0 | 0 | 0 |
| ENSDARG000 | 1:1:1:1:1:1:1: | 184915:1849 | 185422:1854 | -:-:-:-:-:-:-:-: | 1723 | grtp1a | 69 | 31 | 36 | 12 | 26 | 25 |
| ENSDARG000 | 1:1:1:1:1:1:1: | 6243906:624 | 6244089:624 | +:+:+:+:+:+:+: | 2932 | gtf3c3 | 215 | 223 | 342 | 146 | 271 | 341 |
| ENSDARG000 | 1 | 44507134 | 44507253 | + | 120 | 5S_rRNA | 0 | 0 | 0 | 0 | 0 | 0 |
| ENSDARG000 | 1 | 49590701 | 49590823 | + | 123 | 5S_rRNA | 0 | 0 | 0 | 0 | 0 | 0 |
| ENSDARG000 | 1 | 34668236 | 34668351 | - | 116 | 5S_rRNA | 0 | 0 | 0 | 0 | 0 | 0 |
| ENSDARG000 | 1 | 3297848 | 3297936 | - | 89 | dre-mir-19a | 0 | 0 | 0 | 0 | 0 | 0 |
| ENSDARG000 | 1 | 46722622 | 46722705 | - | 84 | dre-mir-15a-1 | 0 | 0 | 2 | 0 | 0 | 4 |
| ENSDARG000 | 1:1:1:1:1:1:1: | 3100244:310 | 3100332:310 | -;-;-;-;-;-;-;- | 1635 | gpc5a | 2100 | 1603 | 2173 | 2231 | 2854 | 1492 |
| ENSDARG000 | 1 | 16458525 | 16458740 | + | 216 | U3 | 0 | 0 | 0 | 0 | 0 | 0 |
| ENSDARG000 | 1:1:1:1:1:1:1: | 53568296:53 | 53568966:53 | -:-:-:-:-:-:-:-: | 8418 | USP34 | 814 | 687 | 1559 | 777 | 1064 | 920 |
| ENSDARG000 | 1 | 55128465 | 55128579 | + | 115 | 5S_rRNA | 0 | 0 | 0 | 0 | 0 | 0 |
| ENSDARG000 | 1:1:1:1:1:1:1: | 56421754:56 | 56423884:56 | -:-:-:-:-:-:-:-: | 2890 | slc25a10b | 178 | 249 | 876 | 370 | 616 | 717 |
| ENSDARG000 | 1;1;1;1;1 | 58885602:58 | 58886434:58 | -;-;-;-;- | 1651 | MFAP4 | 0 | 2 | 0 | 0 | 0 | 0 |
| ENSDARG000 | 1:1:1:1:1:1:1: | 59338175:59 | 59338732:59 | -:-:-:-:-:-:-:-: | 2020 | cyp3a65 | 28 | 62 | 26 | 38 | 34 | 18 |
| ENSDARG000 | 1 | 53387765 | 53387881 | + | 117 | 5S_rRNA | 0 | 0 | 0 | 0 | 0 | 0 |
| ENSDARG000 | 1 | 14553221 | 14553315 | - | 95 | NA | 0 | 0 | 0 | 0 | 0 | 0 |
| ENSDARG000 | 1 | 25585 | 27255 | + | 1671 | NA | 0 | 0 | 0 | 0 | 0 | 2 |
| ENSDARG000 | 1;1 | 552275:5582 | 553355:5582 | -;- | 1110 | NA | 20 | 60 | 55 | 18 | 69 | 22 |
| ENSDARG000 | 1;1;1;1;1 | 59328030:59 | 59328339:59 | +;+;+;+;+ | 2088 | NA | 161 | 180 | 313 | 157 | 334 | 156 |
| ENSDARG000 | 1;1;1 | 54411669:54 | 54411715:54 | -;-;- | 240 | NA | 0 | 0 | 0 | 0 | 0 | 0 |
| ENSDARG000 | 1 | 56519163 | 56519279 | - | 117 | 5S_rRNA | 0 | 0 | 0 | 0 | 0 | 0 |
| ENSDARG000 | 1 | 51465468 | 51465582 | + | 115 | 5S_rRNA | 2 | 0 | 0 | 0 | 0 | 0 |
| ENSDARG000 | 1 | 41269773 | 41269887 | + | 115 | 5S_rRNA | 0 | 0 | 0 | 0 | 0 | 0 |
| ENSDARG000 | 1 | 3297484 | 3297570 | - | 87 | dre-mir-19b | 0 | 0 | 0 | 0 | 0 | 0 |
| ENSDARG000 | 1:1:1:1:1:1:1: | 55215787:55 | 55215825:55 | -:-:-:-:-:-:-:-: | 3003 | CC2D1A | 585 | 267 | 508 | 269 | 358 | 467 |
| ENSDARG000 | 1:1:1:1:1:1:1: | 45958904:45 | 45959183:45 | +:+:+:+:+:+:+: | 3636 | arhgef7b | 663 | 490 | 597 | 574 | 677 | 675 |
| ENSDARG000 | 1:1:1:1:1:1:1: | 50694971:50 | 50695094:50 | -:-:-:-:-:-:-:-: | 1425 | FAM13A | 562 | 1286 | 3826 | 4439 | 1411 | 618 |
| ENSDARG000 | 1:1:1:1:1:1:1: | 960209:9625 | 960225:9625 | -:-:-:-:-:-:-:-: | 3095 | gart | 1039 | 1430 | 2728 | 1672 | 1430 | 1797 |
| ENSDARG000 | 1:1:1:1:1:1:1: | 59007536:59 | 59007643:59 | +:+:+:+:+:+:+: | 1095 | cdc37 | 1103 | 999 | 1081 | 542 | 1122 | 511 |
| ENSDARG000 | 1:1:1:1:1:1:1: | 58639497:58 | 58639671:58 | -:-:-:-:-:-:-:-: | 2151 | NA | 472 | 694 | 1719 | 752 | 684 | 323 |
| ENSDARG000 | 1;1 | 28594809:28 | 28599046:28 | -;- | 5856 | NA | 588 | 222 | 712 | 1350 | 517 | 1146 |
| ENSDARG000 | 1 | 37363872 | 37363990 | - | 119 | 5S_rRNA | 0 | 0 | 0 | 0 | 0 | 0 |
| ENSDARG000 | 1 | 34711138 | 34711271 | - | 134 | NA | 0 | 0 | 0 | 0 | 0 | 0 |
| ENSDARG000 | 1 | 16930156 | 16930372 | - | 217 | NA | 0 | 0 | 0 | 0 | 0 | 0 |
| ENSDARG000 | 1:1:1:1:1:1:1: | 59172744:59 | 59173046:59 | -:-:-:-:-:-:-:-: | 3045 | CAMSAP3 | 4 | 2 | 0 | 0 | 14 | 0 |
| ENSDARG000 | 1:1:1:1:1:1:1: | 14283692:14 | 14283885:14 | +:+:+:+:+:+:+: | 2428 | ppp2r2ca | 0 | 0 | 0 | 0 | 1 | 0 |
| ENSDARG000 | 1 | 5078371 | 5078488 | - | 118 | 5S_rRNA | 0 | 0 | 0 | 0 | 0 | 0 |
| ENSDARG000 | 1 | 30244531 | 30244645 | - | 115 | 5S_rRNA | 0 | 0 | 0 | 0 | 0 | 0 |
| ENSDARG000 | 1 | 14730296 | 14730390 | + | 95 | NA | 0 | 0 | 0 | 0 | 0 | 0 |
| ENSDARG000 | 1;1 | 59125767:59 | 59125865:59 | -;- | 303 | NA | 0 | 0 | 0 | 0 | 0 | 1 |
| ENSDARG000 | 1 | 16937417 | 16937633 | - | 217 | NA | 0 | 0 | 0 | 0 | 0 | 0 |
| ENSDARG000 | 1;1;1;1;1 | 1350111:135 | 1350921:135 | -;-;-;-;- | 1364 | NA | 16 | 70 | 78 | 26 | 36 | 22 |
| ENSDARG000 | 1;1;1;1 | 3305549:330 | 3306074:330 | -;-;-;- | 869 | NA | 0 | 0 | 2 | 0 | 0 | 0 |
| ENSDARG000 | 1;1;1 | 4598060:466 | 4598270:466 | +;+;+ | 1302 | NA | 0 | 0 | 0 | 0 | 0 | 0 |
| ENSDARG000 | 1;1;1;1;1;1;1 | 7092273:709 | 7092383:709 | +;+;+;+;+;+;+ | 719 | NA | 0 | 0 | 0 | 0 | 0 | 0 |
| ENSDARG000 | 1;1;1 | 7771557:784 | 7773138:784 | -;-;- | 1805 | NA | 0 | 3 | 0 | 0 | 0 | 0 |
| ENSDARG000 | 1;1;1;1 | 9308271:931 | 9309210:931 | -;-;-;- | 1326 | NA | 0 | 0 | 0 | 0 | 0 | 0 |
| ENSDARG000 | 1;1;1;1 | 12552300:12 | 12554407:12 | -;-;-;- | 2608 | NA | 0 | 0 | 4 | 0 | 0 | 4 |
| ENSDARG000 | 1:1:1:1:1:1:1: | 13373950:13 | 13374638:13 | -;-;-;-;-;-;-;- | 1290 | NA | 0 | 0 | 0 | 0 | 0 | 0 |
| ENSDARG000 | 1:1:1:1:1:1:1: | 13596298:13 | 13599244:13 | -:-:-:-:-:-:-:-: | 4379 | NA | 2 | 0 | 0 | 0 | 4 | 89 |
| ENSDARG000 | 1:1:1:1:1:1:1: | 14565965:14 | 14566761:14 | -;-;-;-;-;-;-;- | 1657 | NA | 4 | 0 | 18 | 2 | 4 | 6 |
| ENSDARG000 | 1:1:1:1:1:1:1: | 15355857:15 | 15357668:15 | -:-:-:-:-:-:-:-: | 5715 | NA | 286 | 117 | 232 | 388 | 130 | 335 |
| ENSDARG000 | 1;1;1;1;1;1;1 | 18002567:18 | 18002660:18 | -;-;-;-;-;-;- | 975 | NA | 0 | 0 | 0 | 1 | 2 | 0 |
| ENSDARG000 | 1:1:1:1:1:1:1: | 18061682:18 | 18062557:18 | -;-;-;-;-;-;-;- | 1591 | NA | 2 | 2 | 0 | 0 | 0 | 2 |
| ENSDARG000 | 1;1;1;1 | 38263693:38 | 38263953:38 | +;+;+;+ | 1755 | NA | 0 | 0 | 0 | 0 | 0 | 0 |
| ENSDARG000 | 1:1:1:1:1:1:1: | 38928636:38 | 38929077:38 | +:+:+:+:+:+:+: | 1603 | NA | 0 | 0 | 0 | 0 | 0 | 0 |
| ENSDARG000 | 1:1:1:1:1:1:1: | 40663536:40 | 40666902:40 | +:+:+:+:+:+:+: | 5778 | NA | 8 | 0 | 0 | 0 | 0 | 0 |
| ENSDARG000 | 1:1:1:1:1:1:1: | 40683491:40 | 40683608:40 | +:+:+:+:+:+:+: | 4754 | NA | 0 | 0 | 0 | 16 | 0 | 2 |
| ENSDARG000 | 1:1:1:1:1:1:1: | 40693323:40 | 40693499:40 | +:+:+:+:+:+:+: | 6146 | NA | 20 | 0 | 2 | 6 | 12 | 0 |
| ENSDARG000 | 1;1;1;1 | 40716379:40 | 40716861:40 | +;+;+;+ | 3660 | NA | 0 | 0 | 0 | 0 | 2 | 0 |
| ENSDARG000 | 1;1;1;1;1;1 | 43940902:43 | 43941303:43 | +;+;+;+;+;+ | 667 | NA | 0 | 0 | 0 | 0 | 2 | 0 |
| ENSDARG000 | 1;1;1 | 43952298:43 | 43952631:43 | -;-;- | 703 | NA | 0 | 0 | 0 | 0 | 0 | 0 |
| ENSDARG000 | 1;1;1;1 | 44968462:44 | 44968537:44 | +;+;+;+ | 1453 | NA | 6 | 19 | 0 | 1 | 8 | 4 |
| ENSDARG000 | 1:1:1:1:1:1:1: | 45307161:45 | 45307528:45 | -:-:-:-:-:-:-:-: | 2147 | NA | 0 | 0 | 0 | 0 | 0 | 2 |
| ENSDARG000 | 1;1;1;1 | 46039237:46 | 46039448:46 | +;+;+;+ | 1288 | NA | 0 | 0 | 0 | 0 | 0 | 0 |
| ENSDARG000 | 1;1;1;1;1 | 50737700:50 | 50737745:50 | -;-;-;-;- | 1201 | NA | 9 | 6 | 9 | 6 | 16 | 18 |
| ENSDARG000 | 1;1;1;1;1;1 | 52353162:52 | 52353347:52 | +;+;+;+;+;+ | 5639 | NA | 0 | 0 | 0 | 0 | 0 | 0 |
| ENSDARG000 | 1;1;1 | 54159959:54 | 54161905:54 | -;-;- | 2661 | NA | 0 | 0 | 4 | 0 | 6 | 0 |
| ENSDARG000 | 1;1;1;1 | 56638671:56 | 56640736:56 | -;-;-;- | 2536 | NA | 0 | 0 | 0 | 0 | 0 | 4 |
| ENSDARG000 | 1:1:1:1:1:1:1: | 56671781:56 | 56671959:56 | -;-;-;-;-;-;- | 1894 | NA | 0 | 2 | 0 | 0 | 2 | 0 |
| ENSDARG000 | 1;1;1;1;1 | 59395119:59 | 59396879:59 | -;-;-;-;- | 3298 | NA | 2 | 0 | 0 | 0 | 0 | 0 |
| ENSDARG000 | 1;1;1;1;1;1 | 59423277:59 | 59423517:59 | -;-;-;-;-;- | 1748 | NA | 5 | 0 | 1 | 0 | 4 | 0 |
| ENSDARG000 | 1;1;1;1;1 | 59430489:59 | 59432860:59 | -;-;-;-;- | 3217 | NA | 16 | 0 | 0 | 0 | 1 | 0 |
| ENSDARG000 | 1;1;1;1;1 | 59436255:59 | 59436434:59 | -;-;-;-;- | 1194 | NA | 0 | 0 | 0 | 0 | 1 | 0 |
| ENSDARG000 | 1;1;1;1 | 59451692:59 | 59452659:59 | -;-;-;- | 2028 | NA | 3 | 0 | 1 | 0 | 0 | 0 |
| ENSDARG000 | 1:1:1:1:1:1:1: | 59454067:59 | 59454552:59 | -:-:-:-:-:-:-:-: | 2338 | NA | 4 | 0 | 0 | 0 | 16 | 0 |
| ENSDARG000 | 9;9;9 | 22075528:22 | 22075859:22 | -;-;- | 641 | crygm2a | 0 | 0 | 0 | 0 | 0 | 0 |
| ENSDARG000 | 9:9:9:9:9:9:9: | 38029475:38 | 38030849:38 | -:-:-:-:-:-:-:-: | 2482 | hacd2 | 2686 | 4185 | 3171 | 6934 | 4090 | 3822 |
| ENSDARG000 | 9:9:9:9:9:9:9: | 18605460:18 | 18605610:18 | +:+:+:+:+:+:+: | 1359 | NA | 0 | 0 | 0 | 0 | 0 | 0 |
| ENSDARG000 | 9;9;9;9;9 | 22080122:22 | 22080133:22 | +;+;+;+;+ | 663 | NA | 0 | 0 | 0 | 0 | 0 | 0 |
| ENSDARG000 | 9;9;9 | 22091161:22 | 22091466:22 | -;-;- | 642 | crygm2d13 | 0 | 0 | 0 | 0 | 0 | 0 |
| ENSDARG000 | 9;9;9;9;9 | 22098575:22 | 22098993:22 | -;-;-;-;- | 860 | crygm2d11 | 0 | 0 | 0 | 0 | 0 | 0 |
| ENSDARG000 | 9:9:9:9:9:9:9: | 21030930:21 | 21036588:21 | -:-:-:-:-:-:-:-: | 8686 | tbx15 | 18 | 0 | 2 | 2 | 2 | 2 |
| ENSDARG000 | 9;9;9 | 22116600:22 | 22116907:22 | -;-;- | 623 | crygm2d15 | 0 | 0 | 0 | 0 | 0 | 0 |
| ENSDARG000 | 9:9:9:9:9:9:9: | 30585552:30 | 30585953:30 | +:+:+:+:+:+:+: | 4325 | TBC1D4 | 340 | 343 | 1072 | 547 | 955 | 639 |
| ENSDARG000 | 9;9;9 | 48085039:48 | 48085454:48 | +;+;+ | 4127 | NA | 79 | 26 | 76 | 42 | 62 | 70 |
| ENSDARG000 | 9:9:9:9:9:9:9: | 32797816:32 | 32798382:32 | -:-:-:-:-:-:-:-: | 1729 | gatd3 | 289 | 344 | 475 | 246 | 357 | 309 |
| ENSDARG000 | 9 | 48108174 | 48109197 | + | 1024 | CXCR1 | 0 | 0 | 2 | 2 | 2 | 14 |
| ENSDARG000 | 9:9:9:9:9:9:9: | 38043337:38 | 38043501:38 | +:+:+:+:+:+:+: | 2962 | stam2 | 2742 | 2178 | 2060 | 3358 | 2537 | 2641 |
| ENSDARG000 | 9;9;9;9;9;9;9 | 48134394:48 | 48134704:48 | -;-;-;-;-;-;- | 2346 | KLHL23 | 0 | 0 | 0 | 0 | 0 | 0 |
| ENSDARG000 | 9:9:9:9:9:9:9: | 48123002:48 | 48123307:48 | +:+:+:+:+:+:+: | 6462 | mettl5 | 230 | 251 | 405 | 130 | 268 | 206 |
| ENSDARG000 | 9;9;9;9;9 | 22120874:22 | 22122063:22 | -;-;-;-;- | 1654 | crygm2d18 | 0 | 0 | 0 | 0 | 0 | 0 |
| ENSDARG000 | 9:9:9:9:9:9:9: | 8288268:828 | 8291067:829 | -:-:-:-:-:-:-:-: | 4081 | NA | 2149 | 661 | 1671 | 2982 | 2058 | 3749 |
| ENSDARG000 | 9;9;9;9;9;9 | 22929675:22 | 22929973:22 | +;+;+;+;+;+ | 2359 | tsn | 625 | 607 | 759 | 510 | 819 | 532 |
| ENSDARG000 | 9;9;9;9;9 | 22127117:22 | 22128815:22 | -;-;-;-;- | 2177 | crygm2d17 | 0 | 0 | 0 | 0 | 0 | 0 |
| ENSDARG000 | 9;9;9;9;9;9 | 21072359:21 | 21072938:21 | -;-;-;-;-;- | 1246 | wars2 | 82 | 60 | 106 | 44 | 128 | 82 |
| ENSDARG000 | 9;9;9;9;9 | 22134731:22 | 22135040:22 | -;-;-;-;- | 639 | crygm2d8 | 0 | 0 | 0 | 0 | 0 | 0 |
| ENSDARG000 | 9:9:9:9:9:9:9: | 24065855:24 | 24066491:24 | +:+:+:+:+:+:+: | 7510 | NA | 1243 | 817 | 923 | 1750 | 977 | 1814 |
| ENSDARG000 | 9;9;9;9;9;9 | 22940962:22 | 22941766:22 | -;-;-;-;-;- | 3282 | si:dkev-91i10. | 1322 | 2001 | 2088 | 1039 | 4123 | 1524 |
| ENSDARG000 | 9;9;9;9 | 30677470:30 | 30677815:30 | -;-;-;- | 951 | NA | 10 | 0 | 7 | 0 | 10 | 0 |
| ENSDARG000 | 9;9;9;9;9;9 | 22146301:22 | 22146846:22 | -;-;-;-;-;- | 1534 | crygm2d14 | 0 | 0 | 0 | 0 | 0 | 0 |
| ENSDARG000 | 9:9:9:9:9:9:9: | 42894845:42 | 42896172:42 | -:-:-:-:-:-:-:-: | 98586 | TTN | 8 | 8 | 0 | 18 | 2 | 18 |
| ENSDARG000 | 9;9;9;9;9 | 22157282:22 | 22157778:22 | -;-;-;-;- | 950 | crygm2d16 | 1 | 0 | 0 | 0 | 0 | 0 |
| ENSDARG000 | 9:9:9:9:9:9:9: | 38074082:38 | 38074266:38 | +:+:+:+:+:+:+: | 2563 | cacnb4a | 0 | 0 | 8 | 4 | 0 | 10 |
| ENSDARG000 | 9:9:9:9:9:9:9: | 27804027:27 | 27804290:27 | -:-:-:-:-:-:-:-: | 1711 | NA | 0 | 4 | 10 | 1 | 6 | 0 |
| ENSDARG000 | 9;9;9 | 22181416:22 | 22181782:22 | -;-;- | 685 | crygm2d6 | 0 | 0 | 0 | 0 | 0 | 0 |
| ENSDARG000 | 9:9:9:9:9:9:9: | 32805688:32 | 32806473:32 | -:-:-:-:-:-:-:-: | 2436 | mxc | 9560 | 1724 | 1042 | 195 | 1167 | 498 |
| ENSDARG000 | 9;9;9;9;9;9 | 30720048:30 | 30720185:30 | +;+;+;+;+;+ | 1826 | klf12b | 150 | 97 | 176 | 76 | 164 | 108 |
| ENSDARG000 | 9;9;9;9;9;9;9 | 41040294:41 | 41040596:41 | +;+;+;+;+;+;+ | 1386 | osgepl1 | 76 | 62 | 73 | 67 | 70 | 74 |
| ENSDARG000 | 9:9:9:9:9:9:9: | 8364553:836 | 8365449:836 | +:+:+:+:+:+:+: | 2417 | NA | 10 | 2 | 4 | 1 | 10 | 0 |
| ENSDARG000 | 9:9:9:9:9:9:9: | 41045874:41 | 41046114:41 | -:-:-:-:-:-:-:-: | 4335 | ankar | 4 | 9 | 2 | 0 | 8 | 18 |

| | | | | | | | | | | | | |
|---|---|---|---|---|---|---|---|---|---|---|---|---|
| ENSDARG000 | 9;9;9;9 | 41035925:41 | 41037097:41 | -;-;-;- | 1485 | adat3 | 76 | 48 | 138 | 58 | 110 | 72 |
| ENSDARG000 | 9;9;9;9 | 30800876:30 | 30801054:30 | -;-;-;- | 1278 | KLF5 | 0 | 0 | 0 | 0 | 2 | 0 |
| ENSDARG000 | 9;9;9;9;9 | 22187031:22 | 22187508:22 | -;-;-;-;- | 1002 | NA | 0 | 2 | 0 | 0 | 2 | 0 |
| ENSDARG000 | 9;9;9;9;9;9;9; | 32824639:32 | 32825094:32 | -;-;-;-;-;-;-; | 2115 | mxe | 326 | 236 | 29 | 16 | 82 | 4 |
| ENSDARG000 | 9;9;9;9;9;9;9; | 38140911:38 | 38141022:38 | -;-;-;-;-;-;-; | 1542 | NA | 0 | 0 | 0 | 0 | 0 | 0 |
| ENSDARG000 | 9;9;9;9;9;9;9; | 4250918:425 | 4250987:425 | +;+;+;+;+;+;+; | 8245 | KALRN | 5088 | 4321 | 5059 | 6524 | 5321 | 2729 |
| ENSDARG000 | 9;9;9;9;9;9;9; | 30820290:30 | 30820431:30 | +;+;+;+;+;+;+; | 2754 | dachd | 24 | 10 | 135 | 16 | 65 | 88 |
| ENSDARG000 | 9;9;9;9;9;9;9; | 1313418:133 | 1313523:133 | +;+;+;+;+;+;+; | 2251 | NA | 143 | 102 | 134 | 40 | 127 | 121 |
| ENSDARG000 | 9;9;9;9;9;9;9; | 21146862:21 | 21146937:21 | +;+;+;+;+;+;+; | 2022 | hao2 | 0 | 0 | 8 | 10 | 0 | 0 |
| ENSDARG000 | 9;9;9 | 22204677:22 | 22205080:22 | -;-;- | 819 | crygm2d12 | 0 | 2 | 0 | 0 | 1 | 13 |
| ENSDARG000 | 9;9;9;9;9;9;9; | 23003208:23 | 23003415:23 | +;+;+;+;+;+;+; | 2208 | eaf2 | 660 | 680 | 830 | 382 | 803 | 608 |
| ENSDARG000 | 9;9;9;9;9;9;9; | 32840114:32 | 32841488:32 | -;-;-;-;-;-;-; | 3594 | hpxa | 0 | 0 | 24 | 0 | 6 | 0 |
| ENSDARG000 | 9;9;9;9;9;9;9; | 48185936:48 | 48186817:48 | -;-;-;-;-;-;-; | 2374 | phgdh | 1934 | 2393 | 3676 | 3189 | 1238 | 2043 |
| ENSDARG000 | 9;9;9;9;9;9;9; | 1365747:136 | 1365793:136 | +;+;+;+;+;+;+; | 1628 | prkra | 387 | 804 | 673 | 268 | 559 | 262 |
| ENSDARG000 | 9;9;9 | 18716485:18 | 18716751:18 | +;+;+ | 730 | serp2 | 0 | 0 | 0 | 8 | 0 | 2 |
| ENSDARG000 | 9;9;9;9;9;9;9; | 29581662:29 | 29582016:29 | +;+;+;+;+;+;+; | 5890 | MCF2L | 60 | 16 | 44 | 10 | 41 | 12 |
| ENSDARG000 | 9;9;9;9 | 27817925:27 | 27818185:27 | +;+;+;+ | 638 | NA | 0 | 0 | 0 | 0 | 2 | 0 |
| ENSDARG000 | 9;9;9;9;9;9;9; | 38158570:38 | 38158805:38 | +;+;+;+;+;+;+; | 1559 | nifk | 529 | 755 | 1229 | 522 | 752 | 754 |
| ENSDARG000 | 9;9;9;9;9;9;9; | 27837062:27 | 27837120:27 | +;+;+;+;+;+;+; | 807 | NA | 0 | 0 | 0 | 0 | 0 | 0 |
| ENSDARG000 | 9;9;9;9 | 27834445:27 | 27834706:27 | +;+;+;+ | 639 | NA | 0 | 0 | 0 | 0 | 0 | 3 |
| ENSDARG000 | 9;9;9;9;9;9;9; | 48219111:48 | 48219376:48 | +;+;+;+;+;+;+; | 2682 | NA | 0 | 0 | 0 | 0 | 0 | 8 |
| ENSDARG000 | 9;9;9;9;9;9;9; | 32859967:32 | 32860412:32 | +;+;+;+;+;+;+; | 6026 | NA | 150 | 127 | 122 | 68 | 472 | 42 |
| ENSDARG000 | 9;9;9;9;9 | 22212241:22 | 22212656:22 | -;-;-;-;- | 950 | crygm2d20 | 0 | 0 | 0 | 0 | 0 | 2 |
| ENSDARG000 | 9;9;9;9;9;9;9; | 34433053:34 | 34433138:34 | +;+;+;+;+;+;+; | 3940 | NA | 905 | 624 | 1148 | 792 | 1074 | 796 |
| ENSDARG000 | 9;9;9;9 | 4443918:444 | 4444848:444 | -;-;-;- | 1587 | NA | 0 | 1 | 0 | 5 | 4 | 0 |
| ENSDARG000 | 9;9 | 1372946:137 | 1373486:137 | +;+ | 1993 | NA | 179 | 231 | 369 | 113 | 255 | 183 |
| ENSDARG000 | 9;9;9;9;9;9;9; | 21165017:21 | 21165134:21 | +;+;+;+;+;+;+; | 3437 | NA | 50 | 68 | 67 | 24 | 147 | 164 |
| ENSDARG000 | 9;9 | 32883291:32 | 32883429:32 | -;- | 300 | NA | 0 | 0 | 0 | 0 | 0 | 0 |
| ENSDARG000 | 9;9;9;9;9 | 22231878:22 | 22232279:22 | -;-;-;-;- | 926 | crygm2d5 | 0 | 0 | 0 | 0 | 0 | 0 |
| ENSDARG000 | 9 | 31089649 | 31090593 | - | 945 | GPR18 | 4 | 3 | 6 | 2 | 22 | 12 |
| ENSDARG000 | 9;9;9;9;9;9;9; | 31075896:31 | 31076335:31 | +;+;+;+;+;+;+; | 2539 | clybl | 140 | 136 | 180 | 113 | 262 | 115 |
| ENSDARG000 | 9;9;9;9;9;9;9; | 41080029:41 | 41080101:41 | +;+;+;+;+;+;+; | 1326 | zgc:136439 | 0 | 0 | 0 | 0 | 0 | 0 |
| ENSDARG000 | 9;9;9;9;9;9;9; | 32895003:32 | 32895313:32 | -;-;-;-;-;-;-; | 3460 | fhip1b | 310 | 268 | 282 | 217 | 353 | 224 |
| ENSDARG000 | 9;9 | 31105389:31 | 31106714:31 | -;- | 1448 | gpr183a | 26 | 34 | 20 | 22 | 48 | 80 |
| ENSDARG000 | 9;9;9 | 31134907:31 | 31136004:31 | -;-;- | 1102 | si:ch211-184 | 0 | 0 | 0 | 0 | 0 | 2 |
| ENSDARG000 | 9;9;9 | 31076056:31 | 31076335:31 | +;+;+ | 879 | NA | 0 | 0 | 0 | 0 | 0 | 0 |
| ENSDARG000 | 9;9;9;9;9;9;9; | 41079669:41 | 41083547:41 | -;-;-;-;-;-;-; | 6343 | asnsd1 | 598 | 644 | 824 | 460 | 645 | 618 |
| ENSDARG000 | 9;9;9;9;9 | 22239013:22 | 22239424:22 | -;-;-;-;- | 938 | crygm2d9 | 0 | 0 | 0 | 0 | 0 | 0 |
| ENSDARG000 | 9;9;9;9;9;9;9; | 23009608:23 | 23010016:23 | +;+;+;+;+;+;+; | 2176 | cyp27a3 | 34 | 6 | 8 | 10 | 6 | 6 |
| ENSDARG000 | 9;9;9 | 22244537:22 | 22244944:22 | -;-;- | 831 | crygm2d4 | 0 | 0 | 0 | 0 | 0 | 1 |
| ENSDARG000 | 9;9;9;9;9 | 22271056:22 | 22271558:22 | -;-;-;-;- | 1022 | crygm2d7 | 8 | 0 | 0 | 4 | 0 | 2 |
| ENSDARG000 | 9;9;9;9;9;9;9; | 18732505:18 | 18733813:18 | -;-;-;-;-;-;-; | 4389 | tsc22d1 | 993 | 1004 | 1224 | 862 | 1262 | 1092 |
| ENSDARG000 | 9;9;9;9;9 | 22280471:22 | 22280993:22 | -;-;-;-;- | 1279 | crygm2d3 | 8 | 0 | 0 | 0 | 0 | 3 |
| ENSDARG000 | 9;9;9;9;9;9;9 | 4478199:447 | 4478791:447 | -;-;-;-;-;-;- | 2189 | KCNJ3 | 0 | 1 | 2 | 0 | 0 | 0 |
| ENSDARG000 | 9;9;9;9;9;9;9; | 4538527:456 | 4542306:456 | -;-;-;-;-;-;-; | 5923 | GALNT13 | 42 | 40 | 74 | 52 | 69 | 64 |
| ENSDARG000 | 9;9;9;9;9;9;9; | 48232571:48 | 48232806:48 | +;+;+;+;+;+;+; | 5141 | fastkd1 | 346 | 224 | 504 | 131 | 299 | 258 |
| ENSDARG000 | 9;9;9;9;9;9;9; | 27860332:27 | 27861423:27 | -;-;-;-;-;-;-;- | 2169 | dbr1 | 380 | 385 | 457 | 206 | 388 | 201 |
| ENSDARG000 | 9;9;9;9;9;9;9; | 32930622:32 | 32930740:32 | +;+;+;+;+;+;+; | 2244 | NA | 0 | 0 | 10 | 0 | 6 | 0 |
| ENSDARG000 | 9;9;9;9;9;9;9; | 27855684:27 | 27855947:27 | -;-;-;-;-;-;-; | 1497 | NA | 8 | 1 | 6 | 3 | 1 | 1 |
| ENSDARG000 | 9;9;9;9;9;9;9; | 8372167:837 | 8372287:837 | +;+;+;+;+;+;+; | 2565 | NA | 0 | 2 | 0 | 13 | 0 | 4 |
| ENSDARG000 | 9;9;9;9;9;9;9; | 29616854:29 | 29617041:29 | +;+;+;+;+;+;+; | 2143 | NA | 535 | 755 | 505 | 331 | 463 | 351 |
| ENSDARG000 | 9;9;9;9;9;9;9; | 38163876:38 | 38163984:38 | +;+;+;+;+;+;+; | 7238 | clasp1a | 1018 | 903 | 1070 | 1097 | 1132 | 970 |
| ENSDARG000 | 9;9;9;9;9;9;9; | 1391535:139 | 1391643:139 | -;-;-;-;-;-;-; | 3063 | OSBPL6 | 344 | 190 | 311 | 274 | 725 | 273 |
| ENSDARG000 | 9 | 4609746 | 4610777 | + | 1032 | rprma | 0 | 0 | 0 | 0 | 0 | 0 |
| ENSDARG000 | 9;9;9 | 21194445:21 | 21194694:21 | +;+;+ | 1535 | NA | 21 | 8 | 33 | 41 | 10 | 12 |
| ENSDARG000 | 9;9;9;9;9;9;9; | 23026324:23 | 23027025:23 | -;-;-;-;-;-;-;- | 1473 | rnd3b | 95 | 90 | 42 | 14 | 122 | 116 |
| ENSDARG000 | 9;9;9;9;9 | 22298246:22 | 22298783:22 | -;-;-;-;- | 1062 | crygm2d21 | 2 | 0 | 0 | 0 | 0 | 0 |
| ENSDARG000 | 9;9;9;9;9;9;9; | 41103127:41 | 41103420:41 | +;+;+;+;+;+;+; | 3780 | slc40a1 | 588 | 274 | 348 | 264 | 1505 | 484 |
| ENSDARG000 | 9;9;9;9;9 | 22309838:22 | 22310291:22 | -;-;-;-;- | 978 | crygm2d10 | 2 | 0 | 0 | 2 | 0 | 0 |
| ENSDARG000 | 9;9;9;9 | 31266192:31 | 31268921:31 | -;-;-;- | 3142 | NA | 0 | 0 | 0 | 0 | 2 | 0 |
| ENSDARG000 | 9;9 | 32978302:32 | 32978746:32 | +;+ | 2340 | nhlh2 | 0 | 0 | 0 | 0 | 0 | 0 |
| ENSDARG000 | 9;9;9;9;9 | 22317479:22 | 22317908:22 | -;-;-;-;- | 878 | crygm2d2 | 6 | 0 | 0 | 0 | 0 | 0 |
| ENSDARG000 | 9;9;9;9;9;9;9; | 33009284:33 | 33009354:33 | +;+;+;+;+;+;+; | 1672 | vangl1 | 528 | 418 | 550 | 335 | 668 | 463 |
| ENSDARG000 | 9;9;9;9;9;9 | 48270379:48 | 48271003:48 | -;-;-;-;-;- | 2313 | klhl41a | 2 | 0 | 2 | 6 | 3 | 0 |
| ENSDARG000 | 9;9;9 | 31282161:31 | 31283548:31 | +;+;+ | 3177 | zic2a | 26 | 0 | 0 | 0 | 6 | 6 |
| ENSDARG000 | 9;9;9;9;9;9;9 | 31274108:31 | 31275508:31 | -;-;-;-;-;-;- | 3073 | zic5 | 6 | 0 | 0 | 0 | 4 | 0 |
| ENSDARG000 | 9;9;9;9;9;9;9; | 24116275:24 | 24117175:24 | -;-;-;-;-;-;-; | 1962 | tyw5 | 205 | 171 | 182 | 155 | 172 | 165 |
| ENSDARG000 | 9;9;9;9;9;9;9; | 34484454:34 | 34485189:34 | -;-;-;-;-;-;-; | 3074 | NA | 134 | 135 | 147 | 111 | 142 | 156 |
| ENSDARG000 | 9;9;9;9;9;9;9; | 21203832:21 | 21204294:21 | -;-;-;-;-;-;-; | 821 | NA | 95 | 32 | 63 | 167 | 131 | 183 |
| ENSDARG000 | 9;9;9 | 22338578:22 | 22338973:22 | -;-;- | 815 | crygm2d1 | 4 | 0 | 0 | 0 | 0 | 0 |
| ENSDARG000 | 9;9;9;9;9;9;9; | 4675193:468 | 4675398:468 | +;+;+;+;+;+;+; | 3285 | prpf40a | 1287 | 1503 | 2330 | 1043 | 1822 | 1246 |
| ENSDARG000 | 9;9;9;9;9;9;9; | 48284606:48 | 48285024:48 | -;-;-;-;-;-;-; | 1492 | bbs5 | 108 | 190 | 70 | 46 | 41 | 75 |
| ENSDARG000 | 9;9;9;9;9 | 22359919:22 | 22359974:22 | +;+;+;+;+ | 720 | crygs4 | 60 | 0 | 6 | 24 | 0 | 0 |
| ENSDARG000 | 9;9;9;9;9;9;9 | 22345047:22 | 22345284:22 | -;-;-;-;-;-;- | 1309 | crygm5 | 26 | 0 | 0 | 10 | 0 | 0 |
| ENSDARG000 | 9;9;9;9;9;9;9 | 22354281:22 | 22354930:22 | -;-;-;-;-;-;- | 1212 | crygm3 | 62 | 0 | 0 | 3 | 0 | 0 |
| ENSDARG000 | 9;9;9;9;9;9;9; | 22351443:22 | 22351511:22 | +;+;+;+;+;+;+; | 943 | crygs3 | 15 | 0 | 0 | 7 | 0 | 0 |
| ENSDARG000 | 9;9;9 | 22344595:22 | 22344902:22 | -;-;- | 588 | crygm2f | 5 | 0 | 0 | 0 | 0 | 0 |
| ENSDARG000 | 9;9;9;9;9;9;9; | 18829360:18 | 18829521:18 | +;+;+;+;+;+;+; | 1050 | gtf2f2b | 0 | 26 | 73 | 90 | 103 | 28 |
| ENSDARG000 | 9;9;9;9;9;9;9 | 31347037:31 | 31347247:31 | +;+;+;+;+;+;+ | 1230 | NA | 0 | 0 | 0 | 0 | 0 | 0 |
| ENSDARG000 | 9 | 18876036 | 18877597 | - | 1562 | kctd4 | 0 | 80 | 6 | 0 | 18 | 0 |
| ENSDARG000 | 9;9;9;9;9;9;9; | 1463052:146 | 1463330:146 | -;-;-;-;-;-;-; | 2255 | rbm45 | 20 | 19 | 8 | 8 | 6 | 15 |
| ENSDARG000 | 9;9;9;9;9;9;9; | 23049500:23 | 23049590:23 | +;+;+;+;+;+;+; | 1464 | mmadhcb | 400 | 378 | 522 | 328 | 284 | 264 |
| ENSDARG000 | 9;9;9;9;9;9;9; | 8380728:838 | 8380782:838 | +;+;+;+;+;+;+; | 684 | NA | 0 | 0 | 0 | 0 | 0 | 0 |
| ENSDARG000 | 9;9;9;9;9;9;9; | 43074531:43 | 43074914:43 | -;-;-;-;-;-;-; | 6677 | NA | 12 | 12 | 0 | 20 | 10 | 20 |
| ENSDARG000 | 9;9;9;9;9;9;9; | 41120158:41 | 41120554:41 | -;-;-;-;-;-;-; | 3145 | wdr75 | 467 | 862 | 1166 | 379 | 824 | 586 |
| ENSDARG000 | 9;9;9;9;9;9;9; | 41156287:41 | 41157101:41 | +;+;+;+;+;+;+; | 6267 | STAT4 | 76 | 50 | 18 | 22 | 58 | 76 |
| ENSDARG000 | 9;9;9;9;9;9;9; | 24121036:24 | 24121407:24 | +;+;+;+;+;+;+; | 1432 | maip1 | 174 | 316 | 329 | 192 | 361 | 198 |
| ENSDARG000 | 9;9;9;9;9 | 29652041:29 | 29652716:29 | -;-;-;-;- | 2609 | NA | 16 | 6 | 51 | 11 | 21 | 11 |
| ENSDARG000 | 9;9;9;9;9 | 29636620:29 | 29637890:29 | -;-;-;-;- | 1783 | sprydd7b | 461 | 494 | 573 | 759 | 503 | 682 |
| ENSDARG000 | 9;9;9;9;9;9;9; | 33061564:33 | 33061795:33 | -;-;-;-;-;-;-; | 694 | NA | 4 | 2 | 0 | 4 | 4 | 2 |
| ENSDARG000 | 9;9;9;9;9;9;9; | 29642857:29 | 29643156:29 | +;+;+;+;+;+;+; | 3489 | trim13 | 287 | 322 | 469 | 300 | 447 | 249 |
| ENSDARG000 | 9;9;9 | 18911697:18 | 18911747:18 | +;+;+ | 218 | NA | 0 | 0 | 0 | 0 | 2 | 1 |
| ENSDARG000 | 9;9;9;9;9;9 | 18908721:18 | 18909282:18 | -;-;-;-;-;- | 1257 | NA | 78 | 255 | 617 | 296 | 368 | 298 |
| ENSDARG000 | 9;9;9;9;9;9;9; | 24125445:24 | 24126354:24 | +;+;+;+;+;+;+; | 3701 | NA | 143 | 132 | 199 | 80 | 278 | 162 |
| ENSDARG000 | 9;9;9;9;9;9;9; | 48297412:48 | 48298562:48 | -;-;-;-;-;-;-; | 5671 | col28a2a | 2 | 0 | 0 | 0 | 8 | 8 |
| ENSDARG000 | 9;9;9;9;9;9 | 1485234:148 | 1486385:148 | -;-;-;-;-;- | 2240 | si:ch211-222 | 6 | 14 | 23 | 46 | 11 | 118 |
| ENSDARG000 | 9;9;9;9;9;9;9; | 21222428:21 | 21227814:21 | -;-;-;-;-;-;-; | 6568 | popdc2 | 0 | 0 | 0 | 0 | 0 | 0 |
| ENSDARG000 | 9;9;9;9;9;9;9; | 21214940:21 | 21215239:21 | +;+;+;+;+;+;+; | 3370 | pla1a | 0 | 0 | 4 | 0 | 6 | 3 |
| ENSDARG000 | 9;9;9;9;9;9;9; | 33080438:33 | 33080670:33 | -;-;-;-;-;-;-; | 700 | zgc:172053 | 8 | 22 | 2 | 20 | 36 | 16 |
| ENSDARG000 | 9;9;9;9 | 33074680:33 | 33074840:33 | +;+;+;+ | 790 | NA | 0 | 0 | 0 | 0 | 0 | 0 |
| ENSDARG000 | 9;9;9;9;9;9;9; | 24159280:24 | 24160031:24 | +;+;+;+;+;+;+; | 9885 | HECW2 | 0 | 0 | 2 | 0 | 11 | 0 |
| ENSDARG000 | 9;9;9;9;9;9;9; | 31494941:31 | 31498126:31 | -;-;-;-;-;-;-; | 5905 | tmtc4 | 570 | 566 | 584 | 313 | 614 | 491 |
| ENSDARG000 | 9;9;9;9;9;9;9; | 24192370:24 | 24193173:24 | +;+;+;+;+;+;+; | 2284 | stk17b | 356 | 477 | 302 | 130 | 520 | 302 |
| ENSDARG000 | 9;9;9;9;9;9;9; | 48386928:48 | 48387142:48 | -;-;-;-;-;-;-; | 1276 | zgc:172182 | 0 | 0 | 0 | 0 | 0 | 0 |
| ENSDARG000 | 9;9;9;9;9;9 | 19039608:19 | 19039641:19 | +;+;+;+;+;+ | 663 | chmp2ba | 40 | 15 | 40 | 22 | 45 | 40 |
| ENSDARG000 | 9;9 | 19027432:19 | 19027813:19 | -;- | 538 | NA | 0 | 0 | 0 | 0 | 0 | 0 |
| ENSDARG000 | 9;9;9;9;9 | 23071149:23 | 23071838:23 | -;-;-;-;- | 1237 | lypd6 | 28 | 16 | 21 | 0 | 4 | 44 |
| ENSDARG000 | 9;9;9;9;9;9;9; | 38292919:38 | 38293189:38 | +;+;+;+;+;+;+; | 2862 | tfcp2l1 | 0 | 0 | 2 | 2 | 4 | 0 |
| ENSDARG000 | 9;9;9;9;9;9;9; | 1505206:150 | 1505967:150 | +;+;+;+;+;+;+; | 2685 | NA | 0 | 4 | 0 | 0 | 0 | 2 |
| ENSDARG000 | 9;9;9;9;9;9;9; | 33097289:33 | 33098826:33 | -;-;-;-;-;-;-; | 2858 | casq2 | 0 | 0 | 0 | 0 | 6 | 0 |
| ENSDARG000 | 9;9;9;9;9;9;9; | 23142170:23 | 23143200:23 | -;-;-;-;-;-;-; | 3029 | lypd6b | 20 | 2 | 4 | 0 | 0 | 8 |
| ENSDARG000 | 9;9;9;9;9;9;9; | 43149854:43 | 43150961:43 | -;-;-;-;-;-;-; | 3586 | sestd1 | 640 | 1370 | 890 | 608 | 793 | 906 |
| ENSDARG000 | 9;9;9;9;9;9;9; | 33115926:33 | 33116423:33 | -;-;-;-;-;-;-; | 1668 | si:ch211-125 | 33 | 163 | 137 | 99 | 49 | 48 |
| ENSDARG000 | 9;9;9;9;9;9;9; | 8390737:839 | 8390877:839 | +;+;+;+;+;+;+; | 1612 | NA | 0 | 0 | 0 | 0 | 0 | 0 |
| ENSDARG000 | 9;9;9;9;9;9;9; | 29671553:29 | 29671677:29 | +;+;+;+;+;+;+; | 1122 | rnaseh2b | 100 | 247 | 347 | 157 | 337 | 211 |
| ENSDARG000 | 9 | 33134882 | 33136990 | - | 2109 | SLC5A3 | 153 | 217 | 318 | 207 | 253 | 169 |
| ENSDARG000 | 9;9;9;9;9;9;9; | 27876146:27 | 27876201:27 | +;+;+;+;+;+;+; | 2900 | armc8 | 303 | 219 | 299 | 245 | 308 | 250 |
| ENSDARG000 | 9;9 | 24204180:24 | 24206098:24 | -;- | 2264 | C9H2orf69 | 178 | 100 | 146 | 135 | 186 | 110 |
| ENSDARG000 | 9;9;9;9;9;9;9; | 24208533:24 | 24208573:24 | +;+;+;+;+;+;+; | 1673 | ftcdnl1 | 43 | 104 | 67 | 70 | 223 | 210 |
| ENSDARG000 | 9;9 | 22391566:22 | 22391702:22 | +;+ | 363 | NA | 0 | 0 | 0 | 0 | 0 | 0 |
| ENSDARG000 | 9;9;9;9;9;9;9; | 4730398:473 | 4733645:473 | -;-;-;-;-;-;-; | 7172 | FMNL2 | 1530 | 1240 | 2121 | 1146 | 1724 | 1142 |
| ENSDARG000 | 9;9;9;9;9;9;9; | 22375331:22 | 22375382:22 | +;+;+;+;+;+;+; | 6381 | DGKG | 0 | 0 | 0 | 4 | 3 | 6 |
| ENSDARG000 | 9;9;9;9;9;9;9; | 21235476:21 | 21236356:21 | -;-;-;-;-;-;-; | 1697 | cox17 | 186 | 322 | 253 | 168 | 273 | 283 |
| ENSDARG000 | 9;9;9;9;9;9;9; | 33145522:33 | 33145613:33 | +;+;+;+;+;+;+; | 1410 | atp5po | 3216 | 3513 | 4568 | 3279 | 3648 | 3554 |
| ENSDARG000 | 9;9;9;9;9;9;9; | 19073556:19 | 19075173:19 | -;-;-;-;-;-;-; | 2899 | crfb2 | 250 | 258 | 226 | 192 | 320 | 270 |
| ENSDARG000 | 9;9;9;9;9;9;9; | 21243335:21 | 21244145:21 | +;+;+;+;+;+;+; | 3341 | NA | 0 | 0 | 0 | 0 | 0 | 0 |
| ENSDARG000 | 9;9;9;9;9;9;9; | 48415043:48 | 48415100:48 | +;+;+;+;+;+;+; | 17318 | lrp2a | 2 | 8 | 276 | 212 | 594 | 0 |
| ENSDARG000 | 9;9 | 1575420:157 | 1578699:158 | +;+ | 4378 | NA | 10 | 3 | 0 | 10 | 2 | 0 |
| ENSDARG000 | 9;9;9;9;9;9;9; | 23185615:23 | 23185820:23 | -;-;-;-;-;-;-; | 3304 | kif5c | 0 | 0 | 0 | 0 | 6 | 14 |
| ENSDARG000 | 9;9;9;9;9;9;9; | 38314466:38 | 38315190:38 | +;+;+;+;+;+;+; | 2780 | ddx18 | 1065 | 1340 | 2474 | 952 | 1396 | 1080 |
| ENSDARG000 | 9;9;9;9;9;9;9; | 1589672:158 | 1589878:158 | -;-;-;-;-;-;-; | 2058 | agps | 650 | 467 | 579 | 235 | 511 | 410 |
| ENSDARG000 | 9;9 | 29690678:29 | 29691441:29 | +;+ | 1601 | NA | 4 | 1 | 0 | 17 | 0 | 0 |
| ENSDARG000 | 9;9;9;9 | 29682197:29 | 29684215:29 | -;-;-;- | 2749 | NA | 4 | 4 | 2 | 4 | 12 | 0 |
| ENSDARG000 | 9;9;9;9;9;9;9; | 43243212:43 | 43243553:43 | -;-;-;-;-;-;-; | 1918 | znf385b | 33 | 10 | 199 | 20 | 19 | 45 |
| ENSDARG000 | 9;9;9;9;9;9;9; | 8396755:839 | 8396806:839 | +;+;+;+;+;+;+; | 878 | zgc:171776 | 0 | 0 | 0 | 0 | 0 | 0 |
| ENSDARG000 | 9;9;9;9;9;9;9; | 24214660:24 | 24215388:24 | -;-;-;-;-;-;-; | 2323 | nabp1a | 379 | 436 | 604 | 243 | 708 | 399 |
| ENSDARG000 | 9;9;9;9;9;9;9; | 19095023:19 | 19095430:19 | +;+;+;+;+;+;+; | 2395 | crfb1 | 123 | 108 | 107 | 127 | 140 | 162 |
| ENSDARG000 | 9;9;9;9;9;9;9; | 31534601:31 | 31534606:31 | +;+;+;+;+;+;+; | 3178 | NA | 20 | 8 | 11 | 27 | 16 | 18 |
| ENSDARG000 | 9;9;9;9;9 | 34641237:34 | 34641603:34 | +;+;+;+;+ | 1072 | shox | 4 | 0 | 0 | 0 | 0 | 0 |
| ENSDARG000 | 9;9;9;9;9;9;9 | 19132885:19 | 19133177:19 | -;-;-;-;-;-;- | 1170 | pou1f1 | 6 | 1 | 0 | 0 | 1 | 4 |
| ENSDARG000 | 9;9;9;9;9;9;9; | 33154841:33 | 33154926:33 | +;+;+;+;+;+;+; | 7435 | DOP1B | 730 | 607 | 619 | 500 | 743 | 891 |
| ENSDARG000 | 9;9;9;9;9;9;9; | 29985010:29 | 29985082:29 | +;+;+;+;+;+;+; | 2534 | cmss1 | 232 | 249 | 340 | 112 | 296 | 231 |
| ENSDARG000 | 9;9;9;9;9 | 24220315:24 | 24224452:24 | -;-;-;-;- | 5425 | cavin2a | 177 | 6 | 6 | 2 | 34 | 4 |
| ENSDARG000 | 9;9;9;9;9;9;9; | 41218967:41 | 41220078:41 | +;+;+;+;+;+;+; | 4445 | stat1b | 1811 | 2477 | 280 | 66 | 310 | 279 |
| ENSDARG000 | 9;9;9;9;9;9;9 | 29994439:29 | 29994757:29 | +;+;+;+;+;+;+ | 4583 | tmem30c | 0 | 2 | 4 | 24 | 14 | 16 |
| ENSDARG000 | 9;9;9;9;9;9;9; | 14569211:14 | 14569564:14 | -;-;-;-;-;-;-; | 3540 | NA | 303 | 154 | 289 | 141 | 331 | 276 |
| ENSDARG000 | 9;9;9;9;9;9;9; | 29716380:29 | 29718744:29 | -;-;-;-;-;-;-; | 4080 | il1rapl1a | 575 | 1047 | 889 | 756 | 912 | 874 |
| ENSDARG000 | 9;9;9;9;9;9;9; | 27909927:27 | 27910011:27 | -;-;-;-;-;-;-; | 2743 | NA | 0 | 10 | 0 | 2 | 0 | 0 |
| ENSDARG000 | 9;9;9;9;9;9;9; | 8401796:840 | 8401946:840 | +;+;+;+;+;+;+; | 1461 | si:ch1073-75 | 0 | 0 | 0 | 0 | 0 | 0 |
| ENSDARG000 | 9;9 | 1642605:164 | 1644800:164 | -;- | 2441 | NA | 54 | 32 | 150 | 67 | 80 | 58 |
| ENSDARG000 | 9;9 | 34848860:34 | 34849062:34 | -;- | 272 | NA | 0 | 0 | 0 | 0 | 0 | 0 |
| ENSDARG000 | 9;9;9;9;9 | 1654284:166 | 1654481:166 | +;+;+;+;+ | 4739 | nfe2l2a | 6858 | 4736 | 6126 | 7323 | 7047 | 11388 |
| ENSDARG000 | 9;9;9;9;9;9;9 | 21266684:21 | 21267163:21 | -;-;-;-;-;-;- | 1736 | sap18 | 0 | 0 | 0 | 0 | 0 | 0 |
| ENSDARG000 | 9;9;9;9 | 34866576:34 | 34867074:34 | -;-;-;- | 1238 | slc25a6 | 25 | 67 | 147 | 64 | 48 | 72 |
| ENSDARG000 | 9;9;9;9;9;9;9; | 21259657:21 | 21259926:21 | +;+;+;+;+;+;+; | 5619 | ska3 | 132 | 285 | 272 | 120 | 209 | 160 |
| ENSDARG000 | 9;9;9;9;9;9;9; | 38353452:38 | 38353841:38 | -;-;-;-;-;-;-; | 2256 | ccdc93 | 1018 | 1139 | 1437 | 876 | 1235 | 1110 |
| ENSDARG000 | 9;9;9;9;9;9;9; | 34872733:34 | 34873175:34 | -;-;-;-;-;-;-; | 2245 | asmtl | 203 | 210 | 200 | 158 | 231 | 230 |
| ENSDARG000 | 9;9;9;9;9;9;9; | 24249184:24 | 24249581:24 | -;-;-;-;-;-;-; | 1905 | tmeff2a | 12 | 37 | 111 | 2 | 255 | 409 |
| ENSDARG000 | 9;9;9;9;9;9 | 24390887:24 | 24391119:24 | -;-;-;-;-;- | 647 | NA | 0 | 0 | 0 | 2 | 0 | 0 |
| ENSDARG000 | 9;9;9;9;9;9 | 24384134:24 | 24384178:24 | +;+;+;+;+;+ | 621 | NA | 5 | 0 | 7 | 0 | 0 | 2 |
| ENSDARG000 | 9;9;9;9;9;9;9; | 22631672:22 | 22631970:22 | +;+;+;+;+;+;+; | 3845 | etv5a | 2968 | 1476 | 2240 | 1800 | 2779 | 2175 |
| ENSDARG000 | 9;9;9;9;9;9;9; | 5040985:504 | 5042539:504 | -;-;-;-;-;-;-; | 3938 | nr4a2a | 0 | 0 | 0 | 0 | 0 | 0 |

| | | | | | | | | | | | | |
|---|---|---|---|---|---|---|---|---|---|---|---|---|
| ENSDARG000 | 9 | 28232522 | 28234444 | + | 1923 | fzd5 | 2 | 6 | 2 | 0 | 2 | 8 |
| ENSDARG000 | 9;9;9;9;9;9;9; | 23224761;232 | 23224814;232 | +;+;+;+;+;+;+; | 1618 | NA | 2 | 0 | 0 | 6 | 2 | 0 |
| ENSDARG000 | 9;9;9;9;9;9;9; | 43552515;435 | 43553368;435 | -;-;-;-;-;-;-; | 3788 | cwc22 | 863 | 741 | 1311 | 489 | 1112 | 714 |
| ENSDARG000 | 9;9;9;9;9;9;9; | 23234233;232 | 23236833;232 | -;-;-;-;-;-;-; | 3429 | ccnt2a | 705 | 765 | 1273 | 548 | 1389 | 853 |
| ENSDARG000 | 9;9;9 | 28243624;282 | 28244411;282 | +;+;+ | 1440 | NA | 2 | 4 | 2 | 2 | 0 | 2 |
| ENSDARG000 | 9;9;9;9;9;9;9; | 28237378;282 | 28237774;282 | -;-;-;-;-;-;-; | 1691 | ccnyl1 | 183 | 269 | 295 | 207 | 339 | 237 |
| ENSDARG000 | 9 | 34884933 | 34885902 | - | 970 | P2RY8 | 8 | 18 | 7 | 28 | 6 | 11 |
| ENSDARG000 | 9;9;9;9;9;9;9; | 5862040;5868 | 5862093;5868 | +;+;+;+;+;+;+; | 2311 | pdzk1 | 0 | 0 | 116 | 34 | 136 | 0 |
| ENSDARG000 | 9;9;9;9;9;9 | 24423792;244 | 24423862;244 | +;+;+;+;+;+ | 642 | NA | 2 | 0 | 0 | 1 | 0 | 0 |
| ENSDARG000 | 9;9;9;9;9;9;9; | 34902971;349 | 34903430;349 | -;-;-;-;-;-;-; | 2435 | upf3a | 0 | 0 | 6 | 2 | 0 | 0 |
| ENSDARG000 | 9;9;9;9;9;9;9; | 1692926;1693 | 1693596;1693 | -;-;-;-;-;-;-; | 1688 | hnrnpa3 | 38 | 97 | 92 | 182 | 76 | 68 |
| ENSDARG000 | 9;9;9;9;9 | 24752713;247 | 24753423;247 | -;-;-;-;- | 1240 | NA | 2 | 0 | 0 | 6 | 8 | 0 |
| ENSDARG000 | 9;9;9;9;9;9;9; | 8407778;8407 | 8408105;8408 | +;+;+;+;+;+;+; | 2517 | zgc:153499 | 0 | 0 | 0 | 6 | 0 | 0 |
| ENSDARG000 | 9;9;9;9;9;9;9; | 33207574;332 | 33207621;332 | +;+;+;+;+;+;+; | 692 | si:ch211-125e | 0 | 0 | 0 | 0 | 0 | 0 |
| ENSDARG000 | 9;9;9;9;9;9;9; | 28263908;282 | 28264052;282 | -;-;-;-;-;-;-; | 2288 | creb1b | 124 | 140 | 161 | 184 | 208 | 128 |
| ENSDARG000 | 9;9;9;9;9;9;9; | 24920677;249 | 24920761;249 | +;+;+;+;+;+;+; | 5848 | slc39a10 | 236714 | 145977 | 51501 | 83734 | 130761 | 194758 |
| ENSDARG000 | 9;9;9;9;9;9;9; | 24951129;249 | 24951589;249 | -;-;-;-;-;-;-; | 12296 | DNAH7 | 163 | 44 | 50 | 132 | 88 | 134 |
| ENSDARG000 | 9;9;9;9;9;9;9; | 38385561;383 | 38390580;383 | -;-;-;-;-;-;-; | 9486 | ZNF142 | 279 | 156 | 260 | 289 | 212 | 383 |
| ENSDARG000 | 9;9;9;9;9;9;9; | 38369872;383 | 38369894;383 | +;+;+;+;+;+;+; | 2865 | PLCD4 | 176 | 141 | 106 | 140 | 110 | 125 |
| ENSDARG000 | 9;9;9;9;9;9;9; | 23244292;232 | 23244823;232 | -;-;-;-;-;-;-; | 1571 | acmsd | 4 | 2 | 2 | 10 | 33 | 0 |
| ENSDARG000 | 9;9;9;9;9;9;9; | 21277846;212 | 21278023;212 | +;+;+;+;+;+;+; | 3924 | lats2 | 1120 | 755 | 1135 | 1240 | 1322 | 1599 |
| ENSDARG000 | 9;9;9;9;9;9;9; | 41249457;412 | 41249563;412 | +;+;+;+;+;+;+; | 1454 | NA | 8 | 0 | 2 | 12 | 2 | 13 |
| ENSDARG000 | 9;9;9;9;9;9;9; | 15362300;153 | 15363611;153 | -;-;-;-;-;-;-; | 9001 | fn1a | 1208 | 334 | 444 | 491 | 1010 | 1166 |
| ENSDARG000 | 9;9;9;9;9;9;9; | 43797902;437 | 43798248;437 | +;+;+;+;+;+;+; | 5156 | ube2e3 | 378 | 430 | 880 | 625 | 1497 | 974 |
| ENSDARG000 | 9;9;9;9;9;9;9 | 28341109;283 | 28345928;283 | -;-;-;-;-;-;- | 6011 | klf7b | 504 | 1881 | 1364 | 970 | 286 | 1630 |
| ENSDARG000 | 9;9;9;9;9;9;9; | 33216945;332 | 33216973;332 | +;+;+;+;+;+;+; | 687 | si:ch211-125e | 0 | 0 | 0 | 0 | 0 | 0 |
| ENSDARG000 | 9;9;9;9 | 19248124;192 | 19248846;192 | +;+;+;+ | 4633 | NA | 2 | 0 | 0 | 0 | 0 | 0 |
| ENSDARG000 | 9;9;9;9;9;9;9; | 23255410;232 | 23255762;232 | +;+;+;+;+;+;+; | 1235 | tmem163a | 2001 | 1146 | 915 | 1460 | 551 | 809 |
| ENSDARG000 | 9;9;9;9;9;9;9; | 31589425;315 | 31589681;315 | -;-;-;-;-;-;-; | 7335 | nalcn | 5 | 0 | 16 | 28 | 124 | 40 |
| ENSDARG000 | 9;9;9;9;9;9;9; | 44034790;440 | 44035305;440 | +;+;+;+;+;+;+; | 3388 | ITGA4 | 1067 | 537 | 2649 | 465 | 3203 | 1344 |
| ENSDARG000 | 9;9;9;9;9;9;9; | 33220342;332 | 33220546;332 | +;+;+;+;+;+;+; | 770 | si:ch211-125e | 0 | 0 | 0 | 0 | 0 | 0 |
| ENSDARG000 | 9;9 | 1867386;1883 | 1867431;1883 | +;+ | 278 | NA | 0 | 0 | 0 | 0 | 0 | 0 |
| ENSDARG000 | 9;9 | 1931440;1933 | 1932268;1934 | -;- | 1312 | NA | 708 | 370 | 819 | 635 | 1076 | 1524 |
| ENSDARG000 | 9;9 | 28399361;284 | 28399397;284 | +;+ | 943 | NA | 0 | 0 | 1 | 0 | 1 | 0 |
| ENSDARG000 | 9;9;9;9;9;9;9; | 5893608;5898 | 5893919;5898 | -;-;-;-;-;-;-; | 1774 | GPR89B | 79 | 134 | 112 | 73 | 123 | 115 |
| ENSDARG000 | 9;9;9;9;9;9;9; | 19340216;193 | 19340646;193 | -;-;-;-;-;-;-;- | 1694 | zgc:152951 | 40 | 46 | 42 | 36 | 38 | 16 |
| ENSDARG000 | 9;9 | 1953021;1955 | 1953456;1955 | -;- | 746 | NA | 0 | 0 | 7 | 0 | 2 | 2 |
| ENSDARG000 | 9;9 | 1948465;1950 | 1949915;1951 | -;- | 2288 | hoxd4a | 284 | 127 | 554 | 322 | 481 | 489 |
| ENSDARG000 | 9;9;9;9;9;9;9; | 19367459;193 | 19367534;193 | +;+;+;+;+;+;+; | 6030 | slc37a1 | 604 | 633 | 500 | 210 | 488 | 329 |
| ENSDARG000 | 9;9;9;9;9;9;9; | 1937049;1937 | 1938434;1938 | -;-;-;-;-;-;-; | 3171 | hoxd3a | 1262 | 848 | 1891 | 1007 | 2146 | 1539 |
| ENSDARG000 | 9;9;9;9;9;9;9; | 28598577;285 | 28598620;285 | +;+;+;+;+;+;+; | 1317 | NA | 0 | 0 | 0 | 0 | 0 | 0 |
| ENSDARG000 | 9;9 | 8429239;8450 | 8433244;8454 | -;- | 7379 | IRS2 | 5846 | 3903 | 5431 | 4904 | 8480 | 4835 |
| ENSDARG000 | 9;9;9;9;9;9;9; | 34920001;349 | 34920468;349 | -;-;-;-;-;-;-; | 2343 | cdc16 | 178 | 220 | 360 | 116 | 290 | 197 |
| ENSDARG000 | 9;9 | 1964005;1965 | 1964773;1965 | -;- | 1407 | hoxd9a | 1588 | 1665 | 1936 | 744 | 2505 | 484 |
| ENSDARG000 | 9 | 23356774 | 23358597 | - | 1824 | arl4cb | 140 | 98 | 186 | 86 | 50 | 40 |
| ENSDARG000 | 9;9;9;9;9;9;9; | 22656976;226 | 22657025;226 | +;+;+;+;+;+;+; | 3655 | rbm44 | 0 | 0 | 0 | 4 | 0 | 4 |
| ENSDARG000 | 9;9;9;9;9;9;9; | 21306902;213 | 21306999;213 | +;+;+;+;+;+;+; | 4687 | xpo4 | 492 | 619 | 713 | 343 | 745 | 570 |
| ENSDARG000 | 9;9 | 1967707;1969 | 1968494;1970 | -;- | 1650 | hoxd10a | 529 | 160 | 185 | 56 | 591 | 52 |
| ENSDARG000 | 9;9;9;9;9;9;9; | 38399912;383 | 38400562;384 | +;+;+;+;+;+;+; | 1694 | bcs1l | 230 | 265 | 373 | 142 | 231 | 176 |
| ENSDARG000 | 9;9;9;9;9;9;9; | 44094311;440 | 44094612;440 | -;-;-;-;-;-;-; | 2043 | cerkl | 62 | 22 | 102 | 28 | 69 | 63 |
| ENSDARG000 | 9;9;9;9;9;9;9; | 30013667;300 | 30014007;300 | -;-;-;-;-;-;-; | 2158 | dcbld2 | 408 | 397 | 442 | 217 | 549 | 293 |
| ENSDARG000 | 9;9;9;9;9;9;9; | 28610052;286 | 28610436;286 | -;-;-;-;-;-;-; | 2112 | NA | 2 | 0 | 4 | 18 | 0 | 0 |
| ENSDARG000 | 9;9;9;9 | 33222911;332 | 33223005;332 | +;+;+;+ | 560 | NA | 0 | 0 | 0 | 0 | 0 | 0 |
| ENSDARG000 | 9;9;9;9;9;9;9; | 8634229;8635 | 8635519;8635 | -;-;-;-;-;-;-; | 6085 | COL4A1 | 19970 | 5060 | 12577 | 6160 | 16375 | 10479 |
| ENSDARG000 | 9;9;9 | 48682775;486 | 48683085;486 | +;+;+ | 979 | NA | 0 | 0 | 0 | 0 | 0 | 0 |
| ENSDARG000 | 9;9;9;9;9;9;9; | 48673093;486 | 48675146;486 | -;-;-;-;-;-;-; | 4099 | rdh1 | 0 | 0 | 0 | 0 | 10 | 0 |
| ENSDARG000 | 9;9 | 48713222;487 | 48713312;487 | +;+ | 176 | NA | 0 | 0 | 4 | 0 | 0 | 1 |
| ENSDARG000 | 9;9;9 | 2002701;2004 | 2003465;2004 | +;+;+ | 3314 | evx2 | 2 | 0 | 2 | 0 | 0 | 0 |
| ENSDARG000 | 9;9;9;9;9 | 48706038;487 | 48706357;487 | -;-;-;-;- | 1351 | dhrs9 | 42 | 78 | 46 | 156 | 267 | 154 |
| ENSDARG000 | 9;9 | 1986816;1989 | 1987881;1990 | -;- | 1687 | hoxd13a | 220 | 0 | 0 | 0 | 0 | 0 |
| ENSDARG000 | 9;9 | 48750426;487 | 48750531;487 | -;- | 144 | NA | 0 | 0 | 0 | 0 | 0 | 0 |
| ENSDARG000 | 9;9;9 | 48740524;487 | 48740577;487 | +;+;+ | 320 | NA | 0 | 0 | 0 | 0 | 0 | 0 |
| ENSDARG000 | 9;9;9;9;9 | 1981882;1981 | 1982571;1982 | -;-;-;-;- | 1441 | hoxd12a | 133 | 0 | 0 | 6 | 0 | 0 |
| ENSDARG000 | 9;9;9;9;9;9;9; | 48755638;487 | 48755698;487 | +;+;+;+;+;+;+; | 5051 | NA | 0 | 0 | 0 | 0 | 17 | 0 |
| ENSDARG000 | 9;9 | 1975818;1977 | 1976532;1978 | -;- | 1410 | hoxd11a | 630 | 4 | 0 | 0 | 0 | 0 |
| ENSDARG000 | 9;9;9;9;9;9;9; | 61770;63283; | 62558;63440; | -;-;-;-;-;-;-; | 1689 | il10rb | 921 | 700 | 927 | 743 | 874 | 712 |
| ENSDARG000 | 9;9;9;9;9;9;9; | 38408991;384 | 38409254;384 | +;+;+;+;+;+;+; | 2479 | NA | 220 | 55 | 82 | 60 | 76 | 118 |
| ENSDARG000 | 9;9 | 44291721;442 | 44293814;442 | -;- | 2422 | neurod1 | 0 | 0 | 0 | 0 | 0 | 4 |
| ENSDARG000 | 9;9;9;9;9;9;9; | 48835357;488 | 48835408;488 | -;-;-;-;-;-;-; | 2737 | nostrin | 0 | 0 | 0 | 0 | 0 | 0 |
| ENSDARG000 | 9;9;9;9;9;9;9; | 70157;70280; | 70351;70351; | +;+;+;+;+;+;+; | 9313 | RANBP2 | 2103 | 2207 | 3771 | 1431 | 3216 | 1476 |
| ENSDARG000 | 9;9 | 48845578;488 | 48845620;488 | +;+ | 542 | NA | 0 | 0 | 0 | 0 | 0 | 0 |
| ENSDARG000 | 9;9;9;9;9;9;9; | 48819280;488 | 48819391;488 | +;+;+;+;+;+;+; | 5193 | spc25 | 78 | 342 | 379 | 150 | 264 | 129 |
| ENSDARG000 | 9;9;9;9;9;9;9; | 23364242;233 | 23364654;233 | +;+;+;+;+;+;+; | 2030 | NA | 2 | 0 | 4 | 4 | 0 | 4 |
| ENSDARG000 | 9;9;9;9;9;9;9; | 33236844;332 | 33236869;332 | +;+;+;+;+;+;+; | 770 | lectin | 0 | 0 | 0 | 0 | 0 | 6 |
| ENSDARG000 | 9;9;9;9 | 28685853;286 | 28686140;286 | -;-;-;- | 624 | NA | 0 | 0 | 4 | 0 | 5 | 4 |
| ENSDARG000 | 9;9;9;9;9;9;9; | 34938316;349 | 34939211;349 | -;-;-;-;-;-;-; | 2009 | dcun1d2a | 298 | 223 | 288 | 236 | 294 | 214 |
| ENSDARG000 | 9;9;9;9;9;9 | 5993095;5993 | 5993121;5993 | +;+;+;+;+;+ | 1070 | NA | 10 | 25 | 16 | 10 | 9 | 0 |
| ENSDARG000 | 9;9;9;9;9;9;9; | 28688574;286 | 28689156;286 | +;+;+;+;+;+;+; | 1560 | zgc:162396 | 306 | 345 | 292 | 288 | 319 | 371 |
| ENSDARG000 | 9;9;9;9;9;9;9; | 6009077;6079 | 6009379;6079 | +;+;+;+;+;+;+; | 6453 | st6gal2a | 475 | 100 | 94 | 418 | 168 | 330 |
| ENSDARG000 | 9;9;9 | 15570343;155 | 15570861;155 | -;-;- | 777 | NA | 0 | 0 | 0 | 0 | 0 | 0 |
| ENSDARG000 | 9;9 | 6009202;6012 | 6009379;6012 | +;+ | 439 | NA | 0 | 0 | 0 | 0 | 0 | 0 |
| ENSDARG000 | 9;9;9 | 15567364;155 | 15567604;155 | +;+;+ | 867 | NA | 0 | 0 | 0 | 0 | 0 | 0 |
| ENSDARG000 | 9;9;9;9 | 6118362;6130 | 6118895;6130 | -;-;-;- | 1930 | tmtops2b | 2 | 2 | 0 | 0 | 4 | 0 |
| ENSDARG000 | 9;9;9 | 33244198;332 | 33244312;332 | +;+;+ | 285 | NA | 0 | 0 | 0 | 0 | 0 | 0 |
| ENSDARG000 | 9;9 | 15685746;156 | 15686100;156 | -;- | 485 | NA | 0 | 0 | 0 | 0 | 0 | 0 |
| ENSDARG000 | 9;9;9;9;9;9;9; | 28693592;286 | 28693782;286 | +;+;+;+;+;+;+; | 1555 | zgc:162780 | 0 | 46 | 70 | 10 | 84 | 20 |
| ENSDARG000 | 9;9;9;9;9;9;9; | 15782048;157 | 15782128;157 | -;-;-;-;-;-;-; | 1931 | NA | 18 | 12 | 27 | 39 | 14 | 12 |
| ENSDARG000 | 9;9;9 | 30090656;301 | 30090772;301 | +;+;+ | 2256 | col8a1a | 16 | 0 | 0 | 8 | 0 | 0 |
| ENSDARG000 | 9;9 | 25066661;250 | 25066741;250 | +;+ | 676 | NA | 9 | 0 | 21 | 5 | 0 | 0 |
| ENSDARG000 | 9;9;9;9;9;9;9; | 44304980;443 | 44305077;443 | +;+;+;+;+;+;+; | 5580 | NA | 392 | 291 | 385 | 437 | 501 | 423 |
| ENSDARG000 | 9;9;9 | 31755081;317 | 31755749;317 | -;-;- | 959 | NA | 0 | 0 | 0 | 0 | 0 | 0 |
| ENSDARG000 | 9;9;9;9;9;9 | 31758670;317 | 31759262;317 | -;-;-;-;-;- | 2970 | NA | 12 | 0 | 4 | 6 | 8 | 2 |
| ENSDARG000 | 9;9 | 25073539;250 | 25073619;250 | +;+ | 267 | NA | 0 | 0 | 14 | 13 | 0 | 0 |
| ENSDARG000 | 9;9;9;9;9;9;9; | 48870122;488 | 48877412;488 | -;-;-;-;-;-;-; | 8502 | CERS6 | 20 | 28 | 21 | 18 | 16 | 10 |
| ENSDARG000 | 9;9;9;9;9;9;9; | 31752628;317 | 31752850;317 | +;+;+;+;+;+;+; | 2457 | itgbl1 | 38 | 0 | 3 | 2 | 28 | 10 |
| ENSDARG000 | 9;9;9;9;9;9;9; | 33247202;332 | 33247596;332 | -;-;-;-;-;-;-; | 1677 | NA | 0 | 4 | 0 | 0 | 0 | 0 |
| ENSDARG000 | 9;9;9;9 | 25079209;250 | 25079313;250 | +;+;+;+ | 265 | NA | 4 | 0 | 6 | 0 | 0 | 0 |
| ENSDARG000 | 9;9;9;9;9;9;9; | 28722038;287 | 28722148;287 | +;+;+;+;+;+;+; | 1314 | NA | 2 | 0 | 3 | 12 | 0 | 0 |
| ENSDARG000 | 9;9;9;9;9;9;9; | 41273231;412 | 41273387;412 | -;-;-;-;-;-;-; | 5796 | GLS | 2306 | 2313 | 2050 | 4551 | 2465 | 2255 |
| ENSDARG000 | 9;9;9;9;9;9;9; | 28778649;288 | 28782412;288 | -;-;-;-;-;-;-; | 5832 | ramp1 | 0 | 0 | 2 | 0 | 4 | 0 |
| ENSDARG000 | 9;9;9;9;9;9;9; | 30104920;301 | 30106265;301 | -;-;-;-;-;-;-; | 1618 | NA | 31 | 20 | 18 | 10 | 32 | 23 |
| ENSDARG000 | 9;9;9;9;9;9;9; | 38420012;384 | 38420281;384 | +;+;+;+;+;+;+; | 2255 | CYP27A1 | 0 | 0 | 0 | 0 | 0 | 0 |
| ENSDARG000 | 9;9;9;9;9;9;9; | 33261330;332 | 33261525;332 | +;+;+;+;+;+;+; | 9655 | usp9 | 6950 | 5364 | 7219 | 4568 | 6645 | 5225 |
| ENSDARG000 | 9;9;9;9;9;9;9; | 49448752;494 | 49449008;494 | -;-;-;-;-;-;-; | 14127 | XIRP2 | 4 | 4 | 20 | 4 | 2 | 0 |
| ENSDARG000 | 9;9;9;9;9;9;9; | 25083160;250 | 25083263;250 | -;-;-;-;-;-;-;- | 1595 | NA | 15 | 0 | 0 | 0 | 3 | 12 |
| ENSDARG000 | 9;9;9;9;9 | 6371919;6371 | 6372535;6372 | -;-;-;-;- | 926 | ecrg4a | 200 | 280 | 1073 | 131 | 753 | 150 |
| ENSDARG000 | 9;9;9;9;9;9;9; | 6255682;6255 | 6255783;6256 | +;+;+;+;+;+;+; | 2534 | uxs1 | 1316 | 945 | 1240 | 1031 | 1490 | 1015 |
| ENSDARG000 | 9;9;9 | 8730515;8731 | 8730711;8731 | +;+;+ | 363 | NA | 368 | 115 | 263 | 74 | 187 | 143 |
| ENSDARG000 | 9;9;9;9;9;9;9; | 94884;94885; | 95015;95015; | -;-;-;-;-;-;-; | 1291 | lims2 | 52 | 16 | 18 | 8 | 24 | 10 |
| ENSDARG000 | 9;9;9;9;9;9;9; | 38427572;384 | 38427826;384 | +;+;+;+;+;+;+; | 1818 | CYP27A1 | 18 | 0 | 6 | 0 | 2 | 2 |
| ENSDARG000 | 9;9;9;9;9;9;9; | 21358941;213 | 21359050;213 | +;+;+;+;+;+;+; | 1348 | eef1akmt1 | 224 | 178 | 281 | 134 | 217 | 130 |
| ENSDARG000 | 9;9;9 | 30108641;301 | 30108799;301 | +;+;+ | 1248 | jagn1a | 188 | 184 | 258 | 72 | 152 | 90 |
| ENSDARG000 | 9;9;9;9;9;9;9; | 8761099;8769 | 8761107;8769 | +;+;+;+;+;+;+; | 5022 | COL4A2 | 15378 | 3266 | 8745 | 4433 | 11217 | 7192 |
| ENSDARG000 | 9;9 | 15837398;158 | 15837774;158 | +;+ | 3365 | znfl1b | 0 | 0 | 8 | 0 | 6 | 2 |
| ENSDARG000 | 9;9;9;9;9;9;9; | 38446504;384 | 38446758;384 | +;+;+;+;+;+;+; | 1819 | cyp27a1.4 | 8 | 7 | 36 | 16 | 16 | 21 |
| ENSDARG000 | 9;9;9;9;9;9;9; | 15890558;158 | 15890741;158 | +;+;+;+;+;+;+; | 1119 | NA | 0 | 0 | 0 | 0 | 0 | 0 |
| ENSDARG000 | 9;9;9;9;9;9;9; | 44430705;444 | 44431343;444 | +;+;+;+;+;+;+; | 3629 | ppp1r1c | 869 | 93 | 400 | 668 | 2297 | 56 |
| ENSDARG000 | 9;9 | 15898473;158 | 15899106;158 | -;- | 787 | NA | 0 | 1 | 6 | 6 | 0 | 2 |
| ENSDARG000 | 9;9;9;9;9;9;9; | 22677503;226 | 22677847;226 | +;+;+;+;+;+;+; | 3619 | itgb5 | 286 | 532 | 435 | 211 | 1392 | 2106 |
| ENSDARG000 | 9;9;9;9;9;9 | 23369294;233 | 23369518;233 | -;-;-;-;-;- | 3392 | sh3bp4 | 35 | 30 | 56 | 24 | 4 | 16 |
| ENSDARG000 | 9;9;9;9;9;9;9; | 30112423;301 | 30112553;301 | +;+;+;+;+;+;+; | 1872 | tfg | 4027 | 3775 | 4218 | 3044 | 3918 | 2669 |
| ENSDARG000 | 9;9 | 15901526;159 | 15901725;159 | +;+ | 865 | NA | 0 | 2 | 0 | 4 | 0 | 2 |
| ENSDARG000 | 9;9;9;9;9;9;9; | 34950942;349 | 34951028;349 | +;+;+;+;+;+;+; | 2701 | tfdp1a | 856 | 1216 | 1381 | 788 | 1321 | 1154 |
| ENSDARG000 | 9;9 | 15907033;159 | 15907232;159 | +;+ | 639 | NA | 0 | 0 | 0 | 0 | 0 | 0 |
| ENSDARG000 | 9;9;9;9;9;9;9; | 49808483;498 | 49808750;498 | -;-;-;-;-;-;-; | 4300 | ttc21b | 352 | 341 | 433 | 298 | 374 | 319 |
| ENSDARG000 | 9;9;9;9;9;9;9; | 34970418;349 | 34970928;349 | -;-;-;-;-;-;-; | 1319 | mettl21cb | 0 | 2 | 0 | 0 | 0 | 0 |
| ENSDARG000 | 9;9;9;9;9;9;9; | 8894788;8898 | 8895040;8898 | +;+;+;+;+;+;+; | 3711 | naxd | 886 | 890 | 1152 | 1012 | 1049 | 1095 |
| ENSDARG000 | 9;9;9;9;9 | 31838044;318 | 31838183;318 | -;-;-;-;- | 747 | fgf14 | 2 | 11 | 3 | 0 | 0 | 0 |
| ENSDARG000 | 9;9;9;9;9;9;9; | 6393368;6399 | 6396563;6400 | -;-;-;-;-;-;-;- | 5356 | nck2a | 144 | 68 | 112 | 104 | 100 | 191 |
| ENSDARG000 | 9;9;9;9;9;9;9; | 25096500;250 | 25096864;250 | +;+;+;+;+;+;+; | 6643 | LRCH1 | 438 | 280 | 508 | 590 | 470 | 519 |
| ENSDARG000 | 9;9;9;9;9;9;9; | 19421841;194 | 19421903;194 | +;+;+;+;+;+;+; | 1939 | PDE9A | 0 | 0 | 2 | 0 | 0 | 0 |
| ENSDARG000 | 9;9;9 | 2277317;2283 | 2277646;2284 | +;+;+ | 1070 | NA | 0 | 0 | 2 | 0 | 0 | 0 |
| ENSDARG000 | 9;9;9;9 | 2226463;2228 | 2226731;2228 | -;-;-;- | 812 | NA | 0 | 0 | 0 | 0 | 0 | 0 |
| ENSDARG000 | 9;9 | 19468769;194 | 19469300;194 | -;- | 749 | NA | 0 | 0 | 0 | 0 | 0 | 0 |
| ENSDARG000 | 9;9;9;9;9;9;9; | 19477687;194 | 19477983;194 | -;-;-;-;-;-;-; | 1388 | wdr4 | 92 | 121 | 200 | 44 | 142 | 84 |
| ENSDARG000 | 9;9 | 21363623;213 | 21363953;213 | -;- | 639 | il17d | 5 | 16 | 8 | 8 | 1 | 2 |
| ENSDARG000 | 9;9;9 | 23383414;233 | 23383473;233 | -;-;- | 234 | NA | 0 | 0 | 0 | 3 | 0 | 0 |
| ENSDARG000 | 9;9;9;9;9;9;9; | 21365043;213 | 21366027;213 | -;-;-;-;-;-;-; | 3624 | ift88 | 270 | 185 | 364 | 202 | 377 | 205 |
| ENSDARG000 | 9;9;9;9;9;9;9; | 44498182;444 | 44498435;444 | -;-;-;-;-;-;-; | 2238 | PDE1A | 6 | 23 | 95 | 79 | 26 | 19 |
| ENSDARG000 | 9;9;9;9;9;9;9; | 32069957;320 | 32070135;320 | +;+;+;+;+;+;+; | 1075 | NA | 31 | 45 | 70 | 6 | 221 | 16 |
| ENSDARG000 | 9;9;9;9;9 | 8887528;8887 | 8888464;8888 | -;-;-;-;- | 1412 | rab20 | 751 | 565 | 669 | 708 | 733 | 957 |
| ENSDARG000 | 9;9;9;9;9;9;9; | 6578580;6587 | 6578713;6587 | +;+;+;+;+;+;+; | 2137 | fhl2a | 621 | 557 | 1453 | 94 | 468 | 460 |
| ENSDARG000 | 9;9;9;9;9;9;9; | 23511913;235 | 23512023;235 | +;+;+;+;+;+;+; | 5104 | CNTNAP5 | 11 | 28 | 43 | 18 | 8 | 0 |
| ENSDARG000 | 9;9;9;9 | 6607514;6611 | 6608262;6611 | -;-;-;- | 1434 | C9H2orf49 | 216 | 183 | 223 | 137 | 235 | 194 |
| ENSDARG000 | 9;9;9;9 | 44713033;447 | 44713528;447 | -;-;-;- | 1056 | dusp19a | 0 | 0 | 0 | 0 | 0 | 0 |
| ENSDARG000 | 9;9;9 | 2330861;2334 | 2330978;2334 | -;-;- | 249 | NA | 0 | 0 | 0 | 0 | 0 | 0 |
| ENSDARG000 | 9;9;9;9;9;9;9; | 2330798;2333 | 2331086;2333 | +;+;+;+;+;+;+; | 1811 | atp5mc3a | 1106 | 1630 | 1441 | 632 | 1512 | 1398 |
| ENSDARG000 | 9;9;9;9;9;9;9; | 15982344;159 | 15982521;159 | +;+;+;+;+;+;+; | 2478 | NA | 0 | 0 | 2 | 0 | 0 | 0 |
| ENSDARG000 | 9;9;9;9;9;9;9 | 15936216;159 | 15937071;159 | -;-;-;-;-;-;- | 1809 | crfb4 | 250 | 206 | 208 | 190 | 196 | 242 |
| ENSDARG000 | 9 | 6623536 | 6624663 | - | 1128 | GPR45 | 0 | 6 | 13 | 20 | 6 | 5 |
| ENSDARG000 | 9;9;9;9;9 | 6658594;6658 | 6659621;6661 | -;-;-;-;- | 3073 | pou3f3a | 38 | 22 | 12 | 66 | 54 | 28 |
| ENSDARG000 | 9;9;9;9;9;9;9; | 28870357;288 | 28872353;288 | -;-;-;-;-;-;-; | 5481 | epb41l5 | 486 | 90 | 353 | 282 | 461 | 108 |
| ENSDARG000 | 9;9;9;9;9;9;9 | 34978867;349 | 34979404;349 | -;-;-;-;-;-;- | 1506 | NA | 0 | 0 | 0 | 0 | 0 | 0 |
| ENSDARG000 | 9;9;9;9;9 | 6915611;6918 | 6916603;6918 | -;-;-;-;- | 1771 | tmem182a | 239 | 45 | 19 | 21 | 60 | 166 |
| ENSDARG000 | 9;9;9;9;9;9;9; | 30132498;301 | 30132613;301 | -;-;-;-;-;-;-; | 2805 | NA | 40 | 8 | 20 | 12 | 152 | 54 |
| ENSDARG000 | 9;9;9;9;9;9;9; | 2343096;2343 | 2343187;2343 | +;+;+;+;+;+;+; | 1663 | ATF2 | 76 | 27 | 77 | 51 | 66 | 82 |
| ENSDARG000 | 9;9;9;9;9;9;9; | 38457806;384 | 38457924;384 | +;+;+;+;+;+;+; | 9371 | NA | 894 | 613 | 780 | 304 | 751 | 686 |
| ENSDARG000 | 9;9;9;9;9;9 | 30188764;301 | 30188898;301 | -;-;-;-;-;- | 840 | NA | 0 | 0 | 0 | 0 | 0 | 0 |
| ENSDARG000 | 9;9;9;9;9;9;9; | 38481780;384 | 38481901;384 | +;+;+;+;+;+;+; | 2565 | lss | 179 | 134 | 182 | 34 | 63 | 69 |
| ENSDARG000 | 9;9;9;9;9;9;9; | 33323104;333 | 33323296;333 | -;-;-;-;-;-;-; | 1703 | rpl8 | 23033 | 19974 | 34776 | 30748 | 22406 | 29902 |
| ENSDARG000 | 9;9;9;9;9;9;9; | 25171662;251 | 25172395;251 | -;-;-;-;-;-;-; | 2062 | esd | 1126 | 1349 | 1618 | 1264 | 1297 | 803 |
| ENSDARG000 | 9;9;9;9 | 23696924;236 | 23697282;236 | +;+;+;+ | 765 | NA | 0 | 0 | 4 | 0 | 0 | 0 |

| | | | | | | | | | | | |
|---|---|---|---|---|---|---|---|---|---|---|---|
| ENSDARG000 | 9:9:9:9:9:9:9: | 44721808:447 | 44722528:447 | +:+:+:+:+:+:+: | 4941 | nckap1 | 5232 | 4127 | 5068 | 3913 | 3874 | 4405 |
| ENSDARG000 | 9;9;9;9 | 30204415:302 | 30204541:302 | +;+;+;+ | 1746 | NA | 0 | 6 | 0 | 0 | 0 | 0 |
| ENSDARG000 | 9;9;9;9;9;9 | 23665777:236 | 23665975:236 | +;+;+;+;+;+ | 1473 | gypc | 5015 | 3578 | 7413 | 5733 | 3560 | 3118 |
| ENSDARG000 | 9:9:9:9:9:9:9: | 6934491:6937 | 6934559:6937 | +:+:+:+:+:+:+: | 1423 | MFSD9 | 369 | 91 | 138 | 67 | 274 | 163 |
| ENSDARG000 | 9;9;9;9;9 | 30211038:302 | 30211039:302 | +;+;+;+;+ | 1734 | NA | 0 | 2 | 0 | 0 | 4 | 3 |
| ENSDARG000 | 9:9:9:9:9:9:9: | 2499627:2507 | 2499673:2507 | +:+:+:+:+:+:+: | 1898 | wipf1a | 22 | 44 | 20 | 42 | 178 | 214 |
| ENSDARG000 | 9:9:9:9:9:9:9: | 34996245:349 | 34996357:349 | +:+:+:+:+:+:+: | 1723 | jam2b | 4 | 4 | 4 | 0 | 6 | 0 |
| ENSDARG000 | 9;9;9;9;9;9;9 | 41367848:413 | 41367930:413 | -;-;-;-;-;-;- | 1503 | nab1b | 36 | 31 | 135 | 104 | 32 | 33 |
| ENSDARG000 | 9:9:9:9:9:9:9: | 16097812:161 | 16098724:161 | -:-:-:-:-:-:-:-: | 2023 | upp2 | 0 | 0 | 2 | 0 | 0 | 0 |
| ENSDARG000 | 9;9 | 41445791:414 | 41446234:414 | -;- | 705 | NA | 0 | 0 | 0 | 0 | 0 | 0 |
| ENSDARG000 | 9:9:9:9:9:9:9: | 23740506:237 | 23740730:237 | -:-:-:-:-:-:-:-: | 1454 | ndufa10 | 1379 | 1522 | 1838 | 1039 | 1321 | 1076 |
| ENSDARG000 | 9:9:9:9:9:9:9: | 49906037:499 | 49907428:499 | -:-:-:-:-:-:-:-: | 6854 | scn1laa | 20 | 0 | 0 | 10 | 0 | 20 |
| ENSDARG000 | 9:9:9:9:9:9:9: | 6963039:6966 | 6963540:6966 | -:-:-:-:-:-:-:-: | 2736 | slc9a2 | 24 | 9 | 0 | 8 | 9 | 10 |
| ENSDARG000 | 9;9;9;9;9 | 30245637:302 | 30246195:302 | -;-;-;-;- | 827 | NA | 1092 | 814 | 939 | 871 | 814 | 1463 |
| ENSDARG000 | 9:9:9:9:9:9:9: | 30217828:302 | 30218158:302 | -:-:-:-:-:-:-:-: | 3482 | NA | 394 | 291 | 534 | 267 | 530 | 485 |
| ENSDARG000 | 9:9:9:9:9:9:9: | 41459759:414 | 41460180:414 | +:+:+:+:+:+:+: | 4112 | nemp2 | 694 | 531 | 806 | 550 | 446 | 493 |
| ENSDARG000 | 9;9;9 | 21401189:214 | 21401302:214 | +;+;+ | 2318 | gjb8 | 18 | 4 | 13 | 10 | 4 | 0 |
| ENSDARG000 | 9:9:9:9:9:9:9: | 21383620:213 | 21383778:213 | +:+:+:+:+:+:+: | 3085 | cryl1 | 243 | 358 | 273 | 210 | 318 | 456 |
| ENSDARG000 | 9:9:9:9:9:9:9: | 50000504:500 | 50000775:500 | +:+:+:+:+:+:+: | 1959 | slc38a11 | 1005 | 1048 | 1276 | 243 | 1007 | 944 |
| ENSDARG000 | 9;9;9;9;9;9;9 | 41482831:414 | 41483511:414 | -;-;-;-;-;-;- | 3141 | mfsd6b | 18 | 4 | 22 | 7 | 20 | 22 |
| ENSDARG000 | 9:9:9:9:9:9:9: | 33334501:333 | 33334837:333 | +:+:+:+:+:+:+: | 5068 | ddx3xa | 2655 | 2981 | 2725 | 1409 | 3187 | 2285 |
| ENSDARG000 | 9;9;9;9;9;9 | 22764235:227 | 22764397:227 | +;+;+;+;+;+ | 1247 | tnfaip6 | 227 | 1776 | 498 | 693 | 5562 | 4764 |
| ENSDARG000 | 9;9;9 | 25198249:252 | 25199054:252 | -;-;- | 1208 | HTR2A | 32 | 14 | 45 | 14 | 0 | 0 |
| ENSDARG000 | 9:9:9:9:9:9:9: | 25218834:252 | 25218887:252 | -:-:-:-:-:-:-:-: | 1195 | NA | 9 | 2 | 14 | 32 | 4 | 24 |
| ENSDARG000 | 9;9;9;9;9;9;9 | 35077546:350 | 35077615:350 | +;+;+;+;+;+;+ | 1866 | NA | 90 | 34 | 46 | 34 | 142 | 64 |
| ENSDARG000 | 9;9;9 | 41559355:415 | 41559683:415 | -;-;- | 558 | NA | 0 | 0 | 0 | 0 | 0 | 0 |
| ENSDARG000 | 9:9:9:9:9:9:9: | 35026415:350 | 35028398:350 | -:-:-:-:-:-:-:-: | 4496 | appb | 6695 | 3716 | 4281 | 3371 | 5369 | 4500 |
| ENSDARG000 | 9:9:9:9:9:9:9: | 19489304:194 | 19489368:194 | +:+:+:+:+:+:+: | 2851 | si:ch211-140 | 425 | 276 | 625 | 552 | 683 | 741 |
| ENSDARG000 | 9:9:9:9:9:9:9: | 23748133:237 | 23749114:237 | +:+:+:+:+:+:+: | 1744 | faima | 18 | 16 | 13 | 20 | 19 | 29 |
| ENSDARG000 | 9:9:9:9:9:9:9: | 23754275:237 | 23758866:237 | -:-:-:-:-:-:-:-: | 7162 | NA | 839 | 341 | 538 | 430 | 4 | 60 |
| ENSDARG000 | 9;9;9;9;9;9;9 | 44832032:448 | 44832750:448 | +;+;+;+;+;+;+ | 2994 | frzb | 198 | 10 | 24 | 20 | 43 | 1498 |
| ENSDARG000 | 9:9:9:9:9:9:9: | 28939912:289 | 28942734:289 | -:-:-:-:-:-:-:-: | 6116 | ptpn4a | 1382 | 2032 | 2425 | 1702 | 2478 | 1615 |
| ENSDARG000 | 9;9;9 | 35026304:350 | 35026608:350 | +;+;+ | 2776 | NA | 4 | 0 | 1 | 0 | 0 | 2 |
| ENSDARG000 | 9:9:9:9:9:9:9: | 35014513:350 | 35014655:350 | +:+:+:+:+:+:+: | 3922 | gabpa | 461 | 564 | 588 | 380 | 455 | 424 |
| ENSDARG000 | 9;9;9;9 | 44849158:448 | 44849245:448 | -;-;-;- | 459 | NA | 0 | 0 | 0 | 8 | 0 | 0 |
| ENSDARG000 | 9:9:9:9:9:9:9: | 32073606:320 | 32073807:320 | +:+:+:+:+:+:+: | 8908 | pikfyve | 4945 | 3718 | 5855 | 7327 | 4861 | 5841 |
| ENSDARG000 | 9;9 | 33357011:333 | 33357035:333 | +;+ | 2226 | nyx | 0 | 4 | 0 | 0 | 6 | 4 |
| ENSDARG000 | 9:9:9:9:9:9:9: | 16127209:161 | 16129168:161 | -:-:-:-:-:-:-:-: | 6329 | MYO1B | 5671 | 4869 | 3999 | 4823 | 3592 | 8243 |
| ENSDARG000 | 9;9;9;9;9;9;9 | 25320680:253 | 25321310:253 | -;-;-;-;-;-;- | 1290 | med4 | 170 | 114 | 192 | 108 | 110 | 70 |
| ENSDARG000 | 9:9:9:9:9:9:9: | 2522797:2522 | 2523040:2523 | +:+:+:+:+:+:+: | 7571 | GPR155 | 317 | 219 | 383 | 132 | 362 | 303 |
| ENSDARG000 | 9:9:9:9:9:9:9: | 6997861:6998 | 6998196:6998 | +:+:+:+:+:+:+: | 5064 | inpp4aa | 1063 | 668 | 543 | 639 | 817 | 695 |
| ENSDARG000 | 9:9:9:9:9:9:9: | 44869965:448 | 44870829:448 | -:-:-:-:-:-:-:-: | 3384 | dnajc10 | 1030 | 819 | 1028 | 550 | 1378 | 673 |
| ENSDARG000 | 9:9:9:9:9:9:9: | 25330905:253 | 25331083:253 | +:+:+:+:+:+:+: | 1628 | itm2bb | 1488 | 1182 | 1266 | 1411 | 1229 | 1272 |
| ENSDARG000 | 9 | 21407987 | 21409466 | + | 1480 | gja3 | 0 | 0 | 0 | 0 | 0 | 0 |
| ENSDARG000 | 9:9:9:9:9:9:9: | 23765587:237 | 23765772:237 | +:+:+:+:+:+:+: | 5755 | parp14rs1 | 2157 | 1605 | 1429 | 675 | 1020 | 1002 |
| ENSDARG000 | 9;9 | 33423170:334 | 33423280:334 | +;+ | 1329 | NA | 10 | 0 | 0 | 6 | 306 | 4 |
| ENSDARG000 | 9 | 33417969 | 33419321 | + | 1353 | gpr34b | 0 | 6 | 0 | 0 | 0 | 15 |
| ENSDARG000 | 9:9:9:9:9:9:9: | 33366683:333 | 33367058:333 | -:-:-:-:-:-:-:-: | 3537 | caska | 550 | 598 | 1336 | 934 | 1652 | 488 |
| ENSDARG000 | 9;9;9;9;9;9;9 | 32096331:320 | 32096537:320 | -;-;-;-;-;-;- | 774 | NA | 0 | 3 | 4 | 8 | 4 | 10 |
| ENSDARG000 | 9:9:9:9:9:9:9: | 44897806:448 | 44899951:448 | -:-:-:-:-:-:-:-: | 3914 | zgc:66484 | 226 | 172 | 233 | 150 | 272 | 192 |
| ENSDARG000 | 9:9:9:9:9:9:9: | 8929764:8930 | 8930119:8930 | +:+:+:+:+:+:+: | 2190 | ankrd10b | 351 | 202 | 263 | 221 | 379 | 337 |
| ENSDARG000 | 9:9:9:9:9:9:9: | 2560219:2560 | 2561709:2561 | -:-:-:-:-:-:-:-: | 3593 | scrn3 | 284 | 273 | 130 | 108 | 156 | 148 |
| ENSDARG000 | 9:9:9:9:9:9:9: | 21430311:214 | 21431060:214 | -:-:-:-:-:-:-:-: | 4774 | ZMYM2 | 995 | 1068 | 1759 | 949 | 1280 | 834 |
| ENSDARG000 | 9:9:9:9:9:9:9: | 22780901:227 | 22781136:227 | +:+:+:+:+:+:+: | 7795 | rif1 | 1112 | 1804 | 2605 | 1556 | 2518 | 2534 |
| ENSDARG000 | 9:9:9:9:9:9:9: | 25347814:253 | 25348281:253 | -:-:-:-:-:-:-:-: | 3111 | epc2 | 625 | 514 | 770 | 479 | 585 | 525 |
| ENSDARG000 | 9:9:9:9:9:9:9: | 41612642:416 | 41614003:416 | +:+:+:+:+:+:+: | 11941 | NA | 0 | 10 | 47 | 12 | 0 | 0 |
| ENSDARG000 | 9:9:9:9:9:9:9: | 38502524:385 | 38502820:385 | +:+:+:+:+:+:+: | 2159 | LMLN | 427 | 295 | 483 | 120 | 464 | 292 |
| ENSDARG000 | 9:9:9:9:9:9:9: | 2574122:2574 | 2574303:2574 | +:+:+:+:+:+:+: | 1139 | CIR1 | 244 | 168 | 200 | 136 | 198 | 198 |
| ENSDARG000 | 9:9:9:9:9:9:9: | 23783856:237 | 23784086:237 | -:-:-:-:-:-:-:-: | 3184 | esyt3 | 132 | 6 | 6 | 18 | 40 | 14 |
| ENSDARG000 | 9:9:9:9:9:9:9: | 44907768:449 | 44911541:449 | -:-:-:-:-:-:-:-: | 6769 | vil1 | 36 | 12 | 20 | 0 | 54 | 14 |
| ENSDARG000 | 9:9:9:9:9:9:9: | 25380758:253 | 25384588:253 | -:-:-:-:-:-:-:-: | 6420 | acvr2aa | 2017 | 1208 | 1800 | 1476 | 1764 | 2634 |
| ENSDARG000 | 9;9;9;9 | 2591968:2592 | 2594410:2594 | -;-;-;- | 2497 | sp9 | 56 | 4 | 0 | 0 | 10 | 0 |
| ENSDARG000 | 9:9:9:9:9:9:9: | 2762270:2762 | 2762421:2762 | +:+:+:+:+:+:+: | 2996 | sp3a | 185 | 131 | 292 | 78 | 183 | 43 |
| ENSDARG000 | 9;9;9 | 29034221:290 | 29036259:290 | -;-;- | 2103 | tmem177 | 200 | 242 | 219 | 190 | 248 | 198 |
| ENSDARG000 | 9;9;9 | 16241656:162 | 16241884:162 | +;+;+ | 885 | C9H3orf38 | 94 | 60 | 70 | 55 | 112 | 82 |
| ENSDARG000 | 9:9:9:9:9:9:9: | 38524881:385 | 38525304:385 | +:+:+:+:+:+:+: | 2424 | OSBPL11 | 249 | 169 | 305 | 91 | 276 | 199 |
| ENSDARG000 | 9:9:9:9:9:9:9: | 19518674:195 | 19518742:195 | +:+:+:+:+:+:+: | 1316 | NA | 910 | 832 | 796 | 754 | 767 | 717 |
| ENSDARG000 | 9:9:9:9:9:9:9: | 33660964:336 | 33663905:336 | -:-:-:-:-:-:-:-: | 4420 | mao | 1893 | 3386 | 3398 | 3086 | 5515 | 4794 |
| ENSDARG000 | 9;9;9;9 | 25567605:255 | 25567666:255 | +;+;+;+ | 549 | NA | 0 | 0 | 0 | 0 | 0 | 0 |
| ENSDARG000 | 9;9;9;9;9;9;9 | 41690153:417 | 41690548:417 | +;+;+;+;+;+;+ | 2942 | NA | 1161 | 924 | 1107 | 1241 | 1242 | 1172 |
| ENSDARG000 | 9:9:9:9:9:9:9: | 25568839:257 | 25569056:257 | +:+:+:+:+:+:+: | 6765 | zeb2a | 4582 | 7097 | 6088 | 4647 | 8437 | 5616 |
| ENSDARG000 | 9:9:9:9:9:9:9: | 7072504:7072 | 7072903:7072 | +:+:+:+:+:+:+: | 1582 | si:dkey-4e7.3 | 150 | 212 | 310 | 224 | 217 | 121 |
| ENSDARG000 | 9;9;9;9;9 | 33734850:337 | 33735192:337 | -;-;-;-;- | 811 | NA | 0 | 0 | 0 | 0 | 0 | 0 |
| ENSDARG000 | 9;9;9 | 7081577:7087 | 7081773:7087 | -;-;- | 504 | coa5 | 116 | 122 | 182 | 106 | 84 | 120 |
| ENSDARG000 | 9:9:9:9:9:9:9: | 41744865:417 | 41746062:417 | -:-:-:-:-:-:-:-: | 11904 | NA | 2 | 2 | 0 | 0 | 0 | 0 |
| ENSDARG000 | 9:9:9:9:9:9:9: | 8968702:8968 | 8969102:8969 | +:+:+:+:+:+:+: | 1820 | ube2al | 858 | 955 | 669 | 613 | 925 | 656 |
| ENSDARG000 | 9:9:9:9:9:9:9: | 38564900:385 | 38565169:385 | -:-:-:-:-:-:-:-: | 2019 | NA | 93 | 44 | 31 | 16 | 43 | 70 |
| ENSDARG000 | 9:9:9:9:9:9:9: | 21462467:214 | 21463607:214 | -:-:-:-:-:-:-:-: | 4109 | mphosph8 | 1484 | 1466 | 1645 | 947 | 1635 | 1242 |
| ENSDARG000 | 9:9:9:9:9:9:9: | 33788389:337 | 33788792:337 | +:+:+:+:+:+:+: | 5275 | KDM6A | 1093 | 828 | 2156 | 1101 | 1348 | 1016 |
| ENSDARG000 | 9:9:9:9:9:9:9: | 30248571:302 | 30249150:302 | -:-:-:-:-:-:-:-: | 1907 | NA | 332 | 264 | 480 | 331 | 284 | 326 |
| ENSDARG000 | 9:9:9:9:9:9:9: | 29040386:290 | 29040581:290 | +:+:+:+:+:+:+: | 973 | MRAS | 49 | 25 | 90 | 24 | 62 | 52 |
| ENSDARG000 | 9:9:9:9:9:9:9: | 38588081:385 | 38588185:385 | +:+:+:+:+:+:+: | 2881 | snx4 | 1924 | 1586 | 1487 | 1400 | 1917 | 1498 |
| ENSDARG000 | 9;9 | 2861224:2866 | 2861584:2866 | -;- | 474 | NA | 0 | 0 | 0 | 0 | 0 | 0 |
| ENSDARG000 | 9:9:9:9:9:9:9: | 33783549:337 | 33784072:337 | -:-:-:-:-:-:-:-: | 1308 | fundc1 | 1093 | 1332 | 952 | 1242 | 1450 | 980 |
| ENSDARG000 | 9:9:9:9:9:9:9: | 2879763:2882 | 2880130:2882 | -:-:-:-:-:-:-:-: | 1741 | cdca7a | 270 | 1005 | 1248 | 559 | 1458 | 589 |
| ENSDARG000 | 9:9:9:9:9:9:9: | 44961138:449 | 44962347:449 | -:-:-:-:-:-:-:-: | 3489 | NA | 507 | 543 | 765 | 358 | 1081 | 1335 |
| ENSDARG000 | 9:9:9:9:9:9:9: | 33758720:337 | 33758924:337 | -:-:-:-:-:-:-:-: | 2483 | efhc2 | 0 | 6 | 12 | 0 | 5 | 0 |
| ENSDARG000 | 9;9;9;9;9 | 29064147:290 | 29064311:290 | -;-;-;-;- | 413 | NA | 0 | 0 | 0 | 0 | 0 | 1 |
| ENSDARG000 | 9;9 | 50089000:500 | 50089568:500 | +;+ | 675 | NA | 8 | 0 | 0 | 0 | 0 | 0 |
| ENSDARG000 | 9:9:9:9:9:9:9: | 50038347:500 | 50038895:500 | +:+:+:+:+:+:+: | 4396 | NA | 4758 | 2474 | 3046 | 4248 | 2293 | 3635 |
| ENSDARG000 | 9:9:9:9:9:9:9: | 41815312:418 | 41816089:418 | -:-:-:-:-:-:-:-: | 2791 | NA | 2 | 0 | 0 | 6 | 0 | 12 |
| ENSDARG000 | 9:9:9:9:9:9:9: | 35170335:351 | 35170882:351 | -:-:-:-:-:-:-:-: | 3026 | ncam2 | 6 | 156 | 4 | 44 | 12 | 2 |
| ENSDARG000 | 9;9;9;9;9;9 | 8973080:8974 | 8974591:8974 | -;-;-;-;-;- | 2057 | ing1 | 330 | 286 | 462 | 312 | 324 | 323 |
| ENSDARG000 | 9:9:9:9:9:9:9: | 19550367:195 | 19550482:195 | -:-:-:-:-:-:-:-: | 2020 | NA | 31 | 15 | 25 | 2 | 84 | 20 |
| ENSDARG000 | 9:9:9:9:9:9:9: | 23808546:238 | 23808669:238 | +:+:+:+:+:+:+: | 1877 | dhrs12 | 610 | 759 | 654 | 435 | 532 | 728 |
| ENSDARG000 | 9:9:9:9:9:9:9: | 19529951:195 | 19529974:195 | +:+:+:+:+:+:+: | 2337 | pknox1.1 | 198 | 245 | 456 | 351 | 442 | 294 |
| ENSDARG000 | 9:9:9:9:9:9:9: | 29122039:291 | 29123013:291 | -:-:-:-:-:-:-:-: | 2871 | pth2ra | 0 | 0 | 0 | 0 | 0 | 6 |
| ENSDARG000 | 9:9:9:9:9:9:9: | 2896009:2896 | 2897573:2897 | -:-:-:-:-:-:-:-: | 3466 | map3k20a | 718 | 441 | 569 | 229 | 922 | 169 |
| ENSDARG000 | 9;9;9;9 | 29323163:293 | 29323204:293 | +;+;+;+ | 1058 | OXGR1 | 2 | 22 | 22 | 4 | 9 | 0 |
| ENSDARG000 | 9;9 | 29338986:294 | 29339700:294 | -;- | 1260 | hs6st3b | 8 | 0 | 0 | 0 | 0 | 0 |
| ENSDARG000 | 9:9:9:9:9:9:9: | 9112159:9112 | 9112447:9112 | +:+:+:+:+:+:+: | 5907 | sik1 | 38084 | 6595 | 8923 | 16765 | 11290 | 12205 |
| ENSDARG000 | 9:9:9:9:9:9:9: | 23813593:238 | 23814251:238 | -:-:-:-:-:-:-:-: | 2191 | wdfy2 | 738 | 629 | 550 | 615 | 534 | 834 |
| ENSDARG000 | 9:9:9:9:9:9:9: | 41821613:418 | 41821742:418 | +:+:+:+:+:+:+: | 6566 | col18a1a | 168 | 128 | 94 | 10 | 148 | 64 |
| ENSDARG000 | 9:9:9:9:9:9:9: | 25840720:258 | 25840763:258 | +:+:+:+:+:+:+: | 4311 | gtdc1 | 283 | 179 | 291 | 260 | 234 | 202 |
| ENSDARG000 | 9:9:9:9:9:9:9: | 44994214:449 | 44994535:449 | +:+:+:+:+:+:+: | 2374 | retsatl | 4 | 0 | 0 | 0 | 0 | 0 |
| ENSDARG000 | 9;9 | 21495873:214 | 21495913:214 | +;+ | 716 | NA | 0 | 0 | 0 | 0 | 0 | 0 |
| ENSDARG000 | 9:9:9:9:9:9:9: | 21490822:214 | 21491704:214 | -:-:-:-:-:-:-:-: | 4411 | NA | 0 | 10 | 12 | 8 | 6 | 0 |
| ENSDARG000 | 9:9:9:9:9:9:9: | 32109911:321 | 32110155:321 | -:-:-:-:-:-:-:-: | 3207 | ANKRD44 | 1039 | 662 | 783 | 597 | 749 | 767 |
| ENSDARG000 | 9:9:9:9:9:9:9: | 7091028:7091 | 7095978:7095 | -:-:-:-:-:-:-:-: | 7428 | MGAT4A | 1922 | 2509 | 2677 | 2060 | 3094 | 1708 |
| ENSDARG000 | 9:9:9:9:9:9:9: | 32161623:321 | 32162423:321 | -:-:-:-:-:-:-:-: | 4900 | SF3B1 | 7928 | 5283 | 11679 | 12648 | 9880 | 12286 |
| ENSDARG000 | 9:9:9:9:9:9:9: | 25924953:259 | 25925364:259 | -:-:-:-:-:-:-:-: | 1680 | arhgap15 | 12 | 59 | 50 | 54 | 83 | 84 |
| ENSDARG000 | 9:9:9:9:9:9:9: | 35631440:356 | 35631571:356 | -:-:-:-:-:-:-:-: | 1537 | zp2l1 | 0 | 0 | 0 | 0 | 0 | 0 |
| ENSDARG000 | 9:9:9:9:9:9:9: | 17045875:170 | 17047203:170 | -:-:-:-:-:-:-:-: | 15337 | mycbp2 | 6345 | 3222 | 4445 | 3296 | 4327 | 4491 |
| ENSDARG000 | 9:9:9:9:9:9:9: | 51147380:511 | 51148270:511 | +:+:+:+:+:+:+: | 3744 | ifih1 | 291 | 305 | 151 | 45 | 283 | 188 |
| ENSDARG000 | 9:9:9:9:9:9:9: | 9180620:9183 | 9184681:9184 | -:-:-:-:-:-:-:-: | 5778 | cbsb | 706 | 804 | 1235 | 1494 | 332 | 766 |
| ENSDARG000 | 9;9;9;9 | 51265283:512 | 51265381:512 | +;+;+;+ | 592 | gcgb | 6 | 0 | 0 | 0 | 0 | 0 |
| ENSDARG000 | 9:9:9:9:9:9:9: | 51225345:512 | 51225600:512 | +:+:+:+:+:+:+: | 2927 | NA | 8 | 112 | 30 | 37 | 154 | 121 |
| ENSDARG000 | 9:9:9:9:9:9:9: | 29430432:294 | 29430500:294 | +:+:+:+:+:+:+: | 5765 | UGGT2 | 1118 | 1069 | 1641 | 598 | 1731 | 1007 |
| ENSDARG000 | 9:9:9:9:9:9:9: | 2949272:2949 | 2949731:2949 | -:-:-:-:-:-:-:-: | 3456 | RAPGEF4 | 2 | 4 | 2 | 0 | 6 | 4 |
| ENSDARG000 | 9;9;9;9;9 | 51192135:511 | 51192408:511 | -;-;-;-;- | 921 | NA | 0 | 0 | 0 | 0 | 0 | 0 |
| ENSDARG000 | 9:9:9:9:9:9:9: | 38624478:386 | 38624647:386 | +:+:+:+:+:+:+: | 6970 | znf148 | 850 | 570 | 888 | 482 | 923 | 702 |
| ENSDARG000 | 9:9:9:9:9:9:9: | 23825440:238 | 23825880:238 | +:+:+:+:+:+:+: | 4069 | ints6 | 323 | 317 | 430 | 207 | 398 | 443 |
| ENSDARG000 | 9;9;9;9;9 | 45112537:451 | 45112598:451 | +;+;+;+;+ | 807 | NA | 0 | 0 | 0 | 0 | 0 | 0 |
| ENSDARG000 | 9 | 30263188 | 30264415 | - | 1228 | mid1ip1a | 94 | 12 | 0 | 8 | 46 | 4 |
| ENSDARG000 | 9;9;9;9 | 7228083:7232 | 7229667:7232 | -;-;-;- | 2321 | creg2 | 933 | 638 | 899 | 724 | 1023 | 950 |
| ENSDARG000 | 9:9:9:9:9:9:9: | 45227028:453 | 45227254:453 | +:+:+:+:+:+:+: | 2877 | ADARB1 | 52 | 17 | 26 | 7 | 37 | 21 |
| ENSDARG000 | 9:9:9:9:9:9:9: | 21535885:215 | 21536286:215 | +:+:+:+:+:+:+: | 8448 | arhgef7a | 156 | 104 | 164 | 144 | 85 | 242 |
| ENSDARG000 | 9;9 | 45227101:453 | 45227254:453 | +;+ | 236 | NA | 0 | 0 | 0 | 0 | 0 | 0 |
| ENSDARG000 | 9;9;9;9;9;9 | 7240896:7248 | 7245929:7249 | -;-;-;-;-;- | 7915 | NA | 6 | 2 | 4 | 0 | 18 | 2 |
| ENSDARG000 | 9:9:9:9:9:9:9: | 38644976:386 | 38645053:386 | +:+:+:+:+:+:+: | 2447 | slc12a8 | 4 | 2 | 0 | 2 | 2 | 2 |
| ENSDARG000 | 9:9:9:9:9:9:9: | 307863:31033 | 308050:31044 | +:+:+:+:+:+:+: | 4809 | stac3 | 0 | 0 | 0 | 0 | 0 | 0 |
| ENSDARG000 | 9;9;9;9;9;9;9 | 26086135:260 | 26086698:260 | +;+;+;+;+;+;+ | 5736 | arglu1a | 3866 | 1477 | 3795 | 2627 | 3664 | 3681 |
| ENSDARG000 | 9;9 | 3035128:3037 | 3035355:3037 | +;+ | 435 | NA | 0 | 0 | 0 | 0 | 0 | 0 |
| ENSDARG000 | 9:9:9:9:9:9:9: | 3055566:3055 | 3057346:3057 | +:+:+:+:+:+:+: | 4121 | NA | 147 | 70 | 120 | 75 | 120 | 72 |
| ENSDARG000 | 9:9:9:9:9:9:9: | 30268040:302 | 30268182:302 | -:-:-:-:-:-:-:-: | 1796 | OTC | 0 | 16 | 26 | 5 | 17 | 6 |
| ENSDARG000 | 9;9 | 3055023:3055 | 3055108:3055 | +;+ | 320 | NA | 0 | 0 | 0 | 0 | 0 | 0 |
| ENSDARG000 | 9:9:9:9:9:9:9: | 9230446:9235 | 9230789:9235 | -;-;-;-;-;-;-;- | 1004 | u2af1 | 1021 | 1243 | 1256 | 840 | 996 | 1002 |
| ENSDARG000 | 9:9:9:9:9:9:9: | 26103814:261 | 26103926:261 | +:+:+:+:+:+:+: | 4989 | efnb2a | 778 | 650 | 667 | 709 | 1921 | 820 |
| ENSDARG000 | 9:9:9:9:9:9:9: | 7275185:7275 | 7276394:7275 | -:-:-:-:-:-:-:-: | 2122 | mitd1 | 236 | 153 | 170 | 222 | 184 | 199 |
| ENSDARG000 | 9;9;9;9 | 41990086:419 | 41990186:419 | +;+;+;+ | 449 | NA | 0 | 0 | 0 | 2 | 0 | 0 |
| ENSDARG000 | 9:9:9:9:9:9:9: | 35832294:358 | 35832449:358 | +:+:+:+:+:+:+: | 5294 | sona | 1219 | 1290 | 2203 | 1048 | 1321 | 1388 |
| ENSDARG000 | 9:9:9:9:9:9:9: | 38684871:386 | 38685292:386 | +:+:+:+:+:+:+: | 4550 | heg1 | 537 | 259 | 261 | 229 | 246 | 554 |
| ENSDARG000 | 9:9:9:9:9:9:9: | 33863523:338 | 33863971:338 | -:-:-:-:-:-:-:-: | 1496 | DIPK2B | 32 | 7 | 26 | 10 | 26 | 28 |
| ENSDARG000 | 9:9:9:9:9:9:9: | 22805138:228 | 22805958:228 | -:-:-:-:-:-:-:-: | 19942 | NEB | 12 | 26 | 24 | 0 | 36 | 30 |
| ENSDARG000 | 9 | 7357819 | 7359484 | - | 1666 | NA | 0 | 0 | 2 | 0 | 0 | 0 |
| ENSDARG000 | 9;9;9 | 7358749:7361 | 7359365:7362 | +;+;+ | 1600 | ihha | 2 | 0 | 0 | 0 | 0 | 0 |
| ENSDARG000 | 9;9;9 | 19604867:196 | 19605058:196 | -;-;- | 428 | NA | 2 | 0 | 0 | 0 | 1 | 0 |
| ENSDARG000 | 9:9:9:9:9:9:9: | 42040557:420 | 42040642:420 | +:+:+:+:+:+:+: | 4328 | pcbp3 | 2 | 0 | 0 | 0 | 0 | 2 |
| ENSDARG000 | 9:9:9:9:9:9:9: | 23842884:238 | 23843213:238 | -:-:-:-:-:-:-:-: | 2720 | serpine3 | 2 | 10 | 4 | 0 | 26 | 0 |
| ENSDARG000 | 9:9:9:9:9:9:9: | 7371962:7374 | 7372277:7374 | -;-;-;-;-;-;-;- | 1211 | NA | 0 | 0 | 2 | 10 | 0 | 0 |
| ENSDARG000 | 9:9:9:9:9:9:9: | 35843732:358 | 35844135:358 | +:+:+:+:+:+:+: | 3813 | rcan1a | 1340 | 1532 | 1040 | 971 | 1451 | 1295 |
| ENSDARG000 | 9:9:9:9:9:9:9: | 30274637:302 | 30274671:302 | +:+:+:+:+:+:+: | 5116 | NA | 0 | 0 | 18 | 6 | 0 | 2 |
| ENSDARG000 | 9:9:9:9:9:9:9: | 17306162:173 | 17306236:173 | +:+:+:+:+:+:+: | 2733 | scel | 24 | 0 | 0 | 52 | 24 | 24 |
| ENSDARG000 | 9;9;9;9 | 323460:32541 | 324371:32547 | -;-;-;- | 1164 | NDUFA4L2 | 0 | 0 | 0 | 4 | 0 | 0 |
| ENSDARG000 | 9;9 | 35872698:358 | 35873956:358 | -;- | 1375 | dnajc28 | 38 | 52 | 48 | 26 | 46 | 66 |
| ENSDARG000 | 9;9;9;9;9;9;9 | 35876927:358 | 35877111:358 | +;+;+;+;+;+;+ | 1284 | mab21l3 | 2 | 119 | 1 | 44 | 5 | 33 |
| ENSDARG000 | 9;9 | 26410982:264 | 26411069:264 | -;- | 196 | NA | 0 | 0 | 0 | 0 | 0 | 0 |
| ENSDARG000 | 9:9:9:9:9:9:9: | 26199624:261 | 26199834:261 | +:+:+:+:+:+:+: | 1466 | NA | 0 | 0 | 0 | 0 | 0 | 0 |
| ENSDARG000 | 9;9;9;9;9 | 9251466:9259 | 9251665:9259 | -;-;-;-;- | 1063 | NA | 0 | 0 | 4 | 0 | 18 | 0 |
| ENSDARG000 | 9;9 | 45551068:455 | 45551286:455 | +;+ | 429 | NA | 0 | 0 | 0 | 0 | 0 | 0 |
| ENSDARG000 | 9;9 | 45551053:455 | 45551383:455 | -;- | 1477 | NA | 49 | 39 | 88 | 16 | 41 | 50 |
| ENSDARG000 | 9;9;9;9 | 21571433:215 | 21571493:215 | +;+;+;+ | 960 | NA | 2 | 0 | 1 | 0 | 0 | 0 |

| | | | | | | | | | | | | |
|---|---|---|---|---|---|---|---|---|---|---|---|---|
| ENSDARG000 | 9;9;9;9;9;9;9; | 3135062:3135 | 3135203:3135 | -;-;-;-;-;-;-;-; | 1684 | PDK1 | 191 | 166 | 182 | 135 | 180 | 222 |
| ENSDARG000 | 9;9;9;9;9;9;9; | 51289904:512 | 51289940:512 | -;-;-;-;-;-;-;-; | 2987 | SLC4A10 | 2 | 0 | 0 | 0 | 0 | 0 |
| ENSDARG000 | 9;9;9;9;9;9;9; | 9286419:9286 | 9286862:9286 | -;-;-;-;-;-;-;-; | 1050 | NA | 0 | 0 | 0 | 0 | 0 | 0 |
| ENSDARG000 | 9;9 | 45567910:455 | 45567948:455 | -;- | 498 | NA | 0 | 0 | 0 | 0 | 0 | 0 |
| ENSDARG000 | 9;9;9;9;9;9;9; | 45591239:455 | 45591518:455 | -;-;-;-;-;-;-;-; | 877 | agr1 | 52 | 0 | 0 | 12 | 2 | 6 |
| ENSDARG000 | 9;9;9;9;9;9;9; | 29484043:294 | 29484494:294 | -;-;-;-;-;-;-;-; | 2077 | dnajc3a | 523 | 737 | 1573 | 552 | 915 | 455 |
| ENSDARG000 | 9;9;9 | 33997310:340 | 33997538:340 | +;+;+ | 783 | NA | 0 | 0 | 0 | 0 | 0 | 0 |
| ENSDARG000 | 9;9;9;9;9;9;9; | 19623363:196 | 19623506:196 | +;+;+;+;+;+;+;+; | 1875 | pdxka | 270 | 703 | 160 | 179 | 1096 | 1150 |
| ENSDARG000 | 9;9;9;9;9 | 34070996:340 | 34071100:340 | -;-;-;-;- | 453 | NA | 0 | 0 | 0 | 0 | 0 | 0 |
| ENSDARG000 | 9;9;9;9;9;9;9; | 35901554:359 | 35901680:359 | +;+;+;+;+;+;+;+; | 3398 | atp1a1b | 5467 | 3459 | 3156 | 5651 | 3548 | 977 |
| ENSDARG000 | 9;9;9;9;9;9;9 | 34089156:340 | 34090811:340 | +;+;+;+;+;+;+ | 2604 | ccdc80 | 2561 | 1564 | 2381 | 2726 | 1938 | 1319 |
| ENSDARG000 | 9;9;9;9;9;9;9; | 23859317:238 | 23863078:238 | -;-;-;-;-;-;-;- | 5105 | FAM124A | 4 | 4 | 14 | 0 | 0 | 10 |
| ENSDARG000 | 9;9;9;9;9;9;9; | 34112067:341 | 34113129:341 | -;-;-;-;-;-;-;- | 2476 | slc35a5 | 274 | 109 | 186 | 295 | 202 | 172 |
| ENSDARG000 | 9;9;9;9;9;9;9; | 9295428:9296 | 9296401:9296 | +;+;+;+;+;+;+;+; | 3842 | NA | 0 | 0 | 0 | 0 | 14 | 6 |
| ENSDARG000 | 9 | 21722733 | 21724661 | + | 1929 | sox1a | 0 | 0 | 0 | 0 | 0 | 0 |
| ENSDARG000 | 9;9;9;9;9;9;9; | 32178050:321 | 32178715:321 | +;+;+;+;+;+;+;+; | 2641 | coq10b | 529 | 408 | 437 | 378 | 532 | 368 |
| ENSDARG000 | 9;9;9;9;9;9;9 | 23885441:238 | 23888486:238 | -;-;-;-;-;-;- | 4238 | NA | 0 | 0 | 0 | 3 | 1 | 0 |
| ENSDARG000 | 9;9;9 | 23885643:238 | 23885795:238 | +;+;+ | 428 | NA | 0 | 0 | 0 | 0 | 0 | 0 |
| ENSDARG000 | 9;9;9;9;9;9;9; | 38712474:387 | 38712611:387 | +;+;+;+;+;+;+;+; | 5481 | NA | 0 | 0 | 5 | 0 | 0 | 243 |
| ENSDARG000 | 9;9;9;9;9;9;9; | 30294073:302 | 30294215:302 | +;+;+;+;+;+;+;+; | 2072 | srpx | 22 | 0 | 2 | 0 | 6 | 16 |
| ENSDARG000 | 9;9;9;9;9;9;9; | 7410798:7410 | 7410899:7410 | -;-;-;-;-;-;-;-; | 2110 | BIN1 | 2 | 18 | 6 | 0 | 18 | 0 |
| ENSDARG000 | 9;9;9 | 32190670:321 | 32190788:321 | -;-;- | 546 | NA | 0 | 0 | 0 | 0 | 0 | 0 |
| ENSDARG000 | 9;9 | 32189139:321 | 32189954:321 | +;+ | 2749 | NA | 0 | 0 | 1 | 0 | 0 | 0 |
| ENSDARG000 | 9;9;9 | 26561884:265 | 26562052:265 | -;-;- | 317 | NA | 0 | 0 | 0 | 0 | 0 | 0 |
| ENSDARG000 | 9;9;9;9;9;9;9; | 45605410:456 | 45605662:456 | +;+;+;+;+;+;+;+; | 2293 | traf3ip1 | 86 | 74 | 97 | 53 | 114 | 92 |
| ENSDARG000 | 9;9 | 3153726:3154 | 3154030:3154 | -;- | 355 | NA | 0 | 0 | 0 | 0 | 0 | 0 |
| ENSDARG000 | 9;9;9;9;9;9;9; | 3153466:3153 | 3154305:3154 | +;+;+;+;+;+;+;+; | 2404 | ftr52p | 169 | 53 | 117 | 55 | 96 | 200 |
| ENSDARG000 | 9;9;9;9;9 | 45643838:456 | 45644113:456 | -;-;-;-;- | 954 | NA | 2 | 6 | 12 | 0 | 8 | 6 |
| ENSDARG000 | 9;9;9;9;9;9;9; | 9322424:9323 | 9322830:9323 | -;-;-;-;-;-;-;-;- | 1751 | zgc:113337 | 0 | 0 | 2 | 0 | 10 | 0 |
| ENSDARG000 | 9;9;9;9;9;9;9; | 23895711:238 | 23895856:238 | +;+;+;+;+;+;+;+; | 1684 | cops8 | 1075 | 869 | 1083 | 837 | 836 | 905 |
| ENSDARG000 | 9;9;9;9 | 26694869:266 | 26694917:266 | +;+;+;+ | 838 | NA | 0 | 0 | 0 | 0 | 0 | 0 |
| ENSDARG000 | 9;9;9;9;9;9;9; | 21793565:217 | 21793884:217 | +;+;+;+;+;+;+;+; | 5101 | rev1 | 266 | 219 | 333 | 120 | 495 | 357 |
| ENSDARG000 | 9 | 23900703 | 23902215 | + | 1513 | trim63b | 26 | 14 | 12 | 42 | 86 | 40 |
| ENSDARG000 | 9;9;9;9;9;9;9; | 331425:33262 | 331876:33286 | +;+;+;+;+;+;+;+; | 1994 | actr5 | 122 | 162 | 214 | 110 | 180 | 80 |
| ENSDARG000 | 9;9;9;9;9;9;9; | 343062:34307 | 343219:34321 | +;+;+;+;+;+;+;+; | 1918 | NA | 56 | 42 | 500 | 53 | 172 | 322 |
| ENSDARG000 | 9;9;9;9;9;9 | 27329640:273 | 27330482:273 | +;+;+;+;+;+ | 1882 | gtpbp8 | 116 | 96 | 194 | 137 | 86 | 100 |
| ENSDARG000 | 9;9;9;9;9;9;9; | 51521248:515 | 51521678:515 | -;-;-;-;-;-;-;-; | 1567 | tank | 130 | 300 | 293 | 308 | 388 | 257 |
| ENSDARG000 | 9;9;9;9 | 32200650:322 | 32200741:322 | -;-;-;- | 619 | NA | 0 | 0 | 0 | 0 | 0 | 1 |
| ENSDARG000 | 9;9 | 32217548:322 | 32217685:322 | -;- | 239 | NA | 0 | 0 | 0 | 0 | 0 | 0 |
| ENSDARG000 | 9;9;9;9 | 9371502:9375 | 9371771:9375 | +;+;+;+ | 1909 | NA | 2 | 2 | 0 | 14 | 6 | 30 |
| ENSDARG000 | 9;9;9;9;9;9;9; | 34127005:341 | 34127199:341 | +;+;+;+;+;+;+;+; | 6439 | f5 | 212 | 88 | 44 | 146 | 158 | 142 |
| ENSDARG000 | 9;9;9;9;9;9;9 | 9411009:9414 | 9411557:9414 | -;-;-;-;-;-;- | 1654 | NA | 199 | 715 | 743 | 749 | 1586 | 793 |
| ENSDARG000 | 9;9 | 32231102:322 | 32231239:322 | -;- | 239 | NA | 0 | 0 | 0 | 0 | 0 | 0 |
| ENSDARG000 | 9;9;9 | 32244525:322 | 32244662:322 | -;-;- | 545 | NA | 0 | 0 | 0 | 0 | 0 | 0 |
| ENSDARG000 | 9;9;9;9;9;9;9; | 17348745:173 | 17349300:173 | +;+;+;+;+;+;+;+; | 2379 | slain1a | 0 | 0 | 0 | 0 | 2 | 0 |
| ENSDARG000 | 9;9;9;9;9;9;9; | 38883388:388 | 38883411:388 | +;+;+;+;+;+;+;+; | 5543 | NA | 15 | 14 | 22 | 0 | 10 | 4 |
| ENSDARG000 | 9;9;9;9 | 32259814:322 | 32259905:322 | -;-;-;- | 638 | NA | 0 | 0 | 0 | 1 | 0 | 14 |
| ENSDARG000 | 9;9 | 42157578:421 | 42157620:421 | +;+ | 1198 | lrrc3 | 0 | 0 | 0 | 0 | 0 | 0 |
| ENSDARG000 | 9;9;9;9 | 32274308:322 | 32274399:322 | -;-;-;- | 638 | NA | 0 | 0 | 4 | 3 | 2 | 8 |
| ENSDARG000 | 9;9;9;9;9 | 45680802:456 | 45680956:456 | +;+;+;+;+ | 1314 | asb1 | 87 | 30 | 54 | 34 | 58 | 24 |
| ENSDARG000 | 9;9;9;9;9;9;9; | 29529401:295 | 29530925:295 | -;-;-;-;-;-;-;-; | 3726 | NA | 10 | 3 | 16 | 0 | 8 | 3 |
| ENSDARG000 | 9;9;9;9;9;9 | 27339505:273 | 27340053:273 | +;+;+;+;+;+ | 1828 | slc10a2 | 8 | 10 | 25 | 6 | 63 | 5 |
| ENSDARG000 | 9;9;9;9;9;9;9; | 29520696:295 | 29520989:295 | +;+;+;+;+;+;+;+; | 1427 | NA | 320 | 308 | 428 | 332 | 348 | 288 |
| ENSDARG000 | 9;9 | 45789887:457 | 45790335:457 | +;+ | 488 | NA | 0 | 2 | 0 | 0 | 0 | 0 |
| ENSDARG000 | 9;9 | 45839260:458 | 45839972:458 | +;+ | 1227 | twist2 | 82 | 12 | 12 | 0 | 8 | 4 |
| ENSDARG000 | 9;9;9;9;9;9;9; | 42269059:422 | 42269372:422 | +;+;+;+;+;+;+;+; | 1039 | NA | 215 | 227 | 251 | 161 | 216 | 169 |
| ENSDARG000 | 9;9;9;9;9;9;9; | 3170101:3170 | 3170162:3170 | +;+;+;+;+;+;+;+; | 5486 | dync1i2a | 8396 | 7038 | 7289 | 7900 | 6944 | 8150 |
| ENSDARG000 | 9;9;9;9;9;9 | 36118960:361 | 36119048:361 | -;-;-;-;-;- | 387 | NA | 2 | 0 | 3 | 3 | 0 | 0 |
| ENSDARG000 | 9;9;9;9;9;9;9; | 7456076:7456 | 7456517:7456 | +;+;+;+;+;+;+;+; | 1952 | TMEM198 | 135 | 123 | 259 | 50 | 343 | 169 |
| ENSDARG000 | 9;9;9;9;9;9;9; | 19750351:197 | 19750559:197 | -;-;-;-;-;-;-;- | 659 | NA | 0 | 4 | 0 | 0 | 5 | 22 |
| ENSDARG000 | 9 | 27348809 | 27349361 | + | 553 | NA | 0 | 0 | 0 | 0 | 0 | 0 |
| ENSDARG000 | 9;9;9 | 27352298:273 | 27355281:273 | +;+;+ | 3036 | tlr20.2 | 0 | 2 | 0 | 0 | 0 | 8 |
| ENSDARG000 | 9;9;9 | 19715582:197 | 19715985:197 | -;-;- | 714 | NA | 0 | 28 | 0 | 0 | 24 | 93 |
| ENSDARG000 | 9 | 27371240 | 27371879 | + | 640 | NA | 0 | 0 | 0 | 0 | 6 | 0 |
| ENSDARG000 | 9;9;9;9;9;9;9; | 19681569:196 | 19681603:196 | -;-;-;-;-;-;-;-; | 1431 | NA | 0 | 60 | 0 | 0 | 130 | 121 |
| ENSDARG000 | 9;9;9;9;9;9;9 | 30364667:303 | 30365280:303 | -;-;-;-;-;-;- | 1440 | asb11 | 4 | 2 | 23 | 0 | 18 | 0 |
| ENSDARG000 | 9;9;9;9;9;9;9; | 30323462:303 | 30323607:303 | -;-;-;-;-;-;-;-; | 2453 | SYTL5 | 0 | 26 | 93 | 4 | 4 | 0 |
| ENSDARG000 | 9;9 | 27379014:273 | 27379721:273 | +;+ | 726 | NA | 0 | 0 | 0 | 0 | 0 | 0 |
| ENSDARG000 | 9;9;9;9;9;9;9; | 45884031:458 | 45884877:458 | -;-;-;-;-;-;-;-; | 4848 | hdac4 | 164 | 113 | 200 | 159 | 181 | 128 |
| ENSDARG000 | 9;9;9;9;9;9;9; | 42360679:423 | 42360955:423 | -;-;-;-;-;-;-;-; | 1684 | calcrla | 16 | 2 | 22 | 10 | 0 | 32 |
| ENSDARG000 | 9;9;9;9;9;9;9; | 32301017:323 | 32301069:323 | +;+;+;+;+;+;+;+; | 1811 | hspe1 | 794 | 1431 | 1533 | 860 | 888 | 938 |
| ENSDARG000 | 9;9;9;9;9;9;9; | 21819082:218 | 21819208:218 | +;+;+;+;+;+;+;+; | 1814 | txndc9 | 1378 | 1333 | 1922 | 1366 | 1424 | 1631 |
| ENSDARG000 | 9;9;9 | 354868:35574 | 355578:35588 | -;-;- | 1219 | zgc:158320 | 90 | 66 | 58 | 76 | 42 | 38 |
| ENSDARG000 | 9;9;9;9;9;9;9; | 17415241:174 | 17415658:174 | -;-;-;-;-;-;-;-;- | 2155 | ftr53 | 4 | 40 | 22 | 20 | 12 | 0 |
| ENSDARG000 | 9;9;9;9;9;9;9; | 32286605:322 | 32287714:322 | -;-;-;-;-;-;-;-; | 2873 | hspd1 | 3403 | 5688 | 7406 | 1897 | 5113 | 3404 |
| ENSDARG000 | 9;9;9;9;9 | 21824763:218 | 21824961:218 | -;-;-;-;- | 814 | mrpl30 | 607 | 647 | 992 | 625 | 559 | 408 |
| ENSDARG000 | 9;9;9;9;9;9;9; | 27381318:273 | 27381961:273 | -;-;-;-;-;-;-;-; | 2215 | nepro | 69 | 100 | 175 | 54 | 100 | 92 |
| ENSDARG000 | 9;9;9;9;9;9;9; | 7515864:7515 | 7516222:7516 | -;-;-;-;-;-;-;-; | 1854 | desma | 2 | 6 | 14 | 0 | 6 | 4 |
| ENSDARG000 | 9;9;9;9;9;9;9; | 360252:36072 | 360504:36092 | -;-;-;-;-;-;-;-; | 4193 | NA | 76 | 53 | 115 | 136 | 201 | 85 |
| ENSDARG000 | 9;9;9;9;9;9;9; | 27398323:273 | 27398491:273 | -;-;-;-;-;-;-;-; | 1886 | poglut2 | 447 | 709 | 530 | 356 | 1118 | 660 |
| ENSDARG000 | 9;9;9;9;9;9;9; | 27393468:273 | 27393703:273 | -;-;-;-;-;-;-;-; | 875 | NA | 30 | 148 | 114 | 18 | 91 | 94 |
| ENSDARG000 | 9;9;9;9;9;9;9; | 9441453:9441 | 9441671:9441 | +;+;+;+;+;+;+;+; | 5854 | CFAP91 | 0 | 0 | 0 | 0 | 0 | 2 |
| ENSDARG000 | 9;9 | 17423941:174 | 17425867:174 | +;+ | 1941 | NA | 57 | 35 | 80 | 30 | 0 | 22 |
| ENSDARG000 | 9;9 | 52608045:526 | 52608661:526 | -;- | 939 | TMEM169 | 60 | 48 | 44 | 34 | 43 | 59 |
| ENSDARG000 | 9;9 | 17429170:174 | 17429450:174 | +;+ | 3200 | zgc:101559 | 62 | 62 | 43 | 30 | 98 | 100 |
| ENSDARG000 | 9;9;9;9;9;9 | 38994232:389 | 38994745:389 | -;-;-;-;-;- | 1106 | myl1 | 2 | 2 | 0 | 4 | 10 | 0 |
| ENSDARG000 | 9;9;9;9;9;9;9 | 17438765:174 | 17438912:174 | +;+;+;+;+;+;+ | 1661 | rgcc | 149 | 44 | 68 | 52 | 50 | 54 |
| ENSDARG000 | 9;9;9;9;9;9;9; | 52613820:526 | 52614061:526 | +;+;+;+;+;+;+;+; | 1854 | NA | 0 | 0 | 0 | 0 | 0 | 0 |
| ENSDARG000 | 9;9;9;9;9;9;9 | 21826724:218 | 21826959:218 | +;+;+;+;+;+;+ | 1108 | zgc:162944 | 2666 | 2702 | 2516 | 2100 | 1892 | 2986 |
| ENSDARG000 | 9;9;9;9;9;9;9; | 39010432:390 | 39010491:390 | +;+;+;+;+;+;+;+; | 859 | NA | 2 | 0 | 0 | 0 | 3 | 0 |
| ENSDARG000 | 9;9;9;9;9;9;9 | 19821457:198 | 19821518:198 | +;+;+;+;+;+;+ | 428 | NA | 0 | 0 | 0 | 0 | 0 | 2 |
| ENSDARG000 | 9;9;9;9;9;9;9; | 9502610:9502 | 9502895:9502 | +;+;+;+;+;+;+;+; | 2502 | nr1i2 | 129 | 126 | 166 | 107 | 103 | 204 |
| ENSDARG000 | 9;9;9 | 19837383:198 | 19837583:198 | +;+;+ | 750 | NA | 0 | 0 | 0 | 1 | 0 | 0 |
| ENSDARG000 | 9;9;9;9 | 19861310:198 | 19861856:198 | -;-;-;- | 777 | NA | 0 | 11 | 0 | 0 | 0 | 2 |
| ENSDARG000 | 9;9;9;9;9 | 21833628:218 | 21834597:218 | -;-;-;-;- | 1511 | acod1 | 526 | 697 | 375 | 361 | 304 | 259 |
| ENSDARG000 | 9;9;9;9 | 19945634:199 | 19945842:199 | -;-;-;- | 559 | NA | 2 | 6 | 0 | 0 | 0 | 0 |
| ENSDARG000 | 9;9;9;9;9;9;9; | 20076556:200 | 20077070:200 | -;-;-;-;-;-;-;-;- | 3524 | igsf3 | 340 | 170 | 576 | 162 | 357 | 63 |
| ENSDARG000 | 9;9;9;9;9;9;9; | 30374181:303 | 30375844:303 | -;-;-;-;-;-;-;-; | 3353 | piga | 283 | 417 | 413 | 259 | 489 | 301 |
| ENSDARG000 | 9;9;9;9;9;9;9; | 9577743:9583 | 9580267:9583 | -;-;-;-;-;-;-;-; | 4420 | gsk3ba | 3340 | 2149 | 2147 | 2119 | 2451 | 2290 |
| ENSDARG000 | 9;9;9;9;9;9;9; | 7548533:7559 | 7548844:7559 | +;+;+;+;+;+;+;+; | 3711 | ptprna | 4 | 2 | 4 | 6 | 2 | 2 |
| ENSDARG000 | 9;9;9;9;9;9;9 | 23983866:239 | 23983918:239 | -;-;-;-;-;-;- | 2697 | NA | 24 | 19 | 33 | 0 | 49 | 33 |
| ENSDARG000 | 9;9;9;9;9;9;9; | 23914317:239 | 23915105:239 | -;-;-;-;-;-;-;-; | 14531 | COL6A3 | 1037 | 631 | 2775 | 234 | 2427 | 1615 |
| ENSDARG000 | 9;9;9;9;9;9;9; | 46369932:463 | 46370155:463 | -;-;-;-;-;-;-;-; | 7234 | LCT | 0 | 0 | 0 | 0 | 0 | 0 |
| ENSDARG000 | 9;9;9;9;9;9;9; | 426392:42639 | 426484:42648 | +;+;+;+;+;+;+;+; | 2494 | bzw1b | 1964 | 2881 | 2171 | 1188 | 3564 | 1355 |
| ENSDARG000 | 9;9;9;9;9;9;9; | 46387419:463 | 46388962:463 | -;-;-;-;-;-;-;-; | 7897 | LCT | 0 | 0 | 0 | 0 | 0 | 0 |
| ENSDARG000 | 9 | 21843915 | 21845783 | + | 1869 | kctd12.1 | 38 | 4 | 62 | 22 | 42 | 12 |
| ENSDARG000 | 9;9;9;9;9;9;9; | 42420636:424 | 42422016:424 | -;-;-;-;-;-;-;-; | 2441 | tfpia | 778 | 74 | 644 | 320 | 52 | 344 |
| ENSDARG000 | 9;9;9;9;9;9;9; | 17540619:175 | 17540859:175 | -;-;-;-;-;-;-;-; | 5977 | vwa8 | 403 | 289 | 489 | 210 | 521 | 401 |
| ENSDARG000 | 9;9;9;9;9;9;9; | 39026311:390 | 39026420:390 | +;+;+;+;+;+;+;+; | 5015 | CPS1 | 0 | 0 | 0 | 0 | 0 | 0 |
| ENSDARG000 | 9;9;9;9;9;9;9; | 9675926:9679 | 9680787:9680 | -;-;-;-;-;-;-;-; | 7019 | GPR156 | 24 | 0 | 10 | 0 | 14 | 4 |
| ENSDARG000 | 9;9;9;9;9;9;9; | 52628899:526 | 52629161:526 | -;-;-;-;-;-;-;-; | 3462 | PFKL | 236 | 166 | 222 | 274 | 528 | 276 |
| ENSDARG000 | 9;9 | 20386745:203 | 20386773:203 | +;+ | 809 | NA | 0 | 0 | 0 | 0 | 0 | 0 |
| ENSDARG000 | 9;9;9;9;9;9;9; | 32306862:323 | 32306959:323 | +;+;+;+;+;+;+;+; | 1111 | mob4 | 1014 | 955 | 1223 | 1049 | 1093 | 847 |
| ENSDARG000 | 9;9;9;9;9;9;9; | 34148714:341 | 34148991:341 | +;+;+;+;+;+;+;+; | 2354 | gpr161b | 1532 | 712 | 830 | 214 | 476 | 917 |
| ENSDARG000 | 9;9;9;9;9;9;9; | 7621622:7627 | 7621974:7627 | -;-;-;-;-;-;-;-; | 2646 | dnajb2 | 222 | 168 | 159 | 129 | 164 | 107 |
| ENSDARG000 | 9;9;9;9;9;9;9; | 30387916:303 | 30388002:303 | +;+;+;+;+;+;+;+; | 1738 | si:dkey-18p12 | 31 | 20 | 15 | 16 | 158 | 77 |
| ENSDARG000 | 9;9;9;9;9;9;9; | 36314867:363 | 36315026:363 | +;+;+;+;+;+;+;+; | 13847 | LRP1B | 2909 | 663 | 425 | 426 | 499 | 114 |
| ENSDARG000 | 9;9;9;9;9;9;9; | 32312920:323 | 32314240:323 | -;-;-;-;-;-;-;-; | 3025 | rftn2 | 750 | 807 | 1306 | 666 | 1645 | 822 |
| ENSDARG000 | 9;9;9;9;9;9;9; | 27415754:274 | 27416223:274 | -;-;-;-;-;-;-;-; | 4941 | stxbp5l | 14 | 2 | 25 | 13 | 6 | 8 |
| ENSDARG000 | 9;9;9;9 | 20433812:204 | 20434225:204 | +;+;+;+ | 1107 | NA | 28 | 0 | 0 | 0 | 2 | 12 |
| ENSDARG000 | 9;9;9 | 27411502:274 | 27411638:274 | +;+;+ | 429 | NA | 0 | 3 | 6 | 7 | 2 | 4 |
| ENSDARG000 | 9;9;9;9;9;9;9; | 42546504:426 | 42546828:426 | +;+;+;+;+;+;+;+; | 3356 | gulp1a | 3149 | 158 | 712 | 176 | 10 | 202 |
| ENSDARG000 | 9;9;9;9;9;9;9; | 52694971:526 | 52695235:526 | -;-;-;-;-;-;-;-; | 3409 | MAP3K13 | 18 | 6 | 49 | 20 | 24 | 28 |
| ENSDARG000 | 9;9;9;9 | 9719225:9720 | 9719848:9720 | -;-;-;- | 1690 | LRRC58 | 611 | 180 | 715 | 400 | 781 | 1087 |
| ENSDARG000 | 9;9;9;9;9;9;9; | 32358514:324 | 32358710:324 | +;+;+;+;+;+;+;+; | 3604 | PLCL1 | 76 | 45 | 44 | 28 | 47 | 72 |
| ENSDARG000 | 9;9;9;9;9;9;9; | 3429662:3429 | 3429925:3429 | +;+;+;+;+;+;+;+; | 4173 | ITGA6 | 799 | 1228 | 2113 | 3288 | 489 | 326 |
| ENSDARG000 | 9;9 | 32355235:323 | 32355751:323 | -;- | 661 | NA | 0 | 0 | 0 | 0 | 0 | 0 |
| ENSDARG000 | 9;9;9;9;9;9;9; | 29569375:295 | 29572098:295 | -;-;-;-;-;-;-;-; | 6076 | NA | 82 | 120 | 196 | 106 | 163 | 46 |
| ENSDARG000 | 9;9;9;9;9;9;9 | 3429662:3448 | 3429925:3448 | +;+;+;+;+;+;+ | 1274 | NA | 0 | 0 | 0 | 0 | 0 | 0 |
| ENSDARG000 | 9;9;9;9;9;9;9; | 29548195:295 | 29548727:295 | +;+;+;+;+;+;+;+; | 5340 | NA | 2 | 2 | 4 | 8 | 0 | 0 |
| ENSDARG000 | 9;9;9;9;9;9;9; | 34166064:341 | 34166392:341 | -;-;-;-;-;-;-;-; | 3599 | dcaf6 | 787 | 437 | 547 | 408 | 441 | 410 |
| ENSDARG000 | 9;9;9;9;9;9;9; | 9735493:9742 | 9739205:9742 | -;-;-;-;-;-;-;-; | 5642 | fstl1b | 5293 | 3091 | 10278 | 4280 | 13052 | 5415 |
| ENSDARG000 | 9;9;9;9;9;9;9; | 52838336:528 | 52838465:528 | -;-;-;-;-;-;-;-; | 4445 | trpm2 | 16 | 8 | 8 | 32 | 50 | 23 |
| ENSDARG000 | 9 | 53276356 | 53277722 | + | 1367 | sox21b | 0 | 0 | 0 | 8 | 0 | 0 |
| ENSDARG000 | 9;9;9;9;9;9;9; | 32560399:325 | 32561019:325 | -;-;-;-;-;-;-;-; | 2592 | satb2 | 6 | 14 | 19 | 2 | 20 | 16 |
| ENSDARG000 | 9;9;9;9;9;9;9; | 42653188:426 | 42653832:426 | -;-;-;-;-;-;-;-; | 5063 | col5a2a | 946 | 156 | 209 | 161 | 1169 | 221 |
| ENSDARG000 | 9;9;9;9;9;9;9; | 30421489:304 | 30421635:304 | +;+;+;+;+;+;+;+; | 1851 | zgc:113314 | 337 | 461 | 556 | 493 | 423 | 601 |
| ENSDARG000 | 9;9;9 | 46405587:464 | 46406627:464 | -;-;- | 2391 | gjc4b | 312 | 205 | 726 | 418 | 475 | 242 |
| ENSDARG000 | 9;9 | 39230526:392 | 39231108:392 | -;- | 733 | NA | 0 | 0 | 0 | 0 | 0 | 0 |
| ENSDARG000 | 9;9;9;9;9;9;9; | 53327332:533 | 53328039:533 | +;+;+;+;+;+;+;+; | 2180 | dct | 17679 | 13238 | 5315 | 24791 | 5309 | 12037 |
| ENSDARG000 | 9;9;9;9 | 32670526:326 | 32670675:326 | -;-;-;- | 756 | gzm3.4 | 10 | 3 | 12 | 4 | 6 | 20 |
| ENSDARG000 | 9;9;9;9 | 46421027:464 | 46421255:464 | +;+;+;+ | 623 | NA | 0 | 0 | 0 | 0 | 0 | 0 |
| ENSDARG000 | 9;9;9;9;9;9;9; | 32635546:326 | 32635588:326 | +;+;+;+;+;+;+;+; | 2061 | rabl3 | 187 | 152 | 291 | 180 | 257 | 122 |
| ENSDARG000 | 9;9;9;9;9 | 32667169:326 | 32667318:326 | -;-;-;-;- | 814 | gzm3.3 | 8 | 6 | 10 | 0 | 10 | 9 |
| ENSDARG000 | 9;9;9;9;9 | 32654554:326 | 32654700:326 | -;-;-;-;- | 802 | gzm3 | 5 | 8 | 0 | 0 | 4 | 25 |
| ENSDARG000 | 9;9;9;9 | 7662638:7668 | 7663763:7668 | -;-;-;- | 1556 | tuba8l3 | 3984 | 9268 | 5799 | 6550 | 6070 | 4040 |
| ENSDARG000 | 9;9;9;9 | 32662314:326 | 32662376:326 | -;-;-;- | 334 | NA | 0 | 0 | 0 | 0 | 0 | 0 |
| ENSDARG000 | 9;9;9;9;9;9;9; | 32657695:326 | 32657844:326 | -;-;-;-;-;-;-;-;- | 750 | NA | 0 | 0 | 0 | 0 | 0 | 0 |
| ENSDARG000 | 9;9;9;9;9;9;9; | 32641026:326 | 32642174:326 | -;-;-;-;-;-;-;-; | 2295 | dnajb11 | 1029 | 1638 | 2073 | 417 | 1597 | 1005 |
| ENSDARG000 | 9;9 | 30430188:304 | 30432330:304 | -;- | 2327 | gja8a | 15 | 35 | 22 | 33 | 33 | 60 |
| ENSDARG000 | 9;9;9 | 46692166:467 | 46694410:467 | -;-;- | 2565 | NA | 178 | 51 | 24 | 14 | 6 | 126 |
| ENSDARG000 | 9;9;9;9;9;9 | 9858935:9861 | 9859033:9862 | +;+;+;+;+;+ | 759 | NA | 0 | 0 | 0 | 0 | 0 | 3 |
| ENSDARG000 | 9;9;9 | 30464641:304 | 30464900:304 | +;+;+ | 2009 | gja5a | 4 | 0 | 10 | 0 | 0 | 0 |
| ENSDARG000 | 9;9;9;9;9;9;9; | 17787864:178 | 17788035:178 | +;+;+;+;+;+;+;+; | 3835 | DGKH | 330 | 162 | 350 | 133 | 472 | 308 |
| ENSDARG000 | 9;9 | 42710633:427 | 42710736:427 | -;- | 470 | NA | 0 | 0 | 0 | 0 | 0 | 0 |
| ENSDARG000 | 9;9;9;9;9;9 | 42728269:427 | 42728635:427 | -;-;-;-;-;- | 1189 | fkbp7 | 35 | 2 | 8 | 0 | 8 | 4 |
| ENSDARG000 | 9;9;9 | 440196:44197 | 440444:44226 | -;-;- | 823 | NA | 2 | 0 | 0 | 0 | 0 | 0 |
| ENSDARG000 | 9 | 53535419 | 53537989 | - | 2571 | NA | 14 | 54 | 77 | 2 | 0 | 65 |
| ENSDARG000 | 9;9;9;9 | 53707240:537 | 53709642:537 | +;+;+;+ | 2913 | PCDH8 | 0 | 0 | 2 | 0 | 0 | 0 |
| ENSDARG000 | 9;9;9;9 | 46745348:467 | 46745693:467 | +;+;+;+ | 1208 | igfbp2b | 4 | 0 | 8 | 0 | 0 | 0 |
| ENSDARG000 | 9;9;9;9;9;9;9; | 53725128:537 | 53725330:537 | +;+;+;+;+;+;+;+; | 2208 | cnmd | 8 | 0 | 0 | 0 | 0 | 0 |
| ENSDARG000 | 9;9;9;9;9;9;9; | 30478768:304 | 30478821:304 | +;+;+;+;+;+;+;+; | 1770 | acp6 | 114 | 227 | 171 | 60 | 320 | 95 |

| | | | | | | | | | | | | |
|---|---|---|---|---|---|---|---|---|---|---|---|---|
| ENSDARG000 | 9:9:9:9:9:9:9: | 7678136:767 | 7679111:767 | -;-;-;-;-;-;-;- | 1523 | tuba8l5 | 433 | 497 | 569 | 553 | 324 | 333 |
| ENSDARG000 | 9;9;9;9 | 46820636:46 | 46825661:46 | -;-;-;- | 6049 | igfbp5b | 280 | 472 | 62 | 96 | 222 | 128 |
| ENSDARG000 | 9:9:9:9:9:9:9: | 39313944:39 | 39315439:39 | -:-:-:-:-:-:-:-: | 4902 | ERBB4 | 0 | 0 | 2 | 2 | 0 | 0 |
| ENSDARG000 | 9:9:9:9:9:9:9: | 445471:4484 | 447281:4488 | -;-;-;-;-;-;-;- | 4633 | igsf3l | 72 | 0 | 2 | 10 | 6 | 0 |
| ENSDARG000 | 9:9:9:9:9:9:9: | 47436010:47 | 47436427:47 | -:-:-:-:-:-:-:-: | 6043 | TNS1 | 3309 | 1223 | 2402 | 2050 | 2073 | 2479 |
| ENSDARG000 | 9:9:9:9:9:9:9: | 20483846:20 | 20484380:20 | +:+:+:+:+:+:+: | 7761 | NA | 1633 | 1251 | 1419 | 1200 | 1175 | 1322 |
| ENSDARG000 | 9:9:9:9:9:9:9: | 3490824:349 | 3491128:349 | -;-;-;-;-;-;- | 940 | cybrd1 | 77 | 77 | 68 | 138 | 99 | 191 |
| ENSDARG000 | 9:9:9:9:9:9:9: | 53799614:53 | 53800036:53 | -:-:-:-:-:-:-:-: | 1515 | EDNRB | 4 | 6 | 0 | 8 | 23 | 0 |
| ENSDARG000 | 9;9;9 | 7724152:772 | 7724598:772 | +;+;+ | 999 | mnx2a | 0 | 0 | 0 | 0 | 0 | 0 |
| ENSDARG000 | 9:9:9:9:9:9:9: | 30490504:30 | 30491386:30 | -:-:-:-:-:-:-:-: | 3832 | NA | 701 | 565 | 649 | 526 | 698 | 600 |
| ENSDARG000 | 9:9:9:9:9:9:9: | 34221997:34 | 34222359:34 | +:+:+:+:+:+:+: | 1361 | mpc2b | 922 | 781 | 998 | 952 | 974 | 932 |
| ENSDARG000 | 9:9:9:9:9:9:9: | 9927516:992 | 9927739:992 | +:+:+:+:+:+:+: | 4428 | NA | 262 | 634 | 721 | 466 | 402 | 293 |
| ENSDARG000 | 9:9:9:9:9:9:9: | 21893757:21 | 21894451:21 | -:-:-:-:-:-:-:-: | 7151 | lmo7a | 912 | 108 | 44 | 268 | 118 | 312 |
| ENSDARG000 | 9:9:9:9:9:9:9: | 53842705:53 | 53843971:53 | -:-:-:-:-:-:-:-: | 3712 | cog3 | 537 | 403 | 618 | 476 | 572 | 650 |
| ENSDARG000 | 9:9:9:9:9:9:9: | 458751:4587 | 459217:4592 | -:-:-:-:-:-:-:-: | 2347 | NA | 56 | 47 | 52 | 64 | 63 | 75 |
| ENSDARG000 | 9;9;9;9;9;9;9 | 47961482:47 | 47961831:47 | +;+;+;+;+;+;+ | 2270 | NA | 8 | 0 | 16 | 0 | 4 | 6 |
| ENSDARG000 | 9:9:9:9:9:9:9: | 3498377:349 | 3501343:349 | -:-:-:-:-:-:-:-: | 5111 | dcaf17 | 102 | 64 | 67 | 81 | 71 | 66 |
| ENSDARG000 | 9:9:9:9:9:9:9: | 3519191:351 | 3519342:352 | +:+:+:+:+:+:+: | 1341 | mettl8 | 22 | 15 | 32 | 18 | 31 | 29 |
| ENSDARG000 | 9;9;9;9;9 | 27733334:27 | 27733946:27 | -;-;-;-;- | 1105 | crygs2 | 0 | 0 | 0 | 0 | 0 | 0 |
| ENSDARG000 | 9:9:9:9:9:9:9: | 27720267:27 | 27720511:27 | +:+:+:+:+:+:+: | 2798 | lcmt2 | 53 | 26 | 93 | 30 | 88 | 68 |
| ENSDARG000 | 9:9:9:9:9:9:9: | 34230067:34 | 34230131:34 | +:+:+:+:+:+:+: | 2583 | nxpe3 | 4055 | 4680 | 3051 | 3380 | 5202 | 5776 |
| ENSDARG000 | 9:9:9:9:9:9:9: | 27688274:27 | 27692883:27 | -:-:-:-:-:-:-:-: | 6403 | gtf2e1 | 570 | 484 | 631 | 391 | 542 | 460 |
| ENSDARG000 | 9:9:9:9:9:9:9: | 21971260:21 | 21971693:21 | -:-:-:-:-:-:-:-: | 1314 | uchl3 | 1279 | 1115 | 979 | 711 | 975 | 992 |
| ENSDARG000 | 9:9:9:9:9:9:9: | 27740968:27 | 27741729:27 | -:-:-:-:-:-:-:-: | 2656 | tbccd1 | 85 | 96 | 203 | 143 | 160 | 112 |
| ENSDARG000 | 9:9:9:9:9:9:9: | 7732714:773 | 7732737:773 | +:+:+:+:+:+:+: | 4816 | NA | 0 | 8 | 4 | 0 | 0 | 0 |
| ENSDARG000 | 9:9:9:9:9:9:9: | 53909566:53 | 53909914:53 | -:-:-:-:-:-:-:-: | 2363 | MID1 | 9 | 6 | 6 | 4 | 24 | 8 |
| ENSDARG000 | 9:9:9:9:9:9:9: | 17971935:17 | 17972044:17 | +:+:+:+:+:+:+: | 14219 | AKAP11 | 1631 | 1135 | 1626 | 2333 | 1406 | 1356 |
| ENSDARG000 | 9:9:9:9:9:9:9: | 48007081:48 | 48007504:48 | +:+:+:+:+:+:+: | 1712 | zgc:92380 | 42 | 12 | 8 | 6 | 28 | 28 |
| ENSDARG000 | 9:9:9:9:9:9:9: | 39960118:39 | 39961126:39 | -:-:-:-:-:-:-:-: | 2647 | IKZF2 | 14 | 8 | 17 | 4 | 34 | 10 |
| ENSDARG000 | 9:9:9:9:9:9:9: | 30543139:30 | 30543853:30 | -:-:-:-:-:-:-:-: | 2548 | chaf1b | 154 | 571 | 619 | 294 | 553 | 332 |
| ENSDARG000 | 9:9:9:9:9:9:9: | 9953063:995 | 9953407:995 | -;-;-;-;-;-;-;- | 1596 | prmt2 | 34 | 44 | 73 | 84 | 88 | 24 |
| ENSDARG000 | 9;9;9;9 | 18023288:18 | 18023618:18 | +;+;+;+ | 1548 | NA | 9 | 25 | 10 | 22 | 17 | 38 |
| ENSDARG000 | 9:9:9:9:9:9:9: | 473334:4734 | 473580:4735 | +:+:+:+:+:+:+: | 4734 | NA | 8 | 0 | 12 | 2 | 6 | 4 |
| ENSDARG000 | 9;9;9;9;9 | 40081524:40 | 40082172:40 | -;-;-;-;- | 1119 | NA | 0 | 4 | 0 | 0 | 8 | 1 |
| ENSDARG000 | 9:9:9:9:9:9:9: | 18215940:18 | 18216074:18 | +:+:+:+:+:+:+: | 1675 | dnajc15 | 241 | 227 | 307 | 226 | 149 | 124 |
| ENSDARG000 | 9:9:9:9:9:9:9: | 34246384:34 | 34247370:34 | -:-:-:-:-:-:-:-: | 2829 | me3 | 3604 | 2710 | 3128 | 2448 | 3091 | 2369 |
| ENSDARG000 | 9;9 | 18163629:18 | 18163748:18 | +;+ | 301 | NA | 0 | 0 | 0 | 0 | 0 | 0 |
| ENSDARG000 | 9:9:9:9:9:9:9: | 42732992:42 | 42734315:42 | -:-:-:-:-:-:-:-: | 88557 | TTN | 6 | 0 | 4 | 6 | 24 | 0 |
| ENSDARG000 | 9:9:9:9:9:9:9: | 18162115:18 | 18162245:18 | -:-:-:-:-:-:-:-: | 1101 | EPSTI1 | 158 | 114 | 67 | 61 | 73 | 89 |
| ENSDARG000 | 9;9 | 844221:8485 | 844400:8490 | -;- | 747 | C1QL2 | 0 | 2 | 2 | 10 | 0 | 2 |
| ENSDARG000 | 9;9;9;9;9 | 881589:8831 | 882332:8831 | -;-;-;-;- | 1288 | NA | 12 | 23 | 40 | 51 | 36 | 24 |
| ENSDARG000 | 9:9:9:9:9:9:9: | 20519846:20 | 20520186:20 | +:+:+:+:+:+:+: | 2081 | NA | 0 | 8 | 0 | 0 | 4 | 0 |
| ENSDARG000 | 9:9:9:9:9:9:9: | 48043162:48 | 48043515:48 | -:-:-:-:-:-:-:-: | 1504 | arpc2 | 4145 | 4089 | 4329 | 3384 | 3980 | 3632 |
| ENSDARG000 | 9;9;9;9;9 | 890536:8933 | 890815:8938 | +;+;+;+;+ | 2155 | STEAP3 | 10 | 52 | 48 | 14 | 4 | 2 |
| ENSDARG000 | 9;9;9;9;9 | 900392:9010 | 900760:9011 | -;-;-;-;- | 826 | C9H2orf76 | 82 | 97 | 120 | 52 | 65 | 53 |
| ENSDARG000 | 9:9:9:9:9:9:9: | 40617215:40 | 40617396:40 | -:-:-:-:-:-:-:-: | 2575 | BARD1 | 24 | 111 | 65 | 8 | 64 | 40 |
| ENSDARG000 | 9:9:9:9:9:9:9: | 20526472:20 | 20526920:20 | +:+:+:+:+:+:+: | 4616 | TRIM45 | 58 | 90 | 80 | 50 | 102 | 80 |
| ENSDARG000 | 9;9;9;9 | 40546677:40 | 40546817:40 | +;+;+;+ | 896 | vwc2l | 0 | 0 | 0 | 0 | 0 | 0 |
| ENSDARG000 | 9:9:9:9:9:9:9: | 3548529:354 | 3548727:354 | +:+:+:+:+:+:+: | 3075 | tlk1a | 828 | 837 | 1064 | 686 | 894 | 825 |
| ENSDARG000 | 9:9:9:9:9:9:9: | 34260604:34 | 34261460:34 | -;-;-;-;-;-;- | 2701 | ildr1b | 22 | 12 | 0 | 0 | 8 | 12 |
| ENSDARG000 | 9:9:9:9:9:9:9: | 907459:9075 | 907585:9075 | +:+:+:+:+:+:+: | 1912 | dbi | 3524 | 6399 | 5775 | 4626 | 4066 | 4367 |
| ENSDARG000 | 9:9:9:9:9:9:9: | 9969987:996 | 9970553:997 | -:-:-:-:-:-:-:-: | 7969 | ugt1a2 | 6 | 20 | 38 | 46 | 24 | 52 |
| ENSDARG000 | 9:9:9:9:9:9:9: | 24008879:24 | 24009085:24 | +:+:+:+:+:+:+: | 2124 | NA | 251 | 161 | 1025 | 143 | 280 | 692 |
| ENSDARG000 | 9:9:9:9:9:9:9: | 18310228:18 | 18310359:18 | -:-:-:-:-:-:-:-: | 2267 | ENOX1 | 10 | 3 | 1 | 0 | 2 | 0 |
| ENSDARG000 | 9:9:9:9:9:9:9: | 7909490:790 | 7909715:790 | +:+:+:+:+:+:+: | 5503 | MYO16 | 510 | 928 | 773 | 472 | 542 | 860 |
| ENSDARG000 | 9:9:9:9:9:9:9: | 30558508:30 | 30559144:30 | -:-:-:-:-:-:-:-: | 4001 | morc3a | 256 | 514 | 434 | 406 | 355 | 371 |
| ENSDARG000 | 9:9:9:9:9:9:9: | 32674676:32 | 32674942:32 | -;-;-;-;-;-;-;- | 1384 | ercc1 | 90 | 135 | 247 | 108 | 159 | 160 |
| ENSDARG000 | 9:9:9:9:9:9:9: | 34274501:34 | 34275026:34 | -:-:-:-:-:-:-:-: | 2471 | ildr2 | 36 | 24 | 10 | 16 | 15 | 24 |
| ENSDARG000 | 9;9 | 32728653:32 | 32729927:32 | -;- | 1477 | olig1 | 0 | 0 | 0 | 0 | 0 | 0 |
| ENSDARG000 | 9:9:9:9:9:9:9: | 20554178:20 | 20554389:20 | +:+:+:+:+:+:+: | 6975 | man1a2 | 1345 | 1154 | 1183 | 884 | 1560 | 1138 |
| ENSDARG000 | 9:9:9:9:9:9:9: | 32745088:32 | 32746121:32 | -:-:-:-:-:-:-:-: | 1306 | NA | 0 | 2 | 0 | 18 | 2 | 4 |
| ENSDARG000 | 9;9 | 36760889:36 | 36761062:36 | -;- | 300 | NA | 0 | 0 | 1 | 0 | 1 | 0 |
| ENSDARG000 | 9:9:9:9:9:9:9: | 18572970:18 | 18573513:18 | +:+:+:+:+:+:+: | 4486 | LACC1 | 104 | 78 | 34 | 38 | 45 | 65 |
| ENSDARG000 | 9;9 | 36765623:36 | 36765796:36 | -;- | 300 | NA | 0 | 0 | 1 | 0 | 0 | 0 |
| ENSDARG000 | 9;9 | 36770357:36 | 36770530:36 | -;- | 201 | NA | 0 | 0 | 0 | 0 | 0 | 0 |
| ENSDARG000 | 9;9;9 | 34307140:34 | 34307250:34 | -;-;- | 493 | NA | 0 | 0 | 0 | 0 | 2 | 0 |
| ENSDARG000 | 9;9 | 42889385:42 | 42889680:42 | -;- | 511 | NA | 0 | 0 | 0 | 0 | 0 | 0 |
| ENSDARG000 | 9:9:9:9:9:9:9: | 36794384:36 | 36794443:36 | +:+:+:+:+:+:+: | 1135 | NA | 0 | 0 | 0 | 0 | 0 | 0 |
| ENSDARG000 | 9;9;9;9 | 20780813:20 | 20780920:20 | +;+;+;+ | 1936 | tent5c | 31 | 10 | 68 | 10 | 86 | 97 |
| ENSDARG000 | 9;9;9;9;9 | 36896257:36 | 36896917:36 | -;-;-;-;- | 1351 | ralba | 373 | 412 | 341 | 182 | 323 | 351 |
| ENSDARG000 | 9:9:9:9:9:9:9: | 24025444:24 | 24026471:24 | -:-:-:-:-:-:-:-: | 1834 | rpe | 749 | 762 | 912 | 425 | 770 | 647 |
| ENSDARG000 | 9;9 | 32751428:32 | 32753017:32 | -;- | 1684 | olig2 | 0 | 10 | 0 | 0 | 4 | 4 |
| ENSDARG000 | 9;9;9;9;9 | 10014514:10 | 10015510:10 | +;+;+;+;+ | 1756 | NXPH2 | 0 | 0 | 0 | 2 | 5 | 0 |
| ENSDARG000 | 9:9:9:9:9:9:9: | 34319333:34 | 34319481:34 | +:+:+:+:+:+:+: | 5415 | pou2f1b | 347 | 218 | 305 | 244 | 428 | 238 |
| ENSDARG000 | 9:9:9:9:9:9:9: | 54629993:54 | 54630106:54 | +:+:+:+:+:+:+: | 4379 | egfl6 | 47 | 2 | 10 | 10 | 4 | 6 |
| ENSDARG000 | 9;9;9 | 24043780:24 | 24046287:24 | -;-;- | 2556 | ackr3a | 0 | 0 | 0 | 0 | 0 | 0 |
| ENSDARG000 | 9;9;9;9;9;9;9 | 54686686:54 | 54686838:54 | +;+;+;+;+;+;+ | 944 | rab9a | 915 | 1195 | 958 | 781 | 1043 | 786 |
| ENSDARG000 | 9;9;9;9;9;9;9 | 36946340:36 | 36946434:36 | +;+;+;+;+;+;+ | 662 | NA | 128 | 89 | 99 | 113 | 102 | 111 |
| ENSDARG000 | 9:9:9:9:9:9:9: | 34356895:34 | 34357354:34 | -;-;-;-;-;-;- | 899 | cd247l | 14 | 15 | 4 | 0 | 14 | 39 |
| ENSDARG000 | 9;9;9;9;9;9;9 | 34326430:34 | 34327078:34 | -;-;-;-;-;-;- | 1101 | NA | 0 | 0 | 0 | 0 | 0 | 4 |
| ENSDARG000 | 9:9:9:9:9:9:9: | 3621968:362 | 3622141:362 | -:-:-:-:-:-:-:-: | 1806 | GAD1 | 59 | 46 | 26 | 42 | 64 | 22 |
| ENSDARG000 | 9;9;9;9;9;9 | 22902865:22 | 22903339:22 | -;-;-;-;-;- | 1326 | arl5a | 352 | 358 | 413 | 196 | 423 | 356 |
| ENSDARG000 | 9:9:9:9:9:9:9: | 27750490:27 | 27750605:27 | +:+:+:+:+:+:+: | 4500 | rab5b | 822 | 564 | 806 | 668 | 510 | 423 |
| ENSDARG000 | 9;9;9;9;9;9;9 | 30577925:30 | 30578137:30 | +;+;+;+;+;+;+ | 833 | COMMD6 | 198 | 258 | 262 | 152 | 218 | 268 |
| ENSDARG000 | 9:9:9:9:9:9:9: | 27769526:27 | 27769789:27 | -:-:-:-:-:-:-:-: | 1562 | NA | 0 | 3 | 2 | 0 | 0 | 1 |
| ENSDARG000 | 9:9:9:9:9:9:9: | 20806334:20 | 20807320:20 | -:-:-:-:-:-:-:-: | 2684 | gdap2 | 2858 | 2894 | 3287 | 3157 | 3078 | 2470 |
| ENSDARG000 | 9;9 | 36967294:36 | 36968148:36 | +;+ | 3022 | inhbb | 142 | 48 | 81 | 64 | 32 | 42 |
| ENSDARG000 | 9:9:9:9:9:9:9: | 10050890:10 | 10052098:10 | -:-:-:-:-:-:-:-: | 2666 | spopla | 559 | 478 | 595 | 393 | 773 | 426 |
| ENSDARG000 | 9;9;9;9 | 27782504:27 | 27782762:27 | +;+;+;+ | 633 | NA | 0 | 0 | 0 | 0 | 0 | 1 |
| ENSDARG000 | 9:9:9:9:9:9:9: | 40685613:40 | 40686471:40 | -:-:-:-:-:-:-:-: | 6192 | ABCA12 | 30 | 11 | 0 | 24 | 20 | 14 |
| ENSDARG000 | 9:9:9:9:9:9:9: | 37141836:37 | 37142189:37 | +:+:+:+:+:+:+: | 6621 | gli2a | 1662 | 923 | 749 | 495 | 1453 | 224 |
| ENSDARG000 | 9:9:9:9:9:9:9: | 54751201:54 | 54752110:54 | -:-:-:-:-:-:-:-: | 2380 | gpm6bb | 5683 | 4162 | 7284 | 2872 | 9119 | 3104 |
| ENSDARG000 | 9:9:9:9:9:9:9: | 21977383:21 | 21977527:21 | +:+:+:+:+:+:+: | 3138 | sgo2 | 73 | 261 | 285 | 86 | 192 | 128 |
| ENSDARG000 | 9:9:9:9:9:9:9: | 37320222:37 | 37320326:37 | +:+:+:+:+:+:+: | 3271 | slc15a2 | 134 | 50 | 257 | 156 | 493 | 241 |
| ENSDARG000 | 9:9:9:9:9:9:9: | 20853894:20 | 20854003:20 | +:+:+:+:+:+:+: | 3560 | wdr3 | 1611 | 1825 | 3396 | 2170 | 2043 | 1857 |
| ENSDARG000 | 9:9:9:9:9:9:9: | 37366973:37 | 37367404:37 | +:+:+:+:+:+:+: | 3063 | slc49a4 | 538 | 446 | 359 | 415 | 503 | 466 |
| ENSDARG000 | 9:9:9:9:9:9:9: | 21984109:21 | 21984234:21 | +:+:+:+:+:+:+: | 723 | NA | 13 | 17 | 20 | 9 | 4 | 4 |
| ENSDARG000 | 9:9:9:9:9:9:9: | 37341240:37 | 37341906:37 | -:-:-:-:-:-:-:-: | 2523 | hspbap1 | 147 | 102 | 135 | 53 | 72 | 141 |
| ENSDARG000 | 9:9:9:9:9:9:9: | 10138196:10 | 10138564:10 | -:-:-:-:-:-:-:-: | 1042 | hnmt | 524 | 485 | 175 | 687 | 193 | 847 |
| ENSDARG000 | 9;9 | 3666401:367 | 3669312:367 | -;- | 3084 | sp5a | 0 | 0 | 0 | 0 | 0 | 0 |
| ENSDARG000 | 9:9:9:9:9:9:9: | 54891962:54 | 54892146:54 | +:+:+:+:+:+:+: | 1813 | glra2 | 4 | 0 | 0 | 0 | 0 | 0 |
| ENSDARG000 | 9:9:9:9:9:9:9: | 55056310:55 | 55056559:55 | +:+:+:+:+:+:+: | 1688 | GPR143 | 17258 | 9459 | 9981 | 15324 | 8435 | 10585 |
| ENSDARG000 | 9:9:9:9:9:9:9: | 21990058:21 | 21990144:21 | +:+:+:+:+:+:+: | 1034 | NA | 0 | 0 | 0 | 0 | 0 | 0 |
| ENSDARG000 | 9:9:9:9:9:9:9: | 55093078:55 | 55095486:55 | -:-:-:-:-:-:-:-: | 4177 | tbl1x | 236 | 384 | 479 | 178 | 454 | 204 |
| ENSDARG000 | 9:9:9:9:9:9:9: | 3699397:370 | 3699532:370 | -:-:-:-:-:-:-:-: | 3835 | MYO3B | 0 | 0 | 8 | 0 | 4 | 2 |
| ENSDARG000 | 9;9;9;9;9;9 | 34380299:34 | 34380575:34 | +;+;+;+;+;+ | 2333 | lamp1b | 172 | 172 | 345 | 288 | 130 | 266 |
| ENSDARG000 | 9;9;9 | 40825065:40 | 40825222:40 | +;+;+ | 656 | NA | 6 | 8 | 50 | 17 | 3 | 2 |
| ENSDARG000 | 9;9;9;9 | 40829469:40 | 40829644:40 | -;-;-;- | 610 | NA | 0 | 0 | 0 | 0 | 0 | 0 |
| ENSDARG000 | 9:9:9:9:9:9:9: | 40874194:40 | 40874288:40 | +:+:+:+:+:+:+: | 2193 | hibch | 478 | 582 | 486 | 436 | 598 | 553 |
| ENSDARG000 | 9;9;9;9 | 40895649:40 | 40895916:40 | +;+;+;+ | 1003 | NA | 0 | 0 | 0 | 0 | 0 | 0 |
| ENSDARG000 | 9:9:9:9:9:9:9: | 40851088:40 | 40851241:40 | -;-;-;-;-;-;-;- | 1354 | pofut2 | 195 | 244 | 212 | 93 | 280 | 220 |
| ENSDARG000 | 9:9:9:9:9:9:9: | 21993092:21 | 21993162:21 | +:+:+:+:+:+:+: | 860 | crygm7 | 0 | 0 | 0 | 0 | 0 | 0 |
| ENSDARG000 | 9:9:9:9:9:9:9: | 34389976:34 | 34390593:34 | -;-;-;-;-;-;- | 1642 | grtp1b | 0 | 3 | 6 | 2 | 4 | 4 |
| ENSDARG000 | 9:9:9:9:9:9:9: | 34397516:34 | 34398219:34 | +:+:+:+:+:+:+: | 9676 | med14 | 1118 | 1112 | 1616 | 680 | 1417 | 1070 |
| ENSDARG000 | 9;9;9;9;9 | 21996315:21 | 21996354:21 | +;+;+;+;+ | 559 | NA | 0 | 0 | 0 | 0 | 0 | 0 |
| ENSDARG000 | 9:9:9:9:9:9:9: | 34425728:34 | 34426008:34 | +:+:+:+:+:+:+: | 1514 | NA | 0 | 0 | 0 | 0 | 0 | 0 |
| ENSDARG000 | 9;9;9 | 22000812:22 | 22000850:22 | +;+;+ | 613 | crygm6 | 0 | 0 | 0 | 0 | 0 | 0 |
| ENSDARG000 | 9;9;9 | 22003942:22 | 22003965:22 | +;+;+ | 680 | si:dkev-57a22 | 0 | 0 | 0 | 0 | 0 | 0 |
| ENSDARG000 | 9;9;9;9;9 | 37636957:37 | 37637110:37 | -;-;-;-;- | 1035 | NA | 0 | 0 | 0 | 0 | 0 | 0 |
| ENSDARG000 | 9;9;9 | 22011912:22 | 22011949:22 | +;+;+ | 606 | crygm1 | 0 | 0 | 0 | 0 | 0 | 0 |
| ENSDARG000 | 9:9:9:9:9:9:9: | 37481434:37 | 37481601:37 | -:-:-:-:-:-:-:-: | 4313 | SEMA5B | 47 | 18 | 26 | 25 | 26 | 2 |
| ENSDARG000 | 9;9;9 | 22017368:22 | 22017425:22 | +;+;+ | 628 | zgc:153846 | 0 | 0 | 0 | 0 | 0 | 0 |
| ENSDARG000 | 9:9:9:9:9:9:9: | 3898132:390 | 3898328:390 | -:-:-:-:-:-:-:-: | 7354 | ubr3 | 671 | 573 | 759 | 379 | 982 | 697 |
| ENSDARG000 | 9;9;9 | 40927277:40 | 40931856:40 | -;-;- | 5060 | NA | 52 | 36 | 62 | 32 | 105 | 84 |
| ENSDARG000 | 9;9;9 | 22022105:22 | 22022417:22 | -;-;- | 600 | crygm2c | 0 | 0 | 0 | 0 | 0 | 0 |
| ENSDARG000 | 9;9;9 | 40939336:40 | 40939790:40 | +;+;+ | 2178 | mstnb | 6 | 2 | 0 | 1 | 8 | 0 |
| ENSDARG000 | 9;9;9;9;9;9;9 | 41024973:41 | 41025094:41 | +;+;+;+;+;+;+ | 1783 | ormdl1 | 344 | 493 | 442 | 296 | 359 | 503 |
| ENSDARG000 | 9;9;9 | 40943119:40 | 40943608:40 | -;-;- | 770 | NA | 0 | 2 | 0 | 0 | 1 | 0 |
| ENSDARG000 | 9:9:9:9:9:9:9: | 40948569:40 | 40948609:40 | -:-:-:-:-:-:-:-: | 3036 | pms1 | 82 | 118 | 176 | 106 | 186 | 117 |
| ENSDARG000 | 9;9 | 37817266:37 | 37817733:37 | -;- | 533 | NA | 2 | 0 | 0 | 4 | 0 | 0 |
| ENSDARG000 | 9:9:9:9:9:9:9: | 37754845:37 | 37754946:37 | +:+:+:+:+:+:+: | 3085 | pdia5 | 1178 | 1139 | 2112 | 818 | 1365 | 783 |
| ENSDARG000 | 9;9;9 | 22056172:22 | 22056777:22 | -;-;- | 887 | crygmxl1 | 0 | 0 | 0 | 0 | 0 | 0 |
| ENSDARG000 | 9;9;9 | 22068412:22 | 22068724:22 | -;-;- | 600 | crygm2b | 0 | 0 | 4 | 0 | 0 | 0 |
| ENSDARG000 | 9;9;9;9;9;9;9 | 12428226:12 | 12429568:12 | -;-;-;-;-;-;- | 2480 | ehhadh | 174 | 364 | 256 | 228 | 284 | 188 |
| ENSDARG000 | 9 | 13352308 | 13355071 | - | 2764 | fzd7a | 86 | 86 | 888 | 164 | 80 | 76 |
| ENSDARG000 | 9:9:9:9:9:9:9: | 12664373:12 | 12664477:12 | -:-:-:-:-:-:-:-: | 7573 | NA | 1858 | 2528 | 3290 | 2154 | 1522 | 2098 |
| ENSDARG000 | 9:9:9:9:9:9:9: | 12905591:12 | 12905678:12 | +:+:+:+:+:+:+: | 8566 | NA | 255 | 300 | 364 | 251 | 363 | 285 |
| ENSDARG000 | 9:9:9:9:9:9:9: | 12894578:12 | 12894724:12 | +:+:+:+:+:+:+: | 3431 | NA | 0 | 0 | 0 | 0 | 0 | 0 |
| ENSDARG000 | 9:9:9:9:9:9:9: | 12468500:12 | 12469616:12 | -:-:-:-:-:-:-:-: | 3221 | igf2bp2a | 94 | 48 | 136 | 12 | 35 | 72 |
| ENSDARG000 | 9:9:9:9:9:9:9: | 37850360:37 | 37851230:37 | +:+:+:+:+:+:+: | 4181 | sec22a | 692 | 609 | 760 | 687 | 578 | 714 |
| ENSDARG000 | 9:9:9:9:9:9:9: | 12959760:13 | 12960633:13 | +:+:+:+:+:+:+: | 4183 | bmpr2b | 309 | 193 | 301 | 84 | 246 | 161 |
| ENSDARG000 | 9:9:9:9:9:9:9: | 14023386:14 | 14024074:14 | +:+:+:+:+:+:+: | 3090 | NA | 300 | 294 | 429 | 199 | 286 | 266 |
| ENSDARG000 | 9:9:9:9:9:9:9: | 14044310:14 | 14044490:14 | -:-:-:-:-:-:-:-: | 5387 | FER1L6 | 0 | 4 | 2 | 14 | 4 | 0 |
| ENSDARG000 | 9:9:9:9:9:9:9: | 11822379:11 | 11823081:11 | -;-;-;-;-;-;-;- | 1227 | NA | 0 | 0 | 0 | 0 | 0 | 0 |
| ENSDARG000 | 9:9:9:9:9:9:9: | 12847040:12 | 12847350:12 | +:+:+:+:+:+:+: | 5292 | glb1l | 693 | 1882 | 532 | 460 | 2731 | 2147 |
| ENSDARG000 | 9:9:9:9:9:9:9: | 14202686:14 | 14202730:14 | -:-:-:-:-:-:-:-: | 2052 | ABCB6 | 104 | 75 | 100 | 42 | 95 | 90 |
| ENSDARG000 | 9;9;9 | 14143073:14 | 14143256:14 | +;+;+ | 695 | NA | 0 | 2 | 0 | 0 | 0 | 0 |
| ENSDARG000 | 9:9:9:9:9:9:9: | 13796845:13 | 13797759:13 | -:-:-:-:-:-:-:-: | 1598 | NA | 19 | 28 | 36 | 21 | 25 | 28 |
| ENSDARG000 | 9 | 11532025 | 11534622 | + | 2598 | cdk5r2b | 0 | 0 | 0 | 0 | 0 | 0 |
| ENSDARG000 | 9:9:9:9:9:9:9: | 14367570:14 | 14369313:14 | -:-:-:-:-:-:-:-: | 4784 | nrp2b | 10246 | 6355 | 6463 | 5063 | 11639 | 6095 |
| ENSDARG000 | 9:9:9:9:9:9:9: | 12579077:12 | 12579421:12 | -:-:-:-:-:-:-:-: | 2045 | tra2b | 747 | 838 | 1214 | 847 | 1175 | 1101 |
| ENSDARG000 | 9:9:9:9:9:9:9: | 13999620:13 | 13999719:13 | +:+:+:+:+:+:+: | 1042 | NA | 41 | 9 | 0 | 6 | 12 | 11 |
| ENSDARG000 | 9;9 | 10692905:10 | 10693008:10 | +;+ | 1674 | cxcr4b | 154 | 135 | 67 | 106 | 402 | 310 |
| ENSDARG000 | 9:9:9:9:9:9:9: | 12403289:12 | 12405827:12 | -;-;-;-;-;-;-;- | 3652 | C9H3orf70 | 8 | 2 | 0 | 7 | 0 | 0 |
| ENSDARG000 | 9:9:9:9:9:9:9: | 10152820:10 | 10157463:10 | -:-:-:-:-:-:-:-: | 10462 | THSD7B | 0 | 0 | 0 | 0 | 0 | 6 |
| ENSDARG000 | 9;9;9;9;9;9 | 11576502:11 | 11576842:11 | -;-;-;-;-;- | 1696 | umps | 216 | 535 | 653 | 272 | 433 | 257 |
| ENSDARG000 | 9:9:9:9:9:9:9: | 11548233:11 | 11549438:11 | -;-;-;-;-;-;-;- | 1613 | fev | 0 | 0 | 2 | 0 | 0 | 0 |
| ENSDARG000 | 9;9;9;9 | 12648705:12 | 12649612:12 | -;-;-;- | 1289 | sumo3b | 437 | 1286 | 1809 | 552 | 1331 | 994 |
| ENSDARG000 | 9:9:9:9:9:9:9: | 37872577:37 | 37872850:37 | -:-:-:-:-:-:-:-: | 3814 | adcy5 | 4041 | 2949 | 3333 | 2934 | 3628 | 2886 |
| ENSDARG000 | 9:9:9:9:9:9:9: | 11554089:11 | 11554608:11 | -:-:-:-:-:-:-:-: | 1859 | cryba2b | 0 | 0 | 0 | 0 | 0 | 0 |
| ENSDARG000 | 9;9;9;9 | 11906770:11 | 11909782:11 | -;-;-;- | 3657 | znf804a | 0 | 0 | 0 | 0 | 4 | 8 |
| ENSDARG000 | 9:9:9:9:9:9:9: | 12887491:12 | 12888490:12 | +:+:+:+:+:+:+: | 2037 | NA | 0 | 0 | 0 | 0 | 2 | 0 |

| | | | | | | | | | | | | |
|---|---|---|---|---|---|---|---|---|---|---|---|---|
| ENSDARG000 | 9:9:9:9:9:9:9: | 12444494:124 | 12444721:124 | +:+:+:+:+:+:+: | 1574 | tmem41aa | 165 | 199 | 209 | 112 | 241 | 212 |
| ENSDARG000 | 9:9:9:9:9:9:9: | 14111561:141 | 14113156:141 | -:-:-:-:-:-:-:-: | 4891 | NA | 290 | 499 | 577 | 176 | 424 | 357 |
| ENSDARG000 | 9;9;9;9;9;9;9 | 14152211:141 | 14152833:141 | +;+;+;+;+;+;+ | 4188 | NA | 184 | 0 | 6 | 0 | 30 | 2 |
| ENSDARG000 | 9:9:9:9:9:9:9: | 13229585:132 | 13229627:132 | +:+:+:+:+:+:+: | 3326 | NA | 221 | 326 | 319 | 155 | 498 | 374 |
| ENSDARG000 | 9:9:9:9:9:9:9: | 11597500:116 | 11598530:116 | -:-:-:-:-:-:-:-: | 5012 | itgav | 7197 | 4688 | 4929 | 3130 | 5276 | 4177 |
| ENSDARG000 | 9:9:9:9:9:9:9: | 14010823:140 | 14010947:140 | +:+:+:+:+:+:+: | 1022 | NA | 10 | 15 | 2 | 4 | 28 | 20 |
| ENSDARG000 | 9:9:9:9:9:9:9: | 12259469:122 | 12259560:122 | -:-:-:-:-:-:-:-: | 3479 | nup35 | 217 | 426 | 399 | 117 | 387 | 246 |
| ENSDARG000 | 9:9:9:9:9:9:9: | 11664797:116 | 11665945:116 | -:-:-:-:-:-:-:-: | 2379 | zc3h15 | 1628 | 1793 | 2141 | 988 | 1853 | 1607 |
| ENSDARG000 | 9:9:9:9:9:9:9: | 14090329:140 | 14090504:140 | -:-:-:-:-:-:-:-: | 1361 | prkag3b | 0 | 0 | 6 | 6 | 0 | 0 |
| ENSDARG000 | 9;9;9;9 | 11245962:112 | 11247244:112 | -;-;-;- | 2466 | NA | 376 | 338 | 480 | 252 | 430 | 379 |
| ENSDARG000 | 9:9:9:9:9:9:9: | 12834414:128 | 12834576:128 | -:-:-:-:-:-:-:-: | 3672 | NA | 149 | 259 | 184 | 171 | 395 | 169 |
| ENSDARG000 | 9:9:9:9:9:9:9: | 13120419:131 | 13121212:131 | +:+:+:+:+:+:+: | 1950 | fam117bb | 476 | 152 | 386 | 374 | 425 | 180 |
| ENSDARG000 | 9;9;9;9;9;9;9 | 11281969:112 | 11282941:112 | +;+;+;+;+;+;+ | 1961 | WNT6 | 4 | 0 | 0 | 0 | 0 | 10 |
| ENSDARG000 | 9:9:9:9:9:9:9: | 10781687:107 | 10782610:107 | -:-:-:-:-:-:-:-: | 3353 | NA | 2 | 0 | 0 | 0 | 2 | 2 |
| ENSDARG000 | 9:9:9:9:9:9:9: | 12596038:125 | 12598234:125 | -:-:-:-:-:-:-:-: | 5043 | TSPEAR | 0 | 0 | 2 | 0 | 0 | 0 |
| ENSDARG000 | 9:9:9:9:9:9:9: | 13733468:137 | 13733708:137 | +:+:+:+:+:+:+: | 2257 | abi2a | 254 | 169 | 207 | 209 | 239 | 258 |
| ENSDARG000 | 9:9:9:9:9:9:9: | 13814963:138 | 13817056:138 | -:-:-:-:-:-:-:-: | 4950 | RAPH1 | 646 | 576 | 447 | 323 | 1020 | 753 |
| ENSDARG000 | 9:9:9:9:9:9:9: | 13628635:136 | 13628746:136 | +:+:+:+:+:+:+: | 5151 | ALS2 | 171 | 72 | 160 | 70 | 159 | 105 |
| ENSDARG000 | 9;9 | 10845918:108 | 10846023:108 | -;- | 471 | NA | 0 | 0 | 0 | 0 | 0 | 0 |
| ENSDARG000 | 9:9:9:9:9:9:9: | 12885746:128 | 12885902:128 | -:-:-:-:-:-:-:-: | 944 | NA | 0 | 0 | 0 | 0 | 0 | 0 |
| ENSDARG000 | 9:9:9:9:9:9:9: | 11034314:110 | 11035777:110 | +:+:+:+:+:+:+: | 4114 | asic4a | 0 | 0 | 0 | 2 | 2 | 0 |
| ENSDARG000 | 9:9:9:9:9:9:9: | 11587629:115 | 11587862:115 | +:+:+:+:+:+:+: | 1156 | si:dkev-69o16 | 154 | 160 | 228 | 206 | 145 | 232 |
| ENSDARG000 | 9;9 | 10936773:109 | 10936888:109 | +;+ | 459 | si:dkev-22p6.: | 0 | 0 | 0 | 0 | 0 | 0 |
| ENSDARG000 | 9;9;9;9;9;9 | 13985567:139 | 13985914:139 | +;+;+;+;+;+ | 2891 | NA | 57 | 105 | 10 | 63 | 98 | 156 |
| ENSDARG000 | 9:9:9:9:9:9:9: | 13682133:136 | 13682516:136 | +:+:+:+:+:+:+: | 2312 | MPP4 | 8 | 0 | 2 | 0 | 6 | 10 |
| ENSDARG000 | 9:9:9:9:9:9:9: | 12655278:126 | 12655838:126 | -:-:-:-:-:-:-:-: | 1336 | pttg1ipb | 6499 | 5729 | 5165 | 8919 | 5373 | 8649 |
| ENSDARG000 | 9:9:9:9:9:9:9: | 13714379:137 | 13714466:137 | +:+:+:+:+:+:+: | 1490 | tmem237a | 2 | 2 | 5 | 0 | 16 | 0 |
| ENSDARG000 | 9;9;9;9;9;9;9 | 11379782:113 | 11380105:113 | +;+;+;+;+;+;+ | 1775 | wnt10a | 2 | 0 | 0 | 0 | 0 | 0 |
| ENSDARG000 | 9;9;9;9 | 14304447:143 | 14304878:143 | -;-;-;- | 825 | NA | 0 | 0 | 2 | 0 | 0 | 0 |
| ENSDARG000 | 9;9;9;9;9;9;9 | 707086:70788 | 708183:70818 | -;-;-;-;-;-;- | 1483 | ndufb3 | 852 | 1175 | 999 | 666 | 835 | 647 |
| ENSDARG000 | 9;9;9;9 | 993477:99463 | 993643:99470 | +;+;+;+ | 1473 | NA | 1 | 2 | 0 | 6 | 4 | 1 |
| ENSDARG000 | 9;9;9;9 | 701452:70191 | 701580:70202 | -;-;-;- | 610 | NA | 159 | 118 | 70 | 94 | 159 | 71 |
| ENSDARG000 | 9;9;9;9;9;9 | 51427826:514 | 51432816:514 | -;-;-;-;-;- | 6172 | tbr1b | 0 | 0 | 2 | 0 | 2 | 0 |
| ENSDARG000 | 9:9:9:9:9:9:9: | 2020667:2026 | 2020920:2026 | +:+:+:+:+:+:+: | 1067 | lnpa | 223 | 523 | 379 | 352 | 308 | 286 |
| ENSDARG000 | 9 | 52742196 | 52744063 | + | 1868 | NA | 0 | 0 | 0 | 0 | 0 | 0 |
| ENSDARG000 | 9:9:9:9:9:9:9: | 53011309:530 | 53011706:530 | -:-:-:-:-:-:-:-: | 1989 | NA | 45 | 20 | 27 | 34 | 101 | 47 |
| ENSDARG000 | 9:9:9:9:9:9:9: | 55893789:558 | 55893957:558 | -:-:-:-:-:-:-:-: | 1838 | SH3RF3 | 238 | 105 | 155 | 94 | 196 | 62 |
| ENSDARG000 | 9;9;9;9 | 16854121:168 | 16854224:168 | +;+;+;+ | 990 | CLN5 | 223 | 252 | 228 | 217 | 339 | 423 |
| ENSDARG000 | 9:9:9:9:9:9:9: | 295576:29916 | 299450:29945 | -:-:-:-:-:-:-:-: | 5653 | NA | 319 | 288 | 353 | 178 | 409 | 238 |
| ENSDARG000 | 9 | 49785320 | 49785437 | + | 118 | 5S_rRNA | 0 | 0 | 0 | 0 | 0 | 0 |
| ENSDARG000 | 9 | 38161263 | 38161389 | + | 127 | NA | 0 | 0 | 3 | 0 | 0 | 0 |
| ENSDARG000 | 9:9:9:9:9:9:9: | 55857193:558 | 55857305:558 | +:+:+:+:+:+:+: | 2069 | septin10 | 1768 | 2052 | 2127 | 2230 | 1679 | 1663 |
| ENSDARG000 | 9;9;9;9;9;9;9 | 33783061:337 | 33783076:337 | +;+;+;+;+;+;+ | 141 | NA | 0 | 0 | 0 | 0 | 0 | 0 |
| ENSDARG000 | 9:9:9:9:9:9:9: | 1162216:1165 | 1162273:1165 | +:+:+:+:+:+:+: | 1488 | stk24a | 0 | 1 | 4 | 0 | 0 | 0 |
| ENSDARG000 | 9;9;9;9;9;9 | 56422311:564 | 56423282:564 | +;+;+;+;+;+ | 2597 | gpr39 | 8 | 10 | 8 | 58 | 48 | 12 |
| ENSDARG000 | 9 | 38160446 | 38160572 | + | 127 | NA | 0 | 0 | 0 | 0 | 0 | 0 |
| ENSDARG000 | 9;9;9;9 | 56457544:564 | 56457639:564 | -;-;-;- | 381 | ubxn4 | 0 | 0 | 0 | 0 | 0 | 0 |
| ENSDARG000 | 9;9 | 778745:78446 | 782215:78544 | -;- | 4451 | en1a | 0 | 0 | 2 | 0 | 0 | 0 |
| ENSDARG000 | 9 | 6848989 | 6849104 | + | 116 | 5S_rRNA | 0 | 0 | 0 | 0 | 0 | 0 |
| ENSDARG000 | 9:9:9:9:9:9:9: | 49761929:497 | 49762042:497 | -:-:-:-:-:-:-:-: | 2204 | GALNT3 | 6 | 0 | 0 | 0 | 0 | 0 |
| ENSDARG000 | 9:9:9:9:9:9:9: | 54946499:549 | 54946531:549 | -:-:-:-:-:-:-:-: | 2391 | NA | 13 | 63 | 83 | 38 | 134 | 99 |
| ENSDARG000 | 9;9;9;9;9;9 | 51784922:517 | 51785030:517 | +;+;+;+;+;+ | 756 | NA | 737 | 514 | 767 | 690 | 828 | 768 |
| ENSDARG000 | 9 | 53436689 | 53436772 | - | 84 | MIR19A | 0 | 0 | 0 | 0 | 0 | 0 |
| ENSDARG000 | 9:9:9:9:9:9:9: | 219124:22520 | 219298:22569 | +:+:+:+:+:+:+: | 3532 | map3k12 | 0 | 0 | 0 | 0 | 0 | 0 |
| ENSDARG000 | 9 | 34790750 | 34790887 | + | 138 | 5S_rRNA | 0 | 0 | 0 | 0 | 0 | 0 |
| ENSDARG000 | 9:9:9:9:9:9:9: | 54237100:542 | 54237158:542 | +:+:+:+:+:+:+: | 3096 | RBM27 | 245 | 192 | 347 | 200 | 356 | 326 |
| ENSDARG000 | 9;9;9;9;9;9;9 | 102422:10282 | 102495:10302 | -;-;-;-;-;-;- | 1152 | NA | 326 | 225 | 314 | 276 | 375 | 368 |
| ENSDARG000 | 9 | 53436831 | 53436917 | - | 87 | dre-mir-18b | 0 | 0 | 0 | 0 | 0 | 0 |
| ENSDARG000 | 9 | 27938302 | 27938418 | + | 117 | 5S_rRNA | 0 | 0 | 0 | 0 | 0 | 0 |
| ENSDARG000 | 9 | 358669 | 358783 | + | 115 | 5S_rRNA | 1 | 0 | 16 | 9 | 15 | 5 |
| ENSDARG000 | 9:9:9:9:9:9:9: | 2393764:2403 | 2393782:2403 | +:+:+:+:+:+:+: | 5580 | chn1 | 1241 | 1269 | 1161 | 951 | 1898 | 854 |
| ENSDARG000 | 9 | 19348124 | 19348239 | - | 116 | 5S_rRNA | 0 | 0 | 0 | 0 | 0 | 0 |
| ENSDARG000 | 9 | 29807947 | 29808064 | - | 118 | 5S_rRNA | 0 | 0 | 0 | 0 | 0 | 0 |
| ENSDARG000 | 9 | 54966621 | 54966735 | - | 115 | 5S_rRNA | 0 | 0 | 0 | 0 | 0 | 0 |
| ENSDARG000 | 9 | 7612376 | 7612526 | + | 151 | dre-mir-153c | 0 | 0 | 0 | 0 | 0 | 0 |
| ENSDARG000 | 9;9;9 | 2041535:2041 | 2041607:2042 | +;+;+ | 1146 | NA | 235 | 420 | 392 | 522 | 487 | 387 |
| ENSDARG000 | 9 | 26435073 | 26435187 | - | 115 | 5S_rRNA | 0 | 0 | 0 | 0 | 0 | 0 |
| ENSDARG000 | 9:9:9:9:9:9:9: | 492459:50005 | 492504:50048 | +:+:+:+:+:+:+: | 7106 | NA | 915 | 1107 | 1241 | 845 | 1441 | 1200 |
| ENSDARG000 | 9 | 29652511 | 29652612 | - | 102 | dre-mir-15a-2 | 0 | 0 | 0 | 0 | 0 | 0 |
| ENSDARG000 | 9;9 | 21268739:212 | 21268916:212 | +;+ | 5194 | NA | 246 | 125 | 331 | 166 | 186 | 138 |
| ENSDARG000 | 9;9;9;9;9;9;9 | 52479142:524 | 52479292:524 | -;-;-;-;-;-;- | 1753 | NA | 11 | 28 | 101 | 29 | 61 | 30 |
| ENSDARG000 | 9:9:9:9:9:9:9: | 55839522:558 | 55839536:558 | -:-:-:-:-:-:-:-: | 858 | NA | 0 | 0 | 0 | 0 | 0 | 0 |
| ENSDARG000 | 9:9:9:9:9:9:9: | 10758339:107 | 10763017:107 | -:-:-:-:-:-:-:-: | 5822 | NA | 169 | 121 | 322 | 441 | 141 | 183 |
| ENSDARG000 | 9:9:9:9:9:9:9: | 5253137:5257 | 5255194:5257 | -;-;-;-;-;-;-;- | 3226 | CYTIP | 12 | 10 | 10 | 2 | 6 | 22 |
| ENSDARG000 | 9:9:9:9:9:9:9: | 1039215:1068 | 1039331:1068 | +:+:+:+:+:+:+: | 5964 | DOCK9 | 311 | 432 | 144 | 529 | 109 | 143 |
| ENSDARG000 | 9;9;9 | 10778227:107 | 10778320:107 | -;-;- | 273 | NA | 0 | 0 | 0 | 0 | 0 | 0 |
| ENSDARG000 | 9:9:9:9:9:9:9: | 50316921:503 | 50317128:503 | +:+:+:+:+:+:+: | 2115 | GRB14 | 376 | 144 | 143 | 155 | 345 | 105 |
| ENSDARG000 | 9 | 40137497 | 40137612 | + | 116 | 5S_rRNA | 0 | 0 | 0 | 0 | 0 | 0 |
| ENSDARG000 | 9;9 | 24025027:240 | 24025130:240 | +;+ | 280 | prlh | 29 | 17 | 43 | 10 | 21 | 39 |
| ENSDARG000 | 9;9;9;9 | 7068832:7073 | 7068842:7073 | +;+;+;+ | 87 | NA | 0 | 0 | 0 | 0 | 2 | 0 |
| ENSDARG000 | 9 | 6717677 | 6717801 | - | 125 | 5S_rRNA | 0 | 0 | 0 | 0 | 0 | 0 |
| ENSDARG000 | 9 | 9377863 | 9377978 | + | 116 | 5S_rRNA | 0 | 0 | 0 | 0 | 0 | 0 |
| ENSDARG000 | 9 | 41733533 | 41733610 | + | 78 | dre-mir-26a-1 | 0 | 0 | 0 | 0 | 0 | 0 |
| ENSDARG000 | 9:9:9:9:9:9:9: | 250724:25072 | 253633:25456 | -:-:-:-:-:-:-:-: | 5193 | pcbp2 | 3786 | 3005 | 4624 | 3577 | 3636 | 3246 |
| ENSDARG000 | 9 | 134489 | 135774 | - | 1286 | NA | 414 | 402 | 498 | 386 | 636 | 352 |
| ENSDARG000 | 9:9:9:9:9:9:9: | 54312035:543 | 54312221:543 | -:-:-:-:-:-:-:-: | 4286 | RAD50 | 302 | 636 | 613 | 338 | 712 | 447 |
| ENSDARG000 | 9 | 54143997 | 54147134 | + | 3138 | TLR7 | 0 | 0 | 0 | 0 | 0 | 0 |
| ENSDARG000 | 9 | 56091026 | 56091141 | - | 116 | 5S_rRNA | 0 | 0 | 0 | 0 | 0 | 0 |
| ENSDARG000 | 9;9;9;9;9;9 | 747612:74910 | 748052:74935 | +;+;+;+;+;+ | 1119 | insig2 | 110 | 130 | 70 | 66 | 117 | 100 |
| ENSDARG000 | 9:9:9:9:9:9:9: | 5325015:5326 | 5327620:5327 | -:-:-:-:-:-:-:-: | 3448 | tnfsf13b | 28 | 10 | 23 | 2 | 20 | 20 |
| ENSDARG000 | 9;9;9 | 6802641:6806 | 6802744:6806 | +;+;+ | 300 | NA | 0 | 3 | 24 | 9 | 5 | 10 |
| ENSDARG000 | 9 | 31495648 | 31495765 | + | 118 | 5S_rRNA | 0 | 0 | 0 | 0 | 0 | 0 |
| ENSDARG000 | 9 | 55916413 | 55916544 | + | 132 | 5S_rRNA | 0 | 0 | 0 | 0 | 0 | 0 |
| ENSDARG000 | 9 | 1317931 | 1318095 | + | 165 | U1 | 0 | 0 | 0 | 0 | 0 | 0 |
| ENSDARG000 | 9:9:9:9:9:9:9: | 28140089:281 | 28140219:281 | +:+:+:+:+:+:+: | 7613 | PLEKHM3 | 304 | 264 | 340 | 184 | 245 | 240 |
| ENSDARG000 | 9 | 52082311 | 52082437 | + | 127 | 5S_rRNA | 0 | 0 | 0 | 0 | 0 | 0 |
| ENSDARG000 | 9 | 41593496 | 41593610 | + | 115 | 5S_rRNA | 0 | 0 | 0 | 0 | 0 | 0 |
| ENSDARG000 | 9;9 | 5345236:5350 | 5346413:5351 | -;- | 1325 | abhd13 | 124 | 101 | 188 | 45 | 146 | 114 |
| ENSDARG000 | 9 | 6957256 | 6957373 | + | 118 | 5S_rRNA | 1 | 13 | 2 | 0 | 20 | 0 |
| ENSDARG000 | 9:9:9:9:9:9:9: | 55576475:555 | 55576839:555 | -:-:-:-:-:-:-:-: | 1971 | ARSD | 64 | 64 | 78 | 84 | 239 | 70 |
| ENSDARG000 | 9 | 38786928 | 38787044 | + | 117 | 5S_rRNA | 0 | 0 | 0 | 0 | 0 | 0 |
| ENSDARG000 | 9 | 1325704 | 1325868 | + | 165 | U1 | 1 | 0 | 0 | 0 | 0 | 0 |
| ENSDARG000 | 9 | 1952956 | 1953033 | - | 78 | dre-mir-10b-1 | 0 | 0 | 0 | 0 | 0 | 0 |
| ENSDARG000 | 9:9:9:9:9:9:9: | 711638:71163 | 716230:71623 | -:-:-:-:-:-:-:-: | 7659 | usp37 | 396 | 567 | 759 | 315 | 820 | 537 |
| ENSDARG000 | 9:9:9:9:9:9:9: | 10267:11746: | 10406:11879: | -:-:-:-:-:-:-:-: | 1293 | NA | 30 | 47 | 48 | 52 | 61 | 51 |
| ENSDARG000 | 9:9:9:9:9:9:9: | 19007959:190 | 19007999:190 | -;-;-;-;-;-;-;- | 2214 | VGLL3 | 0 | 0 | 0 | 0 | 10 | 6 |
| ENSDARG000 | 9 | 21204616 | 21204722 | + | 107 | U6 | 0 | 0 | 0 | 0 | 0 | 0 |
| ENSDARG000 | 9:9:9:9:9:9:9: | 537297:53829 | 537684:53871 | +:+:+:+:+:+:+: | 2949 | NA | 32 | 15 | 0 | 4 | 19 | 11 |
| ENSDARG000 | 9;9 | 16823285:168 | 16824085:168 | -;- | 855 | NA | 446 | 221 | 304 | 426 | 694 | 410 |
| ENSDARG000 | 9:9:9:9:9:9:9: | 5271524:5272 | 5272562:5273 | -;-;-;-;-;-;-;- | 2522 | ACVR1C | 0 | 0 | 0 | 0 | 0 | 0 |
| ENSDARG000 | 9:9:9:9:9:9:9: | 205248:20557 | 205478:20565 | -:-:-:-:-:-:-:-: | 1617 | aaas | 282 | 598 | 585 | 282 | 627 | 322 |
| ENSDARG000 | 9 | 45143942 | 45149695 | - | 5754 | NA | 4 | 0 | 6 | 0 | 10 | 4 |
| ENSDARG000 | 9:9:9:9:9:9:9: | 1187805:1188 | 1188658:1189 | -:-:-:-:-:-:-:-: | 2798 | INO80D | 146 | 56 | 177 | 166 | 195 | 108 |
| ENSDARG000 | 9:9:9:9:9:9:9: | 52558181:525 | 52558494:525 | -:-:-:-:-:-:-:-: | 3174 | xrcc5 | 425 | 685 | 786 | 733 | 938 | 534 |
| ENSDARG000 | 9:9:9:9:9:9:9: | 54710916:547 | 54713086:547 | -;-;-;-;-;-;-;- | 2766 | trappc2 | 558 | 338 | 429 | 354 | 361 | 499 |
| ENSDARG000 | 9;9;9;9;9 | 156142:15733 | 156434:15749 | +;+;+;+;+ | 1722 | cldn10b | 0 | 4 | 13 | 2 | 24 | 0 |
| ENSDARG000 | 9;9;9;9;9 | 54158387:541 | 54160966:541 | +;+;+;+;+ | 3030 | NA | 8 | 0 | 4 | 0 | 4 | 6 |
| ENSDARG000 | 9 | 1235778 | 1236848 | - | 1071 | CMKLR2 | 36 | 8 | 34 | 6 | 29 | 38 |
| ENSDARG000 | 9:9:9:9:9:9:9: | 54219304:542 | 54219532:542 | -:-:-:-:-:-:-:-: | 1071 | NA | 343 | 308 | 495 | 269 | 259 | 284 |
| ENSDARG000 | 9;9 | 54290896:542 | 54290991:542 | +;+ | 996 | pou4f3 | 0 | 0 | 0 | 0 | 0 | 0 |
| ENSDARG000 | 9 | 24708653 | 24710074 | + | 1422 | NA | 0 | 0 | 0 | 0 | 0 | 0 |
| ENSDARG000 | 9;9;9;9;9;9 | 34836068:348 | 34836307:348 | -;-;-;-;-;- | 780 | crlf2 | 0 | 10 | 0 | 12 | 17 | 4 |
| ENSDARG000 | 9 | 55231078 | 55231198 | - | 121 | 5S_rRNA | 0 | 0 | 0 | 0 | 0 | 0 |
| ENSDARG000 | 9 | 36782547 | 36782665 | - | 119 | 5S_rRNA | 0 | 0 | 0 | 0 | 0 | 0 |
| ENSDARG000 | 9:9:9:9:9:9:9: | 1137430:1138 | 1137459:1138 | +:+:+:+:+:+:+: | 2870 | SLC15A1 | 0 | 0 | 0 | 0 | 0 | 0 |
| ENSDARG000 | 9;9;9;9;9;9 | 123593:12452 | 124237:12474 | -;-;-;-;-;- | 1443 | SLC5A7 | 10 | 6 | 6 | 0 | 2 | 0 |
| ENSDARG000 | 9 | 43855495 | 43855610 | + | 116 | 5S_rRNA | 0 | 0 | 0 | 0 | 0 | 0 |
| ENSDARG000 | 9:9:9:9:9:9:9: | 48943471:489 | 48943760:489 | +:+:+:+:+:+:+: | 4784 | STK39 | 8 | 20 | 6 | 5 | 22 | 0 |
| ENSDARG000 | 9 | 44796392 | 44796506 | - | 115 | 5S_rRNA | 0 | 0 | 0 | 0 | 0 | 0 |
| ENSDARG000 | 9 | 19029353 | 19029467 | - | 115 | 5S_rRNA | 0 | 0 | 0 | 0 | 0 | 0 |
| ENSDARG000 | 9:9:9:9:9:9:9: | 55455801:554 | 55456118:554 | +:+:+:+:+:+:+: | 8828 | MXRA5 | 4 | 4 | 2 | 0 | 2 | 0 |
| ENSDARG000 | 9:9:9:9:9:9:9: | 23787:23787: | 23870:23870: | -:-:-:-:-:-:-:-: | 3952 | MAP4K4 | 3177 | 2246 | 3109 | 1966 | 2985 | 2739 |
| ENSDARG000 | 9:9:9:9:9:9:9: | 5776359:5776 | 5776888:5776 | -:-:-:-:-:-:-:-: | 1221 | CCKBR | 2 | 0 | 0 | 0 | 0 | 0 |
| ENSDARG000 | 9 | 6184235 | 6184349 | + | 115 | 5S_rRNA | 0 | 0 | 0 | 0 | 0 | 0 |
| ENSDARG000 | 9 | 40509529 | 40509643 | - | 115 | 5S_rRNA | 0 | 0 | 0 | 0 | 0 | 0 |
| ENSDARG000 | 9:9:9:9:9:9:9: | 55730059:557 | 55730221:557 | -;-;-;-;-;-;-;- | 1229 | asmt | 74 | 25 | 52 | 6 | 6 | 46 |
| ENSDARG000 | 9 | 38469446 | 38469566 | - | 121 | 5S_rRNA | 0 | 0 | 0 | 0 | 0 | 0 |
| ENSDARG000 | 9 | 28632385 | 28632500 | + | 116 | 5S_rRNA | 0 | 0 | 0 | 0 | 0 | 0 |
| ENSDARG000 | 9 | 34969450 | 34969567 | + | 118 | 5S_rRNA | 0 | 0 | 0 | 0 | 0 | 0 |
| ENSDARG000 | 9:9:9:9:9:9:9: | 711638:71182 | 712193:71219 | +:+:+:+:+:+:+: | 972 | ybey | 0 | 0 | 0 | 0 | 0 | 0 |
| ENSDARG000 | 9;9;9 | 55783686:557 | 55784426:557 | -;-;- | 1518 | SOWAHC | 6 | 6 | 0 | 14 | 18 | 6 |
| ENSDARG000 | 9;9;9 | 52398531:523 | 52398622:523 | +;+;+ | 273 | NA | 626 | 192 | 231 | 156 | 64 | 533 |
| ENSDARG000 | 9:9:9:9:9:9:9: | 563547:56487 | 563575:56496 | +:+:+:+:+:+:+: | 12519 | NA | 4596 | 5902 | 884 | 390 | 658 | 721 |
| ENSDARG000 | 9;9 | 963627:97003 | 963940:97027 | +;+ | 552 | NA | 114 | 110 | 116 | 105 | 148 | 93 |
| ENSDARG000 | 9 | 10343864 | 10343980 | + | 117 | 5S_rRNA | 0 | 0 | 0 | 0 | 0 | 0 |
| ENSDARG000 | 9;9;9;9 | 54301736:543 | 54302225:543 | -;-;-;- | 892 | il13 | 14 | 8 | 12 | 15 | 12 | 20 |
| ENSDARG000 | 9;9;9;9;9 | 50599362:506 | 50599557:506 | +;+;+;+;+ | 6511 | fign | 76 | 55 | 150 | 50 | 132 | 90 |
| ENSDARG000 | 9 | 22576945 | 22577059 | + | 115 | 5S_rRNA | 0 | 0 | 0 | 0 | 0 | 0 |
| ENSDARG000 | 9 | 55248200 | 55248314 | + | 115 | 5S_rRNA | 0 | 0 | 0 | 0 | 0 | 0 |
| ENSDARG000 | 9;9;9;9;9;9 | 52949626:529 | 52950673:529 | -;-;-;-;-;- | 2101 | NA | 8 | 2 | 4 | 2 | 0 | 0 |
| ENSDARG000 | 9:9:9:9:9:9:9: | 16835900:168 | 16835922:168 | -;-;-;-;-;-;-;- | 699 | RSPH1 | 55 | 121 | 73 | 26 | 123 | 126 |
| ENSDARG000 | 9 | 31187738 | 31187859 | - | 122 | 5S_rRNA | 0 | 0 | 0 | 0 | 0 | 0 |
| ENSDARG000 | 9:9:9:9:9:9:9: | 49659114:496 | 49659299:496 | +:+:+:+:+:+:+: | 3265 | CSRNP3 | 271 | 91 | 55 | 104 | 102 | 40 |
| ENSDARG000 | 9 | 41682926 | 41683010 | + | 85 | dre-mir-732 | 0 | 0 | 0 | 0 | 0 | 0 |
| ENSDARG000 | 9:9:9:9:9:9:9: | 56194410:561 | 56194575:561 | +:+:+:+:+:+:+: | 3093 | TBC1D8 | 512 | 338 | 372 | 263 | 402 | 265 |
| ENSDARG000 | 9:9:9:9:9:9:9: | 54716436:547 | 54716714:547 | +:+:+:+:+:+:+: | 2951 | ofd1 | 367 | 360 | 474 | 245 | 434 | 259 |
| ENSDARG000 | 9 | 47420391 | 47420507 | - | 117 | 5S_rRNA | 0 | 0 | 0 | 0 | 0 | 0 |
| ENSDARG000 | 9 | 52172483 | 52172599 | - | 117 | 5S_rRNA | 0 | 0 | 0 | 0 | 0 | 0 |
| ENSDARG000 | 9;9;9;9 | 52378484:523 | 52378956:523 | -;-;-;- | 740 | dap1b | 184 | 43 | 111 | 472 | 327 | 685 |
| ENSDARG000 | 9 | 38204407 | 38204528 | - | 122 | NA | 0 | 0 | 0 | 0 | 0 | 0 |
| ENSDARG000 | 9:9:9:9:9:9:9: | 16449398:164 | 16449458:164 | +:+:+:+:+:+:+: | 2892 | EPHA3 | 24 | 6 | 40 | 4 | 14 | 4 |
| ENSDARG000 | 9:9:9:9:9:9:9: | 10699038:107 | 10699428:107 | +:+:+:+:+:+:+: | 1964 | dars1 | 1381 | 1401 | 2511 | 3112 | 930 | 1614 |

| | | | | | | | | | | | | |
|---|---|---|---|---|---|---|---|---|---|---|---|---|
| ENSDARG000 | 9 | 37181440 | 37181557 | + | 118 | 5S_rRNA | 0 | 0 | 0 | 0 | 0 | 0 |
| ENSDARG000 | 9 | 6401679 | 6401794 | + | 116 | 5S_rRNA | 0 | 0 | 0 | 0 | 0 | 0 |
| ENSDARG000 | 9 | 17662950 | 17663065 | + | 116 | 5S_rRNA | 0 | 0 | 0 | 0 | 0 | 0 |
| ENSDARG000 | 9;9;9;9;9;9;9;9 | 56225410;562 | 56226271;562 | -;-;-;-;-;-;-;-;- | 1228 | rpl31 | 15403 | 16600 | 24213 | 22466 | 15743 | 20763 |
| ENSDARG000 | 9 | 35429202 | 35429316 | + | 115 | 5S_rRNA | 0 | 0 | 0 | 0 | 0 | 0 |
| ENSDARG000 | 9 | 53436280 | 53436374 | - | 95 | dre-mir-92a-2 | 0 | 0 | 0 | 0 | 0 | 0 |
| ENSDARG000 | 9;9 | 917060;92003 | 917136;92325 | +;+ | 3294 | tmem37 | 6 | 198 | 592 | 1290 | 48 | 84 |
| ENSDARG000 | 9 | 41072820 | 41072935 | - | 116 | U5 | 0 | 0 | 0 | 0 | 0 | 0 |
| ENSDARG000 | 9 | 18350367 | 18350483 | + | 117 | 5S_rRNA | 0 | 0 | 0 | 0 | 0 | 0 |
| ENSDARG000 | 9 | 27282322 | 27282436 | + | 115 | 5S_rRNA | 0 | 0 | 0 | 0 | 0 | 0 |
| ENSDARG000 | 9 | 48134214 | 48134285 | - | 72 | NA | 0 | 0 | 0 | 0 | 0 | 0 |
| ENSDARG000 | 9 | 40385130 | 40385247 | - | 118 | 5S_rRNA | 0 | 0 | 0 | 0 | 0 | 0 |
| ENSDARG000 | 9;9;9;9 | 163662;16520 | 164475;16536 | +;+;+;+ | 1212 | NA | 0 | 2 | 4 | 0 | 0 | 0 |
| ENSDARG000 | 9;9;9;9;9;9 | 56413986;564 | 56414886;564 | -;-;-;-;-;- | 1817 | actr3 | 2645 | 2701 | 2501 | 1842 | 2371 | 1978 |
| ENSDARG000 | 9;9;9;9;9;9;9;9 | 52411530;524 | 52411660;524 | +;+;+;+;+;+;+; | 2035 | nme9 | 1 | 7 | 2 | 0 | 9 | 3 |
| ENSDARG000 | 9;9;9;9;9;9;9;9 | 56443416;564 | 56443441;564 | +;+;+;+;+;+;+; | 6304 | NA | 1137 | 762 | 817 | 746 | 956 | 641 |
| ENSDARG000 | 9 | 25856613 | 25856729 | + | 117 | 5S_rRNA | 0 | 0 | 0 | 0 | 0 | 0 |
| ENSDARG000 | 9 | 44932175 | 44932293 | + | 119 | 5S_rRNA | 0 | 0 | 0 | 0 | 0 | 0 |
| ENSDARG000 | 9 | 45540477 | 45540593 | + | 117 | 5S_rRNA | 0 | 0 | 0 | 0 | 0 | 0 |
| ENSDARG000 | 9 | 23911891 | 23912005 | - | 115 | 5S_rRNA | 0 | 0 | 0 | 0 | 0 | 0 |
| ENSDARG000 | 9 | 29652354 | 29652463 | - | 110 | mir16c | 0 | 0 | 0 | 0 | 0 | 0 |
| ENSDARG000 | 9 | 50800357 | 50800473 | + | 117 | 5S_rRNA | 0 | 0 | 0 | 0 | 0 | 0 |
| ENSDARG000 | 9 | 44618159 | 44618275 | - | 117 | 5S_rRNA | 0 | 0 | 3 | 0 | 3 | 0 |
| ENSDARG000 | 9;9;9;9;9;9;9;9 | 190438;19108 | 190607;19117 | -;-;-;-;-;-;-;- | 1449 | NA | 6 | 6 | 16 | 8 | 4 | 10 |
| ENSDARG000 | 9 | 51882534 | 51884024 | + | 1491 | NA | 0 | 0 | 0 | 0 | 0 | 0 |
| ENSDARG000 | 9;9;9;9;9;9;9;9 | 584940;58494 | 585192;58519 | -;-;-;-;-;-;-;- | 4833 | DIP2A | 2940 | 1321 | 2937 | 1639 | 2601 | 1424 |
| ENSDARG000 | 9 | 30296675 | 30296791 | - | 117 | 5S_rRNA | 0 | 0 | 0 | 0 | 0 | 0 |
| ENSDARG000 | 9;9;9 | 33201906;332 | 33202028;332 | -;-;- | 198 | NA | 0 | 0 | 0 | 0 | 0 | 0 |
| ENSDARG000 | 9;9;9;9;9;9;9;9 | 56251000;562 | 56251473;562 | -;-;-;-;-;-;-;- | 2283 | lcp1 | 404 | 653 | 532 | 319 | 1358 | 1414 |
| ENSDARG000 | 9;9;9;9;9;9;9;9 | 54039006;540 | 54039556;540 | +;+;+;+;+;+;+; | 4011 | FRMPD4 | 147 | 214 | 196 | 72 | 68 | 160 |
| ENSDARG000 | 9;9;9;9;9;9;9;9 | 56103173;561 | 56103280;561 | +;+;+;+;+;+;+; | 1515 | edar | 0 | 2 | 6 | 2 | 4 | 2 |
| ENSDARG000 | 9;9;9;9;9;9;9;9 | 168442;16976 | 169465;16992 | -;-;-;-;-;-;-;- | 2599 | plbd2 | 1407 | 976 | 1241 | 1408 | 1463 | 1164 |
| ENSDARG000 | 9 | 29442493 | 29442609 | + | 117 | 5S_rRNA | 0 | 0 | 0 | 0 | 0 | 0 |
| ENSDARG000 | 9 | 24547879 | 24547996 | - | 118 | 5S_rRNA | 0 | 0 | 0 | 0 | 0 | 0 |
| ENSDARG000 | 9;9;9;9 | 53654866;536 | 53658203;536 | -;-;-;- | 6292 | pcdh8 | 0 | 2 | 0 | 2 | 0 | 8 |
| ENSDARG000 | 9 | 8078632 | 8078755 | + | 124 | 5S_rRNA | 0 | 0 | 0 | 0 | 0 | 0 |
| ENSDARG000 | 9 | 1644683 | 1644761 | - | 79 | dre-mir-148 | 0 | 0 | 0 | 0 | 0 | 0 |
| ENSDARG000 | 9 | 53325716 | 53325832 | - | 117 | 5S_rRNA | 0 | 0 | 0 | 0 | 0 | 0 |
| ENSDARG000 | 9 | 29518239 | 29518355 | + | 117 | 5S_rRNA | 0 | 0 | 0 | 0 | 0 | 0 |
| ENSDARG000 | 9 | 13448529 | 13448648 | + | 120 | 5S_rRNA | 0 | 0 | 0 | 0 | 0 | 0 |
| ENSDARG000 | 9 | 1748919 | 1749036 | - | 118 | 5S_rRNA | 0 | 0 | 0 | 0 | 0 | 0 |
| ENSDARG000 | 9 | 33921464 | 33921571 | - | 108 | dre-mir-222a | 0 | 0 | 0 | 0 | 0 | 0 |
| ENSDARG000 | 9 | 45821469 | 45821585 | + | 117 | 5S_rRNA | 0 | 0 | 0 | 0 | 0 | 0 |
| ENSDARG000 | 9;9;9;9;9;9;9;9 | 145093;14573 | 145216;14581 | +;+;+;+;+;+;+; | 2022 | NA | 492 | 660 | 958 | 366 | 777 | 813 |
| ENSDARG000 | 9 | 32525255 | 32525371 | - | 117 | 5S_rRNA | 0 | 0 | 0 | 0 | 0 | 0 |
| ENSDARG000 | 9;9 | 159720;16159 | 159951;16183 | +;+ | 474 | NA | 0 | 0 | 5 | 0 | 0 | 0 |
| ENSDARG000 | 9;9;9;9;9;9;9;9 | 53364472;533 | 53364668;533 | -;-;-;-;-;-;-;-;- | 1632 | gpc6b | 2 | 176 | 101 | 182 | 160 | 70 |
| ENSDARG000 | 9;9;9;9;9;9;9;9 | 46644633;466 | 46644769;466 | +;+;+;+;+;+;+; | 3300 | slc4a3 | 2 | 2 | 1 | 0 | 2 | 0 |
| ENSDARG000 | 9;9;9;9;9;9 | 51655636;516 | 51656633;516 | +;+;+;+;+;+ | 3303 | NA | 50 | 60 | 172 | 92 | 230 | 96 |
| ENSDARG000 | 9 | 14291932 | 14297132 | + | 5201 | NA | 0 | 0 | 0 | 0 | 0 | 0 |
| ENSDARG000 | 9 | 43331395 | 43331511 | + | 117 | 5S_rRNA | 0 | 0 | 0 | 0 | 0 | 0 |
| ENSDARG000 | 9 | 11551604 | 11551704 | - | 101 | dre-mir-1788 | 0 | 0 | 0 | 0 | 0 | 0 |
| ENSDARG000 | 9;9;9;9 | 5243476;5243 | 5243799;5243 | +;+;+;+ | 2410 | NA | 22 | 8 | 26 | 26 | 30 | 38 |
| ENSDARG000 | 9;9;9;9;9;9;9;9 | 10557144;106 | 10557161;106 | +;+;+;+;+;+;+; | 303 | NA | 1 | 0 | 0 | 0 | 0 | 0 |
| ENSDARG000 | 9 | 48632860 | 48632996 | - | 137 | 5S_rRNA | 0 | 0 | 0 | 0 | 0 | 0 |
| ENSDARG000 | 9 | 37411870 | 37411986 | + | 117 | 5S_rRNA | 0 | 0 | 0 | 0 | 0 | 0 |
| ENSDARG000 | 9 | 2845199 | 2845314 | + | 116 | 5S_rRNA | 0 | 0 | 0 | 0 | 0 | 0 |
| ENSDARG000 | 9 | 49938008 | 49938124 | - | 117 | 5S_rRNA | 0 | 0 | 0 | 0 | 0 | 0 |
| ENSDARG000 | 9 | 53436944 | 53437026 | - | 83 | dre-mir-17a-2 | 0 | 0 | 0 | 0 | 0 | 0 |
| ENSDARG000 | 9;9;9;9 | 4429593;4429 | 4429648;4429 | +;+;+;+ | 942 | NA | 706 | 642 | 743 | 1017 | 657 | 375 |
| ENSDARG000 | 9 | 33920932 | 33921024 | - | 93 | dre-mir-221 | 0 | 0 | 0 | 0 | 0 | 0 |
| ENSDARG000 | 9 | 11571973 | 11572055 | - | 83 | dre-mir-375-2 | 0 | 0 | 0 | 0 | 0 | 0 |
| ENSDARG000 | 9 | 6409157 | 6409271 | - | 115 | 5S_rRNA | 0 | 0 | 0 | 0 | 0 | 0 |
| ENSDARG000 | 9 | 26228449 | 26228563 | - | 115 | 5S_rRNA | 0 | 0 | 0 | 0 | 0 | 0 |
| ENSDARG000 | 9;9;9;9;9;9;9;9 | 10770880;107 | 10770919;107 | +;+;+;+;+;+;+; | 3142 | gmppaa | 139 | 180 | 147 | 144 | 172 | 142 |
| ENSDARG000 | 9;9;9;9 | 52492639;525 | 52493463;525 | +;+;+;+ | 1611 | marchf4 | 0 | 0 | 0 | 0 | 0 | 0 |
| ENSDARG000 | 9;9;9;9;9 | 54178475;541 | 54178518;541 | +;+;+;+;+ | 677 | tmsb4x | 18451 | 26848 | 25930 | 27794 | 23501 | 26776 |
| ENSDARG000 | 9;9;9;9;9;9;9;9 | 28103735;281 | 28104747;281 | +;+;+;+;+;+;+; | 4322 | idh1 | 2732 | 3099 | 2244 | 1201 | 3254 | 2136 |
| ENSDARG000 | 9;9;9;9;9;9 | 3397407;3397 | 3398441;3398 | -;-;-;-;-;- | 1704 | dlx2a | 4755 | 5442 | 5587 | 4696 | 8632 | 5743 |
| ENSDARG000 | 9;9;9;9;9;9;9;9 | 51891;52044; | 51938;52145; | +;+;+;+;+;+;+; | 1302 | hycc2 | 403 | 236 | 548 | 392 | 452 | 199 |
| ENSDARG000 | 9;9;9 | 3388099;3388 | 3388545;3388 | +;+;+ | 1214 | dlx1a | 18 | 56 | 2197 | 1428 | 3050 | 918 |
| ENSDARG000 | 9;9;9;9;9;9;9;9 | 265988;26880 | 267296;26889 | -;-;-;-;-;-;-;- | 4604 | kif5aa | 32 | 22 | 20 | 0 | 24 | 6 |
| ENSDARG000 | 9;9;9;9;9;9 | 55748544;557 | 55751093;557 | -;-;-;-;-;- | 3790 | akap17a | 211 | 228 | 396 | 335 | 220 | 240 |
| ENSDARG000 | 9;9;9;9;9;9;9;9 | 795609;79571 | 796127;79612 | -;-;-;-;-;-;-;- | 2064 | NA | 594 | 580 | 1030 | 557 | 3491 | 673 |
| ENSDARG000 | 9;9;9;9;9;9;9;9 | 56232548;562 | 56232727;562 | +;+;+;+;+;+;+; | 1872 | cnot11 | 392 | 416 | 643 | 340 | 476 | 233 |
| ENSDARG000 | 9 | 44122969 | 44123093 | - | 125 | 5S_rRNA | 0 | 0 | 0 | 0 | 0 | 0 |
| ENSDARG000 | 9 | 21432545 | 21432661 | - | 117 | 5S_rRNA | 0 | 0 | 4 | 0 | 0 | 0 |
| ENSDARG000 | 9 | 35701655 | 35701772 | - | 118 | 5S_rRNA | 0 | 0 | 0 | 0 | 0 | 0 |
| ENSDARG000 | 9 | 1003441 | 1003555 | - | 115 | 5S_rRNA | 0 | 0 | 0 | 0 | 0 | 0 |
| ENSDARG000 | 9 | 3981374 | 3981489 | + | 116 | 5S_rRNA | 0 | 0 | 0 | 0 | 0 | 0 |
| ENSDARG000 | 9 | 34753044 | 34753159 | - | 116 | 5S_rRNA | 0 | 0 | 0 | 0 | 0 | 0 |
| ENSDARG000 | 9;9;9;9;9;9;9 | 19052374;190 | 19052402;190 | -;-;-;-;-;-;- | 222 | NA | 3 | 2 | 7 | 10 | 12 | 4 |
| ENSDARG000 | 9;9;9;9;9;9;9 | 55154414;551 | 55154744;551 | +;+;+;+;+;+;+ | 2392 | ANOS1 | 4 | 4 | 0 | 0 | 0 | 0 |
| ENSDARG000 | 9 | 45064155 | 45064269 | + | 115 | 5S_rRNA | 0 | 0 | 0 | 0 | 0 | 0 |
| ENSDARG000 | 9 | 54885574 | 54885690 | - | 117 | 5S_rRNA | 0 | 0 | 0 | 0 | 0 | 0 |
| ENSDARG000 | 9 | 32785417 | 32785537 | - | 121 | 5S_rRNA | 0 | 0 | 0 | 0 | 0 | 0 |
| ENSDARG000 | 9;9;9;9;9;9;9;9 | 56336873;563 | 56337103;563 | -;-;-;-;-;-;-;- | 2907 | rab3gap1 | 1144 | 832 | 937 | 757 | 802 | 735 |
| ENSDARG000 | 9;9;9;9;9;9;9;9 | 52971097;529 | 52974294;529 | -;-;-;-;-;-;-;- | 5008 | NA | 48 | 140 | 171 | 61 | 236 | 164 |
| ENSDARG000 | 9;9;9;9;9;9;9;9 | 51995481;520 | 52001250;520 | -;-;-;-;-;-;-;- | 10460 | TANC1 | 98 | 22 | 63 | 60 | 127 | 24 |
| ENSDARG000 | 9;9;9;9;9;9;9;9 | 55321322;553 | 55321426;553 | +;+;+;+;+;+;+; | 3206 | nlgn4xb | 0 | 0 | 0 | 0 | 0 | 0 |
| ENSDARG000 | 9;9;9;9;9;9;9;9 | 1251514;1251 | 1251828;1251 | +;+;+;+;+;+;+; | 2177 | NA | 0 | 0 | 7 | 0 | 0 | 4 |
| ENSDARG000 | 9 | 18316403 | 18316520 | + | 118 | 5S_rRNA | 0 | 0 | 0 | 0 | 0 | 0 |
| ENSDARG000 | 9;9;9;9;9;9;9;9 | 925969;92630 | 926121;92634 | -;-;-;-;-;-;-;- | 1338 | sctr | 2 | 0 | 0 | 0 | 0 | 0 |
| ENSDARG000 | 9;9;9;9;9;9;9;9 | 3282369;3283 | 3282539;3284 | +;+;+;+;+;+;+; | 2123 | hat1 | 1560 | 2276 | 2751 | 1578 | 2512 | 1883 |
| ENSDARG000 | 9 | 46795447 | 46795567 | - | 121 | 5S_rRNA | 0 | 0 | 0 | 0 | 0 | 0 |
| ENSDARG000 | 9 | 40278147 | 40278263 | + | 117 | 5S_rRNA | 0 | 0 | 0 | 0 | 0 | 0 |
| ENSDARG000 | 9 | 53487060 | 53487175 | - | 116 | 5S_rRNA | 0 | 0 | 0 | 0 | 0 | 0 |
| ENSDARG000 | 9;9;9;9;9;9;9;9 | 16867458;168 | 16868531;168 | -;-;-;-;-;-;-;- | 2030 | fbxl3a | 321 | 299 | 354 | 411 | 232 | 347 |
| ENSDARG000 | 9;9;9;9;9;9;9;9 | 54984537;549 | 54985089;549 | +;+;+;+;+;+;+; | 2605 | mospd2 | 808 | 599 | 675 | 614 | 639 | 494 |
| ENSDARG000 | 9 | 24335163 | 24335279 | - | 117 | 5S_rRNA | 0 | 0 | 0 | 0 | 0 | 0 |
| ENSDARG000 | 9 | 6430416 | 6430530 | + | 115 | 5S_rRNA | 0 | 0 | 0 | 0 | 0 | 0 |
| ENSDARG000 | 9;9 | 54417141;544 | 54417194;544 | +;+ | 2042 | drd1b | 6 | 0 | 12 | 0 | 0 | 12 |
| ENSDARG000 | 9;9;9;9;9 | 23124252;231 | 23124312;231 | +;+;+;+;+ | 159 | NA | 0 | 0 | 0 | 0 | 0 | 0 |
| ENSDARG000 | 9;9;9;9;9;9;9;9 | 4024250;4026 | 4024994;4026 | -;-;-;-;-;-;-;- | 3464 | MYLK | 54 | 16 | 22 | 32 | 78 | 54 |
| ENSDARG000 | 9 | 53637932 | 53646223 | + | 8292 | NA | 0 | 0 | 0 | 0 | 0 | 0 |
| ENSDARG000 | 9 | 1379376 | 1379490 | - | 115 | 5S_rRNA | 6 | 2 | 0 | 0 | 4 | 1 |
| ENSDARG000 | 9 | 31993311 | 31993433 | + | 123 | 5S_rRNA | 0 | 0 | 0 | 0 | 0 | 0 |
| ENSDARG000 | 9 | 12828005 | 12828120 | + | 116 | 5S_rRNA | 0 | 0 | 0 | 0 | 0 | 0 |
| ENSDARG000 | 9 | 30707839 | 30707954 | - | 116 | 5S_rRNA | 0 | 0 | 0 | 0 | 0 | 0 |
| ENSDARG000 | 9 | 43775251 | 43775365 | + | 115 | 5S_rRNA | 0 | 0 | 0 | 0 | 0 | 0 |
| ENSDARG000 | 9 | 23046267 | 23046385 | + | 119 | 5S_rRNA | 0 | 0 | 0 | 0 | 0 | 0 |
| ENSDARG000 | 9 | 37877434 | 37877550 | + | 117 | 5S_rRNA | 0 | 0 | 0 | 0 | 0 | 0 |
| ENSDARG000 | 9;9;9;9 | 54282494;542 | 54282500;542 | -;-;-;- | 462 | NA | 19 | 14 | 15 | 29 | 8 | 32 |
| ENSDARG000 | 9;9;9;9;9;9;9;9 | 19326099;193 | 19326109;193 | +;+;+;+;+;+;+; | 384 | NA | 0 | 1 | 0 | 0 | 0 | 0 |
| ENSDARG000 | 9 | 11447777 | 11447909 | + | 133 | 5S_rRNA | 0 | 0 | 0 | 0 | 0 | 0 |
| ENSDARG000 | 9;9;9;9;9;9 | 407678;40897 | 408406;40926 | -;-;-;-;-;- | 1644 | NA | 9 | 2 | 17 | 33 | 39 | 14 |
| ENSDARG000 | 9;9;9;9;9;9;9 | 9009123;9009 | 9009590;9009 | -;-;-;-;-;-;- | 1243 | NA | 0 | 0 | 0 | 0 | 0 | 0 |
| ENSDARG000 | 9;9;9;9;9 | 9355372;9359 | 9357087;9360 | -;-;-;-;- | 2377 | NA | 0 | 0 | 0 | 0 | 0 | 4 |
| ENSDARG000 | 9;9;9;9;9;9 | 13316919;133 | 13317206;133 | -;-;-;-;-;- | 1630 | NA | 0 | 0 | 0 | 0 | 0 | 0 |
| ENSDARG000 | 9;9;9 | 26604063;267 | 26604475;267 | +;+;+ | 734 | NA | 0 | 0 | 0 | 0 | 0 | 0 |
| ENSDARG000 | 9;9;9;9;9;9;9 | 26896785;268 | 26896952;268 | +;+;+;+;+;+;+ | 5953 | NA | 0 | 2 | 2 | 0 | 0 | 0 |
| ENSDARG000 | 9;9;9 | 30695283;306 | 30695503;306 | +;+;+ | 1879 | NA | 0 | 0 | 10 | 0 | 0 | 0 |
| ENSDARG000 | 9;9;9;9;9;9;9;9 | 34701942;347 | 34702009;347 | -;-;-;-;-;-;-;- | 441 | NA | 0 | 0 | 0 | 0 | 0 | 0 |
| ENSDARG000 | 9;9;9;9;9 | 34822849;348 | 34823004;348 | +;+;+;+;+ | 1455 | NA | 0 | 0 | 0 | 0 | 0 | 0 |
| ENSDARG000 | 9;9;9;9 | 36937694;369 | 36937763;369 | -;-;-;- | 376 | NA | 4 | 0 | 0 | 2 | 0 | 0 |
| ENSDARG000 | 9;9;9 | 40388789;404 | 40388998;404 | -;-;- | 433 | NA | 0 | 0 | 0 | 0 | 0 | 0 |
| ENSDARG000 | 9;9;9;9;9;9 | 40767055;407 | 40768089;407 | -;-;-;-;-;- | 1587 | NA | 6 | 0 | 0 | 0 | 5 | 0 |
| ENSDARG000 | 9;9;9;9;9;9 | 41509948;415 | 41510070;415 | +;+;+;+;+;+ | 1462 | NA | 0 | 0 | 19 | 0 | 0 | 1 |
| ENSDARG000 | 9;9;9;9 | 41788217;417 | 41788440;417 | +;+;+;+ | 1358 | NA | 0 | 0 | 0 | 0 | 0 | 0 |
| ENSDARG000 | 9;9;9;9;9;9;9 | 46489846;465 | 46490227;465 | +;+;+;+;+;+;+ | 2559 | NA | 0 | 0 | 0 | 0 | 0 | 0 |
| ENSDARG000 | 9;9;9;9;9;9;9 | 49869692;498 | 49869834;498 | +;+;+;+;+;+;+ | 1218 | NA | 0 | 0 | 0 | 0 | 0 | 0 |
| ENSDARG000 | 9;9;9 | 51043455;510 | 51043662;510 | +;+;+ | 708 | NA | 0 | 0 | 0 | 0 | 0 | 0 |
| ENSDARG000 | 9;9;9;9;9 | 53436297;534 | 53436353;534 | -;-;-;-;- | 726 | NA | 0 | 0 | 0 | 0 | 0 | 0 |
| ENSDARG000 | 9;9;9;9;9;9;9 | 55666315;556 | 55666422;556 | +;+;+;+;+;+;+ | 1965 | NA | 53 | 18 | 53 | 53 | 67 | 31 |
| ENSDARG000 | 16;16;16;16;1 | 22022533;220 | 22022571;220 | +;+;+;+;+;+;+ | 550 | NA | 0 | 0 | 0 | 0 | 0 | 0 |
| ENSDARG000 | 16;16 | 45087810;450 | 45088222;450 | -;- | 772 | NA | 6 | 0 | 0 | 32 | 0 | 15 |
| ENSDARG000 | 16;16;16 | 22061003;220 | 22061136;220 | -;-;- | 375 | NA | 0 | 0 | 0 | 0 | 0 | 0 |
| ENSDARG000 | 16;16;16;16;1 | 7242610;7248 | 7243019;7249 | +;+;+;+;+;+;+; | 1577 | sri | 1291 | 1323 | 1184 | 1317 | 1344 | 1402 |
| ENSDARG000 | 16;16;16 | 7227584;7293 | 7227687;7293 | +;+;+ | 663 | NA | 0 | 0 | 0 | 0 | 0 | 0 |
| ENSDARG000 | 16;16 | 7226676;7231 | 7227236;7231 | +;+ | 907 | NA | 0 | 0 | 2 | 0 | 3 | 2 |
| ENSDARG000 | 16;16;16;16;1 | 7214005;7216 | 7217533;7217 | -;-;-;-;-;-;-;- | 4750 | nt5c3a | 467 | 598 | 564 | 403 | 680 | 403 |
| ENSDARG000 | 16;16 | 42471455;424 | 42471528;424 | +;+ | 1217 | NA | 56 | 20 | 108 | 58 | 52 | 18 |
| ENSDARG000 | 16;16;16;16;1 | 5196226;5199 | 5196849;5199 | +;+;+;+;+;+;+; | 4816 | si:ch211-151 | 0 | 4 | 3 | 2 | 1 | 7 |
| ENSDARG000 | 16;16;16;16;1 | 7202082;7208 | 7203643;7208 | -;-;-;-;-;-;- | 2824 | NA | 110 | 59 | 72 | 74 | 77 | 88 |
| ENSDARG000 | 16;16;16;16;1 | 5184402;5187 | 5184508;5188 | +;+;+;+;+;+;+; | 2579 | SOGA3 | 0 | 0 | 0 | 0 | 0 | 0 |
| ENSDARG000 | 16;16;16;16;1 | 5184402;5187 | 5184508;5188 | +;+;+;+;+;+;+; | 6035 | SOGA3 | 0 | 0 | 0 | 0 | 0 | 0 |
| ENSDARG000 | 16;16;16;16;1 | 9433738;9437 | 9434293;9437 | -;-;-;-;-;-;-;- | 3093 | prpf3 | 567 | 641 | 709 | 323 | 592 | 460 |
| ENSDARG000 | 16;16;16;16;1 | 35535171;355 | 35535368;355 | +;+;+;+;+ | 1819 | cited4b | 5128 | 3932 | 5101 | 5189 | 5380 | 6056 |
| ENSDARG000 | 16;16 | 45104613;451 | 45104949;451 | -;- | 619 | NA | 31 | 0 | 8 | 66 | 4 | 13 |
| ENSDARG000 | 16;16 | 45130921;451 | 45131013;451 | -;- | 423 | NA | 0 | 0 | 0 | 0 | 0 | 0 |
| ENSDARG000 | 16;16 | 45140858;451 | 45141320;451 | -;- | 792 | NA | 6 | 28 | 0 | 4 | 14 | 43 |
| ENSDARG000 | 16;16;16;16;1 | 26132951;261 | 26133021;261 | -;-;-;-;- | 687 | cd79a | 4 | 1 | 6 | 18 | 4 | 5 |
| ENSDARG000 | 16;16;16 | 42477155;424 | 42477555;424 | +;+;+ | 642 | NA | 0 | 0 | 0 | 0 | 0 | 0 |
| ENSDARG000 | 16;16;16;16;1 | 29555801;295 | 29555856;295 | +;+;+;+;+;+;+ | 711 | ensab | 161 | 107 | 130 | 73 | 141 | 93 |
| ENSDARG000 | 16;16;16;16;1 | 378766;37980 | 379853;37985 | -;-;-;-;-;-;-;- | 2302 | irx4a | 0 | 0 | 0 | 0 | 0 | 0 |
| ENSDARG000 | 16;16;16;16;1 | 26140895;261 | 26143720;261 | -;-;-;-;-;-;-;- | 5965 | arhgef1b | 2715 | 1334 | 2790 | 2015 | 1522 | 2719 |
| ENSDARG000 | 16;16;16;16;1 | 29551023;295 | 29551490;295 | -;-;-;-;-;-;-;- | 1664 | hormad1 | 2 | 0 | 0 | 4 | 12 | 0 |
| ENSDARG000 | 16;16;16;16;1 | 11029762;110 | 11030301;110 | +;+;+;+;+ | 7377 | ERF | 520 | 592 | 626 | 354 | 592 | 532 |
| ENSDARG000 | 16;16;16;16;1 | 14093781;140 | 14095777;140 | -;-;-;-;-;-;-;- | 8354 | NA | 220 | 64 | 224 | 246 | 141 | 172 |
| ENSDARG000 | 16;16;16;16;1 | 38556836;385 | 38557024;385 | -;-;-;-;-;-;-;- | 1787 | eif3ea | 5802 | 5762 | 8977 | 7429 | 6429 | 7870 |
| ENSDARG000 | 16;16;16;16;1 | 12474861;124 | 12475816;124 | -;-;-;-;-;-;-;- | 4303 | NA | 283 | 1266 | 4632 | 2290 | 671 | 691 |
| ENSDARG000 | 16;16;16;16;1 | 9495583;9495 | 9495849;9495 | +;+;+;+;+;+;+; | 1485 | NA | 6 | 0 | 0 | 0 | 7 | 0 |

| | | | | | | | | | | | | |
|---|---|---|---|---|---|---|---|---|---|---|---|---|
| ENSDARG000 | 16;16 | 45149355:451 | 45149899:451 | -;- | 927 | si:ch73-343g1 | 0 | 12 | 2 | 0 | 2 | 2 |
| ENSDARG000 | 16;16;16;16;1 | 26081481:260 | 26081974:260 | -;-;-;-;-;-;-;-; | 2735 | NA | 681 | 555 | 734 | 314 | 1020 | 1071 |
| ENSDARG000 | 16;16;16;16;1 | 22236961:222 | 22237249:222 | -;-;-;-;-;-;-;-; | 1311 | NA | 0 | 0 | 0 | 0 | 2 | 0 |
| ENSDARG000 | 16;16;16;16;1 | 14201401:142 | 14201605:142 | +;+;+;+;+;+;+;+; | 1867 | dap3 | 420 | 514 | 809 | 361 | 524 | 348 |
| ENSDARG000 | 16;16;16;16;1 | 42481447:424 | 42481543:424 | +;+;+;+;+;+;+;+; | 2987 | herpud2 | 684 | 766 | 685 | 686 | 584 | 595 |
| ENSDARG000 | 16;16;16;16;1 | 16968682:169 | 16968758:169 | +;+;+;+;+;+;+;+; | 1920 | si:ch211-120k | 2 | 0 | 4 | 1 | 0 | 0 |
| ENSDARG000 | 16;16 | 45172793:451 | 45173324:451 | -;- | 958 | si:dkev-33i11. | 24 | 33 | 36 | 20 | 118 | 20 |
| ENSDARG000 | 16;16;16;16 | 27561114:275 | 27563816:275 | -;-;-;- | 2974 | rbm12ba | 367 | 420 | 624 | 441 | 504 | 808 |
| ENSDARG000 | 16;16;16;16;1 | 555746:55868 | 560946:55946 | -;-;-;-;-;-;-;-; | 6850 | irx2a | 0 | 0 | 0 | 0 | 12 | 2 |
| ENSDARG000 | 16;16;16;16;1 | 27564270:275 | 27564490:275 | +;+;+;+;+;+;+;+; | 3274 | TMEM67 | 418 | 356 | 445 | 435 | 308 | 528 |
| ENSDARG000 | 16;16;16;16;1 | 16969086:169 | 16969088:169 | +;+;+;+;+;+;+;+; | 745 | rpl18 | 15107 | 14382 | 23423 | 24252 | 16088 | 23875 |
| ENSDARG000 | 16;16 | 9526107:9527 | 9526269:9527 | +;+ | 344 | NA | 0 | 0 | 0 | 0 | 2 | 0 |
| ENSDARG000 | 16;16;16;16;1 | 42509011:425 | 42509944:425 | -;-;-;-;-;-;-;- | 2126 | tbx20 | 0 | 0 | 0 | 0 | 0 | 0 |
| ENSDARG000 | 16;16;16;16;1 | 52926139:529 | 52926539:529 | -;-;-;-;-;-;-;-; | 2339 | brd9 | 494 | 495 | 641 | 384 | 643 | 596 |
| ENSDARG000 | 16;16;16;16;1 | 24646600:246 | 24649424:246 | -;-;-;-;-;-;-;-;- | 3791 | pon1 | 180 | 110 | 86 | 224 | 107 | 173 |
| ENSDARG000 | 16;16;16;16;1 | 22288777:222 | 22289455:222 | -;-;-;-;-;- | 1508 | aqp10a | 0 | 0 | 2 | 0 | 0 | 0 |
| ENSDARG000 | 16;16 | 45186673:451 | 45187292:451 | -;- | 1023 | NA | 0 | 0 | 0 | 0 | 0 | 0 |
| ENSDARG000 | 16;16;16;16;1 | 52966812:529 | 52966892:529 | +;+;+;+;+;+;+;+; | 1755 | trip13 | 30 | 246 | 231 | 96 | 142 | 85 |
| ENSDARG000 | 16;16;16;16;1 | 9540033:9540 | 9540252:9540 | +;+;+;+;+;+;+;+; | 1324 | ca14 | 0 | 0 | 0 | 0 | 0 | 0 |
| ENSDARG000 | 16;16;16;16;1 | 35555029:355 | 35557035:355 | -;-;-;-;-;-;-;-; | 5672 | scmh1 | 393 | 693 | 855 | 424 | 962 | 716 |
| ENSDARG000 | 16;16;16;16;1 | 11151699:111 | 11151972:111 | +;+;+;+;+;+;+;+; | 11698 | NA | 1400 | 969 | 1270 | 781 | 1358 | 1101 |
| ENSDARG000 | 16;16;16;16;1 | 29586004:295 | 29586446:295 | +;+;+;+;+ | 1838 | mcl1b | 4329 | 4425 | 4721 | 6182 | 4267 | 3547 |
| ENSDARG000 | 16;16;16;16;1 | 1334431:1338 | 1334609:1338 | +;+;+;+;+;+;+;+; | 1419 | CELF3 | 38 | 40 | 132 | 44 | 67 | 105 |
| ENSDARG000 | 16;16;16;16;1 | 32676517:326 | 32677107:326 | -;-;-;-;-;-;-;-; | 3164 | usp45 | 323 | 297 | 379 | 190 | 353 | 325 |
| ENSDARG000 | 16;16;16;16;1 | 22295662:222 | 22295932:222 | -;-;-;-;-;- | 1098 | NA | 22 | 6 | 39 | 73 | 13 | 121 |
| ENSDARG000 | 16;16;16;16;1 | 1362250:1383 | 1362551:1384 | +;+;+;+;+;+;+;+; | 5984 | CERS2 | 3258 | 3820 | 4926 | 3912 | 3494 | 3399 |
| ENSDARG000 | 16;16;16;16;1 | 41495670:414 | 41496243:414 | -;-;-;-;- | 1181 | cmtm7 | 1729 | 1780 | 2399 | 2370 | 2430 | 2733 |
| ENSDARG000 | 16;16;16;16;1 | 29630965:296 | 29631081:296 | +;+;+;+;+;+;+;+; | 1304 | VPS72 | 112 | 67 | 117 | 696 | 62 | 141 |
| ENSDARG000 | 16;16;16;16;1 | 31601181:316 | 31602202:316 | -;-;-;-;-;-;-;-; | 4556 | NA | 677 | 526 | 724 | 416 | 777 | 539 |
| ENSDARG000 | 16;16;16;16;1 | 22345513:223 | 22345731:223 | +;+;+;+;+;+;+;+ | 982 | C16H1orf43 | 178 | 215 | 202 | 235 | 235 | 229 |
| ENSDARG000 | 16;16;16;16;1 | 14216581:142 | 14216838:142 | +;+;+;+;+;+;+;+; | 4227 | GBA1 | 2334 | 1298 | 1188 | 1339 | 2325 | 2918 |
| ENSDARG000 | 16;16;16;16;1 | 22363700:223 | 22365796:223 | -;-;-;-;-;-;-;-; | 3237 | tuft1a | 1673 | 362 | 526 | 272 | 201 | 511 |
| ENSDARG000 | 16;16;16;16;1 | 5251768:5252 | 5251804:5253 | +;+;+;+;+;+;+;+; | 16427 | NA | 2003 | 881 | 1429 | 916 | 1099 | 1642 |
| ENSDARG000 | 16;16;16;16;1 | 11227246:112 | 11228107:112 | -;-;-;-;-;-;-;-; | 4627 | NA | 204 | 244 | 242 | 222 | 314 | 260 |
| ENSDARG000 | 16;16;16;16;1 | 45249055:452 | 45253090:452 | -;-;-;-;-;-;-;-; | 5510 | hpn | 26 | 0 | 8 | 0 | 6 | 4 |
| ENSDARG000 | 16;16;16;16;1 | 24656958:246 | 24657162:246 | -;-;-;-;-;-;-;-; | 1250 | pon3.2 | 310 | 222 | 182 | 308 | 343 | 526 |
| ENSDARG000 | 16;16;16;16;1 | 45202602:452 | 45203248:452 | -;-;-;-;-;-;-;-; | 1214 | fxyd3 | 3922 | 2990 | 3097 | 4185 | 2843 | 5185 |
| ENSDARG000 | 16;16;16 | 45242381:452 | 45242477:452 | -;-;- | 468 | NA | 0 | 0 | 0 | 0 | 0 | 0 |
| ENSDARG000 | 16;16;16;16;1 | 38659475:386 | 38659589:386 | +;+;+;+;+;+;+;+; | 1475 | emc2 | 1276 | 1222 | 1585 | 1186 | 1329 | 1090 |
| ENSDARG000 | 16;16;16;16;1 | 45228101:452 | 45228508:452 | -;-;-;-;-;-;-;-; | 875 | fxyd11 | 0 | 0 | 0 | 0 | 0 | 0 |
| ENSDARG000 | 16;16;16;16;1 | 19637384:196 | 19637566:196 | +;+;+;+;+;+;+;+; | 6194 | macc1 | 98 | 12 | 28 | 28 | 30 | 10 |
| ENSDARG000 | 16;16 | 45203090:452 | 45203216:452 | +;+ | 392 | NA | 5 | 2 | 2 | 0 | 5 | 0 |
| ENSDARG000 | 16;16;16;16 | 45217307:452 | 45217356:452 | +;+;+;+ | 539 | NA | 0 | 4 | 0 | 3 | 9 | 4 |
| ENSDARG000 | 16;16;16;16;1 | 41505255:415 | 41505401:415 | -;-;-;-;-;-;-;-; | 1243 | siglec15l | 0 | 0 | 0 | 0 | 0 | 0 |
| ENSDARG000 | 16;16;16;16;1 | 12504929:125 | 12505508:125 | -;-;-;-;-;-;-;-; | 1583 | cox6b2 | 2382 | 2701 | 2846 | 2407 | 2556 | 2914 |
| ENSDARG000 | 16;16;16;16;1 | 22478161:224 | 22478511:224 | -;-;-;-;- | 678 | NA | 0 | 2 | 0 | 0 | 0 | 2 |
| ENSDARG000 | 16;16;16;16;1 | 14231406:142 | 14232294:142 | -;-;-;-;-;-;-;-;- | 1769 | rnf115a | 766 | 571 | 668 | 548 | 680 | 546 |
| ENSDARG000 | 16;16;16;16;1 | 48282321:482 | 48282571:482 | -;-;-;-;-;-;-;- | 1220 | eif3ha | 2036 | 1866 | 3470 | 2075 | 2152 | 2499 |
| ENSDARG000 | 16;16;16;16;1 | 1465624:1470 | 1471104:1471 | -;-;-;-;-;-;-;-; | 7449 | sim1a | 52 | 36 | 84 | 62 | 112 | 56 |
| ENSDARG000 | 16;16;16 | 41517188:415 | 41517257:415 | +;+;+ | 1062 | si:dkev-11p23 | 0 | 16 | 14 | 0 | 0 | 26 |
| ENSDARG000 | 16;16;16;16;1 | 42667560:426 | 42667884:426 | +;+;+;+;+;+;+;+; | 2339 | dpy19l1l | 240 | 191 | 333 | 148 | 428 | 217 |
| ENSDARG000 | 16;16 | 41524790:415 | 41525217:415 | -;- | 517 | NA | 2 | 0 | 0 | 0 | 2 | 0 |
| ENSDARG000 | 16;16;16 | 19677484:196 | 19677557:196 | +;+;+ | 521 | NA | 0 | 2 | 0 | 0 | 2 | 0 |
| ENSDARG000 | 16;16;16;16;1 | 24662252:246 | 24662458:246 | -;-;-;-;-;-;-;-;- | 1132 | pon3.1 | 219 | 204 | 118 | 396 | 220 | 294 |
| ENSDARG000 | 16;16;16;16 | 32729223:327 | 32729495:327 | +;+;+;+ | 826 | tstd3 | 244 | 252 | 196 | 225 | 186 | 267 |
| ENSDARG000 | 16;16;16;16;1 | 32736588:327 | 32736821:327 | +;+;+;+;+;+;+;+; | 3344 | zgc:172323 | 0 | 0 | 10 | 6 | 4 | 0 |
| ENSDARG000 | 16;16;16;16;1 | 17007089:170 | 17007716:170 | -;-;-;-;-;-;-;-; | 3643 | NA | 10 | 0 | 0 | 10 | 2 | 2 |
| ENSDARG000 | 16;16;16;16 | 22555855:225 | 22556260:225 | -;-;-;- | 808 | NA | 0 | 0 | 0 | 0 | 0 | 0 |
| ENSDARG000 | 16;16;16;16;1 | 22520712:225 | 22525341:225 | -;-;-;-;-;-;-;-; | 8499 | NA | 23 | 4 | 16 | 2 | 10 | 10 |
| ENSDARG000 | 16;16;16;16 | 41532616:415 | 41533923:415 | -;-;-;- | 1570 | rpp25l | 64 | 170 | 264 | 80 | 244 | 132 |
| ENSDARG000 | 16;16;16;16;1 | 29650698:296 | 29650992:296 | +;+;+;+;+;+;+;+; | 2382 | tmod4 | 14 | 12 | 20 | 14 | 34 | 22 |
| ENSDARG000 | 16;16;16;16 | 19732543:197 | 19733282:197 | +;+;+;+ | 1215 | twist1b | 6 | 2 | 18 | 24 | 2 | 0 |
| ENSDARG000 | 16;16;16;16;1 | 11242443:112 | 11242750:112 | +;+;+;+;+;+;+;+; | 3332 | gsk3ab | 2239 | 2320 | 2652 | 1906 | 2452 | 2039 |
| ENSDARG000 | 16;16;16;16;1 | 17060758:170 | 17063549:170 | -;-;-;-;-;-;-;-; | 4373 | tnfrsf1a | 1076 | 1211 | 1345 | 1268 | 1169 | 1158 |
| ENSDARG000 | 16;16;16;16 | 9564503:9565 | 9564910:9565 | -;-;-;- | 865 | NA | 0 | 0 | 0 | 0 | 0 | 0 |
| ENSDARG000 | 16;16;16;16;1 | 38818864:388 | 38818881:388 | +;+;+;+;+;+;+;+; | 1452 | TRHR | 0 | 0 | 0 | 0 | 2 | 0 |
| ENSDARG000 | 16;16;16;16;1 | 1504280:1508 | 1504402:1509 | -;-;-;-;-;-;-;-; | 6086 | ASCC3 | 1072 | 848 | 1264 | 453 | 1362 | 632 |
| ENSDARG000 | 16;16 | 38920128:389 | 38920833:389 | -;- | 792 | NA | 0 | 0 | 0 | 0 | 0 | 0 |
| ENSDARG000 | 16;16;16;16;1 | 24669883:246 | 24670097:246 | -;-;-;-;-;-;-;-; | 1336 | pon2 | 247 | 232 | 309 | 246 | 518 | 324 |
| ENSDARG000 | 16;16;16;16;1 | 14249546:142 | 14249622:142 | +;+;+;+;+;+;+;+; | 2027 | polr3c | 245 | 118 | 198 | 66 | 194 | 166 |
| ENSDARG000 | 16;16;16;16;1 | 14276364:142 | 14281204:142 | -;-;-;-;-;-;-;-; | 8704 | ITGA10 | 22 | 8 | 0 | 0 | 22 | 37 |
| ENSDARG000 | 16;16;16;16;1 | 48409066:484 | 48409775:484 | -;-;-;-;-;-;-;-; | 3111 | rad21a | 4102 | 4203 | 6039 | 3579 | 4598 | 3392 |
| ENSDARG000 | 16;16;16;16;1 | 2637478:2637 | 2637823:2637 | -;-;-;-;-;-;-;-; | 1413 | lyplal1 | 76 | 110 | 111 | 106 | 104 | 116 |
| ENSDARG000 | 16;16;16;16;1 | 35593785:355 | 35595059:355 | +;+;+;+;+;+;+;+; | 5015 | NA | 50 | 57 | 75 | 42 | 87 | 54 |
| ENSDARG000 | 16;16 | 41542079:415 | 41542662:415 | -;- | 643 | NA | 0 | 0 | 0 | 0 | 0 | 0 |
| ENSDARG000 | 16;16 | 41546763:415 | 41547093:415 | +;+ | 459 | NA | 0 | 0 | 0 | 0 | 0 | 0 |
| ENSDARG000 | 16;16;16;16;1 | 19840146:198 | 19843181:198 | -;-;-;-;-;-;-;-; | 6955 | hdac9b | 1802 | 1506 | 1101 | 958 | 1271 | 1118 |
| ENSDARG000 | 16;16;16;16 | 32749591:327 | 32749932:327 | +;+;+;+ | 2671 | prdm13 | 0 | 0 | 0 | 0 | 4 | 0 |
| ENSDARG000 | 16;16;16;16;1 | 22587661:225 | 22588641:225 | +;+;+;+;+;+;+;+; | 1994 | NA | 74 | 9 | 16 | 10 | 8 | 43 |
| ENSDARG000 | 16;16;16;16;1 | 17089114:170 | 17089731:170 | +;+;+;+;+;+;+;+ | 1739 | NA | 6 | 8 | 4 | 38 | 16 | 54 |
| ENSDARG000 | 16;16;16;16;1 | 41570653:415 | 41570681:415 | +;+;+;+;+ | 2708 | aste1a | 514 | 6 | 52 | 57 | 0 | 0 |
| ENSDARG000 | 16;16 | 27593698:276 | 27593762:276 | +;+ | 2249 | NA | 14 | 172 | 90 | 54 | 64 | 146 |
| ENSDARG000 | 16;16;16;16;1 | 27595562:275 | 27596479:275 | -;-;-;-;-;-;-;-; | 2172 | NA | 2 | 4 | 8 | 2 | 4 | 0 |
| ENSDARG000 | 16;16;16;16;1 | 31623975:316 | 31626954:316 | -;-;-;-;-;-;-;-; | 3977 | NA | 1964 | 134 | 36 | 43 | 14 | 182 |
| ENSDARG000 | 16;16;16;16;1 | 17102103:171 | 17103172:171 | -;-;-;-;-;-;-;-;- | 2446 | NA | 14 | 23 | 0 | 0 | 22 | 44 |
| ENSDARG000 | 16;16;16;16;1 | 12517535:125 | 12517797:125 | +;+;+;+;+;+;+;+; | 3613 | rasip1 | 91 | 10 | 22 | 6 | 22 | 58 |
| ENSDARG000 | 16;16;16;16;1 | 11297703:113 | 11297775:113 | +;+;+;+;+;+;+;+ | 3102 | znf574 | 202 | 243 | 255 | 104 | 228 | 181 |
| ENSDARG000 | 16;16;16;16;1 | 2745006:2746 | 2745164:2747 | -;-;-;-;-;-;- | 1242 | tgfb2 | 24 | 8 | 8 | 0 | 6 | 6 |
| ENSDARG000 | 16;16 | 12524732:125 | 12525322:125 | -;- | 829 | NA | 2 | 0 | 0 | 0 | 0 | 2 |
| ENSDARG000 | 16;16;16;16;1 | 32820327:328 | 32823995:328 | -;-;-;-;-;-;-;- | 4346 | MCHR2 | 0 | 0 | 0 | 0 | 0 | 0 |
| ENSDARG000 | 16;16;16;16;1 | 22618620:226 | 22618665:226 | +;+;+;+;+;+;+;+; | 7101 | CHRNB2 | 14 | 2 | 4 | 0 | 3 | 0 |
| ENSDARG000 | 16;16;16;16;1 | 32832213:328 | 32832422:328 | -;-;-;-;-;-;-;-; | 2475 | me1 | 14 | 6 | 36 | 6 | 40 | 12 |
| ENSDARG000 | 16;16;16;16;1 | 26234059:262 | 26237775:262 | -;-;-;-;-;-;-;-; | 4821 | NA | 0 | 4 | 12 | 0 | 10 | 2 |
| ENSDARG000 | 16;16;16;16;1 | 9580699:9580 | 9580876:9581 | +;+;+;+;+;+;+;+; | 4401 | taf2 | 481 | 538 | 728 | 322 | 734 | 583 |
| ENSDARG000 | 16;16 | 32995882:330 | 32996073:330 | +;+ | 2329 | prss35 | 82 | 0 | 6 | 0 | 18 | 8 |
| ENSDARG000 | 16;16;16;16;1 | 11483811:114 | 11483975:114 | +;+;+;+;+;+;+;+; | 3924 | grik5 | 0 | 2 | 2 | 6 | 12 | 14 |
| ENSDARG000 | 16;16;16 | 2814233:2815 | 2815331:2815 | -;-;- | 1837 | acot22 | 0 | 0 | 0 | 0 | 0 | 0 |
| ENSDARG000 | 16;16;16;16;1 | 24681177:246 | 24681315:246 | +;+;+;+;+;+;+;+; | 1885 | ywhabl | 10108 | 11821 | 11104 | 11399 | 9081 | 9476 |
| ENSDARG000 | 16;16;16 | 11437869:115 | 11438276:115 | -;-;- | 661 | NA | 0 | 0 | 0 | 0 | 0 | 0 |
| ENSDARG000 | 16;16;16;16;1 | 38940758:389 | 38940823:389 | +;+;+;+;+;+;+;+; | 802 | eny2 | 218 | 351 | 280 | 122 | 262 | 182 |
| ENSDARG000 | 16;16;16;16;1 | 22681818:226 | 22682320:226 | -;-;-;-;- | 1190 | s100t | 8 | 4 | 2 | 0 | 8 | 2 |
| ENSDARG000 | 16;16;16;16;1 | 31646390:316 | 31646410:316 | -;-;-;-;-;- | 651 | NA | 0 | 0 | 0 | 0 | 0 | 2 |
| ENSDARG000 | 16;16;16;16;1 | 48460873:485 | 48462355:485 | +;+;+;+;+;+;+;+; | 3820 | ext1a | 146 | 134 | 156 | 152 | 185 | 166 |
| ENSDARG000 | 16;16 | 7315144:7361 | 7317006:7362 | -;- | 2671 | NA | 0 | 0 | 0 | 0 | 0 | 0 |
| ENSDARG000 | 16;16;16;16;1 | 52999778:530 | 53000004:530 | +;+;+;+;+;+;+;+; | 1432 | nkd2b | 14 | 2 | 4 | 16 | 6 | 8 |
| ENSDARG000 | 16;16;16;16;1 | 17121986:171 | 17122236:171 | -;-;-;-;-;-;-;-; | 2714 | IFFO1 | 12 | 4 | 10 | 3 | 60 | 48 |
| ENSDARG000 | 16;16;16;16;1 | 31720042:317 | 31720444:317 | -;-;-;-;-;-;-;-; | 3214 | ptpn6 | 308 | 124 | 151 | 92 | 399 | 357 |
| ENSDARG000 | 16;16;16;16;1 | 2820340:2820 | 2820676:2820 | +;+;+;+;+;+;+;+; | 1331 | FEZ2 | 143 | 65 | 75 | 152 | 103 | 155 |
| ENSDARG000 | 16;16;16;16;1 | 22695211:226 | 22697076:226 | -;-;-;-;-;-;- | 2115 | si:ch211-105c | 404 | 12 | 22 | 22 | 31 | 70 |
| ENSDARG000 | 16;16 | 26378297:263 | 26378801:263 | -;- | 764 | NA | 36 | 12 | 70 | 15 | 30 | 40 |
| ENSDARG000 | 16;16 | 7376471:7378 | 7377171:7379 | -;- | 1093 | med18 | 114 | 118 | 148 | 60 | 136 | 84 |
| ENSDARG000 | 16;16;16;16;1 | 48616220:486 | 48616556:486 | +;+;+;+;+;+;+;+; | 3732 | pbx2 | 1089 | 616 | 1008 | 754 | 1107 | 945 |
| ENSDARG000 | 16;16;16;16 | 39000605:390 | 39000788:390 | -;-;-;- | 468 | NA | 2 | 0 | 4 | 0 | 10 | 0 |
| ENSDARG000 | 16;16;16;16;1 | 20012190:200 | 20012280:200 | -;-;-;-;-;-;-;-; | 2851 | ahr1a | 0 | 0 | 0 | 0 | 1 | 0 |
| ENSDARG000 | 16;16;16;16;1 | 24698849:246 | 24699234:246 | -;-;-;-;-;-;-;-;- | 1176 | NA | 0 | 0 | 0 | 0 | 0 | 0 |
| ENSDARG000 | 16;16 | 39125589:391 | 39127774:391 | -;- | 2649 | gdf6a | 7085 | 911 | 2095 | 1491 | 3728 | 68 |
| ENSDARG000 | 16;16;16;16;1 | 42693247:426 | 42693349:426 | -;-;-;-;-;-;-;-; | 3524 | SLC4A7 | 215 | 251 | 504 | 134 | 275 | 154 |
| ENSDARG000 | 16;16;16;16;1 | 11558868:115 | 11558956:115 | +;+;+;+;+;+;+;+; | 2017 | zgc:198329 | 0 | 0 | 0 | 0 | 0 | 0 |
| ENSDARG000 | 16;16;16;16;1 | 17163193:171 | 17163903:171 | -;-;-;-;-;-;-;-; | 2166 | opn9 | 0 | 0 | 2 | 0 | 0 | 0 |
| ENSDARG000 | 16;16;16;16;1 | 12660477:126 | 12660540:126 | +;+;+;+;+ | 869 | aicda | 0 | 0 | 0 | 0 | 0 | 0 |
| ENSDARG000 | 16;16;16;16;1 | 7380463:7380 | 7380549:7380 | +;+;+;+;+;+;+;+; | 2311 | atg5 | 599 | 506 | 647 | 574 | 670 | 607 |
| ENSDARG000 | 16;16;16;16;1 | 29685557:296 | 29686483:296 | -;-;-;-;-;-;-;-; | 1682 | scnm1 | 158 | 122 | 136 | 130 | 146 | 233 |
| ENSDARG000 | 16;16;16;16;1 | 39146696:391 | 39147392:391 | +;+;+;+;+;+;+;+; | 4896 | sybu | 16 | 6 | 0 | 4 | 4 | 6 |
| ENSDARG000 | 16;16;16;16;1 | 48645217:486 | 48646159:486 | -;-;-;-;-;-;-;-; | 2939 | NA | 123 | 26 | 56 | 2 | 34 | 34 |
| ENSDARG000 | 16;16;16;16;1 | 31761825:317 | 31764107:317 | -;-;-;-;-;-;-;-; | 8252 | NA | 4616 | 3390 | 5716 | 2643 | 6107 | 4109 |
| ENSDARG000 | 16;16;16;16;1 | 41585645:415 | 41586311:415 | -;-;-;-;-;-;-;-; | 3533 | ATP2C1 | 693 | 507 | 544 | 243 | 698 | 516 |
| ENSDARG000 | 16;16;16;16;1 | 48671460:486 | 48671561:486 | +;+;+;+;+;+;+;+; | 977 | NA | 15 | 28 | 26 | 10 | 82 | 48 |
| ENSDARG000 | 16;16;16;16;1 | 24721556:247 | 24722060:247 | +;+;+;+;+;+;+;+; | 2346 | SMG9 | 272 | 371 | 366 | 246 | 376 | 284 |
| ENSDARG000 | 16;16;16;16;1 | 26380461:263 | 26384422:263 | -;-;-;-;-;-;-;-; | 11676 | MEGF8 | 1619 | 890 | 1561 | 864 | 1808 | 1506 |
| ENSDARG000 | 16;16;16;16;1 | 45303096:453 | 45303344:453 | -;-;-;-;-;-;-;-; | 2379 | NA | 10 | 2 | 0 | 0 | 14 | 0 |
| ENSDARG000 | 16;16;16;16;1 | 39172842:391 | 39173008:391 | -;-;-;-;-;-;-;-; | 1887 | NA | 0 | 0 | 0 | 0 | 0 | 0 |
| ENSDARG000 | 16;16;16;16;1 | 53203370:532 | 53203467:532 | +;+;+;+;+;+;+;+; | 1638 | NA | 27 | 24 | 179 | 88 | 14 | 17 |
| ENSDARG000 | 16;16;16;16;1 | 2840775:2843 | 2840871:2843 | +;+;+;+;+ | 744 | CLEC3B | 110 | 95 | 244 | 227 | 317 | 331 |
| ENSDARG000 | 16;16;16;16;1 | 11573471:115 | 11573529:115 | +;+;+;+;+ | 811 | NA | 0 | 0 | 0 | 0 | 0 | 0 |
| ENSDARG000 | 16;16;16 | 24733741:247 | 24733931:247 | +;+;+ | 2602 | NA | 93 | 54 | 77 | 38 | 76 | 140 |
| ENSDARG000 | 16;16;16;16;1 | 22722352:227 | 22722432:227 | -;-;-;-;- | 3036 | NA | 4 | 6 | 2 | 0 | 7 | 0 |
| ENSDARG000 | 16;16;16;16;1 | 12669765:126 | 12670274:126 | -;-;-;-;-;-;-;-; | 1586 | mfap5 | 34 | 2 | 6 | 18 | 34 | 46 |
| ENSDARG000 | 16;16;16;16;1 | 2847565:2850 | 2848065:2850 | -;-;-;-;-;-;-;-;- | 2614 | cdcp1a | 114 | 68 | 90 | 104 | 48 | 49 |
| ENSDARG000 | 16;16;16;16;1 | 5579744:5580 | 5579909:5580 | +;+;+;+;+ | 725 | NA | 0 | 2 | 1 | 0 | 6 | 0 |
| ENSDARG000 | 16;16;16;16;1 | 39185103:391 | 39185685:391 | -;-;-;-;-;-;- | 1368 | ebag9 | 560 | 740 | 923 | 678 | 617 | 700 |
| ENSDARG000 | 16;16;16;16;1 | 11585576:115 | 11585674:115 | +;+;+;+;+ | 820 | NA | 0 | 0 | 0 | 0 | 0 | 0 |
| ENSDARG000 | 16;16;16;16;1 | 7430998:7433 | 7432819:7433 | -;-;-;-;-;-;- | 3694 | prdm1a | 226 | 90 | 42 | 68 | 122 | 110 |
| ENSDARG000 | 16;16;16;16 | 5597600:5600 | 5597849:5601 | +;+;+;+ | 2071 | zgc:91890 | 630 | 1080 | 1208 | 890 | 649 | 625 |
| ENSDARG000 | 16;16;16;16;1 | 35661331:356 | 35662177:356 | +;+;+;+;+;+;+;+; | 6453 | map7d1a | 90 | 266 | 312 | 284 | 256 | 166 |
| ENSDARG000 | 16;16;16;16;1 | 22737481:227 | 22737503:227 | -;-;-;-;-;-;- | 2892 | NA | 1 | 0 | 0 | 6 | 1 | 7 |
| ENSDARG000 | 16;16;16;16;1 | 24819118:248 | 24821940:248 | -;-;-;-;- | 3551 | NA | 1159 | 1210 | 3711 | 3566 | 976 | 913 |
| ENSDARG000 | 16;16;16;16;1 | 24743871:247 | 24743967:247 | -;-;-;-;-;-;-;-; | 1778 | NA | 78 | 0 | 6 | 24 | 6 | 80 |
| ENSDARG000 | 16;16;16;16;1 | 7552084:7552 | 7552210:7552 | -;-;-;-;- | 600 | NA | 3 | 0 | 0 | 0 | 0 | 0 |
| ENSDARG000 | 16;16 | 7564527:7565 | 7564955:7566 | +;+ | 866 | NA | 0 | 0 | 4 | 0 | 0 | 0 |
| ENSDARG000 | 16;16;16 | 11597386:115 | 11597612:115 | +;+;+ | 720 | NA | 0 | 0 | 0 | 0 | 0 | 0 |
| ENSDARG000 | 16;16;16;16;1 | 45337663:453 | 45337794:453 | +;+;+;+;+;+;+;+; | 3476 | dnase2 | 396 | 406 | 404 | 552 | 1005 | 400 |
| ENSDARG000 | 16;16;16;16;1 | 20055878:200 | 20056056:200 | +;+;+;+;+;+;+;+; | 6717 | ANKRD28 | 1317 | 1246 | 1708 | 771 | 1480 | 1073 |
| ENSDARG000 | 16;16;16;16;1 | 27603739:276 | 27603814:276 | +;+;+;+;+ | 2066 | si:ch211-197h | 602 | 777 | 947 | 472 | 1091 | 1002 |
| ENSDARG000 | 16;16;16;16;1 | 14383479:143 | 14385348:143 | -;-;-;-;-;-;-;- | 2575 | crabp2a | 137 | 416 | 945 | 26 | 126 | 82 |
| ENSDARG000 | 16;16;16;16;1 | 9609721:9610 | 9609799:9610 | +;+;+;+;+;+;+;+; | 3119 | enpp2 | 12 | 12 | 14 | 12 | 26 | 18 |
| ENSDARG000 | 16;16;16;16;1 | 53247831:532 | 53248192:532 | -;-;-;-;-;-;-;-; | 927 | NA | 10 | 38 | 14 | 8 | 35 | 60 |
| ENSDARG000 | 16;16;16;16;1 | 48678655:486 | 48678844:486 | +;+;+;+;+;+;+;+; | 2603 | TAP1 | 504 | 398 | 556 | 197 | 499 | 428 |
| ENSDARG000 | 16;16;16;16;1 | 53252951:532 | 53253074:532 | +;+;+;+;+;+;+;+; | 1205 | si:ch211-269k | 14 | 26 | 18 | 2 | 41 | 28 |
| ENSDARG000 | 16;16;16;16;1 | 12710904:127 | 12711906:127 | -;-;-;-;-;-;-;- | 1832 | sbk3 | 0 | 0 | 0 | 0 | 2 | 0 |
| ENSDARG000 | 16;16;16;16;1 | 27614989:276 | 27615294:276 | +;+;+;+;+ | 1345 | glipr2l | 32 | 79 | 12 | 13 | 22 | 24 |
| ENSDARG000 | 16;16;16 | 29690708:296 | 29691400:296 | +;+;+ | 2033 | lysmd1 | 417 | 381 | 252 | 356 | 237 | 382 |
| ENSDARG000 | 16;16 | 22779229:227 | 22779441:227 | -;- | 719 | lenep | 0 | 0 | 0 | 6 | 4 | 4 |

| | | | | | | | | | | | | |
|---|---|---|---|---|---|---|---|---|---|---|---|---|
| ENSDARG000 | 16:16:16:16:1 | 39196727:392 | 39196994:392 | +;+;+;+;+;+;+ | 1975 | zdhhc3a | 376 | 372 | 271 | 278 | 655 | 436 |
| ENSDARG000 | 16:16:16:16:1 | 22781854:227 | 22784532:227 | -:-:-:-:-:-:-: | 4374 | SHC1 | 702 | 722 | 727 | 406 | 874 | 605 |
| ENSDARG000 | 16:16:16:16:1 | 2899611:2905 | 2899778:2905 | +:+:+:+:+:+:+: | 2684 | lars2 | 188 | 190 | 134 | 69 | 166 | 66 |
| ENSDARG000 | 16:16:16:16:1 | 11603096:116 | 11603166:116 | +:+:+:+:+:+:+: | 1479 | NA | 0 | 0 | 0 | 0 | 0 | 0 |
| ENSDARG000 | 16:16:16:16:1 | 22750174:227 | 22754410:227 | -:-:-:-:-:-:-: | 5661 | pbxip1b | 318 | 323 | 257 | 256 | 485 | 310 |
| ENSDARG000 | 16:16:16:16:1 | 29698565:296 | 29699453:296 | -;-;-;-;- | 2089 | tnfaip8l2b | 50 | 6 | 20 | 2 | 28 | 88 |
| ENSDARG000 | 16;16;16 | 39237768:392 | 39238778:392 | -;-;- | 1478 | tmem42a | 152 | 148 | 240 | 170 | 252 | 124 |
| ENSDARG000 | 16:16:16:16:1 | 27624395:276 | 27624508:276 | -:-:-:-:-:-:-: | 5876 | NA | 22 | 22 | 6 | 18 | 12 | 10 |
| ENSDARG000 | 16;16 | 14416346:144 | 14417075:144 | -;- | 940 | NA | 22 | 38 | 86 | 8 | 14 | 18 |
| ENSDARG000 | 16:16:16:16:1 | 42772678:427 | 42772799:427 | +:+:+:+:+:+:+: | 1507 | si:ch211-135r | 20 | 0 | 0 | 0 | 0 | 0 |
| ENSDARG000 | 16:16:16:16:1 | 7626535:7626 | 7626593:7627 | +:+:+:+:+:+:+: | 4106 | STX12 | 2695 | 2114 | 1743 | 2612 | 1905 | 3511 |
| ENSDARG000 | 16:16:16:16:1 | 17189839:171 | 17192972:171 | -:-:-:-:-:-:-: | 4973 | gapdh | 16 | 12 | 126 | 32 | 200 | 46 |
| ENSDARG000 | 16:16:16:16:1 | 5612547:5612 | 5612725:5612 | +:+:+:+:+:+:+: | 1366 | CYTH2 | 244 | 141 | 168 | 148 | 232 | 198 |
| ENSDARG000 | 16;16;16 | 39242339:392 | 39242517:392 | +;+;+ | 1055 | zgc:77056 | 210 | 190 | 179 | 68 | 178 | 156 |
| ENSDARG000 | 16;16 | 29903644:299 | 29903928:299 | +;+ | 475 | NA | 0 | 0 | 0 | 0 | 0 | 0 |
| ENSDARG000 | 16:16:16:16:1 | 11623956:116 | 11624023:116 | +;+;+;+;+ | 1038 | NA | 0 | 0 | 0 | 0 | 0 | 0 |
| ENSDARG000 | 16;16 | 14479160:144 | 14479291:144 | +;+ | 454 | NA | 0 | 0 | 0 | 0 | 0 | 0 |
| ENSDARG000 | 16:16:16:16:1 | 45364153:453 | 45364680:453 | -:-:-:-:-:-:-: | 2458 | rgl2 | 1024 | 441 | 455 | 517 | 630 | 702 |
| ENSDARG000 | 16:16:16:16:1 | 35821321:358 | 35821622:358 | -:-:-:-:-:-:-: | 1324 | NA | 134 | 54 | 177 | 33 | 86 | 57 |
| ENSDARG000 | 16:16:16:16:1 | 30002605:301 | 30002993:301 | +:+:+:+:+:+:+: | 7536 | sema6e | 5944 | 2679 | 2770 | 3620 | 2837 | 7376 |
| ENSDARG000 | 16;16 | 24842662:248 | 24842919:248 | +;+ | 603 | NA | 0 | 0 | 0 | 0 | 0 | 0 |
| ENSDARG000 | 16;16 | 24896017:248 | 24896541:248 | +;+ | 958 | NA | 0 | 0 | 0 | 0 | 0 | 0 |
| ENSDARG000 | 16:16:16:16:1 | 24905556:249 | 24905777:249 | +:+:+:+:+:+:+: | 1415 | NA | 0 | 0 | 0 | 0 | 0 | 0 |
| ENSDARG000 | 16:16:16:16:1 | 39248644:392 | 39250295:392 | -:-:-:-:-:-:-: | 2741 | gpd1l | 274 | 243 | 671 | 299 | 278 | 363 |
| ENSDARG000 | 16;16 | 30002752:300 | 30002993:300 | +;+ | 792 | NA | 34 | 8 | 0 | 2 | 0 | 4 |
| ENSDARG000 | 16:16:16:16:1 | 27658977:276 | 27661289:276 | -:-:-:-:-:-:-: | 4221 | NA | 192 | 481 | 483 | 80 | 446 | 198 |
| ENSDARG000 | 16;16;16;16 | 14528374:145 | 14529069:145 | -;-;-;- | 935 | NA | 0 | 0 | 2 | 4 | 6 | 6 |
| ENSDARG000 | 16;16;16;16 | 9672126:9673 | 9672617:9673 | -;-;-;- | 977 | mal2 | 0 | 2 | 2 | 0 | 0 | 0 |
| ENSDARG000 | 16;16;16 | 11652591:116 | 11652866:116 | +;+;+ | 643 | NA | 0 | 0 | 0 | 0 | 0 | 0 |
| ENSDARG000 | 16:16:16:16:1 | 14572008:145 | 14572204:145 | -;-;-;-;-;-;-;- | 1372 | dscc1 | 56 | 286 | 221 | 194 | 181 | 97 |
| ENSDARG000 | 16:16:16:16:1 | 35870456:358 | 35870791:358 | +:+:+:+:+:+:+: | 3246 | THRAP3 | 52 | 44 | 205 | 430 | 111 | 68 |
| ENSDARG000 | 16:16:16:16:1 | 45424907:454 | 45425215:454 | +:+:+:+:+:+:+: | 1385 | PHF1 | 3 | 0 | 1 | 4 | 4 | 1 |
| ENSDARG000 | 16:16:16:16:1 | 9680908:9683 | 9681805:9683 | -;-;-;-;-;- | 1376 | COLEC10 | 0 | 35 | 30 | 18 | 22 | 4 |
| ENSDARG000 | 16;16;16 | 7653905:7656 | 7654513:7656 | +;+;+ | 2873 | popdc3 | 18 | 6 | 38 | 40 | 14 | 24 |
| ENSDARG000 | 16:16:16:16:1 | 9713850:9714 | 9714161:9714 | +:+:+:+:+:+:+: | 2768 | NA | 363 | 604 | 499 | 351 | 540 | 293 |
| ENSDARG000 | 16:16:16:16:1 | 42829735:428 | 42829914:428 | +:+:+:+:+:+:+: | 1249 | polr3glb | 243 | 281 | 335 | 240 | 314 | 176 |
| ENSDARG000 | 16:16:16:16:1 | 39385483:393 | 39387132:393 | -:-:-:-:-:-:-: | 7122 | stt3b | 4967 | 5292 | 5840 | 3110 | 6347 | 3783 |
| ENSDARG000 | 16:16:16:16:1 | 39271123:392 | 39271609:392 | +:+:+:+:+:+:+: | 6568 | OSBPL10 | 2212 | 1853 | 1258 | 853 | 1713 | 1861 |
| ENSDARG000 | 16:16:16:16:1 | 35905031:359 | 35905258:359 | +:+:+:+:+:+:+: | 2623 | sh3d21 | 1087 | 1685 | 1088 | 1772 | 1526 | 1404 |
| ENSDARG000 | 16:16:16:16:1 | 5650495:5652 | 5650696:5652 | -;-;-;-;-;-;- | 2038 | NA | 6 | 4 | 0 | 0 | 0 | 0 |
| ENSDARG000 | 16:16:16:16:1 | 12730311:127 | 12730726:127 | +:+:+:+:+:+:+: | 4294 | epn1a | 1957 | 1547 | 2328 | 1784 | 2096 | 2037 |
| ENSDARG000 | 16:16:16:16:1 | 53278406:532 | 53278726:532 | +:+:+:+:+:+:+: | 1996 | ptdss1a | 714 | 390 | 524 | 317 | 662 | 348 |
| ENSDARG000 | 16:16:16:16:1 | 26439518:264 | 26440133:264 | +:+:+:+:+:+:+: | 3993 | trim35-28 | 916 | 835 | 1095 | 703 | 939 | 836 |
| ENSDARG000 | 16:16:16:16:1 | 14588141:145 | 14588457:145 | +:+:+:+:+:+:+: | 3245 | deptor | 1866 | 849 | 908 | 879 | 2141 | 1465 |
| ENSDARG000 | 16:16:16:16:1 | 31802203:318 | 31802635:318 | +:+:+:+:+:+:+: | 2212 | wnt4b | 6 | 6 | 0 | 15 | 13 | 8 |
| ENSDARG000 | 16:16:16:16:1 | 27734872:277 | 27737363:277 | -;-;-;-;- | 3949 | steap4 | 3729 | 2669 | 5923 | 1974 | 11696 | 1540 |
| ENSDARG000 | 16:16:16:16:1 | 11660839:116 | 11660890:116 | +:+:+:+:+:+:+: | 1225 | NA | 0 | 0 | 0 | 0 | 0 | 0 |
| ENSDARG000 | 16:16:16:16:1 | 48714048:487 | 48714208:487 | +:+:+:+:+:+:+: | 4608 | brd2b | 348 | 100 | 242 | 150 | 164 | 172 |
| ENSDARG000 | 16:16:16:16:1 | 41668474:416 | 41668578:416 | +:+:+:+:+:+:+: | 1899 | NA | 2 | 2 | 0 | 1 | 6 | 0 |
| ENSDARG000 | 16:16:16:16:1 | 33005688:330 | 33007163:330 | -:-:-:-:-:-:-: | 4417 | SNAP91 | 20 | 4 | 0 | 0 | 14 | 3 |
| ENSDARG000 | 16;16 | 27957808:279 | 27957951:279 | +;+ | 268 | NA | 0 | 0 | 4 | 0 | 0 | 0 |
| ENSDARG000 | 16;16;16;16 | 24940377:249 | 24941052:249 | -;-;-;- | 1716 | has1 | 4 | 0 | 2 | 2 | 12 | 0 |
| ENSDARG000 | 16:16:16:16:1 | 11683548:116 | 11683630:116 | +:+:+:+:+:+:+: | 2101 | NA | 0 | 0 | 0 | 0 | 0 | 2 |
| ENSDARG000 | 16:16:16:16:1 | 45456066:455 | 45456405:455 | +:+:+:+:+:+:+: | 10829 | NA | 119 | 161 | 190 | 289 | 900 | 235 |
| ENSDARG000 | 16:16:16:16:1 | 28112364:281 | 28113103:281 | -:-:-:-:-:-:-: | 1877 | NA | 642 | 648 | 810 | 368 | 595 | 654 |
| ENSDARG000 | 16:16:16:16:1 | 53325585:533 | 53325639:533 | +;+;+;+;+ | 354 | NA | 34 | 15 | 28 | 22 | 19 | 34 |
| ENSDARG000 | 16:16:16:16:1 | 7662609:7662 | 7662781:7662 | +:+:+:+:+:+:+: | 3049 | bves | 16 | 12 | 20 | 24 | 10 | 36 |
| ENSDARG000 | 16;16;16 | 24971717:249 | 24971749:249 | +;+;+ | 1921 | NA | 16 | 28 | 21 | 28 | 50 | 30 |
| ENSDARG000 | 16:16:16:16:1 | 5678071:5678 | 5678224:5678 | +:+:+:+:+:+:+: | 2491 | zgc:158689 | 150 | 38 | 52 | 6 | 16 | 22 |
| ENSDARG000 | 16:16:16:16:1 | 35944222:359 | 35945437:359 | -:-:-:-:-:-:-: | 1688 | eva1ba | 6118 | 7594 | 8050 | 7016 | 12021 | 8850 |
| ENSDARG000 | 16:16:16:16:1 | 9726038:9726 | 9726163:9726 | +:+:+:+:+:+:+: | 5599 | pip5k1ab | 794 | 754 | 842 | 755 | 913 | 938 |
| ENSDARG000 | 16:16:16:16:1 | 26449615:264 | 26449909:264 | +:+:+:+:+:+:+: | 5516 | epb41b | 66 | 28 | 99 | 70 | 240 | 115 |
| ENSDARG000 | 16;16;16 | 48753664:487 | 48753741:487 | +;+;+ | 475 | NA | 8 | 0 | 0 | 0 | 0 | 0 |
| ENSDARG000 | 16:16:16:16:1 | 31815244:318 | 31816978:318 | -;-;-;-;-;- | 2372 | cdc42l | 4398 | 5140 | 5195 | 3771 | 4630 | 4815 |
| ENSDARG000 | 16:16:16:16:1 | 42869054:428 | 42869669:428 | -:-:-:-:-:-:-: | 2018 | txnipb | 2 | 0 | 2 | 0 | 0 | 0 |
| ENSDARG000 | 16:16:16:16:1 | 28270037:282 | 28270756:282 | +:+:+:+:+:+:+: | 2796 | mindy3 | 156 | 298 | 284 | 223 | 289 | 234 |
| ENSDARG000 | 16;16;16 | 36582486:365 | 36582576:365 | +;+;+ | 383 | NA | 0 | 0 | 5 | 0 | 0 | 0 |
| ENSDARG000 | 16:16:16:16:1 | 36608041:366 | 36608110:366 | -;-;-;-;-;-;- | 690 | NA | 2 | 0 | 0 | 2 | 0 | 0 |
| ENSDARG000 | 16:16:16:16:1 | 41723048:417 | 41723258:417 | -;-;-;-;-;-;- | 604 | NA | 0 | 0 | 0 | 0 | 0 | 0 |
| ENSDARG000 | 16:16:16:16:1 | 41712634:417 | 41712689:417 | -;-;-;-;- | 556 | NA | 0 | 0 | 0 | 0 | 0 | 0 |
| ENSDARG000 | 16:16:16:16:1 | 41717313:417 | 41717770:417 | +:+:+:+:+:+:+: | 2226 | NA | 268 | 688 | 912 | 327 | 909 | 500 |
| ENSDARG000 | 16;16 | 42873740:428 | 42874358:428 | -;- | 817 | NA | 0 | 0 | 2 | 0 | 0 | 0 |
| ENSDARG000 | 16:16:16:16:1 | 41685001:416 | 41686617:416 | -:-:-:-:-:-:-: | 4418 | CEP85 | 47 | 134 | 187 | 133 | 148 | 102 |
| ENSDARG000 | 16:16:16:16:1 | 11724230:117 | 11724402:117 | +:+:+:+:+:+:+: | 3793 | ceacam1 | 0 | 0 | 0 | 0 | 0 | 0 |
| ENSDARG000 | 16:16:16:16:1 | 9779680:9780 | 9779865:9780 | +;+;+;+;+ | 1060 | vamp1b | 4 | 2 | 6 | 0 | 6 | 0 |
| ENSDARG000 | 16:16:16:16:1 | 45693413:456 | 45693799:456 | +:+:+:+:+:+:+: | 3764 | smg5 | 1134 | 993 | 1354 | 533 | 1374 | 901 |
| ENSDARG000 | 16:16:16:16:1 | 9762261:9766 | 9762317:9766 | +:+:+:+:+:+:+: | 1530 | psmd4b | 1696 | 1638 | 1687 | 970 | 1859 | 1500 |
| ENSDARG000 | 16;16 | 42876446:428 | 42876874:428 | +;+ | 866 | NA | 0 | 0 | 5 | 0 | 0 | 0 |
| ENSDARG000 | 16:16:16:16:1 | 20167811:201 | 20167941:201 | +:+:+:+:+:+:+: | 2520 | hacl1 | 301 | 296 | 271 | 210 | 354 | 214 |
| ENSDARG000 | 16:16:16:16:1 | 42883799:428 | 42884699:428 | -:-:-:-:-:-:-: | 1891 | hjv | 0 | 0 | 0 | 0 | 0 | 0 |
| ENSDARG000 | 16:16:16:16:1 | 53387085:533 | 53387148:533 | +:+:+:+:+:+:+: | 6053 | KIF13A | 2461 | 1262 | 1473 | 1732 | 1862 | 1430 |
| ENSDARG000 | 16;16;16 | 24978203:249 | 24978461:249 | +;+;+ | 4037 | si:dkeyp-84f3. | 615 | 642 | 932 | 502 | 884 | 802 |
| ENSDARG000 | 16:16:16:16:1 | 28383758:283 | 28384103:283 | +:+:+:+:+:+:+: | 4544 | ITGA8 | 2408 | 6 | 4 | 34 | 42 | 67 |
| ENSDARG000 | 16:16:16:16:1 | 49601838:496 | 49602257:496 | +:+:+:+:+:+:+: | 865 | si:dkey-82o10 | 93 | 167 | 115 | 104 | 139 | 123 |
| ENSDARG000 | 16:16:16:16:1 | 12758494:127 | 12758597:127 | -:-:-:-:-:-:-: | 1804 | NA | 287 | 267 | 343 | 332 | 287 | 215 |
| ENSDARG000 | 16:16:16:16:1 | 9787383:9791 | 9790366:9791 | -:-:-:-:-:-:-: | 4800 | NA | 797 | 256 | 288 | 42 | 545 | 322 |
| ENSDARG000 | 16:16:16:16:1 | 7688368:7689 | 7688464:7689 | -;-;-;-;-;- | 882 | NA | 0 | 3 | 11 | 0 | 8 | 0 |
| ENSDARG000 | 16;16;16 | 17204251:172 | 17204644:172 | -;-;- | 764 | nppcl | 2 | 0 | 0 | 0 | 0 | 0 |
| ENSDARG000 | 16;16 | 24999679:250 | 25000140:250 | -;- | 1145 | NA | 0 | 4 | 5 | 6 | 0 | 4 |
| ENSDARG000 | 16;16;16;16 | 25011994:250 | 25012181:250 | +;+;+;+ | 9208 | znf1035 | 1806 | 2166 | 3000 | 944 | 3256 | 2344 |
| ENSDARG000 | 16:16:16:16:1 | 22864313:228 | 22865114:228 | +:+:+:+:+:+:+: | 5449 | flad1 | 414 | 401 | 464 | 237 | 460 | 354 |
| ENSDARG000 | 16;16;16;16 | 42897582:428 | 42897756:428 | +;+;+;+ | 576 | NA | 0 | 0 | 0 | 0 | 0 | 0 |
| ENSDARG000 | 16:16:16:16:1 | 42900238:429 | 42900602:429 | -:-:-:-:-:-:-: | 3279 | thbs3a | 100 | 24 | 16 | 0 | 62 | 18 |
| ENSDARG000 | 16:16:16:16:1 | 31827502:318 | 31827817:318 | +:+:+:+:+:+:+: | 5856 | mlf2 | 1130 | 1044 | 1388 | 1232 | 1044 | 1258 |
| ENSDARG000 | 16;16 | 26494756:264 | 26495021:264 | +;+ | 658 | NA | 0 | 0 | 0 | 0 | 0 | 2 |
| ENSDARG000 | 16:16:16:16:1 | 12793911:127 | 12794142:127 | -:-:-:-:-:-:-: | 1108 | isoc2 | 222 | 231 | 279 | 280 | 385 | 264 |
| ENSDARG000 | 16:16:16:16:1 | 36650984:366 | 36651153:366 | +:+:+:+:+:+:+: | 8943 | NA | 84 | 32 | 140 | 32 | 184 | 14 |
| ENSDARG000 | 16;16 | 26503093:265 | 26503430:265 | -;- | 416 | NA | 0 | 0 | 2 | 0 | 1 | 8 |
| ENSDARG000 | 16;16;16;16 | 7693881:7694 | 7693980:7694 | -;-;-;- | 541 | NA | 3 | 10 | 13 | 4 | 18 | 4 |
| ENSDARG000 | 16:16:16:16:1 | 30408697:304 | 30409187:304 | -:-:-:-:-:-:-: | 2396 | cct3 | 7099 | 8454 | 10489 | 6582 | 6887 | 7441 |
| ENSDARG000 | 16:16:16:16:1 | 17234540:172 | 17235326:172 | -:-:-:-:-:-:-: | 2931 | nop2 | 554 | 885 | 1229 | 339 | 924 | 713 |
| ENSDARG000 | 16:16:16:16:1 | 49629886:496 | 49630054:496 | -:-:-:-:-:-:-: | 1861 | efhb | 4 | 0 | 0 | 0 | 0 | 0 |
| ENSDARG000 | 16:16:16:16:1 | 26510630:265 | 26511116:265 | -:-:-:-:-:-:-: | 2786 | NA | 249 | 422 | 684 | 542 | 570 | 283 |
| ENSDARG000 | 16;16 | 3809616:3814 | 3809912:3815 | +;+ | 871 | NA | 0 | 0 | 0 | 0 | 0 | 0 |
| ENSDARG000 | 16:16:16:16:1 | 45739193:457 | 45739590:457 | +:+:+:+:+:+:+: | 2550 | PAQR6 | 1855 | 1016 | 1231 | 929 | 1243 | 1152 |
| ENSDARG000 | 16:16:16:16:1 | 11760919:117 | 11760942:117 | -;-;-;-;-;-;-;- | 784 | PAFAH1B3 | 0 | 0 | 0 | 0 | 0 | 0 |
| ENSDARG000 | 16;16;16;16 | 11760602:117 | 11760942:117 | -;-;-;- | 613 | zgc:158701 | 0 | 0 | 0 | 0 | 0 | 0 |
| ENSDARG000 | 16:16:16:16:1 | 11779534:117 | 11779736:117 | +;+;+;+;+;+;+ | 3037 | NA | 38 | 28 | 52 | 36 | 24 | 70 |
| ENSDARG000 | 16:16:16:16:1 | 45796602:457 | 45797366:457 | -:-:-:-:-:-:-: | 3114 | ntrk1 | 0 | 0 | 2 | 4 | 0 | 0 |
| ENSDARG000 | 16:16:16:16:1 | 11773395:117 | 11775739:117 | -:-:-:-:-:-:-: | 3169 | pafah1b3 | 1182 | 1028 | 981 | 1179 | 889 | 1656 |
| ENSDARG000 | 16:16:16:16:1 | 14684916:146 | 14685009:146 | +:+:+:+:+:+:+: | 8308 | COL14A1 | 24 | 4 | 0 | 0 | 10 | 6 |
| ENSDARG000 | 16:16:16:16:1 | 39505163:395 | 39505790:395 | -:-:-:-:-:-:-: | 2390 | tgfbr2b | 122 | 125 | 164 | 82 | 347 | 182 |
| ENSDARG000 | 16;16 | 3928067:3931 | 3928155:3931 | +;+ | 331 | NA | 0 | 0 | 0 | 0 | 0 | 0 |
| ENSDARG000 | 16;16 | 7697878:7698 | 7698915:7699 | +;+ | 1330 | NA | 4 | 0 | 0 | 0 | 0 | 0 |
| ENSDARG000 | 16;16;16 | 25066880:250 | 25067004:250 | +;+;+ | 4460 | NA | 462 | 414 | 710 | 180 | 556 | 490 |
| ENSDARG000 | 16:16:16:16:1 | 3982590:3985 | 3982640:3985 | +;+;+;+;+;+ | 1100 | ZC3H12A | 56 | 8 | 0 | 0 | 6 | 0 |
| ENSDARG000 | 16:16:16:16:1 | 9807263:9807 | 9807568:9807 | -:-:-:-:-:-:-: | 2181 | grhl2a | 141 | 16 | 8 | 4 | 28 | 30 |
| ENSDARG000 | 16;16;16;16 | 5717760:5720 | 5718705:5720 | -;-;-;- | 1155 | ndufa3 | 507 | 614 | 426 | 218 | 340 | 378 |
| ENSDARG000 | 16 | 17252487 | 17253389 | + | 903 | gtf3c6 | 40 | 67 | 43 | 73 | 80 | 44 |
| ENSDARG000 | 16;16 | 25074029:250 | 25074083:250 | +;+ | 4381 | NA | 202 | 226 | 346 | 100 | 297 | 285 |
| ENSDARG000 | 16:16:16:16:1 | 42938859:429 | 42941206:429 | -;-;-;-;-;-;- | 3102 | mtx1a | 375 | 309 | 444 | 217 | 564 | 340 |
| ENSDARG000 | 16:16:16:16:1 | 25080764:250 | 25080995:250 | -;-;-;-;- | 871 | prss1 | 2 | 0 | 4 | 10 | 6 | 12 |
| ENSDARG000 | 16:16:16:16:1 | 31853919:318 | 31854045:318 | +:+:+:+:+:+:+: | 5812 | NA | 10 | 0 | 8 | 2 | 22 | 12 |
| ENSDARG000 | 16:16:16:16:1 | 17254489:172 | 17254902:172 | -;-;-;-;- | 1084 | emg1 | 264 | 190 | 284 | 198 | 238 | 226 |
| ENSDARG000 | 16:16:16:16:1 | 53455638:534 | 53455964:534 | +:+:+:+:+:+:+: | 1357 | rbm24b | 0 | 0 | 10 | 0 | 2 | 0 |
| ENSDARG000 | 16:16:16:16:1 | 22896453:228 | 22897161:228 | -;-;-;-;-;-;- | 1267 | NA | 16 | 2 | 4 | 9 | 6 | 0 |
| ENSDARG000 | 16:16:16:16:1 | 12812214:128 | 12812955:128 | +:+:+:+:+:+:+: | 3064 | u2af2a | 1174 | 1060 | 1825 | 1083 | 1385 | 1198 |
| ENSDARG000 | 16:16:16:16:1 | 33062570:330 | 33063158:330 | -:-:-:-:-:-:-: | 8213 | DOP1A | 703 | 393 | 690 | 585 | 637 | 595 |
| ENSDARG000 | 16:16:16:16:1 | 23017031:230 | 23017573:230 | -;-;-;-;- | 873 | NA | 0 | 2 | 0 | 0 | 0 | 4 |
| ENSDARG000 | 16:16:16:16:1 | 4010967:4011 | 4013931:4013 | -:-:-:-:-:-:-: | 4558 | NA | 0 | 0 | 0 | 0 | 0 | 0 |
| ENSDARG000 | 16;16;16 | 5774977:5776 | 5775071:5776 | +;+;+ | 770 | ccka | 0 | 0 | 24 | 4 | 256 | 44 |
| ENSDARG000 | 16:16:16:16:1 | 25095483:250 | 25096573:250 | +;+;+;+;+ | 2679 | NA | 345 | 486 | 624 | 406 | 537 | 583 |
| ENSDARG000 | 16:16:16:16:1 | 28578352:285 | 28578528:285 | +:+:+:+:+:+:+: | 2478 | nmt2 | 648 | 646 | 712 | 482 | 550 | 549 |
| ENSDARG000 | 16:16:16:16:1 | 9833256:9835 | 9835289:9835 | -:-:-:-:-:-:-: | 3106 | ncalda | 1028 | 1078 | 1621 | 840 | 1525 | 1148 |
| ENSDARG000 | 16:16:16:16:1 | 31879167:318 | 31879246:318 | -;-;-;-;-;- | 452 | NA | 0 | 0 | 0 | 0 | 0 | 0 |
| ENSDARG000 | 16:16:16:16:1 | 7703534:7703 | 7703623:7703 | +:+:+:+:+:+:+: | 3052 | cnot10 | 282 | 524 | 527 | 224 | 376 | 343 |
| ENSDARG000 | 16:16:16:16:1 | 4078240:4078 | 4078664:4078 | +:+:+:+:+:+:+: | 4532 | INPP5B | 787 | 775 | 837 | 968 | 913 | 820 |
| ENSDARG000 | 16;16 | 25107344:251 | 25107734:251 | +;+ | 4260 | zgc:66448 | 452 | 448 | 749 | 244 | 710 | 512 |
| ENSDARG000 | 16;16;16;16 | 42968097:429 | 42968330:429 | -;-;-;- | 1256 | NA | 0 | 0 | 0 | 0 | 0 | 0 |
| ENSDARG000 | 16:16:16:16:1 | 49647402:496 | 49647471:496 | +:+:+:+:+:+:+: | 1279 | rab5ab | 604 | 428 | 761 | 306 | 544 | 473 |
| ENSDARG000 | 16;16;16 | 42975366:429 | 42975930:429 | -;-;- | 866 | NA | 0 | 4 | 7 | 8 | 20 | 0 |
| ENSDARG000 | 16:16:16:16:1 | 23087326:230 | 23087981:230 | +:+:+:+:+:+:+: | 2273 | efna3b | 262 | 650 | 493 | 393 | 826 | 437 |
| ENSDARG000 | 16:16:16:16:1 | 20198064:202 | 20199032:202 | +:+:+:+:+:+:+: | 6677 | COLQ | 10 | 0 | 0 | 2 | 12 | 31 |
| ENSDARG000 | 16;16 | 25116827:251 | 25117362:251 | +;+ | 6876 | NA | 222 | 207 | 366 | 203 | 280 | 258 |
| ENSDARG000 | 16:16:16:16:1 | 17302452:173 | 17302627:173 | -:-:-:-:-:-:-: | 2195 | zyx | 772 | 751 | 684 | 479 | 821 | 628 |
| ENSDARG000 | 16:16:16:16:1 | 43000828:430 | 43001452:430 | -:-:-:-:-:-:-: | 1682 | nudt17 | 474 | 465 | 385 | 388 | 396 | 456 |
| ENSDARG000 | 16:16:16:16:1 | 25126935:251 | 25127014:251 | +;+;+;+;+ | 861 | zgc:92590 | 2 | 0 | 2 | 0 | 10 | 4 |
| ENSDARG000 | 16:16:16:16:1 | 49729339:497 | 49729569:497 | -:-:-:-:-:-:-: | 6139 | ZNF385D | 16 | 0 | 33 | 0 | 0 | 0 |
| ENSDARG000 | 16:16:16:16:1 | 17263027:172 | 17263561:172 | -:-:-:-:-:-:-: | 2890 | NA | 2 | 0 | 12 | 12 | 28 | 14 |
| ENSDARG000 | 16:16:16:16:1 | 30422481:304 | 30424015:304 | -:-:-:-:-:-:-: | 3015 | pdia7 | 1156 | 931 | 1144 | 619 | 1123 | 916 |
| ENSDARG000 | 16;16;16;16 | 12836143:128 | 12836904:128 | +;+;+;+ | 1415 | NA | 0 | 0 | 4 | 0 | 0 | 4 |
| ENSDARG000 | 16:16:16:16:1 | 36721230:367 | 36721797:367 | -:-:-:-:-:-:-: | 4662 | PIK3R4 | 1180 | 693 | 1243 | 779 | 1184 | 918 |
| ENSDARG000 | 16:16:16:16:1 | 20265499:202 | 20268547:202 | -:-:-:-:-:-:-: | 4589 | plekha8 | 571 | 579 | 794 | 477 | 982 | 761 |
| ENSDARG000 | 16:16:16:16:1 | 53489676:534 | 53489810:534 | +;+;+;+;+;+;+ | 1657 | grinab | 1674 | 1468 | 2000 | 1634 | 1559 | 1817 |
| ENSDARG000 | 16:16:16:16:1 | 25137483:251 | 25137609:251 | +:+:+:+:+:+:+: | 8122 | NA | 424 | 264 | 414 | 389 | 472 | 357 |
| ENSDARG000 | 16;16;16;16 | 45904153:459 | 45904739:459 | -;-;-;- | 858 | NA | 0 | 0 | 0 | 0 | 0 | 0 |
| ENSDARG000 | 16:16:16:16:1 | 50006145:500 | 50007034:500 | +:+:+:+:+:+:+: | 4419 | ube2e2 | 246 | 1417 | 352 | 180 | 368 | 296 |
| ENSDARG000 | 16:16:16:16:1 | 4178949:4181 | 4179854:4181 | +:+:+:+:+:+:+: | 3419 | yrdc | 135 | 373 | 465 | 144 | 278 | 203 |
| ENSDARG000 | 16:16:16:16:1 | 23282655:232 | 23282935:232 | +:+:+:+:+:+:+: | 1911 | efna1b | 66 | 4 | 2 | 18 | 23 | 0 |
| ENSDARG000 | 16;16 | 11785967:117 | 11788044:117 | -;- | 3142 | tlr21 | 14 | 0 | 8 | 8 | 32 | 8 |
| ENSDARG000 | 16:16:16:16:1 | 20294976:202 | 20295337:202 | +;+;+;+;+;+;+ | 1883 | fkbp14 | 674 | 526 | 788 | 1223 | 1178 | 932 |
| ENSDARG000 | 16:16:16:16:1 | 26530426:265 | 26531098:265 | -:-:-:-:-:-:-: | 1959 | sgk2b | 19 | 20 | 34 | 2 | 48 | 16 |

| | | | | | | | | | | | |
|---|---|---|---|---|---|---|---|---|---|---|---|
| ENSDARG000 | 16;16;16;16 | 11790640:117 | 11791057:117 | -;-;-;- | 743 | cnfn | 2 | 0 | 0 | 0 | 0 | 0 |
| ENSDARG000 | 16:16:16:16:1 | 12882381:128 | 12884529:128 | -:-:-:-:-:-:-:-: | 3099 | CACNG8 | 0 | 14 | 124 | 28 | 226 | 6 |
| ENSDARG000 | 16:16:16:16:1 | 33101851:331 | 33104591:331 | -;-;-;-;-;-;- | 3146 | pnrc2 | 26571 | 13412 | 17469 | 21888 | 12508 | 21156 |
| ENSDARG000 | 16;16;16 | 28592495:285 | 28593964:285 | -;-;- | 1570 | rpp38 | 123 | 182 | 242 | 84 | 226 | 158 |
| ENSDARG000 | 16;16;16 | 11818126:118 | 11818182:118 | +;+;+ | 1893 | cxcr3.2 | 68 | 80 | 120 | 42 | 304 | 232 |
| ENSDARG000 | 16:16:16:16:1 | 5786770:5795 | 5789412:5795 | -:-:-:-:-:-:-:-: | 5170 | TRAK1 | 764 | 811 | 1123 | 646 | 1153 | 572 |
| ENSDARG000 | 16:16:16:16:1 | 31900127:319 | 31900940:319 | -:-:-:-:-:-:-:-: | 2232 | rbfox1l | 0 | 0 | 4 | 2 | 2 | 0 |
| ENSDARG000 | 16:16:16:16:1 | 39858672:398 | 39858950:398 | +;+;+;+;+;+ | 2973 | NA | 94 | 17 | 66 | 47 | 120 | 106 |
| ENSDARG000 | 16:16:16:16:1 | 39643268:396 | 39643574:396 | -:-:-:-:-:-:-:-: | 2176 | rbms3 | 251 | 71 | 125 | 45 | 177 | 199 |
| ENSDARG000 | 16:16:16:16:1 | 4196932:4200 | 4197315:4200 | -;-;-;-;- | 1007 | rrp15 | 356 | 721 | 847 | 322 | 488 | 380 |
| ENSDARG000 | 16:16:16:16:1 | 33109496:331 | 33109897:331 | +;+;+;+;+ | 6194 | cnr2 | 6 | 0 | 6 | 4 | 42 | 14 |
| ENSDARG000 | 16:16:16:16:1 | 45912647:459 | 45914801:459 | -;-;-;-;-;-;- | 2927 | afp4 | 0 | 0 | 20 | 28 | 24 | 0 |
| ENSDARG000 | 16:16:16:16:1 | 28596555:285 | 28596640:285 | +;+;+;+;+;+;+ | 1056 | acbd7 | 155 | 44 | 121 | 182 | 120 | 28 |
| ENSDARG000 | 16:16:16:16:1 | 43014417:430 | 43015600:430 | -:-:-:-:-:-:-:-: | 2518 | si:dkev-7i14.5 | 10 | 29 | 31 | 52 | 32 | 109 |
| ENSDARG000 | 16:16:16:16:1 | 28601974:286 | 28602145:286 | +:+:+:+:+:+:+:+: | 3327 | crot | 525 | 258 | 619 | 496 | 610 | 650 |
| ENSDARG000 | 16:16:16:16:1 | 12963020:129 | 12964079:129 | -:-:-:-:-:-:-:-: | 1710 | CACNG7 | 0 | 4 | 2 | 4 | 4 | 6 |
| ENSDARG000 | 16;16 | 20309371:203 | 20311200:203 | -;- | 1972 | NA | 16 | 14 | 0 | 34 | 47 | 6 |
| ENSDARG000 | 16:16:16:16:1 | 45922175:459 | 45922311:459 | +;+;+;+;+;+ | 937 | rbm8a | 1779 | 2508 | 2209 | 1130 | 1773 | 1380 |
| ENSDARG000 | 16:16:16:16:1 | 23303859:233 | 23304104:233 | +;+;+;+;+;+ | 1276 | slc50a1 | 75 | 64 | 55 | 56 | 106 | 51 |
| ENSDARG000 | 16;16 | 11834516:118 | 11835708:118 | +;+ | 4038 | cxcr3.3 | 422 | 418 | 580 | 732 | 320 | 474 |
| ENSDARG000 | 16:16:16:16:1 | 50089417:501 | 50089669:501 | +:+:+:+:+:+:+:+: | 4079 | nr1d2a | 5254 | 6332 | 4637 | 5561 | 4420 | 3651 |
| ENSDARG000 | 16;16 | 30505279:305 | 30505621:305 | -;- | 430 | NA | 0 | 0 | 0 | 0 | 0 | 0 |
| ENSDARG000 | 16:16:16:16:1 | 30438041:304 | 30438243:304 | +:+:+:+:+:+:+:+: | 5659 | NA | 46 | 0 | 20 | 2 | 10 | 32 |
| ENSDARG000 | 16;16;16 | 9977858:9979 | 9978451:9979 | -;-;- | 1191 | id4 | 20 | 18 | 11 | 25 | 53 | 17 |
| ENSDARG000 | 16:16:16:16:1 | 11853212:118 | 11855648:118 | -;-;-;-;- | 2860 | cxcr3.1 | 18 | 12 | 2 | 2 | 57 | 54 |
| ENSDARG000 | 16;16 | 4210477:4211 | 4211017:4211 | -;- | 684 | NA | 0 | 0 | 2 | 1 | 0 | 0 |
| ENSDARG000 | 16:16:16:16:1 | 10025896:100 | 10026435:100 | -:-:-:-:-:-:-:-: | 2533 | cep76 | 34 | 81 | 51 | 90 | 82 | 90 |
| ENSDARG000 | 16:16:16:16:1 | 23339062:233 | 23339725:233 | -;-;-;-;-;- | 975 | NA | 112 | 8 | 0 | 14 | 13 | 0 |
| ENSDARG000 | 16:16:16:16:1 | 4287542:4293 | 4289066:4293 | -;-;-;-;-;-;- | 2658 | vegfaa | 84 | 707 | 2825 | 2802 | 299 | 115 |
| ENSDARG000 | 16:16:16:16:1 | 11911897:119 | 11912374:119 | -:-:-:-:-:-:-:-: | 2052 | cd4-1 | 34 | 22 | 12 | 4 | 106 | 73 |
| ENSDARG000 | 16:16:16:16:1 | 23353541:233 | 23353931:233 | -:-:-:-:-:-:-:-: | 2562 | TRIM46 | 0 | 0 | 0 | 0 | 0 | 0 |
| ENSDARG000 | 16:16:16:16:1 | 36748538:367 | 36748712:367 | +:+:+:+:+:+:+:+: | 1785 | decr1 | 138 | 144 | 170 | 174 | 228 | 288 |
| ENSDARG000 | 16:16:16:16:1 | 23335528:233 | 23335977:233 | -;-;-;-;-;-;-;- | 1130 | NA | 0 | 0 | 0 | 0 | 0 | 0 |
| ENSDARG000 | 16:16:16:16:1 | 41740365:417 | 41743525:417 | -:-:-:-:-:-:-:-: | 5041 | NA | 1329 | 1009 | 1217 | 700 | 1383 | 1456 |
| ENSDARG000 | 16:16:16:16:1 | 43028079:430 | 43028634:430 | -:-:-:-:-:-:-:-: | 1352 | si:dkev-7i14.6 | 4070 | 3245 | 3710 | 2984 | 3233 | 3687 |
| ENSDARG000 | 16:16:16:16:1 | 23326085:233 | 23326391:233 | +:+:+:+:+:+:+:+: | 4967 | krtcap2 | 705 | 966 | 1294 | 578 | 1016 | 619 |
| ENSDARG000 | 16:16:16:16:1 | 25163443:251 | 25163564:251 | +;+;+;+;+ | 2011 | znf576.2 | 474 | 519 | 676 | 528 | 590 | 531 |
| ENSDARG000 | 16:16:16:16:1 | 33121106:331 | 33121655:331 | +:+:+:+:+:+:+:+: | 2755 | akirin1 | 767 | 860 | 821 | 589 | 936 | 697 |
| ENSDARG000 | 16;16;16;16 | 7742658:7748 | 7748585:7748 | -;-;-;- | 7688 | trim71 | 2055 | 973 | 1510 | 1855 | 2162 | 2196 |
| ENSDARG000 | 16:16:16:16:1 | 45930962:459 | 45931150:459 | +:+:+:+:+:+:+:+: | 3736 | otud7b | 319 | 236 | 384 | 180 | 476 | 213 |
| ENSDARG000 | 16;16;16;16 | 25171832:251 | 25172056:251 | +;+;+;+ | 3216 | NA | 288 | 194 | 337 | 222 | 377 | 319 |
| ENSDARG000 | 16:16:16:16:1 | 25184207:251 | 25184276:251 | +;+;+;+;+ | 606 | hcst | 5 | 6 | 0 | 2 | 6 | 22 |
| ENSDARG000 | 16:16:16:16:1 | 25196572:251 | 25196684:251 | +;+;+;+;+ | 796 | tyrobp | 4 | 0 | 1 | 16 | 8 | 6 |
| ENSDARG000 | 16:16:16:16:1 | 15052513:150 | 15052635:150 | -;-;-;-;-;- | 693 | mrpl13 | 311 | 351 | 503 | 214 | 322 | 379 |
| ENSDARG000 | 16:16:16:16:1 | 4497302:4497 | 4497324:4497 | +:+:+:+:+:+:+:+: | 1646 | ttc29 | 0 | 1 | 0 | 0 | 0 | 0 |
| ENSDARG000 | 16:16:16:16:1 | 53519048:535 | 53519227:535 | +:+:+:+:+:+:+:+: | 2357 | smpd5 | 64 | 61 | 76 | 80 | 49 | 58 |
| ENSDARG000 | 16:16:16:16:1 | 20318400:203 | 20318571:203 | -:-:-:-:-:-:-:-: | 2170 | CHN2 | 554 | 640 | 427 | 974 | 1133 | 399 |
| ENSDARG000 | 16:16:16:16:1 | 13020776:130 | 13020933:130 | -:-:-:-:-:-:-:-: | 1964 | NA | 0 | 0 | 0 | 2 | 0 | 0 |
| ENSDARG000 | 16:16:16:16:1 | 30520357:305 | 30525553:305 | -:-:-:-:-:-:-:-: | 8044 | lmna | 7099 | 11964 | 7856 | 7876 | 10454 | 6811 |
| ENSDARG000 | 16:16:16:16:1 | 6021908:6035 | 6021996:6035 | +:+:+:+:+:+:+:+: | 893 | NA | 0 | 0 | 0 | 0 | 0 | 0 |
| ENSDARG000 | 16:16:16:16:1 | 5898878:5901 | 5899597:5902 | +:+:+:+:+:+:+:+: | 5259 | ULK4 | 50 | 66 | 77 | 20 | 131 | 46 |
| ENSDARG000 | 16:16:16:16:1 | 31921812:319 | 31922902:319 | +:+:+:+:+:+:+:+: | 4959 | rps9 | 16242 | 17879 | 26137 | 21993 | 16888 | 23367 |
| ENSDARG000 | 16:16:16:16:1 | 33142734:331 | 33142786:331 | +:+:+:+:+:+:+:+: | 4586 | rhbdl2 | 232 | 268 | 443 | 571 | 412 | 483 |
| ENSDARG000 | 16:16:16:16:1 | 41794715:417 | 41795126:417 | -:-:-:-:-:-:-:-: | 1980 | psmc4 | 2747 | 2678 | 3184 | 1588 | 2857 | 2023 |
| ENSDARG000 | 16:16:16:16:1 | 31924934:319 | 31925314:319 | -:-:-:-:-:-:-:-: | 1924 | NA | 2 | 5 | 0 | 0 | 4 | 20 |
| ENSDARG000 | 16:16:16:16:1 | 43047972:430 | 43048260:430 | -:-:-:-:-:-:-:-: | 1971 | NA | 485 | 139 | 297 | 1687 | 729 | 1340 |
| ENSDARG000 | 16:16:16:16:1 | 50114739:501 | 50114834:501 | -:-:-:-:-:-:-:-: | 2943 | NEK10 | 0 | 0 | 3 | 2 | 0 | 0 |
| ENSDARG000 | 16:16:16:16:1 | 4513597:4517 | 4517752:4517 | -;-;-;-;-;-;- | 4915 | ppp1r8b | 606 | 737 | 1043 | 394 | 824 | 582 |
| ENSDARG000 | 16 | 25231963 | 25233515 | - | 1553 | lratd2b | 20 | 29 | 28 | 10 | 12 | 26 |
| ENSDARG000 | 16:16:16:16:1 | 10055492:100 | 10056133:100 | -:-:-:-:-:-:-:-: | 2364 | ptpn2a | 281 | 166 | 160 | 116 | 278 | 167 |
| ENSDARG000 | 16;16;16 | 11960417:119 | 11960856:119 | -;-;- | 759 | NA | 0 | 1 | 0 | 4 | 1 | 0 |
| ENSDARG000 | 16:16:16:16:1 | 7802730:7805 | 7805689:7805 | -:-:-:-:-:-:-:-: | 4567 | TCAIM | 296 | 234 | 264 | 271 | 196 | 300 |
| ENSDARG000 | 16:16:16:16:1 | 11960588:119 | 11960856:119 | -:-:-:-:-:-:-:-: | 1570 | NA | 16 | 10 | 12 | 13 | 19 | 14 |
| ENSDARG000 | 16:16:16:16:1 | 43077909:430 | 43078121:430 | +:+:+:+:+:+:+:+: | 1819 | rundc3b | 4 | 8 | 4 | 14 | 0 | 10 |
| ENSDARG000 | 16;16 | 7856500:7881 | 7857249:7881 | -;- | 976 | NA | 0 | 0 | 4 | 6 | 2 | 0 |
| ENSDARG000 | 16;16;16;16 | 25245857:252 | 25246199:252 | +;+;+;+ | 2093 | KLHL38 | 2 | 0 | 0 | 0 | 0 | 2 |
| ENSDARG000 | 16:16:16:16:1 | 7968020:7968 | 7968310:7968 | -:-:-:-:-:-:-:-: | 1236 | NA | 12 | 13 | 6 | 2 | 30 | 17 |
| ENSDARG000 | 16 | 30575901 | 30577136 | + | 1236 | mc2r | 2 | 2 | 4 | 0 | 0 | 4 |
| ENSDARG000 | 16:16:16:16:1 | 36768674:367 | 36768853:367 | +:+:+:+:+:+:+:+: | 3758 | NA | 469 | 188 | 152 | 164 | 180 | 754 |
| ENSDARG000 | 16:16:16:16:1 | 36765934:367 | 36766623:367 | -:-:-:-:-:-:-:-: | 1544 | CALB1 | 0 | 0 | 281 | 22 | 236 | 0 |
| ENSDARG000 | 16:16:16:16:1 | 28629242:286 | 28629527:286 | -:-:-:-:-:-:-:-: | 4032 | abcb4 | 123 | 175 | 176 | 92 | 143 | 105 |
| ENSDARG000 | 16;16 | 50163352:501 | 50164218:501 | +;+ | 930 | NA | 0 | 0 | 0 | 0 | 0 | 0 |
| ENSDARG000 | 16:16:16:16:1 | 46000956:460 | 46001317:460 | +:+:+:+:+:+:+:+: | 9157 | mtmr11 | 186 | 165 | 145 | 137 | 307 | 73 |
| ENSDARG000 | 16:16:16:16:1 | 36807924:368 | 36808425:368 | -:-:-:-:-:-:-:-: | 1654 | pnp4b | 0 | 6 | 26 | 0 | 0 | 0 |
| ENSDARG000 | 16:16:16:16:1 | 20496691:204 | 20497336:204 | +:+:+:+:+:+:+:+: | 2989 | cpvl | 2628 | 1798 | 1761 | 2270 | 2758 | 2351 |
| ENSDARG000 | 16:16:16:16:1 | 17389116:173 | 17389268:173 | +:+:+:+:+:+:+:+: | 2810 | FAM131B | 0 | 0 | 6 | 0 | 0 | 0 |
| ENSDARG000 | 16:16:16:16:1 | 53569071:535 | 53569206:535 | +;+;+;+;+ | 570 | NA | 5 | 2 | 0 | 0 | 0 | 0 |
| ENSDARG000 | 16:16:16:16:1 | 53574817:535 | 53574911:535 | +;+;+;+;+ | 966 | NA | 2 | 0 | 0 | 0 | 0 | 0 |
| ENSDARG000 | 16 | 53585707 | 53586356 | - | 650 | NA | 8 | 0 | 6 | 0 | 1 | 0 |
| ENSDARG000 | 16 | 30582989 | 30584279 | - | 1291 | mc5rb | 0 | 0 | 0 | 0 | 0 | 0 |
| ENSDARG000 | 16:16:16:16:1 | 50166421:501 | 50166824:501 | -;-;-;-;-;-;-;- | 1253 | lim2.5 | 0 | 2 | 12 | 0 | 6 | 0 |
| ENSDARG000 | 16:16:16:16:1 | 25259313:252 | 25259551:252 | +:+:+:+:+:+:+:+: | 1812 | fbxo32 | 20 | 6 | 5 | 13 | 28 | 62 |
| ENSDARG000 | 16;16;16;16 | 30604387:306 | 30604530:306 | +;+;+;+ | 1554 | fam210ab | 224 | 213 | 357 | 133 | 335 | 316 |
| ENSDARG000 | 16:16:16:16:1 | 15097763:150 | 15097917:150 | +:+:+:+:+:+:+:+: | 2852 | MTBP | 34 | 46 | 67 | 42 | 74 | 106 |
| ENSDARG000 | 16:16:16:16:1 | 4531730:4533 | 4531996:4533 | +;+;+;+;+;+ | 1032 | tfpt | 199 | 162 | 197 | 144 | 129 | 195 |
| ENSDARG000 | 16;16 | 46093254:460 | 46093688:460 | -;- | 840 | NA | 18 | 3 | 12 | 0 | 3 | 14 |
| ENSDARG000 | 16;16 | 41806740:418 | 41806898:418 | -;- | 569 | si:dkev-199f5 | 0 | 0 | 1 | 0 | 0 | 0 |
| ENSDARG000 | 16:16:16:16:1 | 50183190:501 | 50183249:501 | -:-:-:-:-:-:-:-: | 1991 | NA | 0 | 0 | 0 | 0 | 0 | 0 |
| ENSDARG000 | 16;16;16;16 | 30621684:306 | 30622204:306 | -;-;-;- | 996 | ldlrad4b | 58 | 38 | 75 | 34 | 92 | 33 |
| ENSDARG000 | 16:16:16:16:1 | 17477152:174 | 17478426:174 | -:-:-:-:-:-:-:-: | 3869 | CLCN1 | 2 | 0 | 0 | 0 | 0 | 0 |
| ENSDARG000 | 16;16;16 | 41826584:418 | 41826759:418 | +;+;+ | 663 | NA | 8 | 0 | 19 | 40 | 0 | 1 |
| ENSDARG000 | 16:16:16:16:1 | 53603945:536 | 53604013:536 | +:+:+:+:+:+:+:+: | 2122 | sacm1lb | 295 | 311 | 555 | 246 | 523 | 321 |
| ENSDARG000 | 16:16:16:16:1 | 20529200:205 | 20530676:205 | -:-:-:-:-:-:-:-: | 2912 | CREB5 | 6 | 26 | 30 | 16 | 28 | 9 |
| ENSDARG000 | 16:16:16:16:1 | 33163858:331 | 33164170:331 | +;+;+;+;+;+;+ | 2324 | rragca | 627 | 706 | 851 | 288 | 897 | 733 |
| ENSDARG000 | 16:16:16:16:1 | 23397785:233 | 23397856:233 | +:+:+:+:+:+:+:+: | 872 | s100a10b | 5228 | 10608 | 4700 | 11812 | 6460 | 5842 |
| ENSDARG000 | 16:16:16:16:1 | 26547152:265 | 26547322:265 | +:+:+:+:+:+:+:+: | 3850 | PTPN3 | 98 | 28 | 46 | 32 | 46 | 54 |
| ENSDARG000 | 16:16:16:16:1 | 10097122:100 | 10097514:100 | -:-:-:-:-:-:-:-: | 8923 | NA | 142 | 327 | 397 | 156 | 383 | 258 |
| ENSDARG000 | 16:16:16:16:1 | 43126252:431 | 43127005:431 | -:-:-:-:-:-:-:-: | 1910 | slc25a40 | 243 | 234 | 340 | 228 | 247 | 234 |
| ENSDARG000 | 16:16:16:16:1 | 23403602:234 | 23403715:234 | +;+;+;+;+ | 1563 | s100w | 811 | 671 | 259 | 587 | 304 | 703 |
| ENSDARG000 | 16;16 | 33515766:335 | 33516171:335 | -;- | 742 | NA | 0 | 0 | 0 | 0 | 0 | 0 |
| ENSDARG000 | 16:16:16:16:1 | 26601364:266 | 26601618:266 | +:+:+:+:+:+:+:+: | 2076 | EPB41L4B | 25 | 0 | 2 | 6 | 2 | 8 |
| ENSDARG000 | 16;16 | 41840011:418 | 41840169:418 | -;- | 569 | NA | 0 | 0 | 0 | 0 | 0 | 0 |
| ENSDARG000 | 16:16:16:16:1 | 17548695:175 | 17551418:175 | -:-:-:-:-:-:-:-: | 4705 | casp2 | 328 | 460 | 523 | 314 | 340 | 407 |
| ENSDARG000 | 16;16 | 26601000:266 | 26601041:266 | -;- | 350 | NA | 2 | 0 | 0 | 0 | 0 | 0 |
| ENSDARG000 | 16;16 | 26619613:266 | 26619833:266 | -;- | 356 | NA | 3 | 0 | 0 | 0 | 0 | 0 |
| ENSDARG000 | 16:16:16:16:1 | 25275767:252 | 25276395:252 | -:-:-:-:-:-:-:-: | 1338 | PRELID3A | 16 | 45 | 20 | 12 | 24 | 6 |
| ENSDARG000 | 16:16:16:16:1 | 33566997:335 | 33567108:335 | -:-:-:-:-:-:-:-: | 1254 | NA | 0 | 0 | 0 | 0 | 0 | 0 |
| ENSDARG000 | 16:16:16:16:1 | 4540981:4542 | 4542639:4542 | -;-;-;-;-;-;- | 2374 | mrpl51 | 477 | 591 | 742 | 374 | 614 | 433 |
| ENSDARG000 | 16:16:16:16:1 | 30735426:307 | 30738002:307 | -:-:-:-:-:-:-:-: | 8799 | ptk2ab | 248 | 289 | 410 | 273 | 420 | 303 |
| ENSDARG000 | 16:16:16:16:1 | 46111849:461 | 46112140:461 | +:+:+:+:+:+:+:+: | 8959 | SV2A | 6 | 20 | 20 | 8 | 12 | 2 |
| ENSDARG000 | 16:16:16:16:1 | 50214819:502 | 50215319:502 | -:-:-:-:-:-:-:-: | 1388 | etfb | 925 | 1135 | 1262 | 1136 | 1140 | 828 |
| ENSDARG000 | 16:16:16:16:1 | 20738740:207 | 20738934:207 | +;+;+;+;+ | 819 | JAZF1 | 55 | 26 | 38 | 33 | 36 | 12 |
| ENSDARG000 | 16:16:16:16:1 | 13179146:131 | 13182340:131 | -:-:-:-:-:-:-:-: | 6134 | NA | 56 | 0 | 0 | 0 | 0 | 6 |
| ENSDARG000 | 16:16:16:16:1 | 40024883:400 | 40025052:400 | +;+;+;+;+ | 823 | hint3 | 59 | 49 | 85 | 51 | 60 | 40 |
| ENSDARG000 | 16;16 | 41853715:418 | 41853873:418 | -;- | 1205 | NA | 0 | 0 | 0 | 0 | 2 | 0 |
| ENSDARG000 | 16;16 | 40036711:400 | 40036959:400 | -;- | 352 | NA | 0 | 2 | 1 | 0 | 0 | 0 |
| ENSDARG000 | 16;16 | 40040813:400 | 40041457:400 | -;- | 747 | NA | 0 | 0 | 0 | 0 | 0 | 0 |
| ENSDARG000 | 16:16:16:16:1 | 15164983:151 | 15165271:151 | -;-;-;-;-;- | 1849 | sntb1 | 23 | 26 | 22 | 12 | 23 | 12 |
| ENSDARG000 | 16:16:16:16:1 | 17578607:175 | 17581339:175 | -:-:-:-:-:-:-:-: | 3737 | m6pr | 1929 | 2112 | 2433 | 1758 | 2544 | 2754 |
| ENSDARG000 | 16;16 | 41867176:418 | 41867334:418 | -;- | 569 | NA | 0 | 0 | 0 | 0 | 0 | 0 |
| ENSDARG000 | 16 | 33593116 | 33595985 | + | 2870 | pou3f1 | 2 | 0 | 4 | 0 | 12 | 0 |
| ENSDARG000 | 16;16;16 | 23431189:234 | 23431257:234 | +;+;+ | 408 | icn | 31731 | 38255 | 20733 | 45481 | 21177 | 30250 |
| ENSDARG000 | 16:16:16:16:1 | 28663184:286 | 28667059:286 | -:-:-:-:-:-:-:-: | 7540 | ADAM15 | 566 | 325 | 502 | 499 | 340 | 1497 |
| ENSDARG000 | 16:16:16:16:1 | 20934353:209 | 20934556:209 | +:+:+:+:+:+:+:+: | 1524 | skap2 | 273 | 565 | 195 | 200 | 528 | 194 |
| ENSDARG000 | 16:16:16:16:1 | 4570417:4570 | 4570672:4570 | -:-:-:-:-:-:-:-: | 3628 | arnt | 428 | 109 | 368 | 328 | 405 | 254 |
| ENSDARG000 | 16:16:16:16:1 | 31942527:319 | 31942740:319 | +:+:+:+:+:+:+:+: | 4515 | phc1 | 566 | 559 | 638 | 460 | 656 | 663 |
| ENSDARG000 | 16;16 | 20904754:209 | 20905502:209 | +;+ | 1878 | hoxa11b | 1391 | 2 | 0 | 0 | 8 | 8 |
| ENSDARG000 | 16:16:16:16:1 | 50240323:502 | 50240430:502 | +;+;+;+;+ | 497 | NA | 47 | 75 | 73 | 89 | 40 | 138 |
| ENSDARG000 | 16:16:16:16:1 | 20931585:209 | 20932758:209 | -:-:-:-:-:-:-:-: | 2694 | NA | 1 | 4 | 4 | 12 | 17 | 0 |
| ENSDARG000 | 16;16 | 20926673:209 | 20927486:209 | +;+ | 1914 | hoxa2b | 87 | 26 | 2 | 0 | 8 | 0 |
| ENSDARG000 | 16:16:16:16:1 | 41873708:418 | 41874052:418 | +:+:+:+:+:+:+:+: | 1828 | scn1ba | 32 | 12 | 16 | 15 | 3 | 43 |
| ENSDARG000 | 16:16:16:16:1 | 20805322:208 | 20806798:208 | -:-:-:-:-:-:-:-: | 3922 | tax1bp1b | 2406 | 2882 | 3839 | 2219 | 3484 | 1732 |
| ENSDARG000 | 16:16:16:16:1 | 25285998:252 | 25286109:252 | +;+;+;+;+ | 1999 | NA | 78 | 39 | 58 | 38 | 38 | 37 |
| ENSDARG000 | 16;16 | 20910186:209 | 20910938:209 | +;+ | 1649 | hoxa10b | 1725 | 22 | 0 | 0 | 0 | 0 |
| ENSDARG000 | 16;16 | 20895904:208 | 20896580:208 | +;+ | 1546 | hoxa13b | 62 | 0 | 0 | 0 | 4 | 0 |
| ENSDARG000 | 16;16;16 | 23487051:234 | 23487129:234 | +;+;+ | 434 | icn2 | 104 | 2 | 1 | 10 | 6 | 4 |
| ENSDARG000 | 16:16:16:16:1 | 43139504:431 | 43139792:431 | +:+:+:+:+:+:+:+: | 2556 | dbf4 | 73 | 119 | 122 | 97 | 148 | 94 |
| ENSDARG000 | 16:16:16:16:1 | 20871015:208 | 20871144:208 | +:+:+:+:+:+:+:+: | 2317 | hibadhb | 1133 | 1145 | 1515 | 800 | 1118 | 957 |
| ENSDARG000 | 16:16:16:16:1 | 33645721:336 | 33645998:336 | -;-;-;-;-;-;-;- | 1095 | utp11 | 253 | 458 | 438 | 152 | 411 | 300 |
| ENSDARG000 | 16;16 | 20915319:209 | 20915963:209 | +;+ | 1495 | hoxa9b | 789 | 358 | 0 | 0 | 76 | 0 |
| ENSDARG000 | 16:16:16:16:1 | 50289916:502 | 50290093:502 | +;+;+;+;+;+ | 675 | hamp | 40 | 5 | 13 | 57 | 24 | 68 |
| ENSDARG000 | 16:16:16:16:1 | 10287588:102 | 10289053:102 | -:-:-:-:-:-:-:-: | 2831 | flot1b | 4345 | 3163 | 4054 | 4317 | 4598 | 5081 |
| ENSDARG000 | 16:16:16:16:1 | 33655890:337 | 33656168:337 | +:+:+:+:+:+:+:+: | 2678 | FHL3 | 1475 | 1187 | 2786 | 429 | 1459 | 1419 |
| ENSDARG000 | 16:16:16:16:1 | 11965371:119 | 11966061:119 | -:-:-:-:-:-:-:-: | 3324 | usp5 | 3291 | 3397 | 3605 | 2346 | 3746 | 2117 |
| ENSDARG000 | 16:16:16:16:1 | 23495600:235 | 23496035:235 | +:+:+:+:+:+:+:+: | 2140 | snx27b | 329 | 227 | 294 | 262 | 328 | 168 |
| ENSDARG000 | 16:16:16:16:1 | 13424534:134 | 13424906:134 | +:+:+:+:+:+:+:+: | 2098 | pitpnc1b | 112 | 30 | 38 | 56 | 54 | 132 |
| ENSDARG000 | 16:16:16:16:1 | 10264286:102 | 10264766:102 | +:+:+:+:+:+:+:+: | 2237 | MRS2 | 211 | 210 | 337 | 190 | 159 | 180 |
| ENSDARG000 | 16;16;16;16 | 28728347:287 | 28728417:287 | +;+;+;+ | 1074 | NA | 2 | 0 | 0 | 4 | 0 | 0 |
| ENSDARG000 | 16:16:16:16:1 | 28727383:287 | 28727499:287 | +:+:+:+:+:+:+:+: | 2329 | dcst2 | 20 | 22 | 23 | 44 | 14 | 58 |
| ENSDARG000 | 16:16:16:16:1 | 46205693:462 | 46206204:462 | +;+;+;+;+ | 1320 | bola1 | 190 | 159 | 249 | 107 | 146 | 190 |
| ENSDARG000 | 16:16:16:16:1 | 28711556:287 | 28711910:287 | -:-:-:-:-:-:-:-: | 2264 | NA | 6 | 2 | 6 | 30 | 0 | 12 |
| ENSDARG000 | 16;16;16;16 | 46294337:463 | 46295352:463 | +;+;+;+ | 3077 | nr2f5 | 74 | 388 | 164 | 28 | 80 | 112 |
| ENSDARG000 | 16:16:16:16:1 | 50434668:507 | 50434725:507 | +:+:+:+:+:+:+:+: | 2359 | iglon5 | 0 | 14 | 0 | 8 | 16 | 4 |
| ENSDARG000 | 16;16;16;16 | 33752102:337 | 33752517:337 | +;+;+;+ | 1453 | NA | 0 | 0 | 0 | 0 | 0 | 0 |
| ENSDARG000 | 16;16;16 | 33750067:337 | 33750253:337 | -;-;- | 475 | NA | 0 | 0 | 0 | 0 | 0 | 0 |
| ENSDARG000 | 16;16;16;16 | 15369634:153 | 15372994:153 | -;-;-;- | 4614 | has2 | 20 | 4 | 0 | 0 | 6 | 4 |
| ENSDARG000 | 16:16:16:16:1 | 40043673:400 | 40043794:400 | +:+:+:+:+:+:+:+: | 2131 | trmt11 | 124 | 128 | 199 | 74 | 136 | 128 |
| ENSDARG000 | 16;16 | 15708204:157 | 15708673:157 | -;- | 573 | NA | 0 | 0 | 0 | 0 | 0 | 0 |
| ENSDARG000 | 16;16 | 31960916:319 | 31961015:319 | +;+ | 265 | NA | 0 | 0 | 0 | 0 | 0 | 0 |
| ENSDARG000 | 16:16:16:16:1 | 31965340:319 | 31969161:319 | -:-:-:-:-:-:-:-: | 5150 | styk1b | 11 | 4 | 4 | 2 | 2 | 2 |
| ENSDARG000 | 16;16 | 15709411:157 | 15709447:157 | +;+ | 333 | NA | 0 | 0 | 0 | 0 | 0 | 0 |
| ENSDARG000 | 16:16:16:16:1 | 46350620:463 | 46351280:463 | -;-;-;-;-;-;-;- | 1055 | NA | 0 | 2 | 2 | 0 | 2 | 4 |

| | | | | | | | | | | | | |
|---|---|---|---|---|---|---|---|---|---|---|---|---|
| ENSDARG000 | 16:16:16:16:1 | 53642820:536 | 53643095:536 | -:-:-:-:-:-:-:-: | 1973 | NA | 7 | 0 | 1 | 0 | 2 | 0 |
| ENSDARG000 | 16;16;16;16 | 15779159:157 | 15779246:157 | -;-;-;- | 462 | NA | 0 | 0 | 0 | 0 | 0 | 0 |
| ENSDARG000 | 16;16 | 15813506:158 | 15813658:158 | -;- | 195 | NA | 0 | 0 | 0 | 0 | 0 | 0 |
| ENSDARG000 | 16;16;16 | 15866224:158 | 15866267:158 | +;+;+ | 293 | NA | 0 | 0 | 0 | 0 | 1 | 0 |
| ENSDARG000 | 16:16:16:16:1 | 46401756:464 | 46401801:464 | +:+:+:+:+:+:+: | 1432 | rpz2 | 268 | 234 | 321 | 224 | 208 | 300 |
| ENSDARG000 | 16;16;16 | 26640677:266 | 26640783:266 | +;+;+ | 544 | NA | 0 | 0 | 0 | 0 | 0 | 0 |
| ENSDARG000 | 16;16;16 | 46397661:464 | 46402195:464 | -;-;- | 5181 | NA | 0 | 14 | 0 | 8 | 6 | 26 |
| ENSDARG000 | 16;16;16 | 46401576:464 | 46401801:464 | +;+;+ | 1127 | rpz | 567 | 590 | 453 | 417 | 756 | 417 |
| ENSDARG000 | 16:16:16:16:1 | 13505064:135 | 13506378:135 | -;-;-;-;-;-;-;- | 2756 | NA | 2 | 2 | 0 | 4 | 0 | 0 |
| ENSDARG000 | 16:16:16:16:1 | 33772902:337 | 33775237:337 | -:-:-:-:-:-:-:-: | 3588 | rspo1 | 12 | 10 | 0 | 4 | 0 | 0 |
| ENSDARG000 | 16:16:16:16:1 | 41902949:419 | 41906080:419 | -:-:-:-:-:-:-:-: | 6897 | NA | 366 | 321 | 375 | 198 | 360 | 320 |
| ENSDARG000 | 16:16:16:16:1 | 23531583:235 | 23531668:235 | +:+:+:+:+:+:+: | 5421 | adar | 2253 | 1692 | 1817 | 1260 | 1686 | 1206 |
| ENSDARG000 | 16:16:16:16:1 | 13522299:135 | 13522348:135 | -;-;-;-;-;- | 876 | NA | 2 | 0 | 0 | 0 | 0 | 2 |
| ENSDARG000 | 16:16:16:16:1 | 43152727:431 | 43153220:431 | +:+:+:+:+:+:+: | 5702 | ADAM22 | 110 | 170 | 343 | 342 | 471 | 82 |
| ENSDARG000 | 16;16 | 41948075:419 | 41948488:419 | -;- | 464 | NA | 0 | 0 | 0 | 0 | 0 | 0 |
| ENSDARG000 | 16;16;16 | 41955855:419 | 41956159:419 | -;-;- | 649 | NA | 2 | 0 | 0 | 0 | 1 | 0 |
| ENSDARG000 | 16;16;16 | 28754403:287 | 28754537:287 | +;+;+ | 995 | s100v1 | 36 | 2 | 8 | 5 | 2 | 26 |
| ENSDARG000 | 16:16:16:16:1 | 26645663:266 | 26647070:266 | -:-:-:-:-:-:-:-: | 3669 | esrp1 | 110 | 4 | 4 | 4 | 29 | 14 |
| ENSDARG000 | 16:16:16:16:1 | 8258203:826 | 8258905:826 | -:-:-:-:-:-:-:-: | 2097 | entpd3 | 4 | 196 | 178 | 120 | 26 | 4 |
| ENSDARG000 | 16;16;16 | 43227724:432 | 43230805:432 | -;-;- | 3544 | cldn12 | 1968 | 2002 | 2647 | 2696 | 3190 | 1704 |
| ENSDARG000 | 16:16:16:16:1 | 33902006:339 | 33902218:339 | +:+:+:+:+:+:+: | 2776 | gnl2 | 1262 | 1008 | 1939 | 799 | 1049 | 1162 |
| ENSDARG000 | 16:16:16:16:1 | 28764017:287 | 28764343:287 | +:+:+:+:+:+:+: | 4150 | NA | 490 | 417 | 442 | 677 | 592 | 732 |
| ENSDARG000 | 16:16:16:16:1 | 33925743:339 | 33925853:339 | -:-:-:-:-:-:-:-: | 2117 | dnali1 | 14 | 21 | 39 | 32 | 49 | 34 |
| ENSDARG000 | 16:16:16:16:1 | 25296389:252 | 25296683:252 | +:+:+:+:+:+:+: | 3611 | TBC1D31 | 129 | 166 | 235 | 162 | 216 | 136 |
| ENSDARG000 | 16;16;16;16 | 46453176:464 | 46453374:464 | +;+;+;+ | 1690 | rpz3 | 51 | 98 | 59 | 59 | 82 | 51 |
| ENSDARG000 | 16:16:16:16:1 | 16041643:160 | 16043331:160 | -:-:-:-:-:-:-:-: | 4345 | ANKIB1 | 1348 | 716 | 898 | 936 | 1194 | 1094 |
| ENSDARG000 | 16:16:16:16:1 | 23598908:235 | 23599368:236 | +:+:+:+:+:+:+: | 3836 | KCNN3 | 2 | 4 | 0 | 0 | 2 | 0 |
| ENSDARG000 | 16:16:16:16:1 | 25316973:253 | 25317125:253 | +:+:+:+:+:+:+: | 2539 | dync1i1 | 4 | 8 | 16 | 27 | 13 | 4 |
| ENSDARG000 | 16;16 | 16055484:160 | 16055612:160 | +;+ | 1834 | NA | 47 | 12 | 9 | 42 | 61 | 61 |
| ENSDARG000 | 16;16 | 8287771:828 | 8287914:828 | -;- | 280 | NA | 0 | 4 | 0 | 0 | 0 | 0 |
| ENSDARG000 | 16:16:16:16:1 | 46492756:464 | 46493059:464 | +;+;+;+;+ | 2125 | rpz5 | 343 | 185 | 321 | 159 | 402 | 180 |
| ENSDARG000 | 16;16 | 50889118:508 | 50889978:508 | -;- | 906 | NA | 63 | 37 | 30 | 40 | 40 | 22 |
| ENSDARG000 | 16:16:16:16:1 | 50926455:509 | 50926829:509 | -;-;-;-;-;-;-;- | 3331 | NA | 4 | 5 | 0 | 0 | 0 | 4 |
| ENSDARG000 | 16:16:16:16:1 | 28792158:287 | 28792312:287 | +:+:+:+:+:+:+: | 2027 | zgc:171704 | 1558 | 4878 | 9381 | 16994 | 5534 | 1560 |
| ENSDARG000 | 16:16:16:16:1 | 50953547:509 | 50953690:509 | +:+:+:+:+:+:+: | 893 | NA | 9 | 0 | 7 | 19 | 34 | 2 |
| ENSDARG000 | 16;16;16;16 | 40131526:401 | 40131711:401 | +;+;+;+ | 480 | NA | 0 | 0 | 0 | 0 | 0 | 0 |
| ENSDARG000 | 16;16 | 40338584:403 | 40338994:403 | -;- | 668 | NA | 0 | 0 | 0 | 0 | 0 | 0 |
| ENSDARG000 | 16:16:16:16:1 | 40131473:401 | 40131711:401 | +;+;+;+;+;+;+ | 1112 | cenpw | 57 | 86 | 60 | 104 | 80 | 56 |
| ENSDARG000 | 16:16:16:16:1 | 46497149:464 | 46497249:464 | +;+;+;+;+;+ | 1663 | rpz4 | 427 | 684 | 554 | 531 | 559 | 518 |
| ENSDARG000 | 16:16:16:16:1 | 40340222:403 | 40340719:403 | +:+:+:+:+:+:+: | 2739 | mettl6 | 146 | 106 | 146 | 128 | 182 | 160 |
| ENSDARG000 | 16;16 | 13559511:135 | 13559805:135 | -;- | 335 | NA | 22 | 12 | 19 | 9 | 18 | 4 |
| ENSDARG000 | 16:16:16:16:1 | 23788616:237 | 23788893:237 | -:-:-:-:-:-:-:-: | 1105 | rps27.2 | 5170 | 5618 | 7765 | 7118 | 4606 | 8307 |
| ENSDARG000 | 16:16:16:16:1 | 40301056:403 | 40301748:403 | +:+:+:+:+:+:+: | 2254 | rspo3 | 7 | 0 | 1 | 1 | 5 | 18 |
| ENSDARG000 | 16:16:16:16:1 | 36934183:369 | 36934771:369 | -:-:-:-:-:-:-:-: | 2968 | aldh5a1 | 575 | 385 | 485 | 194 | 580 | 393 |
| ENSDARG000 | 16:16:16:16:1 | 8298542:830 | 8301306:830 | -:-:-:-:-:-:-:-: | 4346 | ddc | 47 | 146 | 48 | 63 | 94 | 10 |
| ENSDARG000 | 16:16:16:16:1 | 23796612:237 | 23796702:238 | +:+:+:+:+:+:+: | 1915 | rab13 | 654 | 368 | 342 | 398 | 412 | 280 |
| ENSDARG000 | 16:16:16:16:1 | 28797703:287 | 28799332:287 | -:-:-:-:-:-:-:-: | 2970 | plin6 | 18 | 18 | 36 | 52 | 30 | 14 |
| ENSDARG000 | 16:16:16:16:1 | 50969248:509 | 50969482:509 | +:+:+:+:+:+:+: | 1856 | NA | 18 | 4 | 14 | 67 | 70 | 10 |
| ENSDARG000 | 16:16:16:16:1 | 23811554:238 | 23811781:238 | +:+:+:+:+:+:+: | 6031 | NA | 0 | 0 | 0 | 0 | 28 | 0 |
| ENSDARG000 | 16:16:16:16:1 | 32009203:320 | 32009353:320 | -:-:-:-:-:-:-:-: | 1554 | gstk1 | 300 | 253 | 612 | 648 | 373 | 602 |
| ENSDARG000 | 16:16:16:16:1 | 26680508:266 | 26680573:266 | +:+:+:+:+:+:+: | 5715 | VIRMA | 1411 | 1175 | 1698 | 922 | 1697 | 1334 |
| ENSDARG000 | 16:16:16:16:1 | 31982083:319 | 31982180:319 | -:-:-:-:-:-:-:-: | 865 | NA | 0 | 0 | 1 | 4 | 0 | 0 |
| ENSDARG000 | 16:16:16:16:1 | 31982149:319 | 31982180:319 | -:-:-:-:-:-:-:-: | 2192 | gstk4 | 24 | 26 | 30 | 21 | 22 | 17 |
| ENSDARG000 | 16:16:16:16:1 | 13568697:135 | 13569451:135 | -;-;-;-;-;-;-;- | 1788 | ntd5 | 12 | 0 | 12 | 0 | 10 | 6 |
| ENSDARG000 | 16:16:16:16:1 | 41984032:419 | 41984548:419 | -;-;-;-;- | 916 | pycard | 44 | 20 | 26 | 28 | 68 | 30 |
| ENSDARG000 | 16:16:16:16:1 | 41974438:419 | 41974746:419 | +;+;+;+;+ | 3619 | si:ch211-225 | 3 | 0 | 0 | 0 | 0 | 0 |
| ENSDARG000 | 16:16:16:16:1 | 13599693:136 | 13603395:136 | -;-;-;-;-;-;-;- | 5177 | dbpb | 1602 | 919 | 2582 | 1538 | 3623 | 2301 |
| ENSDARG000 | 16:16:16:16:1 | 51084472:510 | 51084650:510 | -:-:-:-:-:-:-:-: | 3026 | ago1 | 62 | 44 | 71 | 50 | 114 | 44 |
| ENSDARG000 | 16;16;16;16 | 28842414:288 | 28842881:288 | -;-;-;- | 1717 | syt11b | 14 | 18 | 49 | 73 | 10 | 10 |
| ENSDARG000 | 16;16 | 17704917:177 | 17706020:177 | -;- | 1224 | NA | 0 | 0 | 0 | 0 | 0 | 0 |
| ENSDARG000 | 16:16:16:16:1 | 4615661:461 | 4616113:461 | -:-:-:-:-:-:-:-: | 2381 | cyp4t8 | 6 | 6 | 0 | 2 | 12 | 191 |
| ENSDARG000 | 16;16;16;16 | 10318893:103 | 10319050:103 | +;+;+;+ | 2319 | tubb5 | 440 | 1121 | 505 | 418 | 534 | 366 |
| ENSDARG000 | 16:16:16:16:1 | 8364085:836 | 8367873:836 | -:-:-:-:-:-:-:-: | 6077 | GRB10 | 242 | 395 | 967 | 504 | 503 | 251 |
| ENSDARG000 | 16;16;16;16 | 17711215:177 | 17712136:177 | -;-;-;- | 1690 | zgc:174935 | 8 | 0 | 5 | 8 | 0 | 0 |
| ENSDARG000 | 16;16 | 25339936:253 | 25341818:253 | -;- | 1993 | NA | 4 | 0 | 0 | 10 | 0 | 0 |
| ENSDARG000 | 16:16:16:16:1 | 13616239:136 | 13619899:136 | -;-;-;-;-;-;- | 6744 | NA | 1047 | 632 | 1323 | 607 | 1269 | 1193 |
| ENSDARG000 | 16;16;16;16 | 16148761:161 | 16148964:161 | -;-;-;- | 636 | fabp4b | 66 | 2 | 0 | 0 | 0 | 0 |
| ENSDARG000 | 16:16:16:16:1 | 43291709:432 | 43291927:432 | -:-:-:-:-:-:-:-: | 3228 | washc5 | 2151 | 2403 | 2187 | 2103 | 2448 | 2565 |
| ENSDARG000 | 16:16:16:16:1 | 25342152:253 | 25342319:253 | +:+:+:+:+:+:+: | 3788 | ZFAT | 519 | 180 | 692 | 660 | 563 | 644 |
| ENSDARG000 | 16:16:16:16:1 | 6512481:653 | 6512842:653 | -:-:-:-:-:-:-:-: | 1322 | NA | 2 | 0 | 2 | 0 | 0 | 0 |
| ENSDARG000 | 16;16;16;16 | 21033633:210 | 21035498:210 | -;-;-;- | 2241 | snx10b | 28 | 17 | 14 | 8 | 9 | 16 |
| ENSDARG000 | 16:16:16:16:1 | 41993551:419 | 41993964:419 | -;-;-;-;-;- | 1542 | caspa | 72 | 40 | 70 | 24 | 78 | 46 |
| ENSDARG000 | 16:16:16:16:1 | 21038520:210 | 21040497:210 | -:-:-:-:-:-:-:-: | 2854 | cbx3b | 863 | 850 | 979 | 962 | 1098 | 829 |
| ENSDARG000 | 16:16:16:16:1 | 46518114:465 | 46518849:465 | -:-:-:-:-:-:-:-: | 2287 | NA | 1954 | 2713 | 2943 | 2535 | 3383 | 2633 |
| ENSDARG000 | 16:16:16:16:1 | 33930864:339 | 33931096:339 | +:+:+:+:+:+:+: | 1776 | snip1 | 466 | 456 | 465 | 295 | 499 | 402 |
| ENSDARG000 | 16:16:16:16:1 | 54209504:542 | 54209576:542 | +:+:+:+:+:+:+: | 4871 | xrcc1 | 250 | 667 | 743 | 350 | 570 | 398 |
| ENSDARG000 | 16:16:16:16:1 | 11992498:119 | 11992895:119 | +:+:+:+:+:+:+: | 2536 | P3H3 | 562 | 891 | 528 | 894 | 886 | 373 |
| ENSDARG000 | 16:16:16:16:1 | 10329701:103 | 10329764:103 | +:+:+:+:+:+:+: | 5977 | mdc1 | 564 | 553 | 822 | 296 | 652 | 637 |
| ENSDARG000 | 16:16:16:16:1 | 17714664:177 | 17714888:177 | +:+:+:+:+:+:+: | 1197 | si:dkev-87o1. | 67 | 8 | 12 | 2 | 0 | 0 |
| ENSDARG000 | 16:16:16:16:1 | 25364313:253 | 25364767:253 | -;-;-;-;-;- | 1198 | rbfa | 110 | 96 | 150 | 67 | 130 | 118 |
| ENSDARG000 | 16:16:16:16:1 | 28861342:288 | 28861813:288 | -:-:-:-:-:-:-:-: | 2049 | NA | 62 | 161 | 75 | 131 | 81 | 120 |
| ENSDARG000 | 16:16:16:16:1 | 8529772:852 | 8532442:853 | -:-:-:-:-:-:-:-: | 7274 | cobl | 2 | 6 | 12 | 26 | 54 | 8 |
| ENSDARG000 | 16:16:16:16:1 | 16165029:161 | 16166531:161 | -;-;-;-;- | 2102 | stmn2a | 5 | 6 | 6 | 16 | 6 | 42 |
| ENSDARG000 | 16:16:16:16:1 | 36974523:369 | 36974742:369 | -:-:-:-:-:-:-:-: | 1528 | snrnp48 | 144 | 167 | 171 | 202 | 228 | 214 |
| ENSDARG000 | 16:16:16:16:1 | 30871614:308 | 30876017:308 | -:-:-:-:-:-:-:-: | 7323 | NA | 14 | 6 | 8 | 10 | 33 | 26 |
| ENSDARG000 | 16:16:16:16:1 | 4654333:465 | 4654543:465 | +;+;+;+;+;+;+ | 1102 | LIN28A | 0 | 0 | 0 | 0 | 0 | 0 |
| ENSDARG000 | 16:16:16:16:1 | 25371653:253 | 25376975:253 | -;-;-;-;-;- | 6175 | adnp2a | 284 | 389 | 338 | 336 | 489 | 320 |
| ENSDARG000 | 16;16 | 4685665:469 | 4685871:469 | -;- | 306 | NA | 0 | 0 | 0 | 0 | 0 | 0 |
| ENSDARG000 | 16:16:16:16:1 | 4695075:469 | 4695394:469 | +:+:+:+:+:+:+: | 1791 | mecr | 181 | 125 | 185 | 108 | 165 | 153 |
| ENSDARG000 | 16:16:16:16:1 | 42007090:420 | 42009383:420 | -:-:-:-:-:-:-:-: | 3590 | etsrp | 50 | 0 | 6 | 12 | 0 | 36 |
| ENSDARG000 | 16;16 | 8668519:866 | 8668576:867 | +;+ | 1507 | NA | 6 | 4 | 19 | 18 | 26 | 0 |
| ENSDARG000 | 16:16:16:16:1 | 51154918:511 | 51154965:511 | +:+:+:+:+:+:+: | 1758 | dhdds | 260 | 365 | 678 | 434 | 327 | 278 |
| ENSDARG000 | 16:16:16:16:1 | 33938227:339 | 33938282:339 | +:+:+:+:+:+:+: | 1264 | gpn2 | 218 | 240 | 274 | 274 | 190 | 185 |
| ENSDARG000 | 16:16:16:16:1 | 6588022:658 | 6588938:658 | -:-:-:-:-:-:-:-: | 2215 | NA | 0 | 0 | 2 | 0 | 0 | 0 |
| ENSDARG000 | 16:16:16:16:1 | 46553980:465 | 46556551:465 | -;-;-;-;- | 2671 | NA | 0 | 0 | 0 | 0 | 0 | 0 |
| ENSDARG000 | 16;16;16;16 | 8673636:867 | 8673690:867 | +;+;+;+ | 2467 | NA | 0 | 14 | 15 | 1 | 21 | 48 |
| ENSDARG000 | 16:16:16:16:1 | 46530244:465 | 46531042:465 | -;-;-;-;-;- | 2100 | NA | 0 | 10 | 19 | 16 | 4 | 0 |
| ENSDARG000 | 16;16;16 | 25388503:253 | 25389443:253 | -;-;- | 1638 | pard6ga | 0 | 2 | 0 | 0 | 0 | 0 |
| ENSDARG000 | 16:16:16:16:1 | 46567408:465 | 46569330:465 | -;-;-;-;-;- | 2496 | zgc:174938 | 0 | 0 | 1 | 0 | 0 | 0 |
| ENSDARG000 | 16:16:16:16:1 | 26720738:267 | 26721416:267 | -:-:-:-:-:-:-:-: | 1528 | NA | 0 | 4 | 0 | 0 | 0 | 0 |
| ENSDARG000 | 16:16:16:16:1 | 25394763:253 | 25395314:253 | -;-;-;-;-;- | 1606 | zgc:136493 | 52 | 60 | 86 | 88 | 82 | 98 |
| ENSDARG000 | 16:16:16:16:1 | 42018041:420 | 42018163:420 | +:+:+:+:+:+:+: | 2830 | fli1rs | 112 | 47 | 82 | 40 | 154 | 203 |
| ENSDARG000 | 16:16:16:16:1 | 43417632:434 | 43417767:434 | -:-:-:-:-:-:-:-: | 3123 | oxr1a | 273 | 186 | 279 | 194 | 279 | 224 |
| ENSDARG000 | 16:16:16:16:1 | 30905551:309 | 30909824:309 | -:-:-:-:-:-:-:-: | 7307 | SLC45A4 | 209 | 201 | 279 | 199 | 392 | 206 |
| ENSDARG000 | 16:16:16:16:1 | 54248706:542 | 54248900:542 | +:+:+:+:+:+:+: | 1535 | drd2l | 0 | 0 | 0 | 0 | 1 | 0 |
| ENSDARG000 | 16:16:16:16:1 | 40346447:403 | 40346798:403 | +:+:+:+:+:+:+: | 848 | NA | 89 | 114 | 113 | 79 | 69 | 57 |
| ENSDARG000 | 16:16:16:16:1 | 33953644:339 | 33954543:339 | +:+:+:+:+:+:+: | 8160 | ARID1A | 2503 | 1515 | 2789 | 1800 | 2762 | 2205 |
| ENSDARG000 | 16:16:16:16:1 | 28881235:288 | 28881365:288 | +;+;+;+;+;+;+ | 1697 | chtopb | 31 | 124 | 151 | 539 | 31 | 64 |
| ENSDARG000 | 16:16:16:16:1 | 10346277:103 | 10346458:103 | +:+:+:+:+:+:+: | 3602 | NA | 6 | 2 | 0 | 0 | 24 | 8 |
| ENSDARG000 | 16:16:16:16:1 | 23921610:239 | 23921658:239 | +;+;+;+;+;+;+ | 960 | apoa4b.3 | 0 | 0 | 0 | 0 | 0 | 0 |
| ENSDARG000 | 16:16:16:16:1 | 23880004:238 | 23881162:238 | -;-;-;-;-;- | 1735 | OSGIN2 | 10 | 2 | 0 | 0 | 5 | 1 |
| ENSDARG000 | 16;16 | 6634106:663 | 6634508:663 | -;- | 1053 | NA | 0 | 0 | 0 | 0 | 0 | 0 |
| ENSDARG000 | 16;16;16;16 | 6632969:664 | 6635296:665 | +;+;+;+ | 7273 | SALL3 | 7 | 0 | 10 | 0 | 0 | 18 |
| ENSDARG000 | 16:16:16:16:1 | 23975930:239 | 23976135:239 | +:+:+:+:+:+:+: | 1096 | apoc4 | 0 | 0 | 2 | 0 | 0 | 0 |
| ENSDARG000 | 16;16;16 | 10387719:103 | 10387850:103 | +;+;+ | 337 | NA | 0 | 0 | 0 | 0 | 0 | 0 |
| ENSDARG000 | 16:16:16:16:1 | 16210208:162 | 16210557:162 | -;-;-;-;-;- | 922 | upp1 | 143 | 128 | 137 | 88 | 206 | 109 |
| ENSDARG000 | 16;16;16;16 | 13636704:136 | 13636896:136 | -;-;-;- | 2930 | NA | 0 | 0 | 0 | 0 | 0 | 0 |
| ENSDARG000 | 16:16:16:16:1 | 23923746:239 | 23923807:239 | +:+:+:+:+:+:+: | 769 | NA | 0 | 0 | 4 | 0 | 0 | 0 |
| ENSDARG000 | 16:16:16:16:1 | 30933044:309 | 30933064:309 | +:+:+:+:+:+:+: | 1516 | NA | 0 | 0 | 2 | 32 | 0 | 0 |
| ENSDARG000 | 16:16:16:16:1 | 23960744:239 | 23960920:239 | +:+:+:+:+:+:+: | 2116 | apoeb | 672 | 40 | 40 | 62 | 160 | 176 |
| ENSDARG000 | 16:16:16:16:1 | 42048362:420 | 42048694:420 | -;-;-;-;-;-;- | 1576 | zp3d.2 | 0 | 0 | 2 | 0 | 4 | 0 |
| ENSDARG000 | 16:16:16:16:1 | 46576702:465 | 46577762:465 | -:-:-:-:-:-:-:-: | 1313 | NA | 0 | 0 | 0 | 0 | 0 | 0 |
| ENSDARG000 | 16:16:16:16:1 | 37166572:371 | 37168898:371 | -:-:-:-:-:-:-:-: | 7914 | NA | 119 | 97 | 75 | 73 | 79 | 118 |
| ENSDARG000 | 16:16:16:16:1 | 30948451:309 | 30949621:309 | -:-:-:-:-:-:-:-: | 2812 | dgat1b | 140 | 93 | 82 | 60 | 43 | 153 |
| ENSDARG000 | 16:16:16:16:1 | 12022543:120 | 12022707:120 | +:+:+:+:+:+:+: | 1958 | gnb3a | 0 | 0 | 0 | 0 | 0 | 0 |
| ENSDARG000 | 16:16:16:16:1 | 10397440:103 | 10398317:103 | -;-;-;-;-;- | 1626 | NA | 4 | 8 | 20 | 28 | 12 | 0 |
| ENSDARG000 | 16:16:16:16:1 | 23837847:238 | 23838021:238 | +:+:+:+:+:+:+: | 3993 | nbn | 156 | 202 | 305 | 111 | 286 | 196 |
| ENSDARG000 | 16:16:16:16:1 | 21131015:211 | 21131461:211 | -;-;-;-;-;-;-;- | 889 | NA | 10 | 6 | 41 | 12 | 10 | 14 |
| ENSDARG000 | 16;16 | 30970501:309 | 30970589:309 | +;+ | 468 | NA | 0 | 0 | 0 | 0 | 0 | 0 |
| ENSDARG000 | 16:16:16:16:1 | 26732086:267 | 26732140:267 | +:+:+:+:+:+:+: | 3751 | RAD54B | 8 | 71 | 90 | 50 | 102 | 27 |
| ENSDARG000 | 16;16 | 28903774:289 | 28903989:289 | +;+ | 764 | NA | 5 | 0 | 4 | 1 | 2 | 0 |
| ENSDARG000 | 16:16:16:16:1 | 28908478:289 | 28909237:289 | +:+:+:+:+:+:+: | 4108 | NA | 20 | 51 | 124 | 117 | 130 | 42 |
| ENSDARG000 | 16;16 | 42066582:420 | 42066674:420 | +;+ | 787 | NA | 0 | 0 | 0 | 0 | 0 | 0 |
| ENSDARG000 | 16:16:16:16:1 | 42071696:420 | 42073769:420 | -:-:-:-:-:-:-:-: | 4403 | dyrk1b | 512 | 455 | 578 | 341 | 545 | 399 |
| ENSDARG000 | 16;16 | 13668166:136 | 13668367:136 | -;- | 410 | NA | 0 | 1 | 7 | 0 | 4 | 0 |
| ENSDARG000 | 16:16:16:16:1 | 42057918:420 | 42058945:420 | -:-:-:-:-:-:-:-: | 2320 | zp3d.1 | 0 | 0 | 0 | 0 | 0 | 0 |
| ENSDARG000 | 16:16:16:16:1 | 23891443:238 | 23891579:238 | +:+:+:+:+:+:+: | 2143 | gtpbp10 | 289 | 201 | 403 | 218 | 258 | 276 |
| ENSDARG000 | 16;16;16 | 26747753:267 | 26748182:267 | -;-;- | 978 | NA | 0 | 0 | 0 | 0 | 0 | 0 |
| ENSDARG000 | 16:16:16:16:1 | 23947196:239 | 23947244:239 | +:+:+:+:+:+:+: | 1111 | apoa4b.2 | 0 | 0 | 7 | 0 | 4 | 0 |
| ENSDARG000 | 16;16;16;16 | 13672768:136 | 13674865:136 | -;-;-;- | 2968 | ube2s | 590 | 1326 | 1242 | 698 | 1278 | 812 |
| ENSDARG000 | 16:16:16:16:1 | 12052010:120 | 12052567:120 | -:-:-:-:-:-:-:-: | 2211 | cdca3 | 487 | 1198 | 1322 | 574 | 813 | 440 |
| ENSDARG000 | 16:16:16:16:1 | 23972126:239 | 23972382:239 | +:+:+:+:+:+:+: | 783 | apoc1 | 0 | 6 | 122 | 4 | 20 | 10 |
| ENSDARG000 | 16:16:16:16:1 | 17763848:177 | 17764096:177 | +:+:+:+:+:+:+: | 1516 | them4 | 88 | 184 | 108 | 20 | 127 | 66 |
| ENSDARG000 | 16:16:16:16:1 | 23913943:239 | 23914272:239 | +:+:+:+:+:+:+: | 1796 | apoa4b.1 | 6 | 0 | 12 | 4 | 0 | 0 |
| ENSDARG000 | 16:16:16:16:1 | 46596206:465 | 46599705:465 | -:-:-:-:-:-:-:-: | 4854 | NA | 617 | 479 | 543 | 210 | 666 | 805 |
| ENSDARG000 | 16:16:16:16:1 | 25407021:254 | 25407627:254 | +:+:+:+:+:+:+: | 4647 | JARID2 | 1885 | 981 | 1323 | 953 | 1071 | 1095 |
| ENSDARG000 | 16:16:16:16:1 | 33992319:339 | 33992420:339 | +:+:+:+:+:+:+: | 3699 | zdhhc18a | 404 | 281 | 421 | 334 | 334 | 487 |
| ENSDARG000 | 16:16:16:16:1 | 10413551:104 | 10413984:104 | +:+:+:+:+:+:+: | 1128 | ino80e | 124 | 121 | 198 | 128 | 180 | 164 |
| ENSDARG000 | 16:16:16:16:1 | 43765993:437 | 43766154:437 | +:+:+:+:+:+:+: | 1761 | NA | 327 | 333 | 396 | 181 | 360 | 302 |
| ENSDARG000 | 16:16:16:16:1 | 51180938:511 | 51181174:511 | +:+:+:+:+:+:+: | 1270 | hmgn2 | 11627 | 15386 | 21119 | 13629 | 14511 | 10894 |
| ENSDARG000 | 16:16:16:16:1 | 16222394:162 | 16223836:162 | -:-:-:-:-:-:-:-: | 1933 | gra | 463 | 653 | 513 | 358 | 622 | 664 |
| ENSDARG000 | 16:16:16:16:1 | 4710977:471 | 4711711:471 | -:-:-:-:-:-:-:-: | 1767 | smpdl3b | 8 | 61 | 42 | 10 | 48 | 16 |
| ENSDARG000 | 16:16:16:16:1 | 6727195:673 | 6727263:673 | +:+:+:+:+:+:+: | 2941 | NA | 199 | 141 | 234 | 139 | 204 | 265 |
| ENSDARG000 | 16:16:16:16:1 | 40353816:403 | 40354232:403 | -:-:-:-:-:-:-:-: | 1851 | NA | 27 | 19 | 13 | 8 | 9 | 25 |
| ENSDARG000 | 16:16:16:16:1 | 54568513:545 | 54568829:545 | +:+:+:+:+:+:+: | 5104 | DENND4B | 547 | 690 | 1119 | 561 | 963 | 526 |
| ENSDARG000 | 16:16:16:16:1 | 13683807:136 | 13683858:136 | +:+:+:+:+:+:+: | 1520 | josd2 | 186 | 241 | 210 | 134 | 180 | 184 |
| ENSDARG000 | 16:16:16:16:1 | 32014552:320 | 32014689:320 | +:+:+:+:+:+:+: | 2251 | mboat7 | 727 | 871 | 855 | 739 | 1128 | 1142 |
| ENSDARG000 | 16:16:16:16:1 | 17913936:179 | 17914011:179 | +:+:+:+:+:+:+: | 2596 | NA | 0 | 3 | 8 | 10 | 24 | 4 |
| ENSDARG000 | 16;16 | 31186083:311 | 31187401:311 | -;- | 1833 | scrt1b | 0 | 0 | 0 | 0 | 0 | 0 |
| ENSDARG000 | 16:16:16:16:1 | 37470717:374 | 37471257:374 | +:+:+:+:+:+:+: | 9542 | ADGRB1 | 6 | 10 | 15 | 0 | 16 | 48 |
| ENSDARG000 | 16;16 | 34046733:340 | 34047146:340 | +;+ | 3433 | tent5ba | 20 | 39 | 144 | 96 | 294 | 82 |
| ENSDARG000 | 16;16;16 | 40425873:404 | 40426840:404 | -;-;- | 1210 | plekhf2 | 272 | 260 | 340 | 285 | 298 | 226 |

| | | | | | | | | | | | | |
|---|---|---|---|---|---|---|---|---|---|---|---|---|
| ENSDARG000 | 16:16:16:16:1 | 46639772:466 | 46640083:466 | -:-:-:-:-:-:-:-: | 1149 | tmem176l.2 | 18 | 0 | 2 | 0 | 0 | 0 |
| ENSDARG000 | 16:16:16:16:1 | 40487732:404 | 40487933:404 | -;-;-;-;-;-;-;- | 1108 | NDUFAF6 | 174 | 124 | 188 | 116 | 196 | 148 |
| ENSDARG000 | 16:16:16:16:1 | 31258283:312 | 31261387:312 | -:-:-:-:-:-:-:-: | 8375 | NA | 8216 | 9889 | 10836 | 8195 | 7816 | 9692 |
| ENSDARG000 | 16:16:16:16:1 | 16232621:162 | 16234086:162 | -;-;-;-;- | 2718 | rbm12bb | 645 | 491 | 865 | 502 | 847 | 719 |
| ENSDARG000 | 16;16 | 10422836:104 | 10423253:104 | +;+ | 3569 | NA | 32 | 2 | 15 | 8 | 17 | 41 |
| ENSDARG000 | 16:16:16:16:1 | 51192655:511 | 51192772:511 | +:+:+:+:+:+:+: | 1523 | NA | 602 | 677 | 400 | 1580 | 1919 | 210 |
| ENSDARG000 | 16:16:16:16:1 | 32029090:320 | 32029228:320 | +:+:+:+:+:+:+: | 2810 | NA | 6 | 0 | 2 | 6 | 6 | 0 |
| ENSDARG000 | 16:16:16:16:1 | 40560622:405 | 40560856:405 | +;+;+;+;+;+;+ | 1587 | tp53inp1 | 120 | 75 | 72 | 101 | 86 | 47 |
| ENSDARG000 | 16:16:16:16:1 | 16253710:162 | 16254073:162 | +:+:+:+:+:+:+: | 11279 | NA | 1821 | 1298 | 1969 | 1282 | 1971 | 1696 |
| ENSDARG000 | 16;16;16 | 32028079:320 | 32028337:320 | -;-;- | 541 | NA | 0 | 0 | 0 | 0 | 0 | 0 |
| ENSDARG000 | 16:16:16:16:1 | 40508882:405 | 40509022:405 | +:+:+:+:+:+:+: | 2464 | nagpa | 834 | 639 | 878 | 442 | 733 | 627 |
| ENSDARG000 | 16:16:16:16:1 | 21144669:211 | 21145160:211 | -:-:-:-:-:-:-:-: | 3085 | NA | 8 | 0 | 0 | 0 | 8 | 0 |
| ENSDARG000 | 16:16:16:16:1 | 40575742:405 | 40575760:405 | +:+:+:+:+:+:+: | 4644 | ccne2 | 74 | 534 | 711 | 190 | 408 | 191 |
| ENSDARG000 | 16;16 | 37876779:378 | 37876875:378 | +;+ | 920 | si:ch211-198 | 16 | 2 | 8 | 0 | 0 | 10 |
| ENSDARG000 | 16:16:16:16:1 | 4725829:4725 | 4725925:4725 | +:+:+:+:+:+:+: | 2679 | NA | 12 | 21 | 24 | 3 | 56 | 40 |
| ENSDARG000 | 16:16:16:16:1 | 43818896:438 | 43822377:438 | -:-:-:-:-:-:-:-: | 5232 | zfpm2a | 26 | 34 | 37 | 18 | 22 | 21 |
| ENSDARG000 | 16;16;16 | 40521738:405 | 40521820:405 | -;-;- | 576 | NA | 0 | 0 | 1 | 0 | 1 | 0 |
| ENSDARG000 | 16:16:16:16:1 | 46649469:466 | 46652352:466 | -:-:-:-:-:-:-:-: | 3726 | tmem176l.4 | 484 | 68 | 90 | 58 | 188 | 135 |
| ENSDARG000 | 16:16:16:16:1 | 12064576:120 | 12065458:120 | -:-:-:-:-:-:-:-: | 2986 | pex5 | 380 | 287 | 308 | 145 | 290 | 229 |
| ENSDARG000 | 16:16:16:16:1 | 18145893:181 | 18146458:181 | -:-:-:-:-:-:-:-: | 1143 | NA | 17 | 0 | 0 | 18 | 8 | 38 |
| ENSDARG000 | 16:16:16:16:1 | 18142127:181 | 18142404:181 | +:+:+:+:+:+:+: | 1924 | NA | 97 | 0 | 0 | 40 | 36 | 85 |
| ENSDARG000 | 16:16:16:16:1 | 51244297:512 | 51245096:512 | -:-:-:-:-:-:-:-: | 1895 | NA | 708 | 318 | 279 | 190 | 271 | 235 |
| ENSDARG000 | 16:16:16:16:1 | 34111919:341 | 34112125:341 | +:+:+:+:+:+:+: | 2108 | tcea3 | 6 | 2 | 36 | 20 | 34 | 32 |
| ENSDARG000 | 16:16:16:16:1 | 6750289:6750 | 6750827:6750 | +:+:+:+:+:+:+: | 6350 | ZNF236 | 342 | 235 | 402 | 188 | 374 | 405 |
| ENSDARG000 | 16:16:16:16:1 | 51266502:512 | 51266585:512 | -:-:-:-:-:-:-:-: | 1527 | serpinb1l1 | 1188 | 1406 | 839 | 821 | 1133 | 1728 |
| ENSDARG000 | 16:16:16:16:1 | 16318028:163 | 16318390:163 | -;-;-;-;-;- | 1405 | NA | 397 | 166 | 343 | 725 | 526 | 645 |
| ENSDARG000 | 16;16;16;16 | 26766423:267 | 26766996:267 | +;+;+;+ | 1538 | gem | 2 | 14 | 9 | 14 | 28 | 0 |
| ENSDARG000 | 16:16:16:16:1 | 13714058:137 | 13714215:137 | -:-:-:-:-:-:-:-: | 1892 | NA | 12 | 1 | 9 | 6 | 0 | 14 |
| ENSDARG000 | 16:16:16:16:1 | 4746588:4746 | 4746876:4746 | -:-:-:-:-:-:-:-: | 1640 | rpa2 | 168 | 918 | 988 | 288 | 660 | 432 |
| ENSDARG000 | 16:16:16:16:1 | 25504109:255 | 25505006:255 | -:-:-:-:-:-:-:-: | 2662 | dtnbp1a | 2631 | 2859 | 2611 | 3012 | 2960 | 3143 |
| ENSDARG000 | 16:16:16:16:1 | 51281959:512 | 51282444:512 | -;-;-;-;-;-;-;- | 1492 | zgc:173729 | 18 | 0 | 2 | 0 | 8 | 2 |
| ENSDARG000 | 16:16:16:16:1 | 51292624:512 | 51293073:512 | -;-;-;-;-;-;-;- | 1419 | serpinb1l4 | 71 | 0 | 2 | 0 | 4 | 0 |
| ENSDARG000 | 16:16:16:16:1 | 44298902:443 | 44299257:443 | +:+:+:+:+:+:+: | 2772 | dpys | 0 | 4 | 30 | 10 | 28 | 0 |
| ENSDARG000 | 16;16 | 51370106:514 | 51373047:514 | -;- | 3003 | fam110d | 339 | 51 | 191 | 110 | 426 | 8 |
| ENSDARG000 | 16:16:16:16:1 | 37891735:378 | 37892294:378 | +:+:+:+:+:+:+: | 4770 | cep162 | 128 | 143 | 225 | 105 | 167 | 176 |
| ENSDARG000 | 16:16:16:16:1 | 26774182:267 | 26774254:267 | +:+:+:+:+:+:+: | 3663 | cdh17 | 0 | 12 | 268 | 36 | 624 | 4 |
| ENSDARG000 | 16;16 | 31385083:313 | 31385297:313 | -;- | 306 | NA | 0 | 0 | 0 | 0 | 0 | 0 |
| ENSDARG000 | 16;16 | 31396647:314 | 31397213:314 | +;+ | 607 | SCX | 0 | 0 | 1 | 0 | 0 | 0 |
| ENSDARG000 | 16;16 | 51941830:519 | 51942456:519 | -;- | 679 | NA | 153 | 82 | 230 | 80 | 264 | 175 |
| ENSDARG000 | 16:16:16:16:1 | 42177659:421 | 42178059:421 | -:-:-:-:-:-:-:-: | 1436 | fbl | 2214 | 2515 | 4227 | 1462 | 2478 | 1867 |
| ENSDARG000 | 16:16:16:16:1 | 28994709:289 | 28994779:289 | +:+:+:+:+:+:+: | 6756 | gon4lb | 471 | 434 | 639 | 254 | 584 | 440 |
| ENSDARG000 | 16:16:16:16:1 | 16358348:163 | 16358487:163 | -:-:-:-:-:-:-:-: | 9094 | NBEAL2 | 54 | 18 | 50 | 42 | 111 | 98 |
| ENSDARG000 | 16:16:16:16:1 | 42156064:421 | 42156742:421 | -:-:-:-:-:-:-:-: | 2164 | ALKBH8 | 78 | 176 | 213 | 54 | 218 | 102 |
| ENSDARG000 | 16;16;16 | 18368914:183 | 18369011:183 | -;-;- | 282 | NA | 0 | 0 | 0 | 0 | 0 | 0 |
| ENSDARG000 | 16:16:16:16:1 | 10429770:104 | 10429961:104 | +:+:+:+:+:+:+: | 4150 | vars1 | 2104 | 2434 | 3702 | 1152 | 2427 | 1482 |
| ENSDARG000 | 16:16:16:16:1 | 46666074:466 | 46666336:466 | +:+:+:+:+:+:+: | 3328 | ubqln4 | 2896 | 3154 | 4354 | 2157 | 3788 | 2397 |
| ENSDARG000 | 16:16:16:16:1 | 18495540:184 | 18497873:184 | -;-;-;-;-;-;-;- | 2891 | NA | 4 | 0 | 0 | 0 | 0 | 0 |
| ENSDARG000 | 16:16:16:16:1 | 31428488:314 | 31428911:314 | -:-:-:-:-:-:-:-: | 1426 | zgc:194210 | 0 | 12 | 5 | 8 | 4 | 0 |
| ENSDARG000 | 16:16:16:16:1 | 31434524:314 | 31438007:314 | -:-:-:-:-:-:-:-: | 5264 | csnk2a1 | 3581 | 4407 | 5190 | 4025 | 3998 | 3288 |
| ENSDARG000 | 16:16:16:16:1 | 37947342:379 | 37948795:379 | -;-;-;-;- | 2001 | mrap2a | 0 | 16 | 2 | 0 | 8 | 10 |
| ENSDARG000 | 16:16:16:16:1 | 37948330:379 | 37948795:379 | -:-:-:-:-:-:-:-: | 3470 | NA | 178 | 70 | 34 | 8 | 165 | 62 |
| ENSDARG000 | 16:16:16:16:1 | 13810018:138 | 13811126:138 | -:-:-:-:-:-:-:-: | 1988 | leng9 | 76 | 102 | 112 | 74 | 102 | 41 |
| ENSDARG000 | 16:16:16:16:1 | 44338805:443 | 44339519:443 | -:-:-:-:-:-:-:-: | 1774 | NA | 0 | 0 | 0 | 0 | 3 | 0 |
| ENSDARG000 | 16:16:16:16:1 | 32059785:320 | 32060276:320 | +:+:+:+:+:+:+: | 10049 | NA | 41 | 4 | 20 | 86 | 110 | 4 |
| ENSDARG000 | 16:16:16:16:1 | 51950510:519 | 51953209:519 | -;-;-;-;- | 3439 | pdcl | 1839 | 1725 | 2197 | 1907 | 1722 | 1492 |
| ENSDARG000 | 16;16;16;16 | 32051572:320 | 32052019:320 | +;+;+;+ | 1696 | leng1 | 273 | 209 | 275 | 261 | 224 | 180 |
| ENSDARG000 | 16:16:16:16:1 | 6803342:6803 | 6803933:6803 | -:-:-:-:-:-:-:-: | 2792 | mbpb | 1108 | 700 | 786 | 754 | 1253 | 852 |
| ENSDARG000 | 16;16;16 | 34160835:341 | 34161203:341 | +;+;+ | 2362 | oprd1b | 4 | 5 | 44 | 130 | 14 | 22 |
| ENSDARG000 | 16;16;16;16 | 46684855:466 | 46685095:466 | +;+;+;+ | 785 | lamtor2 | 588 | 666 | 595 | 594 | 551 | 558 |
| ENSDARG000 | 16:16:16:16:1 | 18534834:185 | 18535729:185 | +:+:+:+:+:+:+: | 6470 | rxrbb | 1429 | 1394 | 1489 | 1298 | 1569 | 1557 |
| ENSDARG000 | 16;16 | 6828552:6829 | 6828737:6829 | -;- | 526 | NA | 0 | 2 | 0 | 7 | 4 | 0 |
| ENSDARG000 | 16;16 | 26816753:268 | 26818728:268 | -;- | 2072 | pdp1 | 212 | 203 | 132 | 130 | 229 | 278 |
| ENSDARG000 | 16:16:16:16:1 | 26846495:268 | 26847082:268 | +;+;+;+;+;+ | 1873 | trim35-29 | 2 | 6 | 10 | 2 | 4 | 0 |
| ENSDARG000 | 16:16:16:16:1 | 4770266:4770 | 4770497:4770 | +:+:+:+:+:+:+: | 1896 | ube3d | 58 | 107 | 168 | 44 | 124 | 56 |
| ENSDARG000 | 16;16;16;16 | 42193163:422 | 42193634:422 | -;-;-;- | 1736 | kcnj21 | 0 | 0 | 0 | 0 | 2 | 0 |
| ENSDARG000 | 16:16:16:16:1 | 26850481:268 | 26851274:268 | -:-:-:-:-:-:-:-: | 1402 | ino80c | 434 | 479 | 713 | 769 | 425 | 557 |
| ENSDARG000 | 16:16:16:16:1 | 21242491:212 | 21242879:212 | +:+:+:+:+:+:+: | 4638 | OSBPL3 | 362 | 323 | 473 | 324 | 546 | 252 |
| ENSDARG000 | 16;16;16 | 54672219:546 | 54672322:546 | +;+;+ | 274 | NA | 0 | 0 | 0 | 0 | 0 | 0 |
| ENSDARG000 | 16:16:16:16:1 | 46725087:467 | 46725483:467 | +;+;+;+;+ | 1647 | rab11al | 46 | 14 | 35 | 60 | 16 | 27 |
| ENSDARG000 | 16:16:16:16:1 | 12102336:121 | 12106281:121 | -:-:-:-:-:-:-:-: | 7049 | CLSTN3 | 34 | 10 | 9 | 0 | 26 | 10 |
| ENSDARG000 | 16:16:16:16:1 | 52105227:521 | 52105343:521 | +:+:+:+:+:+:+: | 1266 | MAN1C1 | 4 | 8 | 17 | 10 | 62 | 12 |
| ENSDARG000 | 16:16:16:16:1 | 34185613:341 | 34186368:341 | -:-:-:-:-:-:-:-: | 2049 | rcc1 | 448 | 734 | 796 | 316 | 805 | 350 |
| ENSDARG000 | 16:16:16:16:1 | 25535993:255 | 25536222:255 | +;+;+;+;+;+;+ | 2031 | mylipb | 233 | 120 | 164 | 82 | 222 | 136 |
| ENSDARG000 | 16:16:16:16:1 | 40594128:405 | 40594595:405 | -:-:-:-:-:-:-:-: | 8554 | NA | 20 | 32 | 38 | 38 | 24 | 44 |
| ENSDARG000 | 16;16 | 40681928:406 | 40682022:406 | -;- | 247 | NA | 0 | 0 | 0 | 0 | 0 | 0 |
| ENSDARG000 | 16:16:16:16:1 | 38015350:380 | 38020585:380 | -;-;-;-;-;- | 6060 | plekho1b | 2157 | 1854 | 1650 | 1418 | 2532 | 1441 |
| ENSDARG000 | 16:16:16:16:1 | 32082359:320 | 32082756:320 | +:+:+:+:+:+:+: | 1984 | prpf31 | 464 | 429 | 373 | 218 | 396 | 340 |
| ENSDARG000 | 16:16:16:16:1 | 42319438:423 | 42322673:423 | -:-:-:-:-:-:-:-: | 5226 | NA | 12693 | 7824 | 6113 | 12422 | 9937 | 10614 |
| ENSDARG000 | 16:16:16:16:1 | 54674556:546 | 54674691:546 | +:+:+:+:+:+:+: | 1502 | NA | 40 | 112 | 177 | 18 | 172 | 122 |
| ENSDARG000 | 16:16:16:16:1 | 26863414:269 | 26863763:269 | +:+:+:+:+:+:+: | 4221 | GALNT1 | 512 | 948 | 989 | 528 | 660 | 789 |
| ENSDARG000 | 16:16:16:16:1 | 16529748:165 | 16529914:165 | +;+;+;+;+;+;+ | 766 | ccdc12 | 109 | 54 | 81 | 11 | 200 | 80 |
| ENSDARG000 | 16:16:16:16:1 | 16554687:165 | 16556657:165 | -:-:-:-:-:-:-:-: | 5010 | NA | 58 | 2 | 30 | 12 | 42 | 60 |
| ENSDARG000 | 16:16:16:16:1 | 25555309:255 | 25561324:255 | -;-;-;-;- | 8697 | ATXN1 | 212 | 295 | 850 | 657 | 728 | 159 |
| ENSDARG000 | 16;16 | 16526975:165 | 16527107:165 | -;- | 407 | NA | 0 | 0 | 0 | 0 | 0 | 0 |
| ENSDARG000 | 16:16:16:16:1 | 21425114:214 | 21425209:214 | +:+:+:+:+:+:+: | 2384 | gsdmeb | 52 | 44 | 81 | 36 | 129 | 83 |
| ENSDARG000 | 16 | 4834527 | 4836218 | - | 1692 | tpbgb | 738 | 710 | 714 | 475 | 766 | 699 |
| ENSDARG000 | 16;16;16;16 | 32092866:320 | 32093057:320 | +;+;+;+ | 895 | NA | 0 | 0 | 0 | 0 | 0 | 0 |
| ENSDARG000 | 16:16:16:16:1 | 34199297:342 | 34199493:342 | -:-:-:-:-:-:-:-: | 3155 | phactr4b | 436 | 503 | 345 | 266 | 502 | 457 |
| ENSDARG000 | 16:16:16:16:1 | 13818500:138 | 13818698:138 | +:+:+:+:+:+:+: | 4287 | flcn | 1298 | 927 | 1405 | 1476 | 1130 | 1196 |
| ENSDARG000 | 16:16:16:16:1 | 25593339:255 | 25594225:255 | -;-;-;-;- | 1782 | fam8a1b | 330 | 193 | 356 | 302 | 271 | 297 |
| ENSDARG000 | 16 | 40954481 | 40955839 | + | 1359 | gbp | 376 | 453 | 583 | 312 | 756 | 356 |
| ENSDARG000 | 16:16:16:16:1 | 4838808:4839 | 4839228:4839 | +:+:+:+:+:+:+: | 4947 | IBTK | 903 | 756 | 1478 | 531 | 1102 | 616 |
| ENSDARG000 | 16;16 | 40991911:410 | 40992728:410 | -;- | 1970 | KCNV1 | 0 | 0 | 0 | 0 | 0 | 4 |
| ENSDARG000 | 16:16:16:16:1 | 44653266:446 | 44653599:446 | -:-:-:-:-:-:-:-: | 2495 | dcaf13 | 920 | 843 | 1154 | 635 | 883 | 744 |
| ENSDARG000 | 16:16:16:16:1 | 54761897:547 | 54761990:547 | -;-;-;-;- | 457 | NA | 0 | 0 | 0 | 0 | 0 | 0 |
| ENSDARG000 | 16:16:16:16:1 | 46869010:469 | 46869464:469 | +:+:+:+:+:+:+: | 5529 | THSD7A | 282 | 101 | 76 | 176 | 219 | 212 |
| ENSDARG000 | 16:16:16:16:1 | 29025292:290 | 29025561:290 | +:+:+:+:+:+:+: | 3723 | gpatch4 | 160 | 349 | 338 | 70 | 298 | 93 |
| ENSDARG000 | 16;16;16 | 16597539:166 | 16598516:166 | -;-;- | 1256 | NA | 160 | 104 | 118 | 128 | 146 | 270 |
| ENSDARG000 | 16:16:16:16:1 | 18580724:185 | 18580756:185 | -;-;-;-;-;- | 757 | NA | 2 | 0 | 4 | 0 | 0 | 0 |
| ENSDARG000 | 16:16:16:16:1 | 42423530:424 | 42424923:424 | -:-:-:-:-:-:-:-: | 4601 | SMARCC1 | 2548 | 2465 | 3940 | 1435 | 3960 | 2107 |
| ENSDARG000 | 16:16:16:16:1 | 6864244:6864 | 6864883:6864 | +:+:+:+:+:+:+: | 4447 | ARHGEF2 | 398 | 416 | 610 | 220 | 482 | 367 |
| ENSDARG000 | 16:16:16:16:1 | 12236339:122 | 12236433:122 | +:+:+:+:+:+:+: | 1974 | tpi1b | 3934 | 9406 | 15337 | 12211 | 7633 | 6727 |
| ENSDARG000 | 16:16:16:16:1 | 27126473:271 | 27127975:271 | -:-:-:-:-:-:-:-: | 3281 | TMEM245 | 1267 | 1386 | 1398 | 869 | 1802 | 1381 |
| ENSDARG000 | 16:16:16:16:1 | 5013868:5013 | 5014598:5015 | -:-:-:-:-:-:-:-: | 2342 | bckdhb | 1012 | 957 | 1052 | 824 | 1112 | 1108 |
| ENSDARG000 | 16:16:16:16:1 | 12174437:121 | 12175868:121 | -:-:-:-:-:-:-:-: | 3042 | lpcat3 | 1190 | 1129 | 1131 | 631 | 1312 | 988 |
| ENSDARG000 | 16:16:16:16:1 | 41015163:410 | 41015256:410 | +:+:+:+:+:+:+: | 2865 | dek | 282 | 754 | 942 | 487 | 678 | 587 |
| ENSDARG000 | 16:16:16:16:1 | 34300279:343 | 34302064:343 | -;-;-;-;-;-;-;- | 3019 | NA | 0 | 2 | 0 | 0 | 0 | 8 |
| ENSDARG000 | 16:16:16:16:1 | 16605517:166 | 16607140:166 | -:-:-:-:-:-:-:-: | 3226 | NA | 169 | 18 | 34 | 4 | 28 | 6 |
| ENSDARG000 | 16:16:16:16:1 | 47359754:473 | 47363931:473 | -:-:-:-:-:-:-:-: | 6183 | NA | 167 | 128 | 198 | 132 | 254 | 218 |
| ENSDARG000 | 16:16:16:16:1 | 38109941:381 | 38110360:381 | -:-:-:-:-:-:-:-: | 1439 | NA | 0 | 0 | 0 | 0 | 0 | 0 |
| ENSDARG000 | 16;16 | 34400806:344 | 34401271:344 | -;- | 595 | NA | 0 | 0 | 0 | 0 | 0 | 0 |
| ENSDARG000 | 16:16:16:16:1 | 18608978:186 | 18609060:186 | -:-:-:-:-:-:-:-: | 4940 | FHOD3 | 0 | 0 | 0 | 0 | 0 | 2 |
| ENSDARG000 | 16:16:16:16:1 | 34321394:343 | 34323007:343 | -:-:-:-:-:-:-:-: | 7547 | NA | 96 | 80 | 61 | 28 | 38 | 106 |
| ENSDARG000 | 16:16:16:16:1 | 25599379:255 | 25599887:255 | -:-:-:-:-:-:-:-: | 2399 | zgc:110410 | 2 | 6 | 5 | 2 | 5 | 2 |
| ENSDARG000 | 16;16;16;16 | 54770206:547 | 54770408:547 | +;+;+;+ | 623 | NA | 0 | 4 | 1 | 0 | 0 | 4 |
| ENSDARG000 | 16;16;16;16 | 13857913:138 | 13858495:138 | -;-;-;- | 952 | NA | 0 | 0 | 0 | 0 | 0 | 0 |
| ENSDARG000 | 16:16:16:16:1 | 54780544:547 | 54780895:547 | +:+:+:+:+:+:+: | 1597 | malsu1 | 482 | 300 | 470 | 380 | 400 | 412 |
| ENSDARG000 | 16:16:16:16:1 | 13855039:138 | 13855559:138 | +:+:+:+:+:+:+: | 2604 | zgc:174888 | 480 | 349 | 542 | 359 | 422 | 412 |
| ENSDARG000 | 16:16:16:16:1 | 34459066:344 | 34459198:344 | -:-:-:-:-:-:-:-: | 1405 | SERINC2 | 169 | 70 | 174 | 156 | 159 | 84 |
| ENSDARG000 | 16:16:16:16:1 | 13860299:138 | 13860561:138 | +;+;+;+;+;+;+ | 1646 | grwd1 | 456 | 712 | 733 | 256 | 690 | 497 |
| ENSDARG000 | 16;16;16;16 | 44679659:446 | 44679782:446 | -;-;-;- | 474 | NA | 4 | 0 | 0 | 3 | 1 | 0 |
| ENSDARG000 | 16:16:16:16:1 | 32110195:321 | 32111622:321 | -:-:-:-:-:-:-:-: | 4220 | cnot3a | 1470 | 1222 | 1664 | 1192 | 1785 | 1348 |
| ENSDARG000 | 16;16 | 44704372:447 | 44704415:447 | +;+ | 490 | NA | 0 | 0 | 1 | 4 | 1 | 6 |
| ENSDARG000 | 16:16:16:16:1 | 44678302:446 | 44678586:446 | +:+:+:+:+:+:+: | 1371 | zgc:101716 | 25 | 44 | 48 | 33 | 40 | 26 |
| ENSDARG000 | 16:16:16:16:1 | 41032961:410 | 41034556:410 | -;-;-;-;- | 3450 | NA | 0 | 0 | 0 | 0 | 0 | 0 |
| ENSDARG000 | 16:16:16:16:1 | 41042310:410 | 41042439:410 | +;+;+;+;+;+ | 678 | NA | 0 | 0 | 0 | 3 | 0 | 0 |
| ENSDARG000 | 16;16;16 | 41034317:410 | 41034556:410 | -;-;- | 530 | NA | 0 | 0 | 0 | 0 | 0 | 0 |
| ENSDARG000 | 16:16:16:16:1 | 27224068:272 | 27224464:272 | +:+:+:+:+:+:+: | 1614 | sec61b | 1275 | 1751 | 2551 | 1914 | 1372 | 1303 |
| ENSDARG000 | 16:16:16:16:1 | 47428721:474 | 47429154:474 | +:+:+:+:+:+:+: | 1361 | slc25a32b | 57 | 121 | 194 | 12 | 76 | 104 |
| ENSDARG000 | 16:16:16:16:1 | 27215938:272 | 27217399:272 | -;-;-;-;-;- | 2070 | alg2 | 468 | 720 | 879 | 726 | 599 | 438 |
| ENSDARG000 | 16:16:16:16:1 | 5110455:5110 | 5110754:5110 | -:-:-:-:-:-:-:-: | 3330 | ttk | 606 | 1581 | 1598 | 846 | 1242 | 565 |
| ENSDARG000 | 16:16:16:16:1 | 38119004:381 | 38119172:381 | +:+:+:+:+:+:+: | 7266 | pogzb | 548 | 711 | 857 | 416 | 913 | 694 |
| ENSDARG000 | 16:16:16:16:1 | 42464009:424 | 42464057:424 | +:+:+:+:+:+:+: | 1342 | NA | 0 | 0 | 0 | 0 | 0 | 0 |
| ENSDARG000 | 16:16:16:16:1 | 47389539:473 | 47391067:473 | -:-:-:-:-:-:-:-: | 3675 | septin7b | 1993 | 1796 | 2481 | 1781 | 2292 | 1868 |
| ENSDARG000 | 16:16:16:16:1 | 6937186:6938 | 6938622:6938 | -;-;-;-;- | 2762 | pmvk | 16 | 42 | 19 | 4 | 51 | 22 |
| ENSDARG000 | 16:16:16:16:1 | 6944311:6944 | 6944374:6944 | +:+:+:+:+:+:+: | 1838 | eaf1 | 343 | 309 | 410 | 329 | 377 | 389 |
| ENSDARG000 | 16:16:16:16:1 | 16621242:166 | 16621751:166 | -:-:-:-:-:-:-:-: | 6359 | NA | 6 | 2 | 82 | 2 | 236 | 14 |
| ENSDARG000 | 16:16:16:16:1 | 21442362:214 | 21446310:214 | -:-:-:-:-:-:-:-: | 6148 | pals2a | 302 | 253 | 449 | 148 | 367 | 404 |
| ENSDARG000 | 16;16;16;16 | 29030937:290 | 29031003:290 | -;-;-;- | 634 | NA | 22 | 11 | 16 | 20 | 21 | 13 |
| ENSDARG000 | 16;16;16;16 | 25608778:256 | 25608954:256 | +;+;+;+ | 3779 | zhx2a | 1265 | 880 | 1048 | 1257 | 1083 | 860 |
| ENSDARG000 | 16:16:16:16:1 | 25643445:256 | 25644608:256 | -:-:-:-:-:-:-:-: | 2193 | derl1 | 1039 | 1128 | 1491 | 947 | 1574 | 1254 |
| ENSDARG000 | 16:16:16:16:1 | 10477533:104 | 10478132:104 | -:-:-:-:-:-:-:-: | 2174 | abhd16a | 742 | 976 | 826 | 507 | 913 | 835 |
| ENSDARG000 | 16:16:16:16:1 | 29072874:290 | 29076690:290 | -:-:-:-:-:-:-:-: | 7228 | mef2d | 1132 | 884 | 1379 | 1069 | 1385 | 1246 |
| ENSDARG000 | 16:16:16:16:1 | 54785264:547 | 54785683:547 | -:-:-:-:-:-:-:-: | 2989 | khdc4 | 996 | 652 | 1066 | 885 | 1262 | 1186 |
| ENSDARG000 | 16:16:16:16:1 | 34479064:344 | 34479237:344 | +;+;+;+;+;+ | 5087 | NA | 0 | 2 | 0 | 0 | 12 | 2 |
| ENSDARG000 | 16:16:16:16:1 | 44768361:447 | 44768387:447 | +:+:+:+:+:+:+: | 2331 | upk1a | 70 | 0 | 0 | 12 | 20 | 16 |
| ENSDARG000 | 16:16:16:16:1 | 34523515:345 | 34523602:345 | +;+;+;+;+ | 1687 | stmn1b | 16 | 4 | 10 | 2 | 6 | 22 |
| ENSDARG000 | 16:16:16:16:1 | 18974064:189 | 18974301:189 | +:+:+:+:+:+:+: | 2538 | slc6a19b | 2 | 15 | 54 | 60 | 108 | 2 |
| ENSDARG000 | 16:16:16:16:1 | 34522977:345 | 34523058:345 | +;+;+;+;+ | 1599 | paqr7b | 114 | 119 | 167 | 20 | 286 | 225 |
| ENSDARG000 | 16:16:16:16:1 | 9400661:9403 | 9400892:9403 | +:+:+:+:+:+:+: | 4811 | ice1 | 399 | 291 | 632 | 299 | 411 | 334 |
| ENSDARG000 | 16:16:16:16:1 | 38138246:381 | 38138450:381 | +:+:+:+:+:+:+: | 1825 | selenbp1 | 74 | 83 | 265 | 86 | 192 | 83 |
| ENSDARG000 | 16;16;16 | 54804815:548 | 54807003:548 | -;-;- | 3083 | zbtb7b | 80 | 94 | 54 | 86 | 135 | 396 |
| ENSDARG000 | 16:16:16:16:1 | 9421002:9422 | 9423497:9423 | -;-;-;-;- | 2683 | NA | 211 | 212 | 200 | 154 | 221 | 200 |
| ENSDARG000 | 16:16:16:16:1 | 32131568:321 | 32131705:321 | +:+:+:+:+:+:+: | 6009 | NA | 458 | 243 | 375 | 348 | 349 | 337 |
| ENSDARG000 | 16:16:16:16:1 | 23978978:239 | 23979043:239 | +:+:+:+:+:+:+: | 1020 | apoa2 | 0 | 0 | 468 | 6 | 0 | 4 |
| ENSDARG000 | 16;16;16;16 | 29212421:292 | 29212719:292 | +;+;+;+ | 2424 | NA | 0 | 0 | 0 | 0 | 0 | 0 |
| ENSDARG000 | 16:16:16:16:1 | 34557016:345 | 34560690:345 | -:-:-:-:-:-:-:-: | 4540 | NA | 2 | 0 | 6 | 0 | 10 | 0 |
| ENSDARG000 | 16:16:16:16:1 | 16665203:166 | 16665507:166 | -:-:-:-:-:-:-:-: | 5979 | c4b | 364 | 158 | 166 | 72 | 181 | 210 |
| ENSDARG000 | 16;16 | 34875441:348 | 34875905:348 | -;- | 691 | NA | 0 | 0 | 0 | 0 | 0 | 0 |
| ENSDARG000 | 16:16:16:16:1 | 19013018:190 | 19013125:190 | +:+:+:+:+:+:+: | 4777 | NA | 0 | 0 | 0 | 48 | 0 | 16 |
| ENSDARG000 | 16:16:16:16:1 | 44788331:447 | 44788880:447 | -:-:-:-:-:-:-:-: | 7072 | NA | 128 | 122 | 137 | 38 | 227 | 154 |
| ENSDARG000 | 16:16:16:16:1 | 13875867:138 | 13877342:138 | -;-;-;-;-;-;-;- | 1709 | cdc42ep5 | 356 | 325 | 395 | 329 | 284 | 392 |

| | | | | | | | | | | | | |
|---|---|---|---|---|---|---|---|---|---|---|---|---|
| ENSDARG000 | 16:16:16:16:1 | 25664078:256 | 25664097:256 | +:+:+:+:+:+:+: | 878 | nsmce2 | 43 | 112 | 126 | 194 | 128 | 92 |
| ENSDARG000 | 16:16:16:16:1 | 29265796:292 | 29268360:292 | -:-:-:-:-:-:-:-: | 3933 | rhbg | 42 | 0 | 8 | 24 | 42 | 44 |
| ENSDARG000 | 16;16 | 32152199:321 | 32152868:321 | -;- | 748 | NA | 0 | 0 | 0 | 0 | 0 | 0 |
| ENSDARG000 | 16:16:16:16:1 | 32152612:321 | 32152838:321 | +;+;+;+;+;+ | 2694 | gprc6a | 0 | 0 | 0 | 0 | 0 | 0 |
| ENSDARG000 | 16:16:16:16:1 | 19025023:190 | 19025125:190 | -:-:-:-:-:-:-:-: | 913 | NA | 0 | 0 | 0 | 0 | 0 | 0 |
| ENSDARG000 | 16:16:16:16:1 | 5145132:5145 | 5145744:5145 | -:-:-:-:-:-:-:-: | 1588 | dctn3 | 1913 | 1821 | 1854 | 2498 | 1624 | 2135 |
| ENSDARG000 | 16:16:16:16:1 | 10522596:105 | 10522627:105 | -:-:-:-:-:-:-:-: | 1624 | g6fl | 0 | 0 | 8 | 0 | 0 | 0 |
| ENSDARG000 | 16:16:16:16:1 | 6957582:6957 | 6957714:6957 | +:+:+:+:+:+:+: | 2790 | BBS9 | 62 | 68 | 45 | 36 | 120 | 34 |
| ENSDARG000 | 16:16:16:16:1 | 38159758:381 | 38159880:381 | +:+:+:+:+:+:+: | 5108 | pi4kb | 1251 | 1605 | 1946 | 972 | 1620 | 1187 |
| ENSDARG000 | 16:16:16:16:1 | 23984179:239 | 23985179:239 | +:+:+:+:+:+:+: | 5467 | apoc2 | 0 | 0 | 8 | 0 | 6 | 2 |
| ENSDARG000 | 16:16:16:16:1 | 41060161:410 | 41060263:410 | +:+:+:+:+:+:+: | 3947 | SCAP | 567 | 379 | 816 | 263 | 776 | 491 |
| ENSDARG000 | 16:16:16:16:1 | 34916162:349 | 34916359:349 | -:-:-:-:-:-:-:-: | 4794 | ptprub | 83 | 4 | 10 | 22 | 22 | 4 |
| ENSDARG000 | 16:16:16:16:1 | 29303971:293 | 29304043:293 | +;+;+;+;+;+ | 1098 | hapln2 | 4 | 0 | 0 | 14 | 8 | 22 |
| ENSDARG000 | 16:16:16:16:1 | 10557504:106 | 10557967:106 | +;+;+;+;+;+;+ | 6267 | NA | 1013 | 404 | 1171 | 446 | 382 | 902 |
| ENSDARG000 | 16;16 | 35206855:352 | 35207243:352 | +;+ | 493 | NA | 0 | 0 | 0 | 0 | 0 | 0 |
| ENSDARG000 | 16:16:16:16:1 | 54829284:548 | 54829795:548 | +:+:+:+:+:+:+: | 2831 | pabpc1a | 36328 | 34327 | 54072 | 43146 | 37713 | 39566 |
| ENSDARG000 | 16:16:16:16:1 | 35343493:353 | 35344192:353 | +:+:+:+:+:+:+: | 5357 | NA | 62 | 72 | 40 | 72 | 66 | 52 |
| ENSDARG000 | 16:16:16:16:1 | 21547583:215 | 21552425:215 | -:-:-:-:-:-:-:-: | 8131 | DDR1 | 180 | 48 | 335 | 57 | 439 | 248 |
| ENSDARG000 | 16;16;16 | 35379720:353 | 35379865:353 | +;+;+ | 1883 | si:dkev-34d22 | 0 | 4 | 0 | 8 | 0 | 0 |
| ENSDARG000 | 16;16;16 | 10724406:107 | 10725063:107 | +;+;+ | 3859 | zgc:165409 | 136 | 140 | 208 | 106 | 212 | 187 |
| ENSDARG000 | 16:16:16:16:1 | 29310145:293 | 29311728:293 | -:-:-:-:-:-:-:-: | 5529 | bcan | 14 | 60 | 224 | 60 | 96 | 33 |
| ENSDARG000 | 16:16:16:16:1 | 5156420:5157 | 5156618:5157 | +;+;+;+;+;+;+ | 2172 | elovl4a | 50 | 0 | 2 | 39 | 2 | 45 |
| ENSDARG000 | 16:16:16:16:1 | 25673107:256 | 25674253:256 | -:-:-:-:-:-:-:-: | 2607 | tomm40 | 1383 | 1528 | 1871 | 759 | 1571 | 1193 |
| ENSDARG000 | 16:16:16:16:1 | 13883872:138 | 13884036:138 | +:+:+:+:+:+:+: | 784 | atg12 | 74 | 79 | 93 | 136 | 121 | 130 |
| ENSDARG000 | 16:16:16:16:1 | 27345383:273 | 27346222:273 | +:+:+:+:+:+:+: | 3771 | nr4a3 | 571 | 233 | 200 | 296 | 409 | 373 |
| ENSDARG000 | 16:16:16:16:1 | 10776688:107 | 10776992:107 | +:+:+:+:+:+:+: | 5084 | atp1a3b | 268 | 32 | 24 | 22 | 56 | 24 |
| ENSDARG000 | 16:16:16:16:1 | 27355441:273 | 27356219:273 | -;-;-;-;-;- | 1614 | NA | 3 | 0 | 0 | 0 | 17 | 10 |
| ENSDARG000 | 16:16:16:16:1 | 21651712:216 | 21652078:216 | -:-:-:-:-:-:-:-: | 2051 | gnl1 | 1444 | 923 | 1046 | 699 | 1207 | 1118 |
| ENSDARG000 | 16:16:16:16:1 | 24030272:240 | 24030365:240 | -:-:-:-:-:-:-:-: | 2222 | bcam | 34 | 2 | 16 | 24 | 26 | 19 |
| ENSDARG000 | 16:16:16:16:1 | 31453980:314 | 31455683:314 | -:-:-:-:-:-:-:-: | 2788 | NA | 1020 | 1277 | 887 | 1779 | 881 | 1196 |
| ENSDARG000 | 16:16:16:16:1 | 7154266:7154 | 7154843:7154 | +:+:+:+:+:+:+: | 2491 | bmper | 0 | 4 | 0 | 2 | 8 | 2 |
| ENSDARG000 | 16;16 | 16757219:167 | 16758895:167 | -;- | 1906 | NA | 0 | 0 | 0 | 0 | 0 | 0 |
| ENSDARG000 | 16:16:16:16:1 | 16824678:168 | 16824749:168 | +:+:+:+:+:+:+: | 4104 | NA | 0 | 1 | 4 | 0 | 0 | 0 |
| ENSDARG000 | 16:16:16:16:1 | 29366446:293 | 29371287:293 | -;-;-;-;-;-;- | 5807 | isg20l2 | 141 | 136 | 189 | 126 | 182 | 116 |
| ENSDARG000 | 16:16:16:16:1 | 12294284:122 | 12294835:122 | -:-:-:-:-:-:-:-: | 2557 | trpv6 | 0 | 3 | 0 | 0 | 0 | 0 |
| ENSDARG000 | 16:16:16:16:1 | 13898399:139 | 13898495:139 | +:+:+:+:+:+:+: | 1377 | NA | 2 | 14 | 28 | 29 | 20 | 6 |
| ENSDARG000 | 16:16:16:16:1 | 35385690:353 | 35387048:353 | -:-:-:-:-:-:-:-: | 3179 | NA | 0 | 0 | 0 | 0 | 0 | 0 |
| ENSDARG000 | 16:16:16:16:1 | 47454586:474 | 47454876:474 | -:-:-:-:-:-:-:-: | 1066 | CTHRC1 | 12 | 0 | 15 | 3 | 4 | 7 |
| ENSDARG000 | 16:16:16:16:1 | 12256527:122 | 12260363:122 | -:-:-:-:-:-:-:-: | 6406 | leng8 | 4321 | 2597 | 4666 | 3994 | 5480 | 4837 |
| ENSDARG000 | 16:16:16:16:1 | 12281032:122 | 12281306:122 | +:+:+:+:+:+:+: | 1469 | si:dkev-26c10 | 10 | 18 | 18 | 0 | 3 | 4 |
| ENSDARG000 | 16;16;16 | 13911692:139 | 13913002:139 | -;-;- | 2059 | rcvrn2 | 0 | 0 | 2 | 0 | 0 | 0 |
| ENSDARG000 | 16:16:16:16:1 | 27383717:273 | 27383832:273 | +:+:+:+:+:+:+: | 1573 | stx17 | 297 | 170 | 163 | 131 | 239 | 194 |
| ENSDARG000 | 16:16:16:16:1 | 12265158:122 | 12265266:122 | +:+:+:+:+:+:+: | 1075 | zgc:92606 | 1385 | 1447 | 1795 | 3947 | 1042 | 1414 |
| ENSDARG000 | 16:16:16:16:1 | 38201840:382 | 38202112:382 | +:+:+:+:+:+:+: | 4020 | myo1eb | 136 | 30 | 16 | 32 | 32 | 30 |
| ENSDARG000 | 16;16 | 31508827:315 | 31509477:315 | -;- | 733 | NA | 0 | 0 | 0 | 0 | 0 | 0 |
| ENSDARG000 | 16:16:16:16:1 | 19029297:190 | 19029507:190 | +:+:+:+:+:+:+: | 5134 | rapgef5b | 4 | 0 | 23 | 0 | 14 | 8 |
| ENSDARG000 | 16:16:16:16:1 | 31511739:315 | 31511952:315 | +:+:+:+:+:+:+: | 1778 | ndrg1b | 8 | 12 | 14 | 54 | 0 | 0 |
| ENSDARG000 | 16:16:16:16:1 | 21698980:216 | 21699301:216 | +:+:+:+:+:+:+: | 6800 | si:ch211-154 | 324 | 308 | 555 | 285 | 467 | 389 |
| ENSDARG000 | 16;16;16 | 19145147:191 | 19145679:191 | -;-;- | 803 | NA | 0 | 0 | 0 | 2 | 0 | 0 |
| ENSDARG000 | 16:16:16:16:1 | 32167430:321 | 32169208:321 | -:-:-:-:-:-:-:-: | 3312 | kpna5 | 904 | 749 | 1071 | 689 | 896 | 831 |
| ENSDARG000 | 16:16:16:16:1 | 10828787:108 | 10830504:108 | -:-:-:-:-:-:-:-: | 2325 | rabac1 | 1646 | 838 | 974 | 826 | 907 | 2049 |
| ENSDARG000 | 16;16 | 13957756:139 | 13958039:139 | -;- | 418 | NA | 0 | 0 | 0 | 0 | 0 | 0 |
| ENSDARG000 | 16:16:16:16:1 | 44881829:448 | 44882356:448 | -:-:-:-:-:-:-:-: | 1804 | rbm42 | 346 | 397 | 568 | 292 | 424 | 381 |
| ENSDARG000 | 16;16;16;16 | 13955586:139 | 13956148:139 | -;-;-;- | 1214 | NA | 6 | 0 | 0 | 0 | 0 | 0 |
| ENSDARG000 | 16:16:16:16:1 | 24188114:241 | 24190081:241 | -:-:-:-:-:-:-:-: | 3579 | rps19 | 10622 | 11470 | 16557 | 13157 | 10447 | 13621 |
| ENSDARG000 | 16;16 | 35395597:353 | 35395930:353 | +;+ | 693 | NA | 4 | 4 | 5 | 0 | 13 | 8 |
| ENSDARG000 | 16:16:16:16:1 | 29376751:293 | 29376927:293 | +:+:+:+:+:+:+: | 2604 | mettl25b | 116 | 54 | 80 | 60 | 66 | 108 |
| ENSDARG000 | 16;16;16;16 | 35401543:354 | 35401951:354 | +;+;+;+ | 1715 | rab42b | 24 | 8 | 15 | 10 | 15 | 16 |
| ENSDARG000 | 16:16:16:16:1 | 31518873:315 | 31519514:315 | -;-;-;-;- | 1600 | ccn4b | 2 | 4 | 10 | 6 | 30 | 14 |
| ENSDARG000 | 16:16:16:16:1 | 32184485:321 | 32184571:321 | +:+:+:+:+:+:+: | 2055 | zufsp | 64 | 56 | 58 | 24 | 98 | 81 |
| ENSDARG000 | 16:16:16:16:1 | 41112346:411 | 41113580:411 | -:-:-:-:-:-:-:-: | 6762 | ptpn23a | 1427 | 986 | 1607 | 821 | 1717 | 1350 |
| ENSDARG000 | 16:16:16:16:1 | 10841163:108 | 10841216:108 | +:+:+:+:+:+:+: | 3983 | dedd1 | 2816 | 682 | 1969 | 1635 | 2770 | 3827 |
| ENSDARG000 | 16;16 | 13965923:139 | 13966177:139 | +;+ | 2653 | zgc:162509 | 16 | 15 | 13 | 6 | 40 | 16 |
| ENSDARG000 | 16:16:16:16:1 | 44906324:449 | 44906621:449 | +:+:+:+:+:+:+: | 1834 | cd22 | 31 | 40 | 24 | 20 | 42 | 16 |
| ENSDARG000 | 16:16:16:16:1 | 21751922:217 | 21752275:217 | -;-;-;-;-;-;-;- | 1196 | cops7a | 795 | 687 | 901 | 442 | 691 | 662 |
| ENSDARG000 | 16;16;16 | 13987735:139 | 13988082:139 | -;-;- | 869 | NA | 36 | 22 | 161 | 116 | 12 | 22 |
| ENSDARG000 | 16:16:16:16:1 | 32197554:321 | 32197856:321 | -:-:-:-:-:-:-:-: | 1185 | rwdd1 | 1306 | 1400 | 1651 | 1024 | 1438 | 1349 |
| ENSDARG000 | 16;16;16 | 54875530:548 | 54875894:548 | +;+;+ | 3063 | nr0b2a | 16 | 22 | 8 | 4 | 18 | 4 |
| ENSDARG000 | 16;16;16 | 29385565:293 | 29385928:293 | -;-;- | 628 | s100a1 | 0 | 0 | 0 | 0 | 0 | 0 |
| ENSDARG000 | 16:16:16:16:1 | 21779920:217 | 21780167:217 | -;-;-;-;-;- | 1054 | cldc1 | 64 | 142 | 98 | 82 | 4 | 6 |
| ENSDARG000 | 16:16:16:16:1 | 27442549:274 | 27442706:274 | +:+:+:+:+:+:+: | 3827 | invs | 248 | 292 | 533 | 456 | 294 | 236 |
| ENSDARG000 | 16:16:16:16:1 | 44920012:449 | 44920066:449 | -:-:-:-:-:-:-:-: | 2178 | NA | 0 | 10 | 11 | 14 | 12 | 10 |
| ENSDARG000 | 16:16:16:16:1 | 27417200:274 | 27418075:274 | -:-:-:-:-:-:-:-: | 2083 | erp44 | 2315 | 2867 | 4644 | 3176 | 3349 | 2557 |
| ENSDARG000 | 16:16:16:16:1 | 54890835:548 | 54895299:548 | -;-;-;-;-;-;- | 6003 | NA | 695 | 545 | 344 | 379 | 484 | 716 |
| ENSDARG000 | 16:16:16:16:1 | 13993351:140 | 13993406:140 | +:+:+:+:+:+:+: | 5435 | fdps | 692 | 492 | 575 | 254 | 316 | 309 |
| ENSDARG000 | 16:16:16:16:1 | 54930747:549 | 54933350:549 | -:-:-:-:-:-:-:-: | 3139 | tmem222a | 297 | 287 | 380 | 236 | 336 | 344 |
| ENSDARG000 | 16;16;16 | 13993285:140 | 13993406:140 | +;+;+ | 2202 | NA | 0 | 3 | 28 | 8 | 6 | 0 |
| ENSDARG000 | 16:16:16:16:1 | 47531516:475 | 47531675:475 | -:-:-:-:-:-:-:-: | 11025 | CSMD3 | 42 | 0 | 12 | 2 | 8 | 20 |
| ENSDARG000 | 16:16:16:16:1 | 38240027:382 | 38240196:382 | +:+:+:+:+:+:+: | 2197 | prune | 186 | 177 | 287 | 152 | 164 | 180 |
| ENSDARG000 | 16:16:16:16:1 | 10918252:109 | 10919534:109 | +:+:+:+:+:+:+: | 7182 | NA | 11 | 10 | 13 | 11 | 23 | 0 |
| ENSDARG000 | 16:16:16:16:1 | 16849220:168 | 16849433:168 | +:+:+:+:+:+:+: | 13151 | NA | 392 | 323 | 666 | 326 | 680 | 684 |
| ENSDARG000 | 16:16:16:16:1 | 24213078:242 | 24214593:242 | -:-:-:-:-:-:-:-: | 2826 | cadm4 | 70 | 40 | 112 | 14 | 168 | 76 |
| ENSDARG000 | 16:16:16:16:1 | 35408856:354 | 35409661:354 | -:-:-:-:-:-:-:-: | 1612 | taf12 | 1178 | 1110 | 1408 | 1282 | 1180 | 1110 |
| ENSDARG000 | 16:16:16:16:1 | 29387675:293 | 29392372:293 | -:-:-:-:-:-:-:-: | 6986 | tlr18 | 10 | 0 | 2 | 0 | 0 | 0 |
| ENSDARG000 | 16;16 | 24516092:245 | 24516136:245 | +;+ | 854 | NA | 0 | 0 | 0 | 0 | 0 | 0 |
| ENSDARG000 | 16:16:16:16:1 | 25682577:256 | 25683958:256 | -:-:-:-:-:-:-:-: | 3835 | NA | 1969 | 2138 | 2974 | 1636 | 3117 | 2161 |
| ENSDARG000 | 16:16:16:16:1 | 35415575:354 | 35415740:354 | -:-:-:-:-:-:-:-: | 2886 | ctps1b | 158 | 330 | 264 | 232 | 308 | 234 |
| ENSDARG000 | 16;16 | 21807422:218 | 21807908:218 | -;- | 805 | NA | 0 | 0 | 0 | 0 | 10 | 0 |
| ENSDARG000 | 16;16 | 21800385:218 | 21801746:218 | -;- | 1478 | NA | 0 | 0 | 0 | 0 | 0 | 0 |
| ENSDARG000 | 16;16;16;16 | 21808418:218 | 21808731:218 | -;-;-;- | 995 | dicp3.3 | 46 | 21 | 70 | 21 | 110 | 46 |
| ENSDARG000 | 16:16:16:16:1 | 21789703:217 | 21789832:217 | +:+:+:+:+:+:+: | 2491 | trim108 | 4 | 4 | 0 | 0 | 0 | 0 |
| ENSDARG000 | 16;16;16;16 | 41135018:411 | 41135300:411 | +;+;+;+ | 566 | NA | 0 | 0 | 0 | 0 | 0 | 0 |
| ENSDARG000 | 16:16:16:16:1 | 29407666:294 | 29414701:294 | -:-:-:-:-:-:-:-: | 8713 | si:ch211-113 | 0 | 4 | 16 | 10 | 10 | 8 |
| ENSDARG000 | 16:16:16:16:1 | 52706059:527 | 52706176:527 | -:-:-:-:-:-:-:-: | 3196 | azin1a | 222 | 266 | 376 | 274 | 305 | 196 |
| ENSDARG000 | 16;16 | 32205660:322 | 32206514:322 | -;- | 1418 | CALHM5 | 15 | 80 | 42 | 71 | 77 | 100 |
| ENSDARG000 | 16:16:16:16:1 | 31530762:315 | 31531025:315 | -:-:-:-:-:-:-:-: | 8356 | tg | 0 | 18 | 2 | 0 | 6 | 10 |
| ENSDARG000 | 16:16:16:16:1 | 24522824:245 | 24523568:245 | -;-;-;-;-;-;- | 1840 | si:ch211-79k1 | 52 | 96 | 68 | 16 | 220 | 90 |
| ENSDARG000 | 16;16 | 32216552:322 | 32217745:322 | -;- | 1794 | calhm5.1 | 19 | 12 | 12 | 23 | 29 | 13 |
| ENSDARG000 | 16;16 | 32226664:322 | 32227384:322 | -;- | 1503 | calhm6 | 10 | 6 | 26 | 0 | 50 | 46 |
| ENSDARG000 | 16:16:16:16:1 | 54947768:549 | 54949378:549 | -:-:-:-:-:-:-:-: | 3679 | wdtc1 | 619 | 430 | 732 | 558 | 471 | 627 |
| ENSDARG000 | 16:16:16:16:1 | 41162854:411 | 41162932:411 | +;+;+;+;+;+;+ | 1323 | sc:d156 | 0 | 0 | 0 | 0 | 0 | 0 |
| ENSDARG000 | 16:16:16:16:1 | 24551898:245 | 24554648:245 | -;-;-;-;-;- | 3211 | si:ch211-79k1 | 375 | 263 | 448 | 523 | 1233 | 507 |
| ENSDARG000 | 16:16:16:16:1 | 44983613:449 | 44985358:449 | -;-;-;-;-;-;- | 2777 | sult2st3 | 114 | 62 | 49 | 40 | 66 | 170 |
| ENSDARG000 | 16:16:16:16:1 | 38277339:382 | 38277481:382 | +:+:+:+:+:+:+: | 1812 | bnipl | 64 | 12 | 8 | 6 | 36 | 24 |
| ENSDARG000 | 16:16:16:16:1 | 41171949:411 | 41172130:411 | +:+:+:+:+:+:+: | 2062 | NA | 0 | 0 | 2 | 0 | 0 | 0 |
| ENSDARG000 | 16:16:16:16:1 | 19181163:191 | 19182959:191 | -:-:-:-:-:-:-:-: | 10741 | DNAH11 | 38 | 18 | 44 | 126 | 179 | 90 |
| ENSDARG000 | 16:16:16:16:1 | 38311887:383 | 38313421:383 | -;-;-;-;- | 2158 | cdc42se1 | 2429 | 2366 | 2115 | 2385 | 1396 | 1927 |
| ENSDARG000 | 16:16:16:16:1 | 19306522:193 | 19306602:193 | -:-:-:-:-:-:-:-: | 6084 | NA | 0 | 0 | 3 | 6 | 18 | 8 |
| ENSDARG000 | 16;16;16;16 | 29455974:294 | 29458765:294 | -;-;-;- | 3072 | lingo4b | 593 | 606 | 861 | 522 | 1278 | 1088 |
| ENSDARG000 | 16:16:16:16:1 | 32238020:322 | 32238163:322 | +:+:+:+:+:+:+: | 3295 | klhl32 | 667 | 498 | 468 | 700 | 436 | 445 |
| ENSDARG000 | 16;16;16 | 41465511:414 | 41465552:414 | +;+;+ | 609 | NA | 0 | 2 | 1 | 1 | 4 | 0 |
| ENSDARG000 | 16:16:16:16:1 | 41467188:414 | 41470573:414 | -;-;-;-;-;-;-;- | 3994 | cmtm6 | 4411 | 5665 | 7845 | 7190 | 7860 | 8007 |
| ENSDARG000 | 16:16:16:16:1 | 41345477:413 | 41346285:413 | -:-:-:-:-:-:-:-: | 3010 | CPNE4 | 12 | 36 | 26 | 8 | 36 | 12 |
| ENSDARG000 | 16:16:16:16:1 | 38350704:383 | 38351106:383 | +:+:+:+:+:+:+: | 4913 | elp6 | 192 | 138 | 226 | 161 | 181 | 114 |
| ENSDARG000 | 16:16:16:16:1 | 38337783:383 | 38337955:383 | +:+:+:+:+:+:+: | 1935 | gabpb2b | 115 | 132 | 205 | 132 | 171 | 167 |
| ENSDARG000 | 16:16:16:16:1 | 12335274:123 | 12335868:123 | -:-:-:-:-:-:-:-: | 3518 | NA | 36 | 16 | 14 | 6 | 10 | 22 |
| ENSDARG000 | 16:16:16:16:1 | 52748263:527 | 52748417:527 | +:+:+:+:+:+:+: | 1869 | atp6v1c1a | 1718 | 1698 | 2119 | 2004 | 1990 | 1896 |
| ENSDARG000 | 16:16:16:16:1 | 45019449:450 | 45019890:450 | -;-;-;-;-;-;- | 1113 | tmem147 | 900 | 1213 | 1418 | 862 | 1148 | 1150 |
| ENSDARG000 | 16:16:16:16:1 | 32281200:322 | 32281605:322 | -:-:-:-:-:-:-:-: | 4401 | mms22l | 82 | 350 | 354 | 244 | 276 | 170 |
| ENSDARG000 | 16:16:16:16:1 | 29492749:294 | 29493533:294 | +:+:+:+:+:+:+: | 2397 | ctsk | 1027 | 282 | 326 | 489 | 1674 | 377 |
| ENSDARG000 | 16:16:16:16:1 | 24590538:245 | 24591806:245 | -:-:-:-:-:-:-:-: | 2037 | fxyd7 | 27 | 17 | 26 | 52 | 59 | 76 |
| ENSDARG000 | 16:16:16:16:1 | 25701985:257 | 25703009:257 | -:-:-:-:-:-:-:-: | 4376 | NA | 531 | 725 | 653 | 1714 | 678 | 391 |
| ENSDARG000 | 16:16:16:16:1 | 54983349:549 | 54983418:549 | +;+;+;+;+;+ | 286 | NA | 0 | 0 | 4 | 3 | 10 | 0 |
| ENSDARG000 | 16;16 | 21822734:218 | 21823040:218 | -;- | 391 | NA | 2 | 0 | 0 | 0 | 0 | 10 |
| ENSDARG000 | 16:16:16:16:1 | 27508381:275 | 27508587:275 | -:-:-:-:-:-:-:-: | 3130 | tex10 | 533 | 549 | 1043 | 460 | 753 | 537 |
| ENSDARG000 | 16:16:16:16:1 | 21827332:218 | 21827423:218 | -;-;-;-;-;-;- | 835 | NA | 0 | 0 | 2 | 0 | 0 | 4 |
| ENSDARG000 | 16:16:16:16:1 | 14029283:140 | 14029501:140 | +:+:+:+:+:+:+: | 2254 | NA | 13 | 11 | 10 | 12 | 36 | 7 |
| ENSDARG000 | 16:16:16:16:1 | 45044642:450 | 45045033:450 | -:-:-:-:-:-:-:-: | 1452 | gapdhs | 17435 | 38436 | 65255 | 38527 | 29752 | 19608 |
| ENSDARG000 | 16:16:16:16:1 | 32305253:323 | 32305440:323 | +;+;+;+;+;+;+ | 2242 | NA | 2 | 0 | 0 | 0 | 0 | 0 |
| ENSDARG000 | 16;16;16 | 21851856:218 | 21852170:218 | -;-;- | 692 | NA | 0 | 2 | 1 | 0 | 28 | 8 |
| ENSDARG000 | 16 | 32559821 | 32563092 | + | 3272 | pou3f2b | 4 | 0 | 0 | 0 | 0 | 0 |
| ENSDARG000 | 16;16 | 45077593:450 | 45077810:450 | -;- | 578 | NA | 0 | 0 | 0 | 4 | 0 | 4 |
| ENSDARG000 | 16:16:16:16:1 | 25761101:257 | 25761711:257 | +:+:+:+:+:+:+: | 2191 | cblc | 38 | 3 | 12 | 0 | 5 | 12 |
| ENSDARG000 | 16:16:16:16:1 | 24626448:246 | 24626590:246 | +:+:+:+:+:+:+: | 1781 | NA | 1106 | 71 | 52 | 63 | 112 | 112 |
| ENSDARG000 | 16;16;16 | 25822455:258 | 25824199:258 | -;-;- | 1865 | irge4 | 694 | 522 | 382 | 318 | 245 | 303 |
| ENSDARG000 | 16:16:16:16:1 | 25806400:258 | 25806603:258 | +;+;+;+;+ | 5409 | NA | 334 | 228 | 310 | 242 | 204 | 236 |
| ENSDARG000 | 16;16 | 25989943:259 | 25990137:259 | +;+ | 594 | NA | 0 | 0 | 0 | 2 | 0 | 0 |
| ENSDARG000 | 16:16:16:16:1 | 21881694:218 | 21882547:218 | -:-:-:-:-:-:-:-: | 1886 | dicp3.1 | 19 | 25 | 66 | 23 | 134 | 88 |
| ENSDARG000 | 16:16:16:16:1 | 24601140:246 | 24601553:246 | -:-:-:-:-:-:-:-: | 1275 | fxyd6l | 1875 | 960 | 1837 | 3168 | 2635 | 3548 |
| ENSDARG000 | 16:16:16:16:1 | 32566218:325 | 32567462:325 | -;-;-;-;-;-;- | 3130 | fbxl4 | 333 | 177 | 218 | 263 | 304 | 174 |
| ENSDARG000 | 16:16:16:16:1 | 55003121:550 | 55004823:550 | -:-:-:-:-:-:-:-: | 3368 | cndp1 | 0 | 0 | 0 | 3 | 2 | 0 |
| ENSDARG000 | 16:16:16:16:1 | 38360002:383 | 38360195:383 | +:+:+:+:+:+:+: | 2467 | zgc:113232 | 8 | 2 | 24 | 7 | 20 | 10 |
| ENSDARG000 | 16:16:16:16:1 | 29509133:295 | 29509195:295 | +:+:+:+:+:+:+: | 1416 | ctss2.1 | 192 | 237 | 315 | 258 | 1030 | 488 |
| ENSDARG000 | 16:16:16:16:1 | 26012569:260 | 26012632:260 | +;+;+;+;+;+ | 850 | prss59.1 | 6 | 0 | 0 | 0 | 0 | 2 |
| ENSDARG000 | 16:16:16:16:1 | 32630074:326 | 32630342:326 | -:-:-:-:-:-:-:-: | 1493 | FAXC | 16 | 4 | 4 | 0 | 14 | 10 |
| ENSDARG000 | 16:16:16:16:1 | 14064397:140 | 14065093:140 | -;-;-;-;-;- | 1765 | trim109 | 0 | 0 | 0 | 0 | 0 | 0 |
| ENSDARG000 | 16:16:16:16:1 | 21891811:218 | 21892465:218 | -:-:-:-:-:-:-:-: | 5190 | setdb1b | 1026 | 817 | 1305 | 771 | 1032 | 756 |
| ENSDARG000 | 16:16:16:16:1 | 14078748:140 | 14078886:140 | +;+;+;+;+;+;+ | 962 | NA | 0 | 0 | 1 | 0 | 2 | 0 |
| ENSDARG000 | 16:16:16:16:1 | 26017360:260 | 26017443:260 | +;+;+;+;+;+ | 879 | prss59.2 | 4 | 0 | 0 | 12 | 2 | 4 |
| ENSDARG000 | 16:16:16:16:1 | 27543308:275 | 27543589:275 | +:+:+:+:+:+:+: | 2694 | NA | 516 | 546 | 666 | 426 | 556 | 389 |
| ENSDARG000 | 16:16:16:16:1 | 38394371:383 | 38394442:383 | +:+:+:+:+:+:+: | 1311 | cd83 | 150 | 74 | 175 | 120 | 536 | 324 |
| ENSDARG000 | 16:16:16:16:1 | 32662469:326 | 32664706:326 | -:-:-:-:-:-:-:-: | 6778 | pnisr | 2201 | 1296 | 2803 | 1839 | 2599 | 2265 |
| ENSDARG000 | 16:16:16:16:1 | 32652350:326 | 32652895:326 | -;-;-;-;-;-;-;- | 1834 | coq3 | 242 | 203 | 210 | 165 | 170 | 175 |
| ENSDARG000 | 16:16:16:16:1 | 19422295:194 | 19422459:194 | -;-;-;-;- | 537 | NA | 0 | 1 | 0 | 6 | 0 | 0 |
| ENSDARG000 | 16:16:16:16:1 | 29514473:295 | 29514530:295 | +:+:+:+:+:+:+: | 1445 | ctss2.2 | 247 | 108 | 195 | 404 | 570 | 321 |
| ENSDARG000 | 16:16:16:16:1 | 29521736:295 | 29523467:295 | -;-;-;-;- | 3353 | onecutl | 0 | 4 | 0 | 0 | 0 | 0 |
| ENSDARG000 | 16;16;16 | 29515971:295 | 29516184:295 | -;-;- | 428 | NA | 6 | 0 | 0 | 0 | 5 | 0 |
| ENSDARG000 | 16;16;16 | 26020886:260 | 26021282:260 | +;+;+ | 918 | mrpl17 | 274 | 356 | 401 | 175 | 341 | 340 |
| ENSDARG000 | 16:16:16:16:1 | 38453876:384 | 38455547:384 | -;-;-;-;-;-;-;- | 2299 | rspo2 | 10 | 4 | 0 | 0 | 0 | 0 |
| ENSDARG000 | 16:16:16:16:1 | 19537038:195 | 19541635:195 | -:-:-:-:-:-:-:-: | 8897 | NA | 41 | 2 | 0 | 10 | 16 | 45 |
| ENSDARG000 | 16:16:16:16:1 | 19536614:195 | 19536959:195 | +;+;+;+;+;+;+ | 2050 | sp8b | 0 | 0 | 0 | 0 | 0 | 0 |
| ENSDARG000 | 16:16:16:16:1 | 21917673:219 | 21918314:219 | +:+:+:+:+:+:+: | 4536 | znf687a | 294 | 208 | 273 | 217 | 296 | 305 |

| | | | | | | | | | | | |
|---|---|---|---|---|---|---|---|---|---|---|---|
| ENSDARG000 | 16:16:16:16:1 | 26027011:26( | 26027269:26( | -:-:-:-:-:-:-:-: | 1756 | TMEM145 | 0 | 10 | 29 | 23 | 50 | 65 |
| ENSDARG000 | 16:16:16:16:1 | 21946091:21 | 21947298:21 | -:-:-:-:-:-:-:-: | 5927 | NA | 0 | 10 | 0 | 0 | 4 | 8 |
| ENSDARG000 | 16;16;16;16 | 22009140:22( | 22009249:22( | -;-;-;- | 335 | NA | 0 | 0 | 0 | 0 | 0 | 0 |
| ENSDARG000 | 16:16:16:16:1 | 22015648:22( | 22016276:22( | -;-;-;-;-;-;-;- | 1174 | NA | 0 | 0 | 0 | 0 | 0 | 0 |
| ENSDARG000 | 16:16:16:16:1 | 1100559:1101 | 1100952:1101 | +:+:+:+:+:+:+: | 3973 | ADAMTS16 | 24 | 33 | 17 | 0 | 32 | 0 |
| ENSDARG000 | 16:16:16:16:1 | 6395311:6395 | 6395944:6396 | -:-:-:-:-:-:-:-: | 2243 | mboat1 | 8 | 20 | 60 | 0 | 39 | 12 |
| ENSDARG000 | 16:16:16:16:1 | 8716800:8716 | 8717076:8717 | +:+:+:+:+:+:+: | 5437 | CCR8 | 2 | 8 | 17 | 2 | 94 | 50 |
| ENSDARG000 | 16:16:16:16:1 | 55059026:55( | 55059117:55( | +;+;+;+;+;+;+ | 1172 | NA | 57 | 55 | 2 | 0 | 8 | 22 |
| ENSDARG000 | 16:16:16:16:1 | 21016820:21( | 21016839:21( | +:+:+:+:+:+:+: | 309 | NA | 2 | 4 | 0 | 13 | 8 | 8 |
| ENSDARG000 | 16;16;16 | 33987892:339 | 33988218:339 | +;+;+ | 1942 | pigv | 98 | 116 | 219 | 190 | 205 | 191 |
| ENSDARG000 | 16;16 | 28087866:28( | 28088271:28( | -;- | 1333 | NA | 0 | 0 | 0 | 2 | 0 | 0 |
| ENSDARG000 | 16;16;16;16 | 6386941:6387 | 6387172:6387 | -;-;-;- | 637 | ndufb9 | 1327 | 1003 | 1237 | 924 | 1042 | 903 |
| ENSDARG000 | 16:16:16:16:1 | 2565433:2565 | 2565626:2565 | +:+:+:+:+:+:+: | 5909 | preb | 614 | 803 | 890 | 466 | 859 | 538 |
| ENSDARG000 | 16:16:16:16:1 | 1802307:1803 | 1802586:1804 | +:+:+:+:+:+:+: | 7921 | GRIK2 | 0 | 0 | 4 | 0 | 6 | 0 |
| ENSDARG000 | 16:16:16:16:1 | 49088698:49( | 49088724:49( | +:+:+:+:+:+:+: | 3598 | PLCL2 | 233 | 493 | 333 | 175 | 610 | 265 |
| ENSDARG000 | 16:16:16:16:1 | 31676281:316 | 31678579:316 | -;-;-;-;-;-;-;- | 3736 | c1s.1 | 32 | 24 | 31 | 38 | 48 | 56 |
| ENSDARG000 | 16:16:16:16:1 | 36072329:36( | 36076192:36( | -:-:-:-:-:-:-:-: | 4887 | OSCP1 | 68 | 92 | 79 | 51 | 85 | 94 |
| ENSDARG000 | 16;16;16 | 20161805:201 | 20161857:201 | +;+;+ | 443 | c16h2orf66 | 0 | 0 | 0 | 0 | 0 | 0 |
| ENSDARG000 | 16 | 48158309 | 48158423 | + | 115 | 5S_rRNA | 0 | 0 | 0 | 0 | 0 | 0 |
| ENSDARG000 | 16 | 50961246 | 50961362 | - | 117 | 5S_rRNA | 0 | 0 | 0 | 0 | 0 | 0 |
| ENSDARG000 | 16 | 12250268 | 12250323 | - | 56 | U7 | 0 | 0 | 7 | 1 | 11 | 0 |
| ENSDARG000 | 16;16 | 46583604:465 | 46585843:465 | -;- | 2294 | NA | 0 | 0 | 8 | 0 | 8 | 104 |
| ENSDARG000 | 16:16:16:16:1 | 36095582:361 | 36100182:361 | -;-;-;-;-;-;-;- | 5598 | mrps15 | 360 | 454 | 593 | 428 | 388 | 356 |
| ENSDARG000 | 16:16:16:16:1 | 24705852:247 | 24706398:247 | -;-;-;-;- | 965 | NA | 0 | 0 | 0 | 0 | 0 | 0 |
| ENSDARG000 | 16:16:16:16:1 | 54402310:544 | 54402603:544 | -;-;-;-;-;-;- | 1125 | osr2 | 6 | 0 | 6 | 0 | 14 | 0 |
| ENSDARG000 | 16;16;16 | 51326237:513 | 51326281:513 | +;+;+ | 483 | C16H1orf232 | 4 | 0 | 0 | 0 | 0 | 0 |
| ENSDARG000 | 16;16;16;16 | 27157683:271 | 27158775:271 | -;-;-;- | 1979 | NA | 335 | 493 | 392 | 232 | 493 | 374 |
| ENSDARG000 | 16;16 | 46807214:468 | 46807461:468 | +;+ | 261 | NA | 0 | 0 | 0 | 0 | 0 | 0 |
| ENSDARG000 | 16;16;16;16 | 54925437:549 | 54926948:549 | -;-;-;- | 2136 | tpbgl | 6 | 4 | 0 | 0 | 0 | 6 |
| ENSDARG000 | 16:16:16:16:1 | 36038130:36( | 36038404:36( | -:-:-:-:-:-:-:-: | 1429 | STK40 | 1032 | 953 | 1427 | 1116 | 1178 | 900 |
| ENSDARG000 | 16;16;16;16 | 30316177:303 | 30316794:303 | +;+;+;+ | 4344 | NA | 38 | 0 | 8 | 8 | 12 | 0 |
| ENSDARG000 | 16 | 48023293 | 48023408 | + | 116 | 5S_rRNA | 0 | 0 | 0 | 0 | 0 | 0 |
| ENSDARG000 | 16 | 49995478 | 49995594 | - | 117 | 5S_rRNA | 0 | 0 | 0 | 0 | 0 | 0 |
| ENSDARG000 | 16 | 24394346 | 24394463 | - | 118 | 5S_rRNA | 0 | 0 | 0 | 0 | 0 | 0 |
| ENSDARG000 | 16:16:16:16:1 | 22177254:221 | 22179278:221 | -:-:-:-:-:-:-:-: | 4005 | il6r | 46 | 28 | 64 | 18 | 100 | 212 |
| ENSDARG000 | 16:16:16:16:1 | 49393200:493 | 49393390:493 | -:-:-:-:-:-:-:-: | 2457 | NA | 75 | 14 | 34 | 7 | 36 | 44 |
| ENSDARG000 | 16 | 40183743 | 40183859 | + | 117 | 5S_rRNA | 0 | 0 | 0 | 0 | 0 | 0 |
| ENSDARG000 | 16 | 23466094 | 23471387 | - | 5294 | NA | 0 | 0 | 0 | 0 | 4 | 0 |
| ENSDARG000 | 16:16:16:16:1 | 2515223:2517 | 2515405:2517 | -:-:-:-:-:-:-:-: | 3360 | ADCY3 | 302 | 103 | 143 | 110 | 108 | 183 |
| ENSDARG000 | 16 | 25328531 | 25328647 | - | 117 | 5S_rRNA | 0 | 0 | 0 | 5 | 0 | 0 |
| ENSDARG000 | 16 | 32574110 | 32574236 | - | 127 | 5S_rRNA | 0 | 0 | 0 | 0 | 0 | 0 |
| ENSDARG000 | 16 | 5107298 | 5107417 | - | 120 | 5S_rRNA | 0 | 0 | 0 | 0 | 0 | 0 |
| ENSDARG000 | 16 | 53542355 | 53542469 | + | 115 | 5S_rRNA | 0 | 0 | 0 | 0 | 0 | 0 |
| ENSDARG000 | 16 | 44605326 | 44605440 | + | 115 | 5S_rRNA | 0 | 0 | 0 | 0 | 0 | 0 |
| ENSDARG000 | 16 | 26411646 | 26411778 | + | 133 | 5S_rRNA | 0 | 0 | 0 | 0 | 0 | 0 |
| ENSDARG000 | 16 | 24898171 | 24898253 | + | 83 | dre-mir-125a- | 0 | 0 | 0 | 0 | 0 | 0 |
| ENSDARG000 | 16:16:16:16:1 | 6182501:6183 | 6183293:6183 | -:-:-:-:-:-:-:-: | 3376 | ctnnb1 | 12590 | 10369 | 12981 | 8084 | 11481 | 7586 |
| ENSDARG000 | 16 | 4527481 | 4527647 | - | 167 | NA | 0 | 0 | 1 | 0 | 11 | 0 |
| ENSDARG000 | 16:16:16:16:1 | 36901549:369 | 36902052:369 | -;-;-;-;-;- | 1613 | NA | 8 | 8 | 0 | 4 | 8 | 8 |
| ENSDARG000 | 16 | 27415697 | 27415812 | + | 116 | 5S_rRNA | 0 | 0 | 0 | 0 | 0 | 0 |
| ENSDARG000 | 16;16;16 | 43347966:433 | 43348031:433 | +;+;+ | 1568 | ccdc127a | 73 | 52 | 102 | 69 | 153 | 30 |
| ENSDARG000 | 16 | 34724568 | 34724682 | + | 115 | 5S_rRNA | 0 | 0 | 0 | 0 | 0 | 0 |
| ENSDARG000 | 16:16:16:16:1 | 47198112:472 | 47198140:472 | -:-:-:-:-:-:-:-: | 438 | NA | 0 | 0 | 1 | 0 | 7 | 3 |
| ENSDARG000 | 16 | 2729373 | 2729489 | + | 117 | 5S_rRNA | 0 | 0 | 0 | 0 | 0 | 0 |
| ENSDARG000 | 16 | 12239407 | 12239460 | - | 54 | U7 | 0 | 0 | 1 | 2 | 2 | 0 |
| ENSDARG000 | 16;16 | 221739:22412 | 221945:22948 | +;+ | 5561 | nrsn1 | 22 | 63 | 78 | 4 | 34 | 0 |
| ENSDARG000 | 16;16 | 44754799:447 | 44759382:447 | -;- | 6152 | NA | 4 | 0 | 0 | 0 | 0 | 0 |
| ENSDARG000 | 16 | 50507176 | 50507295 | + | 120 | 5S_rRNA | 0 | 0 | 0 | 0 | 0 | 0 |
| ENSDARG000 | 16;16 | 50917913:509 | 50917972:509 | -;- | 936 | NA | 0 | 1 | 2 | 0 | 12 | 14 |
| ENSDARG000 | 16;16;16;16 | 29043813:29( | 29044628:29( | +;+;+;+ | 8377 | nes | 22 | 4 | 28 | 86 | 0 | 22 |
| ENSDARG000 | 16 | 45666735 | 45666852 | + | 118 | 5S_rRNA | 0 | 3 | 1 | 0 | 1 | 4 |
| ENSDARG000 | 16 | 38021019 | 38021134 | + | 116 | 5S_rRNA | 2 | 0 | 1 | 0 | 0 | 0 |
| ENSDARG000 | 16;16;16;16 | 6336952:6342 | 6337186:6342 | +;+;+;+ | 2376 | cmtm8b | 32 | 8 | 24 | 17 | 10 | 8 |
| ENSDARG000 | 16:16:16:16:1 | 52901490:529 | 52901695:529 | -;-;-;-;-;-;-;- | 954 | ZDHHC11B | 0 | 8 | 48 | 31 | 16 | 22 |
| ENSDARG000 | 16:16:16:16:1 | 248239:2496( | 248262:24967 | -:-:-:-:-:-:-:-: | 1890 | slc6a3 | 0 | 0 | 0 | 4 | 10 | 0 |
| ENSDARG000 | 16 | 40463365 | 40464426 | + | 1062 | NA | 0 | 0 | 0 | 0 | 10 | 2 |
| ENSDARG000 | 16:16:16:16:1 | 43401005:434 | 43401560:434 | +:+:+:+:+:+:+: | 2093 | sqlea | 99 | 80 | 245 | 27 | 31 | 32 |
| ENSDARG000 | 16 | 43555464 | 43555579 | - | 116 | 5S_rRNA | 0 | 0 | 0 | 0 | 0 | 0 |
| ENSDARG000 | 16 | 43679267 | 43679611 | - | 345 | NA | 0 | 6 | 4 | 0 | 0 | 0 |
| ENSDARG000 | 16:16:16:16:1 | 22199510:222 | 22199552:222 | -:-:-:-:-:-:-:-: | 1578 | NA | 0 | 0 | 6 | 0 | 20 | 2 |
| ENSDARG000 | 16:16:16:16:1 | 53590586:535 | 53590966:535 | +:+:+:+:+:+:+: | 2166 | gpatch3 | 238 | 168 | 235 | 90 | 178 | 170 |
| ENSDARG000 | 16:16:16:16:1 | 52847079:528 | 52847164:528 | +:+:+:+:+:+:+: | 1820 | cep72 | 49 | 76 | 66 | 52 | 122 | 34 |
| ENSDARG000 | 16:16:16:16:1 | 26824691:268 | 26825280:268 | +;+;+;+;+;+ | 3255 | NA | 126 | 68 | 80 | 84 | 52 | 68 |
| ENSDARG000 | 16 | 30648387 | 30648504 | - | 118 | 5S_rRNA | 0 | 0 | 0 | 0 | 0 | 0 |
| ENSDARG000 | 16 | 10916290 | 10916404 | - | 115 | 5S_rRNA | 0 | 0 | 0 | 0 | 0 | 0 |
| ENSDARG000 | 16:16:16:16:1 | 17639896:176 | 17640763:176 | -;-;-;-;-;-;-;- | 2168 | ccdc106a | 487 | 430 | 553 | 549 | 428 | 521 |
| ENSDARG000 | 16:16:16:16:1 | 54459135:544 | 54459400:544 | -:-:-:-:-:-:-:-: | 2489 | cyp11c1 | 4 | 4 | 6 | 6 | 2 | 4 |
| ENSDARG000 | 16:16:16:16:1 | 54633238:546 | 54633282:546 | -:-:-:-:-:-:-:-: | 1713 | spag1b | 4 | 0 | 6 | 2 | 0 | 0 |
| ENSDARG000 | 16 | 53917415 | 53919115 | - | 1701 | fzd1 | 187 | 72 | 162 | 33 | 168 | 121 |
| ENSDARG000 | 16 | 8478880 | 8479009 | - | 130 | 5S_rRNA | 0 | 0 | 0 | 0 | 0 | 0 |
| ENSDARG000 | 16 | 46340543 | 46340657 | - | 115 | 5S_rRNA | 0 | 0 | 0 | 0 | 0 | 0 |
| ENSDARG000 | 16:16:16:16:1 | 26113457:261 | 26113472:261 | +:+:+:+:+:+:+: | 258 | NA | 2 | 1 | 4 | 5 | 4 | 8 |
| ENSDARG000 | 16:16:16:16:1 | 36126310:361 | 36127032:361 | +:+:+:+:+:+:+: | 3758 | sh3bp5b | 184 | 24 | 380 | 72 | 106 | 114 |
| ENSDARG000 | 16:16:16:16:1 | 21673723:216 | 21674043:216 | -;-;-;-;-;-;-;- | 1311 | znf384b | 243 | 237 | 243 | 129 | 206 | 162 |
| ENSDARG000 | 16 | 17516948 | 17517082 | + | 135 | 5S_rRNA | 0 | 0 | 0 | 0 | 0 | 0 |
| ENSDARG000 | 16:16:16:16:1 | 39498:50370: | 39646:50550: | +;+;+;+;+;+ | 1032 | CDKAL1 | 356 | 360 | 480 | 188 | 298 | 272 |
| ENSDARG000 | 16 | 47012341 | 47012457 | + | 117 | 5S_rRNA | 0 | 0 | 0 | 0 | 0 | 0 |
| ENSDARG000 | 16 | 29155738 | 29155852 | - | 115 | 5S_rRNA | 0 | 0 | 0 | 0 | 0 | 0 |
| ENSDARG000 | 16:16:16:16:1 | 54166932:541 | 54166981:541 | -:-:-:-:-:-:-:-: | 951 | NA | 702 | 252 | 431 | 343 | 405 | 418 |
| ENSDARG000 | 16:16:16:16:1 | 43368572:433 | 43368689:433 | +:+:+:+:+:+:+: | 1769 | rnf144b | 207 | 223 | 293 | 189 | 229 | 291 |
| ENSDARG000 | 16 | 9302079 | 9302194 | - | 116 | 5S_rRNA | 0 | 0 | 0 | 0 | 0 | 0 |
| ENSDARG000 | 16:16:16:16:1 | 2394193:2403 | 2394841:2403 | -;-;-;-;-;-;- | 1397 | NA | 21 | 7 | 32 | 2 | 9 | 0 |
| ENSDARG000 | 16:16:16:16:1 | 12632428:126 | 12632681:126 | +;+;+;+;+ | 4210 | nat14 | 126 | 88 | 125 | 36 | 110 | 180 |
| ENSDARG000 | 16 | 16154324 | 16154458 | + | 135 | 5S_rRNA | 0 | 0 | 0 | 0 | 0 | 0 |
| ENSDARG000 | 16 | 35726018 | 35726132 | + | 115 | 5S_rRNA | 0 | 0 | 0 | 0 | 0 | 0 |
| ENSDARG000 | 16;16;16 | 43353106:433 | 43353198:433 | -;-;- | 492 | NA | 2 | 58 | 151 | 177 | 25 | 34 |
| ENSDARG000 | 16 | 4874364 | 4874481 | + | 118 | 5S_rRNA | 0 | 0 | 0 | 0 | 0 | 0 |
| ENSDARG000 | 16 | 17705898 | 17705987 | - | 90 | dre-mir-190b | 0 | 0 | 0 | 0 | 0 | 0 |
| ENSDARG000 | 16:16:16:16:1 | 813780:81378 | 814016:81401 | +;+;+;+;+;+;+ | 1730 | irx1a | 2 | 26 | 91 | 208 | 445 | 0 |
| ENSDARG000 | 16:16:16:16:1 | 31977693:319 | 31977736:319 | -;-;-;-;-;-;- | 648 | NA | 0 | 0 | 0 | 0 | 0 | 0 |
| ENSDARG000 | 16:16:16:16:1 | 3185541:3186 | 3185990:3186 | +:+:+:+:+:+:+: | 1616 | NA | 136 | 83 | 57 | 55 | 90 | 77 |
| ENSDARG000 | 16:16:16:16:1 | 47272825:472 | 47273729:472 | -:-:-:-:-:-:-:-: | 3540 | mios | 586 | 506 | 573 | 322 | 678 | 591 |
| ENSDARG000 | 16 | 5607138 | 5612480 | - | 5343 | NA | 14 | 10 | 4 | 22 | 10 | 16 |
| ENSDARG000 | 16:16:16:16:1 | 25174628:251 | 25174673:251 | -;-;-;-;-;-;-;- | 162 | NA | 5 | 2 | 1 | 5 | 4 | 6 |
| ENSDARG000 | 16:16:16:16:1 | 55208398:552 | 55208511:552 | -:-:-:-:-:-:-:-: | 4920 | iqgap3 | 250 | 1299 | 1578 | 510 | 1022 | 431 |
| ENSDARG000 | 16:16:16:16:1 | 21790604:217 | 21790621:217 | -;-;-;-;-;-;-;- | 984 | NA | 0 | 0 | 0 | 0 | 0 | 0 |
| ENSDARG000 | 16 | 17247222 | 17247363 | - | 142 | NA | 0 | 0 | 1 | 3 | 0 | 5 |
| ENSDARG000 | 16 | 24881122 | 24881218 | + | 97 | dre-let-7d-1 | 0 | 0 | 0 | 0 | 0 | 0 |
| ENSDARG000 | 16 | 51755392 | 51755506 | - | 115 | 5S_rRNA | 0 | 0 | 0 | 0 | 0 | 0 |
| ENSDARG000 | 16:16:16:16:1 | 6356460:6358 | 6357296:6359 | -:-:-:-:-:-:-:-: | 1605 | scgn | 6 | 22 | 10 | 8 | 14 | 24 |
| ENSDARG000 | 16 | 38562527 | 38562644 | - | 118 | 5S_rRNA | 4 | 0 | 11 | 0 | 4 | 12 |
| ENSDARG000 | 16 | 40217043 | 40220920 | + | 3878 | NA | 80 | 44 | 0 | 0 | 0 | 98 |
| ENSDARG000 | 16:16:16:16:1 | 47308856:473 | 47308976:473 | +:+:+:+:+:+:+: | 3330 | col28a1b | 10 | 4 | 12 | 26 | 30 | 4 |
| ENSDARG000 | 16 | 16573414 | 16573530 | - | 117 | 5S_rRNA | 0 | 0 | 0 | 0 | 0 | 0 |
| ENSDARG000 | 16;16 | 46430627:464 | 46430702:464 | +;+ | 1260 | rpz6 | 0 | 0 | 0 | 0 | 3 | 2 |
| ENSDARG000 | 16:16:16:16:1 | 3103182:3106 | 3103634:3106 | -:-:-:-:-:-:-:-: | 2687 | csf3r | 320 | 128 | 84 | 176 | 408 | 670 |
| ENSDARG000 | 16:16:16:16:1 | 1228073:1229 | 1228168:1229 | +:+:+:+:+:+:+: | 1275 | NA | 304 | 261 | 360 | 174 | 415 | 227 |
| ENSDARG000 | 16:16:16:16:1 | 54513267:545 | 54513454:545 | +;+;+;+;+;+;+ | 2851 | fzd6 | 1429 | 889 | 1464 | 1062 | 1678 | 1036 |
| ENSDARG000 | 16:16:16:16:1 | 28599138:286 | 28599165:286 | -;-;-;-;-;- | 120 | NA | 0 | 0 | 0 | 0 | 0 | 0 |
| ENSDARG000 | 16:16:16:16:1 | 8188188:8188 | 8189698:8189 | -;-;-;-;-;-;-;- | 3675 | GASK1A | 6 | 0 | 2 | 0 | 6 | 8 |
| ENSDARG000 | 16:16:16:16:1 | 31542645:315 | 31542723:315 | +:+:+:+:+:+:+: | 819 | SLA | 2 | 8 | 16 | 0 | 8 | 24 |
| ENSDARG000 | 16:16:16:16:1 | 9275689:9275 | 9275841:9275 | -:-:-:-:-:-:-:-: | 3511 | ADCY2 | 0 | 0 | 0 | 0 | 0 | 0 |
| ENSDARG000 | 16 | 37771483 | 37771600 | + | 118 | 5S_rRNA | 0 | 0 | 0 | 0 | 0 | 0 |
| ENSDARG000 | 16:16:16:16:1 | 52786644:527 | 52786902:527 | -:-:-:-:-:-:-:-: | 1449 | NA | 6 | 8 | 13 | 2 | 24 | 0 |
| ENSDARG000 | 16 | 23592464 | 23592599 | - | 136 | 5S_rRNA | 0 | 0 | 0 | 0 | 0 | 0 |
| ENSDARG000 | 16;16;16 | 2469889:2472 | 2470031:2472 | +;+;+ | 312 | CNIH4 | 130 | 141 | 112 | 46 | 181 | 94 |
| ENSDARG000 | 16;16 | 26738404:267 | 26738801:267 | +;+ | 984 | FSBP | 7 | 3 | 11 | 1 | 16 | 6 |
| ENSDARG000 | 16;16 | 2433696:2434 | 2433888:2434 | +;+ | 510 | NA | 2 | 0 | 0 | 2 | 0 | 0 |
| ENSDARG000 | 16 | 32672820 | 32672936 | + | 117 | 5S_rRNA | 0 | 0 | 0 | 0 | 0 | 0 |
| ENSDARG000 | 16:16:16:16:1 | 277789:27778 | 277991:27799 | -:-:-:-:-:-:-:-: | 1646 | NA | 138 | 182 | 189 | 36 | 258 | 126 |
| ENSDARG000 | 16 | 25341416 | 25341543 | + | 128 | dre-mir-30d | 0 | 0 | 0 | 0 | 0 | 0 |
| ENSDARG000 | 16:16:16:16:1 | 31706841:317 | 31709940:317 | -;-;-;-;-;-;-;- | 3889 | rbp5 | 2 | 16 | 14 | 0 | 10 | 10 |
| ENSDARG000 | 16:16:16:16:1 | 42270499:422 | 42270512:422 | +:+:+:+:+:+:+: | 471 | NA | 16 | 11 | 12 | 15 | 11 | 8 |
| ENSDARG000 | 16;16;16 | 47348766:473 | 47348939:473 | -;-;- | 1026 | NA | 0 | 0 | 2 | 0 | 0 | 0 |
| ENSDARG000 | 16 | 17244908 | 17245047 | - | 140 | NA | 2 | 0 | 0 | 1 | 0 | 11 |
| ENSDARG000 | 16:16:16:16:1 | 53685812:536 | 53686480:536 | +;+;+;+;+ | 2050 | ggctb | 0 | 28 | 10 | 0 | 26 | 42 |
| ENSDARG000 | 16;16;16 | 43344475:433 | 43345334:433 | +;+;+ | 1294 | mrpl32 | 268 | 333 | 390 | 282 | 342 | 410 |
| ENSDARG000 | 16;16;16;16 | 5539189:554( | 5539746:554( | +;+;+;+ | 1020 | NA | 0 | 0 | 0 | 0 | 0 | 0 |
| ENSDARG000 | 16 | 18864270 | 18864386 | - | 117 | 5S_rRNA | 0 | 0 | 0 | 0 | 0 | 0 |
| ENSDARG000 | 16;16;16;16 | 1471:2019:24 | 1479:2176:25 | +;+;+;+ | 321 | NA | 72 | 56 | 77 | 18 | 56 | 44 |
| ENSDARG000 | 16 | 42898429 | 42898510 | + | 82 | dre-mir-92b | 0 | 0 | 0 | 0 | 0 | 0 |
| ENSDARG000 | 16;16 | 21867161:218 | 21867476:218 | +;+ | 627 | NA | 0 | 0 | 14 | 0 | 2 | 2 |
| ENSDARG000 | 16;16 | 30301539:303 | 30301586:303 | +;+ | 231 | NA | 4 | 0 | 0 | 4 | 0 | 2 |
| ENSDARG000 | 16 | 23607231 | 23607352 | + | 122 | 5S_rRNA | 0 | 0 | 0 | 0 | 0 | 0 |
| ENSDARG000 | 16 | 25341724 | 25341811 | + | 88 | dre-mir-30b | 0 | 0 | 0 | 0 | 0 | 0 |
| ENSDARG000 | 16 | 54644604 | 54644720 | + | 117 | NA | 0 | 0 | 0 | 0 | 0 | 0 |
| ENSDARG000 | 16;16;16;16 | 12611362:126 | 12611529:126 | +;+;+;+ | 660 | NA | 18 | 54 | 35 | 64 | 2 | 2 |
| ENSDARG000 | 16:16:16:16:1 | 21738194:217 | 21738245:217 | +;+;+;+;+ | 661 | NA | 2 | 0 | 7 | 17 | 7 | 4 |
| ENSDARG000 | 16:16:16:16:1 | 48791226:488 | 48791606:488 | -;-;-;-;-;-;- | 1554 | NA | 60 | 12 | 26 | 22 | 21 | 6 |
| ENSDARG000 | 16:16:16:16:1 | 31658196:316 | 31659159:316 | -;-;-;-;-;- | 1624 | c1s.2 | 0 | 0 | 0 | 0 | 0 | 0 |
| ENSDARG000 | 16:16:16:16:1 | 54450407:544 | 54450651:544 | -:-:-:-:-:-:-:-: | 1787 | pklr | 0 | 10 | 78 | 18 | 38 | 8 |
| ENSDARG000 | 16:16:16:16:1 | 52542550:525 | 52542762:525 | -:-:-:-:-:-:-:-: | 8388 | ubr5 | 4517 | 4660 | 7010 | 2803 | 7018 | 3417 |
| ENSDARG000 | 16:16:16:16:1 | 52343905:523 | 52344265:523 | +;+;+;+;+;+;+ | 2279 | ifnlr1 | 6 | 0 | 0 | 0 | 0 | 0 |
| ENSDARG000 | 16:16:16:16:1 | 47207691:472 | 47207707:472 | +:+:+:+:+:+:+: | 1353 | ICA1 | 4 | 4 | 8 | 2 | 15 | 17 |
| ENSDARG000 | 16:16:16:16:1 | 8047896:8051 | 8047924:8055 | -:-:-:-:-:-:-:-: | 5262 | snrka | 258 | 253 | 358 | 191 | 410 | 302 |
| ENSDARG000 | 16:16:16:16:1 | 44435453:444 | 44435575:444 | -:-:-:-:-:-:-:-: | 3361 | RIMS2 | 14 | 4 | 4 | 5 | 8 | 4 |
| ENSDARG000 | 16 | 48257013 | 48257130 | + | 118 | 5S_rRNA | 0 | 0 | 0 | 0 | 0 | 0 |
| ENSDARG000 | 16 | 1740937 | 1741058 | + | 122 | 5S_rRNA | 0 | 0 | 0 | 0 | 0 | 0 |
| ENSDARG000 | 16:16:16:16:1 | 152182:15873 | 152773:15881 | +:+:+:+:+:+:+: | 5367 | NA | 408 | 262 | 374 | 358 | 354 | 496 |
| ENSDARG000 | 16 | 37157005 | 37157121 | - | 117 | 5S_rRNA | 0 | 0 | 0 | 0 | 1 | 0 |
| ENSDARG000 | 16:16:16:16:1 | 54019456:54( | 54019524:54( | -;-;-;-;-;-;-;- | 972 | cdk14 | 187 | 197 | 295 | 230 | 259 | 201 |
| ENSDARG000 | 16:16:16:16:1 | 55078477:55( | 55078583:55( | +:+:+:+:+:+:+: | 1856 | azi2 | 166 | 88 | 102 | 82 | 118 | 62 |

| | | | | | | | | | | | |
|---|---|---|---|---|---|---|---|---|---|---|---|
| ENSDARG000 | 16 | 3353501 | 3353617 | + | 117 | 5S_rRNA | 0 | 0 | 0 | 0 | 0 | 0 |
| ENSDARG000 | 16 | 5870573 | 5870689 | - | 117 | 5S_rRNA | 0 | 0 | 0 | 0 | 1 | 0 |
| ENSDARG000 | 16:16:16:16:1 | 25975426:259 | 25977354:259 | -;-;-;-;- | 2050 | NA | 4 | 0 | 0 | 0 | 0 | 0 |
| ENSDARG000 | 16;16;16 | 54355842:543 | 54356107:543 | -;-;- | 911 | NA | 118 | 51 | 115 | 40 | 154 | 109 |
| ENSDARG000 | 16:16:16:16:1 | 8139515:8139 | 8139810:8139 | +;+;+;+;+;+ | 2617 | pomgnt2 | 88 | 132 | 142 | 66 | 423 | 106 |
| ENSDARG000 | 16;16;16 | 52771199:527 | 52771417:527 | +;+;+ | 648 | baalca | 2 | 20 | 12 | 0 | 15 | 0 |
| ENSDARG000 | 16;16;16;16 | 53125918:531 | 53126082:531 | +;+;+;+ | 1479 | NA | 16 | 10 | 57 | 80 | 2 | 0 |
| ENSDARG000 | 16:16:16:16:1 | 43319342:433 | 43319503:433 | -:-:-:-:-:-:-:-:-: | 4056 | ATAD2 | 330 | 1212 | 1615 | 501 | 981 | 679 |
| ENSDARG000 | 16 | 17120064 | 17120186 | - | 123 | 5S_rRNA | 3 | 6 | 0 | 0 | 0 | 0 |
| ENSDARG000 | 16:16:16:16:1 | 36167907:361 | 36169904:361 | -:-:-:-:-:-:-:-:-: | 4593 | capn7 | 726 | 490 | 802 | 473 | 758 | 720 |
| ENSDARG000 | 16:16:16:16:1 | 15942887:159 | 15943641:159 | -:-:-:-:-:-:-:-:-: | 3308 | UBE2O | 340 | 105 | 170 | 189 | 97 | 438 |
| ENSDARG000 | 16:16:16:16:1 | 55168245:551 | 55168440:551 | +;+;+;+;+;+;+ | 2609 | eomesb | 0 | 2 | 0 | 0 | 0 | 0 |
| ENSDARG000 | 16 | 20895112 | 20895175 | - | 64 | NA | 0 | 0 | 0 | 1 | 0 | 0 |
| ENSDARG000 | 16:16:16:16:1 | 12266366:122 | 12266374:122 | -;-;-;-;-;-;- | 177 | NA | 4 | 0 | 3 | 11 | 9 | 0 |
| ENSDARG000 | 16 | 12380730 | 12380846 | + | 117 | 5S_rRNA | 0 | 0 | 0 | 0 | 0 | 0 |
| ENSDARG000 | 16 | 47943143 | 47943257 | - | 115 | 5S_rRNA | 0 | 0 | 0 | 0 | 0 | 0 |
| ENSDARG000 | 16 | 50638670 | 50638786 | - | 117 | 5S_rRNA | 0 | 0 | 0 | 0 | 0 | 0 |
| ENSDARG000 | 16 | 27206669 | 27206797 | + | 129 | 5S_rRNA | 0 | 0 | 0 | 0 | 0 | 0 |
| ENSDARG000 | 16 | 17570810 | 17570934 | + | 125 | 5S_rRNA | 0 | 0 | 0 | 0 | 0 | 0 |
| ENSDARG000 | 16:16:16:16:1 | 49005321:490 | 49005419:490 | +;+;+;+;+;+ | 645 | NA | 6 | 10 | 2 | 12 | 0 | 6 |
| ENSDARG000 | 16:16:16:16:1 | 51001957:510 | 51002308:510 | -:-:-:-:-:-:-:-:-: | 3073 | NA | 197 | 92 | 309 | 102 | 294 | 186 |
| ENSDARG000 | 16;16;16;16 | 52512025:525 | 52512103:525 | +;+;+;+ | 433 | fabp10a | 0 | 0 | 121 | 0 | 2 | 0 |
| ENSDARG000 | 16;16;16;16 | 36095663:360 | 36097559:360 | -;-;-;- | 2225 | NA | 0 | 0 | 0 | 0 | 0 | 0 |
| ENSDARG000 | 16 | 1212674 | 1212788 | + | 115 | 5S_rRNA | 0 | 0 | 3 | 0 | 0 | 4 |
| ENSDARG000 | 16:16:16:16:1 | 54641230:546 | 54641387:546 | +:+:+:+:+:+:+: | 2195 | fbxo43 | 0 | 0 | 0 | 0 | 0 | 0 |
| ENSDARG000 | 16:16:16:16:1 | 43335696:433 | 43336860:433 | -;-;-;-;-;-;-;- | 1753 | psma2a | 1224 | 1392 | 1203 | 898 | 1137 | 1321 |
| ENSDARG000 | 16 | 50935190 | 50935306 | - | 117 | 5S_rRNA | 0 | 0 | 0 | 0 | 0 | 0 |
| ENSDARG000 | 16 | 12300169 | 12300286 | + | 118 | 5S_rRNA | 0 | 0 | 0 | 0 | 0 | 0 |
| ENSDARG000 | 16:16:16:16:1 | 31662256:316 | 31663270:316 | -:-:-:-:-:-:-:-:-: | 3267 | c1r | 0 | 0 | 0 | 2 | 0 | 0 |
| ENSDARG000 | 16:16:16:16:1 | 17671301:176 | 17671410:176 | -:-:-:-:-:-:-:-:-: | 3477 | NA | 123 | 440 | 604 | 182 | 270 | 79 |
| ENSDARG000 | 16 | 12012074 | 12012188 | - | 115 | 5S_rRNA | 0 | 0 | 0 | 0 | 0 | 0 |
| ENSDARG000 | 16 | 19704424 | 19704538 | + | 115 | 5S_rRNA | 0 | 0 | 0 | 0 | 0 | 0 |
| ENSDARG000 | 16 | 21451155 | 21451284 | - | 130 | 5S_rRNA | 0 | 0 | 0 | 0 | 0 | 0 |
| ENSDARG000 | 16;16 | 46760625:467 | 46761721:467 | -;- | 1509 | NA | 36 | 18 | 66 | 65 | 20 | 24 |
| ENSDARG000 | 16:16:16:16:1 | 8751815:8759 | 8752360:8760 | -:-:-:-:-:-:-:-:-: | 8958 | NA | 1335 | 782 | 956 | 786 | 2466 | 1109 |
| ENSDARG000 | 16 | 28388046 | 28388163 | + | 118 | 5S_rRNA | 0 | 0 | 0 | 0 | 0 | 0 |
| ENSDARG000 | 16:16:16:16:1 | 40138033:401 | 40138047:401 | -;-;-;-;-;- | 144 | NA | 0 | 0 | 1 | 0 | 3 | 5 |
| ENSDARG000 | 16:16:16:16:1 | 51883725:518 | 51885518:518 | -:-:-:-:-:-:-:-:-: | 2188 | rpl28 | 8562 | 7506 | 12460 | 11314 | 8335 | 10728 |
| ENSDARG000 | 16 | 2735676 | 2735799 | - | 124 | 5S_rRNA | 0 | 0 | 0 | 0 | 0 | 0 |
| ENSDARG000 | 16:16:16:16:1 | 4055331:4059 | 4055587:4059 | +:+:+:+:+:+:+: | 1514 | NA | 0 | 4 | 4 | 0 | 0 | 0 |
| ENSDARG000 | 16 | 49547771 | 49547886 | + | 116 | 5S_rRNA | 0 | 0 | 0 | 0 | 0 | 0 |
| ENSDARG000 | 16 | 20913413 | 20913484 | + | 72 | mir196d | 0 | 0 | 0 | 0 | 0 | 0 |
| ENSDARG000 | 16:16:16:16:1 | 4132537:4133 | 4132649:4133 | +:+:+:+:+:+:+: | 5402 | mtf1 | 164 | 105 | 271 | 213 | 246 | 148 |
| ENSDARG000 | 16:16:16:16:1 | 1254390:1262 | 1255349:1263 | +:+:+:+:+:+:+: | 4207 | NA | 2391 | 1334 | 4501 | 1089 | 4057 | 4197 |
| ENSDARG000 | 16 | 45588214 | 45588330 | + | 117 | 5S_rRNA | 0 | 0 | 0 | 0 | 0 | 0 |
| ENSDARG000 | 16:16:16:16:1 | 25133335:251 | 25133792:251 | +;+;+;+;+ | 1446 | NA | 0 | 2 | 8 | 22 | 6 | 0 |
| ENSDARG000 | 16;16;16 | 68069:68124: | 69930:69570: | +;+;+ | 2943 | sox4b | 60 | 30 | 74 | 18 | 128 | 100 |
| ENSDARG000 | 16 | 47203422 | 47204301 | + | 880 | NA | 17 | 0 | 36 | 0 | 29 | 58 |
| ENSDARG000 | 16 | 5061499 | 5061629 | - | 131 | 5S_rRNA | 0 | 0 | 0 | 0 | 0 | 0 |
| ENSDARG000 | 16 | 46216808 | 46216909 | - | 102 | U6 | 3 | 0 | 0 | 0 | 0 | 0 |
| ENSDARG000 | 16:16:16:16:1 | 2482498:2484 | 2482563:2484 | -:-:-:-:-:-:-:-:-: | 1432 | WDR26 | 360 | 196 | 190 | 142 | 296 | 274 |
| ENSDARG000 | 16 | 2284056 | 2284174 | + | 119 | 5S_rRNA | 0 | 0 | 0 | 0 | 0 | 0 |
| ENSDARG000 | 16 | 33343151 | 33343265 | + | 115 | 5S_rRNA | 0 | 0 | 0 | 0 | 0 | 0 |
| ENSDARG000 | 16 | 9237132 | 9237246 | + | 115 | 5S_rRNA | 0 | 0 | 0 | 0 | 0 | 0 |
| ENSDARG000 | 16:16:16:16:1 | 3067134:3068 | 3067343:3068 | +:+:+:+:+:+:+: | 2656 | cyb5r4 | 288 | 344 | 234 | 203 | 240 | 243 |
| ENSDARG000 | 16:16:16:16:1 | 9053275:9056 | 9053307:9057 | +:+:+:+:+:+:+: | 13653 | DNAH5 | 26 | 4 | 4 | 0 | 16 | 14 |
| ENSDARG000 | 16:16:16:16:1 | 3004422:3014 | 3004467:3014 | +;+;+;+;+;+;+ | 564 | NA | 730 | 514 | 557 | 515 | 770 | 366 |
| ENSDARG000 | 16:16:16:16:1 | 54691594:546 | 54691716:546 | +:+:+:+:+:+:+: | 1005 | NA | 0 | 0 | 0 | 0 | 0 | 0 |
| ENSDARG000 | 16;16 | 52536999:525 | 52537116:525 | -;- | 183 | NA | 83 | 98 | 96 | 59 | 129 | 57 |
| ENSDARG000 | 16;16 | 8695595:8699 | 8695768:8701 | +;+ | 1985 | NA | 6 | 0 | 8 | 0 | 4 | 0 |
| ENSDARG000 | 16;16 | 4907690:4913 | 4908091:4913 | +;+ | 1182 | TENT5A | 0 | 8 | 74 | 15 | 22 | 0 |
| ENSDARG000 | 16:16:16:16:1 | 21491891:214 | 21492739:214 | -:-:-:-:-:-:-:-:-: | 4526 | NA | 248 | 145 | 296 | 302 | 316 | 376 |
| ENSDARG000 | 16:16:16:16:1 | 53710416:537 | 53710920:537 | +:+:+:+:+:+:+: | 3097 | NOD1 | 329 | 249 | 361 | 176 | 464 | 239 |
| ENSDARG000 | 16 | 30080543 | 30080661 | + | 119 | 5S_rRNA | 6 | 3 | 10 | 5 | 3 | 2 |
| ENSDARG000 | 16:16:16:16:1 | 53733714:537 | 53734553:537 | -;-;-;-;- | 1611 | znrf2b | 107 | 70 | 122 | 54 | 114 | 92 |
| ENSDARG000 | 16 | 40011869 | 40011991 | + | 123 | 5S_rRNA | 0 | 0 | 0 | 0 | 0 | 0 |
| ENSDARG000 | 16:16:16:16:1 | 31639110:316 | 31640542:316 | -;-;-;-;-;- | 2185 | NA | 18 | 0 | 0 | 2 | 4 | 0 |
| ENSDARG000 | 16 | 10324549 | 10324665 | + | 117 | 5S_rRNA | 0 | 1 | 2 | 0 | 0 | 0 |
| ENSDARG000 | 16 | 20393693 | 20393809 | - | 117 | 5S_rRNA | 0 | 0 | 0 | 0 | 0 | 0 |
| ENSDARG000 | 16:16:16:16:1 | 55105545:551 | 55106855:551 | -;-;-;-;-;-;-;- | 1645 | cmc1 | 276 | 227 | 202 | 198 | 222 | 238 |
| ENSDARG000 | 16:16:16:16:1 | 2351520:2358 | 2355559:2358 | -:-:-:-:-:-:-:-:-: | 6912 | HACE1 | 268 | 305 | 649 | 673 | 370 | 191 |
| ENSDARG000 | 16:16:16:16:1 | 46695777:467 | 46695947:467 | +;+;+;+;+ | 1289 | rab25b | 10 | 2 | 2 | 2 | 0 | 0 |
| ENSDARG000 | 16 | 30933249 | 30934319 | - | 1071 | GPR20 | 2 | 0 | 0 | 0 | 0 | 4 |
| ENSDARG000 | 16 | 45018424 | 45018540 | - | 117 | 5S_rRNA | 0 | 0 | 0 | 0 | 0 | 0 |
| ENSDARG000 | 16 | 51776536 | 51776640 | + | 105 | dre-mir-455-2 | 0 | 0 | 0 | 0 | 0 | 0 |
| ENSDARG000 | 16 | 39885134 | 39885251 | - | 118 | 5S_rRNA | 0 | 0 | 0 | 0 | 0 | 0 |
| ENSDARG000 | 16:16:16:16:1 | 39485608:394 | 39485628:394 | +:+:+:+:+:+:+: | 1146 | NA | 0 | 0 | 0 | 0 | 0 | 0 |
| ENSDARG000 | 16 | 34351945 | 34352061 | - | 117 | 5S_rRNA | 0 | 0 | 0 | 0 | 0 | 0 |
| ENSDARG000 | 16:16:16:16:1 | 51462620:514 | 51462860:515 | +:+:+:+:+:+:+: | 1794 | SLC9A1 | 20 | 4 | 20 | 12 | 32 | 15 |
| ENSDARG000 | 16;16 | 49249669:492 | 49249889:492 | +;+ | 264 | NA | 0 | 0 | 0 | 0 | 0 | 0 |
| ENSDARG000 | 16 | 1220446 | 1220562 | + | 117 | 5S_rRNA | 0 | 2 | 0 | 0 | 0 | 0 |
| ENSDARG000 | 16 | 41882445 | 41882560 | - | 116 | 5S_rRNA | 0 | 0 | 0 | 0 | 0 | 0 |
| ENSDARG000 | 16;16 | 52875093:528 | 52875496:528 | -;- | 1031 | TPPP | 8 | 6 | 10 | 2 | 5 | 2 |
| ENSDARG000 | 16:16:16:16:1 | 54474621:544 | 54475132:544 | -:-:-:-:-:-:-:-:-: | 1914 | clk2b | 130 | 134 | 202 | 62 | 203 | 147 |
| ENSDARG000 | 16 | 12247862 | 12247920 | - | 59 | U7 | 0 | 0 | 4 | 0 | 2 | 0 |
| ENSDARG000 | 16:16:16:16:1 | 27166157:271 | 27166329:271 | -;-;-;-;-;- | 900 | FRRS1L | 6 | 8 | 10 | 0 | 4 | 2 |
| ENSDARG000 | 16:16:16:16:1 | 22761846:227 | 22761852:227 | +:+:+:+:+:+:+: | 192 | NA | 2 | 0 | 0 | 3 | 3 | 2 |
| ENSDARG000 | 16 | 35094846 | 35094960 | - | 115 | 5S_rRNA | 0 | 0 | 0 | 0 | 0 | 0 |
| ENSDARG000 | 16 | 12253415 | 12253471 | - | 57 | U7 | 0 | 3 | 13 | 6 | 12 | 1 |
| ENSDARG000 | 16:16:16:16:1 | 1199202:1199 | 1199359:1199 | +:+:+:+:+:+:+: | 3288 | NA | 588 | 344 | 558 | 408 | 493 | 492 |
| ENSDARG000 | 16:16:16:16:1 | 42289076:422 | 42289675:422 | -:-:-:-:-:-:-:-:-: | 2013 | ppox | 94 | 119 | 120 | 40 | 144 | 79 |
| ENSDARG000 | 16:16:16:16:1 | 3463727:3470 | 3464155:3470 | -:-:-:-:-:-:-:-:-: | 3759 | GRIK3 | 30 | 0 | 4 | 22 | 0 | 30 |
| ENSDARG000 | 16:16:16:16:1 | 7985886:7991 | 7985997:7991 | +:+:+:+:+:+:+: | 2658 | ano10a | 2026 | 1979 | 1846 | 1488 | 2039 | 2157 |
| ENSDARG000 | 16;16;16 | 25831165:258 | 25831206:258 | -;-;- | 1116 | NA | 5 | 6 | 0 | 16 | 10 | 0 |
| ENSDARG000 | 16:16:16:16:1 | 28547157:285 | 28547385:285 | +:+:+:+:+:+:+: | 2580 | FAM171A1 | 236 | 56 | 220 | 64 | 147 | 154 |
| ENSDARG000 | 16 | 29216274 | 29216360 | + | 87 | dre-mir-9-1 | 0 | 0 | 0 | 0 | 0 | 0 |
| ENSDARG000 | 16:16:16:16:1 | 54303491:543 | 54303834:543 | +;+;+;+;+;+;+ | 1394 | ethe1 | 82 | 124 | 146 | 221 | 144 | 104 |
| ENSDARG000 | 16 | 31634470 | 31634595 | + | 126 | 5S_rRNA | 0 | 0 | 0 | 0 | 0 | 0 |
| ENSDARG000 | 16 | 19190514 | 19190631 | - | 118 | 5S_rRNA | 0 | 0 | 0 | 0 | 0 | 0 |
| ENSDARG000 | 16 | 23523261 | 23523375 | - | 115 | 5S_rRNA | 3 | 11 | 19 | 17 | 4 | 6 |
| ENSDARG000 | 16 | 45174446 | 45174560 | - | 115 | 5S_rRNA | 0 | 0 | 0 | 0 | 1 | 0 |
| ENSDARG000 | 16;16;16;16 | 420122:48189 | 420225:48207 | +;+;+;+ | 1851 | NA | 0 | 0 | 0 | 0 | 0 | 0 |
| ENSDARG000 | 16:16:16:16:1 | 4234727:4234 | 4235716:4235 | -;-;-;-;- | 1588 | NA | 0 | 0 | 0 | 0 | 0 | 0 |
| ENSDARG000 | 16;16;16;16 | 14421800:144 | 14423095:144 | -;-;-;- | 2898 | NA | 0 | 0 | 0 | 0 | 0 | 0 |
| ENSDARG000 | 16:16:16:16:1 | 14431515:144 | 14432614:144 | -;-;-;-;-;- | 2114 | NA | 0 | 0 | 0 | 0 | 0 | 0 |
| ENSDARG000 | 16;16;16;16 | 14479551:145 | 14479830:145 | +;+;+;+ | 1758 | NA | 0 | 0 | 0 | 0 | 0 | 0 |
| ENSDARG000 | 16:16:16:16:1 | 19617088:196 | 19617213:196 | -;-;-;-;- | 780 | NA | 0 | 0 | 0 | 2 | 0 | 0 |
| ENSDARG000 | 16;16;16 | 19626526:196 | 19627008:196 | -;-;- | 622 | NA | 4 | 0 | 0 | 0 | 0 | 0 |
| ENSDARG000 | 16:16:16:16:1 | 19631414:196 | 19631439:196 | -;-;-;-;-;-;-;- | 1588 | NA | 3 | 0 | 2 | 5 | 0 | 14 |
| ENSDARG000 | 16;16;16 | 19686713:197 | 19686929:197 | +;+;+ | 653 | NA | 0 | 0 | 0 | 0 | 0 | 0 |
| ENSDARG000 | 16;16;16 | 19805021:198 | 19805857:198 | +;+;+ | 1039 | NA | 0 | 2 | 9 | 0 | 3 | 0 |
| ENSDARG000 | 16;16;16;16 | 19807943:198 | 19808320:198 | +;+;+;+ | 915 | NA | 2 | 0 | 16 | 0 | 0 | 0 |
| ENSDARG000 | 16;16;16 | 19828376:198 | 19830356:198 | -;-;- | 2336 | NA | 2 | 0 | 1 | 2 | 0 | 0 |
| ENSDARG000 | 16;16;16 | 20041857:200 | 20042586:200 | -;-;- | 1038 | NA | 2 | 0 | 0 | 0 | 0 | 4 |
| ENSDARG000 | 16;16;16 | 22060569:220 | 22061136:220 | -;-;- | 745 | NA | 0 | 0 | 1 | 2 | 4 | 4 |
| ENSDARG000 | 16;16;16 | 32397507:324 | 32397663:324 | +;+;+ | 362 | NA | 0 | 0 | 0 | 0 | 0 | 0 |
| ENSDARG000 | 16:16:16:16:1 | 33254300:332 | 33254603:332 | +;+;+;+;+;+ | 914 | NA | 0 | 0 | 0 | 0 | 0 | 0 |
| ENSDARG000 | 16:16:16:16:1 | 39029883:390 | 39030189:390 | -;-;-;-;-;- | 1069 | NA | 28 | 0 | 0 | 0 | 0 | 0 |
| ENSDARG000 | 16;16;16 | 40075545:400 | 40075727:400 | +;+;+ | 343 | NA | 4 | 0 | 0 | 4 | 0 | 2 |
| ENSDARG000 | 16:16:16:16:1 | 44187324:441 | 44188464:441 | -;-;-;-;-;-;-;-;- | 1536 | NA | 0 | 0 | 2 | 0 | 2 | 4 |
| ENSDARG000 | 16:16:16:16:1 | 44224076:442 | 44224267:442 | -;-;-;-;- | 891 | NA | 4 | 0 | 2 | 0 | 0 | 0 |
| ENSDARG000 | 16;16;16;16 | 51559132:515 | 51559680:515 | +;+;+;+ | 3250 | NA | 0 | 3 | 5 | 5 | 15 | 12 |
| ENSDARG000 | 16;16;16 | 52385594:523 | 52385851:523 | +;+;+ | 1009 | NA | 0 | 0 | 0 | 0 | 0 | 0 |
| ENSDARG000 | 16;16;16 | 53186147:531 | 53186302:531 | +;+;+ | 3322 | NA | 0 | 0 | 33 | 26 | 10 | 0 |
| ENSDARG000 | 16:16:16:16:1 | 54341485:543 | 54341571:543 | +;+;+;+;+ | 3733 | NA | 17 | 2 | 1 | 14 | 2 | 0 |
| ENSDARG000 | 16;16;16 | 55192189:551 | 55192804:551 | -;-;- | 850 | NA | 4 | 0 | 0 | 0 | 2 | 0 |
| ENSDARG000 | 20:20:20:20:2 | 23657860:236 | 23657915:236 | +;+;+;+;+;+;+ | 3631 | NA | 209 | 201 | 191 | 138 | 216 | 134 |
| ENSDARG000 | 20:20:20:20:2 | 9355912:9355 | 9355998:9355 | +:+:+:+:+:+:+: | 3256 | si:dkev-174m | 8 | 6 | 4 | 8 | 16 | 12 |
| ENSDARG000 | 20:20:20:20:2 | 26572456:265 | 26572764:265 | -:-:-:-:-:-:-:-:-: | 3194 | exoc2 | 1348 | 1738 | 2076 | 1556 | 2024 | 1590 |
| ENSDARG000 | 20:20:20:20:2 | 2669017:2669 | 2669053:2670 | +:+:+:+:+:+:+: | 1198 | slc35a1 | 481 | 434 | 468 | 332 | 626 | 421 |
| ENSDARG000 | 20:20:20:20:2 | 812012:81216 | 812363:81236 | +:+:+:+:+:+:+: | 1566 | mei4 | 14 | 0 | 6 | 0 | 20 | 8 |
| ENSDARG000 | 20:20:20:20:2 | 18209895:182 | 18210362:182 | +:+:+:+:+:+:+: | 3087 | NA | 824 | 973 | 868 | 591 | 962 | 680 |
| ENSDARG000 | 20:20:20:20:2 | 32406011:324 | 32406259:324 | +:+:+:+:+:+:+: | 1903 | snx3 | 1634 | 1932 | 1647 | 1748 | 1621 | 2320 |
| ENSDARG000 | 20;20;20;20 | 13956106:139 | 13956275:139 | +;+;+;+ | 1147 | NA | 0 | 0 | 0 | 0 | 0 | 0 |
| ENSDARG000 | 20:20:20:20:2 | 23670494:236 | 23670610:236 | +:+:+:+:+:+:+: | 5420 | NA | 291 | 200 | 370 | 194 | 448 | 356 |
| ENSDARG000 | 20;20;20 | 13969414:139 | 13969601:139 | +;+;+ | 1414 | rd3 | 12 | 4 | 8 | 13 | 27 | 60 |
| ENSDARG000 | 20;20 | 9421448:9427 | 9422838:9428 | -;- | 1774 | rdh14b | 236 | 270 | 364 | 122 | 216 | 230 |
| ENSDARG000 | 20:20:20:20:2 | 25639470:256 | 25639851:256 | -:-:-:-:-:-:-:-:-: | 2666 | si:dkeyp-117h | 28 | 105 | 173 | 36 | 90 | 56 |
| ENSDARG000 | 20:20:20:20:2 | 826459:83402 | 826807:83424 | +:+:+:+:+:+:+: | 2104 | nt5e | 64 | 12 | 22 | 0 | 74 | 22 |
| ENSDARG000 | 20:20:20:20:2 | 14003060:140 | 14003757:140 | -:-:-:-:-:-:-:-:-: | 1378 | si:ch211-22i1 | 430 | 427 | 454 | 386 | 350 | 495 |
| ENSDARG000 | 20:20:20:20:2 | 6533260:6533 | 6533345:6533 | +:+:+:+:+:+:+: | 1592 | si:ch211-191a | 56 | 8 | 12 | 44 | 16 | 6 |
| ENSDARG000 | 20:20:20:20:2 | 32437396:324 | 32438653:324 | -;-;-;-;-;-;-;- | 2710 | nr2e1 | 0 | 2 | 0 | 0 | 0 | 12 |
| ENSDARG000 | 20:20:20:20:2 | 28650025:286 | 28652742:286 | -:-:-:-:-:-:-:-:-: | 8493 | sipa1l1 | 76 | 8 | 358 | 20 | 102 | 46 |
| ENSDARG000 | 20;20 | 25643044:256 | 25643199:256 | +;+ | 518 | NA | 0 | 0 | 0 | 0 | 0 | 0 |
| ENSDARG000 | 20:20:20:20:2 | 25625872:256 | 25626686:256 | +:+:+:+:+:+:+: | 2680 | ppat | 1899 | 2842 | 3594 | 2750 | 4210 | 2680 |
| ENSDARG000 | 20:20:20:20:2 | 6543674:6551 | 6543879:6551 | +:+:+:+:+:+:+: | 1265 | NA | 264 | 207 | 264 | 232 | 440 | 459 |
| ENSDARG000 | 20:20:20:20:2 | 34157308:341 | 34158998:341 | -:-:-:-:-:-:-:-:-: | 18497 | NA | 64 | 14 | 616 | 378 | 1892 | 229 |
| ENSDARG000 | 20:20:20:20:2 | 32466886:324 | 32466956:324 | +;+;+;+;+;+ | 642 | NA | 2 | 2 | 0 | 0 | 0 | 0 |
| ENSDARG000 | 20:20:20:20:2 | 25618114:256 | 25618433:256 | -:-:-:-:-:-:-:-:-: | 4098 | paics | 13845 | 23152 | 19784 | 25034 | 27779 | 17344 |
| ENSDARG000 | 20:20:20:20:2 | 32473584:324 | 32473641:324 | +:+:+:+:+:+:+: | 4622 | OSTM1 | 3998 | 3085 | 2392 | 3205 | 4733 | 3656 |
| ENSDARG000 | 20:20:20:20:2 | 29676874:296 | 29677180:296 | -:-:-:-:-:-:-:-:-: | 1923 | e2f6 | 347 | 315 | 448 | 286 | 442 | 254 |
| ENSDARG000 | 20:20:20:20:2 | 9428965:9428 | 9429475:9429 | -:-:-:-:-:-:-:-:-: | 1201 | crip2l | 267 | 148 | 1036 | 38 | 400 | 307 |
| ENSDARG000 | 20;20 | 29680929:296 | 29681075:296 | +;+ | 367 | NA | 0 | 2 | 0 | 0 | 4 | 0 |
| ENSDARG000 | 20:20:20:20:2 | 18260358:182 | 18261380:182 | +:+:+:+:+:+:+: | 3059 | NA | 6 | 2 | 0 | 2 | 6 | 2 |
| ENSDARG000 | 20:20:20:20:2 | 14014813:140 | 14016743:140 | -:-:-:-:-:-:-:-:-: | 3699 | rhag | 54 | 17 | 86 | 30 | 228 | 162 |
| ENSDARG000 | 20;20 | 26683933:266 | 26685395:266 | +;+ | 1463 | foxq1b | 8 | 3 | 0 | 6 | 5 | 6 |
| ENSDARG000 | 20;20 | 26690036:266 | 26691374:266 | +;+ | 2027 | NA | 0 | 0 | 0 | 0 | 0 | 0 |
| ENSDARG000 | 20:20:20:20:2 | 20816747:208 | 20817150:208 | -:-:-:-:-:-:-:-:-: | 1971 | ckbb | 23808 | 21407 | 18710 | 32884 | 37940 | 14963 |
| ENSDARG000 | 20:20:20:20:2 | 18306122:183 | 18306569:183 | -:-:-:-:-:-:-:-:-: | 1345 | psma8 | 3450 | 4358 | 3999 | 2148 | 3828 | 2493 |
| ENSDARG000 | 20:20:20:20:2 | 32497260:324 | 32497388:324 | +:+:+:+:+:+:+: | 1315 | NA | 94 | 97 | 128 | 144 | 147 | 135 |
| ENSDARG000 | 20:20:20:20:2 | 41021054:410 | 41021743:410 | +:+:+:+:+:+:+: | 7969 | MAN1A1 | 2408 | 1910 | 1540 | 1747 | 1534 | 2391 |
| ENSDARG000 | 20:20:20:20:2 | 2681992:2682 | 2682114:2682 | -:-:-:-:-:-:-:-:-: | 2171 | rars2 | 62 | 92 | 84 | 70 | 78 | 58 |

| | | | | | | | | | | | | |
|---|---|---|---|---|---|---|---|---|---|---|---|---|
| ENSDARG000 | 20;20;20 | 18325556:183 | 18325750:183 | +;+;+ | 2765 | NA | 94 | 136 | 160 | 181 | 71 | 162 |
| ENSDARG000 | 20 | 26702377 | 26704331 | + | 1955 | foxc1b | 56 | 48 | 32 | 22 | 40 | 6 |
| ENSDARG000 | 20;20;20;20;2 | 14096858:140 | 14098394:140 | -;-;-;-;-;-;-;-; | 4287 | zpax2 | 0 | 0 | 0 | 0 | 0 | 0 |
| ENSDARG000 | 20;20 | 6581799:6583 | 6582296:6583 | -;- | 792 | NA | 26 | 10 | 8 | 10 | 14 | 20 |
| ENSDARG000 | 20;20;20;20 | 41248480:413 | 41248660:413 | +;+;+;+ | 1064 | NA | 0 | 0 | 0 | 0 | 0 | 0 |
| ENSDARG000 | 20;20 | 18345518:183 | 18346898:183 | -;- | 1726 | gpr132b | 8 | 28 | 24 | 4 | 32 | 68 |
| ENSDARG000 | 20;20;20;20;2 | 32501748:325 | 32501968:325 | +;+;+;+;+;+;+; | 4664 | sec63 | 1705 | 2113 | 2893 | 1248 | 2512 | 1959 |
| ENSDARG000 | 20;20;20;20;2 | 29687636:296 | 29687740:296 | -;-;-;-;- | 809 | NA | 0 | 3 | 0 | 0 | 0 | 0 |
| ENSDARG000 | 20;20;20;20;2 | 29690901:296 | 29691288:296 | +;+;+;+;+;+;+; | 2862 | mboat2b | 324 | 318 | 344 | 385 | 298 | 315 |
| ENSDARG000 | 20;20;20;20;2 | 23742574:237 | 23742896:237 | +;+;+;+;+;+;+; | 4714 | sh3rf1 | 2987 | 4581 | 5807 | 2253 | 7754 | 2213 |
| ENSDARG000 | 20;20;20;20;2 | 18352770:183 | 18354783:183 | -;-;-;-;-;-;-;-; | 4121 | vipas39 | 202 | 204 | 209 | 101 | 176 | 214 |
| ENSDARG000 | 20;20;20;20;2 | 9474841:9475 | 9475139:9475 | +;+;+;+;+;+;+; | 3220 | rad51b | 22 | 71 | 103 | 32 | 66 | 56 |
| ENSDARG000 | 20;20;20;20;2 | 28782903:287 | 28784217:287 | -;-;-;-;-;-;- | 1958 | rab15 | 2 | 8 | 12 | 10 | 16 | 8 |
| ENSDARG000 | 20;20;20;20;2 | 41549200:415 | 41549914:415 | +;+;+;+;+;+;+; | 8801 | FAM184A | 0 | 18 | 8 | 42 | 0 | 0 |
| ENSDARG000 | 20;20;20;20;2 | 32523576:325 | 32523909:325 | +;+;+;+;+;+;+; | 1866 | SCML4 | 6 | 0 | 32 | 18 | 100 | 0 |
| ENSDARG000 | 20;20;20;20;2 | 26709009:267 | 26709516:267 | -;-;-;-;-;-;-;-; | 3302 | gmds | 606 | 212 | 247 | 124 | 304 | 211 |
| ENSDARG000 | 20;20;20;20;2 | 851583:85158 | 851943:85194 | -;-;-;-;-;-;-;-; | 3419 | snx14 | 1561 | 1276 | 1777 | 1654 | 1467 | 1624 |
| ENSDARG000 | 20;20;20;20;2 | 6590220:6602 | 6590553:6602 | +;+;+;+;+;+;+; | 3819 | NA | 2396 | 2283 | 1715 | 2375 | 2918 | 3817 |
| ENSDARG000 | 20;20 | 9575374:9580 | 9578206:9580 | -;- | 3021 | zfp36l1b | 10525 | 4746 | 11380 | 9159 | 8570 | 8620 |
| ENSDARG000 | 20;20;20;20;2 | 34320635:343 | 34320741:343 | +;+;+;+;+;+;+; | 3708 | ivns1abpa | 7792 | 9372 | 11852 | 6516 | 19300 | 16564 |
| ENSDARG000 | 20;20;20;20;2 | 9643499:9643 | 9643961:9643 | -;-;-;-;-;-;-;-; | 4086 | NA | 10 | 6 | 4 | 0 | 28 | 2 |
| ENSDARG000 | 20;20;20;20;2 | 41801913:418 | 41802109:418 | +;+;+;+;+;+;+; | 4126 | mcm9 | 66 | 204 | 220 | 78 | 276 | 271 |
| ENSDARG000 | 20;20;20;20;2 | 23838814:238 | 23840590:238 | -;-;-;-;-;-;- | 2723 | si:dkev-15i16. | 2 | 67 | 56 | 43 | 123 | 43 |
| ENSDARG000 | 20 | 34374735 | 34379656 | + | 4922 | NA | 18 | 0 | 10 | 17 | 0 | 32 |
| ENSDARG000 | 20;20;20;20;2 | 34343649:343 | 34344568:343 | -;-;-;-;-;-;-;-; | 3934 | NA | 154 | 124 | 144 | 78 | 168 | 168 |
| ENSDARG000 | 20;20;20;20;2 | 32669475:326 | 32669617:326 | +;+;+;+;+;+;+; | 781 | NA | 0 | 0 | 0 | 0 | 0 | 0 |
| ENSDARG000 | 20;20;20;20 | 18387358:183 | 18387738:183 | +;+;+;+ | 1748 | NA | 0 | 0 | 0 | 0 | 0 | 0 |
| ENSDARG000 | 20;20 | 14114258:141 | 14114350:141 | +;+ | 1610 | kcns3b | 0 | 0 | 0 | 0 | 0 | 0 |
| ENSDARG000 | 20;20;20;20;2 | 32655858:326 | 32656985:326 | -;-;-;-;-;-;-;-; | 1626 | NA | 0 | 0 | 12 | 0 | 2 | 0 |
| ENSDARG000 | 20;20;20;20;2 | 32666384:326 | 32667230:326 | -;-;-;-;- | 1398 | NA | 0 | 0 | 0 | 0 | 0 | 0 |
| ENSDARG000 | 20;20;20;20;2 | 29743904:297 | 29744041:297 | +;+;+;+;+;+;+; | 6130 | kidins220b | 1829 | 963 | 1506 | 1083 | 1697 | 1632 |
| ENSDARG000 | 20;20;20;20 | 14258545:142 | 14258764:142 | -;-;-;- | 1727 | NA | 0 | 0 | 0 | 0 | 0 | 2 |
| ENSDARG000 | 20;20;20;20;2 | 14144289:141 | 14147549:141 | -;-;-;-;-;-;-;-; | 4836 | DLGAP2 | 4 | 0 | 10 | 0 | 0 | 0 |
| ENSDARG000 | 20;20;20;20;2 | 23847380:238 | 23847834:238 | -;-;-;-;-;-;-;- | 1884 | NA | 68 | 49 | 73 | 38 | 142 | 74 |
| ENSDARG000 | 20;20;20;20;2 | 41859277:418 | 41859646:418 | +;+;+;+;+;+;+; | 5828 | CEP85L | 420 | 418 | 461 | 610 | 397 | 500 |
| ENSDARG000 | 20;20;20;20;2 | 41911214:419 | 41913414:419 | -;-;-;-;-;-;- | 3083 | pln | 6 | 2 | 0 | 10 | 0 | 2 |
| ENSDARG000 | 20;20;20;20;2 | 23860858:238 | 23861739:238 | -;-;-;-;-;-;-;-; | 2355 | katna1 | 220 | 186 | 150 | 86 | 112 | 194 |
| ENSDARG000 | 20;20;20;20;2 | 23855025:238 | 23855161:238 | +;+;+;+;+;+;+; | 2103 | ginm1 | 2753 | 1812 | 1721 | 1871 | 1788 | 2254 |
| ENSDARG000 | 20;20;20;20;2 | 34388425:343 | 34388494:343 | +;+;+;+;+;+;+; | 2920 | TRMT1L | 336 | 305 | 402 | 209 | 462 | 310 |
| ENSDARG000 | 20;20;20;20;2 | 28803642:288 | 28804152:288 | +;+;+;+;+;+;+; | 2763 | fntb | 547 | 452 | 900 | 712 | 555 | 776 |
| ENSDARG000 | 20;20 | 32756763:327 | 32756897:327 | +;+ | 1944 | lratd1 | 47 | 44 | 90 | 8 | 50 | 16 |
| ENSDARG000 | 20;20;20;20;2 | 20896622:208 | 20897162:208 | -;-;-;-;-;-;-;-; | 1690 | xrcc3 | 38 | 105 | 110 | 89 | 100 | 74 |
| ENSDARG000 | 20;20;20;20;2 | 26827265:268 | 26827935:268 | -;-;-;-;-;-;-;-; | 3506 | MYLK4 | 8 | 6 | 0 | 0 | 6 | 0 |
| ENSDARG000 | 20;20;20;20;2 | 41967312:419 | 41967933:419 | -;-;-;-;-;-;-;- | 2098 | si:dkeyp-114g | 2 | 0 | 0 | 0 | 0 | 0 |
| ENSDARG000 | 20;20;20;20;2 | 32838467:328 | 32838733:328 | -;-;-;-;-;-;-;-; | 7399 | nbas | 451 | 440 | 768 | 360 | 658 | 433 |
| ENSDARG000 | 20;20;20;20;2 | 18391860:183 | 18393511:183 | -;-;-;-;-;-;-;-; | 7622 | cdc42bpb | 2357 | 1762 | 2424 | 1597 | 3066 | 2268 |
| ENSDARG000 | 20;20;20;20;2 | 29784927:297 | 29785393:297 | -;-;-;-;- | 1464 | id2b | 98 | 12 | 139 | 21 | 58 | 104 |
| ENSDARG000 | 20;20 | 14454908:144 | 14454946:144 | -;- | 238 | NA | 0 | 0 | 0 | 0 | 0 | 0 |
| ENSDARG000 | 20;20 | 9683994:9693 | 9684941:9694 | +;+ | 2330 | ptger2b | 78 | 8 | 20 | 8 | 24 | 22 |
| ENSDARG000 | 20;20 | 9720121:9721 | 9720293:9722 | +;+ | 634 | NA | 0 | 0 | 0 | 0 | 0 | 0 |
| ENSDARG000 | 20;20;20;20;2 | 34400715:344 | 34400812:344 | +;+;+;+;+;+;+; | 4248 | niban1a | 28 | 6 | 6 | 0 | 4 | 8 |
| ENSDARG000 | 20;20;20;20;2 | 29836276:298 | 29837654:298 | -;-;-;-;-;-;-;-; | 2416 | rnf144ab | 402 | 368 | 655 | 589 | 690 | 766 |
| ENSDARG000 | 20;20;20;20;2 | 14458930:144 | 14459800:144 | -;-;-;-;-;-;-;-; | 1954 | C20H12orf57 | 292 | 332 | 537 | 398 | 348 | 356 |
| ENSDARG000 | 20;20;20;20;2 | 41998502:419 | 41999192:419 | -;-;-;-;-;-;-;-; | 2213 | SLC35F1 | 2 | 0 | 60 | 2 | 24 | 6 |
| ENSDARG000 | 20;20;20;20;2 | 26880668:268 | 26880906:268 | +;+;+;+;+;+;+; | 3307 | serpinb1 | 60 | 20 | 44 | 58 | 144 | 44 |
| ENSDARG000 | 20;20;20;20;2 | 9751743:9752 | 9752602:9752 | -;-;-;-;-;-;-;-; | 1959 | si:dkev-63i12. | 12 | 14 | 22 | 14 | 22 | 28 |
| ENSDARG000 | 20;20;20;20;2 | 18559913:185 | 18559972:185 | +;+;+;+;+;+;+; | 1331 | NA | 4 | 0 | 2 | 0 | 8 | 4 |
| ENSDARG000 | 20 | 30032628 | 30035326 | - | 2699 | sox11b | 154 | 245 | 194 | 96 | 404 | 89 |
| ENSDARG000 | 20;20;20;20;2 | 6756247:6773 | 6756300:6774 | +;+;+;+;+;+;+; | 3147 | igfbp3 | 6100 | 5787 | 6609 | 5722 | 15058 | 7244 |
| ENSDARG000 | 20;20;20;20 | 30031578:300 | 30034470:300 | +;+;+;+ | 3455 | NA | 22 | 44 | 32 | 45 | 108 | 35 |
| ENSDARG000 | 20;20;20;20;2 | 18551657:185 | 18551768:185 | +;+;+;+;+;+;+ | 1182 | si:dkeyp-72h1 | 2 | 0 | 12 | 8 | 11 | 0 |
| ENSDARG000 | 20;20;20 | 18565758:185 | 18566369:185 | -;-;- | 1465 | NA | 2 | 2 | 0 | 0 | 0 | 0 |
| ENSDARG000 | 20;20;20;20 | 14465973:144 | 14466162:144 | +;+;+;+ | 981 | NA | 0 | 0 | 0 | 0 | 0 | 0 |
| ENSDARG000 | 20;20;20;20;2 | 34420623:344 | 34421852:344 | -;-;-;-;-;-;-;-; | 3648 | kifap3a | 600 | 465 | 515 | 591 | 511 | 589 |
| ENSDARG000 | 20;20;20;20;2 | 2715949:2715 | 2716465:2716 | -;-;-;-;-;-;-;-; | 1959 | akirin2 | 944 | 1460 | 1637 | 1141 | 1375 | 1059 |
| ENSDARG000 | 20;20;20;20;2 | 6808774:6809 | 6809722:6809 | -;-;-;-;-;-;-;- | 1773 | igfbp1a | 860 | 764 | 4117 | 1714 | 729 | 734 |
| ENSDARG000 | 20;20;20;20;2 | 42221158:422 | 42224212:422 | -;-;-;-;-;-;-;-; | 4126 | nus1 | 812 | 1313 | 1720 | 1155 | 1144 | 1138 |
| ENSDARG000 | 20;20;20;20;2 | 33175190:331 | 33175239:331 | +;+;+;+;+;+;+; | 2283 | ddx1 | 1129 | 1071 | 1358 | 639 | 1015 | 803 |
| ENSDARG000 | 20;20;20;20;2 | 9776390:9777 | 9776973:9777 | -;-;-;-;-;-;-;-; | 1402 | cgrrf1 | 130 | 90 | 122 | 156 | 126 | 82 |
| ENSDARG000 | 20;20;20;20;2 | 30355436:303 | 30356093:303 | -;-;-;-;-;-;-;-; | 2200 | allc | 0 | 0 | 10 | 0 | 2 | 0 |
| ENSDARG000 | 20;20;20 | 26892761:268 | 26893542:268 | +;+;+ | 1683 | NA | 16 | 8 | 26 | 18 | 61 | 48 |
| ENSDARG000 | 20;20;20;20;2 | 9763364:9763 | 9763484:9763 | +;+;+;+;+;+;+; | 1561 | psmc6 | 2061 | 2377 | 2353 | 979 | 2301 | 1619 |
| ENSDARG000 | 20;20;20;20;2 | 42246948:422 | 42247396:422 | +;+;+;+;+;+;+; | 2337 | gopc | 334 | 279 | 310 | 139 | 372 | 276 |
| ENSDARG000 | 20;20;20;20 | 34455645:344 | 34455998:344 | +;+;+;+ | 1037 | ntmt2 | 2 | 0 | 0 | 0 | 0 | 0 |
| ENSDARG000 | 20;20;20;20;2 | 20901791:209 | 20902002:209 | +;+;+;+;+;+;+; | 4529 | brf1b | 122 | 141 | 155 | 155 | 168 | 128 |
| ENSDARG000 | 20;20;20;20;2 | 20927827:209 | 20929268:209 | -;-;-;-;-;-;-;-; | 2554 | btbd6b | 568 | 321 | 280 | 358 | 234 | 364 |
| ENSDARG000 | 20;20;20;20;2 | 902412:90241 | 903183:90318 | -;-;-;-;- | 2954 | cnr1 | 0 | 6 | 0 | 0 | 0 | 0 |
| ENSDARG000 | 20;20;20;20;2 | 26899745:269 | 26899912:269 | +;+;+;+;+;+ | 1155 | NA | 13 | 0 | 0 | 0 | 0 | 6 |
| ENSDARG000 | 20;20;20;20;2 | 6894886:6894 | 6895231:6895 | -;-;-;-;-;-;-;-; | 3822 | adcy1a | 0 | 0 | 0 | 0 | 10 | 0 |
| ENSDARG000 | 20;20;20;20;2 | 34501152:345 | 34501380:345 | +;+;+;+;+;+;+; | 2453 | gorab | 178 | 202 | 289 | 130 | 264 | 158 |
| ENSDARG000 | 20;20;20;20;2 | 34511678:345 | 34512290:345 | +;+;+;+;+;+;+; | 2828 | prrx1b | 166 | 12 | 3 | 0 | 8 | 19 |
| ENSDARG000 | 20;20;20 | 33294428:332 | 33294561:332 | +;+;+ | 2488 | mycn | 3977 | 4475 | 5796 | 4986 | 3383 | 8202 |
| ENSDARG000 | 20;20;20;20;2 | 28832762:288 | 28834147:288 | -;-;-;-;-;-;-;-; | 3237 | max | 2154 | 1946 | 3112 | 2201 | 2305 | 2638 |
| ENSDARG000 | 20;20;20;20;2 | 42296256:423 | 42300253:423 | -;-;-;-;-;-;-;-; | 7037 | dcbld1 | 2759 | 2983 | 2998 | 2096 | 5216 | 1802 |
| ENSDARG000 | 20;20;20;20;2 | 7052870:7053 | 7053525:7053 | -;-;-;-;-;-;-;-; | 1948 | sirt5 | 457 | 402 | 717 | 566 | 482 | 264 |
| ENSDARG000 | 20;20;20;20;2 | 14632492:146 | 14635830:146 | -;-;-;-;-;-;-;-; | 4787 | scrn2 | 362 | 309 | 364 | 256 | 544 | 210 |
| ENSDARG000 | 20;20 | 33461088:334 | 33461383:334 | -;- | 918 | NA | 0 | 0 | 0 | 0 | 0 | 0 |
| ENSDARG000 | 20;20;20 | 33370214:334 | 33370640:334 | -;-;- | 803 | NA | 0 | 0 | 0 | 1 | 0 | 0 |
| ENSDARG000 | 20;20 | 14657141:146 | 14657282:146 | +;+ | 1845 | NA | 15 | 24 | 14 | 0 | 12 | 20 |
| ENSDARG000 | 20;20;20;20 | 42431542:424 | 42431723:424 | -;-;-;- | 1163 | vgll2a | 0 | 0 | 12 | 4 | 0 | 0 |
| ENSDARG000 | 20;20;20;20;2 | 943879:94665 | 944622:94678 | -;-;-;-;-;-;-;-; | 2649 | rngtt | 361 | 267 | 457 | 295 | 461 | 228 |
| ENSDARG000 | 20;20;20;20;2 | 23877299:238 | 23878445:238 | -;-;-;-;-;-;-;-; | 17739 | MDN1 | 1566 | 934 | 2486 | 928 | 1490 | 1061 |
| ENSDARG000 | 20;20;20;20;2 | 42506612:425 | 42506816:425 | -;-;-;-;-;-;-;-; | 2771 | rfx6 | 0 | 0 | 2 | 4 | 2 | 2 |
| ENSDARG000 | 20;20;20;20;2 | 26905158:269 | 26905250:269 | +;+;+;+;+;+;+; | 1511 | rwdd2b | 153 | 108 | 150 | 88 | 161 | 98 |
| ENSDARG000 | 20;20;20;20;2 | 25645459:256 | 25645820:256 | +;+;+;+;+;+;+; | 3802 | AASDH | 38 | 42 | 86 | 58 | 67 | 42 |
| ENSDARG000 | 20;20;20;20;2 | 33467161:334 | 33468552:334 | -;-;-;-;-;-;-;-; | 4640 | smek1 | 2077 | 1950 | 2621 | 1783 | 2615 | 1828 |
| ENSDARG000 | 20;20;20;20;2 | 2731436:2731 | 2731619:2731 | +;+;+;+;+;+;+; | 5150 | syncrip | 8939 | 11123 | 12813 | 6001 | 11932 | 7682 |
| ENSDARG000 | 20;20;20;20;2 | 18661624:186 | 18661724:186 | +;+;+;+;+;+;+; | 2729 | tnfaip2a | 10044 | 6496 | 6479 | 6029 | 6131 | 4807 |
| ENSDARG000 | 20;20;20;20 | 33493924:334 | 33494517:334 | -;-;-;- | 921 | erh | 1276 | 1685 | 1713 | 1387 | 1355 | 1406 |
| ENSDARG000 | 20;20;20;20;2 | 42537768:425 | 42537962:425 | +;+;+;+;+;+;+; | 6462 | tdrd15 | 12 | 12 | 54 | 38 | 14 | 14 |
| ENSDARG000 | 20;20;20;20;2 | 25659839:256 | 25661633:256 | -;-;-;-;-;-;-;-; | 5485 | cracd | 48 | 58 | 88 | 93 | 62 | 134 |
| ENSDARG000 | 20;20;20;20;2 | 34537736:345 | 34537898:345 | +;+;+;+;+;+;+; | 11037 | NA | 3442 | 1866 | 1586 | 3784 | 2111 | 4360 |
| ENSDARG000 | 20;20;20;20;2 | 7070634:7070 | 7070890:7070 | -;-;-;-;-;-;-;-; | 1863 | si:ch211-121a | 0 | 0 | 0 | 0 | 8 | 0 |
| ENSDARG000 | 20;20;20;20;2 | 30372352:303 | 30372463:303 | -;-;-;-;-;-;-;-; | 690 | rps7 | 17056 | 15877 | 25762 | 22122 | 15527 | 25420 |
| ENSDARG000 | 20;20;20;20;2 | 42563618:425 | 42563695:425 | +;+;+;+;+;+;+; | 7969 | igf2r | 1923 | 1008 | 1706 | 1004 | 1614 | 1391 |
| ENSDARG000 | 20;20;20;20;2 | 33499378:334 | 33499892:334 | -;-;-;-;-;-;-;-; | 1975 | paplnb | 28 | 4 | 12 | 6 | 50 | 5 |
| ENSDARG000 | 20;20;20;20;2 | 18702624:187 | 18702839:187 | +;+;+;+;+;+;+; | 5380 | eif5 | 10745 | 10560 | 11195 | 7404 | 14881 | 8130 |
| ENSDARG000 | 20;20 | 25710791:257 | 25711784:257 | -;- | 1280 | NA | 0 | 0 | 0 | 0 | 0 | 0 |
| ENSDARG000 | 20;20;20;20;2 | 25711718:257 | 25711873:257 | +;+;+;+;+;+;+; | 4259 | cep135 | 149 | 279 | 342 | 154 | 322 | 243 |
| ENSDARG000 | 20;20;20;20;2 | 26916639:269 | 26916829:269 | +;+;+;+;+;+ | 1959 | serpinb1l2 | 34 | 88 | 149 | 90 | 99 | 127 |
| ENSDARG000 | 20;20;20;20;2 | 14683699:146 | 14686636:146 | -;-;-;-;-;-;-;-; | 6871 | NA | 5831 | 3782 | 6181 | 8818 | 8546 | 4975 |
| ENSDARG000 | 20;20;20;20;2 | 7084154:7088 | 7084622:7088 | +;+;+;+;+;+ | 1637 | NA | 26 | 32 | 4 | 72 | 44 | 2 |
| ENSDARG000 | 20;20;20;20;2 | 28850233:288 | 28850488:288 | +;+;+;+;+;+;+; | 969 | NA | 0 | 0 | 0 | 0 | 0 | 2 |
| ENSDARG000 | 20;20;20;20;2 | 30378803:303 | 30378888:303 | +;+;+;+;+;+;+; | 1357 | rnaseh1 | 424 | 458 | 540 | 291 | 522 | 396 |
| ENSDARG000 | 20 | 33513664 | 33515890 | - | 2227 | NA | 0 | 2 | 0 | 0 | 0 | 0 |
| ENSDARG000 | 20;20;20;20;2 | 26931496:269 | 26931918:269 | -;-;-;-;-;-;-;-; | 3393 | NA | 334 | 186 | 263 | 228 | 246 | 238 |
| ENSDARG000 | 20;20;20;20;2 | 26920681:269 | 26921318:269 | -;-;-;-;-;-;-;-; | 3118 | NA | 177 | 147 | 273 | 219 | 268 | 230 |
| ENSDARG000 | 20;20;20;20;2 | 33519435:335 | 33519648:335 | +;+;+;+;+;+;+; | 2384 | drc1 | 2 | 6 | 6 | 0 | 2 | 4 |
| ENSDARG000 | 20;20;20;20;2 | 7122527:7122 | 7122989:7122 | -;-;-;-;-;-;-;-; | 829 | NA | 6 | 2 | 0 | 16 | 30 | 2 |
| ENSDARG000 | 20;20 | 1121458:1125 | 1121875:1125 | +;+ | 808 | pnrc1 | 71 | 44 | 69 | 25 | 78 | 84 |
| ENSDARG000 | 20;20;20;20;2 | 42688488:426 | 42689363:426 | -;-;-;-;-;-;-;-; | 3241 | plg | 2 | 0 | 28 | 4 | 0 | 0 |
| ENSDARG000 | 20;20;20;20;2 | 23947004:239 | 23947214:239 | +;+;+;+;+;+;+; | 6363 | casp8ap2 | 410 | 334 | 610 | 206 | 474 | 388 |
| ENSDARG000 | 20;20;20;20;2 | 42668875:426 | 42669323:426 | +;+;+;+;+;+;+; | 2849 | slc22a2 | 1812 | 186 | 1079 | 964 | 1616 | 1848 |
| ENSDARG000 | 20;20;20;20;2 | 9604:10847:1 | 10751:11017: | -;-;-;-;-;-;- | 2956 | zgc:174972 | 10 | 28 | 36 | 25 | 16 | 51 |
| ENSDARG000 | 20;20;20;20;2 | 30445971:304 | 30446178:304 | +;+;+;+;+;+;+; | 4144 | myt1la | 0 | 0 | 0 | 0 | 2 | 2 |
| ENSDARG000 | 20;20;20 | 23960525:239 | 23960640:239 | +;+;+ | 2114 | GJA10 | 0 | 0 | 0 | 0 | 0 | 0 |
| ENSDARG000 | 20;20;20;20;2 | 7149100:7149 | 7149411:7150 | -;-;-;-;-;-;-;-; | 1976 | dhcr24 | 440 | 220 | 134 | 500 | 1697 | 120 |
| ENSDARG000 | 20;20;20;20;2 | 14789148:147 | 14789668:147 | +;+;+;+;+;+;+; | 2394 | tmed5 | 714 | 972 | 1023 | 704 | 1019 | 644 |
| ENSDARG000 | 20;20;20;20;2 | 28861435:288 | 28861521:288 | +;+;+;+;+;+;+; | 2077 | SLC10A1 | 0 | 0 | 0 | 0 | 0 | 0 |
| ENSDARG000 | 20;20;20;20;2 | 18721311:187 | 18724295:187 | -;-;-;-;-;-;- | 4153 | enpp5 | 114 | 36 | 43 | 14 | 74 | 62 |
| ENSDARG000 | 20;20 | 14810886:148 | 14810931:148 | +;+ | 1593 | pigc | 134 | 211 | 196 | 108 | 266 | 77 |
| ENSDARG000 | 20;20;20;20;2 | 23972948:239 | 23975081:239 | -;-;-;-;-;-;-;-; | 5373 | BACH2 | 157 | 67 | 110 | 74 | 137 | 172 |
| ENSDARG000 | 20;20;20;20;2 | 18740518:187 | 18741129:187 | +;+;+;+;+;+;+; | 6015 | fam167ab | 10 | 3 | 4 | 2 | 18 | 0 |
| ENSDARG000 | 20;20;20;20 | 34647500:346 | 34647940:346 | -;-;-;- | 963 | NA | 35 | 275 | 104 | 130 | 149 | 30 |
| ENSDARG000 | 20;20;20;20;2 | 21022641:210 | 21022898:210 | +;+;+;+;+;+;+; | 1243 | NA | 8 | 0 | 4 | 0 | 0 | 0 |
| ENSDARG000 | 20;20;20;20;2 | 9781932:9782 | 9783117:9783 | +;+;+;+;+;+;+; | 3788 | ddhd1b | 622 | 499 | 308 | 284 | 430 | 497 |
| ENSDARG000 | 20;20;20;20;2 | 28869969:288 | 28871999:288 | -;-;-;-;-;-;-;-; | 4709 | srsf5b | 1955 | 2505 | 3392 | 2575 | 3251 | 2672 |
| ENSDARG000 | 20;20;20 | 17667:19253: | 17722:19373: | +;+;+ | 1013 | NA | 28 | 36 | 49 | 52 | 41 | 22 |
| ENSDARG000 | 20;20;20;20;2 | 34653916:346 | 34654696:346 | -;-;-;-;-;-;-;-; | 2461 | slc25a24 | 1257 | 1045 | 972 | 1044 | 1239 | 1166 |
| ENSDARG000 | 20;20;20;20;2 | 18774129:187 | 18774625:187 | -;-;-;-;-;-;-;-; | 2395 | ccm2 | 318 | 246 | 340 | 179 | 217 | 267 |
| ENSDARG000 | 20;20;20 | 21066100:210 | 21066148:210 | +;+;+ | 597 | NA | 2 | 0 | 0 | 0 | 0 | 4 |
| ENSDARG000 | 20;20;20 | 33532296:335 | 33532609:335 | +;+;+ | 3520 | kcnf1a | 2 | 4 | 2 | 0 | 0 | 0 |
| ENSDARG000 | 20;20 | 7180298:7191 | 7180892:7192 | +;+ | 1124 | si:dkeyp-51f1 | 12 | 4 | 4 | 18 | 8 | 0 |
| ENSDARG000 | 20;20;20;20 | 33555381:335 | 33555552:335 | -;-;-;- | 544 | NA | 2 | 0 | 0 | 1 | 2 | 1 |
| ENSDARG000 | 20;20 | 34663322:346 | 34663348:346 | +;+ | 608 | NA | 2 | 0 | 0 | 13 | 10 | 3 |
| ENSDARG000 | 20;20 | 34661555:346 | 34661609:346 | +;+ | 173 | NA | 0 | 0 | 0 | 0 | 0 | 0 |
| ENSDARG000 | 20;20;20;20;2 | 21268795:213 | 21269110:213 | +;+;+;+;+ | 1925 | nudt14 | 673 | 384 | 705 | 189 | 481 | 448 |
| ENSDARG000 | 20;20;20;20;2 | 25370:55937: | 26570:56037: | +;+;+;+;+;+ | 4026 | LRFN2 | 38 | 43 | 63 | 198 | 94 | 4 |
| ENSDARG000 | 20;20;20;20;2 | 1132092:1132 | 1132391:1132 | -;-;-;-;-;-;-;-; | 1548 | gabrr1 | 0 | 0 | 5 | 0 | 0 | 0 |
| ENSDARG000 | 20;20;20;20;2 | 33571312:335 | 33571663:335 | -;-;-;-;-;-;-;- | 1498 | NA | 208 | 4 | 304 | 370 | 5 | 193 |
| ENSDARG000 | 20;20;20;20 | 33560419:335 | 33560670:335 | +;+;+;+ | 659 | si:dkev-65b13 | 0 | 0 | 2 | 0 | 0 | 0 |
| ENSDARG000 | 20;20;20 | 33564425:335 | 33564563:335 | -;-;- | 545 | NA | 0 | 0 | 2 | 0 | 1 | 0 |
| ENSDARG000 | 20;20;20;20;2 | 2879700:2882 | 2879813:2882 | -;-;-;-;-;-;-;-; | 1458 | CDK19 | 52 | 44 | 32 | 20 | 45 | 87 |
| ENSDARG000 | 20;20;20;20;2 | 7209972:7209 | 7210380:7210 | +;+;+;+;+;+ | 1414 | si:dkeyp-51f1 | 24 | 50 | 101 | 26 | 100 | 80 |
| ENSDARG000 | 20;20 | 25739979:257 | 25741281:257 | -;- | 1707 | chst14 | 154 | 77 | 100 | 101 | 188 | 152 |
| ENSDARG000 | 20;20;20 | 18801525:188 | 18803273:188 | -;-;- | 2565 | xkr6b | 0 | 0 | 0 | 0 | 2 | 0 |
| ENSDARG000 | 20;20;20 | 7243879:7248 | 7244074:7248 | +;+;+ | 886 | bsnd | 0 | 4 | 6 | 0 | 10 | 0 |
| ENSDARG000 | 20;20;20;20;2 | 18936455:189 | 18937134:189 | -;-;-;-;-;-;-;-; | 2567 | NA | 0 | 0 | 0 | 0 | 0 | 0 |
| ENSDARG000 | 20;20;20;20;2 | 24132073:241 | 24132109:241 | +;+;+;+;+;+;+; | 1035 | NA | 0 | 0 | 0 | 2 | 0 | 4 |
| ENSDARG000 | 20;20 | 84692;87253 | 85775;88281 | -;- | 2113 | NA | 1 | 0 | 14 | 0 | 34 | 2 |
| ENSDARG000 | 20 | 88168 | 90255 | + | 2088 | zgc:112001 | 0 | 0 | 0 | 0 | 0 | 0 |
| ENSDARG000 | 20;20;20;20;2 | 30578163:305 | 30579101:305 | +;+;+;+;+;+;+; | 3083 | cep43 | 486 | 511 | 467 | 457 | 610 | 561 |

| | | | | | | | | | | | | |
|---|---|---|---|---|---|---|---|---|---|---|---|---|
| ENSDARG000 | 20:20:20:20:2 | 26939742:269 | 26939938:269 | +:+:+:+:+:+:+: | 3044 | cdca4 | 996 | 868 | 1383 | 940 | 1340 | 1232 |
| ENSDARG000 | 20:20:20:20:2 | 21391181:213 | 21391411:213 | +:+:+:+:+:+:+: | 6808 | jag2b | 2754 | 2265 | 2072 | 1362 | 2141 | 2318 |
| ENSDARG000 | 20:20:20:20:2 | 2950005:2961 | 2950351:2961 | +:+:+:+:+:+:+: | 5735 | amd1 | 2845 | 5023 | 4366 | 1182 | 6077 | 4673 |
| ENSDARG000 | 20:20:20:20:2 | 14817630:148 | 14819146:148 | -:-:-:-:-:-:-:-:-: | 4454 | dnm3a | 572 | 390 | 432 | 1090 | 432 | 1060 |
| ENSDARG000 | 20:20:20:20:2 | 25853012:258 | 25853549:258 | -:-:-:-:-:-:-:-:-: | 3411 | exoc1 | 2385 | 1851 | 1812 | 2023 | 1851 | 1988 |
| ENSDARG000 | 20:20:20:20:2 | 19006703:190 | 19006736:190 | +;+;+;+;+;+;+ | 1814 | pinx1 | 512 | 544 | 1104 | 476 | 635 | 654 |
| ENSDARG000 | 20:20:20:20:2 | 7262434:7267 | 7264528:7267 | -:-:-:-:-:-:-:-:-: | 4846 | dsc2l | 299 | 84 | 102 | 84 | 360 | 100 |
| ENSDARG000 | 20:20:20:20:2 | 18992679:189 | 18992782:189 | +:+:+:+:+:+:+: | 2382 | tdh | 6131 | 6517 | 14948 | 15343 | 22291 | 25652 |
| ENSDARG000 | 20 | 14875460 | 14875546 | + | 87 | mir199-3a | 0 | 0 | 0 | 0 | 0 | 0 |
| ENSDARG000 | 20:20:20:20:2 | 14926881:149 | 14928444:149 | -:-:-:-:-:-:-:-:-: | 4004 | mettl13 | 250 | 306 | 363 | 184 | 228 | 220 |
| ENSDARG000 | 20:20:20:20:2 | 9828769:9828 | 9829091:9829 | +:+:+:+:+:+:+: | 3376 | stxbp6l | 73 | 6 | 23 | 109 | 60 | 85 |
| ENSDARG000 | 20 | 14873030 | 14873330 | - | 301 | NA | 0 | 0 | 0 | 0 | 0 | 0 |
| ENSDARG000 | 20 | 14877639 | 14877738 | + | 100 | mir214a | 0 | 0 | 0 | 0 | 0 | 0 |
| ENSDARG000 | 20:20:20:20:2 | 34671386:346 | 34671545:346 | +:+:+:+:+:+:+: | 2365 | elp3 | 264 | 395 | 630 | 389 | 608 | 392 |
| ENSDARG000 | 20:20:20:20:2 | 18943406:189 | 18943932:189 | +:+:+:+:+:+:+: | 2442 | mtmr9 | 150 | 90 | 91 | 94 | 172 | 124 |
| ENSDARG000 | 20:20:20:20:2 | 98179:100229 | 98501:100520 | +;+;+;+;+;+ | 3479 | olfcb1 | 0 | 0 | 0 | 0 | 0 | 0 |
| ENSDARG000 | 20:20:20:20:2 | 24149794:241 | 24152926:241 | -:-:-:-:-:-:-:-:-: | 6021 | map3k7 | 1235 | 910 | 1433 | 718 | 1407 | 1254 |
| ENSDARG000 | 20;20 | 30610547:306 | 30610644:306 | +;+ | 1231 | ccr6a | 39 | 7 | 27 | 9 | 37 | 39 |
| ENSDARG000 | 20;20 | 106481:11295 | 109163:11331 | +;+ | 3050 | NA | 0 | 0 | 0 | 0 | 0 | 0 |
| ENSDARG000 | 20;20 | 42968785:429 | 42969309:429 | -;- | 648 | pomcb | 0 | 0 | 0 | 0 | 0 | 0 |
| ENSDARG000 | 20;20 | 26947800:269 | 26948361:269 | +;+ | 4764 | si:dkev-177p2 | 635 | 445 | 994 | 786 | 1176 | 548 |
| ENSDARG000 | 20;20 | 26959533:269 | 26959724:269 | -;- | 347 | NA | 0 | 0 | 0 | 0 | 0 | 0 |
| ENSDARG000 | 20;20;20 | 113949:11607 | 114818:11622 | -;-;- | 1132 | ypel5 | 509 | 363 | 481 | 382 | 360 | 477 |
| ENSDARG000 | 20;20 | 33679500:336 | 33679751:336 | +;+ | 460 | si:ch211-199r | 0 | 0 | 0 | 6 | 0 | 0 |
| ENSDARG000 | 20:20:20:20:2 | 33601046:336 | 33601710:336 | -:-:-:-:-:-:-:-:-: | 5319 | rock2b | 1421 | 1479 | 3910 | 2944 | 3216 | 1545 |
| ENSDARG000 | 20:20:20:20:2 | 26966725:269 | 26967568:269 | +:+:+:+:+:+:+: | 2375 | ahsa1a | 6 | 0 | 14 | 18 | 2 | 0 |
| ENSDARG000 | 20:20:20:20:2 | 1158393:1158 | 1159043:1159 | -:-:-:-:-:-:-:-:-: | 1887 | GABRR2 | 13 | 6 | 15 | 23 | 14 | 23 |
| ENSDARG000 | 20:20:20:20:2 | 30702531:307 | 30702848:307 | +:+:+:+:+:+:+: | 3350 | acbd3 | 939 | 1405 | 1566 | 831 | 1549 | 1003 |
| ENSDARG000 | 20:20:20:20:2 | 2982381:2986 | 2983688:2986 | -:-:-:-:-:-:-:-:-: | 2532 | ppil4 | 306 | 270 | 417 | 198 | 307 | 412 |
| ENSDARG000 | 20:20:20:20:2 | 1177751:1182 | 1179783:1182 | -;-;-;-;-;-;- | 2891 | ube2j1 | 653 | 562 | 681 | 344 | 560 | 605 |
| ENSDARG000 | 20:20:20:20:2 | 42978499:429 | 42978563:429 | +:+:+:+:+:+:+: | 2269 | NA | 2 | 4 | 0 | 0 | 12 | 0 |
| ENSDARG000 | 20;20 | 9958506:9963 | 9961646:9963 | -;- | 3259 | gjd2b | 0 | 0 | 0 | 0 | 0 | 0 |
| ENSDARG000 | 20:20:20:20:2 | 122123:12448 | 122920:12472 | -:-:-:-:-:-:-:-:-: | 2389 | slc16a10 | 312 | 523 | 459 | 381 | 436 | 655 |
| ENSDARG000 | 20:20:20:20:2 | 7327627:7327 | 7327974:7327 | +:+:+:+:+:+:+: | 4038 | dsg2.1 | 400 | 46 | 38 | 74 | 128 | 43 |
| ENSDARG000 | 20:20:20:20:2 | 30725778:307 | 30726099:307 | +:+:+:+:+:+:+: | 6838 | NHSL1 | 99 | 100 | 44 | 25 | 111 | 58 |
| ENSDARG000 | 20:20:20:20:2 | 9970786:9971 | 9971142:9971 | -;-;-;-;-;-;- | 1570 | actc2 | 4 | 2 | 0 | 0 | 0 | 0 |
| ENSDARG000 | 20:20:20:20:2 | 43016437:430 | 43017260:430 | -:-:-:-:-:-:-:-:-: | 3347 | SNX9 | 6288 | 3252 | 4249 | 4116 | 4594 | 4238 |
| ENSDARG000 | 20:20:20:20:2 | 25879826:258 | 25880579:258 | +;+;+;+;+ | 3179 | zgc:153896 | 62 | 7 | 55 | 62 | 65 | 70 |
| ENSDARG000 | 20;20;20 | 10057750:100 | 10061650:100 | -;-;- | 4195 | NA | 557 | 568 | 672 | 474 | 717 | 626 |
| ENSDARG000 | 20:20:20:20:2 | 25886491:258 | 25887014:258 | -:-:-:-:-:-:-:-:-: | 3961 | mideasa | 556 | 300 | 372 | 322 | 467 | 662 |
| ENSDARG000 | 20:20:20:20:2 | 7443224:7443 | 7443286:7443 | -:-:-:-:-:-:-:-:-: | 8499 | NA | 811 | 878 | 1557 | 438 | 936 | 1052 |
| ENSDARG000 | 20:20:20:20:2 | 30894093:308 | 30895157:308 | -;-;-;-;-;-;-;- | 1682 | hebp2 | 562 | 245 | 186 | 209 | 185 | 496 |
| ENSDARG000 | 20:20:20:20:2 | 34702410:347 | 34703462:347 | -;-;-;-;-;- | 2184 | si:ch211-63o2 | 69 | 221 | 178 | 97 | 189 | 91 |
| ENSDARG000 | 20;20;20 | 34717403:347 | 34717596:347 | +;+;+ | 1260 | pnocb | 0 | 6 | 6 | 2 | 52 | 6 |
| ENSDARG000 | 20:20:20:20:2 | 10098590:100 | 10100308:101 | -:-:-:-:-:-:-:-:-: | 3889 | meis2b | 0 | 0 | 8 | 0 | 0 | 0 |
| ENSDARG000 | 20:20:20:20:2 | 33708318:337 | 33708380:337 | +:+:+:+:+:+:+: | 5560 | NA | 0 | 0 | 9 | 0 | 0 | 0 |
| ENSDARG000 | 20:20:20:20:2 | 150890:15211 | 151917:15225 | -:-:-:-:-:-:-:-:-: | 1808 | rpf2 | 1420 | 2052 | 1971 | 1242 | 1572 | 1916 |
| ENSDARG000 | 20:20:20:20:2 | 34737738:347 | 34738858:347 | -:-:-:-:-:-:-:-:-: | 2392 | znf395b | 494 | 289 | 382 | 435 | 483 | 691 |
| ENSDARG000 | 20:20:20:20:2 | 43083745:430 | 43084045:430 | +:+:+:+:+:+:+: | 2062 | moxd1l | 0 | 0 | 0 | 0 | 0 | 0 |
| ENSDARG000 | 20:20:20:20:2 | 33739059:337 | 33739180:337 | +:+:+:+:+:+:+: | 1075 | ppil6 | 0 | 0 | 0 | 0 | 0 | 0 |
| ENSDARG000 | 20:20:20:20:2 | 30902138:309 | 30902704:309 | -:-:-:-:-:-:-:-:-: | 2293 | KIF25 | 0 | 0 | 0 | 0 | 0 | 0 |
| ENSDARG000 | 20;20 | 30902538:309 | 30902744:309 | +;+ | 728 | NA | 0 | 0 | 0 | 0 | 0 | 0 |
| ENSDARG000 | 20:20:20:20:2 | 3005472:3006 | 3006588:3006 | -:-:-:-:-:-:-:-:-: | 6737 | map3k5 | 176 | 163 | 818 | 315 | 953 | 410 |
| ENSDARG000 | 20;20;20 | 34766183:347 | 34767478:347 | -;-;- | 1450 | paqr8 | 5 | 0 | 0 | 10 | 0 | 0 |
| ENSDARG000 | 20:20:20:20:2 | 26987416:269 | 26988041:269 | +:+:+:+:+:+:+: | 4360 | sel1l | 1606 | 1670 | 1927 | 816 | 1741 | 1437 |
| ENSDARG000 | 20:20:20:20:2 | 33744065:337 | 33744267:337 | +:+:+:+:+:+:+: | 1481 | henmt1 | 0 | 2 | 26 | 6 | 16 | 3 |
| ENSDARG000 | 20;20 | 43113465:431 | 43113867:431 | +;+ | 1121 | dusp23a | 26 | 29 | 0 | 16 | 22 | 10 |
| ENSDARG000 | 20:20:20:20:2 | 28885990:288 | 28889528:288 | -:-:-:-:-:-:-:-:-: | 4981 | susd6 | 1754 | 999 | 1228 | 1064 | 2043 | 1389 |
| ENSDARG000 | 20:20:20:20:2 | 33752967:337 | 33753079:337 | -:-:-:-:-:-:-:-:-: | 1473 | fam102bb | 123 | 156 | 276 | 161 | 271 | 215 |
| ENSDARG000 | 20:20:20:20:2 | 43379029:433 | 43379165:433 | +:+:+:+:+:+:+: | 1523 | unc93a | 0 | 0 | 0 | 0 | 0 | 0 |
| ENSDARG000 | 20:20:20:20:2 | 27003558:270 | 27003680:270 | +:+:+:+:+:+:+: | 3110 | NA | 4 | 18 | 36 | 12 | 12 | 10 |
| ENSDARG000 | 20:20:20:20:2 | 43120132:431 | 43123132:431 | -:-:-:-:-:-:-:-:-: | 8947 | AFDN | 1179 | 485 | 724 | 492 | 927 | 413 |
| ENSDARG000 | 20:20:20:20:2 | 33792156:337 | 33792308:337 | +:+:+:+:+:+:+: | 2652 | olfml2ba | 2 | 9 | 13 | 8 | 74 | 20 |
| ENSDARG000 | 20;20 | 1198079:1198 | 1198339:1198 | -;- | 541 | NA | 0 | 0 | 2 | 0 | 0 | 0 |
| ENSDARG000 | 20;20;20 | 1196386:1197 | 1196465:1198 | +;+;+ | 979 | lyrm2 | 27 | 41 | 92 | 66 | 65 | 108 |
| ENSDARG000 | 20:20:20:20:2 | 10166297:101 | 10166764:101 | +;+;+;+;+;+;+ | 3268 | kcnk10a | 0 | 0 | 0 | 0 | 0 | 0 |
| ENSDARG000 | 20:20:20:20:2 | 43345882:433 | 43345913:433 | -:-:-:-:-:-:-:-:-: | 943 | NA | 0 | 0 | 0 | 2 | 0 | 0 |
| ENSDARG000 | 20:20:20:20:2 | 34773646:347 | 34773974:347 | -:-:-:-:-:-:-:-:-: | 2456 | stmn4 | 16 | 212 | 82 | 77 | 421 | 54 |
| ENSDARG000 | 20:20:20:20:2 | 21583639:215 | 21583694:215 | +:+:+:+:+:+:+: | 3249 | esr2a | 0 | 2 | 8 | 0 | 2 | 0 |
| ENSDARG000 | 20:20:20:20:2 | 3108597:3111 | 3108828:3112 | +:+:+:+:+:+:+: | 6010 | CEP170B | 0 | 0 | 0 | 0 | 0 | 0 |
| ENSDARG000 | 20;20;20 | 34792166:347 | 34792205:347 | +;+;+ | 1221 | il17a/f3 | 0 | 0 | 0 | 0 | 0 | 0 |
| ENSDARG000 | 20:20:20:20:2 | 34770197:347 | 34770387:347 | +:+:+:+:+:+:+: | 3062 | mcm3 | 741 | 2615 | 3131 | 1096 | 3116 | 1700 |
| ENSDARG000 | 20;20;20 | 24365947:243 | 24366317:243 | +;+;+ | 903 | NA | 0 | 0 | 0 | 0 | 0 | 0 |
| ENSDARG000 | 20:20:20:20:2 | 34845672:348 | 34846157:348 | +:+:+:+:+:+:+: | 3882 | emilin1a | 105 | 19 | 60 | 0 | 24 | 32 |
| ENSDARG000 | 20:20:20:20:2 | 43398431:434 | 43398887:434 | -;-;-;-;- | 798 | NA | 0 | 0 | 0 | 0 | 0 | 0 |
| ENSDARG000 | 20:20:20:20:2 | 43396318:433 | 43396656:433 | +;+;+;+;+;+;+ | 4234 | NA | 2 | 0 | 5 | 45 | 8 | 0 |
| ENSDARG000 | 20;20 | 43406707:434 | 43406923:434 | -;- | 655 | NA | 0 | 0 | 0 | 0 | 0 | 0 |
| ENSDARG000 | 20;20;20;20 | 24449637:244 | 24450106:244 | -;-;-;- | 1324 | NA | 2 | 0 | 2 | 0 | 0 | 1 |
| ENSDARG000 | 20:20:20:20:2 | 186277:18627 | 186835:18683 | -;-;-;-;- | 1294 | smim8 | 92 | 154 | 218 | 68 | 142 | 112 |
| ENSDARG000 | 20;20;20;20 | 43424002:434 | 43424130:434 | +;+;+;+ | 350 | NA | 0 | 0 | 4 | 4 | 2 | 0 |
| ENSDARG000 | 20;20 | 24448007:244 | 24448120:244 | +;+ | 547 | NA | 0 | 0 | 0 | 0 | 0 | 0 |
| ENSDARG000 | 20;20 | 24418405:245 | 24418756:245 | +;+ | 1148 | NA | 0 | 0 | 0 | 0 | 0 | 0 |
| ENSDARG000 | 20 | 10287061 | 10288156 | - | 1096 | BDKRB2 | 8 | 0 | 0 | 0 | 0 | 0 |
| ENSDARG000 | 20:20:20:20:2 | 159277:15930 | 160324:16032 | -:-:-:-:-:-:-:-:-: | 6039 | si:ch211-241i | 5312 | 2156 | 1555 | 3436 | 556 | 3621 |
| ENSDARG000 | 20:20:20:20:2 | 43432092:434 | 43432219:434 | -;-;-;-;-;-;- | 1686 | pimr109 | 0 | 0 | 2 | 0 | 0 | 0 |
| ENSDARG000 | 20;20 | 1200366:1201 | 1200494:1202 | +;+ | 742 | NA | 27 | 9 | 13 | 25 | 2 | 39 |
| ENSDARG000 | 20;20;20;20 | 10353201:103 | 10354499:103 | -;-;-;- | 1620 | NA | 306 | 5 | 37 | 2 | 8 | 18 |
| ENSDARG000 | 20:20:20:20:2 | 25904199:259 | 25904416:259 | +;+;+;+;+;+ | 2159 | slc35f6 | 709 | 645 | 763 | 465 | 860 | 861 |
| ENSDARG000 | 20;20 | 24754446:247 | 24754504:247 | +;+ | 279 | NA | 0 | 0 | 0 | 0 | 0 | 0 |
| ENSDARG000 | 20:20:20:20:2 | 25911342:259 | 25911693:259 | +:+:+:+:+:+:+: | 5277 | ttbk2b | 268 | 308 | 315 | 212 | 355 | 321 |
| ENSDARG000 | 20:20:20:20:2 | 30922595:309 | 30923438:309 | -:-:-:-:-:-:-:-:-: | 1945 | acat2 | 886 | 802 | 989 | 553 | 802 | 586 |
| ENSDARG000 | 20:20:20:20:2 | 33875256:339 | 33875341:339 | +:+:+:+:+:+:+: | 2729 | rxrgb | 2 | 10 | 44 | 1 | 34 | 22 |
| ENSDARG000 | 20:20:20:20:2 | 43457425:434 | 43458321:434 | -:-:-:-:-:-:-:-:-: | 2216 | pimr133 | 0 | 0 | 0 | 0 | 0 | 0 |
| ENSDARG000 | 20;20;20;20 | 3145497:3151 | 3146542:3152 | -;-;-;- | 3381 | NA | 920 | 384 | 372 | 277 | 647 | 685 |
| ENSDARG000 | 20:20:20:20:2 | 33809434:338 | 33811758:338 | -:-:-:-:-:-:-:-:-: | 4367 | atf6 | 1484 | 1542 | 3762 | 2639 | 1950 | 1470 |
| ENSDARG000 | 20;20 | 10484015:104 | 10484498:104 | -;- | 892 | glrx5 | 702 | 762 | 811 | 532 | 713 | 494 |
| ENSDARG000 | 20:20:20:20:2 | 7584211:7584 | 7584394:7584 | +:+:+:+:+:+:+: | 1019 | bloc1s2 | 350 | 720 | 776 | 333 | 437 | 497 |
| ENSDARG000 | 20:20:20:20:2 | 43491526:434 | 43491632:434 | -:-:-:-:-:-:-:-:-: | 1511 | pimr132 | 0 | 0 | 0 | 0 | 0 | 0 |
| ENSDARG000 | 20;20;20 | 28935413:289 | 28935930:289 | +;+;+ | 1425 | NA | 172 | 160 | 177 | 96 | 222 | 138 |
| ENSDARG000 | 20:20:20:20:2 | 1203238:1203 | 1203303:1203 | -:-:-:-:-:-:-:-:-: | 3239 | ankrd6b | 37 | 32 | 88 | 80 | 56 | 81 |
| ENSDARG000 | 20:20:20:20:2 | 43527890:435 | 43528101:435 | -;-;-;-;-;-;- | 2166 | NA | 0 | 0 | 2 | 0 | 0 | 0 |
| ENSDARG000 | 20;20 | 10489474:104 | 10489568:104 | +;+ | 208 | NA | 0 | 0 | 0 | 0 | 0 | 0 |
| ENSDARG000 | 20:20:20:20:2 | 14941264:149 | 14941334:149 | +:+:+:+:+:+:+: | 752 | itpa | 116 | 188 | 148 | 70 | 100 | 102 |
| ENSDARG000 | 20;20;20 | 28946709:289 | 28947340:289 | -;-;- | 1356 | NA | 0 | 0 | 0 | 2 | 0 | 0 |
| ENSDARG000 | 20:20:20:20:2 | 43567760:435 | 43567831:435 | -;-;-;-;-;-;-;- | 1649 | NA | 0 | 0 | 0 | 0 | 0 | 0 |
| ENSDARG000 | 20:20:20:20:2 | 43597003:435 | 43597163:435 | -;-;-;-;-;- | 2062 | NA | 0 | 0 | 0 | 0 | 0 | 0 |
| ENSDARG000 | 20:20:20:20:2 | 3162423:3162 | 3162720:3162 | -;-;-;-;-;- | 827 | NA | 84 | 37 | 60 | 68 | 77 | 44 |
| ENSDARG000 | 20:20:20:20:2 | 19066596:190 | 19066661:190 | +:+:+:+:+:+:+: | 5357 | sox7 | 78 | 9 | 42 | 24 | 38 | 34 |
| ENSDARG000 | 20:20:20:20:2 | 14968031:149 | 14968201:149 | +:+:+:+:+:+:+: | 3321 | vamp4 | 265 | 234 | 224 | 170 | 178 | 242 |
| ENSDARG000 | 20:20:20:20:2 | 43602810:436 | 43602949:436 | +:+:+:+:+:+:+: | 5672 | NA | 0 | 4 | 0 | 0 | 0 | 0 |
| ENSDARG000 | 20:20:20:20:2 | 1240461:1244 | 1244663:1244 | -:-:-:-:-:-:-:-:-: | 7634 | lats1 | 454 | 322 | 543 | 266 | 512 | 544 |
| ENSDARG000 | 20;20;20;20 | 25929440:259 | 25930047:259 | -;-;-;- | 901 | churc1 | 409 | 423 | 509 | 294 | 419 | 421 |
| ENSDARG000 | 20;20;20 | 28960340:289 | 28960615:289 | +;+;+ | 1782 | gpr176 | 0 | 0 | 0 | 0 | 0 | 0 |
| ENSDARG000 | 20:20:20:20:2 | 3218893:3223 | 3219763:3223 | -:-:-:-:-:-:-:-:-: | 2436 | spint1b | 100 | 16 | 4 | 2 | 26 | 38 |
| ENSDARG000 | 20:20:20:20:2 | 21638397:216 | 21639719:216 | -:-:-:-:-:-:-:-:-: | 28487 | syne2a | 108 | 24 | 64 | 78 | 94 | 26 |
| ENSDARG000 | 20:20:20:20:2 | 25936370:259 | 25936771:259 | +:+:+:+:+:+:+: | 8432 | NA | 13 | 35 | 91 | 19 | 88 | 33 |
| ENSDARG000 | 20:20:20:20:2 | 25955118:259 | 25955886:259 | -:-:-:-:-:-:-:-:-: | 2994 | CAPN3 | 0 | 0 | 4 | 0 | 2 | 0 |
| ENSDARG000 | 20:20:20:20:2 | 190028:19002 | 190189:19018 | +;+;+;+;+;+ | 2615 | gjb7 | 7 | 2 | 4 | 2 | 4 | 0 |
| ENSDARG000 | 20:20:20:20:2 | 10498704:104 | 10499126:104 | +:+:+:+:+:+:+: | 3350 | clmnb | 48 | 6 | 0 | 6 | 0 | 18 |
| ENSDARG000 | 20;20;20;20 | 19119800:191 | 19119960:191 | +;+;+;+ | 12281 | NA | 8 | 6 | 0 | 0 | 2 | 0 |
| ENSDARG000 | 20:20:20:20:2 | 29036328:290 | 29039238:290 | -:-:-:-:-:-:-:-:-: | 7446 | THBS1 | 280 | 26 | 50 | 32 | 80 | 84 |
| ENSDARG000 | 20:20:20:20:2 | 30937934:309 | 30938075:309 | +:+:+:+:+:+:+: | 1216 | sod2 | 510 | 526 | 720 | 508 | 739 | 466 |
| ENSDARG000 | 20:20:20:20:2 | 24817118:248 | 24821193:248 | -:-:-:-:-:-:-:-:-: | 7150 | epha7 | 35 | 22 | 20 | 4 | 12 | 2 |
| ENSDARG000 | 20:20:20:20:2 | 34868933:348 | 34869002:348 | +:+:+:+:+:+:+: | 3813 | ankef1a | 4 | 4 | 6 | 4 | 16 | 4 |
| ENSDARG000 | 20:20:20:20:2 | 30931414:309 | 30932617:309 | -:-:-:-:-:-:-:-:-: | 2578 | wtap | 1065 | 1147 | 1524 | 1049 | 1698 | 1165 |
| ENSDARG000 | 20;20 | 19170062:191 | 19170133:191 | -;- | 324 | NA | 0 | 0 | 0 | 0 | 0 | 0 |
| ENSDARG000 | 20:20:20:20:2 | 43648369:436 | 43648580:436 | +:+:+:+:+:+:+: | 3871 | parp1 | 1159 | 1338 | 2242 | 824 | 2248 | 1432 |
| ENSDARG000 | 20:20:20:20:2 | 27020201:270 | 27020449:270 | +:+:+:+:+:+:+: | 1740 | chga | 0 | 2 | 0 | 0 | 0 | 12 |
| ENSDARG000 | 20:20:20:20:2 | 34913069:349 | 34913189:349 | +:+:+:+:+:+:+: | 2780 | snap25a | 30 | 36 | 34 | 0 | 40 | 42 |
| ENSDARG000 | 20:20:20:20:2 | 10538025:105 | 10538053:105 | +:+:+:+:+:+:+: | 1658 | serpina1l | 2 | 0 | 135 | 4 | 8 | 0 |
| ENSDARG000 | 20;20 | 43658361:436 | 43659953:436 | -;- | 2117 | NA | 1 | 0 | 1 | 0 | 2 | 4 |
| ENSDARG000 | 20:20:20:20:2 | 3238879:3242 | 3239552:3242 | -:-:-:-:-:-:-:-:-: | 3427 | rps6kc1 | 578 | 525 | 552 | 421 | 582 | 514 |
| ENSDARG000 | 20:20:20:20:2 | 1272526:1272 | 1272634:1272 | +:+:+:+:+:+:+: | 1369 | hsd3b2 | 0 | 0 | 0 | 0 | 0 | 0 |
| ENSDARG000 | 20:20:20:20:2 | 33922339:339 | 33922574:339 | +:+:+:+:+:+:+: | 1496 | lmx1a | 0 | 0 | 0 | 0 | 0 | 0 |
| ENSDARG000 | 20:20:20:20:2 | 19175031:191 | 19175092:191 | +:+:+:+:+:+:+: | 4580 | NA | 4 | 10 | 8 | 0 | 36 | 40 |
| ENSDARG000 | 20;20 | 7612282:7629 | 7612640:7629 | -;- | 449 | NA | 0 | 0 | 0 | 0 | 0 | 2 |
| ENSDARG000 | 20:20:20:20:2 | 10544100:105 | 10544128:105 | +:+:+:+:+:+:+: | 1651 | serpina1 | 0 | 0 | 76 | 0 | 2 | 0 |
| ENSDARG000 | 20:20:20:20:2 | 1278184:1279 | 1279298:1280 | +;+;+;+;+;+ | 1883 | lrp11 | 0 | 1 | 0 | 0 | 10 | 0 |
| ENSDARG000 | 20 | 7799615 | 7801342 | + | 1728 | NA | 16 | 8 | 0 | 16 | 50 | 0 |
| ENSDARG000 | 20;20 | 34985123:349 | 34985413:349 | +;+ | 584 | NA | 0 | 0 | 0 | 0 | 0 | 6 |
| ENSDARG000 | 20:20:20:20:2 | 34967410:349 | 34967677:349 | -:-:-:-:-:-:-:-:-: | 1662 | sccpdhb | 426 | 700 | 689 | 693 | 863 | 841 |
| ENSDARG000 | 20:20:20:20:2 | 27049523:270 | 27050513:270 | -:-:-:-:-:-:-:-:-: | 2231 | itpk1a | 78 | 110 | 185 | 32 | 97 | 28 |
| ENSDARG000 | 20:20:20:20:2 | 196541:20425 | 203898:20439 | -;-;-;-;-;-;-;- | 8412 | znf292b | 1752 | 1528 | 3431 | 1758 | 2280 | 1334 |
| ENSDARG000 | 20:20:20:20:2 | 7964356:7965 | 7967352:7967 | -:-:-:-:-:-:-:-:-: | 4628 | PLPP3 | 335 | 457 | 645 | 74 | 628 | 228 |
| ENSDARG000 | 20:20:20:20:2 | 3277620:3277 | 3277774:3277 | +:+:+:+:+:+:+: | 1647 | ndufaf7 | 87 | 192 | 210 | 49 | 163 | 137 |
| ENSDARG000 | 20:20:20:20:2 | 10723292:107 | 10723324:107 | +:+:+:+:+:+:+: | 3082 | si:ch211-155i | 0 | 0 | 0 | 4 | 0 | 0 |
| ENSDARG000 | 20;20 | 27087539:270 | 27087642:270 | +;+ | 1949 | tmem251 | 714 | 600 | 728 | 568 | 573 | 748 |
| ENSDARG000 | 20;20 | 27090908:270 | 27091310:270 | -;- | 611 | si:dkeyp-55f1 | 84 | 174 | 265 | 190 | 250 | 235 |
| ENSDARG000 | 20:20:20:20:2 | 8081709:8085 | 8082282:8085 | -:-:-:-:-:-:-:-:-: | 1369 | NA | 6 | 0 | 0 | 0 | 0 | 0 |
| ENSDARG000 | 20:20:20:20:2 | 3290231:3290 | 3290600:3290 | -:-:-:-:-:-:-:-:-: | 3366 | cebpz | 660 | 1058 | 1409 | 469 | 1031 | 791 |
| ENSDARG000 | 20;20 | 3290188:3290 | 3290600:3290 | -;- | 495 | NA | 0 | 2 | 2 | 0 | 0 | 0 |
| ENSDARG000 | 20:20:20:20:2 | 1293988:1296 | 1296257:1296 | -:-:-:-:-:-:-:-:-: | 3179 | pcmt | 552 | 629 | 710 | 633 | 567 | 660 |
| ENSDARG000 | 20:20:20:20:2 | 34991620:349 | 34991969:349 | -:-:-:-:-:-:-:-:-: | 1574 | cnstb | 132 | 46 | 50 | 44 | 30 | 70 |
| ENSDARG000 | 20;20 | 11098239:111 | 11106093:111 | -;- | 7961 | flrt2 | 48 | 2 | 18 | 14 | 4 | 2 |
| ENSDARG000 | 20:20:20:20:2 | 27093042:270 | 27093407:270 | +:+:+:+:+:+:+: | 2123 | ubr7 | 583 | 929 | 1202 | 509 | 1041 | 836 |
| ENSDARG000 | 20:20:20:20:2 | 8101070:8101 | 8101141:8101 | -;-;-;-;-;-;- | 3071 | NA | 4 | 0 | 0 | 0 | 0 | 0 |
| ENSDARG000 | 20:20:20:20:2 | 19212962:192 | 19213641:192 | +:+:+:+:+:+:+: | 1457 | FNDC4 | 0 | 0 | 0 | 0 | 0 | 0 |
| ENSDARG000 | 20:20:20:20:2 | 35013434:350 | 35014345:350 | -:-:-:-:-:-:-:-:-: | 6360 | NA | 1 | 18 | 25 | 14 | 2 | 11 |
| ENSDARG000 | 20:20:20:20:2 | 19276932:192 | 19278283:192 | -:-:-:-:-:-:-:-:-: | 2525 | NA | 0 | 2 | 2 | 0 | 12 | 0 |
| ENSDARG000 | 20:20:20:20:2 | 25225112:252 | 25225547:252 | +:+:+:+:+:+:+: | 4399 | moxd1 | 1389 | 402 | 437 | 361 | 737 | 708 |
| ENSDARG000 | 20;20 | 19415093:194 | 19415590:194 | +;+ | 573 | NA | 0 | 0 | 0 | 0 | 0 | 0 |
| ENSDARG000 | 20;20;20 | 19382427:193 | 19382803:193 | -;-;- | 1068 | NA | 0 | 0 | 0 | 0 | 0 | 0 |

| | | | | | | | | | | | |
|---|---|---|---|---|---|---|---|---|---|---|---|
| ENSDARG000 | 20;20;20;20;2 | 19373428:193 | 19375071:193 | -;-;-;-;-;-;-;-; | 3867 | NA | 48 | 119 | 92 | 50 | 41 | 76 |
| ENSDARG000 | 20;20;20;20;2 | 29209615:292 | 29209761:292 | +;+;+;+;+;+;+;+; | 1962 | katnbl1 | 260 | 326 | 387 | 247 | 293 | 282 |
| ENSDARG000 | 20;20;20;20;2 | 31002188:310 | 31003265:310 | -;-;-;-;-;-;-;-; | 6152 | FNDC1 | 62 | 4 | 8 | 14 | 100 | 18 |
| ENSDARG000 | 20;20;20;20;2 | 33991801:339 | 33992371:339 | +;+;+;+;+;+;+;+; | 1377 | zp3a.1 | 0 | 0 | 0 | 0 | 0 | 0 |
| ENSDARG000 | 20;20;20;20;2 | 33969430:339 | 33971267:339 | -;-;-;-;-;-;-;-; | 4616 | sele | 85 | 12 | 303 | 50 | 45 | 148 |
| ENSDARG000 | 20;20 | 11629881:116 | 11629943:116 | -;- | 185 | NA | 0 | 0 | 0 | 0 | 0 | 0 |
| ENSDARG000 | 20;20;20;20;2 | 35058634:350 | 35059220:350 | +;+;+;+;+;+;+;+; | 4447 | hnrnpub | 3225 | 4584 | 5154 | 2237 | 5324 | 3062 |
| ENSDARG000 | 20;20;20;20;2 | 33948642:339 | 33948777:339 | -;-;-;-;-;-;-;-; | 2914 | selp | 2 | 21 | 8 | 19 | 55 | 50 |
| ENSDARG000 | 20;20;20;20;2 | 218886:22188 | 219292:22198 | +;+;+;+;+;+;+;+; | 6221 | lama4 | 18851 | 11110 | 15116 | 15545 | 20589 | 11230 |
| ENSDARG000 | 20;20;20;20;2 | 33994580:339 | 33994845:339 | +;+;+;+;+;+;+;+; | 3187 | si:dkev-97o5.: | 171 | 265 | 308 | 98 | 360 | 221 |
| ENSDARG000 | 20;20;20 | 43715157:437 | 43717425:437 | -;-;- | 2709 | mixl1 | 4 | 4 | 5 | 4 | 10 | 0 |
| ENSDARG000 | 20;20;20;20;2 | 43691208:436 | 43691276:436 | +;+;+;+;+;+;+;+; | 3103 | lin9 | 162 | 245 | 190 | 110 | 120 | 126 |
| ENSDARG000 | 20;20;20;20;2 | 8122693:8122 | 8123388:8123 | -;-;-;-;-;-;-;-; | 2981 | dab1a | 6 | 0 | 0 | 0 | 0 | 0 |
| ENSDARG000 | 20;20;20;20;2 | 11731039:117 | 11731116:117 | +;+;+;+;+;+;+;+; | 1841 | NA | 2 | 2 | 4 | 0 | 3 | 0 |
| ENSDARG000 | 20;20;20;20 | 33962828:339 | 33963007:339 | -;-;-;- | 834 | NA | 0 | 0 | 0 | 0 | 0 | 0 |
| ENSDARG000 | 20;20;20;20;2 | 33987465:339 | 33988084:339 | +;+;+;+;+;+;+;+; | 1429 | zp3a.2 | 0 | 3 | 0 | 0 | 2 | 10 |
| ENSDARG000 | 20;20;20;20;2 | 3312859:3316 | 3313315:3317 | -;-;-;-;- | 1776 | marcksa | 98 | 305 | 220 | 124 | 152 | 193 |
| ENSDARG000 | 20;20;20;20;2 | 25340814:253 | 25341059:253 | +;+;+;+;+ | 2119 | ccn2a | 510 | 88 | 128 | 120 | 220 | 152 |
| ENSDARG000 | 20 | 33961587 | 33962059 | - | 473 | NA | 0 | 0 | 0 | 0 | 0 | 0 |
| ENSDARG000 | 20;20;20;20;2 | 19423823:194 | 19424050:194 | +;+;+;+;+ | 3345 | znf513a | 247 | 213 | 383 | 215 | 361 | 314 |
| ENSDARG000 | 20;20;20;20;2 | 34012571:340 | 34013105:340 | -;-;-;-;-;-;-;-; | 3024 | scyl3 | 402 | 257 | 425 | 136 | 407 | 430 |
| ENSDARG000 | 20;20;20;20;2 | 27114874:271 | 27115725:271 | -;-;-;-;-;-;-;-; | 3905 | btbd7 | 77 | 108 | 192 | 173 | 132 | 44 |
| ENSDARG000 | 20;20;20;20;2 | 34039281:340 | 34039522:340 | -;-;-;-;-;-;-;-; | 7418 | tprb | 1867 | 1466 | 2570 | 883 | 2325 | 1122 |
| ENSDARG000 | 20;20;20;20;2 | 11800063:118 | 11800120:118 | +;+;+;+;+ | 725 | NA | 0 | 0 | 0 | 2 | 1 | 2 |
| ENSDARG000 | 20;20;20;20 | 3339248:3340 | 3339764:3341 | +;+;+;+ | 2517 | mfsd4b | 258 | 126 | 193 | 178 | 274 | 202 |
| ENSDARG000 | 20;20 | 12203762:122 | 12203874:122 | +;+ | 1849 | NA | 18 | 113 | 136 | 97 | 362 | 125 |
| ENSDARG000 | 20;20;20 | 31072892:310 | 31073128:310 | -;-;- | 614 | NA | 0 | 0 | 0 | 0 | 0 | 0 |
| ENSDARG000 | 20;20;20 | 33981946:339 | 33982141:339 | +;+;+ | 749 | si:dkev-51e6.: | 143 | 76 | 171 | 168 | 154 | 243 |
| ENSDARG000 | 20;20 | 27194833:271 | 27195079:271 | +;+ | 973 | NA | 0 | 4 | 13 | 0 | 1 | 0 |
| ENSDARG000 | 20;20;20;20;2 | 34029820:340 | 34030112:340 | +;+;+;+;+;+;+;+; | 2867 | prg4b | 27 | 0 | 12 | 14 | 9 | 4 |
| ENSDARG000 | 20;20;20;20 | 35073981:350 | 35074573:350 | -;-;-;- | 1381 | lft1 | 0 | 0 | 0 | 0 | 0 | 0 |
| ENSDARG000 | 20;20;20;20;2 | 33959570:339 | 33959798:339 | +;+;+;+;+ | 789 | NA | 0 | 0 | 0 | 0 | 0 | 0 |
| ENSDARG000 | 20;20;20;20;2 | 3346242:3346 | 3347084:3347 | -;-;-;-;-;-;-;-; | 10451 | REV3L | 908 | 790 | 1556 | 603 | 1511 | 1470 |
| ENSDARG000 | 20;20;20;20;2 | 43737022:437 | 43737135:437 | -;-;-;-;-;-;-;-; | 2992 | zpax1 | 8 | 4 | 8 | 20 | 5 | 8 |
| ENSDARG000 | 20;20 | 12239930:122 | 12240080:122 | -;- | 262 | NA | 0 | 0 | 0 | 0 | 0 | 0 |
| ENSDARG000 | 20;20;20;20;2 | 19512622:195 | 19517347:195 | -;-;-;-;-;-;-;-; | 6585 | eif2b4 | 365 | 469 | 1039 | 556 | 530 | 421 |
| ENSDARG000 | 20;20;20;20;2 | 1316495:1316 | 1316920:1316 | +;+;+;+;+;+;+;+; | 1632 | nup43 | 159 | 332 | 519 | 134 | 303 | 238 |
| ENSDARG000 | 20;20;20;20;2 | 27222849:272 | 27223595:272 | -;-;-;-;-;-;-;-; | 1443 | NA | 0 | 65 | 37 | 54 | 58 | 17 |
| ENSDARG000 | 20;20;20;20;2 | 12605718:126 | 12605838:126 | -;-;-;-;- | 835 | NA | 6 | 0 | 0 | 0 | 0 | 0 |
| ENSDARG000 | 20;20;20;20;2 | 12598637:125 | 12598705:125 | +;+;+;+;+;+ | 961 | NA | 0 | 0 | 0 | 0 | 0 | 0 |
| ENSDARG000 | 20;20 | 12566101:126 | 12566354:126 | -;- | 282 | NA | 12 | 4 | 0 | 4 | 0 | 0 |
| ENSDARG000 | 20;20;20;20;2 | 19436187:194 | 19438484:194 | -;-;-;-;-;-;-;-; | 4379 | snx17 | 709 | 414 | 491 | 308 | 596 | 454 |
| ENSDARG000 | 20;20;20;20;2 | 35085224:350 | 35086770:350 | +;+;+;+;+;+;+;+; | 9910 | cdc42bpab | 43 | 4 | 10 | 0 | 32 | 8 |
| ENSDARG000 | 20;20;20;20;2 | 31076488:310 | 31076616:310 | +;+;+;+;+;+;+;+; | 7224 | otofa | 24 | 0 | 0 | 10 | 6 | 0 |
| ENSDARG000 | 20;20;20 | 43746523:437 | 43746974:437 | -;-;- | 804 | ttc32 | 90 | 104 | 208 | 113 | 114 | 85 |
| ENSDARG000 | 20;20;20;20;2 | 19512727:195 | 19512866:195 | +;+;+;+;+;+;+;+; | 1757 | atraid | 0 | 0 | 0 | 0 | 0 | 0 |
| ENSDARG000 | 20;20;20;20;2 | 43760059:437 | 43761088:437 | -;-;-;-;-;-;-;-; | 2624 | matn3a | 4 | 11 | 21 | 11 | 166 | 2 |
| ENSDARG000 | 20;20;20;20;2 | 21815613:218 | 21816281:218 | -;-;-;-;-;-;-;-; | 4100 | daam1b | 402 | 204 | 381 | 38 | 188 | 181 |
| ENSDARG000 | 20;20;20;20;2 | 25350435:253 | 25350532:253 | +;+;+;+;+;+;+;+; | 2113 | fam228a | 0 | 0 | 2 | 0 | 2 | 0 |
| ENSDARG000 | 20;20;20;20;2 | 43775793:437 | 43776918:437 | -;-;-;-;-;-;-;- | 2081 | laptm4a | 3820 | 4033 | 4207 | 4557 | 4400 | 4554 |
| ENSDARG000 | 20;20;20;20;2 | 1329509:1329 | 1329802:1329 | +;+;+;+;+;+;+;+; | 3969 | tab2 | 774 | 611 | 860 | 544 | 829 | 726 |
| ENSDARG000 | 20;20;20;20;2 | 35208020:352 | 35208108:352 | +;+;+;+;+;+;+;+; | 1915 | fbxo16 | 0 | 8 | 8 | 0 | 0 | 0 |
| ENSDARG000 | 20;20;20;20 | 12702923:127 | 12703057:127 | +;+;+;+ | 699 | FAM177A1 | 94 | 84 | 155 | 159 | 118 | 155 |
| ENSDARG000 | 20;20;20;20;2 | 27240388:272 | 27240934:272 | +;+;+;+;+;+ | 4459 | NA | 36 | 722 | 483 | 408 | 480 | 303 |
| ENSDARG000 | 20;20;20;20;2 | 43792866:437 | 43794854:437 | -;-;-;-;-;-;-;-; | 4632 | mark3b | 1700 | 1059 | 1691 | 1449 | 2156 | 1532 |
| ENSDARG000 | 20;20;20;20 | 12907233:129 | 12907342:129 | +;+;+;+ | 636 | NA | 0 | 0 | 0 | 0 | 0 | 0 |
| ENSDARG000 | 20;20;20 | 8794502:8833 | 8794874:8833 | -;-;- | 666 | NA | 0 | 0 | 0 | 0 | 0 | 0 |
| ENSDARG000 | 20;20;20;20;2 | 26002534:260 | 26003120:260 | +;+;+;+;+;+;+;+; | 3732 | nrbp1 | 904 | 1183 | 1188 | 774 | 1396 | 1010 |
| ENSDARG000 | 20;20;20;20;2 | 35208855:352 | 35213252:352 | -;-;-;-;-;-;- | 6668 | fzd3a | 32 | 8 | 2 | 0 | 14 | 10 |
| ENSDARG000 | 20;20 | 8993860:8995 | 8994094:8995 | +;+ | 479 | NA | 0 | 0 | 0 | 0 | 0 | 0 |
| ENSDARG000 | 20;20;20;20;2 | 43925266:439 | 43925524:439 | +;+;+;+;+;+;+;+; | 2504 | clic5b | 0 | 0 | 2 | 6 | 8 | 0 |
| ENSDARG000 | 20;20;20;20 | 12936096:129 | 12936289:129 | -;-;-;- | 361 | NA | 0 | 0 | 0 | 0 | 0 | 0 |
| ENSDARG000 | 20;20;20;20;2 | 27316678:273 | 27317212:273 | -;-;-;-;-;-;-;-; | 2655 | NA | 0 | 0 | 0 | 0 | 0 | 0 |
| ENSDARG000 | 20;20;20;20;2 | 27371394:273 | 27372942:273 | -;-;-;-;- | 2750 | serpina10a | 3157 | 2112 | 2799 | 4061 | 3584 | 4195 |
| ENSDARG000 | 20;20;20;20;2 | 27298783:273 | 27298926:273 | +;+;+;+;+;+;+;+; | 3698 | PPP4R4 | 12 | 18 | 10 | 15 | 2 | 14 |
| ENSDARG000 | 20;20;20;20;2 | 27297152:272 | 27297389:272 | +;+;+;+;+;+ | 1677 | NA | 0 | 0 | 0 | 0 | 0 | 0 |
| ENSDARG000 | 20;20;20;20;2 | 27295701:272 | 27298895:272 | -;-;-;-;-;-;-;-; | 5451 | NA | 0 | 0 | 0 | 0 | 0 | 0 |
| ENSDARG000 | 20;20;20;20;2 | 43994848:439 | 43994928:439 | -;-;-;-;-;-;-;-; | 2653 | runx2b | 596 | 6 | 11 | 4 | 20 | 6 |
| ENSDARG000 | 20;20;20;20 | 44311448:443 | 44311821:443 | +;+;+;+ | 1841 | opn8b | 28 | 0 | 24 | 6 | 0 | 12 |
| ENSDARG000 | 20;20;20;20;2 | 9093366:9096 | 9097698:9097 | -;-;-;-;-;-;-;-; | 6758 | mysm1 | 286 | 250 | 444 | 176 | 456 | 221 |
| ENSDARG000 | 20;20;20;20;2 | 44372745:443 | 44373383:443 | -;-;-;-;-;-;-;-; | 2880 | mmut | 965 | 719 | 905 | 601 | 797 | 1219 |
| ENSDARG000 | 20;20;20;20 | 21999156:220 | 21999576:220 | -;-;-;- | 901 | NA | 0 | 0 | 0 | 0 | 0 | 0 |
| ENSDARG000 | 20;20;20;20;2 | 26026518:260 | 26027814:260 | -;-;-;-;-;-;-;-; | 3195 | NA | 14 | 0 | 2 | 4 | 2 | 2 |
| ENSDARG000 | 20;20;20;20;2 | 9058679:9075 | 9059500:9075 | -;-;-;-;-;-;- | 2489 | OMA1 | 28 | 47 | 80 | 38 | 57 | 34 |
| ENSDARG000 | 20;20;20;20;2 | 1354695:1355 | 1355799:1355 | -;-;-;-;-;-;-;-; | 2812 | scara5 | 43 | 0 | 6 | 0 | 38 | 4 |
| ENSDARG000 | 20;20;20;20;2 | 29217835:292 | 29218204:292 | +;+;+;+;+ | 1854 | emc7b | 1860 | 1794 | 2252 | 1603 | 2233 | 1980 |
| ENSDARG000 | 20;20;20;20;2 | 22045089:220 | 22045291:220 | +;+;+;+;+;+;+;+; | 1438 | nmu | 0 | 0 | 8 | 0 | 0 | 0 |
| ENSDARG000 | 20;20;20;20;2 | 1385674:1385 | 1385833:1386 | +;+;+;+;+;+;+;+; | 1503 | mmachc | 22 | 72 | 90 | 8 | 34 | 38 |
| ENSDARG000 | 20;20 | 12968914:129 | 12969591:129 | +;+ | 724 | NA | 0 | 0 | 0 | 0 | 0 | 0 |
| ENSDARG000 | 20;20 | 13001068:130 | 13001260:130 | +;+ | 910 | NA | 0 | 0 | 0 | 0 | 0 | 0 |
| ENSDARG000 | 20;20;20;20;2 | 35255403:352 | 35255594:352 | +;+;+;+;+;+;+;+; | 2972 | fam49a | 1191 | 1195 | 1267 | 1050 | 1399 | 1048 |
| ENSDARG000 | 20;20;20;20 | 44464345:444 | 44465092:444 | -;-;-;- | 1046 | fkbp1b | 425 | 722 | 900 | 435 | 678 | 861 |
| ENSDARG000 | 20;20;20;20;2 | 44507667:445 | 44507697:445 | -;-;-;-;-;-;-;-; | 2148 | mfsd2b | 1 | 2 | 10 | 0 | 7 | 0 |
| ENSDARG000 | 20;20;20;20;2 | 15552657:155 | 15554606:155 | +;+;+;+;+ | 1950 | jun | 2812 | 781 | 1657 | 1425 | 1124 | 1344 |
| ENSDARG000 | 20;20;20;20 | 44498056:445 | 44498123:445 | +;+;+;+ | 2811 | wdcp | 34 | 58 | 72 | 46 | 60 | 60 |
| ENSDARG000 | 20;20;20 | 44516163:445 | 44516246:445 | +;+;+ | 512 | NA | 0 | 0 | 0 | 0 | 4 | 0 |
| ENSDARG000 | 20;20;20;20;2 | 25359064:253 | 25359265:253 | -;-;-;-;-;-;-;-; | 5735 | itsn2a | 20 | 14 | 71 | 16 | 67 | 14 |
| ENSDARG000 | 20;20;20;20;2 | 262995:26476 | 263245:26484 | +;+;+;+;+;+;+;+; | 2388 | TUBE1 | 0 | 0 | 0 | 0 | 0 | 0 |
| ENSDARG000 | 20;20;20;20;2 | 305035:30834 | 305485:30844 | +;+;+;+;+;+;+ | 1622 | si:dkev-119m! | 8 | 6 | 0 | 4 | 2 | 0 |
| ENSDARG000 | 20;20 | 15560141:155 | 15561523:155 | +;+ | 2235 | caiap | 0 | 13 | 0 | 0 | 0 | 0 |
| ENSDARG000 | 20;20;20;20;2 | 276753:27770 | 277539:27790 | -;-;-;-;- | 1561 | ccn6 | 0 | 15 | 23 | 12 | 6 | 1 |
| ENSDARG000 | 20;20;20;20;2 | 263056:26305 | 263245:26324 | +;+;+;+;+;+;+;+; | 2009 | tube1 | 0 | 0 | 0 | 1 | 0 | 0 |
| ENSDARG000 | 20;20 | 29228337:292 | 29229854:292 | -;- | 1518 | chrm5b | 6 | 20 | 2 | 6 | 0 | 0 |
| ENSDARG000 | 20;20;20;20;2 | 44566939:445 | 44567168:445 | -;-;-;-;-;-;-;-; | 915 | UBXN2A | 88 | 34 | 119 | 47 | 133 | 26 |
| ENSDARG000 | 20;20;20;20;2 | 1398564:1400 | 1399037:1401 | +;+;+;+;+;+ | 1383 | C20H6orf58 | 0 | 0 | 7 | 0 | 0 | 0 |
| ENSDARG000 | 20;20;20;20;2 | 15565926:155 | 15566051:155 | +;+;+;+;+ | 923 | si:dkev-86e18 | 96 | 105 | 122 | 52 | 74 | 151 |
| ENSDARG000 | 20;20 | 15725659:157 | 15725764:157 | -;- | 450 | NA | 0 | 0 | 0 | 0 | 0 | 0 |
| ENSDARG000 | 20;20;20;20;2 | 13204066:132 | 13204846:132 | -;-;-;-;-;-;-;-; | 2715 | NA | 0 | 0 | 0 | 0 | 0 | 0 |
| ENSDARG000 | 20;20;20;20;2 | 13175801:132 | 13175957:132 | +;+;+;+;+;+;+;+; | 1609 | NA | 0 | 0 | 0 | 0 | 0 | 0 |
| ENSDARG000 | 20;20;20;20;2 | 13175379:132 | 13175957:132 | +;+;+;+;+;+;+;+; | 4364 | PPP2R5A | 34 | 40 | 43 | 14 | 131 | 48 |
| ENSDARG000 | 20;20;20;20;2 | 44582318:445 | 44582606:445 | +;+;+;+;+;+;+;+; | 4602 | ATAD2B | 553 | 459 | 590 | 155 | 709 | 410 |
| ENSDARG000 | 20;20;20;20;2 | 13141408:131 | 13141501:131 | +;+;+;+;+;+;+;+; | 2368 | dtl | 34 | 183 | 202 | 52 | 108 | 79 |
| ENSDARG000 | 20;20;20;20;2 | 29231556:292 | 29231798:292 | +;+;+;+;+;+;+;+; | 2401 | aven | 202 | 211 | 258 | 202 | 226 | 166 |
| ENSDARG000 | 20;20;20;20;2 | 13020489:130 | 13020966:130 | -;-;-;-;-;-;-;-; | 4195 | ints7 | 324 | 254 | 405 | 185 | 363 | 262 |
| ENSDARG000 | 20;20;20;20;2 | 29268444:292 | 29269878:292 | -;-;-;-;-;-;-;-; | 16422 | RYR3 | 75 | 34 | 20 | 24 | 37 | 27 |
| ENSDARG000 | 20;20;20;20;2 | 26045253:260 | 26045838:260 | -;-;-;-;-;-;-;-; | 957 | gtf2h5 | 403 | 152 | 202 | 186 | 192 | 218 |
| ENSDARG000 | 20;20;20;20;2 | 1412193:1414 | 1412666:1414 | +;+;+;+;+;+ | 1369 | C20H6orf58 | 0 | 0 | 61 | 0 | 0 | 0 |
| ENSDARG000 | 20;20;20;20;2 | 15857041:158 | 15857901:158 | -;-;-;-;-;-;-;-; | 2959 | fam20b | 720 | 659 | 741 | 639 | 1154 | 717 |
| ENSDARG000 | 20;20;20;20;2 | 45643823:456 | 45644307:456 | -;-;-;-;-;-;-;-; | 1612 | napbb | 60 | 124 | 114 | 16 | 28 | 60 |
| ENSDARG000 | 20;20;20 | 15891961:158 | 15892074:158 | +;+;+ | 736 | NA | 0 | 0 | 1 | 0 | 1 | 6 |
| ENSDARG000 | 20 | 35382438 | 35382617 | + | 180 | NA | 0 | 0 | 0 | 0 | 0 | 0 |
| ENSDARG000 | 20;20;20;20 | 35382482:353 | 35382617:353 | +;+;+;+ | 3234 | vsnl1a | 18 | 26 | 9 | 47 | 12 | 6 |
| ENSDARG000 | 20;20;20;20;2 | 1423192:1424 | 1423689:1424 | -;-;-;-;-;- | 2455 | themis | 6 | 8 | 8 | 2 | 23 | 6 |
| ENSDARG000 | 20;20 | 9124369:9124 | 9124613:9125 | +;+ | 1212 | saysd1 | 76 | 46 | 100 | 82 | 80 | 100 |
| ENSDARG000 | 20;20;20;20;2 | 45669176:456 | 45669380:456 | -;-;-;-;-;-;-;-; | 2616 | gpcpd1 | 746 | 1007 | 648 | 488 | 872 | 752 |
| ENSDARG000 | 20;20 | 26062519:260 | 26063584:260 | -;- | 1390 | myct1a | 6 | 6 | 0 | 2 | 2 | 12 |
| ENSDARG000 | 20;20;20 | 27384138:273 | 27385187:273 | -;-;- | 1957 | prph2lb | 18 | 2 | 30 | 3 | 18 | 4 |
| ENSDARG000 | 20;20;20;20;2 | 26048304:260 | 26049038:260 | -;-;-;-;-;-;-;-; | 2769 | serac1 | 181 | 109 | 219 | 185 | 121 | 223 |
| ENSDARG000 | 20;20;20;20;2 | 45741566:457 | 45741702:457 | +;+;+;+;+ | 2778 | chgb | 50 | 29 | 28 | 17 | 8 | 14 |
| ENSDARG000 | 20;20;20 | 26059710:260 | 26060399:260 | +;+;+ | 1828 | NA | 6 | 1 | 1 | 1 | 5 | 4 |
| ENSDARG000 | 20;20;20;20;2 | 45754323:457 | 45754620:457 | -;-;-;-;-;-;-;-; | 1715 | trmt6 | 140 | 227 | 260 | 83 | 123 | 131 |
| ENSDARG000 | 20;20;20;20;2 | 35461736:354 | 35462970:354 | -;-;-;-;-;-;-;-; | 2740 | pla2g7 | 2050 | 4031 | 1628 | 3571 | 1931 | 4065 |
| ENSDARG000 | 20;20 | 19543262:195 | 19543538:195 | -;- | 830 | NA | 0 | 0 | 0 | 0 | 0 | 0 |
| ENSDARG000 | 20;20;20;20;2 | 35433192:354 | 35433368:354 | +;+;+;+;+;+ | 1329 | sf3b6 | 668 | 701 | 729 | 628 | 684 | 640 |
| ENSDARG000 | 20;20;20;20;2 | 35438300:354 | 35438656:354 | +;+;+;+;+;+;+;+; | 7305 | tdrd6 | 20 | 50 | 6 | 54 | 82 | 16 |
| ENSDARG000 | 20 | 22068160 | 22070094 | - | 1935 | NA | 0 | 0 | 0 | 0 | 0 | 0 |
| ENSDARG000 | 20;20;20;20;2 | 22067337:220 | 22069036:220 | +;+;+;+;+;+;+;+; | 6535 | clocka | 87 | 92 | 122 | 68 | 162 | 168 |
| ENSDARG000 | 20;20;20;20;2 | 31217495:312 | 31217632:312 | +;+;+;+;+;+;+;+; | 2058 | pdia6 | 2733 | 1734 | 3676 | 1073 | 2062 | 1566 |
| ENSDARG000 | 20;20;20;20;2 | 19562383:195 | 19562857:195 | +;+;+;+;+ | 1150 | nme6 | 90 | 81 | 172 | 66 | 72 | 107 |
| ENSDARG000 | 20;20;20;20;2 | 31269778:312 | 31269832:312 | +;+;+;+;+;+;+;+; | 12857 | apobb.1 | 0 | 18 | 136 | 14 | 2 | 0 |
| ENSDARG000 | 20;20;20;20;2 | 45775935:457 | 45776652:457 | -;-;-;-;-;-;-;-; | 2618 | FERMT1 | 6 | 0 | 0 | 4 | 20 | 8 |
| ENSDARG000 | 20;20;20;20;2 | 31232923:312 | 31233892:312 | -;-;-;-;-;-;-;-; | 2167 | hpcal1 | 18 | 424 | 459 | 648 | 117 | 26 |
| ENSDARG000 | 20;20;20;20 | 27393668:273 | 27393987:273 | +;+;+;+ | 1154 | tmem179aa | 230 | 145 | 165 | 103 | 208 | 244 |
| ENSDARG000 | 20;20;20;20;2 | 31287356:312 | 31287648:312 | +;+;+;+;+;+;+;+; | 4408 | slc22a16 | 6 | 1 | 20 | 0 | 4 | 8 |
| ENSDARG000 | 20;20;20;20;2 | 9128256:9128 | 9128949:9128 | +;+;+;+;+;+;+;+; | 2155 | bpnt1 | 502 | 436 | 346 | 375 | 527 | 278 |
| ENSDARG000 | 20;20;20 | 27416520:274 | 27416656:274 | +;+;+ | 368 | NA | 0 | 0 | 0 | 0 | 0 | 0 |
| ENSDARG000 | 20;20;20;20;2 | 1441548:1442 | 1442687:1442 | -;-;-;-;-;-;-;-; | 4901 | PTPRK | 2102 | 1006 | 2335 | 1584 | 4019 | 1138 |
| ENSDARG000 | 20;20;20;20 | 19579835:195 | 19580731:195 | -;-;-;- | 1614 | NA | 4 | 307 | 10 | 4 | 8 | 1372 |
| ENSDARG000 | 20;20;20 | 45893173:458 | 45893522:458 | +;+;+ | 1961 | bmp2b | 16 | 6 | 24 | 2 | 36 | 2 |
| ENSDARG000 | 20;20;20;20;2 | 4712776:4716 | 4713999:4716 | -;-;-;-;-;-;-;-; | 4217 | papola | 3499 | 2616 | 3790 | 1966 | 4186 | 2854 |
| ENSDARG000 | 20;20;20;20;2 | 45907764:459 | 45908513:459 | -;-;-;-;-;-;-;-; | 1795 | NA | 0 | 0 | 0 | 0 | 0 | 0 |
| ENSDARG000 | 20;20 | 45985461:459 | 45985819:459 | +;+ | 1019 | NA | 0 | 0 | 0 | 0 | 0 | 0 |
| ENSDARG000 | 20;20;20;20 | 45989179:459 | 45989954:459 | +;+;+;+ | 1170 | NA | 22 | 4 | 0 | 14 | 6 | 8 |
| ENSDARG000 | 20;20;20 | 19779299:197 | 19779573:197 | +;+;+ | 1250 | NA | 3158 | 4327 | 1664 | 409 | 1740 | 2977 |
| ENSDARG000 | 20;20;20;20;2 | 19793620:198 | 19794188:198 | +;+;+;+;+;+;+;+; | 3735 | CHRNA2 | 8 | 406 | 2 | 0 | 2 | 282 |
| ENSDARG000 | 20;20;20;20;2 | 27426507:274 | 27427428:274 | +;+;+;+;+;+;+;+; | 10621 | KIF26A | 5248 | 1509 | 748 | 1489 | 1124 | 2220 |
| ENSDARG000 | 20;20;20;20;2 | 27427112:274 | 27427428:274 | +;+;+;+;+ | 958 | NA | 0 | 0 | 0 | 0 | 0 | 0 |
| ENSDARG000 | 20;20;20;20;2 | 15982482:159 | 15983028:159 | +;+;+;+;+ | 3683 | angptl1a | 100 | 50 | 140 | 14 | 175 | 338 |
| ENSDARG000 | 20;20;20;20;2 | 46039591:460 | 46039724:460 | +;+;+;+;+;+;+;+; | 2474 | si:dkev-7c18.: | 391 | 0 | 2 | 48 | 19 | 4 |
| ENSDARG000 | 20;20;20;20;2 | 15933028:159 | 15938303:159 | -;-;-;-;-;-;-;-; | 7284 | ralgps2 | 346 | 378 | 390 | 430 | 525 | 530 |
| ENSDARG000 | 20 | 46045735 | 46046758 | - | 1024 | taar12j | 0 | 0 | 0 | 0 | 0 | 0 |
| ENSDARG000 | 20 | 46052685 | 46053720 | - | 1036 | taar12h | 0 | 0 | 0 | 0 | 0 | 0 |
| ENSDARG000 | 20 | 46060635 | 46061650 | - | 1016 | taar12e | 0 | 0 | 0 | 0 | 0 | 0 |
| ENSDARG000 | 20 | 46068598 | 46069609 | - | 1012 | taar12g | 0 | 0 | 0 | 0 | 0 | 0 |
| ENSDARG000 | 20;20;20;20;2 | 25440514:254 | 25441126:254 | -;-;-;-;-;-;-;-; | 1253 | mrpl19 | 553 | 620 | 759 | 328 | 488 | 558 |
| ENSDARG000 | 20;20;20 | 320415:32101 | 320524:32122 | +;+;+ | 680 | NA | 10 | 5 | 0 | 0 | 8 | 2 |
| ENSDARG000 | 20 | 46073680 | 46074694 | - | 1015 | taar12f | 0 | 0 | 0 | 0 | 0 | 0 |
| ENSDARG000 | 20 | 46084808 | 46085840 | - | 1033 | taar12a | 0 | 0 | 0 | 0 | 0 | 0 |
| ENSDARG000 | 20 | 46100634 | 46101648 | - | 1015 | taar12b | 0 | 0 | 0 | 0 | 0 | 0 |
| ENSDARG000 | 20 | 46104102 | 46105119 | - | 1018 | taar12c | 0 | 0 | 0 | 0 | 0 | 0 |
| ENSDARG000 | 20 | 46113445 | 46114467 | - | 1023 | taar12i | 0 | 0 | 0 | 0 | 0 | 0 |
| ENSDARG000 | 20 | 22179916 | 22180508 | + | 593 | NA | 11 | 35 | 25 | 28 | 26 | 34 |

| | | | | | | | | | | | | |
|---|---|---|---|---|---|---|---|---|---|---|---|---|
| ENSDARG000 | 20 | 46127591 | 46128590 | - | 1000 | taar1b | 0 | 0 | 0 | 0 | 0 | 0 |
| ENSDARG000 | 20 | 46131812 | 46132911 | + | 1100 | taar11 | 0 | 0 | 0 | 0 | 0 | 0 |
| ENSDARG000 | 20 | 46137100 | 46138110 | - | 1011 | NA | 0 | 0 | 0 | 0 | 0 | 0 |
| ENSDARG000 | 20 | 46145718 | 46146729 | - | 1012 | NA | 0 | 0 | 0 | 0 | 0 | 0 |
| ENSDARG000 | 20 | 46154142 | 46155153 | - | 1012 | taar10c | 0 | 0 | 0 | 0 | 0 | 0 |
| ENSDARG000 | 20 | 46157976 | 46158987 | - | 1012 | taar10a | 0 | 0 | 0 | 0 | 0 | 0 |
| ENSDARG000 | 20:20:20:20:2 | 22186297:221 | 22187413:221 | -;-;-;-;-;- | 2163 | tmem165 | 1856 | 1598 | 1895 | 1158 | 2439 | 1441 |
| ENSDARG000 | 20:20:20:20:2 | 19827379:198 | 19828258:198 | -:-:-:-:-:-:-:-: | 3790 | ptk2bb | 40 | 8 | 0 | 6 | 39 | 10 |
| ENSDARG000 | 20:20:20:20:2 | 46213553:462 | 46213613:462 | +:+:+:+:+:+:+: | 2408 | STX7 | 34 | 2 | 50 | 4 | 40 | 28 |
| ENSDARG000 | 20 | 46183505 | 46184530 | + | 1026 | taar13b | 0 | 0 | 0 | 0 | 0 | 0 |
| ENSDARG000 | 20 | 46206998 | 46207984 | + | 987 | taar15 | 0 | 0 | 0 | 0 | 0 | 0 |
| ENSDARG000 | 20 | 46172649 | 46173674 | + | 1026 | taar13e | 0 | 0 | 0 | 0 | 0 | 0 |
| ENSDARG000 | 20 | 46245450 | 46246407 | + | 958 | taar14l | 0 | 0 | 0 | 0 | 0 | 0 |
| ENSDARG000 | 20 | 46176565 | 46177592 | + | 1028 | taar13a | 0 | 0 | 0 | 0 | 0 | 0 |
| ENSDARG000 | 20 | 46240675 | 46241809 | + | 1135 | NA | 0 | 0 | 0 | 0 | 0 | 0 |
| ENSDARG000 | 20 | 46191630 | 46192655 | + | 1026 | taar13d | 0 | 0 | 0 | 0 | 0 | 0 |
| ENSDARG000 | 20:20:20:20:2 | 31309364:313 | 31311233:313 | -;-;-;-;-;-;-;- | 2904 | usta | 36 | 35 | 30 | 14 | 54 | 12 |
| ENSDARG000 | 20:20:20:20:2 | 35473288:354 | 35473366:354 | +:+:+:+:+:+:+: | 2340 | mep1a.1 | 62 | 79 | 16 | 58 | 170 | 162 |
| ENSDARG000 | 20:20:20:20:2 | 46164338:461 | 46165438:461 | -;-;-;-;-;-;- | 1600 | taar10 | 0 | 0 | 0 | 0 | 0 | 0 |
| ENSDARG000 | 20;20 | 22194193:221 | 22194494:221 | +;+ | 978 | NA | 4 | 8 | 42 | 34 | 41 | 17 |
| ENSDARG000 | 20:20:20:20:2 | 22194318:221 | 22194624:221 | -;-;-;-;- | 1015 | srd5a3 | 102 | 125 | 133 | 85 | 148 | 51 |
| ENSDARG000 | 20 | 46237000 | 46237963 | + | 964 | taar14a | 0 | 0 | 0 | 0 | 0 | 0 |
| ENSDARG000 | 20 | 46202188 | 46203215 | + | 1028 | taar13c | 0 | 0 | 0 | 0 | 0 | 0 |
| ENSDARG000 | 20;20 | 46250944:462 | 46252080:462 | +;+ | 1137 | taar14h | 0 | 0 | 0 | 0 | 0 | 0 |
| ENSDARG000 | 20 | 46255057 | 46256016 | + | 960 | taar14i | 0 | 0 | 0 | 0 | 0 | 0 |
| ENSDARG000 | 20 | 46260580 | 46261653 | + | 1074 | taar14j | 0 | 0 | 0 | 0 | 0 | 0 |
| ENSDARG000 | 20 | 46268906 | 46270063 | + | 1158 | taar14e | 0 | 0 | 0 | 0 | 0 | 0 |
| ENSDARG000 | 20;20 | 46277906:462 | 46278334:462 | +;+ | 866 | NA | 0 | 0 | 0 | 4 | 0 | 0 |
| ENSDARG000 | 20;20 | 46274118:462 | 46274876:462 | +;+ | 996 | taar14g | 0 | 0 | 0 | 0 | 0 | 0 |
| ENSDARG000 | 20;20 | 13360656:133 | 13361545:133 | -;- | 1173 | kcnk3a | 0 | 2 | 0 | 0 | 0 | 0 |
| ENSDARG000 | 20:20:20:20:2 | 22220988:222 | 22221279:222 | +:+:+:+:+:+:+: | 5381 | kdr | 319 | 52 | 183 | 68 | 61 | 90 |
| ENSDARG000 | 20:20:20:20:2 | 25473056:254 | 25473887:254 | -:-:-:-:-:-:-:-: | 1342 | NA | 0 | 0 | 6 | 2 | 0 | 2 |
| ENSDARG000 | 20:20:20:20:2 | 25445826:254 | 25445904:254 | +:+:+:+:+:+:+: | 5174 | pfas | 896 | 1387 | 3695 | 805 | 1876 | 3909 |
| ENSDARG000 | 20 | 46286082 | 46287334 | + | 1253 | taar14f | 0 | 0 | 0 | 0 | 0 | 0 |
| ENSDARG000 | 20 | 46291135 | 46292527 | + | 1393 | taar14d | 0 | 0 | 0 | 0 | 0 | 0 |
| ENSDARG000 | 20 | 46303923 | 46305446 | + | 1524 | taar14b | 0 | 0 | 0 | 0 | 0 | 0 |
| ENSDARG000 | 20:20:20:20:2 | 46311242:463 | 46312249:463 | +:+:+:+:+:+:+: | 2580 | FLVCR2 | 183 | 300 | 169 | 108 | 194 | 137 |
| ENSDARG000 | 20:20:20:20:2 | 16170848:161 | 16170943:161 | +:+:+:+:+:+:+: | 3801 | zyg11 | 578 | 393 | 338 | 217 | 419 | 458 |
| ENSDARG000 | 20:20:20:20:2 | 27646772:276 | 27647341:276 | +:+:+:+:+:+:+: | 8241 | MTHFD1 | 213 | 228 | 295 | 146 | 271 | 208 |
| ENSDARG000 | 20:20:20:20:2 | 34084331:340 | 34085295:340 | -;-;-;-;-;-;-;- | 2335 | pdcb | 0 | 64 | 291 | 70 | 0 | 0 |
| ENSDARG000 | 20:20:20:20:2 | 25486021:254 | 25486250:254 | +:+:+:+:+:+:+: | 5930 | hook1 | 82 | 14 | 221 | 227 | 162 | 197 |
| ENSDARG000 | 20:20:20:20:2 | 16205080:162 | 16205450:162 | -:-:-:-:-:-:-:-: | 2070 | echdc2 | 278 | 284 | 220 | 187 | 320 | 221 |
| ENSDARG000 | 20:20:20:20:2 | 46351606:463 | 46353870:463 | -;-;-;-;-;- | 3112 | bmf2 | 94 | 136 | 136 | 243 | 26 | 62 |
| ENSDARG000 | 20;20;20 | 16163142:161 | 16163682:161 | -;-;- | 1055 | coa7 | 58 | 108 | 124 | 55 | 85 | 67 |
| ENSDARG000 | 20:20:20:20:2 | 46371458:463 | 46371635:463 | +:+:+:+:+:+:+: | 3732 | NA | 26 | 4 | 8 | 4 | 28 | 20 |
| ENSDARG000 | 20;20 | 31722252:317 | 31722277:317 | +;+ | 742 | NA | 0 | 0 | 2 | 0 | 1 | 1 |
| ENSDARG000 | 20:20:20:20:2 | 31428819:314 | 31432194:314 | -:-:-:-:-:-:-:-: | 9014 | sash1a | 2115 | 4216 | 3222 | 3464 | 2920 | 2810 |
| ENSDARG000 | 20;20 | 27691307:276 | 27691431:276 | +;+ | 1964 | NA | 2 | 4 | 9 | 6 | 21 | 2 |
| ENSDARG000 | 20:20:20:20:2 | 9172301:9173 | 9172632:9173 | -:-:-:-:-:-:-:-: | 5919 | NA | 1977 | 1413 | 2189 | 731 | 2325 | 1320 |
| ENSDARG000 | 20;20;20 | 27707766:277 | 27709006:277 | -;-;- | 1639 | NA | 2 | 0 | 4 | 0 | 10 | 0 |
| ENSDARG000 | 20:20:20:20:2 | 35495687:354 | 35496549:354 | -:-:-:-:-:-:-:-: | 4020 | NA | 127 | 313 | 340 | 313 | 329 | 277 |
| ENSDARG000 | 20;20;20;20 | 27712714:277 | 27712884:277 | +;+;+;+ | 4742 | zbtb1 | 666 | 344 | 712 | 401 | 717 | 662 |
| ENSDARG000 | 20:20:20:20:2 | 35484070:354 | 35484152:354 | +:+:+:+:+:+:+: | 2719 | mep1a.2 | 0 | 6 | 1 | 0 | 2 | 7 |
| ENSDARG000 | 20:20:20:20:2 | 13403281:134 | 13403460:134 | +:+:+:+:+:+:+: | 4508 | enpp1 | 2084 | 466 | 254 | 301 | 485 | 890 |
| ENSDARG000 | 20:20:20:20:2 | 26095530:261 | 26095837:261 | +:+:+:+:+:+:+: | 26957 | SYNE1 | 1324 | 941 | 1344 | 1155 | 925 | 848 |
| ENSDARG000 | 20:20:20:20:2 | 34121818:341 | 34122907:341 | -:-:-:-:-:-:-:-: | 2556 | ptgs2b | 189 | 674 | 5432 | 3427 | 410 | 74 |
| ENSDARG000 | 20:20:20:20:2 | 46442580:464 | 46442738:464 | +:+:+:+:+:+:+: | 2927 | zc3h14 | 668 | 427 | 627 | 547 | 650 | 484 |
| ENSDARG000 | 20;20 | 27732578:277 | 27733791:277 | -;- | 1214 | zgc:153157 | 8 | 2 | 206 | 8 | 15 | 0 |
| ENSDARG000 | 20:20:20:20:2 | 328412:32903 | 329122:32912 | +:+:+:+:+:+:+: | 2766 | fynb | 3715 | 2918 | 2460 | 2006 | 3610 | 2956 |
| ENSDARG000 | 20:20:20:20:2 | 46427963:464 | 46428040:464 | +:+:+:+:+:+:+: | 2131 | rad51 | 65 | 254 | 282 | 138 | 212 | 102 |
| ENSDARG000 | 20;20 | 27749133:277 | 27749272:277 | +;+ | 3354 | vrtn | 0 | 0 | 44 | 0 | 2 | 0 |
| ENSDARG000 | 20;20;20 | 27755471:277 | 27755544:277 | +;+;+ | 441 | NA | 0 | 0 | 0 | 0 | 0 | 0 |
| ENSDARG000 | 20:20:20:20:2 | 2118356:2121 | 2118675:2121 | -;-;-;-;-;- | 867 | tmem244 | 2 | 2 | 8 | 0 | 2 | 2 |
| ENSDARG000 | 20:20:20:20:2 | 2134816:2139 | 2134827:2140 | +:+:+:+:+:+:+: | 3318 | l3mbtl3 | 169 | 153 | 430 | 185 | 279 | 417 |
| ENSDARG000 | 20:20:20:20:2 | 46442282:464 | 46442533:464 | -:-:-:-:-:-:-:-: | 1825 | rmdn3 | 214 | 222 | 305 | 275 | 162 | 207 |
| ENSDARG000 | 20:20:20:20:2 | 35517572:355 | 35517773:355 | -:-:-:-:-:-:-:-: | 2850 | NA | 0 | 0 | 0 | 0 | 14 | 0 |
| ENSDARG000 | 20:20:20:20:2 | 20030968:200 | 20031078:200 | +:+:+:+:+:+:+: | 1647 | NA | 0 | 0 | 0 | 0 | 0 | 0 |
| ENSDARG000 | 20:20:20:20:2 | 19964854:200 | 19966089:200 | -:-:-:-:-:-:-:-: | 3452 | kcnh5a | 50 | 10 | 118 | 54 | 204 | 26 |
| ENSDARG000 | 20:20:20:20:2 | 22335511:223 | 22336232:223 | -:-:-:-:-:-:-:-: | 3558 | kita | 7298 | 4398 | 2882 | 2097 | 3916 | 5299 |
| ENSDARG000 | 20;20;20;20 | 35541282:355 | 35541734:355 | -;-;-;- | 645 | NA | 0 | 0 | 0 | 2 | 0 | 0 |
| ENSDARG000 | 20;20;20 | 400487:40268 | 400683:40277 | +;+;+ | 798 | NDUFAF4 | 68 | 104 | 87 | 72 | 74 | 50 |
| ENSDARG000 | 20;20;20 | 34133914:341 | 34134525:341 | -;-;- | 892 | NA | 0 | 0 | 0 | 1 | 0 | 0 |
| ENSDARG000 | 20:20:20:20:2 | 34129465:341 | 34129479:341 | +:+:+:+:+:+:+: | 2812 | PLA2G4A | 0 | 23 | 257 | 102 | 28 | 6 |
| ENSDARG000 | 20:20:20:20:2 | 27822044:278 | 27822287:278 | -;-;-;-;-;-;- | 1615 | syndig1l | 26 | 0 | 22 | 8 | 6 | 16 |
| ENSDARG000 | 20:20:20:20:2 | 35551636:355 | 35553139:355 | -;-;-;-;-;-;-;-;- | 2111 | NA | 0 | 4 | 5 | 2 | 0 | 4 |
| ENSDARG000 | 20;20 | 405811:40923 | 406298:41113 | +;+ | 2386 | GPR63 | 136 | 90 | 86 | 54 | 98 | 110 |
| ENSDARG000 | 20;20 | 27919945:279 | 27920136:279 | +;+ | 234 | NA | 0 | 0 | 0 | 0 | 0 | 0 |
| ENSDARG000 | 20:20:20:20:2 | 27904245:279 | 27904437:279 | +;+;+;+;+ | 1225 | NA | 0 | 0 | 0 | 0 | 0 | 0 |
| ENSDARG000 | 20:20:20:20:2 | 418513:42009 | 418772:42027 | -:-:-:-:-:-:-:-: | 2872 | ufl1 | 666 | 732 | 1397 | 1202 | 990 | 757 |
| ENSDARG000 | 20:20:20:20:2 | 35576739:355 | 35576893:355 | -:-:-:-:-:-:-:-: | 29476 | NA | 66 | 16 | 63 | 27 | 86 | 20 |
| ENSDARG000 | 20:20:20:20:2 | 4742400:4742 | 4742487:4742 | -:-:-:-:-:-:-:-: | 2424 | ak7a | 906 | 54 | 283 | 1841 | 650 | 630 |
| ENSDARG000 | 20:20:20:20:2 | 28245164:282 | 28245640:282 | +:+:+:+:+:+:+: | 7391 | dll4 | 124 | 12 | 38 | 0 | 27 | 76 |
| ENSDARG000 | 20:20:20:20:2 | 441942:44261 | 442830:44283 | -:-:-:-:-:-:-:-: | 2213 | FRK | 32 | 18 | 37 | 8 | 86 | 20 |
| ENSDARG000 | 20:20:20:20:2 | 46523078:465 | 46523308:465 | -;-;-;-;-;-;-;- | 1247 | eif2b2 | 575 | 518 | 895 | 643 | 565 | 444 |
| ENSDARG000 | 20:20:20:20:2 | 46536598:465 | 46537766:465 | -:-:-:-:-:-:-:-: | 1755 | tmed10 | 4348 | 4455 | 5898 | 4384 | 5479 | 3465 |
| ENSDARG000 | 20;20;20;20 | 9211237:9211 | 9211437:9211 | +;+;+;+ | 656 | NA | 0 | 0 | 0 | 0 | 0 | 0 |
| ENSDARG000 | 20:20:20:20:2 | 35568552:355 | 35569849:355 | -:-:-:-:-:-:-:-: | 4045 | ADGRF1 | 44 | 65 | 59 | 62 | 41 | 35 |
| ENSDARG000 | 20:20:20:20:2 | 9223514:9227 | 9223766:9227 | +;+;+;+;+ | 2515 | kcnk5b | 273 | 76 | 195 | 200 | 154 | 184 |
| ENSDARG000 | 20:20:20:20:2 | 16279963:162 | 16280399:162 | +:+:+:+:+:+:+: | 1839 | NA | 0 | 6 | 0 | 0 | 0 | 0 |
| ENSDARG000 | 20;20;20 | 2227860:2240 | 2228402:2241 | +;+;+ | 2220 | tmem200a | 0 | 18 | 12 | 4 | 0 | 0 |
| ENSDARG000 | 20;20 | 2261933:2266 | 2262203:2266 | +;+ | 405 | NA | 0 | 0 | 0 | 0 | 0 | 0 |
| ENSDARG000 | 20:20:20:20:2 | 474288:47916 | 474676:47925 | +:+:+:+:+:+:+: | 1958 | nt5dc1 | 189 | 310 | 319 | 130 | 435 | 197 |
| ENSDARG000 | 20:20:20:20:2 | 28266327:282 | 28270881:282 | -:-:-:-:-:-:-:-: | 9499 | ino80 | 649 | 577 | 654 | 324 | 639 | 567 |
| ENSDARG000 | 20:20:20:20:2 | 31745949:317 | 31749486:317 | -:-:-:-:-:-:-:-: | 7466 | stxbp5a | 16 | 39 | 36 | 14 | 80 | 38 |
| ENSDARG000 | 20;20 | 2281933:2284 | 2282136:2284 | +;+ | 522 | NA | 0 | 0 | 0 | 0 | 0 | 0 |
| ENSDARG000 | 20;20 | 46553223:465 | 46553564:465 | +;+ | 723 | NA | 19 | 15 | 21 | 40 | 30 | 44 |
| ENSDARG000 | 20:20:20:20:2 | 46544052:465 | 46544061:465 | +:+:+:+:+:+:+: | 1303 | NA | 10 | 12 | 17 | 23 | 2 | 10 |
| ENSDARG000 | 20:20:20:20:2 | 46552311:465 | 46553459:465 | -;-;-;-;-;-;- | 1699 | fosab | 21995 | 7291 | 16896 | 16230 | 6633 | 15378 |
| ENSDARG000 | 20:20:20:20:2 | 28266892:282 | 28267576:282 | +;+;+;+;+ | 1705 | chac1 | 0 | 0 | 0 | 0 | 0 | 0 |
| ENSDARG000 | 20:20:20:20:2 | 22435180:224 | 22437003:224 | -:-:-:-:-:-:-:-: | 6274 | pdgfra | 122 | 8 | 52 | 10 | 60 | 18 |
| ENSDARG000 | 20:20:20:20:2 | 535458:53981 | 535594:54028 | +:+:+:+:+:+:+: | 4405 | dse | 766 | 527 | 661 | 629 | 527 | 623 |
| ENSDARG000 | 20:20:20:20:2 | 2287298:2290 | 2287573:2290 | -:-:-:-:-:-:-:-: | 3797 | EPB41L2 | 2229 | 1851 | 3802 | 1638 | 2268 | 1633 |
| ENSDARG000 | 20:20:20:20:2 | 46586678:465 | 46586806:465 | +:+:+:+:+:+:+: | 2366 | jdp2b | 4661 | 1988 | 3174 | 1765 | 922 | 2842 |
| ENSDARG000 | 20:20:20:20:2 | 46572550:465 | 46572593:465 | +:+:+:+:+:+:+: | 1006 | batf | 2 | 12 | 0 | 10 | 24 | 10 |
| ENSDARG000 | 20:20:20:20:2 | 34151670:341 | 34151774:341 | +:+:+:+:+:+:+: | 1660 | arpc5b | 1350 | 1101 | 1901 | 1575 | 1768 | 1704 |
| ENSDARG000 | 20:20:20:20:2 | 46651916:466 | 46652141:466 | +:+:+:+:+:+:+: | 3851 | NA | 0 | 0 | 0 | 0 | 14 | 0 |
| ENSDARG000 | 20;20 | 34151733:341 | 34151976:341 | -;- | 502 | NA | 0 | 11 | 2 | 2 | 5 | 7 |
| ENSDARG000 | 20;20 | 22483048:224 | 22483835:224 | -;- | 1273 | gsx2 | 0 | 0 | 0 | 0 | 0 | 0 |
| ENSDARG000 | 20:20:20:20:2 | 13564931:135 | 13569083:135 | -:-:-:-:-:-:-:-: | 6685 | ezrb | 5307 | 3586 | 4385 | 3150 | 4073 | 5090 |
| ENSDARG000 | 20:20:20:20:2 | 4769762:4770 | 4770479:4770 | -:-:-:-:-:-:-:-: | 2809 | galca | 1149 | 253 | 683 | 1168 | 1053 | 1113 |
| ENSDARG000 | 20;20 | 16356267:163 | 16356624:163 | -;- | 488 | NA | 0 | 0 | 0 | 0 | 0 | 0 |
| ENSDARG000 | 20;20 | 16332325:164 | 16333586:164 | -;- | 2025 | NA | 4 | 0 | 0 | 0 | 6 | 2 |
| ENSDARG000 | 20;20 | 2460864:2479 | 2461041:2479 | +;+ | 585 | NA | 542 | 967 | 886 | 881 | 1117 | 1022 |
| ENSDARG000 | 20:20:20:20:2 | 22739792:227 | 22740665:227 | -:-:-:-:-:-:-:-: | 2650 | NA | 0 | 0 | 4 | 0 | 10 | 0 |
| ENSDARG000 | 20;20;20;20 | 560805:56214 | 561055:56230 | +;+;+;+ | 1846 | NA | 96 | 61 | 179 | 46 | 149 | 116 |
| ENSDARG000 | 20:20:20:20:2 | 16437741:164 | 16439264:164 | -:-:-:-:-:-:-:-: | 2819 | gtf2a1 | 1241 | 1254 | 1313 | 1060 | 1222 | 843 |
| ENSDARG000 | 20:20:20:20:2 | 22666548:226 | 22666820:226 | +:+:+:+:+:+:+: | 3415 | lnx1 | 231 | 580 | 441 | 876 | 1060 | 155 |
| ENSDARG000 | 20:20:20:20:2 | 13533544:135 | 13533766:135 | +:+:+:+:+:+:+: | 2130 | SYTL3 | 13 | 4 | 0 | 13 | 8 | 23 |
| ENSDARG000 | 20:20:20:20:2 | 29436601:294 | 29438582:294 | +:+:+:+:+:+:+: | 4244 | FMN1 | 0 | 0 | 0 | 0 | 0 | 0 |
| ENSDARG000 | 20:20:20:20:2 | 46699021:466 | 46699568:466 | +:+:+:+:+:+:+: | 5478 | wdr26b | 1494 | 1179 | 1568 | 927 | 1542 | 1386 |
| ENSDARG000 | 20:20:20:20:2 | 20186706:202 | 20187292:202 | +;+;+;+;+ | 2760 | rhoj | 538 | 344 | 672 | 300 | 668 | 548 |
| ENSDARG000 | 20;20 | 2497521:2498 | 2497538:2498 | -;- | 628 | NA | 0 | 4 | 2 | 2 | 0 | 0 |
| ENSDARG000 | 20:20:20:20:2 | 46736682:467 | 46738810:467 | -;-;-;-;-;-;-;- | 2453 | NA | 88 | 46 | 54 | 114 | 52 | 62 |
| ENSDARG000 | 20:20:20:20:2 | 16477570:164 | 16478636:164 | -:-:-:-:-:-:-:-: | 2160 | ches1 | 69 | 69 | 86 | 47 | 69 | 58 |
| ENSDARG000 | 20:20:20:20:2 | 46741074:467 | 46741330:467 | +:+:+:+:+:+:+: | 1877 | fez2b | 128 | 62 | 111 | 70 | 110 | 92 |
| ENSDARG000 | 20:20:20:20:2 | 31929377:319 | 31930078:319 | -:-:-:-:-:-:-:-: | 5313 | ADGB | 62 | 10 | 38 | 28 | 16 | 26 |
| ENSDARG000 | 20:20:20:20:2 | 29471151:294 | 29472146:294 | -:-:-:-:-:-:-:-: | 2290 | SCG5 | 0 | 0 | 0 | 0 | 0 | 0 |
| ENSDARG000 | 20:20:20:20:2 | 572037:57215 | 572344:57234 | +:+:+:+:+:+:+: | 5723 | smyd2b | 23 | 26 | 146 | 12 | 36 | 48 |
| ENSDARG000 | 20:20:20:20:2 | 9241794:9241 | 9242630:9242 | -:-:-:-:-:-:-:-: | 2776 | syt14b | 10 | 8 | 24 | 7 | 14 | 27 |
| ENSDARG000 | 20;20 | 29469753:294 | 29470985:294 | -;- | 1297 | grem1a | 0 | 0 | 0 | 0 | 0 | 0 |
| ENSDARG000 | 20:20:20:20:2 | 16639848:166 | 16640274:166 | +:+:+:+:+:+:+: | 1988 | tmem30ab | 2041 | 1710 | 1982 | 1763 | 2091 | 2091 |
| ENSDARG000 | 20:20:20:20:2 | 9277299:9278 | 9277972:9278 | +;+;+;+;+;+ | 1912 | NA | 186 | 169 | 99 | 122 | 100 | 160 |
| ENSDARG000 | 20:20:20:20:2 | 591505:59361 | 591665:59392 | +;+;+;+;+;+;+ | 1470 | kcnk2b | 0 | 6 | 2 | 0 | 2 | 0 |
| ENSDARG000 | 20:20:20:20:2 | 20229586:202 | 20233252:202 | -:-:-:-:-:-:-:-: | 5442 | ppp2r5ea | 1339 | 1074 | 1306 | 1068 | 1347 | 1108 |
| ENSDARG000 | 20:20:20:20:2 | 46765733:467 | 46766137:467 | -;-;-;-;- | 1042 | NA | 0 | 0 | 0 | 0 | 0 | 0 |
| ENSDARG000 | 20:20:20:20:2 | 46790056:467 | 46790634:467 | -:-:-:-:-:-:-:-: | 1840 | esrrgb | 45 | 6 | 0 | 8 | 0 | 2 |
| ENSDARG000 | 20:20:20:20:2 | 46830828:468 | 46835781:468 | -:-:-:-:-:-:-:-: | 6332 | GPATCH2 | 942 | 610 | 798 | 584 | 807 | 899 |
| ENSDARG000 | 20:20:20:20:2 | 25515259:255 | 25515619:255 | -:-:-:-:-:-:-:-: | 1828 | cyp2n13 | 172 | 280 | 660 | 184 | 2023 | 1564 |
| ENSDARG000 | 20:20:20:20:2 | 2499622:2499 | 2500104:2500 | -:-:-:-:-:-:-:-: | 4659 | med23 | 1699 | 1188 | 2265 | 1458 | 1604 | 882 |
| ENSDARG000 | 20:20:20:20:2 | 46867816:468 | 46868054:468 | +:+:+:+:+:+:+: | 2701 | NA | 14 | 17 | 91 | 27 | 11 | 4 |
| ENSDARG000 | 20:20:20:20:2 | 32009271:320 | 32010300:320 | -:-:-:-:-:-:-:-: | 2501 | rab32a | 3058 | 4061 | 2834 | 2398 | 3085 | 2914 |
| ENSDARG000 | 20;20 | 16703050:167 | 16703389:167 | +;+ | 1239 | kcnk13b | 0 | 0 | 0 | 0 | 0 | 0 |
| ENSDARG000 | 20:20:20:20:2 | 4793516:4793 | 4793828:4793 | +:+:+:+:+:+:+: | 1522 | LGALS8 | 887 | 254 | 445 | 400 | 855 | 395 |
| ENSDARG000 | 20:20:20:20:2 | 46924390:469 | 46924467:469 | +:+:+:+:+:+:+: | 2416 | GNPAT | 26 | 4 | 13 | 4 | 17 | 11 |
| ENSDARG000 | 20;20;20;20 | 16719769:167 | 16719821:167 | +;+;+;+ | 460 | NA | 1 | 0 | 0 | 8 | 0 | 2 |
| ENSDARG000 | 20:20:20:20:2 | 28354269:283 | 28355289:283 | -;-;-;-;- | 3544 | vps18 | 1595 | 1344 | 1474 | 1254 | 1665 | 1411 |
| ENSDARG000 | 20:20:20:20:2 | 16721933:167 | 16721971:167 | +:+:+:+:+:+:+: | 1501 | psmc1b | 1534 | 1257 | 1420 | 766 | 1369 | 899 |
| ENSDARG000 | 20 | 46957091 | 46959768 | - | 2678 | NA | 964 | 999 | 1195 | 1247 | 1324 | 837 |
| ENSDARG000 | 20:20:20:20:2 | 28364742:283 | 28365072:283 | +;+;+;+;+;+ | 1933 | rhov | 151 | 102 | 132 | 176 | 298 | 82 |
| ENSDARG000 | 20;20 | 48803248:488 | 48803624:488 | +;+ | 1151 | nkx2.4b | 0 | 0 | 0 | 0 | 0 | 0 |
| ENSDARG000 | 20:20:20:20:2 | 32047385:320 | 32049467:320 | -:-:-:-:-:-:-:-: | 5021 | GRM1 | 14 | 14 | 19 | 30 | 8 | 11 |
| ENSDARG000 | 20:20:20:20:2 | 38322444:383 | 38322727:383 | +:+:+:+:+:+:+: | 1556 | stum | 2 | 8 | 3 | 20 | 18 | 4 |
| ENSDARG000 | 20;20 | 48886477:488 | 48886545:488 | +;+ | 664 | NA | 0 | 0 | 0 | 0 | 0 | 0 |
| ENSDARG000 | 20;20 | 48893342:488 | 48893422:488 | +;+ | 267 | NA | 0 | 0 | 0 | 0 | 0 | 0 |
| ENSDARG000 | 20;20;20 | 20275001:202 | 20279285:202 | -;-;- | 5481 | NA | 1524 | 776 | 989 | 715 | 1242 | 828 |
| ENSDARG000 | 20:20:20:20:2 | 48810901:488 | 48811546:488 | -:-:-:-:-:-:-:-: | 3758 | xrn2 | 1633 | 1431 | 1827 | 1013 | 1510 | 1349 |
| ENSDARG000 | 20:20:20:20:2 | 46967269:469 | 46967723:469 | -:-:-:-:-:-:-:-: | 3186 | dnmt3aa | 491 | 505 | 621 | 206 | 834 | 322 |
| ENSDARG000 | 20:20:20:20:2 | 4827735:4827 | 4828355:4828 | -:-:-:-:-:-:-:-: | 1753 | zdhhc14 | 838 | 141 | 443 | 449 | 901 | 313 |
| ENSDARG000 | 20:20:20:20:2 | 26359756:263 | 26361553:263 | -:-:-:-:-:-:-:-: | 3663 | esr1 | 6 | 4 | 8 | 4 | 4 | 8 |
| ENSDARG000 | 20;20 | 47104702:471 | 47104717:471 | -;- | 194 | NA | 1 | 0 | 0 | 0 | 0 | 0 |
| ENSDARG000 | 20:20:20:20:2 | 25533064:255 | 25533132:255 | +:+:+:+:+:+:+: | 2399 | cyp2v1 | 580 | 408 | 450 | 467 | 601 | 747 |
| ENSDARG000 | 20;20 | 47110856:471 | 47111175:471 | +;+ | 1216 | NA | 0 | 0 | 0 | 0 | 0 | 0 |

| | | | | | | | | | | | |
|---|---|---|---|---|---|---|---|---|---|---|---|
| ENSDARG000 | 20:20:20:20:2 | 38386282:383 | 38387393:383 | -:-:-:-:-:-:-:-: | 3363 | ITPKB | 196 | 100 | 177 | 46 | 215 | 137 |
| ENSDARG000 | 20:20:20:20:2 | 26387179:263 | 26388361:263 | -;-;-;-;-;-;-;- | 1861 | armt1 | 307 | 420 | 587 | 436 | 614 | 485 |
| ENSDARG000 | 20;20;20 | 47120504:471 | 47120612:471 | -;-;- | 786 | NA | 0 | 0 | 0 | 0 | 0 | 0 |
| ENSDARG000 | 20;20;20 | 47143900:471 | 47143952:471 | +;+;+ | 1791 | NA | 2 | 23 | 22 | 2 | 10 | 6 |
| ENSDARG000 | 20:20:20:20:2 | 25534669:255 | 25535216:255 | -:-:-:-:-:-:-:-: | 1941 | cyp2ad2 | 0 | 4 | 62 | 19 | 54 | 30 |
| ENSDARG000 | 20:20:20:20:2 | 32118559:321 | 32118759:321 | +;+;+;+;+;+;+ | 3732 | cd164 | 10506 | 7688 | 7359 | 8324 | 10338 | 11549 |
| ENSDARG000 | 20:20:20:20:2 | 616114:61675 | 616839:61683 | -:-:-:-:-:-:-:-: | 2373 | NA | 0 | 6 | 8 | 0 | 10 | 0 |
| ENSDARG000 | 20:20:20:20:2 | 29476852:294 | 29477396:294 | -:-:-:-:-:-:-:-: | 1600 | actc1a | 2 | 0 | 2 | 0 | 16 | 0 |
| ENSDARG000 | 20:20:20:20:2 | 20326611:203 | 20328249:203 | -:-:-:-:-:-:-:-: | 4410 | hif1ab | 4535 | 3803 | 6112 | 6345 | 4320 | 2672 |
| ENSDARG000 | 20:20:20:20:2 | 47153293:471 | 47153510:471 | -;-;-;-;-;- | 1316 | NA | 0 | 0 | 19 | 0 | 10 | 4 |
| ENSDARG000 | 20:20:20:20:2 | 25525802:255 | 25526107:255 | -;-;-;-;-;-;-;- | 1709 | cyp2ad6 | 4 | 0 | 0 | 0 | 0 | 20 |
| ENSDARG000 | 20 | 47187917 | 47188966 | - | 1050 | NA | 0 | 0 | 0 | 0 | 0 | 0 |
| ENSDARG000 | 20:20:20:20:2 | 25548425:255 | 25548913:255 | -:-:-:-:-:-:-:-: | 2060 | cyp2ad3 | 795 | 67 | 261 | 127 | 167 | 948 |
| ENSDARG000 | 20:20:20:20:2 | 20314574:203 | 20314818:203 | -:-:-:-:-:-:-:-: | 1437 | snapc1a | 6 | 8 | 14 | 10 | 0 | 0 |
| ENSDARG000 | 20;20;20 | 47196506:471 | 47196566:471 | +;+;+ | 1042 | NA | 1 | 4 | 2 | 0 | 5 | 0 |
| ENSDARG000 | 20;20;20;20 | 4885298:4887 | 4885403:4888 | +;+;+;+ | 921 | tmem242 | 144 | 246 | 169 | 146 | 184 | 138 |
| ENSDARG000 | 20;20;20 | 47264962:472 | 47265817:472 | -;-;- | 1098 | NA | 0 | 0 | 0 | 0 | 0 | 28 |
| ENSDARG000 | 20;20 | 23026378:230 | 23026570:230 | +;+ | 686 | NA | 0 | 0 | 0 | 0 | 0 | 0 |
| ENSDARG000 | 20:20:20:20:2 | 48918512:489 | 48918752:489 | +:+:+:+:+:+:+: | 1953 | NA | 0 | 0 | 7 | 0 | 0 | 0 |
| ENSDARG000 | 20;20;20;20 | 23022985:230 | 23024263:230 | -;-;-;- | 1713 | rasl11b | 24 | 32 | 61 | 26 | 216 | 42 |
| ENSDARG000 | 20:20:20:20:2 | 25558878:255 | 25559067:255 | +;+;+;+;+ | 1280 | NA | 2 | 11 | 9 | 23 | 5 | 12 |
| ENSDARG000 | 20:20:20:20:2 | 47290004:472 | 47291218:472 | -:-:-:-:-:-:-:-: | 3371 | dtnba | 1061 | 1112 | 1206 | 1211 | 1284 | 738 |
| ENSDARG000 | 20:20:20:20:2 | 23007470:230 | 23007604:230 | +;+;+;+;+ | 623 | NA | 10 | 5 | 1 | 0 | 0 | 4 |
| ENSDARG000 | 20:20:20:20:2 | 22799641:227 | 22800787:228 | +:+:+:+:+:+:+: | 2379 | scfd2 | 302 | 131 | 166 | 61 | 110 | 110 |
| ENSDARG000 | 20:20:20:20:2 | 49105429:491 | 49105691:491 | -;-;-;-;-;- | 779 | naa20 | 441 | 482 | 691 | 508 | 465 | 458 |
| ENSDARG000 | 20;20;20 | 26394324:264 | 26394797:264 | +;+;+ | 3863 | zbtb2b | 666 | 603 | 900 | 520 | 842 | 763 |
| ENSDARG000 | 20;20 | 16742089:167 | 16742306:167 | -;- | 339 | NA | 1 | 0 | 0 | 0 | 0 | 0 |
| ENSDARG000 | 20:20:20:20:2 | 4897996:4897 | 4901595:4901 | -:-:-:-:-:-:-:-: | 8905 | arid1b | 1246 | 966 | 1287 | 1151 | 1416 | 874 |
| ENSDARG000 | 20;20 | 4906078:4907 | 4906555:4907 | -;- | 646 | NA | 2 | 0 | 0 | 0 | 6 | 0 |
| ENSDARG000 | 20;20 | 5051189:5057 | 5051347:5057 | +;+ | 285 | NA | 0 | 0 | 0 | 3 | 0 | 8 |
| ENSDARG000 | 20:20:20:20:2 | 49119633:491 | 49119724:491 | +:+:+:+:+:+:+: | 4036 | CD109 | 10 | 0 | 0 | 2 | 0 | 4 |
| ENSDARG000 | 20:20:20:20:2 | 48097252:480 | 48097266:480 | -:-:-:-:-:-:-:-: | 2466 | efhc1 | 34 | 31 | 27 | 17 | 27 | 13 |
| ENSDARG000 | 20:20:20:20:2 | 2589414:2589 | 2589591:2589 | +:+:+:+:+:+:+: | 2876 | il20ra | 43 | 4 | 6 | 6 | 6 | 4 |
| ENSDARG000 | 20:20:20:20:2 | 48116476:481 | 48116678:481 | +:+:+:+:+:+:+: | 1723 | tram2 | 398 | 398 | 430 | 347 | 446 | 325 |
| ENSDARG000 | 20:20:20:20:2 | 16743056:167 | 16743203:167 | +:+:+:+:+:+:+: | 1517 | calm1b | 492 | 456 | 412 | 544 | 391 | 546 |
| ENSDARG000 | 20:20:20:20:2 | 29492469:294 | 29493348:294 | -:-:-:-:-:-:-:-: | 2448 | rrm2 | 145 | 2385 | 1911 | 546 | 1599 | 471 |
| ENSDARG000 | 20:20:20:20:2 | 26407758:264 | 26410439:264 | -:-:-:-:-:-:-:-: | 9966 | akap12b | 469 | 366 | 1314 | 106 | 378 | 691 |
| ENSDARG000 | 20:20:20:20:2 | 38458084:384 | 38458197:384 | +:+:+:+:+:+:+: | 3256 | coq8aa | 135 | 84 | 137 | 138 | 214 | 244 |
| ENSDARG000 | 20:20:20:20:2 | 38524827:385 | 38524923:385 | +:+:+:+:+:+:+: | 2603 | NA | 44 | 76 | 48 | 40 | 57 | 63 |
| ENSDARG000 | 20:20:20:20:2 | 32141562:321 | 32141996:321 | -:-:-:-:-:-:-:-: | 1909 | cep57l1 | 102 | 134 | 211 | 138 | 110 | 161 |
| ENSDARG000 | 20:20:20:20:2 | 16830415:168 | 16835169:168 | -:-:-:-:-:-:-:-: | 5758 | NA | 130 | 190 | 133 | 109 | 176 | 80 |
| ENSDARG000 | 20;20 | 38504733:385 | 38505703:385 | -;- | 1291 | NA | 1 | 0 | 0 | 10 | 0 | 4 |
| ENSDARG000 | 20:20:20:20:2 | 38517441:385 | 38518973:385 | -:-:-:-:-:-:-:-: | 2832 | si:ch211-245I | 78 | 140 | 98 | 77 | 87 | 75 |
| ENSDARG000 | 20:20:20:20:2 | 49451764:494 | 49451885:494 | -:-:-:-:-:-:-:-: | 11059 | COL12A1 | 489 | 12 | 66 | 4 | 199 | 58 |
| ENSDARG000 | 20:20:20:20:2 | 700616:70078 | 700701:70101 | +:+:+:+:+:+:+: | 1960 | NA | 507 | 446 | 770 | 511 | 562 | 394 |
| ENSDARG000 | 20:20:20:20:2 | 23083800:230 | 23084139:230 | +:+:+:+:+:+:+: | 1417 | usp46 | 36 | 19 | 27 | 24 | 53 | 60 |
| ENSDARG000 | 20:20:20:20:2 | 16884386:168 | 16884533:168 | -:-:-:-:-:-:-:-: | 1210 | psma6b | 631 | 815 | 803 | 603 | 870 | 718 |
| ENSDARG000 | 20;20 | 16883555:168 | 16883589:168 | -;- | 251 | NA | 0 | 0 | 0 | 0 | 0 | 0 |
| ENSDARG000 | 20:20:20:20:2 | 16881883:168 | 16882166:168 | +:+:+:+:+:+:+: | 1553 | nfkbiaa | 902 | 475 | 534 | 440 | 815 | 925 |
| ENSDARG000 | 20:20:20:20:2 | 49940818:499 | 49941438:499 | -:-:-:-:-:-:-:-: | 3704 | extl3 | 450 | 138 | 1406 | 337 | 898 | 211 |
| ENSDARG000 | 20;20 | 48482925:484 | 48485354:484 | -;- | 2430 | insm1a | 2 | 0 | 0 | 0 | 0 | 0 |
| ENSDARG000 | 20:20:20:20:2 | 48488954:484 | 48489312:484 | -;-;-;-;-;-;-;- | 1775 | NA | 4 | 8 | 2 | 8 | 4 | 16 |
| ENSDARG000 | 20;20;20 | 5065268:5067 | 5066066:5067 | +;+;+ | 2439 | NA | 142 | 130 | 108 | 83 | 130 | 150 |
| ENSDARG000 | 20:20:20:20:2 | 23160512:231 | 23161689:231 | -:-:-:-:-:-:-:-: | 3175 | spata18 | 0 | 8 | 25 | 4 | 42 | 6 |
| ENSDARG000 | 20:20:20:20:2 | 13632526:136 | 13633100:136 | -:-:-:-:-:-:-:-: | 1692 | rsph3 | 0 | 4 | 6 | 0 | 0 | 0 |
| ENSDARG000 | 20:20:20:20:2 | 35719375:357 | 35719886:357 | -:-:-:-:-:-:-:-: | 5836 | NA | 21 | 0 | 0 | 6 | 2 | 6 |
| ENSDARG000 | 20:20:20:20:2 | 26478197:264 | 26481034:264 | -:-:-:-:-:-:-:-: | 6304 | mthfd1l | 887 | 270 | 767 | 350 | 221 | 441 |
| ENSDARG000 | 20:20:20:20:2 | 35712884:357 | 35712943:357 | +;+;+;+;+ | 398 | NA | 0 | 0 | 0 | 0 | 0 | 0 |
| ENSDARG000 | 20:20:20:20:2 | 48588679:485 | 48588931:485 | -:-:-:-:-:-:-:-: | 1076 | mgst3a | 180 | 120 | 256 | 129 | 157 | 124 |
| ENSDARG000 | 20:20:20:20:2 | 50015470:500 | 50015805:500 | +:+:+:+:+:+:+: | 1937 | riox1 | 274 | 377 | 662 | 268 | 384 | 323 |
| ENSDARG000 | 20:20:20:20:2 | 50052116:500 | 50052189:500 | +:+:+:+:+:+:+: | 6480 | cpsf2 | 802 | 934 | 991 | 594 | 988 | 472 |
| ENSDARG000 | 20:20:20:20:2 | 48694056:486 | 48695703:486 | -:-:-:-:-:-:-:-: | 4995 | PAX1 | 0 | 6 | 0 | 0 | 4 | 0 |
| ENSDARG000 | 20:20:20:20:2 | 50115335:501 | 50115793:501 | +:+:+:+:+:+:+: | 2084 | SLC24A4 | 4 | 0 | 0 | 0 | 0 | 11 |
| ENSDARG000 | 20 | 50238453 | 50239406 | - | 954 | tmem121ab | 0 | 0 | 4 | 0 | 0 | 0 |
| ENSDARG000 | 20;20 | 50248092:502 | 50248641:502 | +;+ | 710 | NA | 0 | 0 | 0 | 0 | 0 | 0 |
| ENSDARG000 | 20:20:20:20:2 | 2610550:2610 | 2610823:2611 | -:-:-:-:-:-:-:-: | 3892 | NA | 490 | 899 | 1100 | 360 | 791 | 448 |
| ENSDARG000 | 20:20:20:20:2 | 48738887:487 | 48739145:487 | -;-;-;-;- | 770 | NA | 0 | 0 | 0 | 0 | 0 | 0 |
| ENSDARG000 | 20;20 | 48782068:487 | 48782581:487 | +;+ | 1634 | nkx2.2b | 0 | 0 | 0 | 0 | 0 | 0 |
| ENSDARG000 | 20:20:20:20:2 | 5074118:5074 | 5074222:5074 | +:+:+:+:+:+:+: | 2363 | ccnk | 425 | 419 | 673 | 263 | 492 | 365 |
| ENSDARG000 | 20:20:20:20:2 | 48739154:487 | 48739267:487 | +:+:+:+:+:+:+: | 1550 | gfus.3 | 2 | 0 | 0 | 0 | 0 | 0 |
| ENSDARG000 | 20:20:20:20:2 | 715739:73350 | 715819:73363 | +:+:+:+:+:+:+: | 5181 | myo6a | 1413 | 849 | 1197 | 714 | 1286 | 943 |
| ENSDARG000 | 20:20:20:20:2 | 35797998:358 | 35800849:358 | -;-;-;-;-;- | 4854 | tnfrsf21 | 6539 | 3778 | 3504 | 2831 | 6217 | 6119 |
| ENSDARG000 | 20;20 | 28611272:286 | 28611693:286 | -;- | 472 | NA | 0 | 2 | 0 | 0 | 0 | 0 |
| ENSDARG000 | 20:20:20:20:2 | 28641979:286 | 28641990:286 | +;+;+;+;+ | 855 | NA | 0 | 0 | 0 | 0 | 0 | 0 |
| ENSDARG000 | 20:20:20:20:2 | 50852356:508 | 50852419:508 | +:+:+:+:+:+:+: | 3544 | gphnb | 0 | 0 | 0 | 2 | 4 | 4 |
| ENSDARG000 | 20:20:20:20:2 | 23173710:231 | 23173746:231 | +;+;+;+;+;+ | 1080 | sgcb | 452 | 308 | 341 | 270 | 528 | 470 |
| ENSDARG000 | 20:20:20:20:2 | 35857399:359 | 35857642:359 | +:+:+:+:+:+:+: | 3764 | cd2ap | 964 | 1047 | 1022 | 589 | 1139 | 855 |
| ENSDARG000 | 20:20:20:20:2 | 16946132:169 | 16949951:169 | -:-:-:-:-:-:-:-: | 6695 | NA | 8 | 0 | 0 | 0 | 0 | 0 |
| ENSDARG000 | 20;20;20;20 | 13673384:136 | 13673566:136 | -;-;-;- | 622 | NA | 0 | 0 | 0 | 0 | 0 | 0 |
| ENSDARG000 | 20;20;20;20 | 32231609:322 | 32231762:322 | +;+;+;+ | 678 | NA | 4 | 0 | 0 | 0 | 0 | 0 |
| ENSDARG000 | 20:20:20:20:2 | 13647987:136 | 13650924:136 | -:-:-:-:-:-:-:-: | 5155 | tagapb | 77 | 98 | 43 | 34 | 272 | 200 |
| ENSDARG000 | 20:20:20:20:2 | 32183829:321 | 32184218:321 | -:-:-:-:-:-:-:-: | 1261 | si:ch211-51a1 | 17 | 24 | 23 | 38 | 18 | 22 |
| ENSDARG000 | 20:20:20:20:2 | 32152355:321 | 32152460:321 | +:+:+:+:+:+:+: | 4992 | sesn1 | 277 | 193 | 256 | 259 | 369 | 652 |
| ENSDARG000 | 20:20:20:20:2 | 5106568:5106 | 5106774:5106 | +:+:+:+:+:+:+: | 2163 | cyp46a1.1 | 4 | 10 | 11 | 6 | 12 | 0 |
| ENSDARG000 | 20;20 | 23185530:231 | 23185762:231 | +;+ | 403 | NA | 0 | 0 | 0 | 6 | 0 | 0 |
| ENSDARG000 | 20:20:20:20:2 | 51006362:510 | 51006471:510 | +:+:+:+:+:+:+: | 2097 | pals1b | 0 | 2 | 4 | 0 | 0 | 0 |
| ENSDARG000 | 20:20:20:20:2 | 32228210:322 | 32231529:322 | -:-:-:-:-:-:-:-: | 5802 | ARMC2 | 0 | 2 | 2 | 4 | 0 | 2 |
| ENSDARG000 | 20:20:20:20:2 | 51088606:510 | 51090006:510 | -;-;-;-;-;-;-;- | 2181 | atp6v1d | 2604 | 2400 | 2155 | 2345 | 2222 | 2102 |
| ENSDARG000 | 20:20:20:20:2 | 51168912:511 | 51169281:511 | +;+;+;+;+ | 2499 | NA | 6 | 0 | 12 | 18 | 0 | 30 |
| ENSDARG000 | 20;20;20 | 51062867:510 | 51063217:510 | -;-;- | 529 | NA | 0 | 0 | 0 | 0 | 0 | 0 |
| ENSDARG000 | 20:20:20:20:2 | 51104367:511 | 51104513:511 | +:+:+:+:+:+:+: | 1345 | eif2s1b | 3366 | 3546 | 5256 | 4374 | 3563 | 4518 |
| ENSDARG000 | 20;20 | 17038903:170 | 17039190:170 | -;- | 810 | si:dkey-5n18.1 | 132 | 63 | 132 | 82 | 438 | 210 |
| ENSDARG000 | 20;20 | 51062874:511 | 51063016:511 | +;+ | 812 | NA | 0 | 0 | 0 | 0 | 0 | 0 |
| ENSDARG000 | 20;20;20 | 51061695:510 | 51061925:510 | +;+;+ | 5446 | NA | 226 | 293 | 375 | 222 | 485 | 268 |
| ENSDARG000 | 20:20:20:20:2 | 51131933:511 | 51132084:511 | -:-:-:-:-:-:-:-: | 3331 | rbm25b | 1404 | 1484 | 1971 | 1071 | 1769 | 1208 |
| ENSDARG000 | 20;20;20;20 | 29502065:295 | 29502789:295 | -;-;-;- | 2061 | klf11a | 266 | 454 | 902 | 72 | 850 | 58 |
| ENSDARG000 | 20;20;20 | 26535342:265 | 26536581:265 | -;-;- | 1368 | stx11b.2 | 2 | 0 | 1 | 6 | 8 | 0 |
| ENSDARG000 | 20:20:20:20:2 | 35998073:360 | 35998117:360 | +:+:+:+:+:+:+: | 1516 | opn5 | 439 | 622 | 562 | 248 | 60 | 1164 |
| ENSDARG000 | 20:20:20:20:2 | 23245620:232 | 23246266:232 | -:-:-:-:-:-:-:-: | 1682 | ociad1 | 555 | 659 | 748 | 709 | 775 | 732 |
| ENSDARG000 | 20:20:20:20:2 | 51197758:511 | 51198571:511 | +:+:+:+:+:+:+: | 5522 | hsp90ab1 | 86563 | 78644 | 138736 | 120965 | 115487 | 131580 |
| ENSDARG000 | 20:20:20:20:2 | 17581881:177 | 17582055:177 | +:+:+:+:+:+:+: | 5028 | cdh2 | 134 | 390 | 94 | 6 | 165 | 70 |
| ENSDARG000 | 20;20 | 23254477:232 | 23254726:232 | +;+ | 755 | NA | 17 | 8 | 15 | 1 | 12 | 7 |
| ENSDARG000 | 20:20:20:20:2 | 13750537:137 | 13755286:137 | -;-;-;-;-;-;- | 5863 | opn8c | 4 | 0 | 0 | 0 | 2 | 0 |
| ENSDARG000 | 20:20:20:20:2 | 23203946:232 | 23206189:232 | -:-:-:-:-:-:-:-: | 3416 | dcun1d4 | 243 | 171 | 247 | 188 | 256 | 242 |
| ENSDARG000 | 20:20:20:20:2 | 29513470:295 | 29514704:295 | -:-:-:-:-:-:-:-: | 3362 | taf1b | 137 | 181 | 282 | 83 | 256 | 217 |
| ENSDARG000 | 20;20;20 | 36066408:361 | 36069785:361 | -;-;- | 4604 | ptchd4 | 0 | 0 | 0 | 0 | 2 | 0 |
| ENSDARG000 | 20:20:20:20:2 | 23238833:232 | 23238938:232 | +;+;+;+;+;+;+ | 1483 | ociad2 | 16 | 89 | 22 | 116 | 10 | 44 |
| ENSDARG000 | 20:20:20:20:2 | 36233873:362 | 36234412:362 | +:+:+:+:+:+:+: | 1851 | CNIH3 | 0 | 0 | 0 | 0 | 0 | 0 |
| ENSDARG000 | 20;20 | 26538137:265 | 26538213:265 | +;+ | 1556 | stx11b.1 | 22 | 1 | 2 | 4 | 16 | 0 |
| ENSDARG000 | 20:20:20:20:2 | 51237646:512 | 51240138:512 | -:-:-:-:-:-:-:-: | 3321 | slc35b2 | 220 | 309 | 330 | 210 | 349 | 253 |
| ENSDARG000 | 20:20:20:20:2 | 36380266:363 | 36380905:363 | -;-;-;-;- | 1769 | NA | 0 | 0 | 2 | 0 | 0 | 0 |
| ENSDARG000 | 20:20:20:20:2 | 5563042:5564 | 5564738:5564 | +:+:+:+:+:+:+: | 6226 | nrxn3b | 0 | 0 | 0 | 0 | 2 | 0 |
| ENSDARG000 | 20:20:20:20:2 | 26543683:265 | 26544455:265 | -:-:-:-:-:-:-:-: | 2247 | si:dkev-25e12 | 343 | 750 | 461 | 367 | 323 | 367 |
| ENSDARG000 | 20:20:20:20:2 | 13781779:137 | 13782056:137 | +:+:+:+:+:+:+: | 3715 | lpgat1 | 1101 | 1266 | 1480 | 989 | 1332 | 1413 |
| ENSDARG000 | 20;20;20 | 51312883:513 | 51312941:513 | +;+;+ | 2515 | tmem151ba | 0 | 0 | 0 | 0 | 0 | 2 |
| ENSDARG000 | 20 | 51341381 | 51341517 | - | 137 | NA | 0 | 0 | 0 | 0 | 0 | 0 |
| ENSDARG000 | 20;20 | 51342164:513 | 51342234:513 | -;- | 412 | NA | 0 | 0 | 0 | 0 | 0 | 0 |
| ENSDARG000 | 20:20:20:20:2 | 51344305:513 | 51344543:513 | -:-:-:-:-:-:-:-: | 1630 | TCTE1 | 2 | 0 | 14 | 4 | 10 | 0 |
| ENSDARG000 | 20:20:20:20:2 | 20360337:203 | 20363698:203 | -:-:-:-:-:-:-:-: | 6165 | prkchb | 66 | 50 | 54 | 33 | 46 | 48 |
| ENSDARG000 | 20:20:20:20:2 | 38543314:385 | 38543504:385 | +:+:+:+:+:+:+: | 3527 | gtf3c2 | 100 | 122 | 164 | 48 | 147 | 84 |
| ENSDARG000 | 20:20:20:20:2 | 18163355:181 | 18163433:181 | +:+:+:+:+:+:+: | 1695 | aqp4 | 0 | 0 | 2 | 0 | 0 | 2 |
| ENSDARG000 | 20;20;20;20 | 32270907:322 | 32271136:322 | +;+;+;+ | 498 | NA | 0 | 0 | 0 | 0 | 0 | 0 |
| ENSDARG000 | 20:20:20:20:2 | 29565906:295 | 29565959:295 | +;+;+;+;+;+ | 1959 | ywhaqa | 5006 | 6749 | 5864 | 4326 | 5839 | 3857 |
| ENSDARG000 | 20;20;20;20 | 32286765:322 | 32288642:322 | -;-;-;- | 4540 | foxo3b | 494 | 884 | 598 | 494 | 750 | 630 |
| ENSDARG000 | 20:20:20:20:2 | 32337522:323 | 32337935:323 | -:-:-:-:-:-:-:-: | 1871 | afg1lb | 48 | 62 | 130 | 60 | 87 | 82 |
| ENSDARG000 | 20:20:20:20:2 | 51363686:513 | 51364017:513 | -:-:-:-:-:-:-:-: | 1953 | rnf8 | 326 | 364 | 445 | 440 | 416 | 320 |
| ENSDARG000 | 20;20 | 51389901:513 | 51390559:513 | +;+ | 1013 | NA | 213 | 123 | 226 | 149 | 304 | 282 |
| ENSDARG000 | 20:20:20:20:2 | 36393912:363 | 36396929:363 | -:-:-:-:-:-:-:-: | 4833 | lbr | 482 | 933 | 1185 | 464 | 839 | 613 |
| ENSDARG000 | 20;20 | 51396223:513 | 51398880:513 | -;- | 2768 | NA | 206 | 214 | 408 | 186 | 408 | 124 |
| ENSDARG000 | 20;20;20;20 | 51407080:514 | 51407470:514 | +;+;+;+ | 671 | NA | 0 | 0 | 0 | 0 | 0 | 9 |
| ENSDARG000 | 20 | 51464670 | 51466071 | + | 1402 | NA | 9 | 26 | 39 | 4 | 14 | 20 |
| ENSDARG000 | 20;20;20;20 | 51466046:514 | 51466068:514 | +;+;+;+ | 3513 | tlr5b | 216 | 164 | 308 | 169 | 122 | 203 |
| ENSDARG000 | 20:20:20:20:2 | 51478939:514 | 51478980:514 | +;+;+;+;+;+ | 2893 | NA | 86 | 84 | 62 | 56 | 70 | 175 |
| ENSDARG000 | 20:20:20:20:2 | 51489195:514 | 51492741:514 | -:-:-:-:-:-:-:-: | 4623 | disp1 | 165 | 182 | 282 | 166 | 218 | 169 |
| ENSDARG000 | 20;20;20 | 51680006:516 | 51680692:516 | -;-;- | 1054 | NA | 2 | 0 | 0 | 2 | 0 | 2 |
| ENSDARG000 | 20:20:20:20:2 | 26556174:265 | 26556244:265 | +:+:+:+:+:+:+: | 1695 | NA | 11 | 10 | 10 | 20 | 46 | 36 |
| ENSDARG000 | 20;20;20 | 51695479:516 | 51696075:516 | -;-;- | 963 | NA | 1 | 0 | 0 | 3 | 4 | 0 |
| ENSDARG000 | 20:20:20:20:2 | 51700633:517 | 51701087:517 | -:-:-:-:-:-:-:-: | 2299 | brox | 1104 | 882 | 1166 | 1227 | 1042 | 872 |
| ENSDARG000 | 20:20:20:20:2 | 38595935:385 | 38596162:385 | -:-:-:-:-:-:-:-: | 1535 | slc30a2 | 2 | 2 | 0 | 0 | 0 | 0 |
| ENSDARG000 | 20:20:20:20:2 | 51730658:517 | 51730912:517 | +:+:+:+:+:+:+: | 1587 | aida | 558 | 669 | 581 | 385 | 567 | 816 |
| ENSDARG000 | 20;20 | 51748497:517 | 51748959:517 | -;- | 592 | NA | 0 | 0 | 0 | 0 | 2 | 0 |
| ENSDARG000 | 20:20:20:20:2 | 51813432:518 | 51813760:518 | +:+:+:+:+:+:+: | 1964 | taf1a | 126 | 128 | 132 | 66 | 152 | 116 |
| ENSDARG000 | 20:20:20:20:2 | 6089507:6089 | 6089638:6089 | -:-:-:-:-:-:-:-: | 3902 | pum2 | 381 | 248 | 428 | 297 | 383 | 317 |
| ENSDARG000 | 20:20:20:20:2 | 29586529:295 | 29586598:295 | +:+:+:+:+:+:+: | 3219 | ADAM17 | 204 | 198 | 206 | 132 | 387 | 193 |
| ENSDARG000 | 20:20:20:20:2 | 36419947:364 | 36422336:364 | -:-:-:-:-:-:-:-: | 6335 | enah | 725 | 1319 | 1803 | 754 | 2995 | 544 |
| ENSDARG000 | 20:20:20:20:2 | 51758426:517 | 51759669:517 | -:-:-:-:-:-:-:-: | 7182 | NA | 1901 | 1606 | 2657 | 1220 | 2401 | 1668 |
| ENSDARG000 | 20:20:20:20:2 | 778608:78030 | 778755:78038 | -;-;-;-;-;-;- | 1233 | NA | 0 | 0 | 0 | 0 | 0 | 0 |
| ENSDARG000 | 20:20:20:20:2 | 6142433:6142 | 6142533:6142 | +:+:+:+:+:+:+: | 843 | ttr | 0 | 0 | 3 | 0 | 0 | 0 |
| ENSDARG000 | 20;20;20 | 2642855:2644 | 2643029:2645 | +;+;+ | 1346 | zgc:101562 | 20 | 32 | 30 | 30 | 32 | 48 |
| ENSDARG000 | 20:20:20:20:2 | 38620666:386 | 38620795:386 | +:+:+:+:+:+:+: | 3164 | NA | 871 | 844 | 1022 | 961 | 1473 | 1288 |
| ENSDARG000 | 20:20:20:20:2 | 51917506:519 | 51917771:519 | -;-;-;-;- | 1769 | DUSP10 | 28 | 8 | 35 | 24 | 8 | 2 |
| ENSDARG000 | 20;20 | 13894123:139 | 13894916:139 | +;+ | 2996 | slc30a1a | 1129 | 1892 | 1347 | 854 | 2123 | 1168 |
| ENSDARG000 | 20:20:20:20:2 | 13883131:138 | 13883264:138 | +:+:+:+:+:+:+: | 2999 | nek2 | 36 | 140 | 167 | 76 | 98 | 32 |
| ENSDARG000 | 20:20:20:20:2 | 52233981:522 | 52234200:522 | -:-:-:-:-:-:-:-: | 2266 | RHPN1 | 2 | 1 | 0 | 2 | 4 | 0 |
| ENSDARG000 | 20;20 | 20499869:205 | 20500627:205 | +;+ | 5417 | six1b | 0 | 0 | 0 | 0 | 0 | 2 |
| ENSDARG000 | 20;20 | 52152792:522 | 52152912:522 | +;+ | 1565 | NA | 0 | 0 | 0 | 0 | 0 | 0 |
| ENSDARG000 | 20;20 | 13894918:138 | 13895157:138 | -;- | 693 | NA | 2 | 3 | 2 | 2 | 0 | 2 |
| ENSDARG000 | 20:20:20:20:2 | 52195525:521 | 52196146:521 | -;-;-;-;-;- | 1489 | hlx1 | 50 | 28 | 24 | 6 | 48 | 46 |
| ENSDARG000 | 20;20 | 20531002:205 | 20531758:205 | -;- | 1615 | six6b | 0 | 0 | 0 | 0 | 0 | 0 |
| ENSDARG000 | 20;20;20 | 20484827:204 | 20485667:204 | +;+;+ | 2406 | six4b | 12 | 2 | 4 | 2 | 12 | 10 |
| ENSDARG000 | 20;20;20;20 | 20428328:204 | 20428369:204 | -;-;-;- | 756 | NA | 0 | 0 | 0 | 0 | 0 | 0 |
| ENSDARG000 | 20:20:20:20:2 | 52273446:522 | 52273793:522 | -:-:-:-:-:-:-:-: | 2014 | naprt | 1271 | 1242 | 1222 | 699 | 1367 | 678 |

| | | | | | | | | | | | | |
|---|---|---|---|---|---|---|---|---|---|---|---|---|
| ENSDARG000 | 20;20;20 | 52299192:523 | 52299410:523 | +;+;+ | 1345 | NA | 0 | 0 | 0 | 0 | 0 | 0 |
| ENSDARG000 | 20;20;20;20;2 | 52332972:523 | 52333500:523 | -;-;-;-;-;- | 855 | si:ch1073-287 | 70 | 46 | 75 | 63 | 83 | 89 |
| ENSDARG000 | 20;20;20;20;2 | 52389858:523 | 52390159:523 | +;+;+;+;+;+;+; | 3081 | ARHGAP39 | 65 | 159 | 253 | 106 | 120 | 61 |
| ENSDARG000 | 20;20;20;20;2 | 52458765:524 | 52459016:524 | +;+;+;+;+;+;+; | 1622 | gfus.1 | 830 | 953 | 947 | 373 | 973 | 686 |
| ENSDARG000 | 20;20;20;20;2 | 6163932:616 | 6167503:616 | -;-;-;-;-;-;-; | 4926 | b4galt6 | 1080 | 1522 | 1548 | 1009 | 1212 | 1245 |
| ENSDARG000 | 20;20;20;20;2 | 52492192:525 | 52492321:525 | +;+;+;+;+;+;+; | 4404 | gfus.2 | 163 | 40 | 83 | 69 | 74 | 113 |
| ENSDARG000 | 20;20;20;20;2 | 38671894:386 | 38672082:386 | +;+;+;+;+;+;+; | 875 | mpv17 | 52 | 33 | 88 | 30 | 50 | 18 |
| ENSDARG000 | 20;20;20;20;2 | 20600971:206 | 20605959:206 | -;-;-;-;-;- | 6539 | ppm1ab | 644 | 557 | 966 | 899 | 925 | 766 |
| ENSDARG000 | 20;20;20;20;2 | 52529044:525 | 52529187:525 | +;+;+;+;+;+;+; | 2427 | pycr3 | 118 | 98 | 153 | 55 | 156 | 89 |
| ENSDARG000 | 20;20;20;20;2 | 36623807:366 | 36623974:366 | +;+;+;+;+;+ | 725 | srp9 | 218 | 393 | 390 | 165 | 322 | 268 |
| ENSDARG000 | 20;20;20;20;2 | 25563105:255 | 25563406:255 | +;+;+;+;+;+;+; | 2498 | cyp2p6 | 2 | 4 | 18 | 0 | 24 | 24 |
| ENSDARG000 | 20;20;20;20;2 | 52546186:525 | 52546268:525 | +;+;+;+;+;+;+; | 1946 | eef1db | 4371 | 4691 | 6556 | 5974 | 3958 | 5853 |
| ENSDARG000 | 20;20;20;20;2 | 36628059:366 | 36628325:366 | +;+;+;+;+;+;+; | 2325 | ephx1 | 214 | 471 | 528 | 432 | 592 | 413 |
| ENSDARG000 | 20;20;20;20;2 | 6493648:649 | 6493747:649 | +;+;+;+;+;+;+; | 2320 | mep1ba | 2 | 2 | 0 | 0 | 6 | 0 |
| ENSDARG000 | 20;20;20;20;2 | 6516523:651 | 6516844:651 | -;-;-;-;-;-;-; | 2666 | mcm3l | 30 | 18 | 4 | 22 | 8 | 0 |
| ENSDARG000 | 20;20;20;20;2 | 29619820:296 | 29620309:296 | -;-;-;-;-;-;-; | 2913 | cpsf3 | 1105 | 1326 | 1689 | 1003 | 1526 | 1306 |
| ENSDARG000 | 20;20;20;20;2 | 52686229:526 | 52686825:526 | -;-;-;-;-;-;- | 962 | NA | 0 | 0 | 0 | 0 | 0 | 0 |
| ENSDARG000 | 20;20;20;20;2 | 52613021:526 | 52613789:526 | -;-;-;-;-;- | 1141 | NA | 0 | 0 | 0 | 0 | 0 | 0 |
| ENSDARG000 | 20;20;20;20;2 | 52584532:525 | 52584600:525 | +;+;+;+;+;+;+; | 6272 | NA | 8 | 22 | 10 | 8 | 30 | 4 |
| ENSDARG000 | 20;20;20;20;2 | 38724575:387 | 38724620:387 | +;+;+;+;+ | 1909 | uts1 | 0 | 0 | 3 | 6 | 0 | 0 |
| ENSDARG000 | 20;20;20;20;2 | 23315242:233 | 23315314:233 | +;+;+;+;+;+;+; | 11147 | fryl | 2816 | 1864 | 3941 | 2355 | 3923 | 3004 |
| ENSDARG000 | 20;20;20;20;2 | 52702012:527 | 52702786:527 | -;-;-;-;-;- | 1141 | NA | 1 | 0 | 0 | 0 | 0 | 0 |
| ENSDARG000 | 20;20;20;20;2 | 52663031:526 | 52663627:526 | -;-;-;-;-;- | 969 | NA | 1 | 0 | 0 | 0 | 0 | 0 |
| ENSDARG000 | 20;20;20;20;2 | 25597527:256 | 25597725:256 | +;+;+;+;+;+;+; | 1573 | NA | 0 | 28 | 22 | 44 | 48 | 9 |
| ENSDARG000 | 20;20;20;20;2 | 52599959:525 | 52600555:526 | -;-;-;-;-;- | 969 | NA | 0 | 0 | 0 | 0 | 0 | 0 |
| ENSDARG000 | 20;20;20;20;2 | 38730291:387 | 38730602:387 | -;-;-;-;-;-;-; | 1642 | trim54 | 10 | 0 | 0 | 12 | 2 | 0 |
| ENSDARG000 | 20;20;20;20;2 | 25581627:255 | 25581886:255 | +;+;+;+;+;+;+; | 2328 | cyp2p9 | 0 | 0 | 0 | 0 | 0 | 0 |
| ENSDARG000 | 20;20;20;20;2 | 52639377:526 | 52639975:526 | -;-;-;-;-;- | 965 | NA | 0 | 0 | 0 | 0 | 0 | 0 |
| ENSDARG000 | 20;20;20;20;2 | 25600105:256 | 25600384:256 | -;-;-;-;-;-;-;- | 812 | NA | 0 | 0 | 30 | 2 | 0 | 2 |
| ENSDARG000 | 20;20;20;20;2 | 52628875:526 | 52629477:526 | -;-;-;-;-;- | 969 | NA | 0 | 0 | 0 | 0 | 0 | 0 |
| ENSDARG000 | 20;20;20 | 25580906:255 | 25581061:255 | +;+;+ | 379 | NA | 0 | 0 | 0 | 0 | 0 | 0 |
| ENSDARG000 | 20;20;20;20;2 | 25568694:255 | 25568980:255 | +;+;+;+;+;+;+; | 2194 | cyp2p7 | 5 | 0 | 0 | 0 | 0 | 34 |
| ENSDARG000 | 20;20;20;20;2 | 36646761:366 | 36647238:366 | -;-;-;-;-;-;-; | 2320 | slc5a6a | 16 | 43 | 103 | 30 | 20 | 55 |
| ENSDARG000 | 20;20;20;20;2 | 25586099:255 | 25586511:255 | +;+;+;+;+;+;+; | 1806 | cyp2p10 | 244 | 0 | 0 | 0 | 0 | 165 |
| ENSDARG000 | 20;20;20;20;2 | 25575391:255 | 25575611:255 | +;+;+;+;+;+;+; | 1867 | cyp2p8 | 0 | 0 | 0 | 0 | 0 | 0 |
| ENSDARG000 | 20;20 | 20611896:206 | 20612288:206 | -;- | 588 | NA | 0 | 0 | 0 | 0 | 0 | 2 |
| ENSDARG000 | 20;20;20;20;2 | 52907397:529 | 52907756:529 | -;-;-;-;-;-;-; | 1939 | fdft1 | 321 | 187 | 345 | 96 | 138 | 171 |
| ENSDARG000 | 20;20;20;20;2 | 52886927:528 | 52887983:528 | -;-;-;-;-;-;-; | 2097 | ctsbb | 4 | 6 | 9 | 0 | 20 | 0 |
| ENSDARG000 | 20;20;20;20;2 | 38763953:387 | 38765908:387 | -;-;-;-;-;-;-; | 2953 | dnajc5ga | 1016 | 1180 | 1144 | 925 | 1273 | 1025 |
| ENSDARG000 | 20;20;20;20;2 | 52952116:529 | 52952285:529 | -;-;-;-;-;- | 1214 | gata4 | 2 | 4 | 9 | 0 | 0 | 4 |
| ENSDARG000 | 20;20;20;20;2 | 53300588:533 | 53300745:533 | -;-;-;-;-;-;-; | 2088 | WASF1 | 172 | 64 | 94 | 88 | 76 | 40 |
| ENSDARG000 | 20;20;20;20;2 | 36675083:366 | 36675538:366 | -;-;-;-;-;-;-; | 1863 | rpl7l1 | 649 | 1155 | 1586 | 827 | 1081 | 886 |
| ENSDARG000 | 20;20;20;20;2 | 53368611:533 | 53368839:533 | +;+;+;+;+;+;+; | 2008 | cdc40 | 578 | 486 | 775 | 543 | 542 | 546 |
| ENSDARG000 | 20;20;20;20;2 | 53408284:534 | 53408616:534 | -;-;-;-;-;-;-; | 1220 | METTL24 | 58 | 65 | 79 | 64 | 66 | 36 |
| ENSDARG000 | 20;20;20;20;2 | 36682051:367 | 36682578:367 | +;+;+;+;+;+;+; | 5929 | NCOA1 | 486 | 244 | 554 | 294 | 446 | 416 |
| ENSDARG000 | 20;20;20;20;2 | 20637866:206 | 20638256:206 | +;+;+;+;+;+;+; | 6055 | rtn1b | 9252 | 9471 | 5617 | 8442 | 9883 | 8564 |
| ENSDARG000 | 20;20;20;20;2 | 53439665:534 | 53439729:534 | +;+;+;+;+;+;+; | 8203 | NA | 4 | 0 | 36 | 2 | 0 | 0 |
| ENSDARG000 | 20;20;20;20;2 | 53474963:534 | 53475069:534 | +;+;+;+;+;+;+; | 2518 | NA | 121 | 249 | 366 | 138 | 264 | 90 |
| ENSDARG000 | 20;20;20;20;2 | 23423814:234 | 23424147:234 | -;-;-;-;-;-;- | 1321 | zar1 | 0 | 8 | 25 | 0 | 18 | 4 |
| ENSDARG000 | 20;20;20 | 20713385:207 | 20713651:207 | +;+;+ | 1167 | NA | 2 | 10 | 2 | 32 | 2 | 0 |
| ENSDARG000 | 20;20;20;20;2 | 29634653:296 | 29635017:296 | +;+;+;+;+;+;+; | 4876 | asap2b | 264 | 142 | 86 | 110 | 102 | 140 |
| ENSDARG000 | 20;20;20;20;2 | 23431485:234 | 23432252:234 | -;-;-;-;-;- | 1617 | slc10a4 | 0 | 2 | 12 | 2 | 12 | 0 |
| ENSDARG000 | 20;20;20;20;2 | 23440632:234 | 23441406:234 | +;+;+;+;+ | 1547 | si:dkev-90m5. | 0 | 0 | 0 | 0 | 0 | 0 |
| ENSDARG000 | 20;20;20;20;2 | 53521648:535 | 53521849:535 | +;+;+;+;+;+;+; | 2269 | pak6b | 57 | 0 | 1 | 0 | 0 | 31 |
| ENSDARG000 | 20;20 | 36797612:367 | 36798420:368 | -;- | 1056 | ptrhd1 | 94 | 138 | 223 | 98 | 206 | 158 |
| ENSDARG000 | 20;20;20;20;2 | 23446243:234 | 23447883:234 | -;-;-;-;-;-;- | 3481 | slain2 | 955 | 869 | 747 | 658 | 876 | 989 |
| ENSDARG000 | 20 | 53577502 | 53583436 | + | 5935 | myh6 | 0 | 0 | 4 | 0 | 2 | 0 |
| ENSDARG000 | 20;20;20;20;2 | 36801804:368 | 36802048:368 | -;-;-;-;-;-;-;- | 1472 | slc25a27 | 47 | 34 | 34 | 29 | 48 | 15 |
| ENSDARG000 | 20;20;20;20;2 | 53600638:536 | 53602186:536 | -;-;-;-;-;- | 1964 | slc25a29 | 46 | 41 | 72 | 96 | 57 | 62 |
| ENSDARG000 | 20;20;20;20;2 | 38788319:387 | 38788993:387 | -;-;-;-;-;-;-; | 8054 | cad | 1223 | 1392 | 2720 | 847 | 1934 | 1202 |
| ENSDARG000 | 20;20;20;20;2 | 36806398:368 | 36806528:368 | +;+;+;+;+;+;+; | 2271 | abracl | 3149 | 3979 | 2926 | 2218 | 3167 | 3006 |
| ENSDARG000 | 20;20;20;20;2 | 53628534:536 | 53628648:536 | +;+;+;+;+;+;+; | 3907 | slc25a47a | 159 | 76 | 234 | 188 | 80 | 198 |
| ENSDARG000 | 20;20 | 53802521:538 | 53804728:538 | +;+ | 2720 | dio3b | 135 | 328 | 624 | 219 | 151 | 248 |
| ENSDARG000 | 20;20;20;20;2 | 53865133:538 | 53865198:538 | +;+;+;+;+;+;+; | 2514 | NA | 0 | 0 | 0 | 0 | 0 | 0 |
| ENSDARG000 | 20;20;20;20;2 | 20726231:207 | 20726305:207 | +;+;+;+;+;+;+ | 900 | atp5mj | 1142 | 1480 | 1231 | 1130 | 1084 | 1274 |
| ENSDARG000 | 20;20;20;20;2 | 53898193:539 | 53904066:539 | -;-;-;-;-;-;-; | 7723 | ppp2r5cb | 1390 | 1082 | 1483 | 888 | 1658 | 1370 |
| ENSDARG000 | 20;20;20;20;2 | 36820965:368 | 36821259:368 | +;+;+;+;+;+ | 2055 | heca | 154 | 14 | 42 | 92 | 51 | 57 |
| ENSDARG000 | 20;20;20;20;2 | 53967350:539 | 53967730:539 | -;-;-;-;-;-;-; | 3062 | hsp90aa1.2 | 3958 | 4097 | 5906 | 11344 | 1886 | 1171 |
| ENSDARG000 | 20;20;20;20;2 | 36845787:368 | 36849969:368 | -;-;-;-;-;-;-; | 5810 | TXLNB | 12 | 0 | 0 | 6 | 23 | 4 |
| ENSDARG000 | 20;20;20;20;2 | 36860807:368 | 36860944:368 | +;+;+;+;+;+;+; | 1168 | NA | 0 | 0 | 0 | 0 | 0 | 0 |
| ENSDARG000 | 20;20;20;20;2 | 53982317:539 | 53982630:539 | -;-;-;-;-;-;-; | 2804 | hsp90aa1.1 | 14 | 34 | 22 | 4 | 7 | 14 |
| ENSDARG000 | 20;20;20;20;2 | 23475381:234 | 23475760:234 | -;-;-;-;-;-;-; | 2855 | anapc4 | 152 | 350 | 380 | 104 | 429 | 238 |
| ENSDARG000 | 20;20;20;20;2 | 20731052:207 | 20731105:207 | +;+;+;+;+;+;+; | 7140 | ppp1r13bb | 1078 | 642 | 1061 | 767 | 1029 | 874 |
| ENSDARG000 | 20;20;20;20;2 | 54013212:540 | 54013474:540 | -;-;-;-;-;-;-; | 931 | NA | 0 | 0 | 0 | 0 | 0 | 0 |
| ENSDARG000 | 20;20;20;20;2 | 54018005:540 | 54018119:540 | +;+;+;+;+;+;+; | 700 | NA | 0 | 0 | 0 | 0 | 0 | 0 |
| ENSDARG000 | 20;20;20;20;2 | 54027778:540 | 54027992:540 | +;+;+;+;+;+;+; | 825 | NA | 0 | 0 | 0 | 0 | 0 | 0 |
| ENSDARG000 | 20;20;20;20;2 | 54024559:540 | 54024773:540 | +;+;+;+;+;+;+; | 891 | si:dkev-241l7. | 0 | 0 | 0 | 0 | 0 | 0 |
| ENSDARG000 | 20;20;20;20;2 | 54034512:540 | 54034726:540 | +;+;+;+;+;+;+; | 835 | si:dkev-241l7. | 0 | 0 | 0 | 0 | 0 | 0 |
| ENSDARG000 | 20;20 | 36915143:369 | 36917304:369 | -;- | 2345 | cited2 | 124 | 179 | 707 | 181 | 1473 | 254 |
| ENSDARG000 | 20;20;20;20;2 | 36925954:369 | 36926134:369 | +;+;+;+;+;+;+; | 1246 | NA | 0 | 0 | 0 | 0 | 0 | 0 |
| ENSDARG000 | 20;20;20 | 37256654:372 | 37256721:372 | +;+;+ | 234 | NA | 0 | 0 | 0 | 0 | 0 | 0 |
| ENSDARG000 | 20;20;20;20;2 | 37263663:372 | 37263698:372 | -;-;-;-;-;-;- | 1922 | NMBR | 0 | 0 | 0 | 0 | 0 | 2 |
| ENSDARG000 | 20;20;20;20;2 | 38831588:388 | 38831803:388 | -;-;-;-;-;- | 1164 | si:dkev-221h1 | 0 | 1 | 12 | 18 | 12 | 0 |
| ENSDARG000 | 20;20;20;20;2 | 54036981:540 | 54037457:540 | +;+;+;+;+;+;+ | 2723 | WDR20 | 178 | 122 | 170 | 142 | 190 | 160 |
| ENSDARG000 | 20;20 | 54067041:540 | 54068748:540 | -;- | 1987 | mgat2 | 500 | 367 | 501 | 354 | 547 | 366 |
| ENSDARG000 | 20;20;20 | 54079341:540 | 54079465:540 | +;+;+ | 346 | rps29 | 5767 | 5438 | 4782 | 3168 | 2306 | 3684 |
| ENSDARG000 | 20;20;20 | 54088980:541 | 54089044:541 | +;+;+ | 239 | NA | 0 | 0 | 0 | 0 | 0 | 0 |
| ENSDARG000 | 20;20;20;20;2 | 54274431:542 | 54274623:542 | +;+;+;+;+;+;+; | 2457 | actr10 | 3749 | 2820 | 3466 | 3813 | 3461 | 2717 |
| ENSDARG000 | 20;20;20;20;2 | 37294112:372 | 37294173:372 | +;+;+;+;+;+;+; | 1997 | gje1a | 0 | 0 | 0 | 0 | 0 | 0 |
| ENSDARG000 | 20;20;20;20;2 | 23498255:235 | 23498489:235 | +;+;+;+;+;+;+; | 6187 | PALLD | 135 | 280 | 197 | 126 | 1172 | 141 |
| ENSDARG000 | 20;20;20;20;2 | 39190854:391 | 39191758:391 | -;-;-;-;-;-;-; | 2274 | cyp39a1 | 8 | 14 | 8 | 16 | 19 | 11 |
| ENSDARG000 | 20;20;20;20;2 | 54290356:542 | 54290721:542 | +;+;+;+;+;+;+; | 1327 | zp2.2 | 2 | 0 | 0 | 0 | 0 | 0 |
| ENSDARG000 | 20;20;20;20;2 | 39055432:390 | 39056280:390 | -;-;-;-;-;-;-;- | 1713 | rcan2 | 16 | 7 | 295 | 320 | 34 | 10 |
| ENSDARG000 | 20;20;20;20;2 | 54295213:542 | 54295594:542 | +;+;+;+;+;+;+; | 1352 | zp2.3 | 0 | 0 | 0 | 0 | 0 | 0 |
| ENSDARG000 | 20;20;20;20;2 | 38837238:388 | 38837396:388 | +;+;+;+;+;+;+; | 5655 | ift172 | 372 | 315 | 409 | 226 | 340 | 326 |
| ENSDARG000 | 20;20;20;20;2 | 54299419:542 | 54299798:542 | +;+;+;+;+;+;+; | 1357 | NA | 0 | 0 | 0 | 0 | 0 | 0 |
| ENSDARG000 | 20;20;20;20;2 | 39048114:390 | 39048527:390 | +;+;+;+;+;+;+; | 1389 | NA | 18 | 4 | 249 | 407 | 28 | 8 |
| ENSDARG000 | 20;20;20;20;2 | 54304800:543 | 54305180:543 | +;+;+;+;+;+;+; | 1352 | zp2.6 | 0 | 0 | 0 | 0 | 0 | 0 |
| ENSDARG000 | 20;20;20;20;2 | 38910214:389 | 38910412:389 | +;+;+;+;+;+;+; | 1030 | msraa | 195 | 201 | 263 | 240 | 316 | 224 |
| ENSDARG000 | 20;20;20;20;2 | 37383034:373 | 37383287:373 | +;+;+;+;+;+;+; | 4287 | adgrg6 | 127 | 2 | 24 | 10 | 171 | 6 |
| ENSDARG000 | 20;20;20;20;2 | 37300296:373 | 37300902:373 | +;+;+;+;+;+;+; | 4874 | vta1 | 942 | 856 | 1080 | 874 | 843 | 1048 |
| ENSDARG000 | 20;20;20;20 | 37375914:373 | 37376216:373 | -;-;-;- | 547 | NA | 1 | 0 | 0 | 0 | 0 | 0 |
| ENSDARG000 | 20;20;20;20;2 | 54309148:543 | 54309527:543 | +;+;+;+;+;+;+; | 1351 | zp2.1 | 0 | 1 | 0 | 0 | 0 | 0 |
| ENSDARG000 | 20;20;20;20;2 | 54312970:543 | 54313342:543 | +;+;+;+;+;+;+; | 1415 | zp2.5 | 0 | 0 | 0 | 0 | 0 | 0 |
| ENSDARG000 | 20;20;20;20;2 | 54316884:543 | 54316968:543 | +;+;+;+;+;+ | 807 | NA | 0 | 0 | 0 | 0 | 0 | 2 |
| ENSDARG000 | 20;20;20;20;2 | 54333774:543 | 54334542:543 | +;+;+;+;+;+ | 7925 | NA | 933 | 964 | 1278 | 672 | 1942 | 992 |
| ENSDARG000 | 20;20;20;20;2 | 23650832:236 | 23651546:236 | -;-;-;-;-;-;-; | 1365 | cbr4 | 161 | 143 | 127 | 89 | 219 | 208 |
| ENSDARG000 | 20;20;20;20;2 | 54349333:543 | 54349524:543 | +;+;+;+;+;+;+; | 2991 | znf410 | 647 | 410 | 670 | 545 | 585 | 571 |
| ENSDARG000 | 20;20;20;20;2 | 37475836:374 | 37476284:374 | -;-;-;-;-;-;- | 772 | si:ch211-202r | 0 | 0 | 0 | 0 | 0 | 0 |
| ENSDARG000 | 20;20;20;20;2 | 54360268:543 | 54361067:543 | -;-;-;-;-;-;-; | 2549 | NA | 32 | 18 | 22 | 22 | 8 | 12 |
| ENSDARG000 | 20;20 | 37610051:376 | 37610126:376 | -;- | 399 | NA | 0 | 0 | 0 | 0 | 1 | 0 |
| ENSDARG000 | 20;20;20;20;2 | 37495157:374 | 37495281:374 | +;+;+;+;+;+;+ | 1362 | NA | 2 | 0 | 0 | 0 | 0 | 0 |
| ENSDARG000 | 20;20;20;20 | 54383838:543 | 54384930:543 | +;+;+;+ | 3592 | NA | 0 | 0 | 0 | 2 | 0 | 0 |
| ENSDARG000 | 20;20;20;20;2 | 37511067:375 | 37513938:375 | -;-;-;-;-;-;-; | 10325 | hivep2a | 403 | 408 | 990 | 823 | 792 | 420 |
| ENSDARG000 | 20;20;20;20;2 | 54685143:546 | 54685148:546 | +;+;+;+;+;+;+; | 5580 | pak5 | 2 | 6 | 4 | 2 | 0 | 12 |
| ENSDARG000 | 20;20;20;20;2 | 54874121:549 | 54874238:549 | -;-;-;-;- | 499 | NA | 1 | 0 | 1 | 6 | 1 | 3 |
| ENSDARG000 | 20;20;20;20;2 | 54993357:549 | 54993475:549 | -;-;-;-;-;-;-;- | 1478 | NA | 0 | 0 | 0 | 0 | 0 | 0 |
| ENSDARG000 | 20 | 55169000 | 55169889 | - | 890 | NA | 11 | 17 | 54 | 13 | 3 | 45 |
| ENSDARG000 | 20;20;20;20;2 | 37661229:376 | 37661687:376 | +;+;+;+;+;+;+; | 1881 | AIG1 | 15 | 6 | 4 | 0 | 0 | 4 |
| ENSDARG000 | 20;20;20;20;2 | 37733181:377 | 37733931:377 | -;-;-;-;-;-;-; | 3055 | adat2 | 68 | 135 | 204 | 104 | 216 | 190 |
| ENSDARG000 | 20;20;20;20;2 | 39223235:392 | 39223403:392 | +;+;+;+;+;+;+; | 6020 | reps1 | 1718 | 1504 | 2726 | 2214 | 1940 | 1743 |
| ENSDARG000 | 20;20;20;20;2 | 39260475:392 | 39260896:392 | -;-;-;-;-;-;-; | 1955 | clu | 571 | 555 | 297 | 302 | 543 | 800 |
| ENSDARG000 | 20;20;20;20;2 | 37760493:377 | 37760711:377 | +;+;+;+;+;+;+; | 2074 | pex3 | 270 | 302 | 438 | 224 | 534 | 422 |
| ENSDARG000 | 20;20;20;20;2 | 39283849:392 | 39284113:392 | +;+;+;+;+;+;+; | 3240 | SCARA3 | 62 | 6 | 8 | 2 | 62 | 12 |
| ENSDARG000 | 20;20;20;20;2 | 39332579:393 | 39332924:393 | -;-;-;-;-;-;- | 679 | ccl38.1 | 0 | 20 | 0 | 0 | 16 | 84 |
| ENSDARG000 | 20;20;20;20;2 | 39338700:393 | 39339177:393 | -;-;-;-;-;-;-; | 1390 | ccdc25 | 815 | 781 | 896 | 507 | 785 | 788 |
| ENSDARG000 | 20;20;20;20 | 37794633:377 | 37794768:377 | +;+;+;+ | 2701 | atf3 | 2522 | 1773 | 4198 | 4513 | 751 | 1919 |
| ENSDARG000 | 20;20;20 | 37812699:378 | 37813035:378 | -;-;- | 716 | batf3 | 16 | 18 | 34 | 16 | 15 | 38 |
| ENSDARG000 | 20;20;20;20;2 | 37814345:378 | 37815014:378 | -;-;-;-;-;- | 1201 | nsl1 | 6 | 168 | 114 | 64 | 74 | 80 |
| ENSDARG000 | 20;20;20;20;2 | 37820990:378 | 37821180:378 | +;+;+;+;+;+;+; | 1352 | tatdn3 | 124 | 102 | 109 | 21 | 112 | 109 |
| ENSDARG000 | 20;20;20;20;2 | 39344889:393 | 39345024:393 | +;+;+;+;+;+;+; | 2339 | esco2 | 110 | 554 | 447 | 111 | 354 | 168 |
| ENSDARG000 | 20;20;20;20;2 | 39359729:393 | 39360576:393 | -;-;-;-;-;-;-; | 1960 | pbk | 236 | 571 | 466 | 252 | 388 | 286 |
| ENSDARG000 | 20;20;20;20;2 | 39373480:393 | 39373705:393 | +;+;+;+;+;+;+; | 1638 | NA | 0 | 2 | 0 | 2 | 0 | 0 |
| ENSDARG000 | 20;20;20;20;2 | 39389777:393 | 39389838:393 | -;-;-;-;-;-;-; | 685 | si:dkev-217m5 | 0 | 0 | 0 | 0 | 0 | 0 |
| ENSDARG000 | 20;20;20;20;2 | 39394308:393 | 39394536:393 | +;+;+;+;+;+ | 1676 | pimr136 | 3 | 1 | 0 | 0 | 0 | 0 |
| ENSDARG000 | 20;20;20;20;2 | 39410844:394 | 39411040:394 | -;-;-;-;-;-;-; | 977 | NA | 0 | 0 | 0 | 0 | 0 | 2 |
| ENSDARG000 | 20;20;20;20;2 | 37831352:378 | 37831497:378 | -;-;-;-;- | 501 | NA | 21 | 0 | 33 | 44 | 13 | 10 |
| ENSDARG000 | 20;20;20 | 39419083:394 | 39419174:394 | -;-;- | 279 | NA | 9 | 6 | 0 | 2 | 3 | 4 |
| ENSDARG000 | 20;20;20;20;2 | 37844035:378 | 37844679:378 | +;+;+;+;+;+;+; | 2644 | flvcr1 | 153 | 238 | 311 | 346 | 65 | 100 |
| ENSDARG000 | 20;20;20;20;2 | 39416729:394 | 39416957:394 | +;+;+;+;+;+;+; | 1964 | NA | 64 | 8 | 0 | 10 | 30 | 12 |
| ENSDARG000 | 20;20;20;20;2 | 39457598:394 | 39457841:394 | +;+;+;+;+;+;+; | 2841 | pimr128 | 24 | 2 | 3 | 3 | 8 | 10 |
| ENSDARG000 | 20;20;20;20;2 | 39458596:394 | 39460017:394 | -;-;-;-;- | 2004 | NA | 25 | 2 | 8 | 0 | 14 | 5 |
| ENSDARG000 | 20;20;20 | 39493111:394 | 39493165:394 | -;-;- | 242 | NA | 3 | 0 | 1 | 0 | 0 | 0 |
| ENSDARG000 | 20;20;20;20;2 | 39489931:394 | 39490062:394 | +;+;+;+;+;+;+; | 2296 | NA | 26 | 4 | 1 | 10 | 4 | 4 |
| ENSDARG000 | 20;20;20;20;2 | 37866171:378 | 37867000:378 | +;+;+;+;+;+;+; | 5557 | VASH2 | 1777 | 2203 | 1341 | 1744 | 960 | 2333 |
| ENSDARG000 | 20;20;20;20;2 | 39520628:395 | 39521030:395 | -;-;-;-;-;-;-; | 1516 | pex7 | 411 | 208 | 202 | 124 | 187 | 190 |
| ENSDARG000 | 20;20;20;20;2 | 39588897:395 | 39590212:395 | -;-;-;-;- | 1816 | hey2 | 6 | 0 | 0 | 0 | 6 | 0 |
| ENSDARG000 | 20;20;20;20;2 | 39685572:396 | 39685640:396 | +;+;+;+;+;+;+; | 884 | hddc2 | 87 | 78 | 122 | 54 | 131 | 78 |
| ENSDARG000 | 20;20;20;20;2 | 39601856:396 | 39601977:396 | -;-;-;-;-;-;-; | 1500 | NA | 0 | 0 | 4 | 0 | 2 | 0 |
| ENSDARG000 | 20;20;20;20;2 | 37920096:379 | 37920835:379 | -;-;-;-;-;-;-; | 2853 | angel2 | 186 | 187 | 176 | 169 | 126 | 185 |
| ENSDARG000 | 20;20;20;20;2 | 39691793:396 | 39693275:396 | -;-;-;-;-;-;-; | 2184 | tpd52l1 | 481 | 700 | 452 | 312 | 374 | 648 |
| ENSDARG000 | 20;20;20;20;2 | 39749448:397 | 39750052:397 | -;-;-;-;-;-;-; | 3392 | rnf217 | 103 | 45 | 55 | 59 | 125 | 104 |
| ENSDARG000 | 20;20;20;20;2 | 38031439:380 | 38031576:380 | +;+;+;+;+;+;+; | 2763 | galnt14 | 32 | 14 | 153 | 9 | 40 | 34 |
| ENSDARG000 | 20;20;20;20;2 | 39806370:398 | 39807610:398 | -;-;-;-;-;-;-; | 2640 | nkain2 | 52 | 38 | 16 | 26 | 2 | 32 |
| ENSDARG000 | 20;20 | 39919232:399 | 39919703:399 | -;- | 575 | NA | 2 | 0 | 0 | 0 | 0 | 0 |
| ENSDARG000 | 20;20 | 39920443:399 | 39920476:399 | +;+ | 326 | NA | 0 | 0 | 0 | 0 | 0 | 0 |
| ENSDARG000 | 20;20;20;20;2 | 38201644:382 | 38202389:382 | +;+;+;+;+;+ | 2680 | ehd3 | 122 | 119 | 162 | 42 | 205 | 311 |
| ENSDARG000 | 20;20;20;20;2 | 40150612:401 | 40150946:401 | +;+;+;+;+;+;+; | 3186 | NA | 0 | 0 | 0 | 0 | 0 | 0 |
| ENSDARG000 | 20;20;20;20;2 | 38257766:382 | 38258029:382 | +;+;+;+;+;+;+; | 1716 | ccl38a.3 | 16 | 12 | 0 | 29 | 10 | 6 |
| ENSDARG000 | 20;20;20 | 38279523:382 | 38279612:382 | +;+;+ | 532 | ccl38a.5 | 2 | 0 | 4 | 8 | 28 | 0 |
| ENSDARG000 | 20;20;20 | 38285671:382 | 38285877:382 | +;+;+ | 643 | ccl38a.4 | 0 | 0 | 0 | 3 | 14 | 4 |
| ENSDARG000 | 20;20;20;20;2 | 40237441:402 | 40237502:402 | +;+;+;+;+ | 6120 | NA | 0 | 0 | 0 | 0 | 0 | 0 |

| | | | | | | | | | | | | |
|---|---|---|---|---|---|---|---|---|---|---|---|---|
| ENSDARG000 | 20;20;20;20;2 | 40269891:402 | 40270080:402 | -;-;-;-;- | 1514 | clvs2 | 0 | 2 | 0 | 10 | 0 | 0 |
| ENSDARG000 | 20;20;20;20;2 | 40337423:403 | 40337659:403 | -;-;-;-;-;-;-;-;-; | 1639 | smpdl3a | 191 | 74 | 43 | 16 | 124 | 95 |
| ENSDARG000 | 20;20;20;20 | 40369257:403 | 40369521:403 | -;-;-;- | 673 | fabp7b | 183 | 34 | 8 | 6 | 110 | 24 |
| ENSDARG000 | 20;20;20 | 40371897:403 | 40373001:403 | -;-;- | 1409 | pkib | 425 | 36 | 22 | 66 | 0 | 12 |
| ENSDARG000 | 20;20;20;20;2 | 40457599:404 | 40457724:404 | +;+;+;+;+;+;+; | 2885 | serinc1 | 9541 | 4951 | 4257 | 5167 | 5495 | 6193 |
| ENSDARG000 | 20;20;20;20;2 | 40475576:404 | 40476290:404 | -;-;-;-;-;-;-;-;-; | 2239 | hsf2 | 1258 | 766 | 757 | 498 | 1301 | 934 |
| ENSDARG000 | 20;20;20 | 40488776:404 | 40488852:404 | +;+;+ | 370 | NA | 0 | 0 | 0 | 0 | 0 | 0 |
| ENSDARG000 | 20;20;20;20;2 | 40715163:407 | 40717741:407 | -;-;-;-;-;-;-;-;-; | 2926 | gja1b | 409 | 206 | 642 | 183 | 629 | 277 |
| ENSDARG000 | 20;20;20 | 40724207:407 | 40725142:407 | -;-;- | 1146 | gja13.2 | 127 | 133 | 200 | 88 | 250 | 274 |
| ENSDARG000 | 20;20;20 | 40757141:407 | 40758404:407 | -;-;- | 1334 | gja11 | 0 | 0 | 14 | 34 | 0 | 0 |
| ENSDARG000 | 20;20;20;20;2 | 40766645:407 | 40766769:407 | +;+;+;+;+;+;+; | 4193 | tbc1d32 | 129 | 116 | 246 | 178 | 184 | 188 |
| ENSDARG000 | 20;20;20 | 40753518:407 | 40754791:407 | -;-;- | 1343 | gja13.1 | 0 | 6 | 73 | 16 | 66 | 0 |
| ENSDARG000 | 20;20;20;20;2 | 40739964:407 | 40740707:407 | +;+;+;+;+ | 805 | NA | 16 | 16 | 50 | 14 | 59 | 30 |
| ENSDARG000 | 20;20;20 | 40744754:407 | 40747114:407 | -;-;- | 2480 | gja12.1 | 0 | 0 | 14 | 12 | 56 | 0 |
| ENSDARG000 | 20;20;20;20;2 | 40761717:407 | 40762217:407 | -;-;-;-;-;-;-;-;-; | 1740 | hsdl1 | 179 | 210 | 276 | 207 | 267 | 193 |
| ENSDARG000 | 20;20;20 | 172162:18403 | 172212:18404 | +;+;+ | 93 | NA | 0 | 1 | 0 | 0 | 0 | 0 |
| ENSDARG000 | 20;20;20;20;2 | 15083987:150 | 15085434:150 | -;-;-;-;-;-;-;-;-; | 2768 | si:dkev-239i2( | 13 | 9 | 15 | 13 | 23 | 10 |
| ENSDARG000 | 20;20;20;20;2 | 50650143:506 | 50650345:506 | +;+;+;+;+;+;+; | 1740 | FUT8 | 0 | 0 | 0 | 0 | 0 | 0 |
| ENSDARG000 | 20;20;20;20 | 49719141:497 | 49719314:497 | -;-;-;- | 3258 | FILIP1 | 15 | 4 | 6 | 0 | 26 | 6 |
| ENSDARG000 | 20;20;20;20;2 | 47994741:479 | 47995503:479 | -;-;-;-;-;- | 2433 | GAREM2 | 0 | 0 | 6 | 0 | 0 | 0 |
| ENSDARG000 | 20 | 19543379 | 19543462 | - | 84 | dre-mir-133b | 0 | 0 | 0 | 0 | 0 | 0 |
| ENSDARG000 | 20;20;20;20;2 | 48452655:484 | 48453022:484 | -;-;-;-;-;-;-;-;-; | 1658 | NA | 24 | 48 | 112 | 41 | 68 | 58 |
| ENSDARG000 | 20 | 4051752 | 4054045 | + | 2294 | exoc8 | 542 | 908 | 791 | 358 | 942 | 837 |
| ENSDARG000 | 20;20;20;20;2 | 42780162:427 | 42780297:427 | +;+;+;+;+ | 1848 | pimr213 | 0 | 0 | 0 | 0 | 0 | 0 |
| ENSDARG000 | 20;20;20;20;2 | 5302186:5306 | 5302703:5306 | -;-;-;-;-;- | 1314 | ism2b | 4 | 0 | 0 | 4 | 0 | 0 |
| ENSDARG000 | 20;20;20;20;2 | 28511941:285 | 28511992:285 | -;-;-;-;-;-;-;-;-; | 1945 | rgs6 | 4 | 0 | 0 | 0 | 0 | 0 |
| ENSDARG000 | 20 | 26119339 | 26119459 | - | 121 | 5S_rRNA | 0 | 0 | 0 | 0 | 0 | 0 |
| ENSDARG000 | 20;20;20;20;2 | 5330540:5336 | 5333710:5337 | -;-;-;-;-;-;-;-;-; | 4767 | sptlc2b | 1292 | 1813 | 2230 | 929 | 1827 | 944 |
| ENSDARG000 | 20;20;20;20;2 | 28418484:284 | 28423446:284 | -;-;-;-;-;-;-;-;-; | 6397 | wdr21 | 393 | 271 | 392 | 361 | 330 | 437 |
| ENSDARG000 | 20 | 35422698 | 35422827 | + | 130 | 5S_rRNA | 0 | 0 | 0 | 0 | 0 | 0 |
| ENSDARG000 | 20 | 10489621 | 10489890 | + | 270 | NA | 2 | 1 | 9 | 0 | 1 | 6 |
| ENSDARG000 | 20 | 3322604 | 3322820 | - | 217 | NA | 0 | 0 | 0 | 0 | 0 | 0 |
| ENSDARG000 | 20 | 8936903 | 8937004 | - | 102 | U6 | 0 | 0 | 0 | 0 | 0 | 0 |
| ENSDARG000 | 20;20;20;20;2 | 53160860:531 | 53160871:531 | +;+;+;+;+;+;+ | 132 | NA | 0 | 0 | 0 | 0 | 0 | 0 |
| ENSDARG000 | 20;20;20;20 | 39513435:395 | 39513707:395 | -;-;-;- | 1218 | SLC35D3 | 4033 | 36 | 83 | 58 | 35 | 64 |
| ENSDARG000 | 20 | 41352846 | 41352960 | - | 115 | 5S_rRNA | 0 | 0 | 0 | 0 | 0 | 0 |
| ENSDARG000 | 20;20 | 45404656:454 | 45409320:454 | -;- | 4991 | TRIB2 | 245 | 111 | 239 | 74 | 200 | 81 |
| ENSDARG000 | 20 | 47104710 | 47104873 | + | 164 | U1 | 0 | 0 | 0 | 0 | 0 | 0 |
| ENSDARG000 | 20 | 28708907 | 28709023 | + | 117 | 5S_rRNA | 0 | 0 | 0 | 0 | 0 | 0 |
| ENSDARG000 | 20;20;20;20;2 | 54512067:545 | 54512425:545 | -;-;-;-;-;- | 1059 | ppp1r35 | 43 | 53 | 82 | 42 | 106 | 56 |
| ENSDARG000 | 20;20 | 54196543:541 | 54197897:541 | -;- | 1443 | arf6a | 1348 | 1841 | 1989 | 1265 | 1483 | 1147 |
| ENSDARG000 | 20;20 | 54525404:545 | 54525446:545 | -;- | 699 | zgc:153012 | 450 | 216 | 427 | 506 | 415 | 330 |
| ENSDARG000 | 20;20;20;20;2 | 47434709:474 | 47435298:474 | +;+;+;+;+;+ | 1294 | rab10 | 1532 | 2371 | 2085 | 1400 | 1641 | 1755 |
| ENSDARG000 | 20 | 27338006 | 27338121 | + | 116 | 5S_rRNA | 0 | 0 | 0 | 0 | 0 | 0 |
| ENSDARG000 | 20 | 47407150 | 47407264 | + | 115 | 5S_rRNA | 0 | 0 | 0 | 0 | 0 | 0 |
| ENSDARG000 | 20 | 16427790 | 16427905 | + | 116 | U5 | 0 | 0 | 0 | 0 | 0 | 0 |
| ENSDARG000 | 20 | 42405098 | 42405212 | - | 115 | 5S_rRNA | 0 | 0 | 0 | 0 | 0 | 0 |
| ENSDARG000 | 20 | 48613321 | 48613436 | + | 116 | U5 | 0 | 0 | 0 | 0 | 0 | 0 |
| ENSDARG000 | 20 | 17784447 | 17784568 | - | 122 | 5S_rRNA | 0 | 0 | 0 | 0 | 0 | 0 |
| ENSDARG000 | 20;20;20;20;2 | 19494273:194 | 19494282:194 | -;-;-;-;-;-;-;-;-; | 339 | NA | 4 | 0 | 3 | 0 | 0 | 2 |
| ENSDARG000 | 20;20;20;20;2 | 6308564:6312 | 6308798:6312 | -;-;-;-;-;-;-;-;-; | 4437 | TRAPPC8 | 1120 | 704 | 981 | 498 | 1039 | 770 |
| ENSDARG000 | 20;20;20;20;2 | 54174978:541 | 54175242:541 | -;-;-;-;-;-;- | 1239 | NA | 1569 | 682 | 1974 | 910 | 1524 | 796 |
| ENSDARG000 | 20;20;20;20 | 18974990:189 | 18975023:189 | -;-;-;- | 87 | NA | 0 | 3 | 2 | 6 | 5 | 30 |
| ENSDARG000 | 20 | 6919748 | 6919864 | + | 117 | 5S_rRNA | 0 | 0 | 0 | 0 | 0 | 0 |
| ENSDARG000 | 20;20;20;20;2 | 48413712:484 | 48414067:484 | +;+;+;+;+ | 1257 | NA | 1438 | 1791 | 1425 | 1210 | 2691 | 1307 |
| ENSDARG000 | 20 | 9299434 | 9299549 | + | 116 | 5S_rRNA | 0 | 0 | 0 | 0 | 0 | 0 |
| ENSDARG000 | 20 | 3322605 | 3322820 | - | 216 | U3 | 0 | 0 | 0 | 0 | 0 | 0 |
| ENSDARG000 | 20 | 28478203 | 28478318 | + | 116 | 5S_rRNA | 0 | 0 | 2 | 0 | 0 | 0 |
| ENSDARG000 | 20 | 9906268 | 9906384 | - | 117 | 5S_rRNA | 0 | 0 | 0 | 0 | 0 | 0 |
| ENSDARG000 | 20 | 16671441 | 16671555 | + | 115 | 5S_rRNA | 0 | 0 | 0 | 0 | 0 | 0 |
| ENSDARG000 | 20 | 35510865 | 35510980 | - | 116 | 5S_rRNA | 0 | 0 | 0 | 0 | 0 | 0 |
| ENSDARG000 | 20 | 5074924 | 5075054 | + | 131 | dre-mir-7146 | 0 | 0 | 0 | 0 | 0 | 0 |
| ENSDARG000 | 20;20;20;20;2 | 22869608:228 | 22869647:228 | -;-;-;-;-;-;-;-;-; | 1248 | NA | 2 | 1 | 5 | 3 | 0 | 1 |
| ENSDARG000 | 20;20;20;20;2 | 42761881:427 | 42762016:427 | +;+;+;+;+;+ | 1722 | NA | 0 | 0 | 0 | 0 | 0 | 0 |
| ENSDARG000 | 20;20;20;20;2 | 15164482:151 | 15164606:151 | +;+;+;+;+;+;+; | 1050 | prdx6 | 4389 | 7590 | 6831 | 5590 | 7317 | 6136 |
| ENSDARG000 | 20;20;20;20;2 | 5237286:5239 | 5237682:5239 | -;-;-;-;-;-;-;-;-; | 1829 | cyp46a1.2 | 72 | 75 | 117 | 111 | 156 | 224 |
| ENSDARG000 | 20 | 7341078 | 7341194 | + | 117 | 5S_rRNA | 7 | 0 | 0 | 0 | 0 | 0 |
| ENSDARG000 | 20;20;20 | 53169969:531 | 53172077:531 | -;-;- | 2383 | NA | 398 | 430 | 562 | 224 | 332 | 238 |
| ENSDARG000 | 20 | 1846081 | 1847433 | - | 1353 | NA | 0 | 0 | 0 | 0 | 0 | 0 |
| ENSDARG000 | 20;20;20;20 | 44250691:442 | 44250760:442 | +;+;+;+ | 1008 | opn8a | 0 | 0 | 0 | 0 | 0 | 0 |
| ENSDARG000 | 20;20;20;20;2 | 29514914:295 | 29514919:295 | +;+;+;+;+;+;+; | 1716 | NA | 2 | 4 | 10 | 0 | 7 | 0 |
| ENSDARG000 | 20;20;20;20;2 | 54213688:542 | 54215312:542 | -;-;-;-;-;-;-;-;-; | 3797 | prpf39 | 2494 | 2981 | 4683 | 2539 | 4075 | 2730 |
| ENSDARG000 | 20;20;20;20;2 | 3828237:3832 | 3828651:3832 | -;-;-;-;-;-;-;-;-; | 1911 | cnksr3 | 32 | 12 | 14 | 10 | 22 | 24 |
| ENSDARG000 | 20 | 50351539 | 50351653 | - | 115 | 5S_rRNA | 0 | 0 | 0 | 0 | 0 | 0 |
| ENSDARG000 | 20 | 23232690 | 23232804 | + | 115 | U5 | 0 | 0 | 0 | 0 | 0 | 0 |
| ENSDARG000 | 20;20;20 | 1607457:1607 | 1607467:1607 | -;-;- | 414 | NA | 191 | 72 | 224 | 149 | 286 | 118 |
| ENSDARG000 | 20;20 | 54506428:545 | 54506949:545 | -;- | 795 | NA | 2 | 4 | 10 | 7 | 8 | 2 |
| ENSDARG000 | 20 | 44139745 | 44139861 | + | 117 | 5S_rRNA | 0 | 0 | 0 | 0 | 0 | 0 |
| ENSDARG000 | 20;20;20;20;2 | 47927720:479 | 47932548:479 | -;-;-;-;-;-;-;-;-; | 6981 | hadhb | 1041 | 1206 | 1562 | 898 | 1400 | 1054 |
| ENSDARG000 | 20;20;20;20;2 | 47686629:477 | 47690602:477 | -;-;-;-;-;-;- | 4999 | tfap2b | 12 | 4 | 12 | 2 | 2 | 0 |
| ENSDARG000 | 20;20;20;20;2 | 15093609:150 | 15094449:150 | -;-;-;-;-;-;-;-;-; | 2207 | fmo5 | 147 | 39 | 78 | 115 | 142 | 130 |
| ENSDARG000 | 20 | 23076201 | 23076317 | + | 117 | 5S_rRNA | 0 | 0 | 0 | 0 | 0 | 0 |
| ENSDARG000 | 20 | 41228064 | 41228179 | - | 116 | 5S_rRNA | 0 | 0 | 0 | 0 | 0 | 0 |
| ENSDARG000 | 20;20;20;20;2 | 5985329:5987 | 5986233:5987 | +;+;+;+;+;+;+; | 2587 | cep128 | 4 | 18 | 12 | 2 | 24 | 8 |
| ENSDARG000 | 20 | 44935039 | 44935155 | + | 117 | 5S_rRNA | 0 | 0 | 0 | 0 | 0 | 0 |
| ENSDARG000 | 20 | 23081660 | 23081796 | - | 137 | NA | 0 | 0 | 5 | 2 | 0 | 0 |
| ENSDARG000 | 20 | 49199293 | 49199484 | - | 192 | NA | 0 | 0 | 0 | 0 | 0 | 0 |
| ENSDARG000 | 20;20 | 50048836:500 | 50048872:500 | -;- | 177 | NDUFB1 | 214 | 319 | 146 | 138 | 124 | 218 |
| ENSDARG000 | 20;20;20;20;2 | 2099243:2099 | 2099271:2099 | +;+;+;+;+;+;+ | 147 | NA | 0 | 0 | 0 | 0 | 0 | 0 |
| ENSDARG000 | 20 | 42967274 | 42967388 | - | 115 | 5S_rRNA | 0 | 0 | 0 | 0 | 0 | 0 |
| ENSDARG000 | 20 | 3323566 | 3323782 | - | 217 | NA | 0 | 0 | 0 | 0 | 0 | 0 |
| ENSDARG000 | 20;20;20;20;2 | 5378841:5382 | 5381223:5382 | -;-;-;-;-;-;-;-;-; | 4406 | snw1 | 1414 | 1356 | 1845 | 1157 | 1608 | 1079 |
| ENSDARG000 | 20;20;20;20;2 | 42918755:429 | 42918764:429 | +;+;+;+;+;+;+; | 2454 | EFR3B | 140 | 97 | 120 | 40 | 225 | 116 |
| ENSDARG000 | 20;20;20 | 45620076:456 | 45620127:456 | +;+;+ | 853 | NA | 33 | 54 | 59 | 36 | 60 | 65 |
| ENSDARG000 | 20 | 3327390 | 3327605 | - | 216 | U3 | 0 | 0 | 0 | 0 | 0 | 0 |
| ENSDARG000 | 20;20;20;20;2 | 54419592:544 | 54423440:544 | -;-;-;-;- | 4731 | yy1b | 475 | 590 | 654 | 402 | 625 | 559 |
| ENSDARG000 | 20;20;20;20;2 | 28434196:284 | 28434701:284 | +;+;+;+;+;+;+; | 1795 | dpf3 | 89 | 57 | 109 | 56 | 154 | 88 |
| ENSDARG000 | 20 | 47106094 | 47106257 | + | 164 | U1 | 0 | 0 | 0 | 0 | 0 | 0 |
| ENSDARG000 | 20 | 22910198 | 22910312 | - | 115 | 5S_rRNA | 0 | 0 | 0 | 0 | 0 | 0 |
| ENSDARG000 | 20 | 30373865 | 30374084 | - | 220 | SNORA73 | 18 | 2 | 3 | 20 | 3 | 4 |
| ENSDARG000 | 20;20 | 31985461:319 | 31985645:319 | +;+ | 234 | NA | 0 | 0 | 0 | 0 | 0 | 0 |
| ENSDARG000 | 20;20;20;20;2 | 2648405:2652 | 2648761:2652 | -;-;-;-;-;-;-;-;-; | 2079 | cfap206 | 0 | 0 | 0 | 14 | 6 | 0 |
| ENSDARG000 | 20 | 52827010 | 52827126 | - | 117 | 5S_rRNA | 0 | 0 | 0 | 0 | 0 | 0 |
| ENSDARG000 | 20 | 38905531 | 38905613 | - | 83 | dre-mir-124-2 | 0 | 0 | 0 | 0 | 0 | 0 |
| ENSDARG000 | 20 | 3325446 | 3325661 | - | 216 | U3 | 0 | 0 | 0 | 0 | 0 | 0 |
| ENSDARG000 | 20 | 27054095 | 27054220 | - | 126 | 5S_rRNA | 0 | 0 | 2 | 0 | 0 | 0 |
| ENSDARG000 | 20;20;20;20;2 | 15023263:150 | 15023511:150 | -;-;-;-;-;-;-;-;-; | 8721 | PRRC2C | 2351 | 1824 | 4133 | 1294 | 3399 | 1766 |
| ENSDARG000 | 20;20;20;20;2 | 28399593:284 | 28401938:284 | -;-;-;-;-;-;-;-;-; | 3101 | pigh | 246 | 282 | 324 | 371 | 362 | 370 |
| ENSDARG000 | 20;20;20;20 | 15600167:156 | 15600763:156 | +;+;+;+ | 1570 | faslg | 0 | 18 | 0 | 0 | 0 | 18 |
| ENSDARG000 | 20 | 21332014 | 21332128 | - | 115 | 5S_rRNA | 0 | 0 | 0 | 0 | 0 | 0 |
| ENSDARG000 | 20;20 | 4026408:4030 | 4026683:4031 | -;- | 1011 | fam89a | 149 | 217 | 130 | 93 | 151 | 119 |
| ENSDARG000 | 20 | 3329513 | 3329727 | - | 215 | U3 | 0 | 0 | 0 | 0 | 0 | 0 |
| ENSDARG000 | 20;20;20;20;2 | 48100261:481 | 48100486:481 | +;+;+;+;+ | 1464 | xkr5a | 29 | 19 | 35 | 13 | 16 | 29 |
| ENSDARG000 | 20 | 53278746 | 53279747 | + | 1002 | GPR6 | 0 | 12 | 0 | 0 | 0 | 2 |
| ENSDARG000 | 20;20;20;20 | 3916406:3918 | 3916741:3918 | +;+;+;+ | 4934 | C20H1orf198 | 1415 | 1411 | 1630 | 1182 | 1879 | 1281 |
| ENSDARG000 | 20 | 32460275 | 32460391 | - | 117 | 5S_rRNA | 0 | 0 | 0 | 0 | 0 | 0 |
| ENSDARG000 | 20 | 30372663 | 30372883 | - | 221 | SNORA73 | 0 | 7 | 19 | 12 | 30 | 14 |
| ENSDARG000 | 20;20 | 484256:48766 | 486487:48778 | -;- | 2356 | COL10A1 | 398 | 66 | 89 | 18 | 386 | 10 |
| ENSDARG000 | 20;20;20;20;2 | 661037:66682 | 661075:66753 | +;+;+;+;+ | 1464 | nrl | 0 | 0 | 0 | 0 | 0 | 0 |
| ENSDARG000 | 20 | 31045075 | 31045205 | - | 131 | 5S_rRNA | 0 | 0 | 0 | 0 | 0 | 0 |
| ENSDARG000 | 20 | 31387055 | 31387171 | + | 117 | 5S_rRNA | 0 | 0 | 0 | 0 | 0 | 0 |
| ENSDARG000 | 20 | 24435671 | 24443680 | - | 8010 | NA | 0 | 0 | 6 | 18 | 12 | 4 |
| ENSDARG000 | 20;20;20;20;2 | 4334008:4337 | 4334133:4337 | +;+;+;+;+;+;+; | 1386 | NA | 50 | 6 | 35 | 4 | 101 | 12 |
| ENSDARG000 | 20 | 3334543 | 3334758 | - | 216 | U3 | 0 | 0 | 0 | 0 | 0 | 0 |
| ENSDARG000 | 20 | 4405937 | 4407228 | - | 1292 | NA | 0 | 0 | 0 | 0 | 0 | 0 |
| ENSDARG000 | 20;20;20;20;2 | 2039518:2042 | 2039653:2042 | +;+;+;+;+;+;+; | 1581 | NA | 141 | 166 | 131 | 67 | 118 | 118 |
| ENSDARG000 | 20;20 | 51647431:516 | 51650044:516 | -;- | 2852 | NA | 172 | 184 | 198 | 176 | 174 | 172 |
| ENSDARG000 | 20;20;20;20;2 | 47383841:473 | 47383955:473 | -;-;-;-;-;-;- | 2265 | KIF3C | 10 | 2 | 9 | 9 | 8 | 4 |
| ENSDARG000 | 20;20;20;20;2 | 5132138:5132 | 5132344:5132 | +;+;+;+;+;+;+; | 1931 | cyp46a1.5 | 4 | 7 | 7 | 5 | 6 | 0 |
| ENSDARG000 | 20 | 39331969 | 39332092 | + | 124 | 5S_rRNA | 0 | 0 | 0 | 0 | 0 | 0 |
| ENSDARG000 | 20 | 2908551 | 2908665 | + | 115 | 5S_rRNA | 0 | 0 | 0 | 0 | 0 | 0 |
| ENSDARG000 | 20 | 3328354 | 3328569 | - | 216 | U3 | 0 | 0 | 0 | 0 | 0 | 0 |
| ENSDARG000 | 20 | 48612257 | 48612370 | + | 114 | U5 | 0 | 0 | 0 | 0 | 0 | 0 |
| ENSDARG000 | 20;20;20;20;2 | 47717389:477 | 47718687:477 | -;-;-;-;-;-;-;-;-; | 2701 | tfap2d | 54 | 0 | 3 | 54 | 51 | 68 |
| ENSDARG000 | 20 | 5613879 | 5613996 | - | 118 | 5S_rRNA | 0 | 0 | 0 | 0 | 0 | 0 |
| ENSDARG000 | 20 | 51516202 | 51516318 | + | 117 | 5S_rRNA | 0 | 0 | 0 | 0 | 0 | 0 |
| ENSDARG000 | 20 | 50361059 | 50361175 | - | 117 | 5S_rRNA | 0 | 0 | 0 | 0 | 0 | 0 |
| ENSDARG000 | 20 | 4126140 | 4126254 | - | 115 | 5S_rRNA | 0 | 0 | 0 | 0 | 0 | 0 |
| ENSDARG000 | 20;20;20;20;2 | 53181017:531 | 53181144:531 | +;+;+;+;+;+;+; | 1587 | NA | 357 | 452 | 417 | 259 | 437 | 276 |
| ENSDARG000 | 20;20;20;20;2 | 5158419:5158 | 5159066:5159 | -;-;-;-;-;-;-;-;-; | 2136 | ccdc85cb | 92 | 54 | 54 | 92 | 121 | 66 |
| ENSDARG000 | 20;20;20;20;2 | 54437053:544 | 54437090:544 | -;-;-;-;-;-;-;-;-; | 1111 | evlb | 12 | 18 | 23 | 24 | 7 | 35 |
| ENSDARG000 | 20;20;20 | 4392687:4413 | 4393748:4414 | +;+;+ | 3383 | NA | 59 | 6 | 0 | 0 | 10 | 26 |
| ENSDARG000 | 20;20;20;20;2 | 48292634:482 | 48292730:482 | -;-;-;-;-;-;-;-;-; | 2862 | IPO13 | 350 | 276 | 589 | 137 | 550 | 214 |
| ENSDARG000 | 20 | 48610352 | 48610467 | + | 116 | U5 | 0 | 0 | 0 | 0 | 0 | 0 |
| ENSDARG000 | 20;20;20 | 47491247:475 | 47491417:475 | +;+;+ | 633 | lin28b | 0 | 4 | 0 | 4 | 0 | 4 |
| ENSDARG000 | 20 | 48607137 | 48607244 | - | 108 | U5 | 0 | 0 | 0 | 0 | 0 | 0 |
| ENSDARG000 | 20 | 3332612 | 3332826 | - | 215 | U3 | 0 | 0 | 0 | 0 | 0 | 0 |
| ENSDARG000 | 20 | 33701117 | 33701233 | - | 117 | 5S_rRNA | 0 | 0 | 0 | 0 | 0 | 0 |
| ENSDARG000 | 20 | 21194649 | 21194764 | + | 116 | 5S_rRNA | 0 | 0 | 0 | 0 | 0 | 0 |
| ENSDARG000 | 20 | 16651125 | 16651223 | + | 99 | 5S_rRNA | 0 | 0 | 0 | 0 | 4 | 0 |
| ENSDARG000 | 20 | 17257101 | 17259573 | - | 2473 | NA | 15 | 0 | 22 | 2 | 4 | 22 |
| ENSDARG000 | 20 | 47077979 | 47078093 | - | 115 | 5S_rRNA | 0 | 0 | 0 | 0 | 0 | 0 |
| ENSDARG000 | 20 | 47982137 | 47982271 | + | 135 | 5S_rRNA | 0 | 0 | 0 | 0 | 0 | 0 |
| ENSDARG000 | 20 | 19544648 | 19544746 | - | 99 | dre-mir-206-1 | 0 | 0 | 0 | 0 | 0 | 0 |
| ENSDARG000 | 20 | 51211932 | 51212048 | + | 117 | 5S_rRNA | 0 | 0 | 0 | 0 | 0 | 0 |
| ENSDARG000 | 20 | 5321215 | 5321329 | - | 115 | 5S_rRNA | 0 | 0 | 0 | 0 | 0 | 0 |
| ENSDARG000 | 20 | 52144191 | 52144305 | - | 115 | 5S_rRNA | 0 | 0 | 0 | 0 | 0 | 0 |
| ENSDARG000 | 20;20;20 | 54323218:543 | 54323457:543 | +;+;+ | 381 | NA | 4 | 15 | 13 | 15 | 0 | 16 |
| ENSDARG000 | 20 | 3332611 | 3332826 | - | 216 | NA | 0 | 0 | 0 | 0 | 0 | 0 |
| ENSDARG000 | 20;20;20;20;2 | 8028613:8040 | 8028899:8040 | +;+;+;+;+;+;+; | 3579 | PRKAA2 | 40 | 12 | 20 | 14 | 30 | 8 |
| ENSDARG000 | 20;20;20;20;2 | 48543492:485 | 48543558:485 | +;+;+;+;+;+;+ | 1008 | NA | 1 | 0 | 0 | 2 | 0 | 0 |
| ENSDARG000 | 20;20;20;20 | 54201459:542 | 54201643:542 | -;-;-;- | 471 | faua | 7782 | 7929 | 12095 | 11009 | 7896 | 9023 |

| | | | | | | | | | | | |
|---|---|---|---|---|---|---|---|---|---|---|---|
| ENSDARG000 | 20 | 10053405 | 10053701 | - | 297 | Metazoa_SRP | 0 | 0 | 0 | 0 | 0 | 0 |
| ENSDARG000 | 20 | 23766953 | 23767069 | + | 117 | 5S_rRNA | 0 | 4 | 0 | 0 | 0 | 0 |
| ENSDARG000 | 20:20:20:20:2 | 3954579:3956 | 3955107:3956 | -:-:-:-:-:-:-:-:-: | 2924 | ttc13 | 515 | 336 | 430 | 330 | 697 | 413 |
| ENSDARG000 | 20 | 39032821 | 39032936 | - | 116 | 5S_rRNA | 0 | 0 | 0 | 0 | 0 | 0 |
| ENSDARG000 | 20:20:20:20:2 | 3997684:4000 | 3997779:4000 | +;+;+;+;+ | 3335 | arv1 | 210 | 266 | 287 | 215 | 293 | 441 |
| ENSDARG000 | 20 | 42748295 | 42750271 | - | 1977 | ttll2 | 0 | 0 | 0 | 0 | 0 | 0 |
| ENSDARG000 | 20:20:20:20:2 | 49787584:497 | 49787603:497 | +:+:+:+:+:+:+:+: | 1497 | HMBOX1 | 171 | 61 | 154 | 109 | 188 | 107 |
| ENSDARG000 | 20:20:20:20:2 | 48037019:480 | 48037791:480 | -:-:-:-:-:-:-:-:-: | 2696 | prep | 301 | 146 | 348 | 105 | 379 | 210 |
| ENSDARG000 | 20 | 38774972 | 38775087 | + | 116 | 5S_rRNA | 0 | 0 | 0 | 0 | 0 | 0 |
| ENSDARG000 | 20:20:20:20:2 | 3943237:3945 | 3943362:3945 | -:-:-:-:-:-:-:-:-: | 903 | selenol | 30 | 76 | 60 | 30 | 93 | 73 |
| ENSDARG000 | 20:20:20:20:2 | 4157815:4158 | 4157993:4158 | +:+:+:+:+:+:+:+: | 2162 | GNPAT | 312 | 347 | 357 | 224 | 377 | 366 |
| ENSDARG000 | 20 | 26594764 | 26594879 | + | 116 | 5S_rRNA | 0 | 0 | 0 | 0 | 0 | 0 |
| ENSDARG000 | 20:20:20:20:2 | 4221978:4222 | 4222092:4222 | +:+:+:+:+:+:+:+: | 3706 | ipcef1 | 170 | 160 | 113 | 156 | 212 | 144 |
| ENSDARG000 | 20 | 30374686 | 30374906 | - | 221 | SNORA73 | 2 | 12 | 24 | 6 | 24 | 11 |
| ENSDARG000 | 20;20;20 | 1960092:1961 | 1960232:1961 | +;+;+ | 399 | NA | 2 | 0 | 0 | 0 | 0 | 0 |
| ENSDARG000 | 20 | 19773360 | 19773475 | - | 116 | 5S_rRNA | 0 | 0 | 0 | 0 | 0 | 0 |
| ENSDARG000 | 20 | 5423962 | 5424096 | + | 135 | 5S_rRNA | 0 | 0 | 0 | 0 | 0 | 0 |
| ENSDARG000 | 20;20 | 48168809:481 | 48169185:481 | -;- | 423 | NA | 41 | 8 | 73 | 24 | 67 | 28 |
| ENSDARG000 | 20;20 | 3913646:3915 | 3913858:3915 | -;- | 402 | NA | 0 | 2 | 0 | 0 | 0 | 0 |
| ENSDARG000 | 20 | 47549572 | 47550577 | - | 1006 | NA | 0 | 0 | 0 | 0 | 0 | 0 |
| ENSDARG000 | 20:20:20:20:2 | 51274297:512 | 51274836:512 | -;-;-;-;-;- | 1755 | nfkbie | 262 | 250 | 214 | 88 | 247 | 236 |
| ENSDARG000 | 20 | 44198698 | 44198815 | + | 118 | 5S_rRNA | 0 | 0 | 0 | 0 | 0 | 0 |
| ENSDARG000 | 20;20 | 33610271:336 | 33610522:336 | +;+ | 460 | NA | 0 | 0 | 0 | 0 | 0 | 0 |
| ENSDARG000 | 20:20:20:20:2 | 6241855:6248 | 6242511:6249 | -;-;-;-;- | 1368 | opn7a | 0 | 0 | 0 | 0 | 0 | 0 |
| ENSDARG000 | 20 | 3324483 | 3324698 | - | 216 | U3 | 0 | 0 | 0 | 0 | 0 | 0 |
| ENSDARG000 | 20 | 27405878 | 27405962 | + | 85 | dre-mir-203b | 0 | 0 | 0 | 0 | 0 | 0 |
| ENSDARG000 | 20:20:20:20:2 | 47953047:479 | 47953129:479 | +:+:+:+:+:+:+:+: | 2758 | hadhaa | 49 | 148 | 287 | 128 | 186 | 96 |
| ENSDARG000 | 20 | 43082949 | 43083067 | - | 119 | 5S_rRNA | 0 | 0 | 0 | 0 | 0 | 0 |
| ENSDARG000 | 20 | 29495300 | 29495435 | - | 136 | NA | 0 | 0 | 2 | 0 | 1 | 0 |
| ENSDARG000 | 20 | 3323567 | 3323782 | - | 216 | U3 | 0 | 0 | 0 | 0 | 0 | 0 |
| ENSDARG000 | 20 | 13369227 | 13369343 | - | 117 | 5S_rRNA | 0 | 0 | 0 | 0 | 0 | 0 |
| ENSDARG000 | 20 | 32649843 | 32649959 | - | 117 | 5S_rRNA | 0 | 0 | 0 | 0 | 0 | 0 |
| ENSDARG000 | 20:20:20:20:2 | 4149523:4149 | 4149674:4150 | -;-;-;-;-;-;- | 827 | zgc:194224 | 86 | 99 | 122 | 82 | 146 | 128 |
| ENSDARG000 | 20:20:20:20:2 | 44799375:447 | 44799558:447 | -:-:-:-:-:-:-:-:-: | 2745 | KLHL29 | 28 | 72 | 59 | 12 | 14 | 4 |
| ENSDARG000 | 20;20;20 | 3934516:3935 | 3934657:3935 | +;+;+ | 732 | NA | 0 | 2 | 0 | 0 | 8 | 3 |
| ENSDARG000 | 20:20:20:20:2 | 1834262:1837 | 1834304:1837 | +:+:+:+:+:+:+:+: | 1938 | NA | 14 | 0 | 2 | 2 | 10 | 8 |
| ENSDARG000 | 20 | 17642809 | 17642923 | - | 115 | 5S_rRNA | 0 | 0 | 0 | 0 | 0 | 0 |
| ENSDARG000 | 20:20:20:20:2 | 4033193:4043 | 4038330:4043 | -;-;-;-;- | 5983 | sprtn | 220 | 178 | 189 | 155 | 256 | 218 |
| ENSDARG000 | 20 | 19217898 | 19218029 | + | 132 | 5S_rRNA | 0 | 0 | 0 | 0 | 0 | 0 |
| ENSDARG000 | 20:20:20:20:2 | 49816798:498 | 49817030:498 | -:-:-:-:-:-:-:-:-: | 6276 | KIF13B | 63 | 30 | 10 | 2 | 3 | 4 |
| ENSDARG000 | 20:20:20:20:2 | 53069721:530 | 53069853:530 | -:-:-:-:-:-:-:-:-: | 666 | NA | 41 | 13 | 70 | 86 | 63 | 78 |
| ENSDARG000 | 20:20:20:20:2 | 52825215:528 | 52825316:528 | -;-;-;-;-;-;-;-;- | 1303 | tbc1d7 | 338 | 355 | 448 | 375 | 444 | 489 |
| ENSDARG000 | 20;20;20;20 | 49697463:497 | 49697521:497 | -;-;-;- | 252 | COX7A2 | 188 | 235 | 477 | 398 | 252 | 272 |
| ENSDARG000 | 20:20:20:20:2 | 2064400:2066 | 2064528:2067 | -:-:-:-:-:-:-:-:-: | 1644 | ARHGAP18 | 66 | 24 | 42 | 23 | 44 | 47 |
| ENSDARG000 | 20:20:20:20:2 | 49895220:498 | 49895330:498 | +:+:+:+:+:+:+:+: | 2299 | ints9 | 240 | 195 | 350 | 166 | 281 | 194 |
| ENSDARG000 | 20:20:20:20:2 | 54666222:546 | 54666281:546 | +:+:+:+:+:+:+:+: | 3100 | NA | 317 | 165 | 276 | 250 | 416 | 234 |
| ENSDARG000 | 20 | 44912213 | 44912331 | - | 119 | 5S_rRNA | 0 | 0 | 0 | 0 | 0 | 0 |
| ENSDARG000 | 20 | 23081660 | 23081781 | - | 122 | NA | 0 | 0 | 0 | 0 | 0 | 0 |
| ENSDARG000 | 20 | 13818395 | 13818510 | - | 116 | 5S_rRNA | 0 | 0 | 0 | 0 | 0 | 0 |
| ENSDARG000 | 20 | 48611195 | 48611310 | + | 116 | U5 | 0 | 0 | 0 | 0 | 0 | 0 |
| ENSDARG000 | 20:20:20:20:2 | 28407720:284 | 28408205:284 | +:+:+:+:+:+:+:+: | 2340 | ZFYVE1 | 355 | 195 | 229 | 182 | 225 | 237 |
| ENSDARG000 | 20;20;20 | 46614805:466 | 46614922:466 | -;-;- | 315 | NA | 0 | 0 | 0 | 0 | 0 | 0 |
| ENSDARG000 | 20 | 42632985 | 42633105 | + | 121 | 5S_rRNA | 0 | 0 | 0 | 0 | 0 | 0 |
| ENSDARG000 | 20:20:20:20:2 | 15142972:151 | 15142977:151 | -;-;-;-;-;-;-;- | 1714 | plpp6 | 34 | 36 | 74 | 2 | 52 | 34 |
| ENSDARG000 | 20:20:20:20:2 | 4060839:4087 | 4061594:4087 | +:+:+:+:+:+:+:+: | 2094 | TRIM67 | 2 | 0 | 0 | 8 | 4 | 0 |
| ENSDARG000 | 20:20:20:20:2 | 44082532:440 | 44082578:440 | +;+;+;+;+;+ | 162 | NA | 0 | 0 | 0 | 0 | 0 | 0 |
| ENSDARG000 | 20:20:20:20:2 | 1996202:1999 | 1996314:1999 | +:+:+:+:+:+:+:+: | 2706 | NA | 165 | 2 | 18 | 59 | 88 | 12 |
| ENSDARG000 | 20:20:20:20:2 | 38892235:388 | 38892251:388 | -:-:-:-:-:-:-:-:-: | 219 | NA | 3 | 0 | 0 | 0 | 0 | 2 |
| ENSDARG000 | 20 | 19357658 | 19357772 | + | 115 | 5S_rRNA | 0 | 0 | 0 | 0 | 0 | 0 |
| ENSDARG000 | 20 | 26194726 | 26194841 | - | 116 | 5S_rRNA | 0 | 0 | 0 | 0 | 0 | 0 |
| ENSDARG000 | 20 | 3334542 | 3334758 | - | 217 | NA | 0 | 0 | 0 | 0 | 0 | 0 |
| ENSDARG000 | 20 | 36280048 | 36280164 | + | 117 | 5S_rRNA | 0 | 0 | 0 | 0 | 0 | 0 |
| ENSDARG000 | 20 | 7646762 | 7648325 | - | 1564 | NA | 0 | 0 | 0 | 0 | 0 | 0 |
| ENSDARG000 | 20:20:20:20:2 | 7398090:7399 | 7398269:7399 | +:+:+:+:+:+:+:+: | 2004 | PCSK9 | 60 | 37 | 10 | 8 | 34 | 25 |
| ENSDARG000 | 20:20:20:20:2 | 42881104:428 | 42881425:428 | +;+;+;+;+;+ | 2518 | pimr110 | 0 | 0 | 6 | 0 | 2 | 6 |
| ENSDARG000 | 20 | 44348942 | 44349057 | - | 116 | 5S_rRNA | 1 | 0 | 4 | 0 | 0 | 0 |
| ENSDARG000 | 20 | 6335296 | 6335411 | + | 116 | 5S_rRNA | 0 | 0 | 0 | 0 | 0 | 0 |
| ENSDARG000 | 20:20:20:20:2 | 49081967:490 | 49082065:490 | +:+:+:+:+:+:+:+: | 2371 | crnkl1 | 587 | 643 | 754 | 516 | 710 | 418 |
| ENSDARG000 | 20 | 23082069 | 23082190 | - | 122 | NA | 0 | 1 | 26 | 0 | 4 | 10 |
| ENSDARG000 | 20:20:20:20:2 | 5256275:5257 | 5256702:5257 | -:-:-:-:-:-:-:-:-: | 1843 | cyp46a1.4 | 10 | 7 | 35 | 10 | 42 | 93 |
| ENSDARG000 | 20;20 | 45853154:458 | 45853170:458 | +;+ | 399 | NA | 0 | 0 | 0 | 0 | 0 | 0 |
| ENSDARG000 | 20:20:20:20:2 | 5272655:5274 | 5272768:5274 | -:-:-:-:-:-:-:-:-: | 1476 | cyp46a1.3 | 10 | 37 | 19 | 22 | 42 | 26 |
| ENSDARG000 | 20:20:20:20:2 | 15114957:151 | 15116496:151 | -:-:-:-:-:-:-:-:-: | 3370 | si:dkev-239i2( | 0 | 2 | 2 | 8 | 44 | 0 |
| ENSDARG000 | 20:20:20:20:2 | 36220847:362 | 36220977:362 | -;-;-;-;-;-;- | 706 | NA | 0 | 0 | 0 | 0 | 0 | 0 |
| ENSDARG000 | 20 | 52919856 | 52919986 | + | 131 | 5S_rRNA | 0 | 0 | 0 | 0 | 0 | 0 |
| ENSDARG000 | 20 | 3330476 | 3330691 | - | 216 | U3 | 0 | 0 | 0 | 0 | 0 | 0 |
| ENSDARG000 | 20:20:20:20:2 | 52774730:527 | 52775060:527 | +:+:+:+:+:+:+:+: | 1629 | phactr1 | 179 | 245 | 361 | 241 | 330 | 154 |
| ENSDARG000 | 20:20:20:20:2 | 49319834:493 | 49320112:493 | -;-;-;-;- | 662 | NA | 0 | 0 | 0 | 0 | 0 | 0 |
| ENSDARG000 | 20;20;20 | 492529:49319 | 492595:49324 | +;+;+ | 497 | NA | 0 | 10 | 15 | 14 | 0 | 7 |
| ENSDARG000 | 20:20:20:20:2 | 1903481:1905 | 1903597:1906 | +:+:+:+:+:+:+:+: | 2535 | NA | 14 | 2 | 14 | 4 | 25 | 10 |
| ENSDARG000 | 20:20:20:20:2 | 54829098:548 | 54829257:548 | -;-;-;-;-;- | 627 | NA | 2 | 0 | 11 | 0 | 16 | 2 |
| ENSDARG000 | 20 | 1088658 | 1089603 | + | 946 | NA | 0 | 0 | 0 | 0 | 0 | 0 |
| ENSDARG000 | 20 | 49033004 | 49033118 | + | 115 | 5S_rRNA | 0 | 0 | 0 | 0 | 0 | 0 |
| ENSDARG000 | 20:20:20:20:2 | 54249339:542 | 54249393:542 | -:-:-:-:-:-:-:-:-: | 669 | FKBP3 | 354 | 370 | 290 | 215 | 255 | 229 |
| ENSDARG000 | 20;20;20;20 | 15015557:150 | 15016330:150 | +;+;+;+ | 2495 | myoc | 20 | 0 | 2 | 0 | 15 | 10 |
| ENSDARG000 | 20:20:20:20:2 | 6035427:6048 | 6035788:6048 | +:+:+:+:+:+:+:+: | 2561 | tshr | 0 | 0 | 2 | 0 | 2 | 2 |
| ENSDARG000 | 20 | 5995589 | 5995703 | - | 115 | 5S_rRNA | 0 | 0 | 0 | 0 | 0 | 0 |
| ENSDARG000 | 20 | 3333576 | 3333791 | - | 216 | U3 | 0 | 0 | 0 | 0 | 0 | 0 |
| ENSDARG000 | 20 | 49813144 | 49813254 | - | 111 | 5S_rRNA | 1 | 10 | 9 | 0 | 2 | 8 |
| ENSDARG000 | 20 | 8934408 | 8934523 | + | 116 | 5S_rRNA | 0 | 0 | 0 | 0 | 0 | 0 |
| ENSDARG000 | 20 | 14998956 | 14999072 | + | 117 | 5S_rRNA | 2 | 0 | 0 | 0 | 0 | 0 |
| ENSDARG000 | 20 | 30373207 | 30373427 | - | 221 | SNORA73 | 42 | 62 | 126 | 29 | 76 | 51 |
| ENSDARG000 | 20 | 12830448 | 12834749 | + | 4302 | NA | 4 | 10 | 8 | 0 | 6 | 18 |
| ENSDARG000 | 20 | 47759786 | 47759908 | + | 123 | 5S_rRNA | 2 | 0 | 0 | 0 | 0 | 0 |
| ENSDARG000 | 20 | 25237009 | 25237124 | + | 116 | 5S_rRNA | 0 | 0 | 0 | 0 | 0 | 0 |
| ENSDARG000 | 20:20:20:20:2 | 52851793:528 | 52855059:528 | -:-:-:-:-:-:-:-:-: | 6624 | DSP | 158 | 74 | 26 | 14 | 66 | 68 |
| ENSDARG000 | 20 | 46883360 | 46883487 | - | 128 | 5S_rRNA | 0 | 0 | 0 | 0 | 0 | 0 |
| ENSDARG000 | 20:20:20:20:2 | 3403647:3404 | 3403925:3404 | +:+:+:+:+:+:+:+: | 2940 | NA | 10 | 2 | 1 | 1 | 7 | 0 |
| ENSDARG000 | 20;20;20 | 4304457:4306 | 4305569:4306 | +;+;+ | 1934 | NA | 31 | 2 | 31 | 2 | 78 | 6 |
| ENSDARG000 | 20:20:20:20:2 | 4309521:4311 | 4310031:4312 | +;+;+;+;+ | 2758 | NA | 14 | 0 | 4 | 0 | 20 | 2 |
| ENSDARG000 | 20:20:20:20:2 | 6012451:6014 | 6012909:6014 | -;-;-;-;-;- | 1185 | NA | 0 | 8 | 10 | 0 | 22 | 12 |
| ENSDARG000 | 20:20:20:20:2 | 6028201:6033 | 6028394:6034 | -:-:-:-:-:-:-:-:-: | 1301 | NA | 4 | 8 | 11 | 2 | 8 | 4 |
| ENSDARG000 | 20;20;20 | 21478028:214 | 21478520:214 | +;+;+ | 893 | NA | 0 | 0 | 0 | 0 | 0 | 0 |
| ENSDARG000 | 20;20;20;20 | 27579772:275 | 27579840:275 | +;+;+;+ | 554 | NA | 0 | 0 | 0 | 0 | 0 | 0 |
| ENSDARG000 | 20;20;20 | 28995857:289 | 28996090:289 | +;+;+ | 949 | NA | 2 | 0 | 0 | 0 | 0 | 0 |
| ENSDARG000 | 20:20:20:20:2 | 52064027:520 | 52064316:520 | +;+;+;+;+ | 372 | NA | 0 | 0 | 0 | 0 | 0 | 0 |
| ENSDARG000 | 8;8;8 | 10561922:106 | 10562635:106 | +;+;+ | 992 | NA | 0 | 2 | 4 | 4 | 0 | 8 |
| ENSDARG000 | 8:8:8:8:8:8:8: | 43814821:438 | 43817003:438 | -:-:-:-:-:-:-:-:-: | 3995 | ADGRD1 | 458 | 2 | 258 | 67 | 732 | 57 |
| ENSDARG000 | 8:8:8:8:8:8:8: | 24745041:247 | 24745334:247 | +:+:+:+:+:+:+:+: | 3100 | slc16a4 | 127 | 59 | 105 | 72 | 279 | 94 |
| ENSDARG000 | 8:8:8:8:8:8:8: | 27630692:276 | 27630753:276 | -:-:-:-:-:-:-:-:-: | 3131 | mov10b.2 | 1020 | 529 | 120 | 310 | 138 | 731 |
| ENSDARG000 | 8:8:8:8:8:8:8: | 53080515:530 | 53081199:530 | +:+:+:+:+:+:+:+: | 2478 | wdr46 | 362 | 582 | 645 | 243 | 541 | 391 |
| ENSDARG000 | 8:8:8:8:8:8:8: | 24730120:247 | 24730183:247 | +:+:+:+:+:+:+:+: | 1322 | lamtor5 | 628 | 548 | 588 | 631 | 582 | 715 |
| ENSDARG000 | 8;8;8 | 24717002:247 | 24717077:247 | +;+;+ | 787 | NA | 0 | 0 | 0 | 0 | 0 | 0 |
| ENSDARG000 | 8;8 | 33132502:331 | 33134063:331 | -;- | 1657 | zbtb34 | 66 | 38 | 103 | 35 | 60 | 62 |
| ENSDARG000 | 8;8;8 | 24714733:247 | 24715264:247 | -;-;- | 896 | prok1 | 0 | 0 | 2 | 0 | 0 | 2 |
| ENSDARG000 | 8:8:8:8:8:8:8: | 48629495:486 | 48630043:486 | -:-:-:-:-:-:-:-:-: | 3677 | CLCN6 | 458 | 290 | 322 | 216 | 363 | 273 |
| ENSDARG000 | 8:8:8:8:8:8:8: | 18460807:184 | 18460845:184 | +:+:+:+:+:+:+:+: | 2680 | alkal1 | 0 | 0 | 0 | 0 | 0 | 10 |
| ENSDARG000 | 8:8:8:8:8:8:8: | 14211746:142 | 14212780:142 | -:-:-:-:-:-:-:-:-: | 4705 | sin3b | 572 | 577 | 796 | 355 | 835 | 526 |
| ENSDARG000 | 8:8:8:8:8:8:8: | 10823069:108 | 10823425:108 | +:+:+:+:+:+:+:+: | 1556 | NA | 427 | 356 | 346 | 341 | 253 | 462 |
| ENSDARG000 | 8:8:8:8:8:8:8: | 33258580:332 | 33261375:332 | -:-:-:-:-:-:-:-:-: | 4631 | lmx1bb | 4 | 9 | 6 | 168 | 39 | 26 |
| ENSDARG000 | 8:8:8:8:8:8:8: | 7571095:7571 | 7571247:7571 | -:-:-:-:-:-:-:-:-: | 2362 | NA | 127 | 139 | 99 | 85 | 136 | 103 |
| ENSDARG000 | 8;8;8;8 | 50190742:502 | 50190974:502 | +;+;+;+ | 2613 | slc25a37 | 156 | 41 | 208 | 88 | 221 | 154 |
| ENSDARG000 | 8;8;8;8;8 | 50256930:502 | 50257376:502 | -;-;-;-;- | 1170 | nkx3-1 | 0 | 4 | 2 | 0 | 0 | 0 |
| ENSDARG000 | 8;8;8;8;8;8 | 33382787:333 | 33383232:333 | -;-;-;-;-;- | 900 | NA | 0 | 0 | 0 | 0 | 0 | 0 |
| ENSDARG000 | 8;8 | 43934595:439 | 43936673:439 | -;- | 2190 | NA | 4 | 0 | 2 | 64 | 2 | 6 |
| ENSDARG000 | 8:8:8:8:8:8:8: | 16688370:166 | 16688834:166 | -:-:-:-:-:-:-:-:-: | 1956 | rpe65b | 104 | 60 | 193 | 37 | 72 | 425 |
| ENSDARG000 | 8:8:8:8:8:8:8: | 391590:39159 | 391824:39230 | -:-:-:-:-:-:-:-:-: | 3906 | trim36 | 38 | 37 | 20 | 32 | 24 | 18 |
| ENSDARG000 | 8:8:8:8:8:8:8: | 41256612:412 | 41261401:412 | -:-:-:-:-:-:-:-:-: | 7540 | rnf10 | 6845 | 5844 | 7332 | 6010 | 7067 | 5975 |
| ENSDARG000 | 8:8:8:8:8:8:8: | 16676894:166 | 16676992:166 | +:+:+:+:+:+:+:+: | 2856 | si:ch211-198r | 174 | 146 | 169 | 127 | 149 | 227 |
| ENSDARG000 | 8 | 53107380 | 53108207 | - | 828 | NA | 4 | 0 | 4 | 0 | 0 | 14 |
| ENSDARG000 | 8:8:8:8:8:8:8: | 13106760:131 | 13107211:131 | +:+:+:+:+:+:+:+: | 9247 | itgb4 | 284 | 28 | 32 | 16 | 56 | 26 |
| ENSDARG000 | 8:8:8:8:8:8:8: | 27659244:276 | 27659471:276 | -:-:-:-:-:-:-:-:-: | 3234 | mov10b.1 | 16904 | 5254 | 1731 | 1521 | 807 | 12505 |
| ENSDARG000 | 8;8 | 412312:41424 | 412540:41459 | +;+ | 578 | NA | 56 | 22 | 19 | 33 | 51 | 20 |
| ENSDARG000 | 8:8:8:8:8:8:8: | 411977:41327 | 412690:41338 | -:-:-:-:-:-:-:-:-: | 1596 | pggt1b | 657 | 584 | 662 | 449 | 728 | 424 |
| ENSDARG000 | 8;8 | 44011114:440 | 44011164:440 | -;- | 153 | NA | 0 | 0 | 0 | 0 | 0 | 0 |
| ENSDARG000 | 8:8:8:8:8:8:8: | 43941313:439 | 43941339:439 | -:-:-:-:-:-:-:-:-: | 4519 | RIMBP2 | 2 | 0 | 3 | 26 | 10 | 2 |
| ENSDARG000 | 8:8:8:8:8:8:8: | 14244476:142 | 14244533:142 | -:-:-:-:-:-:-:-:-: | 2435 | xpr1a | 4 | 0 | 3 | 0 | 10 | 4 |
| ENSDARG000 | 8;8 | 423393:42394 | 425255:42525 | -;- | 1867 | fem1c | 482 | 397 | 484 | 476 | 572 | 281 |
| ENSDARG000 | 8:8:8:8:8:8:8: | 33420677:334 | 33421275:334 | -:-:-:-:-:-:-:-:-: | 1420 | mvb12bb | 0 | 0 | 0 | 0 | 2 | 0 |
| ENSDARG000 | 8:8:8:8:8:8:8: | 10862353:108 | 10862854:108 | +:+:+:+:+:+:+:+: | 4264 | BRPF3 | 694 | 550 | 743 | 216 | 895 | 535 |
| ENSDARG000 | 8:8:8:8:8:8:8: | 23165749:231 | 23165873:231 | +:+:+:+:+:+:+:+: | 1019 | dnajc5aa | 125 | 159 | 142 | 102 | 126 | 132 |
| ENSDARG000 | 8;8;8;8 | 20488322:204 | 20488503:204 | +;+;+;+ | 1858 | zgc:101100 | 1 | 0 | 0 | 0 | 6 | 0 |
| ENSDARG000 | 8:8:8:8:8:8:8: | 18506141:185 | 18506382:185 | -:-:-:-:-:-:-:-:-: | 2546 | NA | 114 | 85 | 79 | 62 | 108 | 150 |
| ENSDARG000 | 8:8:8:8:8:8:8: | 21353878:213 | 21353967:213 | +:+:+:+:+:+:+:+: | 2959 | alas2 | 96 | 60 | 273 | 35 | 634 | 148 |
| ENSDARG000 | 8;8 | 50284859:502 | 50285549:502 | -;- | 1176 | nkx2.7 | 0 | 0 | 0 | 0 | 0 | 0 |
| ENSDARG000 | 8;8;8;8;8;8 | 7616573:7616 | 7617841:7617 | -;-;-;-;-;- | 1687 | mecp2 | 92 | 70 | 158 | 130 | 181 | 86 |
| ENSDARG000 | 8:8:8:8:8:8:8: | 18515711:185 | 18518251:185 | -:-:-:-:-:-:-:-:-: | 5373 | nexn | 4 | 6 | 10 | 2 | 9 | 6 |
| ENSDARG000 | 8:8:8:8:8:8:8: | 21366771:213 | 21366943:213 | -:-:-:-:-:-:-:-:-: | 1407 | ela2l | 10 | 0 | 0 | 4 | 0 | 0 |
| ENSDARG000 | 8;8;8;8;8 | 38415374:384 | 38415791:384 | +;+;+;+;+ | 2418 | nkx6.3 | 0 | 1 | 0 | 0 | 0 | 4 |
| ENSDARG000 | 8:8:8:8:8:8:8: | 53120278:531 | 53120433:531 | +:+:+:+:+:+:+:+: | 4310 | nr2c2 | 649 | 435 | 732 | 495 | 422 | 606 |
| ENSDARG000 | 8;8;8;8 | 50322769:503 | 50323120:503 | -;-;-;- | 990 | stc1 | 30 | 16 | 114 | 50 | 203 | 6 |
| ENSDARG000 | 8:8:8:8:8:8:8: | 10956468:109 | 10957191:109 | -:-:-:-:-:-:-:-:-: | 4049 | trim33 | 811 | 825 | 1724 | 604 | 934 | 1120 |
| ENSDARG000 | 8:8:8:8:8:8:8: | 33622404:336 | 33623090:336 | -;-;-;-;-;-;-;- | 1141 | NA | 0 | 0 | 0 | 0 | 0 | 0 |
| ENSDARG000 | 8;8;8;8 | 18515040:185 | 18515271:185 | -;-;-;- | 907 | NA | 0 | 0 | 0 | 0 | 0 | 0 |
| ENSDARG000 | 8 | 23190986 | 23194408 | - | 3423 | NA | 257 | 368 | 414 | 256 | 376 | 50 |
| ENSDARG000 | 8:8:8:8:8:8:8: | 23194212:231 | 23194410:231 | +:+:+:+:+:+:+:+: | 3850 | tpd52l2a | 1010 | 767 | 783 | 1166 | 826 | 1168 |
| ENSDARG000 | 8:8:8:8:8:8:8: | 10937523:109 | 10938455:109 | -:-:-:-:-:-:-:-:-: | 1825 | slc66a1 | 178 | 81 | 152 | 93 | 96 | 106 |
| ENSDARG000 | 8:8:8:8:8:8:8: | 24854600:248 | 24854672:248 | +:+:+:+:+:+:+:+: | 4043 | slc6a17 | 900 | 673 | 1324 | 1744 | 640 | 517 |
| ENSDARG000 | 8:8:8:8:8:8:8: | 50463227:504 | 50463672:504 | -:-:-:-:-:-:-:-:-: | 1379 | ido1 | 0 | 2 | 18 | 0 | 13 | 0 |
| ENSDARG000 | 8:8:8:8:8:8:8: | 10903325:109 | 10903726:109 | -:-:-:-:-:-:-:-:-: | 2047 | nr1h5 | 5 | 12 | 18 | 14 | 0 | 0 |
| ENSDARG000 | 8;8;8;8 | 24866985:248 | 24867419:248 | -;-;-;- | 669 | NA | 0 | 0 | 0 | 0 | 0 | 0 |
| ENSDARG000 | 8:8:8:8:8:8:8: | 14381272:143 | 14381663:143 | +:+:+:+:+:+:+:+: | 3161 | acbd6 | 614 | 317 | 424 | 218 | 458 | 478 |

| | | | | | | | | | | | | |
|---|---|---|---|---|---|---|---|---|---|---|---|---|
| ENSDARG000 | 8;8;8;8;8;8;8; | 668184:6942 | 668244:6945 | +;+;+;+;+;+;+; | 1146 | RNF165 | 118 | 71 | 133 | 148 | 168 | 110 |
| ENSDARG000 | 8;8;8;8;8;8 | 14414353:144 | 14414910:144 | -;-;-;-;-;- | 1085 | NA | 0 | 0 | 0 | 0 | 2 | 0 |
| ENSDARG000 | 8;8;8;8;8;8 | 50493095:504 | 50493286:504 | +;+;+;+;+;+ | 1511 | RHOBTB2 | 2 | 0 | 2 | 0 | 12 | 0 |
| ENSDARG000 | 8;8;8;8;8 | 27743550:277 | 27743968:277 | +;+;+;+;+ | 1413 | wnt2bb | 0 | 0 | 0 | 0 | 0 | 0 |
| ENSDARG000 | 8;8;8;8;8;8;8; | 48673596:486 | 48673740:486 | -;-;-;-;-;-;-; | 1112 | NA | 4 | 0 | 0 | 0 | 0 | 0 |
| ENSDARG000 | 8;8;8;8;8;8;8; | 20495889:204 | 20495948:204 | +;+;+;+;+;+;+; | 3119 | csnk1g2b | 1186 | 1122 | 986 | 678 | 1233 | 753 |
| ENSDARG000 | 8;8 | 50727220:507 | 50727550:507 | +;+ | 3258 | egr3 | 250 | 106 | 102 | 64 | 543 | 374 |
| ENSDARG000 | 8;8;8;8;8;8;8; | 2438638:2442 | 2438702:2442 | +;+;+;+;+;+;+; | 1155 | enkd1 | 0 | 4 | 0 | 4 | 0 | 0 |
| ENSDARG000 | 8;8;8;8;8;8;8; | 50742975:507 | 50743161:507 | +;+;+;+;+;+;+; | 4681 | bin3 | 1399 | 1024 | 1487 | 1439 | 1485 | 1712 |
| ENSDARG000 | 8;8;8;8;8;8;8; | 38421333:384 | 38423646:384 | -;-;-;-;-;-;-; | 3892 | inpp5l | 578 | 714 | 538 | 332 | 613 | 788 |
| ENSDARG000 | 8;8;8;8;8;8;8; | 30787111:307 | 30788032:307 | -;-;-;-;-;-;-; | 1674 | gstt1a | 3958 | 4624 | 2915 | 5759 | 3467 | 4267 |
| ENSDARG000 | 8;8;8;8;8;8;8; | 44211374:442 | 44211492:442 | -;-;-;-;-;-;-; | 1589 | STX2 | 40 | 31 | 24 | 126 | 59 | 56 |
| ENSDARG000 | 8;8;8;8;8 | 41274211:412 | 41274658:412 | -;-;-;-;- | 916 | pop5 | 192 | 174 | 118 | 200 | 142 | 276 |
| ENSDARG000 | 8;8;8;8;8;8;8; | 33895773:338 | 33897409:338 | -;-;-;-;-;-;-; | 3986 | pbx3b | 4 | 12 | 68 | 0 | 44 | 6 |
| ENSDARG000 | 8;8;8;8;8;8;8 | 24928632:249 | 24928699:249 | -;-;-;-;-;-;- | 1812 | kcnc4 | 30 | 592 | 579 | 688 | 1555 | 351 |
| ENSDARG000 | 8;8;8 | 48691469:486 | 48691706:486 | +;+;+ | 440 | NA | 0 | 0 | 0 | 1 | 2 | 0 |
| ENSDARG000 | 8;8;8;8;8;8;8; | 53150972:531 | 53151178:531 | -;-;-;-;-;-;-; | 2452 | RBSN | 196 | 118 | 207 | 101 | 180 | 166 |
| ENSDARG000 | 8;8;8;8;8;8;8; | 25811892:258 | 25812166:258 | -;-;-;-;-;-;-;- | 1201 | taf10 | 552 | 380 | 537 | 290 | 392 | 545 |
| ENSDARG000 | 8;8;8;8 | 53143211:531 | 53143434:531 | +;+;+;+ | 6188 | mrps25 | 882 | 648 | 1344 | 1020 | 994 | 913 |
| ENSDARG000 | 8;8;8;8;8;8;8 | 50942240:509 | 50942429:509 | +;+;+;+;+;+;+ | 3657 | b2ml | 7268 | 5077 | 7870 | 5141 | 7010 | 7171 |
| ENSDARG000 | 8;8;8;8;8;8;8; | 714875:7151 | 715592:7155 | -;-;-;-;-;-;-; | 7102 | LOXHD1 | 2 | 6 | 20 | 33 | 36 | 0 |
| ENSDARG000 | 8;8;8;8;8;8;8; | 48703005:487 | 48703157:487 | -;-;-;-;-;-;-; | 1101 | NA | 4 | 0 | 6 | 0 | 0 | 0 |
| ENSDARG000 | 8;8;8 | 38505383:385 | 38505879:385 | -;-;- | 957 | FOXB2 | 0 | 0 | 0 | 0 | 0 | 0 |
| ENSDARG000 | 8;8;8;8;8;8;8; | 27823012:278 | 27824590:278 | -;-;-;-;-;-;-; | 2455 | cttnbp2nlb | 203 | 55 | 60 | 52 | 22 | 423 |
| ENSDARG000 | 8;8;8;8;8;8;8; | 27807266:278 | 27808084:278 | +;+;+;+;+;+;+; | 2708 | capza1b | 4348 | 5426 | 4664 | 4620 | 4227 | 4598 |
| ENSDARG000 | 8;8;8;8 | 50953776:509 | 50953943:509 | +;+;+;+ | 845 | UBL4A | 266 | 268 | 401 | 235 | 314 | 206 |
| ENSDARG000 | 8;8;8;8;8;8;8; | 30797363:308 | 30797639:308 | +;+;+;+;+;+;+; | 1618 | MMP11 | 10 | 63 | 31 | 10 | 101 | 45 |
| ENSDARG000 | 8;8;8;8;8;8;8; | 41281801:413 | 41282039:413 | +;+;+;+;+;+;+; | 2099 | olfml2a | 12 | 4 | 67 | 22 | 8 | 0 |
| ENSDARG000 | 8 | 48732153 | 48732858 | + | 706 | NA | 1 | 0 | 4 | 29 | 8 | 0 |
| ENSDARG000 | 8;8;8;8;8;8;8; | 25818622:258 | 25819629:258 | -;-;-;-;-;-;-;- | 3101 | efhd2 | 1755 | 2260 | 2038 | 1965 | 2137 | 1304 |
| ENSDARG000 | 8;8;8;8;8;8 | 14453933:144 | 14455605:144 | -;-;-;-;-;- | 2627 | lhx4 | 156 | 24 | 84 | 59 | 17 | 14 |
| ENSDARG000 | 8;8 | 24968098:249 | 24970036:249 | -;- | 2020 | hspa1b | 20 | 66 | 66 | 64 | 58 | 48 |
| ENSDARG000 | 8;8;8;8 | 7750953:775 | 7751473:775 | -;-;-;- | 934 | NA | 0 | 0 | 2 | 0 | 0 | 0 |
| ENSDARG000 | 8;8;8;8;8;8;8; | 7737062:774 | 7737126:774 | +;+;+;+;+;+;+; | 2645 | FGD1 | 333 | 238 | 250 | 191 | 289 | 104 |
| ENSDARG000 | 8;8 | 50966806:509 | 50966980:509 | +;+ | 637 | NA | 619 | 667 | 658 | 654 | 526 | 542 |
| ENSDARG000 | 8;8;8;8 | 41413919:414 | 41414037:414 | +;+;+;+ | 759 | NA | 0 | 0 | 0 | 0 | 0 | 0 |
| ENSDARG000 | 8;8;8;8;8;8;8 | 25858224:258 | 25858812:258 | -;-;-;-;-;-;- | 1481 | opn7c | 3693 | 2746 | 3800 | 2386 | 6194 | 3086 |
| ENSDARG000 | 8;8;8;8 | 38564942:385 | 38565328:385 | +;+;+;+ | 957 | rfk | 146 | 161 | 144 | 47 | 184 | 121 |
| ENSDARG000 | 8;8;8;8;8;8;8; | 13161680:131 | 13162399:131 | -;-;-;-;-;-;-; | 1529 | zgc:194990 | 0 | 0 | 0 | 2 | 0 | 0 |
| ENSDARG000 | 8;8;8;8;8;8;8; | 20564621:205 | 20565305:205 | +;+;+;+;+;+;+; | 2303 | NA | 10 | 20 | 14 | 10 | 46 | 23 |
| ENSDARG000 | 8;8;8;8 | 41453691:414 | 41454077:414 | +;+;+;+ | 1425 | arpc5la | 482 | 935 | 741 | 436 | 648 | 485 |
| ENSDARG000 | 8;8;8;8;8;8;8; | 50979018:509 | 50979111:509 | +;+;+;+;+;+;+; | 3329 | NA | 335 | 185 | 287 | 206 | 375 | 178 |
| ENSDARG000 | 8;8;8;8;8;8;8 | 48739990:487 | 48740142:487 | -;-;-;-;-;-;- | 808 | NA | 0 | 0 | 2 | 7 | 0 | 0 |
| ENSDARG000 | 8;8;8;8;8;8;8; | 50968990:509 | 50969534:509 | -;-;-;-;-;-;-; | 1260 | rab7b | 1904 | 1514 | 2152 | 1891 | 1669 | 1368 |
| ENSDARG000 | 8;8;8;8;8;8;8; | 21376290:213 | 21376342:213 | +;+;+;+;+;+;+; | 890 | ela2 | 0 | 0 | 2 | 0 | 6 | 0 |
| ENSDARG000 | 8;8;8;8;8;8;8; | 20624510:206 | 20624765:206 | +;+;+;+;+;+;+; | 1891 | NA | 28 | 20 | 86 | 60 | 104 | 36 |
| ENSDARG000 | 8;8;8;8;8;8;8; | 2456854:2456 | 2457049:2457 | +;+;+;+;+;+;+; | 1710 | polb | 198 | 296 | 340 | 189 | 282 | 314 |
| ENSDARG000 | 8;8 | 902750:9120 | 903291:9128 | -;- | 1352 | ptger4a | 1 | 0 | 0 | 0 | 0 | 0 |
| ENSDARG000 | 8;8;8;8 | 48754895:487 | 48755047:487 | -;-;-;- | 864 | NA | 0 | 0 | 3 | 5 | 0 | 0 |
| ENSDARG000 | 8;8;8;8;8;8;8; | 14503610:145 | 14504778:145 | -;-;-;-;-;-;-; | 2786 | qsox1 | 1259 | 131 | 345 | 102 | 377 | 142 |
| ENSDARG000 | 8;8;8;8 | 38593904:386 | 38594198:386 | -;-;-;- | 685 | NA | 4 | 0 | 0 | 31 | 6 | 0 |
| ENSDARG000 | 8;8;8;8;8;8;8 | 30921403:309 | 30921706:309 | +;+;+;+;+;+;+ | 1741 | derl3 | 68 | 215 | 404 | 146 | 230 | 178 |
| ENSDARG000 | 8;8;8;8;8;8;8; | 53171002:531 | 53171394:531 | -;-;-;-;-;-;-; | 1362 | sult1st4 | 0 | 0 | 4 | 23 | 4 | 0 |
| ENSDARG000 | 8;8;8;8;8;8;8; | 926673:9282 | 927405:9284 | -;-;-;-;-;-;-; | 1780 | mrps27 | 314 | 304 | 512 | 224 | 375 | 270 |
| ENSDARG000 | 8;8;8;8;8;8;8; | 48768873:487 | 48769180:487 | -;-;-;-;-;-;-; | 1277 | NA | 0 | 0 | 0 | 0 | 0 | 0 |
| ENSDARG000 | 8;8;8;8;8;8;8; | 7778770:778 | 7779269:778 | +;+;+;+;+;+;+; | 2256 | tfe3a | 185 | 107 | 223 | 134 | 125 | 135 |
| ENSDARG000 | 8;8;8;8;8 | 944232:9492 | 944418:9493 | +;+;+;+;+ | 803 | NA | 0 | 0 | 0 | 0 | 0 | 0 |
| ENSDARG000 | 8;8;8;8;8;8;8; | 25900049:259 | 25900351:259 | +;+;+;+;+;+;+; | 1917 | rhoab | 1888 | 2792 | 2463 | 1595 | 4115 | 2071 |
| ENSDARG000 | 8;8;8;8;8;8;8; | 955162:9560 | 955220:9568 | -;-;-;-;-;-;-; | 2363 | ZNF366 | 18 | 0 | 6 | 2 | 0 | 16 |
| ENSDARG000 | 8;8 | 38624155:386 | 38624315:386 | -;- | 536 | NA | 0 | 2 | 2 | 0 | 0 | 0 |
| ENSDARG000 | 8;8;8;8;8;8;8; | 46327350:463 | 46327473:463 | +;+;+;+;+;+;+; | 2716 | si:dkev-75a21 | 364 | 559 | 339 | 233 | 333 | 520 |
| ENSDARG000 | 8 | 44295983 | 44298964 | - | 2982 | fzd10 | 92 | 54 | 100 | 54 | 100 | 75 |
| ENSDARG000 | 8;8;8;8 | 28207354:282 | 28207519:282 | -;-;-;- | 1353 | NA | 0 | 2 | 0 | 0 | 0 | 0 |
| ENSDARG000 | 8;8;8;8;8;8;8; | 44224693:442 | 44225419:442 | -;-;-;-;-;-;-; | 4042 | piwil1 | 8 | 4 | 18 | 26 | 28 | 3 |
| ENSDARG000 | 8;8;8;8;8;8;8; | 28065803:280 | 28066358:280 | +;+;+;+;+;+;+; | 3361 | kcnd3 | 174 | 22 | 2 | 6 | 14 | 0 |
| ENSDARG000 | 8;8;8;8;8;8;8 | 11008763:110 | 11009270:110 | -;-;-;-;-;-;- | 1110 | bcas2 | 943 | 1054 | 1484 | 774 | 939 | 897 |
| ENSDARG000 | 8;8;8;8;8;8;8; | 46386601:463 | 46386647:463 | +;+;+;+;+;+;+; | 2225 | ogg1 | 190 | 101 | 120 | 145 | 80 | 173 |
| ENSDARG000 | 8;8;8;8;8;8;8; | 25892319:258 | 25893013:258 | +;+;+;+;+;+;+; | 5470 | tmem115 | 1136 | 859 | 898 | 1075 | 859 | 1146 |
| ENSDARG000 | 8;8;8;8;8;8;8; | 13190314:131 | 13190702:131 | -;-;-;-;-;-;-; | 1889 | NA | 6 | 4 | 0 | 0 | 18 | 0 |
| ENSDARG000 | 8;8;8;8;8;8;8; | 24971890:249 | 24973509:249 | -;-;-;-;-;-;-; | 3627 | ahcyl1 | 140 | 124 | 442 | 242 | 703 | 201 |
| ENSDARG000 | 8;8;8;8 | 20825987:208 | 20826194:208 | +;+;+;+ | 569 | NA | 0 | 0 | 0 | 0 | 0 | 0 |
| ENSDARG000 | 8;8;8;8;8 | 20852995:208 | 20853429:208 | -;-;-;-;- | 1208 | dohh | 481 | 473 | 833 | 591 | 577 | 413 |
| ENSDARG000 | 8;8;8;8;8;8;8 | 20832584:208 | 20833488:208 | -;-;-;-;-;-;- | 1757 | smim24 | 0 | 2 | 4 | 0 | 0 | 2 |
| ENSDARG000 | 8;8;8;8;8;8;8; | 46353203:463 | 46353466:463 | -;-;-;-;-;-;-; | 2600 | qars1 | 1280 | 1723 | 2276 | 738 | 1974 | 1104 |
| ENSDARG000 | 8;8;8 | 20842134:208 | 20842263:208 | -;-;- | 362 | NA | 0 | 0 | 0 | 0 | 2 | 0 |
| ENSDARG000 | 8;8;8;8;8;8;8 | 20846257:208 | 20847004:208 | -;-;-;-;-;-;- | 1140 | NA | 0 | 0 | 0 | 0 | 0 | 1 |
| ENSDARG000 | 8;8;8;8;8;8;8; | 21383347:213 | 21383418:213 | +;+;+;+;+;+;+; | 1215 | fhad1 | 0 | 0 | 0 | 0 | 0 | 0 |
| ENSDARG000 | 8;8;8;8;8;8;8; | 53185079:531 | 53185400:531 | -;-;-;-;-;-;-; | 1737 | GABRD | 0 | 0 | 0 | 0 | 0 | 0 |
| ENSDARG000 | 8;8;8;8;8;8;8; | 18545185:185 | 18545686:185 | +;+;+;+;+;+;+; | 3293 | tab3 | 437 | 547 | 579 | 484 | 655 | 381 |
| ENSDARG000 | 8;8;8;8;8;8 | 48778961:487 | 48779123:487 | -;-;-;-;-;- | 2485 | NA | 2 | 2 | 4 | 0 | 10 | 15 |
| ENSDARG000 | 8;8;8;8;8;8;8; | 14559378:145 | 14561383:145 | -;-;-;-;-;-;-; | 11400 | CEP350 | 1982 | 1457 | 1654 | 1021 | 2157 | 1344 |
| ENSDARG000 | 8;8;8;8;8;8;8; | 7828414:782 | 7828491:782 | +;+;+;+;+;+;+; | 1418 | NA | 414 | 185 | 189 | 296 | 584 | 308 |
| ENSDARG000 | 8;8;8;8;8;8;8; | 7839208:784 | 7840914:784 | -;-;-;-;-;-;-;- | 2302 | zgc:113363 | 65 | 35 | 47 | 38 | 53 | 53 |
| ENSDARG000 | 8;8;8;8;8;8 | 48801402:488 | 48801726:488 | +;+;+;+;+;+ | 1197 | TPRG1L | 88 | 40 | 62 | 71 | 69 | 161 |
| ENSDARG000 | 8;8;8;8;8;8;8; | 13244371:132 | 13244835:132 | -;-;-;-;-;-;-; | 2073 | kcnn1b | 0 | 2 | 0 | 0 | 0 | 0 |
| ENSDARG000 | 8;8;8;8;8;8;8 | 21406769:214 | 21406908:214 | +;+;+;+;+;+;+ | 1060 | NA | 0 | 0 | 0 | 0 | 0 | 0 |
| ENSDARG000 | 8;8;8;8;8;8;8; | 7854130:785 | 7854334:785 | +;+;+;+;+;+;+; | 2732 | cxxc1a | 626 | 559 | 736 | 578 | 718 | 628 |
| ENSDARG000 | 8;8;8;8;8;8;8; | 48848200:488 | 48848464:488 | +;+;+;+;+;+;+; | 2666 | tp73 | 14 | 2 | 0 | 0 | 0 | 4 |
| ENSDARG000 | 8;8;8;8 | 999421:1001 | 999539:1001 | +;+;+;+ | 499 | fabp1b.1 | 0 | 0 | 0 | 0 | 0 | 0 |
| ENSDARG000 | 8;8;8;8;8;8;8; | 41467922:414 | 41469314:414 | -;-;-;-;-;-;-; | 4510 | golga1 | 685 | 647 | 1050 | 815 | 854 | 657 |
| ENSDARG000 | 8;8;8;8;8;8;8; | 23213320:232 | 23213468:232 | +;+;+;+;+;+;+; | 1309 | ppdpfa | 758 | 1401 | 144 | 666 | 304 | 314 |
| ENSDARG000 | 8;8;8;8;8;8;8; | 30929704:309 | 30930038:309 | -;-;-;-;-;-;-; | 1512 | smarcb1a | 703 | 756 | 996 | 596 | 998 | 787 |
| ENSDARG000 | 8;8 | 4798158:479 | 4798262:480 | +;+ | 2302 | hsp70l | 883 | 574 | 3051 | 7354 | 36 | 48 |
| ENSDARG000 | 8;8;8 | 2478862:248 | 2479230:248 | +;+;+ | 893 | NA | 18 | 0 | 0 | 0 | 2 | 2 |
| ENSDARG000 | 8;8;8;8;8;8;8; | 48816082:488 | 48816665:488 | -;-;-;-;-;-;-; | 1995 | wrap73 | 154 | 107 | 62 | 69 | 127 | 62 |
| ENSDARG000 | 8;8;8;8;8;8;8; | 16710634:167 | 16713730:167 | -;-;-;-;-;-;-; | 5961 | depdc1a | 88 | 388 | 506 | 262 | 353 | 189 |
| ENSDARG000 | 8;8;8;8;8;8 | 4803906:480 | 4804193:480 | +;+;+;+;+;+ | 1549 | tmem127 | 596 | 475 | 629 | 362 | 643 | 439 |
| ENSDARG000 | 8;8 | 41505590:415 | 41505697:415 | +;+ | 394 | NA | 0 | 0 | 0 | 0 | 0 | 0 |
| ENSDARG000 | 8;8;8;8;8;8;8; | 16702106:167 | 16702289:167 | -;-;-;-;-;-;-; | 1672 | rpe65c | 63 | 73 | 195 | 1 | 77 | 251 |
| ENSDARG000 | 8;8;8;8;8 | 2487250:248 | 2487338:248 | +;+;+;+;+ | 1000 | dynll1 | 3986 | 4489 | 3759 | 2475 | 4749 | 3627 |
| ENSDARG000 | 8;8;8;8;8;8;8 | 4838114:484 | 4838356:484 | +;+;+;+;+;+;+ | 1057 | gatd3l | 0 | 0 | 0 | 0 | 0 | 0 |
| ENSDARG000 | 8;8;8;8;8;8;8; | 11019051:110 | 11019379:110 | -;-;-;-;-;-;-; | 3699 | DENND2C | 380 | 305 | 720 | 267 | 397 | 596 |
| ENSDARG000 | 8;8 | 28259347:282 | 28259594:282 | +;+ | 2356 | INKA2 | 85 | 249 | 149 | 69 | 140 | 47 |
| ENSDARG000 | 8;8;8;8;8;8;8; | 23227100:232 | 23227795:232 | -;-;-;-;-;-;-;- | 2040 | NA | 0 | 2 | 0 | 0 | 0 | 50 |
| ENSDARG000 | 8;8;8;8;8;8;8; | 28241100:282 | 28244377:282 | -;-;-;-;-;-;-; | 5634 | ddx20 | 219 | 281 | 354 | 200 | 273 | 238 |
| ENSDARG000 | 8;8;8 | 18565904:185 | 18570516:185 | -;-;- | 5127 | tmem47 | 64 | 78 | 243 | 164 | 88 | 946 |
| ENSDARG000 | 8;8;8;8;8;8;8; | 51050554:510 | 51050812:510 | +;+;+;+;+;+;+; | 4430 | DISP3 | 3 | 0 | 2 | 0 | 5 | 2 |
| ENSDARG000 | 8;8;8;8;8;8;8; | 28267546:282 | 28268590:282 | -;-;-;-;-;-;-; | 1843 | rap1aa | 1240 | 1319 | 1581 | 904 | 1585 | 1162 |
| ENSDARG000 | 8;8;8;8 | 1009831:1011 | 1009964:1011 | +;+;+;+ | 559 | NA | 26 | 32 | 2 | 0 | 28 | 0 |
| ENSDARG000 | 8;8 | 21426001:214 | 21426183:214 | +;+ | 247 | NA | 0 | 0 | 0 | 0 | 0 | 0 |
| ENSDARG000 | 8;8;8;8;8 | 21437908:214 | 21437989:214 | +;+;+;+;+ | 738 | NA | 0 | 0 | 0 | 0 | 0 | 0 |
| ENSDARG000 | 8;8;8;8;8;8;8; | 7875110:787 | 7875226:787 | +;+;+;+;+;+;+; | 8264 | mbd1a | 510 | 332 | 654 | 542 | 568 | 868 |
| ENSDARG000 | 8;8;8;8;8;8;8; | 18588551:185 | 18588646:185 | +;+;+;+;+;+;+; | 4391 | prrg1 | 243 | 251 | 484 | 435 | 275 | 454 |
| ENSDARG000 | 8;8;8;8;8;8;8; | 21588067:216 | 21589003:216 | +;+;+;+;+;+;+; | 6988 | AJAP1 | 2 | 14 | 18 | 62 | 119 | 8 |
| ENSDARG000 | 8;8;8;8;8;8;8; | 13351698:133 | 13352605:133 | -;-;-;-;-;-;-;- | 1411 | ccdc124 | 922 | 989 | 1232 | 1028 | 1006 | 992 |
| ENSDARG000 | 8;8;8;8;8;8;8; | 34341885:343 | 34342676:343 | -;-;-;-;-;-;-; | 5428 | gapvd1 | 503 | 423 | 767 | 330 | 635 | 383 |
| ENSDARG000 | 8;8;8;8;8;8;8; | 25019542:250 | 25022580:250 | -;-;-;-;-;-;-; | 4130 | nfyal | 59 | 52 | 118 | 90 | 83 | 47 |
| ENSDARG000 | 8;8;8;8;8;8;8; | 7975745:801 | 7976031:801 | +;+;+;+;+;+;+; | 2273 | NA | 33 | 18 | 36 | 26 | 11 | 27 |
| ENSDARG000 | 8;8;8;8;8;8;8; | 2493794:249 | 2493977:249 | -;-;-;-;-;-;-; | 978 | rpl6 | 26315 | 22700 | 38872 | 36645 | 28176 | 37195 |
| ENSDARG000 | 8;8;8;8;8;8;8; | 18830759:188 | 18831041:188 | +;+;+;+;+;+;+; | 2140 | mpnd | 435 | 303 | 364 | 360 | 391 | 533 |
| ENSDARG000 | 8;8;8;8;8;8;8; | 1015243:101 | 1015667:101 | -;-;-;-;-;-;-; | 1819 | smyd1b | 87 | 14 | 110 | 55 | 36 | 12 |
| ENSDARG000 | 8;8;8;8;8;8;8; | 4856440:485 | 4856630:485 | -;-;-;-;-;-;-; | 5760 | NA | 18 | 78 | 80 | 22 | 118 | 56 |
| ENSDARG000 | 8;8 | 20861358:208 | 20861950:208 | -;- | 899 | NA | 29 | 36 | 53 | 23 | 32 | 75 |
| ENSDARG000 | 8;8;8 | 1065458:106 | 1065819:106 | +;+;+ | 1409 | sprb | 12 | 2 | 132 | 0 | 33 | 38 |
| ENSDARG000 | 8;8;8;8;8;8;8; | 11080571:110 | 11080933:110 | -;-;-;-;-;-;-; | 2500 | ampd1 | 2 | 3 | 8 | 26 | 22 | 10 |
| ENSDARG000 | 8;8;8;8;8;8;8; | 18842278:188 | 18842836:188 | -;-;-;-;-;-;-; | 2318 | sh3gl1b | 469 | 455 | 566 | 464 | 716 | 440 |
| ENSDARG000 | 8;8;8;8;8;8;8; | 8046086:804 | 8047024:804 | +;+;+;+;+;+;+; | 2426 | abcd1 | 300 | 152 | 319 | 290 | 255 | 334 |
| ENSDARG000 | 8;8;8;8;8;8;8 | 44340504:443 | 44341184:443 | -;-;-;-;-;-;- | 1440 | NA | 130 | 309 | 108 | 54 | 174 | 298 |
| ENSDARG000 | 8;8;8 | 46418996:464 | 46419211:464 | +;+;+ | 1271 | NA | 23 | 2 | 5 | 0 | 17 | 15 |
| ENSDARG000 | 8;8;8;8;8;8;8; | 2509625:251 | 2510338:251 | -;-;-;-;-;-;-; | 1900 | acaa2 | 421 | 424 | 748 | 548 | 603 | 784 |
| ENSDARG000 | 8;8;8;8;8;8;8; | 13364950:133 | 13365018:133 | +;+;+;+;+;+;+; | 2029 | slc5a5 | 0 | 0 | 0 | 0 | 0 | 0 |
| ENSDARG000 | 8;8 | 23245596:232 | 23245972:232 | +;+ | 3357 | NA | 2 | 4 | 6 | 14 | 0 | 6 |
| ENSDARG000 | 8;8;8;8;8;8;8; | 1082100:109 | 1082416:109 | +;+;+;+;+;+;+; | 2880 | LZTS3 | 16 | 42 | 16 | 32 | 31 | 97 |
| ENSDARG000 | 8;8;8;8;8;8;8; | 53204017:532 | 53204094:532 | +;+;+;+;+;+;+; | 5711 | NA | 2 | 2 | 6 | 6 | 10 | 0 |
| ENSDARG000 | 8;8;8;8;8;8;8; | 44373764:443 | 44373903:443 | -;-;-;-;-;-;-; | 1321 | mhc2d8.45b1 | 10 | 0 | 4 | 0 | 0 | 20 |
| ENSDARG000 | 8;8;8;8;8;8;8; | 14710542:147 | 14710807:147 | +;+;+;+;+;+;+; | 7025 | CACNA1E | 2 | 2 | 4 | 4 | 12 | 0 |
| ENSDARG000 | 8;8;8;8;8;8;8; | 51304099:513 | 51309192:513 | -;-;-;-;-;-;-; | 8224 | kansl3 | 1170 | 1118 | 1372 | 839 | 1078 | 1028 |
| ENSDARG000 | 8;8 | 23260411:232 | 23261096:232 | +;+ | 3308 | NA | 0 | 0 | 0 | 0 | 0 | 0 |
| ENSDARG000 | 8;8;8;8;8;8;8; | 23252850:232 | 23254079:232 | -;-;-;-;-;-;-; | 3985 | NA | 0 | 0 | 3 | 2 | 4 | 0 |
| ENSDARG000 | 8;8 | 38922317:389 | 38922406:389 | +;+ | 431 | NA | 0 | 0 | 0 | 0 | 0 | 0 |
| ENSDARG000 | 8 | 44420108 | 44421418 | + | 1311 | NA | 0 | 0 | 0 | 0 | 0 | 0 |
| ENSDARG000 | 8;8;8;8;8;8;8; | 46447061:464 | 46447173:464 | -;-;-;-;-;-;-; | 943 | sult1st7 | 0 | 0 | 0 | 0 | 0 | 0 |
| ENSDARG000 | 8;8;8;8 | 2530065:253 | 2530231:253 | +;+;+;+ | 883 | mrpl40 | 198 | 225 | 329 | 162 | 254 | 232 |
| ENSDARG000 | 8;8;8;8 | 41571030:415 | 41571076:415 | +;+;+;+ | 371 | NA | 0 | 0 | 0 | 0 | 0 | 0 |
| ENSDARG000 | 8;8;8;8;8;8 | 41533268:415 | 41533881:415 | +;+;+;+;+;+ | 1816 | NA | 0 | 0 | 0 | 0 | 0 | 0 |
| ENSDARG000 | 8;8 | 16726386:167 | 16726494:167 | +;+ | 738 | smim15 | 464 | 528 | 657 | 342 | 534 | 430 |
| ENSDARG000 | 8;8;8;8;8;8;8; | 18885108:188 | 18885584:188 | -;-;-;-;-;-;-; | 2051 | rorca | 1182 | 569 | 716 | 462 | 653 | 1433 |
| ENSDARG000 | 8;8;8 | 2536280:253 | 2536934:253 | -;-;- | 912 | si:ch211-51h | 60 | 53 | 104 | 36 | 24 | 64 |
| ENSDARG000 | 8;8 | 38938290:389 | 38938553:389 | -;- | 450 | NA | 0 | 0 | 0 | 0 | 0 | 0 |
| ENSDARG000 | 8;8;8;8;8;8;8 | 44442220:444 | 44442440:444 | +;+;+;+;+;+;+ | 1797 | mhc1laa | 224 | 207 | 217 | 99 | 269 | 165 |
| ENSDARG000 | 8;8;8;8;8;8 | 1118328:112 | 1118418:112 | +;+;+;+;+;+ | 1602 | ubox5 | 122 | 93 | 158 | 85 | 108 | 148 |
| ENSDARG000 | 8;8;8 | 28294476:282 | 28294636:282 | -;-;- | 578 | NA | 0 | 0 | 0 | 0 | 0 | 1 |
| ENSDARG000 | 8;8;8 | 28297066:283 | 28297764:283 | -;-;- | 2610 | LMOD1 | 2 | 7 | 0 | 4 | 2 | 0 |
| ENSDARG000 | 8;8;8;8;8;8;8; | 48942470:489 | 48942598:489 | +;+;+;+;+;+;+; | 4150 | rer1 | 894 | 1245 | 1426 | 602 | 1250 | 953 |
| ENSDARG000 | 8;8;8;8;8;8;8; | 39511932:395 | 39512120:395 | +;+;+;+;+;+;+; | 5543 | LZTS1 | 170 | 16 | 16 | 4 | 164 | 4 |
| ENSDARG000 | 8;8;8;8 | 16772174:167 | 16772222:167 | +;+;+;+ | 661 | NA | 0 | 0 | 0 | 0 | 0 | 0 |
| ENSDARG000 | 8;8;8;8;8;8;8; | 16738282:167 | 16738605:167 | +;+;+;+;+;+;+; | 2506 | ercc8 | 212 | 174 | 254 | 110 | 236 | 182 |
| ENSDARG000 | 8;8;8;8;8;8;8; | 25913787:259 | 25913955:259 | +;+;+;+;+;+;+; | 3035 | sema3h | 11 | 0 | 2 | 2 | 9 | 8 |
| ENSDARG000 | 8;8;8;8;8 | 11114754:111 | 11116646:111 | -;-;-;-;- | 2636 | nras | 338 | 475 | 501 | 206 | 323 | 359 |
| ENSDARG000 | 8;8;8;8;8;8;8; | 41647539:416 | 41647875:416 | +;+;+;+;+;+;+; | 1048 | si:ch211-158 | 7 | 11 | 21 | 10 | 25 | 103 |
| ENSDARG000 | 8;8;8;8;8;8;8; | 16758304:167 | 16758508:167 | +;+;+;+;+;+;+; | 1894 | elovl7a | 30 | 0 | 2 | 10 | 10 | 2 |
| ENSDARG000 | 8;8;8;8;8;8;8; | 21978057:219 | 21978807:219 | -;-;-;-;-;-;-; | 5759 | NA | 90 | 180 | 206 | 243 | 209 | 123 |

| | | | | | | | | | | | | |
|---|---|---|---|---|---|---|---|---|---|---|---|---|
| ENSDARG000 | 8;8;8;8 | 16732834:167 | 16734328:167 | -;-;-;- | 1835 | ndufaf2 | 112 | 68 | 111 | 40 | 108 | 82 |
| ENSDARG000 | 8;8;8 | 1148876:1151 | 1149005:1151 | +;+;+ | 517 | avp | 0 | 0 | 0 | 0 | 0 | 0 |
| ENSDARG000 | 8;8;8;8;8;8;8; | 28305213:283 | 28307448:283 | -;-;-;-;-;-;-;-; | 7019 | kdm5bb | 2745 | 2481 | 4190 | 2700 | 3557 | 2223 |
| ENSDARG000 | 8;8;8;8;8;8;8 | 25034529:250 | 25034580:250 | +;+;+;+;+;+;+ | 1583 | ngrn | 240 | 100 | 208 | 184 | 162 | 465 |
| ENSDARG000 | 8;8;8;8;8;8;8; | 20863025:208 | 20863247:208 | +;+;+;+;+;+;+; | 2808 | fzr1a | 867 | 703 | 1002 | 632 | 721 | 932 |
| ENSDARG000 | 8;8;8 | 41710359:417 | 41710452:417 | +;+;+ | 823 | NA | 0 | 0 | 2 | 0 | 0 | 0 |
| ENSDARG000 | 8;8 | 41712799:417 | 41713029:417 | +;+ | 384 | NA | 10 | 0 | 2 | 0 | 0 | 2 |
| ENSDARG000 | 8;8;8;8;8;8;8; | 11136384:111 | 11136982:111 | -;-;-;-;-;-;-;-; | 1469 | ASB13 | 92 | 100 | 66 | 124 | 112 | 108 |
| ENSDARG000 | 8;8;8;8;8;8;8; | 39607466:396 | 39607547:396 | +;+;+;+;+;+;+; | 2631 | msi1b | 4 | 2 | 6 | 12 | 2 | 8 |
| ENSDARG000 | 8;8;8;8;8;8;8; | 30949968:309 | 30950143:309 | -;-;-;-;-;-;-;-; | 2442 | ralgds | 139 | 45 | 126 | 58 | 86 | 97 |
| ENSDARG000 | 8;8;8;8;8;8;8; | 23355484:233 | 23356298:233 | +;+;+;+;+;+;+; | 7594 | dnmt3ba | 230 | 608 | 808 | 196 | 520 | 350 |
| ENSDARG000 | 8;8;8;8;8;8;8; | 8166285:8196 | 8166408:8197 | +;+;+;+;+;+;+; | 6490 | NA | 2166 | 1713 | 1680 | 1476 | 1596 | 2381 |
| ENSDARG000 | 8;8;8;8;8;8;8; | 46471875:464 | 46471981:464 | -;-;-;-;-;-;-;-; | 1200 | NA | 0 | 0 | 0 | 0 | 0 | 0 |
| ENSDARG000 | 8;8;8;8;8;8 | 1170015:1170 | 1170105:1170 | +;+;+;+;+;+ | 505 | ccl27a | 30 | 0 | 0 | 0 | 0 | 0 |
| ENSDARG000 | 8;8;8;8;8;8;8; | 18936603:189 | 18936818:189 | -;-;-;-;-;-;-;-; | 1947 | NA | 59 | 33 | 20 | 4 | 76 | 26 |
| ENSDARG000 | 8;8;8;8;8;8;8; | 48965767:489 | 48966318:489 | +;+;+;+;+;+;+; | 3038 | AAK1 | 176 | 72 | 183 | 89 | 279 | 75 |
| ENSDARG000 | 8 | 41987350 | 41988113 | + | 764 | NA | 0 | 0 | 0 | 0 | 0 | 0 |
| ENSDARG000 | 8;8;8;8;8;8;8; | 25041713:250 | 25043702:250 | -;-;-;-;-;-;-;-; | 4584 | sort1b | 3673 | 2637 | 2922 | 2920 | 4218 | 1689 |
| ENSDARG000 | 8;8;8;8;8;8;8; | 42059495:420 | 42059611:420 | -;-;-;-;-;-;-;- | 1488 | gfra2a | 30 | 22 | 101 | 0 | 69 | 22 |
| ENSDARG000 | 8;8;8;8;8;8;8; | 2543525:2543 | 2543724:2543 | +;+;+;+;+;+;+; | 2824 | gle1 | 357 | 318 | 517 | 184 | 577 | 283 |
| ENSDARG000 | 8;8;8;8;8;8;8; | 44475793:444 | 44475838:444 | +;+;+;+;+;+;+; | 1592 | NA | 0 | 0 | 0 | 0 | 0 | 0 |
| ENSDARG000 | 8;8;8;8;8;8;8; | 31030613:310 | 31031802:310 | -;-;-;-;-;-;-;-; | 7697 | snrnp200 | 3039 | 2926 | 4839 | 1520 | 3931 | 2252 |
| ENSDARG000 | 8;8;8;8;8;8;8; | 1187928:1188 | 1188022:1188 | +;+;+;+;+;+;+; | 1313 | SLC35D2 | 65 | 36 | 28 | 4 | 53 | 46 |
| ENSDARG000 | 8;8;8;8;8;8 | 44549290:445 | 44549493:445 | +;+;+;+;+;+ | 860 | NA | 0 | 0 | 0 | 0 | 0 | 0 |
| ENSDARG000 | 8;8;8;8;8;8;8; | 35212233:353 | 35213274:353 | +;+;+;+;+;+;+; | 5667 | TMEM132C | 2 | 1 | 0 | 0 | 0 | 3 |
| ENSDARG000 | 8;8;8;8;8;8;8; | 46499098:464 | 46499501:464 | -;-;-;-;-;-;-;- | 1614 | sult1st1 | 14 | 261 | 485 | 0 | 10 | 81 |
| ENSDARG000 | 8;8;8;8;8;8;8; | 31016180:310 | 31016236:310 | +;+;+;+;+;+;+; | 3516 | odf2b | 84 | 146 | 169 | 69 | 176 | 64 |
| ENSDARG000 | 8;8;8;8;8;8;8; | 11151805:111 | 11152110:111 | -;-;-;-;-;-;-;-; | 1340 | plekha2 | 290 | 418 | 348 | 269 | 273 | 397 |
| ENSDARG000 | 8;8;8;8;8;8;8; | 6424721:6425 | 6425179:6425 | -;-;-;-;-;-;-;- | 1015 | NA | 0 | 0 | 0 | 0 | 0 | 7 |
| ENSDARG000 | 8;8;8;8;8;8;8; | 31008287:310 | 31008633:310 | +;+;+;+;+;+;+; | 2195 | ptgesl | 332 | 591 | 511 | 190 | 566 | 370 |
| ENSDARG000 | 8;8;8;8;8;8;8; | 13389115:133 | 13389301:133 | +;+;+;+;+;+;+; | 3638 | NA | 38 | 66 | 73 | 94 | 120 | 168 |
| ENSDARG000 | 8;8 | 23387884:233 | 23389988:233 | -;- | 2247 | NA | 0 | 1 | 0 | 2 | 1 | 0 |
| ENSDARG000 | 8;8;8;8;8;8;8; | 23381892:233 | 23382694:233 | +;+;+;+;+;+;+; | 3528 | mapre1a | 109 | 128 | 174 | 81 | 229 | 171 |
| ENSDARG000 | 8;8;8;8;8;8;8; | 51341612:513 | 51341827:513 | -;-;-;-;-;-;-;-; | 1394 | si:ch211-198 | 0 | 0 | 0 | 0 | 0 | 0 |
| ENSDARG000 | 8 | 53464216 | 53465017 | + | 802 | NA | 0 | 0 | 0 | 0 | 0 | 0 |
| ENSDARG000 | 8;8;8;8;8;8;8; | 44566708:445 | 44567191:445 | -;-;-;-;-;-;-;-; | 1367 | rsph14 | 0 | 0 | 0 | 0 | 0 | 0 |
| ENSDARG000 | 8;8;8;8;8;8;8; | 42545757:425 | 42549786:425 | -;-;-;-;-;-;-;-; | 5181 | NA | 632 | 971 | 156 | 701 | 252 | 1587 |
| ENSDARG000 | 8;8;8;8;8;8;8; | 35809081:358 | 35810236:358 | -;-;-;-;-;-;-;- | 2953 | slc15a4 | 954 | 613 | 972 | 595 | 961 | 1018 |
| ENSDARG000 | 8;8;8;8;8;8;8; | 53474755:534 | 53475250:534 | -;-;-;-;-;-;-;-; | 2157 | CHDH | 94 | 116 | 90 | 82 | 152 | 128 |
| ENSDARG000 | 8;8;8;8;8;8;8; | 49065348:490 | 49065496:490 | +;+;+;+;+;+;+; | 2217 | ddx51 | 269 | 328 | 526 | 222 | 428 | 241 |
| ENSDARG000 | 8;8;8;8;8;8;8; | 46513743:465 | 46514329:465 | -;-;-;-;-;-;-;- | 1622 | sult1st2 | 0 | 0 | 8 | 0 | 0 | 0 |
| ENSDARG000 | 8;8;8;8;8 | 44599405:446 | 44600342:446 | -;-;-;-;- | 1878 | bag4 | 345 | 278 | 319 | 204 | 329 | 256 |
| ENSDARG000 | 8;8;8;8;8;8;8; | 1210468:1210 | 1210507:1210 | -;-;-;-;-;-;-;-; | 1400 | znf367 | 144 | 512 | 519 | 267 | 473 | 267 |
| ENSDARG000 | 8;8 | 1221881:1226 | 1222813:1227 | +;+ | 1356 | NA | 10 | 7 | 18 | 7 | 14 | 17 |
| ENSDARG000 | 8;8;8;8;8;8;8; | 1222132:1222 | 1222616:1222 | -;-;-;-;-;-;-;-; | 2697 | cdc14b | 508 | 752 | 839 | 434 | 737 | 565 |
| ENSDARG000 | 8;8;8;8;8;8;8; | 25959940:259 | 25959988:259 | +;+;+;+;+;+;+; | 3658 | borcs6 | 496 | 402 | 458 | 320 | 467 | 667 |
| ENSDARG000 | 8;8;8;8 | 44613135:446 | 44613311:446 | +;+;+;+ | 919 | lsm1 | 86 | 209 | 182 | 119 | 278 | 94 |
| ENSDARG000 | 8;8;8;8;8;8;8; | 11173794:111 | 11174848:111 | -;-;-;-;-;-;-;-; | 11505 | NA | 1680 | 1077 | 1822 | 1175 | 1476 | 1372 |
| ENSDARG000 | 8;8;8;8;8;8 | 46536893:465 | 46536926:465 | +;+;+;+;+;+ | 1248 | NA | 0 | 2 | 4 | 0 | 4 | 0 |
| ENSDARG000 | 8;8;8;8;8;8;8; | 51350453:513 | 51351256:513 | -;-;-;-;-;-;-;-; | 3104 | chmp7 | 930 | 933 | 897 | 675 | 809 | 723 |
| ENSDARG000 | 8;8 | 6589486:6608 | 6589744:6609 | -;- | 540 | NA | 0 | 0 | 0 | 0 | 0 | 0 |
| ENSDARG000 | 8;8 | 28593707:285 | 28594407:285 | +;+ | 1041 | tcf15 | 16 | 0 | 14 | 0 | 4 | 4 |
| ENSDARG000 | 8;8;8;8;8;8;8; | 6576940:6581 | 6577018:6582 | +;+;+;+;+;+;+; | 1344 | vsig8b | 3 | 1 | 0 | 2 | 8 | 6 |
| ENSDARG000 | 8;8;8;8;8;8;8; | 20880848:208 | 20880933:208 | +;+;+;+;+;+;+; | 1987 | si:ch73-196i1 | 2 | 0 | 0 | 0 | 0 | 8 |
| ENSDARG000 | 8;8;8;8;8;8;8; | 44623540:446 | 44623645:446 | +;+;+;+;+;+;+; | 2140 | NA | 221 | 163 | 304 | 120 | 236 | 324 |
| ENSDARG000 | 8;8;8;8;8;8;8; | 8291492:8291 | 8291562:8292 | +;+;+;+;+;+;+; | 2557 | SRPK3 | 164 | 132 | 148 | 59 | 169 | 88 |
| ENSDARG000 | 8;8 | 53496201:534 | 53496295:534 | +;+ | 233 | NA | 0 | 0 | 0 | 0 | 0 | 0 |
| ENSDARG000 | 8;8;8;8;8;8;8; | 35964482:359 | 35964769:359 | +;+;+;+;+;+;+; | 1254 | GLT1D1 | 4 | 4 | 10 | 2 | 11 | 3 |
| ENSDARG000 | 8;8;8;8;8;8;8; | 14855471:148 | 14855524:148 | +;+;+;+;+;+;+; | 3378 | NA | 4 | 0 | 0 | 0 | 0 | 0 |
| ENSDARG000 | 8;8;8;8;8;8 | 46552778:465 | 46552811:465 | +;+;+;+;+;+ | 1249 | NA | 6 | 0 | 0 | 0 | 2 | 0 |
| ENSDARG000 | 8;8;8;8;8;8;8; | 16990120:170 | 16990178:170 | +;+;+;+;+;+;+; | 2112 | PDE4D | 57 | 41 | 95 | 18 | 48 | 30 |
| ENSDARG000 | 8;8;8;8 | 36354125:363 | 36354220:363 | +;+;+;+ | 939 | NA | 0 | 0 | 0 | 0 | 0 | 0 |
| ENSDARG000 | 8;8;8;8;8;8;8; | 17055292:170 | 17055920:170 | -;-;-;-;-;-;-;- | 1680 | rab3c | 10 | 26 | 66 | 102 | 0 | 116 |
| ENSDARG000 | 8;8;8;8 | 28627060:286 | 28627123:286 | +;+;+;+ | 885 | NA | 126 | 117 | 56 | 53 | 146 | 129 |
| ENSDARG000 | 8;8;8;8;8;8;8 | 22277198:222 | 22277358:222 | +;+;+;+;+;+;+ | 1326 | dffb | 204 | 182 | 209 | 160 | 216 | 164 |
| ENSDARG000 | 8;8;8;8;8 | 28629741:286 | 28630141:286 | +;+;+;+;+ | 1865 | NA | 10 | 11 | 0 | 2 | 24 | 38 |
| ENSDARG000 | 8 | 28654289 | 28655338 | - | 1050 | mc3r | 0 | 0 | 0 | 0 | 0 | 0 |
| ENSDARG000 | 8;8;8;8;8;8;8; | 51375308:513 | 51376044:513 | -;-;-;-;-;-;-;- | 1769 | lgi3 | 4 | 0 | 0 | 8 | 2 | 0 |
| ENSDARG000 | 8;8;8;8;8;8;8; | 44703854:447 | 44703966:447 | +;+;+;+;+;+;+; | 1378 | star | 6 | 4 | 10 | 0 | 18 | 12 |
| ENSDARG000 | 8;8;8;8;8 | 25973151:259 | 25974147:259 | -;-;-;-;- | 2255 | si:dkev-72l14. | 0 | 2 | 0 | 0 | 0 | 0 |
| ENSDARG000 | 8;8;8;8;8;8;8; | 49092039:490 | 49092085:490 | +;+;+;+;+;+;+; | 5540 | golm1 | 1801 | 1232 | 1759 | 1454 | 1820 | 1582 |
| ENSDARG000 | 8;8;8;8;8;8 | 23395469:233 | 23395533:233 | +;+;+;+;+;+ | 1456 | suv39h1a | 0 | 6 | 0 | 4 | 4 | 1 |
| ENSDARG000 | 8;8;8;8;8;8;8; | 46559332:465 | 46559782:465 | -;-;-;-;-;-;-;-; | 1625 | sult1st3 | 0 | 4 | 4 | 10 | 114 | 0 |
| ENSDARG000 | 8;8;8;8;8;8;8; | 53517998:535 | 53518400:535 | -;-;-;-;-;-;-;-; | 2281 | actr8 | 240 | 261 | 328 | 157 | 270 | 188 |
| ENSDARG000 | 8;8;8;8 | 42609777:426 | 42610309:426 | -;-;-;- | 1450 | kazald3 | 8 | 14 | 12 | 20 | 30 | 90 |
| ENSDARG000 | 8;8;8;8;8;8;8; | 23390844:233 | 23391451:233 | +;+;+;+;+;+;+; | 3104 | NA | 10 | 0 | 0 | 0 | 0 | 0 |
| ENSDARG000 | 8;8;8;8;8;8 | 25079470:250 | 25079564:250 | +;+;+;+;+;+ | 1966 | sypl2b | 5241 | 6139 | 6528 | 6115 | 7573 | 7077 |
| ENSDARG000 | 8;8;8;8;8;8;8; | 11204059:112 | 11205237:112 | -;-;-;-;-;-;-;-; | 3834 | unc45b | 2 | 18 | 62 | 12 | 2 | 10 |
| ENSDARG000 | 8;8;8;8;8;8;8; | 17069577:170 | 17069646:170 | +;+;+;+;+;+;+; | 996 | si:dkev-190g1 | 33 | 34 | 88 | 110 | 28 | 171 |
| ENSDARG000 | 8;8;8;8;8;8;8; | 8341082:8341 | 8341645:8341 | -;-;-;-;-;-;-;-; | 1132 | tsr2 | 497 | 504 | 697 | 325 | 522 | 410 |
| ENSDARG000 | 8;8;8;8 | 13417986:134 | 13418127:134 | -;-;-;- | 767 | NA | 0 | 0 | 0 | 0 | 0 | 0 |
| ENSDARG000 | 8;8;8;8;8;8;8; | 6651253:6656 | 6651405:6656 | -;-;-;-;-;-;-;-; | 5682 | dock5 | 94 | 41 | 40 | 51 | 76 | 91 |
| ENSDARG000 | 8;8;8;8;8;8;8; | 26007308:260 | 26008269:260 | +;+;+;+;+;+;+; | 5039 | xpc | 131 | 167 | 156 | 101 | 136 | 85 |
| ENSDARG000 | 8;8;8;8;8;8 | 39650940:396 | 39651206:396 | -;-;-;-;-;- | 1230 | NA | 0 | 0 | 0 | 0 | 0 | 2 |
| ENSDARG000 | 8;8;8;8;8;8;8; | 22283564:222 | 22284659:222 | -;-;-;-;-;-;-;-; | 2235 | si:ch211-147a | 214 | 186 | 283 | 195 | 156 | 188 |
| ENSDARG000 | 8;8;8;8;8;8;8; | 28667021:286 | 28667387:286 | +;+;+;+;+;+;+; | 4866 | slc13a3 | 0 | 2 | 26 | 2 | 62 | 0 |
| ENSDARG000 | 8;8;8;8 | 13427678:134 | 13427819:134 | -;-;-;- | 766 | NA | 0 | 0 | 0 | 0 | 0 | 0 |
| ENSDARG000 | 8;8;8;8;8;8;8; | 36362898:363 | 36365664:363 | -;-;-;-;-;-;-;- | 3549 | NA | 0 | 0 | 0 | 0 | 0 | 0 |
| ENSDARG000 | 8;8;8;8;8;8;8; | 25091407:250 | 25091578:250 | +;+;+;+;+;+;+; | 2337 | atxn7l2b | 170 | 144 | 252 | 258 | 255 | 158 |
| ENSDARG000 | 8;8;8;8;8;8;8; | 42629748:426 | 42629895:426 | +;+;+;+;+;+;+; | 1474 | crybb2 | 0 | 0 | 0 | 0 | 0 | 0 |
| ENSDARG000 | 8;8;8;8 | 13441808:134 | 13441949:134 | -;-;-;- | 767 | NA | 0 | 0 | 0 | 0 | 0 | 0 |
| ENSDARG000 | 8;8;8;8;8;8;8; | 39674707:396 | 39674834:396 | +;+;+;+;+;+;+; | 1390 | prkab1b | 1001 | 1019 | 927 | 924 | 952 | 1028 |
| ENSDARG000 | 8;8;8;8;8;8 | 46587593:465 | 46587774:465 | +;+;+;+;+;+ | 2478 | tas1r2.1 | 0 | 0 | 0 | 0 | 0 | 0 |
| ENSDARG000 | 8;8;8;8;8;8;8; | 19214207:192 | 19215006:192 | -;-;-;-;-;-;-;-; | 1074 | NA | 0 | 0 | 0 | 0 | 0 | 0 |
| ENSDARG000 | 8;8;8;8;8;8;8; | 19213867:192 | 19215006:192 | -;-;-;-;-;-;-;-; | 1412 | NA | 0 | 0 | 0 | 0 | 0 | 0 |
| ENSDARG000 | 8;8;8;8;8;8;8; | 44714336:447 | 44714541:447 | +;+;+;+;+;+;+; | 1506 | ELMOD3 | 59 | 29 | 56 | 44 | 45 | 40 |
| ENSDARG000 | 8;8;8;8;8;8;8; | 36377006:363 | 36378321:363 | -;-;-;-;-;-;-;- | 1941 | mhc2d8.37a2 | 10 | 0 | 4 | 187 | 4 | 0 |
| ENSDARG000 | 8;8;8;8;8;8;8; | 17078692:170 | 17078908:170 | +;+;+;+;+;+;+; | 2415 | plk2b | 8975 | 3846 | 6416 | 4229 | 3470 | 6109 |
| ENSDARG000 | 8;8;8;8 | 13453219:134 | 13453360:134 | -;-;-;- | 766 | NA | 0 | 0 | 0 | 0 | 0 | 0 |
| ENSDARG000 | 8;8;8 | 28695914:287 | 28696054:287 | +;+;+ | 1629 | ocstamp | 34 | 41 | 34 | 26 | 58 | 16 |
| ENSDARG000 | 8;8;8;8;8;8;8; | 11234168:112 | 11235058:112 | -;-;-;-;-;-;-;-; | 2743 | pip5k1bb | 1413 | 74 | 74 | 186 | 250 | 110 |
| ENSDARG000 | 8;8;8;8;8;8;8; | 8362441:8372 | 8363154:8372 | -;-;-;-;-;-;-;-; | 2972 | CDK16 | 320 | 370 | 383 | 229 | 334 | 290 |
| ENSDARG000 | 8 | 23415023 | 23416362 | - | 1340 | gpr173 | 0 | 0 | 8 | 0 | 0 | 0 |
| ENSDARG000 | 8;8;8;8;8;8 | 46610020:466 | 46610184:466 | +;+;+;+;+;+ | 2473 | tas1r2.2 | 2 | 0 | 0 | 0 | 0 | 0 |
| ENSDARG000 | 8 | 22289320 | 22290886 | + | 1567 | b3gnt7l | 131 | 8 | 10 | 6 | 10 | 189 |
| ENSDARG000 | 8;8 | 25117390:251 | 25118621:251 | -;- | 1366 | AMIGO1 | 289 | 103 | 301 | 331 | 242 | 66 |
| ENSDARG000 | 8;8;8;8;8;8;8; | 36397818:363 | 36399164:363 | -;-;-;-;-;-;-;- | 2396 | NA | 0 | 0 | 0 | 0 | 0 | 0 |
| ENSDARG000 | 8;8;8;8;8;8;8; | 49117506:491 | 49117554:491 | +;+;+;+;+;+;+; | 2291 | rad21l1 | 0 | 0 | 0 | 0 | 0 | 0 |
| ENSDARG000 | 8;8 | 2575993:2578 | 2576559:2579 | +;+ | 786 | si:ch211-51h9 | 75 | 106 | 237 | 88 | 116 | 150 |
| ENSDARG000 | 8;8;8;8;8;8;8; | 20897849:208 | 20898157:208 | -;-;-;-;-;-;-;-; | 2244 | haus5 | 30 | 192 | 202 | 62 | 258 | 50 |
| ENSDARG000 | 8;8;8 | 6863462:6865 | 6864488:6865 | +;+;+ | 2713 | nefmb | 0 | 0 | 0 | 0 | 0 | 0 |
| ENSDARG000 | 8;8;8;8;8;8;8; | 28724692:287 | 28724902:287 | +;+;+;+;+;+;+; | 2493 | CCSER1 | 32 | 28 | 18 | 50 | 43 | 72 |
| ENSDARG000 | 8;8 | 25173317:251 | 25173459:251 | +;+ | 1744 | GPR61 | 4388 | 1664 | 1358 | 2672 | 1544 | 2872 |
| ENSDARG000 | 8;8;8;8;8;8;8; | 25145464:251 | 25145660:251 | +;+;+;+;+;+;+; | 1995 | gnai3 | 2607 | 1381 | 1079 | 1097 | 1388 | 1955 |
| ENSDARG000 | 8;8;8;8;8;8;8; | 14886452:148 | 14886514:148 | +;+;+;+;+;+;+; | 2088 | soat1 | 7876 | 4735 | 4213 | 3394 | 7025 | 4258 |
| ENSDARG000 | 8;8;8;8;8 | 36466771:364 | 36467445:364 | -;-;-;-;- | 1428 | si:busm1-266 | 0 | 0 | 0 | 0 | 0 | 0 |
| ENSDARG000 | 8;8;8;8;8;8;8; | 53537103:535 | 53537525:535 | -;-;-;-;-;-;-;-; | 1348 | selenok | 695 | 452 | 552 | 380 | 486 | 501 |
| ENSDARG000 | 8;8;8;8;8;8;8; | 23436411:234 | 23436445:234 | +;+;+;+;+;+;+; | 2174 | NA | 0 | 0 | 0 | 0 | 4 | 8 |
| ENSDARG000 | 8;8;8;8 | 46641314:466 | 46641438:466 | +;+;+;+ | 905 | her3 | 0 | 0 | 0 | 0 | 0 | 0 |
| ENSDARG000 | 8;8;8;8 | 13470855:134 | 13470996:134 | -;-;-;- | 765 | NA | 0 | 0 | 0 | 0 | 0 | 0 |
| ENSDARG000 | 8;8;8;8 | 36472444:364 | 36473389:364 | -;-;-;- | 1590 | mhc2daa | 2 | 18 | 4 | 6 | 22 | 16 |
| ENSDARG000 | 8;8;8;8;8;8;8 | 6873853:6874 | 6875839:6875 | -;-;-;-;-;-;- | 3209 | neflb | 50 | 128 | 89 | 164 | 62 | 71 |
| ENSDARG000 | 8;8;8;8;8;8;8; | 2585256:2586 | 2586104:2586 | -;-;-;-;-;-;-;-; | 2043 | seta | 2670 | 3809 | 5934 | 2552 | 4675 | 3270 |
| ENSDARG000 | 8;8;8;8 | 13485196:134 | 13485337:134 | -;-;-;- | 766 | NA | 0 | 0 | 0 | 0 | 0 | 0 |
| ENSDARG000 | 8;8;8 | 17087060:170 | 17087084:170 | +;+;+ | 798 | NA | 0 | 0 | 10 | 2 | 0 | 6 |
| ENSDARG000 | 8;8;8;8;8;8 | 11325310:113 | 11325472:113 | +;+;+;+;+;+ | 2780 | fxn | 106 | 116 | 140 | 82 | 126 | 64 |
| ENSDARG000 | 8;8;8;8;8;8;8; | 22297813:222 | 22297979:222 | -;-;-;-;-;-;-;-; | 3601 | CEP104 | 245 | 180 | 229 | 99 | 217 | 202 |
| ENSDARG000 | 8;8;8;8;8;8;8 | 6880262:6881 | 6880857:6881 | -;-;-;-;-;-;- | 2995 | NA | 158 | 242 | 299 | 89 | 321 | 207 |
| ENSDARG000 | 8;8;8;8;8;8;8; | 17143501:171 | 17143589:171 | +;+;+;+;+;+;+; | 1785 | mier3b | 171 | 83 | 219 | 109 | 147 | 175 |
| ENSDARG000 | 8;8;8;8;8 | 19232238:192 | 19232621:192 | -;-;-;-;- | 1500 | ABHD17A | 237 | 143 | 198 | 70 | 185 | 172 |
| ENSDARG000 | 8;8;8;8;8;8;8; | 26034623:260 | 26034727:260 | +;+;+;+;+;+;+; | 2394 | arih2 | 1374 | 1329 | 1233 | 1078 | 1351 | 1106 |
| ENSDARG000 | 8;8;8 | 29581495:296 | 29582959:296 | -;-;- | 1852 | NA | 0 | 0 | 0 | 0 | 0 | 0 |
| ENSDARG000 | 8 | 29593986 | 29595615 | + | 1630 | atoh1a | 4 | 6 | 0 | 0 | 0 | 0 |
| ENSDARG000 | 8;8;8;8;8;8;8; | 28900689:289 | 28901328:289 | +;+;+;+;+;+;+; | 3799 | grid2 | 6 | 0 | 6 | 0 | 0 | 2 |
| ENSDARG000 | 8;8;8;8;8;8;8; | 44759576:447 | 44759798:447 | +;+;+;+;+;+;+; | 4181 | NA | 32 | 19 | 10 | 0 | 86 | 14 |
| ENSDARG000 | 8;8;8;8;8;8;8; | 51430597:514 | 51434739:514 | -;-;-;-;-;-;-;-; | 7424 | fgfr1a | 46 | 0 | 344 | 142 | 305 | 12 |
| ENSDARG000 | 8;8;8;8;8;8;8; | 23485079:235 | 23485781:235 | +;+;+;+;+;+;+; | 3057 | sema3gb | 0 | 0 | 2 | 0 | 2 | 0 |
| ENSDARG000 | 8;8;8;8;8;8;8; | 6903327:6903 | 6904145:6904 | -;-;-;-;-;-;-;-; | 1714 | asb6 | 94 | 160 | 219 | 79 | 215 | 185 |
| ENSDARG000 | 8;8;8;8;8;8;8; | 14915332:149 | 14915572:149 | +;+;+;+;+;+;+; | 8723 | abl2 | 1043 | 790 | 977 | 661 | 1013 | 812 |
| ENSDARG000 | 8;8;8;8;8;8;8; | 25247245:252 | 25247469:252 | +;+;+;+;+;+;+; | 4908 | AMPD2 | 475 | 215 | 173 | 293 | 259 | 334 |
| ENSDARG000 | 8;8;8;8 | 49149907:491 | 49150010:491 | +;+;+;+ | 819 | NA | 0 | 0 | 2 | 0 | 0 | 0 |
| ENSDARG000 | 8;8 | 49149904:491 | 49150253:491 | -;- | 426 | NA | 0 | 0 | 0 | 0 | 0 | 0 |
| ENSDARG000 | 8;8;8;8;8;8;8; | 42641812:426 | 42643224:426 | -;-;-;-;-;-;-;-; | 6915 | NA | 662 | 353 | 268 | 212 | 385 | 386 |
| ENSDARG000 | 8;8;8;8;8 | 13503377:135 | 13503546:135 | +;+;+;+;+ | 1932 | fut9d | 0 | 0 | 0 | 0 | 0 | 0 |
| ENSDARG000 | 8;8;8;8;8;8;8; | 36500061:365 | 36500338:365 | +;+;+;+;+;+;+; | 2859 | slc7a4 | 36 | 16 | 125 | 6 | 35 | 65 |
| ENSDARG000 | 8;8;8;8;8 | 2595424:2597 | 2595764:2598 | -;-;-;-;- | 934 | zdhhc12a | 186 | 205 | 230 | 167 | 279 | 252 |
| ENSDARG000 | 8;8;8;8;8;8;8; | 25225590:252 | 25226416:252 | -;-;-;-;-;-;-;-; | 2586 | gnat2 | 95 | 34 | 39 | 92 | 38 | 81 |
| ENSDARG000 | 8;8;8;8;8;8;8 | 13518768:135 | 13518873:135 | +;+;+;+;+;+;+ | 942 | NA | 0 | 0 | 0 | 2 | 0 | 0 |
| ENSDARG000 | 8;8 | 13534385:135 | 13535729:135 | -;- | 1454 | b3gnt3.1 | 0 | 0 | 0 | 0 | 0 | 0 |
| ENSDARG000 | 8;8;8 | 46896445:468 | 46897234:468 | -;-;- | 1090 | hes2.2 | 0 | 0 | 0 | 0 | 0 | 2 |
| ENSDARG000 | 8;8;8;8;8;8;8; | 39742459:397 | 39743271:397 | -;-;-;-;-;-;-;-; | 1861 | NA | 39 | 0 | 26 | 32 | 29 | 12 |
| ENSDARG000 | 8;8;8;8;8 | 39760258:397 | 39760413:397 | +;+;+;+;+ | 1080 | cox6a1 | 1987 | 2365 | 1974 | 1606 | 2138 | 1805 |
| ENSDARG000 | 8;8;8;8 | 39724138:397 | 39724201:397 | +;+;+;+ | 2101 | pla2g1b | 0 | 0 | 0 | 0 | 0 | 0 |
| ENSDARG000 | 8;8;8;8;8;8;8; | 39733278:397 | 39733978:397 | -;-;-;-;-;-;-;-; | 1493 | asgrl1 | 2 | 0 | 6 | 6 | 6 | 0 |
| ENSDARG000 | 8;8;8;8;8;8;8; | 46921333:469 | 46921738:469 | -;-;-;-;-;-;-;-; | 857 | hes2.1 | 0 | 0 | 0 | 0 | 0 | 0 |
| ENSDARG000 | 8;8;8;8;8;8;8; | 11425048:114 | 11425098:114 | +;+;+;+;+;+;+; | 4325 | tjp2b | 4023 | 1589 | 1499 | 4516 | 1871 | 714 |
| ENSDARG000 | 8;8;8;8;8;8;8; | 42712747:427 | 42712898:427 | +;+;+;+;+;+;+; | 2132 | zgc:194007 | 22 | 2 | 0 | 0 | 2 | 0 |
| ENSDARG000 | 8;8;8;8;8;8;8; | 2606068:2606 | 2607417:2606 | -;-;-;-;-;-;-;-; | 3220 | slc25a25a | 1976 | 1713 | 1971 | 2080 | 1563 | 1114 |
| ENSDARG000 | 8;8;8;8;8;8;8; | 49169768:491 | 49170108:491 | -;-;-;-;-;-;-;-; | 1070 | fkbp1aa | 6070 | 7315 | 7817 | 7233 | 6991 | 5538 |
| ENSDARG000 | 8 | 36529691 | 36530048 | + | 358 | NA | 0 | 0 | 0 | 0 | 0 | 0 |
| ENSDARG000 | 8;8 | 13572531:135 | 13573702:135 | -;- | 1261 | b3gnt3.2 | 14 | 0 | 2 | 0 | 18 | 0 |
| ENSDARG000 | 8;8;8;8;8;8;8; | 1269941:1269 | 1270281:1270 | -;-;-;-;-;-;-;-; | 905 | PRXL2C | 119 | 108 | 63 | 82 | 84 | 108 |

| | | | | | | | | | | | | |
|---|---|---|---|---|---|---|---|---|---|---|---|---|
| ENSDARG000 | 8:8:8:8:8:8:8: | 17163494:171 | 17163879:171 | -:-:-:-:-:-:-:-: | 866 | mrps36 | 474 | 661 | 686 | 309 | 494 | 405 |
| ENSDARG000 | 8:8:8:8:8:8:8: | 19253205:192 | 19253809:192 | -:-:-:-:-:-:-:-: | 2477 | zgc:77486 | 474 | 361 | 577 | 421 | 393 | 282 |
| ENSDARG000 | 8:8:8:8:8:8:8: | 36541949:365 | 36543882:365 | -:-:-:-:-:-:-:-: | 3743 | NA | 6 | 0 | 14 | 2 | 8 | 2 |
| ENSDARG000 | 8;8 | 13597405:136 | 13598591:136 | -;- | 1275 | b3gnt3.3 | 0 | 2 | 2 | 0 | 0 | 0 |
| ENSDARG000 | 8:8:8:8:8:8:8: | 46939391:469 | 46939859:469 | +:+:+:+:+:+:+: | 5250 | espn | 6 | 0 | 0 | 0 | 0 | 0 |
| ENSDARG000 | 8;8;8;8 | 20918207:209 | 20918315:209 | +;+;+;+ | 611 | NA | 0 | 0 | 0 | 0 | 0 | 0 |
| ENSDARG000 | 8;8;8;8;8;8;8 | 22327251:223 | 22327860:223 | +;+;+;+;+;+;+ | 2177 | lrrc47 | 338 | 447 | 900 | 281 | 448 | 364 |
| ENSDARG000 | 8:8:8:8:8:8:8: | 44850710:448 | 44850773:448 | -:-:-:-:-:-:-:-: | 2015 | NA | 0 | 0 | 0 | 0 | 0 | 0 |
| ENSDARG000 | 8;8;8 | 13630492:136 | 13631660:136 | -;-;- | 1427 | b3gnt3.4 | 454 | 372 | 156 | 287 | 328 | 105 |
| ENSDARG000 | 8:8:8:8:8:8:8: | 29635968:296 | 29636335:296 | +:+:+:+:+:+:+: | 4722 | smarcad1a | 389 | 793 | 1154 | 308 | 1136 | 524 |
| ENSDARG000 | 8;8;8 | 2642384:2645 | 2642765:2645 | +;+;+ | 961 | naif1 | 40 | 3 | 16 | 22 | 28 | 12 |
| ENSDARG000 | 8:8:8:8:8:8:8: | 1284784:1290 | 1284831:1290 | +:+:+:+:+:+:+: | 6258 | NA | 192 | 144 | 186 | 122 | 213 | 112 |
| ENSDARG000 | 8;8;8;8 | 13694950:136 | 13695307:136 | +;+;+;+ | 766 | NA | 5 | 0 | 0 | 0 | 0 | 0 |
| ENSDARG000 | 8:8:8:8:8:8:8: | 8459192:8459 | 8459253:8459 | +:+:+:+:+:+:+: | 1431 | comta | 21 | 55 | 73 | 20 | 45 | 16 |
| ENSDARG000 | 8:8:8:8:8:8:8: | 6920834:6921 | 6921708:6921 | -:-:-:-:-:-:-:-: | 2497 | wdr13 | 878 | 979 | 721 | 653 | 993 | 656 |
| ENSDARG000 | 8;8;8;8 | 13643799:136 | 13644242:136 | -;-;-;- | 761 | NA | 68 | 89 | 111 | 96 | 89 | 73 |
| ENSDARG000 | 8:8:8:8:8:8:8: | 13700605:137 | 13701288:137 | +:+:+:+:+:+:+: | 4572 | lonrf1l | 528 | 753 | 432 | 402 | 1158 | 517 |
| ENSDARG000 | 8;8;8;8 | 20940459:209 | 20940517:209 | -;-;-;- | 582 | NA | 0 | 0 | 0 | 0 | 0 | 4 |
| ENSDARG000 | 8;8;8;8 | 20937980:209 | 20938062:209 | +;+;+;+ | 324 | NA | 2 | 0 | 0 | 0 | 0 | 0 |
| ENSDARG000 | 8:8:8:8:8:8:8: | 26059677:260 | 26059941:260 | +:+:+:+:+:+:+: | 2335 | impdh2 | 182 | 312 | 384 | 146 | 316 | 200 |
| ENSDARG000 | 8:8:8:8:8:8:8: | 13653699:136 | 13653822:136 | -:-:-:-:-:-:-:-: | 2082 | NA | 32 | 81 | 90 | 10 | 60 | 52 |
| ENSDARG000 | 8:8:8:8:8:8:8: | 2656231:2656 | 2656413:2656 | +:+:+:+:+:+:+: | 1681 | EEIG1 | 89 | 59 | 103 | 74 | 85 | 112 |
| ENSDARG000 | 8;8;8;8;8 | 13722656:137 | 13722779:137 | -;-;-;-;- | 704 | NA | 6 | 6 | 5 | 11 | 4 | 2 |
| ENSDARG000 | 8;8;8;8 | 6954984:6955 | 6955045:6955 | +;+;+;+ | 519 | si:ch211-255 | 0 | 0 | 0 | 0 | 0 | 0 |
| ENSDARG000 | 8;8;8 | 44893628:448 | 44894789:448 | -;-;- | 1594 | praf2 | 487 | 468 | 839 | 799 | 818 | 464 |
| ENSDARG000 | 8:8:8:8:8:8:8: | 23560183:235 | 23560411:235 | -:-:-:-:-:-:-:-: | 1735 | wasb | 116 | 101 | 170 | 170 | 298 | 512 |
| ENSDARG000 | 8:8:8:8:8:8:8: | 8483664:8483 | 8484364:8484 | -;-;-;-;-;-;-;- | 1731 | abt1 | 217 | 239 | 436 | 128 | 265 | 234 |
| ENSDARG000 | 8:8:8:8:8:8:8: | 49173131:491 | 49173496:491 | -:-:-:-:-:-:-:-: | 2009 | mfsd2al2 | 28 | 11 | 3 | 14 | 14 | 36 |
| ENSDARG000 | 8:8:8:8:8:8:8: | 1632622:1636 | 1637196:1637 | -:-:-:-:-:-:-:-: | 5969 | dgcr6 | 146 | 180 | 224 | 151 | 173 | 172 |
| ENSDARG000 | 8:8:8:8:8:8:8: | 47099033:471 | 47099147:471 | +:+:+:+:+:+:+: | 2680 | arhgef16 | 40 | 14 | 10 | 4 | 20 | 13 |
| ENSDARG000 | 8:8:8:8:8:8:8: | 20950344:209 | 20950494:209 | +:+:+:+:+:+:+: | 1388 | NA | 28 | 4 | 12 | 64 | 22 | 26 |
| ENSDARG000 | 8:8:8:8:8:8:8: | 14986833:149 | 14987220:149 | +:+:+:+:+:+:+: | 2781 | fnbp1l | 6620 | 6853 | 5357 | 4680 | 7608 | 6307 |
| ENSDARG000 | 8:8:8:8:8:8:8: | 19268474:192 | 19271360:192 | -:-:-:-:-:-:-:-: | 4130 | plin3 | 1358 | 2406 | 1910 | 3249 | 1624 | 1422 |
| ENSDARG000 | 8:8:8:8:8:8:8: | 1534237:1571 | 1534663:1572 | -;-;-;-;-;-;-;- | 1529 | NA | 0 | 2 | 0 | 0 | 0 | 0 |
| ENSDARG000 | 8:8:8:8:8:8:8: | 42764234:428 | 42764319:428 | +:+:+:+:+:+:+: | 1375 | kcnip3a | 2 | 2 | 0 | 0 | 0 | 0 |
| ENSDARG000 | 8:8:8:8:8:8:8: | 17167876:171 | 17167959:171 | +:+:+:+:+:+:+: | 1066 | cenph | 15 | 162 | 131 | 60 | 105 | 98 |
| ENSDARG000 | 8:8:8:8:8:8:8: | 36554816:365 | 36554969:365 | +:+:+:+:+:+:+: | 3610 | sf3a1 | 1660 | 2564 | 2235 | 1080 | 1874 | 2236 |
| ENSDARG000 | 8:8:8:8:8:8:8: | 6967108:6969 | 6967329:6970 | +:+:+:+:+:+:+: | 3385 | asic1a | 6 | 2 | 4 | 0 | 16 | 8 |
| ENSDARG000 | 8;8;8;8;8 | 42860031:428 | 42860283:428 | -;-;-;-;- | 565 | NA | 0 | 0 | 0 | 0 | 0 | 0 |
| ENSDARG000 | 8:8:8:8:8:8:8: | 26083808:260 | 26084164:260 | +:+:+:+:+:+:+: | 2599 | dalrd3 | 546 | 327 | 716 | 403 | 498 | 472 |
| ENSDARG000 | 8:8:8:8:8:8:8: | 36570508:365 | 36570665:365 | +:+:+:+:+:+:+: | 1700 | pold2 | 75 | 200 | 371 | 112 | 298 | 123 |
| ENSDARG000 | 8;8 | 49206057:492 | 49206427:492 | -;- | 921 | NA | 107 | 99 | 120 | 67 | 205 | 118 |
| ENSDARG000 | 8:8:8:8:8:8:8: | 22341606:223 | 22342089:223 | -;-;-;-;-;-;-;- | 685 | smim1 | 0 | 0 | 2 | 0 | 4 | 0 |
| ENSDARG000 | 8:8:8:8:8:8:8: | 8532407:8532 | 8532969:8532 | +:+:+:+:+:+:+: | 2745 | grm6a | 0 | 0 | 0 | 0 | 0 | 0 |
| ENSDARG000 | 8:8:8:8:8:8:8: | 17174796:171 | 17175054:171 | -:-:-:-:-:-:-:-: | 2016 | pola2 | 90 | 544 | 600 | 325 | 462 | 212 |
| ENSDARG000 | 8:8:8:8:8:8:8: | 2757821:2757 | 2757968:2757 | +:+:+:+:+:+:+: | 2293 | sh3glb2a | 147 | 76 | 160 | 129 | 125 | 80 |
| ENSDARG000 | 8:8:8:8:8:8:8: | 7033049:7033 | 7033156:7033 | +:+:+:+:+:+:+: | 1418 | gpd1a | 116 | 119 | 116 | 147 | 141 | 260 |
| ENSDARG000 | 8;8;8;8;8;8 | 44921123:449 | 44921495:449 | -;-;-;-;-;- | 825 | timm17b | 62 | 95 | 74 | 34 | 46 | 75 |
| ENSDARG000 | 8:8:8:8:8:8:8: | 22345282:223 | 22345606:223 | +:+:+:+:+:+:+: | 1230 | mad2l2 | 104 | 132 | 186 | 72 | 156 | 130 |
| ENSDARG000 | 8:8:8:8:8:8:8: | 49230948:492 | 49231345:492 | -:-:-:-:-:-:-:-: | 1762 | PRICKLE3 | 17 | 16 | 63 | 31 | 10 | 0 |
| ENSDARG000 | 8;8 | 29655948:296 | 29657498:296 | -;- | 2001 | mpeg1.3 | 0 | 0 | 0 | 0 | 0 | 0 |
| ENSDARG000 | 8:8:8:8:8:8:8: | 15084386:150 | 15084561:150 | -:-:-:-:-:-:-:-: | 3360 | bcar3 | 276 | 62 | 251 | 64 | 672 | 124 |
| ENSDARG000 | 8:8:8:8:8:8:8: | 23579974:235 | 23582000:235 | -:-:-:-:-:-:-:-: | 4065 | slc38a5b | 1730 | 2975 | 6209 | 5284 | 2933 | 3537 |
| ENSDARG000 | 8:8:8:8:8:8:8: | 44926629:449 | 44927035:449 | +:+:+:+:+:+:+: | 3269 | CRAT | 58 | 38 | 6 | 2 | 2 | 131 |
| ENSDARG000 | 8:8:8:8:8:8:8: | 11542852:115 | 11543515:115 | -;-;-;-;-;-;-;- | 1793 | si:ch211-248 | 15 | 0 | 0 | 0 | 2 | 0 |
| ENSDARG000 | 8:8:8:8:8:8:8: | 7055619:7059 | 7055703:7059 | -;-;-;-;-;-;-;- | 3075 | NA | 86 | 77 | 183 | 93 | 185 | 178 |
| ENSDARG000 | 8;8;8;8;8 | 29670527:296 | 29673062:296 | -;-;-;-;- | 3295 | prf1.6 | 0 | 12 | 10 | 14 | 18 | 32 |
| ENSDARG000 | 8:8:8:8:8:8:8: | 11578318:115 | 11578496:115 | -:-:-:-:-:-:-:-: | 2070 | fnbp1a | 163 | 118 | 224 | 295 | 60 | 60 |
| ENSDARG000 | 8:8:8:8:8:8:8: | 31057115:310 | 31057737:310 | -:-:-:-:-:-:-:-: | 3872 | slc20a1a | 9037 | 4502 | 5957 | 17698 | 7856 | 7521 |
| ENSDARG000 | 8:8:8:8:8:8:8: | 19283143:192 | 19285395:192 | -:-:-:-:-:-:-:-: | 4786 | rgl1 | 16825 | 12036 | 12658 | 10645 | 16559 | 11823 |
| ENSDARG000 | 8:8:8:8:8:8:8: | 47143829:471 | 47144170:471 | -:-:-:-:-:-:-:-: | 2192 | ybx1 | 34423 | 31306 | 55291 | 37927 | 45774 | 31055 |
| ENSDARG000 | 8;8 | 29688511:296 | 29689860:296 | -;- | 1804 | NA | 0 | 4 | 0 | 0 | 0 | 6 |
| ENSDARG000 | 8:8:8:8:8:8:8: | 17184602:171 | 17184774:171 | +:+:+:+:+:+:+: | 2287 | dimt1l | 328 | 406 | 396 | 176 | 402 | 329 |
| ENSDARG000 | 8;8;8 | 2798109:2800 | 2798234:2800 | -;-;- | 701 | NA | 0 | 0 | 0 | 0 | 0 | 0 |
| ENSDARG000 | 8;8;8;8;8 | 49310785:493 | 49311415:493 | -;-;-;-;- | 1299 | plp2b | 919 | 1564 | 886 | 989 | 987 | 1180 |
| ENSDARG000 | 8;8 | 2800179:2806 | 2800264:2806 | +;+ | 315 | NA | 0 | 0 | 0 | 0 | 0 | 0 |
| ENSDARG000 | 8:8:8:8:8:8:8: | 26125218:261 | 26128110:261 | +:+:+:+:+:+:+: | 9001 | CELSR3 | 28 | 44 | 391 | 165 | 132 | 18 |
| ENSDARG000 | 8:8:8:8:8:8:8: | 39767915:397 | 39768149:397 | +:+:+:+:+:+:+: | 3413 | hps4 | 1076 | 604 | 912 | 747 | 747 | 835 |
| ENSDARG000 | 8:8:8:8:8:8:8: | 51573124:515 | 51573471:515 | -;-;-;-;-;-;-;- | 820 | ankrd39 | 27 | 31 | 44 | 12 | 22 | 20 |
| ENSDARG000 | 8;8;8;8;8 | 20956078:209 | 20956145:209 | +;+;+;+;+ | 782 | NA | 2 | 5 | 6 | 1 | 0 | 0 |
| ENSDARG000 | 8:8:8:8:8:8:8: | 3085120:3085 | 3086089:3085 | +:+:+:+:+:+:+: | 3677 | lhx2b | 6 | 0 | 0 | 0 | 0 | 0 |
| ENSDARG000 | 8;8 | 29701428:297 | 29702531:297 | -;- | 1643 | prf1.5 | 2 | 6 | 0 | 0 | 10 | 22 |
| ENSDARG000 | 8;8;8 | 7086916:7089 | 7087057:7090 | +;+;+ | 991 | NA | 0 | 1 | 2 | 0 | 0 | 0 |
| ENSDARG000 | 8:8:8:8:8:8:8: | 7084974:7086 | 7085282:7086 | -:-:-:-:-:-:-:-: | 1538 | NA | 0 | 0 | 0 | 0 | 10 | 4 |
| ENSDARG000 | 8:8:8:8:8:8:8: | 22355909:223 | 22355950:223 | +:+:+:+:+:+:+: | 2469 | zgc:153631 | 805 | 675 | 692 | 351 | 464 | 266 |
| ENSDARG000 | 8:8:8:8:8:8:8: | 8643901:8643 | 8644052:8644 | +:+:+:+:+:+:+: | 3412 | NA | 299 | 285 | 225 | 116 | 264 | 191 |
| ENSDARG000 | 8;8 | 3389168:3413 | 3390429:3413 | -;- | 1343 | fut9b.5 | 0 | 0 | 0 | 0 | 0 | 0 |
| ENSDARG000 | 8:8:8:8:8:8:8: | 42917515:429 | 42917825:429 | +:+:+:+:+:+:+: | 4114 | SLC23A2 | 160 | 56 | 204 | 172 | 403 | 52 |
| ENSDARG000 | 8;8 | 3405612:3406 | 3405710:3407 | +;+ | 1434 | fut9b.6 | 14 | 0 | 0 | 4 | 10 | 0 |
| ENSDARG000 | 8;8 | 3379815:3380 | 3379866:3380 | +;+ | 849 | fut9b.4 | 4 | 0 | 0 | 0 | 1 | 0 |
| ENSDARG000 | 8:8:8:8:8:8:8: | 3345133:3345 | 3346401:3346 | -;-;-;-;-;-;-;- | 1489 | fut9b.3 | 0 | 0 | 0 | 0 | 0 | 0 |
| ENSDARG000 | 8;8;8;8;8;8;8 | 20982389:209 | 20982471:209 | +;+;+;+;+;+;+ | 809 | NA | 0 | 0 | 0 | 0 | 0 | 0 |
| ENSDARG000 | 8:8:8:8:8:8:8: | 25295160:252 | 25295289:252 | +:+:+:+:+:+:+: | 1114 | gstm.1 | 3068 | 2796 | 3801 | 2230 | 4612 | 2787 |
| ENSDARG000 | 8;8 | 3334708:3336 | 3335920:3336 | -;- | 1379 | fut9b.2 | 10 | 0 | 0 | 0 | 0 | 0 |
| ENSDARG000 | 8:8:8:8:8:8:8: | 20994433:209 | 20994512:209 | +:+:+:+:+:+:+: | 1297 | NA | 0 | 4 | 0 | 2 | 0 | 0 |
| ENSDARG000 | 8;8;8;8;8 | 3307765:3307 | 3309193:3309 | -;-;-;-;- | 1708 | fut9b.1 | 6 | 4 | 0 | 0 | 23 | 12 |
| ENSDARG000 | 8:8:8:8:8:8:8: | 45003659:450 | 45003691:450 | +:+:+:+:+:+:+: | 4203 | si:ch211-163 | 6 | 2 | 8 | 2 | 6 | 0 |
| ENSDARG000 | 8;8;8 | 45035418:450 | 45035754:450 | -;-;- | 597 | NA | 8 | 20 | 4 | 18 | 6 | 18 |
| ENSDARG000 | 8;8;8;8;8 | 45186989:451 | 45187097:451 | +;+;+;+;+ | 565 | NA | 0 | 0 | 0 | 0 | 0 | 0 |
| ENSDARG000 | 8:8:8:8:8:8:8: | 23615132:236 | 23615530:236 | +:+:+:+:+:+:+: | 3491 | ccdc22 | 1441 | 2174 | 1877 | 1922 | 1677 | 1881 |
| ENSDARG000 | 8;8;8;8;8;8;8 | 31099264:310 | 31100219:311 | -;-;-;-;-;-;- | 1775 | vgll4l | 78 | 16 | 32 | 156 | 24 | 2 |
| ENSDARG000 | 8:8:8:8:8:8:8: | 49358744:493 | 49359316:493 | -:-:-:-:-:-:-:-: | 4661 | NA | 0 | 0 | 0 | 0 | 0 | 38 |
| ENSDARG000 | 8;8 | 47188154:471 | 47188244:471 | +;+ | 744 | NA | 2 | 0 | 0 | 0 | 0 | 0 |
| ENSDARG000 | 8;8 | 47154662:471 | 47155518:471 | +;+ | 1911 | NA | 0 | 4 | 0 | 0 | 0 | 0 |
| ENSDARG000 | 8:8:8:8:8:8:8: | 15239549:152 | 15239616:152 | +:+:+:+:+:+:+: | 2182 | paox1 | 95 | 118 | 189 | 88 | 258 | 158 |
| ENSDARG000 | 8:8:8:8:8:8:8: | 47152704:471 | 47154310:471 | -:-:-:-:-:-:-:-: | 2889 | ppih | 230 | 459 | 302 | 118 | 148 | 284 |
| ENSDARG000 | 8;8;8;8;8 | 29710110:297 | 29712255:297 | -;-;-;-;- | 2634 | mpeg1.1 | 198 | 74 | 209 | 60 | 572 | 226 |
| ENSDARG000 | 8;8 | 49378890:493 | 49379069:493 | +;+ | 729 | NA | 0 | 0 | 0 | 0 | 0 | 8 |
| ENSDARG000 | 8;8;8 | 39795918:397 | 39796484:397 | +;+;+ | 1388 | si:ch211-170 | 182 | 63 | 27 | 121 | 47 | 20 |
| ENSDARG000 | 8:8:8:8:8:8:8: | 1651821:1651 | 1651947:1651 | +:+:+:+:+:+:+: | 869 | NA | 0 | 0 | 0 | 0 | 0 | 0 |
| ENSDARG000 | 8 | 29718692 | 29719393 | - | 702 | si:ch211-103 | 18 | 8 | 236 | 78 | 587 | 50 |
| ENSDARG000 | 8;8;8;8;8 | 21010850:210 | 21010915:210 | +;+;+;+;+ | 780 | NA | 6 | 0 | 2 | 0 | 2 | 0 |
| ENSDARG000 | 8;8;8;8;8 | 49384263:493 | 49384905:493 | -;-;-;-;- | 2794 | NA | 0 | 0 | 0 | 0 | 0 | 92 |
| ENSDARG000 | 8;8;8;8 | 45200860:452 | 45201054:452 | -;-;-;- | 645 | NA | 0 | 4 | 0 | 6 | 0 | 2 |
| ENSDARG000 | 8:8:8:8:8:8:8: | 51586483:515 | 51586527:515 | -:-:-:-:-:-:-:-: | 1468 | kctd9a | 141 | 58 | 153 | 117 | 146 | 115 |
| ENSDARG000 | 8;8;8;8;8;8;8 | 29726620:297 | 29726917:297 | +;+;+;+;+;+;+ | 1496 | tspan36 | 28284 | 19740 | 12608 | 21351 | 12072 | 22571 |
| ENSDARG000 | 8:8:8:8:8:8:8: | 29742237:297 | 29742485:297 | +:+:+:+:+:+:+: | 7452 | mapk4 | 234 | 172 | 100 | 190 | 220 | 36 |
| ENSDARG000 | 8 | 3426522 | 3427574 | + | 1053 | NA | 6 | 10 | 6 | 12 | 7 | 20 |
| ENSDARG000 | 8;8;8;8;8;8 | 47209659:472 | 47210177:472 | -;-;-;-;-;- | 1894 | NA | 0 | 0 | 0 | 0 | 0 | 0 |
| ENSDARG000 | 8:8:8:8:8:8:8: | 11642070:116 | 11642486:116 | +:+:+:+:+:+:+: | 3840 | ift81 | 151 | 156 | 136 | 74 | 158 | 142 |
| ENSDARG000 | 8:8:8:8:8:8:8: | 49403668:494 | 49406761:494 | -:-:-:-:-:-:-:-: | 4078 | sypb | 0 | 7 | 8 | 0 | 4 | 6 |
| ENSDARG000 | 8;8;8 | 15251448:152 | 15251660:152 | +;+;+ | 863 | zgc:171480 | 170 | 166 | 195 | 122 | 176 | 104 |
| ENSDARG000 | 8;8 | 19356072:193 | 19356577:193 | +;+ | 2638 | mpeg1.2 | 958 | 178 | 90 | 104 | 104 | 56 |
| ENSDARG000 | 8;8;8;8;8;8 | 39816523:398 | 39817082:398 | -;-;-;-;-;- | 1181 | zgc:162025 | 72 | 43 | 57 | 46 | 72 | 59 |
| ENSDARG000 | 8:8:8:8:8:8:8: | 42998944:430 | 42999325:430 | +:+:+:+:+:+:+: | 4042 | rassf2a | 1880 | 723 | 1581 | 1264 | 1512 | 1570 |
| ENSDARG000 | 8:8:8:8:8:8:8: | 15254564:152 | 15254695:152 | +:+:+:+:+:+:+: | 2201 | slc5a9 | 2 | 2 | 12 | 6 | 35 | 0 |
| ENSDARG000 | 8:8:8:8:8:8:8: | 39802506:398 | 39803074:398 | +:+:+:+:+:+:+: | 2961 | hnf1a | 2 | 4 | 0 | 12 | 1 | 2 |
| ENSDARG000 | 8;8;8;8;8 | 3431671:3434 | 3431940:3434 | +;+;+;+;+ | 1587 | ctu1 | 117 | 128 | 205 | 100 | 183 | 163 |
| ENSDARG000 | 8:8:8:8:8:8:8: | 31119548:311 | 31119758:311 | +:+:+:+:+:+:+: | 4981 | syn1 | 0 | 0 | 14 | 2 | 0 | 0 |
| ENSDARG000 | 8:8:8:8:8:8:8: | 21030142:210 | 21030235:210 | +:+:+:+:+:+:+: | 1530 | NA | 1128 | 185 | 187 | 333 | 307 | 261 |
| ENSDARG000 | 8;8;8;8;8;8;8 | 17395929:174 | 17399707:174 | -;-;-;-;-;-;- | 6147 | skia | 934 | 888 | 928 | 851 | 1197 | 834 |
| ENSDARG000 | 8:8:8:8:8:8:8: | 13794785:137 | 13794960:137 | -:-:-:-:-:-:-:-: | 940 | NA | 0 | 0 | 0 | 0 | 0 | 0 |
| ENSDARG000 | 8:8:8:8:8:8:8: | 25299069:252 | 25299169:252 | +:+:+:+:+:+:+: | 1547 | gstm.2 | 527 | 66 | 70 | 62 | 144 | 82 |
| ENSDARG000 | 8:8:8:8:8:8:8: | 23639124:236 | 23639599:236 | +:+:+:+:+:+:+: | 3123 | nt5dc2 | 515 | 431 | 642 | 534 | 471 | 497 |
| ENSDARG000 | 8;8;8;8;8;8 | 3649036:3659 | 3649290:3659 | +;+;+;+;+;+ | 1421 | rab35b | 287 | 314 | 336 | 151 | 241 | 233 |
| ENSDARG000 | 8;8 | 3688979:3690 | 3689056:3691 | -;- | 498 | NA | 0 | 0 | 0 | 0 | 0 | 0 |
| ENSDARG000 | 8;8;8 | 17529151:175 | 17529350:175 | +;+;+ | 528 | NA | 10 | 13 | 33 | 11 | 15 | 4 |
| ENSDARG000 | 8:8:8:8:8:8:8: | 47219107:472 | 47219206:472 | +:+:+:+:+:+:+: | 3826 | mthfr | 420 | 509 | 1412 | 631 | 734 | 296 |
| ENSDARG000 | 8:8:8:8:8:8:8: | 29770098:297 | 29775580:297 | -:-:-:-:-:-:-:-: | 8352 | slc20a2 | 3436 | 2144 | 3636 | 2241 | 3578 | 2561 |
| ENSDARG000 | 8:8:8:8:8:8:8: | 3701042:3701 | 3701284:3701 | -:-:-:-:-:-:-:-: | 2066 | BICDL1 | 0 | 0 | 0 | 0 | 0 | 0 |
| ENSDARG000 | 8:8:8:8:8:8:8: | 19364568:193 | 19364841:193 | -:-:-:-:-:-:-:-: | 1880 | colgalt2b | 0 | 0 | 0 | 0 | 2 | 4 |
| ENSDARG000 | 8:8:8:8:8:8:8: | 45226586:452 | 45227222:452 | -:-:-:-:-:-:-:-: | 5260 | NA | 0 | 0 | 0 | 0 | 0 | 0 |
| ENSDARG000 | 8:8:8:8:8:8:8: | 21047321:210 | 21048468:210 | -;-;-;-;-;-;-;- | 2134 | NA | 580 | 417 | 508 | 294 | 551 | 519 |
| ENSDARG000 | 8:8:8:8:8:8:8: | 8686130:8691 | 8687130:8691 | -;-;-;-;-;-;- | 1824 | zgc:86609 | 436 | 451 | 553 | 548 | 637 | 566 |
| ENSDARG000 | 8:8:8:8:8:8:8: | 26226044:262 | 26226097:262 | +:+:+:+:+:+:+: | 4655 | wdr6 | 302 | 252 | 461 | 150 | 400 | 484 |
| ENSDARG000 | 8:8:8:8:8:8:8: | 11687254:116 | 11687429:116 | +:+:+:+:+:+:+: | 4051 | atp2a2a | 116 | 253 | 279 | 72 | 267 | 65 |
| ENSDARG000 | 8;8;8;8;8 | 26220175:262 | 26221443:262 | -;-;-;-;- | 2175 | NA | 8 | 13 | 40 | 8 | 29 | 8 |
| ENSDARG000 | 8:8:8:8:8:8:8: | 7097740:7097 | 7097984:7097 | +:+:+:+:+:+:+: | 6557 | ABTB1 | 513 | 293 | 582 | 615 | 593 | 876 |
| ENSDARG000 | 8:8:8:8:8:8:8: | 1741938:1742 | 1742622:1742 | -:-:-:-:-:-:-:-: | 4048 | pi4kab | 86 | 84 | 89 | 38 | 133 | 68 |
| ENSDARG000 | 8:8:8:8:8:8:8: | 36591791:365 | 36592386:365 | -:-:-:-:-:-:-:-: | 1997 | gpkow | 242 | 252 | 288 | 176 | 316 | 228 |
| ENSDARG000 | 8:8:8:8:8:8:8: | 22377770:223 | 22378062:223 | +:+:+:+:+:+:+: | 886 | NA | 590 | 375 | 438 | 429 | 471 | 484 |
| ENSDARG000 | 8 | 51634803 | 51635930 | - | 1128 | npy8br | 0 | 0 | 20 | 0 | 2 | 0 |
| ENSDARG000 | 8:8:8:8:8:8:8: | 17554313:175 | 17554400:175 | -:-:-:-:-:-:-:-: | 1800 | prkcz | 906 | 355 | 494 | 445 | 470 | 585 |
| ENSDARG000 | 8:8:8:8:8:8:8: | 36582728:365 | 36582808:365 | +:+:+:+:+:+:+: | 1122 | pqbp1 | 347 | 466 | 531 | 192 | 414 | 349 |
| ENSDARG000 | 8:8:8:8:8:8:8: | 3932793:3944 | 3933579:3944 | -:-:-:-:-:-:-:-: | 4695 | mtmr3 | 375 | 272 | 351 | 302 | 374 | 371 |
| ENSDARG000 | 8:8:8:8:8:8:8: | 49433663:494 | 49433739:494 | +:+:+:+:+:+:+: | 2261 | nfu1 | 292 | 306 | 431 | 538 | 475 | 399 |
| ENSDARG000 | 8;8;8;8;8;8;8 | 51707781:517 | 51707915:517 | +;+;+;+;+;+;+ | 7145 | KLHL22 | 702 | 265 | 511 | 340 | 463 | 282 |
| ENSDARG000 | 8:8:8:8:8:8:8: | 1766206:1769 | 1766254:1770 | +:+:+:+:+:+:+: | 2471 | serpind1 | 0 | 0 | 2 | 16 | 6 | 0 |
| ENSDARG000 | 8;8;8 | 17660185:176 | 17660290:176 | -;-;- | 524 | NA | 0 | 0 | 0 | 0 | 0 | 0 |
| ENSDARG000 | 8:8:8:8:8:8:8: | 3944108:3947 | 3944197:3947 | -;-;-;-;-;-;- | 1218 | NA | 0 | 0 | 0 | 0 | 0 | 0 |
| ENSDARG000 | 8:8:8:8:8:8:8: | 26253252:262 | 26253356:262 | +:+:+:+:+:+:+: | 2403 | NA | 4 | 0 | 0 | 0 | 2 | 0 |
| ENSDARG000 | 8:8:8:8:8:8:8: | 15269423:152 | 15269632:152 | +:+:+:+:+:+:+: | 2016 | gclm | 662 | 1152 | 754 | 741 | 948 | 543 |
| ENSDARG000 | 8:8:8:8:8:8:8: | 31375119:313 | 31376523:313 | -:-:-:-:-:-:-:-: | 2699 | creb3l3l | 1348 | 1132 | 1671 | 1589 | 949 | 1231 |
| ENSDARG000 | 8;8;8;8 | 22393908:223 | 22394008:223 | -;-;-;- | 468 | zgc:158862 | 109 | 36 | 79 | 58 | 84 | 0 |
| ENSDARG000 | 8:8:8:8:8:8:8: | 51733907:517 | 51737720:517 | -;-;-;-;-;-;-;- | 4949 | tbx16 | 4 | 10 | 2 | 3 | 4 | 2 |
| ENSDARG000 | 8;8;8;8;8;8 | 31367667:313 | 31368353:313 | -;-;-;-;-;- | 1393 | gadd45ga | 4773 | 1667 | 3815 | 3118 | 1599 | 2156 |
| ENSDARG000 | 8:8:8:8:8:8:8: | 25302172:253 | 25302254:253 | +:+:+:+:+:+:+: | 2728 | gstm.3 | 129 | 60 | 128 | 55 | 145 | 20 |
| ENSDARG000 | 8:8:8:8:8:8:8: | 49478026:494 | 49478523:494 | -:-:-:-:-:-:-:-: | 1212 | opn7d | 8 | 149 | 256 | 37 | 246 | 45 |
| ENSDARG000 | 8:8:8:8:8:8:8: | 22438398:224 | 22438635:224 | +:+:+:+:+:+:+: | 1374 | zgc:153759 | 1030 | 648 | 788 | 1279 | 862 | 0 |
| ENSDARG000 | 8:8:8:8:8:8:8: | 22404981:224 | 22405614:224 | +:+:+:+:+:+:+: | 4590 | NA | 48 | 28 | 37 | 38 | 44 | 0 |
| ENSDARG000 | 8:8:8:8:8:8:8: | 22404981:224 | 22405614:224 | +:+:+:+:+:+:+: | 1969 | NA | 0 | 0 | 0 | 0 | 0 | 0 |
| ENSDARG000 | 8:8:8:8:8:8:8: | 13849999:138 | 13850357:138 | +:+:+:+:+:+:+: | 2030 | doc2d | 115 | 0 | 2 | 4 | 0 | 0 |

| | | | | | | | | | | | |
|---|---|---|---|---|---|---|---|---|---|---|---|
| ENSDARG000 | 8;8;8;8;8;8;8 | 23658626:23 | 23658952:23 | +;+;+;+;+;+;+ | 4139 | tbc1d25 | 174 | 158 | 195 | 138 | 163 | 110 |
| ENSDARG000 | 8;8;8 | 43053519:43 | 43053586:43 | +;+;+ | 898 | prnpa | 1 | 0 | 17 | 1 | 10 | 8 |
| ENSDARG000 | 8;8;8;8;8;8;8 | 8699085:869 | 8699385:869 | +;+;+;+;+;+;+ | 1016 | uxt | 214 | 124 | 237 | 154 | 132 | 163 |
| ENSDARG000 | 8;8;8;8;8;8;8 | 45293743:45 | 45293821:45 | +;+;+;+;+;+;+ | 3918 | ubap2b | 2590 | 2385 | 4574 | 2363 | 3617 | 2327 |
| ENSDARG000 | 8;8;8;8;8;8 | 39827198:39 | 39827353:39 | -;-;-;-;-;- | 1580 | NA | 22 | 35 | 5 | 0 | 10 | 17 |
| ENSDARG000 | 8;8;8;8;8;8 | 43094799:43 | 43097791:43 | -;-;-;-;-;- | 3943 | ccdc92 | 4 | 1 | 16 | 6 | 6 | 2 |
| ENSDARG000 | 8;8;8;8;8;8 | 15277874:15 | 15278002:15 | +;+;+;+;+;+ | 3154 | dnttip2 | 942 | 1223 | 2559 | 979 | 1509 | 1281 |
| ENSDARG000 | 8;8;8;8;8;8;8 | 36621598:36 | 36621982:36 | +;+;+;+;+;+;+ | 3887 | NA | 0 | 0 | 0 | 0 | 6 | 4 |
| ENSDARG000 | 8;8;8 | 17778259:17 | 17778780:17 | -;-;- | 4269 | NA | 15 | 3 | 2 | 0 | 8 | 13 |
| ENSDARG000 | 8;8;8;8;8;8;8 | 17775247:17 | 17775789:17 | +;+;+;+;+;+;+ | 2785 | NA | 21 | 8 | 15 | 29 | 12 | 12 |
| ENSDARG000 | 8;8;8;8;8;8;8 | 11746483:11 | 11746960:11 | -;-;-;-;-;-;- | 2091 | anapc7 | 135 | 112 | 148 | 170 | 192 | 128 |
| ENSDARG000 | 8;8;8;8;8;8;8 | 49570884:49 | 49570899:49 | +;+;+;+;+;+;+ | 2299 | NA | 632 | 4945 | 4653 | 5728 | 5442 | 3509 |
| ENSDARG000 | 8;8;8;8 | 43340995:43 | 43341223:43 | +;+;+;+ | 1070 | rflna | 40 | 2 | 8 | 6 | 10 | 20 |
| ENSDARG000 | 8;8;8;8;8;8;8 | 39839543:39 | 39840983:39 | -;-;-;-;-;-;- | 2774 | zgc:174931 | 153 | 144 | 105 | 109 | 65 | 328 |
| ENSDARG000 | 8;8;8;8;8 | 36815070:36 | 36815222:36 | -;-;-;-;- | 478 | NA | 2 | 0 | 0 | 0 | 0 | 1 |
| ENSDARG000 | 8;8;8;8;8;8;8 | 36803415:36 | 36803857:36 | +;+;+;+;+;+;+ | 8710 | IQSEC2 | 1037 | 655 | 369 | 469 | 801 | 1173 |
| ENSDARG000 | 8;8;8;8;8;8;8 | 26268100:26 | 26268856:26 | +;+;+;+;+;+;+ | 3694 | NA | 10 | 6 | 18 | 20 | 20 | 8 |
| ENSDARG000 | 8;8;8;8;8;8;8 | 7144066:716 | 7144119:716 | +;+;+;+;+;+;+ | 2611 | slc6a6a | 4 | 2 | 4 | 0 | 6 | 2 |
| ENSDARG000 | 8;8;8;8;8;8 | 51779734:51 | 51779877:51 | -;-;-;-;-;- | 953 | NA | 0 | 0 | 0 | 0 | 0 | 0 |
| ENSDARG000 | 8;8;8;8;8;8;8 | 15287424:15 | 15288033:15 | -;-;-;-;-;-;- | 1937 | spata6 | 98 | 54 | 210 | 92 | 370 | 108 |
| ENSDARG000 | 8;8;8;8;8;8;8 | 51779734:51 | 51779877:51 | -;-;-;-;-;-;- | 6923 | CABIN1 | 200 | 87 | 237 | 143 | 213 | 132 |
| ENSDARG000 | 8;8;8;8;8;8;8 | 29852353:29 | 29852396:29 | +;+;+;+;+;+;+ | 2439 | hsd17b3 | 2 | 14 | 35 | 2 | 41 | 8 |
| ENSDARG000 | 8;8;8;8;8;8;8 | 23689992:23 | 23692221:23 | -;-;-;-;-;-;- | 3659 | si:ch73-237c | 105 | 44 | 48 | 39 | 170 | 60 |
| ENSDARG000 | 8;8;8;8;8;8;8 | 23672254:23 | 23675914:23 | -;-;-;-;-;-;- | 5216 | cfp | 98 | 62 | 132 | 96 | 267 | 274 |
| ENSDARG000 | 8;8;8;8;8;8;8 | 26292560:26 | 26292719:26 | +;+;+;+;+;+;+ | 4532 | mgll | 290 | 298 | 626 | 234 | 438 | 292 |
| ENSDARG000 | 8;8;8;8;8;8;8 | 21052473:21 | 21053603:21 | -;-;-;-;-;-;- | 3171 | zgc:112962 | 641 | 304 | 393 | 306 | 364 | 679 |
| ENSDARG000 | 8;8;8;8;8;8;8 | 39848504:39 | 39853079:39 | -;-;-;-;-;-;- | 5447 | unc119.1 | 546 | 323 | 652 | 303 | 379 | 246 |
| ENSDARG000 | 8;8;8;8;8;8;8 | 4044578:404 | 4045232:404 | -;-;-;-;-;-;- | 4372 | CUX2 | 10 | 2 | 28 | 2 | 8 | 20 |
| ENSDARG000 | 8;8;8;8;8;8;8 | 17787307:17 | 17788073:17 | -;-;-;-;-;-;- | 1815 | si:ch211-150 | 138 | 72 | 152 | 118 | 124 | 184 |
| ENSDARG000 | 8;8;8;8;8;8;8 | 8712446:872 | 8712620:872 | +;+;+;+;+;+;+ | 1902 | ELK1 | 145 | 46 | 93 | 48 | 151 | 121 |
| ENSDARG000 | 8;8;8;8;8;8;8 | 14003232:14 | 14003729:14 | -;-;-;-;-;-;- | 5052 | atp2b3a | 0 | 3 | 6 | 4 | 0 | 2 |
| ENSDARG000 | 8;8;8;8;8;8 | 47319804:47 | 47320213:47 | -;-;-;-;-;- | 1526 | pex10 | 94 | 150 | 106 | 54 | 69 | 132 |
| ENSDARG000 | 8;8;8;8;8;8 | 39870714:39 | 39871339:39 | -;-;-;-;-;- | 1626 | mlec | 391 | 354 | 688 | 191 | 597 | 242 |
| ENSDARG000 | 8;8;8;8;8;8;8 | 49617946:49 | 49617976:49 | -;-;-;-;-;-;- | 1683 | FRMD3 | 8 | 0 | 11 | 33 | 11 | 9 |
| ENSDARG000 | 8;8;8;8;8;8;8 | 22477922:22 | 22478130:22 | +;+;+;+;+;+;+ | 2062 | NA | 0 | 0 | 0 | 0 | 0 | 0 |
| ENSDARG000 | 8;8;8;8;8;8;8 | 31387563:31 | 31388318:31 | -;-;-;-;-;-;- | 2140 | NA | 6 | 6 | 6 | 0 | 0 | 0 |
| ENSDARG000 | 8;8;8;8;8;8;8 | 1839695:183 | 1839764:183 | +;+;+;+;+;+;+ | 5867 | snap29 | 1121 | 410 | 433 | 375 | 1063 | 426 |
| ENSDARG000 | 8 | 8845932 | 8848166 | + | 2235 | NA | 6 | 0 | 4 | 0 | 0 | 4 |
| ENSDARG000 | 8;8;8;8;8;8;8 | 43379436:43 | 43382137:43 | -;-;-;-;-;-;- | 10270 | ncor2 | 2718 | 2219 | 2670 | 1595 | 3593 | 2198 |
| ENSDARG000 | 8;8;8;8;8;8;8 | 45334255:45 | 45334396:45 | +;+;+;+;+;+;+ | 2715 | pabpc1l | 0 | 0 | 0 | 0 | 0 | 0 |
| ENSDARG000 | 8;8;8;8;8;8;8 | 25308003:25 | 25308636:25 | -;-;-;-;-;-;- | 3266 | eps8l3b | 2 | 4 | 12 | 12 | 25 | 2 |
| ENSDARG000 | 8;8;8;8;8;8;8 | 13960655:13 | 13961010:13 | -;-;-;-;-;-;- | 2654 | serping1 | 4 | 0 | 44 | 0 | 0 | 0 |
| ENSDARG000 | 8;8;8;8;8 | 17800380:17 | 17801457:17 | -;-;-;-;- | 1923 | st6galnac5b | 4 | 16 | 38 | 24 | 34 | 32 |
| ENSDARG000 | 8;8;8;8;8;8;8 | 47342586:47 | 47342767:47 | +;+;+;+;+;+;+ | 6172 | plch2a | 22 | 8 | 12 | 32 | 10 | 22 |
| ENSDARG000 | 8;8;8;8;8 | 13991297:13 | 13991967:13 | -;-;-;-;- | 1656 | si:dkeyp-110 | 63 | 73 | 92 | 52 | 71 | 85 |
| ENSDARG000 | 8;8;8;8;8;8;8 | 13971417:13 | 13971620:13 | +;+;+;+;+;+;+ | 1061 | NA | 0 | 0 | 0 | 0 | 0 | 0 |
| ENSDARG000 | 8;8;8;8 | 13978486:13 | 13978758:13 | -;-;-;- | 685 | igf3 | 5 | 0 | 16 | 9 | 13 | 6 |
| ENSDARG000 | 8;8;8;8;8;8;8 | 25329032:25 | 25330594:25 | -;-;-;-;-;-;- | 3309 | atp5pb | 4193 | 4629 | 4962 | 3444 | 4428 | 3406 |
| ENSDARG000 | 8;8;8;8;8;8;8 | 4333914:433 | 4333938:433 | +;+;+;+;+;+;+ | 1833 | myl2b | 0 | 0 | 0 | 0 | 0 | 0 |
| ENSDARG000 | 8;8;8;8;8;8;8 | 23703464:23 | 23703591:23 | +;+;+;+;+;+;+ | 4833 | ppardb | 621 | 451 | 534 | 356 | 589 | 567 |
| ENSDARG000 | 8;8 | 26329818:26 | 26329935:26 | -;- | 451 | NA | 0 | 0 | 0 | 0 | 0 | 0 |
| ENSDARG000 | 8;8;8;8 | 26373149:26 | 26373199:26 | +;+;+;+ | 1604 | hyal3 | 4 | 5 | 25 | 12 | 27 | 14 |
| ENSDARG000 | 8;8;8;8;8;8;8 | 26329482:26 | 26330923:26 | +;+;+;+;+;+;+ | 3514 | prkar2aa | 4261 | 3481 | 3796 | 2975 | 3928 | 2878 |
| ENSDARG000 | 8;8;8;8;8;8;8 | 7205524:720 | 7206537:720 | -;-;-;-;-;-;- | 4292 | grip2a | 3 | 32 | 106 | 70 | 90 | 29 |
| ENSDARG000 | 8;8;8;8;8;8;8 | 11779547:11 | 11779863:11 | -;-;-;-;-;-;- | 3962 | RAPGEF1 | 214 | 189 | 274 | 185 | 295 | 285 |
| ENSDARG000 | 8;8;8;8;8;8;8 | 23725957:23 | 23726040:23 | +;+;+;+;+;+;+ | 2085 | mkrn4 | 446 | 225 | 334 | 248 | 268 | 362 |
| ENSDARG000 | 8;8 | 31438472:31 | 31438692:31 | +;+ | 613 | NA | 0 | 31 | 0 | 0 | 0 | 16 |
| ENSDARG000 | 8;8;8;8;8;8;8 | 31435452:31 | 31435648:31 | +;+;+;+;+;+;+ | 2665 | selenop | 244 | 110 | 2018 | 1172 | 3000 | 826 |
| ENSDARG000 | 8;8;8;8;8;8;8 | 22490069:22 | 22490104:22 | +;+;+;+;+;+;+ | 880 | NA | 0 | 0 | 0 | 0 | 0 | 0 |
| ENSDARG000 | 8;8;8;8;8;8;8 | 47633438:47 | 47633470:47 | +;+;+;+;+;+;+ | 2261 | si:ch211-251 | 72 | 60 | 685 | 422 | 341 | 391 |
| ENSDARG000 | 8;8;8 | 22495431:22 | 22495843:22 | +;+;+ | 637 | NA | 0 | 0 | 0 | 0 | 0 | 0 |
| ENSDARG000 | 8;8;8;8 | 22499362:22 | 22500744:22 | -;-;-;- | 3338 | NA | 374 | 353 | 521 | 386 | 585 | 349 |
| ENSDARG000 | 8;8;8;8;8;8;8 | 39890186:39 | 39890387:39 | -;-;-;-;-;-;- | 2309 | cabp1a | 0 | 0 | 0 | 0 | 4 | 0 |
| ENSDARG000 | 8;8;8;8;8;8;8 | 36983559:36 | 36983979:36 | +;+;+;+;+;+;+ | 6384 | kdm5c | 3807 | 3095 | 5518 | 3521 | 5306 | 3156 |
| ENSDARG000 | 8;8;8;8;8;8;8 | 17838924:17 | 17839147:17 | +;+;+;+;+;+;+ | 3992 | SLC44A5 | 119 | 26 | 219 | 88 | 302 | 173 |
| ENSDARG000 | 8;8;8;8;8;8;8 | 8860902:886 | 8861693:886 | +;+;+;+;+;+;+ | 2349 | otud5a | 596 | 604 | 611 | 312 | 529 | 438 |
| ENSDARG000 | 8;8;8;8;8;8;8 | 49706387:49 | 49706734:49 | -;-;-;-;-;-;- | 1957 | gkap1 | 573 | 540 | 631 | 492 | 570 | 608 |
| ENSDARG000 | 8;8;8;8;8;8;8 | 19504011:19 | 19504226:19 | +;+;+;+;+;+;+ | 1789 | sec22bb | 398 | 374 | 403 | 284 | 288 | 394 |
| ENSDARG000 | 8;8;8;8;8;8 | 4368534:438 | 4369058:438 | +;+;+;+;+;+ | 4955 | pptc7a | 772 | 613 | 679 | 961 | 984 | 847 |
| ENSDARG000 | 8;8;8;8;8;8;8 | 51933983:51 | 51936453:51 | -;-;-;-;-;-;- | 4354 | CEP78 | 38 | 64 | 58 | 42 | 52 | 56 |
| ENSDARG000 | 8;8;8;8;8;8;8 | 19514294:19 | 19514603:19 | +;+;+;+;+;+;+ | 2292 | NA | 0 | 8 | 10 | 10 | 14 | 2 |
| ENSDARG000 | 8;8;8;8;8;8;8 | 1873345:187 | 1873750:187 | -;-;-;-;-;-;- | 1036 | NA | 24 | 2 | 8 | 16 | 6 | 5 |
| ENSDARG000 | 8;8;8;8;8;8;8 | 22509508:22 | 22511986:22 | -;-;-;-;-;-;- | 3390 | apex2 | 158 | 228 | 325 | 248 | 486 | 124 |
| ENSDARG000 | 8;8;8;8;8;8;8 | 31465222:31 | 31466352:31 | -;-;-;-;-;-;- | 2359 | ghra | 4 | 19 | 62 | 2 | 36 | 43 |
| ENSDARG000 | 8;8;8;8;8;8;8 | 1926178:192 | 1926304:192 | +;+;+;+;+;+;+ | 1174 | NA | 2 | 0 | 0 | 0 | 0 | 2 |
| ENSDARG000 | 8;8;8;8;8;8 | 26382604:26 | 26383198:26 | -;-;-;-;-;- | 1460 | si:dkey-20d21 | 6 | 3 | 12 | 12 | 4 | 16 |
| ENSDARG000 | 8;8;8 | 29894444:29 | 29894689:29 | +;+;+ | 1280 | NA | 6 | 1 | 6 | 36 | 1 | 0 |
| ENSDARG000 | 8;8;8;8;8;8;8 | 29898715:29 | 29899582:29 | -;-;-;-;-;-;- | 4165 | ercc6l2 | 237 | 191 | 275 | 168 | 242 | 244 |
| ENSDARG000 | 8;8;8;8 | 15476625:15 | 15477099:15 | +;+;+;+ | 1587 | NA | 2 | 0 | 0 | 0 | 0 | 0 |
| ENSDARG000 | 8;8;8;8;8;8;8 | 31646031:31 | 31646087:31 | -;-;-;-;-;-;- | 1227 | FBXO4 | 102 | 121 | 134 | 88 | 100 | 173 |
| ENSDARG000 | 8;8;8;8;8;8 | 31704624:31 | 31705412:31 | -;-;-;-;-;- | 1610 | rimoc1 | 712 | 590 | 798 | 739 | 648 | 822 |
| ENSDARG000 | 8;8;8;8;8;8;8 | 11884671:11 | 11885090:11 | -;-;-;-;-;-;- | 1440 | med27 | 174 | 244 | 364 | 266 | 296 | 321 |
| ENSDARG000 | 8;8;8;8;8;8;8 | 4407994:440 | 4408478:440 | -;-;-;-;-;-;- | 1840 | rad9b | 0 | 4 | 0 | 0 | 0 | 0 |
| ENSDARG000 | 8;8;8;8;8;8;8 | 8927870:892 | 8928113:892 | +;+;+;+;+;+;+ | 2535 | pim2 | 354 | 384 | 746 | 2072 | 428 | 1166 |
| ENSDARG000 | 8;8;8;8;8;8 | 15502785:15 | 15504169:15 | -;-;-;-;-;- | 2720 | bend5 | 278 | 157 | 244 | 148 | 318 | 216 |
| ENSDARG000 | 8;8;8;8;8;8;8 | 47677208:47 | 47677341:47 | +;+;+;+;+;+;+ | 5007 | dpp9 | 1850 | 1835 | 2361 | 1280 | 2019 | 1616 |
| ENSDARG000 | 8;8;8;8;8;8;8 | 17921429:17 | 17921489:17 | -;-;-;-;-;-;- | 735 | NA | 0 | 0 | 7 | 0 | 5 | 0 |
| ENSDARG000 | 8;8;8;8;8;8;8 | 39965117:39 | 39966345:39 | -;-;-;-;-;-;- | 2947 | asphd2 | 192 | 247 | 122 | 48 | 240 | 336 |
| ENSDARG000 | 8;8;8 | 22516728:22 | 22516789:22 | +;+;+ | 646 | NA | 2 | 10 | 0 | 6 | 0 | 9 |
| ENSDARG000 | 8;8;8;8;8;8;8 | 51958968:51 | 51959088:51 | +;+;+;+;+;+;+ | 1098 | NA | 0 | 0 | 0 | 0 | 12 | 0 |
| ENSDARG000 | 8;8;8;8;8;8;8 | 26396552:26 | 26396794:26 | +;+;+;+;+;+;+ | 2007 | amt | 166 | 180 | 339 | 78 | 287 | 249 |
| ENSDARG000 | 8;8;8;8;8;8;8 | 17933475:17 | 17933649:17 | +;+;+;+;+;+;+ | 2239 | erich3 | 0 | 8 | 0 | 0 | 0 | 0 |
| ENSDARG000 | 8;8;8;8;8;8;8 | 11993150:11 | 11993363:11 | +;+;+;+;+;+;+ | 1861 | NTNG2 | 45 | 1240 | 522 | 288 | 2375 | 444 |
| ENSDARG000 | 8;8;8;8 | 8936912:893 | 8937128:893 | +;+;+;+ | 2252 | NA | 10 | 26 | 80 | 118 | 66 | 86 |
| ENSDARG000 | 8;8;8;8;8;8;8 | 19548985:19 | 19549312:19 | +;+;+;+;+;+;+ | 10095 | notch2 | 5071 | 5020 | 3453 | 3979 | 4496 | 5212 |
| ENSDARG000 | 8;8;8;8;8;8;8 | 39998467:39 | 39998888:39 | +;+;+;+;+;+;+ | 4782 | ggt5a | 18 | 0 | 4 | 0 | 4 | 2 |
| ENSDARG000 | 8;8;8;8;8;8;8 | 1946994:194 | 1947108:194 | -;-;-;-;-;-;- | 825 | SLC25A1 | 0 | 0 | 0 | 0 | 0 | 2 |
| ENSDARG000 | 8;8;8;8;8;8;8 | 49730163:49 | 49730454:49 | -;-;-;-;-;-;- | 1180 | C8H9orf64 | 163 | 142 | 466 | 291 | 294 | 120 |
| ENSDARG000 | 8;8;8;8;8;8;8 | 45361735:45 | 45362037:45 | +;+;+;+;+;+;+ | 1783 | chmp4bb | 2587 | 1984 | 2129 | 1623 | 1978 | 2144 |
| ENSDARG000 | 8;8;8;8;8;8;8 | 37030974:37 | 37031580:37 | -;-;-;-;-;-;- | 1889 | renbp | 242 | 109 | 152 | 45 | 237 | 137 |
| ENSDARG000 | 8;8;8;8 | 8947623:895 | 8947831:895 | +;+;+;+ | 1252 | slc35a2 | 91 | 98 | 178 | 125 | 144 | 112 |
| ENSDARG000 | 8;8;8;8;8;8;8 | 29962635:29 | 29963191:29 | +;+;+;+;+;+;+ | 4834 | ptch1 | 323 | 285 | 378 | 311 | 402 | 465 |
| ENSDARG000 | 8;8;8 | 12118097:12 | 12118746:12 | +;+;+ | 1349 | endog | 70 | 235 | 264 | 70 | 238 | 127 |
| ENSDARG000 | 8;8;8;8;8;8;8 | 16004154:16 | 16004684:16 | +;+;+;+;+;+;+ | 7728 | elavl4 | 0 | 0 | 2 | 0 | 10 | 12 |
| ENSDARG000 | 8;8 | 29963024:29 | 29963191:29 | +;+ | 361 | NA | 0 | 0 | 0 | 0 | 0 | 0 |
| ENSDARG000 | 8;8;8;8;8;8;8 | 47726919:47 | 47726930:47 | +;+;+;+;+;+;+ | 4850 | NA | 40 | 114 | 278 | 76 | 99 | 166 |
| ENSDARG000 | 8;8;8;8;8;8;8 | 31716872:31 | 31717409:31 | +;+;+;+;+;+;+ | 3808 | oxct1a | 730 | 549 | 433 | 484 | 339 | 1034 |
| ENSDARG000 | 8;8;8;8;8;8;8 | 2116429:212 | 2116545:212 | -;-;-;-;-;-;- | 1221 | NA | 18 | 8 | 2 | 0 | 10 | 4 |
| ENSDARG000 | 8;8 | 16062172:16 | 16062291:16 | -;- | 470 | NA | 0 | 0 | 0 | 0 | 0 | 0 |
| ENSDARG000 | 8 | 7315769 | 7316206 | + | 438 | NA | 0 | 0 | 0 | 0 | 0 | 0 |
| ENSDARG000 | 8;8;8;8 | 7315625:731 | 7316250:731 | +;+;+;+ | 1820 | selenoh | 1244 | 2109 | 2079 | 1528 | 1896 | 1834 |
| ENSDARG000 | 8;8;8;8;8;8;8 | 7311478:731 | 7311691:731 | +;+;+;+;+;+;+ | 5700 | ssuh2rs1 | 970 | 836 | 975 | 1057 | 1200 | 1962 |
| ENSDARG000 | 8;8;8;8;8;8;8 | 45418985:45 | 45419626:45 | -;-;-;-;-;-;- | 1525 | ywhabb | 4022 | 4852 | 4662 | 2885 | 4447 | 3431 |
| ENSDARG000 | 8;8;8;8;8;8;8 | 4434422:443 | 4434503:443 | +;+;+;+;+;+;+ | 1392 | srrd | 30 | 78 | 74 | 80 | 70 | 75 |
| ENSDARG000 | 8;8;8;8;8;8;8 | 45381298:45 | 45381331:45 | +;+;+;+;+;+;+ | 1858 | NA | 133 | 27 | 99 | 187 | 243 | 49 |
| ENSDARG000 | 8;8;8;8;8;8;8 | 23731483:23 | 23731804:23 | +;+;+;+;+;+;+ | 1833 | fance | 132 | 100 | 92 | 105 | 144 | 136 |
| ENSDARG000 | 8;8;8;8;8;8;8 | 45394914:45 | 45396070:45 | -;-;-;-;-;-;- | 2166 | mak16 | 343 | 480 | 778 | 370 | 413 | 455 |
| ENSDARG000 | 8;8;8;8;8;8;8 | 8973425:897 | 8973512:897 | +;+;+;+;+;+;+ | 1543 | bcap31 | 1893 | 1684 | 2105 | 1402 | 2237 | 1879 |
| ENSDARG000 | 8;8;8;8 | 47804284:47 | 47804446:47 | +;+;+;+ | 482 | ndufa7 | 211 | 335 | 272 | 200 | 270 | 193 |
| ENSDARG000 | 8;8;8;8;8;8 | 47817454:47 | 47817548:47 | +;+;+;+;+;+ | 1009 | NA | 9 | 0 | 1 | 0 | 2 | 8 |
| ENSDARG000 | 8;8 | 4435799:444 | 4436150:444 | -;- | 755 | NA | 0 | 0 | 0 | 0 | 0 | 0 |
| ENSDARG000 | 8;8;8;8;8;8;8 | 47758621:47 | 47759115:47 | -;-;-;-;-;-;- | 4557 | kank3 | 1350 | 775 | 1286 | 700 | 1100 | 1176 |
| ENSDARG000 | 8;8;8;8;8;8;8 | 12134127:12 | 12134294:12 | -;-;-;-;-;-;- | 3559 | DAB2IP | 1081 | 693 | 1020 | 641 | 1176 | 759 |
| ENSDARG000 | 8;8;8;8;8;8;8 | 17958419:17 | 17958495:17 | -;-;-;-;-;-;- | 2508 | tnni3k | 0 | 0 | 0 | 2 | 0 | 16 |
| ENSDARG000 | 8;8;8;8;8;8;8 | 37046010:37 | 37047265:37 | -;-;-;-;-;-;- | 2580 | NA | 0 | 0 | 4 | 0 | 0 | 0 |
| ENSDARG000 | 8;8;8;8;8;8;8 | 43668476:43 | 43668817:43 | -;-;-;-;-;-;- | 9217 | EP400 | 1560 | 978 | 1925 | 730 | 2032 | 1264 |
| ENSDARG000 | 8;8 | 26427549:26 | 26427629:26 | +;+ | 197 | NA | 0 | 5 | 3 | 0 | 0 | 0 |
| ENSDARG000 | 8;8;8;8;8;8;8 | 26410197:26 | 26410587:26 | +;+;+;+;+;+;+ | 2507 | ifrd2 | 792 | 1034 | 1532 | 498 | 1327 | 915 |
| ENSDARG000 | 8;8;8;8;8;8;8 | 37058899:37 | 37060891:37 | -;-;-;-;-;-;- | 4787 | zgc:162200 | 588 | 1078 | 1101 | 420 | 1031 | 704 |
| ENSDARG000 | 8;8;8;8;8;8;8 | 40042632:40 | 40043025:40 | -;-;-;-;-;-;- | 2192 | ggt1a | 197 | 2 | 10 | 12 | 0 | 0 |
| ENSDARG000 | 8;8;8;8 | 25339517:25 | 25339602:25 | +;+;+;+ | 734 | NA | 12 | 4 | 3 | 0 | 15 | 10 |
| ENSDARG000 | 8;8;8;8;8;8;8 | 51992344:51 | 51992911:51 | -;-;-;-;-;-;- | 10224 | vps13a | 3124 | 2291 | 4147 | 3338 | 2524 | 2261 |
| ENSDARG000 | 8;8;8;8;8;8;8 | 49744801:49 | 49746512:49 | -;-;-;-;-;-;- | 4477 | hnrnpk | 2443 | 1823 | 3165 | 2701 | 2688 | 2237 |
| ENSDARG000 | 8;8;8 | 31821396:31 | 31821586:31 | +;+;+ | 2378 | plcxd3 | 8 | 2 | 10 | 0 | 0 | 0 |
| ENSDARG000 | 8;8;8;8;8;8;8 | 25351863:25 | 25352029:25 | +;+;+;+;+;+;+ | 2816 | dnase1l1l | 0 | 0 | 2 | 0 | 0 | 0 |
| ENSDARG000 | 8;8;8;8;8;8;8 | 22521729:22 | 22523176:22 | -;-;-;-;-;-;- | 4966 | csde1 | 12357 | 12788 | 16047 | 10990 | 14130 | 13505 |
| ENSDARG000 | 8;8;8;8;8;8;8 | 21082361:21 | 21083196:21 | -;-;-;-;-;-;- | 2291 | aldh9a1a.2 | 59 | 46 | 123 | 137 | 80 | 7 |
| ENSDARG000 | 8;8;8;8;8;8 | 2215708:221 | 2216131:221 | -;-;-;-;-;- | 1198 | NA | 102 | 75 | 102 | 65 | 99 | 77 |
| ENSDARG000 | 8;8;8;8;8;8;8 | 9052123:905 | 9052484:905 | -;-;-;-;-;-;- | 2126 | slc6a8 | 824 | 955 | 3281 | 2083 | 1738 | 518 |
| ENSDARG000 | 8;8;8;8;8;8;8 | 23738122:23 | 23738169:23 | +;+;+;+;+;+;+ | 1353 | rpl10a | 17432 | 16086 | 26684 | 22689 | 18117 | 25571 |
| ENSDARG000 | 8;8;8;8;8;8;8 | 17987215:17 | 17987235:17 | +;+;+;+;+;+;+ | 974 | NA | 0 | 0 | 0 | 0 | 0 | 0 |
| ENSDARG000 | 8;8;8;8;8;8;8 | 30112624:30 | 30112869:30 | +;+;+;+;+;+;+ | 2739 | fancc | 16 | 136 | 75 | 20 | 104 | 27 |
| ENSDARG000 | 8;8;8;8;8;8;8 | 17983369:17 | 17985014:17 | -;-;-;-;-;-;- | 2164 | fpgt | 160 | 142 | 141 | 114 | 134 | 93 |
| ENSDARG000 | 8;8;8 | 17990195:17 | 17990263:17 | +;+;+ | 795 | NA | 0 | 0 | 0 | 0 | 0 | 0 |
| ENSDARG000 | 8;8;8 | 7359294:737 | 7359449:737 | +;+;+ | 3759 | pcsk1nl | 0 | 0 | 0 | 4 | 0 | 2 |
| ENSDARG000 | 8;8;8;8;8;8;8 | 40107885:40 | 40108273:40 | -;-;-;-;-;-;- | 2122 | NA | 4 | 1 | 0 | 0 | 0 | 0 |
| ENSDARG000 | 8;8;8;8;8;8;8 | 40080728:40 | 40081910:40 | +;+;+;+;+;+;+ | 1999 | NA | 25 | 0 | 60 | 22 | 39 | 0 |
| ENSDARG000 | 8;8;8;8;8;8;8 | 37111643:37 | 37111748:37 | +;+;+;+;+;+;+ | 1733 | ren | 0 | 0 | 0 | 0 | 0 | 0 |
| ENSDARG000 | 8;8;8;8;8;8;8 | 9149436:915 | 9149509:915 | +;+;+;+;+;+;+ | 949 | NA | 0 | 11 | 723 | 47 | 298 | 177 |
| ENSDARG000 | 8;8;8;8;8;8;8 | 25359394:25 | 25359661:25 | +;+;+;+;+;+;+ | 3441 | NA | 4 | 12 | 12 | 8 | 22 | 72 |
| ENSDARG000 | 8;8;8;8;8;8;8 | 7382006:738 | 7382037:738 | -;-;-;-;-;-;- | 862 | NA | 0 | 0 | 0 | 0 | 2 | 0 |
| ENSDARG000 | 8;8 | 16257220:16 | 16258150:16 | -;- | 1419 | dmrta2 | 0 | 0 | 0 | 0 | 0 | 0 |
| ENSDARG000 | 8;8;8;8;8;8;8 | 31872992:31 | 31873053:31 | +;+;+;+;+;+;+ | 3087 | NA | 113 | 42 | 44 | 40 | 90 | 76 |
| ENSDARG000 | 8;8;8 | 2233329:223 | 2233668:223 | -;-;- | 575 | NA | 14 | 16 | 18 | 5 | 33 | 12 |
| ENSDARG000 | 8;8;8;8;8;8;8 | 19621511:19 | 19622010:19 | +;+;+;+;+;+;+ | 1610 | slc35a3a | 98 | 75 | 129 | 78 | 101 | 166 |
| ENSDARG000 | 8;8;8;8;8;8;8 | 12340247:12 | 12342512:12 | -;-;-;-;-;-;- | 4073 | phf19 | 263 | 771 | 743 | 450 | 780 | 596 |
| ENSDARG000 | 8;8;8;8;8;8;8 | 4445232:444 | 4446464:444 | -;-;-;-;-;-;- | 5068 | NA | 4470 | 3185 | 3335 | 3656 | 2699 | 3621 |
| ENSDARG000 | 8;8 | 45444493:45 | 45444616:45 | +;+ | 181 | NA | 0 | 0 | 0 | 0 | 0 | 0 |
| ENSDARG000 | 8;8;8;8;8;8;8 | 37168570:37 | 37168849:37 | +;+;+;+;+;+;+ | 1452 | csf1b | 12 | 2 | 22 | 0 | 22 | 0 |
| ENSDARG000 | 8;8;8;8;8;8;8 | 25368034:25 | 25368747:25 | -;-;-;-;-;-;- | 2894 | kcnq2a | 13 | 6 | 8 | 0 | 0 | 11 |
| ENSDARG000 | 8;8;8;8 | 45608540:45 | 45608735:45 | +;+;+;+ | 742 | NA | 0 | 4 | 0 | 0 | 0 | 0 |
| ENSDARG000 | 8;8;8 | 47829361:47 | 47830206:47 | -;-;- | 2649 | si:dkeyp-104 | 566 | 450 | 481 | 419 | 506 | 607 |

| | | | | | | | | | | | | |
|---|---|---|---|---|---|---|---|---|---|---|---|---|
| ENSDARG000 | 8:8:8:8:8:8:8: | 23745167:237 | 23745286:237 | +:+:+:+:+:+:+: | 834 | NA | 2 | 4 | 0 | 0 | 1 | 0 |
| ENSDARG000 | 8 | 19668654 | 19670305 | + | 1652 | foxe3 | 6 | 0 | 14 | 4 | 0 | 2 |
| ENSDARG000 | 8:8:8:8:8:8:8: | 23776570:237 | 23777047:237 | -:-:-:-:-:-:-: | 1012 | zgc:195245 | 0 | 0 | 0 | 0 | 0 | 0 |
| ENSDARG000 | 8:8:8:8:8:8:8: | 23753273:237 | 23753990:237 | -:-:-:-:-:-:-: | 3314 | NA | 218 | 2 | 8 | 0 | 0 | 18 |
| ENSDARG000 | 8:8:8:8:8:8:8: | 16274778:162 | 16275527:162 | -:-:-:-:-:-:-: | 2898 | faf1 | 833 | 807 | 940 | 374 | 795 | 582 |
| ENSDARG000 | 8 | 19674369 | 19676640 | + | 2272 | foxd2 | 14 | 4 | 58 | 260 | 166 | 12 |
| ENSDARG000 | 8;8;8 | 40275830:402 | 40276572:402 | +;+;+ | 3994 | ORAI1 | 73 | 94 | 81 | 72 | 124 | 128 |
| ENSDARG000 | 8;8;8;8;8;8;8 | 19801865:198 | 19802682:198 | -;-;-;-;-;-;- | 2327 | trabd2b | 2 | 0 | 0 | 8 | 10 | 6 |
| ENSDARG000 | 8:8:8:8:8:8:8: | 26432639:264 | 26433123:264 | +:+:+:+:+:+:+: | 3175 | zgc:136971 | 64 | 68 | 51 | 106 | 34 | 68 |
| ENSDARG000 | 8 | 40325690 | 40327397 | - | 1708 | aplnra | 18 | 0 | 16 | 0 | 0 | 18 |
| ENSDARG000 | 8;8;8;8;8;8 | 19739127:197 | 19739415:197 | +;+;+;+;+;+ | 1418 | NA | 0 | 0 | 0 | 0 | 1 | 26 |
| ENSDARG000 | 8:8:8:8:8:8:8: | 21094884:210 | 21095629:210 | -:-:-:-:-:-:-: | 2504 | aldh9a1a.1 | 5106 | 5168 | 5930 | 8095 | 5459 | 9627 |
| ENSDARG000 | 8:8:8:8:8:8:8: | 2260912:2261 | 2261212:2261 | -:-:-:-:-:-:-: | 1995 | PLAT | 71 | 26 | 6 | 4 | 16 | 16 |
| ENSDARG000 | 8:8:8:8:8:8:8: | 37228611:372 | 37232569:372 | -:-:-:-:-:-:-: | 7232 | rbm39b | 1644 | 1485 | 1866 | 1662 | 1801 | 1757 |
| ENSDARG000 | 8:8:8:8:8:8:8: | 45762484:457 | 45765200:457 | -:-:-:-:-:-:-: | 6410 | ogdha | 4711 | 3831 | 5004 | 4249 | 4596 | 5004 |
| ENSDARG000 | 8:8:8:8:8:8:8: | 30159329:301 | 30159577:301 | -:-:-:-:-:-:-: | 3616 | aopep | 49 | 70 | 76 | 42 | 38 | 90 |
| ENSDARG000 | 8:8:8:8:8:8:8: | 40426766:404 | 40431336:404 | -:-:-:-:-:-:-: | 5447 | NA | 1840 | 2207 | 1522 | 1082 | 2925 | 2143 |
| ENSDARG000 | 8;8 | 45767653:457 | 45768269:457 | +;+ | 821 | NA | 0 | 0 | 0 | 0 | 0 | 0 |
| ENSDARG000 | 8:8:8:8:8:8:8: | 45754871:457 | 45756009:457 | -:-:-:-:-:-:-: | 1815 | ppiaa | 30769 | 34027 | 44683 | 53289 | 37003 | 34116 |
| ENSDARG000 | 8;8;8;8;8 | 16407884:164 | 16408010:164 | +;+;+;+;+ | 4540 | cdkn2c | 270 | 738 | 531 | 519 | 866 | 544 |
| ENSDARG000 | 8:8:8:8:8:8:8: | 7393435:7394 | 7394136:7394 | -:-:-:-:-:-:-: | 4234 | HDAC6 | 597 | 750 | 725 | 490 | 832 | 725 |
| ENSDARG000 | 8:8:8:8:8:8:8: | 19948606:199 | 19948872:199 | -:-:-:-:-:-:-: | 1273 | NA | 28 | 42 | 26 | 10 | 52 | 53 |
| ENSDARG000 | 8:8:8:8:8:8:8: | 12408840:124 | 12409245:124 | -:-:-:-:-:-:-: | 2220 | traf1 | 42 | 26 | 36 | 18 | 47 | 68 |
| ENSDARG000 | 8:8:8:8:8:8:8: | 43736072:437 | 43736461:437 | -:-:-:-:-:-:-: | 3654 | ulk1a | 74 | 56 | 182 | 103 | 132 | 76 |
| ENSDARG000 | 8;8;8;8 | 19958485:199 | 19958556:199 | +;+;+;+ | 215 | NA | 0 | 0 | 0 | 0 | 0 | 0 |
| ENSDARG000 | 8;8;8;8 | 19965223:199 | 19965294:199 | +;+;+;+ | 222 | NA | 0 | 0 | 0 | 0 | 0 | 0 |
| ENSDARG000 | 8;8;8 | 16424722:164 | 16425698:164 | -;-;- | 1698 | rnf11b | 236 | 118 | 131 | 98 | 230 | 142 |
| ENSDARG000 | 8;8;8;8 | 19951733:199 | 19951804:199 | +;+;+;+ | 215 | NA | 0 | 0 | 0 | 0 | 0 | 0 |
| ENSDARG000 | 8:8:8:8:8:8:8: | 17993788:179 | 17993967:179 | -:-:-:-:-:-:-: | 3261 | acot11b | 4 | 0 | 6 | 0 | 0 | 4 |
| ENSDARG000 | 8:8:8:8:8:8:8: | 18010568:180 | 18011133:180 | +:+:+:+:+:+:+: | 3218 | ssbp3b | 498 | 437 | 657 | 411 | 718 | 532 |
| ENSDARG000 | 8:8:8:8:8:8:8: | 31901318:319 | 31902307:319 | -;-;-;-;-;-;-;- | 2526 | rgs7bpa | 4 | 0 | 0 | 0 | 0 | 0 |
| ENSDARG000 | 8:8:8:8:8:8:8: | 40457107:404 | 40457540:404 | -:-:-:-:-:-:-: | 1337 | myl7 | 0 | 0 | 0 | 0 | 0 | 2 |
| ENSDARG000 | 8;8 | 25569903:255 | 25569979:255 | +;+ | 889 | NA | 2 | 0 | 0 | 0 | 8 | 0 |
| ENSDARG000 | 8:8:8:8:8:8:8: | 25423133:254 | 25426626:254 | -:-:-:-:-:-:-: | 8930 | PREX1 | 2766 | 1421 | 2305 | 1411 | 2470 | 2393 |
| ENSDARG000 | 8;8;8;8;8;8;8 | 23781028:237 | 23781654:237 | -;-;-;-;-;-;- | 1628 | si:ch211-163l | 0 | 0 | 5 | 0 | 0 | 5 |
| ENSDARG000 | 8:8:8:8:8:8:8: | 23783066:237 | 23783107:237 | +:+:+:+:+:+:+: | 4468 | ELAPOR1 | 0 | 0 | 0 | 0 | 4 | 10 |
| ENSDARG000 | 8:8:8:8:8:8:8: | 9222684:9226 | 9223425:9226 | -:-:-:-:-:-:-: | 6597 | plxna3 | 825 | 565 | 3151 | 615 | 2145 | 1491 |
| ENSDARG000 | 8:8:8:8:8:8:8: | 40476811:404 | 40477394:404 | +:+:+:+:+:+:+: | 2973 | gck | 2 | 2 | 8 | 5 | 10 | 4 |
| ENSDARG000 | 8:8:8:8:8:8:8: | 16514830:165 | 16519829:165 | -:-:-:-:-:-:-: | 8039 | EPS15 | 1081 | 799 | 1301 | 1131 | 1206 | 1080 |
| ENSDARG000 | 8;8;8 | 49766338:497 | 49766571:497 | +;+;+ | 2087 | rmi1 | 51 | 84 | 112 | 43 | 112 | 86 |
| ENSDARG000 | 8:8:8:8:8:8:8: | 2319091:2319 | 2319262:2319 | +:+:+:+:+:+:+: | 2858 | ikbkb | 460 | 331 | 422 | 237 | 422 | 415 |
| ENSDARG000 | 8:8:8:8:8:8:8: | 16470358:164 | 16471109:164 | -:-:-:-:-:-:-: | 2850 | TTC39A | 2897 | 2791 | 7197 | 3617 | 6592 | 7772 |
| ENSDARG000 | 8;8 | 12451739:124 | 12452348:124 | -;- | 790 | FIBCD1 | 39 | 70 | 47 | 52 | 84 | 39 |
| ENSDARG000 | 8:8:8:8:8:8:8: | 7463911:7464 | 7464300:7464 | -:-:-:-:-:-:-: | 1242 | NA | 4 | 4 | 0 | 0 | 2 | 0 |
| ENSDARG000 | 8:8:8:8:8:8:8: | 12452086:124 | 12452348:124 | -:-:-:-:-:-:-: | 5243 | NA | 224 | 256 | 107 | 173 | 220 | 62 |
| ENSDARG000 | 8:8:8:8:8:8:8: | 47850073:478 | 47850275:478 | +:+:+:+:+:+:+: | 3050 | plod1a | 1045 | 2487 | 5279 | 3867 | 2492 | 1102 |
| ENSDARG000 | 8:8:8:8:8:8:8: | 21104507:211 | 21105068:211 | -:-:-:-:-:-:-: | 1692 | tmco1 | 555 | 740 | 912 | 462 | 812 | 762 |
| ENSDARG000 | 8;8;8;8;8 | 31923858:319 | 31924060:319 | -;-;-;-;- | 1090 | RNF180 | 0 | 0 | 0 | 0 | 1 | 0 |
| ENSDARG000 | 8:8:8:8:8:8:8: | 4516831:4516 | 4517114:4517 | -:-:-:-:-:-:-: | 3717 | dhx37 | 512 | 644 | 810 | 280 | 516 | 372 |
| ENSDARG000 | 8:8:8:8:8:8:8: | 45850139:458 | 45850261:458 | -:-:-:-:-:-:-: | 3823 | adam9 | 1359 | 1120 | 1308 | 889 | 1186 | 1218 |
| ENSDARG000 | 8:8:8:8:8:8:8: | 7494941:7495 | 7495356:7495 | -:-:-:-:-:-:-: | 1573 | cdk20 | 41 | 77 | 35 | 80 | 62 | 48 |
| ENSDARG000 | 8:8:8:8:8:8:8: | 49778486:497 | 49778946:497 | +:+:+:+:+:+:+: | 4135 | NTRK2 | 153 | 77 | 126 | 32 | 301 | 33 |
| ENSDARG000 | 8 | 31941650 | 31943435 | + | 1786 | htr1aa | 0 | 0 | 0 | 0 | 0 | 0 |
| ENSDARG000 | 8:8:8:8:8:8:8: | 9699111:9699 | 9699345:9699 | +:+:+:+:+:+:+: | 3937 | gripap1 | 675 | 440 | 580 | 362 | 467 | 549 |
| ENSDARG000 | 8:8:8:8:8:8:8: | 45941942:459 | 45942516:459 | +:+:+:+:+:+:+: | 1709 | nudt18 | 68 | 44 | 88 | 74 | 98 | 94 |
| ENSDARG000 | 8:8:8:8:8:8:8: | 26586484:265 | 26587451:265 | -:-:-:-:-:-:-: | 2989 | sema3b | 12 | 18 | 6 | 6 | 14 | 9 |
| ENSDARG000 | 8;8;8 | 12843300:128 | 12843558:128 | -;-;- | 604 | NA | 6 | 6 | 6 | 8 | 4 | 2 |
| ENSDARG000 | 8:8:8:8:8:8:8: | 26522013:265 | 26522338:265 | +:+:+:+:+:+:+: | 3253 | sema3bl | 1192 | 530 | 166 | 1176 | 264 | 160 |
| ENSDARG000 | 8:8:8:8:8:8:8: | 19977712:199 | 19977939:199 | +:+:+:+:+:+:+: | 2466 | znf692 | 85 | 80 | 78 | 30 | 151 | 35 |
| ENSDARG000 | 8:8:8:8:8:8:8: | 9641252:9644 | 9641997:9644 | -:-:-:-:-:-:-: | 2453 | ptpn18 | 498 | 910 | 2454 | 1155 | 5366 | 600 |
| ENSDARG000 | 8:8:8:8:8:8:8: | 21114338:211 | 21114480:211 | +:+:+:+:+:+:+: | 3918 | uck2a | 184 | 439 | 586 | 214 | 320 | 420 |
| ENSDARG000 | 8:8:8:8:8:8:8: | 9684479:9686 | 9684688:9686 | +:+:+:+:+:+:+: | 1923 | MON1B | 4435 | 5286 | 6211 | 4673 | 7492 | 4976 |
| ENSDARG000 | 8:8:8:8:8:8:8: | 41035230:410 | 41035289:410 | +:+:+:+:+:+:+: | 3277 | gpat2 | 14 | 18 | 4 | 19 | 30 | 12 |
| ENSDARG000 | 8:8:8:8:8:8:8: | 22544290:225 | 22544855:225 | -:-:-:-:-:-:-: | 2089 | porcnl | 422 | 371 | 398 | 144 | 485 | 346 |
| ENSDARG000 | 8;8 | 37259398:372 | 37260687:372 | -;- | 1749 | rh50 | 0 | 0 | 0 | 0 | 0 | 0 |
| ENSDARG000 | 8:8:8:8:8:8:8: | 30325902:303 | 30326920:303 | -:-:-:-:-:-:-: | 7240 | DOCK8 | 3403 | 3291 | 3112 | 2858 | 2825 | 2895 |
| ENSDARG000 | 8:8:8:8:8:8:8: | 32028367:320 | 32031008:320 | -:-:-:-:-:-:-: | 6479 | ipo11 | 953 | 760 | 1150 | 402 | 885 | 721 |
| ENSDARG000 | 8:8:8:8:8:8:8: | 23809843:238 | 23809937:238 | +:+:+:+:+:+:+: | 1821 | si:ch211-163l | 2 | 0 | 0 | 0 | 2 | 0 |
| ENSDARG000 | 8;8 | 30368731:303 | 30368754:303 | +;+ | 243 | NA | 0 | 0 | 0 | 0 | 0 | 0 |
| ENSDARG000 | 8;8;8;8;8;8 | 4576132:4577 | 4578079:4578 | -;-;-;-;-;- | 2374 | gp1bb | 12 | 26 | 10 | 6 | 56 | 14 |
| ENSDARG000 | 8;8;8;8;8 | 21133307:211 | 21133559:211 | +;+;+;+;+ | 734 | magoh | 574 | 745 | 963 | 464 | 676 | 516 |
| ENSDARG000 | 8;8 | 49928535:499 | 49928601:499 | -;- | 122 | NA | 0 | 0 | 0 | 0 | 0 | 0 |
| ENSDARG000 | 8:8:8:8:8:8:8: | 45961617:459 | 45962510:459 | -:-:-:-:-:-:-: | 3131 | fhip2b | 497 | 272 | 478 | 373 | 327 | 389 |
| ENSDARG000 | 8:8:8:8:8:8:8: | 49885638:498 | 49885939:498 | -:-:-:-:-:-:-: | 3686 | agtpbp1 | 379 | 283 | 266 | 144 | 310 | 341 |
| ENSDARG000 | 8:8:8:8:8:8:8: | 4588398:4588 | 4589505:4589 | -:-:-:-:-:-:-: | 2278 | septin5a | 257 | 166 | 74 | 90 | 86 | 56 |
| ENSDARG000 | 8:8:8:8:8:8:8: | 12854105:128 | 12857275:128 | -:-:-:-:-:-:-: | 4613 | SLC2A6 | 294 | 285 | 200 | 97 | 391 | 243 |
| ENSDARG000 | 8 | 4697764 | 4699320 | + | 1557 | cldn5a | 34 | 0 | 27 | 0 | 5 | 6 |
| ENSDARG000 | 8:8:8:8:8:8:8: | 37280532:372 | 37284527:372 | -:-:-:-:-:-:-: | 7871 | NA | 0 | 0 | 0 | 0 | 0 | 0 |
| ENSDARG000 | 8:8:8:8:8:8:8: | 7506827:7508 | 7508488:7508 | -:-:-:-:-:-:-: | 8027 | hcfc1b | 1942 | 1969 | 3471 | 1354 | 3222 | 2078 |
| ENSDARG000 | 8;8;8;8;8 | 26701038:267 | 26702459:267 | -;-;-;-;- | 2311 | tmem51a | 152 | 99 | 112 | 125 | 65 | 58 |
| ENSDARG000 | 8:8:8:8:8:8:8: | 25590841:255 | 25591815:255 | -:-:-:-:-:-:-: | 2315 | stk38a | 550 | 646 | 682 | 434 | 714 | 548 |
| ENSDARG000 | 8;8;8 | 26702003:267 | 26702029:267 | -;-;- | 584 | NA | 2 | 6 | 7 | 13 | 0 | 0 |
| ENSDARG000 | 8:8:8:8:8:8:8: | 22761854:227 | 22762647:227 | -:-:-:-:-:-:-: | 5111 | NA | 18 | 22 | 33 | 8 | 27 | 38 |
| ENSDARG000 | 8;8;8;8 | 37370860:373 | 37370947:373 | +;+;+;+ | 438 | NA | 0 | 0 | 0 | 0 | 0 | 0 |
| ENSDARG000 | 8:8:8:8:8:8:8: | 20037368:200 | 20037394:200 | +:+:+:+:+:+:+: | 3833 | NA | 2716 | 1373 | 2445 | 1251 | 1320 | 2803 |
| ENSDARG000 | 8:8:8:8:8:8:8: | 12975963:129 | 12976093:129 | +:+:+:+:+:+:+: | 851 | DRAM2 | 2766 | 2398 | 1959 | 3296 | 3047 | 1368 |
| ENSDARG000 | 8;8 | 52161861:521 | 52162325:521 | +;+ | 523 | NA | 0 | 0 | 0 | 0 | 0 | 0 |
| ENSDARG000 | 8:8:8:8:8:8:8: | 52106818:521 | 52109924:521 | -:-:-:-:-:-:-: | 5114 | tcf7l1b | 304 | 128 | 209 | 100 | 195 | 207 |
| ENSDARG000 | 8:8:8:8:8:8:8: | 47897734:479 | 47897771:479 | +:+:+:+:+:+:+: | 5369 | mfn2 | 618 | 933 | 1096 | 555 | 920 | 664 |
| ENSDARG000 | 8:8:8:8:8:8:8: | 21137446:211 | 21138164:211 | -:-:-:-:-:-:-: | 3191 | cpt2 | 368 | 564 | 618 | 256 | 742 | 344 |
| ENSDARG000 | 8;8;8;8;8;8 | 4779744:4781 | 4780664:4781 | -;-;-;-;-;- | 1840 | ciao1 | 857 | 820 | 879 | 628 | 876 | 655 |
| ENSDARG000 | 8:8:8:8:8:8:8: | 26714532:267 | 26714746:267 | -:-:-:-:-:-:-: | 2910 | kazna | 19 | 25 | 26 | 16 | 65 | 22 |
| ENSDARG000 | 8;8;8 | 46010838:460 | 46011202:460 | +;+;+ | 683 | NA | 59 | 71 | 71 | 52 | 56 | 42 |
| ENSDARG000 | 8;8;8;8;8;8;8 | 52284075:522 | 52284331:522 | +;+;+;+;+;+;+ | 1740 | ube2d4 | 89 | 53 | 148 | 24 | 101 | 46 |
| ENSDARG000 | 8:8:8:8:8:8:8: | 4734005:4737 | 4734434:4737 | -:-:-:-:-:-:-: | 2211 | cdc45 | 74 | 225 | 250 | 70 | 170 | 137 |
| ENSDARG000 | 8;8;8;8 | 18039570:180 | 18040179:180 | -;-;-;- | 1474 | dio1 | 14 | 22 | 64 | 22 | 10 | 55 |
| ENSDARG000 | 8:8:8:8:8:8:8: | 25615781:256 | 25615842:256 | +:+:+:+:+:+:+: | 3840 | SLC38A5 | 532 | 1724 | 1635 | 1027 | 742 | 692 |
| ENSDARG000 | 8:8:8:8:8:8:8: | 2412583:2414 | 2413287:2415 | -:-:-:-:-:-:-: | 1823 | vdac3 | 3630 | 3226 | 4831 | 5582 | 3595 | 4445 |
| ENSDARG000 | 8;8;8 | 22844700:228 | 22845447:228 | -;-;- | 876 | NA | 0 | 6 | 24 | 4 | 3 | 4 |
| ENSDARG000 | 8:8:8:8:8:8:8: | 4745340:4761 | 4745342:4761 | +:+:+:+:+:+:+: | 1888 | ufd1l | 1135 | 860 | 1097 | 838 | 1118 | 1072 |
| ENSDARG000 | 8;8;8 | 16565837:165 | 16566023:165 | -;-;- | 418 | NA | 1 | 0 | 0 | 0 | 0 | 0 |
| ENSDARG000 | 8:8:8:8:8:8:8: | 23826985:238 | 23827930:238 | +:+:+:+:+:+:+: | 3255 | mapk14a | 1139 | 1300 | 1296 | 987 | 1242 | 1092 |
| ENSDARG000 | 8:8:8:8:8:8:8: | 26818446:268 | 26818657:268 | +:+:+:+:+:+:+: | 1835 | si:ch211-156j | 188 | 602 | 1515 | 1108 | 1111 | 198 |
| ENSDARG000 | 8:8:8:8:8:8:8: | 18044257:180 | 18044552:180 | +:+:+:+:+:+:+: | 2760 | NA | 28 | 0 | 0 | 0 | 4 | 1 |
| ENSDARG000 | 8:8:8:8:8:8:8: | 16582339:165 | 16583689:165 | -;-;-;-;-;-;-;- | 2513 | calr | 2236 | 3853 | 4711 | 1557 | 4179 | 2905 |
| ENSDARG000 | 8;8;8 | 46025122:460 | 46025441:460 | -;-;- | 675 | NA | 14 | 22 | 46 | 35 | 27 | 51 |
| ENSDARG000 | 8:8:8:8:8:8:8: | 37447606:374 | 37447770:374 | -:-:-:-:-:-:-: | 2049 | eif2d | 280 | 288 | 559 | 240 | 476 | 369 |
| ENSDARG000 | 8;8;8;8;8;8 | 16597139:166 | 16598203:166 | -;-;-;-;-;- | 2028 | tor3a | 464 | 123 | 344 | 65 | 307 | 228 |
| ENSDARG000 | 8:8:8:8:8:8:8: | 18200331:182 | 18200774:182 | -:-:-:-:-:-:-: | 1610 | elovl8b | 48 | 84 | 112 | 49 | 60 | 36 |
| ENSDARG000 | 8:8:8:8:8:8:8: | 20072987:200 | 20074548:200 | -:-:-:-:-:-:-: | 4114 | rfx2 | 310 | 134 | 197 | 133 | 136 | 134 |
| ENSDARG000 | 8;8;8;8;8;8 | 18193116:181 | 18193816:181 | -;-;-;-;-;- | 1245 | rps8b | 50 | 6 | 4 | 2 | 0 | 24 |
| ENSDARG000 | 8;8;8 | 46058144:460 | 46058197:460 | +;+;+ | 337 | NA | 0 | 0 | 0 | 0 | 0 | 0 |
| ENSDARG000 | 8;8;8;8;8;8 | 46057696:460 | 46058169:460 | -;-;-;-;-;- | 1849 | NA | 0 | 0 | 0 | 10 | 2 | 0 |
| ENSDARG000 | 8:8:8:8:8:8:8: | 21146262:211 | 21146397:211 | +:+:+:+:+:+:+: | 1329 | porcn | 4 | 0 | 0 | 2 | 0 | 0 |
| ENSDARG000 | 8;8 | 48095911:482 | 48095998:482 | +;+ | 438 | NA | 0 | 0 | 0 | 0 | 0 | 0 |
| ENSDARG000 | 8;8 | 48092911:480 | 48093191:480 | -;- | 677 | NA | 2 | 0 | 0 | 0 | 0 | 0 |
| ENSDARG000 | 8:8:8:8:8:8:8: | 30425700:304 | 30425825:304 | +:+:+:+:+:+:+: | 1653 | cbwd | 287 | 242 | 386 | 175 | 296 | 241 |
| ENSDARG000 | 8:8:8:8:8:8:8: | 49936585:499 | 49936633:499 | +:+:+:+:+:+:+: | 2400 | naa35 | 868 | 796 | 939 | 618 | 874 | 793 |
| ENSDARG000 | 8:8:8:8:8:8:8: | 22851472:228 | 22852003:228 | +:+:+:+:+:+:+: | 7710 | cacna1fb | 0 | 0 | 0 | 4 | 0 | 10 |
| ENSDARG000 | 8;8 | 26859639:268 | 26859688:268 | +;+ | 4243 | NA | 4 | 4 | 33 | 39 | 35 | 4 |
| ENSDARG000 | 8:8:8:8:8:8:8: | 32376936:323 | 32377578:323 | -:-:-:-:-:-:-: | 2817 | lipg | 538 | 609 | 1431 | 461 | 121 | 322 |
| ENSDARG000 | 8:8:8:8:8:8:8: | 12989372:129 | 12989766:129 | -:-:-:-:-:-:-: | 2538 | dennd2da | 1390 | 1233 | 979 | 1629 | 1528 | 992 |
| ENSDARG000 | 8:8:8:8:8:8:8: | 48514551:485 | 48514674:485 | +:+:+:+:+:+:+: | 1006 | NA | 4 | 4 | 13 | 0 | 8 | 2 |
| ENSDARG000 | 8:8:8:8:8:8:8: | 52491436:525 | 52491797:525 | +:+:+:+:+:+:+: | 1585 | RNF122 | 43 | 67 | 134 | 130 | 71 | 73 |
| ENSDARG000 | 8;8 | 13027524:130 | 13027797:130 | +;+ | 337 | NA | 0 | 0 | 0 | 0 | 0 | 0 |
| ENSDARG000 | 8:8:8:8:8:8:8: | 48480529:484 | 48480553:484 | +:+:+:+:+:+:+: | 3372 | NA | 14 | 4 | 5 | 0 | 6 | 0 |
| ENSDARG000 | 8:8:8:8:8:8:8: | 25629722:256 | 25630045:256 | +:+:+:+:+:+:+: | 2535 | NA | 36 | 62 | 34 | 26 | 72 | 62 |
| ENSDARG000 | 8:8:8:8:8:8:8: | 23861461:238 | 23861513:238 | +:+:+:+:+:+:+: | 2614 | srpk1a | 461 | 329 | 387 | 162 | 472 | 336 |
| ENSDARG000 | 8:8:8:8:8:8:8: | 23876928:238 | 23881197:238 | -;-;-;-;-;-;-;- | 5769 | lhfpl5b | 47 | 24 | 26 | 14 | 43 | 53 |
| ENSDARG000 | 8:8:8:8:8:8:8: | 26868105:268 | 26868691:268 | +:+:+:+:+:+:+: | 4577 | rimkla | 3235 | 4914 | 5687 | 5526 | 3694 | 3122 |
| ENSDARG000 | 8;8 | 21159122:211 | 21159319:211 | +;+ | 1020 | spryd4 | 128 | 212 | 397 | 334 | 212 | 256 |
| ENSDARG000 | 8;8;8;8 | 52515188:525 | 52515337:525 | +;+;+;+ | 3552 | NA | 3786 | 2280 | 3004 | 2891 | 2916 | 2891 |
| ENSDARG000 | 8:8:8:8:8:8:8: | 16615798:166 | 16616913:166 | -:-:-:-:-:-:-: | 4721 | osbpl9 | 1958 | 1573 | 1890 | 2109 | 3238 | 2580 |
| ENSDARG000 | 8:8:8:8:8:8:8: | 9866351:9867 | 9866526:9868 | +:+:+:+:+:+:+: | 2656 | kcnd1 | 2 | 0 | 4 | 0 | 0 | 0 |
| ENSDARG000 | 8:8:8:8:8:8:8: | 41186027:411 | 41186588:411 | -:-:-:-:-:-:-: | 1648 | fahd2a | 430 | 234 | 492 | 447 | 640 | 166 |
| ENSDARG000 | 8:8:8:8:8:8:8: | 16609666:166 | 16610180:166 | -:-:-:-:-:-:-: | 3458 | NA | 291 | 375 | 686 | 323 | 873 | 434 |
| ENSDARG000 | 8:8:8:8:8:8:8: | 37465380:374 | 37465659:374 | +:+:+:+:+:+:+: | 1270 | mapkapk2b | 0 | 2 | 0 | 0 | 0 | 0 |
| ENSDARG000 | 8:8:8:8:8:8:8: | 21166053:211 | 21166216:211 | -:-:-:-:-:-:-: | 1858 | GLS2 | 0 | 0 | 22 | 6 | 80 | 10 |
| ENSDARG000 | 8 | 14058646 | 14060244 | + | 1599 | ugt5e1 | 0 | 0 | 0 | 0 | 0 | 0 |
| ENSDARG000 | 8:8:8:8:8:8:8: | 14058185:140 | 14062928:140 | -:-:-:-:-:-:-: | 8258 | dedd | 604 | 633 | 810 | 629 | 644 | 615 |
| ENSDARG000 | 8:8:8:8:8:8:8: | 49975160:499 | 49975208:499 | +:+:+:+:+:+:+: | 2092 | gfpt1 | 1893 | 1929 | 1172 | 563 | 1822 | 996 |
| ENSDARG000 | 8;8 | 37489495:374 | 37490431:374 | +;+ | 1089 | FMOD | 4 | 4 | 0 | 0 | 0 | 4 |
| ENSDARG000 | 8:8:8:8:8:8:8: | 26902358:269 | 26903018:269 | -;-;-;-;-;-;-;- | 1754 | zmynd12 | 0 | 4 | 17 | 8 | 2 | 12 |
| ENSDARG000 | 8:8:8:8:8:8:8: | 52530889:525 | 52531029:525 | +:+:+:+:+:+:+: | 2895 | stambpb | 421 | 421 | 637 | 1006 | 532 | 463 |
| ENSDARG000 | 8:8:8:8:8:8:8: | 50066802:500 | 50066972:500 | -:-:-:-:-:-:-: | 1764 | ANTXR1 | 14 | 6 | 2 | 0 | 6 | 10 |
| ENSDARG000 | 8:8:8:8:8:8:8: | 20140321:201 | 20140913:201 | +:+:+:+:+:+:+: | 4717 | acsbg2 | 2672 | 3611 | 2305 | 1527 | 2011 | 2854 |
| ENSDARG000 | 8:8:8:8:8:8:8: | 10001805:100 | 10002100:100 | +:+:+:+:+:+:+: | 2750 | NA | 4 | 2 | 0 | 2 | 17 | 4 |
| ENSDARG000 | 8:8:8:8:8:8:8: | 32389838:323 | 32390017:323 | +:+:+:+:+:+:+: | 1830 | mthfd2l | 0 | 12 | 7 | 14 | 4 | 0 |
| ENSDARG000 | 8:8:8:8:8:8:8: | 18204117:182 | 18204492:182 | -:-:-:-:-:-:-: | 1827 | NA | 26 | 48 | 70 | 14 | 30 | 83 |
| ENSDARG000 | 8:8:8:8:8:8:8: | 32402441:324 | 32403032:324 | +:+:+:+:+:+:+: | 1852 | NA | 236 | 14 | 0 | 6 | 8 | 20 |
| ENSDARG000 | 8;8;8;8 | 23917588:239 | 23918019:239 | +;+;+;+ | 1526 | NA | 2 | 3 | 0 | 0 | 2 | 0 |
| ENSDARG000 | 8:8:8:8:8:8:8: | 23916647:239 | 23916741:239 | +:+:+:+:+:+:+: | 1692 | cpne5a | 0 | 0 | 0 | 0 | 2 | 0 |
| ENSDARG000 | 8;8;8;8;8;8 | 10033326:100 | 10033970:100 | -;-;-;-;-;- | 1115 | NA | 0 | 22 | 4 | 0 | 6 | 2 |
| ENSDARG000 | 8;8;8;8;8 | 48558100:485 | 48558234:485 | +;+;+;+;+ | 534 | NA | 0 | 0 | 0 | 0 | 0 | 0 |
| ENSDARG000 | 8 | 37514822 | 37515754 | + | 933 | NA | 0 | 0 | 0 | 0 | 0 | 0 |
| ENSDARG000 | 8 | 37527575 | 37531221 | + | 3647 | or90j1 | 0 | 0 | 0 | 0 | 0 | 0 |
| ENSDARG000 | 8:8:8:8:8:8:8: | 26923613:269 | 26924459:269 | -;-;-;-;-;-;-;- | 2205 | slc16a1b | 1727 | 2297 | 2896 | 2350 | 2301 | 2020 |
| ENSDARG000 | 8;8;8;8 | 10068272:100 | 10068405:100 | +;+;+;+ | 771 | NA | 0 | 2 | 0 | 0 | 0 | 2 |
| ENSDARG000 | 8;8;8 | 37700090:377 | 37701406:377 | +;+;+ | 2127 | adrb3a | 0 | 0 | 2 | 0 | 8 | 2 |
| ENSDARG000 | 8;8 | 26977429:269 | 26977557:269 | -;- | 282 | NA | 0 | 0 | 0 | 0 | 0 | 0 |
| ENSDARG000 | 8:8:8:8:8:8:8: | 10304981:103 | 10305348:103 | +:+:+:+:+:+:+: | 2669 | pim1 | 13516 | 9595 | 11107 | 11398 | 9832 | 14389 |
| ENSDARG000 | 8:8:8:8:8:8:8: | 52550241:525 | 52550643:525 | -:-:-:-:-:-:-: | 3933 | NA | 2155 | 1900 | 2043 | 1757 | 2342 | 1613 |

| | | | | | | | | | | | | |
|---|---|---|---|---|---|---|---|---|---|---|---|---|
| ENSDARG000 | 8:8:8:8:8:8:8: | 27203326:272 | 27203559:272 | -:-:-:-:-:-:-:-:-: | 1515 | NA | 2 | 0 | 0 | 0 | 4 | 0 |
| ENSDARG000 | 8:8:8:8:8:8:8: | 18214081:182 | 18214201:182 | -:-:-:-:-:-:-:-:-: | 2056 | NA | 0 | 0 | 4 | 0 | 4 | 4 |
| ENSDARG000 | 8;8;8;8 | 14085894:140 | 14086211:140 | -;-;-;- | 798 | si:ch211-229r | 4 | 0 | 0 | 0 | 0 | 0 |
| ENSDARG000 | 8;8 | 48603398:486 | 48603564:486 | +;+ | 886 | nppal | 0 | 0 | 0 | 0 | 36 | 0 |
| ENSDARG000 | 8 | 30452945 | 30454096 | + | 1152 | foxd5 | 8 | 8 | 13 | 24 | 4 | 0 |
| ENSDARG000 | 8:8:8:8:8:8:8: | 25650229:256 | 25650591:256 | -:-:-:-:-:-:-:-:-: | 3416 | sema3ga | 594 | 190 | 141 | 501 | 322 | 192 |
| ENSDARG000 | 8:8:8:8:8:8:8: | 50134007:501 | 50134197:501 | -:-:-:-:-:-:-:-:-: | 463 | NA | 0 | 0 | 0 | 0 | 0 | 0 |
| ENSDARG000 | 8:8:8:8:8:8:8: | 37749263:377 | 37749320:377 | +:+:+:+:+:+:+:+: | 999 | npm2a | 0 | 2 | 0 | 0 | 0 | 0 |
| ENSDARG000 | 8:8:8:8:8:8:8: | 13036221:130 | 13036456:130 | -:-:-:-:-:-:-:-:-: | 3976 | NA | 2 | 2 | 10 | 39 | 2 | 38 |
| ENSDARG000 | 8:8:8:8:8:8:8: | 24107030:241 | 24109961:241 | -;-;-;-;-;-;-;-;- | 4037 | dclre1b | 160 | 323 | 424 | 144 | 358 | 353 |
| ENSDARG000 | 8:8:8:8:8:8:8: | 20194155:201 | 20197397:201 | -:-:-:-:-:-:-:-:-: | 5109 | MLLT1 | 612 | 486 | 486 | 466 | 479 | 545 |
| ENSDARG000 | 8;8 | 20229705:202 | 20230811:202 | +;+ | 1773 | NA | 0 | 0 | 0 | 0 | 0 | 4 |
| ENSDARG000 | 8;8 | 24122498:241 | 24123182:241 | -;- | 1026 | adora1b | 14 | 4 | 0 | 0 | 10 | 0 |
| ENSDARG000 | 8:8:8:8:8:8:8: | 20232421:202 | 20233191:202 | -:-:-:-:-:-:-:-:-: | 1574 | acer1 | 2 | 0 | 0 | 0 | 4 | 2 |
| ENSDARG000 | 8;8 | 10323443:103 | 10323842:103 | -;- | 539 | NA | 4 | 0 | 0 | 0 | 10 | 8 |
| ENSDARG000 | 8;8;8;8 | 14092794:140 | 14092857:140 | -;-;-;- | 482 | NA | 0 | 0 | 0 | 0 | 0 | 0 |
| ENSDARG000 | 8;8;8;8;8;8;8 | 27361687:273 | 27362330:273 | -;-;-;-;-;-;- | 1951 | tafa3b | 158 | 55 | 48 | 98 | 20 | 24 |
| ENSDARG000 | 8;8;8;8;8 | 52619365:526 | 52619649:526 | +;+;+;+;+ | 4195 | ccn1l2 | 56 | 0 | 7 | 0 | 2 | 2 |
| ENSDARG000 | 8:8:8:8:8:8:8: | 10339869:103 | 10340122:103 | +:+:+:+:+:+:+:+: | 1952 | TBC1D22B | 144 | 70 | 127 | 74 | 110 | 40 |
| ENSDARG000 | 8:8:8:8:8:8:8: | 21195420:211 | 21195666:211 | +:+:+:+:+:+:+:+: | 5937 | col2a1a | 34 | 34 | 72 | 14 | 30 | 36 |
| ENSDARG000 | 8;8;8;8;8 | 48609521:486 | 48609713:486 | +;+;+;+;+ | 947 | nppb | 0 | 0 | 0 | 0 | 0 | 0 |
| ENSDARG000 | 8;8;8;8 | 14096167:140 | 14096751:140 | -;-;-;- | 1107 | si:ch211-229r | 2 | 0 | 0 | 0 | 0 | 0 |
| ENSDARG000 | 8:8:8:8:8:8:8: | 41229233:412 | 41229243:412 | +:+:+:+:+:+:+:+: | 2770 | zgc:152830 | 2234 | 2714 | 2894 | 1711 | 3134 | 2167 |
| ENSDARG000 | 8:8:8:8:8:8:8: | 30456161:304 | 30456418:304 | +:+:+:+:+:+:+:+: | 1704 | pgm5 | 6 | 4 | 0 | 0 | 2 | 0 |
| ENSDARG000 | 8;8 | 48613040:486 | 48613184:486 | +;+ | 1624 | nppa | 0 | 2 | 1 | 0 | 0 | 4 |
| ENSDARG000 | 8:8:8:8:8:8:8: | 22930627:229 | 22931490:229 | +:+:+:+:+:+:+:+: | 5219 | sypa | 338 | 697 | 1029 | 3832 | 1457 | 5206 |
| ENSDARG000 | 8;8;8;8;8 | 46217861:462 | 46218417:462 | +;+;+;+;+ | 1626 | angptl7 | 50 | 24 | 32 | 10 | 184 | 14 |
| ENSDARG000 | 8:8:8:8:8:8:8: | 46077609:460 | 46078493:460 | -:-:-:-:-:-:-:-:-: | 8700 | mtor | 2470 | 1832 | 2927 | 1513 | 2550 | 1687 |
| ENSDARG000 | 8:8:8:8:8:8:8: | 52637507:526 | 52637754:526 | +:+:+:+:+:+:+:+: | 736 | NA | 0 | 4 | 4 | 0 | 4 | 1 |
| ENSDARG000 | 8;8;8;8;8;8 | 24246659:242 | 24246956:242 | -;-;-;-;-;- | 1290 | zgc:110353 | 2 | 0 | 0 | 0 | 2 | 0 |
| ENSDARG000 | 8:8:8:8:8:8:8: | 14104496:141 | 14105098:141 | -:-:-:-:-:-:-:-:-: | 1723 | bgna | 14 | 10 | 6 | 6 | 26 | 2 |
| ENSDARG000 | 8;8;8;8 | 50143194:501 | 50143583:501 | -;-;-;- | 631 | NA | 30 | 124 | 102 | 48 | 168 | 294 |
| ENSDARG000 | 8:8:8:8:8:8:8: | 18230948:182 | 18231509:182 | -:-:-:-:-:-:-:-:-: | 1578 | guk1b | 0 | 0 | 0 | 0 | 0 | 0 |
| ENSDARG000 | 8:8:8:8:8:8:8: | 22955262:229 | 22955963:229 | +:+:+:+:+:+:+:+: | 1341 | plp2a | 2 | 18 | 0 | 6 | 0 | 8 |
| ENSDARG000 | 8;8;8;8;8 | 27596435:276 | 27596550:276 | -;-;-;-;- | 565 | NA | 0 | 0 | 0 | 0 | 0 | 0 |
| ENSDARG000 | 8:8:8:8:8:8:8: | 27555314:275 | 27555432:275 | +:+:+:+:+:+:+:+: | 1085 | rhocb | 2260 | 1542 | 1123 | 1786 | 1005 | 1690 |
| ENSDARG000 | 8:8:8:8:8:8:8: | 32481577:324 | 32483171:324 | -:-:-:-:-:-:-:-:-: | 3394 | NA | 12 | 8 | 18 | 0 | 6 | 48 |
| ENSDARG000 | 8:8:8:8:8:8:8: | 24281512:242 | 24281748:242 | +:+:+:+:+:+:+:+: | 3134 | mmp9 | 1823 | 1349 | 714 | 1328 | 1202 | 1304 |
| ENSDARG000 | 8:8:8:8:8:8:8: | 21254192:212 | 21254206:212 | +:+:+:+:+:+:+:+: | 9170 | itpr3 | 299 | 48 | 528 | 166 | 601 | 256 |
| ENSDARG000 | 8:8:8:8:8:8:8: | 37755099:377 | 37755300:377 | +:+:+:+:+:+:+:+: | 2564 | got1l1 | 0 | 0 | 0 | 0 | 0 | 0 |
| ENSDARG000 | 8;8 | 21265263:212 | 21266105:212 | -;- | 1531 | GPR37L1 | 0 | 0 | 0 | 0 | 0 | 0 |
| ENSDARG000 | 8:8:8:8:8:8:8: | 30501170:305 | 30502529:305 | +:+:+:+:+:+:+:+: | 4003 | BCR | 1942 | 1031 | 1956 | 1700 | 2358 | 1301 |
| ENSDARG000 | 8:8:8:8:8:8:8: | 21225064:212 | 21225239:212 | +:+:+:+:+:+:+:+: | 4975 | cry3a | 749 | 816 | 950 | 829 | 1133 | 1599 |
| ENSDARG000 | 8;8;8;8;8 | 22962690:229 | 22964486:229 | -;-;-;-;- | 2593 | NA | 0 | 0 | 0 | 0 | 0 | 0 |
| ENSDARG000 | 8;8;8;8 | 52713232:527 | 52714437:527 | -;-;-;- | 1568 | tubb2b | 0 | 0 | 0 | 0 | 0 | 0 |
| ENSDARG000 | 8:8:8:8:8:8:8: | 50150834:501 | 50151027:501 | +:+:+:+:+:+:+:+: | 2673 | entpd4 | 447 | 418 | 600 | 250 | 645 | 265 |
| ENSDARG000 | 8;8;8;8;8;8;8 | 25708067:257 | 25709725:257 | -;-;-;-;-;-;- | 3406 | tspy | 708 | 526 | 672 | 544 | 645 | 740 |
| ENSDARG000 | 8:8:8:8:8:8:8: | 20250484:202 | 20250941:202 | -:-:-:-:-:-:-:-:-: | 3684 | myo1f | 122 | 155 | 112 | 106 | 420 | 498 |
| ENSDARG000 | 8:8:8:8:8:8:8: | 18242505:182 | 18245178:182 | -:-:-:-:-:-:-:-:-: | 5189 | RNF220 | 0 | 0 | 16 | 2 | 5 | 2 |
| ENSDARG000 | 8;8;8;8;8 | 22977304:229 | 22977423:229 | +;+;+;+;+ | 751 | samd10a | 0 | 0 | 0 | 6 | 4 | 0 |
| ENSDARG000 | 8;8;8;8;8 | 37905666:379 | 37905792:379 | +;+;+;+;+ | 2676 | prlhr2a | 0 | 0 | 0 | 0 | 0 | 0 |
| ENSDARG000 | 8:8:8:8:8:8:8: | 13064750:130 | 13064893:130 | +:+:+:+:+:+:+:+: | 1657 | sap30bp | 486 | 356 | 444 | 380 | 492 | 432 |
| ENSDARG000 | 8:8:8:8:8:8:8: | 18354485:183 | 18356811:183 | -:-:-:-:-:-:-:-:-: | 3769 | dmap1 | 457 | 567 | 610 | 289 | 555 | 467 |
| ENSDARG000 | 8:8:8:8:8:8:8: | 52732107:527 | 52733571:527 | -:-:-:-:-:-:-:-:-: | 2011 | fgf17 | 0 | 3 | 0 | 0 | 0 | 0 |
| ENSDARG000 | 8:8:8:8:8:8:8: | 32516160:325 | 32516779:325 | +:+:+:+:+:+:+:+: | 2070 | ncs1b | 26 | 18 | 105 | 10 | 501 | 108 |
| ENSDARG000 | 8:8:8:8:8:8:8: | 25716308:257 | 25716780:257 | -:-:-:-:-:-:-:-:-: | 1784 | foxp3a | 6 | 12 | 2 | 4 | 20 | 38 |
| ENSDARG000 | 8:8:8:8:8:8:8: | 14130245:141 | 14130471:141 | -:-:-:-:-:-:-:-:-: | 2111 | si:dkey-6n6.1 | 22 | 51 | 8 | 16 | 20 | 50 |
| ENSDARG000 | 8;8;8 | 14180064:141 | 14180471:141 | -;-;- | 816 | ndufb11 | 149 | 118 | 167 | 136 | 142 | 113 |
| ENSDARG000 | 8:8:8:8:8:8:8: | 38026683:380 | 38028123:380 | -:-:-:-:-:-:-:-:-: | 4782 | adgra2 | 24 | 24 | 18 | 0 | 144 | 6 |
| ENSDARG000 | 8:8:8:8:8:8:8: | 14170567:141 | 14172297:141 | -:-:-:-:-:-:-:-:-: | 3426 | rhoaa | 3900 | 4375 | 4414 | 3469 | 4100 | 4657 |
| ENSDARG000 | 8:8:8:8:8:8:8: | 14158021:141 | 14158232:141 | +:+:+:+:+:+:+:+: | 1449 | tpk2 | 221 | 254 | 238 | 275 | 185 | 331 |
| ENSDARG000 | 8;8;8;8;8 | 30664077:306 | 30664411:306 | +;+;+;+;+ | 5165 | adora2aa | 2 | 4 | 0 | 8 | 4 | 10 |
| ENSDARG000 | 8:8:8:8:8:8:8: | 30600386:306 | 30600684:306 | +:+:+:+:+:+:+:+: | 7167 | specc1la | 1050 | 683 | 1009 | 528 | 947 | 916 |
| ENSDARG000 | 8:8:8:8:8:8:8: | 18369177:183 | 18369665:183 | -:-:-:-:-:-:-:-:-: | 6339 | NA | 2 | 4 | 2 | 0 | 0 | 0 |
| ENSDARG000 | 8;8;8;8 | 25733872:257 | 25734965:257 | +;+;+;+ | 4439 | NA | 199 | 163 | 186 | 90 | 264 | 186 |
| ENSDARG000 | 8;8 | 25731012:257 | 25731151:257 | -;- | 577 | NA | 0 | 0 | 0 | 0 | 0 | 0 |
| ENSDARG000 | 8:8:8:8:8:8:8: | 23009662:230 | 23009773:230 | +:+:+:+:+:+:+:+: | 2043 | uckl1a | 0 | 0 | 2 | 2 | 0 | 0 |
| ENSDARG000 | 8:8:8:8:8:8:8: | 24296639:242 | 24297670:242 | +:+:+:+:+:+:+:+: | 13889 | ZNF335 | 424 | 305 | 581 | 295 | 323 | 300 |
| ENSDARG000 | 8:8:8:8:8:8:8: | 52788997:527 | 52789169:527 | +:+:+:+:+:+:+:+: | 3376 | ADGRD2 | 0 | 0 | 0 | 0 | 0 | 0 |
| ENSDARG000 | 8:8:8:8:8:8:8: | 32613680:326 | 32614171:326 | +:+:+:+:+:+:+:+: | 2585 | NA | 0 | 0 | 0 | 0 | 6 | 0 |
| ENSDARG000 | 8:8:8:8:8:8:8: | 32613916:326 | 32614171:326 | +:+:+:+:+:+:+:+: | 13402 | NA | 12 | 14 | 14 | 4 | 16 | 0 |
| ENSDARG000 | 8;8;8;8;8;8;8 | 24329433:243 | 24329530:243 | -;-;-;-;-;-;- | 1352 | NA | 0 | 1 | 0 | 0 | 0 | 0 |
| ENSDARG000 | 8:8:8:8:8:8:8: | 38190233:381 | 38191768:381 | -:-:-:-:-:-:-:-:-: | 2865 | pdlim2 | 92 | 26 | 35 | 9 | 109 | 30 |
| ENSDARG000 | 8:8:8:8:8:8:8: | 23040727:230 | 23041523:230 | -:-:-:-:-:-:-:-:-: | 2524 | si:dkey-70p6. | 14 | 2 | 75 | 2 | 40 | 98 |
| ENSDARG000 | 8:8:8:8:8:8:8: | 23034718:231 | 23034772:231 | +:+:+:+:+:+:+:+: | 4651 | ythdf1 | 1069 | 1122 | 1422 | 707 | 1203 | 904 |
| ENSDARG000 | 8;8;8;8;8;8 | 23093155:230 | 23093370:230 | +;+;+;+;+;+ | 1502 | nkain5 | 4 | 2 | 14 | 0 | 0 | 16 |
| ENSDARG000 | 8:8:8:8:8:8:8: | 25767610:257 | 25768024:257 | +:+:+:+:+:+:+:+: | 6096 | cacna1sb | 0 | 4 | 3 | 0 | 26 | 0 |
| ENSDARG000 | 8:8:8:8:8:8:8: | 14185295:141 | 14185752:141 | -:-:-:-:-:-:-:-:-: | 2205 | def6a | 34 | 54 | 18 | 42 | 38 | 52 |
| ENSDARG000 | 8;8;8 | 25771171:257 | 25771474:257 | -;-;- | 561 | NA | 0 | 0 | 0 | 0 | 0 | 0 |
| ENSDARG000 | 8:8:8:8:8:8:8: | 25750963:257 | 25752601:257 | -:-:-:-:-:-:-:-:-: | 3500 | suv39h1b | 80 | 246 | 303 | 68 | 169 | 140 |
| ENSDARG000 | 8:8:8:8:8:8:8: | 25761654:257 | 25761691:257 | +:+:+:+:+:+:+:+: | 2096 | tmem9 | 241 | 288 | 392 | 334 | 414 | 409 |
| ENSDARG000 | 8:8:8:8:8:8:8: | 52833443:528 | 52834756:528 | -:-:-:-:-:-:-:-:-: | 2852 | nr5a1a | 0 | 0 | 0 | 0 | 0 | 0 |
| ENSDARG000 | 8 | 30784020 | 30784982 | + | 963 | tas2r200.1 | 0 | 10 | 0 | 0 | 0 | 49 |
| ENSDARG000 | 8;8 | 30780985:307 | 30782089:307 | +;+ | 1105 | tas2r200.2 | 0 | 4 | 0 | 0 | 0 | 19 |
| ENSDARG000 | 8:8:8:8:8:8:8: | 18450052:184 | 18450249:184 | -:-:-:-:-:-:-:-:-: | 3521 | zgc:153738 | 0 | 0 | 0 | 0 | 6 | 6 |
| ENSDARG000 | 8;8 | 30778205:307 | 30778317:307 | +;+ | 206 | NA | 0 | 2 | 0 | 0 | 0 | 5 |
| ENSDARG000 | 8;8;8;8;8;8 | 30730628:307 | 30731869:307 | -;-;-;-;-;- | 2150 | gucd1 | 259 | 251 | 200 | 132 | 200 | 169 |
| ENSDARG000 | 8;8;8;8 | 30742898:307 | 30743014:307 | +;+;+;+ | 658 | snrpd3 | 0 | 0 | 0 | 0 | 0 | 0 |
| ENSDARG000 | 8:8:8:8:8:8:8: | 30758169:307 | 30758790:307 | -:-:-:-:-:-:-:-:-: | 1877 | p2rx7 | 764 | 577 | 687 | 916 | 999 | 1244 |
| ENSDARG000 | 8:8:8:8:8:8:8: | 30699429:307 | 30699574:307 | +:+:+:+:+:+:+:+: | 1467 | upb1 | 0 | 6 | 4 | 0 | 28 | 0 |
| ENSDARG000 | 8:8:8:8:8:8:8: | 30743263:307 | 30743408:307 | +:+:+:+:+:+:+:+: | 4366 | p2rx4b | 20 | 2 | 18 | 2 | 35 | 20 |
| ENSDARG000 | 8:8:8:8:8:8:8: | 20298369:203 | 20298922:203 | -:-:-:-:-:-:-:-:-: | 3978 | adamts10 | 392 | 58 | 512 | 95 | 353 | 235 |
| ENSDARG000 | 8:8:8:8:8:8:8: | 24706551:247 | 24706769:247 | +:+:+:+:+:+:+:+: | 1145 | NA | 0 | 0 | 2 | 0 | 0 | 1 |
| ENSDARG000 | 8;8 | 32787850:327 | 32788042:327 | -;- | 430 | NA | 0 | 0 | 0 | 0 | 0 | 0 |
| ENSDARG000 | 8;8 | 53040551:530 | 53040592:530 | -;- | 235 | NA | 0 | 0 | 0 | 0 | 0 | 0 |
| ENSDARG000 | 8:8:8:8:8:8:8: | 52888295:528 | 52888564:528 | -:-:-:-:-:-:-:-:-: | 1986 | nr6a1a | 0 | 0 | 0 | 0 | 0 | 0 |
| ENSDARG000 | 8;8;8 | 32801818:328 | 32803227:328 | -;-;- | 1884 | zgc:194839 | 6 | 2 | 0 | 0 | 12 | 0 |
| ENSDARG000 | 8;8;8;8 | 32851241:328 | 32851592:328 | -;-;-;- | 2329 | si:dkey-56i24. | 4 | 10 | 34 | 0 | 33 | 90 |
| ENSDARG000 | 8;8;8;8 | 53051701:530 | 53051912:530 | +;+;+;+ | 605 | NA | 24 | 52 | 38 | 4 | 38 | 12 |
| ENSDARG000 | 8:8:8:8:8:8:8: | 38318357:383 | 38321573:383 | -:-:-:-:-:-:-:-:-: | 7206 | sorbs3 | 164 | 140 | 130 | 26 | 371 | 193 |
| ENSDARG000 | 8 | 23122151 | 23122372 | - | 222 | NA | 0 | 0 | 0 | 0 | 0 | 0 |
| ENSDARG000 | 8;8;8;8 | 23128233:231 | 23128261:231 | +;+;+;+ | 355 | NA | 13 | 0 | 1 | 0 | 0 | 0 |
| ENSDARG000 | 8 | 23142946 | 23143712 | + | 767 | NA | 0 | 0 | 0 | 0 | 0 | 0 |
| ENSDARG000 | 8;8;8;8;8 | 53064920:530 | 53065061:530 | +;+;+;+;+ | 733 | nadka | 65 | 94 | 60 | 38 | 135 | 77 |
| ENSDARG000 | 8:8:8:8:8:8:8: | 20393250:203 | 20393400:203 | +:+:+:+:+:+:+:+: | 2580 | zap70 | 4 | 56 | 10 | 0 | 30 | 82 |
| ENSDARG000 | 8;8 | 33105974:331 | 33106032:331 | +;+ | 840 | si:dkey-191c1 | 0 | 0 | 0 | 0 | 0 | 0 |
| ENSDARG000 | 8:8:8:8:8:8:8: | 32914626:329 | 32915210:329 | -:-:-:-:-:-:-:-:-: | 2776 | ralgps1 | 8 | 6 | 0 | 2 | 0 | 2 |
| ENSDARG000 | 8;8;8;8 | 33035709:330 | 33036676:330 | +;+;+;+ | 3472 | angptl2b | 94 | 0 | 14 | 0 | 0 | 2 |
| ENSDARG000 | 8:8:8:8:8:8:8: | 23147218:231 | 23147276:231 | +:+:+:+:+:+:+:+: | 4945 | gid8a | 190 | 148 | 223 | 154 | 270 | 258 |
| ENSDARG000 | 8:8:8:8:8:8:8: | 23152244:231 | 23152457:231 | +:+:+:+:+:+:+:+: | 4291 | NA | 0 | 0 | 0 | 2 | 0 | 0 |
| ENSDARG000 | 8:8:8:8:8:8:8: | 20415824:204 | 20416342:204 | +:+:+:+:+:+:+:+: | 4889 | mob3a | 958 | 1257 | 1329 | 1238 | 1265 | 1156 |
| ENSDARG000 | 8:8:8:8:8:8:8: | 20438884:204 | 20438996:204 | +:+:+:+:+:+:+:+: | 3385 | mknk2b | 5616 | 8020 | 26514 | 44725 | 6367 | 6140 |
| ENSDARG000 | 8;8 | 20448121:204 | 20448282:204 | -;- | 496 | NA | 0 | 0 | 0 | 0 | 0 | 0 |
| ENSDARG000 | 8:8:8:8:8:8:8: | 20455134:204 | 20455465:204 | +:+:+:+:+:+:+:+: | 4124 | rexo1 | 327 | 223 | 387 | 167 | 295 | 342 |
| ENSDARG000 | 8:8:8:8:8:8:8: | 144154:14899 | 144252:14909 | +:+:+:+:+:+:+:+: | 1521 | SNX2 | 1607 | 1667 | 1556 | 1078 | 1586 | 1353 |
| ENSDARG000 | 8:8:8:8:8:8:8: | 3820134:3821 | 3820277:3821 | +:+:+:+:+:+:+:+: | 4749 | CIT | 4 | 2 | 0 | 0 | 0 | 0 |
| ENSDARG000 | 8;8;8;8;8;8 | 28257219:282 | 28257231:282 | +;+;+;+;+;+ | 162 | NA | 0 | 6 | 3 | 17 | 5 | 6 |
| ENSDARG000 | 8:8:8:8:8:8:8: | 34731982:347 | 34732027:347 | +:+:+:+:+:+:+:+: | 1621 | hpdb | 54 | 6 | 2 | 2 | 24 | 28 |
| ENSDARG000 | 8;8;8;8;8 | 15311289:153 | 15311382:153 | -;-;-;-;- | 689 | AGBL4 | 0 | 0 | 23 | 8 | 68 | 1 |
| ENSDARG000 | 8:8:8:8:8:8:8: | 36057318:360 | 36063010:360 | -:-:-:-:-:-:-:-:-: | 9484 | TMEM132D | 6 | 22 | 8 | 10 | 22 | 4 |
| ENSDARG000 | 8:8:8:8:8:8:8: | 34486656:344 | 34487041:344 | +:+:+:+:+:+:+:+: | 1894 | NA | 0 | 0 | 0 | 0 | 0 | 0 |
| ENSDARG000 | 8:8:8:8:8:8:8: | 22567363:225 | 22567597:225 | -:-:-:-:-:-:-:-:-: | 6622 | IQSEC2 | 16 | 62 | 39 | 18 | 45 | 54 |
| ENSDARG000 | 8:8:8:8:8:8:8: | 635704:63614 | 635893:63648 | +:+:+:+:+:+:+:+: | 1467 | NA | 4 | 4 | 0 | 0 | 6 | 1 |
| ENSDARG000 | 8:8:8:8:8:8:8: | 650342:65284 | 650399:65284 | +:+:+:+:+:+:+:+: | 895 | PLPP1 | 159 | 218 | 182 | 85 | 119 | 180 |
| ENSDARG000 | 8;8;8;8;8 | 537809:53990 | 537908:53998 | +;+;+;+;+ | 2008 | srp19 | 432 | 432 | 512 | 244 | 417 | 282 |
| ENSDARG000 | 8;8 | 53794022:538 | 53800714:538 | -;- | 7068 | BSN | 0 | 2 | 0 | 0 | 0 | 0 |
| ENSDARG000 | 8:8:8:8:8:8:8: | 54188978:541 | 54189281:541 | +:+:+:+:+:+:+:+: | 4844 | NA | 40 | 6 | 2 | 0 | 6 | 4 |
| ENSDARG000 | 8:8:8:8:8:8:8: | 52479026:524 | 52479175:524 | +:+:+:+:+:+:+:+: | 1395 | NA | 204 | 84 | 60 | 40 | 30 | 30 |
| ENSDARG000 | 8;8 | 47698098:476 | 47698269:476 | +;+ | 354 | NA | 0 | 0 | 0 | 0 | 0 | 0 |
| ENSDARG000 | 8;8 | 547317:55441 | 547471:55454 | -;- | 285 | NA | 0 | 0 | 0 | 0 | 0 | 1 |
| ENSDARG000 | 8;8;8 | 39663612:396 | 39664098:396 | +;+;+ | 1988 | TMEM233 | 0 | 0 | 0 | 0 | 0 | 0 |
| ENSDARG000 | 8;8 | 28547687:285 | 28547973:285 | +;+ | 497 | SRXN1 | 34 | 22 | 26 | 2 | 22 | 52 |
| ENSDARG000 | 8:8:8:8:8:8:8: | 50840565:508 | 50842255:508 | -:-:-:-:-:-:-:-:-: | 3981 | NA | 12 | 0 | 4 | 4 | 20 | 0 |
| ENSDARG000 | 8 | 19023261 | 19023377 | + | 117 | 5S_rRNA | 0 | 0 | 0 | 0 | 0 | 0 |
| ENSDARG000 | 8 | 438111 | 438226 | + | 116 | 5S_rRNA | 0 | 0 | 0 | 0 | 0 | 0 |
| ENSDARG000 | 8 | 18584693 | 18584807 | - | 115 | 5S_rRNA | 0 | 0 | 0 | 0 | 0 | 0 |
| ENSDARG000 | 8;8;8;8;8;8 | 28452738:284 | 28453642:284 | +;+;+;+;+;+ | 6005 | peds1 | 1317 | 1502 | 1288 | 752 | 1064 | 1423 |
| ENSDARG000 | 8:8:8:8:8:8:8: | 54081819:540 | 54081932:540 | +:+:+:+:+:+:+:+: | 4885 | prickle2a | 292 | 88 | 98 | 321 | 30 | 180 |
| ENSDARG000 | 8 | 42508390 | 42508506 | - | 117 | 5S_rRNA | 0 | 0 | 0 | 0 | 0 | 0 |
| ENSDARG000 | 8 | 4875198 | 4875317 | - | 120 | 5S_rRNA | 0 | 0 | 0 | 0 | 0 | 0 |
| ENSDARG000 | 8 | 54076195 | 54077740 | - | 1546 | rho | 18 | 0 | 4 | 8 | 4 | 0 |
| ENSDARG000 | 8 | 42134003 | 42134119 | - | 117 | 5S_rRNA | 0 | 0 | 0 | 0 | 0 | 0 |
| ENSDARG000 | 8 | 17252239 | 17252354 | - | 116 | 5S_rRNA | 0 | 0 | 0 | 0 | 0 | 0 |
| ENSDARG000 | 8 | 44659813 | 44659927 | + | 115 | 5S_rRNA | 0 | 0 | 0 | 0 | 0 | 0 |
| ENSDARG000 | 8 | 20414846 | 20414961 | - | 116 | 5S_rRNA | 0 | 0 | 0 | 0 | 0 | 0 |
| ENSDARG000 | 8:8:8:8:8:8:8: | 5233450:5236 | 5233729:5236 | -:-:-:-:-:-:-:-:-: | 1341 | NA | 408 | 315 | 369 | 189 | 539 | 252 |
| ENSDARG000 | 8:8:8:8:8:8:8: | 34572359:345 | 34572744:345 | +:+:+:+:+:+:+:+: | 2199 | NA | 0 | 0 | 0 | 0 | 0 | 0 |
| ENSDARG000 | 8 | 12612772 | 12612888 | - | 117 | 5S_rRNA | 0 | 0 | 0 | 0 | 0 | 0 |
| ENSDARG000 | 8;8;8 | 53730368:537 | 53733659:537 | -;-;- | 3996 | tlr9 | 4 | 7 | 1 | 2 | 1 | 25 |
| ENSDARG000 | 8:8:8:8:8:8:8: | 3530761:3537 | 3530768:3537 | +:+:+:+:+:+:+:+: | 7695 | GCN1 | 7293 | 4669 | 10350 | 5067 | 7726 | 5637 |
| ENSDARG000 | 8;8;8;8;8;8;8 | 18601252:186 | 18602344:186 | -;-;-;-;-;-;- | 2358 | cpox | 499 | 934 | 1244 | 654 | 1017 | 878 |
| ENSDARG000 | 8:8:8:8:8:8:8: | 40238647:402 | 40243230:402 | -:-:-:-:-:-:-:-:-: | 7548 | kdm2ba | 864 | 953 | 769 | 442 | 950 | 989 |
| ENSDARG000 | 8:8:8:8:8:8:8: | 34476532:344 | 34476814:344 | +:+:+:+:+:+:+:+: | 1530 | NA | 0 | 0 | 0 | 0 | 0 | 0 |
| ENSDARG000 | 8;8;8;8;8;8 | 36321336:363 | 36325691:363 | -;-;-;-;-;- | 5086 | NA | 0 | 380 | 0 | 0 | 0 | 886 |
| ENSDARG000 | 8;8;8;8 | 44362452:443 | 44362570:443 | -;-;-;- | 735 | mhc2dcb | 0 | 0 | 0 | 0 | 0 | 0 |
| ENSDARG000 | 8;8;8 | 53639625:536 | 53640603:536 | -;-;- | 1873 | lrtm1 | 263 | 100 | 153 | 64 | 457 | 108 |
| ENSDARG000 | 8 | 24064210 | 24064295 | + | 86 | NA | 0 | 0 | 0 | 0 | 0 | 0 |
| ENSDARG000 | 8 | 10216 | 10324 | + | 109 | dre-mir-7a-3 | 0 | 0 | 0 | 0 | 0 | 0 |
| ENSDARG000 | 8;8;8 | 1433647:1433 | 1433719:1433 | +;+;+ | 907 | efna5b | 29 | 36 | 528 | 124 | 346 | 205 |
| ENSDARG000 | 8;8;8;8 | 52701811:527 | 52702124:527 | +;+;+;+ | 5997 | bcl2l16 | 5 | 9 | 21 | 23 | 6 | 12 |
| ENSDARG000 | 8 | 40644838 | 40646418 | + | 1581 | adra2b | 3354 | 830 | 803 | 1888 | 3657 | 737 |
| ENSDARG000 | 8 | 6138368 | 6138484 | - | 117 | 5S_rRNA | 0 | 0 | 0 | 0 | 0 | 0 |

| | | | | | | | | | | | |
|---|---|---|---|---|---|---|---|---|---|---|---|
| ENSDARG000 | 8:8:8:8:8:8:8: | 51165392:511 | 51166332:511 | -:-:-:-:-:-:-:-:-: | 3800 | bmp1a | 4127 | 851 | 1365 | 1110 | 1351 | 547 |
| ENSDARG000 | 8;8;8;8 | 50520139:505 | 50520341:505 | -;-;-;- | 654 | NA | 0 | 0 | 8 | 0 | 0 | 0 |
| ENSDARG000 | 8 | 39806314 | 39806430 | + | 117 | 5S_rRNA | 0 | 0 | 0 | 0 | 0 | 0 |
| ENSDARG000 | 8;8;8 | 532228:53436 | 533035:53462 | -;-;- | 1175 | zgc:101664 | 72 | 28 | 56 | 18 | 49 | 62 |
| ENSDARG000 | 8 | 51027839 | 51027953 | - | 115 | 5S_rRNA | 16 | 2 | 4 | 10 | 23 | 16 |
| ENSDARG000 | 8;8;8;8;8;8 | 35032633:350 | 35032959:350 | +;+;+;+;+;+ | 1574 | NA | 0 | 0 | 0 | 0 | 0 | 0 |
| ENSDARG000 | 8;8;8;8 | 40628926:406 | 40629779:406 | +;+;+;+ | 3241 | dusp2 | 12779 | 3993 | 3469 | 7638 | 2969 | 5372 |
| ENSDARG000 | 8:8:8:8:8:8:8: | 50531709:505 | 50531889:505 | +:+:+:+:+:+:+: | 1646 | NA | 0 | 0 | 0 | 0 | 2 | 0 |
| ENSDARG000 | 8 | 37383101 | 37383231 | + | 131 | 5S_rRNA | 0 | 0 | 0 | 0 | 0 | 0 |
| ENSDARG000 | 8:8:8:8:8:8:8: | 24422393:244 | 24422490:244 | -:-:-:-:-:-:-:-:-: | 1380 | NA | 0 | 0 | 0 | 0 | 0 | 1 |
| ENSDARG000 | 8 | 13238973 | 13239092 | - | 120 | 5S_rRNA | 0 | 0 | 0 | 0 | 0 | 0 |
| ENSDARG000 | 8 | 26082308 | 26082409 | + | 102 | dre-mir-462 | 0 | 0 | 0 | 0 | 0 | 0 |
| ENSDARG000 | 8;8 | 54284961:542 | 54285843:542 | +;+ | 1194 | NA | 810 | 433 | 780 | 721 | 890 | 504 |
| ENSDARG000 | 8 | 6199315 | 6199429 | + | 115 | 5S_rRNA | 0 | 0 | 0 | 0 | 0 | 0 |
| ENSDARG000 | 8;8 | 54302700:543 | 54303701:543 | -;- | 1242 | exorh | 6 | 12 | 8 | 22 | 24 | 48 |
| ENSDARG000 | 8:8:8:8:8:8:8: | 54263946:542 | 54264400:542 | +:+:+:+:+:+:+: | 1847 | TMCC1 | 78 | 19 | 18 | 49 | 6 | 28 |
| ENSDARG000 | 8:8:8:8:8:8:8: | 34562983:345 | 34563368:345 | +:+:+:+:+:+:+: | 2187 | NA | 0 | 0 | 0 | 0 | 0 | 0 |
| ENSDARG000 | 8:8:8:8:8:8:8: | 54013199:540 | 54013660:540 | +:+:+:+:+:+:+: | 1854 | NA | 432 | 192 | 188 | 161 | 250 | 280 |
| ENSDARG000 | 8;8 | 34624025:346 | 34624845:346 | +;+ | 2929 | NA | 0 | 0 | 0 | 0 | 0 | 0 |
| ENSDARG000 | 8;8;8;8;8;8;8 | 4963390:4964 | 4963617:4964 | -;-;-;-;-;-;- | 1020 | tmem230b | 298 | 398 | 398 | 288 | 322 | 368 |
| ENSDARG000 | 8 | 50517782 | 50517903 | - | 122 | 5S_rRNA | 2 | 0 | 0 | 0 | 0 | 0 |
| ENSDARG000 | 8 | 53356779 | 53356896 | - | 118 | 5S_rRNA | 0 | 0 | 0 | 0 | 0 | 0 |
| ENSDARG000 | 8;8;8;8;8 | 2738518:2744 | 2738928:2745 | -;-;-;-;- | 1299 | NA | 5 | 11 | 16 | 4 | 8 | 12 |
| ENSDARG000 | 8 | 26142533 | 26142647 | + | 115 | 5S_rRNA | 0 | 0 | 0 | 0 | 0 | 0 |
| ENSDARG000 | 8:8:8:8:8:8:8: | 34439052:344 | 34439437:344 | +:+:+:+:+:+:+: | 1635 | NA | 0 | 0 | 0 | 0 | 0 | 0 |
| ENSDARG000 | 8 | 30176805 | 30176922 | + | 118 | 5S_rRNA | 0 | 0 | 0 | 0 | 0 | 0 |
| ENSDARG000 | 8:8:8:8:8:8:8: | 22582146:225 | 22582205:225 | +:+:+:+:+:+:+: | 2697 | PDZD4 | 1 | 0 | 0 | 0 | 4 | 0 |
| ENSDARG000 | 8:8:8:8:8:8:8: | 426828:42705 | 426973:42716 | -:-:-:-:-:-:-:-:-: | 1158 | NA | 0 | 0 | 0 | 0 | 2 | 0 |
| ENSDARG000 | 8:8:8:8:8:8:8: | 5844174:5844 | 5844367:5844 | -;-;-;-;-;-;-;- | 1790 | LRRTM4 | 0 | 0 | 0 | 0 | 2 | 0 |
| ENSDARG000 | 8;8;8;8 | 36283548:362 | 36283842:362 | -;-;-;- | 928 | mhc2dga | 0 | 55 | 0 | 0 | 0 | 67 |
| ENSDARG000 | 8:8:8:8:8:8:8: | 19489854:194 | 19490251:194 | +:+:+:+:+:+:+: | 2285 | npl | 419 | 219 | 267 | 316 | 424 | 410 |
| ENSDARG000 | 8:8:8:8:8:8:8: | 247163:24805 | 247323:24817 | +:+:+:+:+:+:+: | 2790 | CEP120 | 178 | 370 | 414 | 210 | 456 | 200 |
| ENSDARG000 | 8 | 15916210 | 15916327 | - | 118 | 5S_rRNA | 0 | 0 | 0 | 0 | 0 | 0 |
| ENSDARG000 | 8 | 28448267 | 28449782 | - | 1516 | cebpb | 2628 | 1802 | 1268 | 2599 | 2891 | 2406 |
| ENSDARG000 | 8:8:8:8:8:8:8: | 44828432:448 | 44828547:448 | -;-;-;-;-;-;-;- | 837 | NA | 0 | 0 | 0 | 0 | 0 | 0 |
| ENSDARG000 | 8 | 16611666 | 16611747 | - | 82 | SNORD79 | 0 | 0 | 0 | 0 | 0 | 0 |
| ENSDARG000 | 8 | 23342883 | 23342998 | + | 116 | 5S_rRNA | 0 | 0 | 0 | 0 | 0 | 0 |
| ENSDARG000 | 8;8;8;8;8;8;8 | 28467893:284 | 28468016:284 | +;+;+;+;+;+;+ | 3670 | slc52a3 | 0 | 2 | 2 | 8 | 0 | 2 |
| ENSDARG000 | 8:8:8:8:8:8:8: | 53747925:537 | 53748030:537 | -:-:-:-:-:-:-:-:-: | 2238 | NA | 636 | 503 | 612 | 422 | 772 | 631 |
| ENSDARG000 | 8;8;8;8;8;8;8 | 24500116:245 | 24500213:245 | -;-;-;-;-;-;- | 1380 | NA | 0 | 0 | 0 | 0 | 0 | 1 |
| ENSDARG000 | 8:8:8:8:8:8:8: | 4621243:4630 | 4622117:4630 | -;-;-;-;-;-;-;- | 1669 | zgc:63587 | 1773 | 2090 | 1950 | 1788 | 2277 | 2063 |
| ENSDARG000 | 8;8;8;8 | 36128477:361 | 36132116:361 | -;-;-;- | 4532 | NA | 0 | 0 | 0 | 8 | 0 | 0 |
| ENSDARG000 | 8 | 48175308 | 48175423 | + | 116 | 5S_rRNA | 0 | 0 | 0 | 0 | 0 | 0 |
| ENSDARG000 | 8 | 9111072 | 9111190 | + | 119 | 5S_rRNA | 0 | 0 | 0 | 0 | 0 | 0 |
| ENSDARG000 | 8;8;8;8;8 | 28345352:283 | 28345988:283 | -;-;-;-;- | 1803 | tuba5 | 6589 | 12399 | 11163 | 7612 | 10770 | 6620 |
| ENSDARG000 | 8:8:8:8:8:8:8: | 52415603:524 | 52415847:524 | +:+:+:+:+:+:+: | 1480 | gins4 | 52 | 194 | 144 | 28 | 121 | 92 |
| ENSDARG000 | 8 | 16611935 | 16612017 | - | 83 | SNORD79 | 0 | 0 | 3 | 0 | 1 | 0 |
| ENSDARG000 | 8:8:8:8:8:8:8: | 34443759:344 | 34444144:344 | +:+:+:+:+:+:+: | 1635 | NA | 0 | 0 | 0 | 0 | 0 | 0 |
| ENSDARG000 | 8;8;8;8;8;8 | 18615938:186 | 18616102:186 | +;+;+;+;+;+ | 1744 | NA | 2 | 0 | 0 | 0 | 0 | 0 |
| ENSDARG000 | 8 | 8591671 | 8591788 | - | 118 | 5S_rRNA | 0 | 0 | 0 | 0 | 0 | 0 |
| ENSDARG000 | 8;8;8 | 24799688:248 | 24800056:248 | -;-;- | 1281 | NA | 12 | 12 | 16 | 55 | 4 | 39 |
| ENSDARG000 | 8;8;8;8;8;8 | 5202589:5204 | 5202798:5204 | -;-;-;-;-;- | 871 | bnip3la | 362 | 270 | 200 | 173 | 471 | 248 |
| ENSDARG000 | 8 | 35376243 | 35376357 | - | 115 | 5S_rRNA | 0 | 0 | 0 | 0 | 0 | 0 |
| ENSDARG000 | 8 | 23122762 | 23122874 | - | 113 | dre-mir-124-6 | 0 | 2 | 0 | 0 | 0 | 0 |
| ENSDARG000 | 8:8:8:8:8:8:8: | 31406731:314 | 31407335:314 | -:-:-:-:-:-:-:-:-: | 2272 | znf131 | 246 | 309 | 346 | 278 | 296 | 320 |
| ENSDARG000 | 8:8:8:8:8:8:8: | 36268747:362 | 36268755:362 | +:+:+:+:+:+:+: | 4142 | mhc2dgb | 0 | 85 | 0 | 0 | 0 | 86 |
| ENSDARG000 | 8:8:8:8:8:8:8: | 595122:59963 | 595180:59971 | -:-:-:-:-:-:-:-:-: | 1775 | NA | 335 | 244 | 263 | 183 | 420 | 315 |
| ENSDARG000 | 8:8:8:8:8:8:8: | 471342:47375 | 471394:47395 | +:+:+:+:+:+:+: | 1617 | nudt12 | 256 | 234 | 294 | 217 | 194 | 204 |
| ENSDARG000 | 8;8 | 26372115:263 | 26372251:263 | +;+ | 1067 | NAA80 | 69 | 87 | 38 | 42 | 41 | 46 |
| ENSDARG000 | 8;8;8 | 1697194:1697 | 1697287:1698 | -;-;- | 288 | NA | 13 | 6 | 8 | 3 | 18 | 3 |
| ENSDARG000 | 8:8:8:8:8:8:8: | 34932397:349 | 34932770:349 | +:+:+:+:+:+:+: | 3632 | NA | 6 | 2 | 0 | 0 | 4 | 0 |
| ENSDARG000 | 8:8:8:8:8:8:8: | 16742091:167 | 16742150:167 | -:-:-:-:-:-:-:-:-: | 252 | NA | 2 | 0 | 4 | 0 | 3 | 1 |
| ENSDARG000 | 8:8:8:8:8:8:8: | 34448496:344 | 34448855:344 | +:+:+:+:+:+:+: | 1609 | NA | 0 | 0 | 0 | 0 | 0 | 0 |
| ENSDARG000 | 8;8;8;8;8;8 | 39708674:397 | 39708693:397 | -;-;-;-;-;- | 132 | NA | 0 | 1 | 0 | 0 | 0 | 4 |
| ENSDARG000 | 8:8:8:8:8:8:8: | 25988617:259 | 25988626:259 | -:-:-:-:-:-:-:-:-: | 294 | NA | 5 | 0 | 0 | 0 | 1 | 0 |
| ENSDARG000 | 8;8;8 | 68864:75048: | 69125:75686: | +;+;+ | 1398 | prr16 | 0 | 6 | 20 | 6 | 6 | 8 |
| ENSDARG000 | 8 | 11860028 | 11860144 | + | 117 | 5S_rRNA | 0 | 0 | 0 | 0 | 0 | 0 |
| ENSDARG000 | 8 | 6194470 | 6194590 | + | 121 | 5S_rRNA | 0 | 0 | 0 | 0 | 0 | 0 |
| ENSDARG000 | 8 | 24979920 | 24980036 | + | 117 | 5S_rRNA | 0 | 0 | 0 | 0 | 2 | 0 |
| ENSDARG000 | 8:8:8:8:8:8:8: | 46758264:467 | 46758434:467 | -;-;-;-;-;-;-;- | 1057 | acot7 | 242 | 147 | 254 | 74 | 215 | 130 |
| ENSDARG000 | 8 | 14806174 | 14806290 | + | 117 | 5S_rRNA | 0 | 0 | 0 | 0 | 0 | 0 |
| ENSDARG000 | 8 | 16613029 | 16613109 | - | 81 | NA | 0 | 0 | 0 | 0 | 0 | 0 |
| ENSDARG000 | 8:8:8:8:8:8:8: | 15633157:156 | 15633751:156 | -:-:-:-:-:-:-:-:-: | 1325 | agbl4 | 0 | 0 | 0 | 0 | 0 | 0 |
| ENSDARG000 | 8;8;8;8;8;8 | 53452681:534 | 53452755:534 | +;+;+;+;+;+ | 624 | NA | 0 | 0 | 0 | 0 | 0 | 0 |
| ENSDARG000 | 8:8:8:8:8:8:8: | 12925385:129 | 12925586:129 | +:+:+:+:+:+:+: | 3733 | kif2a | 912 | 1175 | 1507 | 1221 | 1078 | 1098 |
| ENSDARG000 | 8;8 | 38963323:389 | 38964660:389 | -;- | 1920 | NA | 4 | 0 | 0 | 0 | 0 | 4 |
| ENSDARG000 | 8:8:8:8:8:8:8: | 34540801:345 | 34541186:345 | +:+:+:+:+:+:+: | 1696 | NA | 0 | 0 | 0 | 0 | 0 | 0 |
| ENSDARG000 | 8;8 | 31248917:312 | 31252137:312 | +;+ | 4167 | AJM1 | 262 | 377 | 817 | 716 | 1637 | 507 |
| ENSDARG000 | 8:8:8:8:8:8:8: | 53862745:538 | 53862986:538 | -:-:-:-:-:-:-:-:-: | 2670 | mib2 | 509 | 334 | 811 | 670 | 578 | 458 |
| ENSDARG000 | 8 | 1783164 | 1783280 | - | 117 | 5S_rRNA | 0 | 0 | 0 | 0 | 3 | 0 |
| ENSDARG000 | 8 | 32188146 | 32188273 | + | 128 | 5S_rRNA | 0 | 0 | 0 | 0 | 0 | 0 |
| ENSDARG000 | 8 | 26005588 | 26005669 | + | 82 | dre-mir-135b | 0 | 0 | 0 | 0 | 0 | 0 |
| ENSDARG000 | 8:8:8:8:8:8:8: | 34531954:345 | 34532339:345 | +:+:+:+:+:+:+: | 1635 | NA | 0 | 0 | 0 | 0 | 0 | 0 |
| ENSDARG000 | 8;8;8 | 54220551:542 | 54222078:542 | -;-;- | 1768 | trh | 8 | 8 | 4 | 0 | 8 | 0 |
| ENSDARG000 | 8;8;8;8;8;8 | 28358161:283 | 28358280:283 | +;+;+;+;+;+ | 1596 | adipor1b | 293 | 324 | 335 | 240 | 351 | 422 |
| ENSDARG000 | 8;8;8 | 53918234:539 | 53919714:539 | -;-;- | 2232 | ubiad1 | 549 | 498 | 704 | 443 | 750 | 534 |
| ENSDARG000 | 8 | 8321482 | 8321596 | + | 115 | 5S_rRNA | 0 | 0 | 0 | 0 | 2 | 0 |
| ENSDARG000 | 8 | 26082534 | 26082623 | + | 90 | dre-mir-731 | 0 | 0 | 0 | 0 | 0 | 0 |
| ENSDARG000 | 8 | 16611114 | 16611194 | - | 81 | NA | 0 | 0 | 0 | 0 | 0 | 0 |
| ENSDARG000 | 8 | 16611327 | 16611411 | - | 85 | NA | 0 | 0 | 0 | 0 | 0 | 0 |
| ENSDARG000 | 8;8;8;8 | 53544094:535 | 53544253:535 | -;-;-;- | 1077 | parapinopsina | 0 | 0 | 0 | 0 | 0 | 0 |
| ENSDARG000 | 8:8:8:8:8:8:8: | 53423408:534 | 53423464:534 | +:+:+:+:+:+:+: | 1242 | NA | 0 | 2 | 0 | 0 | 0 | 0 |
| ENSDARG000 | 8 | 16611328 | 16611408 | - | 81 | NA | 0 | 0 | 0 | 0 | 0 | 0 |
| ENSDARG000 | 8;8;8;8;8 | 28350921:283 | 28351809:283 | -;-;-;-;- | 1530 | KLHL12 | 119 | 78 | 58 | 94 | 149 | 114 |
| ENSDARG000 | 8:8:8:8:8:8:8: | 54202554:542 | 54202670:542 | +:+:+:+:+:+:+: | 1221 | CRELD1 | 429 | 428 | 277 | 116 | 473 | 335 |
| ENSDARG000 | 8:8:8:8:8:8:8: | 34749288:347 | 34749531:347 | -:-:-:-:-:-:-:-:-: | 4745 | NA | 0 | 0 | 0 | 0 | 0 | 0 |
| ENSDARG000 | 8;8;8;8;8 | 622640:62567 | 622788:62572 | +;+;+;+;+ | 1256 | spinb | 117 | 160 | 155 | 82 | 89 | 128 |
| ENSDARG000 | 8:8:8:8:8:8:8: | 53388005:533 | 53388194:533 | +:+:+:+:+:+:+: | 5086 | dcp1a | 175 | 159 | 315 | 81 | 255 | 172 |
| ENSDARG000 | 8;8 | 19642199:196 | 19642828:196 | -;- | 893 | fam78bb | 0 | 0 | 5 | 0 | 0 | 0 |
| ENSDARG000 | 8 | 7158483 | 7158597 | - | 115 | 5S_rRNA | 0 | 0 | 0 | 0 | 0 | 0 |
| ENSDARG000 | 8;8;8;8;8;8;8 | 34567717:345 | 34568048:345 | +;+;+;+;+;+;+ | 1003 | NA | 0 | 0 | 0 | 0 | 0 | 0 |
| ENSDARG000 | 8 | 4705665 | 4705828 | + | 164 | U1 | 0 | 0 | 0 | 0 | 0 | 0 |
| ENSDARG000 | 8 | 48390562 | 48390678 | + | 117 | 5S_rRNA | 0 | 0 | 0 | 0 | 0 | 0 |
| ENSDARG000 | 8:8:8:8:8:8:8: | 34922981:349 | 34923340:349 | +:+:+:+:+:+:+: | 1609 | NA | 0 | 0 | 0 | 0 | 0 | 0 |
| ENSDARG000 | 8 | 42352201 | 42352317 | - | 117 | 5S_rRNA | 0 | 0 | 0 | 0 | 0 | 0 |
| ENSDARG000 | 8 | 35172594 | 35173496 | + | 903 | NA | 4 | 1 | 0 | 0 | 4 | 4 |
| ENSDARG000 | 8;8;8 | 40177361:401 | 40178747:401 | -;-;- | 1971 | gpx8 | 98 | 70 | 124 | 90 | 94 | 68 |
| ENSDARG000 | 8:8:8:8:8:8:8: | 52314542:523 | 52314778:523 | +:+:+:+:+:+:+: | 6431 | dbnlb | 1811 | 1596 | 2938 | 2953 | 1957 | 1625 |
| ENSDARG000 | 8:8:8:8:8:8:8: | 53563949:535 | 53563999:535 | +:+:+:+:+:+:+: | 3472 | CACNA2D3 | 4 | 8 | 9 | 0 | 14 | 4 |
| ENSDARG000 | 8:8:8:8:8:8:8: | 38649989:386 | 38650465:386 | -:-:-:-:-:-:-:-:-: | 2473 | pcsk5b | 86 | 70 | 97 | 66 | 121 | 107 |
| ENSDARG000 | 8:8:8:8:8:8:8: | 10920205:109 | 10920219:109 | +:+:+:+:+:+:+: | 264 | NA | 2 | 1 | 0 | 0 | 4 | 1 |
| ENSDARG000 | 8:8:8:8:8:8:8: | 44358443:443 | 44358680:443 | +:+:+:+:+:+:+: | 429 | NA | 0 | 0 | 0 | 0 | 0 | 0 |
| ENSDARG000 | 8:8:8:8:8:8:8: | 6410933:6411 | 6411027:6411 | +:+:+:+:+:+:+: | 1823 | LRRTM4 | 0 | 0 | 0 | 0 | 0 | 0 |
| ENSDARG000 | 8:8:8:8:8:8:8: | 53979248:539 | 53979488:539 | -;-;-;-;-;-;-;- | 1065 | SLC35E2B | 419 | 278 | 283 | 246 | 260 | 267 |
| ENSDARG000 | 8;8;8;8;8 | 52442622:524 | 52442688:524 | +;+;+;+;+ | 2619 | ppp1r3c2b | 88 | 72 | 40 | 40 | 33 | 132 |
| ENSDARG000 | 8:8:8:8:8:8:8: | 53269657:533 | 53269956:533 | +:+:+:+:+:+:+: | 6252 | gnb1a | 8873 | 9660 | 10079 | 12225 | 8296 | 8774 |
| ENSDARG000 | 8;8;8;8;8;8 | 28398780:284 | 28401061:284 | -;-;-;-;-;- | 3064 | ptpn1 | 330 | 402 | 365 | 350 | 409 | 476 |
| ENSDARG000 | 8 | 24788499 | 24791060 | - | 2562 | RBM15 | 660 | 330 | 619 | 322 | 702 | 625 |
| ENSDARG000 | 8:8:8:8:8:8:8: | 40573137:405 | 40573403:405 | +:+:+:+:+:+:+: | 1233 | izumo1 | 0 | 2 | 0 | 0 | 0 | 0 |
| ENSDARG000 | 8 | 47060401 | 47060515 | + | 115 | 5S_rRNA | 0 | 0 | 0 | 0 | 0 | 0 |
| ENSDARG000 | 8:8:8:8:8:8:8: | 45741044:457 | 45741064:457 | +:+:+:+:+:+:+: | 209 | NA | 0 | 2 | 0 | 0 | 0 | 0 |
| ENSDARG000 | 8;8;8;8;8;8;8 | 52377516:523 | 52377602:523 | +;+;+;+;+;+;+ | 2115 | arid5a | 32 | 52 | 40 | 41 | 163 | 135 |
| ENSDARG000 | 8;8;8;8;8;8 | 34766211:347 | 34766310:347 | -;-;-;-;-;- | 507 | NA | 0 | 0 | 0 | 2 | 2 | 0 |
| ENSDARG000 | 8 | 6778332 | 6778447 | + | 116 | 5S_rRNA | 0 | 0 | 0 | 0 | 0 | 0 |
| ENSDARG000 | 8 | 30167630 | 30167704 | - | 75 | NA | 0 | 0 | 0 | 0 | 0 | 0 |
| ENSDARG000 | 8 | 22789531 | 22789649 | + | 119 | 5S_rRNA | 0 | 0 | 0 | 0 | 0 | 0 |
| ENSDARG000 | 8 | 2868953 | 2869069 | + | 117 | 5S_rRNA | 0 | 0 | 0 | 0 | 0 | 0 |
| ENSDARG000 | 8 | 10052019 | 10052133 | - | 115 | 5S_rRNA | 0 | 0 | 0 | 0 | 0 | 0 |
| ENSDARG000 | 8:8:8:8:8:8:8: | 40210398:402 | 40210635:402 | +:+:+:+:+:+:+: | 5094 | rnf34a | 293 | 201 | 269 | 162 | 414 | 156 |
| ENSDARG000 | 8 | 5067728 | 5067849 | - | 122 | 5S_rRNA | 0 | 0 | 0 | 0 | 0 | 0 |
| ENSDARG000 | 8;8;8;8 | 32501405:325 | 32501686:325 | -;-;-;- | 1010 | RFESD | 24 | 71 | 39 | 28 | 26 | 92 |
| ENSDARG000 | 8 | 46766429 | 46766545 | + | 117 | 5S_rRNA | 0 | 0 | 0 | 0 | 0 | 0 |
| ENSDARG000 | 8 | 1767905 | 1768021 | + | 117 | 5S_rRNA | 0 | 0 | 0 | 0 | 0 | 0 |
| ENSDARG000 | 8:8:8:8:8:8:8: | 40185600:401 | 40186835:401 | -:-:-:-:-:-:-:-:-: | 4174 | anapc5 | 232 | 321 | 411 | 266 | 372 | 258 |
| ENSDARG000 | 8 | 37126124 | 37126240 | + | 117 | 5S_rRNA | 0 | 0 | 0 | 0 | 0 | 0 |
| ENSDARG000 | 8 | 41335070 | 41335184 | + | 115 | 5S_rRNA | 0 | 0 | 0 | 0 | 0 | 0 |
| ENSDARG000 | 8 | 53023679 | 53023764 | + | 86 | dre-mir-181b- | 0 | 0 | 0 | 0 | 0 | 0 |
| ENSDARG000 | 8 | 11548419 | 11548519 | - | 101 | dre-mir-126a | 0 | 0 | 0 | 0 | 0 | 0 |
| ENSDARG000 | 8:8:8:8:8:8:8: | 24110045:241 | 24110080:241 | +:+:+:+:+:+:+: | 357 | NA | 0 | 0 | 0 | 1 | 0 | 0 |
| ENSDARG000 | 8:8:8:8:8:8:8: | 34434345:344 | 34434730:344 | +:+:+:+:+:+:+: | 1635 | NA | 0 | 0 | 0 | 2 | 0 | 0 |
| ENSDARG000 | 8:8:8:8:8:8:8: | 54211861:542 | 54212095:542 | -:-:-:-:-:-:-:-:-: | 1505 | NA | 180 | 120 | 264 | 267 | 184 | 235 |
| ENSDARG000 | 8 | 15632732 | 15632847 | - | 116 | 5S_rRNA | 0 | 0 | 0 | 0 | 0 | 0 |
| ENSDARG000 | 8 | 16610248 | 16610319 | - | 72 | NA | 0 | 0 | 0 | 0 | 0 | 0 |
| ENSDARG000 | 8:8:8:8:8:8:8: | 45708567:457 | 45708725:457 | -:-:-:-:-:-:-:-:-: | 1933 | SLC18A1 | 4 | 0 | 5 | 6 | 0 | 0 |
| ENSDARG000 | 8 | 13877728 | 13877845 | - | 118 | 5S_rRNA | 0 | 0 | 0 | 0 | 0 | 0 |
| ENSDARG000 | 8 | 32931643 | 32931759 | - | 117 | 5S_rRNA | 0 | 0 | 0 | 0 | 0 | 0 |
| ENSDARG000 | 8:8:8:8:8:8:8: | 34527239:345 | 34527624:345 | +:+:+:+:+:+:+: | 2040 | NA | 0 | 0 | 0 | 0 | 0 | 0 |
| ENSDARG000 | 8 | 16612808 | 16612881 | - | 74 | NA | 0 | 0 | 0 | 0 | 0 | 0 |
| ENSDARG000 | 8 | 30999914 | 31000028 | - | 115 | 5S_rRNA | 0 | 0 | 0 | 0 | 0 | 0 |
| ENSDARG000 | 8:8:8:8:8:8:8: | 34988481:349 | 34988826:349 | +:+:+:+:+:+:+: | 1593 | NA | 0 | 0 | 0 | 0 | 0 | 0 |
| ENSDARG000 | 8;8;8;8;8;8 | 46656170:466 | 46656267:466 | -;-;-;-;-;- | 1866 | GPR153 | 56 | 30 | 23 | 12 | 10 | 21 |
| ENSDARG000 | 8:8:8:8:8:8:8: | 2017428:2019 | 2017763:2019 | -:-:-:-:-:-:-:-:-: | 4917 | NA | 48 | 22 | 77 | 82 | 38 | 69 |
| ENSDARG000 | 8;8;8 | 554531:57203 | 554555:57204 | +;+;+ | 117 | NA | 0 | 0 | 0 | 0 | 0 | 0 |
| ENSDARG000 | 8 | 1979823 | 1979937 | + | 115 | 5S_rRNA | 0 | 0 | 0 | 0 | 0 | 0 |
| ENSDARG000 | 8 | 23781005 | 23781394 | + | 390 | NA | 0 | 0 | 0 | 0 | 0 | 0 |
| ENSDARG000 | 8;8;8 | 1725968:1726 | 1726079:1726 | -;-;- | 420 | NA | 32 | 12 | 29 | 2 | 44 | 10 |
| ENSDARG000 | 8 | 30167435 | 30167525 | - | 91 | dre-mir-27b | 0 | 0 | 0 | 0 | 0 | 0 |
| ENSDARG000 | 8:8:8:8:8:8:8: | 36142734:361 | 36142848:361 | +:+:+:+:+:+:+: | 1044 | mhc2d8.35b2 | 0 | 0 | 0 | 0 | 0 | 0 |
| ENSDARG000 | 8;8;8;8;8 | 35735234:357 | 35735266:357 | -;-;-;-;- | 795 | NA | 0 | 0 | 0 | 0 | 0 | 0 |
| ENSDARG000 | 8 | 6266442 | 6266559 | - | 118 | 5S_rRNA | 0 | 0 | 0 | 0 | 0 | 0 |
| ENSDARG000 | 8:8:8:8:8:8:8: | 18624658:186 | 18624733:186 | +:+:+:+:+:+:+: | 2124 | fsd1 | 7 | 2 | 6 | 10 | 4 | 18 |
| ENSDARG000 | 8;8;8;8;8;8 | 28371496:283 | 28371734:283 | +;+;+;+;+;+ | 1156 | klhl12 | 61 | 25 | 66 | 26 | 67 | 70 |

| | | | | | | | | | | | | |
|---|---|---|---|---|---|---|---|---|---|---|---|---|
| ENSDARG000 | 8;8;8 | 5024468:5049 | 5024800:5051 | +;+;+ | 2781 | adra1aa | 0 | 0 | 0 | 0 | 0 | 0 |
| ENSDARG000 | 8;8 | 28527776:285 | 28528319:285 | +;+ | 2661 | scrt2 | 0 | 4 | 2 | 0 | 0 | 0 |
| ENSDARG000 | 8;8;8;8;8 | 41634225:416 | 41634241:416 | -;-;-;-;- | 126 | NA | 0 | 0 | 0 | 0 | 0 | 0 |
| ENSDARG000 | 8:8:8:8:8:8:8: | 1239:1239:13 | 1323:1323:14 | +:+:+:+:+:+:+: | 4062 | tmed7 | 2564 | 2096 | 2388 | 2211 | 2849 | 1736 |
| ENSDARG000 | 8 | 10439725 | 10439844 | - | 120 | 5S_rRNA | 0 | 0 | 0 | 0 | 0 | 0 |
| ENSDARG000 | 8 | 39864840 | 39864958 | - | 119 | 5S_rRNA | 0 | 0 | 0 | 0 | 2 | 0 |
| ENSDARG000 | 8;8;8 | 37998387:380 | 37998991:380 | -;-;- | 1466 | NA | 16 | 12 | 6 | 0 | 12 | 0 |
| ENSDARG000 | 8;8 | 34998570:349 | 34998852:350 | +;+ | 1530 | NA | 0 | 0 | 0 | 0 | 0 | 0 |
| ENSDARG000 | 8 | 40932722 | 40932838 | + | 117 | 5S_rRNA | 0 | 0 | 0 | 0 | 0 | 0 |
| ENSDARG000 | 8 | 28512492 | 28512607 | + | 116 | 5S_rRNA | 0 | 0 | 0 | 0 | 0 | 0 |
| ENSDARG000 | 8 | 10843952 | 10844068 | + | 117 | 5S_rRNA | 2 | 0 | 2 | 0 | 2 | 7 |
| ENSDARG000 | 8 | 31221039 | 31221157 | + | 119 | 5S_rRNA | 0 | 0 | 0 | 0 | 0 | 0 |
| ENSDARG000 | 8 | 1024042 | 1024158 | - | 117 | 5S_rRNA | 0 | 0 | 0 | 0 | 0 | 0 |
| ENSDARG000 | 8:8:8:8:8:8:8: | 34720244:347 | 34720603:347 | +:+:+:+:+:+:+: | 1609 | NA | 0 | 0 | 0 | 0 | 0 | 0 |
| ENSDARG000 | 8:8:8:8:8:8:8: | 5119360:5122 | 5119451:5122 | -:-:-:-:-:-:-:-: | 1728 | NA | 0 | 19 | 33 | 2 | 8 | 36 |
| ENSDARG000 | 8:8:8:8:8:8:8: | 31349522:313 | 31349532:313 | +:+:+:+:+:+:+: | 183 | NA | 0 | 0 | 6 | 1 | 1 | 0 |
| ENSDARG000 | 8;8;8;8;8 | 53670395:536 | 53671110:536 | -;-;-;-;- | 1856 | wnt5a | 558 | 20 | 90 | 108 | 172 | 56 |
| ENSDARG000 | 8;8;8;8 | 52408583:524 | 52408695:524 | -;-;-;- | 384 | COX5B | 76 | 137 | 88 | 88 | 94 | 90 |
| ENSDARG000 | 8:8:8:8:8:8:8: | 34927665:349 | 34928050:349 | +:+:+:+:+:+:+: | 1635 | NA | 0 | 0 | 0 | 0 | 0 | 0 |
| ENSDARG000 | 8:8:8:8:8:8:8: | 40532304:405 | 40533057:405 | -:-:-:-:-:-:-:-: | 4066 | NPC1L1 | 2 | 2 | 0 | 0 | 0 | 1 |
| ENSDARG000 | 8:8:8:8:8:8:8: | 19398335:194 | 19402523:194 | -:-:-:-:-:-:-:-: | 5095 | zgc:92140 | 1088 | 919 | 973 | 1075 | 898 | 1223 |
| ENSDARG000 | 8:8:8:8:8:8:8: | 53928472:539 | 53928561:539 | +:+:+:+:+:+:+: | 1116 | MMP23B | 8 | 72 | 8 | 34 | 46 | 60 |
| ENSDARG000 | 8:8:8:8:8:8:8: | 3496204:3498 | 3496211:3498 | +:+:+:+:+:+:+: | 1578 | PXN | 341 | 100 | 286 | 200 | 263 | 168 |
| ENSDARG000 | 8:8:8:8:8:8:8: | 53948194:539 | 53948830:539 | -:-:-:-:-:-:-:-: | 3007 | cdk11b | 1134 | 999 | 1518 | 577 | 1407 | 714 |
| ENSDARG000 | 8 | 14415698 | 14415823 | + | 126 | 5S_rRNA | 0 | 0 | 0 | 0 | 0 | 0 |
| ENSDARG000 | 8:8:8:8:8:8:8: | 18650414:186 | 18650941:186 | -:-:-:-:-:-:-:-: | 1861 | stap2b | 8 | 0 | 0 | 6 | 0 | 10 |
| ENSDARG000 | 8;8 | 36339470:363 | 36339814:363 | +;+ | 2771 | NA | 0 | 0 | 0 | 0 | 0 | 0 |
| ENSDARG000 | 8;8;8 | 387622:38862 | 388080:38871 | +;+;+ | 1267 | pym1 | 189 | 225 | 277 | 198 | 206 | 100 |
| ENSDARG000 | 8:8:8:8:8:8:8: | 53344726:533 | 53345185:533 | +:+:+:+:+:+:+: | 2569 | BICD2 | 422 | 397 | 589 | 489 | 571 | 456 |
| ENSDARG000 | 8:8:8:8:8:8:8: | 34629184:346 | 34629557:346 | +:+:+:+:+:+:+: | 1623 | NA | 0 | 0 | 0 | 0 | 0 | 0 |
| ENSDARG000 | 8:8:8:8:8:8:8: | 2878756:2890 | 2879000:2891 | +:+:+:+:+:+:+: | 3375 | NA | 2 | 8 | 0 | 0 | 0 | 0 |
| ENSDARG000 | 8:8:8:8:8:8:8: | 41688:41882: | 41777:42003: | -:-:-:-:-:-:-:-: | 2178 | hsd17b4 | 901 | 756 | 835 | 440 | 1060 | 758 |
| ENSDARG000 | 8;8;8;8;8;8 | 43852743:438 | 43852760:438 | +;+;+;+;+;+ | 810 | NA | 0 | 0 | 0 | 0 | 1 | 0 |
| ENSDARG000 | 8:8:8:8:8:8:8: | 24352719:243 | 24352816:243 | -:-:-:-:-:-:-:-: | 1652 | NA | 0 | 0 | 0 | 0 | 0 | 0 |
| ENSDARG000 | 8:8:8:8:8:8:8: | 34685407:346 | 34685792:346 | +:+:+:+:+:+:+: | 1696 | NA | 0 | 0 | 0 | 0 | 0 | 0 |
| ENSDARG000 | 8:8:8:8:8:8:8: | 54135642:541 | 54135757:541 | +:+:+:+:+:+:+: | 3906 | BRPF1 | 327 | 249 | 307 | 158 | 317 | 178 |
| ENSDARG000 | 8 | 8520042 | 8520173 | - | 132 | 5S_rRNA | 0 | 0 | 0 | 0 | 0 | 0 |
| ENSDARG000 | 8 | 27727970 | 27728086 | - | 117 | 5S_rRNA | 0 | 0 | 0 | 0 | 0 | 0 |
| ENSDARG000 | 8 | 44040796 | 44040913 | - | 118 | 5S_rRNA | 0 | 0 | 0 | 0 | 0 | 0 |
| ENSDARG000 | 8 | 47072220 | 47072337 | + | 118 | 5S_rRNA | 0 | 0 | 0 | 0 | 0 | 0 |
| ENSDARG000 | 8:8:8:8:8:8:8: | 19469678:194 | 19469958:194 | -:-:-:-:-:-:-:-: | 2613 | EDEM3 | 966 | 618 | 840 | 588 | 1187 | 730 |
| ENSDARG000 | 8 | 38430563 | 38430679 | + | 117 | 5S_rRNA | 0 | 0 | 1 | 0 | 1 | 0 |
| ENSDARG000 | 8 | 29912634 | 29912750 | + | 117 | 5S_rRNA | 0 | 0 | 0 | 0 | 0 | 0 |
| ENSDARG000 | 8:8:8:8:8:8:8: | 104114:10483 | 104186:10490 | +:+:+:+:+:+:+: | 2442 | sncaip | 190 | 158 | 262 | 72 | 264 | 243 |
| ENSDARG000 | 8 | 47275296 | 47275410 | - | 115 | 5S_rRNA | 0 | 0 | 0 | 0 | 0 | 0 |
| ENSDARG000 | 8;8;8;8 | 44462599:444 | 44462723:444 | -;-;-;- | 936 | NA | 0 | 0 | 0 | 0 | 0 | 0 |
| ENSDARG000 | 8 | 47505885 | 47506001 | - | 117 | 5S_rRNA | 0 | 0 | 0 | 0 | 0 | 0 |
| ENSDARG000 | 8:8:8:8:8:8:8: | 12951155:129 | 12951412:129 | +:+:+:+:+:+:+: | 1706 | CEPT1 | 20 | 3 | 35 | 18 | 14 | 38 |
| ENSDARG000 | 8;8;8;8 | 36416285:364 | 36416380:364 | +;+;+;+ | 1873 | si:busm1-194 | 0 | 0 | 0 | 0 | 0 | 0 |
| ENSDARG000 | 8 | 37620554 | 37620670 | + | 117 | 5S_rRNA | 0 | 0 | 0 | 0 | 0 | 0 |
| ENSDARG000 | 8 | 9893672 | 9893788 | + | 117 | 5S_rRNA | 0 | 0 | 0 | 0 | 0 | 0 |
| ENSDARG000 | 8 | 45411581 | 45411745 | - | 165 | U1 | 0 | 0 | 0 | 0 | 0 | 0 |
| ENSDARG000 | 8;8;8;8;8 | 52453146:524 | 52453867:524 | -;-;-;-;- | 1337 | phyhip | 0 | 12 | 0 | 0 | 0 | 0 |
| ENSDARG000 | 8;8;8;8;8;8;8 | 25342896:253 | 25343095:253 | +;+;+;+;+;+;+ | 3509 | NA | 10 | 14 | 31 | 6 | 75 | 66 |
| ENSDARG000 | 8:8:8:8:8:8:8: | 43078044:430 | 43078242:430 | +:+:+:+:+:+:+: | 1338 | NA | 0 | 0 | 0 | 0 | 0 | 0 |
| ENSDARG000 | 8 | 53023539 | 53023617 | + | 79 | dre-mir-181a- | 0 | 0 | 0 | 0 | 0 | 0 |
| ENSDARG000 | 8 | 30162492 | 30162633 | - | 142 | dre-mir-24-3 | 8 | 0 | 0 | 0 | 0 | 0 |
| ENSDARG000 | 8 | 18670237 | 18670362 | + | 126 | 5S_rRNA | 0 | 0 | 0 | 0 | 0 | 0 |
| ENSDARG000 | 8 | 33783438 | 33783554 | - | 117 | 5S_rRNA | 0 | 0 | 0 | 0 | 0 | 0 |
| ENSDARG000 | 8 | 4245784 | 4245902 | + | 119 | 5S_rRNA | 0 | 0 | 0 | 0 | 0 | 0 |
| ENSDARG000 | 8:8:8:8:8:8:8: | 54055390:540 | 54055516:540 | +:+:+:+:+:+:+: | 3381 | MAGI1 | 210 | 313 | 330 | 327 | 188 | 22 |
| ENSDARG000 | 8;8;8;8 | 7644265:7646 | 7644771:7647 | +;+;+;+ | 1858 | NA | 0 | 2 | 7 | 0 | 0 | 0 |
| ENSDARG000 | 8;8;8;8;8 | 21395015:213 | 21395702:213 | +;+;+;+;+ | 1467 | NA | 0 | 0 | 0 | 0 | 0 | 0 |
| ENSDARG000 | 8:8:8:8:8:8:8: | 21723326:218 | 21723463:218 | -;-;-;-;-;-;-;- | 1054 | NA | 0 | 0 | 0 | 0 | 0 | 0 |
| ENSDARG000 | 8;8;8 | 27168264:271 | 27168464:271 | +;+;+ | 1428 | NA | 0 | 0 | 0 | 0 | 0 | 0 |
| ENSDARG000 | 8:8:8:8:8:8:8: | 27179689:271 | 27179873:271 | +:+:+:+:+:+:+: | 1129 | NA | 0 | 4 | 4 | 2 | 4 | 12 |
| ENSDARG000 | 8:8:8:8:8:8:8: | 27183274:271 | 27183871:271 | -;-;-;-;-;-;-;- | 1239 | NA | 0 | 0 | 0 | 0 | 0 | 0 |
| ENSDARG000 | 8;8;8;8 | 27190786:271 | 27191622:271 | +;+;+;+ | 1648 | NA | 8 | 0 | 0 | 0 | 0 | 0 |
| ENSDARG000 | 8:8:8:8:8:8:8: | 27197082:271 | 27197412:271 | -;-;-;-;-;-;-;- | 1079 | NA | 2 | 0 | 0 | 4 | 2 | 0 |
| ENSDARG000 | 8;8;8 | 32427579:324 | 32428436:324 | -;-;- | 2073 | NA | 0 | 0 | 0 | 0 | 0 | 0 |
| ENSDARG000 | 8:8:8:8:8:8:8: | 32435738:324 | 32435853:324 | -:-:-:-:-:-:-:-: | 2465 | NA | 0 | 0 | 0 | 0 | 0 | 0 |
| ENSDARG000 | 8;8;8;8;8;8 | 32451956:324 | 32452194:324 | -;-;-;-;-;- | 1357 | NA | 0 | 0 | 3 | 0 | 0 | 0 |
| ENSDARG000 | 8;8;8;8 | 32453652:324 | 32454096:324 | -;-;-;- | 932 | NA | 0 | 0 | 3 | 0 | 0 | 0 |
| ENSDARG000 | 8;8;8;8 | 32456407:324 | 32456567:324 | -;-;-;- | 874 | NA | 0 | 0 | 0 | 0 | 0 | 0 |
| ENSDARG000 | 8:8:8:8:8:8:8: | 32460395:324 | 32460586:324 | -:-:-:-:-:-:-:-: | 2088 | NA | 0 | 0 | 0 | 0 | 0 | 0 |
| ENSDARG000 | 8;8;8 | 32472628:324 | 32473291:324 | -;-;- | 996 | NA | 0 | 0 | 0 | 0 | 0 | 0 |
| ENSDARG000 | 8;8;8 | 33593650:336 | 33594024:336 | -;-;- | 615 | NA | 0 | 0 | 0 | 0 | 0 | 0 |
| ENSDARG000 | 8;8;8 | 34271072:342 | 34271228:342 | -;-;- | 334 | NA | 0 | 0 | 0 | 0 | 0 | 0 |
| ENSDARG000 | 8;8;8;8;8 | 34461160:344 | 34461487:344 | -;-;-;-;- | 521 | NA | 32 | 14 | 40 | 32 | 18 | 32 |
| ENSDARG000 | 8:8:8:8:8:8:8: | 36528233:365 | 36528919:365 | -:-:-:-:-:-:-:-: | 1679 | NA | 2 | 4 | 4 | 0 | 0 | 8 |
| ENSDARG000 | 8;8;8;8 | 38991828:389 | 38992089:389 | +;+;+;+ | 902 | NA | 0 | 2 | 0 | 0 | 0 | 0 |
| ENSDARG000 | 8;8;8;8;8 | 39332416:393 | 39332582:393 | +;+;+;+;+ | 447 | NA | 0 | 0 | 0 | 0 | 0 | 0 |
| ENSDARG000 | 8;8;8;8;8 | 40502765:405 | 40502976:405 | +;+;+;+;+ | 5298 | NA | 44 | 651 | 2 | 6 | 46 | 644 |
| ENSDARG000 | 8;8;8 | 50889589:508 | 50890439:508 | -;-;- | 920 | NA | 0 | 0 | 0 | 0 | 1 | 0 |
| ENSDARG000 | 8;8;8;8 | 52601058:526 | 52601377:526 | +;+;+;+ | 771 | NA | 0 | 0 | 0 | 0 | 0 | 0 |
| ENSDARG000 | 8;8;8;8;8 | 53841302:538 | 53843880:538 | -;-;-;-;- | 3118 | NA | 52 | 22 | 51 | 8 | 37 | 48 |
| ENSDARG000 | 8;8;8 | 54027682:540 | 54027936:540 | +;+;+ | 809 | NA | 12 | 6 | 10 | 22 | 8 | 18 |
| ENSDARG000 | 8:8:8:8:8:8:8: | 54115872:541 | 54116186:541 | +:+:+:+:+:+:+: | 6183 | NA | 0 | 8 | 0 | 2 | 2 | 0 |
| ENSDARG000 | 17:17:17:17:1 | 20605013:206 | 20605379:206 | +:+:+:+:+:+:+: | 1222 | NA | 8 | 0 | 8 | 8 | 0 | 177 |
| ENSDARG000 | 17;17;17 | 27971144:279 | 27972332:279 | -;-;- | 1305 | PTAFR | 48 | 65 | 53 | 18 | 46 | 12 |
| ENSDARG000 | 17:17:17:17:1 | 23820450:238 | 23824862:238 | -;-;-;-;- | 6037 | PDZD8 | 1375 | 1300 | 1121 | 700 | 1934 | 794 |
| ENSDARG000 | 17:17:17:17:1 | 24855384:248 | 24856699:248 | -:-:-:-:-:-:-:-: | 2690 | hmgcl | 313 | 296 | 295 | 459 | 403 | 407 |
| ENSDARG000 | 17;17;17;17 | 23926796:239 | 23927002:239 | +;+;+;+ | 2893 | pex13 | 258 | 382 | 425 | 250 | 293 | 278 |
| ENSDARG000 | 17;17;17;17 | 28005763:280 | 28006179:280 | +;+;+;+ | 8327 | LUZP1 | 730 | 1010 | 459 | 334 | 1357 | 220 |
| ENSDARG000 | 17:17:17:17:1 | 45501378:455 | 45502933:455 | -;-;-;-;-;-;-;- | 3756 | HHIPL2 | 0 | 0 | 5 | 9 | 34 | 0 |
| ENSDARG000 | 17:17:17:17:1 | 45472608:454 | 45472945:454 | +:+:+:+:+:+:+: | 4632 | marc1 | 190 | 156 | 106 | 189 | 148 | 165 |
| ENSDARG000 | 17:17:17:17:1 | 35225614:352 | 35225833:352 | -:-:-:-:-:-:-:-: | 1394 | itgb1bp1 | 977 | 1099 | 1099 | 1304 | 1073 | 975 |
| ENSDARG000 | 17:17:17:17:1 | 41081512:410 | 41082078:410 | +:+:+:+:+:+:+: | 4221 | babam2 | 800 | 596 | 773 | 598 | 770 | 573 |
| ENSDARG000 | 17:17:17:17:1 | 52600506:526 | 52600631:526 | -:-:-:-:-:-:-:-: | 1239 | NA | 0 | 0 | 0 | 0 | 0 | 0 |
| ENSDARG000 | 17;17;17;17 | 8879812:8880 | 8880175:8880 | -;-;-;- | 630 | nkl.3 | 4 | 4 | 5 | 0 | 8 | 4 |
| ENSDARG000 | 17:17:17:17:1 | 40999835:410 | 41000219:410 | -;-;-;-;-;-;-;- | 1250 | rbks | 66 | 77 | 114 | 95 | 118 | 120 |
| ENSDARG000 | 17:17:17:17:1 | 23937262:239 | 23937918:239 | +:+:+:+:+:+:+: | 3628 | SANBR | 26 | 19 | 24 | 22 | 14 | 44 |
| ENSDARG000 | 17:17:17:17:1 | 18773961:187 | 18774236:187 | -:-:-:-:-:-:-:-: | 1620 | vrk1 | 298 | 486 | 597 | 340 | 453 | 379 |
| ENSDARG000 | 17:17:17:17:1 | 43889371:438 | 43889556:438 | +:+:+:+:+:+:+: | 3920 | MSH4 | 34 | 140 | 32 | 162 | 59 | 181 |
| ENSDARG000 | 17:17:17:17:1 | 31151215:311 | 31153079:311 | -;-;-;-;-;-;- | 3207 | NA | 174 | 78 | 190 | 309 | 135 | 236 |
| ENSDARG000 | 17:17:17:17:1 | 38030327:380 | 38030384:380 | +:+:+:+:+:+:+: | 1954 | slc25a21 | 108 | 110 | 133 | 56 | 102 | 90 |
| ENSDARG000 | 17 | 29276544 | 29277731 | + | 1188 | ankrd9 | 0 | 0 | 0 | 0 | 0 | 0 |
| ENSDARG000 | 17:17:17:17:1 | 5829851:5836 | 5830091:5836 | -;-;-;-;-;-;-;- | 1538 | NA | 0 | 0 | 0 | 0 | 0 | 0 |
| ENSDARG000 | 17:17:17:17:1 | 5863150:5867 | 5863655:5868 | -;-;-;-;-;-;- | 1211 | NA | 0 | 0 | 4 | 0 | 2 | 0 |
| ENSDARG000 | 17;17 | 5837308:5837 | 5837536:5837 | +;+ | 296 | NA | 0 | 0 | 0 | 0 | 0 | 0 |
| ENSDARG000 | 17 | 11419083 | 11419622 | + | 540 | NA | 4 | 7 | 21 | 5 | 1 | 2 |
| ENSDARG000 | 17:17:17:17:1 | 52612866:526 | 52613099:526 | +:+:+:+:+:+:+: | 4336 | NA | 390 | 275 | 345 | 518 | 408 | 580 |
| ENSDARG000 | 17:17:17:17:1 | 8899016:8907 | 8899153:8907 | +:+:+:+:+:+:+: | 2484 | psmc1a | 2586 | 2503 | 2690 | 1637 | 2447 | 1794 |
| ENSDARG000 | 17:17:17:17:1 | 52632094:526 | 52632445:526 | -;-;-;-;-;-;- | 1328 | fam98b | 134 | 213 | 223 | 186 | 128 | 168 |
| ENSDARG000 | 17:17:17:17:1 | 31181950:311 | 31182037:311 | -:-:-:-:-:-:-:-: | 1240 | NA | 0 | 0 | 0 | 0 | 0 | 0 |
| ENSDARG000 | 17:17:17:17:1 | 52639492:526 | 52641906:526 | -;-;-;-;-;-;- | 3438 | spred1 | 2017 | 1603 | 2450 | 1838 | 2389 | 1687 |
| ENSDARG000 | 17:17:17:17:1 | 8896383:8896 | 8896516:8896 | -:-:-:-:-:-:-:-: | 1016 | nkl.2 | 108 | 470 | 137 | 210 | 429 | 2502 |
| ENSDARG000 | 17:17:17:17:1 | 21994043:219 | 21995313:219 | -:-:-:-:-:-:-:-: | 3511 | NA | 0 | 0 | 10 | 0 | 74 | 0 |
| ENSDARG000 | 17:17:17:17:1 | 29282863:292 | 29283987:292 | -:-:-:-:-:-:-:-: | 5257 | tecpr2 | 210 | 174 | 262 | 187 | 250 | 323 |
| ENSDARG000 | 17:17:17:17:1 | 25856671:258 | 25856931:258 | +:+:+:+:+:+:+: | 4699 | WAPL | 405 | 347 | 689 | 254 | 538 | 386 |
| ENSDARG000 | 17:17:17:17:1 | 20616016:206 | 20616255:206 | +:+:+:+:+:+:+: | 2215 | NA | 0 | 0 | 0 | 0 | 2 | 0 |
| ENSDARG000 | 17:17:17:17:1 | 35243753:352 | 35243879:352 | +:+:+:+:+:+:+: | 1205 | iah1 | 764 | 735 | 870 | 711 | 1072 | 512 |
| ENSDARG000 | 17:17:17:17:1 | 14467015:144 | 14468657:144 | -:-:-:-:-:-:-:-: | 5924 | DAAM1 | 1270 | 1430 | 2062 | 2008 | 1496 | 2100 |
| ENSDARG000 | 17:17:17:17:1 | 11421643:114 | 11421868:114 | -:-:-:-:-:-:-:-: | 1271 | psma3 | 2979 | 3226 | 3360 | 2112 | 3107 | 2289 |
| ENSDARG000 | 17:17:17:17:1 | 28082441:280 | 28085363:280 | -:-:-:-:-:-:-:-: | 6157 | kdm1a | 1391 | 1258 | 1898 | 1088 | 1751 | 1251 |
| ENSDARG000 | 17:17:17:17:1 | 22002857:220 | 22002970:220 | -:-:-:-:-:-:-:-: | 1914 | slc22a7b.1 | 8 | 4 | 6 | 14 | 68 | 0 |
| ENSDARG000 | 17;17;17;17 | 58211:67514: | 58423:67663: | +;+;+;+ | 1165 | NA | 8 | 4 | 4 | 0 | 0 | 5 |
| ENSDARG000 | 17:17:17:17:1 | 49785495:497 | 49786942:497 | -:-:-:-:-:-:-:-: | 10798 | COL12A1 | 1980 | 407 | 209 | 294 | 1066 | 1021 |
| ENSDARG000 | 17;17 | 80167;86280 | 82404;86548 | -;- | 2507 | GREM1 | 4 | 0 | 0 | 8 | 0 | 0 |
| ENSDARG000 | 17;17;17;17 | 24869371:248 | 24870315:248 | -;-;-;- | 2314 | zbtb8a | 71 | 58 | 48 | 64 | 77 | 67 |
| ENSDARG000 | 17:17:17:17:1 | 24877392:248 | 24880695:248 | -:-:-:-:-:-:-:-: | 4558 | gale | 215 | 232 | 198 | 98 | 266 | 136 |
| ENSDARG000 | 17:17:17:17:1 | 29319587:293 | 29319666:293 | +:+:+:+:+:+:+: | 3254 | ptpreb | 61 | 22 | 29 | 14 | 58 | 56 |
| ENSDARG000 | 17:17:17:17:1 | 18810904:188 | 18811122:188 | +;+;+;+;+;+;+ | 2869 | NA | 6 | 0 | 0 | 0 | 10 | 4 |
| ENSDARG000 | 17;17 | 112829:11373 | 112901:11462 | +;+ | 960 | NA | 0 | 0 | 0 | 0 | 0 | 0 |
| ENSDARG000 | 17:17:17:17:1 | 92784:94610: | 94881:94881: | -:-:-:-:-:-:-:-: | 4867 | arhgap11a | 109 | 425 | 428 | 170 | 286 | 158 |
| ENSDARG000 | 17:17:17:17:1 | 15502964:155 | 15504509:155 | -:-:-:-:-:-:-:-: | 3215 | fyna | 16 | 16 | 50 | 20 | 56 | 20 |
| ENSDARG000 | 17;17 | 25893485:258 | 25894345:258 | -;- | 938 | NA | 0 | 0 | 0 | 0 | 0 | 0 |
| ENSDARG000 | 17:17:17:17:1 | 35247497:352 | 35250506:352 | -:-:-:-:-:-:-:-: | 6109 | adam17a | 7761 | 4964 | 4653 | 5418 | 5000 | 5706 |
| ENSDARG000 | 17:17:17:17:1 | 31185276:311 | 31185503:311 | +:+:+:+:+:+:+: | 4581 | disp2 | 4 | 0 | 0 | 0 | 24 | 4 |
| ENSDARG000 | 17:17:17:17:1 | 18877918:188 | 18878291:188 | -:-:-:-:-:-:-:-: | 2453 | ak7b | 4 | 0 | 2 | 0 | 0 | 0 |
| ENSDARG000 | 17:17:17:17:1 | 38236469:382 | 38237450:382 | -:-:-:-:-:-:-:-: | 2084 | pax9 | 8 | 2 | 0 | 0 | 2 | 0 |
| ENSDARG000 | 17;17 | 35278788:352 | 35279019:352 | +;+ | 1779 | NA | 0 | 2 | 0 | 3 | 0 | 0 |
| ENSDARG000 | 17:17:17:17:1 | 35285876:352 | 35287662:352 | -;-;-;-;-;-;- | 2755 | rnf144aa | 131 | 280 | 291 | 458 | 356 | 279 |
| ENSDARG000 | 17:17:17:17:1 | 6016243:6016 | 6016750:6016 | -:-:-:-:-:-:-:-: | 2516 | ephx2 | 273 | 603 | 548 | 357 | 497 | 483 |
| ENSDARG000 | 17;17 | 38255105:382 | 38255362:382 | +;+ | 1166 | nkx2.9 | 0 | 0 | 0 | 0 | 0 | 0 |
| ENSDARG000 | 17;17 | 38262408:382 | 38262936:382 | +;+ | 1808 | nkx2.1 | 0 | 2 | 2 | 0 | 0 | 0 |
| ENSDARG000 | 17:17:17:17:1 | 45529453:455 | 45530243:455 | -:-:-:-:-:-:-:-: | 2816 | susd4 | 0 | 0 | 4 | 0 | 2 | 0 |
| ENSDARG000 | 17:17:17:17:1 | 18889637:188 | 18890037:188 | -:-:-:-:-:-:-:-: | 2320 | galcb | 1723 | 998 | 741 | 1038 | 1150 | 1157 |
| ENSDARG000 | 17:17:17:17:1 | 22032296:220 | 22033574:220 | -:-:-:-:-:-:-:-: | 5200 | NA | 48 | 15 | 123 | 96 | 161 | 64 |
| ENSDARG000 | 17:17:17:17:1 | 32657202:326 | 32662365:326 | -:-:-:-:-:-:-:-: | 6426 | KCNK2 | 6 | 0 | 0 | 0 | 0 | 0 |
| ENSDARG000 | 17:17:17:17:1 | 23964132:239 | 23966641:239 | +:+:+:+:+:+:+: | 7994 | XPO1 | 1759 | 2237 | 3590 | 2684 | 3343 | 2458 |
| ENSDARG000 | 17:17:17:17:1 | 41337246:413 | 41337546:413 | +:+:+:+:+:+:+: | 2486 | mocs1 | 599 | 348 | 919 | 376 | 1842 | 332 |
| ENSDARG000 | 17:17:17:17:1 | 11445671:114 | 11448723:114 | -;-;-;-;-;- | 4145 | NA | 115 | 85 | 100 | 53 | 119 | 105 |
| ENSDARG000 | 17:17:17:17:1 | 25955003:259 | 25955206:259 | +:+:+:+:+:+:+: | 6367 | GRID1 | 140 | 431 | 240 | 407 | 351 | 76 |
| ENSDARG000 | 17;17;17 | 31205064:312 | 31206071:312 | -;-;- | 1995 | rpusd2 | 123 | 155 | 246 | 96 | 242 | 147 |
| ENSDARG000 | 17 | 20639180 | 20640505 | + | 1326 | npy4r | 0 | 0 | 0 | 0 | 0 | 0 |
| ENSDARG000 | 17;17;17 | 376891:37774 | 377110:37778 | +;+;+ | 755 | NA | 0 | 0 | 2 | 0 | 0 | 0 |
| ENSDARG000 | 17;17;17 | 378485:37914 | 379037:37922 | -;-;- | 863 | NA | 0 | 0 | 2 | 0 | 3 | 0 |
| ENSDARG000 | 17:17:17:17:1 | 45577326:455 | 45577425:455 | +:+:+:+:+:+:+: | 3043 | NA | 70 | 8 | 6 | 0 | 8 | 12 |
| ENSDARG000 | 17;17 | 52841830:528 | 52842073:528 | +;+ | 466 | NA | 0 | 0 | 0 | 0 | 0 | 0 |
| ENSDARG000 | 17:17:17:17:1 | 52822422:528 | 52822978:528 | +:+:+:+:+:+:+: | 3468 | meis2a | 8 | 12 | 12 | 2 | 13 | 10 |
| ENSDARG000 | 17:17:17:17:1 | 11470572:114 | 11474111:114 | -:-:-:-:-:-:-:-: | 5924 | HNRNPU | 1810 | 1307 | 1708 | 994 | 1875 | 1626 |

| | | | | | | | | | | | | |
|---|---|---|---|---|---|---|---|---|---|---|---|---|
| ENSDARG000 | 17;17;17;17;1 | 24891756:248 | 24892958:248 | -;-;-;-;-;-;-;-; | 2915 | NA | 42 | 2 | 0 | 6 | 12 | 8 |
| ENSDARG000 | 17;17;17;17 | 389218:3952 | 392205:3958 | +;+;+;+ | 3809 | NA | 24 | 1 | 11 | 0 | 0 | 0 |
| ENSDARG000 | 17;17;17;17;1 | 14605794:146 | 14606429:146 | -;-;-;-;-;-;-;- | 1511 | sdsl | 0 | 0 | 0 | 8 | 4 | 0 |
| ENSDARG000 | 17;17;17;17;1 | 41383756:413 | 41384374:413 | -;-;-;-;-;-;-;-; | 2335 | NA | 0 | 2 | 2 | 0 | 0 | 0 |
| ENSDARG000 | 17;17;17;17;1 | 20653773:206 | 20655221:206 | -;-;-;-;-;-;-;- | 3053 | slc16a9a | 289 | 442 | 377 | 713 | 599 | 379 |
| ENSDARG000 | 17;17;17;17;1 | 15534815:155 | 15535845:155 | +;+;+;+;+;+;+; | 4105 | marcksb | 2580 | 5088 | 4258 | 2395 | 2912 | 2811 |
| ENSDARG000 | 17;17;17;17;1 | 24917350:249 | 24918422:249 | -;-;-;-;-;-;-;-; | 3394 | NA | 79 | 52 | 78 | 50 | 81 | 69 |
| ENSDARG000 | 17;17;17;17;1 | 38295847:382 | 38296046:382 | +;+;+;+;+;+;+; | 1393 | mbip | 103 | 137 | 191 | 116 | 152 | 94 |
| ENSDARG000 | 17;17;17;17;1 | 32720413:327 | 32723290:327 | -;-;-;-;-;-;-;-; | 11212 | CENPF | 1006 | 1991 | 3117 | 1265 | 2114 | 1124 |
| ENSDARG000 | 17;17;17;17;1 | 24936059:249 | 24937758:249 | +;+;+;+;+;+;+; | 6997 | DLGAP2 | 8 | 0 | 0 | 0 | 1 | 4 |
| ENSDARG000 | 17 | 41463942 | 41466145 | + | 2204 | insm1b | 0 | 0 | 2 | 0 | 6 | 4 |
| ENSDARG000 | 17;17;17;17;1 | 35357451:353 | 35358149:353 | -;-;-;-;-;- | 1623 | rsad2 | 113 | 276 | 26 | 4 | 6 | 0 |
| ENSDARG000 | 17;17 | 24917350:249 | 24918422:249 | -;- | 1260 | zgc:193593 | 2 | 4 | 2 | 4 | 0 | 0 |
| ENSDARG000 | 17;17;17;17;1 | 22072337:220 | 22072544:220 | +;+;+;+;+;+;+; | 2413 | nphp1 | 86 | 46 | 112 | 22 | 68 | 56 |
| ENSDARG000 | 17;17;17;17;1 | 29345606:293 | 29346007:293 | +;+;+;+;+;+;+; | 4054 | KCTD3 | 365 | 172 | 360 | 309 | 392 | 219 |
| ENSDARG000 | 17;17;17;17;1 | 20673963:206 | 20675178:206 | -;-;-;-;-;-;-;- | 2598 | ccdc6a | 1434 | 1476 | 1797 | 1708 | 1355 | 1502 |
| ENSDARG000 | 17;17;17;17;1 | 52939748:529 | 52941603:529 | -;-;-;-;-;-;-;-; | 3125 | cdin1 | 46 | 67 | 62 | 42 | 92 | 48 |
| ENSDARG000 | 17;17;17 | 15542077:155 | 15544481:155 | -;-;- | 2639 | col10a1a | 337 | 58 | 84 | 0 | 532 | 18 |
| ENSDARG000 | 17;17;17;17;1 | 7844927:7845 | 7845829:7845 | -;-;-;-;-;-;-;-; | 27036 | NA | 154 | 117 | 154 | 212 | 444 | 270 |
| ENSDARG000 | 17;17;17;17;1 | 43926523:439 | 43926662:439 | +;+;+;+;+;+;+; | 4518 | ktn1 | 2226 | 2335 | 2797 | 1778 | 2532 | 2251 |
| ENSDARG000 | 17;17;17;17;1 | 14614145:146 | 14615173:146 | -;-;-;-;-;-;-;-; | 4978 | ppp1r13ba | 1150 | 648 | 419 | 1297 | 585 | 1146 |
| ENSDARG000 | 17;17;17;17;1 | 6079881:6088 | 6080041:6088 | +;+;+;+;+;+ | 2368 | chrna2a | 0 | 0 | 0 | 0 | 0 | 2 |
| ENSDARG000 | 17;17;17;17;1 | 41475606:414 | 41477491:414 | -;-;-;-;-;-;-;-; | 8039 | RALGAPA2 | 1147 | 789 | 1230 | 917 | 1882 | 894 |
| ENSDARG000 | 17;17;17;17;1 | 11500628:115 | 11500665:115 | +;+;+;+;+;+;+ | 595 | efcab2 | 0 | 0 | 0 | 0 | 6 | 0 |
| ENSDARG000 | 17;17;17;17;1 | 28102487:281 | 28102561:281 | +;+;+;+;+;+;+; | 4112 | tmem30b | 60 | 2 | 0 | 6 | 2 | 8 |
| ENSDARG000 | 17;17;17;17;1 | 15559046:156 | 15560013:156 | +;+;+;+;+;+;+; | 8559 | NA | 16 | 6 | 46 | 18 | 33 | 14 |
| ENSDARG000 | 17;17;17;17;1 | 15658867:156 | 15661499:156 | -;-;-;-;-;-;-;-; | 3390 | manea | 452 | 320 | 644 | 341 | 534 | 362 |
| ENSDARG000 | 17;17;17;17 | 31221761:312 | 31221878:312 | +;+;+;+ | 1169 | ccdc32 | 684 | 604 | 699 | 672 | 709 | 492 |
| ENSDARG000 | 17;17;17;17;1 | 31214167:312 | 31216810:312 | -;-;-;-;-;-;-;-; | 3921 | pcmtl | 170 | 194 | 363 | 360 | 252 | 238 |
| ENSDARG000 | 17;17;17;17 | 41762283:417 | 41762329:417 | +;+;+;+ | 371 | NA | 0 | 6 | 0 | 0 | 0 | 0 |
| ENSDARG000 | 17;17;17;17;1 | 15605572:156 | 15606029:156 | -;-;-;-;-;- | 1255 | fhl5 | 0 | 0 | 0 | 0 | 0 | 0 |
| ENSDARG000 | 17;17;17;17 | 15631619:156 | 15640446:156 | -;-;-;- | 9401 | fut9a | 4 | 2 | 26 | 4 | 8 | 0 |
| ENSDARG000 | 17;17;17;17;1 | 15580665:155 | 15581192:155 | -;-;-;-;-;-;-;-; | 2992 | NA | 2 | 0 | 8 | 2 | 28 | 0 |
| ENSDARG000 | 17;17;17;17;1 | 45602640:456 | 45602896:456 | +;+;+;+;+;+;+; | 2542 | NA | 0 | 2 | 0 | 0 | 0 | 0 |
| ENSDARG000 | 17;17;17;17;1 | 35362851:353 | 35363457:353 | +;+;+;+;+ | 1254 | CMPK2 | 184 | 185 | 48 | 12 | 22 | 22 |
| ENSDARG000 | 17;17;17;17;1 | 53156530:531 | 53156758:531 | +;+;+;+;+;+;+; | 2629 | dph6 | 339 | 236 | 360 | 179 | 242 | 394 |
| ENSDARG000 | 17;17;17;17;1 | 11507131:115 | 11507625:115 | +;+;+;+;+;+;+; | 11088 | KIF26B | 12 | 26 | 56 | 8 | 40 | 186 |
| ENSDARG000 | 17;17;17;17;1 | 38393189:383 | 38395588:383 | -;-;-;-;-;-;-;-; | 4743 | RPS6KA5 | 42 | 22 | 46 | 14 | 66 | 64 |
| ENSDARG000 | 17;17;17 | 19017394:190 | 19020067:190 | -;-;- | 3145 | FLRT2 | 6 | 6 | 2 | 2 | 12 | 0 |
| ENSDARG000 | 17;17;17;17 | 22102791:221 | 22102912:221 | +;+;+;+ | 886 | malb | 4 | 0 | 2 | 0 | 0 | 0 |
| ENSDARG000 | 17 | 35878544 | 35881841 | - | 3298 | sox11a | 298 | 310 | 587 | 150 | 1348 | 198 |
| ENSDARG000 | 17;17;17;17 | 36446514:364 | 36446719:364 | -;-;-;- | 624 | NA | 0 | 0 | 0 | 0 | 6 | 0 |
| ENSDARG000 | 17;17;17;17;1 | 49983157:499 | 49983364:499 | -;-;-;-;-;-;-;-; | 662 | cox7a2a | 601 | 890 | 682 | 493 | 569 | 600 |
| ENSDARG000 | 17;17;17;17;1 | 36483287:364 | 36483572:364 | -;-;-;-;-;-;-;-; | 1493 | NA | 0 | 0 | 0 | 0 | 0 | 0 |
| ENSDARG000 | 17;17;17;17;1 | 6113680:6114 | 6113709:6114 | -;-;-;-;-;-;-;-; | 3413 | PTK2B | 35 | 59 | 30 | 40 | 100 | 111 |
| ENSDARG000 | 17;17;17;17;1 | 8951449:8954 | 8951875:8954 | -;-;-;-;-;-;-;- | 2364 | zranb1b | 779 | 387 | 484 | 427 | 512 | 608 |
| ENSDARG000 | 17;17;17;17;1 | 53292215:532 | 53292914:532 | +;+;+;+;+;+ | 1354 | NA | 6 | 15 | 31 | 16 | 33 | 4 |
| ENSDARG000 | 17;17;17;17 | 23994633:239 | 23994741:239 | +;+;+;+ | 646 | NA | 0 | 0 | 0 | 0 | 0 | 0 |
| ENSDARG000 | 17;17;17;17;1 | 8988374:9009 | 8988484:9009 | +;+;+;+;+;+;+; | 9232 | AKAP6 | 74 | 24 | 26 | 24 | 42 | 44 |
| ENSDARG000 | 17;17;17;17;1 | 45639247:456 | 45639603:456 | +;+;+;+;+;+;+; | 1666 | vcpkmt | 84 | 63 | 102 | 48 | 72 | 82 |
| ENSDARG000 | 17;17;17;17;1 | 24006792:240 | 24006949:240 | +;+;+;+;+;+;+; | 1822 | NA | 0 | 2 | 0 | 0 | 2 | 0 |
| ENSDARG000 | 17;17;17;17;1 | 36516695:365 | 36518878:365 | -;-;-;-;-;-;-;-; | 3606 | colec11 | 0 | 10 | 0 | 0 | 2 | 0 |
| ENSDARG000 | 17;17;17;17;1 | 20701582:207 | 20702067:207 | -;-;-;-;-;-;-;-; | 14067 | ANK3 | 70 | 41 | 28 | 43 | 31 | 24 |
| ENSDARG000 | 17;17;17;17;1 | 11728920:117 | 11729035:117 | -;-;-;-;-;-;-;-; | 1358 | smyd3 | 56 | 20 | 82 | 34 | 80 | 39 |
| ENSDARG000 | 17;17 | 29722085:297 | 29722402:297 | -;- | 525 | NA | 0 | 0 | 0 | 0 | 0 | 0 |
| ENSDARG000 | 17;17 | 29723351:297 | 29723766:297 | +;+ | 1511 | NA | 0 | 0 | 2 | 6 | 0 | 0 |
| ENSDARG000 | 17;17;17;17;1 | 29376854:293 | 29376943:293 | -;-;-;-;-;-;-;-; | 15939 | USH2A | 10 | 16 | 63 | 28 | 175 | 13 |
| ENSDARG000 | 17;17;17;17;1 | 12058509:120 | 12058579:120 | +;+;+;+;+;+;+; | 1894 | tfb2m | 184 | 253 | 272 | 168 | 160 | 280 |
| ENSDARG000 | 17;17;17 | 6276559:6279 | 6276779:6280 | +;+;+ | 789 | dusp23b | 96 | 136 | 98 | 74 | 86 | 146 |
| ENSDARG000 | 17;17;17;17;1 | 22107702:221 | 22108212:221 | -;-;-;-;-;-;-;-; | 1793 | mrps5 | 438 | 573 | 805 | 391 | 585 | 422 |
| ENSDARG000 | 17;17;17;17;1 | 26477611:264 | 26481873:264 | -;-;-;-;-;-;-;-; | 7460 | CCSER2 | 108 | 73 | 155 | 278 | 227 | 136 |
| ENSDARG000 | 17;17;17;17;1 | 19343015:193 | 19343555:193 | -;-;-;-;- | 1628 | gsc | 0 | 0 | 0 | 0 | 0 | 0 |
| ENSDARG000 | 17;17;17;17;1 | 14692808:146 | 14694868:146 | -;-;-;-;-;-;-;-; | 3482 | ptp4a2a | 1286 | 1148 | 1293 | 955 | 1673 | 1205 |
| ENSDARG000 | 17;17;17;17;1 | 32745763:327 | 32746475:327 | -;-;-;-;-;-;-;-; | 2647 | smyd2a | 624 | 819 | 986 | 492 | 763 | 1204 |
| ENSDARG000 | 17;17 | 28166859:281 | 28169155:281 | -;- | 2633 | htr1d | 0 | 0 | 0 | 0 | 0 | 0 |
| ENSDARG000 | 17;17 | 9308875:9310 | 9310219:9310 | -;- | 1381 | NA | 4 | 7 | 20 | 7 | 5 | 0 |
| ENSDARG000 | 17;17;17;17;1 | 24036791:240 | 24036979:240 | +;+;+;+;+;+;+; | 4807 | b3gnt2b | 2266 | 1522 | 1273 | 1445 | 1884 | 2446 |
| ENSDARG000 | 17;17;17;17;1 | 6358160:6358 | 6358750:6358 | -;-;-;-;-;-;-;-; | 1813 | txlnbb | 0 | 8 | 4 | 4 | 8 | 6 |
| ENSDARG000 | 17;17;17;17;1 | 53296851:532 | 53296931:532 | +;+;+;+;+;+;+; | 3409 | DDX24 | 512 | 514 | 1178 | 462 | 624 | 388 |
| ENSDARG000 | 17;17;17;17;1 | 49998159:500 | 49999239:500 | -;-;-;-;-;-;- | 2146 | tmem30aa | 3250 | 1924 | 1729 | 1529 | 2559 | 2562 |
| ENSDARG000 | 17;17;17;17;1 | 53304723:533 | 53305087:533 | -;-;-;-;-;-;-;-; | 1473 | NA | 12 | 15 | 53 | 33 | 29 | 16 |
| ENSDARG000 | 17;17;17;17;1 | 9912036:9913 | 9912952:9913 | -;-;-;-;-;-;- | 2315 | egln3 | 681 | 3251 | 8631 | 9769 | 2672 | 630 |
| ENSDARG000 | 17;17;17;17;1 | 25188694:251 | 25189125:251 | -;-;-;-;-;- | 1058 | NA | 0 | 0 | 0 | 0 | 0 | 0 |
| ENSDARG000 | 17;17;17;17;1 | 25187222:251 | 25187342:251 | +;+;+;+;+;+;+; | 1856 | cln8 | 231 | 266 | 264 | 109 | 279 | 242 |
| ENSDARG000 | 17;17;17;17;1 | 25197180:251 | 25197324:251 | +;+;+;+;+;+;+; | 4102 | ARHGEF10 | 260 | 229 | 257 | 335 | 183 | 182 |
| ENSDARG000 | 17;17;17;17;1 | 19419989:194 | 19423969:194 | -;-;-;-;-;-;-;-; | 9940 | dicer1 | 983 | 739 | 1201 | 720 | 1022 | 881 |
| ENSDARG000 | 17;17 | 8175998:8178 | 8176097:8179 | +;+ | 811 | NA | 4 | 0 | 10 | 0 | 0 | 4 |
| ENSDARG000 | 17;17;17;17;1 | 44005888:440 | 44007361:440 | -;-;-;-;-;-;-;-; | 1690 | NA | 124 | 117 | 226 | 122 | 174 | 174 |
| ENSDARG000 | 17 | 36588281 | 36589605 | + | 1325 | htr1b | 0 | 0 | 0 | 0 | 0 | 0 |
| ENSDARG000 | 17;17;17;17;1 | 22311413:223 | 22311580:223 | +;+;+;+;+;+;+; | 4357 | slc8a1b | 4 | 4 | 8 | 14 | 2 | 12 |
| ENSDARG000 | 17;17 | 9944495:9950 | 9945358:9950 | -;- | 1589 | sptssa | 520 | 687 | 609 | 453 | 502 | 616 |
| ENSDARG000 | 17;17;17;17;1 | 6438051:6440 | 6438680:6441 | -;-;-;-;-;-;- | 1983 | fuca2 | 2957 | 1724 | 2102 | 2793 | 3025 | 2808 |
| ENSDARG000 | 17;17;17;17;1 | 26543690:265 | 26544170:265 | +;+;+;+;+;+;+; | 1267 | NA | 2 | 0 | 0 | 0 | 2 | 0 |
| ENSDARG000 | 17;17;17;17;1 | 44030692:440 | 44030985:440 | +;+;+;+;+;+ | 2448 | peli2 | 2111 | 892 | 1366 | 1259 | 2649 | 1982 |
| ENSDARG000 | 17;17;17;17;1 | 8183393:8184 | 8183578:8184 | +;+;+;+;+;+;+; | 7649 | TULP4 | 98 | 96 | 248 | 66 | 125 | 158 |
| ENSDARG000 | 17;17;17;17;1 | 24109012:241 | 24109546:241 | +;+;+;+;+;+;+; | 6249 | EHBP1 | 1553 | 932 | 1348 | 1060 | 1243 | 1093 |
| ENSDARG000 | 17;17 | 36673484:366 | 36673593:366 | -;- | 557 | NA | 0 | 0 | 0 | 0 | 0 | 0 |
| ENSDARG000 | 17;17;17;17;1 | 36627099:366 | 36627608:366 | +;+;+;+;+;+;+; | 3833 | NA | 82 | 0 | 0 | 0 | 0 | 0 |
| ENSDARG000 | 17;17;17;17 | 26569601:265 | 26569793:265 | +;+;+;+ | 2547 | NA | 6 | 0 | 4 | 13 | 8 | 6 |
| ENSDARG000 | 17;17;17;17;1 | 28340138:284 | 28340948:284 | +;+;+;+;+;+;+; | 4214 | MDGA2 | 2 | 10 | 12 | 14 | 23 | 4 |
| ENSDARG000 | 17;17;17;17 | 15744090:157 | 15744528:157 | -;-;-;- | 2153 | gja10a | 12 | 0 | 0 | 0 | 0 | 0 |
| ENSDARG000 | 17;17;17;17;1 | 38447473:384 | 38447555:384 | +;+;+;+;+;+;+; | 4392 | NA | 70 | 176 | 212 | 76 | 178 | 134 |
| ENSDARG000 | 17;17;17;17;1 | 45648836:456 | 45648986:456 | +;+;+;+;+;+;+; | 2417 | zgc:162184 | 1193 | 246 | 152 | 480 | 253 | 625 |
| ENSDARG000 | 17;17;17;17;1 | 41992054:419 | 41992321:419 | +;+;+;+;+;+;+; | 2597 | kiz | 73 | 102 | 107 | 57 | 85 | 110 |
| ENSDARG000 | 17 | 14711765 | 14712599 | + | 835 | gjb9a | 0 | 0 | 0 | 0 | 0 | 0 |
| ENSDARG000 | 17;17;17;17;1 | 9951618:9955 | 9952261:9955 | -;-;-;-;-;-;-;- | 1912 | eapp | 158 | 216 | 278 | 122 | 237 | 194 |
| ENSDARG000 | 17;17;17;17;1 | 15756828:157 | 15757882:157 | -;-;-;-;-;-;-;-; | 2531 | ANKRD6 | 19 | 13 | 4 | 3 | 2 | 22 |
| ENSDARG000 | 17;17;17;17;1 | 50013594:500 | 50014729:500 | -;-;-;-;-;-;-;- | 6674 | FILIP1 | 4 | 14 | 16 | 2 | 0 | 0 |
| ENSDARG000 | 17;17;17;17;1 | 2479303:2482 | 2479634:2482 | -;-;-;-;-;-;- | 1072 | NA | 4 | 33 | 30 | 18 | 38 | 22 |
| ENSDARG000 | 17;17;17 | 44109203:441 | 44109239:441 | -;-;- | 275 | NA | 0 | 0 | 0 | 1 | 1 | 0 |
| ENSDARG000 | 17;17;17;17;1 | 44115441:441 | 44115707:441 | +;+;+;+;+;+;+; | 6510 | TMEM260 | 607 | 480 | 495 | 300 | 596 | 436 |
| ENSDARG000 | 17;17;17;17;1 | 6467398:6469 | 6467620:6469 | +;+;+;+;+;+;+; | 2218 | COQ8A | 54 | 83 | 111 | 88 | 158 | 96 |
| ENSDARG000 | 17;17;17;17;1 | 32822588:328 | 32823319:328 | -;-;-;-;-;-;-;- | 3008 | prox1a | 38 | 6 | 6 | 0 | 32 | 12 |
| ENSDARG000 | 17;17;17;17;1 | 22530820:225 | 22531123:225 | +;+;+;+;+;+ | 1887 | NA | 12 | 8 | 12 | 8 | 48 | 38 |
| ENSDARG000 | 17;17;17;17;1 | 26611929:266 | 26612158:266 | +;+;+;+;+;+;+; | 6117 | ttc7b | 985 | 937 | 1042 | 849 | 954 | 904 |
| ENSDARG000 | 17;17;17;17;1 | 22544648:225 | 22549615:225 | -;-;-;-;- | 5341 | NA | 202 | 2 | 2 | 34 | 56 | 22 |
| ENSDARG000 | 17 | 14722171 | 14726824 | - | 4654 | NA | 18 | 20 | 20 | 12 | 14 | 4 |
| ENSDARG000 | 17;17;17;17;1 | 45731865:457 | 45733964:457 | -;-;-;-;-;- | 2312 | arf6b | 1438 | 1730 | 1643 | 982 | 1803 | 1109 |
| ENSDARG000 | 17;17;17;17;1 | 20923691:209 | 20923799:209 | +;+;+;+;+;+;+; | 1367 | cdk1 | 634 | 1715 | 1656 | 777 | 1214 | 874 |
| ENSDARG000 | 17;17;17;17;1 | 15788100:157 | 15788458:157 | +;+;+;+;+;+;+ | 2669 | rragd | 136 | 170 | 78 | 79 | 148 | 128 |
| ENSDARG000 | 17;17;17;17;1 | 26589879:265 | 26590722:265 | -;-;-;-;-;-;-;-; | 3189 | fam149b1 | 322 | 178 | 271 | 289 | 296 | 220 |
| ENSDARG000 | 17;17;17;17;1 | 26608510:266 | 26608978:266 | -;-;-;-;-;- | 1394 | mrpl57 | 223 | 325 | 302 | 236 | 280 | 203 |
| ENSDARG000 | 17;17;17;17;1 | 50074372:500 | 50074542:500 | +;+;+;+;+;+;+; | 6808 | vps39 | 4557 | 2450 | 4060 | 4318 | 3657 | 4524 |
| ENSDARG000 | 17 | 31237866 | 31238986 | + | 1121 | prlh2r | 0 | 0 | 0 | 0 | 0 | 2 |
| ENSDARG000 | 17;17;17;17;1 | 24309951:243 | 24311053:243 | -;-;-;-;- | 2709 | NA | 7 | 3 | 31 | 12 | 17 | 31 |
| ENSDARG000 | 17;17;17;17;1 | 12075805:120 | 12075935:120 | +;+;+;+;+;+;+; | 2570 | NA | 301 | 120 | 140 | 88 | 136 | 258 |
| ENSDARG000 | 17;17;17;17;1 | 33158294:331 | 33158609:331 | +;+;+;+;+;+;+; | 3267 | snx9a | 274 | 234 | 241 | 121 | 367 | 325 |
| ENSDARG000 | 17;17;17;17;1 | 38476300:384 | 38476562:384 | +;+;+;+;+;+;+; | 13029 | NA | 420 | 339 | 614 | 753 | 372 | 704 |
| ENSDARG000 | 17;17;17;17;1 | 45737992:457 | 45738377:457 | +;+;+;+;+;+;+; | 3376 | aspg | 271 | 172 | 239 | 114 | 345 | 199 |
| ENSDARG000 | 17;17;17;17;1 | 9963436:9965 | 9965376:9965 | -;-;-;-;-;-;-;-; | 3134 | snx6 | 352 | 459 | 493 | 337 | 575 | 658 |
| ENSDARG000 | 17;17;17;17;1 | 53311243:533 | 53311511:533 | +;+;+;+;+;+;+; | 2933 | asb2b | 4 | 23 | 20 | 12 | 14 | 18 |
| ENSDARG000 | 17;17;17;17;1 | 14731315:147 | 14731831:147 | -;-;-;-;-;-;-;-; | 4895 | NA | 581 | 667 | 1042 | 486 | 846 | 682 |
| ENSDARG000 | 17;17;17;17;1 | 15845765:158 | 15846023:158 | +;+;+;+;+;+;+; | 1693 | gabrr2a | 0 | 0 | 0 | 0 | 0 | 0 |
| ENSDARG000 | 17;17;17;17;1 | 6490089:6491 | 6491947:6491 | -;-;-;-;-;-;-;-; | 3403 | dnajc5gb | 915 | 754 | 562 | 675 | 775 | 805 |
| ENSDARG000 | 17;17;17;17;1 | 31346560:313 | 31347381:313 | -;-;-;-;-;-;-;-; | 2431 | ivd | 871 | 810 | 868 | 795 | 1024 | 1305 |
| ENSDARG000 | 17;17;17 | 15882066:158 | 15882753:158 | -;-;- | 874 | NA | 0 | 0 | 0 | 0 | 1 | 0 |
| ENSDARG000 | 17;17;17;17;1 | 22559055:225 | 22559134:225 | +;+;+;+;+;+ | 628 | NA | 0 | 0 | 0 | 0 | 0 | 0 |
| ENSDARG000 | 17;17;17;17;1 | 15899039:159 | 15899187:159 | +;+;+;+;+;+;+; | 7057 | syne3 | 918 | 793 | 858 | 652 | 961 | 1092 |
| ENSDARG000 | 17;17;17;17;1 | 22563836:225 | 22565962:225 | -;-;-;-;-;-;-;-; | 5079 | exo1 | 31 | 134 | 149 | 32 | 100 | 75 |
| ENSDARG000 | 17;17;17;17;1 | 8211992:8212 | 8212044:8212 | +;+;+;+;+;+;+; | 1534 | slc18b1 | 17 | 6 | 18 | 2 | 20 | 82 |
| ENSDARG000 | 17;17;17;17;1 | 20942058:209 | 20942641:209 | -;-;-;-;-;- | 1335 | TMEM26 | 56 | 1 | 10 | 4 | 10 | 10 |
| ENSDARG000 | 17;17;17;17;1 | 31276437:312 | 31279168:312 | -;-;-;-;-;-;-;-; | 6224 | BAHD1 | 328 | 151 | 376 | 410 | 310 | 428 |
| ENSDARG000 | 17;17;17;17;1 | 19499157:195 | 19499371:195 | +;+;+;+;+;+;+; | 2736 | slc22a15 | 148 | 130 | 102 | 92 | 172 | 210 |
| ENSDARG000 | 17;17;17;17;1 | 2549503:2550 | 2549529:2550 | +;+;+;+;+;+;+; | 394 | NA | 360 | 6 | 6 | 10 | 12 | 2 |
| ENSDARG000 | 17;17;17;17;1 | 29797876:297 | 29798460:297 | -;-;-;-;-;-;-;-; | 1991 | esrrga | 0 | 0 | 4 | 0 | 10 | 2 |
| ENSDARG000 | 17;17;17;17;1 | 19481049:194 | 19481189:194 | +;+;+;+;+;+;+; | 3689 | kif11 | 899 | 1889 | 1804 | 1054 | 1506 | 739 |
| ENSDARG000 | 17;17;17 | 44249608:442 | 44250576:442 | +;+;+ | 1571 | NA | 0 | 2 | 4 | 6 | 18 | 14 |
| ENSDARG000 | 17;17;17;17;1 | 44244659:442 | 44246364:442 | -;-;-;-;-;-;-;-; | 2452 | otx2b | 24 | 0 | 4 | 0 | 12 | 0 |
| ENSDARG000 | 17;17;17;17 | 42095247:420 | 42096378:420 | -;-;-;- | 1684 | nkx2.4a | 0 | 0 | 0 | 0 | 0 | 0 |
| ENSDARG000 | 17;17 | 25289431:252 | 25291530:252 | +;+ | 2100 | KBTBD11 | 137 | 56 | 48 | 36 | 56 | 61 |
| ENSDARG000 | 17;17;17;17;1 | 12138715:121 | 12138953:121 | +;+;+;+;+;+;+; | 1859 | sccpdha.2 | 1728 | 97 | 66 | 126 | 271 | 515 |
| ENSDARG000 | 17;17;17;17;1 | 45815353:458 | 45815779:458 | +;+;+;+;+;+;+; | 9527 | KIF26A | 136 | 52 | 136 | 38 | 278 | 18 |
| ENSDARG000 | 17;17;17 | 42469888:424 | 42469922:424 | +;+;+ | 246 | NA | 0 | 0 | 0 | 0 | 0 | 0 |
| ENSDARG000 | 17;17 | 42561960:425 | 42562631:425 | -;- | 718 | linc.alien | 0 | 0 | 0 | 0 | 0 | 0 |
| ENSDARG000 | 17;17;17;17;1 | 36687094:366 | 36687372:366 | -;-;-;-;-;-;-;-; | 4325 | myo6b | 22 | 24 | 71 | 0 | 41 | 41 |
| ENSDARG000 | 17;17;17;17;1 | 20963477:209 | 20965311:209 | -;-;-;-;-;-;-;- | 3459 | phyhipla | 18 | 0 | 0 | 8 | 0 | 6 |
| ENSDARG000 | 17;17;17;17;1 | 30205258:302 | 30205459:302 | +;+;+;+;+;+;+; | 1255 | spata17 | 0 | 0 | 0 | 0 | 0 | 0 |
| ENSDARG000 | 17;17 | 42565998:425 | 42567458:425 | -;- | 1772 | foxa2 | 0 | 0 | 0 | 0 | 0 | 0 |
| ENSDARG000 | 17;17;17 | 33226955:332 | 33227191:332 | +;+;+ | 1224 | pomca | 0 | 0 | 0 | 0 | 0 | 0 |
| ENSDARG000 | 17;17;17;17;1 | 30140099:301 | 30140417:301 | -;-;-;-;-;-;- | 1486 | NA | 0 | 0 | 0 | 0 | 0 | 0 |
| ENSDARG000 | 17;17;17;17;1 | 33206780:332 | 33207299:332 | -;-;-;-;-;-;- | 4462 | NA | 140 | 99 | 93 | 64 | 226 | 72 |
| ENSDARG000 | 17;17;17;17 | 10007071:100 | 10009380:100 | -;-;-;- | 2768 | cfl2 | 1232 | 1685 | 1679 | 1348 | 1870 | 1128 |
| ENSDARG000 | 17;17;17;17;1 | 24318753:243 | 24319099:243 | +;+;+;+;+;+;+ | 2513 | otx1 | 8 | 16 | 10 | 4 | 12 | 8 |
| ENSDARG000 | 17;17;17;17;1 | 8229868:8229 | 8230295:8230 | -;-;-;-;-;-;-;-; | 4133 | ahi1 | 161 | 90 | 205 | 62 | 129 | 159 |
| ENSDARG000 | 17;17;17;17;1 | 25326054:253 | 25326449:253 | +;+;+;+;+;+;+; | 2247 | tmem54a | 89 | 50 | 72 | 62 | 68 | 78 |
| ENSDARG000 | 17;17;17;17;1 | 2569521:2569 | 2569608:2569 | -;-;-;-;-;-;-;-; | 4807 | zgc:173556 | 0 | 3 | 0 | 0 | 6 | 0 |
| ENSDARG000 | 17;17;17;17;1 | 44413770:444 | 44414185:444 | -;-;-;-;-;-;-;-; | 2481 | exoc5 | 446 | 494 | 588 | 159 | 519 | 336 |
| ENSDARG000 | 17;17;17;17;1 | 46387086:463 | 46387562:463 | +;+;+;+;+;+;+; | 1078 | NA | 824 | 1256 | 1397 | 525 | 3944 | 933 |
| ENSDARG000 | 17;17;17;17;1 | 42610094:426 | 42615092:426 | -;-;-;-;-;-;-;-; | 8283 | PRKD3 | 2867 | 1694 | 1800 | 2627 | 2598 | 2591 |
| ENSDARG000 | 17;17;17;17;1 | 44355293:444 | 44355586:444 | -;-;-;-;- | 761 | NA | 71 | 0 | 5 | 4 | 14 | 0 |
| ENSDARG000 | 17;17;17;17;1 | 53324006:533 | 53324276:533 | -;-;-;-;-;-;-;-; | 1254 | exd1 | 0 | 4 | 6 | 10 | 0 | 0 |
| ENSDARG000 | 17;17;17;17;1 | 25328351:253 | 25328459:253 | -;-;-;-;-;-;-;-; | 1935 | zpcx | 4 | 2 | 17 | 6 | 29 | 16 |

| | | | | | | | | | | | | |
|---|---|---|---|---|---|---|---|---|---|---|---|---|
| ENSDARG000 | 17:17:17:17:1 | 10037543:10 | 10037764:10 | -:-:-:-:-:-:-:-: | 3929 | NA | 371 | 419 | 687 | 201 | 494 | 387 |
| ENSDARG000 | 17:17:17:17:1 | 50127648:50 | 50127969:50 | +:+:+:+:+:+:+: | 2677 | GALNT16 | 0 | 2 | 0 | 2 | 2 | 9 |
| ENSDARG000 | 17:17:17:17:1 | 10031791:10 | 10032824:10 | -:-:-:-:-:-:-:-: | 2609 | NA | 297 | 421 | 530 | 326 | 502 | 349 |
| ENSDARG000 | 17;17 | 50122243:50 | 50122304:50 | +;+ | 552 | NA | 5 | 6 | 0 | 4 | 0 | 1 |
| ENSDARG000 | 17:17:17:17:1 | 25303671:25 | 25305973:25 | -:-:-:-:-:-:-:-: | 4609 | pabpc4 | 338 | 301 | 696 | 333 | 269 | 362 |
| ENSDARG000 | 17:17:17:17:1 | 38566717:38 | 38566920:38 | +:+:+:+:+:+:+: | 7639 | sptb | 23 | 12 | 114 | 21 | 341 | 88 |
| ENSDARG000 | 17:17:17:17:1 | 22577472:22 | 22577809:22 | +:+:+:+:+:+:+: | 1944 | yipf4 | 1142 | 1164 | 1371 | 1390 | 1231 | 1467 |
| ENSDARG000 | 17:17:17:17:1 | 14784258:14 | 14784439:14 | +:+:+:+:+:+:+: | 2301 | rtraf | 659 | 885 | 1014 | 534 | 834 | 700 |
| ENSDARG000 | 17:17:17:17:1 | 33232748:33 | 33233761:33 | -:-:-:-:-:-:-:-: | 4383 | EFR3B | 189 | 71 | 163 | 132 | 291 | 91 |
| ENSDARG000 | 17:17:17:17:1 | 6563307:657 | 6563428:657 | +:+:+:+:+:+:+: | 3827 | NA | 2 | 0 | 0 | 8 | 6 | 0 |
| ENSDARG000 | 17:17:17:17:1 | 12159529:12 | 12159787:12 | +:+:+:+:+:+:+: | 1827 | sccpdha.1 | 3381 | 495 | 480 | 1162 | 1128 | 1552 |
| ENSDARG000 | 17:17:17:17:1 | 6535694:653 | 6536316:653 | +:+:+:+:+:+:+: | 5235 | slc5a6b | 36 | 40 | 83 | 38 | 45 | 39 |
| ENSDARG000 | 17:17:17:17:1 | 24439532:24 | 24439668:24 | +:+:+:+:+:+:+: | 1724 | mdh1ab | 7 | 0 | 12 | 2 | 14 | 14 |
| ENSDARG000 | 17:17:17:17:1 | 24331048:24 | 24331105:24 | -:-:-:-:-:-:-:-: | 3210 | WDPCP | 44 | 44 | 51 | 57 | 112 | 30 |
| ENSDARG000 | 17:17:17:17:1 | 6528067:652 | 6528391:652 | -:-:-:-:-:-:-:-: | 1947 | cenpo | 7 | 48 | 59 | 53 | 26 | 11 |
| ENSDARG000 | 17:17:17:17:1 | 53418445:53 | 53418633:53 | +:+:+:+:+:+:+: | 2234 | slc9a1b | 20 | 0 | 11 | 10 | 49 | 45 |
| ENSDARG000 | 17;17;17;17 | 28607417:28 | 28607981:28 | -;-;-;- | 3728 | NA | 116 | 149 | 485 | 133 | 278 | 260 |
| ENSDARG000 | 17:17:17:17:1 | 28611746:28 | 28611825:28 | +:+:+:+:+:+:+: | 4034 | mis18bp1 | 71 | 266 | 301 | 150 | 161 | 117 |
| ENSDARG000 | 17:17:17:17:1 | 6553795:659 | 6554022:659 | -:-:-:-:-:-:-:-: | 1165 | ANKRD66 | 0 | 0 | 0 | 0 | 0 | 0 |
| ENSDARG000 | 17;17;17 | 19513571:19 | 19513745:19 | +;+;+ | 497 | NA | 0 | 0 | 0 | 0 | 0 | 0 |
| ENSDARG000 | 17:17:17:17:1 | 19522856:19 | 19525391:19 | -:-:-:-:-:-:-:-: | 4032 | cyp26c1 | 2 | 0 | 0 | 0 | 10 | 3 |
| ENSDARG000 | 17:17:17:17:1 | 12177901:12 | 12178490:12 | -:-:-:-:-:-:-:-: | 2062 | NA | 4 | 0 | 0 | 0 | 0 | 4 |
| ENSDARG000 | 17;17;17 | 30307454:30 | 30307804:30 | -;-;- | 530 | NA | 0 | 0 | 0 | 0 | 0 | 0 |
| ENSDARG000 | 17:17:17:17:1 | 53425037:53 | 53425444:53 | +;+;+;+;+;+;+ | 963 | NA | 5 | 8 | 7 | 2 | 8 | 13 |
| ENSDARG000 | 17:17:17:17:1 | 26713111:26 | 26714164:26 | -:-:-:-:-:-:-:-: | 1833 | calm1a | 1410 | 660 | 845 | 1299 | 1027 | 796 |
| ENSDARG000 | 17:17:17:17:1 | 12196980:12 | 12197038:12 | -:-:-:-:-:-:-:-: | 1410 | NA | 0 | 0 | 5 | 0 | 0 | 0 |
| ENSDARG000 | 17:17:17:17:1 | 26718648:26 | 26721090:26 | +:+:+:+:+:+:+: | 6775 | NRDE2 | 164 | 93 | 136 | 71 | 194 | 138 |
| ENSDARG000 | 17;17;17;17 | 50195267:50 | 50195753:50 | -;-;-;- | 1371 | JDP2 | 95 | 65 | 56 | 43 | 70 | 84 |
| ENSDARG000 | 17:17:17:17:1 | 42812265:42 | 42812488:42 | -:-:-:-:-:-:-:-: | 1726 | mocs3 | 363 | 174 | 311 | 288 | 285 | 358 |
| ENSDARG000 | 17:17:17:17:1 | 42842632:42 | 42842890:42 | +:+:+:+:+:+:+: | 1261 | QPCT | 633 | 14 | 0 | 136 | 108 | 499 |
| ENSDARG000 | 17:17:17:17:1 | 50228998:50 | 50229726:50 | -;-;-;-;-;-;- | 1315 | NA | 249 | 94 | 610 | 808 | 123 | 20 |
| ENSDARG000 | 17:17:17:17:1 | 30369107:30 | 30369521:30 | +:+:+:+:+:+:+: | 7734 | NA | 58 | 50 | 10 | 14 | 0 | 2 |
| ENSDARG000 | 17:17:17:17:1 | 53427931:53 | 53428156:53 | +:+:+:+:+:+:+: | 1216 | srsf10b | 154 | 172 | 289 | 113 | 167 | 133 |
| ENSDARG000 | 17:17:17:17:1 | 31363000:31 | 31363767:31 | +;+;+;+;+;+;+ | 3108 | itpka | 242 | 211 | 274 | 210 | 220 | 362 |
| ENSDARG000 | 17:17:17:17:1 | 53438795:53 | 53438968:53 | -:-:-:-:-:-:-:-: | 869 | mycbp | 76 | 84 | 159 | 81 | 107 | 46 |
| ENSDARG000 | 17:17:17:17:1 | 44441042:44 | 44441321:44 | +:+:+:+:+:+:+: | 1895 | ap5m1 | 320 | 433 | 350 | 173 | 451 | 345 |
| ENSDARG000 | 17:17:17:17:1 | 8270864:827 | 8271085:827 | +:+:+:+:+:+:+: | 6653 | NA | 213 | 72 | 135 | 118 | 141 | 145 |
| ENSDARG000 | 17 | 53435279 | 53437459 | + | 2181 | gja9b | 4 | 12 | 12 | 14 | 6 | 4 |
| ENSDARG000 | 17:17:17:17:1 | 20983798:20 | 20986349:20 | -:-:-:-:-:-:-:-: | 5662 | BICC1 | 174 | 28 | 79 | 208 | 90 | 284 |
| ENSDARG000 | 17:17:17:17:1 | 53439606:53 | 53439766:53 | +;+;+;+;+ | 908 | NA | 4 | 7 | 1 | 0 | 0 | 2 |
| ENSDARG000 | 17:17:17:17:1 | 53431272:53 | 53431400:53 | +;+;+;+;+ | 750 | NA | 0 | 0 | 0 | 0 | 0 | 0 |
| ENSDARG000 | 17:17:17:17:1 | 8291561:829 | 8292575:829 | +:+:+:+:+:+:+: | 5628 | NA | 113 | 0 | 0 | 120 | 94 | 4 |
| ENSDARG000 | 17;17;17 | 50244443:50 | 50245780:50 | -;-;- | 1569 | acyp1 | 190 | 170 | 178 | 178 | 130 | 212 |
| ENSDARG000 | 17:17:17:17:1 | 24445818:24 | 24445898:24 | +:+:+:+:+:+:+: | 7012 | ugp2b | 4046 | 4538 | 4418 | 5800 | 4997 | 4572 |
| ENSDARG000 | 17;17;17;17 | 8308182:831 | 8308705:831 | -;-;-;- | 1318 | lft2 | 10 | 6 | 0 | 0 | 0 | 0 |
| ENSDARG000 | 17:17:17:17:1 | 12220781:12 | 12222340:12 | -:-:-:-:-:-:-:-: | 9380 | AHCTF1 | 2128 | 2380 | 2819 | 1230 | 2717 | 2209 |
| ENSDARG000 | 17:17:17:17:1 | 38602602:38 | 38603327:38 | +:+:+:+:+:+:+: | 9374 | ccdc88c | 1499 | 158 | 573 | 706 | 1922 | 489 |
| ENSDARG000 | 17:17:17:17:1 | 15983247:15 | 15983641:15 | +:+:+:+:+:+:+: | 3525 | NA | 92 | 105 | 243 | 0 | 56 | 83 |
| ENSDARG000 | 17;17 | 14835107:14 | 14835130:14 | +;+ | 214 | NA | 0 | 0 | 0 | 0 | 0 | 0 |
| ENSDARG000 | 17:17:17:17:1 | 14796926:14 | 14797397:14 | -:-:-:-:-:-:-:-: | 5686 | NID2 | 1452 | 1079 | 2234 | 1457 | 2022 | 1119 |
| ENSDARG000 | 17:17:17:17:1 | 33296436:33 | 33297000:33 | +:+:+:+:+:+:+: | 6000 | dnajc27 | 432 | 305 | 387 | 219 | 390 | 368 |
| ENSDARG000 | 17:17:17:17:1 | 19607387:19 | 19610275:19 | -;-;-;-;-;-;- | 3865 | reep3a | 198 | 12 | 235 | 62 | 146 | 162 |
| ENSDARG000 | 17:17:17:17:1 | 10062045:10 | 10062635:10 | -:-:-:-:-:-:-:-: | 1877 | zgc:109986 | 155 | 168 | 372 | 332 | 351 | 299 |
| ENSDARG000 | 17:17:17:17:1 | 46739693:46 | 46739777:46 | +;+;+;+;+ | 784 | NA | 0 | 0 | 0 | 0 | 0 | 0 |
| ENSDARG000 | 17:17:17:17:1 | 2684992:269 | 2685026:269 | -:-:-:-:-:-:-:-: | 6146 | NA | 48 | 76 | 60 | 22 | 242 | 40 |
| ENSDARG000 | 17:17:17:17:1 | 2606697:261 | 2610829:261 | -:-:-:-:-:-:-:-: | 7905 | PTPN21 | 841 | 462 | 770 | 587 | 726 | 675 |
| ENSDARG000 | 17:17:17:17:1 | 22687170:22 | 22688133:22 | -;-;-;-;- | 1257 | birc6-as2 | 0 | 2 | 18 | 72 | 0 | 2 |
| ENSDARG000 | 17:17:17:17:1 | 22587356:22 | 22587787:22 | +:+:+:+:+:+:+: | 15893 | BIRC6 | 3615 | 2331 | 4553 | 2302 | 4417 | 2707 |
| ENSDARG000 | 17:17:17:17:1 | 8323348:832 | 8324522:832 | +:+:+:+:+:+:+: | 6093 | CDC42BPA | 218 | 145 | 363 | 140 | 282 | 238 |
| ENSDARG000 | 17:17:17:17:1 | 46815648:46 | 46815733:46 | -;-;-;-;-;- | 991 | NA | 0 | 0 | 0 | 0 | 0 | 0 |
| ENSDARG000 | 17:17:17:17:1 | 31400584:31 | 31401678:31 | -:-:-:-:-:-:-:-: | 4775 | ltk | 240 | 65 | 2184 | 72 | 517 | 944 |
| ENSDARG000 | 17;17;17 | 50248152:50 | 50248223:50 | +;+;+ | 1902 | NA | 12 | 5 | 6 | 16 | 4 | 2 |
| ENSDARG000 | 17;17;17;17 | 44463230:44 | 44464158:44 | +;+;+;+ | 4019 | naa30 | 712 | 699 | 899 | 534 | 927 | 794 |
| ENSDARG000 | 17:17:17:17:1 | 21051022:21 | 21051430:21 | -:-:-:-:-:-:-:-: | 1023 | ube2d1a | 276 | 116 | 204 | 133 | 158 | 156 |
| ENSDARG000 | 17:17:17:17:1 | 12255642:12 | 12256149:12 | +:+:+:+:+:+:+: | 1485 | NA | 0 | 0 | 0 | 0 | 0 | 0 |
| ENSDARG000 | 17:17:17:17:1 | 46818521:46 | 46818813:46 | +;+;+;+;+;+;+ | 1252 | NA | 0 | 0 | 0 | 0 | 0 | 0 |
| ENSDARG000 | 17:17:17:17:1 | 44502707:44 | 44502948:44 | -:-:-:-:-:-:-:-: | 1651 | SLC35F4 | 2 | 10 | 0 | 0 | 34 | 0 |
| ENSDARG000 | 17:17:17:17:1 | 12265095:12 | 12265194:12 | +;+;+;+;+;+;+ | 1016 | NA | 0 | 0 | 0 | 0 | 0 | 0 |
| ENSDARG000 | 17:17:17:17:1 | 42983983:42 | 42984212:42 | -:-:-:-:-:-:-:-: | 1667 | amy2al2 | 0 | 0 | 2 | 4 | 4 | 0 |
| ENSDARG000 | 17:17:17:17:1 | 16051448:16 | 16052622:16 | -:-:-:-:-:-:-:-: | 3588 | NA | 1476 | 1580 | 2100 | 3371 | 2494 | 1798 |
| ENSDARG000 | 17:17:17:17:1 | 16046132:16 | 16046455:16 | +:+:+:+:+:+:+: | 2505 | NA | 44 | 44 | 182 | 206 | 52 | 24 |
| ENSDARG000 | 17:17:17:17:1 | 46827818:46 | 46829173:46 | -;-;-;-;- | 1907 | NA | 4 | 0 | 9 | 16 | 4 | 2 |
| ENSDARG000 | 17:17:17:17:1 | 44697604:44 | 44698091:44 | +;+;+;+;+;+;+ | 1608 | NA | 6 | 55 | 186 | 34 | 10 | 6 |
| ENSDARG000 | 17;17 | 28643458:28 | 28643792:28 | -;- | 522 | NA | 2 | 0 | 0 | 0 | 0 | 0 |
| ENSDARG000 | 17:17:17:17:1 | 28624321:28 | 28624499:28 | +:+:+:+:+:+:+: | 7764 | HEATR5A | 1230 | 614 | 765 | 1064 | 872 | 880 |
| ENSDARG000 | 17:17:17:17:1 | 10074360:10 | 10074417:10 | +:+:+:+:+:+:+: | 2816 | srp54 | 2217 | 1905 | 2766 | 1461 | 2553 | 1761 |
| ENSDARG000 | 17:17:17:17:1 | 46848659:46 | 46848765:46 | -;-;-;-;-;- | 903 | NA | 9 | 0 | 0 | 0 | 0 | 4 |
| ENSDARG000 | 17:17:17:17:1 | 24479880:24 | 24480665:24 | -;-;-;-;-;- | 2052 | peli1b | 133 | 66 | 102 | 91 | 274 | 204 |
| ENSDARG000 | 17:17:17:17:1 | 12274016:12 | 12274115:12 | +;+;+;+;+;+;+ | 1014 | NA | 0 | 0 | 0 | 0 | 0 | 2 |
| ENSDARG000 | 17:17:17:17:1 | 33311784:33 | 33312502:33 | +:+:+:+:+:+:+: | 5566 | NA | 4 | 8 | 16 | 19 | 30 | 8 |
| ENSDARG000 | 17:17:17:17:1 | 8498782:850 | 8499067:850 | -:-:-:-:-:-:-:-: | 3552 | fzd3b | 361 | 198 | 112 | 46 | 62 | 22 |
| ENSDARG000 | 17:17:17:17:1 | 26753967:26 | 26754701:26 | +:+:+:+:+:+:+: | 5153 | NA | 1922 | 2417 | 4021 | 2141 | 2605 | 2548 |
| ENSDARG000 | 17;17 | 26783926:26 | 26784034:26 | -;- | 569 | NA | 0 | 0 | 0 | 0 | 0 | 0 |
| ENSDARG000 | 17:17:17:17:1 | 38738840:38 | 38740870:38 | -:-:-:-:-:-:-:-: | 4025 | DGLUCY | 194 | 232 | 573 | 549 | 710 | 330 |
| ENSDARG000 | 17;17;17;17 | 44756247:44 | 44756408:44 | +;+;+;+ | 1767 | cipca | 363 | 193 | 434 | 240 | 336 | 213 |
| ENSDARG000 | 17:17:17:17:1 | 2727807:273 | 2727879:273 | +:+:+:+:+:+:+: | 1757 | kcnk10b | 0 | 2 | 0 | 0 | 0 | 0 |
| ENSDARG000 | 17:17:17:17:1 | 12285285:12 | 12285387:12 | +;+;+;+;+;+;+ | 1017 | NA | 0 | 0 | 0 | 0 | 0 | 0 |
| ENSDARG000 | 17:17:17:17:1 | 50261603:50 | 50261727:50 | +:+:+:+:+:+:+: | 2415 | syncripl | 1528 | 2459 | 3232 | 975 | 2446 | 1201 |
| ENSDARG000 | 17:17:17:17:1 | 36848786:36 | 36850395:36 | -:-:-:-:-:-:-:-: | 3920 | NA | 0 | 0 | 0 | 0 | 0 | 0 |
| ENSDARG000 | 17;17;17 | 14886828:14 | 14887854:14 | +;+;+ | 2681 | ptger2a | 42 | 8 | 42 | 26 | 39 | 14 |
| ENSDARG000 | 17;17;17 | 44767725:44 | 44769441:44 | -;-;- | 2607 | zdhhc22 | 0 | 2 | 0 | 0 | 0 | 0 |
| ENSDARG000 | 17:17:17:17:1 | 6603691:660 | 6604427:660 | -:-:-:-:-:-:-:-: | 6457 | NA | 2 | 0 | 0 | 0 | 0 | 0 |
| ENSDARG000 | 17;17;17;17 | 46851026:46 | 46851439:46 | +;+;+;+ | 1509 | NA | 0 | 2 | 0 | 6 | 0 | 2 |
| ENSDARG000 | 17:17:17:17:1 | 19626479:19 | 19626554:19 | +:+:+:+:+:+:+: | 6845 | rgs7a | 6 | 0 | 30 | 0 | 8 | 4 |
| ENSDARG000 | 17;17;17 | 2810776:281 | 2812002:281 | -;-;- | 1331 | GPR65 | 4 | 6 | 0 | 2 | 20 | 10 |
| ENSDARG000 | 17 | 2833685 | 2834764 | - | 1080 | bdkrb1 | 0 | 0 | 0 | 0 | 0 | 0 |
| ENSDARG000 | 17:17:17:17:1 | 30469744:30 | 30470441:30 | -;-;-;-;-;- | 1648 | pfn4 | 0 | 0 | 0 | 0 | 0 | 0 |
| ENSDARG000 | 17:17:17:17:1 | 21063403:21 | 21064161:21 | -:-:-:-:-:-:-:-: | 1594 | vsx1 | 0 | 0 | 0 | 0 | 0 | 0 |
| ENSDARG000 | 17:17:17:17:1 | 30442518:30 | 30442873:30 | +:+:+:+:+:+:+: | 5050 | lpin1a | 97 | 48 | 67 | 137 | 55 | 925 |
| ENSDARG000 | 17:17:17:17:1 | 24537185:24 | 24537574:24 | -:-:-:-:-:-:-:-: | 2439 | NA | 2 | 2 | 6 | 4 | 28 | 4 |
| ENSDARG000 | 17;17;17 | 26803470:26 | 26803898:26 | +;+;+ | 2389 | pgrmc2 | 2130 | 2002 | 2026 | 2004 | 1973 | 1857 |
| ENSDARG000 | 17:17:17:17:1 | 12297984:12 | 12298854:12 | -:-:-:-:-:-:-:-: | 1828 | snap25b | 147 | 78 | 30 | 20 | 32 | 240 |
| ENSDARG000 | 17;17 | 19730001:19 | 19731406:19 | +;+ | 2427 | GREM2 | 0 | 0 | 2 | 0 | 0 | 0 |
| ENSDARG000 | 17:17:17:17:1 | 14965570:15 | 14966244:15 | +;+;+;+;+;+;+ | 1775 | gpr137c | 1172 | 556 | 824 | 1202 | 1130 | 648 |
| ENSDARG000 | 17:17:17:17:1 | 46864116:46 | 46864478:46 | +;+;+;+;+;+;+ | 1328 | NA | 0 | 0 | 0 | 0 | 0 | 0 |
| ENSDARG000 | 17:17:17:17:1 | 14918101:14 | 14919367:14 | -:-:-:-:-:-:-:-: | 5452 | TXNDC16 | 286 | 278 | 471 | 322 | 439 | 359 |
| ENSDARG000 | 17:17:17:17:1 | 44780166:44 | 44780424:44 | +:+:+:+:+:+:+: | 4553 | tmem63c | 0 | 0 | 0 | 0 | 0 | 0 |
| ENSDARG000 | 17:17:17:17:1 | 6667441:666 | 6668071:666 | +:+:+:+:+:+:+: | 4098 | NA | 43 | 15 | 14 | 14 | 60 | 49 |
| ENSDARG000 | 17:17:17:17:1 | 50289039:50 | 50289983:50 | -:-:-:-:-:-:-:-: | 3862 | NA | 19 | 0 | 10 | 4 | 0 | 0 |
| ENSDARG000 | 17;17;17 | 46885146:46 | 46885251:46 | -;-;- | 243 | NA | 0 | 0 | 0 | 0 | 0 | 0 |
| ENSDARG000 | 17:17:17:17:1 | 3124129:312 | 3124217:312 | +:+:+:+:+:+:+: | 2550 | zgc:136872 | 4 | 0 | 0 | 0 | 0 | 0 |
| ENSDARG000 | 17:17:17:17:1 | 6715575:671 | 6716558:671 | -:-:-:-:-:-:-:-: | 2398 | vsnl1b | 0 | 0 | 4 | 0 | 11 | 14 |
| ENSDARG000 | 17;17 | 46911890:46 | 46912126:46 | +;+ | 368 | NA | 0 | 0 | 0 | 0 | 0 | 0 |
| ENSDARG000 | 17:17:17:17:1 | 46931919:46 | 46932007:46 | -;-;-;-;-;- | 992 | NA | 0 | 0 | 0 | 0 | 0 | 0 |
| ENSDARG000 | 17:17:17:17:1 | 43005428:43 | 43005810:43 | -;-;-;-;-;- | 1558 | NA | 28 | 34 | 56 | 38 | 54 | 28 |
| ENSDARG000 | 17:17:17:17:1 | 21126495:21 | 21126820:21 | +:+:+:+:+:+:+: | 2860 | pygb | 1726 | 1139 | 696 | 925 | 1104 | 1072 |
| ENSDARG000 | 17:17:17:17:1 | 26815021:26 | 26815168:26 | +:+:+:+:+:+:+: | 1685 | asmt2 | 382 | 180 | 142 | 393 | 285 | 207 |
| ENSDARG000 | 17:17:17:17:1 | 33340675:33 | 33340952:33 | +:+:+:+:+:+:+: | 5906 | XDH | 2 | 16 | 106 | 8 | 28 | 36 |
| ENSDARG000 | 17:17:17:17:1 | 12408188:12 | 12408551:12 | +:+:+:+:+:+:+: | 2267 | khk | 133 | 133 | 170 | 213 | 197 | 225 |
| ENSDARG000 | 17:17:17:17:1 | 21102301:21 | 21102425:21 | +:+:+:+:+:+:+: | 1677 | entpd6 | 376 | 218 | 304 | 204 | 255 | 301 |
| ENSDARG000 | 17:17:17:17:1 | 25344734:25 | 25345314:25 | -:-:-:-:-:-:-:-: | 2118 | lck | 8 | 54 | 0 | 10 | 16 | 58 |
| ENSDARG000 | 17:17:17:17:1 | 22760126:22 | 22760470:22 | +:+:+:+:+:+:+: | 3939 | ttc27 | 532 | 451 | 793 | 276 | 434 | 362 |
| ENSDARG000 | 17;17;17 | 43013171:43 | 43013244:43 | +;+;+ | 1901 | gskip | 1554 | 1914 | 2191 | 1943 | 1951 | 2022 |
| ENSDARG000 | 17:17:17:17:1 | 12397033:12 | 12397154:12 | -:-:-:-:-:-:-:-: | 2186 | ANKEF1 | 121 | 83 | 11 | 0 | 11 | 171 |
| ENSDARG000 | 17;17;17;17 | 43021662:43 | 43021781:43 | -;-;-;- | 516 | npc2.2 | 52 | 95 | 283 | 56 | 482 | 185 |
| ENSDARG000 | 17:17:17:17:1 | 26850124:26 | 26850624:26 | -:-:-:-:-:-:-:-: | 1620 | si:dkev-221l4. | 0 | 0 | 0 | 0 | 0 | 0 |
| ENSDARG000 | 17;17 | 26848590:26 | 26848999:26 | +;+ | 488 | NA | 0 | 0 | 0 | 0 | 0 | 0 |
| ENSDARG000 | 17:17:17:17:1 | 26828027:26 | 26828128:26 | +:+:+:+:+:+:+: | 7287 | jade1 | 692 | 575 | 844 | 592 | 805 | 752 |
| ENSDARG000 | 17:17:17:17:1 | 43030317:43 | 43030899:43 | -:-:-:-:-:-:-:-: | 1320 | npc2.1 | 2086 | 2525 | 2748 | 2461 | 2794 | 2865 |
| ENSDARG000 | 17:17:17:17:1 | 46935061:46 | 46935079:46 | +;+;+;+;+;+;+ | 984 | NA | 0 | 0 | 0 | 0 | 0 | 0 |
| ENSDARG000 | 17;17;17 | 16110285:16 | 16111498:16 | -;-;- | 1461 | NA | 14 | 6 | 0 | 0 | 2 | 10 |
| ENSDARG000 | 17;17 | 16110177:16 | 16110265:16 | +;+ | 383 | NA | 0 | 0 | 0 | 0 | 3 | 0 |
| ENSDARG000 | 17:17:17:17:1 | 16090160:16 | 16090470:16 | +:+:+:+:+:+:+: | 3020 | znf395a | 1330 | 1732 | 3776 | 5417 | 1584 | 2055 |
| ENSDARG000 | 17:17:17:17:1 | 19753559:19 | 19758200:19 | -:-:-:-:-:-:-:-: | 9725 | NA | 10144 | 6097 | 7442 | 7138 | 9510 | 7827 |
| ENSDARG000 | 17;17;17 | 46989113:46 | 46989341:46 | -;-;- | 590 | NA | 0 | 0 | 0 | 0 | 0 | 0 |
| ENSDARG000 | 17:17:17:17:1 | 36867281:36 | 36867959:36 | -;-;-;-;-;- | 1496 | mapre3a | 42 | 35 | 118 | 21 | 62 | 62 |
| ENSDARG000 | 17:17:17:17:1 | 10090638:10 | 10090747:10 | +:+:+:+:+:+:+: | 5564 | sec23a | 1793 | 1707 | 2570 | 1825 | 2199 | 1758 |
| ENSDARG000 | 17;17 | 3183276:318 | 3184830:319 | -;- | 2095 | TMEM151B | 0 | 0 | 8 | 0 | 0 | 0 |
| ENSDARG000 | 17:17:17:17:1 | 38784362:38 | 38784714:38 | -:-:-:-:-:-:-:-: | 2224 | slc24a4a | 1719 | 948 | 648 | 1058 | 1006 | 1538 |
| ENSDARG000 | 17:17:17:17:1 | 24569988:24 | 24570573:24 | -:-:-:-:-:-:-:-: | 3166 | NA | 247 | 200 | 269 | 170 | 294 | 216 |
| ENSDARG000 | 17:17:17:17:1 | 31535764:31 | 31536050:31 | -:-:-:-:-:-:-:-: | 4546 | RPAP1 | 587 | 468 | 758 | 290 | 629 | 436 |
| ENSDARG000 | 17:17:17:17:1 | 50313650:50 | 50313747:50 | -:-:-:-:-:-:-:-: | 8425 | OTOF | 4 | 4 | 4 | 0 | 0 | 0 |
| ENSDARG000 | 17:17:17:17:1 | 15018219:15 | 15018580:15 | -:-:-:-:-:-:-:-: | 2078 | ero1a | 941 | 3929 | 11950 | 10934 | 3210 | 1097 |
| ENSDARG000 | 17:17:17:17:1 | 47116500:47 | 47116593:47 | +:+:+:+:+:+:+: | 3330 | NA | 148 | 75 | 96 | 38 | 151 | 119 |
| ENSDARG000 | 17:17:17:17:1 | 47238338:47 | 47238703:47 | +;+;+;+;+ | 885 | NA | 0 | 2 | 0 | 0 | 0 | 1 |
| ENSDARG000 | 17:17:17:17:1 | 47090497:47 | 47090549:47 | +:+:+:+:+:+:+: | 1173 | zgc:103755 | 18 | 7 | 8 | 11 | 17 | 4 |
| ENSDARG000 | 17:17:17:17:1 | 43032529:43 | 43032622:43 | +:+:+:+:+:+:+: | 981 | isca2 | 236 | 282 | 283 | 337 | 268 | 285 |
| ENSDARG000 | 17;17;17 | 19831676:19 | 19834599:19 | -;-;- | 3277 | CHRM3 | 0 | 0 | 12 | 0 | 10 | 0 |
| ENSDARG000 | 17:17:17:17:1 | 44961908:44 | 44963998:44 | -;-;-;-;-;-;-;- | 3093 | ngb | 0 | 0 | 2 | 0 | 6 | 0 |
| ENSDARG000 | 17:17:17:17:1 | 47187870:47 | 47194168:47 | -:-:-:-:-:-:-:-: | 8882 | ALK | 36 | 2 | 130 | 0 | 46 | 47 |
| ENSDARG000 | 17:17:17:17:1 | 6765621:676 | 6765765:676 | +:+:+:+:+:+:+: | 1986 | afg1la | 134 | 131 | 203 | 108 | 244 | 108 |
| ENSDARG000 | 17:17:17:17:1 | 8595588:859 | 8598369:859 | -:-:-:-:-:-:-:-: | 8005 | ctbp2a | 118 | 73 | 359 | 33 | 118 | 76 |
| ENSDARG000 | 17:17:17:17:1 | 3379673:339 | 3379778:339 | +:+:+:+:+:+:+: | 4267 | sntg2 | 44 | 78 | 208 | 50 | 391 | 8 |
| ENSDARG000 | 17:17:17:17:1 | 44980837:44 | 44981262:44 | -;-;-;-;-;-;-;- | 2339 | FAM161B | 38 | 70 | 80 | 40 | 86 | 66 |
| ENSDARG000 | 17:17:17:17:1 | 33375469:33 | 33375870:33 | +:+:+:+:+:+:+: | 1701 | zgc:162964 | 60 | 57 | 36 | 32 | 76 | 34 |
| ENSDARG000 | 17:17:17:17:1 | 28670132:28 | 28670486:28 | +:+:+:+:+:+:+: | 9401 | hectd1 | 7276 | 4833 | 7961 | 5925 | 6676 | 5619 |
| ENSDARG000 | 17:17:17:17:1 | 25397070:25 | 25397451:25 | +:+:+:+:+:+:+: | 2379 | zgc:154055 | 54 | 6 | 13 | 18 | 50 | 77 |
| ENSDARG000 | 17:17:17:17:1 | 36898874:36 | 36901305:36 | -:-:-:-:-:-:-:-: | 4529 | dpysl5a | 116 | 358 | 376 | 162 | 182 | 242 |
| ENSDARG000 | 17:17:17:17:1 | 45009868:45 | 45010437:45 | +:+:+:+:+:+:+: | 5629 | coq6 | 657 | 473 | 664 | 521 | 891 | 662 |
| ENSDARG000 | 17:17:17:17:1 | 25392524:25 | 25393105:25 | -;-;-;-;- | 973 | FAM167B | 37 | 58 | 42 | 28 | 7 | 67 |
| ENSDARG000 | 17:17:17:17:1 | 24618640:24 | 24618732:24 | +:+:+:+:+:+:+: | 1014 | commd9 | 278 | 231 | 335 | 156 | 257 | 214 |
| ENSDARG000 | 17;17;17 | 24613255:24 | 24613607:24 | +;+;+ | 819 | atp5if1b | 482 | 558 | 425 | 566 | 494 | 614 |

| | | | | | | | | | | | | |
|---|---|---|---|---|---|---|---|---|---|---|---|---|
| ENSDARG000 | 17:17:17:17:1 | 24615091:246 | 24615214:246 | +:+:+:+:+:+:+: | 2464 | rpl13a | 8319 | 8755 | 15607 | 13743 | 10677 | 13679 |
| ENSDARG000 | 17:17:17:17:1 | 45030407:450 | 45032014:450 | -:-:-:-:-:-:-:-: | 3522 | ENTPD5 | 12 | 2 | 24 | 0 | 24 | 12 |
| ENSDARG000 | 17:17:17:17:1 | 24602633:246 | 24602792:246 | -;-;-;-;-;-;-;- | 457 | DYNLT1 | 0 | 5 | 1 | 2 | 0 | 0 |
| ENSDARG000 | 17:17:17:17:1 | 24604373:246 | 24604719:246 | +:+:+:+:+:+:+: | 2722 | zmpste24 | 1081 | 1235 | 1472 | 662 | 1516 | 1052 |
| ENSDARG000 | 17:17:17:17:1 | 50389823:503 | 50389858:503 | +:+:+:+:+:+:+: | 2603 | NA | 94 | 28 | 106 | 42 | 120 | 80 |
| ENSDARG000 | 17:17:17:17:1 | 24590177:245 | 24590381:245 | +:+:+:+:+:+:+: | 6342 | NA | 1168 | 1204 | 2417 | 1164 | 1847 | 1560 |
| ENSDARG000 | 17:17:17:17:1 | 25414033:254 | 25414335:254 | +:+:+:+:+:+:+: | 3152 | tdh2 | 18 | 0 | 6 | 2 | 6 | 24 |
| ENSDARG000 | 17:17:17:17:1 | 24623561:246 | 24623684:246 | +:+:+:+:+:+:+: | 1528 | NA | 78 | 85 | 83 | 80 | 99 | 109 |
| ENSDARG000 | 17;17;17 | 31580960:315 | 31581271:315 | +;+;+ | 6150 | NA | 2 | 16 | 8 | 0 | 14 | 2 |
| ENSDARG000 | 17:17:17:17:1 | 38927889:389 | 38929469:389 | -:-:-:-:-:-:-:-: | 5458 | crim1 | 254 | 128 | 243 | 168 | 309 | 370 |
| ENSDARG000 | 17;17 | 50433850:504 | 50435047:504 | -;- | 1374 | NA | 24 | 28 | 142 | 71 | 78 | 80 |
| ENSDARG000 | 17:17:17:17:1 | 3841012:3846 | 3841242:3847 | -:-:-:-:-:-:-:-: | 3021 | PLCB1 | 0 | 0 | 0 | 0 | 0 | 0 |
| ENSDARG000 | 17:17:17:17:1 | 50416634:504 | 50417670:504 | -:-:-:-:-:-:-:-: | 2557 | NA | 48 | 39 | 43 | 44 | 104 | 51 |
| ENSDARG000 | 17;17;17 | 50413084:504 | 50417670:504 | -;-;- | 4978 | NA | 57 | 32 | 15 | 11 | 56 | 27 |
| ENSDARG000 | 17;17;17 | 50424407:504 | 50426894:504 | -;-;- | 2924 | NA | 90 | 59 | 355 | 136 | 348 | 375 |
| ENSDARG000 | 17:17:17:17:1 | 31592191:315 | 31592360:315 | +:+:+:+:+:+:+: | 2342 | NA | 6 | 8 | 8 | 4 | 5 | 2 |
| ENSDARG000 | 17:17:17:17:1 | 33388899:333 | 33389879:333 | -:-:-:-:-:-:-:-: | 8507 | ECT2L | 4 | 4 | 0 | 2 | 0 | 0 |
| ENSDARG000 | 17 | 10242166 | 10243888 | + | 1723 | clec14a | 186 | 14 | 44 | 24 | 28 | 52 |
| ENSDARG000 | 17:17:17:17:1 | 26879725:268 | 26882360:268 | -;-;-;-;-;-;-;- | 3564 | rcan3 | 0 | 0 | 1 | 0 | 0 | 4 |
| ENSDARG000 | 17 | 10247563 | 10249095 | - | 1533 | SSTR1 | 0 | 0 | 0 | 0 | 0 | 2 |
| ENSDARG000 | 17:17:17:17:1 | 3993772:3993 | 3993967:3993 | +:+:+:+:+:+:+: | 1883 | tmx4 | 281 | 478 | 336 | 235 | 624 | 356 |
| ENSDARG000 | 17:17:17:17:1 | 33453689:334 | 33454307:334 | +:+:+:+:+:+:+: | 5791 | NA | 113 | 114 | 173 | 71 | 351 | 196 |
| ENSDARG000 | 17:17:17:17:1 | 33433566:334 | 33433703:334 | +:+:+:+:+:+:+: | 1474 | snap23.2 | 76 | 54 | 44 | 17 | 6 | 14 |
| ENSDARG000 | 17:17:17:17:1 | 33409038:334 | 33409673:334 | -:-:-:-:-:-:-:-: | 1869 | ccdc28a | 12 | 4 | 3 | 10 | 15 | 9 |
| ENSDARG000 | 17;17 | 6793001:6806 | 6794145:6810 | +;+ | 5649 | foxo3a | 784 | 308 | 475 | 525 | 587 | 1063 |
| ENSDARG000 | 17:17:17:17:1 | 30473507:304 | 30474672:304 | -:-:-:-:-:-:-:-: | 6492 | itsn2b | 3289 | 2364 | 3759 | 3287 | 2550 | 2350 |
| ENSDARG000 | 17:17:17:17:1 | 10258376:102 | 10258924:102 | -:-:-:-:-:-:-:-: | 6359 | TTC6 | 55 | 17 | 56 | 48 | 135 | 28 |
| ENSDARG000 | 17;17 | 33418262:334 | 33418557:334 | -;- | 393 | NA | 6 | 1 | 1 | 6 | 3 | 12 |
| ENSDARG000 | 17:17:17:17:1 | 33415319:334 | 33415669:334 | +:+:+:+:+:+:+: | 4305 | snap23.1 | 1252 | 913 | 997 | 736 | 1210 | 897 |
| ENSDARG000 | 17:17:17:17:1 | 22956427:229 | 22957132:229 | +:+:+:+:+:+:+: | 4997 | LTBP1 | 474 | 385 | 1249 | 121 | 751 | 234 |
| ENSDARG000 | 17;17;17;17 | 6822337:6824 | 6822634:6824 | +;+;+;+ | 752 | NA | 0 | 0 | 0 | 0 | 0 | 0 |
| ENSDARG000 | 17:17:17:17:1 | 26928418:269 | 26929945:269 | -:-:-:-:-:-:-:-: | 2650 | nipal3 | 926 | 749 | 725 | 396 | 1116 | 905 |
| ENSDARG000 | 17:17:17:17:1 | 43039463:430 | 43040588:430 | -:-:-:-:-:-:-:-: | 7695 | EML5 | 28 | 16 | 74 | 41 | 44 | 30 |
| ENSDARG000 | 17:17:17:17:1 | 36936918:369 | 36941861:369 | -:-:-:-:-:-:-:-: | 8188 | dnmt3ab | 1312 | 1393 | 1238 | 271 | 2286 | 575 |
| ENSDARG000 | 17:17:17:17:1 | 12439966:124 | 12440741:124 | -:-:-:-:-:-:-:-: | 4153 | NA | 20459 | 20724 | 22585 | 16240 | 25192 | 20325 |
| ENSDARG000 | 17:17:17:17:1 | 15033822:150 | 15034472:150 | +:+:+:+:+:+:+: | 8997 | SAMD4A | 733 | 462 | 943 | 861 | 821 | 666 |
| ENSDARG000 | 17:17:17:17:1 | 15157213:151 | 15157324:151 | +:+:+:+:+:+:+: | 976 | NA | 0 | 0 | 2 | 4 | 0 | 1 |
| ENSDARG000 | 17:17:17:17:1 | 15121977:151 | 15122180:151 | -:-:-:-:-:-:-:-: | 5485 | gch1 | 605 | 215 | 2844 | 767 | 1135 | 4937 |
| ENSDARG000 | 17:17:17:17:1 | 31606635:316 | 31606973:316 | -:-:-:-:-:-:-:-: | 1479 | haus2 | 32 | 105 | 119 | 38 | 88 | 68 |
| ENSDARG000 | 17;17 | 15040992:150 | 15041817:150 | -;- | 969 | NA | 0 | 0 | 1 | 0 | 0 | 0 |
| ENSDARG000 | 17:17:17:17:1 | 15148797:151 | 15150341:151 | -:-:-:-:-:-:-:-: | 5073 | wdhd1 | 125 | 423 | 542 | 228 | 471 | 274 |
| ENSDARG000 | 17:17:17:17:1 | 4030493:4037 | 4030657:4037 | +:+:+:+:+:+:+: | 1311 | hao1 | 0 | 0 | 6 | 21 | 32 | 1 |
| ENSDARG000 | 17:17:17:17:1 | 48126944:481 | 48127618:481 | +;+;+;+;+;+;+ | 1933 | slc39a9 | 382 | 428 | 475 | 197 | 519 | 369 |
| ENSDARG000 | 17:17:17:17:1 | 20009132:200 | 20014331:200 | -:-:-:-:-:-:-:-: | 20050 | RYR2 | 4 | 6 | 14 | 0 | 0 | 8 |
| ENSDARG000 | 17:17:17:17:1 | 16192486:162 | 16193622:162 | +:+:+:+:+:+:+: | 10225 | KIF13B | 2109 | 1309 | 2105 | 788 | 2612 | 3037 |
| ENSDARG000 | 17:17:17:17:1 | 50460321:504 | 50462044:504 | -;-;-;-;-;-;- | 2986 | NA | 49 | 65 | 82 | 34 | 51 | 69 |
| ENSDARG000 | 17:17:17:17:1 | 12658411:126 | 12658638:126 | +:+:+:+:+:+:+: | 1849 | gpn1 | 391 | 348 | 492 | 250 | 394 | 343 |
| ENSDARG000 | 17;17 | 10318071:103 | 10318279:103 | +;+ | 1891 | foxa1 | 8 | 10 | 4 | 2 | 50 | 14 |
| ENSDARG000 | 17:17:17:17:1 | 48164536:481 | 48164589:481 | +:+:+:+:+:+:+: | 2003 | PLEKHD1 | 0 | 0 | 0 | 0 | 0 | 0 |
| ENSDARG000 | 17:17:17:17:1 | 4239437:4239 | 4240414:4240 | -;-;-;-;-;-;-;- | 2155 | gdf3 | 6 | 18 | 2 | 12 | 2 | 8 |
| ENSDARG000 | 17;17 | 4227254:4231 | 4228109:4231 | -;- | 1325 | bmp2a | 39 | 32 | 13 | 21 | 10 | 7 |
| ENSDARG000 | 17:17:17:17:1 | 39242437:392 | 39242685:392 | +:+:+:+:+:+:+: | 1833 | zgc:174356 | 2 | 0 | 8 | 8 | 14 | 16 |
| ENSDARG000 | 17:17:17:17:1 | 50478455:504 | 50478983:504 | -;-;-;-;-;-;- | 1745 | NA | 46 | 20 | 31 | 37 | 35 | 31 |
| ENSDARG000 | 17:17:17:17:1 | 6828550:6829 | 6828639:6829 | +:+:+:+:+:+:+: | 6284 | NA | 12 | 6 | 16 | 12 | 8 | 28 |
| ENSDARG000 | 17:17:17:17:1 | 28706946:287 | 28707418:287 | +:+:+:+:+:+:+: | 4060 | strn3 | 1298 | 1401 | 1831 | 1136 | 1794 | 1401 |
| ENSDARG000 | 17:17:17:17:1 | 10325632:103 | 10326386:103 | -:-:-:-:-:-:-:-: | 2364 | mipol1 | 76 | 66 | 60 | 92 | 68 | 36 |
| ENSDARG000 | 17:17:17:17:1 | 28703971:287 | 28704399:287 | -:-:-:-:-:-:-:-: | 939 | ap4s1 | 258 | 263 | 206 | 165 | 353 | 313 |
| ENSDARG000 | 17:17:17:17:1 | 48486308:484 | 48489206:484 | -:-:-:-:-:-:-:-: | 4846 | NA | 5 | 35 | 0 | 4 | 41 | 61 |
| ENSDARG000 | 17:17:17:17:1 | 31611854:316 | 31612403:316 | +;+;+;+;+;+ | 1615 | adprs | 246 | 351 | 387 | 177 | 395 | 241 |
| ENSDARG000 | 17:17:17:17:1 | 21154576:211 | 21158951:211 | -:-:-:-:-:-:-:-: | 6079 | abhd12 | 702 | 580 | 639 | 349 | 1161 | 1142 |
| ENSDARG000 | 17:17:17:17:1 | 6935280:6937 | 6938811:6938 | -:-:-:-:-:-:-:-: | 4832 | zbtb24 | 30 | 16 | 42 | 28 | 47 | 24 |
| ENSDARG000 | 17:17:17:17:1 | 26965351:269 | 26965461:269 | +:+:+:+:+:+:+: | 3019 | NA | 104 | 0 | 2 | 0 | 16 | 4 |
| ENSDARG000 | 17;17;17;17 | 50564070:505 | 50564301:505 | +;+;+;+ | 2991 | bmp4 | 20 | 0 | 4 | 0 | 1 | 0 |
| ENSDARG000 | 17:17:17:17:1 | 6956696:6957 | 6957069:6957 | +:+:+:+:+:+:+: | 3841 | ddo | 80 | 76 | 90 | 53 | 94 | 64 |
| ENSDARG000 | 17:17:17:17:1 | 12698532:126 | 12698600:126 | +:+:+:+:+:+:+: | 3657 | stmn4l | 2 | 0 | 2 | 2 | 2 | 16 |
| ENSDARG000 | 17;17 | 48574770:485 | 48575161:485 | -;- | 523 | NA | 0 | 0 | 0 | 0 | 2 | 0 |
| ENSDARG000 | 17:17:17:17:1 | 21220934:212 | 21221910:212 | -:-:-:-:-:-:-:-: | 3236 | hspa12a | 56 | 107 | 215 | 8 | 28 | 8 |
| ENSDARG000 | 17;17;17 | 12710430:127 | 12711017:127 | -;-;- | 875 | il17a/f1 | 0 | 0 | 0 | 0 | 0 | 0 |
| ENSDARG000 | 17:17:17:17:1 | 50655619:506 | 50656518:506 | +:+:+:+:+:+:+: | 7350 | DDHD1 | 18 | 10 | 50 | 26 | 50 | 8 |
| ENSDARG000 | 17:17:17:17:1 | 48683769:486 | 48684802:486 | -;-;-;-;- | 1949 | kcnk5a | 79 | 10 | 24 | 2 | 14 | 28 |
| ENSDARG000 | 17:17:17:17:1 | 37070096:370 | 37070572:370 | -:-:-:-:-:-:-:-: | 2483 | dtnbb | 20 | 6 | 6 | 0 | 0 | 2 |
| ENSDARG000 | 17:17:17:17:1 | 25426091:254 | 25426155:254 | +:+:+:+:+:+:+: | 4499 | srrm1 | 4633 | 2985 | 6703 | 5934 | 5534 | 6928 |
| ENSDARG000 | 17:17:17:17:1 | 4297076:4297 | 4297265:4298 | -:-:-:-:-:-:-:-: | 2635 | napba | 0 | 6 | 0 | 0 | 0 | 0 |
| ENSDARG000 | 17:17:17:17:1 | 48736160:487 | 48736897:487 | -;-;-;-;- | 1423 | NA | 0 | 0 | 0 | 0 | 0 | 0 |
| ENSDARG000 | 17:17:17:17:1 | 21311401:213 | 21311577:213 | -:-:-:-:-:-:-:-: | 1945 | shtn1 | 14 | 0 | 8 | 0 | 8 | 2 |
| ENSDARG000 | 17:17:17:17:1 | 21295132:212 | 21295399:212 | +:+:+:+:+:+:+: | 2066 | eno4 | 0 | 0 | 0 | 0 | 0 | 0 |
| ENSDARG000 | 17:17:17:17:1 | 50701748:507 | 50702088:507 | +:+:+:+:+:+:+: | 2908 | fermt2 | 1126 | 772 | 1075 | 1495 | 961 | 939 |
| ENSDARG000 | 17:17:17:17:1 | 7380550:7389 | 7381390:7390 | -:-:-:-:-:-:-:-: | 3645 | GRM1 | 0 | 0 | 0 | 0 | 0 | 0 |
| ENSDARG000 | 17:17:17:17:1 | 31621788:316 | 31622239:316 | +:+:+:+:+:+:+: | 1299 | cinp | 132 | 101 | 165 | 64 | 116 | 98 |
| ENSDARG000 | 17:17:17:17:1 | 7471987:7472 | 7472542:7472 | -;-;-;-;-;-;- | 2899 | NA | 2 | 0 | 0 | 0 | 0 | 0 |
| ENSDARG000 | 17:17:17:17:1 | 7440458:7443 | 7444326:7444 | -:-:-:-:-:-:-:-: | 7629 | NA | 6 | 0 | 2 | 0 | 0 | 0 |
| ENSDARG000 | 17:17:17:17:1 | 10501647:105 | 10501900:105 | +:+:+:+:+:+:+: | 3493 | tyro3 | 26 | 6 | 20 | 6 | 4 | 0 |
| ENSDARG000 | 17:17:17:17:1 | 43297456:432 | 43298156:432 | -:-:-:-:-:-:-:-: | 2420 | ITPK1 | 210 | 125 | 122 | 160 | 184 | 207 |
| ENSDARG000 | 17:17:17:17:1 | 39697621:396 | 39697759:396 | +:+:+:+:+:+:+: | 5433 | eprs1 | 4451 | 4986 | 10010 | 6273 | 4575 | 4079 |
| ENSDARG000 | 17:17:17:17:1 | 48745740:487 | 48746257:487 | -:-:-:-:-:-:-:-: | 2918 | kif6 | 0 | 0 | 2 | 6 | 0 | 1 |
| ENSDARG000 | 17:17:17:17:1 | 16298122:163 | 16298580:163 | -;-;-;-;-;-;-;- | 1860 | hmbox1a | 109 | 26 | 86 | 42 | 50 | 126 |
| ENSDARG000 | 17:17:17:17:1 | 37162571:371 | 37164931:371 | -:-:-:-:-:-:-:-: | 5052 | ASXL2 | 276 | 207 | 287 | 137 | 254 | 324 |
| ENSDARG000 | 17:17:17:17:1 | 7489235:7490 | 7490807:7490 | -;-;-;-;-;-;-;- | 4003 | NA | 12 | 6 | 30 | 4 | 36 | 9 |
| ENSDARG000 | 17 | 7504109 | 7505978 | - | 1870 | NA | 7 | 4 | 5 | 6 | 6 | 2 |
| ENSDARG000 | 17:17:17:17:1 | 31623356:316 | 31624921:316 | -;-;-;-;-;-;- | 4066 | NA | 623 | 468 | 772 | 583 | 657 | 568 |
| ENSDARG000 | 17;17 | 16336278:163 | 16337735:163 | -;- | 1989 | kcnk13a | 0 | 0 | 0 | 2 | 2 | 4 |
| ENSDARG000 | 17;17 | 26982035:269 | 26982562:269 | -;- | 1701 | NA | 14 | 10 | 11 | 34 | 14 | 13 |
| ENSDARG000 | 17;17 | 26981132:269 | 26981602:269 | +;+ | 1319 | NA | 1 | 2 | 2 | 0 | 6 | 5 |
| ENSDARG000 | 17:17:17:17:1 | 4325693:4328 | 4325765:4328 | +:+:+:+:+:+:+: | 2954 | MCM8 | 28 | 176 | 164 | 78 | 116 | 88 |
| ENSDARG000 | 17;17;17 | 12730025:127 | 12730178:127 | +;+;+ | 882 | il17a/f2 | 0 | 0 | 0 | 0 | 0 | 0 |
| ENSDARG000 | 17:17:17:17:1 | 7510220:7510 | 7510277:7511 | +:+:+:+:+:+:+: | 1999 | KLHL10 | 0 | 0 | 2 | 0 | 6 | 0 |
| ENSDARG000 | 17:17:17:17:1 | 43410753:434 | 43411134:434 | -:-:-:-:-:-:-:-: | 3297 | plk4 | 72 | 402 | 367 | 176 | 311 | 245 |
| ENSDARG000 | 17:17:17:17:1 | 16348830:163 | 16349149:163 | -:-:-:-:-:-:-:-: | 2349 | TDP1 | 124 | 144 | 215 | 107 | 233 | 208 |
| ENSDARG000 | 17:17:17:17:1 | 26990531:269 | 26990946:269 | -:-:-:-:-:-:-:-: | 2640 | cnksr1 | 1036 | 688 | 397 | 570 | 841 | 736 |
| ENSDARG000 | 17:17:17:17:1 | 51138092:511 | 51138251:511 | -:-:-:-:-:-:-:-: | 3236 | NA | 52 | 5 | 10 | 4 | 30 | 18 |
| ENSDARG000 | 17:17:17:17:1 | 31652640:316 | 31653467:316 | -;-;-;-;- | 1720 | vsx2 | 0 | 16 | 6 | 8 | 0 | 2 |
| ENSDARG000 | 17:17:17:17:1 | 28743661:287 | 28744199:287 | -:-:-:-:-:-:-:-: | 2303 | coch | 57 | 12 | 284 | 300 | 1732 | 489 |
| ENSDARG000 | 17:17:17:17:1 | 30522764:305 | 30523973:305 | +:+:+:+:+:+:+: | 8652 | NA | 190 | 168 | 380 | 195 | 513 | 364 |
| ENSDARG000 | 17:17:17:17:1 | 12737389:127 | 12737652:127 | +:+:+:+:+:+:+: | 1422 | lrrc57 | 311 | 234 | 363 | 207 | 215 | 485 |
| ENSDARG000 | 17:17:17:17:1 | 45136353:451 | 45140515:451 | -:-:-:-:-:-:-:-: | 5677 | tmed8 | 668 | 644 | 524 | 474 | 510 | 668 |
| ENSDARG000 | 17:17:17:17:1 | 37204229:372 | 37204583:372 | -;-;-;-;-;-;-;- | 2814 | kif3cb | 4 | 4 | 0 | 0 | 0 | 6 |
| ENSDARG000 | 17:17:17:17:1 | 48831579:488 | 48831746:488 | +:+:+:+:+:+:+: | 3359 | DAAM2 | 46 | 16 | 30 | 26 | 6 | 22 |
| ENSDARG000 | 17:17:17:17:1 | 45154497:451 | 45154875:451 | -;-;-;-;-;- | 1499 | NOXRED1 | 0 | 0 | 0 | 0 | 0 | 0 |
| ENSDARG000 | 17:17:17:17:1 | 10566490:105 | 10566610:105 | +:+:+:+:+:+:+: | 12354 | mgaa | 1445 | 1031 | 2787 | 2361 | 1907 | 1977 |
| ENSDARG000 | 17 | 51201217 | 51202339 | - | 1123 | NA | 0 | 2 | 0 | 4 | 0 | 0 |
| ENSDARG000 | 17:17:17:17:1 | 31667625:316 | 31668970:316 | -:-:-:-:-:-:-:-: | 1750 | lin52 | 56 | 332 | 190 | 124 | 164 | 110 |
| ENSDARG000 | 17:17:17:17:1 | 20122019:201 | 20122293:201 | -:-:-:-:-:-:-:-: | 2986 | actn2b | 385 | 536 | 956 | 491 | 321 | 1047 |
| ENSDARG000 | 17:17:17:17:1 | 7522777:7524 | 7522810:7524 | +:+:+:+:+:+:+: | 1906 | KLHL10 | 0 | 0 | 0 | 0 | 0 | 0 |
| ENSDARG000 | 17;17;17 | 48943204:489 | 48943590:489 | -;-;- | 540 | NA | 0 | 4 | 0 | 4 | 0 | 0 |
| ENSDARG000 | 17;17;17;17 | 51224421:512 | 51224595:512 | +;+;+;+ | 1382 | adi1 | 846 | 1001 | 1029 | 1085 | 944 | 972 |
| ENSDARG000 | 17:17:17:17:1 | 43430365:434 | 43431274:434 | -:-:-:-:-:-:-:-: | 3476 | HSPA4L | 0 | 0 | 8 | 0 | 4 | 6 |
| ENSDARG000 | 17:17:17:17:1 | 51206225:512 | 51207307:512 | -:-:-:-:-:-:-:-: | 3182 | psen1 | 987 | 1042 | 489 | 688 | 728 | 728 |
| ENSDARG000 | 17:17:17:17:1 | 10593398:105 | 10593929:105 | +:+:+:+:+:+:+: | 5991 | NA | 403 | 192 | 129 | 106 | 191 | 229 |
| ENSDARG000 | 17:17:17:17:1 | 48947762:489 | 48947844:489 | +:+:+:+:+:+:+: | 1719 | hs1bp3 | 894 | 834 | 1222 | 1018 | 1222 | 1216 |
| ENSDARG000 | 17:17:17:17:1 | 12753579:127 | 12754399:127 | -:-:-:-:-:-:-:-: | 2311 | brms1la | 536 | 298 | 385 | 378 | 359 | 665 |
| ENSDARG000 | 17;17 | 37215820:372 | 37216632:372 | +;+ | 1824 | slc30a1b | 398 | 192 | 88 | 36 | 102 | 87 |
| ENSDARG000 | 17:17:17:17:1 | 30591287:305 | 30591715:305 | +:+:+:+:+:+:+: | 6705 | NA | 32 | 206 | 121 | 65 | 890 | 183 |
| ENSDARG000 | 17:17:17:17:1 | 30604819:306 | 30605667:306 | -:-:-:-:-:-:-:-: | 1986 | SH3YL1 | 59 | 53 | 34 | 45 | 67 | 35 |
| ENSDARG000 | 17:17:17:17:1 | 4368859:4372 | 4369439:4372 | +;+;+;+;+;+;+ | 1439 | crls1 | 126 | 238 | 250 | 125 | 192 | 164 |
| ENSDARG000 | 17;17;17 | 45199552:451 | 45199621:451 | +;+;+ | 206 | NA | 0 | 0 | 0 | 0 | 0 | 0 |
| ENSDARG000 | 17:17:17:17:1 | 16423721:164 | 16423887:164 | +:+:+:+:+:+:+: | 1562 | efcab11 | 2 | 10 | 8 | 8 | 6 | 0 |
| ENSDARG000 | 17:17:17:17:1 | 28760586:287 | 28762432:287 | -;-;-;-;-;- | 2699 | opn6b | 8 | 4 | 10 | 6 | 12 | 6 |
| ENSDARG000 | 17:17:17:17:1 | 28771028:287 | 28771235:287 | -:-:-:-:-:-:-:-: | 2124 | scfd1 | 1009 | 934 | 1392 | 835 | 1206 | 968 |
| ENSDARG000 | 17:17:17:17:1 | 16564921:165 | 16565365:165 | +:+:+:+:+:+:+: | 4008 | foxn3 | 32 | 0 | 46 | 0 | 44 | 27 |
| ENSDARG000 | 17:17:17:17:1 | 48964632:489 | 48965425:489 | +:+:+:+:+:+:+: | 5286 | NA | 1178 | 1120 | 1719 | 1008 | 1615 | 1115 |
| ENSDARG000 | 17:17:17:17:1 | 21372291:213 | 21373284:213 | -:-:-:-:-:-:-:-: | 2547 | atp6v1ba | 4987 | 6763 | 5728 | 3895 | 5073 | 3950 |
| ENSDARG000 | 17:17:17:17:1 | 10711641:107 | 10711868:107 | -:-:-:-:-:-:-:-: | 1324 | jmjd7 | 160 | 132 | 203 | 125 | 132 | 117 |
| ENSDARG000 | 17:17:17:17:1 | 45216084:452 | 45216969:452 | -:-:-:-:-:-:-:-: | 4521 | NA | 146 | 71 | 114 | 83 | 110 | 176 |
| ENSDARG000 | 17:17:17:17:1 | 10738126:107 | 10738495:107 | +;+;+;+;+;+;+ | 1581 | tbpl2 | 0 | 0 | 0 | 0 | 0 | 0 |
| ENSDARG000 | 17:17:17:17:1 | 31696657:316 | 31697233:316 | -:-:-:-:-:-:-:-: | 3318 | pomt2 | 445 | 275 | 558 | 238 | 397 | 465 |
| ENSDARG000 | 17:17:17:17:1 | 16755287:167 | 16755325:167 | +:+:+:+:+:+:+: | 7598 | ston2 | 174 | 95 | 118 | 42 | 182 | 32 |
| ENSDARG000 | 17:17:17:17:1 | 43468732:434 | 43468882:434 | +:+:+:+:+:+:+: | 1454 | chmp3 | 366 | 379 | 353 | 404 | 268 | 417 |
| ENSDARG000 | 17:17:17:17:1 | 37253706:372 | 37254029:372 | +:+:+:+:+:+:+: | 2324 | TMEM62 | 94 | 263 | 261 | 100 | 322 | 112 |
| ENSDARG000 | 17:17:17:17:1 | 37227936:372 | 37228039:372 | +:+:+:+:+:+:+: | 3133 | hadhab | 2461 | 2274 | 2514 | 1418 | 2338 | 1764 |
| ENSDARG000 | 17;17;17;17 | 8748655:8749 | 8749239:8749 | -;-;-;- | 946 | tex36 | 0 | 0 | 0 | 0 | 0 | 0 |
| ENSDARG000 | 17:17:17:17:1 | 7534180:7534 | 7534492:7534 | +:+:+:+:+:+:+: | 6056 | SHPRH | 352 | 205 | 263 | 218 | 320 | 288 |
| ENSDARG000 | 17:17:17:17:1 | 51231210:512 | 51232848:512 | -:-:-:-:-:-:-:-: | 3767 | trappc12 | 373 | 254 | 328 | 164 | 338 | 326 |
| ENSDARG000 | 17:17:17:17:1 | 4386649:4386 | 4386793:4386 | -:-:-:-:-:-:-:-: | 1881 | KLHL10 | 0 | 0 | 2 | 0 | 0 | 2 |
| ENSDARG000 | 17:17:17:17:1 | 43486436:434 | 43486467:434 | +:+:+:+:+:+:+: | 1687 | NA | 99 | 2 | 34 | 58 | 18 | 188 |
| ENSDARG000 | 17:17:17:17:1 | 39741926:397 | 39742060:397 | +:+:+:+:+:+:+: | 1187 | NA | 4 | 0 | 6 | 0 | 0 | 6 |
| ENSDARG000 | 17;17;17;17 | 43512876:435 | 43513183:435 | -;-;-;- | 686 | mrpl35 | 236 | 295 | 312 | 198 | 187 | 169 |
| ENSDARG000 | 17:17:17:17:1 | 12767640:127 | 12767808:127 | +:+:+:+:+:+:+: | 8101 | RALGAPA1 | 1187 | 785 | 1307 | 787 | 1206 | 989 |
| ENSDARG000 | 17:17:17:17:1 | 20144626:201 | 20147999:201 | -:-:-:-:-:-:-:-: | 5249 | p4ha1b | 662 | 1837 | 6588 | 4322 | 1859 | 804 |
| ENSDARG000 | 17;17 | 16873417:168 | 16873904:168 | +;+ | 2166 | dio2 | 0 | 2 | 0 | 8 | 6 | 0 |
| ENSDARG000 | 17 | 39760145 | 39761086 | - | 942 | NA | 0 | 0 | 0 | 0 | 4 | 0 |
| ENSDARG000 | 17:17:17:17:1 | 25444323:254 | 25444774:254 | +:+:+:+:+:+:+: | 3222 | clic4 | 3166 | 1614 | 1861 | 4358 | 1648 | 1620 |
| ENSDARG000 | 17:17:17:17:1 | 45259259:452 | 45259333:452 | +:+:+:+:+:+:+: | 3189 | GANC | 323 | 146 | 220 | 186 | 225 | 182 |
| ENSDARG000 | 17:17:17:17:1 | 27064158:270 | 27064964:270 | +:+:+:+:+:+:+: | 6756 | rps6ka1 | 636 | 397 | 276 | 314 | 620 | 626 |
| ENSDARG000 | 17:17:17:17:1 | 49081828:490 | 49082436:490 | +:+:+:+:+:+:+: | 7160 | TIAM2 | 1435 | 752 | 762 | 969 | 444 | 476 |
| ENSDARG000 | 17:17:17:17:1 | 33495194:334 | 33495456:334 | +;+;+;+;+;+ | 1173 | pth2 | 0 | 0 | 0 | 0 | 0 | 0 |
| ENSDARG000 | 17:17:17:17:1 | 43519290:435 | 43520306:435 | -:-:-:-:-:-:-:-: | 2471 | NA | 19404 | 13998 | 12213 | 12779 | 16808 | 18505 |
| ENSDARG000 | 17:17:17:17:1 | 39768680:397 | 39769290:397 | -;-;-;-;-;-;- | 1844 | NA | 0 | 0 | 0 | 1 | 4 | 0 |
| ENSDARG000 | 17;17;17 | 21432693:214 | 21434610:214 | -;-;- | 2555 | vax1 | 0 | 0 | 0 | 0 | 0 | 0 |
| ENSDARG000 | 17;17;17 | 23298928:233 | 23298968:233 | +;+;+ | 1536 | NA | 4 | 0 | 0 | 0 | 0 | 0 |
| ENSDARG000 | 17;17 | 33501766:335 | 33504807:335 | -;- | 3281 | TMEM121 | 0 | 6 | 0 | 12 | 2 | 2 |
| ENSDARG000 | 17;17 | 15196727:152 | 15198439:152 | -;- | 1943 | socs4 | 0 | 8 | 12 | 18 | 2 | 6 |
| ENSDARG000 | 17:17:17:17:1 | 51627209:516 | 51627311:516 | +;+;+;+;+;+ | 2138 | zgc:113142 | 10 | 0 | 184 | 2 | 98 | 0 |

| | | | | | | | | | | | | |
|---|---|---|---|---|---|---|---|---|---|---|---|---|
| ENSDARG000 | 17:17:17:17:1 | 4398102:440 | 4398319:440 | +;+;+;+;+;+;+; | 8240 | NA | 30 | 33 | 28 | 19 | 34 | 24 |
| ENSDARG000 | 17;17 | 23311377:233 | 23311565:233 | +;+ | 2170 | ppp1r3ca | 0 | 4 | 8 | 6 | 0 | 0 |
| ENSDARG000 | 17;17;17;17 | 8854108:885 | 8854490:885 | -;-;-;- | 612 | nkl.4 | 2 | 13 | 0 | 1 | 25 | 58 |
| ENSDARG000 | 17:17:17:17:1 | 39776296:397 | 39776449:397 | -;-;-;-;-;-;- | 1135 | NA | 0 | 0 | 0 | 0 | 0 | 0 |
| ENSDARG000 | 17:17:17:17:1 | 24632323:246 | 24632535:246 | +;+;+;+;+;+;+; | 3389 | MAP4K3 | 304 | 217 | 381 | 344 | 304 | 444 |
| ENSDARG000 | 17:17:17:17:1 | 8799451:879 | 8799910:879 | +;+;+;+;+;+;+; | 6823 | tonsl | 48 | 241 | 282 | 66 | 183 | 168 |
| ENSDARG000 | 17:17:17:17:1 | 33708682:337 | 33709737:337 | -;-;-;-;-;-;-;- | 1615 | dnal1 | 126 | 106 | 120 | 183 | 117 | 130 |
| ENSDARG000 | 17;17;17;17 | 39783268:397 | 39783617:397 | -;-;-;- | 954 | NA | 0 | 0 | 0 | 0 | 0 | 0 |
| ENSDARG000 | 17:17:17:17:1 | 43534115:435 | 43534164:435 | -;-;-;-;-;-;-; | 1858 | NT5C1A | 64 | 22 | 39 | 50 | 12 | 28 |
| ENSDARG000 | 17:17:17:17:1 | 33688220:336 | 33688851:336 | -;-;-;-;-;-;-; | 1963 | NA | 59 | 70 | 49 | 38 | 106 | 65 |
| ENSDARG000 | 17;17 | 51642687:516 | 51643974:516 | -;- | 1373 | CCR6 | 2 | 2 | 4 | 4 | 0 | 0 |
| ENSDARG000 | 17:17:17:17:1 | 33699379:337 | 33699507:337 | -;-;-;-;-;-;-; | 1111 | dnal1 | 0 | 0 | 0 | 0 | 0 | 0 |
| ENSDARG000 | 17:17:17:17:1 | 4427463:443 | 4427580:443 | +;+;+;+;+ | 2508 | NA | 104 | 161 | 153 | 107 | 251 | 201 |
| ENSDARG000 | 17;17 | 23411202:234 | 23416897:234 | -;- | 5696 | NA | 4 | 0 | 0 | 0 | 0 | 0 |
| ENSDARG000 | 17;17;17 | 43543898:435 | 43544044:435 | +;+;+ | 413 | NA | 7 | 283 | 67 | 73 | 31 | 81 |
| ENSDARG000 | 17:17:17:17:1 | 8760793:876 | 8760955:876 | -;-;-;-;- | 920 | NA | 0 | 2 | 0 | 0 | 0 | 0 |
| ENSDARG000 | 17:17:17:17:1 | 8754020:875 | 8754515:875 | +;+;+;+;+;+;+; | 8987 | EDRF1 | 324 | 201 | 339 | 174 | 328 | 275 |
| ENSDARG000 | 17:17:17:17:1 | 51655229:516 | 51655424:516 | -;-;-;-;-;-;-; | 1868 | ATP6V1C2 | 0 | 4 | 2 | 2 | 4 | 0 |
| ENSDARG000 | 17:17:17:17:1 | 25481466:255 | 25481984:255 | +;+;+;+;+;+;+; | 10552 | NA | 8151 | 5773 | 7202 | 7287 | 9110 | 7447 |
| ENSDARG000 | 17:17:17:17:1 | 15213496:152 | 15213666:152 | +;+;+;+;+;+;+; | 1658 | gnpnat1 | 214 | 96 | 221 | 116 | 158 | 156 |
| ENSDARG000 | 17:17:17:17:1 | 39790388:398 | 39790456:398 | +;+;+;+;+;+;+; | 1276 | NA | 110 | 70 | 130 | 36 | 120 | 149 |
| ENSDARG000 | 17:17:17:17:1 | 21452258:214 | 21452474:214 | +;+;+;+;+;+;+; | 3366 | NA | 65 | 4 | 18 | 0 | 50 | 8 |
| ENSDARG000 | 17:17:17:17:1 | 16906523:169 | 16908448:169 | -;-;-;-;-;-;-; | 10405 | nrxn3a | 2 | 4 | 58 | 11 | 71 | 341 |
| ENSDARG000 | 17:17:17:17:1 | 15220042:152 | 15220547:152 | -;-;-;-;-;-;-; | 1459 | styx | 295 | 421 | 679 | 613 | 351 | 369 |
| ENSDARG000 | 17:17:17:17:1 | 25550235:255 | 25550452:255 | -;-;-;-;-;-;-;- | 1370 | rtn4ip1 | 126 | 189 | 119 | 84 | 196 | 137 |
| ENSDARG000 | 17;17;17;17 | 25512924:255 | 25513116:255 | -;-;-;- | 828 | NA | 0 | 0 | 0 | 4 | 0 | 0 |
| ENSDARG000 | 17:17:17:17:1 | 23434211:234 | 23434440:234 | -;-;-;-;-;-;-;- | 1166 | pcgf5a | 66 | 46 | 84 | 36 | 65 | 72 |
| ENSDARG000 | 17;17;17 | 31717192:317 | 31717655:317 | -;-;- | 839 | dtd2 | 57 | 72 | 78 | 62 | 58 | 42 |
| ENSDARG000 | 17:17:17:17:1 | 31719034:317 | 31719383:317 | +;+;+;+;+;+;+; | 1468 | nubpl | 152 | 125 | 174 | 133 | 193 | 167 |
| ENSDARG000 | 17:17:17:17:1 | 5061135:507 | 5061338:507 | +;+;+;+;+;+;+; | 2786 | cdc5l | 2095 | 2416 | 5065 | 2193 | 2297 | 2341 |
| ENSDARG000 | 17:17:17:17:1 | 30661348:306 | 30662115:306 | -;-;-;-;-;- | 2141 | alkal2b | 890 | 136 | 14 | 146 | 20 | 132 |
| ENSDARG000 | 17:17:17:17:1 | 20173882:201 | 20174078:201 | +;+;+;+;+;+;+; | 3997 | NA | 10 | 20 | 27 | 30 | 29 | 4 |
| ENSDARG000 | 17:17:17:17:1 | 20188704:201 | 20189175:201 | -;-;-;-;-;-;-; | 2258 | ecd | 199 | 175 | 257 | 144 | 280 | 232 |
| ENSDARG000 | 17:17:17:17:1 | 23462972:234 | 23463148:234 | +;+;+;+;+;+;+; | 1682 | ANKRD1 | 2 | 0 | 0 | 0 | 0 | 2 |
| ENSDARG000 | 17;17 | 30700627:307 | 30701586:307 | -;- | 1328 | zgc:194392 | 0 | 0 | 4 | 0 | 18 | 0 |
| ENSDARG000 | 17:17:17:17:1 | 23470967:234 | 23471619:234 | +;+;+;+;+;+;+; | 7861 | NA | 136 | 602 | 720 | 247 | 580 | 251 |
| ENSDARG000 | 17:17:17:17:1 | 21477892:214 | 21478049:214 | +;+;+;+;+;+;+; | 2944 | NA | 0 | 0 | 0 | 0 | 0 | 0 |
| ENSDARG000 | 17;17 | 40021476:400 | 40021943:400 | -;- | 507 | NA | 0 | 0 | 0 | 0 | 0 | 0 |
| ENSDARG000 | 17:17:17:17:1 | 49276061:492 | 49276069:492 | +;+;+;+;+;+;+; | 3145 | zgc:113176 | 60 | 64 | 84 | 37 | 98 | 96 |
| ENSDARG000 | 17:17:17:17:1 | 10748366:107 | 10748857:107 | +;+;+;+;+;+;+; | 2864 | atg14 | 503 | 301 | 357 | 363 | 449 | 448 |
| ENSDARG000 | 17:17:17:17:1 | 49220126:492 | 49220985:492 | -;-;-;-;-;-;-; | 1976 | tfb1m | 113 | 95 | 175 | 76 | 91 | 152 |
| ENSDARG000 | 17:17:17:17:1 | 28798725:287 | 28799838:288 | -;-;-;-;-;-;-; | 3360 | g2e3 | 144 | 763 | 722 | 366 | 422 | 262 |
| ENSDARG000 | 17:17:17:17:1 | 43565628:435 | 43565868:435 | -;-;-;-;-;-;-; | 3365 | zfyve28 | 527 | 257 | 446 | 267 | 394 | 415 |
| ENSDARG000 | 17:17:17:17:1 | 12942021:129 | 12942913:129 | +;+;+;+;+;+;+; | 5342 | nfkbiab | 3339 | 2137 | 3115 | 2470 | 2226 | 2795 |
| ENSDARG000 | 17 | 40109059 | 40110782 | - | 1724 | NA | 0 | 0 | 0 | 0 | 0 | 0 |
| ENSDARG000 | 17:17:17:17:1 | 10783599:107 | 10786754:107 | -;-;-;-;- | 3729 | fbxo34 | 1066 | 812 | 902 | 770 | 1293 | 972 |
| ENSDARG000 | 17;17;17;17 | 17763167:177 | 17763439:177 | -;-;-;- | 548 | slirp | 276 | 386 | 236 | 96 | 206 | 345 |
| ENSDARG000 | 17:17:17:17:1 | 17491226:174 | 17493086:174 | -;-;-;-;-;-;-; | 3764 | adck1 | 74 | 92 | 104 | 28 | 120 | 84 |
| ENSDARG000 | 17:17:17:17:1 | 40860016:408 | 40860324:408 | -;-;-;-;-;-;-;- | 1453 | supt7l | 76 | 80 | 124 | 100 | 173 | 106 |
| ENSDARG000 | 17:17:17:17:1 | 51682429:516 | 51682641:516 | +;+;+;+;+;+;+; | 3003 | nol10 | 794 | 589 | 784 | 335 | 506 | 506 |
| ENSDARG000 | 17:17:17:17:1 | 30704068:307 | 30704140:307 | +;+;+;+;+;+;+; | 13486 | APOB | 16 | 2 | 30 | 6 | 0 | 0 |
| ENSDARG000 | 17:17:17:17:1 | 40879331:408 | 40880183:408 | +;+;+;+;+;+;+; | 6884 | NA | 396 | 311 | 570 | 204 | 388 | 354 |
| ENSDARG000 | 17:17:17:17:1 | 37287852:372 | 37288072:372 | -;-;-;-;-;-;-; | 1386 | ptgr2 | 380 | 292 | 283 | 291 | 396 | 207 |
| ENSDARG000 | 17:17:17:17:1 | 5351922:542 | 5352135:542 | +;+;+;+;+;+;+; | 1712 | runx2a | 8 | 0 | 5 | 0 | 0 | 0 |
| ENSDARG000 | 17:17:17:17:1 | 31739418:317 | 31739639:317 | +;+;+;+;+;+;+; | 7488 | ARHGAP5 | 1362 | 1841 | 2243 | 1297 | 2944 | 1263 |
| ENSDARG000 | 17:17:17:17:1 | 20204122:202 | 20204141:202 | +;+;+;+;+;+;+ | 590 | gnrh3 | 0 | 0 | 2 | 0 | 0 | 0 |
| ENSDARG000 | 17:17:17:17:1 | 5158439:516 | 5162183:516 | -;-;-;-;-;-;-; | 5552 | supt3h | 465 | 221 | 422 | 358 | 442 | 264 |
| ENSDARG000 | 17:17:17:17:1 | 25563979:255 | 25564044:255 | +;+;+;+;+;+;+; | 4758 | qrsl1 | 105 | 129 | 164 | 119 | 190 | 126 |
| ENSDARG000 | 17;17 | 25585487:255 | 25586009:255 | -;- | 644 | si:ch211-214 | 0 | 4 | 1 | 0 | 17 | 41 |
| ENSDARG000 | 17;17 | 25587625:255 | 25588115:255 | -;- | 612 | si:ch211-214 | 0 | 0 | 0 | 0 | 0 | 0 |
| ENSDARG000 | 17;17;17;17 | 37310662:373 | 37311512:373 | -;-;-;- | 1178 | NA | 0 | 0 | 0 | 4 | 0 | 0 |
| ENSDARG000 | 17:17:17:17:1 | 37301860:373 | 37302268:373 | +;+;+;+;+;+;+; | 9643 | NA | 1489 | 1650 | 2879 | 3391 | 1592 | 1949 |
| ENSDARG000 | 17;17 | 28822396:288 | 28822479:288 | +;+ | 896 | NA | 0 | 0 | 0 | 0 | 0 | 0 |
| ENSDARG000 | 17:17:17:17:1 | 43595692:435 | 43596007:435 | +;+;+;+;+;+;+; | 5377 | NA | 0 | 0 | 6 | 2 | 6 | 4 |
| ENSDARG000 | 17:17:17:17:1 | 7595356:760 | 7595617:760 | +;+;+;+;+;+;+; | 1425 | NA | 0 | 0 | 6 | 0 | 0 | 0 |
| ENSDARG000 | 17:17:17:17:1 | 43610894:436 | 43612423:436 | -;-;-;-;-;-;-; | 2957 | rtkn2 | 2 | 64 | 48 | 26 | 39 | 21 |
| ENSDARG000 | 17:17:17:17:1 | 10817675:108 | 10818333:108 | +;+;+;+;+;+;+; | 1663 | NA | 0 | 0 | 3 | 2 | 0 | 0 |
| ENSDARG000 | 17:17:17:17:1 | 10824659:108 | 10825738:108 | -;-;-;-;-;- | 1678 | lgals3b | 2709 | 2010 | 2124 | 3475 | 1851 | 2542 |
| ENSDARG000 | 17:17:17:17:1 | 49313504:493 | 49313762:493 | -;-;-;-;-;-;-; | 2506 | orc3 | 210 | 660 | 782 | 174 | 468 | 303 |
| ENSDARG000 | 17:17:17:17:1 | 15247086:152 | 15247425:152 | -;-;-;-;-;-;-; | 4056 | NA | 6 | 1 | 14 | 6 | 20 | 6 |
| ENSDARG000 | 17:17:17:17:1 | 21514777:215 | 21515008:215 | +;+;+;+;+;+;+; | 5690 | CHST15 | 16 | 92 | 26 | 2 | 64 | 48 |
| ENSDARG000 | 17 | 27176243 | 27183448 | + | 7206 | NA | 0 | 0 | 0 | 0 | 0 | 0 |
| ENSDARG000 | 17;17;17 | 7620220:762 | 7620302:762 | +;+;+ | 2473 | fbxo30a | 267 | 241 | 247 | 113 | 274 | 188 |
| ENSDARG000 | 17:17:17:17:1 | 24668528:246 | 24669049:246 | -;-;-;-;-;-;-; | 2736 | NA | 7260 | 4636 | 8081 | 9257 | 5333 | 8478 |
| ENSDARG000 | 17:17:17:17:1 | 40912071:409 | 40912575:409 | -;-;-;-;- | 1756 | NA | 60 | 26 | 60 | 80 | 52 | 38 |
| ENSDARG000 | 17:17:17:17:1 | 7643531:764 | 7643573:764 | -;-;-;-;- | 532 | NA | 10 | 2 | 2 | 0 | 0 | 23 |
| ENSDARG000 | 17:17:17:17:1 | 33719415:337 | 33719504:337 | +;+;+;+;+;+;+; | 3470 | fut8a | 753 | 562 | 367 | 258 | 525 | 438 |
| ENSDARG000 | 17:17:17:17:1 | 31805775:318 | 31807089:318 | -;-;-;-;-;-;-;- | 2127 | abraxas2 | 133 | 199 | 156 | 60 | 182 | 123 |
| ENSDARG000 | 17:17:17:17:1 | 27186948:271 | 27191657:271 | -;-;-;-;-;-;-; | 9558 | asap3 | 136 | 66 | 249 | 52 | 96 | 120 |
| ENSDARG000 | 17:17:17:17:1 | 12955390:129 | 12955588:129 | -;-;-;-;-;-;-; | 1043 | psma6a | 924 | 1128 | 1341 | 683 | 1186 | 752 |
| ENSDARG000 | 17:17:17:17:1 | 24681280:246 | 24681868:246 | -;-;-;-;- | 954 | morn2 | 13 | 14 | 26 | 102 | 2 | 15 |
| ENSDARG000 | 17;17 | 7727464:773 | 7728737:773 | -;- | 1526 | stx11a | 54 | 151 | 91 | 120 | 113 | 70 |
| ENSDARG000 | 17;17;17;17 | 7763630:777 | 7764171:777 | +;+;+;+ | 10675 | NA | 1940 | 882 | 4730 | 519 | 2166 | 2348 |
| ENSDARG000 | 17:17:17:17:1 | 20214213:202 | 20214783:202 | -;-;-;-;-;-;-; | 1236 | mgmt | 123 | 78 | 101 | 149 | 104 | 136 |
| ENSDARG000 | 17;17 | 15297398:153 | 15297468:153 | +;+ | 2620 | NA | 820 | 292 | 773 | 1149 | 1189 | 1349 |
| ENSDARG000 | 17;17;17;17 | 11007091:110 | 11008335:110 | -;-;-;- | 2148 | dact1 | 199 | 82 | 229 | 166 | 167 | 68 |
| ENSDARG000 | 17:17:17:17:1 | 28882977:288 | 28883517:288 | +;+;+;+;+;+;+; | 3886 | PRKD1 | 0 | 8 | 0 | 0 | 2 | 30 |
| ENSDARG000 | 17;17;17;17 | 28944581:289 | 28945272:289 | -;-;-;- | 1419 | NA | 0 | 0 | 0 | 0 | 0 | 0 |
| ENSDARG000 | 17:17:17:17:1 | 43623598:436 | 43624027:436 | +;+;+;+;+;+;+; | 2047 | NA | 2 | 8 | 10 | 4 | 10 | 4 |
| ENSDARG000 | 17:17:17:17:1 | 15309432:153 | 15309761:153 | -;-;-;-;-;-;-; | 3814 | fam184aa | 30 | 18 | 79 | 81 | 98 | 48 |
| ENSDARG000 | 17;17;17 | 21607239:216 | 21607832:216 | -;-;- | 1674 | GPR26 | 0 | 0 | 0 | 0 | 0 | 0 |
| ENSDARG000 | 17;17 | 21564607:216 | 21564769:216 | +;+ | 357 | NA | 0 | 0 | 0 | 0 | 0 | 0 |
| ENSDARG000 | 17:17:17:17:1 | 31820401:318 | 31820652:318 | +;+;+;+;+;+;+ | 1246 | eef1akmt2 | 66 | 130 | 172 | 68 | 158 | 51 |
| ENSDARG000 | 17;17 | 23542115:235 | 23543448:235 | -;- | 2020 | NA | 10 | 0 | 0 | 0 | 0 | 0 |
| ENSDARG000 | 17 | 11327164 | 11329959 | - | 2796 | irf2bpl | 690 | 351 | 394 | 198 | 434 | 228 |
| ENSDARG000 | 17;17;17 | 7786246:778 | 7787573:778 | -;-;- | 1727 | zbtb2a | 77 | 47 | 86 | 25 | 44 | 52 |
| ENSDARG000 | 17:17:17:17:1 | 11072942:110 | 11073635:110 | -;-;-;-;-;-;-; | 5868 | si:ch211-185 | 65 | 85 | 137 | 30 | 104 | 48 |
| ENSDARG000 | 17:17:17:17:1 | 17764979:177 | 17765126:177 | +;+;+;+;+;+;+; | 1259 | alkbh1 | 42 | 75 | 167 | 57 | 75 | 104 |
| ENSDARG000 | 17:17:17:17:1 | 12971304:129 | 12971476:129 | -;-;-;-;-;-;-; | 2394 | PRORP | 122 | 76 | 135 | 76 | 87 | 92 |
| ENSDARG000 | 17;17 | 43659940:436 | 43661745:436 | +;+ | 1806 | adob | 373 | 610 | 890 | 760 | 606 | 467 |
| ENSDARG000 | 17:17:17:17:1 | 30749251:307 | 30749365:307 | +;+;+;+;+;+;+; | 2084 | ldah | 118 | 139 | 96 | 70 | 125 | 96 |
| ENSDARG000 | 17:17:17:17:1 | 13007288:130 | 13007482:130 | +;+;+;+;+;+;+; | 1778 | ppp2r3c | 361 | 192 | 308 | 362 | 250 | 346 |
| ENSDARG000 | 17:17:17:17:1 | 24684778:246 | 24684848:246 | +;+;+;+;+;+;+; | 951 | znf593 | 260 | 379 | 590 | 448 | 255 | 297 |
| ENSDARG000 | 17;17;17;17 | 11073258:110 | 11073635:110 | -;-;-;- | 615 | NA | 0 | 0 | 0 | 0 | 0 | 0 |
| ENSDARG000 | 17:17:17:17:1 | 11364199:113 | 11364358:113 | -;-;-;-;-;- | 786 | NA | 0 | 0 | 0 | 0 | 0 | 0 |
| ENSDARG000 | 17:17:17:17:1 | 51743908:517 | 51744043:517 | +;+;+;+;+;+;+; | 5194 | odc1 | 1474 | 2729 | 2081 | 1031 | 3969 | 1260 |
| ENSDARG000 | 17:17:17:17:1 | 37385811:373 | 37386166:373 | -;-;-;-;-;-;-; | 1171 | crip1 | 286 | 224 | 202 | 104 | 274 | 225 |
| ENSDARG000 | 17;17;17 | 23549117:235 | 23549526:235 | +;+;+ | 1275 | zgc:65997 | 102 | 56 | 53 | 72 | 54 | 114 |
| ENSDARG000 | 17:17:17:17:1 | 25609826:256 | 25610028:256 | -;-;-;-;-;-;-; | 5360 | rab3gap2 | 1082 | 1032 | 1332 | 1179 | 1168 | 1036 |
| ENSDARG000 | 17:17:17:17:1 | 43663504:436 | 43665366:436 | -;-;-;-;- | 2325 | egr2a | 364 | 152 | 313 | 561 | 96 | 230 |
| ENSDARG000 | 17:17:17:17:1 | 5566138:556 | 5567347:556 | -;-;-;-;-;- | 1906 | clic5a | 4 | 12 | 6 | 10 | 6 | 2 |
| ENSDARG000 | 17:17:17:17:1 | 21760032:218 | 21760181:218 | +;+;+;+;+;+;+; | 3745 | ikzf5 | 330 | 302 | 412 | 358 | 346 | 436 |
| ENSDARG000 | 17:17:17:17:1 | 25590127:255 | 25590421:255 | +;+;+;+;+;+;+; | 8372 | iars2 | 708 | 845 | 1093 | 557 | 863 | 741 |
| ENSDARG000 | 17 | 29117027 | 29119362 | - | 2336 | foxg1a | 4 | 0 | 0 | 0 | 0 | 0 |
| ENSDARG000 | 17;17 | 21782043:217 | 21783049:217 | -;- | 1331 | hmx2 | 0 | 0 | 0 | 0 | 0 | 0 |
| ENSDARG000 | 17;17 | 21790606:217 | 21792110:217 | -;- | 2121 | hmx3a | 2 | 0 | 8 | 0 | 0 | 0 |
| ENSDARG000 | 17:17:17:17:1 | 23554932:235 | 23555443:235 | +;+;+;+;+;+;+; | 2842 | pank1a | 207 | 126 | 113 | 61 | 184 | 226 |
| ENSDARG000 | 17;17 | 30835076:308 | 30836034:308 | -;- | 1189 | GDF7 | 0 | 4 | 10 | 0 | 0 | 16 |
| ENSDARG000 | 17:17:17:17:1 | 43850516:438 | 43851119:438 | -;-;-;-;-;-;-; | 1609 | ahsa1b | 423 | 601 | 856 | 451 | 757 | 401 |
| ENSDARG000 | 17:17:17:17:1 | 27243447:272 | 27244713:272 | -;-;-;-;-;- | 2871 | NA | 46 | 102 | 175 | 61 | 112 | 82 |
| ENSDARG000 | 17:17:17:17:1 | 24687338:246 | 24687708:246 | +;+;+;+;+;+;+; | 3253 | selenon | 1425 | 1609 | 3198 | 2237 | 1744 | 1406 |
| ENSDARG000 | 17:17:17:17:1 | 32158951:321 | 32159441:321 | +;+;+;+;+;+;+; | 5044 | ADAM12 | 389 | 182 | 89 | 195 | 125 | 136 |
| ENSDARG000 | 17:17:17:17:1 | 45347726:453 | 45351075:453 | -;-;-;-;-;-;-; | 8375 | ZNF106 | 606 | 643 | 812 | 564 | 1342 | 751 |
| ENSDARG000 | 17:17:17:17:1 | 7795559:779 | 7795748:779 | -;-;-;-;-;-;-; | 1731 | rmnd1 | 219 | 138 | 170 | 79 | 117 | 156 |
| ENSDARG000 | 17:17:17:17:1 | 29124190:291 | 29124621:291 | -;-;-;-;-;-;-; | 13082 | NA | 10 | 20 | 46 | 36 | 80 | 8 |
| ENSDARG000 | 17;17 | 49345771:493 | 49346422:493 | -;- | 974 | gpx2 | 0 | 0 | 0 | 0 | 0 | 0 |
| ENSDARG000 | 17;17;17;17 | 5599014:560 | 5599750:560 | -;-;-;- | 1938 | enpp4 | 130 | 88 | 198 | 102 | 256 | 160 |
| ENSDARG000 | 17:17:17:17:1 | 11372531:113 | 11372631:113 | +;+;+;+;+;+;+ | 1212 | timm9 | 164 | 227 | 310 | 145 | 170 | 200 |
| ENSDARG000 | 17:17:17:17:1 | 29195962:291 | 29196728:291 | -;-;-;-;-;- | 2052 | ehd4 | 109 | 83 | 78 | 16 | 252 | 141 |
| ENSDARG000 | 17;17;17 | 27271399:272 | 27272063:272 | -;-;- | 1169 | id3 | 458 | 253 | 631 | 287 | 948 | 254 |
| ENSDARG000 | 17:17:17:17:1 | 15388479:153 | 15388931:153 | +;+;+;+;+;+;+; | 3137 | NA | 825 | 1619 | 3107 | 1805 | 8206 | 1270 |
| ENSDARG000 | 17:17:17:17:1 | 5611166:561 | 5611194:561 | +;+;+;+;+ | 4124 | fam167aa | 0 | 0 | 17 | 0 | 0 | 20 |
| ENSDARG000 | 17:17:17:17:1 | 20219153:202 | 20220405:202 | -;-;-;-;-;-;-; | 3508 | NA | 0 | 0 | 0 | 2 | 2 | 32 |
| ENSDARG000 | 17:17:17:17:1 | 17804727:178 | 17804963:178 | +;+;+;+;+;+;+; | 2211 | sptlc2a | 1825 | 2549 | 2378 | 1387 | 3476 | 3775 |
| ENSDARG000 | 17:17:17:17:1 | 20231801:202 | 20232348:202 | -;-;-;-;-;-;-; | 1401 | bnip4 | 70 | 98 | 158 | 130 | 94 | 96 |
| ENSDARG000 | 17:17:17:17:1 | 5662379:567 | 5663327:567 | -;-;-;-;-;- | 2427 | XKR6 | 0 | 5 | 10 | 0 | 4 | 0 |
| ENSDARG000 | 17:17:17:17:1 | 30843881:308 | 30843971:308 | +;+;+;+;+;+;+; | 2409 | tpp1 | 4924 | 3520 | 3338 | 3800 | 5103 | 4128 |
| ENSDARG000 | 17:17:17:17:1 | 5768608:579 | 5768998:579 | +;+;+;+;+;+;+; | 10186 | NA | 0 | 0 | 2 | 10 | 2 | 0 |
| ENSDARG000 | 17;17;17;17 | 5756547:576 | 5756807:576 | -;-;-;- | 938 | NA | 0 | 0 | 1 | 0 | 0 | 0 |
| ENSDARG000 | 17;17;17 | 32360673:323 | 32361356:323 | +;+;+ | 1797 | NA | 10 | 6 | 12 | 35 | 6 | 27 |
| ENSDARG000 | 17:17:17:17:1 | 23578316:235 | 23578679:235 | +;+;+;+;+;+;+; | 4006 | SLC16A12 | 12 | 0 | 18 | 4 | 4 | 0 |
| ENSDARG000 | 17:17:17:17:1 | 32343121:323 | 32343565:323 | +;+;+;+;+;+;+; | 3289 | DHX32 | 0 | 20 | 24 | 46 | 63 | 12 |
| ENSDARG000 | 17:17:17:17:1 | 27314088:273 | 27314136:273 | -;-;-;-;-;-;-; | 3851 | NA | 380 | 210 | 308 | 394 | 506 | 462 |
| ENSDARG000 | 17:17:17:17:1 | 37445525:374 | 37446903:374 | -;-;-;-;-;-;-;- | 2072 | crip2 | 564 | 1512 | 2499 | 1486 | 2708 | 1060 |
| ENSDARG000 | 17:17:17:17:1 | 51764310:517 | 51764526:517 | +;+;+;+;+;+;+; | 4220 | NA | 0 | 0 | 10 | 2 | 0 | 0 |
| ENSDARG000 | 17:17:17:17:1 | 49354343:493 | 49354542:493 | -;-;-;-;-;-;-; | 3671 | mthfd1b | 1286 | 2334 | 3773 | 1420 | 2596 | 2298 |
| ENSDARG000 | 17;17 | 15413412:154 | 15413987:154 | +;+ | 1959 | GJA1 | 1529 | 1857 | 1455 | 1150 | 1529 | 1686 |
| ENSDARG000 | 17;17;17 | 15422670:154 | 15423327:154 | -;-;- | 832 | NA | 60 | 57 | 43 | 106 | 28 | 65 |
| ENSDARG000 | 17:17:17:17:1 | 21826825:218 | 21827565:218 | -;-;-;-;-;-;-; | 1883 | NA | 6 | 10 | 7 | 0 | 0 | 24 |
| ENSDARG000 | 17:17:17:17:1 | 7821890:782 | 7822001:782 | +;+;+;+;+;+;+; | 3313 | NA | 0 | 0 | 0 | 0 | 0 | 0 |
| ENSDARG000 | 17:17:17:17:1 | 11377970:113 | 11379686:113 | -;-;-;-;-;-;-; | 6046 | arid4a | 1261 | 1374 | 1879 | 1297 | 1700 | 1063 |
| ENSDARG000 | 17:17:17:17:1 | 15433518:154 | 15433654:154 | +;+;+;+;+;+;+; | 858 | fabp7a | 216 | 1566 | 1081 | 121 | 567 | 64 |
| ENSDARG000 | 17;17 | 23598205:236 | 23600384:236 | -;- | 2264 | NA | 292 | 14 | 0 | 80 | 2 | 330 |
| ENSDARG000 | 17:17:17:17:1 | 32374876:325 | 32374953:325 | +;+;+;+;+;+;+; | 4237 | ywhaqb | 10403 | 14270 | 16012 | 12372 | 12689 | 11777 |
| ENSDARG000 | 17:17:17:17:1 | 37510808:375 | 37511119:375 | +;+;+;+;+;+;+; | 3717 | MTA1 | 508 | 392 | 362 | 340 | 390 | 426 |
| ENSDARG000 | 17:17:17:17:1 | 51796668:518 | 51797917:518 | -;-;-;-;-;-;-; | 3011 | exd2 | 360 | 264 | 490 | 244 | 419 | 346 |
| ENSDARG000 | 17:17:17:17:1 | 32365941:323 | 32366634:323 | -;-;-;-;-;- | 2064 | klf11b | 1400 | 1102 | 1630 | 1182 | 768 | 941 |
| ENSDARG000 | 17:17:17:17:1 | 32391056:323 | 32391315:323 | -;-;-;-;-;-;-; | 3057 | grhl1 | 60 | 3 | 10 | 17 | 8 | 20 |
| ENSDARG000 | 17;17 | 24714816:247 | 24714968:247 | +;+ | 340 | NA | 0 | 0 | 0 | 0 | 0 | 0 |
| ENSDARG000 | 17;17 | 24701071:247 | 24701813:247 | -;- | 1676 | NA | 82 | 64 | 69 | 106 | 66 | 65 |
| ENSDARG000 | 17:17:17:17:1 | 24695594:246 | 24696909:246 | -;-;-;-;-;-;-; | 3588 | NA | 310 | 202 | 339 | 378 | 372 | 494 |
| ENSDARG000 | 17;17 | 23607195:236 | 23608993:236 | -;- | 1879 | ifit15 | 0 | 0 | 0 | 2 | 0 | 0 |

| | | | | | | | | | | | |
|---|---|---|---|---|---|---|---|---|---|---|---|
| ENSDARG000 | 17;17;17;17 | 32365256:323 | 32365334:323 | +;+;+;+ | 716 | NA | 0 | 0 | 0 | 4 | 3 | 7 |
| ENSDARG000 | 17;17;17;17;1 | 45378908:453 | 45379650:453 | -;-;-;-;-;-;-;-;- | 2199 | pacc1 | 309 | 390 | 306 | 243 | 500 | 389 |
| ENSDARG000 | 17 | 27400703 | 27401354 | + | 652 | NA | 0 | 0 | 0 | 0 | 0 | 0 |
| ENSDARG000 | 17;17;17;17;1 | 21833719:218 | 21833885:218 | -;-;-;-;-;-;-;-;- | 2612 | NA | 0 | 0 | 0 | 0 | 1 | 6 |
| ENSDARG000 | 17 | 27402261 | 27402343 | + | 83 | dre-mir-124-4 | 0 | 0 | 0 | 0 | 0 | 0 |
| ENSDARG000 | 17;17;17;17;1 | 17861681:178 | 17862370:178 | +;+;+;+;+;+ | 3920 | NA | 6 | 0 | 8 | 0 | 8 | 0 |
| ENSDARG000 | 17;17;17;17;1 | 27407918:274 | 27408912:274 | -;-;-;-;-;-;-;-;- | 3365 | ythdf2 | 792 | 927 | 1160 | 639 | 1328 | 821 |
| ENSDARG000 | 17;17 | 23612616:236 | 23613905:236 | -;- | 1354 | ifit14 | 18 | 19 | 27 | 13 | 18 | 25 |
| ENSDARG000 | 17;17;17;17;1 | 45395846:453 | 45396289:453 | +;+;+;+;+;+;+ | 1470 | nenf | 655 | 609 | 621 | 538 | 526 | 692 |
| ENSDARG000 | 17;17;17;17;1 | 23619924:236 | 23620241:236 | -;-;-;-;-;-;-;-;- | 1383 | NA | 207 | 159 | 86 | 126 | 38 | 133 |
| ENSDARG000 | 17;17;17;17;1 | 34186632:342 | 34186777:342 | +;+;+;+;+;+;+;+ | 3052 | pals1a | 535 | 250 | 253 | 167 | 285 | 157 |
| ENSDARG000 | 17;17;17;17;1 | 43863708:438 | 43863796:438 | +;+;+;+;+;+;+;+ | 3600 | amy2al1 | 20 | 101 | 172 | 104 | 88 | 94 |
| ENSDARG000 | 17;17;17;17;1 | 34244345:342 | 34244502:342 | +;+;+;+;+;+;+ | 1523 | eif2s1a | 1067 | 1155 | 1503 | 964 | 1041 | 1151 |
| ENSDARG000 | 17;17;17;17;1 | 30857008:308 | 30858041:308 | -;-;-;-;- | 2041 | yy1a | 1062 | 1140 | 1387 | 975 | 1176 | 1072 |
| ENSDARG000 | 17;17;17;17;1 | 29216367:292 | 29216858:292 | -;-;-;-;-;-;-;- | 937 | NA | 5 | 3 | 20 | 10 | 4 | 26 |
| ENSDARG000 | 17;17;17;17 | 34706888:347 | 34707304:347 | -;-;-;- | 657 | NA | 5 | 7 | 17 | 16 | 3 | 14 |
| ENSDARG000 | 17;17;17;17;1 | 34805897:348 | 34806362:348 | +;+;+;+;+ | 1854 | id2a | 1512 | 1050 | 1072 | 1027 | 1863 | 2090 |
| ENSDARG000 | 17;17;17 | 30894431:308 | 30894730:308 | +;+;+ | 1793 | degs2 | 2 | 0 | 0 | 0 | 0 | 0 |
| ENSDARG000 | 17;17;17;17;1 | 30897292:309 | 30901365:309 | -;-;-;-;-;-;-;-;- | 6367 | evla | 1470 | 1133 | 1149 | 1338 | 1114 | 1100 |
| ENSDARG000 | 17;17;17;17;1 | 17939095:179 | 17940190:179 | -;-;-;-;-;-;-;- | 3000 | hhipl1 | 2 | 48 | 61 | 9 | 39 | 10 |
| ENSDARG000 | 17;17;17;17;1 | 17914369:179 | 17914583:179 | +;+;+;+;+;+;+;+ | 2367 | ccdc85ca | 0 | 0 | 10 | 40 | 40 | 6 |
| ENSDARG000 | 17;17;17;17;1 | 20237727:202 | 20238717:202 | +;+;+;+;+;+;+;+ | 2152 | smndc1 | 738 | 854 | 896 | 560 | 774 | 950 |
| ENSDARG000 | 17;17;17;17;1 | 13044473:130 | 13046286:130 | -;-;-;-;-;-;-;-;- | 4255 | NA | 688 | 611 | 873 | 376 | 996 | 588 |
| ENSDARG000 | 17;17;17;17;1 | 45404758:454 | 45404892:454 | +;+;+;+;+;+;+;+ | 3030 | tagapa | 175 | 129 | 199 | 168 | 423 | 769 |
| ENSDARG000 | 17;17;17;17;1 | 25635953:256 | 25637273:256 | -;-;-;-;-;-;-;-;- | 2350 | ppp1cb | 2660 | 2300 | 2904 | 3551 | 2225 | 2687 |
| ENSDARG000 | 17;17;17;17;1 | 37624704:376 | 37624960:376 | +;+;+;+;+;+;+;+ | 1963 | tmem229b | 122 | 77 | 48 | 26 | 83 | 20 |
| ENSDARG000 | 17;17;17;17;1 | 51821906:518 | 51826243:518 | -;-;-;-;-;-;-;-;- | 5815 | numb | 592 | 386 | 699 | 719 | 838 | 512 |
| ENSDARG000 | 17;17;17;17 | 27434626:274 | 27434902:274 | +;+;+;+ | 1551 | vgll2b | 4 | 10 | 1 | 2 | 0 | 0 |
| ENSDARG000 | 17;17 | 37792617:377 | 37794319:377 | -;- | 1899 | zfp36l1a | 3360 | 1403 | 2966 | 1155 | 2147 | 1900 |
| ENSDARG000 | 17;17;17;17;1 | 49415354:494 | 49421575:494 | -;-;-;-;-;-;- | 7395 | znf292a | 1034 | 867 | 1354 | 682 | 1425 | 1129 |
| ENSDARG000 | 17;17;17;17;1 | 13634412:136 | 13634572:136 | -;-;-;-;-;-;-;-;- | 1853 | NA | 0 | 0 | 0 | 4 | 0 | 10 |
| ENSDARG000 | 17;17;17;17;1 | 45413324:454 | 45413731:454 | +;+;+;+;+;+;+;+ | 5403 | ezra | 38 | 10 | 24 | 4 | 30 | 6 |
| ENSDARG000 | 17;17;17;17;1 | 24718272:247 | 24718361:247 | +;+;+;+;+;+;+;+ | 3084 | mtfr1l | 768 | 727 | 830 | 308 | 858 | 520 |
| ENSDARG000 | 17;17;17;17;1 | 49439219:494 | 49440695:494 | -;-;-;-;-;-;- | 1794 | cga | 0 | 4 | 0 | 0 | 2 | 0 |
| ENSDARG000 | 17;17;17;17;1 | 29227502:292 | 29228556:292 | -;-;-;-;-;-;-;-;- | 2465 | traf3 | 233 | 196 | 310 | 149 | 245 | 136 |
| ENSDARG000 | 17;17;17;17;1 | 29251662:292 | 29253062:292 | -;-;-;-;-;-;-;-;- | 3082 | rcor1 | 1115 | 1490 | 1795 | 1110 | 1628 | 1213 |
| ENSDARG000 | 17;17;17;17;1 | 23650569:236 | 23651706:236 | -;-;-;-;-;-;-;-;- | 3316 | ptena | 2132 | 1583 | 2073 | 1553 | 1878 | 1865 |
| ENSDARG000 | 17;17 | 49587635:495 | 49587968:495 | -;- | 720 | NA | 0 | 0 | 0 | 0 | 0 | 0 |
| ENSDARG000 | 17;17;17;17;1 | 49597112:495 | 49597186:495 | +;+;+;+;+ | 936 | NA | 0 | 0 | 0 | 0 | 0 | 0 |
| ENSDARG000 | 17;17 | 45454943:454 | 45455408:454 | +;+ | 1593 | NA | 0 | 0 | 0 | 0 | 0 | 0 |
| ENSDARG000 | 17;17;17;17;1 | 37932433:379 | 37932939:379 | +;+;+;+;+;+;+;+ | 6513 | plekhh1 | 28 | 36 | 26 | 4 | 70 | 12 |
| ENSDARG000 | 17;17;17;17;1 | 20321323:203 | 20321825:203 | -;-;-;-;-;-;-;-;- | 4225 | NA | 0 | 2 | 0 | 0 | 0 | 0 |
| ENSDARG000 | 17;17;17;17;1 | 27456804:274 | 27457000:274 | +;+;+;+;+;+;+;+ | 2094 | ctsl.1 | 100 | 14 | 3 | 0 | 8 | 6 |
| ENSDARG000 | 17;17;17;17;1 | 49637257:496 | 49637369:496 | +;+;+;+;+;+;+;+ | 1538 | zgc:113372 | 91 | 103 | 107 | 81 | 108 | 114 |
| ENSDARG000 | 17;17;17 | 49631844:496 | 49633159:496 | -;-;- | 1889 | zgc:171599 | 108 | 37 | 103 | 74 | 94 | 72 |
| ENSDARG000 | 17;17;17;17;1 | 20248177:202 | 20248546:202 | -;-;-;-;-;-;-;-;- | 2490 | add3b | 170 | 58 | 923 | 180 | 716 | 324 |
| ENSDARG000 | 17;17;17;17;1 | 34809643:348 | 34815337:348 | -;-;-;-;-;-;-;-;- | 10220 | KIDINS220 | 160 | 132 | 154 | 195 | 194 | 341 |
| ENSDARG000 | 17;17;17;17;1 | 20251491:202 | 20251564:202 | +;+;+;+;+;+ | 787 | NA | 0 | 1 | 0 | 0 | 0 | 0 |
| ENSDARG000 | 17;17;17;17;1 | 24747209:247 | 24747551:247 | +;+;+;+;+;+;+;+ | 1977 | STAC | 30 | 0 | 2 | 4 | 2 | 0 |
| ENSDARG000 | 17;17;17;17;1 | 18031304:180 | 18031978:180 | +;+;+;+;+;+;+;+ | 4411 | setd3 | 833 | 843 | 1118 | 488 | 1079 | 932 |
| ENSDARG000 | 17;17;17;17;1 | 25677458:256 | 25677570:256 | +;+;+;+;+;+;+;+ | 3019 | kcnh1a | 6 | 9 | 57 | 0 | 227 | 5 |
| ENSDARG000 | 17;17;17;17;1 | 25649939:256 | 25650738:256 | -;-;-;-;-;-;-;-;- | 4251 | NA | 2 | 2 | 2 | 0 | 0 | 6 |
| ENSDARG000 | 17;17;17;17;1 | 27545183:275 | 27545418:275 | +;+;+;+;+;+;+ | 1402 | pacrg | 4 | 0 | 4 | 0 | 2 | 0 |
| ENSDARG000 | 17;17;17;17;1 | 15449401:154 | 15449559:154 | -;-;-;-;-;-;-;-;- | 7313 | NA | 2 | 0 | 0 | 0 | 0 | 2 |
| ENSDARG000 | 17;17;17;17;1 | 13664442:136 | 13664523:136 | +;+;+;+;+;+ | 3083 | LRFN5 | 0 | 0 | 10 | 0 | 0 | 0 |
| ENSDARG000 | 17;17;17;17 | 14005588:140 | 14005753:140 | -;-;-;- | 478 | NA | 0 | 0 | 0 | 0 | 0 | 0 |
| ENSDARG000 | 17;17;17;17 | 24800332:248 | 24800440:248 | +;+;+;+ | 1055 | NA | 0 | 0 | 0 | 0 | 6 | 0 |
| ENSDARG000 | 17;17;17;17;1 | 51906053:519 | 51906562:519 | +;+;+;+;+;+;+;+ | 1446 | FLVCR2 | 0 | 0 | 0 | 0 | 0 | 0 |
| ENSDARG000 | 17;17;17;17;1 | 14425219:144 | 14425357:144 | +;+;+;+;+;+;+;+ | 2387 | NA | 0 | 0 | 8 | 0 | 2 | 0 |
| ENSDARG000 | 17;17;17;17 | 24808035:248 | 24808292:248 | +;+;+;+ | 783 | NA | 0 | 2 | 0 | 0 | 8 | 2 |
| ENSDARG000 | 17;17;17;17;1 | 21870044:218 | 21870218:218 | -;-;-;-;-;-;-;-;- | 1934 | NA | 0 | 0 | 0 | 0 | 2 | 6 |
| ENSDARG000 | 17;17;17;17;1 | 18117029:181 | 18118206:181 | +;+;+;+;+;+;+;+ | 8460 | bcl11ba | 36 | 6 | 2 | 4 | 44 | 34 |
| ENSDARG000 | 17;17;17 | 18237272:182 | 18237447:182 | +;+;+ | 543 | NA | 0 | 0 | 0 | 0 | 0 | 0 |
| ENSDARG000 | 17;17;17;17;1 | 30976484:309 | 30980192:309 | -;-;-;-;-;-;-;-;- | 7830 | eml1 | 2957 | 1986 | 1192 | 2287 | 1475 | 2429 |
| ENSDARG000 | 17;17;17;17;1 | 34969123:349 | 34970373:349 | -;-;-;-;-;-;-;-;- | 3033 | MBOAT2 | 305 | 314 | 208 | 381 | 293 | 379 |
| ENSDARG000 | 17;17;17;17;1 | 32531854:325 | 32532261:325 | +;+;+;+;+;+;+;+ | 2620 | gcfc2 | 229 | 213 | 361 | 232 | 369 | 209 |
| ENSDARG000 | 17;17;17;17;1 | 24815186:248 | 24815385:248 | -;-;-;-;-;-;- | 1781 | trmt61b | 111 | 84 | 124 | 54 | 181 | 100 |
| ENSDARG000 | 17;17;17;17;1 | 14443406:144 | 14444400:144 | -;-;-;-;-;-;-;-;- | 2113 | jkamp | 419 | 466 | 410 | 397 | 575 | 539 |
| ENSDARG000 | 17;17;17;17;1 | 24809221:248 | 24809277:248 | +;+;+;+;+;+;+;+ | 2033 | spdya | 10 | 69 | 0 | 32 | 0 | 1 |
| ENSDARG000 | 17;17;17;17;1 | 23681086:236 | 23682063:236 | -;-;-;-;-;-;-;-;- | 2817 | papss2a | 52 | 26 | 26 | 28 | 30 | 18 |
| ENSDARG000 | 17;17;17;17;1 | 27723490:277 | 27723840:277 | +;+;+;+;+;+;+;+ | 8502 | qkia | 7926 | 7006 | 13376 | 8600 | 10759 | 7137 |
| ENSDARG000 | 17;17;17;17;1 | 23710659:237 | 23711886:237 | -;-;-;-;- | 2385 | minpp1a | 456 | 275 | 355 | 314 | 289 | 298 |
| ENSDARG000 | 17;17;17;17;1 | 23729717:237 | 23729950:237 | +;+;+;+;+;+;+;+ | 4107 | sgms1a | 923 | 726 | 985 | 1401 | 654 | 969 |
| ENSDARG000 | 17;17;17;17;1 | 21887823:218 | 21887940:218 | +;+;+;+;+;+;+;+ | 2551 | plekha1a | 1702 | 2678 | 2121 | 2677 | 2247 | 2644 |
| ENSDARG000 | 17;17;17 | 23770848:237 | 23771134:237 | +;+;+ | 1241 | kcnk18 | 0 | 0 | 0 | 2 | 0 | 1 |
| ENSDARG000 | 17;17;17;17;1 | 20500229:205 | 20500465:205 | +;+;+;+;+;+;+;+ | 2553 | neurl1ab | 4 | 6 | 0 | 22 | 6 | 16 |
| ENSDARG000 | 17;17;17;17 | 52493499:524 | 52494296:524 | -;-;-;- | 2432 | prox2 | 8 | 2 | 0 | 0 | 0 | 0 |
| ENSDARG000 | 17;17;17;17;1 | 25777330:257 | 25777547:257 | -;-;-;-;-;-;-;-;- | 1726 | hhat | 72 | 68 | 142 | 86 | 127 | 53 |
| ENSDARG000 | 17;17;17;17;1 | 25833947:258 | 25834364:258 | +;+;+;+;+;+;+;+ | 4728 | acss1 | 1250 | 1133 | 1126 | 761 | 1140 | 1223 |
| ENSDARG000 | 17;17;17;17;1 | 20522081:205 | 20524193:205 | -;-;-;-;-;-;- | 4132 | NA | 44 | 12 | 2 | 4 | 4 | 2 |
| ENSDARG000 | 17;17;17;17;1 | 35097024:351 | 35097567:351 | +;+;+;+;+;+;+;+ | 4258 | asap2a | 103 | 135 | 318 | 145 | 298 | 114 |
| ENSDARG000 | 17;17;17;17;1 | 52524355:525 | 52524415:525 | +;+;+;+;+;+;+;+ | 2551 | dlst | 3663 | 3813 | 4216 | 3153 | 3238 | 3045 |
| ENSDARG000 | 17;17;17;17;1 | 32608010:326 | 32611182:326 | -;-;-;-;-;-;- | 4243 | NA | 58 | 226 | 270 | 76 | 210 | 142 |
| ENSDARG000 | 17;17;17;17 | 32571584:326 | 32571916:326 | +;+;+;+ | 3098 | eva1a | 14 | 11 | 29 | 10 | 35 | 22 |
| ENSDARG000 | 17;17;17;17;1 | 23782324:237 | 23782417:237 | +;+;+;+;+;+;+;+ | 1762 | slc18a2 | 27 | 20 | 28 | 24 | 16 | 43 |
| ENSDARG000 | 17;17;17;17;1 | 24821627:248 | 24821834:248 | +;+;+;+;+;+;+;+ | 2724 | wdr43 | 1416 | 2162 | 4043 | 2113 | 2193 | 2235 |
| ENSDARG000 | 17;17;17;17;1 | 20585328:205 | 20585378:205 | +;+;+;+;+;+;+ | 569 | NA | 0 | 0 | 0 | 0 | 0 | 0 |
| ENSDARG000 | 17;17;17;17;1 | 20569806:205 | 20569921:205 | +;+;+;+;+;+;+;+ | 4740 | samd8 | 72 | 48 | 116 | 69 | 50 | 124 |
| ENSDARG000 | 17;17;17;17;1 | 31096326:310 | 31096351:310 | -;-;-;-;-;-;- | 563 | NA | 48 | 4 | 42 | 11 | 12 | 25 |
| ENSDARG000 | 17;17;17;17;1 | 31125155:311 | 31125330:311 | +;+;+;+;+;+;+;+ | 1096 | NA | 2 | 0 | 0 | 0 | 0 | 0 |
| ENSDARG000 | 17;17;17;17;1 | 32622931:326 | 32623020:326 | +;+;+;+;+;+;+;+ | 2376 | ctsba | 9211 | 14 | 0 | 0 | 0 | 8475 |
| ENSDARG000 | 17;17;17;17;1 | 52559952:525 | 52560214:525 | -;-;-;-;-;-;-;-;- | 2577 | RPS6KL1 | 36 | 18 | 14 | 28 | 0 | 20 |
| ENSDARG000 | 17;17;17;17;1 | 32630789:326 | 32631617:326 | -;-;-;-;-;-;-;-;- | 1760 | NA | 40 | 38 | 72 | 36 | 35 | 41 |
| ENSDARG000 | 17;17;17;17;1 | 52581012:525 | 52582792:525 | -;-;-;-;- | 2404 | NA | 0 | 0 | 0 | 0 | 0 | 10 |
| ENSDARG000 | 17;17;17;17;1 | 21919131:219 | 21919734:219 | +;+;+;+;+;+;+;+ | 2078 | htra1a | 106 | 611 | 254 | 176 | 168 | 172 |
| ENSDARG000 | 17;17;17;17;1 | 52589913:525 | 52590887:525 | -;-;-;-;-;-;-;- | 1627 | NA | 0 | 0 | 0 | 0 | 0 | 0 |
| ENSDARG000 | 17 | 24843401 | 24844543 | + | 1143 | gjb10 | 32 | 6 | 21 | 6 | 20 | 12 |
| ENSDARG000 | 17;17;17 | 24848976:248 | 24849025:248 | +;+;+ | 2102 | gjb3 | 122 | 46 | 36 | 32 | 22 | 26 |
| ENSDARG000 | 17;17;17;17;1 | 21964472:219 | 21964641:219 | +;+;+;+;+;+;+;+ | 1595 | NA | 2 | 8 | 0 | 0 | 10 | 2 |
| ENSDARG000 | 17;17;17;17;1 | 21982617:219 | 21982796:219 | -;-;-;-;-;-;-;-;- | 1964 | NA | 0 | 0 | 0 | 0 | 0 | 0 |
| ENSDARG000 | 17;17;17;17;1 | 1514711:1515 | 1514839:1516 | +;+;+;+;+;+ | 744 | AKT1 | 359 | 144 | 333 | 149 | 341 | 219 |
| ENSDARG000 | 17;17;17;17;1 | 51517750:515 | 51517784:515 | +;+;+;+;+;+;+;+ | 4541 | PXDN | 3156 | 1917 | 2180 | 1442 | 2227 | 1313 |
| ENSDARG000 | 17;17;17;17;1 | 1544903:1544 | 1545251:1545 | +;+;+;+;+;+;+;+ | 5065 | CEP170B | 215 | 61 | 196 | 118 | 195 | 92 |
| ENSDARG000 | 17;17;17;17;1 | 584369:58585 | 584416:58614 | +;+;+;+;+ | 750 | C17H14orf28 | 103 | 26 | 56 | 38 | 22 | 40 |
| ENSDARG000 | 17 | 48162951 | 48163065 | - | 115 | 5S_rRNA | 0 | 0 | 0 | 0 | 0 | 0 |
| ENSDARG000 | 17;17;17;17;1 | 117024:11711 | 117351:11735 | -;-;-;-;-;-;-;-;- | 1773 | actc1b | 68 | 108 | 52 | 122 | 164 | 46 |
| ENSDARG000 | 17;17;17;17;1 | 53250802:532 | 53251128:532 | +;+;+;+;+;+;+ | 1113 | VASH1 | 258 | 172 | 145 | 106 | 121 | 123 |
| ENSDARG000 | 17;17;17;17;1 | 654759:65612 | 655080:65630 | +;+;+;+;+;+;+ | 894 | NA | 10 | 45 | 26 | 11 | 47 | 25 |
| ENSDARG000 | 17 | 24837682 | 24838927 | - | 1246 | NA | 0 | 0 | 0 | 0 | 0 | 0 |
| ENSDARG000 | 17;17;17;17;1 | 11808031:118 | 11808039:118 | +;+;+;+;+;+;+;+ | 354 | NA | 1 | 4 | 0 | 2 | 2 | 0 |
| ENSDARG000 | 17;17;17 | 7364911:7369 | 7365418:7369 | -;-;- | 1009 | RAB32 | 0 | 0 | 2 | 0 | 0 | 0 |
| ENSDARG000 | 17;17;17;17;1 | 53344385:533 | 53344456:533 | -;-;-;-;-;-;-;- | 921 | NA | 84 | 200 | 120 | 81 | 73 | 123 |
| ENSDARG000 | 17 | 23573130 | 23573215 | + | 86 | dre-mir-107b | 0 | 0 | 0 | 0 | 0 | 0 |
| ENSDARG000 | 17;17;17;17 | 2063693:2091 | 2064219:2091 | +;+;+;+ | 1432 | ppp1r14d | 0 | 12 | 10 | 0 | 48 | 0 |
| ENSDARG000 | 17 | 620561 | 620677 | + | 117 | 5S_rRNA | 1 | 0 | 0 | 0 | 0 | 0 |
| ENSDARG000 | 17 | 8090357 | 8090474 | + | 118 | 5S_rRNA | 0 | 0 | 0 | 0 | 0 | 0 |
| ENSDARG000 | 17 | 3085649 | 3085765 | + | 117 | 5S_rRNA | 0 | 0 | 0 | 0 | 0 | 0 |
| ENSDARG000 | 17 | 44483490 | 44483617 | - | 128 | 5S_rRNA | 0 | 0 | 0 | 0 | 0 | 0 |
| ENSDARG000 | 17 | 24615707 | 24615794 | + | 88 | SNORD33 | 0 | 0 | 0 | 0 | 0 | 0 |
| ENSDARG000 | 17;17;17;17;1 | 599084:60149 | 600233:60170 | -;-;-;-;- | 2555 | KLHL28 | 216 | 176 | 203 | 116 | 204 | 145 |
| ENSDARG000 | 17 | 25063171 | 25063287 | + | 117 | 5S_rRNA | 0 | 0 | 0 | 0 | 0 | 0 |
| ENSDARG000 | 17;17;17 | 50048491:500 | 50049457:500 | -;-;- | 1489 | zgc:100951 | 0 | 2 | 0 | 0 | 0 | 0 |
| ENSDARG000 | 17 | 28624048 | 28624176 | - | 129 | 5S_rRNA | 0 | 0 | 0 | 0 | 0 | 0 |
| ENSDARG000 | 17 | 11400583 | 11400665 | - | 83 | NA | 0 | 0 | 0 | 0 | 0 | 0 |
| ENSDARG000 | 17 | 47391517 | 47391634 | + | 118 | 5S_rRNA | 0 | 0 | 0 | 0 | 0 | 0 |
| ENSDARG000 | 17 | 18283949 | 18284063 | - | 115 | 5S_rRNA | 0 | 0 | 0 | 0 | 0 | 0 |
| ENSDARG000 | 17 | 37390003 | 37390117 | - | 115 | 5S_rRNA | 4 | 0 | 0 | 0 | 0 | 4 |
| ENSDARG000 | 17 | 15517118 | 15517232 | + | 115 | 5S_rRNA | 0 | 0 | 0 | 0 | 0 | 0 |
| ENSDARG000 | 17;17;17;17;1 | 42274240:422 | 42274579:422 | +;+;+;+;+;+;+;+ | 1941 | pax1a | 8 | 8 | 31 | 20 | 20 | 22 |
| ENSDARG000 | 17;17;17 | 14452329:144 | 14452345:144 | +;+;+ | 1275 | GPR135 | 57 | 18 | 19 | 2 | 16 | 8 |
| ENSDARG000 | 17;17;17;17;1 | 23146976:231 | 23147047:231 | +;+;+;+;+;+;+;+ | 3375 | rasgrp3 | 382 | 304 | 281 | 323 | 316 | 238 |
| ENSDARG000 | 17;17;17 | 10111655:101 | 10111850:101 | -;-;- | 1661 | NA | 0 | 0 | 2 | 0 | 0 | 0 |
| ENSDARG000 | 17;17 | 50973017:509 | 50973024:509 | +;+ | 867 | NA | 4 | 0 | 12 | 0 | 30 | 6 |
| ENSDARG000 | 17 | 32701705 | 32701821 | + | 117 | 5S_rRNA | 0 | 0 | 0 | 0 | 0 | 0 |
| ENSDARG000 | 17;17;17;17;1 | 45114015:451 | 45114521:451 | -;-;-;-;-;-;-;- | 1300 | zgc:163014 | 874 | 448 | 245 | 349 | 177 | 524 |
| ENSDARG000 | 17;17 | 11401947:114 | 11401949:114 | +;+ | 12 | NA | 0 | 0 | 0 | 0 | 0 | 0 |
| ENSDARG000 | 17;17;17;17 | 36901790:369 | 36901855:369 | -;-;-;- | 303 | NA | 1 | 0 | 0 | 0 | 0 | 0 |
| ENSDARG000 | 17 | 29272963 | 29273081 | - | 119 | 5S_rRNA | 0 | 0 | 0 | 0 | 0 | 0 |
| ENSDARG000 | 17;17;17;17;1 | 13088594:130 | 13089006:130 | +;+;+;+;+;+;+;+ | 2257 | gemin2 | 248 | 347 | 500 | 196 | 385 | 330 |
| ENSDARG000 | 17 | 6958610 | 6958730 | - | 121 | 5S_rRNA | 0 | 0 | 0 | 0 | 0 | 0 |
| ENSDARG000 | 17;17;17;17;1 | 6969846:6970 | 6970057:6970 | -;-;-;-;-;-;-;- | 1146 | UST | 0 | 0 | 1 | 2 | 1 | 0 |
| ENSDARG000 | 17 | 205194 | 205310 | - | 117 | 5S_rRNA | 0 | 0 | 0 | 0 | 0 | 0 |
| ENSDARG000 | 17 | 42486740 | 42492668 | - | 5929 | NA | 0 | 0 | 0 | 0 | 0 | 0 |
| ENSDARG000 | 17 | 21917448 | 21917564 | + | 117 | 5S_rRNA | 0 | 0 | 0 | 0 | 0 | 0 |
| ENSDARG000 | 17;17;17;17;1 | 6217704:6225 | 6217828:6226 | +;+;+;+;+;+;+;+ | 3407 | NA | 14 | 32 | 14 | 2 | 50 | 44 |
| ENSDARG000 | 17;17;17;17 | 53353419:533 | 53353653:533 | -;-;-;- | 549 | NA | 71 | 41 | 60 | 47 | 40 | 41 |
| ENSDARG000 | 17;17;17;17;1 | 9308425:9308 | 9308579:9308 | +;+;+;+;+;+;+;+ | 7063 | NPAS3 | 32 | 26 | 128 | 47 | 202 | 22 |
| ENSDARG000 | 17;17;17;17;1 | 4207278:4207 | 4207289:4207 | +;+;+;+;+;+;+;+ | 537 | NA | 1 | 0 | 0 | 0 | 1 | 0 |
| ENSDARG000 | 17 | 39531815 | 39531929 | - | 115 | 5S_rRNA | 0 | 0 | 0 | 0 | 0 | 0 |
| ENSDARG000 | 17 | 23348329 | 23348469 | + | 141 | U4 | 0 | 0 | 0 | 0 | 0 | 0 |
| ENSDARG000 | 17 | 39641681 | 39641798 | - | 118 | 5S_rRNA | 0 | 0 | 0 | 0 | 0 | 0 |
| ENSDARG000 | 17 | 14262422 | 14262538 | - | 117 | 5S_rRNA | 0 | 0 | 0 | 0 | 0 | 0 |
| ENSDARG000 | 17;17;17;17;1 | 2348540:2354 | 2349465:2354 | -;-;-;-;-;-;-;-;- | 3490 | NA | 4 | 42 | 49 | 56 | 136 | 64 |
| ENSDARG000 | 17;17 | 50068460:500 | 50069603:500 | -;- | 1340 | NA | 0 | 4 | 12 | 34 | 2 | 12 |
| ENSDARG000 | 17 | 5962975 | 5963065 | - | 91 | dre-mir-133c | 0 | 0 | 0 | 0 | 0 | 0 |
| ENSDARG000 | 17;17;17;17;1 | 7229384:7236 | 7232842:7236 | -;-;-;-;-;-;-;-;- | 7083 | STXBP5 | 988 | 654 | 933 | 982 | 735 | 1094 |
| ENSDARG000 | 17 | 21130261 | 21130377 | - | 117 | 5S_rRNA | 0 | 0 | 0 | 0 | 0 | 0 |
| ENSDARG000 | 17 | 8522906 | 8523030 | - | 125 | 5S_rRNA | 0 | 0 | 0 | 0 | 0 | 0 |
| ENSDARG000 | 17;17;17;17;1 | 25296982:252 | 25297104:252 | -;-;-;-;-;-;-;-;- | 1255 | ppie | 149 | 149 | 229 | 182 | 135 | 116 |
| ENSDARG000 | 17 | 30496724 | 30496840 | + | 117 | 5S_rRNA | 1 | 0 | 0 | 0 | 0 | 3 |
| ENSDARG000 | 17;17;17;17;1 | 734133:73436 | 734291:73450 | -;-;-;-;-;-;-;- | 915 | WARS1 | 189 | 205 | 297 | 86 | 194 | 172 |
| ENSDARG000 | 17 | 16538205 | 16538320 | + | 116 | 5S_rRNA | 0 | 0 | 0 | 0 | 0 | 0 |

| | | | | | | | | | | | | |
|---|---|---|---|---|---|---|---|---|---|---|---|---|
| ENSDARG000 | 17;17;17;17;1 | 6452511;6452 | 6452623;6452 | +;+;+;+;+;+;+; | 1140 | TATDN3 | 36 | 63 | 32 | 46 | 38 | 72 |
| ENSDARG000 | 17;17;17;17;1 | 52059020;520 | 52059178;520 | -;-;-;-;-;-;- | 2160 | tgfb3 | 165 | 12 | 28 | 6 | 28 | 18 |
| ENSDARG000 | 17;17;17;17;1 | 25821388;258 | 25821410;258 | -;-;-;-;-;-;-;- | 162 | NA | 6 | 0 | 0 | 1 | 5 | 0 |
| ENSDARG000 | 17 | 12801464 | 12801579 | + | 116 | 5S_rRNA | 0 | 0 | 0 | 0 | 0 | 0 |
| ENSDARG000 | 17;17;17;17;1 | 51020114;510 | 51020395;510 | -;-;-;-;-;-;-;-; | 4575 | aqr | 1211 | 922 | 1721 | 807 | 1526 | 812 |
| ENSDARG000 | 17 | 15901425 | 15901547 | + | 123 | 5S_rRNA | 0 | 0 | 0 | 0 | 0 | 0 |
| ENSDARG000 | 17 | 44291405 | 44291520 | + | 116 | 5S_rRNA | 0 | 0 | 0 | 0 | 0 | 0 |
| ENSDARG000 | 17;17;17;17;1 | 1632294;1641 | 1632839;1641 | +;+;+;+;+ | 855 | srp14 | 327 | 636 | 735 | 593 | 824 | 430 |
| ENSDARG000 | 17;17;17 | 28073062;280 | 28073115;280 | +;+;+ | 429 | NA | 0 | 0 | 0 | 2 | 0 | 0 |
| ENSDARG000 | 17;17;17;17;1 | 5931530;5931 | 5931883;5932 | +;+;+;+;+;+;+; | 2393 | ZNF513 | 78 | 48 | 148 | 54 | 84 | 66 |
| ENSDARG000 | 17;17;17;17;1 | 53284040;532 | 53284063;532 | +;+;+;+;+;+;+; | 936 | atxn3 | 241 | 198 | 228 | 198 | 168 | 185 |
| ENSDARG000 | 17 | 36614315 | 36614432 | + | 118 | 5S_rRNA | 0 | 0 | 0 | 0 | 0 | 0 |
| ENSDARG000 | 17 | 5967249 | 5967346 | - | 98 | mir206-2 | 0 | 0 | 0 | 0 | 0 | 0 |
| ENSDARG000 | 17;17;17;17;1 | 49484640;494 | 49485514;494 | +;+;+;+;+;+;+; | 7238 | AREL1 | 694 | 445 | 744 | 366 | 719 | 614 |
| ENSDARG000 | 17 | 39692582 | 39692696 | + | 115 | 5S_rRNA | 0 | 0 | 0 | 0 | 0 | 0 |
| ENSDARG000 | 17 | 2165656 | 2165770 | - | 115 | 5S_rRNA | 0 | 0 | 0 | 0 | 0 | 0 |
| ENSDARG000 | 17;17 | 7124059;7217 | 7124085;7218 | -;- | 1196 | SAMD5 | 0 | 3 | 22 | 7 | 15 | 0 |
| ENSDARG000 | 17;17;17;17;1 | 5976683;5985 | 5976908;5985 | +;+;+;+;+;+;+; | 1341 | plaat1l | 0 | 0 | 8 | 2 | 22 | 0 |
| ENSDARG000 | 17;17;17;17 | 46451371;464 | 46451553;464 | -;-;-;- | 774 | TMEM179 | 0 | 8 | 2 | 12 | 0 | 0 |
| ENSDARG000 | 17 | 24617464 | 24617542 | + | 79 | SNORD33 | 2 | 1 | 7 | 4 | 5 | 4 |
| ENSDARG000 | 17;17;17;17;1 | 12687687;126 | 12687718;126 | -;-;-;-;-;-;- | 171 | NA | 0 | 0 | 0 | 0 | 0 | 0 |
| ENSDARG000 | 17 | 49451177 | 49451298 | - | 122 | 5S_rRNA | 0 | 0 | 0 | 0 | 0 | 0 |
| ENSDARG000 | 17;17;17;17;1 | 6393719;6394 | 6394563;6394 | -;-;-;-;- | 5816 | NA | 10 | 20 | 6 | 2 | 42 | 2 |
| ENSDARG000 | 17;17;17;17;1 | 48314724;483 | 48314919;483 | +;+;+;+;+;+;+; | 1386 | smoc1 | 16 | 0 | 0 | 0 | 4 | 0 |
| ENSDARG000 | 17 | 51908071 | 51908186 | + | 116 | 5S_rRNA | 0 | 0 | 0 | 0 | 0 | 0 |
| ENSDARG000 | 17 | 37309329 | 37309446 | - | 118 | 5S_rRNA | 0 | 0 | 0 | 0 | 0 | 0 |
| ENSDARG000 | 17 | 39778647 | 39778744 | + | 98 | dre-mir-2185- | 0 | 0 | 0 | 0 | 0 | 0 |
| ENSDARG000 | 17 | 265173 | 265287 | + | 115 | 5S_rRNA | 0 | 0 | 0 | 0 | 0 | 0 |
| ENSDARG000 | 17 | 23383599 | 23383739 | + | 141 | U4 | 0 | 0 | 0 | 0 | 0 | 0 |
| ENSDARG000 | 17;17;17;17 | 13031497;130 | 13032681;130 | +;+;+;+ | 2230 | FBXO33 | 446 | 237 | 340 | 320 | 302 | 326 |
| ENSDARG000 | 17;17;17;17;1 | 8574646;8581 | 8578906;8581 | -;-;-;-;-;- | 5229 | NA | 2 | 2 | 0 | 0 | 2 | 2 |
| ENSDARG000 | 17;17;17;17;1 | 22014345;220 | 22014390;220 | -;-;-;-;-;-;-;-; | 1674 | NA | 0 | 0 | 0 | 0 | 0 | 0 |
| ENSDARG000 | 17 | 11374774 | 11374896 | + | 123 | 5S_rRNA | 0 | 2 | 0 | 0 | 2 | 0 |
| ENSDARG000 | 17 | 10954241 | 10954356 | - | 116 | 5S_rRNA | 0 | 0 | 0 | 0 | 0 | 0 |
| ENSDARG000 | 17;17;17;17;1 | 3717384;3717 | 3717473;3717 | -;-;-;-;-;-;-;-; | 3429 | PLCB4 | 4 | 0 | 0 | 0 | 2 | 0 |
| ENSDARG000 | 17;17;17;17 | 23255365;232 | 23255371;232 | +;+;+;+ | 1146 | NA | 8 | 1 | 0 | 1 | 0 | 0 |
| ENSDARG000 | 17 | 836949 | 837063 | + | 115 | 5S_rRNA | 0 | 0 | 0 | 0 | 0 | 0 |
| ENSDARG000 | 17 | 36950438 | 36950555 | - | 118 | 5S_rRNA | 0 | 0 | 0 | 0 | 0 | 0 |
| ENSDARG000 | 17 | 41759808 | 41759925 | + | 118 | 5S_rRNA | 0 | 0 | 0 | 0 | 0 | 0 |
| ENSDARG000 | 17 | 23356350 | 23356490 | + | 141 | U4 | 0 | 0 | 0 | 0 | 0 | 0 |
| ENSDARG000 | 17;17;17;17;1 | 47086682;470 | 47086712;470 | -;-;-;-;-;-;-;-; | 213 | NA | 0 | 0 | 4 | 0 | 0 | 0 |
| ENSDARG000 | 17;17;17;17;1 | 3281440;3281 | 3283329;3283 | -;-;-;-;-;-;-;-; | 3255 | WSCD1 | 6 | 2 | 0 | 0 | 2 | 8 |
| ENSDARG000 | 17;17;17;17 | 1360192;1362 | 1360312;1362 | +;+;+;+ | 2401 | siva1 | 54 | 132 | 110 | 86 | 142 | 122 |
| ENSDARG000 | 17;17;17 | 741424;74658 | 744299;74672 | -;-;- | 3160 | NA | 0 | 2 | 14 | 2 | 12 | 12 |
| ENSDARG000 | 17;17 | 47083446;470 | 47085431;470 | -;- | 5679 | NA | 4 | 0 | 0 | 0 | 0 | 0 |
| ENSDARG000 | 17;17;17;17;1 | 5846202;5895 | 5847234;5896 | +;+;+;+;+;+;+; | 4879 | FNDC4 | 6 | 0 | 0 | 0 | 0 | 0 |
| ENSDARG000 | 17;17;17;17;1 | 918005;91825 | 918061;91844 | -;-;-;-;-;-;-;-; | 5295 | DYNC1H1 | 11092 | 3789 | 6893 | 5736 | 7981 | 4978 |
| ENSDARG000 | 17;17;17 | 49566461;495 | 49567413;495 | -;-;- | 1257 | NA | 0 | 0 | 0 | 0 | 0 | 0 |
| ENSDARG000 | 17;17;17;17;1 | 50798484;508 | 50798645;508 | +;+;+;+;+;+ | 2215 | stxbp6 | 1054 | 1441 | 1804 | 1271 | 1876 | 879 |
| ENSDARG000 | 17;17;17;17;1 | 1998041;1998 | 2000333;2000 | -;-;-;-;-;-;-;-; | 3855 | spint1a | 3079 | 183 | 115 | 152 | 118 | 352 |
| ENSDARG000 | 17;17;17;17;1 | 13099476;131 | 13099617;131 | +;+;+;+;+;+;+; | 2592 | pnn | 1761 | 1295 | 2480 | 1364 | 1915 | 1675 |
| ENSDARG000 | 17 | 46740206 | 46740303 | - | 98 | NA | 0 | 0 | 0 | 0 | 0 | 0 |
| ENSDARG000 | 17 | 2673793 | 2673907 | + | 115 | 5S_rRNA | 0 | 0 | 0 | 0 | 0 | 0 |
| ENSDARG000 | 17;17;17;17;1 | 1193815;1256 | 1194063;1256 | +;+;+;+;+;+ | 5253 | NA | 217 | 89 | 82 | 90 | 161 | 248 |
| ENSDARG000 | 17;17;17;17;1 | 1664632;1667 | 1665083;1668 | -;-;-;-;-;- | 1087 | xgb | 7 | 109 | 141 | 521 | 1560 | 6 |
| ENSDARG000 | 17;17;17;17;1 | 51940768;519 | 51940900;519 | +;+;+;+;+;+;+; | 3771 | NA | 69 | 56 | 96 | 42 | 58 | 36 |
| ENSDARG000 | 17 | 31384434 | 31384548 | - | 115 | 5S_rRNA | 0 | 0 | 0 | 0 | 0 | 0 |
| ENSDARG000 | 17 | 8542203 | 8545189 | + | 2987 | NA | 0 | 0 | 0 | 0 | 0 | 0 |
| ENSDARG000 | 17;17;17;17;1 | 53273684;532 | 53273980;532 | -;-;-;-;-;-;-;-; | 1541 | GABRR1 | 229 | 158 | 343 | 207 | 254 | 224 |
| ENSDARG000 | 17;17;17;17;1 | 1323699;1330 | 1323902;1330 | +;+;+;+;+;+;+; | 4028 | adss1 | 3276 | 3455 | 1019 | 3889 | 432 | 1750 |
| ENSDARG000 | 17;17;17;17 | 40989973;409 | 40990107;409 | +;+;+;+ | 749 | mrpl33 | 152 | 70 | 37 | 28 | 24 | 177 |
| ENSDARG000 | 17;17;17;17;1 | 39834947;398 | 39834976;398 | +;+;+;+;+;+;+; | 3177 | UBR1 | 465 | 298 | 532 | 295 | 413 | 335 |
| ENSDARG000 | 17 | 39426092 | 39426212 | + | 121 | 5S_rRNA | 0 | 0 | 0 | 0 | 0 | 0 |
| ENSDARG000 | 17;17;17;17 | 51934501;519 | 51934589;519 | -;-;-;- | 1202 | ERG28 | 10 | 22 | 27 | 14 | 0 | 21 |
| ENSDARG000 | 17;17;17;17;1 | 6511423;6512 | 6511605;6512 | -;-;-;-;-;-;-;-; | 1571 | TP53I3 | 10 | 18 | 23 | 21 | 25 | 8 |
| ENSDARG000 | 17 | 8472101 | 8472217 | + | 117 | 5S_rRNA | 0 | 0 | 8 | 0 | 0 | 0 |
| ENSDARG000 | 17 | 32246129 | 32246255 | - | 127 | 5S_rRNA | 0 | 0 | 0 | 0 | 0 | 0 |
| ENSDARG000 | 17;17;17;17;1 | 2580541;2580 | 2580635;2580 | -;-;-;-;-;-;-;- | 1332 | zp3 | 0 | 1 | 0 | 4 | 0 | 0 |
| ENSDARG000 | 17 | 11682 | 11797 | + | 116 | 5S_rRNA | 0 | 0 | 0 | 0 | 0 | 0 |
| ENSDARG000 | 17;17;17;17 | 450956;45095 | 451103;45110 | +;+;+;+ | 886 | zgc:194887 | 0 | 0 | 0 | 0 | 0 | 0 |
| ENSDARG000 | 17 | 463746 | 465285 | - | 1540 | chrm5a | 0 | 0 | 0 | 0 | 0 | 0 |
| ENSDARG000 | 17 | 996509 | 998695 | + | 2187 | cyp1c2 | 4 | 0 | 0 | 0 | 6 | 10 |
| ENSDARG000 | 17;17;17;17;1 | 6985428;6985 | 6988440;6988 | -;-;-;-;-;-;-;-; | 6507 | SASH1 | 4815 | 3279 | 5629 | 3696 | 6283 | 3814 |
| ENSDARG000 | 17 | 18798995 | 18799131 | + | 137 | 5S_rRNA | 0 | 0 | 0 | 0 | 0 | 0 |
| ENSDARG000 | 17 | 22786174 | 22786291 | - | 118 | 5S_rRNA | 0 | 0 | 1 | 0 | 0 | 0 |
| ENSDARG000 | 17 | 33585346 | 33585462 | + | 117 | 5S_rRNA | 0 | 0 | 0 | 0 | 0 | 0 |
| ENSDARG000 | 17 | 4660919 | 4661043 | + | 125 | 5S_rRNA | 0 | 0 | 0 | 0 | 0 | 0 |
| ENSDARG000 | 17 | 37453676 | 37453807 | + | 132 | 5S_rRNA | 0 | 0 | 2 | 0 | 3 | 0 |
| ENSDARG000 | 17 | 4870403 | 4870522 | + | 120 | 5S_rRNA | 0 | 0 | 0 | 0 | 0 | 0 |
| ENSDARG000 | 17 | 8300390 | 8300504 | + | 115 | 5S_rRNA | 0 | 0 | 0 | 0 | 0 | 0 |
| ENSDARG000 | 17 | 23341874 | 23342014 | + | 141 | U4 | 0 | 0 | 0 | 0 | 0 | 0 |
| ENSDARG000 | 17 | 38724473 | 38725534 | + | 1062 | GPR68 | 8 | 2 | 0 | 0 | 2 | 0 |
| ENSDARG000 | 17;17;17;17;1 | 12924930;129 | 12925312;129 | -;-;-;-;- | 1350 | INSM2 | 12 | 10 | 6 | 2 | 10 | 32 |
| ENSDARG000 | 17;17;17;17;1 | 45071531;450 | 45073681;450 | -;-;-;-;-;-;-;-; | 3880 | aldh6a1 | 2145 | 2030 | 3606 | 2426 | 2995 | 3514 |
| ENSDARG000 | 17 | 52619473 | 52619590 | - | 118 | 5S_rRNA | 0 | 0 | 0 | 0 | 0 | 0 |
| ENSDARG000 | 17 | 46670007 | 46670103 | - | 97 | NA | 0 | 0 | 0 | 0 | 0 | 0 |
| ENSDARG000 | 17 | 22925209 | 22925337 | - | 129 | 5S_rRNA | 0 | 0 | 0 | 0 | 0 | 0 |
| ENSDARG000 | 17;17;17;17;1 | 49529197;495 | 49529427;495 | -;-;-;-;-;-;- | 1428 | NA | 0 | 0 | 0 | 0 | 8 | 0 |
| ENSDARG000 | 17;17;17;17;1 | 23212727;232 | 23213434;232 | -;-;-;-;-;-;-;- | 1584 | FAM98A | 342 | 584 | 514 | 228 | 610 | 296 |
| ENSDARG000 | 17;17;17;17;1 | 48920799;489 | 48922318;489 | -;-;-;-;-;- | 2564 | gnmt | 487 | 415 | 400 | 695 | 318 | 1283 |
| ENSDARG000 | 17 | 35915010 | 35915126 | + | 117 | 5S_rRNA | 0 | 0 | 0 | 0 | 0 | 0 |
| ENSDARG000 | 17 | 22319080 | 22324727 | - | 5648 | NA | 0 | 0 | 0 | 0 | 0 | 4 |
| ENSDARG000 | 17;17;17;17 | 6625717;6625 | 6626151;6626 | +;+;+;+ | 4612 | NA | 0 | 0 | 0 | 0 | 0 | 0 |
| ENSDARG000 | 17;17;17;17;1 | 4578552;4578 | 4579723;4579 | +;+;+;+;+;+;+; | 5665 | KLHL29 | 0 | 0 | 28 | 10 | 16 | 40 |
| ENSDARG000 | 17;17;17;17 | 890988;89152 | 890989;89174 | +;+;+;+ | 297 | NA | 1613 | 858 | 995 | 921 | 1006 | 746 |
| ENSDARG000 | 17;17;17;17;1 | 48433817;484 | 48434081;484 | +;+;+;+;+;+;+; | 5631 | NA | 2 | 0 | 0 | 0 | 0 | 0 |
| ENSDARG000 | 17;17;17;17 | 443264;44367 | 443391;44380 | +;+;+;+ | 628 | emc7a | 2 | 8 | 2 | 2 | 6 | 0 |
| ENSDARG000 | 17;17;17;17;1 | 23237239;232 | 23237331;232 | -;-;-;-;- | 1055 | NA | 466 | 387 | 363 | 373 | 390 | 284 |
| ENSDARG000 | 17;17;17;17;1 | 42208623;422 | 42212100;422 | -;-;-;-;-;-;-;-; | 4649 | nkx2.2a | 0 | 0 | 0 | 0 | 0 | 0 |
| ENSDARG000 | 17 | 50524573 | 50525944 | + | 1372 | NA | 0 | 0 | 0 | 0 | 0 | 0 |
| ENSDARG000 | 17 | 38987296 | 38987415 | - | 120 | 5S_rRNA | 0 | 0 | 0 | 0 | 0 | 0 |
| ENSDARG000 | 17 | 23331419 | 23331559 | + | 141 | U4 | 0 | 0 | 0 | 0 | 0 | 0 |
| ENSDARG000 | 17;17;17;17 | 40211155;402 | 40216791;402 | -;-;-;- | 7246 | LRFN2 | 0 | 0 | 0 | 0 | 0 | 0 |
| ENSDARG000 | 17;17;17 | 1627379;1627 | 1627406;1628 | +;+;+ | 213 | NA | 18 | 13 | 1 | 0 | 13 | 13 |
| ENSDARG000 | 17;17;17;17 | 729423;72979 | 729605;73005 | +;+;+;+ | 699 | NA | 10 | 8 | 16 | 4 | 4 | 9 |
| ENSDARG000 | 17;17;17;17;1 | 45305645;453 | 45305928;453 | +;+;+;+;+;+;+; | 3218 | capn3a | 40 | 23 | 19 | 67 | 63 | 21 |
| ENSDARG000 | 17 | 33298348 | 33298470 | + | 123 | 5S_rRNA | 2 | 0 | 0 | 0 | 0 | 0 |
| ENSDARG000 | 17;17;17;17;1 | 135590;13615 | 136016;13622 | +;+;+;+;+;+ | 890 | NA | 196 | 198 | 138 | 101 | 167 | 104 |
| ENSDARG000 | 17 | 22827063 | 22827147 | + | 85 | dre-mir-456 | 0 | 0 | 0 | 0 | 0 | 0 |
| ENSDARG000 | 17 | 44058811 | 44058927 | - | 117 | 5S_rRNA | 0 | 0 | 0 | 0 | 0 | 0 |
| ENSDARG000 | 17 | 37272019 | 37272133 | + | 115 | 5S_rRNA | 4 | 0 | 0 | 0 | 1 | 0 |
| ENSDARG000 | 17;17;17;17;1 | 2162916;2214 | 2163346;2214 | +;+;+;+;+;+;+; | 2564 | PAK6 | 2 | 0 | 8 | 0 | 0 | 0 |
| ENSDARG000 | 17;17;17;17;1 | 51262556;512 | 51262705;512 | +;+;+;+;+;+;+; | 10037 | eipr1 | 543 | 486 | 632 | 429 | 732 | 530 |
| ENSDARG000 | 17;17;17;17 | 50044389;500 | 50044454;500 | -;-;-;- | 513 | NA | 0 | 0 | 0 | 0 | 1 | 0 |
| ENSDARG000 | 17 | 19900754 | 19900857 | - | 104 | dre-mir-2187 | 0 | 0 | 0 | 0 | 0 | 0 |
| ENSDARG000 | 17;17;17 | 50052174;500 | 50053137;500 | -;-;- | 1489 | NA | 0 | 0 | 0 | 0 | 0 | 0 |
| ENSDARG000 | 17 | 39149567 | 39149683 | - | 117 | 5S_rRNA | 0 | 0 | 0 | 0 | 0 | 0 |
| ENSDARG000 | 17;17 | 50037644;500 | 50040553;500 | -;- | 2983 | NA | 0 | 2 | 0 | 0 | 0 | 0 |
| ENSDARG000 | 17;17;17;17 | 31914877;319 | 31915660;319 | +;+;+;+ | 2124 | INSYN2A | 2 | 0 | 4 | 0 | 0 | 0 |
| ENSDARG000 | 17;17;17;17;1 | 50052747;500 | 50052934;500 | -;-;-;-;- | 1182 | NA | 0 | 0 | 2 | 0 | 1 | 0 |
| ENSDARG000 | 17 | 11386859 | 11386940 | - | 82 | NA | 0 | 1 | 4 | 0 | 10 | 2 |
| ENSDARG000 | 17 | 25861193 | 25861787 | - | 595 | NA | 0 | 0 | 0 | 0 | 0 | 0 |
| ENSDARG000 | 17;17;17;17 | 53356451;533 | 53356491;533 | +;+;+;+ | 513 | NA | 124 | 84 | 117 | 65 | 84 | 33 |
| ENSDARG000 | 17 | 46849249 | 46849346 | + | 98 | NA | 0 | 0 | 0 | 0 | 0 | 0 |
| ENSDARG000 | 17 | 46819748 | 46819845 | - | 98 | NA | 0 | 0 | 0 | 0 | 0 | 0 |
| ENSDARG000 | 17 | 11398076 | 11398157 | - | 82 | NA | 0 | 0 | 0 | 0 | 0 | 0 |
| ENSDARG000 | 17 | 39771488 | 39771585 | + | 98 | dre-mir-2185- | 0 | 0 | 0 | 0 | 0 | 0 |
| ENSDARG000 | 17;17;17 | 262803;26722 | 262961;26732 | -;-;- | 381 | NA | 0 | 2 | 0 | 0 | 0 | 0 |
| ENSDARG000 | 17 | 2928074 | 2928192 | + | 119 | 5S_rRNA | 0 | 0 | 0 | 0 | 0 | 0 |
| ENSDARG000 | 17;17;17;17;1 | 1393122;1393 | 1393548;1393 | -;-;-;-;-;- | 1433 | NA | 45 | 60 | 61 | 34 | 45 | 28 |
| ENSDARG000 | 17;17;17 | 51499789;515 | 51499792;515 | +;+;+ | 411 | NA | 4 | 0 | 0 | 0 | 0 | 0 |
| ENSDARG000 | 17 | 26619840 | 26619957 | + | 118 | 5S_rRNA | 0 | 0 | 0 | 0 | 0 | 0 |
| ENSDARG000 | 17 | 23350420 | 23350560 | + | 141 | U4 | 0 | 0 | 0 | 0 | 0 | 0 |
| ENSDARG000 | 17;17;17;17;1 | 956821;96775 | 957395;96795 | +;+;+;+;+;+;+; | 3438 | NA | 981 | 759 | 906 | 974 | 1059 | 819 |
| ENSDARG000 | 17 | 7320042 | 7320165 | + | 124 | 5S_rRNA | 0 | 0 | 0 | 0 | 0 | 0 |
| ENSDARG000 | 17 | 32069258 | 32069372 | + | 115 | 5S_rRNA | 0 | 0 | 0 | 0 | 0 | 0 |
| ENSDARG000 | 17 | 1499230 | 1499349 | + | 120 | 5S_rRNA | 0 | 0 | 0 | 0 | 0 | 0 |
| ENSDARG000 | 17;17;17;17;1 | 52300018;523 | 52300443;523 | +;+;+;+;+;+ | 2426 | esrrb | 0 | 6 | 14 | 0 | 16 | 0 |
| ENSDARG000 | 17;17;17;17;1 | 1074503;1120 | 1074749;1120 | -;-;-;-;-;-;-;-; | 1087 | dnajc17 | 54 | 96 | 92 | 74 | 164 | 57 |
| ENSDARG000 | 17;17;17;17;1 | 2130018;2130 | 2130169;2130 | +;+;+;+;+;+;+; | 6078 | NA | 170 | 440 | 476 | 160 | 375 | 255 |
| ENSDARG000 | 17;17;17;17;1 | 53349626;533 | 53350071;533 | -;-;-;-;-;-;-;-; | 1839 | NA | 972 | 498 | 587 | 666 | 546 | 460 |
| ENSDARG000 | 17;17;17;17;1 | 1992495;1993 | 1992513;1993 | +;+;+;+;+;+;+; | 345 | NA | 0 | 0 | 0 | 0 | 0 | 2 |
| ENSDARG000 | 17 | 23223804 | 23223920 | + | 117 | 5S_rRNA | 0 | 1 | 0 | 0 | 0 | 0 |
| ENSDARG000 | 17;17;17;17;1 | 15111319;151 | 15111335;151 | +;+;+;+;+;+;+; | 216 | NA | 2 | 5 | 6 | 2 | 0 | 8 |
| ENSDARG000 | 17 | 28530467 | 28530582 | - | 116 | 5S_rRNA | 0 | 0 | 0 | 0 | 0 | 0 |
| ENSDARG000 | 17 | 7661208 | 7661324 | + | 117 | 5S_rRNA | 0 | 0 | 0 | 0 | 0 | 0 |
| ENSDARG000 | 17;17;17;17;1 | 2587132;2587 | 2587536;2587 | -;-;-;-;-;-;-;- | 1618 | zp3.2 | 0 | 0 | 0 | 3 | 0 | 0 |
| ENSDARG000 | 17 | 32456803 | 32456919 | + | 117 | 5S_rRNA | 0 | 0 | 0 | 0 | 0 | 0 |
| ENSDARG000 | 17 | 496611 | 496725 | + | 115 | 5S_rRNA | 0 | 0 | 0 | 0 | 0 | 0 |
| ENSDARG000 | 17 | 17879594 | 17879709 | - | 116 | 5S_rRNA | 0 | 0 | 0 | 0 | 0 | 0 |
| ENSDARG000 | 17 | 12868830 | 12868944 | + | 115 | 5S_rRNA | 0 | 0 | 0 | 0 | 0 | 0 |
| ENSDARG000 | 17 | 35617986 | 35618104 | - | 119 | 5S_rRNA | 0 | 0 | 0 | 0 | 0 | 0 |
| ENSDARG000 | 17 | 47197876 | 47197992 | + | 117 | 5S_rRNA | 0 | 0 | 0 | 0 | 0 | 0 |
| ENSDARG000 | 17 | 13340933 | 13341049 | - | 117 | 5S_rRNA | 0 | 0 | 0 | 0 | 0 | 0 |
| ENSDARG000 | 17 | 29637205 | 29637321 | - | 117 | 5S_rRNA | 0 | 0 | 0 | 0 | 0 | 0 |
| ENSDARG000 | 17;17;17;17 | 706630;71313 | 707373;71322 | -;-;-;- | 906 | SLC25A29 | 0 | 0 | 0 | 2 | 0 | 0 |
| ENSDARG000 | 17;17;17;17;1 | 189276;18933 | 189300;18935 | -;-;-;-;-;-;- | 942 | NA | 0 | 0 | 2 | 0 | 6 | 4 |
| ENSDARG000 | 17;17;17;17;1 | 41302660;413 | 41303244;413 | +;+;+;+;+;+ | 3632 | fosl2 | 1863 | 672 | 920 | 1100 | 721 | 955 |
| ENSDARG000 | 17;17;17;17;1 | 2108984;2110 | 2109065;2110 | -;-;-;-;-;-;-;-; | 1293 | zfyve19 | 541 | 344 | 675 | 467 | 559 | 482 |
| ENSDARG000 | 17;17;17 | 49452032;494 | 49452125;494 | +;+;+ | 363 | CGA | 2 | 0 | 0 | 0 | 0 | 0 |
| ENSDARG000 | 17;17;17;17 | 856681;86003 | 856734;86132 | -;-;-;- | 1791 | WDR20 | 369 | 353 | 407 | 221 | 396 | 204 |
| ENSDARG000 | 17;17;17;17;1 | 691453;69207 | 691615;69218 | +;+;+;+;+;+;+; | 1380 | NA | 16 | 48 | 77 | 98 | 86 | 34 |

| | | | | | | | | | | | | |
|---|---|---|---|---|---|---|---|---|---|---|---|---|
| ENSDARG000 | 17:17:17:17:1 | 48906344:489 | 48906493:489 | -;-;-;-;-;-;- | 876 | LGALS8 | 0 | 0 | 0 | 0 | 0 | 0 |
| ENSDARG000 | 17 | 6013555 | 6013671 | - | 117 | 5S_rRNA | 0 | 0 | 0 | 0 | 0 | 0 |
| ENSDARG000 | 17;17;17 | 53356403:533 | 53356422:533 | +;+;+ | 291 | NA | 15 | 5 | 6 | 4 | 8 | 3 |
| ENSDARG000 | 17 | 23335601 | 23335741 | + | 141 | U4 | 0 | 0 | 0 | 0 | 0 | 0 |
| ENSDARG000 | 17 | 3291340 | 3291456 | - | 117 | 5S_rRNA | 0 | 0 | 0 | 0 | 0 | 0 |
| ENSDARG000 | 17 | 25864305 | 25864419 | - | 115 | 5S_rRNA | 0 | 0 | 0 | 0 | 0 | 0 |
| ENSDARG000 | 17 | 17778968 | 17779083 | - | 116 | 5S_rRNA | 0 | 0 | 0 | 0 | 0 | 0 |
| ENSDARG000 | 17 | 49510866 | 49511000 | + | 135 | 5S_rRNA | 0 | 0 | 0 | 0 | 0 | 0 |
| ENSDARG000 | 17 | 8366705 | 8366816 | - | 112 | NA | 0 | 0 | 0 | 0 | 0 | 0 |
| ENSDARG000 | 17 | 1704442 | 1705013 | - | 572 | NA | 34 | 13 | 14 | 20 | 27 | 6 |
| ENSDARG000 | 17 | 23346238 | 23346378 | + | 141 | U4 | 0 | 0 | 0 | 0 | 0 | 0 |
| ENSDARG000 | 17;17 | 6763586:676 | 6763772:676 | -;- | 219 | NA | 0 | 0 | 0 | 0 | 0 | 0 |
| ENSDARG000 | 17:17:17:17:1 | 660126:6655 | 660851:6657 | -;-;-;-;- | 1455 | soul3 | 1173 | 721 | 1406 | 1793 | 777 | 800 |
| ENSDARG000 | 17:17:17:17:1 | 45106424:451 | 45106465:451 | +;+;+;+;+;+;+ | 1574 | gstz1 | 391 | 215 | 275 | 449 | 448 | 452 |
| ENSDARG000 | 17:17:17:17:1 | 26593562:265 | 26593588:265 | +;+;+;+;+;+;+ | 213 | NA | 0 | 0 | 1 | 0 | 0 | 0 |
| ENSDARG000 | 17;17;17 | 1063988:106 | 1064123:106 | +;+;+ | 778 | gchfr | 32 | 48 | 40 | 58 | 72 | 76 |
| ENSDARG000 | 17;17 | 2902113:294 | 2906417:294 | -;- | 4781 | C17H14orf13 | 0 | 0 | 0 | 0 | 0 | 0 |
| ENSDARG000 | 17 | 20703179 | 20703297 | - | 119 | 5S_rRNA | 0 | 0 | 0 | 0 | 0 | 0 |
| ENSDARG000 | 17;17 | 37249736:372 | 37250317:372 | +;+ | 654 | NA | 1 | 0 | 0 | 0 | 8 | 0 |
| ENSDARG000 | 17 | 39785815 | 39785912 | + | 98 | dre-mir-2185- | 0 | 0 | 0 | 0 | 0 | 0 |
| ENSDARG000 | 17;17;17;17 | 50057704:500 | 50058667:500 | -;-;-;- | 1486 | NA | 0 | 2 | 2 | 0 | 0 | 0 |
| ENSDARG000 | 17 | 23387806 | 23387946 | + | 141 | U4 | 0 | 0 | 0 | 0 | 0 | 0 |
| ENSDARG000 | 17 | 32652381 | 32652505 | + | 125 | 5S_rRNA | 0 | 0 | 0 | 0 | 0 | 0 |
| ENSDARG000 | 17 | 40684 | 41722 | + | 1039 | NA | 0 | 0 | 0 | 0 | 0 | 0 |
| ENSDARG000 | 17 | 803591 | 803706 | + | 116 | 5S_rRNA | 0 | 0 | 0 | 0 | 0 | 0 |
| ENSDARG000 | 17:17:17:17:1 | 1496107:149 | 1496153:149 | +;+;+;+;+;+;+ | 784 | NA | 291 | 142 | 206 | 107 | 228 | 168 |
| ENSDARG000 | 17:17:17:17:1 | 39873584:398 | 39873850:398 | -;-;-;-;-;-;- | 1984 | NA | 0 | 0 | 0 | 0 | 4 | 0 |
| ENSDARG000 | 17:17:17:17:1 | 43843536:438 | 43843734:438 | +;+;+;+;+;+ | 647 | NA | 1 | 0 | 0 | 0 | 2 | 0 |
| ENSDARG000 | 17:17:17:17:1 | 5623514:562 | 5623834:562 | +;+;+;+;+ | 1666 | NA | 0 | 8 | 4 | 0 | 0 | 8 |
| ENSDARG000 | 17 | 16852272 | 16852405 | - | 134 | 5S_rRNA | 0 | 0 | 0 | 0 | 0 | 0 |
| ENSDARG000 | 17:17:17:17:1 | 24588834:245 | 24588847:245 | -;-;-;-;-;-;-;-;- | 947 | NA | 7 | 6 | 15 | 2 | 11 | 6 |
| ENSDARG000 | 17 | 49227204 | 49227319 | - | 116 | 5S_rRNA | 0 | 0 | 0 | 2 | 0 | 0 |
| ENSDARG000 | 17;17 | 6382275:638 | 6382674:638 | +;+ | 1892 | GJE1 | 0 | 8 | 6 | 6 | 6 | 4 |
| ENSDARG000 | 17:17:17:17:1 | 33999630:339 | 33999693:339 | +;+;+;+;+;+;+ | 5582 | gphna | 1904 | 2531 | 4192 | 3389 | 3019 | 4444 |
| ENSDARG000 | 17:17:17:17:1 | 6331994:633 | 6332084:633 | -;-;-;-;-;-;-;-;- | 1996 | slc22a3 | 0 | 0 | 0 | 0 | 24 | 0 |
| ENSDARG000 | 17 | 45808153 | 45808248 | + | 96 | dre-mir-203a | 0 | 0 | 0 | 0 | 0 | 0 |
| ENSDARG000 | 17 | 18350726 | 18350845 | - | 120 | 5S_rRNA | 0 | 0 | 0 | 0 | 0 | 0 |
| ENSDARG000 | 17:17:17:17:1 | 13019499:130 | 13020792:130 | -;-;-;-;- | 1756 | fam177a1 | 1012 | 717 | 952 | 868 | 895 | 1236 |
| ENSDARG000 | 17 | 43941937 | 43942051 | + | 115 | 5S_rRNA | 0 | 0 | 0 | 0 | 0 | 0 |
| ENSDARG000 | 17;17;17;17 | 6275206:627 | 6275235:627 | -;-;-;- | 129 | NA | 0 | 0 | 0 | 0 | 0 | 0 |
| ENSDARG000 | 17;17;17 | 132555:1390 | 133297:1393 | +;+;+ | 1259 | lrr1 | 86 | 126 | 186 | 70 | 208 | 92 |
| ENSDARG000 | 17 | 23330171 | 23330311 | + | 141 | U4 | 0 | 0 | 0 | 0 | 0 | 0 |
| ENSDARG000 | 17 | 35431724 | 35437222 | + | 5499 | NA | 0 | 0 | 0 | 0 | 0 | 0 |
| ENSDARG000 | 17 | 36635072 | 36635189 | - | 118 | 5S_rRNA | 0 | 0 | 0 | 0 | 0 | 0 |
| ENSDARG000 | 17:17:17:17:1 | 49457158:494 | 49457642:494 | -;-;-;-;-;-;-;- | 1096 | fcf1 | 670 | 497 | 728 | 353 | 562 | 430 |
| ENSDARG000 | 17;17;17;17 | 14695446:146 | 14695499:146 | -;-;-;- | 384 | NA | 16 | 1 | 6 | 2 | 2 | 2 |
| ENSDARG000 | 17 | 47510280 | 47510396 | - | 117 | 5S_rRNA | 0 | 0 | 0 | 0 | 0 | 0 |
| ENSDARG000 | 17:17:17:17:1 | 44399360:444 | 44399392:444 | -;-;-;-;-;-;-;- | 756 | NA | 10 | 0 | 0 | 8 | 0 | 0 |
| ENSDARG000 | 17;17;17;17 | 337801:3464 | 338567:3472 | +;+;+;+ | 1762 | NA | 0 | 14 | 16 | 0 | 0 | 6 |
| ENSDARG000 | 17:17:17:17:1 | 634641:6352 | 634706:6354 | +;+;+;+;+;+;+ | 5925 | NA | 97 | 72 | 120 | 36 | 75 | 53 |
| ENSDARG000 | 17;17;17 | 811182:8118 | 811403:8119 | -;-;- | 494 | NA | 45 | 67 | 98 | 23 | 52 | 34 |
| ENSDARG000 | 17;17;17 | 1915788:192 | 1915934:192 | -;-;- | 580 | NA | 0 | 0 | 0 | 0 | 1 | 0 |
| ENSDARG000 | 17;17;17;17 | 2401258:241 | 2405367:241 | -;-;-;- | 4408 | NA | 2 | 2 | 2 | 0 | 2 | 2 |
| ENSDARG000 | 17;17;17 | 7676631:767 | 7676750:767 | -;-;- | 406 | NA | 0 | 0 | 0 | 0 | 0 | 0 |
| ENSDARG000 | 17;17;17;17 | 7741616:774 | 7741927:774 | +;+;+;+ | 2107 | NA | 1 | 1 | 7 | 6 | 6 | 0 |
| ENSDARG000 | 17;17;17;17 | 10920738:109 | 10920875:109 | -;-;-;- | 823 | NA | 0 | 0 | 0 | 0 | 0 | 0 |
| ENSDARG000 | 17:17:17:17:1 | 13221920:133 | 13222302:133 | +;+;+;+;+;+ | 732 | NA | 0 | 0 | 0 | 0 | 0 | 0 |
| ENSDARG000 | 17;17;17;17 | 13813423:138 | 13813580:138 | +;+;+;+ | 4907 | NA | 0 | 0 | 0 | 0 | 0 | 0 |
| ENSDARG000 | 17;17;17 | 18660353:186 | 18660836:186 | +;+;+ | 675 | NA | 0 | 0 | 0 | 0 | 0 | 0 |
| ENSDARG000 | 17;17;17;17 | 19023661:190 | 19024080:190 | +;+;+;+ | 837 | NA | 0 | 1 | 0 | 4 | 6 | 0 |
| ENSDARG000 | 17;17;17;17 | 19349313:193 | 19349640:193 | -;-;-;- | 1537 | NA | 0 | 0 | 0 | 0 | 0 | 0 |
| ENSDARG000 | 17:17:17:17:1 | 22521636:225 | 22524272:225 | -;-;-;-;- | 3552 | NA | 0 | 0 | 0 | 0 | 0 | 0 |
| ENSDARG000 | 17:17:17:17:1 | 23222177:232 | 23222602:232 | +;+;+;+;+;+;+ | 2394 | NA | 697 | 823 | 1127 | 651 | 982 | 545 |
| ENSDARG000 | 17:17:17:17:1 | 24803274:248 | 24803854:248 | +;+;+;+;+;+;+ | 1137 | NA | 0 | 2 | 0 | 0 | 0 | 0 |
| ENSDARG000 | 17;17;17 | 29976009:300 | 29976355:300 | -;-;- | 668 | NA | 0 | 0 | 0 | 0 | 0 | 0 |
| ENSDARG000 | 17;17;17 | 35554571:355 | 35558522:355 | -;-;- | 4356 | NA | 0 | 0 | 0 | 0 | 0 | 0 |
| ENSDARG000 | 17:17:17:17:1 | 35901065:359 | 35901651:359 | -;-;-;-;-;- | 1782 | NA | 0 | 0 | 0 | 0 | 0 | 0 |
| ENSDARG000 | 17;17;17 | 37900421:379 | 37900600:379 | +;+;+ | 1537 | NA | 0 | 0 | 0 | 0 | 0 | 1 |
| ENSDARG000 | 17:17:17:17:1 | 43708399:437 | 43708767:437 | -;-;-;-;-;-;-;-;- | 4298 | NA | 839 | 412 | 1077 | 896 | 964 | 678 |
| ENSDARG000 | 17;17;17;17 | 45177626:451 | 45177717:451 | +;+;+;+ | 1916 | NA | 0 | 0 | 0 | 0 | 0 | 0 |
| ENSDARG000 | 17:17:17:17:1 | 46561140:466 | 46561173:466 | -;-;-;-;-;- | 254 | NA | 0 | 0 | 0 | 0 | 0 | 0 |
| ENSDARG000 | 17;17;17 | 46978855:469 | 46978896:469 | -;-;- | 141 | NA | 0 | 0 | 0 | 0 | 0 | 0 |
| ENSDARG000 | 17:17:17:17:1 | 52476873:524 | 52477193:524 | -;-;-;-;-;-;- | 1199 | NA | 0 | 0 | 0 | 0 | 0 | 0 |
| ENSDARG000 | 17:17:17:17:1 | 53388239:533 | 53392736:533 | -;-;-;-;-;-;-;-;- | 5242 | NA | 0 | 4 | 18 | 9 | 1 | 8 |
| ENSDARG000 | 14:14:14:14:1 | 26379484:263 | 26382228:263 | -;-;-;-;-;-;-;-;- | 4425 | b4galt7 | 194 | 319 | 490 | 208 | 468 | 278 |
| ENSDARG000 | 14:14:14:14:1 | 4084120:408 | 4085056:408 | -;-;-;-;-;-;-;-;- | 2727 | irf2 | 468 | 418 | 355 | 324 | 314 | 296 |
| ENSDARG000 | 14:14:14:14:1 | 36886950:368 | 36886974:368 | +;+;+;+;+;+;+ | 1128 | LECT2 | 1161 | 377 | 426 | 158 | 260 | 611 |
| ENSDARG000 | 14:14:14:14:1 | 24215046:242 | 24215256:242 | +;+;+;+;+;+ | 2555 | stc2a | 62 | 12 | 125 | 38 | 58 | 66 |
| ENSDARG000 | 14:14:14:14:1 | 33264303:332 | 33264350:332 | +;+;+;+;+;+;+ | 1540 | pdzd11 | 189 | 189 | 209 | 210 | 353 | 174 |
| ENSDARG000 | 14:14:14:14:1 | 34501245:345 | 34501459:345 | +;+;+;+;+;+;+ | 1497 | NA | 9 | 10 | 0 | 2 | 2 | 20 |
| ENSDARG000 | 14:14:14:14:1 | 36885524:368 | 36885680:368 | +;+;+;+;+ | 729 | lect2.1 | 72 | 0 | 42 | 60 | 417 | 30 |
| ENSDARG000 | 14 | 7305686 | 7306983 | - | 1298 | NA | 0 | 0 | 0 | 0 | 0 | 0 |
| ENSDARG000 | 14:14:14:14:1 | 17329895:173 | 17330053:173 | +;+;+;+;+;+;+ | 5254 | WFS1 | 668 | 506 | 431 | 421 | 664 | 518 |
| ENSDARG000 | 14:14:14:14:1 | 33219370:332 | 33221360:332 | -;-;-;-;-;-;-;-;- | 4023 | GDPD2 | 88 | 0 | 8 | 2 | 30 | 34 |
| ENSDARG000 | 14:14:14:14:1 | 31668510:316 | 31670839:316 | -;-;-;-;-;-;-;-;- | 4931 | MAP7D3 | 3294 | 2778 | 3022 | 2932 | 3529 | 2864 |
| ENSDARG000 | 14:14:14:14:1 | 26395314:263 | 26395428:263 | -;-;-;-;-;-;-;-;- | 2091 | pimr212 | 0 | 0 | 0 | 0 | 0 | 0 |
| ENSDARG000 | 14;14 | 24241241:242 | 24241684:242 | +;+ | 1657 | nkx2.5 | 26 | 57 | 235 | 158 | 71 | 124 |
| ENSDARG000 | 14:14:14:14:1 | 7320948:732 | 7321273:732 | -;-;-;-;-;-;-;-;- | 3843 | trappc11 | 1864 | 1966 | 1863 | 1380 | 1545 | 1816 |
| ENSDARG000 | 14:14:14:14:1 | 17376940:173 | 17377047:173 | +;+;+;+;+;+;+ | 2218 | spon2b | 83 | 0 | 11 | 0 | 6 | 0 |
| ENSDARG000 | 14:14:14:14:1 | 12066695:120 | 12067025:120 | -;-;-;-;- | 1053 | tmsb1 | 262 | 26 | 54 | 22 | 94 | 71 |
| ENSDARG000 | 14:14:14:14:1 | 46199941:462 | 46200877:462 | -;-;-;-;-;-;-;-;- | 2725 | setd7 | 561 | 397 | 585 | 399 | 572 | 639 |
| ENSDARG000 | 14:14:14:14:1 | 21820034:218 | 21820147:218 | +;+;+;+;+;+;+ | 5010 | ctbp1 | 3946 | 6190 | 5842 | 4794 | 4745 | 6124 |
| ENSDARG000 | 14;14;14;14 | 52418247:524 | 52423474:524 | -;-;-;- | 6301 | rest | 912 | 700 | 1082 | 717 | 881 | 859 |
| ENSDARG000 | 14:14:14:14:1 | 9077314:908 | 9077671:908 | -;-;-;-;- | 667 | polr1d | 166 | 225 | 208 | 79 | 86 | 120 |
| ENSDARG000 | 14:14:14:14:1 | 24248391:242 | 24248814:242 | -;-;-;-;-;- | 928 | BNIP1 | 73 | 99 | 139 | 67 | 139 | 86 |
| ENSDARG000 | 14:14:14:14:1 | 12076821:120 | 12077318:120 | -;-;-;-;-;-;- | 1537 | prps1b | 414 | 829 | 1024 | 565 | 570 | 649 |
| ENSDARG000 | 14:14:14:14:1 | 4125363:413 | 4130161:413 | -;-;-;-;-;-;- | 5614 | CASP3 | 52 | 10 | 38 | 31 | 16 | 6 |
| ENSDARG000 | 14:14:14:14:1 | 26404640:264 | 26406022:264 | -;-;-;-;- | 2010 | tmed9 | 2932 | 3432 | 4102 | 2350 | 3874 | 2988 |
| ENSDARG000 | 14:14:14:14:1 | 17382803:173 | 17383027:173 | +;+;+;+;+ | 896 | NA | 0 | 0 | 0 | 0 | 0 | 0 |
| ENSDARG000 | 14:14:14:14:1 | 4151374:415 | 4151541:415 | +;+;+;+;+;+;+ | 1776 | dhrs13l1 | 444 | 348 | 352 | 339 | 490 | 490 |
| ENSDARG000 | 14;14 | 4160855:416 | 4161200:416 | -;- | 459 | NA | 0 | 6 | 0 | 0 | 1 | 1 |
| ENSDARG000 | 14:14:14:14:1 | 52440161:524 | 52440810:524 | +;+;+;+;+;+ | 1665 | b4galt1l | 558 | 674 | 743 | 409 | 1110 | 640 |
| ENSDARG000 | 14:14:14:14:1 | 34512011:345 | 34512333:345 | -;-;-;-;- | 976 | zgc:194246 | 2 | 72 | 0 | 0 | 0 | 98 |
| ENSDARG000 | 14;14;14 | 34510700:345 | 34510967:345 | -;-;- | 442 | NA | 0 | 6 | 0 | 0 | 0 | 10 |
| ENSDARG000 | 14:14:14:14:1 | 12109535:121 | 12109887:121 | +;+;+;+;+ | 2988 | kctd12b | 0 | 0 | 0 | 8 | 0 | 0 |
| ENSDARG000 | 14;14;14 | 9117226:912 | 9117777:912 | -;-;- | 795 | NA | 0 | 8 | 0 | 0 | 0 | 4 |
| ENSDARG000 | 14:14:14:14:1 | 17397534:173 | 17398140:173 | +;+;+;+;+;+;+ | 1831 | NA | 0 | 0 | 0 | 0 | 0 | 0 |
| ENSDARG000 | 14:14:14:14:1 | 52476520:524 | 52476895:524 | -;-;-;-;-;-;- | 1281 | exosc3 | 75 | 39 | 100 | 41 | 41 | 76 |
| ENSDARG000 | 14:14:14:14:1 | 4163773:416 | 4164238:416 | -;-;-;-;-;-;-;-;- | 2713 | NA | 36 | 151 | 134 | 116 | 134 | 86 |
| ENSDARG000 | 14;14;14;14 | 12169979:121 | 12170030:121 | +;+;+;+ | 3587 | rhogd | 260 | 308 | 256 | 196 | 363 | 276 |
| ENSDARG000 | 14:14:14:14:1 | 12142813:121 | 12143042:121 | +;+;+;+;+;+;+ | 4655 | hspa9 | 2979 | 4389 | 6612 | 2385 | 3523 | 3784 |
| ENSDARG000 | 14:14:14:14:1 | 46216703:462 | 46216890:462 | +;+;+;+;+;+;+ | 1106 | mgst2 | 32 | 50 | 34 | 48 | 14 | 53 |
| ENSDARG000 | 14:14:14:14:1 | 36896641:368 | 36897162:368 | +;+;+;+;+;+;+ | 983 | NA | 0 | 0 | 0 | 0 | 0 | 0 |
| ENSDARG000 | 14:14:14:14:1 | 9142162:915 | 9145096:915 | -;-;-;-;-;-;-;-;- | 4616 | arhgef9b | 55 | 24 | 4 | 14 | 16 | 48 |
| ENSDARG000 | 14:14:14:14:1 | 30773764:307 | 30774120:307 | +;+;+;+;+;+;+ | 4852 | atl3 | 8234 | 5917 | 6722 | 5375 | 8259 | 4897 |
| ENSDARG000 | 14;14 | 34514336:345 | 34515098:345 | +;+ | 1865 | foxi3b | 0 | 0 | 0 | 0 | 0 | 0 |
| ENSDARG000 | 14:14:14:14:1 | 80685:81919 | 81309:82086 | +;+;+;+;+;+;+ | 2052 | NA | 3 | 0 | 0 | 0 | 0 | 0 |
| ENSDARG000 | 14:14:14:14:1 | 14841685:148 | 14841837:148 | +;+;+;+;+;+;+ | 1393 | slbp | 159 | 707 | 565 | 314 | 494 | 362 |
| ENSDARG000 | 14:14:14:14:1 | 26413417:264 | 26414308:264 | -;-;-;-;-;-;-;- | 1890 | lman2 | 1864 | 2673 | 3229 | 2518 | 3374 | 2348 |
| ENSDARG000 | 14:14:14:14:1 | 52481288:524 | 52481347:524 | +;+;+;+;+;+;+ | 3950 | tcerg1a | 891 | 804 | 1442 | 804 | 1050 | 765 |
| ENSDARG000 | 14:14:14:14:1 | 37545341:375 | 37546254:375 | +;+;+;+;+;+;+ | 7425 | PCDH1 | 175 | 8 | 42 | 36 | 25 | 54 |
| ENSDARG000 | 14:14:14:14:1 | 38654300:386 | 38654329:386 | +;+;+;+;+ | 1914 | rnasel3 | 0 | 10 | 16 | 0 | 0 | 0 |
| ENSDARG000 | 14:14:14:14:1 | 94603:94946 | 94684:95326 | +;+;+;+;+;+;+ | 2332 | mcm7 | 755 | 2605 | 3490 | 1021 | 2614 | 1253 |
| ENSDARG000 | 14:14:14:14:1 | 38656776:386 | 38657653:386 | +;+;+;+;+;+;+ | 6521 | FBXO38 | 640 | 369 | 384 | 256 | 545 | 470 |
| ENSDARG000 | 14:14:14:14:1 | 29200772:293 | 29201249:293 | +;+;+;+;+;+;+ | 11579 | TENM2 | 2557 | 1391 | 2183 | 4503 | 2566 | 4789 |
| ENSDARG000 | 14;14;14 | 46250864:462 | 46251330:462 | -;-;- | 739 | NA | 109 | 54 | 20 | 28 | 12 | 6 |
| ENSDARG000 | 14;14;14 | 46233704:462 | 46234589:462 | -;-;- | 1095 | NA | 0 | 0 | 0 | 0 | 0 | 0 |
| ENSDARG000 | 14:14:14:14:1 | 46223458:462 | 46225630:462 | +;+;+;+;+;+;+ | 5392 | NA | 0 | 0 | 0 | 0 | 2 | 14 |
| ENSDARG000 | 14:14:14:14:1 | 695863:6958 | 696189:6961 | +;+;+;+;+;+;+ | 851 | NA | 0 | 0 | 66 | 71 | 4 | 0 |
| ENSDARG000 | 14;14;14 | 46221004:462 | 46223850:462 | -;-;- | 3087 | NA | 6 | 0 | 12 | 0 | 2 | 0 |
| ENSDARG000 | 14:14:14:14:1 | 9265741:926 | 9266490:926 | -;-;-;-;- | 1651 | asb12b | 0 | 0 | 0 | 0 | 0 | 0 |
| ENSDARG000 | 14:14:14:14:1 | 41887070:418 | 41887343:418 | +;+;+;+;+;+;+ | 6838 | NA | 90 | 90 | 60 | 38 | 62 | 139 |
| ENSDARG000 | 14:14:14:14:1 | 9287683:930 | 9287752:930 | +;+;+;+;+;+;+ | 2104 | msnb | 10 | 0 | 0 | 0 | 0 | 0 |
| ENSDARG000 | 14;14;14;14 | 699229:7015 | 699764:7016 | -;-;-;- | 1108 | si:zfos-741a1 | 8 | 21 | 76 | 44 | 29 | 22 |
| ENSDARG000 | 14:14:14:14:1 | 34547554:345 | 34547776:345 | +;+;+;+;+;+;+ | 1592 | NA | 0 | 4 | 0 | 0 | 0 | 0 |
| ENSDARG000 | 14:14:14:14:1 | 17421668:174 | 17421763:174 | +;+;+;+;+ | 1449 | NA | 0 | 0 | 0 | 0 | 2 | 0 |
| ENSDARG000 | 14;14;14;14 | 42110473:421 | 42110778:421 | -;-;-;- | 549 | NA | 0 | 0 | 0 | 0 | 0 | 0 |
| ENSDARG000 | 14:14:14:14:1 | 17442913:174 | 17444523:174 | -;-;-;-;-;-;- | 3132 | fgfrl1a | 307 | 453 | 585 | 728 | 1258 | 488 |
| ENSDARG000 | 14:14:14:14:1 | 29769332:297 | 29769410:297 | +;+;+;+;+;+ | 918 | TSTD1 | 98 | 234 | 163 | 102 | 158 | 119 |
| ENSDARG000 | 14:14:14:14:1 | 21861099:218 | 21861145:218 | +;+;+;+;+;+ | 4724 | spon2a | 3983 | 6519 | 1229 | 4746 | 6470 | 13330 |
| ENSDARG000 | 14:14:14:14:1 | 33270387:332 | 33271717:332 | -;-;-;-;-;-;-;-;- | 2386 | stard14 | 99 | 2 | 6 | 33 | 50 | 10 |
| ENSDARG000 | 14:14:14:14:1 | 42172257:421 | 42175081:421 | +;+;+;+;+;+;+ | 4405 | pcdh18b | 2323 | 2099 | 2999 | 2050 | 2771 | 3450 |
| ENSDARG000 | 14:14:14:14:1 | 26426085:264 | 26427576:264 | -;-;-;-;-;-;-;-;- | 3817 | NA | 362 | 348 | 318 | 196 | 394 | 211 |
| ENSDARG000 | 14:14:14:14:1 | 31721171:317 | 31721268:317 | +;+;+;+;+;+;+ | 4586 | NA | 0 | 0 | 0 | 0 | 0 | 0 |
| ENSDARG000 | 14:14:14:14:1 | 21905492:219 | 21905608:219 | +;+;+;+;+;+;+ | 4554 | NA | 20 | 18 | 21 | 18 | 100 | 100 |
| ENSDARG000 | 14:14:14:14:1 | 7377552:737 | 7377742:737 | +;+;+;+;+;+;+ | 2126 | hars | 2170 | 2715 | 3600 | 2271 | 2239 | 2279 |
| ENSDARG000 | 14;14 | 4178236:418 | 4178358:418 | +;+ | 2898 | NA | 411 | 359 | 565 | 450 | 517 | 367 |
| ENSDARG000 | 14:14:14:14:1 | 4191136:419 | 4197312:419 | -;-;-;-;-;-;-;-;- | 9566 | NA | 1101 | 1249 | 1345 | 1511 | 596 | 975 |
| ENSDARG000 | 14:14:14:14:1 | 34558480:345 | 34558582:345 | +;+;+;+;+;+;+ | 1087 | pttg1 | 150 | 192 | 200 | 199 | 198 | 180 |
| ENSDARG000 | 14:14:14:14:1 | 29782508:297 | 29782997:297 | -;-;-;-;-;-;-;-;- | 1592 | pdlim3b | 14 | 27 | 29 | 0 | 9 | 4 |
| ENSDARG000 | 14:14:14:14:1 | 52522717:525 | 52525866:525 | -;-;-;-;-;-;-;-;- | 4346 | PPP2R2B | 116 | 130 | 54 | 21 | 34 | 44 |
| ENSDARG000 | 14:14:14:14:1 | 29775402:297 | 29775485:297 | +;+;+;+;+;+;+ | 1972 | C14H4orf47 | 10 | 12 | 4 | 0 | 4 | 2 |
| ENSDARG000 | 14:14:14:14:1 | 24258562:242 | 24259289:242 | +;+;+;+;+;+;+ | 1477 | CHRM1 | 0 | 0 | 0 | 0 | 0 | 0 |
| ENSDARG000 | 14:14:14:14:1 | 42978695:429 | 42979904:429 | -;-;-;-;-;-;-;- | 4672 | pcdh10b | 1116 | 320 | 4746 | 395 | 1236 | 2889 |
| ENSDARG000 | 14:14:14:14:1 | 12178915:121 | 12179184:121 | +;+;+;+;+;+;+ | 2140 | hdac3 | 364 | 485 | 597 | 252 | 398 | 504 |
| ENSDARG000 | 14 | 24277556 | 24279957 | + | 2402 | hnrnpa0a | 2446 | 2644 | 2785 | 1498 | 2993 | 2627 |
| ENSDARG000 | 14:14:14:14:1 | 24278047:242 | 24278478:242 | +;+;+;+;+;+;+ | 1633 | NA | 393 | 286 | 515 | 372 | 687 | 700 |
| ENSDARG000 | 14:14:14:14:1 | 31751260:317 | 31751434:317 | +;+;+;+;+;+ | 1757 | si:dkeyp-11e3 | 12 | 0 | 0 | 0 | 16 | 8 |
| ENSDARG000 | 14:14:14:14:1 | 24267892:242 | 24268889:242 | -;-;-;-;-;-;-;-;- | 2620 | dpf2l | 418 | 543 | 530 | 280 | 407 | 418 |
| ENSDARG000 | 14:14:14:14:1 | 4276387:427 | 4276460:427 | +;+;+;+;+;+;+ | 4901 | NA | 1695 | 1463 | 1560 | 1723 | 1300 | 1249 |
| ENSDARG000 | 14:14:14:14:1 | 21923741:219 | 21924131:219 | -;-;-;-;-;-;-;-;- | 1884 | rad9a | 8 | 78 | 85 | 36 | 26 | 30 |
| ENSDARG000 | 14;14 | 43554490:435 | 43555565:435 | +;+ | 1830 | adra1bb | 0 | 0 | 4 | 0 | 1 | 0 |

| | | | | | | | | | | | | |
|---|---|---|---|---|---|---|---|---|---|---|---|---|
| ENSDARG000 | 14:14:14:14:1 | 52565660:525 | 52565832:525 | +;+;+;+;+;+;+ | 4684 | atp6v0e1 | 1507 | 1698 | 1026 | 1481 | 1164 | 1036 |
| ENSDARG000 | 14:14:14:14:1 | 43400776:434 | 43400983:434 | -;-;-;-;-;-;- | 1070 | NA | 0 | 0 | 0 | 0 | 0 | 0 |
| ENSDARG000 | 14;14;14;14 | 52563794:525 | 52563899:525 | +;+;+;+ | 588 | rpl26 | 1 | 2 | 0 | 0 | 0 | 0 |
| ENSDARG000 | 14:14:14:14:1 | 26437891:264 | 26438321:264 | +;+;+;+;+;+;+; | 2333 | gpr137 | 87 | 70 | 78 | 61 | 93 | 112 |
| ENSDARG000 | 14:14:14:14:1 | 17566383:175 | 17568365:175 | -;-;-;-;-;-;-; | 3336 | rnf4 | 215 | 261 | 267 | 193 | 204 | 291 |
| ENSDARG000 | 14:14:14:14:1 | 26452740:264 | 26453435:264 | -;-;-;-;-;-;-; | 2545 | syvn1 | 966 | 836 | 1327 | 665 | 1068 | 812 |
| ENSDARG000 | 14:14:14:14:1 | 17580571:175 | 17581470:175 | -;-;-;-;-;-;-; | 2002 | selenot2 | 2489 | 2545 | 2062 | 1720 | 2675 | 2044 |
| ENSDARG000 | 14:14:14:14:1 | 43577164:435 | 43577932:435 | -;-;-;-;-;-;-; | 1682 | eif4ea | 417 | 503 | 537 | 238 | 454 | 435 |
| ENSDARG000 | 14:14:14:14:1 | 712115:71211 | 712380:71238 | +;+;+;+;+;+;+; | 2649 | ints10 | 274 | 321 | 373 | 154 | 331 | 224 |
| ENSDARG000 | 14:14:14:14:1 | 38762977:387 | 38763840:387 | -;-;-;-;-;-;-; | 2732 | csnk1a1 | 2916 | 3075 | 3992 | 3270 | 3840 | 3182 |
| ENSDARG000 | 14:14:14:14:1 | 30795559:307 | 30795666:307 | +;+;+;+;+;+;+ | 1766 | cfl1 | 23690 | 27702 | 29690 | 22280 | 25136 | 24111 |
| ENSDARG000 | 14;14;14;14 | 31766144:317 | 31766525:317 | +;+;+;+ | 2065 | tmtops3a | 11 | 12 | 0 | 9 | 2 | 0 |
| ENSDARG000 | 14:14:14:14:1 | 21942383:219 | 21944777:219 | -;-;-;-;-;-;-; | 3586 | p2rx3a | 0 | 2 | 0 | 0 | 0 | 0 |
| ENSDARG000 | 14:14:14:14:1 | 34562426:345 | 34563088:345 | -;-;-;-;-;-;-; | 3225 | afap1l1a | 136 | 176 | 196 | 44 | 224 | 157 |
| ENSDARG000 | 14:14:14:14:1 | 21934331:219 | 21934468:219 | +;+;+;+;+;+ | 4166 | zgc:113229 | 2 | 3 | 0 | 0 | 2 | 14 |
| ENSDARG000 | 14:14:14:14:1 | 31788733:317 | 31789164:317 | +;+;+;+;+ | 1791 | cd40lg | 29 | 16 | 7 | 3 | 23 | 49 |
| ENSDARG000 | 14:14:14:14:1 | 46301973:463 | 46305546:463 | -;-;-;-;-;-;-; | 4211 | NA | 0 | 1 | 1 | 0 | 6 | 2 |
| ENSDARG000 | 14:14:14:14:1 | 14847304:148 | 14847623:148 | +;+;+;+;+;+;+; | 4130 | NA | 199 | 146 | 245 | 187 | 262 | 251 |
| ENSDARG000 | 14;14;14;14 | 30802465:308 | 30805059:308 | -;-;-;- | 2906 | NA | 210 | 96 | 197 | 389 | 306 | 181 |
| ENSDARG000 | 14:14:14:14:1 | 4295933:4298 | 4296549:4298 | -;-;-;-;-;-;-; | 2703 | GUF1 | 202 | 268 | 296 | 84 | 274 | 314 |
| ENSDARG000 | 14:14:14:14:1 | 31855592:318 | 31855695:318 | -;-;-;-;-;-;-; | 2793 | rbmx | 2911 | 2090 | 3697 | 2050 | 3295 | 2468 |
| ENSDARG000 | 14:14:14:14:1 | 43590816:435 | 43591006:435 | -;-;-;-;-;-;-;- | 1325 | adh5 | 3636 | 3578 | 6680 | 5652 | 5729 | 4325 |
| ENSDARG000 | 14;14;14 | 22022441:220 | 22022831:220 | +;+;+ | 4183 | CLCF1 | 20 | 40 | 22 | 0 | 2 | 0 |
| ENSDARG000 | 14:14:14:14:1 | 21980446:219 | 21980936:219 | -;-;-;-;-;-;-; | 2925 | ssrp1a | 449 | 1256 | 1879 | 456 | 1294 | 673 |
| ENSDARG000 | 14:14:14:14:1 | 22076596:220 | 22076903:220 | +;+;+;+;+;+;+; | 2967 | slc43a1a | 282 | 767 | 400 | 326 | 246 | 778 |
| ENSDARG000 | 14:14:14:14:1 | 12222543:122 | 12226853:122 | -;-;-;-;-;-;-; | 7690 | NA | 0 | 0 | 0 | 2 | 18 | 0 |
| ENSDARG000 | 14:14:14:14:1 | 4796168:4798 | 4797011:4798 | +;+;+;+;+;+;+; | 8196 | ap1ar | 454 | 452 | 392 | 253 | 479 | 425 |
| ENSDARG000 | 14:14:14:14:1 | 31799787:318 | 31803468:318 | -;-;-;-;-;-;-; | 6858 | NA | 8605 | 4535 | 6215 | 4356 | 9196 | 11530 |
| ENSDARG000 | 14;14 | 15064870:150 | 15065472:150 | +;+ | 1829 | nkx1.2lb | 0 | 0 | 0 | 0 | 0 | 0 |
| ENSDARG000 | 14:14:14:14:1 | 15070494:150 | 15070918:150 | -;-;-;-;-;-;-; | 3042 | uvssa | 182 | 144 | 176 | 106 | 125 | 194 |
| ENSDARG000 | 14;14;14 | 731864:73260 | 733565:73510 | -;-;- | 3259 | tlr1 | 10 | 12 | 6 | 2 | 30 | 13 |
| ENSDARG000 | 14:14:14:14:1 | 5383060:5385 | 5383406:5386 | +;+;+;+;+ | 4545 | lbx2 | 27 | 0 | 0 | 0 | 4 | 8 |
| ENSDARG000 | 14:14:14:14:1 | 740032:74334 | 740730:74385 | +;+;+;+;+;+;+; | 2988 | NA | 11 | 6 | 1 | 15 | 7 | 5 |
| ENSDARG000 | 14:14:14:14:1 | 24283915:242 | 24284234:242 | +;+;+;+;+;+;+; | 5889 | KLHL3 | 337 | 372 | 369 | 371 | 455 | 402 |
| ENSDARG000 | 14:14:14:14:1 | 5394630:5394 | 5394809:5394 | -;-;-;-;-;-;-; | 1419 | pcgf1 | 141 | 76 | 104 | 46 | 92 | 118 |
| ENSDARG000 | 14;14 | 43614697:436 | 43615025:436 | -;- | 386 | NA | 0 | 0 | 0 | 0 | 0 | 3 |
| ENSDARG000 | 14:14:14:14:1 | 43603993:436 | 43605571:436 | -;-;-;-;-;-;- | 2348 | gar1 | 1370 | 1453 | 1707 | 540 | 1083 | 1089 |
| ENSDARG000 | 14:14:14:14:1 | 30854553:308 | 30856187:308 | -;-;-;-;-;-;-; | 2853 | ubl3b | 726 | 634 | 884 | 701 | 797 | 657 |
| ENSDARG000 | 14;14 | 17593284:175 | 17596674:175 | -;- | 3722 | rab33a | 2 | 2 | 0 | 0 | 0 | 0 |
| ENSDARG000 | 14;14;14;14 | 12316581:123 | 12316706:123 | +;+;+;+ | 1189 | NA | 23 | 10 | 18 | 24 | 21 | 26 |
| ENSDARG000 | 14;14 | 12316606:123 | 12316706:123 | +;+ | 1377 | GJB1 | 17 | 0 | 0 | 12 | 8 | 10 |
| ENSDARG000 | 14:14:14:14:1 | 757435:75851 | 758221:75892 | -;-;-;-;-;-;- | 2122 | trim35-27 | 0 | 4 | 10 | 0 | 6 | 0 |
| ENSDARG000 | 14;14;14;14 | 7397804:7401 | 7398979:7402 | -;-;-;- | 2193 | dnd1 | 1 | 0 | 4 | 0 | 0 | 0 |
| ENSDARG000 | 14:14:14:14:1 | 29807097:298 | 29808822:298 | -;-;-;-;-;-;-; | 8250 | SORBS2 | 12 | 2 | 0 | 12 | 8 | 10 |
| ENSDARG000 | 14:14:14:14:1 | 33280718:332 | 33282560:332 | -;-;-;-;-;-;-; | 2713 | rab41 | 1219 | 982 | 1313 | 875 | 1092 | 859 |
| ENSDARG000 | 14:14:14:14:1 | 9343860:9343 | 9344438:9344 | -;-;-;-;-;- | 1313 | TENT5D | 0 | 0 | 0 | 0 | 0 | 0 |
| ENSDARG000 | 14;14;14 | 44013419:441 | 44014244:441 | +;+;+ | 1266 | NA | 0 | 0 | 0 | 0 | 0 | 0 |
| ENSDARG000 | 14:14:14:14:1 | 9390850:9391 | 9391656:9391 | -;-;-;-;-;-;-; | 1969 | tbx22 | 2 | 0 | 0 | 0 | 0 | 0 |
| ENSDARG000 | 14:14:14:14:1 | 46375241:463 | 46377153:463 | -;-;-;-;-;-;-; | 4644 | bbs7 | 170 | 234 | 216 | 136 | 178 | 229 |
| ENSDARG000 | 14;14;14 | 44545092:445 | 44545433:445 | +;+;+ | 3714 | LINGO2 | 32 | 4 | 26 | 4 | 0 | 8 |
| ENSDARG000 | 14:14:14:14:1 | 7452302:7480 | 7452829:7480 | +;+;+;+;+;+;+; | 3619 | gfra3 | 2 | 0 | 0 | 0 | 0 | 0 |
| ENSDARG000 | 14;14;14;14 | 44769364:447 | 44771143:447 | -;-;-;- | 2428 | si:dkev-109l4. | 182 | 181 | 456 | 305 | 430 | 349 |
| ENSDARG000 | 14;14;14;14 | 9405099:9412 | 9408325:9412 | -;-;-;- | 3655 | sh3bgrl | 906 | 537 | 1195 | 637 | 1516 | 622 |
| ENSDARG000 | 14:14:14:14:1 | 17599875:176 | 17603866:176 | -;-;-;-;-;-;-; | 9588 | NA | 12 | 6 | 0 | 14 | 24 | 4 |
| ENSDARG000 | 14:14:14:14:1 | 46410766:464 | 46410796:464 | +;+;+;+;+;+;+; | 1202 | anxa5a | 0 | 0 | 0 | 2 | 0 | 0 |
| ENSDARG000 | 14;14;14;14 | 15155684:151 | 15155959:151 | +;+;+;+ | 711 | spink2.1 | 10 | 6 | 8 | 18 | 28 | 2 |
| ENSDARG000 | 14:14:14:14:1 | 12370375:123 | 12370945:123 | -;-;-;-;-;-;-; | 2160 | magt1 | 1417 | 1681 | 1927 | 1615 | 1579 | 1834 |
| ENSDARG000 | 14:14:14:14:1 | 44777478:447 | 44777872:447 | -;-;-;-;-;-;-;- | 1413 | grhpra | 304 | 260 | 375 | 340 | 442 | 316 |
| ENSDARG000 | 14:14:14:14:1 | 38786258:387 | 38786393:387 | +;+;+;+;+;+;+ | 1030 | si:ch211-195l | 42 | 3 | 33 | 57 | 16 | 3 |
| ENSDARG000 | 14:14:14:14:1 | 5506983:5512 | 5508384:5512 | -;-;-;-;- | 2376 | fgf24 | 6 | 0 | 0 | 0 | 0 | 0 |
| ENSDARG000 | 14:14:14:14:1 | 22109241:221 | 22109441:221 | -;-;-;-;-;-;-; | 869 | si:dkev-6i22.5 | 50 | 144 | 182 | 54 | 116 | 72 |
| ENSDARG000 | 14:14:14:14:1 | 44790904:447 | 44790924:447 | +;+;+;+;+;+;+; | 1864 | zgc:195212 | 469 | 492 | 1307 | 791 | 626 | 504 |
| ENSDARG000 | 14:14:14:14:1 | 22102203:221 | 22102461:221 | -;-;-;-;-;-;-;- | 1420 | med19a | 500 | 585 | 610 | 442 | 600 | 576 |
| ENSDARG000 | 14;14 | 44774089:447 | 44774327:447 | +;+ | 650 | tomm5 | 726 | 869 | 1458 | 1280 | 928 | 1128 |
| ENSDARG000 | 14:14:14:14:1 | 22113331:221 | 22113947:221 | +;+;+;+;+;+;+; | 1607 | tmx2a | 0 | 0 | 2 | 2 | 0 | 0 |
| ENSDARG000 | 14:14:14:14:1 | 34634145:346 | 34635672:346 | -;-;-;-;-;-;-; | 4550 | ablim3 | 7293 | 4143 | 3107 | 4294 | 3718 | 6345 |
| ENSDARG000 | 14:14:14:14:1 | 15170667:151 | 15170850:151 | -;-;-;-;-;-;- | 411 | spink2.2 | 2 | 12 | 2 | 6 | 76 | 14 |
| ENSDARG000 | 14:14:14:14:1 | 26467120:264 | 26467915:264 | -;-;-;-;-;-;-; | 2834 | smad5 | 1370 | 1593 | 1944 | 1441 | 1841 | 1232 |
| ENSDARG000 | 14:14:14:14:1 | 9421510:9422 | 9421961:9422 | +;+;+;+;+;+;+; | 1582 | hmgn6 | 4375 | 4866 | 5257 | 4258 | 5738 | 5209 |
| ENSDARG000 | 14 | 790166 | 792959 | + | 2794 | adra2da | 0 | 0 | 20 | 0 | 7 | 0 |
| ENSDARG000 | 14:14:14:14:1 | 30877962:308 | 30881367:308 | -;-;-;-;-;-;-; | 6012 | SLC7A3 | 374 | 285 | 433 | 394 | 192 | 442 |
| ENSDARG000 | 14:14:14:14:1 | 7724876:7739 | 7725309:7739 | -;-;-;-;- | 1531 | si:ch211-39i2 | 421 | 641 | 1147 | 1226 | 387 | 404 |
| ENSDARG000 | 14:14:14:14:1 | 18667247:186 | 18671373:186 | -;-;-;-;-;- | 4819 | SLITRK4 | 0 | 48 | 6 | 0 | 0 | 2 |
| ENSDARG000 | 14:14:14:14:1 | 15257685:153 | 15257789:153 | +;+;+;+;+;+;+; | 840 | NA | 0 | 8 | 12 | 0 | 0 | 24 |
| ENSDARG000 | 14:14:14:14:1 | 7697694:7697 | 7697778:7697 | +;+;+;+;+;+;+; | 3357 | brd8b | 762 | 438 | 641 | 332 | 713 | 472 |
| ENSDARG000 | 14;14;14;14 | 15331486:153 | 15331615:153 | +;+;+;+ | 505 | NA | 0 | 6 | 2 | 0 | 0 | 37 |
| ENSDARG000 | 14:14:14:14:1 | 24316770:243 | 24319119:243 | -;-;-;-;-;-;-; | 6216 | fam13b | 2091 | 2577 | 3826 | 2103 | 6333 | 2233 |
| ENSDARG000 | 14;14;14;14 | 15231097:152 | 15231231:152 | +;+;+;+ | 533 | NA | 2 | 1 | 6 | 2 | 0 | 16 |
| ENSDARG000 | 14;14;14;14 | 15204935:152 | 15204959:152 | +;+;+;+ | 794 | NA | 0 | 0 | 0 | 0 | 0 | 0 |
| ENSDARG000 | 14:14:14:14:1 | 15191176:151 | 15191417:151 | +;+;+;+;+;+;+; | 1621 | NA | 2 | 6 | 14 | 18 | 72 | 80 |
| ENSDARG000 | 14;14;14;14 | 15257658:152 | 15257789:152 | +;+;+;+ | 545 | NA | 0 | 0 | 0 | 0 | 0 | 0 |
| ENSDARG000 | 14;14 | 5560755:5562 | 5561064:5563 | -;- | 588 | NA | 0 | 0 | 0 | 0 | 0 | 0 |
| ENSDARG000 | 14:14:14:14:1 | 38795518:387 | 38796161:387 | -;-;-;-;-;-;-;- | 1922 | SIL1 | 696 | 1124 | 2151 | 835 | 979 | 834 |
| ENSDARG000 | 14:14:14:14:1 | 5588029:5590 | 5590514:5590 | -;-;-;-;-;-;-; | 3431 | npm1b | 352 | 371 | 644 | 388 | 503 | 377 |
| ENSDARG000 | 14:14:14:14:1 | 18781082:187 | 18781439:187 | +;+;+;+;+;+;+; | 3869 | NA | 0 | 0 | 0 | 0 | 0 | 0 |
| ENSDARG000 | 14;14;14 | 19258702:192 | 19261733:192 | +;+;+ | 3032 | SLITRK2 | 2 | 4 | 0 | 28 | 92 | 4 |
| ENSDARG000 | 14:14:14:14:1 | 834619:83648 | 837425:83742 | -;-;-;-;-;-;-; | 5044 | SLC34A1 | 0 | 4 | 347 | 44 | 562 | 0 |
| ENSDARG000 | 14:14:14:14:1 | 872016:87303 | 872899:87311 | -;-;-;-;-;-;-; | 2715 | rgs14a | 21 | 32 | 9 | 23 | 84 | 60 |
| ENSDARG000 | 14:14:14:14:1 | 20156477:201 | 20156729:201 | +;+;+;+;+;+;+; | 4413 | fmr1 | 1375 | 1558 | 1447 | 1206 | 1719 | 2185 |
| ENSDARG000 | 14;14 | 865625:87270 | 865671:87289 | -;- | 238 | NA | 0 | 0 | 0 | 0 | 0 | 0 |
| ENSDARG000 | 14:14:14:14:1 | 863031:86500 | 864328:86506 | -;-;-;-;- | 1580 | NA | 0 | 0 | 0 | 0 | 0 | 0 |
| ENSDARG000 | 14 | 46441617 | 46442459 | - | 843 | NA | 0 | 0 | 0 | 0 | 2 | 4 |
| ENSDARG000 | 14:14:14:14:1 | 5656510:5656 | 5659536:5657 | -;-;-;-;-;-;-; | 4265 | tlx2 | 0 | 0 | 0 | 0 | 0 | 0 |
| ENSDARG000 | 14;14 | 46449070:464 | 46449900:464 | -;- | 1497 | fgfbp2a | 123 | 2 | 10 | 6 | 22 | 8 |
| ENSDARG000 | 14:14:14:14:1 | 38810385:388 | 38812174:388 | -;-;-;-;-;-;-; | 4408 | spdl1 | 252 | 655 | 589 | 360 | 429 | 308 |
| ENSDARG000 | 14:14:14:14:1 | 9432627:9437 | 9433068:9437 | +;+;+;+;+;+ | 960 | NA | 0 | 0 | 0 | 0 | 0 | 0 |
| ENSDARG000 | 14:14:14:14:1 | 34780886:347 | 34781190:347 | +;+;+;+;+;+;+; | 5865 | SH3TC2 | 11489 | 5447 | 6726 | 6838 | 9681 | 11050 |
| ENSDARG000 | 14;14 | 24396829:243 | 24397232:243 | -;- | 807 | NA | 4 | 0 | 2 | 26 | 0 | 0 |
| ENSDARG000 | 14:14:14:14:1 | 9456675:9462 | 9456836:9463 | +;+;+;+;+;+;+; | 2076 | rell2 | 0 | 10 | 26 | 22 | 30 | 22 |
| ENSDARG000 | 14:14:14:14:1 | 26511041:265 | 26512074:265 | -;-;-;-;-;-;-; | 3037 | tgfbi | 2477 | 3618 | 5592 | 1962 | 5168 | 1203 |
| ENSDARG000 | 14:14:14:14:1 | 20342553:203 | 20342791:203 | +;+;+;+;+;+;+; | 12720 | AFF2 | 56 | 39 | 82 | 2 | 20 | 8 |
| ENSDARG000 | 14;14;14;14 | 5690650:5690 | 5690831:5691 | +;+;+;+ | 550 | NA | 0 | 0 | 0 | 0 | 0 | 0 |
| ENSDARG000 | 14:14:14:14:1 | 38845674:388 | 38848166:388 | +;+;+;+;+;+ | 2493 | hnrnpa0b | 206 | 123 | 198 | 223 | 273 | 270 |
| ENSDARG000 | 14 | 38845504 | 38847375 | - | 1872 | NA | 14 | 13 | 0 | 0 | 5 | 3 |
| ENSDARG000 | 14;14;14;14 | 24400799:244 | 24401999:244 | -;-;-;- | 1732 | cxcl14 | 296 | 1170 | 1168 | 1327 | 3209 | 733 |
| ENSDARG000 | 14;14 | 5816332:5820 | 5816792:5820 | +;+ | 860 | NA | 0 | 0 | 0 | 0 | 0 | 0 |
| ENSDARG000 | 14:14:14:14:1 | 5804928:5810 | 5805225:5810 | -;-;-;-;-;- | 1396 | kazald2 | 0 | 0 | 0 | 0 | 0 | 0 |
| ENSDARG000 | 14 | 38849432 | 38850933 | + | 1502 | hnrnpa0l | 10223 | 11264 | 12372 | 5719 | 10468 | 7776 |
| ENSDARG000 | 14:14:14:14:1 | 30898255:308 | 30899240:308 | -;-;-;-;-;-;-; | 4649 | NA | 150 | 563 | 556 | 178 | 401 | 214 |
| ENSDARG000 | 14:14:14:14:1 | 12391871:123 | 12391989:123 | +;+;+;+;+;+;+; | 7886 | thoc2 | 2825 | 2643 | 4447 | 2654 | 3400 | 3489 |
| ENSDARG000 | 14:14:14:14:1 | 15495088:154 | 15495165:154 | +;+;+;+;+;+;+; | 409 | NA | 0 | 0 | 16 | 2 | 23 | 48 |
| ENSDARG000 | 14:14:14:14:1 | 29941266:299 | 29941650:299 | +;+;+;+;+;+;+; | 5105 | FAM149A | 0 | 0 | 0 | 2 | 0 | 0 |
| ENSDARG000 | 14;14;14;14 | 15543331:155 | 15543456:155 | +;+;+;+ | 521 | NA | 0 | 2 | 0 | 10 | 58 | 34 |
| ENSDARG000 | 14:14:14:14:1 | 29975268:299 | 29975619:299 | +;+;+;+;+;+;+; | 5017 | cyp4v2b | 48 | 81 | 79 | 83 | 143 | 0 |
| ENSDARG000 | 14;14;14 | 901847:90335 | 901955:90341 | +;+;+ | 228 | NA | 0 | 0 | 0 | 0 | 0 | 0 |
| ENSDARG000 | 14:14:14:14:1 | 33297781:332 | 33299144:332 | -;-;-;-;-;-;-; | 3524 | septin6 | 2475 | 2866 | 4647 | 3343 | 4141 | 2655 |
| ENSDARG000 | 14;14;14;14 | 15597049:155 | 15597126:155 | +;+;+;+ | 410 | spink2.5 | 0 | 6 | 6 | 8 | 13 | 104 |
| ENSDARG000 | 14;14;14 | 904850:90675 | 905228:90685 | +;+;+ | 736 | NA | 0 | 0 | 0 | 0 | 0 | 4 |
| ENSDARG000 | 14:14:14:14:1 | 44804326:448 | 44804467:448 | +;+;+;+;+;+;+; | 5054 | slc30a9 | 1187 | 1351 | 1883 | 761 | 1906 | 1513 |
| ENSDARG000 | 14:14:14:14:1 | 7764331:7764 | 7764958:7764 | -;-;-;-;-;-;-; | 1699 | sting1 | 40 | 2 | 2 | 6 | 34 | 10 |
| ENSDARG000 | 14;14 | 34825465:348 | 34830268:348 | -;- | 5042 | adrb2a | 720 | 88 | 10 | 12 | 40 | 198 |
| ENSDARG000 | 14:14:14:14:1 | 30910114:309 | 30910545:309 | +;+;+;+;+ | 5206 | FOXO4 | 2044 | 956 | 1705 | 2004 | 1015 | 1933 |
| ENSDARG000 | 14:14:14:14:1 | 34951202:349 | 34951264:349 | +;+;+;+;+;+;+; | 1470 | il12ba | 10 | 6 | 4 | 18 | 0 | 37 |
| ENSDARG000 | 14;14 | 918287:91931 | 918375:91989 | +;+ | 668 | NA | 0 | 0 | 0 | 0 | 0 | 0 |
| ENSDARG000 | 14:14:14:14:1 | 909769:90979 | 909893:90989 | +;+;+;+;+;+;+; | 1274 | arl3l2 | 0 | 0 | 0 | 2 | 4 | 0 |
| ENSDARG000 | 14:14:14:14:1 | 20869618:208 | 20870367:208 | -;-;-;-;-;-;-; | 2291 | ids | 247 | 151 | 119 | 112 | 204 | 215 |
| ENSDARG000 | 14:14:14:14:1 | 20893065:208 | 20893159:208 | +;+;+;+;+ | 786 | lygl1 | 773 | 399 | 356 | 366 | 561 | 566 |
| ENSDARG000 | 14:14:14:14:1 | 20911310:209 | 20911411:209 | +;+;+;+;+ | 761 | lygl2 | 1 | 0 | 0 | 0 | 0 | 4 |
| ENSDARG000 | 14:14:14:14:1 | 20900989:209 | 20901042:209 | +;+;+;+;+ | 847 | zgc:162941 | 13 | 2 | 0 | 0 | 2 | 4 |
| ENSDARG000 | 14:14:14:14:1 | 9481443:9481 | 9482184:9481 | +;+;+;+;+;+;+; | 5050 | st3gal5 | 818 | 800 | 1146 | 669 | 1068 | 1013 |
| ENSDARG000 | 14:14:14:14:1 | 15620640:156 | 15620663:156 | +;+;+;+;+;+;+ | 430 | NA | 0 | 0 | 0 | 0 | 0 | 4 |
| ENSDARG000 | 14;14;14 | 9507291:9516 | 9508316:9516 | -;-;- | 2086 | atoh8 | 40 | 12 | 24 | 14 | 18 | 6 |
| ENSDARG000 | 14:14:14:14:1 | 22129096:221 | 22129131:221 | +;+;+;+;+;+;+; | 2421 | ccng1 | 9970 | 19514 | 20380 | 25548 | 23435 | 13856 |
| ENSDARG000 | 14:14:14:14:1 | 26546175:265 | 26546353:265 | +;+;+;+;+;+ | 3620 | NA | 0 | 0 | 0 | 0 | 0 | 0 |
| ENSDARG000 | 14:14:14:14:1 | 46530976:465 | 46531557:465 | -;-;-;-;- | 910 | NA | 0 | 0 | 0 | 0 | 0 | 0 |
| ENSDARG000 | 14:14:14:14:1 | 5835134:5835 | 5835332:5835 | +;+;+;+;+;+;+; | 2124 | aup1 | 1120 | 1234 | 1798 | 1389 | 1099 | 1330 |
| ENSDARG000 | 14:14:14:14:1 | 46456091:464 | 46459309:464 | -;-;-;-;-;-;-; | 7296 | prom1a | 3561 | 5405 | 3494 | 5934 | 5361 | 4218 |
| ENSDARG000 | 14:14:14:14:1 | 38852969:388 | 38853618:388 | -;-;-;-;-;-;-; | 2954 | gsr | 532 | 551 | 1027 | 776 | 1239 | 738 |
| ENSDARG000 | 14:14:14:14:1 | 15662563:156 | 15664982:156 | -;-;-;-;-;-;-; | 4301 | neurl1b | 3098 | 2193 | 924 | 2105 | 1751 | 3495 |
| ENSDARG000 | 14:14:14:14:1 | 31865099:318 | 31865637:318 | +;+;+;+;+;+;+; | 4835 | NA | 1395 | 1071 | 1257 | 920 | 1687 | 1266 |
| ENSDARG000 | 14:14:14:14:1 | 24491818:244 | 24493226:244 | -;-;-;-;-;-;-; | 6432 | slit3 | 164 | 8 | 84 | 57 | 160 | 24 |
| ENSDARG000 | 14:14:14:14:1 | 7789093:7798 | 7790798:7798 | -;-;-;-;-;-;-; | 3631 | ppp3cb | 54 | 181 | 120 | 306 | 124 | 307 |
| ENSDARG000 | 14:14:14:14:1 | 26566936:265 | 26567112:265 | +;+;+;+;+;+;+; | 8860 | NA | 134 | 170 | 271 | 46 | 176 | 128 |
| ENSDARG000 | 14:14:14:14:1 | 923522:92380 | 923919:92391 | +;+;+;+;+;+;+; | 4549 | NA | 0 | 0 | 0 | 0 | 0 | 0 |
| ENSDARG000 | 14:14:14:14:1 | 34966598:349 | 34966790:349 | +;+;+;+;+;+;+; | 5711 | RNF145 | 408 | 349 | 198 | 76 | 453 | 256 |
| ENSDARG000 | 14;14;14;14 | 22140258:221 | 22140589:221 | -;-;-;- | 912 | nudcd2 | 207 | 231 | 243 | 125 | 323 | 165 |
| ENSDARG000 | 14:14:14:14:1 | 22152060:221 | 22152581:221 | +;+;+;+;+;+;+; | 2874 | gabra6a | 0 | 10 | 0 | 32 | 8 | 28 |
| ENSDARG000 | 14;14 | 31890001:318 | 31891412:318 | -;- | 1762 | GPR101 | 0 | 0 | 0 | 0 | 0 | 0 |
| ENSDARG000 | 14:14:14:14:1 | 44826592:448 | 44830357:448 | -;-;-;-;-;-;- | 4646 | GRXCR1 | 4 | 12 | 53 | 16 | 36 | 18 |
| ENSDARG000 | 14:14:14:14:1 | 9581896:9583 | 9582112:9584 | +;+;+;+;+;+;+ | 6298 | tmem129 | 253 | 143 | 351 | 303 | 349 | 234 |
| ENSDARG000 | 14:14:14:14:1 | 9592921:9601 | 9596785:9601 | -;-;-;-;-;-;-; | 7065 | NA | 1820 | 1121 | 1586 | 1405 | 1836 | 1262 |
| ENSDARG000 | 14:14:14:14:1 | 44860335:448 | 44860702:448 | +;+;+;+;+;+;+; | 2247 | atp8a1 | 87 | 104 | 434 | 263 | 262 | 142 |
| ENSDARG000 | 14;14 | 15740152:157 | 15740196:157 | +;+ | 754 | NA | 0 | 12 | 0 | 10 | 2 | 8 |
| ENSDARG000 | 14;14 | 26703331:267 | 26704610:267 | -;- | 1393 | neurog1 | 0 | 0 | 0 | 0 | 0 | 0 |
| ENSDARG000 | 14:14:14:14:1 | 33329420:333 | 33329729:333 | +;+;+;+;+;+ | 1765 | NA | 8 | 32 | 42 | 12 | 52 | 72 |
| ENSDARG000 | 14:14:14:14:1 | 35013656:350 | 35013748:350 | +;+;+;+;+;+;+; | 3494 | ebf3a | 26 | 2 | 18 | 0 | 14 | 22 |
| ENSDARG000 | 14:14:14:14:1 | 5866894:5866 | 5866967:5866 | -;-;-;-;-;-;-; | 2529 | loxl3a | 11331 | 2 | 9657 | 4416 | 4331 | 5902 |
| ENSDARG000 | 14;14 | 30105611:301 | 30110907:301 | -;- | 5496 | MTNR1A | 0 | 8 | 24 | 8 | 19 | 0 |
| ENSDARG000 | 14:14:14:14:1 | 5861435:5861 | 5861624:5861 | +;+;+;+;+;+;+ | 1833 | tubb4b | 21210 | 25229 | 37114 | 36060 | 23623 | 21525 |
| ENSDARG000 | 14:14:14:14:1 | 26715626:267 | 26715767:267 | +;+;+;+;+;+;+; | 1587 | polr2g | 329 | 633 | 769 | 375 | 617 | 450 |
| ENSDARG000 | 14:14:14:14:1 | 31905235:319 | 31905276:319 | +;+;+;+;+;+;+; | 1760 | NA | 0 | 0 | 0 | 0 | 0 | 0 |
| ENSDARG000 | 14:14:14:14:1 | 26719691:267 | 26719760:267 | +;+;+;+;+;+;+; | 1520 | eef1g | 23476 | 23387 | 39172 | 34251 | 29137 | 37021 |

| | | | | | | | | | | | | |
|---|---|---|---|---|---|---|---|---|---|---|---|---|
| ENSDARG000 | 14;14 | 32022272:32 | 32023277:32 | +;+ | 2423 | zic6 | 0 | 0 | 0 | 0 | 0 | 0 |
| ENSDARG000 | 14;14;14;14;1 | 33331773:33 | 33331986:33 | -;-;-;-;-;-;-;-; | 1211 | rpl39 | 6857 | 8221 | 8536 | 6618 | 4652 | 8580 |
| ENSDARG000 | 14;14 | 26715192:26 | 26715374:26 | -;- | 287 | NA | 0 | 0 | 0 | 0 | 0 | 0 |
| ENSDARG000 | 14;14;14 | 32011689:32 | 32013986:32 | -;-;- | 3711 | zic3 | 8 | 0 | 0 | 0 | 4 | 2 |
| ENSDARG000 | 14;14;14;14;1 | 33339322:33 | 33340287:33 | -;-;-;-;-;-;-;-; | 2580 | upf3b | 1067 | 730 | 877 | 670 | 968 | 762 |
| ENSDARG000 | 14;14;14;14;1 | 35279343:35 | 35279708:35 | +;+;+;+;+;+;+;+; | 5535 | clint1a | 2017 | 1379 | 2205 | 1160 | 1605 | 1563 |
| ENSDARG000 | 14;14;14;14;1 | 937194:9421 | 942483:9424 | -;-;-;-;-;-;-;-; | 7443 | acsl1b | 370 | 298 | 436 | 155 | 330 | 180 |
| ENSDARG000 | 14;14;14;14;1 | 12533560:12 | 12536437:12 | -;-;-;-;-;-;-;-; | 6083 | gria3b | 2 | 6 | 0 | 0 | 10 | 0 |
| ENSDARG000 | 14;14;14 | 35263591:35 | 35263663:35 | +;+;+ | 1070 | NA | 0 | 0 | 0 | 0 | 0 | 0 |
| ENSDARG000 | 14;14;14;14;1 | 15760253:15 | 15761632:15 | -;-;-;-;-;-;-;- | 2716 | dusp1 | 10682 | 2159 | 3826 | 5533 | 3983 | 6074 |
| ENSDARG000 | 14;14;14;14;1 | 20918701:20 | 20918788:20 | +;+;+;+;+;+;+;+; | 6431 | zgc:66433 | 575 | 975 | 2047 | 160 | 152 | 2073 |
| ENSDARG000 | 14;14;14 | 13036560:13 | 13040551:13 | -;-;- | 4261 | NA | 0 | 0 | 0 | 0 | 0 | 0 |
| ENSDARG000 | 14;14;14;14;1 | 7892021:789 | 7892406:789 | +;+;+;+;+;+;+;+; | 2354 | ube2d2 | 2636 | 3142 | 3559 | 2595 | 3489 | 3185 |
| ENSDARG000 | 14;14;14;14;1 | 30156449:30 | 30159741:30 | -;-;-;-;-;-;-;-; | 16650 | FAT1 | 306 | 14 | 22 | 46 | 22 | 88 |
| ENSDARG000 | 14;14;14 | 7902374:790 | 7902530:790 | +;+;+ | 2991 | zgc:110843 | 184 | 483 | 174 | 287 | 249 | 510 |
| ENSDARG000 | 14;14 | 13353701:13 | 13353990:13 | +;+ | 764 | si:dkey-48g21 | 0 | 0 | 3 | 0 | 4 | 3 |
| ENSDARG000 | 14;14 | 973103:9765 | 975009:9769 | -;- | 2247 | ARSI | 48 | 24 | 36 | 26 | 30 | 8 |
| ENSDARG000 | 14;14;14;14;1 | 7932973:793 | 7933099:794 | +;+;+;+;+;+;+;+ | 3824 | NA | 1281 | 1351 | 1166 | 1358 | 1002 | 941 |
| ENSDARG000 | 14;14;14;14;1 | 33355163:33 | 33355271:33 | -;-;-;-;-;-;- | 2031 | NA | 0 | 0 | 0 | 0 | 0 | 0 |
| ENSDARG000 | 14;14;14;14;1 | 24840669:24 | 24841131:24 | +;+;+;+;+;+;+;+; | 3486 | NA | 22 | 28 | 6 | 0 | 18 | 4 |
| ENSDARG000 | 14;14;14;14;1 | 8043441:805 | 8043962:805 | -;-;-;-;-;- | 1459 | zgc:92242 | 1725 | 1303 | 1144 | 1466 | 1249 | 835 |
| ENSDARG000 | 14;14;14;14;1 | 989733:9897 | 989972:9899 | +;+;+;+;+;+;+;+; | 2429 | NA | 324 | 498 | 658 | 273 | 407 | 401 |
| ENSDARG000 | 14;14;14;14;1 | 32074541:32 | 32074617:32 | +;+;+;+;+;+;+;+; | 2475 | NA | 0 | 0 | 0 | 0 | 0 | 0 |
| ENSDARG000 | 14;14;14 | 5936996:593 | 5938248:593 | +;+;+ | 10205 | kctd8 | 0 | 2 | 10 | 6 | 8 | 0 |
| ENSDARG000 | 14;14 | 32071775:32 | 32077983:32 | -;- | 6439 | NA | 4 | 0 | 0 | 0 | 0 | 0 |
| ENSDARG000 | 14;14;14;14;1 | 24934736:24 | 24935207:24 | +;+;+;+;+;+;+;+; | 7415 | NA | 926 | 1132 | 821 | 507 | 1104 | 649 |
| ENSDARG000 | 14;14;14;14;1 | 33365130:33 | 33365238:33 | -;-;-;-;-;-;- | 2031 | NA | 0 | 0 | 0 | 0 | 0 | 0 |
| ENSDARG000 | 14;14;14;14;1 | 26759332:26 | 26759600:26 | +;+;+;+;+ | 20569 | NA | 49275 | 45963 | 45152 | 64098 | 43174 | 41355 |
| ENSDARG000 | 14;14;14;14;1 | 9717678:971 | 9718363:971 | -;-;-;-;-;-;-;-; | 2440 | il1rapl2 | 92 | 2 | 16 | 2 | 6 | 0 |
| ENSDARG000 | 14;14;14;14;1 | 22172047:22 | 22172236:22 | +;+;+;+;+;+;+;+; | 3400 | gabrb2a | 6 | 0 | 0 | 0 | 2 | 0 |
| ENSDARG000 | 14;14;14;14;1 | 30272891:30 | 30272995:30 | +;+;+;+;+;+;+;+; | 2134 | asah1a | 943 | 691 | 1088 | 858 | 896 | 941 |
| ENSDARG000 | 14;14;14;14;1 | 15768942:15 | 15769095:15 | +;+;+;+;+;+;+;+; | 3418 | ergic1 | 175 | 263 | 240 | 139 | 375 | 255 |
| ENSDARG000 | 14;14;14;14;1 | 6159162:615 | 6159635:615 | +;+;+;+;+;+;+;+; | 1877 | bscl2l | 0 | 2 | 199 | 22 | 84 | 96 |
| ENSDARG000 | 14;14;14;14;1 | 30279391:30 | 30279557:30 | +;+;+;+;+;+;+ | 1192 | NA | 11 | 15 | 17 | 21 | 13 | 6 |
| ENSDARG000 | 14;14;14;14;1 | 33382032:33 | 33382147:33 | -;-;-;-;-;-;- | 1998 | NA | 0 | 0 | 0 | 0 | 0 | 0 |
| ENSDARG000 | 14;14;14;14;1 | 30922330:30 | 30922491:30 | -;-;-;-;-;-;-;-; | 1728 | NA | 398 | 366 | 323 | 232 | 457 | 268 |
| ENSDARG000 | 14;14;14;14;1 | 13453130:13 | 13453309:13 | +;+;+;+;+;+;+;+; | 3427 | pls3 | 3901 | 3313 | 2677 | 2285 | 4051 | 3230 |
| ENSDARG000 | 14;14 | 6155374:615 | 6156091:615 | -;- | 763 | NA | 0 | 0 | 0 | 0 | 0 | 0 |
| ENSDARG000 | 14;14;14;14;1 | 30955683:30 | 30956003:30 | -;-;-;-;-;- | 751 | C14HXorf65 | 63 | 33 | 41 | 35 | 65 | 106 |
| ENSDARG000 | 14;14;14;14;1 | 26796684:26 | 26796979:26 | +;+;+;+;+;+;+;+; | 2577 | klhl4 | 8 | 0 | 2 | 4 | 14 | 13 |
| ENSDARG000 | 14;14;14;14;1 | 30961549:30 | 30962614:30 | -;-;-;-;-;-;-;- | 1864 | il2rgb | 26 | 18 | 22 | 14 | 35 | 53 |
| ENSDARG000 | 14;14;14;14;1 | 8127893:815 | 8128277:815 | +;+;+;+;+;+;+;+; | 6606 | PSD2 | 293 | 110 | 559 | 96 | 560 | 68 |
| ENSDARG000 | 14;14;14;14;1 | 33391851:33 | 33391966:33 | -;-;-;-;-;-;- | 1881 | NA | 0 | 0 | 0 | 0 | 0 | 0 |
| ENSDARG000 | 14;14;14;14;1 | 21005372:21 | 21005580:21 | +;+;+;+;+;+;+ | 1411 | NA | 0 | 0 | 2 | 0 | 0 | 0 |
| ENSDARG000 | 14;14;14;14;1 | 21007957:21 | 21008051:21 | +;+;+;+;+ | 812 | NA | 0 | 0 | 0 | 0 | 0 | 1 |
| ENSDARG000 | 14;14;14;14;1 | 38875078:38 | 38875245:38 | -;-;-;-;-;-;-;- | 1117 | uqcrq | 686 | 722 | 786 | 436 | 513 | 626 |
| ENSDARG000 | 14;14;14;14;1 | 10374530:10 | 10375100:10 | -;-;-;-;-;-;- | 1133 | commd5 | 1275 | 979 | 1370 | 845 | 1134 | 1231 |
| ENSDARG000 | 14;14;14;14;1 | 10338417:10 | 10344465:10 | -;-;-;-;-;-;- | 7134 | fam199x | 1896 | 904 | 848 | 1432 | 2159 | 862 |
| ENSDARG000 | 14;14;14;14;1 | 35392736:35 | 35393150:35 | -;-;-;-;-;-;-;- | 1435 | LSM11 | 23 | 82 | 56 | 8 | 20 | 46 |
| ENSDARG000 | 14;14 | 35405518:35 | 35405875:35 | +;+ | 3107 | ZBTB3 | 204 | 183 | 207 | 128 | 152 | 182 |
| ENSDARG000 | 14 | 21025026 | 21025388 | - | 363 | NA | 1 | 0 | 0 | 0 | 1 | 0 |
| ENSDARG000 | 14;14;14;14;1 | 35409323:35 | 35410249:35 | -;-;-;-;-;-;-;-; | 1900 | rnaseh2c | 346 | 610 | 365 | 212 | 412 | 555 |
| ENSDARG000 | 14;14 | 38878482:38 | 38878712:38 | +;+ | 1057 | timm8a | 196 | 164 | 160 | 98 | 158 | 240 |
| ENSDARG000 | 14;14;14;14;1 | 45306073:45 | 45306240:45 | +;+;+;+;+;+;+;+; | 4276 | sfxn5b | 54 | 22 | 68 | 110 | 104 | 106 |
| ENSDARG000 | 14;14;14;14;1 | 30967442:30 | 30967829:30 | -;-;-;-;-;- | 959 | zgc:92907 | 153 | 136 | 173 | 120 | 150 | 195 |
| ENSDARG000 | 14;14;14;14;1 | 21038462:21 | 21038480:21 | +;+;+;+;+;+;+;+; | 1450 | NA | 0 | 0 | 0 | 0 | 0 | 0 |
| ENSDARG000 | 14;14;14;14;1 | 22293787:22 | 22293975:22 | +;+;+;+;+;+;+;+; | 10978 | ATP10B | 18 | 3 | 0 | 28 | 0 | 2 |
| ENSDARG000 | 14;14;14;14;1 | 25034750:25 | 25035010:25 | -;-;-;-;-;-;-;-; | 1138 | tmem216 | 28 | 33 | 14 | 76 | 17 | 28 |
| ENSDARG000 | 14;14;14;14;1 | 25005403:25 | 25005838:25 | -;-;-;-;-;-;-;-; | 3984 | pde6a | 9 | 26 | 27 | 67 | 14 | 28 |
| ENSDARG000 | 14;14;14;14;1 | 1007169:100 | 1007333:100 | +;+;+;+;+;+;+;+; | 5516 | NA | 117 | 100 | 335 | 255 | 89 | 88 |
| ENSDARG000 | 14;14;14;14;1 | 22367361:22 | 22372398:22 | -;-;-;-;-;-;-;-; | 5832 | NA | 110 | 110 | 323 | 765 | 54 | 107 |
| ENSDARG000 | 14;14;14;14;1 | 32183978:32 | 32185598:32 | -;-;-;-;-;-;-;-; | 4100 | fgf13a | 0 | 0 | 0 | 8 | 6 | 6 |
| ENSDARG000 | 14;14;14 | 30975163:30 | 30975877:30 | -;-;- | 1862 | rap2c | 2210 | 3330 | 4214 | 3158 | 4048 | 4498 |
| ENSDARG000 | 14;14 | 6193458:619 | 6193705:619 | -;- | 460 | NA | 0 | 0 | 0 | 0 | 0 | 0 |
| ENSDARG000 | 14;14;14;14;1 | 6182346:618 | 6182397:618 | +;+;+;+;+;+;+;+; | 1969 | NA | 87 | 76 | 132 | 134 | 33 | 51 |
| ENSDARG000 | 14 | 8261417 | 8271686 | - | 10270 | PURA | 1324 | 902 | 1189 | 630 | 1810 | 1176 |
| ENSDARG000 | 14;14;14;14;1 | 46629941:46 | 46634280:46 | -;-;-;-;-;-;-;-; | 5893 | TAPT1 | 159 | 104 | 124 | 122 | 200 | 165 |
| ENSDARG000 | 14;14;14;14 | 45339986:45 | 45343540:45 | -;-;-;- | 4248 | zgc:153018 | 611 | 857 | 2012 | 1844 | 1309 | 1139 |
| ENSDARG000 | 14;14;14;14;1 | 27120406:27 | 27121863:27 | -;-;-;-;-;-;- | 4786 | PCDH11Y | 24 | 2 | 52 | 10 | 25 | 6 |
| ENSDARG000 | 14;14 | 26995114:27 | 27000630:27 | -;- | 5710 | NA | 4 | 0 | 28 | 4 | 8 | 0 |
| ENSDARG000 | 14;14;14;14;1 | 38882652:38 | 38886332:38 | -;-;-;-;-;-;-;-; | 4188 | zgc:101583 | 968 | 1048 | 1402 | 954 | 2225 | 1288 |
| ENSDARG000 | 14;14;14;14;1 | 46678711:46 | 46684872:46 | -;-;-;-;-;-;-;- | 7351 | ldb2a | 142 | 243 | 227 | 201 | 739 | 374 |
| ENSDARG000 | 14;14;14;14;1 | 13752432:13 | 13752538:13 | +;+;+;+;+;+ | 2105 | cnga2a | 0 | 0 | 0 | 0 | 0 | 0 |
| ENSDARG000 | 14;14;14;14 | 27322650:27 | 27323164:27 | -;-;-;- | 745 | NA | 0 | 0 | 0 | 0 | 0 | 0 |
| ENSDARG000 | 14;14;14;14;1 | 15816509:15 | 15817529:15 | -;-;-;-;-;-;-;-; | 5248 | flt4 | 96 | 4 | 78 | 58 | 50 | 122 |
| ENSDARG000 | 14;14;14;14;1 | 21042178:21 | 21042272:21 | +;+;+;+;+ | 812 | NA | 0 | 0 | 3 | 0 | 0 | 0 |
| ENSDARG000 | 14;14;14 | 27998756:28 | 28000086:28 | -;-;- | 1684 | NA | 2301 | 2819 | 1404 | 1386 | 3330 | 1358 |
| ENSDARG000 | 14;14 | 28013626:28 | 28016056:28 | -;- | 2933 | NA | 42 | 13 | 9 | 8 | 7 | 3 |
| ENSDARG000 | 14;14;14 | 27999805:28 | 28000086:28 | -;-;- | 759 | TSC22D3 | 0 | 0 | 0 | 0 | 0 | 0 |
| ENSDARG000 | 14;14;14;14;1 | 33402642:33 | 33402750:33 | -;-;-;-;-;-;- | 2070 | NA | 0 | 0 | 0 | 0 | 0 | 0 |
| ENSDARG000 | 14;14;14;14;1 | 8275115:832 | 8275593:832 | +;+;+;+;+;+;+;+; | 4329 | nrg2b | 165 | 0 | 59 | 22 | 245 | 26 |
| ENSDARG000 | 14;14;14;14;1 | 6209810:620 | 6210125:621 | -;-;-;-;-;-;-;-; | 2528 | hdx | 156 | 100 | 152 | 108 | 120 | 101 |
| ENSDARG000 | 14;14;14;14;1 | 6228996:622 | 6229163:622 | +;+;+;+;+;+;+;+; | 2223 | smu1a | 755 | 785 | 990 | 605 | 902 | 499 |
| ENSDARG000 | 14;14;14;14 | 45350008:45 | 45350178:45 | +;+;+;+ | 1071 | ttc9c | 160 | 176 | 190 | 165 | 152 | 204 |
| ENSDARG000 | 14 | 8383549 | 8383923 | - | 375 | NA | 2 | 0 | 0 | 0 | 0 | 0 |
| ENSDARG000 | 14;14;14;14;1 | 14027427:14 | 14027430:14 | +;+;+;+;+;+;+;+; | 1987 | rraga | 1901 | 1084 | 1054 | 1569 | 2440 | 1332 |
| ENSDARG000 | 14;14;14 | 10450216:10 | 10450280:10 | +;+;+ | 1560 | cysltr1 | 6 | 2 | 0 | 0 | 12 | 6 |
| ENSDARG000 | 14;14;14;14;1 | 38891722:38 | 38892468:38 | -;-;-;-;-;-;-;-; | 2634 | BTK | 86 | 1953 | 3058 | 901 | 10949 | 591 |
| ENSDARG000 | 14;14;14;14;1 | 45365157:45 | 45367827:45 | -;-;-;-;-;- | 3692 | NA | 56 | 80 | 651 | 378 | 758 | 291 |
| ENSDARG000 | 14;14;14;14 | 10487990:10 | 10492247:10 | -;-;-;- | 4483 | LPAR4 | 0 | 0 | 0 | 0 | 0 | 0 |
| ENSDARG000 | 14;14;14 | 10505464:10 | 10505549:10 | +;+;+ | 1445 | p2ry10 | 4 | 2 | 4 | 0 | 12 | 10 |
| ENSDARG000 | 14;14;14;14;1 | 22397251:22 | 22397691:22 | +;+;+;+;+;+;+;+; | 9286 | atp7a | 3092 | 2831 | 3914 | 3855 | 3763 | 2583 |
| ENSDARG000 | 14;14 | 1111332:111 | 1111406:112 | +;+ | 2532 | NA | 66 | 80 | 115 | 38 | 56 | 56 |
| ENSDARG000 | 14;14 | 31081862:31 | 31083515:31 | -;- | 2476 | hs6st2 | 526 | 583 | 688 | 622 | 720 | 694 |
| ENSDARG000 | 14;14;14;14 | 10567104:10 | 10567388:10 | +;+;+;+ | 3037 | NA | 98 | 36 | 12 | 46 | 124 | 161 |
| ENSDARG000 | 14;14;14;14;1 | 21052504:21 | 21053125:21 | -;-;-;-;-;-;-;-; | 1709 | NA | 22 | 6 | 12 | 0 | 15 | 4 |
| ENSDARG000 | 14;14;14;14;1 | 30986754:30 | 30993327:30 | -;-;-;-;-;-;-;-; | 9667 | mbnl3 | 1751 | 1098 | 3195 | 3162 | 2007 | 2628 |
| ENSDARG000 | 14;14;14;14;1 | 45406299:45 | 45406476:45 | +;+;+;+;+;+;+;+; | 1274 | NA | 2 | 0 | 17 | 18 | 11 | 6 |
| ENSDARG000 | 14;14;14;14;1 | 14224730:14 | 14225333:14 | +;+;+;+;+;+;+;+; | 4963 | nlgn3a | 0 | 8 | 264 | 22 | 156 | 42 |
| ENSDARG000 | 14;14;14;14;1 | 46873229:46 | 46877212:46 | -;-;-;-;-;-;- | 4656 | qdpra | 5773 | 6928 | 8247 | 8179 | 13457 | 6245 |
| ENSDARG000 | 14;14;14;14;1 | 38932441:38 | 38932876:38 | -;-;-;-;-;-;- | 1483 | gla | 30 | 22 | 64 | 81 | 76 | 14 |
| ENSDARG000 | 14;14;14;14;1 | 10607446:10 | 10608046:10 | -;-;-;-;-;-;-;-; | 1685 | NA | 134 | 175 | 210 | 227 | 356 | 206 |
| ENSDARG000 | 14;14;14 | 8442248:844 | 8442760:844 | +;+;+ | 2715 | NA | 6 | 4 | 0 | 2 | 2 | 0 |
| ENSDARG000 | 14;14;14;14;1 | 15972124:15 | 15974719:15 | -;-;-;-;-;-;-;- | 3641 | trim105 | 129 | 76 | 128 | 155 | 187 | 134 |
| ENSDARG000 | 14;14;14;14;1 | 30285613:30 | 30285717:30 | +;+;+;+;+;+;+;+; | 6092 | mtus1a | 427 | 224 | 461 | 556 | 285 | 214 |
| ENSDARG000 | 14;14;14;14;1 | 35414632:35 | 35414759:35 | +;+;+;+;+;+;+;+; | 1451 | trmt12 | 35 | 48 | 84 | 72 | 87 | 109 |
| ENSDARG000 | 14;14;14;14;1 | 45434680:45 | 45436708:45 | -;-;-;-;- | 3220 | ehd1a | 3329 | 2833 | 3311 | 3029 | 4565 | 3359 |
| ENSDARG000 | 14;14;14;14;1 | 1124409:112 | 1124684:112 | +;+;+;+;+;+;+;+; | 6813 | SEC24B | 1647 | 1580 | 1971 | 1162 | 2151 | 1144 |
| ENSDARG000 | 14;14;14;14;1 | 33413980:33 | 33414053:33 | +;+;+;+;+;+;+;+; | 1234 | ndufa1 | 726 | 634 | 681 | 519 | 694 | 559 |
| ENSDARG000 | 14;14;14;14;1 | 14560932:14 | 14561316:14 | -;-;-;-;- | 706 | si:dkey-27i16. | 83 | 82 | 52 | 69 | 281 | 217 |
| ENSDARG000 | 14 | 22447662 | 22454885 | + | 7224 | NA | 62 | 144 | 70 | 86 | 68 | 178 |
| ENSDARG000 | 14;14;14;14;1 | 32430982:32 | 32431120:32 | +;+;+;+;+;+;+;+; | 1719 | f9a | 0 | 0 | 6 | 0 | 0 | 0 |
| ENSDARG000 | 14;14;14;14;1 | 8450841:845 | 8451284:845 | -;-;-;-;-;-;-;-; | 1375 | eif1ad | 280 | 305 | 423 | 339 | 289 | 350 |
| ENSDARG000 | 14 | 16022157 | 16023337 | + | 1181 | npy7r | 0 | 0 | 0 | 0 | 0 | 0 |
| ENSDARG000 | 14;14;14;14;1 | 45471642:45 | 45471759:45 | +;+;+;+;+;+;+;+; | 1529 | ubxn1 | 743 | 767 | 927 | 633 | 840 | 632 |
| ENSDARG000 | 14;14 | 14568437:14 | 14568531:14 | +;+ | 3121 | pcdh20 | 26 | 0 | 2 | 0 | 2 | 16 |
| ENSDARG000 | 14;14;14;14;1 | 35442511:35 | 35443062:35 | -;-;-;-;-;-;-;-; | 1942 | SRPX2 | 26 | 4 | 0 | 13 | 4 | 17 |
| ENSDARG000 | 14;14;14;14;1 | 35424539:35 | 35424729:35 | +;+;+;+;+;+;+;+; | 2957 | sytl4 | 13 | 2 | 38 | 20 | 28 | 10 |
| ENSDARG000 | 14;14;14;14;1 | 16034447:16 | 16034568:16 | +;+;+;+;+;+;+;+; | 1864 | prelid1a | 1165 | 2160 | 4076 | 3078 | 1998 | 1265 |
| ENSDARG000 | 14;14;14;14;1 | 21065434:21 | 21067976:21 | -;-;-;-;-;-;-;-; | 5682 | rnf20 | 874 | 1017 | 1169 | 432 | 1009 | 691 |
| ENSDARG000 | 14;14;14;14;1 | 16068318:16 | 16069652:16 | -;-;-;-;-;- | 2090 | mxd3 | 131 | 260 | 281 | 126 | 121 | 263 |
| ENSDARG000 | 14 | 22455575 | 22455834 | - | 260 | NA | 0 | 0 | 0 | 0 | 0 | 0 |
| ENSDARG000 | 14;14;14;14 | 25043810:25 | 25043915:25 | +;+;+;+ | 478 | NA | 0 | 0 | 0 | 0 | 0 | 0 |
| ENSDARG000 | 14;14 | 22457230:22 | 22457674:22 | +;+ | 2099 | gdf9 | 0 | 0 | 0 | 0 | 0 | 0 |
| ENSDARG000 | 14;14;14;14;1 | 45488922:45 | 45489153:45 | -;-;-;-;-;-;-;-; | 2287 | NA | 98 | 44 | 122 | 66 | 135 | 115 |
| ENSDARG000 | 14;14;14;14;1 | 2119162:214 | 2123968:214 | -;-;-;-;- | 6856 | NA | 0 | 0 | 0 | 0 | 0 | 0 |
| ENSDARG000 | 14;14;14;14;1 | 2123123:212 | 2123968:212 | -;-;-;-;-;-;-;-; | 4343 | NA | 0 | 0 | 0 | 0 | 0 | 0 |
| ENSDARG000 | 14;14;14;14;1 | 2123123:214 | 2123968:214 | -;-;-;-;- | 3434 | pcdh2ab3 | 0 | 0 | 0 | 0 | 0 | 0 |
| ENSDARG000 | 14;14;14;14 | 2123123:214 | 2123968:214 | -;-;-;- | 3577 | pcdh2ac | 0 | 0 | 0 | 0 | 0 | 0 |
| ENSDARG000 | 14;14;14;14;1 | 2123564:212 | 2123968:212 | -;-;-;-;-;-;-;- | 3033 | pcdh2ab2 | 0 | 0 | 0 | 0 | 0 | 0 |
| ENSDARG000 | 14;14;14;14 | 2123249:214 | 2123968:214 | -;-;-;- | 3308 | NA | 0 | 0 | 0 | 0 | 0 | 0 |
| ENSDARG000 | 14;14;14;14 | 2123249:214 | 2123968:214 | -;-;-;- | 3317 | pcdh2ab1 | 8 | 0 | 6 | 0 | 8 | 0 |
| ENSDARG000 | 14;14;14;14;1 | 2123123:212 | 2123968:212 | -;-;-;-;-;-;-;-; | 8298 | pcdh2ab8 | 0 | 0 | 14 | 0 | 0 | 0 |
| ENSDARG000 | 14;14;14;14 | 16083818:16 | 16084460:16 | +;+;+;+ | 3116 | rnf103 | 1010 | 682 | 688 | 979 | 700 | 950 |
| ENSDARG000 | 14;14;14;14;1 | 14577246:14 | 14577699:14 | +;+;+;+;+;+;+ | 4964 | plp1a | 40 | 42 | 40 | 60 | 62 | 100 |
| ENSDARG000 | 14;14;14;14;1 | 22467672:22 | 22467693:22 | +;+;+;+;+;+;+;+; | 2519 | nudt22 | 328 | 565 | 361 | 412 | 390 | 416 |
| ENSDARG000 | 14;14;14;14 | 2123123:214 | 2123968:214 | -;-;-;- | 3398 | pcdh2ab12 | 0 | 0 | 0 | 0 | 0 | 0 |
| ENSDARG000 | 14;14;14;14 | 2123123:214 | 2123968:214 | -;-;-;- | 3395 | pcdh2ab6 | 13 | 0 | 8 | 3 | 2 | 0 |
| ENSDARG000 | 14 | 2187511 | 2189889 | - | 2379 | NA | 0 | 0 | 0 | 0 | 0 | 0 |
| ENSDARG000 | 14 | 2320220 | 2322484 | - | 2265 | NA | 0 | 4 | 0 | 0 | 0 | 0 |
| ENSDARG000 | 14 | 2200749 | 2203124 | - | 2376 | NA | 9 | 4 | 5 | 3 | 0 | 0 |
| ENSDARG000 | 14;14;14;14 | 2123123:214 | 2123968:214 | -;-;-;- | 3373 | pcdh2ab5 | 0 | 0 | 0 | 0 | 0 | 0 |
| ENSDARG000 | 14;14;14;14;1 | 2118775:211 | 2123968:212 | -;-;-;-;-;-;-;-; | 20230 | pcdh2ab9 | 0 | 0 | 0 | 0 | 3 | 0 |
| ENSDARG000 | 14;14;14;14;1 | 2123323:214 | 2123968:214 | -;-;-;-;- | 3183 | pcdh2aa1 | 0 | 0 | 0 | 0 | 0 | 0 |
| ENSDARG000 | 14;14;14;14;1 | 39002116:39 | 39003047:39 | -;-;-;-;-;-;-;-; | 2971 | glra4a | 0 | 0 | 1 | 0 | 0 | 6 |
| ENSDARG000 | 14;14;14;14 | 2123123:214 | 2123968:214 | -;-;-;- | 3470 | pcdh2ab10 | 0 | 0 | 7 | 0 | 12 | 0 |
| ENSDARG000 | 14;14;14;14;1 | 2123123:212 | 2123968:212 | -;-;-;-;-;-;-;- | 3379 | pcdh2aa15 | 0 | 0 | 0 | 0 | 0 | 0 |
| ENSDARG000 | 14;14;14;14;1 | 33418500:33 | 33418950:33 | -;-;-;-;-;-;-;-; | 1730 | nkap | 858 | 974 | 1053 | 943 | 918 | 865 |
| ENSDARG000 | 14;14;14;14;1 | 30359124:30 | 30359455:30 | -;-;-;-;-;- | 1402 | pdgfrl | 108 | 100 | 153 | 443 | 196 | 108 |
| ENSDARG000 | 14;14;14;14;1 | 6245942:624 | 6246100:624 | -;-;-;-;-;-;-;-; | 4318 | ELP1 | 638 | 638 | 1235 | 534 | 716 | 649 |
| ENSDARG000 | 14;14;14;14;1 | 25052810:25 | 25052838:25 | +;+;+;+;+;+;+;+; | 1431 | NA | 8 | 54 | 48 | 43 | 54 | 16 |
| ENSDARG000 | 14;14;14;14;1 | 31090151:31 | 31092606:31 | -;-;-;-;-;-;-;- | 3998 | gpc4 | 2202 | 2056 | 4377 | 2851 | 3869 | 2274 |
| ENSDARG000 | 14;14;14;14;1 | 28426602:28 | 28426940:28 | -;-;-;-;-;-;- | 1900 | nek12 | 6 | 9 | 32 | 16 | 26 | 0 |
| ENSDARG000 | 14;14 | 45551993:45 | 45555820:45 | -;- | 3899 | INTS5 | 212 | 266 | 428 | 146 | 332 | 295 |
| ENSDARG000 | 14;14;14;14;1 | 45514469:45 | 45515730:45 | -;-;-;-;-;-;-;-; | 3980 | GANAB | 4811 | 4210 | 6350 | 3977 | 5265 | 3930 |
| ENSDARG000 | 14;14;14;14 | 28432737:28 | 28433015:28 | +;+;+;+ | 785 | pin4 | 378 | 509 | 547 | 298 | 433 | 390 |
| ENSDARG000 | 14;14;14;14;1 | 8456870:845 | 8457135:845 | +;+;+;+;+;+;+;+; | 1836 | tmco6 | 98 | 118 | 136 | 111 | 70 | 146 |
| ENSDARG000 | 14;14;14;14;1 | 22481833:22 | 22482391:22 | -;-;-;-;-;-;-;-; | 2924 | NA | 74 | 173 | 141 | 91 | 216 | 130 |
| ENSDARG000 | 14;14;14;14;1 | 10624911:10 | 10625178:10 | +;+;+;+;+;+;+;+; | 1977 | nup62l | 873 | 989 | 1344 | 584 | 1000 | 842 |
| ENSDARG000 | 14;14;14;14;1 | 30367602:30 | 30369685:30 | -;-;-;-;-;-;-;-; | 4199 | slc7a2 | 760 | 677 | 794 | 562 | 994 | 737 |
| ENSDARG000 | 14;14;14;14;1 | 32440639:32 | 32444719:32 | -;-;-;-;-;-;-;-; | 8207 | MCF2 | 48 | 14 | 22 | 0 | 2 | 44 |
| ENSDARG000 | 14;14;14;14 | 14603731:14 | 14604522:14 | -;-;-;- | 878 | RAB9B | 138 | 57 | 115 | 62 | 108 | 115 |

| | | | | | | | | | | | | |
|---|---|---|---|---|---|---|---|---|---|---|---|---|
| ENSDARG000 | 14;14;14;14;1 | 39255437:392 | 39255469:392 | +;+;+;+;+;+;+; | 7767 | DIAPH2 | 1510 | 1018 | 1266 | 1099 | 1307 | 1320 |
| ENSDARG000 | 14;14;14;14;1 | 16093889:160 | 16094111:160 | +;+;+;+;+;+;+; | 2950 | immt | 2440 | 2818 | 3190 | 2259 | 2451 | 2498 |
| ENSDARG000 | 14;14;14;14;1 | 47000351:470 | 47005022:470 | -;-;-;-;-;-;-;-; | 7649 | fstl5 | 0 | 0 | 0 | 0 | 4 | 0 |
| ENSDARG000 | 14;14;14;14;1 | 28438947:284 | 28439011:284 | +;+;+;+;+;+;+; | 3852 | acsl4a | 1667 | 4614 | 1813 | 1141 | 1703 | 1133 |
| ENSDARG000 | 14;14;14 | 47085362:473 | 47085633:473 | +;+;+ | 489 | NA | 0 | 0 | 0 | 0 | 0 | 0 |
| ENSDARG000 | 14;14;14;14;1 | 28459439:284 | 28459746:284 | -;-;-;-;-;-;-;-; | 1284 | nxt2 | 112 | 133 | 148 | 120 | 96 | 136 |
| ENSDARG000 | 14;14;14;14;1 | 31176952:312 | 31177884:312 | -;-;-;-;-;-;-;- | 2558 | gpc3 | 0 | 2 | 0 | 0 | 2 | 0 |
| ENSDARG000 | 14;14 | 33427837:334 | 33427943:334 | +;+ | 4913 | zbtb33 | 503 | 334 | 497 | 288 | 413 | 512 |
| ENSDARG000 | 14;14;14;14;1 | 33435855:334 | 33437479:334 | -;-;-;-;-;-;-;-; | 2793 | TMEM255A | 26 | 50 | 29 | 27 | 66 | 92 |
| ENSDARG000 | 14;14;14;14;1 | 14620830:146 | 14623471:146 | -;-;-;-;-;-;-;-; | 5902 | znf185 | 101 | 60 | 44 | 65 | 30 | 44 |
| ENSDARG000 | 14;14;14;14;1 | 8475007:8476 | 8475122:8476 | +;+;+;+;+;+;+; | 2913 | dnajc4 | 287 | 118 | 272 | 180 | 268 | 222 |
| ENSDARG000 | 14;14;14;14;1 | 28464736:284 | 28464914:284 | +;+;+;+;+ | 1287 | psmd10 | 343 | 470 | 411 | 315 | 455 | 329 |
| ENSDARG000 | 14;14;14;14;1 | 35464994:354 | 35465229:354 | +;+;+;+;+;+;+; | 6358 | NA | 243 | 147 | 118 | 62 | 260 | 175 |
| ENSDARG000 | 14;14;14;14;1 | 28473173:284 | 28473583:284 | +;+;+;+;+;+ | 2196 | xiap | 1002 | 1070 | 1154 | 2319 | 952 | 974 |
| ENSDARG000 | 14;14;14;14;1 | 45559268:455 | 45559316:455 | +;+;+;+;+ | 1298 | rarres3 | 30 | 26 | 10 | 6 | 22 | 16 |
| ENSDARG000 | 14;14;14;14 | 14608099:146 | 14608209:146 | +;+;+;+ | 2262 | fut11 | 94 | 72 | 145 | 94 | 102 | 69 |
| ENSDARG000 | 14;14;14;14;1 | 16126983:161 | 16127218:161 | -;-;-;-;-;-;-;-; | 2320 | ptcd3 | 540 | 617 | 723 | 614 | 715 | 620 |
| ENSDARG000 | 14 | 31473866 | 31475391 | + | 1526 | NA | 77 | 44 | 62 | 76 | 40 | 100 |
| ENSDARG000 | 14;14;14;14;1 | 21140612:211 | 21140731:211 | +;+;+;+;+;+;+; | 3317 | CTBP1 | 888 | 680 | 793 | 735 | 513 | 645 |
| ENSDARG000 | 14;14;14;14;1 | 35499678:354 | 35499823:354 | -;-;-;-;-;-;- | 839 | mrpl18 | 377 | 581 | 640 | 255 | 491 | 390 |
| ENSDARG000 | 14;14;14;14;1 | 31476973:314 | 31480265:314 | -;-;-;-;-;-;-;-; | 4424 | cab39l1 | 1174 | 724 | 1579 | 1111 | 1640 | 929 |
| ENSDARG000 | 14;14;14;14;1 | 33458294:334 | 33458765:334 | +;+;+;+;+;+;+; | 2484 | ATP1B4 | 4 | 4 | 0 | 0 | 4 | 4 |
| ENSDARG000 | 14;14;14;14;1 | 21123867:211 | 21125358:211 | -;-;-;-;-;-;-;- | 2669 | maea | 724 | 697 | 679 | 553 | 763 | 702 |
| ENSDARG000 | 14;14;14;14;1 | 45565891:455 | 45565959:455 | +;+;+;+;+;+;+; | 3115 | rarres3l | 718 | 409 | 338 | 498 | 543 | 452 |
| ENSDARG000 | 14;14;14;14;1 | 21106444:211 | 21106653:211 | +;+;+;+;+;+;+; | 5534 | aldob | 157 | 4 | 2981 | 87 | 331 | 130 |
| ENSDARG000 | 14;14;14;14;1 | 31488579:314 | 31488636:314 | +;+;+;+;+;+;+; | 2355 | phf6 | 294 | 245 | 410 | 359 | 329 | 279 |
| ENSDARG000 | 14;14;14;14;1 | 8504410:8530 | 8504978:8530 | +;+;+;+;+;+;+; | 7452 | NA | 4 | 0 | 4 | 0 | 16 | 0 |
| ENSDARG000 | 14;14;14;14;1 | 40108638:401 | 40108976:401 | +;+;+;+;+;+;+; | 2804 | NA | 4 | 0 | 0 | 0 | 0 | 0 |
| ENSDARG000 | 14;14;14;14;1 | 22498757:224 | 22499008:224 | +;+;+;+;+;+;+; | 1621 | smyd5 | 183 | 355 | 349 | 46 | 230 | 226 |
| ENSDARG000 | 14;14;14;14;1 | 21115029:211 | 21115420:211 | -;-;-;-;-;-;-;-; | 1262 | NA | 2 | 6 | 2 | 2 | 0 | 21 |
| ENSDARG000 | 14;14;14;14;1 | 25063318:250 | 25063639:250 | -;-;-;-;-;-;-;-; | 5193 | NA | 1633 | 1239 | 2117 | 1430 | 1893 | 1683 |
| ENSDARG000 | 14;14;14;14;1 | 47779846:477 | 47784509:477 | -;-;-;-;-;-;-;-; | 10701 | RAPGEF2 | 2136 | 2136 | 1644 | 1229 | 1915 | 1278 |
| ENSDARG000 | 14;14;14;14;1 | 8594367:8603 | 8594457:8603 | +;+;+;+;+;+;+; | 3477 | stx5a | 653 | 558 | 813 | 490 | 784 | 527 |
| ENSDARG000 | 14;14;14;14;1 | 2337973:2341 | 2338005:2341 | -;-;-;-;- | 2529 | NA | 0 | 0 | 0 | 0 | 1 | 0 |
| ENSDARG000 | 14 | 2359230 | 2361692 | - | 2463 | NA | 0 | 0 | 0 | 0 | 0 | 0 |
| ENSDARG000 | 14 | 2356243 | 2358774 | - | 2532 | NA | 0 | 0 | 0 | 0 | 0 | 0 |
| ENSDARG000 | 14 | 2349946 | 2352384 | - | 2439 | NA | 0 | 0 | 0 | 0 | 0 | 0 |
| ENSDARG000 | 14 | 2353398 | 2355833 | - | 2436 | NA | 0 | 0 | 0 | 0 | 0 | 0 |
| ENSDARG000 | 14 | 2346509 | 2348917 | - | 2409 | NA | 0 | 0 | 0 | 0 | 0 | 0 |
| ENSDARG000 | 14 | 2362347 | 2364761 | - | 2415 | NA | 0 | 0 | 0 | 0 | 0 | 0 |
| ENSDARG000 | 14;14;14;14;1 | 33473804:334 | 33473994:334 | -;-;-;-;-;-;-;-; | 3237 | lamp2 | 1110 | 1186 | 1237 | 1292 | 1714 | 1486 |
| ENSDARG000 | 14;14;14;14;1 | 35691889:356 | 35692162:356 | +;+;+;+;+;+;+; | 2573 | glrbb | 18 | 11 | 0 | 0 | 0 | 4 |
| ENSDARG000 | 14;14 | 33478661:334 | 33479354:334 | +;+ | 766 | NA | 0 | 0 | 0 | 0 | 0 | 0 |
| ENSDARG000 | 14;14;14;14;1 | 10656975:106 | 10657200:106 | +;+;+;+;+;+;+; | 8686 | atrx | 5089 | 4714 | 6914 | 2521 | 6901 | 3773 |
| ENSDARG000 | 14;14;14;14;1 | 35748206:357 | 35748622:357 | +;+;+;+;+;+;+; | 5242 | gria2b | 0 | 2 | 0 | 0 | 0 | 2 |
| ENSDARG000 | 14;14;14;14;1 | 32507811:325 | 32508080:325 | -;-;-;-;-;-;-;-; | 4558 | ATP11C | 144 | 117 | 92 | 53 | 191 | 149 |
| ENSDARG000 | 14;14;14;14;1 | 45575035:455 | 45575825:455 | -;-;-;-;-;-;-;-; | 3352 | NA | 924 | 913 | 1173 | 724 | 988 | 746 |
| ENSDARG000 | 14;14;14;14;1 | 35567321:355 | 35568430:355 | -;-;-;-;-;-;-;-; | 2736 | PDGFC | 2 | 6 | 6 | 0 | 34 | 0 |
| ENSDARG000 | 14;14;14;14;1 | 2382690:2387 | 2387410:2387 | -;-;-;-;-;-;-;-; | 9016 | NA | 0 | 0 | 2 | 4 | 8 | 0 |
| ENSDARG000 | 14;14;14 | 8619126:8622 | 8619956:8623 | -;-;- | 1299 | zgc:162144 | 22 | 2 | 23 | 4 | 6 | 0 |
| ENSDARG000 | 14;14;14;14;1 | 22591624:226 | 22592011:226 | +;+;+;+;+;+;+; | 2924 | NA | 0 | 0 | 0 | 0 | 0 | 0 |
| ENSDARG000 | 14;14;14;14;1 | 8638353:8646 | 8638533:8646 | +;+;+;+;+;+;+; | 10645 | NA | 1208 | 620 | 702 | 703 | 754 | 520 |
| ENSDARG000 | 14;14;14;14;1 | 30398546:303 | 30398741:303 | +;+;+;+;+;+;+; | 2826 | mtmr7a | 64 | 18 | 30 | 185 | 28 | 8 |
| ENSDARG000 | 14;14;14;14;1 | 16151368:161 | 16151824:161 | +;+;+;+;+;+;+; | 7583 | POLR1A | 1047 | 917 | 1748 | 1157 | 1376 | 903 |
| ENSDARG000 | 14;14 | 45596583:456 | 45597176:456 | +;+ | 974 | zgc:194285 | 31 | 29 | 43 | 29 | 59 | 37 |
| ENSDARG000 | 14;14;14 | 22841462:228 | 22841557:228 | +;+;+ | 545 | NA | 0 | 0 | 0 | 0 | 0 | 0 |
| ENSDARG000 | 14;14;14;14;1 | 23076127:230 | 23076360:230 | +;+;+;+;+;+;+; | 4315 | ENOX2 | 72 | 46 | 74 | 36 | 48 | 66 |
| ENSDARG000 | 14;14;14;14;1 | 2487672:2487 | 2487703:2487 | +;+;+;+;+;+;+; | 4893 | fgf18a | 6 | 0 | 5 | 0 | 8 | 17 |
| ENSDARG000 | 14;14;14;14;1 | 6287871:6289 | 6288064:6289 | +;+;+;+;+;+;+; | 5004 | TBC1D2 | 767 | 2039 | 1002 | 1355 | 1010 | 918 |
| ENSDARG000 | 14;14;14;14;1 | 2661646:2661 | 2661839:2661 | -;-;-;-;-;-;-;-; | 1934 | SPOCK1 | 0 | 0 | 0 | 0 | 0 | 0 |
| ENSDARG000 | 14;14;14;14;1 | 48045193:480 | 48045366:480 | +;+;+;+;+;+;+; | 1790 | ppid | 1388 | 1450 | 2466 | 1493 | 1603 | 1551 |
| ENSDARG000 | 14;14;14;14;1 | 8710568:8715 | 8710709:8715 | +;+;+;+;+;+;+; | 9914 | NA | 0 | 2 | 0 | 0 | 0 | 0 |
| ENSDARG000 | 14;14;14;14;1 | 8722152:8722 | 8722243:8722 | -;-;-;-;-;- | 1408 | NA | 0 | 0 | 0 | 0 | 0 | 0 |
| ENSDARG000 | 14 | 45603739 | 45604667 | - | 929 | si:ch211-276i | 368 | 353 | 578 | 368 | 482 | 341 |
| ENSDARG000 | 14;14;14;14;1 | 8725216:8725 | 8725572:8726 | +;+;+;+;+;+;+ | 1394 | NA | 0 | 0 | 0 | 0 | 0 | 0 |
| ENSDARG000 | 14;14;14;14;1 | 25079782:250 | 25080108:250 | -;-;-;-;-;-;-;-; | 3991 | NA | 2526 | 2213 | 3574 | 2051 | 2820 | 2453 |
| ENSDARG000 | 14;14 | 45608576:456 | 45609037:456 | +;+ | 874 | NA | 206 | 248 | 343 | 192 | 340 | 222 |
| ENSDARG000 | 14;14 | 45636960:456 | 45637262:456 | +;+ | 638 | NA | 0 | 0 | 0 | 0 | 0 | 0 |
| ENSDARG000 | 14 | 45631236 | 45638712 | - | 7477 | NA | 21245 | 20390 | 23924 | 18815 | 27804 | 25835 |
| ENSDARG000 | 14;14;14;14;1 | 48070872:480 | 48071454:480 | -;-;-;-;-;-;-;-; | 2498 | etfdh | 448 | 434 | 530 | 373 | 438 | 476 |
| ENSDARG000 | 14;14;14;14;1 | 2941797:2942 | 2942324:2942 | -;-;-;-;-;-;-;-; | 2976 | bicc2 | 0 | 4 | 0 | 0 | 0 | 2 |
| ENSDARG000 | 14;14;14;14;1 | 40321004:403 | 40321432:403 | +;+;+;+;+;+ | 3794 | NA | 0 | 0 | 0 | 0 | 0 | 0 |
| ENSDARG000 | 14;14;14;14;1 | 40300961:403 | 40305738:403 | -;-;-;-;-;-;-;-; | 8561 | pcdh19 | 76 | 10 | 62 | 0 | 119 | 12 |
| ENSDARG000 | 14;14;14;14;1 | 30413312:304 | 30414182:304 | +;+;+;+;+;+;+; | 4465 | cnot7 | 127 | 95 | 195 | 124 | 148 | 168 |
| ENSDARG000 | 14;14;14;14;1 | 2986680:2986 | 2991641:2991 | -;-;-;-;-;-;-;-; | 8446 | ndst1a | 12 | 6 | 57 | 20 | 4 | 88 |
| ENSDARG000 | 14;14;14;14;1 | 48107216:481 | 48111030:481 | -;-;-;-;-;-;-;-; | 6084 | rxfp1 | 0 | 2 | 0 | 0 | 0 | 2 |
| ENSDARG000 | 14;14;14;14;1 | 45645024:456 | 45645350:456 | +;+;+;+;+;+;+ | 4061 | NA | 0 | 1 | 2 | 0 | 0 | 0 |
| ENSDARG000 | 14;14;14;14;1 | 14646717:146 | 14647335:146 | -;-;-;-;-;-;-;-; | 1686 | nsdhl | 270 | 282 | 222 | 178 | 81 | 192 |
| ENSDARG000 | 14;14;14;14;1 | 40641350:406 | 40641856:406 | +;+;+;+;+;+;+; | 4992 | tnmd | 5591 | 4277 | 5856 | 4691 | 9950 | 1640 |
| ENSDARG000 | 14;14;14;14;1 | 40745833:407 | 40748526:407 | -;-;-;-;-;-;-;-; | 4781 | elf1 | 2985 | 2395 | 2477 | 3969 | 3540 | 2949 |
| ENSDARG000 | 14;14 | 48580369:485 | 48580760:485 | -;- | 579 | NA | 0 | 0 | 0 | 0 | 0 | 0 |
| ENSDARG000 | 14;14;14;14;1 | 48734794:487 | 48741545:487 | -;-;-;-;-;-;-;-; | 8351 | cnot6b | 623 | 379 | 309 | 96 | 390 | 592 |
| ENSDARG000 | 14;14;14 | 10751249:107 | 10752740:107 | -;-;- | 1898 | fgf16 | 6 | 0 | 0 | 0 | 0 | 0 |
| ENSDARG000 | 14;14;14;14;1 | 45702521:457 | 45702735:457 | +;+;+;+;+;+;+; | 5068 | ccdc88b | 44 | 98 | 62 | 57 | 312 | 224 |
| ENSDARG000 | 14;14;14;14;1 | 45675306:456 | 45675403:456 | +;+;+;+;+;+;+; | 2820 | rom1b | 0 | 2 | 0 | 7 | 0 | 0 |
| ENSDARG000 | 14;14;14;14;1 | 6331499:6350 | 6344024:6350 | -;-;-;-;-;-;-;-; | 14740 | SLC44A1 | 454 | 555 | 543 | 580 | 899 | 745 |
| ENSDARG000 | 14;14;14 | 45698879:456 | 45699338:456 | -;-;- | 683 | NA | 0 | 0 | 0 | 0 | 0 | 0 |
| ENSDARG000 | 14;14;14;14;1 | 23518110:235 | 23518397:235 | +;+;+;+;+;+;+; | 4779 | NA | 52 | 24 | 10 | 16 | 86 | 14 |
| ENSDARG000 | 14;14;14;14;1 | 3038473:3038 | 3038723:3038 | +;+;+;+;+;+;+; | 1273 | cd74a | 644 | 1106 | 2732 | 740 | 5264 | 904 |
| ENSDARG000 | 14;14;14;14;1 | 40874608:408 | 40874703:408 | +;+;+;+;+;+;+; | 1749 | NA | 2 | 0 | 6 | 0 | 2 | 0 |
| ENSDARG000 | 14;14;14;14;1 | 40852497:408 | 40852782:408 | +;+;+;+;+;+;+; | 2373 | taf7 | 654 | 787 | 891 | 775 | 831 | 819 |
| ENSDARG000 | 14;14;14;14;1 | 33486179:334 | 33487053:334 | +;+;+;+;+;+;+; | 6508 | cul4b | 2245 | 2131 | 3234 | 2384 | 3245 | 3066 |
| ENSDARG000 | 14;14;14 | 16218769:162 | 16218914:162 | +;+;+ | 349 | NA | 0 | 0 | 0 | 0 | 0 | 0 |
| ENSDARG000 | 14;14;14;14;1 | 48766996:487 | 48767658:487 | -;-;-;-;-;-;-;-; | 2924 | NA | 236 | 168 | 220 | 160 | 195 | 316 |
| ENSDARG000 | 14;14;14;14;1 | 10954345:109 | 10954489:109 | +;+;+;+;+;+;+; | 2893 | zdhhc15b | 442 | 256 | 398 | 212 | 432 | 367 |
| ENSDARG000 | 14;14;14;14;1 | 8770152:8781 | 8773255:8782 | -;-;-;-;-;-;-;- | 4853 | NA | 14 | 8 | 2 | 4 | 25 | 24 |
| ENSDARG000 | 14;14;14;14;1 | 32664833:326 | 32665661:326 | -;-;-;-;-;-;-;-; | 2661 | NA | 18 | 6 | 205 | 137 | 124 | 4 |
| ENSDARG000 | 14 | 32742701 | 32744464 | - | 1764 | sox3 | 10 | 0 | 70 | 38 | 42 | 8 |
| ENSDARG000 | 14;14;14;14 | 32751464:327 | 32751633:327 | +;+;+;+ | 740 | NA | 0 | 0 | 3 | 2 | 0 | 0 |
| ENSDARG000 | 14;14;14;14;1 | 30426956:304 | 30428355:304 | -;-;-;-;-;-;-;-; | 2825 | zdhhc2 | 5082 | 1981 | 782 | 3273 | 552 | 2742 |
| ENSDARG000 | 14;14;14;14;1 | 25100267:251 | 25102363:251 | -;-;-;-;- | 5222 | NA | 109 | 159 | 176 | 78 | 146 | 208 |
| ENSDARG000 | 14;14;14;14;1 | 25133764:251 | 25134651:251 | -;-;-;-;-;-;-;-; | 2543 | HTR4 | 14 | 6 | 4 | 0 | 4 | 2 |
| ENSDARG000 | 14;14;14;14;1 | 48812189:488 | 48812708:488 | +;+;+;+;+;+;+; | 6987 | NA | 4 | 0 | 2 | 0 | 0 | 0 |
| ENSDARG000 | 14;14;14;14;1 | 23593989:235 | 23594163:235 | +;+;+;+;+;+;+; | 6924 | med12 | 1733 | 1282 | 1939 | 950 | 1667 | 1458 |
| ENSDARG000 | 14;14;14;14;1 | 48862987:488 | 48863171:488 | +;+;+;+;+;+;+; | 2760 | zgc:154054 | 14 | 2 | 0 | 0 | 4 | 4 |
| ENSDARG000 | 14;14;14;14 | 8806227:8809 | 8806772:8809 | -;-;-;- | 1009 | trmt112 | 350 | 377 | 743 | 370 | 384 | 345 |
| ENSDARG000 | 14;14;14;14;1 | 3056713:3057 | 3056882:3057 | +;+;+;+;+ | 880 | NA | 0 | 0 | 0 | 0 | 2 | 0 |
| ENSDARG000 | 14;14;14;14;1 | 33507608:335 | 33507820:335 | +;+;+;+;+;+ | 894 | mcts1 | 876 | 1055 | 1775 | 994 | 1578 | 1059 |
| ENSDARG000 | 14;14;14;14;1 | 25438015:254 | 25441361:254 | -;-;-;-;-;-;- | 5011 | SLC26A2 | 311 | 821 | 2020 | 563 | 3393 | 1110 |
| ENSDARG000 | 14;14;14 | 23679049:236 | 23680454:236 | -;-;- | 2067 | MARS2 | 76 | 46 | 99 | 36 | 67 | 48 |
| ENSDARG000 | 14;14;14;14;1 | 25464681:254 | 25465582:254 | +;+;+;+;+;+;+; | 6742 | NA | 2 | 20 | 40 | 0 | 24 | 8 |
| ENSDARG000 | 14;14;14;14 | 48932235:489 | 48935054:489 | -;-;-;- | 3160 | scocb | 1094 | 1039 | 1123 | 600 | 905 | 1000 |
| ENSDARG000 | 14;14 | 6462273:6468 | 6462643:6468 | -;- | 609 | NA | 0 | 0 | 0 | 0 | 0 | 0 |
| ENSDARG000 | 14;14;14;14;1 | 6423973:6429 | 6424142:6429 | +;+;+;+;+;+;+; | 9432 | abca1b | 265 | 163 | 923 | 108 | 724 | 332 |
| ENSDARG000 | 14 | 8815868 | 8816933 | - | 1066 | NA | 962 | 831 | 514 | 1457 | 514 | 775 |
| ENSDARG000 | 14;14;14;14;1 | 32773672:327 | 32774125:327 | -;-;-;-;- | 1329 | NA | 18 | 0 | 19 | 11 | 17 | 6 |
| ENSDARG000 | 14;14;14;14;1 | 32783972:327 | 32784296:327 | +;+;+;+;+ | 1914 | crhr2 | 4 | 10 | 10 | 4 | 30 | 0 |
| ENSDARG000 | 14;14;14;14;1 | 16287968:162 | 16288255:162 | +;+;+;+;+;+;+; | 2419 | prpf19 | 1130 | 1416 | 2044 | 966 | 1899 | 982 |
| ENSDARG000 | 14;14;14;14;1 | 14662116:146 | 14662140:146 | +;+;+;+;+ | 2045 | CETN1 | 22 | 37 | 30 | 32 | 12 | 20 |
| ENSDARG000 | 14;14;14;14;1 | 3071110:3074 | 3071603:3074 | +;+;+;+;+;+;+; | 5791 | HMGXB3 | 136 | 85 | 172 | 28 | 114 | 122 |
| ENSDARG000 | 14;14;14;14 | 45883687:459 | 45883836:459 | +;+;+;+ | 7287 | FLRT1 | 448 | 99 | 101 | 56 | 32 | 194 |
| ENSDARG000 | 14;14;14;14;1 | 45789454:457 | 45790456:457 | -;-;-;-;-;-;-;-; | 2534 | macrod1 | 210 | 96 | 122 | 162 | 146 | 179 |
| ENSDARG000 | 14;14 | 33515515:335 | 33518735:335 | -;- | 3458 | c1galt1c1 | 378 | 405 | 710 | 470 | 1014 | 470 |
| ENSDARG000 | 14;14;14;14;1 | 3062608:3065 | 3065513:3065 | -;-;-;-;-;-;-;-; | 3789 | slc35a4 | 458 | 488 | 696 | 341 | 710 | 479 |
| ENSDARG000 | 14;14;14;14;1 | 32793272:327 | 32793713:327 | -;-;-;-;-;-;-;-; | 4105 | inppl1b | 620 | 197 | 200 | 704 | 131 | 859 |
| ENSDARG000 | 14;14;14;14;1 | 48960078:489 | 48960707:489 | +;+;+;+;+;+;+ | 1290 | mif | 832 | 1000 | 1202 | 846 | 1111 | 886 |
| ENSDARG000 | 14;14;14;14;1 | 28486213:284 | 28486413:284 | +;+;+;+;+;+;+; | 6693 | STAG2 | 2020 | 2541 | 3394 | 2022 | 3149 | 2485 |
| ENSDARG000 | 14;14;14;14;1 | 48944320:489 | 48945377:489 | -;-;-;-;- | 2184 | NA | 74 | 78 | 84 | 53 | 85 | 122 |
| ENSDARG000 | 14;14;14;14;1 | 14667268:146 | 14667903:146 | -;-;-;-;-;-;-;-; | 2300 | gcna | 17 | 59 | 91 | 20 | 42 | 38 |
| ENSDARG000 | 14;14;14;14 | 35876876:358 | 35877498:358 | -;-;-;- | 1988 | GASK1B | 224 | 559 | 323 | 106 | 319 | 98 |
| ENSDARG000 | 14;14;14;14;1 | 31509922:315 | 31510094:315 | +;+;+;+;+;+;+; | 3577 | hprt1 | 1547 | 1799 | 4376 | 2083 | 2444 | 2078 |
| ENSDARG000 | 14;14;14;14;1 | 25505468:255 | 25505665:255 | +;+;+;+;+;+ | 1477 | thoc3 | 228 | 402 | 406 | 138 | 344 | 240 |
| ENSDARG000 | 14;14;14;14;1 | 30453342:304 | 30456469:304 | -;-;-;-;-;-;-;-; | 4964 | MICU3 | 13 | 20 | 23 | 18 | 34 | 33 |
| ENSDARG000 | 14;14;14;14;1 | 35884853:358 | 35885177:358 | -;-;-;-;-;-;-;-; | 1505 | tmem144b | 0 | 4 | 0 | 0 | 0 | 0 |
| ENSDARG000 | 14;14;14;14;1 | 8893529:8894 | 8893855:8894 | -;-;-;-;-;-;-;-; | 2437 | nxf1a | 6 | 2 | 14 | 22 | 4 | 2 |
| ENSDARG000 | 14;14;14;14;1 | 35892802:358 | 35892955:358 | +;+;+;+;+;+;+; | 1741 | adh5l | 233 | 225 | 41 | 89 | 51 | 231 |
| ENSDARG000 | 14;14;14;14;1 | 33568838:335 | 33569396:335 | -;-;-;-;-;-;- | 1558 | SASH3 | 50 | 84 | 78 | 30 | 205 | 131 |
| ENSDARG000 | 14;14;14;14;1 | 8940326:8947 | 8940397:8947 | +;+;+;+;+;+;+; | 3776 | rps6kal | 1675 | 1321 | 2119 | 3118 | 1603 | 1837 |
| ENSDARG000 | 14;14;14;14;1 | 45970033:459 | 45970285:459 | +;+;+;+;+;+;+; | 2577 | fermt3b | 70 | 123 | 140 | 72 | 160 | 112 |
| ENSDARG000 | 14;14;14;14;1 | 48982236:489 | 48982503:489 | +;+;+;+;+;+;+ | 4292 | clp1 | 396 | 416 | 513 | 176 | 480 | 420 |
| ENSDARG000 | 14;14;14;14;1 | 33525196:335 | 33525517:335 | +;+;+;+;+;+;+; | 3587 | zdhhc9 | 339 | 333 | 260 | 204 | 166 | 370 |
| ENSDARG000 | 14;14;14;14;1 | 14687726:146 | 14688473:146 | -;-;-;-;-;-;-;-; | 4354 | ogt.2 | 2573 | 1214 | 2991 | 1727 | 3340 | 2573 |
| ENSDARG000 | 14;14;14;14;1 | 10981703:109 | 10981885:109 | +;+;+;+;+;+;+; | 2744 | abcb7 | 822 | 950 | 1260 | 340 | 1032 | 775 |
| ENSDARG000 | 14;14;14;14 | 25534604:255 | 25540007:255 | -;-;-;- | 5772 | cplx2 | 16 | 10 | 1003 | 18 | 2422 | 28 |
| ENSDARG000 | 14;14;14 | 30491890:304 | 30492166:304 | +;+;+ | 1289 | fgf20b | 2 | 0 | 0 | 0 | 0 | 0 |
| ENSDARG000 | 14;14 | 48998812:490 | 48999091:490 | +;+ | 1062 | NA | 153 | 95 | 161 | 50 | 151 | 214 |
| ENSDARG000 | 14;14;14;14;1 | 32837914:328 | 32838195:328 | +;+;+;+;+;+;+; | 4138 | arr3b | 8 | 4 | 10 | 0 | 8 | 2 |
| ENSDARG000 | 14;14;14;14;1 | 25590603:255 | 25591199:255 | -;-;-;-;-;-;-;-; | 2085 | slc25a48 | 32 | 2 | 4 | 35 | 16 | 21 |
| ENSDARG000 | 14;14;14;14;1 | 49007501:490 | 49007637:490 | +;+;+;+;+;+;+; | 5372 | zdhhc5b | 1205 | 699 | 1032 | 481 | 1281 | 1048 |
| ENSDARG000 | 14;14;14;14;1 | 40919736:409 | 40919905:409 | -;-;-;-;-;-;-;-; | 3604 | drp2 | 0 | 0 | 6 | 0 | 2 | 0 |
| ENSDARG000 | 14;14;14;14;1 | 3123517:3124 | 3123637:3124 | -;-;-;-;-;-;-;-; | 1446 | NA | 2 | 1 | 70 | 92 | 31 | 0 |
| ENSDARG000 | 14;14;14;14 | 30580681:305 | 30582076:305 | -;-;-;- | 1899 | NA | 0 | 2 | 14 | 30 | 0 | 0 |
| ENSDARG000 | 14;14;14;14;1 | 23687678:236 | 23687967:236 | +;+;+;+;+;+;+; | 3010 | hspa4b | 3245 | 3436 | 4532 | 2363 | 3521 | 2698 |
| ENSDARG000 | 14;14;14;14;1 | 30557361:305 | 30557495:305 | +;+;+;+;+;+;+; | 1646 | mrpl11 | 255 | 405 | 495 | 234 | 352 | 247 |
| ENSDARG000 | 14;14;14;14 | 16308840:163 | 16309525:163 | -;-;-;- | 3822 | PCDH12 | 80 | 53 | 204 | 66 | 188 | 595 |
| ENSDARG000 | 14;14;14;14;1 | 41244877:412 | 41245244:412 | -;-;-;-;-;-;-;-; | 2508 | CENPI | 60 | 353 | 425 | 170 | 316 | 172 |
| ENSDARG000 | 14;14;14;14 | 30538482:305 | 30539292:305 | -;-;-;- | 1166 | her11 | 0 | 0 | 0 | 8 | 0 | 2 |
| ENSDARG000 | 14;14;14;14;1 | 49040190:490 | 49040806:490 | -;-;-;-;-;-;-;-; | 1924 | septin8b | 2 | 8 | 449 | 525 | 893 | 32 |
| ENSDARG000 | 14;14;14 | 30543008:305 | 30543092:305 | +;+;+ | 1326 | her5 | 0 | 0 | 0 | 4 | 0 | 0 |
| ENSDARG000 | 14;14;14;14;1 | 30550397:305 | 30550528:305 | -;-;-;-;-;-;-;- | 1326 | pfdn6 | 430 | 544 | 744 | 484 | 445 | 474 |
| ENSDARG000 | 14;14;14;14;1 | 21204541:212 | 21205705:212 | -;-;-;-;-;-;-;-; | 2559 | NA | 0 | 0 | 6 | 0 | 0 | 2 |
| ENSDARG000 | 14;14;14;14;1 | 46003973:460 | 46004132:460 | +;+;+;+;+;+;+; | 2551 | slu7 | 967 | 995 | 1060 | 905 | 998 | 958 |

| | | | | | | | | | | | | |
|---|---|---|---|---|---|---|---|---|---|---|---|---|
| ENSDARG000 | 14;14;14;14;1 | 16345003;163 | 16345119;163 | +;+;+;+;+;+;+; | 1356 | itln3 | 8 | 0 | 18 | 0 | 0 | 6 |
| ENSDARG000 | 14;14;14;14;1 | 6520310;6521 | 6520901;6521 | -;-;-;-;-;-;-;- | 1719 | nipsnap3a | 8 | 122 | 120 | 134 | 122 | 66 |
| ENSDARG000 | 14;14;14;14 | 28530153;285 | 28530535;285 | -;-;-;- | 1075 | zgc:113364 | 0 | 0 | 0 | 0 | 0 | 0 |
| ENSDARG000 | 14;14;14;14;1 | 3143337;3143 | 3144064;3144 | -;-;-;-;-;-;-;-; | 3685 | csf1ra | 4434 | 1905 | 7951 | 6172 | 3765 | 7031 |
| ENSDARG000 | 14;14;14 | 11103718;112 | 11103893;112 | +;+;+ | 8055 | NEXMIF | 22 | 16 | 26 | 26 | 4 | 2 |
| ENSDARG000 | 14;14;14;14;1 | 6529630;6529 | 6529782;6529 | +;+;+;+;+;+ | 2045 | NA | 0 | 0 | 0 | 0 | 1 | 0 |
| ENSDARG000 | 14;14;14;14;1 | 35901249;359 | 35901294;359 | +;+;+;+;+;+;+; | 1469 | zgc:77938 | 35 | 29 | 68 | 101 | 94 | 93 |
| ENSDARG000 | 14;14;14;14;1 | 23709134;237 | 23709285;237 | +;+;+;+;+;+;+; | 3353 | gnpda1 | 716 | 732 | 554 | 406 | 466 | 498 |
| ENSDARG000 | 14;14;14;14;1 | 25816874;258 | 25817746;258 | +;+;+;+;+;+;+; | 8779 | glra1 | 98 | 118 | 431 | 312 | 700 | 332 |
| ENSDARG000 | 14;14;14;14;1 | 31529958;315 | 31530459;315 | +;+;+;+;+;+;+; | 3111 | pabir2 | 463 | 472 | 715 | 568 | 631 | 653 |
| ENSDARG000 | 14;14;14;14;1 | 33591626;335 | 33592523;335 | -;-;-;-;-;-;-;-; | 2837 | xpnpep2 | 28 | 20 | 36 | 18 | 140 | 54 |
| ENSDARG000 | 14;14;14;14;1 | 16370063;163 | 16372147;163 | -;-;-;-;- | 2433 | LTC4S | 8 | 6 | 0 | 0 | 28 | 6 |
| ENSDARG000 | 14;14;14;14;1 | 21222287;212 | 21222344;212 | +;+;+;+;+;+;+; | 1161 | NA | 10 | 2 | 2 | 0 | 0 | 0 |
| ENSDARG000 | 14;14;14;14;1 | 28563081;285 | 28563361;285 | -;-;-;-;-;-;-;-; | 1057 | insb | 0 | 0 | 0 | 0 | 0 | 0 |
| ENSDARG000 | 14;14;14;14;1 | 21232586;212 | 21233261;212 | -;-;-;-;- | 1210 | si:ch211-175r | 40 | 42 | 24 | 38 | 38 | 80 |
| ENSDARG000 | 14;14;14;14;1 | 11289851;112 | 11289995;112 | +;+;+;+;+;+;+ | 5016 | RLIM | 958 | 910 | 1042 | 492 | 978 | 715 |
| ENSDARG000 | 14;14;14;14;1 | 23717165;237 | 23717406;237 | +;+;+;+;+;+;+ | 1820 | ndfip1 | 3508 | 3603 | 3596 | 3442 | 3477 | 4227 |
| ENSDARG000 | 14;14;14;14;1 | 28545198;285 | 28545371;285 | +;+;+;+;+;+;+; | 3313 | hmmr | 282 | 733 | 874 | 331 | 557 | 398 |
| ENSDARG000 | 14;14;14;14;1 | 46020282;460 | 46020734;460 | +;+;+;+;+;+;+; | 4933 | C1QTNF2 | 8 | 6 | 8 | 4 | 16 | 0 |
| ENSDARG000 | 14;14;14;14;1 | 6535426;6536 | 6535897;6536 | +;+;+;+;+;+ | 1481 | thg1l | 138 | 209 | 236 | 126 | 167 | 158 |
| ENSDARG000 | 14;14;14;14;1 | 21238682;212 | 21238937;212 | +;+;+;+;+;+;+; | 3055 | smim19 | 497 | 353 | 474 | 518 | 459 | 586 |
| ENSDARG000 | 14;14;14 | 28567635;286 | 28568130;286 | +;+;+ | 1793 | NA | 0 | 0 | 0 | 0 | 0 | 0 |
| ENSDARG000 | 14;14;14;14;1 | 16382507;163 | 16385853;163 | -;-;-;-;-;-;-;-; | 5484 | MAML1 | 532 | 272 | 583 | 400 | 696 | 436 |
| ENSDARG000 | 14;14;14 | 23731949;237 | 23733330;237 | -;-;- | 1805 | fgf1a | 2 | 0 | 4 | 0 | 2 | 0 |
| ENSDARG000 | 14;14;14;14;1 | 35932410;361 | 35933454;361 | -;-;-;-;-;-;-;-; | 2356 | NA | 0 | 0 | 0 | 0 | 0 | 0 |
| ENSDARG000 | 14;14;14;14;1 | 36231126;362 | 36231211;362 | +;+;+;+;+;+;+; | 2062 | elovl6 | 111 | 252 | 271 | 70 | 135 | 168 |
| ENSDARG000 | 14;14;14;14;1 | 49078606;490 | 49079390;490 | -;-;-;-;-;-;-;-; | 1730 | sfxn1 | 253 | 254 | 639 | 227 | 528 | 215 |
| ENSDARG000 | 14;14;14;14;1 | 36218072;362 | 36218749;362 | +;+;+;+;+;+;+; | 4684 | pitx2 | 0 | 0 | 0 | 0 | 0 | 2 |
| ENSDARG000 | 14;14;14 | 32852388;328 | 32852699;328 | +;+;+ | 3082 | nkrf | 714 | 758 | 1038 | 382 | 855 | 725 |
| ENSDARG000 | 14;14;14;14;1 | 6546516;6546 | 6546754;6547 | +;+;+;+;+;+;+; | 4561 | NA | 58 | 146 | 116 | 127 | 115 | 198 |
| ENSDARG000 | 14;14;14;14;1 | 11307214;113 | 11307766;113 | -;-;-;-;-;- | 1048 | NA | 0 | 3 | 15 | 16 | 4 | 10 |
| ENSDARG000 | 14;14;14;14;1 | 33722950;337 | 33723416;337 | +;+;+;+;+ | 5000 | apln | 184 | 240 | 932 | 206 | 294 | 164 |
| ENSDARG000 | 14;14;14;14;1 | 3411771;3423 | 3411931;3424 | +;+;+;+;+;+;+; | 2709 | trpc3 | 0 | 0 | 0 | 0 | 2 | 0 |
| ENSDARG000 | 14;14;14 | 46028003;460 | 46028573;460 | +;+;+ | 3390 | NOCT | 898 | 253 | 306 | 188 | 341 | 289 |
| ENSDARG000 | 14;14;14;14;1 | 31550759;315 | 31551305;315 | -;-;-;-;-;-;-;-; | 1409 | mospd1 | 317 | 237 | 266 | 214 | 214 | 244 |
| ENSDARG000 | 14;14;14;14;1 | 32859430;328 | 32860584;328 | -;-;-;-;-;-;-;-; | 1990 | ube2a | 968 | 1155 | 1062 | 640 | 1194 | 776 |
| ENSDARG000 | 14;14;14;14;1 | 11317070;113 | 11317619;113 | -;-;-;-;-;- | 1053 | NA | 0 | 6 | 1 | 0 | 1 | 7 |
| ENSDARG000 | 14;14;14;14;1 | 25923373;259 | 25926374;259 | -;-;-;-;-;-;-;-; | 4408 | g3bp1 | 2950 | 6064 | 6456 | 2003 | 6850 | 2886 |
| ENSDARG000 | 14;14;14;14;1 | 46047761;460 | 46049031;460 | -;-;-;-;-;-;-;- | 2661 | elf2a | 594 | 324 | 538 | 369 | 448 | 499 |
| ENSDARG000 | 14;14;14;14;1 | 11338375;113 | 11338919;113 | -;-;-;-;-;- | 1357 | NA | 0 | 4 | 6 | 0 | 1 | 0 |
| ENSDARG000 | 14;14 | 21332522;213 | 21335044;213 | -;- | 3158 | egr1 | 52435 | 22433 | 41803 | 67834 | 15269 | 54413 |
| ENSDARG000 | 14;14;14;14;1 | 3495542;3497 | 3495596;3497 | +;+;+;+;+;+;+ | 873 | gstp1.1 | 0 | 0 | 1 | 0 | 0 | 4 |
| ENSDARG000 | 14;14;14;14;1 | 23746710;237 | 23748719;237 | -;-;-;-;-;-;-;-; | 5363 | nr3c1 | 1422 | 1307 | 1514 | 1751 | 1540 | 1550 |
| ENSDARG000 | 14;14;14;14 | 11366695;113 | 11367247;113 | -;-;-;- | 1049 | NA | 0 | 0 | 2 | 0 | 1 | 2 |
| ENSDARG000 | 14;14;14;14;1 | 32866748;328 | 32867330;328 | -;-;-;-;-;-;-;-; | 3245 | NA | 1184 | 636 | 1440 | 2095 | 768 | 1085 |
| ENSDARG000 | 14;14;14;14;1 | 21596716;216 | 21598933;216 | -;-;-;-;-;-;- | 3083 | reep2 | 2 | 4 | 32 | 4 | 0 | 12 |
| ENSDARG000 | 14;14;14;14;1 | 49118860;491 | 49119750;491 | -;-;-;-;-;-;- | 2097 | ppp2caa | 744 | 592 | 800 | 368 | 903 | 575 |
| ENSDARG000 | 14;14;14;14 | 11378295;113 | 11378847;113 | -;-;-;- | 1048 | NA | 0 | 0 | 0 | 0 | 0 | 0 |
| ENSDARG000 | 14;14 | 30625386;306 | 30625638;306 | +;+ | 336 | NA | 0 | 1 | 0 | 0 | 0 | 0 |
| ENSDARG000 | 14;14;14;14;1 | 11388813;113 | 11389366;113 | -;-;-;-;-;- | 1047 | NA | 14 | 6 | 4 | 0 | 15 | 6 |
| ENSDARG000 | 14;14;14;14;1 | 46074311;460 | 46078538;460 | -;-;-;-;-;-;-;-; | 9472 | mgarpa | 230 | 290 | 423 | 128 | 389 | 288 |
| ENSDARG000 | 14;14 | 41276307;412 | 41280524;412 | -;- | 4467 | tmem35 | 86 | 90 | 599 | 166 | 482 | 116 |
| ENSDARG000 | 14;14;14;14;1 | 9009600;9009 | 9009743;9009 | +;+;+;+;+;+;+; | 1941 | NA | 0 | 0 | 6 | 2 | 8 | 5 |
| ENSDARG000 | 14;14;14;14;1 | 31566517;315 | 31566791;315 | +;+;+;+;+;+;+; | 3149 | ints6l | 252 | 175 | 227 | 119 | 273 | 246 |
| ENSDARG000 | 14;14;14;14;1 | 41293110;412 | 41293765;412 | -;-;-;-;-;-;-;-; | 2072 | arl13a | 0 | 0 | 0 | 0 | 0 | 4 |
| ENSDARG000 | 14;14;14;14;1 | 14721300;147 | 14725535;147 | -;-;-;-;-;-;-;-; | 9250 | ogt.1 | 5530 | 2490 | 6902 | 4707 | 7278 | 5807 |
| ENSDARG000 | 14;14;14;14;1 | 33788835;337 | 33791366;337 | -;-;-;-;-;-;-;-; | 4699 | NA | 0 | 6 | 20 | 10 | 10 | 8 |
| ENSDARG000 | 14;14;14;14;1 | 16432738;164 | 16435164;164 | -;-;-;-;-;-;-;-; | 4225 | canx | 25929 | 23562 | 28103 | 13088 | 28226 | 16418 |
| ENSDARG000 | 14;14 | 49135264;491 | 49135485;491 | +;+ | 1333 | si:ch1073-44 | 206 | 208 | 255 | 264 | 221 | 230 |
| ENSDARG000 | 14;14;14;14;1 | 32881625;328 | 32882084;328 | -;-;-;-;-;- | 1415 | slc25a5 | 37279 | 38478 | 46345 | 46933 | 36328 | 43449 |
| ENSDARG000 | 14;14;14;14;1 | 30636955;306 | 30638616;306 | -;-;-;-;-;-;-;- | 4736 | npas4a | 44 | 8 | 32 | 0 | 10 | 0 |
| ENSDARG000 | 14;14;14;14;1 | 3507326;3507 | 3507380;3508 | +;+;+;+;+;+;+;+ | 893 | gstp1.2 | 6790 | 7179 | 5363 | 5665 | 8402 | 5509 |
| ENSDARG000 | 14 | 49138952 | 49139795 | + | 844 | NA | 0 | 0 | 2 | 0 | 8 | 0 |
| ENSDARG000 | 14;14;14;14;1 | 33825482;338 | 33829201;338 | -;-;-;-;-;-;-;-; | 7475 | ocrl | 2190 | 1457 | 2332 | 1236 | 2767 | 4489 |
| ENSDARG000 | 14;14;14;14 | 11395383;113 | 11395551;113 | +;+;+;+ | 864 | glod5 | 406 | 347 | 327 | 262 | 316 | 304 |
| ENSDARG000 | 14;14;14;14 | 6615564;6618 | 6615754;6618 | +;+;+;+ | 3280 | NA | 1068 | 2458 | 1053 | 2291 | 754 | 4486 |
| ENSDARG000 | 14;14;14;14;1 | 32884731;328 | 32885673;328 | -;-;-;-;- | 1879 | slc25a43 | 599 | 477 | 236 | 288 | 208 | 471 |
| ENSDARG000 | 14;14;14 | 32887808;328 | 32887965;328 | +;+;+ | 780 | NA | 13 | 12 | 0 | 0 | 3 | 0 |
| ENSDARG000 | 14;14;14;14;1 | 41318412;413 | 41318531;413 | +;+;+;+;+;+;+; | 3050 | xkrx | 292 | 288 | 214 | 244 | 232 | 352 |
| ENSDARG000 | 14;14;14;14 | 46114659;461 | 46114764;461 | -;-;-;- | 428 | ndufc1 | 214 | 274 | 231 | 124 | 202 | 164 |
| ENSDARG000 | 14;14;14;14;1 | 11427052;114 | 11428344;114 | -;-;-;-;-;-;-;-; | 2509 | irg1l | 12 | 6 | 0 | 0 | 0 | 0 |
| ENSDARG000 | 14;14;14;14;1 | 49152341;491 | 49152693;491 | +;+;+;+;+;+;+; | 10686 | NA | 917 | 767 | 1002 | 377 | 1014 | 834 |
| ENSDARG000 | 14;14;14;14;1 | 16483926;164 | 16485364;164 | -;-;-;-;-;-;-;-; | 3199 | mgat4b | 471 | 34 | 22 | 102 | 45 | 247 |
| ENSDARG000 | 14;14;14;14;1 | 32918172;329 | 32918689;329 | +;+;+;+;+;+;+; | 3906 | lnx2b | 84 | 2 | 21 | 12 | 56 | 22 |
| ENSDARG000 | 14;14;14;14;1 | 41345175;413 | 41345253;413 | +;+;+;+;+;+;+; | 3098 | nox1 | 14 | 1 | 2 | 28 | 4 | 6 |
| ENSDARG000 | 14 | 23811808 | 23812725 | + | 918 | kctd16a | 0 | 0 | 0 | 2 | 3 | 0 |
| ENSDARG000 | 14;14;14 | 32913680;329 | 32914404;329 | -;-;- | 1192 | NA | 0 | 0 | 0 | 0 | 0 | 0 |
| ENSDARG000 | 14;14;14;14;1 | 3516567;3516 | 3516780;3516 | +;+;+;+;+;+;+; | 5957 | fnta | 637 | 681 | 699 | 627 | 810 | 670 |
| ENSDARG000 | 14;14;14;14;1 | 21622154;216 | 21623772;216 | -;-;-;-;-;-;-;-; | 7927 | NA | 2854 | 2607 | 5707 | 4019 | 3489 | 2582 |
| ENSDARG000 | 14;14;14;14;1 | 30669662;306 | 30673264;306 | -;-;-;-;-;-;-;-; | 4993 | efemp2a | 594 | 990 | 2167 | 1551 | 3791 | 912 |
| ENSDARG000 | 14;14;14;14;1 | 23874062;239 | 23874383;239 | +;+;+;+;+;+;+; | 6665 | NA | 8 | 0 | 16 | 1 | 10 | 4 |
| ENSDARG000 | 14;14;14;14;1 | 16765791;167 | 16770438;167 | -;-;-;-;-;-;-;-; | 6147 | mrnip | 68 | 83 | 76 | 122 | 116 | 54 |
| ENSDARG000 | 14;14;14;14;1 | 11430796;114 | 11430873;114 | +;+;+;+;+;+;+; | 3256 | NA | 6 | 0 | 8 | 2 | 0 | 0 |
| ENSDARG000 | 14;14;14;14;1 | 31618982;316 | 31619384;316 | +;+;+;+;+;+;+; | 8262 | slc9a6a | 774 | 507 | 1069 | 628 | 1228 | 675 |
| ENSDARG000 | 14;14;14;14;1 | 16765992;167 | 16766290;167 | +;+;+;+;+;+;+; | 2131 | sqstm1 | 836 | 593 | 648 | 1421 | 421 | 682 |
| ENSDARG000 | 14;14;14;14;1 | 31615577;316 | 31616393;316 | -;-;-;-;-;-;-;-; | 1327 | mmgt1 | 747 | 824 | 1012 | 731 | 927 | 819 |
| ENSDARG000 | 14;14;14;14;1 | 46118834;461 | 46119088;461 | +;+;+;+;+;+;+; | 3537 | naa15a | 1756 | 1941 | 3015 | 967 | 1960 | 1585 |
| ENSDARG000 | 14;14 | 31633146;316 | 31633373;316 | -;- | 384 | NA | 0 | 0 | 0 | 0 | 0 | 0 |
| ENSDARG000 | 14;14;14;14;1 | 6628863;6635 | 6635771;6635 | -;-;-;-;-;-;-;-; | 8604 | NA | 4 | 2 | 4 | 0 | 0 | 4 |
| ENSDARG000 | 14;14;14;14;1 | 9033315;9036 | 9037033;9036 | -;-;-;-;-;-;-;-; | 4516 | sybl1 | 756 | 722 | 691 | 572 | 692 | 985 |
| ENSDARG000 | 14;14;14;14;1 | 33859695;338 | 33860619;338 | -;-;-;-;-;-;-;-; | 2393 | NA | 23 | 14 | 113 | 186 | 440 | 10 |
| ENSDARG000 | 14;14;14;14;1 | 23970818;239 | 23970961;239 | +;+;+;+;+;+;+; | 3310 | kif3a | 585 | 740 | 756 | 721 | 690 | 654 |
| ENSDARG000 | 14;14;14;14;1 | 41361858;413 | 41362010;413 | -;-;-;-;-;-;-;-; | 1993 | cstf2 | 1102 | 1034 | 1680 | 824 | 1394 | 952 |
| ENSDARG000 | 14;14;14;14 | 23999568;240 | 23999815;240 | -;-;-;- | 587 | il4 | 0 | 2 | 0 | 6 | 6 | 20 |
| ENSDARG000 | 14;14;14;14;1 | 36279826;362 | 36280070;362 | -;-;-;-;-;-;-;-; | 3930 | egf | 0 | 0 | 0 | 0 | 0 | 0 |
| ENSDARG000 | 14 | 3571256 | 3576820 | + | 5565 | NA | 0 | 0 | 0 | 0 | 0 | 0 |
| ENSDARG000 | 14;14;14;14;1 | 32951842;329 | 32953086;329 | -;-;-;-;-;- | 1741 | chic1 | 346 | 374 | 322 | 269 | 377 | 387 |
| ENSDARG000 | 14;14;14;14;1 | 30713736;307 | 30714044;307 | -;-;-;-;-;-;-;-; | 1678 | fibpa | 796 | 596 | 737 | 617 | 813 | 554 |
| ENSDARG000 | 14;14;14 | 32961718;329 | 32962573;329 | -;-;- | 1638 | cdx4 | 0 | 0 | 0 | 0 | 0 | 0 |
| ENSDARG000 | 14;14;14;14;1 | 11450498;114 | 11451106;114 | -;-;-;-;-;-;-;- | 3128 | si:ch211-153l | 92 | 2 | 0 | 4 | 4 | 6 |
| ENSDARG000 | 14;14;14;14;1 | 25937948;259 | 25939654;259 | -;-;-;-;-;-;-;-; | 3618 | sparc | 6495 | 12554 | 15718 | 9697 | 32598 | 15138 |
| ENSDARG000 | 14;14;14;14;1 | 3717745;3732 | 3719081;3732 | -;-;-;-;-;-;-;-; | 4242 | PIGG | 166 | 211 | 236 | 96 | 266 | 134 |
| ENSDARG000 | 14;14 | 33875058;338 | 33875715;338 | -;- | 721 | NA | 0 | 0 | 0 | 0 | 0 | 0 |
| ENSDARG000 | 14;14;14;14;1 | 33892300;338 | 33892533;338 | -;-;-;-;- | 640 | urp1 | 20 | 27 | 12 | 33 | 37 | 27 |
| ENSDARG000 | 14;14;14;14;1 | 33882973;338 | 33883221;338 | +;+;+;+;+;+ | 1431 | clic2 | 494 | 762 | 494 | 579 | 618 | 698 |
| ENSDARG000 | 14;14 | 6928500;6931 | 6928798;6931 | -;- | 924 | NA | 1 | 0 | 0 | 0 | 0 | 0 |
| ENSDARG000 | 14;14;14;14;1 | 6933400;6933 | 6934438;6934 | -;-;-;-;-;-;-;-; | 2383 | NA | 3319 | 1662 | 4869 | 4009 | 4358 | 3773 |
| ENSDARG000 | 14;14;14;14;1 | 49220026;492 | 49220121;492 | +;+;+;+;+;+;+; | 2761 | rmnd5b | 1143 | 998 | 994 | 556 | 894 | 1120 |
| ENSDARG000 | 14;14 | 46177524;461 | 46178016;461 | +;+ | 8108 | rab33ba | 212 | 110 | 395 | 258 | 314 | 180 |
| ENSDARG000 | 14;14;14;14;1 | 21671107;216 | 21671138;216 | +;+;+;+;+;+;+; | 1002 | ndufs8b | 0 | 0 | 0 | 0 | 0 | 0 |
| ENSDARG000 | 14;14;14;14;1 | 41406321;414 | 41406598;414 | +;+;+;+;+;+;+; | 7709 | NA | 142 | 127 | 218 | 170 | 228 | 118 |
| ENSDARG000 | 14;14;14;14;1 | 21685537;216 | 21685643;216 | +;+;+;+;+;+;+; | 1539 | ran | 8275 | 11824 | 12156 | 9172 | 9473 | 9810 |
| ENSDARG000 | 14;14;14;14;1 | 11457500;114 | 11457581;114 | +;+;+;+;+;+;+; | 2296 | si:ch211-153l | 336 | 10 | 0 | 10 | 6 | 26 |
| ENSDARG000 | 14;14 | 30730749;307 | 30731447;307 | +;+ | 4078 | ccdc85b | 2345 | 2329 | 3419 | 3154 | 3618 | 3104 |
| ENSDARG000 | 14;14;14;14;1 | 3944826;3951 | 3945189;3951 | +;+;+;+;+;+;+; | 1857 | lrp2bp | 0 | 0 | 0 | 0 | 0 | 0 |
| ENSDARG000 | 14;14;14;14;1 | 6946482;6947 | 6946839;6947 | +;+;+;+;+;+;+; | 1331 | rack1 | 27422 | 26551 | 43887 | 44009 | 30567 | 43377 |
| ENSDARG000 | 14;14;14 | 21684475;216 | 21684654;216 | -;-;- | 769 | NA | 0 | 10 | 0 | 3 | 0 | 24 |
| ENSDARG000 | 14;14;14;14;1 | 3960541;3960 | 3960982;3960 | -;-;-;-;-;-;-;-; | 3989 | snx25 | 700 | 458 | 497 | 582 | 559 | 482 |
| ENSDARG000 | 14;14;14;14;1 | 9056071;9059 | 9056137;9059 | +;+;+;+;+;+;+; | 1336 | NA | 0 | 0 | 0 | 0 | 6 | 0 |
| ENSDARG000 | 14;14;14;14;1 | 46190807;461 | 46191148;461 | -;-;-;-;-;-;-;-; | 1168 | zgc:113425 | 6 | 2 | 0 | 2 | 0 | 0 |
| ENSDARG000 | 14;14;14;14;1 | 6954579;6955 | 6954920;6955 | +;+;+;+;+ | 969 | nme5 | 1 | 21 | 5 | 6 | 1 | 27 |
| ENSDARG000 | 14;14;14;14;1 | 11504504;115 | 11505377;115 | -;-;-;-;-;-;-;-; | 2365 | zgc:174917 | 2 | 0 | 0 | 14 | 10 | 2 |
| ENSDARG000 | 14;14;14 | 24033718;240 | 24033775;240 | +;+;+ | 643 | NA | 8 | 12 | 14 | 8 | 49 | 90 |
| ENSDARG000 | 14;14;14;14;1 | 32983769;329 | 32985424;329 | -;-;-;-;-;-;-;-; | 5790 | kdrl | 380 | 70 | 210 | 30 | 135 | 250 |
| ENSDARG000 | 14 | 24042760 | 24043851 | + | 1092 | NA | 0 | 0 | 2 | 0 | 0 | 0 |
| ENSDARG000 | 14;14;14;14;1 | 33898976;339 | 33902114;339 | -;-;-;-;-;-;-;-; | 4997 | NA | 19 | 6 | 0 | 2 | 2 | 6 |
| ENSDARG000 | 14;14;14;14;1 | 30739476;307 | 30742502;307 | -;-;-;-;-;- | 4005 | fosl1a | 7311 | 4417 | 8965 | 10232 | 2089 | 5709 |
| ENSDARG000 | 14;14;14;14;1 | 41469478;414 | 41470533;414 | -;-;-;-;-;-;-;-; | 2013 | tspan7 | 305 | 273 | 493 | 330 | 566 | 282 |
| ENSDARG000 | 14;14;14;14;1 | 24092094;240 | 24092644;240 | -;-;-;-;-;- | 1303 | NA | 12 | 4 | 8 | 8 | 7 | 2 |
| ENSDARG000 | 14;14;14;14;1 | 11509638;115 | 11510589;115 | -;-;-;-;-;- | 1911 | NA | 146 | 198 | 258 | 136 | 255 | 227 |
| ENSDARG000 | 14;14;14;14;1 | 24096588;240 | 24099599;240 | -;-;-;-;-;-;-;-; | 6456 | cpeb4a | 2854 | 2310 | 2267 | 2422 | 3754 | 2291 |
| ENSDARG000 | 14;14;14;14;1 | 31651533;316 | 31651591;316 | +;+;+;+;+;+;+; | 1757 | fhl1a | 14 | 2 | 0 | 13 | 10 | 1 |
| ENSDARG000 | 14;14;14;14;1 | 41466368;414 | 41467500;414 | -;-;-;-;- | 1373 | mid1ip1l | 198 | 72 | 216 | 152 | 248 | 99 |
| ENSDARG000 | 14;14;14;14 | 36328425;363 | 36329778;363 | -;-;-;- | 2533 | lrit3a | 0 | 0 | 0 | 0 | 6 | 0 |
| ENSDARG000 | 14;14;14;14;1 | 16777308;167 | 16778131;167 | -;-;-;-;-;-;-;-; | 3783 | tcirg1b | 2569 | 2023 | 2270 | 1272 | 2726 | 1737 |
| ENSDARG000 | 14;14 | 24080077;240 | 24081014;240 | -;- | 1300 | msx2a | 2 | 0 | 2 | 4 | 1 | 0 |
| ENSDARG000 | 14;14;14;14;1 | 36348549;363 | 36352267;363 | -;-;-;-;-;-;-;- | 5148 | gpm6aa | 88 | 2 | 2 | 0 | 40 | 36 |
| ENSDARG000 | 14;14;14;14;1 | 4047215;4051 | 4048894;4051 | -;-;-;-;-;-;-;-; | 3171 | enpp6 | 16 | 8 | 12 | 18 | 36 | 0 |
| ENSDARG000 | 14;14;14;14;1 | 49251331;492 | 49251483;492 | +;+;+;+;+;+;+; | 2178 | anxa6 | 4 | 4 | 82 | 78 | 76 | 6 |
| ENSDARG000 | 14;14;14;14;1 | 25982962;259 | 25983255;259 | -;-;-;-;-;- | 690 | atox1 | 846 | 795 | 1468 | 827 | 1285 | 1726 |
| ENSDARG000 | 14;14;14;14;1 | 25986895;259 | 25987039;259 | +;+;+;+;+;+;+; | 4313 | SLC36A1 | 1618 | 1704 | 2239 | 2274 | 3507 | 4082 |
| ENSDARG000 | 14;14;14;14;1 | 33939213;339 | 33941580;339 | -;-;-;-;-;- | 3503 | zdhhc24 | 362 | 409 | 335 | 212 | 397 | 339 |
| ENSDARG000 | 14;14 | 33048519;330 | 33048857;330 | -;- | 539 | NA | 0 | 0 | 0 | 0 | 0 | 0 |
| ENSDARG000 | 14;14;14;14;1 | 33049579;330 | 33049709;330 | +;+;+;+;+;+;+; | 3197 | TEX11 | 0 | 6 | 1 | 0 | 2 | 4 |
| ENSDARG000 | 14;14;14;14;1 | 11762991;118 | 11763384;118 | +;+;+;+;+;+;+; | 11212 | FRMPD3 | 0 | 6 | 0 | 60 | 12 | 221 |
| ENSDARG000 | 14;14;14;14 | 49264184;492 | 49264533;492 | -;-;-;- | 1435 | NA | 0 | 0 | 0 | 0 | 4 | 0 |
| ENSDARG000 | 14;14;14;14;1 | 6962271;6962 | 6962403;6962 | +;+;+;+;+;+;+; | 4736 | hnrnpaba | 4507 | 7702 | 9201 | 3522 | 6719 | 4720 |
| ENSDARG000 | 14;14;14;14;1 | 21699129;216 | 21699450;216 | +;+;+;+;+;+;+; | 2679 | stx3b | 48 | 74 | 70 | 55 | 18 | 152 |
| ENSDARG000 | 14;14;14;14;1 | 41490262;414 | 41490872;414 | +;+;+;+;+;+;+; | 6178 | NA | 0 | 0 | 6 | 0 | 2 | 0 |
| ENSDARG000 | 14;14;14;14;1 | 33955807;339 | 33955900;339 | +;+;+;+;+ | 1551 | NA | 0 | 0 | 0 | 0 | 0 | 0 |
| ENSDARG000 | 14;14;14;14;1 | 30753666;307 | 30753751;307 | +;+;+;+;+;+;+; | 1987 | yif1a | 407 | 681 | 702 | 515 | 603 | 474 |
| ENSDARG000 | 14;14 | 26075799;260 | 26076626;260 | -;- | 1101 | mtnr1al | 69 | 150 | 531 | 153 | 1672 | 463 |
| ENSDARG000 | 14;14;14;14;1 | 41528635;415 | 41533123;415 | -;-;-;-;-;-;-;-; | 7598 | NA | 0 | 0 | 0 | 0 | 0 | 0 |
| ENSDARG000 | 14;14;14;14;1 | 30762131;307 | 30762270;307 | +;+;+;+;+ | 1258 | NA | 10 | 2 | 16 | 0 | 20 | 0 |
| ENSDARG000 | 14;14;14;14;1 | 41518257;415 | 41518475;415 | +;+;+;+;+;+;+; | 2531 | chrnb3b | 0 | 0 | 0 | 0 | 0 | 0 |
| ENSDARG000 | 14;14;14;14;1 | 41576849;415 | 41576923;415 | +;+;+;+;+;+;+; | 1730 | nitr13 | 2 | 0 | 0 | 0 | 0 | 0 |
| ENSDARG000 | 14;14;14;14 | 33973146;339 | 33974928;339 | -;-;-;- | 2069 | foxa | 0 | 0 | 0 | 0 | 0 | 0 |
| ENSDARG000 | 14;14;14;14;1 | 41567403;415 | 41568062;415 | -;-;-;-;-;- | 1381 | NA | 0 | 0 | 0 | 0 | 0 | 0 |
| ENSDARG000 | 14;14;14;14;1 | 16813816;168 | 16814073;168 | +;+;+;+;+;+;+; | 6348 | NA | 339 | 104 | 588 | 351 | 343 | 579 |
| ENSDARG000 | 14;14;14 | 41587813;415 | 41588469;415 | -;-;- | 809 | NA | 0 | 0 | 0 | 0 | 4 | 0 |
| ENSDARG000 | 14;14;14;14;1 | 49376011;493 | 49376325;493 | +;+;+;+;+;+;+; | 7783 | tnip1 | 5377 | 2611 | 2451 | 1927 | 2490 | 3252 |
| ENSDARG000 | 14;14;14;14;1 | 41600340;416 | 41602627;416 | -;-;-;-;-;-;-;-; | 3577 | fgfrl1b | 36 | 12 | 38 | 0 | 48 | 4 |

| | | | | | | | | | | | | |
|---|---|---|---|---|---|---|---|---|---|---|---|---|
| ENSDARG000 | 14:14:14:14:1 | 11975344:119 | 11979594:119 | -;-;-;-;-;-;-;-;- | 5656 | znf711 | 560 | 338 | 563 | 313 | 376 | 509 |
| ENSDARG000 | 14:14:14:14:1 | 21754521:217 | 21754704:217 | +;+;+;+;+;+;+; | 7129 | KDM2A | 1622 | 1791 | 1715 | 1504 | 1524 | 1840 |
| ENSDARG000 | 14:14:14:14:1 | 12020936:120 | 12021797:120 | -;-;-;-;-;-;-;-;-; | 2711 | zgc:66447 | 314 | 219 | 297 | 248 | 289 | 278 |
| ENSDARG000 | 14:14:14:14:1 | 36393854:363 | 36394025:363 | -;-;-;-;-;-;-;-;-; | 989 | spata4 | 0 | 0 | 0 | 0 | 2 | 0 |
| ENSDARG000 | 14:14:14:14:1 | 33982007:339 | 33984998:339 | -;-;-;-;-;-;-;-;-; | 6800 | cyfip2 | 406 | 582 | 371 | 268 | 478 | 716 |
| ENSDARG000 | 14:14:14:14:1 | 36398282:363 | 36398597:363 | -;-;-;-;-;-;-;-;-; | 1700 | asb5a | 0 | 0 | 6 | 3 | 0 | 4 |
| ENSDARG000 | 14:14:14:14:1 | 26103330:261 | 26106920:261 | -;-;-;-;-;-;-;-;-; | 16505 | FAT2 | 363 | 18 | 20 | 26 | 60 | 28 |
| ENSDARG000 | 14;14;14 | 26164878:261 | 26165283:261 | +;+;+ | 1610 | NA | 0 | 0 | 0 | 0 | 0 | 0 |
| ENSDARG000 | 14:14:14:14:1 | 21783229:217 | 21783483:217 | +;+;+;+;+;+;+; | 2454 | ANKRD13D | 90 | 96 | 128 | 135 | 123 | 132 |
| ENSDARG000 | 14:14:14:14:1 | 49439887:494 | 49444581:494 | -;-;-;-;- | 5189 | gpx3 | 7690 | 29130 | 37313 | 9757 | 57790 | 36625 |
| ENSDARG000 | 14:14:14:14:1 | 36428441:364 | 36428769:364 | -;-;-;-;-;-;-;-;-; | 1286 | aga | 496 | 461 | 366 | 344 | 499 | 453 |
| ENSDARG000 | 14:14:14:14:1 | 36414856:364 | 36415104:364 | +;+;+;+;+;+;+; | 2301 | neil3 | 88 | 395 | 318 | 173 | 186 | 137 |
| ENSDARG000 | 14:14:14:14:1 | 6972868:6972 | 6973458:6973 | -;-;-;-;-;- | 1325 | eif4ebp3l | 788 | 794 | 1536 | 1222 | 592 | 934 |
| ENSDARG000 | 14;14;14 | 26363537:263 | 26365616:263 | -;-;- | 2987 | emx3 | 64 | 0 | 2 | 0 | 2 | 6 |
| ENSDARG000 | 14:14:14:14:1 | 49462855:494 | 49463005:494 | +;+;+;+;+;+;+; | 3736 | dctn4 | 5969 | 3263 | 3618 | 2158 | 2860 | 4569 |
| ENSDARG000 | 14:14:14:14:1 | 17029813:170 | 17029937:170 | -;-;-;-;-;- | 949 | dicp2.1 | 17 | 17 | 17 | 18 | 12 | 36 |
| ENSDARG000 | 14:14:14:14:1 | 17040484:170 | 17040821:170 | -;-;-;-;-;-;-;-;-; | 1277 | dicp2.2 | 22 | 66 | 28 | 9 | 18 | 22 |
| ENSDARG000 | 14:14:14:14:1 | 17064126:170 | 17066791:170 | -;-;-;-;-;-;-;-;-; | 4422 | phox2bb | 0 | 0 | 0 | 0 | 0 | 0 |
| ENSDARG000 | 14:14:14:14:1 | 6991142:6991 | 6991375:6991 | +;+;+;+;+;+;+; | 2088 | hnrnph1 | 358 | 295 | 389 | 390 | 322 | 290 |
| ENSDARG000 | 14:14:14:14:1 | 14806692:148 | 14807059:148 | +;+;+;+;+;+;+; | 5412 | NA | 8 | 0 | 2 | 0 | 4 | 0 |
| ENSDARG000 | 14:14:14:14:1 | 33070284:330 | 33071666:330 | -;-;-;-;-;-;-;-;-; | 4604 | DLG3 | 159 | 215 | 168 | 80 | 172 | 212 |
| ENSDARG000 | 14;14 | 17087349:170 | 17089243:170 | -;- | 2038 | NA | 0 | 0 | 0 | 0 | 0 | 0 |
| ENSDARG000 | 14:14:14:14:1 | 49811926:498 | 49812087:498 | -;-;-;-;-;-;-;-;-; | 1959 | NA | 182 | 102 | 169 | 78 | 164 | 108 |
| ENSDARG000 | 14;14 | 34064249:340 | 34064409:340 | +;+ | 1073 | med7 | 255 | 291 | 279 | 152 | 328 | 244 |
| ENSDARG000 | 14:14:14:14:1 | 34068075:340 | 34068896:340 | +;+;+;+;+;+;+; | 5581 | NA | 956 | 658 | 492 | 696 | 1100 | 1254 |
| ENSDARG000 | 14:14:14:14:1 | 34045284:340 | 34046095:340 | -;-;-;-;-;-;-;-;-; | 2752 | itk | 4 | 10 | 12 | 24 | 4 | 32 |
| ENSDARG000 | 14:14:14:14:1 | 17108003:171 | 17108369:171 | -;-;-;-;-;-;-;-;-; | 1359 | smtnl1 | 0 | 0 | 0 | 10 | 0 | 18 |
| ENSDARG000 | 14:14:14:14:1 | 36454087:364 | 36454257:364 | +;+;+;+;+;+;+; | 15514 | NA | 4 | 4 | 23 | 14 | 28 | 0 |
| ENSDARG000 | 14:14:14:14:1 | 49865082:498 | 49865907:498 | -;-;-;-;-;-;- | 1525 | ube2d2l | 738 | 818 | 747 | 738 | 728 | 694 |
| ENSDARG000 | 14:14:14:14:1 | 36752033:367 | 36752563:367 | -;-;-;-;-;-;-;- | 1594 | CTSO | 284 | 189 | 345 | 311 | 424 | 266 |
| ENSDARG000 | 14:14:14:14:1 | 36738063:367 | 36738138:367 | +;+;+;+;+;+;+; | 2337 | tdo2a | 9 | 9 | 4 | 0 | 6 | 24 |
| ENSDARG000 | 14:14:14:14:1 | 52323322:523 | 52323620:523 | +;+;+;+;+;+;+; | 4038 | CPXM1 | 75 | 12 | 260 | 35 | 175 | 196 |
| ENSDARG000 | 14:14:14:14:1 | 17125428:171 | 17125694:171 | +;+;+;+;+;+;+; | 3077 | slc43a3b | 302 | 364 | 928 | 256 | 1262 | 360 |
| ENSDARG000 | 14:14:14:14:1 | 52356897:523 | 52357050:523 | +;+;+;+;+;+;+; | 1940 | COX18 | 140 | 145 | 208 | 106 | 169 | 99 |
| ENSDARG000 | 14;14;14 | 17197132:171 | 17197256:171 | +;+;+ | 2535 | rtn4rl2b | 0 | 0 | 22 | 8 | 6 | 0 |
| ENSDARG000 | 14:14:14:14:1 | 14836468:148 | 14836698:148 | +;+;+;+;+;+;+; | 1738 | si:dkev-102m | 142 | 187 | 4 | 63 | 153 | 232 |
| ENSDARG000 | 14:14:14:14:1 | 52377557:523 | 52380629:523 | -;-;-;-;-;-;-;-;-; | 6981 | polr2b | 1869 | 1810 | 2224 | 651 | 2142 | 1077 |
| ENSDARG000 | 14:14:14:14:1 | 52408619:524 | 52409818:524 | +;+;+;+;+;+;+; | 3467 | noa1 | 115 | 108 | 224 | 82 | 280 | 180 |
| ENSDARG000 | 14:14:14:14:1 | 52369262:523 | 52369729:523 | +;+;+;+;+;+;+; | 908 | igfbp7 | 174 | 23 | 196 | 39 | 398 | 194 |
| ENSDARG000 | 14:14:14:14:1 | 34125855:341 | 34131114:341 | -;-;-;-;-;-;-;-;-; | 8014 | gria1a | 38 | 20 | 222 | 4 | 134 | 6 |
| ENSDARG000 | 14;14 | 34154456:341 | 34154879:341 | +;+ | 1217 | NA | 0 | 0 | 0 | 0 | 0 | 0 |
| ENSDARG000 | 14;14;14;14 | 33195619:332 | 33195883:332 | -;-;-;- | 1070 | NA | 0 | 0 | 1 | 0 | 0 | 0 |
| ENSDARG000 | 14:14:14:14:1 | 36765612:367 | 36766756:367 | -;-;-;-;-;-;-;-;-; | 3054 | rnf130 | 3100 | 3061 | 2406 | 1500 | 4353 | 2614 |
| ENSDARG000 | 14;14 | 36869273:368 | 36869524:368 | +;+ | 4682 | F2RL3 | 38 | 116 | 28 | 32 | 36 | 74 |
| ENSDARG000 | 14;14;14 | 34486629:344 | 34486818:344 | +;+;+ | 1321 | NA | 18 | 65 | 16 | 32 | 10 | 52 |
| ENSDARG000 | 14:14:14:14:1 | 17230283:172 | 17230727:172 | -;-;-;-;-;-;-;-;-; | 4216 | JAKMIP1 | 4 | 22 | 30 | 18 | 16 | 32 |
| ENSDARG000 | 14:14:14:14:1 | 34490445:344 | 34490544:344 | +;+;+;+;+;+;+; | 1710 | wnt8a | 0 | 0 | 0 | 0 | 0 | 0 |
| ENSDARG000 | 14:14:14:14:1 | 34490445:344 | 34490544:344 | +;+;+;+;+;+;+; | 3285 | NA | 0 | 0 | 0 | 0 | 0 | 0 |
| ENSDARG000 | 14:14:14:14:1 | 34495216:344 | 34495330:344 | +;+;+;+;+;+ | 1208 | wnt8-2 | 0 | 0 | 0 | 0 | 0 | 0 |
| ENSDARG000 | 14:14:14:14:1 | 7016934:7017 | 7017362:7017 | -;-;-;-;-;-;-;-;-; | 2448 | RUFY1 | 264 | 308 | 200 | 179 | 213 | 201 |
| ENSDARG000 | 14:14:14:14:1 | 7048930:7049 | 7049148:7049 | +;+;+;+;+;+;+; | 1891 | hbegfa | 254 | 75 | 92 | 150 | 42 | 106 |
| ENSDARG000 | 14;14 | 7120620:7128 | 7128239:7128 | -;- | 8108 | NA | 80 | 20 | 26 | 6 | 241 | 12 |
| ENSDARG000 | 14:14:14:14:1 | 7129816:7131 | 7131565:7131 | -;-;-;-;-;-;-;-;-; | 3112 | trpt1 | 362 | 313 | 347 | 328 | 759 | 183 |
| ENSDARG000 | 14:14:14:14:1 | 7140997:7140 | 7141359:7141 | +;+;+;+;+;+;+; | 3357 | ctsf | 6426 | 2645 | 4362 | 3933 | 7128 | 5717 |
| ENSDARG000 | 14:14:14:14:1 | 7168447:7183 | 7176872:7187 | -;-;-;-;-;-;-;-;-; | 12712 | STOX2 | 795 | 613 | 912 | 753 | 900 | 784 |
| ENSDARG000 | 14 | 5117072 | 5119497 | + | 2426 | nanos1 | 8314 | 2761 | 10188 | 5071 | 7068 | 9308 |
| ENSDARG000 | 14;14;14 | 567216:57021 | 567820:57031 | -;-;- | 926 | fgf2 | 205 | 34 | 73 | 134 | 92 | 8 |
| ENSDARG000 | 14:14:14:14:1 | 51023954:510 | 51024125:510 | -;-;-;-;-;- | 1509 | CLTB | 327 | 315 | 282 | 98 | 433 | 243 |
| ENSDARG000 | 14:14:14:14:1 | 49602:52148: | 49986:52294: | +;+;+;+;+;+;+; | 1764 | OTOP1 | 2 | 0 | 2 | 0 | 0 | 0 |
| ENSDARG000 | 14:14:14:14:1 | 240487:24191 | 241802:24209 | -;-;-;-;-;-;-;-;- | 2105 | aurkb | 239 | 696 | 822 | 482 | 572 | 304 |
| ENSDARG000 | 14:14:14:14:1 | 146857:14712 | 147034:14729 | +;+;+;+;+;+;+; | 1145 | NA | 0 | 8 | 2 | 2 | 25 | 26 |
| ENSDARG000 | 14:14:14:14:1 | 109016:10933 | 109102:10949 | +;+;+;+;+;+;+;+ | 1116 | gpc2 | 6 | 12 | 39 | 2 | 24 | 6 |
| ENSDARG000 | 14:14:14:14:1 | 9497032:9510 | 9497045:9510 | +;+;+;+;+;+;+; | 204 | NA | 0 | 0 | 2 | 2 | 4 | 1 |
| ENSDARG000 | 14 | 41066998 | 41067114 | - | 117 | 5S_rRNA | 0 | 0 | 0 | 0 | 0 | 0 |
| ENSDARG000 | 14:14:14:14:1 | 51164056:511 | 51164691:511 | -;-;-;-;-;-;-;-;-; | 3452 | unc5a | 2 | 0 | 6 | 0 | 26 | 0 |
| ENSDARG000 | 14:14:14:14:1 | 51056605:510 | 51056700:510 | +;+;+;+;+;+;+; | 1260 | pdlim7 | 26 | 0 | 10 | 3 | 16 | 15 |
| ENSDARG000 | 14 | 11634865 | 11635080 | - | 216 | U3 | 0 | 0 | 0 | 0 | 0 | 0 |
| ENSDARG000 | 14:14:14:14:1 | 1719367:1721 | 1720119:1721 | +;+;+;+;+;+;+; | 2526 | TRPC7 | 0 | 0 | 0 | 0 | 2 | 0 |
| ENSDARG000 | 14:14:14:14:1 | 46313135:463 | 46313439:463 | +;+;+;+;+;+;+; | 1883 | cryba1l1 | 5 | 8 | 2 | 8 | 12 | 3 |
| ENSDARG000 | 14:14:14:14:1 | 52573476:525 | 52576456:525 | -;-;-;-;-;-;-;-;- | 4269 | NA | 114 | 49 | 320 | 208 | 179 | 128 |
| ENSDARG000 | 14:14:14:14:1 | 26909:27544: | 27207:27705: | -;-;-;-;-;-;-;- | 1554 | zbtb49 | 171 | 116 | 162 | 126 | 122 | 148 |
| ENSDARG000 | 14:14:14:14:1 | 50994138:509 | 50994821:509 | -;-;-;-;-;-;-;-;-; | 1859 | faf2 | 1100 | 1116 | 1451 | 773 | 1267 | 941 |
| ENSDARG000 | 14:14:14:14:1 | 32964:33159: | 33077:33242: | +;+;+;+;+;+ | 1000 | lyar | 266 | 413 | 499 | 286 | 368 | 297 |
| ENSDARG000 | 14:14:14:14:1 | 28281744:283 | 28282651:283 | +;+;+;+;+;+;+; | 2306 | MID2 | 280 | 228 | 136 | 116 | 270 | 201 |
| ENSDARG000 | 14:14:14:14:1 | 1351214:1355 | 1351234:1355 | +;+;+;+;+;+ | 5997 | NA | 42 | 36 | 62 | 50 | 50 | 46 |
| ENSDARG000 | 14 | 30740988 | 30741116 | + | 129 | 5S_rRNA | 0 | 0 | 0 | 0 | 0 | 0 |
| ENSDARG000 | 14 | 28128158 | 28128278 | - | 121 | 5S_rRNA | 0 | 0 | 0 | 0 | 0 | 0 |
| ENSDARG000 | 14 | 11637354 | 11637570 | - | 217 | NA | 0 | 0 | 0 | 0 | 0 | 0 |
| ENSDARG000 | 14 | 100660 | 100746 | + | 87 | dre-mir-19d | 0 | 0 | 0 | 0 | 0 | 0 |
| ENSDARG000 | 14:14:14:14:1 | 1402558:1405 | 1402729:1405 | -;-;-;-;-;-;-;-;-; | 2821 | zgc:152774 | 27 | 18 | 33 | 34 | 37 | 59 |
| ENSDARG000 | 14 | 14250880 | 14250995 | - | 116 | 5S_rRNA | 0 | 0 | 0 | 0 | 0 | 0 |
| ENSDARG000 | 14 | 11614550 | 11614765 | - | 216 | U3 | 0 | 0 | 0 | 0 | 0 | 0 |
| ENSDARG000 | 14 | 11629832 | 11630048 | - | 217 | NA | 0 | 0 | 0 | 0 | 0 | 0 |
| ENSDARG000 | 14:14:14:14:1 | 1365097:1365 | 1365214:1365 | +;+;+;+;+;+;+; | 3661 | TBC1D9B | 271 | 255 | 416 | 244 | 269 | 274 |
| ENSDARG000 | 14 | 31326601 | 31326717 | - | 117 | 5S_rRNA | 0 | 0 | 0 | 0 | 0 | 0 |
| ENSDARG000 | 14:14:14:14:1 | 1348065:1350 | 1348154:1350 | -;-;-;-;- | 516 | NA | 121 | 57 | 154 | 201 | 77 | 138 |
| ENSDARG000 | 14 | 10816297 | 10816413 | + | 117 | 5S_rRNA | 0 | 0 | 0 | 0 | 0 | 0 |
| ENSDARG000 | 14;14 | 45200278:452 | 45203793:452 | -;- | 4159 | shisa3 | 2 | 0 | 0 | 0 | 2 | 10 |
| ENSDARG000 | 14 | 11634864 | 11635080 | - | 217 | NA | 0 | 0 | 0 | 0 | 0 | 0 |
| ENSDARG000 | 14:14:14:14:1 | 4478416:4486 | 4479148:4487 | -;-;-;-;-;-;-;-;- | 2138 | NA | 0 | 0 | 0 | 0 | 0 | 0 |
| ENSDARG000 | 14;14;14 | 1644798:1645 | 1644914:1645 | -;-;- | 252 | NA | 2 | 4 | 5 | 1 | 6 | 28 |
| ENSDARG000 | 14:14:14:14:1 | 683947:68639 | 684267:68649 | -;-;-;-;-;- | 817 | ube2ka | 262 | 239 | 309 | 228 | 200 | 217 |
| ENSDARG000 | 14;14 | 51015205:510 | 51015405:510 | +;+ | 741 | higd2a | 235 | 185 | 308 | 208 | 239 | 218 |
| ENSDARG000 | 14 | 42211828 | 42211961 | - | 134 | 5S_rRNA | 0 | 0 | 0 | 0 | 0 | 0 |
| ENSDARG000 | 14 | 14837073 | 14837270 | + | 198 | SNORA74 | 0 | 0 | 0 | 0 | 0 | 0 |
| ENSDARG000 | 14 | 2407328 | 2407431 | - | 104 | Vault | 0 | 0 | 0 | 0 | 0 | 0 |
| ENSDARG000 | 14:14:14:14:1 | 353099:35309 | 358443:35844 | -;-;-;-;-;-;-;-;-; | 18523 | FAT4 | 181 | 34 | 24 | 12 | 68 | 45 |
| ENSDARG000 | 14 | 31521930 | 31522007 | - | 78 | dre-mir-457b | 6 | 2 | 6 | 3 | 0 | 0 |
| ENSDARG000 | 14:14:14:14:1 | 50839282:508 | 50839437:508 | -;-;-;-;-;-;-;-;-; | 3929 | NA | 0 | 0 | 2 | 0 | 0 | 0 |
| ENSDARG000 | 14 | 47579542 | 47579667 | + | 126 | 5S_rRNA | 0 | 0 | 0 | 0 | 0 | 0 |
| ENSDARG000 | 14 | 3757655 | 3757772 | + | 118 | 5S_rRNA | 0 | 0 | 0 | 0 | 0 | 0 |
| ENSDARG000 | 14 | 51753080 | 51753197 | - | 118 | 5S_rRNA | 0 | 0 | 0 | 0 | 0 | 0 |
| ENSDARG000 | 14;14 | 212867:21474 | 214520:21505 | -;- | 1964 | nkx3-2 | 10 | 12 | 2 | 14 | 2 | 4 |
| ENSDARG000 | 14 | 45468955 | 45469031 | - | 77 | dre-mir-194b | 0 | 0 | 0 | 0 | 0 | 0 |
| ENSDARG000 | 14:14:14:14:1 | 9246361:9248 | 9246374:9248 | +;+;+;+;+;+;+; | 798 | NA | 0 | 4 | 0 | 0 | 0 | 8 |
| ENSDARG000 | 14:14:14:14:1 | 26247319:262 | 26247833:262 | +;+;+;+;+;+;+; | 3929 | NA | 656 | 614 | 163 | 407 | 518 | 223 |
| ENSDARG000 | 14;14;14;14 | 2243:2540:27 | 2392:2584:29 | +;+;+;+ | 438 | CYTL1 | 12 | 13 | 5 | 8 | 22 | 30 |
| ENSDARG000 | 14 | 18799434 | 18799558 | + | 125 | 5S_rRNA | 0 | 0 | 0 | 0 | 0 | 0 |
| ENSDARG000 | 14:14:14:14:1 | 555674:55650 | 556785:55678 | +;+;+;+;+;+;+; | 1999 | nudt6 | 407 | 277 | 187 | 221 | 331 | 78 |
| ENSDARG000 | 14 | 37625722 | 37625837 | + | 116 | 5S_rRNA | 0 | 0 | 0 | 0 | 0 | 0 |
| ENSDARG000 | 14 | 11640626 | 11640841 | - | 216 | U3 | 0 | 0 | 0 | 0 | 0 | 0 |
| ENSDARG000 | 14:14:14:14:1 | 1887677:1889 | 1888952:1890 | -;-;-;-;-;-;-;-;-; | 30619 | pcdh2g13 | 9623 | 6259 | 10738 | 8406 | 12399 | 7789 |
| ENSDARG000 | 14 | 33644656 | 33644771 | + | 116 | 5S_rRNA | 0 | 0 | 0 | 0 | 0 | 0 |
| ENSDARG000 | 14 | 40542910 | 40543026 | - | 117 | 5S_rRNA | 0 | 0 | 0 | 0 | 0 | 0 |
| ENSDARG000 | 14:14:14:14:1 | 1303930:1308 | 1304390:1308 | -;-;-;-;-;- | 1220 | il21 | 2 | 16 | 4 | 0 | 6 | 6 |
| ENSDARG000 | 14 | 31467376 | 31467460 | - | 85 | dre-mir-18c | 0 | 0 | 0 | 0 | 0 | 0 |
| ENSDARG000 | 14 | 6186714 | 6186833 | - | 120 | 5S_rRNA | 2 | 0 | 2 | 7 | 0 | 0 |
| ENSDARG000 | 14:14:14:14:1 | 220639:22116 | 220738:22123 | -;-;-;-;-;-;-;-;-; | 5234 | NA | 724 | 620 | 780 | 422 | 767 | 655 |
| ENSDARG000 | 14 | 36938443 | 36938576 | + | 134 | 5S_rRNA | 0 | 0 | 0 | 0 | 0 | 0 |
| ENSDARG000 | 14;14;14;14 | 15484544:154 | 15484942:154 | +;+;+;+ | 779 | NA | 0 | 0 | 0 | 0 | 0 | 0 |
| ENSDARG000 | 14 | 1563985 | 1565317 | - | 1333 | NA | 0 | 0 | 0 | 0 | 0 | 0 |
| ENSDARG000 | 14 | 24357400 | 24357517 | + | 118 | 5S_rRNA | 0 | 0 | 2 | 0 | 5 | 0 |
| ENSDARG000 | 14:14:14:14:1 | 69716:71924: | 69955:72051: | -;-;-;-;-;-;-;-;- | 1257 | NA | 2 | 20 | 70 | 2 | 39 | 39 |
| ENSDARG000 | 14:14:14:14:1 | 45028062:450 | 45029131:450 | +;+;+;+;+;+;+; | 3100 | ATP8A1 | 218 | 146 | 611 | 519 | 635 | 272 |
| ENSDARG000 | 14:14:14:14:1 | 1595725:1605 | 1597432:1605 | -;-;-;-;-;-;- | 2566 | NA | 8 | 8 | 12 | 2 | 30 | 28 |
| ENSDARG000 | 14 | 467104 | 470505 | - | 3402 | ANKRD50 | 414 | 355 | 386 | 382 | 428 | 298 |
| ENSDARG000 | 14;14 | 2046786:2047 | 2047064:2050 | -;- | 2370 | NA | 1 | 2 | 2 | 3 | 0 | 6 |
| ENSDARG000 | 14 | 1687357 | 1688712 | + | 1356 | hrh2a | 0 | 0 | 0 | 0 | 0 | 0 |
| ENSDARG000 | 14:14:14:14:1 | 1070028:1070 | 1070248:1070 | -;-;-;-;-;-;-;-;-; | 1787 | NA | 91 | 13 | 89 | 7 | 73 | 58 |
| ENSDARG000 | 14;14;14;14 | 51098036:511 | 51098085:511 | +;+;+;+ | 480 | NA | 0 | 0 | 0 | 0 | 4 | 0 |
| ENSDARG000 | 14:14:14:14:1 | 14992909:149 | 14993030:149 | +;+;+;+;+;+ | 1874 | NA | 6 | 0 | 6 | 2 | 7 | 0 |
| ENSDARG000 | 14 | 48576584 | 48576700 | - | 117 | 5S_rRNA | 0 | 0 | 0 | 0 | 0 | 0 |
| ENSDARG000 | 14:14:14:14:1 | 49944368:499 | 49948766:499 | -;-;-;-;-;-;-;-;-; | 7466 | FAM193B | 307 | 238 | 410 | 141 | 237 | 237 |
| ENSDARG000 | 14:14:14:14:1 | 247456:24897 | 248393:24907 | -;-;-;-;-;- | 1419 | tmem107 | 48 | 25 | 56 | 107 | 47 | 16 |
| ENSDARG000 | 14 | 2478994 | 2479918 | + | 925 | NA | 0 | 0 | 0 | 0 | 0 | 0 |
| ENSDARG000 | 14 | 11609562 | 11609778 | - | 217 | NA | 0 | 0 | 0 | 0 | 0 | 0 |
| ENSDARG000 | 14:14:14:14:1 | 46342882:463 | 46343106:463 | +;+;+;+;+;+;+; | 4129 | tmem33 | 1307 | 1475 | 1705 | 768 | 1940 | 1294 |
| ENSDARG000 | 14:14:14:14:1 | 4610439:4627 | 4611610:4627 | -;-;-;-;-;-;-;-;- | 2659 | gabra4 | 17 | 4 | 16 | 98 | 10 | 22 |
| ENSDARG000 | 14 | 31540443 | 31540598 | + | 156 | NA | 2 | 2 | 4 | 2 | 3 | 1 |
| ENSDARG000 | 14 | 33934997 | 33935113 | + | 117 | 5S_rRNA | 0 | 0 | 0 | 0 | 0 | 0 |
| ENSDARG000 | 14 | 49754834 | 49754950 | - | 117 | 5S_rRNA | 0 | 0 | 0 | 0 | 0 | 0 |
| ENSDARG000 | 14 | 51273185 | 51273300 | - | 116 | 5S_rRNA | 0 | 0 | 0 | 0 | 0 | 0 |
| ENSDARG000 | 14;14;14;14 | 591216:59153 | 591386:59162 | +;+;+;+ | 1163 | NA | 547 | 221 | 327 | 490 | 348 | 294 |
| ENSDARG000 | 14 | 40827926 | 40828042 | - | 117 | 5S_rRNA | 0 | 0 | 0 | 0 | 2 | 0 |
| ENSDARG000 | 14:14:14:14:1 | 50918769:509 | 50918804:509 | +;+;+;+;+;+;+; | 5138 | rnf44 | 253 | 150 | 314 | 72 | 238 | 180 |
| ENSDARG000 | 14 | 31860750 | 31860827 | - | 78 | NA | 0 | 2 | 0 | 4 | 8 | 1 |
| ENSDARG000 | 14 | 24562801 | 24562883 | - | 83 | dre-mir-218a- | 0 | 0 | 0 | 0 | 0 | 0 |
| ENSDARG000 | 14:14:14:14:1 | 597532:60314 | 597885:60419 | +;+;+;+;+;+;+; | 3775 | NA | 88 | 52 | 41 | 71 | 79 | 47 |
| ENSDARG000 | 14:14:14:14:1 | 3186485:3190 | 3190167:3190 | -;-;-;-;-;-;-;-;-; | 7594 | pdgfrb | 2049 | 449 | 1993 | 1044 | 931 | 1497 |
| ENSDARG000 | 14:14:14:14:1 | 2549313:2549 | 2550686:2550 | -;-;-;-;-;-;-;-;-; | 3194 | fbxw11a | 378 | 407 | 380 | 348 | 494 | 367 |
| ENSDARG000 | 14 | 42224893 | 42231293 | - | 6401 | NA | 0 | 0 | 0 | 0 | 0 | 0 |
| ENSDARG000 | 14:14:14:14:1 | 26224541:262 | 26224661:262 | +;+;+;+;+;+;+; | 1128 | gm2a | 3558 | 3372 | 4097 | 4934 | 7049 | 5382 |
| ENSDARG000 | 14:14:14:14:1 | 49683767:497 | 49684211:497 | +;+;+;+;+;+;+; | 5618 | tspan17 | 342 | 132 | 167 | 58 | 261 | 260 |
| ENSDARG000 | 14:14:14:14:1 | 46354648:463 | 46358255:463 | -;-;-;-;-;-;-;-;-; | 5478 | ccna2 | 182 | 1270 | 1171 | 304 | 688 | 430 |
| ENSDARG000 | 14 | 39154371 | 39154489 | + | 119 | 5S_rRNA | 0 | 0 | 0 | 0 | 0 | 0 |
| ENSDARG000 | 14;14;14 | 46722493:467 | 46722530:467 | +;+;+ | 414 | NA | 0 | 0 | 0 | 0 | 0 | 0 |
| ENSDARG000 | 14 | 38742313 | 38742411 | + | 99 | dre-mir-143 | 0 | 0 | 0 | 0 | 0 | 0 |
| ENSDARG000 | 14:14:14:14:1 | 164556:16668 | 164571:16680 | +;+;+;+;+;+;+; | 1821 | WDR1 | 4905 | 5754 | 4993 | 4354 | 4725 | 3249 |
| ENSDARG000 | 14;14;14;14 | 10909:10909: | 11719:11693: | -;-;-;- | 1162 | msx1a | 0 | 0 | 2 | 0 | 0 | 0 |
| ENSDARG000 | 14:14:14:14:1 | 2589465:2592 | 2589939:2592 | -;-;-;-;-;-;-;-;- | 1664 | etf1a | 0 | 0 | 0 | 0 | 0 | 0 |
| ENSDARG000 | 14 | 100767 | 100848 | + | 82 | dre-mir-25 | 0 | 0 | 0 | 0 | 0 | 0 |
| ENSDARG000 | 14;14;14 | 484389:48936 | 484503:49002 | -;-;- | 863 | ankrd50 | 80 | 54 | 91 | 48 | 78 | 40 |

| | | | | | | | | | | | | |
|---|---|---|---|---|---|---|---|---|---|---|---|---|
| ENSDARG000 | 14:14:14:14:1 | 49604362:496 | 49604435:496 | +;+;+;+;+ | 3617 | NA | 549 | 12 | 47 | 0 | 57 | 28 |
| ENSDARG000 | 14 | 11627338 | 11627553 | - | 216 | U3 | 0 | 0 | 0 | 0 | 0 | 0 |
| ENSDARG000 | 14 | 2407736 | 2407840 | - | 105 | Vault | 0 | 0 | 0 | 0 | 0 | 0 |
| ENSDARG000 | 14;14;14 | 9590540:9652 | 9590794:9652 | -;-;- | 378 | NA | 0 | 0 | 0 | 0 | 0 | 0 |
| ENSDARG000 | 14 | 31533083 | 31533220 | + | 138 | NA | 3 | 2 | 0 | 2 | 0 | 0 |
| ENSDARG000 | 14:14:14:14:1 | 24075254:240 | 24075268:240 | +;+;+;+;+;+;+; | 300 | NA | 20 | 25 | 31 | 8 | 26 | 17 |
| ENSDARG000 | 14 | 33333448 | 33333581 | - | 134 | NA | 0 | 0 | 0 | 0 | 0 | 0 |
| ENSDARG000 | 14:14:14:14:1 | 1170968:1170 | 1170992:1171 | +;+;+;+;+ | 1214 | hopx | 72 | 86 | 100 | 30 | 314 | 108 |
| ENSDARG000 | 14 | 2406978 | 2407072 | - | 95 | dre-mir-733 | 0 | 0 | 0 | 0 | 0 | 0 |
| ENSDARG000 | 14:14:14:14:1 | 39712697:397 | 39712772:397 | -;-;-;-;-;-;-;-; | 1857 | NA | 10 | 0 | 0 | 34 | 2 | 40 |
| ENSDARG000 | 14:14:14:14:1 | 46157379:461 | 46157951:461 | -;-;-;-;-;-;-;-; | 2097 | mlsl | 18 | 16 | 20 | 12 | 12 | 4 |
| ENSDARG000 | 14 | 52049158 | 52049274 | - | 117 | 5S_rRNA | 0 | 0 | 0 | 0 | 0 | 0 |
| ENSDARG000 | 14;14;14 | 50807558:508 | 50808269:508 | +;+;+ | 2517 | si:ch211-59p2 | 0 | 0 | 0 | 0 | 0 | 0 |
| ENSDARG000 | 14 | 25159129 | 25159206 | - | 78 | dre-mir-458 | 0 | 0 | 0 | 0 | 0 | 0 |
| ENSDARG000 | 14:14:14:14:1 | 50770537:507 | 50770573:507 | +;+;+;+;+;+ | 506 | sncb | 0 | 0 | 0 | 0 | 1 | 0 |
| ENSDARG000 | 14:14:14:14:1 | 540157:54048 | 540234:54053 | -;-;-;-;-;-;-;-; | 2133 | NA | 166 | 100 | 258 | 89 | 130 | 78 |
| ENSDARG000 | 14:14:14:14:1 | 48305439:483 | 48308674:483 | -;-;-;-;-;- | 4302 | NA | 8 | 2 | 0 | 0 | 2 | 0 |
| ENSDARG000 | 14 | 11612057 | 11612272 | - | 216 | U3 | 0 | 0 | 0 | 0 | 0 | 0 |
| ENSDARG000 | 14;14;14;14 | 1274301:1277 | 1275034:1277 | -;-;-;- | 1632 | NA | 2 | 16 | 0 | 6 | 24 | 32 |
| ENSDARG000 | 14:14:14:14:1 | 3363368:3367 | 3367851:3367 | -;-;-;-;-;-;-;-;- | 5840 | NA | 0 | 0 | 0 | 2 | 0 | 2 |
| ENSDARG000 | 14;14;14;14 | 1192314:1194 | 1192713:1194 | -;-;-;- | 1205 | arl9 | 450 | 588 | 598 | 422 | 661 | 393 |
| ENSDARG000 | 14:14:14:14:1 | 1658827:1659 | 1659033:1659 | -;-;-;-;- | 495 | NA | 11 | 7 | 6 | 0 | 15 | 23 |
| ENSDARG000 | 14;14;14 | 3287740:3305 | 3288753:3306 | +;+;+ | 4042 | cdx1a | 20 | 0 | 0 | 0 | 7 | 2 |
| ENSDARG000 | 14 | 7361749 | 7361865 | - | 117 | 5S_rRNA | 0 | 0 | 0 | 0 | 0 | 0 |
| ENSDARG000 | 14 | 45631180 | 45631235 | - | 56 | NA | 0 | 0 | 4 | 2 | 2 | 0 |
| ENSDARG000 | 14 | 48349487 | 48349602 | - | 116 | 5S_rRNA | 0 | 0 | 0 | 0 | 0 | 0 |
| ENSDARG000 | 14 | 6107039 | 6107155 | + | 117 | 5S_rRNA | 0 | 0 | 0 | 0 | 0 | 0 |
| ENSDARG000 | 14 | 48409215 | 48409319 | + | 105 | NA | 0 | 0 | 0 | 0 | 0 | 0 |
| ENSDARG000 | 14 | 11629833 | 11630048 | - | 216 | U3 | 0 | 0 | 0 | 0 | 0 | 0 |
| ENSDARG000 | 14 | 11637355 | 11637570 | - | 216 | U3 | 0 | 0 | 0 | 0 | 0 | 0 |
| ENSDARG000 | 14 | 24958280 | 24958396 | - | 117 | 5S_rRNA | 0 | 0 | 0 | 0 | 0 | 0 |
| ENSDARG000 | 14 | 48661503 | 48661619 | + | 117 | 5S_rRNA | 0 | 0 | 0 | 0 | 0 | 0 |
| ENSDARG000 | 14;14;14 | 51829317:518 | 51829511:518 | -;-;- | 402 | CPLX1 | 153 | 0 | 4 | 0 | 0 | 27 |
| ENSDARG000 | 14 | 2502276 | 2502390 | + | 115 | NA | 0 | 0 | 0 | 0 | 0 | 0 |
| ENSDARG000 | 14:14:14:14:1 | 1524220:1528 | 1526058:1528 | -;-;-;-;-;-;-;-; | 3078 | NA | 6 | 27 | 23 | 11 | 91 | 166 |
| ENSDARG000 | 14 | 44276488 | 44276605 | - | 118 | 5S_rRNA | 0 | 0 | 0 | 0 | 0 | 0 |
| ENSDARG000 | 14:14:14:14:1 | 39156:39572: | 39210:39713: | +;+;+;+;+;+;+; | 459 | TMEM128 | 8 | 7 | 4 | 6 | 10 | 0 |
| ENSDARG000 | 14 | 31859430 | 31859506 | - | 77 | NA | 0 | 0 | 0 | 0 | 3 | 0 |
| ENSDARG000 | 14 | 4304052 | 4304170 | + | 119 | 5S_rRNA | 0 | 1 | 0 | 0 | 0 | 0 |
| ENSDARG000 | 14:14:14:14:1 | 1240235:1240 | 1240468:1240 | +;+;+;+;+;+;+; | 4966 | adad1 | 0 | 2 | 4 | 0 | 4 | 0 |
| ENSDARG000 | 14:14:14:14:1 | 45259644:452 | 45260536:452 | +;+;+;+;+ | 5007 | NA | 281 | 190 | 307 | 253 | 410 | 439 |
| ENSDARG000 | 14:14:14:14:1 | 10362077:103 | 10362090:103 | +;+;+;+;+;+;+; | 210 | NA | 5 | 0 | 11 | 0 | 6 | 0 |
| ENSDARG000 | 14 | 40447368 | 40447484 | - | 117 | 5S_rRNA | 0 | 0 | 0 | 0 | 0 | 0 |
| ENSDARG000 | 14 | 2406945 | 2407048 | - | 104 | Vault | 0 | 0 | 0 | 0 | 0 | 0 |
| ENSDARG000 | 14 | 14909553 | 14909670 | + | 118 | 5S_rRNA | 0 | 0 | 0 | 0 | 0 | 0 |
| ENSDARG000 | 14 | 38743941 | 38744053 | + | 113 | dre-mir-145 | 0 | 0 | 0 | 0 | 0 | 0 |
| ENSDARG000 | 14:14:14:14:1 | 1203608:1204 | 1203791:1204 | -;-;-;-;-;-;-;-; | 2373 | SRP72 | 1856 | 2384 | 4118 | 1572 | 2963 | 1511 |
| ENSDARG000 | 14 | 29353007 | 29353123 | - | 117 | 5S_rRNA | 0 | 0 | 0 | 0 | 0 | 0 |
| ENSDARG000 | 14 | 40252289 | 40252412 | + | 124 | 5S_rRNA | 0 | 0 | 0 | 0 | 0 | 0 |
| ENSDARG000 | 14 | 11614549 | 11614765 | - | 217 | NA | 0 | 0 | 0 | 0 | 0 | 0 |
| ENSDARG000 | 14 | 100502 | 100575 | + | 74 | dre-mir-93 | 0 | 0 | 0 | 0 | 0 | 0 |
| ENSDARG000 | 14 | 11609563 | 11609778 | - | 216 | U3 | 0 | 0 | 0 | 0 | 0 | 0 |
| ENSDARG000 | 14 | 31467167 | 31467260 | - | 94 | dre-mir-20b | 0 | 0 | 0 | 0 | 0 | 0 |
| ENSDARG000 | 14;14 | 2076215:2076 | 2076216:2076 | +;+ | 183 | NA | 0 | 4 | 1 | 0 | 0 | 1 |
| ENSDARG000 | 14 | 2035749 | 2036604 | - | 856 | NA | 0 | 2 | 0 | 0 | 0 | 3 |
| ENSDARG000 | 14;14;14;14 | 26229056:262 | 26229423:262 | +;+;+;+ | 884 | GM2A | 2 | 0 | 0 | 0 | 0 | 7 |
| ENSDARG000 | 14 | 11632363 | 11632578 | - | 216 | U3 | 0 | 0 | 0 | 0 | 0 | 0 |
| ENSDARG000 | 14 | 48409093 | 48409183 | + | 91 | NA | 0 | 0 | 0 | 0 | 0 | 0 |
| ENSDARG000 | 14 | 43343468 | 43343585 | - | 118 | 5S_rRNA | 0 | 0 | 0 | 0 | 0 | 0 |
| ENSDARG000 | 14:14:14:14:1 | 45213093:452 | 45214294:452 | +;+;+;+;+ | 3582 | BEND4 | 0 | 0 | 6 | 0 | 12 | 0 |
| ENSDARG000 | 14 | 52520204 | 52521460 | - | 1257 | GPR151 | 0 | 0 | 10 | 0 | 2 | 0 |
| ENSDARG000 | 14:14:14:14:1 | 1537536:1538 | 1537621:1538 | -;-;-;-;-;-;- | 738 | NA | 7 | 8 | 9 | 5 | 66 | 27 |
| ENSDARG000 | 14 | 1695724 | 1695842 | - | 119 | 5S_rRNA | 0 | 0 | 0 | 0 | 0 | 0 |
| ENSDARG000 | 14:14:14:14:1 | 8819064:8819 | 8819300:8819 | -;-;-;-;-;-;-;-; | 6069 | ATG2A | 3520 | 2513 | 1802 | 2756 | 1778 | 1561 |
| ENSDARG000 | 14 | 49086765 | 49086881 | + | 117 | 5S_rRNA | 1 | 0 | 0 | 0 | 0 | 0 |
| ENSDARG000 | 14;14 | 2095394:2097 | 2095421:2097 | +;+ | 186 | NA | 1 | 0 | 6 | 1 | 0 | 0 |
| ENSDARG000 | 14 | 27175726 | 27175841 | - | 116 | 5S_rRNA | 0 | 0 | 0 | 0 | 0 | 0 |
| ENSDARG000 | 14:14:14:14:1 | 50034196:500 | 50034233:500 | +;+;+;+;+;+;+; | 1230 | NA | 0 | 0 | 0 | 0 | 0 | 0 |
| ENSDARG000 | 14 | 44517029 | 44517148 | - | 120 | 5S_rRNA | 0 | 0 | 0 | 0 | 0 | 0 |
| ENSDARG000 | 14:14:14:14:1 | 49994501:499 | 49997465:499 | -;-;-;-;-;-;-;-; | 4416 | abhd18 | 258 | 263 | 308 | 204 | 317 | 256 |
| ENSDARG000 | 14:14:14:14:1 | 20351:22895: | 20540:22962: | +;+;+;+;+;+;+; | 1103 | stx18 | 192 | 238 | 256 | 192 | 214 | 190 |
| ENSDARG000 | 14:14:14:14:1 | 4090432:4094 | 4090439:4094 | +;+;+;+;+;+;+; | 162 | NA | 0 | 2 | 3 | 0 | 0 | 1 |
| ENSDARG000 | 14;14 | 497695:49887 | 498845:49897 | -;- | 1254 | spry1 | 1160 | 920 | 1156 | 927 | 1426 | 1210 |
| ENSDARG000 | 14:14:14:14:1 | 1431077:1431 | 1435576:1435 | -;-;-;-;-;-;-;-; | 6946 | pmt | 40 | 35 | 222 | 41 | 34 | 22 |
| ENSDARG000 | 14 | 2747924 | 2748045 | + | 122 | 5S_rRNA | 0 | 0 | 0 | 0 | 0 | 0 |
| ENSDARG000 | 14 | 31860408 | 31860470 | - | 63 | NA | 1 | 0 | 4 | 0 | 4 | 0 |
| ENSDARG000 | 14 | 31466671 | 31466782 | - | 112 | mir363 | 0 | 0 | 3 | 0 | 0 | 0 |
| ENSDARG000 | 14 | 31467026 | 31467130 | - | 105 | dre-mir-19c | 0 | 0 | 1 | 0 | 0 | 0 |
| ENSDARG000 | 14 | 10678393 | 10678511 | - | 119 | 5S_rRNA | 0 | 0 | 0 | 0 | 0 | 0 |
| ENSDARG000 | 14:14:14:14:1 | 669982:67209 | 670392:67225 | -;-;-;-;-;-;- | 1437 | NA | 12 | 3 | 22 | 2 | 0 | 2 |
| ENSDARG000 | 14 | 27556486 | 27556622 | - | 137 | 5S_rRNA | 0 | 0 | 0 | 0 | 0 | 0 |
| ENSDARG000 | 14 | 5455707 | 5455823 | + | 117 | 5S_rRNA | 0 | 0 | 0 | 0 | 0 | 0 |
| ENSDARG000 | 14 | 18367834 | 18367949 | - | 116 | 5S_rRNA | 0 | 0 | 0 | 0 | 0 | 0 |
| ENSDARG000 | 14:14:14:14:1 | 3325867:3330 | 3327620:3330 | -;-;-;-;-;-;-;-; | 6207 | NA | 46 | 170 | 106 | 24 | 186 | 206 |
| ENSDARG000 | 14;14;14;14 | 252650:25265 | 252785:25278 | +;+;+;+ | 2283 | NA | 0 | 0 | 0 | 0 | 0 | 0 |
| ENSDARG000 | 14;14;14;14 | 1816294:1816 | 1816572:1816 | -;-;-;- | 711 | NA | 0 | 0 | 0 | 0 | 0 | 0 |
| ENSDARG000 | 14;14;14;14 | 3540872:3542 | 3541153:3542 | +;+;+;+ | 681 | NA | 0 | 0 | 0 | 0 | 0 | 0 |
| ENSDARG000 | 14;14;14;14 | 4692749:4702 | 4692913:4702 | -;-;-;- | 329 | NA | 0 | 0 | 0 | 0 | 0 | 0 |
| ENSDARG000 | 14:14:14:14:1 | 4952114:4952 | 4952912:4952 | -;-;-;-;-;-;-;-;- | 1523 | NA | 0 | 11 | 0 | 47 | 3 | 5 |
| ENSDARG000 | 14;14;14 | 5128881:5129 | 5129099:5129 | +;+;+ | 1138 | NA | 0 | 0 | 0 | 0 | 0 | 0 |
| ENSDARG000 | 14;14;14 | 7582453:7588 | 7582600:7589 | +;+;+ | 1914 | NA | 0 | 0 | 0 | 0 | 0 | 0 |
| ENSDARG000 | 14:14:14:14:1 | 14161512:141 | 14161720:141 | -;-;-;-;-;- | 627 | NA | 0 | 0 | 3 | 0 | 0 | 0 |
| ENSDARG000 | 14:14:14:14:1 | 18850749:188 | 18851178:188 | -;-;-;-;- | 825 | NA | 0 | 0 | 1 | 0 | 0 | 0 |
| ENSDARG000 | 14;14;14 | 19553010:195 | 19553200:195 | -;-;- | 1139 | NA | 0 | 4 | 2 | 0 | 0 | 0 |
| ENSDARG000 | 14:14:14:14:1 | 19806641:198 | 19806802:198 | +;+;+;+;+;+;+ | 3318 | NA | 38 | 80 | 72 | 12 | 26 | 62 |
| ENSDARG000 | 14;14;14;14 | 22807498:228 | 22807837:228 | +;+;+;+ | 586 | NA | 0 | 0 | 0 | 0 | 0 | 0 |
| ENSDARG000 | 14:14:14:14:1 | 26272035:262 | 26272549:262 | +;+;+;+;+;+;+; | 2812 | NA | 2 | 0 | 0 | 4 | 0 | 0 |
| ENSDARG000 | 14:14:14:14:1 | 30511481:305 | 30512039:305 | -;-;-;-;-;-;- | 1653 | NA | 0 | 0 | 0 | 0 | 0 | 0 |
| ENSDARG000 | 14;14;14 | 34529884:345 | 34530098:345 | -;-;- | 471 | NA | 0 | 0 | 4 | 0 | 0 | 0 |
| ENSDARG000 | 14;14;14 | 37027924:370 | 37028274:370 | -;-;- | 809 | NA | 0 | 0 | 0 | 0 | 2 | 0 |
| ENSDARG000 | 14:14:14:14:1 | 37166528:371 | 37166601:371 | -;-;-;-;-;-;-;-; | 2451 | NA | 0 | 0 | 0 | 0 | 0 | 0 |
| ENSDARG000 | 14:14:14:14:1 | 37417722:374 | 37418303:374 | +;+;+;+;+;+;+; | 2651 | NA | 0 | 0 | 0 | 0 | 0 | 0 |
| ENSDARG000 | 14:14:14:14:1 | 37846691:378 | 37846859:378 | +;+;+;+;+;+;+; | 2171 | NA | 0 | 0 | 0 | 0 | 0 | 0 |
| ENSDARG000 | 14:14:14:14:1 | 38064232:380 | 38064443:380 | +;+;+;+;+;+;+ | 846 | NA | 0 | 0 | 0 | 0 | 0 | 0 |
| ENSDARG000 | 14:14:14:14:1 | 38235594:382 | 38235823:382 | +;+;+;+;+ | 1273 | NA | 0 | 0 | 0 | 0 | 0 | 0 |
| ENSDARG000 | 14;14;14 | 38715673:387 | 38715850:387 | +;+;+ | 5154 | NA | 2 | 4 | 2 | 0 | 3 | 6 |
| ENSDARG000 | 14;14;14;14 | 39192439:391 | 39192818:391 | +;+;+;+ | 2581 | NA | 0 | 1 | 0 | 0 | 1 | 0 |
| ENSDARG000 | 14;14;14 | 40123046:401 | 40123233:401 | +;+;+ | 640 | NA | 0 | 0 | 0 | 0 | 0 | 0 |
| ENSDARG000 | 14;14;14;14 | 43178078:431 | 43178663:431 | -;-;-;- | 1490 | NA | 0 | 0 | 0 | 0 | 0 | 0 |
| ENSDARG000 | 14;14;14 | 43428851:434 | 43428904:434 | -;-;- | 270 | NA | 0 | 0 | 0 | 0 | 0 | 0 |
| ENSDARG000 | 14;14;14;14 | 44613241:447 | 44613408:447 | +;+;+;+ | 654 | NA | 51 | 45 | 64 | 62 | 30 | 39 |
| ENSDARG000 | 14;14;14;14 | 48673253:486 | 48673314:486 | -;-;-;- | 451 | NA | 0 | 0 | 0 | 0 | 0 | 0 |
| ENSDARG000 | 14;14;14 | 49496163:495 | 49496416:495 | +;+;+ | 1392 | NA | 0 | 0 | 0 | 1 | 0 | 0 |
| ENSDARG000 | 14:14:14:14:1 | 49931223:499 | 49934372:499 | -;-;-;-;- | 4159 | NA | 135 | 100 | 112 | 46 | 94 | 118 |
| ENSDARG000 | 14:14:14:14:1 | 50426351:504 | 50426648:504 | +;+;+;+;+ | 1454 | NA | 0 | 0 | 0 | 0 | 0 | 0 |
| ENSDARG000 | 13;13 | 11674764:116 | 11678572:116 | -;- | 4172 | NA | 528 | 538 | 649 | 494 | 676 | 960 |
| ENSDARG000 | 13:13:13:13:1 | 29925397:299 | 29925596:299 | +;+;+;+;+;+;+; | 2136 | cuedc2 | 150 | 188 | 218 | 232 | 206 | 130 |
| ENSDARG000 | 13:13:13:13:1 | 269822:27052 | 270316:27059 | -;-;-;-;-;-;-;-; | 1624 | slc30a6 | 351 | 398 | 549 | 375 | 478 | 190 |
| ENSDARG000 | 13:13:13:13:1 | 281214:28205 | 281464:28205 | +;+;+;+;+ | 781 | mpc1 | 520 | 514 | 510 | 459 | 556 | 452 |
| ENSDARG000 | 13:13:13:13:1 | 285874:28587 | 286177:28617 | -;-;-;-;-;-;-;-; | 4811 | NA | 0 | 8 | 11 | 0 | 10 | 2 |
| ENSDARG000 | 13:13:13:13:1 | 6086730:6088 | 6086823:6090 | +;+;+;+;+;+;+; | 2828 | fam120b | 101 | 104 | 92 | 52 | 139 | 129 |
| ENSDARG000 | 13:13:13:13:1 | 9819194:9819 | 9819609:9820 | +;+;+;+;+;+;+; | 3599 | dennd10 | 762 | 1329 | 1604 | 998 | 886 | 1593 |
| ENSDARG000 | 13:13:13:13:1 | 36868365:368 | 36869762:368 | -;-;-;-;-;-;-;-; | 3516 | trim9 | 10 | 0 | 26 | 0 | 2 | 0 |
| ENSDARG000 | 13:13:13:13:1 | 11690277:116 | 11691030:116 | -;-;-;-;-;-;- | 1446 | SLC39A8 | 0 | 0 | 0 | 0 | 0 | 4 |
| ENSDARG000 | 13:13:13:13:1 | 11700107:117 | 11700188:117 | +;+;+;+;+;+ | 698 | NA | 0 | 0 | 0 | 0 | 0 | 0 |
| ENSDARG000 | 13;13 | 2851139:2852 | 2851350:2853 | -;- | 346 | NA | 0 | 0 | 0 | 0 | 0 | 0 |
| ENSDARG000 | 13:13:13:13:1 | 23037585:230 | 23038183:230 | -;-;-;-;-;-;-;-; | 2792 | supv3l1 | 190 | 308 | 397 | 84 | 335 | 264 |
| ENSDARG000 | 13:13:13:13:1 | 311380:31259 | 311589:31268 | +;+;+;+;+;+;+; | 1439 | irf1a | 12 | 10 | 19 | 1 | 8 | 11 |
| ENSDARG000 | 13;13 | 2854325:2856 | 2854553:2856 | -;- | 363 | NA | 0 | 0 | 0 | 0 | 0 | 0 |
| ENSDARG000 | 13:13:13:13:1 | 319575:31957 | 319996:31999 | -;-;-;-;-;-;-;-; | 2545 | ddx21 | 5311 | 3287 | 7434 | 5296 | 4267 | 6771 |
| ENSDARG000 | 13 | 44818447 | 44819254 | - | 808 | NA | 58 | 30 | 44 | 30 | 34 | 36 |
| ENSDARG000 | 13:13:13:13:1 | 40284872:402 | 40287120:402 | -;-;-;-;-;-;-;-; | 4030 | pyroxd2 | 147 | 268 | 513 | 422 | 286 | 128 |
| ENSDARG000 | 13:13:13:13:1 | 353175:35358 | 353762:35376 | -;-;-;-;-;-;-;-; | 2697 | VIT | 712 | 496 | 536 | 312 | 2247 | 103 |
| ENSDARG000 | 13;13 | 2857461:2859 | 2857685:2859 | -;- | 359 | NA | 0 | 0 | 0 | 0 | 0 | 0 |
| ENSDARG000 | 13:13:13:13:1 | 337814:33893 | 338430:33904 | -;-;-;-;-;-;-;-; | 2681 | NA | 74 | 16 | 50 | 6 | 23 | 16 |
| ENSDARG000 | 13:13:13:13:1 | 2861265:2861 | 2861366:2861 | +;+;+;+;+;+;+; | 2907 | XPO5 | 160 | 210 | 347 | 72 | 348 | 170 |
| ENSDARG000 | 13;13;13 | 6142124:6144 | 6142211:6144 | +;+;+ | 303 | NA | 0 | 0 | 0 | 0 | 0 | 0 |
| ENSDARG000 | 13:13:13:13:1 | 11828516:118 | 11829398:118 | +;+;+;+;+;+;+; | 5739 | sufu | 1011 | 838 | 1146 | 848 | 1105 | 1001 |
| ENSDARG000 | 13:13:13:13:1 | 3109064:3110 | 3109300:3110 | -;-;-;-;-;-;-;-; | 1698 | pkdcca | 10 | 18 | 102 | 26 | 34 | 6 |
| ENSDARG000 | 13;13 | 13578552:135 | 13579221:135 | +;+ | 1342 | foxi2 | 0 | 0 | 0 | 0 | 0 | 0 |
| ENSDARG000 | 13:13:13:13:1 | 36958086:369 | 36958199:369 | +;+;+;+;+;+;+; | 4840 | frmd6 | 140 | 115 | 42 | 66 | 88 | 37 |
| ENSDARG000 | 13;13;13;13 | 44827349:448 | 44828282:448 | -;-;-;- | 1013 | si:dkeyp-2e4.6 | 4 | 4 | 4 | 0 | 11 | 23 |
| ENSDARG000 | 13;13 | 8987957:8993 | 8988044:8994 | +;+ | 1029 | OXER1 | 6 | 4 | 5 | 8 | 0 | 34 |
| ENSDARG000 | 13 | 44835774 | 44836727 | - | 954 | si:dkeyp-2e4.1 | 17 | 34 | 32 | 31 | 11 | 15 |
| ENSDARG000 | 13:13:13:13:1 | 44827514:448 | 44828282:448 | -;-;-;-;-;-;- | 2772 | si:dkeyp-2e4.3 | 26 | 27 | 73 | 68 | 54 | 81 |
| ENSDARG000 | 13;13;13;13 | 40327562:403 | 40327612:403 | +;+;+;+ | 370 | NA | 0 | 0 | 0 | 1 | 0 | 4 |
| ENSDARG000 | 13;13 | 44831575:448 | 44832526:448 | -;- | 1055 | NA | 25 | 63 | 43 | 30 | 38 | 52 |
| ENSDARG000 | 13;13 | 6160581:6161 | 6160676:6161 | -;- | 270 | NA | 0 | 0 | 0 | 0 | 0 | 0 |
| ENSDARG000 | 13;13;13;13 | 40334727:403 | 40335249:403 | +;+;+;+ | 3417 | slc25a28 | 845 | 703 | 816 | 534 | 1187 | 903 |
| ENSDARG000 | 13:13:13:13:1 | 29952633:299 | 29953614:299 | -;-;-;-;-;-;-;-; | 2249 | hif1an | 828 | 1279 | 2013 | 1730 | 1188 | 754 |
| ENSDARG000 | 13:13:13:13:1 | 23058268:230 | 23059607:230 | -;-;-;-;-;-;-;-; | 2678 | vps26a | 2834 | 3420 | 2599 | 2204 | 2989 | 2257 |
| ENSDARG000 | 13;13 | 13587296:135 | 13587827:135 | -;- | 579 | NA | 0 | 0 | 0 | 0 | 0 | 0 |
| ENSDARG000 | 13;13;13 | 3198438:3199 | 3198615:3199 | -;-;- | 435 | NA | 0 | 0 | 0 | 0 | 0 | 0 |
| ENSDARG000 | 13:13:13:13:1 | 450002:45089 | 450309:45110 | -;-;-;-;-;-;-;-; | 6083 | HEATR5B | 779 | 603 | 841 | 391 | 1017 | 468 |
| ENSDARG000 | 13:13:13:13:1 | 21888214:219 | 21888344:219 | +;+;+;+;+;+;+; | 4761 | NDST2 | 602 | 520 | 748 | 228 | 943 | 614 |
| ENSDARG000 | 13;13 | 3259543:3260 | 3260166:3260 | +;+ | 720 | NA | 0 | 0 | 0 | 0 | 0 | 0 |
| ENSDARG000 | 13;13;13 | 3252950:3254 | 3253878:3255 | +;+;+ | 2345 | prph2b | 2 | 0 | 0 | 1 | 0 | 0 |
| ENSDARG000 | 13:13:13:13:1 | 29991090:299 | 29994216:299 | -;-;-;-;-;- | 3669 | wnt8b | 8 | 0 | 0 | 0 | 0 | 0 |
| ENSDARG000 | 13:13:13:13:1 | 8999261:9000 | 8999590:9000 | +;+;+;+;+;+;+ | 1641 | NA | 0 | 0 | 0 | 0 | 0 | 0 |
| ENSDARG000 | 13:13:13:13:1 | 9831010:9832 | 9831178:9832 | -;-;-;-;-;-;-;-; | 1182 | sfxn4 | 162 | 110 | 204 | 134 | 199 | 138 |
| ENSDARG000 | 13:13:13:13:1 | 482911:48482 | 483111:48493 | +;+;+;+;+ | 1041 | fbxo28 | 517 | 345 | 604 | 267 | 458 | 341 |
| ENSDARG000 | 13;13;13 | 40360173:403 | 40360198:403 | +;+;+ | 854 | NA | 2 | 0 | 0 | 0 | 0 | 0 |

| | | | | | | | | | | | | |
|---|---|---|---|---|---|---|---|---|---|---|---|---|
| ENSDARG000 | 13:13:13:13:1 | 11963789:119 | 11964259:119 | -:-:-:-:-:-:-:-:-: | 1075 | arl3a | 26 | 8 | 22 | 2 | 36 | 18 |
| ENSDARG000 | 13:13:13:13:1 | 13660564:136 | 13660648:136 | -;-;-;-;-;-;-;-;- | 1421 | NA | 0 | 0 | 0 | 0 | 0 | 0 |
| ENSDARG000 | 13:13:13:13:1 | 40409215:404 | 40410222:404 | -;-;-;-;- | 1550 | nkx2.3 | 6 | 0 | 0 | 0 | 0 | 0 |
| ENSDARG000 | 13:13:13:13:1 | 6188759:6189 | 6189313:6189 | +:+:+:+:+:+:+: | 2949 | ppm1g | 1190 | 2197 | 2824 | 923 | 2121 | 1350 |
| ENSDARG000 | 13:13:13:13:1 | 44857087:448 | 44857200:448 | +;+;+;+;+;+;+ | 1126 | zbtb8os | 445 | 246 | 385 | 187 | 284 | 208 |
| ENSDARG000 | 13:13:13:13:1 | 23070581:230 | 23071527:230 | -;-;-;-;- | 1961 | srgn | 610 | 678 | 634 | 520 | 1655 | 1454 |
| ENSDARG000 | 13;13 | 44848343:448 | 44849277:448 | -;- | 1019 | si:dkeyp-2e4.1 | 13 | 12 | 5 | 15 | 14 | 20 |
| ENSDARG000 | 13:13:13:13:1 | 16922705:169 | 16923446:169 | -:-:-:-:-:-:-:-:-: | 2578 | lrmda | 404 | 315 | 394 | 328 | 483 | 517 |
| ENSDARG000 | 13;13;13;13 | 17464402:174 | 17464464:174 | +;+;+;+ | 591 | NA | 2 | 6 | 6 | 14 | 19 | 22 |
| ENSDARG000 | 13:13:13:13:1 | 33256847:332 | 33257160:332 | +;+;+;+;+ | 758 | oip5 | 20 | 70 | 78 | 20 | 30 | 30 |
| ENSDARG000 | 13:13:13:13:1 | 502230:50231 | 502503:50250 | +;+;+;+;+;+ | 1639 | degs1 | 2368 | 3078 | 2633 | 1492 | 2879 | 2544 |
| ENSDARG000 | 13:13:13:13:1 | 27000382:270 | 27000759:270 | -;-;-;-;-;-;- | 2547 | ccdc85a | 0 | 2 | 4 | 0 | 3 | 0 |
| ENSDARG000 | 13:13:13:13:1 | 11975954:119 | 11976679:119 | -;-;-;-;-;-;-;- | 3051 | npm3 | 609 | 712 | 883 | 470 | 706 | 673 |
| ENSDARG000 | 13:13:13:13:1 | 23083858:230 | 23085052:230 | -;-;-;-;-;- | 2231 | kifbp | 815 | 856 | 719 | 652 | 1001 | 650 |
| ENSDARG000 | 13:13:13:13:1 | 23093743:230 | 23095024:230 | +:+:+:+:+:+:+: | 8106 | NA | 220 | 271 | 346 | 196 | 400 | 309 |
| ENSDARG000 | 13:13:13:13:1 | 35856463:358 | 35856905:358 | +;+;+;+;+ | 1870 | kcnk1b | 6 | 14 | 4 | 0 | 4 | 6 |
| ENSDARG000 | 13;13 | 17468161:174 | 17468964:174 | +;+ | 2492 | znf503 | 1678 | 719 | 3126 | 1048 | 2576 | 2825 |
| ENSDARG000 | 13:13:13:13:1 | 534453:53445 | 534564:53456 | +:+:+:+:+:+:+: | 1384 | NA | 65 | 47 | 94 | 79 | 89 | 107 |
| ENSDARG000 | 13:13:13:13:1 | 516258:51684 | 516423:51692 | -:-:-:-:-:-:-:-:-: | 3054 | nvl | 364 | 410 | 745 | 475 | 542 | 336 |
| ENSDARG000 | 13:13:13:13:1 | 9091958:9091 | 9092290:9092 | +:+:+:+:+:+:+: | 2414 | NA | 0 | 0 | 0 | 0 | 0 | 0 |
| ENSDARG000 | 13:13:13:13:1 | 30007554:300 | 30010641:300 | -:-:-:-:-:-:-:-:-: | 4671 | scdb | 862 | 18 | 4 | 217 | 20 | 38 |
| ENSDARG000 | 13;13;13 | 22104298:221 | 22104808:221 | +;+;+ | 9735 | NA | 6 | 0 | 2 | 0 | 1 | 6 |
| ENSDARG000 | 13:13:13:13:1 | 9046473:9047 | 9046607:9047 | +;+;+;+;+;+;+ | 1446 | NA | 0 | 4 | 0 | 0 | 0 | 0 |
| ENSDARG000 | 13:13:13:13:1 | 9002575:9002 | 9003471:9003 | -:-:-:-:-:-:-:-:-: | 3366 | NA | 0 | 2 | 0 | 0 | 0 | 0 |
| ENSDARG000 | 13:13:13:13:1 | 24781188:247 | 24782710:247 | -:-:-:-:-:-:-:-:-: | 6377 | slka | 2193 | 2210 | 2896 | 2727 | 2203 | 2691 |
| ENSDARG000 | 13:13:13:13:1 | 37022601:370 | 37022672:370 | +:+:+:+:+:+:+: | 3420 | esr2b | 8 | 2 | 0 | 4 | 14 | 0 |
| ENSDARG000 | 13:13:13:13:1 | 9162465:9163 | 9162769:9163 | +;+;+;+;+;+;+ | 1537 | NA | 0 | 0 | 0 | 0 | 0 | 0 |
| ENSDARG000 | 13:13:13:13:1 | 33298338:332 | 33298460:332 | +:+:+:+:+:+:+: | 1332 | iqcc | 4 | 34 | 12 | 14 | 14 | 30 |
| ENSDARG000 | 13:13:13:13:1 | 40437550:404 | 40437796:404 | +:+:+:+:+:+:+: | 3737 | got1 | 1057 | 1357 | 1159 | 687 | 1340 | 853 |
| ENSDARG000 | 13:13:13:13:1 | 9071364:9072 | 9071694:9072 | +;+;+;+;+;+;+ | 1642 | pimr151 | 0 | 0 | 0 | 0 | 0 | 0 |
| ENSDARG000 | 13:13:13:13:1 | 13668991:136 | 13669376:136 | +;+;+;+;+;+;+ | 1335 | NA | 0 | 0 | 0 | 0 | 0 | 0 |
| ENSDARG000 | 13:13:13:13:1 | 9016867:9017 | 9017197:9018 | +;+;+;+;+;+;+ | 1642 | NA | 0 | 0 | 0 | 0 | 0 | 0 |
| ENSDARG000 | 13:13:13:13:1 | 9184801:9184 | 9185130:9185 | +:+:+:+:+:+:+: | 1641 | NA | 0 | 0 | 0 | 2 | 0 | 0 |
| ENSDARG000 | 13:13:13:13:1 | 33261131:332 | 33265491:332 | -:-:-:-:-:-:-:-:-: | 5060 | oip5-as1 | 2262 | 967 | 1389 | 760 | 1116 | 1461 |
| ENSDARG000 | 13:13:13:13:1 | 9115978:9116 | 9116579:9116 | -:-:-:-:-:-:-:-:-: | 1131 | NA | 0 | 0 | 0 | 0 | 0 | 0 |
| ENSDARG000 | 13:13:13:13:1 | 9020301:9020 | 9020909:9020 | -:-:-:-:-:-:-:-:-: | 1459 | NA | 20 | 11 | 19 | 33 | 36 | 2 |
| ENSDARG000 | 13:13:13:13:1 | 33282095:332 | 33282262:332 | +;+;+;+;+;+ | 981 | ccdc28b | 12 | 99 | 120 | 140 | 121 | 190 |
| ENSDARG000 | 13:13:13:13:1 | 33268657:332 | 33268838:332 | +:+:+:+:+:+:+: | 1857 | tmem39b | 308 | 264 | 337 | 257 | 296 | 322 |
| ENSDARG000 | 13:13:13:13:1 | 9165861:9165 | 9166454:9166 | -:-:-:-:-:-:-:-:-: | 1438 | HTRA2 | 0 | 0 | 0 | 0 | 0 | 0 |
| ENSDARG000 | 13:13:13:13:1 | 17672527:176 | 17672705:176 | +:+:+:+:+:+:+: | 955 | comtd1 | 0 | 0 | 16 | 0 | 0 | 0 |
| ENSDARG000 | 13:13:13:13:1 | 13681681:136 | 13681759:136 | +;+;+;+;+ | 900 | cfd | 395 | 101 | 108 | 96 | 202 | 177 |
| ENSDARG000 | 13:13:13:13:1 | 48617974:486 | 48620215:486 | +:+:+:+:+:+:+: | 3166 | NA | 56 | 0 | 6 | 2 | 18 | 2 |
| ENSDARG000 | 13;13;13;13 | 9003127:9003 | 9003290:9003 | -;-;-;- | 535 | NA | 0 | 0 | 0 | 0 | 0 | 0 |
| ENSDARG000 | 13:13:13:13:1 | 9217235:9217 | 9217839:9217 | -:-:-:-:-:-:-:-:-: | 1243 | HTRA2 | 0 | 0 | 1 | 0 | 0 | 1 |
| ENSDARG000 | 13:13:13:13:1 | 9213805:9214 | 9214134:9215 | +;+;+;+;+;+;+ | 1641 | NA | 0 | 0 | 0 | 0 | 0 | 0 |
| ENSDARG000 | 13:13:13:13:1 | 9115984:9116 | 9116579:9116 | -:-:-:-:-:-:-:-:-: | 2361 | HTRA2 | 0 | 21 | 30 | 14 | 1 | 10 |
| ENSDARG000 | 13;13;13;13 | 27040887:270 | 27041037:270 | +;+;+;+ | 1494 | soul2 | 16 | 8 | 2 | 4 | 10 | 0 |
| ENSDARG000 | 13:13:13:13:1 | 45009957:450 | 45010123:450 | -:-:-:-:-:-:-:-:-: | 2441 | khdrbs1a | 16815 | 17497 | 25342 | 12901 | 21064 | 15448 |
| ENSDARG000 | 13;13 | 30033036:300 | 30034754:300 | -;- | 1719 | trmt2b | 62 | 108 | 92 | 28 | 104 | 76 |
| ENSDARG000 | 13:13:13:13:1 | 9188209:9188 | 9188806:9188 | -:-:-:-:-:-:-:-:-: | 1236 | NA | 5 | 0 | 1 | 1 | 1 | 0 |
| ENSDARG000 | 13:13:13:13:1 | 22119798:221 | 22119940:221 | +:+:+:+:+:+:+: | 5275 | NA | 1221 | 932 | 1707 | 2091 | 4059 | 2002 |
| ENSDARG000 | 13:13:13:13:1 | 9074797:9074 | 9075398:9075 | -:-:-:-:-:-:-:-:-: | 1238 | HTRA2 | 0 | 0 | 0 | 0 | 0 | 0 |
| ENSDARG000 | 13:13:13:13:1 | 27073901:271 | 27073923:271 | +:+:+:+:+:+:+: | 1839 | SLC24A3 | 23 | 4 | 23 | 14 | 50 | 24 |
| ENSDARG000 | 13:13:13:13:1 | 9112544:9112 | 9112874:9112 | +:+:+:+:+:+:+: | 1917 | NA | 0 | 0 | 0 | 0 | 0 | 0 |
| ENSDARG000 | 13:13:13:13:1 | 40501455:405 | 40501457:405 | +:+:+:+:+:+:+: | 1683 | HPSE2 | 0 | 0 | 0 | 0 | 0 | 0 |
| ENSDARG000 | 13:13:13:13:1 | 9842276:9842 | 9842512:9842 | +:+:+:+:+:+:+: | 1523 | prdx3 | 928 | 1051 | 1494 | 815 | 1201 | 883 |
| ENSDARG000 | 13;13;13 | 22191361:221 | 22191874:221 | -;-;- | 845 | NA | 0 | 0 | 0 | 0 | 0 | 0 |
| ENSDARG000 | 13:13:13:13:1 | 9137116:9137 | 9137447:9138 | +;+;+;+;+;+;+ | 1643 | NA | 0 | 0 | 0 | 2 | 0 | 0 |
| ENSDARG000 | 13:13:13:13:1 | 9095390:9095 | 9095986:9095 | -:-:-:-:-:-:-:-:-: | 1443 | NA | 0 | 4 | 2 | 2 | 0 | 2 |
| ENSDARG000 | 13;13 | 17694845:176 | 17695361:176 | +;+ | 2136 | ifit8 | 130 | 124 | 18 | 0 | 0 | 0 |
| ENSDARG000 | 13:13:13:13:1 | 48747449:487 | 48748360:487 | -:-:-:-:-:-:-:-:-: | 1711 | ntpcr | 216 | 175 | 218 | 220 | 289 | 307 |
| ENSDARG000 | 13:13:13:13:1 | 11988464:119 | 11990754:119 | -:-:-:-:-:-:-:-:-: | 5516 | oga | 2757 | 2014 | 2559 | 3705 | 3068 | 4029 |
| ENSDARG000 | 13:13:13:13:1 | 17702522:177 | 17703692:177 | +;+;+;+;+ | 2216 | NA | 0 | 4 | 1 | 0 | 0 | 0 |
| ENSDARG000 | 13:13:13:13:1 | 30035253:300 | 30035594:300 | +:+:+:+:+:+:+: | 4644 | dnajb12a | 859 | 659 | 680 | 595 | 1057 | 661 |
| ENSDARG000 | 13;13;13 | 24829996:248 | 24830135:248 | +;+;+ | 561 | si:ch211-223 | 0 | 0 | 0 | 0 | 0 | 0 |
| ENSDARG000 | 13:13:13:13:1 | 35880573:358 | 35880735:358 | -:-:-:-:-:-:-:-:-: | 3337 | tacc3 | 1064 | 2154 | 2319 | 1286 | 1892 | 786 |
| ENSDARG000 | 13;13;13;13 | 24842857:248 | 24842901:248 | +;+;+;+ | 1136 | dusp13a | 0 | 0 | 0 | 0 | 0 | 0 |
| ENSDARG000 | 13;13;13 | 24834199:248 | 24834492:248 | +;+;+ | 1760 | zgc:153981 | 12 | 0 | 2 | 6 | 10 | 0 |
| ENSDARG000 | 13;13;13;13 | 24851102:248 | 24851208:248 | +;+;+;+ | 1192 | NA | 0 | 0 | 0 | 2 | 0 | 0 |
| ENSDARG000 | 13:13:13:13:1 | 27216641:272 | 27217015:272 | +;+;+;+;+;+ | 1900 | NA | 80 | 128 | 231 | 172 | 121 | 184 |
| ENSDARG000 | 13:13:13:13:1 | 40716319:407 | 40717979:407 | -;-;-;-;-;- | 2714 | st3gal7 | 6 | 4 | 14 | 35 | 21 | 12 |
| ENSDARG000 | 13:13:13:13:1 | 27232694:272 | 27232998:272 | +:+:+:+:+:+:+: | 6831 | RIN2 | 826 | 397 | 504 | 362 | 650 | 664 |
| ENSDARG000 | 13:13:13:13:1 | 40686133:406 | 40686192:406 | +:+:+:+:+:+:+: | 4984 | hps1 | 10735 | 7040 | 6660 | 7579 | 4544 | 7852 |
| ENSDARG000 | 13:13:13:13:1 | 17714296:177 | 17715208:177 | -:-:-:-:-:-:-:-:-: | 1890 | vdac2 | 6743 | 8193 | 12117 | 9757 | 7909 | 9825 |
| ENSDARG000 | 13:13:13:13:1 | 22249636:222 | 22250304:222 | +:+:+:+:+:+:+: | 6606 | NA | 10 | 0 | 2 | 2 | 20 | 0 |
| ENSDARG000 | 13:13:13:13:1 | 3263853:3264 | 3264783:3264 | -:-:-:-:-:-:-:-:-: | 6295 | UBR2 | 1500 | 947 | 1187 | 1006 | 1340 | 981 |
| ENSDARG000 | 13:13:13:13:1 | 555321:55564 | 555351:55717 | -:-:-:-:-:-:-:-:-: | 6039 | slit1a | 16 | 14 | 335 | 11 | 198 | 38 |
| ENSDARG000 | 13;13 | 48757751:487 | 48760984:487 | -;- | 4263 | NA | 4 | 4 | 6 | 34 | 30 | 8 |
| ENSDARG000 | 13;13 | 48765887:487 | 48766371:487 | +;+ | 678 | NA | 0 | 0 | 0 | 0 | 4 | 1 |
| ENSDARG000 | 13:13:13:13:1 | 22264914:222 | 22265130:222 | +;+;+;+;+;+ | 1919 | myoz1a | 0 | 0 | 0 | 0 | 4 | 0 |
| ENSDARG000 | 13:13:13:13:1 | 31473190:314 | 31473261:314 | +:+:+:+:+:+:+: | 4088 | NA | 28 | 23 | 18 | 5 | 6 | 22 |
| ENSDARG000 | 13:13:13:13:1 | 49057081:490 | 49057408:490 | -:-:-:-:-:-:-:-:-: | 3225 | disc1 | 2164 | 1319 | 2453 | 1640 | 2064 | 1576 |
| ENSDARG000 | 13:13:13:13:1 | 45035756:450 | 45036625:450 | -:-:-:-:-:-:-:-:-: | 1862 | chp1 | 495 | 416 | 546 | 244 | 767 | 307 |
| ENSDARG000 | 13:13:13:13:1 | 13693722:136 | 13693863:136 | +:+:+:+:+:+:+: | 3826 | dnaaf9 | 949 | 781 | 1238 | 1596 | 1356 | 1262 |
| ENSDARG000 | 13:13:13:13:1 | 6208365:6210 | 6208528:6210 | -;-;-;-;-;-;-;- | 723 | si:zfos-1056e | 2 | 0 | 0 | 0 | 0 | 0 |
| ENSDARG000 | 13:13:13:13:1 | 27314795:273 | 27314851:273 | +:+:+:+:+:+:+: | 3647 | eef1a1a | 462 | 247 | 1054 | 507 | 202 | 464 |
| ENSDARG000 | 13:13:13:13:1 | 27298673:273 | 27298825:273 | +:+:+:+:+:+:+: | 2202 | slc17a5 | 144 | 54 | 83 | 68 | 125 | 76 |
| ENSDARG000 | 13:13:13:13:1 | 49175624:491 | 49176552:491 | +:+:+:+:+:+:+: | 1403 | EGLN1 | 257 | 119 | 0 | 257 | 198 | 112 |
| ENSDARG000 | 13:13:13:13:1 | 9855380:9857 | 9857278:9857 | -:-:-:-:-:-:-:-:-: | 3689 | tm9sf3 | 2850 | 2311 | 2833 | 1857 | 3193 | 2696 |
| ENSDARG000 | 13:13:13:13:1 | 24871050:248 | 24873661:248 | -:-:-:-:-:-:-:-:-: | 6104 | KAT6B | 1649 | 665 | 1352 | 1012 | 2000 | 940 |
| ENSDARG000 | 13:13:13:13:1 | 23132666:231 | 23132748:231 | +:+:+:+:+:+:+: | 6848 | NA | 1781 | 1166 | 2751 | 2789 | 2186 | 2249 |
| ENSDARG000 | 13:13:13:13:1 | 22280983:222 | 22281147:222 | +:+:+:+:+:+:+: | 7256 | usp54a | 82 | 16 | 16 | 32 | 40 | 9 |
| ENSDARG000 | 13:13:13:13:1 | 17751866:177 | 17752635:177 | -:-:-:-:-:-:-:-:-: | 7482 | marchf8 | 288 | 256 | 496 | 280 | 474 | 258 |
| ENSDARG000 | 13:13:13:13:1 | 30054996:300 | 30055400:300 | +:+:+:+:+:+:+: | 2296 | SPOCK2 | 62 | 15 | 41 | 12 | 6 | 12 |
| ENSDARG000 | 13:13:13:13:1 | 6219551:6219 | 6219820:6220 | +:+:+:+:+:+:+: | 2292 | apmap | 717 | 446 | 615 | 513 | 566 | 477 |
| ENSDARG000 | 13:13:13:13:1 | 37171976:371 | 37172150:371 | -;-;-;-;- | 539 | NA | 36 | 81 | 27 | 62 | 98 | 90 |
| ENSDARG000 | 13:13:13:13:1 | 37075322:370 | 37078192:370 | -:-:-:-:-:-:-:-:-: | 7312 | NA | 138 | 162 | 212 | 162 | 275 | 292 |
| ENSDARG000 | 13;13 | 49443226:494 | 49443258:494 | -;- | 1001 | irf2bp2a | 2437 | 1823 | 2778 | 1794 | 1580 | 1237 |
| ENSDARG000 | 13:13:13:13:1 | 12007904:120 | 12008484:120 | -:-:-:-:-:-:-:-:-: | 4265 | NA | 573 | 1079 | 1494 | 257 | 1202 | 616 |
| ENSDARG000 | 13;13;13 | 37227050:372 | 37227200:372 | -;-;- | 409 | NA | 0 | 0 | 2 | 7 | 0 | 0 |
| ENSDARG000 | 13;13;13 | 37186303:371 | 37186457:371 | -;-;- | 343 | NA | 14 | 41 | 18 | 19 | 60 | 39 |
| ENSDARG000 | 13;13;13 | 37208460:372 | 37208673:372 | -;-;- | 434 | NA | 22 | 9 | 12 | 0 | 20 | 12 |
| ENSDARG000 | 13;13;13 | 37199298:371 | 37199521:371 | -;-;- | 538 | NA | 17 | 18 | 24 | 15 | 31 | 12 |
| ENSDARG000 | 13:13:13:13:1 | 646700:64886 | 646980:64901 | +:+:+:+:+:+:+: | 4869 | tp53bp2a | 2268 | 1538 | 3091 | 2152 | 1945 | 2617 |
| ENSDARG000 | 13:13:13:13:1 | 22421151:224 | 22421379:224 | +:+:+:+:+:+:+: | 1945 | opn4a | 0 | 0 | 0 | 0 | 0 | 0 |
| ENSDARG000 | 13:13:13:13:1 | 49597258:495 | 49597394:495 | -:-:-:-:-:-:-:-:-: | 3778 | ARID4B | 251 | 143 | 288 | 149 | 154 | 176 |
| ENSDARG000 | 13:13:13:13:1 | 24918096:249 | 24920153:249 | -:-:-:-:-:-:-:-:-: | 3507 | adka | 681 | 1146 | 1758 | 822 | 1740 | 886 |
| ENSDARG000 | 13:13:13:13:1 | 6245219:6247 | 6248274:6248 | -:-:-:-:-:-:-:-:-: | 3593 | tuba4l | 69 | 216 | 105 | 165 | 54 | 114 |
| ENSDARG000 | 13:13:13:13:1 | 33304187:333 | 33304343:333 | +:+:+:+:+:+:+: | 1738 | dcdc2b | 10 | 6 | 13 | 17 | 6 | 19 |
| ENSDARG000 | 13:13:13:13:1 | 13743202:137 | 13744526:137 | -:-:-:-:-:-:-:-:-: | 2100 | NA | 0 | 4 | 2 | 12 | 6 | 0 |
| ENSDARG000 | 13:13:13:13:1 | 33314673:333 | 33315122:333 | -:-:-:-:-:-:-:-:-: | 1053 | tmem234 | 26 | 92 | 96 | 104 | 80 | 73 |
| ENSDARG000 | 13:13:13:13:1 | 6259392:6266 | 6259906:6266 | -;-;-;-;-;-;-;- | 1947 | agpat5 | 1294 | 1151 | 1503 | 779 | 1770 | 1261 |
| ENSDARG000 | 13:13:13:13:1 | 35904335:359 | 35905416:359 | -;-;-;-;- | 2174 | mycla | 798 | 284 | 602 | 132 | 472 | 273 |
| ENSDARG000 | 13:13:13:13:1 | 40729618:407 | 40729687:407 | +:+:+:+:+:+:+: | 888 | ANKRD2 | 0 | 0 | 0 | 6 | 0 | 6 |
| ENSDARG000 | 13:13:13:13:1 | 27328098:273 | 27328991:273 | +:+:+:+:+:+:+: | 1830 | CGAS | 46 | 38 | 9 | 0 | 26 | 0 |
| ENSDARG000 | 13:13:13:13:1 | 27332656:273 | 27332916:273 | -:-:-:-:-:-:-:-:-: | 2518 | ddx43 | 0 | 0 | 2 | 0 | 0 | 0 |
| ENSDARG000 | 13:13:13:13:1 | 45092766:450 | 45098415:450 | -:-:-:-:-:-:-:-:-: | 7946 | runx3 | 11575 | 3460 | 8254 | 10982 | 9011 | 11461 |
| ENSDARG000 | 13:13:13:13:1 | 23227531:232 | 23227717:232 | +;+;+;+;+ | 974 | NA | 0 | 7 | 0 | 2 | 4 | 0 |
| ENSDARG000 | 13:13:13:13:1 | 23233956:232 | 23235109:232 | -:-:-:-:-:-:-:-:-: | 2069 | NA | 0 | 0 | 4 | 0 | 0 | 0 |
| ENSDARG000 | 13:13:13:13:1 | 13770980:137 | 13771131:137 | +:+:+:+:+:+:+: | 2871 | slc4a11 | 22 | 2 | 4 | 17 | 12 | 10 |
| ENSDARG000 | 13:13:13:13:1 | 31520537:315 | 31520649:315 | +:+:+:+:+:+:+: | 4652 | pcnx4 | 492 | 366 | 412 | 407 | 575 | 548 |
| ENSDARG000 | 13:13:13:13:1 | 30135675:301 | 30136228:301 | -;-;-;-;-;-;-;- | 2369 | tysnd1 | 124 | 84 | 131 | 70 | 120 | 152 |
| ENSDARG000 | 13:13:13:13:1 | 9878544:9878 | 9878706:9878 | -:-:-:-:-:-:-:-:-: | 1720 | si:ch211-117 | 0 | 0 | 0 | 2 | 10 | 0 |
| ENSDARG000 | 13;13;13;13 | 45332203:453 | 45332580:453 | +;+;+;+ | 2417 | znfl1g | 2 | 6 | 4 | 0 | 0 | 5 |
| ENSDARG000 | 13:13:13:13:1 | 6295173:6295 | 6295599:6295 | +:+:+:+:+:+:+: | 1630 | angpt2a | 26 | 43 | 6 | 56 | 58 | 50 |
| ENSDARG000 | 13:13:13:13:1 | 35925490:359 | 35925779:359 | +:+:+:+:+:+:+: | 1943 | mfsd2aa | 664 | 836 | 364 | 184 | 294 | 432 |
| ENSDARG000 | 13:13:13:13:1 | 23282549:233 | 23282762:233 | +:+:+:+:+:+:+: | 2070 | khdrbs2 | 0 | 6 | 6 | 0 | 0 | 14 |
| ENSDARG000 | 13:13:13:13:1 | 49774315:497 | 49775200:497 | -:-:-:-:-:-:-:-:-: | 2399 | b3galnt2 | 146 | 264 | 232 | 142 | 258 | 112 |
| ENSDARG000 | 13;13;13 | 45321119:453 | 45324505:453 | -;-;- | 3611 | NA | 0 | 0 | 0 | 0 | 0 | 0 |
| ENSDARG000 | 13:13:13:13:1 | 6286594:6288 | 6286786:6288 | -:-:-:-:-:-:-:-:-: | 2517 | mcph1 | 4 | 32 | 52 | 20 | 9 | 15 |
| ENSDARG000 | 13:13:13:13:1 | 681628:68198 | 681798:68212 | +:+:+:+:+:+:+: | 4843 | abcc2 | 49 | 40 | 148 | 123 | 128 | 86 |
| ENSDARG000 | 13:13:13:13:1 | 18019709:180 | 18019903:180 | -:-:-:-:-:-:-:-:-: | 2748 | zfand4 | 120 | 67 | 109 | 78 | 96 | 45 |
| ENSDARG000 | 13:13:13:13:1 | 25199849:251 | 25199916:251 | +:+:+:+:+:+:+: | 2256 | ap3m1 | 281 | 396 | 470 | 318 | 513 | 396 |
| ENSDARG000 | 13;13;13;13 | 35955562:359 | 35956149:359 | +;+;+;+ | 1130 | si:ch211-67f1 | 22 | 0 | 0 | 6 | 0 | 0 |
| ENSDARG000 | 13:13:13:13:1 | 31536709:315 | 31536911:315 | -;-;-;-;-;- | 1299 | dhrs7 | 586 | 398 | 439 | 331 | 517 | 494 |
| ENSDARG000 | 13:13:13:13:1 | 40746281:407 | 40749701:407 | -:-:-:-:-:-:-:-:-: | 3862 | morn4 | 58 | 62 | 32 | 26 | 88 | 28 |
| ENSDARG000 | 13:13:13:13:1 | 30144937:301 | 30145989:301 | -:-:-:-:-:-:-:-:-: | 1750 | sar1ab | 16 | 6 | 28 | 38 | 0 | 48 |
| ENSDARG000 | 13:13:13:13:1 | 31545530:315 | 31546127:315 | +:+:+:+:+:+:+: | 5804 | ppm1aa | 356 | 366 | 529 | 193 | 326 | 440 |
| ENSDARG000 | 13;13;13;13 | 35974362:359 | 35975212:359 | -;-;-;- | 1683 | zgc:153911 | 769 | 59 | 58 | 156 | 111 | 183 |
| ENSDARG000 | 13:13:13:13:1 | 13930263:139 | 13930582:139 | +:+:+:+:+:+:+: | 3165 | rpia | 1212 | 1110 | 1815 | 795 | 1581 | 1532 |
| ENSDARG000 | 13:13:13:13:1 | 12045475:120 | 12045613:120 | +:+:+:+:+:+:+: | 1526 | gng2 | 76 | 168 | 193 | 810 | 194 | 499 |
| ENSDARG000 | 13:13:13:13:1 | 733027:74046 | 734333:74049 | +:+:+:+:+:+:+: | 11063 | nrxn1b | 2 | 26 | 27 | 8 | 2 | 12 |
| ENSDARG000 | 13:13:13:13:1 | 45431660:454 | 45431721:454 | +:+:+:+:+:+:+: | 1622 | syf2 | 1732 | 1161 | 1467 | 1236 | 1344 | 1424 |
| ENSDARG000 | 13:13:13:13:1 | 40754580:407 | 40754621:407 | +:+:+:+:+:+:+: | 1264 | NA | 0 | 2 | 0 | 0 | 0 | 0 |
| ENSDARG000 | 13;13 | 40770628:407 | 40770947:407 | +;+ | 3319 | nkx1.2la | 0 | 0 | 0 | 0 | 0 | 0 |
| ENSDARG000 | 13:13:13:13:1 | 3327417:3328 | 3327462:3328 | -:-:-:-:-:-:-:-:-: | 2410 | NA | 0 | 0 | 0 | 0 | 0 | 0 |
| ENSDARG000 | 13:13:13:13:1 | 31636732:316 | 31637250:316 | -;-;-;-;-;- | 2363 | six4a | 18 | 0 | 0 | 0 | 0 | 0 |
| ENSDARG000 | 13;13 | 31619961:316 | 31620737:316 | -;- | 1573 | six1a | 0 | 0 | 0 | 0 | 0 | 0 |
| ENSDARG000 | 13:13:13:13:1 | 31583034:316 | 31583126:316 | +;+;+;+;+ | 1348 | six6a | 0 | 8 | 6 | 4 | 8 | 12 |
| ENSDARG000 | 13:13:13:13:1 | 6342410:6418 | 6342689:6418 | +;+;+;+;+ | 1449 | csmd1a | 0 | 0 | 0 | 0 | 6 | 0 |
| ENSDARG000 | 13:13:13:13:1 | 37273010:372 | 37273118:372 | +;+;+;+;+;+;+ | 623 | gphb5 | 0 | 0 | 0 | 0 | 0 | 0 |
| ENSDARG000 | 13:13:13:13:1 | 23488282:234 | 23488658:234 | -:-:-:-:-:-:-:-:-: | 2231 | prim2 | 92 | 407 | 448 | 221 | 391 | 306 |
| ENSDARG000 | 13:13:13:13:1 | 3354895:3354 | 3355055:3355 | -:-:-:-:-:-:-:-:-: | 1500 | prkn | 76 | 37 | 58 | 72 | 53 | 54 |
| ENSDARG000 | 13:13:13:13:1 | 27378857:273 | 27379897:273 | -:-:-:-:-:-:-:-:-: | 3606 | KCNQ5 | 65 | 0 | 232 | 22 | 73 | 32 |
| ENSDARG000 | 13:13:13:13:1 | 7322245:7333 | 7322565:7333 | +;+;+;+;+;+ | 1392 | NA | 10 | 0 | 0 | 6 | 9 | 0 |
| ENSDARG000 | 13:13:13:13:1 | 35983997:359 | 35984239:359 | -;-;-;-;- | 887 | NA | 27 | 1 | 0 | 0 | 1 | 7 |
| ENSDARG000 | 13;13;13 | 27369309:273 | 27369399:273 | +;+;+ | 558 | NA | 3 | 0 | 15 | 2 | 5 | 0 |
| ENSDARG000 | 13:13:13:13:1 | 813579:81572 | 814932:81590 | -:-:-:-:-:-:-:-:-: | 2928 | polh | 247 | 493 | 499 | 342 | 432 | 319 |
| ENSDARG000 | 13:13:13:13:1 | 35995238:359 | 35995316:359 | -;-;-;-;- | 663 | NA | 8 | 0 | 0 | 0 | 2 | 0 |
| ENSDARG000 | 13:13:13:13:1 | 13945218:139 | 13945656:139 | +:+:+:+:+:+:+: | 4402 | EIF2AK3 | 639 | 695 | 1043 | 686 | 957 | 578 |
| ENSDARG000 | 13:13:13:13:1 | 45468403:454 | 45468838:454 | -:-:-:-:-:-:-:-:-: | 3151 | rsrp1 | 4639 | 1442 | 3093 | 3639 | 5300 | 4543 |
| ENSDARG000 | 13:13:13:13:1 | 18073211:180 | 18073761:180 | -:-:-:-:-:-:-:-:-: | 4787 | NA | 4042 | 2981 | 4064 | 4855 | 5856 | 3018 |

| | | | | | | | | | | | | |
|---|---|---|---|---|---|---|---|---|---|---|---|---|
| ENSDARG000 | 13:13:13:13:1 | 3667230:3667 | 3667374:3667 | +:+:+:+:+:+:+: | 1681 | QKI | 384 | 224 | 542 | 224 | 694 | 352 |
| ENSDARG000 | 13 | 7346601 | 7347745 | + | 1145 | NA | 10 | 2 | 2 | 2 | 12 | 4 |
| ENSDARG000 | 13:13:13:13:1 | 9889676:989( | 9889820:989( | -:-:-:-:-:-:-:-:-: | 1296 | NA | 0 | 0 | 29 | 0 | 6 | 0 |
| ENSDARG000 | 13:13:13:13:1 | 30152217:301 | 30152545:301 | -:-:-:-:-:-:-:-:-: | 1788 | ppa1b | 1368 | 1336 | 1906 | 820 | 1273 | 962 |
| ENSDARG000 | 13:13:13:13:1 | 40815012:408 | 40815089:408 | +:+:+:+:+:+:+: | 3095 | prkg1a | 24 | 13 | 9 | 37 | 38 | 24 |
| ENSDARG000 | 13:13:13:13:1 | 36037970:36( | 36040674:36( | -:-:-:-:-:-:-:-:-: | 4103 | lgmn | 7806 | 8680 | 10395 | 9520 | 16506 | 10171 |
| ENSDARG000 | 13:13:13:13:1 | 7387822:7389 | 7388033:739( | +:+:+:+:+:+:+: | 2697 | exoc3l4 | 20 | 2 | 18 | 0 | 8 | 0 |
| ENSDARG000 | 13;13;13;13 | 45476181:454 | 45476511:454 | +;+;+;+ | 1158 | mgst3b | 1575 | 1504 | 1843 | 1779 | 2159 | 1976 |
| ENSDARG000 | 13:13:13:13:1 | 7368994:7372 | 7373090:7372 | -:-:-:-:-:-:-:-:-: | 6388 | NA | 18 | 0 | 7 | 0 | 14 | 0 |
| ENSDARG000 | 13:13:13:13:1 | 37302575:373 | 37302961:373 | -:-:-:-:-:-:-:-:-: | 2955 | KCNH5 | 2 | 0 | 4 | 0 | 0 | 0 |
| ENSDARG000 | 13:13:13:13:1 | 12120664:121 | 12120747:121 | +:+:+:+:+:+:+: | 1674 | NA | 0 | 0 | 0 | 0 | 0 | 0 |
| ENSDARG000 | 13:13:13:13:1 | 31648271:316 | 31648490:316 | +:+:+:+:+:+:+: | 1243 | mnat1 | 126 | 102 | 224 | 108 | 130 | 138 |
| ENSDARG000 | 13:13:13:13:1 | 25211939:252 | 25212557:252 | -:-:-:-:-:-:-:-:-: | 4491 | vcla | 2744 | 1786 | 2017 | 1169 | 1946 | 1726 |
| ENSDARG000 | 13;13;13 | 30163515:301 | 30163833:301 | +;+;+ | 2841 | eif4ebp2 | 5144 | 3865 | 4527 | 4089 | 5175 | 4410 |
| ENSDARG000 | 13:13:13:13:1 | 9896368:9896 | 9896924:9896 | +:+:+:+:+:+:+: | 2952 | SRBD1 | 92 | 105 | 175 | 63 | 148 | 97 |
| ENSDARG000 | 13:13:13:13:1 | 27631926:276 | 27634923:276 | -:-:-:-:-:-:-:-:-: | 7877 | RIMS1 | 28 | 20 | 20 | 6 | 16 | 2 |
| ENSDARG000 | 13:13:13:13:1 | 45485185:454 | 45485284:454 | +:+:+:+:+:+:+: | 1979 | tmem50a | 4364 | 4465 | 4132 | 4328 | 5216 | 5118 |
| ENSDARG000 | 13:13:13:13:1 | 7442023:7444 | 7442171:7444 | +:+:+:+:+:+:+: | 2181 | NA | 151 | 39 | 328 | 50 | 164 | 218 |
| ENSDARG000 | 13;13;13;13 | 41022502:41( | 41022719:41( | +;+;+;+ | 1286 | dkk1a | 1259 | 5294 | 7785 | 4835 | 11777 | 6506 |
| ENSDARG000 | 13;13;13;13 | 3874415:3875 | 3875221:3875 | -;-;-;- | 1276 | NA | 99 | 97 | 67 | 52 | 99 | 105 |
| ENSDARG000 | 13:13:13:13:1 | 12174937:121 | 12175016:121 | +:+:+:+:+:+:+: | 2168 | GABRG1 | 0 | 0 | 0 | 0 | 0 | 0 |
| ENSDARG000 | 13:13:13:13:1 | 50064464:50( | 50067925:50( | -:-:-:-:-:-:-:-:-: | 7214 | NID1 | 3452 | 1128 | 3124 | 2129 | 2846 | 1648 |
| ENSDARG000 | 13:13:13:13:1 | 829585:83024 | 829724:83029 | +;+;+;+;+;+ | 1053 | gsta.2 | 63 | 42 | 72 | 54 | 86 | 41 |
| ENSDARG000 | 13:13:13:13:1 | 7462715:7462 | 7462912:7463 | -:-:-:-:-:-:-:-:-: | 1720 | NA | 6 | 0 | 0 | 0 | 7 | 8 |
| ENSDARG000 | 13:13:13:13:1 | 23623346:236 | 23623427:236 | +;+;+;+;+;+ | 996 | il22ra2 | 0 | 0 | 0 | 0 | 0 | 2 |
| ENSDARG000 | 13:13:13:13:1 | 30169233:301 | 30169813:301 | +:+:+:+:+:+:+: | 5342 | pald1b | 9 | 1 | 20 | 21 | 23 | 20 |
| ENSDARG000 | 13:13:13:13:1 | 37421333:374 | 37422839:374 | -:-:-:-:-:-:-:-:-: | 3154 | ppp2r5eb | 1370 | 1393 | 1701 | 1276 | 1587 | 1337 |
| ENSDARG000 | 13:13:13:13:1 | 844143:84415 | 844261:84426 | +:+:+:+:+:+:+: | 1044 | NA | 984 | 1099 | 252 | 848 | 862 | 162 |
| ENSDARG000 | 13:13:13:13:1 | 852289:85229 | 852403:8524( | -;-;-;-;-;-;-;-;- | 611 | tmem14a | 67 | 159 | 107 | 108 | 157 | 124 |
| ENSDARG000 | 13;13 | 12206937:122 | 12206980:122 | +;+ | 699 | NA | 0 | 4 | 0 | 0 | 0 | 0 |
| ENSDARG000 | 13:13:13:13:1 | 30225262:302 | 30225615:302 | +:+:+:+:+:+:+: | 3569 | rps24 | 11296 | 11867 | 17818 | 14780 | 10643 | 18326 |
| ENSDARG000 | 13:13:13:13:1 | 7477500:7477 | 7477540:7477 | +:+:+:+:+:+:+: | 7161 | NA | 0 | 0 | 0 | 0 | 0 | 0 |
| ENSDARG000 | 13:13:13:13:1 | 41125358:411 | 41125918:411 | -:-:-:-:-:-:-:-:-: | 7300 | pcdh15a | 0 | 0 | 17 | 0 | 2 | 6 |
| ENSDARG000 | 13:13:13:13:1 | 23634817:236 | 23635433:236 | -:-:-:-:-:-:-:-:-: | 4213 | MAP3K21 | 759 | 418 | 737 | 561 | 1037 | 243 |
| ENSDARG000 | 13:13:13:13:1 | 22476742:224 | 22476884:224 | +:+:+:+:+:+:+: | 6806 | ldb3a | 0 | 6 | 8 | 1 | 0 | 0 |
| ENSDARG000 | 13;13 | 12362447:123 | 12362494:123 | +;+ | 601 | NA | 0 | 5 | 0 | 0 | 1 | 4 |
| ENSDARG000 | 13:13:13:13:1 | 12299997:123 | 12300531:123 | +:+:+:+:+:+:+: | 6580 | NA | 0 | 2 | 0 | 0 | 0 | 14 |
| ENSDARG000 | 13:13:13:13:1 | 33320063:333 | 33320400:333 | -;-;-;-;-;- | 777 | NA | 6 | 0 | 0 | 0 | 0 | 0 |
| ENSDARG000 | 13:13:13:13:1 | 30215765:302 | 30216305:302 | +;+;+;+;+;+;+ | 1434 | QRFPR | 0 | 0 | 0 | 0 | 0 | 0 |
| ENSDARG000 | 13;13;13;13 | 41124594:411 | 41125918:411 | -;-;-;- | 1555 | NA | 0 | 0 | 2 | 0 | 0 | 0 |
| ENSDARG000 | 13;13;13 | 37469093:374 | 37471026:374 | -;-;- | 2639 | wdr89 | 154 | 201 | 246 | 156 | 132 | 192 |
| ENSDARG000 | 13:13:13:13:1 | 3915260:3916 | 3916269:3916 | -:-:-:-:-:-:-:-:-: | 2863 | ncoa4 | 1845 | 1617 | 2413 | 1523 | 1932 | 727 |
| ENSDARG000 | 13:13:13:13:1 | 3895363:3895 | 3895462:3895 | +:+:+:+:+:+:+: | 972 | NA | 8 | 1 | 8 | 23 | 2 | 0 |
| ENSDARG000 | 13:13:13:13:1 | 36053985:36( | 36055069:36( | -:-:-:-:-:-:-:-:-: | 8196 | PCNX1 | 851 | 610 | 1050 | 620 | 828 | 790 |
| ENSDARG000 | 13;13;13;13 | 33342455:333 | 33343047:333 | -;-;-;- | 1366 | zgc:172120 | 42 | 0 | 0 | 2 | 2 | 0 |
| ENSDARG000 | 13:13:13:13:1 | 50518815:505 | 50520889:505 | -:-:-:-:-:-:-:-:-: | 3517 | blnk | 41 | 55 | 137 | 58 | 121 | 123 |
| ENSDARG000 | 13:13:13:13:1 | 3880882:3881 | 3881310:3881 | +:+:+:+:+:+:+: | 1089 | NA | 0 | 0 | 4 | 0 | 10 | 10 |
| ENSDARG000 | 13:13:13:13:1 | 25297257:252 | 25298518:252 | -:-:-:-:-:-:-:-:-: | 2408 | plaua | 151 | 50 | 278 | 104 | 237 | 173 |
| ENSDARG000 | 13;13;13 | 50520668:505 | 50520723:505 | +;+;+ | 336 | NA | 0 | 0 | 0 | 0 | 0 | 0 |
| ENSDARG000 | 13:13:13:13:1 | 37491804:374 | 37494280:374 | -;-;-;-;-;-;- | 3239 | SGPP1 | 362 | 433 | 292 | 114 | 583 | 161 |
| ENSDARG000 | 13:13:13:13:1 | 25286538:253 | 25286647:253 | +:+:+:+:+:+:+: | 4412 | chuk | 2794 | 2823 | 2698 | 2477 | 2821 | 2792 |
| ENSDARG000 | 13:13:13:13:1 | 45496766:455 | 45497420:455 | -:-:-:-:-:-:-:-:-: | 2073 | rhd | 10 | 0 | 5 | 77 | 34 | 34 |
| ENSDARG000 | 13:13:13:13:1 | 31687973:316 | 31688203:316 | +:+:+:+:+:+:+: | 2039 | slc38a6 | 184 | 130 | 156 | 138 | 261 | 209 |
| ENSDARG000 | 13:13:13:13:1 | 33368140:333 | 33368666:333 | +:+:+:+:+:+:+: | 5076 | brf1a | 68 | 78 | 192 | 18 | 52 | 80 |
| ENSDARG000 | 13:13:13:13:1 | 37528497:375 | 37529035:375 | -;-;-;-;-;-;-;- | 1069 | NA | 0 | 2 | 0 | 0 | 0 | 0 |
| ENSDARG000 | 13:13:13:13:1 | 18128275:181 | 18128454:181 | -:-:-:-:-:-:-:-:-: | 2121 | alox5a | 38 | 20 | 59 | 27 | 248 | 70 |
| ENSDARG000 | 13:13:13:13:1 | 31681988:316 | 31682151:316 | -;-;-;-;- | 1586 | trmt5 | 126 | 132 | 267 | 123 | 247 | 261 |
| ENSDARG000 | 13:13:13:13:1 | 1070705:1073 | 1071648:1073 | -:-:-:-:-:-:-:-:-: | 1914 | plek | 66 | 126 | 109 | 52 | 247 | 122 |
| ENSDARG000 | 13;13;13 | 1089942:1095 | 1090278:1095 | +;+;+ | 1015 | cnrip1b | 0 | 0 | 0 | 2 | 8 | 0 |
| ENSDARG000 | 13:13:13:13:1 | 27768898:277 | 27769004:277 | +:+:+:+:+:+:+: | 1623 | asrgl1 | 160 | 210 | 212 | 321 | 185 | 366 |
| ENSDARG000 | 13:13:13:13:1 | 12378074:123 | 12378424:123 | -:-:-:-:-:-:-:-:-: | 1173 | commd8 | 610 | 518 | 572 | 399 | 512 | 446 |
| ENSDARG000 | 13:13:13:13:1 | 14006118:14( | 14006585:14( | +:+:+:+:+:+:+: | 4974 | atrn | 2344 | 2043 | 4699 | 2655 | 8139 | 1884 |
| ENSDARG000 | 13:13:13:13:1 | 45524475:455 | 45524680:455 | +:+:+:+:+:+:+: | 2717 | maco1b | 572 | 475 | 659 | 293 | 594 | 565 |
| ENSDARG000 | 13:13:13:13:1 | 23669659:236 | 23670128:236 | +:+:+:+:+:+:+: | 10873 | NA | 417 | 89 | 177 | 126 | 357 | 90 |
| ENSDARG000 | 13:13:13:13:1 | 41887575:418 | 41888026:418 | -:-:-:-:-:-:-:-:-: | 982 | NA | 118 | 51 | 106 | 45 | 86 | 82 |
| ENSDARG000 | 13:13:13:13:1 | 7555720:7556 | 7557634:7557 | -:-:-:-:-:-:-:-:-: | 2504 | pitx3 | 0 | 0 | 2 | 0 | 0 | 0 |
| ENSDARG000 | 13:13:13:13:1 | 1100197:1106 | 1100350:1107 | +;+;+;+;+;+ | 1203 | ppp3r1a | 702 | 499 | 719 | 490 | 917 | 690 |
| ENSDARG000 | 13:13:13:13:1 | 33611703:336 | 33612418:336 | -:-:-:-:-:-:-:-:-: | 4890 | rrbp1a | 718 | 716 | 1241 | 616 | 1413 | 610 |
| ENSDARG000 | 13:13:13:13:1 | 3938526:3941 | 3938777:3942 | +:+:+:+:+:+:+: | 1948 | yipf3 | 1187 | 1131 | 1448 | 933 | 1238 | 1145 |
| ENSDARG000 | 13:13:13:13:1 | 23728176:237 | 23728537:237 | -;-;-;-;-;- | 1185 | rgs17 | 245 | 262 | 275 | 220 | 313 | 232 |
| ENSDARG000 | 13;13;13;13 | 33606739:336 | 33607041:336 | +;+;+;+ | 1098 | cfl1l | 558 | 32 | 12 | 62 | 100 | 140 |
| ENSDARG000 | 13:13:13:13:1 | 18179096:181 | 18179928:181 | -:-:-:-:-:-:-:-:-: | 1767 | slc25a16 | 275 | 260 | 308 | 58 | 324 | 264 |
| ENSDARG000 | 13;13 | 37543498:375 | 37543811:375 | -;- | 494 | NA | 4 | 2 | 12 | 23 | 17 | 8 |
| ENSDARG000 | 13;13 | 27785329:278 | 27785868:278 | +;+ | 2855 | NA | 12 | 14 | 26 | 16 | 36 | 2 |
| ENSDARG000 | 13:13:13:13:1 | 45709380:457 | 45710149:457 | +:+:+:+:+:+:+: | 5585 | NA | 4 | 4 | 0 | 8 | 8 | 2 |
| ENSDARG000 | 13:13:13:13:1 | 27900446:279 | 27902261:279 | -:-:-:-:-:-:-:-:-: | 2595 | ogfrl1 | 8 | 38 | 22 | 8 | 2 | 49 |
| ENSDARG000 | 13;13;13 | 41910342:419 | 41911985:419 | -;-;- | 2001 | cisd1 | 714 | 715 | 862 | 850 | 747 | 881 |
| ENSDARG000 | 13:13:13:13:1 | 37586491:375 | 37587370:375 | -:-:-:-:-:-:-:-:-: | 2163 | snapc1b | 169 | 172 | 279 | 200 | 235 | 182 |
| ENSDARG000 | 13;13;13;13 | 27951688:279 | 27952513:279 | +;+;+;+ | 1674 | b3gat2 | 14 | 50 | 78 | 9 | 35 | 567 |
| ENSDARG000 | 13:13:13:13:1 | 7575563:7578 | 7575718:7578 | +:+:+:+:+:+:+: | 9570 | GBF1 | 1599 | 1176 | 2070 | 941 | 2105 | 1467 |
| ENSDARG000 | 13:13:13:13:1 | 36135122:361 | 36135618:361 | -:-:-:-:-:-:-:-:-: | 1222 | med6 | 284 | 286 | 306 | 208 | 320 | 314 |
| ENSDARG000 | 13:13:13:13:1 | 1125185:1126 | 1125420:1126 | -:-:-:-:-:-:-:-:-: | 1222 | pno1 | 2340 | 1928 | 3142 | 2698 | 1865 | 2952 |
| ENSDARG000 | 13:13:13:13:1 | 18202703:182 | 18204199:182 | +:+:+:+:+:+:+: | 5568 | TET1 | 474 | 262 | 687 | 409 | 696 | 975 |
| ENSDARG000 | 13:13:13:13:1 | 28009727:28( | 28010786:28( | -:-:-:-:-:-:-:-:-: | 2547 | smap1 | 1797 | 1587 | 1831 | 1290 | 1791 | 1401 |
| ENSDARG000 | 13:13:13:13:1 | 3954540:3954 | 3954986:3954 | +:+:+:+:+:+:+: | 1504 | LRRC73 | 4 | 0 | 0 | 0 | 2 | 1 |
| ENSDARG000 | 13:13:13:13:1 | 41917606:419 | 41917847:419 | +:+:+:+:+:+:+: | 3529 | polr1b | 301 | 369 | 639 | 168 | 579 | 414 |
| ENSDARG000 | 13:13:13:13:1 | 31716820:317 | 31717328:317 | +:+:+:+:+:+:+: | 3399 | prkcha | 163 | 40 | 34 | 84 | 125 | 186 |
| ENSDARG000 | 13:13:13:13:1 | 37605896:376 | 37606112:376 | -;-;-;-;-;-;-;- | 1222 | ifi27.1 | 1253 | 1144 | 673 | 298 | 658 | 428 |
| ENSDARG000 | 13:13:13:13:1 | 25380432:253 | 25380531:253 | +;+;+;+;+;+;+ | 1486 | gsto1 | 813 | 1464 | 1835 | 2267 | 1565 | 1776 |
| ENSDARG000 | 13:13:13:13:1 | 12389849:123 | 12390320:123 | +:+:+:+:+:+:+: | 6288 | atp10d | 1749 | 1223 | 1136 | 898 | 1384 | 1310 |
| ENSDARG000 | 13:13:13:13:1 | 36144341:361 | 36144357:361 | +:+:+:+:+:+:+: | 3064 | TTC9 | 48 | 117 | 292 | 327 | 118 | 150 |
| ENSDARG000 | 13;13;13;13 | 14186985:141 | 14187044:141 | +;+;+;+ | 540 | NA | 0 | 0 | 1 | 1 | 0 | 0 |
| ENSDARG000 | 13:13:13:13:1 | 14191293:141 | 14192712:141 | -:-:-:-:-:-:-:-:-: | 4143 | gfra4a | 0 | 2 | 0 | 0 | 0 | 0 |
| ENSDARG000 | 13:13:13:13:1 | 3979341:3982 | 3984377:3984 | -:-:-:-:-:-:-:-:-: | 6429 | NA | 4819 | 3096 | 4142 | 3092 | 5325 | 2265 |
| ENSDARG000 | 13:13:13:13:1 | 45786113:457 | 45786280:457 | -;-;-;-;-;-;-;- | 1119 | FNDC5 | 0 | 0 | 0 | 0 | 0 | 0 |
| ENSDARG000 | 13;13;13 | 28264834:282 | 28265253:282 | -;-;- | 688 | sdhaf4 | 140 | 199 | 200 | 208 | 244 | 194 |
| ENSDARG000 | 13:13:13:13:1 | 45840427:458 | 45840490:458 | +:+:+:+:+:+:+: | 1269 | NA | 0 | 0 | 0 | 0 | 0 | 0 |
| ENSDARG000 | 13;13;13 | 28267997:282 | 28269412:282 | -;-;- | 2223 | tlx1 | 12 | 40 | 28 | 38 | 50 | 191 |
| ENSDARG000 | 13;13;13 | 23843712:238 | 23843987:238 | +;+;+ | 1378 | oprm1 | 105 | 44 | 223 | 51 | 189 | 78 |
| ENSDARG000 | 13;13 | 23897975:239 | 23898075:239 | +;+ | 564 | sf3b5 | 399 | 456 | 619 | 374 | 546 | 348 |
| ENSDARG000 | 13:13:13:13:1 | 7753049:7755 | 7753739:7755 | -:-:-:-:-:-:-:-:-: | 2372 | macroh2a2 | 2463 | 3320 | 4269 | 2437 | 3748 | 3106 |
| ENSDARG000 | 13;13;13;13 | 4205724:4205 | 4205789:4206 | +;+;+;+ | 1036 | DLK2 | 175 | 62 | 78 | 355 | 71 | 61 |
| ENSDARG000 | 13:13:13:13:1 | 37609458:376 | 37609777:376 | -:-:-:-:-:-:-:-:-: | 1380 | ifi27.3 | 189 | 347 | 36 | 0 | 0 | 2 |
| ENSDARG000 | 13:13:13:13:1 | 31757331:317 | 31757359:317 | +:+:+:+:+:+:+: | 2233 | hif1aa | 542 | 1080 | 1175 | 787 | 1208 | 452 |
| ENSDARG000 | 13:13:13:13:1 | 1142724:1143 | 1143028:1143 | -;-;-;-;-;- | 942 | psmb1 | 3015 | 3857 | 3582 | 1976 | 3110 | 2346 |
| ENSDARG000 | 13;13;13 | 1155536:1163 | 1155786:1163 | +;+;+ | 3265 | perp | 1287 | 1034 | 954 | 871 | 1045 | 590 |
| ENSDARG000 | 13:13:13:13:1 | 28417297:284 | 28417434:284 | +:+:+:+:+:+:+: | 1662 | slc2a15a | 213 | 300 | 1225 | 136 | 718 | 1203 |
| ENSDARG000 | 13:13:13:13:1 | 1131748:1132 | 1132023:1132 | +:+:+:+:+:+:+: | 1825 | dnaaf10 | 179 | 300 | 178 | 174 | 274 | 104 |
| ENSDARG000 | 13;13 | 28305798:283 | 28306419:283 | -;- | 1459 | lbx1a | 22 | 72 | 52 | 39 | 50 | 118 |
| ENSDARG000 | 13:13:13:13:1 | 28406913:284 | 28407100:284 | +:+:+:+:+:+:+: | 3033 | NA | 20 | 18 | 26 | 36 | 42 | 68 |
| ENSDARG000 | 13:13:13:13:1 | 28276898:282 | 28277272:282 | -;-;-;-;-;-;-;- | 1032 | NA | 2 | 25 | 23 | 16 | 23 | 92 |
| ENSDARG000 | 13:13:13:13:1 | 45931232:459 | 45931723:459 | -;-;-;-;-;- | 1161 | NA | 4 | 6 | 0 | 1 | 16 | 10 |
| ENSDARG000 | 13 | 45967179 | 45968256 | + | 1078 | olig4 | 0 | 0 | 0 | 2 | 2 | 0 |
| ENSDARG000 | 13:13:13:13:1 | 14902318:149 | 14902705:149 | -;-;-;-;-;-;-;- | 1085 | ap5s1 | 481 | 608 | 565 | 380 | 540 | 469 |
| ENSDARG000 | 13;13 | 41944138:419 | 41945971:419 | -;- | 2026 | nanp | 58 | 56 | 118 | 46 | 64 | 62 |
| ENSDARG000 | 13:13:13:13:1 | 4218329:4218 | 4218595:4218 | -;-;-;-;-;-;-;- | 667 | dnph1 | 54 | 144 | 329 | 88 | 154 | 120 |
| ENSDARG000 | 13:13:13:13:1 | 36164639:361 | 36164759:361 | -:-:-:-:-:-:-:-:-: | 1967 | MAP3K9 | 0 | 0 | 1 | 0 | 0 | 0 |
| ENSDARG000 | 13:13:13:13:1 | 23919480:239 | 23919556:239 | -:-:-:-:-:-:-:-:-: | 2356 | NA | 1191 | 955 | 1057 | 972 | 978 | 1161 |
| ENSDARG000 | 13:13:13:13:1 | 45980163:459 | 45980340:459 | +:+:+:+:+:+:+: | 2139 | bsdc1 | 1756 | 1308 | 1656 | 1569 | 1676 | 1504 |
| ENSDARG000 | 13:13:13:13:1 | 42011285:42( | 42012456:42( | +;+;+;+;+ | 3386 | cyp1b1 | 2 | 0 | 4 | 0 | 4 | 8 |
| ENSDARG000 | 13:13:13:13:1 | 25396896:253 | 25396972:253 | +:+:+:+:+:+:+: | 1088 | gsto2 | 593 | 535 | 537 | 778 | 524 | 802 |
| ENSDARG000 | 13:13:13:13:1 | 36196778:362 | 36197360:362 | -;-;-;-;-;- | 2960 | slc8a3 | 0 | 4 | 4 | 0 | 10 | 6 |
| ENSDARG000 | 13:13:13:13:1 | 18311410:183 | 18311538:183 | +:+:+:+:+:+:+: | 1994 | eif4e1c | 522 | 1286 | 1168 | 390 | 934 | 608 |
| ENSDARG000 | 13;13;13;13 | 42245221:422 | 42245551:422 | +;+;+;+ | 1187 | opn3 | 4 | 0 | 2 | 24 | 0 | 6 |
| ENSDARG000 | 13:13:13:13:1 | 4225173:4225 | 4225250:4225 | +:+:+:+:+:+:+: | 1917 | mea1 | 388 | 291 | 262 | 344 | 312 | 178 |
| ENSDARG000 | 13:13:13:13:1 | 4232568:4238 | 4234875:4238 | -:-:-:-:-:-:-:-:-: | 4065 | ppp2r5d | 2833 | 2139 | 1870 | 1653 | 2154 | 1505 |
| ENSDARG000 | 13:13:13:13:1 | 23996070:239 | 23996775:239 | -:-:-:-:-:-:-:-:-: | 2926 | cog2 | 466 | 482 | 664 | 414 | 768 | 455 |
| ENSDARG000 | 13:13:13:13:1 | 23988442:239 | 23988487:239 | +;+;+;+;+ | 1676 | agt | 34 | 38 | 88 | 82 | 28 | 39 |
| ENSDARG000 | 13;13;13;13 | 36311894:363 | 36312705:363 | -;-;-;- | 1187 | cox16 | 140 | 203 | 180 | 106 | 142 | 156 |
| ENSDARG000 | 13:13:13:13:1 | 14915344:149 | 14916512:149 | -:-:-:-:-:-:-:-:-: | 3191 | cdc25b | 186 | 449 | 643 | 301 | 378 | 320 |
| ENSDARG000 | 13:13:13:13:1 | 37617518:376 | 37618152:376 | -:-:-:-:-:-:-:-:-: | 991 | ifi27.2 | 1237 | 2091 | 110 | 48 | 40 | 60 |
| ENSDARG000 | 13:13:13:13:1 | 30421472:304 | 30421525:304 | +:+:+:+:+:+:+: | 6486 | zmiz1a | 664 | 683 | 589 | 436 | 563 | 871 |
| ENSDARG000 | 13:13:13:13:1 | 1182257:1182 | 1182563:1182 | +:+:+:+:+:+:+: | 2333 | TNFAIP3 | 1725 | 915 | 2430 | 2597 | 1737 | 1184 |
| ENSDARG000 | 13:13:13:13:1 | 8131622:8135 | 8131648:8135 | +:+:+:+:+:+:+: | 3680 | NA | 0 | 50 | 71 | 18 | 3 | 45 |
| ENSDARG000 | 13:13:13:13:1 | 8168712:8175 | 8172815:8175 | -;-;-;-;-;-;-;- | 5892 | NA | 50 | 8 | 134 | 70 | 124 | 56 |
| ENSDARG000 | 13:13:13:13:1 | 9237493:9238 | 9237822:9238 | +;+;+;+;+;+;+ | 1641 | NA | 0 | 0 | 0 | 0 | 0 | 0 |
| ENSDARG000 | 13:13:13:13:1 | 8255106:8255 | 8255427:8255 | +:+:+:+:+:+:+: | 3287 | hells | 244 | 980 | 1276 | 302 | 1087 | 556 |
| ENSDARG000 | 13:13:13:13:1 | 9240925:924( | 9241526:9241 | -:-:-:-:-:-:-:-:-: | 1443 | HTRA2 | 0 | 0 | 0 | 1 | 1 | 0 |
| ENSDARG000 | 13:13:13:13:1 | 31805824:318 | 31810631:318 | -;-;-;-;-;-;- | 5579 | SERTAD4 | 1098 | 1642 | 863 | 1446 | 647 | 1576 |
| ENSDARG000 | 13;13;13;13 | 33655404:336 | 33656113:336 | +;+;+;+ | 1304 | mgme1 | 28 | 100 | 74 | 43 | 88 | 49 |
| ENSDARG000 | 13:13:13:13:1 | 33669936:336 | 33670499:336 | -;-;-;-;- | 826 | zgc:163030 | 37 | 4 | 19 | 97 | 81 | 0 |
| ENSDARG000 | 13;13;13 | 1345276:1347 | 1345910:1347 | -;-;- | 884 | si:ch73-52p7. | 52 | 54 | 80 | 4 | 164 | 63 |
| ENSDARG000 | 13:13:13:13:1 | 33665305:336 | 33665525:336 | -;-;-;-;- | 896 | NA | 92 | 48 | 90 | 31 | 58 | 52 |
| ENSDARG000 | 13:13:13:13:1 | 46014087:46( | 46015606:46( | -:-:-:-:-:-:-:-:-: | 2523 | NA | 0 | 0 | 2 | 12 | 4 | 2 |
| ENSDARG000 | 13:13:13:13:1 | 33675556:336 | 33675746:336 | -:-:-:-:-:-:-:-:-: | 1678 | cst3 | 2342 | 2049 | 1953 | 2626 | 2753 | 3717 |
| ENSDARG000 | 13:13:13:13:1 | 33658871:336 | 33659748:336 | -:-:-:-:-:-:-:-:-: | 1671 | si:dkev-76k16 | 4 | 33 | 87 | 65 | 93 | 101 |
| ENSDARG000 | 13:13:13:13:1 | 33645413:336 | 33646735:336 | -:-:-:-:-:-:-:-:-: | 2582 | snx5 | 1366 | 1519 | 1895 | 1425 | 1826 | 2039 |
| ENSDARG000 | 13;13;13 | 14976108:149 | 14976511:149 | +;+;+ | 991 | noto | 0 | 0 | 0 | 0 | 0 | 0 |
| ENSDARG000 | 13;13 | 25403413:254 | 25405250:254 | -;- | 1988 | itprip | 453 | 427 | 599 | 654 | 412 | 600 |
| ENSDARG000 | 13:13:13:13:1 | 31844011:318 | 31844324:318 | -:-:-:-:-:-:-:-:-: | 1835 | SYT14 | 31 | 2 | 14 | 10 | 8 | 2 |
| ENSDARG000 | 13;13;13;13 | 37623276:376 | 37623815:376 | -;-;-;- | 1082 | ifi27.4 | 526 | 593 | 60 | 102 | 44 | 252 |
| ENSDARG000 | 13:13:13:13:1 | 42309688:423 | 42309932:423 | +:+:+:+:+:+:+: | 3973 | ide | 1524 | 1641 | 1752 | 808 | 1601 | 1420 |
| ENSDARG000 | 13;13;13 | 25410594:254 | 25410880:254 | +;+;+ | 996 | CALHM3 | 0 | 0 | 0 | 0 | 0 | 3 |
| ENSDARG000 | 13;13;13 | 15004398:15( | 15004956:15( | +;+;+ | 1235 | emx1 | 0 | 0 | 0 | 0 | 4 | 0 |
| ENSDARG000 | 13:13:13:13:1 | 24022963:24( | 24023386:24( | +;+;+;+;+;+;+ | 1671 | pgbd5 | 1016 | 865 | 487 | 408 | 559 | 483 |
| ENSDARG000 | 13:13:13:13:1 | 10023256:10( | 10023329:10( | +;+;+;+;+ | 848 | NA | 22 | 35 | 44 | 18 | 25 | 6 |
| ENSDARG000 | 13;13 | 46177478:461 | 46177764:461 | +;+ | 1694 | marcksl1a | 1902 | 3468 | 6036 | 6146 | 4932 | 7282 |
| ENSDARG000 | 13;13 | 8272043:8277 | 8272212:8279 | -;- | 2335 | NA | 14 | 17 | 16 | 0 | 25 | 6 |
| ENSDARG000 | 13:13:13:13:1 | 12456412:124 | 12456543:124 | +:+:+:+:+:+:+: | 1517 | dapp1 | 2 | 8 | 9 | 0 | 18 | 24 |
| ENSDARG000 | 13:13:13:13:1 | 25412307:254 | 25412565:254 | +;+;+;+;+ | 3309 | CALHM1 | 0 | 0 | 0 | 0 | 0 | 0 |
| ENSDARG000 | 13:13:13:13:1 | 36321088:363 | 36321718:363 | -:-:-:-:-:-:-:-:-: | 4390 | actn1 | 1693 | 1599 | 1128 | 1414 | 1680 | 1488 |
| ENSDARG000 | 13:13:13:13:1 | 25421531:254 | 25421960:254 | +;+;+;+;+ | 5279 | calhm2.1 | 767 | 751 | 390 | 598 | 819 | 715 |

| | | | | | | | | | | | | |
|---|---|---|---|---|---|---|---|---|---|---|---|---|
| ENSDARG000 | 13:13:13:13:1 | 25418561:254 | 25419106:254 | +:+:+:+:+:+:+: | 1183 | CALHM2 | 0 | 0 | 0 | 0 | 0 | 0 |
| ENSDARG000 | 13:13:13:13:1 | 1357552:1368 | 1357746:1368 | +:+:+:+:+:+:+: | 1497 | ifngr1l | 569 | 326 | 372 | 425 | 720 | 432 |
| ENSDARG000 | 13;13;13 | 37633904:376 | 37634654:376 | -;-;- | 851 | NA | 2 | 9 | 12 | 0 | 4 | 6 |
| ENSDARG000 | 13:13:13:13:1 | 15020822:150 | 15021281:150 | -:-:-:-:-:-:-:-: | 1425 | sfxn5a | 66 | 3 | 58 | 30 | 148 | 2 |
| ENSDARG000 | 13:13:13:13:1 | 10080027:100 | 10080613:100 | -;-;-;-;-;-;-;- | 1443 | NA | 0 | 0 | 0 | 0 | 0 | 0 |
| ENSDARG000 | 13:13:13:13:1 | 8281223:8285 | 8282126:8285 | -:-:-:-:-:-:-:-: | 2808 | atl2 | 799 | 708 | 763 | 396 | 932 | 535 |
| ENSDARG000 | 13:13:13:13:1 | 10287702:103 | 10290603:103 | -;-;-;-;-;- | 3954 | NA | 0 | 0 | 0 | 0 | 0 | 0 |
| ENSDARG000 | 13;13 | 10258007:102 | 10259087:102 | -;- | 1889 | six3a | 0 | 0 | 0 | 0 | 0 | 0 |
| ENSDARG000 | 13;13;13;13 | 10164234:101 | 10164365:101 | +;+;+;+ | 648 | NA | 0 | 0 | 0 | 0 | 0 | 0 |
| ENSDARG000 | 13;13 | 10232695:102 | 10233507:102 | +;+ | 1609 | six2a | 10 | 0 | 4 | 2 | 10 | 0 |
| ENSDARG000 | 13:13:13:13:1 | 1381953:1386 | 1382310:1386 | +;+;+;+;+;+;+ | 1521 | rab23 | 249 | 234 | 328 | 92 | 254 | 137 |
| ENSDARG000 | 13;13 | 4332783:4335 | 4333055:4335 | +;+ | 589 | NA | 0 | 0 | 0 | 0 | 0 | 0 |
| ENSDARG000 | 13:13:13:13:1 | 4307583:4307 | 4311941:4307 | -:-:-:-:-:-:-:-: | 7641 | NA | 1820 | 1569 | 2588 | 1713 | 2496 | 2011 |
| ENSDARG000 | 13:13:13:13:1 | 30572172:305 | 30572432:305 | +:+:+:+:+:+:+: | 1032 | ppifa | 99 | 61 | 99 | 90 | 88 | 76 |
| ENSDARG000 | 13:13:13:13:1 | 31925254:319 | 31925905:319 | -:-:-:-:-:-:-:-: | 2951 | utp25 | 587 | 552 | 926 | 266 | 642 | 541 |
| ENSDARG000 | 13:13:13:13:1 | 9258379:9259 | 9258708:9259 | +;+;+;+;+;+;+ | 1641 | NA | 0 | 0 | 0 | 0 | 0 | 0 |
| ENSDARG000 | 13:13:13:13:1 | 28495419:284 | 28496014:284 | +:+:+:+:+:+:+: | 2238 | fgf8a | 0 | 0 | 0 | 0 | 2 | 0 |
| ENSDARG000 | 13;13;13 | 1404924:1406 | 1405436:1406 | -;-;- | 844 | bag2 | 31 | 10 | 11 | 0 | 44 | 2 |
| ENSDARG000 | 13;13 | 28500327:285 | 28501801:285 | -;- | 1697 | NA | 0 | 0 | 0 | 0 | 0 | 0 |
| ENSDARG000 | 13:13:13:13:1 | 28512530:285 | 28513291:285 | +:+:+:+:+:+:+: | 2418 | fbxw4 | 299 | 188 | 179 | 154 | 296 | 155 |
| ENSDARG000 | 13:13:13:13:1 | 32144370:321 | 32144629:321 | +;+;+;+;+ | 1738 | osr1 | 54 | 2 | 8 | 2 | 38 | 14 |
| ENSDARG000 | 13:13:13:13:1 | 15091264:150 | 15091521:150 | -;-;-;-;- | 2100 | NA | 0 | 8 | 38 | 0 | 20 | 14 |
| ENSDARG000 | 13:13:13:13:1 | 37638758:376 | 37639176:376 | -:-:-:-:-:-:-:-: | 693 | ifi27.6 | 254 | 551 | 46 | 12 | 46 | 20 |
| ENSDARG000 | 13:13:13:13:1 | 4338882:4342 | 4339762:4342 | -:-:-:-:-:-:-:-: | 6875 | NA | 1070 | 833 | 1623 | 810 | 1404 | 1248 |
| ENSDARG000 | 13:13:13:13:1 | 24068577:240 | 24070988:240 | -:-:-:-:-:-:-:-: | 4104 | galnt2 | 1148 | 853 | 1261 | 529 | 990 | 1119 |
| ENSDARG000 | 13:13:13:13:1 | 4368504:4370 | 4368629:4370 | +:+:+:+:+:+:+: | 2548 | sft2d1 | 2087 | 2188 | 2298 | 2417 | 2115 | 2598 |
| ENSDARG000 | 13:13:13:13:1 | 25428677:254 | 25429411:254 | +:+:+:+:+:+:+: | 4165 | NA | 933 | 311 | 378 | 550 | 427 | 625 |
| ENSDARG000 | 13:13:13:13:1 | 9261815:9261 | 9262416:9262 | -:-:-:-:-:-:-:-: | 1446 | HTRA2 | 5 | 1 | 4 | 5 | 9 | 6 |
| ENSDARG000 | 13:13:13:13:1 | 37645572:376 | 37646163:376 | -;-;-;-;-;-;- | 860 | ifi27.7 | 505 | 597 | 282 | 270 | 212 | 346 |
| ENSDARG000 | 13;13;13;13 | 25443984:254 | 25444043:254 | +;+;+;+ | 211 | NA | 0 | 0 | 0 | 0 | 0 | 0 |
| ENSDARG000 | 13:13:13:13:1 | 18331509:183 | 18331550:183 | +:+:+:+:+:+:+: | 3280 | zgc:110319 | 276 | 248 | 314 | 255 | 311 | 400 |
| ENSDARG000 | 13:13:13:13:1 | 12479264:124 | 12479572:124 | -:-:-:-:-:-:-:-: | 1537 | lamtor3 | 662 | 446 | 612 | 423 | 531 | 586 |
| ENSDARG000 | 13:13:13:13:1 | 32446169:324 | 32446375:324 | +;+;+;+;+;+;+ | 2207 | nt5c1ba | 0 | 0 | 2 | 0 | 0 | 7 |
| ENSDARG000 | 13:13:13:13:1 | 46298560:462 | 46300912:462 | -:-:-:-:-:-:-:-: | 5768 | fgfr2 | 225 | 32 | 312 | 130 | 778 | 160 |
| ENSDARG000 | 13:13:13:13:1 | 18332685:183 | 18333650:183 | -;-;-;-;-;-;- | 1668 | si:dkev-228d1 | 8 | 0 | 0 | 0 | 2 | 0 |
| ENSDARG000 | 13;13 | 32454262:324 | 32454640:324 | +;+ | 1638 | rdh14a | 505 | 377 | 337 | 382 | 342 | 654 |
| ENSDARG000 | 13:13:13:13:1 | 15182200:152 | 15182240:152 | +:+:+:+:+:+:+: | 2570 | pank2 | 1148 | 716 | 1081 | 1240 | 1178 | 1857 |
| ENSDARG000 | 13 | 25449681 | 25450725 | + | 1045 | atoh7 | 0 | 2 | 0 | 0 | 0 | 0 |
| ENSDARG000 | 13:13:13:13:1 | 15182149:151 | 15182240:151 | +:+:+:+:+:+:+: | 4766 | mavs | 1392 | 1310 | 1363 | 1117 | 1394 | 1108 |
| ENSDARG000 | 13:13:13:13:1 | 1409786:1409 | 1409940:1409 | -:-:-:-:-:-:-:-: | 3769 | NA | 811 | 677 | 836 | 809 | 918 | 926 |
| ENSDARG000 | 13:13:13:13:1 | 37653851:376 | 37654007:376 | +:+:+:+:+:+:+: | 5878 | PHF3 | 1059 | 978 | 1789 | 896 | 1456 | 1281 |
| ENSDARG000 | 13:13:13:13:1 | 30579260:305 | 30584478:305 | -;-;-;-;-;- | 6405 | zcchc24 | 934 | 443 | 611 | 751 | 600 | 1037 |
| ENSDARG000 | 13:13:13:13:1 | 37650673:376 | 37650850:376 | -;-;-;-;-;-;-;-;- | 904 | ifi27.8 | 488 | 380 | 443 | 575 | 519 | 415 |
| ENSDARG000 | 13 | 4405282 | 4407074 | + | 1793 | PRR18 | 6 | 4 | 8 | 8 | 4 | 10 |
| ENSDARG000 | 13:13:13:13:1 | 4409294:4409 | 4409605:4409 | +:+:+:+:+:+:+: | 1720 | NA | 0 | 4 | 0 | 2 | 2 | 0 |
| ENSDARG000 | 13:13:13:13:1 | 42361599:423 | 42361960:423 | -;-;-;-;-;- | 1387 | marchf5 | 349 | 229 | 296 | 218 | 354 | 382 |
| ENSDARG000 | 13:13:13:13:1 | 4439074:4440 | 4439106:4441 | +:+:+:+:+:+:+: | 1616 | TBXT | 4 | 0 | 20 | 0 | 5 | 6 |
| ENSDARG000 | 13:13:13:13:1 | 42406883:424 | 42407696:424 | +:+:+:+:+:+:+: | 1882 | cpeb3 | 470 | 145 | 270 | 293 | 293 | 332 |
| ENSDARG000 | 13:13:13:13:1 | 4445553:4445 | 4445965:4445 | -;-;-;-;- | 927 | NA | 2 | 2 | 0 | 6 | 2 | 4 |
| ENSDARG000 | 13 | 32575777 | 32577386 | - | 1610 | kcns3a | 2 | 0 | 0 | 0 | 0 | 0 |
| ENSDARG000 | 13:13:13:13:1 | 10360444:103 | 10363465:103 | -:-:-:-:-:-:-:-: | 4247 | CAMKMT | 89 | 132 | 168 | 40 | 156 | 71 |
| ENSDARG000 | 13;13;13;13 | 10494920:106 | 10495167:106 | -;-;-;- | 808 | NA | 0 | 0 | 0 | 0 | 0 | 0 |
| ENSDARG000 | 13:13:13:13:1 | 9285112:9285 | 9285349:9286 | +;+;+;+;+;+;+ | 1479 | NA | 0 | 0 | 0 | 0 | 0 | 0 |
| ENSDARG000 | 13;13 | 10409762:104 | 10410045:104 | +;+ | 786 | NA | 0 | 0 | 0 | 0 | 0 | 0 |
| ENSDARG000 | 13:13:13:13:1 | 30655172:306 | 30656490:306 | -:-:-:-:-:-:-:-: | 3456 | paox | 18 | 14 | 12 | 24 | 14 | 6 |
| ENSDARG000 | 13;13;13 | 33811842:338 | 33815043:338 | -;-;- | 4225 | flrt3 | 76 | 50 | 68 | 14 | 66 | 36 |
| ENSDARG000 | 13;13;13;13 | 28580357:285 | 28580563:285 | +;+;+;+ | 1185 | wbp1la | 159 | 68 | 94 | 38 | 153 | 82 |
| ENSDARG000 | 13:13:13:13:1 | 8306399:8306 | 8306699:8306 | +:+:+:+:+:+:+: | 1506 | galm | 462 | 463 | 546 | 571 | 565 | 382 |
| ENSDARG000 | 13:13:13:13:1 | 4505079:4505 | 4505451:4505 | +:+:+:+:+:+:+: | 3135 | pde10a | 2322 | 924 | 1857 | 2118 | 2478 | 1310 |
| ENSDARG000 | 13:13:13:13:1 | 24231327:242 | 24234871:242 | -:-:-:-:-:-:-:-: | 9263 | urb2 | 280 | 565 | 856 | 219 | 625 | 483 |
| ENSDARG000 | 13:13:13:13:1 | 25472005:254 | 25473337:254 | -:-:-:-:-:-:-:-: | 2941 | tial1 | 2128 | 1871 | 2846 | 1770 | 2781 | 2885 |
| ENSDARG000 | 13:13:13:13:1 | 32580382:325 | 32580780:325 | -:-:-:-:-:-:-:-: | 4824 | wdr35 | 144 | 93 | 226 | 143 | 156 | 242 |
| ENSDARG000 | 13:13:13:13:1 | 9288383:9288 | 9288987:9288 | -:-:-:-:-:-:-:-: | 2290 | HTRA2 | 191 | 328 | 596 | 374 | 257 | 136 |
| ENSDARG000 | 13;13;13;13 | 12488863:124 | 12489563:124 | -;-;-;- | 1289 | si:dkev-20i10. | 116 | 105 | 131 | 60 | 143 | 143 |
| ENSDARG000 | 13:13:13:13:1 | 24263049:242 | 24263548:242 | +:+:+:+:+:+:+: | 3882 | ABCB10 | 332 | 374 | 729 | 279 | 567 | 891 |
| ENSDARG000 | 13:13:13:13:1 | 10621257:106 | 10621513:106 | +:+:+:+:+:+:+: | 2379 | prepl | 30 | 11 | 28 | 14 | 72 | 20 |
| ENSDARG000 | 13:13:13:13:1 | 42480852:424 | 42481622:424 | -:-:-:-:-:-:-:-: | 7078 | BTAF1 | 1491 | 1240 | 2009 | 864 | 1814 | 1424 |
| ENSDARG000 | 13:13:13:13:1 | 15234521:152 | 15234695:152 | +:+:+:+:+:+:+: | 3901 | loxl3b | 748 | 894 | 2316 | 1378 | 1607 | 923 |
| ENSDARG000 | 13:13:13:13:1 | 1430094:1430 | 1430274:1430 | +:+:+:+:+:+:+: | 2433 | NA | 132 | 36 | 126 | 65 | 134 | 138 |
| ENSDARG000 | 13:13:13:13:1 | 25455319:254 | 25455529:254 | +:+:+:+:+:+:+: | 2718 | PKD2L1 | 0 | 2 | 0 | 4 | 0 | 0 |
| ENSDARG000 | 13:13:13:13:1 | 36406274:364 | 36410775:364 | -:-:-:-:-:-:-:-: | 5898 | dcaf5 | 308 | 282 | 224 | 182 | 358 | 216 |
| ENSDARG000 | 13:13:13:13:1 | 30664819:306 | 30666205:306 | -:-:-:-:-:-:-:-: | 2680 | rassf4a | 8 | 26 | 26 | 4 | 26 | 0 |
| ENSDARG000 | 13:13:13:13:1 | 4671698:4671 | 4672134:4672 | +:+:+:+:+:+:+: | 2292 | pla2g12b | 0 | 0 | 6 | 0 | 2 | 0 |
| ENSDARG000 | 13:13:13:13:1 | 34627550:346 | 34629956:346 | -:-:-:-:-:-:-:-: | 4484 | KIF16B | 4 | 2 | 27 | 0 | 14 | 10 |
| ENSDARG000 | 13:13:13:13:1 | 10649069:106 | 10651092:106 | -:-:-:-:-:-:-:-: | 3754 | SLC3A1 | 14 | 3 | 13 | 24 | 72 | 6 |
| ENSDARG000 | 13:13:13:13:1 | 36446187:364 | 36446305:364 | -:-:-:-:-:-:-:-: | 2637 | pacs2 | 298 | 142 | 273 | 252 | 400 | 154 |
| ENSDARG000 | 13:13:13:13:1 | 28601195:286 | 28602376:286 | -;-;-;-;-;-;- | 2799 | cyp17a1 | 0 | 0 | 0 | 0 | 0 | 6 |
| ENSDARG000 | 13:13:13:13:1 | 1928361:1940 | 1928475:1940 | +:+:+:+:+:+:+: | 1188 | hmgcll1 | 0 | 0 | 4 | 0 | 0 | 0 |
| ENSDARG000 | 13;13;13;13 | 9303554:9307 | 9304166:9307 | -;-;-;- | 1295 | mrps26 | 202 | 384 | 582 | 265 | 237 | 458 |
| ENSDARG000 | 13:13:13:13:1 | 8320455:8320 | 8321021:8321 | -:-:-:-:-:-:-:-: | 5200 | DHX57 | 595 | 480 | 848 | 258 | 785 | 417 |
| ENSDARG000 | 13:13:13:13:1 | 4685309:4685 | 4685666:4685 | -:-:-:-:-:-:-:-: | 2003 | OIT3 | 0 | 1 | 56 | 0 | 4 | 0 |
| ENSDARG000 | 13:13:13:13:1 | 10682427:106 | 10684121:106 | -:-:-:-:-:-:-:-: | 3342 | ppm1ba | 213 | 102 | 228 | 86 | 261 | 216 |
| ENSDARG000 | 13:13:13:13:1 | 1990970:1992 | 1991822:1992 | -:-:-:-:-:-:-:-: | 2162 | NA | 10 | 8 | 0 | 0 | 0 | 2 |
| ENSDARG000 | 13 | 4712421 | 4714106 | - | 1686 | NA | 220 | 161 | 150 | 160 | 138 | 228 |
| ENSDARG000 | 13:13:13:13:1 | 37992158:379 | 37992203:379 | -;-;-;-;- | 700 | NA | 0 | 0 | 0 | 0 | 0 | 0 |
| ENSDARG000 | 13:13:13:13:1 | 12528043:125 | 12528282:125 | +;+;+;+;+;+;+ | 1361 | rrh | 0 | 0 | 0 | 0 | 0 | 0 |
| ENSDARG000 | 13:13:13:13:1 | 4722200:4723 | 4722420:4724 | -;-;-;-;-;-;-;- | 1323 | mcu | 40 | 36 | 61 | 54 | 42 | 32 |
| ENSDARG000 | 13;13;13 | 9313777:9315 | 9315460:9315 | -;-;- | 2930 | si:dkey-33c12 | 48 | 25 | 54 | 26 | 42 | 35 |
| ENSDARG000 | 13;13;13;13 | 15575137:155 | 15575913:155 | -;-;-;- | 2561 | NA | 2 | 6 | 3 | 0 | 0 | 3 |
| ENSDARG000 | 13;13 | 38190598:381 | 38190670:381 | -;- | 420 | NA | 0 | 0 | 0 | 0 | 0 | 0 |
| ENSDARG000 | 13;13 | 38213694:382 | 38213784:382 | +;+ | 2110 | zgc:171579 | 14 | 6 | 8 | 6 | 20 | 32 |
| ENSDARG000 | 13:13:13:13:1 | 28612313:286 | 28612452:286 | +;+;+;+;+ | 754 | borcs7 | 100 | 126 | 109 | 74 | 64 | 160 |
| ENSDARG000 | 13:13:13:13:1 | 2020215:2026 | 2020384:2026 | -;-;-;-;-;-;- | 1317 | hcrtr2 | 0 | 0 | 0 | 0 | 1 | 6 |
| ENSDARG000 | 13:13:13:13:1 | 18366299:183 | 18366360:183 | +:+:+:+:+:+:+: | 4917 | ccar1 | 835 | 619 | 1012 | 602 | 1125 | 717 |
| ENSDARG000 | 13;13 | 2010287:2029 | 2010386:2029 | +;+ | 363 | NA | 0 | 0 | 0 | 0 | 0 | 0 |
| ENSDARG000 | 13:13:13:13:1 | 24287093:242 | 24287157:242 | +;+;+;+;+;+ | 2542 | ccsapb | 428 | 533 | 522 | 475 | 587 | 644 |
| ENSDARG000 | 13:13:13:13:1 | 4871886:4883 | 4872149:4883 | +:+:+:+:+:+:+: | 2969 | micu1 | 894 | 879 | 949 | 1058 | 1051 | 1076 |
| ENSDARG000 | 13;13;13 | 24283670:242 | 24284151:242 | -;-;- | 692 | NA | 0 | 0 | 0 | 0 | 0 | 0 |
| ENSDARG000 | 13:13:13:13:1 | 24279021:242 | 24279104:242 | +:+:+:+:+:+:+: | 1569 | acta1b | 2 | 12 | 2 | 9 | 18 | 0 |
| ENSDARG000 | 13:13:13:13:1 | 38226157:382 | 38226263:382 | +:+:+:+:+:+:+: | 796 | si:dkeyp-4c7.: | 0 | 0 | 0 | 0 | 0 | 0 |
| ENSDARG000 | 13;13;13;13 | 42544009:425 | 42544104:425 | +;+;+;+ | 1949 | si:dkey-221i1 | 12 | 0 | 0 | 2 | 0 | 0 |
| ENSDARG000 | 13:13:13:13:1 | 28634862:286 | 28636230:286 | -:-:-:-:-:-:-:-: | 4586 | nt5c2a | 480 | 424 | 612 | 786 | 651 | 599 |
| ENSDARG000 | 13:13:13:13:1 | 36526806:365 | 36530871:365 | -:-:-:-:-:-:-:-: | 6315 | srsf5a | 5220 | 5276 | 8903 | 7085 | 7477 | 7458 |
| ENSDARG000 | 13:13:13:13:1 | 28618086:286 | 28619616:286 | +:+:+:+:+:+:+: | 2817 | cnnm2a | 60 | 55 | 62 | 12 | 122 | 72 |
| ENSDARG000 | 13:13:13:13:1 | 2070368:2075 | 2077060:2077 | -:-:-:-:-:-:-:-: | 7962 | FAM83B | 107 | 2 | 4 | 8 | 36 | 16 |
| ENSDARG000 | 13:13:13:13:1 | 4973211:4973 | 4974187:4974 | -:-:-:-:-:-:-:-: | 3839 | nolc1 | 1566 | 2203 | 4106 | 917 | 2529 | 2030 |
| ENSDARG000 | 13:13:13:13:1 | 32627444:326 | 32627664:326 | -:-:-:-:-:-:-:-: | 1681 | matn3b | 0 | 0 | 0 | 0 | 2 | 10 |
| ENSDARG000 | 13:13:13:13:1 | 30704361:307 | 30704382:307 | -:-:-:-:-:-:-:-: | 907 | tmem72 | 0 | 0 | 0 | 0 | 1 | 0 |
| ENSDARG000 | 13:13:13:13:1 | 8375612:8381 | 8381547:8381 | -:-:-:-:-:-:-:-: | 8581 | epas1b | 10 | 12 | 8 | 4 | 14 | 8 |
| ENSDARG000 | 13:13:13:13:1 | 42564666:425 | 42564674:425 | -;-;-;-;-;- | 7192 | NA | 595 | 198 | 290 | 407 | 250 | 441 |
| ENSDARG000 | 13;13;13;13 | 32636197:326 | 32636289:326 | +;+;+;+ | 1283 | mtres1 | 148 | 206 | 135 | 197 | 212 | 212 |
| ENSDARG000 | 13:13:13:13:1 | 34689663:346 | 34689865:346 | +:+:+:+:+:+:+: | 6738 | tasp1 | 233 | 193 | 318 | 188 | 295 | 292 |
| ENSDARG000 | 13:13:13:13:1 | 10799380:108 | 10799652:108 | +:+:+:+:+:+:+: | 4829 | lrpprc | 1961 | 1835 | 2737 | 1054 | 1986 | 1233 |
| ENSDARG000 | 13:13:13:13:1 | 9339919:9343 | 9340761:9343 | -:-:-:-:-:-:-:-: | 3098 | si:dkev-33c12 | 1626 | 1602 | 1709 | 929 | 2336 | 1348 |
| ENSDARG000 | 13:13:13:13:1 | 15580758:155 | 15581323:155 | +:+:+:+:+:+:+: | 4505 | mark3a | 1643 | 1200 | 1579 | 1400 | 1667 | 1550 |
| ENSDARG000 | 13:13:13:13:1 | 42602406:426 | 42602444:426 | +:+:+:+:+:+:+: | 2403 | mlh1 | 110 | 164 | 180 | 106 | 219 | 166 |
| ENSDARG000 | 13:13:13:13:1 | 38302665:383 | 38302705:383 | +:+:+:+:+:+:+: | 5110 | ADGRB3 | 34 | 207 | 146 | 432 | 369 | 261 |
| ENSDARG000 | 13;13;13;13 | 30804367:308 | 30804530:308 | +;+;+;+ | 3041 | cxcl12a | 256 | 30 | 383 | 80 | 267 | 344 |
| ENSDARG000 | 13;13 | 42597760:426 | 42597894:426 | -;- | 246 | NA | 0 | 0 | 0 | 0 | 0 | 0 |
| ENSDARG000 | 13:13:13:13:1 | 18471546:184 | 18472022:184 | +:+:+:+:+:+:+: | 3598 | stox1 | 244 | 66 | 574 | 89 | 392 | 86 |
| ENSDARG000 | 13;13;13;13 | 25486608:254 | 25486880:254 | +;+;+;+ | 2239 | bag3 | 1004 | 606 | 629 | 641 | 505 | 901 |
| ENSDARG000 | 13:13:13:13:1 | 8452354:8464 | 8454045:8465 | -:-:-:-:-:-:-:-: | 4757 | PRKCE | 26 | 12 | 30 | 32 | 16 | 58 |
| ENSDARG000 | 13;13 | 4995433:5000 | 4995940:5000 | +;+ | 771 | mrps6 | 172 | 267 | 340 | 118 | 132 | 254 |
| ENSDARG000 | 13:13:13:13:1 | 12543634:125 | 12544014:125 | -:-:-:-:-:-:-:-: | 3090 | enpep | 18 | 4 | 6 | 0 | 30 | 4 |
| ENSDARG000 | 13:13:13:13:1 | 2150518:2152 | 2152920:2152 | -:-:-:-:-:-:-:-: | 3405 | NA | 22 | 6 | 8 | 22 | 22 | 10 |
| ENSDARG000 | 13;13 | 18507592:185 | 18507737:185 | +;+ | 3579 | si:ch211-198a | 2395 | 3031 | 3501 | 2260 | 3074 | 2759 |
| ENSDARG000 | 13;13;13 | 30855319:308 | 30855872:308 | -;-;- | 740 | NA | 0 | 0 | 0 | 0 | 0 | 0 |
| ENSDARG000 | 13;13;13 | 18514788:185 | 18515116:185 | +;+;+ | 2657 | tlr4ba | 12 | 10 | 4 | 4 | 46 | 4 |
| ENSDARG000 | 13:13:13:13:1 | 32641540:326 | 32645348:326 | -;-;-;-;-;-;- | 4176 | bend3 | 893 | 437 | 568 | 586 | 755 | 513 |
| ENSDARG000 | 13:13:13:13:1 | 46927350:469 | 46927912:469 | +:+:+:+:+:+:+: | 1487 | MTRF1L | 25 | 49 | 63 | 42 | 44 | 24 |
| ENSDARG000 | 13:13:13:13:1 | 25531004:255 | 25535471:255 | -:-:-:-:-:-:-:-: | 7010 | mcmbp | 1659 | 2097 | 2379 | 1216 | 2372 | 2306 |
| ENSDARG000 | 13:13:13:13:1 | 25505580:255 | 25505676:255 | +:+:+:+:+:+:+: | 3363 | inpp5f | 874 | 710 | 552 | 199 | 838 | 687 |
| ENSDARG000 | 13:13:13:13:1 | 5004924:5005 | 5005072:5005 | +:+:+:+:+:+:+: | 2417 | psap | 28870 | 23637 | 28705 | 28504 | 27496 | 35865 |
| ENSDARG000 | 13:13:13:13:1 | 32660544:326 | 32660941:326 | -;-;-;-;-;-;- | 1669 | pdss2 | 210 | 338 | 259 | 142 | 300 | 153 |
| ENSDARG000 | 13:13:13:13:1 | 18533005:185 | 18533401:185 | +;+;+;+;+;+ | 1257 | NA | 3 | 0 | 0 | 0 | 0 | 0 |
| ENSDARG000 | 13;13;13 | 34752314:347 | 34752351:347 | +;+;+ | 576 | NA | 0 | 0 | 0 | 0 | 0 | 0 |
| ENSDARG000 | 13:13:13:13:1 | 18545819:185 | 18546070:185 | +:+:+:+:+:+:+: | 2301 | zgc:154058 | 241 | 59 | 182 | 142 | 614 | 128 |
| ENSDARG000 | 13:13:13:13:1 | 34765141:347 | 34766299:347 | -:-:-:-:-:-:-:-: | 4750 | ism1 | 67 | 0 | 4 | 0 | 18 | 0 |
| ENSDARG000 | 13:13:13:13:1 | 22571943:225 | 22572029:225 | +;+;+;+;+ | 563 | NA | 2 | 0 | 4 | 0 | 0 | 0 |
| ENSDARG000 | 13:13:13:13:1 | 22619684:226 | 22620603:226 | -:-:-:-:-:-:-:-: | 3868 | mmrn2a | 204 | 65 | 72 | 37 | 62 | 42 |
| ENSDARG000 | 13;13;13;13 | 36557205:365 | 36558063:365 | -;-;-;- | 1273 | synj2bp | 202 | 338 | 332 | 207 | 425 | 183 |
| ENSDARG000 | 13:13:13:13:1 | 22555342:225 | 22555977:225 | +:+:+:+:+:+:+: | 5007 | bmpr1aa | 899 | 675 | 1076 | 914 | 810 | 912 |
| ENSDARG000 | 13;13;13 | 18523661:185 | 18523936:185 | +;+;+ | 2736 | tlr4bb | 4 | 1 | 0 | 0 | 2 | 0 |
| ENSDARG000 | 13:13:13:13:1 | 30874153:308 | 30874303:308 | +:+:+:+:+:+:+: | 5362 | ERCC6 | 241 | 165 | 255 | 129 | 172 | 212 |
| ENSDARG000 | 13;13;13 | 18520738:185 | 18520828:185 | +;+;+ | 2473 | tlr4al | 1 | 4 | 4 | 0 | 1 | 7 |
| ENSDARG000 | 13:13:13:13:1 | 32740509:327 | 32740604:327 | +:+:+:+:+:+:+: | 2823 | sobpa | 23 | 22 | 52 | 20 | 48 | 35 |
| ENSDARG000 | 13;13;13 | 32793864:327 | 32794977:327 | -;-;- | 1557 | NA | 0 | 0 | 0 | 0 | 0 | 0 |
| ENSDARG000 | 13:13:13:13:1 | 32807419:328 | 32807677:328 | +;+;+;+;+;+;+ | 1400 | slc66a3 | 224 | 197 | 277 | 249 | 416 | 286 |
| ENSDARG000 | 13:13:13:13:1 | 34804243:348 | 34804990:348 | -:-:-:-:-:-:-:-: | 2738 | sptlc3 | 278 | 222 | 60 | 150 | 40 | 60 |
| ENSDARG000 | 13;13;13;13 | 12591508:125 | 12591938:125 | -;-;-;- | 1317 | lrit3b | 0 | 0 | 0 | 0 | 0 | 0 |
| ENSDARG000 | 13:13:13:13:1 | 36568635:365 | 36569353:365 | -:-:-:-:-:-:-:-: | 2336 | lgals3a | 4200 | 3431 | 3794 | 4109 | 4317 | 4171 |
| ENSDARG000 | 13;13 | 8604710:8621 | 8605191:8628 | +;+ | 7308 | socs5b | 878 | 721 | 1589 | 509 | 1371 | 588 |
| ENSDARG000 | 13;13 | 8649711:8649 | 8649842:8650 | -;- | 242 | NA | 1 | 1 | 0 | 0 | 1 | 0 |
| ENSDARG000 | 13:13:13:13:1 | 46941930:469 | 46942100:469 | +:+:+:+:+:+:+: | 5331 | fbxo5 | 28 | 140 | 202 | 75 | 128 | 116 |
| ENSDARG000 | 13:13:13:13:1 | 24295896:243 | 24296781:243 | -;-;-;-;-;-;-;- | 1687 | rab4a | 558 | 399 | 410 | 400 | 597 | 594 |
| ENSDARG000 | 13;13 | 8655119:8655 | 8655238:8655 | -;- | 230 | NA | 0 | 0 | 0 | 0 | 0 | 0 |
| ENSDARG000 | 13:13:13:13:1 | 8682176:8683 | 8685755:8685 | -;-;-;-;-;-;- | 4023 | mcfd2 | 783 | 560 | 971 | 574 | 905 | 672 |
| ENSDARG000 | 13:13:13:13:1 | 2442841:2442 | 2442920:2442 | +:+:+:+:+:+:+: | 6689 | ARFGEF3 | 80 | 32 | 21 | 59 | 22 | 33 |
| ENSDARG000 | 13:13:13:13:1 | 8677166:8677 | 8677403:8677 | +:+:+:+:+:+:+: | 979 | prop1 | 0 | 0 | 2 | 0 | 0 | 0 |
| ENSDARG000 | 13:13:13:13:1 | 46987955:469 | 46988459:469 | -;-;-;-;-;- | 1109 | vip | 18 | 0 | 4 | 4 | 10 | 2 |
| ENSDARG000 | 13;13;13;13 | 35041892:350 | 35042924:350 | -;-;-;- | 1606 | BTBD3 | 0 | 0 | 0 | 0 | 0 | 0 |
| ENSDARG000 | 13:13:13:13:1 | 10926528:109 | 10927276:109 | -:-:-:-:-:-:-:-: | 3013 | abcg8 | 0 | 0 | 2 | 0 | 2 | 0 |

| | | | | | | | | | | | | |
|---|---|---|---|---|---|---|---|---|---|---|---|---|
| ENSDARG000 | 13:13:13:13:1 | 10945337:109 | 10945666:109 | +:+:+:+:+:+:+: | 2471 | abcg5 | 0 | 0 | 1 | 2 | 0 | 0 |
| ENSDARG000 | 13:13:13:13:1 | 24325413:243 | 24326800:243 | -;-;-;-;-;-;- | 3516 | NA | 4 | 0 | 0 | 3 | 0 | 0 |
| ENSDARG000 | 13:13:13:13:1 | 28705143:287 | 28705170:287 | +:+:+:+:+:+:+: | 6999 | ldb1a | 2075 | 1583 | 1960 | 2350 | 1840 | 2832 |
| ENSDARG000 | 13:13:13:13:1 | 38621517:386 | 38622264:386 | -:-:-:-:-:-:-:-: | 2808 | lmbrd1 | 888 | 685 | 837 | 1024 | 939 | 1115 |
| ENSDARG000 | 13:13:13:13:1 | 8693410:8696 | 8693535:8696 | +:+:+:+:+:+:+: | 3099 | TTC7A | 4 | 24 | 90 | 0 | 34 | 16 |
| ENSDARG000 | 13:13:13:13:1 | 15682075:156 | 15683006:156 | -:-:-:-:-:-:-:-: | 2586 | ckba | 235 | 261 | 154 | 189 | 64 | 593 |
| ENSDARG000 | 13:13:13:13:1 | 28682614:286 | 28683352:286 | -:-:-:-:-:-:-:-: | 1978 | pcgf6 | 40 | 42 | 76 | 54 | 68 | 68 |
| ENSDARG000 | 13;13;13 | 28675686:286 | 28676745:286 | +;+;+ | 1844 | inaa | 0 | 2 | 0 | 0 | 4 | 0 |
| ENSDARG000 | 13:13:13:13:1 | 28690355:286 | 28690475:286 | +:+:+:+:+:+:+: | 1612 | polr1c | 212 | 205 | 520 | 241 | 345 | 276 |
| ENSDARG000 | 13:13:13:13:1 | 30912117:309 | 30912250:309 | +:+:+:+:+:+:+: | 3750 | drgx | 0 | 2 | 0 | 0 | 0 | 0 |
| ENSDARG000 | 13:13:13:13:1 | 15701596:157 | 15702145:157 | +:+:+:+:+:+:+: | 2760 | trmt61a | 250 | 208 | 419 | 103 | 234 | 178 |
| ENSDARG000 | 13:13:13:13:1 | 12606821:126 | 12607003:126 | +:+:+:+:+:+:+: | 2308 | metap1 | 1632 | 2288 | 3012 | 1558 | 1979 | 2121 |
| ENSDARG000 | 13:13:13:13:1 | 9368619:9368 | 9368667:9368 | +:+:+:+:+:+:+: | 7106 | NA | 455 | 230 | 553 | 410 | 529 | 379 |
| ENSDARG000 | 13:13:13:13:1 | 32815241:328 | 32816455:328 | -:-:-:-:-:-:-:-: | 5766 | rock2a | 1657 | 1856 | 2018 | 1976 | 1935 | 1606 |
| ENSDARG000 | 13:13:13:13:1 | 28747934:287 | 28748033:287 | +:+:+:+:+:+:+: | 3945 | NA | 49 | 119 | 120 | 155 | 76 | 441 |
| ENSDARG000 | 13:13:13:13:1 | 36585399:365 | 36585652:365 | +:+:+:+:+:+:+: | 2179 | gmfb | 3330 | 4058 | 4455 | 3860 | 5384 | 4369 |
| ENSDARG000 | 13;13 | 35182994:351 | 35183024:351 | -;- | 97 | NA | 0 | 0 | 0 | 0 | 0 | 0 |
| ENSDARG000 | 13:13:13:13:1 | 36592194:365 | 36592281:365 | +:+:+:+:+:+:+: | 2373 | cnih1 | 1891 | 1880 | 1882 | 1277 | 2022 | 1769 |
| ENSDARG000 | 13;13;13 | 36587770:366 | 36587885:366 | -;-;- | 337 | NA | 0 | 0 | 2 | 1 | 9 | 0 |
| ENSDARG000 | 13;13;13;13 | 28701233:287 | 28701297:287 | +;+;+;+ | 523 | NA | 8 | 1 | 5 | 7 | 4 | 10 |
| ENSDARG000 | 13:13:13:13:1 | 30951155:309 | 30951327:309 | +:+:+:+:+:+:+: | 2442 | vstm4a | 24 | 3 | 22 | 1 | 48 | 24 |
| ENSDARG000 | 13;13;13;13 | 28677581:286 | 28677748:286 | -;-;-;- | 566 | NA | 0 | 0 | 0 | 4 | 3 | 4 |
| ENSDARG000 | 13;13;13 | 35206835:352 | 35206888:352 | -;-;- | 352 | NA | 0 | 0 | 0 | 0 | 0 | 0 |
| ENSDARG000 | 13;13;13 | 24371815:243 | 24373254:243 | -;-;- | 2143 | rhoua | 362 | 276 | 222 | 313 | 368 | 479 |
| ENSDARG000 | 13:13:13:13:1 | 38814521:388 | 38814644:388 | +:+:+:+:+:+:+: | 1193 | col19a1 | 0 | 0 | 0 | 0 | 0 | 0 |
| ENSDARG000 | 13:13:13:13:1 | 24380132:243 | 24380548:243 | +;+;+;+;+;+;+ | 1410 | pdcd2 | 11 | 16 | 15 | 20 | 34 | 26 |
| ENSDARG000 | 13:13:13:13:1 | 18628920:186 | 18630081:186 | -;-;-;-;-;-;-;- | 2517 | mat1a | 164 | 97 | 317 | 88 | 369 | 450 |
| ENSDARG000 | 13:13:13:13:1 | 24381344:243 | 24384915:243 | -:-:-:-:-:-:-:-: | 5236 | tbp | 246 | 443 | 380 | 239 | 417 | 364 |
| ENSDARG000 | 13:13:13:13:1 | 30985047:309 | 30985143:309 | -:-:-:-:-:-:-:-: | 2964 | NA | 584 | 422 | 449 | 535 | 958 | 574 |
| ENSDARG000 | 13:13:13:13:1 | 5031197:5031 | 5031529:5031 | -:-:-:-:-:-:-:-: | 2230 | abcg2c | 309 | 365 | 527 | 339 | 261 | 358 |
| ENSDARG000 | 13;13 | 31002658:310 | 31002727:310 | +;+ | 827 | LRRC18 | 6 | 0 | 6 | 0 | 8 | 8 |
| ENSDARG000 | 13:13:13:13:1 | 25549425:255 | 25549607:255 | +:+:+:+:+:+:+: | 3382 | sec23ip | 2196 | 1581 | 2229 | 1194 | 2009 | 1614 |
| ENSDARG000 | 13:13:13:13:1 | 42621205:426 | 42622212:426 | -:-:-:-:-:-:-:-: | 3781 | lrrfip2 | 611 | 743 | 820 | 566 | 852 | 602 |
| ENSDARG000 | 13:13:13:13:1 | 22659153:226 | 22659376:226 | +:+:+:+:+:+:+: | 1298 | sncga | 5006 | 1280 | 580 | 3032 | 2761 | 427 |
| ENSDARG000 | 13:13:13:13:1 | 35339182:353 | 35339516:353 | +:+:+:+:+:+:+: | 6754 | jag1b | 82 | 12 | 25 | 14 | 86 | 44 |
| ENSDARG000 | 13:13:13:13:1 | 8760695:8760 | 8761008:8761 | -:-:-:-:-:-:-:-: | 949 | si:ch211-93n2 | 2 | 0 | 0 | 2 | 3 | 0 |
| ENSDARG000 | 13:13:13:13:1 | 8777633:8781 | 8778597:8781 | -:-:-:-:-:-:-:-: | 2630 | calm2a | 2355 | 3691 | 3306 | 1568 | 3196 | 2860 |
| ENSDARG000 | 13:13:13:13:1 | 18659325:186 | 18659432:186 | +:+:+:+:+:+:+: | 1762 | selenou1a | 886 | 708 | 846 | 898 | 1009 | 737 |
| ENSDARG000 | 13:13:13:13:1 | 2426993:2429 | 2427819:2429 | -:-:-:-:-:-:-:-: | 2496 | fbxo9 | 1395 | 1041 | 1424 | 1175 | 1500 | 1180 |
| ENSDARG000 | 13:13:13:13:1 | 10970614:109 | 10970768:109 | +:+:+:+:+:+:+: | 1423 | dync2li1 | 24 | 13 | 16 | 3 | 19 | 40 |
| ENSDARG000 | 13:13:13:13:1 | 10980755:109 | 10984281:109 | -:-:-:-:-:-:-:-: | 7933 | NA | 2421 | 4411 | 4487 | 3257 | 2898 | 3670 |
| ENSDARG000 | 13:13:13:13:1 | 32901358:329 | 32902902:329 | -:-:-:-:-:-:-:-: | 2047 | si:dkev-18i18. | 0 | 0 | 0 | 0 | 0 | 2 |
| ENSDARG000 | 13:13:13:13:1 | 38990939:390 | 38990983:390 | +:+:+:+:+:+:+: | 6802 | NA | 4 | 0 | 0 | 0 | 0 | 0 |
| ENSDARG000 | 13;13;13;13 | 32993911:329 | 32994008:329 | -;-;-;- | 1437 | KCNF1 | 0 | 0 | 0 | 2 | 4 | 0 |
| ENSDARG000 | 13:13:13:13:1 | 47007075:470 | 47007316:470 | +:+:+:+:+:+:+: | 9431 | ANAPC1 | 405 | 455 | 778 | 182 | 628 | 310 |
| ENSDARG000 | 13:13:13:13:1 | 24396666:243 | 24396870:243 | +:+:+:+:+:+:+: | 1663 | zgc:153169 | 62 | 38 | 82 | 49 | 64 | 47 |
| ENSDARG000 | 13:13:13:13:1 | 8799168:8800 | 8799471:8801 | -:-:-:-:-:-:-:-: | 2453 | NA | 2 | 10 | 4 | 1 | 0 | 2 |
| ENSDARG000 | 13:13:13:13:1 | 9408101:9408 | 9408180:9408 | +:+:+:+:+:+:+: | 1665 | bbof1 | 0 | 0 | 2 | 0 | 0 | 2 |
| ENSDARG000 | 13:13:13:13:1 | 15762278:157 | 15763805:157 | -:-:-:-:-:-:-:-: | 2896 | bag5 | 148 | 144 | 189 | 139 | 183 | 218 |
| ENSDARG000 | 13:13:13:13:1 | 22679084:226 | 22679938:226 | -:-:-:-:-:-:-:-: | 2482 | glud1a | 6483 | 7165 | 5008 | 5605 | 10641 | 4677 |
| ENSDARG000 | 13:13:13:13:1 | 22698215:227 | 22698411:227 | +:+:+:+:+:+:+: | 4574 | shld2 | 132 | 154 | 181 | 170 | 171 | 153 |
| ENSDARG000 | 13:13:13:13:1 | 22675802:226 | 22675986:226 | +;+;+;+;+;+ | 794 | zgc:193505 | 88 | 36 | 2 | 94 | 48 | 11 |
| ENSDARG000 | 13;13;13 | 8825192:8827 | 8827104:8827 | -;-;- | 3943 | NA | 0 | 0 | 0 | 0 | 0 | 0 |
| ENSDARG000 | 13:13:13:13:1 | 25569831:255 | 25571515:255 | -;-;-;-;-;-;- | 3015 | csgalnact2 | 1324 | 725 | 755 | 928 | 772 | 975 |
| ENSDARG000 | 13:13:13:13:1 | 9432501:9432 | 9432543:9432 | +;+;+;+;+;+;+ | 878 | zgc:123321 | 0 | 2 | 0 | 0 | 0 | 0 |
| ENSDARG000 | 13:13:13:13:1 | 32997020:329 | 32997767:329 | -:-:-:-:-:-:-:-: | 3395 | NA | 1076 | 913 | 1874 | 2003 | 1846 | 2672 |
| ENSDARG000 | 13:13:13:13:1 | 8840772:8841 | 8841002:8841 | +:+:+:+:+:+:+: | 1476 | epcam | 1061 | 113 | 54 | 76 | 179 | 286 |
| ENSDARG000 | 13;13;13 | 5069552:5076 | 5073343:5077 | -;-;- | 4182 | SLC29A3 | 414 | 727 | 849 | 687 | 507 | 1975 |
| ENSDARG000 | 13:13:13:13:1 | 24402406:244 | 24402591:244 | +;+;+;+;+;+;+ | 1810 | rab1ab | 4527 | 6324 | 6087 | 3684 | 6182 | 4587 |
| ENSDARG000 | 13:13:13:13:1 | 12633223:126 | 12633475:126 | -:-:-:-:-:-:-:-: | 2206 | adh8b | 2232 | 1194 | 1844 | 2704 | 1680 | 464 |
| ENSDARG000 | 13:13:13:13:1 | 36613359:366 | 36613544:366 | +:+:+:+:+:+:+: | 1050 | NA | 0 | 0 | 2 | 0 | 0 | 0 |
| ENSDARG000 | 13:13:13:13:1 | 36612930:366 | 36613270:366 | -:-:-:-:-:-:-:-: | 984 | cdkn3 | 154 | 102 | 165 | 120 | 132 | 98 |
| ENSDARG000 | 13:13:13:13:1 | 18677428:186 | 18678002:186 | -:-:-:-:-:-:-:-: | 1811 | sfxn3 | 1159 | 842 | 750 | 601 | 1002 | 1016 |
| ENSDARG000 | 13:13:13:13:1 | 31070181:310 | 31070582:310 | +:+:+:+:+:+:+: | 1931 | NA | 103 | 75 | 94 | 61 | 78 | 31 |
| ENSDARG000 | 13:13:13:13:1 | 24421515:244 | 24421838:244 | -:-:-:-:-:-:-:-: | 2501 | cep68 | 151 | 94 | 170 | 140 | 243 | 194 |
| ENSDARG000 | 13:13:13:13:1 | 5243167:5247 | 5247513:5247 | -;-;-;-;-;-;- | 6232 | NA | 233 | 196 | 270 | 220 | 399 | 296 |
| ENSDARG000 | 13;13 | 5397541:5400 | 5398405:5401 | +;+ | 1409 | NA | 0 | 0 | 0 | 0 | 0 | 0 |
| ENSDARG000 | 13;13;13;13 | 9440326:9440 | 9440726:9440 | -;-;-;- | 989 | NA | 0 | 0 | 2 | 0 | 4 | 0 |
| ENSDARG000 | 13:13:13:13:1 | 35391609:353 | 35393056:353 | -:-:-:-:-:-:-:-: | 3205 | slx4ip | 38 | 105 | 101 | 29 | 112 | 16 |
| ENSDARG000 | 13:13:13:13:1 | 2394264:2394 | 2394419:2394 | +:+:+:+:+:+:+: | 2696 | elovl5 | 306 | 323 | 221 | 156 | 268 | 255 |
| ENSDARG000 | 13:13:13:13:1 | 35472803:354 | 35472863:354 | +:+:+:+:+:+:+: | 1815 | mkks | 8 | 34 | 45 | 12 | 6 | 20 |
| ENSDARG000 | 13:13:13:13:1 | 15800742:158 | 15801182:158 | +;+;+;+;+;+ | 1249 | coa8 | 155 | 227 | 234 | 143 | 188 | 199 |
| ENSDARG000 | 13:13:13:13:1 | 35510893:355 | 35511907:355 | -:-:-:-:-:-:-:-: | 2162 | memo1 | 723 | 702 | 680 | 297 | 937 | 645 |
| ENSDARG000 | 13:13:13:13:1 | 47361104:473 | 47361211:473 | -;-;-;-;-;-;- | 848 | bcl2l11 | 55 | 40 | 140 | 145 | 31 | 23 |
| ENSDARG000 | 13;13 | 31126449:311 | 31126768:311 | -;- | 436 | NA | 2 | 0 | 6 | 2 | 0 | 0 |
| ENSDARG000 | 13:13:13:13:1 | 31108334:311 | 31108531:311 | +:+:+:+:+:+:+: | 2087 | ARHGAP22 | 289 | 135 | 174 | 60 | 275 | 70 |
| ENSDARG000 | 13:13:13:13:1 | 35483618:354 | 35483732:354 | +;+;+;+;+;+ | 807 | NA | 0 | 0 | 0 | 0 | 0 | 0 |
| ENSDARG000 | 13;13;13;13 | 35484229:354 | 35484918:354 | -;-;-;- | 1277 | srd5a2b | 0 | 0 | 0 | 0 | 0 | 0 |
| ENSDARG000 | 13:13:13:13:1 | 18701906:187 | 18702089:187 | +:+:+:+:+:+:+: | 2865 | pdzd7a | 0 | 0 | 0 | 0 | 0 | 0 |
| ENSDARG000 | 13:13:13:13:1 | 33024547:330 | 33024970:330 | +:+:+:+:+:+:+: | 2257 | arg2 | 179 | 198 | 498 | 553 | 77 | 322 |
| ENSDARG000 | 13;13;13 | 42676202:426 | 42676266:426 | +;+;+ | 624 | mrpl14 | 203 | 217 | 280 | 176 | 289 | 187 |
| ENSDARG000 | 13:13:13:13:1 | 24429918:244 | 24432573:244 | -:-:-:-:-:-:-:-: | 4957 | slc1a4 | 220 | 1719 | 2526 | 1084 | 700 | 1111 |
| ENSDARG000 | 13;13;13 | 24500963:245 | 24501148:245 | +;+;+ | 5557 | sertad2b | 1519 | 699 | 968 | 1187 | 1592 | 1499 |
| ENSDARG000 | 13:13:13:13:1 | 18801094:188 | 18801873:188 | -:-:-:-:-:-:-:-: | 3011 | lzts2a | 366 | 195 | 260 | 122 | 582 | 249 |
| ENSDARG000 | 13:13:13:13:1 | 47427160:474 | 47427336:474 | -:-:-:-:-:-:-:-: | 2085 | ACOXL | 73 | 51 | 72 | 56 | 130 | 49 |
| ENSDARG000 | 13:13:13:13:1 | 42679767:426 | 42682159:426 | -:-:-:-:-:-:-:-: | 4609 | capn1a | 126 | 62 | 434 | 283 | 420 | 104 |
| ENSDARG000 | 13:13:13:13:1 | 36633355:366 | 36633524:366 | +;+;+;+;+;+ | 1398 | NA | 8 | 6 | 0 | 0 | 12 | 6 |
| ENSDARG000 | 13:13:13:13:1 | 36638955:366 | 36640863:366 | -:-:-:-:-:-:-:-: | 5462 | sos2 | 1314 | 960 | 1520 | 728 | 1586 | 1021 |
| ENSDARG000 | 13:13:13:13:1 | 25584556:255 | 25586525:255 | -:-:-:-:-:-:-:-: | 5094 | ret | 30 | 74 | 424 | 58 | 6 | 571 |
| ENSDARG000 | 13:13:13:13:1 | 12653975:126 | 12654255:126 | -:-:-:-:-:-:-:-: | 1810 | adh8a | 79 | 14 | 49 | 18 | 150 | 24 |
| ENSDARG000 | 13:13:13:13:1 | 33040436:330 | 33041491:330 | -:-:-:-:-:-:-:-: | 1813 | vti1b | 370 | 387 | 357 | 349 | 368 | 324 |
| ENSDARG000 | 13:13:13:13:1 | 8847421:8848 | 8848611:8848 | -:-:-:-:-:-:-:-: | 5411 | PLEKHH2 | 574 | 187 | 318 | 209 | 718 | 194 |
| ENSDARG000 | 13;13 | 24552254:245 | 24552547:245 | +;+ | 438 | lgalslb | 24 | 21 | 36 | 6 | 24 | 44 |
| ENSDARG000 | 13:13:13:13:1 | 15816573:158 | 15816932:158 | +:+:+:+:+:+:+: | 2183 | klc1a | 361 | 524 | 564 | 264 | 578 | 600 |
| ENSDARG000 | 13:13:13:13:1 | 5490551:5491 | 5491358:5491 | -:-:-:-:-:-:-:-: | 3053 | meis1b | 862 | 985 | 1911 | 1312 | 2452 | 769 |
| ENSDARG000 | 13:13:13:13:1 | 15853677:158 | 15854769:158 | -:-:-:-:-:-:-:-: | 3569 | NA | 26 | 0 | 8 | 8 | 19 | 0 |
| ENSDARG000 | 13:13:13:13:1 | 9447020:9447 | 9447190:9447 | -;-;-;-;-;-;- | 1642 | NA | 0 | 0 | 0 | 0 | 0 | 0 |
| ENSDARG000 | 13:13:13:13:1 | 22717366:227 | 22717545:227 | +:+:+:+:+:+:+: | 4210 | nfkb2 | 1495 | 996 | 1751 | 852 | 1082 | 826 |
| ENSDARG000 | 13:13:13:13:1 | 24579108:245 | 24579198:245 | +:+:+:+:+:+:+: | 1421 | echs1 | 939 | 909 | 1100 | 768 | 977 | 1074 |
| ENSDARG000 | 13:13:13:13:1 | 33066605:330 | 33067847:330 | -:-:-:-:-:-:-:-: | 8850 | zfyve26 | 708 | 488 | 622 | 364 | 828 | 672 |
| ENSDARG000 | 13:13:13:13:1 | 24584401:245 | 24584541:245 | +;+;+;+;+;+;+ | 1049 | fuom | 12 | 4 | 11 | 7 | 20 | 19 |
| ENSDARG000 | 13:13:13:13:1 | 2357637:2366 | 2358110:2366 | +:+:+:+:+:+:+: | 3045 | gclc | 527 | 665 | 771 | 318 | 777 | 585 |
| ENSDARG000 | 13:13:13:13:1 | 9464758:9465 | 9464928:9465 | -;-;-;-;-;-;- | 1643 | NA | 0 | 0 | 0 | 0 | 0 | 0 |
| ENSDARG000 | 13;13;13;13 | 24585224:245 | 24587042:245 | -;-;-;- | 2500 | NA | 10 | 4 | 23 | 0 | 19 | 29 |
| ENSDARG000 | 13:13:13:13:1 | 9450826:9458 | 9450958:9458 | +;+;+;+;+;+;+;+ | 1237 | NA | 0 | 0 | 0 | 0 | 0 | 0 |
| ENSDARG000 | 13:13:13:13:1 | 39119460:391 | 39120345:391 | -:-:-:-:-:-:-:-: | 3013 | col9a1b | 0 | 10 | 4 | 16 | 4 | 69 |
| ENSDARG000 | 13;13 | 12664866:126 | 12666569:126 | -;- | 1965 | fam241a | 256 | 120 | 271 | 150 | 216 | 296 |
| ENSDARG000 | 13:13:13:13:1 | 33055548:330 | 33055715:330 | +;+;+;+;+;+;+;+ | 1422 | rdh12 | 155 | 258 | 294 | 116 | 133 | 104 |
| ENSDARG000 | 13:13:13:13:1 | 19283936:192 | 19284140:192 | +:+:+:+:+:+:+: | 4667 | eif3s10 | 10949 | 9593 | 17192 | 10600 | 11481 | 12058 |
| ENSDARG000 | 13;13;13;13 | 31193485:311 | 31193650:311 | -;-;-;- | 530 | NA | 9 | 6 | 8 | 0 | 5 | 4 |
| ENSDARG000 | 13:13:13:13:1 | 31177934:311 | 31178003:311 | +:+:+:+:+:+:+: | 4044 | NA | 351 | 148 | 261 | 111 | 270 | 110 |
| ENSDARG000 | 13;13 | 5570952:5572 | 5571110:5574 | +;+ | 2137 | NA | 0 | 0 | 0 | 6 | 0 | 0 |
| ENSDARG000 | 13:13:13:13:1 | 31147518:311 | 31148382:311 | -:-:-:-:-:-:-:-: | 3201 | mapk8a | 262 | 153 | 288 | 100 | 301 | 242 |
| ENSDARG000 | 13:13:13:13:1 | 35528607:355 | 35528674:355 | +:+:+:+:+:+:+: | 3191 | NA | 0 | 0 | 0 | 0 | 0 | 0 |
| ENSDARG000 | 13:13:13:13:1 | 47573405:475 | 47573828:475 | +:+:+:+:+:+:+: | 3001 | lhcgr | 209 | 57 | 80 | 211 | 53 | 111 |
| ENSDARG000 | 13;13;13;13 | 5579097:5590 | 5579390:5590 | -;-;-;- | 525 | NA | 0 | 0 | 0 | 0 | 0 | 0 |
| ENSDARG000 | 13:13:13:13:1 | 47623916:476 | 47624438:476 | +:+:+:+:+:+:+: | 5697 | NA | 30 | 32 | 54 | 4 | 94 | 48 |
| ENSDARG000 | 13:13:13:13:1 | 11073282:110 | 11073516:110 | +:+:+:+:+:+:+: | 2707 | sdccag8 | 102 | 101 | 106 | 82 | 118 | 54 |
| ENSDARG000 | 13:13:13:13:1 | 42725712:427 | 42728815:427 | -:-:-:-:-:-:-:-: | 5407 | capn2a | 3426 | 7037 | 4666 | 5088 | 5491 | 8101 |
| ENSDARG000 | 13 | 2352289 | 2352995 | + | 707 | NA | 0 | 0 | 0 | 0 | 0 | 0 |
| ENSDARG000 | 13:13:13:13:1 | 9468535:9468 | 9468670:9468 | +:+:+:+:+:+:+: | 1451 | HTRA2 | 6 | 2 | 0 | 10 | 2 | 0 |
| ENSDARG000 | 13:13:13:13:1 | 12671513:126 | 12671560:126 | +;+;+;+;+;+;+;+ | 1563 | eif4eb | 1363 | 1688 | 990 | 948 | 1399 | 1423 |
| ENSDARG000 | 13:13:13:13:1 | 48358467:483 | 48358870:483 | +:+:+:+:+:+:+: | 4519 | msh6 | 653 | 1714 | 2479 | 1042 | 1878 | 1208 |
| ENSDARG000 | 13 | 12739283 | 12741158 | + | 1876 | lingo2b | 8 | 6 | 2 | 0 | 0 | 22 |
| ENSDARG000 | 13;13;13 | 19322686:193 | 19323279:193 | +;+;+ | 1986 | emx2 | 2 | 4 | 3 | 2 | 6 | 86 |
| ENSDARG000 | 13:13:13:13:1 | 48381035:483 | 48382148:483 | -:-:-:-:-:-:-:-: | 4146 | fbxo11a | 1130 | 1033 | 1339 | 1121 | 1386 | 1086 |
| ENSDARG000 | 13:13:13:13:1 | 5870644:5874 | 5872725:5874 | -;-;-;-;-;-;- | 3030 | spred2b | 1575 | 1538 | 3409 | 2005 | 2687 | 1828 |
| ENSDARG000 | 13:13:13:13:1 | 42762226:427 | 42762379:427 | +:+:+:+:+:+:+: | 1813 | NA | 0 | 0 | 0 | 0 | 0 | 0 |
| ENSDARG000 | 13:13:13:13:1 | 2338761:2338 | 2338813:2338 | +:+:+:+:+:+:+: | 4628 | klhl31 | 0 | 0 | 4 | 0 | 4 | 6 |
| ENSDARG000 | 13:13:13:13:1 | 19610562:196 | 19610792:196 | -:-:-:-:-:-:-:-: | 1948 | RAB11FIP2 | 267 | 176 | 302 | 134 | 241 | 179 |
| ENSDARG000 | 13;13 | 2349076:2349 | 2349191:2350 | -;- | 888 | NA | 0 | 0 | 0 | 0 | 0 | 0 |
| ENSDARG000 | 13:13:13:13:1 | 12761707:127 | 12761797:127 | +:+:+:+:+:+:+: | 2427 | C13H9orf72 | 364 | 491 | 383 | 280 | 350 | 444 |
| ENSDARG000 | 13;13 | 48358433:483 | 48358574:483 | -;- | 263 | NA | 0 | 0 | 0 | 0 | 0 | 0 |
| ENSDARG000 | 13;13;13;13 | 24597711:246 | 24597780:246 | +;+;+;+ | 650 | NA | 0 | 0 | 0 | 0 | 0 | 0 |
| ENSDARG000 | 13:13:13:13:1 | 24602410:246 | 24602834:246 | -;-;-;-;-;-;- | 1063 | NA | 1 | 2 | 0 | 2 | 0 | 0 |
| ENSDARG000 | 13:13:13:13:1 | 19766896:197 | 19767130:197 | -:-:-:-:-:-:-:-: | 1304 | fam204a | 356 | 212 | 381 | 289 | 417 | 310 |
| ENSDARG000 | 13:13:13:13:1 | 8892784:8892 | 8892889:8892 | +:+:+:+:+:+:+: | 9748 | thada | 478 | 174 | 470 | 252 | 593 | 344 |
| ENSDARG000 | 13:13:13:13:1 | 28785814:287 | 28785890:287 | +:+:+:+:+:+:+: | 1352 | nsmce4a | 73 | 329 | 243 | 167 | 306 | 148 |
| ENSDARG000 | 13;13 | 43099336:431 | 43100141:431 | -;- | 1101 | NA | 0 | 0 | 0 | 0 | 0 | 0 |
| ENSDARG000 | 13;13 | 43395269:434 | 43395321:434 | -;- | 190 | NA | 0 | 0 | 0 | 0 | 0 | 0 |
| ENSDARG000 | 13;13;13;13 | 43400443:434 | 43400768:434 | +;+;+;+ | 3021 | dact2 | 10 | 0 | 0 | 0 | 0 | 0 |
| ENSDARG000 | 13;13;13 | 2523032:2523 | 2523120:2523 | +;+;+ | 1378 | lhb | 2 | 0 | 0 | 12 | 0 | 2 |
| ENSDARG000 | 13:13:13:13:1 | 2538636:2617 | 2538718:2617 | +:+:+:+:+:+:+: | 903 | PLPP4 | 0 | 0 | 0 | 0 | 0 | 0 |
| ENSDARG000 | 13;13;13 | 28903959:289 | 28904279:289 | +;+;+ | 3248 | ly6pge | 0 | 0 | 0 | 0 | 0 | 0 |
| ENSDARG000 | 13:13:13:13:1 | 5946607:5953 | 5946630:5954 | -:-:-:-:-:-:-:-: | 1381 | actr2b | 574 | 914 | 1744 | 1253 | 2203 | 821 |
| ENSDARG000 | 13;13 | 24662238:246 | 24662754:246 | +;+ | 2028 | msx3 | 42 | 0 | 2 | 0 | 58 | 26 |
| ENSDARG000 | 13:13:13:13:1 | 35635672:356 | 35635859:356 | +:+:+:+:+:+:+: | 5054 | NA | 26 | 12 | 14 | 0 | 26 | 2 |
| ENSDARG000 | 13:13:13:13:1 | 36666662:366 | 36667163:366 | -:-:-:-:-:-:-:-: | 1809 | l2hgdh | 62 | 170 | 164 | 85 | 146 | 106 |
| ENSDARG000 | 13:13:13:13:1 | 9492954:9493 | 9493124:9493 | -;-;-;-;-;-;- | 1644 | NA | 0 | 0 | 0 | 0 | 0 | 0 |
| ENSDARG000 | 13:13:13:13:1 | 15918449:159 | 15918732:159 | -:-:-:-:-:-:-:-: | 1880 | ttl | 208 | 199 | 214 | 189 | 186 | 104 |
| ENSDARG000 | 13:13:13:13:1 | 22740038:227 | 22740292:227 | -:-:-:-:-:-:-:-: | 2475 | PSD | 656 | 23 | 335 | 14 | 417 | 15 |
| ENSDARG000 | 13:13:13:13:1 | 24665334:246 | 24665500:246 | -;-;-;-;-;- | 951 | znf511 | 74 | 41 | 72 | 68 | 58 | 100 |
| ENSDARG000 | 13:13:13:13:1 | 36680564:366 | 36680645:366 | +:+:+:+:+:+:+: | 940 | dmac2l | 166 | 55 | 99 | 110 | 133 | 124 |
| ENSDARG000 | 13:13:13:13:1 | 25642050:256 | 25642288:256 | -:-:-:-:-:-:-:-: | 1987 | si:dkey-192p2 | 0 | 0 | 6 | 0 | 0 | 2 |
| ENSDARG000 | 13:13:13:13:1 | 20227045:202 | 20227415:202 | -;-;-;-;-;- | 1108 | NA | 0 | 6 | 50 | 8 | 18 | 2 |
| ENSDARG000 | 13:13:13:13:1 | 19884631:199 | 19885393:199 | +:+:+:+:+:+:+: | 4733 | ATRNL1 | 12 | 18 | 25 | 43 | 34 | 27 |
| ENSDARG000 | 13 | 22819485 | 22819667 | - | 183 | NA | 0 | 0 | 1 | 0 | 3 | 0 |
| ENSDARG000 | 13:13:13:13:1 | 20217490:202 | 20217836:202 | -;-;-;-;-;- | 1068 | NA | 0 | 4 | 19 | 0 | 8 | 8 |
| ENSDARG000 | 13:13:13:13:1 | 43507383:435 | 43509916:435 | -:-:-:-:-:-:-:-: | 4549 | ablim1a | 12 | 8 | 6 | 6 | 24 | 30 |
| ENSDARG000 | 13:13:13:13:1 | 2732951:2733 | 2733183:2733 | +:+:+:+:+:+:+: | 5855 | wdr11 | 1041 | 833 | 948 | 736 | 1020 | 790 |
| ENSDARG000 | 13:13:13:13:1 | 9496748:9496 | 9496875:9496 | +:+:+:+:+:+:+: | 1368 | HTRA2 | 4 | 0 | 2 | 0 | 4 | 0 |
| ENSDARG000 | 13;13;13;13 | 33117528:331 | 33117710:331 | +;+;+;+ | 1951 | si:ch211-10a2 | 0 | 0 | 0 | 0 | 0 | 0 |

| | | | | | | | | | | | |
|---|---|---|---|---|---|---|---|---|---|---|---|
| ENSDARG000 | 13:13:13:13:1 | 9517816:9518 | 9517986:9518 | -;-;-;-;-;-;- | 1643 | NA | 0 | 0 | 0 | 0 | 0 | 0 |
| ENSDARG000 | 13:13:13:13:1 | 12801092:128 | 12801182:128 | +;+;+;+;+;+;+; | 16205 | kiaa1109 | 2315 | 1344 | 2121 | 1382 | 2521 | 1882 |
| ENSDARG000 | 13:13:13:13:1 | 24671481:246 | 24671568:246 | +;+;+;+;+;+;+; | 3337 | adam8a | 1562 | 1331 | 1125 | 452 | 1703 | 977 |
| ENSDARG000 | 13:13:13:13:1 | 8954667:8955 | 8954741:8955 | +;+;+;+;+;+;+; | 1895 | haao | 18 | 16 | 34 | 17 | 100 | 46 |
| ENSDARG000 | 13:13:13:13:1 | 36693152:366 | 36693448:366 | -;-;-;-;-;-;-;-; | 1510 | cdkl1 | 0 | 8 | 39 | 24 | 4 | 2 |
| ENSDARG000 | 13:13:13:13:1 | 33123484:331 | 33125514:331 | -;-;-;-;-;-;-;- | 2995 | plek2 | 26 | 14 | 0 | 0 | 2 | 14 |
| ENSDARG000 | 13;13;13 | 15933168:159 | 15933230:159 | +;+;+ | 2118 | fignl1 | 84 | 72 | 149 | 106 | 142 | 76 |
| ENSDARG000 | 13;13 | 20379392:203 | 20380130:203 | -;- | 838 | apcs | 0 | 0 | 34 | 0 | 0 | 0 |
| ENSDARG000 | 13:13:13:13:1 | 22821647:228 | 22822038:228 | +;+;+;+;+;+;+; | 4880 | NA | 40 | 164 | 271 | 41 | 110 | 122 |
| ENSDARG000 | 13:13:13:13:1 | 9521629:9521 | 9521764:9521 | +;+;+;+;+;+;+; | 1243 | HTRA2 | 0 | 4 | 3 | 3 | 12 | 8 |
| ENSDARG000 | 13:13:13:13:1 | 33155428:331 | 33156243:331 | -;-;-;-;-;-;-;-; | 2428 | fbln5 | 211 | 245 | 133 | 60 | 88 | 27 |
| ENSDARG000 | 13:13:13:13:1 | 20392656:203 | 20392799:203 | -;-;-;-;-;-;-;-; | 3206 | gfra1a | 8 | 2 | 2 | 0 | 6 | 0 |
| ENSDARG000 | 13:13:13:13:1 | 33170350:331 | 33175483:331 | -;-;-;-;-;-;-;-; | 10865 | trip11 | 1424 | 1217 | 1776 | 812 | 1836 | 1255 |
| ENSDARG000 | 13:13:13:13:1 | 29127960:291 | 29128747:291 | +;+;+;+;+;+;+; | 7294 | unc5b | 161 | 10 | 62 | 24 | 48 | 31 |
| ENSDARG000 | 13:13:13:13:1 | 24679674:246 | 24679909:246 | +;+;+;+;+;+;+; | 1888 | spef1 | 2 | 0 | 0 | 0 | 2 | 0 |
| ENSDARG000 | 13:13:13:13:1 | 9543180:9543 | 9543350:9543 | -;-;-;-;-;-;- | 1643 | NA | 0 | 0 | 0 | 0 | 0 | 10 |
| ENSDARG000 | 13:13:13:13:1 | 15957508:159 | 15959208:159 | -;-;-;-;-;-;-;-; | 2685 | ikzf1 | 30 | 26 | 6 | 12 | 82 | 34 |
| ENSDARG000 | 13:13:13:13:1 | 25725547:257 | 25729638:257 | -;-;-;-;-;-;-;-; | 5884 | sgpl1 | 1561 | 1616 | 2142 | 1650 | 1364 | 1703 |
| ENSDARG000 | 13;13;13 | 9548849:9550 | 9549033:9551 | +;+;+ | 4584 | pomk | 250 | 132 | 116 | 196 | 214 | 147 |
| ENSDARG000 | 13;13;13;13 | 25720969:257 | 25721116:257 | +;+;+;+ | 820 | pcbd1 | 428 | 550 | 397 | 443 | 516 | 412 |
| ENSDARG000 | 13:13:13:13:1 | 20524921:205 | 20525133:205 | +;+;+;+;+;+;+; | 1061 | CCDC172 | 4 | 0 | 0 | 0 | 0 | 0 |
| ENSDARG000 | 13:13:13:13:1 | 11175208:111 | 11176528:111 | -;-;-;-;-;-;-;-; | 4574 | akt3a | 300 | 400 | 376 | 150 | 471 | 266 |
| ENSDARG000 | 13:13:13:13:1 | 31285660:312 | 31286143:312 | +;+;+;+;+ | 2065 | GDF10 | 172 | 242 | 274 | 607 | 321 | 181 |
| ENSDARG000 | 13:13:13:13:1 | 5978809:5978 | 5978895:5978 | +;+;+;+;+;+;+; | 2258 | phf10 | 343 | 389 | 697 | 289 | 517 | 453 |
| ENSDARG000 | 13:13:13:13:1 | 9559461:9560 | 9560045:9560 | +;+;+;+;+;+;+ | 3051 | wdr32 | 270 | 414 | 415 | 186 | 316 | 268 |
| ENSDARG000 | 13:13:13:13:1 | 2830574:2832 | 2830742:2832 | +;+;+;+;+;+;+; | 4254 | NA | 606 | 634 | 60 | 490 | 22 | 24 |
| ENSDARG000 | 13:13:13:13:1 | 43610350:436 | 43610461:436 | -;-;-;-;-;-;-;-; | 2625 | fhip2a | 221 | 184 | 198 | 120 | 235 | 166 |
| ENSDARG000 | 13;13;13 | 29404218:294 | 29406045:294 | -;-;- | 2126 | C13H10orf53 | 0 | 0 | 0 | 0 | 0 | 0 |
| ENSDARG000 | 13:13:13:13:1 | 9575516:9581 | 9576020:9581 | -;-;-;-;-;-;-;-; | 2741 | CPXM1 | 0 | 6 | 10 | 8 | 2 | 0 |
| ENSDARG000 | 13:13:13:13:1 | 39182099:391 | 39182373:391 | +;+;+;+;+;+;+; | 5433 | fam135a | 867 | 703 | 1114 | 673 | 1115 | 935 |
| ENSDARG000 | 13:13:13:13:1 | 29406814:294 | 29409637:294 | -;-;-;-;-;-;-;-; | 4650 | chata | 0 | 0 | 0 | 0 | 0 | 0 |
| ENSDARG000 | 13:13:13:13:1 | 36705595:367 | 36706472:367 | -;-;-;-;-;-;-;-; | 3942 | map4k5 | 1347 | 1642 | 2181 | 1939 | 1946 | 1508 |
| ENSDARG000 | 13:13:13:13:1 | 6075572:6076 | 6076303:6076 | -;-;-;-;-;-;-;-; | 3339 | dld | 48 | 6 | 2 | 6 | 0 | 6 |
| ENSDARG000 | 13;13;13;13 | 16150654:161 | 16150805:161 | -;-;-;- | 1226 | vwc2 | 0 | 0 | 9 | 50 | 0 | 0 |
| ENSDARG000 | 13:13:13:13:1 | 35689749:356 | 35690279:356 | +;+;+;+;+;+;+; | 7207 | PSME4 | 34 | 40 | 48 | 36 | 79 | 14 |
| ENSDARG000 | 13:13:13:13:1 | 22839821:228 | 22840064:228 | -;-;-;-;-;-;-;-; | 1422 | pbld | 2 | 0 | 8 | 2 | 18 | 14 |
| ENSDARG000 | 13 | 29422049 | 29424454 | - | 2406 | slc18a3a | 0 | 2 | 0 | 0 | 0 | 0 |
| ENSDARG000 | 13:13:13:13:1 | 20535159:205 | 20536062:205 | -;-;-;-;-;-;-;-; | 1884 | si:ch1073-126 | 0 | 0 | 10 | 0 | 0 | 0 |
| ENSDARG000 | 13:13:13:13:1 | 29446226:294 | 29446388:294 | -;-;-;-;-;-;- | 1197 | rgra | 2 | 0 | 1 | 0 | 2 | 0 |
| ENSDARG000 | 13:13:13:13:1 | 16261245:162 | 16261374:162 | +;+;+;+;+;+;+; | 4515 | mocos | 4170 | 1328 | 5122 | 3785 | 9820 | 2872 |
| ENSDARG000 | 13:13:13:13:1 | 22843830:228 | 22844139:228 | -;-;-;-;-;-;-;-; | 1257 | NA | 0 | 0 | 22 | 4 | 46 | 0 |
| ENSDARG000 | 13:13:13:13:1 | 25762908:257 | 25763566:257 | -;-;-;-;-;-;-;-; | 2308 | pdlim1 | 873 | 906 | 544 | 1050 | 491 | 841 |
| ENSDARG000 | 13:13:13:13:1 | 24701406:247 | 24704538:247 | -;-;-;-;-;-;-;-; | 4419 | erlin1 | 1492 | 1571 | 1915 | 1095 | 2055 | 1571 |
| ENSDARG000 | 13;13;13 | 25761279:257 | 25762124:257 | +;+;+ | 1113 | neurog3 | 0 | 0 | 0 | 0 | 0 | 6 |
| ENSDARG000 | 13;13;13;13 | 31292700:312 | 31294440:312 | -;-;-;- | 2351 | prdm8 | 4 | 6 | 8 | 2 | 4 | 2 |
| ENSDARG000 | 13:13:13:13:1 | 43639867:436 | 43640084:436 | +;+;+;+;+;+;+; | 2048 | zfyve21 | 972 | 935 | 1466 | 1398 | 1395 | 1389 |
| ENSDARG000 | 13:13:13:13:1 | 11436130:114 | 11438483:114 | +;+;+;+;+;+ | 7068 | zbtb18 | 565 | 304 | 327 | 242 | 473 | 314 |
| ENSDARG000 | 13:13:13:13:1 | 9612395:9612 | 9612506:9612 | +;+;+;+;+;+;+; | 5531 | NA | 255 | 279 | 296 | 119 | 176 | 333 |
| ENSDARG000 | 13;13;13;13 | 29462249:294 | 29462670:294 | +;+;+;+ | 3427 | lrit1a | 0 | 4 | 0 | 0 | 0 | 0 |
| ENSDARG000 | 13;13 | 11445400:114 | 11445768:114 | +;+ | 745 | NA | 0 | 0 | 0 | 0 | 0 | 0 |
| ENSDARG000 | 13;13;13 | 29470442:294 | 29470593:294 | +;+;+ | 2116 | lrit2 | 0 | 0 | 0 | 0 | 0 | 0 |
| ENSDARG000 | 13:13:13:13:1 | 31297016:312 | 31297125:312 | +;+;+;+;+;+;+; | 2015 | NA | 700 | 366 | 623 | 452 | 574 | 234 |
| ENSDARG000 | 13:13:13:13:1 | 11489953:114 | 11490750:114 | -;-;-;-;-;-;-;-; | 2187 | adss2 | 107 | 126 | 110 | 144 | 88 | 159 |
| ENSDARG000 | 13:13:13:13:1 | 22849796:228 | 22850048:228 | -;-;-;-;-;-;-;-; | 1178 | pbld2 | 0 | 0 | 2 | 0 | 0 | 4 |
| ENSDARG000 | 13:13:13:13:1 | 16279890:162 | 16279960:162 | +;+;+;+;+;+;+; | 2564 | anxa11a | 3081 | 2898 | 3943 | 4816 | 4655 | 2892 |
| ENSDARG000 | 13:13:13:13:1 | 29480010:294 | 29482376:294 | -;-;-;-;-;-;-;-; | 4850 | cdhr1a | 0 | 0 | 2 | 0 | 12 | 0 |
| ENSDARG000 | 13:13:13:13:1 | 39236141:392 | 39236175:392 | +;+;+;+;+;+;+; | 2985 | FRMD1 | 104 | 48 | 22 | 23 | 28 | 18 |
| ENSDARG000 | 13 | 20546967 | 20547498 | - | 532 | NA | 0 | 0 | 0 | 0 | 0 | 0 |
| ENSDARG000 | 13:13:13:13:1 | 33211014:332 | 33211543:332 | -;-;-;-;-;-;-;-; | 2933 | golga5 | 1237 | 843 | 1506 | 1225 | 1193 | 790 |
| ENSDARG000 | 13;13;13 | 20856611:208 | 20856758:208 | +;+;+ | 358 | NA | 0 | 0 | 0 | 0 | 0 | 0 |
| ENSDARG000 | 13:13:13:13:1 | 21600946:216 | 21601314:216 | +;+;+;+;+;+;+; | 1708 | SH2D4B | 0 | 0 | 0 | 0 | 0 | 0 |
| ENSDARG000 | 13:13:13:13:1 | 12993851:129 | 12994853:129 | -;-;-;-;-;-;-;-; | 5357 | nsd2 | 409 | 1240 | 1570 | 655 | 1327 | 730 |
| ENSDARG000 | 13:13:13:13:1 | 11550454:115 | 11550973:115 | +;+;+;+;+;+;+; | 1764 | desi2 | 396 | 436 | 557 | 547 | 362 | 504 |
| ENSDARG000 | 13:13:13:13:1 | 21638454:216 | 21639061:216 | -;-;-;-;-;-;-;- | 1442 | tspan14 | 1014 | 1203 | 1500 | 893 | 1116 | 1449 |
| ENSDARG000 | 13:13:13:13:1 | 33246150:332 | 33246429:332 | +;+;+;+;+;+;+; | 1977 | ndufaf1 | 198 | 344 | 336 | 184 | 438 | 283 |
| ENSDARG000 | 13:13:13:13:1 | 22863516:228 | 22863589:228 | +;+;+;+;+;+;+; | 2565 | hnrnph3 | 276 | 258 | 313 | 363 | 251 | 254 |
| ENSDARG000 | 13:13:13:13:1 | 36764715:367 | 36764872:367 | +;+;+;+;+;+;+; | 2198 | atl1 | 254 | 249 | 138 | 147 | 84 | 150 |
| ENSDARG000 | 13 | 11577407 | 11578304 | - | 898 | NA | 0 | 0 | 0 | 2 | 2 | 1 |
| ENSDARG000 | 13:13:13:13:1 | 29506660:295 | 29506961:295 | -;-;-;-;-;-;- | 1015 | NA | 0 | 0 | 0 | 0 | 0 | 0 |
| ENSDARG000 | 13:13:13:13:1 | 9678262:9678 | 9678562:9678 | +;+;+;+;+;+;+; | 4126 | SEMA4G | 14 | 0 | 0 | 0 | 0 | 0 |
| ENSDARG000 | 13:13:13:13:1 | 29510023:295 | 29510472:295 | +;+;+;+;+;+ | 1909 | chst3a | 0 | 0 | 0 | 0 | 0 | 0 |
| ENSDARG000 | 13:13:13:13:1 | 33251721:332 | 33252284:332 | -;-;-;-;-;-;-;-; | 1923 | nusap1 | 310 | 1049 | 1202 | 657 | 796 | 498 |
| ENSDARG000 | 13:13:13:13:1 | 33229446:332 | 33230570:332 | -;-;-;-;-;-;-;-; | 3122 | rtf1 | 1112 | 1166 | 1784 | 1030 | 1488 | 1645 |
| ENSDARG000 | 13:13:13:13:1 | 44702196:447 | 44703591:447 | -;-;-;-;-;-;-;-; | 2639 | zfand3 | 2412 | 1390 | 1797 | 1764 | 1708 | 1362 |
| ENSDARG000 | 13:13:13:13:1 | 25795511:257 | 25796537:257 | -;-;-;-;-;-;-;-; | 2137 | rel | 404 | 382 | 326 | 308 | 367 | 303 |
| ENSDARG000 | 13;13;13 | 21656713:216 | 21658301:216 | -;-;- | 1867 | mxtx1 | 0 | 0 | 0 | 0 | 0 | 0 |
| ENSDARG000 | 13:13:13:13:1 | 21666521:216 | 21668398:216 | -;-;-;-;-;- | 3013 | elovl6l | 26 | 4 | 0 | 0 | 10 | 0 |
| ENSDARG000 | 13:13:13:13:1 | 31343326:313 | 31343697:313 | -;-;-;-;-;-;-;-; | 4231 | rrp12 | 2055 | 2045 | 3710 | 1227 | 2633 | 2282 |
| ENSDARG000 | 13;13;13;13 | 39315881:393 | 39316034:393 | +;+;+;+ | 894 | NA | 0 | 0 | 0 | 0 | 0 | 0 |
| ENSDARG000 | 13:13:13:13:1 | 39277178:392 | 39278054:392 | +;+;+;+;+;+;+; | 5452 | NA | 188 | 92 | 266 | 105 | 261 | 231 |
| ENSDARG000 | 13:13:13:13:1 | 11581942:115 | 11586776:115 | -;-;-;-;-;-;-;-; | 8643 | DCTN1 | 0 | 34 | 36 | 80 | 22 | 18 |
| ENSDARG000 | 13:13:13:13:1 | 16342926:163 | 16343226:163 | +;+;+;+;+;+;+; | 9454 | dlg5a | 880 | 496 | 917 | 656 | 1219 | 782 |
| ENSDARG000 | 13:13:13:13:1 | 35745572:357 | 35745813:357 | +;+;+;+;+ | 2594 | GPR75 | 0 | 0 | 0 | 0 | 0 | 0 |
| ENSDARG000 | 13:13:13:13:1 | 24717880:247 | 24717945:247 | +;+;+;+;+;+;+; | 5120 | CFAP43 | 0 | 0 | 0 | 0 | 6 | 8 |
| ENSDARG000 | 13;13;13 | 24742724:247 | 24743344:247 | -;-;- | 1221 | sfr1 | 12 | 40 | 27 | 12 | 8 | 23 |
| ENSDARG000 | 13:13:13:13:1 | 44763835:447 | 44764230:447 | -;-;-;-;-;-;-;-; | 2274 | btbd9 | 352 | 269 | 367 | 224 | 360 | 322 |
| ENSDARG000 | 13:13:13:13:1 | 44788077:447 | 44788976:447 | -;-;-;-;-;- | 1523 | glo1 | 816 | 718 | 1128 | 1251 | 717 | 904 |
| ENSDARG000 | 13:13:13:13:1 | 36787077:367 | 36788054:367 | -;-;-;-;- | 2282 | sav1 | 892 | 969 | 1083 | 932 | 869 | 1033 |
| ENSDARG000 | 13:13:13:13:1 | 22876467:228 | 22877278:228 | -;-;-;-;-;-;-;-; | 3103 | rufy2 | 1097 | 531 | 816 | 743 | 954 | 718 |
| ENSDARG000 | 13:13:13:13:1 | 39346298:393 | 39346536:393 | -;-;-;-;-;-;-;-; | 1287 | SLC35F3 | 0 | 0 | 2 | 0 | 0 | 0 |
| ENSDARG000 | 13:13:13:13:1 | 21674445:216 | 21674539:216 | +;+;+;+;+;+;+; | 1455 | mtg1 | 106 | 186 | 214 | 132 | 183 | 141 |
| ENSDARG000 | 13;13 | 39684492:397 | 39684691:397 | +;+ | 825 | NA | 0 | 0 | 0 | 0 | 0 | 0 |
| ENSDARG000 | 13:13:13:13:1 | 39656610:397 | 39657739:397 | -;-;-;-;-;-;-;-; | 2859 | zgc:171482 | 0 | 0 | 4 | 0 | 0 | 3 |
| ENSDARG000 | 13:13:13:13:1 | 29770837:297 | 29771505:297 | +;+;+;+;+;+;+; | 5138 | pax2a | 0 | 0 | 23 | 0 | 32 | 0 |
| ENSDARG000 | 13;13 | 21682770:216 | 21684091:216 | -;- | 1454 | sprn | 0 | 0 | 5 | 0 | 4 | 1 |
| ENSDARG000 | 13:13:13:13:1 | 25824345:258 | 25825470:258 | -;-;-;-;-;-;-;-; | 3725 | papolg | 1248 | 997 | 1843 | 1042 | 1354 | 1255 |
| ENSDARG000 | 13:13:13:13:1 | 35750690:357 | 35751325:357 | -;-;-;-;-;-;-;-; | 2955 | erlec1 | 1285 | 1123 | 1337 | 725 | 1581 | 1006 |
| ENSDARG000 | 13:13:13:13:1 | 13033424:130 | 13034134:130 | +;+;+;+;+;+;+; | 3397 | letm1 | 601 | 486 | 477 | 234 | 457 | 285 |
| ENSDARG000 | 13;13;13 | 21689267:216 | 21693582:216 | -;-;- | 4934 | NA | 4 | 0 | 0 | 0 | 0 | 12 |
| ENSDARG000 | 13;13;13 | 39911551:399 | 39912722:399 | -;-;- | 1976 | sfrp5 | 0 | 0 | 4 | 0 | 8 | 2 |
| ENSDARG000 | 13:13:13:13:1 | 36800334:368 | 36803771:368 | -;-;-;-;-;-;-;-; | 9712 | NIN | 125 | 156 | 184 | 120 | 197 | 190 |
| ENSDARG000 | 13:13:13:13:1 | 16521898:165 | 16522895:165 | +;+;+;+;+;+;+; | 5793 | kcnma1a | 0 | 6 | 9 | 2 | 2 | 16 |
| ENSDARG000 | 13:13:13:13:1 | 21712284:217 | 21712584:217 | -;-;-;-;-;-;-;-; | 1997 | NA | 6 | 0 | 0 | 0 | 0 | 0 |
| ENSDARG000 | 13:13:13:13:1 | 40019001:400 | 40019243:400 | +;+;+;+;+;+;+; | 2191 | GOLGA7B | 0 | 0 | 0 | 0 | 0 | 0 |
| ENSDARG000 | 13:13:13:13:1 | 31371536:313 | 31372905:313 | -;-;-;-;-;- | 2272 | ubtd1a | 28 | 28 | 20 | 25 | 0 | 6 |
| ENSDARG000 | 13:13:13:13:1 | 31379784:313 | 31381470:313 | -;-;-;-;-;-;-;-; | 2906 | zdhhc16a | 169 | 106 | 223 | 126 | 120 | 152 |
| ENSDARG000 | 13:13:13:13:1 | 22924024:229 | 22926314:229 | -;-;-;-;-;-;-;-; | 3247 | tspan15 | 26 | 14 | 14 | 42 | 20 | 2 |
| ENSDARG000 | 13;13;13 | 21768447:217 | 21768759:217 | +;+;+ | 1695 | chchd1 | 308 | 587 | 522 | 292 | 316 | 404 |
| ENSDARG000 | 13:13:13:13:1 | 21779975:217 | 21780040:217 | +;+;+;+;+;+;+; | 7404 | NA | 6 | 8 | 20 | 0 | 0 | 16 |
| ENSDARG000 | 13:13:13:13:1 | 40064481:400 | 40065277:400 | -;-;-;-;-;-;-;-; | 2838 | CRTAC1 | 8 | 0 | 6 | 2 | 2 | 0 |
| ENSDARG000 | 13:13:13:13:1 | 35765317:357 | 35765484:357 | +;+;+;+;+;+;+; | 1935 | agpat4 | 233 | 327 | 250 | 198 | 181 | 220 |
| ENSDARG000 | 13:13:13:13:1 | 35787873:357 | 35788638:357 | -;-;-;-;-;-;-;-; | 5742 | NA | 937 | 1091 | 1350 | 793 | 1416 | 1015 |
| ENSDARG000 | 13:13:13:13:1 | 22964868:229 | 22965462:229 | +;+;+;+;+;+;+ | 2736 | tacr2 | 4 | 0 | 12 | 2 | 2 | 0 |
| ENSDARG000 | 13:13:13:13:1 | 31390313:313 | 31390487:313 | +;+;+;+;+ | 2294 | mettl18 | 144 | 248 | 249 | 114 | 236 | 196 |
| ENSDARG000 | 13:13:13:13:1 | 31393192:313 | 31393701:313 | -;-;-;-;-;- | 1774 | pgam1a | 3417 | 6785 | 11815 | 9583 | 5411 | 1936 |
| ENSDARG000 | 13:13:13:13:1 | 24750078:247 | 24750428:247 | +;+;+;+;+;+;+; | 5454 | NA | 191 | 58 | 13 | 10 | 30 | 15 |
| ENSDARG000 | 13:13:13:13:1 | 25846528:258 | 25853366:258 | +;+;+;+;+ | 10033 | bcl11aa | 676 | 784 | 322 | 999 | 447 | 579 |
| ENSDARG000 | 13:13:13:13:1 | 40182487:401 | 40186180:401 | -;-;-;-;-;-;-;-; | 6382 | LOXL4 | 66 | 62 | 82 | 70 | 72 | 54 |
| ENSDARG000 | 13:13:13:13:1 | 21826096:218 | 21826546:218 | +;+;+;+;+;+;+; | 8217 | zswim8 | 2083 | 1322 | 2256 | 1317 | 2458 | 2018 |
| ENSDARG000 | 13:13:13:13:1 | 26780004:267 | 26780564:267 | -;-;-;-;-;-;-;-; | 2733 | vrk2 | 426 | 373 | 337 | 450 | 609 | 399 |
| ENSDARG000 | 13:13:13:13:1 | 26703897:267 | 26704185:267 | +;+;+;+;+;+;+; | 2029 | fancl | 30 | 40 | 69 | 38 | 91 | 44 |
| ENSDARG000 | 13:13:13:13:1 | 22972888:229 | 22974135:229 | -;-;-;-;-;-;-;-; | 4061 | hk1 | 1385 | 3784 | 10415 | 6113 | 3451 | 1094 |
| ENSDARG000 | 13:13:13:13:1 | 26226410:263 | 26226566:263 | -;-;-;-;-;-;-;-; | 1558 | si:rp71-1h3.1 | 0 | 8 | 2 | 0 | 12 | 1 |
| ENSDARG000 | 13:13:13:13:1 | 36846789:368 | 36847110:368 | -;-;-;-;-;-;-;-; | 2797 | pygl | 0 | 2 | 44 | 0 | 8 | 0 |
| ENSDARG000 | 13:13:13:13:1 | 40247379:402 | 40250439:402 | -;-;-;-;-;-;-;-; | 4414 | zgc:123010 | 1436 | 916 | 987 | 917 | 912 | 854 |
| ENSDARG000 | 13:13:13:13:1 | 31424586:314 | 31425496:314 | -;-;-;-;-;-;-;-; | 3972 | rtn1a | 5924 | 6881 | 6952 | 4810 | 6064 | 4392 |
| ENSDARG000 | 13:13:13:13:1 | 31402067:314 | 31402289:314 | +;+;+;+;+;+;+; | 4524 | tdrd9 | 2 | 16 | 10 | 34 | 20 | 4 |
| ENSDARG000 | 13:13:13:13:1 | 23017498:230 | 23018067:230 | -;-;-;-;-;-;-;-; | 3280 | hkdc1 | 71 | 8 | 20 | 0 | 24 | 10 |
| ENSDARG000 | 13:13:13:13:1 | 18521493:185 | 18521501:185 | -;-;-;-;-;-;-;-; | 1467 | NA | 6 | 0 | 1 | 0 | 0 | 0 |
| ENSDARG000 | 13 | 44495063 | 44496070 | + | 1008 | NA | 0 | 0 | 0 | 0 | 0 | 0 |
| ENSDARG000 | 13:13:13:13:1 | 2515900:2515 | 2516135:2516 | -;-;-;-;-;-;-;- | 578 | cox20 | 94 | 127 | 88 | 62 | 80 | 88 |
| ENSDARG000 | 13:13:13:13:1 | 33488032:334 | 33488097:334 | +;+;+;+;+;+;+; | 3621 | NA | 54 | 12 | 47 | 0 | 46 | 18 |
| ENSDARG000 | 13:13:13:13:1 | 33703784:337 | 33703820:337 | +;+;+;+;+;+;+; | 1359 | macrod2 | 434 | 347 | 311 | 306 | 314 | 373 |
| ENSDARG000 | 13:13:13:13:1 | 2315110:2319 | 2315398:2319 | -;-;-;-;-;-;-;-; | 2033 | eloal | 0 | 0 | 0 | 0 | 0 | 0 |
| ENSDARG000 | 13;13;13 | 42046770:420 | 42047415:420 | -;-;- | 1243 | NA | 18 | 10 | 24 | 6 | 48 | 16 |
| ENSDARG000 | 13:13:13:13:1 | 47710434:477 | 47710494:477 | +;+;+;+;+;+;+; | 687 | TMEM87B | 80 | 130 | 124 | 47 | 156 | 75 |
| ENSDARG000 | 13:13:13:13:1 | 44299676:443 | 44300005:443 | -;-;-;-;-;-;-;-; | 3359 | MDGA1 | 532 | 2 | 8 | 72 | 0 | 62 |
| ENSDARG000 | 13:13:13:13:1 | 7049210:7049 | 7049410:7049 | +;+;+;+;+;+;+; | 5426 | rnaset2 | 3297 | 2401 | 2998 | 2887 | 3095 | 4577 |
| ENSDARG000 | 13:13:13:13:1 | 42552495:425 | 42552811:425 | -;-;-;-;-;- | 890 | NA | 6 | 0 | 20 | 0 | 10 | 4 |
| ENSDARG000 | 13;13;13 | 669177:67040 | 669278:67046 | +;+;+ | 213 | NA | 4 | 1 | 0 | 0 | 0 | 0 |
| ENSDARG000 | 13:13:13:13:1 | 51966573:519 | 51966860:519 | -;-;-;-;-;- | 2067 | slc30a10 | 110 | 12 | 19 | 24 | 88 | 4 |
| ENSDARG000 | 13;13;13 | 15682803:156 | 15682804:156 | +;+;+ | 252 | NA | 0 | 0 | 0 | 0 | 0 | 0 |
| ENSDARG000 | 13;13;13 | 51949171:519 | 51949277:519 | -;-;- | 483 | NA | 26 | 24 | 32 | 5 | 13 | 18 |
| ENSDARG000 | 13;13;13;13 | 50621840:506 | 50622341:506 | -;-;-;- | 2904 | vox | 12 | 0 | 2 | 16 | 4 | 18 |
| ENSDARG000 | 13:13:13:13:1 | 45582391:456 | 45582661:456 | +;+;+;+;+;+;+; | 1615 | ldlrap1b | 112 | 110 | 106 | 73 | 118 | 65 |
| ENSDARG000 | 13:13:13:13:1 | 44849564:448 | 44849582:448 | -;-;-;-;-;-;-;-; | 168 | NA | 0 | 0 | 0 | 0 | 0 | 0 |
| ENSDARG000 | 13:13:13:13:1 | 51140464:511 | 51140808:511 | -;-;-;-;-;-;-;-; | 1893 | NA | 0 | 0 | 0 | 0 | 47 | 0 |
| ENSDARG000 | 13;13;13 | 40397517:403 | 40398564:403 | -;-;- | 1530 | nkx3.3 | 0 | 0 | 0 | 0 | 0 | 0 |
| ENSDARG000 | 13;13 | 38084077:380 | 38084234:380 | -;- | 1044 | NA | 0 | 0 | 0 | 0 | 0 | 0 |
| ENSDARG000 | 13:13:13:13:1 | 47821524:478 | 47821609:478 | +;+;+;+;+;+;+; | 3120 | NA | 153 | 117 | 320 | 235 | 328 | 206 |
| ENSDARG000 | 13:13:13:13:1 | 33141279:331 | 33141588:331 | -;-;-;-;-;-;-;-; | 1637 | TC2N | 0 | 2 | 0 | 0 | 0 | 0 |
| ENSDARG000 | 13 | 38150517 | 38150637 | + | 121 | 5S_rRNA | 0 | 0 | 0 | 0 | 0 | 0 |
| ENSDARG000 | 13 | 47049912 | 47050026 | + | 115 | 5S_rRNA | 0 | 0 | 0 | 0 | 0 | 0 |
| ENSDARG000 | 13 | 41844953 | 41845067 | - | 115 | 5S_rRNA | 0 | 0 | 0 | 0 | 0 | 0 |
| ENSDARG000 | 13 | 18649492 | 18649682 | - | 191 | U2 | 0 | 0 | 0 | 0 | 0 | 0 |
| ENSDARG000 | 13:13:13:13:1 | 1542493:1542 | 1542678:1542 | +;+;+;+;+;+ | 5863 | NA | 99 | 23 | 42 | 13 | 57 | 44 |
| ENSDARG000 | 13 | 13844755 | 13844873 | + | 119 | 5S_rRNA | 0 | 0 | 0 | 0 | 0 | 0 |
| ENSDARG000 | 13:13:13:13:1 | 40462366:404 | 40463845:404 | -;-;-;-;-;-;-;-; | 4311 | CNNM1 | 0 | 0 | 0 | 0 | 0 | 0 |
| ENSDARG000 | 13:13:13:13:1 | 33394116:333 | 33395541:333 | -;-;-;-;-;-;-;- | 2130 | btbd6a | 315 | 175 | 228 | 146 | 179 | 301 |
| ENSDARG000 | 13 | 22123090 | 22123205 | + | 116 | 5S_rRNA | 0 | 0 | 0 | 0 | 0 | 0 |
| ENSDARG000 | 13 | 26567066 | 26567184 | - | 119 | 5S_rRNA | 0 | 0 | 0 | 0 | 0 | 0 |
| ENSDARG000 | 13:13:13:13:1 | 49867268:498 | 49867406:498 | -;-;-;-;-;-;-;-; | 8001 | LYST | 12839 | 6894 | 9578 | 8345 | 8353 | 9026 |

| | | | | | | | | | | | |
|---|---|---|---|---|---|---|---|---|---|---|---|
| ENSDARG000 | 13:13:13:13:1 | 1136462:113 | 1136482:113 | -:-:-:-:-:-:-:-:-: | 339 | NA | 9 | 4 | 5 | 3 | 9 | 0 |
| ENSDARG000 | 13 | 51001059 | 51001178 | + | 120 | 5S_rRNA | 0 | 0 | 0 | 0 | 0 | 0 |
| ENSDARG000 | 13:13:13:13:1 | 50387365:503 | 50387573:503 | -:-:-:-:-:-:-:-:-: | 3647 | HERC4 | 256 | 398 | 433 | 383 | 360 | 266 |
| ENSDARG000 | 13:13:13:13:1 | 50151407:501 | 50152186:501 | +;+;+;+;+;+;+ | 4733 | gpr137ba | 244 | 130 | 206 | 42 | 267 | 191 |
| ENSDARG000 | 13 | 825904 | 826225 | - | 322 | rn7sk | 4 | 8 | 22 | 4 | 0 | 24 |
| ENSDARG000 | 13 | 18382289 | 18382405 | + | 117 | 5S_rRNA | 0 | 0 | 0 | 0 | 0 | 0 |
| ENSDARG000 | 13 | 39242555 | 39242672 | - | 118 | 5S_rRNA | 0 | 0 | 0 | 0 | 0 | 0 |
| ENSDARG000 | 13 | 42910630 | 42910746 | - | 117 | 5S_rRNA | 0 | 0 | 0 | 0 | 0 | 0 |
| ENSDARG000 | 13:13:13:13:1 | 52006253:520 | 52007190:520 | +:+:+:+:+:+:+: | 7653 | npas4l | 10 | 8 | 2 | 12 | 16 | 22 |
| ENSDARG000 | 13:13:13:13:1 | 51853716:518 | 51854193:518 | +:+:+:+:+:+:+: | 2457 | NA | 10 | 4 | 12 | 16 | 6 | 12 |
| ENSDARG000 | 13;13;13 | 42124566:421 | 42124647:421 | +;+;+ | 1734 | cdc42ep3 | 20 | 0 | 6 | 2 | 6 | 2 |
| ENSDARG000 | 13;13;13 | 51710725:517 | 51710864:517 | +;+;+ | 4945 | PWWP2B | 805 | 712 | 864 | 624 | 1149 | 555 |
| ENSDARG000 | 13:13:13:13:1 | 11876145:118 | 11877009:118 | +:+:+:+:+:+:+: | 4253 | trim8a | 662 | 675 | 753 | 548 | 702 | 600 |
| ENSDARG000 | 13:13:13:13:1 | 33651416:336 | 33651430:336 | +:+:+:+:+:+:+: | 387 | NA | 0 | 0 | 0 | 1 | 0 | 0 |
| ENSDARG000 | 13:13:13:13:1 | 47750933:477 | 47752799:477 | -;-;-;-;-;-;-;- | 3198 | NA | 176 | 557 | 3042 | 1242 | 3423 | 1644 |
| ENSDARG000 | 13;13;13;13 | 47803499:478 | 47806202:478 | -;-;-;- | 3701 | zmp:0000001 | 16 | 6 | 16 | 14 | 15 | 18 |
| ENSDARG000 | 13;13;13 | 48224541:482 | 48225433:482 | -;-;- | 1257 | KCNK12 | 0 | 0 | 0 | 0 | 0 | 0 |
| ENSDARG000 | 13 | 35962350 | 35962466 | - | 117 | 5S_rRNA | 0 | 0 | 0 | 0 | 0 | 0 |
| ENSDARG000 | 13:13:13:13:1 | 28819768:288 | 28820513:288 | +:+:+:+:+:+:+: | 7511 | NA | 220 | 105 | 159 | 90 | 120 | 156 |
| ENSDARG000 | 13;13;13 | 2908764:2911 | 2908876:2911 | +;+;+ | 399 | wu:fj16a03 | 18 | 9 | 11 | 4 | 2 | 12 |
| ENSDARG000 | 13;13;13;13 | 47810211:478 | 47810307:478 | +;+;+;+ | 956 | zmp:0000001 | 20 | 13 | 15 | 0 | 8 | 14 |
| ENSDARG000 | 13;13;13;13 | 44897170:449 | 44897309:449 | -;-;-;- | 1590 | NA | 138 | 35 | 236 | 22 | 96 | 35 |
| ENSDARG000 | 13:13:13:13:1 | 674351:6746 | 674420:6746 | +:+:+:+:+:+:+: | 756 | cutc | 328 | 300 | 449 | 267 | 409 | 347 |
| ENSDARG000 | 13;13 | 129128:1333 | 129275:1335 | -;- | 297 | NA | 12 | 9 | 9 | 16 | 4 | 4 |
| ENSDARG000 | 13;13 | 11670967:116 | 11671432:116 | -;- | 664 | NA | 2 | 0 | 0 | 0 | 0 | 0 |
| ENSDARG000 | 13:13:13:13:1 | 1726953:172 | 1727676:172 | +:+:+:+:+:+:+: | 7585 | NA | 8 | 12 | 0 | 10 | 9 | 2 |
| ENSDARG000 | 13;13;13;13 | 33688474:336 | 33688517:336 | +;+;+;+ | 654 | NA | 0 | 2 | 0 | 0 | 1 | 1 |
| ENSDARG000 | 13 | 50602639 | 50602714 | + | 76 | dre-mir-202 | 0 | 0 | 0 | 0 | 0 | 0 |
| ENSDARG000 | 13 | 7381518 | 7381633 | + | 116 | 5S_rRNA | 0 | 0 | 0 | 0 | 0 | 0 |
| ENSDARG000 | 13 | 10264741 | 10264905 | - | 165 | NA | 0 | 0 | 0 | 0 | 0 | 0 |
| ENSDARG000 | 13 | 403103 | 404113 | + | 1011 | NA | 0 | 0 | 0 | 0 | 0 | 0 |
| ENSDARG000 | 13:13:13:13:1 | 29238850:292 | 29238937:292 | +:+:+:+:+:+:+: | 6012 | MYOF | 424 | 99 | 73 | 70 | 93 | 102 |
| ENSDARG000 | 13:13:13:13:1 | 29352842:293 | 29353712:293 | +:+:+:+:+:+:+: | 6619 | OGDHL | 0 | 0 | 0 | 1 | 0 | 6 |
| ENSDARG000 | 13:13:13:13:1 | 83010:83838: | 83075:84014: | -:-:-:-:-:-:-:-:-: | 972 | pole2 | 62 | 164 | 214 | 98 | 168 | 64 |
| ENSDARG000 | 13 | 20026967 | 20027084 | + | 118 | 5S_rRNA | 0 | 0 | 0 | 0 | 0 | 0 |
| ENSDARG000 | 13 | 10804713 | 10804835 | + | 123 | 5S_rRNA | 0 | 0 | 0 | 0 | 0 | 0 |
| ENSDARG000 | 13;13 | 228045:2286 | 228179:2290 | +;+ | 555 | prorsd1 | 54 | 138 | 92 | 62 | 132 | 22 |
| ENSDARG000 | 13 | 51102744 | 51102859 | + | 116 | 5S_rRNA | 0 | 0 | 0 | 0 | 0 | 0 |
| ENSDARG000 | 13 | 34969519 | 34969645 | + | 127 | 5S_rRNA | 0 | 0 | 0 | 0 | 0 | 0 |
| ENSDARG000 | 13 | 24127063 | 24127179 | - | 117 | 5S_rRNA | 0 | 0 | 0 | 0 | 0 | 0 |
| ENSDARG000 | 13 | 37674178 | 37674294 | + | 117 | 5S_rRNA | 0 | 0 | 1 | 0 | 0 | 0 |
| ENSDARG000 | 13 | 29590994 | 29591115 | + | 122 | 5S_rRNA | 0 | 0 | 0 | 0 | 0 | 0 |
| ENSDARG000 | 13 | 13210610 | 13210724 | + | 115 | 5S_rRNA | 0 | 0 | 0 | 0 | 0 | 0 |
| ENSDARG000 | 13;13;13;13 | 30692669:306 | 30692768:306 | +;+;+;+ | 2258 | NA | 94 | 64 | 123 | 16 | 24 | 26 |
| ENSDARG000 | 13 | 36399260 | 36399375 | - | 116 | 5S_rRNA | 0 | 0 | 0 | 0 | 0 | 0 |
| ENSDARG000 | 13:13:13:13:1 | 159504:1612 | 159519:1613 | +:+:+:+:+:+:+: | 1311 | NA | 42 | 34 | 95 | 0 | 0 | 42 |
| ENSDARG000 | 13;13;13 | 50607664:506 | 50610605:506 | -;-;- | 3263 | vent | 24 | 0 | 0 | 0 | 2 | 2 |
| ENSDARG000 | 13 | 20248893 | 20249009 | + | 117 | 5S_rRNA | 0 | 0 | 0 | 0 | 0 | 0 |
| ENSDARG000 | 13 | 39532050 | 39533365 | + | 1316 | marveld1 | 1488 | 1125 | 2020 | 1275 | 1663 | 1418 |
| ENSDARG000 | 13;13 | 866293:8697 | 866517:8698 | -;- | 279 | NA | 31 | 31 | 28 | 26 | 22 | 15 |
| ENSDARG000 | 13:13:13:13:1 | 51981764:519 | 51981917:519 | +;+;+;+;+;+ | 681 | KLHDC3 | 320 | 287 | 477 | 364 | 329 | 288 |
| ENSDARG000 | 13:13:13:13:1 | 36537426:365 | 36537674:365 | -;-;-;-;- | 597 | NA | 48 | 20 | 14 | 1 | 16 | 13 |
| ENSDARG000 | 13;13;13;13 | 19871646:198 | 19872243:198 | +;+;+;+ | 2042 | NA | 0 | 4 | 8 | 2 | 0 | 0 |
| ENSDARG000 | 13 | 32984654 | 32984769 | + | 116 | 5S_rRNA | 0 | 0 | 0 | 0 | 0 | 0 |
| ENSDARG000 | 13:13:13:13:1 | 36923052:369 | 36923298:369 | +:+:+:+:+:+:+: | 3558 | tmx1 | 793 | 925 | 831 | 474 | 758 | 765 |
| ENSDARG000 | 13 | 11565736 | 11565863 | - | 128 | 5S_rRNA | 0 | 0 | 0 | 4 | 0 | 1 |
| ENSDARG000 | 13 | 47904068 | 47904182 | + | 115 | 5S_rRNA | 2 | 0 | 0 | 0 | 0 | 0 |
| ENSDARG000 | 13 | 29755293 | 29755427 | + | 135 | 5S_rRNA | 0 | 0 | 0 | 0 | 0 | 0 |
| ENSDARG000 | 13:13:13:13:1 | 50778187:508 | 50778362:508 | +:+:+:+:+:+:+: | 1461 | dntt | 0 | 0 | 0 | 0 | 0 | 0 |
| ENSDARG000 | 13 | 48265070 | 48265188 | - | 119 | 5S_rRNA | 0 | 0 | 0 | 0 | 0 | 0 |
| ENSDARG000 | 13 | 42042226 | 42042342 | + | 117 | 5S_rRNA | 0 | 0 | 0 | 0 | 0 | 0 |
| ENSDARG000 | 13 | 31075833 | 31075947 | - | 115 | 5S_rRNA | 0 | 0 | 0 | 0 | 0 | 0 |
| ENSDARG000 | 13 | 416648 | 417628 | + | 981 | NA | 0 | 0 | 0 | 0 | 0 | 0 |
| ENSDARG000 | 13:13:13:13:1 | 4656504:4656 | 4657864:4657 | -;-;-;-;-;-;-;-;- | 2525 | c1d | 154 | 241 | 403 | 154 | 216 | 294 |
| ENSDARG000 | 13 | 11974452 | 11974568 | - | 117 | 5S_rRNA | 4 | 2 | 7 | 0 | 0 | 7 |
| ENSDARG000 | 13 | 31579448 | 31579564 | - | 117 | 5S_rRNA | 0 | 0 | 0 | 0 | 0 | 0 |
| ENSDARG000 | 13:13:13:13:1 | 48482256:485 | 48482593:485 | +;+;+;+;+ | 1255 | FOXN2 | 10 | 0 | 6 | 0 | 4 | 0 |
| ENSDARG000 | 13 | 48046112 | 48046228 | + | 117 | 5S_rRNA | 0 | 0 | 6 | 0 | 0 | 3 |
| ENSDARG000 | 13 | 52101368 | 52101482 | - | 115 | 5S_rRNA | 12 | 0 | 12 | 12 | 29 | 10 |
| ENSDARG000 | 13 | 438347 | 438462 | + | 116 | 5S_rRNA | 0 | 0 | 1 | 0 | 0 | 0 |
| ENSDARG000 | 13;13;13 | 44233720:442 | 44233998:442 | -;-;- | 554 | NA | 123 | 2 | 2 | 8 | 0 | 13 |
| ENSDARG000 | 13:13:13:13:1 | 40508:41146: | 41047:41348: | -;-;-;-;- | 2515 | ston1 | 551 | 354 | 573 | 493 | 553 | 533 |
| ENSDARG000 | 13 | 34883246 | 34883370 | + | 125 | NA | 0 | 0 | 0 | 0 | 0 | 0 |
| ENSDARG000 | 13:13:13:13:1 | 31172833:311 | 31173007:311 | +;+;+;+;+ | 732 | NA | 35 | 14 | 37 | 5 | 27 | 18 |
| ENSDARG000 | 13:13:13:13:1 | 49727333:497 | 49727625:497 | +;+;+;+;+;+;+ | 4132 | ggps1 | 558 | 556 | 678 | 319 | 606 | 626 |
| ENSDARG000 | 13:13:13:13:1 | 6960782:6961 | 6960824:6961 | -:-:-:-:-:-:-:-:-: | 3282 | NA | 299 | 101 | 205 | 539 | 274 | 213 |
| ENSDARG000 | 13:13:13:13:1 | 50450190:504 | 50454700:504 | -;-;-;-;-;- | 5455 | CCNJ | 6 | 32 | 29 | 12 | 16 | 16 |
| ENSDARG000 | 13:13:13:13:1 | 51956397:519 | 51956544:519 | +;+;+;+;+;+ | 2789 | lclat1 | 1812 | 1112 | 1159 | 790 | 1726 | 985 |
| ENSDARG000 | 13;13 | 50544625:505 | 50544768:505 | -;- | 726 | NA | 0 | 0 | 2 | 0 | 0 | 0 |
| ENSDARG000 | 13:13:13:13:1 | 33462232:334 | 33462423:334 | +:+:+:+:+:+:+: | 2955 | ftr81 | 0 | 0 | 4 | 0 | 0 | 0 |
| ENSDARG000 | 13;13;13;13 | 50375800:503 | 50375953:503 | +;+;+;+ | 522 | cox5b2 | 6 | 14 | 52 | 10 | 14 | 0 |
| ENSDARG000 | 13 | 34638629 | 34638746 | - | 118 | 5S_rRNA | 0 | 0 | 0 | 0 | 0 | 0 |
| ENSDARG000 | 13;13;13;13 | 52082648:520 | 52082791:520 | -;-;-;- | 398 | tmem254 | 8 | 15 | 99 | 31 | 73 | 58 |
| ENSDARG000 | 13 | 14384421 | 14384538 | - | 118 | 5S_rRNA | 0 | 0 | 0 | 0 | 0 | 0 |
| ENSDARG000 | 13 | 22050183 | 22050309 | + | 127 | 5S_rRNA | 0 | 0 | 0 | 0 | 0 | 0 |
| ENSDARG000 | 13 | 48902945 | 48903059 | + | 115 | 5S_rRNA | 0 | 0 | 0 | 0 | 0 | 0 |
| ENSDARG000 | 13:13:13:13:1 | 46803979:468 | 46804300:468 | +;+;+;+;+;+;+ | 1156 | NA | 875 | 977 | 712 | 1060 | 709 | 575 |
| ENSDARG000 | 13 | 44100845 | 44100965 | + | 121 | 5S_rRNA | 0 | 0 | 0 | 0 | 0 | 0 |
| ENSDARG000 | 13 | 6032844 | 6038477 | - | 5634 | NA | 0 | 0 | 0 | 0 | 0 | 0 |
| ENSDARG000 | 13 | 52094098 | 52094212 | - | 115 | 5S_rRNA | 0 | 0 | 0 | 0 | 0 | 0 |
| ENSDARG000 | 13 | 22203524 | 22203638 | + | 115 | 5S_rRNA | 0 | 0 | 0 | 0 | 0 | 0 |
| ENSDARG000 | 13:13:13:13:1 | 2894536:2896 | 2894574:2897 | +;+;+;+;+;+;+ | 699 | NA | 99 | 78 | 184 | 38 | 130 | 64 |
| ENSDARG000 | 13 | 24828753 | 24828869 | + | 117 | 5S_rRNA | 0 | 0 | 0 | 0 | 0 | 0 |
| ENSDARG000 | 13:13:13:13:1 | 50190664:501 | 50190943:501 | -:-:-:-:-:-:-:-:-: | 1847 | pkz | 302 | 298 | 137 | 50 | 112 | 61 |
| ENSDARG000 | 13 | 49580507 | 49580639 | - | 133 | NA | 4 | 0 | 0 | 0 | 0 | 0 |
| ENSDARG000 | 13;13;13 | 330611:3354 | 331506:3365 | -;-;- | 2469 | zgc:171534 | 0 | 0 | 0 | 0 | 0 | 0 |
| ENSDARG000 | 13 | 11822170 | 11822283 | + | 114 | dre-mir-146a | 5 | 8 | 28 | 34 | 1 | 10 |
| ENSDARG000 | 13;13 | 861730:8651 | 861891:8651 | -;- | 189 | NA | 43 | 23 | 22 | 3 | 44 | 2 |
| ENSDARG000 | 13 | 9524382 | 9525527 | - | 1146 | NA | 0 | 0 | 0 | 0 | 0 | 0 |
| ENSDARG000 | 13 | 16307767 | 16307883 | - | 117 | 5S_rRNA | 2 | 0 | 0 | 0 | 0 | 0 |
| ENSDARG000 | 13:13:13:13:1 | 51893676:518 | 51894471:518 | -;-;-;-;-;-;- | 1609 | mrpl2 | 1476 | 403 | 671 | 857 | 1076 | 970 |
| ENSDARG000 | 13:13:13:13:1 | 50207943:502 | 50208072:502 | -:-:-:-:-:-:-:-:-: | 2075 | eif2ak2 | 211 | 165 | 122 | 62 | 134 | 62 |
| ENSDARG000 | 13 | 6414284 | 6414400 | + | 117 | 5S_rRNA | 0 | 0 | 0 | 0 | 0 | 0 |
| ENSDARG000 | 13 | 37178602 | 37178716 | - | 115 | 5S_rRNA | 0 | 0 | 0 | 0 | 0 | 0 |
| ENSDARG000 | 13 | 406635 | 407645 | + | 1011 | NA | 0 | 0 | 0 | 0 | 0 | 0 |
| ENSDARG000 | 13:13:13:13:1 | 330555:3330 | 330566:3330 | +:+:+:+:+:+:+: | 183 | NA | 34 | 30 | 13 | 25 | 114 | 0 |
| ENSDARG000 | 13:13:13:13:1 | 43223491:432 | 43223757:432 | -:-:-:-:-:-:-:-:-: | 2000 | eif3s6ip | 4464 | 3690 | 7797 | 5569 | 5036 | 5574 |
| ENSDARG000 | 13:13:13:13:1 | 2254238:2257 | 2255440:2257 | -:-:-:-:-:-:-:-:-: | 2361 | lrrc1 | 612 | 608 | 612 | 553 | 503 | 518 |
| ENSDARG000 | 13 | 23821186 | 23821300 | + | 115 | 5S_rRNA | 0 | 0 | 0 | 0 | 0 | 0 |
| ENSDARG000 | 13:13:13:13:1 | 46718518:467 | 46718977:467 | +:+:+:+:+:+:+: | 2965 | TMEM63B | 905 | 563 | 810 | 455 | 972 | 724 |
| ENSDARG000 | 13 | 35370569 | 35370684 | - | 116 | 5S_rRNA | 0 | 0 | 0 | 0 | 0 | 0 |
| ENSDARG000 | 13 | 27804067 | 27804145 | + | 79 | dre-mir-30c | 0 | 0 | 0 | 0 | 0 | 0 |
| ENSDARG000 | 13 | 49123570 | 49123687 | - | 118 | 5S_rRNA | 0 | 0 | 0 | 0 | 0 | 0 |
| ENSDARG000 | 13:13:13:13:1 | 17776301:177 | 17776323:177 | +:+:+:+:+:+:+: | 318 | NA | 0 | 0 | 0 | 0 | 0 | 1 |
| ENSDARG000 | 13:13:13:13:1 | 50235706:502 | 50236494:502 | -;-;-;-;-;-;-;- | 1658 | gpatch11 | 264 | 221 | 270 | 138 | 225 | 204 |
| ENSDARG000 | 13:13:13:13:1 | 50345345:503 | 50349638:503 | -;-;-;-;-;-;-;-;- | 6340 | SIRT1 | 387 | 289 | 296 | 138 | 302 | 225 |
| ENSDARG000 | 13;13 | 49827015:498 | 49827143:498 | -;- | 228 | GNG4 | 41 | 21 | 10 | 7 | 15 | 31 |
| ENSDARG000 | 13:13:13:13:1 | 49153761:491 | 49154690:491 | -;-;-;-;-;- | 1616 | tsnax | 145 | 122 | 134 | 70 | 96 | 74 |
| ENSDARG000 | 13:13:13:13:1 | 2927637:2927 | 2927872:2927 | -:-:-:-:-:-:-:-:-: | 2871 | EML4 | 689 | 692 | 1068 | 556 | 1100 | 554 |
| ENSDARG000 | 13;13 | 41819817:418 | 41820004:418 | +;+ | 312 | NA | 0 | 0 | 0 | 0 | 0 | 0 |
| ENSDARG000 | 13:13:13:13:1 | 7171066:7177 | 7171257:7177 | -:-:-:-:-:-:-:-:-: | 3912 | ninl | 81 | 160 | 179 | 76 | 135 | 120 |
| ENSDARG000 | 13:13:13:13:1 | 42272515:422 | 42273188:422 | -:-:-:-:-:-:-:-:-: | 2124 | kmo | 4 | 2 | 6 | 8 | 48 | 0 |
| ENSDARG000 | 13;13 | 50670744:506 | 50670972:506 | -;- | 357 | NA | 862 | 4 | 14 | 205 | 282 | 333 |
| ENSDARG000 | 13:13:13:13:1 | 46830736:468 | 46835511:468 | -:-:-:-:-:-:-:-:-: | 6503 | slc29a1a | 3925 | 3414 | 4631 | 3370 | 3311 | 6698 |
| ENSDARG000 | 13 | 30632105 | 30632219 | - | 115 | 5S_rRNA | 0 | 0 | 1 | 0 | 0 | 0 |
| ENSDARG000 | 13 | 27795130 | 27795209 | + | 80 | dre-mir-30e-2 | 0 | 0 | 0 | 0 | 0 | 0 |
| ENSDARG000 | 13:13:13:13:1 | 1535511:1535 | 1535651:1535 | +:+:+:+:+:+:+: | 1975 | NA | 169 | 50 | 135 | 60 | 168 | 86 |
| ENSDARG000 | 13;13;13 | 31144305:311 | 31144325:311 | +;+;+ | 360 | NA | 84 | 29 | 39 | 32 | 63 | 40 |
| ENSDARG000 | 13;13;13;13 | 9333317:9334 | 9333354:9335 | -;-;-;- | 810 | NA | 3 | 0 | 0 | 0 | 12 | 0 |
| ENSDARG000 | 13:13:13:13:1 | 912123:91692 | 912314:91702 | +:+:+:+:+:+:+: | 1941 | pcsk2 | 4 | 2 | 0 | 0 | 26 | 0 |
| ENSDARG000 | 13:13:13:13:1 | 9100:9317:96 | 9229:9372:97 | +;+;+;+;+;+;+ | 1722 | ppp4r3b | 632 | 627 | 910 | 314 | 933 | 288 |
| ENSDARG000 | 13 | 12105778 | 12105894 | - | 117 | 5S_rRNA | 0 | 0 | 0 | 0 | 0 | 0 |
| ENSDARG000 | 13:13:13:13:1 | 30684186:306 | 30684355:306 | -;-;-;-;-;-;- | 1050 | NA | 0 | 0 | 0 | 0 | 0 | 0 |
| ENSDARG000 | 13;13;13 | 33695678:336 | 33696471:336 | -;-;- | 1466 | mad2l1bp | 6 | 38 | 28 | 14 | 40 | 38 |
| ENSDARG000 | 13 | 10912486 | 10912602 | - | 117 | 5S_rRNA | 0 | 0 | 1 | 0 | 0 | 0 |
| ENSDARG000 | 13 | 8539267 | 8539381 | + | 115 | 5S_rRNA | 0 | 0 | 0 | 0 | 0 | 0 |
| ENSDARG000 | 13:13:13:13:1 | 216320:2166 | 216673:2166 | -:-:-:-:-:-:-:-:-: | 3226 | rtn4b | 329 | 273 | 445 | 256 | 471 | 158 |
| ENSDARG000 | 13 | 27690160 | 27690276 | - | 117 | 5S_rRNA | 0 | 0 | 0 | 0 | 0 | 0 |
| ENSDARG000 | 13:13:13:13:1 | 52061034:520 | 52061137:520 | +;+;+;+;+;+ | 1155 | NA | 0 | 0 | 1 | 0 | 0 | 0 |
| ENSDARG000 | 13:13:13:13:1 | 43130895:431 | 43131909:431 | -;-;-;-;-;-;- | 1986 | vsir | 212 | 314 | 447 | 336 | 471 | 668 |
| ENSDARG000 | 13:13:13:13:1 | 51502950:515 | 51503652:515 | -;-;-;-;-;-;-;-;- | 1869 | valopa | 2 | 0 | 4 | 0 | 8 | 0 |
| ENSDARG000 | 13:13:13:13:1 | 432790:4333 | 432951:4334 | -;-;-;-;-;- | 842 | EFEMP1 | 0 | 0 | 3 | 2 | 14 | 10 |
| ENSDARG000 | 13;13;13 | 51579851:515 | 51580265:515 | +;+;+ | 2865 | nkx6.2 | 153 | 36 | 187 | 37 | 11 | 154 |
| ENSDARG000 | 13:13:13:13:1 | 43247936:432 | 43247948:432 | +:+:+:+:+:+:+: | 1396 | SMOC2 | 8 | 2 | 0 | 2 | 18 | 2 |
| ENSDARG000 | 13 | 13644445 | 13644562 | - | 118 | 5S_rRNA | 0 | 0 | 0 | 0 | 0 | 0 |
| ENSDARG000 | 13:13:13:13:1 | 47863695:478 | 47864526:478 | -:-:-:-:-:-:-:-:-: | 4387 | itga9 | 964 | 366 | 1560 | 545 | 1728 | 564 |
| ENSDARG000 | 13 | 46227713 | 46227829 | + | 117 | 5S_rRNA | 0 | 0 | 0 | 0 | 0 | 0 |
| ENSDARG000 | 13:13:13:13:1 | 16230556:162 | 16231109:162 | -:-:-:-:-:-:-:-:-: | 1408 | zgc:110045 | 42 | 28 | 105 | 249 | 110 | 27 |
| ENSDARG000 | 13 | 30836656 | 30836776 | + | 121 | 5S_rRNA | 0 | 0 | 0 | 0 | 0 | 0 |
| ENSDARG000 | 13 | 18715310 | 18715426 | - | 117 | 5S_rRNA | 0 | 0 | 0 | 0 | 0 | 0 |
| ENSDARG000 | 13 | 48982686 | 48982800 | + | 115 | 5S_rRNA | 0 | 0 | 0 | 0 | 0 | 0 |
| ENSDARG000 | 13 | 50793756 | 50793872 | + | 117 | 5S_rRNA | 0 | 0 | 0 | 0 | 0 | 0 |
| ENSDARG000 | 13:13:13:13:1 | 835390:8378 | 835476:8379 | +;+;+;+;+;+ | 672 | NA | 42 | 18 | 57 | 21 | 19 | 26 |
| ENSDARG000 | 13 | 9873894 | 9874010 | + | 117 | 5S_rRNA | 0 | 0 | 0 | 0 | 0 | 0 |
| ENSDARG000 | 13 | 28924846 | 28924965 | + | 120 | 5S_rRNA | 0 | 0 | 0 | 0 | 0 | 0 |
| ENSDARG000 | 13;13;13 | 51909828:519 | 51909905:519 | -;-;- | 780 | srfb | 6 | 18 | 14 | 0 | 6 | 4 |
| ENSDARG000 | 13:13:13:13:1 | 33878:34712: | 34358:34801: | -;-;-;-;-;-;-;-;- | 1575 | gtf2a1l | 18 | 20 | 25 | 23 | 78 | 52 |
| ENSDARG000 | 13 | 411607 | 412617 | + | 1011 | NA | 0 | 0 | 0 | 0 | 0 | 0 |
| ENSDARG000 | 13:13:13:13:1 | 49541003:495 | 49545050:495 | -:-:-:-:-:-:-:-:-: | 5366 | tomm20a | 1156 | 1284 | 1933 | 721 | 1426 | 844 |
| ENSDARG000 | 13 | 30251741 | 30251857 | + | 117 | 5S_rRNA | 0 | 0 | 0 | 0 | 0 | 0 |
| ENSDARG000 | 13 | 25530534 | 25530648 | - | 115 | 5S_rRNA | 0 | 0 | 0 | 0 | 0 | 0 |
| ENSDARG000 | 13 | 32392383 | 32392499 | + | 117 | 5S_rRNA | 0 | 0 | 0 | 0 | 0 | 0 |

| | | | | | | | | | | | |
|---|---|---|---|---|---|---|---|---|---|---|---|
| ENSDARG000 | 13 | 32846493 | 32846609 | + | 117 | 5S_rRNA | 0 | 0 | 2 | 0 | 0 | 0 |
| ENSDARG000 | 13 | 15223296 | 15223383 | + | 88 | dre-mir-103 | 0 | 0 | 0 | 0 | 0 | 0 |
| ENSDARG000 | 13 | 2386413 | 2386533 | + | 121 | NA | 0 | 0 | 0 | 0 | 0 | 0 |
| ENSDARG000 | 13 | 47154419 | 47154533 | - | 115 | 5S_rRNA | 0 | 0 | 0 | 0 | 0 | 0 |
| ENSDARG000 | 13 | 41662972 | 41663088 | - | 117 | 5S_rRNA | 0 | 0 | 0 | 0 | 0 | 0 |
| ENSDARG000 | 13:13:13:13:1 | 46520804:465 | 46521290:465 | -:-:-:-:-:-:-:-:-: | 1937 | ATE1 | 394 | 336 | 384 | 170 | 1301 | 366 |
| ENSDARG000 | 13 | 18709249 | 18709363 | + | 115 | 5S_rRNA | 0 | 0 | 0 | 0 | 0 | 0 |
| ENSDARG000 | 13:13:13:13:1 | 24830737:248 | 24830748:248 | -:-:-:-:-:-:-:-:-: | 564 | NA | 54 | 31 | 40 | 61 | 45 | 33 |
| ENSDARG000 | 13;13;13;13 | 30696286:306 | 30696401:306 | +;+;+;+ | 800 | cxcl18a.1 | 26 | 2 | 12 | 0 | 8 | 10 |
| ENSDARG000 | 13 | 44865594 | 44865710 | + | 117 | 5S_rRNA | 0 | 0 | 0 | 0 | 0 | 0 |
| ENSDARG000 | 13:13:13:13:1 | 1872767:1884 | 1873440:1884 | +;+;+;+;+;+;+ | 2203 | bmp5 | 6 | 4 | 8 | 0 | 2 | 0 |
| ENSDARG000 | 13 | 13670440 | 13670537 | - | 98 | NA | 0 | 0 | 0 | 0 | 0 | 0 |
| ENSDARG000 | 13:13:13:13:1 | 11714869:117 | 11720083:117 | -:-:-:-:-:-:-:-:-: | 7789 | manba | 751 | 506 | 657 | 479 | 828 | 594 |
| ENSDARG000 | 13:13:13:13:1 | 11746412:117 | 11746870:117 | -:-:-:-:-:-:-:-:-: | 1192 | ube2d3 | 563 | 660 | 840 | 391 | 580 | 632 |
| ENSDARG000 | 13:13:13:13:1 | 7241170:7242 | 7241278:7243 | +:+:+:+:+:+:+:+: | 4061 | AIFM2 | 1024 | 1406 | 900 | 555 | 1181 | 1336 |
| ENSDARG000 | 13 | 41230293 | 41230409 | - | 117 | 5S_rRNA | 0 | 0 | 0 | 0 | 0 | 0 |
| ENSDARG000 | 13 | 28740235 | 28740369 | + | 135 | NA | 3 | 0 | 0 | 10 | 3 | 3 |
| ENSDARG000 | 13 | 46489534 | 46489649 | - | 116 | 5S_rRNA | 0 | 0 | 0 | 0 | 0 | 0 |
| ENSDARG000 | 13 | 38853137 | 38853252 | - | 116 | 5S_rRNA | 0 | 0 | 0 | 0 | 0 | 0 |
| ENSDARG000 | 13 | 2833092 | 2833207 | + | 116 | 5S_rRNA | 0 | 0 | 0 | 0 | 0 | 0 |
| ENSDARG000 | 13:13:13:13:1 | 43050562:430 | 43050638:430 | +:+:+:+:+:+:+:+: | 9179 | cdh23 | 8 | 17 | 12 | 5 | 17 | 10 |
| ENSDARG000 | 13:13:13:13:1 | 29292011:292 | 29292278:292 | +:+:+:+:+:+:+:+: | 4995 | PARG | 990 | 932 | 1136 | 504 | 994 | 698 |
| ENSDARG000 | 13:13:13:13:1 | 45087:45608: | 45116:45736: | -:-:-:-:-:-:-:-:-: | 2010 | ppp1r21 | 1454 | 1150 | 1330 | 1039 | 1217 | 1195 |
| ENSDARG000 | 13:13:13:13:1 | 16219082:162 | 16219442:162 | -;-;-;-;-;-;- | 748 | NA | 11 | 5 | 60 | 132 | 76 | 15 |
| ENSDARG000 | 13:13:13:13:1 | 1799003:1799 | 1799012:1800 | +:+:+:+:+:+:+:+: | 2154 | NA | 28 | 0 | 4 | 0 | 26 | 4 |
| ENSDARG000 | 13:13:13:13:1 | 7292061:7302 | 7292163:7302 | +;+;+;+;+;+;+ | 12209 | NA | 146 | 60 | 375 | 16 | 508 | 676 |
| ENSDARG000 | 13:13:13:13:1 | 33452096:334 | 33452701:334 | +;+;+;+;+ | 1620 | NA | 0 | 0 | 0 | 0 | 0 | 0 |
| ENSDARG000 | 13 | 3054823 | 3054937 | - | 115 | 5S_rRNA | 0 | 0 | 0 | 0 | 0 | 0 |
| ENSDARG000 | 13 | 23961292 | 23961406 | - | 115 | 5S_rRNA | 0 | 0 | 0 | 0 | 0 | 0 |
| ENSDARG000 | 13 | 51493927 | 51494042 | - | 116 | 5S_rRNA | 0 | 0 | 0 | 0 | 0 | 0 |
| ENSDARG000 | 13;13 | 1015749:1024 | 1016488:1025 | +;+ | 1461 | prokr1b | 0 | 0 | 0 | 0 | 0 | 0 |
| ENSDARG000 | 13:13:13:13:1 | 1575276:1577 | 1575458:1578 | +:+:+:+:+:+:+:+: | 11550 | NA | 6514 | 2704 | 3417 | 3599 | 4907 | 3729 |
| ENSDARG000 | 13:13:13:13:1 | 50328815:503 | 50328957:503 | -;-;-;-;-;- | 678 | CACUL1 | 8 | 8 | 0 | 2 | 4 | 3 |
| ENSDARG000 | 13:13:13:13:1 | 46183639:461 | 46183698:461 | -:-:-:-:-:-:-:-:-: | 1411 | NA | 6 | 0 | 0 | 0 | 6 | 4 |
| ENSDARG000 | 13 | 30511607 | 30511723 | + | 117 | 5S_rRNA | 0 | 0 | 0 | 0 | 0 | 0 |
| ENSDARG000 | 13 | 18644890 | 18645005 | - | 116 | 5S_rRNA | 0 | 0 | 0 | 0 | 0 | 0 |
| ENSDARG000 | 13:13:13:13:1 | 48064335:480 | 48064349:480 | -:-:-:-:-:-:-:-:-: | 6694 | GOLGA4 | 1147 | 909 | 1275 | 864 | 1391 | 1148 |
| ENSDARG000 | 13:13:13:13:1 | 33689740:336 | 33689954:336 | +:+:+:+:+:+:+:+: | 1568 | EPHX1 | 0 | 20 | 24 | 38 | 25 | 20 |
| ENSDARG000 | 13:13:13:13:1 | 16027559:160 | 16030497:160 | -;-;-;-;-;-;-;- | 4143 | SPATA48 | 0 | 0 | 0 | 0 | 0 | 0 |
| ENSDARG000 | 13;13;13 | 39324865:393 | 39325018:393 | +;+;+ | 1521 | NA | 0 | 0 | 0 | 0 | 0 | 0 |
| ENSDARG000 | 13:13:13:13:1 | 7164345:7164 | 7164519:7164 | +;+;+;+;+;+;+ | 1163 | gins1 | 62 | 273 | 219 | 64 | 175 | 149 |
| ENSDARG000 | 13 | 1765507 | 1765622 | + | 116 | 5S_rRNA | 0 | 0 | 0 | 0 | 0 | 0 |
| ENSDARG000 | 13:13:13:13:1 | 985442:98653 | 986259:98662 | -;-;-;-;- | 1269 | BFSP1 | 0 | 0 | 0 | 0 | 0 | 0 |
| ENSDARG000 | 13 | 34644150 | 34644281 | - | 132 | 5S_rRNA | 0 | 0 | 0 | 0 | 0 | 0 |
| ENSDARG000 | 13:13:13:13:1 | 52071971:520 | 52071994:520 | +:+:+:+:+:+:+:+: | 165 | NA | 18 | 0 | 14 | 21 | 34 | 24 |
| ENSDARG000 | 13 | 13661130 | 13661227 | + | 98 | NA | 0 | 0 | 0 | 0 | 0 | 0 |
| ENSDARG000 | 13:13:13:13:1 | 11767603:117 | 11768094:117 | -:-:-:-:-:-:-:-:-: | 2523 | NA | 349 | 385 | 380 | 201 | 335 | 293 |
| ENSDARG000 | 13:13:13:13:1 | 111212:11191 | 111877:11201 | +:+:+:+:+:+:+:+: | 1701 | dnaaf2 | 24 | 27 | 97 | 33 | 154 | 50 |
| ENSDARG000 | 13:13:13:13:1 | 50368668:503 | 50368823:503 | +;+;+;+;+ | 706 | dnajc12 | 115 | 124 | 142 | 216 | 90 | 140 |
| ENSDARG000 | 13 | 31694931 | 31695047 | - | 117 | 5S_rRNA | 0 | 0 | 0 | 0 | 0 | 0 |
| ENSDARG000 | 13 | 34426248 | 34426365 | + | 118 | 5S_rRNA | 0 | 0 | 0 | 0 | 0 | 0 |
| ENSDARG000 | 13:13:13:13:1 | 7086735:7107 | 7086833:7107 | +:+:+:+:+:+:+:+: | 2202 | RPS6KA2 | 44 | 12 | 4 | 22 | 38 | 24 |
| ENSDARG000 | 13:13:13:13:1 | 38798045:388 | 38798072:388 | -:-:-:-:-:-:-:-:-: | 321 | NA | 0 | 0 | 0 | 0 | 0 | 0 |
| ENSDARG000 | 13 | 35592242 | 35592377 | - | 136 | 5S_rRNA | 0 | 0 | 0 | 0 | 0 | 0 |
| ENSDARG000 | 13:13:13:13:1 | 46196354:461 | 46196818:461 | -;-;-;-;- | 1507 | NA | 8 | 0 | 0 | 2 | 0 | 0 |
| ENSDARG000 | 13:13:13:13:1 | 233482:23369 | 233584:23375 | +:+:+:+:+:+:+:+: | 1053 | cfap36 | 211 | 333 | 377 | 323 | 224 | 184 |
| ENSDARG000 | 13:13:13:13:1 | 713941:71579 | 714139:71597 | +:+:+:+:+:+:+:+: | 1583 | cox15 | 509 | 643 | 494 | 432 | 404 | 333 |
| ENSDARG000 | 13 | 51063727 | 51065048 | - | 1322 | NA | 0 | 0 | 0 | 0 | 0 | 0 |
| ENSDARG000 | 13 | 28179434 | 28179550 | - | 117 | 5S_rRNA | 0 | 0 | 0 | 0 | 0 | 0 |
| ENSDARG000 | 13 | 255067 | 257821 | + | 2755 | foxg1d | 0 | 0 | 0 | 0 | 0 | 0 |
| ENSDARG000 | 13;13 | 51844058:518 | 51844264:518 | -;- | 285 | lbh | 62 | 15 | 14 | 33 | 34 | 27 |
| ENSDARG000 | 13 | 44088148 | 44088251 | - | 104 | 5S_rRNA | 0 | 0 | 0 | 0 | 0 | 0 |
| ENSDARG000 | 13:13:13:13:1 | 13213567:132 | 13218872:132 | -:-:-:-:-:-:-:-:-: | 8112 | fgfr3 | 205 | 8 | 150 | 36 | 477 | 40 |
| ENSDARG000 | 13 | 44384277 | 44384393 | + | 117 | 5S_rRNA | 0 | 0 | 0 | 0 | 0 | 0 |
| ENSDARG000 | 13 | 13863530 | 13863645 | + | 116 | 5S_rRNA | 0 | 0 | 0 | 0 | 0 | 0 |
| ENSDARG000 | 13 | 39969349 | 39969465 | + | 117 | 5S_rRNA | 0 | 0 | 0 | 0 | 0 | 0 |
| ENSDARG000 | 13:13:13:13:1 | 421231:42126 | 421571:42157 | +:+:+:+:+:+:+:+: | 2505 | lgi1a | 116 | 164 | 252 | 259 | 437 | 80 |
| ENSDARG000 | 13:13:13:13:1 | 50287009:502 | 50287218:502 | -;-;-;-;-;- | 2052 | NA | 0 | 1 | 0 | 0 | 0 | 1 |
| ENSDARG000 | 13;13 | 1305363:1307 | 1307864:1308 | -;- | 2617 | olig3 | 0 | 0 | 0 | 0 | 0 | 0 |
| ENSDARG000 | 13;13;13;13 | 8011303:8011 | 8011392:8011 | +;+;+;+ | 365 | NA | 0 | 0 | 1 | 4 | 0 | 0 |
| ENSDARG000 | 13:13:13:13:1 | 10160856:101 | 10160925:101 | +;+;+;+;+ | 685 | NA | 0 | 0 | 0 | 0 | 0 | 0 |
| ENSDARG000 | 13:13:13:13:1 | 12498462:124 | 12499030:124 | +;+;+;+;+;+ | 4274 | NA | 4 | 10 | 8 | 4 | 0 | 0 |
| ENSDARG000 | 13:13:13:13:1 | 19151268:191 | 19152290:191 | +:+:+:+:+:+:+:+: | 3550 | NA | 0 | 0 | 0 | 0 | 0 | 0 |
| ENSDARG000 | 13;13;13 | 20707546:207 | 20707842:207 | +;+;+ | 998 | NA | 0 | 0 | 0 | 0 | 6 | 0 |
| ENSDARG000 | 13;13;13 | 22778060:227 | 22778134:227 | -;-;- | 2931 | NA | 47 | 0 | 22 | 0 | 46 | 0 |
| ENSDARG000 | 13;13;13;13 | 26051785:261 | 26052628:261 | -;-;-;- | 1215 | NA | 0 | 0 | 0 | 0 | 4 | 0 |
| ENSDARG000 | 13;13;13;13 | 27363949:273 | 27364304:273 | +;+;+;+ | 2141 | NA | 0 | 0 | 0 | 0 | 0 | 0 |
| ENSDARG000 | 13;13;13 | 29749474:297 | 29749726:297 | -;-;- | 1283 | NA | 0 | 0 | 0 | 1 | 0 | 4 |
| ENSDARG000 | 13;13;13;13 | 31042514:310 | 31043946:310 | -;-;-;- | 2539 | NA | 106 | 59 | 113 | 94 | 200 | 114 |
| ENSDARG000 | 13:13:13:13:1 | 31052536:310 | 31053446:310 | -:-:-:-:-:-:-:-:-: | 3262 | NA | 68 | 81 | 119 | 64 | 158 | 101 |
| ENSDARG000 | 13:13:13:13:1 | 31983061:319 | 31984178:319 | -;-;-;-;- | 1461 | NA | 0 | 0 | 2 | 12 | 6 | 3 |
| ENSDARG000 | 13;13;13 | 32023488:320 | 32023842:320 | -;-;- | 644 | NA | 0 | 0 | 2 | 0 | 6 | 0 |
| ENSDARG000 | 13;13;13;13 | 43921833:439 | 43921950:439 | +;+;+;+ | 620 | NA | 0 | 0 | 0 | 0 | 0 | 0 |
| ENSDARG000 | 13:13:13:13:1 | 45942158:459 | 45942523:459 | +;+;+;+;+ | 1026 | NA | 0 | 0 | 0 | 0 | 0 | 0 |
| ENSDARG000 | 13;13;13 | 46202830:462 | 46202970:462 | -;-;- | 374 | NA | 0 | 0 | 0 | 0 | 0 | 0 |
| ENSDARG000 | 13;13;13;13 | 50481817:504 | 50482242:505 | +;+;+;+ | 7630 | NA | 12 | 0 | 16 | 22 | 18 | 0 |
| ENSDARG000 | 13;13;13 | 50709317:507 | 50712176:507 | -;-;- | 3424 | NA | 4 | 5 | 2 | 0 | 4 | 8 |
| ENSDARG000 | 13:13:13:13:1 | 50730055:507 | 50730869:507 | -;-;-;-;- | 1438 | NA | 1 | 2 | 9 | 0 | 0 | 5 |
| ENSDARG000 | 13:13:13:13:1 | 51297553:513 | 51297672:513 | +:+:+:+:+:+:+:+: | 2381 | NA | 0 | 0 | 2 | 0 | 0 | 0 |
| ENSDARG000 | 13:13:13:13:1 | 51428398:514 | 51428993:514 | +;+;+;+;+ | 835 | NA | 0 | 0 | 0 | 0 | 0 | 0 |
| ENSDARG000 | 13:13:13:13:1 | 51445949:514 | 51446048:514 | +:+:+:+:+:+:+:+: | 5466 | NA | 0 | 4 | 0 | 0 | 5 | 6 |
| ENSDARG000 | 13;13;13 | 52107945:521 | 52109004:521 | -;-;- | 1317 | NA | 44 | 2 | 2 | 0 | 12 | 2 |
| ENSDARG000 | 18:18:18:18:1 | 22739732:227 | 22740319:227 | -:-:-:-:-:-:-:-:-: | 2751 | hsf4 | 2 | 0 | 2 | 0 | 0 | 0 |
| ENSDARG000 | 18:18:18:18:1 | 27732285:277 | 27732504:277 | +:+:+:+:+:+:+:+: | 1880 | tspan18b | 1231 | 778 | 220 | 491 | 504 | 757 |
| ENSDARG000 | 18:18:18:18:1 | 32916351:329 | 32916550:329 | +:+:+:+:+:+:+:+: | 6444 | olfcg2 | 0 | 0 | 0 | 0 | 0 | 0 |
| ENSDARG000 | 18:18:18:18:1 | 18995247:189 | 18995854:189 | -:-:-:-:-:-:-:-:-: | 2854 | ints14 | 330 | 371 | 431 | 273 | 326 | 336 |
| ENSDARG000 | 18;18 | 22739732:227 | 22740315:227 | -;- | 730 | NA | 0 | 0 | 0 | 0 | 0 | 0 |
| ENSDARG000 | 18;18 | 27821856:278 | 27821888:278 | +;+ | 594 | NA | 0 | 0 | 0 | 0 | 0 | 0 |
| ENSDARG000 | 18:18:18:18:1 | 18982077:189 | 18982296:189 | +:+:+:+:+:+:+:+: | 1694 | hacd3 | 270 | 370 | 258 | 196 | 354 | 462 |
| ENSDARG000 | 18:18:18:18:1 | 7073130:7073 | 7073211:7073 | +:+:+:+:+:+:+:+: | 897 | NA | 10 | 10 | 8 | 4 | 2 | 17 |
| ENSDARG000 | 18;18 | 19025989:190 | 19026451:190 | +;+ | 714 | NA | 0 | 5 | 2 | 0 | 0 | 0 |
| ENSDARG000 | 18:18:18:18:1 | 20825375:208 | 20825654:208 | +:+:+:+:+:+:+:+: | 1981 | TTC23 | 133 | 189 | 517 | 200 | 495 | 245 |
| ENSDARG000 | 18:18:18:18:1 | 19006063:190 | 19006487:190 | +:+:+:+:+:+:+:+: | 3753 | SLC24A1 | 0 | 2 | 0 | 0 | 0 | 0 |
| ENSDARG000 | 18:18:18:18:1 | 32950480:329 | 32950747:329 | +:+:+:+:+:+:+:+: | 4233 | olfcg3 | 0 | 6 | 0 | 0 | 0 | 0 |
| ENSDARG000 | 18:18:18:18:1 | 32979166:329 | 32979374:329 | +;+;+;+;+;+ | 2508 | olfcg4 | 0 | 40 | 0 | 0 | 0 | 0 |
| ENSDARG000 | 18:18:18:18:1 | 3572314:3572 | 3572568:3572 | +;+;+;+;+ | 1118 | serp1 | 596 | 1386 | 1750 | 1333 | 1063 | 856 |
| ENSDARG000 | 18:18:18:18:1 | 27843776:278 | 27844015:278 | -:-:-:-:-:-:-:-:-: | 1313 | iqcg | 0 | 0 | 0 | 0 | 0 | 0 |
| ENSDARG000 | 18:18:18:18:1 | 39060687:390 | 39060852:390 | +:+:+:+:+:+:+:+: | 1241 | zgc:171509 | 94 | 12 | 6 | 46 | 54 | 136 |
| ENSDARG000 | 18:18:18:18:1 | 8889888:8889 | 8889990:8889 | +:+:+:+:+:+:+:+: | 1059 | NA | 0 | 0 | 4 | 0 | 0 | 0 |
| ENSDARG000 | 18:18:18:18:1 | 7090920:7092 | 7091263:7092 | -;-;-;-;-;-;-;- | 1254 | cfap161 | 1 | 0 | 0 | 0 | 0 | 0 |
| ENSDARG000 | 18;18 | 15770461:157 | 15771554:157 | -;- | 1094 | NA | 0 | 0 | 0 | 0 | 0 | 0 |
| ENSDARG000 | 18:18:18:18:1 | 7099162:7099 | 7103969:7103 | -;-;-;-;-;-;- | 7945 | NA | 0 | 0 | 0 | 0 | 0 | 4 |
| ENSDARG000 | 18;18;18;18 | 20859661:208 | 20861888:208 | -;-;-;- | 3222 | synm | 30 | 30 | 78 | 152 | 62 | 58 |
| ENSDARG000 | 18:18:18:18:1 | 17474989:174 | 17475247:174 | +;+;+;+;+;+;+ | 871 | NA | 0 | 0 | 0 | 0 | 0 | 0 |
| ENSDARG000 | 18:18:18:18:1 | 8901846:8901 | 8901917:8901 | +:+:+:+:+:+:+:+: | 1293 | si:dkey-5i3.5 | 85 | 124 | 126 | 77 | 137 | 111 |
| ENSDARG000 | 18;18;18;18 | 15778110:157 | 15778245:157 | +;+;+;+ | 1795 | ube2na | 2578 | 3715 | 3656 | 2306 | 3782 | 2757 |
| ENSDARG000 | 18:18:18:18:1 | 3579829:3579 | 3580184:3580 | +:+:+:+:+:+:+:+: | 5163 | LRCH3 | 483 | 332 | 472 | 499 | 488 | 449 |
| ENSDARG000 | 18:18:18:18:1 | 27904515:279 | 27904843:279 | +:+:+:+:+:+:+:+: | 4358 | hipk3b | 699 | 328 | 818 | 390 | 682 | 443 |
| ENSDARG000 | 18:18:18:18:1 | 41527877:415 | 41528117:415 | +;+;+;+;+;+ | 1287 | selenot1b | 440 | 551 | 566 | 402 | 433 | 530 |
| ENSDARG000 | 18:18:18:18:1 | 32992646:329 | 32992854:329 | +;+;+;+;+;+ | 2502 | olfcg5 | 0 | 0 | 0 | 0 | 0 | 0 |
| ENSDARG000 | 18:18:18:18:1 | 20869923:208 | 20870002:208 | +;+;+;+;+;+ | 4784 | PGPEP1L | 22 | 24 | 59 | 83 | 49 | 35 |
| ENSDARG000 | 18:18:18:18:1 | 15795566:157 | 15796115:157 | +:+:+:+:+:+:+:+: | 1679 | nudt4a | 116 | 98 | 79 | 111 | 54 | 118 |
| ENSDARG000 | 18:18:18:18:1 | 22793465:227 | 22793872:227 | +:+:+:+:+:+:+:+: | 2755 | gnao1a | 647 | 1233 | 1053 | 1392 | 1372 | 1110 |
| ENSDARG000 | 18 | 17483557 | 17485419 | - | 1863 | foxl1 | 0 | 6 | 8 | 0 | 0 | 0 |
| ENSDARG000 | 18:18:18:18:1 | 22859479:228 | 22859793:228 | -;-;-;-;-;- | 971 | NA | 0 | 0 | 0 | 0 | 0 | 0 |
| ENSDARG000 | 18:18:18:18:1 | 7122577:7122 | 7123059:7123 | -:-:-:-:-:-:-:-:-: | 1719 | cd9a | 46 | 46 | 132 | 38 | 108 | 84 |
| ENSDARG000 | 18;18;18;18 | 41542542:415 | 41543249:415 | +;+;+;+ | 1311 | tsen34 | 230 | 158 | 235 | 134 | 212 | 240 |
| ENSDARG000 | 18:18:18:18:1 | 45645357:456 | 45645491:456 | +:+:+:+:+:+:+:+: | 1275 | eif3m | 3749 | 3034 | 5953 | 4939 | 3778 | 4883 |
| ENSDARG000 | 18:18:18:18:1 | 32996837:329 | 32997039:329 | +;+;+;+;+;+ | 2562 | olfch1 | 0 | 0 | 0 | 0 | 0 | 0 |
| ENSDARG000 | 18:18:18:18:1 | 45619558:456 | 45619721:456 | -;-;-;-;-;- | 577 | NA | 0 | 0 | 0 | 0 | 0 | 0 |
| ENSDARG000 | 18:18:18:18:1 | 39067575:390 | 39067604:390 | +:+:+:+:+:+:+:+: | 2101 | gnb5b | 0 | 0 | 0 | 4 | 0 | 0 |
| ENSDARG000 | 18;18;18 | 4771136:4771 | 4771209:4771 | +;+;+ | 490 | NA | 2 | 0 | 0 | 0 | 0 | 0 |
| ENSDARG000 | 18;18;18 | 39106722:391 | 39106797:391 | +;+;+ | 1349 | bcl2l10 | 444 | 588 | 726 | 354 | 801 | 484 |
| ENSDARG000 | 18;18 | 4783878:4804 | 4783983:4804 | -;- | 299 | NA | 0 | 0 | 1 | 0 | 0 | 0 |
| ENSDARG000 | 18:18:18:18:1 | 33009828:330 | 33010033:330 | +;+;+;+;+;+ | 2574 | olfcj1 | 0 | 0 | 0 | 0 | 0 | 0 |
| ENSDARG000 | 18:18:18:18:1 | 28102620:281 | 28102839:281 | +:+:+:+:+:+:+:+: | 2178 | NA | 0 | 0 | 2 | 0 | 0 | 0 |
| ENSDARG000 | 18:18:18:18:1 | 22994113:229 | 22994512:229 | +:+:+:+:+:+:+:+: | 4168 | cbfb | 8238 | 10462 | 9667 | 4372 | 9214 | 9144 |
| ENSDARG000 | 18:18:18:18:1 | 17493859:174 | 17493922:174 | +:+:+:+:+:+:+:+: | 1307 | phaf1 | 328 | 204 | 382 | 324 | 173 | 249 |
| ENSDARG000 | 18:18:18:18:1 | 12760802:127 | 12760943:127 | -;-;-;-;-;-;-;- | 1100 | NA | 0 | 0 | 2 | 0 | 0 | 0 |
| ENSDARG000 | 18:18:18:18:1 | 12655766:126 | 12655791:126 | +:+:+:+:+:+:+:+: | 2272 | tbxas1 | 253 | 109 | 246 | 449 | 246 | 505 |
| ENSDARG000 | 18;18 | 21021699:210 | 21021907:210 | +;+ | 534 | NA | 0 | 0 | 0 | 0 | 0 | 0 |
| ENSDARG000 | 18:18:18:18:1 | 20881462:208 | 20883234:208 | -:-:-:-:-:-:-:-:-: | 6460 | igf1ra | 445 | 304 | 610 | 324 | 534 | 394 |
| ENSDARG000 | 18;18 | 28303891:283 | 28304111:283 | +;+ | 348 | NA | 0 | 0 | 0 | 0 | 0 | 0 |
| ENSDARG000 | 18;18;18 | 4845053:4848 | 4845415:4848 | -;-;- | 606 | NA | 0 | 0 | 1 | 0 | 0 | 0 |
| ENSDARG000 | 18:18:18:18:1 | 8912113:8912 | 8912683:8912 | +:+:+:+:+:+:+:+: | 1573 | tmem243a | 1421 | 1021 | 908 | 830 | 965 | 690 |
| ENSDARG000 | 18:18:18:18:1 | 50627085:506 | 50627172:506 | +;+;+;+;+;+ | 2651 | CHRNB4 | 0 | 2 | 0 | 0 | 0 | 0 |
| ENSDARG000 | 18:18:18:18:1 | 28320730:283 | 28321932:283 | -;-;-;-;-;- | 2180 | NA | 0 | 0 | 14 | 6 | 0 | 6 |
| ENSDARG000 | 18 | 50706476 | 50707287 | - | 812 | NA | 47 | 20 | 34 | 16 | 28 | 62 |
| ENSDARG000 | 18:18:18:18:1 | 50650512:506 | 50650769:506 | +:+:+:+:+:+:+:+: | 9155 | NA | 8 | 12 | 4 | 8 | 22 | 0 |
| ENSDARG000 | 18:18:18:18:1 | 33090443:330 | 33092053:330 | -;-;-;-;-;-;-;- | 2946 | NA | 0 | 0 | 0 | 0 | 0 | 0 |
| ENSDARG000 | 18:18:18:18:1 | 15841449:158 | 15841613:158 | +:+:+:+:+:+:+:+: | 4338 | EEA1 | 518 | 383 | 506 | 288 | 653 | 369 |
| ENSDARG000 | 18:18:18:18:1 | 4902498:4902 | 4902760:4902 | -:-:-:-:-:-:-:-:-: | 1222 | NA | 0 | 0 | 0 | 0 | 0 | 0 |
| ENSDARG000 | 18:18:18:18:1 | 41549764:415 | 41549910:415 | +;+;+;+;+;+ | 1359 | bcl7bb | 138 | 124 | 120 | 122 | 136 | 192 |
| ENSDARG000 | 18:18:18:18:1 | 33100606:331 | 33100820:331 | +;+;+;+;+;+ | 2538 | NA | 0 | 0 | 0 | 0 | 0 | 0 |
| ENSDARG000 | 18:18:18:18:1 | 35742838:357 | 35743100:357 | +:+:+:+:+:+:+:+: | 3445 | rasgrp4 | 143 | 64 | 73 | 38 | 97 | 121 |
| ENSDARG000 | 18;18;18;18 | 39185907:391 | 39186880:391 | -;-;-;- | 1205 | NA | 0 | 0 | 0 | 0 | 0 | 0 |
| ENSDARG000 | 18:18:18:18:1 | 39170127:391 | 39174388:391 | -:-:-:-:-:-:-:-:-: | 6557 | mapk6 | 1207 | 1298 | 976 | 593 | 1147 | 966 |
| ENSDARG000 | 18:18:18:18:1 | 28564640:285 | 28565029:285 | +:+:+:+:+:+:+:+: | 3128 | spon1a | 0 | 0 | 0 | 0 | 0 | 0 |
| ENSDARG000 | 18:18:18:18:1 | 50638006:506 | 50638174:506 | +;+;+;+;+;+ | 3410 | CHRNA3 | 0 | 2 | 0 | 0 | 0 | 0 |
| ENSDARG000 | 18 | 17511995 | 17513426 | - | 1432 | B3GNT9 | 134 | 182 | 182 | 98 | 202 | 119 |
| ENSDARG000 | 18 | 50707179 | 50709715 | + | 2537 | rap2b | 6921 | 7484 | 7243 | 7259 | 9434 | 7584 |
| ENSDARG000 | 18:18:18:18:1 | 50600989:506 | 50601710:506 | -:-:-:-:-:-:-:-:-: | 2312 | ube2q1 | 468 | 366 | 372 | 274 | 493 | 332 |

| | | | | | | | | | | | | |
|---|---|---|---|---|---|---|---|---|---|---|---|---|
| ENSDARG000 | 18:18:18:18:1 | 12839126:128 | 12839712:128 | -:-:-:-:-:-:-:-: | 2359 | parp12a | 511 | 480 | 397 | 387 | 447 | 281 |
| ENSDARG000 | 18:18:18:18:1 | 33132266:331 | 33132479:331 | +;+;+;+;+;+ | 2540 | NA | 0 | 0 | 0 | 0 | 0 | 0 |
| ENSDARG000 | 18:18:18:18:1 | 33171765:331 | 33172863:331 | +:+:+:+:+:+:+: | 4533 | NA | 0 | 0 | 0 | 0 | 0 | 0 |
| ENSDARG000 | 18;18;18 | 50684059:506 | 50684149:506 | +;+;+ | 4512 | P2RY1 | 68 | 130 | 185 | 68 | 164 | 34 |
| ENSDARG000 | 18:18:18:18:1 | 50536202:505 | 50539558:505 | -:-:-:-:-:-:-:-: | 5350 | zgc:158464 | 50 | 8 | 40 | 0 | 17 | 13 |
| ENSDARG000 | 18:18:18:18:1 | 41560822:415 | 41561403:415 | +:+:+:+:+:+:+: | 5167 | baz1b | 666 | 776 | 1064 | 264 | 1021 | 665 |
| ENSDARG000 | 18:18:18:18:1 | 33212589:332 | 33212778:332 | +;+;+;+;+;+ | 2512 | NA | 0 | 0 | 0 | 0 | 0 | 0 |
| ENSDARG000 | 18:18:18:18:1 | 21061216:210 | 21061318:210 | +;+;+;+;+ | 660 | NA | 0 | 0 | 5 | 0 | 0 | 0 |
| ENSDARG000 | 18:18:18:18:1 | 33237127:332 | 33237264:332 | +;+;+;+;+;+ | 2468 | NA | 0 | 0 | 0 | 0 | 0 | 0 |
| ENSDARG000 | 18:18:18:18:1 | 33248996:332 | 33251216:332 | -:-:-:-:-:-:-:-: | 4196 | NA | 0 | 0 | 0 | 0 | 0 | 0 |
| ENSDARG000 | 18;18 | 28937061:289 | 28937425:289 | -;- | 888 | NA | 0 | 0 | 0 | 0 | 0 | 0 |
| ENSDARG000 | 18:18:18:18:1 | 33276609:332 | 33276925:332 | +;+;+;+;+ | 2444 | NA | 0 | 0 | 0 | 0 | 0 | 0 |
| ENSDARG000 | 18:18:18:18:1 | 33265514:332 | 33265802:332 | +;+;+;+;+ | 2323 | NA | 0 | 0 | 0 | 0 | 0 | 0 |
| ENSDARG000 | 18:18:18:18:1 | 33185819:331 | 33186134:331 | +:+:+:+:+:+:+: | 8483 | NA | 0 | 0 | 0 | 0 | 0 | 0 |
| ENSDARG000 | 18:18:18:18:1 | 33264609:332 | 33265344:332 | +:+:+:+:+:+:+: | 3375 | NA | 0 | 0 | 0 | 0 | 0 | 0 |
| ENSDARG000 | 18:18:18:18:1 | 35807588:358 | 35808467:358 | -:-:-:-:-:-:-:-: | 3219 | hnrnpul1 | 2518 | 2262 | 2935 | 1998 | 2245 | 2240 |
| ENSDARG000 | 18:18:18:18:1 | 17516569:175 | 17517002:175 | -:-:-:-:-:-:-:-: | 2578 | bbs2 | 347 | 433 | 329 | 235 | 235 | 214 |
| ENSDARG000 | 18;18 | 23187128:231 | 23187495:231 | -;- | 398 | NA | 0 | 0 | 0 | 0 | 0 | 1 |
| ENSDARG000 | 18:18:18:18:1 | 23187419:231 | 23187671:231 | +:+:+:+:+:+:+: | 3175 | mef2aa | 657 | 455 | 668 | 326 | 551 | 671 |
| ENSDARG000 | 18:18:18:18:1 | 39292911:392 | 39293338:392 | -:-:-:-:-:-:-:-: | 2284 | leo1 | 806 | 664 | 927 | 431 | 723 | 535 |
| ENSDARG000 | 18;18 | 35838752:358 | 35839845:358 | -;- | 1258 | opa3 | 98 | 138 | 214 | 76 | 183 | 117 |
| ENSDARG000 | 18:18:18:18:1 | 15887830:158 | 15888411:158 | -:-:-:-:-:-:-:-: | 3713 | plekhg7 | 4 | 4 | 3 | 9 | 18 | 8 |
| ENSDARG000 | 18:18:18:18:1 | 35842933:358 | 35843084:358 | +:+:+:+:+:+:+: | 4225 | ppp1r13l | 210 | 18 | 85 | 68 | 80 | 67 |
| ENSDARG000 | 18:18:18:18:1 | 29145681:291 | 29146085:291 | +:+:+:+:+:+:+: | 4993 | NA | 90 | 10 | 10 | 2 | 10 | 0 |
| ENSDARG000 | 18:18:18:18:1 | 12867400:128 | 12868208:128 | -:-:-:-:-:-:-:-: | 3747 | SRGAP1 | 374 | 180 | 1092 | 206 | 763 | 62 |
| ENSDARG000 | 18:18:18:18:1 | 21071023:210 | 21071130:210 | +;+;+;+;+;+ | 1001 | arpin | 236 | 215 | 193 | 264 | 279 | 339 |
| ENSDARG000 | 18:18:18:18:1 | 7150144:7151 | 7150198:7151 | +:+:+:+:+:+:+: | 8829 | vwf | 127 | 21 | 46 | 12 | 35 | 46 |
| ENSDARG000 | 18:18:18:18:1 | 39327010:393 | 39327104:393 | +:+:+:+:+:+:+: | 1476 | tmod2 | 433 | 332 | 370 | 144 | 385 | 295 |
| ENSDARG000 | 18:18:18:18:1 | 35725:36068: | 36213:36213: | -:-:-:-:-:-:-:-: | 2103 | gatm | 1253 | 192 | 1090 | 1024 | 1664 | 212 |
| ENSDARG000 | 18;18 | 7196132:7201 | 7196284:7201 | -;- | 448 | NA | 0 | 0 | 0 | 0 | 0 | 0 |
| ENSDARG000 | 18;18 | 19101679:191 | 19101854:191 | +;+ | 521 | NA | 0 | 0 | 0 | 0 | 0 | 0 |
| ENSDARG000 | 18:18:18:18:1 | 19044839:190 | 19045262:190 | -:-:-:-:-:-:-:-: | 6006 | dennd4a | 128 | 344 | 179 | 172 | 281 | 169 |
| ENSDARG000 | 18:18:18:18:1 | 39416357:394 | 39416607:394 | +:+:+:+:+:+:+: | 1527 | lysmd2 | 302 | 184 | 219 | 146 | 217 | 130 |
| ENSDARG000 | 18:18:18:18:1 | 8917766:8922 | 8917906:8922 | +:+:+:+:+:+:+: | 3272 | elapor2a | 43 | 14 | 2 | 4 | 8 | 0 |
| ENSDARG000 | 18;18;18 | 8937718:8942 | 8937787:8943 | +;+;+ | 296 | NA | 0 | 0 | 0 | 0 | 0 | 0 |
| ENSDARG000 | 18:18:18:18:1 | 21077528:210 | 21078292:210 | -:-:-:-:-:-:-:-: | 3456 | anpeplb | 8 | 5 | 20 | 19 | 14 | 40 |
| ENSDARG000 | 18:18:18:18:1 | 19110047:191 | 19110350:191 | -;-;-;-;-;- | 825 | NA | 0 | 2 | 0 | 0 | 0 | 0 |
| ENSDARG000 | 18 | 41648812 | 41650648 | - | 1837 | fzd9b | 253 | 33 | 164 | 137 | 261 | 63 |
| ENSDARG000 | 18;18;18 | 41657487:416 | 41657582:416 | +;+;+ | 458 | NA | 0 | 0 | 0 | 0 | 0 | 0 |
| ENSDARG000 | 18;18;18;18 | 41740742:417 | 41741197:417 | -;-;-;- | 1820 | grk7b | 36 | 14 | 10 | 38 | 17 | 22 |
| ENSDARG000 | 18:18:18:18:1 | 19120984:191 | 19121013:191 | +:+:+:+:+:+:+: | 1107 | NA | 0 | 0 | 0 | 0 | 0 | 0 |
| ENSDARG000 | 18;18 | 36037223:360 | 36037369:360 | +;+ | 303 | TMEM91 | 0 | 0 | 0 | 4 | 0 | 0 |
| ENSDARG000 | 18:18:18:18:1 | 13195745:131 | 13196360:131 | -:-:-:-:-:-:-:-: | 2138 | klhl36 | 159 | 58 | 245 | 159 | 244 | 96 |
| ENSDARG000 | 18;18;18 | 17534627:175 | 17534735:175 | +;+;+ | 594 | mt2 | 2091 | 7739 | 3345 | 2119 | 1970 | 622 |
| ENSDARG000 | 18:18:18:18:1 | 15937610:159 | 15937701:159 | +:+:+:+:+:+:+: | 6030 | ITPR2 | 380 | 192 | 491 | 287 | 514 | 296 |
| ENSDARG000 | 18;18;18 | 29402623:294 | 29405645:294 | +;+;+ | 3086 | mafa | 54 | 10 | 48 | 26 | 54 | 30 |
| ENSDARG000 | 18:18:18:18:1 | 19131773:191 | 19131946:191 | +;+;+;+;+ | 3232 | rab11a | 7484 | 7803 | 7161 | 5574 | 7388 | 6792 |
| ENSDARG000 | 18:18:18:18:1 | 39434589:394 | 39435418:394 | -:-:-:-:-:-:-:-: | 2367 | scg3 | 26 | 2 | 2 | 2 | 10 | 0 |
| ENSDARG000 | 18;18;18 | 41757263:417 | 41757682:417 | -;-;- | 725 | rnf7 | 770 | 1005 | 943 | 587 | 907 | 766 |
| ENSDARG000 | 18;18 | 21101835:211 | 21103477:211 | -;- | 2021 | si:dkev-12e7.: | 286 | 168 | 132 | 112 | 257 | 213 |
| ENSDARG000 | 18:18:18:18:1 | 41768357:417 | 41768445:417 | +:+:+:+:+:+:+: | 3063 | NA | 10 | 11 | 15 | 11 | 5 | 12 |
| ENSDARG000 | 18:18:18:18:1 | 36060295:360 | 36060413:360 | -;-;-;-;-;-;-;- | 836 | exosc5 | 114 | 189 | 134 | 80 | 84 | 128 |
| ENSDARG000 | 18:18:18:18:1 | 41820039:418 | 41820577:418 | +:+:+:+:+:+:+: | 1662 | pgr | 0 | 0 | 0 | 0 | 0 | 0 |
| ENSDARG000 | 18:18:18:18:1 | 29885190:298 | 29889739:298 | -:-:-:-:-:-:-:-: | 5648 | cmc2 | 130 | 145 | 232 | 174 | 192 | 202 |
| ENSDARG000 | 18;18;18;18 | 13203831:132 | 13204164:132 | +;+;+;+ | 1398 | cotl1 | 8575 | 12865 | 9500 | 12508 | 10073 | 12857 |
| ENSDARG000 | 18:18:18:18:1 | 33284202:332 | 33284387:332 | +;+;+;+;+;+ | 2570 | NA | 0 | 0 | 0 | 0 | 0 | 0 |
| ENSDARG000 | 18:18:18:18:1 | 19143437:191 | 19143612:191 | -:-:-:-:-:-:-:-: | 3748 | NA | 270 | 0 | 15 | 0 | 0 | 0 |
| ENSDARG000 | 18:18:18:18:1 | 7283283:7284 | 7283360:7284 | +:+:+:+:+:+:+: | 1649 | ANO2 | 0 | 0 | 0 | 0 | 0 | 0 |
| ENSDARG000 | 18:18:18:18:1 | 21113216:211 | 21113549:211 | +:+:+:+:+:+:+: | 4073 | kmt5b | 1053 | 828 | 1291 | 729 | 1528 | 976 |
| ENSDARG000 | 18:18:18:18:1 | 33310811:333 | 33311015:333 | +;+;+;+;+;+ | 2583 | NA | 0 | 0 | 0 | 0 | 0 | 0 |
| ENSDARG000 | 18:18:18:18:1 | 21122818:211 | 21123195:211 | +:+:+:+:+:+:+: | 3333 | chka | 365 | 1046 | 2308 | 1558 | 1009 | 1300 |
| ENSDARG000 | 18:18:18:18:1 | 54354:55186: | 54841:55389: | +:+:+:+:+:+:+: | 1967 | zgc:158482 | 1532 | 453 | 1534 | 1171 | 1982 | 1684 |
| ENSDARG000 | 18:18:18:18:1 | 13235714:132 | 13236632:132 | +:+:+:+:+:+:+: | 9979 | plcg2 | 112 | 97 | 101 | 144 | 269 | 148 |
| ENSDARG000 | 18:18:18:18:1 | 33333411:333 | 33334315:333 | -;-;-;-;-;- | 2607 | olfcu1 | 0 | 0 | 0 | 0 | 0 | 0 |
| ENSDARG000 | 18:18:18:18:1 | 36066389:360 | 36066522:360 | +:+:+:+:+:+:+: | 2368 | bckdha | 946 | 953 | 1194 | 867 | 1304 | 1194 |
| ENSDARG000 | 18:18:18:18:1 | 29898955:299 | 29899226:299 | +:+:+:+:+:+:+: | 1808 | cenpn | 62 | 192 | 300 | 93 | 168 | 90 |
| ENSDARG000 | 18:18:18:18:1 | 17537311:175 | 17537383:175 | +:+:+:+:+:+:+: | 6116 | nup93 | 783 | 1164 | 1712 | 642 | 1365 | 1066 |
| ENSDARG000 | 18:18:18:18:1 | 33322515:333 | 33322708:333 | +;+;+;+;+;+ | 2535 | NA | 0 | 0 | 0 | 0 | 0 | 0 |
| ENSDARG000 | 18:18:18:18:1 | 23373683:233 | 23374350:233 | +:+:+:+:+:+:+: | 2828 | NA | 12 | 8 | 6 | 8 | 10 | 2 |
| ENSDARG000 | 18:18:18:18:1 | 42150947:421 | 42151865:421 | -:-:-:-:-:-:-:-: | 4039 | cntn5 | 22 | 0 | 2 | 10 | 4 | 2 |
| ENSDARG000 | 18:18:18:18:1 | 13236255:132 | 13236279:132 | +:+:+:+:+:+:+: | 1403 | meak7 | 0 | 0 | 0 | 0 | 0 | 0 |
| ENSDARG000 | 18;18 | 13281654:132 | 13281875:132 | -;- | 277 | NA | 0 | 0 | 0 | 0 | 0 | 0 |
| ENSDARG000 | 18:18:18:18:1 | 62932:66153: | 63465:66362: | +:+:+:+:+:+:+: | 3019 | slc27a2a | 139 | 122 | 168 | 152 | 283 | 190 |
| ENSDARG000 | 18:18:18:18:1 | 81867:82027: | 82218:82218: | -:-:-:-:-:-:-:-: | 1322 | gabpb1 | 76 | 23 | 62 | 46 | 70 | 42 |
| ENSDARG000 | 18:18:18:18:1 | 73434:75027: | 75237:75237: | -:-:-:-:-:-:-:-: | 3100 | hdc | 397 | 62 | 110 | 61 | 60 | 24 |
| ENSDARG000 | 18:18:18:18:1 | 7309150:7310 | 7309269:7310 | +:+:+:+:+:+:+: | 1827 | cers3b | 39 | 7 | 5 | 0 | 19 | 15 |
| ENSDARG000 | 18:18:18:18:1 | 39487486:394 | 39487679:394 | +:+:+:+:+:+:+: | 1753 | acadl | 618 | 490 | 636 | 356 | 504 | 520 |
| ENSDARG000 | 18:18:18:18:1 | 7317545:7327 | 7317627:7328 | -:-:-:-:-:-:-:-: | 1848 | NA | 2 | 16 | 31 | 0 | 16 | 12 |
| ENSDARG000 | 18;18 | 23830265:238 | 23831161:238 | -;- | 1090 | NA | 0 | 0 | 0 | 0 | 0 | 0 |
| ENSDARG000 | 18:18:18:18:1 | 23868650:238 | 23871170:238 | -:-:-:-:-:-:-:-: | 4270 | nr2f2 | 53 | 2 | 26 | 10 | 44 | 26 |
| ENSDARG000 | 18:18:18:18:1 | 33340868:333 | 33340977:333 | +;+;+;+;+;+ | 2463 | olfcq1 | 0 | 0 | 0 | 0 | 0 | 0 |
| ENSDARG000 | 18;18 | 23877624:238 | 23877963:238 | +;+ | 5146 | NA | 0 | 0 | 0 | 0 | 0 | 0 |
| ENSDARG000 | 18:18:18:18:1 | 19419120:194 | 19419503:194 | +:+:+:+:+:+:+: | 2093 | map2k1 | 599 | 522 | 406 | 642 | 380 | 507 |
| ENSDARG000 | 18:18:18:18:1 | 8972227:8972 | 8972303:8972 | -:-:-:-:-:-:-:-: | 2932 | grm3 | 0 | 0 | 0 | 0 | 0 | 0 |
| ENSDARG000 | 18:18:18:18:1 | 36105930:361 | 36106133:361 | -;-;-;-;- | 1226 | b3gat1a | 0 | 2 | 4 | 0 | 0 | 1 |
| ENSDARG000 | 18:18:18:18:1 | 19451070:194 | 19451320:194 | -:-:-:-:-:-:-:-: | 2521 | rpl4 | 35028 | 32108 | 54289 | 49580 | 38892 | 50275 |
| ENSDARG000 | 18;18;18 | 29918062:299 | 29918175:299 | -;-;- | 645 | NA | 0 | 0 | 0 | 0 | 0 | 0 |
| ENSDARG000 | 18:18:18:18:1 | 33353271:333 | 33353529:333 | +:+:+:+:+:+:+: | 2969 | NA | 0 | 0 | 0 | 0 | 0 | 0 |
| ENSDARG000 | 18:18:18:18:1 | 29922939:299 | 29923494:299 | -;-;-;-;- | 1299 | mhc2dbb | 0 | 0 | 2 | 0 | 2 | 0 |
| ENSDARG000 | 18;18 | 36141233:361 | 36141452:361 | +;+ | 404 | NA | 0 | 0 | 0 | 0 | 0 | 0 |
| ENSDARG000 | 18;18 | 42773594:427 | 42775429:427 | +;+ | 2045 | NA | 0 | 0 | 0 | 0 | 0 | 0 |
| ENSDARG000 | 18:18:18:18:1 | 42781622:427 | 42781996:427 | -;-;-;-;-;- | 783 | ttc36 | 8 | 0 | 0 | 0 | 2 | 0 |
| ENSDARG000 | 18:18:18:18:1 | 9171778:9186 | 9172014:9186 | +:+:+:+:+:+:+: | 2814 | sema3d | 38 | 4 | 8 | 4 | 42 | 16 |
| ENSDARG000 | 18:18:18:18:1 | 36455307:364 | 36455568:364 | -:-:-:-:-:-:-:-: | 1222 | NA | 5 | 0 | 15 | 2 | 7 | 17 |
| ENSDARG000 | 18:18:18:18:1 | 33373811:333 | 33374133:333 | +:+:+:+:+:+:+: | 2682 | olfcq3 | 0 | 0 | 0 | 0 | 0 | 0 |
| ENSDARG000 | 18:18:18:18:1 | 36582815:365 | 36582912:365 | +:+:+:+:+:+:+: | 2275 | siae | 820 | 620 | 621 | 368 | 863 | 644 |
| ENSDARG000 | 18;18 | 39522468:395 | 39524039:395 | -;- | 1795 | tnfaip8l3 | 10 | 8 | 18 | 2 | 34 | 0 |
| ENSDARG000 | 18:18:18:18:1 | 42970208:429 | 42970219:429 | +:+:+:+:+:+:+: | 2403 | NECTIN1 | 91 | 99 | 123 | 57 | 155 | 92 |
| ENSDARG000 | 18:18:18:18:1 | 24273428:242 | 24273754:242 | -:-:-:-:-:-:-:-: | 1682 | si:ch211-147r | 0 | 0 | 0 | 0 | 0 | 0 |
| ENSDARG000 | 18:18:18:18:1 | 7345417:7348 | 7345826:7349 | +;+;+;+;+ | 1787 | glipr1b | 0 | 2 | 0 | 0 | 0 | 2 |
| ENSDARG000 | 18;18;18 | 43365890:433 | 43366423:433 | +;+;+ | 1841 | NA | 0 | 4 | 0 | 0 | 0 | 0 |
| ENSDARG000 | 18:18:18:18:1 | 33386372:333 | 33386592:333 | +;+;+;+;+;+ | 2604 | NA | 0 | 0 | 0 | 0 | 0 | 0 |
| ENSDARG000 | 18:18:18:18:1 | 21178721:211 | 21180041:211 | -:-:-:-:-:-:-:-: | 2573 | got2a | 865 | 806 | 865 | 797 | 1146 | 811 |
| ENSDARG000 | 18:18:18:18:1 | 7363242:7367 | 7363521:7367 | +:+:+:+:+:+:+: | 2351 | NA | 8 | 32 | 70 | 6 | 15 | 300 |
| ENSDARG000 | 18:18:18:18:1 | 21171465:211 | 21171982:211 | -;-;-;-;-;- | 1238 | si:dkev-12e7.4 | 12 | 4 | 0 | 0 | 0 | 16 |
| ENSDARG000 | 18:18:18:18:1 | 7375923:7376 | 7377718:7377 | -;-;-;-;-;-;-;- | 2187 | bbs10 | 10 | 18 | 22 | 0 | 10 | 27 |
| ENSDARG000 | 18;18 | 7381624:7381 | 7381837:7381 | -;- | 275 | NA | 0 | 0 | 0 | 0 | 0 | 0 |
| ENSDARG000 | 18:18:18:18:1 | 43846936:438 | 43847295:438 | -:-:-:-:-:-:-:-: | 2208 | TREH | 0 | 0 | 0 | 5 | 0 | 0 |
| ENSDARG000 | 18:18:18:18:1 | 9323211:9323 | 9323380:9323 | +:+:+:+:+:+:+: | 3096 | sema3ab | 873 | 390 | 617 | 125 | 522 | 470 |
| ENSDARG000 | 18:18:18:18:1 | 24477526:244 | 24477750:244 | +:+:+:+:+:+:+: | 1147 | NA | 0 | 0 | 0 | 0 | 0 | 0 |
| ENSDARG000 | 18;18;18;18 | 29950233:299 | 29950448:299 | +;+;+;+ | 2271 | ATMIN | 74 | 122 | 88 | 110 | 99 | 90 |
| ENSDARG000 | 18:18:18:18:1 | 29958653:299 | 29959612:299 | -;-;-;-;-;-;-;- | 1324 | si:ch73-103b9 | 22 | 4 | 16 | 34 | 4 | 12 |
| ENSDARG000 | 18:18:18:18:1 | 43865901:438 | 43870785:438 | -:-:-:-:-:-:-:-: | 6897 | DDX6 | 466 | 372 | 395 | 346 | 489 | 352 |
| ENSDARG000 | 18:18:18:18:1 | 33408250:334 | 33408470:334 | +;+;+;+;+;+ | 2604 | NA | 0 | 0 | 0 | 0 | 0 | 0 |
| ENSDARG000 | 18:18:18:18:1 | 21198762:211 | 21199373:211 | -:-:-:-:-:-:-:-: | 1365 | calb2a | 0 | 0 | 0 | 8 | 0 | 0 |
| ENSDARG000 | 18:18:18:18:1 | 33420995:334 | 33421160:334 | +;+;+;+;+;+ | 2528 | NA | 0 | 0 | 0 | 0 | 0 | 0 |
| ENSDARG000 | 18:18:18:18:1 | 36631923:366 | 36632084:366 | +;+;+;+;+;+ | 3904 | znf296 | 68 | 20 | 45 | 52 | 92 | 46 |
| ENSDARG000 | 18:18:18:18:1 | 36601403:366 | 36601598:366 | +:+:+:+:+:+:+: | 5421 | clasrp | 779 | 667 | 966 | 707 | 1034 | 939 |
| ENSDARG000 | 18 | 33443264 | 33444257 | + | 994 | NA | 0 | 0 | 0 | 0 | 0 | 0 |
| ENSDARG000 | 18:18:18:18:1 | 39620535:396 | 39620988:396 | -;-;-;-;-;-;-;- | 1862 | cyp19a1a | 16 | 4 | 40 | 12 | 5 | 0 |
| ENSDARG000 | 18:18:18:18:1 | 33464345:334 | 33464417:334 | +:+:+:+:+:+:+: | 2669 | NA | 0 | 0 | 0 | 0 | 0 | 0 |
| ENSDARG000 | 18:18:18:18:1 | 29968205:299 | 29972310:299 | -:-:-:-:-:-:-:-: | 10037 | NA | 1071 | 659 | 1209 | 862 | 1066 | 1107 |
| ENSDARG000 | 18;18 | 24560187:245 | 24560244:245 | -;- | 134 | NA | 0 | 4 | 4 | 0 | 2 | 6 |
| ENSDARG000 | 18:18:18:18:1 | 9493720:9524 | 9493865:9524 | +:+:+:+:+:+:+: | 3044 | sema3e | 2075 | 439 | 323 | 418 | 382 | 736 |
| ENSDARG000 | 18:18:18:18:1 | 7392147:7395 | 7395743:7395 | -:-:-:-:-:-:-:-: | 4312 | si:dkev-30c15 | 0 | 0 | 2 | 0 | 12 | 6 |
| ENSDARG000 | 18:18:18:18:1 | 13300768:133 | 13300979:133 | +:+:+:+:+:+:+: | 3048 | NA | 51 | 22 | 137 | 14 | 105 | 26 |
| ENSDARG000 | 18:18:18:18:1 | 7387090:7387 | 7387276:7387 | +:+:+:+:+:+:+: | 1169 | NA | 0 | 0 | 0 | 0 | 0 | 0 |
| ENSDARG000 | 18;18;18 | 24562188:245 | 24562280:245 | +;+;+ | 1649 | lysmd4 | 154 | 114 | 138 | 94 | 78 | 112 |
| ENSDARG000 | 18:18:18:18:1 | 33490315:334 | 33490496:334 | +:+:+:+:+:+:+: | 2568 | NA | 0 | 0 | 0 | 0 | 0 | 0 |
| ENSDARG000 | 18:18:18:18:1 | 21257261:212 | 21258120:212 | -;-;-;-;-;-;-;- | 1984 | pnp6 | 7 | 10 | 41 | 22 | 173 | 0 |
| ENSDARG000 | 18:18:18:18:1 | 5213338:5216 | 5213554:5216 | +:+:+:+:+:+:+: | 2003 | slc24a5 | 4401 | 4152 | 785 | 4950 | 1461 | 4077 |
| ENSDARG000 | 18:18:18:18:1 | 95770:96279: | 95936:96423: | +:+:+:+:+:+:+: | 4181 | USP8 | 3626 | 2336 | 2928 | 2207 | 2908 | 2359 |
| ENSDARG000 | 18;18;18 | 16111185:161 | 16111544:161 | -;-;- | 687 | sspn | 279 | 485 | 213 | 915 | 140 | 432 |
| ENSDARG000 | 18:18:18:18:1 | 16125852:161 | 16126008:161 | +;+;+;+;+ | 3155 | bhlhe41 | 5904 | 3662 | 3654 | 3575 | 3750 | 3655 |
| ENSDARG000 | 18:18:18:18:1 | 33521609:335 | 33521772:335 | +;+;+;+;+;+ | 2499 | olfcq10 | 0 | 0 | 0 | 0 | 0 | 0 |
| ENSDARG000 | 18:18:18:18:1 | 30028135:300 | 30028249:300 | +:+:+:+:+:+:+: | 1409 | NA | 4 | 0 | 4 | 10 | 2 | 6 |
| ENSDARG000 | 18:18:18:18:1 | 45792035:457 | 45792063:457 | +:+:+:+:+:+:+: | 4755 | abcc5 | 11981 | 3110 | 7595 | 9260 | 16920 | 9277 |
| ENSDARG000 | 18:18:18:18:1 | 24737234:247 | 24737419:247 | +:+:+:+:+:+:+: | 1660 | NA | 10 | 8 | 0 | 0 | 6 | 0 |
| ENSDARG000 | 18:18:18:18:1 | 17583479:175 | 17583731:175 | +:+:+:+:+:+:+: | 4358 | slc12a3 | 0 | 0 | 86 | 0 | 116 | 0 |
| ENSDARG000 | 18:18:18:18:1 | 33550667:335 | 33550869:335 | +:+:+:+:+:+:+: | 2574 | NA | 0 | 0 | 0 | 0 | 0 | 0 |
| ENSDARG000 | 18:18:18:18:1 | 13324140:133 | 13326777:133 | -:-:-:-:-:-:-:-: | 4855 | CMIP | 4201 | 2995 | 3515 | 2508 | 3157 | 3196 |
| ENSDARG000 | 18:18:18:18:1 | 39649660:396 | 39650274:396 | +:+:+:+:+:+:+: | 2056 | gldn | 20 | 0 | 0 | 4 | 14 | 6 |
| ENSDARG000 | 18:18:18:18:1 | 19456648:194 | 19456764:194 | +:+:+:+:+:+:+: | 2026 | zwilch | 73 | 183 | 198 | 70 | 251 | 87 |
| ENSDARG000 | 18:18:18:18:1 | 7400929:7402 | 7401138:7402 | -:-:-:-:-:-:-:-: | 2213 | NA | 0 | 2 | 3 | 8 | 8 | 8 |
| ENSDARG000 | 18:18:18:18:1 | 45955042:459 | 45955177:459 | +:+:+:+:+:+:+: | 1799 | ap2m1b | 1981 | 1845 | 2316 | 1512 | 1929 | 1697 |
| ENSDARG000 | 18:18:18:18:1 | 9615147:9617 | 9615421:9618 | +:+:+:+:+:+:+: | 31976 | NA | 50 | 2 | 20 | 6 | 16 | 14 |
| ENSDARG000 | 18:18:18:18:1 | 16133595:161 | 16133855:161 | +:+:+:+:+:+:+: | 2091 | ctsd | 5503 | 3855 | 5468 | 4168 | 6120 | 6086 |
| ENSDARG000 | 18;18 | 13477102:134 | 13477527:134 | -;- | 551 | NA | 0 | 0 | 0 | 0 | 0 | 0 |
| ENSDARG000 | 18:18:18:18:1 | 5233354:5233 | 5233737:5233 | -:-:-:-:-:-:-:-: | 2776 | myef2 | 1651 | 2120 | 2150 | 1405 | 2196 | 1426 |
| ENSDARG000 | 18:18:18:18:1 | 113928:11460 | 114738:11473 | -:-:-:-:-:-:-:-: | 6335 | trpm7 | 2243 | 1760 | 2298 | 1215 | 2598 | 1975 |
| ENSDARG000 | 18:18:18:18:1 | 36664654:366 | 36664897:366 | +:+:+:+:+:+:+: | 2677 | NA | 1352 | 2997 | 6252 | 2111 | 10107 | 4112 |
| ENSDARG000 | 18:18:18:18:1 | 33557125:335 | 33557350:335 | +;+;+;+;+;+ | 2594 | NA | 0 | 0 | 0 | 0 | 0 | 0 |
| ENSDARG000 | 18:18:18:18:1 | 39669255:396 | 39670863:396 | -:-:-:-:-:-:-:-: | 10832 | dmxl2 | 2523 | 1804 | 1844 | 2178 | 1752 | 1396 |
| ENSDARG000 | 18:18:18:18:1 | 30129771:301 | 30129867:301 | +:+:+:+:+:+:+: | 588 | NA | 0 | 0 | 0 | 0 | 0 | 0 |
| ENSDARG000 | 18:18:18:18:1 | 13792490:138 | 13792701:138 | +:+:+:+:+:+:+: | 4088 | CDH13 | 0 | 12 | 23 | 28 | 16 | 34 |
| ENSDARG000 | 18;18;18;18 | 19487702:194 | 19488205:194 | -;-;-;- | 830 | snapc5 | 412 | 522 | 628 | 601 | 638 | 530 |
| ENSDARG000 | 18:18:18:18:1 | 7425222:7425 | 7428335:7428 | -:-:-:-:-:-:-:-: | 6457 | NA | 284 | 1222 | 723 | 232 | 481 | 498 |
| ENSDARG000 | 18:18:18:18:1 | 9749375:9751 | 9749410:9751 | +:+:+:+:+:+:+: | 3111 | CACNA2D1 | 0 | 0 | 7 | 0 | 0 | 0 |
| ENSDARG000 | 18;18;18;18 | 19648275:196 | 19649710:196 | +;+;+;+ | 2113 | smad6b | 240 | 81 | 207 | 133 | 198 | 79 |
| ENSDARG000 | 18:18:18:18:1 | 173603:18474 | 173971:18494 | +;+;+;+;+ | 2246 | larp6a | 389 | 501 | 678 | 340 | 743 | 402 |
| ENSDARG000 | 18:18:18:18:1 | 16151774:161 | 16152163:161 | -:-:-:-:-:-:-:-: | 2517 | slc6a15 | 1272 | 941 | 384 | 1345 | 27 | 2193 |
| ENSDARG000 | 18 | 10296228 | 10298162 | - | 1935 | lrrc4.2 | 0 | 0 | 0 | 0 | 0 | 0 |

| | | | | | | | | | | | |
|---|---|---|---|---|---|---|---|---|---|---|---|
| ENSDARG000 | 18;18 | 10689772:106 | 10689915:106 | +;+ | 501 | lepa | 4 | 6 | 0 | 0 | 0 | 0 |
| ENSDARG000 | 18:18:18:18:1 | 46099223:461 | 46099313:461 | -:-:-:-:-:-:-:-:-: | 3825 | SPTBN4 | 2 | 0 | 0 | 0 | 6 | 6 |
| ENSDARG000 | 18:18:18:18:1 | 33570170:335 | 33570390:335 | +;+;+;+;+;+ | 2588 | NA | 0 | 0 | 0 | 0 | 0 | 0 |
| ENSDARG000 | 18:18:18:18:1 | 36769758:367 | 36770017:367 | +:+:+:+:+:+:+: | 2802 | fosb | 4512 | 510 | 1928 | 1182 | 667 | 1373 |
| ENSDARG000 | 18:18:18:18:1 | 21273749:212 | 21274125:212 | +:+:+:+:+:+:+: | 15817 | hydin | 6 | 20 | 40 | 50 | 36 | 4 |
| ENSDARG000 | 18:18:18:18:1 | 204608:20460 | 205336:20533 | -:-:-:-:-:-:-:-:-: | 1539 | tm2d3 | 602 | 544 | 501 | 368 | 620 | 420 |
| ENSDARG000 | 18:18:18:18:1 | 43896153:438 | 43897571:438 | -:-:-:-:-:-:-:-:-: | 5202 | bcl9l | 512 | 321 | 628 | 329 | 482 | 400 |
| ENSDARG000 | 18:18:18:18:1 | 14233553:142 | 14234372:142 | -;-;-;-;- | 1839 | mlycd | 64 | 68 | 62 | 32 | 54 | 56 |
| ENSDARG000 | 18:18:18:18:1 | 24919614:249 | 24920061:249 | +:+:+:+:+:+:+: | 5298 | rgma | 366 | 124 | 318 | 90 | 142 | 188 |
| ENSDARG000 | 18:18:18:18:1 | 30157034:301 | 30159676:301 | +:+:+:+:+:+:+: | 10949 | gse1b | 4936 | 5029 | 6865 | 6868 | 9133 | 4347 |
| ENSDARG000 | 18:18:18:18:1 | 210838:21229 | 212444:21244 | -:-:-:-:-:-:-:-:-: | 3093 | NA | 54 | 16 | 82 | 40 | 88 | 26 |
| ENSDARG000 | 18;18 | 39937225:399 | 39937582:399 | +;+ | 1020 | NA | 8 | 24 | 40 | 36 | 4 | 30 |
| ENSDARG000 | 18:18:18:18:1 | 33580391:335 | 33580880:335 | +:+:+:+:+:+:+: | 5652 | NA | 0 | 0 | 0 | 0 | 0 | 0 |
| ENSDARG000 | 18:18:18:18:1 | 36782930:367 | 36782987:367 | +:+:+:+:+:+:+: | 1742 | C18H19orf47 | 194 | 152 | 243 | 123 | 195 | 106 |
| ENSDARG000 | 18:18:18:18:1 | 24933473:249 | 24935388:249 | -:-:-:-:-:-:-:-:-: | 7837 | chd2 | 1058 | 1139 | 2624 | 1710 | 2082 | 1901 |
| ENSDARG000 | 18:18:18:18:1 | 14277003:142 | 14277052:142 | +:+:+:+:+:+:+: | 2616 | NA | 0 | 4 | 6 | 2 | 37 | 6 |
| ENSDARG000 | 18:18:18:18:1 | 302796:30282 | 303140:30314 | +:+:+:+:+:+:+: | 3966 | lsm14ab | 2381 | 2211 | 3094 | 2155 | 2703 | 2150 |
| ENSDARG000 | 18:18:18:18:1 | 16192083:161 | 16192114:161 | +:+:+:+:+:+:+: | 4035 | NA | 10 | 6 | 0 | 4 | 0 | 4 |
| ENSDARG000 | 18;18 | 40364630:403 | 40364854:403 | -;- | 291 | NA | 0 | 4 | 1 | 1 | 1 | 5 |
| ENSDARG000 | 18:18:18:18:1 | 40354998:403 | 40355046:403 | +:+:+:+:+:+:+: | 4101 | sema6dl | 490 | 207 | 234 | 448 | 75 | 753 |
| ENSDARG000 | 18:18:18:18:1 | 19772874:197 | 19773223:197 | +:+:+:+:+:+:+: | 2167 | smad3b | 213 | 191 | 276 | 180 | 202 | 220 |
| ENSDARG000 | 18:18:18:18:1 | 44053052:440 | 44053636:440 | -:-:-:-:-:-:-:-:-: | 4423 | cdon | 431 | 281 | 1754 | 264 | 626 | 425 |
| ENSDARG000 | 18:18:18:18:1 | 10692945:106 | 10693480:106 | -:-:-:-:-:-:-:-:-: | 3150 | rbm28 | 826 | 840 | 1468 | 667 | 786 | 955 |
| ENSDARG000 | 18;18;18 | 7452809:7452 | 7454422:7454 | -;-;- | 1728 | pdp2 | 169 | 196 | 209 | 44 | 225 | 153 |
| ENSDARG000 | 18:18:18:18:1 | 30481030:304 | 30481388:304 | -;-;-;-;- | 853 | gins2 | 22 | 217 | 306 | 112 | 219 | 98 |
| ENSDARG000 | 18:18:18:18:1 | 14334123:143 | 14334559:143 | -:-:-:-:-:-:-:-:-: | 1166 | hsbp1b | 502 | 623 | 621 | 542 | 595 | 492 |
| ENSDARG000 | 18;18;18 | 36806545:368 | 36807217:368 | +;+;+ | 951 | ttc9b | 0 | 2 | 0 | 0 | 0 | 0 |
| ENSDARG000 | 18:18:18:18:1 | 46151505:461 | 46151757:461 | +:+:+:+:+:+:+: | 1652 | blvrb | 358 | 798 | 319 | 287 | 458 | 434 |
| ENSDARG000 | 18;18;18;18 | 16246806:162 | 16247007:162 | +;+;+;+ | 963 | alx1 | 6 | 0 | 2 | 6 | 0 | 14 |
| ENSDARG000 | 18:18:18:18:1 | 30500690:305 | 30501879:305 | -:-:-:-:-:-:-:-:-: | 1821 | emc8 | 699 | 708 | 1047 | 541 | 881 | 803 |
| ENSDARG000 | 18;18 | 16258787:163 | 16259057:163 | -;- | 306 | NA | 0 | 0 | 1 | 0 | 0 | 0 |
| ENSDARG000 | 18:18:18:18:1 | 7456597:7456 | 7456729:7456 | +:+:+:+:+:+:+: | 3472 | terb1 | 4 | 13 | 14 | 13 | 0 | 75 |
| ENSDARG000 | 18:18:18:18:1 | 33591704:335 | 33591935:335 | +:+:+:+:+:+:+: | 6125 | NA | 0 | 0 | 0 | 0 | 0 | 0 |
| ENSDARG000 | 18;18;18;18 | 16314591:163 | 16315822:163 | -;-;-;- | 1293 | RASSF9 | 16 | 0 | 4 | 0 | 8 | 8 |
| ENSDARG000 | 18:18:18:18:1 | 328689:32873 | 328934:32883 | +;+;+;+;+;+ | 919 | ss18l2 | 162 | 125 | 230 | 170 | 250 | 136 |
| ENSDARG000 | 18:18:18:18:1 | 19812608:198 | 19813168:198 | -:-:-:-:-:-:-:-:-: | 1562 | aagab | 327 | 409 | 280 | 206 | 414 | 313 |
| ENSDARG000 | 18:18:18:18:1 | 333679:33647 | 336789:33678 | -:-:-:-:-:-:-:-:-: | 6938 | GARRE1 | 897 | 853 | 937 | 816 | 1179 | 923 |
| ENSDARG000 | 18:18:18:18:1 | 17600570:176 | 17601011:176 | +:+:+:+:+:+:+: | 5657 | herpud1 | 1108 | 1338 | 2465 | 984 | 1150 | 795 |
| ENSDARG000 | 18;18;18;18 | 336443:33817 | 336789:33825 | -;-;-;- | 730 | NA | 0 | 0 | 0 | 0 | 0 | 0 |
| ENSDARG000 | 18:18:18:18:1 | 46162204:461 | 46162369:461 | +;+;+;+;+;+ | 3692 | sertad3 | 16 | 230 | 166 | 54 | 144 | 74 |
| ENSDARG000 | 18:18:18:18:1 | 16330025:163 | 16330061:163 | +;+;+;+;+;+;+ | 941 | NTS | 0 | 0 | 0 | 0 | 0 | 0 |
| ENSDARG000 | 18;18 | 44153674:441 | 44154009:441 | +;+ | 440 | NA | 0 | 0 | 0 | 0 | 0 | 0 |
| ENSDARG000 | 18:18:18:18:1 | 5510251:5514 | 5510330:5514 | +:+:+:+:+:+:+: | 3295 | slc30a4 | 1873 | 987 | 1509 | 1401 | 1278 | 1174 |
| ENSDARG000 | 18:18:18:18:1 | 17611627:176 | 17611776:176 | +:+:+:+:+:+:+: | 1789 | cetp | 2 | 2 | 2 | 12 | 18 | 0 |
| ENSDARG000 | 18:18:18:18:1 | 16426266:164 | 16426284:164 | +;+;+;+;+;+;+ | 793 | NA | 0 | 0 | 0 | 0 | 2 | 0 |
| ENSDARG000 | 18:18:18:18:1 | 16387745:163 | 16388901:163 | -:-:-:-:-:-:-:-:-: | 2017 | MGAT4C | 0 | 0 | 0 | 0 | 2 | 0 |
| ENSDARG000 | 18:18:18:18:1 | 14342326:143 | 14342525:143 | +:+:+:+:+:+:+: | 748 | NA | 8 | 4 | 0 | 4 | 2 | 2 |
| ENSDARG000 | 18:18:18:18:1 | 5506025:5506 | 5506392:5506 | -;-;-;-;- | 870 | bloc1s6 | 1496 | 963 | 968 | 1632 | 1106 | 1352 |
| ENSDARG000 | 18:18:18:18:1 | 33622917:336 | 33623355:336 | +;+;+;+;+;+;+ | 2692 | NA | 0 | 0 | 0 | 0 | 0 | 8 |
| ENSDARG000 | 18;18;18;18 | 14477740:144 | 14477823:144 | +;+;+;+ | 2630 | kcng4a | 0 | 4 | 4 | 0 | 0 | 0 |
| ENSDARG000 | 18:18:18:18:1 | 46183136:461 | 46183438:461 | +;+;+;+;+;+;+ | 1086 | NA | 0 | 0 | 1 | 0 | 0 | 0 |
| ENSDARG000 | 18:18:18:18:1 | 5528981:5530 | 5529009:5530 | +:+:+:+:+:+:+: | 694 | NA | 0 | 0 | 0 | 3 | 0 | 0 |
| ENSDARG000 | 18:18:18:18:1 | 46194037:461 | 46194416:461 | -;-;-;-;-;- | 3260 | NA | 8 | 9 | 2 | 8 | 0 | 4 |
| ENSDARG000 | 18:18:18:18:1 | 5530083:5530 | 5530258:5530 | -:-:-:-:-:-:-:-:-: | 2333 | spata5l1 | 179 | 244 | 204 | 322 | 235 | 281 |
| ENSDARG000 | 18;18;18 | 46171104:461 | 46174479:461 | -;-;- | 4113 | kcnk6 | 585 | 1071 | 236 | 216 | 104 | 674 |
| ENSDARG000 | 18:18:18:18:1 | 5523918:5524 | 5525043:5525 | -:-:-:-:-:-:-:-:-: | 2209 | C18H15orf48 | 6409 | 8694 | 16479 | 21762 | 8141 | 10714 |
| ENSDARG000 | 18:18:18:18:1 | 36868951:368 | 36870708:368 | -;-;-;-;-;-;- | 2369 | NA | 0 | 2 | 0 | 0 | 0 | 0 |
| ENSDARG000 | 18;18 | 7479201:7480 | 7479921:7481 | -;- | 783 | si:dkev-238c7 | 180 | 251 | 215 | 128 | 122 | 457 |
| ENSDARG000 | 18:18:18:18:1 | 14564085:145 | 14564185:145 | +:+:+:+:+:+:+: | 3509 | NA | 0 | 0 | 42 | 20 | 64 | 0 |
| ENSDARG000 | 18:18:18:18:1 | 19820008:198 | 19820111:198 | +:+:+:+:+:+:+: | 4309 | iqch | 4 | 2 | 0 | 0 | 0 | 0 |
| ENSDARG000 | 18:18:18:18:1 | 24990268:249 | 24990527:249 | -:-:-:-:-:-:-:-:-: | 1508 | NA | 909 | 817 | 1153 | 1391 | 1193 | 932 |
| ENSDARG000 | 18:18:18:18:1 | 44281623:442 | 44281747:442 | -;-;-;-;-;- | 1264 | NA | 0 | 0 | 0 | 0 | 0 | 0 |
| ENSDARG000 | 18:18:18:18:1 | 44303153:443 | 44303651:443 | -;-;-;-;-;-;-;- | 1525 | si:ch211-151h | 2 | 0 | 0 | 3 | 0 | 6 |
| ENSDARG000 | 18:18:18:18:1 | 402048:40354 | 402259:40363 | +:+:+:+:+:+:+: | 1908 | gpib | 198 | 102 | 74 | 134 | 132 | 20 |
| ENSDARG000 | 18:18:18:18:1 | 14595546:145 | 14595944:145 | +:+:+:+:+:+:+: | 2306 | wfdc1 | 36 | 134 | 184 | 112 | 672 | 12 |
| ENSDARG000 | 18:18:18:18:1 | 21408794:214 | 21409125:214 | +:+:+:+:+:+:+: | 4530 | necab2 | 2 | 0 | 0 | 0 | 18 | 4 |
| ENSDARG000 | 18:18:18:18:1 | 438263:44225 | 438415:44232 | -:-:-:-:-:-:-:-:-: | 2231 | wtip | 169 | 110 | 226 | 107 | 173 | 105 |
| ENSDARG000 | 18:18:18:18:1 | 30511230:305 | 30511427:305 | +;+;+;+;+;+;+ | 1308 | dusp22a | 104 | 48 | 81 | 102 | 67 | 136 |
| ENSDARG000 | 18:18:18:18:1 | 36903456:369 | 36904781:369 | -;-;-;-;-;-;-;- | 3352 | NA | 0 | 2 | 0 | 0 | 0 | 0 |
| ENSDARG000 | 18:18:18:18:1 | 30507839:305 | 30507871:305 | +:+:+:+:+:+:+: | 709 | cox4i1 | 3269 | 4016 | 3426 | 2929 | 2963 | 3366 |
| ENSDARG000 | 18:18:18:18:1 | 44320014:443 | 44322582:443 | -:-:-:-:-:-:-:-:-: | 6446 | prdm10 | 379 | 285 | 449 | 231 | 440 | 416 |
| ENSDARG000 | 18;18;18;18 | 501116:50111 | 502713:50272 | -;-;-;- | 1727 | sdr42e1 | 128 | 106 | 134 | 50 | 136 | 87 |
| ENSDARG000 | 18:18:18:18:1 | 25003207:250 | 25003876:250 | +;+;+;+;+ | 1275 | fam174b | 250 | 193 | 299 | 227 | 244 | 258 |
| ENSDARG000 | 18:18:18:18:1 | 7500434:7500 | 7500490:7500 | +:+:+:+:+:+:+: | 1728 | NA | 0 | 2 | 0 | 0 | 0 | 0 |
| ENSDARG000 | 18 | 40460130 | 40461725 | + | 1596 | ugt5c1 | 96 | 38 | 79 | 63 | 41 | 103 |
| ENSDARG000 | 18;18;18 | 40462445:404 | 40462502:404 | +;+;+ | 1872 | ugt5c2 | 164 | 143 | 253 | 165 | 117 | 466 |
| ENSDARG000 | 18:18:18:18:1 | 19972853:199 | 19973073:199 | +:+:+:+:+:+:+: | 5627 | skor1b | 0 | 2 | 0 | 0 | 0 | 2 |
| ENSDARG000 | 18:18:18:18:1 | 25027495:250 | 25028346:250 | -;-;-;-;-;- | 2321 | st8sia2 | 0 | 0 | 0 | 0 | 0 | 0 |
| ENSDARG000 | 18;18 | 40471826:404 | 40471883:404 | +;+ | 2844 | ugt5c3 | 1010 | 833 | 896 | 413 | 886 | 1082 |
| ENSDARG000 | 18:18:18:18:1 | 21645309:216 | 21646005:216 | -:-:-:-:-:-:-:-:-: | 4150 | mbtps1 | 1879 | 1286 | 1605 | 1068 | 1720 | 1772 |
| ENSDARG000 | 18:18:18:18:1 | 19990412:199 | 19990435:199 | +:+:+:+:+:+:+: | 2217 | PIAS1 | 121 | 95 | 196 | 114 | 155 | 92 |
| ENSDARG000 | 18:18:18:18:1 | 36914221:369 | 36914354:369 | +:+:+:+:+:+:+: | 3339 | NA | 0 | 10 | 5 | 10 | 0 | 0 |
| ENSDARG000 | 18:18:18:18:1 | 25073984:250 | 25074360:250 | -:-:-:-:-:-:-:-:-: | 2467 | slco3a1a | 1557 | 923 | 1040 | 868 | 1954 | 774 |
| ENSDARG000 | 18:18:18:18:1 | 46211210:462 | 46223407:462 | -:-:-:-:-:-:-:-:-: | 13523 | NA | 36 | 31 | 99 | 4 | 18 | 22 |
| ENSDARG000 | 18:18:18:18:1 | 10884996:108 | 10885157:108 | +:+:+:+:+:+:+: | 3710 | NA | 87 | 63 | 117 | 103 | 75 | 84 |
| ENSDARG000 | 18:18:18:18:1 | 14619544:146 | 14619651:146 | +:+:+:+:+:+:+: | 2302 | utp4 | 735 | 771 | 3241 | 1215 | 1479 | 1688 |
| ENSDARG000 | 18:18:18:18:1 | 10784730:108 | 10785133:108 | +:+:+:+:+:+:+: | 6522 | mical3a | 280 | 229 | 580 | 219 | 307 | 219 |
| ENSDARG000 | 18;18;18;18 | 46253933:462 | 46254226:462 | -;-;-;- | 1794 | NA | 26 | 37 | 1 | 24 | 7 | 31 |
| ENSDARG000 | 18 | 40479684 | 40481028 | - | 1345 | zgc:101040 | 0 | 2 | 8 | 3 | 32 | 0 |
| ENSDARG000 | 18:18:18:18:1 | 33652976:336 | 33653188:336 | +;+;+;+;+;+ | 2599 | NA | 0 | 0 | 0 | 0 | 0 | 0 |
| ENSDARG000 | 18:18:18:18:1 | 21686115:216 | 21688912:216 | -:-:-:-:-:-:-:-:-: | 6203 | fan1 | 219 | 251 | 316 | 218 | 179 | 163 |
| ENSDARG000 | 18:18:18:18:1 | 36948879:369 | 36949288:369 | -:-:-:-:-:-:-:-:-: | 3189 | MAP3K10 | 12 | 4 | 22 | 0 | 6 | 2 |
| ENSDARG000 | 18:18:18:18:1 | 507618:50783 | 507885:50788 | +:+:+:+:+:+:+: | 7636 | nedd4a | 885 | 658 | 790 | 1299 | 709 | 1292 |
| ENSDARG000 | 18:18:18:18:1 | 30567945:305 | 30568205:305 | +:+:+:+:+:+:+: | 1767 | irf8 | 196 | 171 | 353 | 128 | 282 | 267 |
| ENSDARG000 | 18:18:18:18:1 | 44364530:443 | 44367521:443 | -:-:-:-:-:-:-:-:-: | 5599 | aplp2 | 7352 | 3345 | 4526 | 3546 | 7660 | 4346 |
| ENSDARG000 | 18:18:18:18:1 | 21712296:217 | 21712580:217 | -;-;-;-;-;-;- | 690 | NA | 9 | 7 | 15 | 9 | 17 | 7 |
| ENSDARG000 | 18:18:18:18:1 | 40486457:404 | 40486818:404 | -:-:-:-:-:-:-:-:-: | 1752 | chrna5 | 7 | 5 | 281 | 4 | 30 | 8 |
| ENSDARG000 | 18:18:18:18:1 | 7520493:7524 | 7522489:7524 | -:-:-:-:-:-:-:-:-: | 3277 | si:dkey-30c15 | 13 | 11 | 8 | 0 | 14 | 10 |
| ENSDARG000 | 18:18:18:18:1 | 46263169:462 | 46263475:462 | -:-:-:-:-:-:-:-:-: | 1984 | PLD3 | 388 | 247 | 28 | 121 | 96 | 276 |
| ENSDARG000 | 18;18;18 | 25180927:251 | 25181286:251 | -;-;- | 473 | NA | 0 | 0 | 2 | 0 | 0 | 0 |
| ENSDARG000 | 18:18:18:18:1 | 16715864:167 | 16716104:167 | +:+:+:+:+:+:+: | 4669 | eif4g2b | 7199 | 6621 | 7684 | 5794 | 6638 | 5779 |
| ENSDARG000 | 18:18:18:18:1 | 33675664:336 | 33675823:336 | +;+;+;+;+;+ | 2550 | NA | 0 | 0 | 0 | 0 | 0 | 0 |
| ENSDARG000 | 18:18:18:18:1 | 37015185:371 | 37015501:371 | +:+:+:+:+:+:+: | 5782 | SIPA1L3 | 864 | 435 | 656 | 557 | 617 | 719 |
| ENSDARG000 | 18:18:18:18:1 | 7553950:7554 | 7554216:7554 | +:+:+:+:+:+:+: | 4788 | ARF5 | 756 | 948 | 966 | 691 | 838 | 707 |
| ENSDARG000 | 18:18:18:18:1 | 21728194:217 | 21732050:217 | -;-;-;-;-;-;-;- | 5538 | pskh1 | 368 | 24 | 99 | 68 | 48 | 40 |
| ENSDARG000 | 18;18;18 | 7566007:7567 | 7566105:7567 | +;+;+ | 561 | NA | 5012 | 17730 | 10269 | 16040 | 10728 | 10529 |
| ENSDARG000 | 18;18 | 30847237:308 | 30848455:308 | +;+ | 1383 | foxf1 | 2 | 0 | 0 | 0 | 0 | 0 |
| ENSDARG000 | 18;18 | 30854496:308 | 30854907:308 | -;- | 462 | NA | 0 | 0 | 0 | 0 | 0 | 0 |
| ENSDARG000 | 18:18:18:18:1 | 30878196:308 | 30878407:308 | +:+:+:+:+:+:+: | 4147 | mthfsd | 18 | 32 | 106 | 20 | 51 | 105 |
| ENSDARG000 | 18:18:18:18:1 | 40525408:405 | 40525772:405 | +:+:+:+:+:+:+: | 1904 | NA | 4 | 4 | 0 | 0 | 20 | 9 |
| ENSDARG000 | 18:18:18:18:1 | 31016379:310 | 31016439:310 | +:+:+:+:+:+:+: | 587 | uraha | 824 | 895 | 669 | 2584 | 518 | 2632 |
| ENSDARG000 | 18 | 20031046 | 20032089 | + | 1044 | NA | 1 | 0 | 8 | 6 | 0 | 2 |
| ENSDARG000 | 18:18:18:18:1 | 21913685:219 | 21914442:219 | -:-:-:-:-:-:-:-:-: | 1327 | nutf2 | 1262 | 1990 | 1758 | 978 | 1519 | 1204 |
| ENSDARG000 | 18:18:18:18:1 | 30998472:309 | 30998802:309 | +:+:+:+:+:+:+: | 3616 | cd151l | 288 | 580 | 426 | 556 | 374 | 392 |
| ENSDARG000 | 18:18:18:18:1 | 21782074:217 | 21784639:217 | -;-;-;-;-;- | 3016 | nrn1la | 4 | 2 | 0 | 0 | 0 | 0 |
| ENSDARG000 | 18;18 | 46350574:463 | 46352938:463 | -;- | 2763 | foxa3 | 22 | 4 | 2 | 4 | 6 | 0 |
| ENSDARG000 | 18:18:18:18:1 | 21858744:218 | 21862901:218 | -:-:-:-:-:-:-:-:-: | 8173 | edc4 | 935 | 1012 | 1303 | 489 | 1169 | 814 |
| ENSDARG000 | 18;18;18;18 | 25225524:252 | 25225596:252 | +;+;+;+ | 633 | si:dkeyp-59c1 | 2 | 16 | 32 | 35 | 6 | 28 |
| ENSDARG000 | 18;18 | 46361544:463 | 46362889:463 | -;- | 3178 | irf2bp1 | 1337 | 1532 | 1970 | 1058 | 1801 | 1355 |
| ENSDARG000 | 18:18:18:18:1 | 44532199:445 | 44532381:445 | +:+:+:+:+:+:+: | 5214 | st14a | 206 | 28 | 10 | 26 | 14 | 42 |
| ENSDARG000 | 18:18:18:18:1 | 25229634:252 | 25231008:252 | -:-:-:-:-:-:-:-:-: | 3201 | NA | 0 | 0 | 0 | 0 | 0 | 0 |
| ENSDARG000 | 18:18:18:18:1 | 33685119:336 | 33685368:336 | +:+:+:+:+:+:+: | 3449 | NA | 0 | 0 | 0 | 0 | 0 | 0 |
| ENSDARG000 | 18:18:18:18:1 | 20047374:200 | 20047445:200 | +:+:+:+:+:+:+: | 2322 | NA | 489 | 387 | 1461 | 1161 | 2413 | 998 |
| ENSDARG000 | 18:18:18:18:1 | 20034023:200 | 20034207:200 | +:+:+:+:+:+:+: | 1662 | morf4l1 | 1013 | 1348 | 1614 | 1132 | 1433 | 1361 |
| ENSDARG000 | 18:18:18:18:1 | 25391171:253 | 25391643:253 | -;-;-;-;-;-;- | 1762 | gnrhr4 | 0 | 0 | 0 | 0 | 0 | 0 |
| ENSDARG000 | 18:18:18:18:1 | 40552620:405 | 40552778:405 | +:+:+:+:+:+:+: | 1937 | si:ch211-132b | 0 | 0 | 0 | 0 | 4 | 0 |
| ENSDARG000 | 18:18:18:18:1 | 37237714:372 | 37237906:372 | -;-;-;-;-;- | 833 | six9 | 8 | 4 | 1 | 4 | 12 | 0 |
| ENSDARG000 | 18:18:18:18:1 | 37219140:372 | 37220018:372 | +;+;+;+;+;+;+ | 2464 | qpctla | 379 | 500 | 517 | 292 | 499 | 252 |
| ENSDARG000 | 18;18 | 25515893:255 | 25516035:255 | -;- | 235 | NA | 0 | 0 | 0 | 0 | 0 | 0 |
| ENSDARG000 | 18;18;18 | 25546227:255 | 25546282:255 | +;+;+ | 741 | pex11a | 24 | 59 | 19 | 28 | 41 | 51 |
| ENSDARG000 | 18:18:18:18:1 | 14633974:146 | 14634122:146 | +:+:+:+:+:+:+: | 4454 | vps9d1 | 369 | 110 | 1213 | 399 | 883 | 653 |
| ENSDARG000 | 18;18;18;18 | 618005:61961 | 618041:61973 | +;+;+;+ | 517 | NA | 0 | 0 | 0 | 0 | 0 | 0 |
| ENSDARG000 | 18:18:18:18:1 | 46370385:463 | 46370714:463 | -:-:-:-:-:-:-:-:-: | 1629 | slc19a3b | 9 | 0 | 1 | 0 | 0 | 0 |
| ENSDARG000 | 18:18:18:18:1 | 7569329:7569 | 7569549:7569 | +;+;+;+;+;+ | 1364 | NA | 9478 | 2224 | 574 | 86 | 1147 | 2193 |
| ENSDARG000 | 18:18:18:18:1 | 25563292:255 | 25563402:255 | -:-:-:-:-:-:-:-:-: | 2660 | NA | 24 | 8 | 28 | 2 | 8 | 14 |
| ENSDARG000 | 18:18:18:18:1 | 16744307:167 | 16744442:167 | +:+:+:+:+:+:+: | 1229 | lyve1b | 2 | 14 | 14 | 0 | 14 | 44 |
| ENSDARG000 | 18:18:18:18:1 | 33714357:337 | 33714462:337 | -;-;-;-;- | 678 | NA | 1 | 0 | 0 | 0 | 0 | 0 |
| ENSDARG000 | 18:18:18:18:1 | 31033122:310 | 31033307:310 | -:-:-:-:-:-:-:-:-: | 1795 | gas8 | 197 | 191 | 262 | 238 | 251 | 209 |
| ENSDARG000 | 18:18:18:18:1 | 46382484:463 | 46382565:463 | +:+:+:+:+:+:+: | 1748 | daw1 | 0 | 0 | 0 | 4 | 6 | 0 |
| ENSDARG000 | 18:18:18:18:1 | 16749091:167 | 16749152:167 | +:+:+:+:+:+:+: | 925 | rnf141 | 168 | 188 | 239 | 146 | 146 | 246 |
| ENSDARG000 | 18:18:18:18:1 | 5543677:5543 | 5543726:5543 | +:+:+:+:+:+:+: | 2665 | NA | 14 | 4 | 4 | 20 | 12 | 0 |
| ENSDARG000 | 18:18:18:18:1 | 17624295:176 | 17624799:176 | +:+:+:+:+:+:+: | 6586 | nlrc5 | 407 | 570 | 335 | 251 | 361 | 352 |
| ENSDARG000 | 18;18;18;18 | 33725576:337 | 33725642:337 | +;+;+;+ | 2370 | NA | 214 | 0 | 31 | 430 | 88 | 218 |
| ENSDARG000 | 18:18:18:18:1 | 10926197:109 | 10926226:109 | +:+:+:+:+:+:+: | 1626 | ttc38 | 3575 | 528 | 674 | 1628 | 820 | 2018 |
| ENSDARG000 | 18;18;18 | 37245331:372 | 37245564:372 | -;-;- | 2454 | six5 | 132 | 66 | 119 | 90 | 152 | 101 |
| ENSDARG000 | 18;18 | 7575886:7579 | 7575940:7581 | +;+ | 1646 | NA | 398 | 0 | 12 | 0 | 26 | 4 |
| ENSDARG000 | 18:18:18:18:1 | 7584386:7584 | 7584591:7584 | +;+;+;+;+ | 821 | NA | 1443 | 1106 | 82 | 18 | 38 | 84 |
| ENSDARG000 | 18:18:18:18:1 | 887122:88788 | 887979:88797 | -:-:-:-:-:-:-:-:-: | 1723 | parp6a | 34 | 24 | 33 | 40 | 16 | 88 |
| ENSDARG000 | 18:18:18:18:1 | 46395909:464 | 46396064:464 | -:-:-:-:-:-:-:-:-: | 4791 | sphkap | 18 | 0 | 0 | 0 | 0 | 2 |
| ENSDARG000 | 18:18:18:18:1 | 33762090:337 | 33762147:337 | +;+;+;+;+;+ | 902 | NA | 0 | 0 | 0 | 0 | 2 | 0 |
| ENSDARG000 | 18:18:18:18:1 | 47313715:473 | 47314082:473 | +:+:+:+:+:+:+: | 1057 | BARX2 | 28 | 0 | 0 | 0 | 0 | 0 |
| ENSDARG000 | 18;18;18 | 47283187:473 | 47283356:473 | -;-;- | 610 | NA | 0 | 0 | 0 | 0 | 0 | 0 |
| ENSDARG000 | 18:18:18:18:1 | 47301086:473 | 47301313:473 | +;+;+;+;+ | 1462 | rbm7 | 208 | 340 | 319 | 295 | 251 | 238 |
| ENSDARG000 | 18;18 | 7591381:7591 | 7591431:7593 | +;+ | 1676 | NA | 68 | 12 | 0 | 6 | 8 | 6 |
| ENSDARG000 | 18:18:18:18:1 | 37259800:372 | 37260000:372 | +:+:+:+:+:+:+: | 2204 | tbcb | 1863 | 1816 | 2014 | 1626 | 1803 | 1953 |
| ENSDARG000 | 18:18:18:18:1 | 10967461:109 | 10967637:109 | -;-;-;-;-;-;- | 726 | tspan33b | 27 | 0 | 0 | 7 | 3 | 0 |
| ENSDARG000 | 18;18;18;18 | 37272568:372 | 37272745:372 | +;+;+;+ | 862 | NA | 2 | 6 | 4 | 4 | 2 | 0 |
| ENSDARG000 | 18:18:18:18:1 | 33779440:337 | 33779804:337 | +:+:+:+:+:+:+: | 4250 | NA | 0 | 0 | 6 | 17 | 20 | 0 |
| ENSDARG000 | 18:18:18:18:1 | 11016203:110 | 11016274:110 | -:-:-:-:-:-:-:-:-: | 1400 | TSPAN9 | 83 | 5 | 27 | 19 | 24 | 0 |
| ENSDARG000 | 18:18:18:18:1 | 33788340:337 | 33788401:337 | +:+:+:+:+:+:+: | 3350 | NAMPT | 2 | 19 | 4 | 16 | 6 | 0 |
| ENSDARG000 | 18;18 | 11281593:113 | 11281668:113 | -;- | 157 | NA | 0 | 0 | 0 | 0 | 0 | 0 |
| ENSDARG000 | 18:18:18:18:1 | 16762806:167 | 16763264:167 | -:-:-:-:-:-:-:-:-: | 3081 | ampd3b | 99 | 36 | 18 | 126 | 80 | 60 |

| | | | | | | | | | | | |
|---|---|---|---|---|---|---|---|---|---|---|---|
| ENSDARG000 | 18:18:18:18:1 | 31056645:31 | 31057330:31 | +:+:+:+:+:+:+: | 1813 | mvda | 274 | 354 | 335 | 95 | 160 | 101 |
| ENSDARG000 | 18:18:18:18:1 | 951092:9555 | 955826:9558 | -:-:-:-:-:-:-:-: | 5650 | dhdh.2 | 959 | 480 | 891 | 773 | 666 | 444 |
| ENSDARG000 | 18:18:18:18:1 | 907266:9109 | 907370:9110 | +:+:+:+:+:+:+: | 2722 | pkma | 7511 | 16275 | 26778 | 26189 | 9569 | 7371 |
| ENSDARG000 | 18:18:18:18:1 | 7594012:759 | 7594130:759 | +:+:+:+:+:+:+: | 2258 | zgc:77752 | 2 | 2 | 0 | 0 | 0 | 0 |
| ENSDARG000 | 18:18:18:18:1 | 40572294:405 | 40572439:405 | +:+:+:+:+:+:+: | 1756 | si:ch211-132 | 0 | 10 | 16 | 24 | 26 | 0 |
| ENSDARG000 | 18:18:18:18:1 | 25653599:25 | 25653927:25 | +:+:+:+:+:+:+: | 1509 | NA | 0 | 0 | 0 | 0 | 0 | 4 |
| ENSDARG000 | 18:18:18:18:1 | 48423973:48 | 48424104:48 | +:+:+:+:+:+:+: | 3245 | fli1 | 157 | 32 | 86 | 10 | 118 | 146 |
| ENSDARG000 | 18:18:18:18:1 | 16876833:16 | 16877699:16 | -:-:-:-:-:-:-:-: | 2777 | swap70b | 92 | 90 | 92 | 68 | 146 | 64 |
| ENSDARG000 | 18:18:18:18:1 | 16918426:16 | 16919600:16 | -:-:-:-:-:-:-:-: | 3590 | wee1 | 300 | 791 | 881 | 307 | 642 | 464 |
| ENSDARG000 | 18:18:18:18:1 | 44566863:44 | 44567599:44 | +:+:+:+:+:+:+: | 4295 | dhx34 | 150 | 219 | 291 | 143 | 257 | 139 |
| ENSDARG000 | 18:18:18:18:1 | 5588068:558 | 5589321:558 | -:-:-:-:-:-:-:-: | 2614 | cyp1a | 12 | 2 | 24 | 2 | 44 | 4 |
| ENSDARG000 | 18:18:18:18:1 | 20225961:20 | 20226292:20 | +:+:+:+:+:+:+: | 4159 | tle3a | 229 | 162 | 425 | 158 | 206 | 169 |
| ENSDARG000 | 18:18:18:18:1 | 11858397:11 | 11859152:11 | +:+:+:+:+:+:+: | 1784 | TMTC2 | 227 | 7 | 30 | 6 | 147 | 58 |
| ENSDARG000 | 18:18:18:18:1 | 17660158:17 | 17660590:17 | +:+:+:+:+:+:+: | 3633 | cpne2 | 252 | 145 | 215 | 124 | 66 | 82 |
| ENSDARG000 | 18:18:18:18:1 | 33818577:33 | 33818797:33 | +;+;+;+;+;+ | 2607 | olfcq19 | 0 | 0 | 0 | 0 | 0 | 0 |
| ENSDARG000 | 18;18;18;18 | 11970987:11 | 11971210:11 | +;+;+;+ | 1026 | NA | 0 | 0 | 2 | 6 | 0 | 0 |
| ENSDARG000 | 18:18:18:18:1 | 970246:9708 | 970447:9710 | -;-;-;-;-;-;- | 1247 | dhdh.1 | 565 | 490 | 692 | 787 | 703 | 512 |
| ENSDARG000 | 18:18:18:18:1 | 7622040:762 | 7622575:762 | -;-;-;-;-;- | 1413 | trim35-30 | 0 | 0 | 0 | 0 | 2 | 0 |
| ENSDARG000 | 18:18:18:18:1 | 7611298:761 | 7611776:761 | +:+:+:+:+:+:+: | 5217 | odf3b | 2 | 0 | 2 | 0 | 0 | 0 |
| ENSDARG000 | 18:18:18:18:1 | 25754838:25 | 25755314:25 | -:-:-:-:-:-:-:-: | 2737 | zgc:162879 | 23 | 0 | 4 | 9 | 4 | 20 |
| ENSDARG000 | 18:18:18:18:1 | 33914168:33 | 33914277:33 | +;+;+;+;+;+ | 2490 | NA | 0 | 0 | 0 | 0 | 0 | 0 |
| ENSDARG000 | 18;18;18 | 25752592:25 | 25752729:25 | +;+;+ | 750 | serf2a | 58 | 81 | 85 | 183 | 104 | 100 |
| ENSDARG000 | 18:18:18:18:1 | 37283765:37 | 37286212:37 | -;-;-;-;-;-;-;- | 3940 | yap1 | 2850 | 1873 | 1972 | 2378 | 2330 | 2015 |
| ENSDARG000 | 18:18:18:18:1 | 21931741:21 | 21932420:21 | -:-:-:-:-:-:-:-: | 3073 | tsnaxip1 | 56 | 56 | 82 | 40 | 46 | 59 |
| ENSDARG000 | 18:18:18:18:1 | 12034985:12 | 12035649:12 | -:-:-:-:-:-:-:-: | 2309 | slc23a4 | 8794 | 7270 | 12011 | 8573 | 13056 | 10683 |
| ENSDARG000 | 18;18;18 | 5677508:568 | 5677760:568 | -;-;- | 601 | cplx3b | 4 | 18 | 17 | 0 | 3 | 2 |
| ENSDARG000 | 18:18:18:18:1 | 1004814:100 | 1005538:100 | -:-:-:-:-:-:-:-: | 3450 | HCN4 | 20 | 6 | 14 | 26 | 2 | 6 |
| ENSDARG000 | 18:18:18:18:1 | 33970385:33 | 33973732:33 | -:-:-:-:-:-:-:-: | 5414 | NA | 0 | 0 | 0 | 0 | 0 | 0 |
| ENSDARG000 | 18:18:18:18:1 | 14671789:14 | 14672370:14 | -;-;-;-;-;- | 1202 | si:dkev-238o1 | 47 | 0 | 26 | 225 | 12 | 0 |
| ENSDARG000 | 18:18:18:18:1 | 40584288:40 | 40584492:40 | +:+:+:+:+:+:+: | 2008 | si:ch211-132 | 0 | 1 | 13 | 0 | 8 | 0 |
| ENSDARG000 | 18:18:18:18:1 | 12058403:12 | 12058618:12 | +:+:+:+:+:+:+: | 2674 | BICD1 | 32 | 10 | 47 | 34 | 71 | 60 |
| ENSDARG000 | 18:18:18:18:1 | 6039141:605 | 6040031:605 | +:+:+:+:+:+:+: | 14510 | NA | 636 | 546 | 633 | 482 | 652 | 860 |
| ENSDARG000 | 18:18:18:18:1 | 37380125:37 | 37380443:37 | -:-:-:-:-:-:-:-: | 3415 | NA | 0 | 2 | 0 | 0 | 4 | 0 |
| ENSDARG000 | 18:18:18:18:1 | 7639401:763 | 7639969:763 | +:+:+:+:+:+:+: | 1554 | rabl2 | 46 | 68 | 79 | 67 | 87 | 63 |
| ENSDARG000 | 18:18:18:18:1 | 33981474:33 | 33987236:33 | -:-:-:-:-:-:-:-: | 9767 | PLCH1 | 0 | 8 | 4 | 4 | 0 | 0 |
| ENSDARG000 | 18:18:18:18:1 | 44596787:44 | 44599875:44 | -:-:-:-:-:-:-:-: | 3948 | spred3 | 0 | 0 | 0 | 0 | 0 | 0 |
| ENSDARG000 | 18;18;18 | 37409823:37 | 37409864:37 | -;-;- | 435 | NA | 0 | 0 | 0 | 0 | 0 | 0 |
| ENSDARG000 | 18;18 | 48499029:48 | 48500149:48 | -;- | 1293 | kcnj1a.5 | 0 | 0 | 0 | 0 | 0 | 0 |
| ENSDARG000 | 18;18 | 48515579:48 | 48516693:48 | -;- | 1282 | kcnj1a.3 | 0 | 0 | 0 | 0 | 0 | 0 |
| ENSDARG000 | 18;18 | 48507075:48 | 48508189:48 | -;- | 1287 | kcnj1a.4 | 0 | 0 | 0 | 0 | 0 | 0 |
| ENSDARG000 | 18:18:18:18:1 | 48498366:48 | 48498466:48 | -;-;-;-;-;-;- | 2097 | kcnj1a.2 | 0 | 0 | 0 | 0 | 0 | 0 |
| ENSDARG000 | 18:18:18:18:1 | 48491007:48 | 48492653:48 | -;-;-;-;- | 1952 | kcnj1a.6 | 0 | 0 | 0 | 0 | 0 | 0 |
| ENSDARG000 | 18:18:18:18:1 | 12147971:12 | 12148058:12 | +:+:+:+:+:+:+: | 3445 | fgd4a | 54 | 92 | 181 | 110 | 203 | 53 |
| ENSDARG000 | 18:18:18:18:1 | 1158197:116 | 1161624:116 | -:-:-:-:-:-:-:-: | 4670 | nptnb | 1 | 0 | 2 | 4 | 2 | 4 |
| ENSDARG000 | 18;18;18 | 14684115:14 | 14684204:14 | +;+;+ | 2307 | NA | 130 | 73 | 190 | 231 | 196 | 60 |
| ENSDARG000 | 18:18:18:18:1 | 1154189:115 | 1154374:115 | +:+:+:+:+:+:+: | 785 | NA | 0 | 0 | 0 | 0 | 0 | 0 |
| ENSDARG000 | 18;18;18;18 | 1145571:114 | 1145931:114 | +;+;+;+ | 821 | NA | 0 | 0 | 0 | 0 | 0 | 0 |
| ENSDARG000 | 18:18:18:18:1 | 6126506:612 | 6126757:612 | +:+:+:+:+:+:+: | 3283 | NA | 0 | 4 | 22 | 6 | 14 | 6 |
| ENSDARG000 | 18:18:18:18:1 | 1151453:115 | 1152539:115 | -;-;-;-;-;-;- | 1376 | NA | 2 | 1 | 6 | 7 | 0 | 18 |
| ENSDARG000 | 18:18:18:18:1 | 6145307:614 | 6145849:614 | -;-;-;-;-;-;-;- | 1152 | gcsha | 560 | 542 | 658 | 346 | 523 | 412 |
| ENSDARG000 | 18:18:18:18:1 | 17707688:17 | 17708028:17 | -:-:-:-:-:-:-:-: | 1428 | psme3ip1 | 849 | 758 | 903 | 702 | 860 | 682 |
| ENSDARG000 | 18:18:18:18:1 | 44615591:44 | 44616334:44 | -:-:-:-:-:-:-:-: | 1481 | psmd8 | 1703 | 1809 | 2103 | 1074 | 1977 | 1278 |
| ENSDARG000 | 18;18;18 | 14688643:14 | 14688816:14 | -;-;- | 931 | pdf | 53 | 130 | 192 | 126 | 126 | 139 |
| ENSDARG000 | 18:18:18:18:1 | 37836960:37 | 37838748:37 | -:-:-:-:-:-:-:-: | 2970 | luzp2 | 12 | 0 | 4 | 0 | 2 | 0 |
| ENSDARG000 | 18:18:18:18:1 | 14693682:14 | 14694086:14 | +:+:+:+:+:+:+: | 2756 | uri1 | 1380 | 1450 | 2440 | 1615 | 1625 | 1476 |
| ENSDARG000 | 18;18;18;18 | 48541660:48 | 48541750:48 | +;+;+;+ | 296 | NA | 2 | 8 | 4 | 0 | 5 | 0 |
| ENSDARG000 | 18:18:18:18:1 | 48542900:48 | 48544551:48 | -;-;-;-;-;-;- | 2076 | kcnj1a.1 | 23 | 1 | 14 | 0 | 12 | 1 |
| ENSDARG000 | 18:18:18:18:1 | 21951551:21 | 21951903:21 | +:+:+:+:+:+:+: | 2573 | ranbp10 | 532 | 363 | 547 | 443 | 485 | 393 |
| ENSDARG000 | 18:18:18:18:1 | 20435821:20 | 20436789:20 | -:-:-:-:-:-:-:-: | 4904 | kif23 | 257 | 1065 | 976 | 455 | 791 | 323 |
| ENSDARG000 | 18:18:18:18:1 | 16924865:16 | 16925199:16 | -:-:-:-:-:-:-:-: | 2785 | znf143b | 364 | 371 | 515 | 334 | 574 | 443 |
| ENSDARG000 | 18:18:18:18:1 | 16940440:16 | 16940692:16 | +:+:+:+:+:+:+: | 2299 | NA | 61 | 28 | 102 | 30 | 60 | 43 |
| ENSDARG000 | 18:18:18:18:1 | 25783446:25 | 25784212:25 | -:-:-:-:-:-:-:-: | 3100 | sema4ba | 26 | 31 | 23 | 2 | 11 | 3 |
| ENSDARG000 | 18:18:18:18:1 | 16950724:16 | 16951108:16 | -;-;-;-;- | 898 | akip1 | 142 | 46 | 113 | 85 | 108 | 103 |
| ENSDARG000 | 18:18:18:18:1 | 48576527:48 | 48576662:48 | +;+;+;+;+;+;+ | 4877 | KCNJ5 | 0 | 0 | 0 | 0 | 8 | 4 |
| ENSDARG000 | 18:18:18:18:1 | 12236044:12 | 12236373:12 | -:-:-:-:-:-:-:-: | 6199 | nup205 | 1592 | 1650 | 3540 | 1451 | 2303 | 1472 |
| ENSDARG000 | 18;18;18 | 40609153:40 | 40609305:40 | +;+;+ | 429 | NA | 0 | 0 | 0 | 0 | 0 | 0 |
| ENSDARG000 | 18;18;18 | 14731784:14 | 14734809:14 | -;-;- | 3705 | NA | 2178 | 732 | 857 | 2138 | 986 | 2071 |
| ENSDARG000 | 18:18:18:18:1 | 7663802:766 | 7664351:766 | -:-:-:-:-:-:-:-: | 6426 | SHANK3 | 195 | 86 | 268 | 139 | 163 | 43 |
| ENSDARG000 | 18:18:18:18:1 | 44630415:44 | 44630460:44 | +:+:+:+:+:+:+: | 9577 | BICRA | 815 | 564 | 1238 | 715 | 990 | 833 |
| ENSDARG000 | 18 | 31226288 | 31227615 | - | 1328 | mc1r | 4191 | 3665 | 4030 | 3672 | 6214 | 6156 |
| ENSDARG000 | 18:18:18:18:1 | 44649804:44 | 44650095:44 | +;+;+;+;+;+;+ | 3004 | ehd2b | 6809 | 10866 | 5944 | 10251 | 12538 | 8610 |
| ENSDARG000 | 18:18:18:18:1 | 6155258:616 | 6155329:616 | +:+:+:+:+:+:+: | 8545 | OTOGL | 0 | 0 | 2 | 0 | 0 | 0 |
| ENSDARG000 | 18:18:18:18:1 | 17725410:17 | 17725712:17 | +:+:+:+:+:+:+: | 3186 | rspry1 | 3786 | 2456 | 2592 | 3033 | 1898 | 3342 |
| ENSDARG000 | 18:18:18:18:1 | 31280984:31 | 31281271:31 | +:+:+:+:+:+:+: | 1713 | def8 | 750 | 1060 | 966 | 1242 | 1339 | 1250 |
| ENSDARG000 | 18:18:18:18:1 | 40617717:40 | 40617921:40 | +:+:+:+:+:+:+: | 3043 | zgc:158427 | 0 | 0 | 0 | 0 | 12 | 0 |
| ENSDARG000 | 18:18:18:18:1 | 48864367:48 | 48865018:48 | +:+:+:+:+:+:+: | 4089 | ppp1r37 | 1716 | 1130 | 1579 | 1124 | 1943 | 1240 |
| ENSDARG000 | 18:18:18:18:1 | 16960966:16 | 16961076:16 | +:+:+:+:+:+:+: | 3558 | NA | 320 | 2 | 46 | 70 | 276 | 63 |
| ENSDARG000 | 18:18:18:18:1 | 38191346:38 | 38191470:38 | +:+:+:+:+:+:+: | 3316 | nucb2b | 224 | 387 | 883 | 269 | 368 | 196 |
| ENSDARG000 | 18:18:18:18:1 | 16978628:16 | 16978764:16 | +:+:+:+:+:+:+: | 978 | NA | 4 | 0 | 2 | 2 | 0 | 0 |
| ENSDARG000 | 18:18:18:18:1 | 20459888:20 | 20460521:20 | -;-;-;-;-;-;-;- | 1682 | paqr5a | 10 | 2 | 3 | 0 | 0 | 10 |
| ENSDARG000 | 18:18:18:18:1 | 16986495:16 | 16986600:16 | +;+;+;+;+;+ | 952 | NA | 82 | 98 | 36 | 9 | 15 | 84 |
| ENSDARG000 | 18:18:18:18:1 | 38209055:38 | 38210357:38 | -;-;-;-;-;-;- | 2461 | si:dkev-10o6. | 3 | 1 | 19 | 17 | 2 | 7 |
| ENSDARG000 | 18:18:18:18:1 | 8231138:823 | 8231530:823 | +:+:+:+:+:+:+: | 1892 | arsa | 236 | 134 | 147 | 96 | 193 | 173 |
| ENSDARG000 | 18:18:18:18:1 | 14772474:14 | 14773237:14 | -:-:-:-:-:-:-:-: | 2545 | mtss1la | 883 | 24 | 25 | 29 | 907 | 40 |
| ENSDARG000 | 18:18:18:18:1 | 38213778:38 | 38217033:38 | -:-:-:-:-:-:-:-: | 6693 | nat10 | 823 | 1134 | 2477 | 1166 | 1220 | 1289 |
| ENSDARG000 | 18:18:18:18:1 | 50880096:50 | 50880282:50 | +:+:+:+:+:+:+: | 5736 | NA | 28 | 26 | 91 | 88 | 30 | 24 |
| ENSDARG000 | 18:18:18:18:1 | 16989764:16 | 16990433:16 | -:-:-:-:-:-:-:-: | 2224 | tph2 | 52 | 22 | 56 | 34 | 40 | 21 |
| ENSDARG000 | 18:18:18:18:1 | 22078144:22 | 22078216:22 | +;+;+;+;+;+ | 1504 | NA | 0 | 4 | 0 | 0 | 0 | 0 |
| ENSDARG000 | 18;18;18 | 22028674:22 | 22032156:22 | -;-;- | 3890 | pard6a | 46 | 16 | 44 | 14 | 28 | 38 |
| ENSDARG000 | 18:18:18:18:1 | 17754510:17 | 17754707:17 | +:+:+:+:+:+:+: | 2891 | si:ch211-216 | 800 | 503 | 1033 | 1224 | 739 | 859 |
| ENSDARG000 | 18:18:18:18:1 | 49001425:49 | 49002912:49 | -:-:-:-:-:-:-:-: | 7758 | NA | 173 | 290 | 970 | 962 | 703 | 74 |
| ENSDARG000 | 18:18:18:18:1 | 20468157:20 | 20468305:20 | +:+:+:+:+:+:+: | 3815 | ddb2 | 38 | 50 | 90 | 72 | 197 | 174 |
| ENSDARG000 | 18;18 | 31845504:31 | 31845918:31 | +;+ | 545 | NA | 0 | 0 | 0 | 0 | 0 | 0 |
| ENSDARG000 | 18:18:18:18:1 | 6300608:630 | 6300653:630 | +:+:+:+:+:+:+: | 1131 | NA | 4 | 1 | 2 | 1 | 0 | 0 |
| ENSDARG000 | 18:18:18:18:1 | 20481982:20 | 20483369:20 | +:+:+:+:+:+:+: | 5249 | kbtbd4 | 198 | 165 | 240 | 167 | 219 | 234 |
| ENSDARG000 | 18:18:18:18:1 | 34225520:34 | 34225761:34 | +;+;+;+;+;+ | 2790 | NA | 0 | 0 | 0 | 0 | 0 | 0 |
| ENSDARG000 | 18;18;18;18 | 26296165:26 | 26296787:26 | -;-;-;- | 827 | NA | 0 | 0 | 0 | 0 | 2 | 0 |
| ENSDARG000 | 18:18:18:18:1 | 31912759:31 | 31913656:31 | -;-;-;-;-;- | 2645 | olfcs2 | 0 | 0 | 0 | 0 | 0 | 0 |
| ENSDARG000 | 18:18:18:18:1 | 8255265:825 | 8255337:825 | -:-:-:-:-:-:-:-: | 5080 | MAPK8IP2 | 0 | 0 | 0 | 0 | 0 | 0 |
| ENSDARG000 | 18:18:18:18:1 | 17002164:17 | 17002733:17 | -:-:-:-:-:-:-:-: | 2356 | tbc1d15 | 427 | 389 | 630 | 465 | 396 | 477 |
| ENSDARG000 | 18;18;18 | 2554666:256 | 2554741:256 | +;+;+ | 1087 | P2RY2 | 4 | 0 | 2 | 0 | 10 | 0 |
| ENSDARG000 | 18;18 | 2585298:258 | 2586290:258 | +;+ | 993 | P2RY2 | 0 | 0 | 0 | 0 | 0 | 0 |
| ENSDARG000 | 18:18:18:18:1 | 6319447:632 | 6319555:632 | +:+:+:+:+:+:+: | 7179 | NA | 6 | 0 | 0 | 0 | 7 | 0 |
| ENSDARG000 | 18 | 2593756 | 2594781 | + | 1026 | P2RY2 | 0 | 0 | 0 | 0 | 0 | 10 |
| ENSDARG000 | 18:18:18:18:1 | 26337869:26 | 26338519:26 | +:+:+:+:+:+:+: | 4177 | RASGRF1 | 0 | 4 | 2 | 0 | 10 | 0 |
| ENSDARG000 | 18:18:18:18:1 | 44673990:44 | 44674158:44 | +:+:+:+:+:+:+: | 1588 | napab | 1409 | 1437 | 1699 | 1243 | 1752 | 1311 |
| ENSDARG000 | 18:18:18:18:1 | 2600959:260 | 2603885:260 | -:-:-:-:-:-:-:-: | 5015 | relt | 540 | 618 | 511 | 236 | 897 | 344 |
| ENSDARG000 | 18:18:18:18:1 | 40643660:40 | 40643864:40 | +:+:+:+:+:+:+: | 1975 | si:ch211-132 | 0 | 0 | 0 | 0 | 4 | 0 |
| ENSDARG000 | 18:18:18:18:1 | 34362608:34 | 34363140:34 | +:+:+:+:+:+:+: | 2850 | kcnab1a | 70 | 24 | 110 | 12 | 401 | 4 |
| ENSDARG000 | 18:18:18:18:1 | 31965347:31 | 31966242:31 | -;-;-;-;-;- | 2535 | olfcs1 | 0 | 0 | 0 | 0 | 0 | 0 |
| ENSDARG000 | 18:18:18:18:1 | 34542525:34 | 34542997:34 | -;-;-;-;-;-;-;- | 940 | ssr3 | 2798 | 4259 | 6469 | 3225 | 4041 | 2150 |
| ENSDARG000 | 18;18 | 49124810:49 | 49127338:49 | +;+ | 6029 | NA | 0 | 0 | 0 | 0 | 4 | 0 |
| ENSDARG000 | 18;18;18 | 49193824:49 | 49193893:49 | +;+;+ | 1742 | NA | 0 | 2 | 8 | 13 | 12 | 8 |
| ENSDARG000 | 18:18:18:18:1 | 14844669:14 | 14845811:14 | -:-:-:-:-:-:-:-: | 2224 | mapk12a | 230 | 181 | 70 | 75 | 164 | 172 |
| ENSDARG000 | 18:18:18:18:1 | 22109379:22 | 22109650:22 | +;+;+;+;+;+ | 987 | zgc:158868 | 6 | 0 | 6 | 16 | 4 | 12 |
| ENSDARG000 | 18:18:18:18:1 | 17786548:17 | 17786749:17 | +:+:+:+:+:+:+: | 9744 | znf423 | 459 | 380 | 585 | 498 | 598 | 326 |
| ENSDARG000 | 18:18:18:18:1 | 49225552:49 | 49225749:49 | +:+:+:+:+:+:+: | 4470 | NA | 2149 | 3211 | 7378 | 9602 | 3943 | 1979 |
| ENSDARG000 | 18:18:18:18:1 | 34599315:34 | 34599455:34 | +:+:+:+:+:+:+: | 2842 | tiparp | 594 | 845 | 1746 | 1997 | 876 | 456 |
| ENSDARG000 | 18;18 | 35072976:35 | 35073274:35 | +;+ | 1199 | NA | 0 | 0 | 0 | 0 | 0 | 0 |
| ENSDARG000 | 18:18:18:18:1 | 17959681:17 | 17959812:17 | +;+;+;+;+ | 1598 | NA | 0 | 0 | 0 | 0 | 0 | 0 |
| ENSDARG000 | 18:18:18:18:1 | 6422528:642 | 6422950:642 | -:-:-:-:-:-:-:-: | 3943 | IQSEC3 | 0 | 0 | 0 | 0 | 0 | 0 |
| ENSDARG000 | 18:18:18:18:1 | 14863407:14 | 14864251:14 | -:-:-:-:-:-:-:-: | 2886 | selenoo1 | 444 | 402 | 382 | 346 | 419 | 354 |
| ENSDARG000 | 18:18:18:18:1 | 17020967:17 | 17021431:17 | +;+;+;+;+;+;+ | 1898 | chtf8 | 66 | 137 | 154 | 104 | 112 | 142 |
| ENSDARG000 | 18:18:18:18:1 | 22123072:22 | 22123421:22 | +;+;+;+;+;+ | 1055 | NA | 0 | 0 | 0 | 0 | 0 | 0 |
| ENSDARG000 | 18:18:18:18:1 | 31999208:31 | 31999393:31 | +:+:+:+:+:+:+: | 1720 | NA | 2 | 6 | 6 | 30 | 0 | 0 |
| ENSDARG000 | 18:18:18:18:1 | 2681241:268 | 2685619:268 | -:-:-:-:-:-:-:-: | 7343 | ARHGEF17 | 197 | 176 | 314 | 653 | 247 | 356 |
| ENSDARG000 | 18:18:18:18:1 | 22138924:22 | 22139380:22 | +:+:+:+:+:+:+: | 5115 | ripor1 | 1686 | 1755 | 3235 | 1868 | 3960 | 1868 |
| ENSDARG000 | 18;18;18 | 17027874:17 | 17028803:17 | -;-;- | 1668 | has3 | 28 | 6 | 15 | 29 | 28 | 8 |
| ENSDARG000 | 18:18:18:18:1 | 8340886:834 | 8341611:834 | +:+:+:+:+:+:+: | 3674 | cpt1b | 1 | 9 | 22 | 12 | 40 | 56 |
| ENSDARG000 | 18;18 | 22149561:22 | 22150041:22 | -;- | 531 | NA | 1 | 0 | 0 | 0 | 0 | 5 |
| ENSDARG000 | 18:18:18:18:1 | 6479963:647 | 6480054:648 | +:+:+:+:+:+:+: | 1730 | wash1 | 650 | 511 | 588 | 472 | 526 | 511 |
| ENSDARG000 | 18:18:18:18:1 | 35128685:35 | 35128780:35 | +:+:+:+:+:+:+: | 2916 | NA | 434 | 238 | 780 | 846 | 204 | 957 |
| ENSDARG000 | 18:18:18:18:1 | 8320165:832 | 8320831:832 | +:+:+:+:+:+:+: | 3775 | chkb | 435 | 844 | 908 | 428 | 862 | 725 |
| ENSDARG000 | 18:18:18:18:1 | 20494413:20 | 20494676:20 | +:+:+:+:+:+:+: | 1377 | rapsn | 0 | 0 | 4 | 0 | 2 | 0 |
| ENSDARG000 | 18:18:18:18:1 | 17032312:17 | 17034268:17 | -:-:-:-:-:-:-:-: | 9123 | TANGO6 | 169 | 234 | 214 | 132 | 174 | 161 |
| ENSDARG000 | 18;18 | 44703343:44 | 44703451:44 | +;+ | 1628 | NA | 52 | 12 | 5 | 14 | 76 | 12 |
| ENSDARG000 | 18:18:18:18:1 | 14885545:14 | 14886373:14 | -:-:-:-:-:-:-:-: | 2128 | trabd | 409 | 657 | 727 | 427 | 550 | 345 |
| ENSDARG000 | 18;18 | 17072075:17 | 17072112:17 | +;+ | 525 | NA | 0 | 0 | 0 | 0 | 0 | 0 |
| ENSDARG000 | 18:18:18:18:1 | 40668615:40 | 40669443:40 | -;-;-;-;-;-;-;- | 3236 | si:ch211-132 | 3627 | 617 | 2369 | 1187 | 5147 | 1289 |
| ENSDARG000 | 18:18:18:18:1 | 44746594:44 | 44746621:44 | +:+:+:+:+:+:+: | 1845 | NA | 73 | 101 | 105 | 195 | 192 | 134 |
| ENSDARG000 | 18:18:18:18:1 | 26422124:26 | 26422278:26 | +:+:+:+:+:+:+: | 1533 | ctsh | 2073 | 1580 | 1316 | 1488 | 1957 | 1556 |
| ENSDARG000 | 18:18:18:18:1 | 49248389:49 | 49248630:49 | +:+:+:+:+:+:+: | 4115 | yif1b | 386 | 1024 | 791 | 620 | 747 | 436 |
| ENSDARG000 | 18:18:18:18:1 | 12276567:12 | 12277682:12 | -:-:-:-:-:-:-:-: | 1735 | fam107b | 2701 | 3766 | 3506 | 8031 | 3785 | 2668 |
| ENSDARG000 | 18:18:18:18:1 | 18000695:18 | 18000894:18 | +:+:+:+:+:+:+: | 2441 | si:ch211-212 | 735 | 628 | 515 | 517 | 728 | 780 |
| ENSDARG000 | 18:18:18:18:1 | 20516527:20 | 20517068:20 | -:-:-:-:-:-:-:-: | 3450 | ighmbp2 | 154 | 134 | 210 | 68 | 152 | 146 |
| ENSDARG000 | 18:18:18:18:1 | 6502441:650 | 6503377:650 | -:-:-:-:-:-:-:-: | 3733 | ddx11 | 259 | 240 | 320 | 252 | 254 | 297 |
| ENSDARG000 | 18:18:18:18:1 | 35173683:35 | 35174046:35 | +:+:+:+:+:+:+: | 4273 | cfap45 | 86 | 24 | 52 | 54 | 12 | 52 |
| ENSDARG000 | 18:18:18:18:1 | 40698550:40 | 40699187:40 | -:-:-:-:-:-:-:-: | 1764 | ccnp | 243 | 107 | 240 | 74 | 541 | 70 |
| ENSDARG000 | 18:18:18:18:1 | 22285992:22 | 22286139:22 | +:+:+:+:+:+:+: | 6551 | ctcf | 1705 | 2161 | 2806 | 1497 | 3051 | 2058 |
| ENSDARG000 | 18;18;18;18 | 18113318:18 | 18113369:18 | +;+;+;+ | 822 | NA | 0 | 0 | 0 | 0 | 0 | 0 |
| ENSDARG000 | 18;18;18 | 18104235:18 | 18104764:18 | +;+;+ | 1715 | cbln1 | 10 | 27 | 36 | 21 | 44 | 17 |
| ENSDARG000 | 18:18:18:18:1 | 49273989:49 | 49275444:49 | -;-;-;-;-;-;- | 2116 | NA | 48 | 42 | 68 | 88 | 96 | 114 |
| ENSDARG000 | 18;18;18 | 49309953:49 | 49311491:49 | -;-;- | 2447 | NA | 446 | 569 | 265 | 176 | 920 | 202 |
| ENSDARG000 | 18;18;18;18 | 49282063:49 | 49282620:49 | +;+;+;+ | 1159 | NA | 0 | 0 | 0 | 2 | 0 | 0 |
| ENSDARG000 | 18:18:18:18:1 | 44768829:44 | 44769029:44 | +:+:+:+:+:+:+: | 5632 | ilvbl | 570 | 503 | 625 | 455 | 548 | 480 |
| ENSDARG000 | 18:18:18:18:1 | 17076680:17 | 17080206:17 | -:-:-:-:-:-:-:-: | 7173 | zc3h18 | 1184 | 1145 | 1436 | 753 | 1395 | 882 |
| ENSDARG000 | 18:18:18:18:1 | 14906570:14 | 14906935:14 | -:-:-:-:-:-:-:-: | 3231 | panx2 | 8 | 0 | 4 | 6 | 10 | 0 |
| ENSDARG000 | 18:18:18:18:1 | 35402759:35 | 35403601:35 | -:-:-:-:-:-:-:-: | 1907 | snrpa | 1140 | 1308 | 2066 | 1041 | 1591 | 918 |
| ENSDARG000 | 18:18:18:18:1 | 32005058:32 | 32005633:32 | -:-:-:-:-:-:-:-: | 1721 | NA | 0 | 0 | 0 | 0 | 0 | 0 |
| ENSDARG000 | 18:18:18:18:1 | 18405992:18 | 18406541:18 | +;+;+;+;+;+;+ | 3990 | n4bp1 | 587 | 396 | 510 | 420 | 698 | 588 |
| ENSDARG000 | 18:18:18:18:1 | 17075985:17 | 17076426:17 | -;-;-;-;- | 714 | il17c | 6 | 0 | 5 | 2 | 0 | 0 |
| ENSDARG000 | 18:18:18:18:1 | 49411417:49 | 49412813:49 | +;+;+;+;+ | 3104 | si:cabz01090 | 0 | 0 | 0 | 0 | 2 | 10 |

| | | | | | | | | | | | | |
|---|---|---|---|---|---|---|---|---|---|---|---|---|
| ENSDARG000 | 18:18:18:18:1 | 14928040:149 | 14928133:149 | -:-:-:-:-:-:-:-: | 1392 | mlc1 | 2 | 12 | 10 | 0 | 2 | 0 |
| ENSDARG000 | 18:18:18:18:1 | 26428829:264 | 26429761:264 | +:+:+:+:+:+:+:+: | 5188 | NA | 2505 | 2902 | 1254 | 2034 | 2634 | 2514 |
| ENSDARG000 | 18:18:18:18:1 | 40711188:407 | 40712712:407 | -:-:-:-:-:-:-:-: | 3363 | akt2 | 2159 | 2046 | 2431 | 1606 | 2287 | 1867 |
| ENSDARG000 | 18:18:18:18:1 | 18439332:184 | 18439497:184 | +;+;+;+;+;+ | 3151 | siah1 | 307 | 296 | 408 | 374 | 365 | 371 |
| ENSDARG000 | 18:18:18:18:1 | 12364733:123 | 12364930:123 | -:-:-:-:-:-:-:-: | 5181 | NA | 378 | 236 | 632 | 467 | 488 | 476 |
| ENSDARG000 | 18;18 | 12451709:124 | 12451840:124 | -;- | 329 | NA | 8 | 5 | 3 | 16 | 6 | 3 |
| ENSDARG000 | 18;18;18;18 | 32018101:320 | 32019990:320 | +;+;+;+ | 3450 | NA | 0 | 0 | 0 | 0 | 0 | 0 |
| ENSDARG000 | 18:18:18:18:1 | 32084939:320 | 32085223:320 | +:+:+:+:+:+:+:+: | 2107 | NA | 4 | 0 | 0 | 18 | 0 | 18 |
| ENSDARG000 | 18:18:18:18:1 | 6536293:6536 | 6536710:6536 | +:+:+:+:+:+:+:+: | 2329 | fkbp4 | 2080 | 2246 | 2397 | 1255 | 2465 | 1897 |
| ENSDARG000 | 18:18:18:18:1 | 20532618:205 | 20532847:205 | +:+:+:+:+:+:+:+: | 2480 | agk | 399 | 390 | 602 | 228 | 587 | 443 |
| ENSDARG000 | 18:18:18:18:1 | 44795711:447 | 44796118:447 | +:+:+:+:+:+:+:+: | 2305 | FAM118B | 53 | 82 | 181 | 62 | 231 | 66 |
| ENSDARG000 | 18:18:18:18:1 | 35407921:354 | 35408182:354 | +;+;+;+;+;+;+ | 1375 | ACTMAP | 16 | 40 | 64 | 41 | 118 | 30 |
| ENSDARG000 | 18:18:18:18:1 | 18475674:184 | 18476293:184 | -:-:-:-:-:-:-:-: | 3018 | lonp2 | 639 | 514 | 477 | 555 | 562 | 508 |
| ENSDARG000 | 18:18:18:18:1 | 17151916:171 | 17152088:171 | +;+;+;+;+;+;+ | 1963 | trhr2 | 0 | 0 | 0 | 0 | 0 | 0 |
| ENSDARG000 | 18:18:18:18:1 | 26649860:266 | 26649883:266 | -:-:-:-:-:-:-:-: | 2529 | NA | 141 | 103 | 94 | 62 | 152 | 124 |
| ENSDARG000 | 18:18:18:18:1 | 18527170:185 | 18528007:185 | -;-;-;-;-;- | 1758 | il34 | 323 | 420 | 218 | 330 | 180 | 362 |
| ENSDARG000 | 18:18:18:18:1 | 26460444:264 | 26461036:264 | -:-:-:-:-:-:-:-: | 1731 | FRMD5 | 10 | 6 | 0 | 9 | 2 | 1 |
| ENSDARG000 | 18;18 | 14989079:149 | 14989240:149 | +;+ | 234 | NA | 0 | 0 | 0 | 0 | 0 | 0 |
| ENSDARG000 | 18:18:18:18:1 | 14956165:149 | 14956374:149 | -:-:-:-:-:-:-:-: | 3129 | ABTB3 | 38 | 41 | 25 | 95 | 43 | 24 |
| ENSDARG000 | 18:18:18:18:1 | 32135370:321 | 32136979:321 | -;-;-;-;-;- | 3348 | olfcw1 | 0 | 0 | 0 | 0 | 0 | 0 |
| ENSDARG000 | 18:18:18:18:1 | 35429333:354 | 35431849:354 | -:-:-:-:-:-:-:-: | 3844 | itpkcb | 1201 | 1203 | 2366 | 1540 | 1638 | 1081 |
| ENSDARG000 | 18:18:18:18:1 | 20541316:205 | 20543532:205 | -:-:-:-:-:-:-:-: | 3829 | dennd11 | 535 | 611 | 409 | 432 | 408 | 454 |
| ENSDARG000 | 18:18:18:18:1 | 12464461:124 | 12465397:124 | -:-:-:-:-:-:-:-: | 4508 | hipk2 | 228 | 110 | 330 | 114 | 474 | 190 |
| ENSDARG000 | 18:18:18:18:1 | 18547023:185 | 18551240:185 | -:-:-:-:-:-:-:-: | 8254 | sf3b3 | 2143 | 2898 | 3885 | 1346 | 2875 | 2198 |
| ENSDARG000 | 18;18;18 | 32146708:321 | 32146934:321 | -;-;- | 699 | NA | 4 | 2 | 0 | 0 | 0 | 8 |
| ENSDARG000 | 18:18:18:18:1 | 44826173:448 | 44827204:448 | -:-:-:-:-:-:-:-: | 3061 | SRPRA | 1387 | 1538 | 2353 | 1066 | 2274 | 1119 |
| ENSDARG000 | 18:18:18:18:1 | 17270427:172 | 17273784:172 | -:-:-:-:-:-:-:-: | 5050 | zfpm1 | 961 | 148 | 273 | 520 | 128 | 933 |
| ENSDARG000 | 18:18:18:18:1 | 32444636:324 | 32444901:324 | +;+;+;+;+;+ | 2618 | NA | 0 | 0 | 0 | 0 | 0 | 0 |
| ENSDARG000 | 18:18:18:18:1 | 15106518:151 | 15107044:151 | +:+:+:+:+:+:+:+: | 3019 | cry1b | 201 | 130 | 88 | 237 | 286 | 181 |
| ENSDARG000 | 18:18:18:18:1 | 15175478:151 | 15175746:151 | +;+;+;+;+ | 1567 | si:ch73-62b1 | 0 | 0 | 0 | 0 | 0 | 0 |
| ENSDARG000 | 18:18:18:18:1 | 6558146:6558 | 6558695:6558 | +:+:+:+:+:+:+:+: | 7661 | B4GALNT3 | 36 | 2 | 4 | 2 | 14 | 30 |
| ENSDARG000 | 18:18:18:18:1 | 26719787:267 | 26719815:267 | +:+:+:+:+:+:+:+: | 7064 | NA | 0 | 0 | 10 | 0 | 7 | 0 |
| ENSDARG000 | 18:18:18:18:1 | 44849809:448 | 44849939:448 | +:+:+:+:+:+:+:+: | 2098 | arfgap2 | 1087 | 974 | 1424 | 458 | 1091 | 620 |
| ENSDARG000 | 18:18:18:18:1 | 22302635:223 | 22302758:223 | +:+:+:+:+:+:+:+: | 5241 | carmil2 | 1523 | 86 | 26 | 4 | 24 | 151 |
| ENSDARG000 | 18;18 | 32456046:324 | 32456308:324 | +;+ | 1227 | NA | 2 | 5 | 9 | 8 | 16 | 11 |
| ENSDARG000 | 18:18:18:18:1 | 15287065:152 | 15287833:152 | -:-:-:-:-:-:-:-: | 2937 | ric8b | 718 | 769 | 695 | 280 | 744 | 661 |
| ENSDARG000 | 18:18:18:18:1 | 40761913:407 | 40763074:407 | -:-:-:-:-:-:-:-: | 2964 | NA | 584 | 505 | 608 | 552 | 565 | 525 |
| ENSDARG000 | 18;18 | 32525411:325 | 32525718:325 | -;- | 630 | NA | 0 | 0 | 0 | 0 | 0 | 0 |
| ENSDARG000 | 18:18:18:18:1 | 44886077:448 | 44886523:448 | -:-:-:-:-:-:-:-: | 2520 | NA | 35 | 42 | 63 | 51 | 38 | 28 |
| ENSDARG000 | 18:18:18:18:1 | 26770967:267 | 26771981:267 | -:-:-:-:-:-:-:-: | 1886 | kti12 | 95 | 131 | 232 | 106 | 141 | 126 |
| ENSDARG000 | 18:18:18:18:1 | 18815654:188 | 18815804:188 | -;-;-;-;- | 778 | NA | 0 | 2 | 3 | 0 | 4 | 0 |
| ENSDARG000 | 18:18:18:18:1 | 18636273:186 | 18637080:186 | -:-:-:-:-:-:-:-: | 4471 | aars1 | 5152 | 6149 | 14504 | 8148 | 4370 | 5900 |
| ENSDARG000 | 18:18:18:18:1 | 17408470:174 | 17408700:174 | -:-:-:-:-:-:-:-: | 1826 | ces2a | 74 | 0 | 20 | 0 | 0 | 0 |
| ENSDARG000 | 18:18:18:18:1 | 32674236:327 | 32675149:327 | -:-:-:-:-:-:-:-: | 3476 | NA | 0 | 0 | 0 | 0 | 0 | 0 |
| ENSDARG000 | 18:18:18:18:1 | 32674212:326 | 32675149:326 | -;-;-;-;-;- | 2573 | NA | 0 | 0 | 0 | 0 | 0 | 0 |
| ENSDARG000 | 18:18:18:18:1 | 6625348:6625 | 6625539:6625 | -:-:-:-:-:-:-:-: | 794 | tnni1c | 0 | 0 | 0 | 0 | 0 | 0 |
| ENSDARG000 | 18;18;18;18 | 18744462:187 | 18744612:187 | -;-;-;- | 534 | NA | 0 | 0 | 0 | 0 | 0 | 0 |
| ENSDARG000 | 18;18;18 | 18799118:187 | 18799683:187 | +;+;+ | 685 | NA | 0 | 0 | 0 | 0 | 0 | 0 |
| ENSDARG000 | 18;18;18 | 18720679:187 | 18720829:187 | -;-;- | 487 | NA | 0 | 0 | 0 | 0 | 0 | 0 |
| ENSDARG000 | 18:18:18:18:1 | 18696207:186 | 18696357:186 | -;-;-;-;- | 759 | NA | 7 | 4 | 4 | 4 | 1 | 10 |
| ENSDARG000 | 18;18;18 | 18675702:186 | 18676516:186 | +;+;+ | 934 | NA | 0 | 0 | 0 | 0 | 0 | 0 |
| ENSDARG000 | 18;18;18 | 18827249:188 | 18828116:188 | +;+;+ | 987 | NA | 0 | 0 | 0 | 0 | 0 | 0 |
| ENSDARG000 | 18:18:18:18:1 | 18803539:188 | 18804510:188 | -:-:-:-:-:-:-:-: | 4038 | NA | 146 | 233 | 69 | 34 | 14 | 48 |
| ENSDARG000 | 18;18;18 | 18709049:187 | 18709786:187 | +;+;+ | 857 | NA | 0 | 0 | 0 | 0 | 0 | 0 |
| ENSDARG000 | 18;18;18 | 18770075:187 | 18770525:187 | +;+;+ | 570 | NA | 0 | 0 | 0 | 0 | 0 | 0 |
| ENSDARG000 | 18:18:18:18:1 | 20566817:205 | 20566897:205 | +:+:+:+:+:+:+:+: | 1743 | bida | 66 | 94 | 106 | 76 | 142 | 58 |
| ENSDARG000 | 18;18;18 | 18733425:187 | 18733990:187 | +;+;+ | 685 | NA | 0 | 0 | 0 | 0 | 0 | 0 |
| ENSDARG000 | 18:18:18:18:1 | 18787384:187 | 18787534:187 | -;-;-;-;- | 768 | NA | 0 | 0 | 0 | 5 | 0 | 0 |
| ENSDARG000 | 18:18:18:18:1 | 20560442:205 | 20560557:205 | +:+:+:+:+:+:+:+: | 2245 | wee2 | 4 | 2 | 10 | 10 | 6 | 6 |
| ENSDARG000 | 18:18:18:18:1 | 44913422:449 | 44913832:449 | -:-:-:-:-:-:-:-: | 1451 | pex16 | 312 | 311 | 286 | 208 | 296 | 358 |
| ENSDARG000 | 18:18:18:18:1 | 44935173:449 | 44935265:449 | +:+:+:+:+:+:+:+: | 932 | NA | 0 | 0 | 0 | 0 | 4 | 0 |
| ENSDARG000 | 18:18:18:18:1 | 15334117:153 | 15335089:153 | -:-:-:-:-:-:-:-: | 3461 | rfx4 | 10 | 8 | 2 | 0 | 0 | 0 |
| ENSDARG000 | 18:18:18:18:1 | 26788083:267 | 26788444:267 | -;-;-;-;-;- | 992 | sec11a | 1094 | 1473 | 1881 | 803 | 1491 | 1131 |
| ENSDARG000 | 18;18 | 26784968:267 | 26785601:267 | -;- | 818 | nmba | 0 | 0 | 2 | 0 | 6 | 0 |
| ENSDARG000 | 18:18:18:18:1 | 32780450:327 | 32780672:327 | +;+;+;+;+;+ | 2540 | NA | 0 | 0 | 0 | 0 | 0 | 0 |
| ENSDARG000 | 18:18:18:18:1 | 32801603:328 | 32801829:328 | +;+;+;+;+;+ | 2580 | olfcg6 | 0 | 2 | 0 | 0 | 0 | 2 |
| ENSDARG000 | 18;18;18 | 22405707:224 | 22407393:224 | -;-;- | 2235 | tmem170a | 328 | 271 | 291 | 232 | 357 | 350 |
| ENSDARG000 | 18:18:18:18:1 | 8367602:8368 | 8368636:8368 | -:-:-:-:-:-:-:-: | 2186 | sephs1 | 458 | 802 | 894 | 282 | 1016 | 462 |
| ENSDARG000 | 18:18:18:18:1 | 50089000:500 | 50089166:500 | +:+:+:+:+:+:+:+: | 2681 | SCAMP5 | 0 | 0 | 0 | 0 | 0 | 0 |
| ENSDARG000 | 18:18:18:18:1 | 26800061:268 | 26800525:268 | -;-;-;-;-;-;-;- | 3420 | znf592 | 453 | 270 | 494 | 351 | 479 | 562 |
| ENSDARG000 | 18:18:18:18:1 | 22416051:224 | 22416257:224 | +;+;+;+;+;+;+ | 1741 | cfdp1 | 300 | 281 | 535 | 303 | 335 | 321 |
| ENSDARG000 | 18:18:18:18:1 | 32844439:328 | 32844647:328 | +;+;+;+;+;+ | 2523 | olfcg1 | 0 | 0 | 0 | 0 | 0 | 0 |
| ENSDARG000 | 18:18:18:18:1 | 45114392:451 | 45115023:451 | +:+:+:+:+:+:+:+: | 7656 | large2 | 24 | 18 | 58 | 86 | 42 | 2 |
| ENSDARG000 | 18:18:18:18:1 | 38246940:382 | 38248311:382 | -:-:-:-:-:-:-:-: | 4498 | caprin1b | 10648 | 9237 | 20167 | 17279 | 8621 | 16803 |
| ENSDARG000 | 18:18:18:18:1 | 50124297:501 | 50128763:501 | -:-:-:-:-:-:-:-: | 6763 | loxl1 | 244 | 190 | 1290 | 957 | 1636 | 1232 |
| ENSDARG000 | 18:18:18:18:1 | 26829086:268 | 26829110:268 | +:+:+:+:+:+:+:+: | 2287 | slc28a1 | 2 | 2 | 2 | 0 | 0 | 1 |
| ENSDARG000 | 18:18:18:18:1 | 32906619:329 | 32906694:329 | +;+;+;+;+;+ | 699 | NA | 0 | 0 | 0 | 0 | 0 | 0 |
| ENSDARG000 | 18:18:18:18:1 | 17418254:174 | 17418359:174 | +:+:+:+:+:+:+:+: | 1748 | ces3 | 14 | 0 | 10 | 0 | 0 | 0 |
| ENSDARG000 | 18:18:18:18:1 | 45483282:454 | 45483317:454 | +:+:+:+:+:+:+:+: | 2523 | NA | 0 | 2 | 5 | 2 | 0 | 0 |
| ENSDARG000 | 18:18:18:18:1 | 15569627:155 | 15571729:155 | -:-:-:-:-:-:-:-: | 4771 | arntl2 | 104 | 132 | 264 | 135 | 151 | 86 |
| ENSDARG000 | 18:18:18:18:1 | 22550595:225 | 22550897:225 | -;-;-;-;-;- | 822 | NA | 0 | 2 | 0 | 2 | 6 | 2 |
| ENSDARG000 | 18:18:18:18:1 | 22504200:225 | 22504211:225 | +:+:+:+:+:+:+:+: | 5442 | bcar1 | 532 | 501 | 730 | 584 | 706 | 386 |
| ENSDARG000 | 18;18;18 | 40875873:408 | 40876341:408 | -;-;- | 745 | snrpd2 | 636 | 923 | 1158 | 434 | 798 | 697 |
| ENSDARG000 | 18 | 26853891 | 26854959 | - | 1069 | ch25hl1.1 | 14 | 2 | 0 | 0 | 0 | 2 |
| ENSDARG000 | 18 | 40900568 | 40901707 | - | 1140 | foxg1c | 32 | 0 | 6 | 0 | 8 | 9 |
| ENSDARG000 | 18:18:18:18:1 | 8382131:8385 | 8384395:8385 | -:-:-:-:-:-:-:-: | 3478 | BEND7 | 49 | 0 | 25 | 7 | 48 | 18 |
| ENSDARG000 | 18;18;18;18 | 40902636:409 | 40903327:409 | -;-;-;- | 1209 | pglyrp5 | 22 | 18 | 4 | 4 | 6 | 35 |
| ENSDARG000 | 18;18 | 22695407:226 | 22696357:226 | -;- | 951 | NA | 0 | 0 | 0 | 0 | 0 | 6 |
| ENSDARG000 | 18;18 | 22701030:227 | 22701983:227 | -;- | 954 | NA | 0 | 0 | 0 | 0 | 0 | 0 |
| ENSDARG000 | 18:18:18:18:1 | 40909821:409 | 40910170:409 | -:-:-:-:-:-:-:-: | 1230 | polr2i | 215 | 164 | 275 | 95 | 155 | 165 |
| ENSDARG000 | 18:18:18:18:1 | 6706140:6708 | 6706384:6708 | +:+:+:+:+:+:+:+: | 2430 | lmf2a | 215 | 174 | 464 | 224 | 326 | 236 |
| ENSDARG000 | 18;18 | 22712638:227 | 22713621:227 | -;- | 984 | NA | 0 | 4 | 0 | 0 | 0 | 0 |
| ENSDARG000 | 18:18:18:18:1 | 50275933:502 | 50276489:502 | +:+:+:+:+:+:+:+: | 2392 | insyn1 | 0 | 0 | 0 | 0 | 0 | 18 |
| ENSDARG000 | 18;18 | 22716708:227 | 22717673:227 | -;- | 966 | NA | 0 | 0 | 0 | 0 | 0 | 0 |
| ENSDARG000 | 18:18:18:18:1 | 8402129:8402 | 8402308:8402 | +:+:+:+:+:+:+:+: | 1557 | prpf18 | 437 | 353 | 451 | 222 | 466 | 460 |
| ENSDARG000 | 18:18:18:18:1 | 50523870:505 | 50524149:505 | +:+:+:+:+:+:+:+: | 3480 | ubl7b | 667 | 451 | 595 | 367 | 490 | 366 |
| ENSDARG000 | 18:18:18:18:1 | 50433722:504 | 50435298:504 | -:-:-:-:-:-:-:-: | 3814 | cd276 | 515 | 665 | 654 | 398 | 453 | 469 |
| ENSDARG000 | 18:18:18:18:1 | 50353005:503 | 50353735:503 | -;-;-;-;-;-;-;- | 1311 | NA | 2 | 0 | 0 | 0 | 0 | 0 |
| ENSDARG000 | 18:18:18:18:1 | 6732195:6732 | 6732711:6732 | -:-:-:-:-:-:-:-: | 1311 | miox | 0 | 2 | 6 | 0 | 5 | 5 |
| ENSDARG000 | 18:18:18:18:1 | 18870625:188 | 18871510:188 | -:-:-:-:-:-:-:-: | 1399 | arl2bp | 64 | 34 | 14 | 38 | 44 | 62 |
| ENSDARG000 | 18;18;18;18 | 26880472:268 | 26880783:268 | -;-;-;- | 1139 | NA | 0 | 0 | 0 | 0 | 0 | 0 |
| ENSDARG000 | 18:18:18:18:1 | 26863139:268 | 26863717:268 | -:-:-:-:-:-:-:-: | 2741 | si:dkey-24l11. | 270 | 239 | 181 | 232 | 241 | 180 |
| ENSDARG000 | 18:18:18:18:1 | 18861359:188 | 18861421:188 | +:+:+:+:+:+:+:+: | 1180 | pllp | 8 | 18 | 5 | 3 | 7 | 4 |
| ENSDARG000 | 18:18:18:18:1 | 22721551:227 | 22721921:227 | -;-;-;-;-;- | 1201 | nudt21 | 686 | 762 | 776 | 440 | 774 | 504 |
| ENSDARG000 | 18;18;18 | 38288877:382 | 38289409:382 | +;+;+ | 1805 | lmo2 | 12 | 6 | 26 | 12 | 34 | 18 |
| ENSDARG000 | 18 | 26895994 | 26896826 | + | 833 | ch25hl1.2 | 2 | 10 | 2 | 33 | 0 | 5 |
| ENSDARG000 | 18;18 | 6753677:6766 | 6754423:6766 | -;- | 786 | ADM2 | 0 | 6 | 29 | 13 | 0 | 0 |
| ENSDARG000 | 18:18:18:18:1 | 8503370:8504 | 8503433:8504 | -:-:-:-:-:-:-:-: | 3012 | frmd4a | 1330 | 866 | 1556 | 1186 | 1744 | 958 |
| ENSDARG000 | 18;18;18;18 | 40944144:409 | 40944284:409 | -;-;-;- | 716 | NA | 0 | 0 | 0 | 0 | 0 | 0 |
| ENSDARG000 | 18:18:18:18:1 | 15616167:156 | 15616264:156 | +:+:+:+:+:+:+:+: | 1773 | slc17a8 | 0 | 0 | 0 | 0 | 6 | 0 |
| ENSDARG000 | 18:18:18:18:1 | 38321039:383 | 38321644:383 | +:+:+:+:+:+:+:+: | 2033 | alx4b | 273 | 88 | 974 | 92 | 213 | 319 |
| ENSDARG000 | 18:18:18:18:1 | 40993196:409 | 40993279:409 | +:+:+:+:+:+:+:+: | 929 | NA | 2 | 0 | 0 | 0 | 0 | 0 |
| ENSDARG000 | 18;18 | 38749547:387 | 38749850:387 | +;+ | 1136 | NA | 2 | 0 | 2 | 0 | 8 | 0 |
| ENSDARG000 | 18:18:18:18:1 | 26899316:269 | 26899479:269 | +;+;+;+;+;+;+ | 1803 | tspan3a | 953 | 878 | 1149 | 1665 | 1379 | 1320 |
| ENSDARG000 | 18:18:18:18:1 | 41198559:411 | 41202236:412 | -;-;-;-;-;- | 5038 | NA | 175 | 230 | 251 | 359 | 186 | 149 |
| ENSDARG000 | 18;18;18;18 | 38755023:387 | 38756353:387 | +;+;+;+ | 1858 | onecut1 | 52 | 74 | 57 | 95 | 112 | 36 |
| ENSDARG000 | 18:18:18:18:1 | 20581685:205 | 20585908:205 | -:-:-:-:-:-:-:-: | 5497 | bcl2l13 | 220 | 1133 | 962 | 498 | 1263 | 709 |
| ENSDARG000 | 18:18:18:18:1 | 45594085:455 | 45594206:455 | -;-;-;-;-;-;-;- | 1389 | wt1b | 14 | 28 | 73 | 20 | 20 | 14 |
| ENSDARG000 | 18;18;18;18 | 41226444:412 | 41226966:412 | -;-;-;- | 931 | fbxo36a | 2 | 2 | 8 | 0 | 2 | 2 |
| ENSDARG000 | 18:18:18:18:1 | 45526483:455 | 45526702:455 | +:+:+:+:+:+:+:+: | 7216 | KIFC3 | 637 | 578 | 826 | 478 | 662 | 350 |
| ENSDARG000 | 18:18:18:18:1 | 6785640:6790 | 6785703:6790 | -;-;-;-;- | 727 | NA | 2 | 2 | 2 | 0 | 0 | 4 |
| ENSDARG000 | 18:18:18:18:1 | 18879733:188 | 18880197:188 | +:+:+:+:+:+:+:+: | 2456 | arih1 | 636 | 545 | 600 | 447 | 513 | 561 |
| ENSDARG000 | 18:18:18:18:1 | 17428258:174 | 17428562:174 | +:+:+:+:+:+:+:+: | 3234 | kiaa0513 | 462 | 325 | 349 | 366 | 285 | 348 |
| ENSDARG000 | 18:18:18:18:1 | 38774584:387 | 38775388:387 | +:+:+:+:+:+:+:+: | 7463 | atosa | 1127 | 713 | 395 | 664 | 769 | 926 |
| ENSDARG000 | 18:18:18:18:1 | 3475688:3476 | 3476895:3476 | -:-:-:-:-:-:-:-: | 3698 | capn5a | 5092 | 2249 | 2547 | 2459 | 1783 | 3660 |
| ENSDARG000 | 18;18 | 3498989:3501 | 3499073:3502 | +;+ | 1019 | ompa | 16 | 56 | 16 | 12 | 48 | 9 |
| ENSDARG000 | 18;18 | 18910366:189 | 18910455:189 | +;+ | 128 | NA | 0 | 0 | 1 | 0 | 0 | 0 |
| ENSDARG000 | 18:18:18:18:1 | 8805575:8812 | 8805649:8812 | +:+:+:+:+:+:+:+: | 2766 | impdh1a | 317 | 212 | 156 | 130 | 260 | 234 |
| ENSDARG000 | 18:18:18:18:1 | 26930956:269 | 26931092:269 | -:-:-:-:-:-:-:-: | 1262 | pstpip1a | 25 | 18 | 20 | 34 | 25 | 93 |
| ENSDARG000 | 18:18:18:18:1 | 18931484:189 | 18932767:189 | -;-;-;-;-;- | 4428 | NA | 277 | 143 | 417 | 182 | 112 | 299 |
| ENSDARG000 | 18:18:18:18:1 | 15644559:156 | 15644647:156 | +:+:+:+:+:+:+:+: | 2044 | nr1h4 | 0 | 0 | 6 | 0 | 9 | 0 |
| ENSDARG000 | 18:18:18:18:1 | 6815744:6815 | 6815998:6815 | -:-:-:-:-:-:-:-: | 3570 | PPP6R2 | 314 | 246 | 407 | 270 | 604 | 327 |
| ENSDARG000 | 18;18;18 | 38885309:388 | 38885407:388 | +;+;+ | 4425 | arpp19a | 2183 | 2330 | 3021 | 2493 | 2612 | 2252 |
| ENSDARG000 | 18:18:18:18:1 | 8869465:8869 | 8873278:8873 | -:-:-:-:-:-:-:-: | 6497 | NA | 0 | 0 | 0 | 0 | 0 | 4 |
| ENSDARG000 | 18;18;18;18 | 8872926:8873 | 8873278:8873 | -;-;-;- | 671 | NA | 0 | 0 | 0 | 0 | 0 | 0 |
| ENSDARG000 | 18:18:18:18:1 | 26967414:269 | 26968581:269 | -;-;-;-;-;-;-;- | 2070 | rcn2 | 184 | 160 | 212 | 186 | 240 | 164 |
| ENSDARG000 | 18:18:18:18:1 | 15683157:156 | 15683215:156 | +:+:+:+:+:+:+:+: | 4101 | NA | 0 | 0 | 0 | 0 | 0 | 0 |
| ENSDARG000 | 18;18;18 | 3530769:3531 | 3530808:3531 | +;+;+ | 923 | NA | 0 | 0 | 7 | 0 | 14 | 6 |
| ENSDARG000 | 18:18:18:18:1 | 41232719:412 | 41232799:412 | +:+:+:+:+:+:+:+: | 7834 | trip12 | 3010 | 2100 | 2969 | 1695 | 3794 | 2923 |
| ENSDARG000 | 18:18:18:18:1 | 6868826:6869 | 6869467:6869 | -:-:-:-:-:-:-:-: | 1826 | NA | 1 | 12 | 10 | 2 | 3 | 2 |
| ENSDARG000 | 18:18:18:18:1 | 6857071:6857 | 6857280:6857 | +:+:+:+:+:+:+:+: | 2721 | dnaja2b | 1808 | 1003 | 1502 | 986 | 1338 | 1399 |
| ENSDARG000 | 18:18:18:18:1 | 20613241:206 | 20613715:206 | -;-;-;-;- | 1509 | wnt2 | 0 | 0 | 0 | 0 | 0 | 0 |
| ENSDARG000 | 18:18:18:18:1 | 18975539:189 | 18976678:189 | -;-;-;-;- | 1886 | ppcdc | 128 | 106 | 115 | 92 | 140 | 144 |
| ENSDARG000 | 18:18:18:18:1 | 15706160:157 | 15706295:157 | +:+:+:+:+:+:+:+: | 3493 | NA | 0 | 0 | 0 | 0 | 0 | 0 |
| ENSDARG000 | 18:18:18:18:1 | 20642378:206 | 20642743:206 | -:-:-:-:-:-:-:-: | 1725 | asz1 | 0 | 0 | 6 | 0 | 0 | 0 |
| ENSDARG000 | 18:18:18:18:1 | 38900092:389 | 38900160:389 | +:+:+:+:+:+:+:+: | 6200 | MYO5A | 5531 | 1900 | 4592 | 4312 | 3331 | 2753 |
| ENSDARG000 | 18:18:18:18:1 | 27077853:270 | 27078219:270 | +:+:+:+:+:+:+:+: | 2378 | ppp1r15b | 90 | 56 | 150 | 223 | 14 | 14 |
| ENSDARG000 | 18:18:18:18:1 | 3533787:3535 | 3536256:3536 | -:-:-:-:-:-:-:-: | 3643 | dcun1d5 | 316 | 410 | 357 | 114 | 466 | 193 |
| ENSDARG000 | 18:18:18:18:1 | 27000850:270 | 27001170:270 | +:+:+:+:+:+:+:+: | 10221 | PIK3C2A | 912 | 853 | 1290 | 1040 | 1206 | 811 |
| ENSDARG000 | 18:18:18:18:1 | 6900622:6903 | 6900839:6904 | -:-:-:-:-:-:-:-: | 7974 | NA | 865 | 762 | 878 | 457 | 1380 | 703 |
| ENSDARG000 | 18:18:18:18:1 | 15758375:157 | 15758386:157 | +:+:+:+:+:+:+:+: | 4017 | NA | 0 | 0 | 0 | 0 | 0 | 0 |
| ENSDARG000 | 18:18:18:18:1 | 41313999:413 | 41314104:413 | +:+:+:+:+:+:+:+: | 2642 | veph1 | 0 | 9 | 56 | 34 | 20 | 3 |
| ENSDARG000 | 18;18;18 | 41364437:413 | 41365783:413 | -;-;- | 2164 | PTX3 | 54 | 22 | 0 | 8 | 34 | 32 |
| ENSDARG000 | 18:18:18:18:1 | 6985876:6985 | 6986009:6986 | +:+:+:+:+:+:+:+: | 948 | NA | 0 | 0 | 0 | 0 | 4 | 0 |
| ENSDARG000 | 18:18:18:18:1 | 20677090:206 | 20677236:206 | +:+:+:+:+:+:+:+: | 4814 | cftr | 2 | 0 | 16 | 0 | 2 | 10 |
| ENSDARG000 | 18:18:18:18:1 | 41495841:414 | 41495926:414 | +;+;+;+;+ | 1696 | ankrd67 | 2 | 0 | 4 | 0 | 0 | 0 |
| ENSDARG000 | 18:18:18:18:1 | 7015283:7016 | 7016856:7016 | -:-:-:-:-:-:-:-: | 3325 | calub | 900 | 528 | 1289 | 478 | 1656 | 619 |
| ENSDARG000 | 18:18:18:18:1 | 7037349:7037 | 7037773:7037 | -:-:-:-:-:-:-:-: | 1974 | utp15 | 377 | 432 | 723 | 333 | 424 | 302 |
| ENSDARG000 | 18:18:18:18:1 | 41512102:415 | 41512436:415 | +:+:+:+:+:+:+:+: | 4159 | ccnl1a | 2544 | 1903 | 3158 | 3664 | 2230 | 1745 |
| ENSDARG000 | 18:18:18:18:1 | 20726149:207 | 20726394:207 | -:-:-:-:-:-:-:-: | 5222 | CTTNBP2 | 2162 | 1493 | 2124 | 1890 | 3052 | 2584 |
| ENSDARG000 | 18:18:18:18:1 | 3552641:3553 | 3552811:3553 | -:-:-:-:-:-:-:-: | 1870 | eif2a | 1121 | 1080 | 1608 | 874 | 1278 | 860 |
| ENSDARG000 | 18:18:18:18:1 | 27101096:271 | 27101611:271 | +:+:+:+:+:+:+:+: | 6200 | plekha7a | 4 | 4 | 2 | 6 | 13 | 0 |

| | | | | | | | | | | | | |
|---|---|---|---|---|---|---|---|---|---|---|---|---|
| ENSDARG000 | 18:18:18:18:1 | 27309404:273 | 27309957:273 | -;-;-;-;- | 1003 | zgc:56106 | 114 | 102 | 113 | 105 | 150 | 98 |
| ENSDARG000 | 18:18:18:18:1 | 27318589:273 | 27318731:273 | +:+:+:+:+:+:+: | 1365 | cd81b | 8 | 16 | 22 | 10 | 30 | 4 |
| ENSDARG000 | 18:18:18:18:1 | 27337994:273 | 27338273:273 | +;+;+;+;+;+;+ | 2208 | NA | 235 | 246 | 256 | 138 | 210 | 247 |
| ENSDARG000 | 18:18:18:18:1 | 27439680:274 | 27439985:274 | +:+:+:+:+:+:+: | 4934 | tp53i11b | 6356 | 3978 | 2568 | 2771 | 2659 | 3886 |
| ENSDARG000 | 18:18:18:18:1 | 27571448:275 | 27571562:275 | +:+:+:+:+:+:+: | 3342 | cd82b | 428 | 252 | 323 | 194 | 349 | 184 |
| ENSDARG000 | 18:18:18:18:1 | 5192343:5195 | 5195333:5195 | -;-;-;-;-;-;- | 3795 | golim4a | 393 | 234 | 373 | 347 | 513 | 226 |
| ENSDARG000 | 18;18;18;18 | 3004572:3006 | 3004597:3006 | +;+;+;+ | 315 | NA | 44 | 8 | 34 | 32 | 31 | 24 |
| ENSDARG000 | 18;18;18 | 783936:78985 | 784089:79002 | +;+;+ | 1801 | rpp25b | 0 | 0 | 0 | 0 | 0 | 0 |
| ENSDARG000 | 18;18;18 | 13123723:131 | 13125872:131 | -;-;- | 2586 | C7H19orf12 | 500 | 473 | 654 | 580 | 568 | 414 |
| ENSDARG000 | 18;18;18 | 5428472:5436 | 5428765:5436 | +;+;+ | 663 | fgf7 | 72 | 5 | 18 | 23 | 8 | 1 |
| ENSDARG000 | 18:18:18:18:1 | 46699731:467 | 46699842:467 | -:-:-:-:-:-:-:-:- | 2235 | dner | 0 | 0 | 0 | 0 | 0 | 0 |
| ENSDARG000 | 18;18;18 | 48979123:489 | 48979367:489 | -;-;- | 1791 | NA | 6 | 6 | 10 | 6 | 11 | 11 |
| ENSDARG000 | 18;18 | 3052408:3059 | 3052590:3063 | +;+ | 4465 | KCTD14 | 0 | 0 | 4 | 0 | 6 | 0 |
| ENSDARG000 | 18:18:18:18:1 | 2189211:2190 | 2189365:2190 | +;+;+;+;+;+ | 853 | ccpg1 | 452 | 161 | 281 | 273 | 297 | 271 |
| ENSDARG000 | 18:18:18:18:1 | 2436987:2443 | 2441723:2444 | -;-;-;-;- | 5344 | NA | 34 | 18 | 35 | 20 | 35 | 20 |
| ENSDARG000 | 18:18:18:18:1 | 5850837:5853 | 5851175:5853 | +;+;+;+;+ | 2210 | cog8 | 267 | 246 | 318 | 188 | 246 | 244 |
| ENSDARG000 | 18:18:18:18:1 | 3037998:3037 | 3038229:3038 | +:+:+:+:+:+:+: | 3754 | rps3 | 21323 | 22081 | 35903 | 31855 | 23406 | 30012 |
| ENSDARG000 | 18:18:18:18:1 | 34667486:346 | 34667498:346 | +;+;+;+;+;+;+ | 1330 | NA | 0 | 0 | 0 | 0 | 0 | 0 |
| ENSDARG000 | 18 | 10016508 | 10016626 | + | 119 | 5S_rRNA | 0 | 0 | 0 | 0 | 0 | 0 |
| ENSDARG000 | 18 | 13948810 | 13948924 | + | 115 | 5S_rRNA | 0 | 0 | 0 | 0 | 0 | 0 |
| ENSDARG000 | 18 | 19454800 | 19454867 | - | 68 | SNORD18 | 0 | 0 | 0 | 1 | 0 | 0 |
| ENSDARG000 | 18:18:18:18:1 | 33053133:330 | 33053430:330 | +:+:+:+:+:+:+: | 4600 | olfcd2 | 0 | 0 | 0 | 0 | 0 | 0 |
| ENSDARG000 | 18:18:18:18:1 | 31117136:311 | 31117517:311 | +:+:+:+:+:+:+: | 2412 | tcf25 | 798 | 966 | 939 | 576 | 910 | 752 |
| ENSDARG000 | 18 | 12532046 | 12532160 | - | 115 | 5S_rRNA | 0 | 0 | 0 | 0 | 0 | 0 |
| ENSDARG000 | 18 | 1703984 | 1706599 | + | 2616 | SLITRK3 | 4 | 2 | 2 | 2 | 4 | 20 |
| ENSDARG000 | 18 | 22083772 | 22083886 | - | 115 | 5S_rRNA | 0 | 0 | 0 | 0 | 0 | 0 |
| ENSDARG000 | 18 | 15036151 | 15036269 | - | 119 | 5S_rRNA | 0 | 0 | 0 | 0 | 0 | 0 |
| ENSDARG000 | 18;18 | 47094334:471 | 47095727:471 | -;- | 1699 | NA | 12 | 0 | 0 | 0 | 7 | 0 |
| ENSDARG000 | 18 | 18577474 | 18577549 | - | 76 | NA | 0 | 0 | 0 | 0 | 0 | 0 |
| ENSDARG000 | 18:18:18:18:1 | 31610744:316 | 31611007:316 | +:+:+:+:+:+:+: | 3114 | dhx36 | 312 | 329 | 424 | 245 | 283 | 266 |
| ENSDARG000 | 18:18:18:18:1 | 45708744:457 | 45708863:457 | +:+:+:+:+:+:+: | 1855 | depdc7a | 6 | 0 | 2 | 10 | 2 | 3 |
| ENSDARG000 | 18 | 10441343 | 10441458 | - | 116 | 5S_rRNA | 0 | 0 | 0 | 0 | 0 | 0 |
| ENSDARG000 | 18 | 3062162 | 3062278 | - | 117 | 5S_rRNA | 0 | 0 | 0 | 0 | 0 | 0 |
| ENSDARG000 | 18 | 3047003 | 3047149 | + | 147 | SNORD15 | 0 | 2 | 6 | 3 | 0 | 12 |
| ENSDARG000 | 18 | 21191665 | 21191784 | + | 120 | 5S_rRNA | 0 | 0 | 0 | 0 | 0 | 0 |
| ENSDARG000 | 18 | 9267063 | 9267178 | - | 116 | 5S_rRNA | 0 | 0 | 0 | 0 | 0 | 0 |
| ENSDARG000 | 18:18:18:18:1 | 44080948:440 | 44080970:440 | +:+:+:+:+:+:+: | 1050 | NA | 0 | 0 | 3 | 4 | 0 | 1 |
| ENSDARG000 | 18 | 37962778 | 37962897 | + | 120 | 5S_rRNA | 0 | 0 | 0 | 0 | 0 | 0 |
| ENSDARG000 | 18:18:18:18:1 | 14710:17213: | 14805:17345: | -:-:-:-:-:-:-:-:- | 1932 | pde8a | 429 | 177 | 423 | 115 | 356 | 114 |
| ENSDARG000 | 18:18:18:18:1 | 2377067:2377 | 2379590:2379 | -:-:-:-:-:-:-:-:- | 5226 | NA | 15 | 2 | 58 | 22 | 4 | 12 |
| ENSDARG000 | 18;18;18 | 15259388:152 | 15263662:152 | -;-;- | 4719 | tmem263 | 712 | 903 | 1393 | 923 | 1114 | 589 |
| ENSDARG000 | 18 | 47503777 | 47503894 | - | 118 | 5S_rRNA | 0 | 0 | 0 | 0 | 0 | 0 |
| ENSDARG000 | 18 | 19423011 | 19423128 | - | 118 | 5S_rRNA | 0 | 0 | 0 | 0 | 0 | 0 |
| ENSDARG000 | 18:18:18:18:1 | 11556620:115 | 11559132:115 | -:-:-:-:-:-:-:-:- | 5003 | CRACR2A | 0 | 0 | 4 | 2 | 16 | 36 |
| ENSDARG000 | 18 | 27334941 | 27335059 | - | 119 | 5S_rRNA | 0 | 0 | 0 | 0 | 2 | 0 |
| ENSDARG000 | 18 | 8522895 | 8523009 | - | 115 | 5S_rRNA | 0 | 0 | 2 | 0 | 0 | 0 |
| ENSDARG000 | 18 | 5003269 | 5003384 | + | 116 | 5S_rRNA | 0 | 0 | 0 | 0 | 0 | 0 |
| ENSDARG000 | 18:18:18:18:1 | 2203945:2205 | 2204474:2205 | -:-:-:-:-:-:-:-:- | 1994 | pigb | 835 | 504 | 745 | 733 | 613 | 556 |
| ENSDARG000 | 18:18:18:18:1 | 45862414:458 | 45863560:458 | +:+:+:+:+:+:+: | 6032 | rnpepl1 | 1439 | 1745 | 1134 | 1182 | 1789 | 1586 |
| ENSDARG000 | 18 | 3040274 | 3040422 | + | 149 | SNORD15 | 0 | 3 | 5 | 0 | 5 | 0 |
| ENSDARG000 | 18 | 41668184 | 41668309 | + | 126 | 5S_rRNA | 3 | 1 | 2 | 0 | 0 | 3 |
| ENSDARG000 | 18:18:18:18:1 | 3332999:3334 | 3333284:3334 | +:+:+:+:+:+:+: | 1944 | GDPD4 | 2 | 0 | 0 | 0 | 2 | 0 |
| ENSDARG000 | 18:18:18:18:1 | 7543347:7543 | 7543413:7543 | +;+;+;+;+;+ | 2201 | arf5 | 2994 | 3377 | 3959 | 2103 | 3128 | 2565 |
| ENSDARG000 | 18:18:18:18:1 | 5917625:5945 | 5917962:5945 | +:+:+:+:+:+:+: | 2390 | glg1b | 238 | 273 | 266 | 233 | 433 | 103 |
| ENSDARG000 | 18:18:18:18:1 | 50040849:500 | 50040986:500 | -:-:-:-:-:-:-:-:- | 4962 | ABCC8 | 0 | 0 | 4 | 4 | 0 | 0 |
| ENSDARG000 | 18:18:18:18:1 | 5137241:5139 | 5137694:5140 | +:+:+:+:+:+:+: | 4689 | serpini1 | 14 | 0 | 8 | 0 | 18 | 2 |
| ENSDARG000 | 18:18:18:18:1 | 5086318:5087 | 5088174:5088 | -:-:-:-:-:-:-:-:- | 2587 | pdcd10a | 1161 | 1047 | 1369 | 1234 | 1187 | 1086 |
| ENSDARG000 | 18 | 9811020 | 9811152 | + | 133 | 5S_rRNA | 0 | 0 | 0 | 0 | 0 | 0 |
| ENSDARG000 | 18:18:18:18:1 | 31089957:310 | 31090330:310 | -;-;-;-;-;- | 959 | pdcd5 | 700 | 989 | 1376 | 731 | 906 | 1028 |
| ENSDARG000 | 18:18:18:18:1 | 15382029:153 | 15382158:153 | -:-:-:-:-:-:-:-:- | 3393 | POLR3B | 635 | 766 | 853 | 383 | 818 | 565 |
| ENSDARG000 | 18 | 50461981 | 50463268 | + | 1288 | NA | 0 | 0 | 0 | 0 | 0 | 0 |
| ENSDARG000 | 18;18;18;18 | 48943875:489 | 48944233:489 | +;+;+;+ | 3126 | NA | 18 | 0 | 0 | 0 | 6 | 0 |
| ENSDARG000 | 18 | 8129622 | 8129736 | - | 115 | 5S_rRNA | 0 | 0 | 0 | 0 | 0 | 0 |
| ENSDARG000 | 18 | 19374839 | 19374955 | - | 117 | 5S_rRNA | 0 | 0 | 0 | 0 | 0 | 0 |
| ENSDARG000 | 18:18:18:18:1 | 48279202:482 | 48279464:482 | -;-;-;-;-;-;- | 798 | ets1 | 56 | 63 | 32 | 28 | 16 | 58 |
| ENSDARG000 | 18 | 19454486 | 19454583 | - | 98 | SNORD16 | 0 | 0 | 0 | 0 | 0 | 0 |
| ENSDARG000 | 18;18;18;18 | 48983506:489 | 48983690:489 | -;-;-;- | 1654 | NA | 0 | 0 | 0 | 0 | 1 | 1 |
| ENSDARG000 | 18 | 16721799 | 16721982 | + | 184 | SNORD97 | 0 | 0 | 0 | 0 | 0 | 0 |
| ENSDARG000 | 18;18 | 48753162:487 | 48753742:487 | -;- | 612 | TIRAP | 2 | 0 | 0 | 0 | 8 | 0 |
| ENSDARG000 | 18 | 29175093 | 29175208 | + | 116 | 5S_rRNA | 0 | 0 | 0 | 0 | 0 | 0 |
| ENSDARG000 | 18:18:18:18:1 | 46840797:468 | 46840886:468 | -:-:-:-:-:-:-:-:- | 2855 | GRAMD1B | 137 | 54 | 133 | 26 | 66 | 123 |
| ENSDARG000 | 18 | 46577021 | 46577139 | + | 119 | 5S_rRNA | 0 | 0 | 0 | 0 | 0 | 0 |
| ENSDARG000 | 18 | 38017901 | 38018018 | + | 118 | 5S_rRNA | 0 | 0 | 0 | 0 | 0 | 0 |
| ENSDARG000 | 18:18:18:18:1 | 26096814:260 | 26097195:260 | -;-;-;-;-;- | 975 | idh2 | 5285 | 5700 | 7761 | 5089 | 4893 | 4785 |
| ENSDARG000 | 18:18:18:18:1 | 8339596:8339 | 8339874:8339 | +;+;+;+;+;+;+ | 5726 | NA | 2 | 0 | 8 | 2 | 22 | 31 |
| ENSDARG000 | 18:18:18:18:1 | 50826764:508 | 50826847:508 | -:-:-:-:-:-:-:-:- | 3429 | ddb1 | 2718 | 2335 | 3464 | 2128 | 2739 | 1978 |
| ENSDARG000 | 18 | 24642987 | 24643102 | + | 116 | 5S_rRNA | 0 | 0 | 0 | 0 | 0 | 0 |
| ENSDARG000 | 18 | 8397209 | 8397325 | + | 117 | 5S_rRNA | 0 | 0 | 3 | 0 | 2 | 2 |
| ENSDARG000 | 18 | 27306924 | 27307040 | - | 117 | 5S_rRNA | 0 | 0 | 0 | 0 | 0 | 0 |
| ENSDARG000 | 18:18:18:18:1 | 2228737:2249 | 2228918:2249 | +;+;+;+;+;+ | 1474 | rab27a | 5535 | 3631 | 4513 | 4078 | 5508 | 5421 |
| ENSDARG000 | 18:18:18:18:1 | 814328:81442 | 814491:81449 | +:+:+:+:+:+:+: | 5043 | fam219b | 168 | 145 | 190 | 135 | 130 | 313 |
| ENSDARG000 | 18:18:18:18:1 | 3367779:3368 | 3368734:3368 | -:-:-:-:-:-:-:-:- | 7690 | myo7aa | 24 | 9 | 10 | 0 | 10 | 16 |
| ENSDARG000 | 18:18:18:18:1 | 11506561:115 | 11507353:115 | +:+:+:+:+:+:+: | 1957 | PRMT8 | 0 | 0 | 0 | 0 | 0 | 0 |
| ENSDARG000 | 18 | 3044053 | 3044201 | + | 149 | SNORD15 | 0 | 3 | 2 | 0 | 0 | 0 |
| ENSDARG000 | 18:18:18:18:1 | 27590442:275 | 27590473:275 | -:-:-:-:-:-:-:-:- | 216 | NA | 0 | 0 | 0 | 0 | 0 | 0 |
| ENSDARG000 | 18:18:18:18:1 | 41857403:418 | 41857491:418 | -:-:-:-:-:-:-:-:- | 2622 | ARHGAP42 | 128 | 42 | 72 | 22 | 62 | 14 |
| ENSDARG000 | 18:18:18:18:1 | 13077800:130 | 13077990:130 | +:+:+:+:+:+:+: | 1578 | GAN | 81 | 59 | 68 | 67 | 73 | 54 |
| ENSDARG000 | 18:18:18:18:1 | 35491473:354 | 35491568:354 | -:-:-:-:-:-:-:-:- | 15237 | ryr1b | 0 | 4 | 18 | 6 | 2 | 6 |
| ENSDARG000 | 18 | 9111816 | 9111932 | + | 117 | 5S_rRNA | 0 | 0 | 0 | 0 | 0 | 0 |
| ENSDARG000 | 18 | 18251499 | 18251615 | + | 117 | 5S_rRNA | 0 | 0 | 0 | 0 | 0 | 0 |
| ENSDARG000 | 18:18:18:18:1 | 31073265:310 | 31073336:310 | +;+;+;+;+;+ | 1033 | cyba | 768 | 728 | 498 | 560 | 844 | 728 |
| ENSDARG000 | 18 | 27136865 | 27136982 | - | 118 | 5S_rRNA | 0 | 0 | 0 | 0 | 0 | 0 |
| ENSDARG000 | 18 | 15712429 | 15712545 | + | 117 | 5S_rRNA | 0 | 0 | 0 | 0 | 0 | 0 |
| ENSDARG000 | 18:18:18:18:1 | 21164591:211 | 21165514:211 | -;-;-;-;-;-;- | 2122 | NA | 0 | 0 | 0 | 0 | 2 | 2 |
| ENSDARG000 | 18:18:18:18:1 | 15459318:154 | 15459990:154 | -;-;-;-;-;-;- | 1554 | endouc | 86 | 26 | 128 | 30 | 104 | 30 |
| ENSDARG000 | 18 | 9607916 | 9608042 | + | 127 | 5S_rRNA | 0 | 0 | 0 | 0 | 0 | 0 |
| ENSDARG000 | 18 | 1148960 | 1149076 | + | 117 | 5S_rRNA | 0 | 0 | 5 | 0 | 0 | 0 |
| ENSDARG000 | 18;18;18;18 | 3096612:3100 | 3097295:3100 | -;-;-;- | 980 | aamdc | 147 | 270 | 120 | 164 | 66 | 292 |
| ENSDARG000 | 18;18;18 | 3558381:3562 | 3558409:3562 | -;-;- | 66 | NA | 0 | 2 | 4 | 0 | 2 | 1 |
| ENSDARG000 | 18;18 | 5781194:5781 | 5781288:5781 | -;- | 699 | RGS9BP | 0 | 0 | 0 | 0 | 0 | 0 |
| ENSDARG000 | 18;18;18 | 46634371:466 | 46635932:466 | -;-;- | 1905 | pid1 | 37 | 2 | 43 | 12 | 16 | 38 |
| ENSDARG000 | 18:18:18:18:1 | 3064237:3066 | 3064290:3066 | +:+:+:+:+:+:+: | 2940 | INTS4 | 339 | 249 | 411 | 245 | 365 | 354 |
| ENSDARG000 | 18:18:18:18:1 | 50839633:508 | 50839823:508 | -:-:-:-:-:-:-:-:- | 3092 | PDPR | 1158 | 622 | 908 | 660 | 1129 | 1133 |
| ENSDARG000 | 18:18:18:18:1 | 50795174:507 | 50795394:507 | -;-;-;-;-;- | 926 | taldo1 | 5961 | 5222 | 5596 | 3350 | 5820 | 2590 |
| ENSDARG000 | 18 | 7897120 | 7897234 | - | 115 | 5S_rRNA | 0 | 0 | 0 | 0 | 0 | 0 |
| ENSDARG000 | 18 | 46798159 | 46799199 | + | 1041 | NA | 31 | 28 | 55 | 8 | 142 | 69 |
| ENSDARG000 | 18 | 15875662 | 15875780 | - | 119 | 5S_rRNA | 0 | 0 | 0 | 0 | 0 | 1 |
| ENSDARG000 | 18 | 35819449 | 35819563 | + | 115 | 5S_rRNA | 6 | 0 | 0 | 3 | 0 | 0 |
| ENSDARG000 | 18:18:18:18:1 | 21622414:216 | 21622516:216 | -:-:-:-:-:-:-:-:- | 1480 | slc38a8a | 0 | 0 | 0 | 0 | 0 | 0 |
| ENSDARG000 | 18:18:18:18:1 | 16053455:160 | 16053464:160 | +:+:+:+:+:+:+: | 1475 | NA | 193 | 139 | 223 | 233 | 243 | 183 |
| ENSDARG000 | 18:18:18:18:1 | 50978940:509 | 50979288:509 | -:-:-:-:-:-:-:-:- | 1228 | NA | 0 | 0 | 0 | 0 | 4 | 0 |
| ENSDARG000 | 18:18:18:18:1 | 13162728:131 | 13162789:131 | +:+:+:+:+:+:+: | 2023 | tat | 40 | 8 | 553 | 38 | 351 | 97 |
| ENSDARG000 | 18:18:18:18:1 | 15469015:154 | 15469981:154 | -:-:-:-:-:-:-:-:- | 4094 | ppfibp1b | 1621 | 1233 | 1764 | 1273 | 934 | 1079 |
| ENSDARG000 | 18 | 28795559 | 28795675 | - | 117 | 5S_rRNA | 0 | 0 | 0 | 0 | 0 | 0 |
| ENSDARG000 | 18;18;18 | 3157911:3163 | 3160654:3163 | -;-;- | 3544 | aqp11 | 208 | 365 | 328 | 122 | 403 | 472 |
| ENSDARG000 | 18:18:18:18:1 | 1291434:1292 | 1292339:1292 | -:-:-:-:-:-:-:-:- | 2360 | pepd | 2012 | 3235 | 1262 | 1809 | 2456 | 2014 |
| ENSDARG000 | 18 | 7874631 | 7874748 | + | 118 | 5S_rRNA | 0 | 0 | 0 | 0 | 0 | 0 |
| ENSDARG000 | 18;18;18;18 | 5454341:5455 | 5454529:5455 | +;+;+;+ | 5221 | dtwd1 | 70 | 83 | 62 | 72 | 66 | 59 |
| ENSDARG000 | 18:18:18:18:1 | 48168755:481 | 48168770:481 | +;+;+;+;+ | 702 | smtla | 0 | 0 | 0 | 0 | 0 | 0 |
| ENSDARG000 | 18 | 44119137 | 44119262 | + | 126 | 5S_rRNA | 0 | 0 | 0 | 0 | 0 | 0 |
| ENSDARG000 | 18:18:18:18:1 | 45676788:456 | 45677564:456 | +:+:+:+:+:+:+: | 6881 | QSER1 | 928 | 555 | 1044 | 597 | 1147 | 1002 |
| ENSDARG000 | 18 | 20633789 | 20633923 | - | 135 | 5S_rRNA | 0 | 0 | 0 | 0 | 0 | 0 |
| ENSDARG000 | 18 | 39615775 | 39615905 | - | 131 | 5S_rRNA | 0 | 0 | 0 | 0 | 0 | 0 |
| ENSDARG000 | 18 | 19451870 | 19451937 | - | 68 | SNORD18 | 3 | 1 | 14 | 3 | 11 | 3 |
| ENSDARG000 | 18:18:18:18:1 | 26048128:260 | 26048168:260 | +:+:+:+:+:+:+: | 2452 | znf710a | 48 | 30 | 20 | 70 | 70 | 16 |
| ENSDARG000 | 18:18:18:18:1 | 9996813:9999 | 9996846:9999 | +:+:+:+:+:+:+: | 2058 | HGF | 28 | 0 | 12 | 2 | 13 | 0 |
| ENSDARG000 | 18 | 40023772 | 40023888 | + | 117 | 5S_rRNA | 0 | 0 | 0 | 0 | 0 | 0 |
| ENSDARG000 | 18;18 | 35375229:353 | 35375839:353 | -;- | 1168 | NA | 4 | 10 | 14 | 14 | 12 | 0 |
| ENSDARG000 | 18 | 409018 | 409134 | - | 117 | 5S_rRNA | 0 | 0 | 0 | 0 | 0 | 0 |
| ENSDARG000 | 18 | 3049515 | 3049659 | + | 145 | SNORD15 | 0 | 0 | 0 | 0 | 0 | 0 |
| ENSDARG000 | 18 | 26124663 | 26126258 | - | 1596 | NA | 0 | 0 | 0 | 0 | 4 | 0 |
| ENSDARG000 | 18;18;18;18 | 46773198:467 | 46773319:467 | +;+;+;+ | 549 | NA | 0 | 0 | 0 | 0 | 0 | 0 |
| ENSDARG000 | 18 | 45050835 | 45050950 | + | 116 | 5S_rRNA | 0 | 0 | 0 | 0 | 0 | 0 |
| ENSDARG000 | 18:18:18:18:1 | 660578:66967 | 660990:66975 | +:+:+:+:+:+:+: | 1543 | CELF6 | 0 | 0 | 0 | 0 | 0 | 0 |
| ENSDARG000 | 18 | 9224413 | 9224528 | + | 116 | 5S_rRNA | 0 | 0 | 0 | 0 | 0 | 0 |
| ENSDARG000 | 18:18:18:18:1 | 5813996:5817 | 5814149:5818 | +:+:+:+:+:+:+: | 1487 | zdhhc7 | 164 | 136 | 218 | 136 | 153 | 177 |
| ENSDARG000 | 18;18;18;18 | 16798854:168 | 16800029:168 | -;-;-;- | 1512 | NA | 98 | 30 | 28 | 118 | 150 | 24 |
| ENSDARG000 | 18 | 18569241 | 18569322 | - | 82 | NA | 0 | 0 | 0 | 0 | 0 | 0 |
| ENSDARG000 | 18:18:18:18:1 | 1837668:1837 | 1837752:1837 | +:+:+:+:+:+:+: | 3987 | NA | 2324 | 1784 | 3909 | 2846 | 1770 | 3146 |
| ENSDARG000 | 18:18:18:18:1 | 18087260:181 | 18087311:181 | -;-;-;-;-;- | 126 | NA | 0 | 0 | 6 | 0 | 1 | 0 |
| ENSDARG000 | 18:18:18:18:1 | 33032203:330 | 33032510:330 | +;+;+;+;+;+ | 2658 | NA | 0 | 0 | 0 | 0 | 0 | 0 |
| ENSDARG000 | 18:18:18:18:1 | 45889496:459 | 45889656:459 | +:+:+:+:+:+:+: | 3666 | DVL3 | 586 | 667 | 837 | 738 | 828 | 702 |
| ENSDARG000 | 18 | 45258981 | 45259097 | + | 117 | 5S_rRNA | 0 | 0 | 0 | 0 | 0 | 0 |
| ENSDARG000 | 18 | 7882779 | 7882893 | + | 115 | 5S_rRNA | 0 | 0 | 0 | 0 | 0 | 0 |
| ENSDARG000 | 18:18:18:18:1 | 1249617:1249 | 1251256:1251 | -;-;-;-;-;- | 1847 | ugt5f1 | 66 | 106 | 66 | 116 | 60 | 116 |
| ENSDARG000 | 18 | 41931719 | 41931833 | + | 115 | 5S_rRNA | 0 | 0 | 0 | 0 | 0 | 0 |
| ENSDARG000 | 18:18:18:18:1 | 48696094:486 | 48696641:486 | -:-:-:-:-:-:-:-:- | 1902 | st3gal4 | 288 | 403 | 436 | 524 | 393 | 615 |
| ENSDARG000 | 18 | 3942020 | 3942136 | + | 117 | 5S_rRNA | 0 | 0 | 0 | 0 | 0 | 0 |
| ENSDARG000 | 18 | 48438497 | 48438613 | + | 117 | 5S_rRNA | 0 | 0 | 0 | 0 | 0 | 0 |
| ENSDARG000 | 18:18:18:18:1 | 48793536:488 | 48794078:488 | +;+;+;+;+ | 7599 | bmp16 | 14 | 4 | 24 | 2 | 38 | 30 |
| ENSDARG000 | 18:18:18:18:1 | 31299076:313 | 31299090:313 | -:-:-:-:-:-:-:-:- | 492 | NA | 8 | 1 | 6 | 3 | 4 | 8 |
| ENSDARG000 | 18:18:18:18:1 | 5273953:5276 | 5273998:5277 | +:+:+:+:+:+:+: | 2046 | SLC12A1 | 0 | 0 | 48 | 0 | 42 | 0 |
| ENSDARG000 | 18 | 19542102 | 19542231 | - | 130 | 7SK | 0 | 0 | 0 | 0 | 0 | 0 |
| ENSDARG000 | 18:18:18:18:1 | 50801307:508 | 50801361:508 | +:+:+:+:+:+:+: | 1229 | tspan4a | 190 | 140 | 194 | 179 | 263 | 252 |
| ENSDARG000 | 18:18:18:18:1 | 5875268:5875 | 5875321:5875 | +:+:+:+:+:+:+: | 5489 | WDR59 | 80 | 110 | 129 | 128 | 138 | 125 |
| ENSDARG000 | 18 | 5277875 | 5277991 | + | 117 | 5S_rRNA | 0 | 0 | 0 | 0 | 0 | 0 |
| ENSDARG000 | 18 | 44006572 | 44006688 | + | 117 | 5S_rRNA | 0 | 0 | 0 | 0 | 0 | 0 |
| ENSDARG000 | 18 | 19451582 | 19451651 | - | 70 | SNORD18 | 0 | 1 | 3 | 0 | 3 | 0 |
| ENSDARG000 | 18:18:18:18:1 | 18612388:186 | 18612798:186 | +;+;+;+;+;+ | 1125 | st3gal2 | 18 | 14 | 27 | 12 | 42 | 20 |
| ENSDARG000 | 18 | 15588087 | 15588201 | - | 115 | 5S_rRNA | 0 | 0 | 0 | 0 | 0 | 0 |
| ENSDARG000 | 18:18:18:18:1 | 3022188:3024 | 3022907:3025 | +;+;+;+;+;+;+ | 7550 | NA | 16 | 6 | 8 | 12 | 12 | 16 |
| ENSDARG000 | 18;18 | 43891051:438 | 43891107:438 | +;+ | 1137 | CXCR5 | 6 | 2 | 0 | 0 | 2 | 0 |

| | | | | | | | | | | | | |
|---|---|---|---|---|---|---|---|---|---|---|---|---|
| ENSDARG000 | 18:18:18:18:1 | 50942208:509 | 50942514:509 | -:-:-:-:-:-:-:-: | 1651 | st7 | 138 | 148 | 349 | 144 | 193 | 263 |
| ENSDARG000 | 18:18:18:18:1 | 32563554:325 | 32563733:325 | +;+;+;+;+;+;+ | 2538 | NA | 0 | 0 | 0 | 0 | 0 | 0 |
| ENSDARG000 | 18:18:18:18:1 | 5618368:5619 | 5618520:5619 | +:+:+:+:+:+:+: | 1224 | ulk3 | 120 | 87 | 120 | 104 | 146 | 121 |
| ENSDARG000 | 18 | 20846190 | 20846313 | - | 124 | 5S_rRNA | 0 | 1 | 4 | 0 | 0 | 0 |
| ENSDARG000 | 18 | 18725978 | 18726093 | + | 116 | 5S_rRNA | 0 | 0 | 0 | 0 | 0 | 0 |
| ENSDARG000 | 18 | 17675340 | 17675455 | + | 116 | 5S_rRNA | 0 | 0 | 0 | 0 | 0 | 0 |
| ENSDARG000 | 18:18:18:18:1 | 50961953:509 | 50962213:509 | +:+:+:+:+:+:+: | 1605 | ppfia1 | 628 | 418 | 714 | 422 | 652 | 426 |
| ENSDARG000 | 18 | 27736724 | 27736840 | + | 117 | 5S_rRNA | 0 | 0 | 0 | 0 | 0 | 0 |
| ENSDARG000 | 18:18:18:18:1 | 47534181:475 | 47534199:475 | -:-:-:-:-:-:-:-: | 1725 | KIRREL3 | 225 | 152 | 202 | 158 | 178 | 198 |
| ENSDARG000 | 18:18:18:18:1 | 34158178:341 | 34158395:341 | -:-:-:-:-:-:-:-: | 2284 | slc33a1 | 419 | 459 | 572 | 297 | 711 | 318 |
| ENSDARG000 | 18:18:18:18:1 | 15271993:152 | 15272158:152 | +;+;+;+;+;+;+ | 1296 | NA | 21 | 35 | 62 | 29 | 39 | 27 |
| ENSDARG000 | 18:18:18:18:1 | 46056907:460 | 46057209:460 | -;-;-;-;-;- | 1230 | NA | 0 | 0 | 3 | 2 | 0 | 0 |
| ENSDARG000 | 18;18 | 48758916:487 | 48758957:487 | +;+ | 590 | gemin7 | 130 | 148 | 149 | 164 | 112 | 128 |
| ENSDARG000 | 18 | 17580159 | 17580245 | + | 87 | dre-mir-138-1 | 0 | 0 | 0 | 0 | 0 | 0 |
| ENSDARG000 | 18 | 378947 | 379061 | - | 115 | 5S_rRNA | 0 | 0 | 0 | 0 | 0 | 0 |
| ENSDARG000 | 18:18:18:18:1 | 45990394:459 | 45991290:459 | +:+:+:+:+:+:+: | 1947 | mmp23bb | 15 | 25 | 37 | 27 | 23 | 37 |
| ENSDARG000 | 18 | 29307525 | 29307641 | - | 117 | 5S_rRNA | 0 | 0 | 0 | 0 | 0 | 0 |
| ENSDARG000 | 18:18:18:18:1 | 6638726:6638 | 6639257:6639 | +:+:+:+:+:+:+: | 4785 | c2cd5 | 2474 | 2205 | 2038 | 1921 | 2016 | 2686 |
| ENSDARG000 | 18:18:18:18:1 | 5861755:5863 | 5862021:5863 | -;-;-;-;-;-;-;- | 1577 | NOB1 | 290 | 507 | 550 | 395 | 357 | 261 |
| ENSDARG000 | 18 | 10742049 | 10742163 | + | 115 | 5S_rRNA | 0 | 0 | 0 | 0 | 0 | 0 |
| ENSDARG000 | 18 | 11830905 | 11831020 | + | 116 | 5S_rRNA | 0 | 0 | 0 | 0 | 0 | 0 |
| ENSDARG000 | 18 | 18577814 | 18577894 | - | 81 | NA | 0 | 0 | 0 | 0 | 2 | 0 |
| ENSDARG000 | 18 | 19452751 | 19452881 | - | 131 | NA | 1 | 8 | 10 | 0 | 0 | 6 |
| ENSDARG000 | 18:18:18:18:1 | 13063210:130 | 13063765:130 | -;-;-;-;-;- | 1547 | rxylt1 | 188 | 162 | 216 | 106 | 204 | 186 |
| ENSDARG000 | 18 | 35573181 | 35573295 | - | 115 | 5S_rRNA | 0 | 0 | 0 | 0 | 0 | 0 |
| ENSDARG000 | 18;18;18 | 7264961:7268 | 7265025:7268 | +;+;+ | 390 | NA | 0 | 0 | 0 | 0 | 0 | 0 |
| ENSDARG000 | 18:18:18:18:1 | 15554381:155 | 15554578:155 | -;-;-;-;- | 684 | si:ch211-245i | 0 | 0 | 0 | 0 | 0 | 0 |
| ENSDARG000 | 18 | 27196210 | 27196326 | + | 117 | 5S_rRNA | 0 | 0 | 0 | 0 | 0 | 0 |
| ENSDARG000 | 18 | 5337658 | 5337772 | - | 115 | 5S_rRNA | 0 | 0 | 0 | 0 | 0 | 0 |
| ENSDARG000 | 18:18:18:18:1 | 5172848:5178 | 5172858:5178 | +;+;+;+;+;+;+ | 5164 | NA | 0 | 0 | 8 | 0 | 8 | 0 |
| ENSDARG000 | 18:18:18:18:1 | 3140682:3142 | 3141049:3142 | +;+;+;+;+;+;+ | 1260 | clns1a | 422 | 632 | 686 | 414 | 660 | 470 |
| ENSDARG000 | 18 | 29085058 | 29085174 | - | 117 | 5S_rRNA | 0 | 0 | 0 | 0 | 0 | 0 |
| ENSDARG000 | 18 | 3535058 | 3535174 | - | 117 | 5S_rRNA | 0 | 0 | 0 | 0 | 0 | 0 |
| ENSDARG000 | 18;18 | 15147931:151 | 15150463:151 | -;- | 2583 | mterf2 | 56 | 94 | 89 | 38 | 64 | 63 |
| ENSDARG000 | 18:18:18:18:1 | 46010081:460 | 46010272:460 | -:-:-:-:-:-:-:-: | 1080 | pcyt1ab | 397 | 394 | 547 | 692 | 509 | 436 |
| ENSDARG000 | 18:18:18:18:1 | 32615075:326 | 32615299:326 | +;+;+;+;+;+ | 5354 | NA | 0 | 0 | 0 | 0 | 0 | 0 |
| ENSDARG000 | 18:18:18:18:1 | 47522973:475 | 47523440:475 | -;-;-;-;- | 849 | NA | 79 | 93 | 77 | 76 | 103 | 136 |
| ENSDARG000 | 18 | 3260143 | 3260258 | + | 116 | 5S_rRNA | 0 | 0 | 0 | 0 | 0 | 0 |
| ENSDARG000 | 18 | 8174254 | 8174368 | - | 115 | 5S_rRNA | 0 | 0 | 0 | 0 | 0 | 0 |
| ENSDARG000 | 18 | 13544250 | 13544367 | - | 118 | 5S_rRNA | 0 | 0 | 0 | 0 | 0 | 0 |
| ENSDARG000 | 18 | 29213133 | 29213249 | - | 117 | 5S_rRNA | 0 | 0 | 0 | 0 | 0 | 0 |
| ENSDARG000 | 18 | 41713076 | 41713190 | + | 115 | 5S_rRNA | 0 | 0 | 0 | 0 | 0 | 0 |
| ENSDARG000 | 18;18;18;18 | 33074718:330 | 33079332:330 | -;-;-;- | 5462 | NA | 0 | 0 | 0 | 0 | 0 | 0 |
| ENSDARG000 | 18:18:18:18:1 | 50914231:509 | 50914375:509 | -:-:-:-:-:-:-:-: | 2567 | CTTN | 3359 | 3131 | 3229 | 2975 | 3663 | 2666 |
| ENSDARG000 | 18 | 9804348 | 9804464 | - | 117 | 5S_rRNA | 0 | 0 | 0 | 0 | 0 | 0 |
| ENSDARG000 | 18 | 6524776 | 6524890 | + | 115 | 5S_rRNA | 0 | 0 | 0 | 0 | 0 | 0 |
| ENSDARG000 | 18:18:18:18:1 | 34181655:341 | 34181703:341 | +:+:+:+:+:+:+: | 6143 | gmps | 4672 | 3759 | 6286 | 2360 | 6861 | 5055 |
| ENSDARG000 | 18:18:18:18:1 | 5843583:5844 | 5844630:5844 | -;-;-;-;- | 1471 | nip7 | 260 | 352 | 518 | 186 | 270 | 241 |
| ENSDARG000 | 18:18:18:18:1 | 45781115:457 | 45781258:457 | +:+:+:+:+:+:+: | 3574 | rpl35a | 9147 | 11002 | 14267 | 12816 | 8788 | 12789 |
| ENSDARG000 | 18 | 1495837 | 1495952 | + | 116 | 5S_rRNA | 0 | 0 | 0 | 0 | 0 | 0 |
| ENSDARG000 | 18 | 10670661 | 10670738 | + | 78 | NA | 0 | 0 | 0 | 0 | 0 | 0 |
| ENSDARG000 | 18:18:18:18:1 | 19463425:194 | 19468326:194 | -:-:-:-:-:-:-:-: | 6913 | LCTL | 0 | 2 | 17 | 2 | 20 | 7 |
| ENSDARG000 | 18;18;18 | 2153530:2156 | 2153712:2156 | +;+;+ | 1298 | pygo1 | 49 | 46 | 74 | 20 | 59 | 47 |
| ENSDARG000 | 18;18 | 19883325:198 | 19883667:198 | +;+ | 3334 | C18H15orf61 | 23 | 14 | 0 | 0 | 0 | 20 |
| ENSDARG000 | 18:18:18:18:1 | 50924556:509 | 50924597:509 | +:+:+:+:+:+:+: | 3042 | NA | 52 | 4 | 0 | 8 | 12 | 10 |
| ENSDARG000 | 18 | 4445571 | 4445680 | + | 110 | U6 | 0 | 0 | 0 | 0 | 0 | 0 |
| ENSDARG000 | 18:18:18:18:1 | 45734297:457 | 45734849:457 | -:-:-:-:-:-:-:-: | 2683 | cstf3 | 1751 | 1959 | 2699 | 1736 | 2241 | 1986 |
| ENSDARG000 | 18 | 24316181 | 24316299 | - | 119 | 5S_rRNA | 0 | 0 | 0 | 0 | 0 | 0 |
| ENSDARG000 | 18 | 10080090 | 10080204 | - | 115 | 5S_rRNA | 0 | 0 | 0 | 0 | 0 | 0 |
| ENSDARG000 | 18 | 40466790 | 40466904 | + | 115 | 5S_rRNA | 0 | 0 | 0 | 2 | 0 | 0 |
| ENSDARG000 | 18:18:18:18:1 | 5490668:5491 | 5490693:5491 | +:+:+:+:+:+:+: | 2225 | mibp2 | 12468 | 11580 | 26363 | 26572 | 13661 | 20274 |
| ENSDARG000 | 18:18:18:18:1 | 41097406:410 | 41097485:410 | -:-:-:-:-:-:-:-: | 2990 | RASA2 | 130 | 127 | 263 | 192 | 215 | 122 |
| ENSDARG000 | 18 | 7812401 | 7812516 | - | 116 | 5S_rRNA | 0 | 0 | 0 | 0 | 0 | 0 |
| ENSDARG000 | 18 | 12945617 | 12945731 | - | 115 | 5S_rRNA | 0 | 0 | 0 | 0 | 0 | 0 |
| ENSDARG000 | 18 | 11457213 | 11457330 | + | 118 | 5S_rRNA | 0 | 0 | 0 | 0 | 0 | 0 |
| ENSDARG000 | 18:18:18:18:1 | 13182528:131 | 13182702:131 | +:+:+:+:+:+:+: | 1511 | zgc:56622 | 77 | 43 | 147 | 105 | 81 | 86 |
| ENSDARG000 | 18:18:18:18:1 | 5308392:5309 | 5308464:5310 | +;+;+;+;+;+;+ | 4979 | dut | 738 | 836 | 1253 | 662 | 927 | 640 |
| ENSDARG000 | 18:18:18:18:1 | 28988373:290 | 28988828:290 | +;+;+;+;+;+;+ | 4089 | SYT9 | 0 | 0 | 0 | 0 | 4 | 0 |
| ENSDARG000 | 18:18:18:18:1 | 2837563:2837 | 2837735:2837 | +:+:+:+:+:+:+: | 1981 | fam168a | 232 | 239 | 281 | 163 | 294 | 328 |
| ENSDARG000 | 18 | 23472146 | 23472260 | + | 115 | 5S_rRNA | 0 | 0 | 0 | 0 | 0 | 0 |
| ENSDARG000 | 18;18;18;18 | 48670741:486 | 48670754:486 | +;+;+;+ | 300 | NA | 0 | 0 | 1 | 0 | 2 | 1 |
| ENSDARG000 | 18 | 2506032 | 2506169 | - | 138 | 5S_rRNA | 0 | 0 | 0 | 0 | 0 | 0 |
| ENSDARG000 | 18 | 5020563 | 5020679 | - | 117 | 5S_rRNA | 0 | 0 | 0 | 0 | 0 | 0 |
| ENSDARG000 | 18 | 35806848 | 35806966 | + | 119 | 5S_rRNA | 0 | 0 | 0 | 0 | 0 | 1 |
| ENSDARG000 | 18;18;18;18 | 642889:64311 | 642966:64317 | +;+;+;+ | 330 | NA | 22 | 33 | 22 | 54 | 55 | 6 |
| ENSDARG000 | 18:18:18:18:1 | 5362559:5364 | 5362640:5364 | -:-:-:-:-:-:-:-: | 1503 | NA | 619 | 379 | 571 | 386 | 459 | 327 |
| ENSDARG000 | 18:18:18:18:1 | 35229115:352 | 35229245:352 | +:+:+:+:+:+:+: | 1828 | tbrg1 | 221 | 283 | 554 | 322 | 266 | 296 |
| ENSDARG000 | 18;18;18 | 8845376:8849 | 8847467:8849 | -;-;- | 3272 | NA | 2 | 0 | 0 | 0 | 2 | 0 |
| ENSDARG000 | 18;18;18 | 6860822:6862 | 6860916:6862 | -;-;- | 225 | NA | 0 | 0 | 0 | 0 | 0 | 0 |
| ENSDARG000 | 18;18 | 2222447:2225 | 2222592:2226 | +;+ | 528 | pigbos1 | 93 | 106 | 160 | 178 | 113 | 147 |
| ENSDARG000 | 18:18:18:18:1 | 1615:1902:20 | 1761:2012:21 | +;+;+;+;+ | 528 | homer2 | 10 | 18 | 8 | 0 | 6 | 6 |
| ENSDARG000 | 18:18:18:18:1 | 1768804:1769 | 1768881:1769 | +:+:+:+:+:+:+: | 1344 | NA | 3 | 0 | 6 | 0 | 23 | 8 |
| ENSDARG000 | 18 | 37458446 | 37458562 | + | 117 | 5S_rRNA | 0 | 0 | 0 | 0 | 0 | 0 |
| ENSDARG000 | 18:18:18:18:1 | 45666489:456 | 45666823:456 | +:+:+:+:+:+:+: | 4113 | prrg4 | 104 | 82 | 87 | 58 | 74 | 32 |
| ENSDARG000 | 18 | 39736389 | 39736503 | + | 115 | 5S_rRNA | 0 | 0 | 0 | 0 | 0 | 0 |
| ENSDARG000 | 18 | 19452210 | 19452278 | - | 69 | SNORD18 | 3 | 2 | 1 | 1 | 4 | 5 |
| ENSDARG000 | 18:18:18:18:1 | 808911:81056 | 809066:81068 | +;+;+;+;+ | 673 | cox5ab | 713 | 794 | 1054 | 654 | 758 | 614 |
| ENSDARG000 | 18:18:18:18:1 | 2168695:2169 | 2169284:2169 | +:+:+:+:+:+:+: | 1737 | dnaaf4 | 0 | 6 | 11 | 0 | 39 | 22 |
| ENSDARG000 | 18:18:18:18:1 | 49969568:500 | 49969966:500 | +;+;+;+;+ | 3847 | mob2b | 396 | 337 | 340 | 418 | 446 | 167 |
| ENSDARG000 | 18 | 19455144 | 19455241 | - | 98 | SNORD16 | 0 | 0 | 0 | 0 | 0 | 0 |
| ENSDARG000 | 18:18:18:18:1 | 48953963:489 | 48954009:489 | +:+:+:+:+:+:+: | 2526 | SHKBP1 | 510 | 312 | 520 | 289 | 505 | 483 |
| ENSDARG000 | 18 | 9090995 | 9091111 | - | 117 | 5S_rRNA | 0 | 0 | 0 | 0 | 0 | 0 |
| ENSDARG000 | 18:18:18:18:1 | 50852359:508 | 50852371:508 | -;-;-;-;-;-;- | 828 | NA | 2 | 2 | 0 | 0 | 0 | 0 |
| ENSDARG000 | 18:18:18:18:1 | 34861568:348 | 34862055:348 | +;+;+;+;+;+ | 1620 | NA | 4 | 0 | 4 | 15 | 0 | 12 |
| ENSDARG000 | 18:18:18:18:1 | 50948003:509 | 50948098:509 | -:-:-:-:-:-:-:-: | 1314 | ST7 | 90 | 55 | 146 | 122 | 79 | 51 |
| ENSDARG000 | 18 | 26747811 | 26747927 | + | 117 | 5S_rRNA | 0 | 0 | 0 | 0 | 0 | 0 |
| ENSDARG000 | 18 | 7998576 | 7998697 | + | 122 | 5S_rRNA | 0 | 0 | 0 | 0 | 0 | 0 |
| ENSDARG000 | 18:18:18:18:1 | 15132112:151 | 15132550:151 | +;+;+;+;+;+ | 3094 | zgc:153031 | 0 | 40 | 38 | 68 | 38 | 156 |
| ENSDARG000 | 18:18:18:18:1 | 34146007:341 | 34147260:341 | -;-;-;-;- | 1747 | C18H3orf33 | 32 | 38 | 26 | 52 | 22 | 60 |
| ENSDARG000 | 18:18:18:18:1 | 33080843:330 | 33082231:330 | -;-;-;-;-;- | 3147 | NA | 0 | 0 | 0 | 0 | 0 | 0 |
| ENSDARG000 | 18 | 39586546 | 39586665 | - | 120 | 5S_rRNA | 0 | 0 | 0 | 0 | 0 | 0 |
| ENSDARG000 | 18 | 5631378 | 5631492 | - | 115 | 5S_rRNA | 0 | 0 | 0 | 0 | 0 | 0 |
| ENSDARG000 | 18:18:18:18:1 | 6587:7473:87 | 7275:7632:89 | -;-;-;-;-;- | 1866 | NA | 358 | 214 | 371 | 228 | 261 | 182 |
| ENSDARG000 | 18 | 26203303 | 26203389 | - | 87 | dre-mir-184-1 | 0 | 0 | 0 | 0 | 0 | 0 |
| ENSDARG000 | 18:18:18:18:1 | 3169579:3169 | 3170008:3170 | +:+:+:+:+:+:+: | 6210 | pak1 | 210 | 146 | 246 | 110 | 331 | 118 |
| ENSDARG000 | 18 | 25158661 | 25158777 | - | 117 | 5S_rRNA | 0 | 0 | 0 | 0 | 0 | 0 |
| ENSDARG000 | 18:18:18:18:1 | 4101719:4156 | 4102233:4156 | +;+;+;+;+;+ | 1678 | NA | 0 | 0 | 0 | 8 | 0 | 0 |
| ENSDARG000 | 18:18:18:18:1 | 160101:16134 | 160256:16155 | -;-;-;-;-;- | 1551 | NA | 0 | 0 | 0 | 0 | 0 | 0 |
| ENSDARG000 | 18;18;18;18 | 272590:27948 | 275637:27959 | -;-;-;- | 4176 | NA | 12 | 8 | 22 | 16 | 24 | 0 |
| ENSDARG000 | 18:18:18:18:1 | 830866:83086 | 831249:83124 | +:+:+:+:+:+:+: | 7143 | NA | 0 | 0 | 0 | 0 | 0 | 0 |
| ENSDARG000 | 18:18:18:18:1 | 1270387:1270 | 1271286:1271 | -;-;-;-;- | 1470 | NA | 49 | 4 | 0 | 0 | 10 | 0 |
| ENSDARG000 | 18;18;18;18 | 1447840:1450 | 1448017:1450 | +;+;+;+ | 879 | NA | 0 | 0 | 0 | 0 | 0 | 0 |
| ENSDARG000 | 18;18;18 | 2139002:2139 | 2139527:2139 | +;+;+ | 1005 | NA | 0 | 0 | 0 | 0 | 0 | 0 |
| ENSDARG000 | 18:18:18:18:1 | 2314106:2363 | 2314723:2363 | -;-;-;-;- | 1867 | NA | 0 | 0 | 0 | 0 | 0 | 0 |
| ENSDARG000 | 18;18;18 | 3307765:3323 | 3308013:3323 | +;+;+ | 1920 | NA | 0 | 0 | 0 | 0 | 6 | 0 |
| ENSDARG000 | 18;18;18;18 | 4351636:4443 | 4351729:4444 | +;+;+;+ | 1004 | NA | 0 | 2 | 2 | 0 | 0 | 0 |
| ENSDARG000 | 18;18;18;18 | 4632843:4636 | 4634250:4636 | -;-;-;- | 1929 | NA | 12 | 0 | 0 | 8 | 2 | 4 |
| ENSDARG000 | 18:18:18:18:1 | 7000194:7000 | 7000898:7000 | +:+:+:+:+:+:+: | 1940 | NA | 2 | 0 | 3 | 2 | 22 | 4 |
| ENSDARG000 | 18:18:18:18:1 | 8090216:8090 | 8091193:8091 | -;-;-;-;- | 1617 | NA | 0 | 0 | 0 | 0 | 0 | 0 |
| ENSDARG000 | 18;18;18;18 | 8100905:8101 | 8101390:8101 | +;+;+;+ | 1912 | NA | 0 | 0 | 0 | 0 | 0 | 0 |
| ENSDARG000 | 18:18:18:18:1 | 8897821:8897 | 8898906:8898 | -;-;-;-;- | 1549 | NA | 0 | 0 | 0 | 0 | 0 | 2 |
| ENSDARG000 | 18;18;18 | 9980878:9982 | 9981174:9983 | +;+;+ | 1535 | NA | 8 | 2 | 2 | 0 | 2 | 0 |
| ENSDARG000 | 18:18:18:18:1 | 10106306:101 | 10106460:101 | +:+:+:+:+:+:+: | 2001 | NA | 0 | 0 | 0 | 0 | 0 | 0 |
| ENSDARG000 | 18;18;18 | 20089381:200 | 20089797:200 | +;+;+ | 813 | NA | 0 | 0 | 0 | 0 | 0 | 0 |
| ENSDARG000 | 18:18:18:18:1 | 23538522:235 | 23538820:235 | -;-;-;-;-;- | 1849 | NA | 0 | 0 | 0 | 0 | 0 | 0 |
| ENSDARG000 | 18:18:18:18:1 | 24741499:247 | 24743774:247 | -;-;-;-;-;- | 4034 | NA | 4 | 0 | 0 | 0 | 0 | 0 |
| ENSDARG000 | 18;18;18;18 | 33043295:330 | 33044691:330 | -;-;-;- | 1826 | NA | 0 | 0 | 0 | 0 | 0 | 0 |
| ENSDARG000 | 18;18;18 | 33735335:337 | 33735433:337 | +;+;+ | 490 | NA | 0 | 0 | 0 | 0 | 5 | 0 |
| ENSDARG000 | 18;18;18 | 38859119:388 | 38862186:388 | -;-;- | 3559 | NA | 0 | 0 | 0 | 0 | 0 | 0 |
| ENSDARG000 | 18:18:18:18:1 | 43638947:436 | 43639232:436 | +;+;+;+;+;+ | 1653 | NA | 0 | 0 | 0 | 0 | 0 | 0 |
| ENSDARG000 | 18;18;18 | 44036074:440 | 44036158:440 | +;+;+ | 1366 | NA | 0 | 0 | 0 | 0 | 0 | 0 |
| ENSDARG000 | 18;18;18 | 45313968:453 | 45314134:453 | +;+;+ | 416 | NA | 0 | 0 | 0 | 0 | 0 | 0 |
| ENSDARG000 | 18:18:18:18:1 | 45763546:457 | 45763914:457 | +;+;+;+;+;+;+ | 3437 | NA | 0 | 0 | 0 | 0 | 2 | 0 |
| ENSDARG000 | 18;18;18;18 | 47990791:480 | 47990861:480 | +;+;+;+ | 451 | NA | 5 | 0 | 0 | 3 | 0 | 0 |
| ENSDARG000 | 18:18:18:18:1 | 50782247:507 | 50783247:507 | +:+:+:+:+:+:+: | 2377 | NA | 0 | 0 | 0 | 0 | 0 | 0 |
| ENSDARG000 | 18;18;18 | 50786633:507 | 50787295:507 | +;+;+ | 2580 | NA | 1 | 0 | 2 | 0 | 0 | 0 |
| ENSDARG000 | 18:18:18:18:1 | 50866261:508 | 50868872:508 | -;-;-;-;-;- | 3476 | NA | 2 | 0 | 1 | 1 | 2 | 0 |
| ENSDARG000 | 12:12:12:12:1 | 4396308:4398 | 4396324:4398 | -:-:-:-:-:-:-:-: | 3680 | trpm4b3 | 168 | 4 | 13 | 25 | 9 | 23 |
| ENSDARG000 | 12:12:12:12:1 | 33790121:337 | 33790313:337 | -;-;-;-;-;-;-;- | 1529 | galk1 | 146 | 270 | 163 | 230 | 183 | 182 |
| ENSDARG000 | 12:12:12:12:1 | 16332416:163 | 16333133:163 | -:-:-:-:-:-:-:-: | 3623 | HECTD2 | 167 | 90 | 274 | 166 | 139 | 162 |
| ENSDARG000 | 12:12:12:12:1 | 22404108:224 | 22404171:224 | +:+:+:+:+:+:+: | 4972 | NA | 800 | 408 | 950 | 1013 | 874 | 971 |
| ENSDARG000 | 12:12:12:12:1 | 31537604:315 | 31537770:315 | +:+:+:+:+:+:+: | 1331 | NA | 0 | 0 | 0 | 0 | 0 | 0 |
| ENSDARG000 | 12:12:12:12:1 | 33805850:338 | 33807182:338 | -:-:-:-:-:-:-:-: | 3828 | twnk | 134 | 196 | 296 | 124 | 176 | 194 |
| ENSDARG000 | 12:12:12:12:1 | 25775734:257 | 25776057:257 | +:+:+:+:+:+:+: | 7075 | EPAS1 | 8 | 0 | 6 | 0 | 16 | 22 |
| ENSDARG000 | 12;12;12 | 33817327:338 | 33817613:338 | +;+;+ | 1082 | mrpl43 | 142 | 164 | 206 | 139 | 207 | 158 |
| ENSDARG000 | 12:12:12:12:1 | 48841182:488 | 48841996:488 | +:+:+:+:+:+:+: | 7696 | DLG5 | 2322 | 1018 | 1770 | 1127 | 1791 | 1574 |
| ENSDARG000 | 12:12:12:12:1 | 22450548:224 | 22451991:224 | -:-:-:-:-:-:-:-: | 6050 | NEURL4 | 522 | 463 | 633 | 431 | 448 | 415 |
| ENSDARG000 | 12:12:12:12:1 | 7254112:7256 | 7254356:7256 | +;+;+;+;+;+;+ | 1356 | tfam | 406 | 520 | 709 | 436 | 578 | 579 |
| ENSDARG000 | 12:12:12:12:1 | 2505519:2506 | 2506279:2506 | -:-:-:-:-:-:-:-: | 2240 | mapk8b | 321 | 154 | 204 | 189 | 242 | 224 |
| ENSDARG000 | 12:12:12:12:1 | 16405598:164 | 16408483:164 | -:-:-:-:-:-:-:-: | 4605 | pcgf5b | 118 | 84 | 68 | 74 | 152 | 156 |
| ENSDARG000 | 12:12:12:12:1 | 7399403:7406 | 7399459:7406 | +:+:+:+:+:+:+: | 1823 | BICC1 | 39 | 5 | 6 | 0 | 13 | 4 |
| ENSDARG000 | 12:12:12:12:1 | 31608905:316 | 31608953:316 | +;+;+;+;+;+;+ | 531 | NA | 3 | 18 | 3 | 18 | 14 | 3 |
| ENSDARG000 | 12:12:12:12:1 | 22511015:225 | 22511469:225 | -;-;-;-;-;-;-;- | 1573 | shbg | 0 | 2 | 6 | 0 | 0 | 0 |
| ENSDARG000 | 12:12:12:12:1 | 31616412:316 | 31616753:316 | +:+:+:+:+:+:+: | 2250 | cpn1 | 0 | 1 | 3 | 8 | 6 | 0 |
| ENSDARG000 | 12:12:12:12:1 | 49135755:491 | 49135891:491 | +:+:+:+:+:+:+: | 1276 | NA | 0 | 0 | 0 | 0 | 0 | 0 |
| ENSDARG000 | 12:12:12:12:1 | 31608908:316 | 31608953:316 | +:+:+:+:+:+:+: | 1996 | CPN1 | 0 | 0 | 0 | 0 | 0 | 0 |
| ENSDARG000 | 12:12:12:12:1 | 18234557:182 | 18235923:182 | +:+:+:+:+:+:+: | 2523 | FAM20C | 36 | 107 | 84 | 62 | 29 | 108 |
| ENSDARG000 | 12;12;12;12 | 4573696:4573 | 4573791:4574 | +;+;+;+ | 651 | NA | 0 | 0 | 6 | 0 | 0 | 0 |
| ENSDARG000 | 12:12:12:12:1 | 30046320:300 | 30046549:300 | +:+:+:+:+:+:+: | 8669 | ablim1b | 61 | 112 | 138 | 26 | 64 | 18 |
| ENSDARG000 | 12:12:12:12:1 | 49146715:491 | 49146899:491 | -:-:-:-:-:-:-:-: | 1557 | bub3 | 323 | 718 | 819 | 470 | 576 | 419 |
| ENSDARG000 | 12:12:12:12:1 | 33831515:338 | 33832777:338 | -:-:-:-:-:-:-:-: | 3315 | SEMA4G | 0 | 0 | 0 | 6 | 0 | 0 |
| ENSDARG000 | 12:12:12:12:1 | 4580372:4580 | 4580485:4580 | -:-:-:-:-:-:-:-: | 1536 | irf3 | 201 | 356 | 482 | 291 | 152 | 159 |

| | | | | | | | | | | | | |
|---|---|---|---|---|---|---|---|---|---|---|---|---|
| ENSDARG000 | 12;12;12 | 13164706:131 | 13164871:131 | +;+;+ | 875 | PAQR4 | 0 | 0 | 0 | 21 | 0 | 0 |
| ENSDARG000 | 12;12 | 49153888:491 | 49154470:491 | -;- | 838 | HMX3 | 0 | 0 | 0 | 0 | 0 | 0 |
| ENSDARG000 | 12:12:12:12:1 | 13188616:131 | 13189412:131 | -:-:-:-:-:-:-:-: | 2260 | pelo | 644 | 959 | 1129 | 476 | 869 | 645 |
| ENSDARG000 | 12;12;12 | 18358082:183 | 18358297:183 | +;+;+ | 1039 | foxl3 | 0 | 0 | 0 | 0 | 0 | 2 |
| ENSDARG000 | 12:12:12:12:1 | 19362335:193 | 19362712:193 | +:+:+:+:+:+:+: | 4853 | gspt1 | 620 | 597 | 816 | 492 | 692 | 512 |
| ENSDARG000 | 12:12:12:12:1 | 25845494:258 | 25846106:258 | +:+:+:+:+:+:+: | 2659 | glud1b | 1994 | 2408 | 4588 | 1711 | 5685 | 4795 |
| ENSDARG000 | 12:12:12:12:1 | 16440708:164 | 16440805:164 | +:+:+:+:+:+:+: | 1348 | ankrd1b | 0 | 1 | 2 | 0 | 0 | 0 |
| ENSDARG000 | 12:12:12:12:1 | 18381982:183 | 18382442:183 | -:-:-:-:-:-:-:-: | 1904 | tom1l2 | 224 | 88 | 197 | 186 | 234 | 180 |
| ENSDARG000 | 12:12:12:12:1 | 16446228:164 | 16446612:164 | -:-:-:-:-:-:-:-: | 1619 | rpp30 | 80 | 105 | 201 | 85 | 123 | 105 |
| ENSDARG000 | 12:12:12:12:1 | 4601330:4601 | 4602607:4602 | -:-:-:-:-:-:-:-: | 9923 | PRR12 | 2803 | 1964 | 3337 | 2178 | 2968 | 2758 |
| ENSDARG000 | 12:12:12:12:1 | 18410608:184 | 18411027:184 | -:-:-:-:-:-:-:-: | 1963 | atpaf2 | 114 | 192 | 316 | 114 | 204 | 156 |
| ENSDARG000 | 12:12:12:12:1 | 49156997:491 | 49158651:491 | -:-:-:-:-:-:-:-: | 3787 | ACADSB | 736 | 609 | 797 | 821 | 562 | 906 |
| ENSDARG000 | 12:12:12:12:1 | 7445595:7445 | 7445979:7445 | +:+:+:+:+:+:+: | 3120 | phyhiplb | 10 | 32 | 4 | 14 | 2 | 6 |
| ENSDARG000 | 12;12 | 22528750:225 | 22537588:225 | -;- | 8887 | NA | 1309 | 1166 | 1516 | 2126 | 1563 | 1932 |
| ENSDARG000 | 12:12:12:12:1 | 31638045:316 | 31638429:316 | +:+:+:+:+:+:+: | 9923 | dnmbp | 884 | 432 | 582 | 570 | 628 | 550 |
| ENSDARG000 | 12:12:12:12:1 | 33894396:338 | 33894611:338 | +:+:+:+:+:+:+: | 4257 | mfsd13a | 1426 | 1482 | 1862 | 915 | 1179 | 807 |
| ENSDARG000 | 12:12:12:12:1 | 10381957:103 | 10382640:103 | -:-:-:-:-:-:-:-: | 3084 | eef2k | 3656 | 3644 | 7027 | 9828 | 4533 | 4910 |
| ENSDARG000 | 12:12:12:12:1 | 33916939:339 | 33917101:339 | +:+:+:+:+:+:+: | 4200 | trim8b | 1128 | 981 | 1749 | 978 | 1249 | 692 |
| ENSDARG000 | 12:12:12:12:1 | 22560067:225 | 22560470:225 | +:+:+:+:+:+:+: | 8182 | POLR2A | 4173 | 3037 | 5046 | 2509 | 5612 | 3613 |
| ENSDARG000 | 12:12:12:12:1 | 4643996:4645 | 4644111:4645 | -:-:-:-:-:-:-:-: | 2670 | NA | 1 | 0 | 0 | 0 | 2 | 2 |
| ENSDARG000 | 12:12:12:12:1 | 19384615:193 | 19384713:193 | +:+:+:+:+:+:+: | 1495 | rsl1d1 | 1028 | 1183 | 2471 | 1284 | 1126 | 1654 |
| ENSDARG000 | 12:12:12:12:1 | 27589607:275 | 27589859:275 | +:+:+:+:+:+:+: | 2943 | SPATA20 | 795 | 654 | 1064 | 935 | 922 | 987 |
| ENSDARG000 | 12:12:12:12:1 | 31713239:317 | 31713378:317 | +:+:+:+:+:+:+: | 2217 | habp2 | 0 | 0 | 2 | 0 | 0 | 0 |
| ENSDARG000 | 12:12:12:12:1 | 13205955:132 | 13206277:132 | +;+;+;+;+;+;+ | 2233 | ppp1cab | 726 | 931 | 1396 | 807 | 984 | 950 |
| ENSDARG000 | 12;12;12 | 25885910:258 | 25886451:258 | -;-;- | 696 | si:dkey-193p1 | 176 | 18 | 16 | 8 | 32 | 6 |
| ENSDARG000 | 12:12:12:12:1 | 25950834:259 | 25952165:259 | -:-:-:-:-:-:-:-: | 3258 | bmpr1ab | 336 | 495 | 538 | 316 | 503 | 412 |
| ENSDARG000 | 12;12;12 | 4677193:4677 | 4677451:4677 | -;-;- | 483 | NA | 0 | 0 | 0 | 4 | 2 | 0 |
| ENSDARG000 | 12:12:12:12:1 | 16452575:164 | 16452648:164 | +:+:+:+:+:+:+: | 4950 | NA | 131 | 423 | 478 | 200 | 432 | 178 |
| ENSDARG000 | 12:12:12:12:1 | 18445604:184 | 18445842:184 | +:+:+:+:+:+:+: | 2169 | noxo1b | 8 | 0 | 2 | 0 | 0 | 8 |
| ENSDARG000 | 12:12:12:12:1 | 2522642:2524 | 2522791:2524 | +:+:+:+:+:+:+: | 3113 | FRMPD2 | 0 | 0 | 0 | 13 | 0 | 0 |
| ENSDARG000 | 12;12 | 19401854:194 | 19402423:194 | +;+ | 964 | NA | 0 | 2 | 0 | 0 | 0 | 0 |
| ENSDARG000 | 12:12:12:12:1 | 19408373:194 | 19408701:194 | +:+:+:+:+:+:+: | 3765 | SNX29 | 585 | 664 | 643 | 469 | 805 | 534 |
| ENSDARG000 | 12:12:12:12:1 | 33990947:339 | 33993334:339 | -;-;-;-;- | 3127 | timp2a | 3845 | 3860 | 3863 | 4583 | 2792 | 3462 |
| ENSDARG000 | 12:12:12:12:1 | 7757663:7757 | 7757860:7757 | -:-:-:-:-:-:-:-: | 2587 | NA | 90 | 78 | 91 | 39 | 132 | 24 |
| ENSDARG000 | 12:12:12:12:1 | 33966414:339 | 33968252:339 | -;-;-;-;-;- | 2396 | arl3b | 856 | 600 | 711 | 858 | 680 | 722 |
| ENSDARG000 | 12:12:12:12:1 | 33975065:339 | 33975251:339 | +:+:+:+:+:+:+: | 2092 | sfxn2 | 581 | 491 | 836 | 696 | 629 | 696 |
| ENSDARG000 | 12:12:12:12:1 | 31720673:317 | 31721066:317 | -:-:-:-:-:-:-:-: | 2944 | srsf2a | 3134 | 3414 | 5178 | 2658 | 4268 | 3505 |
| ENSDARG000 | 12:12:12:12:1 | 13244149:132 | 13244682:132 | +:+:+:+:+:+:+: | 5649 | atp2a1l | 10 | 18 | 12 | 30 | 42 | 4 |
| ENSDARG000 | 12;12;12;12 | 18458502:184 | 18458830:184 | +;+;+;+ | 1242 | NA | 0 | 0 | 0 | 0 | 0 | 0 |
| ENSDARG000 | 12;12;12 | 36669128:367 | 36672311:367 | -;-;- | 4621 | NA | 4 | 6 | 0 | 0 | 0 | 0 |
| ENSDARG000 | 12:12:12:12:1 | 22607761:226 | 22607815:226 | +:+:+:+:+:+:+: | 1926 | NA | 22 | 28 | 6 | 90 | 0 | 12 |
| ENSDARG000 | 12:12:12:12:1 | 27704015:277 | 27704661:277 | +:+:+:+:+:+:+: | 7751 | CACNA1G | 26 | 2 | 6 | 0 | 24 | 2 |
| ENSDARG000 | 12:12:12:12:1 | 22576404:225 | 22576679:225 | +:+:+:+:+:+:+: | 6088 | capgb | 3435 | 2377 | 376 | 4535 | 1170 | 1474 |
| ENSDARG000 | 12:12:12:12:1 | 18468066:184 | 18468839:184 | -:-:-:-:-:-:-:-: | 3879 | NA | 0 | 0 | 0 | 0 | 0 | 0 |
| ENSDARG000 | 12:12:12:12:1 | 4686145:4690 | 4686586:4691 | +:+:+:+:+:+:+: | 4714 | NA | 1161 | 1078 | 1307 | 645 | 1462 | 1113 |
| ENSDARG000 | 12:12:12:12:1 | 26023948:260 | 26025867:260 | -:-:-:-:-:-:-:-: | 4051 | ldb3b | 4 | 0 | 0 | 0 | 6 | 0 |
| ENSDARG000 | 12:12:12:12:1 | 16592408:165 | 16592914:165 | -:-:-:-:-:-:-:-: | 1455 | si:dkev-239i1 | 0 | 0 | 0 | 0 | 0 | 0 |
| ENSDARG000 | 12:12:12:12:1 | 36783721:367 | 36785050:367 | -:-:-:-:-:-:-:-: | 2404 | NA | 10 | 12 | 50 | 36 | 73 | 0 |
| ENSDARG000 | 12;12 | 2579237:2617 | 2579395:2617 | -;- | 312 | NA | 0 | 0 | 0 | 0 | 0 | 0 |
| ENSDARG000 | 12:12:12:12:1 | 2578020:2579 | 2578066:2579 | +;+;+;+;+ | 970 | NA | 0 | 0 | 0 | 0 | 0 | 0 |
| ENSDARG000 | 12;12;12 | 2631711:2638 | 2632041:2639 | +;+;+ | 1362 | gdf10b | 4 | 0 | 1 | 0 | 0 | 0 |
| ENSDARG000 | 12;12 | 2648043:2650 | 2648367:2651 | +;+ | 1170 | gdf2 | 0 | 0 | 6 | 0 | 0 | 0 |
| ENSDARG000 | 12 | 2660733 | 2663471 | + | 2739 | RBP3 | 0 | 2 | 0 | 0 | 0 | 0 |
| ENSDARG000 | 12:12:12:12:1 | 30233977:302 | 30234387:302 | +:+:+:+:+:+:+: | 4204 | AFAP1L2 | 46 | 22 | 48 | 6 | 8 | 28 |
| ENSDARG000 | 12:12:12:12:1 | 4733892:4739 | 4735020:4739 | -:-:-:-:-:-:-:-: | 2513 | NA | 142 | 473 | 553 | 198 | 437 | 1602 |
| ENSDARG000 | 12;12 | 18524953:185 | 18525853:185 | +;+ | 2185 | neurl2 | 0 | 0 | 0 | 0 | 0 | 0 |
| ENSDARG000 | 12:12:12:12:1 | 28346722:283 | 28351399:283 | -:-:-:-:-:-:-:-: | 5914 | psmd11b | 646 | 1121 | 967 | 352 | 978 | 536 |
| ENSDARG000 | 12:12:12:12:1 | 28310647:283 | 28311109:283 | -:-:-:-:-:-:-:-: | 1345 | ogfod3 | 44 | 47 | 57 | 11 | 71 | 31 |
| ENSDARG000 | 12:12:12:12:1 | 8168272:8227 | 8168494:8227 | +:+:+:+:+:+:+: | 4536 | NA | 526 | 413 | 446 | 307 | 498 | 358 |
| ENSDARG000 | 12:12:12:12:1 | 16617138:166 | 16617474:166 | -:-:-:-:-:-:-:-: | 1351 | NA | 0 | 0 | 0 | 0 | 0 | 0 |
| ENSDARG000 | 12;12;12;12 | 2665081:2666 | 2666314:2666 | +;+;+;+ | 2354 | rbp3 | 2 | 2 | 9 | 0 | 5 | 0 |
| ENSDARG000 | 12 | 8474868 | 8476513 | + | 1646 | adoa | 264 | 242 | 464 | 243 | 302 | 242 |
| ENSDARG000 | 12;12;12 | 10412009:104 | 10413021:104 | -;-;- | 1918 | mosmob | 44 | 24 | 74 | 80 | 60 | 74 |
| ENSDARG000 | 12:12:12:12:1 | 18542954:185 | 18543900:185 | +:+:+:+:+:+:+: | 4154 | mlst8 | 412 | 347 | 461 | 288 | 415 | 396 |
| ENSDARG000 | 12:12:12:12:1 | 19792281:197 | 19793706:197 | -;-;-;-;-;-;-;- | 2502 | shisa9a | 4 | 8 | 0 | 0 | 4 | 0 |
| ENSDARG000 | 12;12;12;12 | 8483838:8484 | 8485565:8485 | -;-;-;- | 2180 | egr2b | 82 | 8 | 32 | 14 | 43 | 80 |
| ENSDARG000 | 12:12:12:12:1 | 31729075:317 | 31729585:317 | +:+:+:+:+:+:+: | 4296 | RNF157 | 4 | 6 | 10 | 2 | 66 | 2 |
| ENSDARG000 | 12:12:12:12:1 | 16634442:166 | 16634665:166 | -;-;-;-;-;-;- | 1150 | NA | 0 | 0 | 0 | 0 | 0 | 0 |
| ENSDARG000 | 12;12 | 18565553:185 | 18566443:185 | -;- | 1684 | pgp | 346 | 313 | 312 | 70 | 289 | 154 |
| ENSDARG000 | 12:12:12:12:1 | 18551597:185 | 18552671:185 | -;-;-;-;-;- | 1733 | bricd5 | 2 | 0 | 0 | 0 | 0 | 0 |
| ENSDARG000 | 12:12:12:12:1 | 18578597:185 | 18578788:185 | +:+:+:+:+:+:+: | 4318 | grid2ipb | 34 | 14 | 13 | 40 | 38 | 24 |
| ENSDARG000 | 12:12:12:12:1 | 22628704:226 | 22628995:226 | +:+:+:+:+:+:+: | 2907 | mfsd8 | 923 | 708 | 1187 | 833 | 1211 | 1124 |
| ENSDARG000 | 12:12:12:12:1 | 4817962:4823 | 4818701:4827 | -;-;-;-;-;-;- | 6247 | NA | 8 | 0 | 0 | 0 | 0 | 4 |
| ENSDARG000 | 12:12:12:12:1 | 30312308:303 | 30312447:303 | -:-:-:-:-:-:-:-: | 2470 | VWA2 | 248 | 75 | 9 | 304 | 10 | 0 |
| ENSDARG000 | 12:12:12:12:1 | 26221999:262 | 26222178:262 | -:-:-:-:-:-:-:-: | 2942 | USP54 | 6 | 2 | 0 | 0 | 6 | 2 |
| ENSDARG000 | 12:12:12:12:1 | 18529228:185 | 18529484:185 | +:+:+:+:+:+:+: | 5168 | MEIOB | 9 | 10 | 36 | 50 | 49 | 43 |
| ENSDARG000 | 12:12:12:12:1 | 30344076:303 | 30346467:303 | -:-:-:-:-:-:-:-: | 6183 | tdrd1 | 10 | 0 | 18 | 0 | 12 | 2 |
| ENSDARG000 | 12:12:12:12:1 | 18571455:185 | 18571830:185 | -:-:-:-:-:-:-:-: | 1698 | zdhhc4 | 223 | 197 | 399 | 203 | 220 | 281 |
| ENSDARG000 | 12;12;12;12 | 18529311:185 | 18529484:185 | +;+;+;+ | 967 | msrb1b | 0 | 0 | 0 | 0 | 0 | 0 |
| ENSDARG000 | 12:12:12:12:1 | 16691586:166 | 16691921:166 | -:-:-:-:-:-:-:-: | 1802 | MGC174155 | 0 | 0 | 0 | 11 | 0 | 5 |
| ENSDARG000 | 12:12:12:12:1 | 16691584:166 | 16691921:166 | -;-;-;-;-;-;-;- | 1289 | wu:fa26c03 | 0 | 0 | 0 | 12 | 0 | 0 |
| ENSDARG000 | 12:12:12:12:1 | 13282797:132 | 13282843:132 | +:+:+:+:+:+:+: | 1832 | irf9 | 957 | 1239 | 1017 | 975 | 931 | 772 |
| ENSDARG000 | 12:12:12:12:1 | 2676878:2677 | 2676890:2677 | +:+:+:+:+:+:+: | 6734 | NA | 469 | 318 | 878 | 172 | 739 | 190 |
| ENSDARG000 | 12:12:12:12:1 | 10424676:104 | 10424994:104 | -:-:-:-:-:-:-:-: | 1974 | uqcrc2b | 2260 | 2639 | 2492 | 1417 | 2244 | 1596 |
| ENSDARG000 | 12:12:12:12:1 | 8569685:8569 | 8569876:8569 | +;+;+;+;+;+;+ | 2140 | nrbf2b | 407 | 274 | 454 | 369 | 371 | 463 |
| ENSDARG000 | 12:12:12:12:1 | 19866865:198 | 19867081:198 | +:+:+:+:+:+:+: | 2583 | cpped1 | 564 | 725 | 422 | 268 | 340 | 356 |
| ENSDARG000 | 12:12:12:12:1 | 22645324:226 | 22645769:226 | -:-:-:-:-:-:-:-: | 1887 | zcchc4 | 168 | 112 | 221 | 132 | 236 | 248 |
| ENSDARG000 | 12:12:12:12:1 | 4836856:4837 | 4837636:4837 | -;-;-;-;-;-;- | 1484 | zgc:163073 | 0 | 0 | 0 | 0 | 0 | 0 |
| ENSDARG000 | 12;12;12 | 31795031:318 | 31795303:318 | +;+;+ | 723 | NA | 0 | 4 | 0 | 0 | 0 | 0 |
| ENSDARG000 | 12:12:12:12:1 | 31773435:317 | 31773480:317 | +;+;+;+;+ | 682 | NA | 0 | 0 | 0 | 0 | 0 | 0 |
| ENSDARG000 | 12:12:12:12:1 | 22657923:226 | 22658088:226 | +:+:+:+:+:+:+: | 1319 | NA | 5 | 2 | 6 | 4 | 16 | 16 |
| ENSDARG000 | 12:12:12:12:1 | 31775182:317 | 31775215:317 | +:+:+:+:+:+:+: | 2327 | lrrc59 | 400 | 456 | 886 | 545 | 580 | 369 |
| ENSDARG000 | 12:12:12:12:1 | 31789379:317 | 31789976:317 | -;-;-;-;-;-;-;- | 1384 | nat9 | 376 | 352 | 371 | 298 | 496 | 356 |
| ENSDARG000 | 12;12 | 28367557:283 | 28367833:283 | +;+ | 1385 | cdk5r1b | 0 | 0 | 2 | 4 | 0 | 2 |
| ENSDARG000 | 12:12:12:12:1 | 19958845:199 | 19959679:199 | +:+:+:+:+:+:+: | 4504 | ercc4 | 354 | 272 | 440 | 218 | 496 | 292 |
| ENSDARG000 | 12:12:12:12:1 | 28377745:284 | 28381144:284 | -:-:-:-:-:-:-:-: | 6701 | MYO1D | 1818 | 1385 | 1097 | 716 | 1482 | 1098 |
| ENSDARG000 | 12:12:12:12:1 | 34051434:340 | 34051995:340 | +:+:+:+:+:+:+: | 3646 | cyth1b | 3800 | 3011 | 3048 | 5369 | 2495 | 2983 |
| ENSDARG000 | 12:12:12:12:1 | 26385096:263 | 26385838:263 | -:-:-:-:-:-:-:-: | 2299 | myoz1b | 0 | 0 | 0 | 0 | 0 | 0 |
| ENSDARG000 | 12:12:12:12:1 | 36891483:368 | 36891654:368 | +;+;+;+;+;+;+ | 2240 | cox10 | 504 | 473 | 556 | 304 | 591 | 428 |
| ENSDARG000 | 12:12:12:12:1 | 16743052:167 | 16743151:167 | -:-:-:-:-:-:-:-: | 1044 | NA | 0 | 1 | 0 | 0 | 0 | 0 |
| ENSDARG000 | 12:12:12:12:1 | 2780190:2780 | 2781182:2781 | -;-;-;-;-;-;- | 1757 | ubtd1b | 94 | 77 | 283 | 79 | 198 | 118 |
| ENSDARG000 | 12:12:12:12:1 | 26408351:264 | 26413725:264 | -;-;-;-;-;-;-;- | 7244 | SYNPO2L | 10 | 4 | 4 | 0 | 8 | 0 |
| ENSDARG000 | 12:12:12:12:1 | 19976400:199 | 19976624:199 | +:+:+:+:+:+:+: | 4134 | MRTFB | 260 | 108 | 286 | 102 | 326 | 374 |
| ENSDARG000 | 12:12:12:12:1 | 16758972:167 | 16759313:167 | -;-;-;-;-;-;-;- | 1288 | zgc:174154 | 0 | 0 | 0 | 0 | 0 | 0 |
| ENSDARG000 | 12:12:12:12:1 | 4845448:4845 | 4845505:4845 | +:+:+:+:+:+:+: | 1240 | ghrhr2 | 0 | 0 | 6 | 0 | 0 | 0 |
| ENSDARG000 | 12:12:12:12:1 | 8581875:8582 | 8582844:8582 | -:-:-:-:-:-:-:-: | 9138 | JMJD1C | 3364 | 2365 | 3970 | 2501 | 3841 | 3659 |
| ENSDARG000 | 12:12:12:12:1 | 2849584:2852 | 2851577:2853 | -:-:-:-:-:-:-:-: | 3221 | zdhhc16b | 183 | 326 | 624 | 288 | 416 | 162 |
| ENSDARG000 | 12:12:12:12:1 | 2804505:2806 | 2804640:2806 | +:+:+:+:+:+:+: | 3499 | mms19 | 759 | 715 | 1268 | 653 | 1085 | 494 |
| ENSDARG000 | 12:12:12:12:1 | 20059859:200 | 20060169:200 | -:-:-:-:-:-:-:-: | 2717 | ubald1a | 580 | 404 | 453 | 356 | 506 | 457 |
| ENSDARG000 | 12:12:12:12:1 | 13311939:133 | 13312300:133 | -:-:-:-:-:-:-:-: | 996 | emc9 | 261 | 293 | 189 | 177 | 178 | 152 |
| ENSDARG000 | 12;12 | 36971952:370 | 36972554:370 | +;+ | 5335 | hs3st3b1b | 36 | 4 | 8 | 4 | 44 | 22 |
| ENSDARG000 | 12:12:12:12:1 | 37214518:372 | 37214623:372 | +;+;+;+;+;+;+ | 1007 | NA | 0 | 0 | 0 | 0 | 0 | 0 |
| ENSDARG000 | 12:12:12:12:1 | 22680115:227 | 22680435:227 | +:+:+:+:+:+:+: | 3507 | ABLIM2 | 275 | 210 | 136 | 106 | 127 | 167 |
| ENSDARG000 | 12:12:12:12:1 | 31909346:319 | 31911609:319 | -:-:-:-:-:-:-:-: | 4940 | GRIN2C | 0 | 0 | 0 | 0 | 0 | 0 |
| ENSDARG000 | 12:12:12:12:1 | 16874645:168 | 16874968:168 | -;-;-;-;-;-;-;- | 1275 | si:dkev-269i1. | 10 | 4 | 0 | 11 | 0 | 0 |
| ENSDARG000 | 12:12:12:12:1 | 13331526:133 | 13331937:133 | -:-:-:-:-:-:-:-: | 1366 | lsm5 | 110 | 121 | 207 | 98 | 120 | 121 |
| ENSDARG000 | 12:12:12:12:1 | 4869248:4869 | 4869499:4869 | +:+:+:+:+:+:+: | 5559 | scn4aa | 0 | 0 | 0 | 0 | 0 | 0 |
| ENSDARG000 | 12;12;12;12 | 28543064:285 | 28543608:285 | -;-;-;- | 1202 | NA | 0 | 0 | 2 | 0 | 14 | 8 |
| ENSDARG000 | 12:12:12:12:1 | 28559408:285 | 28561922:285 | -:-:-:-:-:-:-:-: | 3727 | PDK2 | 20 | 16 | 94 | 80 | 18 | 74 |
| ENSDARG000 | 12:12:12:12:1 | 13319132:133 | 13319560:133 | +:+:+:+:+:+:+: | 1706 | psme2 | 2046 | 2048 | 1656 | 1037 | 1527 | 1228 |
| ENSDARG000 | 12;12;12 | 10443785:104 | 10444014:104 | +;+;+ | 761 | snu13b | 624 | 725 | 1084 | 344 | 706 | 592 |
| ENSDARG000 | 12:12:12:12:1 | 26434478:264 | 26434583:264 | +:+:+:+:+:+:+: | 3063 | itgb3b | 44 | 28 | 8 | 8 | 22 | 10 |
| ENSDARG000 | 12:12:12:12:1 | 32073660:320 | 32073686:320 | +:+:+:+:+:+:+: | 3347 | STXBP4 | 10 | 12 | 10 | 24 | 13 | 6 |
| ENSDARG000 | 12:12:12:12:1 | 22672323:226 | 22672352:226 | +:+:+:+:+:+:+: | 1224 | cdca9 | 0 | 2 | 0 | 0 | 1 | 0 |
| ENSDARG000 | 12;12;12;12 | 34258139:342 | 34258247:342 | +;+;+;+ | 2264 | socs3b | 10159 | 7208 | 12695 | 10553 | 4272 | 5863 |
| ENSDARG000 | 12:12:12:12:1 | 34378733:343 | 34379106:343 | +;+;+;+;+;+;+ | 1469 | tha1 | 207 | 363 | 261 | 159 | 222 | 180 |
| ENSDARG000 | 12:12:12:12:1 | 32068294:320 | 32068892:320 | -:-:-:-:-:-:-:-: | 1342 | cox11 | 199 | 166 | 237 | 170 | 225 | 201 |
| ENSDARG000 | 12:12:12:12:1 | 10448226:104 | 10448436:104 | +:+:+:+:+:+:+: | 1578 | crym | 4 | 8 | 4 | 4 | 6 | 4 |
| ENSDARG000 | 12:12:12:12:1 | 32039177:320 | 32040214:320 | -:-:-:-:-:-:-:-: | 2047 | rab40b | 83 | 53 | 51 | 105 | 95 | 104 |
| ENSDARG000 | 12:12:12:12:1 | 16895644:168 | 16895973:168 | -;-;-;-;-;-;-;- | 1278 | NA | 0 | 0 | 0 | 11 | 0 | 1 |
| ENSDARG000 | 12:12:12:12:1 | 30360184:303 | 30360640:303 | +:+:+:+:+:+:+: | 2882 | NA | 4 | 2 | 0 | 0 | 2 | 8 |
| ENSDARG000 | 12:12:12:12:1 | 20122256:201 | 20122600:201 | +:+:+:+:+:+:+: | 4944 | mgrn1a | 1062 | 691 | 678 | 643 | 526 | 638 |
| ENSDARG000 | 12:12:12:12:1 | 37275336:372 | 37278057:372 | -;-;-;-;-;-;-;- | 3161 | pmp22b | 337 | 797 | 574 | 566 | 282 | 714 |
| ENSDARG000 | 12:12:12:12:1 | 37327867:373 | 37328318:373 | -:-:-:-:-:-:-:-: | 1783 | TEKT3 | 4 | 2 | 0 | 2 | 4 | 0 |
| ENSDARG000 | 12:12:12:12:1 | 20149305:201 | 20149558:201 | +;+;+;+;+;+;+ | 2316 | foxj1b | 2 | 0 | 0 | 2 | 0 | 4 |
| ENSDARG000 | 12:12:12:12:1 | 8822717:8851 | 8823035:8851 | +:+:+:+:+:+:+: | 1816 | reep3b | 1146 | 1077 | 1143 | 1005 | 1013 | 819 |
| ENSDARG000 | 12:12:12:12:1 | 2870671:2875 | 2870770:2875 | +:+:+:+:+:+:+: | 1829 | prkar1ab | 735 | 428 | 634 | 269 | 583 | 582 |
| ENSDARG000 | 12:12:12:12:1 | 16919411:169 | 16919752:169 | -;-;-;-;-;-;-;- | 1288 | si:dkev-26g8.5 | 2 | 1 | 0 | 0 | 0 | 3 |
| ENSDARG000 | 12:12:12:12:1 | 4900981:4903 | 4901197:4904 | +;+;+;+;+ | 843 | NA | 0 | 6 | 2 | 0 | 6 | 2 |
| ENSDARG000 | 12:12:12:12:1 | 10464897:104 | 10466001:104 | -;-;-;-;-;-;-;- | 2091 | rac1a | 2241 | 3612 | 2252 | 2100 | 1987 | 1968 |
| ENSDARG000 | 12:12:12:12:1 | 8954761:8956 | 8955379:8956 | -:-:-:-:-:-:-:-: | 1879 | cyp26a1 | 0 | 0 | 6 | 12 | 4 | 12 |
| ENSDARG000 | 12:12:12:12:1 | 16938807:169 | 16939149:169 | -;-;-;-;-;-;- | 1289 | zgc:174855 | 2 | 0 | 6 | 2 | 0 | 5 |
| ENSDARG000 | 12:12:12:12:1 | 4911907:4913 | 4912018:4913 | -;-;-;-;- | 553 | ifnphi4 | 0 | 0 | 0 | 0 | 0 | 0 |
| ENSDARG000 | 12:12:12:12:1 | 16949391:169 | 16949415:169 | +:+:+:+:+:+:+: | 1237 | zgc:174153 | 20 | 14 | 0 | 132 | 0 | 10 |
| ENSDARG000 | 12;12;12;12 | 34429702:344 | 34430434:344 | -;-;-;- | 1471 | birc5a | 232 | 461 | 488 | 199 | 143 | 214 |
| ENSDARG000 | 12;12;12;12 | 34416175:344 | 34416756:344 | -;-;-;- | 1279 | TMEM235 | 402 | 362 | 718 | 14 | 130 | 216 |
| ENSDARG000 | 12:12:12:12:1 | 16953415:169 | 16953935:169 | +;+;+;+;+;+;+ | 2959 | pank1b | 1162 | 1060 | 939 | 290 | 1212 | 552 |
| ENSDARG000 | 12:12:12:12:1 | 34438277:344 | 34439459:344 | -;-;-;-;- | 3389 | fscn2b | 4 | 4 | 2 | 0 | 0 | 4 |
| ENSDARG000 | 12:12:12:12:1 | 2906610:2907 | 2907002:2907 | -:-:-:-:-:-:-:-: | 1781 | fam20a | 4 | 4 | 0 | 0 | 6 | 8 |
| ENSDARG000 | 12:12:12:12:1 | 34481728:345 | 34481911:345 | -;-;-;-;- | 566 | NA | 0 | 0 | 0 | 0 | 0 | 0 |
| ENSDARG000 | 12:12:12:12:1 | 26467847:264 | 26467942:264 | +:+:+:+:+:+:+: | 2025 | ndel1a | 112 | 108 | 142 | 250 | 65 | 222 |
| ENSDARG000 | 12:12:12:12:1 | 4920451:4920 | 4920567:4920 | +:+:+:+:+:+:+: | 6274 | plekhm1 | 2189 | 3248 | 3897 | 5156 | 8673 | 6402 |
| ENSDARG000 | 12:12:12:12:1 | 16967715:169 | 16967918:169 | +;+;+;+;+;+;+ | 2322 | slc16a12b | 22 | 1 | 25 | 0 | 44 | 3 |
| ENSDARG000 | 12:12:12:12:1 | 28574863:286 | 28575065:286 | +:+:+:+:+:+:+: | 4324 | TBKBP1 | 0 | 6 | 0 | 0 | 4 | 66 |
| ENSDARG000 | 12:12:12:12:1 | 28749189:287 | 28749757:287 | +;+;+;+;+;+ | 2280 | tbx21 | 24 | 16 | 3 | 4 | 13 | 34 |
| ENSDARG000 | 12;12;12;12 | 37401331:374 | 37401390:374 | +;+;+;+ | 1598 | NA | 4 | 0 | 0 | 0 | 0 | 3 |
| ENSDARG000 | 12;12 | 37407105:374 | 37411172:374 | -;- | 4334 | CDC42EP4 | 2556 | 1584 | 1951 | 1433 | 1642 | 1506 |
| ENSDARG000 | 12:12:12:12:1 | 30367079:303 | 30367489:303 | +:+:+:+:+:+:+: | 7807 | CCDC186 | 334 | 249 | 286 | 224 | 256 | 312 |
| ENSDARG000 | 12:12:12:12:1 | 20227395:202 | 20227849:202 | -:-:-:-:-:-:-:-: | 2897 | NA | 205 | 199 | 641 | 532 | 814 | 919 |
| ENSDARG000 | 12;12 | 16989120:169 | 16990511:169 | -;- | 1440 | ifit12 | 136 | 114 | 189 | 247 | 153 | 175 |
| ENSDARG000 | 12 | 16997206 | 16998442 | - | 1237 | NA | 14 | 65 | 2 | 2 | 18 | 26 |
| ENSDARG000 | 12;12;12 | 20349865:203 | 20350063:203 | -;-;- | 590 | hbbe2 | 0 | 0 | 0 | 0 | 0 | 2 |

| | | | | | | | | | | | | |
|---|---|---|---|---|---|---|---|---|---|---|---|---|
| ENSDARG000 | 12;12;12;12;1 | 2951186:2953 | 2951532:2953 | -;-;-;-;-;-;-;-; | 3300 | NA | 0 | 2 | 0 | 0 | 0 | 4 |
| ENSDARG000 | 12;12;12;12;1 | 20342451:203 | 20342504:203 | +;+;+;+;+;+;+;+; | 1262 | hbae5 | 0 | 2 | 0 | 0 | 4 | 0 |
| ENSDARG000 | 12;12;12;12;1 | 8968275:8976 | 8969799:8976 | -;-;-;-;-;-;-;-; | 6519 | exoc6 | 430 | 482 | 663 | 329 | 860 | 525 |
| ENSDARG000 | 12;12;12;12;1 | 34561304:345 | 34565533:345 | -;-;-;-;-;-;-;-; | 12727 | BAHCC1 | 1711 | 985 | 1121 | 942 | 1063 | 2195 |
| ENSDARG000 | 12;12 | 26489370:264 | 26490011:264 | -;- | 867 | NA | 54 | 42 | 102 | 66 | 112 | 64 |
| ENSDARG000 | 12 | 34732262 | 34733438 | + | 1177 | NA | 24 | 50 | 94 | 24 | 44 | 40 |
| ENSDARG000 | 12;12;12;12;1 | 20354691:203 | 20355966:203 | -;-;-;-;-;-;-;- | 1892 | aqp8a.2 | 4 | 0 | 0 | 0 | 4 | 0 |
| ENSDARG000 | 12;12;12 | 20339413:203 | 20339610:203 | -;-;- | 525 | hbbe3 | 0 | 0 | 0 | 0 | 0 | 0 |
| ENSDARG000 | 12;12;12 | 20336070:203 | 20336404:203 | +;+;+ | 760 | hbae4 | 0 | 0 | 14 | 7 | 28 | 0 |
| ENSDARG000 | 12;12;12;12;1 | 28773589:287 | 28774524:287 | -;-;-;-;-;-;-;-; | 3888 | osbpl7 | 86 | 2 | 2 | 5 | 16 | 44 |
| ENSDARG000 | 12;12;12;12;1 | 10478666:104 | 10479048:104 | -;-;-;-;-;-;-;-; | 3282 | NA | 0 | 0 | 0 | 2 | 6 | 0 |
| ENSDARG000 | 12;12;12;12 | 26491637:264 | 26491786:264 | +;+;+;+ | 1580 | armc7 | 48 | 48 | 87 | 27 | 66 | 46 |
| ENSDARG000 | 12;12;12;12;1 | 30429833:304 | 30429873:304 | +;+;+;+;+;+;+;+; | 940 | NA | 0 | 0 | 0 | 0 | 0 | 0 |
| ENSDARG000 | 12;12 | 17019905:170 | 17022210:170 | -;- | 2412 | ifit11 | 250 | 276 | 122 | 102 | 317 | 365 |
| ENSDARG000 | 12;12;12;12 | 26512569:265 | 26513427:265 | +;+;+;+ | 2600 | chad | 60 | 4 | 0 | 0 | 10 | 12 |
| ENSDARG000 | 12;12 | 17025726:170 | 17027571:170 | -;- | 1979 | ifit10 | 361 | 1401 | 74 | 14 | 126 | 158 |
| ENSDARG000 | 12;12;12;12;1 | 26497167:264 | 26497824:264 | -;-;-;-;-;-;-;-; | 3405 | acsf2 | 33 | 37 | 52 | 12 | 42 | 122 |
| ENSDARG000 | 12;12 | 17033494:170 | 17034955:170 | -;- | 1546 | ifit9 | 112 | 86 | 62 | 36 | 80 | 37 |
| ENSDARG000 | 12;12;12;12;1 | 4971515:4984 | 4972022:4985 | +;+;+;+;+;+;+;+; | 4401 | NA | 355 | 214 | 161 | 580 | 796 | 530 |
| ENSDARG000 | 12;12;12;12;1 | 2993523:2995 | 2993594:2996 | +;+;+;+;+;+;+;+; | 2639 | LRRC45 | 485 | 440 | 422 | 319 | 338 | 539 |
| ENSDARG000 | 12 | 17042754 | 17043874 | + | 1121 | ch25h | 182 | 94 | 361 | 362 | 336 | 230 |
| ENSDARG000 | 12;12;12;12;1 | 13344896:133 | 13345050:133 | +;+;+;+;+;+;+;+; | 5388 | drosha | 484 | 530 | 676 | 197 | 768 | 336 |
| ENSDARG000 | 12;12;12;12;1 | 18663154:186 | 18663214:186 | +;+;+;+;+;+;+ | 2206 | NA | 0 | 0 | 0 | 0 | 0 | 0 |
| ENSDARG000 | 12;12;12;12;1 | 17100021:171 | 17100522:171 | +;+;+;+;+;+;+;+; | 2794 | acta2 | 36 | 30 | 8 | 2 | 14 | 8 |
| ENSDARG000 | 12;12;12;12;1 | 22823219:228 | 22823333:228 | +;+;+;+;+;+;+;+; | 3573 | AFAP1 | 275 | 246 | 285 | 42 | 164 | 97 |
| ENSDARG000 | 12;12;12;12;1 | 17117405:171 | 17118988:171 | -;-;-;-;-;-;-;-; | 3851 | STAMBPL1 | 48 | 118 | 59 | 18 | 96 | 64 |
| ENSDARG000 | 12;12;12;12;1 | 37463513:374 | 37464013:374 | -;-;-;-;-;-;-;-; | 6802 | SDK2 | 82 | 84 | 354 | 88 | 573 | 58 |
| ENSDARG000 | 12;12;12;12;1 | 34763795:347 | 34763993:347 | +;+;+;+;+;+;+;+; | 5870 | SLC38A10 | 2281 | 1718 | 1829 | 1649 | 2221 | 2431 |
| ENSDARG000 | 12;12;12;12;1 | 5048044:5048 | 5048201:5048 | +;+;+;+;+;+;+;+; | 2317 | kif22 | 337 | 969 | 802 | 436 | 588 | 532 |
| ENSDARG000 | 12;12;12;12;1 | 3022882:3026 | 3023021:3026 | +;+;+;+;+;+;+ | 4241 | rac3b | 3284 | 3367 | 2006 | 3860 | 2151 | 4598 |
| ENSDARG000 | 12;12;12;12;1 | 30435416:304 | 30435711:304 | -;-;-;-;-;-;-;-; | 2534 | NA | 0 | 0 | 0 | 0 | 2 | 0 |
| ENSDARG000 | 12;12;12;12;1 | 5081759:5084 | 5082028:5084 | +;+;+;+;+;+ | 1952 | NA | 66 | 56 | 136 | 14 | 36 | 32 |
| ENSDARG000 | 12;12;12;12;1 | 18681423:186 | 18681598:186 | +;+;+;+;+;+;+;+; | 4909 | rgs9b | 0 | 0 | 0 | 0 | 6 | 0 |
| ENSDARG000 | 12;12;12;12;1 | 13404784:134 | 13405520:134 | +;+;+;+;+;+;+;+; | 8034 | KCNH4 | 2 | 4 | 2 | 16 | 0 | 0 |
| ENSDARG000 | 12;12;12;12;1 | 17154655:171 | 17154929:171 | +;+;+;+;+;+;+ | 1106 | ankrd22 | 27 | 16 | 30 | 46 | 44 | 33 |
| ENSDARG000 | 12;12;12;12 | 32159272:321 | 32159519:321 | +;+;+;+ | 1039 | hlfb | 30 | 123 | 79 | 18 | 88 | 15 |
| ENSDARG000 | 12;12;12;12;1 | 20366109:203 | 20366364:203 | -;-;-;-;- | 1029 | aqp8a.1 | 652 | 62 | 32 | 61 | 96 | 132 |
| ENSDARG000 | 12;12;12;12;1 | 28796622:287 | 28796930:287 | -;-;-;-;-;-;- | 1488 | mrpl10 | 218 | 252 | 233 | 130 | 292 | 184 |
| ENSDARG000 | 12;12;12;12 | 32179211:321 | 32179587:321 | -;-;-;- | 884 | SMIM36 | 0 | 0 | 2 | 0 | 0 | 0 |
| ENSDARG000 | 12;12;12 | 34825062:348 | 34825225:348 | -;-;- | 532 | ndufaf8 | 34 | 40 | 52 | 28 | 52 | 46 |
| ENSDARG000 | 12;12;12;12;1 | 3043308:3045 | 3043672:3045 | -;-;-;-;-;-;-;-; | 1548 | dcxr | 208 | 272 | 191 | 300 | 236 | 293 |
| ENSDARG000 | 12;12;12;12;1 | 5102670:5102 | 5102780:5103 | +;+;+;+;+;+;+;+; | 2200 | cep55l | 153 | 252 | 270 | 202 | 225 | 120 |
| ENSDARG000 | 12;12;12;12;1 | 281166:28154 | 281402:28159 | -;-;-;-;-;-;-;-; | 1578 | map2k4b | 1219 | 1071 | 1708 | 1487 | 1250 | 1112 |
| ENSDARG000 | 12;12;12;12;1 | 32186095:321 | 32186573:321 | -;-;-;-;- | 621 | NA | 0 | 0 | 0 | 0 | 0 | 0 |
| ENSDARG000 | 12;12;12;12;1 | 17158642:171 | 17159405:171 | -;-;-;-;-;-;-;-; | 1839 | lipf | 736 | 564 | 948 | 737 | 1201 | 721 |
| ENSDARG000 | 12;12;12;12;1 | 45661810:456 | 45662668:456 | -;-;-;-;- | 1903 | fbxl15 | 162 | 108 | 96 | 60 | 186 | 114 |
| ENSDARG000 | 12;12;12;12 | 32199651:322 | 32199756:322 | +;+;+;+ | 609 | NA | 0 | 4 | 6 | 2 | 14 | 4 |
| ENSDARG000 | 12;12;12;12;1 | 13527422:135 | 13528279:135 | -;-;-;-;-;-;-;-; | 2531 | ghdc | 276 | 256 | 242 | 148 | 329 | 212 |
| ENSDARG000 | 12;12;12;12;1 | 32292564:323 | 32292741:323 | +;+;+;+;+;+;+;+; | 5373 | ANKFN1 | 13 | 23 | 41 | 61 | 86 | 84 |
| ENSDARG000 | 12;12;12;12;1 | 9096963:9099 | 9097961:9099 | -;-;-;-;-;-;-;-; | 3768 | NA | 550 | 666 | 873 | 643 | 536 | 414 |
| ENSDARG000 | 12;12;12;12;1 | 45676667:456 | 45677470:456 | +;+;+;+;+;+;+;+; | 2695 | gpx9 | 0 | 0 | 0 | 0 | 0 | 0 |
| ENSDARG000 | 12;12;12;12;1 | 34827808:348 | 34827958:348 | +;+;+;+;+;+;+;+; | 2113 | TEPSIN | 136 | 122 | 199 | 191 | 198 | 159 |
| ENSDARG000 | 12 | 45748117 | 45748861 | - | 745 | NA | 0 | 0 | 0 | 0 | 0 | 0 |
| ENSDARG000 | 12;12;12;12;1 | 45799982:458 | 45801077:458 | -;-;-;-;-;-;-;-; | 2650 | pax2b | 0 | 0 | 0 | 0 | 0 | 0 |
| ENSDARG000 | 12;12;12;12;1 | 28799988:288 | 28800203:288 | +;+;+;+;+;+;+;+ | 2155 | pnpo | 228 | 234 | 322 | 372 | 264 | 209 |
| ENSDARG000 | 12;12;12;12;1 | 22921477:229 | 22921664:229 | +;+;+;+;+;+;+;+; | 2350 | rab18a | 761 | 517 | 622 | 533 | 613 | 601 |
| ENSDARG000 | 12;12;12;12;1 | 26546820:265 | 26547497:265 | -;-;-;-;-;-;-;-; | 2846 | NA | 639 | 600 | 648 | 301 | 462 | 408 |
| ENSDARG000 | 12;12;12;12;1 | 20378376:203 | 20379663:203 | -;-;-;-;-;-;-;-; | 2767 | lcmt1 | 326 | 446 | 462 | 253 | 370 | 330 |
| ENSDARG000 | 12 | 32388860 | 32390444 | + | 1585 | nog3 | 34 | 0 | 0 | 0 | 0 | 8 |
| ENSDARG000 | 12;12;12;12;1 | 26538861:265 | 26539174:265 | +;+;+;+;+;+;+;+; | 2913 | NA | 79 | 128 | 163 | 69 | 107 | 69 |
| ENSDARG000 | 12;12;12;12 | 34844927:348 | 34848071:348 | -;-;-;- | 3506 | si:dkev-21c1.1 | 420 | 360 | 458 | 346 | 500 | 436 |
| ENSDARG000 | 12;12;12;12;1 | 5115929:5116 | 5116221:5116 | -;-;-;-;-;-;-;-; | 1574 | rbp4 | 182 | 17 | 34 | 38 | 160 | 98 |
| ENSDARG000 | 12;12;12;12;1 | 288108:28876 | 288948:28894 | -;-;-;-;-;-;-;- | 1865 | adprm | 156 | 308 | 346 | 331 | 181 | 88 |
| ENSDARG000 | 12;12;12;12;1 | 17201522:172 | 17201743:172 | +;+;+;+;+;+;+;+; | 2038 | rnls | 102 | 161 | 202 | 106 | 100 | 178 |
| ENSDARG000 | 12 | 30442230 | 30443562 | - | 1333 | adrb1 | 4 | 21 | 34 | 0 | 52 | 8 |
| ENSDARG000 | 12;12;12;12;1 | 3054442:3058 | 3054803:3061 | -;-;-;-;-;-;-;-; | 4124 | rfng | 1074 | 630 | 269 | 877 | 493 | 1174 |
| ENSDARG000 | 12;12;12;12;1 | 30517386:305 | 30517746:305 | -;-;-;-;-;-;-;-; | 1514 | plekhs1.1 | 4 | 2 | 0 | 0 | 4 | 8 |
| ENSDARG000 | 12;12;12;12;1 | 30458409:304 | 30461513:304 | -;-;-;-;-;-;-;-; | 5008 | NHLRC2 | 751 | 1054 | 864 | 423 | 829 | 573 |
| ENSDARG000 | 12;12;12;12;1 | 30528623:305 | 30531240:305 | -;-;-;-;-;-;-;-; | 3928 | plekhs1.2 | 2 | 0 | 0 | 0 | 0 | 0 |
| ENSDARG000 | 12;12;12;12;1 | 5129245:5133 | 5129823:5133 | +;+;+;+;+;+;+;+; | 4527 | pde6c | 12 | 0 | 4 | 18 | 8 | 2 |
| ENSDARG000 | 12;12;12;12;1 | 30501044:305 | 30501637:305 | +;+;+;+;+;+;+;+; | 3316 | dclre1a | 60 | 183 | 177 | 66 | 102 | 103 |
| ENSDARG000 | 12;12;12;12;1 | 13560082:135 | 13560152:135 | -;-;-;-;-;-;-;-; | 3775 | stat5b | 604 | 508 | 794 | 400 | 738 | 657 |
| ENSDARG000 | 12;12;12;12 | 32402084:324 | 32402192:324 | +;+;+;+ | 542 | NA | 0 | 0 | 0 | 0 | 0 | 0 |
| ENSDARG000 | 12;12;12;12;1 | 32413524:324 | 32413851:324 | -;-;-;-;-;- | 1803 | enpp7.1 | 0 | 0 | 0 | 0 | 0 | 0 |
| ENSDARG000 | 12;12;12;12;1 | 34854562:348 | 34854816:348 | +;+;+;+;+;+ | 1850 | NA | 70 | 50 | 80 | 52 | 83 | 72 |
| ENSDARG000 | 12;12;12;12;1 | 3078221:3078 | 3078267:3078 | +;+;+;+;+;+;+;+; | 1779 | NA | 2 | 6 | 0 | 30 | 3 | 0 |
| ENSDARG000 | 12;12;12;12;1 | 46231498:462 | 46231895:462 | -;-;-;-;-;-;-;-; | 3705 | NA | 10 | 2 | 0 | 0 | 0 | 0 |
| ENSDARG000 | 12;12;12;12;1 | 10517473:105 | 10517921:105 | -;-;-;-;-;-;-;-; | 6668 | myof | 664 | 186 | 2483 | 156 | 715 | 1704 |
| ENSDARG000 | 12;12;12;12;1 | 46264190:462 | 46264296:462 | -;-;-;-;-;-;-;-; | 3328 | NA | 12 | 0 | 0 | 3 | 0 | 0 |
| ENSDARG000 | 12;12;12;12;1 | 46188692:461 | 46188796:461 | -;-;-;-;-;-;-;-; | 5064 | NA | 1 | 2 | 0 | 0 | 12 | 0 |
| ENSDARG000 | 12;12;12;12;1 | 46152983:461 | 46153303:461 | -;-;-;-;-;-;-;-; | 4827 | NA | 4 | 0 | 0 | 0 | 2 | 0 |
| ENSDARG000 | 12;12;12;12;1 | 46209859:462 | 46210410:462 | -;-;-;-;-;-;-;-; | 3755 | NA | 5 | 0 | 0 | 4 | 0 | 0 |
| ENSDARG000 | 12;12;12;12;1 | 18718065:187 | 18718229:187 | +;+;+;+;+;+;+;+; | 4958 | mrtfab | 912 | 841 | 900 | 607 | 1312 | 1024 |
| ENSDARG000 | 12;12;12;12;1 | 34876125:348 | 34877222:348 | -;-;-;-;-;-;-;-; | 3520 | BICRAL | 302 | 345 | 421 | 380 | 446 | 420 |
| ENSDARG000 | 12;12;12;12;1 | 46283843:462 | 46283982:462 | -;-;-;-;-;-;-;-; | 3419 | NA | 7 | 1 | 1 | 3 | 0 | 0 |
| ENSDARG000 | 12;12;12;12;1 | 46120836:461 | 46120942:461 | -;-;-;-;-;-;-;-; | 3441 | zgc:153932 | 4 | 0 | 0 | 3 | 0 | 0 |
| ENSDARG000 | 12;12;12;12;1 | 32423375:324 | 32423613:324 | -;-;-;-;-;- | 1705 | enpp7.2 | 0 | 0 | 0 | 0 | 0 | 0 |
| ENSDARG000 | 12;12;12;12;1 | 28808645:288 | 28809310:288 | -;-;-;-;- | 1452 | prr15lb | 0 | 2 | 6 | 12 | 14 | 0 |
| ENSDARG000 | 12;12;12;12;1 | 26566422:265 | 26566832:265 | -;-;-;-;-;-;-;- | 2157 | SLC16A5 | 82 | 56 | 60 | 38 | 39 | 80 |
| ENSDARG000 | 12;12;12;12;1 | 32431703:324 | 32431805:324 | +;+;+;+;+;+;+;+; | 752 | NA | 0 | 0 | 0 | 0 | 0 | 0 |
| ENSDARG000 | 12;12;12;12;1 | 5156332:5157 | 5156740:5157 | -;-;-;-;-;-;-;-; | 1424 | fra10ac1 | 142 | 121 | 140 | 94 | 149 | 146 |
| ENSDARG000 | 12;12;12;12;1 | 22961894:229 | 22963777:229 | -;-;-;-;-;-;-;-; | 3852 | mkxa | 122 | 38 | 66 | 2 | 101 | 0 |
| ENSDARG000 | 12;12;12;12 | 20492884:204 | 20493835:204 | +;+;+;+ | 3819 | NA | 0 | 2 | 0 | 0 | 12 | 18 |
| ENSDARG000 | 12;12;12;12;1 | 9183626:9184 | 9183725:9184 | +;+;+;+;+;+;+;+; | 3003 | tubgcp2 | 394 | 500 | 490 | 378 | 494 | 310 |
| ENSDARG000 | 12;12;12;12;1 | 38546885:385 | 38547207:385 | -;-;-;-;- | 660 | NA | 6 | 7 | 7 | 16 | 20 | 17 |
| ENSDARG000 | 12;12 | 38545890:385 | 38546595:385 | +;+ | 1048 | NA | 12 | 9 | 4 | 11 | 10 | 4 |
| ENSDARG000 | 12;12;12;12;1 | 32729470:328 | 32729879:328 | +;+;+;+;+;+;+;+; | 7945 | RBFOX3 | 18 | 10 | 10 | 2 | 18 | 0 |
| ENSDARG000 | 12;12 | 26608883:266 | 26609112:266 | +;+ | 514 | NA | 0 | 0 | 0 | 0 | 0 | 0 |
| ENSDARG000 | 12;12;12;12;1 | 26611435:266 | 26611817:266 | -;-;-;-;-;-;-;-; | 1879 | hexdc | 110 | 49 | 72 | 107 | 82 | 81 |
| ENSDARG000 | 12;12;12;12;1 | 26594013:265 | 26594322:265 | -;-;-;-;-;- | 1337 | kctd2 | 44 | 36 | 84 | 35 | 54 | 47 |
| ENSDARG000 | 12;12;12;12;1 | 17399852:174 | 17401611:174 | -;-;-;-;-;-;-;-; | 3461 | ptenb | 913 | 809 | 1528 | 695 | 1388 | 787 |
| ENSDARG000 | 12;12 | 10599604:106 | 10599747:106 | +;+ | 1282 | NA | 0 | 0 | 0 | 0 | 0 | 0 |
| ENSDARG000 | 12;12;12;12;1 | 20506197:205 | 20506323:205 | +;+;+;+;+;+;+;+ | 1342 | NA | 2 | 0 | 0 | 0 | 0 | 12 |
| ENSDARG000 | 12;12;12;12;1 | 28825735:288 | 28825933:288 | +;+;+;+;+;+;+;+; | 4644 | cdk5rap3 | 424 | 522 | 979 | 395 | 576 | 460 |
| ENSDARG000 | 12;12;12;12 | 20552190:205 | 20552478:205 | +;+;+;+ | 1669 | gna13a | 431 | 542 | 645 | 404 | 554 | 700 |
| ENSDARG000 | 12;12;12;12;1 | 5189776:5190 | 5189970:5190 | +;+;+;+;+;+;+;+; | 2070 | lgi1b | 2 | 8 | 4 | 2 | 0 | 6 |
| ENSDARG000 | 12;12;12;12;1 | 9250107:9250 | 9250287:9250 | +;+;+;+;+;+;+;+ | 1194 | NA | 0 | 0 | 0 | 6 | 0 | 0 |
| ENSDARG000 | 12 | 34891529 | 34892823 | + | 1295 | tbcc | 214 | 258 | 302 | 201 | 188 | 202 |
| ENSDARG000 | 12;12;12;12;1 | 38556462:385 | 38556515:385 | +;+;+;+;+;+;+;+; | 1060 | rpl38 | 3035 | 2819 | 3120 | 2344 | 1657 | 2618 |
| ENSDARG000 | 12;12;12 | 34896956:348 | 34897843:348 | +;+;+ | 2484 | prph2a | 8 | 0 | 8 | 10 | 0 | 0 |
| ENSDARG000 | 12;12;12;12;1 | 23026760:230 | 23027150:230 | -;-;-;-;-;-;-;-; | 3622 | ODAD2 | 1 | 0 | 0 | 2 | 1 | 0 |
| ENSDARG000 | 12;12;12;12;1 | 20583552:205 | 20583605:205 | +;+;+;+;+;+;+;+; | 2310 | arsg | 16 | 16 | 24 | 20 | 4 | 55 |
| ENSDARG000 | 12;12;12;12;1 | 20575640:205 | 20577403:205 | -;-;-;-;-;-;-;-; | 3670 | SLC16A6 | 0 | 0 | 0 | 0 | 0 | 0 |
| ENSDARG000 | 12;12;12;12;1 | 34953038:349 | 34953856:349 | +;+;+;+;+;+;+;+; | 2664 | qki2 | 1175 | 929 | 1631 | 874 | 1212 | 1303 |
| ENSDARG000 | 12;12;12;12 | 33298838:333 | 33299778:333 | -;-;-;- | 1362 | c1qtnf1 | 2 | 6 | 0 | 4 | 0 | 6 |
| ENSDARG000 | 12;12;12;12;1 | 28839839:288 | 28840491:288 | -;-;-;-;-;-;- | 1759 | NA | 419 | 386 | 533 | 298 | 341 | 549 |
| ENSDARG000 | 12;12;12;12;1 | 3095699:3098 | 3096984:3099 | -;-;-;-;-;-;-;-; | 5610 | col1a1b | 3697 | 984 | 1781 | 670 | 8756 | 535 |
| ENSDARG000 | 12;12;12;12;1 | 10706772:107 | 10707002:107 | +;+;+;+;+;+;+;+; | 5539 | top2a | 1514 | 4298 | 4318 | 1701 | 3403 | 1676 |
| ENSDARG000 | 12;12;12;12;1 | 33320504:333 | 33320984:333 | +;+;+;+;+;+;+;+; | 3081 | csnk1db | 3464 | 3428 | 4843 | 4389 | 5228 | 3630 |
| ENSDARG000 | 12;12;12;12;1 | 30540911:305 | 30543474:305 | -;-;-;-;-;-;-;-; | 4487 | plekhs1.3 | 30 | 0 | 0 | 0 | 0 | 0 |
| ENSDARG000 | 12;12;12;12;1 | 17436904:174 | 17437057:174 | +;+;+;+;+;+;+;+; | 1289 | atad1b | 491 | 638 | 583 | 418 | 711 | 544 |
| ENSDARG000 | 12;12;12;12;1 | 18792315:187 | 18796204:187 | -;-;-;-;-;-;-;-; | 5622 | NA | 10 | 6 | 2 | 0 | 0 | 0 |
| ENSDARG000 | 12;12;12;12;1 | 5209822:5215 | 5210275:5215 | +;+;+;+;+;+;+ | 3971 | SLC35G1 | 34 | 22 | 22 | 10 | 34 | 18 |
| ENSDARG000 | 12;12;12;12;1 | 18895657:188 | 18896321:188 | -;-;-;-;-;-;-;-; | 1358 | desi1b | 615 | 408 | 260 | 192 | 486 | 226 |
| ENSDARG000 | 12;12;12;12;1 | 33350256:333 | 33352042:333 | -;-;-;-;-;-;-;-; | 3126 | slc16a3b | 58 | 631 | 1865 | 360 | 178 | 54 |
| ENSDARG000 | 12;12;12;12;1 | 13652361:136 | 13652558:136 | +;+;+;+;+;+;+;+; | 4865 | oplah | 162 | 162 | 222 | 103 | 294 | 159 |
| ENSDARG000 | 12;12;12;12;1 | 5251647:5307 | 5252738:5307 | +;+;+;+;+;+;+;+; | 7267 | plce1 | 79 | 0 | 4 | 0 | 12 | 28 |
| ENSDARG000 | 12;12;12;12;1 | 30550721:305 | 30550834:305 | -;-;-;-;-;-;-;- | 776 | NA | 0 | 0 | 0 | 0 | 0 | 0 |
| ENSDARG000 | 12;12;12;12;1 | 28854410:288 | 28854620:288 | +;+;+;+;+;+;+;+; | 5677 | nfe2l1b | 2623 | 2976 | 2235 | 2345 | 3959 | 2901 |
| ENSDARG000 | 12;12;12;12;1 | 17458949:174 | 17459849:174 | -;-;-;-;-;-;-;-; | 2674 | papss2b | 1166 | 1140 | 1072 | 893 | 1256 | 763 |
| ENSDARG000 | 12;12;12;12;1 | 9257457:9258 | 9257993:9258 | -;-;-;-;-;-;-;-; | 2143 | pth3r | 0 | 0 | 0 | 0 | 0 | 0 |
| ENSDARG000 | 12;12;12;12;1 | 30563550:305 | 30563828:305 | +;+;+;+;+;+;+ | 1350 | NA | 4 | 0 | 0 | 0 | 0 | 0 |
| ENSDARG000 | 12;12;12;12;1 | 30569621:305 | 30571299:305 | -;-;-;-;-;-;-;-; | 2801 | casp7 | 524 | 717 | 558 | 451 | 342 | 464 |
| ENSDARG000 | 12;12;12;12;1 | 38563073:385 | 38563501:385 | +;+;+;+;+;+;+;+; | 3415 | ttyh2 | 206 | 201 | 263 | 159 | 202 | 214 |
| ENSDARG000 | 12;12;12;12;1 | 26621906:266 | 26622243:266 | +;+;+;+;+;+;+;+; | 5223 | arhgap12b | 49 | 74 | 277 | 34 | 106 | 485 |
| ENSDARG000 | 12;12;12;12;1 | 23175962:231 | 23178577:231 | -;-;-;-;-;-;-;-; | 4365 | mpp7a | 50 | 34 | 12 | 2 | 12 | 38 |
| ENSDARG000 | 12;12;12;12;1 | 9305832:9307 | 9306350:9308 | +;+;+;+;+;+ | 1325 | taco1 | 136 | 266 | 347 | 194 | 314 | 280 |
| ENSDARG000 | 12;12;12;12;1 | 18899396:188 | 18899666:188 | +;+;+;+;+;+;+;+; | 2228 | xrcc6 | 619 | 749 | 867 | 481 | 888 | 568 |
| ENSDARG000 | 12;12;12;12;1 | 35046704:350 | 35047109:350 | +;+;+;+;+;+;+;+; | 1394 | timm23a | 420 | 591 | 780 | 352 | 611 | 481 |
| ENSDARG000 | 12;12;12;12;1 | 17482607:174 | 17483286:174 | -;-;-;-;-;-;-;-; | 1811 | minpp1b | 114 | 94 | 138 | 80 | 152 | 128 |
| ENSDARG000 | 12;12;12;12;1 | 28870462:288 | 28871541:288 | -;-;-;-;-;-;-;-; | 2267 | cbx1b | 535 | 643 | 657 | 512 | 598 | 631 |
| ENSDARG000 | 12;12;12;12;1 | 30586599:305 | 30586929:305 | +;+;+;+;+;+;+;+; | 7232 | nrap | 0 | 2 | 0 | 0 | 2 | 0 |
| ENSDARG000 | 12;12;12;12;1 | 20605191:206 | 20608568:206 | -;-;-;-;-;-;- | 4211 | snx11 | 202 | 217 | 184 | 145 | 272 | 254 |
| ENSDARG000 | 12;12;12;12;1 | 33452915:334 | 33453889:334 | -;-;-;-;-;-;- | 1607 | cybc1 | 216 | 188 | 306 | 266 | 240 | 250 |
| ENSDARG000 | 12;12;12;12;1 | 13722385:137 | 13726999:137 | -;-;-;-;-;-;- | 5377 | foxh1 | 6 | 0 | 5 | 10 | 38 | 2 |
| ENSDARG000 | 12;12;12;12;1 | 33460794:334 | 33461059:334 | +;+;+;+;+;+;+;+; | 2316 | narf | 252 | 278 | 483 | 477 | 381 | 201 |
| ENSDARG000 | 12;12;12;12;1 | 13742778:137 | 13743116:137 | +;+;+;+;+;+;+;+; | 4177 | PPP1R16A | 31 | 4 | 28 | 14 | 184 | 6 |
| ENSDARG000 | 12;12;12;12;1 | 35051636:350 | 35052758:350 | -;-;-;-;-;-;-;-; | 2189 | zgc:112285 | 0 | 1 | 2 | 1 | 4 | 0 |
| ENSDARG000 | 12;12;12;12;1 | 5399680:5399 | 5399862:5399 | -;-;-;-;-;-;-;-; | 3002 | noc3l | 676 | 736 | 1284 | 574 | 836 | 512 |
| ENSDARG000 | 12;12;12;12;1 | 10952794:110 | 10952846:110 | +;+;+;+;+;+;+;+; | 3029 | raraa | 170 | 104 | 120 | 168 | 264 | 123 |
| ENSDARG000 | 12;12;12;12;1 | 35067863:350 | 35068043:350 | +;+;+;+;+;+;+;+; | 1579 | SYT15 | 0 | 4 | 14 | 19 | 12 | 2 |
| ENSDARG000 | 12;12;12;12;1 | 33361948:333 | 33362351:333 | +;+;+;+;+;+;+;+; | 18655 | ccdc57 | 3547 | 3962 | 3545 | 1583 | 2709 | 3004 |
| ENSDARG000 | 12;12;12;12;1 | 23371542:233 | 23371815:233 | +;+;+;+;+;+;+;+; | 3062 | waca | 1453 | 1195 | 1482 | 1032 | 1281 | 1153 |
| ENSDARG000 | 12;12;12;12;1 | 9320446:9320 | 9320494:9320 | +;+;+;+;+;+;+;+; | 2299 | NA | 0 | 0 | 10 | 0 | 0 | 0 |
| ENSDARG000 | 12;12;12;12;1 | 18906407:189 | 18906546:189 | +;+;+;+;+;+;+;+; | 4059 | josd1 | 462 | 440 | 436 | 422 | 415 | 459 |
| ENSDARG000 | 12;12;12 | 23424108:234 | 23424520:234 | +;+;+ | 1569 | bambia | 124 | 34 | 22 | 34 | 14 | 52 |
| ENSDARG000 | 12;12;12;12;1 | 3714076:3716 | 3715950:3716 | -;-;-;-;-;- | 2328 | vps25 | 875 | 692 | 1182 | 756 | 1216 | 675 |
| ENSDARG000 | 12;12;12;12;1 | 38770654:387 | 38770765:387 | +;+;+;+;+;+;+;+; | 5711 | KIF19 | 6 | 0 | 0 | 0 | 0 | 6 |
| ENSDARG000 | 12;12;12;12;1 | 3723650:3723 | 3723840:3724 | +;+;+;+;+;+;+;+; | 1278 | pagr1 | 220 | 358 | 512 | 150 | 408 | 256 |
| ENSDARG000 | 12;12;12 | 30630073:306 | 30630133:306 | +;+;+ | 617 | NA | 0 | 0 | 0 | 0 | 0 | 0 |
| ENSDARG000 | 12;12;12;12;1 | 9360316:9360 | 9360743:9360 | -;-;-;-;-;-;-;-; | 4098 | erbb2 | 373 | 487 | 604 | 264 | 432 | 418 |
| ENSDARG000 | 12;12;12;12;1 | 13811631:138 | 13812577:138 | -;-;-;-;-;-;-;-; | 3479 | ADAM11 | 22 | 12 | 30 | 2 | 58 | 2 |

| | | | | | | | | | | | |
|---|---|---|---|---|---|---|---|---|---|---|---|
| ENSDARG000 | 12:12:12:12:1 | 35084809:350 | 35088103:350 | -;-;-;-;- | 3851 | si:dkev-21e13 | 0 | 3 | 2 | 12 | 0 | 8 |
| ENSDARG000 | 12:12:12:12:1 | 17504559:175 | 17504866:175 | +:+:+:+:+:+:+: | 3275 | CYTH3 | 54 | 46 | 20 | 22 | 134 | 68 |
| ENSDARG000 | 12:12:12:12:1 | 18916285:189 | 18916597:189 | +;+;+;+;+;+ | 1678 | CBX7 | 34 | 28 | 20 | 63 | 6 | 64 |
| ENSDARG000 | 12:12:12:12:1 | 20618336:206 | 20618427:206 | +:+:+:+:+:+:+: | 1911 | NA | 6 | 18 | 4 | 0 | 0 | 26 |
| ENSDARG000 | 12:12:12:12:1 | 11167870:111 | 11167907:111 | +:+:+:+:+:+:+: | 3098 | vps35l | 1521 | 1226 | 1757 | 969 | 1553 | 1146 |
| ENSDARG000 | 12:12:12:12:1 | 38807604:388 | 38807699:388 | +:+:+:+:+:+:+: | 6473 | abca5 | 494 | 387 | 965 | 529 | 612 | 517 |
| ENSDARG000 | 12:12:12:12:1 | 18927065:189 | 18931974:189 | -:-:-:-:-:-:-: | 10242 | EP300 | 1404 | 941 | 1513 | 950 | 1399 | 1203 |
| ENSDARG000 | 12:12:12:12:1 | 3740605:3740 | 3740997:3740 | -:-:-:-:-:-:-: | 1374 | TLCD3B | 122 | 33 | 127 | 36 | 109 | 149 |
| ENSDARG000 | 12:12:12:12:1 | 3766557:3767 | 3767403:3767 | -:-:-:-:-:-:-: | 2527 | NA | 282 | 195 | 466 | 226 | 317 | 314 |
| ENSDARG000 | 12;12;12;12 | 35098915:351 | 35099925:351 | -;-;-;- | 1914 | si:ch73-127m | 144 | 167 | 184 | 88 | 212 | 156 |
| ENSDARG000 | 12:12:12:12:1 | 30653047:306 | 30653359:306 | +:+:+:+:+:+:+: | 4897 | NA | 93 | 5 | 16 | 2 | 72 | 10 |
| ENSDARG000 | 12;12;12 | 35106575:351 | 35106615:351 | +;+;+ | 429 | NA | 0 | 0 | 0 | 0 | 0 | 0 |
| ENSDARG000 | 12:12:12:12:1 | 3789114:3789 | 3789935:3789 | -:-:-:-:-:-:-: | 3977 | NA | 261 | 300 | 427 | 218 | 482 | 279 |
| ENSDARG000 | 12:12:12:12:1 | 26750274:267 | 26752983:267 | -;-;-;-;-;-;- | 5299 | zeb1b | 4753 | 3604 | 5788 | 5121 | 6285 | 5223 |
| ENSDARG000 | 12;12 | 11244914:112 | 11247293:112 | -;- | 2644 | NA | 6 | 0 | 0 | 0 | 0 | 0 |
| ENSDARG000 | 12:12:12:12:1 | 13905286:139 | 13905498:139 | +:+:+:+:+:+:+: | 2282 | fkbp10b | 99 | 299 | 314 | 52 | 439 | 106 |
| ENSDARG000 | 12:12:12:12:1 | 23454729:234 | 23454807:234 | +;+;+;+;+ | 545 | NA | 0 | 0 | 0 | 0 | 0 | 0 |
| ENSDARG000 | 12:12:12:12:1 | 26894321:268 | 26894360:268 | +;+;+;+;+ | 840 | NA | 0 | 0 | 0 | 0 | 0 | 0 |
| ENSDARG000 | 12:12:12:12:1 | 26877381:269 | 26877447:269 | +;+;+;+;+ | 3166 | znf438 | 284 | 207 | 247 | 224 | 324 | 188 |
| ENSDARG000 | 12:12:12:12:1 | 13888670:138 | 13891515:138 | -:-:-:-:-:-:-: | 4029 | NA | 38 | 208 | 319 | 81 | 130 | 97 |
| ENSDARG000 | 12:12:12:12:1 | 23647546:236 | 23647975:236 | -;-;-;-;-;-;- | 1724 | MAP3K8 | 29 | 82 | 116 | 98 | 168 | 120 |
| ENSDARG000 | 12:12:12:12:1 | 28888975:288 | 28889165:288 | +:+:+:+:+:+:+: | 2373 | phkg2 | 223 | 241 | 180 | 108 | 134 | 280 |
| ENSDARG000 | 12:12:12:12:1 | 11262954:112 | 11263771:112 | -:-:-:-:-:-:-: | 1760 | get4 | 735 | 877 | 1082 | 762 | 967 | 834 |
| ENSDARG000 | 12:12:12:12:1 | 5424108:5424 | 5424808:5424 | -:-:-:-:-:-:-: | 3349 | NA | 288 | 242 | 330 | 195 | 382 | 142 |
| ENSDARG000 | 12:12:12:12:1 | 28899760:289 | 28900983:289 | -:-:-:-:-:-:-: | 1930 | ccdc189 | 0 | 1 | 2 | 0 | 0 | 0 |
| ENSDARG000 | 12:12:12:12:1 | 17570675:175 | 17571252:175 | -:-:-:-:-:-:-: | 5061 | NA | 330 | 392 | 470 | 281 | 468 | 330 |
| ENSDARG000 | 12:12:12:12:1 | 17591786:175 | 17593030:175 | -:-:-:-:-:-:-: | 3324 | eif2ak1 | 467 | 515 | 853 | 426 | 475 | 528 |
| ENSDARG000 | 12:12:12:12:1 | 23664335:236 | 23664548:236 | +:+:+:+:+:+:+: | 2356 | MTPAP | 130 | 166 | 154 | 66 | 127 | 155 |
| ENSDARG000 | 12:12:12:12:1 | 27021979:270 | 27022589:270 | +:+:+:+:+:+:+: | 5303 | NA | 577 | 532 | 646 | 476 | 719 | 585 |
| ENSDARG000 | 12:12:12:12:1 | 27027311:270 | 27028283:270 | -:-:-:-:-:-:-: | 1920 | chmp2a | 1029 | 909 | 763 | 768 | 730 | 785 |
| ENSDARG000 | 12:12:12:12:1 | 20627164:206 | 20627834:206 | +:+:+:+:+:+:+: | 4413 | stx4 | 621 | 927 | 1013 | 692 | 881 | 954 |
| ENSDARG000 | 12:12:12:12:1 | 387850:39005 | 387858:39039 | +:+:+:+:+:+:+: | 1367 | NA | 0 | 0 | 0 | 0 | 0 | 2 |
| ENSDARG000 | 12:12:12:12:1 | 20641102:206 | 20641176:206 | +:+:+:+:+:+:+: | 3338 | calcoco2 | 334 | 195 | 160 | 114 | 186 | 239 |
| ENSDARG000 | 12:12:12:12:1 | 23762966:237 | 23763256:237 | +:+:+:+:+:+:+: | 7745 | NA | 26 | 6 | 12 | 0 | 14 | 8 |
| ENSDARG000 | 12:12:12:12:1 | 35119762:351 | 35120006:351 | +:+:+:+:+:+:+: | 2693 | NA | 2 | 0 | 2 | 4 | 3 | 10 |
| ENSDARG000 | 12:12:12:12:1 | 20652957:206 | 20653327:206 | -;-;-;-;-;-;- | 1265 | snf8 | 248 | 231 | 342 | 257 | 276 | 263 |
| ENSDARG000 | 12;12 | 415711:43076 | 416422:43124 | -;- | 1198 | hs3st3l | 0 | 0 | 0 | 0 | 0 | 0 |
| ENSDARG000 | 12;12 | 410038:45590 | 410561:45651 | +;+ | 1139 | NA | 0 | 0 | 1 | 0 | 0 | 0 |
| ENSDARG000 | 12:12:12:12:1 | 455552:45619 | 456103:45638 | -;-;-;-;-;-;-;- | 1655 | dhrs7cb | 4 | 4 | 11 | 0 | 48 | 0 |
| ENSDARG000 | 12;12;12 | 5458462:5458 | 5459742:5459 | +;+;+ | 1300 | NA | 0 | 0 | 0 | 0 | 0 | 0 |
| ENSDARG000 | 12:12:12:12:1 | 5492804:5495 | 5493347:5495 | -;-;-;-;-;- | 1553 | NA | 24 | 4 | 4 | 0 | 6 | 2 |
| ENSDARG000 | 12:12:12:12:1 | 18970790:189 | 18970870:189 | +:+:+:+:+:+:+: | 5929 | l3mbtl2 | 248 | 288 | 343 | 138 | 406 | 222 |
| ENSDARG000 | 12:12:12:12:1 | 30705769:307 | 30706637:307 | +:+:+:+:+:+:+: | 1647 | kcnk1a | 4 | 0 | 0 | 0 | 8 | 0 |
| ENSDARG000 | 12:12:12:12:1 | 11315849:113 | 11316409:113 | -:-:-:-:-:-:-: | 2959 | zgc:174164 | 22 | 6 | 4 | 6 | 6 | 2 |
| ENSDARG000 | 12:12:12:12:1 | 18989486:189 | 18989771:189 | -;-;-;-;-;-;-;- | 2681 | chadlb | 0 | 8 | 0 | 0 | 0 | 0 |
| ENSDARG000 | 12:12:12:12:1 | 9449665:9449 | 9450158:9450 | -:-:-:-:-:-:-: | 1522 | pgap3 | 236 | 182 | 242 | 81 | 342 | 158 |
| ENSDARG000 | 12:12:12:12:1 | 35203091:352 | 35203276:352 | +:+:+:+:+:+:+: | 3960 | NA | 50 | 129 | 68 | 36 | 185 | 91 |
| ENSDARG000 | 12:12:12:12:1 | 9482875:9490 | 9483975:9490 | -;-;-;-;- | 1808 | si:ch211-207i | 59 | 84 | 74 | 60 | 77 | 46 |
| ENSDARG000 | 12:12:12:12:1 | 9497646:9497 | 9497784:9497 | +:+:+:+:+:+:+: | 1368 | dnajc9 | 143 | 436 | 579 | 233 | 471 | 303 |
| ENSDARG000 | 12:12:12:12:1 | 30723178:307 | 30723838:307 | +:+:+:+:+:+:+: | 3385 | slc35f3a | 0 | 0 | 0 | 0 | 0 | 0 |
| ENSDARG000 | 12:12:12:12:1 | 3848109:3848 | 3848180:3848 | +:+:+:+:+:+:+: | 1554 | cabp5a | 0 | 0 | 0 | 0 | 2 | 0 |
| ENSDARG000 | 12:12:12:12:1 | 30740797:307 | 30741764:307 | -:-:-:-:-:-:-: | 2659 | entpd1 | 86 | 113 | 113 | 101 | 464 | 385 |
| ENSDARG000 | 12:12:12:12:1 | 33484458:334 | 33485048:334 | +:+:+:+:+:+:+: | 2397 | nherf1a | 982 | 283 | 321 | 440 | 691 | 304 |
| ENSDARG000 | 12:12:12:12:1 | 38878830:388 | 38878876:388 | +:+:+:+:+:+:+: | 10411 | NA | 115 | 8 | 0 | 25 | 1 | 10 |
| ENSDARG000 | 12:12:12:12:1 | 5530247:5545 | 5530589:5545 | +:+:+:+:+:+:+: | 4260 | NA | 226 | 102 | 100 | 72 | 273 | 66 |
| ENSDARG000 | 12:12:12:12:1 | 11352630:113 | 11353092:113 | +:+:+:+:+:+:+: | 2583 | NA | 422 | 410 | 431 | 426 | 404 | 346 |
| ENSDARG000 | 12:12:12:12:1 | 17603528:176 | 17603808:176 | +:+:+:+:+:+:+: | 2940 | PMS2 | 89 | 209 | 274 | 116 | 197 | 162 |
| ENSDARG000 | 12;12 | 13961268:139 | 13963051:139 | -;- | 2350 | klhl11 | 230 | 228 | 278 | 279 | 163 | 217 |
| ENSDARG000 | 12:12:12:12:1 | 23812530:238 | 23812684:238 | +:+:+:+:+:+:+: | 10218 | svila | 973 | 1313 | 1348 | 3309 | 812 | 1424 |
| ENSDARG000 | 12:12:12:12:1 | 3871452:3875 | 3871516:3875 | +:+:+:+:+:+:+: | 1748 | nif3l1 | 202 | 192 | 366 | 166 | 252 | 169 |
| ENSDARG000 | 12:12:12:12:1 | 27034594:270 | 27035254:270 | +:+:+:+:+:+:+: | 5907 | NA | 1521 | 1142 | 1493 | 1185 | 1748 | 1431 |
| ENSDARG000 | 12:12:12:12:1 | 3893032:3893 | 3893169:3893 | -:-:-:-:-:-:-: | 1088 | NA | 8 | 4 | 18 | 0 | 2 | 8 |
| ENSDARG000 | 12:12:12:12:1 | 13973058:139 | 13974494:139 | -:-:-:-:-:-:-: | 4823 | ACLY | 101 | 32 | 114 | 117 | 101 | 134 |
| ENSDARG000 | 12:12:12:12:1 | 33541600:335 | 33541718:335 | -:-:-:-:-:-:-: | 2270 | MBTD1 | 316 | 176 | 284 | 122 | 271 | 172 |
| ENSDARG000 | 12:12:12:12:1 | 465701:46632 | 466446:46644 | +:+:+:+:+:+:+: | 4539 | NA | 1549 | 1258 | 1764 | 920 | 1645 | 1012 |
| ENSDARG000 | 12:12:12:12:1 | 19007926:190 | 19008632:190 | -:-:-:-:-:-:-: | 2514 | rangap1b | 84 | 149 | 170 | 22 | 64 | 78 |
| ENSDARG000 | 12:12:12:12:1 | 28910762:289 | 28910773:289 | +:+:+:+:+:+:+: | 3815 | RNF40 | 1762 | 1522 | 2355 | 1355 | 2068 | 1752 |
| ENSDARG000 | 12:12:12:12:1 | 27061720:270 | 27062019:270 | +:+:+:+:+:+:+: | 14407 | NA | 2443 | 2000 | 3183 | 1262 | 3229 | 1991 |
| ENSDARG000 | 12:12:12:12:1 | 17620067:176 | 17620381:176 | +:+:+:+:+:+:+: | 2636 | rsph10b | 0 | 18 | 10 | 4 | 8 | 2 |
| ENSDARG000 | 12:12:12:12:1 | 14079097:140 | 14079264:140 | +:+:+:+:+:+:+: | 2518 | GPT | 0 | 1 | 14 | 10 | 26 | 2 |
| ENSDARG000 | 12:12:12:12:1 | 35345830:353 | 35348467:353 | -:-:-:-:-:-:-: | 5112 | camk2g1 | 1564 | 1333 | 1873 | 1611 | 1633 | 1970 |
| ENSDARG000 | 12:12:12:12:1 | 17633354:176 | 17633844:176 | -:-:-:-:-:-:-: | 3267 | dlgap5 | 183 | 507 | 661 | 363 | 455 | 265 |
| ENSDARG000 | 12:12:12:12:1 | 30788912:307 | 30789022:307 | +:+:+:+:+:+:+: | 5654 | aldh18a1 | 980 | 1273 | 2567 | 1064 | 1624 | 1093 |
| ENSDARG000 | 12:12:12:12:1 | 20662426:206 | 20662791:206 | -;-;-;-;-;-;-;- | 1536 | gip | 0 | 0 | 0 | 0 | 0 | 0 |
| ENSDARG000 | 12;12;12;12 | 20667301:206 | 20667786:206 | +;+;+;+ | 2450 | NA | 162 | 125 | 153 | 138 | 164 | 274 |
| ENSDARG000 | 12:12:12:12:1 | 3912544:3917 | 3912749:3917 | +:+:+:+:+:+:+: | 3049 | tbx6 | 88 | 58 | 93 | 217 | 88 | 96 |
| ENSDARG000 | 12:12:12:12:1 | 14138340:141 | 14138504:141 | -;-;-;-;- | 3141 | NA | 0 | 0 | 2 | 6 | 0 | 0 |
| ENSDARG000 | 12:12:12:12:1 | 9510477:9510 | 9510925:9510 | -;-;-;-;-;-;- | 960 | NA | 57 | 74 | 62 | 18 | 66 | 62 |
| ENSDARG000 | 12;12;12 | 35510152:355 | 35510247:355 | -;-;- | 322 | NA | 0 | 0 | 0 | 0 | 0 | 0 |
| ENSDARG000 | 12:12:12:12:1 | 3926168:3929 | 3927499:3929 | -:-:-:-:-:-:-: | 2803 | zgc:92040 | 0 | 2 | 0 | 6 | 14 | 0 |
| ENSDARG000 | 12:12:12:12:1 | 19029285:190 | 19029509:190 | +:+:+:+:+:+:+: | 1716 | PLBD1 | 2 | 0 | 0 | 0 | 0 | 0 |
| ENSDARG000 | 12:12:12:12:1 | 11388724:113 | 11391111:113 | -;-;-;-;-;-;-;- | 3775 | htra1b | 584 | 317 | 347 | 182 | 183 | 360 |
| ENSDARG000 | 12:12:12:12:1 | 19036380:190 | 19036830:190 | +:+:+:+:+:+:+: | 4907 | NA | 292 | 269 | 311 | 158 | 374 | 381 |
| ENSDARG000 | 12:12:12:12:1 | 46543572:465 | 46543689:465 | +:+:+:+:+:+:+: | 2696 | hid1b | 0 | 0 | 0 | 0 | 4 | 4 |
| ENSDARG000 | 12:12:12:12:1 | 38929663:389 | 38929715:389 | +:+:+:+:+:+:+: | 16486 | NA | 409 | 6 | 6 | 53 | 69 | 52 |
| ENSDARG000 | 12:12:12:12:1 | 9519250:9519 | 9519453:9519 | -:-:-:-:-:-:-: | 1511 | nudt13 | 34 | 32 | 8 | 6 | 16 | 24 |
| ENSDARG000 | 12:12:12:12:1 | 20678187:206 | 20678316:206 | +:+:+:+:+:+:+: | 1256 | st6galnac1.1 | 44 | 11 | 11 | 24 | 6 | 17 |
| ENSDARG000 | 12:12:12:12:1 | 35543160:355 | 35545057:355 | -:-:-:-:-:-:-: | 6237 | sec24c | 1785 | 1493 | 2402 | 1241 | 2035 | 1269 |
| ENSDARG000 | 12:12:12:12:1 | 27096835:270 | 27096975:271 | +:+:+:+:+:+:+: | 3115 | ttll6 | 6 | 8 | 12 | 4 | 12 | 0 |
| ENSDARG000 | 12:12:12:12:1 | 46582168:465 | 46582902:465 | +;+;+;+;+;+;+ | 2038 | USH1G | 0 | 0 | 0 | 0 | 0 | 0 |
| ENSDARG000 | 12;12;12;12 | 28937092:289 | 28937147:289 | +;+;+;+ | 2100 | NA | 391 | 422 | 581 | 757 | 540 | 609 |
| ENSDARG000 | 12:12:12:12:1 | 46634736:466 | 46634992:466 | +;+;+;+;+;+;+ | 2230 | trim16 | 157 | 74 | 32 | 62 | 42 | 144 |
| ENSDARG000 | 12;12;12 | 28962590:289 | 28963476:289 | +;+;+ | 887 | NA | 13 | 2 | 5 | 9 | 27 | 19 |
| ENSDARG000 | 12;12;12;12 | 28964034:289 | 28964286:289 | -;-;-;- | 496 | NA | 0 | 0 | 4 | 0 | 0 | 0 |
| ENSDARG000 | 12:12:12:12:1 | 28946708:289 | 28949148:289 | -:-:-:-:-:-:-: | 7201 | NA | 1145 | 1015 | 1357 | 808 | 1491 | 1188 |
| ENSDARG000 | 12:12:12:12:1 | 3944229:3945 | 3944586:3945 | -:-:-:-:-:-:-: | 3207 | eif3c | 11022 | 9684 | 18378 | 11526 | 10903 | 12686 |
| ENSDARG000 | 12;12;12 | 30838167:308 | 30838711:308 | -;-;- | 848 | crygmx | 0 | 0 | 0 | 0 | 0 | 0 |
| ENSDARG000 | 12:12:12:12:1 | 23991276:239 | 23991835:239 | +:+:+:+:+:+:+: | 7392 | psme4b | 1384 | 1480 | 1873 | 781 | 1473 | 1320 |
| ENSDARG000 | 12:12:12:12:1 | 28955766:289 | 28955876:289 | +:+:+:+:+:+:+: | 2533 | znf668 | 198 | 175 | 272 | 126 | 290 | 254 |
| ENSDARG000 | 12:12:12:12:1 | 19065679:190 | 19065877:190 | -:-:-:-:-:-:-: | 3445 | NA | 20 | 66 | 4 | 10 | 4 | 6 |
| ENSDARG000 | 12:12:12:12:1 | 30842677:308 | 30843137:308 | -;-;-;-;-;-;- | 1419 | crygmxl2 | 0 | 0 | 0 | 0 | 0 | 0 |
| ENSDARG000 | 12:12:12:12:1 | 28971580:289 | 28971845:289 | -:-:-:-:-:-:-: | 1102 | cd37 | 2 | 0 | 11 | 8 | 26 | 14 |
| ENSDARG000 | 12:12:12:12:1 | 9542124:9551 | 9542228:9551 | +:+:+:+:+:+:+: | 2310 | p4ha1a | 478 | 309 | 313 | 293 | 418 | 206 |
| ENSDARG000 | 12:12:12:12:1 | 33561418:335 | 33561894:335 | -:-:-:-:-:-:-: | 3321 | tdrkh | 260 | 291 | 489 | 207 | 605 | 414 |
| ENSDARG000 | 12:12:12:12:1 | 5587347:5591 | 5587700:5591 | -:-:-:-:-:-:-: | 3529 | itga3b | 2 | 2 | 30 | 0 | 10 | 14 |
| ENSDARG000 | 12;12;12;12 | 14149686:141 | 14150148:141 | +;+;+;+ | 2910 | kbtbd2 | 439 | 603 | 639 | 358 | 446 | 439 |
| ENSDARG000 | 12;12 | 27120117:271 | 27120367:271 | -;- | 343 | NA | 0 | 0 | 0 | 13 | 97 | 0 |
| ENSDARG000 | 12:12:12:12:1 | 532019:53343 | 532110:53546 | +;+;+;+;+;+ | 2545 | NA | 210 | 167 | 344 | 97 | 285 | 183 |
| ENSDARG000 | 12;12 | 27117609:271 | 27118239:271 | +;+ | 1089 | hoxb8b | 0 | 0 | 2 | 24 | 324 | 0 |
| ENSDARG000 | 12:12:12:12:1 | 46683114:466 | 46683269:466 | +:+:+:+:+:+:+: | 1152 | tvp23b | 510 | 589 | 634 | 498 | 626 | 531 |
| ENSDARG000 | 12:12:12:12:1 | 11496740:115 | 11497143:115 | -:-:-:-:-:-:-: | 1513 | PLEKHA1 | 149 | 136 | 142 | 79 | 191 | 142 |
| ENSDARG000 | 12:12:12:12:1 | 17657224:176 | 17657715:176 | -:-:-:-:-:-:-: | 2080 | ccz1 | 1123 | 857 | 921 | 637 | 1064 | 791 |
| ENSDARG000 | 12:12:12:12:1 | 549598:54959 | 550241:55156 | -;-;-;-;-;-;-;- | 2516 | bsk146 | 60 | 16 | 38 | 8 | 24 | 18 |
| ENSDARG000 | 12:12:12:12:1 | 20699534:207 | 20700308:207 | +:+:+:+:+:+:+: | 5034 | CASKIN2 | 12 | 4 | 15 | 14 | 10 | 17 |
| ENSDARG000 | 12;12;12 | 5708400:5709 | 5708881:5709 | +;+;+ | 1591 | dlx3b | 110 | 56 | 6 | 18 | 30 | 98 |
| ENSDARG000 | 12:12:12:12:1 | 3964351:3967 | 3965246:3967 | -:-:-:-:-:-:-: | 1979 | ppp4cb | 1031 | 1267 | 1289 | 474 | 1292 | 895 |
| ENSDARG000 | 12:12:12:12:1 | 20691310:206 | 20691694:206 | +:+:+:+:+:+:+: | 1937 | st6galnac1.2 | 0 | 0 | 2 | 0 | 0 | 0 |
| ENSDARG000 | 12:12:12:12:1 | 14166036:141 | 14167110:141 | -:-:-:-:-:-:-: | 3471 | avl9 | 650 | 572 | 522 | 505 | 580 | 791 |
| ENSDARG000 | 12;12;12 | 30933851:309 | 30934324:309 | +;+;+ | 757 | NA | 0 | 0 | 0 | 0 | 0 | 0 |
| ENSDARG000 | 12;12;12 | 5719616:5726 | 5720098:5726 | -;-;- | 1291 | dlx4b | 4 | 0 | 2 | 0 | 8 | 12 |
| ENSDARG000 | 12:12:12:12:1 | 19099264:190 | 19100005:191 | -;-;-;-;-;-;- | 1348 | csdc2a | 10 | 42 | 26 | 138 | 24 | 90 |
| ENSDARG000 | 12:12:12:12:1 | 35587971:355 | 35588223:355 | +:+:+:+:+:+:+: | 2547 | chmp6b | 1796 | 1651 | 2089 | 1336 | 2052 | 1919 |
| ENSDARG000 | 12;12;12 | 5842639:5904 | 5842918:5905 | -;-;- | 689 | NA | 0 | 0 | 0 | 0 | 0 | 0 |
| ENSDARG000 | 12:12:12:12:1 | 38983989:389 | 38984051:389 | +:+:+:+:+:+:+: | 7264 | NA | 32 | 0 | 0 | 2 | 0 | 0 |
| ENSDARG000 | 12:12:12:12:1 | 5977777:5978 | 5977890:5978 | +;+;+;+;+;+ | 538 | NA | 7 | 5 | 11 | 7 | 11 | 10 |
| ENSDARG000 | 12:12:12:12:1 | 17695986:176 | 17696338:176 | -:-:-:-:-:-:-: | 1872 | pvalb9 | 30 | 0 | 32 | 1 | 75 | 0 |
| ENSDARG000 | 12:12:12:12:1 | 5981333:5984 | 5982037:5984 | -:-:-:-:-:-:-: | 2516 | kat7b | 437 | 526 | 675 | 419 | 562 | 399 |
| ENSDARG000 | 12;12;12 | 24056000:240 | 24057406:240 | -;-;- | 1962 | chac2 | 78 | 62 | 112 | 62 | 60 | 90 |
| ENSDARG000 | 12:12:12:12:1 | 33583497:335 | 33584831:335 | -;-;-;-;- | 1697 | NA | 0 | 0 | 0 | 0 | 0 | 0 |
| ENSDARG000 | 12;12 | 27127139:271 | 27127604:271 | +;+ | 1194 | hoxb6b | 0 | 0 | 0 | 1 | 49 | 0 |
| ENSDARG000 | 12 | 36259018 | 36260532 | - | 1515 | kcnj2a | 274 | 11 | 32 | 23 | 132 | 8 |
| ENSDARG000 | 12 | 36267761 | 36268723 | - | 963 | KCNJ16 | 0 | 0 | 4 | 10 | 8 | 0 |
| ENSDARG000 | 12:12:12:12:1 | 3981346:3982 | 3981957:3982 | -;-;-;-;-;- | 6759 | NA | 114 | 128 | 190 | 64 | 284 | 232 |
| ENSDARG000 | 12:12:12:12:1 | 33602429:336 | 33606590:336 | -:-:-:-:-:-:-: | 8674 | TMEM94 | 483 | 560 | 954 | 287 | 597 | 586 |
| ENSDARG000 | 12:12:12:12:1 | 560512:56110 | 561472:56147 | -:-:-:-:-:-:-: | 9787 | NA | 3325 | 3717 | 1360 | 618 | 878 | 622 |
| ENSDARG000 | 12:12:12:12:1 | 39034225:390 | 39034376:390 | +:+:+:+:+:+:+: | 4601 | NA | 0 | 0 | 0 | 0 | 0 | 0 |
| ENSDARG000 | 12:12:12:12:1 | 19106571:191 | 19107827:191 | -:-:-:-:-:-:-: | 3495 | aco2 | 5244 | 4537 | 5663 | 2639 | 5045 | 3962 |
| ENSDARG000 | 12:12:12:12:1 | 619669:62050 | 619690:62074 | -;-;-;-;-;-;- | 1050 | NA | 0 | 7 | 12 | 2 | 14 | 3 |
| ENSDARG000 | 12:12:12:12:1 | 24060894:240 | 24061343:240 | +:+:+:+:+:+:+: | 1890 | asb3 | 190 | 180 | 273 | 98 | 224 | 160 |
| ENSDARG000 | 12;12 | 39203745:392 | 39204103:392 | +;+ | 1020 | NA | 0 | 3 | 4 | 0 | 0 | 0 |
| ENSDARG000 | 12:12:12:12:1 | 17705218:177 | 17705486:177 | -:-:-:-:-:-:-: | 826 | pvalb3 | 2 | 4 | 0 | 1 | 0 | 2 |
| ENSDARG000 | 12:12:12:12:1 | 3998764:4002 | 3999177:4002 | -:-:-:-:-:-:-: | 3143 | NA | 2 | 0 | 0 | 0 | 11 | 0 |
| ENSDARG000 | 12:12:12:12:1 | 36390422:363 | 36390436:363 | +;+;+;+;+;+ | 1146 | galr2a | 0 | 0 | 0 | 0 | 0 | 0 |
| ENSDARG000 | 12:12:12:12:1 | 17710425:177 | 17710684:177 | -;-;-;-;-;-;-;- | 1181 | pvalb2 | 16 | 24 | 16 | 20 | 46 | 10 |
| ENSDARG000 | 12:12:12:12:1 | 39409601:394 | 39409779:394 | -;-;-;-;- | 499 | NA | 0 | 0 | 0 | 0 | 0 | 0 |
| ENSDARG000 | 12;12;12 | 17716512:177 | 17716592:177 | +;+;+ | 786 | NA | 8 | 0 | 0 | 0 | 0 | 0 |
| ENSDARG000 | 12:12:12:12:1 | 28986308:289 | 28986413:289 | +;+;+;+;+;+ | 625 | gall | 0 | 0 | 0 | 0 | 0 | 0 |
| ENSDARG000 | 12:12:12:12:1 | 14214755:142 | 14214861:142 | +:+:+:+:+:+:+: | 2185 | mto1 | 425 | 342 | 551 | 484 | 524 | 435 |
| ENSDARG000 | 12:12:12:12:1 | 46696867:466 | 46697051:466 | +:+:+:+:+:+:+: | 3149 | exoc7 | 1431 | 1024 | 1256 | 1489 | 1273 | 1161 |
| ENSDARG000 | 12:12:12:12:1 | 11601559:116 | 11601708:116 | -:-:-:-:-:-:-: | 2047 | NA | 0 | 0 | 1 | 0 | 0 | 0 |
| ENSDARG000 | 12:12:12:12:1 | 36413886:364 | 36413974:364 | +:+:+:+:+:+:+: | 1569 | NA | 0 | 0 | 0 | 0 | 0 | 0 |
| ENSDARG000 | 12:12:12:12:1 | 36416173:364 | 36416736:364 | +:+:+:+:+:+:+: | 8207 | unk | 164 | 137 | 330 | 92 | 250 | 164 |
| ENSDARG000 | 12;12 | 27129659:271 | 27130338:271 | +;+ | 1626 | hoxb5b | 0 | 0 | 0 | 10 | 40 | 8 |
| ENSDARG000 | 12:12:12:12:1 | 39444747:394 | 39445103:394 | -;-;-;-;-;- | 924 | NA | 67 | 33 | 19 | 182 | 66 | 13 |
| ENSDARG000 | 12:12:12:12:1 | 14556092:145 | 14556130:145 | +:+:+:+:+:+:+: | 1932 | becn1 | 750 | 530 | 637 | 414 | 598 | 663 |
| ENSDARG000 | 12:12:12:12:1 | 4036409:4036 | 4036476:4036 | +;+;+;+;+;+ | 1250 | zgc:123217 | 0 | 0 | 0 | 0 | 0 | 0 |
| ENSDARG000 | 12:12:12:12:1 | 30977736:309 | 30982183:309 | -:-:-:-:-:-:-: | 7466 | tcf7l2 | 137 | 160 | 244 | 86 | 156 | 196 |
| ENSDARG000 | 12;12 | 14920088:149 | 14921518:149 | -;- | 1604 | neurod2 | 0 | 0 | 1 | 0 | 0 | 0 |
| ENSDARG000 | 12:12:12:12:1 | 27156943:271 | 27157205:271 | +:+:+:+:+:+:+: | 3252 | skap1 | 0 | 15 | 5 | 0 | 40 | 16 |
| ENSDARG000 | 12:12:12:12:1 | 4052180:4057 | 4052523:4057 | -;-;-;-;-;-;- | 1596 | aldoab | 4 | 6 | 6 | 4 | 6 | 12 |
| ENSDARG000 | 12:12:12:12:1 | 40205995:402 | 40206759:402 | +:+:+:+:+:+:+: | 3375 | NA | 2 | 0 | 2 | 0 | 0 | 0 |
| ENSDARG000 | 12:12:12:12:1 | 28995942:289 | 28996372:289 | +;+;+;+;+ | 1298 | valopb | 0 | 2 | 0 | 0 | 0 | 0 |

| | | | | | | | | | | | | |
|---|---|---|---|---|---|---|---|---|---|---|---|---|
| ENSDARG000 | 12:12:12:12:1 | 27144696:271 | 27144903:271 | -;-;-;-;-;- | 782 | NA | 0 | 0 | 0 | 0 | 0 | 0 |
| ENSDARG000 | 12:12:12:12:1 | 27206736:272 | 27207519:272 | -;-;-;-;-;-;-;-; | 1922 | psme3 | 667 | 1078 | 1483 | 482 | 1007 | 719 |
| ENSDARG000 | 12:12:12:12:1 | 9597151:9601 | 9597314:9601 | -;-;-;-;-;-;-;- | 1041 | arg1 | 0 | 0 | 14 | 0 | 46 | 0 |
| ENSDARG000 | 12;12 | 27141140:271 | 27142102:271 | +;+ | 1606 | hoxb1b | 0 | 0 | 0 | 14 | 252 | 0 |
| ENSDARG000 | 12:12:12:12:1 | 24342303:243 | 24342705:243 | +;+;+;+;+;+;+;+; | 11699 | nrxn1a | 53 | 202 | 64 | 377 | 132 | 235 |
| ENSDARG000 | 12:12:12:12:1 | 630295:63055 | 630668:63066 | -;-;-;-;-;-;-;-; | 1423 | sult2st1 | 405 | 34 | 60 | 24 | 119 | 68 |
| ENSDARG000 | 12;12 | 638435:63955 | 638613:63994 | +;+ | 572 | NA | 0 | 0 | 4 | 0 | 0 | 0 |
| ENSDARG000 | 12:12:12:12:1 | 33661781:336 | 33662378:336 | -;-;-;-;-;-;-;-; | 5445 | NA | 0 | 12 | 8 | 8 | 20 | 4 |
| ENSDARG000 | 12:12:12:12:1 | 14965587:149 | 14966240:149 | -;-;-;-;-;-;-;-; | 2347 | pkmyt1 | 164 | 174 | 197 | 162 | 240 | 212 |
| ENSDARG000 | 12;12 | 17754859:177 | 17754964:177 | +;+ | 1060 | bhlha15 | 0 | 0 | 4 | 0 | 2 | 6 |
| ENSDARG000 | 12:12:12:12:1 | 17889465:178 | 17889692:178 | -;-;-;-;- | 1281 | nptx2b | 0 | 0 | 0 | 0 | 6 | 0 |
| ENSDARG000 | 12:12:12:12:1 | 6002715:6003 | 6002904:6003 | +;+;+;+;+;+;+ | 959 | ptges3l | 0 | 4 | 1 | 2 | 12 | 4 |
| ENSDARG000 | 12:12:12:12:1 | 15002757:150 | 15002801:150 | +;+;+;+;+;+;+;+; | 839 | mylpfb | 6 | 8 | 6 | 2 | 11 | 12 |
| ENSDARG000 | 12:12:12:12:1 | 19138452:191 | 19138590:191 | +;+;+;+;+;+;+;+; | 1103 | phf5a | 140 | 208 | 209 | 87 | 136 | 148 |
| ENSDARG000 | 12:12:12:12:1 | 29054907:290 | 29055332:290 | +;+;+;+;+;+;+;+; | 2758 | gabrz | 0 | 0 | 0 | 0 | 0 | 0 |
| ENSDARG000 | 12:12:12:12:1 | 6023490:6023 | 6024139:6024 | -;-;-;-;-;-;-;-; | 1372 | g6pc1a.1 | 2 | 0 | 2 | 0 | 0 | 0 |
| ENSDARG000 | 12:12:12:12:1 | 19144833:191 | 19146697:191 | -;-;-;-;-;-;-;-;- | 2932 | tefa | 3102 | 4155 | 2979 | 3792 | 6959 | 3401 |
| ENSDARG000 | 12:12:12:12:1 | 29075171:290 | 29075584:290 | -;-;-;-;-;-;-;- | 869 | NA | 0 | 4 | 0 | 0 | 0 | 0 |
| ENSDARG000 | 12:12:12:12:1 | 20788243:207 | 20788487:207 | -;-;-;-;-;-;-;-; | 1509 | nme2a | 0 | 0 | 0 | 0 | 0 | 0 |
| ENSDARG000 | 12:12:12:12:1 | 24738962:247 | 24739196:247 | +;+;+;+;+;+;+;+; | 2543 | fshr | 0 | 0 | 0 | 0 | 0 | 0 |
| ENSDARG000 | 12:12:12:12:1 | 29131689:291 | 29132385:291 | +;+;+;+;+;+;+;+; | 6209 | NA | 0 | 0 | 0 | 0 | 0 | 2 |
| ENSDARG000 | 12:12:12:12:1 | 36474565:364 | 36474832:364 | -;-;-;-;-;-;-;-; | 4219 | NA | 44 | 94 | 53 | 14 | 164 | 116 |
| ENSDARG000 | 12;12;12 | 9669927:9670 | 9670038:9670 | -;-;- | 326 | NA | 0 | 0 | 0 | 0 | 0 | 0 |
| ENSDARG000 | 12:12:12:12:1 | 9619909:9621 | 9620333:9621 | -;-;-;-;-;-;-;-; | 6867 | heatr1 | 1333 | 1038 | 2211 | 706 | 1673 | 1060 |
| ENSDARG000 | 12:12:12:12:1 | 4114401:4116 | 4114639:4116 | +;+;+;+;+;+;+;+; | 5121 | NA | 171 | 199 | 279 | 174 | 532 | 604 |
| ENSDARG000 | 12;12 | 29223278:292 | 29229713:292 | -;- | 6896 | NA | 624 | 561 | 610 | 456 | 748 | 586 |
| ENSDARG000 | 12:12:12:12:1 | 648334:65037 | 648483:65055 | -;-;-;-;-;- | 1043 | sult2st2 | 26 | 0 | 0 | 0 | 0 | 2 |
| ENSDARG000 | 12:12:12:12:1 | 11650146:116 | 11650403:116 | +;+;+;+;+;+;+;+; | 5475 | WAPL | 1232 | 895 | 1522 | 877 | 1210 | 1155 |
| ENSDARG000 | 12:12:12:12:1 | 6041575:6041 | 6041837:6041 | +;+;+;+;+;+;+;+; | 2976 | g6pc1a.2 | 48 | 58 | 156 | 102 | 305 | 39 |
| ENSDARG000 | 12;12;12 | 29236274:292 | 29236375:292 | +;+;+ | 1327 | mxtx2 | 0 | 0 | 0 | 0 | 0 | 0 |
| ENSDARG000 | 12;12;12;12 | 20807012:208 | 20807118:208 | +;+;+;+ | 943 | NA | 0 | 0 | 0 | 0 | 0 | 0 |
| ENSDARG000 | 12:12:12:12:1 | 24796517:247 | 24798629:247 | -;-;-;-;-;-;-;-; | 3965 | foxn2b | 668 | 747 | 1014 | 1044 | 1015 | 1063 |
| ENSDARG000 | 12:12:12:12:1 | 657704:66015 | 658721:66029 | -;-;-;-;-;-;-;-; | 4000 | NA | 28 | 6 | 2 | 38 | 24 | 4 |
| ENSDARG000 | 12:12:12:12:1 | 21298317:212 | 21298705:212 | +;+;+;+;+;+;+;+; | 6343 | ca10a | 108 | 30 | 30 | 26 | 149 | 6 |
| ENSDARG000 | 12:12:12:12:1 | 46740584:467 | 46740613:467 | +;+;+;+;+;+;+;+; | 5289 | plaub | 10 | 24 | 112 | 10 | 39 | 22 |
| ENSDARG000 | 12:12:12:12:1 | 19158832:191 | 19159560:191 | -;-;-;-;-;-;-;-; | 3660 | ZC3H7B | 2033 | 1511 | 2429 | 1323 | 2060 | 1959 |
| ENSDARG000 | 12:12:12:12:1 | 46841918:468 | 46842503:468 | +;+;+;+;+;+;+;+; | 2937 | adkb | 351 | 290 | 409 | 235 | 420 | 413 |
| ENSDARG000 | 12:12:12:12:1 | 4147714:4149 | 4147756:4149 | +;+;+;+;+;+;+;+; | 3037 | NA | 15 | 16 | 11 | 6 | 28 | 42 |
| ENSDARG000 | 12:12:12:12:1 | 33734849:337 | 33739314:337 | -;-;-;-;-;-;-;-; | 8368 | llgl2 | 818 | 827 | 970 | 584 | 943 | 470 |
| ENSDARG000 | 12:12:12:12:1 | 17927871:179 | 17927912:179 | +;+;+;+;+;+;+;+; | 12749 | trrap | 1264 | 1068 | 1732 | 709 | 1612 | 1115 |
| ENSDARG000 | 12:12:12:12:1 | 4170921:4171 | 4171277:4171 | -;-;-;-;-;-;-;-; | 1627 | NA | 0 | 2 | 2 | 10 | 0 | 0 |
| ENSDARG000 | 12:12:12:12:1 | 6049355:6050 | 6049763:6050 | -;-;-;-;-;-;-;-; | 1589 | aarsd1 | 215 | 180 | 228 | 124 | 194 | 156 |
| ENSDARG000 | 12:12:12:12:1 | 15197857:151 | 15198305:151 | -;-;-;-;-;-;- | 1600 | sult1st6 | 10 | 2 | 12 | 3 | 10 | 12 |
| ENSDARG000 | 12:12:12:12:1 | 46869271:468 | 46869657:468 | +;+;+;+;+;+;+ | 2182 | hpdl | 14 | 46 | 16 | 10 | 14 | 28 |
| ENSDARG000 | 12:12:12:12:1 | 15148618:151 | 15148928:151 | +;+;+;+;+;+;+;+; | 1882 | adprh | 469 | 453 | 146 | 176 | 263 | 150 |
| ENSDARG000 | 12;12;12;12 | 21684239:216 | 21684333:216 | +;+;+;+ | 661 | mrpl27 | 240 | 237 | 326 | 146 | 254 | 240 |
| ENSDARG000 | 12:12:12:12:1 | 21669385:216 | 21673031:216 | -;-;-;-;-;-;-;-; | 5331 | NA | 34 | 112 | 128 | 122 | 88 | 60 |
| ENSDARG000 | 12:12:12:12:1 | 29240124:292 | 29240262:292 | +;+;+;+;+;+;+;+; | 4662 | bms1 | 1307 | 881 | 1497 | 541 | 1099 | 862 |
| ENSDARG000 | 12:12:12:12:1 | 27213733:272 | 27213954:272 | +;+;+;+;+;+;+;+; | 4386 | nbr1a | 805 | 427 | 751 | 940 | 803 | 1037 |
| ENSDARG000 | 12;12;12 | 4196493:4196 | 4196582:4196 | -;-;- | 357 | NA | 10 | 0 | 0 | 0 | 0 | 0 |
| ENSDARG000 | 12;12;12;12 | 4201826:4203 | 4202056:4203 | -;-;-;- | 846 | NA | 0 | 0 | 4 | 0 | 0 | 0 |
| ENSDARG000 | 12 | 11768114 | 11768889 | - | 776 | NA | 0 | 0 | 1 | 2 | 4 | 0 |
| ENSDARG000 | 12:12:12:12:1 | 46883785:468 | 46883888:468 | +;+;+;+;+;+;+;+; | 1700 | fam53b | 730 | 710 | 1130 | 881 | 793 | 630 |
| ENSDARG000 | 12:12:12:12:1 | 695619:72542 | 695654:72559 | +;+;+;+;+;+;+;+; | 5771 | NA | 719 | 577 | 911 | 632 | 1087 | 712 |
| ENSDARG000 | 12:12:12:12:1 | 12112384:121 | 12112721:121 | +;+;+;+;+;+;+;+; | 4474 | GRID1 | 0 | 0 | 0 | 0 | 0 | 0 |
| ENSDARG000 | 12:12:12:12:1 | 6065661:6098 | 6065887:6098 | +;+;+;+;+;+;+;+; | 2753 | sgms1b | 385 | 258 | 509 | 364 | 282 | 386 |
| ENSDARG000 | 12:12:12:12:1 | 19188542:191 | 19188763:191 | +;+;+;+;+;+;+;+; | 2286 | cby1 | 114 | 56 | 170 | 113 | 154 | 134 |
| ENSDARG000 | 12:12:12:12:1 | 46950584:469 | 46951103:469 | -;-;-;-;-;-;-;-;- | 1442 | lhpp | 211 | 192 | 252 | 115 | 167 | 76 |
| ENSDARG000 | 12;12;12 | 4217178:4218 | 4217634:4218 | -;-;- | 863 | vkorc1 | 2 | 18 | 21 | 8 | 20 | 7 |
| ENSDARG000 | 12:12:12:12:1 | 4220353:4222 | 4220616:4222 | +;+;+;+;+;+;+;+; | 4213 | mapk7 | 324 | 318 | 372 | 297 | 306 | 442 |
| ENSDARG000 | 12:12:12:12:1 | 36524084:365 | 36524264:365 | +;+;+;+;+;+;+;+; | 3247 | NA | 12 | 0 | 0 | 0 | 6 | 0 |
| ENSDARG000 | 12:12:12:12:1 | 24841719:248 | 24843451:248 | -;-;-;-;-;-;-;-; | 4766 | NA | 748 | 640 | 767 | 540 | 777 | 716 |
| ENSDARG000 | 12:12:12:12:1 | 27231212:272 | 27231403:272 | +;+;+;+;+;+;+;+; | 2087 | tmem106a | 777 | 572 | 830 | 849 | 614 | 1095 |
| ENSDARG000 | 12;12 | 804314:80678 | 805027:80844 | +;+ | 2378 | NA | 44 | 24 | 0 | 23 | 53 | 10 |
| ENSDARG000 | 12:12:12:12:1 | 9703172:9744 | 9704497:9744 | +;+;+;+;+;+;+;+; | 2779 | PPP1R9B | 424 | 144 | 239 | 157 | 216 | 71 |
| ENSDARG000 | 12:12:12:12:1 | 12954938:129 | 12955806:129 | -;-;-;-;-;-;-;-; | 6355 | NA | 315 | 161 | 64 | 145 | 132 | 182 |
| ENSDARG000 | 12:12:12:12:1 | 27236871:272 | 27237095:272 | -;-;-;-;-;-;-;-;- | 566 | NA | 38 | 22 | 11 | 35 | 19 | 46 |
| ENSDARG000 | 12;12 | 847187:85028 | 849289:85045 | -;- | 2273 | tob1b | 1184 | 549 | 773 | 973 | 864 | 1146 |
| ENSDARG000 | 12:12:12:12:1 | 19191787:191 | 19192156:191 | +;+;+;+;+;+;+;+; | 2582 | slc16a8 | 68 | 47 | 228 | 186 | 252 | 137 |
| ENSDARG000 | 12:12:12:12:1 | 46960579:469 | 46960616:469 | +;+;+;+;+;+;+;+; | 5693 | oat | 62 | 94 | 289 | 112 | 122 | 79 |
| ENSDARG000 | 12:12:12:12:1 | 15273304:152 | 15273412:152 | +;+;+;+;+;+;+;+; | 5464 | MED1 | 900 | 582 | 837 | 370 | 780 | 631 |
| ENSDARG000 | 12:12:12:12:1 | 6117682:6119 | 6117783:6119 | +;+;+;+;+;+;+;+; | 2805 | asah2 | 16 | 16 | 117 | 70 | 42 | 36 |
| ENSDARG000 | 12:12:12:12:1 | 982796:98485 | 983436:98511 | -;-;-;-;- | 1431 | kdelr2b | 702 | 1023 | 1028 | 647 | 991 | 682 |
| ENSDARG000 | 12:12:12:12:1 | 19199735:192 | 19199973:192 | +;+;+;+;+;+;+ | 2073 | pdap1a | 2689 | 2820 | 2939 | 1686 | 2775 | 1924 |
| ENSDARG000 | 12:12:12:12:1 | 9781524:9786 | 9783650:9786 | -;-;-;-;-;-;-;-; | 3133 | prdm9 | 358 | 261 | 349 | 302 | 382 | 312 |
| ENSDARG000 | 12:12:12:12:1 | 15363463:153 | 15363634:153 | +;+;+;+;+;+;+;+; | 1903 | plxdc1 | 234 | 4 | 64 | 0 | 94 | 18 |
| ENSDARG000 | 12:12:12:12:1 | 29289387:292 | 29290106:292 | -;-;-;-;-;-;-;-; | 2385 | SH2D4B | 0 | 4 | 0 | 0 | 2 | 0 |
| ENSDARG000 | 12:12:12:12:1 | 29350698:293 | 29351906:293 | -;-;-;-;-;-;-;-;- | 2837 | NA | 68 | 16 | 30 | 16 | 26 | 1 |
| ENSDARG000 | 12;12;12 | 29312652:293 | 29312799:293 | +;+;+ | 1019 | NA | 0 | 0 | 0 | 0 | 0 | 0 |
| ENSDARG000 | 12:12:12:12:1 | 4241458:4241 | 4242146:4242 | -;-;-;-;-;-;-;-;- | 1538 | zgc:92313 | 32 | 2 | 0 | 0 | 40 | 2 |
| ENSDARG000 | 12;12;12 | 42364157:423 | 42367090:423 | -;-;- | 3099 | NA | 16 | 1 | 10 | 16 | 6 | 20 |
| ENSDARG000 | 12:12:12:12:1 | 47446158:474 | 47446346:474 | +;+;+;+;+;+;+;+; | 5384 | NA | 1188 | 755 | 1005 | 331 | 1269 | 501 |
| ENSDARG000 | 12:12:12:12:1 | 13091842:130 | 13091999:130 | +;+;+;+;+;+;+;+; | 3913 | NA | 0 | 0 | 0 | 0 | 0 | 0 |
| ENSDARG000 | 12;12 | 47424427:474 | 47424501:474 | +;+ | 1583 | CHRM3 | 0 | 0 | 0 | 0 | 0 | 0 |
| ENSDARG000 | 12:12:12:12:1 | 19205086:192 | 19205125:192 | +;+;+;+;+;+;+;+; | 2859 | NA | 2 | 17 | 18 | 28 | 66 | 34 |
| ENSDARG000 | 12:12:12:12:1 | 31161564:311 | 31161657:311 | -;-;-;-;-;-;-;-; | 1058 | vti1a | 217 | 196 | 156 | 179 | 205 | 125 |
| ENSDARG000 | 12 | 6158225 | 6159545 | - | 1321 | rltgr | 8 | 0 | 8 | 0 | 14 | 2 |
| ENSDARG000 | 12:12:12:12:1 | 29763546:297 | 29763716:297 | +;+;+;+;+;+;+;+; | 1698 | gfra1b | 69 | 15 | 99 | 75 | 106 | 26 |
| ENSDARG000 | 12:12:12:12:1 | 24906106:249 | 24906491:249 | -;-;-;-;-;-;-;-; | 3054 | msh2 | 292 | 1086 | 1193 | 442 | 1120 | 686 |
| ENSDARG000 | 12:12:12:12:1 | 9817440:9817 | 9817650:9817 | +;+;+;+;+;+;+;+; | 2747 | rundc3ab | 0 | 0 | 28 | 0 | 0 | 52 |
| ENSDARG000 | 12:12:12:12:1 | 42369566:423 | 42369992:423 | -;-;-;-;-;-;-;-; | 1525 | glrx3 | 1302 | 1434 | 1792 | 819 | 1490 | 1286 |
| ENSDARG000 | 12:12:12:12:1 | 6160446:6161 | 6161260:6161 | -;-;-;-;-;-;-;-; | 2584 | A1CF | 2 | 6 | 10 | 6 | 4 | 2 |
| ENSDARG000 | 12:12:12:12:1 | 992442:99310 | 993526:99352 | -;-;-;-;-;-;-;-; | 2207 | mettl9 | 696 | 657 | 902 | 468 | 846 | 696 |
| ENSDARG000 | 12:12:12:12:1 | 4294923:4295 | 4295821:4295 | -;-;-;-;-;-;-;-; | 1998 | ca15b | 0 | 0 | 0 | 0 | 0 | 0 |
| ENSDARG000 | 12:12:12:12:1 | 15477111:154 | 15477478:154 | +;+;+;+;+;+;+;+; | 4870 | NA | 216 | 363 | 521 | 441 | 1083 | 729 |
| ENSDARG000 | 12;12;12;12 | 19240610:192 | 19241425:192 | -;-;-;- | 2145 | NA | 14 | 2 | 8 | 0 | 10 | 12 |
| ENSDARG000 | 12:12:12:12:1 | 42400277:424 | 42400595:424 | +;+;+;+;+;+;+ | 3606 | NA | 94 | 104 | 128 | 62 | 94 | 66 |
| ENSDARG000 | 12;12 | 15565344:155 | 15566567:155 | -;- | 1546 | hexim1 | 898 | 1038 | 1218 | 812 | 1132 | 833 |
| ENSDARG000 | 12:12:12:12:1 | 6195191:6195 | 6195450:6195 | +;+;+;+;+;+;+;+; | 3776 | PRKG1 | 0 | 2 | 6 | 0 | 0 | 4 |
| ENSDARG000 | 12:12:12:12:1 | 29842836:298 | 29844976:298 | -;-;-;-;-;-;-;-; | 7028 | ATRNL1 | 162 | 132 | 550 | 130 | 445 | 382 |
| ENSDARG000 | 12;12 | 47543832:475 | 47544579:475 | -;- | 866 | grem2b | 0 | 1 | 0 | 0 | 0 | 0 |
| ENSDARG000 | 12;12 | 6195199:6217 | 6195450:6217 | +;+ | 539 | NA | 0 | 0 | 0 | 0 | 0 | 0 |
| ENSDARG000 | 12:12:12:12:1 | 19281189:192 | 19281431:192 | +;+;+;+;+;+;+;+; | 7867 | NA | 2459 | 1643 | 2438 | 1406 | 2080 | 1746 |
| ENSDARG000 | 12:12:12:12:1 | 6465557:6465 | 6465806:6465 | +;+;+;+;+;+;+;+; | 1150 | dkk1b | 0 | 0 | 0 | 0 | 0 | 0 |
| ENSDARG000 | 12:12:12:12:1 | 15568761:155 | 15569165:155 | -;-;-;-;-;-;-;-; | 1754 | acbd4 | 189 | 206 | 494 | 438 | 428 | 471 |
| ENSDARG000 | 12;12;12;12 | 6391032:6430 | 6391178:6430 | -;-;-;- | 592 | NA | 0 | 4 | 4 | 0 | 0 | 0 |
| ENSDARG000 | 12:12:12:12:1 | 19272963:192 | 19275530:192 | -;-;-;-;-;-;-;-; | 4081 | NA | 10 | 12 | 50 | 30 | 26 | 20 |
| ENSDARG000 | 12:12:12:12:1 | 9849476:9854 | 9849826:9854 | +;+;+;+;+;+;+;+; | 2156 | fam117ab | 6 | 28 | 42 | 4 | 30 | 34 |
| ENSDARG000 | 12:12:12:12:1 | 31422557:314 | 31422765:314 | +;+;+;+;+;+;+;+; | 2255 | zdhhc6 | 786 | 870 | 906 | 588 | 845 | 696 |
| ENSDARG000 | 12:12:12:12:1 | 31460877:314 | 31461230:314 | +;+;+;+;+;+;+;+; | 3384 | NA | 4 | 4 | 2 | 0 | 0 | 0 |
| ENSDARG000 | 12:12:12:12:1 | 21697540:216 | 21697701:216 | +;+;+;+;+;+;+;+; | 11674 | NA | 12 | 2 | 8 | 8 | 7 | 3 |
| ENSDARG000 | 12;12 | 27243059:272 | 27243301:272 | +;+ | 1601 | arl4d | 3518 | 4779 | 2475 | 3991 | 5331 | 2137 |
| ENSDARG000 | 12:12:12:12:1 | 31438579:314 | 31439913:314 | -;-;-;-;-;-;-;-; | 3793 | acsl5 | 2 | 4 | 6 | 2 | 7 | 0 |
| ENSDARG000 | 12;12;12 | 27285994:272 | 27286176:272 | +;+;+ | 3720 | NA | 4 | 8 | 33 | 46 | 4 | 12 |
| ENSDARG000 | 12;12;12;12 | 9880493:9883 | 9880862:9883 | +;+;+;+ | 1403 | ndufa4b | 108 | 384 | 1313 | 180 | 256 | 269 |
| ENSDARG000 | 12;12 | 27331324:273 | 27331650:273 | +;+ | 2238 | SOST | 24 | 0 | 2 | 16 | 10 | 10 |
| ENSDARG000 | 12:12:12:12:1 | 9949941:9953 | 9950720:9953 | -;-;-;-;-;- | 1974 | ngfrb | 436 | 174 | 79 | 281 | 531 | 36 |
| ENSDARG000 | 12:12:12:12:1 | 47573586:475 | 47573620:475 | -;-;-;-;-;-;-;-; | 1245 | RGS7 | 4 | 7 | 11 | 2 | 4 | 0 |
| ENSDARG000 | 12:12:12:12:1 | 15622621:156 | 15623220:156 | +;+;+;+;+;+;+;+; | 3137 | PLCD3 | 32 | 32 | 34 | 32 | 43 | 30 |
| ENSDARG000 | 12:12:12:12:1 | 1000323:1000 | 1000604:1000 | +;+;+;+;+;+;+;+; | 2211 | NA | 6 | 6 | 0 | 2 | 20 | 1 |
| ENSDARG000 | 12:12:12:12:1 | 42436328:424 | 42437053:424 | +;+;+;+;+;+;+;+; | 5271 | ebf3a | 97 | 0 | 2 | 10 | 28 | 42 |
| ENSDARG000 | 12:12:12:12:1 | 13153184:131 | 13153390:131 | -;-;-;-;- | 534 | NA | 18 | 0 | 0 | 10 | 0 | 2 |
| ENSDARG000 | 12:12:12:12:1 | 13118540:131 | 13118706:131 | +;+;+;+;+;+;+;+; | 4111 | cmn | 2 | 0 | 0 | 50 | 8 | 0 |
| ENSDARG000 | 12;12;12 | 10060636:100 | 10061156:100 | -;-;- | 1213 | NA | 0 | 0 | 0 | 0 | 0 | 0 |
| ENSDARG000 | 12;12 | 25043156:250 | 25045329:250 | -;- | 2554 | socs5a | 991 | 779 | 1157 | 590 | 1549 | 1003 |
| ENSDARG000 | 12:12:12:12:1 | 10053852:100 | 10053900:100 | +;+;+;+;+;+;+;+; | 3054 | slc4a1b | 0 | 0 | 0 | 0 | 0 | 0 |
| ENSDARG000 | 12:12:12:12:1 | 47638430:476 | 47638746:476 | -;-;-;-;-;-;-;-; | 1741 | fh | 542 | 756 | 795 | 721 | 855 | 766 |
| ENSDARG000 | 12:12:12:12:1 | 4308522:4310 | 4308794:4310 | -;-;-;-;-;-;-;- | 1163 | zgc:153760 | 0 | 0 | 0 | 0 | 0 | 0 |
| ENSDARG000 | 12:12:12:12:1 | 25085751:250 | 25086283:250 | +;+;+;+;+;+;+ | 1932 | gch2 | 10887 | 20903 | 12947 | 11347 | 24157 | 22096 |
| ENSDARG000 | 12:12:12:12:1 | 43376436:433 | 43380038:433 | -;-;-;-;-;-;-;-; | 6428 | PTPRE | 2656 | 20474 | 13425 | 8483 | 17988 | 2724 |
| ENSDARG000 | 12;12;12 | 27462225:274 | 27462908:274 | +;+;+ | 2771 | meox1 | 46 | 2 | 8 | 0 | 6 | 28 |
| ENSDARG000 | 12:12:12:12:1 | 4320041:4322 | 4320765:4322 | -;-;-;-;-;-;-;-;- | 1529 | ca15a | 0 | 0 | 0 | 0 | 0 | 0 |
| ENSDARG000 | 12:12:12:12:1 | 48340133:483 | 48340268:483 | +;+;+;+;+;+;+;+; | 1551 | ddit4 | 92 | 148 | 423 | 479 | 198 | 59 |
| ENSDARG000 | 12:12:12:12:1 | 1014418:1015 | 1015339:1015 | -;-;-;-;-;-;-;-; | 3359 | polr3e | 745 | 707 | 1449 | 517 | 929 | 653 |
| ENSDARG000 | 12:12:12:12:1 | 25097754:250 | 25097980:250 | +;+;+;+;+;+;+ | 1214 | pigf | 305 | 421 | 369 | 318 | 383 | 390 |
| ENSDARG000 | 12:12:12:12:1 | 6542789:6544 | 6544006:6545 | -;-;-;-;- | 2794 | NA | 0 | 0 | 0 | 0 | 0 | 0 |
| ENSDARG000 | 12:12:12:12:1 | 25110016:251 | 25111478:251 | -;-;-;-;-;-;- | 2296 | rhoq | 390 | 231 | 102 | 169 | 236 | 180 |
| ENSDARG000 | 12:12:12:12:1 | 21799130:217 | 21799390:217 | -;-;-;-;-;-;-;-; | 1869 | THRA | 989 | 707 | 768 | 444 | 1242 | 451 |
| ENSDARG000 | 12:12:12:12:1 | 27536095:275 | 27536390:275 | +;+;+;+;+;+;+;+; | 1974 | etv4 | 4773 | 2892 | 5057 | 4106 | 5717 | 4237 |
| ENSDARG000 | 12:12:12:12:1 | 10115964:101 | 10116160:101 | +;+;+;+;+;+;+;+; | 1906 | atxn7l3b | 167 | 149 | 252 | 117 | 211 | 272 |
| ENSDARG000 | 12;12;12;12 | 27494784:275 | 27495105:275 | -;-;-;- | 915 | NA | 3 | 0 | 5 | 0 | 2 | 0 |
| ENSDARG000 | 12:12:12:12:1 | 27569125:275 | 27571434:275 | -;-;-;-;-;-;-;-; | 5849 | DHX8 | 1221 | 1166 | 1509 | 706 | 1281 | 1204 |
| ENSDARG000 | 12:12:12:12:1 | 48346693:483 | 48346919:483 | -;-;-;-;-;-;-;-; | 2100 | dnajb12b | 216 | 293 | 368 | 296 | 296 | 171 |
| ENSDARG000 | 12:12:12:12:1 | 6566233:6566 | 6567694:6567 | -;-;-;-;-;-;-;-; | 6041 | pcdh15b | 0 | 0 | 4 | 0 | 0 | 0 |
| ENSDARG000 | 12:12:12:12:1 | 10134366:101 | 10134733:101 | +;+;+;+;+;+;+;+; | 3116 | SMARCD2 | 514 | 474 | 801 | 535 | 550 | 633 |
| ENSDARG000 | 12:12:12:12:1 | 4338298:4340 | 4339049:4340 | -;-;-;-;-;-;-;- | 1675 | ca15c | 0 | 0 | 0 | 0 | 0 | 2 |
| ENSDARG000 | 12:12:12:12:1 | 15666949:156 | 15667054:156 | +;+;+;+;+;+;+;+; | 1577 | nmt1b | 8 | 0 | 56 | 8 | 54 | 12 |
| ENSDARG000 | 12;12 | 25209369:252 | 25210655:252 | -;- | 1910 | KCNG3 | 8 | 16 | 21 | 28 | 24 | 21 |
| ENSDARG000 | 12;12;12 | 25197383:251 | 25197734:252 | -;-;- | 662 | cox7a2l | 544 | 799 | 1143 | 609 | 774 | 683 |
| ENSDARG000 | 12;12;12;12 | 25201543:252 | 25201722:252 | +;+;+;+ | 1095 | NA | 2 | 2 | 18 | 14 | 8 | 10 |
| ENSDARG000 | 12;12;12 | 19340786:193 | 19342033:193 | -;-;- | 1504 | maff | 805 | 681 | 886 | 1072 | 498 | 582 |
| ENSDARG000 | 12:12:12:12:1 | 19320657:193 | 19321015:193 | +;+;+;+;+;+;+;+; | 3794 | tmem184ba | 1249 | 1204 | 1302 | 988 | 1548 | 1045 |
| ENSDARG000 | 12:12:12:12:1 | 22164410:221 | 22164550:221 | -;-;-;-;- | 466 | NA | 2 | 0 | 2 | 0 | 0 | 0 |
| ENSDARG000 | 12:12:12:12:1 | 19305390:193 | 19306280:193 | +;+;+;+;+;+;+;+; | 5557 | csnk1e | 560 | 988 | 380 | 817 | 372 | 696 |
| ENSDARG000 | 12:12:12:12:1 | 22230412:222 | 22231246:222 | -;-;-;-;-;-;-;-;- | 1521 | ormdl3 | 28 | 2 | 18 | 7 | 6 | 12 |
| ENSDARG000 | 12:12:12:12:1 | 1286642:1288 | 1286719:1288 | +;+;+;+;+;+;+;+; | 2452 | NA | 140 | 180 | 134 | 228 | 264 | 221 |
| ENSDARG000 | 12:12:12:12:1 | 4360817:4361 | 4360855:4361 | -;-;-;-;-;-;- | 810 | NA | 0 | 8 | 0 | 4 | 6 | 12 |
| ENSDARG000 | 12:12:12:12:1 | 48390715:483 | 48390931:483 | +;+;+;+;+;+;+;+; | 1537 | scd | 866 | 869 | 397 | 308 | 369 | 284 |
| ENSDARG000 | 12:12:12:12:1 | 4377624:4378 | 4378048:4378 | -;-;-;-;-;- | 1753 | NA | 26 | 1 | 0 | 6 | 0 | 0 |
| ENSDARG000 | 12:12:12:12:1 | 7149448:7149 | 7149577:7149 | -;-;-;-;- | 1970 | NA | 2 | 0 | 6 | 2 | 34 | 6 |
| ENSDARG000 | 12:12:12:12:1 | 30036425:300 | 30036959:300 | -;-;-;-;-;-;-;-; | 1205 | trub1 | 74 | 66 | 133 | 44 | 110 | 170 |
| ENSDARG000 | 12;12;12;12 | 22258962:222 | 22259398:222 | +;+;+;+ | 1498 | wnt3 | 2 | 0 | 0 | 0 | 0 | 0 |

| | | | | | | | | | | | | |
|---|---|---|---|---|---|---|---|---|---|---|---|---|
| ENSDARG000 | 12:12:12:12:1 | 10163585:101 | 10163675:101 | +:+:+:+:+:+:+: | 1557 | psmc5 | 3189 | 3143 | 3315 | 1961 | 3254 | 2181 |
| ENSDARG000 | 12:12:12:12:1 | 7193779:720 | 7199763:720 | -;-;-;-;-;-;-;- | 6935 | ipmkb | 424 | 447 | 571 | 696 | 516 | 491 |
| ENSDARG000 | 12:12:12:12:1 | 48419226:484 | 48419989:484 | -:-:-:-:-:-:-:-: | 4867 | SEC31B | 797 | 454 | 927 | 491 | 828 | 682 |
| ENSDARG000 | 12:12:12:12:1 | 19356623:193 | 19356696:193 | +:+:+:+:+:+:+: | 1350 | dmc1 | 0 | 0 | 1 | 4 | 0 | 0 |
| ENSDARG000 | 12;12;12;12 | 19348538:193 | 19348781:193 | +;+;+;+ | 1241 | tomm22 | 813 | 977 | 837 | 567 | 730 | 631 |
| ENSDARG000 | 12:12:12:12:1 | 7237666:724 | 7238173:724 | -:-:-:-:-:-:-:-: | 1580 | ube2d1b | 615 | 625 | 781 | 518 | 605 | 577 |
| ENSDARG000 | 12;12 | 1398404:139 | 1398710:140 | +;+ | 1312 | rasd1 | 441 | 100 | 138 | 110 | 156 | 206 |
| ENSDARG000 | 12:12:12:12:1 | 25223799:252 | 25223907:252 | +:+:+:+:+:+:+: | 2685 | mta3 | 1131 | 1226 | 1824 | 1252 | 1394 | 1089 |
| ENSDARG000 | 12;12;12;12 | 1403806:140 | 1404557:140 | -;-;-;- | 1067 | MED9 | 200 | 136 | 99 | 76 | 128 | 116 |
| ENSDARG000 | 12:12:12:12:1 | 15690363:156 | 15690678:156 | +:+:+:+:+:+:+: | 8702 | NA | 3529 | 2865 | 2460 | 2857 | 3702 | 3148 |
| ENSDARG000 | 12:12:12:12:1 | 48469755:484 | 48469917:484 | -;-;-;-;-;-;- | 1034 | ndufb8 | 1159 | 1452 | 1490 | 980 | 1289 | 1172 |
| ENSDARG000 | 12:12:12:12:1 | 1469090:146 | 1469456:146 | +:+:+:+:+:+:+: | 3858 | usp22 | 417 | 357 | 434 | 198 | 396 | 212 |
| ENSDARG000 | 12:12:12:12:1 | 31476169:314 | 31476597:314 | -:-:-:-:-:-:-:-: | 1663 | tectb | 0 | 0 | 0 | 0 | 2 | 0 |
| ENSDARG000 | 12:12:12:12:1 | 22310365:223 | 22311508:223 | -:-:-:-:-:-:-:-: | 3593 | nsfb | 1171 | 1261 | 1278 | 890 | 1280 | 944 |
| ENSDARG000 | 12;12;12;12 | 10179452:101 | 10179485:101 | +;+;+;+ | 786 | NA | 2 | 0 | 0 | 2 | 0 | 0 |
| ENSDARG000 | 12:12:12:12:1 | 1947593:194 | 1948528:194 | -;-;-;-;-;- | 1835 | sox9a | 54 | 4 | 18 | 8 | 30 | 24 |
| ENSDARG000 | 12:12:12:12:1 | 10199219:101 | 10200141:102 | -:-:-:-:-:-:-:-: | 3000 | mpp2b | 0 | 2 | 0 | 2 | 0 | 0 |
| ENSDARG000 | 12:12:12:12:1 | 31494830:315 | 31494932:315 | +:+:+:+:+:+:+: | 4763 | GPAM | 70 | 109 | 126 | 36 | 139 | 140 |
| ENSDARG000 | 12:12:12:12:1 | 2446837:245 | 2446957:245 | +:+:+:+:+:+:+: | 2261 | ARHGAP22 | 18 | 8 | 24 | 12 | 16 | 4 |
| ENSDARG000 | 12;12;12 | 2428300:247 | 2428402:247 | -;-;- | 337 | NA | 0 | 0 | 2 | 0 | 0 | 0 |
| ENSDARG000 | 12;12 | 2428247:242 | 2428424:243 | +;+ | 951 | LRRC18 | 0 | 0 | 4 | 0 | 0 | 0 |
| ENSDARG000 | 12:12:12:12:1 | 48480632:485 | 48480842:485 | +:+:+:+:+:+:+: | 2372 | ARHGAP44 | 8 | 8 | 8 | 0 | 10 | 6 |
| ENSDARG000 | 12;12;12 | 16062560:160 | 16063757:160 | -;-;- | 2419 | NA | 56 | 4 | 20 | 0 | 62 | 68 |
| ENSDARG000 | 12:12:12:12:1 | 16087077:160 | 16087300:160 | +;+;+;+;+;+;+ | 4782 | znf281b | 613 | 430 | 741 | 535 | 586 | 736 |
| ENSDARG000 | 12;12;12 | 25292876:252 | 25294122:252 | -;-;- | 1470 | OXER1 | 791 | 1277 | 1089 | 1285 | 791 | 1115 |
| ENSDARG000 | 12;12;12;12 | 10304435:103 | 10304745:103 | -;-;-;- | 642 | pyyb | 0 | 2 | 0 | 0 | 0 | 0 |
| ENSDARG000 | 12;12 | 25376639:253 | 25379064:253 | -;- | 2656 | zfp36l2 | 686 | 469 | 823 | 628 | 394 | 430 |
| ENSDARG000 | 12:12:12:12:1 | 25312018:253 | 25312141:253 | +:+:+:+:+:+:+: | 1306 | NA | 3 | 0 | 10 | 0 | 19 | 7 |
| ENSDARG000 | 12:12:12:12:1 | 48784731:487 | 48784822:487 | +:+:+:+:+:+:+: | 2602 | ZMIZ1 | 81 | 22 | 82 | 57 | 83 | 58 |
| ENSDARG000 | 12:12:12:12:1 | 10337674:103 | 10339095:103 | -:-:-:-:-:-:-:-: | 2774 | tut1 | 122 | 204 | 209 | 94 | 120 | 110 |
| ENSDARG000 | 12:12:12:12:1 | 25432626:254 | 25433054:254 | +:+:+:+:+:+:+: | 6314 | ppm1bb | 1714 | 1482 | 1958 | 1353 | 1964 | 1789 |
| ENSDARG000 | 12:12:12:12:1 | 48803098:488 | 48803301:488 | +;+;+;+;+;+ | 1135 | ppifb | 559 | 689 | 884 | 534 | 838 | 724 |
| ENSDARG000 | 12;12 | 25600685:256 | 25601576:256 | +;+ | 1759 | six3b | 0 | 0 | 0 | 0 | 0 | 0 |
| ENSDARG000 | 12;12 | 25610019:256 | 25610554:256 | -;- | 1334 | six2b | 2 | 0 | 2 | 0 | 0 | 0 |
| ENSDARG000 | 12:12:12:12:1 | 22359700:223 | 22362163:223 | -:-:-:-:-:-:-:-: | 5591 | NA | 114 | 16 | 48 | 16 | 20 | 30 |
| ENSDARG000 | 12:12:12:12:1 | 10353333:103 | 10353611:103 | -:-:-:-:-:-:-:-: | 2310 | ndc80 | 407 | 813 | 831 | 206 | 510 | 350 |
| ENSDARG000 | 12:12:12:12:1 | 48815988:488 | 48816227:488 | +:+:+:+:+:+:+: | 2919 | anxa11b | 6320 | 5236 | 5747 | 5370 | 5913 | 7249 |
| ENSDARG000 | 12:12:12:12:1 | 25640480:256 | 25641130:256 | +:+:+:+:+:+:+: | 4572 | PRKCE | 2156 | 1389 | 1763 | 2904 | 1808 | 2189 |
| ENSDARG000 | 12:12:12:12:1 | 10366617:103 | 10368381:103 | -:-:-:-:-:-:-:-: | 8187 | mki67 | 730 | 2929 | 4012 | 1011 | 2811 | 1435 |
| ENSDARG000 | 12:12:12:12:1 | 16106459:161 | 16106624:161 | +:+:+:+:+:+:+: | 2220 | muc13b | 0 | 0 | 18 | 6 | 53 | 0 |
| ENSDARG000 | 12:12:12:12:1 | 16132612:161 | 16132775:161 | +:+:+:+:+:+:+: | 14302 | NA | 35 | 2 | 45 | 16 | 121 | 36 |
| ENSDARG000 | 12:12:12:12:1 | 16233077:162 | 16233252:162 | +:+:+:+:+:+:+: | 2097 | MPP3 | 0 | 0 | 0 | 0 | 0 | 0 |
| ENSDARG000 | 12:12:12:12:1 | 16281312:162 | 16281363:162 | +;+;+;+;+;+ | 2695 | ppp1r3cb | 122 | 136 | 264 | 156 | 264 | 254 |
| ENSDARG000 | 12:12:12:12:1 | 48898806:488 | 48898919:488 | -:-:-:-:-:-:-:-: | 1389 | KCNMA1 | 2 | 0 | 0 | 0 | 0 | 0 |
| ENSDARG000 | 12:12:12:12:1 | 7609928:761 | 7613163:761 | -:-:-:-:-:-:-:-: | 4446 | ccdc6b | 397 | 501 | 810 | 578 | 652 | 474 |
| ENSDARG000 | 12:12:12:12:1 | 46483618:464 | 46483665:464 | +:+:+:+:+:+:+: | 1271 | NA | 4 | 0 | 0 | 0 | 0 | 20 |
| ENSDARG000 | 12:12:12:12:1 | 41420355:414 | 41420507:414 | +:+:+:+:+:+:+: | 2391 | EPC1 | 757 | 489 | 705 | 362 | 648 | 574 |
| ENSDARG000 | 12:12:12:12:1 | 45225166:452 | 45225567:452 | -;-;-;-;-;-;-;- | 2212 | NA | 40 | 72 | 66 | 52 | 153 | 50 |
| ENSDARG000 | 12:12:12:12:1 | 41697664:417 | 41698291:417 | +;+;+;+;+ | 1137 | bnip3 | 43 | 11 | 14 | 39 | 50 | 22 |
| ENSDARG000 | 12:12:12:12:1 | 204095:2052 | 204443:2053 | -:-:-:-:-:-:-:-: | 2234 | NA | 3134 | 4435 | 901 | 10479 | 3434 | 5943 |
| ENSDARG000 | 12:12:12:12:1 | 48581588:486 | 48581966:486 | +:+:+:+:+:+:+: | 7241 | SYDE1 | 4144 | 1123 | 2366 | 2585 | 3049 | 3711 |
| ENSDARG000 | 12:12:12:12:1 | 48241841:482 | 48241901:482 | +:+:+:+:+:+:+: | 1369 | ppa1a | 2 | 0 | 0 | 6 | 0 | 6 |
| ENSDARG000 | 12;12 | 43662980:436 | 43663857:436 | -;- | 1571 | foxi1 | 0 | 0 | 4 | 0 | 6 | 4 |
| ENSDARG000 | 12:12:12:12:1 | 47783814:477 | 47784002:477 | -;-;-;-;-;-;- | 997 | dydc2 | 0 | 0 | 0 | 0 | 0 | 0 |
| ENSDARG000 | 12:12:12:12:1 | 1349418:134 | 1349436:134 | -:-:-:-:-:-:-:-: | 861 | RASSF4 | 6 | 0 | 0 | 2 | 2 | 0 |
| ENSDARG000 | 12:12:12:12:1 | 47794089:477 | 47795073:477 | +:+:+:+:+:+:+: | 5493 | polr3a | 581 | 474 | 1027 | 260 | 569 | 562 |
| ENSDARG000 | 12 | 1556147 | 1557286 | - | 1140 | SSTR2 | 0 | 0 | 0 | 0 | 0 | 0 |
| ENSDARG000 | 12:12:12:12:1 | 8003872:800 | 8003873:800 | +:+:+:+:+:+:+: | 2098 | RHOBTB1 | 0 | 0 | 0 | 0 | 0 | 0 |
| ENSDARG000 | 12:12:12:12:1 | 16555603:165 | 16555937:165 | -;-;-;-;-;-;-;- | 1284 | NA | 0 | 0 | 0 | 12 | 0 | 0 |
| ENSDARG000 | 12;12 | 15078124:150 | 15078330:150 | +;+ | 379 | NA | 0 | 0 | 0 | 0 | 0 | 0 |
| ENSDARG000 | 12;12;12;12 | 52468:53325 | 53209:53608 | -;-;-;- | 1372 | si:ch1073-357 | 233 | 296 | 207 | 176 | 227 | 113 |
| ENSDARG000 | 12 | 37698602 | 37698717 | + | 116 | 5S_rRNA | 0 | 0 | 0 | 0 | 0 | 0 |
| ENSDARG000 | 12:12:12:12:1 | 47967761:479 | 47968101:479 | -:-:-:-:-:-:-:-: | 2403 | ADAMTS14 | 20 | 2 | 0 | 14 | 6 | 38 |
| ENSDARG000 | 12 | 41764153 | 41764276 | - | 124 | 5S_rRNA | 0 | 0 | 0 | 0 | 0 | 0 |
| ENSDARG000 | 12:12:12:12:1 | 1087263:109 | 1087482:109 | +:+:+:+:+:+:+: | 1728 | LMF1 | 264 | 242 | 312 | 136 | 327 | 256 |
| ENSDARG000 | 12 | 26140205 | 26140321 | - | 117 | 5S_rRNA | 0 | 0 | 0 | 0 | 0 | 0 |
| ENSDARG000 | 12 | 16654681 | 16654795 | - | 115 | 5S_rRNA | 0 | 0 | 0 | 0 | 0 | 0 |
| ENSDARG000 | 12 | 37959814 | 37959930 | - | 117 | 5S_rRNA | 0 | 0 | 0 | 0 | 0 | 0 |
| ENSDARG000 | 12 | 37792016 | 37792131 | - | 116 | 5S_rRNA | 0 | 0 | 0 | 0 | 0 | 0 |
| ENSDARG000 | 12:12:12:12:1 | 197424:1976 | 197429:1977 | +;+;+;+;+ | 384 | NA | 161 | 184 | 99 | 258 | 101 | 188 |
| ENSDARG000 | 12:12:12:12:1 | 964760:9651 | 964952:9653 | -:-:-:-:-:-:-:-: | 2007 | DAGLB | 779 | 739 | 782 | 610 | 863 | 689 |
| ENSDARG000 | 12;12;12 | 118704:1222 | 118913:1224 | -;-;- | 543 | NA | 0 | 2 | 0 | 6 | 3 | 0 |
| ENSDARG000 | 12 | 5052443 | 5052578 | + | 136 | NA | 0 | 0 | 0 | 0 | 0 | 0 |
| ENSDARG000 | 12 | 7420032 | 7420148 | + | 117 | 5S_rRNA | 0 | 0 | 0 | 0 | 0 | 0 |
| ENSDARG000 | 12:12:12:12:1 | 312448:3126 | 312571:3127 | -;-;-;-;-;-;-;- | 420 | PTS | 126 | 60 | 464 | 389 | 333 | 246 |
| ENSDARG000 | 12 | 43738618 | 43738733 | - | 116 | 5S_rRNA | 0 | 0 | 0 | 0 | 0 | 0 |
| ENSDARG000 | 12:12:12:12:1 | 32020121:320 | 32020154:320 | +:+:+:+:+:+:+: | 399 | NA | 2 | 0 | 0 | 3 | 0 | 2 |
| ENSDARG000 | 12 | 5061567 | 5061697 | + | 131 | NA | 0 | 0 | 4 | 0 | 0 | 0 |
| ENSDARG000 | 12 | 8443904 | 8444021 | - | 118 | 5S_rRNA | 0 | 0 | 0 | 0 | 0 | 0 |
| ENSDARG000 | 12:12:12:12:1 | 3314504:331 | 3316580:331 | -:-:-:-:-:-:-:-: | 6231 | TANC2 | 173 | 128 | 242 | 202 | 191 | 125 |
| ENSDARG000 | 12 | 24565478 | 24565592 | + | 115 | 5S_rRNA | 0 | 0 | 0 | 0 | 0 | 0 |
| ENSDARG000 | 12:12:12:12:1 | 46404307:464 | 46404354:464 | +:+:+:+:+:+:+: | 1271 | NA | 0 | 0 | 0 | 0 | 0 | 6 |
| ENSDARG000 | 12:12:12:12:1 | 36834901:368 | 36835152:368 | +:+:+:+:+:+:+: | 2661 | NA | 20 | 6 | 0 | 6 | 8 | 6 |
| ENSDARG000 | 12:12:12:12:1 | 25094099:250 | 25095050:250 | -;-;-;-;- | 1241 | cript | 589 | 535 | 440 | 459 | 491 | 543 |
| ENSDARG000 | 12:12:12:12:1 | 34259742:342 | 34259771:342 | -;-;-;-;-;-;- | 177 | NA | 22 | 2 | 5 | 3 | 17 | 14 |
| ENSDARG000 | 12:12:12:12:1 | 44064620:440 | 44064820:440 | -:-:-:-:-:-:-:-: | 2487 | NA | 12 | 0 | 0 | 0 | 1 | 0 |
| ENSDARG000 | 12 | 35021516 | 35021614 | - | 99 | 5S_rRNA | 106 | 67 | 79 | 141 | 111 | 124 |
| ENSDARG000 | 12;12 | 48117443:481 | 48122280:481 | -;- | 5081 | NPFFR1 | 1543 | 10 | 14 | 0 | 14 | 10 |
| ENSDARG000 | 12 | 18598850 | 18598964 | + | 115 | 5S_rRNA | 0 | 0 | 0 | 0 | 0 | 0 |
| ENSDARG000 | 12:12:12:12:1 | 17794284:177 | 17794623:177 | -;-;-;-;-;-;-;- | 1497 | NA | 24 | 22 | 24 | 18 | 27 | 8 |
| ENSDARG000 | 12:12:12:12:1 | 45132906:451 | 45133173:451 | -:-:-:-:-:-:-:-: | 1311 | FANK1 | 62 | 36 | 60 | 72 | 97 | 71 |
| ENSDARG000 | 12:12:12:12:1 | 8074343:807 | 8074501:807 | +:+:+:+:+:+:+: | 1012 | cabcoco1 | 2 | 2 | 24 | 0 | 6 | 0 |
| ENSDARG000 | 12:12:12:12:1 | 3652015:365 | 3652218:365 | -:-:-:-:-:-:-:-: | 1419 | NA | 10 | 4 | 16 | 8 | 24 | 0 |
| ENSDARG000 | 12:12:12:12:1 | 48979060:489 | 48979683:489 | -:-:-:-:-:-:-:-: | 2361 | NA | 4 | 6 | 214 | 22 | 140 | 16 |
| ENSDARG000 | 12;12;12;12 | 2381213:238 | 2381230:238 | +;+;+;+ | 339 | NA | 0 | 0 | 0 | 0 | 0 | 0 |
| ENSDARG000 | 12:12:12:12:1 | 47907277:479 | 47907299:479 | +:+:+:+:+:+:+: | 2117 | TBATA | 2 | 0 | 0 | 0 | 8 | 0 |
| ENSDARG000 | 12:12:12:12:1 | 45339637:453 | 45340139:453 | -;-;-;-;-;- | 1086 | NA | 11316 | 4099 | 1499 | 4103 | 5152 | 1158 |
| ENSDARG000 | 12;12;12;12 | 17701625:177 | 17701643:177 | +;+;+;+ | 93 | NA | 0 | 0 | 0 | 0 | 0 | 0 |
| ENSDARG000 | 12;12 | 25945560:259 | 25947719:259 | +;+ | 2579 | NA | 37 | 22 | 12 | 5 | 6 | 0 |
| ENSDARG000 | 12 | 24418224 | 24418340 | + | 117 | 5S_rRNA | 0 | 0 | 0 | 0 | 0 | 0 |
| ENSDARG000 | 12:12:12:12:1 | 46386983:463 | 46387030:463 | +:+:+:+:+:+:+: | 1271 | NA | 24 | 2 | 0 | 14 | 74 | 156 |
| ENSDARG000 | 12;12;12 | 36092218:360 | 36092872:360 | +;+;+ | 5851 | BTBD17 | 6 | 4 | 0 | 0 | 0 | 0 |
| ENSDARG000 | 12:12:12:12:1 | 41723040:417 | 41724061:417 | -:-:-:-:-:-:-:-: | 2222 | ppp2r2d | 676 | 704 | 929 | 471 | 621 | 800 |
| ENSDARG000 | 12:12:12:12:1 | 25894539:258 | 25894919:258 | -;-;-;-;- | 856 | sncgb | 6 | 12 | 6 | 0 | 78 | 0 |
| ENSDARG000 | 12:12:12:12:1 | 4460210:446 | 4460341:446 | -:-:-:-:-:-:-:-: | 1581 | NA | 15 | 0 | 0 | 1 | 0 | 4 |
| ENSDARG000 | 12 | 4286366 | 4286480 | - | 115 | 5S_rRNA | 0 | 0 | 2 | 0 | 0 | 4 |
| ENSDARG000 | 12 | 12335216 | 12335331 | - | 116 | 5S_rRNA | 0 | 0 | 0 | 0 | 0 | 0 |
| ENSDARG000 | 12;12;12;12 | 35947865:359 | 35948120:359 | -;-;-;- | 462 | NA | 0 | 0 | 0 | 0 | 0 | 0 |
| ENSDARG000 | 12 | 23110163 | 23110277 | + | 115 | 5S_rRNA | 0 | 0 | 0 | 0 | 0 | 0 |
| ENSDARG000 | 12:12:12:12:1 | 42592:43389 | 43292:43510 | -:-:-:-:-:-:-:-: | 2074 | wdr45b | 923 | 802 | 1224 | 895 | 1038 | 883 |
| ENSDARG000 | 12;12;12;12 | 35980541:359 | 35981948:359 | -;-;-;- | 1704 | pde6gb | 22 | 4 | 6 | 108 | 4 | 47 |
| ENSDARG000 | 12:12:12:12:1 | 3287783:329 | 3288477:329 | -;-;-;-;- | 1198 | lsm12b | 1300 | 1469 | 2214 | 1618 | 1327 | 1501 |
| ENSDARG000 | 12:12:12:12:1 | 17815725:178 | 17816948:178 | -:-:-:-:-:-:-:-: | 2933 | baiap2l1a | 14 | 14 | 18 | 2 | 4 | 0 |
| ENSDARG000 | 12 | 5057221 | 5057356 | + | 136 | NA | 0 | 0 | 0 | 0 | 0 | 0 |
| ENSDARG000 | 12:12:12:12:1 | 47439:47714 | 47624:47741 | +;+;+;+;+;+ | 654 | mrpl58 | 134 | 100 | 140 | 143 | 153 | 109 |
| ENSDARG000 | 12:12:12:12:1 | 44162747:441 | 44162827:441 | -:-:-:-:-:-:-:-: | 1293 | NA | 0 | 0 | 0 | 0 | 0 | 0 |
| ENSDARG000 | 12 | 1693014 | 1694053 | - | 1040 | NA | 0 | 0 | 0 | 0 | 0 | 0 |
| ENSDARG000 | 12 | 29837882 | 29837998 | - | 117 | 5S_rRNA | 0 | 0 | 0 | 0 | 0 | 0 |
| ENSDARG000 | 12 | 18125658 | 18125774 | + | 117 | 5S_rRNA | 0 | 0 | 0 | 0 | 0 | 0 |
| ENSDARG000 | 12:12:12:12:1 | 4260779:426 | 4260886:426 | +:+:+:+:+:+:+: | 1722 | NA | 54 | 87 | 74 | 56 | 177 | 89 |
| ENSDARG000 | 12 | 15381694 | 15381814 | + | 121 | 5S_rRNA | 0 | 0 | 0 | 0 | 0 | 0 |
| ENSDARG000 | 12:12:12:12:1 | 45186533:451 | 45189303:451 | -;-;-;-;-;-;- | 4209 | bccip | 1403 | 1125 | 1569 | 893 | 1327 | 1248 |
| ENSDARG000 | 12:12:12:12:1 | 44119432:441 | 44121071:441 | -:-:-:-:-:-:-:-: | 7634 | NA | 0 | 2 | 0 | 0 | 0 | 0 |
| ENSDARG000 | 12 | 29811844 | 29811960 | + | 117 | 5S_rRNA | 0 | 0 | 0 | 0 | 0 | 0 |
| ENSDARG000 | 12 | 17706792 | 17706901 | - | 110 | dre-mir-2192 | 0 | 0 | 0 | 0 | 0 | 0 |
| ENSDARG000 | 12:12:12:12:1 | 35809503:358 | 35813971:358 | -;-;-;-;-;-;-;- | 5869 | NA | 92 | 86 | 79 | 16 | 50 | 4 |
| ENSDARG000 | 12:12:12:12:1 | 39685485:397 | 39685823:397 | +:+:+:+:+:+:+: | 1992 | NA | 12 | 20 | 133 | 8 | 62 | 30 |
| ENSDARG000 | 12:12:12:12:1 | 292452:2941 | 292464:2941 | +:+:+:+:+:+:+: | 417 | NA | 853 | 909 | 497 | 984 | 1162 | 2435 |
| ENSDARG000 | 12:12:12:12:1 | 47773964:477 | 47773989:477 | +;+;+;+;+;+ | 126 | NA | 1 | 2 | 0 | 2 | 0 | 2 |
| ENSDARG000 | 12:12:12:12:1 | 17865374:178 | 17866148:178 | +:+:+:+:+:+:+: | 2695 | tmem130 | 707 | 744 | 2365 | 2064 | 1932 | 4647 |
| ENSDARG000 | 12 | 46836271 | 46838062 | - | 1792 | NA | 636 | 847 | 830 | 344 | 1035 | 495 |
| ENSDARG000 | 12:12:12:12:1 | 10679026:106 | 10679303:106 | -;-;-;-;-;-;-;- | 1432 | NA | 8 | 2 | 14 | 4 | 2 | 8 |
| ENSDARG000 | 12 | 32095170 | 32095284 | - | 115 | 5S_rRNA | 0 | 0 | 0 | 0 | 0 | 0 |
| ENSDARG000 | 12 | 8860193 | 8860307 | - | 115 | 5S_rRNA | 6 | 1 | 0 | 9 | 0 | 0 |
| ENSDARG000 | 12 | 21437346 | 21437458 | + | 113 | 5S_rRNA | 0 | 0 | 0 | 0 | 0 | 0 |
| ENSDARG000 | 12 | 37938792 | 37941733 | - | 2942 | NA | 0 | 0 | 0 | 0 | 0 | 0 |
| ENSDARG000 | 12 | 17727542 | 17727651 | + | 110 | dre-mir-338-2 | 0 | 0 | 0 | 0 | 0 | 0 |
| ENSDARG000 | 12:12:12:12:1 | 44041915:440 | 44041995:440 | -;-;-;-;-;- | 924 | NA | 6 | 2 | 0 | 0 | 8 | 0 |
| ENSDARG000 | 12:12:12:12:1 | 48965049:489 | 48965177:489 | -;-;-;-;-;-;-;- | 873 | rgrb | 731 | 217 | 412 | 540 | 362 | 210 |
| ENSDARG000 | 12:12:12:12:1 | 44009628:440 | 44009815:440 | -:-:-:-:-:-:-:-: | 1937 | NA | 0 | 6 | 0 | 0 | 0 | 0 |
| ENSDARG000 | 12:12:12:12:1 | 43848634:440 | 43850183:440 | -:-:-:-:-:-:-:-: | 6565 | NA | 7 | 2 | 0 | 0 | 0 | 0 |
| ENSDARG000 | 12:12:12:12:1 | 46098784:460 | 46099205:460 | -:-:-:-:-:-:-:-: | 2755 | NA | 35 | 4 | 0 | 2 | 4 | 0 |
| ENSDARG000 | 12;12 | 7699632:771 | 7699759:771 | -;- | 210 | NA | 31 | 12 | 64 | 15 | 63 | 5 |
| ENSDARG000 | 12:12:12:12:1 | 46425800:464 | 46425847:464 | +:+:+:+:+:+:+: | 1795 | NA | 1 | 0 | 0 | 0 | 0 | 4 |
| ENSDARG000 | 12;12;12 | 48744016:487 | 48745441:487 | +;+;+ | 1593 | NA | 0 | 0 | 1 | 0 | 4 | 5 |
| ENSDARG000 | 12;12;12;12 | 4533921:453 | 4534306:453 | -;-;-;- | 847 | NA | 117 | 16 | 5 | 20 | 36 | 29 |
| ENSDARG000 | 12 | 35710521 | 35710605 | - | 85 | dre-mir-338-1 | 0 | 0 | 0 | 0 | 0 | 0 |
| ENSDARG000 | 12;12;12;12 | 48972915:489 | 48973813:489 | +;+;+;+ | 2742 | lrit1b | 4 | 0 | 1 | 0 | 8 | 0 |
| ENSDARG000 | 12 | 13553730 | 13556536 | - | 2807 | NA | 0 | 2 | 0 | 0 | 0 | 0 |
| ENSDARG000 | 12;12 | 48958913:489 | 48959739:489 | -;- | 1296 | slc18a3b | 0 | 0 | 0 | 0 | 0 | 0 |
| ENSDARG000 | 12;12;12 | 41348969:413 | 41350120:413 | +;+;+ | 2569 | NA | 0 | 0 | 0 | 0 | 0 | 0 |
| ENSDARG000 | 12 | 9117052 | 9117178 | - | 127 | 5S_rRNA | 0 | 0 | 0 | 0 | 0 | 0 |
| ENSDARG000 | 12:12:12:12:1 | 32370:33853 | 33741:34020 | -;-;-;-;-;-;-;- | 2962 | foxk2a | 1317 | 1097 | 1646 | 1092 | 1360 | 888 |
| ENSDARG000 | 12:12:12:12:1 | 48634927:486 | 48634957:486 | +:+:+:+:+:+:+: | 1015 | zgc:165653 | 7 | 8 | 4 | 49 | 14 | 8 |
| ENSDARG000 | 12 | 20189169 | 20189286 | - | 118 | 5S_rRNA | 0 | 0 | 0 | 0 | 0 | 0 |
| ENSDARG000 | 12:12:12:12:1 | 34206753:342 | 34206914:342 | -:-:-:-:-:-:-:-: | 1966 | PGS1 | 139 | 198 | 225 | 36 | 218 | 101 |
| ENSDARG000 | 12 | 35109719 | 35109833 | - | 115 | 5S_rRNA | 0 | 0 | 0 | 0 | 0 | 0 |
| ENSDARG000 | 12 | 30202191 | 30202307 | - | 117 | 5S_rRNA | 0 | 0 | 0 | 0 | 0 | 0 |
| ENSDARG000 | 12:12:12:12:1 | 35956245:359 | 35956402:359 | -:-:-:-:-:-:-:-: | 2811 | nploc4 | 927 | 772 | 1120 | 516 | 959 | 840 |
| ENSDARG000 | 12 | 39503564 | 39503678 | + | 115 | 5S_rRNA | 0 | 0 | 0 | 0 | 0 | 0 |
| ENSDARG000 | 12 | 35973856 | 35973973 | - | 118 | 5S_rRNA | 0 | 0 | 0 | 0 | 0 | 0 |
| ENSDARG000 | 12;12;12;12 | 47663419:476 | 47663762:476 | +;+;+;+ | 531 | NA | 8 | 0 | 0 | 8 | 8 | 0 |

| | | | | | | | | | | | | |
|---|---|---|---|---|---|---|---|---|---|---|---|---|
| ENSDARG000 | 12 | 30509283 | 30509399 | + | 117 | 5S_rRNA | 0 | 0 | 0 | 0 | 0 | 0 |
| ENSDARG000 | 12;12;12;12 | 3262564:3264 | 3262718:3264 | +;+;+;+ | 3533 | tmem101 | 294 | 310 | 747 | 342 | 364 | 346 |
| ENSDARG000 | 12;12;12;12;1 | 8064029:8064 | 8064510:8064 | -;-;-;-;-;- | 1196 | tmem26b | 0 | 2 | 0 | 4 | 2 | 6 |
| ENSDARG000 | 12;12;12;12;1 | 45512415:455 | 45512696:455 | -;-;-;-;-;- | 894 | NA | 1 | 0 | 9 | 4 | 6 | 0 |
| ENSDARG000 | 12;12;12;12;1 | 35654749:356 | 35654886:356 | +;+;+;+;+;+;+; | 1227 | BAIAP2 | 12 | 6 | 0 | 2 | 0 | 0 |
| ENSDARG000 | 12 | 40411566 | 40411682 | + | 117 | 5S_rRNA | 0 | 0 | 0 | 0 | 0 | 0 |
| ENSDARG000 | 12 | 28500121 | 28500239 | - | 119 | 5S_rRNA | 0 | 0 | 0 | 0 | 0 | 0 |
| ENSDARG000 | 12;12;12;12;1 | 45060789:450 | 45060922:450 | +;+;+;+;+;+;+; | 1794 | ADAM12 | 149 | 110 | 244 | 115 | 260 | 169 |
| ENSDARG000 | 12 | 7865470 | 7870791 | + | 5322 | NA | 0 | 0 | 0 | 1 | 3 | 0 |
| ENSDARG000 | 12;12;12;12;1 | 43701777:437 | 43706402:437 | -;-;-;-;-;- | 5779 | NA | 0 | 0 | 0 | 0 | 0 | 0 |
| ENSDARG000 | 12;12;12;12;1 | 1368645:1368 | 1368814:1369 | -;-;-;-;-;-;- | 1050 | NA | 0 | 0 | 0 | 0 | 0 | 0 |
| ENSDARG000 | 12;12;12;12;1 | 45493054:454 | 45493672:454 | -;-;-;-;-;-;-;-;-; | 1294 | NA | 1155 | 346 | 402 | 240 | 554 | 706 |
| ENSDARG000 | 12;12;12;12;1 | 49166074:491 | 49166354:491 | -;-;-;-;-;- | 1098 | PSTK | 147 | 120 | 252 | 107 | 167 | 203 |
| ENSDARG000 | 12;12;12 | 9059:10291:1 | 9538:10996:1 | -;-;- | 1287 | shisa6 | 0 | 0 | 20 | 38 | 0 | 2 |
| ENSDARG000 | 12 | 19388143 | 19388273 | + | 131 | NA | 0 | 0 | 0 | 0 | 0 | 0 |
| ENSDARG000 | 12;12 | 47774120:477 | 47774227:477 | -;- | 273 | NA | 2 | 25 | 2 | 14 | 0 | 6 |
| ENSDARG000 | 12;12;12;12;1 | 137149:13741 | 137337:13756 | -;-;-;-;-;-;-;-;-; | 1964 | dnah9 | 0 | 4 | 0 | 0 | 2 | 2 |
| ENSDARG000 | 12 | 40104436 | 40104551 | - | 116 | 5S_rRNA | 0 | 0 | 0 | 0 | 0 | 0 |
| ENSDARG000 | 12;12 | 16522472:165 | 16523225:165 | -;- | 1501 | HTR7 | 0 | 0 | 0 | 0 | 0 | 0 |
| ENSDARG000 | 12 | 12613335 | 12613451 | + | 117 | 5S_rRNA | 0 | 0 | 0 | 0 | 0 | 0 |
| ENSDARG000 | 12 | 29013573 | 29013690 | + | 118 | 5S_rRNA | 0 | 0 | 0 | 0 | 4 | 0 |
| ENSDARG000 | 12;12;12;12;1 | 35914103:359 | 35914289:359 | -;-;-;-;-;-;-;-;-; | 12211 | NA | 2060 | 1093 | 466 | 250 | 881 | 496 |
| ENSDARG000 | 12 | 23577266 | 23577391 | + | 126 | 5S_rRNA | 0 | 0 | 0 | 0 | 0 | 0 |
| ENSDARG000 | 12 | 7502962 | 7503098 | - | 137 | 5S_rRNA | 0 | 0 | 0 | 0 | 0 | 0 |
| ENSDARG000 | 12;12;12;12;1 | 46462090:464 | 46462137:464 | +;+;+;+;+;+;+; | 1271 | NA | 1 | 1 | 0 | 4 | 0 | 2 |
| ENSDARG000 | 12 | 35611698 | 35611814 | + | 117 | 5S_rRNA | 0 | 0 | 0 | 0 | 0 | 0 |
| ENSDARG000 | 12 | 21569783 | 21569899 | - | 117 | 5S_rRNA | 0 | 0 | 0 | 0 | 0 | 0 |
| ENSDARG000 | 12;12;12;12;1 | 3301149:3303 | 3301469:3304 | +;+;+;+;+;+;+ | 1822 | g6pc3 | 121 | 217 | 213 | 60 | 165 | 136 |
| ENSDARG000 | 12 | 16493352 | 16493468 | - | 117 | 5S_rRNA | 0 | 2 | 0 | 0 | 0 | 0 |
| ENSDARG000 | 12 | 13457237 | 13457356 | + | 120 | 5S_rRNA | 0 | 0 | 0 | 0 | 0 | 0 |
| ENSDARG000 | 12;12;12;12;1 | 48094278:480 | 48094427:480 | -;-;-;-;-;-;-;-;-; | 4011 | pald1a | 2833 | 3505 | 4767 | 7409 | 4197 | 6416 |
| ENSDARG000 | 12 | 29959940 | 29960054 | + | 115 | 5S_rRNA | 0 | 0 | 0 | 0 | 0 | 0 |
| ENSDARG000 | 12 | 41460518 | 41460634 | + | 117 | 5S_rRNA | 0 | 0 | 0 | 0 | 0 | 0 |
| ENSDARG000 | 12;12;12;12;1 | 41510492:415 | 41510959:415 | +;+;+;+;+;+;+; | 3227 | KIF5B | 1587 | 803 | 896 | 873 | 1136 | 727 |
| ENSDARG000 | 12 | 29432075 | 29432191 | + | 117 | 5S_rRNA | 0 | 0 | 0 | 0 | 0 | 0 |
| ENSDARG000 | 12;12;12;12;1 | 43669294:436 | 43670252:436 | -;-;-;-;-;-;-;- | 1900 | zgc:112964 | 0 | 0 | 5 | 0 | 2 | 0 |
| ENSDARG000 | 12;12;12;12;1 | 20412293:204 | 20412970:204 | +;+;+;+;+;+;+; | 3323 | arhgap17a | 374 | 296 | 550 | 238 | 537 | 417 |
| ENSDARG000 | 12;12;12;12 | 48319221:483 | 48319256:483 | +;+;+;+ | 561 | anapc16 | 318 | 276 | 329 | 258 | 292 | 248 |
| ENSDARG000 | 12;12;12 | 43585527:435 | 43586586:435 | +;+;+ | 5888 | CLRN3 | 0 | 0 | 3 | 0 | 4 | 0 |
| ENSDARG000 | 12;12;12;12;1 | 41631521:416 | 41632142:416 | -;-;-;-;-;-;-;-;-; | 5526 | NA | 64 | 8 | 177 | 34 | 55 | 254 |
| ENSDARG000 | 12;12;12;12;1 | 47698356:477 | 47698834:477 | +;+;+;+;+ | 2333 | LZTS2 | 308 | 117 | 188 | 275 | 116 | 226 |
| ENSDARG000 | 12 | 16901645 | 16901759 | - | 115 | 5S_rRNA | 0 | 0 | 0 | 0 | 0 | 0 |
| ENSDARG000 | 12;12;12;12;1 | 44362897:443 | 44362992:443 | -;-;-;-;-;-;-;-;-; | 5523 | dock1 | 1175 | 1039 | 1647 | 1069 | 1452 | 1021 |
| ENSDARG000 | 12;12;12;12;1 | 30498412:304 | 30498419:304 | +;+;+;+;+;+;+; | 276 | NA | 0 | 0 | 0 | 0 | 0 | 0 |
| ENSDARG000 | 12;12;12;12;1 | 38695808:386 | 38695990:386 | +;+;+;+;+;+;+; | 1894 | NA | 0 | 0 | 3 | 0 | 0 | 0 |
| ENSDARG000 | 12;12;12;12;1 | 24952902:249 | 24953570:249 | +;+;+;+;+;+;+; | 2354 | calm2b | 6260 | 9406 | 8244 | 7722 | 9759 | 7329 |
| ENSDARG000 | 12 | 14545475 | 14551077 | - | 5603 | NA | 0 | 0 | 0 | 0 | 0 | 0 |
| ENSDARG000 | 12;12;12;12;1 | 45263874:452 | 45264004:452 | -;-;-;-;-;-;-;-;-; | 2199 | FDXR | 1414 | 415 | 470 | 302 | 430 | 334 |
| ENSDARG000 | 12;12;12;12;1 | 47162456:471 | 47162503:471 | +;+;+;+;+;+;+; | 16743 | NA | 6 | 2 | 4 | 0 | 4 | 0 |
| ENSDARG000 | 12;12;12;12;1 | 48216662:482 | 48216803:482 | +;+;+;+;+;+;+; | 1238 | lrrc20 | 107 | 186 | 257 | 162 | 238 | 143 |
| ENSDARG000 | 12;12;12;12 | 1072486:1078 | 1073122:1078 | -;-;-;- | 1982 | sox8a | 6 | 4 | 54 | 4 | 10 | 22 |
| ENSDARG000 | 12;12;12;12;1 | 44610912:446 | 44612112:446 | +;+;+;+;+ | 3802 | INSYN2A | 0 | 6 | 0 | 0 | 6 | 15 |
| ENSDARG000 | 12 | 11643551 | 11643671 | + | 121 | 5S_rRNA | 0 | 0 | 0 | 0 | 0 | 0 |
| ENSDARG000 | 12 | 5051562 | 5051697 | + | 136 | NA | 0 | 0 | 0 | 0 | 0 | 0 |
| ENSDARG000 | 12 | 17345550 | 17345667 | + | 118 | 5S_rRNA | 0 | 0 | 0 | 0 | 0 | 0 |
| ENSDARG000 | 12;12;12;12;1 | 45238292:452 | 45238377:452 | +;+;+;+;+;+;+; | 1248 | mrpl38 | 445 | 581 | 693 | 368 | 441 | 355 |
| ENSDARG000 | 12;12;12 | 36054721:360 | 36055676:360 | -;-;- | 1567 | GPR142 | 0 | 0 | 0 | 0 | 0 | 0 |
| ENSDARG000 | 12;12;12;12;1 | 47081783:470 | 47081813:470 | +;+;+;+;+;+;+; | 3792 | mtr | 358 | 619 | 1427 | 282 | 653 | 313 |
| ENSDARG000 | 12;12;12 | 28117365:281 | 28117883:281 | +;+;+ | 5881 | NA | 8 | 8 | 0 | 0 | 0 | 2 |
| ENSDARG000 | 12;12;12;12;1 | 26084882:260 | 26085046:260 | -;-;-;-;-;-;-;-;-; | 1698 | opn4b | 8 | 2 | 0 | 0 | 0 | 2 |
| ENSDARG000 | 12;12;12;12;1 | 47778786:477 | 47778950:477 | -;-;-;-;-;- | 939 | selenou1b | 68 | 0 | 0 | 34 | 16 | 6 |
| ENSDARG000 | 12 | 22578404 | 22578530 | + | 127 | 5S_rRNA | 0 | 2 | 0 | 0 | 0 | 0 |
| ENSDARG000 | 12 | 19385426 | 19385555 | + | 130 | NA | 0 | 0 | 0 | 0 | 0 | 0 |
| ENSDARG000 | 12 | 1490472 | 1490588 | - | 117 | 5S_rRNA | 0 | 0 | 0 | 0 | 0 | 0 |
| ENSDARG000 | 12 | 28945532 | 28945646 | + | 115 | 5S_rRNA | 0 | 0 | 0 | 0 | 0 | 0 |
| ENSDARG000 | 12 | 7732534 | 7732650 | + | 117 | 5S_rRNA | 0 | 0 | 0 | 0 | 0 | 0 |
| ENSDARG000 | 12;12;12 | 316238:31887 | 316618:31898 | +;+;+ | 612 | rcvrnb | 4 | 6 | 1 | 87 | 2 | 26 |
| ENSDARG000 | 12;12;12;12;1 | 47945664:479 | 47945783:479 | +;+;+;+;+;+;+; | 911 | NA | 0 | 0 | 0 | 0 | 0 | 0 |
| ENSDARG000 | 12;12;12;12;1 | 30119165:301 | 30119181:301 | -;-;-;-;-;-;-;-;-; | 318 | NA | 0 | 0 | 0 | 0 | 0 | 0 |
| ENSDARG000 | 12 | 44626890 | 44627007 | + | 118 | 5S_rRNA | 0 | 0 | 0 | 0 | 0 | 0 |
| ENSDARG000 | 12;12;12;12;1 | 48559480:485 | 48559808:485 | -;-;-;-;-;-;- | 1099 | cyp4f3 | 51 | 30 | 98 | 34 | 76 | 67 |
| ENSDARG000 | 12;12;12;12;1 | 4487561:4490 | 4487864:4490 | -;-;-;-;-;-;-;-;-; | 3990 | trpm4b.2 | 712 | 99 | 48 | 70 | 115 | 204 |
| ENSDARG000 | 12;12;12;12 | 43692048:436 | 43692275:436 | -;-;-;- | 894 | NA | 0 | 0 | 0 | 0 | 0 | 0 |
| ENSDARG000 | 12;12 | 47122104:471 | 47122214:471 | +;+ | 1067 | NA | 4 | 0 | 0 | 0 | 0 | 0 |
| ENSDARG000 | 12 | 29970513 | 29970629 | + | 117 | 5S_rRNA | 0 | 0 | 0 | 0 | 0 | 0 |
| ENSDARG000 | 12 | 5089815 | 5089930 | + | 116 | 5S_rRNA | 0 | 0 | 0 | 0 | 0 | 0 |
| ENSDARG000 | 12;12;12;12 | 5775853:5776 | 5775882:5776 | +;+;+;+ | 507 | NA | 0 | 0 | 0 | 0 | 0 | 0 |
| ENSDARG000 | 12 | 27134869 | 27134967 | + | 99 | dre-mir-10a | 0 | 0 | 0 | 0 | 0 | 0 |
| ENSDARG000 | 12 | 24214401 | 24214517 | + | 117 | 5S_rRNA | 0 | 0 | 0 | 0 | 0 | 0 |
| ENSDARG000 | 12 | 13658022 | 13658144 | + | 123 | 5S_rRNA | 0 | 0 | 0 | 0 | 0 | 0 |
| ENSDARG000 | 12 | 6708129 | 6708243 | + | 115 | 5S_rRNA | 0 | 0 | 0 | 0 | 0 | 0 |
| ENSDARG000 | 12 | 31499300 | 31499434 | - | 135 | 5S_rRNA | 0 | 0 | 0 | 0 | 0 | 0 |
| ENSDARG000 | 12 | 9648228 | 9648345 | - | 118 | 5S_rRNA | 0 | 0 | 0 | 0 | 0 | 0 |
| ENSDARG000 | 12;12;12;12;1 | 1592146:1592 | 1592253:1592 | +;+;+;+;+;+;+; | 1051 | SLC39A11 | 155 | 83 | 153 | 92 | 95 | 113 |
| ENSDARG000 | 12;12;12 | 49006048:490 | 49006606:490 | +;+;+ | 3225 | chst3b | 10 | 24 | 20 | 30 | 11 | 10 |
| ENSDARG000 | 12;12;12;12;1 | 48204891:482 | 48205229:482 | +;+;+;+;+;+;+ | 3918 | ndr2 | 0 | 0 | 0 | 11 | 0 | 14 |
| ENSDARG000 | 12 | 28298859 | 28299929 | + | 1071 | NA | 0 | 1 | 0 | 0 | 0 | 0 |
| ENSDARG000 | 12;12;12;12;1 | 1338341:1340 | 1339653:1340 | -;-;-;-;-;-;- | 2756 | PAOX | 0 | 6 | 0 | 0 | 2 | 2 |
| ENSDARG000 | 12;12;12;12;1 | 36027746:360 | 36028100:360 | -;-;-;-;-;- | 1938 | gprc5c | 99 | 22 | 16 | 8 | 24 | 0 |
| ENSDARG000 | 12 | 31195826 | 31195950 | + | 125 | 5S_rRNA | 0 | 0 | 0 | 0 | 0 | 0 |
| ENSDARG000 | 12 | 36159664 | 36159753 | - | 90 | NA | 0 | 0 | 0 | 0 | 0 | 0 |
| ENSDARG000 | 12;12;12;12;1 | 48674381:486 | 48674418:486 | +;+;+;+;+;+;+; | 1868 | uros | 151 | 108 | 127 | 104 | 164 | 250 |
| ENSDARG000 | 12;12;12;12;1 | 10644074:106 | 10644190:106 | -;-;-;-;-;-;-;-;-; | 2970 | med24 | 478 | 552 | 682 | 377 | 720 | 517 |
| ENSDARG000 | 12;12;12 | 8509640:8510 | 8509913:8510 | -;-;- | 414 | NA | 2 | 0 | 0 | 0 | 0 | 0 |
| ENSDARG000 | 12;12;12;12;1 | 33960406:339 | 33960414:339 | -;-;-;-;-;-;-;-;-; | 297 | NA | 1 | 4 | 0 | 3 | 5 | 3 |
| ENSDARG000 | 12 | 16572195 | 16572309 | - | 115 | 5S_rRNA | 0 | 0 | 0 | 0 | 0 | 0 |
| ENSDARG000 | 12;12;12;12;1 | 48683208:486 | 48683412:486 | +;+;+;+;+;+;+; | 1908 | mmp21 | 2 | 1 | 1 | 0 | 2 | 3 |
| ENSDARG000 | 12;12;12;12 | 43852970:439 | 43853321:439 | -;-;-;- | 703 | NA | 3 | 2 | 0 | 2 | 0 | 2 |
| ENSDARG000 | 12;12;12;12 | 49003489:490 | 49003630:490 | -;-;-;- | 471 | NA | 481 | 280 | 379 | 368 | 361 | 335 |
| ENSDARG000 | 12;12;12;12;1 | 20567382:205 | 20567669:205 | +;+;+;+;+;+;+; | 954 | AMZ2 | 2 | 0 | 0 | 2 | 2 | 2 |
| ENSDARG000 | 12;12;12;12 | 3571770:3571 | 3572041:3572 | +;+;+;+ | 772 | coa3a | 266 | 306 | 431 | 187 | 333 | 294 |
| ENSDARG000 | 12 | 33334674 | 33334789 | - | 116 | 5S_rRNA | 8 | 2 | 0 | 0 | 7 | 0 |
| ENSDARG000 | 12;12;12;12;1 | 3220285:3220 | 3220454:3221 | -;-;-;-;-;-;-;-;-; | 1845 | NA | 0 | 0 | 0 | 0 | 0 | 0 |
| ENSDARG000 | 12 | 49100365 | 49102223 | + | 1859 | NA | 2 | 0 | 4 | 0 | 0 | 0 |
| ENSDARG000 | 12;12;12;12;1 | 47044707:470 | 47044987:470 | +;+;+;+;+;+;+; | 2661 | ZRANB1 | 887 | 557 | 1096 | 787 | 743 | 699 |
| ENSDARG000 | 12 | 37553703 | 37553818 | - | 116 | 5S_rRNA | 0 | 0 | 0 | 0 | 0 | 0 |
| ENSDARG000 | 12 | 7561065 | 7561202 | + | 138 | 5S_rRNA | 0 | 0 | 0 | 0 | 0 | 0 |
| ENSDARG000 | 12;12;12;12;1 | 45150709:451 | 45151139:451 | +;+;+;+;+;+;+; | 2513 | dhx32a | 18 | 29 | 20 | 10 | 16 | 29 |
| ENSDARG000 | 12;12;12 | 35650321:356 | 35650327:356 | +;+;+ | 153 | NA | 0 | 0 | 0 | 1 | 0 | 1 |
| ENSDARG000 | 12;12;12 | 44018826:440 | 44019197:440 | -;-;- | 531 | NA | 0 | 0 | 0 | 0 | 0 | 0 |
| ENSDARG000 | 12 | 31916943 | 31917052 | + | 110 | dre-mir-2196 | 0 | 0 | 0 | 0 | 0 | 0 |
| ENSDARG000 | 12;12;12;12;1 | 46512881:465 | 46512928:465 | +;+;+;+;+;+;+; | 1272 | NA | 6 | 0 | 0 | 2 | 9 | 39 |
| ENSDARG000 | 12;12;12;12;1 | 46605600:466 | 46605606:466 | +;+;+;+;+;+;+; | 5563 | FADS6 | 0 | 0 | 0 | 0 | 0 | 0 |
| ENSDARG000 | 12;12;12;12;1 | 7268392:7269 | 7269301:7269 | -;-;-;-;-;-;-;- | 2013 | zgc:171971 | 181 | 112 | 271 | 90 | 178 | 221 |
| ENSDARG000 | 12;12;12;12;1 | 10631266:106 | 10631347:106 | +;+;+;+;+ | 738 | csf3a | 54 | 67 | 44 | 24 | 85 | 61 |
| ENSDARG000 | 12;12;12;12;1 | 45200744:452 | 45200875:452 | +;+;+;+;+;+;+; | 3502 | wbp2 | 633 | 626 | 527 | 1025 | 640 | 579 |
| ENSDARG000 | 12 | 9345364 | 9345485 | - | 122 | 5S_rRNA | 0 | 0 | 0 | 0 | 0 | 0 |
| ENSDARG000 | 12 | 21794002 | 21794116 | + | 115 | 5S_rRNA | 156 | 105 | 118 | 129 | 215 | 130 |
| ENSDARG000 | 12 | 46603441 | 46603555 | + | 115 | 5S_rRNA | 0 | 0 | 0 | 0 | 0 | 0 |
| ENSDARG000 | 12 | 18738413 | 18738529 | - | 117 | 5S_rRNA | 0 | 0 | 0 | 0 | 0 | 0 |
| ENSDARG000 | 12 | 31047320 | 31047435 | - | 116 | 5S_rRNA | 0 | 0 | 0 | 0 | 0 | 0 |
| ENSDARG000 | 12 | 31208436 | 31208551 | + | 116 | 5S_rRNA | 0 | 0 | 0 | 0 | 0 | 0 |
| ENSDARG000 | 12;12;12;12;1 | 35936447:359 | 35937207:359 | -;-;-;-;-;-;-;-;-; | 3308 | dnai2a | 2 | 1 | 4 | 0 | 0 | 0 |
| ENSDARG000 | 12;12;12;12;1 | 2399130:2402 | 2399181:2402 | +;+;+;+;+;+;+; | 1074 | VSTM4 | 62 | 200 | 337 | 98 | 557 | 10 |
| ENSDARG000 | 12;12;12;12;1 | 1139690:1187 | 1139911:1187 | +;+;+;+;+;+;+; | 687 | RAB26 | 0 | 0 | 0 | 0 | 0 | 0 |
| ENSDARG000 | 12;12;12;12;1 | 48542998:485 | 48543514:485 | -;-;-;-;-;-;- | 1248 | NA | 0 | 4 | 4 | 0 | 2 | 12 |
| ENSDARG000 | 12;12;12;12;1 | 41580783:415 | 41581287:415 | -;-;-;-;-;-;-;-;-; | 2675 | dpysl4 | 69 | 32 | 205 | 18 | 10 | 129 |
| ENSDARG000 | 12;12;12;12;1 | 41991635:419 | 41992182:419 | +;+;+;+;+;+;+; | 6063 | TCERG1L | 0 | 0 | 0 | 0 | 0 | 0 |
| ENSDARG000 | 12;12;12;12;1 | 40810418:409 | 40810509:409 | +;+;+;+;+;+;+; | 7487 | CDH18 | 314 | 94 | 42 | 150 | 74 | 6 |
| ENSDARG000 | 12 | 39430202 | 39430317 | + | 116 | 5S_rRNA | 0 | 0 | 0 | 0 | 0 | 0 |
| ENSDARG000 | 12;12;12;12;1 | 899223:89922 | 899341:89934 | -;-;-;-;-;-;-;-;-; | 4222 | SPAG9 | 30239 | 16292 | 21622 | 29578 | 24251 | 29841 |
| ENSDARG000 | 12 | 10210932 | 10211048 | + | 117 | 5S_rRNA | 0 | 0 | 0 | 0 | 0 | 0 |
| ENSDARG000 | 12;12;12;12;1 | 36109071:361 | 36109600:361 | +;+;+;+;+;+;+; | 4689 | map2k6 | 843 | 941 | 1187 | 655 | 1343 | 880 |
| ENSDARG000 | 12;12;12;12 | 48613795:486 | 48614924:486 | -;-;-;- | 1494 | zgc:92749 | 8 | 14 | 4 | 34 | 20 | 24 |
| ENSDARG000 | 12 | 16960693 | 16960781 | + | 89 | dre-mir-107a | 0 | 0 | 0 | 0 | 0 | 0 |
| ENSDARG000 | 12;12;12 | 3249054:3249 | 3249388:3249 | -;-;- | 580 | gngt2b | 4 | 4 | 0 | 0 | 0 | 0 |
| ENSDARG000 | 12 | 32951782 | 32951896 | + | 115 | 5S_rRNA | 0 | 0 | 0 | 0 | 0 | 0 |
| ENSDARG000 | 12 | 27659061 | 27659175 | - | 115 | 5S_rRNA | 0 | 0 | 0 | 0 | 1 | 0 |
| ENSDARG000 | 12;12;12 | 7653863:7654 | 7654422:7654 | -;-;- | 980 | NA | 56 | 37 | 136 | 47 | 148 | 28 |
| ENSDARG000 | 12 | 20498983 | 20499080 | + | 98 | NA | 0 | 0 | 0 | 0 | 0 | 0 |
| ENSDARG000 | 12;12 | 49114483:491 | 49115435:491 | +;+ | 2502 | NA | 26 | 4 | 9 | 4 | 2 | 8 |
| ENSDARG000 | 12 | 34881777 | 34881895 | + | 119 | 5S_rRNA | 0 | 0 | 0 | 0 | 0 | 0 |
| ENSDARG000 | 12;12;12;12;1 | 46791907:467 | 46792059:467 | +;+;+;+;+;+;+; | 3124 | vclb | 1452 | 1835 | 1850 | 615 | 1917 | 973 |
| ENSDARG000 | 12;12;12;12;1 | 35702809:357 | 35702837:357 | -;-;-;-;-;-;-;-;-; | 4974 | AATK | 196 | 768 | 674 | 880 | 310 | 455 |
| ENSDARG000 | 12;12;12;12 | 49125510:491 | 49125774:491 | +;+;+;+ | 660 | NA | 0 | 0 | 0 | 0 | 0 | 0 |
| ENSDARG000 | 12;12;12;12;1 | 47871878:478 | 47872260:478 | -;-;-;-;-;-;- | 1287 | QRFPR | 30 | 0 | 0 | 2 | 0 | 12 |
| ENSDARG000 | 12 | 23910832 | 23910949 | + | 118 | 5S_rRNA | 0 | 0 | 0 | 0 | 0 | 0 |
| ENSDARG000 | 12;12;12;12;1 | 43730276:437 | 43730448:437 | -;-;-;-;-;-;-;-;-; | 2503 | NA | 82 | 0 | 2 | 4 | 14 | 2 |
| ENSDARG000 | 12 | 16598743 | 16598857 | - | 115 | 5S_rRNA | 0 | 0 | 0 | 0 | 0 | 0 |
| ENSDARG000 | 12;12;12;12;1 | 7589193:7594 | 7590252:7595 | -;-;-;-;-;- | 2553 | slc16a9b | 658 | 765 | 928 | 655 | 1303 | 790 |
| ENSDARG000 | 12;12;12;12;1 | 18556929:185 | 18556962:185 | +;+;+;+;+;+;+; | 222 | NA | 1 | 2 | 2 | 5 | 0 | 4 |
| ENSDARG000 | 12;12;12;12;1 | 48307111:483 | 48307332:483 | -;-;-;-;-;- | 855 | ascc1 | 246 | 240 | 247 | 138 | 246 | 134 |
| ENSDARG000 | 12;12;12;12;1 | 35836610:358 | 35841884:358 | -;-;-;-;-;-;-;-;-; | 8669 | cep131 | 402 | 253 | 373 | 150 | 380 | 414 |
| ENSDARG000 | 12 | 36809059 | 36809175 | - | 117 | 5S_rRNA | 0 | 0 | 0 | 0 | 0 | 0 |
| ENSDARG000 | 12;12;12;12;1 | 1455147:1455 | 1455201:1455 | +;+;+;+;+;+;+; | 1275 | COPS3 | 676 | 767 | 851 | 633 | 738 | 774 |
| ENSDARG000 | 12 | 25648127 | 25648243 | + | 117 | 5S_rRNA | 0 | 0 | 0 | 0 | 0 | 2 |
| ENSDARG000 | 12 | 4497364 | 4497479 | + | 116 | 5S_rRNA | 0 | 0 | 0 | 0 | 0 | 0 |
| ENSDARG000 | 12 | 39345392 | 39345506 | - | 115 | 5S_rRNA | 0 | 0 | 0 | 0 | 0 | 0 |
| ENSDARG000 | 12;12;12;12 | 13740216:137 | 13740970:137 | +;+;+;+ | 1897 | NA | 0 | 0 | 0 | 2 | 0 | 1 |
| ENSDARG000 | 12;12;12 | 25073305:250 | 25073662:250 | +;+;+ | 1510 | NA | 0 | 2 | 0 | 0 | 0 | 2 |
| ENSDARG000 | 12;12;12;12 | 26200885:262 | 26201470:262 | -;-;-;- | 1718 | NA | 0 | 0 | 10 | 14 | 0 | 0 |
| ENSDARG000 | 12;12;12 | 28686992:286 | 28687065:286 | -;-;- | 326 | NA | 0 | 0 | 0 | 0 | 0 | 0 |

| | | | | | | | | | | | | |
|---|---|---|---|---|---|---|---|---|---|---|---|---|
| ENSDARG000 | 12:12:12:12:1 | 35889699:358 | 35890155:358 | -;-;-;-;-;- | 2246 | NA | 8 | 0 | 12 | 2 | 0 | 2 |
| ENSDARG000 | 12;12;12;12 | 38146860:381 | 38147558:381 | -;-;-;- | 1275 | NA | 0 | 0 | 0 | 0 | 0 | 0 |
| ENSDARG000 | 12:12:12:12:1 | 39883711:399 | 39884348:399 | -;-;-;-;-;-;-;-; | 2881 | NA | 6 | 0 | 0 | 0 | 0 | 0 |
| ENSDARG000 | 12:12:12:12:1 | 40529226:405 | 40529493:405 | +;+;+;+;+;+;+; | 1181 | NA | 0 | 0 | 0 | 0 | 0 | 0 |
| ENSDARG000 | 12;12;12;12 | 42273544:423 | 42275126:423 | -;-;-;- | 1911 | NA | 0 | 0 | 0 | 10 | 0 | 0 |
| ENSDARG000 | 12:12:12:12:1 | 42582803:426 | 42583431:426 | -;-;-;-;-;- | 2476 | NA | 0 | 0 | 0 | 0 | 0 | 0 |
| ENSDARG000 | 12;12;12;12 | 42802988:428 | 42803104:428 | +;+;+;+ | 465 | NA | 0 | 0 | 0 | 0 | 0 | 0 |
| ENSDARG000 | 12:12:12:12:1 | 43108661:431 | 43108781:431 | +;+;+;+;+ | 1261 | NA | 2 | 0 | 0 | 6 | 0 | 2 |
| ENSDARG000 | 12;12;12;12 | 43201606:432 | 43201758:432 | -;-;-;- | 718 | NA | 0 | 0 | 0 | 0 | 0 | 0 |
| ENSDARG000 | 12;12;12 | 43306014:433 | 43306075:433 | -;-;- | 360 | NA | 0 | 0 | 0 | 0 | 0 | 0 |
| ENSDARG000 | 12;12;12 | 44330624:443 | 44331029:443 | -;-;- | 864 | NA | 0 | 0 | 0 | 0 | 0 | 0 |
| ENSDARG000 | 12;12;12;12 | 45368547:453 | 45368755:453 | -;-;-;- | 616 | NA | 1402 | 624 | 494 | 370 | 371 | 350 |
| ENSDARG000 | 12;12;12 | 45419756:454 | 45419991:454 | -;-;- | 778 | NA | 263 | 76 | 7 | 4 | 32 | 25 |
| ENSDARG000 | 12;12;12;12 | 45467926:454 | 45468125:454 | -;-;-;- | 768 | NA | 69 | 2 | 0 | 0 | 0 | 18 |
| ENSDARG000 | 12:12:12:12:1 | 46046268:460 | 46048595:460 | +;+;+;+;+;+;+; | 5265 | NA | 0 | 0 | 0 | 0 | 1 | 0 |
| ENSDARG000 | 12:12:12:12:1 | 46046268:460 | 46049015:460 | -;-;-;-;-;-;-;-; | 4467 | NA | 0 | 0 | 0 | 0 | 0 | 0 |
| ENSDARG000 | 12:12:12:12:1 | 46071521:460 | 46074047:460 | -;-;-;-;-;-;- | 4094 | NA | 0 | 0 | 0 | 0 | 0 | 0 |
| ENSDARG000 | 12;12;12;12 | 47749126:477 | 47749395:477 | -;-;-;- | 782 | NA | 0 | 0 | 0 | 0 | 0 | 0 |
| ENSDARG000 | 12:12:12:12:1 | 49033723:490 | 49034669:490 | +;+;+;+;+;+ | 3568 | NA | 6 | 0 | 2 | 0 | 4 | 0 |
| ENSDARG000 | 19:19:19:19:1 | 44009473:440 | 44009655:440 | +;+;+;+;+;+;+; | 1583 | NA | 175 | 61 | 121 | 20 | 128 | 47 |
| ENSDARG000 | 19:19:19:19:1 | 7249041:7249 | 7249979:7249 | -;-;-;-;-;-;-;-; | 2885 | rxrba | 994 | 841 | 1285 | 735 | 1223 | 944 |
| ENSDARG000 | 19:19:19:19:1 | 33836439:338 | 33836832:338 | -;-;-;-;-;-;-;-; | 1730 | otud6b | 264 | 286 | 384 | 263 | 289 | 241 |
| ENSDARG000 | 19:19:19:19:1 | 33828976:338 | 33829087:338 | -;-;-;-;-;-;-;-; | 1094 | NA | 0 | 0 | 0 | 0 | 0 | 0 |
| ENSDARG000 | 19:19:19:19:1 | 9460432:9460 | 9461888:9461 | -;-;-;-;-;-;-;-; | 2384 | vamp1a | 0 | 0 | 0 | 9 | 2 | 4 |
| ENSDARG000 | 19:19:19:19:1 | 798589:80111 | 799241:80138 | -;-;-;-;-;-;-;- | 2177 | glmp | 1993 | 1577 | 1514 | 1700 | 1761 | 1707 |
| ENSDARG000 | 19;19;19 | 9455218:9457 | 9455358:9460 | +;+;+ | 2618 | NA | 6176 | 7111 | 8773 | 5260 | 7505 | 4646 |
| ENSDARG000 | 19:19:19:19:1 | 7938121:7938 | 7938290:7938 | +;+;+;+;+;+;+; | 3507 | si:dkev-266f7. | 38 | 8 | 8 | 8 | 36 | 30 |
| ENSDARG000 | 19:19:19:19:1 | 17514631:175 | 17518974:175 | -;-;-;-;-;-;-;-; | 6264 | thrb | 321 | 398 | 351 | 420 | 667 | 1945 |
| ENSDARG000 | 19:19:19:19:1 | 40115977:401 | 40116324:401 | +;+;+;+;+;+;+; | 6405 | zmym4.2 | 1652 | 1172 | 1926 | 1361 | 1669 | 1657 |
| ENSDARG000 | 19;19;19;19 | 7242117:7242 | 7242164:7242 | +;+;+;+ | 577 | NA | 0 | 0 | 0 | 0 | 0 | 0 |
| ENSDARG000 | 19;19 | 23308419:233 | 23308934:233 | +;+ | 1136 | NA | 0 | 0 | 0 | 0 | 0 | 0 |
| ENSDARG000 | 19:19:19:19:1 | 23300120:233 | 23300235:233 | +;+;+;+;+ | 2001 | NA | 4 | 0 | 2 | 0 | 8 | 0 |
| ENSDARG000 | 19:19:19:19:1 | 23296616:232 | 23296748:232 | +;+;+;+;+ | 3975 | irgf1 | 10 | 12 | 22 | 0 | 24 | 18 |
| ENSDARG000 | 19 | 47574615 | 47574823 | + | 209 | NA | 0 | 0 | 0 | 0 | 0 | 0 |
| ENSDARG000 | 19:19:19:19:1 | 29834440:298 | 29838129:298 | -;-;-;-;-;-;-;-; | 5602 | txlna | 1290 | 1424 | 1355 | 928 | 1545 | 1178 |
| ENSDARG000 | 19;19;19 | 44039849:440 | 44040859:440 | +;+;+ | 1687 | lrrc14b | 30 | 8 | 15 | 8 | 48 | 13 |
| ENSDARG000 | 19:19:19:19:1 | 19786117:197 | 19786852:197 | +;+;+;+;+ | 1441 | hoxa1a | 5 | 0 | 0 | 41 | 0 | 81 |
| ENSDARG000 | 19;19;19;19 | 816208:81720 | 816364:81740 | +;+;+;+ | 3850 | nrm | 234 | 154 | 218 | 222 | 202 | 244 |
| ENSDARG000 | 19:19:19:19:1 | 19734769:197 | 19735484:197 | -;-;-;-;-;-;-;- | 1284 | NA | 9 | 0 | 0 | 3 | 0 | 0 |
| ENSDARG000 | 19:19:19:19:1 | 19717971:197 | 19719602:197 | -;-;-;-;-;-;-;-; | 3183 | evx1 | 3 | 0 | 0 | 0 | 0 | 0 |
| ENSDARG000 | 19:19:19:19:1 | 823945:82578 | 823988:82929 | +;+;+;+;+ | 4779 | NA | 58 | 62 | 50 | 32 | 114 | 102 |
| ENSDARG000 | 19:19:19:19:1 | 19747430:197 | 19747504:197 | +;+;+;+;+ | 1827 | hoxa9a | 42 | 0 | 0 | 0 | 0 | 0 |
| ENSDARG000 | 19:19:19:19:1 | 27366932:273 | 27372244:273 | -;-;-;-;-;- | 6648 | mgat1b | 256 | 233 | 363 | 146 | 655 | 272 |
| ENSDARG000 | 19:19:19:19:1 | 25656812:256 | 25658539:256 | -;-;-;-;-;-;-;-; | 3284 | asns | 3554 | 3558 | 4923 | 2368 | 4618 | 3607 |
| ENSDARG000 | 19;19;19;19 | 19717971:197 | 19719602:197 | -;-;-;- | 2035 | EVX1 | 0 | 0 | 0 | 0 | 0 | 0 |
| ENSDARG000 | 19:19:19:19:1 | 6213680:6217 | 6217770:6217 | -;-;-;-;-;-;-;-; | 6351 | POU2F2 | 30 | 2 | 2 | 2 | 0 | 14 |
| ENSDARG000 | 19;19 | 19750223:197 | 19750691:197 | +;+ | 498 | NA | 0 | 0 | 0 | 0 | 0 | 0 |
| ENSDARG000 | 19;19;19;19 | 19782413:197 | 19782740:197 | +;+;+;+ | 836 | NA | 2 | 0 | 3 | 62 | 0 | 115 |
| ENSDARG000 | 19:19:19:19:1 | 1658698:1662 | 1658925:1662 | +;+;+;+;+;+;+; | 2624 | klhl7 | 184 | 163 | 195 | 142 | 284 | 163 |
| ENSDARG000 | 19:19:19:19:1 | 19754950:197 | 19755109:197 | +;+;+;+;+;+;+; | 5234 | hoxa3a | 210 | 2 | 2 | 91 | 12 | 226 |
| ENSDARG000 | 19;19 | 7954679:7956 | 7954733:7957 | +;+ | 1810 | NA | 12 | 0 | 0 | 0 | 2 | 29 |
| ENSDARG000 | 19 | 7966116 | 7968667 | + | 2552 | NA | 1383 | 120 | 350 | 380 | 982 | 1808 |
| ENSDARG000 | 19:19:19:19:1 | 13193215:131 | 13196617:131 | -;-;-;-;-;-;-;-; | 4615 | zfpm2b | 32 | 2 | 174 | 56 | 210 | 76 |
| ENSDARG000 | 19:19:19:19:1 | 44049501:440 | 44049619:440 | -;-;-;-;-;-;-;-; | 692 | uqcrb | 0 | 0 | 1 | 1 | 0 | 2 |
| ENSDARG000 | 19;19 | 19759577:197 | 19760128:197 | +;+ | 888 | hoxa5a | 14 | 0 | 4 | 0 | 0 | 0 |
| ENSDARG000 | 19;19 | 19729506:197 | 19730292:197 | +;+ | 1669 | hoxa13a | 8 | 0 | 0 | 0 | 0 | 0 |
| ENSDARG000 | 19;19;19;19 | 19737214:197 | 19737921:197 | +;+;+;+ | 1090 | hoxa11a | 10 | 0 | 0 | 0 | 0 | 0 |
| ENSDARG000 | 19:19:19:19:1 | 44058557:440 | 44058804:440 | -;-;-;-;-;-;-;-; | 1471 | mterf3 | 49 | 90 | 76 | 101 | 85 | 68 |
| ENSDARG000 | 19:19:19:19:1 | 23368001:233 | 23368082:233 | -;-;-;-;- | 550 | NA | 0 | 0 | 0 | 0 | 0 | 0 |
| ENSDARG000 | 19;19;19;19 | 8086491:8088 | 8086954:8089 | -;-;-;- | 1186 | si:dkev-266f7. | 0 | 0 | 0 | 0 | 0 | 0 |
| ENSDARG000 | 19:19:19:19:1 | 23381331:233 | 23381419:233 | -;-;-;-;- | 575 | NA | 0 | 0 | 0 | 0 | 0 | 0 |
| ENSDARG000 | 19;19;19;19 | 44049647:440 | 44049839:440 | -;-;-;- | 668 | NA | 0 | 0 | 0 | 0 | 0 | 0 |
| ENSDARG000 | 19:19:19:19:1 | 4968947:4969 | 4969032:4969 | +;+;+;+;+;+;+; | 2855 | stard3 | 1836 | 1443 | 1182 | 1922 | 1578 | 2086 |
| ENSDARG000 | 19:19:19:19:1 | 23418078:234 | 23418166:234 | -;-;-;-;- | 557 | NA | 0 | 0 | 0 | 0 | 0 | 0 |
| ENSDARG000 | 19:19:19:19:1 | 1688727:1688 | 1689115:1689 | +;+;+;+;+;+;+; | 6987 | DENND3 | 394 | 148 | 265 | 239 | 219 | 220 |
| ENSDARG000 | 19:19:19:19:1 | 42047427:420 | 42047584:420 | +;+;+;+;+;+;+; | 3718 | si:ch211-13c6 | 1138 | 942 | 1563 | 1271 | 1616 | 1276 |
| ENSDARG000 | 19:19:19:19:1 | 8144556:8144 | 8144918:8144 | +;+;+;+;+;+;+; | 4728 | NA | 0 | 2 | 0 | 0 | 0 | 0 |
| ENSDARG000 | 19:19:19:19:1 | 23436133:234 | 23436221:234 | -;-;-;-;- | 557 | NA | 0 | 0 | 0 | 0 | 0 | 0 |
| ENSDARG000 | 19:19:19:19:1 | 33850705:338 | 33850923:338 | +;+;+;+;+;+;+; | 4088 | pex1 | 328 | 220 | 306 | 148 | 389 | 236 |
| ENSDARG000 | 19:19:19:19:1 | 21814047:218 | 21815462:218 | -;-;-;-;-;-;-;-; | 2678 | NA | 46 | 31 | 134 | 67 | 398 | 66 |
| ENSDARG000 | 19:19:19:19:1 | 13375838:133 | 13376764:133 | +;+;+;+;+;+;+ | 4700 | lrp12 | 1863 | 1134 | 1402 | 1434 | 1485 | 1482 |
| ENSDARG000 | 19:19:19:19:1 | 17688350:177 | 17688869:177 | -;-;-;-;-;-;-;-; | 5653 | top2b | 3060 | 2516 | 3699 | 1830 | 3973 | 2925 |
| ENSDARG000 | 19:19:19:19:1 | 1790797:1795 | 1790991:1795 | +;+;+;+;+;+;+; | 3637 | PTK2 | 403 | 395 | 662 | 489 | 671 | 369 |
| ENSDARG000 | 19:19:19:19:1 | 29854223:298 | 29854373:298 | +;+;+;+;+;+;+; | 1282 | si:ch73-130a3 | 62 | 39 | 70 | 28 | 66 | 85 |
| ENSDARG000 | 19:19:19:19:1 | 23450493:234 | 23450761:234 | -;-;-;-;- | 896 | NA | 0 | 0 | 0 | 0 | 0 | 0 |
| ENSDARG000 | 19:19:19:19:1 | 8481802:8496 | 8482083:8496 | +;+;+;+;+ | 2545 | efna1a | 100 | 2 | 102 | 78 | 72 | 10 |
| ENSDARG000 | 19;19 | 3237790:3240 | 3238984:3240 | -;- | 1595 | si:ch211-133n | 1548 | 735 | 1838 | 1165 | 2471 | 1346 |
| ENSDARG000 | 19:19:19:19:1 | 9481698:9483 | 9482446:9483 | -;-;-;-;-;-;-;- | 2829 | IFFO1 | 6 | 2 | 22 | 4 | 27 | 6 |
| ENSDARG000 | 19:19:19:19:1 | 25679538:256 | 25680313:256 | -;-;-;-;- | 1380 | bloc1s4 | 1572 | 1073 | 2271 | 1482 | 1763 | 1646 |
| ENSDARG000 | 19:19:19:19:1 | 42919279:429 | 42919871:429 | -;-;-;-;-;- | 2020 | cyhr1 | 723 | 442 | 778 | 473 | 866 | 686 |
| ENSDARG000 | 19:19:19:19:1 | 27399168:273 | 27399339:273 | -;-;-;-;-;-;-;-; | 3798 | vars2 | 338 | 308 | 407 | 172 | 395 | 293 |
| ENSDARG000 | 19;19;19 | 35357196:353 | 35357440:353 | -;-;- | 542 | ndufs5 | 768 | 899 | 965 | 518 | 749 | 557 |
| ENSDARG000 | 19;19;19;19 | 42938754:429 | 42938835:429 | -;-;-;- | 1646 | DCLK3 | 0 | 0 | 2 | 0 | 0 | 0 |
| ENSDARG000 | 19:19:19:19:1 | 8506178:8587 | 8506469:8587 | +;+;+;+;+;+;+; | 1567 | s100a10a | 0 | 0 | 6 | 0 | 0 | 2 |
| ENSDARG000 | 19:19:19:19:1 | 8582685:8582 | 8583356:8583 | -;-;-;-;-;-;-;-; | 2768 | trim46b | 0 | 0 | 0 | 0 | 0 | 1 |
| ENSDARG000 | 19;19;19 | 8533989:8534 | 8535012:8534 | -;-;- | 1082 | dpm3 | 304 | 128 | 274 | 241 | 362 | 242 |
| ENSDARG000 | 19;19 | 3242926:3243 | 3243756:3244 | -;- | 1200 | zgc:158445 | 0 | 0 | 12 | 0 | 18 | 0 |
| ENSDARG000 | 19:19:19:19:1 | 10892465:109 | 10893398:109 | -;-;-;-;-;-;-;-; | 2908 | pip5k1aa | 897 | 639 | 469 | 397 | 639 | 694 |
| ENSDARG000 | 19:19:19:19:1 | 42955280:429 | 42955333:429 | +;+;+;+;+;+;+; | 835 | fabp3 | 4777 | 5843 | 6089 | 6159 | 3737 | 5461 |
| ENSDARG000 | 19:19:19:19:1 | 6328234:6332 | 6330258:6332 | -;-;-;-;-;-;-;-; | 5379 | atp1a3a | 54 | 62 | 104 | 44 | 115 | 184 |
| ENSDARG000 | 19;19 | 42962095:429 | 42962572:429 | -;- | 505 | NA | 0 | 2 | 3 | 0 | 2 | 1 |
| ENSDARG000 | 19:19:19:19:1 | 13410903:134 | 13411090:134 | +;+;+;+;+;+;+; | 3103 | srsf4 | 1684 | 1572 | 2231 | 1771 | 2117 | 2022 |
| ENSDARG000 | 19;19;19;19 | 63567:63594: | 63643:63643: | +;+;+;+ | 961 | NA | 2 | 30 | 47 | 67 | 34 | 3 |
| ENSDARG000 | 19 | 35368602 | 35369837 | + | 1236 | NA | 28 | 6 | 81 | 51 | 93 | 100 |
| ENSDARG000 | 19:19:19:19:1 | 49721:49992: | 49870:50121: | +;+;+;+;+;+;+; | 3689 | NA | 113 | 69 | 90 | 30 | 140 | 58 |
| ENSDARG000 | 19;19 | 13420884:134 | 13421526:134 | +;+ | 3067 | NA | 12 | 8 | 14 | 6 | 82 | 4 |
| ENSDARG000 | 19:19:19:19:1 | 33887589:338 | 33888748:338 | -;-;-;-;-;-;-;-; | 3881 | elmo1 | 510 | 1313 | 2660 | 1632 | 2241 | 1106 |
| ENSDARG000 | 19:19:19:19:1 | 23468771:234 | 23469039:234 | -;-;-;-;- | 896 | NA | 0 | 0 | 0 | 0 | 0 | 0 |
| ENSDARG000 | 19:19:19:19:1 | 9512982:9516 | 9514356:9516 | -;-;-;-;-;-;- | 2116 | ing4 | 882 | 816 | 1265 | 1017 | 1108 | 1036 |
| ENSDARG000 | 19:19:19:19:1 | 35379899:353 | 35380316:353 | -;-;-;-;-;-;-;- | 2109 | rnf19b | 269 | 261 | 408 | 431 | 522 | 195 |
| ENSDARG000 | 19;19 | 60878;64007 | 61070;64189 | -;- | 376 | NA | 0 | 0 | 0 | 0 | 0 | 0 |
| ENSDARG000 | 19:19:19:19:1 | 1852078:1852 | 1852387:1852 | -;-;-;-;-;-;-;-; | 1222 | rplp1 | 15443 | 16252 | 21815 | 27860 | 14673 | 17851 |
| ENSDARG000 | 19:19:19:19:1 | 29861859:298 | 29862013:298 | -;-;-;-;-;-;-;-; | 2128 | KPNA6 | 347 | 341 | 599 | 220 | 375 | 321 |
| ENSDARG000 | 19:19:19:19:1 | 31593453:315 | 31594087:315 | -;-;-;-;-;-;-;-; | 4031 | ripor2 | 472 | 605 | 396 | 593 | 572 | 835 |
| ENSDARG000 | 19:19:19:19:1 | 42235739:422 | 42236185:422 | -;-;-;-;- | 1023 | si:ch211-191i | 0 | 0 | 0 | 0 | 0 | 1 |
| ENSDARG000 | 19;19;19 | 42142381:422 | 42143707:422 | +;+;+ | 1544 | NA | 2 | 0 | 0 | 0 | 4 | 0 |
| ENSDARG000 | 19:19:19:19:1 | 42227400:422 | 42227800:422 | +;+;+;+;+;+;+; | 3181 | jtb | 288 | 248 | 371 | 368 | 318 | 273 |
| ENSDARG000 | 19;19;19 | 42241530:422 | 42241932:422 | -;-;- | 727 | cart4 | 0 | 0 | 0 | 0 | 0 | 0 |
| ENSDARG000 | 19:19:19:19:1 | 23488241:234 | 23488329:234 | -;-;-;-;- | 522 | NA | 0 | 0 | 0 | 0 | 0 | 0 |
| ENSDARG000 | 19;19;19 | 25699329:257 | 25702093:257 | -;-;- | 3213 | pard6gb | 822 | 628 | 631 | 691 | 554 | 823 |
| ENSDARG000 | 19:19:19:19:1 | 40182641:401 | 40182768:401 | -;-;-;-;-;-;-;-; | 771 | efcab1 | 2 | 7 | 6 | 4 | 4 | 8 |
| ENSDARG000 | 19:19:19:19:1 | 23525358:235 | 23525446:235 | -;-;-;-;- | 555 | NA | 0 | 0 | 0 | 0 | 0 | 0 |
| ENSDARG000 | 19:19:19:19:1 | 40196462:401 | 40196658:401 | -;-;-;-;-;-;-;-; | 783 | grna.2 | 19 | 29 | 6 | 21 | 54 | 19 |
| ENSDARG000 | 19:19:19:19:1 | 42996968:429 | 42997596:429 | -;-;-;-;-;-;- | 1432 | zcchc17 | 292 | 269 | 270 | 220 | 308 | 235 |
| ENSDARG000 | 19:19:19:19:1 | 40186385:401 | 40186495:401 | +;+;+;+;+;+;+ | 2093 | grnas | 0 | 0 | 0 | 0 | 0 | 0 |
| ENSDARG000 | 19:19:19:19:1 | 40167363:401 | 40167702:401 | -;-;-;-;-;- | 986 | illr4 | 50 | 13 | 3 | 8 | 4 | 43 |
| ENSDARG000 | 19:19:19:19:1 | 40177246:401 | 40177448:401 | +;+;+;+;+ | 1203 | rbm48 | 150 | 126 | 263 | 215 | 170 | 322 |
| ENSDARG000 | 19:19:19:19:1 | 40186677:401 | 40187697:401 | -;-;-;-;-;-;-;- | 1618 | grna.1 | 2 | 11 | 37 | 4 | 46 | 146 |
| ENSDARG000 | 19:19:19:19:1 | 9533008:9533 | 9533368:9533 | +;+;+;+;+;+;+; | 3154 | NA | 0 | 4 | 4 | 2 | 2 | 0 |
| ENSDARG000 | 19:19:19:19:1 | 35418683:354 | 35419557:354 | -;-;-;-;-;-;-;-; | 1575 | ak2 | 1445 | 1518 | 2842 | 2156 | 1531 | 1548 |
| ENSDARG000 | 19;19;19 | 10917602:109 | 10917951:109 | -;-;- | 729 | NA | 0 | 0 | 0 | 0 | 0 | 0 |
| ENSDARG000 | 19:19:19:19:1 | 23548906:235 | 23548994:235 | -;-;-;-;- | 543 | NA | 0 | 0 | 0 | 0 | 0 | 0 |
| ENSDARG000 | 19:19:19:19:1 | 17834769:178 | 17838181:178 | -;-;-;-;-;-;-;-; | 7304 | ints8 | 546 | 452 | 712 | 314 | 750 | 591 |
| ENSDARG000 | 19:19:19:19:1 | 23573725:235 | 23573810:235 | -;-;-;-;-;- | 1266 | NA | 0 | 0 | 0 | 0 | 0 | 0 |
| ENSDARG000 | 19:19:19:19:1 | 7276060:7276 | 7276663:7276 | -;-;-;-;-;-;-;-; | 2844 | sdha | 2563 | 2346 | 2740 | 1918 | 2449 | 1849 |
| ENSDARG000 | 19:19:19:19:1 | 34073877:340 | 34074452:340 | -;-;-;-;-;-;-;-; | 1299 | rp9 | 350 | 267 | 386 | 313 | 374 | 363 |
| ENSDARG000 | 19:19:19:19:1 | 8612839:8628 | 8613169:8628 | +;+;+;+;+;+;+; | 2099 | snx27a | 807 | 272 | 492 | 273 | 613 | 574 |
| ENSDARG000 | 19:19:19:19:1 | 10937932:109 | 10938531:109 | +;+;+;+;+ | 1817 | NA | 26 | 10 | 0 | 0 | 0 | 0 |
| ENSDARG000 | 19;19 | 17834204:178 | 17834677:178 | -;- | 531 | NA | 8 | 5 | 17 | 15 | 4 | 13 |
| ENSDARG000 | 19:19:19:19:1 | 9612163:9612 | 9613428:9613 | -;-;-;-;-;-;-;-; | 3951 | clcn1a | 0 | 0 | 0 | 0 | 0 | 0 |
| ENSDARG000 | 19;19;19 | 27448625:274 | 27449063:274 | +;+;+ | 1289 | ppp1r11 | 200 | 191 | 263 | 128 | 168 | 144 |
| ENSDARG000 | 19:19:19:19:1 | 23592248:235 | 23592336:235 | -;-;-;-;-;- | 1269 | NA | 0 | 0 | 0 | 0 | 0 | 0 |
| ENSDARG000 | 19;19 | 67006;67615 | 67220;68058 | +;+ | 659 | NA | 8 | 13 | 36 | 53 | 13 | 4 |
| ENSDARG000 | 19:19:19:19:1 | 3247555:3248 | 3248230:3248 | -;-;-;-;-;- | 1309 | illr1 | 0 | 6 | 20 | 12 | 38 | 0 |
| ENSDARG000 | 19:19:19:19:1 | 31743104:317 | 31746060:317 | -;-;-;-;-;-;-;-; | 4542 | si:dkeyp-120h | 397 | 498 | 816 | 414 | 617 | 575 |
| ENSDARG000 | 19:19:19:19:1 | 23625263:236 | 23625351:236 | -;-;-;-;-;- | 1269 | NA | 0 | 0 | 0 | 0 | 0 | 0 |
| ENSDARG000 | 19:19:19:19:1 | 69238:69651: | 69553:69726: | -;-;-;-;-;-;-;-; | 1031 | dnajc8 | 932 | 868 | 854 | 322 | 593 | 555 |
| ENSDARG000 | 19:19:19:19:1 | 13639486:136 | 13639584:136 | -;-;-;-;-;-;-;-; | 3131 | EPB41 | 0 | 4 | 4 | 2 | 22 | 0 |
| ENSDARG000 | 19:19:19:19:1 | 25971000:260 | 25971505:260 | +;+;+;+;+;+;+; | 4421 | jarid2b | 228 | 167 | 163 | 153 | 235 | 215 |
| ENSDARG000 | 19;19 | 11013622:110 | 11013654:110 | +;+ | 232 | NA | 0 | 0 | 0 | 0 | 0 | 2 |
| ENSDARG000 | 19:19:19:19:1 | 10985369:109 | 10989777:109 | -;-;-;-;-;-;-;-; | 6957 | tpm3 | 5797 | 8079 | 6357 | 5780 | 5539 | 5832 |
| ENSDARG000 | 19:19:19:19:1 | 44068250:440 | 44068605:440 | -;-;-;-;-;-;-;-; | 2230 | rad21b | 200 | 250 | 227 | 261 | 306 | 186 |
| ENSDARG000 | 19:19:19:19:1 | 31771270:317 | 31771523:317 | +;+;+;+;+ | 1822 | stmn2b | 0 | 6 | 0 | 0 | 0 | 2 |
| ENSDARG000 | 19;19;19;19 | 26161709:261 | 26162014:261 | -;-;-;- | 843 | dtnbp1b | 0 | 0 | 2 | 0 | 12 | 0 |
| ENSDARG000 | 19:19:19:19:1 | 6606197:6606 | 6606788:6606 | -;-;-;-;-;-;-;-; | 1439 | pvrl2l | 16 | 17 | 34 | 13 | 49 | 68 |
| ENSDARG000 | 19;19;19;19 | 10979983:109 | 10980038:109 | +;+;+;+ | 772 | NA | 0 | 0 | 0 | 0 | 0 | 0 |
| ENSDARG000 | 19:19:19:19:1 | 34091552:340 | 34093168:340 | -;-;-;-;-;-;- | 2313 | si:dkey-184p1 | 173 | 87 | 174 | 103 | 101 | 106 |
| ENSDARG000 | 19:19:19:19:1 | 4990320:4990 | 4990384:4990 | +;+;+;+;+;+;+; | 1865 | zgc:91968 | 54065 | 34431 | 24146 | 58579 | 37796 | 37152 |
| ENSDARG000 | 19:19:19:19:1 | 3257869:3258 | 3258016:3258 | -;-;-;-;-;- | 749 | NA | 0 | 0 | 0 | 0 | 0 | 0 |
| ENSDARG000 | 19:19:19:19:1 | 31797567:318 | 31800494:318 | -;-;-;-;- | 3322 | hey1 | 43 | 2 | 18 | 0 | 36 | 28 |
| ENSDARG000 | 19:19:19:19:1 | 1856250:1856 | 1856740:1856 | -;-;-;-;-;-;-;-; | 2324 | clptm1l | 1107 | 1123 | 1283 | 1023 | 1222 | 1036 |
| ENSDARG000 | 19:19:19:19:1 | 26340736:263 | 26340957:263 | +;+;+;+;+;+;+ | 2600 | mylipa | 326 | 210 | 217 | 122 | 402 | 498 |
| ENSDARG000 | 19:19:19:19:1 | 13803155:138 | 13803895:138 | -;-;-;-;-;-;-;-; | 1579 | ccn2b | 27 | 2 | 2 | 7 | 11 | 16 |
| ENSDARG000 | 19:19:19:19:1 | 48018464:480 | 48018987:480 | +;+;+;+;+;+;+; | 2093 | UBE2M | 1967 | 1749 | 2127 | 1858 | 1961 | 1641 |
| ENSDARG000 | 19;19;19 | 3263724:3264 | 3263847:3265 | -;-;- | 337 | NA | 0 | 0 | 0 | 0 | 0 | 0 |
| ENSDARG000 | 19;19;19 | 13796657:138 | 13796755:138 | +;+;+ | 494 | NA | 0 | 0 | 0 | 0 | 0 | 0 |
| ENSDARG000 | 19:19:19:19:1 | 6896184:6896 | 6896385:6896 | +;+;+;+;+ | 1661 | NA | 40 | 38 | 56 | 89 | 66 | 45 |
| ENSDARG000 | 19:19:19:19:1 | 43004408:430 | 43004672:430 | +;+;+;+;+;+;+; | 1431 | snrnp40 | 231 | 355 | 374 | 178 | 367 | 307 |
| ENSDARG000 | 19:19:19:19:1 | 3266914:3266 | 3267521:3267 | -;-;-;-;-;-;-;-; | 1390 | illr3 | 0 | 0 | 0 | 0 | 0 | 0 |
| ENSDARG000 | 19;19 | 6904314:6906 | 6904677:6908 | +;+ | 2206 | NA | 22 | 96 | 53 | 70 | 49 | 28 |
| ENSDARG000 | 19:19:19:19:1 | 31873308:318 | 31873457:318 | +;+;+;+;+;+;+ | 943 | NA | 0 | 0 | 4 | 1 | 2 | 0 |
| ENSDARG000 | 19;19;19;19 | 31878454:318 | 31879204:318 | -;-;-;- | 2267 | zbtb10 | 663 | 431 | 600 | 426 | 613 | 499 |

| | | | | | | | | | | | | |
|---|---|---|---|---|---|---|---|---|---|---|---|---|
| ENSDARG000 | 19;19;19 | 48017892:48 | 48018342:48 | -;-;- | 612 | NA | 1 | 0 | 1 | 0 | 2 | 0 |
| ENSDARG000 | 19 | 48018006 | 48018346 | + | 341 | NA | 0 | 0 | 0 | 0 | 0 | 0 |
| ENSDARG000 | 19;19;19;19;1 | 9666961:9666 | 9667137:9667 | +;+;+;+;+;+;+; | 3880 | NA | 12 | 2 | 0 | 0 | 4 | 2 |
| ENSDARG000 | 19;19 | 9676517:9683 | 9676734:9683 | -;- | 586 | NA | 0 | 0 | 0 | 0 | 0 | 0 |
| ENSDARG000 | 19;19;19;19 | 6910384:6914 | 6910563:6916 | +;+;+;+ | 2348 | NA | 167 | 311 | 300 | 152 | 240 | 219 |
| ENSDARG000 | 19;19;19;19 | 27454227:274 | 27454509:274 | -;-;-;- | 3445 | NA | 353 | 319 | 410 | 442 | 392 | 304 |
| ENSDARG000 | 19;19;19;19;1 | 8680580:8689 | 8681664:8689 | -;-;-;-;-;-;-; | 2513 | chrna11 | 0 | 0 | 6 | 0 | 14 | 0 |
| ENSDARG000 | 19;19;19;19;1 | 23637789:236 | 23637900:236 | +;+;+;+;+ | 994 | NA | 0 | 0 | 0 | 0 | 0 | 0 |
| ENSDARG000 | 19;19;19;19;1 | 42263073:422 | 42264039:422 | -;-;-;-;-;-;-;- | 1461 | si:ch211-191i | 589 | 673 | 131 | 431 | 444 | 323 |
| ENSDARG000 | 19;19 | 6918135:6921 | 6918342:6923 | +;+ | 2431 | NA | 25 | 81 | 76 | 20 | 65 | 38 |
| ENSDARG000 | 19;19;19;19;1 | 23652399:236 | 23652452:236 | +;+;+;+;+;+;+; | 532 | NA | 0 | 0 | 0 | 0 | 0 | 0 |
| ENSDARG000 | 19;19;19;19;1 | 31904836:319 | 31905057:319 | +;+;+;+;+;+;+; | 1078 | tpd52 | 4234 | 3244 | 3941 | 4090 | 5278 | 5609 |
| ENSDARG000 | 19;19;19;19;1 | 3275361:3275 | 3275538:3275 | -;-;-;-;-;-;- | 799 | illr2 | 0 | 0 | 0 | 0 | 0 | 0 |
| ENSDARG000 | 19;19;19;19;1 | 30348676:303 | 30349638:303 | -;-;-;-;-;-;-; | 3995 | khdrbs1b | 2533 | 2377 | 3608 | 1656 | 4634 | 2509 |
| ENSDARG000 | 19;19 | 6928229:6932 | 6928550:6934 | +;+ | 2191 | NA | 25 | 18 | 36 | 30 | 35 | 49 |
| ENSDARG000 | 19;19;19;19;1 | 34169055:341 | 34169294:341 | +;+;+;+;+;+;+; | 2805 | poc1bl | 331 | 282 | 277 | 142 | 269 | 233 |
| ENSDARG000 | 19;19;19;19;1 | 23660324:236 | 23660358:236 | -;-;-;-;- | 501 | NA | 0 | 0 | 0 | 0 | 0 | 0 |
| ENSDARG000 | 19;19;19;19;1 | 35433155:354 | 35434849:354 | -;-;-;-;-;-;-; | 5436 | anln | 578 | 2382 | 2140 | 1001 | 1615 | 870 |
| ENSDARG000 | 19;19;19;19;1 | 7292445:7292 | 7292671:7292 | +;+;+;+;+;+ | 1149 | ccdc127b | 58 | 48 | 38 | 42 | 90 | 116 |
| ENSDARG000 | 19;19;19;19;1 | 23708263:237 | 23708351:237 | -;-;-;-;- | 557 | NA | 0 | 0 | 0 | 0 | 0 | 0 |
| ENSDARG000 | 19;19;19;19;1 | 23745832:237 | 23745913:237 | -;-;-;-;-;- | 1045 | NA | 0 | 0 | 0 | 0 | 0 | 0 |
| ENSDARG000 | 19;19 | 3285305:3287 | 3285826:3287 | -;- | 849 | NA | 0 | 0 | 0 | 0 | 0 | 0 |
| ENSDARG000 | 19;19;19;19;1 | 23767577:237 | 23767665:237 | -;-;-;-;- | 557 | NA | 0 | 0 | 0 | 0 | 0 | 0 |
| ENSDARG000 | 19;19;19;19;1 | 43017628:430 | 43018489:430 | +;+;+;+;+;+;+; | 6663 | nkain1 | 638 | 725 | 685 | 828 | 660 | 874 |
| ENSDARG000 | 19;19;19;19;1 | 44123618:441 | 44124412:441 | -;-;-;-;-;-;-; | 11366 | CSMD3 | 0 | 0 | 16 | 0 | 4 | 0 |
| ENSDARG000 | 19;19;19;19;1 | 5000885:5003 | 5001154:5003 | -;-;-;-;-;-;-; | 6679 | chd4a | 3342 | 3448 | 5298 | 1615 | 5166 | 2200 |
| ENSDARG000 | 19;19;19;19;1 | 3290098:3290 | 3290227:3290 | -;-;-;-;-;-;-;- | 828 | NA | 6 | 0 | 2 | 0 | 2 | 4 |
| ENSDARG000 | 19;19;19;19;1 | 6938289:6938 | 6938413:6938 | +;+;+;+;+;+;+; | 3796 | FLOT1 | 32 | 63 | 85 | 254 | 49 | 308 |
| ENSDARG000 | 19;19 | 43029382:430 | 43029512:430 | -;- | 436 | NA | 0 | 0 | 0 | 0 | 0 | 0 |
| ENSDARG000 | 19;19;19;19;1 | 1872794:1873 | 1872967:1873 | +;+;+;+;+;+ | 1231 | snrpd1 | 736 | 1120 | 1170 | 694 | 746 | 662 |
| ENSDARG000 | 19;19;19;19;1 | 72661:74327: | 75519:75519: | -;-;-;-;-;-;-; | 4882 | hnrnpr | 2239 | 2896 | 3848 | 2073 | 3152 | 2129 |
| ENSDARG000 | 19;19;19;19;1 | 1879611:1881 | 1881280:1881 | -;-;-;-;-;-;-; | 6736 | NA | 2 | 3 | 15 | 31 | 26 | 1 |
| ENSDARG000 | 19;19;19;19;1 | 13905068:139 | 13906040:139 | -;-;-;-;-;-;-; | 1954 | stx12 | 2207 | 2573 | 2333 | 1785 | 2465 | 2268 |
| ENSDARG000 | 19;19;19 | 19976990:199 | 19977143:199 | +;+;+ | 1233 | npvf | 0 | 0 | 0 | 0 | 0 | 0 |
| ENSDARG000 | 19;19;19;19 | 1878500:1879 | 1878678:1879 | +;+;+;+ | 758 | ggcta | 0 | 0 | 2 | 0 | 1 | 0 |
| ENSDARG000 | 19;19;19;19;1 | 42328423:423 | 42328591:423 | +;+;+;+;+;+;+; | 2456 | anp32e | 2951 | 2792 | 3625 | 2034 | 3521 | 2621 |
| ENSDARG000 | 19;19 | 3300795:3303 | 3301864:3303 | -;- | 1467 | NA | 1 | 0 | 30 | 0 | 1 | 0 |
| ENSDARG000 | 19;19;19;19 | 8743260:8747 | 8743556:8747 | -;-;-;- | 588 | rps27.1 | 8140 | 8108 | 12148 | 13210 | 7866 | 14104 |
| ENSDARG000 | 19;19;19;19;1 | 23794835:237 | 23794880:237 | -;-;-;-;-;-;-; | 1392 | NA | 0 | 0 | 0 | 0 | 0 | 0 |
| ENSDARG000 | 19;19;19;19;1 | 9685165:9685 | 9687572:9685 | -;-;-;-;-;-;-; | 4017 | slc2a3a | 10 | 248 | 2080 | 917 | 56 | 36 |
| ENSDARG000 | 19;19;19;19;1 | 7299865:7300 | 7299902:7300 | +;+;+;+;+;+;+; | 2553 | SQLE | 0 | 0 | 0 | 0 | 0 | 0 |
| ENSDARG000 | 19;19;19;19 | 8760795:8761 | 8761190:8761 | -;-;-;- | 888 | NA | 0 | 0 | 0 | 0 | 0 | 2 |
| ENSDARG000 | 19;19;19;19;1 | 26423590:264 | 26423723:264 | +;+;+;+;+;+;+; | 5171 | npr1a | 20 | 2 | 26 | 0 | 14 | 15 |
| ENSDARG000 | 19;19;19;19;1 | 8758336:8775 | 8761459:8775 | -;-;-;-;-;-;-; | 5196 | cers2a | 308 | 556 | 439 | 355 | 388 | 513 |
| ENSDARG000 | 19;19;19;19;1 | 3376430:3379 | 3377106:3379 | -;-;-;-;- | 1220 | edn1 | 0 | 10 | 2 | 0 | 0 | 6 |
| ENSDARG000 | 19;19;19;19;1 | 23870661:238 | 23870741:238 | -;-;-;-;- | 594 | NA | 0 | 0 | 0 | 0 | 0 | 0 |
| ENSDARG000 | 19;19;19;19;1 | 40248697:402 | 40249257:402 | +;+;+;+;+;+;+; | 5907 | cdk6 | 482 | 341 | 1251 | 486 | 1813 | 842 |
| ENSDARG000 | 19;19;19;19;1 | 3383025:3384 | 3387949:3387 | -;-;-;-;-;-;-; | 13276 | HIVEP1 | 652 | 565 | 881 | 512 | 1090 | 605 |
| ENSDARG000 | 19;19;19;19;1 | 27463934:274 | 27464116:274 | -;-;-;-;-;-;-; | 1526 | NA | 182 | 91 | 366 | 238 | 294 | 297 |
| ENSDARG000 | 19;19;19;19;1 | 30387999:303 | 30388248:303 | +;+;+;+;+;+;+; | 2295 | tspan13b | 10 | 2 | 0 | 0 | 12 | 0 |
| ENSDARG000 | 19;19;19;19;1 | 21919856:219 | 21920947:219 | +;+;+;+;+ | 2535 | GALR1 | 0 | 0 | 0 | 0 | 0 | 0 |
| ENSDARG000 | 19;19 | 19984181:199 | 19984357:199 | +;+ | 277 | NA | 0 | 0 | 0 | 0 | 0 | 0 |
| ENSDARG000 | 19;19;19;19;1 | 34230108:342 | 34230292:342 | +;+;+;+;+;+;+; | 1566 | GALNT12 | 166 | 74 | 111 | 88 | 100 | 125 |
| ENSDARG000 | 19;19 | 22016891:220 | 22017423:220 | +;+ | 1006 | NA | 0 | 0 | 3 | 1 | 0 | 0 |
| ENSDARG000 | 19;19;19;19;1 | 30400372:304 | 30402138:304 | -;-;-;-;-;-;-; | 2300 | agr2 | 20 | 36 | 22 | 28 | 58 | 50 |
| ENSDARG000 | 19;19;19;19 | 22062202:220 | 22062888:220 | +;+;+;+ | 6395 | sall3b | 308 | 62 | 158 | 34 | 84 | 16 |
| ENSDARG000 | 19;19;19;19;1 | 42367286:423 | 42368785:423 | -;-;-;-;-;-;-; | 2135 | plekho1a | 20 | 8 | 34 | 4 | 14 | 20 |
| ENSDARG000 | 19;19;19;19;1 | 13923600:139 | 13925131:139 | -;-;-;-;-;-;-; | 2551 | trnau1apa | 140 | 150 | 272 | 172 | 298 | 228 |
| ENSDARG000 | 19;19;19;19;1 | 31956534:319 | 31956877:319 | -;-;-;-;-;-;-;- | 2044 | znf704 | 119 | 27 | 53 | 39 | 128 | 103 |
| ENSDARG000 | 19;19 | 3573901:3577 | 3574010:3578 | +;+ | 748 | NA | 2 | 0 | 0 | 0 | 0 | 0 |
| ENSDARG000 | 19;19;19 | 3539601:3542 | 3540485:3542 | -;-;- | 1343 | TMEM170B | 68 | 21 | 19 | 22 | 52 | 19 |
| ENSDARG000 | 19;19 | 34274504:342 | 34274570:342 | +;+ | 560 | NA | 0 | 0 | 0 | 0 | 0 | 0 |
| ENSDARG000 | 19;19;19;19;1 | 23875382:238 | 23877640:238 | -;-;-;-;-;-;-; | 4461 | MATCAP2 | 0 | 0 | 4 | 0 | 18 | 6 |
| ENSDARG000 | 19;19;19;19;1 | 34311374:344 | 34311685:344 | +;+;+;+;+;+;+; | 2808 | gabbr2 | 4 | 0 | 2 | 4 | 2 | 0 |
| ENSDARG000 | 19;19;19;19;1 | 35455172:354 | 35459950:354 | -;-;-;-;-;-;-; | 6240 | ptp4a2b | 3634 | 4955 | 6291 | 5233 | 5100 | 2631 |
| ENSDARG000 | 19;19;19;19;1 | 7309050:7309 | 7310517:7310 | -;-;-;-;-;-;-; | 4211 | oxr1b | 120 | 4 | 42 | 28 | 16 | 138 |
| ENSDARG000 | 19;19;19;19;1 | 9732084:9732 | 9732559:9732 | -;-;-;-;-;-;-; | 2929 | NA | 0 | 0 | 0 | 0 | 0 | 0 |
| ENSDARG000 | 19;19;19;19;1 | 40335271:403 | 40335416:403 | +;+;+;+;+;+;+; | 1699 | fam133b | 679 | 623 | 706 | 853 | 714 | 844 |
| ENSDARG000 | 19;19;19;19;1 | 3653976:3680 | 3654129:3681 | +;+;+;+;+;+;+ | 2909 | nedd9 | 50 | 4 | 30 | 34 | 44 | 32 |
| ENSDARG000 | 19;19 | 9751075:9756 | 9751311:9757 | +;+ | 720 | NA | 0 | 0 | 0 | 0 | 0 | 0 |
| ENSDARG000 | 19 | 9771417 | 9776075 | - | 4659 | NA | 2 | 0 | 2 | 0 | 0 | 10 |
| ENSDARG000 | 19;19;19;19;1 | 9786722:9790 | 9787133:9791 | +;+;+;+;+;+;+; | 2014 | cacng6a | 0 | 0 | 0 | 0 | 0 | 0 |
| ENSDARG000 | 19;19;19;19;1 | 6960388:6966 | 6966432:6966 | -;-;-;-;-;-;-; | 7660 | znf384a | 588 | 979 | 1120 | 691 | 1179 | 1072 |
| ENSDARG000 | 19;19;19;19;1 | 84052:84054: | 84335:84335: | -;-;-;-;-;- | 534 | atp5if1a | 980 | 1262 | 1261 | 794 | 899 | 640 |
| ENSDARG000 | 19;19;19;19;1 | 30408317:304 | 30411514:304 | -;-;-;-;-;-;-; | 5870 | ppp1r10 | 2283 | 1939 | 1977 | 1957 | 2512 | 1753 |
| ENSDARG000 | 19;19;19;19;1 | 11039517:110 | 11040853:110 | -;-;-;-;-;-;-; | 4238 | arhgef1a | 1435 | 965 | 841 | 732 | 974 | 994 |
| ENSDARG000 | 19;19;19;19;1 | 32048379:320 | 32051574:320 | -;-;-;-;-;-;-; | 4666 | NA | 1295 | 1182 | 1769 | 1290 | 1516 | 1992 |
| ENSDARG000 | 19;19;19;19;1 | 19988869:199 | 19988974:199 | +;+;+;+;+;+;+; | 3694 | osbpl3a | 52 | 4 | 23 | 10 | 19 | 28 |
| ENSDARG000 | 19;19;19;19;1 | 42397287:424 | 42398058:424 | -;-;-;-;-;-;-; | 2596 | vps45 | 603 | 410 | 517 | 332 | 553 | 508 |
| ENSDARG000 | 19;19;19;19;1 | 23897666:238 | 23898501:238 | -;-;-;-;-;-;-; | 3671 | chtopa | 2177 | 2052 | 2300 | 1408 | 2371 | 1588 |
| ENSDARG000 | 19;19;19;19;1 | 8816657:8816 | 8817852:8817 | -;-;-;-;-;-;-; | 2907 | celf3a | 148 | 95 | 196 | 188 | 130 | 126 |
| ENSDARG000 | 19 | 3726561 | 3726781 | - | 221 | NA | 0 | 0 | 0 | 0 | 0 | 0 |
| ENSDARG000 | 19;19;19;19;1 | 45375669:453 | 45377097:453 | -;-;-;-;-;-;-; | 4616 | trps1 | 216 | 87 | 235 | 190 | 309 | 170 |
| ENSDARG000 | 19;19;19;19;1 | 3776563:3777 | 3777068:3777 | -;-;-;-;-;-;-; | 1740 | NA | 4 | 4 | 6 | 0 | 2 | 3 |
| ENSDARG000 | 19;19;19;19;1 | 1934433:1934 | 1934926:1934 | -;-;-;-;-;-;-; | 2157 | znrf2a | 20 | 8 | 22 | 4 | 14 | 0 |
| ENSDARG000 | 19;19;19;19;1 | 3729008:3729 | 3729785:3729 | -;-;-;-;-;-;-; | 2043 | NA | 3 | 0 | 0 | 0 | 2 | 3 |
| ENSDARG000 | 19;19;19;19;1 | 3762885:3764 | 3763680:3764 | -;-;-;-;-;-;- | 1990 | NA | 1 | 0 | 2 | 0 | 0 | 0 |
| ENSDARG000 | 19;19;19;19;1 | 43037657:430 | 43037746:430 | +;+;+;+;+;+;+; | 4007 | pum1 | 2923 | 3186 | 3677 | 2519 | 3433 | 2449 |
| ENSDARG000 | 19;19;19;19;1 | 22074468:220 | 22074575:220 | +;+;+;+;+;+;+; | 3631 | atp9b | 575 | 430 | 602 | 376 | 660 | 473 |
| ENSDARG000 | 19;19 | 3760178:3761 | 3760636:3761 | -;- | 503 | NA | 0 | 0 | 0 | 0 | 0 | 0 |
| ENSDARG000 | 19;19;19;19;1 | 3747060:3747 | 3747837:3747 | -;-;-;-;-;-;-; | 2085 | NA | 0 | 0 | 0 | 0 | 0 | 0 |
| ENSDARG000 | 19;19;19;19;1 | 3717988:3718 | 3718904:3718 | -;-;-;-;- | 1215 | smim13 | 606 | 556 | 635 | 469 | 752 | 642 |
| ENSDARG000 | 19;19;19;19;1 | 3729013:3737 | 3729785:3737 | -;-;-;-;-;-;-; | 3074 | btr18 | 13 | 16 | 17 | 12 | 19 | 11 |
| ENSDARG000 | 19;19 | 3775114:3775 | 3775341:3775 | -;- | 266 | NA | 0 | 0 | 0 | 0 | 0 | 0 |
| ENSDARG000 | 19;19 | 3776589:3777 | 3777068:3777 | -;- | 507 | NA | 0 | 0 | 0 | 0 | 0 | 0 |
| ENSDARG000 | 19;19;19;19;1 | 45856171:458 | 45856520:458 | -;-;-;-;-;-;-; | 2165 | eif3hb | 385 | 220 | 398 | 296 | 310 | 236 |
| ENSDARG000 | 19;19;19;19;1 | 34742706:347 | 34743080:347 | +;+;+;+;+;+;+; | 964 | abitram | 98 | 122 | 146 | 67 | 154 | 78 |
| ENSDARG000 | 19;19;19 | 13939750:139 | 13941739:139 | -;-;- | 2307 | sh3bgrl3 | 2274 | 2918 | 3065 | 3010 | 2486 | 3609 |
| ENSDARG000 | 19;19;19 | 45962016:459 | 45962384:459 | +;+;+ | 1052 | utp23 | 194 | 151 | 259 | 108 | 164 | 208 |
| ENSDARG000 | 19;19;19;19;1 | 34691645:346 | 34692041:346 | -;-;-;-;-;-;-; | 2749 | elp2 | 182 | 308 | 408 | 141 | 277 | 256 |
| ENSDARG000 | 19;19;19;19;1 | 92570:94288: | 94206:94338: | -;-;-;-;-;-;-; | 5262 | ADGRB1 | 22 | 6 | 2 | 26 | 36 | 12 |
| ENSDARG000 | 19;19;19 | 32158010:321 | 32158112:321 | +;+;+ | 396 | mrpl53 | 132 | 141 | 200 | 71 | 163 | 159 |
| ENSDARG000 | 19 | 13961396 | 13962469 | - | 1074 | paqr7a | 2 | 4 | 0 | 8 | 0 | 9 |
| ENSDARG000 | 19;19;19;19;1 | 9813287:9817 | 9815099:9817 | -;-;-;-;-;-;-; | 3117 | cacng8a | 0 | 0 | 0 | 0 | 0 | 0 |
| ENSDARG000 | 19;19;19;19 | 32166702:321 | 32166838:321 | +;+;+;+ | 674 | fabp4a | 208 | 58 | 142 | 28 | 152 | 234 |
| ENSDARG000 | 19;19;19;19;1 | 42418227:424 | 42418573:424 | -;-;-;-;-;-;- | 1401 | zgc:153441 | 2 | 2 | 0 | 0 | 4 | 26 |
| ENSDARG000 | 19;19;19;19;1 | 48024457:480 | 48024648:480 | +;+;+;+;+;+;+; | 3774 | kpnb1 | 4775 | 7538 | 9645 | 5410 | 7417 | 4928 |
| ENSDARG000 | 19;19 | 5089416:5091 | 5089823:5092 | -;- | 1601 | NA | 2 | 11 | 7 | 0 | 0 | 0 |
| ENSDARG000 | 19;19;19;19;1 | 5072918:5073 | 5073014:5073 | +;+;+;+;+;+;+; | 2759 | eno2 | 6 | 14 | 4 | 6 | 8 | 12 |
| ENSDARG000 | 19;19;19;19;1 | 32201991:322 | 32202393:322 | +;+;+;+;+ | 894 | NA | 56 | 53 | 38 | 70 | 58 | 48 |
| ENSDARG000 | 19;19;19;19;1 | 13994563:140 | 13994805:140 | +;+;+;+;+;+ | 944 | TMEM222 | 107 | 45 | 44 | 47 | 18 | 47 |
| ENSDARG000 | 19;19;19;19;1 | 7408235:7409 | 7408435:7409 | +;+;+;+;+;+;+ | 1307 | naxe | 210 | 388 | 348 | 158 | 290 | 290 |
| ENSDARG000 | 19;19 | 14059349:140 | 14059785:140 | +;+ | 1740 | tpbga | 2 | 6 | 6 | 5 | 12 | 0 |
| ENSDARG000 | 19;19 | 45970692:459 | 45970909:459 | +;+ | 3264 | NA | 0 | 0 | 0 | 0 | 0 | 0 |
| ENSDARG000 | 19;19;19;19;1 | 9840771:9842 | 9842476:9842 | -;-;-;-;-;-;-; | 2509 | cacng7a | 0 | 0 | 0 | 0 | 0 | 0 |
| ENSDARG000 | 19;19 | 45983647:459 | 45983888:459 | -;- | 391 | NA | 10 | 10 | 1 | 6 | 3 | 5 |
| ENSDARG000 | 19;19;19 | 35496282:355 | 35498772:355 | -;-;- | 3045 | col8a2 | 11 | 12 | 5 | 0 | 51 | 12 |
| ENSDARG000 | 19;19;19;19;1 | 26640096:266 | 26640444:266 | +;+;+;+;+;+;+; | 4173 | ints3 | 538 | 482 | 827 | 381 | 656 | 566 |
| ENSDARG000 | 19;19;19;19;1 | 142044:14227 | 142172:14263 | +;+;+;+;+;+ | 4629 | NA | 158 | 214 | 208 | 322 | 192 | 131 |
| ENSDARG000 | 19;19;19;19;1 | 18310386:183 | 18310808:183 | -;-;-;-;-;-;-; | 1577 | 42sp43 | 0 | 0 | 0 | 0 | 8 | 0 |
| ENSDARG000 | 19;19;19;19;1 | 14109348:141 | 14109439:141 | +;+;+;+;+;+;+; | 2184 | zgc:175136 | 0 | 0 | 0 | 0 | 0 | 0 |
| ENSDARG000 | 19;19;19;19;1 | 6990970:6990 | 6991095:6991 | +;+;+;+;+;+;+; | 2560 | kifc1 | 402 | 1419 | 1493 | 726 | 1166 | 698 |
| ENSDARG000 | 19;19;19;19;1 | 32257472:323 | 32257707:323 | +;+;+;+;+;+;+; | 3735 | atxn1a | 255 | 153 | 308 | 150 | 472 | 137 |
| ENSDARG000 | 19;19;19;19;1 | 14113886:141 | 14114307:141 | +;+;+;+;+;+;+ | 2439 | NA | 4 | 2 | 2 | 0 | 0 | 0 |
| ENSDARG000 | 19;19;19;19;1 | 11129318:111 | 11129449:111 | -;-;-;-;-;- | 961 | NA | 0 | 2 | 4 | 0 | 2 | 0 |
| ENSDARG000 | 19;19;19;19;1 | 3803437:3806 | 3804497:3806 | -;-;-;-;-;- | 2209 | zc3h12ab | 284 | 176 | 224 | 238 | 286 | 394 |
| ENSDARG000 | 19;19;19;19;1 | 9909962:9914 | 9917685:9917 | -;-;-;-;-;-;-; | 10684 | NA | 20 | 6 | 11 | 4 | 36 | 24 |
| ENSDARG000 | 19;19;19;19;1 | 35722902:357 | 35723579:357 | -;-;-;-;- | 1223 | trappc3 | 1050 | 838 | 819 | 817 | 904 | 678 |
| ENSDARG000 | 19;19;19;19;1 | 8893952:8897 | 8896517:8897 | -;-;-;-;-;-;-; | 4262 | rprd2a | 608 | 498 | 918 | 413 | 640 | 573 |
| ENSDARG000 | 19;19;19;19;1 | 40350468:403 | 40350588:403 | +;+;+;+;+;+;+; | 1950 | hepacam2 | 4 | 4 | 8 | 4 | 16 | 0 |
| ENSDARG000 | 19;19;19;19;1 | 45991054:459 | 45991383:459 | -;-;-;-;-;-;-; | 3071 | krit1 | 520 | 392 | 480 | 333 | 484 | 434 |
| ENSDARG000 | 19;19;19;19;1 | 7413942:7416 | 7415099:7416 | -;-;-;-;- | 1822 | rab25a | 19 | 0 | 0 | 0 | 2 | 4 |
| ENSDARG000 | 19;19;19;19;1 | 43314700:433 | 43314851:433 | +;+;+;+;+;+;+; | 2066 | matn1 | 3 | 0 | 1 | 0 | 14 | 4 |
| ENSDARG000 | 19;19;19;19 | 3840955:3842 | 3841065:3843 | +;+;+;+ | 921 | LSM10 | 0 | 0 | 0 | 0 | 0 | 0 |
| ENSDARG000 | 19;19;19;19;1 | 22216778:222 | 22217121:222 | +;+;+;+;+;+;+; | 3905 | nfatc1 | 637 | 559 | 850 | 691 | 1396 | 505 |
| ENSDARG000 | 19;19;19;19;1 | 3826782:3827 | 3827005:3827 | +;+;+;+;+;+;+; | 1733 | oscp1a | 0 | 0 | 0 | 0 | 4 | 0 |
| ENSDARG000 | 19;19;19;19;1 | 156757:15914 | 156961:15924 | +;+;+;+;+;+;+; | 730 | NA | 20 | 16 | 4 | 22 | 2 | 20 |
| ENSDARG000 | 19;19;19;19;1 | 32488572:324 | 32488947:324 | -;-;-;-;-;-;- | 1774 | fuca1.2 | 697 | 756 | 2114 | 883 | 1916 | 1124 |
| ENSDARG000 | 19;19;19;19;1 | 32472996:324 | 32474927:324 | -;-;-;-;-;-;-;- | 3052 | gmpr | 101 | 61 | 198 | 120 | 247 | 112 |
| ENSDARG000 | 19;19;19;19;1 | 20037857:200 | 20038086:200 | +;+;+;+;+;+;+; | 2051 | NA | 249 | 225 | 568 | 126 | 361 | 505 |
| ENSDARG000 | 19;19;19;19;1 | 40379771:403 | 40379906:403 | +;+;+;+;+;+;+; | 1045 | VPS50 | 0 | 0 | 0 | 0 | 0 | 0 |
| ENSDARG000 | 19;19;19;19 | 23919096:239 | 23919211:239 | +;+;+;+ | 2221 | snapin | 1562 | 1402 | 1518 | 1853 | 1536 | 2106 |
| ENSDARG000 | 19;19;19;19;1 | 40379874:403 | 40379906:403 | +;+;+;+;+;+;+; | 4161 | VPS50 | 505 | 808 | 710 | 707 | 813 | 464 |
| ENSDARG000 | 19;19;19;19;1 | 34751687:347 | 34751851:347 | -;-;-;-;-;-;-; | 2432 | ctnnal1 | 458 | 223 | 435 | 224 | 420 | 261 |
| ENSDARG000 | 19;19;19;19;1 | 30428913:304 | 30429670:304 | -;-;-;-;-;-;-; | 3704 | abcf1 | 3441 | 4225 | 5547 | 2202 | 3641 | 2414 |
| ENSDARG000 | 19;19;19;19;1 | 35799384:358 | 35799985:358 | +;+;+;+;+;+;+; | 1790 | angpt2b | 116 | 99 | 47 | 30 | 309 | 16 |
| ENSDARG000 | 19;19;19;19;1 | 42432625:424 | 42432744:424 | +;+;+;+;+;+;+; | 4538 | pogza | 242 | 151 | 258 | 90 | 180 | 408 |
| ENSDARG000 | 19;19;19;19 | 10177115:101 | 10178404:101 | -;-;-;- | 2992 | dbpa | 870 | 364 | 701 | 422 | 1141 | 546 |
| ENSDARG000 | 19;19 | 14153486:141 | 14153745:141 | -;- | 726 | NA | 0 | 0 | 0 | 0 | 0 | 26 |
| ENSDARG000 | 19;19;19;19;1 | 3849378:3849 | 3849529:3849 | +;+;+;+;+ | 2809 | eva1bb | 1317 | 1103 | 1531 | 1655 | 1041 | 1023 |
| ENSDARG000 | 19;19;19;19;1 | 48037798:480 | 48038183:480 | -;-;-;-;-;-;-; | 1032 | csf3b | 32 | 18 | 10 | 34 | 4 | 24 |
| ENSDARG000 | 19;19;19;19;1 | 232536:23312 | 232608:23323 | +;+;+;+;+;+;+; | 4493 | SYNGAP1 | 18 | 21 | 49 | 47 | 72 | 28 |
| ENSDARG000 | 19;19;19;19;1 | 30423117:304 | 30423224:304 | +;+;+;+;+;+;+ | 1124 | mrps18b | 306 | 435 | 630 | 256 | 507 | 416 |
| ENSDARG000 | 19;19;19;19;1 | 23932259:239 | 23932495:239 | +;+;+;+;+;+;+; | 1743 | smad10a | 77 | 49 | 60 | 47 | 61 | 20 |
| ENSDARG000 | 19;19;19 | 1953540:1955 | 1953783:1955 | -;-;- | 664 | mturn | 0 | 0 | 0 | 0 | 0 | 0 |
| ENSDARG000 | 19;19;19;19;1 | 5094218:5094 | 5094731:5094 | -;-;-;-;-;-;-; | 1287 | tpi1a | 170 | 139 | 180 | 148 | 200 | 136 |
| ENSDARG000 | 19;19;19;19;1 | 8928434:8932 | 8930228:8932 | -;-;-;-;-;- | 2631 | ciarta | 2736 | 2563 | 3122 | 1969 | 3853 | 2237 |
| ENSDARG000 | 19;19 | 18363317:183 | 18363790:183 | -;- | 956 | NA | 0 | 0 | 4 | 2 | 2 | 2 |
| ENSDARG000 | 19;19;19;19;1 | 10201170:102 | 10207750:102 | -;-;-;-;-;-;- | 7139 | znf865 | 1754 | 1389 | 2316 | 1235 | 1994 | 1806 |
| ENSDARG000 | 19;19;19;19;1 | 7424347:7424 | 7424419:7424 | +;+;+;+;+ | 1391 | sf3b4 | 698 | 613 | 993 | 352 | 789 | 564 |
| ENSDARG000 | 19;19;19 | 2229162:2230 | 2229650:2231 | -;-;- | 1156 | twist1a | 58 | 10 | 8 | 6 | 12 | 2 |
| ENSDARG000 | 19;19;19;19;1 | 18368869:183 | 18368966:183 | +;+;+;+;+;+;+ | 438 | NA | 0 | 0 | 0 | 0 | 0 | 0 |

| | | | | | | | | | | | |
|---|---|---|---|---|---|---|---|---|---|---|---|
| ENSDARG000 | 19:19:19:19:1 | 22318000:223 | 22318609:223 | -;-;-;-;-;-;- | 2561 | NA | 392 | 10 | 62 | 2 | 30 | 378 |
| ENSDARG000 | 19:19:19:19:1 | 3865210:3866 | 3866491:3866 | -;-;-;-;-;-;-;-; | 4744 | thrap3b | 3038 | 3406 | 4477 | 2696 | 3590 | 3238 |
| ENSDARG000 | 19:19:19:19:1 | 14158075:141 | 14158266:141 | +;+;+;+;+;+;+; | 1320 | nudc | 1086 | 1139 | 1400 | 641 | 1050 | 992 |
| ENSDARG000 | 19 | 2235690 | 2235933 | - | 244 | FERD3L | 0 | 0 | 0 | 0 | 0 | 0 |
| ENSDARG000 | 19;19;19 | 295147:29946 | 297073:29953 | -;-;- | 2200 | lingo4a | 2 | 0 | 0 | 0 | 6 | 0 |
| ENSDARG000 | 19:19:19:19:1 | 18417093:184 | 18417243:184 | -;-;-;-;-;-;-;-; | 977 | NA | 9 | 4 | 11 | 4 | 16 | 0 |
| ENSDARG000 | 19:19:19:19:1 | 34913855:349 | 34914408:349 | -;-;-;-;-;-;-;- | 1350 | sla1a | 21 | 20 | 17 | 25 | 114 | 133 |
| ENSDARG000 | 19:19:19:19:1 | 48049202:480 | 48049488:480 | +;+;+;+;+;+;+; | 1851 | psmd3 | 3966 | 3951 | 4612 | 2567 | 4598 | 2935 |
| ENSDARG000 | 19:19:19:19:1 | 20061892:200 | 20062364:200 | -;-;-;-;-;-;-;-; | 2480 | pals2b | 124 | 142 | 178 | 88 | 277 | 163 |
| ENSDARG000 | 19:19:19:19:1 | 317629:31835 | 318009:31845 | -;-;-;-;-;-;-;-; | 1753 | gpd1c | 2897 | 2228 | 1185 | 2354 | 1499 | 2085 |
| ENSDARG000 | 19:19:19:19:1 | 22333321:223 | 22333889:223 | -;-;-;-;- | 1803 | slc52a2 | 170 | 191 | 211 | 122 | 270 | 250 |
| ENSDARG000 | 19;19;19 | 22341811:223 | 22344741:223 | -;-;- | 3189 | NA | 172 | 266 | 215 | 75 | 227 | 224 |
| ENSDARG000 | 19:19:19:19:1 | 310494:31185 | 310572:31197 | +;+;+;+;+;+;+; | 1702 | ctss1 | 56 | 98 | 137 | 69 | 166 | 188 |
| ENSDARG000 | 19;19;19;19 | 7001170:7003 | 7001274:7003 | +;+;+;+ | 2580 | zbtb22b | 93 | 55 | 86 | 76 | 95 | 119 |
| ENSDARG000 | 19:19:19:19:1 | 40716975:407 | 40717825:407 | -;-;-;-;-;-;-;-; | 3096 | CALCR | 0 | 1059 | 103 | 354 | 369 | 40 |
| ENSDARG000 | 19:19:19:19:1 | 42456922:424 | 42457060:424 | -;-;-;-;-;-;- | 967 | psmb4 | 2827 | 2496 | 3020 | 1467 | 2580 | 1853 |
| ENSDARG000 | 19:19:19:19:1 | 11167289:111 | 11167932:111 | -;-;-;-;-;-;-;-; | 1901 | NA | 76 | 12 | 48 | 30 | 48 | 50 |
| ENSDARG000 | 19;19;19;19 | 23982466:240 | 23982634:240 | +;+;+;+ | 4441 | syt11a | 903 | 712 | 1918 | 1667 | 1028 | 2428 |
| ENSDARG000 | 19:19:19:19:1 | 43341115:433 | 43341198:433 | +;+;+;+;+;+;+; | 2194 | sesn2 | 226 | 463 | 544 | 451 | 169 | 199 |
| ENSDARG000 | 19:19:19:19:1 | 10217598:102 | 10220815:102 | -;-;-;-;-;-;-;- | 7243 | NA | 0 | 22 | 6 | 8 | 6 | 6 |
| ENSDARG000 | 19:19:19:19:1 | 36114846:361 | 36115848:361 | -;-;-;-;-;-;-;-; | 12563 | NA | 64 | 24 | 461 | 364 | 89 | 40 |
| ENSDARG000 | 19:19:19:19:1 | 42469058:424 | 42470737:424 | +;+;+;+;+;+;+; | 3014 | NA | 15 | 10 | 12 | 0 | 2 | 27 |
| ENSDARG000 | 19:19:19:19:1 | 8958134:8958 | 8958577:8958 | -;-;-;-;- | 921 | mrps21 | 176 | 280 | 252 | 152 | 202 | 152 |
| ENSDARG000 | 19;19;19;19 | 8973042:8974 | 8973253:8974 | +;+;+;+ | 3535 | crabp2b | 10727 | 4233 | 7916 | 5571 | 8646 | 6542 |
| ENSDARG000 | 19:19:19:19:1 | 34950200:349 | 34950377:349 | +;+;+;+;+;+;+; | 1510 | ccn4a | 2 | 0 | 0 | 2 | 2 | 2 |
| ENSDARG000 | 19:19:19:19:1 | 11214007:112 | 11214151:112 | +;+;+;+;+;+;+; | 2604 | NA | 0 | 8 | 4 | 4 | 16 | 52 |
| ENSDARG000 | 19:19:19:19:1 | 34959967:349 | 34961282:349 | -;-;-;-;-;-;-;-; | 2576 | ndrg1a | 7426 | 12961 | 25487 | 26088 | 10722 | 7322 |
| ENSDARG000 | 19:19:19:19:1 | 27479563:274 | 27480119:274 | +;+;+;+;+;+;+; | 2764 | atat1 | 76 | 84 | 69 | 101 | 123 | 121 |
| ENSDARG000 | 19:19:19:19:1 | 36625275:366 | 36625300:366 | +;+;+;+;+;+;+ | 1233 | NA | 6 | 40 | 0 | 0 | 0 | 47 |
| ENSDARG000 | 19:19:19:19:1 | 20110699:201 | 20112149:201 | -;-;-;-;-;-;- | 1913 | npy | 0 | 0 | 0 | 0 | 0 | 4 |
| ENSDARG000 | 19:19:19:19:1 | 7014455:7014 | 7015305:7015 | -;-;-;-;-;-;-;-; | 3191 | daxx | 210 | 430 | 465 | 210 | 436 | 320 |
| ENSDARG000 | 19:19:19:19:1 | 7432710:7436 | 7435640:7437 | -;-;-;-;-;-;-;-; | 4348 | gabpb2a | 200 | 576 | 646 | 265 | 466 | 444 |
| ENSDARG000 | 19:19:19:19:1 | 26714949:267 | 26715051:267 | +;+;+;+;+;+;+; | 3006 | pdia8 | 30 | 45 | 33 | 13 | 40 | 20 |
| ENSDARG000 | 19:19:19:19:1 | 48060464:480 | 48060657:480 | +;+;+;+;+;+;+; | 3286 | zgc:85936 | 158 | 444 | 557 | 252 | 481 | 279 |
| ENSDARG000 | 19;19 | 18484706:184 | 18485179:184 | -;- | 569 | NA | 6 | 21 | 19 | 14 | 15 | 10 |
| ENSDARG000 | 19:19:19:19:1 | 2275019:2279 | 2275211:2279 | +;+;+;+;+;+;+; | 2639 | NA | 0 | 18 | 18 | 6 | 1 | 0 |
| ENSDARG000 | 19:19:19:19:1 | 18423378:184 | 18423747:184 | -;-;-;-;-;-;-;-; | 1841 | p3h4 | 220 | 251 | 544 | 227 | 602 | 240 |
| ENSDARG000 | 19:19:19:19:1 | 22347627:223 | 22348898:223 | +;+;+;+;+;+;+; | 2701 | fbxl6 | 212 | 95 | 122 | 127 | 138 | 234 |
| ENSDARG000 | 19:19:19:19:1 | 8985197:8985 | 8985638:8985 | +;+;+;+;+;+;+; | 2385 | scamp3 | 1672 | 2548 | 1937 | 1648 | 1967 | 2227 |
| ENSDARG000 | 19:19:19:19:1 | 24039830:240 | 24039927:240 | +;+;+;+;+;+;+; | 1621 | rit1 | 703 | 437 | 844 | 628 | 628 | 525 |
| ENSDARG000 | 19;19 | 22360956:223 | 22380042:223 | -;- | 19233 | eppk1 | 146 | 14 | 28 | 56 | 118 | 90 |
| ENSDARG000 | 19:19:19:19:1 | 7038694:7039 | 7038978:7039 | -;-;-;-;-;-;-;- | 1715 | NA | 1407 | 1558 | 1560 | 920 | 1528 | 989 |
| ENSDARG000 | 19;19 | 8983355:8985 | 8983944:8985 | -;- | 766 | NA | 15 | 4 | 6 | 2 | 5 | 8 |
| ENSDARG000 | 19:19:19:19:1 | 24068223:240 | 24068299:240 | +;+;+;+;+;+;+; | 1443 | pex11b | 156 | 152 | 167 | 144 | 190 | 190 |
| ENSDARG000 | 19:19:19:19:1 | 42490936:424 | 42491298:424 | -;-;-;-;-;-;-;-; | 1963 | zgc:110239 | 5767 | 4324 | 3191 | 5559 | 4304 | 6464 |
| ENSDARG000 | 19:19:19:19:1 | 26681848:266 | 26682056:266 | +;+;+;+;+;+;+; | 3099 | NA | 36 | 86 | 121 | 50 | 142 | 86 |
| ENSDARG000 | 19:19:19:19:1 | 5106226:5106 | 5106989:5106 | -;-;-;-;-;-;-;-; | 2642 | gnb3b | 3 | 0 | 5 | 0 | 0 | 0 |
| ENSDARG000 | 19;19 | 40832493:408 | 40832542:408 | +;+ | 793 | NA | 0 | 0 | 0 | 0 | 0 | 0 |
| ENSDARG000 | 19:19:19:19:1 | 11231083:112 | 11231578:112 | -;-;-;-;-;-;-;-; | 1045 | ssr2 | 2082 | 2783 | 4649 | 2647 | 3423 | 2569 |
| ENSDARG000 | 19:19:19:19:1 | 46019313:460 | 46020574:460 | -;-;-;-;-;-;-;-; | 5699 | nup153 | 1617 | 1860 | 2509 | 960 | 2653 | 1407 |
| ENSDARG000 | 19:19:19:19:1 | 40848109:408 | 40849097:408 | -;-;-;-;- | 1665 | tfpi2 | 79 | 40 | 14 | 40 | 26 | 100 |
| ENSDARG000 | 19;19;19;19 | 20143850:201 | 20146837:201 | -;-;-;- | 3366 | NA | 16 | 0 | 13 | 10 | 8 | 12 |
| ENSDARG000 | 19;19;19 | 2315208:2315 | 2317558:2317 | -;-;- | 2445 | sp8a | 110 | 102 | 64 | 364 | 298 | 2 |
| ENSDARG000 | 19:19:19:19:1 | 3890506:3892 | 3890662:3892 | -;-;-;-;-;-;- | 934 | NA | 10 | 4 | 22 | 64 | 40 | 41 |
| ENSDARG000 | 19:19:19:19:1 | 5134624:5135 | 5135319:5135 | +;+;+;+;+;+;+; | 6345 | NA | 1727 | 1378 | 2285 | 1384 | 1958 | 1980 |
| ENSDARG000 | 19:19:19:19:1 | 32495419:324 | 32495782:324 | -;-;-;-;-;-;-;- | 1775 | fuca1.1 | 1178 | 883 | 1206 | 1051 | 1346 | 1097 |
| ENSDARG000 | 19:19:19:19:1 | 7043634:7044 | 7043658:7045 | +;+;+;+;+;+;+; | 1387 | NA | 0 | 124 | 40 | 55 | 25 | 28 |
| ENSDARG000 | 19:19:19:19:1 | 14187540:141 | 14188667:141 | -;-;-;-;-;-;-;-; | 2216 | tbxta | 0 | 0 | 0 | 0 | 0 | 0 |
| ENSDARG000 | 19:19:19:19:1 | 40856534:408 | 40856752:408 | +;+;+;+;+ | 921 | gngt1 | 0 | 0 | 0 | 0 | 4 | 6 |
| ENSDARG000 | 19:19:19:19:1 | 2364552:2546 | 2364821:2546 | +;+;+;+;+;+;+; | 5629 | sp4 | 521 | 672 | 705 | 627 | 724 | 515 |
| ENSDARG000 | 19;19 | 7445644:7450 | 7446707:7450 | -;- | 1330 | mllt11 | 0 | 0 | 0 | 0 | 4 | 6 |
| ENSDARG000 | 19;19 | 42505974:425 | 42506130:425 | +;+ | 651 | NA | 0 | 0 | 0 | 0 | 0 | 0 |
| ENSDARG000 | 19;19;19;19 | 24123409:241 | 24124618:241 | -;-;-;- | 1519 | zgc:64022 | 2 | 0 | 6 | 2 | 28 | 6 |
| ENSDARG000 | 19:19:19:19:1 | 20158657:201 | 20159277:201 | -;-;-;-;-;-;-;-; | 1373 | fam221a | 4 | 6 | 6 | 16 | 4 | 12 |
| ENSDARG000 | 19:19:19:19:1 | 42547699:425 | 42548102:425 | -;-;-;-;-;-;-;-;- | 1392 | zgc:123103 | 0 | 0 | 106 | 6 | 5 | 0 |
| ENSDARG000 | 19:19:19:19:1 | 10252052:102 | 10255900:102 | -;-;-;-;-;-;-;-; | 6582 | cnot3b | 651 | 506 | 935 | 344 | 668 | 584 |
| ENSDARG000 | 19:19:19:19:1 | 40861853:408 | 40862107:408 | +;+;+;+;+;+;+; | 1634 | zgc:85777 | 546 | 533 | 505 | 497 | 577 | 580 |
| ENSDARG000 | 19:19:19:19:1 | 7065988:7066 | 7066531:7066 | -;-;-;-;-;-;- | 1982 | tapbp.2 | 1318 | 705 | 757 | 438 | 859 | 458 |
| ENSDARG000 | 19:19:19:19:1 | 2569150:2570 | 2569530:2570 | -;-;-;-;-;-;-;-; | 2255 | NA | 260 | 680 | 940 | 426 | 634 | 588 |
| ENSDARG000 | 19:19:19:19:1 | 7453996:7454 | 7454223:7454 | +;+;+;+;+;+;+; | 2628 | mindy1 | 768 | 354 | 525 | 496 | 534 | 412 |
| ENSDARG000 | 19;19;19;19 | 32505747:325 | 32506055:325 | -;-;-;- | 1619 | zbtb8b | 317 | 229 | 300 | 204 | 340 | 265 |
| ENSDARG000 | 19:19:19:19:1 | 9004256:9004 | 9004412:9004 | +;+;+;+;+;+;+; | 1247 | NA | 10 | 4 | 20 | 4 | 30 | 2 |
| ENSDARG000 | 19:19:19:19:1 | 24126775:241 | 24131204:241 | -;-;-;-;-;-;-;-; | 5681 | thap7 | 319 | 200 | 458 | 370 | 280 | 302 |
| ENSDARG000 | 19;19;19 | 11247054:112 | 11247768:112 | -;-;- | 1330 | NA | 1 | 6 | 6 | 0 | 12 | 7 |
| ENSDARG000 | 19:19:19:19:1 | 27522425:275 | 27523299:275 | -;-;-;-;-;-;-;-; | 2111 | NA | 28 | 22 | 14 | 18 | 32 | 34 |
| ENSDARG000 | 19:19:19:19:1 | 9032073:9033 | 9032500:9033 | +;+;+;+;+;+;+; | 12685 | ash1l | 2040 | 1453 | 2690 | 1566 | 2468 | 1961 |
| ENSDARG000 | 19;19 | 18596786:185 | 18596968:185 | -;- | 315 | NA | 0 | 0 | 0 | 0 | 0 | 0 |
| ENSDARG000 | 19:19:19:19:1 | 18493782:185 | 18494120:185 | +;+;+;+;+;+;+; | 1978 | FKBP10 | 385 | 412 | 728 | 618 | 740 | 424 |
| ENSDARG000 | 19;19;19 | 9030019:9030 | 9030581:9030 | -;-;- | 649 | NA | 0 | 0 | 0 | 0 | 0 | 0 |
| ENSDARG000 | 19:19:19:19:1 | 32553874:325 | 32553998:325 | +;+;+;+;+ | 1619 | heyl | 0 | 5 | 0 | 0 | 0 | 0 |
| ENSDARG000 | 19:19:19:19:1 | 11256617:112 | 11256675:112 | +;+;+;+;+ | 3742 | NA | 2 | 0 | 0 | 0 | 0 | 8 |
| ENSDARG000 | 19:19:19:19:1 | 3900129:3900 | 3900443:3900 | -;-;-;-;-;-;-;-; | 1137 | NA | 13 | 97 | 41 | 20 | 86 | 35 |
| ENSDARG000 | 19:19:19:19:1 | 329808:33006 | 332863:33286 | -;-;-;-;-;-;-;-; | 4378 | golph3l | 70 | 10 | 12 | 2 | 8 | 24 |
| ENSDARG000 | 19:19:19:19:1 | 11270847:112 | 11271503:112 | -;-;-;-;-;-;- | 2345 | eepd1 | 30 | 42 | 62 | 26 | 108 | 3 |
| ENSDARG000 | 19:19:19:19:1 | 14351560:143 | 14353228:143 | +;+;+;+;+;+;+; | 10786 | ARID1A | 2304 | 1869 | 2848 | 1555 | 2653 | 2065 |
| ENSDARG000 | 19:19:19:19:1 | 11264584:112 | 11264716:112 | +;+;+;+;+ | 477 | NA | 0 | 0 | 3 | 5 | 5 | 6 |
| ENSDARG000 | 19;19;19 | 24161712:241 | 24163052:241 | -;-;- | 1898 | prf1.2 | 0 | 1 | 2 | 9 | 14 | 9 |
| ENSDARG000 | 19:19:19:19:1 | 30450125:304 | 30450386:304 | +;+;+;+;+;+;+ | 1924 | si:ch211-215a | 158 | 197 | 257 | 155 | 265 | 202 |
| ENSDARG000 | 19 | 24187594 | 24188743 | - | 1150 | NA | 0 | 0 | 0 | 0 | 0 | 0 |
| ENSDARG000 | 19:19:19:19:1 | 42553029:425 | 42553271:425 | -;-;-;-;-;-;-;-; | 1584 | NA | 4 | 6 | 126 | 12 | 86 | 0 |
| ENSDARG000 | 19:19:19:19:1 | 46066889:460 | 46067615:460 | -;-;-;-;-;-;-;-; | 2043 | PTDSS1 | 19 | 12 | 24 | 34 | 28 | 32 |
| ENSDARG000 | 19 | 46091318 | 46091605 | - | 288 | NA | 0 | 0 | 0 | 0 | 0 | 0 |
| ENSDARG000 | 19;19;19;19 | 2590182:2590 | 2590297:2590 | +;+;+;+ | 864 | NA | 2 | 5 | 46 | 0 | 0 | 4 |
| ENSDARG000 | 19;19;19;19 | 40899910:409 | 40900363:409 | -;-;-;- | 766 | bet1 | 136 | 152 | 184 | 196 | 104 | 128 |
| ENSDARG000 | 19:19:19:19:1 | 7072854:7072 | 7072981:7072 | +;+;+;+;+;+;+; | 2551 | mhc1uba | 6 | 22393 | 5826 | 6260 | 6429 | 6206 |
| ENSDARG000 | 19:19:19:19:1 | 7482784:7489 | 7487648:7489 | -;-;-;-;-;-;-;-; | 5783 | NA | 381 | 289 | 406 | 363 | 509 | 391 |
| ENSDARG000 | 19:19:19:19:1 | 24196088:241 | 24197428:241 | -;-;-;-;- | 2018 | PRF1 | 0 | 35 | 26 | 7 | 77 | 141 |
| ENSDARG000 | 19:19:19:19:1 | 22407819:224 | 22416685:224 | -;-;-;-;-;-;-;-; | 19476 | pleca | 34052 | 12540 | 9387 | 14674 | 9417 | 11807 |
| ENSDARG000 | 19:19:19:19:1 | 3909980:3910 | 3910172:3910 | -;-;-;-;-;-;-;-; | 977 | NA | 0 | 0 | 6 | 7 | 1 | 6 |
| ENSDARG000 | 19:19:19:19:1 | 24221868:242 | 24223218:242 | -;-;-;-;- | 2271 | NA | 46 | 309 | 102 | 142 | 226 | 662 |
| ENSDARG000 | 19:19:19:19:1 | 46095210:460 | 46095339:460 | +;+;+;+;+ | 913 | STMND1 | 0 | 6 | 0 | 0 | 0 | 0 |
| ENSDARG000 | 19:19:19:19:1 | 11331822:113 | 11332946:113 | -;-;-;-;-;-;-;-; | 2470 | septin7a | 5326 | 4042 | 5901 | 3689 | 4217 | 4686 |
| ENSDARG000 | 19:19:19:19:1 | 32571727:325 | 32573449:325 | -;-;-;-;-;-;-;-; | 3726 | zgc:91944 | 449 | 371 | 600 | 420 | 553 | 706 |
| ENSDARG000 | 19:19:19:19:1 | 2602903:2619 | 2603011:2619 | +;+;+;+;+;+;+; | 2895 | hycc1 | 1035 | 1010 | 956 | 975 | 916 | 694 |
| ENSDARG000 | 19:19:19:19:1 | 24262263:242 | 24264268:242 | -;-;-;-;-;-;- | 2421 | s100v2 | 170 | 28 | 44 | 6 | 34 | 60 |
| ENSDARG000 | 19:19:19:19:1 | 20163826:201 | 20164118:201 | +;+;+;+;+;+;+ | 1623 | ccdc126 | 173 | 218 | 316 | 236 | 270 | 221 |
| ENSDARG000 | 19:19:19:19:1 | 42559143:425 | 42559881:425 | -;-;-;-;-;-;-;-; | 1903 | si:dkey-228a1 | 53 | 45 | 145 | 39 | 51 | 66 |
| ENSDARG000 | 19:19:19:19:1 | 40983221:409 | 40983468:409 | +;+;+;+;+;+;+; | 5750 | col1a2 | 2683 | 496 | 990 | 184 | 6200 | 434 |
| ENSDARG000 | 19;19 | 14573998:145 | 14574381:145 | +;+ | 2981 | tent5bb | 34 | 12 | 12 | 0 | 58 | 2 |
| ENSDARG000 | 19:19:19:19:1 | 27561664:275 | 27561911:275 | -;-;-;-;-;-;-;- | 972 | NA | 0 | 0 | 0 | 0 | 0 | 0 |
| ENSDARG000 | 19:19:19:19:1 | 42567539:425 | 42568253:425 | -;-;-;-;-;-;-;-; | 1778 | zgc:103438 | 439 | 419 | 532 | 299 | 608 | 568 |
| ENSDARG000 | 19:19:19:19:1 | 18617608:186 | 18617859:186 | -;-;-;-;-;-;-;-; | 1500 | rps18 | 10430 | 10868 | 16677 | 15626 | 10079 | 16937 |
| ENSDARG000 | 19:19:19:19:1 | 27546401:275 | 27548728:275 | -;-;-;-;-;-;-;-; | 4129 | NA | 0 | 0 | 0 | 0 | 0 | 0 |
| ENSDARG000 | 19:19:19:19:1 | 26730298:267 | 26731357:267 | -;-;-;-;-;-;-;-; | 2275 | csnk2b | 3334 | 3202 | 3671 | 2854 | 3248 | 3065 |
| ENSDARG000 | 19;19 | 14850833:148 | 14850904:148 | +;+ | 527 | NA | 0 | 0 | 0 | 0 | 0 | 0 |
| ENSDARG000 | 19:19:19:19:1 | 27568231:275 | 27568476:275 | -;-;-;-;-;-;- | 880 | NA | 0 | 0 | 0 | 0 | 0 | 0 |
| ENSDARG000 | 19:19:19:19:1 | 24324967:243 | 24325745:243 | +;+;+;+;+;+;+; | 6645 | NA | 1672 | 927 | 1305 | 737 | 1269 | 2468 |
| ENSDARG000 | 19:19:19:19:1 | 40941131:409 | 40941710:409 | -;-;-;-;-;-;- | 2048 | NA | 0 | 0 | 0 | 0 | 0 | 0 |
| ENSDARG000 | 19:19:19:19:1 | 42591150:425 | 42591614:425 | -;-;-;-;-;-;-;-; | 2235 | fkbp9 | 401 | 629 | 659 | 355 | 680 | 256 |
| ENSDARG000 | 19;19;19 | 14921000:149 | 14921450:149 | +;+;+ | 1430 | oprd1a | 0 | 0 | 0 | 0 | 0 | 0 |
| ENSDARG000 | 19:19:19:19:1 | 42573724:425 | 42574818:425 | -;-;-;-;-;-;-;-; | 2706 | sytl1 | 8 | 2 | 0 | 0 | 20 | 4 |
| ENSDARG000 | 19:19:19:19:1 | 42560956:425 | 42560997:425 | +;+;+;+;+ | 899 | NA | 0 | 0 | 0 | 3 | 0 | 0 |
| ENSDARG000 | 19:19:19:19:1 | 342094:34213 | 342178:34217 | +;+;+;+;+;+ | 475 | ensaa | 334 | 111 | 85 | 181 | 109 | 200 |
| ENSDARG000 | 19:19:19:19:1 | 30475281:304 | 30475376:304 | -;-;-;-;-;-;-;-; | 4348 | bag6l | 389 | 285 | 523 | 260 | 400 | 379 |
| ENSDARG000 | 19:19:19:19:1 | 46113828:461 | 46114181:461 | +;+;+;+;+;+;+ | 3259 | rbm24a | 40 | 10 | 70 | 16 | 52 | 29 |
| ENSDARG000 | 19 | 37118547 | 37119891 | + | 1345 | gjb9b | 8 | 12 | 30 | 10 | 12 | 0 |
| ENSDARG000 | 19;19;19 | 348729:35121 | 349306:35146 | +;+;+ | 1409 | mcl1a | 15265 | 9021 | 12674 | 20972 | 9364 | 10536 |
| ENSDARG000 | 19:19:19:19:1 | 37120491:371 | 37120530:371 | +;+;+;+;+ | 1450 | pef1 | 1636 | 1286 | 1372 | 1369 | 1504 | 1071 |
| ENSDARG000 | 19:19:19:19:1 | 3919591:3919 | 3919922:3919 | -;-;-;-;-;-;-;-; | 1179 | NA | 2 | 1 | 6 | 6 | 4 | 6 |
| ENSDARG000 | 19:19:19:19:1 | 7495791:7497 | 7495899:7498 | +;+;+;+;+;+ | 874 | mrpl24 | 348 | 497 | 586 | 278 | 479 | 396 |
| ENSDARG000 | 19:19:19:19:1 | 43439401:434 | 43439824:434 | +;+;+;+;+;+;+ | 6603 | AHDC1 | 0 | 12 | 14 | 2 | 8 | 26 |
| ENSDARG000 | 19;19 | 7501070:7501 | 7501288:7501 | -;- | 472 | NA | 1 | 2 | 0 | 0 | 0 | 0 |
| ENSDARG000 | 19;19;19;19 | 43429805:434 | 43430099:434 | -;-;-;- | 805 | NA | 0 | 2 | 4 | 0 | 0 | 1 |
| ENSDARG000 | 19:19:19:19:1 | 43523690:435 | 43523810:435 | +;+;+;+;+;+;+; | 3321 | wasf2 | 2487 | 2362 | 2626 | 2490 | 2614 | 2133 |
| ENSDARG000 | 19:19:19:19:1 | 11632834:117 | 11633017:117 | -;-;-;-;-;-;-;-; | 3022 | ctdp1 | 68 | 216 | 291 | 120 | 262 | 110 |
| ENSDARG000 | 19:19:19:19:1 | 5158306:5163 | 5164018:5164 | -;-;-;-;-;-;-;-; | 6827 | stx1b | 0 | 8 | 8 | 2 | 4 | 2 |
| ENSDARG000 | 19:19:19:19:1 | 2670130:2670 | 2670219:2670 | +;+;+;+;+;+;+; | 3483 | NA | 248 | 213 | 140 | 64 | 158 | 218 |
| ENSDARG000 | 19:19:19:19:1 | 43359075:433 | 43359162:433 | +;+;+;+;+;+;+; | 4404 | yrk | 242 | 128 | 239 | 119 | 372 | 313 |
| ENSDARG000 | 19:19:19:19:1 | 366034:36798 | 366345:36814 | +;+;+;+;+;+ | 2900 | vps72a | 1805 | 1229 | 2671 | 2782 | 2449 | 1858 |
| ENSDARG000 | 19:19:19:19:1 | 7518304:7519 | 7519063:7519 | -;-;-;-;-;- | 1739 | mrpl9 | 234 | 328 | 415 | 182 | 370 | 266 |
| ENSDARG000 | 19:19:19:19:1 | 7124337:7124 | 7124896:7125 | +;+;+;+;+;+;+; | 3190 | tap2a | 239 | 678 | 318 | 100 | 337 | 147 |
| ENSDARG000 | 19:19:19:19:1 | 46158078:461 | 46158249:461 | +;+;+;+;+;+;+; | 3488 | cap2 | 30 | 32 | 69 | 2 | 7 | 88 |
| ENSDARG000 | 19;19;19;19 | 24394560:243 | 24394679:243 | +;+;+;+ | 818 | si:dkev-81h8. | 0 | 0 | 0 | 4 | 0 | 0 |
| ENSDARG000 | 19:19:19:19:1 | 7506380:7507 | 7506411:7507 | +;+;+;+;+;+;+; | 4820 | prcc | 427 | 663 | 787 | 567 | 801 | 693 |
| ENSDARG000 | 19:19:19:19:1 | 7529612:7533 | 7531210:7533 | -;-;-;-;-;- | 2577 | lix1l | 156 | 242 | 232 | 73 | 194 | 208 |
| ENSDARG000 | 19:19:19:19:1 | 7105838:7106 | 7106013:7106 | -;-;-;-;-;-;-;-; | 1620 | psmb8a | 1610 | 2191 | 1469 | 847 | 1115 | 1231 |
| ENSDARG000 | 19:19:19:19:1 | 32603627:326 | 32603742:326 | +;+;+;+;+;+ | 1854 | tmem54b | 0 | 0 | 5 | 2 | 6 | 5 |
| ENSDARG000 | 19;19 | 37135700:371 | 37135778:371 | +;+ | 1174 | smim12 | 238 | 299 | 338 | 347 | 586 | 370 |
| ENSDARG000 | 19 | 37153409 | 37154436 | - | 1028 | gja4 | 2753 | 2371 | 3500 | 2623 | 3225 | 1711 |
| ENSDARG000 | 19:19:19:19:1 | 7111779:7112 | 7112062:7112 | -;-;-;-;-;-;-;-; | 1486 | psmb13a | 445 | 714 | 338 | 198 | 310 | 455 |
| ENSDARG000 | 19:19:19:19:1 | 26739813:267 | 26745869:267 | -;-;-;-;-;-;-;-; | 9335 | prrc2a | 5140 | 3913 | 5910 | 3836 | 5763 | 4280 |
| ENSDARG000 | 19:19:19:19:1 | 9091673:9091 | 9091901:9091 | +;+;+;+;+;+;+; | 1976 | lrrc71 | 0 | 34 | 24 | 60 | 14 | 46 |
| ENSDARG000 | 19:19:19:19:1 | 37458610:374 | 37459668:374 | +;+;+;+;+;+;+; | 3846 | dlgap3 | 34 | 31 | 98 | 78 | 81 | 34 |
| ENSDARG000 | 19:19:19:19:1 | 10299002:102 | 10299871:102 | -;-;-;-;-;-;-;-; | 3685 | u2af2b | 1784 | 2218 | 2918 | 1497 | 2257 | 1734 |
| ENSDARG000 | 19;19;19;19 | 2685779:2685 | 2686014:2686 | +;+;+;+ | 584 | tomm7 | 338 | 491 | 381 | 324 | 362 | 358 |
| ENSDARG000 | 19:19:19:19:1 | 7120380:7120 | 7120744:7121 | +;+;+;+;+;+ | 1153 | psmb9a | 406 | 765 | 478 | 155 | 485 | 384 |
| ENSDARG000 | 19;19 | 373811:37933 | 376392:37975 | -;- | 2999 | tnfaip8l2a | 44 | 104 | 56 | 21 | 68 | 155 |
| ENSDARG000 | 19:19:19:19:1 | 7115223:7117 | 7115357:7117 | +;+;+;+;+;+ | 873 | psmb12 | 943 | 680 | 781 | 520 | 814 | 677 |
| ENSDARG000 | 19:19:19:19:1 | 20177208:201 | 20178060:201 | +;+;+;+;+;+;+; | 3575 | tra2a | 1298 | 1371 | 1857 | 1200 | 1705 | 1469 |
| ENSDARG000 | 19:19:19:19:1 | 37501652:375 | 37502112:375 | -;-;-;-;-;-;-;-; | 1560 | ilf2 | 2744 | 3602 | 4672 | 2507 | 3400 | 2633 |
| ENSDARG000 | 19;19;19 | 2896188:2897 | 2896453:2897 | +;+;+ | 513 | NA | 0 | 0 | 0 | 0 | 0 | 0 |
| ENSDARG000 | 19:19:19:19:1 | 15142527:151 | 15144523:151 | -;-;-;-;-;-;-;-; | 5864 | NA | 178 | 90 | 168 | 170 | 212 | 190 |

| | | | | | | | | | | | |
|---|---|---|---|---|---|---|---|---|---|---|---|
| ENSDARG000 | 19:19:19:19:1 | 30517311:305 | 30522478:305 | -:-:-:-:-:-:-:-: | 6052 | hpcal4 | 2 | 6 | 8 | 0 | 26 | 2 |
| ENSDARG000 | 19:19:19:19:1 | 5259503:5259 | 5259839:5259 | -;-;-;-;-;-;- | 1854 | prf1.3 | 0 | 0 | 0 | 0 | 10 | 2 |
| ENSDARG000 | 19;19;19;19 | 5291935:5299 | 5292409:5300 | +;+;+;+ | 2121 | NA | 50 | 52 | 73 | 65 | 71 | 93 |
| ENSDARG000 | 19:19:19:19:1 | 27571706:275 | 27572607:275 | -:-:-:-:-:-:-:-: | 1895 | NA | 0 | 0 | 0 | 0 | 0 | 0 |
| ENSDARG000 | 19:19:19:19:1 | 2930132:2934 | 2930989:2934 | -:-:-:-:-:-:-:-: | 3259 | bop1 | 700 | 644 | 1207 | 440 | 827 | 580 |
| ENSDARG000 | 19:19:19:19:1 | 32622177:326 | 32624879:326 | -:-:-:-:-:-:-:-: | 3765 | hpca | 47 | 104 | 298 | 177 | 184 | 183 |
| ENSDARG000 | 19:19:19:19:1 | 11855330:118 | 11855464:118 | +:+:+:+:+:+:+:+: | 3913 | adck5 | 296 | 382 | 197 | 153 | 358 | 130 |
| ENSDARG000 | 19:19:19:19:1 | 3056450:3056 | 3056840:3056 | +:+:+:+:+:+:+:+: | 4030 | hsf1 | 282 | 268 | 468 | 294 | 373 | 309 |
| ENSDARG000 | 19;19;19;19 | 37509638:375 | 37509729:375 | +;+;+;+ | 3263 | NA | 2 | 0 | 0 | 0 | 0 | 0 |
| ENSDARG000 | 19;19;19 | 34998029:349 | 34998755:349 | -;-;- | 1182 | zgc:113424 | 0 | 2 | 0 | 0 | 0 | 0 |
| ENSDARG000 | 19;19 | 3001473:3008 | 3002317:3009 | +;+ | 1432 | scxa | 2 | 0 | 0 | 0 | 2 | 4 |
| ENSDARG000 | 19;19;19 | 34994433:349 | 34994975:349 | -;-;- | 1007 | si:rp71-45k5.: | 0 | 0 | 2 | 0 | 0 | 8 |
| ENSDARG000 | 19:19:19:19:1 | 30633453:306 | 30633829:306 | +:+:+:+:+:+:+:+: | 4734 | fam49al | 30 | 48 | 107 | 20 | 248 | 130 |
| ENSDARG000 | 19:19:19:19:1 | 24419693:244 | 24419748:244 | -:-:-:-:-:-:-:-: | 3304 | thbs3b | 12 | 2 | 6 | 4 | 8 | 8 |
| ENSDARG000 | 19:19:19:19:1 | 32766423:327 | 32770970:327 | -:-:-:-:-:-:-:-: | 5346 | nt5c1aa | 0 | 0 | 6 | 0 | 6 | 0 |
| ENSDARG000 | 19:19:19:19:1 | 390298:40678 | 390396:40691 | +:+:+:+:+:+:+:+: | 3378 | sema6d | 106 | 84 | 160 | 190 | 72 | 132 |
| ENSDARG000 | 19 | 46218099 | 46219035 | - | 937 | NA | 15 | 58 | 43 | 38 | 32 | 15 |
| ENSDARG000 | 19;19 | 20191887:201 | 20197474:201 | -;- | 5770 | NA | 66 | 22 | 73 | 32 | 57 | 38 |
| ENSDARG000 | 19:19:19:19:1 | 15276713:152 | 15277978:152 | -;-;-;-;- | 1775 | edn2 | 14 | 0 | 10 | 10 | 8 | 20 |
| ENSDARG000 | 19:19:19:19:1 | 9111550:9113 | 9111834:9114 | +:+:+:+:+:+:+:+: | 8049 | setdb1a | 614 | 598 | 859 | 408 | 768 | 702 |
| ENSDARG000 | 19:19:19:19:1 | 26810208:268 | 26810731:268 | -;-;-;-;-;-;-;-;- | 1636 | neu1 | 724 | 544 | 648 | 543 | 690 | 740 |
| ENSDARG000 | 19;19;19 | 5315987:5318 | 5316481:5320 | +;+;+ | 2517 | si:dkeyp-113d | 714 | 609 | 1023 | 608 | 947 | 658 |
| ENSDARG000 | 19:19:19:19:1 | 27577099:275 | 27577638:275 | -:-:-:-:-:-:-:-: | 1345 | NA | 0 | 0 | 0 | 0 | 0 | 0 |
| ENSDARG000 | 19:19:19:19:1 | 5324653:5324 | 5324858:5325 | -;-;-;-;-;-;-;- | 1371 | NA | 33 | 10 | 4 | 14 | 36 | 0 |
| ENSDARG000 | 19:19:19:19:1 | 46222428:462 | 46222439:462 | +:+:+:+:+:+:+:+: | 1538 | vps28 | 900 | 776 | 1087 | 988 | 1089 | 993 |
| ENSDARG000 | 19:19:19:19:1 | 35001253:350 | 35005398:350 | -:-:-:-:-:-:-:-: | 4890 | psma2b | 1252 | 1474 | 1763 | 885 | 1531 | 1312 |
| ENSDARG000 | 19:19:19:19:1 | 32807236:328 | 32807375:328 | +:+:+:+:+:+:+:+: | 2994 | stk3 | 1203 | 1372 | 1404 | 878 | 1548 | 1246 |
| ENSDARG000 | 19:19:19:19:1 | 35002479:350 | 35002558:350 | +;+;+;+;+;+;+ | 625 | wdyhv1 | 0 | 0 | 0 | 0 | 0 | 0 |
| ENSDARG000 | 19:19:19:19:1 | 20201254:202 | 20201767:202 | +:+:+:+:+:+:+:+: | 6514 | igf2bp3 | 1333 | 989 | 2178 | 1278 | 1863 | 1681 |
| ENSDARG000 | 19:19:19:19:1 | 3929084:3929 | 3929636:3929 | -:-:-:-:-:-:-:-: | 3136 | map7d1b | 1674 | 1298 | 1184 | 1072 | 1226 | 946 |
| ENSDARG000 | 19:19:19:19:1 | 15306968:153 | 15308474:153 | -:-:-:-:-:-:-:-: | 7473 | NA | 18 | 4 | 18 | 0 | 0 | 0 |
| ENSDARG000 | 19:19:19:19:1 | 37620342:377 | 37620861:377 | +:+:+:+:+:+:+:+: | 5424 | thsd7aa | 10 | 0 | 0 | 0 | 5 | 0 |
| ENSDARG000 | 19:19:19:19:1 | 46240171:462 | 46240584:462 | +:+:+:+:+:+:+:+: | 2293 | mapk15 | 0 | 1 | 0 | 0 | 1 | 0 |
| ENSDARG000 | 19:19:19:19:1 | 46243620:462 | 46243647:462 | +:+:+:+:+:+:+:+: | 1428 | NA | 0 | 0 | 0 | 0 | 0 | 0 |
| ENSDARG000 | 19:19:19:19:1 | 46259619:462 | 46259722:462 | +;+;+;+;+;+;+;+ | 1803 | grinaa | 561 | 366 | 625 | 687 | 369 | 293 |
| ENSDARG000 | 19:19:19:19:1 | 24448584:244 | 24449091:244 | -;-;-;-;-;-;-;- | 1265 | mtx1b | 122 | 84 | 118 | 70 | 122 | 110 |
| ENSDARG000 | 19:19:19:19:1 | 41006975:410 | 41007378:410 | +:+:+:+:+:+:+:+: | 2892 | casd1 | 462 | 353 | 497 | 482 | 506 | 584 |
| ENSDARG000 | 19:19:19:19:1 | 5339364:5342 | 5339613:5342 | -:-:-:-:-:-:-:-: | 1578 | krtt1c19e | 0 | 0 | 0 | 6 | 16 | 12 |
| ENSDARG000 | 19:19:19:19:1 | 11897631:118 | 11897909:118 | -:-:-:-:-:-:-:-: | 9408 | cpsf1 | 1057 | 1251 | 1814 | 730 | 1351 | 1031 |
| ENSDARG000 | 19:19:19:19:1 | 30722767:307 | 30723766:307 | -:-:-:-:-:-:-:-: | 2664 | trit1 | 261 | 363 | 441 | 164 | 310 | 261 |
| ENSDARG000 | 19:19:19:19:1 | 15408363:154 | 15409056:154 | -:-:-:-:-:-:-:-: | 1963 | serinc2 | 3391 | 2454 | 2171 | 2124 | 3218 | 2966 |
| ENSDARG000 | 19:19:19:19:1 | 27581750:275 | 27581989:275 | -;-;-;-;-;- | 790 | NA | 0 | 0 | 0 | 0 | 0 | 0 |
| ENSDARG000 | 19:19:19:19:1 | 11952309:119 | 11955786:119 | -:-:-:-:-:-:-:-: | 6785 | NA | 68 | 56 | 52 | 58 | 51 | 71 |
| ENSDARG000 | 19:19:19:19:1 | 18627100:186 | 18627405:186 | +:+:+:+:+:+:+:+: | 2529 | VPS52 | 508 | 443 | 583 | 273 | 627 | 440 |
| ENSDARG000 | 19:19:19:19:1 | 27586823:275 | 27586974:275 | -;-;-;-;-;-;-;-;- | 876 | si:dkeyp-46h3 | 0 | 0 | 0 | 0 | 0 | 0 |
| ENSDARG000 | 19;19;19 | 27589335:275 | 27589502:275 | -;-;- | 593 | NA | 0 | 0 | 0 | 0 | 0 | 0 |
| ENSDARG000 | 19:19:19:19:1 | 46309553:463 | 46309854:463 | +;+;+;+;+;+;+;+ | 1927 | RPRD1A | 412 | 396 | 510 | 379 | 458 | 314 |
| ENSDARG000 | 19:19:19:19:1 | 7559920:7563 | 7560043:7563 | -;-;-;-;- | 861 | NA | 1 | 8 | 10 | 4 | 16 | 0 |
| ENSDARG000 | 19;19 | 7565388:7565 | 7565561:7566 | -;- | 621 | NA | 0 | 0 | 0 | 0 | 0 | 0 |
| ENSDARG000 | 19:19:19:19:1 | 7549854:7549 | 7549947:7549 | +:+:+:+:+:+:+:+: | 3688 | NA | 628 | 644 | 1542 | 633 | 794 | 775 |
| ENSDARG000 | 19:19:19:19:1 | 35013129:350 | 35013193:350 | +;+;+;+;+;+;+;+ | 1083 | zgc:91910 | 1243 | 1507 | 1721 | 1023 | 1721 | 1329 |
| ENSDARG000 | 19:19:19:19:1 | 7575141:7575 | 7575298:7575 | +;+;+;+;+;+;+;+ | 2525 | s100u | 214 | 95 | 113 | 88 | 118 | 28 |
| ENSDARG000 | 19;19;19 | 7567763:7568 | 7567797:7568 | +;+;+ | 462 | s100a11 | 0 | 2 | 11 | 0 | 2 | 0 |
| ENSDARG000 | 19:19:19:19:1 | 30805513:308 | 30807642:308 | -;-;-;-;- | 3062 | myclb | 48 | 54 | 32 | 48 | 54 | 25 |
| ENSDARG000 | 19:19:19:19:1 | 24488403:244 | 24488923:244 | +;+;+;+;+;+;+ | 2753 | txnipa | 4656 | 8393 | 6686 | 9860 | 13596 | 15000 |
| ENSDARG000 | 19:19:19:19:1 | 26828181:268 | 26828415:268 | -;-;-;-;-;-;- | 828 | NA | 139 | 108 | 38 | 84 | 81 | 113 |
| ENSDARG000 | 19:19:19:19:1 | 420350:42044 | 420697:42069 | -:-:-:-:-:-:-:-: | 2331 | dus3l | 756 | 635 | 942 | 1113 | 668 | 1060 |
| ENSDARG000 | 19:19:19:19:1 | 5349656:5349 | 5350098:5350 | -:-:-:-:-:-:-:-: | 1995 | krt92 | 0 | 0 | 8 | 0 | 4 | 0 |
| ENSDARG000 | 19:19:19:19:1 | 10325360:103 | 10326048:103 | -:-:-:-:-:-:-:-: | 1806 | ccdc106b | 0 | 0 | 0 | 0 | 0 | 8 |
| ENSDARG000 | 19:19:19:19:1 | 37848830:378 | 37849331:378 | +:+:+:+:+:+:+:+: | 3494 | nxph1 | 0 | 0 | 2 | 4 | 2 | 2 |
| ENSDARG000 | 19:19:19:19:1 | 27608927:276 | 27610078:276 | -:-:-:-:-:-:-:-: | 5007 | adcy2b | 39 | 3 | 30 | 14 | 16 | 52 |
| ENSDARG000 | 19:19:19:19:1 | 32855139:328 | 32855192:328 | +:+:+:+:+:+:+:+: | 681 | rpl30 | 9409 | 10798 | 15454 | 12841 | 9029 | 13769 |
| ENSDARG000 | 19:19:19:19:1 | 35019304:350 | 35019704:350 | -;-;-;-;-;-;- | 1553 | bmp8a | 77 | 85 | 675 | 826 | 2145 | 224 |
| ENSDARG000 | 19;19;19;19 | 32840143:328 | 32840250:328 | -;-;-;- | 539 | si:dkey-284p5 | 1 | 2 | 2 | 3 | 19 | 6 |
| ENSDARG000 | 19:19:19:19:1 | 32822269:328 | 32822410:328 | +:+:+:+:+:+:+:+: | 1419 | rida | 172 | 263 | 366 | 146 | 350 | 316 |
| ENSDARG000 | 19:19:19:19:1 | 9150014:9150 | 9150149:9150 | +:+:+:+:+:+:+:+: | 2787 | clk2a | 1539 | 723 | 2003 | 1964 | 1752 | 2051 |
| ENSDARG000 | 19:19:19:19:1 | 32864577:328 | 32867360:328 | -;-;-;-;-;-;- | 3579 | laptm4b | 3133 | 2692 | 2968 | 2591 | 3705 | 3589 |
| ENSDARG000 | 19:19:19:19:1 | 24545731:245 | 24546733:245 | -:-:-:-:-:-:-:-: | 1722 | polr3gla | 102 | 154 | 158 | 74 | 102 | 178 |
| ENSDARG000 | 19:19:19:19:1 | 30867845:308 | 30868075:308 | +:+:+:+:+:+:+:+: | 3319 | mfsd2ab | 1624 | 2029 | 1716 | 976 | 1501 | 1364 |
| ENSDARG000 | 19:19:19:19:1 | 20247435:202 | 20252007:202 | -:-:-:-:-:-:-:-: | 6477 | GPNMB | 1122 | 638 | 3287 | 183 | 1137 | 894 |
| ENSDARG000 | 19:19:19:19:1 | 26858039:268 | 26858215:268 | -;-;-;-;-;-;- | 1127 | NA | 0 | 0 | 0 | 0 | 0 | 0 |
| ENSDARG000 | 19:19:19:19:1 | 46343385:463 | 46343471:463 | +:+:+:+:+:+:+:+: | 3839 | SLC30A8 | 1532 | 1248 | 1450 | 784 | 2135 | 2759 |
| ENSDARG000 | 19:19:19:19:1 | 38167468:381 | 38168081:381 | +:+:+:+:+:+:+:+: | 4079 | phf14 | 333 | 361 | 459 | 256 | 403 | 313 |
| ENSDARG000 | 19:19:19:19:1 | 22622355:226 | 22624635:226 | -:-:-:-:-:-:-:-: | 4121 | eif3eb | 712 | 859 | 1715 | 1468 | 1198 | 761 |
| ENSDARG000 | 19:19:19:19:1 | 18657029:186 | 18657394:186 | -:-:-:-:-:-:-:-: | 1202 | NA | 0 | 0 | 0 | 0 | 0 | 0 |
| ENSDARG000 | 19:19:19:19:1 | 46372115:463 | 46372155:463 | +;+;+;+;+ | 979 | med30 | 190 | 229 | 409 | 491 | 372 | 433 |
| ENSDARG000 | 19:19:19:19:1 | 15423057:154 | 15423544:154 | -;-;-;-;-;- | 1593 | NA | 0 | 0 | 0 | 0 | 0 | 0 |
| ENSDARG000 | 19:19:19:19:1 | 24575077:245 | 24575350:245 | +:+:+:+:+:+:+:+: | 1926 | NA | 6 | 0 | 0 | 0 | 0 | 0 |
| ENSDARG000 | 19:19:19:19:1 | 4016702:4017 | 4017807:4018 | -;-;-;-;-;-;-;-;- | 2067 | stpg1 | 0 | 0 | 2 | 0 | 0 | 0 |
| ENSDARG000 | 19:19:19:19:1 | 551963:55311 | 552342:55321 | +:+:+:+:+:+:+:+: | 7619 | NA | 1396 | 1029 | 1254 | 1225 | 1842 | 905 |
| ENSDARG000 | 19 | 24596324 | 24597277 | + | 954 | si:dkeyp-92c9 | 343 | 189 | 181 | 269 | 235 | 173 |
| ENSDARG000 | 19:19:19:19:1 | 5356421:5356 | 5356604:5356 | -:-:-:-:-:-:-:-: | 1561 | cyt1l | 0 | 0 | 0 | 0 | 0 | 6 |
| ENSDARG000 | 19;19 | 43547465:435 | 43549897:435 | -;- | 2590 | gpr3 | 4 | 0 | 0 | 0 | 0 | 0 |
| ENSDARG000 | 19:19:19:19:1 | 42609132:426 | 42609602:426 | +;+;+;+;+;+;+;+ | 1501 | crtap | 869 | 1407 | 1318 | 1070 | 1394 | 742 |
| ENSDARG000 | 19;19 | 10331325:103 | 10331411:103 | +;+ | 1444 | tmem238a | 22 | 0 | 8 | 2 | 18 | 4 |
| ENSDARG000 | 19:19:19:19:1 | 4024443:4024 | 4024562:4024 | +:+:+:+:+:+:+:+: | 1023 | meaf6 | 386 | 411 | 688 | 356 | 507 | 414 |
| ENSDARG000 | 19:19:19:19:1 | 41038668:410 | 41039044:410 | -:-:-:-:-:-:-:-: | 1882 | sgce | 2054 | 1952 | 1669 | 2722 | 1623 | 2839 |
| ENSDARG000 | 19;19 | 46416806:464 | 46417013:464 | -;- | 410 | NA | 11 | 10 | 15 | 15 | 40 | 27 |
| ENSDARG000 | 19:19:19:19:1 | 43563179:435 | 43563266:435 | +;+;+;+;+;+;+;+ | 718 | CD164L2 | 1 | 0 | 0 | 0 | 0 | 4 |
| ENSDARG000 | 19:19:19:19:1 | 10339538:103 | 10339609:103 | +:+:+:+:+:+:+:+: | 3579 | rcvrn3 | 0 | 0 | 0 | 0 | 2 | 0 |
| ENSDARG000 | 19:19:19:19:1 | 4038589:4039 | 4038719:4040 | +;+;+;+;+;+;+;+ | 1660 | NA | 23 | 12 | 4 | 2 | 0 | 2 |
| ENSDARG000 | 19:19:19:19:1 | 46768266:467 | 46773366:467 | -;-;-;-;- | 5829 | abrab | 0 | 0 | 0 | 0 | 0 | 0 |
| ENSDARG000 | 19:19:19:19:1 | 24640467:246 | 24640626:246 | -:-:-:-:-:-:-:-: | 3610 | myo1g | 45 | 63 | 46 | 37 | 58 | 174 |
| ENSDARG000 | 19:19:19:19:1 | 15440841:154 | 15441581:154 | +:+:+:+:+:+:+:+: | 6853 | lin28ab | 0 | 0 | 0 | 0 | 0 | 0 |
| ENSDARG000 | 19:19:19:19:1 | 38387255:383 | 38387380:383 | -:-:-:-:-:-:-:-: | 1257 | smap2 | 297 | 125 | 157 | 128 | 140 | 313 |
| ENSDARG000 | 19:19:19:19:1 | 26869428:268 | 26873100:268 | -:-:-:-:-:-:-:-: | 8300 | skic2 | 614 | 576 | 626 | 360 | 682 | 420 |
| ENSDARG000 | 19:19:19:19:1 | 42629535:426 | 42631016:426 | -;-;-;-;- | 2379 | NA | 2 | 0 | 0 | 0 | 1 | 0 |
| ENSDARG000 | 19:19:19:19:1 | 26922780:269 | 26923307:269 | +:+:+:+:+:+:+:+: | 3168 | nelfe | 567 | 475 | 672 | 449 | 625 | 445 |
| ENSDARG000 | 19:19:19:19:1 | 27807415:278 | 27807609:278 | -:-:-:-:-:-:-:-: | 1838 | TENT4A | 72 | 137 | 231 | 120 | 203 | 98 |
| ENSDARG000 | 19:19:19:19:1 | 4051535:4051 | 4051767:4051 | +:+:+:+:+:+:+:+: | 2067 | NA | 5 | 4 | 2 | 0 | 10 | 1 |
| ENSDARG000 | 19:19:19:19:1 | 41075488:410 | 41075681:410 | +:+:+:+:+:+:+:+: | 4631 | PPP1R9A | 709 | 197 | 214 | 287 | 156 | 387 |
| ENSDARG000 | 19:19:19:19:1 | 48300293:483 | 48300753:483 | -;-;-;-;-;-;-;- | 1035 | NA | 2 | 0 | 0 | 0 | 0 | 0 |
| ENSDARG000 | 19:19:19:19:1 | 10356345:103 | 10356881:103 | -:-:-:-:-:-:-:-: | 1710 | si:ch211-232r | 0 | 0 | 0 | 0 | 0 | 0 |
| ENSDARG000 | 19:19:19:19:1 | 42660158:426 | 42660434:426 | +:+:+:+:+:+:+:+: | 2170 | fbxl2 | 367 | 126 | 324 | 290 | 172 | 218 |
| ENSDARG000 | 19:19:19:19:1 | 20274264:202 | 20274558:202 | +:+:+:+:+:+:+:+: | 3345 | oxnad1 | 154 | 176 | 220 | 167 | 118 | 114 |
| ENSDARG000 | 19:19:19:19:1 | 3086851:3087 | 3087347:3087 | -:-:-:-:-:-:-:-: | 3554 | si:ch211-80h1 | 36 | 92 | 218 | 356 | 48 | 334 |
| ENSDARG000 | 19;19 | 48308750:483 | 48309916:483 | -;- | 1338 | si:ch73-359m | 0 | 6 | 8 | 0 | 48 | 4 |
| ENSDARG000 | 19:19:19:19:1 | 5362420:5362 | 5362747:5362 | -:-:-:-:-:-:-:-: | 1673 | cyt1 | 0 | 0 | 6 | 0 | 26 | 0 |
| ENSDARG000 | 19:19:19:19:1 | 24750466:247 | 24753868:247 | -:-:-:-:-:-:-:-: | 4191 | NA | 76 | 83 | 108 | 32 | 126 | 113 |
| ENSDARG000 | 19;19 | 48328898:483 | 48329611:483 | -;- | 842 | NA | 0 | 0 | 0 | 0 | 0 | 2 |
| ENSDARG000 | 19:19:19:19:1 | 30884706:308 | 30885017:308 | +:+:+:+:+:+:+:+: | 2741 | yars1 | 1587 | 1998 | 3340 | 1614 | 1363 | 1431 |
| ENSDARG000 | 19:19:19:19:1 | 4061712:4062 | 4062263:4062 | +:+:+:+:+:+:+:+: | 2686 | NA | 62 | 165 | 61 | 15 | 84 | 53 |
| ENSDARG000 | 19:19:19:19:1 | 4059200:4059 | 4059308:4059 | +:+:+:+:+:+:+:+: | 2085 | btr25 | 2 | 1 | 8 | 0 | 4 | 1 |
| ENSDARG000 | 19:19:19:19:1 | 46812272:468 | 46812929:468 | -;-;-;-;-;-;-;-;- | 2164 | angpt1 | 6 | 8 | 0 | 0 | 6 | 0 |
| ENSDARG000 | 19;19;19 | 48334573:483 | 48335380:483 | -;-;- | 1094 | NA | 0 | 0 | 3 | 0 | 0 | 0 |
| ENSDARG000 | 19:19:19:19:1 | 27836913:278 | 27837785:278 | -:-:-:-:-:-:-:-: | 2147 | srd5a1 | 100 | 210 | 153 | 96 | 80 | 141 |
| ENSDARG000 | 19:19:19:19:1 | 7595924:7597 | 7596009:7598 | +:+:+:+:+:+:+:+: | 1706 | s100s | 0 | 0 | 2 | 0 | 0 | 0 |
| ENSDARG000 | 19:19:19:19:1 | 41169996:411 | 41170327:411 | +;+;+;+;+;+;+;+ | 1727 | asb4 | 69 | 8 | 29 | 38 | 16 | 34 |
| ENSDARG000 | 19:19:19:19:1 | 43579786:435 | 43579854:435 | +:+:+:+:+:+:+:+: | 4356 | NA | 526 | 857 | 1117 | 1804 | 585 | 1627 |
| ENSDARG000 | 19:19:19:19:1 | 10375782:103 | 10380141:103 | -:-:-:-:-:-:-:-: | 5997 | epn1b | 4 | 10 | 2 | 4 | 0 | 30 |
| ENSDARG000 | 19:19:19:19:1 | 10396042:103 | 10396238:103 | +:+:+:+:+:+:+:+: | 1599 | necap1 | 8 | 14 | 38 | 31 | 34 | 58 |
| ENSDARG000 | 19:19:19:19:1 | 7627070:7627 | 7627133:7627 | +;+;+;+;+;+;+ | 3733 | pygo2 | 614 | 659 | 805 | 263 | 720 | 577 |
| ENSDARG000 | 19:19:19:19:1 | 15485287:154 | 15485731:154 | +:+:+:+:+:+:+:+: | 4811 | pdik1l | 174 | 121 | 151 | 97 | 121 | 177 |
| ENSDARG000 | 19:19:19:19:1 | 9174166:9174 | 9174288:9174 | +:+:+:+:+:+:+:+: | 1120 | si:ch211-81a5 | 12 | 2 | 50 | 0 | 17 | 36 |
| ENSDARG000 | 19;19 | 47123688:471 | 47123840:471 | -;- | 406 | NA | 0 | 0 | 0 | 0 | 0 | 0 |
| ENSDARG000 | 19;19 | 47135677:471 | 47135923:471 | +;+ | 639 | NA | 0 | 0 | 2 | 0 | 0 | 0 |
| ENSDARG000 | 19:19:19:19:1 | 24872159:248 | 24872727:248 | +:+:+:+:+:+:+:+: | 3601 | si:ch211-195l | 15038 | 8126 | 3333 | 4312 | 18772 | 7287 |
| ENSDARG000 | 19:19:19:19:1 | 38422059:384 | 38422338:384 | +:+:+:+:+:+:+:+: | 3866 | col9a2 | 11779 | 155 | 271 | 196 | 28 | 161 |
| ENSDARG000 | 19:19:19:19:1 | 4089494:4092 | 4091212:4092 | -:-:-:-:-:-:-:-: | 2572 | fhl3b | 254 | 194 | 90 | 225 | 106 | 390 |
| ENSDARG000 | 19:19:19:19:1 | 32895379:328 | 32896940:328 | -:-:-:-:-:-:-:-: | 3268 | mtdha | 3930 | 4398 | 5637 | 3169 | 4952 | 4467 |
| ENSDARG000 | 19:19:19:19:1 | 41190258:411 | 41191575:411 | -:-:-:-:-:-:-:-: | 2612 | pdk4 | 85 | 61 | 188 | 210 | 136 | 261 |
| ENSDARG000 | 19:19:19:19:1 | 9186175:9188 | 9186490:9189 | +:+:+:+:+:+:+:+: | 2517 | NA | 0 | 0 | 0 | 0 | 0 | 0 |
| ENSDARG000 | 19:19:19:19:1 | 27858866:278 | 27859029:278 | +:+:+:+:+:+:+:+: | 3196 | nsun2 | 880 | 1078 | 1157 | 693 | 980 | 791 |
| ENSDARG000 | 19:19:19:19:1 | 4126905:4128 | 4127317:4128 | -;-;-;-;-;-;- | 1574 | rragcb | 148 | 110 | 179 | 52 | 128 | 185 |
| ENSDARG000 | 19:19:19:19:1 | 7152966:7154 | 7153125:7154 | +:+:+:+:+:+:+:+: | 5628 | brd2a | 2325 | 2073 | 3010 | 1906 | 2449 | 1883 |
| ENSDARG000 | 19:19:19:19:1 | 5367330:5367 | 5367614:5367 | -:-:-:-:-:-:-:-: | 1706 | krt17 | 791 | 20 | 8 | 42 | 266 | 144 |
| ENSDARG000 | 19:19:19:19:1 | 15521997:155 | 15522069:155 | +;+;+;+;+;+;+;+ | 1282 | ppp1r8a | 74 | 33 | 61 | 39 | 74 | 70 |
| ENSDARG000 | 19:19:19:19:1 | 47191005:471 | 47191653:471 | -;-;-;-;- | 1375 | sdc2 | 3645 | 2296 | 4930 | 2473 | 5275 | 3368 |
| ENSDARG000 | 19;19 | 48339349:483 | 48339802:483 | -;- | 592 | NA | 0 | 0 | 0 | 0 | 0 | 6 |
| ENSDARG000 | 19;19 | 48342937:483 | 48344307:483 | -;- | 1534 | NA | 0 | 0 | 0 | 0 | 0 | 0 |
| ENSDARG000 | 19;19;19 | 591490:59149 | 592467:59266 | -;-;- | 1178 | FUT9 | 0 | 0 | 0 | 0 | 0 | 0 |
| ENSDARG000 | 19:19:19:19:1 | 41290713:412 | 41290868:412 | -:-:-:-:-:-:-:-: | 2047 | SLC25A13 | 0 | 0 | 6 | 0 | 0 | 2 |
| ENSDARG000 | 19:19:19:19:1 | 35039902:350 | 35041483:350 | -:-:-:-:-:-:-:-: | 18720 | NA | 8311 | 4575 | 6324 | 6266 | 7414 | 5470 |
| ENSDARG000 | 19:19:19:19:1 | 598657:60240 | 599685:60257 | -:-:-:-:-:-:-:-: | 2597 | cyp51 | 1182 | 1168 | 1461 | 706 | 581 | 674 |
| ENSDARG000 | 19;19 | 22727940:227 | 22728938:227 | +;+ | 2112 | trhrb | 0 | 0 | 4 | 0 | 1 | 0 |
| ENSDARG000 | 19:19:19:19:1 | 18832318:188 | 18832615:188 | -:-:-:-:-:-:-:-: | 1224 | NA | 0 | 2 | 0 | 0 | 0 | 3 |
| ENSDARG000 | 19:19:19:19:1 | 7636941:7636 | 7637072:7637 | +:+:+:+:+:+:+:+: | 4757 | cgnb | 22 | 20 | 12 | 12 | 14 | 20 |
| ENSDARG000 | 19:19:19:19:1 | 43604256:436 | 43604532:436 | +:+:+:+:+:+:+:+: | 1670 | si:ch211-199 | 268 | 194 | 350 | 279 | 307 | 292 |
| ENSDARG000 | 19;19;19;19 | 10415484:104 | 10416875:104 | -;-;-;- | 1902 | NA | 2 | 0 | 4 | 0 | 0 | 2 |
| ENSDARG000 | 19;19 | 15571290:156 | 15571976:156 | +;+ | 3840 | foxo6b | 12 | 8 | 20 | 2 | 26 | 8 |
| ENSDARG000 | 19:19:19:19:1 | 22759116:227 | 22759647:227 | -:-:-:-:-:-:-:-: | 2162 | nudcd1 | 921 | 755 | 1054 | 689 | 1133 | 653 |
| ENSDARG000 | 19:19:19:19:1 | 48351624:483 | 48351857:483 | +:+:+:+:+:+:+:+: | 2318 | sgo1 | 52 | 255 | 254 | 152 | 170 | 104 |
| ENSDARG000 | 19;19;19;19 | 10429810:104 | 10429980:104 | -;-;-;- | 578 | NA | 18 | 12 | 4 | 2 | 20 | 46 |
| ENSDARG000 | 19;19 | 15851226:158 | 15853533:158 | -;- | 2445 | cited4a | 116 | 79 | 396 | 326 | 1379 | 175 |
| ENSDARG000 | 19:19:19:19:1 | 4139065:4142 | 4139117:4143 | +:+:+:+:+:+:+:+: | 3097 | TRAPPC9 | 405 | 464 | 400 | 342 | 421 | 353 |
| ENSDARG000 | 19:19:19:19:1 | 20301373:203 | 20302050:203 | -:-:-:-:-:-:-:-: | 2598 | RFTN1 | 6 | 2 | 13 | 4 | 3 | 12 |
| ENSDARG000 | 19:19:19:19:1 | 41401133:414 | 41402078:414 | -;-;-;-;-;-;-;-;- | 2035 | sem1 | 966 | 1130 | 1012 | 636 | 967 | 793 |
| ENSDARG000 | 19:19:19:19:1 | 9212031:9212 | 9212079:9212 | +:+:+:+:+:+:+:+: | 2572 | ndufv1 | 2802 | 2289 | 3159 | 2141 | 2660 | 2212 |
| ENSDARG000 | 19:19:19:19:1 | 10454305:104 | 10454737:104 | -;-;-;-;-;-;- | 1511 | NA | 14 | 10 | 22 | 2 | 12 | 14 |
| ENSDARG000 | 19 | 10473612 | 10476024 | + | 2413 | NA | 6 | 2 | 0 | 0 | 0 | 0 |
| ENSDARG000 | 19:19:19:19:1 | 47311020:473 | 47311175:473 | +;+;+;+;+;+;+ | 1496 | NA | 174 | 90 | 224 | 158 | 265 | 112 |
| ENSDARG000 | 19:19:19:19:1 | 47290287:472 | 47290342:472 | +:+:+:+:+:+:+:+: | 947 | tpmt.1 | 240 | 326 | 229 | 202 | 294 | 208 |

| | | | | | | | | | | | | |
|---|---|---|---|---|---|---|---|---|---|---|---|---|
| ENSDARG000 | 19:19:19:19:1 | 10485847:104 | 10486010:104 | +:+:+:+:+:+:+: | 6043 | NA | 10 | 6 | 24 | 2 | 22 | 2 |
| ENSDARG000 | 19:19:19:19:1 | 47299166:472 | 47299227:472 | +:+:+:+:+:+:+: | 993 | tpmt.2 | 51 | 79 | 92 | 46 | 49 | 92 |
| ENSDARG000 | 19;19 | 4443285:4444 | 4443508:4447 | +;+ | 2983 | NA | 205 | 123 | 144 | 235 | 139 | 163 |
| ENSDARG000 | 19;19;19;19 | 27957119:279 | 27957339:279 | -;-;-;- | 929 | UBE2QL1 | 167 | 224 | 288 | 144 | 873 | 96 |
| ENSDARG000 | 19:19:19:19:1 | 43612179:436 | 43612503:436 | -:-:-:-:-:-:-:-: | 4027 | fam83hb | 1268 | 945 | 1547 | 1112 | 1585 | 1219 |
| ENSDARG000 | 19:19:19:19:1 | 5378563:5378 | 5378827:5379 | -:-:-:-:-:-:-:-: | 1754 | krt91 | 4595 | 292 | 153 | 174 | 1393 | 449 |
| ENSDARG000 | 19:19:19:19:1 | 16015881:160 | 16016224:160 | +:+:+:+:+:+:+: | 6863 | ctps1a | 950 | 1636 | 2256 | 821 | 1847 | 1045 |
| ENSDARG000 | 19;19 | 10508323:105 | 10508850:105 | +;+ | 1017 | NA | 2 | 0 | 0 | 0 | 0 | 0 |
| ENSDARG000 | 19:19:19:19:1 | 24891747:248 | 24891848:248 | +:+:+:+:+:+:+: | 4995 | eya3 | 303 | 272 | 472 | 278 | 463 | 406 |
| ENSDARG000 | 19:19:19:19:1 | 48364312:483 | 48364955:483 | -;-;-;-;-;- | 1070 | rpl14 | 8615 | 8677 | 13557 | 11204 | 8485 | 12407 |
| ENSDARG000 | 19;19 | 4463389:4530 | 4464531:4531 | +;+ | 2060 | kcnk9 | 0 | 0 | 0 | 0 | 0 | 0 |
| ENSDARG000 | 19;19;19;19 | 10518448:105 | 10519029:105 | +;+;+;+ | 1074 | NA | 0 | 2 | 0 | 0 | 8 | 0 |
| ENSDARG000 | 19;19 | 10527228:105 | 10527779:105 | +;+ | 1017 | NA | 0 | 0 | 0 | 0 | 0 | 0 |
| ENSDARG000 | 19:19:19:19:1 | 10537443:105 | 10542073:105 | -:-:-:-:-:-:-:-: | 5422 | lim2.4 | 6 | 12 | 24 | 10 | 20 | 78 |
| ENSDARG000 | 19:19:19:19:1 | 619200:61925 | 619428:61942 | +:+:+:+:+:+:+: | 2134 | nup42 | 112 | 119 | 118 | 75 | 137 | 117 |
| ENSDARG000 | 19;19 | 10536093:105 | 10536635:105 | +;+ | 1038 | si:dkey-211g8 | 0 | 0 | 4 | 2 | 11 | 10 |
| ENSDARG000 | 19:19:19:19:1 | 32916372:329 | 32916668:329 | -:-:-:-:-:-:-:-: | 1602 | rrm2b | 154 | 112 | 166 | 90 | 214 | 138 |
| ENSDARG000 | 19:19:19:19:1 | 27970292:279 | 27970602:279 | +:+:+:+:+:+:+: | 1096 | med10 | 402 | 349 | 531 | 417 | 484 | 361 |
| ENSDARG000 | 19;19;19 | 41470216:414 | 41470805:414 | -;-;- | 1302 | dlx5a | 4 | 0 | 6 | 0 | 2 | 6 |
| ENSDARG000 | 19:19:19:19:1 | 3127912:3130 | 3128070:3130 | +;+;+;+;+ | 509 | NA | 0 | 0 | 0 | 0 | 0 | 2 |
| ENSDARG000 | 19:19:19:19:1 | 43654238:436 | 43655535:436 | -;-;-;-;-;- | 2679 | si:ch211-193k | 0 | 0 | 0 | 0 | 0 | 0 |
| ENSDARG000 | 19;19 | 41446936:414 | 41448307:414 | -;- | 1560 | NA | 0 | 0 | 0 | 0 | 0 | 0 |
| ENSDARG000 | 19:19:19:19:1 | 41419519:414 | 41419596:414 | +:+:+:+:+:+:+: | 1823 | dlx6a | 4 | 0 | 0 | 0 | 0 | 0 |
| ENSDARG000 | 19:19:19:19:1 | 4773710:4775 | 4773874:4775 | -:-:-:-:-:-:-:-: | 1119 | NA | 201 | 0 | 111 | 96 | 77 | 117 |
| ENSDARG000 | 19:19:19:19:1 | 48372446:483 | 48373245:483 | +:+:+:+:+:+:+: | 2230 | NA | 0 | 0 | 4 | 0 | 8 | 0 |
| ENSDARG000 | 19:19:19:19:1 | 18846923:188 | 18847966:188 | -:-:-:-:-:-:-:-: | 2181 | ppt2b | 150 | 51 | 118 | 72 | 116 | 112 |
| ENSDARG000 | 19:19:19:19:1 | 5418006:5418 | 5418247:5418 | +:+:+:+:+:+:+: | 1565 | eif1 | 9724 | 8591 | 8529 | 8956 | 8384 | 6411 |
| ENSDARG000 | 19:19:19:19:1 | 7682109:7688 | 7686948:7688 | -;-;-;-;-;- | 5839 | NA | 0 | 0 | 0 | 2 | 0 | 0 |
| ENSDARG000 | 19:19:19:19:1 | 12121944:121 | 12122340:121 | -:-:-:-:-:-:-:-: | 2886 | pabpc1b | 5075 | 4303 | 13791 | 8053 | 9487 | 6260 |
| ENSDARG000 | 19:19:19:19:1 | 7694487:7700 | 7699210:7701 | -;-;-;-;-;- | 5632 | NA | 26 | 2 | 0 | 0 | 0 | 0 |
| ENSDARG000 | 19:19:19:19:1 | 20389104:203 | 20390293:203 | -:-:-:-:-:-:-:-: | 2413 | dazl | 4 | 10 | 6 | 4 | 0 | 4 |
| ENSDARG000 | 19;19 | 18879002:188 | 18879549:188 | -;- | 817 | NA | 0 | 0 | 0 | 0 | 0 | 0 |
| ENSDARG000 | 19:19:19:19:1 | 627899:62929 | 628162:63059 | +:+:+:+:+:+:+: | 3752 | tert | 122 | 458 | 488 | 94 | 443 | 293 |
| ENSDARG000 | 19;19 | 16032383:160 | 16032695:160 | +;+ | 1090 | rab42a | 0 | 0 | 0 | 0 | 0 | 0 |
| ENSDARG000 | 19:19:19:19:1 | 28114503:281 | 28115103:281 | +:+:+:+:+:+:+: | 4096 | irx4b | 0 | 0 | 4 | 0 | 0 | 0 |
| ENSDARG000 | 19;19 | 18881888:188 | 18882369:188 | +;+ | 956 | NA | 2 | 0 | 0 | 0 | 0 | 0 |
| ENSDARG000 | 19;19;19;19 | 28127654:281 | 28127981:281 | -;-;-;- | 1746 | irx1b | 0 | 0 | 7 | 0 | 4 | 0 |
| ENSDARG000 | 19:19:19:19:1 | 42693855:426 | 42694408:426 | +:+:+:+:+:+:+: | 7098 | clasp2 | 1997 | 1720 | 2901 | 1719 | 2803 | 2215 |
| ENSDARG000 | 19:19:19:19:1 | 7717449:7717 | 7717517:7717 | +:+:+:+:+:+:+: | 2712 | tuft1b | 187 | 62 | 106 | 138 | 217 | 254 |
| ENSDARG000 | 19:19:19:19:1 | 7173613:7173 | 7173934:7173 | +:+:+:+:+:+:+: | 1724 | hsd17b8 | 6 | 72 | 50 | 36 | 60 | 56 |
| ENSDARG000 | 19:19:19:19:1 | 43669122:436 | 43669433:436 | +:+:+:+:+:+:+: | 4102 | NA | 320 | 220 | 304 | 301 | 372 | 309 |
| ENSDARG000 | 19:19:19:19:1 | 48382963:483 | 48383070:483 | -:-:-:-:-:-:-:-: | 1490 | si:ch73-359m | 1 | 2 | 8 | 2 | 35 | 12 |
| ENSDARG000 | 19:19:19:19:1 | 9231766:9232 | 9232993:9232 | +:+:+:+:+:+:+: | 11593 | NA | 1411 | 816 | 2035 | 895 | 1454 | 1329 |
| ENSDARG000 | 19:19:19:19:1 | 48389299:483 | 48391481:483 | +:+:+:+:+:+:+: | 7221 | NA | 0 | 6 | 0 | 0 | 0 | 2 |
| ENSDARG000 | 19;19 | 9232539:9233 | 9233145:9234 | -;- | 684 | NA | 0 | 0 | 0 | 0 | 0 | 0 |
| ENSDARG000 | 19;19 | 24922531:249 | 24923047:249 | +;+ | 2376 | NA | 0 | 0 | 0 | 0 | 2 | 2 |
| ENSDARG000 | 19:19:19:19:1 | 3137518:3142 | 3138778:3142 | -:-:-:-:-:-:-:-: | 3344 | stm | 15 | 0 | 6 | 0 | 2 | 30 |
| ENSDARG000 | 19:19:19:19:1 | 32930476:329 | 32931667:329 | -:-:-:-:-:-:-:-: | 2845 | azin1b | 3990 | 3788 | 5349 | 4330 | 3776 | 4219 |
| ENSDARG000 | 19:19:19:19:1 | 38473758:384 | 38473944:384 | -:-:-:-:-:-:-:-: | 6103 | NA | 216 | 37 | 203 | 28 | 435 | 37 |
| ENSDARG000 | 19:19:19:19:1 | 16064439:160 | 16064487:160 | +:+:+:+:+:+:+: | 4532 | gmeb1 | 423 | 334 | 522 | 460 | 502 | 417 |
| ENSDARG000 | 19:19:19:19:1 | 645872:64652 | 646323:64669 | -:-:-:-:-:-:-:-: | 2153 | slc6a18 | 0 | 2 | 4 | 20 | 54 | 0 |
| ENSDARG000 | 19:19:19:19:1 | 3135271:3138 | 3135480:3138 | +:+:+:+:+:+:+: | 4565 | zgc:86598 | 2327 | 2300 | 4315 | 3337 | 3186 | 2247 |
| ENSDARG000 | 19;19;19 | 24938136:249 | 24939059:249 | -;-;- | 1553 | NA | 2 | 0 | 2 | 0 | 0 | 1 |
| ENSDARG000 | 19:19:19:19:1 | 22850657:228 | 22850846:228 | +:+:+:+:+:+:+: | 13133 | pkhd1l1.1 | 2403 | 1176 | 1425 | 2009 | 4237 | 2053 |
| ENSDARG000 | 19:19:19:19:1 | 18903533:189 | 18903742:189 | +:+:+:+:+:+:+: | 1992 | slc39a7 | 2116 | 2517 | 1861 | 1598 | 2303 | 1936 |
| ENSDARG000 | 19:19:19:19:1 | 3186669:3187 | 3187501:3187 | -:-:-:-:-:-:-:-: | 1233 | si:ch211-133r | 0 | 4 | 0 | 0 | 0 | 6 |
| ENSDARG000 | 19;19;19 | 24982672:249 | 24983508:249 | -;-;- | 1396 | NA | 8 | 2 | 0 | 10 | 2 | 4 |
| ENSDARG000 | 19:19:19:19:1 | 12164428:121 | 12165403:121 | -;-;-;-;-;- | 1720 | ywhaz | 14598 | 15735 | 17523 | 16708 | 16931 | 13035 |
| ENSDARG000 | 19;19;19 | 24994270:250 | 24995612:250 | -;-;- | 2000 | NA | 0 | 0 | 0 | 0 | 2 | 0 |
| ENSDARG000 | 19;19;19 | 25056812:250 | 25057542:250 | -;-;- | 1233 | NA | 12 | 12 | 6 | 0 | 38 | 14 |
| ENSDARG000 | 19;19;19 | 25072053:250 | 25072922:250 | -;-;- | 1668 | xkr8.3 | 86 | 63 | 111 | 88 | 131 | 133 |
| ENSDARG000 | 19;19 | 5425832:5428 | 5425962:5428 | +;+ | 338 | NA | 0 | 0 | 0 | 0 | 0 | 0 |
| ENSDARG000 | 19:19:19:19:1 | 28281201:282 | 28281363:282 | -;-;-;-;-;-;- | 818 | ndufs6 | 486 | 545 | 585 | 450 | 412 | 368 |
| ENSDARG000 | 19:19:19:19:1 | 4800068:4800 | 4800321:4800 | +:+:+:+:+:+:+: | 3484 | NA | 652 | 14 | 306 | 490 | 283 | 279 |
| ENSDARG000 | 19:19:19:19:1 | 48410195:484 | 48410271:484 | +:+:+:+:+:+:+: | 2730 | si:ch1073-376 | 0 | 0 | 24 | 22 | 118 | 108 |
| ENSDARG000 | 19:19:19:19:1 | 30897255:309 | 30900790:309 | -;-;-;-;- | 3978 | si:ch211-194e | 1376 | 538 | 791 | 1305 | 367 | 1095 |
| ENSDARG000 | 19:19:19:19:1 | 18989911:189 | 18991589:189 | -:-:-:-:-:-:-:-: | 3218 | dync1li1 | 768 | 822 | 889 | 516 | 837 | 727 |
| ENSDARG000 | 19:19:19:19:1 | 10558535:105 | 10558634:105 | +:+:+:+:+:+:+: | 4071 | usf2 | 2607 | 2889 | 2020 | 2431 | 2604 | 3007 |
| ENSDARG000 | 19:19:19:19:1 | 5480327:5525 | 5480517:5525 | +:+:+:+:+:+:+: | 2560 | JUP | 50 | 12 | 6 | 42 | 18 | 6 |
| ENSDARG000 | 19:19:19:19:1 | 43715911:437 | 43716048:437 | +:+:+:+:+:+:+: | 1943 | cap1 | 383 | 472 | 534 | 428 | 527 | 467 |
| ENSDARG000 | 19:19:19:19:1 | 47394270:474 | 47394416:474 | +:+:+:+:+:+:+: | 1643 | psmb2 | 2015 | 2730 | 2692 | 1611 | 2456 | 1970 |
| ENSDARG000 | 19:19:19:19:1 | 30990129:309 | 30990875:309 | +:+:+:+:+:+:+: | 4309 | NA | 0 | 0 | 0 | 0 | 0 | 0 |
| ENSDARG000 | 19;19 | 28284120:282 | 28284236:282 | +;+ | 2183 | mrpl36 | 470 | 504 | 730 | 302 | 711 | 554 |
| ENSDARG000 | 19:19:19:19:1 | 26967357:269 | 26968064:269 | -:-:-:-:-:-:-:-: | 2576 | sacm1la | 849 | 666 | 669 | 364 | 654 | 563 |
| ENSDARG000 | 19 | 47415590 | 47416405 | - | 816 | NA | 5 | 10 | 0 | 23 | 0 | 28 |
| ENSDARG000 | 19:19:19:19:1 | 4828251:4828 | 4828722:4828 | -:-:-:-:-:-:-:-: | 2303 | FBXL20 | 206 | 49 | 134 | 107 | 164 | 196 |
| ENSDARG000 | 19:19:19:19:1 | 41479990:414 | 41480213:414 | +:+:+:+:+:+:+: | 2966 | ago2 | 182 | 97 | 73 | 158 | 162 | 170 |
| ENSDARG000 | 19:19:19:19:1 | 43750676:437 | 43750853:437 | +:+:+:+:+:+:+: | 1816 | NA | 829 | 387 | 409 | 365 | 480 | 225 |
| ENSDARG000 | 19:19:19:19:1 | 12206158:122 | 12207040:122 | -:-:-:-:-:-:-:-: | 1533 | znf706 | 748 | 817 | 1277 | 949 | 761 | 996 |
| ENSDARG000 | 19:19:19:19:1 | 43736269:437 | 43741096:437 | -:-:-:-:-:-:-:-: | 6514 | ppt1 | 4428 | 2940 | 1914 | 3815 | 3319 | 3817 |
| ENSDARG000 | 19;19 | 5651325:5653 | 5651381:5655 | +;+ | 1304 | ft1 | 0 | 10 | 0 | 0 | 4 | 14 |
| ENSDARG000 | 19:19:19:19:1 | 663796:66752 | 665912:66768 | -:-:-:-:-:-:-:-: | 3908 | slc6a19a.1 | 0 | 2 | 156 | 28 | 160 | 0 |
| ENSDARG000 | 19:19:19:19:1 | 26957425:269 | 26958792:269 | -;-;-;-;- | 2281 | zbtb12.2 | 204 | 215 | 273 | 176 | 239 | 238 |
| ENSDARG000 | 19;19;19;19 | 18983900:189 | 18984175:189 | +;+;+;+ | 1373 | NA | 0 | 0 | 0 | 12 | 8 | 0 |
| ENSDARG000 | 19:19:19:19:1 | 7757682:7757 | 7757831:7757 | +:+:+:+:+:+:+: | 5557 | ubap2l | 2842 | 2411 | 3855 | 1938 | 3698 | 2394 |
| ENSDARG000 | 19:19:19:19:1 | 16429638:164 | 16430971:164 | -:-:-:-:-:-:-:-: | 2559 | NA | 0 | 0 | 0 | 0 | 0 | 0 |
| ENSDARG000 | 19:19:19:19:1 | 16222618:162 | 16222815:162 | +:+:+:+:+:+:+: | 8323 | PTPRU | 20 | 0 | 4 | 6 | 0 | 0 |
| ENSDARG000 | 19:19:19:19:1 | 20404153:204 | 20405463:204 | -:-:-:-:-:-:-:-: | 3790 | tbc1d5 | 2548 | 2200 | 3047 | 2582 | 3440 | 3120 |
| ENSDARG000 | 19:19:19:19:1 | 32947910:329 | 32948145:329 | +:+:+:+:+:+:+: | 1928 | atp6v1c1b | 682 | 815 | 792 | 582 | 639 | 787 |
| ENSDARG000 | 19:19:19:19:1 | 5659526:5662 | 5659709:5662 | -;-;-;-;-;-;- | 1467 | si:ch211-264f | 68 | 122 | 50 | 36 | 108 | 173 |
| ENSDARG000 | 19:19:19:19:1 | 25088150:250 | 25090240:250 | -:-:-:-:-:-:-:-: | 4427 | ptp4a3a | 705 | 1421 | 829 | 616 | 1047 | 739 |
| ENSDARG000 | 19:19:19:19:1 | 41509956:415 | 41510237:415 | +;+;+;+;+;+ | 1081 | NA | 37 | 20 | 142 | 33 | 139 | 123 |
| ENSDARG000 | 19:19:19:19:1 | 47429722:474 | 47431549:474 | -:-:-:-:-:-:-:-: | 3593 | tfap2e | 12701 | 10168 | 10450 | 13949 | 11862 | 7801 |
| ENSDARG000 | 19 | 10587307 | 10587858 | + | 552 | NA | 137 | 0 | 19 | 94 | 16 | 22 |
| ENSDARG000 | 19:19:19:19:1 | 10592778:105 | 10592834:105 | +;+;+;+;+;+;+ | 982 | NA | 79 | 29 | 2 | 0 | 1 | 4 |
| ENSDARG000 | 19:19:19:19:1 | 28291062:282 | 28291445:282 | +:+:+:+:+:+:+: | 2573 | lpcat1 | 411 | 506 | 387 | 321 | 539 | 440 |
| ENSDARG000 | 19:19:19:19:1 | 30999833:309 | 31000484:310 | -:-:-:-:-:-:-:-: | 2220 | rbbp4 | 733 | 2188 | 2506 | 787 | 2172 | 1232 |
| ENSDARG000 | 19:19:19:19:1 | 684502:68870 | 684913:68888 | -:-:-:-:-:-:-:-: | 2404 | NA | 0 | 0 | 40 | 61 | 46 | 0 |
| ENSDARG000 | 19:19:19:19:1 | 47502581:475 | 47502652:475 | +:+:+:+:+:+:+: | 2873 | NA | 0 | 0 | 0 | 0 | 0 | 0 |
| ENSDARG000 | 19:19:19:19:1 | 10603405:106 | 10604061:106 | +;+;+;+;+ | 1814 | si:dkey-211g8 | 357 | 82 | 18 | 13 | 6 | 18 |
| ENSDARG000 | 19:19:19:19:1 | 47523730:475 | 47524362:475 | -:-:-:-:-:-:-:-: | 1628 | scg5 | 0 | 0 | 0 | 0 | 0 | 0 |
| ENSDARG000 | 19:19:19:19:1 | 41515759:415 | 41516366:415 | -;-;-;-;-;- | 936 | chrac1 | 370 | 375 | 527 | 287 | 428 | 439 |
| ENSDARG000 | 19:19:19:19:1 | 32979132:329 | 32979782:329 | +:+:+:+:+:+:+: | 4029 | spire1a | 2904 | 1958 | 2095 | 3468 | 2610 | 1932 |
| ENSDARG000 | 19:19:19:19:1 | 12234975:122 | 12235047:122 | +:+:+:+:+:+:+: | 3699 | grhl2b | 13 | 4 | 12 | 0 | 8 | 0 |
| ENSDARG000 | 19:19:19:19:1 | 4856351:4866 | 4858167:4867 | +:+:+:+:+:+:+: | 6364 | CDK12 | 1772 | 1625 | 2517 | 1541 | 1933 | 1362 |
| ENSDARG000 | 19:19:19:19:1 | 5674907:5676 | 5675134:5676 | +:+:+:+:+:+:+: | 2368 | pdk2b | 110 | 56 | 276 | 162 | 420 | 222 |
| ENSDARG000 | 19 | 10611788 | 10612774 | - | 987 | si:dkey-211g8 | 0 | 4 | 0 | 0 | 0 | 0 |
| ENSDARG000 | 19:19:19:19:1 | 38766613:387 | 38766664:387 | -:-:-:-:-:-:-:-: | 2198 | NA | 4 | 6 | 75 | 4 | 24 | 0 |
| ENSDARG000 | 19:19:19:19:1 | 712127:71212 | 712291:71229 | +:+:+:+:+:+:+: | 7224 | NA | 2 | 14 | 6 | 6 | 8 | 2 |
| ENSDARG000 | 19:19:19:19:1 | 9277327:9277 | 9277376:9277 | +:+:+:+:+:+:+: | 2315 | si:rp71-15k1. | 0 | 0 | 0 | 0 | 0 | 0 |
| ENSDARG000 | 19;19 | 16711856:167 | 16711883:167 | +;+ | 338 | NA | 0 | 0 | 0 | 0 | 0 | 0 |
| ENSDARG000 | 19;19;19 | 10624644:106 | 10626312:106 | -;-;- | 1819 | si:dkey-211g8 | 0 | 0 | 0 | 0 | 0 | 0 |
| ENSDARG000 | 19:19:19:19:1 | 41520892:415 | 41521253:415 | +:+:+:+:+:+:+: | 4641 | NA | 336 | 190 | 310 | 199 | 270 | 317 |
| ENSDARG000 | 19:19:19:19:1 | 4912817:4912 | 4913141:4913 | +:+:+:+:+:+:+: | 4690 | ppp1r1b | 1914 | 825 | 1155 | 1763 | 1594 | 1766 |
| ENSDARG000 | 19 | 10630679 | 10631482 | - | 804 | NA | 0 | 0 | 0 | 0 | 0 | 0 |
| ENSDARG000 | 19;19;19 | 39275973:392 | 39276667:392 | +;+;+ | 1205 | NA | 0 | 0 | 3 | 0 | 4 | 0 |
| ENSDARG000 | 19;19 | 39588596:395 | 39588960:395 | -;- | 457 | NA | 0 | 0 | 0 | 0 | 0 | 0 |
| ENSDARG000 | 19:19:19:19:1 | 7179740:7180 | 7181485:7181 | -:-:-:-:-:-:-:-: | 8059 | col11a2 | 136 | 6 | 295 | 102 | 1939 | 326 |
| ENSDARG000 | 19;19;19 | 33084066:330 | 33084355:330 | -;-;- | 1752 | klhl38a | 26 | 6 | 2 | 0 | 2 | 4 |
| ENSDARG000 | 19;19;19 | 39927406:399 | 39927465:399 | +;+;+ | 922 | NA | 0 | 0 | 0 | 0 | 0 | 0 |
| ENSDARG000 | 19:19:19:19:1 | 28356753:283 | 28357310:283 | -;-;-;-;- | 848 | nrsn1l | 0 | 0 | 0 | 8 | 0 | 4 |
| ENSDARG000 | 19:19:19:19:1 | 10636369:106 | 10636896:106 | -:-:-:-:-:-:-:-: | 1666 | tekt2 | 0 | 0 | 0 | 0 | 0 | 0 |
| ENSDARG000 | 19:19:19:19:1 | 770300:77043 | 770626:77062 | +:+:+:+:+:+:+: | 2216 | gstr | 3730 | 3152 | 2371 | 3236 | 2756 | 3518 |
| ENSDARG000 | 19;19;19 | 25151416:251 | 25151645:251 | +;+;+ | 981 | NA | 0 | 0 | 0 | 0 | 0 | 0 |
| ENSDARG000 | 19:19:19:19:1 | 31007901:310 | 31010071:310 | -:-:-:-:-:-:-:-: | 3883 | bzw2 | 164 | 303 | 391 | 62 | 974 | 497 |
| ENSDARG000 | 19:19:19:19:1 | 5690933:5693 | 5691973:5693 | -;-;-;-;-;- | 1903 | zgc:174904 | 66 | 120 | 188 | 50 | 746 | 112 |
| ENSDARG000 | 19;19;19 | 25154066:251 | 25154235:251 | +;+;+ | 839 | NA | 0 | 0 | 0 | 0 | 0 | 0 |
| ENSDARG000 | 19:19:19:19:1 | 7799036:7800 | 7799237:7801 | +;+;+;+;+;+;+ | 1286 | hax1 | 344 | 296 | 593 | 502 | 216 | 424 |
| ENSDARG000 | 19 | 25155542 | 25157191 | - | 1650 | NA | 2 | 10 | 0 | 4 | 0 | 0 |
| ENSDARG000 | 19:19:19:19:1 | 25163540:251 | 25163752:251 | -:-:-:-:-:-:-:-: | 1342 | rims3 | 0 | 8 | 2 | 0 | 16 | 0 |
| ENSDARG000 | 19:19:19:19:1 | 12297100:122 | 12298150:122 | -:-:-:-:-:-:-:-: | 1818 | ncaldb | 22 | 10 | 6 | 0 | 57 | 63 |
| ENSDARG000 | 19:19:19:19:1 | 17205143:172 | 17208337:172 | -:-:-:-:-:-:-:-: | 4134 | stmn1a | 236 | 513 | 774 | 438 | 591 | 354 |
| ENSDARG000 | 19:19:19:19:1 | 41551335:415 | 41551588:415 | +:+:+:+:+:+:+: | 2422 | NA | 0 | 0 | 0 | 0 | 0 | 0 |
| ENSDARG000 | 19:19:19:19:1 | 33093395:330 | 33093648:330 | +:+:+:+:+:+:+: | 3940 | fam91a1 | 1103 | 960 | 1339 | 884 | 1255 | 985 |
| ENSDARG000 | 19:19:19:19:1 | 28678094:286 | 28679566:286 | -:-:-:-:-:-:-:-: | 2242 | NA | 2 | 6 | 0 | 4 | 0 | 0 |
| ENSDARG000 | 19;19 | 28786149:287 | 28789409:287 | -;- | 3261 | sox4a | 1207 | 937 | 1214 | 727 | 378 | 1219 |
| ENSDARG000 | 19 | 33139164 | 33140403 | + | 1240 | lratd2a | 8 | 12 | 20 | 10 | 14 | 28 |
| ENSDARG000 | 19;19;19 | 5709847:5710 | 5710414:5710 | -;-;- | 874 | NA | 0 | 0 | 0 | 0 | 0 | 0 |
| ENSDARG000 | 19:19:19:19:1 | 20453831:204 | 20454458:204 | -:-:-:-:-:-:-:-: | 2524 | satb1a | 76 | 64 | 98 | 68 | 139 | 85 |
| ENSDARG000 | 19;19 | 28734048:287 | 28734163:287 | -;- | 597 | NA | 0 | 0 | 0 | 0 | 0 | 0 |
| ENSDARG000 | 19:19:19:19:1 | 28788968:287 | 28788996:287 | +:+:+:+:+:+:+: | 4978 | NA | 127 | 74 | 32 | 110 | 27 | 28 |
| ENSDARG000 | 19:19:19:19:1 | 12350069:123 | 12350249:123 | +;+;+;+;+;+;+ | 1075 | psmg2 | 459 | 437 | 743 | 401 | 409 | 692 |
| ENSDARG000 | 19:19:19:19:1 | 41701660:417 | 41702016:417 | +:+:+:+:+:+:+: | 2635 | gatad2b | 509 | 206 | 423 | 171 | 513 | 380 |
| ENSDARG000 | 19:19:19:19:1 | 10649424:106 | 10649875:106 | -:-:-:-:-:-:-:-: | 1652 | olah | 103 | 96 | 138 | 122 | 144 | 107 |
| ENSDARG000 | 19:19:19:19:1 | 40025928:400 | 40026815:400 | +:+:+:+:+:+:+: | 3478 | sfpq | 4323 | 3634 | 4498 | 2956 | 4143 | 4372 |
| ENSDARG000 | 19:19:19:19:1 | 5731849:5733 | 5732030:5733 | -:-:-:-:-:-:-:-: | 2007 | NA | 97 | 81 | 60 | 54 | 69 | 103 |
| ENSDARG000 | 19:19:19:19:1 | 7810028:7810 | 7810114:7810 | +:+:+:+:+:+:+: | 1527 | aqp10b | 18 | 2 | 8 | 18 | 4 | 2 |
| ENSDARG000 | 19:19:19:19:1 | 776208:77954 | 778180:77965 | -:-:-:-:-:-:-:-: | 3982 | msto1 | 607 | 458 | 764 | 405 | 660 | 670 |
| ENSDARG000 | 19:19:19:19:1 | 25352289:253 | 25353410:253 | -;-;-;-;-;-;- | 2733 | glcci1a | 223 | 175 | 230 | 178 | 238 | 182 |
| ENSDARG000 | 19;19;19;19 | 27019432:270 | 27019705:270 | +;+;+;+ | 1048 | tnfa | 30 | 12 | 16 | 24 | 22 | 18 |
| ENSDARG000 | 19:19:19:19:1 | 791538:79157 | 791620:79162 | +:+:+:+:+:+:+: | 1595 | tmem79a | 8 | 4 | 0 | 0 | 2 | 0 |
| ENSDARG000 | 19:19:19:19:1 | 5779084:5785 | 5781495:5785 | -:-:-:-:-:-:-:-: | 3387 | NA | 0 | 0 | 6 | 0 | 10 | 0 |
| ENSDARG000 | 19:19:19:19:1 | 47549568:475 | 47549819:475 | -:-:-:-:-:-:-:-: | 1425 | NA | 0 | 0 | 0 | 0 | 0 | 0 |
| ENSDARG000 | 19;19;19;19 | 17259912:172 | 17261039:172 | +;+;+;+ | 3959 | maneal | 4 | 4 | 8 | 8 | 0 | 2 |
| ENSDARG000 | 19:19:19:19:1 | 6059673:6065 | 6060874:6066 | -:-:-:-:-:-:-:-: | 2727 | gsk3aa | 224 | 302 | 398 | 203 | 397 | 342 |
| ENSDARG000 | 19:19:19:19:1 | 7819480:7819 | 7821126:7821 | -:-:-:-:-:-:-:-: | 5695 | znf687b | 639 | 497 | 712 | 483 | 617 | 653 |
| ENSDARG000 | 19:19:19:19:1 | 41769135:417 | 41769241:417 | +:+:+:+:+:+:+: | 1759 | tinagl1 | 8 | 33 | 81 | 41 | 116 | 14 |
| ENSDARG000 | 19:19:19:19:1 | 20495249:205 | 20495472:205 | +:+:+:+:+:+:+: | 4035 | KCNH8 | 0 | 0 | 2 | 0 | 0 | 10 |
| ENSDARG000 | 19;19;19;19 | 28922941:289 | 28923009:289 | -;-;-;- | 493 | NA | 0 | 0 | 0 | 0 | 0 | 0 |
| ENSDARG000 | 19:19:19:19:1 | 19028633:190 | 19028996:190 | +:+:+:+:+:+:+: | 2271 | CPNE4 | 0 | 0 | 0 | 0 | 0 | 0 |
| ENSDARG000 | 19:19:19:19:1 | 22930177:229 | 22930229:229 | +:+:+:+:+:+:+: | 13516 | pkhd1l1.2 | 12 | 10 | 4 | 0 | 2 | 8 |
| ENSDARG000 | 19:19:19:19:1 | 10661520:106 | 10662089:106 | +:+:+:+:+:+:+: | 4832 | ago3b | 264 | 315 | 280 | 94 | 503 | 138 |

| | | | | | | | | | | | |
|---|---|---|---|---|---|---|---|---|---|---|---|
| ENSDARG000 | 19:19:19:19:1 | 17288529:17: | 17293347:17: | -:-:-:-:-:-:-:-: | 6376 | sf3a3 | 1054 | 1119 | 1211 | 610 | 973 | 660 |
| ENSDARG000 | 19:19:19:19:1 | 12362787:12: | 12363485:12: | -:-:-:-:-:-:-:-: | 2052 | ptpn2b | 744 | 718 | 797 | 482 | 732 | 560 |
| ENSDARG000 | 19;19;19 | 25435150:25 | 25435519:25 | -;-;- | 891 | umad1 | 131 | 71 | 109 | 167 | 81 | 82 |
| ENSDARG000 | 19:19:19:19:1 | 42806812:42 | 42807237:42 | +:+:+:+:+:+:+: | 2506 | ubp1 | 289 | 234 | 223 | 182 | 327 | 251 |
| ENSDARG000 | 19:19:19:19:1 | 29274385:29: | 29276132:29: | -:-:-:-:-:-:-:-: | 3872 | e2f3 | 618 | 717 | 1204 | 366 | 1014 | 828 |
| ENSDARG000 | 19;19;19;19 | 7835025:783 | 7835187:783 | +;+;+;+ | 918 | cks1b | 70 | 350 | 405 | 162 | 248 | 196 |
| ENSDARG000 | 19;19;19;19 | 25465025:25 | 25465223:25 | +;+;+;+ | 643 | rpa3 | 44 | 251 | 224 | 104 | 148 | 162 |
| ENSDARG000 | 19:19:19:19:1 | 33258572:33: | 33259041:33: | -:-:-:-:-:-:-:-: | 1894 | cibar1 | 225 | 184 | 244 | 190 | 319 | 278 |
| ENSDARG000 | 19:19:19:19:1 | 31043002:31( | 31043101:31( | +:+:+:+:+:+:+: | 1109 | ANKMY2 | 0 | 0 | 0 | 0 | 0 | 0 |
| ENSDARG000 | 19:19:19:19:1 | 9295244:929 | 9295340:929 | +:+:+:+:+:+:+: | 2614 | si:ch73-15n2 | 0 | 0 | 2 | 0 | 0 | 0 |
| ENSDARG000 | 19;19 | 31061718:31( | 31061913:31( | +;+ | 615 | sostdc1b | 4 | 0 | 0 | 0 | 0 | 0 |
| ENSDARG000 | 19:19:19:19:1 | 20724347:20 | 20724654:20 | +:+:+:+:+:+:+: | 5267 | kat2b | 230 | 144 | 117 | 74 | 254 | 188 |
| ENSDARG000 | 19:19:19:19:1 | 20708460:20 | 20708625:20 | +:+:+:+:+:+:+: | 2667 | rab5aa | 2831 | 2317 | 2672 | 2616 | 2600 | 2411 |
| ENSDARG000 | 19:19:19:19:1 | 12398544:12: | 12399598:12: | -:-:-:-:-:-:-:-: | 2390 | NA | 144 | 483 | 71 | 162 | 186 | 118 |
| ENSDARG000 | 19:19:19:19:1 | 27040533:27( | 27040545:27( | -:-:-:-:-:-:-:-: | 4253 | NA | 1 | 4 | 2 | 0 | 1 | 2 |
| ENSDARG000 | 19:19:19:19:1 | 47564890:47 | 47565768:47 | -:-:-:-:-:-:-:-: | 2430 | rrm2 | 111 | 24 | 537 | 216 | 10 | 369 |
| ENSDARG000 | 19;19 | 42843109:42 | 42843201:42 | -;- | 796 | NA | 130 | 94 | 58 | 64 | 120 | 188 |
| ENSDARG000 | 19:19:19:19:1 | 29303847:29: | 29304065:29: | +:+:+:+:+:+:+: | 2577 | maco1a | 206 | 187 | 194 | 128 | 212 | 204 |
| ENSDARG000 | 19:19:19:19:1 | 29296191:29: | 29299108:29: | -;-;-;-;- | 3647 | srfbp1 | 303 | 427 | 610 | 209 | 447 | 423 |
| ENSDARG000 | 19:19:19:19:1 | 7838644:784( | 7838771:784 | +:+:+:+:+:+:+: | 4541 | NA | 16 | 6 | 2 | 6 | 29 | 37 |
| ENSDARG000 | 19:19:19:19:1 | 25476228:25 | 25476587:25 | +:+:+:+:+:+:+: | 5314 | col28a1a | 14 | 8 | 2 | 4 | 17 | 2 |
| ENSDARG000 | 19:19:19:19:1 | 31079326:31( | 31079480:31( | -;-;-;-;-;-;-;- | 1101 | NA | 3 | 0 | 5 | 2 | 5 | 10 |
| ENSDARG000 | 19:19:19:19:1 | 43790932:43 | 43791691:43 | -:-:-:-:-:-:-:-: | 2362 | NA | 2 | 0 | 4 | 2 | 2 | 6 |
| ENSDARG000 | 19;19 | 25508793:25 | 25508875:25 | +;+ | 476 | NA | 0 | 0 | 0 | 0 | 0 | 0 |
| ENSDARG000 | 19;19 | 33368152:33: | 33369328:33: | -;- | 1530 | nkd3l | 0 | 0 | 0 | 0 | 0 | 0 |
| ENSDARG000 | 19:19:19:19:1 | 41841147:41 | 41842183:41 | -:-:-:-:-:-:-:-: | 10477 | trioa | 1475 | 1214 | 1715 | 1766 | 2368 | 1447 |
| ENSDARG000 | 19;19;19 | 31183495:31 | 31183969:31 | +;+;+ | 870 | meox2b | 2 | 0 | 0 | 0 | 4 | 0 |
| ENSDARG000 | 19;19 | 33372364:33: | 33372547:33: | -;- | 255 | NA | 0 | 0 | 0 | 0 | 0 | 0 |
| ENSDARG000 | 19:19:19:19:1 | 10692166:10 | 10692614:10 | -:-:-:-:-:-:-:-: | 3816 | slc9a3.1 | 0 | 0 | 4 | 0 | 24 | 4 |
| ENSDARG000 | 19:19:19:19:1 | 33464688:33 | 33464769:33 | +:+:+:+:+:+:+: | 5787 | triqk | 639 | 560 | 656 | 488 | 608 | 586 |
| ENSDARG000 | 19:19:19:19:1 | 6085784:609( | 6088444:609( | -:-:-:-:-:-:-:-: | 10240 | NA | 2934 | 1366 | 1810 | 1627 | 2470 | 1712 |
| ENSDARG000 | 19;19 | 31339305:31: | 31341061:31: | -;- | 1896 | arl4ab | 62 | 84 | 114 | 68 | 180 | 76 |
| ENSDARG000 | 19:19:19:19:1 | 43822968:43 | 43823142:43 | +:+:+:+:+:+:+: | 1352 | eloa | 962 | 894 | 1258 | 1055 | 874 | 888 |
| ENSDARG000 | 19:19:19:19:1 | 42847306:42 | 42847625:42 | +:+:+:+:+:+:+: | 4172 | pdcd6ip | 5469 | 4034 | 5207 | 3797 | 5489 | 4226 |
| ENSDARG000 | 19:19:19:19:1 | 40069524:40( | 40069712:40( | +:+:+:+:+:+:+: | 5315 | zmym4.1 | 547 | 348 | 778 | 264 | 726 | 641 |
| ENSDARG000 | 19;19 | 31342023:31: | 31342064:31: | +;+ | 749 | NA | 7 | 2 | 0 | 1 | 2 | 1 |
| ENSDARG000 | 19:19:19:19:1 | 31347675:31: | 31347847:31: | -:-:-:-:-:-:-:-: | 2249 | scin | 29 | 6 | 6 | 12 | 59 | 102 |
| ENSDARG000 | 19:19:19:19:1 | 19193851:19 | 19194098:19 | +:+:+:+:+:+:+: | 1380 | mrpl3 | 617 | 798 | 896 | 554 | 723 | 768 |
| ENSDARG000 | 19:19:19:19:1 | 27334527:27: | 27334666:27: | -:-:-:-:-:-:-:-: | 981 | polr1h | 24 | 19 | 19 | 30 | 35 | 42 |
| ENSDARG000 | 19:19:19:19:1 | 20761108:20 | 20761985:20 | -:-:-:-:-:-:-:-: | 2844 | ngly1 | 614 | 513 | 638 | 226 | 566 | 440 |
| ENSDARG000 | 19:19:19:19:1 | 27283072:27: | 27287665:27: | -:-:-:-:-:-:-:-: | 10513 | SH3BP5L | 199 | 131 | 235 | 116 | 255 | 255 |
| ENSDARG000 | 19:19:19:19:1 | 12406583:12 | 12406918:12 | +:+:+:+:+:+:+: | 3665 | seh1l | 1386 | 1659 | 2094 | 1080 | 1822 | 1440 |
| ENSDARG000 | 19:19:19:19:1 | 27339848:27: | 27339930:27: | +:+:+:+:+:+:+: | 3006 | ddx39b | 466 | 599 | 519 | 294 | 809 | 336 |
| ENSDARG000 | 19:19:19:19:1 | 27316467:27: | 27316908:27: | -:-:-:-:-:-:-:-: | 2338 | gtf2h4 | 2 | 3 | 8 | 7 | 0 | 0 |
| ENSDARG000 | 19:19:19:19:1 | 33553586:33 | 33553684:33 | +:+:+:+:+:+:+: | 6062 | runx1t1 | 20 | 2 | 8 | 0 | 8 | 56 |
| ENSDARG000 | 19:19:19:19:1 | 7864767:786 | 7864942:786 | +:+:+:+:+:+:+: | 1274 | NA | 31 | 2 | 3 | 2 | 33 | 27 |
| ENSDARG000 | 19:19:19:19:1 | 6141913:614 | 6145027:614 | -:-:-:-:-:-:-:-: | 4905 | erf | 1154 | 1120 | 1271 | 958 | 1410 | 960 |
| ENSDARG000 | 19:19:19:19:1 | 19241372:19 | 19241631:19 | +:+:+:+:+:+:+: | 3989 | ptpn23b | 406 | 220 | 306 | 338 | 307 | 469 |
| ENSDARG000 | 19:19:19:19:1 | 29337789:29: | 29337999:29: | +:+:+:+:+:+:+: | 1490 | ldlrap1a | 139 | 126 | 174 | 99 | 270 | 82 |
| ENSDARG000 | 19:19:19:19:1 | 25512797:25 | 25513060:25 | -;-;-;-;- | 1054 | c1galt1a | 4 | 0 | 0 | 0 | 0 | 0 |
| ENSDARG000 | 19;19 | 25526354:25 | 25526768:25 | +;+ | 1010 | sdhaf3 | 68 | 65 | 132 | 58 | 118 | 88 |
| ENSDARG000 | 19:19:19:19:1 | 7878122:787 | 7878615:787 | +:+:+:+:+:+:+: | 2427 | nuggc.2 | 32 | 15 | 13 | 18 | 54 | 132 |
| ENSDARG000 | 19:19:19:19:1 | 20778011:20 | 20778168:20 | +;+;+;+;+ | 833 | nutf2l | 192 | 729 | 839 | 412 | 631 | 584 |
| ENSDARG000 | 19:19:19:19:1 | 43884120:43 | 43884206:43 | +:+:+:+:+:+:+: | 4296 | lypla2 | 7553 | 6399 | 7140 | 4983 | 6710 | 5073 |
| ENSDARG000 | 19:19:19:19:1 | 9344171:934 | 9344341:934 | +:+:+:+:+:+:+: | 1805 | NA | 0 | 0 | 0 | 0 | 0 | 0 |
| ENSDARG000 | 19:19:19:19:1 | 31392350:31: | 31392818:31: | -;-;-;-;-;-;-;- | 1189 | tmem106bb | 740 | 533 | 699 | 691 | 947 | 850 |
| ENSDARG000 | 19:19:19:19:1 | 12444943:12 | 12445106:12 | +:+:+:+:+:+:+: | 3563 | LDLRAD4 | 506 | 236 | 295 | 266 | 354 | 285 |
| ENSDARG000 | 19:19:19:19:1 | 25649626:25 | 25649913:25 | +;+;+;+;+ | 910 | tac1 | 49 | 25 | 33 | 20 | 55 | 50 |
| ENSDARG000 | 19:19:19:19:1 | 10738150:10 | 10738567:10 | -:-:-:-:-:-:-:-: | 2796 | slc9a3.2 | 0 | 0 | 6 | 12 | 20 | 4 |
| ENSDARG000 | 19;19 | 22999181:23( | 22999725:23( | +;+ | 1610 | gdf6b | 0 | 0 | 4 | 0 | 2 | 0 |
| ENSDARG000 | 19:19:19:19:1 | 42886413:42 | 42886510:42 | +:+:+:+:+:+:+: | 2836 | NA | 344 | 234 | 339 | 371 | 217 | 329 |
| ENSDARG000 | 19;19 | 23042486:23( | 23042842:23( | -;- | 394 | NA | 0 | 0 | 0 | 0 | 0 | 0 |
| ENSDARG000 | 19:19:19:19:1 | 29383486:29: | 29384097:29: | -:-:-:-:-:-:-:-: | 1497 | fndc5b | 278 | 24 | 27 | 42 | 98 | 70 |
| ENSDARG000 | 19;19 | 23043685:23( | 23043719:23( | +;+ | 344 | NA | 0 | 0 | 0 | 0 | 0 | 0 |
| ENSDARG000 | 19:19:19:19:1 | 20785724:20 | 20785902:20 | +:+:+:+:+:+:+: | 4838 | adnp2b | 925 | 1311 | 1747 | 957 | 1770 | 1279 |
| ENSDARG000 | 19:19:19:19:1 | 31404686:31 | 31404843:31 | +;+;+;+;+;+;+ | 826 | pip4p2 | 1196 | 997 | 1005 | 873 | 1171 | 1243 |
| ENSDARG000 | 19;19;19;19 | 17377346:17: | 17378381:17: | -;-;-;- | 1730 | nkiras1 | 180 | 187 | 187 | 98 | 204 | 167 |
| ENSDARG000 | 19:19:19:19:1 | 19278082:19: | 19278814:19: | -:-:-:-:-:-:-:-: | 3332 | cspg5b | 0 | 0 | 6 | 6 | 0 | 0 |
| ENSDARG000 | 19;19 | 23046800:23( | 23047147:23( | -;- | 394 | NA | 0 | 0 | 0 | 0 | 0 | 0 |
| ENSDARG000 | 19:19:19:19:1 | 10794072:10 | 10794254:10 | +:+:+:+:+:+:+: | 2539 | lipea | 267 | 128 | 231 | 145 | 155 | 233 |
| ENSDARG000 | 19:19:19:19:1 | 10805858:10 | 10806119:10 | -:-:-:-:-:-:-:-: | 2125 | NA | 79 | 56 | 72 | 80 | 60 | 93 |
| ENSDARG000 | 19:19:19:19:1 | 29798064:29 | 29798206:29 | +;+;+;+;+ | 1600 | marcksl1b | 4326 | 5398 | 3613 | 5400 | 3306 | 6012 |
| ENSDARG000 | 19:19:19:19:1 | 12630414:12 | 12632415:12 | -;-;-;-;-;-;-;- | 2742 | fam210aa | 494 | 673 | 914 | 920 | 695 | 535 |
| ENSDARG000 | 19:19:19:19:1 | 20793237:20 | 20793470:20 | +;+;+;+;+;+;+ | 1066 | txnl4a | 403 | 379 | 502 | 234 | 430 | 280 |
| ENSDARG000 | 19 | 23047917 | 23048219 | - | 303 | NA | 0 | 0 | 0 | 0 | 0 | 0 |
| ENSDARG000 | 19:19:19:19:1 | 7895086:790( | 7895315:790( | +:+:+:+:+:+:+: | 4185 | nuggc.3 | 132 | 7 | 0 | 0 | 38 | 44 |
| ENSDARG000 | 19 | 23048480 | 23048514 | + | 35 | NA | 0 | 0 | 0 | 0 | 0 | 0 |
| ENSDARG000 | 19 | 23049885 | 23050160 | + | 276 | NA | 0 | 0 | 0 | 0 | 0 | 0 |
| ENSDARG000 | 19 | 23050777 | 23051016 | - | 240 | NA | 0 | 0 | 0 | 0 | 0 | 0 |
| ENSDARG000 | 19:19:19:19:1 | 31437689:31 | 31438070:31 | -:-:-:-:-:-:-:-: | 1580 | necab1 | 6 | 2 | 23 | 0 | 4 | 4 |
| ENSDARG000 | 19;19 | 23051978:23( | 23052271:23( | -;- | 343 | NA | 0 | 0 | 0 | 0 | 0 | 0 |
| ENSDARG000 | 19:19:19:19:1 | 12649691:12 | 12650383:12 | +:+:+:+:+:+:+: | 2539 | rnmt | 458 | 627 | 685 | 403 | 731 | 563 |
| ENSDARG000 | 19:19:19:19:1 | 17385561:17: | 17385610:17: | +:+:+:+:+:+:+: | 3488 | rpl15 | 16923 | 16350 | 30334 | 26217 | 18062 | 25920 |
| ENSDARG000 | 19;19 | 23064025:23( | 23064381:23( | -;- | 406 | NA | 0 | 0 | 0 | 0 | 0 | 0 |
| ENSDARG000 | 19;19 | 23080227:23( | 23080272:23( | +;+ | 420 | NA | 0 | 0 | 0 | 0 | 0 | 0 |
| ENSDARG000 | 19 | 23114196 | 23114482 | - | 287 | NA | 0 | 0 | 0 | 0 | 0 | 0 |
| ENSDARG000 | 19:19:19:19:1 | 20801060:20 | 20801159:20 | +:+:+:+:+:+:+: | 2164 | cndp2 | 4677 | 5550 | 5312 | 4096 | 5935 | 6830 |
| ENSDARG000 | 19:19:19:19:1 | 31568992:31 | 31569565:31 | -;-;-;-;-;-;- | 1727 | tdp2b | 34 | 103 | 102 | 76 | 120 | 54 |
| ENSDARG000 | 19:19:19:19:1 | 31541607:31 | 31541615:31 | +:+:+:+:+:+:+: | 2481 | gmnn | 92 | 191 | 208 | 186 | 155 | 191 |
| ENSDARG000 | 19:19:19:19:1 | 10831362:10 | 10831613:10 | +:+:+:+:+:+:+: | 5326 | tomm40l | 300 | 481 | 739 | 379 | 497 | 295 |
| ENSDARG000 | 19:19:19:19:1 | 7929704:792 | 7929912:793( | +:+:+:+:+:+:+: | 1067 | si:dkey-266f7 | 6 | 0 | 0 | 10 | 4 | 0 |
| ENSDARG000 | 19:19:19:19:1 | 31576849:31 | 31577016:31 | +;+;+;+;+;+ | 589 | acot13 | 78 | 91 | 126 | 40 | 42 | 41 |
| ENSDARG000 | 19;19;19 | 31532043:31 | 31533038:31 | +;+;+ | 3183 | tmem64 | 134 | 30 | 62 | 56 | 72 | 63 |
| ENSDARG000 | 19:19:19:19:1 | 31579391:31 | 31580697:31 | -;-;-;-;-;-;- | 3331 | C19H6orf62 | 4809 | 4738 | 5734 | 4496 | 5986 | 5728 |
| ENSDARG000 | 19:19:19:19:1 | 29808471:29 | 29808796:29 | +:+:+:+:+:+:+: | 6034 | hdac1 | 3534 | 3474 | 5140 | 2377 | 4065 | 2932 |
| ENSDARG000 | 19 | 23115057 | 23115094 | + | 38 | NA | 0 | 0 | 0 | 0 | 0 | 0 |
| ENSDARG000 | 19;19 | 23120398:23 | 23120745:23 | -;- | 397 | NA | 0 | 0 | 0 | 0 | 0 | 0 |
| ENSDARG000 | 19 | 23119713 | 23119939 | + | 227 | NA | 0 | 0 | 0 | 0 | 0 | 0 |
| ENSDARG000 | 19:19:19:19:1 | 43905671:43 | 43905884:43 | +:+:+:+:+:+:+: | 4059 | ankib1a | 675 | 539 | 682 | 332 | 783 | 334 |
| ENSDARG000 | 19:19:19:19:1 | 17400263:17 | 17400711:17 | +:+:+:+:+:+:+: | 3986 | nr1d2b | 4710 | 2811 | 4440 | 3005 | 4978 | 3958 |
| ENSDARG000 | 19;19;19 | 20831079:20 | 20831242:20 | +;+;+ | 767 | NA | 0 | 0 | 0 | 0 | 0 | 1 |
| ENSDARG000 | 19;19 | 23121009:23 | 23121292:23 | +;+ | 829 | NA | 0 | 0 | 0 | 0 | 0 | 0 |
| ENSDARG000 | 19;19;19 | 12762887:12 | 12763018:12 | +;+;+ | 1626 | mc5ra | 42 | 14 | 58 | 0 | 30 | 24 |
| ENSDARG000 | 19:19:19:19:1 | 21266008:21 | 21266230:21 | +;+;+;+;+;+;+ | 12103 | tshz1 | 391 | 494 | 628 | 382 | 1010 | 360 |
| ENSDARG000 | 19:19:19:19:1 | 21358624:21 | 21359757:21 | -;-;-;-;-;-;- | 2741 | NA | 0 | 2 | 4 | 13 | 14 | 2 |
| ENSDARG000 | 19;19;19;19 | 12915498:12 | 12915686:12 | +;+;+;+ | 1262 | cthrc1a | 20 | 12 | 4 | 16 | 14 | 16 |
| ENSDARG000 | 19;19 | 23128663:23 | 23129004:23 | -;- | 391 | NA | 0 | 0 | 0 | 0 | 0 | 0 |
| ENSDARG000 | 19:19:19:19:1 | 10845953:10 | 10846102:10 | +:+:+:+:+:+:+: | 1044 | apoa4a | 0 | 6 | 4 | 1 | 8 | 0 |
| ENSDARG000 | 19:19:19:19:1 | 19511394:19 | 19512629:19 | +:+:+:+:+:+:+: | 2223 | jazf1a | 63 | 16 | 13 | 0 | 18 | 12 |
| ENSDARG000 | 19:19:19:19:1 | 23136170:23 | 23137098:23 | -:-:-:-:-:-:-:-: | 2792 | grb10a | 293 | 276 | 459 | 296 | 370 | 345 |
| ENSDARG000 | 19:19:19:19:1 | 12937066:12 | 12939676:12 | -:-:-:-:-:-:-:-: | 3778 | slc25a32a | 179 | 48 | 80 | 74 | 132 | 301 |
| ENSDARG000 | 19:19:19:19:1 | 21698149:21 | 21698568:21 | -;-;-;-;-;-;-;- | 3972 | znf516 | 350 | 144 | 267 | 214 | 54 | 539 |
| ENSDARG000 | 19:19:19:19:1 | 43969363:43 | 43969762:43 | +;+;+;+;+ | 1328 | gatad1 | 229 | 248 | 368 | 127 | 369 | 249 |
| ENSDARG000 | 19;19 | 43978814:43 | 43979145:43 | +;+ | 1474 | NA | 0 | 1 | 7 | 3 | 5 | 1 |
| ENSDARG000 | 19:19:19:19:1 | 10860485:10 | 10860691:10 | +:+:+:+:+:+:+: | 9186 | NA | 477 | 411 | 673 | 385 | 494 | 653 |
| ENSDARG000 | 19;19 | 43987389:43 | 43987720:43 | +;+ | 1485 | NA | 0 | 4 | 0 | 0 | 2 | 6 |
| ENSDARG000 | 19;19;19;19 | 10859219:10 | 10859354:10 | -;-;-;- | 445 | PSMD4 | 0 | 0 | 0 | 0 | 0 | 0 |
| ENSDARG000 | 19:19:19:19:1 | 19549062:19 | 19551933:19 | -:-:-:-:-:-:-:-: | 5255 | tax1bp1a | 907 | 686 | 634 | 394 | 847 | 735 |
| ENSDARG000 | 19:19:19:19:1 | 10871601:10 | 10871995:10 | -:-:-:-:-:-:-:-: | 1815 | psmd4a | 419 | 371 | 351 | 177 | 347 | 207 |
| ENSDARG000 | 19:19:19:19:1 | 19599528:19 | 19599575:19 | +:+:+:+:+:+:+: | 1727 | hibadha | 166 | 80 | 162 | 152 | 134 | 134 |
| ENSDARG000 | 19:19:19:19:1 | 10859194:10 | 10859354:10 | -;-;-;-;-;-;- | 430 | NA | 0 | 0 | 0 | 0 | 0 | 0 |
| ENSDARG000 | 19:19:19:19:1 | 10855158:10 | 10855294:10 | +;+;+;+;+;+ | 1513 | apoea | 2 | 0 | 6 | 1 | 0 | 4 |
| ENSDARG000 | 19;19 | 43995936:43 | 43996289:43 | +;+ | 1537 | nkd3 | 18 | 18 | 17 | 9 | 38 | 9 |
| ENSDARG000 | 19;19 | 44004348:44( | 44004679:44( | +;+ | 1509 | NA | 21 | 22 | 24 | 15 | 19 | 5 |
| ENSDARG000 | 19:19:19:19:1 | 29822500:29 | 29822753:29 | -:-:-:-:-:-:-:-: | 1539 | eif3i | 5404 | 5918 | 9445 | 7864 | 5786 | 6884 |
| ENSDARG000 | 19:19:19:19:1 | 13099541:13( | 13099612:13( | +:+:+:+:+:+:+: | 3571 | NA | 125 | 104 | 140 | 85 | 108 | 154 |
| ENSDARG000 | 19;19;19;19 | 11978209:11 | 11978288:11 | +;+;+;+ | 384 | polr2k | 80 | 107 | 140 | 52 | 62 | 60 |
| ENSDARG000 | 19;19 | 3726193:373 | 3726220:373 | +;+ | 69 | NA | 0 | 0 | 0 | 0 | 0 | 0 |
| ENSDARG000 | 19:19:19:19:1 | 14903379:14 | 14903394:14 | -:-:-:-:-:-:-:-: | 324 | NA | 0 | 0 | 0 | 0 | 0 | 0 |
| ENSDARG000 | 19:19:19:19:1 | 42214934:42 | 42214939:42 | -;-;-;-;- | 156 | NA | 0 | 0 | 0 | 0 | 0 | 0 |
| ENSDARG000 | 19 | 40636572 | 40636691 | - | 120 | 5S_rRNA | 0 | 0 | 0 | 0 | 0 | 0 |
| ENSDARG000 | 19 | 9429857 | 9429944 | + | 88 | dre-let-7d-2 | 0 | 0 | 0 | 0 | 0 | 0 |
| ENSDARG000 | 19:19:19:19:1 | 30495109:30 | 30495164:30 | +:+:+:+:+:+:+: | 171 | NA | 0 | 0 | 0 | 0 | 0 | 0 |
| ENSDARG000 | 19:19:19:19:1 | 19342177:19: | 19342584:19: | -:-:-:-:-:-:-:-: | 3233 | smarcc1b | 508 | 364 | 567 | 340 | 539 | 413 |
| ENSDARG000 | 19 | 25518420 | 25518540 | - | 121 | 5S_rRNA | 0 | 0 | 0 | 0 | 0 | 0 |
| ENSDARG000 | 19;19 | 7237188:724( | 7238001:724( | -;- | 863 | NA | 0 | 0 | 0 | 0 | 0 | 0 |
| ENSDARG000 | 19:19:19:19:1 | 1964005:196 | 1964085:196 | +:+:+:+:+:+:+: | 5531 | SH3BP5 | 665 | 501 | 583 | 536 | 774 | 291 |
| ENSDARG000 | 19 | 7306467 | 7306563 | + | 97 | NA | 0 | 0 | 0 | 0 | 0 | 0 |
| ENSDARG000 | 19 | 27172714 | 27172830 | - | 117 | 5S_rRNA | 0 | 0 | 0 | 0 | 0 | 0 |
| ENSDARG000 | 19:19:19:19:1 | 18118321:18 | 18119075:18 | -:-:-:-:-:-:-:-: | 2586 | snx10a | 40 | 8 | 22 | 2 | 11 | 2 |
| ENSDARG000 | 19;19 | 23672489:23 | 23672797:23 | -;- | 345 | NA | 0 | 0 | 0 | 0 | 0 | 0 |
| ENSDARG000 | 19 | 34207162 | 34207280 | + | 119 | 5S_rRNA | 0 | 0 | 0 | 0 | 0 | 0 |
| ENSDARG000 | 19 | 7468201 | 7468316 | - | 116 | 5S_rRNA | 0 | 0 | 0 | 0 | 0 | 0 |
| ENSDARG000 | 19 | 28441841 | 28441960 | - | 120 | 5S_rRNA | 0 | 0 | 0 | 0 | 0 | 0 |
| ENSDARG000 | 19 | 42595268 | 42595384 | - | 117 | 5S_rRNA | 0 | 0 | 0 | 0 | 0 | 0 |
| ENSDARG000 | 19:19:19:19:1 | 19496479:19 | 19496612:19 | -:-:-:-:-:-:-:-: | 2013 | CREB5 | 147 | 34 | 49 | 73 | 20 | 26 |
| ENSDARG000 | 19 | 18841413 | 18841543 | - | 131 | NA | 0 | 3 | 0 | 0 | 0 | 6 |
| ENSDARG000 | 19 | 26576594 | 26576708 | + | 115 | 5S_rRNA | 0 | 0 | 0 | 0 | 0 | 0 |
| ENSDARG000 | 19 | 25175272 | 25175392 | + | 121 | 5S_rRNA | 0 | 0 | 0 | 0 | 0 | 0 |
| ENSDARG000 | 19 | 23219477 | 23219595 | - | 119 | 5S_rRNA | 0 | 0 | 0 | 0 | 0 | 0 |
| ENSDARG000 | 19 | 20439142 | 20439260 | - | 119 | 5S_rRNA | 0 | 0 | 0 | 0 | 0 | 0 |
| ENSDARG000 | 19;19;19 | 43213527:43 | 43213933:43 | -;-;- | 1134 | gnrhr1 | 4 | 0 | 0 | 0 | 2 | 0 |
| ENSDARG000 | 19 | 29785095 | 29785212 | - | 118 | 5S_rRNA | 0 | 0 | 0 | 0 | 0 | 0 |
| ENSDARG000 | 19 | 5843091 | 5843205 | - | 115 | 5S_rRNA | 0 | 0 | 0 | 0 | 0 | 0 |
| ENSDARG000 | 19 | 42219165 | 42220434 | + | 1270 | NA | 0 | 0 | 0 | 0 | 0 | 0 |
| ENSDARG000 | 19:19:19:19:1 | 17358850:17: | 17359095:17: | +;+;+;+;+;+ | 1204 | ube2e1 | 726 | 880 | 1356 | 1215 | 831 | 916 |
| ENSDARG000 | 19:19:19:19:1 | 950394:9515 | 950500:9516 | -:-:-:-:-:-:-:-: | 2112 | slc44a4 | 27 | 20 | 63 | 43 | 45 | 38 |
| ENSDARG000 | 19 | 5415799 | 5415915 | - | 117 | 5S_rRNA | 0 | 0 | 0 | 0 | 0 | 0 |
| ENSDARG000 | 19:19:19:19:1 | 19398723:19 | 19398859:19 | +;+;+;+;+ | 3813 | ASTE1 | 315 | 340 | 315 | 202 | 377 | 416 |
| ENSDARG000 | 19:19:19:19:1 | 1370504:137 | 1370765:137 | +:+:+:+:+:+:+: | 2730 | dgat1a | 863 | 220 | 72 | 172 | 72 | 397 |
| ENSDARG000 | 19;19 | 43123331:43 | 43123474:43 | +;+ | 243 | NA | 5 | 0 | 2 | 2 | 0 | 0 |
| ENSDARG000 | 19:19:19:19:1 | 5604241:560 | 5604296:560 | +:+:+:+:+:+:+: | 3031 | wipf2b | 402 | 340 | 486 | 370 | 544 | 420 |

| | | | | | | | | | | | | |
|---|---|---|---|---|---|---|---|---|---|---|---|---|
| ENSDARG000 | 19 | 22646182 | 22646299 | - | 118 | 5S_rRNA | 0 | 0 | 8 | 0 | 1 | 0 |
| ENSDARG000 | 19 | 37669660 | 37669774 | + | 115 | 5S_rRNA | 0 | 0 | 0 | 0 | 0 | 0 |
| ENSDARG000 | 19 | 12136736 | 12136871 | - | 136 | NA | 0 | 0 | 0 | 0 | 0 | 4 |
| ENSDARG000 | 19 | 47567713 | 47567848 | - | 136 | NA | 0 | 0 | 0 | 0 | 0 | 0 |
| ENSDARG000 | 19;19;19;19 | 43517049:435 | 43517058:435 | -;-;-;- | 276 | NA | 0 | 0 | 0 | 0 | 0 | 0 |
| ENSDARG000 | 19 | 4776292 | 4776413 | + | 122 | 5S_rRNA | 8 | 0 | 7 | 6 | 4 | 1 |
| ENSDARG000 | 19 | 13833879 | 13833993 | + | 115 | 5S_rRNA | 0 | 0 | 0 | 0 | 0 | 0 |
| ENSDARG000 | 19 | 42337317 | 42337431 | + | 115 | 5S_rRNA | 0 | 0 | 0 | 0 | 4 | 0 |
| ENSDARG000 | 19;19;19;19;1 | 2726819:2726 | 2727182:2727 | +;+;+;+;+;+;+; | 1767 | NA | 6 | 0 | 0 | 0 | 4 | 0 |
| ENSDARG000 | 19;19;19;19;1 | 1465004:1467 | 1465213:1467 | +;+;+;+;+;+;+; | 3223 | NA | 0 | 20 | 0 | 0 | 3 | 0 |
| ENSDARG000 | 19 | 9443772 | 9443878 | + | 107 | dre-mir-125a- | 0 | 0 | 0 | 0 | 0 | 0 |
| ENSDARG000 | 19 | 40781168 | 40781271 | - | 104 | dre-mir-489 | 0 | 0 | 0 | 0 | 0 | 0 |
| ENSDARG000 | 19 | 10012498 | 10012614 | - | 117 | 5S_rRNA | 0 | 0 | 0 | 0 | 0 | 0 |
| ENSDARG000 | 19 | 36242791 | 36242908 | + | 118 | 5S_rRNA | 0 | 0 | 0 | 0 | 0 | 0 |
| ENSDARG000 | 19 | 19709927 | 19710042 | - | 116 | 5S_rRNA | 0 | 0 | 0 | 0 | 0 | 0 |
| ENSDARG000 | 19;19;19;19;1 | 48164242:481 | 48164587:481 | +;+;+;+;+;+;+ | 1658 | NA | 110 | 45 | 127 | 71 | 79 | 71 |
| ENSDARG000 | 19 | 38028498 | 38028618 | - | 121 | 5S_rRNA | 0 | 0 | 0 | 0 | 0 | 0 |
| ENSDARG000 | 19 | 26805766 | 26805882 | + | 117 | 5S_rRNA | 0 | 0 | 0 | 0 | 0 | 0 |
| ENSDARG000 | 19;19;19;19;1 | 4076259:4077 | 4076331:4078 | +;+;+;+;+;+;+ | 1650 | NA | 11 | 19 | 1 | 7 | 7 | 18 |
| ENSDARG000 | 19 | 11725496 | 11725631 | - | 136 | 5S_rRNA | 0 | 0 | 0 | 0 | 0 | 0 |
| ENSDARG000 | 19;19;19 | 33200266:332 | 33205460:332 | -;-;- | 6164 | NA | 2434 | 1755 | 1609 | 2006 | 1550 | 1548 |
| ENSDARG000 | 19 | 19747619 | 19747694 | + | 76 | mir196a-2 | 0 | 0 | 0 | 0 | 0 | 0 |
| ENSDARG000 | 19;19;19;19;1 | 48003691:480 | 48003860:480 | -;-;-;-;-;-;-;-;- | 3913 | zgc:158376 | 166 | 74 | 125 | 51 | 160 | 83 |
| ENSDARG000 | 19;19;19 | 27589201:275 | 27589502:275 | +;+;+ | 3536 | CFAP90 | 4 | 0 | 4 | 6 | 0 | 0 |
| ENSDARG000 | 19;19 | 31591524:315 | 31591526:315 | +;+ | 15 | NA | 41 | 40 | 16 | 66 | 31 | 79 |
| ENSDARG000 | 19;19 | 46724211:467 | 46724990:467 | -;- | 1491 | SAMD12 | 1 | 2 | 0 | 0 | 0 | 6 |
| ENSDARG000 | 19;19;19;19;1 | 48290786:482 | 48290853:482 | +;+;+;+;+;+;+; | 1682 | anxa14 | 0 | 0 | 0 | 0 | 0 | 2 |
| ENSDARG000 | 19 | 32330513 | 32330630 | - | 118 | 5S_rRNA | 0 | 0 | 0 | 0 | 0 | 0 |
| ENSDARG000 | 19 | 18861739 | 18861880 | - | 142 | dre-mir-219-3 | 0 | 0 | 0 | 0 | 0 | 0 |
| ENSDARG000 | 19;19;19;19;1 | 48117388:481 | 48117422:481 | +;+;+;+;+;+;+; | 626 | nme2b.1 | 14159 | 17554 | 19306 | 17571 | 16115 | 16168 |
| ENSDARG000 | 19;19 | 10705459:107 | 10705467:107 | -;- | 15 | NA | 0 | 0 | 0 | 0 | 0 | 0 |
| ENSDARG000 | 19;19;19;19;1 | 40171960:401 | 40171995:401 | +;+;+;+;+;+ | 123 | NA | 0 | 2 | 0 | 0 | 0 | 0 |
| ENSDARG000 | 19 | 22769111 | 22769227 | + | 117 | 5S_rRNA | 0 | 0 | 0 | 0 | 0 | 0 |
| ENSDARG000 | 19 | 5773921 | 5774035 | - | 115 | 5S_rRNA | 0 | 0 | 0 | 0 | 0 | 0 |
| ENSDARG000 | 19;19;19 | 47578811:475 | 47579485:475 | -;-;- | 768 | NA | 0 | 0 | 0 | 0 | 0 | 0 |
| ENSDARG000 | 19 | 7035206 | 7035320 | + | 115 | 5S_rRNA | 0 | 0 | 0 | 0 | 0 | 0 |
| ENSDARG000 | 19 | 6627173 | 6627290 | + | 118 | 5S_rRNA | 0 | 0 | 0 | 0 | 0 | 0 |
| ENSDARG000 | 19 | 27464416 | 27464482 | - | 67 | NA | 0 | 0 | 0 | 0 | 0 | 0 |
| ENSDARG000 | 19 | 37227910 | 37228025 | - | 116 | 5S_rRNA | 0 | 0 | 0 | 0 | 0 | 0 |
| ENSDARG000 | 19;19;19;19;1 | 6856589:6859 | 6856717:6859 | -;-;-;-;-;-;- | 1335 | NA | 4 | 5 | 5 | 20 | 6 | 21 |
| ENSDARG000 | 19 | 19489588 | 19491864 | + | 2277 | TRIL | 8 | 4 | 32 | 4 | 33 | 6 |
| ENSDARG000 | 19;19;19;19;1 | 2867014:2871 | 2867118:2871 | -;-;-;-;-;-;-;- | 1260 | exosc7 | 90 | 197 | 201 | 93 | 152 | 127 |
| ENSDARG000 | 19;19;19;19;1 | 5849008:5849 | 5849109:5849 | -;-;-;-;-;- | 825 | NA | 0 | 0 | 0 | 0 | 0 | 0 |
| ENSDARG000 | 19 | 38464295 | 38464413 | - | 119 | 5S_rRNA | 0 | 0 | 0 | 0 | 0 | 0 |
| ENSDARG000 | 19 | 26192822 | 26192936 | + | 115 | 5S_rRNA | 0 | 0 | 0 | 0 | 0 | 0 |
| ENSDARG000 | 19 | 43077744 | 43077879 | + | 136 | NA | 0 | 0 | 0 | 0 | 0 | 0 |
| ENSDARG000 | 19 | 12131045 | 12131180 | - | 136 | NA | 0 | 0 | 0 | 0 | 0 | 0 |
| ENSDARG000 | 19 | 43080925 | 43081060 | + | 136 | NA | 0 | 0 | 9 | 4 | 1 | 0 |
| ENSDARG000 | 19 | 27465104 | 27465170 | - | 67 | NA | 0 | 0 | 0 | 0 | 0 | 0 |
| ENSDARG000 | 19;19;19;19;1 | 2835240:2841 | 2835820:2841 | +;+;+;+;+;+;+; | 3088 | CDCP1 | 3 | 0 | 12 | 30 | 2 | 1 |
| ENSDARG000 | 19;19;19 | 1119416:1123 | 1120541:1123 | -;-;- | 1646 | slc39a4 | 0 | 2 | 0 | 0 | 0 | 0 |
| ENSDARG000 | 19 | 27468449 | 27468515 | - | 67 | NA | 0 | 0 | 0 | 0 | 0 | 0 |
| ENSDARG000 | 19;19;19;19;1 | 1097393:1103 | 1097630:1103 | +;+;+;+;+;+;+; | 2542 | NA | 14 | 27 | 75 | 14 | 65 | 79 |
| ENSDARG000 | 19 | 35778842 | 35778957 | + | 116 | 5S_rRNA | 0 | 0 | 0 | 0 | 0 | 0 |
| ENSDARG000 | 19 | 21917323 | 21917453 | + | 131 | 5S_rRNA | 0 | 0 | 0 | 0 | 0 | 0 |
| ENSDARG000 | 19 | 10123233 | 10123347 | - | 115 | 5S_rRNA | 0 | 0 | 0 | 0 | 0 | 0 |
| ENSDARG000 | 19;19;19 | 18739085:187 | 18739488:187 | +;+;+ | 5518 | NA | 222 | 8 | 0 | 7 | 20 | 20 |
| ENSDARG000 | 19;19;19;19;1 | 43297546:433 | 43297680:433 | +;+;+;+;+;+;+ | 875 | NA | 129 | 189 | 254 | 162 | 503 | 465 |
| ENSDARG000 | 19;19;19;19;1 | 43504480:435 | 43504833:435 | +;+;+;+;+ | 1372 | NA | 0 | 2 | 0 | 0 | 0 | 0 |
| ENSDARG000 | 19;19;19;19;1 | 2876106:2877 | 2876155:2877 | +;+;+;+;+;+;+; | 4128 | hgh1 | 175 | 202 | 324 | 94 | 202 | 170 |
| ENSDARG000 | 19;19 | 11508960:115 | 11509796:115 | -;- | 1641 | KCNG2 | 92 | 963 | 24 | 448 | 830 | 18 |
| ENSDARG000 | 19 | 23913182 | 23913300 | + | 119 | 5S_rRNA | 3 | 3 | 0 | 0 | 1 | 0 |
| ENSDARG000 | 19;19;19;19;1 | 935565:94089 | 935764:94098 | +;+;+;+;+ | 579 | RNF5 | 294 | 195 | 253 | 313 | 197 | 212 |
| ENSDARG000 | 19;19;19;19;1 | 11984725:119 | 11986009:119 | +;+;+;+;+;+;+; | 2843 | spag1a | 806 | 942 | 611 | 333 | 587 | 609 |
| ENSDARG000 | 19;19;19;19;1 | 47476908:474 | 47477032:474 | +;+;+;+;+ | 1095 | zgc:114119 | 136 | 247 | 261 | 216 | 279 | 171 |
| ENSDARG000 | 19;19 | 1184878:1186 | 1185714:1186 | +;+ | 1562 | scrt1a | 0 | 0 | 0 | 0 | 0 | 0 |
| ENSDARG000 | 19;19;19;19;1 | 46421349:464 | 46421832:464 | -;-;-;-;-;-;-;-;- | 2972 | ext1b | 139 | 121 | 297 | 75 | 342 | 226 |
| ENSDARG000 | 19 | 33977878 | 33977994 | - | 117 | 5S_rRNA | 0 | 0 | 2 | 0 | 0 | 0 |
| ENSDARG000 | 19 | 30518455 | 30518573 | + | 119 | 5S_rRNA | 0 | 0 | 0 | 0 | 0 | 0 |
| ENSDARG000 | 19;19;19;19 | 2395114:2400 | 2397876:2400 | -;-;-;- | 3924 | polr1f | 96 | 240 | 329 | 122 | 195 | 170 |
| ENSDARG000 | 19 | 19264759 | 19264881 | + | 123 | 5S_rRNA | 0 | 0 | 0 | 0 | 0 | 0 |
| ENSDARG000 | 19 | 11193249 | 11193363 | - | 115 | 5S_rRNA | 0 | 0 | 0 | 0 | 0 | 0 |
| ENSDARG000 | 19 | 24757193 | 24757324 | - | 132 | NA | 0 | 2 | 0 | 1 | 3 | 0 |
| ENSDARG000 | 19 | 11278413 | 11278529 | + | 117 | 5S_rRNA | 0 | 1 | 0 | 0 | 9 | 5 |
| ENSDARG000 | 19;19;19 | 170705:17186 | 170947:17385 | +;+;+ | 2861 | rnf139 | 893 | 866 | 789 | 428 | 931 | 560 |
| ENSDARG000 | 19 | 16068970 | 16069103 | + | 134 | NA | 0 | 0 | 0 | 0 | 0 | 0 |
| ENSDARG000 | 19;19;19;19;1 | 48251273:482 | 48252610:482 | +;+;+;+;+;+;+; | 3534 | NA | 36 | 2 | 2 | 14 | 14 | 20 |
| ENSDARG000 | 19 | 4770919 | 4771035 | + | 117 | 5S_rRNA | 1 | 0 | 0 | 0 | 4 | 0 |
| ENSDARG000 | 19;19;19;19;1 | 47975916:479 | 47976927:479 | -;-;-;-;-;-;-;-;- | 3538 | ctnnb2 | 11959 | 7181 | 10844 | 7322 | 7769 | 5642 |
| ENSDARG000 | 19 | 16999627 | 16999742 | - | 116 | 5S_rRNA | 0 | 0 | 0 | 0 | 0 | 0 |
| ENSDARG000 | 19 | 43665391 | 43665511 | + | 121 | 5S_rRNA | 0 | 0 | 0 | 0 | 0 | 0 |
| ENSDARG000 | 19;19;19 | 1406262:1407 | 1407867:1408 | -;-;- | 2605 | OXSM | 190 | 216 | 350 | 628 | 206 | 208 |
| ENSDARG000 | 19;19;19;19;1 | 12030832:120 | 12033914:120 | -;-;-;-;-;-;-;-;- | 5285 | rnf19a | 1940 | 1432 | 2033 | 1685 | 2010 | 1379 |
| ENSDARG000 | 19;19;19 | 24214225:242 | 24214383:242 | -;-;- | 453 | NA | 2 | 0 | 5 | 0 | 0 | 0 |
| ENSDARG000 | 19;19;19;19;1 | 2699136:2701 | 2699348:2701 | -;-;-;-;- | 696 | il6 | 26 | 67 | 24 | 9 | 28 | 30 |
| ENSDARG000 | 19;19;19;19;1 | 18797623:187 | 18798265:187 | +;+;+;+;+;+;+; | 2012 | ddah2 | 854 | 732 | 636 | 700 | 635 | 1016 |
| ENSDARG000 | 19 | 10567371 | 10567486 | + | 116 | 5S_rRNA | 0 | 2 | 0 | 2 | 0 | 0 |
| ENSDARG000 | 19 | 28143135 | 28143254 | + | 120 | 5S_rRNA | 0 | 0 | 0 | 0 | 0 | 0 |
| ENSDARG000 | 19;19;19;19;1 | 18758014:187 | 18758589:187 | +;+;+;+;+;+ | 2440 | clic1 | 898 | 738 | 715 | 646 | 1091 | 970 |
| ENSDARG000 | 19 | 32058477 | 32058599 | + | 123 | 5S_rRNA | 0 | 0 | 2 | 4 | 0 | 0 |
| ENSDARG000 | 19 | 16027419 | 16027490 | - | 72 | NA | 0 | 0 | 0 | 0 | 0 | 0 |
| ENSDARG000 | 19 | 30630943 | 30631060 | + | 118 | 5S_rRNA | 0 | 0 | 0 | 0 | 0 | 0 |
| ENSDARG000 | 19 | 38256276 | 38256392 | + | 117 | 5S_rRNA | 0 | 0 | 3 | 0 | 0 | 0 |
| ENSDARG000 | 19 | 18834727 | 18834857 | - | 131 | NA | 0 | 0 | 0 | 0 | 0 | 0 |
| ENSDARG000 | 19 | 29203142 | 29203257 | + | 116 | 5S_rRNA | 0 | 0 | 0 | 0 | 0 | 0 |
| ENSDARG000 | 19 | 6095618 | 6095732 | + | 115 | 5S_rRNA | 0 | 0 | 0 | 0 | 0 | 0 |
| ENSDARG000 | 19 | 42188990 | 42189104 | + | 115 | 5S_rRNA | 0 | 0 | 0 | 0 | 0 | 0 |
| ENSDARG000 | 19 | 12633149 | 12633263 | - | 115 | 5S_rRNA | 0 | 0 | 0 | 0 | 0 | 0 |
| ENSDARG000 | 19;19;19;19;1 | 43119014:431 | 43119074:431 | +;+;+;+;+;+;+; | 5863 | eef1a1l1 | 247882 | 233781 | 361993 | 335616 | 280138 | 341207 |
| ENSDARG000 | 19 | 8852540 | 8852655 | - | 116 | 5S_rRNA | 0 | 0 | 0 | 0 | 0 | 0 |
| ENSDARG000 | 19 | 27465336 | 27465402 | - | 67 | NA | 0 | 0 | 0 | 3 | 0 | 6 |
| ENSDARG000 | 19;19;19 | 6834380:6836 | 6834707:6837 | -;-;- | 1551 | NA | 4 | 80 | 97 | 16 | 67 | 36 |
| ENSDARG000 | 19;19;19;19;1 | 18131512:181 | 18133015:181 | -;-;-;-;-;-;-;-;- | 2793 | cbx3a | 1338 | 2396 | 2452 | 1210 | 1897 | 1460 |
| ENSDARG000 | 19;19;19;19;1 | 47285773:472 | 47285795:472 | -;-;-;-;-;-;-;-;- | 243 | NA | 0 | 0 | 0 | 2 | 0 | 0 |
| ENSDARG000 | 19;19;19;19;1 | 43256986:432 | 43257066:432 | +;+;+;+;+;+;+; | 1239 | DGKD | 786 | 207 | 581 | 811 | 667 | 251 |
| ENSDARG000 | 19;19;19 | 1999838:2001 | 2000289:2001 | +;+;+ | 2369 | btd | 102 | 76 | 120 | 60 | 222 | 92 |
| ENSDARG000 | 19;19;19;19;1 | 19437648:194 | 19437688:194 | -;-;-;-;-;-;-;- | 1938 | NA | 26 | 93 | 140 | 178 | 394 | 372 |
| ENSDARG000 | 19 | 12046454 | 12046570 | - | 117 | 5S_rRNA | 0 | 0 | 0 | 4 | 0 | 0 |
| ENSDARG000 | 19 | 42678364 | 42678480 | - | 117 | 5S_rRNA | 0 | 0 | 0 | 0 | 0 | 0 |
| ENSDARG000 | 19 | 19728690 | 19728755 | - | 66 | NA | 0 | 0 | 0 | 0 | 0 | 0 |
| ENSDARG000 | 19;19;19;19;1 | 45844363:458 | 45844374:458 | +;+;+;+;+;+;+; | 576 | NA | 0 | 0 | 0 | 2 | 2 | 4 |
| ENSDARG000 | 19;19;19;19;1 | 18812805:188 | 18813100:188 | +;+;+;+;+;+;+; | 2592 | cratb | 381 | 265 | 272 | 232 | 374 | 452 |
| ENSDARG000 | 19;19;19 | 43878049:438 | 43878152:438 | +;+;+ | 502 | pithd1 | 155 | 111 | 150 | 107 | 137 | 106 |
| ENSDARG000 | 19;19;19;19;1 | 206835:20823 | 207002:20847 | +;+;+;+;+;+ | 1854 | scn1bb | 2 | 4 | 4 | 0 | 9 | 4 |
| ENSDARG000 | 19;19;19;19;1 | 14454306:144 | 14454653:144 | +;+;+;+;+;+;+; | 2480 | zdhhc18b | 8 | 6 | 77 | 24 | 14 | 18 |
| ENSDARG000 | 19;19;19;19;1 | 48141623:481 | 48142219:481 | +;+;+;+;+;+;+; | 1740 | btr30 | 0 | 0 | 0 | 0 | 1 | 0 |
| ENSDARG000 | 19;19;19;19 | 11429263:114 | 11429473:114 | -;-;-;- | 614 | hsbp1l1 | 22 | 49 | 80 | 43 | 67 | 48 |
| ENSDARG000 | 19 | 36680091 | 36680207 | + | 117 | 5S_rRNA | 0 | 0 | 0 | 0 | 0 | 0 |
| ENSDARG000 | 19;19;19;19;1 | 1329539:1344 | 1331488:1344 | -;-;-;-;-;-;-;- | 3167 | zgc:63863 | 2769 | 1629 | 905 | 1522 | 544 | 1020 |
| ENSDARG000 | 19 | 16028456 | 16028590 | - | 135 | NA | 0 | 0 | 0 | 0 | 0 | 0 |
| ENSDARG000 | 19;19;19;19;1 | 177455:18336 | 177674:18340 | +;+;+;+;+;+;+ | 639 | tmem65 | 497 | 333 | 400 | 258 | 434 | 302 |
| ENSDARG000 | 19 | 17373632 | 17373686 | + | 55 | NA | 0 | 2 | 5 | 0 | 4 | 0 |
| ENSDARG000 | 19 | 16029308 | 16029443 | - | 136 | NA | 0 | 0 | 0 | 0 | 0 | 0 |
| ENSDARG000 | 19 | 24756843 | 24756974 | - | 132 | NA | 0 | 3 | 0 | 0 | 12 | 2 |
| ENSDARG000 | 19;19;19;19;1 | 3991835:3991 | 3991910:3992 | -;-;-;-;-;-;-;-;- | 1854 | NA | 0 | 0 | 10 | 14 | 6 | 2 |
| ENSDARG000 | 19;19;19 | 432473:43602 | 434552:43648 | -;-;- | 2637 | NA | 114 | 70 | 98 | 135 | 94 | 87 |
| ENSDARG000 | 19;19;19;19;1 | 14903616:149 | 14903641:149 | -;-;-;-;-;-;-;-;- | 372 | NA | 0 | 0 | 0 | 0 | 0 | 0 |
| ENSDARG000 | 19;19;19 | 2857714:2859 | 2858099:2859 | -;-;- | 603 | CLEC3B | 2 | 0 | 18 | 3 | 16 | 6 |
| ENSDARG000 | 19;19;19;19;1 | 167612:16775 | 167664:16779 | +;+;+;+;+;+;+; | 558 | tatdn1 | 209 | 123 | 262 | 171 | 271 | 181 |
| ENSDARG000 | 19 | 46824723 | 46825683 | + | 961 | NA | 0 | 0 | 0 | 0 | 0 | 0 |
| ENSDARG000 | 19 | 38062000 | 38062124 | - | 125 | 5S_rRNA | 0 | 0 | 0 | 0 | 0 | 0 |
| ENSDARG000 | 19 | 16061826 | 16061961 | + | 136 | snrnau11 | 0 | 0 | 0 | 0 | 0 | 0 |
| ENSDARG000 | 19;19;19;19;1 | 2064173:2064 | 2065026:2065 | -;-;-;-;-;-;-;-;- | 3694 | SNX13 | 368 | 255 | 388 | 225 | 331 | 339 |
| ENSDARG000 | 19;19;19;19 | 192741:19348 | 193372:19412 | -;-;-;- | 1972 | NA | 157 | 75 | 116 | 153 | 94 | 84 |
| ENSDARG000 | 19 | 47306623 | 47306738 | + | 116 | 5S_rRNA | 5 | 0 | 6 | 0 | 1 | 0 |
| ENSDARG000 | 19 | 18838411 | 18838541 | - | 131 | NA | 0 | 0 | 0 | 0 | 0 | 0 |
| ENSDARG000 | 19;19;19;19;1 | 10955337:109 | 10957744:109 | -;-;-;-;-;-;-;-;- | 3971 | NA | 0 | 0 | 5 | 12 | 12 | 2 |
| ENSDARG000 | 19;19;19;19;1 | 26828502:268 | 26828543:268 | +;+;+;+;+;+;+; | 1949 | NA | 1009 | 1046 | 1108 | 995 | 1515 | 977 |
| ENSDARG000 | 19 | 12944850 | 12944966 | + | 117 | 5S_rRNA | 0 | 0 | 0 | 0 | 0 | 0 |
| ENSDARG000 | 19;19;19;19 | 1019044:1020 | 1019214:1020 | -;-;-;- | 824 | NA | 151 | 181 | 261 | 75 | 158 | 187 |
| ENSDARG000 | 19 | 39203618 | 39203735 | - | 118 | 5S_rRNA | 0 | 0 | 0 | 0 | 0 | 0 |
| ENSDARG000 | 19;19;19;19;1 | 42071033:420 | 42071909:420 | +;+;+;+;+;+;+; | 4045 | nfyc | 949 | 975 | 1161 | 666 | 1074 | 672 |
| ENSDARG000 | 19;19;19;19;1 | 2408367:2411 | 2408435:2411 | -;-;-;-;-;- | 1451 | NA | 0 | 4 | 0 | 0 | 0 | 0 |
| ENSDARG000 | 19;19;19;19;1 | 48111285:481 | 48111334:481 | +;+;+;+;+;+ | 811 | nme2b.2 | 14 | 43 | 22 | 46 | 36 | 26 |
| ENSDARG000 | 19;19;19;19;1 | 47460748:474 | 47461028:474 | -;-;-;-;-;-;-;- | 2109 | NCDN | 427 | 408 | 594 | 577 | 536 | 418 |
| ENSDARG000 | 19 | 8298994 | 8299110 | + | 117 | 5S_rRNA | 0 | 0 | 0 | 0 | 0 | 0 |
| ENSDARG000 | 19 | 42173114 | 42173228 | - | 115 | 5S_rRNA | 0 | 0 | 0 | 0 | 0 | 0 |
| ENSDARG000 | 19;19;19;19;1 | 43780970:437 | 43780996:437 | +;+;+;+;+;+ | 611 | rpl11 | 11687 | 13156 | 18357 | 17123 | 12092 | 15557 |
| ENSDARG000 | 19;19;19;19;1 | 19596605:196 | 19596625:196 | -;-;-;-;-;-;-;-;- | 348 | NA | 8 | 0 | 0 | 6 | 8 | 13 |
| ENSDARG000 | 19 | 45508865 | 45508981 | + | 117 | 5S_rRNA | 0 | 0 | 0 | 0 | 0 | 0 |
| ENSDARG000 | 19 | 4481183 | 4481297 | - | 115 | 5S_rRNA | 0 | 0 | 0 | 0 | 0 | 0 |
| ENSDARG000 | 19 | 24182775 | 24182891 | - | 117 | 5S_rRNA | 0 | 0 | 0 | 0 | 0 | 0 |
| ENSDARG000 | 19 | 6891821 | 6893275 | + | 1455 | NA | 15 | 2 | 4 | 15 | 4 | 6 |
| ENSDARG000 | 19 | 47400189 | 47400307 | - | 119 | 5S_rRNA | 0 | 0 | 0 | 4 | 0 | 0 |
| ENSDARG000 | 19 | 335627 | 335741 | + | 115 | 5S_rRNA | 0 | 0 | 0 | 0 | 0 | 0 |
| ENSDARG000 | 19 | 34135607 | 34135695 | + | 89 | dre-mir-187 | 0 | 0 | 0 | 0 | 0 | 0 |
| ENSDARG000 | 19 | 21116411 | 21116543 | - | 133 | 5S_rRNA | 0 | 0 | 0 | 0 | 0 | 0 |
| ENSDARG000 | 19 | 7825395 | 7825533 | - | 139 | NA | 0 | 0 | 0 | 0 | 1 | 0 |
| ENSDARG000 | 19;19;19;19;1 | 2018796:2019 | 2018997:2019 | -;-;-;-;-;-;-;-;- | 1698 | NA | 19 | 16 | 66 | 65 | 45 | 33 |
| ENSDARG000 | 19;19 | 1606185:1618 | 1606360:1619 | +;+ | 1098 | lrrc3b | 0 | 0 | 0 | 0 | 0 | 0 |
| ENSDARG000 | 19;19;19;19 | 11432330:114 | 11432532:114 | +;+;+;+ | 762 | SLC66A2 | 250 | 183 | 236 | 138 | 194 | 202 |
| ENSDARG000 | 19;19;19;19;1 | 19412692:194 | 19413269:194 | +;+;+;+;+;+;+; | 1969 | NA | 0 | 0 | 0 | 1 | 0 | 0 |

| | | | | | | | | | | | | |
|---|---|---|---|---|---|---|---|---|---|---|---|---|
| ENSDARG000 | 19:19:19:19:1 | 1003332:1005 | 1003383:1006 | +:+:+:+:+:+:+: | 3724 | zdhhc3b | 626 | 723 | 937 | 636 | 898 | 631 |
| ENSDARG000 | 19 | 37637374 | 37637493 | - | 120 | 5S_rRNA | 0 | 0 | 0 | 0 | 0 | 0 |
| ENSDARG000 | 19 | 36203078 | 36203195 | + | 118 | 5S_rRNA | 0 | 0 | 0 | 0 | 0 | 0 |
| ENSDARG000 | 19:19:19:19:1 | 47783137:477 | 47783500:477 | +:+:+:+:+:+:+: | 854 | c19h1orf109 | 103 | 121 | 202 | 54 | 150 | 165 |
| ENSDARG000 | 19:19:19:19:1 | 1510971:1538 | 1511068:1538 | +;+;+;+;+ | 2379 | SLC45A4 | 28 | 10 | 8 | 4 | 16 | 10 |
| ENSDARG000 | 19 | 4737576 | 4737664 | + | 89 | dre-mir-30a | 0 | 0 | 0 | 0 | 0 | 0 |
| ENSDARG000 | 19:19:19:19:1 | 47771659:477 | 47774050:477 | -:-:-:-:-:-:-:-: | 3272 | cdca8 | 216 | 644 | 680 | 338 | 495 | 302 |
| ENSDARG000 | 19:19:19:19:1 | 2752933:2788 | 2753873:2792 | -:-:-:-:-:-:-:-: | 9739 | NA | 28 | 190 | 217 | 62 | 156 | 78 |
| ENSDARG000 | 19:19:19:19:1 | 17640090:176 | 17640144:176 | +:+:+:+:+:+:+: | 2156 | NA | 0 | 2 | 0 | 4 | 0 | 4 |
| ENSDARG000 | 19;19 | 47522007:475 | 47523238:475 | -;- | 1296 | NA | 0 | 0 | 0 | 0 | 0 | 0 |
| ENSDARG000 | 19 | 4204245 | 4204359 | - | 115 | 5S_rRNA | 0 | 0 | 0 | 0 | 0 | 0 |
| ENSDARG000 | 19 | 32649667 | 32649781 | - | 115 | 5S_rRNA | 0 | 0 | 0 | 0 | 0 | 0 |
| ENSDARG000 | 19:19:19:19:1 | 1393360:1394 | 1393474:1394 | -;-;-;-;-;- | 737 | cfap20 | 259 | 226 | 317 | 332 | 250 | 198 |
| ENSDARG000 | 19 | 24139780 | 24139894 | - | 115 | 5S_rRNA | 0 | 0 | 0 | 0 | 0 | 0 |
| ENSDARG000 | 19 | 7484217 | 7484331 | + | 115 | 5S_rRNA | 0 | 0 | 0 | 0 | 0 | 0 |
| ENSDARG000 | 19:19:19:19:1 | 917852:91975 | 918012:91996 | +:+:+:+:+:+:+: | 1938 | TGFBR2 | 490 | 214 | 169 | 133 | 136 | 270 |
| ENSDARG000 | 19;19;19 | 26937543:269 | 26938523:269 | -;-;- | 1469 | NA | 0 | 2 | 0 | 2 | 4 | 8 |
| ENSDARG000 | 19 | 35420835 | 35420951 | - | 117 | 5S_rRNA | 0 | 0 | 0 | 1 | 0 | 0 |
| ENSDARG000 | 19:19:19:19:1 | 981321:98279 | 981495:98296 | -:-:-:-:-:-:-:-: | 3504 | ehmt2 | 313 | 367 | 558 | 391 | 480 | 302 |
| ENSDARG000 | 19 | 5640504 | 5641825 | + | 1322 | ft2 | 2 | 2 | 2 | 4 | 4 | 14 |
| ENSDARG000 | 19 | 40958500 | 40958616 | + | 117 | 5S_rRNA | 0 | 0 | 0 | 0 | 0 | 0 |
| ENSDARG000 | 19:19:19:19:1 | 18143593:181 | 18144328:181 | -:-:-:-:-:-:-:-: | 4555 | nfe2l3 | 696 | 747 | 935 | 352 | 802 | 561 |
| ENSDARG000 | 19:19:19:19:1 | 47800550:478 | 47800820:478 | -:-:-:-:-:-:-:-: | 2877 | ago4 | 770 | 335 | 763 | 362 | 813 | 531 |
| ENSDARG000 | 19:19:19:19:1 | 21818460:218 | 21818472:218 | +:+:+:+:+:+:+: | 300 | NA | 0 | 0 | 0 | 0 | 0 | 0 |
| ENSDARG000 | 19:19:19:19:1 | 860830:86083 | 861996:86199 | -:-:-:-:-:-:-:-: | 2875 | eomesa | 15 | 41 | 6 | 22 | 56 | 36 |
| ENSDARG000 | 19:19:19:19:1 | 1414604:1416 | 1414672:1416 | +;+;+;+;+;+;+ | 2239 | btr29 | 268 | 240 | 80 | 48 | 73 | 74 |
| ENSDARG000 | 19 | 38464127 | 38464249 | - | 123 | 5S_rRNA | 2 | 0 | 0 | 0 | 0 | 0 |
| ENSDARG000 | 19 | 38948572 | 38948686 | + | 115 | 5S_rRNA | 0 | 0 | 0 | 0 | 0 | 0 |
| ENSDARG000 | 19 | 42915419 | 42915541 | + | 123 | dre-mir-128-2 | 0 | 0 | 0 | 0 | 0 | 0 |
| ENSDARG000 | 19 | 48033299 | 48033440 | + | 142 | NA | 10 | 1 | 5 | 2 | 4 | 5 |
| ENSDARG000 | 19;19 | 48102560:481 | 48102694:481 | +;+ | 312 | NA | 48 | 57 | 38 | 18 | 78 | 36 |
| ENSDARG000 | 19 | 39874831 | 39874947 | + | 117 | 5S_rRNA | 0 | 0 | 0 | 0 | 0 | 0 |
| ENSDARG000 | 19;19;19;19 | 497953:49889 | 498613:49920 | +;+;+;+ | 3759 | NA | 80 | 64 | 98 | 54 | 76 | 79 |
| ENSDARG000 | 19:19:19:19:1 | 1135052:1135 | 1135310:1135 | -;-;-;-;- | 1558 | NA | 0 | 0 | 0 | 0 | 0 | 0 |
| ENSDARG000 | 19;19;19 | 9381450:9407 | 9381678:9407 | +;+;+ | 474 | NA | 2 | 0 | 0 | 0 | 1 | 0 |
| ENSDARG000 | 19;19;19;19 | 13364536:133 | 13364842:133 | -;-;-;- | 1159 | NA | 0 | 0 | 0 | 0 | 0 | 0 |
| ENSDARG000 | 19;19;19 | 16907709:169 | 16908136:169 | +;+;+ | 608 | NA | 0 | 0 | 0 | 0 | 0 | 0 |
| ENSDARG000 | 19;19;19 | 18314328:183 | 18314686:183 | -;-;- | 578 | NA | 0 | 0 | 0 | 3 | 0 | 0 |
| ENSDARG000 | 19;19;19;19 | 18942726:189 | 18943961:189 | -;-;-;- | 2304 | NA | 4 | 2 | 11 | 0 | 40 | 0 |
| ENSDARG000 | 19:19:19:19:1 | 21176257:211 | 21177308:211 | -;-;-;-;-;-;- | 2123 | NA | 0 | 0 | 0 | 0 | 0 | 0 |
| ENSDARG000 | 19:19:19:19:1 | 24077935:240 | 24078090:240 | +;+;+;+;+;+;+ | 572 | NA | 0 | 0 | 0 | 0 | 0 | 0 |
| ENSDARG000 | 19;19;19 | 24569365:245 | 24569964:245 | +;+;+ | 1857 | NA | 0 | 2 | 0 | 0 | 0 | 0 |
| ENSDARG000 | 19:19:19:19:1 | 25084265:250 | 25085379:250 | -;-;-;-;- | 1676 | NA | 0 | 2 | 0 | 0 | 0 | 0 |
| ENSDARG000 | 19:19:19:19:1 | 28803429:288 | 28803615:288 | +;+;+;+;+;+ | 4147 | NA | 4 | 4 | 0 | 7 | 0 | 2 |
| ENSDARG000 | 19;19;19;19 | 30147370:302 | 30147596:302 | -;-;-;- | 626 | NA | 0 | 0 | 0 | 0 | 0 | 0 |
| ENSDARG000 | 19:19:19:19:1 | 30587866:305 | 30588194:305 | +:+:+:+:+:+:+: | 862 | NA | 4 | 0 | 0 | 2 | 4 | 0 |
| ENSDARG000 | 19;19;19 | 30962723:309 | 30962917:309 | -;-;- | 980 | NA | 0 | 0 | 0 | 0 | 0 | 0 |
| ENSDARG000 | 19;19;19 | 38947629:389 | 38948454:389 | +;+;+ | 1160 | NA | 0 | 0 | 11 | 0 | 0 | 0 |
| ENSDARG000 | 19:19:19:19:1 | 39427180:394 | 39427401:394 | -;-;-;-;-;-;-;- | 963 | NA | 0 | 0 | 0 | 0 | 0 | 0 |
| ENSDARG000 | 19;19;19;19 | 45157502:452 | 45157621:452 | -;-;-;- | 513 | NA | 0 | 0 | 0 | 0 | 0 | 0 |
| ENSDARG000 | 19:19:19:19:1 | 47838603:478 | 47839333:478 | +;+;+;+;+;+ | 2453 | NA | 24 | 102 | 127 | 10 | 86 | 36 |
| ENSDARG000 | 19:19:19:19:1 | 47909148:479 | 47909550:479 | +;+;+;+;+;+;+ | 2279 | NA | 51 | 204 | 251 | 72 | 223 | 128 |
| ENSDARG000 | 15:15:15:15:1 | 16122984:161 | 16124245:161 | -:-:-:-:-:-:-:-: | 8092 | BLTP2 | 1400 | 1029 | 1503 | 1072 | 1489 | 1222 |
| ENSDARG000 | 15:15:15:15:1 | 20830146:208 | 20831123:208 | -:-:-:-:-:-:-:-: | 4704 | aldh3a2a | 234 | 407 | 380 | 258 | 480 | 337 |
| ENSDARG000 | 15:15:15:15:1 | 45994123:460 | 45994238:460 | +:+:+:+:+:+:+: | 2061 | lrfn1 | 6 | 2 | 0 | 4 | 10 | 6 |
| ENSDARG000 | 15;15 | 46198434:462 | 46198626:462 | +;+ | 328 | NA | 0 | 0 | 0 | 0 | 1 | 0 |
| ENSDARG000 | 15;15 | 46313082:463 | 46313402:463 | +;+ | 731 | NA | 207 | 71 | 373 | 64 | 89 | 104 |
| ENSDARG000 | 15;15 | 8314932:8319 | 8316313:8319 | -;- | 1815 | NA | 130 | 164 | 118 | 64 | 158 | 88 |
| ENSDARG000 | 15:15:15:15:1 | 8335600:8335 | 8335936:8335 | +:+:+:+:+:+:+: | 3457 | EGLN2 | 289 | 290 | 851 | 597 | 569 | 118 |
| ENSDARG000 | 15:15:15:15:1 | 46329149:463 | 46329981:463 | +;+;+;+;+;+;+ | 2068 | si:ch1073-340 | 47 | 8 | 4 | 0 | 22 | 0 |
| ENSDARG000 | 15:15:15:15:1 | 18789111:187 | 18789179:187 | +;+;+;+;+;+ | 1833 | NA | 32 | 16 | 8 | 43 | 39 | 59 |
| ENSDARG000 | 15:15:15:15:1 | 27303279:273 | 27303587:273 | -:-:-:-:-:-:-:-: | 2387 | NA | 36 | 8 | 41 | 10 | 12 | 2 |
| ENSDARG000 | 15 | 31435957 | 31438111 | + | 2155 | or30bt2 | 0 | 0 | 0 | 0 | 0 | 0 |
| ENSDARG000 | 15;15;15;15 | 8394819:8395 | 8395098:8395 | +;+;+;+ | 1220 | tmem160 | 100 | 112 | 98 | 26 | 144 | 44 |
| ENSDARG000 | 15:15:15:15:1 | 2475894:2479 | 2476330:2480 | +;+;+;+;+;+ | 1652 | panx1a | 423 | 493 | 392 | 269 | 458 | 468 |
| ENSDARG000 | 15:15:15:15:1 | 46344655:463 | 46344860:463 | +;+;+;+;+;+ | 1319 | NA | 0 | 4 | 0 | 0 | 4 | 8 |
| ENSDARG000 | 15:15:15:15:1 | 46356879:463 | 46356966:463 | +:+:+:+:+:+:+: | 1775 | NA | 24 | 0 | 9 | 0 | 2 | 0 |
| ENSDARG000 | 15:15:15:15:1 | 8424711:8425 | 8427378:8427 | -:-:-:-:-:-:-:-: | 5128 | npas1 | 0 | 2 | 0 | 0 | 0 | 0 |
| ENSDARG000 | 15;15;15;15 | 31438568:314 | 31438601:314 | +;+;+;+ | 1209 | or30bt4 | 0 | 0 | 0 | 0 | 0 | 0 |
| ENSDARG000 | 15;15 | 31438568:314 | 31438601:314 | +;+ | 1026 | or30bt5 | 0 | 0 | 0 | 0 | 0 | 0 |
| ENSDARG000 | 15;15;15 | 26992109:269 | 26992412:269 | -;-;- | 747 | NA | 0 | 0 | 2 | 0 | 0 | 0 |
| ENSDARG000 | 15;15;15;15 | 27296676:272 | 27296716:272 | +;+;+;+ | 836 | NA | 0 | 2 | 0 | 5 | 0 | 0 |
| ENSDARG000 | 15:15:15:15:1 | 26940569:269 | 26941131:269 | +:+:+:+:+:+:+: | 5703 | bcas3 | 821 | 384 | 554 | 532 | 533 | 764 |
| ENSDARG000 | 15:15:15:15:1 | 31454930:314 | 31455093:314 | +:+:+:+:+:+:+: | 1495 | NA | 0 | 0 | 0 | 0 | 0 | 0 |
| ENSDARG000 | 15;15 | 46695492:467 | 46695739:467 | +;+ | 584 | NA | 0 | 0 | 0 | 0 | 0 | 0 |
| ENSDARG000 | 15:15:15:15:1 | 46736794:467 | 46737145:467 | -:-:-:-:-:-:-:-: | 2755 | clcn2c | 2 | 2 | 8 | 6 | 0 | 0 |
| ENSDARG000 | 15:15:15:15:1 | 18863872:188 | 18864021:188 | +:+:+:+:+:+:+: | 2417 | cadm1b | 0 | 0 | 0 | 0 | 2 | 0 |
| ENSDARG000 | 15:15:15:15:1 | 46685434:466 | 46685624:466 | -:-:-:-:-:-:-:-: | 2269 | zgc:162872 | 151 | 153 | 195 | 122 | 163 | 119 |
| ENSDARG000 | 15:15:15:15:1 | 29123031:291 | 29123170:291 | +:+:+:+:+:+:+: | 1700 | zgc:101731 | 0 | 4 | 2 | 0 | 0 | 8 |
| ENSDARG000 | 15:15:15:15:1 | 24563504:245 | 24563864:245 | +:+:+:+:+:+:+: | 1572 | dhrs13b.1 | 204 | 131 | 120 | 140 | 120 | 138 |
| ENSDARG000 | 15:15:15:15:1 | 46722510:467 | 46722704:467 | -:-:-:-:-:-:-:-: | 2037 | ilkap | 172 | 105 | 177 | 70 | 149 | 198 |
| ENSDARG000 | 15:15:15:15:1 | 2490204:2490 | 2491069:2491 | -;-;-;-;-;-;- | 1411 | neu4 | 4 | 14 | 0 | 4 | 10 | 26 |
| ENSDARG000 | 15;15 | 46744442:467 | 46744919:467 | -;- | 717 | NA | 0 | 0 | 0 | 0 | 0 | 0 |
| ENSDARG000 | 15;15;15 | 5628363:5628 | 5628641:5628 | +;+;+ | 555 | NA | 0 | 0 | 0 | 0 | 0 | 0 |
| ENSDARG000 | 15;15;15;15 | 31462664:314 | 31462780:314 | +;+;+;+ | 1385 | or30bx1 | 0 | 0 | 0 | 0 | 0 | 0 |
| ENSDARG000 | 15:15:15:15:1 | 5627877:5627 | 5632838:5632 | -:-:-:-:-:-:-:-: | 11664 | urb1 | 392 | 344 | 884 | 246 | 603 | 555 |
| ENSDARG000 | 15 | 31466978 | 31470101 | + | 3124 | or30bw1 | 0 | 0 | 0 | 0 | 0 | 0 |
| ENSDARG000 | 15 | 31470462 | 31470528 | + | 67 | NA | 0 | 0 | 0 | 0 | 0 | 0 |
| ENSDARG000 | 15:15:15:15:1 | 8604477:8604 | 8607684:8607 | -:-:-:-:-:-:-:-: | 8525 | ARHGAP35 | 788 | 865 | 1056 | 630 | 958 | 925 |
| ENSDARG000 | 15;15 | 8762580:8764 | 8762996:8765 | +;+ | 1214 | NA | 0 | 0 | 0 | 0 | 0 | 0 |
| ENSDARG000 | 15 | 31471808 | 31474129 | + | 2322 | or30by3 | 2 | 0 | 0 | 0 | 0 | 0 |
| ENSDARG000 | 15;15;15 | 31476727:314 | 31477915:314 | +;+;+ | 1189 | NA | 0 | 0 | 0 | 0 | 0 | 0 |
| ENSDARG000 | 15:15:15:15:1 | 2498496:2500 | 2498604:2500 | +;+;+;+;+ | 6052 | WDR53 | 219 | 174 | 82 | 144 | 312 | 242 |
| ENSDARG000 | 15 | 31479657 | 31480566 | + | 910 | NA | 0 | 0 | 0 | 0 | 0 | 2 |
| ENSDARG000 | 15:15:15:15:1 | 29140126:291 | 29140224:291 | +;+;+;+;+;+;+ | 2478 | zgc:113149 | 310 | 189 | 269 | 196 | 254 | 279 |
| ENSDARG000 | 15 | 31481939 | 31482892 | + | 954 | or30by1 | 0 | 0 | 0 | 0 | 0 | 2 |
| ENSDARG000 | 15:15:15:15:1 | 31485877:314 | 31486937:314 | -:-:-:-:-:-:-:-: | 5854 | NA | 18 | 4 | 18 | 2 | 14 | 2 |
| ENSDARG000 | 15:15:15:15:1 | 2510761:2511 | 2511248:2511 | -;-;-;-;-;-;-;- | 1342 | srprb | 845 | 1278 | 2169 | 745 | 1399 | 1179 |
| ENSDARG000 | 15:15:15:15:1 | 21875545:218 | 21876600:218 | -:-:-:-:-:-:-:-: | 1539 | zgc:162608 | 2 | 0 | 6 | 14 | 4 | 0 |
| ENSDARG000 | 15:15:15:15:1 | 31500785:315 | 31500950:315 | -;-;-;-;-;- | 2029 | NA | 4 | 0 | 3 | 0 | 10 | 6 |
| ENSDARG000 | 15:15:15:15:1 | 21862295:218 | 21862497:218 | +:+:+:+:+:+:+: | 1868 | NA | 2 | 0 | 12 | 0 | 0 | 0 |
| ENSDARG000 | 15;15 | 19094511:190 | 19094637:190 | -;- | 317 | NA | 0 | 0 | 0 | 0 | 0 | 1 |
| ENSDARG000 | 15:15:15:15:1 | 19036377:190 | 19040388:190 | -:-:-:-:-:-:-:-: | 7061 | NA | 0 | 0 | 7 | 0 | 2 | 0 |
| ENSDARG000 | 15:15:15:15:1 | 20843161:208 | 20844245:208 | +:+:+:+:+:+:+: | 5992 | ULK2 | 723 | 314 | 603 | 350 | 1371 | 636 |
| ENSDARG000 | 15:15:15:15:1 | 16178775:161 | 16179086:161 | -;-;-;-;-;-;-;- | 1237 | glod4 | 255 | 388 | 244 | 202 | 318 | 262 |
| ENSDARG000 | 15:15:15:15:1 | 31515976:315 | 31516094:315 | +;+;+;+;+;+;+ | 1673 | NA | 0 | 0 | 3 | 0 | 3 | 4 |
| ENSDARG000 | 15:15:15:15:1 | 31513065:315 | 31513644:315 | -:-:-:-:-:-:-:-: | 1724 | hmgb1b | 1116 | 1788 | 2693 | 1160 | 1788 | 1194 |
| ENSDARG000 | 15:15:15:15:1 | 29154547:291 | 29154864:291 | -:-:-:-:-:-:-:-: | 1609 | xaf1 | 695 | 434 | 221 | 175 | 110 | 267 |
| ENSDARG000 | 15:15:15:15:1 | 29161656:291 | 29161879:291 | +;+;+;+;+ | 537 | NA | 0 | 0 | 0 | 0 | 0 | 0 |
| ENSDARG000 | 15;15 | 24583682:245 | 24586937:245 | -;- | 3430 | gemin4 | 2 | 0 | 9 | 0 | 0 | 2 |
| ENSDARG000 | 15:15:15:15:1 | 24592325:245 | 24592821:245 | -;-;-;-;- | 912 | NA | 0 | 0 | 0 | 0 | 1 | 0 |
| ENSDARG000 | 15:15:15:15:1 | 24572926:245 | 24573055:245 | +:+:+:+:+:+:+: | 11637 | NA | 614 | 434 | 191 | 351 | 506 | 289 |
| ENSDARG000 | 15:15:15:15:1 | 14203675:142 | 14206845:142 | -:-:-:-:-:-:-:-: | 8489 | zgc:162193 | 73 | 622 | 394 | 56 | 666 | 268 |
| ENSDARG000 | 15:15:15:15:1 | 24588963:245 | 24589114:245 | +:+:+:+:+:+:+: | 2588 | dhrs13b.2 | 86 | 76 | 43 | 16 | 37 | 65 |
| ENSDARG000 | 15:15:15:15:1 | 8767650:8771 | 8767813:8771 | +;+;+;+;+ | 2805 | ap2s1 | 3456 | 3800 | 4059 | 3627 | 4561 | 3008 |
| ENSDARG000 | 15;15;15 | 14202114:142 | 14202435:142 | +;+;+ | 1203 | NA | 0 | 0 | 0 | 0 | 0 | 0 |
| ENSDARG000 | 15:15:15:15:1 | 2520319:2522 | 2520417:2522 | +:+:+:+:+:+:+: | 2494 | cux1b | 794 | 653 | 732 | 533 | 966 | 512 |
| ENSDARG000 | 15:15:15:15:1 | 21882419:218 | 21882477:218 | +:+:+:+:+:+:+: | 5243 | NA | 252 | 205 | 54 | 326 | 32 | 151 |
| ENSDARG000 | 15;15 | 21900980:219 | 21901669:219 | -;- | 799 | NA | 0 | 0 | 0 | 0 | 0 | 0 |
| ENSDARG000 | 15;15;15;15 | 16185118:161 | 16185513:161 | +;+;+;+ | 1370 | mrm3a | 42 | 37 | 55 | 28 | 60 | 63 |
| ENSDARG000 | 15:15:15:15:1 | 16193809:162 | 16194937:162 | -;-;-;-;-;-;- | 2478 | nxn | 470 | 414 | 659 | 450 | 628 | 399 |
| ENSDARG000 | 15:15:15:15:1 | 19131702:191 | 19136107:191 | -:-:-:-:-:-:-:-: | 9744 | NA | 588 | 85 | 16 | 250 | 48 | 66 |
| ENSDARG000 | 15:15:15:15:1 | 21947117:219 | 21947232:219 | +:+:+:+:+:+:+: | 3534 | ttc12 | 10 | 14 | 8 | 39 | 8 | 2 |
| ENSDARG000 | 15:15:15:15:1 | 14246103:142 | 14246817:142 | -;-;-;-;-;- | 1218 | NA | 0 | 2 | 0 | 0 | 0 | 0 |
| ENSDARG000 | 15;15;15;15 | 46801341:468 | 46801352:468 | +;+;+;+ | 245 | NA | 0 | 0 | 0 | 0 | 0 | 0 |
| ENSDARG000 | 15:15:15:15:1 | 46790676:467 | 46792270:467 | -:-:-:-:-:-:-:-: | 4558 | chrd | 17 | 25 | 18 | 16 | 7 | 9 |
| ENSDARG000 | 15:15:15:15:1 | 8813755:8813 | 8815846:8816 | -:-:-:-:-:-:-:-: | 4645 | rab4b | 488 | 313 | 315 | 270 | 418 | 293 |
| ENSDARG000 | 15;15;15;15 | 16328450:163 | 16328846:163 | -;-;-;- | 830 | txndc17 | 529 | 708 | 670 | 394 | 730 | 470 |
| ENSDARG000 | 15;15;15;15 | 46799469:467 | 46799682:467 | +;+;+;+ | 436 | NA | 0 | 0 | 0 | 2 | 0 | 0 |
| ENSDARG000 | 15:15:15:15:1 | 31526225:315 | 31526267:315 | +:+:+:+:+:+:+: | 3119 | NA | 0 | 0 | 1 | 0 | 6 | 0 |
| ENSDARG000 | 15:15:15:15:1 | 29276508:292 | 29276681:292 | +:+:+:+:+:+:+: | 3393 | rap1gap2a | 18 | 2 | 25 | 0 | 10 | 12 |
| ENSDARG000 | 15;15;15 | 20897103:208 | 20897172:208 | +;+;+ | 791 | NA | 0 | 0 | 0 | 0 | 0 | 0 |
| ENSDARG000 | 15:15:15:15:1 | 20900627:209 | 20901811:209 | -:-:-:-:-:-:-:-: | 3734 | usp2a | 131 | 122 | 129 | 105 | 130 | 80 |
| ENSDARG000 | 15;15 | 47418565:474 | 47418674:474 | +;+ | 3033 | NA | 170 | 96 | 265 | 224 | 177 | 130 |
| ENSDARG000 | 15:15:15:15:1 | 16343073:163 | 16343247:163 | +:+:+:+:+:+:+: | 1217 | p2rx8 | 0 | 0 | 0 | 18 | 0 | 0 |
| ENSDARG000 | 15;15;15 | 47440477:474 | 47440798:474 | +;+;+ | 3891 | phox2a | 0 | 0 | 4 | 0 | 0 | 4 |
| ENSDARG000 | 15:15:15:15:1 | 22014029:220 | 22014242:220 | +:+:+:+:+:+:+: | 2448 | ankk1 | 2 | 2 | 0 | 0 | 6 | 0 |
| ENSDARG000 | 15:15:15:15:1 | 14257232:142 | 14257319:142 | -:-:-:-:-:-:-:-: | 1788 | dpf1 | 34 | 73 | 89 | 16 | 58 | 29 |
| ENSDARG000 | 15:15:15:15:1 | 29322203:293 | 29322554:293 | -;-;-;-;-;- | 1380 | NA | 18 | 6 | 8 | 25 | 14 | 10 |
| ENSDARG000 | 15;15;15 | 8990577:8998 | 8994195:8998 | -;-;- | 4077 | rhoub | 34 | 38 | 18 | 444 | 44 | 14 |
| ENSDARG000 | 15:15:15:15:1 | 22046557:220 | 22047616:220 | -:-:-:-:-:-:-:-: | 2613 | drd2a | 105 | 80 | 43 | 0 | 56 | 22 |
| ENSDARG000 | 15:15:15:15:1 | 47452630:474 | 47454605:474 | -:-:-:-:-:-:-:-: | 4895 | inppl1a | 2206 | 2214 | 2905 | 1702 | 2562 | 2192 |
| ENSDARG000 | 15:15:15:15:1 | 24644016:246 | 24644086:246 | +:+:+:+:+:+:+: | 2159 | smtnl | 316 | 120 | 418 | 14 | 184 | 173 |
| ENSDARG000 | 15;15;15;15 | 29343280:293 | 29345446:293 | -;-;-;- | 2498 | tsku | 288 | 170 | 237 | 262 | 253 | 238 |
| ENSDARG000 | 15;15;15 | 1017437:1020 | 1018353:1020 | -;-;- | 1085 | NA | 126 | 74 | 127 | 35 | 92 | 102 |
| ENSDARG000 | 15;15;15 | 16377263:163 | 16382097:163 | -;-;- | 5186 | FAM222B | 570 | 417 | 558 | 316 | 821 | 465 |
| ENSDARG000 | 15:15:15:15:1 | 9010891:9011 | 9011272:9011 | -;-;-;-;-;- | 981 | rtn2a | 0 | 0 | 0 | 0 | 0 | 0 |
| ENSDARG000 | 15:15:15:15:1 | 16387088:163 | 16387315:163 | +:+:+:+:+:+:+: | 3137 | flot2b | 889 | 1260 | 1516 | 1635 | 847 | 981 |
| ENSDARG000 | 15;15;15 | 1025308:1029 | 1026507:1029 | -;-;- | 1374 | NA | 52 | 53 | 53 | 20 | 58 | 93 |
| ENSDARG000 | 15;15;15 | 2589949:2592 | 2591080:2593 | -;-;- | 1682 | dnajc30b | 64 | 105 | 110 | 24 | 148 | 84 |
| ENSDARG000 | 15:15:15:15:1 | 22133839:221 | 22134065:221 | -;-;-;-;-;-;-;- | 1310 | scn3b | 43 | 50 | 16 | 23 | 38 | 110 |
| ENSDARG000 | 15:15:15:15:1 | 31564137:315 | 31564956:315 | -:-:-:-:-:-:-:-: | 3735 | HSPH1 | 244 | 222 | 353 | 634 | 233 | 314 |
| ENSDARG000 | 15:15:15:15:1 | 24673283:246 | 24673824:246 | -:-:-:-:-:-:-:-: | 882 | tmem199 | 255 | 265 | 277 | 294 | 312 | 208 |
| ENSDARG000 | 15:15:15:15:1 | 29362714:293 | 29363494:293 | +:+:+:+:+:+:+: | 6241 | gucy2f | 0 | 2 | 6 | 32 | 8 | 0 |
| ENSDARG000 | 15:15:15:15:1 | 1033243:1033 | 1034119:1034 | -;-;-;-;- | 1215 | si:dkev-77f5.3 | 146 | 144 | 188 | 130 | 188 | 172 |
| ENSDARG000 | 15;15;15 | 2599146:2606 | 2604282:2607 | -;-;- | 6218 | NA | 0 | 0 | 2 | 0 | 4 | 0 |
| ENSDARG000 | 15:15:15:15:1 | 9053059:9057 | 9053838:9057 | +;+;+;+;+;+ | 3396 | ppm1na | 0 | 0 | 0 | 0 | 0 | 0 |
| ENSDARG000 | 15 | 2625028 | 2625663 | - | 636 | NA | 13 | 6 | 22 | 16 | 20 | 4 |
| ENSDARG000 | 15;15;15 | 19284842:192 | 19285063:192 | +;+;+ | 606 | NA | 2 | 0 | 1 | 14 | 0 | 0 |
| ENSDARG000 | 15 | 2631421 | 2632891 | - | 1471 | cldnj | 3 | 2 | 0 | 0 | 0 | 0 |
| ENSDARG000 | 15:15:15:15:1 | 1045374:1047 | 1046218:1048 | -;-;-;-;- | 1794 | AADAC | 1005 | 768 | 257 | 566 | 363 | 2299 |
| ENSDARG000 | 15 | 2638884 | 2640966 | - | 2083 | cldne | 382 | 4 | 0 | 64 | 20 | 12 |

| | | | | | | | | | | | | |
|---|---|---|---|---|---|---|---|---|---|---|---|---|
| ENSDARG000 | 15;15;15;15 | 47721873:47 | 47722121:47 | +;+;+;+ | 1651 | NA | 2 | 0 | 0 | 0 | 0 | 8 |
| ENSDARG000 | 15;15;15;15 | 27356825:27 | 27357101:27 | -;-;-;- | 904 | NA | 130 | 52 | 129 | 81 | 53 | 197 |
| ENSDARG000 | 15;15;15;15;1 | 5722525:572 | 5724186:572 | -;-;-;-;-;-;-;-; | 2753 | phkg1a | 80 | 58 | 68 | 30 | 76 | 44 |
| ENSDARG000 | 15 | 2650787 | 2652640 | - | 1854 | cldnf | 18 | 0 | 0 | 1 | 6 | 4 |
| ENSDARG000 | 15;15;15;15;1 | 47746176:47 | 47746675:47 | +;+;+;+;+;+;+ | 1539 | stard10 | 310 | 128 | 568 | 986 | 104 | 108 |
| ENSDARG000 | 15;15;15;15;1 | 19287632:19 | 19287682:19 | +;+;+;+;+ | 920 | NA | 86 | 58 | 88 | 53 | 120 | 181 |
| ENSDARG000 | 15;15;15;15;1 | 47850875:47 | 47852571:47 | -;-;-;-;-;-;-;- | 2088 | si:ch1073-42 | 6161 | 4658 | 5648 | 4696 | 6738 | 5554 |
| ENSDARG000 | 15;15;15;15;1 | 47828637:47 | 47834672:47 | -;-;-;-;-;- | 6576 | KBTBD3 | 2 | 33 | 14 | 2 | 32 | 6 |
| ENSDARG000 | 15;15;15;15;1 | 24676905:24 | 24677323:24 | +;+;+;+;+;+;+; | 2329 | poldip2 | 518 | 529 | 691 | 286 | 864 | 493 |
| ENSDARG000 | 15;15;15;15;1 | 47867133:47 | 47867374:47 | -;-;-;-;-;-;-;-; | 1832 | NA | 14 | 15 | 43 | 34 | 252 | 110 |
| ENSDARG000 | 15 | 2655566 | 2657508 | - | 1943 | cldna | 776 | 36 | 0 | 32 | 52 | 32 |
| ENSDARG000 | 15;15;15;15;1 | 47845369:47 | 47845445:47 | -;-;-;-;-;-;-;- | 801 | eif3k | 1536 | 1390 | 2484 | 1932 | 1690 | 1785 |
| ENSDARG000 | 15;15;15;15;1 | 5773119:577 | 5773427:577 | -;-;-;-;-;-;- | 961 | chchd2 | 1091 | 1243 | 1603 | 1015 | 1074 | 746 |
| ENSDARG000 | 15;15;15;15;1 | 22164549:22 | 22164800:22 | -;-;-;-;-;-;-;-; | 13203 | NA | 0 | 0 | 0 | 0 | 28 | 1 |
| ENSDARG000 | 15;15;15;15 | 47860045:47 | 47860760:47 | -;-;-;- | 1089 | h3f3b.1 | 829 | 1124 | 1548 | 833 | 990 | 840 |
| ENSDARG000 | 15;15;15;15;1 | 1086656:108 | 1086845:108 | +;+;+;+;+;+;+; | 11599 | IGSF10 | 18 | 4 | 2 | 0 | 14 | 0 |
| ENSDARG000 | 15;15 | 47828719:47 | 47828830:47 | +;+ | 1194 | NA | 0 | 0 | 0 | 0 | 0 | 0 |
| ENSDARG000 | 15;15;15;15;1 | 31590224:31 | 31590334:31 | +;+;+;+;+;+;+; | 1876 | b3glcta | 189 | 70 | 412 | 182 | 204 | 139 |
| ENSDARG000 | 15;15;15;15;1 | 19293744:19 | 19294008:19 | +;+;+;+;+;+;+; | 6308 | JAM3 | 36 | 32 | 36 | 38 | 4 | 24 |
| ENSDARG000 | 15;15;15;15;1 | 16521785:16 | 16522389:16 | +;+;+;+;+;+;+; | 7042 | GALNT17 | 6 | 0 | 0 | 0 | 0 | 2 |
| ENSDARG000 | 15;15;15;15;1 | 29387910:29 | 29388027:29 | +;+;+;+;+;+;+; | 6529 | GDPD5 | 68 | 24 | 110 | 360 | 154 | 178 |
| ENSDARG000 | 15 | 2673935 | 2674590 | + | 656 | NA | 6 | 0 | 4 | 0 | 0 | 4 |
| ENSDARG000 | 15;15;15;15;1 | 27364394:27 | 27365168:27 | +;+;+;+;+;+;+ | 2616 | tbx2b | 3802 | 3017 | 2088 | 1351 | 1184 | 3327 |
| ENSDARG000 | 15;15;15;15;1 | 22311803:22 | 22311977:22 | +;+;+;+;+;+;+ | 2316 | hepacama | 2 | 0 | 0 | 0 | 6 | 4 |
| ENSDARG000 | 15;15;15;15 | 1116664:111 | 1116711:111 | +;+;+;+ | 1439 | p2ry12 | 0 | 2 | 2 | 0 | 6 | 26 |
| ENSDARG000 | 15;15;15;15;1 | 29382454:29 | 29384101:29 | -;-;-;-;-;-;-;-; | 4212 | serpinh1b | 933 | 1655 | 5985 | 4677 | 5761 | 1497 |
| ENSDARG000 | 15;15;15;15;1 | 1134870:113 | 1134930:113 | +;+;+;+;+ | 2906 | NA | 2217 | 14 | 225 | 1528 | 192 | 12 |
| ENSDARG000 | 15 | 2685055 | 2685706 | + | 652 | NA | 111 | 0 | 0 | 6 | 0 | 2 |
| ENSDARG000 | 15;15;15;15;1 | 9072821:907 | 9072944:907 | +;+;+;+;+;+;+; | 2070 | NA | 0 | 0 | 0 | 0 | 0 | 0 |
| ENSDARG000 | 15;15;15;15;1 | 27384798:27 | 27384954:27 | +;+;+;+;+;+;+; | 2878 | tbx4 | 0 | 0 | 0 | 3 | 0 | 10 |
| ENSDARG000 | 15;15;15;15;1 | 31735931:31 | 31736017:31 | +;+;+;+;+;+;+; | 2640 | rxfp2b | 2 | 0 | 0 | 0 | 0 | 2 |
| ENSDARG000 | 15;15;15;15;1 | 2700646:270 | 2701100:270 | -;-;-;-;-;-;-;-;- | 1913 | NA | 12 | 2 | 16 | 6 | 10 | 6 |
| ENSDARG000 | 15;15;15;15;1 | 1148074:114 | 1148143:114 | +;+;+;+;+;+;+; | 1507 | mlf1 | 121 | 62 | 71 | 166 | 29 | 19 |
| ENSDARG000 | 15;15;15;15;1 | 2727670:272 | 2731030:272 | -;-;-;-;-;-;-;-; | 4832 | ppp5c | 1876 | 1857 | 2201 | 1190 | 2080 | 1785 |
| ENSDARG000 | 15;15;15;15;1 | 16564692:16 | 16565925:16 | -;-;-;-;-;-;-;-; | 2561 | CALN1 | 2 | 0 | 0 | 0 | 2 | 0 |
| ENSDARG000 | 15;15;15;15;1 | 22722682:22 | 22722841:22 | +;+;+;+;+;+;+; | 5067 | GRIK4 | 0 | 2 | 0 | 0 | 34 | 0 |
| ENSDARG000 | 15;15;15 | 47903864:47 | 47904190:47 | +;+;+ | 1747 | otx5 | 0 | 2 | 8 | 0 | 0 | 0 |
| ENSDARG000 | 15;15;15;15;1 | 9117330:912 | 9117666:912 | +;+;+;+;+ | 2001 | PTGIR | 5 | 0 | 6 | 10 | 19 | 2 |
| ENSDARG000 | 15 | 16697515 | 16697916 | - | 402 | NA | 0 | 0 | 0 | 0 | 0 | 0 |
| ENSDARG000 | 15;15;15;15;1 | 19324697:19 | 19325223:19 | +;+;+;+;+;+;+; | 2232 | vps26b | 421 | 452 | 546 | 377 | 626 | 501 |
| ENSDARG000 | 15;15;15;15;1 | 47920629:47 | 47921179:47 | -;-;-;-;-;-;- | 1434 | psma6l | 411 | 440 | 264 | 122 | 284 | 270 |
| ENSDARG000 | 15;15;15;15;1 | 14378829:14 | 14379061:14 | +;+;+;+;+;+;+; | 5157 | NA | 452 | 186 | 258 | 320 | 212 | 280 |
| ENSDARG000 | 15;15;15;15;1 | 9205264:920 | 9205460:920 | +;+;+;+;+;+;+; | 1713 | NA | 0 | 0 | 0 | 0 | 0 | 0 |
| ENSDARG000 | 15;15;15;15;1 | 9234052:923 | 9235637:923 | -;-;-;-;-;-;-;-; | 2148 | calm3a | 1028 | 1948 | 2839 | 1083 | 1592 | 1191 |
| ENSDARG000 | 15;15;15;15;1 | 20940904:20 | 20945087:20 | -;-;-;-;-;-;-;-; | 6479 | tbcela | 882 | 446 | 999 | 975 | 699 | 736 |
| ENSDARG000 | 15;15;15;15;1 | 16739160:16 | 16739652:16 | -;-;-;-;-;-;-;-; | 2677 | tyw1 | 788 | 623 | 860 | 565 | 778 | 655 |
| ENSDARG000 | 15;15;15;15;1 | 47929941:47 | 47931591:47 | -;-;-;-;-;-;-;-; | 4352 | SAMD4B | 617 | 496 | 701 | 502 | 725 | 534 |
| ENSDARG000 | 15;15;15;15;1 | 27415772:27 | 27416221:27 | -;-;-;-;-;-;-;-; | 8776 | brip1 | 94 | 261 | 328 | 199 | 264 | 140 |
| ENSDARG000 | 15;15;15;15;1 | 14436159:14 | 14437403:14 | -;-;-;-;-;-;-;-; | 3120 | numbl | 16 | 62 | 95 | 12 | 53 | 36 |
| ENSDARG000 | 15;15;15;15;1 | 29593547:29 | 29594193:29 | -;-;-;-;-;- | 1357 | si:ch211-207 | 48 | 2 | 0 | 28 | 12 | 2 |
| ENSDARG000 | 15;15;15;15;1 | 3284416:328 | 3285235:328 | +;+;+;+;+;+;+; | 6053 | foxo1a | 1365 | 1414 | 678 | 1828 | 1828 | 1285 |
| ENSDARG000 | 15;15;15;15;1 | 29549115:29 | 29550252:29 | -;-;-;-;-;-;- | 2112 | hspa13 | 424 | 90 | 776 | 174 | 156 | 353 |
| ENSDARG000 | 15;15;15;15;1 | 19331159:19 | 19331371:19 | -;-;-;-;-;-;- | 1073 | thyn1 | 722 | 871 | 815 | 696 | 743 | 683 |
| ENSDARG000 | 15;15;15 | 29553047:29 | 29553432:29 | +;+;+ | 599 | NA | 2 | 0 | 0 | 0 | 0 | 0 |
| ENSDARG000 | 15;15;15;15;1 | 29564364:29 | 29564985:29 | -;-;-;-;-;-;-;-; | 2760 | samsn1a | 121 | 145 | 59 | 81 | 206 | 294 |
| ENSDARG000 | 15;15;15;15;1 | 14554298:14 | 14554816:14 | -;-;-;-;-;-;-;-; | 2161 | coq8b | 333 | 196 | 282 | 168 | 260 | 274 |
| ENSDARG000 | 15;15;15;15;1 | 23200217:23 | 23200545:23 | -;-;-;-;-;-;-;-; | 1127 | ccdc153 | 0 | 14 | 0 | 10 | 8 | 18 |
| ENSDARG000 | 15;15;15 | 24737599:24 | 24738121:24 | +;+;+ | 3346 | crk | 1876 | 2357 | 2582 | 1751 | 2663 | 2163 |
| ENSDARG000 | 15;15;15;15;1 | 5784212:578 | 5784798:578 | -;-;-;-;-;-;-;-; | 2928 | hnrnpl2 | 3202 | 3724 | 4990 | 2505 | 3783 | 3081 |
| ENSDARG000 | 15;15;15;15;1 | 24691088:24 | 24693049:24 | +;+;+;+;+;+;+ | 7047 | LRRC75A | 18 | 10 | 17 | 2 | 22 | 46 |
| ENSDARG000 | 15;15;15;15;1 | 9294620:929 | 9294721:929 | +;+;+;+;+;+;+; | 2368 | slc37a4a | 115 | 101 | 140 | 157 | 146 | 115 |
| ENSDARG000 | 15;15;15;15;1 | 9315177:931 | 9316286:931 | -;-;-;-;-;-;-;- | 1744 | trappc4 | 516 | 766 | 489 | 413 | 633 | 292 |
| ENSDARG000 | 15;15;15;15;1 | 1179876:118 | 1181566:118 | -;-;-;-;-;-;-;-; | 2554 | lxn | 268 | 106 | 215 | 259 | 188 | 275 |
| ENSDARG000 | 15;15;15 | 9278992:928 | 9280046:928 | -;-;- | 1693 | NA | 1 | 12 | 36 | 0 | 4 | 0 |
| ENSDARG000 | 15;15;15;15;1 | 1167119:116 | 1167279:116 | +;+;+;+;+;+;+; | 2776 | gfm1 | 653 | 802 | 1045 | 278 | 922 | 475 |
| ENSDARG000 | 15;15;15;15;1 | 3454529:345 | 3456470:345 | -;-;-;-;-;-;-;-; | 3889 | cog6 | 306 | 219 | 272 | 232 | 315 | 278 |
| ENSDARG000 | 15;15;15;15;1 | 23208042:23 | 23208528:23 | +;+;+;+;+;+;+; | 4069 | cbl | 523 | 547 | 834 | 429 | 1124 | 568 |
| ENSDARG000 | 15;15;15;15;1 | 19335646:19 | 19335787:19 | +;+;+;+;+;+;+; | 1983 | acad8 | 386 | 410 | 448 | 246 | 442 | 349 |
| ENSDARG000 | 15;15;15;15;1 | 31806303:31 | 31806730:31 | +;+;+;+;+;+;+; | 10867 | FRY | 5263 | 80 | 649 | 30 | 78 | 70 |
| ENSDARG000 | 15;15 | 27642233:27 | 27642385:27 | -;- | 218 | NA | 0 | 0 | 0 | 0 | 0 | 0 |
| ENSDARG000 | 15;15;15;15;1 | 27703409:27 | 27704788:27 | -;-;-;-;-;-;-;- | 2464 | lhx1a | 2 | 2 | 10 | 0 | 0 | 1 |
| ENSDARG000 | 15;15;15;15;1 | 23271862:23 | 23273602:23 | -;-;-;-;-;-;-;-; | 3597 | mcamb | 520 | 255 | 598 | 2148 | 1499 | 1897 |
| ENSDARG000 | 15;15 | 14592624:14 | 14593016:14 | +;+ | 3397 | fbxo46 | 827 | 657 | 907 | 756 | 830 | 728 |
| ENSDARG000 | 15;15;15;15;1 | 28006768:28 | 28007030:28 | -;-;-;-;-;-;-;-; | 6322 | dhrs13a.3 | 297 | 256 | 315 | 194 | 314 | 356 |
| ENSDARG000 | 15;15;15;15;1 | 19353693:19 | 19355144:19 | -;-;-;-;-;-;- | 2668 | NA | 0 | 0 | 0 | 0 | 0 | 0 |
| ENSDARG000 | 15;15;15;15;1 | 16873690:16 | 16874233:16 | -;-;-;-;-;-;-;-; | 1641 | tmem120ab | 12 | 2 | 2 | 0 | 20 | 2 |
| ENSDARG000 | 15;15;15;15;1 | 20960069:20 | 20960145:20 | -;-;-;-;-;-;-;-; | 2546 | NA | 10 | 2 | 2 | 6 | 16 | 7 |
| ENSDARG000 | 15;15;15;15;1 | 29649515:29 | 29649738:29 | -;-;-;-;-;-;-;-; | 3105 | NA | 28 | 4 | 4 | 2 | 6 | 0 |
| ENSDARG000 | 15;15;15 | 29662401:29 | 29662650:29 | +;+;+ | 4573 | NRIP1 | 830 | 784 | 1860 | 1243 | 838 | 1804 |
| ENSDARG000 | 15 | 23356600 | 23358807 | + | 2208 | RNF26 | 259 | 304 | 352 | 182 | 306 | 327 |
| ENSDARG000 | 15;15;15;15;1 | 1372343:137 | 1372438:138 | +;+;+;+;+;+;+; | 4496 | schip1 | 1789 | 809 | 676 | 906 | 487 | 817 |
| ENSDARG000 | 15;15;15;15;1 | 19456112:19 | 19456159:19 | +;+;+;+;+;+;+; | 7546 | zgc:77784 | 17 | 24 | 2 | 6 | 14 | 2 |
| ENSDARG000 | 15;15;15;15;1 | 24746444:24 | 24746686:24 | +;+;+;+;+;+;+; | 1818 | ywhae1 | 4963 | 5469 | 6413 | 4455 | 5117 | 4510 |
| ENSDARG000 | 15 | 19581907 | 19583015 | - | 1109 | NA | 0 | 0 | 0 | 0 | 0 | 0 |
| ENSDARG000 | 15;15;15;15;1 | 28086193:28 | 28087139:28 | -;-;-;-;-;-;-;-; | 1989 | cryba1a | 10 | 0 | 0 | 0 | 0 | 0 |
| ENSDARG000 | 15;15;15;15;1 | 14602520:14 | 14603040:14 | -;-;-;-;-;-;-;-; | 2334 | slc5a2 | 28 | 8 | 78 | 232 | 133 | 2 |
| ENSDARG000 | 15;15;15;15;1 | 24756860:24 | 24757625:24 | +;+;+;+;+;+;+; | 7375 | CPD | 3042 | 2917 | 2558 | 2447 | 3884 | 2307 |
| ENSDARG000 | 15;15;15;15;1 | 16886196:16 | 16886254:16 | +;+;+;+;+;+;+; | 2669 | gdpd1 | 215 | 339 | 226 | 111 | 515 | 172 |
| ENSDARG000 | 15;15;15;15;1 | 28096152:28 | 28096227:28 | +;+;+;+;+;+;+; | 1293 | crybb1l3 | 0 | 0 | 0 | 2 | 0 | 4 |
| ENSDARG000 | 15;15;15;15;1 | 20991821:20 | 20991992:20 | -;-;-;-;-;-;-;-; | 4714 | NA | 388 | 247 | 471 | 393 | 453 | 466 |
| ENSDARG000 | 15;15;15;15;1 | 28106498:28 | 28106992:28 | +;+;+;+;+;+;+; | 1440 | UNC119 | 39 | 0 | 2 | 52 | 10 | 12 |
| ENSDARG000 | 15;15;15;15;1 | 21079826:21 | 21079916:21 | -;-;-;-;-;-;-;-; | 4510 | NA | 0 | 0 | 0 | 0 | 4 | 0 |
| ENSDARG000 | 15;15 | 29770611:29 | 29775541:29 | -;- | 5254 | NA | 0 | 0 | 10 | 6 | 5 | 0 |
| ENSDARG000 | 15;15;15;15;1 | 29727799:29 | 29728658:29 | +;+;+;+;+;+;+; | 4225 | zgc:153372 | 61 | 65 | 139 | 208 | 92 | 182 |
| ENSDARG000 | 15;15;15;15 | 5803458:580 | 5803658:580 | -;-;-;- | 669 | rbp2a | 14 | 2 | 20 | 0 | 2 | 14 |
| ENSDARG000 | 15;15;15;15;1 | 16897554:16 | 16897681:16 | +;+;+;+;+;+;+; | 805 | YPEL2 | 0 | 6 | 0 | 0 | 2 | 0 |
| ENSDARG000 | 15;15;15 | 23369020:23 | 23369771:23 | -;-;- | 1084 | c1qtnf5 | 0 | 2 | 0 | 2 | 12 | 4 |
| ENSDARG000 | 15;15 | 16900902:16 | 16901487:16 | -;- | 708 | NA | 0 | 0 | 0 | 0 | 0 | 0 |
| ENSDARG000 | 15;15;15 | 16914731:16 | 16915036:16 | +;+;+ | 885 | NA | 0 | 7 | 0 | 0 | 12 | 0 |
| ENSDARG000 | 15;15 | 29947543:29 | 29947989:29 | +;+ | 939 | NA | 0 | 0 | 1 | 0 | 0 | 2 |
| ENSDARG000 | 15;15;15;15;1 | 1440364:144 | 1440407:144 | +;+;+;+;+;+;+ | 949 | il12a | 0 | 4 | 0 | 0 | 0 | 2 |
| ENSDARG000 | 15;15;15;15;1 | 5828994:583 | 5831590:583 | -;-;-;-;-;-;-;-; | 7895 | NA | 996 | 613 | 1038 | 552 | 1022 | 861 |
| ENSDARG000 | 15;15;15;15;1 | 21113965:21 | 21114215:21 | -;-;-;-;-;-;-;-; | 4558 | a2ml | 0 | 1 | 6 | 0 | 0 | 0 |
| ENSDARG000 | 15;15 | 29949334:29 | 29949434:29 | +;+ | 635 | NA | 1 | 0 | 11 | 0 | 0 | 3 |
| ENSDARG000 | 15;15;15;15;1 | 29950658:29 | 29950790:29 | +;+;+;+;+;+;+; | 1888 | NA | 83 | 0 | 428 | 0 | 0 | 112 |
| ENSDARG000 | 15;15;15;15;1 | 19646902:19 | 19647569:19 | +;+;+;+;+;+;+; | 2789 | lim2.3 | 0 | 0 | 0 | 0 | 0 | 0 |
| ENSDARG000 | 15;15;15;15;1 | 23382199:23 | 23383664:23 | -;-;-;-;-;-;-;-; | 13970 | kmt2a | 1649 | 822 | 1929 | 1265 | 1592 | 1622 |
| ENSDARG000 | 15;15;15;15;1 | 30021652:30 | 30021917:30 | -;-;-;-;-;-;- | 2770 | NA | 2 | 2 | 14 | 0 | 16 | 12 |
| ENSDARG000 | 15;15;15;15;1 | 9327252:933 | 9327685:933 | +;+;+;+;+;+;+; | 7938 | sgcg | 272 | 226 | 300 | 0 | 746 | 150 |
| ENSDARG000 | 15;15;15;15;1 | 21137046:21 | 21137211:21 | -;-;-;-;-;-;-;-; | 4452 | sb:cb37 | 0 | 0 | 62 | 0 | 0 | 0 |
| ENSDARG000 | 15;15;15;15;1 | 30126971:30 | 30127484:30 | +;+;+;+;+;+ | 6050 | NA | 1734 | 2150 | 2154 | 980 | 2488 | 1797 |
| ENSDARG000 | 15;15;15;15;1 | 1452834:145 | 1453555:145 | -;-;-;-;-;-;-;-; | 2839 | NA | 71 | 106 | 118 | 112 | 105 | 117 |
| ENSDARG000 | 15;15;15;15;1 | 14630690:14 | 14634550:14 | -;-;-;-;- | 4849 | si:dkev-260i1 | 582 | 1043 | 1084 | 305 | 1272 | 528 |
| ENSDARG000 | 15;15;15;15;1 | 31911989:31 | 31912130:31 | +;+;+;+;+;+;+; | 9167 | brca2 | 304 | 322 | 587 | 175 | 260 | 410 |
| ENSDARG000 | 15;15;15;15;1 | 30157602:30 | 30158825:30 | +;+;+;+;+;+;+; | 3511 | nlk2 | 902 | 991 | 1351 | 607 | 2945 | 1219 |
| ENSDARG000 | 15;15;15;15 | 9414153:941 | 9414579:941 | +;+;+;+ | 2384 | NA | 26 | 4 | 10 | 7 | 3 | 6 |
| ENSDARG000 | 15;15;15;15;1 | 9367101:937 | 9382539:938 | -;-;-;-;-;-;-;-; | 18317 | SACS | 688 | 427 | 538 | 245 | 621 | 477 |
| ENSDARG000 | 15;15;15;15;1 | 16935390:16 | 16939636:16 | -;-;-;-;-;-;-;-; | 8671 | hip1 | 1130 | 1248 | 1274 | 800 | 1461 | 1220 |
| ENSDARG000 | 15;15;15;15;1 | 28129683:28 | 28130816:28 | -;-;-;-;-;-;-;-; | 3049 | foxn1 | 0 | 0 | 0 | 0 | 0 | 0 |
| ENSDARG000 | 15;15;15 | 30310843:30 | 30311152:30 | +;+;+ | 766 | lyrm9 | 94 | 98 | 76 | 54 | 76 | 76 |
| ENSDARG000 | 15;15;15;15;1 | 28175638:28 | 28176042:28 | +;+;+;+;+;+ | 1940 | slc46a1 | 56 | 62 | 104 | 16 | 95 | 82 |
| ENSDARG000 | 15;15;15;15;1 | 5898410:589 | 5899414:589 | -;-;-;-;-;-;-;-; | 1754 | si:ch73-281n1 | 3080 | 5358 | 6456 | 2306 | 4507 | 3098 |
| ENSDARG000 | 15;15;15;15;1 | 19661498:19 | 19661962:19 | -;-;-;-;-;-;-;-; | 2887 | hspa8b | 6 | 180 | 124 | 187 | 6 | 26 |
| ENSDARG000 | 15;15;15;15;1 | 24795473:24 | 24795554:24 | +;+;+;+;+;+;+; | 1406 | gosr1 | 326 | 395 | 420 | 409 | 316 | 250 |
| ENSDARG000 | 15;15;15;15;1 | 28188362:28 | 28189767:28 | -;-;-;-;-;-;-;- | 4019 | sarm1 | 6 | 2 | 15 | 0 | 0 | 6 |
| ENSDARG000 | 15;15;15;15;1 | 21157585:21 | 21158224:21 | -;-;-;-;-;-;-;-; | 4015 | NA | 0 | 0 | 20 | 0 | 0 | 0 |
| ENSDARG000 | 15;15;15;15;1 | 14642641:14 | 14642961:14 | +;+;+;+;+;+;+; | 1751 | sars2 | 110 | 173 | 293 | 142 | 251 | 80 |
| ENSDARG000 | 15;15;15;15;1 | 4172288:417 | 4172565:417 | -;-;-;-;-;-;-;-; | 3261 | ece2b | 72 | 24 | 18 | 44 | 33 | 27 |
| ENSDARG000 | 15;15;15;15;1 | 28208440:28 | 28211912:28 | -;-;-;-;-;-;-;-; | 5235 | SCARF1 | 28 | 4 | 14 | 0 | 46 | 18 |
| ENSDARG000 | 15;15;15;15;1 | 23443665:23 | 23443819:23 | +;+;+;+;+;+;+; | 1919 | phykpl | 150 | 135 | 251 | 176 | 222 | 192 |
| ENSDARG000 | 15;15;15;15;1 | 28227058:28 | 28227497:28 | -;-;-;-;-;-;-;- | 2004 | NA | 172 | 109 | 144 | 96 | 174 | 243 |
| ENSDARG000 | 15;15;15;15;1 | 1494493:149 | 1494783:149 | -;-;-;-;-;-;-;-; | 3133 | ift80 | 237 | 199 | 252 | 251 | 333 | 200 |
| ENSDARG000 | 15;15;15;15;1 | 28202170:28 | 28202246:28 | +;+;+;+;+;+;+; | 1855 | vtna | 0 | 0 | 6 | 0 | 0 | 0 |
| ENSDARG000 | 15;15;15;15;1 | 23423672:23 | 23423908:23 | -;-;-;-;-;-;-;-; | 3711 | UBE4A | 764 | 447 | 572 | 439 | 648 | 616 |
| ENSDARG000 | 15;15;15;15;1 | 31981885:31 | 31982005:31 | +;+;+;+;+;+;+; | 12407 | NA | 34 | 8 | 0 | 4 | 30 | 16 |
| ENSDARG000 | 15;15;15;15;1 | 1534644:153 | 1534823:153 | +;+;+;+;+;+;+; | 4294 | smc4 | 293 | 1410 | 2110 | 737 | 1372 | 622 |
| ENSDARG000 | 15;15;15;15;1 | 30323491:30 | 30323664:30 | +;+;+;+;+;+;+; | 3428 | nos2b | 80 | 60 | 137 | 15 | 51 | 59 |
| ENSDARG000 | 15;15 | 19678772:19 | 19679142:19 | -;- | 525 | NA | 0 | 0 | 0 | 0 | 0 | 0 |
| ENSDARG000 | 15;15;15;15;1 | 19681718:19 | 19682134:19 | +;+;+;+;+;+;+; | 2445 | NA | 0 | 0 | 2 | 0 | 8 | 0 |
| ENSDARG000 | 15;15;15 | 9451978:945 | 9452178:945 | +;+;+ | 542 | NA | 0 | 0 | 0 | 0 | 0 | 0 |
| ENSDARG000 | 15;15;15;15;1 | 9444982:944 | 9446060:944 | -;-;-;-;-;-;-;-; | 3677 | GAB2 | 387 | 174 | 225 | 182 | 281 | 245 |
| ENSDARG000 | 15;15;15;15;1 | 4353161:435 | 4354181:435 | -;-;-;-;-;-;- | 1874 | atp1b3b | 5551 | 3464 | 2953 | 3008 | 3462 | 3539 |
| ENSDARG000 | 15;15;15;15;1 | 5901970:590 | 5902072:590 | +;+;+;+;+ | 964 | get1 | 306 | 296 | 356 | 303 | 364 | 350 |
| ENSDARG000 | 15;15;15;15;1 | 30350061:30 | 30350243:30 | +;+;+;+;+;+;+; | 1081 | mrps23 | 166 | 196 | 428 | 92 | 198 | 253 |
| ENSDARG000 | 15;15;15;15;1 | 9601637:960 | 9601811:960 | -;-;-;-;-;-;-;-; | 1769 | nars2 | 52 | 71 | 108 | 42 | 80 | 50 |
| ENSDARG000 | 15;15;15;15;1 | 32249062:32 | 32249246:32 | +;+;+;+;+;+;+; | 2418 | arfip2a | 358 | 316 | 586 | 404 | 361 | 321 |
| ENSDARG000 | 15;15;15;15;1 | 21167142:21 | 21167243:21 | +;+;+;+;+;+;+; | 4158 | NA | 0 | 0 | 77 | 0 | 2 | 0 |
| ENSDARG000 | 15;15;15;15;1 | 5923851:592 | 5924036:592 | +;+;+;+;+;+;+; | 832 | sh3bgr | 0 | 0 | 0 | 0 | 0 | 2 |
| ENSDARG000 | 15;15;15;15;1 | 19697246:19 | 19698077:19 | -;-;-;-;-;-;-;-; | 6051 | NA | 936 | 250 | 2987 | 271 | 791 | 3434 |
| ENSDARG000 | 15;15;15 | 19688875:19 | 19689081:19 | +;+;+ | 758 | NA | 0 | 1 | 3 | 0 | 2 | 3 |
| ENSDARG000 | 15;15;15;15;1 | 14674928:14 | 14676190:14 | -;-;-;-;-;-;-;-; | 3347 | P4HA3 | 2 | 0 | 2 | 18 | 2 | 0 |
| ENSDARG000 | 15;15;15;15;1 | 4421797:442 | 4422598:442 | -;-;-;-;-;-;-;-; | 3683 | tfdp2 | 227 | 241 | 404 | 189 | 279 | 230 |
| ENSDARG000 | 15;15;15;15;1 | 21180094:21 | 21180195:21 | +;+;+;+;+;+;+; | 4531 | NA | 0 | 0 | 14 | 0 | 0 | 0 |
| ENSDARG000 | 15;15;15 | 21202820:21 | 21202915:21 | +;+;+ | 441 | NA | 0 | 0 | 0 | 0 | 0 | 0 |
| ENSDARG000 | 15;15;15;15;1 | 21220744:21 | 21220898:21 | -;-;-;-;-;-;-;-; | 4437 | NA | 2 | 2 | 28 | 10 | 2 | 10 |
| ENSDARG000 | 15;15;15;15;1 | 32268790:32 | 32268953:32 | +;+;+;+;+;+;+; | 2683 | NA | 8 | 0 | 10 | 0 | 12 | 6 |
| ENSDARG000 | 15;15;15;15;1 | 9861973:986 | 9862027:986 | +;+;+;+;+;+;+; | 1629 | NA | 0 | 0 | 4 | 6 | 0 | 0 |
| ENSDARG000 | 15;15;15;15;1 | 17030473:17 | 17030978:17 | +;+;+;+;+;+;+; | 2060 | plin2 | 215 | 70 | 438 | 174 | 197 | 209 |
| ENSDARG000 | 15;15;15;15;1 | 28248052:28 | 28249586:28 | -;-;-;-;-;-;-;-; | 8726 | prpf8 | 5583 | 4829 | 9075 | 3152 | 7350 | 4617 |
| ENSDARG000 | 15;15 | 9878848:988 | 9879146:988 | +;+ | 898 | NA | 0 | 0 | 0 | 0 | 0 | 0 |
| ENSDARG000 | 15;15;15;15;1 | 30424645:30 | 30426255:30 | -;-;-;-;-;-;-;-; | 4910 | msi2b | 258 | 283 | 538 | 220 | 270 | 178 |
| ENSDARG000 | 15;15;15;15;1 | 23452862:23 | 23452909:23 | -;-;-;-;-;-;-;-; | 1530 | NHERF4 | 0 | 17 | 12 | 0 | 38 | 6 |

| | | | | | | | | | | | | |
|---|---|---|---|---|---|---|---|---|---|---|---|---|
| ENSDARG000 | 15;15;15;15;1 | 32297441:32 | 32297636:32 | +;+;+;+;+;+;+; | 7714 | TRIM3 | 440 | 531 | 962 | 391 | 414 | 283 |
| ENSDARG000 | 15 | 14776854 | 14780602 | - | 3749 | or95a1 | 0 | 0 | 0 | 0 | 0 | 0 |
| ENSDARG000 | 15;15;15;15;1 | 32351886:32 | 32354009:32 | -;-;-;-;-;-;-; | 7462 | NA | 498 | 250 | 363 | 110 | 189 | 482 |
| ENSDARG000 | 15;15;15;15;1 | 10000553:10 | 10004771:10 | -;-;-;-;-;-;-; | 12785 | tenm4 | 376 | 1185 | 1563 | 467 | 1492 | 110 |
| ENSDARG000 | 15;15;15 | 24845145:24 | 24846736:24 | -;-;- | 2704 | TRARG1 | 0 | 0 | 0 | 0 | 0 | 2 |
| ENSDARG000 | 15;15;15 | 1574356:158 | 1578066:158 | -;-;- | 4443 | NA | 188 | 197 | 196 | 184 | 260 | 194 |
| ENSDARG000 | 15;15;15;15;1 | 24873388:24 | 24874133:24 | -;-;-;-;-;-;-;- | 2784 | AIPL1 | 0 | 2 | 4 | 0 | 0 | 0 |
| ENSDARG000 | 15;15;15;15;1 | 23468025:23 | 23469168:23 | -;-;-;-;-;-;-; | 4119 | NLRX1 | 296 | 297 | 453 | 346 | 622 | 290 |
| ENSDARG000 | 15;15;15;15;1 | 1589813:158 | 1590043:159 | -;-;-;-;-;- | 949 | nnr | 2 | 0 | 0 | 0 | 8 | 0 |
| ENSDARG000 | 15 | 14850820 | 14853579 | + | 2760 | NA | 0 | 0 | 0 | 0 | 0 | 0 |
| ENSDARG000 | 15;15;15 | 14826966:14 | 14828918:14 | +;+;+ | 2494 | or94a1 | 0 | 0 | 0 | 0 | 0 | 0 |
| ENSDARG000 | 15;15;15;15;1 | 17074185:17 | 17074268:17 | +;+;+;+;+;+;+; | 2287 | clptm1 | 2437 | 2080 | 2579 | 1425 | 2708 | 2108 |
| ENSDARG000 | 15;15;15;15;1 | 17059622:17 | 17059909:17 | -;-;-;-;-;-;-; | 1495 | lgals17 | 894 | 1555 | 171 | 31 | 92 | 148 |
| ENSDARG000 | 15;15;15;15;1 | 24946321:24 | 24949103:24 | -;-;-;-;- | 4369 | NA | 74 | 11 | 8 | 12 | 20 | 12 |
| ENSDARG000 | 15;15;15;15;1 | 6138423:613 | 6140071:614 | -;-;-;-;-;-;-; | 7599 | DSCAM | 120 | 54 | 136 | 229 | 238 | 104 |
| ENSDARG000 | 15;15;15;15;1 | 19752881:19 | 19753027:19 | +;+;+;+;+ | 975 | NA | 0 | 0 | 0 | 0 | 0 | 0 |
| ENSDARG000 | 15;15;15;15;1 | 19726229:19 | 19729373:19 | -;-;-;-;-;-;-; | 5649 | picalmb | 639 | 533 | 879 | 520 | 774 | 718 |
| ENSDARG000 | 15;15;15;15;1 | 23487730:23 | 23488732:23 | -;-;-;-;-;-;-; | 3126 | abcg4b | 2 | 18 | 6 | 0 | 4 | 14 |
| ENSDARG000 | 15;15;15;15;1 | 464229:4647 | 464652:4648 | -;-;-;-;-;-;-; | 2389 | MFRP | 17 | 12 | 28 | 27 | 39 | 32 |
| ENSDARG000 | 15;15;15;15;1 | 14854666:14 | 14855088:14 | +;+;+;+;+;+;+; | 2784 | diabloa | 84 | 80 | 104 | 22 | 110 | 78 |
| ENSDARG000 | 15;15;15;15;1 | 25007209:25 | 25007723:25 | -;-;-;-;-;- | 2156 | pimr187 | 4 | 3 | 2 | 15 | 11 | 2 |
| ENSDARG000 | 15;15 | 25021085:25 | 25021155:25 | +;+ | 765 | NA | 0 | 0 | 14 | 0 | 0 | 0 |
| ENSDARG000 | 15;15;15;15;1 | 28268135:28 | 28268259:28 | +;+;+;+;+;+;+; | 6854 | myo1cb | 831 | 1115 | 844 | 1129 | 2056 | 1011 |
| ENSDARG000 | 15;15;15;15;1 | 1596508:159 | 1598406:159 | -;-;-;-;-;-;-; | 3793 | kpna4 | 3280 | 3300 | 3298 | 2230 | 3325 | 2695 |
| ENSDARG000 | 15;15;15;15 | 14875726:14 | 14876532:14 | -;-;-;- | 1072 | NA | 0 | 3 | 0 | 0 | 0 | 0 |
| ENSDARG000 | 15;15 | 14883291:14 | 14883444:14 | -;- | 471 | NA | 0 | 0 | 0 | 0 | 0 | 1 |
| ENSDARG000 | 15;15 | 25028482:25 | 25028552:25 | +;+ | 765 | NA | 0 | 0 | 0 | 3 | 0 | 0 |
| ENSDARG000 | 15;15;15;15;1 | 25035947:25 | 25036445:25 | -;-;-;-;-;- | 2086 | NA | 18 | 0 | 4 | 9 | 4 | 9 |
| ENSDARG000 | 15;15;15;15;1 | 32387063:32 | 32387166:32 | +;+;+;+;+;+;+; | 1587 | NA | 32 | 16 | 6 | 6 | 8 | 10 |
| ENSDARG000 | 15;15;15 | 32411660:32 | 32411863:32 | +;+;+ | 728 | NA | 0 | 0 | 0 | 0 | 0 | 0 |
| ENSDARG000 | 15;15;15;15;1 | 32419303:32 | 32419391:32 | +;+;+;+;+;+;+; | 1780 | NA | 2 | 0 | 0 | 0 | 0 | 0 |
| ENSDARG000 | 15 | 17098351 | 17099560 | - | 1210 | mos | 47 | 101 | 125 | 92 | 64 | 61 |
| ENSDARG000 | 15;15;15;15;1 | 32398921:32 | 32399073:32 | +;+;+;+;+;+;+; | 1535 | NA | 0 | 0 | 0 | 0 | 6 | 0 |
| ENSDARG000 | 15;15;15;15;1 | 4549240:454 | 4549800:454 | -;-;-;-;-;-;-; | 1796 | rffl | 332 | 243 | 232 | 306 | 274 | 177 |
| ENSDARG000 | 15;15;15;15;1 | 19783352:19 | 19783484:19 | +;+;+;+;+;+;+; | 1298 | ZPLD1 | 0 | 2 | 0 | 0 | 0 | 0 |
| ENSDARG000 | 15;15 | 10485534:10 | 10485672:10 | +;+ | 405 | NA | 0 | 0 | 0 | 0 | 0 | 0 |
| ENSDARG000 | 15;15;15;15 | 23534126:23 | 23534389:23 | +;+;+;+ | 2826 | NA | 186 | 86 | 167 | 118 | 112 | 208 |
| ENSDARG000 | 15;15;15;15;1 | 23511407:23 | 23513072:23 | -;-;-;-;-;-;-; | 2850 | hmbsb | 762 | 494 | 860 | 632 | 596 | 628 |
| ENSDARG000 | 15;15;15;15;1 | 10752071:10 | 10752208:10 | +;+;+;+;+;+;+; | 1097 | NA | 10 | 6 | 14 | 16 | 0 | 24 |
| ENSDARG000 | 15 | 25059584 | 25060654 | + | 1071 | NA | 0 | 0 | 0 | 0 | 0 | 0 |
| ENSDARG000 | 15;15 | 25055961:25 | 25056031:25 | +;+ | 764 | NA | 0 | 0 | 0 | 0 | 0 | 0 |
| ENSDARG000 | 15;15;15;15;1 | 10750208:10 | 10750458:10 | +;+;+;+;+;+;+; | 894 | NA | 0 | 0 | 0 | 0 | 0 | 0 |
| ENSDARG000 | 15;15;15 | 10756210:10 | 10756753:10 | -;-;- | 840 | NA | 0 | 0 | 0 | 0 | 0 | 0 |
| ENSDARG000 | 15;15;15;15;1 | 15173611:15 | 15173801:15 | +;+;+;+;+;+;+; | 3419 | NA | 31 | 25 | 5 | 8 | 44 | 12 |
| ENSDARG000 | 15;15 | 25063306:25 | 25063376:25 | +;+ | 765 | NA | 0 | 0 | 0 | 0 | 0 | 0 |
| ENSDARG000 | 15;15;15;15;1 | 23523337:23 | 23523585:23 | +;+;+;+;+;+;+; | 4367 | NA | 36 | 32 | 26 | 48 | 30 | 38 |
| ENSDARG000 | 15;15;15;15;1 | 23528010:23 | 23528356:23 | +;+;+;+;+;+;+; | 5193 | NA | 78 | 90 | 76 | 62 | 50 | 161 |
| ENSDARG000 | 15;15 | 32434738:32 | 32434779:32 | +;+ | 625 | NA | 0 | 0 | 0 | 0 | 0 | 0 |
| ENSDARG000 | 15;15;15;15;1 | 15221123:15 | 15221534:15 | -;-;-;-;-;-;-; | 2357 | NA | 160 | 264 | 387 | 208 | 283 | 239 |
| ENSDARG000 | 15;15;15;15;1 | 32436907:32 | 32437772:32 | -;-;-;-;-;-;-; | 9898 | frem2b | 59 | 27 | 78 | 30 | 227 | 56 |
| ENSDARG000 | 15;15;15;15;1 | 19797918:19 | 19798093:19 | +;+;+;+;+;+;+; | 1208 | NA | 4 | 0 | 0 | 0 | 0 | 0 |
| ENSDARG000 | 15;15;15;15;1 | 6459847:645 | 6460311:646 | +;+;+;+;+;+;+; | 1387 | bace2 | 12723 | 10069 | 8613 | 7799 | 12786 | 8933 |
| ENSDARG000 | 15;15;15;15;1 | 28355023:28 | 28357683:28 | +;+;+;+;+;+;+ | 4163 | NA | 33 | 40 | 43 | 19 | 68 | 38 |
| ENSDARG000 | 15;15 | 28356845:28 | 28357638:28 | -;- | 941 | NA | 0 | 0 | 0 | 0 | 0 | 0 |
| ENSDARG000 | 15;15;15;15;1 | 19838458:19 | 19838673:19 | +;+;+;+;+;+;+; | 3078 | alcamb | 86 | 168 | 50 | 6 | 138 | 78 |
| ENSDARG000 | 15;15;15;15;1 | 21252532:21 | 21252645:21 | +;+;+;+;+;+;+; | 2903 | usf1 | 490 | 250 | 417 | 379 | 425 | 475 |
| ENSDARG000 | 15;15;15;15 | 1630815:163 | 1631571:163 | -;-;-;- | 1716 | ppm1lb | 6 | 4 | 6 | 28 | 8 | 0 |
| ENSDARG000 | 15;15;15;15;1 | 17100412:17 | 17100752:17 | +;+;+;+;+;+;+; | 4681 | NA | 1387 | 1241 | 1462 | 1248 | 1282 | 1357 |
| ENSDARG000 | 15;15;15;15;1 | 28368644:28 | 28368944:28 | +;+;+;+;+;+;+; | 2208 | SLC43A2 | 189 | 204 | 156 | 94 | 318 | 90 |
| ENSDARG000 | 15;15;15;15;1 | 25080041:25 | 25080559:25 | -;-;-;-;-;- | 2106 | NA | 16 | 2 | 2 | 24 | 26 | 9 |
| ENSDARG000 | 15;15;15;15;1 | 1653779:165 | 1653892:165 | +;+;+;+;+;+;+; | 2017 | nmd3 | 817 | 661 | 1379 | 769 | 870 | 807 |
| ENSDARG000 | 15;15;15;15;1 | 25084821:25 | 25085576:25 | -;-;-;-;-;-;-; | 1843 | exo5 | 263 | 168 | 215 | 293 | 235 | 247 |
| ENSDARG000 | 15;15;15;15;1 | 6514964:651 | 6515580:651 | -;-;-;-;-;-;-; | 9714 | ATM | 808 | 838 | 1498 | 599 | 1319 | 910 |
| ENSDARG000 | 15;15;15;15;1 | 32633290:32 | 32633740:32 | -;-;-;-;-;-;- | 1459 | NA | 0 | 6 | 25 | 6 | 12 | 18 |
| ENSDARG000 | 15;15;15;15;1 | 25093291:25 | 25093299:25 | -;-;-;-;-;-;- | 2410 | rflnb | 98 | 39 | 59 | 88 | 84 | 6 |
| ENSDARG000 | 15;15;15;15;1 | 15232855:15 | 15233069:15 | +;+;+;+;+;+;+ | 859 | med29 | 122 | 202 | 254 | 138 | 271 | 163 |
| ENSDARG000 | 15;15;15;15;1 | 28410664:28 | 28410824:28 | +;+;+;+;+;+;+; | 1799 | pitpnaa | 140 | 96 | 42 | 38 | 152 | 94 |
| ENSDARG000 | 15;15;15;15;1 | 32642878:32 | 32643145:32 | +;+;+;+;+;+;+; | 3938 | trpc4b | 10 | 8 | 18 | 0 | 14 | 4 |
| ENSDARG000 | 15;15;15;15;1 | 30836981:30 | 30840845:30 | -;-;-;-;-;-;-; | 6881 | akap1b | 806 | 1314 | 2264 | 877 | 1318 | 1054 |
| ENSDARG000 | 15;15 | 32707770:32 | 32707945:32 | +;+ | 770 | NA | 10 | 0 | 0 | 0 | 0 | 0 |
| ENSDARG000 | 15;15;15;15;1 | 17128282:17 | 17128492:17 | -;-;-;-;-;- | 1262 | mrpl28 | 656 | 789 | 1198 | 553 | 836 | 715 |
| ENSDARG000 | 15;15;15;15;1 | 4585266:458 | 4586756:458 | -;-;-;-;-;-;-; | 2458 | eif4h | 2582 | 2682 | 3449 | 1625 | 2944 | 2012 |
| ENSDARG000 | 15;15;15;15;1 | 17147002:17 | 17147522:17 | -;-;-;-;-;-;-; | 2713 | cul5a | 594 | 540 | 598 | 297 | 583 | 560 |
| ENSDARG000 | 15;15 | 11097869:11 | 11098382:11 | -;- | 752 | NA | 0 | 0 | 0 | 0 | 0 | 0 |
| ENSDARG000 | 15;15;15;15;1 | 11218246:11 | 11219833:11 | -;-;-;-;-;-;-;- | 3070 | prcp | 250 | 170 | 178 | 306 | 400 | 312 |
| ENSDARG000 | 15;15 | 11303325:11 | 11303469:11 | +;+ | 908 | NA | 4 | 34 | 21 | 10 | 13 | 19 |
| ENSDARG000 | 15;15;15;15;1 | 32710094:32 | 32710407:32 | +;+;+;+;+;+;+; | 5848 | postnb | 1736 | 2288 | 1421 | 380 | 2055 | 387 |
| ENSDARG000 | 15;15 | 4632782:463 | 4632945:463 | +;+ | 2587 | NA | 0 | 0 | 0 | 0 | 8 | 0 |
| ENSDARG000 | 15;15;15;15 | 1679962:167 | 1682804:168 | -;-;-;- | 3110 | sptssb | 6 | 36 | 16 | 8 | 6 | 12 |
| ENSDARG000 | 15;15 | 15303731:15 | 15304391:15 | -;- | 745 | NA | 0 | 0 | 2 | 0 | 0 | 0 |
| ENSDARG000 | 15;15;15;15;1 | 15258758:15 | 15258919:15 | +;+;+;+;+;+;+; | 7360 | NA | 185 | 201 | 613 | 261 | 184 | 146 |
| ENSDARG000 | 15;15;15;15 | 1705167:170 | 1705610:170 | +;+;+;+ | 2429 | otol1b | 0 | 0 | 2 | 0 | 0 | 1 |
| ENSDARG000 | 15;15;15;15;1 | 4957784:495 | 4958070:495 | -;-;-;-;-;-;- | 1530 | lipt2 | 45 | 40 | 64 | 32 | 59 | 47 |
| ENSDARG000 | 15;15;15;15;1 | 28431621:28 | 28434167:28 | -;-;-;-;-;-;-; | 5130 | GIT1 | 1571 | 906 | 1692 | 872 | 1324 | 954 |
| ENSDARG000 | 15;15;15;15;1 | 4969128:497 | 4969568:497 | +;+;+;+;+;+ | 5389 | rnf169 | 297 | 297 | 238 | 250 | 484 | 252 |
| ENSDARG000 | 15 | 23544431 | 23546452 | - | 2022 | NA | 0 | 10 | 2 | 4 | 4 | 14 |
| ENSDARG000 | 15;15;15;15;1 | 11308462:11 | 11309817:11 | -;-;-;-;- | 1929 | rab30 | 178 | 149 | 238 | 110 | 230 | 248 |
| ENSDARG000 | 15;15;15;15;1 | 28482862:28 | 28483345:28 | +;+;+;+;+;+;+; | 4911 | ankrd13b | 7 | 0 | 10 | 0 | 0 | 0 |
| ENSDARG000 | 15;15;15;15;1 | 19884242:19 | 19884928:19 | +;+;+;+;+;+;+; | 5987 | dyrk1ab | 1173 | 993 | 1306 | 904 | 1141 | 1110 |
| ENSDARG000 | 15;15;15;15;1 | 1718668:172 | 1719565:172 | -;-;-;-;-;-;-; | 2383 | rabgef1l | 157 | 61 | 112 | 88 | 89 | 117 |
| ENSDARG000 | 15;15;15 | 30896722:30 | 30897091:30 | +;+;+ | 1632 | NA | 0 | 0 | 4 | 0 | 1 | 2 |
| ENSDARG000 | 15;15 | 30917282:30 | 30917825:30 | +;+ | 2701 | OMG | 2 | 0 | 34 | 0 | 12 | 14 |
| ENSDARG000 | 15;15;15;15;1 | 30861647:30 | 30864648:30 | -;-;-;-;-;-;-; | 11440 | NF1 | 2559 | 2863 | 3793 | 3293 | 3392 | 5128 |
| ENSDARG000 | 15;15;15;15;1 | 17177635:17 | 17177891:17 | -;-;-;-;-;-;-; | 3070 | NA | 2629 | 284 | 252 | 1552 | 520 | 541 |
| ENSDARG000 | 15;15;15;15;1 | 11381532:11 | 11381580:11 | +;+;+;+;+;+ | 781 | NA | 0 | 0 | 2 | 0 | 2 | 0 |
| ENSDARG000 | 15;15;15;15;1 | 17196527:17 | 17197318:17 | -;-;-;-;-;-;-; | 2608 | NA | 78 | 24 | 175 | 112 | 128 | 554 |
| ENSDARG000 | 15;15;15;15;1 | 23550890:23 | 23551921:23 | +;+;+;+;+;+;+; | 4360 | MARK4 | 1497 | 1302 | 1283 | 1641 | 1316 | 2115 |
| ENSDARG000 | 15;15;15;15;1 | 32798333:32 | 32798499:32 | +;+;+;+;+;+;+; | 3494 | spartb | 411 | 328 | 464 | 177 | 352 | 218 |
| ENSDARG000 | 15;15 | 30922926:30 | 30922979:30 | +;+ | 935 | NA | 0 | 4 | 0 | 0 | 2 | 20 |
| ENSDARG000 | 15;15;15;15;1 | 19900197:19 | 19900460:19 | +;+;+;+;+ | 2559 | thap12b | 159 | 172 | 219 | 148 | 190 | 198 |
| ENSDARG000 | 15;15;15;15;1 | 32821392:32 | 32821876:32 | +;+;+;+;+;+;+; | 9357 | dclk1b | 20 | 4 | 4 | 0 | 0 | 3 |
| ENSDARG000 | 15;15;15 | 11427620:11 | 11427859:11 | +;+;+ | 538 | ndufc2 | 704 | 906 | 1043 | 619 | 713 | 718 |
| ENSDARG000 | 15;15;15;15;1 | 11683114:11 | 11683259:11 | +;+;+;+;+;+;+; | 1424 | si:dkey-31c13 | 38 | 57 | 76 | 65 | 86 | 44 |
| ENSDARG000 | 15;15;15;15;1 | 19915345:19 | 19916493:19 | +;+;+;+;+;+ | 2298 | NA | 0 | 0 | 0 | 0 | 0 | 0 |
| ENSDARG000 | 15;15 | 19929495:19 | 19929895:19 | -;- | 699 | NA | 0 | 2 | 0 | 0 | 0 | 0 |
| ENSDARG000 | 15;15;15;15;1 | 1734444:173 | 1734521:173 | -;-;-;-;-;-;-; | 944 | STX1A | 0 | 0 | 2 | 1 | 0 | 0 |
| ENSDARG000 | 15;15;15;15;1 | 23643718:23 | 23644178:23 | -;-;-;-;-;-;-; | 1770 | ckmb | 22 | 36 | 12 | 28 | 56 | 8 |
| ENSDARG000 | 15;15;15;15;1 | 19990068:19 | 19990346:19 | +;+;+;+;+;+;+; | 2267 | zgc:112083 | 127 | 187 | 254 | 100 | 212 | 120 |
| ENSDARG000 | 15;15 | 17214062:17 | 17214689:17 | -;- | 932 | NA | 18 | 0 | 0 | 0 | 0 | 0 |
| ENSDARG000 | 15;15;15;15;1 | 17251191:17 | 17251304:17 | +;+;+;+;+ | 704 | NA | 2 | 4 | 0 | 0 | 2 | 6 |
| ENSDARG000 | 15;15;15;15;1 | 4988189:498 | 4988262:498 | +;+;+;+;+;+;+; | 1209 | spcs2 | 1813 | 2671 | 1970 | 1510 | 2270 | 1769 |
| ENSDARG000 | 15;15;15;15;1 | 25100726:25 | 25102121:25 | -;-;-;-;-;-;-; | 3862 | vps53 | 807 | 741 | 944 | 712 | 974 | 612 |
| ENSDARG000 | 15;15;15 | 5011080:501 | 5011429:501 | -;-;- | 541 | defbl3 | 0 | 2 | 4 | 1 | 8 | 0 |
| ENSDARG000 | 15;15;15;15;1 | 5028608:503 | 5028900:503 | +;+;+;+;+;+;+; | 3390 | abcg1 | 0 | 6 | 1 | 0 | 4 | 0 |
| ENSDARG000 | 15;15;15;15;1 | 15314189:15 | 15314425:15 | +;+;+;+;+;+;+; | 2270 | camkk1b | 4 | 0 | 4 | 8 | 0 | 8 |
| ENSDARG000 | 15;15 | 17279984:17 | 17280190:17 | -;- | 338 | NA | 0 | 0 | 0 | 0 | 0 | 0 |
| ENSDARG000 | 15;15;15;15;1 | 17258246:17 | 17258643:17 | +;+;+;+;+;+;+; | 7465 | DHX40 | 323 | 142 | 219 | 260 | 179 | 295 |
| ENSDARG000 | 15;15;15;15;1 | 11825366:11 | 11825750:11 | +;+;+;+;+;+;+; | 7236 | PRKD2 | 2455 | 1217 | 1796 | 1471 | 2181 | 1990 |
| ENSDARG000 | 15;15;15;15;1 | 23657051:23 | 23657272:23 | +;+;+;+;+;+;+; | 1971 | klc3 | 0 | 0 | 4 | 6 | 0 | 0 |
| ENSDARG000 | 15;15 | 15348490:15 | 15350749:15 | -;- | 2518 | YWHAG | 223 | 412 | 254 | 70 | 222 | 146 |
| ENSDARG000 | 15;15;15;15;1 | 1749843:175 | 1750151:175 | -;-;-;-;-;-;-; | 1231 | bud23 | 356 | 386 | 530 | 240 | 403 | 366 |
| ENSDARG000 | 15;15;15;15;1 | 23680018:23 | 23680221:23 | -;-;-;-;-;-;-; | 2581 | ercc2 | 302 | 268 | 429 | 340 | 428 | 343 |
| ENSDARG000 | 15;15;15;15;1 | 5086338:508 | 5086390:508 | +;+;+;+;+;+;+; | 2143 | NA | 259 | 133 | 34 | 46 | 15 | 32 |
| ENSDARG000 | 15 | 33070939 | 33073200 | + | 2262 | mab21l1 | 20 | 6 | 5499 | 2741 | 5144 | 3872 |
| ENSDARG000 | 15;15;15;15;1 | 15375607:15 | 15375895:15 | +;+;+;+;+ | 1020 | NA | 0 | 0 | 0 | 0 | 0 | 0 |
| ENSDARG000 | 15;15;15;15;1 | 32895659:32 | 32897167:32 | -;-;-;-;-;-;-; | 10796 | NBEA | 8 | 2 | 27 | 16 | 61 | 5 |
| ENSDARG000 | 15;15 | 11883804:11 | 11884049:11 | +;+ | 1676 | gpr184 | 6 | 4 | 2 | 4 | 0 | 24 |
| ENSDARG000 | 15;15;15;15;1 | 6652396:665 | 6652659:665 | +;+;+;+;+;+;+; | 1595 | nop53 | 3075 | 2359 | 4801 | 5054 | 2404 | 4450 |
| ENSDARG000 | 15;15;15 | 11952956:11 | 11955491:11 | -;-;- | 2791 | PRLHR | 0 | 0 | 0 | 0 | 0 | 0 |
| ENSDARG000 | 15;15;15;15;1 | 1778773:180 | 1778786:180 | -;-;-;-;-;-;-; | 2252 | MMP28 | 12 | 2 | 2 | 0 | 2 | 2 |
| ENSDARG000 | 15;15;15;15;1 | 1765988:176 | 1766863:176 | +;+;+;+;+;+;+; | 6580 | cul3b | 905 | 683 | 1038 | 805 | 738 | 938 |
| ENSDARG000 | 15;15 | 6678521:672 | 6679032:672 | +;+ | 1488 | NA | 0 | 0 | 0 | 0 | 7 | 0 |
| ENSDARG000 | 15;15 | 23693514:23 | 23696133:23 | -;- | 2733 | zgc:154093 | 429 | 38 | 30 | 410 | 28 | 276 |
| ENSDARG000 | 15;15;15;15;1 | 20004766:20 | 20006024:20 | -;-;-;-;-;-;-; | 3241 | NA | 144 | 64 | 76 | 10 | 25 | 94 |
| ENSDARG000 | 15;15;15;15;1 | 23709329:23 | 23712227:23 | -;-;-;-;-;-;-; | 5568 | ZC3H4 | 1389 | 1060 | 1861 | 1231 | 1682 | 1242 |
| ENSDARG000 | 15;15;15;15;1 | 5116179:512 | 5116460:512 | +;+;+;+;+;+;+; | 3392 | pgm2l1 | 173 | 168 | 175 | 92 | 204 | 156 |
| ENSDARG000 | 15;15;15;15;1 | 11982450:11 | 11982781:11 | +;+;+;+;+;+;+; | 2770 | NA | 4 | 0 | 0 | 0 | 0 | 0 |
| ENSDARG000 | 15;15 | 6770586:677 | 6770693:677 | +;+ | 283 | NA | 0 | 0 | 0 | 0 | 0 | 0 |
| ENSDARG000 | 15;15;15;15;1 | 6773876:677 | 6773945:677 | +;+;+;+;+;+;+; | 1905 | dmac2 | 82 | 62 | 149 | 68 | 106 | 72 |
| ENSDARG000 | 15 | 15383894 | 15384011 | - | 118 | NA | 0 | 0 | 0 | 0 | 0 | 0 |
| ENSDARG000 | 15;15;15;15;1 | 25156121:25 | 25156197:25 | +;+;+;+;+;+;+; | 1250 | SLC35F2 | 11 | 0 | 0 | 0 | 0 | 2 |
| ENSDARG000 | 15;15;15;15;1 | 6786757:678 | 6786906:678 | +;+;+;+;+;+;+; | 2589 | sirt2 | 1861 | 1062 | 1348 | 1709 | 2047 | 1866 |
| ENSDARG000 | 15;15;15;15;1 | 25172671:25 | 25176705:25 | -;-;-;-;-;-;- | 5568 | MNT | 2395 | 2953 | 2992 | 3032 | 2841 | 2150 |
| ENSDARG000 | 15;15;15;15;1 | 23722620:23 | 23722901:23 | +;+;+;+;+;+;+; | 1474 | sae1 | 308 | 592 | 794 | 252 | 524 | 372 |
| ENSDARG000 | 15;15;15;15;1 | 25214171:25 | 25214719:25 | -;-;-;-;-;-;-; | 3400 | mettl16 | 301 | 204 | 218 | 180 | 288 | 237 |
| ENSDARG000 | 15;15;15;15;1 | 15390882:15 | 15390989:15 | +;+;+;+;+;+;+; | 1626 | ca4b | 4 | 0 | 0 | 0 | 0 | 0 |
| ENSDARG000 | 15;15;15;15;1 | 28545143:28 | 28550330:28 | -;-;-;-;-;-;-; | 8889 | SSH2 | 1623 | 1347 | 1402 | 1104 | 1401 | 1154 |
| ENSDARG000 | 15;15;15;15;1 | 30987414:30 | 30987734:30 | -;-;-;-;-;-;-; | 1568 | NA | 0 | 0 | 0 | 0 | 0 | 0 |
| ENSDARG000 | 15;15;15;15;1 | 30994538:30 | 30995199:30 | -;-;-;-;-;-;- | 1437 | lgals9l5 | 2 | 0 | 0 | 0 | 0 | 0 |
| ENSDARG000 | 15;15;15 | 31005981:31 | 31006091:31 | -;-;- | 279 | NA | 0 | 0 | 0 | 0 | 0 | 0 |
| ENSDARG000 | 15;15;15;15;1 | 20060643:20 | 20061504:20 | -;-;-;-;-;-;-; | 8004 | med13b | 1009 | 689 | 1043 | 649 | 1245 | 1094 |
| ENSDARG000 | 15;15;15;15 | 23745217:23 | 23748399:23 | -;-;-;- | 3808 | bbc3 | 31 | 54 | 132 | 146 | 67 | 132 |
| ENSDARG000 | 15;15;15;15;1 | 15403560:15 | 15403722:15 | +;+;+;+;+;+;+ | 1445 | dhrs11b.1 | 52 | 145 | 104 | 68 | 158 | 108 |
| ENSDARG000 | 15;15;15;15;1 | 23776417:23 | 23781332:23 | -;-;-;-;-;-;-; | 5733 | rad1 | 62 | 180 | 232 | 102 | 214 | 148 |
| ENSDARG000 | 15;15;15;15;1 | 31013578:31 | 31013838:31 | +;+;+;+;+;+;+ | 1980 | NA | 0 | 0 | 0 | 0 | 0 | 0 |
| ENSDARG000 | 15;15;15;15;1 | 33467298:33 | 33468330:33 | -;-;-;-;-;-;-; | 6122 | NA | 1816 | 76 | 1094 | 108 | 106 | 388 |
| ENSDARG000 | 15;15 | 28595913:28 | 28596158:28 | -;- | 392 | NA | 36 | 25 | 41 | 46 | 9 | 58 |
| ENSDARG000 | 15;15;15;15;1 | 28598647:28 | 28598751:28 | +;+;+;+;+;+ | 1042 | NA | 0 | 0 | 0 | 0 | 0 | 0 |
| ENSDARG000 | 15;15;15;15;1 | 28609025:28 | 28609243:28 | -;-;-;-;-;-;-; | 2510 | slc6a4a | 31 | 15 | 62 | 11 | 60 | 21 |
| ENSDARG000 | 15;15;15;15;1 | 543744:5437 | 543961:5439 | -;-;-;-;-;-;-; | 822 | tagln | 48 | 30 | 27 | 28 | 12 | 12 |
| ENSDARG000 | 15;15;15;15;1 | 17285989:17 | 17286295:17 | +;+;+;+;+;+;+; | 7218 | cltcb | 4433 | 4260 | 5696 | 2785 | 5702 | 3501 |
| ENSDARG000 | 15;15;15;15;1 | 11989063:11 | 11989662:11 | -;-;-;-;-;-;-; | 1734 | NA | 0 | 0 | 2 | 0 | 0 | 2 |

| | | | | | | | | | | | | |
|---|---|---|---|---|---|---|---|---|---|---|---|---|
| ENSDARG000 | 15:15:15:15:1 | 31022014:31 | 31022334:31 | -;-;-;-;-;-;- | 1065 | lgals9l4 | 0 | 0 | 0 | 0 | 0 | 0 |
| ENSDARG000 | 15:15:15:15:1 | 25273185:252 | 25273888:252 | -:-:-:-:-:-:-:-: | 2290 | pafah1b1a | 2960 | 2612 | 2739 | 2702 | 2378 | 2628 |
| ENSDARG000 | 15:15:15:15:1 | 31032255:31 | 31033605:31 | -:-:-:-:-:-:-:-: | 2201 | lgals9l1 | 53 | 90 | 124 | 36 | 220 | 231 |
| ENSDARG000 | 15:15:15:15:1 | 6838039:683 | 6838998:683 | -:-:-:-:-:-:-:-: | 2373 | meis3 | 0 | 0 | 12 | 6 | 0 | 4 |
| ENSDARG000 | 15:15:15:15:1 | 12004777:12 | 12009681:12 | -:-:-:-:-:-:-:-: | 5724 | NA | 0 | 0 | 0 | 0 | 0 | 0 |
| ENSDARG000 | 15:15:15:15:1 | 31054200:31 | 31055551:31 | -:-:-:-:-:-:-:-: | 2454 | lgals9l3 | 369 | 265 | 276 | 89 | 131 | 311 |
| ENSDARG000 | 15:15:15:15:1 | 15415623:154 | 15415945:154 | +;+;+;+;+;+;+ | 1485 | dhrs11b.2 | 22 | 12 | 15 | 42 | 24 | 0 |
| ENSDARG000 | 15;15 | 5155137:515 | 5157401:515 | -;- | 2325 | or64d1 | 0 | 0 | 0 | 0 | 0 | 0 |
| ENSDARG000 | 15 | 5162274 | 5165013 | - | 2740 | NA | 0 | 0 | 0 | 0 | 0 | 0 |
| ENSDARG000 | 15 | 5167690 | 5172170 | - | 4481 | or62c1 | 0 | 0 | 0 | 0 | 0 | 0 |
| ENSDARG000 | 15;15;15;15 | 15445113:154 | 15447337:154 | -;-;-;- | 3254 | NA | 50 | 196 | 4 | 2 | 2 | 0 |
| ENSDARG000 | 15:15:15:15:1 | 1825172:182 | 1825672:182 | -:-:-:-:-:-:-:-: | 2889 | taf15 | 4245 | 5618 | 5810 | 2347 | 5302 | 4159 |
| ENSDARG000 | 15:15:15:15:1 | 33691455:33 | 33691743:33 | +:+:+:+:+:+:+:+: | 2009 | NA | 21 | 44 | 34 | 32 | 68 | 67 |
| ENSDARG000 | 15:15:15:15:1 | 33702918:33 | 33703127:33 | -:-:-:-:-:-:-:-: | 2775 | trmt9b | 10 | 18 | 20 | 18 | 22 | 10 |
| ENSDARG000 | 15:15:15:15:1 | 25321572:25 | 25323163:25 | -:-:-:-:-:-:-:-: | 6052 | cluha | 755 | 1235 | 1225 | 279 | 1435 | 698 |
| ENSDARG000 | 15 | 5175593 | 5178899 | - | 3307 | or62c4 | 0 | 0 | 0 | 0 | 0 | 0 |
| ENSDARG000 | 15 | 5182854 | 5184186 | - | 1333 | or62c2 | 0 | 0 | 0 | 0 | 0 | 0 |
| ENSDARG000 | 15:15:15:15:1 | 23790057:23 | 23790620:23 | -;-;-;-;- | 1467 | tmem97 | 107 | 81 | 89 | 113 | 131 | 68 |
| ENSDARG000 | 15:15:15:15:1 | 556956:5570 | 557397:5573 | -;-;-;-;-;-;-;- | 1631 | NA | 2 | 0 | 0 | 0 | 0 | 0 |
| ENSDARG000 | 15:15:15:15:1 | 31070603:31 | 31071554:31 | -:-:-:-:-:-:-:-: | 5387 | KSR1 | 455 | 259 | 268 | 307 | 267 | 416 |
| ENSDARG000 | 15:15:15:15:1 | 23784842:23 | 23784905:23 | +:+:+:+:+:+:+:+: | 1186 | ift20 | 134 | 108 | 134 | 88 | 167 | 120 |
| ENSDARG000 | 15:15:15:15:1 | 28685625:28 | 28686596:28 | +:+:+:+:+:+:+:+: | 9248 | nova2 | 192 | 10 | 38 | 12 | 92 | 34 |
| ENSDARG000 | 15:15:15:15:1 | 28782701:28 | 28787837:28 | -:-:-:-:-:-:-:-: | 6745 | polr1g | 198 | 318 | 274 | 168 | 298 | 236 |
| ENSDARG000 | 15:15:15:15:1 | 28634580:28 | 28634858:28 | -:-:-:-:-:-:-:-: | 2007 | blmh | 925 | 898 | 893 | 427 | 942 | 665 |
| ENSDARG000 | 15:15:15:15:1 | 15459974:154 | 15460621:154 | -:-:-:-:-:-:-:-: | 1677 | rab34a | 487 | 393 | 347 | 317 | 381 | 407 |
| ENSDARG000 | 15;15;15;15 | 25375923:25 | 25377366:25 | -;-;-;- | 2062 | CCDC92B | 0 | 0 | 78 | 10 | 0 | 31 |
| ENSDARG000 | 15 | 5189408 | 5191672 | - | 2265 | or62c6 | 0 | 0 | 0 | 0 | 0 | 0 |
| ENSDARG000 | 15 | 5196715 | 5198191 | - | 1477 | or64g2 | 0 | 0 | 0 | 0 | 0 | 0 |
| ENSDARG000 | 15:15:15:15:1 | 6926987:692 | 6927516:692 | -:-:-:-:-:-:-:-: | 2116 | ech1 | 302 | 242 | 323 | 186 | 337 | 384 |
| ENSDARG000 | 15;15;15;15 | 572195:5728 | 572740:5729 | -;-;-;- | 1157 | cbln20 | 2 | 6 | 2 | 6 | 0 | 0 |
| ENSDARG000 | 15 | 5202054 | 5203371 | - | 1318 | or64e1 | 0 | 0 | 0 | 0 | 0 | 0 |
| ENSDARG000 | 15;15;15;15 | 25398127:254 | 25400251:254 | -;-;-;- | 2897 | tlcd2 | 971 | 495 | 2493 | 380 | 592 | 2716 |
| ENSDARG000 | 15 | 5205538 | 5207805 | - | 2268 | or64g1 | 0 | 0 | 0 | 0 | 0 | 0 |
| ENSDARG000 | 15 | 5210468 | 5212930 | - | 2463 | or64f1 | 0 | 0 | 8 | 0 | 0 | 0 |
| ENSDARG000 | 15 | 5215807 | 5217693 | - | 1887 | NA | 0 | 0 | 0 | 0 | 0 | 0 |
| ENSDARG000 | 15:15:15:15:1 | 15456536:154 | 15457059:154 | +:+:+:+:+:+:+:+: | 1850 | zgc:174895 | 8 | 0 | 25 | 67 | 6 | 22 |
| ENSDARG000 | 15:15:15:15:1 | 17343319:17 | 17343388:17 | +:+:+:+:+:+:+:+: | 2205 | vmp1 | 603 | 893 | 778 | 265 | 996 | 494 |
| ENSDARG000 | 15;15;15 | 5220992:522 | 5222631:522 | -;-;- | 1682 | or64b1 | 0 | 0 | 0 | 0 | 0 | 0 |
| ENSDARG000 | 15 | 23799461 | 23801499 | + | 2039 | NA | 4 | 225 | 64 | 0 | 224 | 291 |
| ENSDARG000 | 15:15:15:15:1 | 12071037:12 | 12071569:12 | -:-:-:-:-:-:-:-: | 6840 | DSCAML1 | 17 | 71 | 186 | 26 | 170 | 0 |
| ENSDARG000 | 15;15;15;15 | 20204628:202 | 20205790:202 | -;-;-;- | 1583 | ppp1r14ab | 6 | 0 | 0 | 0 | 4 | 8 |
| ENSDARG000 | 15;15 | 577698:5807 | 577871:5819 | +;+ | 1345 | NA | 72 | 64 | 92 | 42 | 74 | 100 |
| ENSDARG000 | 15:15:15:15:1 | 23807495:23 | 23807782:23 | -;-;-;-;-;-;-;- | 1422 | NA | 8 | 5 | 4 | 14 | 26 | 41 |
| ENSDARG000 | 15;15 | 20238428:202 | 20239229:202 | -;- | 993 | NA | 0 | 0 | 0 | 0 | 0 | 0 |
| ENSDARG000 | 15:15:15:15:1 | 20239141:202 | 20239651:202 | +:+:+:+:+:+:+:+: | 2024 | spint2 | 129 | 7 | 6 | 16 | 26 | 15 |
| ENSDARG000 | 15;15 | 5228580:523 | 5230133:523 | -;- | 1601 | or64b4 | 0 | 0 | 0 | 0 | 0 | 0 |
| ENSDARG000 | 15:15:15:15:1 | 590670:5922 | 592904:5929 | -:-:-:-:-:-:-:-: | 4387 | alox5b.3 | 20 | 0 | 38 | 18 | 121 | 2 |
| ENSDARG000 | 15:15:15:15:1 | 20139031:201 | 20139651:201 | -:-:-:-:-:-:-:-: | 3133 | NA | 25 | 1 | 22 | 26 | 123 | 14 |
| ENSDARG000 | 15:15:15:15:1 | 586605:5870 | 588079:5880 | -;-;-;-;- | 1999 | NA | 75 | 30 | 50 | 29 | 72 | 23 |
| ENSDARG000 | 15:15:15:15:1 | 21262917:21 | 21263129:21 | +:+:+:+:+:+:+:+: | 2440 | gkup | 386 | 256 | 332 | 214 | 243 | 175 |
| ENSDARG000 | 15:15:15:15:1 | 31161783:31 | 31163812:31 | -:-:-:-:-:-:-:-: | 3900 | wsb1 | 11151 | 9731 | 22112 | 26300 | 11907 | 18862 |
| ENSDARG000 | 15;15;15 | 5237596:523 | 5239218:523 | -;-;- | 1682 | or64b3 | 0 | 0 | 0 | 0 | 0 | 0 |
| ENSDARG000 | 15:15:15:15:1 | 25438714:254 | 25438834:254 | +:+:+:+:+:+:+:+: | 2399 | aifm4 | 2 | 0 | 2 | 0 | 2 | 4 |
| ENSDARG000 | 15:15:15:15:1 | 33809864:33 | 33810535:33 | -;-;-;-;-;- | 2025 | NA | 99 | 108 | 122 | 93 | 102 | 166 |
| ENSDARG000 | 15:15:15:15:1 | 31188016:31 | 31188250:31 | -:-:-:-:-:-:-:-: | 2129 | cuedc1b | 160 | 172 | 161 | 210 | 188 | 162 |
| ENSDARG000 | 15;15;15;15 | 33807350:33 | 33807747:33 | +;+;+;+ | 1436 | NA | 0 | 0 | 0 | 0 | 0 | 0 |
| ENSDARG000 | 15:15:15:15:1 | 33744549:33 | 33745371:33 | -:-:-:-:-:-:-:-: | 5401 | PDS5B | 1065 | 1215 | 1583 | 938 | 1528 | 1103 |
| ENSDARG000 | 15;15;15 | 5242840:524 | 5245726:524 | -;-;- | 2913 | or64b2 | 0 | 0 | 0 | 0 | 0 | 0 |
| ENSDARG000 | 15;15 | 5250954:525 | 5252082:525 | -;- | 1162 | NA | 0 | 0 | 0 | 0 | 0 | 0 |
| ENSDARG000 | 15:15:15:15:1 | 23816945:23 | 23819662:23 | -:-:-:-:-:-:-:-: | 7480 | USP6 | 604 | 442 | 630 | 452 | 614 | 497 |
| ENSDARG000 | 15:15:15:15:1 | 31244025:31 | 31244560:31 | -;-;-;-;- | 1661 | vezf1b | 190 | 158 | 303 | 131 | 217 | 179 |
| ENSDARG000 | 15:15:15:15:1 | 12251044:12 | 12251230:12 | -;-;-;-;-;-;-;- | 991 | NA | 0 | 0 | 4 | 0 | 0 | 0 |
| ENSDARG000 | 15:15:15:15:1 | 25452092:254 | 25452231:254 | +:+:+:+:+:+:+:+: | 5449 | pak4 | 1408 | 1117 | 1595 | 1217 | 1673 | 1020 |
| ENSDARG000 | 15 | 31276869 | 31277691 | + | 823 | NA | 0 | 0 | 2 | 0 | 0 | 0 |
| ENSDARG000 | 15 | 5261649 | 5262942 | - | 1294 | or64a1 | 0 | 0 | 0 | 0 | 0 | 0 |
| ENSDARG000 | 15:15:15:15:1 | 6948644:694 | 6948838:694 | -:-:-:-:-:-:-:-: | 1305 | mrps22 | 228 | 194 | 240 | 102 | 240 | 218 |
| ENSDARG000 | 15:15:15:15:1 | 25489406:254 | 25489581:254 | +;+;+;+;+ | 1636 | zgc:152863 | 170 | 143 | 122 | 220 | 240 | 124 |
| ENSDARG000 | 15:15:15:15:1 | 31282220:31 | 31283071:31 | -;-;-;-;-;- | 1373 | zgc:171927 | 0 | 0 | 0 | 0 | 0 | 2 |
| ENSDARG000 | 15 | 5277761 | 5280100 | + | 2340 | NA | 0 | 0 | 4 | 35 | 0 | 0 |
| ENSDARG000 | 15 | 5280858 | 5284248 | + | 3391 | or63a2 | 0 | 0 | 0 | 8 | 0 | 0 |
| ENSDARG000 | 15;15 | 5287703:528 | 5287807:528 | +;+ | 1361 | or63b1 | 0 | 0 | 0 | 0 | 0 | 0 |
| ENSDARG000 | 15:15:15:15:1 | 17376367:17 | 17376545:17 | +;+;+;+;+;+ | 916 | NA | 18 | 0 | 0 | 2 | 0 | 9 |
| ENSDARG000 | 15;15 | 28822736:28 | 28824682:28 | -;- | 2246 | GPR4 | 114 | 0 | 0 | 14 | 0 | 2 |
| ENSDARG000 | 15 | 5293785 | 5295023 | + | 1239 | NA | 0 | 0 | 0 | 0 | 0 | 0 |
| ENSDARG000 | 15:15:15:15:1 | 12273947:12 | 12274549:12 | -:-:-:-:-:-:-:-: | 1317 | fxyd6 | 2 | 2 | 0 | 0 | 6 | 0 |
| ENSDARG000 | 15 | 31296517 | 31298174 | + | 1658 | or71aq1 | 0 | 0 | 0 | 0 | 0 | 0 |
| ENSDARG000 | 15;15 | 31303355:31 | 31303402:31 | +;+ | 2023 | or71aq2 | 0 | 0 | 0 | 0 | 0 | 0 |
| ENSDARG000 | 15;15 | 31307825:31 | 31307876:31 | +;+ | 1951 | or71at1 | 0 | 0 | 0 | 0 | 0 | 0 |
| ENSDARG000 | 15 | 31311716 | 31313932 | - | 2217 | or71as2 | 0 | 0 | 0 | 0 | 0 | 0 |
| ENSDARG000 | 15 | 31315209 | 31315548 | - | 340 | NA | 0 | 0 | 0 | 0 | 0 | 0 |
| ENSDARG000 | 15 | 5299404 | 5300815 | + | 1412 | or63b2 | 0 | 0 | 0 | 0 | 0 | 0 |
| ENSDARG000 | 15:15:15:15:1 | 21276735:21 | 21276975:21 | +:+:+:+:+:+:+:+: | 2249 | ubash3bb | 721 | 788 | 891 | 699 | 745 | 651 |
| ENSDARG000 | 15:15:15:15:1 | 23911655:23 | 23911923:23 | +;+;+;+;+;+ | 1386 | NA | 14 | 2 | 14 | 62 | 17 | 32 |
| ENSDARG000 | 15:15:15:15:1 | 25501864:25 | 25505244:25 | -:-:-:-:-:-:-:-: | 5292 | hif1al | 2612 | 2467 | 2513 | 2556 | 3255 | 1407 |
| ENSDARG000 | 15 | 5307128 | 5309213 | - | 2086 | NA | 0 | 0 | 0 | 8 | 0 | 2 |
| ENSDARG000 | 15 | 5321612 | 5323546 | + | 1935 | or55a1 | 0 | 0 | 0 | 0 | 0 | 0 |
| ENSDARG000 | 15 | 31317358 | 31319276 | - | 1919 | or71ar1 | 0 | 0 | 0 | 0 | 0 | 0 |
| ENSDARG000 | 15 | 5332900 | 5334256 | + | 1357 | or55a2 | 0 | 0 | 0 | 0 | 0 | 0 |
| ENSDARG000 | 15 | 5339056 | 5341379 | + | 2324 | NA | 0 | 0 | 0 | 0 | 0 | 0 |
| ENSDARG000 | 15 | 5343186 | 5344504 | + | 1319 | NA | 0 | 0 | 0 | 0 | 0 | 0 |
| ENSDARG000 | 15:15:15:15:1 | 28874441:28 | 28874877:28 | -:-:-:-:-:-:-:-: | 6077 | eml2 | 1703 | 1237 | 994 | 982 | 2058 | 1959 |
| ENSDARG000 | 15;15;15;15 | 21335097:21 | 21335137:21 | +;+;+;+ | 644 | NA | 0 | 0 | 0 | 0 | 0 | 0 |
| ENSDARG000 | 15;15;15 | 611229:6150 | 612266:6151 | -;-;- | 1194 | NA | 34 | 12 | 51 | 20 | 53 | 22 |
| ENSDARG000 | 15 | 31320017 | 31323932 | - | 3916 | or71as1 | 0 | 0 | 0 | 0 | 0 | 0 |
| ENSDARG000 | 15;15 | 31327455:31 | 31327581:31 | +;+ | 1420 | or15.37 | 0 | 0 | 0 | 0 | 0 | 0 |
| ENSDARG000 | 15:15:15:15:1 | 17379512:17 | 17379826:17 | -:-:-:-:-:-:-:-: | 1419 | NA | 6 | 0 | 0 | 13 | 0 | 17 |
| ENSDARG000 | 15;15;15;15 | 31344472:31 | 31344508:31 | +;+;+;+ | 1414 | or40a1 | 0 | 0 | 0 | 0 | 0 | 0 |
| ENSDARG000 | 15;15 | 31350122:31 | 31351908:31 | -;- | 1925 | or42e1 | 0 | 0 | 0 | 0 | 0 | 0 |
| ENSDARG000 | 15 | 31364709 | 31366742 | - | 2034 | or42d1 | 0 | 0 | 0 | 0 | 0 | 0 |
| ENSDARG000 | 15;15 | 31371190:31 | 31372243:31 | -;- | 1170 | or42d3 | 0 | 0 | 0 | 0 | 0 | 0 |
| ENSDARG000 | 15 | 31382015 | 31383588 | - | 1574 | or42c1 | 0 | 0 | 0 | 0 | 0 | 0 |
| ENSDARG000 | 15;15;15 | 31332552:31 | 31332674:31 | +;+;+ | 1206 | or71al2 | 0 | 0 | 0 | 0 | 0 | 0 |
| ENSDARG000 | 15:15:15:15:1 | 20352123:20 | 20352575:20 | +:+:+:+:+:+:+:+: | 3025 | il15l | 12 | 14 | 8 | 4 | 19 | 18 |
| ENSDARG000 | 15:15:15:15:1 | 31331356:31 | 31331374:31 | -:-:-:-:-:-:-:-: | 1912 | or42e3 | 0 | 0 | 0 | 0 | 0 | 0 |
| ENSDARG000 | 15 | 31361787 | 31362141 | - | 355 | NA | 0 | 0 | 0 | 0 | 0 | 0 |
| ENSDARG000 | 15:15:15:15:1 | 5542068:554 | 5542177:554 | -;-;-;-;-;-;-;- | 1007 | NA | 5 | 2 | 4 | 0 | 8 | 8 |
| ENSDARG000 | 15;15;15 | 618376:6189 | 618426:6191 | +;+;+ | 1099 | NA | 67 | 39 | 38 | 6 | 77 | 17 |
| ENSDARG000 | 15:15:15:15:1 | 20281305:20 | 20281839:20 | +:+:+:+:+:+:+:+: | 9220 | NA | 333 | 68 | 557 | 132 | 485 | 369 |
| ENSDARG000 | 15:15:15:15:1 | 618081:6189 | 618425:6191 | +;+;+;+;+;+ | 5698 | NA | 226 | 123 | 218 | 117 | 341 | 177 |
| ENSDARG000 | 15:15:15:15:1 | 21521803:21 | 21522319:21 | -;-;-;-;- | 1090 | NA | 0 | 1 | 0 | 0 | 0 | 0 |
| ENSDARG000 | 15;15 | 20344912:20 | 20345389:20 | -;- | 650 | NA | 0 | 0 | 0 | 0 | 0 | 0 |
| ENSDARG000 | 15:15:15:15:1 | 20363859:20 | 20364684:20 | +:+:+:+:+:+:+:+: | 4604 | supt5h | 1855 | 1486 | 2486 | 1317 | 1835 | 1610 |
| ENSDARG000 | 15:15:15:15:1 | 25528290:25 | 25528477:25 | +:+:+:+:+:+:+:+: | 4443 | NA | 408 | 468 | 665 | 378 | 496 | 583 |
| ENSDARG000 | 15;15;15 | 638457:6417 | 638578:6422 | +;+;+ | 1654 | NA | 57 | 17 | 37 | 13 | 23 | 65 |
| ENSDARG000 | 15:15:15:15:1 | 6968631:696 | 6970301:697 | -:-:-:-:-:-:-:-: | 2161 | si:ch73-311h1 | 220 | 296 | 303 | 120 | 303 | 276 |
| ENSDARG000 | 15:15:15:15:1 | 656069:6566 | 656557:6570 | -;-;-;-;-;- | 3124 | NA | 4 | 0 | 0 | 0 | 2 | 2 |
| ENSDARG000 | 15:15:15:15:1 | 647854:6483 | 648266:6487 | -;-;-;-;-;-;-;- | 3945 | NA | 20 | 5 | 2 | 11 | 14 | 4 |
| ENSDARG000 | 15:15:15:15:1 | 23922264:23 | 23923308:23 | -:-:-:-:-:-:-:-: | 2895 | appbp2 | 673 | 504 | 763 | 603 | 770 | 645 |
| ENSDARG000 | 15:15:15:15:1 | 665156:6657 | 665640:6660 | -:-:-:-:-:-:-:-: | 3133 | NA | 38 | 24 | 4 | 1 | 36 | 27 |
| ENSDARG000 | 15;15;15 | 7032097:703 | 7032162:703 | -;-;- | 382 | NA | 0 | 0 | 0 | 0 | 0 | 0 |
| ENSDARG000 | 15:15:15:15:1 | 1847135:184 | 1848648:184 | -:-:-:-:-:-:-:-: | 4134 | porb | 2720 | 2099 | 3768 | 1764 | 4766 | 2277 |
| ENSDARG000 | 15:15:15:15:1 | 12326161:12 | 12326343:12 | -:-:-:-:-:-:-:-: | 1795 | tmprss13a | 18 | 8 | 8 | 27 | 3 | 12 |
| ENSDARG000 | 15:15:15:15:1 | 33825750:33 | 33826609:33 | -:-:-:-:-:-:-:-: | 2158 | NA | 82 | 18 | 14 | 24 | 38 | 70 |
| ENSDARG000 | 15:15:15:15:1 | 28940352:28 | 28940784:28 | +:+:+:+:+:+:+:+: | 2987 | NA | 0 | 0 | 14 | 0 | 6 | 4 |
| ENSDARG000 | 15:15:15:15:1 | 672984:6737 | 674764:6747 | -;-;-;-;-;-;-;- | 2817 | NA | 36 | 16 | 28 | 18 | 30 | 50 |
| ENSDARG000 | 15;15;15;15 | 7054754:705 | 7054831:705 | +;+;+;+ | 1135 | foxl2a | 0 | 0 | 0 | 0 | 0 | 5 |
| ENSDARG000 | 15:15:15:15:1 | 33843596:33 | 33843709:33 | +:+:+:+:+:+:+:+: | 2564 | inpp5ka | 787 | 447 | 533 | 602 | 577 | 607 |
| ENSDARG000 | 15:15:15:15:1 | 17397321:17 | 17398761:17 | -;-;-;-;-;-;- | 2891 | tubd1 | 86 | 251 | 134 | 26 | 208 | 63 |
| ENSDARG000 | 15;15;15 | 31389403:31 | 31390762:31 | -;-;- | 1477 | or42c3 | 0 | 0 | 0 | 0 | 0 | 0 |
| ENSDARG000 | 15 | 31399923 | 31401361 | - | 1439 | or42c4 | 0 | 0 | 0 | 0 | 0 | 0 |
| ENSDARG000 | 15:15:15:15:1 | 25545976:25 | 25546514:25 | -:-:-:-:-:-:-:-: | 2007 | NA | 6 | 6 | 6 | 0 | 4 | 0 |
| ENSDARG000 | 15 | 31404806 | 31406093 | - | 1288 | or42e2 | 0 | 0 | 0 | 0 | 0 | 0 |
| ENSDARG000 | 15 | 31407840 | 31409013 | - | 1174 | or42d2 | 0 | 0 | 0 | 0 | 0 | 0 |
| ENSDARG000 | 15;15 | 31412595:31 | 31413998:31 | -;- | 1404 | or42c2 | 0 | 0 | 0 | 0 | 0 | 0 |
| ENSDARG000 | 15:15:15:15:1 | 28989573:28 | 28989722:28 | +:+:+:+:+:+:+:+: | 2879 | CLIP3 | 327 | 121 | 204 | 294 | 350 | 541 |
| ENSDARG000 | 15;15 | 694653:6951 | 694790:6951 | -;- | 186 | NA | 0 | 0 | 0 | 0 | 0 | 0 |
| ENSDARG000 | 15:15:15:15:1 | 685007:6946 | 688839:6947 | -;-;-;-;-;-;-;- | 5465 | NA | 119 | 111 | 100 | 46 | 86 | 117 |
| ENSDARG000 | 15;15 | 31417886:31 | 31419546:31 | -;- | 1808 | or42c5 | 0 | 0 | 0 | 0 | 0 | 0 |
| ENSDARG000 | 15:15:15:15:1 | 12377887:12 | 12378172:12 | +;+;+;+;+;+;+ | 3764 | il10ra | 719 | 261 | 319 | 1018 | 976 | 489 |
| ENSDARG000 | 15 | 31426768 | 31428815 | + | 2048 | or30bt1 | 0 | 0 | 0 | 0 | 0 | 0 |
| ENSDARG000 | 15 | 31428332 | 31430719 | + | 2388 | or30bt3 | 0 | 0 | 0 | 0 | 0 | 0 |
| ENSDARG000 | 15:15:15:15:1 | 33868661:33 | 33868957:33 | -:-:-:-:-:-:-:-: | 1293 | NA | 0 | 0 | 0 | 0 | 0 | 0 |
| ENSDARG000 | 15:15:15:15:1 | 33865312:33 | 33865391:33 | +:+:+:+:+:+:+:+: | 1436 | tekt1 | 5 | 0 | 25 | 0 | 0 | 0 |
| ENSDARG000 | 15:15:15:15:1 | 25556922:25 | 25557733:25 | -:-:-:-:-:-:-:-: | 2249 | NA | 2 | 6 | 0 | 12 | 10 | 0 |
| ENSDARG000 | 15;15;15 | 33885528:33 | 33885585:33 | +;+;+ | 484 | NA | 0 | 0 | 0 | 0 | 0 | 0 |
| ENSDARG000 | 15:15:15:15:1 | 15479442:154 | 15479508:154 | +:+:+:+:+:+:+:+: | 2784 | nek8 | 139 | 79 | 68 | 70 | 156 | 46 |
| ENSDARG000 | 15;15;15;15 | 12411196:124 | 12411846:124 | -;-;-;- | 911 | NA | 0 | 1 | 2 | 0 | 0 | 2 |
| ENSDARG000 | 15:15:15:15:1 | 5565641:556 | 5567073:556 | -:-:-:-:-:-:-:-: | 5827 | wdr62 | 1024 | 884 | 727 | 550 | 733 | 939 |
| ENSDARG000 | 15:15:15:15:1 | 21541010:21 | 21542936:21 | -:-:-:-:-:-:-:-: | 8581 | SORL1 | 993 | 430 | 1184 | 414 | 1263 | 1019 |
| ENSDARG000 | 15:15:15:15:1 | 7064819:706 | 7065154:706 | +:+:+:+:+:+:+:+: | 5316 | PIK3CB | 2054 | 1744 | 2251 | 1434 | 2277 | 1749 |
| ENSDARG000 | 15:15:15:15:1 | 12429206:124 | 12429410:124 | +:+:+:+:+:+:+:+: | 2845 | tmprss4a | 77 | 56 | 80 | 78 | 35 | 37 |
| ENSDARG000 | 15;15;15 | 29011290:29 | 29011510:29 | -;-;- | 770 | dharma | 0 | 0 | 0 | 0 | 0 | 0 |
| ENSDARG000 | 15:15:15:15:1 | 714203:7143 | 714253:7144 | +;+;+;+;+;+;+ | 1564 | NA | 14 | 12 | 94 | 64 | 165 | 66 |
| ENSDARG000 | 15:15:15:15:1 | 25576809:25 | 25577553:25 | -:-:-:-:-:-:-:-: | 2531 | NA | 200 | 138 | 263 | 208 | 228 | 188 |
| ENSDARG000 | 15:15:15:15:1 | 33891808:33 | 33892157:33 | -:-:-:-:-:-:-:-: | 2795 | mag | 26 | 2 | 4 | 0 | 18 | 4 |
| ENSDARG000 | 15:15:15:15:1 | 23947932:23 | 23948038:23 | +:+:+:+:+:+:+:+: | 10822 | MYO18A | 1570 | 1698 | 2587 | 1730 | 2922 | 2194 |
| ENSDARG000 | 15:15:15:15:1 | 1897375:190 | 1900202:190 | -:-:-:-:-:-:-:-: | 9903 | DOCK10 | 985 | 1081 | 1233 | 786 | 1102 | 1083 |
| ENSDARG000 | 15:15:15:15:1 | 34699585:34 | 34699814:34 | +;+;+;+;+;+ | 1764 | tnfb | 189 | 102 | 128 | 148 | 332 | 293 |
| ENSDARG000 | 15:15:15:15:1 | 21673454:21 | 21673899:21 | -:-:-:-:-:-:-:-: | 2050 | bco2b | 799 | 96 | 302 | 79 | 461 | 96 |
| ENSDARG000 | 15:15:15:15:1 | 17406920:17 | 17407149:17 | +:+:+:+:+:+:+:+: | 2346 | rps6kb1b | 713 | 872 | 1186 | 439 | 899 | 1032 |
| ENSDARG000 | 15;15;15;15 | 37992275:37 | 37992651:37 | +;+;+;+ | 811 | NA | 0 | 0 | 0 | 0 | 2 | 2 |
| ENSDARG000 | 15;15;15;15 | 7176182:717 | 7176289:717 | +;+;+;+ | 1046 | her8.2 | 0 | 0 | 28 | 61 | 14 | 0 |

| | | | | | | | | | | | |
|---|---|---|---|---|---|---|---|---|---|---|---|
| ENSDARG000 | 15;15;15;15 | 23999599:24( | 23999707:24( | +;+;+;+ | 565 | NA | 0 | 0 | 0 | 0 | 0 | 1 |
| ENSDARG000 | 15:15:15:15:1 | 15516612:155 | 15517005:155 | +:+:+:+:+:+:+: | 3072 | traf4a | 3460 | 1547 | 4766 | 2828 | 3265 | 2420 |
| ENSDARG000 | 15:15:15:15:1 | 21672281:216 | 21672372:216 | +:+:+:+:+:+:+: | 649 | NA | 15 | 0 | 6 | 0 | 34 | 0 |
| ENSDARG000 | 15:15:15:15:1 | 20395309:203 | 20395606:203 | -:-:-:-:-:-:-:-: | 1008 | rnaset2l | 1 | 26 | 8 | 16 | 68 | 16 |
| ENSDARG000 | 15:15:15:15:1 | 1891689:1891 | 1891957:1891 | +:+:+:+:+:+:+: | 1482 | sbds | 226 | 263 | 258 | 230 | 271 | 307 |
| ENSDARG000 | 15;15 | 38006366:38( | 38006616:38( | +;+ | 537 | NA | 0 | 0 | 0 | 0 | 0 | 0 |
| ENSDARG000 | 15;15 | 897280:8990 | 897447:9000 | +;+ | 1125 | NA | 8 | 27 | 16 | 10 | 10 | 30 |
| ENSDARG000 | 15:15:15:15:1 | 846768:84707 | 846814:84724 | +;+;+;+;+;+ | 1201 | NA | 31 | 11 | 16 | 7 | 5 | 16 |
| ENSDARG000 | 15;15 | 38008502:38( | 38008972:38( | -;- | 750 | NA | 0 | 0 | 0 | 0 | 0 | 0 |
| ENSDARG000 | 15:15:15:15:1 | 863931:86476 | 863996:86492 | +;+;+;+;+ | 4298 | NA | 93 | 61 | 128 | 86 | 155 | 73 |
| ENSDARG000 | 15;15 | 857148:85869 | 857364:85903 | +;+ | 551 | NA | 0 | 0 | 0 | 0 | 0 | 0 |
| ENSDARG000 | 15:15:15:15:1 | 934660:93516 | 934766:93535 | +:+:+:+:+:+:+: | 982 | NA | 0 | 2 | 2 | 0 | 0 | 0 |
| ENSDARG000 | 15:15:15:15:1 | 38018301:38( | 38018362:38( | +:+:+:+:+:+:+: | 1121 | NA | 15 | 10 | 3 | 0 | 79 | 71 |
| ENSDARG000 | 15:15:15:15:1 | 791220:79279 | 791505:79288 | +:+:+:+:+:+:+: | 3720 | NA | 106 | 35 | 60 | 83 | 192 | 99 |
| ENSDARG000 | 15:15:15:15:1 | 846892:10046 | 847241:10048 | +;+;+;+;+;+;+ | 2324 | NA | 96 | 60 | 146 | 88 | 97 | 60 |
| ENSDARG000 | 15:15:15:15:1 | 942531:94262 | 942792:9428( | -:-:-:-:-:-:-:-: | 2345 | ALOX5 | 0 | 0 | 0 | 6 | 10 | 6 |
| ENSDARG000 | 15:15:15:15:1 | 25588357:255 | 25596164:255 | -:-:-:-:-:-:-:-: | 10997 | TAOK1 | 3259 | 2155 | 2632 | 2457 | 2776 | 2259 |
| ENSDARG000 | 15:15:15:15:1 | 34717514:347 | 34717775:347 | -:-:-:-:-:-:-:-: | 3275 | GABBR1 | 61 | 55 | 147 | 40 | 194 | 42 |
| ENSDARG000 | 15:15:15:15:1 | 38025319:38( | 38025751:38( | +:+:+:+:+:+:+: | 1529 | NA | 16 | 8 | 8 | 0 | 30 | 20 |
| ENSDARG000 | 15;15;15;15 | 864761:88075 | 864928:88091 | +;+;+;+ | 5611 | NA | 72 | 26 | 69 | 39 | 71 | 37 |
| ENSDARG000 | 15;15;15 | 998839:10009 | 999500:1001( | -;-;- | 855 | NA | 54 | 41 | 84 | 44 | 79 | 23 |
| ENSDARG000 | 15:15:15:15:1 | 12466852:124 | 12467812:124 | -;-;-;-;- | 2072 | scn4ba | 0 | 0 | 0 | 0 | 0 | 0 |
| ENSDARG000 | 15;15;15 | 758188:76178 | 758734:76192 | -;-;- | 739 | NA | 10 | 1 | 0 | 0 | 0 | 0 |
| ENSDARG000 | 15;15;15 | 823437:82414 | 823486:82431 | +;+;+ | 985 | NA | 51 | 44 | 54 | 30 | 38 | 78 |
| ENSDARG000 | 15;15;15 | 872089:87284 | 872135:8730( | +;+;+ | 828 | NA | 5 | 5 | 8 | 14 | 2 | 6 |
| ENSDARG000 | 15;15;15 | 913798:91544 | 914163:9156( | -;-;- | 585 | NA | 10 | 6 | 10 | 2 | 5 | 6 |
| ENSDARG000 | 15;15;15 | 807897:80892 | 807980:80909 | +;+;+ | 1969 | NA | 102 | 114 | 147 | 118 | 163 | 131 |
| ENSDARG000 | 15:15:15:15:1 | 33937756:339 | 33939776:339 | -:-:-:-:-:-:-:-: | 4794 | lsr | 152 | 20 | 18 | 20 | 28 | 54 |
| ENSDARG000 | 15:15:15:15:1 | 29024895:29( | 29024932:29( | +:+:+:+:+:+:+: | 3486 | si:ch211-137a | 6 | 7 | 11 | 2 | 3 | 10 |
| ENSDARG000 | 15:15:15:15:1 | 722475:77952 | 725149:77969 | -:-:-:-:-:-:-:-: | 4508 | zgc:162936 | 60 | 15 | 65 | 84 | 40 | 46 |
| ENSDARG000 | 15:15:15:15:1 | 872085:87989 | 872135:87998 | +:+:+:+:+:+:+: | 4077 | NA | 86 | 89 | 50 | 30 | 117 | 97 |
| ENSDARG000 | 15;15;15;15 | 7187228:7187 | 7187370:7187 | +;+;+;+ | 1526 | her13 | 2 | 2 | 0 | 12 | 12 | 0 |
| ENSDARG000 | 15;15 | 983741:98614 | 983881:99343 | +;+ | 7427 | NA | 384 | 389 | 444 | 132 | 410 | 336 |
| ENSDARG000 | 15;15 | 834697:83652 | 834838:83706 | +;+ | 684 | NA | 4 | 0 | 6 | 0 | 10 | 27 |
| ENSDARG000 | 15:15:15:15:1 | 722475:72247 | 725149:72514 | -:-:-:-:-:-:-:-: | 4201 | NA | 0 | 0 | 0 | 0 | 0 | 0 |
| ENSDARG000 | 15;15;15 | 742198:74512 | 742566:74529 | -;-;- | 596 | NA | 19 | 12 | 39 | 16 | 32 | 36 |
| ENSDARG000 | 15:15:15:15:1 | 872085:87989 | 872135:87998 | +:+:+:+:+:+:+: | 7135 | NA | 111 | 129 | 168 | 79 | 144 | 195 |
| ENSDARG000 | 15:15:15:15:1 | 963292:96671 | 963361:96686 | +:+:+:+:+:+:+: | 2358 | alox5b.2 | 76 | 72 | 12 | 14 | 32 | 6 |
| ENSDARG000 | 15:15:15:15:1 | 34854025:348 | 34855571:348 | -:-:-:-:-:-:-:-: | 2719 | sh3bp5la | 496 | 419 | 464 | 356 | 569 | 450 |
| ENSDARG000 | 15:15:15:15:1 | 38046435:38( | 38046639:38( | +:+:+:+:+:+:+: | 2254 | NA | 112 | 14 | 60 | 12 | 44 | 117 |
| ENSDARG000 | 15:15:15:15:1 | 850829:85146 | 850860:85163 | +;+;+;+;+ | 3214 | NA | 43 | 41 | 37 | 16 | 58 | 26 |
| ENSDARG000 | 15:15:15:15:1 | 749230:76494 | 749682:76526 | -;-;-;-;-;- | 1422 | NA | 2 | 17 | 19 | 4 | 4 | 2 |
| ENSDARG000 | 15:15:15:15:1 | 722475:75024 | 725149:75113 | -;-;-;-;- | 3826 | NA | 26 | 11 | 5 | 10 | 18 | 8 |
| ENSDARG000 | 15:15:15:15:1 | 34867518:348 | 34868794:348 | -;-;-;-;- | 2473 | mgat1a | 395 | 308 | 340 | 443 | 453 | 342 |
| ENSDARG000 | 15:15:15:15:1 | 781717:78265 | 781772:78277 | +;+;+;+;+;+ | 2124 | NA | 34 | 37 | 20 | 11 | 63 | 27 |
| ENSDARG000 | 15:15:15:15:1 | 38071874:38( | 38071916:38( | +;+;+;+;+ | 1099 | NA | 0 | 4 | 0 | 0 | 2 | 20 |
| ENSDARG000 | 15:15:15:15:1 | 834721:84138 | 834838:84143 | +;+;+;+;+ | 3634 | NA | 10 | 42 | 62 | 20 | 29 | 33 |
| ENSDARG000 | 15:15:15:15:1 | 808647:81655 | 809094:8166( | +;+;+;+;+ | 1855 | zgc:113294 | 14 | 42 | 22 | 2 | 22 | 42 |
| ENSDARG000 | 15;15;15 | 729376:73364 | 729936:73378 | -;-;- | 798 | NA | 0 | 0 | 0 | 0 | 0 | 0 |
| ENSDARG000 | 15;15 | 920298:92292 | 921198:92398 | -;- | 1959 | NA | 20 | 18 | 41 | 40 | 44 | 24 |
| ENSDARG000 | 15;15;15 | 736522:73868 | 737209:73882 | -;-;- | 885 | NA | 0 | 1 | 0 | 0 | 0 | 0 |
| ENSDARG000 | 15;15 | 34881013:348 | 34882709:348 | -;- | 2431 | NA | 8 | 10 | 28 | 68 | 2 | 4 |
| ENSDARG000 | 15:15:15:15:1 | 33966739:339 | 33966824:339 | +:+:+:+:+:+:+: | 1932 | tmem106ba | 800 | 511 | 1104 | 646 | 548 | 1031 |
| ENSDARG000 | 15:15:15:15:1 | 34886480:348 | 34888199:348 | -;-;-;-;- | 1843 | rnf183 | 52 | 18 | 19 | 26 | 54 | 0 |
| ENSDARG000 | 15;15;15 | 15629506:156 | 15629622:156 | +;+;+ | 481 | NA | 0 | 15 | 10 | 0 | 15 | 0 |
| ENSDARG000 | 15:15:15:15:1 | 38101135:381 | 38101658:381 | -:-:-:-:-:-:-:-: | 3426 | si:dkey-24p1.: | 0 | 9 | 4 | 0 | 12 | 20 |
| ENSDARG000 | 15:15:15:15:1 | 33994575:339 | 33994837:339 | -:-:-:-:-:-:-:-: | 6103 | NA | 0 | 8 | 972 | 60 | 34 | 18 |
| ENSDARG000 | 15:15:15:15:1 | 33989181:339 | 33989438:339 | +;+;+;+;+;+;+ | 1409 | seraf | 12 | 8 | 8 | 4 | 24 | 0 |
| ENSDARG000 | 15:15:15:15:1 | 7384520:7409 | 7384637:741( | -:-:-:-:-:-:-:-: | 3122 | GBE1 | 119 | 134 | 106 | 53 | 192 | 83 |
| ENSDARG000 | 15:15:15:15:1 | 17441734:174 | 17441945:174 | +:+:+:+:+:+:+: | 6464 | SNX19 | 376 | 314 | 298 | 266 | 350 | 342 |
| ENSDARG000 | 15;15 | 38150465:381 | 38151731:381 | -;- | 1333 | irgq2 | 1 | 4 | 2 | 0 | 2 | 13 |
| ENSDARG000 | 15:15:15:15:1 | 34933552:349 | 34933624:349 | +:+:+:+:+:+:+: | 1980 | NA | 0 | 0 | 2 | 0 | 0 | 0 |
| ENSDARG000 | 15:15:15:15:1 | 20403084:204 | 20403188:204 | +:+:+:+:+:+:+: | 952 | cox7a1 | 82 | 348 | 117 | 154 | 168 | 160 |
| ENSDARG000 | 15:15:15:15:1 | 12521379:125 | 12522935:125 | -:-:-:-:-:-:-:-: | 2641 | scn2b | 16 | 16 | 28 | 42 | 2 | 31 |
| ENSDARG000 | 15:15:15:15:1 | 34899645:348 | 34899985:348 | -:-:-:-:-:-:-:-: | 3840 | dhx16 | 510 | 555 | 712 | 300 | 484 | 429 |
| ENSDARG000 | 15 | 7805779 | 7805920 | + | 142 | NA | 0 | 0 | 0 | 0 | 0 | 0 |
| ENSDARG000 | 15:15:15:15:1 | 38157487:381 | 38157937:381 | -;-;-;-;-;-;-;- | 2391 | NA | 6 | 20 | 6 | 8 | 56 | 132 |
| ENSDARG000 | 15:15:15:15:1 | 38179904:381 | 38180742:381 | -;-;-;-;-;-;-;- | 1892 | NA | 0 | 0 | 0 | 0 | 0 | 0 |
| ENSDARG000 | 15:15:15:15:1 | 7992906:8043 | 7992932:8043 | +:+:+:+:+:+:+: | 1260 | cadm2b | 2 | 0 | 0 | 0 | 2 | 6 |
| ENSDARG000 | 15:15:15:15:1 | 24150090:241 | 24150959:241 | -;-;-;-;-;-;- | 1963 | pipox | 186 | 14 | 10 | 87 | 8 | 58 |
| ENSDARG000 | 15:15:15:15:1 | 15708065:157 | 15708946:157 | -:-:-:-:-:-:-:-: | 2159 | NA | 0 | 0 | 0 | 0 | 0 | 0 |
| ENSDARG000 | 15;15;15 | 38201964:382 | 38202625:382 | -;-;- | 814 | rhoga | 6 | 4 | 16 | 2 | 10 | 20 |
| ENSDARG000 | 15:15:15:15:1 | 34062460:34( | 34062581:34( | +;+;+;+;+ | 1000 | NA | 0 | 0 | 22 | 0 | 0 | 0 |
| ENSDARG000 | 15;15 | 34069746:34( | 34069870:34( | +;+ | 1090 | arl4aa | 36 | 31 | 180 | 47 | 57 | 22 |
| ENSDARG000 | 15:15:15:15:1 | 29085955:29( | 29086375:29( | +;+;+;+;+ | 2967 | NA | 1554 | 637 | 2813 | 953 | 1036 | 3360 |
| ENSDARG000 | 15;15;15;15 | 21684435:216 | 21685544:216 | -;-;-;- | 1481 | sdhdb | 690 | 845 | 837 | 408 | 736 | 550 |
| ENSDARG000 | 15:15:15:15:1 | 34939906:349 | 34940031:349 | +:+:+:+:+:+:+: | 1967 | ppt2a.1 | 0 | 0 | 0 | 0 | 0 | 0 |
| ENSDARG000 | 15:15:15:15:1 | 17603488:176 | 17604564:176 | -:-:-:-:-:-:-:-: | 4066 | NA | 4 | 4 | 0 | 0 | 8 | 0 |
| ENSDARG000 | 15:15:15:15:1 | 13107342:131 | 13107730:131 | -:-:-:-:-:-:-:-: | 3789 | zgc:172282 | 0 | 0 | 0 | 0 | 0 | 0 |
| ENSDARG000 | 15:15:15:15:1 | 34946928:349 | 34947344:349 | +:+:+:+:+:+:+: | 1438 | ppt2a.2 | 0 | 0 | 0 | 0 | 0 | 0 |
| ENSDARG000 | 15:15:15:15:1 | 34946779:349 | 34947609:349 | +:+:+:+:+:+:+: | 9030 | NA | 39 | 3 | 4 | 12 | 2 | 38 |
| ENSDARG000 | 15:15:15:15:1 | 8170884:817( | 8172754:8172 | -:-:-:-:-:-:-:-: | 4074 | bach1a | 1215 | 648 | 2045 | 1337 | 1065 | 1220 |
| ENSDARG000 | 15:15:15:15:1 | 34215796:342 | 34215871:342 | -:-:-:-:-:-:-:-: | 2896 | DGKB | 0 | 0 | 0 | 0 | 0 | 0 |
| ENSDARG000 | 15:15:15:15:1 | 24212847:243 | 24213262:243 | +:+:+:+:+:+:+: | 4524 | SEZ6 | 0 | 2 | 0 | 0 | 0 | 0 |
| ENSDARG000 | 15:15:15:15:1 | 38299563:382 | 38299610:383 | +:+:+:+:+:+:+: | 5444 | NA | 8 | 97 | 2 | 46 | 74 | 0 |
| ENSDARG000 | 15:15:15:15:1 | 21694388:216 | 21695746:216 | -;-;-;-;-;- | 2181 | NA | 475 | 411 | 144 | 8 | 208 | 26 |
| ENSDARG000 | 15:15:15:15:1 | 38299385:382 | 38299610:383 | +;+;+;+;+ | 1014 | NA | 0 | 0 | 0 | 0 | 0 | 0 |
| ENSDARG000 | 15:15:15:15:1 | 38221038:382 | 38221463:382 | +:+:+:+:+:+:+: | 7939 | stim1a | 2805 | 1511 | 2789 | 1449 | 3023 | 2187 |
| ENSDARG000 | 15:15:15:15:1 | 29113616:291 | 29114201:291 | +:+:+:+:+:+:+: | 2153 | capns1b | 2875 | 2987 | 2314 | 4214 | 2979 | 3143 |
| ENSDARG000 | 15:15:15:15:1 | 17621375:176 | 17621545:176 | +:+:+:+:+:+:+: | 4382 | ADAMTS8 | 22 | 0 | 0 | 0 | 34 | 14 |
| ENSDARG000 | 15;15;15 | 20411292:204 | 20411417:204 | +;+;+ | 570 | NA | 2 | 0 | 6 | 15 | 0 | 4 |
| ENSDARG000 | 15:15:15:15:1 | 20406143:204 | 20407299:204 | -;-;-;-;-;- | 1879 | chp2 | 576 | 524 | 560 | 1237 | 504 | 1043 |
| ENSDARG000 | 15:15:15:15:1 | 17722054:177 | 17722121:177 | +:+:+:+:+:+:+: | 2249 | NA | 2 | 2 | 0 | 0 | 4 | 0 |
| ENSDARG000 | 15;15 | 17751150:177 | 17753161:177 | -;- | 2444 | NA | 0 | 0 | 0 | 0 | 0 | 2 |
| ENSDARG000 | 15:15:15:15:1 | 34988148:349 | 34988506:349 | +:+:+:+:+:+:+: | 1576 | NA | 0 | 0 | 0 | 0 | 0 | 0 |
| ENSDARG000 | 15:15:15:15:1 | 17752915:177 | 17753106:177 | +:+:+:+:+:+:+: | 1629 | NA | 28 | 0 | 8 | 0 | 4 | 0 |
| ENSDARG000 | 15:15:15:15:1 | 25635326:256 | 25635473:256 | +:+:+:+:+:+:+: | 2658 | tsr1 | 1246 | 626 | 1942 | 321 | 750 | 828 |
| ENSDARG000 | 15:15:15:15:1 | 35004547:35( | 35004873:35( | -:-:-:-:-:-:-:-: | 2736 | NA | 0 | 2 | 4 | 0 | 0 | 16 |
| ENSDARG000 | 15;15 | 14034254:14( | 14036750:14( | -;- | 2691 | zgc:114130 | 212 | 288 | 415 | 204 | 360 | 380 |
| ENSDARG000 | 15;15 | 14062617:14( | 14063009:14( | -;- | 572 | NA | 2 | 0 | 0 | 0 | 0 | 0 |
| ENSDARG000 | 15:15:15:15:1 | 29092527:29( | 29093310:29( | -:-:-:-:-:-:-:-: | 2679 | zgc:162698 | 1078 | 884 | 1026 | 841 | 1078 | 891 |
| ENSDARG000 | 15:15:15:15:1 | 34334722:343 | 34335283:343 | -:-:-:-:-:-:-:-: | 2032 | agmo | 18 | 56 | 369 | 112 | 400 | 190 |
| ENSDARG000 | 15:15:15:15:1 | 21707783:217 | 21707806:217 | +:+:+:+:+:+:+: | 5550 | nkapd1 | 120 | 108 | 170 | 99 | 150 | 175 |
| ENSDARG000 | 15:15:15:15:1 | 8192715:8192 | 8193210:8193 | +:+:+:+:+:+:+: | 3134 | MIPEP | 389 | 216 | 486 | 112 | 226 | 344 |
| ENSDARG000 | 15:15:15:15:1 | 17848590:178 | 17848889:178 | +:+:+:+:+:+:+: | 2493 | zgc:113279 | 17 | 0 | 0 | 0 | 0 | 0 |
| ENSDARG000 | 15:15:15:15:1 | 35043007:35( | 35043209:35( | +:+:+:+:+:+:+: | 3720 | sesn3 | 454 | 623 | 1909 | 1617 | 2000 | 1848 |
| ENSDARG000 | 15;15;15 | 17844191:178 | 17844879:178 | -;-;- | 818 | NA | 0 | 0 | 0 | 0 | 2 | 0 |
| ENSDARG000 | 15:15:15:15:1 | 35095196:35( | 35095546:35( | -;-;-;-;-;- | 951 | NA | 0 | 0 | 0 | 0 | 0 | 0 |
| ENSDARG000 | 15:15:15:15:1 | 17862207:178 | 17867364:178 | -;-;-;-;-;-;-;- | 6765 | NA | 711 | 1003 | 2810 | 4121 | 446 | 1238 |
| ENSDARG000 | 15:15:15:15:1 | 35036368:35( | 35039809:35( | -:-:-:-:-:-:-:-: | 5238 | bloc1s3 | 102 | 127 | 155 | 238 | 111 | 220 |
| ENSDARG000 | 15:15:15:15:1 | 15771418:157 | 15771469:157 | +;+;+;+;+ | 752 | NA | 0 | 0 | 0 | 0 | 0 | 0 |
| ENSDARG000 | 15:15:15:15:1 | 35036368:35( | 35039809:35( | -:-:-:-:-:-:-:-: | 6935 | ppt2a.3 | 32 | 14 | 42 | 20 | 37 | 112 |
| ENSDARG000 | 15:15:15:15:1 | 35114784:351 | 35115716:351 | -:-:-:-:-:-:-:-: | 2091 | ppt2a.4 | 41 | 10 | 40 | 45 | 33 | 133 |
| ENSDARG000 | 15:15:15:15:1 | 34428308:344 | 34428937:344 | -;-;-;-;-;- | 2363 | MEOX2 | 10 | 0 | 14 | 0 | 22 | 6 |
| ENSDARG000 | 15;15;15;15 | 15779184:157 | 15779380:157 | +;+;+;+ | 687 | NA | 0 | 14 | 34 | 4 | 10 | 6 |
| ENSDARG000 | 15:15:15:15:1 | 24549054:245 | 24549906:245 | +:+:+:+:+:+:+: | 4889 | PHF12 | 676 | 594 | 740 | 367 | 751 | 472 |
| ENSDARG000 | 15:15:15:15:1 | 20463448:204 | 20464780:204 | -:-:-:-:-:-:-:-: | 3657 | dlc | 86 | 40 | 187 | 26 | 120 | 142 |
| ENSDARG000 | 15:15:15:15:1 | 14076004:14( | 14076832:14( | -:-:-:-:-:-:-:-: | 1563 | trappc6bl | 328 | 245 | 366 | 244 | 174 | 436 |
| ENSDARG000 | 15:15:15:15:1 | 34573311:345 | 34574268:345 | -:-:-:-:-:-:-:-: | 2189 | ankmy2a | 313 | 223 | 390 | 197 | 263 | 272 |
| ENSDARG000 | 15:15:15:15:1 | 34495056:344 | 34495448:344 | -:-:-:-:-:-:-:-: | 2056 | crppa | 392 | 444 | 502 | 302 | 674 | 495 |
| ENSDARG000 | 15:15:15:15:1 | 34568701:345 | 34568802:345 | +:+:+:+:+:+:+: | 852 | NA | 2 | 6 | 10 | 15 | 11 | 9 |
| ENSDARG000 | 15;15 | 34562197:345 | 34563140:345 | -;- | 1344 | sostdc1a | 4 | 2 | 10 | 0 | 8 | 10 |
| ENSDARG000 | 15:15:15:15:1 | 15786160:157 | 15786481:157 | +:+:+:+:+:+:+: | 1891 | tada2a | 222 | 190 | 220 | 200 | 180 | 182 |
| ENSDARG000 | 15:15:15:15:1 | 20475694:204 | 20476403:204 | -:-:-:-:-:-:-:-: | 1807 | timm50 | 212 | 469 | 472 | 163 | 350 | 269 |
| ENSDARG000 | 15;15 | 15856178:158 | 15856347:158 | +;+ | 1344 | dusp14 | 0 | 2 | 4 | 2 | 4 | 8 |
| ENSDARG000 | 15:15:15:15:1 | 38339386:383 | 38339546:383 | -:-:-:-:-:-:-:-: | 2520 | NA | 0 | 0 | 0 | 0 | 0 | 0 |
| ENSDARG000 | 15;15;15;15 | 25681044:256 | 25681229:256 | +;+;+;+ | 2617 | hic1 | 16 | 17 | 85 | 2 | 60 | 46 |
| ENSDARG000 | 15:15:15:15:1 | 35130621:351 | 35130876:351 | -;-;-;-;-;-;- | 1422 | NA | 0 | 0 | 0 | 0 | 0 | 0 |
| ENSDARG000 | 15:15:15:15:1 | 21744680:217 | 21744825:217 | +:+:+:+:+:+:+: | 3026 | ppp2r1bb | 4714 | 5455 | 4326 | 3584 | 4479 | 3518 |
| ENSDARG000 | 15;15;15;15 | 14085187:14( | 14086176:14( | -;-;-;- | 2070 | si:ch211-116( | 266 | 136 | 302 | 228 | 208 | 336 |
| ENSDARG000 | 15:15:15:15:1 | 25718974:257 | 25719503:257 | -:-:-:-:-:-:-:-: | 1534 | NA | 0 | 0 | 0 | 0 | 0 | 0 |
| ENSDARG000 | 15:15:15:15:1 | 17887307:178 | 17887324:178 | +:+:+:+:+:+:+: | 4407 | NA | 104 | 4 | 4 | 15 | 6 | 14 |
| ENSDARG000 | 15:15:15:15:1 | 20543770:205 | 20543827:205 | +:+:+:+:+:+:+: | 3719 | SGSM2 | 39 | 22 | 38 | 30 | 18 | 38 |
| ENSDARG000 | 15 | 2051128 | 2052259 | - | 1132 | NA | 0 | 0 | 0 | 6 | 0 | 0 |
| ENSDARG000 | 15;15;15;15 | 38859648:388 | 38859872:388 | +;+;+;+ | 636 | NA | 0 | 1 | 8 | 0 | 0 | 3 |
| ENSDARG000 | 15:15:15:15:1 | 38948423:389 | 38950226:389 | -:-:-:-:-:-:-:-: | 2849 | NA | 2 | 0 | 0 | 0 | 2 | 2 |
| ENSDARG000 | 15:15:15:15:1 | 35139457:351 | 35139780:351 | +;+;+;+;+;+ | 2115 | NA | 1 | 12 | 9 | 13 | 3 | 15 |
| ENSDARG000 | 15:15:15:15:1 | 35156414:351 | 35157196:351 | -:-:-:-:-:-:-:-: | 2581 | agfg1a | 508 | 708 | 1002 | 867 | 748 | 638 |
| ENSDARG000 | 15:15:15:15:1 | 25910501:259 | 25911016:259 | -:-:-:-:-:-:-:-: | 2138 | dph1 | 316 | 236 | 426 | 196 | 330 | 298 |
| ENSDARG000 | 15:15:15:15:1 | 14109652:141 | 14109814:141 | +:+:+:+:+:+:+: | 3412 | clca1 | 0 | 6 | 0 | 0 | 6 | 2 |
| ENSDARG000 | 15:15:15:15:1 | 2182103:2182 | 2182916:2182 | -:-:-:-:-:-:-:-: | 2019 | shox2 | 0 | 0 | 6 | 249 | 0 | 0 |
| ENSDARG000 | 15:15:15:15:1 | 35214546:352 | 35215021:352 | -:-:-:-:-:-:-:-: | 1439 | mrpl44 | 292 | 393 | 559 | 268 | 423 | 320 |
| ENSDARG000 | 15:15:15:15:1 | 26498339:265 | 26498933:265 | -:-:-:-:-:-:-:-: | 2421 | rpa1 | 755 | 1807 | 2091 | 670 | 1865 | 1160 |
| ENSDARG000 | 15:15:15:15:1 | 20529197:205 | 20529449:205 | +:+:+:+:+:+:+: | 1928 | tnfaip1 | 404 | 304 | 323 | 346 | 345 | 262 |
| ENSDARG000 | 15:15:15:15:1 | 15885849:158 | 15886904:158 | -:-:-:-:-:-:-:-: | 5955 | SYNRG | 1418 | 976 | 1591 | 1217 | 1428 | 1057 |
| ENSDARG000 | 15:15:15:15:1 | 39277099:392 | 39278855:392 | -:-:-:-:-:-:-:-: | 7676 | ROBO1 | 1692 | 2103 | 6493 | 3766 | 6980 | 5605 |
| ENSDARG000 | 15:15:15:15:1 | 35226673:352 | 35227535:352 | -:-:-:-:-:-:-:-: | 1790 | mffa | 561 | 571 | 626 | 968 | 609 | 524 |
| ENSDARG000 | 15:15:15:15:1 | 39847495:398 | 39847823:398 | -;-;-;-;- | 739 | NA | 11 | 0 | 0 | 16 | 5 | 6 |
| ENSDARG000 | 15:15:15:15:1 | 17948397:179 | 17949353:179 | -:-:-:-:-:-:-:-: | 5842 | PHLDB1 | 989 | 280 | 540 | 294 | 334 | 272 |
| ENSDARG000 | 15:15:15:15:1 | 39934295:399 | 39936008:399 | -:-:-:-:-:-:-:-: | 4388 | MSH5 | 8 | 1 | 71 | 4 | 16 | 0 |
| ENSDARG000 | 15:15:15:15:1 | 39964943:399 | 39965063:399 | -:-:-:-:-:-:-:-: | 781 | rps5 | 14376 | 14619 | 24784 | 18899 | 15818 | 20293 |
| ENSDARG000 | 15;15;15 | 39957809:399 | 39958535:399 | -;-;- | 1019 | NA | 126 | 106 | 1417 | 6 | 249 | 14 |
| ENSDARG000 | 15:15:15:15:1 | 35253247:352 | 35253580:352 | +:+:+:+:+:+:+: | 3421 | megf6 | 0 | 2 | 2 | 0 | 14 | 12 |
| ENSDARG000 | 15:15:15:15:1 | 21779681:217 | 21784361:217 | -:-:-:-:-:-:-:-: | 8071 | SIK2 | 1338 | 1057 | 824 | 1157 | 1024 | 671 |
| ENSDARG000 | 15:15:15:15:1 | 8235531:8241 | 8237439:8242 | -:-:-:-:-:-:-:-: | 3353 | tnfrsf19 | 101 | 48 | 98 | 52 | 77 | 43 |
| ENSDARG000 | 15:15:15:15:1 | 21842638:218 | 21842743:218 | -:-:-:-:-:-:-:-: | 1502 | LAYN | 554 | 732 | 802 | 366 | 617 | 1284 |
| ENSDARG000 | 15;15;15;15 | 21780764:217 | 21781184:217 | +;+;+;+ | 1717 | NA | 464 | 654 | 546 | 300 | 417 | 1035 |
| ENSDARG000 | 15:15:15:15:1 | 14145346:141 | 14145483:141 | +:+:+:+:+:+:+: | 4067 | NA | 50 | 242 | 342 | 2825 | 78 | 1362 |
| ENSDARG000 | 15:15:15:15:1 | 2190229:2191 | 2190514:2192 | +:+:+:+:+:+:+: | 2797 | rsrc1 | 164 | 184 | 185 | 151 | 153 | 166 |
| ENSDARG000 | 15:15:15:15:1 | 14141045:141 | 14141137:141 | +:+:+:+:+:+:+: | 2482 | NA | 0 | 9 | 21 | 1 | 20 | 47 |
| ENSDARG000 | 15;15;15 | 2324138:2325 | 2324669:2325 | -;-;- | 713 | NA | 0 | 0 | 0 | 0 | 0 | 0 |
| ENSDARG000 | 15:15:15:15:1 | 14163193:141 | 14163256:141 | -:-:-:-:-:-:-:-: | 3744 | pnkp | 186 | 311 | 292 | 138 | 160 | 284 |

| | | | | | | | | | | | |
|---|---|---|---|---|---|---|---|---|---|---|---|
| ENSDARG000 | 15:15:15:15:1 | 39977461:399 | 39977628:399 | +:+:+:+:+:+:+: | 1759 | cab39 | 195 | 209 | 260 | 100 | 306 | 227 |
| ENSDARG000 | 15:15:15:15:1 | 14154174:141 | 14154382:141 | +:+:+:+:+:+:+: | 1532 | NA | 2 | 22 | 39 | 42 | 21 | 35 |
| ENSDARG000 | 15;15;15 | 35440740:354 | 35440957:354 | -;-;- | 605 | NA | 0 | 0 | 0 | 0 | 2 | 0 |
| ENSDARG000 | 15;15 | 40004732:400 | 40004973:400 | +;+ | 1404 | NA | 185 | 171 | 177 | 170 | 227 | 262 |
| ENSDARG000 | 15;15 | 35780040:357 | 35780133:357 | -;- | 646 | NA | 0 | 0 | 0 | 0 | 0 | 0 |
| ENSDARG000 | 15;15 | 35888549:359 | 35892925:359 | -;- | 8274 | IRS1 | 460 | 515 | 608 | 496 | 536 | 212 |
| ENSDARG000 | 15:15:15:15:1 | 26540981:265 | 26541551:265 | -:-:-:-:-:-:-:-: | 2092 | serpinf2b | 2 | 14 | 8 | 14 | 2 | 10 |
| ENSDARG000 | 15:15:15:15:1 | 35698508:357 | 35698551:357 | +:+:+:+:+:+:+: | 3428 | NYAP2 | 2 | 1 | 2 | 0 | 0 | 0 |
| ENSDARG000 | 15:15:15:15:1 | 40008370:400 | 40008676:400 | +;+;+;+;+;+ | 2694 | itm2ca | 4 | 2 | 0 | 0 | 6 | 10 |
| ENSDARG000 | 15 | 40040155 | 40041455 | - | 1301 | lpar5b | 4 | 13 | 12 | 11 | 18 | 30 |
| ENSDARG000 | 15:15:15:15:1 | 40051218:400 | 40052656:400 | -;-;-;-;- | 1714 | gpr55a | 2 | 4 | 0 | 4 | 0 | 4 |
| ENSDARG000 | 15:15:15:15:1 | 26552948:265 | 26555055:265 | -:-:-:-:-:-:-:-: | 8249 | wdr81 | 2832 | 1208 | 1991 | 1761 | 2042 | 1840 |
| ENSDARG000 | 15:15:15:15:1 | 35933094:359 | 35933251:359 | +:+:+:+:+:+:+: | 1641 | rhbdd1 | 96 | 202 | 746 | 727 | 163 | 129 |
| ENSDARG000 | 15:15:15:15:1 | 18102280:181 | 18103392:181 | -:-:-:-:-:-:-:-: | 2757 | arcn1b | 1723 | 1591 | 2207 | 1553 | 1966 | 1477 |
| ENSDARG000 | 15;15;15;15 | 40125409:401 | 40125608:401 | -;-;-;- | 508 | NA | 1 | 0 | 0 | 0 | 0 | 0 |
| ENSDARG000 | 15:15:15:15:1 | 40074923:400 | 40075600:400 | +:+:+:+:+:+:+: | 7638 | NGEF | 22 | 29 | 1 | 6 | 5 | 7 |
| ENSDARG000 | 15:15:15:15:1 | 2421432:2421 | 2421997:2421 | +:+:+:+:+:+:+: | 6350 | hephl1a | 2037 | 1976 | 1754 | 1130 | 2013 | 1165 |
| ENSDARG000 | 15;15;15 | 18119618:181 | 18120541:181 | -;-;- | 1335 | cryaba | 0 | 0 | 0 | 0 | 0 | 2 |
| ENSDARG000 | 15:15:15:15:1 | 26573676:265 | 26573854:265 | +:+:+:+:+:+:+: | 2943 | NA | 4 | 6 | 14 | 0 | 4 | 10 |
| ENSDARG000 | 15:15:15:15:1 | 36054864:360 | 36055398:360 | +:+:+:+:+:+:+: | 9726 | NA | 0 | 0 | 0 | 0 | 0 | 0 |
| ENSDARG000 | 15:15:15:15:1 | 40137564:401 | 40138300:401 | -:-:-:-:-:-:-:-: | 12217 | NA | 290 | 1666 | 190 | 3279 | 3564 | 438 |
| ENSDARG000 | 15:15:15:15:1 | 35951901:359 | 35952029:359 | +;+;+;+;+;+;+ | 1098 | NA | 0 | 0 | 0 | 0 | 0 | 0 |
| ENSDARG000 | 15:15:15:15:1 | 35951888:359 | 35957108:359 | -:-:-:-:-:-:-:-: | 10607 | COL4A4 | 11 | 0 | 4 | 0 | 0 | 0 |
| ENSDARG000 | 15:15:15:15:1 | 15997149:160 | 15999521:160 | -;-;-;-;-;-;-;- | 4167 | hnf1ba | 0 | 0 | 36 | 28 | 68 | 0 |
| ENSDARG000 | 15:15:15:15:1 | 18130109:181 | 18130600:181 | +:+:+:+:+:+:+: | 2676 | dixdc1a | 24 | 6 | 2 | 2 | 6 | 16 |
| ENSDARG000 | 15:15:15:15:1 | 26600611:266 | 26600885:266 | +:+:+:+:+:+:+: | 2003 | NA | 32 | 57 | 2 | 52 | 40 | 13 |
| ENSDARG000 | 15;15;15;15 | 40188076:402 | 40188380:402 | +;+;+;+ | 1297 | efhd1 | 118 | 129 | 130 | 78 | 246 | 22 |
| ENSDARG000 | 15:15:15:15:1 | 26616004:266 | 26616777:266 | -:-:-:-:-:-:-:-: | 2509 | slc47a4 | 8 | 6 | 43 | 60 | 147 | 16 |
| ENSDARG000 | 15:15:15:15:1 | 40217256:402 | 40221270:402 | -:-:-:-:-:-:-:-: | 6938 | kcnj13 | 401 | 518 | 200 | 118 | 627 | 341 |
| ENSDARG000 | 15;15;15 | 40223437:402 | 40223466:402 | +;+;+ | 1536 | NA | 2 | 0 | 0 | 0 | 11 | 4 |
| ENSDARG000 | 15:15:15:15:1 | 40271177:402 | 40271396:402 | +;+;+;+;+;+;+ | 2913 | GAL3ST2 | 8 | 16 | 83 | 9 | 54 | 4 |
| ENSDARG000 | 15;15 | 36115955:361 | 36116557:361 | +;+ | 968 | NA | 0 | 0 | 0 | 0 | 0 | 0 |
| ENSDARG000 | 15;15 | 40292902:403 | 40293044:403 | +;+ | 270 | NA | 0 | 0 | 0 | 0 | 0 | 0 |
| ENSDARG000 | 15:15:15:15:1 | 26678834:266 | 26680164:266 | -:-:-:-:-:-:-:-: | 4859 | PITPNM3 | 207 | 226 | 342 | 242 | 232 | 155 |
| ENSDARG000 | 15;15 | 36141669:361 | 36141825:361 | +;+ | 1816 | NA | 6 | 6 | 32 | 12 | 45 | 22 |
| ENSDARG000 | 15;15 | 36156986:361 | 36157201:361 | +;+ | 693 | sst1.1 | 0 | 0 | 0 | 0 | 0 | 0 |
| ENSDARG000 | 15 | 20779412 | 20781116 | - | 1705 | NA | 0 | 0 | 0 | 0 | 0 | 0 |
| ENSDARG000 | 15:15:15:15:1 | 20652686:206 | 20653345:206 | -:-:-:-:-:-:-:-: | 2963 | tpst1 | 410 | 278 | 484 | 212 | 664 | 394 |
| ENSDARG000 | 15;15 | 16035869:160 | 16036218:160 | +;+ | 1082 | NA | 0 | 0 | 0 | 0 | 0 | 0 |
| ENSDARG000 | 15;15;15 | 16066091:160 | 16067176:160 | -;-;- | 1383 | dynll2a | 916 | 971 | 993 | 795 | 1128 | 920 |
| ENSDARG000 | 15:15:15:15:1 | 18157665:181 | 18157908:181 | -;-;-;-;- | 1313 | pih1d2 | 0 | 0 | 0 | 0 | 0 | 2 |
| ENSDARG000 | 15:15:15:15:1 | 36187434:361 | 36187597:361 | +:+:+:+:+:+:+: | 4979 | MASP1 | 0 | 0 | 2 | 0 | 0 | 6 |
| ENSDARG000 | 15:15:15:15:1 | 16072080:160 | 16073476:160 | -:-:-:-:-:-:-:-: | 2731 | srsf1a | 2746 | 3341 | 4694 | 2277 | 3760 | 2577 |
| ENSDARG000 | 15:15:15:15:1 | 40356379:403 | 40357708:403 | -:-:-:-:-:-:-:-: | 2525 | NA | 0 | 0 | 0 | 2 | 2 | 0 |
| ENSDARG000 | 15:15:15:15:1 | 26848909:268 | 26850208:268 | -:-:-:-:-:-:-:-: | 4183 | NA | 42 | 112 | 117 | 112 | 138 | 54 |
| ENSDARG000 | 15:15:15:15:1 | 18164054:181 | 18164483:181 | -:-:-:-:-:-:-:-: | 2831 | NA | 10 | 0 | 0 | 0 | 2 | 16 |
| ENSDARG000 | 15:15:15:15:1 | 26903267:269 | 26905370:269 | -:-:-:-:-:-:-:-: | 4278 | ccdc9 | 533 | 530 | 829 | 304 | 512 | 415 |
| ENSDARG000 | 15:15:15:15:1 | 18177941:181 | 18178259:181 | -;-;-;-;-;- | 1282 | NA | 12 | 2 | 0 | 0 | 0 | 8 |
| ENSDARG000 | 15:15:15:15:1 | 36228835:362 | 36229320:362 | -;-;-;-;- | 1961 | ostn | 15 | 0 | 2 | 0 | 6 | 0 |
| ENSDARG000 | 15:15:15:15:1 | 16079586:160 | 16080302:160 | -;-;-;-;-;-;-;- | 2174 | aldoca | 0 | 6 | 2 | 10 | 0 | 0 |
| ENSDARG000 | 15 | 20798128 | 20799830 | - | 1703 | NA | 0 | 0 | 0 | 0 | 0 | 0 |
| ENSDARG000 | 15:15:15:15:1 | 36309070:363 | 36309209:363 | +:+:+:+:+:+:+: | 1866 | NA | 0 | 0 | 0 | 0 | 0 | 0 |
| ENSDARG000 | 15:15:15:15:1 | 36329368:363 | 36330117:363 | -:-:-:-:-:-:-:-: | 4094 | NA | 6 | 0 | 0 | 0 | 0 | 0 |
| ENSDARG000 | 15:15:15:15:1 | 16099458:161 | 16101642:161 | -:-:-:-:-:-:-:-: | 3341 | NA | 0 | 34 | 8 | 10 | 8 | 8 |
| ENSDARG000 | 15:15:15:15:1 | 40665310:406 | 40665529:406 | +:+:+:+:+:+:+: | 16245 | fat3a | 8 | 12 | 127 | 12 | 72 | 18 |
| ENSDARG000 | 15:15:15:15:1 | 36349009:363 | 36352246:363 | -;-;-;-;-;-;-;- | 5843 | NA | 217 | 76 | 84 | 18 | 36 | 16 |
| ENSDARG000 | 15:15:15:15:1 | 18198878:181 | 18200198:182 | -:-:-:-:-:-:-:-: | 2469 | btr16 | 14 | 0 | 4 | 2 | 2 | 0 |
| ENSDARG000 | 15:15:15:15:1 | 26933196:269 | 26933776:269 | +;+;+;+;+;+ | 2474 | ppm1da | 130 | 252 | 209 | 100 | 244 | 162 |
| ENSDARG000 | 15:15:15:15:1 | 18189645:181 | 18190273:181 | -:-:-:-:-:-:-:-: | 1701 | NA | 34 | 8 | 9 | 0 | 0 | 11 |
| ENSDARG000 | 15:15:15:15:1 | 36358144:363 | 36358311:363 | -:-:-:-:-:-:-:-: | 2451 | NA | 516 | 94 | 182 | 54 | 43 | 29 |
| ENSDARG000 | 15:15:15:15:1 | 20799943:208 | 20800067:208 | +:+:+:+:+:+:+: | 4756 | ALDH3A2 | 10 | 2 | 108 | 70 | 14 | 88 |
| ENSDARG000 | 15;15 | 41027466:410 | 41027815:410 | +;+ | 1260 | mtnr1ba | 0 | 0 | 2 | 0 | 0 | 0 |
| ENSDARG000 | 15:15:15:15:1 | 41169211:411 | 41173639:411 | -:-:-:-:-:-:-:-: | 6674 | NA | 764 | 627 | 542 | 342 | 732 | 701 |
| ENSDARG000 | 15:15:15:15:1 | 36445350:364 | 36445443:364 | +:+:+:+:+:+:+: | 9280 | NA | 244 | 78 | 14 | 10 | 154 | 94 |
| ENSDARG000 | 15;15 | 41238247:412 | 41239179:412 | -;- | 1073 | smco4 | 259 | 343 | 369 | 291 | 356 | 404 |
| ENSDARG000 | 15:15:15:15:1 | 41290948:412 | 41292141:412 | -:-:-:-:-:-:-:-: | 3723 | usp16 | 0 | 0 | 0 | 0 | 0 | 0 |
| ENSDARG000 | 15:15:15:15:1 | 18212190:182 | 18212737:182 | -:-:-:-:-:-:-:-: | 1695 | NA | 0 | 0 | 0 | 0 | 0 | 0 |
| ENSDARG000 | 15:15:15:15:1 | 41249772:412 | 41250080:412 | -:-:-:-:-:-:-:-: | 9083 | usp16 | 50 | 54 | 43 | 59 | 74 | 63 |
| ENSDARG000 | 15 | 20809709 | 20811411 | - | 1703 | NA | 0 | 0 | 0 | 0 | 0 | 0 |
| ENSDARG000 | 15 | 20828421 | 20830125 | - | 1705 | NA | 0 | 0 | 0 | 0 | 0 | 0 |
| ENSDARG000 | 15 | 18221859 | 18223769 | - | 1911 | or91a1 | 0 | 0 | 0 | 0 | 0 | 0 |
| ENSDARG000 | 15 | 18230494 | 18232712 | - | 2219 | NA | 0 | 0 | 0 | 0 | 0 | 0 |
| ENSDARG000 | 15:15:15:15:1 | 36506014:365 | 36506626:365 | -:-:-:-:-:-:-:-: | 2043 | NA | 44 | 35 | 32 | 36 | 95 | 158 |
| ENSDARG000 | 15 | 18238419 | 18240199 | - | 1781 | or91a3 | 0 | 0 | 0 | 0 | 0 | 0 |
| ENSDARG000 | 15:15:15:15:1 | 18253542:182 | 18260311:182 | -;-;-;-;-;-;- | 8939 | ZBTB16 | 0 | 0 | 2 | 4 | 14 | 0 |
| ENSDARG000 | 15:15:15:15:1 | 18368809:183 | 18369343:183 | -:-:-:-:-:-:-:-: | 4020 | ncam1b | 72 | 10 | 8 | 12 | 2 | 196 |
| ENSDARG000 | 15:15:15:15:1 | 36553823:365 | 36556653:365 | -:-:-:-:-:-:-:-: | 7238 | nphs1 | 8 | 2 | 0 | 0 | 4 | 0 |
| ENSDARG000 | 15:15:15:15:1 | 41529260:415 | 41529438:415 | -;-;-;-;-;-;-;- | 3002 | NA | 2 | 0 | 4 | 0 | 0 | 0 |
| ENSDARG000 | 15:15:15:15:1 | 41471175:414 | 41471371:414 | -;-;-;-;-;-;-;- | 2984 | NA | 2 | 52 | 15 | 44 | 66 | 50 |
| ENSDARG000 | 15:15:15:15:1 | 41448242:414 | 41448435:414 | -:-:-:-:-:-:-:-: | 4175 | NA | 0 | 141 | 13 | 68 | 117 | 49 |
| ENSDARG000 | 15:15:15:15:1 | 41400520:414 | 41400782:414 | -;-;-;-;-;-;- | 2704 | NA | 8 | 0 | 14 | 3 | 0 | 0 |
| ENSDARG000 | 15:15:15:15:1 | 41422248:414 | 41423539:414 | -:-:-:-:-:-:-:-: | 4552 | NA | 0 | 0 | 0 | 1 | 0 | 0 |
| ENSDARG000 | 15:15:15:15:1 | 41349268:413 | 41353587:413 | -:-:-:-:-:-:-:-: | 7731 | NA | 0 | 1 | 10 | 0 | 6 | 1 |
| ENSDARG000 | 15;15;15;15 | 41393401:413 | 41395213:413 | -;-;-;- | 2083 | NA | 1 | 0 | 0 | 0 | 0 | 0 |
| ENSDARG000 | 15:15:15:15:1 | 41312032:413 | 41312602:413 | -:-:-:-:-:-:-:-: | 4314 | NA | 1 | 4 | 62 | 30 | 8 | 4 |
| ENSDARG000 | 15:15:15:15:1 | 41498490:414 | 41498668:415 | -:-:-:-:-:-:-:-: | 3481 | NA | 3 | 50 | 0 | 30 | 27 | 33 |
| ENSDARG000 | 15:15:15:15:1 | 36941490:369 | 36942159:369 | +:+:+:+:+:+:+: | 7639 | kirrel3l | 84 | 44 | 38 | 30 | 36 | 12 |
| ENSDARG000 | 15:15:15:15:1 | 37105986:371 | 37106377:371 | +:+:+:+:+:+:+: | 6507 | APLP1 | 116 | 60 | 82 | 10 | 24 | 26 |
| ENSDARG000 | 15;15 | 37412883:374 | 37413188:374 | +;+ | 1175 | NA | 6 | 22 | 4 | 38 | 30 | 104 |
| ENSDARG000 | 15;15 | 41643120:416 | 41643342:416 | -;- | 379 | NA | 0 | 0 | 0 | 0 | 0 | 0 |
| ENSDARG000 | 15:15:15:15:1 | 41555593:415 | 41555826:415 | +:+:+:+:+:+:+: | 3652 | CLSTN2 | 4 | 6 | 8 | 0 | 0 | 0 |
| ENSDARG000 | 15:15:15:15:1 | 37419552:374 | 37420295:374 | -;-;-;-;-;- | 2176 | NA | 4 | 0 | 0 | 0 | 0 | 0 |
| ENSDARG000 | 15:15:15:15:1 | 41670742:416 | 41671419:416 | -;-;-;-;-;-;-;- | 2045 | spsb4b | 207 | 188 | 305 | 204 | 218 | 166 |
| ENSDARG000 | 15:15:15:15:1 | 41693635:416 | 41698857:416 | -;-;-;-;- | 6355 | NA | 372 | 372 | 612 | 340 | 615 | 624 |
| ENSDARG000 | 15:15:15:15:1 | 37436430:374 | 37436562:374 | +;+;+;+;+;+;+ | 1248 | NA | 846 | 17 | 188 | 314 | 89 | 99 |
| ENSDARG000 | 15:15:15:15:1 | 41711727:417 | 41711870:417 | -;-;-;-;-;- | 1267 | NA | 7 | 16 | 7 | 0 | 6 | 0 |
| ENSDARG000 | 15:15:15:15:1 | 41711743:417 | 41712254:417 | -:-:-:-:-:-:-:-: | 4045 | NA | 348 | 223 | 87 | 226 | 340 | 296 |
| ENSDARG000 | 15:15:15:15:1 | 41716721:417 | 41718269:417 | -;-;-;-;-;-;-;- | 2792 | NA | 72 | 59 | 14 | 34 | 92 | 78 |
| ENSDARG000 | 15:15:15:15:1 | 41735557:417 | 41738480:417 | -:-:-:-:-:-:-:-: | 4143 | NA | 267 | 144 | 84 | 88 | 166 | 98 |
| ENSDARG000 | 15:15:15:15:1 | 37457009:374 | 37461058:374 | -:-:-:-:-:-:-:-: | 15341 | NA | 2163 | 951 | 2064 | 986 | 1732 | 1653 |
| ENSDARG000 | 15:15:15:15:1 | 41746720:417 | 41747297:417 | -;-;-;-;-;-;- | 1809 | NA | 49 | 40 | 36 | 44 | 35 | 38 |
| ENSDARG000 | 15:15:15:15:1 | 41752335:417 | 41752923:417 | -:-:-:-:-:-:-:-: | 1822 | ftr72 | 41 | 172 | 19 | 74 | 30 | 64 |
| ENSDARG000 | 15:15:15:15:1 | 41757442:417 | 41757962:417 | -;-;-;-;-;-;-;- | 1755 | NA | 22 | 33 | 16 | 6 | 21 | 10 |
| ENSDARG000 | 15:15:15:15:1 | 41767675:417 | 41770533:417 | -;-;-;-;-;-;- | 3994 | SLC25A36 | 352 | 435 | 1279 | 1328 | 498 | 430 |
| ENSDARG000 | 15:15:15:15:1 | 41815703:418 | 41815817:418 | +:+:+:+:+:+:+: | 1891 | pxylp1 | 246 | 174 | 1004 | 267 | 327 | 109 |
| ENSDARG000 | 15:15:15:15:1 | 41901710:419 | 41902031:419 | +:+:+:+:+:+:+: | 3616 | NA | 0 | 3 | 0 | 0 | 0 | 0 |
| ENSDARG000 | 15:15:15:15:1 | 41919484:419 | 41919678:419 | +:+:+:+:+:+:+: | 4000 | NA | 22 | 4 | 0 | 0 | 3 | 4 |
| ENSDARG000 | 15:15:15:15:1 | 37545855:375 | 37545978:375 | +;+;+;+;+;+;+ | 976 | psenen | 1180 | 935 | 974 | 972 | 1014 | 997 |
| ENSDARG000 | 15:15:15:15:1 | 41945310:419 | 41950900:419 | -:-:-:-:-:-:-:-: | 8510 | EPHA4 | 22 | 12 | 8 | 0 | 18 | 10 |
| ENSDARG000 | 15:15:15:15:1 | 41932069:419 | 41932237:419 | +:+:+:+:+:+:+: | 3667 | NA | 5 | 0 | 0 | 0 | 0 | 0 |
| ENSDARG000 | 15;15;15 | 37559570:375 | 37559988:375 | +;+;+ | 2061 | hspb6 | 48 | 18 | 28 | 10 | 20 | 54 |
| ENSDARG000 | 15:15:15:15:1 | 37562582:375 | 37562759:375 | -:-:-:-:-:-:-:-: | 2327 | proser3 | 101 | 120 | 109 | 62 | 78 | 74 |
| ENSDARG000 | 15:15:15:15:1 | 37589698:375 | 37589826:375 | +:+:+:+:+:+:+: | 1375 | lin37 | 114 | 266 | 157 | 120 | 207 | 191 |
| ENSDARG000 | 15;15;15 | 42043102:420 | 42043902:420 | -;-;- | 1005 | NA | 0 | 0 | 10 | 0 | 6 | 0 |
| ENSDARG000 | 15;15;15 | 42064183:420 | 42064691:420 | -;-;- | 698 | NA | 0 | 0 | 0 | 0 | 0 | 0 |
| ENSDARG000 | 15;15;15 | 42080394:420 | 42080988:420 | -;-;- | 763 | NA | 95 | 38 | 4 | 0 | 6 | 28 |
| ENSDARG000 | 15 | 42090770 | 42091109 | + | 340 | NA | 2 | 7 | 3 | 0 | 8 | 0 |
| ENSDARG000 | 15;15;15 | 42262701:422 | 42262851:422 | +;+;+ | 363 | NA | 0 | 0 | 6 | 0 | 0 | 0 |
| ENSDARG000 | 15:15:15:15:1 | 42264891:422 | 42265282:422 | -:-:-:-:-:-:-:-: | 2113 | farsb | 1136 | 1290 | 2742 | 2179 | 1189 | 1168 |
| ENSDARG000 | 15:15:15:15:1 | 42285643:422 | 42285841:422 | +:+:+:+:+:+:+: | 4478 | scaf4b | 642 | 600 | 976 | 486 | 657 | 421 |
| ENSDARG000 | 15;15 | 37608742:376 | 37610022:376 | -;- | 1342 | NA | 6 | 8 | 4 | 6 | 20 | 30 |
| ENSDARG000 | 15;15 | 37619004:376 | 37619760:376 | -;- | 804 | NA | 2 | 0 | 0 | 0 | 9 | 24 |
| ENSDARG000 | 15;15 | 37639184:376 | 37639974:376 | -;- | 1022 | NA | 0 | 0 | 0 | 0 | 0 | 0 |
| ENSDARG000 | 15;15 | 37671873:376 | 37673135:376 | -;- | 1506 | NA | 0 | 10 | 0 | 0 | 58 | 119 |
| ENSDARG000 | 15;15;15;15 | 37649268:376 | 37650523:376 | -;-;-;- | 2163 | NA | 0 | 2 | 0 | 0 | 4 | 10 |
| ENSDARG000 | 15;15 | 37707914:377 | 37708665:377 | -;- | 800 | NA | 0 | 0 | 0 | 0 | 0 | 0 |
| ENSDARG000 | 15;15;15;15 | 37707269:377 | 37708665:377 | -;-;-;- | 3456 | NA | 0 | 4 | 0 | 2 | 12 | 47 |
| ENSDARG000 | 15;15 | 37682403:376 | 37683358:376 | -;- | 1004 | NA | 0 | 0 | 0 | 0 | 0 | 0 |
| ENSDARG000 | 15;15;15 | 37727834:377 | 37728409:377 | -;-;- | 908 | NA | 0 | 0 | 0 | 0 | 0 | 4 |
| ENSDARG000 | 15;15 | 37731055:377 | 37732662:377 | -;- | 1822 | NA | 6 | 0 | 0 | 0 | 0 | 6 |
| ENSDARG000 | 15;15 | 37741382:377 | 37742133:377 | -;- | 801 | NA | 6 | 0 | 0 | 0 | 3 | 3 |
| ENSDARG000 | 15;15 | 37751113:377 | 37752044:377 | -;- | 1020 | NA | 0 | 0 | 0 | 0 | 0 | 0 |
| ENSDARG000 | 15;15 | 37766097:377 | 37767418:377 | -;- | 1443 | NA | 2 | 0 | 0 | 0 | 16 | 16 |
| ENSDARG000 | 15;15 | 37773126:377 | 37773865:377 | -;- | 788 | NA | 0 | 0 | 0 | 0 | 0 | 0 |
| ENSDARG000 | 15:15:15:15:1 | 42397125:424 | 42397442:424 | +:+:+:+:+:+:+: | 5257 | TIAM1 | 20 | 11 | 10 | 0 | 6 | 24 |
| ENSDARG000 | 15;15 | 37778574:377 | 37779319:377 | -;- | 794 | NA | 0 | 0 | 0 | 0 | 0 | 0 |
| ENSDARG000 | 15;15 | 37795286:377 | 37797129:377 | -;- | 1949 | NA | 18 | 23 | 25 | 26 | 96 | 50 |
| ENSDARG000 | 15;15 | 37806111:378 | 37806500:378 | -;- | 727 | NA | 0 | 0 | 0 | 0 | 0 | 0 |
| ENSDARG000 | 15:15:15:15:1 | 37820258:378 | 37821073:378 | -:-:-:-:-:-:-:-: | 1830 | ctrl | 4 | 1 | 0 | 0 | 6 | 0 |
| ENSDARG000 | 15:15:15:15:1 | 37831604:378 | 37832147:378 | -:-:-:-:-:-:-:-: | 1497 | si:dkev-238d1 | 0 | 0 | 0 | 0 | 2 | 0 |
| ENSDARG000 | 15:15:15:15:1 | 37834594:378 | 37836509:378 | -:-:-:-:-:-:-:-: | 3791 | hsc70 | 850 | 30 | 439 | 1033 | 64 | 36 |
| ENSDARG000 | 15:15:15:15:1 | 42599501:426 | 42599618:426 | +:+:+:+:+:+:+: | 2640 | GRIK1 | 14 | 8 | 4 | 0 | 0 | 7 |
| ENSDARG000 | 15:15:15:15:1 | 42693455:426 | 42693826:426 | -:-:-:-:-:-:-:-: | 3503 | NA | 92 | 99 | 98 | 28 | 122 | 96 |
| ENSDARG000 | 15:15:15:15:1 | 37854483:378 | 37857062:378 | -:-:-:-:-:-:-:-: | 4188 | NA | 794 | 458 | 278 | 334 | 473 | 504 |
| ENSDARG000 | 15;15 | 37875077:378 | 37875392:378 | +;+ | 392 | NA | 0 | 0 | 0 | 0 | 0 | 0 |
| ENSDARG000 | 15:15:15:15:1 | 42723126:427 | 42723479:427 | -:-:-:-:-:-:-:-: | 3589 | NA | 197 | 73 | 234 | 203 | 168 | 280 |
| ENSDARG000 | 15:15:15:15:1 | 42741272:427 | 42744557:427 | -:-:-:-:-:-:-:-: | 8431 | NA | 1864 | 1510 | 1999 | 1871 | 1440 | 1967 |
| ENSDARG000 | 15:15:15:15:1 | 42767803:429 | 42767923:429 | +:+:+:+:+:+:+: | 3458 | slc8a2b | 0 | 81 | 10 | 2 | 22 | 2 |
| ENSDARG000 | 15;15 | 43093044:430 | 43093075:431 | +;+ | 3494 | kcne4 | 26 | 14 | 17 | 46 | 32 | 4 |
| ENSDARG000 | 15;15 | 43073923:431 | 43074003:431 | -;- | 190 | NA | 0 | 0 | 0 | 0 | 0 | 0 |
| ENSDARG000 | 15:15:15:15:1 | 42995051:429 | 42995409:429 | +:+:+:+:+:+:+: | 2734 | acsl3a | 147 | 79 | 373 | 149 | 460 | 223 |
| ENSDARG000 | 15:15:15:15:1 | 37891045:378 | 37894496:378 | -:-:-:-:-:-:-:-: | 7395 | NA | 2 | 38 | 16 | 28 | 292 | 86 |
| ENSDARG000 | 15;15 | 37927949:379 | 37928782:379 | -;- | 1039 | NA | 0 | 0 | 0 | 0 | 0 | 0 |
| ENSDARG000 | 15;15;15;15 | 37936458:379 | 37936616:379 | +;+;+;+ | 985 | NA | 1 | 6 | 0 | 0 | 26 | 6 |
| ENSDARG000 | 15;15;15 | 37950556:379 | 37950938:379 | +;+;+ | 971 | NA | 0 | 0 | 0 | 0 | 2 | 4 |
| ENSDARG000 | 15:15:15:15:1 | 37954666:379 | 37955012:379 | +;+;+;+;+;+ | 1019 | NA | 0 | 0 | 0 | 0 | 4 | 2 |
| ENSDARG000 | 15:15:15:15:1 | 37965953:379 | 37966075:379 | +;+;+;+;+;+;+ | 1813 | NA | 0 | 0 | 0 | 0 | 6 | 2 |
| ENSDARG000 | 15:15:15:15:1 | 43208593:432 | 43208652:432 | -:-:-:-:-:-:-:-: | 1794 | wdfy1 | 733 | 521 | 431 | 308 | 668 | 487 |
| ENSDARG000 | 15:15:15:15:1 | 43241915:432 | 43242219:432 | -:-:-:-:-:-:-:-: | 1609 | serpine2 | 1115 | 1351 | 1791 | 2803 | 691 | 1499 |
| ENSDARG000 | 15:15:15:15:1 | 37986069:379 | 37986242:379 | +;+;+;+;+;+ | 1425 | NA | 2 | 0 | 4 | 0 | 0 | 2 |
| ENSDARG000 | 15:15:15:15:1 | 37972671:379 | 37973226:379 | +;+;+;+;+;+;+ | 1108 | NA | 0 | 0 | 0 | 0 | 10 | 1 |
| ENSDARG000 | 15:15:15:15:1 | 43286785:432 | 43287513:432 | -:-:-:-:-:-:-:-: | 2169 | prss16 | 2250 | 1465 | 1051 | 1469 | 1806 | 1659 |

| | | | | | | | | | | | | |
|---|---|---|---|---|---|---|---|---|---|---|---|---|
| ENSDARG000 | 15;15;15;15;1 | 43906043:439 | 43906178:439 | +;+;+;+;+;+;+; | 2395 | naalad2 | 56 | 96 | 33 | 147 | 176 | 100 |
| ENSDARG000 | 15;15;15;15;1 | 43961814:439 | 43962141:439 | -;-;-;-;-;-;-;-; | 1381 | chordc1a | 122 | 125 | 160 | 112 | 190 | 171 |
| ENSDARG000 | 15;15;15;15;1 | 43982088:439 | 43983682:439 | -;-;-;-;-;-;-;-; | 2767 | NA | 618 | 377 | 548 | 86 | 525 | 366 |
| ENSDARG000 | 15;15;15;15;1 | 44440136:444 | 44440398:444 | -;-;-;-;-;- | 1203 | aasdhppt | 123 | 114 | 72 | 78 | 130 | 102 |
| ENSDARG000 | 15;15;15;15;1 | 44472356:444 | 44472946:444 | +;+;+;+;+;+ | 2911 | MSANTD4 | 61 | 32 | 43 | 33 | 58 | 51 |
| ENSDARG000 | 15;15;15;15;1 | 44499187:445 | 44500339:445 | -;-;-;-;-;-;-;-; | 3839 | gria4a | 6 | 18 | 166 | 69 | 12 | 2 |
| ENSDARG000 | 15;15;15;15;1 | 45563491:455 | 45563753:455 | +;+;+;+;+;+;+; | 2204 | cldn15lb | 0 | 0 | 0 | 0 | 0 | 0 |
| ENSDARG000 | 15;15;15;15;1 | 45591669:455 | 45591836:455 | +;+;+;+;+;+;+; | 4785 | atg16l1 | 2386 | 1636 | 2499 | 920 | 2216 | 1644 |
| ENSDARG000 | 15;15;15;15;1 | 45640906:456 | 45641005:456 | +;+;+;+;+;+;+; | 1627 | sagb | 0 | 1 | 1 | 14 | 5 | 2 |
| ENSDARG000 | 15;15;15;15 | 45311936:453 | 45312077:453 | -;-;-;- | 843 | NA | 0 | 0 | 0 | 0 | 0 | 0 |
| ENSDARG000 | 15;15 | 3539044:3651 | 3539643:3652 | +;+ | 1451 | NA | 274 | 111 | 164 | 111 | 186 | 93 |
| ENSDARG000 | 15;15 | 4062000:4066 | 4063410:4067 | -;- | 1791 | NA | 4 | 0 | 0 | 0 | 2 | 4 |
| ENSDARG000 | 15;15;15;15;1 | 46386261:465 | 46386321:465 | +;+;+;+;+;+;+ | 1329 | igsf11 | 578 | 378 | 544 | 298 | 653 | 383 |
| ENSDARG000 | 15;15;15;15;1 | 2559875:2568 | 2560757:2568 | +;+;+;+;+;+;+; | 1969 | SH2B2 | 58 | 28 | 33 | 27 | 88 | 81 |
| ENSDARG000 | 15;15;15;15;1 | 3674765:3676 | 3675142:3676 | +;+;+;+;+;+;+; | 1739 | NA | 84 | 34 | 43 | 47 | 52 | 72 |
| ENSDARG000 | 15;15;15;15;1 | 3957876:3961 | 3957913:3961 | -;-;-;-;-;- | 980 | GPR171 | 4 | 0 | 2 | 0 | 10 | 20 |
| ENSDARG000 | 15;15;15;15;1 | 2931882:2933 | 2933917:2933 | -;-;-;-;-;-;-;-; | 2769 | zgc:153184 | 4373 | 2861 | 3010 | 3696 | 5067 | 4306 |
| ENSDARG000 | 15;15;15 | 449097:45026 | 449228:45058 | -;-;- | 664 | NA | 16 | 0 | 0 | 0 | 0 | 2 |
| ENSDARG000 | 15 | 44017433 | 44017563 | - | 131 | SCARNA14 | 0 | 0 | 0 | 0 | 0 | 0 |
| ENSDARG000 | 15 | 42728470 | 42728586 | - | 117 | 5S_rRNA | 0 | 0 | 0 | 0 | 0 | 0 |
| ENSDARG000 | 15;15;15;15;1 | 47334247:473 | 47334399:473 | -;-;-;-;-;-;-;-; | 1110 | anapc15 | 151 | 149 | 176 | 80 | 142 | 177 |
| ENSDARG000 | 15;15 | 25024685:250 | 25025282:250 | +;+ | 1067 | NA | 0 | 0 | 0 | 0 | 0 | 0 |
| ENSDARG000 | 15 | 15383898 | 15383983 | - | 86 | dre-mir-142b | 0 | 0 | 0 | 0 | 0 | 0 |
| ENSDARG000 | 15;15;15;15;1 | 43154744:431 | 43155127:431 | -;-;-;-;- | 884 | ap1s3a | 0 | 4 | 0 | 0 | 0 | 0 |
| ENSDARG000 | 15;15;15;15;1 | 3246323:3246 | 3246536:3246 | -;-;-;-;-;-;- | 934 | stoml3a | 0 | 0 | 0 | 0 | 0 | 0 |
| ENSDARG000 | 15;15;15;15 | 3656212:3734 | 3656473:3736 | -;-;-;- | 1928 | lpar6a | 620 | 183 | 72 | 97 | 170 | 992 |
| ENSDARG000 | 15;15 | 13984879:139 | 13984978:139 | +;+ | 1775 | zgc:162730 | 7342 | 4677 | 5236 | 5867 | 4492 | 5484 |
| ENSDARG000 | 15;15;15;15 | 45668851:456 | 45669737:456 | -;-;-;- | 1410 | NA | 0 | 0 | 0 | 0 | 10 | 0 |
| ENSDARG000 | 15;15;15;15;1 | 44283723:442 | 44283815:442 | +;+;+;+;+;+;+; | 3666 | cwf19l2 | 229 | 307 | 281 | 146 | 197 | 117 |
| ENSDARG000 | 15 | 8987643 | 8987759 | - | 117 | 5S_rRNA | 0 | 0 | 0 | 0 | 0 | 0 |
| ENSDARG000 | 15;15;15 | 47808159:478 | 47808479:478 | -;-;- | 780 | tomt | 0 | 0 | 0 | 0 | 0 | 0 |
| ENSDARG000 | 15 | 8731752 | 8731866 | + | 115 | 5S_rRNA | 0 | 0 | 0 | 0 | 0 | 0 |
| ENSDARG000 | 15 | 2736383 | 2736498 | - | 116 | 5S_rRNA | 5 | 5 | 0 | 3 | 5 | 13 |
| ENSDARG000 | 15;15;15 | 2976120:2981 | 2976207:2981 | -;-;- | 924 | MLNR | 8 | 0 | 1 | 18 | 4 | 5 |
| ENSDARG000 | 15;15;15;15;1 | 34592215:345 | 34592358:345 | +;+;+;+;+;+ | 3952 | tspan13a | 0 | 0 | 0 | 0 | 0 | 0 |
| ENSDARG000 | 15 | 22955017 | 22955132 | - | 116 | 5S_rRNA | 0 | 0 | 0 | 0 | 0 | 0 |
| ENSDARG000 | 15 | 13824472 | 13824604 | - | 133 | 5S_rRNA | 0 | 0 | 0 | 0 | 0 | 0 |
| ENSDARG000 | 15;15;15 | 47161917:471 | 47162047:471 | +;+;+ | 1056 | gap43 | 20 | 2 | 2 | 0 | 0 | 0 |
| ENSDARG000 | 15 | 1543857 | 1543943 | + | 87 | dre-mir-16a | 0 | 0 | 0 | 0 | 0 | 0 |
| ENSDARG000 | 15 | 29800968 | 29801052 | + | 85 | dre-mir-125c | 0 | 0 | 0 | 0 | 0 | 0 |
| ENSDARG000 | 15 | 27646419 | 27646536 | - | 118 | 5S_rRNA | 0 | 0 | 0 | 0 | 0 | 0 |
| ENSDARG000 | 15 | 26106315 | 26106433 | + | 119 | 5S_rRNA | 0 | 0 | 0 | 0 | 0 | 0 |
| ENSDARG000 | 15 | 27654176 | 27654292 | - | 117 | 5S_rRNA | 0 | 0 | 0 | 0 | 0 | 0 |
| ENSDARG000 | 15 | 44750514 | 44750630 | - | 117 | 5S_rRNA | 0 | 0 | 0 | 0 | 0 | 0 |
| ENSDARG000 | 15;15;15;15;1 | 5390734:5395 | 5390845:5395 | -;-;-;-;-;-;-;-; | 1621 | arrb1 | 12 | 7 | 24 | 4 | 17 | 14 |
| ENSDARG000 | 15;15;15;15;1 | 47386939:473 | 47387037:473 | +;+;+;+;+;+;+; | 915 | CLPB | 100 | 29 | 51 | 30 | 46 | 60 |
| ENSDARG000 | 15 | 44861283 | 44861399 | + | 117 | 5S_rRNA | 0 | 0 | 0 | 0 | 0 | 0 |
| ENSDARG000 | 15 | 40069011 | 40069127 | - | 117 | 5S_rRNA | 0 | 0 | 0 | 0 | 0 | 0 |
| ENSDARG000 | 15;15;15;15;1 | 509126:50984 | 509579:50994 | +;+;+;+;+;+;+; | 2158 | ftr86 | 14 | 2 | 4 | 10 | 4 | 0 |
| ENSDARG000 | 15;15 | 26399538:264 | 26399990:264 | +;+ | 2646 | rtn4rl1b | 28 | 7 | 6 | 2 | 4 | 44 |
| ENSDARG000 | 15;15;15;15;1 | 16175:17652: | 16356:17793: | -;-;-;-;-;-;-;-; | 2619 | cyp2y3 | 4 | 9 | 24 | 4 | 18 | 25 |
| ENSDARG000 | 15;15;15;15;1 | 22448456:224 | 22448616:224 | +;+;+;+;+;+;+; | 6477 | ARHGEF12 | 1383 | 987 | 1685 | 999 | 1604 | 1080 |
| ENSDARG000 | 15;15;15;15 | 93131:93344: | 93211:93560: | -;-;-;- | 663 | bace1 | 203 | 169 | 286 | 167 | 248 | 177 |
| ENSDARG000 | 15;15;15;15;1 | 44559448:445 | 44560464:445 | -;-;-;-;-;-;- | 1581 | fam131ab | 0 | 2 | 0 | 0 | 0 | 0 |
| ENSDARG000 | 15;15;15;15;1 | 2776506:2782 | 2779833:2782 | -;-;-;-;-;-;- | 4414 | tgfb1a | 142 | 162 | 150 | 46 | 97 | 54 |
| ENSDARG000 | 15;15;15;15;1 | 102925:10595 | 106902:10690 | -;-;-;-;-;-;-;- | 4200 | apoa1b | 0 | 2 | 918 | 48 | 15 | 6 |
| ENSDARG000 | 15;15;15;15;1 | 3926095:3929 | 3929850:3930 | -;-;-;-;-;-;-;-;- | 5230 | MINDY4B | 12 | 45 | 8 | 32 | 121 | 62 |
| ENSDARG000 | 15;15;15;15;1 | 436776:43951 | 436822:43976 | +;+;+;+;+;+;+; | 1240 | med17 | 307 | 170 | 390 | 134 | 390 | 214 |
| ENSDARG000 | 15;15;15;15;1 | 20340022:203 | 20340042:203 | -;-;-;-;-;-;-;-; | 363 | NA | 6 | 1 | 2 | 0 | 0 | 6 |
| ENSDARG000 | 15;15 | 38007237:380 | 38007650:380 | +;+ | 1891 | NA | 0 | 0 | 0 | 0 | 4 | 0 |
| ENSDARG000 | 15;15;15;15;1 | 3125136:3128 | 3125372:3128 | +;+;+;+;+;+;+; | 4967 | RCBTB2 | 232 | 284 | 335 | 214 | 302 | 241 |
| ENSDARG000 | 15;15;15;15;1 | 5495003:5514 | 5495222:5514 | -;-;-;-;- | 1376 | NA | 0 | 4 | 0 | 0 | 0 | 2 |
| ENSDARG000 | 15;15;15;15;1 | 44070306:440 | 44070429:440 | -;-;-;-;- | 697 | lamtor1 | 264 | 328 | 283 | 203 | 368 | 186 |
| ENSDARG000 | 15 | 27971298 | 27972474 | - | 1177 | NA | 0 | 0 | 0 | 0 | 0 | 2 |
| ENSDARG000 | 15 | 18086702 | 18086816 | + | 115 | 5S_rRNA | 0 | 0 | 0 | 0 | 0 | 0 |
| ENSDARG000 | 15 | 17373789 | 17373915 | + | 127 | dre-mir-21-2 | 0 | 0 | 0 | 0 | 0 | 0 |
| ENSDARG000 | 15 | 44228917 | 44230508 | + | 1592 | NA | 0 | 0 | 0 | 2 | 6 | 0 |
| ENSDARG000 | 15;15;15;15;1 | 3808996:3809 | 3809146:3809 | +;+;+;+;+;+ | 1311 | NA | 40 | 78 | 141 | 187 | 33 | 49 |
| ENSDARG000 | 15;15;15;15;1 | 2820768:2820 | 2820871:2820 | -;-;-;-;-;-;-;-; | 2958 | NA | 101 | 30 | 118 | 94 | 138 | 100 |
| ENSDARG000 | 15 | 18151665 | 18151780 | - | 116 | 5S_rRNA | 0 | 0 | 0 | 0 | 0 | 0 |
| ENSDARG000 | 15;15;15;15;1 | 3255054:3255 | 3255690:3255 | -;-;-;-;-;-;-;-; | 1602 | slc25a15a | 38 | 6 | 6 | 4 | 22 | 8 |
| ENSDARG000 | 15 | 4365158 | 4365272 | - | 115 | 5S_rRNA | 0 | 0 | 0 | 0 | 0 | 0 |
| ENSDARG000 | 15 | 2265969 | 2266083 | - | 115 | 5S_rRNA | 0 | 0 | 0 | 0 | 0 | 0 |
| ENSDARG000 | 15 | 9895858 | 9895986 | + | 129 | 5S_rRNA | 0 | 0 | 0 | 0 | 0 | 0 |
| ENSDARG000 | 15 | 2358511 | 2358625 | + | 115 | 5S_rRNA | 0 | 0 | 0 | 0 | 0 | 0 |
| ENSDARG000 | 15 | 23378071 | 23378188 | + | 118 | 5S_rRNA | 0 | 0 | 0 | 0 | 0 | 0 |
| ENSDARG000 | 15 | 45617016 | 45617296 | + | 281 | NA | 0 | 0 | 0 | 0 | 2 | 0 |
| ENSDARG000 | 15;15;15;15;1 | 6109861:6113 | 6109869:6113 | +;+;+;+;+;+;+ | 239 | PCP4 | 0 | 0 | 0 | 0 | 0 | 0 |
| ENSDARG000 | 15 | 21447656 | 21447800 | + | 145 | dre-mir-125b- | 0 | 0 | 0 | 0 | 0 | 0 |
| ENSDARG000 | 15 | 1873658 | 1873778 | + | 121 | 5S_rRNA | 0 | 0 | 0 | 0 | 0 | 0 |
| ENSDARG000 | 15;15;15;15;1 | 3825117:3827 | 3825819:3828 | +;+;+;+;+;+ | 7893 | NA | 398 | 398 | 951 | 1151 | 1465 | 1154 |
| ENSDARG000 | 15 | 6559050 | 6559185 | + | 136 | 5S_rRNA | 0 | 0 | 0 | 0 | 4 | 0 |
| ENSDARG000 | 15 | 3506309 | 3506423 | + | 115 | 5S_rRNA | 0 | 0 | 0 | 0 | 0 | 0 |
| ENSDARG000 | 15;15;15;15;1 | 47083163:470 | 47083318:470 | +;+;+;+;+;+;+; | 987 | cep57 | 60 | 35 | 55 | 58 | 80 | 66 |
| ENSDARG000 | 15;15 | 4049222:4052 | 4050698:4053 | -;- | 1830 | NA | 2 | 12 | 0 | 4 | 13 | 12 |
| ENSDARG000 | 15;15;15;15;1 | 404891:40523 | 405024:40532 | +;+;+;+;+;+;+; | 855 | NA | 263 | 198 | 151 | 149 | 168 | 182 |
| ENSDARG000 | 15;15;15;15;1 | 47362728:473 | 47362877:473 | +;+;+;+;+;+;+; | 1487 | NA | 18 | 30 | 0 | 14 | 18 | 26 |
| ENSDARG000 | 15;15;15;15;1 | 44250335:442 | 44250461:442 | +;+;+;+;+;+;+; | 1444 | zgc:162962 | 70 | 54 | 121 | 60 | 57 | 58 |
| ENSDARG000 | 15;15;15 | 9387350:9387 | 9387483:9387 | +;+;+ | 375 | NA | 0 | 0 | 0 | 0 | 0 | 0 |
| ENSDARG000 | 15 | 36874251 | 36874366 | - | 116 | 5S_rRNA | 0 | 0 | 0 | 0 | 0 | 0 |
| ENSDARG000 | 15;15;15;15;1 | 22333778:223 | 22333849:223 | +;+;+;+;+;+;+; | 1761 | NA | 1 | 0 | 0 | 0 | 0 | 0 |
| ENSDARG000 | 15;15;15;15;1 | 11643659:116 | 11645431:116 | +;+;+;+;+;+;+; | 5783 | slc1a5 | 1927 | 2460 | 2755 | 1182 | 1217 | 1894 |
| ENSDARG000 | 15;15;15;15;1 | 119057:12086 | 119327:12102 | -;-;-;-;-;-;-;-; | 1263 | NA | 45 | 32 | 46 | 29 | 155 | 24 |
| ENSDARG000 | 15 | 31547734 | 31547852 | - | 119 | 5S_rRNA | 0 | 0 | 0 | 0 | 0 | 0 |
| ENSDARG000 | 15;15;15;15;1 | 454349:45489 | 454812:45508 | -;-;-;-;-;-;- | 1334 | NA | 2 | 0 | 7 | 25 | 0 | 6 |
| ENSDARG000 | 15 | 43829389 | 43829517 | + | 129 | 5S_rRNA | 0 | 0 | 0 | 0 | 0 | 6 |
| ENSDARG000 | 15;15;15;15;1 | 14861335:149 | 14862077:149 | +;+;+;+;+;+ | 2305 | NA | 0 | 12 | 19 | 6 | 24 | 0 |
| ENSDARG000 | 15;15;15;15;1 | 44618623:446 | 44618933:446 | -;-;-;-;-;-;-;-; | 1605 | alp3 | 0 | 32 | 6 | 18 | 0 | 4 |
| ENSDARG000 | 15;15;15;15;1 | 37022018:370 | 37026982:370 | -;-;-;-;-;-;-;-; | 7645 | NA | 304 | 12 | 18 | 34 | 76 | 58 |
| ENSDARG000 | 15;15;15;15;1 | 47100392:471 | 47100620:471 | -;-;-;-;-;- | 763 | NA | 58 | 13 | 50 | 42 | 46 | 40 |
| ENSDARG000 | 15;15;15;15;1 | 34675660:346 | 34675790:346 | +;+;+;+;+ | 1436 | lta | 4 | 4 | 0 | 0 | 2 | 4 |
| ENSDARG000 | 15;15;15;15;1 | 42163356:421 | 42163408:421 | -;-;-;-;-;-;-;-;- | 1428 | pax3b | 14 | 2 | 0 | 5 | 2 | 0 |
| ENSDARG000 | 15;15;15;15;1 | 478524:47869 | 478594:47891 | +;+;+;+;+;+;+; | 2617 | NA | 0 | 0 | 24 | 6 | 8 | 0 |
| ENSDARG000 | 15;15;15;15;1 | 168002:16872 | 168218:16888 | -;-;-;-;-;-;-;-; | 2024 | limk1a | 1167 | 1068 | 1117 | 1144 | 1317 | 1221 |
| ENSDARG000 | 15 | 11565391 | 11565509 | + | 119 | 5S_rRNA | 0 | 0 | 0 | 0 | 0 | 0 |
| ENSDARG000 | 15 | 14315354 | 14315470 | - | 117 | 5S_rRNA | 0 | 0 | 0 | 0 | 0 | 0 |
| ENSDARG000 | 15 | 42528790 | 42528904 | - | 115 | 5S_rRNA | 0 | 0 | 0 | 0 | 0 | 0 |
| ENSDARG000 | 15;15;15;15;1 | 58207:59417: | 58252:59685: | +;+;+;+;+;+;+; | 2407 | AXL | 55 | 166 | 92 | 43 | 245 | 51 |
| ENSDARG000 | 15 | 29503472 | 29510074 | - | 6603 | NA | 5 | 0 | 0 | 0 | 0 | 1 |
| ENSDARG000 | 15 | 30454452 | 30454566 | + | 115 | 5S_rRNA | 0 | 0 | 0 | 0 | 0 | 0 |
| ENSDARG000 | 15 | 7008822 | 7008938 | - | 117 | 5S_rRNA | 0 | 0 | 0 | 0 | 0 | 0 |
| ENSDARG000 | 15 | 31299563 | 31299678 | - | 116 | 5S_rRNA | 0 | 0 | 0 | 0 | 0 | 0 |
| ENSDARG000 | 15;15;15;15;1 | 44021044:440 | 44021454:440 | -;-;-;-;-;-;-;-; | 7806 | wu:fb44b02 | 419 | 1344 | 1808 | 727 | 1419 | 794 |
| ENSDARG000 | 15 | 33663055 | 33663172 | - | 118 | 5S_rRNA | 0 | 0 | 0 | 0 | 0 | 0 |
| ENSDARG000 | 15 | 24908394 | 24908479 | - | 86 | dre-mir-22a | 0 | 3 | 1 | 1 | 2 | 4 |
| ENSDARG000 | 15 | 29795823 | 29795929 | + | 107 | dre-let-7c-1 | 0 | 0 | 0 | 0 | 0 | 0 |
| ENSDARG000 | 15;15;15;15;1 | 47492838:474 | 47492862:474 | +;+;+;+;+;+;+; | 3144 | SIK3 | 2691 | 1508 | 2352 | 2547 | 1987 | 1824 |
| ENSDARG000 | 15;15;15;15;1 | 34618360:346 | 34619060:346 | -;-;-;-;-;-;-;-; | 4756 | bag6 | 2787 | 1926 | 3236 | 1684 | 2990 | 2112 |
| ENSDARG000 | 15;15;15;15 | 44053244:440 | 44053522:440 | +;+;+;+ | 1039 | lrrc51 | 2 | 0 | 0 | 0 | 2 | 0 |
| ENSDARG000 | 15;15;15;15;1 | 1199407:1201 | 1199563:1201 | +;+;+;+;+;+;+; | 1380 | MFSD1 | 417 | 271 | 277 | 132 | 366 | 236 |
| ENSDARG000 | 15 | 43651646 | 43651761 | - | 116 | 5S_rRNA | 1 | 3 | 0 | 0 | 0 | 0 |
| ENSDARG000 | 15 | 45614290 | 45614569 | + | 280 | NA | 0 | 0 | 0 | 0 | 0 | 0 |
| ENSDARG000 | 15 | 28734927 | 28735041 | - | 115 | 5S_rRNA | 0 | 0 | 0 | 0 | 0 | 0 |
| ENSDARG000 | 15;15;15 | 44238147:442 | 44238156:442 | -;-;- | 879 | NA | 107 | 66 | 63 | 43 | 65 | 63 |
| ENSDARG000 | 15;15 | 22302724:223 | 22302768:223 | -;- | 903 | zgc:194114 | 0 | 0 | 0 | 0 | 0 | 0 |
| ENSDARG000 | 15 | 26075363 | 26075480 | - | 118 | 5S_rRNA | 0 | 0 | 0 | 0 | 0 | 0 |
| ENSDARG000 | 15 | 39320120 | 39320234 | + | 115 | 5S_rRNA | 0 | 0 | 0 | 0 | 0 | 0 |
| ENSDARG000 | 15 | 5383062 | 5383180 | - | 119 | 5S_rRNA | 0 | 0 | 0 | 0 | 0 | 0 |
| ENSDARG000 | 15;15;15;15;1 | 45338623:453 | 45338659:453 | -;-;-;-;-;-;-;-; | 1506 | NA | 66 | 64 | 118 | 60 | 122 | 45 |
| ENSDARG000 | 15 | 37580721 | 37580835 | + | 115 | 5S_rRNA | 0 | 0 | 0 | 0 | 0 | 0 |
| ENSDARG000 | 15 | 4036709 | 4037626 | - | 918 | P2RY14 | 36 | 31 | 22 | 20 | 30 | 32 |
| ENSDARG000 | 15;15 | 47582207:475 | 47582662:475 | +;+ | 1244 | NA | 4 | 8 | 6 | 2 | 6 | 2 |
| ENSDARG000 | 15;15;15;15;1 | 44366556:443 | 44366649:443 | +;+;+;+;+;+;+ | 1896 | GUCY1A2 | 0 | 4 | 0 | 0 | 0 | 0 |
| ENSDARG000 | 15;15;15;15;1 | 413543:41694 | 414288:41709 | -;-;-;-;- | 1385 | psph | 1005 | 1186 | 1288 | 862 | 1026 | 758 |
| ENSDARG000 | 15;15;15;15;1 | 2856696:2857 | 2857633:2857 | +;+;+;+;+;+;+; | 5935 | mre11a | 230 | 309 | 318 | 91 | 240 | 259 |
| ENSDARG000 | 15;15 | 3970797:3975 | 3972878:3976 | -;- | 2549 | NA | 8 | 4 | 0 | 0 | 0 | 4 |
| ENSDARG000 | 15;15;15;15;1 | 37331585:373 | 37331850:373 | +;+;+;+;+;+;+ | 1167 | zgc:171592 | 0 | 4 | 0 | 0 | 0 | 0 |
| ENSDARG000 | 15 | 19357270 | 19357389 | - | 120 | 5S_rRNA | 0 | 0 | 0 | 0 | 0 | 0 |
| ENSDARG000 | 15 | 9039050 | 9039164 | + | 115 | 5S_rRNA | 0 | 0 | 0 | 0 | 0 | 0 |
| ENSDARG000 | 15 | 29877274 | 29877388 | - | 115 | 5S_rRNA | 0 | 0 | 0 | 0 | 0 | 0 |
| ENSDARG000 | 15 | 21437633 | 21437728 | + | 96 | dre-mir-100-1 | 0 | 0 | 0 | 0 | 0 | 0 |
| ENSDARG000 | 15;15;15;15;1 | 34172371:341 | 34177410:341 | -;-;-;-;-;-;-;-; | 6605 | etv1 | 6 | 0 | 40 | 12 | 10 | 8 |
| ENSDARG000 | 15;15;15;15;1 | 3766101:3766 | 3766296:3766 | +;+;+;+;+;+;+; | 9452 | RNF14 | 91 | 149 | 135 | 246 | 205 | 135 |
| ENSDARG000 | 15 | 42314628 | 42314755 | + | 128 | 5S_rRNA | 0 | 0 | 0 | 0 | 0 | 0 |
| ENSDARG000 | 15 | 29527420 | 29534022 | - | 6603 | NA | 6 | 0 | 2 | 0 | 2 | 2 |
| ENSDARG000 | 15;15;15;15;1 | 47180744:471 | 47180841:471 | -;-;-;-;-;-;-;-;- | 624 | NA | 0 | 2 | 10 | 0 | 1 | 6 |
| ENSDARG000 | 15 | 25994980 | 25995095 | - | 116 | 5S_rRNA | 0 | 0 | 0 | 0 | 0 | 0 |
| ENSDARG000 | 15 | 14635669 | 14635785 | - | 117 | 5S_rRNA | 0 | 4 | 7 | 0 | 4 | 0 |
| ENSDARG000 | 15;15;15;15 | 2884711:2891 | 2889646:2891 | -;-;-;- | 5369 | guca1c | 10 | 2 | 5 | 0 | 3 | 3 |
| ENSDARG000 | 15 | 22260881 | 22260997 | - | 117 | 5S_rRNA | 0 | 0 | 0 | 0 | 0 | 0 |
| ENSDARG000 | 15;15;15;15;1 | 47838836:478 | 47838862:478 | +;+;+;+;+;+;+; | 220 | NA | 0 | 0 | 3 | 0 | 0 | 0 |
| ENSDARG000 | 15;15;15 | 44093286:440 | 44093413:440 | +;+;+ | 1407 | zgc:112998 | 20 | 80 | 67 | 25 | 68 | 32 |
| ENSDARG000 | 15;15;15;15;1 | 43824672:438 | 43824792:438 | -;-;-;-;-;-;- | 741 | NOX4 | 114 | 67 | 127 | 80 | 198 | 112 |
| ENSDARG000 | 15;15;15;15;1 | 46605482:466 | 46605542:466 | +;+;+;+;+;+;+; | 1573 | NA | 24 | 8 | 13 | 10 | 15 | 17 |
| ENSDARG000 | 15 | 425695 | 425756 | - | 62 | NA | 1 | 0 | 0 | 1 | 0 | 0 |
| ENSDARG000 | 15 | 28375513 | 28375627 | - | 115 | 5S_rRNA | 0 | 0 | 0 | 0 | 0 | 0 |
| ENSDARG000 | 15 | 26440113 | 26440227 | + | 115 | 5S_rRNA | 0 | 0 | 0 | 0 | 0 | 0 |
| ENSDARG000 | 15 | 27794699 | 27794813 | + | 115 | 5S_rRNA | 0 | 0 | 0 | 0 | 0 | 0 |
| ENSDARG000 | 15;15;15 | 3246475:3246 | 3246518:3246 | +;+;+ | 258 | NA | 0 | 0 | 0 | 0 | 0 | 0 |
| ENSDARG000 | 15;15;15;15;1 | 45544589:455 | 45544783:455 | +;+;+;+;+;+;+; | 1002 | PARL | 229 | 154 | 171 | 131 | 203 | 199 |
| ENSDARG000 | 15;15;15;15;1 | 44201056:442 | 44201547:442 | +;+;+;+;+ | 2237 | NA | 6 | 1 | 0 | 2 | 0 | 3 |
| ENSDARG000 | 15;15;15;15;1 | 2850965:2850 | 2852612:2852 | -;-;-;-;- | 3334 | ankrd49 | 125 | 179 | 97 | 85 | 148 | 172 |

| | | | | | | | | | | | | |
|---|---|---|---|---|---|---|---|---|---|---|---|---|
| ENSDARG000 | 15:15:15:15:1 | 4085623:408 | 4087750:409 | -;-;-;-;-;- | 2952 | NA | 2 | 2 | 0 | 6 | 20 | 12 |
| ENSDARG000 | 15;15 | 4009572:401 | 4010160:401 | -;- | 948 | NA | 0 | 0 | 0 | 0 | 0 | 0 |
| ENSDARG000 | 15;15 | 45527281:45 | 45528545:45 | -;- | 1500 | NA | 115 | 70 | 139 | 55 | 111 | 109 |
| ENSDARG000 | 15;15;15 | 22435460:22 | 22435689:22 | +;+;+ | 3634 | tlcd5a | 128 | 97 | 182 | 148 | 155 | 190 |
| ENSDARG000 | 15 | 25469729 | 25469845 | + | 117 | 5S_rRNA | 0 | 0 | 0 | 0 | 0 | 0 |
| ENSDARG000 | 15 | 22414124 | 22414253 | + | 130 | 5S_rRNA | 0 | 0 | 0 | 0 | 0 | 0 |
| ENSDARG000 | 15 | 8682822 | 8682937 | + | 116 | 5S_rRNA | 0 | 0 | 0 | 0 | 0 | 0 |
| ENSDARG000 | 15:15:15:15:1 | 43398317:43 | 43398603:43 | +:+:+:+:+:+:+: | 4162 | actn4 | 3663 | 3265 | 4823 | 2524 | 4102 | 2656 |
| ENSDARG000 | 15:15:15:15:1 | 4128560:413 | 4128688:413 | -:-:-:-:-:-:-:-: | 2652 | ANO7 | 2 | 10 | 0 | 0 | 6 | 4 |
| ENSDARG000 | 15 | 34536443 | 34536579 | + | 137 | 5S_rRNA | 0 | 0 | 0 | 0 | 0 | 0 |
| ENSDARG000 | 15:15:15:15:1 | 42235449:42 | 42236036:42 | +:+:+:+:+:+:+: | 1584 | SGPP2 | 10 | 2 | 0 | 0 | 0 | 0 |
| ENSDARG000 | 15:15:15:15:1 | 18305:18700 | 18596:18876 | -;-;-;-;-;- | 1114 | NA | 4 | 13 | 2 | 0 | 33 | 20 |
| ENSDARG000 | 15 | 16518967 | 16519083 | + | 117 | 5S_rRNA | 0 | 0 | 0 | 0 | 0 | 0 |
| ENSDARG000 | 15;15;15 | 3947696:395 | 3948506:395 | -;-;- | 1424 | clrn1 | 0 | 0 | 0 | 0 | 0 | 0 |
| ENSDARG000 | 15:15:15:15:1 | 46633275:46 | 46633309:46 | -:-:-:-:-:-:-:-: | 2337 | NA | 1371 | 1345 | 1950 | 994 | 1659 | 895 |
| ENSDARG000 | 15 | 1321790 | 1321906 | - | 117 | 5S_rRNA | 0 | 0 | 0 | 0 | 0 | 0 |
| ENSDARG000 | 15:15:15:15:1 | 44184367:44 | 44184794:44 | +;+;+;+;+ | 2264 | zgc:165514 | 0 | 0 | 0 | 0 | 0 | 3 |
| ENSDARG000 | 15:15:15:15:1 | 47811725:47 | 47815970:47 | -;-;-;-;- | 5195 | NA | 0 | 0 | 0 | 0 | 0 | 0 |
| ENSDARG000 | 15:15:15:15:1 | 7299756:730 | 7302633:730 | -:-:-:-:-:-:-:-: | 5185 | SLC7A1 | 330 | 1234 | 1158 | 530 | 752 | 658 |
| ENSDARG000 | 15:15:15:15:1 | 22267847:22 | 22268000:22 | +;+;+;+;+;+ | 1941 | spa17 | 0 | 0 | 0 | 0 | 0 | 0 |
| ENSDARG000 | 15:15:15:15:1 | 43799258:43 | 43800165:43 | +;+;+;+;+ | 1948 | tyr | 34951 | 20832 | 18032 | 28326 | 17578 | 25830 |
| ENSDARG000 | 15 | 4104408 | 4104526 | - | 119 | 5S_rRNA | 0 | 0 | 0 | 0 | 0 | 0 |
| ENSDARG000 | 15 | 24158953 | 24159067 | - | 115 | 5S_rRNA | 0 | 0 | 0 | 0 | 0 | 0 |
| ENSDARG000 | 15 | 24962136 | 24962252 | + | 117 | 5S_rRNA | 0 | 0 | 0 | 0 | 0 | 0 |
| ENSDARG000 | 15;15 | 44190931:44 | 44191902:44 | +;+ | 2349 | NA | 0 | 0 | 0 | 0 | 0 | 0 |
| ENSDARG000 | 15 | 25897446 | 25897539 | + | 94 | dre-mir-212 | 0 | 0 | 0 | 0 | 0 | 0 |
| ENSDARG000 | 15 | 46595466 | 46595580 | + | 115 | 5S_rRNA | 0 | 0 | 0 | 1 | 1 | 1 |
| ENSDARG000 | 15:15:15:15:1 | 518092:5218 | 518231:5222 | +:+:+:+:+:+:+: | 2913 | pcsk7 | 811 | 522 | 827 | 806 | 671 | 368 |
| ENSDARG000 | 15:15:15:15:1 | 4101254:411 | 4105604:411 | -;-;-;-;-;-;- | 5619 | rnf168 | 281 | 480 | 449 | 259 | 494 | 358 |
| ENSDARG000 | 15:15:15:15:1 | 2987593:299 | 2989949:299 | -:-:-:-:-:-:-:-: | 6357 | FNDC3A | 1074 | 838 | 1320 | 1239 | 1204 | 993 |
| ENSDARG000 | 15:15:15:15:1 | 45414883:45 | 45415013:45 | -;-;-;-;- | 1520 | fgf12b | 14 | 17 | 51 | 5 | 34 | 16 |
| ENSDARG000 | 15:15:15:15:1 | 44646592:44 | 44646773:44 | -:-:-:-:-:-:-:-: | 1287 | NA | 525 | 566 | 350 | 131 | 607 | 356 |
| ENSDARG000 | 15 | 1543686 | 1543768 | + | 83 | dre-mir-15c | 0 | 0 | 0 | 0 | 0 | 0 |
| ENSDARG000 | 15;15;15 | 1796313:179 | 1796691:179 | +;+;+ | 1716 | FAM124B | 4 | 0 | 8 | 0 | 8 | 2 |
| ENSDARG000 | 15;15;15;15 | 12030367:12 | 12030860:12 | +;+;+;+ | 3225 | NA | 6 | 0 | 0 | 0 | 0 | 0 |
| ENSDARG000 | 15 | 29795583 | 29795666 | + | 84 | dre-mir-99-1 | 0 | 0 | 0 | 0 | 0 | 0 |
| ENSDARG000 | 15;15;15 | 18137741:18 | 18138028:18 | -;-;- | 765 | NA | 167 | 34 | 69 | 108 | 203 | 17 |
| ENSDARG000 | 15 | 22003082 | 22003196 | + | 115 | 5S_rRNA | 0 | 0 | 0 | 0 | 0 | 0 |
| ENSDARG000 | 15 | 10186054 | 10186169 | - | 116 | 5S_rRNA | 0 | 0 | 0 | 0 | 0 | 0 |
| ENSDARG000 | 15 | 27475258 | 27475374 | + | 117 | 5S_rRNA | 0 | 2 | 0 | 0 | 0 | 0 |
| ENSDARG000 | 15:15:15:15:1 | 43656241:43 | 43657173:43 | -;-;-;-;-;-;-;- | 3588 | grm5b | 12 | 2 | 0 | 0 | 26 | 4 |
| ENSDARG000 | 15:15:15:15:1 | 3219134:322 | 3219286:322 | +;+;+;+;+ | 675 | NA | 0 | 4 | 2 | 0 | 0 | 0 |
| ENSDARG000 | 15 | 26298629 | 26298747 | - | 119 | 5S_rRNA | 0 | 0 | 0 | 0 | 0 | 0 |
| ENSDARG000 | 15:15:15:15:1 | 47618221:47 | 47618288:47 | +:+:+:+:+:+:+: | 1221 | paf1 | 404 | 331 | 431 | 138 | 287 | 247 |
| ENSDARG000 | 15:15:15:15:1 | 499848:5006 | 500579:5007 | -;-;-;-;- | 1633 | nudt8 | 166 | 86 | 254 | 192 | 138 | 134 |
| ENSDARG000 | 15:15:15:15:1 | 4018362:401 | 4022212:402 | -;-;-;-;- | 4422 | NA | 2 | 0 | 6 | 0 | 0 | 0 |
| ENSDARG000 | 15:15:15:15:1 | 11693624:11 | 11693908:11 | +:+:+:+:+:+:+: | 2396 | STRN4 | 906 | 497 | 992 | 244 | 641 | 1074 |
| ENSDARG000 | 15:15:15:15:1 | 35372907:35 | 35372911:35 | -:-:-:-:-:-:-:-: | 2792 | MECOM | 14 | 4 | 2 | 0 | 3 | 6 |
| ENSDARG000 | 15 | 26274640 | 26274757 | - | 118 | 5S_rRNA | 0 | 0 | 0 | 0 | 0 | 0 |
| ENSDARG000 | 15:15:15:15:1 | 45008585:45 | 45008747:45 | -:-:-:-:-:-:-:-: | 12943 | DYNC2H1 | 612 | 336 | 675 | 306 | 488 | 306 |
| ENSDARG000 | 15 | 34155430 | 34155552 | + | 123 | 5S_rRNA | 0 | 0 | 0 | 0 | 0 | 0 |
| ENSDARG000 | 15 | 29670192 | 29670306 | - | 115 | 5S_rRNA | 0 | 0 | 0 | 0 | 0 | 0 |
| ENSDARG000 | 15:15:15:15:1 | 44133972:44 | 44134079:44 | +;+;+;+;+;+;+ | 3805 | NA | 58 | 80 | 78 | 70 | 64 | 84 |
| ENSDARG000 | 15:15:15:15:1 | 43333121:43 | 43333455:43 | +:+:+:+:+:+:+: | 2339 | zgc:85932 | 4 | 0 | 4 | 4 | 0 | 2 |
| ENSDARG000 | 15;15;15;15 | 42559910:42 | 42560616:42 | +;+;+;+ | 2387 | cldn8.3 | 6 | 4 | 2 | 0 | 2 | 0 |
| ENSDARG000 | 15:15:15:15:1 | 46853252:46 | 46853755:46 | +:+:+:+:+:+:+: | 3402 | zgc:153039 | 5013 | 2285 | 1739 | 995 | 1915 | 2026 |
| ENSDARG000 | 15:15:15:15:1 | 47883585:47 | 47883644:47 | -;-;-;-;-;- | 5272 | DMWD | 90 | 85 | 144 | 110 | 104 | 68 |
| ENSDARG000 | 15 | 22720765 | 22720881 | - | 117 | 5S_rRNA | 0 | 0 | 0 | 0 | 0 | 0 |
| ENSDARG000 | 15 | 13524874 | 13524988 | + | 115 | 5S_rRNA | 0 | 0 | 0 | 0 | 0 | 0 |
| ENSDARG000 | 15:15:15:15:1 | 44106641:44 | 44106820:44 | -;-;-;-;-;-;- | 1484 | NA | 19 | 36 | 16 | 22 | 37 | 22 |
| ENSDARG000 | 15:15:15:15:1 | 39096736:39 | 39096900:39 | +:+:+:+:+:+:+: | 4326 | robo2 | 21 | 28 | 24 | 0 | 26 | 36 |
| ENSDARG000 | 15 | 30893237 | 30893355 | + | 119 | 5S_rRNA | 0 | 0 | 0 | 0 | 1 | 0 |
| ENSDARG000 | 15 | 16829762 | 16829876 | + | 115 | 5S_rRNA | 0 | 0 | 0 | 0 | 0 | 0 |
| ENSDARG000 | 15 | 19542820 | 19542934 | - | 115 | 5S_rRNA | 0 | 0 | 0 | 0 | 0 | 0 |
| ENSDARG000 | 15:15:15:15:1 | 43166511:43 | 43166566:43 | +:+:+:+:+:+:+: | 3341 | dhrs12la | 66 | 157 | 28 | 16 | 111 | 18 |
| ENSDARG000 | 15 | 47322860 | 47322976 | + | 117 | 5S_rRNA | 0 | 0 | 4 | 0 | 0 | 0 |
| ENSDARG000 | 15;15;15;15 | 3089212:308 | 3091838:309 | -;-;-;- | 2685 | SLITRK5 | 0 | 0 | 0 | 0 | 0 | 0 |
| ENSDARG000 | 15;15;15 | 35691783:35 | 35691908:35 | +;+;+ | 501 | NA | 2 | 0 | 0 | 0 | 0 | 0 |
| ENSDARG000 | 15:15:15:15:1 | 43604032:43 | 43606213:43 | -;-;-;-;-;-;- | 3145 | ctsc | 4532 | 4745 | 2474 | 2189 | 6128 | 4310 |
| ENSDARG000 | 15:15:15:15:1 | 47525073:47 | 47525092:47 | +:+:+:+:+:+:+: | 2538 | SIDT2 | 866 | 534 | 790 | 617 | 786 | 736 |
| ENSDARG000 | 15:15:15:15:1 | 5973909:597 | 5974032:597 | +:+:+:+:+:+:+: | 6221 | NA | 5 | 4 | 8 | 18 | 0 | 0 |
| ENSDARG000 | 15 | 1078989 | 1079104 | + | 116 | 5S_rRNA | 0 | 0 | 0 | 0 | 0 | 0 |
| ENSDARG000 | 15 | 349451 | 349567 | - | 117 | 5S_rRNA | 0 | 0 | 0 | 0 | 0 | 0 |
| ENSDARG000 | 15 | 27922978 | 27923093 | - | 116 | 5S_rRNA | 0 | 0 | 0 | 0 | 0 | 0 |
| ENSDARG000 | 15 | 25667587 | 25667701 | + | 115 | 5S_rRNA | 0 | 0 | 0 | 0 | 0 | 0 |
| ENSDARG000 | 15:15:15:15:1 | 17788715:17 | 17792669:17 | -;-;-;-;- | 4786 | NA | 0 | 8 | 16 | 0 | 54 | 16 |
| ENSDARG000 | 15;15;15 | 11814969:11 | 11815110:11 | +;+;+ | 453 | NA | 162 | 61 | 123 | 63 | 159 | 119 |
| ENSDARG000 | 15:15:15:15:1 | 42560354:42 | 42560616:42 | +;+;+;+;+ | 1760 | cldn8.2 | 14 | 0 | 0 | 14 | 10 | 2 |
| ENSDARG000 | 15 | 13957722 | 13957850 | + | 129 | 5S_rRNA | 0 | 0 | 0 | 0 | 0 | 0 |
| ENSDARG000 | 15;15 | 5360407:536 | 5360512:536 | +;+ | 3530 | NA | 0 | 0 | 11 | 0 | 26 | 2 |
| ENSDARG000 | 15 | 27569081 | 27569195 | + | 115 | 5S_rRNA | 0 | 0 | 0 | 0 | 0 | 0 |
| ENSDARG000 | 15:15:15:15:1 | 44206698:44 | 44206800:44 | +;+;+;+;+ | 6785 | NA | 3 | 9 | 7 | 3 | 6 | 17 |
| ENSDARG000 | 15 | 34279421 | 34279535 | - | 115 | 5S_rRNA | 0 | 0 | 0 | 0 | 0 | 0 |
| ENSDARG000 | 15 | 35272927 | 35273041 | + | 115 | 5S_rRNA | 0 | 0 | 0 | 0 | 0 | 0 |
| ENSDARG000 | 15;15;15;15 | 11679759:11 | 11681581:11 | -;-;-;- | 2212 | fkrp | 176 | 153 | 198 | 112 | 197 | 198 |
| ENSDARG000 | 15:15:15:15:1 | 22390076:22 | 22390121:22 | +:+:+:+:+:+:+: | 1738 | OAF | 109 | 117 | 87 | 104 | 160 | 67 |
| ENSDARG000 | 15;15;15 | 3790556:379 | 3791044:379 | +;+;+ | 4405 | RNF14 | 6 | 13 | 10 | 29 | 5 | 35 |
| ENSDARG000 | 15:15:15:15:1 | 47008307:47 | 47008598:47 | +:+:+:+:+:+:+: | 2940 | ECEL1 | 0 | 15 | 4 | 0 | 34 | 0 |
| ENSDARG000 | 15 | 21252191 | 21252282 | + | 92 | Vault | 0 | 0 | 0 | 3 | 0 | 0 |
| ENSDARG000 | 15 | 11976143 | 11976260 | - | 118 | 5S_rRNA | 0 | 0 | 0 | 0 | 0 | 0 |
| ENSDARG000 | 15 | 5855111 | 5855225 | + | 115 | 5S_rRNA | 0 | 0 | 0 | 0 | 0 | 0 |
| ENSDARG000 | 15 | 32063292 | 32063409 | - | 118 | 5S_rRNA | 0 | 0 | 0 | 0 | 0 | 0 |
| ENSDARG000 | 15 | 21437876 | 21437959 | + | 84 | dre-let-7a-2 | 0 | 0 | 0 | 0 | 0 | 0 |
| ENSDARG000 | 15;15;15;15 | 47110884:47 | 47111250:47 | -;-;-;- | 1211 | NA | 28 | 11 | 33 | 32 | 31 | 17 |
| ENSDARG000 | 15 | 25897715 | 25897799 | + | 85 | dre-mir-132-1 | 0 | 0 | 0 | 0 | 0 | 0 |
| ENSDARG000 | 15 | 433035 | 434503 | - | 1469 | NA | 10 | 12 | 8 | 1 | 11 | 6 |
| ENSDARG000 | 15:15:15:15:1 | 47341856:47 | 47342175:47 | +;+;+;+;+ | 2337 | folr | 0 | 0 | 4 | 0 | 10 | 2 |
| ENSDARG000 | 15:15:15:15:1 | 44929497:44 | 44929656:44 | +;+;+;+;+;+;+ | 3338 | PDGFD | 48 | 28 | 31 | 6 | 41 | 30 |
| ENSDARG000 | 15 | 47605529 | 47607480 | - | 1952 | socs9 | 132 | 92 | 151 | 89 | 193 | 123 |
| ENSDARG000 | 15:15:15:15:1 | 282731:2827 | 282873:2828 | -:-:-:-:-:-:-:-: | 2428 | NA | 35 | 6 | 4 | 4 | 68 | 2 |
| ENSDARG000 | 15:15:15:15:1 | 2918257:292 | 2919174:292 | -;-;-;-;-;- | 2307 | NA | 278 | 96 | 69 | 164 | 179 | 332 |
| ENSDARG000 | 15:15:15:15:1 | 3169974:318 | 3170321:318 | +;+;+;+;+;+;+ | 3490 | NA | 0 | 2 | 4 | 0 | 4 | 0 |
| ENSDARG000 | 15;15;15 | 10863850:10 | 10863953:10 | +;+;+ | 2497 | NA | 0 | 4 | 0 | 0 | 0 | 0 |
| ENSDARG000 | 15:15:15:15:1 | 10973717:10 | 10975882:10 | -:-:-:-:-:-:-:-: | 3174 | NA | 0 | 0 | 2 | 0 | 9 | 4 |
| ENSDARG000 | 15;15;15 | 13421514:13 | 13421800:13 | +;+;+ | 911 | NA | 0 | 0 | 0 | 0 | 0 | 0 |
| ENSDARG000 | 15;15;15 | 14045121:14 | 14045272:14 | +;+;+ | 372 | NA | 0 | 0 | 0 | 0 | 0 | 0 |
| ENSDARG000 | 15;15;15;15 | 29886423:29 | 29886906:29 | -;-;-;- | 1430 | NA | 0 | 0 | 5 | 0 | 0 | 0 |
| ENSDARG000 | 15;15;15 | 35414466:35 | 35419417:35 | -;-;- | 6075 | NA | 0 | 0 | 0 | 0 | 0 | 0 |
| ENSDARG000 | 15;15;15 | 37259354:37 | 37263273:37 | -;-;- | 4613 | NA | 0 | 2 | 0 | 0 | 8 | 14 |
| ENSDARG000 | 15:15:15:15:1 | 37262329:37 | 37262520:37 | +:+:+:+:+:+:+: | 5316 | NA | 0 | 0 | 0 | 0 | 10 | 0 |
| ENSDARG000 | 15:15:15:15:1 | 37876716:37 | 37879042:37 | -;-;-;-;- | 3116 | NA | 0 | 0 | 0 | 0 | 0 | 0 |
| ENSDARG000 | 15;15;15;15 | 38372120:38 | 38372366:38 | +;+;+;+ | 769 | NA | 0 | 0 | 0 | 0 | 0 | 0 |
| ENSDARG000 | 15;15;15 | 42136087:42 | 42136618:42 | -;-;- | 800 | NA | 0 | 0 | 0 | 0 | 0 | 0 |
| ENSDARG000 | 15:15:15:15:1 | 43999530:44 | 44000266:44 | -:-:-:-:-:-:-:-: | 3411 | NA | 2 | 0 | 10 | 0 | 0 | 6 |
| ENSDARG000 | 15:15:15:15:1 | 46268826:46 | 46268847:46 | +;+;+;+;+;+;+ | 2870 | NA | 20 | 0 | 0 | 0 | 4 | 0 |
| ENSDARG000 | 15;15;15 | 46288890:46 | 46288943:46 | +;+;+ | 632 | NA | 0 | 0 | 0 | 0 | 0 | 0 |
| ENSDARG000 | 15:15:15:15:1 | 46291409:46 | 46291592:46 | +;+;+;+;+ | 670 | NA | 6 | 0 | 0 | 0 | 2 | 0 |
| ENSDARG000 | 15;15;15;15 | 47585808:47 | 47586227:47 | -;-;-;- | 858 | NA | 16 | 26 | 24 | 27 | 14 | 4 |
| ENSDARG000 | 15:15:15:15:1 | 47609322:47 | 47610093:47 | +;+;+;+;+;+;+ | 2185 | NA | 267 | 281 | 394 | 281 | 286 | 375 |
| ENSDARG000 | 23;23;23;23 | 45611649:45 | 45612397:45 | +;+;+;+ | 1152 | NA | 0 | 0 | 0 | 0 | 8 | 0 |
| ENSDARG000 | 23:23:23:23:2 | 39558508:39 | 39558557:39 | +:+:+:+:+:+:+: | 5192 | camk1gb | 22 | 2 | 22 | 0 | 0 | 12 |
| ENSDARG000 | 23:23:23:23:2 | 30707837:30 | 30707937:30 | +:+:+:+:+:+:+: | 2892 | dnajc11a | 1112 | 1136 | 1310 | 464 | 1209 | 856 |
| ENSDARG000 | 23:23:23:23:2 | 2728095:272 | 2728517:272 | +:+:+:+:+:+:+: | 2063 | zgc:114123 | 0 | 6 | 0 | 0 | 0 | 0 |
| ENSDARG000 | 23:23:23:23:2 | 27251626:27 | 27253452:27 | -:-:-:-:-:-:-:-: | 6816 | scn8aa | 0 | 0 | 12 | 0 | 4 | 0 |
| ENSDARG000 | 23:23:23:23:2 | 43870886:43 | 43871014:43 | +:+:+:+:+:+:+: | 3289 | NFXL1 | 577 | 452 | 941 | 551 | 691 | 794 |
| ENSDARG000 | 23:23:23:23:2 | 36297221:36 | 36299328:36 | -:-:-:-:-:-:-:-: | 3544 | cbx5 | 1351 | 2828 | 3197 | 1826 | 2735 | 1944 |
| ENSDARG000 | 23:23:23:23:2 | 45668597:45 | 45668923:45 | -;-;-;-;-;-;-;- | 1718 | NA | 2 | 2 | 0 | 0 | 0 | 0 |
| ENSDARG000 | 23:23:23:23:2 | 19977120:19 | 19977326:19 | +:+:+:+:+:+:+: | 1751 | cfap126 | 7 | 2 | 2 | 0 | 27 | 4 |
| ENSDARG000 | 23:23:23:23:2 | 19968956:19 | 19970253:19 | -;-;-;-;- | 2298 | ccnq | 252 | 227 | 277 | 157 | 340 | 226 |
| ENSDARG000 | 23:23:23:23:2 | 7471072:747 | 7471210:747 | +:+:+:+:+:+:+: | 4248 | NA | 32 | 8 | 38 | 22 | 30 | 28 |
| ENSDARG000 | 23:23:23:23:2 | 31962383:31 | 31963393:31 | -;-;-;-;-;-;- | 1375 | ormdl2 | 453 | 565 | 548 | 579 | 599 | 730 |
| ENSDARG000 | 23;23 | 16744782:16 | 16745221:16 | -;- | 959 | NA | 0 | 0 | 7 | 0 | 3 | 0 |
| ENSDARG000 | 23:23:23:23:2 | 25354856:25 | 25355534:25 | +:+:+:+:+:+:+: | 5982 | fmnl3 | 226 | 56 | 121 | 80 | 82 | 104 |
| ENSDARG000 | 23:23:23:23:2 | 17993791:17 | 17996228:17 | -:-:-:-:-:-:-:-: | 4134 | pm20d1.2 | 432 | 373 | 366 | 329 | 472 | 445 |
| ENSDARG000 | 23:23:23:23:2 | 10724042:10 | 10726081:10 | -:-:-:-:-:-:-:-: | 2937 | eif4e3 | 866 | 781 | 835 | 614 | 884 | 4057 |
| ENSDARG000 | 23;23;23;23 | 30725716:30 | 30726667:30 | -;-;-;- | 1627 | thap3 | 72 | 60 | 98 | 32 | 94 | 118 |
| ENSDARG000 | 23;23 | 39606108:39 | 39606293:39 | +;+ | 800 | g0s2 | 0 | 26 | 94 | 74 | 12 | 26 |
| ENSDARG000 | 23:23:23:23:2 | 39611688:39 | 39611811:39 | +:+:+:+:+:+:+: | 1610 | otud3 | 212 | 212 | 202 | 168 | 170 | 201 |
| ENSDARG000 | 23:23:23:23:2 | 27345826:27 | 27345946:27 | +;+;+;+;+ | 857 | NA | 0 | 0 | 0 | 0 | 0 | 0 |
| ENSDARG000 | 23:23:23:23:2 | 16748806:16 | 16749099:16 | +:+:+:+:+:+:+: | 1446 | fbxo44.1 | 421 | 337 | 356 | 212 | 342 | 403 |
| ENSDARG000 | 23:23:23:23:2 | 7505943:751 | 7506342:751 | +:+:+:+:+:+:+: | 7266 | HCK | 254 | 330 | 360 | 216 | 598 | 500 |
| ENSDARG000 | 23:23:23:23:2 | 39632397:39 | 39636206:39 | -:-:-:-:-:-:-:-: | 5483 | vwa1 | 1144 | 466 | 787 | 1625 | 3046 | 610 |
| ENSDARG000 | 23:23:23:23:2 | 45694055:45 | 45694556:45 | -;-;-;-;-;-;- | 1649 | ednrab | 2 | 0 | 0 | 0 | 0 | 0 |
| ENSDARG000 | 23;23;23 | 43939124:43 | 43939288:43 | +;+;+ | 388 | NA | 0 | 9 | 0 | 0 | 1 | 0 |
| ENSDARG000 | 23:23:23:23:2 | 27352072:27 | 27352205:27 | +;+;+;+;+ | 985 | NA | 0 | 0 | 4 | 0 | 0 | 0 |
| ENSDARG000 | 23:23:23:23:2 | 19984808:19 | 19985547:19 | -:-:-:-:-:-:-:-: | 2132 | B4GALT3 | 6 | 20 | 37 | 6 | 36 | 40 |
| ENSDARG000 | 23:23:23:23:2 | 30730121:30 | 30730614:30 | +:+:+:+:+:+:+: | 6001 | asxl1 | 996 | 790 | 1241 | 937 | 1238 | 954 |
| ENSDARG000 | 23:23:23:23:2 | 2752645:275 | 2752796:275 | +:+:+:+:+:+:+: | 2342 | TOP1 | 16 | 28 | 46 | 15 | 56 | 42 |
| ENSDARG000 | 23;23;23 | 23254976:23 | 23255111:23 | -;-;- | 1003 | NA | 0 | 1 | 0 | 0 | 0 | 0 |
| ENSDARG000 | 23:23:23:23:2 | 31979602:31 | 31979680:31 | +:+:+:+:+:+:+: | 4146 | PAN2 | 608 | 441 | 708 | 348 | 579 | 512 |
| ENSDARG000 | 23:23:23:23:2 | 21380079:21 | 21380401:21 | +:+:+:+:+:+:+: | 1519 | IFFO2 | 6 | 12 | 5 | 0 | 0 | 0 |
| ENSDARG000 | 23:23:23:23:2 | 10762684:10 | 10763952:10 | -;-;-;-;- | 2262 | rybpa | 300 | 225 | 318 | 109 | 320 | 225 |
| ENSDARG000 | 23:23:23:23:2 | 4709744:471 | 4709852:471 | +;+;+;+;+ | 693 | NA | 0 | 0 | 0 | 0 | 0 | 0 |
| ENSDARG000 | 23:23:23:23:2 | 4709607:470 | 4709852:470 | +:+:+:+:+:+:+: | 2557 | raf1a | 145 | 81 | 143 | 139 | 126 | 98 |
| ENSDARG000 | 23:23:23:23:2 | 19182819:19 | 19183177:19 | +:+:+:+:+:+:+: | 2089 | NA | 0 | 0 | 0 | 0 | 0 | 0 |
| ENSDARG000 | 23:23:23:23:2 | 20014030:20 | 20015709:20 | -:-:-:-:-:-:-:-: | 3042 | bgnb | 38 | 4 | 8 | 0 | 14 | 4 |
| ENSDARG000 | 23:23:23:23:2 | 10805188:10 | 10805351:10 | +:+:+:+:+:+:+: | 3089 | ppp4r2a | 616 | 554 | 645 | 154 | 646 | 334 |
| ENSDARG000 | 23:23:23:23:2 | 43950674:43 | 43950937:43 | +:+:+:+:+:+:+: | 3054 | CORIN | 48 | 37 | 56 | 47 | 90 | 48 |
| ENSDARG000 | 23:23:23:23:2 | 18020928:18 | 18021104:18 | -:-:-:-:-:-:-:-: | 1760 | pm20d1.1 | 0 | 0 | 2 | 4 | 6 | 0 |
| ENSDARG000 | 23:23:23:23:2 | 45845159:45 | 45845805:45 | +:+:+:+:+:+:+: | 6626 | lmnl3 | 973 | 799 | 635 | 635 | 1052 | 906 |
| ENSDARG000 | 23;23;23;23 | 13928346:13 | 13928519:13 | +;+;+;+ | 671 | NA | 4 | 5 | 28 | 0 | 4 | 4 |
| ENSDARG000 | 23:23:23:23:2 | 39695827:39 | 39696157:39 | +:+:+:+:+:+:+: | 2568 | TMCO4 | 197 | 225 | 368 | 463 | 263 | 153 |

| | | | | | | | | | | | |
|---|---|---|---|---|---|---|---|---|---|---|---|
| ENSDARG000 | 23:23:23:23:2 | 23399520:234 | 23400414:234 | -;-;-;-;-;-;- | 1551 | her9 | 342 | 570 | 1567 | 682 | 2435 | 644 |
| ENSDARG000 | 23:23:23:23:2 | 30767310:307 | 30767856:307 | -:-:-:-:-:-:-:-: | 4622 | MYT1 | 2 | 0 | 16 | 0 | 2 | 0 |
| ENSDARG000 | 23:23:23:23:2 | 20068880:200 | 20069113:200 | -;-;-;-;-;-;-;- | 1775 | NA | 0 | 0 | 2 | 0 | 0 | 0 |
| ENSDARG000 | 23:23:23:23:2 | 21424747:214 | 21424907:214 | +:+:+:+:+:+:+: | 2466 | tas1r1 | 0 | 0 | 0 | 0 | 2 | 0 |
| ENSDARG000 | 23:23:23:23:2 | 27358722:273 | 27360358:273 | -:-:-:-:-:-:-:-: | 6269 | dip2ba | 3032 | 2917 | 4334 | 3875 | 3780 | 1761 |
| ENSDARG000 | 23;23;23 | 32011768:320 | 32011986:320 | +;+;+ | 4432 | plagx | 322 | 174 | 327 | 176 | 332 | 272 |
| ENSDARG000 | 23:23:23:23:2 | 16760845:167 | 16761118:167 | +:+:+:+:+:+:+: | 5638 | NA | 43 | 44 | 68 | 54 | 81 | 50 |
| ENSDARG000 | 23;23;23 | 39730965:397 | 39731345:397 | -;-;- | 1440 | HTR6 | 43 | 42 | 108 | 127 | 77 | 43 |
| ENSDARG000 | 23:23:23:23:2 | 44157682:441 | 44157879:441 | +:+:+:+:+:+:+: | 2079 | NA | 0 | 4 | 4 | 0 | 0 | 0 |
| ENSDARG000 | 23:23:23:23:2 | 32021803:320 | 32022099:320 | +;+;+;+;+;+ | 952 | trappc6b | 374 | 437 | 358 | 170 | 527 | 472 |
| ENSDARG000 | 23;23;23;23 | 39744369:397 | 39744989:397 | -;-;-;- | 937 | micos10 | 428 | 779 | 627 | 368 | 550 | 492 |
| ENSDARG000 | 23:23:23:23:2 | 30755804:307 | 30756254:307 | -;-;-;-;-;- | 1439 | pcmtd2a | 244 | 68 | 91 | 173 | 108 | 223 |
| ENSDARG000 | 23;23 | 30764287:307 | 30764475:307 | +;+ | 544 | NA | 0 | 0 | 0 | 0 | 0 | 0 |
| ENSDARG000 | 23:23:23:23:2 | 45860676:458 | 45860970:458 | +:+:+:+:+:+:+: | 2507 | syap1 | 1440 | 1241 | 1597 | 1194 | 1367 | 1206 |
| ENSDARG000 | 23;23 | 21444922:214 | 21445853:214 | -;- | 1050 | her12 | 4 | 0 | 0 | 12 | 4 | 0 |
| ENSDARG000 | 23:23:23:23:2 | 216012:21601 | 216071:21608 | +:+:+:+:+:+:+: | 2701 | PDZD4 | 2 | 1 | 0 | 2 | 0 | 8 |
| ENSDARG000 | 23:23:23:23:2 | 10828716:108 | 10829188:108 | -:-:-:-:-:-:-:-: | 3051 | PDZRN3 | 18 | 31 | 95 | 8 | 38 | 42 |
| ENSDARG000 | 23:23:23:23:2 | 16774589:167 | 16774673:167 | +;+;+;+;+;+;+ | 1412 | fbxo44.3 | 233 | 128 | 227 | 69 | 234 | 222 |
| ENSDARG000 | 23;23 | 21452557:214 | 21453412:214 | -;- | 963 | her4.1 | 0 | 0 | 0 | 0 | 0 | 0 |
| ENSDARG000 | 23;23 | 21462021:214 | 21463586:214 | -;- | 1673 | her4.4 | 0 | 0 | 0 | 0 | 0 | 0 |
| ENSDARG000 | 23;23 | 21459263:214 | 21459369:214 | +;+ | 1783 | her4.3 | 0 | 0 | 0 | 0 | 0 | 0 |
| ENSDARG000 | 23:23:23:23:2 | 21455152:214 | 21455260:214 | +:+:+:+:+:+:+: | 2213 | her4.2 | 2 | 0 | 0 | 0 | 0 | 0 |
| ENSDARG000 | 23 | 250798 | 253229 | - | 2432 | NA | 451 | 305 | 385 | 208 | 397 | 476 |
| ENSDARG000 | 23:23:23:23:2 | 19198244:191 | 19198481:191 | +:+:+:+:+:+:+: | 1005 | ccdc115 | 691 | 757 | 501 | 678 | 709 | 621 |
| ENSDARG000 | 23:23:23:23:2 | 255409:26272 | 255412:26281 | -:-:-:-:-:-:-:-: | 1395 | NA | 237 | 68 | 205 | 69 | 160 | 125 |
| ENSDARG000 | 23:23:23:23:2 | 39755641:397 | 39759537:397 | -:-:-:-:-:-:-:-: | 4769 | SIKE1 | 246 | 182 | 213 | 134 | 245 | 246 |
| ENSDARG000 | 23:23:23:23:2 | 23485858:234 | 23486216:234 | +:+:+:+:+:+:+: | 9055 | agrn | 6625 | 6252 | 9627 | 11944 | 13162 | 6967 |
| ENSDARG000 | 23;23;23 | 33828665:338 | 33828713:338 | +;+;+ | 389 | NA | 2 | 0 | 0 | 0 | 0 | 0 |
| ENSDARG000 | 23;23 | 19202636:192 | 19202961:192 | +;+ | 3354 | NA | 294 | 388 | 418 | 234 | 413 | 250 |
| ENSDARG000 | 23;23 | 21469203:214 | 21470820:214 | -;- | 1725 | her4.5 | 0 | 0 | 0 | 0 | 0 | 0 |
| ENSDARG000 | 23:23:23:23:2 | 36306539:363 | 36306604:363 | +:+:+:+:+:+:+: | 1890 | hnrnpa1b | 3493 | 4154 | 4242 | 1806 | 4234 | 2476 |
| ENSDARG000 | 23:23:23:23:2 | 33907899:339 | 33908053:339 | +:+:+:+:+:+:+: | 3110 | cs | 3357 | 4624 | 5022 | 3227 | 4148 | 3952 |
| ENSDARG000 | 23;23 | 21473103:214 | 21473166:214 | +;+ | 761 | si:ch73-21g5. | 2 | 0 | 0 | 0 | 0 | 0 |
| ENSDARG000 | 23:23:23:23:2 | 25452656:254 | 25453102:254 | -:-:-:-:-:-:-:-: | 14024 | LRP1 | 2748 | 1481 | 1603 | 2185 | 3606 | 1377 |
| ENSDARG000 | 23:23:23:23:2 | 13940958:139 | 13941346:139 | -:-:-:-:-:-:-:-: | 4066 | ADCY6 | 411 | 386 | 768 | 348 | 918 | 545 |
| ENSDARG000 | 23:23:23:23:2 | 20110086:201 | 20110289:201 | +;+;+;+;+;+ | 874 | tnnc1b | 0 | 4 | 6 | 0 | 10 | 0 |
| ENSDARG000 | 23:23:23:23:2 | 20113378:201 | 20114183:201 | -:-:-:-:-:-:-:-: | 3022 | tktb | 6695 | 7977 | 11479 | 6367 | 10044 | 7099 |
| ENSDARG000 | 23:23:23:23:2 | 19217528:192 | 19217654:192 | +;+;+;+;+ | 773 | NA | 6 | 1 | 6 | 2 | 1 | 0 |
| ENSDARG000 | 23:23:23:23:2 | 19220381:192 | 19222476:192 | -;-;-;-;-;- | 2633 | oard1 | 211 | 181 | 181 | 144 | 146 | 153 |
| ENSDARG000 | 23:23:23:23:2 | 2789422:2790 | 2789555:2790 | +:+:+:+:+:+:+: | 5568 | plcg1 | 2069 | 1908 | 2267 | 1520 | 2587 | 1918 |
| ENSDARG000 | 23:23:23:23:2 | 45869010:458 | 45869129:458 | +:+:+:+:+:+:+: | 3446 | fbxl5 | 1395 | 1598 | 3515 | 3820 | 1792 | 1243 |
| ENSDARG000 | 23:23:23:23:2 | 21479958:214 | 21480171:214 | +:+:+:+:+:+:+: | 4724 | NA | 0 | 0 | 0 | 0 | 0 | 0 |
| ENSDARG000 | 23:23:23:23:2 | 309063:30909 | 309223:30922 | +:+:+:+:+:+:+: | 3032 | taf11 | 408 | 267 | 475 | 264 | 439 | 409 |
| ENSDARG000 | 23:23:23:23:2 | 11014693:110 | 11014858:110 | -:-:-:-:-:-:-:-: | 2954 | NA | 0 | 0 | 0 | 0 | 0 | 0 |
| ENSDARG000 | 23:23:23:23:2 | 33934877:339 | 33935584:339 | -;-;-;-;- | 1768 | COQ10A | 95 | 130 | 122 | 20 | 178 | 24 |
| ENSDARG000 | 23:23:23:23:2 | 39774931:397 | 39775355:397 | -;-;-;-;- | 1148 | iyd | 0 | 0 | 0 | 0 | 0 | 8 |
| ENSDARG000 | 23;23;23;23 | 33934228:339 | 33934305:339 | +;+;+;+ | 1788 | NA | 141 | 122 | 170 | 153 | 118 | 119 |
| ENSDARG000 | 23:23:23:23:2 | 4751959:4754 | 4752749:4754 | -:-:-:-:-:-:-:-: | 2178 | syn2a | 43 | 16 | 12 | 32 | 8 | 31 |
| ENSDARG000 | 23;23;23;23 | 19227296:192 | 19227859:192 | -;-;-;- | 1302 | guca1b | 0 | 0 | 0 | 0 | 0 | 0 |
| ENSDARG000 | 23:23:23:23:2 | 32039386:320 | 32039487:320 | +:+:+:+:+:+:+: | 3825 | mylk2 | 0 | 4 | 6 | 0 | 0 | 0 |
| ENSDARG000 | 23:23:23:23:2 | 44190852:441 | 44191126:441 | -:-:-:-:-:-:-:-: | 3847 | zgc:158659 | 16 | 2 | 18 | 4 | 9 | 8 |
| ENSDARG000 | 23:23:23:23:2 | 27446929:274 | 27448029:274 | -:-:-:-:-:-:-:-: | 2383 | atf7a | 328 | 257 | 395 | 265 | 255 | 324 |
| ENSDARG000 | 23:23:23:23:2 | 32028574:320 | 32028739:320 | +:+:+:+:+:+:+: | 4317 | tpx2 | 528 | 1685 | 1841 | 827 | 1210 | 733 |
| ENSDARG000 | 23:23:23:23:2 | 21499585:215 | 21500362:215 | -:-:-:-:-:-:-:-: | 2750 | rnf207b | 0 | 0 | 0 | 0 | 0 | 0 |
| ENSDARG000 | 23;23;23;23 | 39807857:398 | 39812862:398 | -;-;-;- | 5434 | ppp1r14c | 80 | 62 | 306 | 88 | 340 | 188 |
| ENSDARG000 | 23:23:23:23:2 | 16807209:168 | 16807564:168 | +:+:+:+:+:+:+: | 1867 | fbxo6.2 | 25 | 60 | 41 | 17 | 64 | 80 |
| ENSDARG000 | 23:23:23:23:2 | 21492151:214 | 21492440:214 | +;+;+;+;+ | 2210 | icmt | 235 | 313 | 278 | 194 | 326 | 280 |
| ENSDARG000 | 23:23:23:23:2 | 45887202:458 | 45887877:458 | -;-;-;-;-;- | 1476 | NA | 10 | 1 | 4 | 4 | 7 | 16 |
| ENSDARG000 | 23:23:23:23:2 | 28582865:286 | 28582925:286 | +:+:+:+:+:+:+: | 4119 | l1cama | 8 | 2 | 0 | 10 | 4 | 2 |
| ENSDARG000 | 23:23:23:23:2 | 4834447:4837 | 4834921:4837 | -:-:-:-:-:-:-:-: | 2429 | slc6a1a | 2 | 0 | 0 | 2 | 0 | 0 |
| ENSDARG000 | 23:23:23:23:2 | 45906137:459 | 45906170:459 | +:+:+:+:+:+:+: | 1978 | abcg2a | 46 | 45 | 199 | 106 | 322 | 99 |
| ENSDARG000 | 23:23:23:23:2 | 36311581:363 | 36312827:363 | -;-;-;-;-;- | 1640 | nfe2 | 0 | 0 | 4 | 6 | 20 | 8 |
| ENSDARG000 | 23:23:23:23:2 | 23918421:239 | 23918517:239 | +:+:+:+:+:+:+: | 3241 | ptpn11b | 210 | 166 | 318 | 304 | 380 | 199 |
| ENSDARG000 | 23:23:23:23:2 | 4777348:4777 | 4778317:4778 | +:+:+:+:+:+:+: | 3882 | VGLL4 | 331 | 185 | 392 | 190 | 292 | 326 |
| ENSDARG000 | 23:23:23:23:2 | 39854566:398 | 39854653:398 | +:+:+:+:+:+:+: | 3167 | NA | 4 | 0 | 2 | 2 | 0 | 16 |
| ENSDARG000 | 23;23;23;23 | 33957350:339 | 33957505:339 | +;+;+;+ | 589 | snrpg | 776 | 1097 | 1396 | 522 | 1067 | 761 |
| ENSDARG000 | 23:23:23:23:2 | 19249968:192 | 19250720:192 | -;-;-;-;- | 1621 | NA | 0 | 0 | 4 | 8 | 0 | 0 |
| ENSDARG000 | 23:23:23:23:2 | 46157638:461 | 46157686:461 | +;+;+;+;+;+;+ | 2911 | btr32 | 62 | 0 | 8 | 76 | 30 | 24 |
| ENSDARG000 | 23:23:23:23:2 | 28685266:286 | 28685422:286 | -;-;-;-;-;-;- | 1489 | ribc1 | 12 | 0 | 0 | 0 | 0 | 0 |
| ENSDARG000 | 23:23:23:23:2 | 16814968:168 | 16815472:168 | +:+:+:+:+:+:+: | 3084 | fbxo44.5 | 415 | 381 | 454 | 358 | 490 | 411 |
| ENSDARG000 | 23:23:23:23:2 | 7548797:7552 | 7548941:7552 | +:+:+:+:+:+:+: | 2466 | tm9sf4 | 1973 | 1733 | 2450 | 1172 | 2488 | 2160 |
| ENSDARG000 | 23:23:23:23:2 | 16823385:168 | 16823404:168 | +:+:+:+:+:+:+: | 1775 | fbxo6.3 | 217 | 200 | 160 | 313 | 306 | 236 |
| ENSDARG000 | 23:23:23:23:2 | 2872855:2873 | 2873444:2873 | -:-:-:-:-:-:-:-: | 4460 | zhx3b | 476 | 336 | 354 | 390 | 380 | 450 |
| ENSDARG000 | 23:23:23:23:2 | 33963619:339 | 33963745:339 | +:+:+:+:+:+:+: | 1776 | plpbp | 480 | 646 | 855 | 648 | 769 | 509 |
| ENSDARG000 | 23:23:23:23:2 | 24085531:240 | 24085666:240 | +:+:+:+:+:+:+: | 2045 | NA | 0 | 0 | 0 | 0 | 0 | 0 |
| ENSDARG000 | 23:23:23:23:2 | 36340520:363 | 36340609:363 | +:+:+:+:+:+:+: | 1016 | copz1 | 952 | 1090 | 1401 | 959 | 1219 | 1119 |
| ENSDARG000 | 23:23:23:23:2 | 18032569:180 | 18033299:180 | -:-:-:-:-:-:-:-: | 2209 | lemd1 | 6 | 120 | 34 | 32 | 91 | 21 |
| ENSDARG000 | 23:23:23:23:2 | 2914577:2914 | 2914648:2914 | +:+:+:+:+:+:+: | 3116 | dhx35 | 350 | 404 | 566 | 236 | 521 | 374 |
| ENSDARG000 | 23:23:23:23:2 | 44236281:442 | 44236778:442 | +:+:+:+:+:+:+: | 6791 | MEPCE | 653 | 681 | 780 | 391 | 1049 | 724 |
| ENSDARG000 | 23:23:23:23:2 | 11285662:113 | 11285766:113 | +:+:+:+:+:+:+: | 8314 | chl1a | 70 | 25 | 5061 | 4577 | 8412 | 154 |
| ENSDARG000 | 23:23:23:23:2 | 46183410:461 | 46183525:461 | +:+:+:+:+:+:+: | 1439 | btr31 | 616 | 365 | 232 | 532 | 407 | 341 |
| ENSDARG000 | 23;23;23 | 44263986:442 | 44264019:442 | +;+;+ | 225 | NA | 0 | 0 | 0 | 5 | 0 | 0 |
| ENSDARG000 | 23:23:23:23:2 | 19473935:194 | 19475746:194 | -:-:-:-:-:-:-:-: | 5848 | NA | 1550 | 993 | 1700 | 759 | 1569 | 1088 |
| ENSDARG000 | 23;23;23 | 11344619:113 | 11344700:113 | -;-;- | 391 | NA | 0 | 1 | 2 | 0 | 0 | 2 |
| ENSDARG000 | 23:23:23:23:2 | 4890693:4892 | 4890773:4892 | +;+;+;+;+;+ | 1089 | tnnc1a | 0 | 0 | 0 | 0 | 0 | 0 |
| ENSDARG000 | 23:23:23:23:2 | 16864130:168 | 16864252:168 | +;+;+;+;+;+ | 1424 | fbxo44.7 | 223 | 230 | 227 | 115 | 256 | 225 |
| ENSDARG000 | 23:23:23:23:2 | 320624:32081 | 320923:32092 | -:-:-:-:-:-:-:-: | 4339 | BLTP3A | 114 | 68 | 142 | 52 | 187 | 37 |
| ENSDARG000 | 23 | 39944652 | 39945578 | - | 927 | NA | 2 | 0 | 6 | 0 | 28 | 0 |
| ENSDARG000 | 23:23:23:23:2 | 27495091:274 | 27499554:274 | -:-:-:-:-:-:-:-: | 4936 | asb8 | 540 | 381 | 500 | 318 | 574 | 534 |
| ENSDARG000 | 23:23:23:23:2 | 3503596:3504 | 3503748:3504 | -;-;-;-;-;- | 1262 | rnf114 | 535 | 398 | 293 | 199 | 362 | 368 |
| ENSDARG000 | 23;23;23 | 3513201:3517 | 3513317:3518 | +;+;+ | 2542 | spata2 | 497 | 332 | 493 | 389 | 325 | 280 |
| ENSDARG000 | 23 | 39951553 | 39952406 | - | 854 | NA | 6 | 0 | 0 | 0 | 0 | 0 |
| ENSDARG000 | 23;23;23 | 39958166:399 | 39959173:399 | -;-;- | 1204 | XCR1 | 0 | 0 | 2 | 0 | 2 | 0 |
| ENSDARG000 | 23:23:23:23:2 | 14097508:140 | 14097827:140 | +:+:+:+:+:+:+: | 2375 | cacnb3a | 0 | 0 | 0 | 2 | 14 | 0 |
| ENSDARG000 | 23:23:23:23:2 | 4899238:4904 | 4901612:4904 | -:-:-:-:-:-:-:-: | 3578 | atp6ap1a | 82 | 116 | 122 | 36 | 98 | 32 |
| ENSDARG000 | 23:23:23:23:2 | 24124684:241 | 24124834:241 | +;+;+;+;+ | 886 | NA | 0 | 4 | 8 | 8 | 11 | 37 |
| ENSDARG000 | 23:23:23:23:2 | 20127883:201 | 20128185:201 | -:-:-:-:-:-:-:-: | 1636 | lrrc23 | 14 | 30 | 42 | 14 | 34 | 44 |
| ENSDARG000 | 23;23 | 24148055:241 | 24148228:241 | +;+ | 503 | NA | 0 | 0 | 0 | 0 | 0 | 0 |
| ENSDARG000 | 23:23:23:23:2 | 24148338:241 | 24148667:241 | -;-;-;-;- | 836 | NA | 0 | 0 | 0 | 0 | 0 | 0 |
| ENSDARG000 | 23:23:23:23:2 | 24140043:241 | 24140235:241 | -:-:-:-:-:-:-:-: | 2912 | ARHGEF19 | 2 | 0 | 6 | 6 | 6 | 8 |
| ENSDARG000 | 23:23:23:23:2 | 39963599:399 | 39964007:399 | +:+:+:+:+:+:+: | 4952 | FYCO1 | 517 | 321 | 438 | 341 | 511 | 441 |
| ENSDARG000 | 23:23:23:23:2 | 44349252:443 | 44349905:443 | +:+:+:+:+:+:+: | 3816 | ephb4b | 1199 | 502 | 779 | 826 | 714 | 324 |
| ENSDARG000 | 23:23:23:23:2 | 28693278:286 | 28693432:286 | +:+:+:+:+:+:+: | 3745 | smc1a | 0 | 1 | 0 | 0 | 0 | 0 |
| ENSDARG000 | 23:23:23:23:2 | 27511709:275 | 27512106:275 | -:-:-:-:-:-:-:-: | 2782 | NA | 1082 | 1119 | 1563 | 800 | 1387 | 1029 |
| ENSDARG000 | 23:23:23:23:2 | 7592009:7592 | 7593711:7593 | -;-;-;-;-;-;-;- | 2542 | plagl2 | 2 | 2 | 2 | 0 | 8 | 28 |
| ENSDARG000 | 23;23;23;23 | 40109353:401 | 40109444:401 | +;+;+;+ | 512 | NA | 0 | 0 | 0 | 2 | 0 | 0 |
| ENSDARG000 | 23:23:23:23:2 | 33977724:339 | 33978865:339 | -:-:-:-:-:-:-:-: | 1759 | tmed4 | 63 | 99 | 155 | 67 | 183 | 87 |
| ENSDARG000 | 23:23:23:23:2 | 21520769:215 | 21522855:215 | -:-:-:-:-:-:-:-: | 4085 | rcc2 | 1557 | 1566 | 2067 | 1054 | 1957 | 1526 |
| ENSDARG000 | 23:23:23:23:2 | 7692042:7692 | 7692200:7692 | +;+;+;+;+;+;+ | 1549 | pofut1 | 230 | 249 | 311 | 190 | 349 | 242 |
| ENSDARG000 | 23:23:23:23:2 | 354418:35502 | 354497:35543 | +:+:+:+:+:+:+: | 2621 | ap4b1 | 231 | 189 | 304 | 235 | 308 | 162 |
| ENSDARG000 | 23;23 | 33978711:339 | 33978862:339 | +;+ | 308 | NA | 0 | 0 | 0 | 0 | 0 | 0 |
| ENSDARG000 | 23:23:23:23:2 | 19490383:194 | 19490715:194 | -:-:-:-:-:-:-:-: | 2124 | ASB14 | 0 | 0 | 0 | 4 | 2 | 2 |
| ENSDARG000 | 23:23:23:23:2 | 16889152:168 | 16889347:168 | +:+:+:+:+:+:+: | 2997 | NA | 640 | 524 | 624 | 529 | 483 | 570 |
| ENSDARG000 | 23:23:23:23:2 | 383782:38485 | 383906:38496 | +;+;+;+;+;+;+ | 5189 | zgc:101663 | 13 | 13 | 22 | 18 | 24 | 21 |
| ENSDARG000 | 23:23:23:23:2 | 20137520:201 | 20137947:201 | -;-;-;-;- | 1633 | NA | 82 | 123 | 136 | 12 | 147 | 16 |
| ENSDARG000 | 23;23 | 44461493:444 | 44461639:444 | +;+ | 336 | NA | 15 | 6 | 5 | 6 | 17 | 4 |
| ENSDARG000 | 23:23:23:23:2 | 36418152:364 | 36418223:364 | +;+;+;+;+;+;+ | 404 | NA | 0 | 0 | 1 | 6 | 0 | 0 |
| ENSDARG000 | 23:23:23:23:2 | 36401572:364 | 36402798:364 | -:-:-:-:-:-:-:-: | 2981 | znf740b | 208 | 249 | 374 | 291 | 236 | 299 |
| ENSDARG000 | 23:23:23:23:2 | 7710447:7710 | 7710882:7710 | +:+:+:+:+:+:+: | 6932 | kif3b | 352 | 328 | 412 | 338 | 350 | 369 |
| ENSDARG000 | 23;23;23 | 44465523:444 | 44466257:444 | -;-;- | 1542 | SPSB2 | 0 | 4 | 0 | 0 | 4 | 0 |
| ENSDARG000 | 23:23:23:23:2 | 409578:41108 | 409634:41120 | +:+:+:+:+:+:+: | 1666 | slc25a55b | 40 | 38 | 6 | 35 | 8 | 52 |
| ENSDARG000 | 23:23:23:23:2 | 4915595:4915 | 4917030:4917 | -:-:-:-:-:-:-:-: | 3699 | tafazzin | 316 | 294 | 466 | 294 | 362 | 259 |
| ENSDARG000 | 23:23:23:23:2 | 34001803:340 | 34001913:340 | +:+:+:+:+:+:+: | 2764 | NA | 8 | 2 | 2 | 4 | 4 | 2 |
| ENSDARG000 | 23:23:23:23:2 | 7762033:7762 | 7762176:7762 | -:-:-:-:-:-:-:-: | 4175 | MYT1 | 920 | 670 | 1009 | 417 | 1204 | 780 |
| ENSDARG000 | 23:23:23:23:2 | 33987382:339 | 33987664:339 | +:+:+:+:+:+:+: | 1653 | fam50a | 499 | 472 | 465 | 298 | 533 | 386 |
| ENSDARG000 | 23:23:23:23:2 | 423351:42414 | 424205:42420 | -:-:-:-:-:-:-:-: | 1708 | NA | 18 | 52 | 138 | 54 | 14 | 106 |
| ENSDARG000 | 23:23:23:23:2 | 20145300:201 | 20146249:201 | -:-:-:-:-:-:-:-: | 6094 | USP19 | 1146 | 1103 | 1227 | 694 | 1061 | 890 |
| ENSDARG000 | 23:23:23:23:2 | 14193547:141 | 14194411:141 | -:-:-:-:-:-:-:-: | 3139 | ddx23 | 1511 | 1114 | 1475 | 819 | 1287 | 984 |
| ENSDARG000 | 23:23:23:23:2 | 27553786:275 | 27554121:275 | -:-:-:-:-:-:-:-: | 3241 | pfkma | 0 | 0 | 4 | 0 | 2 | 0 |
| ENSDARG000 | 23:23:23:23:2 | 8679216:8687 | 8679356:8687 | +;+;+;+;+;+ | 1077 | rgs19 | 331 | 329 | 443 | 334 | 714 | 319 |
| ENSDARG000 | 23:23:23:23:2 | 44472414:444 | 44472718:444 | -:-:-:-:-:-:-:-: | 1592 | actl6b | 4 | 20 | 6 | 6 | 2 | 8 |
| ENSDARG000 | 23:23:23:23:2 | 11669109:116 | 11669645:116 | +:+:+:+:+:+:+: | 3970 | NA | 0 | 2 | 8 | 0 | 0 | 0 |
| ENSDARG000 | 23:23:23:23:2 | 28718863:287 | 28719119:287 | +:+:+:+:+:+:+: | 2959 | srm | 747 | 1385 | 1263 | 536 | 1139 | 955 |
| ENSDARG000 | 23:23:23:23:2 | 457569:46435 | 462318:46455 | -:-:-:-:-:-:-:-: | 8250 | NA | 392 | 401 | 726 | 306 | 621 | 550 |
| ENSDARG000 | 23:23:23:23:2 | 30827959:308 | 30829053:308 | +;+;+;+;+;+;+ | 2196 | NPBWR2 | 2 | 0 | 0 | 0 | 1 | 49 |
| ENSDARG000 | 23;23;23 | 30859086:308 | 30859628:308 | +;+;+ | 1932 | xkr7a | 0 | 0 | 0 | 0 | 2 | 0 |
| ENSDARG000 | 23;23 | 28731379:287 | 28731594:287 | +;+ | 795 | sst7 | 0 | 0 | 2 | 0 | 2 | 0 |
| ENSDARG000 | 23:23:23:23:2 | 14217508:142 | 14218094:142 | +;+;+;+;+ | 1063 | birc7 | 1751 | 1184 | 916 | 599 | 1099 | 993 |
| ENSDARG000 | 23:23:23:23:2 | 24184719:241 | 24185082:241 | -;-;-;-;-;-;- | 839 | NA | 0 | 10 | 13 | 12 | 2 | 2 |
| ENSDARG000 | 23:23:23:23:2 | 24170671:241 | 24170944:241 | -:-:-:-:-:-:-:-: | 3271 | NA | 0 | 0 | 12 | 4 | 6 | 2 |
| ENSDARG000 | 23:23:23:23:2 | 16928506:169 | 16928639:169 | +:+:+:+:+:+:+: | 1545 | fbxo44.8 | 152 | 212 | 171 | 95 | 123 | 121 |
| ENSDARG000 | 23:23:23:23:2 | 36424768:364 | 36425290:364 | -:-:-:-:-:-:-:-: | 1984 | csad | 22 | 2 | 0 | 4 | 32 | 0 |
| ENSDARG000 | 23:23:23:23:2 | 32066168:320 | 32066350:320 | +:+:+:+:+:+:+: | 1294 | cd63 | 6129 | 5099 | 3236 | 3276 | 3008 | 6449 |
| ENSDARG000 | 23:23:23:23:2 | 8718950:8721 | 8719601:8721 | -:-:-:-:-:-:-:-: | 1730 | tcea2 | 376 | 301 | 294 | 294 | 406 | 338 |
| ENSDARG000 | 23:23:23:23:2 | 32079046:320 | 32079275:320 | -:-:-:-:-:-:-:-: | 1656 | letmd1 | 293 | 328 | 332 | 217 | 365 | 215 |
| ENSDARG000 | 23:23:23:23:2 | 28770225:287 | 28770349:287 | +:+:+:+:+:+:+: | 2765 | masp2 | 6 | 2 | 4 | 0 | 2 | 0 |
| ENSDARG000 | 23:23:23:23:2 | 44497701:445 | 44497780:445 | +:+:+:+:+:+:+: | 7855 | NA | 3366 | 1258 | 1972 | 1252 | 2102 | 1716 |
| ENSDARG000 | 23:23:23:23:2 | 480423:48102 | 481170:48117 | -:-:-:-:-:-:-:-: | 2064 | hm13 | 2543 | 3650 | 4132 | 1499 | 3430 | 2167 |
| ENSDARG000 | 23:23:23:23:2 | 20180159:201 | 20180295:201 | -:-:-:-:-:-:-:-: | 5462 | lamb2 | 16 | 0 | 0 | 10 | 28 | 2 |
| ENSDARG000 | 23:23:23:23:2 | 30898013:309 | 30898542:309 | +:+:+:+:+:+:+: | 1945 | CABLES2 | 181 | 253 | 119 | 122 | 176 | 115 |
| ENSDARG000 | 23:23:23:23:2 | 19506023:195 | 19506290:195 | -:-:-:-:-:-:-:-: | 12261 | DNAH12 | 0 | 0 | 8 | 0 | 6 | 2 |
| ENSDARG000 | 23:23:23:23:2 | 40410644:404 | 40410924:404 | +;+;+;+;+;+;+ | 2533 | elovl4b | 10 | 14 | 21 | 40 | 7 | 13 |
| ENSDARG000 | 23;23 | 8797143:8798 | 8797761:8799 | +;+ | 1530 | NA | 112 | 12 | 44 | 8 | 24 | 30 |
| ENSDARG000 | 23:23:23:23:2 | 14265122:142 | 14265501:142 | -;-;-;-;-;-;- | 1218 | nkain4 | 67 | 21 | 86 | 26 | 58 | 72 |
| ENSDARG000 | 23:23:23:23:2 | 11864223:118 | 11866350:118 | -:-:-:-:-:-:-:-: | 5290 | NA | 610 | 539 | 1760 | 3885 | 771 | 629 |
| ENSDARG000 | 23;23;23 | 8989168:8993 | 8989440:8993 | +;+;+ | 2276 | xkr7b | 6 | 0 | 4 | 4 | 8 | 4 |
| ENSDARG000 | 23:23:23:23:2 | 4929442:4929 | 4930216:4930 | -:-:-:-:-:-:-:-: | 1504 | ngfa | 10 | 0 | 0 | 0 | 2 | 0 |
| ENSDARG000 | 23:23:23:23:2 | 16948049:169 | 16948118:169 | +;+;+;+;+ | 725 | NA | 8 | 2 | 4 | 4 | 11 | 2 |
| ENSDARG000 | 23:23:23:23:2 | 25708787:257 | 25708907:257 | +:+:+:+:+:+:+: | 2308 | rbms2b | 258 | 112 | 200 | 153 | 207 | 116 |
| ENSDARG000 | 23;23 | 40466502:404 | 40466861:404 | +;+ | 556 | NA | 0 | 6 | 10 | 0 | 0 | 16 |
| ENSDARG000 | 23;23;23;23 | 40461051:404 | 40462554:404 | -;-;-;- | 1792 | NA | 0 | 0 | 0 | 0 | 0 | 0 |
| ENSDARG000 | 23;23 | 40477220:404 | 40477317:404 | +;+ | 769 | NA | 6 | 1 | 4 | 2 | 8 | 48 |

| | | | | | | | | | | | | |
|---|---|---|---|---|---|---|---|---|---|---|---|---|
| ENSDARG000 | 23:23:23:23:2 | 40452157:404 | 40452840:404 | +:+:+:+:+:+:+: | 2558 | SOGA3 | 0 | 3 | 5 | 7 | 6 | 24 |
| ENSDARG000 | 23:23:23:23:2 | 5211021:5214 | 5212285:5215 | -:-:-:-:-:-:-:-: | 3063 | NA | 0 | 2 | 0 | 0 | 0 | 0 |
| ENSDARG000 | 23:23:23:23:2 | 3538463:3538 | 3538590:3538 | +:+:+:+:+:+:+: | 47482 | NA | 0 | 0 | 0 | 0 | 0 | 0 |
| ENSDARG000 | 23:23:23:23:2 | 44536926:445 | 44537569:445 | -;-;-;-;-;-;-;- | 1259 | psmb6 | 1714 | 1832 | 1656 | 965 | 1580 | 1036 |
| ENSDARG000 | 23:23:23:23:2 | 9035578:9049 | 9035630:9049 | +:+:+:+:+:+:+: | 5597 | CCM2L | 12 | 2 | 2 | 14 | 30 | 12 |
| ENSDARG000 | 23:23:23:23:2 | 28803007:288 | 28803807:288 | -;-;-;-;-;- | 1684 | dffa | 422 | 294 | 308 | 262 | 305 | 241 |
| ENSDARG000 | 23:23:23:23:2 | 36443022:364 | 36444301:364 | -;-;-;-;-;-;- | 1765 | zgc:174906 | 54 | 40 | 94 | 28 | 42 | 56 |
| ENSDARG000 | 23:23:23:23:2 | 9088191:9112 | 9088705:9112 | +:+:+:+:+:+:+: | 1764 | NA | 23 | 7 | 24 | 10 | 20 | 14 |
| ENSDARG000 | 23:23:23:23:2 | 40525985:405 | 40529768:405 | -:-:-:-:-:-:-:-: | 4041 | rnf146 | 1064 | 802 | 778 | 584 | 868 | 840 |
| ENSDARG000 | 23:23:23:23:2 | 34030550:340 | 34030757:340 | +:+:+:+:+:+:+: | 2371 | sars1 | 2474 | 3492 | 5547 | 2554 | 3382 | 2455 |
| ENSDARG000 | 23;23;23 | 9206350:9210 | 9206723:9210 | +;+;+ | 948 | NA | 0 | 0 | 22 | 12 | 2 | 0 |
| ENSDARG000 | 23:23:23:23:2 | 30937757:309 | 30937859:309 | -:-:-:-:-:-:-:-: | 1543 | NA | 127 | 77 | 128 | 76 | 141 | 139 |
| ENSDARG000 | 23:23:23:23:2 | 21544227:215 | 21544386:215 | +:+:+:+:+:+:+: | 4157 | ARHGEF10L | 25 | 8 | 32 | 18 | 76 | 33 |
| ENSDARG000 | 23:23:23:23:2 | 27575539:275 | 27576288:275 | -:-:-:-:-:-:-:-: | 2994 | NA | 248 | 537 | 577 | 453 | 348 | 319 |
| ENSDARG000 | 23:23:23:23:2 | 16956185:169 | 16957079:169 | -;-;-;-;-;- | 1591 | cdc42l2 | 20 | 56 | 90 | 68 | 139 | 84 |
| ENSDARG000 | 23:23:23:23:2 | 24202559:242 | 24203026:242 | -:-:-:-:-:-:-:-: | 3711 | plekhm2 | 673 | 413 | 539 | 592 | 530 | 400 |
| ENSDARG000 | 23:23:23:23:2 | 28809002:288 | 28809059:288 | +:+:+:+:+:+:+: | 4937 | PEX14 | 490 | 395 | 500 | 353 | 520 | 590 |
| ENSDARG000 | 23:23:23:23:2 | 44546319:445 | 44546369:445 | +:+:+:+:+:+:+: | 757 | trappc1 | 430 | 478 | 576 | 402 | 594 | 409 |
| ENSDARG000 | 23:23:23:23:2 | 25770401:257 | 25770918:257 | -:-:-:-:-:-:-:-: | 3114 | NA | 290 | 415 | 652 | 328 | 514 | 424 |
| ENSDARG000 | 23:23:23:23:2 | 495514:49816 | 496739:49824 | -:-:-:-:-:-:-:-: | 3113 | col9a3 | 118 | 9 | 161 | 14 | 264 | 0 |
| ENSDARG000 | 23:23:23:23:2 | 5465806:5465 | 5465922:5466 | +:+:+:+:+:+:+: | 5478 | NA | 8 | 19 | 19 | 15 | 31 | 4 |
| ENSDARG000 | 23:23:23:23:2 | 40604951:406 | 40605547:406 | +:+:+:+:+:+:+: | 3529 | NA | 12 | 0 | 36 | 7 | 25 | 14 |
| ENSDARG000 | 23:23:23:23:2 | 24227296:242 | 24227858:242 | -:-:-:-:-:-:-:-: | 1884 | ddost | 4146 | 4389 | 6274 | 3386 | 5396 | 4208 |
| ENSDARG000 | 23:23:23:23:2 | 9220436:9220 | 9220489:9220 | +:+:+:+:+:+:+: | 747 | rps21 | 4539 | 5187 | 5643 | 4611 | 3233 | 6849 |
| ENSDARG000 | 23;23;23 | 28944314:289 | 28944860:289 | +;+;+ | 899 | NA | 0 | 0 | 0 | 0 | 0 | 0 |
| ENSDARG000 | 23:23:23:23:2 | 28954137:289 | 28954207:289 | -:-:-:-:-:-:-:-: | 5289 | CASZ1 | 41 | 18 | 0 | 0 | 26 | 12 |
| ENSDARG000 | 23:23:23:23:2 | 34047413:340 | 34047645:340 | +:+:+:+:+:+:+: | 1307 | chchd6a | 152 | 247 | 204 | 248 | 195 | 240 |
| ENSDARG000 | 23:23:23:23:2 | 40755764:407 | 40756126:407 | -;-;-;-;-;- | 865 | mdp1 | 294 | 253 | 393 | 222 | 188 | 261 |
| ENSDARG000 | 23;23;23;23 | 12072980:120 | 12073224:120 | +;+;+;+ | 1267 | edn3b | 0 | 0 | 0 | 0 | 2 | 0 |
| ENSDARG000 | 23:23:23:23:2 | 30967686:309 | 30967910:309 | +:+:+:+:+:+:+: | 3594 | NA | 1639 | 2853 | 3792 | 4005 | 4059 | 3332 |
| ENSDARG000 | 23:23:23:23:2 | 20261570:202 | 20263260:202 | -:-:-:-:-:-:-:-: | 7065 | NA | 8 | 4 | 6 | 0 | 10 | 0 |
| ENSDARG000 | 23:23:23:23:2 | 21638258:216 | 21638512:216 | +:+:+:+:+:+:+: | 2330 | igsf21b | 6 | 8 | 0 | 0 | 28 | 4 |
| ENSDARG000 | 23;23;23;23 | 36771593:367 | 36772117:367 | +;+;+;+ | 1372 | MARCHF9 | 0 | 0 | 0 | 0 | 0 | 0 |
| ENSDARG000 | 23:23:23:23:2 | 14407888:144 | 14409186:144 | -;-;-;-;-;- | 2175 | NA | 44 | 12 | 12 | 24 | 46 | 16 |
| ENSDARG000 | 23;23;23;23 | 20325698:203 | 20325773:203 | -;-;-;- | 286 | NA | 0 | 0 | 0 | 0 | 0 | 0 |
| ENSDARG000 | 23:23:23:23:2 | 36816486:368 | 36816542:368 | -:-:-:-:-:-:-:-: | 3374 | NA | 98 | 59 | 108 | 95 | 134 | 115 |
| ENSDARG000 | 23:23:23:23:2 | 21663631:216 | 21664111:216 | +;+;+;+;+;+;+ | 1723 | dhrs3a | 8 | 2 | 9 | 0 | 8 | 0 |
| ENSDARG000 | 23:23:23:23:2 | 16981929:169 | 16983825:169 | -:-:-:-:-:-:-:-: | 6485 | dnmt3bb.2 | 8 | 54 | 93 | 33 | 78 | 46 |
| ENSDARG000 | 23:23:23:23:2 | 32101202:321 | 32101318:321 | +:+:+:+:+:+:+: | 1392 | zgc:56699 | 16 | 10 | 32 | 4 | 14 | 2 |
| ENSDARG000 | 23;23 | 14468003:144 | 14468219:144 | +;+ | 843 | NA | 0 | 0 | 0 | 0 | 0 | 0 |
| ENSDARG000 | 23:23:23:23:2 | 14590483:145 | 14591159:145 | +:+:+:+:+:+:+: | 2497 | NA | 15 | 49 | 33 | 6 | 75 | 15 |
| ENSDARG000 | 23;23 | 14590767:145 | 14591159:145 | +;+ | 614 | NA | 0 | 0 | 0 | 0 | 0 | 0 |
| ENSDARG000 | 23:23:23:23:2 | 31000243:310 | 31000377:310 | +:+:+:+:+:+:+: | 1239 | selenoi | 478 | 521 | 392 | 167 | 317 | 460 |
| ENSDARG000 | 23:23:23:23:2 | 40788357:407 | 40788729:407 | -;-;-;-;-;-;-;- | 1997 | NA | 0 | 2 | 0 | 0 | 0 | 0 |
| ENSDARG000 | 23:23:23:23:2 | 44553647:445 | 44554063:445 | -:-:-:-:-:-:-:-: | 11886 | NA | 0 | 0 | 0 | 0 | 0 | 0 |
| ENSDARG000 | 23:23:23:23:2 | 40772273:407 | 40772648:407 | -;-;-;-;-;-;-;- | 1959 | NA | 0 | 0 | 0 | 0 | 0 | 0 |
| ENSDARG000 | 23:23:23:23:2 | 21683776:216 | 21684045:216 | -;-;-;-;-;-;-;- | 1108 | NA | 14 | 0 | 0 | 2 | 4 | 1 |
| ENSDARG000 | 23:23:23:23:2 | 16961269:169 | 16961734:169 | -:-:-:-:-:-:-:-: | 5474 | dnmt3bb.3 | 60 | 247 | 248 | 75 | 172 | 175 |
| ENSDARG000 | 23;23;23;23 | 44567210:445 | 44573505:445 | -;-;-;- | 6621 | NA | 0 | 0 | 0 | 0 | 0 | 5 |
| ENSDARG000 | 23;23;23 | 25789132:257 | 25789196:257 | -;-;- | 271 | NA | 0 | 6 | 0 | 0 | 0 | 0 |
| ENSDARG000 | 23:23:23:23:2 | 36870354:368 | 36870771:368 | -:-:-:-:-:-:-:-: | 1340 | ube2j2 | 87 | 53 | 112 | 144 | 94 | 63 |
| ENSDARG000 | 23:23:23:23:2 | 5499307:5524 | 5499547:5524 | +:+:+:+:+:+:+: | 1583 | TEAD3 | 42 | 4 | 25 | 0 | 27 | 0 |
| ENSDARG000 | 23:23:23:23:2 | 9230431:9230 | 9231328:9231 | +:+:+:+:+:+:+: | 7806 | acss2 | 1061 | 984 | 1251 | 944 | 1316 | 1180 |
| ENSDARG000 | 23:23:23:23:2 | 12134839:121 | 12134885:121 | +:+:+:+:+:+:+: | 1795 | ttll9 | 18 | 13 | 22 | 38 | 8 | 4 |
| ENSDARG000 | 23:23:23:23:2 | 24238452:242 | 24239088:242 | -:-:-:-:-:-:-:-: | 2934 | ZBTB17 | 461 | 482 | 655 | 417 | 464 | 458 |
| ENSDARG000 | 23:23:23:23:2 | 27591366:275 | 27592922:275 | -:-:-:-:-:-:-:-: | 5110 | phf8 | 993 | 818 | 1170 | 672 | 1152 | 941 |
| ENSDARG000 | 23;23;23 | 34990693:350 | 34991013:350 | +;+;+ | 1250 | CHST13 | 173 | 131 | 162 | 72 | 199 | 46 |
| ENSDARG000 | 23:23:23:23:2 | 18103080:181 | 18103384:181 | +:+:+:+:+:+:+: | 1966 | mfsd4ab | 0 | 0 | 10 | 12 | 68 | 0 |
| ENSDARG000 | 23;23;23 | 44575669:445 | 44577540:445 | -;-;- | 2055 | NA | 2 | 0 | 4 | 0 | 0 | 5 |
| ENSDARG000 | 23:23:23:23:2 | 31041058:310 | 31041235:310 | -:-:-:-:-:-:-:-: | 1157 | NA | 0 | 0 | 4 | 0 | 0 | 4 |
| ENSDARG000 | 23;23 | 25796348:257 | 25797672:257 | -;- | 1546 | fitm2 | 351 | 380 | 513 | 268 | 475 | 427 |
| ENSDARG000 | 23:23:23:23:2 | 40810286:408 | 40810661:408 | -;-;-;-;-;-;-;- | 2219 | NA | 0 | 0 | 0 | 0 | 0 | 0 |
| ENSDARG000 | 23:23:23:23:2 | 25821055:258 | 25821174:258 | +:+:+:+:+:+:+: | 1294 | R3HDML | 0 | 0 | 0 | 0 | 2 | 0 |
| ENSDARG000 | 23:23:23:23:2 | 35027177:350 | 35027926:350 | -:-:-:-:-:-:-:-: | 3116 | NA | 126 | 151 | 196 | 185 | 245 | 206 |
| ENSDARG000 | 23:23:23:23:2 | 44580254:445 | 44580800:445 | +:+:+:+:+:+:+: | 3658 | pfn1 | 4467 | 5624 | 5549 | 8184 | 7633 | 8949 |
| ENSDARG000 | 23:23:23:23:2 | 19558574:195 | 19558780:195 | +:+:+:+:+:+:+: | 2791 | atp6ap1lb | 0 | 0 | 0 | 0 | 0 | 0 |
| ENSDARG000 | 23;23;23;23 | 5631381:5635 | 5631512:5635 | +;+;+;+ | 643 | NA | 6 | 0 | 0 | 0 | 0 | 0 |
| ENSDARG000 | 23:23:23:23:2 | 526502:52818 | 527800:52831 | -:-:-:-:-:-:-:-: | 3072 | samhd1 | 70 | 147 | 155 | 56 | 155 | 52 |
| ENSDARG000 | 23:23:23:23:2 | 24436273:244 | 24440610:244 | -:-:-:-:-:-:-:-: | 14770 | SPEN | 864 | 455 | 1104 | 782 | 881 | 1149 |
| ENSDARG000 | 23:23:23:23:2 | 31107685:311 | 31108132:311 | +:+:+:+:+:+:+: | 2333 | tbx18 | 12 | 0 | 0 | 0 | 2 | 0 |
| ENSDARG000 | 23:23:23:23:2 | 5653224:5653 | 5653406:5653 | -:-:-:-:-:-:-:-: | 1176 | tnnt2a | 6 | 0 | 0 | 0 | 0 | 0 |
| ENSDARG000 | 23:23:23:23:2 | 27622020:276 | 27623061:276 | -;-;-;-;-;- | 1472 | arf3a | 4 | 0 | 0 | 0 | 0 | 0 |
| ENSDARG000 | 23:23:23:23:2 | 29185852:291 | 29186150:291 | -;-;-;-;-;- | 568 | NA | 0 | 0 | 0 | 0 | 0 | 0 |
| ENSDARG000 | 23:23:23:23:2 | 14743087:147 | 14743578:147 | -:-:-:-:-:-:-:-: | 2055 | gss | 154 | 180 | 272 | 183 | 227 | 156 |
| ENSDARG000 | 23:23:23:23:2 | 40872688:408 | 40873981:408 | -;-;-;-;-;- | 1927 | cltrn | 0 | 4 | 34 | 20 | 122 | 0 |
| ENSDARG000 | 23;23;23;23 | 41629503:416 | 41629600:416 | -;-;-;- | 491 | NA | 1 | 0 | 0 | 0 | 0 | 0 |
| ENSDARG000 | 23:23:23:23:2 | 21696560:216 | 21696922:216 | -:-:-:-:-:-:-:-: | 13611 | VPS13D | 1315 | 608 | 896 | 450 | 1047 | 880 |
| ENSDARG000 | 23:23:23:23:2 | 12155165:121 | 12155646:121 | -;-;-;-;-;- | 970 | NA | 38 | 26 | 55 | 103 | 78 | 94 |
| ENSDARG000 | 23:23:23:23:2 | 20340200:203 | 20340443:203 | -;-;-;-;-;- | 772 | si:rp71-17i16. | 0 | 0 | 2 | 0 | 10 | 0 |
| ENSDARG000 | 23;23;23;23 | 27641471:276 | 27642102:276 | -;-;-;- | 1515 | wnt10b | 20 | 0 | 4 | 0 | 3 | 0 |
| ENSDARG000 | 23;23 | 36891729:368 | 36891793:368 | +;+ | 410 | NA | 13 | 0 | 11 | 12 | 0 | 16 |
| ENSDARG000 | 23:23:23:23:2 | 36888913:368 | 36889337:368 | -:-:-:-:-:-:-:-: | 2745 | ACAP3 | 1261 | 1210 | 1403 | 1238 | 1386 | 899 |
| ENSDARG000 | 23;23 | 9307574:9308 | 9308102:9308 | +;+ | 651 | NA | 0 | 0 | 0 | 0 | 0 | 39 |
| ENSDARG000 | 23;23 | 12160900:121 | 12161202:121 | +;+ | 1675 | ppp1r3da | 170 | 300 | 524 | 695 | 276 | 188 |
| ENSDARG000 | 23;23;23;23 | 27652906:276 | 27653057:276 | +;+;+;+ | 1360 | wnt1 | 2 | 1 | 3 | 0 | 12 | 0 |
| ENSDARG000 | 23:23:23:23:2 | 41739721:417 | 41740234:417 | -;-;-;-;-;-;-;- | 1935 | stk35 | 191 | 226 | 235 | 242 | 361 | 195 |
| ENSDARG000 | 23:23:23:23:2 | 29333851:293 | 29335961:293 | -:-:-:-:-:-:-:-: | 3196 | NA | 6 | 8 | 8 | 6 | 4 | 0 |
| ENSDARG000 | 23;23 | 32128399:321 | 32129569:321 | -;- | 1216 | rprm3 | 2 | 0 | 24 | 4 | 0 | 16 |
| ENSDARG000 | 23:23:23:23:2 | 25879320:258 | 25879947:258 | +:+:+:+:+:+:+: | 2399 | pkig | 78 | 100 | 194 | 123 | 131 | 195 |
| ENSDARG000 | 23:23:23:23:2 | 547107:54814 | 547886:54828 | -;-;-;-;-;- | 1690 | MTG2 | 174 | 208 | 241 | 106 | 311 | 108 |
| ENSDARG000 | 23:23:23:23:2 | 9400425:9401 | 9403364:9401 | -:-:-:-:-:-:-:-: | 7200 | SOGA1 | 935 | 1065 | 975 | 533 | 644 | 790 |
| ENSDARG000 | 23:23:23:23:2 | 25868637:258 | 25868680:258 | +;+;+;+;+;+;+ | 3051 | ttpal | 332 | 238 | 336 | 312 | 347 | 355 |
| ENSDARG000 | 23;23;23;23 | 27675581:276 | 27675613:276 | +;+;+;+ | 436 | rps26 | 2763 | 3367 | 3694 | 3316 | 2595 | 3364 |
| ENSDARG000 | 23 | 24485473 | 24485831 | + | 359 | NA | 1 | 2 | 2 | 0 | 2 | 3 |
| ENSDARG000 | 23:23:23:23:2 | 24485657:244 | 24485801:244 | -;-;-;-;-;-;- | 1160 | tmem82 | 10 | 17 | 26 | 22 | 28 | 9 |
| ENSDARG000 | 23:23:23:23:2 | 25832689:258 | 25832764:258 | +:+:+:+:+:+:+: | 4211 | hnf4a | 0 | 2 | 16 | 8 | 38 | 0 |
| ENSDARG000 | 23;23 | 18129860:181 | 18130352:181 | +;+ | 568 | NA | 0 | 0 | 0 | 0 | 0 | 0 |
| ENSDARG000 | 23:23:23:23:2 | 18118535:181 | 18119958:181 | -:-:-:-:-:-:-:-: | 2466 | nucks1b | 1895 | 2004 | 2768 | 1329 | 2744 | 1992 |
| ENSDARG000 | 23:23:23:23:2 | 31245395:312 | 31245716:312 | +:+:+:+:+:+:+: | 1225 | irak1bp1 | 48 | 12 | 60 | 20 | 44 | 20 |
| ENSDARG000 | 23:23:23:23:2 | 37057603:370 | 37062430:370 | -:-:-:-:-:-:-:-: | 7285 | ints11 | 570 | 491 | 660 | 343 | 733 | 566 |
| ENSDARG000 | 23:23:23:23:2 | 20357247:203 | 20357634:203 | -:-:-:-:-:-:-:-: | 3349 | NA | 1 | 6 | 0 | 0 | 0 | 0 |
| ENSDARG000 | 23:23:23:23:2 | 44611864:446 | 44611963:446 | +:+:+:+:+:+:+: | 4120 | eno3 | 2932 | 2478 | 3944 | 5584 | 1498 | 3978 |
| ENSDARG000 | 23:23:23:23:2 | 29357790:293 | 29358455:293 | +:+:+:+:+:+:+: | 3244 | tardbpa | 2739 | 3016 | 4203 | 2385 | 3653 | 2931 |
| ENSDARG000 | 23:23:23:23:2 | 32149269:321 | 32152709:321 | -:-:-:-:-:-:-:-: | 5776 | nr4a1 | 2135 | 371 | 1236 | 666 | 1722 | 2312 |
| ENSDARG000 | 23:23:23:23:2 | 12198242:121 | 12198310:121 | +:+:+:+:+:+:+: | 3699 | NA | 0 | 6 | 14 | 11 | 0 | 0 |
| ENSDARG000 | 23:23:23:23:2 | 41800406:418 | 41805481:418 | -:-:-:-:-:-:-:-: | 6678 | GMEB2 | 262 | 327 | 381 | 318 | 386 | 284 |
| ENSDARG000 | 23:23:23:23:2 | 9508538:9522 | 9508646:9522 | +:+:+:+:+:+:+: | 2505 | osbpl2b | 550 | 519 | 590 | 384 | 490 | 386 |
| ENSDARG000 | 23:23:23:23:2 | 553396:55581 | 553656:55598 | +:+:+:+:+:+:+: | 1702 | lsm14b | 0 | 0 | 0 | 0 | 0 | 0 |
| ENSDARG000 | 23:23:23:23:2 | 41799748:418 | 41799865:418 | +;+;+;+;+;+ | 1305 | pdyn | 0 | 0 | 0 | 0 | 0 | 0 |
| ENSDARG000 | 23:23:23:23:2 | 5687972:5690 | 5688684:5690 | -:-:-:-:-:-:-:-: | 2125 | lad1 | 0 | 2 | 4 | 4 | 0 | 28 |
| ENSDARG000 | 23;23;23 | 35187683:351 | 35188595:351 | -;-;- | 2321 | klf15 | 481 | 203 | 733 | 425 | 440 | 816 |
| ENSDARG000 | 23;23 | 35195740:352 | 35196165:352 | +;+ | 816 | NA | 0 | 0 | 0 | 0 | 0 | 0 |
| ENSDARG000 | 23:23:23:23:2 | 27683014:276 | 27683423:276 | -:-:-:-:-:-:-:-: | 2078 | ikzf4 | 182 | 90 | 98 | 86 | 109 | 106 |
| ENSDARG000 | 23:23:23:23:2 | 31253297:312 | 31255161:312 | -;-;-;-;-;-;- | 2616 | NA | 0 | 8 | 24 | 4 | 0 | 2 |
| ENSDARG000 | 23:23:23:23:2 | 20408227:204 | 20408308:204 | +;+;+;+;+;+ | 580 | TNNC2 | 0 | 4 | 0 | 0 | 0 | 0 |
| ENSDARG000 | 23:23:23:23:2 | 19590006:195 | 19590208:195 | +:+:+:+:+:+:+: | 2697 | slmapb | 1684 | 1164 | 1538 | 999 | 2290 | 859 |
| ENSDARG000 | 23:23:23:23:2 | 5723090:5725 | 5723194:5725 | -;-;-;-;-;- | 546 | NA | 0 | 0 | 0 | 0 | 0 | 0 |
| ENSDARG000 | 23:23:23:23:2 | 5736226:5736 | 5736809:5736 | +:+:+:+:+:+:+: | 1675 | NA | 12 | 30 | 2 | 10 | 10 | 10 |
| ENSDARG000 | 23:23:23:23:2 | 41831224:418 | 41831358:418 | +;+;+;+;+;+ | 968 | scp2b | 650 | 588 | 820 | 436 | 757 | 544 |
| ENSDARG000 | 23;23 | 5756442:5758 | 5757221:5759 | -;- | 1487 | phlda3 | 268 | 659 | 434 | 243 | 697 | 116 |
| ENSDARG000 | 23:23:23:23:2 | 41889093:418 | 41889332:418 | +:+:+:+:+:+:+: | 5077 | PODN | 14 | 6 | 8 | 0 | 38 | 26 |
| ENSDARG000 | 23:23:23:23:2 | 570517:57090 | 571130:57175 | -;-;-;-;-;- | 1694 | kcnk15 | 14 | 6 | 16 | 0 | 6 | 8 |
| ENSDARG000 | 23:23:23:23:2 | 41949575:419 | 41949870:419 | -:-:-:-:-:-:-:-: | 1785 | slc1a7b | 0 | 0 | 0 | 0 | 5 | 6 |
| ENSDARG000 | 23:23:23:23:2 | 9560797:9560 | 9561776:9561 | +:+:+:+:+:+:+: | 2139 | adrm1 | 1347 | 1861 | 2084 | 1088 | 2009 | 882 |
| ENSDARG000 | 23;23 | 41939138:419 | 41939429:419 | +;+ | 643 | NA | 0 | 0 | 0 | 0 | 0 | 0 |
| ENSDARG000 | 23:23:23:23:2 | 18134313:181 | 18134409:181 | +;+;+;+;+;+ | 420 | NA | 0 | 2 | 1 | 0 | 0 | 0 |
| ENSDARG000 | 23:23:23:23:2 | 18177281:181 | 18177884:181 | -:-:-:-:-:-:-:-: | 1236 | TAFA1 | 10 | 0 | 4 | 4 | 0 | 0 |
| ENSDARG000 | 23;23 | 32194408:322 | 32194526:322 | +;+ | 893 | NA | 1 | 0 | 1 | 0 | 2 | 0 |
| ENSDARG000 | 23:23:23:23:2 | 32199588:322 | 32201190:322 | -;-;-;-;-;-;-;- | 2429 | tamalin | 131 | 57 | 147 | 81 | 271 | 341 |
| ENSDARG000 | 23:23:23:23:2 | 24489077:244 | 24489270:244 | -:-:-:-:-:-:-:-: | 1628 | sult1st5 | 40 | 21 | 41 | 85 | 44 | 48 |
| ENSDARG000 | 23;23 | 37090889:370 | 37090980:370 | +;+ | 1289 | ubxn10 | 16 | 21 | 24 | 20 | 40 | 14 |
| ENSDARG000 | 23:23:23:23:2 | 37086159:370 | 37086436:370 | +;+;+;+;+;+ | 1733 | cptp | 57 | 78 | 100 | 116 | 118 | 70 |
| ENSDARG000 | 23:23:23:23:2 | 27696694:276 | 27697238:276 | -:-:-:-:-:-:-:-: | 1498 | DNAJC22 | 0 | 0 | 6 | 4 | 4 | 0 |
| ENSDARG000 | 23:23:23:23:2 | 20422661:204 | 20422736:204 | +:+:+:+:+:+:+: | 1531 | tnnc2.2 | 6 | 12 | 8 | 10 | 28 | 12 |
| ENSDARG000 | 23:23:23:23:2 | 14771979:147 | 14772163:147 | +:+:+:+:+:+:+: | 2964 | exosc10 | 836 | 759 | 1447 | 666 | 1027 | 680 |
| ENSDARG000 | 23;23 | 20428839:204 | 20429544:204 | -;- | 805 | taf13 | 249 | 180 | 146 | 90 | 187 | 312 |
| ENSDARG000 | 23:23:23:23:2 | 18372048:183 | 18372633:183 | -;-;-;-;-;- | 1254 | hsd17b10 | 610 | 582 | 850 | 353 | 714 | 679 |
| ENSDARG000 | 23:23:23:23:2 | 12232773:122 | 12233306:122 | -:-:-:-:-:-:-:-: | 2403 | phactr3a | 930 | 1106 | 949 | 1041 | 240 | 901 |
| ENSDARG000 | 23:23:23:23:2 | 5765514:5767 | 5767540:5767 | -:-:-:-:-:-:-:-: | 2786 | csrp1a | 184 | 1049 | 94 | 731 | 742 | 85 |
| ENSDARG000 | 23:23:23:23:2 | 642001:64205 | 642284:64256 | +:+:+:+:+:+:+: | 1805 | irf10 | 261 | 316 | 350 | 62 | 252 | 90 |
| ENSDARG000 | 23:23:23:23:2 | 578218:57979 | 578460:57986 | +;+;+;+;+;+;+ | 4949 | ogfr | 116 | 195 | 451 | 214 | 672 | 188 |
| ENSDARG000 | 23:23:23:23:2 | 29375676:293 | 29376139:293 | -;-;-;-;-;- | 701 | sst6 | 4 | 12 | 4 | 0 | 4 | 4 |
| ENSDARG000 | 23:23:23:23:2 | 597082:59793 | 597867:59795 | -:-:-:-:-:-:-:-: | 4724 | l1camb | 240 | 61 | 535 | 59 | 26 | 75 |
| ENSDARG000 | 23:23:23:23:2 | 42292748:422 | 42292970:422 | +:+:+:+:+:+:+: | 2590 | cyp2aa12 | 4 | 0 | 16 | 6 | 25 | 2 |
| ENSDARG000 | 23:23:23:23:2 | 42316065:423 | 42316275:423 | +:+:+:+:+:+:+: | 1804 | cyp2aa6 | 493 | 47 | 179 | 60 | 213 | 14 |
| ENSDARG000 | 23:23:23:23:2 | 25946448:259 | 25947684:259 | -:-:-:-:-:-:-:-: | 2321 | ada | 2342 | 2489 | 2288 | 1935 | 1760 | 2256 |
| ENSDARG000 | 23:23:23:23:2 | 35244378:352 | 35244560:352 | -:-:-:-:-:-:-:-: | 1914 | CPNE9 | 0 | 0 | 0 | 0 | 0 | 0 |
| ENSDARG000 | 23:23:23:23:2 | 18386989:183 | 18392201:183 | -:-:-:-:-:-:-:-: | 8920 | FAM120C | 7827 | 6341 | 12085 | 6780 | 9564 | 6885 |
| ENSDARG000 | 23:23:23:23:2 | 6077503:6077 | 6077845:6077 | +:+:+:+:+:+:+: | 2637 | mybpha | 0 | 0 | 0 | 0 | 24 | 10 |
| ENSDARG000 | 23:23:23:23:2 | 27703749:277 | 27704079:277 | +:+:+:+:+:+:+: | 2673 | lmbr1l | 627 | 72 | 42 | 348 | 87 | 186 |
| ENSDARG000 | 23:23:23:23:2 | 42346799:423 | 42347009:423 | +:+:+:+:+:+:+: | 1668 | NA | 10 | 0 | 1 | 0 | 1 | 0 |
| ENSDARG000 | 23:23:23:23:2 | 42336142:423 | 42336336:423 | +:+:+:+:+:+:+: | 5478 | cyp2aa8 | 0 | 0 | 0 | 0 | 0 | 0 |
| ENSDARG000 | 23:23:23:23:2 | 31276162:312 | 31276381:312 | -:-:-:-:-:-:-:-: | 5979 | PHIP | 312 | 247 | 306 | 133 | 353 | 318 |
| ENSDARG000 | 23:23:23:23:2 | 29381441:293 | 29381899:293 | -:-:-:-:-:-:-:-: | 1914 | pgd | 2026 | 2427 | 2232 | 1299 | 2676 | 1701 |
| ENSDARG000 | 23:23:23:23:2 | 37095792:370 | 37096439:370 | -:-:-:-:-:-:-:-: | 2222 | ddx19a | 8 | 46 | 51 | 32 | 27 | 11 |
| ENSDARG000 | 23:23:23:23:2 | 24501918:245 | 24502058:245 | +:+:+:+:+:+:+: | 2275 | szrd1 | 1408 | 1709 | 2440 | 1266 | 2016 | 1712 |
| ENSDARG000 | 23:23:23:23:2 | 3653386:3655 | 3655333:3655 | -:-:-:-:-:-:-:-: | 4210 | pacsin1a | 0 | 6 | 24 | 24 | 50 | 8 |
| ENSDARG000 | 23:23:23:23:2 | 32238337:322 | 32240346:322 | -:-:-:-:-:-:-:-: | 4524 | dgkaa | 187 | 207 | 269 | 120 | 170 | 347 |
| ENSDARG000 | 23:23:23:23:2 | 44633858:446 | 44634420:446 | +;+;+;+;+;+ | 4531 | NA | 0 | 0 | 0 | 0 | 0 | 2 |
| ENSDARG000 | 23:23:23:23:2 | 42362337:423 | 42362556:423 | +:+:+:+:+:+:+: | 1841 | cyp2aa4 | 0 | 0 | 0 | 0 | 0 | 0 |
| ENSDARG000 | 23:23:23:23:2 | 42336084:423 | 42336336:423 | +:+:+:+:+:+:+: | 1399 | NA | 96 | 24 | 70 | 18 | 16 | 0 |
| ENSDARG000 | 23:23:23:23:2 | 25993686:259 | 25994026:259 | +:+:+:+:+:+:+: | 1410 | ccn5 | 36 | 6 | 91 | 28 | 30 | 34 |

| | | | | | | | | | | | |
|---|---|---|---|---|---|---|---|---|---|---|---|
| ENSDARG000 | 23:23:23:23:2 | 35361724:353 | 35363419:353 | -:-:-:-:-:-:-:-: | 4495 | cmtr1 | 1030 | 923 | 1020 | 580 | 873 | 755 |
| ENSDARG000 | 23:23:23:23:2 | 27740592:277 | 27740734:277 | +;+;+;+;+ | 1957 | dhh | 0 | 2 | 22 | 0 | 4 | 0 |
| ENSDARG000 | 23:23:23:23:2 | 12361899:123 | 12362273:123 | +:+:+:+:+:+:+: | 2018 | PIGT | 61 | 45 | 65 | 38 | 100 | 35 |
| ENSDARG000 | 23:23:23:23:2 | 3710946:3714 | 3713972:3714 | -;-;-;-;- | 3738 | nudt3a | 651 | 316 | 723 | 448 | 502 | 648 |
| ENSDARG000 | 23:23:23:23:2 | 3721041:3721 | 3721597:3721 | +:+:+:+:+:+:+: | 1875 | SMIM29 | 3164 | 1788 | 1842 | 2299 | 1673 | 2627 |
| ENSDARG000 | 23;23;23 | 44644911:446 | 44645124:446 | +;+;+ | 1397 | zgc:85858 | 347 | 412 | 740 | 330 | 558 | 441 |
| ENSDARG000 | 23;23 | 21781577:217 | 21781634:217 | +;+ | 442 | NA | 0 | 4 | 2 | 2 | 6 | 0 |
| ENSDARG000 | 23;23 | 21775436:217 | 21775680:217 | -;- | 1143 | LRRC38 | 22 | 0 | 0 | 0 | 0 | 0 |
| ENSDARG000 | 23:23:23:23:2 | 24511417:245 | 24512913:245 | -:-:-:-:-:-:-:-: | 5064 | atp13a2 | 587 | 609 | 680 | 420 | 843 | 448 |
| ENSDARG000 | 23 | 26009266 | 26010376 | + | 1111 | NA | 0 | 20 | 4 | 6 | 0 | 24 |
| ENSDARG000 | 23:23:23:23:2 | 17011454:170 | 17012593:170 | -:-:-:-:-:-:-:-: | 3678 | dnmt3bb.1 | 86 | 236 | 663 | 146 | 423 | 278 |
| ENSDARG000 | 23;23;23;23 | 37185247:371 | 37185446:371 | +;+;+;+ | 759 | NA | 0 | 0 | 0 | 0 | 0 | 0 |
| ENSDARG000 | 23:23:23:23:2 | 19606291:196 | 19607454:196 | +:+:+:+:+:+:+: | 7979 | NA | 42 | 0 | 0 | 2 | 16 | 8 |
| ENSDARG000 | 23;23 | 17044294:170 | 17044941:170 | -;- | 940 | NA | 0 | 2 | 10 | 16 | 2 | 4 |
| ENSDARG000 | 23:23:23:23:2 | 44655942:446 | 44655975:446 | +:+:+:+:+:+:+: | 1037 | NA | 59 | 50 | 87 | 47 | 82 | 49 |
| ENSDARG000 | 23:23:23:23:2 | 44667616:446 | 44667727:446 | -;-;-;-;-;- | 717 | NA | 196 | 64 | 98 | 18 | 131 | 84 |
| ENSDARG000 | 23:23:23:23:2 | 32308814:323 | 32311816:323 | -;-;-;-;-;-;-;-;- | 4381 | rnf41 | 718 | 582 | 933 | 856 | 714 | 631 |
| ENSDARG000 | 23:23:23:23:2 | 9567334:9567 | 9567853:9568 | -:-:-:-:-:-:-:-: | 11542 | lama5 | 128 | 41 | 79 | 28 | 402 | 23 |
| ENSDARG000 | 23:23:23:23:2 | 21803410:218 | 21806907:218 | -:-:-:-:-:-:-:-: | 8542 | EPHB2 | 12 | 11 | 34 | 1 | 22 | 0 |
| ENSDARG000 | 23:23:23:23:2 | 20431140:204 | 20432161:204 | +:+:+:+:+:+:+: | 11806 | slc35c2 | 210 | 171 | 261 | 68 | 351 | 247 |
| ENSDARG000 | 23:23:23:23:2 | 44732863:447 | 44733118:447 | +:+:+:+:+:+:+: | 2553 | atp1b2a | 950 | 1047 | 1526 | 574 | 1700 | 1207 |
| ENSDARG000 | 23:23:23:23:2 | 37227206:372 | 37227427:372 | +:+:+:+:+:+:+: | 1499 | NA | 0 | 2 | 0 | 0 | 7 | 0 |
| ENSDARG000 | 23;23 | 37277073:372 | 37277302:372 | -;- | 305 | NA | 0 | 0 | 0 | 0 | 0 | 0 |
| ENSDARG000 | 23:23:23:23:2 | 37287542:372 | 37288725:372 | -:-:-:-:-:-:-:-: | 2132 | mul1b | 142 | 137 | 229 | 174 | 180 | 109 |
| ENSDARG000 | 23 | 37323962 | 37325961 | + | 2000 | fam43b | 128 | 30 | 12 | 128 | 44 | 12 |
| ENSDARG000 | 23:23:23:23:2 | 26017227:260 | 26017363:260 | +:+:+:+:+:+:+: | 1836 | pfkfb1 | 30 | 45 | 10 | 6 | 56 | 14 |
| ENSDARG000 | 23;23;23;23 | 12418613:124 | 12418871:124 | -;-;-;- | 653 | wfdc2 | 4 | 0 | 2 | 0 | 0 | 2 |
| ENSDARG000 | 23:23:23:23:2 | 24545239:245 | 24545336:245 | -;-;-;-;- | 553 | NA | 0 | 0 | 0 | 0 | 0 | 0 |
| ENSDARG000 | 23:23:23:23:2 | 32335871:323 | 32335954:323 | +:+:+:+:+:+:+: | 2252 | SLC39A5 | 0 | 0 | 0 | 2 | 0 | 0 |
| ENSDARG000 | 23;23;23 | 19670085:196 | 19670219:196 | +;+;+ | 1329 | kctd6b | 112 | 94 | 100 | 142 | 96 | 123 |
| ENSDARG000 | 23:23:23:23:2 | 3751451:3751 | 3753464:3753 | -:-:-:-:-:-:-:-: | 3564 | hmga1a | 8236 | 9167 | 16419 | 5612 | 8524 | 9277 |
| ENSDARG000 | 23:23:23:23:2 | 19655301:196 | 19655490:196 | +:+:+:+:+:+:+: | 1587 | abhd6b | 0 | 2 | 0 | 2 | 0 | 0 |
| ENSDARG000 | 23:23:23:23:2 | 27756984:277 | 27757063:277 | +:+:+:+:+:+:+: | 11881 | KMT2D | 1080 | 600 | 1119 | 592 | 1252 | 1239 |
| ENSDARG000 | 23:23:23:23:2 | 17102960:171 | 17103118:171 | +:+:+:+:+:+:+: | 876 | commd7 | 333 | 374 | 427 | 340 | 387 | 339 |
| ENSDARG000 | 23:23:23:23:2 | 18527028:185 | 18527411:185 | +:+:+:+:+:+:+: | 1554 | AVPR2 | 1477 | 684 | 988 | 817 | 1583 | 826 |
| ENSDARG000 | 23:23:23:23:2 | 35402697:354 | 35402754:354 | +:+:+:+:+:+:+: | 1982 | NA | 60 | 216 | 108 | 8 | 95 | 292 |
| ENSDARG000 | 23 | 35486037 | 35491587 | - | 5551 | FAM110C | 10 | 0 | 18 | 0 | 0 | 10 |
| ENSDARG000 | 23:23:23:23:2 | 24556455:245 | 24556499:245 | +;+;+;+;+;+ | 867 | NA | 0 | 0 | 0 | 0 | 0 | 0 |
| ENSDARG000 | 23:23:23:23:2 | 35468834:354 | 35469853:354 | -:-:-:-:-:-:-:-: | 2641 | fbxo25 | 495 | 127 | 330 | 250 | 472 | 468 |
| ENSDARG000 | 23;23;23;23 | 12428725:124 | 12429605:124 | -;-;-;- | 1415 | snx21 | 38 | 22 | 42 | 25 | 55 | 40 |
| ENSDARG000 | 23:23:23:23:2 | 24571812:245 | 24571966:245 | -;-;-;-;-;-;- | 1064 | TMEM240 | 0 | 0 | 0 | 0 | 0 | 0 |
| ENSDARG000 | 23:23:23:23:2 | 6232895:6272 | 6233069:6272 | +:+:+:+:+:+:+: | 1998 | SYT2 | 0 | 0 | 2 | 0 | 0 | 0 |
| ENSDARG000 | 23:23:23:23:2 | 14829468:148 | 14833835:148 | -:-:-:-:-:-:-:-: | 6508 | ndrg3b | 284 | 272 | 424 | 212 | 387 | 369 |
| ENSDARG000 | 23:23:23:23:2 | 26039524:260 | 26039755:260 | +:+:+:+:+:+:+: | 2717 | NA | 10 | 0 | 0 | 0 | 10 | 0 |
| ENSDARG000 | 23:23:23:23:2 | 14808221:148 | 14808735:148 | -:-:-:-:-:-:-:-: | 2072 | sla2b | 22 | 44 | 3 | 38 | 66 | 93 |
| ENSDARG000 | 23:23:23:23:2 | 31358088:313 | 31358788:313 | -:-:-:-:-:-:-:-: | 1229 | hmgn3 | 952 | 976 | 1162 | 1095 | 892 | 671 |
| ENSDARG000 | 23;23;23;23 | 44772661:447 | 44774061:447 | -;-;-;- | 1763 | NA | 0 | 2 | 6 | 0 | 11 | 0 |
| ENSDARG000 | 23:23:23:23:2 | 18558209:185 | 18563107:185 | -:-:-:-:-:-:-:-: | 6123 | sephs3 | 1754 | 2083 | 2563 | 1543 | 2793 | 1228 |
| ENSDARG000 | 23:23:23:23:2 | 24589633:245 | 24592195:245 | +:+:+:+:+:+:+: | 3635 | NA | 140 | 147 | 136 | 68 | 234 | 173 |
| ENSDARG000 | 23:23:23:23:2 | 27800519:278 | 27802815:278 | -:-:-:-:-:-:-:-: | 4033 | acvr1ba | 1088 | 777 | 990 | 1062 | 372 | 1804 |
| ENSDARG000 | 23:23:23:23:2 | 42454292:424 | 42454539:424 | +:+:+:+:+:+:+: | 1823 | cyp2aa2 | 0 | 0 | 1 | 8 | 0 | 0 |
| ENSDARG000 | 23:23:23:23:2 | 12454542:124 | 12455449:124 | +:+:+:+:+:+:+: | 5694 | NA | 52 | 244 | 228 | 118 | 157 | 174 |
| ENSDARG000 | 23:23:23:23:2 | 42414652:424 | 42414875:424 | +:+:+:+:+:+:+: | 1557 | cyp2aa9 | 0 | 0 | 0 | 0 | 0 | 0 |
| ENSDARG000 | 23:23:23:23:2 | 799100:80411 | 799342:80418 | +:+:+:+:+:+:+: | 3272 | FRMD4B | 183 | 190 | 183 | 62 | 132 | 144 |
| ENSDARG000 | 23;23 | 42414335:424 | 42414712:424 | -;- | 541 | NA | 0 | 0 | 0 | 0 | 0 | 0 |
| ENSDARG000 | 23:23:23:23:2 | 42434348:424 | 42434570:424 | +:+:+:+:+:+:+: | 1883 | cyp2aa1 | 4 | 8 | 0 | 0 | 14 | 0 |
| ENSDARG000 | 23:23:23:23:2 | 29396182:293 | 29399097:293 | -:-:-:-:-:-:-:-: | 10224 | kif1b | 1284 | 992 | 1345 | 867 | 1470 | 1117 |
| ENSDARG000 | 23:23:23:23:2 | 21955056:219 | 21955477:219 | -;-;-;-;-;-;- | 1362 | NA | 0 | 0 | 0 | 0 | 0 | 0 |
| ENSDARG000 | 23:23:23:23:2 | 827248:82724 | 830137:83013 | -:-:-:-:-:-:-:-: | 9963 | NA | 485 | 267 | 516 | 326 | 302 | 549 |
| ENSDARG000 | 23:23:23:23:2 | 26063149:260 | 26065315:260 | -:-:-:-:-:-:-:-: | 3517 | gdi1 | 7527 | 9283 | 7175 | 7261 | 8052 | 5862 |
| ENSDARG000 | 23:23:23:23:2 | 17146561:171 | 17147275:171 | +;+;+;+;+;+;+ | 1789 | NA | 0 | 0 | 1 | 0 | 4 | 0 |
| ENSDARG000 | 23;23 | 17146315:171 | 17146773:171 | +;+ | 551 | NA | 0 | 0 | 0 | 0 | 0 | 0 |
| ENSDARG000 | 23:23:23:23:2 | 31380607:313 | 31381498:313 | -:-:-:-:-:-:-:-: | 2690 | LCA5 | 101 | 70 | 69 | 76 | 121 | 70 |
| ENSDARG000 | 23:23:23:23:2 | 9794730:9794 | 9796669:9796 | -:-:-:-:-:-:-:-: | 4918 | mapre1b | 5289 | 6346 | 6707 | 4397 | 6273 | 4790 |
| ENSDARG000 | 23:23:23:23:2 | 37738863:377 | 37738986:377 | +:+:+:+:+:+:+: | 1126 | NA | 26 | 12 | 23 | 2 | 10 | 38 |
| ENSDARG000 | 23;23 | 37815645:378 | 37815880:378 | +;+ | 1187 | irx7 | 0 | 0 | 0 | 0 | 4 | 4 |
| ENSDARG000 | 23:23:23:23:2 | 18569291:185 | 18569773:185 | -:-:-:-:-:-:-:-: | 1185 | ssr4 | 1717 | 2519 | 3940 | 2470 | 2203 | 1777 |
| ENSDARG000 | 23:23:23:23:2 | 19679557:196 | 19679818:196 | -:-:-:-:-:-:-:-: | 1958 | ognb | 26 | 6 | 12 | 6 | 14 | 6 |
| ENSDARG000 | 23;23;23;23 | 31405497:314 | 31405739:314 | +;+;+;+ | 2144 | sh3bgrl2 | 34 | 50 | 20 | 70 | 103 | 50 |
| ENSDARG000 | 23:23:23:23:2 | 21964325:219 | 21964415:219 | +:+:+:+:+:+:+: | 2089 | NA | 0 | 4 | 0 | 0 | 6 | 0 |
| ENSDARG000 | 23:23:23:23:2 | 31431956:314 | 31431995:314 | +:+:+:+:+:+:+: | 628 | rps12 | 3772 | 2837 | 5558 | 5433 | 3317 | 5156 |
| ENSDARG000 | 23:23:23:23:2 | 32362085:323 | 32362360:323 | -:-:-:-:-:-:-:-: | 3360 | ankrd52a | 257 | 189 | 372 | 83 | 364 | 224 |
| ENSDARG000 | 23;23 | 31401819:314 | 31402333:314 | -;- | 736 | NA | 0 | 0 | 2 | 2 | 4 | 2 |
| ENSDARG000 | 23:23:23:23:2 | 31426493:314 | 31426676:314 | -;-;-;-;-;-;-;- | 1208 | zgc:153284 | 10 | 0 | 1 | 0 | 0 | 0 |
| ENSDARG000 | 23:23:23:23:2 | 17220986:172 | 17221112:172 | +:+:+:+:+:+:+: | 1852 | nol4lb | 346 | 200 | 613 | 193 | 538 | 346 |
| ENSDARG000 | 23:23:23:23:2 | 37823560:378 | 37823987:378 | -;-;-;-;-;-;-;- | 1175 | cd40 | 543 | 726 | 661 | 323 | 360 | 349 |
| ENSDARG000 | 23:23:23:23:2 | 6421414:6421 | 6421569:6421 | -:-:-:-:-:-:-:-: | 1702 | bmp7b | 6 | 2 | 0 | 0 | 6 | 10 |
| ENSDARG000 | 23:23:23:23:2 | 17354154:173 | 17355587:173 | +:+:+:+:+:+:+: | 3094 | srms | 18 | 60 | 136 | 52 | 172 | 60 |
| ENSDARG000 | 23:23:23:23:2 | 24598910:245 | 24599081:245 | +:+:+:+:+:+:+: | 2596 | kansl2 | 2 | 7 | 0 | 5 | 6 | 1 |
| ENSDARG000 | 23 | 17354153 | 17355702 | - | 1550 | NA | 0 | 0 | 0 | 0 | 0 | 0 |
| ENSDARG000 | 23:23:23:23:2 | 12545114:125 | 12545248:125 | +:+:+:+:+:+:+: | 2012 | NA | 64 | 88 | 62 | 70 | 112 | 104 |
| ENSDARG000 | 23:23:23:23:2 | 24600494:246 | 24600746:246 | +:+:+:+:+:+:+: | 10341 | NA | 1170 | 1270 | 2076 | 1332 | 2510 | 1165 |
| ENSDARG000 | 23;23 | 24608592:246 | 24608974:246 | -;- | 636 | NA | 0 | 0 | 2 | 0 | 0 | 0 |
| ENSDARG000 | 23:23:23:23:2 | 27835647:278 | 27836522:278 | -:-:-:-:-:-:-:-: | 2847 | acvrl1 | 1454 | 351 | 160 | 119 | 282 | 614 |
| ENSDARG000 | 23:23:23:23:2 | 20513104:205 | 20513293:205 | +:+:+:+:+:+:+: | 1433 | dpm1 | 507 | 493 | 657 | 258 | 525 | 508 |
| ENSDARG000 | 23:23:23:23:2 | 17387325:173 | 17387798:173 | +:+:+:+:+:+:+: | 2340 | ptk6b | 0 | 12 | 18 | 14 | 36 | 12 |
| ENSDARG000 | 23:23:23:23:2 | 9809540:9809 | 9810517:9810 | -:-:-:-:-:-:-:-: | 5355 | prkcbp1l | 2380 | 2746 | 5478 | 4969 | 2914 | 2628 |
| ENSDARG000 | 23:23:23:23:2 | 4022620:4053 | 4022947:4053 | +:+:+:+:+:+:+: | 1569 | b4galt5 | 548 | 136 | 349 | 436 | 114 | 325 |
| ENSDARG000 | 23:23:23:23:2 | 6544453:6544 | 6544850:6545 | +:+:+:+:+:+:+: | 1682 | spo11 | 0 | 0 | 0 | 0 | 0 | 0 |
| ENSDARG000 | 23:23:23:23:2 | 18574221:185 | 18574882:185 | -:-:-:-:-:-:-:-: | 2652 | ikbkg | 1212 | 863 | 981 | 715 | 970 | 992 |
| ENSDARG000 | 23;23 | 37851830:378 | 37851968:378 | +;+ | 521 | NA | 0 | 0 | 0 | 0 | 0 | 0 |
| ENSDARG000 | 23:23:23:23:2 | 44797665:448 | 44798688:448 | -;-;-;-;-;- | 1655 | st8sia7.1 | 8 | 4 | 4 | 28 | 0 | 18 |
| ENSDARG000 | 23:23:23:23:2 | 44788132:447 | 44788512:447 | -:-:-:-:-:-:-:-: | 1112 | NA | 4 | 0 | 4 | 0 | 22 | 4 |
| ENSDARG000 | 23:23:23:23:2 | 4216196:4216 | 4216558:4216 | -:-:-:-:-:-:-:-: | 1712 | aar2 | 215 | 232 | 456 | 302 | 303 | 203 |
| ENSDARG000 | 23:23:23:23:2 | 27862807:278 | 27863541:278 | -;-;-;-;- | 1481 | ANKRD33 | 0 | 2 | 0 | 0 | 2 | 6 |
| ENSDARG000 | 23:23:23:23:2 | 19691146:196 | 19691353:196 | +:+:+:+:+:+:+: | 2024 | NA | 0 | 0 | 0 | 0 | 2 | 0 |
| ENSDARG000 | 23:23:23:23:2 | 32400806:324 | 32401491:324 | -:-:-:-:-:-:-:-: | 1103 | NA | 2 | 2 | 2 | 0 | 0 | 2 |
| ENSDARG000 | 23:23:23:23:2 | 934927:93940 | 939267:93949 | -:-:-:-:-:-:-:-: | 6691 | NA | 22 | 6 | 0 | 10 | 24 | 22 |
| ENSDARG000 | 23:23:23:23:2 | 32406009:324 | 32406099:324 | +:+:+:+:+:+:+: | 1236 | NA | 4 | 0 | 0 | 0 | 0 | 0 |
| ENSDARG000 | 23:23:23:23:2 | 17431345:174 | 17432893:174 | -:-:-:-:-:-:-:-: | 3313 | tpd52l2b | 1015 | 1161 | 1910 | 1753 | 1511 | 1293 |
| ENSDARG000 | 23:23:23:23:2 | 6586467:6586 | 6587016:6587 | +:+:+:+:+:+:+: | 2419 | rbm38 | 48 | 8 | 68 | 22 | 36 | 66 |
| ENSDARG000 | 23:23:23:23:2 | 42482137:424 | 42482413:424 | +:+:+:+:+:+:+: | 1704 | cyp2aa3 | 0 | 0 | 0 | 0 | 0 | 0 |
| ENSDARG000 | 23:23:23:23:2 | 6635172:6635 | 6635279:6635 | +;+;+;+;+;+ | 782 | NA | 0 | 0 | 0 | 0 | 0 | 0 |
| ENSDARG000 | 23;23;23;23 | 6637611:6639 | 6638684:6639 | -;-;-;- | 2138 | mipb | 0 | 0 | 0 | 0 | 0 | 0 |
| ENSDARG000 | 23 | 20517696 | 20518184 | - | 489 | NA | 0 | 0 | 0 | 0 | 0 | 0 |
| ENSDARG000 | 23:23:23:23:2 | 32392461:323 | 32392659:323 | +:+:+:+:+:+:+: | 3747 | taf8 | 195 | 199 | 208 | 129 | 208 | 168 |
| ENSDARG000 | 23;23;23 | 37955041:379 | 37955728:379 | +;+;+ | 1441 | cbln4 | 94 | 0 | 0 | 0 | 0 | 7 |
| ENSDARG000 | 23:23:23:23:2 | 32412534:324 | 32412575:324 | +:+:+:+:+:+:+: | 1347 | NA | 10 | 0 | 2 | 0 | 2 | 1 |
| ENSDARG000 | 23:23:23:23:2 | 26079467:260 | 26079619:260 | +:+:+:+:+:+:+: | 2347 | atp6ap1b | 6946 | 6847 | 5832 | 8285 | 6888 | 5120 |
| ENSDARG000 | 23:23:23:23:2 | 12682353:126 | 12682827:126 | -:-:-:-:-:-:-:-: | 2719 | NA | 0 | 0 | 6 | 0 | 0 | 0 |
| ENSDARG000 | 23:23:23:23:2 | 21978584:219 | 21978952:219 | +:+:+:+:+:+:+: | 5921 | EIF4G3 | 1823 | 2133 | 2001 | 834 | 2673 | 1179 |
| ENSDARG000 | 23:23:23:23:2 | 4226341:4231 | 4226608:4231 | +:+:+:+:+:+:+: | 1568 | zgc:113278 | 229 | 207 | 405 | 199 | 426 | 357 |
| ENSDARG000 | 23:23:23:23:2 | 35504824:355 | 35504887:355 | +:+:+:+:+:+:+: | 1357 | acp1 | 584 | 654 | 1017 | 530 | 493 | 512 |
| ENSDARG000 | 23:23:23:23:2 | 26094481:260 | 26095438:260 | -:-:-:-:-:-:-:-: | 2292 | rca2.1 | 1948 | 2322 | 2226 | 2123 | 2756 | 2114 |
| ENSDARG000 | 23:23:23:23:2 | 17454691:174 | 17455129:174 | -;-;-;-;-;-;-;- | 1820 | trim101 | 0 | 0 | 6 | 0 | 0 | 0 |
| ENSDARG000 | 23:23:23:23:2 | 42508734:425 | 42508972:425 | -;-;-;-;- | 708 | NA | 0 | 0 | 0 | 6 | 0 | 0 |
| ENSDARG000 | 23:23:23:23:2 | 17427838:174 | 17428201:174 | -;-;-;-;-;-;- | 1036 | ppdpfb | 2310 | 3649 | 7003 | 5467 | 4972 | 5653 |
| ENSDARG000 | 23:23:23:23:2 | 20518504:205 | 20518851:205 | +;+;+;+;+;+;+ | 5976 | adnpb | 363 | 348 | 419 | 286 | 656 | 363 |
| ENSDARG000 | 23;23 | 26102028:261 | 26102098:261 | +;+ | 570 | NA | 6 | 17 | 12 | 17 | 10 | 13 |
| ENSDARG000 | 23:23:23:23:2 | 28077953:280 | 28078217:280 | +:+:+:+:+:+:+: | 2344 | NA | 21 | 2 | 1 | 0 | 0 | 11 |
| ENSDARG000 | 23:23:23:23:2 | 9867483:9867 | 9867557:9867 | +;+;+;+;+;+;+ | 2019 | slc16a7 | 4 | 4 | 11 | 1 | 0 | 6 |
| ENSDARG000 | 23:23:23:23:2 | 980001:98026 | 980479:98047 | -:-:-:-:-:-:-:-: | 3281 | NA | 47 | 18 | 30 | 35 | 55 | 2 |
| ENSDARG000 | 23;23;23;23 | 18603187:186 | 18603738:186 | -;-;-;- | 1157 | cav4b | 0 | 0 | 0 | 1 | 0 | 0 |
| ENSDARG000 | 23;23 | 42524992:425 | 42525070:425 | +;+ | 262 | NA | 0 | 0 | 0 | 0 | 0 | 0 |
| ENSDARG000 | 23;23;23 | 18610016:186 | 18610083:186 | -;-;- | 243 | NA | 0 | 0 | 0 | 0 | 0 | 0 |
| ENSDARG000 | 23:23:23:23:2 | 42643960:426 | 42644740:426 | +:+:+:+:+:+:+: | 2646 | NA | 54 | 18 | 46 | 30 | 26 | 47 |
| ENSDARG000 | 23;23;23 | 4253957:4255 | 4254233:4255 | +;+;+ | 1281 | arl6ip5b | 1021 | 682 | 737 | 1016 | 937 | 1503 |
| ENSDARG000 | 23:23:23:23:2 | 20563779:206 | 20563991:206 | +:+:+:+:+:+:+: | 2695 | NA | 12 | 2 | 2 | 0 | 18 | 10 |
| ENSDARG000 | 23;23;23 | 18611127:186 | 18611259:186 | -;-;- | 798 | NA | 0 | 0 | 4 | 0 | 0 | 0 |
| ENSDARG000 | 23:23:23:23:2 | 44824083:448 | 44824547:448 | -:-:-:-:-:-:-:-: | 1055 | NA | 4 | 4 | 3 | 0 | 6 | 2 |
| ENSDARG000 | 23:23:23:23:2 | 24669035:246 | 24670177:246 | -:-:-:-:-:-:-:-: | 2666 | ctdsp2 | 800 | 138 | 315 | 479 | 360 | 815 |
| ENSDARG000 | 23:23:23:23:2 | 19701587:197 | 19701718:197 | +:+:+:+:+:+:+: | 1741 | dnase1l1 | 393 | 371 | 340 | 216 | 525 | 227 |
| ENSDARG000 | 23:23:23:23:2 | 35545064:355 | 35545142:355 | -;-;-;-;-;- | 578 | alkal2a | 0 | 0 | 0 | 0 | 0 | 0 |
| ENSDARG000 | 23:23:23:23:2 | 38091978:380 | 38092125:380 | +:+:+:+:+:+:+: | 1106 | NA | 108 | 28 | 126 | 8 | 66 | 16 |
| ENSDARG000 | 23:23:23:23:2 | 6673446:6678 | 6675998:6678 | -:-:-:-:-:-:-:-: | 9874 | baz2a | 2921 | 2154 | 4155 | 2336 | 3781 | 2960 |
| ENSDARG000 | 23:23:23:23:2 | 28092083:281 | 28092165:281 | +:+:+:+:+:+:+: | 2022 | c1galt1la | 14 | 4 | 7 | 2 | 42 | 4 |
| ENSDARG000 | 23:23:23:23:2 | 18615992:186 | 18617377:186 | -:-:-:-:-:-:-:-: | 4083 | ARHGAP4 | 1215 | 1091 | 1987 | 1236 | 2695 | 1554 |
| ENSDARG000 | 23:23:23:23:2 | 4260458:4263 | 4260637:4264 | +;+;+;+;+;+ | 1561 | srsf6a | 2438 | 2306 | 3342 | 1702 | 2458 | 1930 |
| ENSDARG000 | 23:23:23:23:2 | 28097282:280 | 28097913:280 | -:-:-:-:-:-:-:-: | 2140 | b4galnt1a | 68 | 31 | 30 | 22 | 38 | 18 |
| ENSDARG000 | 23:23:23:23:2 | 9908940:9908 | 9909124:9909 | -:-:-:-:-:-:-:-: | 1417 | NA | 0 | 0 | 4 | 0 | 0 | 0 |
| ENSDARG000 | 23:23:23:23:2 | 35642061:356 | 35642524:356 | -;-;-;-;- | 937 | tmem18 | 391 | 403 | 421 | 369 | 324 | 519 |
| ENSDARG000 | 23:23:23:23:2 | 1029450:1031 | 1029467:1031 | +:+:+:+:+:+:+: | 1931 | NA | 21 | 69 | 107 | 87 | 75 | 92 |
| ENSDARG000 | 23:23:23:23:2 | 1042624:1044 | 1043305:1045 | -:-:-:-:-:-:-:-: | 2323 | zgc:113423 | 108 | 88 | 152 | 111 | 92 | 106 |
| ENSDARG000 | 23:23:23:23:2 | 26125676:261 | 26125890:261 | -:-:-:-:-:-:-:-: | 1177 | NA | 44 | 0 | 0 | 0 | 16 | 143 |
| ENSDARG000 | 23:23:23:23:2 | 12840080:128 | 12840498:128 | +:+:+:+:+:+:+: | 5638 | smc1al | 2645 | 2993 | 4014 | 2092 | 3889 | 2635 |
| ENSDARG000 | 23:23:23:23:2 | 26142613:261 | 26142908:261 | +:+:+:+:+:+:+: | 2865 | ptpn22 | 56 | 66 | 36 | 98 | 114 | 117 |
| ENSDARG000 | 23:23:23:23:2 | 4274321:4274 | 4274554:4274 | +;+;+;+;+;+ | 1441 | NA | 88 | 89 | 114 | 52 | 86 | 93 |
| ENSDARG000 | 23:23:23:23:2 | 44843860:448 | 44844251:448 | -:-:-:-:-:-:-:-: | 3009 | plod3 | 1807 | 2502 | 3135 | 1396 | 2749 | 1547 |
| ENSDARG000 | 23:23:23:23:2 | 19712557:197 | 19712780:197 | -:-:-:-:-:-:-:-: | 2173 | rpl10 | 16335 | 16150 | 26769 | 23892 | 17359 | 25088 |
| ENSDARG000 | 23:23:23:23:2 | 35650771:356 | 35650965:356 | +:+:+:+:+:+:+: | 2943 | ccnt1 | 216 | 176 | 329 | 53 | 282 | 164 |
| ENSDARG000 | 23;23;23;23 | 38096820:380 | 38097414:380 | -;-;-;- | 823 | NA | 10 | 0 | 0 | 0 | 0 | 2 |
| ENSDARG000 | 23:23:23:23:2 | 18688354:186 | 18688769:186 | +:+:+:+:+:+:+: | 1716 | NA | 2 | 1 | 6 | 0 | 2 | 0 |
| ENSDARG000 | 23:23:23:23:2 | 24693533:246 | 24694674:246 | -;-;-;-;-;- | 2406 | sdf4 | 1607 | 1233 | 1329 | 1086 | 1554 | 1080 |
| ENSDARG000 | 23:23:23:23:2 | 14992800:149 | 14994392:149 | -:-:-:-:-:-:-:-: | 4679 | phf20b | 741 | 980 | 1084 | 359 | 1060 | 664 |
| ENSDARG000 | 23:23:23:23:2 | 6736116:6739 | 6736546:6739 | -;-;-;-;-;-;- | 991 | ptges3b | 2871 | 3764 | 3365 | 1718 | 3022 | 2210 |
| ENSDARG000 | 23:23:23:23:2 | 29507315:295 | 29508275:295 | -:-:-:-:-:-:-:-: | 5210 | ube4b | 2001 | 1596 | 1999 | 1168 | 2230 | 1552 |
| ENSDARG000 | 23:23:23:23:2 | 38151393:381 | 38151752:381 | -;-;-;-;-;-;- | 1120 | PFDN4 | 592 | 805 | 713 | 373 | 553 | 621 |
| ENSDARG000 | 23:23:23:23:2 | 38159715:381 | 38159851:381 | +:+:+:+:+:+:+: | 2467 | zgc:112994 | 7 | 3 | 21 | 5 | 29 | 4 |
| ENSDARG000 | 23:23:23:23:2 | 6752828:6756 | 6753603:6756 | +:+:+:+:+:+:+: | 2348 | p4htma | 85 | 87 | 226 | 239 | 141 | 81 |
| ENSDARG000 | 23:23:23:23:2 | 42726161:427 | 42726345:427 | -:-:-:-:-:-:-:-: | 2644 | NA | 20 | 0 | 2 | 6 | 44 | 22 |
| ENSDARG000 | 23:23:23:23:2 | 26232185:262 | 26232894:262 | -:-:-:-:-:-:-:-: | 4068 | MAGI3 | 168 | 97 | 79 | 38 | 68 | 36 |
| ENSDARG000 | 23:23:23:23:2 | 1058116:1059 | 1058313:1059 | -;-;-;-;-;- | 1012 | NA | 0 | 0 | 0 | 0 | 0 | 3 |

| | | | | | | | | | | | | |
|---|---|---|---|---|---|---|---|---|---|---|---|---|
| ENSDARG000 | 23;23;23;23 | 4288964:4291 | 4289121:4291 | +;+;+;+ | 600 | NA | 7 | 4 | 6 | 0 | 7 | 1 |
| ENSDARG000 | 23;23;23;23;2 | 20640484:206 | 20640635:206 | +;+;+;+;+;+;+; | 4891 | uba1 | 7011 | 7985 | 8899 | 3914 | 9519 | 5407 |
| ENSDARG000 | 23;23;23;23;2 | 35690657:356 | 35691841:356 | -;-;-;-;-;-;-;- | 1972 | tuba1c | 125 | 902 | 51 | 30 | 361 | 193 |
| ENSDARG000 | 23;23;23;23;2 | 35672542:356 | 35672630:356 | +;+;+;+;+;+;+; | 2196 | pmelb | 24 | 4 | 2 | 0 | 2 | 10 |
| ENSDARG000 | 23;23;23;23;2 | 24705424:247 | 24705655:247 | +;+;+;+;+;+;+; | 4030 | c1qtnf12 | 76 | 8 | 2 | 0 | 4 | 6 |
| ENSDARG000 | 23;23;23;23;2 | 28185883:281 | 28186128:281 | -;-;-;-;-;-;-; | 2409 | ZNF385A | 6 | 0 | 0 | 0 | 4 | 0 |
| ENSDARG000 | 23;23;23;23;2 | 22063484:220 | 22064536:220 | -;-;-;-;-;-;-; | 4007 | phc2a | 332 | 512 | 595 | 368 | 486 | 533 |
| ENSDARG000 | 23;23;23;23;2 | 28139489:281 | 28139676:281 | -;-;-;-;-;-;- | 710 | tac3a | 0 | 0 | 0 | 0 | 0 | 0 |
| ENSDARG000 | 23;23;23;23;2 | 9938236:9942 | 9939846:9942 | -;-;-;-;-;-;-; | 8320 | NA | 134 | 94 | 170 | 42 | 64 | 122 |
| ENSDARG000 | 23;23;23;23;2 | 28139826:281 | 28139885:281 | +;+;+;+;+;+;+; | 559 | NA | 0 | 0 | 0 | 0 | 0 | 0 |
| ENSDARG000 | 23;23;23 | 24789205:247 | 24789444:247 | +;+;+ | 1491 | olfml3a | 18 | 4 | 0 | 0 | 6 | 6 |
| ENSDARG000 | 23;23;23;23 | 32499916:325 | 32500202:325 | +;+;+;+ | 1664 | NA | 301 | 231 | 350 | 139 | 225 | 90 |
| ENSDARG000 | 23;23;23;23;2 | 31439713:314 | 31440420:314 | -;-;-;-;-;-;-; | 3001 | eya4 | 20 | 6 | 84 | 42 | 0 | 156 |
| ENSDARG000 | 23;23;23;23;2 | 18702516:187 | 18702958:187 | -;-;-;-;-;-;-; | 1219 | zgc:103759 | 42 | 78 | 46 | 88 | 48 | 24 |
| ENSDARG000 | 23;23;23;23;2 | 4299887:4300 | 4299970:4300 | +;+;+;+;+;+;+; | 1625 | L3MBTL1 | 19 | 14 | 26 | 28 | 39 | 11 |
| ENSDARG000 | 23;23;23;23;2 | 26491931:264 | 26492231:264 | +;+;+;+;+;+;+; | 5816 | NA | 0 | 0 | 2 | 4 | 0 | 0 |
| ENSDARG000 | 23;23;23;23;2 | 24802899:248 | 24802908:248 | -;-;-;-;-;-;-; | 1804 | SYT6 | 0 | 0 | 0 | 0 | 0 | 0 |
| ENSDARG000 | 23;23;23;23;2 | 1181248:1183 | 1181297:1183 | +;+;+;+;+;+;+; | 10792 | UTRN | 2850 | 1163 | 1928 | 2462 | 2147 | 2347 |
| ENSDARG000 | 23;23;23;23;2 | 12897878:129 | 12900712:129 | -;-;-;-;-;-;-; | 4741 | ndnl2 | 323 | 475 | 568 | 332 | 492 | 407 |
| ENSDARG000 | 23;23;23;23;2 | 6795531:6795 | 6795906:6795 | +;+;+;+;+;+;+; | 2673 | NA | 417 | 1039 | 1654 | 1376 | 636 | 408 |
| ENSDARG000 | 23;23;23;23;2 | 15046544:150 | 15048246:150 | -;-;-;-;-;-;-; | 4535 | NA | 1435 | 963 | 912 | 839 | 1110 | 465 |
| ENSDARG000 | 23;23;23;23;2 | 42765700:427 | 42767560:427 | -;-;-;-;-;-;-; | 3566 | pfkfb2a | 67 | 35 | 83 | 14 | 74 | 43 |
| ENSDARG000 | 23;23;23;23;2 | 42810055:428 | 42810792:428 | +;+;+;+;+;+;+; | 1647 | myl9a | 14 | 12 | 31 | 0 | 12 | 18 |
| ENSDARG000 | 23;23;23;23;2 | 17472702:174 | 17477503:174 | -;-;-;-;-;-;-;- | 5521 | DNAJC5 | 1478 | 1658 | 1855 | 1164 | 1485 | 1280 |
| ENSDARG000 | 23;23;23;23;2 | 44881020:448 | 44881091:448 | +;+;+;+;+;+;+; | 2012 | NA | 2 | 2 | 7 | 0 | 8 | 2 |
| ENSDARG000 | 23;23;23;23;2 | 29580584:295 | 29584016:295 | -;-;-;-;-;-;-; | 6926 | clstn1 | 3437 | 2935 | 5817 | 4327 | 7682 | 5236 |
| ENSDARG000 | 23;23;23;23 | 24880227:248 | 24880740:248 | -;-;-;- | 773 | NA | 0 | 0 | 1 | 0 | 0 | 0 |
| ENSDARG000 | 23;23;23;23;2 | 20669149:206 | 20669333:206 | +;+;+;+;+;+;+; | 2286 | arhgef3l | 70 | 86 | 14 | 18 | 118 | 130 |
| ENSDARG000 | 23;23;23;23;2 | 29569893:295 | 29570043:295 | +;+;+;+;+;+;+; | 2319 | lzic | 414 | 442 | 434 | 521 | 395 | 392 |
| ENSDARG000 | 23;23;23;23;2 | 17522867:175 | 17523319:175 | +;+;+;+;+;+;+; | 3066 | slc17a9b | 194 | 261 | 393 | 109 | 383 | 285 |
| ENSDARG000 | 23;23;23;23;2 | 44901398:449 | 44901625:449 | -;-;-;-;-;-;-; | 1684 | NA | 0 | 0 | 14 | 0 | 4 | 0 |
| ENSDARG000 | 23;23;23;23;2 | 29561918:295 | 29562481:295 | -;-;-;-;-;-;-; | 1245 | nmnat1 | 22 | 21 | 36 | 15 | 53 | 33 |
| ENSDARG000 | 23;23;23;23;2 | 29554591:295 | 29555199:295 | -;-;-;-;-;-;- | 1353 | nmnat1-rbp7a | 0 | 0 | 0 | 0 | 0 | 0 |
| ENSDARG000 | 23;23;23;23;2 | 24926407:249 | 24926520:249 | +;+;+;+;+;+;+; | 4088 | klhl21 | 666 | 413 | 661 | 606 | 707 | 491 |
| ENSDARG000 | 23;23;23;23 | 29554594:295 | 29555199:295 | -;-;-;- | 1016 | rbp7a | 0 | 0 | 0 | 0 | 0 | 0 |
| ENSDARG000 | 23;23;23;23;2 | 17509794:175 | 17509876:175 | +;+;+;+;+;+;+; | 4230 | gid8b | 671 | 771 | 834 | 845 | 742 | 739 |
| ENSDARG000 | 23;23;23;23;2 | 42836125:428 | 42837344:428 | -;-;-;-;-;-;-; | 2911 | NA | 328 | 573 | 763 | 327 | 609 | 340 |
| ENSDARG000 | 23;23;23;23 | 35708730:357 | 35708801:357 | +;+;+;+ | 1537 | tuba1a | 211 | 1197 | 1033 | 656 | 422 | 627 |
| ENSDARG000 | 23;23;23;23;2 | 26503876:265 | 26504376:265 | -;-;-;-;-;-;-; | 2695 | tmem269 | 220 | 154 | 1179 | 1001 | 3039 | 1278 |
| ENSDARG000 | 23;23;23;23;2 | 39281229:392 | 39281441:392 | +;+;+;+;+;+;+; | 3250 | kcng1 | 0 | 0 | 0 | 0 | 2 | 0 |
| ENSDARG000 | 23;23;23;23;2 | 35713557:357 | 35713670:357 | +;+;+;+;+;+;+; | 1591 | tuba1b | 1689 | 3630 | 3560 | 2514 | 1863 | 2094 |
| ENSDARG000 | 23;23;23;23;2 | 4324625:4324 | 4324749:4324 | +;+;+;+;+;+;+; | 2553 | sgk2a | 0 | 0 | 21 | 10 | 40 | 4 |
| ENSDARG000 | 23;23;23;23;2 | 32579495:325 | 32579590:325 | -;-;-;-;-;-;-; | 6664 | PLXNA2 | 7 | 6 | 283 | 46 | 150 | 39 |
| ENSDARG000 | 23;23;23;23 | 44923510:449 | 44923761:449 | +;+;+;+ | 1031 | NA | 6 | 0 | 0 | 0 | 0 | 0 |
| ENSDARG000 | 23;23;23;23;2 | 45077358:450 | 45077626:450 | -;-;-;-;-;-;-; | 4650 | NA | 0 | 452 | 520 | 320 | 322 | 586 |
| ENSDARG000 | 23;23;23;23;2 | 22200467:222 | 22200700:222 | +;+;+;+;+;+;+; | 1792 | slc2a1a | 2 | 2 | 30 | 0 | 0 | 0 |
| ENSDARG000 | 23;23;23;23;2 | 10146542:101 | 10146721:101 | +;+;+;+;+;+;+; | 4005 | zgc:171775 | 18 | 0 | 0 | 0 | 2 | 4 |
| ENSDARG000 | 23;23;23;23;2 | 45304830:453 | 45305338:453 | -;-;-;-;-;-;-; | 1514 | sgms2b | 4 | 4 | 0 | 0 | 6 | 11 |
| ENSDARG000 | 23;23 | 43021194:431 | 43021277:431 | -;- | 396 | NA | 0 | 0 | 1 | 0 | 0 | 0 |
| ENSDARG000 | 23;23;23;23;2 | 6845635:6846 | 6846155:6846 | -;-;-;-;-;- | 1623 | NA | 6 | 19 | 0 | 0 | 6 | 2 |
| ENSDARG000 | 23;23;23;23;2 | 45155156:451 | 45155206:451 | -;-;-;-;-;-;-; | 5554 | NA | 282 | 129 | 250 | 184 | 470 | 73 |
| ENSDARG000 | 23;23;23;23;2 | 39346774:393 | 39347317:393 | +;+;+;+;+;+;+; | 3067 | src | 471 | 311 | 417 | 209 | 280 | 248 |
| ENSDARG000 | 23;23;23;23;2 | 45200481:452 | 45200498:452 | +;+;+;+;+;+;+; | 12526 | NA | 754 | 390 | 628 | 406 | 674 | 552 |
| ENSDARG000 | 23;23;23;23;2 | 24941299:249 | 24941859:249 | -;-;-;-;-;-;-; | 2725 | ZBTB48 | 174 | 142 | 255 | 102 | 204 | 137 |
| ENSDARG000 | 23;23;23;23;2 | 24954951:249 | 24955217:249 | +;+;+;+;+;+;+; | 3831 | NOL9 | 747 | 640 | 881 | 580 | 806 | 658 |
| ENSDARG000 | 23;23;23;23;2 | 43177290:431 | 43177453:431 | +;+;+;+;+;+;+; | 7297 | NA | 0 | 0 | 0 | 0 | 0 | 0 |
| ENSDARG000 | 23;23;23;23;2 | 20687340:206 | 20687492:206 | +;+;+;+;+;+;+; | 2048 | NA | 0 | 2 | 0 | 0 | 0 | 0 |
| ENSDARG000 | 23;23;23;23;2 | 35721607:357 | 35722005:357 | -;-;-;-;-;- | 3105 | jph2 | 2 | 0 | 32 | 25 | 0 | 24 |
| ENSDARG000 | 23;23;23 | 43244091:432 | 43244287:432 | -;-;- | 440 | NA | 0 | 0 | 0 | 0 | 0 | 0 |
| ENSDARG000 | 23;23;23;23;2 | 18722715:187 | 18722757:187 | +;+;+;+;+;+;+; | 6262 | NA | 540 | 5284 | 1851 | 547 | 4270 | 830 |
| ENSDARG000 | 23;23;23;23;2 | 15203822:152 | 15206088:152 | -;-;-;-;-;-;-; | 5177 | sulf2b | 470 | 207 | 477 | 34 | 267 | 246 |
| ENSDARG000 | 23;23;23;23;2 | 1526708:1528 | 1526970:1528 | -;-;-;-;- | 930 | EPM2A | 8 | 17 | 23 | 16 | 16 | 15 |
| ENSDARG000 | 23;23;23;23;2 | 6856469:6856 | 6856877:6857 | -;-;-;-;-;-;-; | 1955 | NA | 8 | 8 | 10 | 0 | 2 | 0 |
| ENSDARG000 | 23;23;23;23;2 | 26523998:265 | 26525954:265 | -;-;-;-;-;-;-; | 4591 | NA | 8 | 49 | 82 | 21 | 209 | 8 |
| ENSDARG000 | 23;23;23 | 1561931:1564 | 1562211:1567 | -;-;- | 2733 | fbxo30b | 131 | 158 | 150 | 75 | 126 | 82 |
| ENSDARG000 | 23;23;23;23;2 | 12916059:129 | 12916271:129 | +;+;+;+;+;+;+; | 2706 | NA | 57 | 5 | 73 | 18 | 64 | 10 |
| ENSDARG000 | 23;23;23;23;2 | 28322986:283 | 28323190:283 | +;+;+;+;+;+;+; | 906 | birc5b | 0 | 0 | 4 | 0 | 0 | 0 |
| ENSDARG000 | 23;23;23;23;2 | 1580294:1582 | 1580363:1582 | -;-;-;-;-;-;-;- | 1568 | NA | 0 | 0 | 0 | 0 | 0 | 0 |
| ENSDARG000 | 23;23;23;23;2 | 22267374:223 | 22267485:223 | +;+;+;+;+;+;+; | 3892 | rap1gapa | 33 | 102 | 108 | 102 | 200 | 331 |
| ENSDARG000 | 23 | 17655790 | 17657348 | - | 1559 | bhlhe23 | 0 | 0 | 0 | 0 | 0 | 0 |
| ENSDARG000 | 23;23;23;23;2 | 17658477:176 | 17658840:176 | -;-;-;-;-;-;- | 1225 | NA | 0 | 0 | 0 | 0 | 0 | 0 |
| ENSDARG000 | 23;23 | 17656836:176 | 17657071:176 | +;+ | 621 | NA | 0 | 0 | 0 | 0 | 0 | 0 |
| ENSDARG000 | 23;23;23;23;2 | 17800717:178 | 17801056:178 | +;+;+;+;+;+;+; | 3940 | rnd1a | 8 | 12 | 14 | 24 | 4 | 4 |
| ENSDARG000 | 23 | 17782508 | 17783044 | + | 537 | NA | 0 | 0 | 0 | 0 | 0 | 0 |
| ENSDARG000 | 23;23;23;23;2 | 35759843:357 | 35760025:357 | +;+;+;+;+;+ | 1104 | GDAP1L1 | 4 | 0 | 61 | 35 | 0 | 11 |
| ENSDARG000 | 23;23;23;23;2 | 6872473:6876 | 6873632:6876 | -;-;-;-;-;- | 2128 | trim35-39 | 166 | 181 | 175 | 224 | 210 | 205 |
| ENSDARG000 | 23;23 | 28343988:283 | 28345728:283 | -;- | 1967 | neurod4 | 0 | 0 | 0 | 0 | 0 | 0 |
| ENSDARG000 | 23;23;23;23;2 | 20705849:207 | 20706299:207 | +;+;+;+;+;+;+; | 5449 | NA | 132 | 120 | 186 | 66 | 143 | 143 |
| ENSDARG000 | 23;23;23;23;2 | 43255328:432 | 43255405:432 | +;+;+;+;+;+;+; | 2730 | tgm2a | 0 | 0 | 0 | 0 | 0 | 0 |
| ENSDARG000 | 23;23;23;23;2 | 10173196:101 | 10174032:101 | -;-;-;-;-;-;-; | 2469 | krt5 | 11036 | 516 | 138 | 323 | 2582 | 510 |
| ENSDARG000 | 23;23;23;23;2 | 6882410:6882 | 6882635:6882 | -;-;-;-;-;-;-; | 2563 | NA | 0 | 0 | 0 | 0 | 0 | 2 |
| ENSDARG000 | 23;23;23;23;2 | 35786942:357 | 35789737:357 | -;-;-;-;-;-;-;- | 3038 | mfsd5 | 721 | 573 | 840 | 629 | 843 | 814 |
| ENSDARG000 | 23;23;23;23;2 | 1661743:1664 | 1661812:1664 | +;+;+;+;+;+;+; | 2909 | stxbp3 | 826 | 1112 | 1275 | 981 | 1276 | 1039 |
| ENSDARG000 | 23;23;23;23;2 | 39461989:394 | 39462311:394 | +;+;+;+;+;+;+; | 5190 | mical1 | 187 | 23 | 80 | 22 | 52 | 53 |
| ENSDARG000 | 23;23;23;23;2 | 4348265:4348 | 4348486:4348 | +;+;+;+;+;+;+; | 2810 | mybl2a | 132 | 120 | 157 | 70 | 112 | 148 |
| ENSDARG000 | 23;23;23;23;2 | 24973773:249 | 24974120:249 | +;+;+;+;+;+;+; | 2090 | casp9 | 268 | 171 | 185 | 212 | 143 | 209 |
| ENSDARG000 | 23;23;23;23;2 | 13124085:132 | 13124268:132 | +;+;+;+;+ | 1549 | SAMD10 | 0 | 0 | 0 | 0 | 4 | 0 |
| ENSDARG000 | 23;23;23;23;2 | 20738382:207 | 20738585:207 | -;-;-;-;-;-;-;- | 1520 | znf362b | 150 | 123 | 142 | 70 | 228 | 113 |
| ENSDARG000 | 23;23;23;23;2 | 10249626:102 | 10250400:102 | -;-;-;-;-;-;-; | 2902 | krt8 | 52282 | 43757 | 45879 | 72993 | 33872 | 72059 |
| ENSDARG000 | 23;23;23;23;2 | 35795759:357 | 35795955:357 | -;-;-;-;-;-;-; | 1993 | nr1d4a | 10 | 9 | 19 | 38 | 66 | 8 |
| ENSDARG000 | 23;23 | 35790808:357 | 35790851:357 | +;+ | 350 | NA | 0 | 0 | 0 | 0 | 0 | 0 |
| ENSDARG000 | 23;23;23;23;2 | 19759128:197 | 19759503:197 | +;+;+;+;+;+;+; | 8719 | FLNA | 4396 | 6341 | 4249 | 3855 | 5705 | 4147 |
| ENSDARG000 | 23;23;23;23;2 | 4362463:4362 | 4362682:4362 | +;+;+;+;+;+ | 1324 | adtrp1 | 405 | 308 | 487 | 171 | 353 | 374 |
| ENSDARG000 | 23;23;23;23 | 43300892:433 | 43301348:433 | -;-;-;- | 1120 | NA | 264 | 222 | 317 | 638 | 121 | 263 |
| ENSDARG000 | 23;23 | 31554269:315 | 31554749:315 | -;- | 1063 | tcf21 | 0 | 0 | 0 | 0 | 0 | 0 |
| ENSDARG000 | 23;23;23;23;2 | 13375359:133 | 13376499:133 | +;+;+;+;+;+;+; | 3953 | ZNF512B | 244 | 140 | 172 | 179 | 249 | 88 |
| ENSDARG000 | 23;23;23;23;2 | 20771899:207 | 20772131:207 | +;+;+;+;+;+;+; | 1712 | NA | 4 | 28 | 0 | 2 | 10 | 32 |
| ENSDARG000 | 23;23;23 | 15750018:157 | 15750066:157 | +;+;+ | 392 | NA | 0 | 0 | 0 | 0 | 0 | 0 |
| ENSDARG000 | 23;23;23 | 22496314:225 | 22497820:225 | -;-;- | 2524 | spsb1 | 425 | 320 | 400 | 348 | 274 | 563 |
| ENSDARG000 | 23;23;23;23;2 | 31596194:315 | 31596716:315 | +;+;+;+;+;+;+; | 2051 | tbpl1 | 130 | 144 | 191 | 126 | 179 | 88 |
| ENSDARG000 | 23;23;23;23;2 | 22616063:226 | 22616156:226 | +;+;+;+;+;+;+; | 2124 | slc2a5 | 0 | 0 | 5 | 0 | 6 | 0 |
| ENSDARG000 | 23;23;23;23;2 | 24989387:249 | 24989415:249 | +;+;+;+;+;+;+; | 2993 | arhgap4a | 28 | 38 | 15 | 28 | 80 | 28 |
| ENSDARG000 | 23;23;23;23;2 | 26543716:265 | 26543888:265 | -;-;-;-;-;-;-; | 11038 | HSPG2 | 1396 | 1161 | 1830 | 955 | 3402 | 925 |
| ENSDARG000 | 23;23;23;23;2 | 43320903:433 | 43323818:433 | -;-;-;-;- | 4772 | zgc:174862 | 4 | 2 | 0 | 0 | 0 | 0 |
| ENSDARG000 | 23;23;23;23;2 | 20795781:207 | 20796004:207 | +;+;+;+;+;+ | 1724 | NA | 0 | 0 | 0 | 0 | 0 | 0 |
| ENSDARG000 | 23;23;23;23;2 | 6926183:6932 | 6926444:6932 | +;+;+;+;+;+;+ | 944 | NA | 0 | 0 | 2 | 0 | 0 | 0 |
| ENSDARG000 | 23;23;23;23 | 22576224:225 | 22576342:225 | +;+;+;+ | 794 | NA | 14 | 16 | 28 | 27 | 21 | 3 |
| ENSDARG000 | 23;23;23;23;2 | 20863145:208 | 20863234:208 | +;+;+;+;+;+;+; | 2187 | pax7b | 0 | 4 | 22 | 2 | 22 | 10 |
| ENSDARG000 | 23;23;23;23 | 22597624:225 | 22598166:226 | +;+;+;+ | 1885 | gpr157 | 321 | 124 | 103 | 168 | 126 | 82 |
| ENSDARG000 | 23;23;23;23 | 22590623:225 | 22590848:225 | -;-;-;- | 593 | NA | 0 | 0 | 0 | 4 | 0 | 3 |
| ENSDARG000 | 23;23;23;23;2 | 17839187:178 | 17839411:178 | +;+;+;+;+;+;+; | 1894 | prim1 | 79 | 372 | 374 | 125 | 405 | 159 |
| ENSDARG000 | 23;23;23;23;2 | 1702624:1706 | 1702683:1706 | +;+;+;+;+;+;+; | 3119 | rabggta | 565 | 643 | 843 | 307 | 811 | 270 |
| ENSDARG000 | 23;23;23;23;2 | 17865554:178 | 17866107:178 | +;+;+;+;+;+;+; | 6590 | naca | 12808 | 11546 | 21350 | 19051 | 13537 | 20836 |
| ENSDARG000 | 23;23;23;23;2 | 4373336:4373 | 4373575:4373 | +;+;+;+;+;+;+; | 2806 | NA | 400 | 34 | 255 | 187 | 49 | 474 |
| ENSDARG000 | 23;23;23;23;2 | 10347851:103 | 10348364:103 | +;+;+;+;+;+;+; | 1851 | krt18a.1 | 1856 | 983 | 762 | 574 | 427 | 552 |
| ENSDARG000 | 23;23;23;23;2 | 21067389:210 | 21067892:210 | +;+;+;+;+;+;+; | 1787 | kcnab2b | 2 | 0 | 0 | 12 | 2 | 5 |
| ENSDARG000 | 23;23;23;23;2 | 17847040:178 | 17849453:178 | -;-;-;-;-;-;-; | 4692 | tfip11 | 624 | 617 | 741 | 368 | 631 | 560 |
| ENSDARG000 | 23;23;23;23;2 | 35847200:358 | 35847289:358 | +;+;+;+;+;+;+; | 2538 | rarga | 1374 | 1153 | 1486 | 1193 | 1449 | 776 |
| ENSDARG000 | 23;23;23;23;2 | 28374458:283 | 28374801:283 | +;+;+;+;+;+;+; | 2191 | zgc:153867 | 4800 | 7407 | 6260 | 7468 | 6367 | 7563 |
| ENSDARG000 | 23;23;23;23;2 | 29827134:298 | 29827893:298 | -;-;-;-;-;-;-; | 2714 | tmem201 | 286 | 422 | 526 | 232 | 497 | 349 |
| ENSDARG000 | 23;23;23;23;2 | 33294633:332 | 33294680:332 | +;+;+;+;+;+;+; | 941 | NA | 0 | 0 | 0 | 0 | 2 | 0 |
| ENSDARG000 | 23 | 29841211 | 29844314 | - | 3104 | NA | 242 | 288 | 534 | 336 | 362 | 287 |
| ENSDARG000 | 23;23;23;23;2 | 15850938:158 | 15851552:158 | -;-;-;-;-;-;-;- | 2030 | eef1a2 | 0 | 2 | 0 | 0 | 0 | 0 |
| ENSDARG000 | 23;23;23;23;2 | 4392147:4392 | 4392711:4392 | -;-;-;-;-;-;-; | 1950 | zgc:171857 | 459 | 374 | 328 | 297 | 349 | 484 |
| ENSDARG000 | 23;23;23;23;2 | 31608259:316 | 31608804:316 | -;-;-;-;-;-;-;- | 2494 | slc2a12 | 226 | 137 | 141 | 110 | 240 | 136 |
| ENSDARG000 | 23;23;23;23;2 | 43405205:434 | 43407838:434 | -;-;-;-;-;-;-; | 4431 | rprd1b | 466 | 537 | 604 | 376 | 636 | 426 |
| ENSDARG000 | 23;23;23;23;2 | 43424518:434 | 43424738:434 | +;+;+;+;+;+;+; | 3631 | TTI1 | 172 | 174 | 359 | 121 | 218 | 154 |
| ENSDARG000 | 23;23;23;23;2 | 25038995:250 | 25039194:250 | -;-;-;-;-;-;-; | 1351 | NA | 4 | 0 | 0 | 3 | 0 | 10 |
| ENSDARG000 | 23;23;23;23;2 | 6948941:6950 | 6951507:6951 | -;-;-;-;-;-;-; | 4128 | edem2 | 268 | 250 | 269 | 135 | 412 | 293 |
| ENSDARG000 | 23;23 | 4689626:4691 | 4689726:4692 | +;+ | 931 | NA | 14 | 4 | 0 | 2 | 2 | 10 |
| ENSDARG000 | 23;23 | 36178104:361 | 36178668:361 | +;+ | 8400 | hoxc1a | 6 | 2 | 2 | 44 | 6 | 0 |
| ENSDARG000 | 23;23;23;23;2 | 36111135:361 | 36111326:361 | -;-;-;-;-;-;-; | 1279 | si:dkey-81p22 | 1 | 1 | 0 | 3 | 5 | 19 |
| ENSDARG000 | 23;23;23;23;2 | 10352921:103 | 10353376:103 | +;+;+;+;+;+;+ | 1528 | krt18a.2 | 0 | 0 | 0 | 0 | 0 | 0 |
| ENSDARG000 | 23;23;23;23;2 | 13528053:135 | 13528685:135 | +;+;+;+;+;+;+; | 4450 | UCKL1 | 410 | 354 | 572 | 364 | 557 | 496 |
| ENSDARG000 | 23;23;23;23;2 | 45402128:454 | 45402354:454 | -;-;-;-;-;-;-; | 1187 | aimp1b | 17 | 0 | 6 | 2 | 4 | 10 |
| ENSDARG000 | 23;23;23;23;2 | 15899693:158 | 15900523:159 | -;-;-;-;-;-;-; | 1507 | mrgbp | 343 | 376 | 341 | 183 | 339 | 266 |
| ENSDARG000 | 23;23 | 36118738:361 | 36119224:361 | +;+ | 956 | hoxc5a | 20 | 0 | 21 | 22 | 24 | 72 |
| ENSDARG000 | 23;23;23;23 | 13564223:135 | 13564369:135 | -;-;-;- | 371 | NA | 0 | 0 | 0 | 0 | 0 | 4 |
| ENSDARG000 | 23;23 | 36052944:360 | 36053641:360 | +;+ | 1162 | hoxc13a | 30 | 0 | 0 | 0 | 0 | 0 |
| ENSDARG000 | 23;23 | 35969429:359 | 35969542:359 | +;+ | 214 | NA | 0 | 0 | 0 | 0 | 0 | 0 |
| ENSDARG000 | 23;23;23;23;2 | 36087219:361 | 36087407:361 | +;+;+;+;+;+;+; | 4159 | hoxc3a | 642 | 191 | 235 | 324 | 301 | 211 |
| ENSDARG000 | 23;23;23;23;2 | 6985212:6990 | 6985457:6990 | -;-;-;-;- | 1212 | NA | 4 | 4 | 4 | 0 | 4 | 2 |
| ENSDARG000 | 23;23;23 | 7171653:7216 | 7172246:7216 | -;-;- | 844 | NA | 0 | 0 | 0 | 0 | 0 | 0 |
| ENSDARG000 | 23;23;23;23;2 | 35981839:359 | 35983075:359 | -;-;-;-;-;-;-; | 3982 | calcoco1a | 2031 | 1925 | 1623 | 1256 | 2241 | 1046 |
| ENSDARG000 | 23;23;23;23;2 | 33337734:333 | 33339302:333 | -;-;-;-;-;-;-; | 2945 | si:ch211-226r | 231 | 204 | 263 | 282 | 258 | 434 |
| ENSDARG000 | 23;23;23 | 7350170:7352 | 7350306:7352 | +;+;+ | 1622 | si:dkeyp-73a2 | 0 | 0 | 0 | 0 | 0 | 0 |
| ENSDARG000 | 23;23;23;23 | 36074798:360 | 36075600:360 | +;+;+;+ | 1971 | hoxc11a | 1137 | 0 | 0 | 0 | 0 | 0 |
| ENSDARG000 | 23;23;23;23;2 | 35969228:359 | 35969542:359 | +;+;+;+;+;+;+; | 2401 | hoxc10a | 527 | 16 | 1 | 14 | 7 | 0 |
| ENSDARG000 | 23;23;23;23;2 | 29861243:298 | 29862118:298 | -;-;-;-;-;-;- | 1702 | slc25a33 | 448 | 422 | 421 | 226 | 726 | 488 |
| ENSDARG000 | 23;23 | 36081425:360 | 36084289:360 | -;- | 3057 | NA | 24 | 2 | 0 | 26 | 0 | 0 |
| ENSDARG000 | 23;23 | 36101185:361 | 36101882:361 | +;+ | 1186 | hoxc8a | 131 | 61 | 108 | 161 | 171 | 113 |
| ENSDARG000 | 23;23 | 36130883:361 | 36131495:361 | +;+ | 1161 | hoxc4a | 49 | 0 | 1 | 11 | 12 | 30 |
| ENSDARG000 | 23;23 | 36115541:361 | 36115874:361 | +;+ | 757 | hoxc6a | 78 | 38 | 33 | 85 | 52 | 145 |
| ENSDARG000 | 23;23;23;23;2 | 1730663:1731 | 1730758:1732 | +;+;+;+;+;+;+; | 3008 | TGM1 | 18 | 0 | 0 | 8 | 2 | 4 |
| ENSDARG000 | 23;23 | 36025538:360 | 36025747:360 | -;- | 471 | NA | 0 | 0 | 0 | 2 | 0 | 0 |
| ENSDARG000 | 23;23 | 36095260:360 | 36095906:360 | +;+ | 1170 | hoxc9a | 210 | 401 | 32 | 258 | 649 | 0 |
| ENSDARG000 | 23;23;23;23;2 | 7379728:7380 | 7380034:7381 | +;+;+;+;+;+;+ | 2310 | gata5 | 0 | 0 | 0 | 0 | 0 | 0 |
| ENSDARG000 | 23;23;23;23;2 | 18759542:187 | 18760539:187 | -;-;-;-;-;-;-; | 3210 | trpc4apb | 289 | 198 | 385 | 319 | 377 | 463 |
| ENSDARG000 | 23;23;23;23;2 | 26733232:267 | 26733619:267 | +;+;+;+;+;+;+; | 2700 | zgc:158263 | 21 | 2 | 30 | 0 | 0 | 6 |
| ENSDARG000 | 23;23;23;23;2 | 36150430:361 | 36150488:361 | +;+;+;+;+;+ | 1002 | NA | 18 | 0 | 9 | 26 | 0 | 16 |
| ENSDARG000 | 23 | 36092346 | 36095368 | - | 3023 | NA | 157 | 22 | 0 | 24 | 45 | 0 |
| ENSDARG000 | 23;23 | 36063599:360 | 36064184:360 | +;+ | 825 | hoxc12a | 44 | 0 | 0 | 0 | 0 | 0 |

| | | | | | | | | | | | | |
|---|---|---|---|---|---|---|---|---|---|---|---|---|
| ENSDARG000 | 23;23;23;23;2 | 29883216:298 | 29883244:298 | +;+;+;+;+;+;+; | 2336 | aurkaip1 | 388 | 489 | 664 | 223 | 458 | 302 |
| ENSDARG000 | 23;23;23;23;2 | 18790658:187 | 18795746:187 | -;-;-;-;-;-;-;-; | 6566 | nsfl1c | 2097 | 2441 | 2382 | 1166 | 2527 | 1806 |
| ENSDARG000 | 23;23;23 | 33357163:333 | 33358260:333 | -;-;- | 1497 | slc48a1a | 174 | 104 | 263 | 102 | 202 | 180 |
| ENSDARG000 | 23;23;23;23;2 | 10396842:103 | 10396955:103 | +;+;+;+;+;+;+; | 6370 | eif4ba | 946 | 928 | 1042 | 672 | 859 | 779 |
| ENSDARG000 | 23;23;23;23;2 | 18779043:187 | 18779248:187 | +;+;+;+;+ | 1094 | emp3b | 536 | 577 | 494 | 520 | 331 | 713 |
| ENSDARG000 | 23;23;23;23;2 | 21181074:211 | 21181307:211 | +;+;+;+;+;+;+ | 1316 | NA | 0 | 2 | 0 | 0 | 0 | 0 |
| ENSDARG000 | 23;23;23;23;2 | 21119592:211 | 21120181:211 | -;-;-;-;-;-;-;-; | 5219 | MEGF6 | 20 | 4 | 60 | 24 | 8 | 6 |
| ENSDARG000 | 23;23;23;23;2 | 33381154:333 | 33381508:333 | +;+;+;+;+;+;+; | 3497 | NA | 714 | 477 | 474 | 718 | 645 | 628 |
| ENSDARG000 | 23;23;23;23;2 | 25095568:250 | 25096175:250 | -;-;-;-;-;-;- | 1352 | NA | 0 | 0 | 0 | 0 | 0 | 0 |
| ENSDARG000 | 23;23;23;23;2 | 43668756:436 | 43669157:436 | +;+;+;+;+;+;+; | 5099 | otud4 | 986 | 693 | 695 | 411 | 762 | 496 |
| ENSDARG000 | 23;23;23;23;2 | 29889089:298 | 29889122:298 | +;+;+;+;+;+;+; | 2383 | mxra8b | 926 | 4 | 99 | 21 | 399 | 20 |
| ENSDARG000 | 23;23 | 29900651:299 | 29901005:299 | -;- | 469 | NA | 2 | 0 | 0 | 0 | 0 | 0 |
| ENSDARG000 | 23;23;23;23;2 | 31640239:316 | 31641374:316 | -;-;-;-;-;-;-;-; | 3430 | sgk1 | 1674 | 553 | 345 | 343 | 320 | 250 |
| ENSDARG000 | 23;23;23;23;2 | 22634442:226 | 22635794:226 | -;-;-;-;-;-;-;-; | 2844 | CA6 | 229 | 13 | 14 | 66 | 82 | 54 |
| ENSDARG000 | 23;23;23;23;2 | 26763338:267 | 26764482:267 | -;-;-;-;-;-;-;-; | 3301 | galnt6 | 70 | 32 | 18 | 32 | 68 | 81 |
| ENSDARG000 | 23;23;23;23;2 | 18869416:188 | 18869810:188 | -;-;-;-;-;-;-;- | 2032 | NA | 83 | 135 | 184 | 246 | 171 | 76 |
| ENSDARG000 | 23;23;23;23;2 | 2703044:2703 | 2703263:2703 | +;+;+;+;+;+;+; | 9249 | NA | 2727 | 1373 | 3211 | 2026 | 2462 | 2440 |
| ENSDARG000 | 23;23;23;23;2 | 4699144:4699 | 4700174:4700 | -;-;-;-;-;-;-;-; | 1932 | cnbpa | 2430 | 4019 | 4845 | 2523 | 3791 | 2961 |
| ENSDARG000 | 23;23 | 15946154:159 | 15946507:159 | -;- | 482 | NA | 0 | 0 | 0 | 0 | 0 | 0 |
| ENSDARG000 | 23;23;23;23;2 | 18904674:189 | 18905486:189 | -;-;-;-;-;-;-;-; | 1447 | NA | 0 | 0 | 0 | 11 | 6 | 3 |
| ENSDARG000 | 23;23 | 16356708:163 | 16356782:163 | +;+ | 491 | NA | 0 | 0 | 0 | 0 | 0 | 0 |
| ENSDARG000 | 23;23;23;23;2 | 16430559:164 | 16431365:164 | +;+;+;+;+;+;+ | 1419 | NTSR1 | 6 | 0 | 0 | 2 | 0 | 0 |
| ENSDARG000 | 23;23 | 16444961:164 | 16445040:164 | +;+ | 276 | NA | 0 | 0 | 0 | 0 | 0 | 0 |
| ENSDARG000 | 23;23;23;23 | 16479489:164 | 16479548:164 | -;-;-;- | 1913 | NA | 4 | 2 | 4 | 2 | 22 | 6 |
| ENSDARG000 | 23;23;23;23;2 | 29908348:299 | 29908517:299 | +;+;+;+;+ | 450 | NA | 9 | 0 | 26 | 2 | 11 | 7 |
| ENSDARG000 | 23;23;23;23;2 | 25121998:251 | 25123208:251 | -;-;-;-;-;-;-;-; | 2545 | idh3g | 1342 | 1193 | 1419 | 813 | 1304 | 1142 |
| ENSDARG000 | 23;23;23;23;2 | 10469955:105 | 10470341:105 | +;+;+;+;+;+;+; | 5746 | NA | 1303 | 944 | 2051 | 1364 | 2179 | 1403 |
| ENSDARG000 | 23;23;23;23 | 13721826:137 | 13722792:137 | +;+;+;+ | 1545 | ZBTB46 | 46 | 18 | 105 | 115 | 50 | 80 |
| ENSDARG000 | 23;23;23;23 | 10601324:106 | 10601378:106 | +;+;+;+ | 393 | NA | 8 | 0 | 20 | 4 | 4 | 0 |
| ENSDARG000 | 23;23;23 | 16561233:165 | 16561301:165 | -;-;- | 315 | NA | 0 | 0 | 0 | 0 | 0 | 0 |
| ENSDARG000 | 23;23;23;23;2 | 43695036:436 | 43696406:436 | -;-;-;-;-;-;-;-; | 3311 | abce1 | 2415 | 2893 | 4358 | 1253 | 3230 | 1937 |
| ENSDARG000 | 23;23;23;23;2 | 33447625:334 | 33449733:334 | -;-;-;-;-;-;-;-; | 5776 | itga5 | 3021 | 2770 | 3564 | 4949 | 6451 | 4218 |
| ENSDARG000 | 23;23;23;23;2 | 45494689:454 | 45494887:454 | -;-;-;-;-;-;-;-; | 1185 | NA | 16 | 1 | 6 | 1 | 736 | 2 |
| ENSDARG000 | 23;23;23;23;2 | 18944388:189 | 18944571:189 | +;+;+;+;+ | 965 | cox4i2 | 1040 | 962 | 630 | 1122 | 1238 | 1084 |
| ENSDARG000 | 23;23;23;23;2 | 16620801:166 | 16621335:166 | +;+;+;+;+;+;+; | 2695 | NA | 10 | 10 | 15 | 24 | 12 | 10 |
| ENSDARG000 | 23;23;23;23;2 | 22656477:226 | 22656607:226 | +;+;+;+;+;+;+; | 3751 | eno1a | 14162 | 33314 | 71669 | 72311 | 20453 | 16147 |
| ENSDARG000 | 23;23;23;23;2 | 25135858:251 | 25136114:251 | +;+;+;+;+;+;+; | 1775 | fam3a | 1030 | 962 | 1331 | 1155 | 1159 | 1367 |
| ENSDARG000 | 23;23;23;23;2 | 45538932:455 | 45539241:455 | +;+;+;+;+ | 2113 | NA | 0 | 0 | 0 | 0 | 8 | 0 |
| ENSDARG000 | 23;23;23;23;2 | 45512825:455 | 45512880:455 | +;+;+;+;+;+ | 1212 | prelid1b | 32 | 34 | 59 | 38 | 36 | 61 |
| ENSDARG000 | 23;23;23;23 | 13814978:138 | 13815316:138 | +;+;+;+ | 2260 | lmod3 | 4 | 4 | 4 | 18 | 0 | 14 |
| ENSDARG000 | 23;23;23;23;2 | 29966466:299 | 29966598:299 | +;+;+;+;+;+;+; | 1747 | DVL1 | 66 | 60 | 65 | 32 | 96 | 70 |
| ENSDARG000 | 23;23;23;23;2 | 26797035:267 | 26797337:267 | -;-;-;-;-;-;-;-; | 4569 | ADCY6 | 284 | 625 | 545 | 206 | 195 | 506 |
| ENSDARG000 | 23;23;23;23;2 | 17878969:178 | 17879137:178 | +;+;+;+;+;+;+; | 926 | NA | 119 | 85 | 110 | 80 | 99 | 88 |
| ENSDARG000 | 23;23;23;23;2 | 18953870:189 | 18954976:189 | -;-;-;-;-;- | 2198 | bcl2l1 | 860 | 724 | 917 | 875 | 1076 | 784 |
| ENSDARG000 | 23;23;23;23;2 | 16648621:166 | 16650086:166 | -;-;-;-;-;-;-;-; | 2568 | sdcbp2 | 9321 | 6080 | 8360 | 8293 | 6259 | 6967 |
| ENSDARG000 | 23;23;23;23;2 | 43718115:437 | 43718262:437 | +;+;+;+;+;+;+; | 3942 | anapc10 | 90 | 131 | 121 | 54 | 141 | 51 |
| ENSDARG000 | 23;23;23;23;2 | 10634837:106 | 10639175:106 | -;-;-;-;-;-;-;-; | 6663 | foxp1a | 202 | 78 | 124 | 128 | 180 | 190 |
| ENSDARG000 | 23;23;23;23;2 | 13832347:138 | 13832702:138 | -;-;-;-;-;-;-;- | 974 | naa10 | 1070 | 1197 | 1484 | 1232 | 915 | 1106 |
| ENSDARG000 | 23;23 | 31693368:317 | 31693764:317 | -;- | 587 | NA | 0 | 0 | 0 | 0 | 0 | 0 |
| ENSDARG000 | 23;23;23;23;2 | 25172682:251 | 25173183:251 | +;+;+;+;+;+;+; | 4450 | NA | 589 | 394 | 1094 | 681 | 1522 | 836 |
| ENSDARG000 | 23;23;23;23;2 | 21261313:212 | 21261433:212 | +;+;+;+;+;+;+; | 4769 | EMC1 | 3290 | 2587 | 3652 | 1966 | 3404 | 2296 |
| ENSDARG000 | 23;23;23;23;2 | 31752604:317 | 31753157:317 | -;-;-;-;-;-;- | 1690 | aldh8a1 | 36 | 10 | 14 | 12 | 32 | 32 |
| ENSDARG000 | 23;23;23;23;2 | 30006206:300 | 30006246:300 | +;+;+;+;+;+;+; | 1514 | tnfrsf9a | 12 | 124 | 76 | 45 | 156 | 150 |
| ENSDARG000 | 23;23;23;23;2 | 28391107:283 | 28391242:283 | +;+;+;+;+;+ | 1026 | NA | 2 | 0 | 0 | 2 | 3 | 0 |
| ENSDARG000 | 23;23;23;23;2 | 28399313:283 | 28399492:283 | -;-;-;-;-;- | 1345 | C1QL4 | 0 | 0 | 0 | 0 | 0 | 0 |
| ENSDARG000 | 23;23;23;23;2 | 45560534:455 | 45560778:455 | -;-;-;-;-;-;-;-; | 1564 | cyp17a2 | 0 | 0 | 0 | 0 | 0 | 0 |
| ENSDARG000 | 23;23 | 45579497:455 | 45579724:455 | +;+ | 1954 | egr4 | 52 | 28 | 44 | 22 | 34 | 20 |
| ENSDARG000 | 23;23;23;23;2 | 45584223:455 | 45584577:455 | +;+;+;+;+;+;+; | 4068 | NA | 0 | 2 | 0 | 0 | 0 | 4 |
| ENSDARG000 | 23;23;23;23;2 | 33654231:336 | 33658256:336 | -;-;-;-;-;-;-;-; | 6012 | TFCP2 | 784 | 617 | 810 | 609 | 825 | 766 |
| ENSDARG000 | 23;23;23 | 43750646:437 | 43750701:437 | +;+;+ | 807 | NA | 2 | 0 | 0 | 0 | 0 | 10 |
| ENSDARG000 | 23;23;23;23;2 | 28482780:284 | 28483346:284 | -;-;-;-;-;-;-;-; | 1064 | TMEM107 | 20 | 54 | 38 | 40 | 24 | 22 |
| ENSDARG000 | 23;23;23;23;2 | 33633241:336 | 33634345:336 | -;-;-;-;-;-;-;-; | 2584 | CSRNP2 | 68 | 37 | 57 | 16 | 100 | 19 |
| ENSDARG000 | 23;23;23;23;2 | 31764061:317 | 31768916:317 | -;-;-;-;-;-;-;-; | 8211 | hbs1l | 1189 | 1016 | 1543 | 850 | 1387 | 1088 |
| ENSDARG000 | 23;23 | 19051485:190 | 19051958:190 | +;+ | 733 | NA | 0 | 0 | 0 | 0 | 2 | 0 |
| ENSDARG000 | 23;23;23 | 19066696:190 | 19067555:190 | -;-;- | 1146 | pard6b | 101 | 32 | 146 | 50 | 84 | 42 |
| ENSDARG000 | 23;23;23;23;2 | 18986015:189 | 18986442:189 | -;-;-;-;-;-;-;-; | 6024 | ARFGEF2 | 1383 | 1382 | 2131 | 1456 | 1534 | 1258 |
| ENSDARG000 | 23;23;23;23;2 | 43770149:437 | 43770866:437 | +;+;+;+;+;+;+ | 4240 | rnf150b | 464 | 295 | 588 | 362 | 540 | 611 |
| ENSDARG000 | 23;23;23;23;2 | 26946429:269 | 26946446:269 | +;+;+;+;+;+;+; | 1190 | CACNB3 | 400 | 159 | 299 | 229 | 287 | 325 |
| ENSDARG000 | 23;23;23;23;2 | 26979174:269 | 26979432:269 | +;+;+;+;+;+;+; | 3115 | rnd1b | 643 | 135 | 472 | 415 | 332 | 255 |
| ENSDARG000 | 23;23;23;23;2 | 25232411:252 | 25232789:252 | +;+;+;+;+;+;+; | 4645 | erbb3b | 113 | 10 | 12 | 3 | 18 | 33 |
| ENSDARG000 | 23;23;23;23 | 16688420:166 | 16688651:166 | -;-;-;- | 487 | fkbp1ab | 1219 | 1181 | 1193 | 1062 | 778 | 1370 |
| ENSDARG000 | 23;23;23;23;2 | 13842982:138 | 13847404:138 | -;-;-;-;-;-;-;-; | 6917 | G6PD | 5696 | 3426 | 4372 | 3653 | 8622 | 2624 |
| ENSDARG000 | 23;23;23;23;2 | 22785375:228 | 22785556:228 | +;+;+;+;+;+;+; | 4926 | RERE | 3872 | 1972 | 3525 | 2627 | 3445 | 2359 |
| ENSDARG000 | 23;23;23;23;2 | 17926279:179 | 17926533:179 | +;+;+;+;+;+;+; | 2289 | chia.4 | 0 | 0 | 0 | 0 | 0 | 0 |
| ENSDARG000 | 23;23;23;23;2 | 19813677:198 | 19813763:198 | +;+;+;+;+;+;+; | 2241 | emd | 960 | 1543 | 1301 | 1194 | 1291 | 1066 |
| ENSDARG000 | 23;23;23;23;2 | 16696410:166 | 16696608:166 | -;-;-;-;-;-;-;-; | 1657 | NA | 387 | 512 | 373 | 440 | 347 | 767 |
| ENSDARG000 | 23;23;23;23;2 | 30013240:300 | 30014187:300 | -;-;-;-;-;-;-;-; | 2833 | ccdc187 | 84 | 100 | 173 | 72 | 150 | 124 |
| ENSDARG000 | 23;23;23;23;2 | 27041111:270 | 27042566:270 | -;-;-;-;-;-;-;-; | 3130 | dtx3 | 241 | 130 | 210 | 118 | 199 | 152 |
| ENSDARG000 | 23;23;23;23;2 | 27051919:270 | 27052160:270 | +;+;+;+;+;+;+; | 1701 | ptges3a | 852 | 1251 | 1299 | 642 | 1058 | 976 |
| ENSDARG000 | 23;23 | 43794687:437 | 43794839:438 | +;+ | 3262 | NA | 2 | 2 | 2 | 4 | 4 | 10 |
| ENSDARG000 | 23;23;23;23;2 | 43809484:438 | 43809582:438 | +;+;+;+;+;+;+; | 4007 | PPM1K | 42 | 22 | 86 | 111 | 128 | 371 |
| ENSDARG000 | 23;23;23;23;2 | 17980875:179 | 17981418:179 | +;+;+;+;+;+;+; | 2187 | chia.6 | 12 | 16 | 18 | 12 | 31 | 32 |
| ENSDARG000 | 23;23;23;23;2 | 31815423:318 | 31815615:318 | +;+;+;+;+;+;+; | 3317 | myb | 101 | 21 | 20 | 34 | 102 | 102 |
| ENSDARG000 | 23;23;23;23;2 | 33681677:336 | 33683830:336 | -;-;-;-;-;-;-;-; | 4090 | pou6f1 | 248 | 320 | 463 | 266 | 474 | 335 |
| ENSDARG000 | 23;23;23;23;2 | 28494189:284 | 28494245:284 | +;+;+;+;+;+;+; | 14538 | HUWE1 | 7114 | 4449 | 9367 | 3903 | 8465 | 5171 |
| ENSDARG000 | 23;23;23;23;2 | 22953379:229 | 22953681:229 | +;+;+;+;+ | 816 | NA | 0 | 10 | 22 | 16 | 1 | 14 |
| ENSDARG000 | 23;23;23;23 | 27068225:270 | 27068970:270 | +;+;+;+ | 2844 | mipa | 14 | 2 | 11 | 6 | 32 | 0 |
| ENSDARG000 | 23;23;23;23;2 | 43834376:438 | 43834478:438 | +;+;+;+;+;+ | 732 | NA | 9 | 2 | 8 | 7 | 29 | 4 |
| ENSDARG000 | 23;23;23;23;2 | 21278948:212 | 21279072:212 | +;+;+;+;+;+;+; | 16386 | UBR4 | 6399 | 3847 | 7877 | 4456 | 7039 | 5080 |
| ENSDARG000 | 23;23;23;23;2 | 23020709:230 | 23021570:230 | +;+;+;+;+;+;+; | 4524 | SAMD11 | 1991 | 2267 | 2995 | 2250 | 3536 | 1774 |
| ENSDARG000 | 23;23;23;23;2 | 25291891:252 | 25292107:252 | +;+;+;+;+;+;+; | 2261 | pa2g4b | 3445 | 4062 | 5768 | 3326 | 3898 | 3145 |
| ENSDARG000 | 23;23;23;23;2 | 43849190:438 | 43849387:438 | +;+;+;+;+;+;+; | 2612 | CNGA1 | 4 | 0 | 8 | 0 | 0 | 2 |
| ENSDARG000 | 23;23;23;23 | 33718602:337 | 33718772:337 | +;+;+;+ | 2241 | dazap2 | 2932 | 2134 | 3218 | 2648 | 2931 | 2769 |
| ENSDARG000 | 23;23;23;23;2 | 16719743:167 | 16720874:167 | -;-;-;-;-;-;-;-; | 3937 | NA | 0 | 0 | 2 | 16 | 13 | 0 |
| ENSDARG000 | 23;23;23;23;2 | 19106056:191 | 19106200:191 | +;+;+;+;+;+;+; | 1908 | NA | 62 | 94 | 154 | 76 | 94 | 94 |
| ENSDARG000 | 23;23;23;23;2 | 27078866:270 | 27079741:270 | -;-;-;-;-;-;-;-; | 3661 | stat6 | 906 | 464 | 747 | 331 | 849 | 749 |
| ENSDARG000 | 23;23;23;23;2 | 33731192:337 | 33732135:337 | -;-;-;-;-;-;-;-; | 1856 | NA | 295 | 236 | 418 | 443 | 325 | 416 |
| ENSDARG000 | 23;23;23;23 | 30048849:300 | 30048988:300 | +;+;+;+ | 728 | uts2a | 0 | 0 | 0 | 0 | 0 | 0 |
| ENSDARG000 | 23;23;23;23;2 | 23126130:231 | 23126830:231 | -;-;-;-;-;-;-;-; | 3189 | noc2l | 855 | 851 | 1892 | 567 | 1105 | 561 |
| ENSDARG000 | 23;23;23;23;2 | 19822434:198 | 19823122:198 | -;-;-;-;-;-;-;-; | 4525 | NA | 108 | 227 | 252 | 120 | 232 | 160 |
| ENSDARG000 | 23;23;23;23;2 | 19135915:191 | 19136037:191 | -;-;-;-;-;-;-;- | 2154 | NA | 0 | 0 | 0 | 0 | 0 | 0 |
| ENSDARG000 | 23;23;23;23;2 | 31912882:319 | 31913016:319 | +;+;+;+;+;+;+; | 1353 | armc1l | 176 | 318 | 409 | 146 | 253 | 243 |
| ENSDARG000 | 23;23;23;23;2 | 31901350:319 | 31901898:319 | -;-;-;-;-;-;-;-; | 1804 | mtfr2 | 143 | 357 | 352 | 206 | 143 | 179 |
| ENSDARG000 | 23;23;23;23 | 25305431:253 | 25305534:253 | +;+;+;+ | 461 | NA | 12636 | 24881 | 20371 | 17820 | 11661 | 16260 |
| ENSDARG000 | 23;23;23;23;2 | 25308018:253 | 25308508:253 | +;+;+;+;+;+;+; | 2852 | NA | 4632 | 4486 | 2942 | 3424 | 4690 | 4538 |
| ENSDARG000 | 23;23;23;23;2 | 19147128:191 | 19148843:191 | -;-;-;-;-;-;-;-; | 3140 | ebp | 185 | 398 | 304 | 204 | 172 | 184 |
| ENSDARG000 | 23;23;23;23;2 | 30061364:300 | 30065776:300 | -;-;-;-;-;-;-;-; | 9552 | CAMTA1 | 12 | 4 | 35 | 10 | 16 | 18 |
| ENSDARG000 | 23;23;23;23;2 | 27134942:271 | 27136971:271 | -;-;-;-;-;-;-;-; | 2882 | fignl2 | 50 | 22 | 12 | 44 | 38 | 59 |
| ENSDARG000 | 23;23;23;23;2 | 31920840:319 | 31920967:319 | -;-;-;-;-;-;-;-; | 3290 | NA | 153 | 36 | 62 | 16 | 151 | 68 |
| ENSDARG000 | 23;23 | 33759747:337 | 33759956:337 | +;+ | 732 | NA | 0 | 0 | 0 | 0 | 0 | 0 |
| ENSDARG000 | 23;23;23;23;2 | 33757173:337 | 33761064:337 | -;-;-;-;-;-;-;-; | 6182 | racgap1 | 374 | 1011 | 1153 | 635 | 712 | 498 |
| ENSDARG000 | 23;23;23;23 | 31933583:319 | 31933960:319 | +;+;+;+ | 1104 | NA | 8 | 2 | 4 | 12 | 0 | 16 |
| ENSDARG000 | 23;23;23;23;2 | 30517199:306 | 30517587:306 | -;-;-;-;-;-;- | 698 | NA | 0 | 0 | 0 | 0 | 0 | 0 |
| ENSDARG000 | 23;23;23;23;2 | 33752275:337 | 33752459:337 | +;+;+;+;+ | 1192 | NA | 152 | 172 | 136 | 175 | 160 | 169 |
| ENSDARG000 | 23;23;23;23;2 | 33739215:337 | 33739574:337 | -;-;-;-;-;-;-;-; | 2119 | bin2a | 0 | 2 | 46 | 2 | 19 | 0 |
| ENSDARG000 | 23;23;23;23;2 | 23182037:231 | 23182215:231 | +;+;+;+;+;+;+; | 4307 | KLHL17 | 130 | 130 | 146 | 88 | 200 | 32 |
| ENSDARG000 | 23;23;23;23;2 | 19177891:191 | 19178726:191 | +;+;+;+;+;+;+; | 3548 | NA | 0 | 6 | 0 | 0 | 0 | 14 |
| ENSDARG000 | 23;23;23;23;2 | 19834215:198 | 19839206:198 | -;-;-;-;-;-;-;-; | 9347 | atp2b3b | 6 | 10 | 6 | 10 | 8 | 4 |
| ENSDARG000 | 23;23;23;23;2 | 31942040:319 | 31942248:319 | +;+;+;+;+;+;+; | 1842 | nemp1 | 68 | 230 | 194 | 74 | 169 | 101 |
| ENSDARG000 | 23;23;23;23;2 | 23232136:232 | 23232286:232 | +;+;+;+;+;+;+; | 2742 | NA | 158 | 14 | 4 | 22 | 10 | 15 |
| ENSDARG000 | 23;23;23;23;2 | 38806861:388 | 38807671:388 | +;+;+;+;+;+ | 3813 | ZFP64 | 138 | 152 | 156 | 82 | 142 | 104 |
| ENSDARG000 | 23;23;23;23;2 | 16796:16844: | 16825:17001: | -;-;-;-;-;-;-;-; | 1293 | HDAC7 | 524 | 250 | 332 | 251 | 261 | 175 |
| ENSDARG000 | 23;23;23 | 16840572:168 | 16840715:168 | -;-;- | 345 | NA | 0 | 0 | 2 | 1 | 1 | 2 |
| ENSDARG000 | 23;23;23;23;2 | 25851913:258 | 25851929:258 | -;-;-;-;-;-;-;-; | 252 | NA | 0 | 0 | 0 | 0 | 0 | 0 |
| ENSDARG000 | 23;23;23;23;2 | 26183772:261 | 26183961:261 | -;-;-;-;-;-;-;-; | 2118 | RSBN1 | 246 | 347 | 984 | 654 | 453 | 197 |
| ENSDARG000 | 23;23;23;23;2 | 5565261:5565 | 5565991:5566 | +;+;+;+;+;+;+; | 2263 | smpd2a | 36 | 26 | 34 | 29 | 125 | 34 |
| ENSDARG000 | 23;23;23;23;2 | 39089574:390 | 39089697:390 | +;+;+;+;+;+;+; | 2685 | NFATC2 | 241 | 128 | 226 | 114 | 261 | 209 |
| ENSDARG000 | 23;23;23;23;2 | 45025909:450 | 45025962:450 | +;+;+;+;+;+;+; | 896 | hmgb2b | 2542 | 4776 | 5771 | 2597 | 3849 | 2291 |
| ENSDARG000 | 23;23 | 404975:40665 | 405130:40689 | +;+ | 396 | NA | 0 | 0 | 0 | 0 | 0 | 0 |
| ENSDARG000 | 23;23;23;23;2 | 4583747:4646 | 4583895:4646 | +;+;+;+;+;+;+; | 3023 | IQSEC1 | 90 | 175 | 232 | 112 | 245 | 211 |
| ENSDARG000 | 23;23;23;23;2 | 38245610:382 | 38245821:382 | +;+;+;+;+;+;+; | 5688 | znf217 | 596 | 438 | 578 | 479 | 445 | 494 |
| ENSDARG000 | 23 | 2446978 | 2448234 | - | 1257 | PRSS35 | 4 | 4 | 10 | 7 | 16 | 0 |
| ENSDARG000 | 23;23;23;23;2 | 44427188:444 | 44428156:444 | +;+;+;+;+;+;+ | 2064 | eif4e2rs1 | 597 | 731 | 786 | 578 | 738 | 480 |
| ENSDARG000 | 23;23;23 | 36660750:366 | 36664222:366 | -;-;- | 5482 | NA | 152 | 102 | 158 | 112 | 144 | 133 |
| ENSDARG000 | 23;23;23;23 | 19660002:196 | 19660035:196 | -;-;-;- | 96 | NA | 0 | 0 | 0 | 0 | 0 | 2 |
| ENSDARG000 | 23;23;23 | 2502181:2507 | 2505584:2507 | -;-;- | 3993 | snai1b | 28 | 2 | 10 | 0 | 10 | 0 |
| ENSDARG000 | 23 | 36718375 | 36718495 | - | 121 | 5S_rRNA | 0 | 0 | 0 | 0 | 0 | 0 |
| ENSDARG000 | 23;23;23;23;2 | 41313084:413 | 41313279:413 | -;-;-;-;-;-;-;-; | 3259 | scube3 | 6 | 0 | 4 | 0 | 22 | 6 |
| ENSDARG000 | 23;23;23;23;2 | 45734011:457 | 45734750:457 | +;+;+;+;+;+;+; | 2029 | NA | 2 | 2 | 8 | 0 | 7 | 9 |
| ENSDARG000 | 23 | 30979385 | 30979499 | + | 115 | 5S_rRNA | 0 | 0 | 0 | 0 | 0 | 0 |
| ENSDARG000 | 23 | 34002757 | 34002873 | + | 117 | 5S_rRNA | 0 | 0 | 0 | 0 | 0 | 0 |
| ENSDARG000 | 23;23;23;23;2 | 20803270:208 | 20803454:208 | +;+;+;+;+;+;+; | 1821 | zgc:154075 | 299 | 142 | 226 | 273 | 267 | 311 |
| ENSDARG000 | 23 | 28536361 | 28536457 | + | 97 | dre-let-7g-2 | 0 | 0 | 0 | 0 | 0 | 0 |
| ENSDARG000 | 23 | 17176104 | 17176218 | - | 115 | 5S_rRNA | 0 | 0 | 0 | 0 | 0 | 0 |
| ENSDARG000 | 23 | 13635793 | 13635907 | - | 115 | 5S_rRNA | 0 | 0 | 0 | 0 | 0 | 0 |
| ENSDARG000 | 23 | 36790223 | 36790332 | + | 110 | dre-mir-2191 | 0 | 0 | 0 | 0 | 0 | 0 |
| ENSDARG000 | 23 | 24074083 | 24074155 | + | 73 | dre-mir-7149 | 0 | 0 | 0 | 0 | 0 | 0 |
| ENSDARG000 | 23 | 12083557 | 12083671 | + | 115 | 5S_rRNA | 0 | 0 | 0 | 0 | 0 | 0 |
| ENSDARG000 | 23;23;23;23 | 41679586:416 | 41679755:416 | +;+;+;+ | 468 | NA | 7 | 4 | 0 | 0 | 0 | 0 |
| ENSDARG000 | 23 | 28903330 | 28903456 | + | 127 | 5S_rRNA | 0 | 0 | 0 | 0 | 0 | 0 |
| ENSDARG000 | 23;23;23;23;2 | 45966436:459 | 45966481:459 | +;+;+;+;+;+;+; | 2640 | DNAI3 | 2 | 7 | 1 | 0 | 0 | 0 |
| ENSDARG000 | 23;23;23;23;2 | 711075:71127 | 711446:71144 | -;-;-;-;-;-;-;-; | 1595 | mitfb | 32 | 34 | 105 | 16 | 93 | 98 |
| ENSDARG000 | 23 | 11587417 | 11587532 | - | 116 | 5S_rRNA | 0 | 0 | 0 | 0 | 0 | 0 |
| ENSDARG000 | 23;23;23;23;2 | 43995820:439 | 43996191:439 | +;+;+;+;+ | 2048 | NA | 0 | 0 | 0 | 0 | 0 | 0 |
| ENSDARG000 | 23 | 33301304 | 33301419 | - | 116 | 5S_rRNA | 0 | 0 | 0 | 0 | 0 | 0 |
| ENSDARG000 | 23 | 42604944 | 42605059 | + | 116 | 5S_rRNA | 0 | 0 | 0 | 0 | 0 | 0 |
| ENSDARG000 | 23 | 22591058 | 22591155 | + | 98 | dre-mir-34a | 0 | 0 | 0 | 0 | 0 | 0 |
| ENSDARG000 | 23;23;23;23;2 | 5104743:5104 | 5104813:5105 | +;+;+;+;+;+;+ | 784 | ube2t | 58 | 224 | 252 | 105 | 155 | 122 |
| ENSDARG000 | 23;23 | 19675780:196 | 19676102:196 | +;+ | 842 | smim4 | 112 | 124 | 142 | 60 | 92 | 74 |
| ENSDARG000 | 23 | 42403564 | 42403680 | - | 117 | 5S_rRNA | 0 | 0 | 0 | 0 | 0 | 0 |
| ENSDARG000 | 23;23;23;23;2 | 17437554:174 | 17437564:174 | +;+;+;+;+;+;+; | 246 | NA | 5 | 6 | 23 | 5 | 3 | 19 |
| ENSDARG000 | 23;23;23;23;2 | 17899226:179 | 17899335:179 | +;+;+;+;+;+;+; | 1174 | NA | 820 | 312 | 599 | 256 | 617 | 261 |

| | | | | | | | | | | | | |
|---|---|---|---|---|---|---|---|---|---|---|---|---|
| ENSDARG000 | 23:23:23:23:2 | 22291440:222 | 22291718:222 | +:+:+:+:+:+:+: | 2174 | NA | 0 | 0 | 1 | 2 | 0 | 0 |
| ENSDARG000 | 23;23 | 19213472:192 | 19213758:192 | +;+ | 822 | apobec2b | 0 | 0 | 0 | 0 | 0 | 0 |
| ENSDARG000 | 23:23:23:23:2 | 1926562:1927 | 1927814:1927 | -:-:-:-:-:-:-:-:-: | 3185 | prdm5 | 26 | 32 | 66 | 36 | 42 | 14 |
| ENSDARG000 | 23 | 30607983 | 30608101 | - | 119 | 5S_rRNA | 0 | 0 | 0 | 0 | 0 | 0 |
| ENSDARG000 | 23;23;23;23 | 46036684:460 | 46036788:460 | -;-;-;- | 453 | ddah1 | 60 | 18 | 27 | 30 | 48 | 14 |
| ENSDARG000 | 23:23:23:23:2 | 36497518:365 | 36497652:365 | -:-:-:-:-:-:-:-:-: | 1504 | spryd3 | 336 | 236 | 303 | 183 | 305 | 190 |
| ENSDARG000 | 23;23 | 2185397:2187 | 2185636:2189 | +;+ | 1641 | C1QTNF7 | 2 | 2 | 8 | 0 | 6 | 4 |
| ENSDARG000 | 23 | 19907828 | 19907944 | - | 117 | 5S_rRNA | 0 | 0 | 0 | 0 | 0 | 0 |
| ENSDARG000 | 23:23:23:23:2 | 42272588:422 | 42272692:422 | +:+:+:+:+:+:+: | 1965 | NA | 406 | 351 | 584 | 289 | 514 | 440 |
| ENSDARG000 | 23 | 5431418 | 5431517 | - | 100 | dre-let-7e | 0 | 0 | 0 | 0 | 0 | 0 |
| ENSDARG000 | 23:23:23:23:2 | 13289083:132 | 13291878:132 | -:-:-:-:-:-:-:-:-: | 4181 | NA | 18 | 4 | 4 | 18 | 13 | 6 |
| ENSDARG000 | 23;23 | 28023091:280 | 28024680:280 | -;- | 2060 | sp5l | 0 | 0 | 0 | 0 | 0 | 0 |
| ENSDARG000 | 23:23:23:23:2 | 42690904:426 | 42691853:426 | +:+:+:+:+:+:+: | 2470 | NA | 3 | 14 | 0 | 2 | 4 | 0 |
| ENSDARG000 | 23 | 35864726 | 35864853 | - | 128 | 5S_rRNA | 0 | 0 | 0 | 0 | 0 | 0 |
| ENSDARG000 | 23;23 | 9727410:9727 | 9727418:9727 | +;+ | 21 | NA | 0 | 0 | 0 | 0 | 0 | 0 |
| ENSDARG000 | 23:23:23:23:2 | 45473286:454 | 45473344:454 | -:-:-:-:-:-:-:-:-: | 1104 | NA | 129 | 161 | 163 | 122 | 115 | 128 |
| ENSDARG000 | 23 | 12926092 | 12931693 | - | 5602 | NA | 0 | 2 | 28 | 34 | 0 | 0 |
| ENSDARG000 | 23 | 20373314 | 20374396 | - | 1083 | NA | 0 | 0 | 0 | 0 | 0 | 0 |
| ENSDARG000 | 23 | 45743796 | 45743917 | + | 122 | 5S_rRNA | 0 | 0 | 0 | 0 | 0 | 0 |
| ENSDARG000 | 23;23 | 2669;8476 | 3621;9339 | +;+ | 1817 | twist3 | 32 | 2 | 4 | 0 | 30 | 12 |
| ENSDARG000 | 23 | 26928743 | 26928858 | + | 116 | 5S_rRNA | 0 | 0 | 0 | 2 | 0 | 0 |
| ENSDARG000 | 23 | 24072160 | 24072245 | + | 86 | dre-mir-200a | 0 | 0 | 0 | 0 | 0 | 0 |
| ENSDARG000 | 23 | 22863457 | 22863571 | - | 115 | 5S_rRNA | 0 | 0 | 0 | 0 | 0 | 0 |
| ENSDARG000 | 23 | 9316578 | 9316694 | - | 117 | 5S_rRNA | 0 | 0 | 0 | 0 | 0 | 0 |
| ENSDARG000 | 23:23:23:23:2 | 8281307:8284 | 8286562:8286 | -;-;-;-;-;-;-;- | 6176 | oprl1 | 6 | 400 | 614 | 314 | 1511 | 194 |
| ENSDARG000 | 23:23:23:23:2 | 4483083:4485 | 4483677:4485 | +:+:+:+:+:+:+: | 7150 | NUP210 | 1047 | 2012 | 3364 | 814 | 2477 | 888 |
| ENSDARG000 | 23 | 18731322 | 18731399 | + | 78 | NA | 0 | 0 | 0 | 0 | 0 | 0 |
| ENSDARG000 | 23:23:23:23:2 | 46002859:460 | 46003487:460 | -;-;-;-;- | 2160 | SYDE2 | 395 | 246 | 347 | 216 | 307 | 214 |
| ENSDARG000 | 23:23:23:23:2 | 7743481:7745 | 7743962:7745 | -;-;-;-;-;- | 1536 | PCMTD2 | 113 | 51 | 60 | 106 | 68 | 156 |
| ENSDARG000 | 23 | 38335760 | 38335884 | + | 125 | 5S_rRNA | 0 | 0 | 0 | 0 | 0 | 0 |
| ENSDARG000 | 23 | 13442608 | 13442722 | + | 115 | 5S_rRNA | 0 | 0 | 0 | 0 | 0 | 0 |
| ENSDARG000 | 23 | 36491114 | 36491234 | + | 121 | 5S_rRNA | 53 | 40 | 34 | 73 | 73 | 47 |
| ENSDARG000 | 23 | 31433273 | 31433348 | + | 76 | NA | 6 | 10 | 26 | 8 | 17 | 2 |
| ENSDARG000 | 23:23:23:23:2 | 3777655:3779 | 3778137:3779 | -;-;-;-;- | 1197 | opn7b | 28 | 0 | 60 | 0 | 11 | 24 |
| ENSDARG000 | 23:23:23:23:2 | 32426954:324 | 32426993:324 | +:+:+:+:+:+:+: | 696 | NA | 0 | 0 | 0 | 0 | 0 | 0 |
| ENSDARG000 | 23;23;23 | 5009964:5010 | 5010467:5012 | -;-;- | 2803 | kcna2b | 0 | 0 | 0 | 0 | 0 | 0 |
| ENSDARG000 | 23 | 17874072 | 17874147 | + | 76 | NA | 0 | 0 | 0 | 0 | 0 | 0 |
| ENSDARG000 | 23 | 42191239 | 42191353 | + | 115 | 5S_rRNA | 0 | 0 | 0 | 0 | 0 | 0 |
| ENSDARG000 | 23 | 28925923 | 28926040 | + | 118 | 5S_rRNA | 0 | 0 | 0 | 0 | 0 | 0 |
| ENSDARG000 | 23:23:23:23:2 | 41200854:412 | 41201603:412 | +;+;+;+;+;+ | 4671 | NHS | 260 | 211 | 374 | 353 | 917 | 373 |
| ENSDARG000 | 23;23;23 | 579893:58346 | 579905:58368 | +;+;+ | 339 | NA | 0 | 0 | 0 | 0 | 2 | 0 |
| ENSDARG000 | 23 | 11935584 | 11935700 | + | 117 | 5S_rRNA | 0 | 0 | 0 | 0 | 0 | 0 |
| ENSDARG000 | 23:23:23:23:2 | 24351304:243 | 24351412:243 | -:-:-:-:-:-:-:-:-: | 2940 | NA | 870 | 1475 | 1022 | 1021 | 1184 | 1268 |
| ENSDARG000 | 23;23;23;23 | 38940700:389 | 38941630:389 | +;+;+;+ | 8687 | sall4 | 0 | 2 | 2 | 0 | 8 | 2 |
| ENSDARG000 | 23 | 20471755 | 20471870 | + | 116 | 5S_rRNA | 0 | 0 | 0 | 0 | 0 | 0 |
| ENSDARG000 | 23 | 31433579 | 31433663 | + | 85 | NA | 0 | 0 | 0 | 0 | 0 | 0 |
| ENSDARG000 | 23:23:23:23:2 | 33777519:338 | 33777800:338 | +:+:+:+:+:+:+: | 2625 | NA | 1 | 0 | 0 | 16 | 0 | 6 |
| ENSDARG000 | 23 | 42502565 | 42502690 | + | 126 | 5S_rRNA | 0 | 0 | 0 | 0 | 0 | 0 |
| ENSDARG000 | 23 | 27580164 | 27580292 | - | 129 | 5S_rRNA | 0 | 1 | 4 | 4 | 0 | 1 |
| ENSDARG000 | 23;23;23;23 | 31972384:319 | 31972545:319 | -;-;-;- | 567 | NA | 0 | 2 | 0 | 0 | 0 | 0 |
| ENSDARG000 | 23:23:23:23:2 | 43535224:435 | 43535902:435 | -:-:-:-:-:-:-:-:-: | 3358 | itchb | 216 | 221 | 349 | 146 | 316 | 198 |
| ENSDARG000 | 23:23:23:23:2 | 37579107:375 | 37579397:375 | +:+:+:+:+:+:+: | 3728 | PLEKHG5 | 219 | 60 | 55 | 32 | 63 | 53 |
| ENSDARG000 | 23 | 17501314 | 17501428 | + | 115 | 5S_rRNA | 0 | 0 | 0 | 0 | 0 | 0 |
| ENSDARG000 | 23:23:23:23:2 | 36460239:364 | 36460283:364 | +:+:+:+:+:+:+: | 2378 | lima1a | 1537 | 1577 | 1144 | 1708 | 1084 | 1372 |
| ENSDARG000 | 23:23:23:23:2 | 1078072:1078 | 1078127:1078 | +:+:+:+:+:+:+: | 2543 | slc34a2b | 102 | 136 | 309 | 110 | 359 | 196 |
| ENSDARG000 | 23 | 24849487 | 24849604 | - | 118 | 5S_rRNA | 0 | 0 | 0 | 0 | 0 | 0 |
| ENSDARG000 | 23:23:23:23:2 | 44940700:449 | 44940942:449 | -:-:-:-:-:-:-:-:-: | 2352 | tfr2 | 18 | 33 | 120 | 32 | 16 | 4 |
| ENSDARG000 | 23 | 18289074 | 18289201 | + | 128 | 5S_rRNA | 0 | 0 | 0 | 0 | 0 | 0 |
| ENSDARG000 | 23 | 17783413 | 17783541 | + | 129 | dre-mir-124-5 | 0 | 0 | 0 | 0 | 0 | 0 |
| ENSDARG000 | 23 | 37438834 | 37438950 | + | 117 | 5S_rRNA | 0 | 0 | 0 | 0 | 0 | 0 |
| ENSDARG000 | 23 | 24606237 | 24606374 | + | 138 | NA | 0 | 0 | 0 | 0 | 0 | 0 |
| ENSDARG000 | 23 | 15353133 | 15353249 | + | 117 | 5S_rRNA | 0 | 0 | 0 | 0 | 0 | 0 |
| ENSDARG000 | 23 | 36509997 | 36510119 | - | 123 | U4 | 0 | 0 | 0 | 0 | 0 | 0 |
| ENSDARG000 | 23 | 13714386 | 13714500 | - | 115 | 5S_rRNA | 0 | 0 | 0 | 0 | 0 | 0 |
| ENSDARG000 | 23;23 | 1140306:1143 | 1141178:1144 | -;- | 1516 | psmb11b | 0 | 0 | 0 | 0 | 2 | 2 |
| ENSDARG000 | 23 | 36322010 | 36322124 | - | 115 | 5S_rRNA | 0 | 0 | 0 | 0 | 0 | 0 |
| ENSDARG000 | 23:23:23:23:2 | 2560005:2564 | 2560194:2564 | +:+:+:+:+:+:+: | 1722 | GGT7 | 34 | 54 | 81 | 20 | 81 | 16 |
| ENSDARG000 | 23:23:23:23:2 | 4414164:4414 | 4415002:4414 | +;+;+;+;+;+;+ | 1594 | WNT7A | 0 | 0 | 0 | 0 | 0 | 0 |
| ENSDARG000 | 23 | 1218382 | 1218501 | - | 120 | 5S_rRNA | 0 | 0 | 0 | 2 | 0 | 0 |
| ENSDARG000 | 23 | 36130001 | 36130092 | + | 92 | dre-mir-10b-2 | 0 | 0 | 0 | 0 | 0 | 0 |
| ENSDARG000 | 23 | 9354822 | 9354931 | - | 110 | dre-mir-2193 | 0 | 0 | 0 | 0 | 0 | 0 |
| ENSDARG000 | 23:23:23:23:2 | 44307996:443 | 44308117:443 | +:+:+:+:+:+:+: | 1667 | DLG4 | 0 | 4 | 10 | 10 | 0 | 6 |
| ENSDARG000 | 23 | 33538051 | 33538169 | + | 119 | 5S_rRNA | 0 | 0 | 0 | 0 | 0 | 0 |
| ENSDARG000 | 23 | 28797426 | 28797548 | + | 123 | 5S_rRNA | 0 | 0 | 0 | 0 | 0 | 0 |
| ENSDARG000 | 23;23 | 36653376:366 | 36653464:366 | +;+ | 1459 | gpr182 | 40 | 10 | 138 | 18 | 22 | 56 |
| ENSDARG000 | 23:23:23:23:2 | 40185381:401 | 40190373:401 | -:-:-:-:-:-:-:-:-: | 7446 | NA | 285 | 190 | 253 | 178 | 262 | 276 |
| ENSDARG000 | 23:23:23:23:2 | 24272421:242 | 24272456:242 | +:+:+:+:+:+:+: | 2352 | clcnk | 0 | 0 | 14 | 0 | 18 | 0 |
| ENSDARG000 | 23:23:23:23:2 | 43638982:436 | 43639093:436 | +:+:+:+:+:+:+: | 1469 | slc10a7 | 273 | 284 | 339 | 144 | 468 | 164 |
| ENSDARG000 | 23:23:23:23:2 | 2666944:2667 | 2667042:2667 | +;+;+;+;+;+;+ | 1001 | NA | 44 | 41 | 94 | 48 | 135 | 52 |
| ENSDARG000 | 23 | 40752704 | 40752818 | + | 115 | 5S_rRNA | 0 | 0 | 0 | 0 | 0 | 0 |
| ENSDARG000 | 23;23;23;23 | 29711569:297 | 29713360:297 | -;-;-;- | 2098 | ctnnbip1 | 784 | 945 | 979 | 752 | 1000 | 912 |
| ENSDARG000 | 23:23:23:23:2 | 37566097:375 | 37566566:375 | -;-;-;-;- | 1308 | tor1l3 | 211 | 204 | 108 | 62 | 182 | 180 |
| ENSDARG000 | 23:23:23:23:2 | 45043301:450 | 45043712:450 | -;-;-;-;- | 935 | sh3gl2b | 0 | 0 | 0 | 0 | 0 | 6 |
| ENSDARG000 | 23:23:23:23:2 | 26966515:269 | 26966529:269 | -;-;-;-;-;-;-;- | 207 | NA | 15 | 4 | 16 | 19 | 20 | 33 |
| ENSDARG000 | 23:23:23:23:2 | 5086317:5087 | 5087044:5088 | -;-;-;-;-;-;-;- | 1911 | etv7 | 56 | 54 | 39 | 12 | 46 | 50 |
| ENSDARG000 | 23:23:23:23:2 | 40257233:402 | 40257266:402 | -:-:-:-:-:-:-:-:-: | 360 | NA | 0 | 0 | 0 | 0 | 0 | 0 |
| ENSDARG000 | 23 | 41345744 | 41345858 | - | 115 | 5S_rRNA | 0 | 0 | 0 | 0 | 0 | 0 |
| ENSDARG000 | 23 | 31667429 | 31667543 | + | 115 | 5S_rRNA | 0 | 0 | 0 | 0 | 0 | 0 |
| ENSDARG000 | 23:23:23:23:2 | 2421313:2421 | 2421759:2421 | +:+:+:+:+:+:+: | 2253 | tcp1 | 7479 | 7319 | 10331 | 5367 | 6659 | 6469 |
| ENSDARG000 | 23 | 17873526 | 17873598 | + | 73 | NA | 0 | 0 | 6 | 0 | 0 | 0 |
| ENSDARG000 | 23 | 24073815 | 24073896 | + | 82 | dre-mir-429a | 0 | 0 | 0 | 0 | 0 | 0 |
| ENSDARG000 | 23 | 40592817 | 40592934 | - | 118 | 5S_rRNA | 0 | 0 | 0 | 0 | 0 | 0 |
| ENSDARG000 | 23 | 33119010 | 33119124 | + | 115 | 5S_rRNA | 0 | 0 | 0 | 0 | 0 | 0 |
| ENSDARG000 | 23 | 5431196 | 5431319 | - | 124 | dre-let-7a-5 | 2 | 5 | 3 | 2 | 2 | 2 |
| ENSDARG000 | 23;23;23;23 | 6754910:6763 | 6759606:6763 | -;-;-;- | 5256 | NA | 12 | 0 | 1 | 0 | 12 | 0 |
| ENSDARG000 | 23 | 13604218 | 13604320 | + | 103 | dre-mir-1388 | 0 | 0 | 0 | 0 | 0 | 0 |
| ENSDARG000 | 23:23:23:23:2 | 2050359:2050 | 2055247:2055 | -;-;-;-;-;-;- | 5957 | ndnf | 36 | 14 | 1 | 0 | 12 | 0 |
| ENSDARG000 | 23:23:23:23:2 | 40133136:401 | 40133287:401 | +:+:+:+:+:+:+: | 2721 | gpsm2l | 217 | 111 | 150 | 82 | 215 | 123 |
| ENSDARG000 | 23;23;23;23 | 45785563:458 | 45785656:458 | +;+;+;+ | 618 | VDR | 67 | 21 | 61 | 63 | 45 | 58 |
| ENSDARG000 | 23:23:23:23:2 | 2136344:2136 | 2136567:2136 | +:+:+:+:+:+:+: | 1353 | CPEB2 | 279 | 316 | 259 | 167 | 315 | 262 |
| ENSDARG000 | 23 | 3055981 | 3056096 | - | 116 | 5S_rRNA | 0 | 0 | 0 | 0 | 0 | 0 |
| ENSDARG000 | 23 | 7172120 | 7172203 | - | 84 | dre-mir-133a- | 0 | 0 | 0 | 0 | 0 | 0 |
| ENSDARG000 | 23 | 17682818 | 17682932 | - | 115 | 5S_rRNA | 0 | 0 | 0 | 0 | 0 | 0 |
| ENSDARG000 | 23 | 7912109 | 7912223 | - | 115 | 5S_rRNA | 0 | 0 | 0 | 0 | 0 | 1 |
| ENSDARG000 | 23:23:23:23:2 | 38952529:389 | 38952596:389 | +:+:+:+:+:+:+: | 8237 | ATP9A | 10 | 9 | 16 | 4 | 54 | 26 |
| ENSDARG000 | 23 | 28536665 | 28536778 | + | 114 | dre-let-7h | 0 | 0 | 0 | 0 | 0 | 0 |
| ENSDARG000 | 23:23:23:23:2 | 900955:90622 | 903901:90635 | -;-;-;-;- | 3617 | rhoac | 2525 | 2230 | 2231 | 2090 | 2111 | 2424 |
| ENSDARG000 | 23 | 37492735 | 37492851 | + | 117 | 5S_rRNA | 10 | 10 | 40 | 5 | 4 | 16 |
| ENSDARG000 | 23:23:23:23:2 | 36730713:367 | 36730863:367 | +;+;+;+;+;+ | 1823 | tspan31 | 1473 | 1597 | 1539 | 1243 | 2118 | 1727 |
| ENSDARG000 | 23:23:23:23:2 | 36711024:367 | 36711029:367 | -;-;-;-;-;-;-;- | 246 | NA | 30 | 29 | 35 | 40 | 36 | 42 |
| ENSDARG000 | 23 | 33290273 | 33290361 | + | 89 | dre-mir-205 | 0 | 0 | 0 | 0 | 0 | 0 |
| ENSDARG000 | 23;23;23;23 | 37444933:374 | 37445060:374 | +;+;+;+ | 681 | NA | 0 | 2 | 0 | 0 | 32 | 0 |
| ENSDARG000 | 23:23:23:23:2 | 37493532:374 | 37493558:374 | -:-:-:-:-:-:-:-:-: | 2349 | DNAJC16 | 171 | 273 | 491 | 210 | 328 | 351 |
| ENSDARG000 | 23;23;23;23 | 42254960:422 | 42255011:422 | +;+;+;+ | 402 | NA | 77 | 63 | 64 | 54 | 139 | 110 |
| ENSDARG000 | 23;23;23 | 42219613:422 | 42219776:422 | -;-;- | 969 | gpx7 | 17 | 70 | 87 | 10 | 23 | 21 |
| ENSDARG000 | 23;23;23 | 24258323:242 | 24258937:242 | -;-;- | 1058 | hspb7 | 0 | 0 | 0 | 0 | 0 | 0 |
| ENSDARG000 | 23:23:23:23:2 | 45282858:452 | 45283023:452 | +:+:+:+:+:+:+: | 1287 | NA | 6 | 24 | 18 | 0 | 56 | 0 |
| ENSDARG000 | 23:23:23:23:2 | 6333406:6347 | 6333492:6347 | -;-;-;-;- | 902 | NA | 2 | 0 | 0 | 0 | 2 | 29 |
| ENSDARG000 | 23:23:23:23:2 | 45422433:454 | 45424421:454 | -:-:-:-:-:-:-:-:-: | 3760 | NA | 1169 | 44 | 628 | 160 | 447 | 36 |
| ENSDARG000 | 23;23 | 3405665:3407 | 3409140:3408 | -;- | 3476 | mafba | 823 | 597 | 1206 | 754 | 2086 | 650 |
| ENSDARG000 | 23:23:23:23:2 | 44049509:440 | 44049702:440 | +;+;+;+;+ | 780 | NA | 2 | 16 | 0 | 0 | 16 | 15 |
| ENSDARG000 | 23:23:23:23:2 | 2361184:2368 | 2361519:2368 | +;+;+;+;+ | 732 | NA | 64 | 71 | 79 | 118 | 130 | 55 |
| ENSDARG000 | 23 | 38127452 | 38127569 | - | 118 | 5S_rRNA | 0 | 0 | 0 | 0 | 0 | 0 |
| ENSDARG000 | 23 | 16858917 | 16859033 | + | 117 | 5S_rRNA | 0 | 0 | 0 | 0 | 0 | 0 |
| ENSDARG000 | 23:23:23:23:2 | 37482727:374 | 37483171:374 | +;+;+;+;+;+;+ | 1433 | agmat | 17 | 2 | 40 | 2 | 31 | 29 |
| ENSDARG000 | 23 | 14269311 | 14270315 | + | 1005 | NA | 0 | 0 | 0 | 0 | 0 | 0 |
| ENSDARG000 | 23:23:23:23:2 | 43597961:436 | 43599700:436 | -;-;-;-;- | 2609 | NA | 0 | 2 | 0 | 0 | 18 | 0 |
| ENSDARG000 | 23:23:23:23:2 | 29756603:297 | 29758383:297 | -:-:-:-:-:-:-:-:-: | 4986 | pik3cd | 28 | 34 | 20 | 20 | 129 | 122 |
| ENSDARG000 | 23:23:23:23:2 | 12062294:120 | 12062316:120 | -;-;-;-;-;-;-;- | 165 | NA | 0 | 0 | 0 | 0 | 0 | 0 |
| ENSDARG000 | 23:23:23:23:2 | 43467799:434 | 43468046:434 | -;-;-;-;-;-;- | 1290 | E2F1 | 72 | 170 | 184 | 66 | 141 | 108 |
| ENSDARG000 | 23 | 26675652 | 26675770 | - | 119 | 5S_rRNA | 0 | 0 | 0 | 0 | 0 | 0 |
| ENSDARG000 | 23 | 21623735 | 21623849 | + | 115 | 5S_rRNA | 0 | 0 | 0 | 0 | 0 | 0 |
| ENSDARG000 | 23 | 19308796 | 19308910 | - | 115 | 5S_rRNA | 0 | 0 | 0 | 0 | 0 | 0 |
| ENSDARG000 | 23;23;23;23 | 46030970:460 | 46031387:460 | -;-;-;- | 1487 | C23H1orf52 | 779 | 712 | 831 | 666 | 747 | 730 |
| ENSDARG000 | 23 | 36828472 | 36828588 | - | 117 | 5S_rRNA | 0 | 0 | 0 | 0 | 0 | 0 |
| ENSDARG000 | 23:23:23:23:2 | 44124138:441 | 44124275:441 | +:+:+:+:+:+:+: | 1866 | tec | 109 | 57 | 137 | 36 | 181 | 72 |
| ENSDARG000 | 23:23:23:23:2 | 36616717:366 | 36617019:366 | +:+:+:+:+:+:+: | 2231 | pip4k2ca | 1340 | 1507 | 1795 | 1042 | 1511 | 1306 |
| ENSDARG000 | 23 | 40306244 | 40306371 | + | 128 | 5S_rRNA | 0 | 0 | 0 | 0 | 0 | 0 |
| ENSDARG000 | 23 | 41287333 | 41287449 | - | 117 | 5S_rRNA | 0 | 0 | 0 | 0 | 0 | 0 |
| ENSDARG000 | 23 | 442237 | 442352 | + | 116 | 5S_rRNA | 0 | 0 | 0 | 0 | 0 | 0 |
| ENSDARG000 | 23:23:23:23:2 | 5977965:5977 | 5978047:5978 | +:+:+:+:+:+:+: | 4458 | NAV1 | 1093 | 481 | 913 | 982 | 1334 | 685 |
| ENSDARG000 | 23 | 32875546 | 32875660 | - | 115 | 5S_rRNA | 0 | 0 | 0 | 0 | 0 | 0 |
| ENSDARG000 | 23 | 24675207 | 24675303 | - | 97 | dre-mir-26b | 0 | 0 | 0 | 0 | 0 | 0 |
| ENSDARG000 | 23 | 11532340 | 11532470 | - | 131 | 5S_rRNA | 0 | 0 | 0 | 0 | 0 | 0 |
| ENSDARG000 | 23 | 7581186 | 7581302 | - | 117 | 5S_rRNA | 2 | 0 | 0 | 0 | 0 | 0 |
| ENSDARG000 | 23 | 20187625 | 20187739 | + | 115 | 5S_rRNA | 0 | 0 | 0 | 0 | 0 | 0 |
| ENSDARG000 | 23 | 412542 | 412659 | + | 118 | 5S_rRNA | 0 | 0 | 0 | 0 | 0 | 0 |
| ENSDARG000 | 23 | 28800103 | 28800219 | + | 117 | 5S_rRNA | 0 | 0 | 0 | 0 | 0 | 0 |
| ENSDARG000 | 23 | 36959791 | 36959912 | - | 122 | 5S_rRNA | 0 | 0 | 0 | 0 | 0 | 0 |
| ENSDARG000 | 23 | 12068120 | 12068234 | - | 115 | 5S_rRNA | 0 | 0 | 0 | 0 | 0 | 0 |
| ENSDARG000 | 23 | 1347806 | 1348933 | - | 1128 | NA | 0 | 0 | 0 | 0 | 0 | 0 |
| ENSDARG000 | 23:23:23:23:2 | 36675207:366 | 36675558:366 | -:-:-:-:-:-:-:-:-: | 2742 | agap2 | 24 | 25 | 24 | 12 | 119 | 76 |
| ENSDARG000 | 23:23:23:23:2 | 44712588:447 | 44712786:447 | -;-;-;-;- | 846 | MOGAT1 | 282 | 252 | 243 | 142 | 132 | 388 |
| ENSDARG000 | 23;23;23 | 2542158:2544 | 2542281:2544 | +;+;+ | 522 | NA | 148 | 49 | 73 | 39 | 86 | 9 |
| ENSDARG000 | 23:23:23:23:2 | 7067196:7068 | 7067357:7068 | -:-:-:-:-:-:-:-:-: | 2691 | SLCO4A1 | 0 | 0 | 0 | 0 | 0 | 0 |
| ENSDARG000 | 23 | 13989364 | 13989482 | - | 119 | 5S_rRNA | 0 | 0 | 0 | 0 | 0 | 0 |
| ENSDARG000 | 23 | 29588444 | 29588558 | + | 115 | 5S_rRNA | 0 | 0 | 4 | 0 | 1 | 0 |
| ENSDARG000 | 23:23:23:23:2 | 37530718:375 | 37531013:375 | -;-;-;-;- | 1056 | tor1l1 | 0 | 0 | 0 | 0 | 0 | 0 |
| ENSDARG000 | 23:23:23:23:2 | 46048218:460 | 46049554:460 | -:-:-:-:-:-:-:-:-: | 3328 | NA | 0 | 0 | 2 | 0 | 755 | 0 |
| ENSDARG000 | 23:23:23:23:2 | 44078929:440 | 44079116:440 | +:+:+:+:+:+:+: | 6073 | g3bp2b | 2209 | 1808 | 2643 | 1382 | 2603 | 1565 |
| ENSDARG000 | 23:23:23:23:2 | 40234602:402 | 40235128:402 | -:-:-:-:-:-:-:-:-: | 2459 | NA | 4 | 10 | 35 | 121 | 41 | 18 |
| ENSDARG000 | 23 | 33219005 | 33219128 | - | 124 | 5S_rRNA | 0 | 0 | 0 | 0 | 0 | 0 |

| | | | | | | | | | | | | |
|---|---|---|---|---|---|---|---|---|---|---|---|---|
| ENSDARG000 | 23;23;23;23 | 36601984:366 | 36602485:366 | +;+;+;+ | 1343 | igfbp6b | 146 | 80 | 86 | 54 | 16 | 32 |
| ENSDARG000 | 23:23:23:23:2 | 40951443:409 | 40951737:409 | +:+:+:+:+:+:+: | 1440 | NA | 18 | 0 | 12 | 6 | 12 | 23 |
| ENSDARG000 | 23 | 6825688 | 6825802 | + | 115 | 5S_rRNA | 0 | 0 | 0 | 0 | 0 | 0 |
| ENSDARG000 | 23 | 44010425 | 44010541 | - | 117 | 5S_rRNA | 0 | 0 | 0 | 0 | 0 | 0 |
| ENSDARG000 | 23 | 33968866 | 33968988 | - | 123 | 5S_rRNA | 0 | 0 | 0 | 0 | 0 | 0 |
| ENSDARG000 | 23;23;23;23 | 45822935:458 | 45823116:458 | +;+;+;+ | 822 | VDR | 64 | 13 | 78 | 145 | 74 | 55 |
| ENSDARG000 | 23 | 28614827 | 28614942 | + | 116 | 5S_rRNA | 0 | 0 | 0 | 0 | 0 | 0 |
| ENSDARG000 | 23 | 19713095 | 19713228 | - | 134 | SNORA70 | 0 | 0 | 2 | 1 | 7 | 0 |
| ENSDARG000 | 23 | 25995027 | 25995141 | + | 115 | 5S_rRNA | 0 | 0 | 0 | 0 | 0 | 0 |
| ENSDARG000 | 23 | 18063173 | 18063299 | - | 127 | dre-mir-135c- | 0 | 0 | 0 | 0 | 0 | 0 |
| ENSDARG000 | 23 | 45271141 | 45271255 | + | 115 | 5S_rRNA | 0 | 0 | 0 | 0 | 0 | 0 |
| ENSDARG000 | 23:23:23:23:2 | 46208304:462 | 46208770:462 | -;-;-;-;- | 764 | zgc:162858 | 243 | 85 | 138 | 206 | 165 | 140 |
| ENSDARG000 | 23:23:23:23:2 | 22578482:225 | 22578510:225 | -:-:-:-:-:-:-:-: | 324 | NA | 2 | 0 | 2 | 5 | 9 | 2 |
| ENSDARG000 | 23:23:23:23:2 | 3949548:3951 | 3952297:3952 | -:-:-:-:-:-:-:-: | 4868 | slc9a8 | 255 | 297 | 452 | 414 | 266 | 272 |
| ENSDARG000 | 23:23:23:23:2 | 46191873:461 | 46192127:461 | -:-:-:-:-:-:-:-: | 2310 | tgm1l4 | 50 | 17 | 51 | 82 | 53 | 58 |
| ENSDARG000 | 23:23:23:23:2 | 1644754:1644 | 1645354:1645 | -:-:-:-:-:-:-:-: | 2793 | NA | 0 | 6 | 0 | 6 | 0 | 0 |
| ENSDARG000 | 23 | 31754752 | 31754868 | - | 117 | 5S_rRNA | 0 | 0 | 0 | 0 | 0 | 0 |
| ENSDARG000 | 23:23:23:23:2 | 6141541:6142 | 6141879:6142 | +:+:+:+:+:+:+: | 2625 | NA | 50 | 32 | 4 | 14 | 42 | 120 |
| ENSDARG000 | 23:23:23:23:2 | 37466967:374 | 37467323:374 | +:+:+:+:+:+:+: | 4010 | pink1 | 251 | 119 | 320 | 169 | 164 | 219 |
| ENSDARG000 | 23:23:23:23:2 | 17939611:179 | 17939678:179 | +:+:+:+:+:+:+: | 1883 | chia.5 | 0 | 0 | 0 | 0 | 0 | 2 |
| ENSDARG000 | 23:23:23:23:2 | 37412800:374 | 37415841:374 | -;-;-;-;- | 3663 | NA | 0 | 0 | 0 | 0 | 2 | 0 |
| ENSDARG000 | 23:23:23:23:2 | 24300651:243 | 24301340:243 | -;-;-;-;-;-;- | 1347 | fam131c | 0 | 0 | 0 | 0 | 0 | 8 |
| ENSDARG000 | 23:23:23:23:2 | 37539068:375 | 37539318:375 | -;-;-;-;- | 1752 | NA | 0 | 0 | 0 | 0 | 0 | 0 |
| ENSDARG000 | 23:23:23:23:2 | 44970214:449 | 44970320:449 | -:-:-:-:-:-:-:-: | 3621 | ctc1 | 490 | 490 | 637 | 551 | 907 | 395 |
| ENSDARG000 | 23:23:23:23:2 | 35451050:354 | 35451058:354 | +:+:+:+:+:+:+: | 309 | NA | 1 | 0 | 22 | 7 | 3 | 13 |
| ENSDARG000 | 23 | 24072029 | 24072110 | + | 82 | dre-mir-200b | 0 | 0 | 0 | 0 | 0 | 0 |
| ENSDARG000 | 23 | 15107848 | 15107962 | - | 115 | 5S_rRNA | 0 | 0 | 0 | 0 | 0 | 0 |
| ENSDARG000 | 23;23 | 38379627:384 | 38382649:384 | -;- | 3643 | TSHZ2 | 3955 | 2610 | 3110 | 4891 | 3800 | 3335 |
| ENSDARG000 | 23:23:23:23:2 | 27912079:279 | 27912172:279 | +:+:+:+:+:+:+: | 1617 | ARHGEF25 | 7 | 4 | 12 | 8 | 17 | 18 |
| ENSDARG000 | 23;23;23 | 5108374:5111 | 5108408:5111 | +;+;+ | 706 | zgc:194242 | 4 | 7 | 6 | 12 | 12 | 24 |
| ENSDARG000 | 23:23:23:23:2 | 43489182:434 | 43489233:434 | +:+:+:+:+:+:+: | 3576 | znf341 | 182 | 112 | 227 | 78 | 204 | 200 |
| ENSDARG000 | 23 | 9353552 | 9357627 | + | 4076 | NA | 82 | 141 | 369 | 119 | 87 | 131 |
| ENSDARG000 | 23:23:23:23:2 | 6998633:7000 | 7000625:7000 | -:-:-:-:-:-:-:-: | 4840 | prpf6 | 2030 | 2578 | 3015 | 1510 | 3181 | 1970 |
| ENSDARG000 | 23 | 38329520 | 38330746 | - | 1227 | NA | 0 | 6 | 0 | 0 | 4 | 0 |
| ENSDARG000 | 23;23 | 40908583:409 | 40908895:409 | +;+ | 1305 | NA | 4 | 0 | 0 | 6 | 0 | 0 |
| ENSDARG000 | 23:23:23:23:2 | 45229198:452 | 45229242:452 | +:+:+:+:+:+:+: | 951 | NA | 730 | 116 | 148 | 144 | 516 | 172 |
| ENSDARG000 | 23:23:23:23:2 | 45520704:455 | 45524591:455 | -:-:-:-:-:-:-:-: | 5625 | NA | 107 | 99 | 74 | 60 | 60 | 94 |
| ENSDARG000 | 23 | 7210157 | 7210262 | - | 106 | mir1-1 | 0 | 0 | 0 | 0 | 0 | 0 |
| ENSDARG000 | 23 | 789493 | 790620 | - | 1128 | NA | 0 | 2 | 0 | 0 | 0 | 0 |
| ENSDARG000 | 23:23:23:23:2 | 17782972:177 | 17782998:177 | -;-;-;-;-;-;-;- | 189 | NA | 0 | 0 | 0 | 0 | 0 | 0 |
| ENSDARG000 | 23:23:23:23:2 | 2191773:2191 | 2192384:2192 | +:+:+:+:+:+:+: | 10045 | CC2D2A | 168 | 178 | 206 | 124 | 225 | 88 |
| ENSDARG000 | 23:23:23:23:2 | 45456490:454 | 45456537:454 | +;+;+;+;+ | 4995 | ccn1 | 1458 | 233 | 448 | 592 | 697 | 385 |
| ENSDARG000 | 23:23:23:23:2 | 6649071:6651 | 6649286:6651 | -;-;-;-;-;- | 834 | NA | 0 | 0 | 0 | 0 | 0 | 0 |
| ENSDARG000 | 23;23 | 42131509:421 | 42132024:421 | +;+ | 1256 | SHISAL2A | 0 | 0 | 0 | 0 | 0 | 0 |
| ENSDARG000 | 23 | 9852896 | 9853010 | - | 115 | 5S_rRNA | 0 | 0 | 0 | 0 | 0 | 0 |
| ENSDARG000 | 23 | 32139843 | 32139958 | - | 116 | 5S_rRNA | 0 | 0 | 0 | 0 | 0 | 0 |
| ENSDARG000 | 23;23;23;23 | 37458602:374 | 37458798:374 | +;+;+;+ | 2292 | cdaa | 1120 | 928 | 180 | 569 | 692 | 405 |
| ENSDARG000 | 23;23;23;23 | 2906031:2910 | 2906389:2910 | +;+;+;+ | 1523 | rab5if | 808 | 1569 | 1615 | 486 | 1305 | 850 |
| ENSDARG000 | 23:23:23:23:2 | 40275400:402 | 40275478:402 | +;+;+;+;+ | 5120 | tent5ab | 6 | 18 | 13 | 11 | 74 | 52 |
| ENSDARG000 | 23:23:23:23:2 | 45937468:459 | 45937516:459 | -:-:-:-:-:-:-:-: | 2452 | MCOLN3 | 3 | 2 | 4 | 10 | 2 | 0 |
| ENSDARG000 | 23 | 18382456 | 18382551 | - | 96 | dre-let-7g-1 | 1 | 0 | 0 | 0 | 0 | 2 |
| ENSDARG000 | 23 | 9618213 | 9618335 | + | 123 | 5S_rRNA | 0 | 0 | 0 | 0 | 0 | 0 |
| ENSDARG000 | 23;23;23;23 | 14432323:145 | 14432555:145 | +;+;+;+ | 575 | NA | 2 | 0 | 2 | 0 | 1 | 0 |
| ENSDARG000 | 23 | 29823286 | 29824146 | - | 861 | foxl2l | 8 | 6 | 10 | 14 | 16 | 16 |
| ENSDARG000 | 23:23:23:23:2 | 34321237:343 | 34321489:343 | +:+:+:+:+:+:+: | 6135 | PLXNA1 | 246 | 324 | 442 | 479 | 848 | 390 |
| ENSDARG000 | 23 | 43530601 | 43530715 | - | 115 | 5S_rRNA | 32 | 8 | 19 | 13 | 33 | 17 |
| ENSDARG000 | 23:23:23:23:2 | 17417539:174 | 17417593:174 | +:+:+:+:+:+:+: | 159 | NA | 18 | 13 | 40 | 22 | 23 | 25 |
| ENSDARG000 | 23;23 | 4086493:4090 | 4090419:4091 | -;- | 4359 | NA | 63 | 16 | 329 | 10 | 46 | 170 |
| ENSDARG000 | 23:23:23:23:2 | 24046032:240 | 24046545:240 | -;-;-;-;- | 966 | NA | 0 | 0 | 0 | 0 | 0 | 0 |
| ENSDARG000 | 23 | 36088091 | 36088196 | + | 106 | mir196a-1 | 0 | 0 | 0 | 0 | 0 | 0 |
| ENSDARG000 | 23 | 43335460 | 43335575 | + | 116 | 5S_rRNA | 0 | 0 | 0 | 0 | 0 | 0 |
| ENSDARG000 | 23;23;23;23 | 1118305:1120 | 1118496:1120 | -;-;-;- | 1569 | NA | 0 | 0 | 0 | 0 | 0 | 0 |
| ENSDARG000 | 23;23;23 | 1124514:1138 | 1124788:1138 | -;-;- | 1287 | NA | 0 | 0 | 0 | 0 | 0 | 0 |
| ENSDARG000 | 23:23:23:23:2 | 1147162:1147 | 1147455:1147 | +;+;+;+;+ | 1213 | NA | 0 | 0 | 2 | 0 | 0 | 0 |
| ENSDARG000 | 23:23:23:23:2 | 3913304:3913 | 3913513:3913 | +;+;+;+;+;+;+ | 2193 | NA | 0 | 0 | 0 | 0 | 0 | 0 |
| ENSDARG000 | 23;23;23;23 | 3939908:3939 | 3940577:3940 | +;+;+;+ | 2684 | NA | 145 | 14 | 39 | 22 | 10 | 80 |
| ENSDARG000 | 23;23;23 | 4559564:4570 | 4559793:4570 | +;+;+ | 647 | NA | 0 | 6 | 7 | 0 | 0 | 0 |
| ENSDARG000 | 23;23;23 | 5119732:5124 | 5120720:5124 | -;-;- | 1669 | NA | 0 | 0 | 0 | 0 | 0 | 0 |
| ENSDARG000 | 23:23:23:23:2 | 5132439:5135 | 5132547:5135 | +;+;+;+;+;+ | 3860 | NA | 0 | 22 | 124 | 0 | 142 | 6 |
| ENSDARG000 | 23:23:23:23:2 | 5595941:5595 | 5596115:5596 | +;+;+;+;+;+;+ | 935 | NA | 0 | 0 | 0 | 6 | 0 | 0 |
| ENSDARG000 | 23;23;23;23 | 5871506:5906 | 5873143:5907 | +;+;+;+ | 6814 | NA | 271 | 103 | 212 | 167 | 214 | 84 |
| ENSDARG000 | 23:23:23:23:2 | 6399839:6403 | 6400258:6403 | -;-;-;-;- | 1796 | NA | 0 | 0 | 0 | 0 | 0 | 0 |
| ENSDARG000 | 23;23;23;23 | 7402655:7405 | 7403131:7406 | +;+;+;+ | 1252 | NA | 0 | 0 | 0 | 4 | 0 | 0 |
| ENSDARG000 | 23;23;23 | 7950085:7969 | 7950352:7970 | -;-;- | 603 | NA | 0 | 0 | 0 | 0 | 0 | 0 |
| ENSDARG000 | 23:23:23:23:2 | 15479081:154 | 15479246:154 | +;+;+;+;+ | 2828 | NA | 2 | 0 | 12 | 0 | 2 | 4 |
| ENSDARG000 | 23;23;23 | 32537814:325 | 32538313:325 | -;-;- | 863 | NA | 2 | 0 | 2 | 0 | 0 | 0 |
| ENSDARG000 | 23;23;23 | 34849715:349 | 34849996:349 | +;+;+ | 1500 | NA | 0 | 0 | 0 | 0 | 0 | 0 |
| ENSDARG000 | 23;23;23;23 | 37393617:373 | 37394964:373 | +;+;+;+ | 2482 | NA | 0 | 0 | 0 | 0 | 0 | 0 |
| ENSDARG000 | 23;23;23 | 38266163:382 | 38266868:382 | +;+;+ | 2023 | NA | 0 | 0 | 0 | 0 | 0 | 0 |
| ENSDARG000 | 23;23;23 | 39183246:391 | 39183485:391 | +;+;+ | 779 | NA | 0 | 0 | 0 | 0 | 0 | 0 |
| ENSDARG000 | 23:23:23:23:2 | 41021511:410 | 41022120:410 | +;+;+;+;+;+ | 5008 | NA | 2 | 2 | 0 | 0 | 17 | 0 |
| ENSDARG000 | 23:23:23:23:2 | 41187230:411 | 41187266:411 | +;+;+;+;+;+ | 949 | NA | 0 | 0 | 5 | 3 | 9 | 0 |
| ENSDARG000 | 23;23;23 | 42626534:426 | 42627248:426 | +;+;+ | 5309 | NA | 0 | 4 | 0 | 0 | 0 | 0 |
| ENSDARG000 | 23:23:23:23:2 | 42884603:428 | 42886263:428 | -;-;-;-;-;-;- | 4263 | NA | 0 | 0 | 0 | 0 | 0 | 0 |
| ENSDARG000 | 23;23;23;23 | 43288994:432 | 43289052:432 | +;+;+;+ | 1372 | NA | 0 | 0 | 0 | 0 | 0 | 0 |
| ENSDARG000 | 23:23:23:23:2 | 44013682:440 | 44015265:440 | -;-;-;-;- | 2235 | NA | 8 | 2 | 0 | 22 | 12 | 2 |
| ENSDARG000 | 23;23;23;23 | 44070920:440 | 44070994:440 | +;+;+;+ | 2112 | NA | 0 | 0 | 0 | 0 | 2 | 1 |
| ENSDARG000 | 21:21:21:21:2 | 44709280:447 | 44709741:447 | -;-;-;-;-;- | 1325 | ammecr1 | 791 | 735 | 1080 | 356 | 1142 | 591 |
| ENSDARG000 | 21:21:21:21:2 | 20742876:207 | 20745556:207 | -:-:-:-:-:-:-:-: | 3937 | ghrb | 38 | 34 | 90 | 56 | 114 | 74 |
| ENSDARG000 | 21:21:21:21:2 | 5993188:6025 | 5993502:6025 | +:+:+:+:+:+:+: | 3967 | SLC4A4 | 126 | 116 | 157 | 74 | 255 | 221 |
| ENSDARG000 | 21:21:21:21:2 | 25421368:254 | 25426078:254 | -:-:-:-:-:-:-:-: | 8677 | CNKSR2 | 957 | 457 | 676 | 408 | 1295 | 691 |
| ENSDARG000 | 21;21;21;21 | 1618181:1622 | 1618890:1622 | -;-;-;- | 1214 | vmo1b | 6 | 1 | 4 | 0 | 8 | 2 |
| ENSDARG000 | 21:21:21:21:2 | 27437867:274 | 27438121:274 | -:-:-:-:-:-:-:-: | 1779 | bfb | 2 | 0 | 16 | 0 | 0 | 0 |
| ENSDARG000 | 21:21:21:21:2 | 1629722:1629 | 1630435:1630 | -:-:-:-:-:-:-:-: | 2488 | zgc:152948 | 30 | 74 | 59 | 37 | 52 | 74 |
| ENSDARG000 | 21:21:21:21:2 | 44735181:447 | 44739346:447 | -:-:-:-:-:-:-:-: | 7976 | kif4 | 306 | 1009 | 1441 | 448 | 1041 | 459 |
| ENSDARG000 | 21:21:21:21:2 | 14806342:148 | 14806425:148 | -:-:-:-:-:-:-:-: | 1667 | phpt1 | 184 | 175 | 415 | 94 | 241 | 249 |
| ENSDARG000 | 21:21:21:21:2 | 27448856:274 | 27448936:274 | +:+:+:+:+:+:+: | 2575 | cfbl | 844 | 660 | 565 | 371 | 860 | 629 |
| ENSDARG000 | 21:21:21:21:2 | 44773237:447 | 44773450:447 | +:+:+:+:+:+:+: | 1704 | rnf121 | 896 | 951 | 1082 | 568 | 1124 | 1008 |
| ENSDARG000 | 21:21:21:21:2 | 20771082:207 | 20771263:207 | +:+:+:+:+:+:+: | 1956 | oxct1b | 0 | 0 | 6 | 0 | 0 | 0 |
| ENSDARG000 | 21:21:21:21:2 | 14817273:148 | 14817472:148 | -:-:-:-:-:-:-:-: | 1701 | noc4l | 382 | 502 | 747 | 314 | 671 | 430 |
| ENSDARG000 | 21;21;21 | 30886127:308 | 30886714:308 | -;-;- | 904 | NA | 52 | 6 | 8 | 10 | 8 | 0 |
| ENSDARG000 | 21:21:21:21:2 | 30841241:308 | 30841649:308 | -:-:-:-:-:-:-:-: | 2575 | stim1b | 312 | 323 | 640 | 739 | 28 | 577 |
| ENSDARG000 | 21:21:21:21:2 | 14826583:148 | 14826720:148 | -;-;-;-;-;-;-;- | 1605 | pus1 | 130 | 229 | 312 | 86 | 255 | 158 |
| ENSDARG000 | 21:21:21:21:2 | 6114305:6114 | 6114881:6114 | +:+:+:+:+:+:+: | 3348 | fpgs | 352 | 763 | 1520 | 581 | 585 | 712 |
| ENSDARG000 | 21:21:21:21:2 | 10577527:105 | 10577762:105 | +:+:+:+:+:+:+: | 1492 | ccbe1 | 2 | 0 | 0 | 0 | 2 | 0 |
| ENSDARG000 | 21:21:21:21:2 | 25533531:255 | 25533601:255 | +:+:+:+:+:+:+: | 3604 | NA | 1813 | 1283 | 2040 | 764 | 1543 | 1790 |
| ENSDARG000 | 21:21:21:21:2 | 27464538:274 | 27464626:274 | +:+:+:+:+:+:+: | 2711 | bbs1 | 61 | 64 | 72 | 38 | 28 | 58 |
| ENSDARG000 | 21:21:21:21:2 | 10701834:107 | 10702252:107 | +:+:+:+:+:+:+: | 3604 | lman1 | 910 | 1290 | 1817 | 772 | 1326 | 711 |
| ENSDARG000 | 21;21;21;21 | 11829836:118 | 11830515:118 | -;-;-;- | 915 | rfesd | 0 | 2 | 2 | 0 | 2 | 4 |
| ENSDARG000 | 21:21:21:21:2 | 1647990:1651 | 1648123:1651 | +:+:+:+:+:+:+: | 1431 | fech | 26 | 26 | 77 | 20 | 107 | 28 |
| ENSDARG000 | 21:21:21:21:2 | 44857293:448 | 44857379:448 | +:+:+:+:+:+:+: | 3392 | fstl4 | 4 | 0 | 16 | 10 | 24 | 14 |
| ENSDARG000 | 21;21;21 | 11834880:118 | 11835026:118 | +;+;+ | 1326 | nudt2 | 237 | 346 | 474 | 232 | 278 | 295 |
| ENSDARG000 | 21;21;21 | 30937690:309 | 30937805:309 | +;+;+ | 1957 | rhogb | 328 | 351 | 503 | 305 | 278 | 467 |
| ENSDARG000 | 21:21:21:21:2 | 25544882:255 | 25545061:255 | -:-:-:-:-:-:-:-: | 2010 | NA | 13 | 6 | 6 | 8 | 4 | 6 |
| ENSDARG000 | 21:21:21:21:2 | 45182305:451 | 45182893:451 | -;-;-;-;-;-;-;- | 2210 | NA | 962 | 698 | 921 | 617 | 1096 | 1039 |
| ENSDARG000 | 21:21:21:21:2 | 14839589:148 | 14839684:148 | -:-:-:-:-:-:-:-: | 3956 | ULK1 | 950 | 322 | 919 | 804 | 869 | 1050 |
| ENSDARG000 | 21:21:21:21:2 | 6136981:6137 | 6137093:6137 | +:+:+:+:+:+:+: | 1788 | cdk9 | 500 | 999 | 930 | 296 | 456 | 669 |
| ENSDARG000 | 21;21;21 | 10739846:107 | 10740373:107 | +;+;+ | 2514 | cplx4a | 10 | 5 | 0 | 2 | 8 | 0 |
| ENSDARG000 | 21:21:21:21:2 | 10738337:107 | 10738440:107 | +;+;+;+;+ | 530 | NA | 0 | 5 | 0 | 0 | 0 | 1 |
| ENSDARG000 | 21:21:21:21:2 | 30973697:309 | 30974411:309 | -;-;-;-;-;-;-;- | 1703 | pgap2 | 272 | 237 | 312 | 121 | 369 | 183 |
| ENSDARG000 | 21:21:21:21:2 | 45190582:451 | 45191282:451 | -:-:-:-:-:-:-:-: | 2058 | vdac1 | 4431 | 4334 | 7766 | 7998 | 5353 | 5381 |
| ENSDARG000 | 21:21:21:21:2 | 27482149:274 | 27482344:274 | -:-:-:-:-:-:-:-: | 2443 | CAPN1 | 13 | 20 | 21 | 16 | 15 | 30 |
| ENSDARG000 | 21:21:21:21:2 | 11845241:118 | 11845883:118 | -:-:-:-:-:-:-:-: | 1975 | ube2r2 | 318 | 188 | 255 | 222 | 306 | 201 |
| ENSDARG000 | 21:21:21:21:2 | 45223194:452 | 45223317:452 | +;+;+;+;+;+ | 927 | NA | 0 | 0 | 0 | 0 | 0 | 0 |
| ENSDARG000 | 21;21 | 1698232:1741 | 1698512:1742 | -;- | 1379 | ONECUT2 | 6 | 0 | 1 | 2 | 0 | 0 |
| ENSDARG000 | 21:21:21:21:2 | 25561284:255 | 25561806:255 | -;-;-;-;- | 927 | mrps12 | 292 | 421 | 287 | 159 | 310 | 345 |
| ENSDARG000 | 21:21:21:21:2 | 30996076:309 | 30996714:309 | -:-:-:-:-:-:-:-: | 2514 | ncbp3 | 565 | 585 | 617 | 278 | 546 | 466 |
| ENSDARG000 | 21;21;21 | 10756154:107 | 10756495:107 | +;+;+ | 1411 | rx3 | 0 | 0 | 0 | 0 | 0 | 0 |
| ENSDARG000 | 21:21:21:21:2 | 14892050:148 | 14892372:148 | -:-:-:-:-:-:-:-: | 1837 | MMP17 | 8 | 8 | 28 | 0 | 7 | 4 |
| ENSDARG000 | 21;21;21;21 | 1780790:1788 | 1781519:1788 | -;-;-;- | 1811 | st8sia3 | 0 | 0 | 0 | 0 | 0 | 0 |
| ENSDARG000 | 21:21:21:21:2 | 25570028:255 | 25570672:255 | -;-;-;-;-;-;-;- | 1808 | NPAS4 | 0 | 0 | 3 | 0 | 0 | 0 |
| ENSDARG000 | 21:21:21:21:2 | 22311964:223 | 22312028:223 | -:-:-:-:-:-:-:-: | 1937 | NA | 76 | 33 | 18 | 28 | 32 | 33 |
| ENSDARG000 | 21:21:21:21:2 | 20795472:207 | 20795827:207 | -:-:-:-:-:-:-:-: | 3002 | NA | 112 | 22 | 39 | 57 | 32 | 42 |
| ENSDARG000 | 21;21;21;21 | 6151953:6157 | 6152029:6157 | +;+;+;+ | 2094 | NA | 0 | 42 | 20 | 0 | 0 | 38 |
| ENSDARG000 | 21:21:21:21:2 | 10773513:107 | 10773594:107 | +:+:+:+:+:+:+: | 4778 | ZNF532 | 342 | 247 | 526 | 439 | 488 | 358 |
| ENSDARG000 | 21:21:21:21:2 | 31020299:310 | 31020548:310 | -:-:-:-:-:-:-:-: | 2405 | rap1gap2b | 48 | 51 | 90 | 87 | 60 | 44 |
| ENSDARG000 | 21;21;21 | 10765886:107 | 10766227:107 | -;-;- | 753 | grp | 18 | 0 | 4 | 22 | 2 | 2 |
| ENSDARG000 | 21:21:21:21:2 | 15125497:151 | 15126187:151 | -:-:-:-:-:-:-:-: | 3660 | sfswap | 2064 | 997 | 1984 | 3203 | 1824 | 3026 |
| ENSDARG000 | 21:21:21:21:2 | 11883336:118 | 11883487:118 | +:+:+:+:+:+:+: | 2239 | dcaf12 | 599 | 424 | 458 | 388 | 313 | 432 |
| ENSDARG000 | 21;21;21;21 | 6197223:6199 | 6197417:6199 | +;+;+;+ | 701 | NA | 0 | 16 | 18 | 0 | 0 | 6 |
| ENSDARG000 | 21:21:21:21:2 | 11865972:118 | 11866134:118 | +;+;+;+;+;+;+ | 4989 | ubap1 | 1235 | 1097 | 1197 | 1049 | 1274 | 1124 |
| ENSDARG000 | 21:21:21:21:2 | 25588436:255 | 25590382:255 | -:-:-:-:-:-:-:-: | 4011 | efemp2b | 92 | 32 | 30 | 125 | 88 | 40 |
| ENSDARG000 | 21:21:21:21:2 | 4364843:4364 | 4365013:4365 | -:-:-:-:-:-:-:-: | 1974 | NA | 0 | 0 | 0 | 0 | 0 | 0 |
| ENSDARG000 | 21:21:21:21:2 | 4384702:4384 | 4384872:4385 | -;-;-;-;-;- | 1642 | NA | 0 | 0 | 0 | 0 | 0 | 0 |
| ENSDARG000 | 21:21:21:21:2 | 45238533:452 | 45238607:452 | +;+;+;+;+;+;+ | 526 | NA | 20 | 0 | 10 | 6 | 1 | 32 |
| ENSDARG000 | 21:21:21:21:2 | 22330005:223 | 22330377:223 | +:+:+:+:+:+:+: | 2115 | nadk2 | 239 | 203 | 328 | 321 | 354 | 236 |
| ENSDARG000 | 21:21:21:21:2 | 1990905:1996 | 1991194:1996 | -:-:-:-:-:-:-:-: | 2319 | add1 | 6796 | 6784 | 7785 | 6512 | 8635 | 7345 |
| ENSDARG000 | 21:21:21:21:2 | 45267589:452 | 45268343:452 | +:+:+:+:+:+:+: | 2046 | tcf7 | 14 | 20 | 0 | 0 | 14 | 35 |
| ENSDARG000 | 21:21:21:21:2 | 10828828:108 | 10828996:108 | +:+:+:+:+:+:+: | 2263 | NA | 4 | 5 | 104 | 10 | 64 | 7 |
| ENSDARG000 | 21:21:21:21:2 | 6206215:6206 | 6206313:6206 | +:+:+:+:+:+:+: | 729 | NA | 0 | 39 | 26 | 0 | 0 | 4 |
| ENSDARG000 | 21:21:21:21:2 | 31150438:311 | 31150507:311 | +:+:+:+:+:+:+: | 2647 | st6gal1 | 0 | 0 | 0 | 0 | 0 | 0 |
| ENSDARG000 | 21:21:21:21:2 | 27513859:275 | 27514234:275 | +:+:+:+:+:+:+: | 3396 | pacs1a | 1217 | 727 | 708 | 962 | 519 | 1353 |
| ENSDARG000 | 21:21:21:21:2 | 20914888:209 | 20915123:209 | -:-:-:-:-:-:-:-: | 3988 | c6 | 207 | 66 | 68 | 141 | 105 | 136 |
| ENSDARG000 | 21;21;21;21 | 6212844:6291 | 6213059:6291 | +;+;+;+ | 531 | NA | 35 | 18 | 27 | 12 | 11 | 15 |
| ENSDARG000 | 21:21:21:21:2 | 6290566:6291 | 6290604:6291 | +:+:+:+:+:+:+: | 2101 | FNBP1 | 1531 | 591 | 820 | 772 | 1024 | 817 |
| ENSDARG000 | 21:21:21:21:2 | 20901505:209 | 20901544:209 | +:+:+:+:+:+:+: | 2612 | c7b | 2731 | 847 | 543 | 627 | 18 | 334 |
| ENSDARG000 | 21:21:21:21:2 | 4388489:4395 | 4388612:4395 | +:+:+:+:+:+:+: | 1433 | NA | 2 | 4 | 16 | 0 | 0 | 3 |
| ENSDARG000 | 21:21:21:21:2 | 45341786:453 | 45342335:453 | -:-:-:-:-:-:-:-: | 1116 | skp1 | 6165 | 5329 | 6509 | 5539 | 6306 | 6138 |
| ENSDARG000 | 21:21:21:21:2 | 11916788:119 | 11916865:119 | +:+:+:+:+:+:+: | 4252 | ubap2a | 7311 | 5590 | 7966 | 7010 | 6775 | 5870 |
| ENSDARG000 | 21:21:21:21:2 | 27558381:275 | 27558516:275 | +:+:+:+:+:+:+: | 3925 | zgc:165604 | 0 | 0 | 0 | 0 | 2 | 0 |
| ENSDARG000 | 21:21:21:21:2 | 10866421:108 | 10866826:108 | +:+:+:+:+:+:+: | 5234 | NA | 8 | 0 | 6 | 6 | 2 | 16 |
| ENSDARG000 | 21;21 | 27558490:275 | 27558546:275 | -;- | 367 | NA | 0 | 0 | 0 | 0 | 0 | 0 |

| | | | | | | | | | | | | |
|---|---|---|---|---|---|---|---|---|---|---|---|---|
| ENSDARG000 | 21:21:21:21:2 | 25605368:256 | 25605680:256 | -:-:-:-:-:-:-:-:- | 1663 | fibpb | 4 | 0 | 29 | 26 | 15 | 21 |
| ENSDARG000 | 21:21:21:21:2 | 22347120:223 | 22348118:223 | -:-:-:-:-:-:-:-:- | 2345 | skp2 | 139 | 373 | 359 | 132 | 214 | 153 |
| ENSDARG000 | 21:21:21:21:2 | 31194980:311 | 31197622:311 | -:-:-:-:-:-:-:-:- | 5040 | zgc:152891 | 22 | 58 | 90 | 58 | 100 | 134 |
| ENSDARG000 | 21:21:21:21:2 | 45351660:453 | 45351770:453 | -:-:-:-:-:-:-:-:- | 1947 | ppp2cab | 2553 | 2354 | 2741 | 2100 | 2720 | 2008 |
| ENSDARG000 | 21:21:21:21:2 | 6394929:6397 | 6395067:6397 | +:+:+:+:+:+:+: | 3264 | NA | 172 | 449 | 572 | 209 | 465 | 307 |
| ENSDARG000 | 21;21;21;21 | 25625026:256 | 25625243:256 | +;+;+;+ | 976 | NA | 6 | 0 | 0 | 0 | 0 | 0 |
| ENSDARG000 | 21:21:21:21:2 | 25620209:256 | 25620597:256 | -;-;-;-;- | 1147 | tmem223 | 110 | 132 | 172 | 48 | 134 | 134 |
| ENSDARG000 | 21;21;21;21 | 25614427:256 | 25614756:256 | -;-;-;- | 928 | NA | 6 | 26 | 15 | 8 | 11 | 4 |
| ENSDARG000 | 21:21:21:21:2 | 27575758:275 | 27576021:275 | -:-:-:-:-:-:-:-:- | 7025 | pcnx3 | 241 | 137 | 291 | 196 | 222 | 213 |
| ENSDARG000 | 21;21;21;21 | 2091895:2094 | 2092063:2094 | +;+;+;+ | 1043 | NA | 6 | 0 | 0 | 0 | 2 | 0 |
| ENSDARG000 | 21:21:21:21:2 | 10883014:108 | 10883755:108 | -;-;-;-;- | 1026 | NA | 0 | 0 | 0 | 0 | 0 | 0 |
| ENSDARG000 | 21:21:21:21:2 | 4410733:4410 | 4410863:4410 | +:+:+:+:+:+:+: | 1819 | NA | 1 | 28 | 20 | 10 | 1 | 7 |
| ENSDARG000 | 21:21:21:21:2 | 4425164:4425 | 4425334:4425 | -;-;-;-;-;-;- | 1639 | NA | 0 | 0 | 0 | 0 | 0 | 0 |
| ENSDARG000 | 21:21:21:21:2 | 4445750:4446 | 4445920:4446 | -;-;-;-;-;-;- | 1642 | NA | 0 | 0 | 0 | 0 | 0 | 0 |
| ENSDARG000 | 21:21:21:21:2 | 4406942:4406 | 4407112:4407 | -:-:-:-:-:-:-:-:- | 1974 | NA | 0 | 0 | 0 | 0 | 0 | 0 |
| ENSDARG000 | 21:21:21:21:2 | 4429001:4439 | 4429139:4439 | +;+;+;+;+;+;+ | 1194 | NA | 0 | 5 | 3 | 3 | 0 | 1 |
| ENSDARG000 | 21:21:21:21:2 | 45366229:453 | 45366374:453 | +;+;+;+;+;+;+ | 2065 | ube2b | 198 | 142 | 261 | 180 | 141 | 325 |
| ENSDARG000 | 21:21:21:21:2 | 31232873:312 | 31233789:312 | -;-;-;-;- | 2309 | tpst1l | 453 | 487 | 812 | 889 | 894 | 404 |
| ENSDARG000 | 21:21:21:21:2 | 25643880:256 | 25645676:256 | +;+;+;+;+;+;+;+ | 1797 | NA | 0 | 0 | 0 | 0 | 3 | 0 |
| ENSDARG000 | 21:21:21:21:2 | 2103164:2103 | 2103216:2103 | -:-:-:-:-:-:-:-:- | 2092 | si:rp71-1h20.! | 73 | 46 | 56 | 46 | 78 | 36 |
| ENSDARG000 | 21:21:21:21:2 | 45377211:453 | 45377745:453 | -:-:-:-:-:-:-:-:- | 1429 | cdkn2aipnl | 210 | 286 | 426 | 202 | 300 | 194 |
| ENSDARG000 | 21;21 | 25660613:256 | 25660697:256 | +;+ | 454 | NA | 36 | 11 | 117 | 8 | 8 | 57 |
| ENSDARG000 | 21:21:21:21:2 | 25664393:256 | 25664548:256 | -:-:-:-:-:-:-:-:- | 821 | NA | 41 | 38 | 183 | 9 | 48 | 122 |
| ENSDARG000 | 21:21:21:21:2 | 31244500:312 | 31245017:312 | -;-;-;-;-;-;-;-;- | 1242 | crcp | 122 | 207 | 198 | 150 | 227 | 150 |
| ENSDARG000 | 21:21:21:21:2 | 22369869:223 | 22369966:223 | +;+;+;+;+;+;+;+ | 1014 | NA | 1 | 2 | 18 | 5 | 0 | 10 |
| ENSDARG000 | 21;21;21;21 | 22367312:223 | 22367477:223 | -;-;-;- | 600 | NA | 0 | 0 | 1 | 0 | 0 | 0 |
| ENSDARG000 | 21:21:21:21:2 | 18313152:183 | 18313403:183 | +:+:+:+:+:+:+: | 5030 | RIMBP2 | 2 | 0 | 0 | 0 | 0 | 0 |
| ENSDARG000 | 21;21 | 2133097:2133 | 2133205:2133 | -;- | 161 | NA | 0 | 0 | 0 | 0 | 1 | 0 |
| ENSDARG000 | 21;21 | 2141985:2142 | 2142099:2142 | -;- | 154 | NA | 9 | 3 | 10 | 2 | 10 | 0 |
| ENSDARG000 | 21:21:21:21:2 | 45386033:453 | 45386270:453 | +:+:+:+:+:+:+: | 3239 | JADE2 | 76 | 32 | 146 | 58 | 189 | 101 |
| ENSDARG000 | 21;21 | 2149976:2150 | 2150090:2150 | -;- | 177 | NA | 33 | 18 | 13 | 11 | 31 | 17 |
| ENSDARG000 | 21;21;21;21 | 45496442:454 | 45496910:454 | +;+;+;+ | 1020 | NA | 47 | 4 | 0 | 40 | 16 | 6 |
| ENSDARG000 | 21:21:21:21:2 | 6556635:6570 | 6556802:6570 | +:+:+:+:+:+:+: | 9656 | COL5A1 | 3656 | 2363 | 5599 | 2439 | 11252 | 1619 |
| ENSDARG000 | 21:21:21:21:2 | 15295685:152 | 15295711:153 | +:+:+:+:+:+:+: | 2985 | NA | 0 | 0 | 0 | 0 | 2 | 0 |
| ENSDARG000 | 21:21:21:21:2 | 20933354:209 | 20934053:209 | -:-:-:-:-:-:-:-:- | 1868 | rgs7bpb | 10 | 12 | 0 | 43 | 2 | 12 |
| ENSDARG000 | 21:21:21:21:2 | 2155695:2156 | 2156833:2156 | -;-;-;-;-;-;- | 1497 | NA | 64 | 51 | 73 | 29 | 51 | 62 |
| ENSDARG000 | 21:21:21:21:2 | 11969603:119 | 11969769:119 | +:+:+:+:+:+:+: | 661 | mlnl | 4 | 6 | 2 | 4 | 0 | 10 |
| ENSDARG000 | 21:21:21:21:2 | 31253048:312 | 31253214:312 | +:+:+:+:+:+:+: | 4782 | asl | 143 | 225 | 152 | 203 | 321 | 156 |
| ENSDARG000 | 21:21:21:21:2 | 25669934:256 | 25670148:256 | +:+:+:+:+:+:+: | 1252 | sumf2 | 88 | 131 | 137 | 150 | 150 | 127 |
| ENSDARG000 | 21:21:21:21:2 | 25676097:256 | 25677121:256 | -:-:-:-:-:-:-:-:- | 2175 | phkg1b | 0 | 0 | 2 | 8 | 0 | 2 |
| ENSDARG000 | 21:21:21:21:2 | 11958758:119 | 11959316:119 | -:-:-:-:-:-:-:-:- | 3316 | nop56 | 2561 | 4439 | 6853 | 2977 | 4269 | 4081 |
| ENSDARG000 | 21:21:21:21:2 | 27636288:276 | 27638306:276 | -:-:-:-:-:-:-:-:- | 8631 | nrxn2a | 6 | 0 | 6 | 0 | 8 | 26 |
| ENSDARG000 | 21:21:21:21:2 | 31275327:312 | 31277245:312 | -:-:-:-:-:-:-:-:- | 3387 | ca4c | 0 | 0 | 0 | 0 | 2 | 0 |
| ENSDARG000 | 21:21:21:21:2 | 45502621:455 | 45502817:455 | +:+:+:+:+:+:+: | 1438 | NA | 0 | 0 | 0 | 0 | 0 | 0 |
| ENSDARG000 | 21:21:21:21:2 | 22385125:223 | 22387845:223 | -:-:-:-:-:-:-:-:- | 5498 | NA | 455 | 190 | 222 | 293 | 414 | 554 |
| ENSDARG000 | 21:21:21:21:2 | 2170798:2170 | 2172680:2172 | -;-;-;-;- | 2154 | zgc:163077 | 105 | 54 | 94 | 92 | 55 | 163 |
| ENSDARG000 | 21:21:21:21:2 | 10890258:108 | 10892058:108 | -:-:-:-:-:-:-:-:- | 5009 | NEDD4L | 843 | 636 | 914 | 417 | 722 | 511 |
| ENSDARG000 | 21:21:21:21:2 | 2178909:2178 | 2180510:2180 | -;-;-;-;- | 1785 | zgc:171220 | 62 | 69 | 95 | 32 | 82 | 49 |
| ENSDARG000 | 21:21:21:21:2 | 18405585:184 | 18406662:184 | +:+:+:+:+:+:+: | 5217 | NA | 0 | 0 | 0 | 0 | 0 | 0 |
| ENSDARG000 | 21:21:21:21:2 | 45510448:455 | 45510709:455 | +:+:+:+:+:+:+: | 6621 | fnip1 | 2748 | 2208 | 4122 | 4322 | 3351 | 2838 |
| ENSDARG000 | 21:21:21:21:2 | 28415888:284 | 28415931:284 | -:-:-:-:-:-:-:-:- | 1804 | RASGRP2 | 10 | 4 | 6 | 6 | 18 | 52 |
| ENSDARG000 | 21:21:21:21:2 | 22400951:224 | 22401001:224 | +:+:+:+:+:+:+: | 3949 | lmbrd2b | 1035 | 1002 | 1196 | 1368 | 1025 | 1448 |
| ENSDARG000 | 21:21:21:21:2 | 18597927:186 | 18603111:186 | -:-:-:-:-:-:-:-:- | 8138 | TMEM132C | 0 | 8 | 0 | 0 | 0 | 0 |
| ENSDARG000 | 21:21:21:21:2 | 20940022:209 | 20940674:209 | -;-;-;-;-;-;-;- | 2522 | rnf180b | 157 | 126 | 112 | 108 | 165 | 154 |
| ENSDARG000 | 21;21;21 | 2235947:2237 | 2236412:2237 | -;-;- | 696 | NA | 40 | 18 | 72 | 26 | 25 | 30 |
| ENSDARG000 | 21:21:21:21:2 | 2202270:2203 | 2203497:2203 | -;-;-;-;- | 1639 | zgc:162971 | 146 | 71 | 105 | 90 | 179 | 88 |
| ENSDARG000 | 21:21:21:21:2 | 2219724:2220 | 2221051:2221 | -:-:-:-:-:-:-:-:- | 1827 | zgc:113343 | 44 | 45 | 48 | 57 | 24 | 66 |
| ENSDARG000 | 21;21 | 2212645:2216 | 2213916:2217 | -;- | 2395 | NA | 231 | 182 | 83 | 142 | 152 | 148 |
| ENSDARG000 | 21:21:21:21:2 | 28445052:284 | 28445361:284 | +:+:+:+:+:+:+: | 2735 | pygma | 0 | 0 | 12 | 0 | 8 | 0 |
| ENSDARG000 | 21:21:21:21:2 | 18877130:188 | 18877180:188 | +:+:+:+:+:+:+: | 1668 | si:dkev-65l23. | 0 | 0 | 0 | 0 | 0 | 0 |
| ENSDARG000 | 21:21:21:21:2 | 18866976:188 | 18867339:188 | -;-;-;-;-;-;-;- | 928 | NA | 0 | 0 | 0 | 0 | 0 | 0 |
| ENSDARG000 | 21:21:21:21:2 | 15312542:153 | 15312567:153 | +:+:+:+:+:+:+: | 3019 | NA | 4 | 0 | 0 | 0 | 0 | 0 |
| ENSDARG000 | 21:21:21:21:2 | 15332508:153 | 15332534:153 | +;+;+;+;+ | 1524 | NA | 0 | 0 | 0 | 0 | 0 | 0 |
| ENSDARG000 | 21:21:21:21:2 | 25688388:256 | 25688775:256 | +:+:+:+:+:+:+: | 1786 | NA | 26 | 26 | 4 | 8 | 7 | 6 |
| ENSDARG000 | 21:21:21:21:2 | 31434251:315 | 31434598:315 | +:+:+:+:+:+:+: | 11720 | TENM2 | 4 | 0 | 2 | 0 | 2 | 12 |
| ENSDARG000 | 21;21 | 28496194:285 | 28496419:285 | +;+ | 599 | cox8a | 1633 | 1645 | 2023 | 1537 | 1439 | 1318 |
| ENSDARG000 | 21;21;21 | 20949976:209 | 20951413:209 | +;+;+ | 1438 | htr1ab | 0 | 2 | 2 | 0 | 0 | 0 |
| ENSDARG000 | 21:21:21:21:2 | 28478663:284 | 28478959:284 | +:+:+:+:+:+:+: | 3108 | slc22a6l | 0 | 6 | 35 | 4 | 24 | 0 |
| ENSDARG000 | 21;21;21 | 31434308:315 | 31434598:315 | +;+;+ | 815 | NA | 0 | 0 | 0 | 0 | 0 | 0 |
| ENSDARG000 | 21:21:21:21:2 | 28493077:284 | 28493609:284 | -;-;-;-;-;-;-;- | 874 | NA | 0 | 0 | 0 | 0 | 0 | 0 |
| ENSDARG000 | 21:21:21:21:2 | 22423286:224 | 22423378:224 | +;+;+;+;+;+ | 1112 | capslb | 0 | 0 | 0 | 0 | 8 | 0 |
| ENSDARG000 | 21:21:21:21:2 | 25703078:257 | 25703320:257 | -;-;-;-;-;-;- | 1310 | abhd11 | 227 | 164 | 185 | 94 | 154 | 96 |
| ENSDARG000 | 21:21:21:21:2 | 21010786:210 | 21011241:210 | +;+;+;+;+ | 1592 | ndr1 | 0 | 0 | 0 | 0 | 0 | 0 |
| ENSDARG000 | 21:21:21:21:2 | 4508959:4508 | 4509106:4509 | +:+:+:+:+:+:+: | 1525 | phyhd1 | 872 | 1167 | 1925 | 808 | 1789 | 1364 |
| ENSDARG000 | 21:21:21:21:2 | 15346363:153 | 15346749:153 | +;+;+;+;+;+;+ | 2572 | NA | 0 | 0 | 0 | 0 | 0 | 0 |
| ENSDARG000 | 21:21:21:21:2 | 21013147:210 | 21013882:210 | -;-;-;-;-;- | 1542 | eif4ebp1 | 1978 | 1970 | 2165 | 2028 | 1838 | 1756 |
| ENSDARG000 | 21:21:21:21:2 | 18929319:189 | 18929618:189 | -:-:-:-:-:-:-:-:- | 2867 | med15 | 1032 | 736 | 1064 | 836 | 857 | 766 |
| ENSDARG000 | 21:21:21:21:2 | 6751405:6751 | 6751873:6751 | +:+:+:+:+:+:+: | 2759 | olfm1b | 0 | 0 | 4 | 0 | 6 | 0 |
| ENSDARG000 | 21;21;21 | 25739295:257 | 25741111:257 | -;-;- | 2018 | cldn3c | 66 | 8 | 32 | 24 | 70 | 24 |
| ENSDARG000 | 21;21 | 25747063:257 | 25747574:257 | -;- | 733 | NA | 22 | 26 | 24 | 32 | 39 | 63 |
| ENSDARG000 | 21:21:21:21:2 | 18907102:189 | 18907255:189 | +:+:+:+:+:+:+: | 4804 | smpd4 | 922 | 1080 | 682 | 1134 | 460 | 667 |
| ENSDARG000 | 21;21;21;21 | 15375785:153 | 15376147:153 | +;+;+;+ | 1461 | NA | 0 | 0 | 0 | 0 | 0 | 0 |
| ENSDARG000 | 21 | 25754268 | 25756119 | - | 1852 | cldn3d | 28 | 0 | 6 | 0 | 20 | 0 |
| ENSDARG000 | 21 | 25765734 | 25766840 | + | 1107 | cldnb | 1283 | 54 | 16 | 178 | 286 | 100 |
| ENSDARG000 | 21 | 25773530 | 25774605 | - | 1076 | cldn29 | 2 | 0 | 10 | 0 | 0 | 2 |
| ENSDARG000 | 21;21;21 | 2246339:2247 | 2247167:2247 | -;-;- | 1034 | NA | 47 | 33 | 56 | 21 | 53 | 60 |
| ENSDARG000 | 21:21:21:21:2 | 11105007:111 | 11105116:111 | +:+:+:+:+:+:+: | 6290 | prlra | 8 | 0 | 21 | 14 | 56 | 6 |
| ENSDARG000 | 21 | 25777425 | 25778201 | + | 777 | cldnd | 0 | 0 | 0 | 0 | 0 | 0 |
| ENSDARG000 | 21;21 | 2259700:2261 | 2260504:2261 | -;- | 899 | NA | 0 | 2 | 8 | 18 | 2 | 0 |
| ENSDARG000 | 21 | 25793970 | 25795968 | + | 1999 | cldn30 | 18 | 4 | 10 | 0 | 18 | 36 |
| ENSDARG000 | 21:21:21:21:2 | 18925318:189 | 18925598:189 | +:+:+:+:+:+:+: | 1233 | si:ch211-222r | 6 | 0 | 0 | 0 | 1 | 0 |
| ENSDARG000 | 21:21:21:21:2 | 25801345:258 | 25801576:258 | +:+:+:+:+:+:+: | 5941 | nf2b | 574 | 412 | 512 | 318 | 434 | 510 |
| ENSDARG000 | 21:21:21:21:2 | 15388351:153 | 15388380:153 | +;+;+;+;+;+ | 1560 | NA | 0 | 0 | 0 | 0 | 0 | 0 |
| ENSDARG000 | 21:21:21:21:2 | 15435946:154 | 15436340:154 | +;+;+;+;+ | 1077 | si:dkey-11o15 | 0 | 0 | 0 | 0 | 0 | 0 |
| ENSDARG000 | 21:21:21:21:2 | 25795875:257 | 25796669:257 | -:-:-:-:-:-:-:-:- | 1426 | mettl27 | 29 | 22 | 12 | 4 | 62 | 24 |
| ENSDARG000 | 21:21:21:21:2 | 28502340:285 | 28502406:285 | +:+:+:+:+:+:+: | 2009 | otub1a | 633 | 556 | 776 | 504 | 583 | 528 |
| ENSDARG000 | 21:21:21:21:2 | 11244068:112 | 11244209:112 | +:+:+:+:+:+:+: | 2211 | arid6 | 81 | 3 | 2 | 8 | 1 | 14 |
| ENSDARG000 | 21;21 | 2269346:2273 | 2270047:2273 | -;- | 808 | NA | 16 | 31 | 57 | 22 | 39 | 10 |
| ENSDARG000 | 21:21:21:21:2 | 22429509:224 | 22430214:224 | -;-;-;-;-;-;- | 1617 | il7r | 14 | 12 | 2 | 4 | 42 | 38 |
| ENSDARG000 | 21;21;21 | 2280950:2284 | 2281592:2284 | -;-;- | 779 | NA | 92 | 80 | 158 | 82 | 93 | 74 |
| ENSDARG000 | 21:21:21:21:2 | 21027375:210 | 21027772:210 | -:-:-:-:-:-:-:-:- | 8086 | ANK1 | 52 | 48 | 45 | 32 | 15 | 52 |
| ENSDARG000 | 21;21;21 | 2289254:2297 | 2289995:2297 | -;-;- | 919 | si:ch73-299h1 | 79 | 66 | 88 | 114 | 100 | 160 |
| ENSDARG000 | 21:21:21:21:2 | 11256425:112 | 11257235:112 | -:-:-:-:-:-:-:-:- | 2506 | rtknb | 1177 | 322 | 479 | 461 | 251 | 452 |
| ENSDARG000 | 21:21:21:21:2 | 2302156:2304 | 2305610:2305 | -;-;-;-;- | 4443 | zgc:110688 | 120 | 108 | 130 | 47 | 106 | 94 |
| ENSDARG000 | 21:21:21:21:2 | 28513057:285 | 28513284:285 | -;-;-;-;-;-;-;- | 840 | NA | 0 | 2 | 42 | 2 | 48 | 0 |
| ENSDARG000 | 21:21:21:21:2 | 11367253:113 | 11367630:113 | +:+:+:+:+:+:+: | 1856 | GTF3C5 | 256 | 206 | 412 | 186 | 255 | 290 |
| ENSDARG000 | 21:21:21:21:2 | 11330234:113 | 11330684:113 | -:-:-:-:-:-:-:-:- | 5371 | zgc:162472 | 494 | 344 | 880 | 544 | 634 | 460 |
| ENSDARG000 | 21;21 | 4534156:4539 | 4535908:4539 | -;- | 1869 | dolk | 90 | 76 | 125 | 29 | 157 | 91 |
| ENSDARG000 | 21:21:21:21:2 | 22446240:224 | 22449147:224 | -:-:-:-:-:-:-:-:- | 8531 | myo5b | 18542 | 6504 | 4542 | 8232 | 5013 | 17707 |
| ENSDARG000 | 21;21;21 | 2316058:2318 | 2316811:2318 | -;-;- | 968 | zgc:66483 | 0 | 0 | 2 | 0 | 0 | 0 |
| ENSDARG000 | 21:21:21:21:2 | 28554716:285 | 28555850:285 | -:-:-:-:-:-:-:-:- | 2952 | nrg2a | 10 | 20 | 11 | 2 | 4 | 2 |
| ENSDARG000 | 21;21;21 | 2327338:2328 | 2328517:2328 | -;-;- | 1415 | zgc:113348 | 236 | 272 | 336 | 186 | 275 | 262 |
| ENSDARG000 | 21;21;21 | 32052080:320 | 32052488:320 | -;-;- | 833 | si:ch211-160i | 16 | 15 | 49 | 20 | 4 | 242 |
| ENSDARG000 | 21;21;21 | 28724099:287 | 28724266:287 | +;+;+ | 1264 | puraa | 257 | 275 | 233 | 89 | 218 | 222 |
| ENSDARG000 | 21:21:21:21:2 | 15592426:156 | 15592606:156 | +:+:+:+:+:+:+: | 5089 | smarcb1b | 1329 | 365 | 398 | 931 | 391 | 485 |
| ENSDARG000 | 21;21;21 | 2336289:2338 | 2337226:2338 | -;-;- | 1095 | zgc:193790 | 4 | 12 | 14 | 8 | 12 | 8 |
| ENSDARG000 | 21:21:21:21:2 | 4647784:4649 | 4647989:4649 | -:-:-:-:-:-:-:-:- | 3835 | cdk5rap2 | 215 | 148 | 305 | 127 | 311 | 192 |
| ENSDARG000 | 21:21:21:21:2 | 45880596:458 | 45880779:458 | -;-;-;-;-;-;- | 1303 | galnt10 | 241 | 112 | 204 | 116 | 215 | 109 |
| ENSDARG000 | 21:21:21:21:2 | 25932107:259 | 25932334:259 | +:+:+:+:+:+:+: | 1452 | NA | 7 | 4 | 2 | 0 | 8 | 30 |
| ENSDARG000 | 21:21:21:21:2 | 45897814:458 | 45898273:458 | +:+:+:+:+:+:+: | 2055 | slc22a4 | 2 | 2 | 26 | 4 | 70 | 20 |
| ENSDARG000 | 21;21;21 | 26021496:260 | 26023295:260 | -;-;- | 2200 | sdf2 | 642 | 557 | 783 | 384 | 552 | 662 |
| ENSDARG000 | 21;21;21;21 | 2342856:2344 | 2343338:2344 | -;-;-;- | 750 | NA | 50 | 34 | 41 | 52 | 37 | 50 |
| ENSDARG000 | 21:21:21:21:2 | 32061725:320 | 32065729:320 | -:-:-:-:-:-:-:-:- | 5259 | mat2b | 238 | 258 | 292 | 254 | 212 | 195 |
| ENSDARG000 | 21:21:21:21:2 | 28747069:287 | 28747325:287 | +:+:+:+:+:+:+: | 2773 | zgc:100829 | 1402 | 1525 | 2276 | 2534 | 1150 | 1658 |
| ENSDARG000 | 21;21 | 2359866:2360 | 2360576:2360 | -;- | 869 | NA | 27 | 47 | 60 | 62 | 57 | 43 |
| ENSDARG000 | 21;21 | 2368531:2375 | 2369431:2376 | -;- | 1047 | si:ch211-241b | 32 | 30 | 23 | 73 | 32 | 24 |
| ENSDARG000 | 21:21:21:21:2 | 45904956:459 | 45905442:459 | +:+:+:+:+:+:+: | 6316 | slc22a21 | 253 | 71 | 143 | 118 | 188 | 320 |
| ENSDARG000 | 21:21:21:21:2 | 2395224:2398 | 2395585:2398 | -;-;-;-;-;-;- | 1396 | NA | 1880 | 2194 | 1815 | 2148 | 1258 | 1770 |
| ENSDARG000 | 21:21:21:21:2 | 28863121:288 | 28864489:288 | -;-;-;-;- | 2670 | CXXC5 | 4011 | 4524 | 4096 | 3855 | 6669 | 4358 |
| ENSDARG000 | 21:21:21:21:2 | 11984805:119 | 11985488:119 | -;-;-;-;-;-;- | 1941 | zgc:64106 | 0 | 0 | 10 | 2 | 25 | 10 |
| ENSDARG000 | 21:21:21:21:2 | 11385031:113 | 11385118:113 | +:+:+:+:+:+:+: | 1825 | CEL | 0 | 10 | 23 | 4 | 0 | 11 |
| ENSDARG000 | 21:21:21:21:2 | 32089491:320 | 32092102:320 | -:-:-:-:-:-:-:-:- | 5757 | NA | 2 | 0 | 5 | 10 | 26 | 0 |
| ENSDARG000 | 21:21:21:21:2 | 15636540:156 | 15636667:156 | -;-;-;-;-;-;- | 1590 | NA | 4 | 0 | 4 | 0 | 4 | 3 |
| ENSDARG000 | 21:21:21:21:2 | 32092771:320 | 32092904:320 | +;+;+;+;+;+;+ | 865 | NA | 0 | 0 | 0 | 0 | 0 | 0 |
| ENSDARG000 | 21:21:21:21:2 | 26028947:260 | 26029078:260 | +:+:+:+:+:+:+: | 6394 | supt6h | 2911 | 2035 | 3373 | 2081 | 3069 | 2612 |
| ENSDARG000 | 21:21:21:21:2 | 12010505:120 | 12010720:120 | +;+;+;+;+;+ | 2461 | aqp7 | 71 | 534 | 359 | 223 | 353 | 1045 |
| ENSDARG000 | 21:21:21:21:2 | 28946015:289 | 28946290:289 | -:-:-:-:-:-:-:-:- | 3281 | NA | 6 | 0 | 0 | 0 | 2 | 0 |
| ENSDARG000 | 21:21:21:21:2 | 4702773:4702 | 4702837:4703 | +;+;+;+;+ | 838 | NA | 92 | 26 | 42 | 63 | 20 | 197 |
| ENSDARG000 | 21:21:21:21:2 | 2506013:2508 | 2506078:2508 | +:+:+:+:+:+:+: | 3424 | hmgcrb | 1354 | 1648 | 2239 | 1963 | 1664 | 1413 |
| ENSDARG000 | 21:21:21:21:2 | 21167837:211 | 21171176:211 | -:-:-:-:-:-:-:-:- | 4505 | ftsj1 | 253 | 191 | 328 | 188 | 238 | 174 |
| ENSDARG000 | 21:21:21:21:2 | 28958471:289 | 28958739:289 | +:+:+:+:+:+:+: | 3509 | ppp3ca | 66 | 175 | 286 | 807 | 18 | 513 |
| ENSDARG000 | 21:21:21:21:2 | 32280175:322 | 32281223:322 | -:-:-:-:-:-:-:-:- | 3968 | clk4b | 546 | 393 | 775 | 785 | 624 | 787 |
| ENSDARG000 | 21:21:21:21:2 | 15704556:157 | 15704779:157 | +;+;+;+;+;+;+ | 857 | chchd10 | 106 | 156 | 233 | 96 | 231 | 124 |
| ENSDARG000 | 21:21:21:21:2 | 15709061:157 | 15709258:157 | +;+;+;+;+ | 1224 | ddt | 457 | 521 | 808 | 240 | 704 | 376 |
| ENSDARG000 | 21:21:21:21:2 | 12023796:120 | 12024143:120 | -:-:-:-:-:-:-:-:- | 1256 | tpgs2 | 64 | 202 | 144 | 87 | 164 | 207 |
| ENSDARG000 | 21:21:21:21:2 | 6953538:6953 | 6953650:6953 | +:+:+:+:+:+:+: | 857 | NA | 6 | 2 | 2 | 0 | 0 | 10 |
| ENSDARG000 | 21;21;21 | 18950575:189 | 18951170:189 | +;+;+ | 2780 | crkl | 862 | 975 | 1104 | 1048 | 1157 | 949 |
| ENSDARG000 | 21:21:21:21:2 | 18970857:189 | 18970950:189 | -;-;-;-;- | 796 | NA | 0 | 0 | 0 | 0 | 0 | 0 |
| ENSDARG000 | 21:21:21:21:2 | 15713097:157 | 15713310:157 | +;+;+;+;+ | 1333 | gstt1b | 62 | 32 | 238 | 228 | 30 | 788 |
| ENSDARG000 | 21:21:21:21:2 | 4712134:4712 | 4713354:4713 | -:-:-:-:-:-:-:-:- | 5746 | camsap1a | 8 | 76 | 26 | 24 | 6 | 11 |
| ENSDARG000 | 21;21;21 | 6961635:6961 | 6961686:6961 | +;+;+ | 423 | NA | 8 | 0 | 13 | 2 | 11 | 1 |
| ENSDARG000 | 21;21;21 | 7026893:7030 | 7027464:7030 | -;-;- | 820 | NA | 0 | 2 | 2 | 8 | 4 | 5 |
| ENSDARG000 | 21:21:21:21:2 | 22589577:225 | 22590226:225 | -:-:-:-:-:-:-:-:- | 1813 | nectin1a | 20 | 7 | 10 | 2 | 1 | 8 |
| ENSDARG000 | 21:21:21:21:2 | 26062452:260 | 26062857:260 | -:-:-:-:-:-:-:-:- | 1903 | NA | 11741 | 8553 | 8324 | 8363 | 9762 | 13168 |
| ENSDARG000 | 21:21:21:21:2 | 22630297:226 | 22630393:226 | +:+:+:+:+:+:+: | 1959 | NA | 44 | 2 | 36 | 18 | 18 | 2 |
| ENSDARG000 | 21:21:21:21:2 | 32332616:323 | 32332791:323 | +:+:+:+:+:+:+: | 6668 | NA | 642 | 630 | 635 | 304 | 885 | 490 |
| ENSDARG000 | 21;21 | 7947630:7953 | 7948822:7953 | -;- | 1255 | F2R | 0 | 0 | 0 | 0 | 0 | 4 |
| ENSDARG000 | 21;21;21;21 | 7921717:7921 | 7923065:7923 | -;-;-;- | 1493 | f2rl1.2 | 131 | 7 | 2 | 4 | 28 | 14 |
| ENSDARG000 | 21:21:21:21:2 | 32349190:323 | 32349537:323 | -:-:-:-:-:-:-:-:- | 1926 | MAPK9 | 279 | 313 | 253 | 105 | 276 | 196 |
| ENSDARG000 | 21:21:21:21:2 | 7934573:7935 | 7936031:7936 | -;-;-;-;- | 1695 | f2rl1.1 | 10 | 0 | 0 | 0 | 0 | 0 |
| ENSDARG000 | 21:21:21:21:2 | 11401247:114 | 11401332:114 | +:+:+:+:+:+:+: | 1994 | cel.1 | 6 | 0 | 0 | 4 | 0 | 0 |
| ENSDARG000 | 21:21:21:21:2 | 29077509:290 | 29077774:290 | +:+:+:+:+:+:+: | 1295 | NA | 42 | 4 | 38 | 120 | 12 | 278 |

| | | | | | | | | | | | | |
|---|---|---|---|---|---|---|---|---|---|---|---|---|
| ENSDARG000 | 21:21:21:21:2 | 15723069:157 | 15723365:157 | +:+:+:+:+:+:+: | 1561 | p2rx4a | 8 | 4 | 48 | 2 | 8 | 28 |
| ENSDARG000 | 21:21:21:21:2 | 29090652:290 | 29091262:290 | -:-:-:-:-:-:-:-:-: | 1781 | timd4 | 18 | 4 | 0 | 12 | 26 | 30 |
| ENSDARG000 | 21;21;21 | 7972805:7976 | 7974158:7976 | -;-;- | 1549 | f2r2.1 | 0 | 0 | 0 | 0 | 0 | 0 |
| ENSDARG000 | 21 | 7982302 | 7983300 | - | 999 | NA | 0 | 0 | 0 | 0 | 0 | 0 |
| ENSDARG000 | 21:21:21:21:2 | 26071874:260 | 26071947:260 | +:+:+:+:+:+:+: | 1312 | rpl23a | 20190 | 17676 | 28193 | 32048 | 16544 | 28702 |
| ENSDARG000 | 21;21 | 7998530:8003 | 7999638:8003 | -;- | 1203 | F2R | 1 | 0 | 8 | 2 | 3 | 11 |
| ENSDARG000 | 21:21:21:21:2 | 2540163:2540 | 2540290:2540 | -:-:-:-:-:-:-:-:-: | 1593 | CERT1 | 5134 | 3922 | 4147 | 4484 | 2624 | 2981 |
| ENSDARG000 | 21;21;21 | 8006536:8008 | 8007709:8009 | -;-;- | 1366 | F2R | 2 | 0 | 2 | 0 | 0 | 0 |
| ENSDARG000 | 21:21:21:21:2 | 12036238:120 | 12036354:120 | +:+:+:+:+:+:+: | 2312 | zgc:162344 | 72 | 51 | 90 | 47 | 109 | 102 |
| ENSDARG000 | 21:21:21:21:2 | 18976623:189 | 18977018:189 | -:-:-:-:-:-:-:-:-: | 1995 | MPP4 | 0 | 0 | 2 | 10 | 0 | 0 |
| ENSDARG000 | 21:21:21:21:2 | 11415224:114 | 11415822:114 | +;+;+;+;+;+;+ | 1755 | zgc:92275 | 0 | 4 | 0 | 0 | 0 | 0 |
| ENSDARG000 | 21:21:21:21:2 | 4766244:4767 | 4768293:4768 | -:-:-:-:-:-:-:-:-: | 8382 | notch1a | 766 | 527 | 1149 | 594 | 1178 | 364 |
| ENSDARG000 | 21:21:21:21:2 | 22635404:226 | 22635706:226 | +;+;+;+;+;+;+ | 1015 | NA | 115 | 154 | 154 | 98 | 62 | 97 |
| ENSDARG000 | 21:21:21:21:2 | 22697059:226 | 22697220:226 | +:+:+:+:+:+:+: | 1782 | bud13 | 382 | 221 | 249 | 112 | 258 | 213 |
| ENSDARG000 | 21:21:21:21:2 | 21195487:211 | 21195535:211 | +:+:+:+:+:+:+: | 7816 | NA | 1046 | 675 | 771 | 728 | 812 | 804 |
| ENSDARG000 | 21 | 8027985 | 8028980 | - | 996 | NA | 0 | 0 | 0 | 0 | 0 | 0 |
| ENSDARG000 | 21;21;21;21 | 8046804:8051 | 8048093:8051 | -;-;-;- | 1589 | F2R | 22 | 0 | 0 | 0 | 0 | 0 |
| ENSDARG000 | 21:21:21:21:2 | 3419459:3421 | 3422635:3422 | -;-;-;-;- | 3757 | smad7 | 474 | 227 | 323 | 263 | 434 | 99 |
| ENSDARG000 | 21:21:21:21:2 | 29100972:291 | 29102691:291 | -:-:-:-:-:-:-:-:-: | 3278 | havcr1 | 26 | 64 | 177 | 438 | 280 | 123 |
| ENSDARG000 | 21 | 8058252 | 8059239 | - | 988 | NA | 0 | 0 | 0 | 0 | 0 | 0 |
| ENSDARG000 | 21;21;21;21 | 26080215:260 | 26080771:260 | -;-;-;- | 1076 | tlcd1 | 13 | 0 | 2 | 27 | 12 | 8 |
| ENSDARG000 | 21:21:21:21:2 | 12079878:120 | 12080191:120 | -:-:-:-:-:-:-:-:-: | 2248 | celf4 | 0 | 0 | 0 | 0 | 0 | 0 |
| ENSDARG000 | 21;21;21;21 | 8076666:8081 | 8077994:8081 | -;-;-;- | 2038 | F2R | 38 | 14 | 20 | 3 | 24 | 14 |
| ENSDARG000 | 21:21:21:21:2 | 32390707:323 | 32390796:323 | -:-:-:-:-:-:-:-:-: | 2175 | gfpt2 | 18 | 6 | 32 | 17 | 8 | 3 |
| ENSDARG000 | 21:21:21:21:2 | 15733880:157 | 15733993:157 | +:+:+:+:+:+:+: | 1616 | idh3b | 1870 | 2302 | 2135 | 1997 | 1583 | 1719 |
| ENSDARG000 | 21:21:21:21:2 | 26097732:260 | 26099841:261 | -;-;-;-;-;- | 2973 | nipal4 | 38 | 2 | 4 | 8 | 6 | 32 |
| ENSDARG000 | 21;21;21 | 12343684:123 | 12343721:123 | +;+;+ | 666 | NA | 0 | 0 | 0 | 0 | 0 | 0 |
| ENSDARG000 | 21;21;21 | 19013850:190 | 19015785:190 | -;-;- | 3166 | nefma | 0 | 0 | 0 | 0 | 0 | 0 |
| ENSDARG000 | 21:21:21:21:2 | 18997503:189 | 18997554:189 | +:+:+:+:+:+:+: | 2264 | rpl17 | 14179 | 12084 | 21584 | 20206 | 13130 | 20884 |
| ENSDARG000 | 21;21 | 12475055:124 | 12475098:124 | -;- | 221 | NA | 0 | 0 | 0 | 0 | 0 | 0 |
| ENSDARG000 | 21;21;21 | 12584756:125 | 12585098:125 | -;-;- | 640 | NA | 0 | 0 | 1 | 0 | 0 | 0 |
| ENSDARG000 | 21:21:21:21:2 | 22706253:227 | 22706520:227 | -:-:-:-:-:-:-:-:-: | 991 | NA | 701 | 38 | 4 | 4 | 264 | 2 |
| ENSDARG000 | 21:21:21:21:2 | 3476207:3477 | 3476344:3478 | -:-:-:-:-:-:-:-:-: | 2286 | DYM | 237 | 220 | 301 | 156 | 324 | 237 |
| ENSDARG000 | 21:21:21:21:2 | 32454896:324 | 32458855:324 | -:-:-:-:-:-:-:-:-: | 5084 | zgc:123105 | 1159 | 695 | 1231 | 910 | 1398 | 693 |
| ENSDARG000 | 21:21:21:21:2 | 32476709:324 | 32478259:324 | -;-;-;-;-;-;-;- | 1836 | NA | 0 | 2 | 0 | 6 | 22 | 2 |
| ENSDARG000 | 21:21:21:21:2 | 13045388:130 | 13051108:130 | -;-;-;-;- | 6273 | MYORG | 472 | 310 | 412 | 380 | 457 | 443 |
| ENSDARG000 | 21:21:21:21:2 | 32508908:325 | 32513768:325 | -:-:-:-:-:-:-:-:-: | 8104 | NA | 44 | 4 | 4 | 0 | 26 | 2 |
| ENSDARG000 | 21:21:21:21:2 | 19001669:190 | 19001907:190 | -;-;-;-;-;- | 1037 | pgam2 | 0 | 8 | 2 | 32 | 4 | 26 |
| ENSDARG000 | 21:21:21:21:2 | 19008168:190 | 19009211:190 | +:+:+:+:+:+:+: | 2798 | NA | 10 | 20 | 30 | 173 | 20 | 91 |
| ENSDARG000 | 21:21:21:21:2 | 4875473:4875 | 4875572:4875 | -:-:-:-:-:-:-:-:-: | 3616 | KCNT1 | 0 | 0 | 0 | 0 | 2 | 0 |
| ENSDARG000 | 21;21;21 | 32464568:324 | 32464747:324 | +;+;+ | 777 | NA | 0 | 0 | 0 | 0 | 0 | 0 |
| ENSDARG000 | 21:21:21:21:2 | 21244046:212 | 21244199:212 | +:+:+:+:+:+:+: | 4040 | lifrb | 448 | 522 | 751 | 478 | 122 | 216 |
| ENSDARG000 | 21:21:21:21:2 | 26152332:261 | 26153251:261 | -:-:-:-:-:-:-:-:-: | 3931 | gria1b | 0 | 0 | 2 | 0 | 6 | 0 |
| ENSDARG000 | 21;21;21 | 22712307:227 | 22712966:227 | -;-;- | 940 | c1qb | 56 | 76 | 141 | 80 | 368 | 232 |
| ENSDARG000 | 21:21:21:21:2 | 13066705:130 | 13066949:130 | -:-:-:-:-:-:-:-:-: | 1961 | zgc:109965 | 4 | 12 | 2 | 10 | 0 | 10 |
| ENSDARG000 | 21;21 | 22717824:227 | 22718504:227 | -;- | 930 | c1qc | 12 | 128 | 192 | 42 | 354 | 234 |
| ENSDARG000 | 21;21;21 | 22721957:227 | 22722806:227 | -;-;- | 1133 | c1qa | 2 | 38 | 120 | 50 | 146 | 128 |
| ENSDARG000 | 21;21;21;21 | 22731566:227 | 22732012:227 | -;-;-;- | 2418 | FBXO40 | 0 | 0 | 0 | 0 | 0 | 0 |
| ENSDARG000 | 21;21;21 | 22726759:227 | 22727091:227 | -;-;- | 2163 | fbxo40.1 | 0 | 0 | 0 | 0 | 0 | 0 |
| ENSDARG000 | 21:21:21:21:2 | 21263988:212 | 21264174:212 | +:+:+:+:+:+:+: | 2591 | ccdc61 | 137 | 234 | 162 | 74 | 150 | 130 |
| ENSDARG000 | 21:21:21:21:2 | 29112053:291 | 29112797:291 | -:-:-:-:-:-:-:-:-: | 1984 | NA | 4 | 6 | 2 | 6 | 26 | 42 |
| ENSDARG000 | 21:21:21:21:2 | 3618519:3619 | 3619313:3619 | -:-:-:-:-:-:-:-:-: | 4543 | ATP8B1 | 102 | 4 | 2 | 8 | 16 | 6 |
| ENSDARG000 | 21:21:21:21:2 | 26389391:263 | 26389483:263 | +;+;+;+;+ | 1895 | tmsb | 2 | 0 | 0 | 0 | 2 | 0 |
| ENSDARG000 | 21;21;21;21 | 26252794:263 | 26253544:263 | -;-;-;- | 1311 | NA | 0 | 0 | 6 | 0 | 0 | 0 |
| ENSDARG000 | 21:21:21:21:2 | 22738939:227 | 22739335:227 | +:+:+:+:+:+:+: | 3990 | arhgap42a | 746 | 400 | 481 | 262 | 252 | 399 |
| ENSDARG000 | 21:21:21:21:2 | 26401332:264 | 26401857:264 | -;-;-;-;- | 1541 | eif4ebp3 | 442 | 378 | 386 | 237 | 447 | 967 |
| ENSDARG000 | 21:21:21:21:2 | 29179887:291 | 29179969:291 | +:+:+:+:+:+:+: | 2182 | NA | 0 | 2 | 2 | 0 | 0 | 0 |
| ENSDARG000 | 21:21:21:21:2 | 15756549:157 | 15756887:157 | +:+:+:+:+:+:+: | 3733 | ptpra | 1309 | 1514 | 1916 | 994 | 1777 | 1685 |
| ENSDARG000 | 21:21:21:21:2 | 4934090:4935 | 4934202:4935 | -;-;-;-;-;-;-;- | 1075 | NA | 2 | 0 | 0 | 0 | 0 | 4 |
| ENSDARG000 | 21:21:21:21:2 | 32757601:327 | 32760319:327 | -:-:-:-:-:-:-:-:-: | 4284 | cnot6a | 1234 | 979 | 1478 | 733 | 1331 | 1049 |
| ENSDARG000 | 21:21:21:21:2 | 13093136:130 | 13093834:130 | -:-:-:-:-:-:-:-:-: | 1493 | fam219aa | 466 | 431 | 506 | 360 | 298 | 415 |
| ENSDARG000 | 21:21:21:21:2 | 22801257:228 | 22801441:228 | -:-:-:-:-:-:-:-:-: | 2706 | trpc6a | 89 | 117 | 190 | 302 | 48 | 90 |
| ENSDARG000 | 21 | 32820175 | 32821422 | + | 1248 | adra2db | 0 | 13 | 22 | 0 | 2 | 2 |
| ENSDARG000 | 21:21:21:21:2 | 11468642:114 | 11469212:114 | +:+:+:+:+:+:+: | 5338 | grin1a | 4 | 2 | 4 | 28 | 4 | 0 |
| ENSDARG000 | 21:21:21:21:2 | 21279159:212 | 21279825:212 | +;+;+;+;+;+;+ | 3331 | NA | 282 | 260 | 253 | 194 | 397 | 240 |
| ENSDARG000 | 21:21:21:21:2 | 3715820:3717 | 3717168:3717 | -:-:-:-:-:-:-:-:-: | 2784 | scamp1 | 630 | 582 | 634 | 539 | 834 | 654 |
| ENSDARG000 | 21:21:21:21:2 | 26485849:264 | 26486509:264 | -;-;-;-;-;- | 1584 | zgc:110540 | 22 | 156 | 335 | 101 | 155 | 144 |
| ENSDARG000 | 21 | 4957685 | 4958755 | + | 1071 | B3GALT9 | 0 | 0 | 0 | 4 | 0 | 0 |
| ENSDARG000 | 21;21 | 26482517:264 | 26483083:264 | +;+ | 2753 | NA | 8 | 4 | 12 | 0 | 5 | 5 |
| ENSDARG000 | 21:21:21:21:2 | 26408016:264 | 26408602:264 | -:-:-:-:-:-:-:-:-: | 8538 | ankhd1 | 4006 | 2942 | 5227 | 2933 | 4326 | 3303 |
| ENSDARG000 | 21:21:21:21:2 | 33006650:330 | 33007322:330 | -:-:-:-:-:-:-:-:-: | 2054 | SLC34A1 | 0 | 0 | 4 | 0 | 6 | 0 |
| ENSDARG000 | 21:21:21:21:2 | 4974256:4974 | 4974373:4974 | -:-:-:-:-:-:-:-:-: | 6186 | RNF165 | 116 | 28 | 132 | 151 | 177 | 125 |
| ENSDARG000 | 21:21:21:21:2 | 29189471:291 | 29189903:291 | -:-:-:-:-:-:-:-:-: | 1317 | NA | 0 | 0 | 0 | 0 | 0 | 0 |
| ENSDARG000 | 21;21;21 | 19070921:190 | 19071780:190 | +;+;+ | 2023 | nkx6.1 | 54 | 106 | 26 | 16 | 6 | 25 |
| ENSDARG000 | 21:21:21:21:2 | 3775551:3778 | 3775865:3778 | +:+:+:+:+:+:+: | 2221 | tor1 | 1001 | 1209 | 1181 | 941 | 1448 | 1157 |
| ENSDARG000 | 21:21:21:21:2 | 19077238:190 | 19077514:190 | +;+;+;+;+;+ | 1022 | NA | 0 | 0 | 0 | 0 | 0 | 0 |
| ENSDARG000 | 21:21:21:21:2 | 22828500:228 | 22828807:228 | +;+;+;+;+;+;+ | 1223 | cfap300 | 6 | 0 | 0 | 0 | 6 | 0 |
| ENSDARG000 | 21:21:21:21:2 | 22820970:228 | 22821525:228 | -:-:-:-:-:-:-:-:-: | 1684 | ANGPTL5 | 25 | 15 | 14 | 14 | 16 | 34 |
| ENSDARG000 | 21;21;21;21 | 15824182:158 | 15824343:158 | +;+;+;+ | 659 | gnrh2 | 0 | 0 | 0 | 0 | 0 | 0 |
| ENSDARG000 | 21;21;21 | 15833222:158 | 15833666:158 | +;+;+ | 1795 | GCNT4 | 26 | 21 | 28 | 7 | 69 | 76 |
| ENSDARG000 | 21 | 26493694 | 26495700 | - | 2007 | NA | 9 | 0 | 2 | 0 | 8 | 2 |
| ENSDARG000 | 21:21:21:21:2 | 21309164:213 | 21309244:213 | +;+;+;+;+ | 450 | NA | 8 | 0 | 12 | 13 | 6 | 0 |
| ENSDARG000 | 21;21 | 15844260:158 | 15844284:158 | +;+ | 467 | NA | 3 | 12 | 4 | 0 | 3 | 34 |
| ENSDARG000 | 21:21:21:21:2 | 19301562:193 | 19302775:193 | -:-:-:-:-:-:-:-:-: | 2637 | gpat3 | 555 | 655 | 571 | 441 | 719 | 614 |
| ENSDARG000 | 21:21:21:21:2 | 21319149:213 | 21319766:213 | -:-:-:-:-:-:-:-:-: | 1742 | NA | 4 | 3 | 0 | 0 | 2 | 0 |
| ENSDARG000 | 21:21:21:21:2 | 15865523:158 | 15865528:158 | +;+;+;+;+ | 1410 | GCNT4 | 0 | 0 | 0 | 2 | 0 | 0 |
| ENSDARG000 | 21:21:21:21:2 | 26514603:265 | 26514826:265 | -;-;-;-;-;-;-;- | 918 | NA | 0 | 0 | 0 | 6 | 0 | 2 |
| ENSDARG000 | 21:21:21:21:2 | 22892836:228 | 22892982:228 | +:+:+:+:+:+:+: | 1840 | alg8 | 225 | 286 | 415 | 257 | 339 | 356 |
| ENSDARG000 | 21:21:21:21:2 | 21325500:213 | 21325796:213 | +:+:+:+:+:+:+: | 1613 | NA | 0 | 0 | 0 | 0 | 0 | 0 |
| ENSDARG000 | 21;21;21;21 | 22872726:228 | 22873022:228 | +;+;+;+ | 2476 | NA | 65 | 42 | 88 | 71 | 103 | 50 |
| ENSDARG000 | 21:21:21:21:2 | 15870752:158 | 15870874:158 | +:+:+:+:+:+:+: | 1725 | NA | 432 | 415 | 561 | 984 | 989 | 766 |
| ENSDARG000 | 21:21:21:21:2 | 22851198:228 | 22851419:228 | -:-:-:-:-:-:-:-:-: | 8467 | POLQ | 54 | 125 | 144 | 119 | 105 | 116 |
| ENSDARG000 | 21:21:21:21:2 | 33637083:336 | 33637363:336 | +:+:+:+:+:+:+: | 1279 | NA | 0 | 0 | 4 | 0 | 2 | 0 |
| ENSDARG000 | 21:21:21:21:2 | 26522571:265 | 26522853:265 | +:+:+:+:+:+:+: | 2365 | adssl | 442 | 910 | 802 | 342 | 779 | 476 |
| ENSDARG000 | 21;21;21 | 13147015:131 | 13147952:131 | -;-;- | 1264 | si:dkey-228b2 | 0 | 0 | 0 | 0 | 0 | 0 |
| ENSDARG000 | 21:21:21:21:2 | 22833905:228 | 22834030:228 | +:+:+:+:+:+:+: | 3108 | birc2 | 2035 | 1705 | 2175 | 1982 | 2402 | 2044 |
| ENSDARG000 | 21:21:21:21:2 | 13127742:131 | 13128273:131 | +:+:+:+:+:+:+: | 5123 | ODF2 | 469 | 310 | 320 | 144 | 300 | 226 |
| ENSDARG000 | 21:21:21:21:2 | 22878834:228 | 22879182:228 | +:+:+:+:+:+:+: | 6243 | pcf11 | 2386 | 1840 | 4005 | 3047 | 4148 | 3409 |
| ENSDARG000 | 21:21:21:21:2 | 21339423:213 | 21339449:213 | +;+;+;+;+;+;+ | 1242 | si:ch211-191i | 416 | 394 | 525 | 328 | 530 | 303 |
| ENSDARG000 | 21;21;21;21 | 21339311:213 | 21339449:213 | +;+;+;+ | 2147 | NA | 75 | 78 | 140 | 59 | 174 | 104 |
| ENSDARG000 | 21:21:21:21:2 | 5023049:5023 | 5023509:5023 | -:-:-:-:-:-:-:-:-: | 2280 | C21H18orf25 | 242 | 213 | 317 | 93 | 277 | 227 |
| ENSDARG000 | 21:21:21:21:2 | 22888889:228 | 22889314:228 | -;-;-;-;-;-;-;- | 1461 | ccdc90b | 155 | 186 | 233 | 221 | 65 | 193 |
| ENSDARG000 | 21:21:21:21:2 | 15913128:159 | 15914004:159 | -;-;-;-;- | 2039 | lhx5 | 0 | 0 | 0 | 0 | 0 | 0 |
| ENSDARG000 | 21;21;21;21 | 15911517:159 | 15911621:159 | +;+;+;+ | 636 | NA | 1 | 0 | 0 | 0 | 0 | 0 |
| ENSDARG000 | 21:21:21:21:2 | 21357309:213 | 21357316:213 | +;+;+;+;+ | 1285 | NA | 0 | 0 | 0 | 0 | 0 | 0 |
| ENSDARG000 | 21:21:21:21:2 | 26532789:265 | 26532818:265 | +:+:+:+:+:+:+: | 485 | NA | 0 | 10 | 13 | 40 | 3 | 0 |
| ENSDARG000 | 21:21:21:21:2 | 21367914:213 | 21368687:213 | -;-;-;-;-;- | 2345 | ppm1nb | 0 | 4 | 10 | 30 | 35 | 0 |
| ENSDARG000 | 21:21:21:21:2 | 3789839:3790 | 3790113:3790 | -;-;-;-;-;- | 1162 | nsa2 | 4277 | 3710 | 6157 | 7609 | 3857 | 7361 |
| ENSDARG000 | 21;21 | 29227224:292 | 29228151:292 | +;+ | 1461 | ADRA1B | 0 | 2 | 3 | 1 | 0 | 0 |
| ENSDARG000 | 21:21:21:21:2 | 3796196:3796 | 3796250:3796 | +:+:+:+:+:+:+: | 1630 | spout1 | 134 | 102 | 226 | 77 | 128 | 137 |
| ENSDARG000 | 21:21:21:21:2 | 8198652:8217 | 8198800:8217 | +:+:+:+:+:+:+: | 1502 | nr6a1b | 20 | 10 | 18 | 4 | 0 | 10 |
| ENSDARG000 | 21:21:21:21:2 | 26536950:265 | 26537412:265 | +:+:+:+:+:+:+: | 2622 | stx5al | 768 | 744 | 928 | 679 | 877 | 606 |
| ENSDARG000 | 21:21:21:21:2 | 5066557:5069 | 5066714:5069 | -:-:-:-:-:-:-:-:-: | 1629 | haus1 | 38 | 190 | 178 | 80 | 106 | 30 |
| ENSDARG000 | 21:21:21:21:2 | 29918658:299 | 29918770:299 | -;-;-;-;- | 2019 | zgc:171310 | 0 | 0 | 2 | 0 | 2 | 0 |
| ENSDARG000 | 21:21:21:21:2 | 19319804:193 | 19320063:193 | +:+:+:+:+:+:+: | 1671 | abraxas1 | 108 | 95 | 178 | 92 | 150 | 83 |
| ENSDARG000 | 21:21:21:21:2 | 19328847:193 | 19329154:193 | -;-;-;-;-;- | 720 | mrps18c | 186 | 197 | 242 | 104 | 140 | 195 |
| ENSDARG000 | 21:21:21:21:2 | 29733436:297 | 29734186:297 | -:-:-:-:-:-:-:-:-: | 3186 | NA | 18 | 18 | 34 | 15 | 16 | 13 |
| ENSDARG000 | 21:21:21:21:2 | 33973095:339 | 33973165:339 | -;-;-;-;-;- | 585 | NA | 0 | 0 | 0 | 0 | 0 | 0 |
| ENSDARG000 | 21:21:21:21:2 | 33756110:337 | 33756304:337 | +:+:+:+:+:+:+: | 751 | NA | 0 | 0 | 0 | 0 | 0 | 0 |
| ENSDARG000 | 21:21:21:21:2 | 33724979:337 | 33725397:337 | -:-:-:-:-:-:-:-:-: | 2761 | EBF1 | 8 | 2 | 0 | 0 | 0 | 3 |
| ENSDARG000 | 21:21:21:21:2 | 21374277:213 | 21374342:213 | +:+:+:+:+:+:+: | 1860 | rtn2b | 0 | 0 | 0 | 0 | 0 | 0 |
| ENSDARG000 | 21:21:21:21:2 | 19345558:193 | 19345671:193 | +:+:+:+:+:+:+: | 5438 | hpse | 2255 | 2073 | 1857 | 2523 | 1618 | 2537 |
| ENSDARG000 | 21:21:21:21:2 | 19330774:193 | 19331247:193 | +:+:+:+:+:+:+: | 3424 | helq | 40 | 88 | 66 | 49 | 77 | 74 |
| ENSDARG000 | 21:21:21:21:2 | 19368720:193 | 19369021:193 | +:+:+:+:+:+:+: | 3567 | btc | 2 | 0 | 2 | 0 | 0 | 0 |
| ENSDARG000 | 21:21:21:21:2 | 8276989:8277 | 8277130:8277 | +;+;+;+;+;+;+ | 2555 | nr5a1b | 0 | 0 | 0 | 0 | 0 | 0 |
| ENSDARG000 | 21:21:21:21:2 | 26606741:266 | 26607118:266 | +:+:+:+:+:+:+: | 2050 | esrra | 76 | 177 | 98 | 89 | 223 | 142 |
| ENSDARG000 | 21:21:21:21:2 | 34001712:340 | 34003560:340 | -:-:-:-:-:-:-:-:-: | 3911 | rnf145b | 1652 | 1542 | 1818 | 1486 | 2237 | 2518 |
| ENSDARG000 | 21;21 | 29998561:300 | 29998717:300 | -;- | 238 | NA | 0 | 0 | 14 | 0 | 1 | 0 |
| ENSDARG000 | 21:21:21:21:2 | 30007566:300 | 30007593:300 | +:+:+:+:+:+:+: | 1853 | ttc1 | 304 | 215 | 329 | 299 | 290 | 254 |
| ENSDARG000 | 21:21:21:21:2 | 19418563:194 | 19419215:194 | +:+:+:+:+:+:+: | 1796 | amacr | 106 | 68 | 84 | 98 | 96 | 98 |
| ENSDARG000 | 21:21:21:21:2 | 21448996:214 | 21449213:214 | -:-:-:-:-:-:-:-:-: | 2620 | nectin3b | 12 | 4 | 8 | 0 | 6 | 0 |
| ENSDARG000 | 21:21:21:21:2 | 30017357:300 | 30020873:300 | -;-;-;-;-;-;- | 6509 | PWWP2A | 399 | 328 | 441 | 285 | 372 | 308 |
| ENSDARG000 | 21:21:21:21:2 | 5080789:5081 | 5080932:5081 | +:+:+:+:+:+:+: | 1954 | atp5fa1 | 16129 | 17348 | 23153 | 12975 | 18838 | 13797 |
| ENSDARG000 | 21:21:21:21:2 | 13217896:132 | 13218217:132 | -:-:-:-:-:-:-:-:-: | 2134 | dync2i2 | 30 | 53 | 91 | 12 | 78 | 33 |
| ENSDARG000 | 21:21:21:21:2 | 13224239:132 | 13228113:132 | -:-:-:-:-:-:-:-:-: | 4930 | setb | 3400 | 5081 | 6838 | 3040 | 5304 | 4358 |
| ENSDARG000 | 21:21:21:21:2 | 8341774:8341 | 8341947:8341 | +:+:+:+:+:+:+: | 1240 | psmb7 | 2279 | 2250 | 2770 | 1638 | 2680 | 1771 |
| ENSDARG000 | 21:21:21:21:2 | 13150937:131 | 13151027:131 | +:+:+:+:+:+:+: | 8195 | sptan1 | 27520 | 19065 | 20172 | 22266 | 17474 | 19609 |
| ENSDARG000 | 21;21;21;21 | 30043054:300 | 30043156:300 | +;+;+;+ | 508 | fabp6 | 0 | 0 | 0 | 0 | 0 | 0 |
| ENSDARG000 | 21:21:21:21:2 | 19445942:194 | 19446435:194 | +;+;+;+;+;+;+ | 1745 | slc45a2 | 6615 | 12725 | 8297 | 15368 | 6416 | 11512 |
| ENSDARG000 | 21:21:21:21:2 | 26665014:266 | 26665423:266 | -:-:-:-:-:-:-:-:-: | 6461 | NA | 2013 | 1318 | 2293 | 1779 | 2219 | 2390 |
| ENSDARG000 | 21 | 19487707 | 19489087 | - | 1381 | rxfp3 | 13 | 49 | 5 | 29 | 26 | 0 |
| ENSDARG000 | 21 | 21529091 | 21530868 | - | 1778 | NA | 0 | 0 | 0 | 0 | 0 | 0 |
| ENSDARG000 | 21 | 21577070 | 21580197 | - | 3128 | NA | 0 | 0 | 0 | 0 | 0 | 0 |
| ENSDARG000 | 21 | 21535295 | 21537452 | - | 2158 | NA | 0 | 0 | 0 | 0 | 0 | 0 |
| ENSDARG000 | 21 | 21568737 | 21570134 | - | 1398 | NA | 0 | 0 | 0 | 0 | 0 | 0 |
| ENSDARG000 | 21:21:21:21:2 | 30054772:300 | 30058451:300 | -:-:-:-:-:-:-:-:-: | 5032 | ccnjl | 374 | 188 | 353 | 251 | 202 | 320 |
| ENSDARG000 | 21 | 21547056 | 21549988 | - | 2933 | NA | 0 | 0 | 0 | 0 | 0 | 0 |
| ENSDARG000 | 21 | 21539617 | 21541124 | - | 1508 | NA | 0 | 0 | 0 | 0 | 0 | 0 |
| ENSDARG000 | 21:21:21:21:2 | 16079124:160 | 16080100:160 | -:-:-:-:-:-:-:-:-: | 1839 | mzt2b | 331 | 287 | 469 | 310 | 341 | 348 |
| ENSDARG000 | 21 | 21562851 | 21564011 | - | 1161 | NA | 0 | 0 | 0 | 0 | 0 | 0 |
| ENSDARG000 | 21 | 21553258 | 21555039 | - | 1782 | NA | 0 | 0 | 0 | 0 | 0 | 0 |
| ENSDARG000 | 21 | 21592253 | 21594841 | - | 2589 | NA | 0 | 0 | 1 | 0 | 0 | 0 |
| ENSDARG000 | 21:21:21:21:2 | 21523082:215 | 21523634:215 | -:-:-:-:-:-:-:-:-: | 1224 | diablob | 220 | 269 | 459 | 333 | 223 | 244 |
| ENSDARG000 | 21:21:21:21:2 | 19517492:195 | 19517616:195 | +:+:+:+:+:+:+: | 2092 | NA | 10 | 18 | 11 | 16 | 14 | 31 |
| ENSDARG000 | 21 | 21599225 | 21600640 | - | 1416 | NA | 0 | 0 | 0 | 0 | 0 | 2 |
| ENSDARG000 | 21:21:21:21:2 | 34033580:340 | 34033853:340 | +:+:+:+:+:+:+: | 3342 | map4k6 | 18 | 15 | 12 | 2 | 28 | 36 |
| ENSDARG000 | 21:21:21:21:2 | 19525141:195 | 19525285:195 | +;+;+;+;+ | 880 | NA | 42 | 20 | 11 | 28 | 20 | 100 |
| ENSDARG000 | 21:21:21:21:2 | 19547806:195 | 19548105:195 | +:+:+:+:+:+:+: | 6080 | rai14 | 284 | 561 | 406 | 222 | 192 | 268 |
| ENSDARG000 | 21;21 | 30084823:300 | 30085759:300 | +;+ | 5894 | NA | 15 | 28 | 19 | 4 | 39 | 14 |
| ENSDARG000 | 21;21;21 | 5192016:5194 | 5192708:5194 | +;+;+ | 1515 | hwa | 10 | 2 | 6 | 4 | 6 | 10 |
| ENSDARG000 | 21;21;21;21 | 5169154:5170 | 5169342:5170 | +;+;+;+ | 1612 | zgc:122979 | 4 | 1 | 0 | 6 | 4 | 14 |
| ENSDARG000 | 21:21:21:21:2 | 16089386:160 | 16089898:160 | -:-:-:-:-:-:-:-:-: | 1656 | zgc:193801 | 90 | 36 | 98 | 46 | 96 | 70 |

| | | | | | | | | | | | |
|---|---|---|---|---|---|---|---|---|---|---|---|
| ENSDARG000 | 21:21:21:21:2 | 5129513:513 | 5129736:513 | +:+:+:+:+:+:+: | 3848 | thbs4b | 186 | 4 | 32 | 36 | 62 | 16 |
| ENSDARG000 | 21:21:21:21:2 | 19635486:19 | 19635687:19 | +;+;+;+;+;+;+ | 2753 | fgf10a | 0 | 0 | 0 | 0 | 12 | 0 |
| ENSDARG000 | 21:21:21:21:2 | 34088110:34 | 34088426:34 | +:+:+:+:+:+:+: | 3499 | mtmr1b | 402 | 318 | 207 | 776 | 323 | 676 |
| ENSDARG000 | 21:21:21:21:2 | 19632251:19 | 19632651:19 | -:-:-:-:-:-:-:-: | 1018 | NA | 0 | 0 | 0 | 0 | 0 | 0 |
| ENSDARG000 | 21:21:21:21:2 | 30091417:30 | 30092193:30 | -:-:-:-:-:-:-:-: | 2419 | slc23a1 | 83 | 67 | 49 | 32 | 52 | 9 |
| ENSDARG000 | 21;21;21 | 5165043:517 | 5165076:517 | -;-;- | 606 | NA | 0 | 0 | 0 | 0 | 0 | 0 |
| ENSDARG000 | 21:21:21:21:2 | 5194602:519 | 5195007:519 | -:-:-:-:-:-:-:-: | 1272 | rpl37 | 4867 | 4975 | 6033 | 4384 | 3142 | 4840 |
| ENSDARG000 | 21;21;21;21 | 30116424:30 | 30116877:30 | -;-;-;- | 964 | pfdn1 | 617 | 961 | 848 | 659 | 802 | 740 |
| ENSDARG000 | 21:21:21:21:2 | 16102938:16 | 16103331:16 | -:-:-:-:-:-:-:-: | 1680 | CYB561A3 | 4 | 4 | 0 | 2 | 14 | 0 |
| ENSDARG000 | 21:21:21:21:2 | 30161295:30 | 30162147:30 | -:-:-:-:-:-:-:-: | 1677 | hbegfb | 130 | 187 | 78 | 156 | 139 | 84 |
| ENSDARG000 | 21:21:21:21:2 | 21611867:21 | 21612366:21 | +:+:+:+:+:+:+: | 1344 | b9d2 | 86 | 82 | 90 | 80 | 58 | 45 |
| ENSDARG000 | 21:21:21:21:2 | 30169610:30 | 30170193:30 | -:-:-:-:-:-:-:-: | 1865 | hnrnph1l | 2048 | 2341 | 2399 | 1195 | 2178 | 1417 |
| ENSDARG000 | 21:21:21:21:2 | 16144014:16 | 16144448:16 | +:+:+:+:+:+:+: | 6272 | NA | 6 | 7 | 12 | 4 | 0 | 8 |
| ENSDARG000 | 21:21:21:21:2 | 30194904:30 | 30194962:30 | +:+:+:+:+:+:+: | 3027 | BRD8 | 824 | 613 | 658 | 629 | 875 | 537 |
| ENSDARG000 | 21:21:21:21:2 | 3820779:382 | 3820965:382 | -:-:-:-:-:-:-:-: | 1025 | ST6GALNAC6 | 58 | 18 | 13 | 36 | 7 | 10 |
| ENSDARG000 | 21:21:21:21:2 | 21621042:21 | 21621378:21 | +;+;+;+;+;+;+ | 1403 | NA | 115 | 116 | 62 | 202 | 130 | 424 |
| ENSDARG000 | 21:21:21:21:2 | 30239549:30 | 30240751:30 | -;-;-;-;-;-;- | 2696 | dnajc18 | 527 | 437 | 633 | 503 | 589 | 438 |
| ENSDARG000 | 21:21:21:21:2 | 16172931:16 | 16173429:16 | -:-:-:-:-:-:-:-: | 1686 | tnfrsfa | 889 | 854 | 1142 | 628 | 1146 | 1363 |
| ENSDARG000 | 21;21 | 11503212:11 | 11503425:11 | +;+ | 1115 | NA | 0 | 2 | 0 | 0 | 0 | 0 |
| ENSDARG000 | 21:21:21:21:2 | 3840955:384 | 3841419:384 | -;-;-;-;-;-;-;- | 1221 | st6galnac4 | 112 | 78 | 112 | 54 | 92 | 90 |
| ENSDARG000 | 21:21:21:21:2 | 11509461:11 | 11509602:11 | +:+:+:+:+:+:+: | 1540 | ENTPD2 | 0 | 0 | 0 | 0 | 0 | 0 |
| ENSDARG000 | 21:21:21:21:2 | 3897680:390 | 3897768:390 | +:+:+:+:+:+:+: | 1341 | dolpp1 | 179 | 390 | 490 | 330 | 320 | 298 |
| ENSDARG000 | 21:21:21:21:2 | 30256952:30 | 30261353:30 | -:-:-:-:-:-:-:-: | 5886 | fam53c | 614 | 522 | 752 | 422 | 763 | 766 |
| ENSDARG000 | 21;21;21 | 19692237:19 | 19692889:19 | -;-;- | 920 | NA | 0 | 0 | 0 | 0 | 0 | 0 |
| ENSDARG000 | 21:21:21:21:2 | 19763546:19 | 19765060:19 | -:-:-:-:-:-:-:-: | 4734 | nnt | 5510 | 4290 | 6822 | 3385 | 6408 | 4898 |
| ENSDARG000 | 21;21;21;21 | 21633213:21 | 21633842:21 | -;-;-;- | 1611 | cbr1l | 212 | 390 | 324 | 646 | 254 | 304 |
| ENSDARG000 | 21:21:21:21:2 | 34132747:34 | 34133266:34 | +:+:+:+:+:+:+: | 3881 | TRPC5 | 193 | 813 | 56 | 280 | 6 | 556 |
| ENSDARG000 | 21:21:21:21:2 | 34119759:34 | 34119892:34 | +:+:+:+:+:+:+: | 1599 | hmgb3b | 1144 | 1191 | 1108 | 569 | 1200 | 1031 |
| ENSDARG000 | 21:21:21:21:2 | 3928947:394 | 3929067:394 | +:+:+:+:+:+:+: | 7330 | NA | 4552 | 2022 | 4265 | 3651 | 2991 | 3606 |
| ENSDARG000 | 21;21 | 21649760:21 | 21649889:21 | +;+ | 631 | NA | 0 | 0 | 0 | 0 | 0 | 0 |
| ENSDARG000 | 21 | 21655755 | 21657308 | + | 1554 | or62a2 | 0 | 0 | 0 | 0 | 0 | 0 |
| ENSDARG000 | 21 | 21663062 | 21664421 | + | 1360 | or62b2 | 0 | 0 | 0 | 0 | 0 | 0 |
| ENSDARG000 | 21;21 | 21670421:21 | 21670496:21 | +;+ | 197 | NA | 0 | 0 | 0 | 0 | 0 | 0 |
| ENSDARG000 | 21:21:21:21:2 | 11521163:11 | 11521389:11 | +;+;+;+;+;+;+ | 2738 | zmat1 | 138 | 154 | 304 | 114 | 264 | 231 |
| ENSDARG000 | 21 | 21673516 | 21674819 | + | 1304 | or62b1 | 0 | 0 | 0 | 0 | 0 | 0 |
| ENSDARG000 | 21:21:21:21:2 | 16273782:16 | 16274022:16 | -:-:-:-:-:-:-:-: | 2904 | unc5da | 4 | 0 | 4 | 2 | 4 | 0 |
| ENSDARG000 | 21:21:21:21:2 | 19844195:19 | 19844265:19 | +:+:+:+:+:+:+: | 3107 | NA | 3423 | 3324 | 3451 | 3011 | 4247 | 1809 |
| ENSDARG000 | 21:21:21:21:2 | 19834072:19 | 19835402:19 | +;+;+;+;+;+;+ | 2514 | NA | 20 | 22 | 10 | 84 | 8 | 10 |
| ENSDARG000 | 21:21:21:21:2 | 11535304:11 | 11535530:11 | +:+:+:+:+:+:+: | 3278 | NA | 0 | 0 | 0 | 4 | 0 | 0 |
| ENSDARG000 | 21:21:21:21:2 | 13233377:13 | 13233527:13 | +:+:+:+:+:+:+: | 3883 | SPECC1L | 468 | 544 | 506 | 336 | 503 | 375 |
| ENSDARG000 | 21:21:21:21:2 | 16980141:16 | 16980291:16 | +;+;+;+;+;+ | 943 | aqp3b | 17039 | 22 | 326 | 10 | 679 | 2 |
| ENSDARG000 | 21:21:21:21:2 | 8357602:835 | 8357724:835 | -:-:-:-:-:-:-:-: | 888 | NEK6 | 633 | 627 | 589 | 743 | 445 | 652 |
| ENSDARG000 | 21;21 | 21678476:21 | 21678656:21 | +;+ | 215 | NA | 0 | 0 | 0 | 0 | 0 | 0 |
| ENSDARG000 | 21:21:21:21:2 | 11560153:11 | 11560267:11 | +;+;+;+;+;+ | 1328 | cd8a | 0 | 4 | 2 | 0 | 0 | 20 |
| ENSDARG000 | 21 | 21679086 | 21681627 | + | 2542 | or62b4 | 0 | 0 | 0 | 0 | 0 | 0 |
| ENSDARG000 | 21;21;21;21 | 17005737:17 | 17005836:17 | +;+;+;+ | 955 | vps29 | 1682 | 1697 | 1992 | 1682 | 1746 | 1592 |
| ENSDARG000 | 21 | 21685228 | 21687545 | + | 2318 | or62b3 | 0 | 0 | 0 | 0 | 0 | 0 |
| ENSDARG000 | 21:21:21:21:2 | 17016337:17 | 17016384:17 | +:+:+:+:+:+:+: | 1007 | gpn3 | 90 | 141 | 152 | 84 | 123 | 130 |
| ENSDARG000 | 21:21:21:21:2 | 8407545:840 | 8407906:840 | -:-:-:-:-:-:-:-: | 1366 | lhx2a | 0 | 0 | 3 | 0 | 1 | 0 |
| ENSDARG000 | 21:21:21:21:2 | 30293564:30 | 30293679:30 | +;+;+;+;+;+ | 1196 | zmat2 | 870 | 794 | 727 | 585 | 860 | 800 |
| ENSDARG000 | 21 | 21692307 | 21694124 | + | 1818 | or62b5 | 0 | 0 | 0 | 0 | 0 | 0 |
| ENSDARG000 | 21:21:21:21:2 | 34233492:34 | 34233893:34 | -:-:-:-:-:-:-:-: | 2939 | alg13 | 0 | 0 | 0 | 0 | 0 | 0 |
| ENSDARG000 | 21:21:21:21:2 | 17024002:17 | 17024099:17 | +;+;+;+;+;+;+ | 828 | arpc3 | 3061 | 3051 | 2599 | 2874 | 2993 | 2980 |
| ENSDARG000 | 21 | 21700241 | 21700559 | + | 319 | NA | 0 | 0 | 0 | 0 | 0 | 0 |
| ENSDARG000 | 21:21:21:21:2 | 30288570:30 | 30289135:30 | -:-:-:-:-:-:-:-: | 1559 | slbp2 | 0 | 2 | 10 | 0 | 0 | 2 |
| ENSDARG000 | 21:21:21:21:2 | 26678484:26 | 26679092:26 | -:-:-:-:-:-:-:-: | 2382 | bscl2 | 280 | 224 | 186 | 189 | 311 | 126 |
| ENSDARG000 | 21;21 | 13301798:13 | 13302212:13 | +;+ | 1528 | adora2ab | 2147 | 441 | 283 | 1390 | 490 | 899 |
| ENSDARG000 | 21 | 21702623 | 21704815 | + | 2193 | or62a1 | 0 | 0 | 0 | 0 | 0 | 0 |
| ENSDARG000 | 21 | 21709204 | 21710481 | + | 1278 | or62a3 | 0 | 0 | 0 | 0 | 0 | 0 |
| ENSDARG000 | 21 | 21713531 | 21716216 | + | 2686 | or61a1 | 0 | 0 | 0 | 0 | 2 | 0 |
| ENSDARG000 | 21 | 21725332 | 21726469 | + | 1138 | or61c2 | 0 | 0 | 0 | 0 | 0 | 0 |
| ENSDARG000 | 21:21:21:21:2 | 30306369:30 | 30306718:30 | +:+:+:+:+:+:+: | 2072 | lcp2a | 42 | 66 | 145 | 14 | 176 | 141 |
| ENSDARG000 | 21;21 | 21727264:21 | 21729268:21 | +;+ | 2005 | or61b1 | 0 | 0 | 0 | 0 | 0 | 0 |
| ENSDARG000 | 21;21;21 | 26697536:26 | 26697758:26 | +;+;+ | 1258 | gng3 | 68 | 14 | 2 | 24 | 8 | 26 |
| ENSDARG000 | 21:21:21:21:2 | 34364456:34 | 34365408:34 | -:-:-:-:-:-:-:-: | 3234 | dacha | 6 | 0 | 0 | 0 | 24 | 0 |
| ENSDARG000 | 21 | 21729607 | 21730961 | + | 1355 | or61c1 | 0 | 0 | 0 | 0 | 0 | 0 |
| ENSDARG000 | 21 | 21736733 | 21737327 | + | 595 | NA | 0 | 0 | 0 | 0 | 0 | 0 |
| ENSDARG000 | 21 | 21738150 | 21740682 | + | 2533 | NA | 0 | 0 | 0 | 0 | 1 | 0 |
| ENSDARG000 | 21:21:21:21:2 | 17031635:17 | 17032688:17 | -;-;-;-;- | 1633 | ube2g1b | 914 | 971 | 1321 | 1155 | 952 | 983 |
| ENSDARG000 | 21:21:21:21:2 | 34686764:34 | 34686865:34 | +:+:+:+:+:+:+: | 2559 | chm | 512 | 569 | 571 | 422 | 501 | 380 |
| ENSDARG000 | 21;21 | 30351256:30 | 30351890:30 | +;+ | 1592 | foxi3a | 0 | 0 | 0 | 4 | 0 | 8 |
| ENSDARG000 | 21:21:21:21:2 | 8427059:843 | 8427171:843 | +:+:+:+:+:+:+: | 2059 | DENND1A | 219 | 154 | 306 | 154 | 298 | 278 |
| ENSDARG000 | 21:21:21:21:2 | 30401784:30 | 30402272:30 | -:-:-:-:-:-:-:-: | 1175 | nhp2 | 498 | 692 | 757 | 306 | 861 | 586 |
| ENSDARG000 | 21:21:21:21:2 | 5209476:520 | 5209846:520 | +:+:+:+:+:+:+: | 1567 | st8sia5 | 0 | 0 | 0 | 0 | 0 | 0 |
| ENSDARG000 | 21:21:21:21:2 | 22906479:22 | 22906705:22 | -;-;-;-;- | 1194 | guca1d | 0 | 0 | 14 | 14 | 22 | 4 |
| ENSDARG000 | 21:21:21:21:2 | 34814444:34 | 34814980:34 | +:+:+:+:+:+:+: | 5284 | wdr55 | 154 | 200 | 272 | 61 | 214 | 141 |
| ENSDARG000 | 21:21:21:21:2 | 34835811:34 | 34837004:34 | -:-:-:-:-:-:-:-: | 2255 | zgc:56585 | 19 | 22 | 38 | 33 | 29 | 0 |
| ENSDARG000 | 21;21;21 | 3975456:398 | 3975651:398 | +;+;+ | 497 | NA | 0 | 0 | 0 | 0 | 0 | 0 |
| ENSDARG000 | 21:21:21:21:2 | 30410334:30 | 30410862:30 | -;-;-;-;-;-;- | 975 | grpel2 | 882 | 982 | 1214 | 1021 | 1096 | 774 |
| ENSDARG000 | 21:21:21:21:2 | 3979107:398 | 3979333:398 | -;-;-;-;- | 739 | NTNG2 | 0 | 10 | 0 | 0 | 20 | 0 |
| ENSDARG000 | 21:21:21:21:2 | 26733333:26 | 26733650:26 | +:+:+:+:+:+:+: | 9043 | pcxa | 2742 | 2374 | 2762 | 2998 | 2781 | 2622 |
| ENSDARG000 | 21;21;21 | 21741804:21 | 21741905:21 | +;+;+ | 866 | defbl1 | 15963 | 5797 | 5288 | 1356 | 6190 | 1816 |
| ENSDARG000 | 21:21:21:21:2 | 13327527:13 | 13327643:13 | +;+;+;+;+;+;+ | 653 | snrpd3l | 332 | 442 | 598 | 431 | 498 | 418 |
| ENSDARG000 | 21:21:21:21:2 | 13353263:13 | 13353321:13 | +:+:+:+:+:+:+: | 1301 | NA | 58 | 40 | 31 | 72 | 12 | 18 |
| ENSDARG000 | 21:21:21:21:2 | 26908627:26 | 26909668:26 | -;-;-;-;- | 3200 | NA | 1691 | 628 | 646 | 1953 | 941 | 532 |
| ENSDARG000 | 21:21:21:21:2 | 13383413:13 | 13383492:13 | +:+:+:+:+:+:+: | 3472 | zgc:113162 | 12 | 2 | 24 | 20 | 2 | 12 |
| ENSDARG000 | 21:21:21:21:2 | 22960735:22 | 22961497:22 | -:-:-:-:-:-:-:-: | 3746 | rb1 | 709 | 1301 | 1198 | 594 | 1150 | 740 |
| ENSDARG000 | 21 | 22985078 | 22986277 | + | 1200 | LPAR6 | 7 | 0 | 8 | 13 | 14 | 33 |
| ENSDARG000 | 21:21:21:21:2 | 19868294:19 | 19869488:19 | -;-;-;-;- | 4779 | MFHAS1 | 317 | 254 | 217 | 304 | 233 | 449 |
| ENSDARG000 | 21:21:21:21:2 | 17051478:17 | 17051613:17 | +:+:+:+:+:+:+: | 5931 | atp2a2b | 2723 | 3784 | 5376 | 1033 | 4825 | 1851 |
| ENSDARG000 | 21:21:21:21:2 | 4116437:411 | 4116697:411 | +:+:+:+:+:+:+: | 4412 | rapgef1b | 336 | 287 | 356 | 194 | 424 | 202 |
| ENSDARG000 | 21:21:21:21:2 | 8478774:847 | 8479568:847 | -:-:-:-:-:-:-:-: | 5153 | crb2a | 0 | 0 | 2 | 12 | 0 | 0 |
| ENSDARG000 | 21:21:21:21:2 | 13455308:13 | 13455405:13 | -:-:-:-:-:-:-:-: | 1990 | NSMF | 4 | 0 | 10 | 0 | 2 | 0 |
| ENSDARG000 | 21:21:21:21:2 | 21743599:21 | 21743794:21 | +:+:+:+:+:+:+: | 1815 | pold3 | 720 | 725 | 907 | 229 | 1064 | 295 |
| ENSDARG000 | 21:21:21:21:2 | 19907759:19 | 19908332:19 | -:-:-:-:-:-:-:-: | 1743 | eri1 | 171 | 297 | 266 | 190 | 274 | 174 |
| ENSDARG000 | 21:21:21:21:2 | 23019372:23 | 23020669:23 | -:-:-:-:-:-:-:-: | 3382 | dlat | 2028 | 1849 | 1899 | 948 | 1969 | 1520 |
| ENSDARG000 | 21:21:21:21:2 | 30418034:30 | 30418427:30 | -:-:-:-:-:-:-:-: | 2997 | afap1l1b | 160 | 160 | 194 | 294 | 102 | 359 |
| ENSDARG000 | 21:21:21:21:2 | 34849406:34 | 34849575:34 | +:+:+:+:+:+:+: | 4254 | cdc23 | 182 | 212 | 367 | 126 | 311 | 258 |
| ENSDARG000 | 21:21:21:21:2 | 21771814:21 | 21772635:21 | -:-:-:-:-:-:-:-: | 2114 | chrdl2 | 558 | 38 | 4 | 34 | 1027 | 16 |
| ENSDARG000 | 21:21:21:21:2 | 13577227:13 | 13577413:13 | +:+:+:+:+:+:+: | 2476 | NA | 2 | 0 | 0 | 0 | 0 | 0 |
| ENSDARG000 | 21:21:21:21:2 | 23036314:23 | 23036688:23 | -:-:-:-:-:-:-:-: | 2719 | zw10 | 472 | 864 | 968 | 591 | 681 | 657 |
| ENSDARG000 | 21:21:21:21:2 | 34879268:34 | 34880245:34 | -:-:-:-:-:-:-:-: | 4017 | kif20a | 255 | 103 | 128 | 22 | 176 | 57 |
| ENSDARG000 | 21:21:21:21:2 | 5257018:525 | 5257070:525 | +:+:+:+:+:+:+: | 6879 | LOXHD1 | 0 | 0 | 0 | 0 | 0 | 4 |
| ENSDARG000 | 21:21:21:21:2 | 30470809:30 | 30471232:30 | +:+:+:+:+:+:+: | 6008 | snx12 | 3448 | 2973 | 3796 | 2428 | 4016 | 3119 |
| ENSDARG000 | 21:21:21:21:2 | 21782331:21 | 21782752:21 | -:-:-:-:-:-:-:-: | 2191 | NA | 10 | 12 | 10 | 8 | 16 | 21 |
| ENSDARG000 | 21:21:21:21:2 | 21791343:21 | 21791565:21 | +;+;+;+;+ | 1823 | neu3.1 | 316 | 142 | 167 | 112 | 231 | 106 |
| ENSDARG000 | 21:21:21:21:2 | 19916456:19 | 19918289:19 | -:-:-:-:-:-:-:-: | 2470 | ppp1r3b | 0 | 2 | 0 | 0 | 0 | 4 |
| ENSDARG000 | 21;21;21 | 21796663:21 | 21796863:21 | +;+;+ | 1505 | neu3.2 | 91 | 2 | 0 | 0 | 114 | 0 |
| ENSDARG000 | 21:21:21:21:2 | 23050310:23 | 23051547:23 | -:-:-:-:-:-:-:-: | 4895 | USP28 | 2 | 0 | 4 | 0 | 20 | 0 |
| ENSDARG000 | 21:21:21:21:2 | 23108420:23 | 23108439:23 | +:+:+:+:+:+:+: | 1375 | HTR3B | 0 | 0 | 0 | 0 | 0 | 0 |
| ENSDARG000 | 21:21:21:21:2 | 21803363:21 | 21803408:21 | +:+:+:+:+:+:+: | 3482 | neu3.3 | 2 | 0 | 0 | 0 | 2 | 0 |
| ENSDARG000 | 21:21:21:21:2 | 9366967:936 | 9367141:936 | -:-:-:-:-:-:-:-: | 2434 | sdad1 | 894 | 866 | 1820 | 902 | 894 | 1123 |
| ENSDARG000 | 21:21:21:21:2 | 34929598:34 | 34929769:34 | +:+:+:+:+:+:+: | 1235 | NA | 0 | 0 | 4 | 0 | 8 | 0 |
| ENSDARG000 | 21:21:21:21:2 | 21811867:21 | 21811924:21 | +;+;+;+;+ | 1375 | neu3.4 | 2 | 0 | 0 | 0 | 0 | 0 |
| ENSDARG000 | 21:21:21:21:2 | 23135322:23 | 23135545:23 | +:+:+:+:+:+:+: | 6659 | HTR3A | 0 | 0 | 0 | 0 | 0 | 8 |
| ENSDARG000 | 21:21:21:21:2 | 23159385:23 | 23164367:23 | -:-:-:-:-:-:-:-: | 8126 | zbtb16a | 306 | 32 | 74 | 361 | 176 | 160 |
| ENSDARG000 | 21:21:21:21:2 | 30502002:30 | 30502081:30 | +:+:+:+:+:+:+: | 4519 | slc7a3a | 3305 | 5008 | 6777 | 4618 | 6761 | 4089 |
| ENSDARG000 | 21:21:21:21:2 | 34945033:34 | 34945384:34 | -:-:-:-:-:-:-:-: | 2002 | lipia | 16 | 1 | 6 | 0 | 8 | 6 |
| ENSDARG000 | 21:21:21:21:2 | 21819878:21 | 21820256:21 | -;-;-;-;-;-;-;- | 1415 | wdr73 | 198 | 116 | 177 | 101 | 171 | 184 |
| ENSDARG000 | 21:21:21:21:2 | 4227260:423 | 4228069:423 | -;-;-;-;-;-;- | 1595 | uck1 | 142 | 65 | 62 | 30 | 144 | 89 |
| ENSDARG000 | 21:21:21:21:2 | 26991198:26 | 26991293:26 | +;+;+;+;+;+ | 850 | fkbp2 | 1405 | 1596 | 1831 | 1305 | 1727 | 1433 |
| ENSDARG000 | 21;21;21;21 | 21817975:21 | 21818226:21 | +;+;+;+ | 1197 | neu3.5 | 10 | 1 | 5 | 2 | 18 | 16 |
| ENSDARG000 | 21;21;21 | 30540465:30 | 30541261:30 | -;-;- | 1367 | rab39ba | 234 | 123 | 192 | 238 | 159 | 105 |
| ENSDARG000 | 21;21;21 | 4245857:424 | 4246670:424 | -;-;- | 1328 | dnlz | 76 | 68 | 212 | 154 | 82 | 120 |
| ENSDARG000 | 21;21;21 | 30549512:30 | 30549704:30 | +;+;+ | 642 | NA | 1657 | 579 | 730 | 1177 | 1033 | 1407 |
| ENSDARG000 | 21:21:21:21:2 | 34976600:34 | 34976864:34 | +:+:+:+:+:+:+: | 3303 | NA | 0 | 0 | 0 | 0 | 0 | 0 |
| ENSDARG000 | 21:21:21:21:2 | 19925910:19 | 19926547:19 | +:+:+:+:+:+:+: | 4855 | NA | 2833 | 1756 | 3064 | 1791 | 3222 | 2387 |
| ENSDARG000 | 21;21;21;21 | 121859:1250 | 123250:1251 | -;-;-;- | 2577 | NA | 0 | 0 | 0 | 0 | 0 | 0 |
| ENSDARG000 | 21:21:21:21:2 | 23338059:23 | 23338460:23 | -:-:-:-:-:-:-:-: | 4796 | ncam1a | 28 | 33 | 63 | 2 | 96 | 40 |
| ENSDARG000 | 21:21:21:21:2 | 26996276:26 | 26996970:26 | -:-:-:-:-:-:-:-: | 1444 | ppp1r14ba | 1619 | 548 | 590 | 1787 | 1831 | 589 |
| ENSDARG000 | 21:21:21:21:2 | 17110598:17 | 17112086:17 | +:+:+:+:+:+:+: | 6840 | BCR | 1266 | 1035 | 1178 | 898 | 1642 | 903 |
| ENSDARG000 | 21:21:21:21:2 | 30600337:30 | 30600510:30 | -:-:-:-:-:-:-:-: | 4914 | NA | 2 | 0 | 2 | 0 | 2 | 0 |
| ENSDARG000 | 21;21;21 | 21826529:21 | 21827185:21 | -;-;- | 860 | defbl2 | 0 | 0 | 0 | 0 | 2 | 0 |
| ENSDARG000 | 21:21:21:21:2 | 13663656:13 | 13664326:13 | -;-;-;-;-;-;- | 1779 | clic3 | 2 | 0 | 9 | 14 | 0 | 4 |
| ENSDARG000 | 21:21:21:21:2 | 13611178:13 | 13614406:13 | -:-:-:-:-:-:-:-: | 12985 | pnpla7a | 1180 | 1183 | 1540 | 1027 | 1485 | 1381 |
| ENSDARG000 | 21:21:21:21:2 | 4256291:428 | 4256444:428 | +;+;+;+;+;+;+ | 3055 | lrrc8aa | 683 | 729 | 1041 | 275 | 1026 | 600 |
| ENSDARG000 | 21:21:21:21:2 | 30651064:30 | 30653280:30 | -;-;-;-;-;- | 2510 | si:dkev-22f5.9 | 6 | 0 | 2 | 4 | 12 | 0 |
| ENSDARG000 | 21:21:21:21:2 | 21894067:21 | 21894180:21 | +:+:+:+:+:+:+: | 3911 | gria4b | 0 | 3 | 0 | 0 | 0 | 0 |
| ENSDARG000 | 21:21:21:21:2 | 30662263:30 | 30662396:30 | +:+:+:+:+:+:+: | 7443 | CLCN5 | 2041 | 1753 | 1977 | 1619 | 3078 | 1745 |
| ENSDARG000 | 21:21:21:21:2 | 11565888:11 | 11567087:11 | -:-:-:-:-:-:-:-: | 4844 | cast | 2694 | 1413 | 1026 | 3924 | 2030 | 2149 |
| ENSDARG000 | 21:21:21:21:2 | 30701902:30 | 30702359:30 | -;-;-;-;- | 980 | tnfsf10l3 | 6266 | 5546 | 4536 | 3856 | 10343 | 3890 |
| ENSDARG000 | 21:21:21:21:2 | 30721733:30 | 30721833:30 | +:+:+:+:+:+:+: | 2075 | xndc1 | 68 | 151 | 204 | 120 | 254 | 106 |
| ENSDARG000 | 21:21:21:21:2 | 5368130:537 | 5368608:537 | -:-:-:-:-:-:-:-: | 2112 | psmd5 | 853 | 632 | 762 | 386 | 618 | 462 |
| ENSDARG000 | 21:21:21:21:2 | 5472330:547 | 5472558:547 | +;+;+;+;+ | 1081 | NA | 0 | 0 | 0 | 0 | 0 | 0 |
| ENSDARG000 | 21:21:21:21:2 | 5494561:549 | 5494623:549 | +;+;+;+;+ | 790 | NA | 2 | 8 | 0 | 0 | 0 | 0 |
| ENSDARG000 | 21:21:21:21:2 | 34992550:34 | 34992708:34 | +:+:+:+:+:+:+: | 4085 | NA | 0 | 0 | 2 | 0 | 0 | 0 |
| ENSDARG000 | 21:21:21:21:2 | 27047333:27 | 27048832:27 | -:-:-:-:-:-:-:-: | 4448 | slc8a4a | 0 | 2 | 0 | 12 | 10 | 2 |
| ENSDARG000 | 21;21;21;21 | 27192763:27 | 27193840:27 | -;-;-;- | 1334 | zgc:110782 | 132 | 117 | 134 | 80 | 144 | 83 |
| ENSDARG000 | 21:21:21:21:2 | 5508167:550 | 5508226:550 | +;+;+;+;+ | 766 | NA | 4 | 0 | 0 | 0 | 0 | 0 |
| ENSDARG000 | 21:21:21:21:2 | 17280309:17 | 17280528:17 | -:-:-:-:-:-:-:-: | 1424 | GFI1B | 4 | 0 | 10 | 0 | 28 | 3 |
| ENSDARG000 | 21:21:21:21:2 | 244503:2445 | 244562:2445 | +:+:+:+:+:+:+: | 1457 | stard4 | 216 | 283 | 211 | 182 | 197 | 134 |
| ENSDARG000 | 21:21:21:21:2 | 35041399:35 | 35041861:35 | +:+:+:+:+:+:+: | 3377 | poglut3 | 382 | 435 | 547 | 180 | 545 | 304 |
| ENSDARG000 | 21;21 | 27197711:27 | 27200276:27 | -;- | 2644 | zgc:77262 | 1603 | 1325 | 1658 | 956 | 1727 | 1283 |
| ENSDARG000 | 21;21 | 35078766:35 | 35082337:35 | -;- | 3764 | adrb2b | 0 | 4 | 0 | 0 | 2 | 0 |
| ENSDARG000 | 21:21:21:21:2 | 30746348:30 | 30746370:30 | +:+:+:+:+:+:+: | 3555 | trpc2b | 14 | 4 | 20 | 0 | 45 | 0 |
| ENSDARG000 | 21;21;21 | 35215810:35 | 35216015:35 | +;+;+ | 2273 | ubtd2 | 613 | 901 | 536 | 656 | 561 | 836 |
| ENSDARG000 | 21:21:21:21:2 | 4324546:432 | 4324716:432 | -;-;-;-;-;-;- | 1644 | NA | 0 | 0 | 0 | 0 | 0 | 0 |
| ENSDARG000 | 21:21:21:21:2 | 4348600:435 | 4348738:435 | +:+:+:+:+:+:+: | 1239 | HTRA2 | 0 | 2 | 2 | 0 | 0 | 0 |
| ENSDARG000 | 21:21:21:21:2 | 22099300:22 | 22103137:22 | -:-:-:-:-:-:-:-: | 5319 | elmod1 | 0 | 0 | 0 | 0 | 0 | 0 |
| ENSDARG000 | 21:21:21:21:2 | 9387264:938 | 9387893:938 | -:-:-:-:-:-:-:-: | 8453 | ptpn13 | 453 | 538 | 1259 | 319 | 814 | 162 |
| ENSDARG000 | 21:21:21:21:2 | 4313039:431 | 4313164:431 | +:+:+:+:+:+:+: | 1056 | HTRA2 | 22 | 2 | 4 | 8 | 12 | 0 |
| ENSDARG000 | 21:21:21:21:2 | 35289469:35 | 35290129:35 | +:+:+:+:+:+:+: | 1663 | NA | 8 | 14 | 5 | 2 | 19 | 0 |

| | | | | | | | | | | | |
|---|---|---|---|---|---|---|---|---|---|---|---|
| ENSDARG000 | 21:21:21:21:2 | 4328348:4328 | 4328486:4328 | +:+:+:+:+:+:+: | 1723 | HTRA2 | 0 | 8 | 7 | 2 | 0 | 2 |
| ENSDARG000 | 21:21:21:21:2 | 22115806:221 | 22116710:221 | -:-:-:-:-:-:-:-:-: | 1992 | slc35f2 | 124 | 147 | 215 | 102 | 196 | 159 |
| ENSDARG000 | 21:21:21:21:2 | 30772043:307 | 30772612:307 | -:-:-:-:-:-:-:-:-: | 2928 | rrm1 | 508 | 2104 | 2527 | 697 | 2135 | 946 |
| ENSDARG000 | 21:21:21:21:2 | 4344795:4345 | 4344965:4345 | -;-;-;-;-;-;- | 1644 | NA | 0 | 0 | 0 | 0 | 0 | 0 |
| ENSDARG000 | 21:21:21:21:2 | 27204382:272 | 27205200:272 | -:-:-:-:-:-:-:-:-: | 3819 | MARK2 | 1882 | 1070 | 1567 | 1381 | 1964 | 2375 |
| ENSDARG000 | 21;21;21;21 | 5518619:5519 | 5518711:5519 | +;+;+;+ | 679 | NA | 0 | 0 | 0 | 0 | 0 | 0 |
| ENSDARG000 | 21:21:21:21:2 | 35304974:353 | 35306213:353 | -:-:-:-:-:-:-:-:-: | 2761 | ublcp1 | 82 | 159 | 265 | 60 | 212 | 122 |
| ENSDARG000 | 21:21:21:21:2 | 30794351:308 | 30794846:308 | +:+:+:+:+:+:+: | 4329 | slc6a7 | 60 | 14 | 4 | 12 | 36 | 38 |
| ENSDARG000 | 21;21 | 13676890:136 | 13676970:136 | +;+ | 1275 | fut7 | 0 | 12 | 20 | 18 | 10 | 39 |
| ENSDARG000 | 21:21:21:21:2 | 23953181:239 | 23953272:239 | +:+:+:+:+:+:+: | 1343 | cadm1a | 53 | 14 | 20 | 12 | 6 | 16 |
| ENSDARG000 | 21:21:21:21:2 | 5531138:5531 | 5531226:5531 | +;+;+;+;+ | 1133 | ly6m6 | 0 | 0 | 10 | 14 | 10 | 4 |
| ENSDARG000 | 21:21:21:21:2 | 17301790:173 | 17301823:173 | +:+:+:+:+:+:+: | 5515 | tsc1b | 2746 | 999 | 2097 | 2543 | 1520 | 3120 |
| ENSDARG000 | 21;21;21;21 | 20113470:201 | 20114389:201 | -;-;-;- | 1816 | dusp4 | 5410 | 6028 | 8780 | 5173 | 8231 | 5794 |
| ENSDARG000 | 21:21:21:21:2 | 5560040:5560 | 5560111:5560 | +;+;+;+;+ | 884 | NA | 0 | 0 | 0 | 0 | 0 | 1 |
| ENSDARG000 | 21;21 | 5575027:5577 | 5575315:5578 | -;- | 819 | NA | 0 | 0 | 0 | 0 | 0 | 0 |
| ENSDARG000 | 21:21:21:21:2 | 5580948:5581 | 5581006:5581 | +;+;+;+;+ | 1031 | NA | 0 | 0 | 0 | 0 | 0 | 0 |
| ENSDARG000 | 21:21:21:21:2 | 249970:25119 | 250106:25126 | +:+:+:+:+:+:+: | 4625 | jak2a | 814 | 600 | 736 | 350 | 942 | 488 |
| ENSDARG000 | 21:21:21:21:2 | 37787805:377 | 37788158:377 | -:-:-:-:-:-:-:-:-: | 1539 | NA | 0 | 0 | 0 | 0 | 0 | 3 |
| ENSDARG000 | 21:21:21:21:2 | 20229926:202 | 20230104:202 | +:+:+:+:+:+:+: | 5033 | NA | 248 | 226 | 92 | 132 | 104 | 822 |
| ENSDARG000 | 21;21 | 5589923:5601 | 5590345:5603 | +;+ | 2714 | NA | 0 | 0 | 0 | 0 | 0 | 0 |
| ENSDARG000 | 21:21:21:21:2 | 24531679:245 | 24532404:245 | -;-;-;-;- | 1291 | sc5d | 242 | 270 | 262 | 57 | 149 | 80 |
| ENSDARG000 | 21:21:21:21:2 | 5803245:5812 | 5803552:5813 | -:-:-:-:-:-:-:-:-: | 3687 | NA | 228 | 370 | 388 | 195 | 466 | 184 |
| ENSDARG000 | 21:21:21:21:2 | 20309085:203 | 20310508:203 | -:-:-:-:-:-:-:-:-: | 3305 | zgc:86764 | 66 | 305 | 379 | 118 | 331 | 142 |
| ENSDARG000 | 21:21:21:21:2 | 278938:27906 | 280050:27919 | -;-;-;-;-;-;-;-;- | 1547 | plgrkt | 430 | 208 | 411 | 267 | 446 | 214 |
| ENSDARG000 | 21;21 | 271957:27514 | 272300:27537 | -;- | 575 | rln1 | 37 | 2 | 6 | 0 | 4 | 0 |
| ENSDARG000 | 21 | 5704393 | 5704593 | + | 201 | NA | 0 | 0 | 0 | 0 | 0 | 0 |
| ENSDARG000 | 21:21:21:21:2 | 5780888:5786 | 5782223:5786 | -;-;-;-;-;-;-;-;- | 2347 | ccni | 6663 | 3915 | 6304 | 10606 | 5060 | 6731 |
| ENSDARG000 | 21:21:21:21:2 | 13685853:136 | 13686940:136 | -;-;-;-;- | 2476 | pou5f3 | 6 | 2 | 6 | 10 | 14 | 7 |
| ENSDARG000 | 21:21:21:21:2 | 5635420:5636 | 5636163:5636 | +:+:+:+:+:+:+: | 5283 | SHROOM3 | 356 | 131 | 144 | 349 | 200 | 482 |
| ENSDARG000 | 21:21:21:21:2 | 5800306:5801 | 5800381:5801 | +:+:+:+:+:+:+: | 3880 | ccng2 | 1494 | 792 | 1089 | 2480 | 686 | 811 |
| ENSDARG000 | 21:21:21:21:2 | 35327328:353 | 35327497:353 | +:+:+:+:+:+:+: | 3083 | rars1 | 1102 | 1473 | 2356 | 952 | 1358 | 886 |
| ENSDARG000 | 21:21:21:21:2 | 24536233:245 | 24536295:245 | +:+:+:+:+:+:+: | 571 | anapc13 | 130 | 114 | 131 | 122 | 130 | 112 |
| ENSDARG000 | 21:21:21:21:2 | 40053666:400 | 40054506:400 | -:-:-:-:-:-:-:-:-: | 2840 | slc13a5a | 0 | 0 | 0 | 0 | 10 | 0 |
| ENSDARG000 | 21:21:21:21:2 | 27278120:272 | 27278428:272 | +:+:+:+:+:+:+: | 3296 | NA | 8 | 0 | 4 | 4 | 16 | 4 |
| ENSDARG000 | 21;21 | 27297145:272 | 27297388:272 | +;+ | 1573 | NA | 12 | 0 | 0 | 0 | 0 | 0 |
| ENSDARG000 | 21:21:21:21:2 | 40092301:400 | 40092402:400 | +:+:+:+:+:+:+: | 1805 | NA | 0 | 0 | 14 | 0 | 2 | 0 |
| ENSDARG000 | 21:21:21:21:2 | 17413613:174 | 17413926:174 | -;-;-;-;- | 2545 | GTF3C4 | 182 | 247 | 178 | 108 | 238 | 140 |
| ENSDARG000 | 21:21:21:21:2 | 24539896:245 | 24542130:245 | -:-:-:-:-:-:-:-:-: | 5853 | ARHGAP32 | 446 | 721 | 897 | 687 | 968 | 429 |
| ENSDARG000 | 21:21:21:21:2 | 20322879:203 | 20322916:203 | +:+:+:+:+:+:+: | 1932 | NA | 8 | 4 | 3 | 2 | 0 | 0 |
| ENSDARG000 | 21:21:21:21:2 | 20338613:203 | 20339884:203 | -:-:-:-:-:-:-:-:-: | 1989 | rbp4l | 0 | 0 | 0 | 2 | 0 | 0 |
| ENSDARG000 | 21:21:21:21:2 | 20324562:203 | 20326613:203 | -;-;-;-;- | 2972 | slc26a1 | 38 | 58 | 271 | 110 | 201 | 110 |
| ENSDARG000 | 21:21:21:21:2 | 22124736:221 | 22124772:221 | +:+:+:+:+:+:+: | 2406 | CUL5 | 14 | 14 | 15 | 15 | 42 | 47 |
| ENSDARG000 | 21:21:21:21:2 | 27302752:273 | 27303243:273 | +:+:+:+:+:+:+: | 2788 | sart1 | 1626 | 1718 | 1946 | 938 | 1800 | 1210 |
| ENSDARG000 | 21:21:21:21:2 | 35382443:353 | 35382686:353 | -:-:-:-:-:-:-:-:-: | 3940 | NA | 33 | 30 | 31 | 40 | 139 | 112 |
| ENSDARG000 | 21:21:21:21:2 | 40106448:401 | 40106507:401 | +:+:+:+:+:+:+: | 1946 | serpinf1 | 48 | 54 | 42 | 14 | 106 | 48 |
| ENSDARG000 | 21:21:21:21:2 | 13702566:137 | 13702802:137 | -;-;-;-;-;-;-;-;- | 1359 | npdc1a | 149 | 32 | 65 | 62 | 77 | 219 |
| ENSDARG000 | 21:21:21:21:2 | 40139036:401 | 40141438:401 | -:-:-:-:-:-:-:-:-: | 5671 | slco2b1 | 324 | 44 | 40 | 12 | 108 | 121 |
| ENSDARG000 | 21:21:21:21:2 | 17478493:174 | 17479023:174 | -;-;-;-;- | 1448 | barhl1b | 0 | 0 | 0 | 0 | 0 | 0 |
| ENSDARG000 | 21:21:21:21:2 | 20356458:203 | 20357348:203 | +:+:+:+:+:+:+: | 5593 | PDE6B | 4 | 10 | 22 | 8 | 14 | 32 |
| ENSDARG000 | 21:21:21:21:2 | 27336550:273 | 27336659:273 | +:+:+:+:+:+:+: | 3318 | dpp3 | 1756 | 1861 | 1754 | 1034 | 1950 | 1212 |
| ENSDARG000 | 21:21:21:21:2 | 24703188:247 | 24704092:247 | -:-:-:-:-:-:-:-:-: | 4215 | IGSF9B | 4 | 10 | 0 | 6 | 20 | 2 |
| ENSDARG000 | 21:21:21:21:2 | 35900917:359 | 35901328:359 | +:+:+:+:+:+:+: | 1543 | NA | 10 | 27 | 11 | 4 | 8 | 2 |
| ENSDARG000 | 21:21:21:21:2 | 27327442:273 | 27328148:273 | -:-:-:-:-:-:-:-:-: | 2522 | hif1al2 | 4 | 2 | 12 | 4 | 42 | 12 |
| ENSDARG000 | 21:21:21:21:2 | 13762409:137 | 13762819:137 | -:-:-:-:-:-:-:-:-: | 2743 | NA | 634 | 442 | 672 | 208 | 667 | 441 |
| ENSDARG000 | 21:21:21:21:2 | 22158744:221 | 22158922:221 | +:+:+:+:+:+:+: | 1943 | CAPN5 | 0 | 0 | 0 | 0 | 0 | 0 |
| ENSDARG000 | 21:21:21:21:2 | 37637679:376 | 37637847:376 | -:-:-:-:-:-:-:-:-: | 1555 | ndfip1l | 982 | 775 | 911 | 922 | 1577 | 1180 |
| ENSDARG000 | 21 | 40195452 | 40197014 | + | 1563 | or70b1 | 0 | 0 | 0 | 0 | 0 | 0 |
| ENSDARG000 | 21;21 | 22188010:221 | 22189719:221 | -;- | 1710 | ompb | 0 | 0 | 0 | 0 | 0 | 0 |
| ENSDARG000 | 21:21:21:21:2 | 17541634:175 | 17542651:175 | +:+:+:+:+:+:+: | 4732 | stom | 5193 | 4726 | 6788 | 4674 | 5605 | 5534 |
| ENSDARG000 | 21:21:21:21:2 | 13785969:137 | 13786174:137 | +:+:+:+:+:+:+: | 2172 | slc31a2 | 252 | 168 | 189 | 176 | 192 | 341 |
| ENSDARG000 | 21;21;21 | 40202617:402 | 40202687:402 | +;+;+ | 1819 | or70a12 | 0 | 0 | 0 | 0 | 0 | 0 |
| ENSDARG000 | 21:21:21:21:2 | 24984970:250 | 24985109:250 | +:+:+:+:+:+:+: | 4865 | jam3b | 10978 | 10921 | 10717 | 14542 | 10641 | 20476 |
| ENSDARG000 | 21:21:21:21:2 | 17572421:175 | 17573202:175 | -:-:-:-:-:-:-:-:-: | 2981 | gsna | 6739 | 7647 | 5326 | 7161 | 5861 | 4727 |
| ENSDARG000 | 21:21:21:21:2 | 9576176:9576 | 9576393:9576 | +:+:+:+:+:+:+: | 2952 | mapk10 | 0 | 3 | 0 | 0 | 6 | 0 |
| ENSDARG000 | 21;21;21 | 40215375:402 | 40215479:402 | +;+;+ | 2184 | or70a2 | 0 | 0 | 0 | 0 | 0 | 0 |
| ENSDARG000 | 21;21 | 40219818:402 | 40219867:402 | +;+ | 1608 | or70a17 | 0 | 0 | 0 | 0 | 0 | 0 |
| ENSDARG000 | 21:21:21:21:2 | 5888641:5891 | 5888961:5891 | +:+:+:+:+:+:+: | 2282 | NA | 1014 | 431 | 665 | 1097 | 438 | 1319 |
| ENSDARG000 | 21;21;21 | 40225915:402 | 40226046:402 | +;+;+ | 1412 | or70a3 | 0 | 0 | 0 | 0 | 0 | 0 |
| ENSDARG000 | 21 | 40232910 | 40234525 | + | 1616 | or70a11 | 0 | 0 | 0 | 0 | 0 | 0 |
| ENSDARG000 | 21:21:21:21:2 | 40235905:402 | 40236039:402 | +;+;+;+;+ | 1464 | or70a4 | 0 | 0 | 0 | 0 | 0 | 0 |
| ENSDARG000 | 21 | 40239192 | 40242287 | + | 3096 | or70a13 | 0 | 0 | 0 | 0 | 0 | 0 |
| ENSDARG000 | 21:21:21:21:2 | 13799448:138 | 13802008:138 | -:-:-:-:-:-:-:-:-: | 4401 | niban2a | 171 | 18 | 83 | 8 | 40 | 68 |
| ENSDARG000 | 21;21;21;21 | 20377752:203 | 20378004:203 | -;-;-;- | 557 | atp5mea | 454 | 439 | 494 | 311 | 358 | 490 |
| ENSDARG000 | 21;21 | 40248120:402 | 40250028:402 | -;- | 2020 | NA | 0 | 0 | 0 | 0 | 0 | 0 |
| ENSDARG000 | 21;21 | 40254429:402 | 40254508:402 | +;+ | 1523 | or70a16 | 0 | 0 | 0 | 0 | 0 | 0 |
| ENSDARG000 | 21 | 40261187 | 40263307 | + | 2121 | NA | 0 | 0 | 0 | 0 | 0 | 0 |
| ENSDARG000 | 21:21:21:21:2 | 37668173:376 | 37671918:376 | -:-:-:-:-:-:-:-:-: | 9777 | mpp1 | 1587 | 987 | 1212 | 982 | 2108 | 1215 |
| ENSDARG000 | 21:21:21:21:2 | 27356768:273 | 27357082:273 | -:-:-:-:-:-:-:-:-: | 2344 | NA | 62 | 27 | 55 | 27 | 195 | 26 |
| ENSDARG000 | 21:21:21:21:2 | 17606383:176 | 17606525:176 | +;+;+;+;+;+ | 1090 | NA | 0 | 0 | 0 | 0 | 0 | 2 |
| ENSDARG000 | 21:21:21:21:2 | 27370671:273 | 27370728:273 | +;+;+;+;+;+;+ | 2982 | rbm14a | 1110 | 989 | 1806 | 1131 | 1467 | 1391 |
| ENSDARG000 | 21 | 702078 | 704318 | + | 2241 | NA | 48 | 20 | 14 | 24 | 54 | 36 |
| ENSDARG000 | 21:21:21:21:2 | 25054420:250 | 25054454:250 | +:+:+:+:+:+:+: | 1421 | RFFL | 50 | 42 | 113 | 26 | 34 | 19 |
| ENSDARG000 | 21:21:21:21:2 | 723998:72861 | 724185:72866 | +;+;+;+;+;+ | 2079 | oaz1b | 2727 | 2385 | 2734 | 903 | 2310 | 892 |
| ENSDARG000 | 21:21:21:21:2 | 25068215:250 | 25068668:250 | +:+:+:+:+:+:+: | 6668 | DIXDC1 | 4465 | 1380 | 378 | 2214 | 726 | 1212 |
| ENSDARG000 | 21:21:21:21:2 | 22237964:222 | 22237981:222 | +:+:+:+:+:+:+: | 6602 | MYO7A | 44 | 24 | 4 | 22 | 27 | 14 |
| ENSDARG000 | 21:21:21:21:2 | 13861589:138 | 13861713:138 | +:+:+:+:+:+:+: | 3629 | stxbp1a | 253 | 12 | 838 | 68 | 22 | 328 |
| ENSDARG000 | 21:21:21:21:2 | 17768174:178 | 17768381:178 | +:+:+:+:+:+:+: | 1851 | rxraa | 604 | 426 | 323 | 213 | 463 | 378 |
| ENSDARG000 | 21:21:21:21:2 | 5926777:5929 | 5927293:5929 | +:+:+:+:+:+:+: | 2964 | NA | 244 | 378 | 649 | 313 | 254 | 328 |
| ENSDARG000 | 21;21;21 | 40268274:402 | 40268291:402 | +;+;+ | 1980 | or70a8 | 0 | 0 | 0 | 0 | 0 | 0 |
| ENSDARG000 | 21 | 40276569 | 40279077 | + | 2509 | NA | 0 | 0 | 0 | 0 | 0 | 0 |
| ENSDARG000 | 21;21 | 40280027:402 | 40280146:402 | +;+ | 1333 | or70c1 | 0 | 0 | 0 | 0 | 0 | 0 |
| ENSDARG000 | 21 | 40287100 | 40289652 | + | 2553 | or70a15 | 0 | 0 | 0 | 0 | 0 | 0 |
| ENSDARG000 | 21;21 | 40292801:402 | 40292844:402 | +;+ | 2600 | NA | 0 | 0 | 0 | 0 | 0 | 0 |
| ENSDARG000 | 21 | 20383837 | 20384878 | + | 1042 | hspb9l | 2 | 0 | 0 | 0 | 0 | 0 |
| ENSDARG000 | 21:21:21:21:2 | 25120546:251 | 25120801:251 | +:+:+:+:+:+:+: | 2775 | NA | 694 | 579 | 778 | 379 | 780 | 530 |
| ENSDARG000 | 21:21:21:21:2 | 20386865:203 | 20387161:203 | +;+;+;+;+;+ | 3899 | NA | 162 | 144 | 192 | 82 | 157 | 163 |
| ENSDARG000 | 21:21:21:21:2 | 27382893:273 | 27383231:273 | +:+:+:+:+:+:+: | 3259 | actn3a | 0 | 6 | 6 | 2 | 2 | 0 |
| ENSDARG000 | 21:21:21:21:2 | 17956024:179 | 17956082:179 | +:+:+:+:+:+:+: | 2390 | dnai1.2 | 4 | 11 | 16 | 0 | 10 | 8 |
| ENSDARG000 | 21:21:21:21:2 | 17951286:179 | 17952371:179 | -:-:-:-:-:-:-:-:-: | 2149 | stx2a | 53 | 124 | 63 | 82 | 48 | 60 |
| ENSDARG000 | 21:21:21:21:2 | 944715:94749 | 944874:94757 | +:+:+:+:+:+:+: | 2449 | NARS1 | 5453 | 4506 | 7960 | 5711 | 4505 | 4587 |
| ENSDARG000 | 21 | 40296480 | 40297710 | + | 1231 | NA | 0 | 0 | 0 | 0 | 0 | 0 |
| ENSDARG000 | 21;21 | 40302747:403 | 40302796:403 | +;+ | 1096 | NA | 0 | 0 | 0 | 0 | 0 | 0 |
| ENSDARG000 | 21;21 | 40307476:403 | 40307547:403 | +;+ | 1149 | NA | 0 | 0 | 0 | 0 | 0 | 0 |
| ENSDARG000 | 21:21:21:21:2 | 9733221:9733 | 9733465:9733 | -:-:-:-:-:-:-:-:-: | 2570 | ARHGAP24 | 14 | 51 | 71 | 12 | 41 | 26 |
| ENSDARG000 | 21;21;21;21 | 40313877:403 | 40316494:403 | -;-;-;- | 2833 | NA | 0 | 0 | 2 | 0 | 0 | 0 |
| ENSDARG000 | 21;21 | 25181003:251 | 25181486:251 | +;+ | 2307 | NA | 0 | 0 | 0 | 0 | 12 | 0 |
| ENSDARG000 | 21:21:21:21:2 | 20396858:203 | 20397272:203 | +:+:+:+:+:+:+: | 1485 | zgc:103482 | 4 | 0 | 8 | 10 | 0 | 2 |
| ENSDARG000 | 21 | 5942484 | 5942957 | - | 474 | NA | 26 | 42 | 29 | 38 | 40 | 34 |
| ENSDARG000 | 21:21:21:21:2 | 40353543:403 | 40356182:403 | -:-:-:-:-:-:-:-:-: | 8600 | nup98 | 2516 | 3493 | 3921 | 2403 | 3474 | 1979 |
| ENSDARG000 | 21:21:21:21:2 | 25187210:251 | 25187794:251 | +:+:+:+:+:+:+: | 2044 | si:dkey-183i3. | 20 | 0 | 0 | 2 | 4 | 2 |
| ENSDARG000 | 21:21:21:21:2 | 5947836:5948 | 5947974:5948 | -;-;-;-;- | 1060 | CACFD1 | 62 | 54 | 55 | 44 | 40 | 41 |
| ENSDARG000 | 21:21:21:21:2 | 27407481:274 | 27408267:274 | -:-:-:-:-:-:-:-:-: | 3762 | slc29a2 | 86 | 424 | 478 | 142 | 363 | 226 |
| ENSDARG000 | 21;21;21 | 9987313:9989 | 9988130:9989 | -;-;- | 1395 | esm1 | 66 | 64 | 4 | 26 | 22 | 94 |
| ENSDARG000 | 21:21:21:21:2 | 20488662:205 | 20488766:205 | +:+:+:+:+:+:+: | 4947 | EFNA5 | 62 | 46 | 230 | 45 | 34 | 93 |
| ENSDARG000 | 21:21:21:21:2 | 917593:91829 | 919196:91896 | -:-:-:-:-:-:-:-:-: | 2495 | txnl1 | 1098 | 1203 | 1536 | 897 | 1331 | 1146 |
| ENSDARG000 | 21:21:21:21:2 | 929444:93491 | 929495:93494 | +;+;+;+;+ | 479 | NA | 0 | 0 | 0 | 0 | 0 | 0 |
| ENSDARG000 | 21:21:21:21:2 | 9997418:9997 | 9997760:9997 | +:+:+:+:+:+:+: | 2827 | NA | 15 | 8 | 306 | 93 | 352 | 58 |
| ENSDARG000 | 21:21:21:21:2 | 27416284:274 | 27416409:274 | +:+:+:+:+:+:+: | 2433 | cfb | 0 | 6 | 52 | 0 | 3 | 0 |
| ENSDARG000 | 21:21:21:21:2 | 40402508:404 | 40403615:404 | -:-:-:-:-:-:-:-:-: | 1781 | nsrp1 | 447 | 382 | 414 | 400 | 316 | 280 |
| ENSDARG000 | 21;21;21 | 20715020:207 | 20715237:207 | +;+;+ | 1317 | gadd45gb.1 | 2 | 0 | 0 | 0 | 0 | 12 |
| ENSDARG000 | 21:21:21:21:2 | 13923872:139 | 13924261:139 | -:-:-:-:-:-:-:-:-: | 5121 | NA | 1563 | 2 | 51 | 15 | 26 | 6 |
| ENSDARG000 | 21:21:21:21:2 | 40417116:404 | 40417275:404 | +:+:+:+:+:+:+: | 4066 | ssh2b | 1542 | 709 | 641 | 915 | 983 | 981 |
| ENSDARG000 | 21:21:21:21:2 | 5960443:5963 | 5960591:5963 | +;+;+;+;+;+ | 2005 | mob1bb | 856 | 865 | 1219 | 881 | 968 | 847 |
| ENSDARG000 | 21:21:21:21:2 | 5974590:5975 | 5974808:5975 | +;+;+;+;+;+;+ | 1126 | dck | 114 | 352 | 430 | 158 | 460 | 193 |
| ENSDARG000 | 21:21:21:21:2 | 25198637:252 | 25198696:252 | +:+:+:+:+:+:+: | 705 | si:dkev-183i3. | 4 | 13 | 5 | 28 | 27 | 16 |
| ENSDARG000 | 21:21:21:21:2 | 25206289:252 | 25206754:252 | -:-:-:-:-:-:-:-:-: | 1615 | rfc2 | 230 | 474 | 562 | 257 | 517 | 298 |
| ENSDARG000 | 21:21:21:21:2 | 10253976:102 | 10254988:102 | -:-:-:-:-:-:-:-:-: | 2911 | HCN1 | 0 | 2 | 0 | 0 | 0 | 0 |
| ENSDARG000 | 21;21 | 40523913:405 | 40525341:405 | +;+ | 2499 | NA | 8 | 10 | 8 | 20 | 17 | 8 |
| ENSDARG000 | 21:21:21:21:2 | 13979033:139 | 13983762:139 | -:-:-:-:-:-:-:-:-: | 8317 | WHRN | 268 | 24 | 30 | 130 | 35 | 70 |
| ENSDARG000 | 21:21:21:21:2 | 40484088:404 | 40484151:404 | +:+:+:+:+:+:+: | 1528 | coro6 | 12 | 4 | 9 | 22 | 17 | 10 |
| ENSDARG000 | 21:21:21:21:2 | 37941347:379 | 37941362:379 | -;-;-;-;-;-;- | 1652 | PRRG3 | 1 | 11 | 23 | 35 | 6 | 0 |
| ENSDARG000 | 21:21:21:21:2 | 20722172:207 | 20722927:207 | -;-;-;-;-;-;- | 1580 | NA | 0 | 0 | 0 | 0 | 0 | 0 |
| ENSDARG000 | 21:21:21:21:2 | 40528788:405 | 40533675:405 | -:-:-:-:-:-:-:-:-: | 7953 | NA | 2102 | 1175 | 1446 | 1866 | 1498 | 1392 |
| ENSDARG000 | 21:21:21:21:2 | 1568044:1568 | 1568045:1568 | +:+:+:+:+:+:+: | 678 | RAB27B | 1677 | 1278 | 1236 | 648 | 1256 | 949 |
| ENSDARG000 | 21:21:21:21:2 | 37966933:379 | 37968085:379 | -:-:-:-:-:-:-:-:-: | 3352 | ripply1 | 0 | 0 | 0 | 8 | 0 | 2 |
| ENSDARG000 | 21;21 | 1608929:1609 | 1609034:1609 | -;- | 446 | NA | 5 | 0 | 4 | 2 | 6 | 1 |
| ENSDARG000 | 21:21:21:21:2 | 1586379:1587 | 1586413:1587 | +:+:+:+:+:+:+: | 2714 | wdr91 | 952 | 617 | 402 | 659 | 355 | 761 |
| ENSDARG000 | 21 | 25221940 | 25222528 | + | 589 | NA | 0 | 0 | 0 | 0 | 0 | 0 |
| ENSDARG000 | 21 | 25226558 | 25227191 | + | 634 | sycn.2 | 0 | 0 | 0 | 0 | 0 | 0 |
| ENSDARG000 | 21:21:21:21:2 | 25231160:252 | 25231281:252 | +;+;+;+;+;+;+ | 1346 | gng8 | 0 | 0 | 0 | 0 | 0 | 0 |
| ENSDARG000 | 21;21;21;21 | 17984451:179 | 17985167:179 | -;-;-;- | 1048 | NA | 0 | 0 | 0 | 0 | 0 | 0 |
| ENSDARG000 | 21;21 | 17984872:179 | 17985684:179 | +;+ | 1259 | NA | 0 | 0 | 0 | 0 | 0 | 0 |
| ENSDARG000 | 21;21 | 38002879:380 | 38003028:380 | +;+ | 1873 | cldn2 | 6 | 10 | 14 | 8 | 4 | 11 |
| ENSDARG000 | 21:21:21:21:2 | 37996822:380 | 37997841:380 | -:-:-:-:-:-:-:-:-: | 3311 | rbm41 | 80 | 40 | 73 | 64 | 118 | 92 |
| ENSDARG000 | 21:21:21:21:2 | 25236297:252 | 25236395:252 | +:+:+:+:+:+:+: | 1700 | tmem45b | 10 | 1 | 16 | 0 | 23 | 14 |
| ENSDARG000 | 21:21:21:21:2 | 25246933:252 | 25247194:252 | -:-:-:-:-:-:-:-:-: | 4608 | nfrkb | 652 | 473 | 724 | 463 | 599 | 617 |
| ENSDARG000 | 21:21:21:21:2 | 14217284:142 | 14217890:142 | -:-:-:-:-:-:-:-:-: | 3243 | MAN1B1 | 707 | 312 | 368 | 535 | 428 | 498 |
| ENSDARG000 | 21:21:21:21:2 | 38033226:380 | 38033610:380 | +:+:+:+:+:+:+: | 2299 | klf8 | 336 | 294 | 273 | 338 | 385 | 340 |
| ENSDARG000 | 21:21:21:21:2 | 40589770:405 | 40590032:405 | +:+:+:+:+:+:+: | 2131 | pdk3b | 62 | 34 | 59 | 50 | 67 | 118 |
| ENSDARG000 | 21:21:21:21:2 | 40618615:406 | 40619315:406 | -;-;-;-;-;-;-;- | 1671 | pcyt1bb | 0 | 0 | 10 | 0 | 0 | 0 |
| ENSDARG000 | 21;21;21;21 | 38128877:381 | 38130139:381 | -;-;-;- | 2361 | klf5l | 88 | 0 | 0 | 26 | 10 | 32 |
| ENSDARG000 | 21;21 | 38312549:383 | 38312859:383 | +;+ | 2339 | pou4f4 | 0 | 0 | 0 | 0 | 0 | 0 |
| ENSDARG000 | 21;21 | 38458072:384 | 38458202:384 | -;- | 721 | NA | 73 | 30 | 137 | 85 | 285 | 100 |
| ENSDARG000 | 21:21:21:21:2 | 40632328:406 | 40632458:406 | +;+;+;+;+;+ | 613 | NA | 0 | 0 | 0 | 0 | 0 | 0 |
| ENSDARG000 | 21:21:21:21:2 | 18021087:180 | 18024725:180 | -:-:-:-:-:-:-:-:-: | 6219 | VAV2 | 555 | 74 | 299 | 118 | 292 | 74 |
| ENSDARG000 | 21:21:21:21:2 | 38509465:385 | 38514148:385 | -:-:-:-:-:-:-:-:-: | 6516 | NA | 144 | 26 | 19 | 4 | 26 | 14 |
| ENSDARG000 | 21:21:21:21:2 | 14253404:142 | 14254824:142 | -:-:-:-:-:-:-:-:-: | 6041 | NA | 92 | 22 | 290 | 204 | 406 | 390 |
| ENSDARG000 | 21:21:21:21:2 | 25275693:252 | 25276023:252 | -:-:-:-:-:-:-:-:-: | 3011 | st14b | 52 | 0 | 2 | 24 | 24 | 16 |
| ENSDARG000 | 21:21:21:21:2 | 40685895:406 | 40686540:406 | +:+:+:+:+:+:+: | 2221 | ccdc82 | 338 | 439 | 594 | 350 | 520 | 416 |
| ENSDARG000 | 21:21:21:21:2 | 40712757:407 | 40713042:407 | -;-;-;-;-;-;- | 716 | pomp | 1301 | 1450 | 1804 | 948 | 1532 | 996 |
| ENSDARG000 | 21 | 25323331 | 25325845 | + | 2515 | or92a1 | 0 | 0 | 0 | 0 | 0 | 0 |
| ENSDARG000 | 21 | 25335807 | 25338687 | + | 2881 | or92a4 | 0 | 0 | 0 | 0 | 0 | 0 |

| | | | | | | | | | | | | |
|---|---|---|---|---|---|---|---|---|---|---|---|---|
| ENSDARG000 | 21 | 25344374 | 25345902 | + | 1529 | NA | 0 | 0 | 0 | 0 | 0 | 0 |
| ENSDARG000 | 21:21:21:21:2 | 18274825:182 | 18275258:182 | +:+:+:+:+:+:+: | 3132 | wdr5 | 967 | 1300 | 1244 | 762 | 1462 | 997 |
| ENSDARG000 | 21 | 25346085 | 25346388 | + | 304 | NA | 0 | 0 | 0 | 0 | 0 | 0 |
| ENSDARG000 | 21:21:21:21:2 | 14381227:143 | 14381762:143 | -:-:-:-:-:-:-:-: | 7027 | CACNA1B | 174 | 1621 | 1074 | 286 | 2206 | 508 |
| ENSDARG000 | 21;21 | 25346781:253 | 25347003:253 | +;+ | 223 | NA | 0 | 0 | 0 | 0 | 0 | 0 |
| ENSDARG000 | 21 | 25354106 | 25356174 | + | 2069 | or92a2 | 0 | 0 | 0 | 0 | 0 | 0 |
| ENSDARG000 | 21 | 25364338 | 25365870 | + | 1533 | or92a3 | 0 | 0 | 0 | 0 | 0 | 0 |
| ENSDARG000 | 21:21:21:21:2 | 18266319:182 | 18266958:182 | -:-:-:-:-:-:-:-: | 3439 | brd3a | 742 | 851 | 1336 | 620 | 1169 | 686 |
| ENSDARG000 | 21 | 25365879 | 25366134 | + | 256 | NA | 0 | 0 | 0 | 0 | 0 | 0 |
| ENSDARG000 | 21:21:21:21:2 | 18292535:182 | 18292665:182 | +:+:+:+:+:+:+: | 2067 | DNAI1 | 0 | 2 | 0 | 0 | 2 | 13 |
| ENSDARG000 | 21:21:21:21:2 | 38585686:385 | 38589772:385 | -:-:-:-:-:-:-:-: | 5451 | slc25a14 | 1147 | 1150 | 1280 | 774 | 1093 | 940 |
| ENSDARG000 | 21:21:21:21:2 | 11684830:116 | 11685182:116 | +:+:+:+:+:+:+: | 2516 | pcsk1 | 0 | 0 | 2 | 0 | 0 | 0 |
| ENSDARG000 | 21;21 | 25375172:253 | 25375250:253 | +;+ | 1441 | NA | 0 | 0 | 0 | 0 | 0 | 0 |
| ENSDARG000 | 21;21 | 25376752:253 | 25377275:253 | -;- | 689 | NA | 0 | 0 | 0 | 0 | 0 | 0 |
| ENSDARG000 | 21 | 25376955 | 25377279 | + | 325 | NA | 0 | 0 | 0 | 0 | 0 | 0 |
| ENSDARG000 | 21:21:21:21:2 | 38638979:386 | 38639052:386 | +;+;+;+;+;+ | 1523 | rbmx2 | 166 | 192 | 216 | 217 | 108 | 118 |
| ENSDARG000 | 21:21:21:21:2 | 25380675:253 | 25381079:253 | -:-:-:-:-:-:-:-: | 1722 | ppme1 | 1045 | 1090 | 1336 | 1584 | 1196 | 1088 |
| ENSDARG000 | 21:21:21:21:2 | 40786804:407 | 40786905:407 | -:-:-:-:-:-:-:-: | 1099 | zgc:162239 | 125 | 12 | 46 | 204 | 111 | 116 |
| ENSDARG000 | 21:21:21:21:2 | 38659302:386 | 38660504:386 | -;-;-;-;- | 1787 | siah2l | 226 | 102 | 130 | 111 | 157 | 97 |
| ENSDARG000 | 21:21:21:21:2 | 40805336:408 | 40805516:408 | -:-:-:-:-:-:-:-: | 2584 | NA | 2908 | 567 | 1056 | 2623 | 915 | 2036 |
| ENSDARG000 | 21:21:21:21:2 | 25399133:254 | 25399972:254 | -:-:-:-:-:-:-:-: | 1997 | sms | 674 | 815 | 1019 | 661 | 1193 | 1134 |
| ENSDARG000 | 21:21:21:21:2 | 40839857:408 | 40839940:408 | -:-:-:-:-:-:-:-: | 6233 | elnb | 50 | 0 | 4 | 4 | 71 | 2 |
| ENSDARG000 | 21:21:21:21:2 | 38724046:387 | 38724386:387 | -:-:-:-:-:-:-:-: | 1473 | taf9 | 664 | 567 | 757 | 629 | 625 | 582 |
| ENSDARG000 | 21:21:21:21:2 | 40930177:409 | 40930896:409 | -;-;-;-;-;- | 1430 | yipf5 | 628 | 801 | 785 | 388 | 971 | 719 |
| ENSDARG000 | 21:21:21:21:2 | 11749097:117 | 11749292:117 | +:+:+:+:+:+:+: | 4266 | ell2 | 1487 | 1798 | 2008 | 1201 | 1862 | 1060 |
| ENSDARG000 | 21;21 | 40944530:409 | 40945403:409 | +;+ | 1353 | kctd16b | 0 | 0 | 0 | 0 | 0 | 0 |
| ENSDARG000 | 21:21:21:21:2 | 11776724:117 | 11781668:117 | -:-:-:-:-:-:-:-: | 7104 | NA | 46 | 34 | 73 | 34 | 109 | 17 |
| ENSDARG000 | 21:21:21:21:2 | 11776328:117 | 11776552:117 | +:+:+:+:+:+:+: | 1610 | glrx | 583 | 200 | 431 | 571 | 819 | 491 |
| ENSDARG000 | 21:21:21:21:2 | 38732945:387 | 38733425:387 | +:+:+:+:+:+:+: | 1973 | rab24 | 267 | 259 | 183 | 219 | 265 | 317 |
| ENSDARG000 | 21:21:21:21:2 | 41026612:410 | 41029599:410 | -:-:-:-:-:-:-:-: | 5363 | LARS1 | 1421 | 1634 | 2147 | 1502 | 1475 | 1332 |
| ENSDARG000 | 21:21:21:21:2 | 41016729:410 | 41016758:410 | -:-:-:-:-:-:-:-: | 1115 | PLAC8L1 | 0 | 4 | 6 | 4 | 11 | 0 |
| ENSDARG000 | 21:21:21:21:2 | 38745094:387 | 38745401:387 | +:+:+:+:+:+:+: | 3836 | heatr6 | 225 | 360 | 264 | 80 | 512 | 253 |
| ENSDARG000 | 21:21:21:21:2 | 14644247:146 | 14645915:146 | -:-:-:-:-:-:-:-: | 6056 | ehmt1b | 1184 | 822 | 1397 | 805 | 1401 | 1123 |
| ENSDARG000 | 21:21:21:21:2 | 38817785:388 | 38818322:388 | +:+:+:+:+:+:+: | 4162 | hnf1bb | 0 | 0 | 28 | 14 | 54 | 0 |
| ENSDARG000 | 21:21:21:21:2 | 39005905:390 | 39006775:390 | -;-;-;-;-;-;- | 1781 | traf4b | 32 | 0 | 14 | 10 | 14 | 12 |
| ENSDARG000 | 21:21:21:21:2 | 41256771:412 | 41257904:412 | -;-;-;-;- | 1576 | nsg2 | 32 | 18 | 38 | 4 | 0 | 70 |
| ENSDARG000 | 21:21:21:21:2 | 41597969:416 | 41601988:416 | -:-:-:-:-:-:-:-: | 5090 | PCYOX1L | 609 | 1156 | 594 | 477 | 967 | 741 |
| ENSDARG000 | 21:21:21:21:2 | 14768411:147 | 14769099:147 | -:-:-:-:-:-:-:-: | 1592 | glulc | 0 | 0 | 0 | 0 | 0 | 0 |
| ENSDARG000 | 21:21:21:21:2 | 39029713:390 | 39029893:390 | -:-:-:-:-:-:-:-: | 2130 | aldh3a2b | 399 | 286 | 288 | 361 | 437 | 255 |
| ENSDARG000 | 21:21:21:21:2 | 14784321:147 | 14785360:147 | -:-:-:-:-:-:-:-: | 3827 | NA | 0 | 2 | 2 | 0 | 4 | 0 |
| ENSDARG000 | 21:21:21:21:2 | 41649336:416 | 41649769:416 | +:+:+:+:+:+:+: | 2394 | ppp2r2bb | 24 | 68 | 58 | 121 | 14 | 38 |
| ENSDARG000 | 21:21:21:21:2 | 41819288:418 | 41820691:418 | -:-:-:-:-:-:-:-: | 4222 | cct6a | 7736 | 9102 | 12529 | 9429 | 9134 | 10314 |
| ENSDARG000 | 21:21:21:21:2 | 41837610:418 | 41838325:418 | +:+:+:+:+:+:+: | 3015 | rnf14 | 588 | 563 | 733 | 533 | 727 | 497 |
| ENSDARG000 | 21:21:21:21:2 | 39067512:390 | 39067917:390 | -:-:-:-:-:-:-:-: | 2133 | vtnb | 0 | 0 | 16 | 0 | 0 | 0 |
| ENSDARG000 | 21:21:21:21:2 | 39100289:391 | 39100514:391 | +:+:+:+:+:+:+: | 2595 | slc13a2 | 2 | 4 | 6 | 0 | 16 | 0 |
| ENSDARG000 | 21:21:21:21:2 | 41863540:418 | 41863871:418 | -:-:-:-:-:-:-:-: | 1562 | endou2 | 0 | 0 | 0 | 0 | 0 | 0 |
| ENSDARG000 | 21:21:21:21:2 | 41903760:419 | 41904041:419 | -:-:-:-:-:-:-:-: | 1909 | GABRG2 | 2 | 8 | 0 | 0 | 38 | 0 |
| ENSDARG000 | 21:21:21:21:2 | 39121453:391 | 39122865:391 | -;-;-;-;- | 2239 | unc119b | 4 | 0 | 10 | 4 | 0 | 0 |
| ENSDARG000 | 21;21 | 42043974:420 | 42045383:420 | -;- | 3181 | NA | 0 | 0 | 0 | 0 | 0 | 0 |
| ENSDARG000 | 21:21:21:21:2 | 42050351:420 | 42051235:420 | -:-:-:-:-:-:-:-: | 2305 | gabra1 | 0 | 0 | 0 | 0 | 0 | 0 |
| ENSDARG000 | 21:21:21:21:2 | 42708621:427 | 42708657:427 | +;+;+;+;+;+;+ | 1080 | NA | 0 | 0 | 0 | 0 | 0 | 0 |
| ENSDARG000 | 21:21:21:21:2 | 39185461:391 | 39186174:391 | +:+:+:+:+:+:+: | 2267 | cryba1b | 0 | 0 | 0 | 0 | 0 | 0 |
| ENSDARG000 | 21:21:21:21:2 | 42717424:427 | 42717685:427 | +:+:+:+:+:+:+: | 4722 | SH3PXD2B | 1397 | 867 | 1721 | 1039 | 1705 | 1157 |
| ENSDARG000 | 21:21:21:21:2 | 39197418:391 | 39198100:391 | +:+:+:+:+:+:+: | 3980 | cpdb | 0 | 0 | 0 | 0 | 0 | 0 |
| ENSDARG000 | 21:21:21:21:2 | 42891618:429 | 42891749:429 | +:+:+:+:+:+:+: | 7398 | NA | 0 | 25 | 4 | 23 | 9 | 8 |
| ENSDARG000 | 21:21:21:21:2 | 42930558:429 | 42930600:429 | +;+;+;+;+;+;+ | 739 | STK32B | 0 | 0 | 0 | 0 | 0 | 0 |
| ENSDARG000 | 21:21:21:21:2 | 42970983:429 | 42974316:429 | -:-:-:-:-:-:-:-: | 5438 | dpysl3 | 378 | 405 | 802 | 554 | 418 | 1192 |
| ENSDARG000 | 21;21;21 | 39240897:392 | 39242471:392 | -;-;- | 2248 | trarg1b | 0 | 0 | 0 | 0 | 0 | 0 |
| ENSDARG000 | 21;21 | 39292266:392 | 39292633:392 | +;+ | 1099 | NA | 2 | 8 | 0 | 0 | 0 | 4 |
| ENSDARG000 | 21:21:21:21:2 | 39302793:393 | 39303196:393 | -:-:-:-:-:-:-:-: | 1996 | NA | 0 | 2 | 14 | 0 | 14 | 0 |
| ENSDARG000 | 21:21:21:21:2 | 39336285:393 | 39336381:393 | +;+;+;+;+ | 1158 | NA | 0 | 10 | 19 | 0 | 13 | 2 |
| ENSDARG000 | 21:21:21:21:2 | 39361040:393 | 39362234:393 | +:+:+:+:+:+:+: | 4278 | CLUH | 11 | 57 | 15 | 2 | 0 | 6 |
| ENSDARG000 | 21:21:21:21:2 | 43029708:430 | 43034137:430 | -:-:-:-:-:-:-:-: | 7419 | JAKMIP2 | 156 | 88 | 236 | 116 | 36 | 214 |
| ENSDARG000 | 21:21:21:21:2 | 43080469:430 | 43080539:430 | -:-:-:-:-:-:-:-: | 2086 | NA | 0 | 4 | 0 | 6 | 0 | 6 |
| ENSDARG000 | 21:21:21:21:2 | 43097798:430 | 43098182:430 | -:-:-:-:-:-:-:-: | 2081 | p4ha2 | 1270 | 1807 | 5039 | 4367 | 2266 | 999 |
| ENSDARG000 | 21:21:21:21:2 | 43137500:431 | 43138212:431 | -:-:-:-:-:-:-:-: | 3219 | hspa4a | 1994 | 3040 | 3082 | 1331 | 2602 | 1418 |
| ENSDARG000 | 21:21:21:21:2 | 43171013:431 | 43171374:431 | +:+:+:+:+:+:+: | 2378 | zcchc10 | 210 | 129 | 298 | 216 | 209 | 184 |
| ENSDARG000 | 21:21:21:21:2 | 39423974:394 | 39424152:394 | +:+:+:+:+:+:+: | 3183 | pafah1b1b | 2501 | 2506 | 2566 | 1964 | 2427 | 2014 |
| ENSDARG000 | 21:21:21:21:2 | 43178817:431 | 43179070:431 | +:+:+:+:+:+:+: | 3695 | AFF4 | 822 | 476 | 861 | 557 | 1212 | 694 |
| ENSDARG000 | 21:21:21:21:2 | 43253538:432 | 43253706:432 | +:+:+:+:+:+:+: | 9145 | NA | 14 | 16 | 70 | 4 | 92 | 38 |
| ENSDARG000 | 21:21:21:21:2 | 39461926:394 | 39462767:394 | +:+:+:+:+:+:+: | 6820 | mntb | 413 | 294 | 413 | 203 | 484 | 475 |
| ENSDARG000 | 21:21:21:21:2 | 39512437:395 | 39512677:395 | -:-:-:-:-:-:-:-: | 1859 | septin4a | 2 | 0 | 0 | 0 | 0 | 0 |
| ENSDARG000 | 21;21;21 | 39557433:395 | 39557982:395 | -;-;- | 807 | dynll2b | 1007 | 803 | 694 | 887 | 660 | 757 |
| ENSDARG000 | 21 | 43326913 | 43328056 | - | 1144 | NA | 20 | 2 | 4 | 18 | 2 | 0 |
| ENSDARG000 | 21:21:21:21:2 | 43328685:433 | 43328834:433 | +:+:+:+:+:+:+: | 2021 | septin8a | 2229 | 2432 | 2396 | 2336 | 2595 | 2202 |
| ENSDARG000 | 21:21:21:21:2 | 43379878:433 | 43380126:433 | -:-:-:-:-:-:-:-: | 1466 | CCNI2 | 25 | 28 | 35 | 104 | 10 | 10 |
| ENSDARG000 | 21;21 | 43390401:433 | 43390794:433 | -;- | 633 | NA | 0 | 4 | 0 | 3 | 0 | 7 |
| ENSDARG000 | 21:21:21:21:2 | 43400351:434 | 43400445:434 | +:+:+:+:+:+:+: | 2229 | frmd7 | 6 | 31 | 32 | 35 | 40 | 22 |
| ENSDARG000 | 21:21:21:21:2 | 43420304:434 | 43422449:434 | -:-:-:-:-:-:-:-: | 3868 | stk26 | 552 | 373 | 740 | 467 | 528 | 528 |
| ENSDARG000 | 21:21:21:21:2 | 39705483:397 | 39705811:397 | +:+:+:+:+:+:+: | 3343 | plrdgb | 38 | 14 | 11 | 52 | 4 | 183 |
| ENSDARG000 | 21:21:21:21:2 | 43460631:434 | 43462231:434 | -:-:-:-:-:-:-:-: | 2709 | tmem185 | 698 | 453 | 817 | 528 | 680 | 595 |
| ENSDARG000 | 21:21:21:21:2 | 43475746:434 | 43478174:434 | -;-;-;-;-;-;- | 2965 | ANKRD46 | 1111 | 807 | 941 | 1057 | 994 | 1230 |
| ENSDARG000 | 21:21:21:21:2 | 39992151:399 | 39992663:399 | -:-:-:-:-:-:-:-: | 2261 | slc47a2.1 | 2 | 12 | 119 | 110 | 136 | 34 |
| ENSDARG000 | 21:21:21:21:2 | 43485512:434 | 43485562:434 | +:+:+:+:+:+:+: | 6199 | atg4a | 589 | 240 | 388 | 260 | 394 | 334 |
| ENSDARG000 | 21:21:21:21:2 | 39941184:399 | 39941271:399 | +:+:+:+:+:+:+: | 5795 | slc47a1 | 0 | 0 | 2 | 0 | 20 | 0 |
| ENSDARG000 | 21;21 | 43515673:435 | 43515963:435 | -;- | 3442 | NA | 232 | 114 | 252 | 182 | 203 | 193 |
| ENSDARG000 | 21:21:21:21:2 | 43536339:435 | 43536628:435 | -;-;-;-;- | 2441 | NA | 8 | 2 | 24 | 30 | 6 | 25 |
| ENSDARG000 | 21;21;21;21 | 43559123:435 | 43559385:435 | +;+;+;+ | 1271 | NA | 0 | 0 | 0 | 0 | 0 | 0 |
| ENSDARG000 | 21:21:21:21:2 | 43581341:435 | 43581748:435 | -;-;-;-;-;-;-;- | 1905 | NA | 538 | 300 | 610 | 635 | 212 | 413 |
| ENSDARG000 | 21;21;21 | 43609452:436 | 43610047:436 | -;-;- | 1068 | NA | 122 | 106 | 239 | 220 | 151 | 137 |
| ENSDARG000 | 21:21:21:21:2 | 43630438:436 | 43633714:436 | -:-:-:-:-:-:-:-: | 4214 | NA | 0 | 0 | 1 | 0 | 2 | 0 |
| ENSDARG000 | 21;21;21;21 | 40045224:400 | 40045767:400 | -;-;-;- | 871 | med31 | 226 | 245 | 268 | 292 | 341 | 342 |
| ENSDARG000 | 21:21:21:21:2 | 43644573:436 | 43648421:436 | -:-:-:-:-:-:-:-: | 4421 | NA | 0 | 1 | 2 | 24 | 0 | 0 |
| ENSDARG000 | 21;21;21 | 43653826:436 | 43654569:436 | -;-;- | 923 | NA | 0 | 0 | 0 | 0 | 0 | 0 |
| ENSDARG000 | 21:21:21:21:2 | 43657216:436 | 43664786:436 | -;-;-;-;- | 7815 | NA | 0 | 0 | 0 | 0 | 0 | 0 |
| ENSDARG000 | 21:21:21:21:2 | 43669943:436 | 43670123:436 | +:+:+:+:+:+:+: | 1572 | TMLHE | 152 | 134 | 229 | 111 | 174 | 155 |
| ENSDARG000 | 21:21:21:21:2 | 43702010:437 | 43702102:437 | +:+:+:+:+:+:+: | 1912 | dkc1 | 685 | 1467 | 2238 | 649 | 1398 | 973 |
| ENSDARG000 | 21:21:21:21:2 | 43738398:437 | 43739154:437 | -:-:-:-:-:-:-:-: | 1865 | pdlim4 | 952 | 816 | 1127 | 350 | 779 | 260 |
| ENSDARG000 | 21;21 | 43841656:438 | 43841804:438 | +;+ | 576 | NA | 8 | 26 | 24 | 66 | 172 | 67 |
| ENSDARG000 | 21:21:21:21:2 | 43854594:438 | 43854653:438 | +:+:+:+:+:+:+: | 2894 | NA | 6 | 18 | 14 | 8 | 48 | 92 |
| ENSDARG000 | 21:21:21:21:2 | 43882274:438 | 43882370:438 | +;+;+;+;+ | 1081 | sra1 | 162 | 217 | 256 | 143 | 242 | 205 |
| ENSDARG000 | 21:21:21:21:2 | 43897128:439 | 43897544:439 | -:-:-:-:-:-:-:-: | 1976 | camk2a | 0 | 4 | 15 | 3 | 0 | 0 |
| ENSDARG000 | 21:21:21:21:2 | 44300689:443 | 44300815:443 | +:+:+:+:+:+:+: | 834 | NA | 23 | 29 | 74 | 2 | 10 | 19 |
| ENSDARG000 | 21:21:21:21:2 | 44505613:445 | 44505773:445 | -;-;-;-;-;-;-;- | 842 | zgc:136410 | 2 | 0 | 2 | 2 | 0 | 12 |
| ENSDARG000 | 21:21:21:21:2 | 44557791:445 | 44559148:445 | -:-:-:-:-:-:-:-: | 2564 | fundc2 | 1748 | 2330 | 2043 | 1810 | 1610 | 1940 |
| ENSDARG000 | 21:21:21:21:2 | 44547450:445 | 44547598:445 | -:-:-:-:-:-:-:-: | 1487 | BRCC3 | 221 | 184 | 132 | 111 | 183 | 262 |
| ENSDARG000 | 21;21 | 44557006:445 | 44557142:445 | +;+ | 423 | cmc4 | 8 | 0 | 5 | 14 | 12 | 0 |
| ENSDARG000 | 21:21:21:21:2 | 520479:52209 | 520667:52223 | +:+:+:+:+:+:+: | 2304 | nop14 | 588 | 491 | 662 | 368 | 592 | 474 |
| ENSDARG000 | 21:21:21:21:2 | 39645762:396 | 39646100:396 | -:-:-:-:-:-:-:-: | 1618 | NA | 12 | 7 | 30 | 8 | 57 | 24 |
| ENSDARG000 | 21:21:21:21:2 | 14699693:147 | 14699760:147 | -;-;-;-;-;-;- | 1332 | ARRDC1 | 528 | 36 | 35 | 99 | 76 | 85 |
| ENSDARG000 | 21:21:21:21:2 | 22558187:225 | 22558416:225 | +:+:+:+:+:+:+: | 1754 | chek1 | 40 | 117 | 108 | 72 | 82 | 64 |
| ENSDARG000 | 21;21 | 10076203:100 | 10076969:100 | +;+ | 1180 | NA | 24 | 20 | 200 | 94 | 151 | 30 |
| ENSDARG000 | 21:21:21:21:2 | 45581985:455 | 45582125:455 | -:-:-:-:-:-:-:-: | 1013 | NA | 15 | 5 | 5 | 0 | 6 | 6 |
| ENSDARG000 | 21;21;21;21 | 419801:42627 | 419970:42699 | -;-;-;- | 1167 | klf4 | 83 | 12 | 16 | 28 | 66 | 16 |
| ENSDARG000 | 21;21 | 22692643:226 | 22693621:226 | -;- | 1003 | gig2d | 20 | 87 | 9 | 10 | 9 | 0 |
| ENSDARG000 | 21:21:21:21:2 | 4505117:4505 | 4505285:4505 | -;-;-;-;-;- | 1642 | pimr166 | 0 | 0 | 0 | 0 | 0 | 0 |
| ENSDARG000 | 21:21:21:21:2 | 37423717:374 | 37423809:374 | -:-:-:-:-:-:-:-: | 1582 | FAM114A2 | 531 | 425 | 642 | 400 | 546 | 442 |
| ENSDARG000 | 21:21:21:21:2 | 974555:97499 | 974677:97515 | +;+;+;+;+;+ | 825 | NA | 423 | 324 | 282 | 264 | 516 | 489 |
| ENSDARG000 | 21;21;21;21 | 39567505:395 | 39570987:395 | -;-;-;- | 4032 | srsf1b | 1624 | 1337 | 1391 | 1252 | 1288 | 1524 |
| ENSDARG000 | 21;21 | 2189257:2190 | 2189954:2190 | -;- | 756 | NA | 47 | 18 | 47 | 27 | 13 | 50 |
| ENSDARG000 | 21;21 | 7131970:7135 | 7132263:7136 | +;+ | 1666 | zgc:113019 | 278 | 212 | 262 | 144 | 276 | 331 |
| ENSDARG000 | 21 | 41251431 | 41251548 | - | 118 | 5S_rRNA | 0 | 0 | 0 | 0 | 0 | 0 |
| ENSDARG000 | 21;21;21 | 2721752:2724 | 2722644:2724 | +;+;+ | 1161 | NA | 70 | 77 | 109 | 86 | 119 | 111 |
| ENSDARG000 | 21;21;21;21 | 45875792:458 | 45876839:458 | -;-;-;- | 1525 | sap30l | 617 | 695 | 808 | 657 | 769 | 573 |
| ENSDARG000 | 21:21:21:21:2 | 30563115:305 | 30563244:305 | +:+:+:+:+:+:+: | 951 | NA | 229 | 83 | 366 | 135 | 386 | 122 |
| ENSDARG000 | 21;21 | 44112914:441 | 44113018:441 | +;+ | 1844 | fgf1b | 0 | 56 | 43 | 38 | 51 | 6 |
| ENSDARG000 | 21:21:21:21:2 | 300647:30100 | 301464:30146 | -;-;-;-;- | 1445 | lhfpl2a | 636 | 553 | 650 | 373 | 389 | 290 |
| ENSDARG000 | 21:21:21:21:2 | 37355716:373 | 37356412:373 | +:+:+:+:+:+:+: | 6061 | NA | 287 | 284 | 329 | 368 | 436 | 176 |
| ENSDARG000 | 21 | 26701821 | 26701954 | - | 134 | 5S_rRNA | 0 | 0 | 0 | 0 | 0 | 0 |
| ENSDARG000 | 21:21:21:21:2 | 226351:22685 | 226670:22698 | -:-:-:-:-:-:-:-: | 1236 | bxdc2 | 789 | 880 | 1396 | 485 | 846 | 563 |
| ENSDARG000 | 21 | 7446598 | 7446715 | + | 118 | 5S_rRNA | 0 | 1 | 0 | 0 | 0 | 0 |
| ENSDARG000 | 21:21:21:21:2 | 7157292:7160 | 7158202:7161 | -;-;-;-;- | 1487 | dipk1b | 89 | 28 | 54 | 16 | 12 | 82 |
| ENSDARG000 | 21 | 44568180 | 44568266 | - | 87 | mir222b | 0 | 0 | 0 | 0 | 0 | 0 |
| ENSDARG000 | 21 | 22362097 | 22362215 | - | 119 | U6 | 0 | 0 | 0 | 0 | 0 | 4 |
| ENSDARG000 | 21:21:21:21:2 | 44091342:440 | 44091435:440 | -:-:-:-:-:-:-:-: | 1671 | NA | 0 | 4 | 0 | 0 | 10 | 0 |
| ENSDARG000 | 21 | 35522702 | 35522834 | + | 133 | 5S_rRNA | 0 | 0 | 0 | 0 | 0 | 0 |
| ENSDARG000 | 21;21 | 43999699:440 | 44001103:440 | -;- | 1707 | NA | 136 | 58 | 86 | 74 | 67 | 174 |
| ENSDARG000 | 21:21:21:21:2 | 33187992:331 | 33188504:331 | +:+:+:+:+:+:+: | 2565 | NA | 9 | 31 | 32 | 27 | 10 | 21 |
| ENSDARG000 | 21:21:21:21:2 | 2203316:2203 | 2203339:2203 | +:+:+:+:+:+:+: | 300 | NA | 11 | 17 | 28 | 13 | 12 | 19 |
| ENSDARG000 | 21 | 28783827 | 28783944 | - | 118 | 5S_rRNA | 1 | 0 | 0 | 5 | 0 | 0 |
| ENSDARG000 | 21 | 38063197 | 38063314 | - | 118 | 5S_rRNA | 0 | 0 | 0 | 0 | 0 | 0 |
| ENSDARG000 | 21:21:21:21:2 | 45673433:456 | 45674797:456 | -;-;-;-;-;- | 2024 | sar1b | 1507 | 1764 | 2368 | 1191 | 2100 | 1384 |
| ENSDARG000 | 21:21:21:21:2 | 44041088:440 | 44041129:440 | -:-:-:-:-:-:-:-: | 2937 | NA | 26 | 4 | 1 | 3 | 4 | 0 |
| ENSDARG000 | 21 | 36191330 | 36191453 | + | 124 | 5S_rRNA | 0 | 0 | 0 | 0 | 0 | 0 |
| ENSDARG000 | 21 | 41639979 | 41640058 | + | 80 | dre-mir-146b | 0 | 0 | 2 | 0 | 0 | 0 |
| ENSDARG000 | 21:21:21:21:2 | 33172526:331 | 33172543:331 | +;+;+;+;+;+ | 3527 | arl3l1 | 0 | 0 | 0 | 0 | 0 | 0 |
| ENSDARG000 | 21;21;21;21 | 3188020:3195 | 3188557:3195 | -;-;-;- | 1815 | ZBTB7C | 2 | 2 | 0 | 0 | 0 | 0 |
| ENSDARG000 | 21 | 19155924 | 19156042 | + | 119 | 5S_rRNA | 0 | 0 | 0 | 0 | 0 | 0 |
| ENSDARG000 | 21 | 36488062 | 36488178 | + | 117 | 5S_rRNA | 0 | 0 | 0 | 0 | 0 | 0 |
| ENSDARG000 | 21;21 | 26555989:265 | 26556313:265 | -;- | 621 | NA | 8 | 4 | 10 | 12 | 2 | 0 |
| ENSDARG000 | 21:21:21:21:2 | 198269:19830 | 198492:19849 | +:+:+:+:+:+:+: | 1343 | gsdf | 16 | 4 | 37 | 4 | 26 | 0 |
| ENSDARG000 | 21;21;21;21 | 43651954:436 | 43652373:436 | -;-;-;- | 948 | NA | 0 | 0 | 2 | 0 | 0 | 0 |
| ENSDARG000 | 21:21:21:21:2 | 45659525:456 | 45659767:456 | +:+:+:+:+:+:+: | 1962 | NA | 0 | 0 | 0 | 0 | 0 | 0 |
| ENSDARG000 | 21:21:21:21:2 | 33249478:333 | 33249711:333 | +:+:+:+:+:+:+: | 6148 | ndst1b | 2119 | 1213 | 2027 | 2141 | 2136 | 3692 |
| ENSDARG000 | 21:21:21:21:2 | 382400:38245 | 382624:38262 | +:+:+:+:+:+:+: | 1982 | rad23b | 3011 | 3617 | 3668 | 3596 | 3213 | 3038 |
| ENSDARG000 | 21 | 7600874 | 7600990 | - | 117 | 5S_rRNA | 0 | 0 | 0 | 0 | 0 | 0 |
| ENSDARG000 | 21:21:21:21:2 | 566286:56874 | 566483:56888 | -:-:-:-:-:-:-:-: | 870 | NA | 211 | 118 | 70 | 129 | 101 | 250 |
| ENSDARG000 | 21:21:21:21:2 | 22549720:225 | 22550012:225 | -;-;-;-;-;-;- | 1716 | cfap53 | 0 | 10 | 8 | 4 | 16 | 6 |
| ENSDARG000 | 21:21:21:21:2 | 7298687:7298 | 7298758:7298 | +;+;+;+;+;+ | 540 | NA | 720 | 804 | 880 | 406 | 690 | 483 |
| ENSDARG000 | 21:21:21:21:2 | 42785516:427 | 42785690:427 | -:-:-:-:-:-:-:-: | 2704 | stk10 | 86 | 57 | 125 | 84 | 111 | 52 |
| ENSDARG000 | 21;21;21 | 20707076:207 | 20711014:207 | -;-;- | 4455 | GADD45G | 2 | 0 | 0 | 0 | 0 | 0 |
| ENSDARG000 | 21:21:21:21:2 | 3244146:3247 | 3244325:3247 | +:+:+:+:+:+:+: | 2019 | CTIF | 113 | 36 | 166 | 61 | 111 | 84 |

| | | | | | | | | | | | | |
|---|---|---|---|---|---|---|---|---|---|---|---|---|
| ENSDARG000 | 21;21 | 5882300:588 | 5882544:588 | +;+ | 1185 | uqcr10 | 512 | 965 | 652 | 494 | 618 | 580 |
| ENSDARG000 | 21:21:21:21:2 | 39608637:39 | 39610457:39 | -:-:-:-:-:-:-:-: | 2889 | aldocb | 3176 | 13045 | 24014 | 14093 | 13044 | 6221 |
| ENSDARG000 | 21 | 2706531 | 2707768 | - | 1238 | NA | 0 | 0 | 0 | 0 | 0 | 0 |
| ENSDARG000 | 21:21:21:21:2 | 45717930:45 | 45717954:45 | +:+:+:+:+:+:+:+: | 3247 | ddx46 | 1856 | 1573 | 2375 | 1492 | 1896 | 1355 |
| ENSDARG000 | 21 | 33136948 | 33137083 | - | 136 | 5S_rRNA | 0 | 0 | 0 | 0 | 0 | 0 |
| ENSDARG000 | 21:21:21:21:2 | 5873227:587 | 5873431:587 | -;-;-;-;-;-;- | 740 | rpl35 | 11293 | 11072 | 15275 | 13224 | 10434 | 14642 |
| ENSDARG000 | 21 | 2957592 | 2958422 | - | 831 | zgc:194215 | 3 | 2 | 0 | 0 | 0 | 6 |
| ENSDARG000 | 21:21:21:21:2 | 15781:17803 | 15851:18319 | -;-;-;-;-;-;-;-;- | 2133 | JMY | 378 | 189 | 393 | 171 | 449 | 224 |
| ENSDARG000 | 21:21:21:21:2 | 2946683:294 | 2950579:295 | -;-;-;-;-;-;-;- | 4212 | znf971 | 8 | 14 | 43 | 17 | 12 | 38 |
| ENSDARG000 | 21:21:21:21:2 | 35652184:35 | 35653332:35 | -:-:-:-:-:-:-:-: | 2144 | sgcd | 630 | 500 | 478 | 157 | 219 | 515 |
| ENSDARG000 | 21;21;21 | 41614408:41 | 41614456:41 | -;-;- | 330 | NA | 24 | 26 | 22 | 20 | 26 | 23 |
| ENSDARG000 | 21:21:21:21:2 | 37183124:37 | 37185601:37 | -:-:-:-:-:-:-:-: | 5936 | fgfr4 | 15115 | 6969 | 9658 | 21379 | 28292 | 17304 |
| ENSDARG000 | 21;21;21 | 26709801:26 | 26710665:26 | -;-;- | 1108 | banf1 | 2902 | 3686 | 3226 | 1668 | 2468 | 2047 |
| ENSDARG000 | 21 | 8071998 | 8072121 | + | 124 | 5S_rRNA | 0 | 1 | 1 | 0 | 0 | 0 |
| ENSDARG000 | 21 | 22371030 | 22371136 | - | 107 | U6 | 0 | 0 | 0 | 0 | 0 | 0 |
| ENSDARG000 | 21:21:21:21:2 | 45856443:45 | 45856628:45 | -:-:-:-:-:-:-:-: | 2910 | LARP1 | 1859 | 1832 | 2904 | 817 | 2421 | 1090 |
| ENSDARG000 | 21 | 37072951 | 37073080 | - | 130 | NA | 0 | 1 | 1 | 0 | 0 | 0 |
| ENSDARG000 | 21:21:21:21:2 | 45816030:45 | 45816206:45 | +:+:+:+:+:+:+:+: | 2295 | pitx1 | 0 | 0 | 4 | 0 | 6 | 0 |
| ENSDARG000 | 21 | 38569285 | 38569401 | + | 117 | 5S_rRNA | 0 | 0 | 0 | 0 | 0 | 0 |
| ENSDARG000 | 21 | 42667915 | 42668034 | + | 120 | 5S_rRNA | 0 | 0 | 0 | 0 | 0 | 0 |
| ENSDARG000 | 21 | 29986796 | 29986896 | + | 101 | 5S_rRNA | 0 | 0 | 0 | 0 | 0 | 0 |
| ENSDARG000 | 21 | 24236151 | 24236272 | + | 122 | 5S_rRNA | 0 | 0 | 0 | 0 | 0 | 0 |
| ENSDARG000 | 21;21;21 | 32021272:32 | 32021725:32 | -;-;- | 872 | zgc:165573 | 794 | 669 | 467 | 690 | 878 | 554 |
| ENSDARG000 | 21;21 | 25216397:25 | 25216739:25 | +;+ | 396 | SYCN | 0 | 0 | 0 | 0 | 0 | 0 |
| ENSDARG000 | 21:21:21:21:2 | 233271:2369 | 233455:2370 | +;+;+;+;+;+ | 873 | NA | 278 | 189 | 335 | 220 | 346 | 171 |
| ENSDARG000 | 21:21:21:21:2 | 40731111:40 | 40731335:40 | -:-:-:-:-:-:-:-: | 1443 | APBB3 | 0 | 0 | 0 | 0 | 6 | 0 |
| ENSDARG000 | 21 | 43713644 | 43713776 | + | 133 | NA | 0 | 3 | 12 | 0 | 2 | 2 |
| ENSDARG000 | 21:21:21:21:2 | 39925379:39 | 39926371:39 | -:-:-:-:-:-:-:-: | 1933 | tmigd1 | 16 | 0 | 8 | 6 | 20 | 8 |
| ENSDARG000 | 21:21:21:21:2 | 7605803:760 | 7606266:760 | +:+:+:+:+:+:+:+: | 2060 | wdr41 | 360 | 362 | 473 | 323 | 482 | 559 |
| ENSDARG000 | 21:21:21:21:2 | 28535203:28 | 28535212:28 | +:+:+:+:+:+:+:+: | 600 | NA | 0 | 0 | 1 | 0 | 0 | 0 |
| ENSDARG000 | 21:21:21:21:2 | 2780907:278 | 2781820:278 | -:-:-:-:-:-:-:-: | 2571 | NA | 518 | 257 | 400 | 441 | 416 | 313 |
| ENSDARG000 | 21:21:21:21:2 | 39583458:39 | 39584009:39 | -:-:-:-:-:-:-:-: | 2290 | pigs | 1007 | 788 | 856 | 707 | 846 | 752 |
| ENSDARG000 | 21 | 44282826 | 44282943 | + | 118 | 5S_rRNA | 0 | 0 | 0 | 0 | 0 | 0 |
| ENSDARG000 | 21;21;21;21 | 7545062:755 | 7545223:755 | +;+;+;+ | 673 | tbca | 1205 | 1331 | 1566 | 1184 | 1237 | 1150 |
| ENSDARG000 | 21:21:21:21:2 | 36442085:36 | 36442927:36 | -:-:-:-:-:-:-:-: | 2516 | cnot8 | 650 | 728 | 832 | 474 | 682 | 547 |
| ENSDARG000 | 21:21:21:21:2 | 36465939:36 | 36466005:36 | -:-:-:-:-:-:-:-: | 4243 | WWC1 | 196 | 138 | 257 | 174 | 199 | 78 |
| ENSDARG000 | 21:21:21:21:2 | 45626136:45 | 45627005:45 | +:+:+:+:+:+:+:+: | 2946 | irf1b | 1220 | 1459 | 2298 | 858 | 1185 | 1389 |
| ENSDARG000 | 21:21:21:21:2 | 7188943:720 | 7189280:720 | +;+;+;+;+;+ | 3275 | agpat2 | 762 | 749 | 1047 | 782 | 810 | 870 |
| ENSDARG000 | 21;21;21 | 29918658:29 | 29918770:29 | -;-;- | 1156 | NA | 0 | 0 | 0 | 0 | 0 | 0 |
| ENSDARG000 | 21:21:21:21:2 | 36801620:36 | 36801672:36 | -:-:-:-:-:-:-:-: | 1662 | NA | 1504 | 1001 | 1960 | 1089 | 2412 | 1456 |
| ENSDARG000 | 21 | 18658115 | 18658230 | - | 116 | 5S_rRNA | 0 | 0 | 0 | 0 | 0 | 0 |
| ENSDARG000 | 21 | 28063206 | 28063322 | - | 117 | 5S_rRNA | 0 | 0 | 0 | 0 | 0 | 0 |
| ENSDARG000 | 21 | 6523654 | 6523780 | + | 127 | 5S_rRNA | 0 | 0 | 0 | 0 | 0 | 0 |
| ENSDARG000 | 21;21 | 5846969:585 | 5848222:585 | -;- | 1382 | F2RL2 | 22 | 0 | 0 | 0 | 18 | 32 |
| ENSDARG000 | 21 | 24668221 | 24668337 | + | 117 | 5S_rRNA | 0 | 0 | 0 | 0 | 0 | 0 |
| ENSDARG000 | 21:21:21:21:2 | 26726936:26 | 26727047:26 | +;+;+;+;+;+ | 2124 | calm3b | 13484 | 18671 | 18910 | 14867 | 16991 | 11946 |
| ENSDARG000 | 21;21 | 2364368:236 | 2365572:236 | -;- | 1467 | NA | 41 | 14 | 29 | 40 | 20 | 41 |
| ENSDARG000 | 21:21:21:21:2 | 41557882:41 | 41558000:41 | -;-;-;-;-;-;-;-;- | 2428 | CREBRF | 269 | 56 | 267 | 304 | 163 | 316 |
| ENSDARG000 | 21 | 42840956 | 42841071 | + | 116 | 5S_rRNA | 0 | 0 | 0 | 0 | 0 | 0 |
| ENSDARG000 | 21:21:21:21:2 | 41310551:41 | 41314257:41 | -:-:-:-:-:-:-:-: | 6316 | cpeb4b | 2410 | 1614 | 1692 | 2495 | 1645 | 2377 |
| ENSDARG000 | 21 | 26074446 | 26074513 | + | 68 | SNORD42 | 0 | 0 | 0 | 0 | 0 | 0 |
| ENSDARG000 | 21 | 6392909 | 6393036 | + | 128 | NA | 0 | 0 | 0 | 0 | 0 | 0 |
| ENSDARG000 | 21;21 | 22675031:22 | 22675881:22 | -;- | 880 | gig2h | 12 | 0 | 4 | 8 | 9 | 0 |
| ENSDARG000 | 21;21;21;21 | 41511469:41 | 41512507:41 | +;+;+;+ | 1856 | STC2 | 0 | 6 | 0 | 0 | 0 | 0 |
| ENSDARG000 | 21 | 22677211 | 22678195 | - | 985 | gig2g | 0 | 0 | 0 | 0 | 0 | 0 |
| ENSDARG000 | 21 | 44318855 | 44318972 | + | 118 | 5S_rRNA | 0 | 0 | 0 | 0 | 0 | 0 |
| ENSDARG000 | 21;21;21 | 38849851:38 | 38852875:38 | -;-;- | 3075 | tlr22 | 26 | 16 | 38 | 12 | 136 | 40 |
| ENSDARG000 | 21;21;21;21 | 39673157:39 | 39673570:39 | +;+;+;+ | 2054 | mrm3b | 54 | 52 | 94 | 28 | 78 | 76 |
| ENSDARG000 | 21;21 | 22652864:22 | 22652957:22 | -;- | 426 | gig2k | 0 | 0 | 0 | 0 | 0 | 0 |
| ENSDARG000 | 21:21:21:21:2 | 33459524:33 | 33459677:33 | +;+;+;+;+;+ | 1071 | cd74b | 490 | 631 | 1589 | 384 | 2512 | 1086 |
| ENSDARG000 | 21;21;21;21 | 45712247:45 | 45712481:45 | +;+;+;+ | 1770 | camlg | 351 | 349 | 527 | 474 | 313 | 254 |
| ENSDARG000 | 21:21:21:21:2 | 36381960:36 | 36387076:36 | -;-;-;-;-;-;- | 5613 | mrpl22 | 158 | 246 | 229 | 91 | 259 | 130 |
| ENSDARG000 | 21:21:21:21:2 | 3854414:385 | 3854466:385 | +:+:+:+:+:+:+:+: | 2233 | miga2 | 111 | 68 | 103 | 49 | 114 | 95 |
| ENSDARG000 | 21:21:21:21:2 | 45685757:45 | 45685811:45 | +:+:+:+:+:+:+:+: | 3291 | SEC24A | 2416 | 1804 | 3600 | 1868 | 2980 | 1664 |
| ENSDARG000 | 21;21;21 | 30025031:30 | 30025414:30 | -;-;- | 585 | NA | 95 | 68 | 186 | 76 | 151 | 108 |
| ENSDARG000 | 21:21:21:21:2 | 4540127:454 | 4540152:454 | +:+:+:+:+:+:+:+: | 5199 | NUP188 | 1476 | 1512 | 2525 | 814 | 1984 | 1157 |
| ENSDARG000 | 21:21:21:21:2 | 37090585:37 | 37090918:37 | +:+:+:+:+:+:+:+: | 5370 | znf346 | 1226 | 1493 | 1993 | 1022 | 1627 | 1536 |
| ENSDARG000 | 21;21 | 29294748:29 | 29295074:29 | -;- | 921 | NA | 3 | 0 | 2 | 0 | 0 | 2 |
| ENSDARG000 | 21:21:21:21:2 | 22917221:22 | 22918415:22 | -:-:-:-:-:-:-:-: | 2303 | dub | 96 | 56 | 434 | 303 | 533 | 298 |
| ENSDARG000 | 21:21:21:21:2 | 37477001:37 | 37477587:37 | +:+:+:+:+:+:+:+: | 7620 | AMOT | 4011 | 913 | 1135 | 526 | 847 | 926 |
| ENSDARG000 | 21 | 21761978 | 21762096 | + | 119 | 5S_rRNA | 3 | 0 | 0 | 0 | 4 | 0 |
| ENSDARG000 | 21 | 2957345 | 2957463 | - | 119 | 5S_rRNA | 1 | 0 | 0 | 0 | 0 | 0 |
| ENSDARG000 | 21;21;21 | 43193623:43 | 43193648:43 | +;+;+ | 354 | NA | 0 | 0 | 2 | 0 | 0 | 0 |
| ENSDARG000 | 21 | 30132668 | 30132783 | - | 116 | 5S_rRNA | 0 | 0 | 0 | 0 | 0 | 0 |
| ENSDARG000 | 21 | 10033294 | 10033410 | - | 117 | 5S_rRNA | 0 | 0 | 0 | 0 | 0 | 0 |
| ENSDARG000 | 21:21:21:21:2 | 8533533:853 | 8533561:853 | +;+;+;+;+;+ | 801 | NA | 38 | 34 | 35 | 6 | 25 | 30 |
| ENSDARG000 | 21 | 7240227 | 7240329 | - | 103 | dre-mir-126b | 0 | 0 | 0 | 0 | 0 | 0 |
| ENSDARG000 | 21 | 29105998 | 29106120 | + | 123 | 5S_rRNA | 0 | 0 | 0 | 0 | 0 | 0 |
| ENSDARG000 | 21;21 | 1143141:114 | 1143236:114 | +;+ | 204 | NA | 0 | 0 | 0 | 0 | 0 | 0 |
| ENSDARG000 | 21;21;21 | 44005521:44 | 44005680:44 | -;-;- | 381 | ndufa2 | 312 | 418 | 405 | 236 | 318 | 331 |
| ENSDARG000 | 21:21:21:21:2 | 17011724:17 | 17011848:17 | -:-:-:-:-:-:-:-: | 2224 | NA | 4 | 8 | 8 | 12 | 5 | 14 |
| ENSDARG000 | 21:21:21:21:2 | 51521:53504 | 51657:53683 | +:+:+:+:+:+:+:+: | 2925 | DMGDH | 59 | 14 | 41 | 53 | 83 | 64 |
| ENSDARG000 | 21:21:21:21:2 | 33503835:33 | 33503926:33 | +:+:+:+:+:+:+:+: | 3470 | CLINT1 | 626 | 541 | 572 | 438 | 563 | 537 |
| ENSDARG000 | 21 | 29416076 | 29416966 | - | 891 | NA | 2 | 0 | 0 | 0 | 0 | 0 |
| ENSDARG000 | 21 | 19276169 | 19276285 | + | 117 | 5S_rRNA | 0 | 0 | 0 | 0 | 0 | 0 |
| ENSDARG000 | 21 | 44594444 | 44594560 | + | 117 | 5S_rRNA | 0 | 0 | 0 | 0 | 0 | 0 |
| ENSDARG000 | 21:21:21:21:2 | 3166519:316 | 3166549:316 | +:+:+:+:+:+:+:+: | 3637 | hnrnpabb | 6090 | 8584 | 9980 | 3446 | 8887 | 4676 |
| ENSDARG000 | 21:21:21:21:2 | 7642403:764 | 7642512:764 | -:-:-:-:-:-:-:-: | 2477 | PDE8B | 224 | 108 | 171 | 177 | 309 | 233 |
| ENSDARG000 | 21 | 22669115 | 22669669 | - | 555 | gig2j | 4 | 0 | 0 | 0 | 6 | 12 |
| ENSDARG000 | 21:21:21:21:2 | 7823146:782 | 7823270:782 | +;+;+;+;+;+;+ | 1378 | crhbp | 18 | 0 | 18 | 0 | 32 | 19 |
| ENSDARG000 | 21 | 19147358 | 19147473 | + | 116 | 5S_rRNA | 0 | 0 | 0 | 0 | 0 | 0 |
| ENSDARG000 | 21:21:21:21:2 | 11179914:11 | 11180435:11 | -:-:-:-:-:-:-:-: | 2247 | dnajc21 | 134 | 260 | 311 | 64 | 255 | 124 |
| ENSDARG000 | 21;21;21 | 29325544:29 | 29325656:29 | -;-;- | 1156 | NA | 0 | 2 | 0 | 0 | 3 | 4 |
| ENSDARG000 | 21:21:21:21:2 | 12745431:12 | 12746063:12 | -;-;-;-;- | 1031 | NA | 10 | 0 | 0 | 0 | 0 | 0 |
| ENSDARG000 | 21:21:21:21:2 | 37442458:37 | 37443236:37 | +;+;+;+;+ | 2481 | cox7b | 1223 | 1511 | 1668 | 1306 | 1148 | 1160 |
| ENSDARG000 | 21 | 26179899 | 26180015 | + | 117 | 5S_rRNA | 0 | 0 | 0 | 0 | 0 | 0 |
| ENSDARG000 | 21 | 9235613 | 9235730 | + | 118 | 5S_rRNA | 0 | 0 | 0 | 0 | 0 | 0 |
| ENSDARG000 | 21;21;21;21 | 40338033:40 | 40338700:40 | -;-;-;- | 1524 | NA | 0 | 0 | 0 | 0 | 0 | 0 |
| ENSDARG000 | 21:21:21:21:2 | 33454147:33 | 33454165:33 | +;+;+;+;+ | 507 | rps14 | 11526 | 13696 | 19564 | 19577 | 11106 | 18513 |
| ENSDARG000 | 21 | 11961238 | 11961304 | - | 67 | NA | 0 | 0 | 3 | 2 | 5 | 2 |
| ENSDARG000 | 21 | 43710610 | 43710685 | + | 76 | SNORD83 | 0 | 0 | 0 | 0 | 0 | 0 |
| ENSDARG000 | 21 | 43710296 | 43710372 | + | 77 | SNORD83 | 1 | 0 | 1 | 0 | 2 | 7 |
| ENSDARG000 | 21:21:21:21:2 | 45590391:45 | 45590486:45 | -:-:-:-:-:-:-:-: | 1278 | NA | 13 | 11 | 9 | 17 | 16 | 16 |
| ENSDARG000 | 21:21:21:21:2 | 36366872:36 | 36366877:36 | +:+:+:+:+:+:+:+: | 4852 | GEMIN5 | 370 | 357 | 549 | 419 | 650 | 293 |
| ENSDARG000 | 21:21:21:21:2 | 216440:2173 | 216596:2174 | -:-:-:-:-:-:-:-: | 1451 | nup54 | 611 | 422 | 964 | 539 | 620 | 450 |
| ENSDARG000 | 21:21:21:21:2 | 38855551:38 | 38855689:38 | +:+:+:+:+:+:+:+: | 1955 | ddx52 | 363 | 452 | 724 | 292 | 502 | 352 |
| ENSDARG000 | 21:21:21:21:2 | 7228467:723 | 7228755:723 | -;-;-;-;-;-;-;- | 1330 | egfl7 | 78 | 18 | 28 | 14 | 11 | 34 |
| ENSDARG000 | 21:21:21:21:2 | 45733871:45 | 45734070:45 | +;+;+;+;+;+;+ | 3053 | C21H5orf24 | 215 | 131 | 285 | 159 | 345 | 127 |
| ENSDARG000 | 21:21:21:21:2 | 40021263:40 | 40021492:40 | -:-:-:-:-:-:-:-: | 1761 | slc47a2.2 | 0 | 6 | 0 | 10 | 11 | 0 |
| ENSDARG000 | 21:21:21:21:2 | 42156949:42 | 42158672:42 | -:-:-:-:-:-:-:-: | 3324 | GABRA6 | 0 | 0 | 0 | 0 | 0 | 0 |
| ENSDARG000 | 21 | 40400714 | 40400788 | + | 75 | NA | 8 | 13 | 16 | 32 | 15 | 27 |
| ENSDARG000 | 21 | 3201555 | 3201692 | + | 138 | 5S_rRNA | 0 | 0 | 0 | 0 | 0 | 0 |
| ENSDARG000 | 21:21:21:21:2 | 349091:3516 | 351429:3516 | +:+:+:+:+:+:+:+: | 4029 | ZNF462 | 379 | 214 | 510 | 298 | 535 | 253 |
| ENSDARG000 | 21 | 10881423 | 10881498 | - | 76 | dre-mir-122 | 0 | 0 | 0 | 0 | 0 | 0 |
| ENSDARG000 | 21:21:21:21:2 | 37436907:37 | 37437195:37 | +;+;+;+;+;+ | 2373 | pgrmc1 | 1460 | 1501 | 1509 | 1282 | 1760 | 1290 |
| ENSDARG000 | 21;21;21 | 1007486:101 | 1007549:101 | -;-;- | 237 | NA | 0 | 0 | 0 | 0 | 0 | 0 |
| ENSDARG000 | 21 | 43705608 | 43705684 | + | 77 | SNORD83 | 0 | 2 | 2 | 4 | 5 | 0 |
| ENSDARG000 | 21;21;21 | 7100442:710 | 7100521:711 | +;+;+ | 3643 | ppp1r26 | 289 | 246 | 361 | 131 | 389 | 183 |
| ENSDARG000 | 21:21:21:21:2 | 45757317:45 | 45757561:45 | +:+:+:+:+:+:+:+: | 3611 | macroh2a1 | 750 | 784 | 699 | 699 | 785 | 750 |
| ENSDARG000 | 21 | 21920077 | 21920209 | + | 133 | 5S_rRNA | 0 | 0 | 0 | 0 | 0 | 0 |
| ENSDARG000 | 21;21 | 22679785:22 | 22679830:22 | -;- | 474 | gig2f | 2 | 0 | 0 | 0 | 0 | 0 |
| ENSDARG000 | 21;21;21 | 3003753:300 | 3004452:300 | -;-;- | 1022 | CKS2 | 40 | 313 | 122 | 106 | 199 | 163 |
| ENSDARG000 | 21;21 | 2166462:216 | 2168296:216 | -;- | 1946 | NA | 78 | 26 | 83 | 71 | 100 | 56 |
| ENSDARG000 | 21;21;21;21 | 36581384:36 | 36581677:36 | +;+;+;+ | 4400 | NA | 108 | 146 | 160 | 170 | 187 | 218 |
| ENSDARG000 | 21 | 19044328 | 19044444 | + | 117 | 5S_rRNA | 0 | 0 | 0 | 0 | 0 | 0 |
| ENSDARG000 | 21;21;21 | 43976075:43 | 43980768:43 | -;-;- | 5384 | cdx1b | 0 | 0 | 4 | 0 | 0 | 16 |
| ENSDARG000 | 21 | 22036061 | 22036177 | + | 117 | 5S_rRNA | 0 | 0 | 0 | 0 | 0 | 0 |
| ENSDARG000 | 21 | 43709559 | 43709635 | + | 77 | SNORD83 | 1 | 0 | 2 | 0 | 3 | 0 |
| ENSDARG000 | 21:21:21:21:2 | 26720803:26 | 26721466:26 | +:+:+:+:+:+:+:+: | 2195 | slc3a2b | 6741 | 10927 | 12688 | 9975 | 12454 | 7218 |
| ENSDARG000 | 21:21:21:21:2 | 45838165:45 | 45838405:45 | -;-;-;-;- | 914 | txndc15 | 578 | 471 | 618 | 377 | 698 | 388 |
| ENSDARG000 | 21:21:21:21:2 | 37049427:37 | 37050279:37 | -:-:-:-:-:-:-:-: | 2370 | NA | 411 | 577 | 552 | 356 | 483 | 462 |
| ENSDARG000 | 21:21:21:21:2 | 40408:40588 | 40812:40812 | -:-:-:-:-:-:-:-: | 1895 | bhmt | 62 | 8 | 96 | 2 | 92 | 22 |
| ENSDARG000 | 21:21:21:21:2 | 45693113:45 | 45693151:45 | -;-;-;-;-;- | 276 | NA | 0 | 0 | 0 | 0 | 0 | 1 |
| ENSDARG000 | 21 | 27946274 | 27946391 | - | 118 | 5S_rRNA | 0 | 0 | 0 | 0 | 0 | 0 |
| ENSDARG000 | 21;21 | 2928964:293 | 2933338:293 | -;- | 6111 | NA | 7 | 3 | 10 | 12 | 20 | 31 |
| ENSDARG000 | 21 | 955909 | 956029 | + | 121 | 5S_rRNA | 0 | 0 | 0 | 0 | 0 | 0 |
| ENSDARG000 | 21:21:21:21:2 | 22685206:22 | 22687113:22 | -;-;-;-;-;- | 2862 | gig2e | 18 | 243 | 2 | 0 | 10 | 2 |
| ENSDARG000 | 21 | 24547524 | 24547600 | + | 77 | 5S_rRNA | 0 | 0 | 1 | 0 | 1 | 0 |
| ENSDARG000 | 21;21;21 | 36774542:36 | 36774922:36 | +;+;+ | 1577 | NA | 20 | 45 | 15 | 10 | 43 | 38 |
| ENSDARG000 | 21 | 43709859 | 43709995 | + | 137 | SNORA36B | 0 | 4 | 1 | 0 | 0 | 0 |
| ENSDARG000 | 21 | 7147698 | 7147835 | + | 138 | NA | 0 | 0 | 0 | 0 | 0 | 0 |
| ENSDARG000 | 21 | 4639890 | 4640006 | - | 117 | 5S_rRNA | 0 | 0 | 0 | 0 | 0 | 0 |
| ENSDARG000 | 21:21:21:21:2 | 44581654:44 | 44582097:44 | +:+:+:+:+:+:+:+: | 987 | TMEM164 | 142 | 76 | 216 | 81 | 219 | 112 |
| ENSDARG000 | 21 | 5528953 | 5529069 | + | 117 | 5S_rRNA | 0 | 0 | 0 | 0 | 0 | 0 |
| ENSDARG000 | 21 | 354195 | 354311 | - | 117 | 5S_rRNA | 0 | 0 | 0 | 0 | 0 | 0 |
| ENSDARG000 | 21 | 43059248 | 43059365 | + | 118 | 5S_rRNA | 0 | 0 | 0 | 0 | 0 | 0 |
| ENSDARG000 | 21:21:21:21:2 | 1119046:112 | 1119141:112 | +;+;+;+;+;+ | 819 | NA | 210 | 90 | 161 | 114 | 183 | 192 |
| ENSDARG000 | 21:21:21:21:2 | 36985209:36 | 36986828:36 | -:-:-:-:-:-:-:-: | 2948 | zgc:77151 | 654 | 526 | 582 | 407 | 685 | 606 |
| ENSDARG000 | 21 | 28491642 | 28491759 | + | 118 | 5S_rRNA | 0 | 0 | 0 | 1 | 4 | 0 |
| ENSDARG000 | 21;21;21 | 33465737:33 | 33465928:33 | -;-;- | 582 | NA | 427 | 529 | 551 | 503 | 394 | 458 |
| ENSDARG000 | 21 | 20931733 | 20931852 | - | 120 | 5S_rRNA | 0 | 0 | 0 | 2 | 0 | 3 |
| ENSDARG000 | 21 | 41351545 | 41351665 | - | 121 | 5S_rRNA | 0 | 0 | 0 | 0 | 0 | 0 |
| ENSDARG000 | 21:21:21:21:2 | 36623162:36 | 36623613:36 | +:+:+:+:+:+:+:+: | 2128 | GRK6 | 247 | 153 | 574 | 145 | 679 | 523 |
| ENSDARG000 | 21:21:21:21:2 | 34081718:34 | 34081744:34 | -:-:-:-:-:-:-:-: | 519 | NA | 17 | 66 | 20 | 45 | 36 | 65 |
| ENSDARG000 | 21;21;21 | 29452759:29 | 29452871:29 | -;-;- | 1156 | NA | 3 | 0 | 2 | 2 | 2 | 0 |
| ENSDARG000 | 21;21;21;21 | 7121942:712 | 7121990:712 | +;+;+;+ | 1276 | mrps2 | 244 | 461 | 425 | 345 | 368 | 230 |
| ENSDARG000 | 21:21:21:21:2 | 33093331:33 | 33095667:33 | -:-:-:-:-:-:-:-: | 4479 | NA | 43 | 44 | 36 | 12 | 28 | 43 |
| ENSDARG000 | 21:21:21:21:2 | 4465118:446 | 4465288:446 | -;-;-;-;-;-;- | 1549 | NA | 0 | 0 | 0 | 0 | 0 | 0 |
| ENSDARG000 | 21:21:21:21:2 | 1885698:188 | 1885739:188 | -:-:-:-:-:-:-:-: | 3904 | WDR7 | 736 | 624 | 841 | 380 | 946 | 466 |
| ENSDARG000 | 21 | 13868662 | 13868778 | - | 117 | 5S_rRNA | 0 | 0 | 0 | 0 | 0 | 0 |
| ENSDARG000 | 21;21 | 29687798:29 | 29687910:29 | -;- | 1156 | NA | 0 | 2 | 0 | 3 | 2 | 4 |
| ENSDARG000 | 21:21:21:21:2 | 7582036:758 | 7582134:758 | +;+;+;+;+;+ | 2222 | otpa | 13 | 5 | 12 | 6 | 14 | 30 |

| | | | | | | | | | | | | |
|---|---|---|---|---|---|---|---|---|---|---|---|---|
| ENSDARG000 | 21;21;21 | 29706179:297 | 29706291:297 | -;-;- | 1156 | NA | 0 | 0 | 2 | 3 | 1 | 4 |
| ENSDARG000 | 21 | 6592935 | 6593054 | + | 120 | 5S_rRNA | 0 | 0 | 0 | 0 | 0 | 0 |
| ENSDARG000 | 21:21:21:21:2 | 45839917:458 | 45839937:458 | +:+:+:+:+:+:+: | 2177 | faxdc2 | 313 | 121 | 142 | 162 | 132 | 208 |
| ENSDARG000 | 21;21 | 2102829:211 | 2103254:211 | +;+ | 2699 | NA | 48 | 12 | 63 | 55 | 44 | 91 |
| ENSDARG000 | 21:21:21:21:2 | 40670996:406 | 40671818:406 | -;-;-;-;- | 1740 | ARX | 0 | 0 | 0 | 0 | 0 | 0 |
| ENSDARG000 | 21 | 12962722 | 12962838 | + | 117 | 5S_rRNA | 0 | 0 | 0 | 0 | 0 | 0 |
| ENSDARG000 | 21 | 15340311 | 15340427 | + | 117 | 5S_rRNA | 0 | 0 | 0 | 0 | 0 | 0 |
| ENSDARG000 | 21:21:21:21:2 | 33211306:332 | 33212004:332 | -:-:-:-:-:-:-:-: | 1874 | rbm22 | 310 | 399 | 359 | 260 | 411 | 334 |
| ENSDARG000 | 21:21:21:21:2 | 1320767:132 | 1320886:132 | +:+:+:+:+:+:+: | 2255 | TCF4 | 146 | 164 | 237 | 93 | 324 | 107 |
| ENSDARG000 | 21:21:21:21:2 | 36612440:366 | 36612823:366 | -;-;-;-;- | 817 | nop16 | 538 | 597 | 986 | 450 | 618 | 543 |
| ENSDARG000 | 21;21;21;21 | 42870101:428 | 42870723:428 | -;-;-;- | 1248 | NA | 8 | 4 | 12 | 0 | 10 | 10 |
| ENSDARG000 | 21:21:21:21:2 | 22642698:226 | 22642980:226 | -;-;-;-;- | 802 | gig2l | 8 | 2 | 4 | 4 | 0 | 0 |
| ENSDARG000 | 21:21:21:21:2 | 7741781:774 | 7742166:774 | -:-:-:-:-:-:-:-: | 2560 | aggf1 | 264 | 341 | 508 | 174 | 343 | 176 |
| ENSDARG000 | 21:21:21:21:2 | 42226113:422 | 42226195:422 | +:+:+:+:+:+:+: | 1515 | GABRB2 | 0 | 0 | 0 | 0 | 5 | 0 |
| ENSDARG000 | 21:21:21:21:2 | 3093419:309 | 3094452:309 | +:+:+:+:+:+:+: | 2042 | SHC3 | 43 | 66 | 123 | 92 | 86 | 120 |
| ENSDARG000 | 21 | 38634296 | 38635108 | + | 813 | NA | 358 | 54 | 27 | 166 | 19 | 39 |
| ENSDARG000 | 21:21:21:21:2 | 5862476:586 | 5862632:586 | +;+;+;+;+;+ | 700 | wdr38 | 0 | 0 | 0 | 0 | 0 | 0 |
| ENSDARG000 | 21;21;21 | 27189490:271 | 27189622:271 | +;+;+ | 438 | NA | 143 | 74 | 69 | 122 | 40 | 151 |
| ENSDARG000 | 21 | 5199981 | 5200059 | - | 79 | NA | 0 | 0 | 0 | 0 | 0 | 0 |
| ENSDARG000 | 21 | 2742823 | 2743026 | + | 204 | NA | 91 | 56 | 72 | 61 | 93 | 55 |
| ENSDARG000 | 21;21 | 44684577:446 | 44684741:446 | +;+ | 3447 | spry4 | 9442 | 5025 | 8384 | 9084 | 8808 | 11199 |
| ENSDARG000 | 21 | 12528120 | 12528236 | + | 117 | 5S_rRNA | 0 | 0 | 0 | 0 | 0 | 0 |
| ENSDARG000 | 21;21;21 | 4368735:437 | 4368827:437 | +;+;+ | 321 | NA | 0 | 0 | 0 | 0 | 0 | 0 |
| ENSDARG000 | 21:21:21:21:2 | 41468210:414 | 41469162:414 | +;+;+;+;+;+;+ | 1690 | bod1 | 390 | 432 | 406 | 354 | 441 | 402 |
| ENSDARG000 | 21:21:21:21:2 | 7332963:733 | 7333124:733 | +:+:+:+:+:+:+: | 2031 | AP3B1 | 2428 | 2385 | 2548 | 1591 | 2178 | 1847 |
| ENSDARG000 | 21:21:21:21:2 | 41782307:417 | 41782508:417 | -:-:-:-:-:-:-:-: | 3146 | TCERG1 | 858 | 782 | 1011 | 583 | 1052 | 814 |
| ENSDARG000 | 21;21;21 | 29836723:298 | 29836835:298 | -;-;- | 1156 | NA | 0 | 4 | 1 | 0 | 0 | 0 |
| ENSDARG000 | 21;21;21 | 29508043:295 | 29508155:295 | -;-;- | 1156 | NA | 0 | 8 | 4 | 0 | 0 | 0 |
| ENSDARG000 | 21:21:21:21:2 | 7917215:791 | 7917226:791 | +:+:+:+:+:+:+: | 330 | NA | 0 | 0 | 0 | 0 | 0 | 0 |
| ENSDARG000 | 21:21:21:21:2 | 26657404:266 | 26657499:266 | +:+:+:+:+:+:+: | 867 | prdx5 | 4632 | 7114 | 5442 | 8402 | 6798 | 7829 |
| ENSDARG000 | 21 | 11252511 | 11252626 | + | 116 | 5S_rRNA | 6 | 2 | 3 | 6 | 1 | 0 |
| ENSDARG000 | 21 | 17808847 | 17808962 | - | 116 | 5S_rRNA | 0 | 0 | 0 | 0 | 0 | 0 |
| ENSDARG000 | 21 | 1595170 | 1595286 | + | 117 | 5S_rRNA | 10 | 3 | 0 | 0 | 0 | 0 |
| ENSDARG000 | 21;21;21 | 37411547:374 | 37411700:374 | +;+;+ | 651 | mrps17 | 93 | 143 | 157 | 40 | 114 | 107 |
| ENSDARG000 | 21;21;21;21 | 41144914:411 | 41145874:411 | -;-;-;- | 1457 | msx2b | 343 | 261 | 648 | 463 | 838 | 299 |
| ENSDARG000 | 21 | 12512607 | 12512722 | - | 116 | 5S_rRNA | 0 | 0 | 0 | 0 | 0 | 0 |
| ENSDARG000 | 21;21;21 | 3021702:302 | 3021736:302 | +;+;+ | 243 | NA | 0 | 0 | 2 | 0 | 2 | 0 |
| ENSDARG000 | 21;21;21 | 7863243:788 | 7864140:788 | -;-;- | 1113 | s100z | 2 | 2 | 0 | 6 | 8 | 0 |
| ENSDARG000 | 21 | 29606327 | 29606450 | + | 124 | 5S_rRNA | 0 | 0 | 0 | 0 | 0 | 0 |
| ENSDARG000 | 21:21:21:21:2 | 37445871:374 | 37446377:374 | +:+:+:+:+:+:+: | 3682 | pgk1 | 8671 | 23813 | 38971 | 37136 | 16413 | 13956 |
| ENSDARG000 | 21 | 31337670 | 31337785 | - | 116 | 5S_rRNA | 0 | 0 | 0 | 0 | 0 | 0 |
| ENSDARG000 | 21:21:21:21:2 | 76739:78944 | 76984:79130 | +;+;+;+;+;+ | 1176 | ARSB | 432 | 326 | 399 | 496 | 585 | 427 |
| ENSDARG000 | 21:21:21:21:2 | 103194:1034 | 103360:1035 | +:+:+:+:+:+:+: | 2325 | zer1 | 764 | 510 | 539 | 440 | 542 | 438 |
| ENSDARG000 | 21:21:21:21:2 | 4484558:448 | 4484726:448 | -;-;-;-;-;-;- | 1642 | NA | 0 | 0 | 0 | 0 | 0 | 0 |
| ENSDARG000 | 21:21:21:21:2 | 170038:1708 | 170093:1710 | +:+:+:+:+:+:+: | 2957 | klhl8 | 638 | 348 | 456 | 416 | 662 | 395 |
| ENSDARG000 | 21:21:21:21:2 | 43987153:44 | 43987177:44 | +:+:+:+:+:+:+: | 2114 | ik | 1800 | 1574 | 1998 | 1443 | 1905 | 1383 |
| ENSDARG000 | 21:21:21:21:2 | 11139175:111 | 11139318:111 | +:+:+:+:+:+:+: | 1947 | agxt2 | 10 | 10 | 16 | 18 | 34 | 0 |
| ENSDARG000 | 21:21:21:21:2 | 215751:2158 | 215783:2158 | -;-;-;-;- | 378 | NA | 0 | 2 | 0 | 0 | 0 | 0 |
| ENSDARG000 | 21 | 7348023 | 7348138 | + | 116 | 5S_rRNA | 0 | 0 | 0 | 0 | 0 | 0 |
| ENSDARG000 | 21:21:21:21:2 | 37915193:379 | 37916435:379 | -:-:-:-:-:-:-:-: | 1975 | NA | 104 | 79 | 128 | 174 | 168 | 116 |
| ENSDARG000 | 21 | 30355767 | 30361679 | + | 5913 | NA | 0 | 0 | 2 | 0 | 0 | 4 |
| ENSDARG000 | 21;21;21 | 8144918:815 | 8145531:815 | -;-;- | 903 | OLFML2A | 23 | 0 | 2 | 5 | 37 | 20 |
| ENSDARG000 | 21:21:21:21:2 | 45041684:45 | 45041703:45 | -:-:-:-:-:-:-:-: | 4499 | RAPGEF6 | 316 | 228 | 374 | 337 | 268 | 271 |
| ENSDARG000 | 21;21 | 22672794:226 | 22673368:226 | -;- | 645 | gig2i | 6 | 12 | 12 | 0 | 12 | 8 |
| ENSDARG000 | 21 | 32030636 | 32030752 | + | 117 | 5S_rRNA | 4 | 2 | 2 | 0 | 0 | 2 |
| ENSDARG000 | 21 | 27335125 | 27335261 | + | 137 | 5S_rRNA | 0 | 0 | 0 | 0 | 9 | 0 |
| ENSDARG000 | 21 | 40400938 | 40401012 | + | 75 | NA | 4 | 3 | 11 | 2 | 9 | 4 |
| ENSDARG000 | 21 | 23684720 | 23684836 | + | 117 | 5S_rRNA | 0 | 0 | 0 | 0 | 0 | 0 |
| ENSDARG000 | 21 | 39267225 | 39267306 | - | 82 | NA | 0 | 0 | 2 | 0 | 0 | 0 |
| ENSDARG000 | 21 | 7900107 | 7901424 | + | 1318 | ch25hl2 | 10 | 2 | 26 | 0 | 98 | 17 |
| ENSDARG000 | 21;21;21 | 14511685:145 | 14511726:145 | +;+;+ | 1454 | NA | 19 | 24 | 32 | 7 | 24 | 54 |
| ENSDARG000 | 21 | 24749662 | 24749779 | + | 118 | 5S_rRNA | 0 | 0 | 0 | 0 | 0 | 0 |
| ENSDARG000 | 21 | 17121361 | 17121477 | - | 117 | 5S_rRNA | 0 | 0 | 0 | 0 | 0 | 6 |
| ENSDARG000 | 21;21;21 | 547098:5491 | 547156:5494 | -;-;- | 711 | NA | 2 | 10 | 21 | 0 | 12 | 8 |
| ENSDARG000 | 21:21:21:21:2 | 41535954:415 | 41538998:415 | -;-;-;-;-;- | 4441 | BNIP1 | 456 | 226 | 434 | 356 | 357 | 273 |
| ENSDARG000 | 21;21;21;21 | 1204370:120 | 1205026:120 | +;+;+;+ | 2593 | NA | 0 | 0 | 1 | 0 | 0 | 0 |
| ENSDARG000 | 21;21;21;21 | 8692022:869 | 8692410:869 | +;+;+;+ | 459 | NA | 0 | 0 | 0 | 0 | 0 | 0 |
| ENSDARG000 | 21:21:21:21:2 | 10064675:10 | 10065350:10 | +;+;+;+;+;+ | 1444 | NA | 39 | 12 | 265 | 111 | 190 | 30 |
| ENSDARG000 | 21:21:21:21:2 | 10092302:10 | 10092493:10 | +;+;+;+;+;+;+ | 2452 | NA | 46 | 42 | 272 | 165 | 317 | 46 |
| ENSDARG000 | 21;21;21 | 12355585:123 | 12356119:123 | -;-;- | 926 | NA | 0 | 0 | 0 | 0 | 0 | 0 |
| ENSDARG000 | 21;21;21 | 12518559:125 | 12518961:125 | -;-;- | 653 | NA | 0 | 0 | 0 | 0 | 0 | 0 |
| ENSDARG000 | 21:21:21:21:2 | 13055886:130 | 13057189:130 | -;-;-;-;-;-;-;- | 2247 | NA | 0 | 0 | 0 | 0 | 0 | 0 |
| ENSDARG000 | 21;21;21 | 16891987:168 | 16892082:168 | -;-;- | 303 | NA | 0 | 0 | 0 | 0 | 0 | 0 |
| ENSDARG000 | 21;21;21 | 17488671:174 | 17488852:174 | -;-;- | 543 | NA | 0 | 0 | 0 | 0 | 0 | 0 |
| ENSDARG000 | 21;21;21 | 19746870:197 | 19749607:197 | -;-;- | 3221 | NA | 0 | 0 | 0 | 0 | 0 | 0 |
| ENSDARG000 | 21;21;21;21 | 23845537:239 | 23845702:239 | -;-;-;- | 1335 | NA | 0 | 0 | 0 | 0 | 0 | 0 |
| ENSDARG000 | 21:21:21:21:2 | 24423198:244 | 24423576:244 | +;+;+;+;+;+;+ | 1236 | NA | 0 | 0 | 0 | 0 | 0 | 0 |
| ENSDARG000 | 21:21:21:21:2 | 33058625:330 | 33059241:330 | -;-;-;-;-;-;-;- | 1669 | zgc:194629 | 9 | 0 | 0 | 0 | 0 | 6 |
| ENSDARG000 | 21:21:21:21:2 | 37595791:375 | 37596170:375 | -;-;-;-;- | 1074 | NA | 0 | 0 | 0 | 0 | 0 | 0 |
| ENSDARG000 | 21:21:21:21:2 | 38960917:389 | 38961036:389 | +:+:+:+:+:+:+: | 2225 | NA | 0 | 4 | 0 | 0 | 3 | 0 |
| ENSDARG000 | 21;21;21 | 38976592:389 | 38977018:389 | +;+;+ | 1058 | NA | 0 | 4 | 0 | 0 | 1 | 0 |
| ENSDARG000 | 21;21;21 | 45445457:454 | 45445977:454 | +;+;+ | 1023 | NA | 0 | 0 | 0 | 0 | 0 | 0 |
| ENSDARG000 | 21:21:21:21:2 | 45449733:454 | 45450117:454 | +:+:+:+:+:+:+: | 2374 | NA | 2 | 0 | 4 | 0 | 11 | 0 |
| ENSDARG000 | 21;21;21;21 | 45454503:454 | 45454669:454 | +;+;+;+ | 2122 | NA | 0 | 0 | 0 | 0 | 0 | 0 |
| ENSDARG000 | 21:21:21:21:2 | 45462359:454 | 45463060:454 | +;+;+;+;+;+ | 6279 | NA | 0 | 2 | 0 | 0 | 0 | 0 |
| ENSDARG000 | 21:21:21:21:2 | 45566178:455 | 45566215:455 | +:+:+:+:+:+:+: | 3573 | NA | 16 | 12 | 43 | 25 | 21 | 32 |
| ENSDARG000 | 11:11:11:11:1 | 1396170:139 | 1396471:139 | -:-:-:-:-:-:-:-: | 1117 | rpl29 | 4735 | 4648 | 4527 | 3561 | 2972 | 5030 |
| ENSDARG000 | 11:11:11:11:1 | 5842632:584 | 5842710:584 | +:+:+:+:+:+:+: | 1446 | ndufs7 | 982 | 1268 | 1297 | 1056 | 1175 | 1252 |
| ENSDARG000 | 11:11:11:11:1 | 18560667:185 | 18562821:185 | -:-:-:-:-:-:-:-: | 6487 | ZMYND8 | 540 | 478 | 663 | 404 | 646 | 524 |
| ENSDARG000 | 11:11:11:11:1 | 34488869:344 | 34489680:344 | -:-:-:-:-:-:-:-: | 5000 | PIK3CA | 2084 | 787 | 1118 | 993 | 1213 | 1404 |
| ENSDARG000 | 11;11;11 | 15613218:157 | 15613908:157 | +;+;+ | 1727 | gdf11 | 0 | 1 | 0 | 0 | 0 | 2 |
| ENSDARG000 | 11;11 | 7324704:733 | 7324969:734 | +;+ | 2475 | diras1a | 4 | 14 | 42 | 42 | 0 | 8 |
| ENSDARG000 | 11:11:11:11:1 | 30744266:307 | 30744503:307 | +:+:+:+:+:+:+: | 824 | wdr83os | 728 | 1064 | 1182 | 870 | 868 | 852 |
| ENSDARG000 | 11:11:11:11:1 | 28127199:281 | 28129163:281 | -:-:-:-:-:-:-:-: | 3692 | LACTBL1 | 0 | 0 | 2 | 0 | 0 | 0 |
| ENSDARG000 | 11:11:11:11:1 | 1403741:140 | 1404687:140 | -;-;-;-;- | 1801 | NA | 0 | 2 | 2 | 0 | 0 | 0 |
| ENSDARG000 | 11;11;11 | 15775452:157 | 15775831:157 | -;-;- | 847 | NA | 0 | 0 | 0 | 0 | 0 | 0 |
| ENSDARG000 | 11:11:11:11:1 | 15776072:157 | 15776118:157 | +;+;+;+;+;+ | 930 | NA | 0 | 0 | 0 | 0 | 0 | 0 |
| ENSDARG000 | 11;11 | 1409622:141 | 1410003:141 | +;+ | 1444 | PPP1R3D | 0 | 0 | 0 | 0 | 0 | 0 |
| ENSDARG000 | 11:11:11:11:1 | 7350926:735 | 7351029:735 | -;-;-;-;-;- | 940 | NA | 126 | 173 | 1051 | 102 | 411 | 478 |
| ENSDARG000 | 11:11:11:11:1 | 1422737:142 | 1423307:142 | -:-:-:-:-:-:-:-: | 2606 | phactr3b | 0 | 0 | 8 | 6 | 0 | 2 |
| ENSDARG000 | 11:11:11:11:1 | 24013769:24 | 24014394:24 | -:-:-:-:-:-:-:-: | 2229 | chia.3 | 0 | 0 | 0 | 0 | 0 | 0 |
| ENSDARG000 | 11:11:11:11:1 | 25504215:255 | 25504331:255 | +:+:+:+:+:+:+: | 1683 | tfe3b | 166 | 94 | 239 | 145 | 246 | 183 |
| ENSDARG000 | 11;11;11;11 | 15796093:157 | 15796665:157 | -;-;-;- | 842 | NA | 0 | 0 | 0 | 0 | 0 | 0 |
| ENSDARG000 | 11:11:11:11:1 | 12811906:128 | 12812177:128 | +:+:+:+:+:+:+: | 3975 | rtel1 | 116 | 180 | 305 | 134 | 303 | 171 |
| ENSDARG000 | 11;11 | 15798258:157 | 15798534:157 | -;- | 454 | NA | 0 | 0 | 0 | 0 | 0 | 0 |
| ENSDARG000 | 11;11 | 15799383:15 | 15799748:15 | +;+ | 803 | NA | 0 | 0 | 0 | 0 | 0 | 0 |
| ENSDARG000 | 11:11:11:11:1 | 3336524:334 | 3339544:334 | -;-;-;-;-;- | 3688 | tuba2 | 56 | 28 | 73 | 122 | 134 | 42 |
| ENSDARG000 | 11:11:11:11:1 | 5855923:585 | 5856106:585 | -;-;-;-;-;-;-;- | 1365 | gamt | 59 | 72 | 115 | 92 | 66 | 58 |
| ENSDARG000 | 11:11:11:11:1 | 37010161:37 | 37011626:37 | -:-:-:-:-:-:-:-: | 2059 | NA | 6 | 0 | 0 | 3 | 22 | 10 |
| ENSDARG000 | 11:11:11:11:1 | 36989696:369 | 36990030:369 | +:+:+:+:+:+:+: | 5862 | TKT | 0 | 6 | 10 | 0 | 42 | 0 |
| ENSDARG000 | 11:11:11:11:1 | 19668:21295 | 20413:21485 | -:-:-:-:-:-:-:-: | 3113 | sp1 | 378 | 323 | 629 | 308 | 497 | 164 |
| ENSDARG000 | 11:11:11:11:1 | 18612166:18 | 18612532:18 | +:+:+:+:+:+:+: | 10379 | NCOA3 | 866 | 701 | 796 | 782 | 940 | 654 |
| ENSDARG000 | 11:11:11:11:1 | 3364193:336 | 3368027:336 | -:-:-:-:-:-:-:-: | 5563 | mcrs1 | 555 | 434 | 669 | 597 | 657 | 581 |
| ENSDARG000 | 11:11:11:11:1 | 3345405:335 | 3348295:335 | -;-;-;-;-;- | 3603 | suox | 120 | 48 | 129 | 46 | 125 | 74 |
| ENSDARG000 | 11;11;11 | 15802450:15 | 15802652:15 | -;-;- | 944 | NA | 0 | 0 | 0 | 0 | 0 | 0 |
| ENSDARG000 | 11:11:11:11:1 | 25528557:255 | 25529361:255 | -:-:-:-:-:-:-:-: | 1391 | si:dkev-245f2 | 0 | 0 | 0 | 0 | 0 | 2 |
| ENSDARG000 | 11:11:11:11:1 | 28166165:281 | 28166482:281 | +:+:+:+:+:+:+: | 8719 | ephb2b | 1899 | 76 | 112 | 106 | 26 | 702 |
| ENSDARG000 | 11:11:11:11:1 | 15806392:15 | 15807085:15 | -;-;-;-;- | 1136 | NA | 18 | 4 | 6 | 0 | 8 | 12 |
| ENSDARG000 | 11;11 | 11332652:113 | 11334029:113 | -;- | 1458 | C11H6orf120 | 200 | 200 | 192 | 144 | 237 | 198 |
| ENSDARG000 | 11:11:11:11:1 | 24027715:24 | 24027974:24 | +:+:+:+:+:+:+: | 1398 | NA | 0 | 0 | 0 | 2 | 0 | 0 |
| ENSDARG000 | 11;11;11;11 | 18702362:187 | 18702451:187 | +;+;+;+ | 1793 | NA | 10 | 0 | 0 | 0 | 0 | 0 |
| ENSDARG000 | 11;11 | 18704191:187 | 18704727:187 | -;- | 1006 | id1 | 132 | 58 | 95 | 21 | 87 | 53 |
| ENSDARG000 | 11:11:11:11:1 | 30817943:30 | 30818217:30 | +:+:+:+:+:+:+: | 8718 | cacna1ab | 0 | 0 | 0 | 0 | 0 | 2 |
| ENSDARG000 | 11;11 | 15845588:15 | 15845842:15 | -;- | 696 | NA | 0 | 2 | 4 | 0 | 4 | 6 |
| ENSDARG000 | 11:11:11:11:1 | 7359496:735 | 7359893:736 | -:-:-:-:-:-:-:-: | 4396 | vtg3 | 2 | 0 | 0 | 0 | 0 | 0 |
| ENSDARG000 | 11:11:11:11:1 | 15833565:15 | 15834494:15 | -:-:-:-:-:-:-:-: | 1942 | rap1ab | 435 | 594 | 549 | 274 | 540 | 447 |
| ENSDARG000 | 11:11:11:11:1 | 11347031:11 | 11347336:11 | +:+:+:+:+:+:+: | 760 | NA | 0 | 0 | 0 | 0 | 0 | 0 |
| ENSDARG000 | 11:11:11:11:1 | 1512571:151 | 1512725:151 | +;+;+;+;+;+ | 1497 | acot8 | 303 | 271 | 287 | 261 | 320 | 362 |
| ENSDARG000 | 11:11:11:11:1 | 25539698:255 | 25539916:255 | +:+:+:+:+:+:+: | 2149 | cxxc1b | 278 | 275 | 377 | 276 | 262 | 253 |
| ENSDARG000 | 11:11:11:11:1 | 11360641:11 | 11360958:11 | +:+:+:+:+:+:+: | 902 | NA | 0 | 0 | 0 | 0 | 0 | 0 |
| ENSDARG000 | 11:11:11:11:1 | 34522554:345 | 34522677:345 | +:+:+:+:+:+:+: | 1319 | ZMAT3 | 166 | 21 | 56 | 99 | 47 | 79 |
| ENSDARG000 | 11;11 | 11353459:11 | 11353775:11 | +;+ | 498 | NA | 0 | 0 | 0 | 0 | 0 | 0 |
| ENSDARG000 | 11:11:11:11:1 | 11374021:11 | 11374353:11 | +:+:+:+:+:+:+: | 1015 | NA | 0 | 5 | 0 | 0 | 0 | 0 |
| ENSDARG000 | 11;11 | 28476298:285 | 28478078:285 | +;+ | 3026 | NA | 10 | 2 | 2 | 0 | 4 | 2 |
| ENSDARG000 | 11:11:11:11:1 | 37038420:37 | 37038722:37 | -;-;-;-;-;-;-;- | 895 | NA | 0 | 0 | 0 | 0 | 0 | 0 |
| ENSDARG000 | 11:11:11:11:1 | 15878343:15 | 15878491:15 | +:+:+:+:+:+:+: | 3216 | PANK4 | 950 | 823 | 1138 | 662 | 998 | 1038 |
| ENSDARG000 | 11:11:11:11:1 | 18799186:187 | 18803135:188 | -:-:-:-:-:-:-:-: | 6530 | phtf1 | 582 | 666 | 701 | 684 | 737 | 702 |
| ENSDARG000 | 11:11:11:11:1 | 28517562:285 | 28518050:285 | +:+:+:+:+:+:+: | 2248 | NA | 0 | 0 | 2 | 0 | 0 | 0 |
| ENSDARG000 | 11:11:11:11:1 | 37049105:37 | 37049855:37 | +:+:+:+:+:+:+: | 3556 | BICD2 | 1185 | 787 | 1168 | 658 | 1107 | 792 |
| ENSDARG000 | 11:11:11:11:1 | 18784847:187 | 18786335:187 | -:-:-:-:-:-:-:-: | 3249 | PFKFB2 | 16 | 18 | 21 | 5 | 14 | 25 |
| ENSDARG000 | 11:11:11:11:1 | 24034345:24 | 24034495:24 | +:+:+:+:+:+:+: | 4190 | zc3h11a | 1927 | 1920 | 2870 | 3055 | 3081 | 2178 |
| ENSDARG000 | 11:11:11:11:1 | 5880562:588 | 5880694:588 | +:+:+:+:+:+:+: | 3910 | dazap1 | 40 | 86 | 64 | 8 | 56 | 66 |
| ENSDARG000 | 11:11:11:11:1 | 11387509:11 | 11387841:11 | +:+:+:+:+:+:+: | 1015 | NA | 0 | 0 | 0 | 0 | 0 | 0 |
| ENSDARG000 | 11:11:11:11:1 | 12860260:128 | 12861421:128 | +:+:+:+:+:+:+: | 5028 | zgpat | 369 | 433 | 405 | 135 | 443 | 333 |
| ENSDARG000 | 11:11:11:11:1 | 1522127:152 | 1522332:152 | +:+:+:+:+:+:+: | 3500 | srsf6b | 3134 | 2939 | 4517 | 3287 | 3735 | 4064 |
| ENSDARG000 | 11:11:11:11:1 | 45153104:451 | 45153278:451 | +:+:+:+:+:+:+: | 2755 | tk1 | 52 | 378 | 372 | 72 | 121 | 197 |
| ENSDARG000 | 11:11:11:11:1 | 11407892:114 | 11408153:114 | +:+:+:+:+:+:+: | 1041 | NA | 0 | 0 | 0 | 0 | 0 | 0 |
| ENSDARG000 | 11:11:11:11:1 | 11419186:114 | 11419518:114 | +:+:+:+:+:+:+: | 1112 | NA | 0 | 1 | 0 | 0 | 0 | 0 |
| ENSDARG000 | 11:11:11:11:1 | 34555036:345 | 34555095:345 | -:-:-:-:-:-:-:-: | 2248 | pfkfb4a | 416 | 453 | 538 | 513 | 478 | 451 |
| ENSDARG000 | 11:11:11:11:1 | 88247:89394 | 89594:89594 | -:-:-:-:-:-:-:-: | 3216 | elmo2 | 3210 | 1780 | 2515 | 2365 | 2498 | 1686 |
| ENSDARG000 | 11:11:11:11:1 | 24046179:24 | 24046537:24 | +;+;+;+;+;+;+ | 3361 | maf1a | 863 | 665 | 744 | 736 | 786 | 1105 |
| ENSDARG000 | 11:11:11:11:1 | 45159353:451 | 45159770:451 | -:-:-:-:-:-:-:-: | 2243 | syngr2b | 880 | 689 | 334 | 567 | 334 | 741 |
| ENSDARG000 | 11:11:11:11:1 | 11430552:114 | 11430884:114 | +:+:+:+:+:+:+: | 1015 | NA | 0 | 7 | 0 | 0 | 0 | 0 |
| ENSDARG000 | 11:11:11:11:1 | 12930157:129 | 12930503:129 | +;+;+;+;+;+ | 1056 | NA | 0 | 0 | 0 | 0 | 0 | 0 |
| ENSDARG000 | 11:11:11:11:1 | 12879635:128 | 12880050:128 | +:+:+:+:+:+:+: | 13908 | NA | 0 | 0 | 0 | 0 | 0 | 0 |
| ENSDARG000 | 11;11;11;11 | 24058530:24 | 24060053:24 | -;-;-;- | 2180 | trib3 | 4135 | 3816 | 5630 | 5693 | 4837 | 5881 |
| ENSDARG000 | 11:11:11:11:1 | 25560072:255 | 25560163:255 | +:+:+:+:+:+:+: | 3802 | mbd1b | 568 | 540 | 833 | 505 | 623 | 580 |
| ENSDARG000 | 11:11:11:11:1 | 37095165:37 | 37095224:37 | +:+:+:+:+:+:+: | 2550 | card19 | 470 | 272 | 366 | 432 | 253 | 732 |
| ENSDARG000 | 11;11;11;11 | 5926850:592 | 5926873:592 | +;+;+;+ | 497 | rps15 | 12275 | 10683 | 16455 | 14987 | 11132 | 14709 |
| ENSDARG000 | 11:11:11:11:1 | 45174520:451 | 45174978:451 | -:-:-:-:-:-:-:-: | 2085 | NA | 14 | 6 | 18 | 0 | 46 | 17 |
| ENSDARG000 | 11;11;11 | 37105640:371 | 37106072:371 | -;-;- | 833 | ninj1 | 82 | 83 | 105 | 118 | 124 | 36 |
| ENSDARG000 | 11:11:11:11:1 | 1533097:153 | 1533653:153 | +:+:+:+:+:+:+: | 1961 | ift52 | 245 | 274 | 284 | 212 | 251 | 273 |

| | | | | | | | | | | | | |
|---|---|---|---|---|---|---|---|---|---|---|---|---|
| ENSDARG000 | 11;11 | 31047854:31 | 31048012:31 | +;+ | 898 | NA | 0 | 0 | 0 | 0 | 0 | 0 |
| ENSDARG000 | 11:11:11:11:1 | 37137196:371 | 37137839:371 | +:+:+:+:+:+:+: | 8453 | NA | 2 | 33 | 26 | 67 | 51 | 6 |
| ENSDARG000 | 11:11:11:11:1 | 12982260:129 | 12982574:129 | -;-;-;-;-;- | 2265 | chrna4b | 0 | 0 | 0 | 0 | 0 | 0 |
| ENSDARG000 | 11:11:11:11:1 | 5933635:5934 | 5934505:5934 | -:-:-:-:-:-:-:-: | 2403 | ABHD8 | 6 | 27 | 8 | 29 | 0 | 16 |
| ENSDARG000 | 11:11:11:11:1 | 11454545:114 | 11454862:114 | -;-;-;-;-;-;-;- | 1622 | krt93 | 14 | 0 | 0 | 0 | 0 | 0 |
| ENSDARG000 | 11:11:11:11:1 | 45199369:451 | 45200681:452 | -:-:-:-:-:-:-:-: | 4177 | TMC8 | 52 | 63 | 42 | 52 | 93 | 133 |
| ENSDARG000 | 11;11 | 13032483:130 | 13033292:130 | +;+ | 1182 | ptgfr | 0 | 0 | 0 | 0 | 0 | 0 |
| ENSDARG000 | 11:11:11:11:1 | 13024002:130 | 13024068:130 | +:+:+:+:+:+:+: | 1077 | btf3l4 | 811 | 849 | 1305 | 901 | 944 | 979 |
| ENSDARG000 | 11;11;11 | 31144720:311 | 31144937:311 | -;-;- | 648 | NA | 0 | 0 | 0 | 0 | 0 | 0 |
| ENSDARG000 | 11:11:11:11:1 | 31168697:311 | 31169551:311 | -;-;-;-;-;- | 2179 | NA | 9 | 10 | 10 | 12 | 15 | 22 |
| ENSDARG000 | 11;11;11 | 24064163:240 | 24064264:240 | +;+;+ | 507 | NA | 3 | 1 | 1 | 2 | 6 | 5 |
| ENSDARG000 | 11 | 18832913 | 18833246 | + | 334 | NA | 0 | 0 | 0 | 0 | 0 | 0 |
| ENSDARG000 | 11:11:11:11:1 | 7383177:7383 | 7383532:7383 | -:-:-:-:-:-:-:-: | 2825 | adgrl4 | 38 | 20 | 16 | 6 | 40 | 24 |
| ENSDARG000 | 11;11 | 1547226:155 | 1548550:155 | -;- | 1364 | NA | 0 | 4 | 14 | 0 | 2 | 0 |
| ENSDARG000 | 11:11:11:11:1 | 31190155:311 | 31190292:311 | +:+:+:+:+:+:+: | 4308 | stx10 | 212 | 143 | 181 | 147 | 191 | 212 |
| ENSDARG000 | 11:11:11:11:1 | 37178271:371 | 37178360:371 | +:+:+:+:+:+:+: | 2918 | itih3b.1 | 0 | 4 | 18 | 0 | 0 | 0 |
| ENSDARG000 | 11;11 | 45234480:452 | 45234770:452 | +;+ | 807 | NA | 2 | 0 | 0 | 0 | 2 | 6 |
| ENSDARG000 | 11:11:11:11:1 | 45219558:452 | 45219613:452 | +:+:+:+:+:+:+: | 2661 | NA | 68 | 44 | 58 | 20 | 82 | 88 |
| ENSDARG000 | 11;11;11 | 24116083:241 | 24116361:241 | +;+;+ | 604 | NA | 0 | 0 | 0 | 0 | 0 | 0 |
| ENSDARG000 | 11:11:11:11:1 | 11468566:114 | 11468814:114 | -;-;-;-;-;-;-;- | 1891 | krt94 | 290 | 4 | 22 | 18 | 78 | 74 |
| ENSDARG000 | 11:11:11:11:1 | 5945186:594 | 5947312:594 | -:-:-:-:-:-:-:-: | 2573 | dda1 | 692 | 1091 | 973 | 941 | 774 | 834 |
| ENSDARG000 | 11:11:11:11:1 | 25583950:255 | 25584175:255 | +:+:+:+:+:+:+: | 6475 | ccdc120b | 10 | 2 | 2 | 0 | 4 | 16 |
| ENSDARG000 | 11:11:11:11:1 | 15931188:159 | 15931253:159 | +:+:+:+:+:+:+: | 8617 | draxina | 80 | 46 | 736 | 208 | 314 | 246 |
| ENSDARG000 | 11:11:11:11:1 | 13058613:130 | 13058679:130 | +:+:+:+:+:+:+: | 3356 | NA | 18 | 29 | 13 | 36 | 30 | 15 |
| ENSDARG000 | 11:11:11:11:1 | 37201483:372 | 37201575:372 | +:+:+:+:+:+:+: | 3314 | itih3b.2 | 0 | 0 | 58 | 36 | 0 | 0 |
| ENSDARG000 | 11:11:11:11:1 | 25619892:256 | 25619995:256 | +:+:+:+:+:+:+: | 2190 | comtb | 1159 | 2304 | 1559 | 1578 | 2025 | 2011 |
| ENSDARG000 | 11:11:11:11:1 | 3523358:352 | 3525099:352 | -:-:-:-:-:-:-:-: | 3355 | ddx19b | 813 | 857 | 890 | 410 | 912 | 528 |
| ENSDARG000 | 11:11:11:11:1 | 1551603:155 | 1551813:155 | +:+:+:+:+:+:+: | 4514 | mybl2b | 152 | 793 | 791 | 316 | 617 | 430 |
| ENSDARG000 | 11:11:11:11:1 | 7432533:752 | 7432758:752 | +:+:+:+:+:+:+: | 7803 | ADGRL2 | 50 | 24 | 1482 | 98 | 1085 | 41 |
| ENSDARG000 | 11:11:11:11:1 | 31218956:312 | 31220930:312 | -;-;-;-;-;-;-;- | 3794 | NACC1 | 54 | 47 | 90 | 38 | 61 | 64 |
| ENSDARG000 | 11;11 | 24187610:241 | 24191324:241 | -;- | 3955 | sox12 | 4 | 24 | 102 | 84 | 140 | 46 |
| ENSDARG000 | 11:11:11:11:1 | 24298432:242 | 24298541:242 | +:+:+:+:+:+:+: | 1231 | NA | 10 | 2 | 1 | 6 | 0 | 0 |
| ENSDARG000 | 11:11:11:11:1 | 24251141:242 | 24251673:242 | +:+:+:+:+:+:+: | 4524 | pnp4a | 18527 | 11668 | 23439 | 19721 | 27847 | 30477 |
| ENSDARG000 | 11:11:11:11:1 | 3499111:349 | 3499398:349 | +:+:+:+:+:+:+: | 5604 | NA | 18 | 27 | 26 | 24 | 22 | 45 |
| ENSDARG000 | 11:11:11:11:1 | 34633392:346 | 34633513:346 | -:-:-:-:-:-:-:-: | 12711 | DNAH1 | 2 | 10 | 2 | 0 | 20 | 0 |
| ENSDARG000 | 11:11:11:11:1 | 11504014:115 | 11504404:115 | +:+:+:+:+:+:+: | 1937 | krt98 | 0 | 0 | 0 | 0 | 0 | 0 |
| ENSDARG000 | 11;11 | 13151919:131 | 13152175:131 | +;+ | 295 | NA | 0 | 0 | 0 | 0 | 0 | 0 |
| ENSDARG000 | 11:11:11:11:1 | 13094179:130 | 13095147:130 | -:-:-:-:-:-:-:-: | 2779 | elovl1b | 7902 | 8719 | 5626 | 9012 | 6739 | 8426 |
| ENSDARG000 | 11:11:11:11:1 | 25634041:256 | 25634276:256 | +:+:+:+:+:+:+: | 4424 | grm6b | 527 | 376 | 797 | 1141 | 816 | 613 |
| ENSDARG000 | 11:11:11:11:1 | 31236001:312 | 31236181:312 | +:+:+:+:+:+:+: | 2903 | TRMT1 | 315 | 302 | 386 | 225 | 355 | 252 |
| ENSDARG000 | 11:11:11:11:1 | 31258116:312 | 31258573:312 | -:-:-:-:-:-:-:-: | 3517 | man2b1 | 4622 | 2697 | 3043 | 2355 | 3493 | 2766 |
| ENSDARG000 | 11:11:11:11:1 | 28522364:285 | 28523272:285 | -:-:-:-:-:-:-:-: | 2178 | NA | 0 | 0 | 0 | 0 | 0 | 0 |
| ENSDARG000 | 11:11:11:11:1 | 18873113:188 | 18873916:188 | +:+:+:+:+:+:+: | 9164 | magi1b | 1676 | 1035 | 1623 | 1725 | 1682 | 1394 |
| ENSDARG000 | 11:11:11:11:1 | 5965168:596 | 5966384:596 | -:-:-:-:-:-:-:-: | 4052 | ANO8 | 182 | 28 | 38 | 16 | 44 | 10 |
| ENSDARG000 | 11 | 37297501 | 37298097 | - | 597 | NA | 0 | 1 | 10 | 2 | 0 | 5 |
| ENSDARG000 | 11:11:11:11:1 | 37275448:372 | 37275539:372 | +:+:+:+:+:+:+: | 1380 | NA | 22 | 22 | 24 | 28 | 18 | 0 |
| ENSDARG000 | 11:11:11:11:1 | 37250839:372 | 37250927:372 | +:+:+:+:+:+:+: | 5353 | NA | 34 | 2 | 6 | 0 | 6 | 14 |
| ENSDARG000 | 11;11;11 | 7782736:778 | 7784759:778 | -;-;- | 2274 | NA | 0 | 0 | 0 | 0 | 0 | 0 |
| ENSDARG000 | 11;11 | 7782514:778 | 7782611:778 | +;+ | 538 | NA | 0 | 0 | 0 | 0 | 0 | 0 |
| ENSDARG000 | 11;11;11 | 40893766:408 | 40893926:408 | -;-;- | 336 | NA | 0 | 2 | 2 | 0 | 0 | 0 |
| ENSDARG000 | 11;11 | 40895694:409 | 40895724:409 | +;+ | 467 | NA | 0 | 0 | 0 | 0 | 0 | 0 |
| ENSDARG000 | 11:11:11:11:1 | 40897326:408 | 40897454:408 | +;+;+;+;+ | 750 | NA | 5 | 2 | 8 | 20 | 9 | 4 |
| ENSDARG000 | 11;11;11 | 7880177:795 | 7880265:795 | +;+;+ | 625 | NA | 0 | 0 | 0 | 0 | 0 | 0 |
| ENSDARG000 | 11:11:11:11:1 | 24313931:243 | 24314287:243 | +:+:+:+:+:+:+: | 3359 | rem1 | 32 | 26 | 4 | 10 | 24 | 10 |
| ENSDARG000 | 11:11:11:11:1 | 16014527:160 | 16021315:160 | -:-:-:-:-:-:-:-: | 6916 | zgc:173544 | 0 | 1 | 0 | 0 | 0 | 0 |
| ENSDARG000 | 11:11:11:11:1 | 6009984:601 | 6010667:601 | +:+:+:+:+:+:+: | 2000 | gtpbp3 | 44 | 35 | 76 | 16 | 32 | 60 |
| ENSDARG000 | 11:11:11:11:1 | 41078191:410 | 41078779:410 | -:-:-:-:-:-:-:-: | 1059 | fbxo2 | 3109 | 2858 | 3024 | 4428 | 2113 | 2950 |
| ENSDARG000 | 11:11:11:11:1 | 41089574:410 | 41089716:410 | +:+:+:+:+:+:+: | 2182 | PHF13 | 179 | 200 | 350 | 332 | 221 | 322 |
| ENSDARG000 | 11:11:11:11:1 | 28590733:285 | 28591144:285 | -:-:-:-:-:-:-:-: | 1773 | dhrs3b | 4 | 9 | 30 | 6 | 34 | 0 |
| ENSDARG000 | 11:11:11:11:1 | 1575435:157 | 1575665:157 | +:+:+:+:+:+:+: | 2257 | NA | 26 | 10 | 12 | 0 | 2 | 6 |
| ENSDARG000 | 11:11:11:11:1 | 41110397:411 | 41114597:411 | -:-:-:-:-:-:-:-: | 6081 | DNAJC11 | 116 | 137 | 136 | 54 | 155 | 105 |
| ENSDARG000 | 11:11:11:11:1 | 41110669:411 | 41110700:411 | +:+:+:+:+:+:+: | 776 | NA | 0 | 0 | 0 | 0 | 0 | 0 |
| ENSDARG000 | 11:11:11:11:1 | 34760628:347 | 34761093:347 | +:+:+:+:+:+:+: | 4711 | NA | 18 | 62 | 48 | 0 | 104 | 24 |
| ENSDARG000 | 11:11:11:11:1 | 34779610:347 | 34780352:347 | -;-;-;-;-;- | 984 | chchd4a | 102 | 193 | 148 | 14 | 214 | 154 |
| ENSDARG000 | 11:11:11:11:1 | 6043742:604 | 6044227:604 | -;-;-;-;-;-;- | 2100 | plvapb | 808 | 98 | 492 | 234 | 278 | 312 |
| ENSDARG000 | 11:11:11:11:1 | 11515778:115 | 11515996:115 | -;-;-;-;-;-;- | 1545 | krt15 | 734 | 10 | 12 | 18 | 182 | 122 |
| ENSDARG000 | 11:11:11:11:1 | 7981243:798 | 7985068:798 | -:-:-:-:-:-:-:-: | 7328 | ttll7 | 102 | 72 | 321 | 170 | 216 | 144 |
| ENSDARG000 | 11:11:11:11:1 | 25681335:256 | 25682106:256 | +:+:+:+:+:+:+: | 2240 | otud5b | 740 | 562 | 684 | 679 | 585 | 648 |
| ENSDARG000 | 11:11:11:11:1 | 37310881:373 | 37310907:373 | -:-:-:-:-:-:-:-: | 3620 | ERC2 | 0 | 10 | 54 | 50 | 12 | 7 |
| ENSDARG000 | 11:11:11:11:1 | 1584747:158 | 1584877:158 | +:+:+:+:+:+:+: | 4898 | NA | 735 | 492 | 428 | 325 | 452 | 308 |
| ENSDARG000 | 11:11:11:11:1 | 1602916:160 | 1602945:160 | +:+:+:+:+:+:+: | 5094 | NA | 8 | 0 | 4 | 8 | 10 | 2 |
| ENSDARG000 | 11 | 19095431 | 19097746 | - | 2316 | rhol | 0 | 0 | 0 | 4 | 0 | 0 |
| ENSDARG000 | 11;11;11;11 | 28617564:286 | 28617753:286 | +;+;+;+ | 1577 | aadacl4 | 0 | 2 | 0 | 16 | 76 | 0 |
| ENSDARG000 | 11:11:11:11:1 | 11565996:115 | 11566443:115 | -:-:-:-:-:-:-:-: | 1894 | krt99 | 0 | 0 | 0 | 0 | 0 | 0 |
| ENSDARG000 | 11:11:11:11:1 | 16040517:160 | 16040712:160 | +:+:+:+:+:+:+: | 2320 | agtrap | 6125 | 5432 | 6139 | 5458 | 7606 | 10145 |
| ENSDARG000 | 11;11;11;11 | 28620683:286 | 28621554:286 | +;+;+;+ | 1727 | NA | 0 | 2 | 12 | 0 | 0 | 0 |
| ENSDARG000 | 11:11:11:11:1 | 37431693:374 | 37432113:374 | -:-:-:-:-:-:-:-: | 11809 | BSN | 20 | 27 | 16 | 70 | 75 | 6 |
| ENSDARG000 | 11:11:11:11:1 | 34824099:348 | 34824405:348 | +:+:+:+:+:+:+: | 10598 | slc38a3a | 14 | 14 | 4 | 0 | 4 | 4 |
| ENSDARG000 | 11:11:11:11:1 | 19157879:191 | 19158231:191 | +:+:+:+:+:+:+: | 10565 | adamts9 | 96 | 44 | 86 | 164 | 168 | 34 |
| ENSDARG000 | 11 | 28633717 | 28639498 | - | 5782 | KLHDC7A | 0 | 0 | 0 | 0 | 0 | 10 |
| ENSDARG000 | 11:11:11:11:1 | 25693395:256 | 25694003:256 | +;+;+;+;+;+ | 3809 | MON1B | 464 | 384 | 488 | 341 | 546 | 440 |
| ENSDARG000 | 11:11:11:11:1 | 11595973:115 | 11598699:115 | -:-:-:-:-:-:-:-: | 5235 | NA | 38 | 10 | 56 | 4 | 42 | 14 |
| ENSDARG000 | 11:11:11:11:1 | 16070438:160 | 16070653:160 | +;+;+;+;+;+;+ | 1568 | ctbs | 612 | 627 | 471 | 579 | 789 | 574 |
| ENSDARG000 | 11;11 | 37601325:376 | 37601793:376 | -;- | 591 | NA | 13 | 14 | 2 | 11 | 13 | 0 |
| ENSDARG000 | 11;11 | 42305089:424 | 42307028:424 | +;+ | 3254 | NA | 0 | 0 | 0 | 0 | 0 | 0 |
| ENSDARG000 | 11:11:11:11:1 | 42304585:42 | 42307869:42 | -:-:-:-:-:-:-:-: | 7212 | slmapa | 903 | 609 | 772 | 551 | 722 | 551 |
| ENSDARG000 | 11:11:11:11:1 | 3563091:356 | 3563282:356 | -;-;-;-;-;-;-;- | 935 | NA | 0 | 0 | 0 | 0 | 0 | 0 |
| ENSDARG000 | 11:11:11:11:1 | 25712772:257 | 25714278:257 | -:-:-:-:-:-:-:-: | 5730 | BRPF3 | 371 | 408 | 459 | 166 | 623 | 391 |
| ENSDARG000 | 11:11:11:11:1 | 13159988:131 | 13160281:131 | +;+;+;+;+;+;+ | 1599 | mob3c | 188 | 104 | 206 | 223 | 164 | 168 |
| ENSDARG000 | 11;11 | 3550370:355 | 3551325:355 | -;- | 1458 | camk2n1 | 6 | 2 | 0 | 3 | 2 | 2 |
| ENSDARG000 | 11:11:11:11:1 | 13176568:131 | 13176709:131 | +:+:+:+:+:+:+: | 2777 | mknk1 | 173 | 118 | 131 | 182 | 112 | 90 |
| ENSDARG000 | 11:11:11:11:1 | 6069875:606 | 6069907:607 | +:+:+:+:+:+:+: | 2897 | NA | 2 | 6 | 14 | 24 | 12 | 16 |
| ENSDARG000 | 11:11:11:11:1 | 6051287:605 | 6052231:605 | -:-:-:-:-:-:-:-: | 2837 | babam1 | 552 | 578 | 738 | 537 | 577 | 553 |
| ENSDARG000 | 11:11:11:11:1 | 19271557:192 | 19271680:192 | +:+:+:+:+:+:+: | 7359 | prickle2b | 634 | 280 | 616 | 653 | 438 | 652 |
| ENSDARG000 | 11:11:11:11:1 | 25735478:257 | 25736226:257 | +:+:+:+:+:+:+: | 2765 | NA | 0 | 8 | 3 | 9 | 22 | 2 |
| ENSDARG000 | 11:11:11:11:1 | 31285127:312 | 31285705:312 | +:+:+:+:+:+:+: | 5153 | NA | 51 | 0 | 2 | 4 | 22 | 32 |
| ENSDARG000 | 11:11:11:11:1 | 37631714:376 | 37631859:376 | +:+:+:+:+:+:+: | 557 | NA | 0 | 0 | 0 | 0 | 0 | 0 |
| ENSDARG000 | 11:11:11:11:1 | 37612214:376 | 37612415:376 | +:+:+:+:+:+:+: | 5368 | hp1bp3 | 4116 | 4484 | 5395 | 3844 | 5877 | 4803 |
| ENSDARG000 | 11 | 13193747 | 13194217 | + | 471 | TMEM275 | 0 | 0 | 0 | 0 | 0 | 0 |
| ENSDARG000 | 11:11:11:11:1 | 8129536:812 | 8129734:813 | +:+:+:+:+:+:+: | 2627 | prkacba | 14 | 8 | 11 | 0 | 36 | 28 |
| ENSDARG000 | 11:11:11:11:1 | 28642913:286 | 28644247:286 | -:-:-:-:-:-:-:-: | 3205 | igsf21a | 10 | 1 | 16 | 8 | 14 | 20 |
| ENSDARG000 | 11;11 | 24345697:243 | 24347340:243 | -;- | 1746 | si:ch211-15p | 0 | 0 | 0 | 0 | 0 | 0 |
| ENSDARG000 | 11:11:11:11:1 | 24339362:243 | 24339412:243 | +:+:+:+:+:+:+: | 8105 | rbm39a | 4549 | 3482 | 5389 | 5222 | 6275 | 6154 |
| ENSDARG000 | 11:11:11:11:1 | 11639483:116 | 11639677:116 | -;-;-;-;-;-;- | 793 | NA | 6 | 2 | 12 | 4 | 8 | 0 |
| ENSDARG000 | 11:11:11:11:1 | 3575963:357 | 3576063:357 | +:+:+:+:+:+:+: | 1640 | NA | 2 | 0 | 0 | 0 | 1 | 0 |
| ENSDARG000 | 11:11:11:11:1 | 19490335:194 | 19490881:194 | -:-:-:-:-:-:-:-: | 2291 | slc2a9l1 | 0 | 2 | 12 | 2 | 31 | 0 |
| ENSDARG000 | 11:11:11:11:1 | 24348425:243 | 24348715:243 | +:+:+:+:+:+:+: | 2194 | nfs1 | 1986 | 1887 | 1135 | 940 | 1962 | 2371 |
| ENSDARG000 | 11:11:11:11:1 | 34921492:350 | 34922135:350 | +:+:+:+:+:+:+: | 2592 | gnai2a | 2312 | 2797 | 2848 | 2503 | 2536 | 2777 |
| ENSDARG000 | 11:11:11:11:1 | 8160894:816 | 8161032:816 | -:-:-:-:-:-:-:-: | 1236 | uox | 0 | 0 | 16 | 0 | 0 | 4 |
| ENSDARG000 | 11:11:11:11:1 | 11719575:117 | 11719695:117 | +:+:+:+:+:+:+: | 6758 | jupa | 2087 | 1827 | 2316 | 1492 | 1547 | 1382 |
| ENSDARG000 | 11:11:11:11:1 | 16094661:160 | 16095113:160 | -:-:-:-:-:-:-:-: | 1617 | rpf1 | 412 | 476 | 587 | 251 | 445 | 452 |
| ENSDARG000 | 11:11:11:11:1 | 25752198:257 | 25752438:257 | -:-:-:-:-:-:-:-: | 4722 | PLCH2 | 4 | 0 | 0 | 0 | 0 | 4 |
| ENSDARG000 | 11:11:11:11:1 | 35048642:350 | 35048787:350 | +;+;+;+;+ | 2022 | inka1a | 3675 | 1251 | 2545 | 1367 | 3532 | 2542 |
| ENSDARG000 | 11:11:11:11:1 | 45286911:452 | 45287650:452 | +:+:+:+:+:+:+: | 6147 | pycr1b | 759 | 757 | 1908 | 740 | 926 | 585 |
| ENSDARG000 | 11:11:11:11:1 | 16084581:160 | 16084652:160 | -:-:-:-:-:-:-:-: | 1407 | SPATA1 | 1 | 0 | 1 | 10 | 2 | 0 |
| ENSDARG000 | 11:11:11:11:1 | 8152872:815 | 8152995:815 | +:+:+:+:+:+:+: | 1501 | samd13 | 209 | 230 | 223 | 228 | 224 | 222 |
| ENSDARG000 | 11:11:11:11:1 | 13207898:132 | 13208082:132 | +:+:+:+:+:+:+: | 1866 | atp5f1b | 19510 | 22440 | 25205 | 16439 | 24225 | 17396 |
| ENSDARG000 | 11:11:11:11:1 | 19502297:195 | 19502599:195 | +:+:+:+:+:+:+: | 1430 | psmd6 | 2474 | 2627 | 3474 | 1681 | 2899 | 2323 |
| ENSDARG000 | 11:11:11:11:1 | 3585934:358 | 3586240:358 | +;+;+;+;+;+;+ | 1473 | cdab | 0 | 0 | 2 | 0 | 0 | 0 |
| ENSDARG000 | 11:11:11:11:1 | 45299447:453 | 45299801:453 | +:+:+:+:+:+:+: | 2889 | mafgb | 149 | 78 | 116 | 134 | 134 | 156 |
| ENSDARG000 | 11:11:11:11:1 | 29123597:291 | 29123773:291 | -:-:-:-:-:-:-:-: | 1686 | NA | 0 | 0 | 0 | 0 | 0 | 0 |
| ENSDARG000 | 11:11:11:11:1 | 31323746:313 | 31324515:313 | +:+:+:+:+:+:+: | 9875 | sipa1l2 | 138 | 148 | 598 | 91 | 226 | 390 |
| ENSDARG000 | 11:11:11:11:1 | 19511642:195 | 19517035:195 | -:-:-:-:-:-:-:-: | 8800 | ATXN7 | 303 | 246 | 358 | 195 | 302 | 386 |
| ENSDARG000 | 11:11:11:11:1 | 37638873:376 | 37639075:376 | +:+:+:+:+:+:+: | 4121 | sh2d5 | 12 | 2 | 0 | 0 | 0 | 0 |
| ENSDARG000 | 11:11:11:11:1 | 19603251:196 | 19603305:196 | +:+:+:+:+:+:+: | 777 | thoc7 | 252 | 292 | 385 | 260 | 282 | 361 |
| ENSDARG000 | 11;11 | 11792408:117 | 11792975:117 | +;+ | 934 | NA | 2 | 0 | 2 | 0 | 0 | 0 |
| ENSDARG000 | 11:11:11:11:1 | 42422371:424 | 42423007:424 | +:+:+:+:+:+:+: | 5208 | dennd6aa | 204 | 143 | 330 | 75 | 278 | 148 |
| ENSDARG000 | 11:11:11:11:1 | 11781973:117 | 11783908:117 | -:-:-:-:-:-:-:-: | 3706 | cdc6 | 160 | 523 | 686 | 295 | 469 | 264 |
| ENSDARG000 | 11:11:11:11:1 | 1628538:165 | 1628965:165 | +:+:+:+:+:+:+: | 14875 | LRP1 | 842 | 94 | 192 | 164 | 402 | 206 |
| ENSDARG000 | 11:11:11:11:1 | 25782721:257 | 25783805:257 | -;-;-;-;-;-;-;- | 3561 | skib | 635 | 763 | 671 | 797 | 427 | 281 |
| ENSDARG000 | 11:11:11:11:1 | 45339906:453 | 45340116:453 | +:+:+:+:+:+:+: | 2897 | NA | 372 | 299 | 330 | 383 | 341 | 397 |
| ENSDARG000 | 11:11:11:11:1 | 19622925:196 | 19628025:196 | -:-:-:-:-:-:-:-: | 5905 | SYNPR | 0 | 10 | 33 | 0 | 6 | 24 |
| ENSDARG000 | 11:11:11:11:1 | 13223625:132 | 13223826:132 | +:+:+:+:+:+:+: | 7871 | ABCB11 | 0 | 0 | 10 | 0 | 0 | 0 |
| ENSDARG000 | 11:11:11:11:1 | 35171406:351 | 35171487:351 | +;+;+;+;+;+ | 4684 | mon1a | 228 | 203 | 301 | 176 | 358 | 289 |
| ENSDARG000 | 11;11;11 | 25833219:258 | 25833951:258 | -;-;- | 1381 | tmem51b | 52 | 71 | 39 | 38 | 40 | 12 |
| ENSDARG000 | 11:11:11:11:1 | 35162684:351 | 35163517:351 | -:-:-:-:-:-:-:-: | 2942 | traip | 8 | 102 | 144 | 36 | 74 | 28 |
| ENSDARG000 | 11:11:11:11:1 | 42474694:424 | 42474782:424 | +:+:+:+:+:+:+: | 2464 | NA | 0 | 2 | 6 | 2 | 12 | 0 |
| ENSDARG000 | 11:11:11:11:1 | 29331142:293 | 29335835:293 | -:-:-:-:-:-:-:-: | 8507 | ARHGEF10L | 18 | 28 | 10 | 22 | 22 | 18 |
| ENSDARG000 | 11:11:11:11:1 | 42478184:424 | 42478322:424 | +;+;+;+;+ | 973 | NA | 8 | 2 | 8 | 8 | 4 | 4 |
| ENSDARG000 | 11:11:11:11:1 | 25857941:258 | 25858398:258 | -:-:-:-:-:-:-:-: | 4797 | kaznb | 329 | 290 | 311 | 201 | 474 | 99 |
| ENSDARG000 | 11:11:11:11:1 | 24360744:243 | 24361261:243 | -:-:-:-:-:-:-:-: | 2260 | dvl1b | 233 | 197 | 357 | 345 | 459 | 702 |
| ENSDARG000 | 11:11:11:11:1 | 3593127:359 | 3595710:359 | -:-:-:-:-:-:-:-: | 5190 | DNAJC16 | 1494 | 1369 | 1400 | 1472 | 1407 | 1399 |
| ENSDARG000 | 11:11:11:11:1 | 11797026:117 | 11798691:117 | +:+:+:+:+:+:+: | 3443 | NA | 6 | 2 | 0 | 0 | 4 | 3 |
| ENSDARG000 | 11:11:11:11:1 | 16139897:161 | 16143907:161 | -:-:-:-:-:-:-:-: | 4722 | arpc4l | 1717 | 2096 | 2026 | 1919 | 1846 | 2148 |
| ENSDARG000 | 11:11:11:11:1 | 42482920:424 | 42483184:424 | +;+;+;+;+;+ | 1609 | ARF5 | 8 | 7 | 12 | 0 | 10 | 11 |
| ENSDARG000 | 11:11:11:11:1 | 42494531:424 | 42494796:424 | +;+;+;+;+;+ | 3317 | arf4a | 668 | 737 | 1271 | 488 | 1499 | 609 |
| ENSDARG000 | 11:11:11:11:1 | 16152316:161 | 16152734:161 | +:+:+:+:+:+:+: | 3223 | tada3l | 553 | 555 | 713 | 310 | 458 | 482 |
| ENSDARG000 | 11 | 31558207 | 31559070 | + | 864 | EGLN1 | 0 | 0 | 0 | 0 | 0 | 0 |
| ENSDARG000 | 11:11:11:11:1 | 31558006:315 | 31559070:315 | +;+;+;+;+ | 2335 | egln1b | 198 | 322 | 421 | 340 | 367 | 270 |
| ENSDARG000 | 11:11:11:11:1 | 16137302:161 | 16137399:161 | +:+:+:+:+:+:+: | 1821 | NA | 0 | 0 | 0 | 0 | 0 | 0 |
| ENSDARG000 | 11:11:11:11:1 | 16166117:161 | 16166211:161 | +:+:+:+:+:+:+: | 1461 | hmces | 30 | 37 | 52 | 43 | 33 | 49 |
| ENSDARG000 | 11:11:11:11:1 | 11842724:118 | 11843979:118 | +:+:+:+:+:+:+: | 3033 | NA | 0 | 0 | 1 | 6 | 1 | 0 |
| ENSDARG000 | 11:11:11:11:1 | 11844123:118 | 11845940:118 | -:-:-:-:-:-:-:-: | 4914 | WIPF2 | 1396 | 979 | 1425 | 1131 | 1178 | 1020 |
| ENSDARG000 | 11:11:11:11:1 | 16176603:161 | 16180351:161 | -:-:-:-:-:-:-:-: | 5080 | rab7a | 7554 | 5527 | 5564 | 5423 | 5018 | 6319 |
| ENSDARG000 | 11:11:11:11:1 | 19729771:197 | 19730273:197 | -:-:-:-:-:-:-:-: | 2020 | nampt2 | 858 | 551 | 710 | 287 | 408 | 412 |
| ENSDARG000 | 11:11:11:11:1 | 24429923:244 | 24432064:244 | -:-:-:-:-:-:-:-: | 3604 | mxra8a | 92 | 24 | 14 | 6 | 52 | 22 |
| ENSDARG000 | 11;11;11;11 | 42506052:425 | 42506784:425 | -;-;-;- | 2114 | pde12 | 251 | 431 | 370 | 117 | 390 | 315 |
| ENSDARG000 | 11:11:11:11:1 | 42532464:425 | 42533362:425 | -;-;-;-;-;-;- | 1660 | atp6ap1la | 2 | 0 | 8 | 0 | 4 | 0 |
| ENSDARG000 | 11:11:11:11:1 | 31609481:316 | 31609621:316 | +:+:+:+:+:+:+: | 2077 | zgc:162816 | 112 | 170 | 273 | 124 | 215 | 110 |
| ENSDARG000 | 11:11:11:11:1 | 24464356:244 | 24466292:244 | -:-:-:-:-:-:-:-: | 7524 | PLEKHG5 | 877 | 615 | 954 | 287 | 850 | 408 |
| ENSDARG000 | 11:11:11:11:1 | 6076646:607 | 6076847:607 | +:+:+:+:+:+:+: | 5867 | NA | 70 | 20 | 50 | 63 | 90 | 105 |
| ENSDARG000 | 11:11:11:11:1 | 19780717:197 | 19780941:197 | +:+:+:+:+:+:+: | 4652 | NA | 10 | 24 | 29 | 1 | 22 | 6 |

| | | | | | | | | | | | |
|---|---|---|---|---|---|---|---|---|---|---|---|
| ENSDARG000 | 11:11:11:11:1 | 45354838:453 | 45356688:453 | -:-:-:-:-:-:-:-: | 5929 | TRAPPC10 | 1640 | 1201 | 1674 | 1041 | 1598 | 1010 |
| ENSDARG000 | 11:11:11:11:1 | 19991633:200 | 19991819:200 | -:-:-:-:-:-:-:-: | 2414 | C11H3orf14 | 54 | 38 | 48 | 18 | 32 | 60 |
| ENSDARG000 | 11:11:11:11:1 | 11891580:119 | 11896806:119 | -;-;-;-;-;-;- | 8992 | NA | 106 | 44 | 76 | 104 | 163 | 44 |
| ENSDARG000 | 11:11:11:11:1 | 20056732:200 | 20056900:200 | +:+:+:+:+:+:+: | 1838 | fezf2 | 0 | 0 | 0 | 0 | 0 | 0 |
| ENSDARG000 | 11:11:11:11:1 | 37652870:376 | 37653278:376 | +:+:+:+:+:+:+: | 3008 | kif17 | 0 | 0 | 0 | 0 | 0 | 0 |
| ENSDARG000 | 11;11 | 45389377:453 | 45389664:453 | +;+ | 558 | NA | 31 | 26 | 56 | 38 | 81 | 108 |
| ENSDARG000 | 11:11:11:11:1 | 45388126:453 | 45388457:453 | +:+:+:+:+:+:+: | 3195 | srsf11 | 3946 | 3721 | 7022 | 4340 | 5830 | 5910 |
| ENSDARG000 | 11:11:11:11:1 | 26189703:261 | 26196141:261 | -:-:-:-:-:-:-:-: | 7204 | PRDM2 | 2269 | 2350 | 2069 | 1836 | 2549 | 1857 |
| ENSDARG000 | 11;11;11;11 | 42556395:425 | 42556498:425 | +;+;+;+ | 582 | rps23 | 9902 | 12633 | 15402 | 14075 | 9509 | 13656 |
| ENSDARG000 | 11:11:11:11:1 | 13280248:132 | 13284113:132 | -:-:-:-:-:-:-:-: | 7605 | NA | 24 | 87 | 74 | 38 | 174 | 48 |
| ENSDARG000 | 11 | 42572242 | 42572982 | + | 741 | NA | 0 | 0 | 0 | 0 | 0 | 0 |
| ENSDARG000 | 11:11:11:11:1 | 29566746:295 | 29566851:295 | +;+;+;+;+;+;+ | 869 | NA | 0 | 0 | 0 | 0 | 0 | 0 |
| ENSDARG000 | 11:11:11:11:1 | 8170948:8171 | 8171171:8171 | +:+:+:+:+:+:+: | 1363 | DNASE2B | 14 | 12 | 30 | 14 | 50 | 49 |
| ENSDARG000 | 11:11:11:11:1 | 37683278:376 | 37686665:376 | -:-:-:-:-:-:-:-: | 5353 | zgc:158258 | 0 | 0 | 0 | 0 | 0 | 0 |
| ENSDARG000 | 11:11:11:11:1 | 11911399:119 | 11911721:119 | -:-:-:-:-:-:-:-: | 5006 | NA | 440 | 281 | 452 | 274 | 458 | 322 |
| ENSDARG000 | 11:11:11:11:1 | 42585138:425 | 42585302:425 | +:+:+:+:+:+:+: | 2784 | ASB14 | 6 | 6 | 254 | 12 | 166 | 14 |
| ENSDARG000 | 11:11:11:11:1 | 26228918:262 | 26234056:262 | -:-:-:-:-:-:-:-: | 7190 | foxj3 | 396 | 759 | 766 | 422 | 899 | 614 |
| ENSDARG000 | 11:11:11:11:1 | 3614326:3614 | 3614673:3614 | -:-:-:-:-:-:-:-: | 3058 | itih3a.1 | 0 | 0 | 12 | 0 | 0 | 0 |
| ENSDARG000 | 11:11:11:11:1 | 37708384:377 | 37711985:377 | -;-;-;-;-;-;- | 4246 | yod1 | 190 | 186 | 227 | 156 | 165 | 252 |
| ENSDARG000 | 11:11:11:11:1 | 31625171:316 | 31625422:316 | +:+:+:+:+:+:+: | 5399 | NA | 224 | 70 | 404 | 139 | 273 | 412 |
| ENSDARG000 | 11:11:11:11:1 | 13366773:133 | 13367071:133 | +:+:+:+:+:+:+: | 2769 | NA | 6 | 2 | 0 | 0 | 0 | 0 |
| ENSDARG000 | 11 | 24580579 | 24581114 | + | 536 | NA | 0 | 0 | 0 | 0 | 0 | 0 |
| ENSDARG000 | 11:11:11:11:1 | 24620742:246 | 24620986:246 | +:+:+:+:+:+:+: | 3118 | SYT6 | 0 | 0 | 0 | 0 | 12 | 0 |
| ENSDARG000 | 11:11:11:11:1 | 13423776:134 | 13424559:134 | +:+:+:+:+:+:+: | 4515 | homer3b | 276 | 458 | 713 | 452 | 566 | 590 |
| ENSDARG000 | 11:11:11:11:1 | 29574743:295 | 29578275:295 | -:-:-:-:-:-:-:-: | 10136 | CHD5 | 1875 | 1232 | 838 | 575 | 1510 | 1046 |
| ENSDARG000 | 11;11;11 | 24671676:246 | 24672740:246 | -;-;- | 1540 | olfml3b | 72 | 14 | 28 | 22 | 52 | 26 |
| ENSDARG000 | 11:11:11:11:1 | 6115621:6115 | 6115865:6115 | +:+:+:+:+:+:+: | 4077 | nr2f6b | 22 | 66 | 98 | 209 | 26 | 159 |
| ENSDARG000 | 11:11:11:11:1 | 37720552:377 | 37720750:377 | +:+:+:+:+:+:+: | 598 | snrpe | 494 | 832 | 1015 | 468 | 716 | 494 |
| ENSDARG000 | 11:11:11:11:1 | 8201842:8201 | 8202127:8202 | -;-;-;-;-;-;-;- | 1869 | NA | 0 | 0 | 0 | 0 | 0 | 0 |
| ENSDARG000 | 11 | 24703108 | 24704437 | + | 1330 | GPR25 | 0 | 0 | 4 | 1 | 6 | 4 |
| ENSDARG000 | 11;11;11;11 | 26363435:263 | 26363565:263 | +;+;+;+ | 1233 | ppcs | 202 | 154 | 237 | 121 | 182 | 220 |
| ENSDARG000 | 11:11:11:11:1 | 35189804:351 | 35189920:351 | +;+;+;+;+ | 1490 | NA | 0 | 6 | 0 | 0 | 0 | 0 |
| ENSDARG000 | 11:11:11:11:1 | 37768298:378 | 37768675:378 | +:+:+:+:+:+:+: | 5557 | sox13 | 86 | 16 | 10 | 14 | 52 | 48 |
| ENSDARG000 | 11:11:11:11:1 | 37848705:378 | 37849065:378 | -;-;-;-;-;-;- | 1553 | etnk2 | 92 | 136 | 71 | 58 | 131 | 66 |
| ENSDARG000 | 11:11:11:11:1 | 31680513:316 | 31681049:316 | +:+:+:+:+:+:+: | 5063 | diaph3 | 91 | 192 | 235 | 58 | 137 | 152 |
| ENSDARG000 | 11;11;11 | 11974708:119 | 11975083:119 | +;+;+ | 2590 | zgc:64002 | 84 | 128 | 120 | 40 | 76 | 100 |
| ENSDARG000 | 11:11:11:11:1 | 37905630:379 | 37907050:379 | +:+:+:+:+:+:+: | 2888 | NA | 0 | 2 | 8 | 8 | 0 | 2 |
| ENSDARG000 | 11:11:11:11:1 | 12052791:120 | 12054577:120 | +:+:+:+:+:+:+: | 6688 | NA | 0 | 0 | 2 | 0 | 0 | 2 |
| ENSDARG000 | 11:11:11:11:1 | 29650011:296 | 29652549:296 | -:-:-:-:-:-:-:-: | 3692 | rpl22 | 5539 | 5572 | 6819 | 6719 | 4502 | 6798 |
| ENSDARG000 | 11:11:11:11:1 | 260763:26128 | 261089:26144 | -:-:-:-:-:-:-:-: | 2031 | alas1 | 1924 | 1546 | 2220 | 1490 | 1309 | 2208 |
| ENSDARG000 | 11:11:11:11:1 | 8221835:8221 | 8222119:8222 | -:-:-:-:-:-:-:-: | 2757 | NA | 0 | 0 | 0 | 0 | 0 | 0 |
| ENSDARG000 | 11:11:11:11:1 | 37948839:379 | 37950108:379 | -:-:-:-:-:-:-:-: | 3144 | slc41a1 | 245 | 58 | 126 | 110 | 149 | 140 |
| ENSDARG000 | 11:11:11:11:1 | 8256292:8259 | 8257194:8259 | -:-:-:-:-:-:-:-: | 3695 | NA | 0 | 0 | 0 | 0 | 0 | 2 |
| ENSDARG000 | 11:11:11:11:1 | 20091255:200 | 20092955:200 | -:-:-:-:-:-:-:-: | 3053 | ogfr2 | 588 | 390 | 329 | 154 | 385 | 472 |
| ENSDARG000 | 11:11:11:11:1 | 8239982:8240 | 8240266:8240 | -;-;-;-;-;- | 1521 | NA | 0 | 0 | 0 | 0 | 0 | 0 |
| ENSDARG000 | 11:11:11:11:1 | 24716837:247 | 24716917:247 | +:+:+:+:+:+:+: | 2468 | inavab | 136 | 73 | 221 | 12 | 212 | 183 |
| ENSDARG000 | 11:11:11:11:1 | 12096199:120 | 12096350:120 | -:-:-:-:-:-:-:-: | 1041 | mrpl45 | 280 | 320 | 478 | 154 | 330 | 299 |
| ENSDARG000 | 11:11:11:11:1 | 6136220:6136 | 6136412:6136 | +:+:+:+:+:+:+: | 1965 | tax1bp3 | 1084 | 1100 | 1258 | 1429 | 879 | 1453 |
| ENSDARG000 | 11:11:11:11:1 | 35196587:351 | 35197864:351 | +:+:+:+:+:+:+: | 4221 | NA | 10 | 0 | 14 | 0 | 2 | 0 |
| ENSDARG000 | 11:11:11:11:1 | 13507053:135 | 13507172:135 | +:+:+:+:+:+:+: | 2075 | NA | 0 | 6 | 12 | 50 | 2 | 0 |
| ENSDARG000 | 11;11 | 38039964:380 | 38041079:380 | -;- | 1298 | NA | 4 | 2 | 2 | 2 | 0 | 0 |
| ENSDARG000 | 11:11:11:11:1 | 38033115:380 | 38033645:380 | -;-;-;-;-;-;-;- | 2265 | NA | 5 | 2 | 6 | 20 | 17 | 9 |
| ENSDARG000 | 11:11:11:11:1 | 13524504:135 | 13525082:135 | +:+:+:+:+:+:+: | 3241 | arrdc2 | 1102 | 1150 | 1533 | 2443 | 1232 | 1549 |
| ENSDARG000 | 11:11:11:11:1 | 8284686:8284 | 8284971:8284 | -;-;-;-;-;-;- | 2058 | NA | 0 | 0 | 0 | 0 | 0 | 0 |
| ENSDARG000 | 11:11:11:11:1 | 26370381:263 | 26370839:263 | -:-:-:-:-:-:-:-: | 2016 | utp3 | 421 | 735 | 1060 | 360 | 704 | 661 |
| ENSDARG000 | 11:11:11:11:1 | 16216909:162 | 16217071:162 | +:+:+:+:+:+:+: | 1608 | slc25a26 | 287 | 395 | 454 | 244 | 329 | 364 |
| ENSDARG000 | 11:11:11:11:1 | 45401608:454 | 45403921:454 | -:-:-:-:-:-:-:-: | 3772 | ankrd13c | 566 | 730 | 861 | 527 | 887 | 536 |
| ENSDARG000 | 11:11:11:11:1 | 276057:27605 | 276763:27676 | -;-;-;-;- | 1009 | npffl | 0 | 0 | 4 | 0 | 0 | 4 |
| ENSDARG000 | 11:11:11:11:1 | 38280454:383 | 38280835:383 | +:+:+:+:+:+:+: | 1798 | CDK18 | 2396 | 672 | 884 | 947 | 1200 | 1177 |
| ENSDARG000 | 11:11:11:11:1 | 12108355:121 | 12110117:121 | -:-:-:-:-:-:-:-: | 4426 | npepps | 2094 | 1958 | 2219 | 1335 | 2131 | 1950 |
| ENSDARG000 | 11:11:11:11:1 | 20371179:203 | 20371397:203 | +:+:+:+:+:+:+: | 4923 | cdh4 | 4 | 2 | 0 | 0 | 0 | 8 |
| ENSDARG000 | 11:11:11:11:1 | 281833:28303 | 282909:28318 | -;-;-;-;-;- | 1663 | slc11a2 | 784 | 951 | 1034 | 595 | 1100 | 1484 |
| ENSDARG000 | 11:11:11:11:1 | 24758967:247 | 24759558:247 | +:+:+:+:+:+:+: | 2492 | rnpep | 336 | 398 | 268 | 232 | 371 | 357 |
| ENSDARG000 | 11:11:11:11:1 | 3630627:3630 | 3630962:3630 | -:-:-:-:-:-:-:-: | 3291 | NA | 770 | 211 | 110 | 344 | 52 | 1206 |
| ENSDARG000 | 11:11:11:11:1 | 6205969:6206 | 6206665:6206 | +:+:+:+:+:+:+: | 3020 | NA | 0 | 2 | 2 | 0 | 4 | 2 |
| ENSDARG000 | 11:11:11:11:1 | 45421761:454 | 45421878:454 | +:+:+:+:+:+:+: | 1425 | plaat1 | 0 | 6 | 4 | 4 | 11 | 2 |
| ENSDARG000 | 11:11:11:11:1 | 38429319:384 | 38429680:384 | +:+:+:+:+:+:+: | 2242 | mfsd4aa | 42 | 21 | 27 | 2 | 14 | 26 |
| ENSDARG000 | 11:11:11:11:1 | 6197276:6197 | 6197511:6197 | -:-:-:-:-:-:-:-: | 2606 | pole4 | 3 | 17 | 20 | 2 | 11 | 24 |
| ENSDARG000 | 11:11:11:11:1 | 8303941:8306 | 8306984:8306 | -:-:-:-:-:-:-:-: | 4732 | NA | 2 | 2 | 4 | 2 | 0 | 8 |
| ENSDARG000 | 11:11:11:11:1 | 6182680:6187 | 6183056:6187 | -;-;-;-;-;- | 1163 | ccl44 | 15 | 2 | 0 | 0 | 8 | 0 |
| ENSDARG000 | 11:11:11:11:1 | 6152643:6159 | 6152695:6159 | +:+:+:+:+:+:+: | 2095 | slc1a6 | 2 | 2 | 10 | 10 | 0 | 12 |
| ENSDARG000 | 11:11:11:11:1 | 3847442:3847 | 3848978:3848 | -:-:-:-:-:-:-:-: | 3208 | gata2a | 60 | 10 | 10 | 10 | 56 | 338 |
| ENSDARG000 | 11:11:11:11:1 | 45426609:454 | 45426853:454 | -:-:-:-:-:-:-:-: | 2139 | rfc4 | 98 | 378 | 483 | 213 | 354 | 180 |
| ENSDARG000 | 11:11:11:11:1 | 35364445:354 | 35364618:354 | +:+:+:+:+:+:+: | 5850 | camkvb | 0 | 0 | 0 | 0 | 2 | 0 |
| ENSDARG000 | 11:11:11:11:1 | 32505375:325 | 32505763:325 | +:+:+:+:+:+:+: | 2167 | NA | 0 | 0 | 0 | 0 | 0 | 0 |
| ENSDARG000 | 11:11:11:11:1 | 12175162:121 | 12176692:121 | +:+:+:+:+:+:+: | 2456 | NA | 0 | 0 | 0 | 0 | 0 | 0 |
| ENSDARG000 | 11;11;11 | 32467342:325 | 32468551:325 | -;-;- | 1602 | NA | 0 | 0 | 0 | 0 | 0 | 0 |
| ENSDARG000 | 11:11:11:11:1 | 38470545:384 | 38471880:384 | -:-:-:-:-:-:-:-: | 3523 | elk4 | 125 | 75 | 109 | 60 | 167 | 92 |
| ENSDARG000 | 11:11:11:11:1 | 38505157:385 | 38509339:385 | -;-;-;-;- | 5340 | NA | 725 | 250 | 451 | 624 | 523 | 1843 |
| ENSDARG000 | 11:11:11:11:1 | 24786378:247 | 24786491:247 | +:+:+:+:+:+:+: | 1730 | cyb5r1 | 531 | 636 | 384 | 811 | 433 | 784 |
| ENSDARG000 | 11:11:11:11:1 | 8635785:8697 | 8636682:8697 | -;-;-;-;-;-;- | 1644 | NA | 3 | 13 | 15 | 28 | 68 | 75 |
| ENSDARG000 | 11:11:11:11:1 | 8565323:8565 | 8566080:8566 | +:+:+:+:+:+:+: | 4634 | tbl1xr1a | 552 | 557 | 842 | 648 | 638 | 579 |
| ENSDARG000 | 11:11:11:11:1 | 20887446:208 | 20888707:208 | -:-:-:-:-:-:-:-: | 4421 | NA | 440 | 344 | 440 | 325 | 384 | 344 |
| ENSDARG000 | 11:11:11:11:1 | 38517255:385 | 38518420:385 | -;-;-;-;-;- | 2614 | slc45a3 | 38 | 53 | 55 | 58 | 100 | 48 |
| ENSDARG000 | 11:11:11:11:1 | 29671661:296 | 29671756:296 | +:+:+:+:+:+:+: | 835 | NA | 0 | 2 | 0 | 4 | 0 | 0 |
| ENSDARG000 | 11;11;11 | 42839868:428 | 42839911:428 | +;+;+ | 539 | NA | 0 | 0 | 0 | 0 | 0 | 0 |
| ENSDARG000 | 11;11 | 8578054:8579 | 8578860:8580 | +;+ | 1674 | NA | 6 | 24 | 15 | 8 | 2 | 3 |
| ENSDARG000 | 11:11:11:11:1 | 12188120:121 | 12188407:121 | -:-:-:-:-:-:-:-: | 2305 | krt95 | 0 | 0 | 0 | 0 | 0 | 0 |
| ENSDARG000 | 11:11:11:11:1 | 3892386:3893 | 3892788:3894 | -:-:-:-:-:-:-:-: | 2908 | rpn1 | 6854 | 8186 | 12038 | 6815 | 8877 | 6232 |
| ENSDARG000 | 11:11:11:11:1 | 35447417:354 | 35451992:354 | -:-:-:-:-:-:-:-: | 6946 | SEMA3F | 28 | 13 | 10 | 8 | 40 | 18 |
| ENSDARG000 | 11:11:11:11:1 | 43135198:431 | 43135428:431 | -:-:-:-:-:-:-:-: | 8121 | SPTBN1 | 61051 | 28931 | 38872 | 31757 | 39922 | 33304 |
| ENSDARG000 | 11:11:11:11:1 | 296437:29651 | 296638:29663 | +:+:+:+:+:+:+: | 1040 | mettl1 | 232 | 282 | 333 | 202 | 187 | 241 |
| ENSDARG000 | 11:11:11:11:1 | 38540138:385 | 38542399:385 | -;-;-;-;-;-;- | 3802 | nucks1a | 3167 | 4252 | 5543 | 2076 | 4440 | 2984 |
| ENSDARG000 | 11:11:11:11:1 | 306007:30600 | 306115:30611 | -:-:-:-:-:-:-:-: | 837 | POC1A | 10 | 40 | 61 | 0 | 10 | 4 |
| ENSDARG000 | 11:11:11:11:1 | 16329776:163 | 16333192:163 | -:-:-:-:-:-:-:-: | 7451 | LRIG1 | 322 | 216 | 364 | 196 | 420 | 255 |
| ENSDARG000 | 11;11 | 29682909:296 | 29683700:296 | -;- | 1457 | nr0b1 | 60 | 50 | 42 | 33 | 57 | 42 |
| ENSDARG000 | 11:11:11:11:1 | 1867613:1871 | 1868096:1871 | +:+:+:+:+:+:+: | 10298 | NA | 2684 | 2247 | 2373 | 2054 | 3077 | 2556 |
| ENSDARG000 | 11;11;11;11 | 43135199:431 | 43135428:431 | -;-;-;- | 608 | NA | 0 | 0 | 0 | 0 | 0 | 0 |
| ENSDARG000 | 11;11 | 29734586:297 | 29735857:297 | -;- | 1502 | NA | 0 | 0 | 0 | 0 | 0 | 4 |
| ENSDARG000 | 11;11;11 | 13605623:136 | 13606751:136 | -;-;- | 1379 | si:ch211-1a19 | 0 | 0 | 0 | 0 | 10 | 0 |
| ENSDARG000 | 11:11:11:11:1 | 45436703:454 | 45436789:454 | +:+:+:+:+:+:+: | 4404 | sos1 | 994 | 620 | 852 | 813 | 754 | 742 |
| ENSDARG000 | 11;11 | 13616015:136 | 13616448:136 | -;- | 563 | sid1 | 0 | 0 | 0 | 0 | 0 | 0 |
| ENSDARG000 | 11:11:11:11:1 | 20994026:209 | 20994285:209 | -:-:-:-:-:-:-:-: | 2000 | slc6a14 | 0 | 0 | 10 | 0 | 4 | 10 |
| ENSDARG000 | 11:11:11:11:1 | 24800156:248 | 24800273:248 | +:+:+:+:+:+:+: | 3187 | adipor1a | 992 | 705 | 925 | 610 | 1224 | 784 |
| ENSDARG000 | 11:11:11:11:1 | 35744918:357 | 35745439:357 | -;-;-;-;-;-;- | 1111 | arl8bb | 384 | 522 | 366 | 239 | 388 | 227 |
| ENSDARG000 | 11:11:11:11:1 | 38614476:386 | 38614622:386 | +:+:+:+:+:+:+: | 4661 | NA | 0 | 0 | 0 | 0 | 0 | 2 |
| ENSDARG000 | 11:11:11:11:1 | 13629528:136 | 13630641:136 | +:+:+:+:+:+:+: | 5735 | NA | 178 | 254 | 12 | 10 | 72 | 48 |
| ENSDARG000 | 11 | 45468730 | 45470370 | - | 1641 | NA | 88 | 86 | 151 | 123 | 113 | 69 |
| ENSDARG000 | 11:11:11:11:1 | 35759652:357 | 35761774:357 | -;-;-;-;- | 2804 | bhlhe40 | 9590 | 6966 | 12319 | 13714 | 12814 | 11099 |
| ENSDARG000 | 11:11:11:11:1 | 12218106:122 | 12218378:122 | -:-:-:-:-:-:-:-: | 1287 | NA | 0 | 2 | 0 | 0 | 0 | 0 |
| ENSDARG000 | 11:11:11:11:1 | 21025895:210 | 21026842:210 | -;-;-;-;- | 1916 | fmoda | 2 | 2 | 14 | 0 | 2 | 6 |
| ENSDARG000 | 11:11:11:11:1 | 13642157:136 | 13642392:136 | +:+:+:+:+:+:+: | 2100 | wdr18 | 246 | 327 | 364 | 88 | 396 | 200 |
| ENSDARG000 | 11:11:11:11:1 | 21050326:210 | 21050457:210 | +;+;+;+;+;+;+ | 1526 | zgc:113307 | 2 | 4 | 4 | 6 | 12 | 2 |
| ENSDARG000 | 11:11:11:11:1 | 29739348:297 | 29739405:297 | +:+:+:+:+:+:+: | 2343 | NA | 0 | 16 | 0 | 0 | 1 | 0 |
| ENSDARG000 | 11:11:11:11:1 | 29756860:297 | 29757851:297 | -:-:-:-:-:-:-:-: | 2404 | plekha3 | 240 | 192 | 222 | 317 | 162 | 324 |
| ENSDARG000 | 11:11:11:11:1 | 12229830:122 | 12230098:122 | -:-:-:-:-:-:-:-: | 1270 | NA | 2 | 122 | 0 | 0 | 0 | 0 |
| ENSDARG000 | 11:11:11:11:1 | 26403334:264 | 26403951:264 | +:+:+:+:+:+:+: | 6941 | NA | 2 | 10 | 4 | 10 | 0 | 12 |
| ENSDARG000 | 11:11:11:11:1 | 26376000:263 | 26376102:263 | +;+;+;+;+ | 3713 | rbm12 | 793 | 860 | 1003 | 373 | 912 | 677 |
| ENSDARG000 | 11:11:11:11:1 | 26375979:263 | 26376102:263 | +:+:+:+:+:+:+: | 3684 | cpne1 | 3651 | 6463 | 5569 | 4542 | 6377 | 4901 |
| ENSDARG000 | 11:11:11:11:1 | 1925804:1926 | 1926254:1926 | -:-:-:-:-:-:-:-: | 1387 | FAIM2 | 0 | 0 | 0 | 0 | 0 | 4 |
| ENSDARG000 | 11:11:11:11:1 | 32576300:326 | 32578254:326 | -;-;-;-;- | 5398 | pcdh17 | 54 | 54 | 52 | 54 | 102 | 36 |
| ENSDARG000 | 11;11 | 9292281:9293 | 9292875:9293 | -;- | 678 | NA | 0 | 0 | 0 | 0 | 0 | 4 |
| ENSDARG000 | 11;11 | 9285823:9286 | 9286000:9286 | -;- | 261 | NA | 0 | 0 | 0 | 0 | 0 | 0 |
| ENSDARG000 | 11:11:11:11:1 | 8984557:8984 | 8984743:8984 | -:-:-:-:-:-:-:-: | 2710 | NA | 0 | 0 | 2 | 2 | 0 | 0 |
| ENSDARG000 | 11:11:11:11:1 | 329687:33096 | 330001:33115 | +:+:+:+:+:+:+: | 2142 | cyp27b1 | 2 | 1 | 2 | 2 | 0 | 10 |
| ENSDARG000 | 11:11:11:11:1 | 13798453:137 | 13798553:137 | -;-;-;-;-;- | 470 | NA | 0 | 0 | 0 | 0 | 0 | 0 |
| ENSDARG000 | 11:11:11:11:1 | 12332626:123 | 12332895:123 | -:-:-:-:-:-:-:-: | 1007 | NA | 0 | 35 | 0 | 2 | 0 | 0 |
| ENSDARG000 | 11;11;11;11 | 16574625:165 | 16574761:165 | +;+;+;+ | 5920 | kbtbd8 | 274 | 336 | 350 | 210 | 338 | 236 |
| ENSDARG000 | 11;11;11 | 32731165:327 | 32731609:327 | -;-;- | 732 | NA | 0 | 0 | 0 | 0 | 1 | 11 |
| ENSDARG000 | 11;11;11 | 16669218:166 | 16669424:166 | -;-;- | 680 | NA | 0 | 0 | 0 | 0 | 0 | 0 |
| ENSDARG000 | 11;11;11 | 32752535:327 | 32752595:327 | +;+;+ | 243 | NA | 0 | 0 | 0 | 0 | 0 | 0 |
| ENSDARG000 | 11:11:11:11:1 | 12348479:123 | 12348748:123 | -:-:-:-:-:-:-:-: | 1017 | NA | 0 | 4 | 0 | 0 | 0 | 0 |
| ENSDARG000 | 11;11;11 | 349113:35143 | 349347:35150 | -;-;- | 372 | NA | 0 | 0 | 0 | 0 | 0 | 0 |
| ENSDARG000 | 11:11:11:11:1 | 24815667:248 | 24815746:248 | +:+:+:+:+:+:+: | 1316 | rabif | 288 | 358 | 598 | 166 | 522 | 354 |
| ENSDARG000 | 11:11:11:11:1 | 9564154:9564 | 9564985:9564 | -:-:-:-:-:-:-:-: | 4251 | nlgn1 | 0 | 0 | 4 | 2 | 0 | 4 |
| ENSDARG000 | 11:11:11:11:1 | 38893819:388 | 38893908:388 | -;-;-;-;-;-;- | 2183 | NA | 0 | 0 | 0 | 0 | 0 | 0 |
| ENSDARG000 | 11:11:11:11:1 | 38908406:389 | 38908468:389 | -;-;-;-;- | 830 | NA | 0 | 0 | 0 | 0 | 0 | 0 |
| ENSDARG000 | 11:11:11:11:1 | 29770966:297 | 29771006:297 | +:+:+:+:+:+:+: | 5322 | rpgrb | 122 | 64 | 48 | 62 | 44 | 92 |
| ENSDARG000 | 11;11 | 38912133:389 | 38912471:389 | -;- | 475 | NA | 0 | 0 | 0 | 0 | 0 | 0 |
| ENSDARG000 | 11:11:11:11:1 | 6259048:6261 | 6262209:6262 | -;-;-;-;-;-;-;- | 3688 | ccl25b | 322 | 28 | 4 | 24 | 64 | 31 |
| ENSDARG000 | 11:11:11:11:1 | 38922744:389 | 38922885:389 | -;-;-;-;- | 909 | NA | 0 | 0 | 0 | 4 | 0 | 0 |
| ENSDARG000 | 11;11;11 | 21076872:210 | 21076915:210 | +;+;+ | 2116 | PRELP | 730 | 621 | 1474 | 953 | 699 | 1269 |
| ENSDARG000 | 11;11 | 38926724:389 | 38927062:389 | -;- | 476 | NA | 0 | 0 | 0 | 0 | 0 | 0 |
| ENSDARG000 | 11;11;11;11 | 38947987:389 | 38948443:389 | -;-;-;- | 960 | NA | 0 | 0 | 0 | 0 | 0 | 0 |
| ENSDARG000 | 11;11;11 | 43228412:432 | 43228799:432 | -;-;- | 503 | NA | 4 | 2 | 18 | 12 | 4 | 2 |
| ENSDARG000 | 11:11:11:11:1 | 16738179:167 | 16739196:167 | -:-:-:-:-:-:-:-: | 2390 | suclg2 | 668 | 540 | 818 | 237 | 702 | 468 |
| ENSDARG000 | 11:11:11:11:1 | 1944729:1945 | 1945251:1945 | -:-:-:-:-:-:-:-: | 2487 | nr1d4b | 565 | 267 | 662 | 429 | 526 | 568 |
| ENSDARG000 | 11:11:11:11:1 | 355305:35530 | 355411:35541 | +;+;+;+;+;+;+ | 2228 | pdrg1 | 194 | 215 | 306 | 222 | 241 | 302 |
| ENSDARG000 | 11:11:11:11:1 | 21096339:210 | 21096519:210 | +;+;+;+;+ | 606 | il19l | 6 | 2 | 24 | 10 | 2 | 4 |
| ENSDARG000 | 11:11:11:11:1 | 12362231:123 | 12362503:123 | -:-:-:-:-:-:-:-: | 1116 | NA | 0 | 4 | 0 | 0 | 0 | 0 |
| ENSDARG000 | 11;11;11;11 | 1935893:1945 | 1936074:1945 | +;+;+;+ | 646 | NA | 0 | 0 | 0 | 0 | 0 | 0 |
| ENSDARG000 | 11:11:11:11:1 | 12377621:123 | 12377893:123 | -:-:-:-:-:-:-:-: | 1117 | NA | 0 | 23 | 0 | 0 | 0 | 0 |
| ENSDARG000 | 11:11:11:11:1 | 33172336:331 | 33172412:331 | +:+:+:+:+:+:+: | 1499 | NA | 0 | 0 | 0 | 0 | 0 | 0 |
| ENSDARG000 | 11;11;11;11 | 39013763:390 | 39014643:390 | -;-;-;- | 1457 | cldn19 | 46 | 10 | 44 | 6 | 6 | 16 |
| ENSDARG000 | 11:11:11:11:1 | 21137948:211 | 21138205:211 | +;+;+;+;+ | 655 | il10 | 0 | 0 | 4 | 2 | 0 | 6 |
| ENSDARG000 | 11:11:11:11:1 | 6281647:6282 | 6281871:6282 | +:+:+:+:+:+:+: | 1548 | ctns | 259 | 161 | 213 | 250 | 198 | 314 |
| ENSDARG000 | 11:11:11:11:1 | 33284837:332 | 33285032:332 | +:+:+:+:+:+:+: | 3291 | NA | 0 | 0 | 4 | 2 | 4 | 0 |
| ENSDARG000 | 11:11:11:11:1 | 10388303:103 | 10389486:103 | -:-:-:-:-:-:-:-: | 9707 | ect2 | 132 | 503 | 572 | 242 | 499 | 208 |
| ENSDARG000 | 11:11:11:11:1 | 39075613:390 | 39076287:390 | -:-:-:-:-:-:-:-: | 2774 | P3H1 | 158 | 279 | 486 | 224 | 329 | 174 |
| ENSDARG000 | 11:11:11:11:1 | 12392715:123 | 12392984:123 | -:-:-:-:-:-:-:-: | 1114 | NA | 0 | 21 | 0 | 0 | 0 | 0 |
| ENSDARG000 | 11:11:11:11:1 | 39105019:391 | 39105217:391 | +:+:+:+:+:+:+: | 1011 | C11H1orf50 | 170 | 157 | 121 | 85 | 224 | 124 |
| ENSDARG000 | 11;11;11;11 | 39077610:390 | 39077776:390 | +;+;+;+ | 482 | NA | 0 | 0 | 0 | 0 | 0 | 0 |

| | | | | | | | | | | | | |
|---|---|---|---|---|---|---|---|---|---|---|---|---|
| ENSDARG000 | 11:11:11:11:1 | 410870:41117 | 411875:41187 | -;-;-;-;-;-;- | 5590 | znf831 | 6 | 23 | 66 | 23 | 41 | 60 |
| ENSDARG000 | 11:11:11:11:1 | 14004236:140 | 14004864:140 | +;+;+;+;+;+;+; | 5604 | GRIN3B | 4 | 0 | 0 | 0 | 0 | 0 |
| ENSDARG000 | 11:11:11:11:1 | 21154678:211 | 21156534:211 | -;-;-;-;-;-;-;-; | 3310 | mapkapk2a | 1060 | 1355 | 1402 | 782 | 1162 | 1190 |
| ENSDARG000 | 11;11 | 21185469:211 | 21185811:211 | +;+ | 674 | NA | 0 | 0 | 0 | 0 | 0 | 0 |
| ENSDARG000 | 11:11:11:11:1 | 3959495:3959 | 3959575:3959 | +;+;+;+;+;+;+; | 2169 | gnl3 | 767 | 1059 | 1492 | 528 | 943 | 667 |
| ENSDARG000 | 11:11:11:11:1 | 24819324:248 | 24819746:248 | +;+;+;+;+;+;+; | 5756 | kdm5ba | 4185 | 2777 | 5624 | 4294 | 4395 | 3431 |
| ENSDARG000 | 11:11:11:11:1 | 3909756:3909 | 3910344:3910 | -;-;-;-;-;-;-;-; | 5825 | PBRM1 | 652 | 555 | 913 | 625 | 1047 | 701 |
| ENSDARG000 | 11:11:11:11:1 | 14071689:140 | 14075826:140 | -;-;-;-;-;-;-;-; | 6357 | TMEM259 | 1006 | 656 | 956 | 889 | 876 | 881 |
| ENSDARG000 | 11;11 | 14081540:140 | 14081719:140 | +;+ | 508 | NA | 2 | 2 | 0 | 2 | 4 | 3 |
| ENSDARG000 | 11 | 24819265 | 24820001 | - | 737 | NA | 0 | 1 | 5 | 4 | 5 | 8 |
| ENSDARG000 | 11:11:11:11:1 | 33312601:334 | 33312981:334 | +;+;+;+;+;+;+; | 9005 | NA | 0 | 0 | 0 | 0 | 0 | 0 |
| ENSDARG000 | 11;11 | 43295275:432 | 43296343:432 | +;+ | 1285 | NA | 2 | 0 | 0 | 0 | 2 | 0 |
| ENSDARG000 | 11:11:11:11:1 | 43375326:433 | 43375646:433 | +;+;+;+;+;+;+ | 1538 | sult6b1 | 222 | 224 | 228 | 216 | 412 | 300 |
| ENSDARG000 | 11:11:11:11:1 | 26476153:265 | 26476519:265 | +;+;+;+;+;+;+; | 4467 | NA | 940 | 637 | 599 | 665 | 665 | 1209 |
| ENSDARG000 | 11;11;11 | 21259356:212 | 21263347:212 | -;-;- | 4871 | dyrk3 | 1366 | 1200 | 1735 | 1396 | 1202 | 1534 |
| ENSDARG000 | 11:11:11:11:1 | 6295370:6300 | 6295407:6300 | +;+;+;+;+;+;+; | 1986 | ranbp3a | 124 | 92 | 170 | 130 | 135 | 145 |
| ENSDARG000 | 11:11:11:11:1 | 35777329:357 | 35778437:357 | -;-;-;-;-;-;-;-; | 10288 | ITPR1 | 48 | 22 | 30 | 59 | 105 | 98 |
| ENSDARG000 | 11:11:11:11:1 | 43401201:434 | 43401679:434 | +;+;+;+;+;+;+; | 1814 | vipb | 0 | 0 | 0 | 0 | 0 | 0 |
| ENSDARG000 | 11:11:11:11:1 | 29790626:297 | 29790736:297 | +;+;+;+;+ | 1148 | dynlt3 | 1400 | 1842 | 1472 | 1935 | 1219 | 1542 |
| ENSDARG000 | 11:11:11:11:1 | 39112341:391 | 39115218:391 | -;-;-;-;- | 3274 | ap5b1 | 304 | 226 | 256 | 206 | 315 | 310 |
| ENSDARG000 | 11:11:11:11:1 | 429776:43013 | 429990:43027 | -;-;-;-;-;-;-;-; | 1652 | nuf2 | 258 | 687 | 917 | 454 | 521 | 286 |
| ENSDARG000 | 11:11:11:11:1 | 17093753:170 | 17094789:170 | -;-;-;-;-;-;-;-; | 1548 | NA | 135 | 95 | 123 | 88 | 58 | 101 |
| ENSDARG000 | 11:11:11:11:1 | 43419809:434 | 43420225:434 | +;+;+;+;+;+;+; | 4805 | SLC29A1 | 35 | 4 | 24 | 2 | 114 | 20 |
| ENSDARG000 | 11:11:11:11:1 | 12416065:124 | 12416337:124 | -;-;-;-;-;-;-;-; | 1016 | NA | 0 | 2 | 0 | 0 | 0 | 0 |
| ENSDARG000 | 11;11;11 | 10541258:105 | 10541370:105 | +;+;+ | 2395 | b3gnt5a | 159 | 246 | 150 | 47 | 190 | 261 |
| ENSDARG000 | 11;11 | 10535456:105 | 10535806:105 | +;+ | 811 | NA | 0 | 0 | 0 | 0 | 0 | 2 |
| ENSDARG000 | 11:11:11:11:1 | 10467299:104 | 10469590:104 | -;-;-;-;-;-;-;-; | 9193 | NA | 1385 | 545 | 666 | 1016 | 1042 | 1011 |
| ENSDARG000 | 11 | 6322998 | 6324302 | + | 1305 | NA | 127 | 152 | 206 | 272 | 171 | 147 |
| ENSDARG000 | 11;11;11 | 29818389:298 | 29819488:298 | -;-;- | 1659 | xk | 24 | 15 | 2 | 10 | 14 | 26 |
| ENSDARG000 | 11:11:11:11:1 | 29795155:297 | 29796987:297 | -;-;-;-;-;-;-;-; | 4284 | cybb | 454 | 238 | 322 | 204 | 400 | 495 |
| ENSDARG000 | 11:11:11:11:1 | 26568492:265 | 26569015:265 | -;-;-;-;-;-;-;-; | 1918 | st3gal8 | 539 | 388 | 427 | 575 | 531 | 560 |
| ENSDARG000 | 11:11:11:11:1 | 39120306:391 | 39120395:391 | +;+;+;+;+;+;+; | 2763 | cdc42 | 4148 | 5539 | 5259 | 3910 | 4266 | 4399 |
| ENSDARG000 | 11:11:11:11:1 | 14104417:141 | 14104944:141 | +;+;+;+;+;+;+; | 3147 | med16 | 324 | 401 | 514 | 231 | 506 | 325 |
| ENSDARG000 | 11:11:11:11:1 | 6349443:6350 | 6350448:6350 | -;-;-;-;- | 2678 | NA | 0 | 0 | 0 | 0 | 0 | 0 |
| ENSDARG000 | 11:11:11:11:1 | 12435301:124 | 12435570:124 | -;-;-;-;-;-;-;-; | 1114 | NA | 0 | 2 | 0 | 0 | 0 | 0 |
| ENSDARG000 | 11:11:11:11:1 | 21278687:212 | 21283216:212 | -;-;-;-;-;-;-;-; | 7044 | RASSF5 | 1038 | 1376 | 1480 | 910 | 1883 | 1330 |
| ENSDARG000 | 11:11:11:11:1 | 12450564:124 | 12450836:124 | -;-;-;-;-;-;-;-; | 1104 | zgc:174354 | 0 | 18 | 0 | 0 | 0 | 0 |
| ENSDARG000 | 11:11:11:11:1 | 44337157:443 | 44337799:443 | -;-;-;-;-;-;-;-; | 2103 | LOXL4 | 188 | 448 | 259 | 159 | 338 | 267 |
| ENSDARG000 | 11:11:11:11:1 | 44324761:443 | 44324859:443 | +;+;+;+;+;+;+; | 1452 | NA | 250 | 126 | 245 | 251 | 226 | 155 |
| ENSDARG000 | 11 | 6443457 | 6444163 | + | 707 | NA | 46 | 70 | 71 | 91 | 106 | 66 |
| ENSDARG000 | 11:11:11:11:1 | 44452233:444 | 44452928:444 | -;-;-;-;-;-;-;-; | 3190 | mfn1b | 927 | 957 | 1273 | 734 | 1265 | 869 |
| ENSDARG000 | 11:11:11:11:1 | 6446235:6446 | 6447504:6447 | -;-;-;-;-;-;-;-; | 1773 | larp6b | 2 | 0 | 0 | 0 | 0 | 0 |
| ENSDARG000 | 11:11:11:11:1 | 44491762:444 | 44492025:444 | -;-;-;-;-;-;- | 3189 | NA | 10 | 0 | 6 | 10 | 58 | 4 |
| ENSDARG000 | 11:11:11:11:1 | 14147913:141 | 14148389:141 | +;+;+;+;+;+;+; | 4446 | PLPPR3 | 6 | 0 | 4 | 0 | 36 | 4 |
| ENSDARG000 | 11:11:11:11:1 | 21364551:213 | 21366734:213 | -;-;-;-;-;-;-;-; | 5066 | ikbke | 86 | 87 | 72 | 14 | 160 | 134 |
| ENSDARG000 | 11;11;11 | 29856032:298 | 29856537:298 | +;+;+ | 1248 | GRPR | 0 | 0 | 0 | 0 | 0 | 0 |
| ENSDARG000 | 11:11:11:11:1 | 17137848:171 | 17138623:171 | +;+;+;+;+;+;+; | 2731 | NA | 35 | 2 | 10 | 4 | 8 | 12 |
| ENSDARG000 | 11:11:11:11:1 | 33598213:335 | 33601378:336 | -;-;-;-;-;-;-;-; | 4540 | lims1 | 3134 | 3095 | 2739 | 3141 | 2506 | 2809 |
| ENSDARG000 | 11:11:11:11:1 | 2252983:2258 | 2255353:2258 | -;-;-;-;-;-;-;-; | 3593 | NA | 578 | 521 | 607 | 403 | 561 | 583 |
| ENSDARG000 | 11:11:11:11:1 | 29936743:299 | 29937068:299 | -;-;-;-;-;-;-;-; | 2031 | zgc:113276 | 38 | 55 | 44 | 144 | 67 | 50 |
| ENSDARG000 | 11:11:11:11:1 | 29917613:299 | 29918421:299 | -;-;-;-;-;-;-;-; | 2276 | pir | 1468 | 2641 | 1362 | 1878 | 2651 | 2089 |
| ENSDARG000 | 11:11:11:11:1 | 26604611:266 | 26604680:266 | +;+;+;+;+;+ | 660 | NA | 0 | 0 | 0 | 0 | 0 | 0 |
| ENSDARG000 | 11;11;11;11 | 26609110:266 | 26609230:266 | +;+;+;+ | 1511 | map1lc3a | 535 | 382 | 373 | 647 | 691 | 715 |
| ENSDARG000 | 11:11:11:11:1 | 26604224:266 | 26604298:266 | +;+;+;+;+;+;+; | 853 | dynlrb1 | 500 | 261 | 268 | 583 | 284 | 526 |
| ENSDARG000 | 11:11:11:11:1 | 29889828:298 | 29890394:298 | -;-;-;-;-;-;-;-; | 2102 | vegfd | 855 | 669 | 434 | 719 | 1038 | 921 |
| ENSDARG000 | 11:11:11:11:1 | 21405697:214 | 21410664:214 | -;-;-;-;-;-;-;-; | 8808 | srgap2 | 665 | 705 | 637 | 535 | 737 | 859 |
| ENSDARG000 | 11:11:11:11:1 | 44502410:445 | 44502818:445 | +;+;+;+;+;+;+; | 4924 | ero1b | 1042 | 943 | 1363 | 982 | 1134 | 962 |
| ENSDARG000 | 11:11:11:11:1 | 21586335:215 | 21586463:215 | +;+;+;+;+;+;+; | 5591 | FAM72B | 286 | 266 | 350 | 286 | 278 | 368 |
| ENSDARG000 | 11:11:11:11:1 | 12470089:124 | 12470361:124 | -;-;-;-;-;-;-;-; | 1104 | NA | 0 | 0 | 0 | 0 | 0 | 0 |
| ENSDARG000 | 11:11:11:11:1 | 24851671:248 | 24851913:248 | +;+;+;+;+;+;+; | 4947 | ipo9 | 758 | 1512 | 2174 | 1085 | 1437 | 943 |
| ENSDARG000 | 11:11:11:11:1 | 10669461:106 | 10672760:106 | -;-;-;-;-;-;-;-; | 7147 | SLC4A10 | 1836 | 1014 | 3003 | 2226 | 2304 | 3304 |
| ENSDARG000 | 11:11:11:11:1 | 6456146:6456 | 6456344:6456 | +;+;+;+;+;+;+; | 1603 | gadd45ba | 4938 | 1274 | 2711 | 2965 | 1530 | 2208 |
| ENSDARG000 | 11:11:11:11:1 | 12481385:124 | 12481657:124 | -;-;-;-;-;-;-;-; | 1020 | NA | 0 | 20 | 0 | 0 | 0 | 0 |
| ENSDARG000 | 11:11:11:11:1 | 44526892:445 | 44529840:445 | -;-;-;-;-;-;- | 4214 | NA | 10 | 2 | 2 | 15 | 26 | 4 |
| ENSDARG000 | 11:11:11:11:1 | 26619133:266 | 26619667:266 | -;-;-;-;-;-;-;-; | 2514 | NA | 4 | 14 | 24 | 18 | 26 | 12 |
| ENSDARG000 | 11:11:11:11:1 | 14199802:141 | 14199962:141 | +;+;+;+;+;+;+; | 2022 | palm1a | 108 | 178 | 82 | 231 | 129 | 123 |
| ENSDARG000 | 11:11:11:11:1 | 12497180:124 | 12497449:124 | -;-;-;-;-;-;-;-; | 1017 | NA | 0 | 0 | 0 | 0 | 0 | 0 |
| ENSDARG000 | 11:11:11:11:1 | 44548209:445 | 44548655:445 | +;+;+;+;+;+;+; | 6349 | NID1 | 7533 | 652 | 1190 | 2822 | 3674 | 76 |
| ENSDARG000 | 11;11;11 | 6460694:6460 | 6460766:6460 | +;+;+ | 713 | NA | 4 | 8 | 0 | 19 | 6 | 2 |
| ENSDARG000 | 11:11:11:11:1 | 6650966:6651 | 6651237:6651 | +;+;+;+;+;+;+ | 1327 | NA | 0 | 0 | 0 | 0 | 0 | 0 |
| ENSDARG000 | 11:11:11:11:1 | 24877820:248 | 24878060:248 | +;+;+;+;+;+;+ | 5176 | shisa4 | 335 | 415 | 487 | 134 | 419 | 401 |
| ENSDARG000 | 11;11;11 | 440305:44473 | 440639:44491 | +;+;+ | 5399 | rab43 | 498 | 355 | 786 | 463 | 456 | 437 |
| ENSDARG000 | 11:11:11:11:1 | 33628104:336 | 33628398:336 | +;+;+;+;+;+;+; | 5651 | NA | 0 | 6 | 16 | 0 | 24 | 0 |
| ENSDARG000 | 11:11:11:11:1 | 21612739:216 | 21612947:216 | +;+;+;+;+ | 1209 | NA | 0 | 0 | 0 | 0 | 0 | 0 |
| ENSDARG000 | 11;11 | 6658656:6678 | 6658703:6678 | +;+ | 127 | NA | 0 | 0 | 0 | 0 | 0 | 0 |
| ENSDARG000 | 11:11:11:11:1 | 12510223:125 | 12510492:125 | -;-;-;-;-;-;-;-; | 1016 | NA | 0 | 7 | 0 | 0 | 0 | 0 |
| ENSDARG000 | 11:11:11:11:1 | 33818179:338 | 33818370:338 | +;+;+;+;+;+;+; | 2560 | spoplb | 1272 | 1081 | 1617 | 1226 | 1822 | 796 |
| ENSDARG000 | 11;11 | 29961114:299 | 29961376:299 | -;- | 331 | NA | 0 | 0 | 0 | 0 | 0 | 0 |
| ENSDARG000 | 11;11 | 6780708:6781 | 6780756:6782 | -;- | 317 | NA | 0 | 0 | 0 | 0 | 0 | 0 |
| ENSDARG000 | 11:11:11:11:1 | 12528381:125 | 12528653:125 | -;-;-;-;-;-;-;-; | 1019 | NA | 0 | 8 | 0 | 0 | 0 | 0 |
| ENSDARG000 | 11:11:11:11:1 | 6684968:6739 | 6685490:6739 | +;+;+;+;+;+;+; | 3317 | NA | 229 | 29 | 164 | 80 | 105 | 214 |
| ENSDARG000 | 11:11:11:11:1 | 29965822:299 | 29965943:299 | +;+;+;+;+;+;+ | 1744 | il1fma | 10 | 28 | 56 | 54 | 84 | 54 |
| ENSDARG000 | 11:11:11:11:1 | 457423:45797 | 457881:45816 | -;-;-;-;-;-;- | 1552 | zgc:77375 | 208 | 183 | 1675 | 77 | 494 | 381 |
| ENSDARG000 | 11:11:11:11:1 | 474433:47671 | 476343:47680 | -;-;-;-;-;-;-;-; | 3563 | isy1 | 500 | 519 | 822 | 464 | 597 | 683 |
| ENSDARG000 | 11:11:11:11:1 | 3971716:3972 | 3972355:3972 | -;-;-;-;-;-;-;-; | 2340 | glt8d1 | 234 | 172 | 232 | 158 | 248 | 250 |
| ENSDARG000 | 11:11:11:11:1 | 44637516:446 | 44638011:446 | -;-;-;-;-;-;- | 999 | tomm20b | 1086 | 1406 | 1575 | 1430 | 1178 | 1133 |
| ENSDARG000 | 11:11:11:11:1 | 44617021:446 | 44617193:446 | +;+;+;+;+;+;+; | 4842 | rbm34 | 284 | 383 | 603 | 314 | 438 | 258 |
| ENSDARG000 | 11;11;11 | 33846440:338 | 33847619:338 | -;-;- | 1700 | nxph2b | 5 | 27 | 8 | 2 | 42 | 6 |
| ENSDARG000 | 11:11:11:11:1 | 26669724:266 | 26669838:266 | +;+;+;+;+;+;+; | 3397 | NA | 77 | 48 | 89 | 81 | 73 | 27 |
| ENSDARG000 | 11;11;11 | 24889170:248 | 24889648:248 | +;+;+ | 2715 | LMOD1 | 15 | 0 | 0 | 20 | 2 | 2 |
| ENSDARG000 | 11:11:11:11:1 | 44605727:446 | 44607413:446 | -;-;-;-;-;-;-;-; | 4309 | tbce | 249 | 221 | 309 | 158 | 388 | 157 |
| ENSDARG000 | 11:11:11:11:1 | 39167846:391 | 39169156:391 | -;-;-;-;- | 2242 | wnt4 | 6 | 0 | 0 | 8 | 2 | 0 |
| ENSDARG000 | 11:11:11:11:1 | 10832560:108 | 10832748:108 | -;-;-;-;-;-;-;-; | 1150 | psmd14 | 1457 | 1497 | 1557 | 895 | 1551 | 1121 |
| ENSDARG000 | 11:11:11:11:1 | 24907469:249 | 24907506:249 | +;+;+;+;+;+;+; | 4121 | sulf2a | 0 | 8 | 0 | 2 | 4 | 0 |
| ENSDARG000 | 11:11:11:11:1 | 10812949:108 | 10813699:108 | -;-;-;-;-;- | 1805 | TBR1 | 32 | 31 | 75 | 4 | 53 | 44 |
| ENSDARG000 | 11:11:11:11:1 | 21910343:219 | 21910824:219 | +;+;+;+;+;+;+; | 7652 | foxp4 | 88 | 4 | 32 | 40 | 34 | 34 |
| ENSDARG000 | 11:11:11:11:1 | 12547191:125 | 12547463:125 | -;-;-;-;-;-;-;-; | 1104 | NA | 0 | 2 | 0 | 0 | 0 | 0 |
| ENSDARG000 | 11:11:11:11:1 | 24900123:249 | 24900292:249 | +;+;+;+;+;+;+; | 1704 | timm17a | 1095 | 1498 | 1401 | 524 | 1268 | 706 |
| ENSDARG000 | 11:11:11:11:1 | 26685982:266 | 26686451:266 | -;-;-;-;-;-;-;-; | 2353 | acad9 | 358 | 504 | 486 | 260 | 382 | 393 |
| ENSDARG000 | 11;11;11;11 | 17144309:171 | 17144902:171 | -;-;-;- | 817 | NA | 0 | 0 | 0 | 0 | 0 | 0 |
| ENSDARG000 | 11:11:11:11:1 | 14280598:142 | 14280976:142 | +;+;+;+;+;+;+; | 6379 | NA | 18 | 4 | 34 | 36 | 65 | 20 |
| ENSDARG000 | 11:11:11:11:1 | 17440591:174 | 17440639:174 | +;+;+;+;+;+;+; | 2636 | NA | 0 | 0 | 4 | 0 | 0 | 2 |
| ENSDARG000 | 11:11:11:11:1 | 29981678:299 | 29982173:299 | -;-;-;-;-;-;-;-; | 2871 | ace2 | 0 | 2 | 0 | 0 | 6 | 0 |
| ENSDARG000 | 11:11:11:11:1 | 29975830:299 | 29975871:299 | +;+;+;+;+;+;+; | 1504 | si:ch73-226l1 | 2 | 2 | 2 | 0 | 8 | 6 |
| ENSDARG000 | 11:11:11:11:1 | 33861177:338 | 33861995:338 | -;-;-;-;-;-;-;-; | 1827 | NA | 66 | 69 | 110 | 81 | 98 | 4 |
| ENSDARG000 | 11;11;11;11 | 14295011:142 | 14295966:142 | +;+;+;+ | 1235 | NA | 0 | 0 | 0 | 0 | 0 | 0 |
| ENSDARG000 | 11:11:11:11:1 | 17596979:176 | 17597569:176 | -;-;-;-;-;- | 1469 | tafa4b | 35 | 8 | 0 | 2 | 32 | 0 |
| ENSDARG000 | 11:11:11:11:1 | 3991911:3994 | 3995179:3995 | -;-;-;-;-;-;-;-; | 5986 | nek4 | 278 | 219 | 248 | 110 | 174 | 198 |
| ENSDARG000 | 11;11;11;11 | 3989403:3991 | 3989529:3991 | +;+;+;+ | 760 | spcs1 | 18 | 58 | 50 | 46 | 27 | 22 |
| ENSDARG000 | 11;11;11 | 14309027:143 | 14309304:143 | -;-;- | 444 | NA | 0 | 4 | 2 | 0 | 0 | 10 |
| ENSDARG000 | 11:11:11:11:1 | 491587:49248 | 492734:49273 | -;-;-;-;-;-;-;-; | 2099 | cnbpb | 2246 | 2928 | 4037 | 2518 | 3095 | 2781 |
| ENSDARG000 | 11:11:11:11:1 | 17726649:177 | 17727102:177 | -;-;-;-;-;-;-;-; | 1993 | EOGT | 62 | 55 | 36 | 18 | 80 | 29 |
| ENSDARG000 | 11:11:11:11:1 | 12688642:126 | 12688911:126 | -;-;-;-;-;-;-;-; | 981 | NA | 0 | 0 | 0 | 0 | 0 | 0 |
| ENSDARG000 | 11;11 | 36037076:360 | 36037883:360 | -;- | 999 | setmar | 28 | 34 | 66 | 20 | 41 | 42 |
| ENSDARG000 | 11;11;11 | 14312260:143 | 14312924:143 | +;+;+ | 1729 | NA | 76 | 82 | 120 | 56 | 166 | 88 |
| ENSDARG000 | 11:11:11:11:1 | 22109887:221 | 22110380:221 | +;+;+;+;+;+;+; | 6423 | MDFI | 78 | 222 | 182 | 50 | 170 | 374 |
| ENSDARG000 | 11:11:11:11:1 | 6819050:6834 | 6819115:6834 | +;+;+;+;+ | 814 | rab3ab | 3 | 2 | 28 | 0 | 102 | 2 |
| ENSDARG000 | 11:11:11:11:1 | 29999414:300 | 30001093:300 | +;+;+;+;+;+;+; | 9917 | NHS | 10 | 2 | 8 | 6 | 4 | 10 |
| ENSDARG000 | 11;11;11;11 | 36052731:360 | 36053288:360 | +;+;+;+ | 1977 | NA | 30 | 31 | 34 | 18 | 36 | 30 |
| ENSDARG000 | 11:11:11:11:1 | 36041617:360 | 36043032:360 | -;-;-;-;-;-;-;-; | 2375 | brk1 | 1432 | 1914 | 1982 | 1650 | 1626 | 1664 |
| ENSDARG000 | 11:11:11:11:1 | 12703887:127 | 12704159:127 | -;-;-;-;-;-;-;-; | 984 | NA | 0 | 2 | 0 | 0 | 0 | 0 |
| ENSDARG000 | 11:11:11:11:1 | 12732247:127 | 12734245:127 | -;-;-;-;-;-;-;-; | 3676 | ZRSR2 | 78 | 103 | 138 | 112 | 188 | 166 |
| ENSDARG000 | 11:11:11:11:1 | 26704181:267 | 26707816:267 | -;-;-;-;-;-;-;-; | 7413 | IQSEC1 | 171 | 82 | 72 | 48 | 162 | 116 |
| ENSDARG000 | 11:11:11:11:1 | 22287954:222 | 22292780:222 | -;-;-;-;-;-;-;-; | 6417 | tfeb | 1877 | 1752 | 2039 | 1406 | 1608 | 1834 |
| ENSDARG000 | 11:11:11:11:1 | 2596667:2600 | 2597210:2601 | +;+;+;+;+;+;+; | 3019 | NA | 146 | 171 | 190 | 84 | 220 | 126 |
| ENSDARG000 | 11:11:11:11:1 | 36048579:360 | 36049217:360 | -;-;-;-;-;- | 1416 | gpx1a | 348 | 290 | 355 | 134 | 534 | 574 |
| ENSDARG000 | 11:11:11:11:1 | 12715064:127 | 12715336:127 | -;-;-;-;-;- | 984 | NA | 0 | 0 | 0 | 0 | 0 | 0 |
| ENSDARG000 | 11:11:11:11:1 | 12719944:127 | 12720059:127 | +;+;+;+;+;+;+; | 1856 | ap1s2 | 1356 | 2156 | 2014 | 1621 | 1767 | 1826 |
| ENSDARG000 | 11;11;11;11 | 30091155:300 | 30091360:300 | +;+;+;+ | 2055 | NA | 6 | 0 | 16 | 8 | 18 | 18 |
| ENSDARG000 | 11;11 | 2615070:2615 | 2615193:2616 | +;+ | 575 | NA | 0 | 0 | 0 | 0 | 0 | 0 |
| ENSDARG000 | 11;11 | 10859024:108 | 10859402:108 | -;- | 933 | NA | 0 | 0 | 0 | 0 | 0 | 0 |
| ENSDARG000 | 11:11:11:11:1 | 30106182:301 | 30107569:301 | -;-;-;-;-;-;-;-; | 4495 | SCML2 | 398 | 306 | 538 | 313 | 322 | 410 |
| ENSDARG000 | 11:11:11:11:1 | 17757342:177 | 17758475:177 | -;-;-;-;-;-;-;-; | 4782 | tmf1 | 1048 | 750 | 1161 | 768 | 1303 | 1030 |
| ENSDARG000 | 11:11:11:11:1 | 6859876:6859 | 6860118:6860 | -;-;-;-;-;-;-;-; | 898 | lsm4 | 972 | 1168 | 1245 | 655 | 985 | 903 |
| ENSDARG000 | 11:11:11:11:1 | 2616816:2621 | 2619873:2621 | -;-;-;-;-;-;-;- | 3939 | NA | 273 | 203 | 226 | 84 | 218 | 146 |
| ENSDARG000 | 11:11:11:11:1 | 4026229:4026 | 4026333:4026 | +;+;+;+;+;+;+; | 5440 | CAMK1 | 116 | 99 | 185 | 58 | 287 | 267 |
| ENSDARG000 | 11:11:11:11:1 | 14321113:143 | 14321225:143 | +;+;+;+;+;+;+; | 4093 | ptbp1b | 3512 | 2312 | 3161 | 3337 | 2931 | 3807 |
| ENSDARG000 | 11:11:11:11:1 | 10909183:109 | 10909479:109 | +;+;+;+;+;+;+; | 3596 | RBMS1 | 1275 | 2000 | 1951 | 2372 | 1489 | 1794 |
| ENSDARG000 | 11;11 | 39637740:396 | 39638395:396 | -;- | 1201 | NA | 0 | 0 | 0 | 0 | 0 | 0 |
| ENSDARG000 | 11:11:11:11:1 | 36158134:361 | 36158934:361 | +;+;+;+;+;+;+; | 5545 | grm2b | 3 | 8 | 2 | 2 | 54 | 0 |
| ENSDARG000 | 11:11:11:11:1 | 36215698:362 | 36216017:362 | -;-;-;-;-;-;-;-; | 1753 | rrp9 | 283 | 458 | 714 | 213 | 415 | 359 |
| ENSDARG000 | 11;11 | 501654:50407 | 501961:50408 | +;+ | 324 | NA | 0 | 0 | 5 | 2 | 1 | 3 |
| ENSDARG000 | 11:11:11:11:1 | 39928828:399 | 39928948:399 | +;+;+;+;+;+;+; | 3783 | vamp3 | 6646 | 7765 | 7149 | 6883 | 5370 | 6557 |
| ENSDARG000 | 11:11:11:11:1 | 2633743:2635 | 2633864:2635 | +;+;+;+;+ | 596 | NA | 0 | 0 | 0 | 0 | 0 | 0 |
| ENSDARG000 | 11:11:11:11:1 | 27274355:272 | 27274465:272 | +;+;+;+;+;+;+; | 5774 | FBLN2 | 46 | 8 | 4 | 11 | 48 | 26 |
| ENSDARG000 | 11:11:11:11:1 | 24994705:249 | 24994911:249 | +;+;+;+;+;+;+; | 6172 | arhgap46a | 850 | 914 | 871 | 370 | 898 | 822 |
| ENSDARG000 | 11:11:11:11:1 | 2633791:2638 | 2633864:2638 | +;+;+;+;+;+;+; | 1646 | NA | 0 | 0 | 0 | 0 | 0 | 0 |
| ENSDARG000 | 11:11:11:11:1 | 22363572:223 | 22364638:223 | -;-;-;-;-;-;-;-; | 2182 | tmem183a | 332 | 436 | 412 | 180 | 396 | 240 |
| ENSDARG000 | 11;11 | 36231248:362 | 36231508:362 | +;+ | 1732 | NA | 0 | 0 | 0 | 0 | 0 | 0 |
| ENSDARG000 | 11:11:11:11:1 | 2642013:2643 | 2642259:2644 | +;+;+;+;+;+;+; | 1577 | zgc:193807 | 0 | 0 | 0 | 0 | 0 | 0 |
| ENSDARG000 | 11:11:11:11:1 | 33870357:338 | 33873794:338 | -;-;-;-;-;-;-;-; | 7231 | PHKA2 | 410 | 224 | 373 | 309 | 408 | 521 |
| ENSDARG000 | 11:11:11:11:1 | 10975012:109 | 10975663:109 | +;+;+;+;+;+;+; | 5168 | ITGB6 | 192 | 42 | 54 | 28 | 88 | 28 |
| ENSDARG000 | 11;11;11 | 36243774:362 | 36244462:362 | +;+;+ | 1435 | cav4a | 0 | 1 | 0 | 0 | 0 | 0 |
| ENSDARG000 | 11:11:11:11:1 | 506465:51732 | 506512:51755 | +;+;+;+;+;+;+; | 2668 | raf1b | 564 | 445 | 740 | 524 | 532 | 497 |
| ENSDARG000 | 11:11:11:11:1 | 39959039:399 | 39959145:399 | +;+;+;+;+;+;+; | 11603 | per3 | 2143 | 856 | 2052 | 1009 | 2910 | 1747 |
| ENSDARG000 | 11:11:11:11:1 | 30161168:301 | 30161277:301 | +;+;+;+;+;+;+; | 4400 | cdkl5 | 182 | 50 | 89 | 50 | 129 | 54 |
| ENSDARG000 | 11:11:11:11:1 | 6881001:6882 | 6881048:6882 | +;+;+;+;+;+;+; | 7538 | klhl26 | 99 | 120 | 107 | 64 | 259 | 139 |
| ENSDARG000 | 11:11:11:11:1 | 25044082:250 | 25044233:250 | +;+;+;+;+;+;+; | 4361 | NA | 78 | 142 | 78 | 130 | 102 | 98 |
| ENSDARG000 | 11:11:11:11:1 | 11021396:110 | 11022082:110 | +;+;+;+;+;+;+; | 4960 | PLA2R1 | 460 | 235 | 451 | 156 | 379 | 509 |
| ENSDARG000 | 11:11:11:11:1 | 6869312:6869 | 6869557:6869 | -;-;-;-;-;-;-;-; | 2000 | ddx49 | 194 | 344 | 342 | 188 | 334 | 222 |
| ENSDARG000 | 11:11:11:11:1 | 543365:56535 | 544181:56660 | -;-;-;-;-;-;-;-; | 3617 | mkrn2 | 284 | 197 | 233 | 114 | 212 | 136 |
| ENSDARG000 | 11:11:11:11:1 | 575665:57709 | 575995:57824 | +;+;+;+;+;+ | 1846 | NA | 55 | 96 | 47 | 169 | 6 | 48 |

| | | | | | | | | | | | | |
|---|---|---|---|---|---|---|---|---|---|---|---|---|
| ENSDARG000 | 11;11 | 545461:5483 | 545552:5483 | -;- | 153 | NA | 0 | 0 | 0 | 0 | 0 | 0 |
| ENSDARG000 | 11 | 543362 | 544187 | - | 826 | NA | 0 | 0 | 0 | 0 | 0 | 0 |
| ENSDARG000 | 11;11;11;11;11;1 | 17785182:177 | 17785658:177 | -;-;-;-;-;-;-;-; | 2097 | uba3 | 510 | 621 | 682 | 421 | 626 | 484 |
| ENSDARG000 | 11;11;11;11;11;1 | 11120532:111 | 11120867:111 | +;+;+;+;+;+;+; | 6738 | LY75 | 1576 | 1147 | 870 | 846 | 1827 | 1130 |
| ENSDARG000 | 11;11;11;11;11;1 | 36254284:362 | 36255236:362 | -;-;-;-;-;-;-;-; | 2491 | nfya | 480 | 639 | 705 | 505 | 695 | 370 |
| ENSDARG000 | 11;11;11;11;11;1 | 14343445:143 | 14343521:143 | +;+;+;+;+;+;+; | 1759 | fam174c | 558 | 474 | 544 | 335 | 539 | 604 |
| ENSDARG000 | 11;11;11 | 27403975:274 | 27404080:274 | +;+;+ | 718 | NA | 0 | 0 | 0 | 0 | 0 | 0 |
| ENSDARG000 | 11;11;11;11;11;1 | 33921358:339 | 33924689:339 | -;-;-;-;-;-;-;-; | 5960 | COL6A2 | 1808 | 248 | 562 | 404 | 1154 | 316 |
| ENSDARG000 | 11;11 | 44723019:447 | 44723954:447 | +;+ | 2117 | irf2bp2b | 2794 | 2614 | 2395 | 1729 | 2717 | 2085 |
| ENSDARG000 | 11;11 | 44750935:447 | 44751147:447 | -;- | 592 | NA | 0 | 5 | 2 | 0 | 2 | 4 |
| ENSDARG000 | 11;11;11;11 | 40024182:400 | 40024450:400 | -;-;-;- | 591 | uts2b | 0 | 0 | 0 | 0 | 0 | 0 |
| ENSDARG000 | 11;11;11 | 30252955:302 | 30253449:302 | +;+;+ | 603 | NA | 0 | 0 | 0 | 0 | 0 | 0 |
| ENSDARG000 | 11;11;11;11;11;1 | 30244356:302 | 30244469:302 | +;+;+;+;+;+;+; | 1373 | rs1a | 0 | 0 | 0 | 0 | 0 | 0 |
| ENSDARG000 | 11;11;11;11;11;1 | 30253968:302 | 30254113:302 | +;+;+;+;+;+;+; | 3326 | ppef1 | 62 | 19 | 145 | 53 | 136 | 112 |
| ENSDARG000 | 11;11;11 | 4148808:4161 | 4150609:4161 | -;-;- | 2568 | ccdc3b | 12 | 8 | 4 | 0 | 32 | 20 |
| ENSDARG000 | 11;11;11;11;11;1 | 40032790:400 | 40033083:400 | +;+;+;+;+;+;+; | 2238 | NA | 6 | 14 | 44 | 6 | 32 | 12 |
| ENSDARG000 | 11;11;11;11 | 4187022:4192 | 4190556:4192 | -;-;-;- | 3811 | NA | 0 | 0 | 0 | 0 | 0 | 0 |
| ENSDARG000 | 11;11;11;11 | 27486584:274 | 27488878:274 | -;-;-;- | 3435 | wnt7aa | 28 | 0 | 2 | 28 | 18 | 16 |
| ENSDARG000 | 11;11;11;11;11;1 | 40088141:400 | 40088886:400 | -;-;-;-;-;-;- | 1630 | tnfrsf9b | 14 | 18 | 18 | 26 | 49 | 44 |
| ENSDARG000 | 11 | 27502509 | 27503132 | - | 624 | NA | 21 | 21 | 23 | 8 | 24 | 44 |
| ENSDARG000 | 11;11;11;11;11;1 | 22374419:223 | 22375425:223 | +;+;+;+;+;+;+; | 7017 | ppfia4 | 336 | 364 | 197 | 282 | 190 | 373 |
| ENSDARG000 | 11;11;11;11;11;1 | 40103720:401 | 40104180:401 | -;-;-;-;-;-;-;-; | 2646 | NA | 4 | 0 | 0 | 8 | 2 | 8 |
| ENSDARG000 | 11;11 | 17906902:179 | 17907024:179 | -;- | 175 | NA | 0 | 0 | 0 | 0 | 0 | 0 |
| ENSDARG000 | 11;11;11;11;11;1 | 40134856:401 | 40135172:401 | -;-;-;-;-;-;-;-; | 2090 | EMP3 | 16 | 11 | 59 | 27 | 82 | 106 |
| ENSDARG000 | 11;11;11 | 17956888:179 | 17957521:179 | +;+;+ | 3889 | NA | 178 | 142 | 334 | 378 | 232 | 295 |
| ENSDARG000 | 11;11;11;11;11;1 | 40148181:401 | 40148572:401 | +;+;+;+;+;+;+; | 2631 | NA | 13 | 18 | 47 | 20 | 30 | 25 |
| ENSDARG000 | 11;11;11 | 17959507:179 | 17960297:179 | -;-;- | 1341 | cishb | 65 | 29 | 25 | 74 | 7 | 24 |
| ENSDARG000 | 11;11;11;11;11;1 | 2649664:2649 | 2650250:2650 | +;+;+;+;+;+;+; | 8130 | NA | 3241 | 2212 | 3818 | 3095 | 3095 | 2847 |
| ENSDARG000 | 11;11;11 | 27503232:275 | 27503320:275 | -;-;- | 251 | dre-let-7a-1 | 0 | 3 | 1 | 0 | 0 | 0 |
| ENSDARG000 | 11;11;11;11;11;1 | 4205567:4205 | 4206012:4205 | -;-;-;-;-;-;-;- | 1265 | FAM107A | 40 | 14 | 12 | 10 | 38 | 16 |
| ENSDARG000 | 11;11;11 | 14622379:146 | 14622671:146 | +;+;+ | 1735 | EFNA2 | 9 | 6 | 8 | 23 | 7 | 6 |
| ENSDARG000 | 11;11;11;11;11;1 | 14431273:144 | 14432237:144 | +;+;+;+;+;+ | 2037 | NA | 10 | 2 | 14 | 4 | 2 | 11 |
| ENSDARG000 | 11;11;11;11;11;1 | 583142:58372 | 583927:58391 | +;+;+;+;+;+;+; | 2245 | MKRN2OS | 2 | 0 | 7 | 6 | 2 | 0 |
| ENSDARG000 | 11;11;11;11;11;1 | 588017:58970 | 588281:58979 | -;-;-;-;-;- | 769 | tsen2 | 146 | 172 | 238 | 96 | 126 | 119 |
| ENSDARG000 | 11;11;11;11;11;1 | 40175161:401 | 40178014:401 | -;-;-;-;-;-;-;- | 4278 | znf362a | 336 | 324 | 396 | 286 | 351 | 433 |
| ENSDARG000 | 11;11;11;11;11;1 | 45092866:450 | 45093027:450 | +;+;+;+;+;+;+; | 4030 | si:dkev-93h22 | 254 | 278 | 458 | 174 | 439 | 244 |
| ENSDARG000 | 11;11;11;11;11;1 | 4286630:4286 | 4286738:4286 | -;-;-;-;-;- | 912 | NA | 2 | 13 | 5 | 62 | 105 | 2 |
| ENSDARG000 | 11;11;11 | 40208239:402 | 40208438:402 | +;+;+ | 1435 | NA | 0 | 0 | 0 | 2 | 8 | 0 |
| ENSDARG000 | 11;11;11;11 | 40227753:402 | 40228476:402 | -;-;-;- | 1198 | TRIM62 | 15 | 0 | 4 | 0 | 10 | 0 |
| ENSDARG000 | 11;11 | 27513874:275 | 27513957:275 | +;+ | 447 | NA | 0 | 0 | 0 | 0 | 0 | 0 |
| ENSDARG000 | 11;11;11;11;11;1 | 27511277:275 | 27512485:275 | -;-;-;-;-;-;-;-; | 3559 | ptpdc1a | 77 | 42 | 20 | 41 | 75 | 52 |
| ENSDARG000 | 11;11 | 14667066:146 | 14667500:146 | -;- | 693 | ADORA1 | 0 | 0 | 0 | 0 | 0 | 0 |
| ENSDARG000 | 11;11;11;11;11;1 | 36280523:362 | 36283734:362 | -;-;-;-;-;-;-;-; | 6381 | sort1a | 875 | 1000 | 1682 | 990 | 1310 | 939 |
| ENSDARG000 | 11;11;11 | 40306579:403 | 40306825:403 | +;+;+ | 1554 | NA | 0 | 0 | 0 | 0 | 2 | 0 |
| ENSDARG000 | 11;11;11;11;11;1 | 4299091:4299 | 4299199:4299 | -;-;-;-;-;- | 912 | NA | 71 | 204 | 141 | 236 | 332 | 12 |
| ENSDARG000 | 11;11;11 | 2685112:2687 | 2685468:2688 | +;+;+ | 3896 | b3galt6 | 191 | 164 | 240 | 190 | 188 | 189 |
| ENSDARG000 | 11;11;11;11;11;1 | 33996526:339 | 33999569:339 | -;-;-;-;-;-;-;-; | 5637 | NA | 5055 | 335 | 593 | 1294 | 850 | 1220 |
| ENSDARG000 | 11;11;11;11;11;1 | 30282141:302 | 30282735:302 | +;+;+;+;+;+;+; | 3348 | NA | 107 | 20 | 375 | 103 | 366 | 277 |
| ENSDARG000 | 11;11;11;11;11;1 | 17997403:179 | 17998375:179 | -;-;-;-;-;-;-;-; | 6055 | qrich1 | 255 | 193 | 403 | 158 | 275 | 202 |
| ENSDARG000 | 11;11;11;11;11;1 | 12744575:127 | 12744774:127 | +;+;+;+;+;+;+; | 4035 | rbb4l | 1224 | 1396 | 1974 | 938 | 1637 | 1103 |
| ENSDARG000 | 11;11;11;11;11;1 | 2691243:2693 | 2692597:2693 | -;-;-;-;-;- | 1961 | NA | 10 | 2 | 9 | 12 | 8 | 10 |
| ENSDARG000 | 11;11;11 | 2699951:2702 | 2700097:2702 | +;+;+ | 2130 | tmem167b | 598 | 569 | 511 | 328 | 614 | 526 |
| ENSDARG000 | 11;11;11;11;11;1 | 6946591:6946 | 6946833:6946 | +;+;+;+;+;+ | 1075 | NA | 0 | 0 | 0 | 0 | 0 | 0 |
| ENSDARG000 | 11;11;11;11;11;1 | 25139495:251 | 25139567:251 | +;+;+;+;+;+;+; | 1372 | NA | 0 | 4 | 0 | 4 | 12 | 6 |
| ENSDARG000 | 11;11 | 14693481:146 | 14694502:146 | -;- | 1279 | NA | 0 | 2 | 0 | 0 | 0 | 0 |
| ENSDARG000 | 11;11;11;11;11;1 | 6902306:6902 | 6903128:6903 | +;+;+;+;+;+;+; | 2545 | NA | 4 | 6 | 9 | 4 | 25 | 0 |
| ENSDARG000 | 11;11;11;11;11;1 | 25064519:250 | 25064637:250 | +;+;+;+;+;+;+; | 4628 | ndrg3a | 562 | 273 | 588 | 318 | 515 | 543 |
| ENSDARG000 | 11;11;11;11;11;1 | 14676236:147 | 14676940:147 | +;+;+;+;+;+;+; | 2361 | NA | 0 | 4 | 0 | 0 | 0 | 0 |
| ENSDARG000 | 11;11;11;11;11;1 | 599440:59975 | 599815:59981 | -;-;-;-;- | 1147 | NA | 0 | 0 | 0 | 0 | 0 | 0 |
| ENSDARG000 | 11;11;11;11;11;1 | 6941389:6944 | 6945095:6945 | -;-;-;-;-;-;-;-; | 6222 | comp | 40 | 58 | 30 | 74 | 59 | 36 |
| ENSDARG000 | 11;11;11;11;11;1 | 18053333:180 | 18053641:180 | +;+;+;+;+;+ | 1421 | zgc:175135 | 0 | 0 | 0 | 0 | 0 | 0 |
| ENSDARG000 | 11;11;11;11;11;1 | 17964689:179 | 17964830:179 | +;+;+;+;+;+;+; | 2594 | rpusd4 | 97 | 242 | 404 | 163 | 195 | 200 |
| ENSDARG000 | 11;11;11;11;11;1 | 17965098:179 | 17966543:179 | -;-;-;-;-;-;-;-; | 2742 | twf2b | 84 | 44 | 43 | 20 | 44 | 58 |
| ENSDARG000 | 11;11;11;11;11;1 | 45110865:451 | 45110973:451 | +;+;+;+;+;+;+; | 2827 | NA | 73 | 34 | 36 | 20 | 49 | 84 |
| ENSDARG000 | 11;11;11;11;11;1 | 4492467:4568 | 4492624:4568 | -;-;-;-;-;- | 1090 | NA | 0 | 0 | 0 | 2 | 0 | 0 |
| ENSDARG000 | 11;11;11;11;11;1 | 40331791:403 | 40332513:403 | -;-;-;-;-;- | 1876 | trim62.1 | 46 | 41 | 26 | 27 | 93 | 32 |
| ENSDARG000 | 11;11;11;11;11;1 | 18037729:180 | 18038057:180 | +;+;+;+;+;+ | 1449 | NA | 0 | 0 | 0 | 0 | 0 | 0 |
| ENSDARG000 | 11;11;11 | 45136052:451 | 45136101:451 | +;+;+ | 385 | NA | 0 | 0 | 0 | 0 | 0 | 0 |
| ENSDARG000 | 11;11;11;11 | 27543093:275 | 27543625:275 | +;+;+;+ | 1430 | barx1 | 0 | 0 | 0 | 0 | 0 | 4 |
| ENSDARG000 | 11;11;11;11 | 18023465:180 | 18023563:180 | +;+;+;+ | 2732 | NA | 0 | 0 | 0 | 0 | 0 | 0 |
| ENSDARG000 | 11;11;11;11;11;1 | 45139300:451 | 45140227:451 | -;-;-;-;-;-;-;-; | 2541 | afmid | 22 | 49 | 50 | 63 | 34 | 43 |
| ENSDARG000 | 11;11;11;11;11;1 | 40431629:404 | 40434441:404 | -;-;-;-;-;-;-;- | 3935 | tnfrsf1b | 182 | 369 | 419 | 186 | 264 | 449 |
| ENSDARG000 | 11;11;11;11;11;1 | 36342997:363 | 36343347:363 | -;-;-;-;-;-;-;-; | 1444 | psma5 | 2321 | 3072 | 2737 | 1174 | 2566 | 1578 |
| ENSDARG000 | 11;11;11;11;11;1 | 40467618:404 | 40468418:404 | -;-;-;-;-;-;-;- | 2022 | NA | 0 | 6 | 17 | 6 | 0 | 8 |
| ENSDARG000 | 11;11 | 27592694:276 | 27593187:276 | -;- | 536 | NA | 0 | 0 | 0 | 0 | 0 | 0 |
| ENSDARG000 | 11;11;11;11;11;1 | 27644524:276 | 27646763:276 | -;-;-;-;-;-;-;-; | 5802 | phf2 | 1215 | 723 | 1001 | 656 | 1228 | 977 |
| ENSDARG000 | 11;11;11;11;11;1 | 40505044:405 | 40505767:405 | -;-;-;-;-;-;-;- | 2016 | NA | 9 | 82 | 32 | 41 | 74 | 76 |
| ENSDARG000 | 11;11;11;11;11;1 | 609516:61008 | 609759:61031 | -;-;-;-;-;-;-;- | 1581 | pparg | 12 | 6 | 28 | 2 | 4 | 18 |
| ENSDARG000 | 11;11;11;11 | 27625076:276 | 27625468:276 | +;+;+;+ | 1005 | NA | 0 | 0 | 0 | 2 | 2 | 0 |
| ENSDARG000 | 11;11;11;11;11;1 | 714386:71438 | 714590:71459 | +;+;+;+;+;+;+; | 1189 | timp4.3 | 2 | 2 | 2 | 8 | 4 | 0 |
| ENSDARG000 | 11;11;11;11;11;1 | 705727:70746 | 705901:70755 | +;+;+;+;+ | 940 | timp4.2 | 0 | 16 | 16 | 18 | 130 | 82 |
| ENSDARG000 | 11;11;11;11;11;1 | 675296:67680 | 675528:67740 | -;-;-;-;-;-;-;-; | 2348 | syn2b | 0 | 0 | 0 | 3 | 0 | 0 |
| ENSDARG000 | 11;11;11;11;11;1 | 45122147:451 | 45127429:451 | -;-;-;-;-;-;-;-; | 8434 | cant1b | 55 | 91 | 22 | 8 | 22 | 47 |
| ENSDARG000 | 11;11;11 | 6977268:6983 | 6981061:6984 | -;-;- | 4713 | si:ch211-253 | 238 | 297 | 390 | 217 | 408 | 308 |
| ENSDARG000 | 11;11;11;11;11;1 | 11200550:112 | 11201260:112 | +;+;+;+;+;+;+; | 7317 | myom2a | 2 | 2 | 8 | 2 | 0 | 8 |
| ENSDARG000 | 11;11;11;11;11;1 | 40530748:405 | 40530948:405 | +;+;+;+;+;+;+; | 3563 | MMEL1 | 2 | 14 | 34 | 188 | 248 | 8 |
| ENSDARG000 | 11;11;11;11;11;1 | 30321116:303 | 30322222:303 | +;+;+;+;+;+ | 1922 | ugt1b1 | 22 | 2 | 12 | 0 | 20 | 14 |
| ENSDARG000 | 11;11;11;11;11;1 | 14771908:147 | 14772445:147 | -;-;-;-;-;-;-;-; | 1633 | sgta | 1230 | 1120 | 1602 | 990 | 1312 | 1036 |
| ENSDARG000 | 11;11;11;11;11;1 | 30303785:303 | 30304948:303 | +;+;+;+;+;+ | 2939 | ugt1b5 | 28 | 0 | 74 | 4 | 69 | 10 |
| ENSDARG000 | 11;11;11;11;11;1 | 30306606:303 | 30307487:303 | +;+;+;+;+;+ | 1887 | ugt1b4 | 0 | 0 | 0 | 0 | 0 | 0 |
| ENSDARG000 | 11;11;11;11;11;1 | 30295582:302 | 30297246:302 | +;+;+;+;+;+ | 2608 | ugt1b7 | 6 | 0 | 0 | 0 | 2 | 0 |
| ENSDARG000 | 11;11;11;11;11;1 | 30314885:303 | 30316335:303 | +;+;+;+;+;+ | 2266 | ugt1b2 | 2 | 2 | 27 | 28 | 19 | 6 |
| ENSDARG000 | 11;11;11;11;11;1 | 30310170:303 | 30311082:303 | +;+;+;+;+ | 1728 | ugt1b3 | 0 | 0 | 0 | 0 | 0 | 0 |
| ENSDARG000 | 11 | 30298706 | 30300053 | + | 1348 | ugt1b6p | 0 | 0 | 0 | 0 | 0 | 0 |
| ENSDARG000 | 11;11;11;11;11;1 | 34084532:340 | 34085174:340 | -;-;-;-;-;-;-;-; | 1800 | tdo2b | 0 | 4 | 9 | 4 | 0 | 0 |
| ENSDARG000 | 11;11;11;11;11;1 | 6986589:6986 | 6987607:6987 | -;-;-;-;-;-;- | 1295 | zgc:173548 | 79 | 83 | 90 | 82 | 66 | 84 |
| ENSDARG000 | 11;11;11 | 14827374:148 | 14827465:148 | -;-;- | 901 | NA | 0 | 0 | 0 | 0 | 0 | 0 |
| ENSDARG000 | 11;11 | 22596726:225 | 22597036:225 | +;+ | 772 | NA | 0 | 0 | 0 | 0 | 0 | 0 |
| ENSDARG000 | 11;11;11;11;11;1 | 22592935:225 | 22597179:225 | -;-;-;-;-;-;-;-; | 5524 | myog | 1 | 0 | 5 | 0 | 4 | 0 |
| ENSDARG000 | 11;11;11;11;11;1 | 40622310:406 | 40623062:406 | -;-;-;-;-;-;-;-; | 1661 | prxl2b | 20 | 412 | 327 | 170 | 198 | 92 |
| ENSDARG000 | 11;11;11 | 14862286:148 | 14862480:148 | +;+;+ | 642 | NA | 0 | 0 | 0 | 0 | 0 | 0 |
| ENSDARG000 | 11;11;11;11;11;1 | 14902709:149 | 14902771:149 | +;+;+;+;+ | 975 | NA | 0 | 0 | 0 | 0 | 0 | 0 |
| ENSDARG000 | 11;11;11;11;11;1 | 779754:78037 | 779919:78053 | +;+;+;+;+;+;+; | 1130 | tamm41 | 180 | 225 | 246 | 198 | 175 | 155 |
| ENSDARG000 | 11;11;11;11;11;1 | 2710530:2710 | 2710881:2710 | +;+;+;+;+;+;+; | 5464 | mapk14b | 486 | 411 | 504 | 462 | 603 | 502 |
| ENSDARG000 | 11;11;11;11;11;1 | 769623:77237 | 769713:77245 | +;+;+;+;+ | 5223 | vgll4b | 1291 | 931 | 1456 | 1302 | 1435 | 1132 |
| ENSDARG000 | 11;11;11;11;11;1 | 14915780:149 | 14916378:149 | -;-;-;-;- | 1158 | NA | 7 | 20 | 2 | 6 | 0 | 8 |
| ENSDARG000 | 11;11;11;11;11;1 | 7001948:7002 | 7002216:7002 | -;-;-;-;-;-;-;-; | 2957 | NA | 18 | 4 | 8 | 3 | 4 | 38 |
| ENSDARG000 | 11;11;11;11;11;1 | 22623284:226 | 22623545:226 | +;+;+;+;+ | 2463 | si:ch211-86h | 181 | 143 | 194 | 86 | 150 | 184 |
| ENSDARG000 | 11;11;11;11;11;1 | 36355348:363 | 36355529:363 | +;+;+;+;+;+;+; | 4630 | sypl2a | 2835 | 2917 | 2557 | 1998 | 2377 | 2706 |
| ENSDARG000 | 11;11;11;11;11;1 | 12782296:127 | 12785658:127 | -;-;-;-;-;-;-;-; | 4627 | txlng | 262 | 284 | 532 | 273 | 346 | 300 |
| ENSDARG000 | 11;11;11;11;11;1 | 36379293:363 | 36379429:363 | +;+;+;+;+;+;+; | 2777 | atxn7l2a | 83 | 96 | 170 | 85 | 117 | 147 |
| ENSDARG000 | 11;11;11;11;11;1 | 30332959:303 | 30333508:303 | -;-;-;-;-;-;-;-; | 2273 | cflara | 742 | 633 | 1016 | 456 | 772 | 710 |
| ENSDARG000 | 11;11;11;11;11;1 | 40649412:406 | 40649580:406 | +;+;+;+;+;+;+; | 3102 | slc45a1 | 12 | 6 | 46 | 22 | 28 | 0 |
| ENSDARG000 | 11;11;11;11;11;1 | 27705030:277 | 27706890:277 | -;-;-;-;-;-;-;-; | 5348 | FAM120A | 5106 | 4112 | 7200 | 4714 | 4711 | 4434 |
| ENSDARG000 | 11 | 27738489 | 27744282 | + | 5794 | NA | 10 | 8 | 19 | 5 | 46 | 34 |
| ENSDARG000 | 11;11;11;11;11;1 | 34116734:341 | 34118559:341 | -;-;-;-;-;-;-;-; | 4653 | NA | 112 | 154 | 278 | 158 | 190 | 128 |
| ENSDARG000 | 11;11;11;11;11;1 | 14955484:149 | 14956867:149 | -;-;-;-;-;-;-;-; | 2282 | elavl1b | 705 | 622 | 738 | 465 | 765 | 570 |
| ENSDARG000 | 11;11;11;11 | 11263451:112 | 11264772:112 | -;-;-;- | 2297 | lgsn | 2 | 6 | 12 | 26 | 6 | 4 |
| ENSDARG000 | 11;11;11 | 30348273:303 | 30349028:303 | -;-;- | 1216 | tmem169a | 2 | 0 | 10 | 0 | 2 | 2 |
| ENSDARG000 | 11;11;11;11;11;1 | 7039259:7040 | 7044333:7044 | -;-;-;-;-;-;-;- | 6226 | NA | 2128 | 607 | 479 | 1687 | 328 | 1160 |
| ENSDARG000 | 11;11;11;11;11;1 | 34219991:342 | 34220450:342 | -;-;-;-;-;-;-;-; | 2738 | lsg1 | 555 | 524 | 879 | 482 | 575 | 586 |
| ENSDARG000 | 11;11;11;11;11;1 | 25157374:251 | 25157510:251 | +;+;+;+;+;+;+; | 3918 | trpc4apa | 1153 | 809 | 982 | 730 | 1083 | 917 |
| ENSDARG000 | 11;11;11;11;11;1 | 34201183:342 | 34201439:342 | -;-;-;-;-;-;-;- | 1538 | tmem44 | 2 | 0 | 18 | 2 | 0 | 7 |
| ENSDARG000 | 11;11;11;11;11;1 | 36408636:364 | 36409592:364 | +;+;+;+;+;+;+ | 2210 | cyb561d1 | 20 | 37 | 25 | 4 | 95 | 28 |
| ENSDARG000 | 11;11;11 | 7095176:7099 | 7095309:7101 | +;+;+ | 2320 | NA | 8 | 10 | 0 | 0 | 0 | 14 |
| ENSDARG000 | 11;11;11;11;11;1 | 30355442:303 | 30355854:303 | -;-;-;-;-;-;-;-; | 1935 | pecr | 171 | 159 | 219 | 85 | 209 | 186 |
| ENSDARG000 | 11;11;11;11;11;1 | 40677616:406 | 40678501:406 | -;-;-;-;-;-;-;-; | 1845 | NA | 412 | 1584 | 3759 | 756 | 1908 | 849 |
| ENSDARG000 | 11;11;11;11;11;1 | 7107798:7112 | 7109855:7112 | -;-;-;-;-;-;-;-; | 3473 | bmp7a | 0 | 3 | 0 | 0 | 0 | 8 |
| ENSDARG000 | 11;11;11;11;11;1 | 36449698:364 | 36453000:364 | -;-;-;-;-;-;-;-; | 6956 | USP48 | 609 | 384 | 601 | 640 | 378 | 604 |
| ENSDARG000 | 11;11;11;11;11;1 | 36422233:364 | 36422515:364 | -;-;-;-;-;-;-;-; | 3610 | NA | 117 | 106 | 125 | 73 | 119 | 109 |
| ENSDARG000 | 11;11;11;11;11;1 | 11267493:112 | 11268003:112 | +;+;+;+;+;+;+; | 2498 | ptp4a1 | 3901 | 4086 | 5198 | 4396 | 3474 | 4645 |
| ENSDARG000 | 11;11;11;11;11;1 | 2829812:2832 | 2830043:2833 | -;-;-;-;-;- | 1410 | LHFPL5 | 0 | 0 | 3 | 6 | 0 | 0 |
| ENSDARG000 | 11;11 | 2827553:2828 | 2827749:2828 | +;+ | 781 | NA | 151 | 238 | 259 | 232 | 246 | 224 |
| ENSDARG000 | 11;11;11;11;11;1 | 2763168:2765 | 2763234:2765 | +;+;+;+;+;+;+; | 2370 | srpk1b | 421 | 550 | 783 | 259 | 693 | 387 |
| ENSDARG000 | 11;11 | 15018138:150 | 15018470:150 | -;- | 1035 | NA | 0 | 0 | 0 | 0 | 3 | 0 |
| ENSDARG000 | 11;11;11;11;11;1 | 15017185:150 | 15017439:150 | +;+;+;+;+ | 744 | NA | 0 | 0 | 0 | 0 | 0 | 0 |
| ENSDARG000 | 11;11;11;11;11;1 | 7149504:7149 | 7149564:7150 | +;+;+;+;+;+;+; | 791 | NA | 54 | 10 | 168 | 16 | 53 | 120 |
| ENSDARG000 | 11;11;11;11;11;1 | 27745618:277 | 27746492:277 | -;-;-;-;-;-;- | 3027 | NA | 26 | 20 | 22 | 21 | 100 | 32 |
| ENSDARG000 | 11;11;11 | 25212540:252 | 25212587:252 | +;+;+ | 481 | NA | 0 | 0 | 0 | 0 | 0 | 0 |
| ENSDARG000 | 11;11;11;11;11;1 | 25181886:251 | 25183021:251 | -;-;-;-;-;-;-;-; | 7314 | myh7ba | 10 | 0 | 2 | 16 | 16 | 18 |
| ENSDARG000 | 11;11;11;11;11;1 | 27783758:277 | 27783910:277 | -;-;-;-;-;-;- | 750 | NA | 0 | 0 | 0 | 0 | 0 | 0 |
| ENSDARG000 | 11;11;11;11;11;1 | 15068307:150 | 15068841:150 | -;-;-;-;-;-;-;-; | 2352 | NA | 0 | 0 | 0 | 0 | 0 | 0 |
| ENSDARG000 | 11;11;11;11;11;1 | 22920587:229 | 22926947:229 | -;-;-;-;-;-;-;-; | 11044 | atp2b4 | 27449 | 17646 | 19551 | 23684 | 24981 | 23676 |
| ENSDARG000 | 11;11;11 | 34235356:342 | 34235717:342 | +;+;+ | 2447 | fam43a | 689 | 1124 | 1228 | 2348 | 956 | 982 |
| ENSDARG000 | 11;11;11;11;11;1 | 816106:84366 | 816982:84381 | -;-;-;-;-;-;-;-; | 3605 | ATG7 | 648 | 403 | 462 | 389 | 434 | 478 |
| ENSDARG000 | 11;11;11;11;11;1 | 7151579:7152 | 7152796:7152 | -;-;-;-;-;-;-;-; | 1621 | smim7 | 228 | 178 | 150 | 144 | 226 | 187 |
| ENSDARG000 | 11;11;11;11;11;1 | 30365668:303 | 30367725:303 | -;-;-;-;-;-;-;-; | 4878 | MAP4K3 | 366 | 312 | 323 | 323 | 463 | 434 |
| ENSDARG000 | 11;11 | 34249456:342 | 34249751:342 | -;- | 446 | NA | 0 | 0 | 0 | 0 | 0 | 0 |
| ENSDARG000 | 11;11;11;11;11;1 | 34401460:344 | 34402812:344 | -;-;-;-;-;-;-;- | 2706 | XXYLT1 | 224 | 152 | 308 | 142 | 195 | 253 |
| ENSDARG000 | 11;11;11;11;11;1 | 27817301:278 | 27818792:278 | -;-;-;-;-;-;-;-; | 2907 | cstf1 | 586 | 664 | 745 | 337 | 748 | 516 |
| ENSDARG000 | 11;11;11;11 | 923795:96906 | 923893:96934 | -;-;-;- | 1674 | NA | 0 | 0 | 1 | 0 | 0 | 0 |
| ENSDARG000 | 11;11;11;11;11;1 | 40713723:407 | 40714221:407 | -;-;-;-;-;-;-;-; | 2204 | odad1 | 0 | 0 | 1 | 0 | 0 | 0 |
| ENSDARG000 | 11;11;11;11;11;1 | 2855430:2902 | 2855787:2902 | +;+;+;+;+;+;+; | 5194 | KIF21B | 338 | 21 | 99 | 30 | 74 | 106 |
| ENSDARG000 | 11;11;11;11;11;1 | 25251495:252 | 25252631:252 | -;-;-;-;-;-;- | 2048 | snai1a | 2499 | 1809 | 2369 | 2165 | 1990 | 2059 |
| ENSDARG000 | 11;11;11;11;11;1 | 25257022:252 | 25257147:252 | +;+;+;+;+;+;+; | 2225 | tp53inp2 | 900 | 1278 | 1641 | 1990 | 1673 | 1242 |
| ENSDARG000 | 11;11;11;11;11;1 | 15225534:152 | 15225558:152 | -;-;-;-;-;-;-;-; | 2596 | rpn2 | 4065 | 3772 | 5773 | 4006 | 4607 | 3950 |
| ENSDARG000 | 11;11;11;11;11;1 | 36657025:366 | 36657117:366 | +;+;+;+;+ | 781 | NA | 0 | 0 | 0 | 0 | 0 | 3 |
| ENSDARG000 | 11;11;11;11;11;1 | 36477481:364 | 36477885:364 | +;+;+;+;+ | 1858 | LDLRAD2 | 0 | 6 | 7 | 0 | 2 | 0 |
| ENSDARG000 | 11;11;11;11;11;1 | 36683859:366 | 36684022:366 | +;+;+;+;+ | 1671 | NA | 92 | 0 | 0 | 0 | 0 | 0 |
| ENSDARG000 | 11;11;11;11;11;1 | 36541551:365 | 36541623:365 | +;+;+;+;+;+;+ | 1103 | NA | 2 | 0 | 0 | 0 | 0 | 0 |
| ENSDARG000 | 11;11;11 | 36600242:366 | 36600330:366 | +;+;+ | 689 | NA | 0 | 0 | 0 | 0 | 0 | 0 |
| ENSDARG000 | 11;11;11;11;11;1 | 40731602:407 | 40732168:407 | -;-;-;-;-;-;-;-; | 2555 | tas1r3 | 0 | 0 | 0 | 0 | 11 | 0 |

| | | | | | | | | | | | | |
|---|---|---|---|---|---|---|---|---|---|---|---|---|
| ENSDARG000 | 11;11;11 | 36519971:365 | 36520056:365 | +;+;+ | 510 | NA | 0 | 0 | 0 | 4 | 0 | 0 |
| ENSDARG000 | 11:11:11:11:1 | 36675599:366 | 36675785:366 | +:+:+:+:+:+:+: | 1138 | NA | 0 | 0 | 0 | 0 | 0 | 4 |
| ENSDARG000 | 11;11;11 | 36484562:366 | 36485145:366 | -;-;- | 1055 | NA | 0 | 0 | 0 | 0 | 0 | 0 |
| ENSDARG000 | 11:11:11:11:1 | 36554936:365 | 36555041:365 | +:+:+:+:+:+:+: | 1371 | NA | 0 | 0 | 0 | 0 | 2 | 0 |
| ENSDARG000 | 11:11:11:11:1 | 7158723:7168 | 7158894:7169 | +;+;+;+;+;+ | 1395 | tmem38a | 0 | 2 | 0 | 6 | 0 | 6 |
| ENSDARG000 | 11:11:11:11:1 | 36653213:366 | 36653255:366 | +;+;+;+;+ | 775 | NA | 0 | 0 | 0 | 0 | 0 | 3 |
| ENSDARG000 | 11:11:11:11:1 | 7183025:7183 | 7183143:7183 | +:+:+:+:+:+:+: | 2428 | thop1 | 1341 | 1416 | 1479 | 605 | 1598 | 1081 |
| ENSDARG000 | 11:11:11:11:1 | 36494094:364 | 36494133:364 | +:+:+:+:+:+:+: | 846 | NA | 0 | 0 | 0 | 0 | 4 | 0 |
| ENSDARG000 | 11;11 | 40760718:407 | 40760823:407 | -;- | 445 | NA | 0 | 0 | 0 | 0 | 0 | 0 |
| ENSDARG000 | 11:11:11:11:1 | 36665359:366 | 36665475:366 | +:+:+:+:+:+:+: | 1632 | NA | 0 | 0 | 2 | 0 | 0 | 0 |
| ENSDARG000 | 11:11:11:11:1 | 40812590:408 | 40812814:408 | +:+:+:+:+:+:+: | 5446 | errfi1a | 148 | 22 | 54 | 32 | 46 | 38 |
| ENSDARG000 | 11:11:11:11:1 | 997100:10051 | 999050:10053 | -:-:-:-:-:-:-:-: | 4291 | slc6a1b | 542 | 1201 | 96 | 204 | 946 | 3458 |
| ENSDARG000 | 11:11:11:11:1 | 25276748:252 | 25277207:252 | +:+:+:+:+:+:+: | 2990 | NA | 2 | 2 | 10 | 4 | 28 | 12 |
| ENSDARG000 | 11:11:11:11:1 | 23205889:232 | 23206421:232 | -;-;-;-;-;-;- | 1427 | optc | 12 | 2 | 0 | 6 | 8 | 12 |
| ENSDARG000 | 11:11:11:11:1 | 11274940:112 | 11275745:112 | -:-:-:-:-:-:-:-: | 2893 | col9a1a | 14 | 0 | 8 | 14 | 14 | 4 |
| ENSDARG000 | 11;11;11 | 40858654:408 | 40858812:408 | -;-;- | 887 | NA | 0 | 0 | 0 | 0 | 0 | 0 |
| ENSDARG000 | 11;11;11;11 | 30513656:305 | 30514509:305 | +;+;+;+ | 2289 | tmem178a | 0 | 0 | 0 | 4 | 0 | 0 |
| ENSDARG000 | 11:11:11:11:1 | 23265157:232 | 23265384:232 | +;+;+;+;+;+;+ | 2441 | csf1a | 3667 | 327 | 92 | 2220 | 24 | 945 |
| ENSDARG000 | 11:11:11:11:1 | 6899:6899:69 | 6998:7276:75 | -:-:-:-:-:-:-:-: | 1468 | myg1 | 1279 | 1167 | 1506 | 860 | 1183 | 877 |
| ENSDARG000 | 11:11:11:11:1 | 30535943:305 | 30536921:305 | -:-:-:-:-:-:-:-: | 4173 | slc8a1a | 16 | 6 | 43 | 14 | 16 | 16 |
| ENSDARG000 | 11 | 3176686 | 3176770 | + | 85 | dre-mir-724 | 0 | 0 | 0 | 0 | 0 | 0 |
| ENSDARG000 | 11:11:11:11:1 | 23311452:233 | 23312694:233 | -;-;-;-;- | 1649 | kiss1 | 6 | 0 | 0 | 4 | 12 | 12 |
| ENSDARG000 | 11;11;11;11 | 30628793:306 | 30629119:306 | -;-;-;- | 753 | mala.1 | 0 | 0 | 0 | 0 | 0 | 0 |
| ENSDARG000 | 11:11:11:11:1 | 30626768:306 | 30631215:306 | -;-;-;-;-;-;-;- | 5047 | mala.2 | 108 | 45 | 124 | 46 | 36 | 52 |
| ENSDARG000 | 11:11:11:11:1 | 23327784:233 | 23327838:233 | -;-;-;-;- | 420 | GOLT1A | 0 | 0 | 0 | 0 | 0 | 0 |
| ENSDARG000 | 11:11:11:11:1 | 7214353:7214 | 7214737:7214 | +:+:+:+:+:+:+: | 5422 | NA | 0 | 0 | 0 | 0 | 0 | 0 |
| ENSDARG000 | 11:11:11:11:1 | 25296366:252 | 25296518:252 | +:+:+:+:+:+:+: | 3795 | top1a | 2622 | 2449 | 3373 | 1575 | 4064 | 2122 |
| ENSDARG000 | 11:11:11:11:1 | 5817202:5818 | 5817526:5818 | +:+:+:+:+:+:+: | 1679 | NA | 318 | 305 | 295 | 246 | 300 | 329 |
| ENSDARG000 | 11;11 | 30636351:306 | 30636632:306 | +;+ | 3591 | pgap4 | 338 | 433 | 499 | 343 | 502 | 512 |
| ENSDARG000 | 11:11:11:11:1 | 1065161:1069 | 1065508:1070 | -;-;-;-;-;- | 1363 | NA | 0 | 0 | 0 | 0 | 0 | 0 |
| ENSDARG000 | 11:11:11:11:1 | 7253077:7255 | 7253399:7255 | -;-;-;-;-;-;- | 1193 | zgc:113223 | 4 | 0 | 0 | 0 | 2 | 0 |
| ENSDARG000 | 11:11:11:11:1 | 23342996:233 | 23345188:233 | -:-:-:-:-:-:-:-: | 6882 | PLEKHA6 | 318 | 224 | 757 | 771 | 614 | 1802 |
| ENSDARG000 | 11:11:11:11:1 | 3211826:3211 | 3212027:3212 | +:+:+:+:+:+:+: | 4526 | timeless | 1278 | 1349 | 2018 | 1327 | 1636 | 1505 |
| ENSDARG000 | 11:11:11:11:1 | 30647545:306 | 30647746:306 | +:+:+:+:+:+:+: | 2523 | tpcn3 | 940 | 620 | 622 | 699 | 873 | 1091 |
| ENSDARG000 | 11:11:11:11:1 | 27828212:278 | 27829791:278 | -:-:-:-:-:-:-:-: | 7389 | EIF4G3 | 1032 | 1056 | 1476 | 632 | 1126 | 802 |
| ENSDARG000 | 11:11:11:11:1 | 30663300:306 | 30663517:306 | +:+:+:+:+:+:+: | 5652 | NA | 18 | 26 | 22 | 26 | 25 | 27 |
| ENSDARG000 | 11:11:11:11:1 | 13179:13205: | 13626:13568: | -;-;-;-;-;-;- | 1085 | NA | 362 | 258 | 235 | 398 | 134 | 254 |
| ENSDARG000 | 11:11:11:11:1 | 36688794:366 | 36689503:366 | -:-:-:-:-:-:-:-: | 4338 | NA | 16 | 6 | 5 | 8 | 2 | 24 |
| ENSDARG000 | 11:11:11:11:1 | 3254252:3254 | 3254671:3254 | +:+:+:+:+:+:+: | 3435 | pmela | 318092 | 176579 | 139736 | 162635 | 251541 | 221567 |
| ENSDARG000 | 11:11:11:11:1 | 7261981:7262 | 7262185:7262 | +:+:+:+:+:+:+: | 1617 | reep6 | 0 | 4 | 0 | 0 | 0 | 0 |
| ENSDARG000 | 11:11:11:11:1 | 7276824:7276 | 7277105:7277 | +:+:+:+:+:+:+: | 1395 | NA | 158 | 268 | 164 | 138 | 178 | 220 |
| ENSDARG000 | 11:11:11:11:1 | 7281240:7285 | 7285616:7285 | -:-:-:-:-:-:-:-: | 6450 | APC2 | 0 | 2 | 18 | 0 | 10 | 8 |
| ENSDARG000 | 11:11:11:11:1 | 1094844:1098 | 1095268:1098 | -:-:-:-:-:-:-:-: | 1982 | SLC6A11 | 20 | 33 | 158 | 10 | 78 | 22 |
| ENSDARG000 | 11;11;11;11 | 15345654:153 | 15346341:153 | -;-;-;- | 997 | ghrh | 9 | 70 | 25 | 59 | 48 | 14 |
| ENSDARG000 | 11:11:11:11:1 | 15480695:154 | 15481001:154 | +:+:+:+:+:+:+: | 3291 | cdk5rap1 | 168 | 254 | 292 | 131 | 338 | 306 |
| ENSDARG000 | 11;11 | 23569869:235 | 23570114:235 | +;+ | 638 | NA | 0 | 2 | 0 | 8 | 9 | 0 |
| ENSDARG000 | 11;11;11 | 25313810:253 | 25314972:253 | -;-;- | 4586 | NA | 729 | 2421 | 2099 | 5198 | 1578 | 769 |
| ENSDARG000 | 11:11:11:11:1 | 36777613:367 | 36777949:367 | +:+:+:+:+:+:+: | 2215 | parapinopsint | 12 | 152 | 177 | 354 | 371 | 189 |
| ENSDARG000 | 11:11:11:11:1 | 30729745:307 | 30730256:307 | +:+:+:+:+:+:+: | 2023 | slc22a7a | 345 | 1061 | 7606 | 460 | 2768 | 2691 |
| ENSDARG000 | 11:11:11:11:1 | 25328199:253 | 25328252:253 | +;+;+;+;+;+;+ | 2328 | fam83d | 40 | 307 | 281 | 266 | 284 | 88 |
| ENSDARG000 | 11:11:11:11:1 | 11303458:113 | 11303575:113 | +:+:+:+:+:+:+: | 2048 | NA | 14 | 50 | 110 | 44 | 48 | 40 |
| ENSDARG000 | 11:11:11:11:1 | 11319141:113 | 11319332:113 | -:-:-:-:-:-:-:-: | 1209 | mrpl4 | 389 | 434 | 568 | 321 | 484 | 458 |
| ENSDARG000 | 11;11;11;11 | 25403561:254 | 25403911:254 | +;+;+;+ | 3637 | adnpa | 0 | 0 | 0 | 0 | 0 | 0 |
| ENSDARG000 | 11 | 25382602 | 25384213 | - | 1612 | mafbb | 36 | 76 | 692 | 236 | 1041 | 186 |
| ENSDARG000 | 11;11;11 | 25368296:253 | 25368777:253 | -;-;- | 6057 | NA | 0 | 4 | 4 | 4 | 10 | 0 |
| ENSDARG000 | 11:11:11:11:1 | 25388747:253 | 25388870:253 | +:+:+:+:+:+:+: | 1809 | NA | 0 | 2 | 0 | 0 | 0 | 0 |
| ENSDARG000 | 11:11:11:11:1 | 27968834:279 | 27968893:279 | +:+:+:+:+:+:+: | 2200 | alpl | 211 | 132 | 477 | 141 | 831 | 124 |
| ENSDARG000 | 11:11:11:11:1 | 27919702:279 | 27920779:279 | -:-:-:-:-:-:-:-: | 3394 | ece1 | 1134 | 1301 | 865 | 941 | 979 | 992 |
| ENSDARG000 | 11:11:11:11:1 | 23704410:237 | 23704725:237 | +:+:+:+:+:+:+: | 10350 | nfasca | 2 | 0 | 6 | 0 | 22 | 4 |
| ENSDARG000 | 11:11:11:11:1 | 25411914:254 | 25412507:254 | -;-;-;-;-;-;- | 2096 | gata1a | 4 | 0 | 34 | 16 | 56 | 6 |
| ENSDARG000 | 11:11:11:11:1 | 25430851:254 | 25430987:254 | +:+:+:+:+:+:+: | 1131 | si:dkev-13a21 | 0 | 0 | 7 | 2 | 6 | 0 |
| ENSDARG000 | 11:11:11:11:1 | 23900000:239 | 23900452:239 | -;-;-;-;- | 777 | NA | 0 | 0 | 0 | 0 | 0 | 0 |
| ENSDARG000 | 11;11;11 | 23895562:239 | 23895656:239 | +;+;+ | 369 | NA | 0 | 0 | 0 | 0 | 0 | 0 |
| ENSDARG000 | 11:11:11:11:1 | 3279039:3279 | 3279195:3279 | +:+:+:+:+:+:+: | 1888 | MMP19 | 0 | 16 | 6 | 10 | 12 | 0 |
| ENSDARG000 | 11:11:11:11:1 | 3301277:3303 | 3301924:3303 | -:-:-:-:-:-:-:-: | 1833 | cdk2 | 158 | 550 | 524 | 272 | 498 | 296 |
| ENSDARG000 | 11:11:11:11:1 | 25459697:254 | 25459757:254 | +:+:+:+:+:+:+: | 1531 | opn1sw2 | 4 | 8 | 14 | 8 | 16 | 8 |
| ENSDARG000 | 11;11 | 25475283:254 | 25475564:254 | -;- | 703 | NA | 0 | 0 | 0 | 0 | 0 | 0 |
| ENSDARG000 | 11:11:11:11:1 | 25436440:254 | 25437946:254 | -:-:-:-:-:-:-:-: | 6964 | hcfc1a | 1577 | 2048 | 2600 | 759 | 2031 | 1394 |
| ENSDARG000 | 11:11:11:11:1 | 23933016:239 | 23933210:239 | +:+:+:+:+:+:+: | 3357 | cntn2 | 2 | 4 | 4 | 12 | 8 | 0 |
| ENSDARG000 | 11:11:11:11:1 | 28078148:280 | 28079010:280 | -;-;-;-;-;-;-;- | 1435 | NA | 0 | 0 | 2 | 0 | 32 | 0 |
| ENSDARG000 | 11;11;11 | 28052699:280 | 28053252:280 | -;-;- | 807 | NA | 0 | 0 | 0 | 0 | 0 | 0 |
| ENSDARG000 | 11:11:11:11:1 | 28107632:281 | 28108050:281 | +;+;+;+;+ | 5448 | NA | 0 | 0 | 2 | 0 | 0 | 0 |
| ENSDARG000 | 11:11:11:11:1 | 25477643:254 | 25477843:254 | +:+:+:+:+:+:+: | 1699 | opn1lw1 | 0 | 0 | 0 | 0 | 0 | 0 |
| ENSDARG000 | 11:11:11:11:1 | 25481046:254 | 25481191:254 | +:+:+:+:+:+:+: | 1809 | opn1lw2 | 4 | 0 | 0 | 0 | 10 | 0 |
| ENSDARG000 | 11:11:11:11:1 | 25485774:254 | 25485874:254 | +:+:+:+:+:+:+: | 1932 | gnl3l | 340 | 510 | 1058 | 375 | 602 | 488 |
| ENSDARG000 | 11;11;11 | 41064822:410 | 41064889:410 | +;+;+ | 1386 | svbp | 312 | 171 | 391 | 351 | 428 | 412 |
| ENSDARG000 | 11:11:11:11:1 | 3320598:3321 | 3321331:3322 | -:-:-:-:-:-:-:-: | 3270 | prph | 18 | 14 | 0 | 0 | 0 | 8 |
| ENSDARG000 | 11:11:11:11:1 | 23993298:239 | 23993398:239 | +:+:+:+:+:+:+: | 2810 | chia.1 | 0 | 0 | 0 | 0 | 0 | 0 |
| ENSDARG000 | 11;11 | 43740949:437 | 43741131:437 | +;+ | 711 | ralbb | 152 | 210 | 216 | 170 | 206 | 256 |
| ENSDARG000 | 11:11:11:11:1 | 18130300:181 | 18130577:181 | +;+;+;+;+;+ | 1401 | NA | 0 | 0 | 0 | 0 | 0 | 0 |
| ENSDARG000 | 11;11;11 | 43102421:431 | 43102534:431 | -;-;- | 264 | NA | 0 | 1 | 6 | 6 | 0 | 0 |
| ENSDARG000 | 11:11:11:11:1 | 31869020:318 | 31869095:318 | -:-:-:-:-:-:-:-: | 1854 | NA | 0 | 0 | 0 | 0 | 0 | 0 |
| ENSDARG000 | 11 | 1546117 | 1546231 | + | 115 | 5S_rRNA | 1 | 0 | 0 | 0 | 0 | 0 |
| ENSDARG000 | 11:11:11:11:1 | 11983465:119 | 11984688:119 | -;-;-;-;-;-;- | 3133 | sp2 | 540 | 379 | 648 | 569 | 541 | 522 |
| ENSDARG000 | 11 | 12502551 | 12502847 | + | 297 | Metazoa_SRP | 0 | 0 | 0 | 0 | 0 | 0 |
| ENSDARG000 | 11:11:11:11:1 | 42466834:424 | 42466920:424 | -;-;-;-;-;- | 1062 | arf4b | 10 | 14 | 66 | 54 | 14 | 8 |
| ENSDARG000 | 11:11:11:11:1 | 4953964:5072 | 4954057:5072 | +:+:+:+:+:+:+: | 4245 | ptprga | 175 | 131 | 275 | 112 | 361 | 106 |
| ENSDARG000 | 11:11:11:11:1 | 41846059:418 | 41846496:418 | -;-;-;-;-;-;- | 1061 | mrto4 | 183 | 151 | 238 | 143 | 119 | 221 |
| ENSDARG000 | 11 | 31038409 | 31039533 | - | 1125 | ier2b | 7912 | 688 | 1609 | 2398 | 832 | 2014 |
| ENSDARG000 | 11 | 37299371 | 37299485 | - | 115 | 5S_rRNA | 0 | 0 | 0 | 0 | 0 | 0 |
| ENSDARG000 | 11:11:11:11:1 | 12286940:122 | 12287039:122 | -;-;-;-;- | 525 | NA | 0 | 16 | 0 | 0 | 0 | 0 |
| ENSDARG000 | 11 | 11380904 | 11381204 | - | 301 | Metazoa_SRP | 0 | 0 | 0 | 0 | 0 | 0 |
| ENSDARG000 | 11 | 16122786 | 16123082 | + | 297 | Metazoa_SRP | 0 | 0 | 0 | 0 | 0 | 0 |
| ENSDARG000 | 11;11 | 6431133:6434 | 6431323:6434 | +;+ | 546 | VMAC | 0 | 0 | 0 | 0 | 1 | 2 |
| ENSDARG000 | 11 | 5481753 | 5481884 | + | 132 | NA | 0 | 0 | 0 | 0 | 0 | 2 |
| ENSDARG000 | 11:11:11:11:1 | 44019844:440 | 44019930:440 | -;-;-;-;-;-;-;- | 1216 | NA | 651 | 230 | 468 | 509 | 781 | 544 |
| ENSDARG000 | 11:11:11:11:1 | 44910615:449 | 44910762:449 | -:-:-:-:-:-:-:-: | 2340 | pfklb | 574 | 657 | 1445 | 758 | 954 | 473 |
| ENSDARG000 | 11;11 | 41782683:417 | 41782780:417 | -;- | 231 | NA | 0 | 2 | 2 | 0 | 6 | 0 |
| ENSDARG000 | 11 | 37364446 | 37364560 | + | 115 | 5S_rRNA | 0 | 0 | 0 | 0 | 0 | 0 |
| ENSDARG000 | 11:11:11:11:1 | 3005536:3006 | 3005604:3006 | +:+:+:+:+:+:+: | 2201 | cpne5b | 0 | 0 | 16 | 2 | 6 | 0 |
| ENSDARG000 | 11:11:11:11:1 | 18283948:182 | 18288445:182 | -:-:-:-:-:-:-:-: | 6997 | SFMBT1 | 308 | 376 | 827 | 810 | 502 | 284 |
| ENSDARG000 | 11 | 12691395 | 12691691 | - | 297 | Metazoa_SRP | 0 | 0 | 0 | 0 | 0 | 0 |
| ENSDARG000 | 11;11 | 43114108:431 | 43116249:431 | +;+ | 2142 | foxg1b | 4 | 1 | 0 | 0 | 6 | 0 |
| ENSDARG000 | 11:11:11:11:1 | 44144612:441 | 44145107:441 | -:-:-:-:-:-:-:-: | 2813 | clcn4 | 484 | 530 | 722 | 612 | 864 | 600 |
| ENSDARG000 | 11 | 7150820 | 7150906 | + | 87 | NA | 0 | 0 | 3 | 1 | 0 | 0 |
| ENSDARG000 | 11 | 6327978 | 6328093 | + | 116 | 5S_rRNA | 2 | 0 | 0 | 0 | 2 | 0 |
| ENSDARG000 | 11 | 12213153 | 12213431 | - | 279 | Metazoa_SRP | 0 | 0 | 0 | 0 | 0 | 0 |
| ENSDARG000 | 11 | 27503017 | 27503132 | - | 116 | dre-let-7f | 0 | 0 | 0 | 0 | 0 | 0 |
| ENSDARG000 | 11 | 11445689 | 11445985 | - | 297 | Metazoa_SRP | 0 | 0 | 0 | 0 | 0 | 0 |
| ENSDARG000 | 11 | 12171295 | 12171591 | + | 297 | Metazoa_SRP | 0 | 0 | 0 | 0 | 0 | 0 |
| ENSDARG000 | 11:11:11:11:1 | 12672330:126 | 12672429:126 | -;-;-;-;- | 525 | NA | 0 | 0 | 0 | 0 | 0 | 0 |
| ENSDARG000 | 11:11:11:11:1 | 44932852:449 | 44932928:449 | +:+:+:+:+:+:+: | 1095 | CFAP410 | 74 | 72 | 106 | 92 | 60 | 112 |
| ENSDARG000 | 11 | 12711700 | 12711996 | + | 297 | Metazoa_SRP | 0 | 0 | 0 | 0 | 0 | 0 |
| ENSDARG000 | 11 | 12220890 | 12221186 | - | 297 | Metazoa_SRP | 0 | 0 | 0 | 0 | 0 | 0 |
| ENSDARG000 | 11 | 12593571 | 12593867 | + | 297 | Metazoa_SRP | 0 | 0 | 0 | 0 | 0 | 0 |
| ENSDARG000 | 11:11:11:11:1 | 44804685:448 | 44805707:448 | +:+:+:+:+:+:+: | 6099 | strn | 911 | 756 | 930 | 479 | 1214 | 871 |
| ENSDARG000 | 11:11:11:11:1 | 42150896:421 | 42151017:421 | -:-:-:-:-:-:-:-: | 5976 | FLNB | 203 | 10 | 67 | 10 | 69 | 12 |
| ENSDARG000 | 11 | 11338804 | 11339100 | - | 297 | Metazoa_SRP | 0 | 1 | 2 | 0 | 0 | 0 |
| ENSDARG000 | 11:11:11:11:1 | 5554163:5554 | 5554733:5554 | -;-;-;-;- | 1269 | pex11g | 14 | 16 | 45 | 29 | 30 | 11 |
| ENSDARG000 | 11:11:11:11:1 | 5499661:5505 | 5499991:5505 | +;+;+;+;+;+ | 5014 | slc35e1 | 424 | 451 | 612 | 361 | 535 | 503 |
| ENSDARG000 | 11:11:11:11:1 | 18205766:182 | 18205948:182 | +:+:+:+:+:+:+: | 3229 | tmcc1b | 638 | 846 | 727 | 440 | 820 | 645 |
| ENSDARG000 | 11:11:11:11:1 | 2571486:2576 | 2571847:2576 | -;-;-;-;-;-;- | 2278 | nab2 | 6 | 12 | 10 | 8 | 10 | 30 |
| ENSDARG000 | 11:11:11:11:1 | 40617896:406 | 40617946:406 | +;+;+;+;+ | 114 | NA | 0 | 0 | 6 | 33 | 51 | 7 |
| ENSDARG000 | 11:11:11:11:1 | 12588634:125 | 12588733:125 | -;-;-;-;- | 525 | NA | 0 | 0 | 0 | 0 | 0 | 0 |
| ENSDARG000 | 11:11:11:11:1 | 41540862:415 | 41540938:415 | +:+:+:+:+:+:+: | 1706 | KCNAB2 | 0 | 8 | 10 | 3 | 10 | 3 |
| ENSDARG000 | 11:11:11:11:1 | 2089461:2090 | 2089634:2091 | +;+;+;+;+;+;+ | 1176 | rargb | 203 | 102 | 127 | 59 | 158 | 105 |
| ENSDARG000 | 11 | 12621518 | 12621767 | - | 250 | Metazoa_SRP | 0 | 0 | 0 | 0 | 0 | 0 |
| ENSDARG000 | 11:11:11:11:1 | 12572306:125 | 12572405:125 | -;-;-;-;- | 525 | NA | 0 | 10 | 0 | 2 | 0 | 0 |
| ENSDARG000 | 11 | 11370455 | 11370653 | - | 199 | Metazoa_SRP | 0 | 0 | 0 | 0 | 0 | 0 |
| ENSDARG000 | 11 | 43948091 | 43948885 | - | 795 | INHBB | 0 | 6 | 36 | 0 | 2 | 0 |
| ENSDARG000 | 11;11;11;11 | 44798056:448 | 44798815:448 | -;-;-;- | 1068 | map1lc3c | 2 | 0 | 0 | 0 | 0 | 4 |
| ENSDARG000 | 11;11 | 41994287:419 | 41995731:419 | -;- | 2343 | her15.2 | 0 | 0 | 0 | 6 | 2 | 4 |
| ENSDARG000 | 11:11:11:11:1 | 12632395:126 | 12632569:126 | -:-:-:-:-:-:-:-: | 837 | NA | 0 | 0 | 0 | 0 | 0 | 0 |
| ENSDARG000 | 11 | 18676442 | 18676557 | + | 116 | 5S_rRNA | 0 | 0 | 0 | 0 | 0 | 0 |
| ENSDARG000 | 11 | 20033066 | 20033185 | + | 120 | 5S_rRNA | 0 | 0 | 0 | 0 | 0 | 0 |
| ENSDARG000 | 11 | 16127382 | 16127678 | - | 297 | Metazoa_SRP | 0 | 0 | 0 | 0 | 0 | 0 |
| ENSDARG000 | 11:11:11:11:1 | 41459408:414 | 41459984:414 | +;+;+;+;+;+ | 3156 | park7 | 692 | 1057 | 777 | 705 | 845 | 774 |
| ENSDARG000 | 11 | 11435447 | 11435743 | - | 297 | Metazoa_SRP | 0 | 0 | 0 | 0 | 0 | 0 |
| ENSDARG000 | 11:11:11:11:1 | 2448781:2448 | 2449004:2449 | -:-:-:-:-:-:-:-: | 2286 | slc2a10 | 34 | 36 | 166 | 12 | 54 | 208 |
| ENSDARG000 | 11;11;11;11 | 40573943:405 | 40574104:405 | +;+;+;+ | 306 | NA | 0 | 0 | 0 | 0 | 0 | 0 |
| ENSDARG000 | 11:11:11:11:1 | 5681762:5717 | 5682535:5718 | +:+:+:+:+:+:+: | 2504 | ARID3A | 59 | 36 | 61 | 38 | 83 | 75 |
| ENSDARG000 | 11:11:11:11:1 | 42641404:426 | 42641671:426 | +:+:+:+:+:+:+: | 6340 | il17rd | 70 | 5 | 37 | 25 | 74 | 4 |
| ENSDARG000 | 11 | 41966117 | 41966854 | - | 738 | her2 | 6 | 4 | 6 | 0 | 12 | 6 |
| ENSDARG000 | 11;11;11;11 | 44898714:449 | 44901448:449 | -;-;-;- | 3216 | zgc:171772 | 11974 | 12614 | 14585 | 12690 | 8890 | 13009 |
| ENSDARG000 | 11:11:11:11:1 | 2391469:2391 | 2392072:2392 | +:+:+:+:+:+:+: | 1583 | igfbp6a | 0 | 0 | 0 | 0 | 0 | 0 |
| ENSDARG000 | 11:11:11:11:1 | 20899029:208 | 20899193:208 | +:+:+:+:+:+:+: | 2378 | NA | 28 | 12 | 12 | 0 | 22 | 10 |
| ENSDARG000 | 11 | 11345803 | 11346099 | + | 297 | Metazoa_SRP | 0 | 0 | 0 | 0 | 0 | 0 |
| ENSDARG000 | 11 | 31840068 | 31840189 | + | 122 | 5S_rRNA | 0 | 0 | 0 | 0 | 0 | 0 |
| ENSDARG000 | 11 | 12679732 | 12680028 | + | 297 | Metazoa_SRP | 0 | 0 | 0 | 0 | 0 | 0 |
| ENSDARG000 | 11 | 25205153 | 25205243 | - | 91 | dre-mir-499 | 0 | 0 | 0 | 0 | 0 | 0 |
| ENSDARG000 | 11:11:11:11:1 | 4293288:4293 | 4293310:4293 | -:-:-:-:-:-:-:-: | 2100 | NA | 111 | 451 | 303 | 622 | 606 | 21 |
| ENSDARG000 | 11 | 12623555 | 12623851 | + | 297 | Metazoa_SRP | 0 | 0 | 0 | 0 | 0 | 0 |
| ENSDARG000 | 11 | 16644397 | 16644513 | - | 117 | 5S_rRNA | 0 | 0 | 0 | 0 | 0 | 0 |
| ENSDARG000 | 11:11:11:11:1 | 23593837:235 | 23597611:235 | -:-:-:-:-:-:-:-: | 9532 | PIK3C2B | 816 | 248 | 285 | 588 | 359 | 420 |
| ENSDARG000 | 11 | 459815 | 459930 | - | 116 | 5S_rRNA | 0 | 0 | 0 | 0 | 0 | 0 |
| ENSDARG000 | 11 | 33966158 | 33966272 | - | 115 | 5S_rRNA | 0 | 0 | 0 | 0 | 0 | 0 |
| ENSDARG000 | 11 | 12168002 | 12168298 | - | 297 | Metazoa_SRP | 0 | 0 | 0 | 0 | 0 | 0 |
| ENSDARG000 | 11 | 11424081 | 11424377 | - | 297 | Metazoa_SRP | 0 | 0 | 0 | 0 | 0 | 0 |
| ENSDARG000 | 11:11:11:11:1 | 2480293:2483 | 2481294:2483 | -:-:-:-:-:-:-:-: | 3118 | DGKA | 2 | 2 | 8 | 2 | 4 | 2 |
| ENSDARG000 | 11 | 32596473 | 32596588 | + | 116 | 5S_rRNA | 0 | 0 | 0 | 0 | 0 | 0 |
| ENSDARG000 | 11:11:11:11:1 | 44226200:442 | 44226644:442 | +:+:+:+:+:+:+: | 4276 | gnb4b | 706 | 770 | 716 | 438 | 654 | 829 |
| ENSDARG000 | 11:11:11:11:1 | 41242644:412 | 41243514:412 | +:+:+:+:+:+:+: | 4882 | pax7a | 51 | 264 | 2285 | 490 | 1888 | 1168 |

| | | | | | | | | | | | | |
|---|---|---|---|---|---|---|---|---|---|---|---|---|
| ENSDARG000 | 11 | 33371251 | 33371367 | - | 117 | 5S_rRNA | 0 | 0 | 0 | 0 | 0 | 0 |
| ENSDARG000 | 11;11;11;11 | 42128552:421 | 42128739:421 | -;-;-;- | 474 | NA | 56 | 4 | 12 | 2 | 8 | 3 |
| ENSDARG000 | 11 | 11412602 | 11412898 | - | 297 | Metazoa_SRP | 0 | 0 | 0 | 0 | 0 | 0 |
| ENSDARG000 | 11:11:11:11:1 | 18395879:183 | 18399650:183 | -:-:-:-:-:-:-:-:-: | 8971 | NA | 140 | 32 | 79 | 60 | 46 | 106 |
| ENSDARG000 | 11 | 12317614 | 12317910 | + | 297 | Metazoa_SRP | 0 | 0 | 0 | 0 | 0 | 0 |
| ENSDARG000 | 11 | 12165728 | 12166007 | - | 280 | Metazoa_SRP | 0 | 0 | 0 | 0 | 0 | 0 |
| ENSDARG000 | 11:11:11:11:1 | 5468629:5469 | 5468791:5469 | +:+:+:+:+:+:+: | 6648 | cse1l | 1143 | 2050 | 2596 | 694 | 1904 | 1188 |
| ENSDARG000 | 11 | 17328565 | 17328681 | + | 117 | 5S_rRNA | 0 | 0 | 0 | 0 | 0 | 0 |
| ENSDARG000 | 11 | 38588451 | 38588566 | + | 116 | 5S_rRNA | 0 | 0 | 0 | 0 | 6 | 0 |
| ENSDARG000 | 11 | 12493892 | 12494179 | + | 288 | Metazoa_SRP | 0 | 0 | 0 | 0 | 0 | 0 |
| ENSDARG000 | 11;11 | 394493:39642 | 394849:39672 | -;- | 661 | NA | 3 | 0 | 0 | 0 | 0 | 0 |
| ENSDARG000 | 11:11:11:11:1 | 45187798:451 | 45190639:451 | -;-;-;-;-;-;-;-;- | 4415 | NA | 8 | 2 | 0 | 2 | 18 | 12 |
| ENSDARG000 | 11 | 12440656 | 12440952 | + | 297 | Metazoa_SRP | 0 | 0 | 0 | 0 | 0 | 0 |
| ENSDARG000 | 11 | 11372886 | 11373182 | + | 297 | Metazoa_SRP | 0 | 0 | 0 | 0 | 0 | 0 |
| ENSDARG000 | 11 | 6862252 | 6862368 | + | 117 | 5S_rRNA | 0 | 0 | 0 | 9 | 1 | 0 |
| ENSDARG000 | 11:11:11:11:1 | 12009735:120 | 12011711:120 | -:-:-:-:-:-:-:-:-: | 4857 | socs7 | 22 | 24 | 38 | 16 | 111 | 104 |
| ENSDARG000 | 11 | 12388547 | 12388847 | + | 301 | Metazoa_SRP | 0 | 0 | 0 | 0 | 0 | 0 |
| ENSDARG000 | 11;11;11;11 | 12251840:122 | 12251932:122 | -;-;-;- | 429 | NA | 0 | 6 | 0 | 0 | 0 | 0 |
| ENSDARG000 | 11:11:11:11:1 | 2180072:2180 | 2180759:2180 | +;+;+;+;+ | 4372 | NA | 327 | 0 | 1 | 0 | 0 | 2 |
| ENSDARG000 | 11 | 7149989 | 7150082 | + | 94 | NA | 0 | 0 | 0 | 0 | 0 | 0 |
| ENSDARG000 | 11 | 12185857 | 12186144 | - | 288 | Metazoa_SRP | 0 | 0 | 0 | 0 | 0 | 0 |
| ENSDARG000 | 11:11:11:11:1 | 41712391:417 | 41712620:417 | -:-:-:-:-:-:-:-:-: | 5492 | MEGF6 | 12 | 17 | 38 | 20 | 58 | 28 |
| ENSDARG000 | 11 | 12512966 | 12513262 | - | 297 | Metazoa_SRP | 0 | 0 | 0 | 0 | 0 | 0 |
| ENSDARG000 | 11 | 24178823 | 24178938 | + | 116 | 5S_rRNA | 0 | 0 | 0 | 0 | 0 | 0 |
| ENSDARG000 | 11 | 35093605 | 35093722 | + | 118 | 5S_rRNA | 0 | 0 | 0 | 0 | 0 | 0 |
| ENSDARG000 | 11;11;11 | 2471382:2472 | 2471874:2472 | -;-;- | 900 | NA | 0 | 2 | 2 | 2 | 12 | 5 |
| ENSDARG000 | 11:11:11:11:1 | 44030729:440 | 44030807:440 | -;-;-;-;- | 579 | NA | 156 | 89 | 120 | 170 | 196 | 101 |
| ENSDARG000 | 11:11:11:11:1 | 34580085:345 | 34584849:345 | -:-:-:-:-:-:-:-:-: | 5836 | mapkapk3 | 450 | 234 | 400 | 243 | 305 | 304 |
| ENSDARG000 | 11;11 | 42750214:427 | 42753821:427 | -;- | 3654 | NA | 2 | 9 | 29 | 0 | 9 | 0 |
| ENSDARG000 | 11:11:11:11:1 | 42720203:427 | 42721701:427 | -;-;-;-;- | 1934 | NA | 13 | 0 | 16 | 21 | 4 | 10 |
| ENSDARG000 | 11 | 26748292 | 26748429 | - | 138 | 5S_rRNA | 0 | 0 | 0 | 0 | 0 | 0 |
| ENSDARG000 | 11:11:11:11:1 | 12300579:123 | 12300678:123 | -;-;-;-;- | 525 | NA | 0 | 2 | 0 | 0 | 0 | 0 |
| ENSDARG000 | 11 | 12291945 | 12292241 | + | 297 | Metazoa_SRP | 0 | 0 | 0 | 0 | 0 | 0 |
| ENSDARG000 | 11:11:11:11:1 | 31121340:311 | 31121469:311 | +:+:+:+:+:+:+: | 3736 | NA | 6 | 1 | 3 | 13 | 0 | 0 |
| ENSDARG000 | 11 | 9898513 | 9898630 | + | 118 | 5S_rRNA | 0 | 0 | 0 | 0 | 0 | 0 |
| ENSDARG000 | 11 | 40889180 | 40889294 | + | 115 | 5S_rRNA | 0 | 2 | 0 | 0 | 0 | 0 |
| ENSDARG000 | 11 | 34466395 | 34466513 | - | 119 | 5S_rRNA | 0 | 0 | 0 | 0 | 0 | 0 |
| ENSDARG000 | 11 | 11450155 | 11450442 | + | 288 | Metazoa_SRP | 0 | 0 | 0 | 0 | 0 | 0 |
| ENSDARG000 | 11;11 | 6417450:6420 | 6417804:6420 | -;- | 546 | VMAC | 0 | 2 | 4 | 0 | 0 | 0 |
| ENSDARG000 | 11;11 | 41981959:419 | 41982591:419 | +;+ | 2201 | NA | 2 | 2 | 16 | 0 | 0 | 4 |
| ENSDARG000 | 11;11;11 | 43043171:430 | 43043208:430 | +;+;+ | 207 | NA | 13 | 5 | 32 | 11 | 19 | 10 |
| ENSDARG000 | 11:11:11:11:1 | 691734:69607 | 692725:69617 | +;+;+;+;+ | 1528 | TIMP4 | 0 | 0 | 0 | 0 | 10 | 0 |
| ENSDARG000 | 11;11;11;11 | 2416064:2419 | 2416207:2419 | +;+;+;+ | 1191 | ube2v1 | 1298 | 1616 | 1495 | 1393 | 1302 | 1307 |
| ENSDARG000 | 11:11:11:11:1 | 18183220:181 | 18183634:181 | +;+;+;+;+;+ | 1992 | NA | 0 | 0 | 0 | 0 | 0 | 0 |
| ENSDARG000 | 11:11:11:11:1 | 1148701:1163 | 1150261:1163 | -:-:-:-:-:-:-:-:-: | 5014 | atp2b2 | 10 | 21 | 20 | 6 | 34 | 84 |
| ENSDARG000 | 11:11:11:11:1 | 5666325:5669 | 5666478:5669 | +;+;+;+;+;+;+ | 1812 | r3hdm4 | 381 | 306 | 271 | 158 | 240 | 212 |
| ENSDARG000 | 11;11;11 | 6374448:6376 | 6374623:6376 | +;+;+ | 498 | NA | 0 | 0 | 0 | 0 | 0 | 0 |
| ENSDARG000 | 11 | 21052228 | 21052344 | + | 117 | 5S_rRNA | 0 | 0 | 0 | 0 | 0 | 0 |
| ENSDARG000 | 11:11:11:11:1 | 24001993:240 | 24002651:240 | +:+:+:+:+:+:+: | 3103 | chia.2 | 0 | 0 | 0 | 0 | 4 | 0 |
| ENSDARG000 | 11 | 12431978 | 12432265 | + | 288 | Metazoa_SRP | 0 | 0 | 0 | 0 | 0 | 0 |
| ENSDARG000 | 11 | 12208909 | 12209205 | + | 297 | Metazoa_SRP | 0 | 0 | 0 | 0 | 0 | 0 |
| ENSDARG000 | 11 | 25476099 | 25476185 | - | 87 | dre-mir-726 | 0 | 0 | 0 | 0 | 0 | 0 |
| ENSDARG000 | 11 | 16058191 | 16058307 | - | 117 | 5S_rRNA | 4 | 6 | 8 | 0 | 4 | 7 |
| ENSDARG000 | 11 | 12323630 | 12323926 | - | 297 | Metazoa_SRP | 0 | 0 | 0 | 0 | 0 | 0 |
| ENSDARG000 | 11 | 12189725 | 12190024 | + | 300 | Metazoa_SRP | 0 | 0 | 0 | 0 | 0 | 0 |
| ENSDARG000 | 11:11:11:11:1 | 45255774:452 | 45255863:452 | +:+:+:+:+:+:+: | 1755 | aspscr1 | 661 | 542 | 633 | 570 | 678 | 577 |
| ENSDARG000 | 11:11:11:11:1 | 44300548:443 | 44300651:443 | +;+;+;+;+;+;+ | 657 | HNRNPLL | 2 | 8 | 12 | 8 | 4 | 7 |
| ENSDARG000 | 11;11;11 | 43751263:437 | 43751544:437 | +;+;+ | 1267 | NA | 8 | 6 | 22 | 0 | 10 | 46 |
| ENSDARG000 | 11 | 11415513 | 11415809 | - | 297 | Metazoa_SRP | 0 | 0 | 0 | 0 | 0 | 0 |
| ENSDARG000 | 11 | 11441460 | 11441756 | + | 297 | Metazoa_SRP | 0 | 0 | 0 | 0 | 0 | 0 |
| ENSDARG000 | 11 | 11383833 | 11384130 | - | 298 | Metazoa_SRP | 0 | 0 | 0 | 0 | 0 | 0 |
| ENSDARG000 | 11;11;11 | 42754776:427 | 42758300:427 | -;-;- | 3622 | NA | 2 | 0 | 2 | 0 | 2 | 8 |
| ENSDARG000 | 11 | 44123327 | 44123458 | - | 132 | NA | 0 | 0 | 0 | 0 | 2 | 0 |
| ENSDARG000 | 11:11:11:11:1 | 42929547:429 | 42933847:429 | -:-:-:-:-:-:-:-:-: | 9963 | EML6 | 460 | 306 | 823 | 820 | 1147 | 20 |
| ENSDARG000 | 11 | 12256431 | 12256727 | + | 297 | Metazoa_SRP | 0 | 0 | 0 | 0 | 0 | 0 |
| ENSDARG000 | 11:11:11:11:1 | 41936435:419 | 41936582:419 | +:+:+:+:+:+:+: | 1954 | aldh4a1 | 548 | 482 | 598 | 451 | 918 | 840 |
| ENSDARG000 | 11:11:11:11:1 | 2198831:2202 | 2198999:2203 | +;+;+;+;+;+;+ | 1507 | hoxc6b | 57 | 8 | 8 | 15 | 443 | 8 |
| ENSDARG000 | 11:11:11:11:1 | 44356504:443 | 44356810:443 | +:+:+:+:+:+:+: | 1162 | srsf7b | 134 | 64 | 87 | 67 | 75 | 80 |
| ENSDARG000 | 11 | 12756143 | 12756275 | - | 133 | 5S_rRNA | 0 | 0 | 0 | 0 | 0 | 0 |
| ENSDARG000 | 11:11:11:11:1 | 11175814:111 | 11175978:111 | +;+;+;+;+ | 351 | NA | 0 | 0 | 0 | 0 | 0 | 0 |
| ENSDARG000 | 11:11:11:11:1 | 2434262:2440 | 2434362:2440 | +:+:+:+:+:+:+: | 1170 | NA | 6 | 0 | 0 | 0 | 0 | 0 |
| ENSDARG000 | 11 | 27520456 | 27520573 | + | 118 | 5S_rRNA | 0 | 1 | 0 | 0 | 0 | 0 |
| ENSDARG000 | 11;11;11;11 | 42730639:427 | 42730701:427 | +;+;+;+ | 504 | zgc:194981 | 181 | 75 | 206 | 160 | 256 | 142 |
| ENSDARG000 | 11;11 | 42910904:429 | 42917671:429 | -;- | 6966 | pcdh20 | 4 | 0 | 6 | 6 | 4 | 0 |
| ENSDARG000 | 11:11:11:11:1 | 44937205:449 | 44937288:449 | -:-:-:-:-:-:-:-:-: | 1659 | NA | 82 | 205 | 289 | 116 | 219 | 82 |
| ENSDARG000 | 11 | 11460806 | 11461102 | + | 297 | Metazoa_SRP | 0 | 0 | 0 | 0 | 0 | 0 |
| ENSDARG000 | 11 | 371322 | 371440 | + | 119 | 5S_rRNA | 0 | 0 | 0 | 0 | 0 | 0 |
| ENSDARG000 | 11 | 21752475 | 21752589 | - | 115 | 5S_rRNA | 0 | 0 | 0 | 0 | 0 | 0 |
| ENSDARG000 | 11:11:11:11:1 | 18270313:182 | 18270429:182 | -;-;-;-;-;-;-;- | 1852 | stimate | 123 | 152 | 168 | 232 | 206 | 127 |
| ENSDARG000 | 11 | 12353834 | 12354130 | + | 297 | Metazoa_SRP | 0 | 0 | 0 | 0 | 0 | 0 |
| ENSDARG000 | 11 | 22714015 | 22714131 | + | 117 | 5S_rRNA | 0 | 0 | 0 | 0 | 0 | 0 |
| ENSDARG000 | 11 | 25669839 | 25669955 | - | 117 | 5S_rRNA | 0 | 0 | 0 | 7 | 0 | 0 |
| ENSDARG000 | 11 | 12405730 | 12406015 | + | 286 | Metazoa_SRP | 0 | 0 | 0 | 0 | 0 | 0 |
| ENSDARG000 | 11 | 730807 | 730923 | + | 117 | 5S_rRNA | 0 | 0 | 0 | 0 | 0 | 0 |
| ENSDARG000 | 11:11:11:11:1 | 12241936:122 | 12242035:122 | -;-;-;-;- | 525 | NA | 0 | 15 | 0 | 0 | 0 | 0 |
| ENSDARG000 | 11;11;11 | 42101661:421 | 42102810:421 | -;-;- | 1795 | NA | 0 | 78 | 6 | 0 | 36 | 0 |
| ENSDARG000 | 11;11 | 6422374:6423 | 6422452:6423 | +;+ | 171 | NA | 1 | 0 | 0 | 0 | 0 | 1 |
| ENSDARG000 | 11:11:11:11:1 | 44878571:448 | 44880384:448 | -:-:-:-:-:-:-:-:-: | 3534 | eif4a3 | 2145 | 2956 | 3239 | 1662 | 2654 | 1661 |
| ENSDARG000 | 11:11:11:11:1 | 41858807:419 | 41859829:419 | +:+:+:+:+:+:+: | 1963 | IFFO2 | 72 | 201 | 126 | 50 | 53 | 27 |
| ENSDARG000 | 11:11:11:11:1 | 5565082:5565 | 5565408:5565 | +:+:+:+:+:+:+: | 3775 | APBA3 | 440 | 234 | 395 | 334 | 319 | 324 |
| ENSDARG000 | 11:11:11:11:1 | 2506516:2506 | 2507064:2507 | +:+:+:+:+:+:+: | 1177 | NABP2 | 195 | 172 | 184 | 93 | 187 | 191 |
| ENSDARG000 | 11 | 5531443 | 5531576 | + | 134 | 5S_rRNA | 0 | 0 | 0 | 0 | 0 | 0 |
| ENSDARG000 | 11 | 12542039 | 12542334 | + | 296 | Metazoa_SRP | 0 | 0 | 0 | 0 | 0 | 0 |
| ENSDARG000 | 11 | 12438084 | 12438380 | - | 297 | Metazoa_SRP | 0 | 0 | 0 | 0 | 0 | 0 |
| ENSDARG000 | 11:11:11:11:1 | 3308656:3312 | 3309285:3312 | +:+:+:+:+:+:+: | 1212 | sarnp | 530 | 618 | 979 | 620 | 754 | 613 |
| ENSDARG000 | 11:11:11:11:1 | 6169312:6171 | 6169359:6171 | +:+:+:+:+:+:+: | 189 | NA | 0 | 0 | 1 | 0 | 0 | 0 |
| ENSDARG000 | 11:11:11:11:1 | 44858979:448 | 44862272:448 | -:-:-:-:-:-:-:-:-: | 5019 | opn6a | 4 | 0 | 0 | 2 | 2 | 15 |
| ENSDARG000 | 11:11:11:11:1 | 2240075:2244 | 2240175:2244 | -:-:-:-:-:-:-:-:-: | 1167 | hnrnpa1a | 4371 | 3508 | 4881 | 2120 | 4251 | 2681 |
| ENSDARG000 | 11:11:11:11:1 | 30614485:306 | 30614540:306 | +;+;+;+;+;+;+ | 186 | NA | 0 | 0 | 0 | 0 | 0 | 0 |
| ENSDARG000 | 11;11;11 | 29537756:295 | 29537784:295 | +;+;+ | 2127 | NA | 0 | 0 | 0 | 0 | 0 | 0 |
| ENSDARG000 | 11 | 33428874 | 33428992 | + | 119 | 5S_rRNA | 0 | 0 | 0 | 0 | 0 | 0 |
| ENSDARG000 | 11:11:11:11:1 | 44400864:444 | 44401594:444 | -;-;-;-;-;- | 1407 | il1rapl1b | 12 | 15 | 41 | 60 | 26 | 78 |
| ENSDARG000 | 11 | 35225022 | 35225139 | - | 118 | 5S_rRNA | 0 | 0 | 0 | 0 | 0 | 0 |
| ENSDARG000 | 11 | 12161658 | 12161955 | - | 298 | Metazoa_SRP | 0 | 0 | 0 | 0 | 0 | 0 |
| ENSDARG000 | 11 | 6071960 | 6072066 | - | 107 | U6 | 0 | 0 | 2 | 0 | 0 | 0 |
| ENSDARG000 | 11 | 12578532 | 12578828 | + | 297 | Metazoa_SRP | 0 | 0 | 0 | 0 | 0 | 0 |
| ENSDARG000 | 11:11:11:11:1 | 44178560:441 | 44181462:441 | -:-:-:-:-:-:-:-:-: | 8026 | SHROOM2 | 0 | 0 | 2 | 0 | 10 | 44 |
| ENSDARG000 | 11;11;11;11 | 20953926:209 | 20953976:209 | -;-;-;- | 582 | NA | 0 | 0 | 0 | 8 | 0 | 0 |
| ENSDARG000 | 11:11:11:11:1 | 5588122:5588 | 5588234:5588 | +:+:+:+:+:+:+: | 7743 | ABCA7 | 97 | 140 | 163 | 52 | 192 | 62 |
| ENSDARG000 | 11 | 12289372 | 12289668 | - | 297 | Metazoa_SRP | 0 | 0 | 0 | 0 | 0 | 0 |
| ENSDARG000 | 11 | 12166502 | 12166786 | + | 285 | Metazoa_SRP | 0 | 0 | 0 | 0 | 0 | 0 |
| ENSDARG000 | 11:11:11:11:1 | 44953598:449 | 44954158:449 | -:-:-:-:-:-:-:-:-: | 2262 | ldb1b | 484 | 286 | 463 | 267 | 428 | 330 |
| ENSDARG000 | 11:11:11:11:1 | 12321345:123 | 12321444:123 | -;-;-;-;- | 525 | NA | 0 | 1 | 0 | 0 | 0 | 0 |
| ENSDARG000 | 11 | 30974771 | 30974892 | + | 122 | 5S_rRNA | 0 | 0 | 0 | 0 | 0 | 0 |
| ENSDARG000 | 11 | 12345092 | 12345388 | + | 297 | Metazoa_SRP | 0 | 0 | 0 | 0 | 0 | 0 |
| ENSDARG000 | 11:11:11:11:1 | 220544:22157 | 223876:22387 | -;-;-;-;-;- | 4368 | dusp7 | 1419 | 983 | 1523 | 949 | 1237 | 1181 |
| ENSDARG000 | 11:11:11:11:1 | 42726712:427 | 42726746:427 | +:+:+:+:+:+:+: | 7165 | tdrd3 | 635 | 496 | 927 | 708 | 867 | 597 |
| ENSDARG000 | 11:11:11:11:1 | 22890997:228 | 22893147:228 | -:-:-:-:-:-:-:-:-: | 3229 | mdm4 | 1294 | 1053 | 1778 | 1693 | 1709 | 1498 |
| ENSDARG000 | 11 | 19416178 | 19416315 | - | 138 | 5S_rRNA | 0 | 0 | 0 | 0 | 0 | 0 |
| ENSDARG000 | 11 | 11437891 | 11438187 | - | 297 | Metazoa_SRP | 0 | 0 | 0 | 0 | 0 | 0 |
| ENSDARG000 | 11 | 34355544 | 34355662 | - | 119 | 5S_rRNA | 0 | 0 | 0 | 0 | 0 | 0 |
| ENSDARG000 | 11 | 21650522 | 21650638 | + | 117 | 5S_rRNA | 0 | 0 | 0 | 0 | 0 | 0 |
| ENSDARG000 | 11 | 41375011 | 41375126 | - | 116 | 5S_rRNA | 0 | 0 | 0 | 0 | 0 | 0 |
| ENSDARG000 | 11 | 44335666 | 44335780 | + | 115 | 5S_rRNA | 0 | 0 | 0 | 0 | 0 | 0 |
| ENSDARG000 | 11 | 37214108 | 37214223 | + | 116 | 5S_rRNA | 0 | 0 | 0 | 0 | 0 | 0 |
| ENSDARG000 | 11 | 11426878 | 11427174 | - | 297 | Metazoa_SRP | 0 | 0 | 0 | 0 | 0 | 0 |
| ENSDARG000 | 11 | 12351259 | 12351555 | - | 297 | Metazoa_SRP | 0 | 0 | 0 | 0 | 0 | 0 |
| ENSDARG000 | 11 | 13817569 | 13817684 | + | 116 | 5S_rRNA | 0 | 0 | 0 | 0 | 0 | 0 |
| ENSDARG000 | 11 | 26077947 | 26078061 | + | 115 | 5S_rRNA | 0 | 0 | 0 | 0 | 0 | 0 |
| ENSDARG000 | 11 | 12169361 | 12169654 | - | 294 | Metazoa_SRP | 0 | 0 | 0 | 0 | 0 | 0 |
| ENSDARG000 | 11 | 38175071 | 38175150 | - | 80 | dre-mir-135c- | 0 | 0 | 0 | 0 | 0 | 0 |
| ENSDARG000 | 11:11:11:11:1 | 45244618:452 | 45246066:452 | -;-;-;-;-;-;-;- | 2128 | notum1b | 12 | 6 | 8 | 6 | 6 | 12 |
| ENSDARG000 | 11:11:11:11:1 | 44265722:442 | 44266272:442 | -:-:-:-:-:-:-:-:-: | 3491 | NA | 28 | 48 | 92 | 28 | 42 | 34 |
| ENSDARG000 | 11:11:11:11:1 | 12703523:127 | 12703565:127 | -:-:-:-:-:-:-:-:-: | 261 | NA | 3 | 3 | 7 | 12 | 7 | 7 |
| ENSDARG000 | 11:11:11:11:1 | 161196:16119 | 161219:16121 | -:-:-:-:-:-:-:-:-: | 772 | TMBIM6 | 5191 | 4148 | 5267 | 4915 | 5281 | 4189 |
| ENSDARG000 | 11:11:11:11:1 | 1337620:1339 | 1337789:1339 | -:-:-:-:-:-:-:-:-: | 3955 | iars1 | 3281 | 2852 | 6834 | 3778 | 3022 | 4311 |
| ENSDARG000 | 11:11:11:11:1 | 2217413:2217 | 2218431:2218 | -;-;-;-;- | 2282 | SMUG1 | 98 | 93 | 105 | 70 | 131 | 129 |
| ENSDARG000 | 11:11:11:11:1 | 2540139:2540 | 2543277:2543 | -:-:-:-:-:-:-:-:-: | 3990 | cnpy2 | 38 | 44 | 88 | 16 | 60 | 40 |
| ENSDARG000 | 11 | 12685304 | 12685599 | + | 296 | Metazoa_SRP | 0 | 0 | 0 | 0 | 0 | 0 |
| ENSDARG000 | 11 | 12590999 | 12591296 | - | 298 | Metazoa_SRP | 0 | 0 | 0 | 0 | 0 | 0 |
| ENSDARG000 | 11;11 | 2458264:2461 | 2458492:2466 | +;+ | 5295 | tp53rk | 282 | 328 | 428 | 156 | 402 | 272 |
| ENSDARG000 | 11:11:11:11:1 | 14937904:149 | 14937944:149 | +;+;+;+;+;+;+ | 171 | NA | 7 | 6 | 8 | 0 | 16 | 0 |
| ENSDARG000 | 11:11:11:11:1 | 2464543:2466 | 2464681:2466 | -;-;-;-;-;-;- | 843 | SERINC3 | 0 | 4 | 2 | 0 | 25 | 3 |
| ENSDARG000 | 11 | 31985928 | 31986044 | + | 117 | 5S_rRNA | 0 | 0 | 0 | 0 | 0 | 0 |
| ENSDARG000 | 11 | 29352101 | 29352221 | + | 121 | 5S_rRNA | 0 | 0 | 0 | 0 | 0 | 0 |
| ENSDARG000 | 11 | 12567987 | 12568283 | + | 297 | Metazoa_SRP | 0 | 0 | 0 | 0 | 0 | 0 |
| ENSDARG000 | 11 | 12373666 | 12373962 | + | 297 | Metazoa_SRP | 0 | 0 | 0 | 0 | 0 | 0 |
| ENSDARG000 | 11 | 36994694 | 36994808 | + | 115 | 5S_rRNA | 0 | 0 | 0 | 0 | 0 | 0 |
| ENSDARG000 | 11:11:11:11:1 | 169394:17107 | 169663:17117 | -:-:-:-:-:-:-:-:-: | 2645 | avil | 18 | 2 | 10 | 2 | 0 | 16 |
| ENSDARG000 | 11 | 12162752 | 12163048 | + | 297 | Metazoa_SRP | 0 | 0 | 0 | 0 | 0 | 0 |
| ENSDARG000 | 11:11:11:11:1 | 122634:12376 | 122886:12384 | -:-:-:-:-:-:-:-:-: | 1179 | arfgap1 | 1108 | 898 | 1211 | 555 | 960 | 406 |
| ENSDARG000 | 11:11:11:11:1 | 135249:13560 | 135418:13574 | -;-;-;-;-;-;-;- | 1079 | cdk4 | 746 | 465 | 802 | 492 | 768 | 425 |
| ENSDARG000 | 11;11 | 2172335:2173 | 2172947:2175 | +;+ | 2426 | hoxc12b | 208 | 22 | 2 | 6 | 2 | 6 |
| ENSDARG000 | 11:11:11:11:1 | 2111582:2113 | 2112808:2113 | -:-:-:-:-:-:-:-:-: | 2957 | calcoco1b | 49 | 20 | 12 | 40 | 28 | 30 |
| ENSDARG000 | 11:11:11:11:1 | 42062067:420 | 42065496:420 | -:-:-:-:-:-:-:-:-: | 4931 | ABHD6 | 233 | 126 | 270 | 10 | 231 | 10 |
| ENSDARG000 | 11;11;11 | 44135351:441 | 44135458:441 | +;+;+ | 324 | NA | 9 | 0 | 0 | 0 | 1 | 0 |
| ENSDARG000 | 11;11;11 | 2276354:2276 | 2276394:2276 | -;-;- | 216 | NA | 9 | 17 | 29 | 13 | 11 | 13 |
| ENSDARG000 | 11;11 | 42731890:427 | 42735880:427 | -;- | 4324 | NA | 39 | 52 | 91 | 19 | 68 | 30 |
| ENSDARG000 | 11 | 43044826 | 43044949 | + | 124 | 5S_rRNA | 0 | 0 | 0 | 0 | 0 | 0 |
| ENSDARG000 | 11 | 23655350 | 23655478 | - | 129 | NA | 0 | 0 | 0 | 0 | 0 | 0 |
| ENSDARG000 | 11 | 39089875 | 39089991 | - | 117 | 5S_rRNA | 0 | 0 | 0 | 0 | 1 | 0 |
| ENSDARG000 | 11 | 12536284 | 12536580 | + | 297 | Metazoa_SRP | 0 | 0 | 0 | 0 | 0 | 0 |

| | | | | | | | | | | | | |
|---|---|---|---|---|---|---|---|---|---|---|---|---|
| ENSDARG000 | 11 | 12516933 | 12517229 | + | 297 | Metazoa_SRP | 0 | 0 | 0 | 0 | 0 | 0 |
| ENSDARG000 | 11 | 12693967 | 12694263 | + | 297 | Metazoa_SRP | 0 | 0 | 0 | 0 | 0 | 0 |
| ENSDARG000 | 11;11;11;11;1 | 20896122;208 | 20897045;208 | +;+;+;+;+;+;+ | 3036 | NA | 44 | 14 | 24 | 38 | 34 | 84 |
| ENSDARG000 | 11 | 12223462 | 12223758 | + | 297 | Metazoa_SRP | 0 | 0 | 0 | 0 | 0 | 0 |
| ENSDARG000 | 11 | 11462306 | 11462602 | + | 297 | Metazoa_SRP | 0 | 0 | 0 | 0 | 0 | 0 |
| ENSDARG000 | 11 | 44024762 | 44024878 | + | 117 | 5S_rRNA | 3 | 0 | 0 | 0 | 0 | 0 |
| ENSDARG000 | 11 | 13864846 | 13864964 | + | 119 | 5S_rRNA | 0 | 0 | 0 | 0 | 0 | 0 |
| ENSDARG000 | 11 | 23397270 | 23397385 | + | 116 | 5S_rRNA | 0 | 0 | 0 | 0 | 0 | 0 |
| ENSDARG000 | 11 | 30101142 | 30101257 | - | 116 | 5S_rRNA | 0 | 0 | 0 | 0 | 0 | 4 |
| ENSDARG000 | 11 | 12395518 | 12395814 | - | 297 | Metazoa_SRP | 0 | 0 | 0 | 0 | 0 | 0 |
| ENSDARG000 | 11;11;11;11;1 | 15821433;158 | 15821450;158 | +;+;+;+;+;+;+; | 306 | NA | 4 | 0 | 6 | 4 | 9 | 10 |
| ENSDARG000 | 11 | 43363075 | 43368382 | + | 5308 | NA | 0 | 0 | 0 | 0 | 0 | 0 |
| ENSDARG000 | 11;11;11;11;1 | 43661735;43 | 43661918;43 | +;+;+;+;+;+;+; | 3988 | gli2b | 26 | 0 | 6 | 10 | 8 | 8 |
| ENSDARG000 | 11;11;11;11;1 | 27133560;27 | 27133694;27 | +;+;+;+;+;+;+; | 1335 | hdac11 | 19 | 4 | 24 | 22 | 38 | 46 |
| ENSDARG000 | 11;11;11;11;1 | 141504;1447 | 141593;1448 | +;+;+;+;+;+;+; | 3372 | nckap1l | 74 | 194 | 231 | 114 | 460 | 403 |
| ENSDARG000 | 11;11;11;11;1 | 32007;33428; | 33276;33670; | -;-;-;-;-;-;-;-; | 5158 | col2a1b | 1066 | 557 | 1238 | 651 | 1451 | 344 |
| ENSDARG000 | 11 | 12718121 | 12718398 | + | 278 | Metazoa_SRP | 0 | 0 | 0 | 0 | 0 | 0 |
| ENSDARG000 | 11;11;11;11;1 | 18175893;181 | 18176079;181 | +;+;+;+;+;+ | 1152 | NA | 0 | 0 | 0 | 0 | 0 | 0 |
| ENSDARG000 | 11;11;11 | 18249809;182 | 18250529;182 | -;-;- | 917 | mustn1b | 0 | 0 | 2 | 2 | 5 | 6 |
| ENSDARG000 | 11 | 23159189 | 23159307 | - | 119 | 5S_rRNA | 0 | 0 | 0 | 0 | 0 | 0 |
| ENSDARG000 | 11;11 | 77526;77901 | 77704;78192 | +;+ | 471 | NA | 0 | 0 | 0 | 0 | 0 | 0 |
| ENSDARG000 | 11;11 | 2156430;215 | 2156988;216 | +;+ | 4736 | hoxc13b | 750 | 28 | 0 | 0 | 0 | 6 |
| ENSDARG000 | 11;11;11;11;1 | 42746369;42 | 42747220;42 | -;-;-;-;- | 1843 | NA | 34 | 64 | 112 | 0 | 12 | 15 |
| ENSDARG000 | 11;11;11;11;1 | 21582560;21 | 21582614;21 | -;-;-;-;-;- | 129 | NA | 0 | 0 | 0 | 0 | 0 | 0 |
| ENSDARG000 | 11;11;11;11;1 | 36793473;36 | 36793888;36 | -;-;-;-;-;-;-;-; | 6674 | cacna1da | 97 | 908 | 1506 | 1711 | 2829 | 1223 |
| ENSDARG000 | 11 | 36582258 | 36582372 | + | 115 | 5S_rRNA | 0 | 0 | 0 | 0 | 0 | 0 |
| ENSDARG000 | 11 | 44851474 | 44851593 | + | 120 | 5S_rRNA | 0 | 0 | 0 | 0 | 0 | 0 |
| ENSDARG000 | 11 | 40182681 | 40182796 | - | 116 | 5S_rRNA | 0 | 2 | 0 | 0 | 0 | 0 |
| ENSDARG000 | 11 | 13482671 | 13482787 | - | 117 | 5S_rRNA | 0 | 0 | 0 | 0 | 0 | 0 |
| ENSDARG000 | 11;11;11;11;1 | 12610576;12 | 12610747;12 | -;-;-;-;-;-;- | 834 | NA | 0 | 4 | 0 | 0 | 0 | 0 |
| ENSDARG000 | 11;11;11;11 | 6411169;641 | 6411341;641 | -;-;-;- | 497 | ndufa11 | 505 | 485 | 611 | 407 | 558 | 539 |
| ENSDARG000 | 11;11;11;11;1 | 2554856;255 | 2554956;255 | +;+;+;+;+ | 1101 | zgc:113184 | 30 | 10 | 14 | 2 | 33 | 4 |
| ENSDARG000 | 11 | 12499966 | 12500262 | - | 297 | Metazoa_SRP | 0 | 0 | 0 | 0 | 0 | 0 |
| ENSDARG000 | 11 | 12607076 | 12607370 | + | 295 | Metazoa_SRP | 0 | 0 | 0 | 0 | 0 | 0 |
| ENSDARG000 | 11 | 11418033 | 11418323 | + | 291 | Metazoa_SRP | 0 | 0 | 0 | 0 | 0 | 0 |
| ENSDARG000 | 11 | 11451670 | 11451935 | - | 266 | Metazoa_SRP | 0 | 0 | 0 | 0 | 0 | 0 |
| ENSDARG000 | 11;11;11;11 | 43470176;43 | 43471545;43 | -;-;-;- | 1740 | tmem63bb | 24 | 6 | 51 | 16 | 36 | 118 |
| ENSDARG000 | 11 | 7150547 | 7150640 | + | 94 | NA | 0 | 0 | 1 | 0 | 7 | 3 |
| ENSDARG000 | 11;11;11;11;1 | 41838036;41 | 41839105;41 | +;+;+;+;+;+;+; | 6099 | akr7a3 | 427 | 564 | 632 | 234 | 542 | 444 |
| ENSDARG000 | 11 | 12247688 | 12247984 | + | 297 | Metazoa_SRP | 0 | 0 | 0 | 0 | 0 | 0 |
| ENSDARG000 | 11 | 7116247 | 7116363 | - | 117 | 5S_rRNA | 0 | 0 | 0 | 0 | 0 | 0 |
| ENSDARG000 | 11 | 3953723 | 3953837 | - | 115 | 5S_rRNA | 0 | 0 | 0 | 0 | 0 | 0 |
| ENSDARG000 | 11 | 32191757 | 32191871 | + | 115 | 5S_rRNA | 0 | 0 | 0 | 0 | 0 | 0 |
| ENSDARG000 | 11 | 42625371 | 42625487 | + | 117 | 5S_rRNA | 0 | 0 | 0 | 0 | 1 | 0 |
| ENSDARG000 | 11 | 12215328 | 12215628 | + | 301 | Metazoa_SRP | 0 | 0 | 0 | 0 | 0 | 0 |
| ENSDARG000 | 11 | 12475442 | 12475702 | + | 261 | Metazoa_SRP | 0 | 0 | 0 | 0 | 0 | 0 |
| ENSDARG000 | 11;11;11;11;1 | 3385106;338 | 3385440;338 | -;-;-;-;-;-;-;-; | 2950 | acap3b | 771 | 381 | 507 | 353 | 488 | 661 |
| ENSDARG000 | 11 | 2190621 | 2190730 | + | 110 | mir196c | 0 | 0 | 0 | 0 | 0 | 0 |
| ENSDARG000 | 11;11;11;11;1 | 18157260;181 | 18157446;181 | +;+;+;+;+;+ | 1173 | zgc:173545 | 0 | 0 | 0 | 0 | 0 | 0 |
| ENSDARG000 | 11;11;11;11;1 | 18531984;18 | 18540001;18 | -;-;-;-;-;-;-;-; | 13499 | dido1 | 2811 | 2620 | 3974 | 2239 | 3086 | 2465 |
| ENSDARG000 | 11;11 | 4283014;428 | 4284033;428 | -;- | 2731 | NA | 196 | 788 | 505 | 1324 | 1148 | 51 |
| ENSDARG000 | 11;11;11;11;1 | 39672874;39 | 39673009;39 | +;+;+;+;+;+;+; | 9538 | CAMTA1 | 74 | 78 | 708 | 259 | 29 | 207 |
| ENSDARG000 | 11;11;11;11;1 | 41216191;41 | 41216882;41 | -;-;-;-;-;- | 1101 | mrps16 | 323 | 254 | 322 | 212 | 220 | 261 |
| ENSDARG000 | 11 | 24967771 | 24967887 | - | 117 | 5S_rRNA | 0 | 0 | 0 | 0 | 0 | 0 |
| ENSDARG000 | 11;11;11 | 3538988;354 | 3539008;354 | +;+;+ | 66 | NA | 0 | 0 | 0 | 0 | 0 | 0 |
| ENSDARG000 | 11 | 11429393 | 11429689 | + | 297 | Metazoa_SRP | 0 | 0 | 0 | 0 | 0 | 0 |
| ENSDARG000 | 11 | 11386362 | 11386658 | + | 297 | Metazoa_SRP | 0 | 0 | 0 | 0 | 0 | 0 |
| ENSDARG000 | 11 | 31923111 | 31923225 | + | 115 | 5S_rRNA | 0 | 0 | 0 | 0 | 0 | 0 |
| ENSDARG000 | 11;11;11;11;1 | 42600731;42 | 42600954;42 | +;+;+;+;+;+;+; | 4427 | appl1 | 823 | 628 | 1293 | 602 | 2413 | 588 |
| ENSDARG000 | 11 | 38803611 | 38803738 | + | 128 | 5S_rRNA | 0 | 0 | 0 | 0 | 0 | 0 |
| ENSDARG000 | 11 | 41465107 | 41465236 | + | 130 | 5S_rRNA | 2 | 0 | 0 | 0 | 0 | 0 |
| ENSDARG000 | 11 | 12253860 | 12254156 | - | 297 | Metazoa_SRP | 0 | 0 | 0 | 0 | 0 | 0 |
| ENSDARG000 | 11 | 7150274 | 7150366 | + | 93 | NA | 0 | 0 | 2 | 0 | 1 | 0 |
| ENSDARG000 | 11 | 11367594 | 11367890 | - | 297 | Metazoa_SRP | 0 | 0 | 0 | 0 | 0 | 0 |
| ENSDARG000 | 11 | 12326206 | 12326502 | + | 297 | Metazoa_SRP | 0 | 0 | 0 | 0 | 0 | 0 |
| ENSDARG000 | 11 | 12187306 | 12187602 | + | 297 | Metazoa_SRP | 0 | 0 | 2 | 0 | 0 | 0 |
| ENSDARG000 | 11;11;11;11;1 | 4038040;403 | 4038074;403 | -;-;-;-;-;-;-;-; | 447 | NA | 1 | 3 | 1 | 0 | 0 | 0 |
| ENSDARG000 | 11 | 19180452 | 19180555 | - | 104 | U6 | 0 | 0 | 0 | 0 | 0 | 0 |
| ENSDARG000 | 11 | 11447189 | 11447485 | - | 297 | Metazoa_SRP | 0 | 0 | 0 | 0 | 0 | 0 |
| ENSDARG000 | 11;11;11;11;1 | 2398843;240 | 2398992;240 | +;+;+;+;+;+;+; | 1251 | PIP4K2C | 0 | 0 | 0 | 0 | 0 | 2 |
| ENSDARG000 | 11;11;11;11;1 | 18343890;18 | 18348208;18 | -;-;-;-;-;-;-;-; | 6370 | PRKCD | 1032 | 1514 | 2048 | 1432 | 2430 | 1074 |
| ENSDARG000 | 11;11;11;11;1 | 61190;61190; | 62906;62906; | -;-;-;-;- | 2556 | NA | 0 | 0 | 0 | 6 | 0 | 2 |
| ENSDARG000 | 11;11;11 | 183570;1901 | 187419;1902 | -;-;- | 3976 | NA | 537 | 288 | 303 | 396 | 287 | 446 |
| ENSDARG000 | 11;11;11;11 | 8476961;848 | 8477315;848 | -;-;-;- | 674 | NA | 0 | 0 | 0 | 0 | 0 | 0 |
| ENSDARG000 | 11;11;11;11;1 | 9511533;951 | 9511713;951 | +;+;+;+;+;+;+ | 2462 | NA | 0 | 0 | 0 | 0 | 0 | 0 |
| ENSDARG000 | 11;11;11 | 11588078;11 | 11588114;11 | +;+;+ | 408 | NA | 0 | 0 | 0 | 0 | 0 | 0 |
| ENSDARG000 | 11;11;11 | 17919143;17 | 17919225;17 | +;+;+ | 223 | NA | 0 | 0 | 0 | 0 | 0 | 0 |
| ENSDARG000 | 11;11;11;11;1 | 18144285;181 | 18145173;181 | -;-;-;-;-;-;- | 2267 | NA | 0 | 0 | 0 | 0 | 0 | 0 |
| ENSDARG000 | 11;11;11;11;1 | 23305755;23 | 23307104;23 | -;-;-;-;-;-;-;-; | 2094 | NA | 5 | 0 | 0 | 0 | 0 | 2 |
| ENSDARG000 | 11;11;11;11 | 24018401;24 | 24018514;24 | +;+;+;+ | 446 | NA | 0 | 0 | 0 | 2 | 0 | 0 |
| ENSDARG000 | 11;11;11;11;1 | 28442373;28 | 28442866;28 | +;+;+;+;+;+ | 1135 | NA | 0 | 0 | 0 | 0 | 0 | 0 |
| ENSDARG000 | 11;11;11;11 | 32401204;32 | 32401528;32 | +;+;+;+ | 529 | NA | 0 | 0 | 0 | 0 | 0 | 0 |
| ENSDARG000 | 11;11;11;11 | 33190658;33 | 33190744;33 | -;-;-;- | 270 | NA | 0 | 0 | 0 | 0 | 0 | 0 |
| ENSDARG000 | 11;11;11;11;1 | 35725245;35 | 35726229;35 | -;-;-;-;-;- | 1496 | NA | 0 | 0 | 0 | 0 | 0 | 0 |
| ENSDARG000 | 11;11;11;11;1 | 36505674;36 | 36505953;36 | +;+;+;+;+;+;+; | 2240 | NA | 0 | 0 | 0 | 0 | 0 | 2 |
| ENSDARG000 | 11;11;11;11 | 36564610;36 | 36564684;36 | +;+;+;+ | 605 | NA | 0 | 0 | 0 | 0 | 0 | 4 |
| ENSDARG000 | 11;11;11;11;1 | 36574375;36 | 36574674;36 | +;+;+;+;+;+;+ | 2143 | NA | 0 | 0 | 0 | 0 | 8 | 0 |
| ENSDARG000 | 11;11;11;11;1 | 36587329;36 | 36587356;36 | +;+;+;+;+ | 878 | NA | 0 | 0 | 0 | 0 | 0 | 0 |
| ENSDARG000 | 11;11;11 | 36612354;36 | 36612811;36 | +;+;+ | 576 | NA | 0 | 0 | 0 | 0 | 0 | 0 |
| ENSDARG000 | 11;11;11;11;1 | 36620717;36 | 36622757;36 | -;-;-;-;-;-;- | 3621 | NA | 0 | 0 | 42 | 0 | 12 | 0 |
| ENSDARG000 | 11;11;11;11 | 41192329;41 | 41192536;41 | +;+;+;+ | 4653 | NA | 0 | 0 | 0 | 0 | 0 | 0 |
| ENSDARG000 | 11;11;11 | 41192329;41 | 41199446;41 | -;-;- | 7996 | NA | 0 | 0 | 0 | 0 | 0 | 0 |
| ENSDARG000 | 11;11;11 | 41223485;41 | 41223518;41 | +;+;+ | 499 | NA | 0 | 0 | 0 | 0 | 0 | 0 |
| ENSDARG000 | 11;11;11 | 43070883;43 | 43070953;43 | +;+;+ | 246 | NA | 0 | 0 | 0 | 0 | 0 | 0 |
| ENSDARG000 | 11;11;11;11;1 | 43643158;43 | 43643695;43 | +;+;+;+;+ | 935 | NA | 18 | 13 | 12 | 4 | 2 | 6 |
| ENSDARG000 | 11;11;11;11;1 | 44100339;44 | 44101659;44 | -;-;-;-;-;-;-;-; | 2199 | NA | 130 | 53 | 98 | 118 | 135 | 91 |
| ENSDARG000 | 10;10;10;10;1 | 6697369;669 | 6697575;669 | +;+;+;+;+;+;+; | 3095 | NA | 0 | 0 | 0 | 0 | 0 | 0 |
| ENSDARG000 | 10;10;10;10;1 | 17747880;17 | 17748078;17 | +;+;+;+;+;+;+; | 4782 | PIGO | 440 | 516 | 533 | 159 | 463 | 406 |
| ENSDARG000 | 10;10;10;10;1 | 44581378;44 | 44581988;44 | +;+;+;+;+;+;+; | 7488 | SEZ6L | 0 | 0 | 0 | 0 | 0 | 2 |
| ENSDARG000 | 10;10;10;10;1 | 24371921;24 | 24374723;24 | -;-;-;-;-;-;-;-; | 4637 | slc43a2b | 111 | 148 | 472 | 148 | 608 | 137 |
| ENSDARG000 | 10;10;10;10;1 | 28029578;28 | 28031213;28 | -;-;-;-;-;-;-;-; | 8990 | med13a | 1754 | 1261 | 1847 | 1204 | 2115 | 1657 |
| ENSDARG000 | 10;10;10 | 1396940;139 | 1397465;139 | +;+;+ | 2258 | gdnfa | 12 | 0 | 0 | 12 | 2 | 4 |
| ENSDARG000 | 10;10;10;10;1 | 16029260;16 | 16029415;16 | +;+;+;+;+ | 1786 | phax | 514 | 572 | 640 | 355 | 696 | 506 |
| ENSDARG000 | 10;10;10;10;1 | 37137464;37 | 37137574;37 | +;+;+;+;+;+;+; | 2499 | cuedc1a | 2 | 4 | 8 | 0 | 0 | 39 |
| ENSDARG000 | 10;10;10;10;1 | 41060739;41 | 41061892;41 | -;-;-;-;-;-;-;-; | 5136 | AAK1 | 258 | 249 | 355 | 240 | 527 | 241 |
| ENSDARG000 | 10;10;10;10;1 | 24302433;24 | 24304602;24 | -;-;-;-;-;-;-;-; | 3835 | INPP5K | 0 | 0 | 0 | 0 | 0 | 0 |
| ENSDARG000 | 10;10;10;10 | 24316853;24 | 24317311;24 | +;+;+;+ | 953 | NA | 0 | 0 | 0 | 0 | 2 | 0 |
| ENSDARG000 | 10;10;10;10;1 | 24323751;24 | 24324109;24 | -;-;-;-;-;-;-;-; | 1369 | PITPNA | 22 | 108 | 81 | 64 | 232 | 101 |
| ENSDARG000 | 10;10 | 25726694;25 | 25726728;25 | +;+ | 1530 | ugt5d1 | 0 | 0 | 6 | 0 | 0 | 0 |
| ENSDARG000 | 10;10;10;10 | 17776981;17 | 17777095;17 | +;+;+;+ | 743 | ccl19b | 10 | 2 | 0 | 0 | 0 | 10 |
| ENSDARG000 | 10;10;10;10;1 | 5135842;513 | 5135877;513 | +;+;+;+;+;+;+; | 1829 | zgc:113274 | 16 | 28 | 69 | 102 | 64 | 35 |
| ENSDARG000 | 10;10;10;10;1 | 41159241;41 | 41159497;41 | +;+;+;+;+;+;+; | 1485 | anxa4 | 4743 | 4177 | 5316 | 5013 | 5029 | 5062 |
| ENSDARG000 | 10;10;10;10;1 | 35059856;35 | 35060052;35 | -;-;-;-;-;-;-;- | 1015 | C10H21orf91 | 39 | 28 | 27 | 4 | 20 | 44 |
| ENSDARG000 | 10;10;10;10;1 | 25762196;25 | 25762793;25 | -;-;-;-;-;-;-;-; | 3994 | POSTN | 138 | 32 | 4229 | 66 | 2034 | 45 |
| ENSDARG000 | 10;10;10;10;1 | 17850934;17 | 17851354;17 | +;+;+;+;+;+;+; | 6360 | PHF24 | 8 | 0 | 0 | 0 | 0 | 0 |
| ENSDARG000 | 10;10;10;10;1 | 8091144;809 | 8091252;809 | +;+;+;+;+;+;+; | 741 | sub1a | 1537 | 2495 | 3135 | 1768 | 2186 | 1500 |
| ENSDARG000 | 10;10;10;10;1 | 35114153;35 | 35118434;35 | -;-;-;-;-;-;-;-; | 6188 | ripk4 | 62 | 2 | 0 | 4 | 34 | 38 |
| ENSDARG000 | 10;10;10;10;1 | 6725657;672 | 6725746;672 | -;-;-;-;-;- | 935 | NA | 0 | 0 | 2 | 0 | 0 | 0 |
| ENSDARG000 | 10;10;10;10;1 | 32683089;32 | 32683326;32 | +;+;+;+;+;+;+ | 2028 | ppm1e | 8 | 30 | 83 | 0 | 68 | 25 |
| ENSDARG000 | 10;10;10;10;1 | 44677632;44 | 44677710;44 | +;+;+;+;+;+ | 2939 | tpst2 | 86 | 56 | 30 | 2 | 152 | 50 |
| ENSDARG000 | 10;10;10;10 | 35160010;35 | 35160551;35 | -;-;-;- | 1046 | NA | 1 | 1 | 2 | 0 | 4 | 9 |
| ENSDARG000 | 10;10;10;10;1 | 35152928;35 | 35153188;35 | +;+;+;+;+;+;+; | 1725 | nsun5 | 75 | 103 | 145 | 60 | 129 | 81 |
| ENSDARG000 | 10;10;10 | 14947003;14 | 14947045;14 | +;+;+ | 516 | NA | 0 | 0 | 0 | 0 | 0 | 0 |
| ENSDARG000 | 10;10 | 44692272;44 | 44692342;44 | +;+ | 1785 | ubc | 17524 | 17055 | 20199 | 19518 | 11043 | 11361 |
| ENSDARG000 | 10;10;10;10;1 | 14963898;14 | 14964463;14 | +;+;+;+;+;+;+; | 7250 | NA | 0 | 0 | 0 | 0 | 0 | 0 |
| ENSDARG000 | 10;10;10;10;1 | 44699734;44 | 44700156;44 | +;+;+;+;+;+;+; | 6651 | scarb1 | 418 | 570 | 1540 | 942 | 1933 | 979 |
| ENSDARG000 | 10;10;10;10;1 | 28160265;28 | 28160439;28 | +;+;+;+;+;+;+; | 2240 | rnft1 | 561 | 516 | 518 | 352 | 600 | 338 |
| ENSDARG000 | 10 | 1590604 | 1591103 | + | 500 | NA | 0 | 0 | 0 | 0 | 0 | 0 |
| ENSDARG000 | 10;10 | 8128374;812 | 8129456;812 | -;- | 1083 | NA | 0 | 10 | 2 | 13 | 0 | 0 |
| ENSDARG000 | 10;10;10;10;1 | 8101729;810 | 8101795;810 | +;+;+;+;+;+;+; | 1128 | NA | 46 | 84 | 69 | 151 | 69 | 85 |
| ENSDARG000 | 10;10;10;10;1 | 21867307;21 | 21867523;21 | +;+;+;+;+;+;+; | 1847 | cbln17 | 0 | 0 | 0 | 0 | 0 | 0 |
| ENSDARG000 | 10;10;10;10;1 | 32809620;32 | 32811243;32 | -;-;-;-;-;-;-;-; | 4732 | TRIM37 | 458 | 296 | 747 | 310 | 529 | 353 |
| ENSDARG000 | 10;10;10 | 21899753;21 | 21899842;21 | +;+;+ | 1679 | hrh2b | 2 | 2 | 0 | 0 | 0 | 3 |
| ENSDARG000 | 10;10;10;10;1 | 5149359;514 | 5149566;514 | +;+;+;+;+;+;+; | 2867 | NA | 10 | 2 | 4 | 12 | 4 | 26 |
| ENSDARG000 | 10;10;10;10;1 | 6729670;673 | 6729876;673 | +;+;+;+;+;+;+; | 2940 | NA | 0 | 0 | 0 | 0 | 0 | 0 |
| ENSDARG000 | 10;10;10;10;1 | 21863066;21 | 21863650;21 | -;-;-;-;-;- | 1442 | NA | 0 | 0 | 3 | 4 | 6 | 2 |
| ENSDARG000 | 10;10;10;10;1 | 1597270;159 | 1597504;159 | -;-;-;-;-;-;-;-; | 4400 | nup155 | 1113 | 1324 | 1991 | 590 | 1672 | 782 |
| ENSDARG000 | 10;10;10;10;1 | 17936441;17 | 17936608;17 | +;+;+;+;+;+;+; | 1857 | prkab1a | 340 | 430 | 509 | 676 | 326 | 400 |
| ENSDARG000 | 10;10;10;10;1 | 37173029;37 | 37173244;37 | +;+;+;+;+;+;+; | 3427 | KSR1 | 29 | 12 | 16 | 14 | 36 | 13 |
| ENSDARG000 | 10;10;10;10;1 | 16036246;16 | 16036288;16 | +;+;+;+;+;+;+; | 3819 | lmnb1 | 1701 | 2565 | 3595 | 1450 | 2758 | 1775 |
| ENSDARG000 | 10;10;10;10;1 | 28172344;28 | 28172690;28 | -;-;-;-;-;-;-;-; | 1862 | NA | 215 | 561 | 571 | 318 | 664 | 134 |
| ENSDARG000 | 10;10;10;10;1 | 5159475;517 | 5159660;517 | +;+;+;+;+;+;+; | 1965 | cdc42se2 | 64 | 121 | 214 | 60 | 152 | 105 |
| ENSDARG000 | 10;10 | 44767388;44 | 44767531;44 | +;+ | 525 | NA | 0 | 2 | 0 | 0 | 0 | 2 |
| ENSDARG000 | 10;10;10;10;1 | 44769050;44 | 44769401;44 | -;-;-;-;- | 1476 | tacr1b | 0 | 6 | 0 | 0 | 30 | 20 |
| ENSDARG000 | 10;10;10;10;1 | 5234327;523 | 5234416;523 | +;+;+;+;+;+;+; | 2110 | sptlc1 | 1320 | 1216 | 1306 | 1258 | 1303 | 1647 |
| ENSDARG000 | 10;10;10;10 | 25819996;25 | 25820953;25 | -;-;-;- | 1828 | ftr54 | 0 | 0 | 0 | 0 | 4 | 0 |
| ENSDARG000 | 10;10;10;10;1 | 6747720;674 | 6747926;674 | +;+;+;+;+;+;+; | 3207 | NA | 0 | 0 | 0 | 0 | 0 | 0 |
| ENSDARG000 | 10;10;10 | 41199660;41 | 41199760;41 | +;+;+ | 1676 | adrb3b | 0 | 0 | 0 | 0 | 0 | 0 |
| ENSDARG000 | 10;10 | 41252525;41 | 41252952;41 | +;+ | 1063 | prlhr2b | 0 | 0 | 0 | 0 | 10 | 0 |
| ENSDARG000 | 10;10;10;10;1 | 8265972;826 | 8266490;826 | -;-;-;-;-;-;-;-; | 2301 | PLPP1 | 1359 | 1878 | 1292 | 1923 | 1568 | 2091 |
| ENSDARG000 | 10;10 | 37352480;37 | 37352769;37 | -;- | 463 | NA | 0 | 4 | 0 | 4 | 3 | 1 |
| ENSDARG000 | 10;10;10;10;1 | 37268854;37 | 37268928;37 | +;+;+;+;+;+;+; | 8675 | NF1 | 3832 | 2186 | 4214 | 1829 | 5530 | 3195 |
| ENSDARG000 | 10;10;10;10;1 | 5268054;536 | 5268476;536 | +;+;+;+;+;+;+; | 3990 | ROR2 | 119 | 24 | 65 | 16 | 18 | 65 |
| ENSDARG000 | 10;10;10;10;1 | 8197827;819 | 8197969;819 | +;+;+;+;+;+;+; | 3542 | mtrex | 638 | 767 | 1071 | 309 | 866 | 675 |
| ENSDARG000 | 10;10;10;10;1 | 41279007;41 | 41280384;41 | -;-;-;-;-;-;-;-; | 2807 | brf2 | 40 | 38 | 61 | 21 | 36 | 14 |
| ENSDARG000 | 10;10 | 37334270;37 | 37334996;37 | -;- | 852 | OMG | 64 | 0 | 118 | 123 | 89 | 48 |
| ENSDARG000 | 10;10;10;10;1 | 10636237;10 | 10637018;10 | +;+;+;+;+;+ | 1894 | FAM163B | 0 | 0 | 0 | 0 | 0 | 0 |
| ENSDARG000 | 10;10;10;10 | 41318583;41 | 41318629;41 | -;-;-;- | 616 | NA | 137 | 72 | 113 | 78 | 224 | 78 |
| ENSDARG000 | 10;10;10;10;1 | 25829262;25 | 25830821;25 | -;-;-;-;-;-;-;-; | 4288 | trpc4a | 0 | 6 | 2 | 4 | 12 | 6 |
| ENSDARG000 | 10;10;10;10;1 | 41362390;41 | 41362988;41 | -;-;-;-;-;-;-;-; | 2587 | gpat4 | 482 | 410 | 718 | 390 | 592 | 433 |
| ENSDARG000 | 10;10;10;10;1 | 15024772;15 | 15024847;15 | +;+;+;+;+;+;+; | 1294 | NA | 0 | 0 | 0 | 0 | 0 | 0 |
| ENSDARG000 | 10;10;10;10;1 | 10712508;10 | 10712669;10 | -;-;-;-;- | 850 | NA | 0 | 2 | 0 | 0 | 0 | 0 |

| | | | | | | | | | | | | |
|---|---|---|---|---|---|---|---|---|---|---|---|---|
| ENSDARG000 | 10;10;10;10;1 | 10728870:107 | 10729218:107 | +;+;+;+;+;+;+; | 1682 | swi5 | 60 | 140 | 132 | 54 | 108 | 60 |
| ENSDARG000 | 10;10;10;10;1 | 44811700:448 | 44811852:448 | +;+;+;+;+;+;+; | 6488 | SLC4A5 | 705 | 94 | 445 | 614 | 282 | 384 |
| ENSDARG000 | 10;10;10;10;1 | 1638876:1638 | 1638938:1638 | +;+;+;+;+;+;+; | 3511 | SGSM1 | 24 | 24 | 57 | 110 | 35 | 145 |
| ENSDARG000 | 10;10;10;10;1 | 28311031:283 | 28312352:283 | -;-;-;-;-;-;-;-; | 6921 | cltca | 10436 | 9398 | 11036 | 9796 | 11358 | 7556 |
| ENSDARG000 | 10;10;10;10;1 | 22003750:220 | 22003951:220 | +;+;+;+;+;+;+; | 2241 | kcnip1b | 62 | 8 | 6 | 12 | 76 | 75 |
| ENSDARG000 | 10;10;10;10;1 | 32854901:328 | 32859031:328 | -;-;-;-;-;-;-;-; | 5444 | rabgef1 | 784 | 720 | 862 | 853 | 829 | 720 |
| ENSDARG000 | 10;10 | 15046837:150 | 15048781:150 | -;- | 1945 | cldn23.1 | 26 | 0 | 0 | 2 | 0 | 4 |
| ENSDARG000 | 10;10 | 15050469:150 | 15053614:150 | -;- | 3146 | CLDN23 | 8 | 0 | 0 | 0 | 0 | 0 |
| ENSDARG000 | 10;10;10;10;1 | 35169644:351 | 35170299:351 | -;-;-;-;-;-;-;-; | 4200 | NA | 627 | 688 | 1039 | 292 | 946 | 494 |
| ENSDARG000 | 10;10 | 6765109:6775 | 6765788:6775 | -;- | 772 | macir | 123 | 31 | 79 | 17 | 127 | 91 |
| ENSDARG000 | 10;10;10;10 | 15088534:150 | 15088587:150 | +;+;+;+ | 800 | NA | 2 | 6 | 0 | 2 | 8 | 0 |
| ENSDARG000 | 10 | 5409508 | 5410985 | + | 1478 | NA | 2 | 4 | 2 | 0 | 12 | 6 |
| ENSDARG000 | 10;10 | 5414224:5417 | 5414384:5419 | +;+ | 1957 | nfil3 | 858 | 672 | 981 | 663 | 487 | 242 |
| ENSDARG000 | 10;10;10;10;1 | 25947946:259 | 25948204:259 | +;+;+;+;+;+;+; | 2168 | ufm1 | 609 | 853 | 1208 | 641 | 913 | 886 |
| ENSDARG000 | 10;10;10 | 28306749:283 | 28306854:283 | +;+;+ | 1048 | ptrh2 | 218 | 376 | 230 | 150 | 302 | 96 |
| ENSDARG000 | 10;10;10;10;1 | 5422575:5423 | 5422822:5423 | +;+;+;+;+;+;+; | 1591 | auh | 268 | 345 | 349 | 314 | 368 | 204 |
| ENSDARG000 | 10;10;10;10;1 | 41409255:414 | 41409875:414 | -;-;-;-;-;-;-;-; | 1390 | cabp1b | 0 | 0 | 0 | 0 | 0 | 0 |
| ENSDARG000 | 10;10;10;10;1 | 8401929:8402 | 8402107:8402 | +;+;+;+;+;+;+; | 1570 | hvcn1 | 308 | 263 | 702 | 618 | 432 | 366 |
| ENSDARG000 | 10;10;10;10;1 | 6781123:6782 | 6782757:6782 | -;-;-;-;-;-;-;-; | 5381 | NA | 762 | 486 | 813 | 582 | 731 | 686 |
| ENSDARG000 | 10;10;10;10;1 | 35201850:352 | 35201969:352 | -;-;-;-;-;-;-;- | 691 | NA | 8 | 4 | 11 | 26 | 5 | 10 |
| ENSDARG000 | 10;10 | 1681518:1684 | 1681787:1684 | +;+ | 990 | triap1 | 88 | 182 | 128 | 54 | 172 | 180 |
| ENSDARG000 | 10;10;10;10;1 | 10738880:107 | 10739169:107 | +;+;+;+;+;+;+; | 2460 | slc27a4 | 148 | 108 | 193 | 46 | 186 | 76 |
| ENSDARG000 | 10;10;10;10;1 | 15106808:151 | 15107002:151 | +;+;+;+;+;+;+; | 1936 | scpp8 | 160 | 32 | 2 | 58 | 38 | 40 |
| ENSDARG000 | 10;10;10;10 | 32881586:328 | 32883128:328 | -;-;-;- | 2331 | kctd7 | 28 | 22 | 20 | 62 | 56 | 24 |
| ENSDARG000 | 10;10;10;10;1 | 17987288:179 | 17988122:179 | -;-;-;-;-;-;-;- | 1380 | NA | 135 | 63 | 17 | 98 | 61 | 57 |
| ENSDARG000 | 10;10;10;10;1 | 41624711:416 | 41625339:416 | -;-;-;-;-;-;-;-; | 2468 | NA | 0 | 0 | 11 | 12 | 34 | 0 |
| ENSDARG000 | 10;10;10;10;1 | 44917351:449 | 44917961:449 | -;-;-;-;-;- | 1775 | tuba7l | 70 | 66 | 39 | 29 | 46 | 30 |
| ENSDARG000 | 10;10;10;10 | 44903676:449 | 44903807:449 | +;+;+;+ | 1056 | vps37bb | 166 | 126 | 125 | 49 | 141 | 120 |
| ENSDARG000 | 10;10;10;10;1 | 15120711:151 | 15121556:151 | -;-;-;-;-;-;-;- | 1503 | spp1 | 152 | 29 | 50 | 10 | 106 | 28 |
| ENSDARG000 | 10;10;10;10;1 | 1686975:1686 | 1687624:1687 | -;-;-;-;-;-;-;-; | 1546 | srsf9 | 838 | 862 | 1146 | 925 | 970 | 848 |
| ENSDARG000 | 10;10 | 24397098:243 | 24397585:243 | +;+ | 563 | NA | 0 | 0 | 0 | 0 | 0 | 0 |
| ENSDARG000 | 10;10;10;10;1 | 24394023:243 | 24394374:243 | -;-;-;-;-;-;-;-; | 3723 | HEPH | 2 | 16 | 0 | 14 | 2 | 1 |
| ENSDARG000 | 10;10;10;10;1 | 16061655:160 | 16062308:160 | -;-;-;-;-;-;-;- | 981 | C10H5orf63 | 69 | 53 | 122 | 27 | 94 | 84 |
| ENSDARG000 | 10;10;10;10 | 35238488:352 | 35238940:352 | -;-;-;- | 2998 | SMG8 | 559 | 510 | 531 | 452 | 603 | 655 |
| ENSDARG000 | 10;10;10;10;1 | 18240798:182 | 18240886:182 | +;+;+;+;+ | 565 | NA | 0 | 0 | 0 | 0 | 0 | 0 |
| ENSDARG000 | 10;10;10;10;1 | 32898533:329 | 32903785:329 | -;-;-;-;-;-;-;-; | 9306 | CUX1 | 1018 | 736 | 1502 | 700 | 1349 | 930 |
| ENSDARG000 | 10;10;10;10;1 | 18291971:182 | 18292309:182 | -;-;-;-;-;-;-;-; | 3163 | LRRTM4 | 4 | 2 | 0 | 0 | 0 | 0 |
| ENSDARG000 | 10;10;10;10;1 | 44924684:449 | 44924776:449 | +;+;+;+;+;+;+; | 3281 | NA | 4 | 2 | 14 | 4 | 4 | 1 |
| ENSDARG000 | 10;10;10 | 44997062:450 | 44997507:450 | -;-;- | 1023 | ved | 2 | 10 | 4 | 33 | 19 | 10 |
| ENSDARG000 | 10;10;10;10;1 | 35250205:352 | 35251026:352 | -;-;-;-;-;-;-;-; | 1894 | NA | 178 | 637 | 484 | 340 | 360 | 301 |
| ENSDARG000 | 10;10;10;10;1 | 37400838:374 | 37401052:374 | +;+;+;+;+;+;+; | 2523 | NA | 12 | 2 | 2 | 0 | 16 | 0 |
| ENSDARG000 | 10;10;10;10;1 | 22034477:220 | 22034666:220 | +;+;+;+;+;+;+; | 1922 | npm1a | 3715 | 6669 | 11810 | 5708 | 5757 | 5123 |
| ENSDARG000 | 10;10;10;10 | 5525393:5525 | 5525626:5525 | -;-;-;- | 367 | NA | 0 | 0 | 0 | 0 | 0 | 0 |
| ENSDARG000 | 10;10;10;10;1 | 25982212:259 | 25987505:259 | +;+;+;+;+;+;+; | 14498 | frem2a | 42 | 0 | 0 | 0 | 4 | 68 |
| ENSDARG000 | 10;10;10;10;1 | 18877362:188 | 18877635:188 | +;+;+;+;+;+;+; | 2347 | ppp2r2ab | 773 | 940 | 1026 | 411 | 1044 | 659 |
| ENSDARG000 | 10;10 | 5529734:5541 | 5529906:5543 | +;+ | 1277 | NA | 0 | 0 | 0 | 0 | 0 | 0 |
| ENSDARG000 | 10;10;10 | 1708117:1713 | 1713439:1713 | -;-;- | 5624 | linc-epb41l4a | 19530 | 14489 | 32590 | 15194 | 45598 | 8991 |
| ENSDARG000 | 10;10;10;10;1 | 5533843:5536 | 5537136:5537 | -;-;-;-;-;-;-;-; | 5325 | syk | 123 | 134 | 148 | 115 | 501 | 368 |
| ENSDARG000 | 10;10;10;10;1 | 8527196:8527 | 8527341:8527 | +;+;+;+;+ | 966 | NA | 19 | 22 | 0 | 14 | 24 | 52 |
| ENSDARG000 | 10;10;10;10;1 | 41826331:418 | 41830586:418 | -;-;-;-;-;-;-;-; | 8593 | KDM2B | 715 | 553 | 798 | 897 | 601 | 954 |
| ENSDARG000 | 10;10;10;10;1 | 10788811:107 | 10788912:107 | +;+;+;+;+;+;+; | 1097 | ptgdsa | 0 | 0 | 4 | 0 | 0 | 0 |
| ENSDARG000 | 10;10;10;10;1 | 15224660:152 | 15224795:152 | +;+;+;+;+;+;+; | 1485 | NA | 16 | 27 | 52 | 23 | 83 | 48 |
| ENSDARG000 | 10;10;10;10;1 | 45015225:450 | 45015507:450 | -;-;-;-;-;-;-;-; | 1777 | polm | 28 | 71 | 73 | 78 | 64 | 56 |
| ENSDARG000 | 10;10;10;10;1 | 35257651:352 | 35257993:352 | +;+;+;+;+;+;+; | 1241 | styxl1 | 0 | 4 | 4 | 4 | 0 | 6 |
| ENSDARG000 | 10;10;10;10;1 | 16069987:160 | 16070119:160 | +;+;+;+;+;+;+; | 4121 | megf10 | 7063 | 12351 | 12267 | 9110 | 20063 | 9916 |
| ENSDARG000 | 10;10;10;10;1 | 1722125:1722 | 1723271:1723 | -;-;-;-;-;-;- | 1491 | GAL3ST1 | 11 | 0 | 14 | 16 | 0 | 2 |
| ENSDARG000 | 10;10;10;10;1 | 8534750:8537 | 8534779:8538 | +;+;+;+;+;+;+; | 1577 | NA | 433 | 182 | 202 | 236 | 364 | 344 |
| ENSDARG000 | 10;10;10;10 | 37485403:374 | 37486145:374 | -;-;-;- | 1002 | NA | 5 | 1 | 0 | 0 | 12 | 11 |
| ENSDARG000 | 10;10;10;10;1 | 15255012:152 | 15255375:152 | +;+;+;+;+;+;+; | 4732 | vldlr | 98 | 24 | 84 | 2 | 40 | 76 |
| ENSDARG000 | 10;10;10;10;1 | 37500234:375 | 37500331:375 | +;+;+;+;+;+;+; | 2522 | msi2a | 555 | 404 | 1085 | 437 | 1551 | 278 |
| ENSDARG000 | 10;10;10;10;1 | 45031398:450 | 45031620:450 | +;+;+;+;+;+;+; | 1934 | gnsb | 918 | 390 | 853 | 470 | 766 | 805 |
| ENSDARG000 | 10;10;10;10;1 | 18911152:189 | 18911253:189 | +;+;+;+;+;+;+; | 2641 | bnip3lb | 2288 | 1725 | 3373 | 3151 | 2051 | 1830 |
| ENSDARG000 | 10;10;10;10;1 | 1745475:1745 | 1745859:1745 | -;-;-;-;-;-;-;-; | 2290 | epb41l4a | 42 | 10 | 35 | 6 | 12 | 22 |
| ENSDARG000 | 10;10 | 8618849:8621 | 8619769:8621 | -;- | 953 | NA | 0 | 1 | 0 | 0 | 0 | 2 |
| ENSDARG000 | 10;10;10;10 | 45053577:450 | 45053988:450 | -;-;-;- | 656 | mrps24 | 144 | 222 | 323 | 121 | 210 | 114 |
| ENSDARG000 | 10;10;10;10;1 | 24445698:244 | 24445787:244 | +;+;+;+;+;+;+; | 2360 | NA | 16 | 5 | 118 | 47 | 439 | 8 |
| ENSDARG000 | 10 | 8629275 | 8631361 | + | 2087 | aplnrb | 316 | 763 | 2215 | 1432 | 3148 | 1408 |
| ENSDARG000 | 10;10;10;10;1 | 10801564:108 | 10801823:108 | +;+;+;+;+;+;+; | 2559 | ambp | 0 | 0 | 37 | 0 | 6 | 0 |
| ENSDARG000 | 10;10 | 24439620:244 | 24439900:244 | -;- | 585 | NA | 0 | 0 | 0 | 0 | 3 | 0 |
| ENSDARG000 | 10;10;10;10;1 | 24422017:244 | 24422369:244 | +;+;+;+;+;+;+; | 3839 | paxbp1 | 1398 | 777 | 1679 | 1237 | 1701 | 1697 |
| ENSDARG000 | 10;10;10;10;1 | 37981471:379 | 37981910:379 | +;+;+;+;+;+;+; | 5449 | NA | 0 | 1 | 10 | 4 | 10 | 18 |
| ENSDARG000 | 10;10;10;10;1 | 45071603:450 | 45071664:450 | +;+;+;+;+;+;+; | 1704 | CAMK2B | 0 | 4 | 0 | 0 | 5 | 0 |
| ENSDARG000 | 10;10 | 41914084:419 | 41914343:419 | +;+ | 1299 | orai1b | 158 | 154 | 193 | 64 | 157 | 149 |
| ENSDARG000 | 10;10;10;10 | 8660654:8662 | 8661147:8662 | -;-;-;- | 1169 | GOLPH3 | 971 | 936 | 1047 | 692 | 1238 | 725 |
| ENSDARG000 | 10;10;10;10;1 | 35265083:352 | 35265290:352 | +;+;+;+;+;+;+; | 1426 | tmem120aa | 52 | 40 | 105 | 86 | 119 | 70 |
| ENSDARG000 | 10;10;10;10;1 | 45059702:450 | 45060086:450 | +;+;+;+;+;+ | 1502 | nudcd3 | 668 | 584 | 719 | 579 | 748 | 694 |
| ENSDARG000 | 10;10;10;10;1 | 16165533:161 | 16165609:161 | +;+;+;+;+;+;+; | 3529 | prrc1 | 557 | 627 | 809 | 683 | 944 | 425 |
| ENSDARG000 | 10;10 | 8675890:8677 | 8676048:8677 | -;- | 546 | NA | 0 | 0 | 0 | 0 | 0 | 0 |
| ENSDARG000 | 10;10;10;10;1 | 38168158:381 | 38171879:381 | -;-;-;-;-;-;-;-; | 7474 | usp25 | 1779 | 1624 | 1766 | 2026 | 1776 | 1772 |
| ENSDARG000 | 10;10;10 | 22051054:220 | 22052580:220 | -;-;- | 2382 | tlx3b | 0 | 0 | 1 | 0 | 2 | 0 |
| ENSDARG000 | 10;10;10;10;1 | 41939963:419 | 41940047:419 | +;+;+;+;+;+;+; | 2035 | tmem120b | 45 | 78 | 83 | 70 | 82 | 42 |
| ENSDARG000 | 10;10;10;10;1 | 8680730:8680 | 8680824:8681 | +;+;+;+;+;+ | 1994 | isl1b | 0 | 0 | 2 | 2 | 0 | 0 |
| ENSDARG000 | 10;10;10;10;1 | 26118122:261 | 26118370:261 | +;+;+;+;+;+;+; | 3239 | trim47 | 516 | 557 | 657 | 257 | 764 | 602 |
| ENSDARG000 | 10;10;10;10;1 | 28370816:283 | 28371211:283 | -;-;-;-;-;-;-;-; | 1895 | btg3 | 417 | 839 | 1449 | 1757 | 682 | 344 |
| ENSDARG000 | 10;10;10;10;1 | 1849874:1849 | 1850008:1850 | +;+;+;+;+;+;+; | 9214 | apc | 4476 | 3984 | 6460 | 4070 | 6311 | 5022 |
| ENSDARG000 | 10;10;10;10;1 | 22082596:220 | 22085198:220 | -;-;-;-;- | 2796 | ugt5g2 | 0 | 0 | 0 | 0 | 0 | 0 |
| ENSDARG000 | 10;10;10;10 | 22088465:220 | 22088931:220 | -;-;-;- | 824 | NA | 4 | 0 | 0 | 0 | 0 | 0 |
| ENSDARG000 | 10;10;10;10;1 | 35275965:352 | 35276025:352 | +;+;+;+;+;+;+; | 3596 | pora | 3493 | 3910 | 4765 | 3859 | 3447 | 2522 |
| ENSDARG000 | 10;10;10;10;1 | 33406819:334 | 33407041:334 | +;+;+;+;+;+;+; | 16015 | RYR1 | 0 | 4 | 12 | 0 | 0 | 0 |
| ENSDARG000 | 10;10;10;10;1 | 16225117:162 | 16226397:162 | +;+;+;+;+;+;+; | 7453 | slc12a2 | 9567 | 3013 | 5436 | 4112 | 5733 | 13454 |
| ENSDARG000 | 10;10;10;10;1 | 22110842:221 | 22111693:221 | -;-;-;-;- | 1674 | NA | 4 | 14 | 22 | 8 | 33 | 6 |
| ENSDARG000 | 10;10;10;10;1 | 6884123:6884 | 6884197:6884 | +;+;+;+;+;+;+; | 2889 | ddx4 | 1 | 0 | 4 | 0 | 2 | 0 |
| ENSDARG000 | 10;10;10;10;1 | 41969428:419 | 41970231:419 | -;-;-;-;-;-;-;- | 1553 | rhof | 32 | 64 | 101 | 159 | 158 | 229 |
| ENSDARG000 | 10;10;10;10 | 22123831:221 | 22124533:221 | -;-;-;- | 1060 | ponzr2 | 12 | 0 | 2 | 0 | 0 | 0 |
| ENSDARG000 | 10;10;10;10;1 | 5689510:5689 | 5689610:5689 | +;+;+;+;+;+;+; | 4136 | PAM | 2218 | 1860 | 2358 | 2773 | 2239 | 2603 |
| ENSDARG000 | 10;10;10;10;1 | 26129830:261 | 26132642:261 | -;-;-;-;-;-;-;-; | 6149 | trim3b | 477 | 559 | 1439 | 1816 | 1591 | 957 |
| ENSDARG000 | 10;10;10;10;1 | 28428222:284 | 28428287:284 | +;+;+;+;+;+;+; | 1684 | NA | 0 | 23 | 30 | 20 | 25 | 12 |
| ENSDARG000 | 10;10;10;10;1 | 18952271:189 | 18952465:189 | +;+;+;+;+;+;+; | 2087 | dpysl2b | 1145 | 1277 | 1243 | 1339 | 1044 | 1490 |
| ENSDARG000 | 10;10;10;10;1 | 8688678:8690 | 8688755:8691 | +;+;+;+;+;+;+; | 795 | NA | 12 | 30 | 42 | 6 | 54 | 8 |
| ENSDARG000 | 10;10;10;10 | 38256556:382 | 38261771:382 | -;-;-;- | 5475 | NA | 191 | 200 | 315 | 196 | 174 | 135 |
| ENSDARG000 | 10;10;10;10;1 | 2175583:2232 | 2175654:2232 | +;+;+;+;+;+;+; | 4407 | NA | 15 | 12 | 2043 | 1432 | 1907 | 1037 |
| ENSDARG000 | 10;10;10 | 10860759:108 | 10864156:108 | -;-;- | 3495 | nrarpa | 316 | 64 | 128 | 42 | 108 | 82 |
| ENSDARG000 | 10;10 | 15310811:153 | 15312124:153 | +;+ | 1821 | KCNV2 | 2 | 2 | 6 | 1 | 2 | 0 |
| ENSDARG000 | 10;10;10;10;1 | 41982998:419 | 41985559:419 | -;-;-;-;-;-;-;-; | 3110 | NA | 197 | 108 | 174 | 218 | 359 | 329 |
| ENSDARG000 | 10;10;10 | 28442740:284 | 28442991:284 | -;-;- | 678 | NA | 18 | 21 | 47 | 6 | 34 | 74 |
| ENSDARG000 | 10;10;10;10;1 | 16313851:163 | 16314202:163 | -;-;-;-;-;-;-;-; | 9198 | FBN2 | 14 | 2 | 2 | 8 | 2 | 4 |
| ENSDARG000 | 10;10;10 | 28447906:284 | 28448048:284 | -;-;- | 639 | NA | 0 | 0 | 0 | 0 | 4 | 2 |
| ENSDARG000 | 10;10 | 22134606:221 | 22135085:221 | +;+ | 733 | NA | 0 | 0 | 0 | 0 | 0 | 0 |
| ENSDARG000 | 10;10;10;10;1 | 38417512:384 | 38417661:384 | +;+;+;+;+;+;+; | 1498 | samsn1b | 16 | 6 | 12 | 2 | 23 | 10 |
| ENSDARG000 | 10;10;10;10 | 35314564:353 | 35315093:353 | -;-;-;- | 1402 | NA | 131 | 106 | 130 | 58 | 152 | 119 |
| ENSDARG000 | 10;10;10;10;1 | 26190807:261 | 26192230:261 | -;-;-;-;-;-;-;-; | 8641 | fhdc3 | 180 | 167 | 178 | 259 | 268 | 215 |
| ENSDARG000 | 10;10 | 28454500:284 | 28454763:284 | -;- | 594 | NA | 0 | 1 | 1 | 0 | 2 | 2 |
| ENSDARG000 | 10;10;10;10;1 | 26205779:262 | 26206266:262 | -;-;-;-;-;-;-;-; | 1812 | arfip2b | 0 | 0 | 0 | 0 | 0 | 0 |
| ENSDARG000 | 10;10;10;10;1 | 8730988:8767 | 8731096:8767 | +;+;+;+;+;+;+; | 3755 | ITGA1 | 4132 | 3433 | 3750 | 3024 | 4638 | 4100 |
| ENSDARG000 | 10;10;10;10;1 | 28470307:284 | 28470388:284 | -;-;-;-;-;-;-;-; | 4043 | BBX | 205 | 136 | 168 | 115 | 256 | 116 |
| ENSDARG000 | 10;10;10;10;1 | 10879537:108 | 10880978:108 | -;-;-;-;-;-;-;-; | 4018 | NA | 282 | 223 | 332 | 136 | 309 | 282 |
| ENSDARG000 | 10;10;10;10;1 | 18999841:189 | 19000217:190 | -;-;-;-;- | 2067 | NA | 0 | 0 | 0 | 2 | 2 | 0 |
| ENSDARG000 | 10;10;10;10;1 | 38430359:384 | 38433206:384 | -;-;-;-;-;-;-;-; | 6078 | GDPD5 | 8 | 22 | 155 | 66 | 98 | 76 |
| ENSDARG000 | 10;10;10;10 | 22144768:221 | 22146091:221 | -;-;-;- | 2067 | cldn7b | 492 | 16 | 8 | 90 | 76 | 90 |
| ENSDARG000 | 10;10;10;10 | 19017146:190 | 19017321:190 | +;+;+;+ | 1663 | tmem230a | 226 | 154 | 116 | 166 | 183 | 253 |
| ENSDARG000 | 10;10;10;10;1 | 24504292:245 | 24504592:245 | +;+;+;+;+;+;+; | 5131 | NA | 0 | 2 | 2 | 0 | 4 | 2 |
| ENSDARG000 | 10;10;10 | 35329751:353 | 35329949:353 | +;+;+ | 794 | NA | 0 | 0 | 4 | 0 | 0 | 0 |
| ENSDARG000 | 10;10;10;10;1 | 45205407:452 | 45206826:452 | -;-;-;-;-;-;-;-; | 3703 | NA | 1395 | 880 | 1396 | 1192 | 919 | 820 |
| ENSDARG000 | 10;10;10;10;1 | 2715548:2725 | 2715792:2725 | +;+;+;+;+;+;+; | 2568 | grk5 | 10 | 6 | 2 | 0 | 2 | 0 |
| ENSDARG000 | 10;10;10;10;1 | 45148005:451 | 45148319:451 | +;+;+;+;+;+;+; | 6563 | OGDH | 1102 | 1035 | 1209 | 624 | 1308 | 769 |
| ENSDARG000 | 10;10;10;10;1 | 22161585:221 | 22166645:221 | -;-;-;-;- | 6893 | NA | 0 | 2 | 0 | 0 | 0 | 0 |
| ENSDARG000 | 10;10;10;10;1 | 38526496:385 | 38526598:385 | +;+;+;+;+;+;+; | 795 | acer3 | 48 | 59 | 140 | 39 | 118 | 48 |
| ENSDARG000 | 10;10;10;10;1 | 5788620:5794 | 5794761:5794 | -;-;-;-;-;-;-;-; | 8398 | il6st | 6999 | 3222 | 4478 | 2803 | 3439 | 4121 |
| ENSDARG000 | 10;10;10;10;1 | 35347993:353 | 35348122:353 | +;+;+;+;+;+;+; | 2166 | NA | 0 | 0 | 0 | 0 | 8 | 0 |
| ENSDARG000 | 10;10;10;10;1 | 38512261:385 | 38512434:385 | -;-;-;-;-;- | 1849 | NA | 1 | 0 | 6 | 0 | 2 | 4 |
| ENSDARG000 | 10;10;10;10;1 | 38512270:385 | 38512405:385 | +;+;+;+;+ | 3823 | serpinh1a | 3407 | 4145 | 3042 | 1512 | 3688 | 4167 |
| ENSDARG000 | 10;10;10;10;1 | 41986685:419 | 41986919:419 | +;+;+;+;+;+;+; | 8180 | setd1ba | 583 | 407 | 582 | 470 | 693 | 767 |
| ENSDARG000 | 10;10;10;10;1 | 28731749:287 | 28733190:287 | -;-;-;-;-;-;-;-; | 3371 | alcama | 140 | 147 | 756 | 885 | 667 | 115 |
| ENSDARG000 | 10;10;10;10;1 | 28587446:285 | 28587763:285 | +;+;+;+;+;+;+; | 6532 | CBLB | 4399 | 5093 | 7918 | 3893 | 10550 | 3662 |
| ENSDARG000 | 10;10;10;10;1 | 19026988:190 | 19027071:190 | +;+;+;+;+;+;+; | 9534 | NA | 2 | 14 | 10 | 6 | 30 | 14 |
| ENSDARG000 | 10;10;10;10;1 | 35367722:353 | 35368042:353 | -;-;-;-;-;-;-;-; | 1669 | gabrr3a | 6 | 0 | 0 | 0 | 0 | 0 |
| ENSDARG000 | 10;10;10;10;1 | 22483822:224 | 22484504:224 | -;-;-;-;-;-;-;- | 1247 | epob | 0 | 0 | 0 | 0 | 0 | 0 |
| ENSDARG000 | 10;10;10;10;1 | 15321522:153 | 15321756:153 | -;-;-;-;-;-;-;-; | 2360 | pum3 | 925 | 1137 | 1650 | 555 | 1003 | 768 |
| ENSDARG000 | 10;10;10;10;1 | 10972795:109 | 10973020:109 | +;+;+;+;+;+;+; | 2549 | cdc37l1 | 128 | 182 | 158 | 104 | 185 | 146 |
| ENSDARG000 | 10;10;10;10;1 | 22510280:225 | 22510935:225 | +;+;+;+;+;+;+; | 7835 | GIGYF1 | 1353 | 947 | 1291 | 915 | 1291 | 1383 |
| ENSDARG000 | 10;10;10;10 | 19310688:193 | 19310991:193 | +;+;+;+ | 2188 | LRRTM4 | 4 | 0 | 0 | 0 | 0 | 0 |
| ENSDARG000 | 10;10;10;10;1 | 19427474:194 | 19431790:194 | -;-;-;-;-;-;-;-; | 5609 | NA | 100 | 63 | 231 | 44 | 110 | 6 |
| ENSDARG000 | 10;10;10;10;1 | 8847033:8848 | 8847142:8848 | +;+;+;+;+;+;+ | 835 | NA | 1 | 4 | 0 | 0 | 4 | 14 |
| ENSDARG000 | 10;10;10;10;1 | 24643079:246 | 24643292:246 | +;+;+;+;+;+;+; | 5662 | vps36 | 549 | 519 | 553 | 268 | 572 | 528 |
| ENSDARG000 | 10;10;10;10;1 | 42027285:420 | 42027295:420 | -;-;-;-;-;-;-;-; | 905 | car15 | 0 | 4 | 0 | 0 | 2 | 0 |
| ENSDARG000 | 10;10;10;10;1 | 16501699:165 | 16502270:165 | +;+;+;+;+;+;+; | 2173 | slc27a6 | 45 | 44 | 110 | 70 | 35 | 96 |
| ENSDARG000 | 10;10;10;10;1 | 8862521:8864 | 8862764:8864 | +;+;+;+;+;+ | 828 | NA | 12 | 13 | 0 | 8 | 0 | 1 |
| ENSDARG000 | 10;10;10;10;1 | 24627683:246 | 24627887:246 | +;+;+;+;+;+;+; | 2530 | SLC46A3 | 2 | 0 | 11 | 4 | 3 | 0 |
| ENSDARG000 | 10;10;10;10;1 | 22536693:225 | 22536841:225 | +;+;+;+;+;+;+ | 2791 | NA | 4 | 2 | 24 | 2 | 14 | 3 |
| ENSDARG000 | 10;10;10 | 24678092:246 | 24678180:246 | +;+;+ | 840 | NA | 0 | 0 | 0 | 0 | 0 | 0 |
| ENSDARG000 | 10;10;10;10;1 | 24641229:246 | 24641727:246 | -;-;-;-;-;-;-;-; | 1193 | stoml3b | 82 | 113 | 343 | 88 | 211 | 73 |
| ENSDARG000 | 10;10;10;10;1 | 2759226:2762 | 2759499:2762 | -;-;-;-;-;-;-;-; | 2451 | ash2l | 740 | 919 | 1283 | 605 | 921 | 703 |
| ENSDARG000 | 10;10;10;10;1 | 11000894:110 | 11002061:110 | -;-;-;-;- | 1959 | ak3 | 214 | 378 | 356 | 444 | 371 | 353 |
| ENSDARG000 | 10;10;10;10;1 | 24682307:246 | 24687358:246 | -;-;-;-;-;-;-;-; | 5928 | fgl1b | 33 | 0 | 24 | 3 | 35 | 26 |
| ENSDARG000 | 10;10;10;10;1 | 8869468:8875 | 8869793:8875 | +;+;+;+;+;+;+; | 1713 | NA | 44 | 18 | 2 | 4 | 8 | 73 |
| ENSDARG000 | 10;10;10;10;1 | 24648903:246 | 24650972:246 | -;-;-;-;-;-;-;-; | 4855 | NA | 626 | 475 | 866 | 393 | 773 | 672 |
| ENSDARG000 | 10;10;10;10;1 | 42044220:420 | 42048632:420 | -;-;-;-;-;-;-;-; | 6027 | dgcr2 | 362 | 235 | 301 | 452 | 410 | 584 |
| ENSDARG000 | 10;10;10;10;1 | 35417099:354 | 35417248:354 | +;+;+;+;+;+;+; | 1855 | hhla2a.1 | 52 | 18 | 17 | 6 | 22 | 16 |
| ENSDARG000 | 10;10;10;10;1 | 19723025:197 | 19723367:197 | -;-;-;-;-;-;- | 1343 | GFRA2 | 0 | 0 | 0 | 0 | 0 | 0 |
| ENSDARG000 | 10;10;10;10;1 | 16547259:165 | 16547538:165 | +;+;+;+;+ | 2380 | isoc1 | 316 | 248 | 297 | 157 | 259 | 308 |
| ENSDARG000 | 10;10;10;10;1 | 33544329:335 | 33544577:335 | -;-;-;-;-;-;-;- | 1135 | il2rga | 52 | 66 | 38 | 20 | 120 | 95 |
| ENSDARG000 | 10;10;10;10;1 | 38775408:387 | 38776001:387 | +;+;+;+;+;+;+; | 7411 | dscama | 6 | 2 | 2 | 0 | 6 | 0 |

| | | | | | | | | | | | | |
|---|---|---|---|---|---|---|---|---|---|---|---|---|
| ENSDARG000 | 10:10:10:10:1 | 20070178:200 | 20070214:200 | +:+:+:+:+:+:+: | 5036 | xpo7 | 1165 | 1239 | 1650 | 972 | 1392 | 1349 |
| ENSDARG000 | 10:10:10:10:1 | 29087810:290 | 29092280:290 | -:-:-:-:-:-:-:-: | 5829 | NA | 0 | 0 | 0 | 0 | 0 | 0 |
| ENSDARG000 | 10:10:10:10:1 | 22724059:227 | 22724209:227 | +:+:+:+:+:+:+: | 5989 | kdm6bb | 864 | 895 | 2610 | 786 | 2015 | 1577 |
| ENSDARG000 | 10:10:10:10:1 | 35439952:354 | 35440253:354 | +;+;+;+;+;+;+ | 1765 | NA | 4 | 0 | 0 | 0 | 0 | 0 |
| ENSDARG000 | 10:10:10:10:1 | 20108557:201 | 20108639:201 | +:+:+:+:+:+:+: | 2483 | dmtn | 673 | 663 | 867 | 1124 | 910 | 942 |
| ENSDARG000 | 10;10;10 | 38969429:389 | 38970062:389 | -;-;- | 794 | pcp4a | 0 | 0 | 0 | 0 | 0 | 0 |
| ENSDARG000 | 10:10:10:10:1 | 5823759:5824 | 5823772:5824 | -:-:-:-:-:-:-:-: | 2578 | ANKRD55 | 0 | 0 | 0 | 0 | 0 | 0 |
| ENSDARG000 | 10:10:10:10:1 | 20103067:201 | 20103945:201 | -:-:-:-:-:-:-:-: | 1407 | pbdc1 | 1156 | 1330 | 1503 | 934 | 1079 | 1377 |
| ENSDARG000 | 10;10;10;10 | 20143797:201 | 20143850:201 | -;-;-;- | 509 | NA | 197 | 123 | 198 | 268 | 144 | 191 |
| ENSDARG000 | 10:10:10:10:1 | 29112756:291 | 29113612:291 | -:-:-:-:-:-:-:-: | 2073 | ZPLD1 | 0 | 0 | 0 | 0 | 0 | 2 |
| ENSDARG000 | 10:10:10:10:1 | 35468978:354 | 35469163:354 | +:+:+:+:+:+:+: | 4897 | PHLDB2 | 243 | 142 | 250 | 280 | 183 | 675 |
| ENSDARG000 | 10:10:10:10:1 | 33561567:335 | 33561619:335 | -:-:-:-:-:-:-:-: | 1168 | NA | 0 | 2 | 6 | 0 | 4 | 6 |
| ENSDARG000 | 10;10;10 | 29128202:291 | 29129068:291 | -;-;- | 1125 | NA | 0 | 0 | 0 | 0 | 0 | 0 |
| ENSDARG000 | 10:10:10:10:1 | 29137482:291 | 29138091:291 | +:+:+:+:+:+:+: | 4582 | picalma | 789 | 700 | 1073 | 585 | 793 | 756 |
| ENSDARG000 | 10:10:10:10:1 | 7029664:7050 | 7029772:7051 | +:+:+:+:+:+:+: | 6258 | NA | 3622 | 2251 | 2467 | 2338 | 1830 | 2567 |
| ENSDARG000 | 10:10:10:10:1 | 8885769:8890 | 8885829:8890 | +:+:+:+:+:+:+: | 3594 | ITGA2 | 53 | 2 | 2 | 2 | 3 | 10 |
| ENSDARG000 | 10:10:10:10:1 | 11038192:110 | 11038442:110 | +:+:+:+:+:+:+: | 3003 | ugcg | 3174 | 3187 | 3718 | 3946 | 4340 | 2759 |
| ENSDARG000 | 10 | 35537974 | 35542071 | - | 4098 | NA | 1634 | 660 | 1088 | 1275 | 593 | 1062 |
| ENSDARG000 | 10:10:10:10:1 | 26268790:262 | 26273155:262 | -:-:-:-:-:-:-:-: | 12141 | NA | 62 | 24 | 288 | 22 | 238 | 14 |
| ENSDARG000 | 10:10:10:10:1 | 5852359:5853 | 5853479:5853 | -;-;-;-;-;-;-;- | 2783 | NA | 0 | 0 | 2 | 0 | 0 | 0 |
| ENSDARG000 | 10:10:10:10:1 | 17077177:170 | 17077596:170 | -:-:-:-:-:-:-:-: | 1419 | fam166b | 0 | 0 | 0 | 0 | 0 | 8 |
| ENSDARG000 | 10:10:10:10:1 | 42089267:420 | 42090372:420 | -:-:-:-:-:-:-:-: | 2510 | ess2 | 751 | 770 | 973 | 709 | 644 | 672 |
| ENSDARG000 | 10:10:10:10:1 | 33573838:335 | 33573911:335 | +:+:+:+:+:+:+: | 1784 | cfap298 | 182 | 250 | 338 | 194 | 215 | 138 |
| ENSDARG000 | 10:10:10:10:1 | 26502334:265 | 26502430:265 | -:-:-:-:-:-:-:-: | 1953 | SLC46A3 | 4 | 26 | 22 | 26 | 24 | 10 |
| ENSDARG000 | 10:10:10:10:1 | 17088261:170 | 17088929:170 | +;+;+;+;+;+ | 1855 | NA | 28 | 0 | 0 | 0 | 0 | 6 |
| ENSDARG000 | 10:10:10:10:1 | 22771176:227 | 22771252:227 | +:+:+:+:+:+:+: | 2575 | tmem88a | 20 | 2 | 20 | 18 | 12 | 7 |
| ENSDARG000 | 10:10:10:10:1 | 35554219:355 | 35554367:355 | +:+:+:+:+:+:+: | 1730 | zdhhc20a | 616 | 320 | 315 | 209 | 282 | 385 |
| ENSDARG000 | 10:10:10:10:1 | 15381972:153 | 15383025:153 | -:-:-:-:-:-:-:-: | 2090 | sgtb | 358 | 345 | 377 | 692 | 379 | 441 |
| ENSDARG000 | 10:10:10:10:1 | 39028331:390 | 39031234:390 | -:-:-:-:-:-:-:-: | 4472 | NA | 0 | 0 | 0 | 0 | 0 | 6 |
| ENSDARG000 | 10:10:10:10:1 | 39091353:390 | 39091823:390 | +:+:+:+:+:+:+: | 2499 | NA | 344 | 371 | 545 | 338 | 502 | 407 |
| ENSDARG000 | 10:10:10:10:1 | 15340768:153 | 15341073:153 | +:+:+:+:+:+:+: | 1939 | trappc13 | 476 | 365 | 356 | 316 | 455 | 376 |
| ENSDARG000 | 10;10 | 17104298:171 | 17104695:171 | -;- | 959 | NA | 97 | 105 | 110 | 121 | 94 | 62 |
| ENSDARG000 | 10;10 | 2799285:2801 | 2799463:2801 | +;+ | 837 | pnx | 0 | 2 | 0 | 0 | 0 | 0 |
| ENSDARG000 | 10:10:10:10:1 | 5954787:5954 | 5954943:5954 | +:+:+:+:+:+:+: | 4190 | MAP3K1 | 1893 | 576 | 701 | 1038 | 1108 | 1565 |
| ENSDARG000 | 10:10:10:10:1 | 39084354:390 | 39084489:390 | +;+;+;+;+;+ | 1398 | im:7159173 | 3684 | 4380 | 7315 | 4112 | 5275 | 4761 |
| ENSDARG000 | 10:10:10:10:1 | 22782522:227 | 22782747:227 | +;+;+;+;+ | 980 | si:ch211-237l | 0 | 0 | 0 | 0 | 0 | 0 |
| ENSDARG000 | 10:10:10:10:1 | 26515946:265 | 26516036:265 | +:+:+:+:+:+:+: | 8377 | synj1 | 573 | 378 | 624 | 256 | 744 | 658 |
| ENSDARG000 | 10:10:10:10:1 | 8947228:8947 | 8947434:8947 | -:-:-:-:-:-:-:-: | 1383 | MOCS2 | 355 | 335 | 471 | 205 | 378 | 330 |
| ENSDARG000 | 10:10:10:10:1 | 22789319:227 | 22789874:227 | -:-:-:-:-:-:-:-: | 2964 | pcolcea | 332 | 6 | 18 | 20 | 140 | 0 |
| ENSDARG000 | 10:10:10:10:1 | 45248326:453 | 45248688:453 | +:+:+:+:+:+:+: | 4918 | ZMIZ2 | 990 | 552 | 928 | 531 | 696 | 596 |
| ENSDARG000 | 10:10:10:10:1 | 42111288:421 | 42112380:421 | -:-:-:-:-:-:-:-: | 2466 | stambpa | 2262 | 1861 | 2151 | 3212 | 1715 | 1926 |
| ENSDARG000 | 10:10:10:10:1 | 2807136:2807 | 2807610:2807 | +:+:+:+:+:+:+: | 4932 | aebp1a | 170 | 19 | 70 | 4 | 44 | 26 |
| ENSDARG000 | 10:10:10:10:1 | 35584647:355 | 35585843:355 | -:-:-:-:-:-:-:-: | 5701 | smg6 | 895 | 762 | 835 | 550 | 661 | 708 |
| ENSDARG000 | 10:10:10:10:1 | 33580285:335 | 33580787:335 | -:-:-:-:-:-:-:-: | 1676 | EVA1C | 8 | 1 | 5 | 4 | 2 | 8 |
| ENSDARG000 | 10:10:10:10:1 | 11146321:111 | 11146621:111 | -:-:-:-:-:-:-:-: | 1971 | PTBP3 | 295 | 243 | 580 | 428 | 406 | 317 |
| ENSDARG000 | 10:10:10:10:1 | 17135631:171 | 17136442:171 | -:-:-:-:-:-:-:-: | 2835 | slc5a1 | 6 | 0 | 2 | 24 | 86 | 0 |
| ENSDARG000 | 10:10:10:10:1 | 11260170:112 | 11261631:112 | +:+:+:+:+:+:+: | 5682 | hsdl2 | 691 | 956 | 1455 | 1144 | 1390 | 914 |
| ENSDARG000 | 10:10:10:10:1 | 42134256:421 | 42138446:421 | -:-:-:-:-:-:-:-: | 4992 | dusp11 | 535 | 384 | 863 | 250 | 756 | 326 |
| ENSDARG000 | 10:10:10:10:1 | 33588715:335 | 33588999:335 | +;+;+;+;+ | 1290 | MIS18A | 30 | 102 | 112 | 28 | 82 | 54 |
| ENSDARG000 | 10:10:10:10:1 | 26571174:265 | 26571463:265 | +:+:+:+:+:+:+: | 3731 | slc9a6b | 161 | 148 | 270 | 170 | 236 | 194 |
| ENSDARG000 | 10:10:10:10:1 | 11282883:112 | 11283065:112 | +:+:+:+:+:+:+: | 3738 | NA | 286 | 201 | 379 | 202 | 447 | 318 |
| ENSDARG000 | 10:10:10:10:1 | 29265463:292 | 29265623:292 | +;+;+;+;+;+ | 2612 | CREBZF | 787 | 627 | 1165 | 1044 | 1113 | 748 |
| ENSDARG000 | 10:10:10:10:1 | 7284121:7284 | 7285549:7285 | -:-:-:-:-:-:-:-: | 4190 | nrg1 | 24 | 0 | 0 | 0 | 8 | 4 |
| ENSDARG000 | 10:10:10:10:1 | 8968203:8968 | 8968374:8968 | +:+:+:+:+:+:+: | 2963 | fstb | 150 | 32 | 78 | 12 | 112 | 34 |
| ENSDARG000 | 10;10 | 22828366:228 | 22828529:228 | -;- | 264 | NA | 0 | 0 | 0 | 0 | 0 | 0 |
| ENSDARG000 | 10:10:10:10:1 | 22809715:228 | 22812721:228 | -:-:-:-:-:-:-:-: | 7377 | per1a | 1438 | 604 | 727 | 807 | 1860 | 1275 |
| ENSDARG000 | 10:10:10:10:1 | 33614377:336 | 33615020:336 | -:-:-:-:-:-:-:-: | 2101 | NA | 82 | 12 | 29 | 20 | 29 | 19 |
| ENSDARG000 | 10:10:10:10:1 | 42195732:421 | 42196663:421 | -:-:-:-:-:-:-:-: | 3091 | tcf7l1a | 928 | 421 | 731 | 598 | 797 | 854 |
| ENSDARG000 | 10;10 | 20158838:201 | 20159591:201 | -;- | 1111 | NA | 0 | 2 | 0 | 0 | 6 | 0 |
| ENSDARG000 | 10:10:10:10:1 | 26597990:266 | 26598107:266 | +:+:+:+:+:+:+: | 2202 | fhl1b | 1160 | 619 | 1228 | 838 | 512 | 1692 |
| ENSDARG000 | 10:10:10:10:1 | 2841205:2841 | 2841457:2841 | +:+:+:+:+:+:+: | 1668 | ykt6 | 543 | 602 | 685 | 469 | 670 | 508 |
| ENSDARG000 | 10:10:10:10:1 | 29431529:294 | 29431684:294 | +:+:+:+:+:+:+: | 2788 | dlg2 | 438 | 2 | 45 | 12 | 10 | 44 |
| ENSDARG000 | 10:10:10:10:1 | 6010570:6010 | 6010698:6010 | +:+:+:+:+:+:+: | 3436 | hmgcs1 | 561 | 674 | 977 | 252 | 456 | 372 |
| ENSDARG000 | 10:10:10:10:1 | 9006574:9037 | 9006726:9037 | -:-:-:-:-:-:-:-: | 1236 | ARL15 | 236 | 135 | 189 | 183 | 284 | 142 |
| ENSDARG000 | 10:10:10:10:1 | 20180163:201 | 20180518:201 | +:+:+:+:+:+:+: | 2233 | ppp3ccb | 899 | 773 | 747 | 614 | 750 | 624 |
| ENSDARG000 | 10:10:10:10:1 | 39125637:391 | 39125692:391 | -:-:-:-:-:-:-:-: | 573 | rps25 | 10251 | 10203 | 15385 | 14097 | 9810 | 12778 |
| ENSDARG000 | 10;10 | 9092815:9097 | 9094446:9097 | +;+ | 2405 | snx18b | 139 | 185 | 140 | 56 | 191 | 64 |
| ENSDARG000 | 10;10 | 17166035:171 | 17168110:171 | -;- | 2482 | ywhah | 976 | 1258 | 2065 | 796 | 3174 | 810 |
| ENSDARG000 | 10:10:10:10:1 | 42358426:423 | 42358623:423 | +:+:+:+:+:+:+: | 2376 | dbnla | 42 | 8 | 0 | 0 | 20 | 2 |
| ENSDARG000 | 10:10:10:10:1 | 26634472:266 | 26634620:266 | +:+:+:+:+:+:+: | 2345 | NA | 0 | 0 | 4 | 0 | 2 | 2 |
| ENSDARG000 | 10:10:10:10:1 | 9105007:9107 | 9107457:9107 | -;-;-;-;-;-;- | 3523 | NA | 244 | 108 | 86 | 63 | 190 | 74 |
| ENSDARG000 | 10;10;10;10 | 29731060:297 | 29731288:297 | -;-;-;- | 568 | NA | 0 | 0 | 2 | 0 | 0 | 6 |
| ENSDARG000 | 10:10:10:10:1 | 33626344:336 | 33626485:336 | +:+:+:+:+:+:+: | 2038 | b3glctb | 0 | 0 | 0 | 0 | 0 | 0 |
| ENSDARG000 | 10:10:10:10:1 | 20270831:202 | 20271004:202 | -:-:-:-:-:-:-:-: | 1308 | ZMAT4 | 0 | 0 | 0 | 0 | 0 | 0 |
| ENSDARG000 | 10:10:10:10:1 | 15408501:154 | 15408729:154 | +:+:+:+:+:+:+: | 6060 | nln | 85 | 679 | 1099 | 1022 | 287 | 215 |
| ENSDARG000 | 10:10:10:10:1 | 42374770:423 | 42374897:423 | +:+:+:+:+:+:+: | 643 | cox5ba | 3129 | 4111 | 3797 | 2536 | 3077 | 3182 |
| ENSDARG000 | 10:10:10:10:1 | 39133410:391 | 39136844:391 | -:-:-:-:-:-:-:-: | 5322 | NA | 22 | 120 | 149 | 132 | 88 | 20 |
| ENSDARG000 | 10:10:10:10:1 | 39199547:392 | 39199692:392 | +:+:+:+:+:+:+: | 2036 | ei24 | 522 | 372 | 375 | 256 | 459 | 492 |
| ENSDARG000 | 10;10 | 35651494:356 | 35651792:356 | -;- | 434 | NA | 4 | 2 | 0 | 2 | 4 | 1 |
| ENSDARG000 | 10:10:10:10:1 | 9159279:9164 | 9159609:9164 | +:+:+:+:+:+:+: | 2173 | rasgef1bb | 306 | 242 | 285 | 270 | 264 | 298 |
| ENSDARG000 | 10:10:10:10:1 | 22838364:228 | 22838718:228 | -;-;-;-;- | 897 | vamp2 | 17 | 3 | 17 | 17 | 8 | 24 |
| ENSDARG000 | 10;10 | 35664129:356 | 35664756:356 | -;- | 1013 | ovca2 | 34 | 76 | 97 | 86 | 136 | 116 |
| ENSDARG000 | 10;10;10 | 20351413:203 | 20352198:203 | -;-;- | 1527 | sfrp1b | 108 | 4 | 20 | 0 | 58 | 28 |
| ENSDARG000 | 10:10:10:10:1 | 2856377:2859 | 2856767:2859 | -:-:-:-:-:-:-:-: | 3067 | ddx56 | 338 | 366 | 663 | 280 | 618 | 435 |
| ENSDARG000 | 10:10:10:10:1 | 29735900:297 | 29736461:297 | -:-:-:-:-:-:-:-: | 1605 | ift46 | 66 | 48 | 144 | 132 | 72 | 72 |
| ENSDARG000 | 10:10:10:10:1 | 7483545:7483 | 7483836:7483 | -:-:-:-:-:-:-:-: | 4735 | WRN | 196 | 130 | 308 | 131 | 344 | 238 |
| ENSDARG000 | 10:10:10:10:1 | 22848556:228 | 22849355:228 | -;-;-;-;- | 1211 | and3 | 8 | 2 | 2 | 10 | 0 | 2 |
| ENSDARG000 | 10:10:10:10:1 | 22854803:228 | 22855097:228 | +:+:+:+:+:+:+: | 1909 | ap4m1 | 178 | 171 | 168 | 84 | 150 | 228 |
| ENSDARG000 | 10:10:10:10:1 | 35952532:359 | 35952553:359 | +;+;+;+;+;+ | 1345 | RTN4RL1 | 0 | 0 | 8 | 0 | 2 | 2 |
| ENSDARG000 | 10;10;10;10 | 36150911:361 | 36150932:361 | +;+;+;+ | 1365 | waif2 | 6 | 0 | 0 | 0 | 0 | 0 |
| ENSDARG000 | 10:10:10:10:1 | 20364009:203 | 20364435:203 | +:+:+:+:+:+:+: | 3707 | golga7 | 1824 | 1692 | 2120 | 2025 | 2037 | 2237 |
| ENSDARG000 | 10;10;10 | 42423318:424 | 42423475:424 | +;+;+ | 1441 | npy8ar | 0 | 0 | 0 | 0 | 0 | 0 |
| ENSDARG000 | 10;10 | 36158967:361 | 36160287:361 | +;+ | 1321 | NA | 0 | 0 | 0 | 0 | 0 | 0 |
| ENSDARG000 | 10:10:10:10:1 | 17171604:171 | 17172037:171 | -:-:-:-:-:-:-:-: | 5636 | DEPDC5 | 1763 | 942 | 1681 | 1153 | 1765 | 1886 |
| ENSDARG000 | 10:10:10:10:1 | 39212601:392 | 39212970:392 | +:+:+:+:+:+:+: | 3511 | stt3a | 5496 | 6049 | 8266 | 5255 | 7409 | 5153 |
| ENSDARG000 | 10:10:10:10:1 | 29751029:297 | 29751667:297 | -:-:-:-:-:-:-:-: | 3308 | vps11 | 1526 | 986 | 968 | 711 | 1178 | 1201 |
| ENSDARG000 | 10:10:10:10:1 | 9186651:9190 | 9187618:9190 | -;-;-;-;- | 2083 | NA | 24 | 44 | 100 | 101 | 41 | 62 |
| ENSDARG000 | 10;10 | 36178713:361 | 36178774:361 | +;+ | 1224 | or41a2 | 0 | 0 | 0 | 0 | 0 | 0 |
| ENSDARG000 | 10:10:10:10:1 | 42500895:425 | 42501493:425 | -;-;-;-;-;-;- | 1520 | marchf5l | 811 | 831 | 920 | 746 | 772 | 656 |
| ENSDARG000 | 10 | 36184751 | 36186346 | + | 1596 | or41a1 | 0 | 0 | 0 | 0 | 0 | 0 |
| ENSDARG000 | 10:10:10:10:1 | 9195190:9195 | 9195594:9195 | +:+:+:+:+:+:+: | 1732 | antxr2b | 300 | 688 | 782 | 869 | 648 | 308 |
| ENSDARG000 | 10 | 36194527 | 36196218 | + | 1692 | or41a3 | 0 | 0 | 0 | 0 | 0 | 0 |
| ENSDARG000 | 10;10 | 36197968:361 | 36198047:362 | +;+ | 510 | coa4 | 46 | 40 | 147 | 130 | 92 | 43 |
| ENSDARG000 | 10:10:10:10:1 | 29770120:297 | 29770435:297 | +:+:+:+:+:+:+: | 4856 | hyou1 | 1575 | 2953 | 6217 | 1086 | 2946 | 1321 |
| ENSDARG000 | 10:10:10:10:1 | 33744098:337 | 33744206:337 | +:+:+:+:+:+:+: | 2988 | rxfp2a | 5 | 4 | 2 | 0 | 197 | 16 |
| ENSDARG000 | 10:10:10:10:1 | 26652859:266 | 26652941:266 | +:+:+:+:+:+:+: | 1624 | htatsf1 | 509 | 593 | 624 | 462 | 446 | 467 |
| ENSDARG000 | 10:10:10:10:1 | 26667475:266 | 26667518:266 | +:+:+:+:+:+:+: | 966 | NA | 27 | 5 | 3 | 9 | 14 | 17 |
| ENSDARG000 | 10:10:10:10:1 | 6121558:6121 | 6121662:6121 | +:+:+:+:+:+:+: | 8753 | tln1 | 6866 | 5202 | 8046 | 4904 | 7468 | 5948 |
| ENSDARG000 | 10;10;10 | 7563755:7564 | 7565074:7564 | +;+;+ | 1320 | PURG | 15 | 14 | 32 | 8 | 104 | 18 |
| ENSDARG000 | 10:10:10:10:1 | 20392083:203 | 20392943:203 | +:+:+:+:+:+:+: | 2859 | NA | 192 | 222 | 248 | 192 | 230 | 262 |
| ENSDARG000 | 10 | 36299541 | 36300976 | + | 1436 | or80a7 | 0 | 0 | 0 | 0 | 0 | 0 |
| ENSDARG000 | 10;10;10;10 | 7573316:7575 | 7573359:7575 | +;+;+;+ | 575 | NA | 0 | 10 | 3 | 10 | 19 | 11 |
| ENSDARG000 | 10;10;10;10 | 11319971:113 | 11320222:113 | -;-;-;- | 618 | inip | 66 | 125 | 185 | 72 | 83 | 108 |
| ENSDARG000 | 10 | 36285568 | 36287107 | + | 1540 | or80a11 | 0 | 0 | 0 | 0 | 0 | 0 |
| ENSDARG000 | 10:10:10:10:1 | 36363318:363 | 36364927:363 | -:-:-:-:-:-:-:-:-: | 2670 | ubl3a | 3426 | 2740 | 5076 | 3330 | 5965 | 3724 |
| ENSDARG000 | 10 | 36309072 | 36310557 | + | 1486 | or80a9 | 0 | 0 | 0 | 0 | 0 | 0 |
| ENSDARG000 | 10 | 36294641 | 36296945 | + | 2305 | or80a5 | 0 | 0 | 0 | 0 | 0 | 0 |
| ENSDARG000 | 10 | 36314279 | 36316758 | + | 2480 | or80a4 | 0 | 0 | 0 | 0 | 0 | 0 |
| ENSDARG000 | 10 | 36221505 | 36223005 | - | 1501 | or109-6 | 0 | 0 | 0 | 0 | 0 | 0 |
| ENSDARG000 | 10 | 36345176 | 36346363 | + | 1188 | or30bv1 | 0 | 0 | 0 | 0 | 0 | 0 |
| ENSDARG000 | 10;10;10 | 36251987:362 | 36253988:362 | -;-;- | 2098 | or42b1 | 0 | 0 | 0 | 0 | 0 | 0 |
| ENSDARG000 | 10 | 36322894 | 36325144 | + | 2251 | or80a8 | 0 | 0 | 0 | 0 | 0 | 0 |
| ENSDARG000 | 10;10 | 36205211:362 | 36206384:362 | -;- | 1367 | NA | 0 | 0 | 0 | 0 | 0 | 0 |
| ENSDARG000 | 10 | 36234985 | 36236610 | - | 1626 | or42a5 | 0 | 0 | 0 | 0 | 0 | 0 |
| ENSDARG000 | 10 | 36302967 | 36304931 | + | 1965 | or80a3 | 0 | 0 | 0 | 0 | 0 | 0 |
| ENSDARG000 | 10 | 36287841 | 36288503 | + | 663 | NA | 0 | 0 | 0 | 0 | 0 | 0 |
| ENSDARG000 | 10 | 36328702 | 36330485 | + | 1784 | or80a1 | 0 | 0 | 0 | 0 | 0 | 0 |
| ENSDARG000 | 10 | 36334502 | 36336537 | + | 2036 | or80a13 | 0 | 0 | 0 | 0 | 0 | 0 |
| ENSDARG000 | 10 | 36230602 | 36231741 | - | 1140 | or42a6 | 0 | 0 | 0 | 0 | 0 | 0 |
| ENSDARG000 | 10:10:10:10:1 | 20448638:204 | 20449085:204 | -;-;-;-;-;-;-;- | 1205 | NA | 0 | 50 | 54 | 189 | 20 | 55 |
| ENSDARG000 | 10 | 36242891 | 36244864 | - | 1974 | or42a1 | 0 | 0 | 0 | 0 | 0 | 0 |
| ENSDARG000 | 10 | 36265164 | 36267439 | + | 2276 | or80a2 | 0 | 0 | 0 | 0 | 0 | 0 |
| ENSDARG000 | 10;10;10 | 7585928:7587 | 7586165:7587 | +;+;+ | 777 | NA | 4 | 17 | 23 | 5 | 55 | 41 |
| ENSDARG000 | 10;10 | 36354863:363 | 36354936:363 | +;+ | 1533 | or30bu1 | 0 | 0 | 0 | 0 | 0 | 0 |
| ENSDARG000 | 10;10;10 | 36349225:363 | 36349294:363 | +;+;+ | 1469 | or30bu2 | 0 | 0 | 0 | 0 | 0 | 0 |
| ENSDARG000 | 10 | 36224470 | 36225729 | - | 1260 | or42a2 | 0 | 0 | 0 | 0 | 0 | 0 |
| ENSDARG000 | 10;10 | 36259048:362 | 36260042:362 | -;- | 1133 | or42b2 | 0 | 0 | 0 | 0 | 0 | 0 |
| ENSDARG000 | 10 | 36217239 | 36220061 | - | 2823 | or42a7 | 0 | 0 | 0 | 0 | 0 | 0 |
| ENSDARG000 | 10 | 36277528 | 36279141 | + | 1614 | or80a10 | 0 | 0 | 0 | 0 | 0 | 0 |
| ENSDARG000 | 10 | 36237981 | 36240806 | - | 2826 | or42a8 | 0 | 0 | 0 | 0 | 0 | 0 |
| ENSDARG000 | 10 | 36212575 | 36214582 | - | 2008 | or42a4 | 0 | 0 | 0 | 0 | 0 | 0 |
| ENSDARG000 | 10:10:10:10:1 | 36201194:362 | 36201226:362 | +:+:+:+:+:+:+: | 1274 | NA | 2 | 0 | 0 | 2 | 0 | 0 |
| ENSDARG000 | 10:10:10:10:1 | 9222476:9222 | 9222645:9222 | +:+:+:+:+:+:+: | 2670 | paqr3b | 172 | 209 | 206 | 126 | 195 | 90 |
| ENSDARG000 | 10:10:10:10:1 | 15454745:154 | 15455290:154 | +:+:+:+:+:+:+: | 10330 | erbin | 1563 | 2121 | 2938 | 2986 | 2203 | 2323 |
| ENSDARG000 | 10 | 36272178 | 36274889 | + | 2712 | or80a6 | 0 | 0 | 0 | 0 | 0 | 0 |
| ENSDARG000 | 10:10:10:10:1 | 26689262:266 | 26689610:266 | +;+;+;+;+;+ | 1074 | tmtops3b | 0 | 0 | 0 | 0 | 0 | 0 |
| ENSDARG000 | 10:10:10:10:1 | 42521943:425 | 42522163:425 | +:+:+:+:+:+:+: | 2540 | actr1b | 8593 | 8897 | 7546 | 8009 | 8269 | 7288 |
| ENSDARG000 | 10:10:10:10:1 | 7593185:7593 | 7593641:7593 | +:+:+:+:+:+:+: | 2021 | ppp2cb | 5431 | 5772 | 5481 | 4502 | 5167 | 4063 |
| ENSDARG000 | 10:10:10:10:1 | 26716362:267 | 26717991:267 | -:-:-:-:-:-:-:-: | 3575 | fgf13b | 4 | 0 | 14 | 4 | 0 | 22 |
| ENSDARG000 | 10 | 29816681 | 29817589 | + | 909 | h2ax1 | 3411 | 1921 | 2446 | 1421 | 2765 | 1708 |
| ENSDARG000 | 10 | 29815644 | 29816467 | - | 824 | hist2h2l | 1286 | 744 | 1043 | 715 | 1642 | 803 |
| ENSDARG000 | 10:10:10:10:1 | 29793817:297 | 29794192:297 | -:-:-:-:-:-:-:-: | 1658 | NA | 2 | 0 | 0 | 0 | 0 | 0 |
| ENSDARG000 | 10:10:10:10:1 | 24690534:246 | 24690855:246 | +:+:+:+:+:+:+: | 2629 | tpte | 1008 | 388 | 552 | 430 | 714 | 873 |
| ENSDARG000 | 10:10:10:10:1 | 20503886:205 | 20503961:205 | -:-:-:-:-:-:-:-: | 2266 | DDHD2 | 75 | 85 | 105 | 69 | 101 | 93 |
| ENSDARG000 | 10:10:10:10:1 | 39264802:392 | 39269245:392 | -:-:-:-:-:-:-:-: | 6084 | cry5 | 54 | 51 | 76 | 29 | 139 | 69 |
| ENSDARG000 | 10;10;10;10 | 22912251:229 | 22912336:229 | +;+;+;+ | 706 | med11 | 153 | 178 | 218 | 171 | 160 | 80 |
| ENSDARG000 | 10:10:10:10:1 | 22890791:228 | 22891285:228 | +:+:+:+:+:+:+: | 2367 | arrb2a | 355 | 301 | 396 | 244 | 325 | 379 |
| ENSDARG000 | 10:10:10:10:1 | 39248911:392 | 39249180:392 | +:+:+:+:+:+:+: | 2753 | foxred1 | 28 | 51 | 46 | 4 | 86 | 20 |
| ENSDARG000 | 10:10:10:10:1 | 29820172:298 | 29820575:298 | -:-:-:-:-:-:-:-: | 2020 | zpr1 | 540 | 718 | 911 | 609 | 813 | 709 |
| ENSDARG000 | 10:10:10:10:1 | 22907351:229 | 22907921:229 | -;-;-;-;-;- | 1342 | si:ch1073-143 | 97 | 32 | 84 | 36 | 80 | 95 |
| ENSDARG000 | 10:10:10:10:1 | 26747755:267 | 26747901:267 | +:+:+:+:+:+:+: | 1819 | f9b | 0 | 2 | 7 | 12 | 0 | 8 |
| ENSDARG000 | 10:10:10:10:1 | 7623698:7624 | 7624236:7624 | -:-:-:-:-:-:-:-: | 1236 | ubxn8 | 173 | 89 | 106 | 90 | 80 | 144 |

| | | | | | | | | | | | | |
|---|---|---|---|---|---|---|---|---|---|---|---|---|
| ENSDARG000 | 10;10;10 | 33876131:338 | 33876567:338 | -;-;- | 948 | NA | 0 | 0 | 0 | 0 | 0 | 0 |
| ENSDARG000 | 10:10:10:10:1 | 2876020:2876 | 2876090:2876 | +:+:+:+:+:+:+: | 3156 | NA | 270 | 233 | 301 | 227 | 326 | 230 |
| ENSDARG000 | 10:10:10:10:1 | 11324515:113 | 11328452:113 | -:-:-:-:-:-:-:-: | 6967 | lin54 | 234 | 519 | 520 | 236 | 526 | 332 |
| ENSDARG000 | 10:10:10:10:1 | 9247593:9255 | 9248183:9255 | -:-:-:-:-:-:-:-: | 1864 | anxa3b | 1194 | 2455 | 1923 | 2265 | 2328 | 1493 |
| ENSDARG000 | 10:10:10:10:1 | 29849497:298 | 29849582:298 | +:+:+:+:+:+:+: | 6877 | hspa8 | 70011 | 50676 | 89790 | 52531 | 65286 | 76789 |
| ENSDARG000 | 10:10:10:10:1 | 29855213:298 | 29855433:298 | +:+:+:+:+:+:+: | 3220 | jhy | 8 | 6 | 8 | 0 | 2 | 13 |
| ENSDARG000 | 10;10;10 | 7636811:7637 | 7636967:7637 | +;+;+ | 564 | hint1 | 1016 | 1373 | 1638 | 1193 | 1342 | 1210 |
| ENSDARG000 | 10:10:10:10:1 | 20527427:205 | 20527949:205 | -:-:-:-:-:-:-:-: | 1468 | plpp5 | 382 | 349 | 595 | 412 | 457 | 390 |
| ENSDARG000 | 10:10:10:10:1 | 39283985:392 | 39284188:392 | +:+:+:+:+:+:+: | 3163 | dcps | 56 | 200 | 110 | 28 | 153 | 128 |
| ENSDARG000 | 10:10:10:10:1 | 33895315:339 | 33895503:339 | +:+:+:+:+:+:+: | 12160 | FRY | 30 | 36 | 208 | 194 | 103 | 6 |
| ENSDARG000 | 10:10:10:10:1 | 26771047:267 | 26771662:267 | -:-:-:-:-:-:-:-: | 3341 | MCF2 | 0 | 0 | 0 | 0 | 0 | 0 |
| ENSDARG000 | 10:10:10:10:1 | 42542517:425 | 42542675:425 | +:+:+:+:+:+:+: | 1621 | kctd9b | 299 | 214 | 375 | 338 | 339 | 286 |
| ENSDARG000 | 10;10;10;10 | 26762793:267 | 26763112:267 | -;-;-;- | 1495 | NA | 0 | 4 | 0 | 0 | 0 | 0 |
| ENSDARG000 | 10;10 | 26766385:267 | 26766485:267 | +;+ | 210 | NA | 0 | 0 | 0 | 0 | 0 | 0 |
| ENSDARG000 | 10;10 | 39304422:393 | 39304489:393 | +;+ | 1527 | kcnj1b | 12 | 15 | 14 | 8 | 14 | 13 |
| ENSDARG000 | 10:10:10:10:1 | 17223681:172 | 17224040:172 | -:-:-:-:-:-:-:-: | 1182 | rab36 | 10 | 22 | 2 | 9 | 17 | 6 |
| ENSDARG000 | 10:10:10:10:1 | 11355841:113 | 11356025:113 | +:+:+:+:+:+:+: | 5900 | cops4 | 1136 | 1505 | 1779 | 1484 | 1602 | 1200 |
| ENSDARG000 | 10:10:10:10:1 | 7703251:7704 | 7703419:7704 | +:+:+:+:+:+:+: | 4191 | GGCX | 718 | 692 | 1317 | 947 | 1008 | 883 |
| ENSDARG000 | 10:10:10:10:1 | 39314578:393 | 39314902:393 | -;-;-;-;- | 857 | smtlb | 2 | 0 | 0 | 0 | 0 | 0 |
| ENSDARG000 | 10:10:10:10:1 | 11371961:113 | 11372532:113 | -;-;-;-;-;-;- | 827 | plac8.2 | 0 | 0 | 0 | 0 | 0 | 0 |
| ENSDARG000 | 10:10:10:10:1 | 39476432:394 | 39476958:394 | +:+:+:+:+:+:+: | 6231 | KIRREL3 | 1900 | 110 | 200 | 174 | 202 | 336 |
| ENSDARG000 | 10:10:10:10:1 | 42589391:425 | 42589860:425 | +:+:+:+:+:+:+: | 2965 | fgfr1b | 39 | 16 | 59 | 16 | 85 | 2 |
| ENSDARG000 | 10;10;10 | 29890535:298 | 29891067:298 | -;-;- | 977 | bsx | 0 | 0 | 0 | 0 | 0 | 0 |
| ENSDARG000 | 10:10:10:10:1 | 24709967:247 | 24711263:247 | -;-;-;-;-;- | 2933 | smpd1 | 2054 | 745 | 785 | 676 | 1185 | 1237 |
| ENSDARG000 | 10:10:10:10:1 | 24722580:247 | 24726934:247 | -:-:-:-:-:-:-:-: | 6630 | ilk | 2167 | 2423 | 3232 | 2984 | 2588 | 3141 |
| ENSDARG000 | 10:10:10:10:1 | 7727281:7730 | 7727723:7730 | -;-;-;-;-;-;- | 1452 | snx24 | 52 | 53 | 65 | 46 | 54 | 52 |
| ENSDARG000 | 10:10:10:10:1 | 29896275:298 | 29896444:298 | -;-;-;-;-;-;-;- | 763 | lim2.1 | 0 | 0 | 0 | 0 | 0 | 0 |
| ENSDARG000 | 10:10:10:10:1 | 17460358:174 | 17460938:174 | -:-:-:-:-:-:-:-: | 2682 | nt5c2l1 | 30 | 32 | 40 | 10 | 111 | 70 |
| ENSDARG000 | 10;10 | 29910622:299 | 29910712:299 | +;+ | 181 | NA | 0 | 0 | 0 | 0 | 0 | 0 |
| ENSDARG000 | 10;10;10 | 9281991:9284 | 9282058:9284 | +;+;+ | 792 | NA | 0 | 6 | 5 | 0 | 0 | 0 |
| ENSDARG000 | 10:10:10:10:1 | 62369:63206: | 63736:63736: | -:-:-:-:-:-:-:-: | 3838 | dyrk1aa | 1490 | 1102 | 1665 | 1051 | 1448 | 1115 |
| ENSDARG000 | 10:10:10:10:1 | 42930353:429 | 42931519:429 | -:-:-:-:-:-:-:-: | 2664 | NA | 21 | 108 | 15 | 20 | 303 | 69 |
| ENSDARG000 | 10:10:10:10:1 | 26800213:268 | 26800292:268 | +:+:+:+:+:+:+: | 1724 | arr3a | 4 | 0 | 0 | 0 | 0 | 0 |
| ENSDARG000 | 10;10;10 | 22916113:229 | 22916784:229 | -;-;- | 943 | rnasekb | 206 | 85 | 126 | 142 | 104 | 256 |
| ENSDARG000 | 10:10:10:10:1 | 22918338:229 | 22918601:229 | +:+:+:+:+:+:+: | 1961 | C10H17orf49 | 234 | 336 | 399 | 115 | 289 | 243 |
| ENSDARG000 | 10:10:10:10:1 | 15603082:156 | 15603193:156 | +:+:+:+:+:+:+: | 2588 | zfand5b | 1126 | 1177 | 1306 | 1312 | 1150 | 724 |
| ENSDARG000 | 10:10:10:10:1 | 39893439:398 | 39893766:398 | +;+;+;+;+;+;+ | 1248 | smfn | 292 | 363 | 435 | 234 | 364 | 234 |
| ENSDARG000 | 10:10:10:10:1 | 7738540:7738 | 7739055:7739 | -:-:-:-:-:-:-:-: | 1930 | loxa | 42 | 0 | 14 | 6 | 14 | 10 |
| ENSDARG000 | 10:10:10:10:1 | 26813710:268 | 26813822:268 | +;+;+;+;+;+ | 878 | NA | 0 | 0 | 0 | 0 | 0 | 0 |
| ENSDARG000 | 10:10:10:10:1 | 2899108:2902 | 2899816:2902 | +:+:+:+:+:+:+: | 3323 | ERAP1 | 940 | 686 | 691 | 448 | 915 | 516 |
| ENSDARG000 | 10:10:10:10:1 | 9372702:9373 | 9373079:9373 | +:+:+:+:+:+:+: | 3278 | lhx6a | 44 | 25 | 31 | 46 | 25 | 4 |
| ENSDARG000 | 10:10:10:10:1 | 20543279:205 | 20543556:205 | -:-:-:-:-:-:-:-: | 5089 | NSD3 | 382 | 181 | 484 | 135 | 337 | 380 |
| ENSDARG000 | 10:10:10:10:1 | 9305050:9308 | 9305605:9308 | -;-;-;-;-;-;-;- | 1446 | mrrf | 184 | 273 | 324 | 272 | 275 | 181 |
| ENSDARG000 | 10:10:10:10:1 | 9346383:9346 | 9346550:9346 | +:+:+:+:+:+:+: | 2367 | rbm18 | 183 | 221 | 190 | 135 | 188 | 174 |
| ENSDARG000 | 10:10:10:10:1 | 29963518:299 | 29963746:299 | +:+:+:+:+:+:+: | 1523 | ntm | 0 | 0 | 0 | 0 | 0 | 0 |
| ENSDARG000 | 10;10 | 23022263:230 | 23022441:230 | +;+ | 3278 | NA | 20 | 16 | 22 | 52 | 13 | 22 |
| ENSDARG000 | 10;10;10;10 | 26827113:268 | 26827197:268 | +;+;+;+ | 955 | znrd2 | 434 | 502 | 714 | 411 | 635 | 357 |
| ENSDARG000 | 10;10;10 | 23034329:230 | 23034628:230 | +;+;+ | 705 | NA | 0 | 0 | 2 | 0 | 11 | 0 |
| ENSDARG000 | 10:10:10:10:1 | 17485945:174 | 17486781:174 | -:-:-:-:-:-:-:-: | 2163 | slc2a11l | 0 | 0 | 2 | 2 | 20 | 0 |
| ENSDARG000 | 10;10 | 26834985:268 | 26835649:268 | +;+ | 959 | FAM89B | 56 | 30 | 24 | 76 | 22 | 29 |
| ENSDARG000 | 10:10:10:10:1 | 40160502:401 | 40160589:401 | +:+:+:+:+:+:+: | 4589 | GRAMD1B | 170 | 213 | 340 | 300 | 268 | 375 |
| ENSDARG000 | 10:10:10:10:1 | 11382365:113 | 11382907:113 | -;-;-;-;-;-;- | 792 | plac8.1 | 0 | 0 | 0 | 0 | 0 | 0 |
| ENSDARG000 | 10:10:10:10:1 | 43037064:430 | 43037141:430 | +:+:+:+:+:+:+: | 4528 | atg10 | 153 | 56 | 129 | 166 | 102 | 125 |
| ENSDARG000 | 10:10:10:10:1 | 34001444:340 | 34001854:340 | +:+:+:+:+:+:+: | 7860 | KL | 10 | 0 | 1 | 18 | 6 | 8 |
| ENSDARG000 | 10 | 7769814 | 7772434 | - | 2621 | NA | 206 | 282 | 282 | 134 | 184 | 252 |
| ENSDARG000 | 10:10:10:10:1 | 34021368:340 | 34024501:340 | -:-:-:-:-:-:-:-: | 4950 | rfc3 | 75 | 228 | 323 | 155 | 169 | 186 |
| ENSDARG000 | 10:10:10:10:1 | 23060391:230 | 23060753:230 | +:+:+:+:+:+:+: | 1909 | slc25a1a | 242 | 231 | 218 | 141 | 148 | 207 |
| ENSDARG000 | 10:10:10:10:1 | 76864:77357: | 76954:77500: | +:+:+:+:+:+:+: | 1297 | vps26c | 1463 | 1210 | 1436 | 1293 | 1407 | 1233 |
| ENSDARG000 | 10:10:10:10:1 | 6318227:6318 | 6318461:6318 | +:+:+:+:+:+:+: | 4909 | tpm2 | 314 | 174 | 311 | 244 | 351 | 266 |
| ENSDARG000 | 10:10:10:10:1 | 26871486:268 | 26872294:268 | +:+:+:+:+:+:+: | 10028 | NA | 2997 | 1049 | 1599 | 1971 | 1397 | 2438 |
| ENSDARG000 | 10:10:10:10:1 | 15617741:156 | 15618497:156 | -:-:-:-:-:-:-:-: | 4412 | smc5 | 831 | 647 | 1054 | 680 | 750 | 687 |
| ENSDARG000 | 10:10:10:10:1 | 11394528:113 | 11394958:113 | -;-;-;-;- | 822 | srek1ip1 | 81 | 88 | 146 | 62 | 115 | 60 |
| ENSDARG000 | 10;10 | 26892567:268 | 26893243:268 | -;- | 766 | NA | 0 | 0 | 0 | 1 | 0 | 0 |
| ENSDARG000 | 10:10:10:10:1 | 43088270:431 | 43088448:431 | +:+:+:+:+:+:+: | 1936 | xrcc4 | 80 | 306 | 280 | 164 | 273 | 192 |
| ENSDARG000 | 10:10:10:10:1 | 11398111:113 | 11398332:113 | +:+:+:+:+:+:+: | 1874 | cwc27 | 272 | 230 | 411 | 160 | 338 | 248 |
| ENSDARG000 | 10:10:10:10:1 | 30617826:306 | 30618448:306 | -;-;-;-;-;-;- | 2044 | opcml | 2 | 0 | 0 | 0 | 0 | 0 |
| ENSDARG000 | 10:10:10:10:1 | 36439293:364 | 36439412:364 | +:+:+:+:+:+:+: | 3847 | uspl1 | 687 | 585 | 1459 | 622 | 1498 | 968 |
| ENSDARG000 | 10:10:10:10:1 | 43110788:431 | 43111649:431 | -:-:-:-:-:-:-:-: | 1109 | tmem167a | 411 | 571 | 591 | 370 | 547 | 486 |
| ENSDARG000 | 10;10;10 | 24762216:247 | 24762914:247 | -;-;- | 999 | timm10b | 96 | 120 | 140 | 56 | 128 | 124 |
| ENSDARG000 | 10:10:10:10:1 | 30997238:309 | 31000094:310 | -:-:-:-:-:-:-:-: | 4563 | PANX3 | 4 | 0 | 0 | 0 | 31 | 0 |
| ENSDARG000 | 10:10:10:10:1 | 7778870:7778 | 7779732:7779 | -:-:-:-:-:-:-:-: | 3754 | mpx | 259 | 184 | 174 | 234 | 1030 | 1587 |
| ENSDARG000 | 10:10:10:10:1 | 40284003:402 | 40284144:402 | +:+:+:+:+:+:+: | 5871 | git2b | 1398 | 699 | 1373 | 1302 | 1299 | 1116 |
| ENSDARG000 | 10:10:10:10:1 | 81897:82303: | 82422:82422: | -:-:-:-:-:-:-:-: | 5634 | NA | 2199 | 1569 | 2228 | 2840 | 2033 | 2155 |
| ENSDARG000 | 10:10:10:10:1 | 20608676:206 | 20608961:206 | +:+:+:+:+:+:+: | 2216 | NA | 2 | 4 | 0 | 0 | 24 | 18 |
| ENSDARG000 | 10:10:10:10:1 | 20589969:205 | 20590595:205 | +:+:+:+:+:+:+: | 6768 | LETM2 | 1430 | 1292 | 1311 | 904 | 1284 | 1207 |
| ENSDARG000 | 10:10:10:10:1 | 2924775:2924 | 2925527:2925 | -:-:-:-:-:-:-:-: | 3018 | oclna | 182 | 24 | 16 | 14 | 48 | 22 |
| ENSDARG000 | 10;10;10 | 24778289:247 | 24779129:247 | -;-;- | 1234 | NA | 4 | 6 | 0 | 0 | 0 | 2 |
| ENSDARG000 | 10:10:10:10:1 | 17509980:175 | 17510077:175 | -;-;-;-;-;-;- | 767 | NA | 0 | 0 | 0 | 0 | 0 | 0 |
| ENSDARG000 | 10;10;10 | 24935765:249 | 24935879:249 | -;-;- | 371 | NA | 0 | 2 | 0 | 0 | 0 | 9 |
| ENSDARG000 | 10:10:10:10:1 | 31019607:310 | 31021854:310 | -:-:-:-:-:-:-:-: | 3792 | pknox2 | 44 | 6 | 49 | 0 | 8 | 20 |
| ENSDARG000 | 10:10:10:10:1 | 40321067:403 | 40321169:403 | +:+:+:+:+:+:+: | 856 | NA | 14 | 18 | 0 | 7 | 18 | 12 |
| ENSDARG000 | 10:10:10:10:1 | 17543276:175 | 17543989:175 | -:-:-:-:-:-:-:-: | 1487 | ska1 | 68 | 204 | 260 | 136 | 125 | 78 |
| ENSDARG000 | 10:10:10:10:1 | 17533610:175 | 17533743:175 | -:-:-:-:-:-:-:-: | 1313 | NA | 0 | 1 | 0 | 0 | 6 | 0 |
| ENSDARG000 | 10;10;10 | 24959246:249 | 24959408:249 | -;-;- | 371 | NA | 0 | 0 | 0 | 0 | 0 | 0 |
| ENSDARG000 | 10:10:10:10:1 | 36458563:364 | 36458695:364 | +;+;+;+;+ | 1304 | alox5ap | 20 | 0 | 31 | 46 | 26 | 36 |
| ENSDARG000 | 10:10:10:10:1 | 31222433:312 | 31223231:312 | +:+:+:+:+:+:+: | 1319 | tmem218 | 40 | 48 | 47 | 12 | 45 | 39 |
| ENSDARG000 | 10:10:10:10:1 | 11593998:115 | 11594107:115 | -:-:-:-:-:-:-:-: | 3351 | ADAMTS6 | 6 | 0 | 2 | 0 | 16 | 0 |
| ENSDARG000 | 10;10;10 | 11761380:117 | 11762231:117 | -;-;- | 1125 | NA | 0 | 4 | 0 | 0 | 1 | 0 |
| ENSDARG000 | 10:10:10:10:1 | 43188678:431 | 43188710:431 | +:+:+:+:+:+:+: | 7027 | vcanb | 70 | 99 | 136 | 56 | 400 | 180 |
| ENSDARG000 | 10:10:10:10:1 | 26926654:269 | 26926924:269 | +:+:+:+:+:+:+: | 2010 | rab1bb | 42 | 92 | 140 | 20 | 264 | 28 |
| ENSDARG000 | 10:10:10:10:1 | 6383270:6386 | 6383746:6386 | +;+;+;+;+;+;+ | 10806 | zgc:114200 | 1417 | 841 | 1325 | 865 | 1199 | 1519 |
| ENSDARG000 | 10:10:10:10:1 | 40331191:403 | 40332589:403 | -;-;-;-;- | 3646 | NA | 0 | 0 | 0 | 0 | 0 | 0 |
| ENSDARG000 | 10 | 40347080 | 40348132 | - | 1053 | NA | 0 | 0 | 0 | 0 | 0 | 0 |
| ENSDARG000 | 10 | 40338250 | 40339310 | - | 1061 | NA | 0 | 0 | 0 | 0 | 0 | 0 |
| ENSDARG000 | 10 | 40332261 | 40333319 | - | 1059 | NA | 0 | 0 | 0 | 0 | 0 | 0 |
| ENSDARG000 | 10 | 40375010 | 40375996 | - | 987 | NA | 0 | 0 | 0 | 0 | 0 | 0 |
| ENSDARG000 | 10:10:10:10:1 | 24971473:249 | 24971685:249 | -;-;-;-;-;-;-;- | 1420 | cd209 | 0 | 0 | 0 | 0 | 0 | 0 |
| ENSDARG000 | 10 | 40359577 | 40360635 | - | 1059 | NA | 0 | 0 | 0 | 0 | 0 | 0 |
| ENSDARG000 | 10 | 40351198 | 40352250 | - | 1053 | NA | 0 | 0 | 0 | 0 | 0 | 0 |
| ENSDARG000 | 10:10:10:10:1 | 23070741:230 | 23071719:230 | -:-:-:-:-:-:-:-: | 2647 | nle1 | 246 | 250 | 506 | 207 | 389 | 349 |
| ENSDARG000 | 10:10:10:10:1 | 34035689:340 | 34035885:340 | +;+;+;+;+;+ | 1099 | NA | 4 | 1 | 0 | 0 | 0 | 0 |
| ENSDARG000 | 10:10:10:10:1 | 23099365:230 | 23099988:230 | +:+:+:+:+:+:+: | 10673 | LTN1 | 880 | 705 | 1064 | 404 | 975 | 641 |
| ENSDARG000 | 10:10:10:10:1 | 9397305:9406 | 9398425:9406 | -;-;-;-;- | 1589 | morn5 | 2 | 1 | 10 | 0 | 0 | 0 |
| ENSDARG000 | 10:10:10:10:1 | 110868:11087 | 110920:11092 | +:+:+:+:+:+:+: | 1532 | pigp | 170 | 184 | 176 | 216 | 254 | 212 |
| ENSDARG000 | 10:10:10:10:1 | 2943676:2947 | 2947847:2947 | -:-:-:-:-:-:-:-: | 5993 | MARVELD2 | 325 | 296 | 128 | 184 | 88 | 232 |
| ENSDARG000 | 10;10;10 | 20628698:206 | 20628764:206 | +;+;+ | 2403 | prnpb | 2120 | 223 | 52 | 1517 | 332 | 916 |
| ENSDARG000 | 10:10:10:10:1 | 34043561:340 | 34043731:340 | -;-;-;-;-;-;- | 1648 | NA | 0 | 0 | 0 | 0 | 0 | 0 |
| ENSDARG000 | 10:10:10:10:1 | 6417356:6417 | 6417543:6417 | -:-:-:-:-:-:-:-: | 1321 | NA | 2 | 21 | 37 | 3 | 10 | 4 |
| ENSDARG000 | 10:10:10:10:1 | 25035459:250 | 25036310:250 | -;-;-;-;- | 1561 | mtnr1bb | 0 | 0 | 0 | 0 | 0 | 0 |
| ENSDARG000 | 10 | 20635960 | 20637098 | - | 1139 | sprn2 | 2 | 0 | 0 | 0 | 0 | 0 |
| ENSDARG000 | 10:10:10:10:1 | 43269918:432 | 43274010:432 | -:-:-:-:-:-:-:-: | 5247 | HAPLN1 | 0 | 0 | 0 | 0 | 16 | 0 |
| ENSDARG000 | 10:10:10:10:1 | 7798087:7800 | 7798201:7800 | +:+:+:+:+:+:+: | 1451 | NA | 0 | 10 | 8 | 20 | 56 | 18 |
| ENSDARG000 | 10:10:10:10:1 | 31244619:312 | 31244854:312 | +:+:+:+:+:+:+: | 12079 | NA | 53 | 48 | 74 | 8 | 111 | 32 |
| ENSDARG000 | 10;10;10;10 | 31263274:312 | 31263377:312 | -;-;-;- | 546 | NA | 0 | 0 | 0 | 0 | 0 | 0 |
| ENSDARG000 | 10;10;10;10 | 9410304:9415 | 9410485:9415 | +;+;+;+ | 805 | ndufa8 | 634 | 1039 | 1095 | 598 | 820 | 580 |
| ENSDARG000 | 10:10:10:10:1 | 2975974:2981 | 2976113:2981 | +:+:+:+:+:+:+: | 4545 | NA | 542 | 479 | 700 | 383 | 676 | 476 |
| ENSDARG000 | 10:10:10:10:1 | 6473092:6476 | 6474090:6476 | -:-:-:-:-:-:-:-: | 2289 | dcp2 | 1119 | 965 | 1269 | 1354 | 1465 | 1425 |
| ENSDARG000 | 10:10:10:10:1 | 43355895:433 | 43356044:433 | -:-:-:-:-:-:-:-: | 1933 | EDIL3 | 0 | 0 | 0 | 0 | 4 | 0 |
| ENSDARG000 | 10:10:10:10:1 | 26944418:269 | 26944561:269 | +:+:+:+:+:+:+: | 4843 | frmd8 | 7045 | 7126 | 6411 | 8427 | 9032 | 5614 |
| ENSDARG000 | 10:10:10:10:1 | 34147484:341 | 34147654:341 | -;-;-;-;-;-;- | 1644 | NA | 0 | 3 | 0 | 0 | 0 | 0 |
| ENSDARG000 | 10:10:10:10:1 | 34120543:341 | 34120713:341 | -:-:-:-:-:-:-:-: | 1882 | NA | 0 | 0 | 0 | 0 | 0 | 0 |
| ENSDARG000 | 10:10:10:10:1 | 34059007:340 | 34059177:340 | -;-;-;-;-;-;- | 1644 | NA | 0 | 0 | 0 | 0 | 0 | 0 |
| ENSDARG000 | 10:10:10:10:1 | 34175476:342 | 34175606:342 | +:+:+:+:+:+:+: | 1444 | NA | 6 | 2 | 0 | 0 | 0 | 2 |
| ENSDARG000 | 10:10:10:10:1 | 43406796:434 | 43406987:434 | +:+:+:+:+:+:+: | 5824 | RASA1 | 891 | 462 | 865 | 794 | 657 | 557 |
| ENSDARG000 | 10:10:10:10:1 | 34104733:341 | 34104903:341 | -;-;-;-;-;-;- | 1644 | NA | 0 | 0 | 0 | 0 | 0 | 0 |
| ENSDARG000 | 10:10:10:10:1 | 9534615:9534 | 9535010:9535 | -:-:-:-:-:-:-:-: | 2121 | NA | 12 | 0 | 4 | 4 | 0 | 8 |
| ENSDARG000 | 10:10:10:10:1 | 34067879:340 | 34068004:340 | +:+:+:+:+:+:+: | 1443 | NA | 0 | 2 | 1 | 0 | 1 | 2 |
| ENSDARG000 | 10:10:10:10:1 | 187760:18776 | 187950:18799 | +:+:+:+:+:+:+: | 3047 | ets2 | 820 | 743 | 802 | 2102 | 495 | 313 |
| ENSDARG000 | 10:10:10:10:1 | 34151286:341 | 34151412:341 | +:+:+:+:+:+:+: | 1430 | NA | 12 | 15 | 15 | 16 | 3 | 24 |
| ENSDARG000 | 10:10:10:10:1 | 34047352:340 | 34047490:340 | +:+:+:+:+:+:+: | 1588 | NA | 0 | 0 | 1 | 0 | 0 | 0 |
| ENSDARG000 | 10:10:10:10:1 | 34171674:341 | 34171844:341 | -:-:-:-:-:-:-:-: | 1882 | NA | 0 | 0 | 0 | 0 | 0 | 0 |
| ENSDARG000 | 10:10:10:10:1 | 11767791:117 | 11767927:117 | +:+:+:+:+:+:+: | 1944 | ppwd1 | 356 | 452 | 570 | 222 | 449 | 407 |
| ENSDARG000 | 10:10:10:10:1 | 34108541:341 | 34108667:341 | +:+:+:+:+:+:+: | 1420 | NA | 2 | 2 | 1 | 0 | 2 | 0 |
| ENSDARG000 | 10:10:10:10:1 | 34124421:341 | 34124504:341 | +;+;+;+;+;+ | 1166 | NA | 0 | 0 | 1 | 1 | 0 | 1 |
| ENSDARG000 | 10:10:10:10:1 | 36475860:364 | 36476932:364 | -:-:-:-:-:-:-:-: | 9756 | nipblb | 1150 | 1000 | 1212 | 614 | 1392 | 974 |
| ENSDARG000 | 10:10:10:10:1 | 34187231:341 | 34187401:341 | -;-;-;-;-;-;- | 1643 | NA | 0 | 3 | 0 | 0 | 0 | 0 |
| ENSDARG000 | 10:10:10:10:1 | 34047360:340 | 34047490:340 | +;+;+;+;+;+;+ | 1197 | NA | 0 | 1 | 0 | 0 | 0 | 0 |
| ENSDARG000 | 10:10:10:10:1 | 34220492:342 | 34220660:342 | -:-:-:-:-:-:-:-: | 2509 | NA | 0 | 0 | 0 | 2 | 0 | 0 |
| ENSDARG000 | 10:10:10:10:1 | 6496185:6496 | 6496432:6496 | +;+;+;+;+ | 1234 | reep5 | 7885 | 9545 | 9462 | 13976 | 11234 | 12296 |
| ENSDARG000 | 10:10:10:10:1 | 20641001:206 | 20641501:206 | -:-:-:-:-:-:-:-: | 1687 | rassf2b | 18 | 9 | 24 | 6 | 8 | 36 |
| ENSDARG000 | 10:10:10:10:1 | 34175468:341 | 34175646:341 | +;+;+;+;+;+ | 1162 | NA | 0 | 0 | 0 | 0 | 0 | 0 |
| ENSDARG000 | 10:10:10:10:1 | 34086591:340 | 34086761:340 | -:-:-:-:-:-:-:-: | 1977 | NA | 0 | 0 | 0 | 0 | 0 | 0 |
| ENSDARG000 | 10:10:10:10:1 | 17552733:175 | 17553072:175 | -:-:-:-:-:-:-:-: | 3464 | smarcad1b | 262 | 283 | 403 | 124 | 468 | 109 |
| ENSDARG000 | 10 | 40378761 | 40380136 | - | 1376 | NA | 0 | 0 | 0 | 0 | 0 | 0 |
| ENSDARG000 | 10;10;10 | 43499821:435 | 43499991:435 | -;-;- | 608 | NA | 0 | 0 | 0 | 0 | 0 | 0 |
| ENSDARG000 | 10 | 40383782 | 40384432 | - | 651 | NA | 0 | 0 | 0 | 0 | 0 | 0 |
| ENSDARG000 | 10 | 40387210 | 40388265 | - | 1056 | NA | 0 | 0 | 0 | 0 | 0 | 0 |
| ENSDARG000 | 10;10 | 40396302:403 | 40397363:403 | -;- | 1062 | NA | 0 | 0 | 1 | 0 | 0 | 12 |
| ENSDARG000 | 10 | 40401858 | 40402904 | - | 1047 | NA | 0 | 0 | 0 | 0 | 0 | 0 |
| ENSDARG000 | 10:10:10:10:1 | 26972755:269 | 26972892:269 | +:+:+:+:+:+:+: | 1712 | tm7sf2 | 278 | 237 | 234 | 100 | 156 | 120 |
| ENSDARG000 | 10 | 40403994 | 40404331 | - | 338 | NA | 0 | 0 | 0 | 0 | 0 | 0 |
| ENSDARG000 | 10;10 | 40405283:404 | 40406278:404 | -;- | 1032 | NA | 0 | 0 | 0 | 0 | 0 | 0 |
| ENSDARG000 | 10:10:10:10:1 | 9550419:9553 | 9550453:9554 | +:+:+:+:+:+:+: | 1971 | si:ch211-243 | 0 | 0 | 0 | 0 | 6 | 10 |
| ENSDARG000 | 10:10:10:10:1 | 26961965:269 | 26962141:269 | +:+:+:+:+:+:+: | 2716 | vps51 | 610 | 464 | 566 | 312 | 520 | 425 |
| ENSDARG000 | 10:10:10:10:1 | 43544814:435 | 43550177:435 | -:-:-:-:-:-:-:-: | 6913 | mef2ca | 111 | 160 | 135 | 86 | 169 | 107 |
| ENSDARG000 | 10:10:10:10:1 | 7810455:7810 | 7810929:7811 | -:-:-:-:-:-:-:-: | 3055 | mat2aa | 2103 | 3550 | 3676 | 1686 | 2996 | 2413 |
| ENSDARG000 | 10 | 40409832 | 40410884 | - | 1053 | NA | 0 | 0 | 0 | 4 | 0 | 0 |
| ENSDARG000 | 10;10 | 40415062:404 | 40416054:404 | -;- | 1023 | NA | 0 | 0 | 0 | 0 | 0 | 0 |
| ENSDARG000 | 10:10:10:10:1 | 15648857:156 | 15649350:156 | -:-:-:-:-:-:-:-: | 2564 | NA | 10 | 0 | 0 | 2 | 2 | 0 |
| ENSDARG000 | 10:10:10:10:1 | 3031636:3031 | 3031781:3033 | -;-;-;-;- | 1635 | ANKRD34B | 0 | 0 | 0 | 0 | 0 | 0 |
| ENSDARG000 | 10:10:10:10:1 | 206996:20708 | 207302:20730 | -:-:-:-:-:-:-:-: | 1330 | psmg1 | 1169 | 1552 | 1793 | 903 | 1627 | 1004 |
| ENSDARG000 | 10:10:10:10:1 | 20653716:206 | 20655080:206 | -:-:-:-:-:-:-:-: | 2726 | kcnip3b | 84 | 44 | 195 | 28 | 158 | 98 |

| | | | | | | | | | | | | |
|---|---|---|---|---|---|---|---|---|---|---|---|---|
| ENSDARG000 | 10 | 40418807 | 40419059 | - | 253 | NA | 0 | 0 | 0 | 0 | 0 | 0 |
| ENSDARG000 | 10;10 | 40420052:404 | 40421059:404 | -;- | 1008 | NA | 0 | 0 | 0 | 0 | 0 | 0 |
| ENSDARG000 | 10 | 40428695 | 40429747 | - | 1053 | NA | 0 | 0 | 2 | 0 | 0 | 2 |
| ENSDARG000 | 10;10 | 40442694:404 | 40443746:404 | -;- | 1053 | NA | 0 | 0 | 2 | 0 | 0 | 0 |
| ENSDARG000 | 10;10 | 40447708:404 | 40448760:404 | -;- | 1053 | NA | 0 | 0 | 0 | 0 | 0 | 0 |
| ENSDARG000 | 10;10;10;10;1 | 43705795:437 | 43709261:437 | -;-;-;-;-;-;-;-; | 4682 | cetn3 | 233 | 368 | 420 | 198 | 287 | 224 |
| ENSDARG000 | 10;10;10;10;1 | 6521376:6522 | 6522870:6522 | -;-;-;-;-;-;-;-; | 7266 | chd1 | 564 | 535 | 734 | 222 | 687 | 679 |
| ENSDARG000 | 10 | 40453660 | 40454685 | - | 1026 | NA | 0 | 0 | 0 | 0 | 0 | 0 |
| ENSDARG000 | 10 | 40468372 | 40469394 | - | 1023 | NA | 0 | 0 | 0 | 0 | 0 | 0 |
| ENSDARG000 | 10 | 40461869 | 40462894 | - | 1026 | NA | 0 | 0 | 0 | 0 | 0 | 0 |
| ENSDARG000 | 10;10;10;10;1 | 213878:21393 | 214061:21406 | +;+;+;+;+;+;+; | 925 | mpzl1l | 767 | 599 | 688 | 597 | 681 | 495 |
| ENSDARG000 | 10;10;10;10;1 | 9595575:9595 | 9595837:9595 | +;+;+;+;+;+;+; | 5891 | NA | 604 | 218 | 420 | 570 | 288 | 378 |
| ENSDARG000 | 10;10;10;10;1 | 11790881:118 | 11791060:118 | -;-;-;-;-;-;-;-; | 1755 | trim23 | 370 | 353 | 269 | 177 | 358 | 264 |
| ENSDARG000 | 10;10;10;10;1 | 6625785:6627 | 6625956:6627 | +;+;+;+;+;+;+; | 2182 | NA | 0 | 0 | 0 | 0 | 0 | 0 |
| ENSDARG000 | 10 | 40471629 | 40471895 | - | 267 | NA | 0 | 0 | 0 | 0 | 0 | 0 |
| ENSDARG000 | 10;10;10;10;1 | 236457:23701 | 236616:23720 | -;-;-;-;-;-;- | 1551 | NA | 0 | 4 | 3 | 6 | 3 | 0 |
| ENSDARG000 | 10;10;10;10;1 | 6613279:6614 | 6613465:6615 | +;+;+;+;+;+;+; | 2856 | NA | 0 | 0 | 0 | 0 | 0 | 0 |
| ENSDARG000 | 10;10;10;10;1 | 6637730:6637 | 6637810:6638 | -;-;-;-;-;- | 912 | NA | 0 | 0 | 1 | 0 | 0 | 0 |
| ENSDARG000 | 10;10;10;10;1 | 6611075:6612 | 6611278:6612 | -;-;-;-;-;-;-;-; | 1562 | NA | 1 | 0 | 1 | 0 | 0 | 0 |
| ENSDARG000 | 10;10;10;10;1 | 15777064:157 | 15777331:157 | +;+;+;+;+;+;+; | 3747 | NA | 0 | 0 | 0 | 0 | 0 | 0 |
| ENSDARG000 | 10;10;10;10;1 | 13000370:130 | 13000938:130 | +;+;+;+;+;+;+; | 2796 | lpar1 | 26 | 70 | 162 | 66 | 190 | 100 |
| ENSDARG000 | 10;10;10;10;1 | 23216678:232 | 23216796:232 | -;-;-;-;-;-;-;-; | 1749 | cadm2a | 49 | 245 | 192 | 116 | 79 | 66 |
| ENSDARG000 | 10;10;10;10;1 | 7831439:7831 | 7831687:7831 | -;-;-;-;-;-;-;-; | 1763 | NA | 2 | 2 | 16 | 5 | 2 | 0 |
| ENSDARG000 | 10;10;10;10;1 | 36558804:365 | 36559025:365 | -;-;-;-;-;-;-;-; | 1735 | slc1a3b | 2 | 0 | 0 | 0 | 0 | 2 |
| ENSDARG000 | 10;10;10;10;1 | 23441488:234 | 23441779:234 | +;+;+;+;+ | 1014 | NA | 0 | 0 | 0 | 0 | 0 | 0 |
| ENSDARG000 | 10;10;10;10;1 | 17592273:175 | 17592714:175 | +;+;+;+;+;+;+; | 1877 | KCNN2 | 48 | 20 | 498 | 126 | 183 | 139 |
| ENSDARG000 | 10;10;10;10;1 | 43761480:437 | 43762970:437 | -;-;-;-;-;-;-;- | 2811 | arrdc3b | 4918 | 4051 | 4637 | 7957 | 4644 | 6213 |
| ENSDARG000 | 10;10;10;10 | 20843353:208 | 20843515:208 | -;-;-;- | 542 | NA | 0 | 0 | 0 | 0 | 0 | 0 |
| ENSDARG000 | 10;10 | 252425:25515 | 252511:25704 | +;+ | 1980 | NA | 69 | 8 | 18 | 16 | 14 | 13 |
| ENSDARG000 | 10;10;10;10;1 | 20920364:209 | 20920737:209 | -;-;-;-;-;-;-;-;- | 7379 | PCDH1 | 8 | 0 | 38 | 12 | 103 | 12 |
| ENSDARG000 | 10;10 | 26982254:269 | 26983906:269 | -;- | 1790 | znhit2 | 104 | 96 | 136 | 104 | 112 | 81 |
| ENSDARG000 | 10;10;10;10;1 | 7875076:7881 | 7876071:7881 | -;-;-;-;-;-;-;-; | 2038 | NA | 2 | 0 | 0 | 0 | 6 | 0 |
| ENSDARG000 | 10;10;10 | 7870627:7881 | 7870860:7881 | -;-;- | 795 | NA | 0 | 0 | 2 | 0 | 0 | 0 |
| ENSDARG000 | 10;10;10;10;1 | 9834896:9835 | 9837415:9837 | -;-;-;-;-;-;-;-; | 5718 | STRBP | 153 | 308 | 517 | 215 | 752 | 551 |
| ENSDARG000 | 10 | 40478928 | 40479911 | - | 984 | NA | 0 | 0 | 0 | 0 | 0 | 0 |
| ENSDARG000 | 10;10;10 | 40472282:404 | 40473835:404 | -;-;- | 2683 | NA | 0 | 0 | 0 | 0 | 0 | 0 |
| ENSDARG000 | 10 | 40473603 | 40474628 | - | 1026 | NA | 0 | 0 | 0 | 0 | 0 | 0 |
| ENSDARG000 | 10;10;10;10;1 | 13080092:130 | 13085659:130 | -;-;-;-;-;-;-;-; | 8990 | musk | 239 | 9 | 2 | 4 | 14 | 9 |
| ENSDARG000 | 10;10;10;10;1 | 284303:28430 | 284567:28456 | -;-;-;-;-;-;-;-; | 3747 | emsy | 1053 | 515 | 1193 | 805 | 933 | 732 |
| ENSDARG000 | 10;10;10;10;1 | 36627741:366 | 36628672:366 | -;-;-;-;-;- | 1670 | NA | 1 | 1 | 0 | 0 | 2 | 0 |
| ENSDARG000 | 10;10;10;10;1 | 36613004:366 | 36613567:366 | -;-;-;-;-;-;-;-; | 5767 | NA | 794 | 700 | 735 | 410 | 928 | 881 |
| ENSDARG000 | 10;10;10;10;1 | 15815865:158 | 15816963:158 | -;-;-;-;-;-;-;-; | 5085 | tjp2a | 114 | 0 | 10 | 20 | 24 | 12 |
| ENSDARG000 | 10;10;10;10 | 23502026:235 | 23502218:235 | +;+;+;+ | 567 | NA | 0 | 0 | 4 | 0 | 0 | 2 |
| ENSDARG000 | 10;10;10;10;1 | 21395298:214 | 21396050:214 | +;+;+;+;+;+;+; | 2529 | trpc7a | 0 | 0 | 0 | 0 | 0 | 0 |
| ENSDARG000 | 10;10;10 | 43797130:437 | 43797801:438 | +;+;+ | 2786 | nr2f1b | 0 | 6 | 0 | 0 | 0 | 4 |
| ENSDARG000 | 10 | 40489658 | 40490647 | - | 990 | NA | 0 | 0 | 0 | 0 | 0 | 0 |
| ENSDARG000 | 10;10;10;10;1 | 26990095:269 | 26990180:269 | +;+;+;+;+ | 894 | faub | 276 | 222 | 290 | 199 | 217 | 291 |
| ENSDARG000 | 10;10 | 26991805:269 | 26992003:269 | -;- | 514 | zgc:193742 | 4 | 8 | 6 | 19 | 2 | 6 |
| ENSDARG000 | 10;10;10;10;1 | 21397367:213 | 21399053:213 | +;+;+;+;+;+;+; | 2547 | NA | 0 | 0 | 0 | 0 | 0 | 0 |
| ENSDARG000 | 10;10;10 | 21397320:213 | 21397395:213 | +;+;+ | 899 | NA | 0 | 0 | 0 | 0 | 0 | 0 |
| ENSDARG000 | 10;10;10 | 43831882:438 | 43832315:438 | -;-;- | 558 | NA | 6 | 1 | 0 | 4 | 0 | 2 |
| ENSDARG000 | 10;10;10;10;1 | 3364650:3366 | 3366225:3366 | -;-;-;-;-;-;-;-; | 4421 | NAA25 | 1300 | 1275 | 1764 | 988 | 1411 | 1618 |
| ENSDARG000 | 10 | 40498165 | 40499185 | - | 1021 | NA | 0 | 0 | 0 | 0 | 0 | 0 |
| ENSDARG000 | 10;10 | 40502370:405 | 40503416:405 | -;- | 1047 | NA | 0 | 0 | 0 | 0 | 0 | 0 |
| ENSDARG000 | 10;10;10;10;1 | 17681074:176 | 17681304:176 | +;+;+;+;+;+;+; | 1569 | drg1 | 853 | 1258 | 1509 | 587 | 1371 | 1236 |
| ENSDARG000 | 10;10;10;10;1 | 43838563:438 | 43840172:438 | -;-;-;-;- | 3314 | tlr8b | 22 | 12 | 20 | 42 | 114 | 44 |
| ENSDARG000 | 10;10;10;10;1 | 43862964:438 | 43866387:438 | -;-;-;-;-;-;-;-; | 5721 | MCTP1 | 34 | 66 | 36 | 4 | 70 | 74 |
| ENSDARG000 | 10;10;10;10;1 | 23933778:240 | 23934153:240 | -;-;-;-;-;-;-;-; | 1825 | NA | 0 | 0 | 0 | 0 | 0 | 0 |
| ENSDARG000 | 10;10 | 23992381:239 | 23992517:239 | +;+ | 233 | NA | 0 | 0 | 0 | 0 | 0 | 0 |
| ENSDARG000 | 10;10;10;10;1 | 24048663:240 | 24048974:240 | +;+;+;+;+;+;+; | 3757 | GBE1 | 0 | 0 | 4 | 4 | 0 | 4 |
| ENSDARG000 | 10;10 | 40507526:405 | 40508527:405 | -;- | 1032 | NA | 0 | 0 | 0 | 0 | 0 | 0 |
| ENSDARG000 | 10;10 | 26998414:269 | 26998619:269 | -;- | 612 | si:dkev-88p24 | 114 | 162 | 184 | 102 | 156 | 134 |
| ENSDARG000 | 10;10;10;10;1 | 43936252:439 | 43936382:439 | -;-;-;-;-;-;-;-; | 1203 | septin5b | 52 | 56 | 74 | 19 | 83 | 83 |
| ENSDARG000 | 10;10;10;10 | 27000562:270 | 27000649:270 | -;-;-;- | 463 | NA | 0 | 0 | 4 | 6 | 0 | 0 |
| ENSDARG000 | 10 | 40513612 | 40514643 | - | 1032 | NA | 0 | 0 | 0 | 0 | 0 | 0 |
| ENSDARG000 | 10;10;10;10 | 375042:37857 | 375248:37865 | +;+;+;+ | 1196 | si:ch1073-303 | 0 | 0 | 0 | 0 | 0 | 0 |
| ENSDARG000 | 10;10;10;10;1 | 27006266:270 | 27006732:270 | -;-;-;-;-;-;-;-; | 1058 | uqcc3 | 106 | 144 | 134 | 218 | 170 | 118 |
| ENSDARG000 | 10;10;10;10;1 | 7901012:7902 | 7902397:7902 | -;-;-;-;-;- | 2297 | slc35e4 | 44 | 32 | 32 | 2 | 36 | 32 |
| ENSDARG000 | 10 | 40516749 | 40517598 | - | 850 | NA | 0 | 0 | 0 | 0 | 0 | 0 |
| ENSDARG000 | 10 | 40520823 | 40521100 | - | 278 | NA | 0 | 0 | 0 | 0 | 0 | 0 |
| ENSDARG000 | 10 | 40523707 | 40524741 | - | 1035 | NA | 0 | 6 | 7 | 2 | 6 | 0 |
| ENSDARG000 | 10;10 | 40527237:405 | 40528229:405 | -;- | 1023 | NA | 0 | 0 | 0 | 0 | 0 | 0 |
| ENSDARG000 | 10;10 | 40531123:405 | 40532127:405 | -;- | 1035 | NA | 0 | 0 | 0 | 0 | 0 | 0 |
| ENSDARG000 | 10;10 | 40535971:405 | 40536963:405 | -;- | 1020 | NA | 0 | 0 | 0 | 0 | 0 | 0 |
| ENSDARG000 | 10 | 40537440 | 40538047 | - | 608 | NA | 0 | 0 | 0 | 0 | 0 | 1 |
| ENSDARG000 | 10;10 | 40542904:405 | 40543896:405 | -;- | 1020 | NA | 0 | 0 | 0 | 0 | 0 | 0 |
| ENSDARG000 | 10 | 40548616 | 40549641 | + | 1026 | NA | 0 | 0 | 0 | 0 | 0 | 0 |
| ENSDARG000 | 10;10;10;10;1 | 7916720:7918 | 7918160:7918 | -;-;-;-;-;-;-;-; | 3735 | NA | 0 | 0 | 0 | 0 | 0 | 0 |
| ENSDARG000 | 10 | 40550730 | 40550954 | + | 225 | NA | 0 | 0 | 0 | 0 | 0 | 0 |
| ENSDARG000 | 10;10 | 40552038:405 | 40553090:405 | -;- | 1053 | NA | 0 | 0 | 0 | 0 | 0 | 0 |
| ENSDARG000 | 10;10;10;10;1 | 34315719:343 | 34315768:343 | +;+;+;+;+;+;+; | 6024 | NA | 1216 | 1274 | 1161 | 1092 | 1920 | 1177 |
| ENSDARG000 | 10;10;10;10;1 | 6641514:6643 | 6641720:6643 | +;+;+;+;+;+;+; | 2816 | NA | 0 | 0 | 0 | 0 | 0 | 0 |
| ENSDARG000 | 10 | 40554917 | 40555131 | + | 215 | NA | 0 | 0 | 0 | 0 | 0 | 0 |
| ENSDARG000 | 10;10;10;10;1 | 15937277:159 | 15937822:159 | -;-;-;-;- | 956 | NA | 2 | 2 | 9 | 0 | 0 | 0 |
| ENSDARG000 | 10;10;10;10;1 | 15895988:158 | 15896058:158 | -;-;-;-;-;-;- | 807 | NA | 0 | 0 | 0 | 0 | 0 | 0 |
| ENSDARG000 | 10;10;10;10;1 | 27028019:270 | 27028750:270 | -;-;-;-;-;-;-;-; | 1955 | cnih2 | 0 | 2 | 2 | 0 | 6 | 4 |
| ENSDARG000 | 10;10;10;10;1 | 15881309:158 | 15882028:158 | -;-;-;-;-;-;-;-; | 2678 | pip5k1ba | 83 | 14 | 55 | 6 | 40 | 101 |
| ENSDARG000 | 10 | 399363 | 400213 | + | 851 | FAM222A | 16 | 6 | 29 | 12 | 25 | 0 |
| ENSDARG000 | 10;10;10 | 36654197:366 | 36654297:366 | -;-;- | 266 | NA | 1 | 0 | 0 | 0 | 3 | 1 |
| ENSDARG000 | 10;10;10;10;1 | 36650222:366 | 36650344:366 | +;+;+;+;+;+;+; | 1676 | ucp3 | 10 | 3 | 0 | 4 | 37 | 12 |
| ENSDARG000 | 10;10;10;10;1 | 3428194:3430 | 3428399:3430 | +;+;+;+;+;+;+; | 7284 | ptpn11a | 1726 | 1672 | 2325 | 1379 | 2130 | 1484 |
| ENSDARG000 | 10;10 | 43993352:440 | 43993555:440 | +;+ | 255 | NA | 0 | 0 | 0 | 0 | 0 | 0 |
| ENSDARG000 | 10;10;10;10;1 | 44021147:440 | 44021574:440 | -;-;-;-;-;-;-;-; | 1010 | crybb1 | 0 | 0 | 2 | 0 | 0 | 0 |
| ENSDARG000 | 10;10;10;10;1 | 3419832:3420 | 3420795:3420 | -;-;-;-;-;-;-;-; | 1709 | tmed2 | 2880 | 4313 | 4923 | 3318 | 3970 | 3560 |
| ENSDARG000 | 10;10;10;10;1 | 43999263:439 | 43999735:439 | -;-;-;-;-;-;-;-; | 1611 | acads | 272 | 307 | 289 | 242 | 454 | 432 |
| ENSDARG000 | 10;10;10;10;1 | 3397871:3400 | 3398319:3400 | -;-;-;-;-;-;-;-; | 2263 | ddx55 | 254 | 357 | 426 | 262 | 272 | 284 |
| ENSDARG000 | 10 | 40556203 | 40557210 | - | 1008 | NA | 0 | 0 | 0 | 0 | 0 | 0 |
| ENSDARG000 | 10;10;10;10;1 | 21444654:214 | 21444731:214 | +;+;+;+;+;+;+; | 2917 | fbxw11b | 772 | 973 | 980 | 492 | 884 | 670 |
| ENSDARG000 | 10;10 | 43994471:439 | 43996117:439 | +;+ | 1647 | cldn5b | 298 | 34 | 82 | 50 | 60 | 76 |
| ENSDARG000 | 10;10;10;10;1 | 406146:41248 | 406419:41260 | +;+;+;+;+;+;+ | 1718 | NA | 1024 | 590 | 669 | 295 | 725 | 488 |
| ENSDARG000 | 10 | 40561692 | 40562717 | + | 1026 | NA | 0 | 0 | 0 | 0 | 0 | 0 |
| ENSDARG000 | 10;10;10;10;1 | 21434649:214 | 21435329:214 | +;+;+;+;+;+;+; | 5146 | etf1b | 4912 | 7048 | 8023 | 3610 | 6925 | 3792 |
| ENSDARG000 | 10;10 | 40568735:405 | 40569766:405 | +;+ | 1044 | NA | 0 | 0 | 0 | 0 | 0 | 0 |
| ENSDARG000 | 10 | 40578332 | 40579363 | + | 1032 | NA | 0 | 0 | 0 | 10 | 0 | 0 |
| ENSDARG000 | 10;10;10;10;1 | 27052642:270 | 27052813:270 | +;+;+;+;+;+;+; | 1512 | wdr74 | 186 | 271 | 281 | 142 | 201 | 202 |
| ENSDARG000 | 10;10;10;10;1 | 36662640:366 | 36662795:366 | +;+;+;+;+;+;+; | 2382 | ucp2 | 4270 | 589 | 448 | 696 | 1253 | 1573 |
| ENSDARG000 | 10 | 40583930 | 40584955 | + | 1026 | NA | 0 | 0 | 0 | 0 | 0 | 0 |
| ENSDARG000 | 10;10;10;10;1 | 6661059:6661 | 6661138:6661 | -;-;-;-;-;-;- | 1014 | NA | 0 | 0 | 0 | 0 | 0 | 0 |
| ENSDARG000 | 10;10;10;10;1 | 36673387:366 | 36673609:366 | -;-;-;-;-;-;-;-; | 1684 | dnajb13 | 0 | 0 | 0 | 0 | 0 | 0 |
| ENSDARG000 | 10;10 | 40588802:405 | 40589776:405 | -;- | 975 | NA | 0 | 0 | 0 | 0 | 0 | 0 |
| ENSDARG000 | 10;10;10;10;1 | 428269:42857 | 428318:42876 | +;+;+;+;+;+;+; | 2144 | NA | 66 | 81 | 128 | 129 | 89 | 105 |
| ENSDARG000 | 10;10 | 40593443:405 | 40594447:405 | -;- | 1005 | NA | 0 | 0 | 0 | 0 | 0 | 0 |
| ENSDARG000 | 10 | 40598791 | 40599801 | + | 1011 | NA | 0 | 0 | 0 | 0 | 0 | 0 |
| ENSDARG000 | 10 | 40599935 | 40600502 | + | 568 | NA | 0 | 0 | 0 | 0 | 0 | 0 |
| ENSDARG000 | 10 | 40606084 | 40607106 | + | 1023 | NA | 0 | 0 | 0 | 0 | 0 | 0 |
| ENSDARG000 | 10;10;10;10;1 | 6664911:6666 | 6665114:6666 | +;+;+;+;+;+;+; | 3086 | NA | 0 | 0 | 0 | 0 | 0 | 0 |
| ENSDARG000 | 10;10;10;10;1 | 31800212:318 | 31800564:318 | -;-;-;-;-;- | 1450 | vps26bl | 180 | 165 | 208 | 177 | 152 | 193 |
| ENSDARG000 | 10 | 40607701 | 40608717 | + | 1017 | NA | 0 | 0 | 0 | 0 | 0 | 0 |
| ENSDARG000 | 10;10;10;10;1 | 17700016:177 | 17702496:177 | -;-;-;-;-;-;-;- | 3406 | STARD7 | 1186 | 1368 | 1690 | 1411 | 1617 | 1321 |
| ENSDARG000 | 10;10;10;10 | 25494173:254 | 25494644:254 | -;-;-;- | 1371 | NA | 166 | 12 | 134 | 38 | 33 | 131 |
| ENSDARG000 | 10;10;10;10;1 | 25402463:254 | 25403447:254 | -;-;-;-;-;-;-;-; | 4477 | GRIK1 | 6 | 0 | 465 | 14 | 396 | 148 |
| ENSDARG000 | 10;10;10 | 31809226:318 | 31809815:318 | +;+;+ | 2022 | foxo1b | 20 | 0 | 20 | 1 | 30 | 42 |
| ENSDARG000 | 10 | 40609461 | 40609953 | + | 493 | NA | 0 | 0 | 0 | 0 | 0 | 0 |
| ENSDARG000 | 10;10 | 40611799:406 | 40612794:406 | -;- | 996 | NA | 0 | 0 | 0 | 0 | 0 | 0 |
| ENSDARG000 | 10;10;10;10;1 | 7975585:7976 | 7976168:7976 | -;-;-;-;-;-;-;-; | 3035 | ewsr1a | 3565 | 4037 | 4272 | 2240 | 4225 | 2766 |
| ENSDARG000 | 10;10 | 40616572:406 | 40617567:406 | -;- | 996 | NA | 0 | 0 | 0 | 0 | 0 | 0 |
| ENSDARG000 | 10;10;10;10;1 | 40623414:406 | 40623582:406 | +;+;+;+;+ | 2554 | zgc:172131 | 14 | 2 | 26 | 17 | 0 | 0 |
| ENSDARG000 | 10;10;10;10;1 | 34426256:344 | 34426852:344 | +;+;+;+;+;+;+; | 10818 | nbeaa | 314 | 296 | 461 | 150 | 248 | 242 |
| ENSDARG000 | 10;10;10;10;1 | 15967341:159 | 15967759:159 | +;+;+;+;+;+;+; | 2999 | NA | 470 | 342 | 672 | 306 | 555 | 399 |
| ENSDARG000 | 10;10;10;10;1 | 6677673:6677 | 6677756:6678 | -;-;-;-;-;-;- | 1033 | NA | 0 | 0 | 0 | 0 | 0 | 0 |
| ENSDARG000 | 10;10;10;10;1 | 31938796:319 | 31939205:319 | +;+;+;+;+;+;+; | 6115 | lhfpl6 | 146 | 67 | 125 | 26 | 88 | 106 |
| ENSDARG000 | 10;10;10;10;1 | 6681523:6683 | 6681729:6683 | +;+;+;+;+;+;+; | 3100 | NA | 0 | 0 | 0 | 0 | 0 | 0 |
| ENSDARG000 | 10 | 40629616 | 40630644 | + | 1029 | NA | 0 | 0 | 2 | 1 | 0 | 0 |
| ENSDARG000 | 10;10;10;10;1 | 27068251:270 | 27069024:270 | +;+;+;+;+ | 2752 | ehd1b | 4917 | 4958 | 3256 | 2706 | 3157 | 4568 |
| ENSDARG000 | 10;10 | 40633990:406 | 40635051:406 | +;+ | 1062 | NA | 0 | 0 | 0 | 0 | 0 | 0 |
| ENSDARG000 | 10 | 40638776 | 40639804 | + | 1029 | NA | 0 | 0 | 0 | 0 | 0 | 0 |
| ENSDARG000 | 10;10;10;10;1 | 17713022:177 | 17713435:177 | +;+;+;+;+;+;+; | 4794 | slc20a1b | 25308 | 20029 | 23997 | 37724 | 23055 | 22123 |
| ENSDARG000 | 10;10 | 40642517:406 | 40643602:406 | +;+ | 1086 | NA | 0 | 0 | 0 | 0 | 0 | 0 |
| ENSDARG000 | 10;10 | 40646764:406 | 40647825:406 | +;+ | 1062 | NA | 0 | 0 | 0 | 0 | 0 | 0 |
| ENSDARG000 | 10;10;10;10;1 | 27096043:270 | 27096270:270 | +;+;+;+;+;+;+; | 1960 | cxadr | 26 | 0 | 6 | 4 | 14 | 10 |
| ENSDARG000 | 10;10;10;10;1 | 466926:47481 | 466989:47492 | +;+;+;+;+;+;+; | 3004 | ARVCF | 51 | 131 | 900 | 249 | 94 | 344 |
| ENSDARG000 | 10;10;10;10;1 | 4940767:4947 | 4941453:4947 | -;-;-;-;-;-;-;-; | 2023 | snx30 | 322 | 284 | 426 | 366 | 262 | 314 |
| ENSDARG000 | 10;10 | 40652399:406 | 40653442:406 | +;+ | 1044 | NA | 0 | 0 | 0 | 0 | 0 | 0 |
| ENSDARG000 | 10;10;10;10;1 | 6693518:6693 | 6693608:6693 | -;-;-;-;-;-;- | 1036 | NA | 0 | 0 | 0 | 0 | 0 | 0 |
| ENSDARG000 | 10;10;10;10;1 | 32050906:320 | 32051060:320 | +;+;+;+;+;+;+ | 2448 | NA | 1479 | 2233 | 2515 | 1124 | 1862 | 1262 |
| ENSDARG000 | 10;10 | 40657949:406 | 40658992:406 | +;+ | 1044 | NA | 0 | 0 | 0 | 0 | 0 | 0 |
| ENSDARG000 | 10 | 40660772 | 40661767 | + | 996 | NA | 0 | 0 | 0 | 0 | 0 | 0 |
| ENSDARG000 | 10;10 | 4962731:4963 | 4962832:4963 | +;+ | 268 | NA | 0 | 0 | 0 | 0 | 4 | 1 |
| ENSDARG000 | 10;10;10;10;1 | 36684000:366 | 36684162:366 | -;-;-;-;-;-;-;- | 820 | mrpl48 | 110 | 138 | 313 | 143 | 110 | 234 |
| ENSDARG000 | 10 | 40665182 | 40666204 | + | 1023 | NA | 0 | 0 | 0 | 0 | 0 | 0 |
| ENSDARG000 | 10;10;10;10;1 | 13209580:132 | 13209804:132 | +;+;+;+;+;+;+; | 1602 | rassf6 | 22 | 0 | 0 | 0 | 2 | 2 |
| ENSDARG000 | 10;10;10;10 | 13236100:132 | 13236525:132 | -;-;-;- | 990 | si:busm1-57f2 | 71 | 6 | 6 | 10 | 4 | 10 |
| ENSDARG000 | 10 | 40670847 | 40672605 | + | 1759 | taar19p | 0 | 0 | 0 | 0 | 0 | 0 |
| ENSDARG000 | 10 | 40677476 | 40678507 | + | 1032 | NA | 0 | 0 | 0 | 0 | 0 | 0 |
| ENSDARG000 | 10;10;10;10;1 | 44057177:440 | 44057614:440 | +;+;+;+;+;+;+; | 2431 | grk3 | 646 | 345 | 603 | 292 | 693 | 654 |
| ENSDARG000 | 10;10;10;10;1 | 4964463:4964 | 4964546:4964 | +;+;+;+;+;+;+ | 900 | lyrm7 | 40 | 58 | 137 | 38 | 80 | 20 |
| ENSDARG000 | 10 | 40684758 | 40685789 | + | 1032 | NA | 0 | 0 | 0 | 0 | 0 | 0 |
| ENSDARG000 | 10 | 40690411 | 40691433 | + | 1023 | NA | 0 | 0 | 0 | 0 | 0 | 0 |
| ENSDARG000 | 10;10;10;10;1 | 32058692:320 | 32058962:320 | +;+;+;+;+ | 2692 | thap12a | 326 | 478 | 599 | 252 | 637 | 400 |
| ENSDARG000 | 10;10;10;10;1 | 10210455:102 | 10210719:102 | +;+;+;+;+;+;+; | 2221 | SH2D3C | 245 | 97 | 180 | 323 | 119 | 361 |
| ENSDARG000 | 10;10;10;10;1 | 27182704:271 | 27185161:271 | -;-;-;-;-;-;-;-; | 9335 | AUTS2 | 482 | 438 | 1183 | 522 | 958 | 416 |
| ENSDARG000 | 10 | 40726399 | 40727430 | + | 1032 | NA | 0 | 0 | 0 | 0 | 0 | 0 |

| | | | | | | | | | | | | |
|---|---|---|---|---|---|---|---|---|---|---|---|---|
| ENSDARG000 | 10 | 40722256 | 40723287 | + | 1032 | NA | 0 | 0 | 0 | 2 | 0 | 0 |
| ENSDARG000 | 10;10;10;10;1 | 36695597:366 | 36695870:366 | +;+;+;+;+;+;+; | 5092 | rab6a | 1375 | 874 | 2174 | 1478 | 1861 | 1428 |
| ENSDARG000 | 10;10;10 | 40700311:407 | 40701071:407 | +;+;+ | 2815 | NA | 0 | 0 | 0 | 0 | 0 | 0 |
| ENSDARG000 | 10 | 40711286 | 40712317 | + | 1032 | NA | 0 | 0 | 0 | 0 | 0 | 0 |
| ENSDARG000 | 10 | 40706100 | 40707131 | + | 1032 | NA | 0 | 0 | 0 | 0 | 0 | 0 |
| ENSDARG000 | 10;10;10;10;1 | 506538:50656 | 506663:50666 | +;+;+;+;+;+;+ | 1443 | NA | 0 | 0 | 19 | 8 | 3 | 0 |
| ENSDARG000 | 10;10 | 27738239:277 | 27738927:277 | -;- | 995 | NA | 0 | 0 | 0 | 0 | 0 | 0 |
| ENSDARG000 | 10;10;10;10 | 518546:52639 | 518655:52675 | +;+;+;+ | 1623 | npffr1l3 | 0 | 0 | 0 | 0 | 0 | 0 |
| ENSDARG000 | 10;10;10;10;1 | 4969634:4972 | 4970112:4973 | -;-;-;-;-;-;-;- | 1753 | mat2al | 144 | 243 | 223 | 240 | 314 | 246 |
| ENSDARG000 | 10;10;10;10;1 | 21471316:214 | 21474729:214 | -;-;-;-;- | 3771 | fgf18b | 4 | 15 | 0 | 12 | 4 | 14 |
| ENSDARG000 | 10;10;10;10;1 | 13279079:132 | 13279128:132 | +;+;+;+;+ | 1398 | tmem267 | 73 | 95 | 176 | 112 | 120 | 82 |
| ENSDARG000 | 10;10;10;10;1 | 32066355:320 | 32066625:320 | +;+;+;+;+;+;+; | 2351 | NA | 369 | 236 | 555 | 132 | 580 | 405 |
| ENSDARG000 | 10;10;10;10;1 | 16006905:160 | 16007140:160 | -;-;-;-;-;-;-;-; | 1899 | aldh7a1 | 1925 | 1943 | 2315 | 1460 | 2579 | 2327 |
| ENSDARG000 | 10;10;10;10;1 | 10313024:103 | 10313099:103 | +;+;+;+;+;+;+; | 962 | urm1 | 234 | 350 | 296 | 260 | 285 | 152 |
| ENSDARG000 | 10;10;10;10;1 | 36720166:367 | 36720334:367 | -;-;-;-;-;-;-;-; | 1399 | NA | 0 | 0 | 0 | 0 | 0 | 2 |
| ENSDARG000 | 10;10;10 | 10319308:103 | 10319651:103 | -;-;- | 624 | NA | 10 | 6 | 7 | 10 | 10 | 5 |
| ENSDARG000 | 10;10;10 | 36726238:367 | 36726521:367 | +;+;+ | 565 | NA | 0 | 0 | 0 | 0 | 0 | 0 |
| ENSDARG000 | 10;10;10;10;1 | 7988746:7989 | 7989607:7989 | -;-;-;-;-;-;-;-; | 3632 | ATP6V0A2 | 8 | 4 | 18 | 0 | 22 | 44 |
| ENSDARG000 | 10;10;10;10;1 | 36731039:367 | 36733001:367 | -;-;-;-;-;-;-;-; | 2507 | plekhb1 | 0 | 0 | 0 | 0 | 0 | 0 |
| ENSDARG000 | 10;10 | 40731834:407 | 40732895:407 | +;+ | 1062 | NA | 0 | 0 | 0 | 0 | 0 | 0 |
| ENSDARG000 | 10 | 40737540 | 40738565 | + | 1026 | NA | 0 | 0 | 0 | 0 | 0 | 0 |
| ENSDARG000 | 10;10;10;10;1 | 25572372:255 | 25574692:255 | -;-;-;-;-;-;-;-; | 7704 | TIAM1 | 158 | 397 | 477 | 252 | 415 | 134 |
| ENSDARG000 | 10;10 | 40742685:407 | 40743746:407 | +;+ | 1062 | NA | 0 | 0 | 0 | 0 | 0 | 0 |
| ENSDARG000 | 10;10;10;10;1 | 32104305:321 | 32104545:321 | +;+;+;+;+;+ | 2171 | wnt11 | 44 | 38 | 1547 | 195 | 436 | 453 |
| ENSDARG000 | 10 | 40747873 | 40748898 | + | 1026 | NA | 0 | 0 | 0 | 0 | 0 | 0 |
| ENSDARG000 | 10;10 | 40756352:407 | 40757389:407 | +;+ | 1038 | NA | 0 | 0 | 0 | 0 | 0 | 0 |
| ENSDARG000 | 10;10 | 40762200:407 | 40763234:407 | +;+ | 1035 | NA | 0 | 0 | 0 | 0 | 0 | 0 |
| ENSDARG000 | 10;10 | 40768203:407 | 40769264:407 | +;+ | 1062 | NA | 0 | 0 | 0 | 0 | 0 | 0 |
| ENSDARG000 | 10 | 40774215 | 40775243 | + | 1029 | NA | 0 | 0 | 0 | 0 | 0 | 0 |
| ENSDARG000 | 10;10 | 40781052:407 | 40782113:407 | +;+ | 1062 | NA | 0 | 0 | 0 | 0 | 0 | 0 |
| ENSDARG000 | 10;10;10;10;1 | 13305610:133 | 13307159:133 | -;-;-;-;-;-;-;-; | 2887 | NA | 144 | 120 | 230 | 185 | 197 | 209 |
| ENSDARG000 | 10;10;10;10;1 | 27977373:279 | 27978532:279 | -;-;-;-;-;-;-;-; | 5266 | ints2 | 419 | 529 | 785 | 531 | 724 | 646 |
| ENSDARG000 | 10;10;10;10;1 | 21541925:215 | 21542540:215 | -;-;-;-;-;-;-;-; | 2378 | zgc:165539 | 0 | 0 | 0 | 0 | 0 | 0 |
| ENSDARG000 | 10;10;10;10 | 40792078:407 | 40792186:407 | +;+;+;+ | 1045 | NA | 6 | 8 | 0 | 0 | 0 | 0 |
| ENSDARG000 | 10;10;10;10;1 | 40821804:408 | 40822301:408 | -;-;-;-;-;- | 1445 | pcna | 1050 | 6055 | 7643 | 2631 | 5749 | 3330 |
| ENSDARG000 | 10;10;10;10;1 | 40786196:407 | 40788105:407 | -;-;-;-;-;-;-;-; | 2394 | zgc:113625 | 4 | 0 | 0 | 0 | 0 | 0 |
| ENSDARG000 | 10;10;10;10;1 | 21559605:215 | 21559642:215 | +;+;+;+;+;+;+; | 5978 | pcdh1a6 | 11 | 15 | 25 | 11 | 31 | 11 |
| ENSDARG000 | 10;10;10;10;1 | 21563986:215 | 21566453:215 | +;+;+;+;+;+;+; | 8178 | pcdh1a4 | 44 | 53 | 46 | 39 | 23 | 53 |
| ENSDARG000 | 10;10;10;10 | 21559605:216 | 21562058:216 | +;+;+;+ | 5392 | pcdh1a3 | 4 | 11 | 11 | 0 | 0 | 2 |
| ENSDARG000 | 10;10;10;10;1 | 21570196:215 | 21570590:215 | -;-;-;-;-;-;-;-; | 1383 | NA | 0 | 0 | 2 | 7 | 0 | 0 |
| ENSDARG000 | 10;10;10;10;1 | 34737322:347 | 34738411:347 | -;-;-;-;-;-;-;-; | 4154 | dclk1a | 65 | 267 | 286 | 134 | 688 | 47 |
| ENSDARG000 | 10;10;10;10;1 | 13460130:134 | 13461275:134 | -;-;-;-;-;-;-;-; | 2485 | cntfr | 3836 | 2815 | 3619 | 3047 | 3193 | 3109 |
| ENSDARG000 | 10;10 | 4986050:4987 | 4986216:4987 | +;+ | 525 | NA | 0 | 0 | 0 | 0 | 0 | 2 |
| ENSDARG000 | 10;10;10 | 4987766:4991 | 4987978:4992 | +;+;+ | 862 | si:ch73-234b2 | 40 | 72 | 34 | 54 | 54 | 80 |
| ENSDARG000 | 10;10;10;10;1 | 32264857:322 | 32265760:322 | -;-;-;-;-;-;-;-; | 2720 | uvrag | 294 | 199 | 328 | 172 | 308 | 219 |
| ENSDARG000 | 10;10;10;10;1 | 17731293:177 | 17731623:177 | -;-;-;-;-;-;-;-; | 1101 | NA | 403 | 27 | 43 | 245 | 78 | 298 |
| ENSDARG000 | 10;10;10;10;1 | 8045364:8045 | 8045582:8045 | -;-;-;-;-;-;-;-; | 1021 | zgc:136254 | 0 | 0 | 0 | 0 | 0 | 0 |
| ENSDARG000 | 10;10;10 | 13856792:138 | 13857123:138 | +;+;+ | 707 | NA | 0 | 0 | 0 | 0 | 0 | 0 |
| ENSDARG000 | 10;10 | 13930848:139 | 13931174:139 | +;+ | 394 | NA | 0 | 0 | 0 | 0 | 0 | 0 |
| ENSDARG000 | 10;10;10 | 14210667:142 | 14210823:142 | +;+;+ | 853 | NA | 3 | 0 | 0 | 0 | 0 | 0 |
| ENSDARG000 | 10;10;10;10;1 | 5000623:5002 | 5001026:5002 | -;-;-;-;-;-;-;-; | 2723 | hnrnpd | 2956 | 2106 | 2060 | 1737 | 2272 | 2204 |
| ENSDARG000 | 10;10;10;10;1 | 21804772:218 | 21805294:218 | +;+;+;+;+;+;+ | 4370 | pcdh1g31 | 8 | 1 | 23 | 29 | 8 | 1 |
| ENSDARG000 | 10;10;10;10;1 | 10351685:103 | 10351975:103 | +;+;+;+;+;+;+; | 1864 | CERCAM | 368 | 378 | 545 | 375 | 394 | 442 |
| ENSDARG000 | 10;10;10;10;1 | 5017216:5017 | 5017514:5017 | -;-;-;-;-;-;-;-; | 1836 | hnrnpdl | 1622 | 1195 | 1681 | 1264 | 1838 | 1391 |
| ENSDARG000 | 10;10;10;10;1 | 21786656:217 | 21786935:217 | +;+;+;+;+;+;+; | 4435 | pcdh1g26 | 0 | 4 | 0 | 0 | 0 | 0 |
| ENSDARG000 | 10;10;10;10;1 | 8051616:8052 | 8052209:8052 | -;-;-;-;-;-;-;-; | 1695 | si:ch211-251f | 0 | 0 | 0 | 0 | 0 | 0 |
| ENSDARG000 | 10;10;10;10 | 21793670:218 | 21796108:218 | +;+;+;+ | 4321 | pcdh1gc6 | 10 | 9 | 23 | 12 | 11 | 6 |
| ENSDARG000 | 10;10;10;10;1 | 21677058:216 | 21679505:216 | +;+;+;+;+;+;+ | 2904 | pcdh1gb2 | 39 | 34 | 46 | 12 | 19 | 21 |
| ENSDARG000 | 10;10;10;10;1 | 21670831:216 | 21670901:216 | +;+;+;+;+;+;+; | 2044 | NA | 0 | 0 | 0 | 0 | 0 | 0 |
| ENSDARG000 | 10;10;10;10;1 | 21730585:217 | 21732999:217 | +;+;+;+;+;+;+; | 6965 | pcdh1g22 | 51 | 35 | 76 | 92 | 26 | 22 |
| ENSDARG000 | 10;10;10;10;1 | 36741502:367 | 36741522:367 | +;+;+;+;+;+;+; | 2267 | FCHSD2 | 0 | 0 | 10 | 0 | 2 | 0 |
| ENSDARG000 | 10;10;10;10 | 21660447:218 | 21662938:218 | +;+;+;+ | 4374 | pcdh1g3 | 98 | 55 | 74 | 87 | 54 | 26 |
| ENSDARG000 | 10;10;10;10;1 | 21708463:217 | 21708867:217 | +;+;+;+;+;+;+; | 6838 | pcdh1g11 | 19 | 1 | 21 | 9 | 17 | 5 |
| ENSDARG000 | 10;10;10;10;1 | 21796477:217 | 21799746:217 | +;+;+;+;+;+;+; | 5282 | pcdh1g29 | 4 | 4 | 2 | 1 | 4 | 6 |
| ENSDARG000 | 10;10;10;10;1 | 21701551:217 | 21703522:217 | +;+;+;+;+;+;+; | 7632 | pcdh1g9 | 21 | 34 | 43 | 18 | 51 | 32 |
| ENSDARG000 | 10;10;10;10;1 | 8077969:8078 | 8078189:8078 | -;-;-;-;-;- | 920 | zgc:173443 | 0 | 0 | 0 | 0 | 0 | 0 |
| ENSDARG000 | 10;10;10;10 | 21650828:218 | 21653280:218 | +;+;+;+ | 4335 | pcdh1g1 | 74 | 89 | 94 | 65 | 55 | 19 |
| ENSDARG000 | 10;10;10;10;1 | 21789954:217 | 21792416:217 | +;+;+;+;+;+;+; | 4475 | pcdh1gc5 | 9 | 1 | 1 | 1 | 14 | 5 |
| ENSDARG000 | 10;10;10;10;1 | 8058850:8059 | 8059417:8059 | -;-;-;-;-;-;-;-; | 1508 | si:ch211-251f | 0 | 0 | 0 | 0 | 0 | 0 |
| ENSDARG000 | 10;10;10;10;1 | 21801030:218 | 21803484:218 | +;+;+;+;+;+;+; | 4467 | pcdh1g30 | 17 | 9 | 18 | 0 | 5 | 2 |
| ENSDARG000 | 10;10;10;10;1 | 34872195:348 | 34872382:348 | -;-;-;-;-;-;-;-; | 998 | ccdc169 | 0 | 0 | 0 | 0 | 6 | 0 |
| ENSDARG000 | 10;10;10;10 | 21656654:218 | 21659158:218 | +;+;+;+ | 4387 | pcdh1g2 | 92 | 95 | 141 | 156 | 91 | 78 |
| ENSDARG000 | 10;10;10;10;1 | 21783213:218 | 21785818:218 | +;+;+;+;+ | 4487 | pcdh1g33 | 0 | 1 | 0 | 1 | 1 | 0 |
| ENSDARG000 | 10;10;10;10 | 21718468:218 | 21720873:218 | +;+;+;+ | 2814 | pcdh1gb9 | 11 | 12 | 40 | 2 | 22 | 0 |
| ENSDARG000 | 10;10;10 | 21814475:218 | 21814800:218 | -;-;- | 534 | NA | 0 | 0 | 3 | 7 | 0 | 0 |
| ENSDARG000 | 10 | 21722892 | 21725312 | + | 2421 | NA | 6 | 28 | 17 | 33 | 22 | 7 |
| ENSDARG000 | 10 | 21726853 | 21729246 | + | 2394 | NA | 5 | 8 | 59 | 42 | 35 | 3 |
| ENSDARG000 | 10;10;10;10 | 21807497:218 | 21810013:218 | +;+;+;+ | 4399 | pcdh1g32 | 0 | 4 | 14 | 52 | 10 | 4 |
| ENSDARG000 | 10;10;10;10;1 | 21737745:217 | 21738167:217 | +;+;+;+;+;+;+; | 4444 | pcdh1g18 | 113 | 84 | 85 | 97 | 87 | 101 |
| ENSDARG000 | 10;10;10;10;1 | 40951697:409 | 40954406:409 | -;-;-;-;-;-;- | 3195 | npffr1l1 | 0 | 0 | 2 | 0 | 4 | 0 |
| ENSDARG000 | 10;10;10;10;1 | 25694395:256 | 25694717:256 | -;-;-;-;-;-;- | 1250 | sod1 | 895 | 998 | 1137 | 762 | 1414 | 883 |
| ENSDARG000 | 10;10;10;10;1 | 40848354:408 | 40848370:408 | -;-;-;-;-;-;-;-; | 5348 | bmp1b | 547 | 49 | 103 | 256 | 358 | 658 |
| ENSDARG000 | 10;10;10;10;1 | 25697636:257 | 25702412:257 | -;-;-;-;-;-;-;-; | 7192 | SCAF4 | 859 | 843 | 1147 | 770 | 1081 | 1007 |
| ENSDARG000 | 10;10 | 40939664:409 | 40939770:409 | +;+ | 496 | NA | 0 | 0 | 0 | 0 | 0 | 0 |
| ENSDARG000 | 10;10;10;10;1 | 36795861:368 | 36796409:368 | -;-;-;-;-;-;- | 1635 | dhrs13a.1 | 618 | 757 | 411 | 342 | 602 | 692 |
| ENSDARG000 | 10;10;10;10;1 | 36950773:369 | 36951722:369 | -;-;-;-;-;-;-;-; | 8656 | NA | 185 | 6 | 606 | 613 | 78 | 195 |
| ENSDARG000 | 10;10;10;10;1 | 34905980:349 | 34908764:349 | -;-;-;-;-;-;-;-; | 4382 | ccna1 | 6 | 0 | 0 | 2 | 0 | 0 |
| ENSDARG000 | 10;10;10;10;1 | 36828069:368 | 36828504:368 | -;-;-;-;-;-;-;-; | 3697 | SEZ6 | 0 | 0 | 6 | 38 | 0 | 4 |
| ENSDARG000 | 10;10;10;10;1 | 14263684:142 | 14264167:142 | -;-;-;-;-;-;-;-; | 2320 | galt | 562 | 422 | 375 | 439 | 483 | 524 |
| ENSDARG000 | 10;10;10;10;1 | 34889328:348 | 34889894:348 | -;-;-;-;-;-;-;- | 2188 | sparta | 6 | 0 | 0 | 0 | 0 | 0 |
| ENSDARG000 | 10;10;10;10;1 | 36805085:368 | 36805457:368 | +;+;+;+;+;+ | 844 | NA | 5 | 0 | 0 | 0 | 0 | 0 |
| ENSDARG000 | 10;10;10;10;1 | 32501961:325 | 32502508:325 | -;-;-;-;-;-;-;-; | 2329 | dgat2 | 0 | 0 | 4 | 0 | 0 | 0 |
| ENSDARG000 | 10;10;10;10;1 | 36811372:368 | 36812242:368 | -;-;-;-;-;-;-;-; | 4082 | PHF12 | 569 | 387 | 480 | 315 | 530 | 629 |
| ENSDARG000 | 10;10 | 37041078:370 | 37041445:370 | -;- | 539 | NA | 0 | 0 | 0 | 0 | 0 | 0 |
| ENSDARG000 | 10;10;10;10;1 | 32560667:325 | 32562311:325 | +;+;+;+;+;+;+; | 2722 | NA | 109 | 40 | 2 | 144 | 8 | 12 |
| ENSDARG000 | 10;10;10;10;1 | 32534814:325 | 32535475:325 | -;-;-;-;-;-;-;-; | 2257 | mogat2 | 4 | 12 | 0 | 0 | 0 | 0 |
| ENSDARG000 | 10;10;10;10;1 | 40973524:409 | 40974080:409 | +;+;+;+;+;+;+; | 2737 | ANTXR1 | 12 | 4 | 5 | 15 | 30 | 4 |
| ENSDARG000 | 10;10;10;10 | 14488625:144 | 14488867:144 | +;+;+;+ | 1154 | sigmar1 | 2426 | 2052 | 2453 | 1330 | 2469 | 2077 |
| ENSDARG000 | 10;10;10;10;1 | 34960751:349 | 34962184:349 | -;-;-;-;-;-;-;-; | 8357 | smad9 | 56 | 38 | 44 | 4 | 38 | 38 |
| ENSDARG000 | 10;10;10;10;1 | 34937548:349 | 34937677:349 | +;+;+;+;+;+;+; | 835 | NA | 0 | 0 | 0 | 0 | 0 | 0 |
| ENSDARG000 | 10;10;10;10 | 34951060:349 | 34951528:349 | -;-;-;- | 1326 | NA | 0 | 0 | 0 | 0 | 0 | 0 |
| ENSDARG000 | 10;10;10;10;1 | 34984617:349 | 34984973:349 | -;-;-;-;-;-;-;-; | 1508 | alg5 | 34 | 48 | 28 | 12 | 30 | 25 |
| ENSDARG000 | 10;10;10 | 34958583:349 | 34959176:349 | +;+;+ | 1442 | rfxap | 0 | 0 | 0 | 0 | 0 | 0 |
| ENSDARG000 | 10;10;10;10;1 | 14499201:145 | 14499327:145 | +;+;+;+;+;+;+; | 1418 | katnal2 | 0 | 8 | 2 | 0 | 4 | 0 |
| ENSDARG000 | 10;10;10;10;1 | 5025696:5026 | 5025964:5026 | +;+;+;+;+;+ | 1947 | enoph1 | 363 | 300 | 316 | 290 | 276 | 306 |
| ENSDARG000 | 10;10;10;10;1 | 10564261:105 | 10564629:105 | -;-;-;-;-;-;-;-; | 2280 | dbh | 0 | 0 | 2 | 0 | 0 | 0 |
| ENSDARG000 | 10;10;10;10;1 | 32613706:326 | 32613825:326 | -;-;-;-;-;-;-;-; | 2960 | umodl1 | 2 | 6 | 4 | 4 | 4 | 4 |
| ENSDARG000 | 10;10;10;10;1 | 10386435:103 | 10386706:103 | +;+;+;+;+;+;+; | 7049 | sardh | 232 | 266 | 557 | 314 | 488 | 448 |
| ENSDARG000 | 10;10 | 32646402:326 | 32646502:326 | +;+ | 491 | NA | 17 | 33 | 48 | 23 | 36 | 26 |
| ENSDARG000 | 10;10;10;10;1 | 32657496:326 | 32658099:326 | -;-;-;-;-;-;-;- | 1327 | atg101 | 391 | 267 | 301 | 360 | 238 | 370 |
| ENSDARG000 | 10;10;10;10;1 | 32662886:326 | 32663657:326 | +;+;+;+;+;+;+; | 1934 | rad51c | 16 | 40 | 59 | 30 | 45 | 19 |
| ENSDARG000 | 10;10;10;10;1 | 35002786:350 | 35002826:350 | +;+;+;+;+;+;+; | 1267 | exosc8 | 169 | 269 | 379 | 169 | 230 | 163 |
| ENSDARG000 | 10;10;10;10;1 | 14871136:148 | 14873313:148 | -;-;-;-;-;-;-;-; | 4051 | smad2 | 1433 | 1491 | 1589 | 1024 | 1829 | 1092 |
| ENSDARG000 | 10;10;10;10;1 | 14936045:149 | 14936541:149 | -;-;-;-;-;-;-;-;- | 1108 | NA | 0 | 0 | 0 | 0 | 0 | 0 |
| ENSDARG000 | 10;10;10;10;1 | 37083654:370 | 37083697:370 | +;+;+;+;+;+;+; | 1473 | VEZF1 | 158 | 131 | 283 | 104 | 159 | 97 |
| ENSDARG000 | 10;10;10;10;1 | 35012094:350 | 35012243:350 | -;-;-;-;-;-;-;-; | 2793 | SUPT20H | 252 | 222 | 349 | 243 | 298 | 237 |
| ENSDARG000 | 10;10;10;10;1 | 5041977:5042 | 5042477:5042 | -;-;-;-;-;-;-;-; | 1791 | TMEM150C | 16 | 10 | 8 | 6 | 3 | 0 |
| ENSDARG000 | 10;10;10;10;1 | 5060191:5060 | 5060420:5060 | +;+;+;+;+;+;+; | 1475 | NA | 6 | 2 | 6 | 8 | 4 | 0 |
| ENSDARG000 | 10;10;10;10;1 | 5092511:5094 | 5094819:5094 | -;-;-;-;-;-;-;-; | 6071 | sec31a | 397 | 429 | 845 | 354 | 756 | 394 |
| ENSDARG000 | 10;10;10;10;1 | 8437930:8450 | 8438131:8450 | +;+;+;+;+;+ | 894 | PPTC7 | 138 | 102 | 101 | 52 | 126 | 108 |
| ENSDARG000 | 10;10;10;10 | 44462447:444 | 44465311:444 | -;-;-;- | 3327 | ccdc62 | 32 | 22 | 34 | 36 | 32 | 20 |
| ENSDARG000 | 10;10;10;10 | 42677891:426 | 42678807:426 | -;-;-;- | 1972 | stc1l | 0 | 0 | 0 | 0 | 213 | 0 |
| ENSDARG000 | 10;10 | 4095917:4112 | 4099022:4112 | +;+ | 3162 | MN1 | 48 | 32 | 143 | 20 | 116 | 8 |
| ENSDARG000 | 10;10;10;10;1 | 8416117:8418 | 8416412:8418 | -;-;-;-;-;-;-;-; | 1810 | NA | 89 | 134 | 121 | 105 | 127 | 57 |
| ENSDARG000 | 10;10;10;10;1 | 21938762:219 | 21939017:219 | -;-;-;-;- | 785 | NA | 256 | 284 | 247 | 315 | 329 | 410 |
| ENSDARG000 | 10 | 37927100 | 37928125 | + | 1026 | bhlha9 | 2 | 0 | 0 | 0 | 0 | 0 |
| ENSDARG000 | 10;10;10;10;1 | 33253952:332 | 33255251:332 | -;-;-;-;-;-;- | 2391 | tbl2 | 738 | 784 | 1129 | 512 | 953 | 816 |
| ENSDARG000 | 10;10;10;10;1 | 23553533:235 | 23554056:235 | +;+;+;+;+ | 2726 | NA | 0 | 4 | 2 | 2 | 26 | 4 |
| ENSDARG000 | 10;10;10;10;1 | 644574:64457 | 644831:64483 | -;-;-;-;-;-;-;-; | 2734 | glis3 | 0 | 2 | 0 | 16 | 6 | 0 |
| ENSDARG000 | 10;10;10;10;1 | 3299829:3302 | 3299981:3302 | +;+;+;+;+;+;+; | 5765 | zgc:56235 | 368 | 333 | 471 | 342 | 501 | 340 |
| ENSDARG000 | 10;10;10;10;1 | 44363195:443 | 44363383:443 | +;+;+;+;+;+;+ | 2602 | kmt5aa | 1711 | 1708 | 2066 | 1251 | 1939 | 1642 |
| ENSDARG000 | 10;10;10;10;1 | 38663917:386 | 38664117:386 | +;+;+;+;+;+;+; | 4887 | acat1 | 935 | 896 | 1368 | 1017 | 1284 | 1469 |
| ENSDARG000 | 10;10;10;10;1 | 42873163:428 | 42876976:428 | -;-;-;-;- | 5361 | NA | 19246 | 8153 | 7056 | 9066 | 8896 | 11072 |
| ENSDARG000 | 10;10;10;10;1 | 2499383:2502 | 2499404:2502 | -;-;-;-;-;-;-;-; | 1981 | NA | 432 | 396 | 616 | 161 | 764 | 351 |
| ENSDARG000 | 10;10;10;10;1 | 8481660:8482 | 8482014:8482 | +;+;+;+;+ | 1694 | NA | 0 | 4 | 0 | 0 | 0 | 12 |
| ENSDARG000 | 10;10 | 2680079:2681 | 2680450:2682 | -;- | 1001 | pdxp | 80 | 160 | 200 | 88 | 141 | 93 |
| ENSDARG000 | 10;10;10 | 23548674:235 | 23549416:235 | +;+;+ | 1503 | NA | 0 | 0 | 0 | 0 | 6 | 0 |
| ENSDARG000 | 10;10;10 | 44373349:443 | 44373370:443 | +;+;+ | 762 | snrnp35 | 101 | 119 | 175 | 144 | 100 | 95 |
| ENSDARG000 | 10;10 | 31344227:313 | 31344479:313 | +;+ | 461 | NA | 2 | 0 | 0 | 0 | 0 | 0 |
| ENSDARG000 | 10 | 805244 | 805359 | + | 116 | 5S_rRNA | 0 | 0 | 0 | 0 | 0 | 0 |
| ENSDARG000 | 10;10;10;10 | 15064433:150 | 15064964:150 | +;+;+;+ | 5716 | NA | 26 | 40 | 26 | 8 | 80 | 40 |
| ENSDARG000 | 10;10;10;10;1 | 4235998:4240 | 4236054:4241 | +;+;+;+;+;+;+; | 999 | PLPPR1 | 119 | 4 | 2 | 50 | 24 | 145 |
| ENSDARG000 | 10;10;10;10;1 | 21913105:219 | 21915825:219 | -;-;-;-;-;-;-;-; | 10536 | NA | 889 | 121 | 97 | 143 | 89 | 659 |
| ENSDARG000 | 10;10;10;10 | 116115:11749 | 117406:11755 | -;-;-;- | 1467 | ripply3 | 0 | 0 | 4 | 0 | 2 | 0 |
| ENSDARG000 | 10;10 | 19569052:195 | 19569977:195 | +;+ | 1149 | NA | 2 | 0 | 4 | 0 | 0 | 0 |
| ENSDARG000 | 10;10 | 589501:59090 | 590367:59184 | +;+ | 1814 | NA | 265 | 317 | 347 | 322 | 308 | 373 |
| ENSDARG000 | 10 | 158590 | 159597 | + | 1008 | KCNJ15 | 2 | 0 | 17 | 0 | 42 | 4 |
| ENSDARG000 | 10 | 3506540 | 3506654 | - | 115 | 5S_rRNA | 0 | 0 | 0 | 0 | 0 | 0 |
| ENSDARG000 | 10 | 8343188 | 8343305 | + | 118 | 5S_rRNA | 0 | 0 | 0 | 0 | 0 | 0 |
| ENSDARG000 | 10;10;10;10;1 | 44309997:443 | 44310139:443 | -;-;-;-;-;-;-;-; | 4158 | sbno1 | 1566 | 1525 | 1939 | 932 | 1638 | 1123 |
| ENSDARG000 | 10;10;10;10;1 | 33240410:332 | 33245304:332 | -;-;-;-;-;- | 5567 | bcl7ba | 248 | 451 | 378 | 198 | 402 | 164 |
| ENSDARG000 | 10;10;10;10;1 | 38560032:385 | 38560142:385 | +;+;+;+;+;+;+; | 1665 | NA | 7 | 4 | 2 | 0 | 2 | 23 |
| ENSDARG000 | 10;10;10;10;1 | 44042033:440 | 44042264:440 | +;+;+;+;+;+;+; | 1807 | cryba4 | 0 | 1 | 2 | 0 | 0 | 0 |
| ENSDARG000 | 10 | 27414246 | 27414362 | - | 117 | 5S_rRNA | 0 | 0 | 0 | 0 | 0 | 0 |
| ENSDARG000 | 10 | 11774405 | 11774519 | + | 115 | 5S_rRNA | 0 | 0 | 0 | 0 | 0 | 0 |
| ENSDARG000 | 10;10;10;10;1 | 25312337:253 | 25313428:253 | -;-;-;-;-;-;-;-; | 2369 | tmem135 | 460 | 453 | 498 | 222 | 417 | 332 |
| ENSDARG000 | 10;10;10;10;1 | 33367355:333 | 33370121:333 | -;-;-;-;-;-;- | 3560 | ska2 | 369 | 289 | 445 | 266 | 323 | 368 |
| ENSDARG000 | 10 | 14732283 | 14732399 | - | 117 | 5S_rRNA | 0 | 0 | 0 | 0 | 0 | 0 |
| ENSDARG000 | 10;10;10;10;1 | 27021164:270 | 27021189:270 | +;+;+;+;+;+;+; | 198 | NA | 0 | 0 | 0 | 0 | 0 | 0 |
| ENSDARG000 | 10;10;10;10;1 | 38593645:385 | 38593797:385 | +;+;+;+;+;+;+; | 1694 | mmp13a | 369 | 431 | 301 | 349 | 347 | 324 |
| ENSDARG000 | 10 | 29333059 | 29333175 | + | 117 | 5S_rRNA | 0 | 0 | 0 | 0 | 0 | 0 |
| ENSDARG000 | 10 | 20908442 | 20908556 | - | 115 | 5S_rRNA | 0 | 0 | 0 | 0 | 0 | 0 |

| | | | | | | | | | | | |
|---|---|---|---|---|---|---|---|---|---|---|---|
| ENSDARG000 | 10;10;10;10;1 | 16592851;165 | 16593527;165 | +;+;+;+;+;+;+; | 5229 | chsy3 | 213 | 168 | 149 | 93 | 149 | 194 |
| ENSDARG000 | 10;10;10 | 2582254;2584 | 2582710;2584 | +;+;+ | 962 | nxnl2 | 37 | 47 | 45 | 56 | 43 | 33 |
| ENSDARG000 | 10;10;10;10 | 3322663;3323 | 3325394;3325 | -;-;-;- | 2819 | tor4aa | 192 | 194 | 292 | 202 | 204 | 222 |
| ENSDARG000 | 10;10;10 | 42159471;421 | 42159623;421 | +;+;+ | 555 | NA | 6 | 0 | 0 | 0 | 0 | 0 |
| ENSDARG000 | 10;10;10;10;1 | 28392426;283 | 28393164;283 | -;-;-;-;-;- | 1733 | NA | 2 | 4 | 6 | 2 | 22 | 18 |
| ENSDARG000 | 10 | 44907768 | 44907883 | + | 116 | 5S_rRNA | 0 | 0 | 0 | 0 | 0 | 0 |
| ENSDARG000 | 10;10;10;10;1 | 1294463;1296 | 1294813;1296 | -;-;-;-;-;-;-;- | 1596 | opn4xa | 14 | 0 | 6 | 0 | 0 | 0 |
| ENSDARG000 | 10;10;10;10;1 | 12517059;125 | 12517436;125 | +;+;+;+;+;+;+; | 3245 | NA | 0 | 0 | 0 | 0 | 0 | 0 |
| ENSDARG000 | 10 | 22752232 | 22752348 | - | 117 | 5S_rRNA | 0 | 0 | 5 | 0 | 5 | 14 |
| ENSDARG000 | 10 | 1899595 | 1900662 | + | 1068 | NA | 18 | 699 | 251 | 169 | 757 | 61 |
| ENSDARG000 | 10;10;10;10;1 | 17235370;172 | 17235441;172 | +;+;+;+;+;+;+; | 3535 | sppl3 | 914 | 658 | 808 | 601 | 848 | 893 |
| ENSDARG000 | 10;10;10;10;1 | 2568793;2568 | 2568863;2568 | +;+;+;+;+;+;+; | 821 | NA | 167 | 133 | 262 | 102 | 255 | 261 |
| ENSDARG000 | 10;10;10;10;1 | 7653184;7653 | 7653973;7653 | -;-;-;-;-;-;-; | 1809 | pcyox1 | 278 | 255 | 571 | 230 | 443 | 204 |
| ENSDARG000 | 10 | 28292843 | 28292959 | - | 117 | 5S_rRNA | 0 | 0 | 0 | 0 | 0 | 0 |
| ENSDARG000 | 10 | 40741330 | 40741447 | - | 118 | 5S_rRNA | 0 | 0 | 0 | 0 | 0 | 0 |
| ENSDARG000 | 10;10;10;10 | 6967293;6970 | 6967406;6970 | -;-;-;- | 594 | NA | 7 | 9 | 10 | 14 | 14 | 6 |
| ENSDARG000 | 10;10;10;10 | 41668483;416 | 41668994;416 | +;+;+;+ | 1328 | lrrc75bb | 7 | 13 | 18 | 35 | 20 | 10 |
| ENSDARG000 | 10;10;10;10;1 | 25108853;251 | 25109365;251 | -;-;-;-;-;-;-; | 13089 | FAT3 | 4 | 0 | 6 | 0 | 2 | 10 |
| ENSDARG000 | 10;10;10;10;1 | 25219728;252 | 25219809;252 | +;+;+;+;+;+;+; | 7623 | grm5a | 0 | 0 | 0 | 0 | 0 | 0 |
| ENSDARG000 | 10;10;10;10;1 | 18557102;185 | 18557174;185 | +;+;+;+;+;+;+; | 780 | NA | 0 | 0 | 0 | 0 | 0 | 2 |
| ENSDARG000 | 10;10;10;10;1 | 41765616;417 | 41766041;417 | +;+;+;+;+;+;+; | 1705 | rnf34b | 2 | 1 | 2 | 2 | 0 | 0 |
| ENSDARG000 | 10;10;10;10;1 | 360237;36106 | 360350;36124 | -;-;-;-;-;-;-; | 1260 | NA | 2 | 0 | 0 | 0 | 10 | 0 |
| ENSDARG000 | 10 | 21236564 | 21236678 | + | 115 | 5S_rRNA | 0 | 0 | 0 | 0 | 0 | 0 |
| ENSDARG000 | 10;10;10;10;1 | 3268581;3271 | 3269824;3271 | -;-;-;-;-;-;-;- | 2242 | slc25a1b | 508 | 279 | 146 | 77 | 151 | 230 |
| ENSDARG000 | 10;10;10;10;1 | 19554604;195 | 19554815;195 | +;+;+;+;+;+;+; | 2384 | atp6v1b2 | 212 | 129 | 241 | 300 | 200 | 176 |
| ENSDARG000 | 10;10;10;10;1 | 45371044;453 | 45371119;453 | -;-;-;-;-;-;- | 804 | NA | 93 | 91 | 83 | 50 | 81 | 57 |
| ENSDARG000 | 10 | 20256454 | 20257175 | + | 722 | tcimb | 54 | 40 | 26 | 15 | 38 | 26 |
| ENSDARG000 | 10;10;10 | 2684958;2688 | 2685246;2688 | +;+;+ | 582 | SETD9 | 54 | 26 | 47 | 36 | 49 | 43 |
| ENSDARG000 | 10;10;10;10;1 | 304596;30518 | 304836;30521 | -;-;-;-;-;-;- | 969 | NHLRC3 | 407 | 250 | 266 | 298 | 295 | 356 |
| ENSDARG000 | 10;10;10;10;1 | 41926888;419 | 41927001;419 | -;-;-;-;-;- | 1393 | morn3 | 14 | 7 | 4 | 4 | 14 | 27 |
| ENSDARG000 | 10;10;10;10;1 | 36037977;360 | 36038059;360 | +;+;+;+;+;+;+; | 2110 | katnal1 | 74 | 162 | 154 | 206 | 152 | 130 |
| ENSDARG000 | 10;10;10;10 | 24865535;248 | 24866554;248 | -;-;-;- | 2385 | NA | 0 | 0 | 0 | 0 | 0 | 2 |
| ENSDARG000 | 10;10;10;10;1 | 42740862;427 | 42745580;427 | -;-;-;-;-;-;-; | 5449 | zgc:100918 | 7227 | 2374 | 7569 | 7170 | 2964 | 4463 |
| ENSDARG000 | 10;10;10;10;1 | 600164;60042 | 600714;60071 | -;-;-;-;-;-;-; | 2780 | rfx3 | 58 | 54 | 144 | 95 | 61 | 84 |
| ENSDARG000 | 10;10;10;10;1 | 4499943;4499 | 4499963;4500 | +;+;+;+;+;+;+; | 2013 | PLK2 | 1166 | 1042 | 1418 | 856 | 2661 | 1471 |
| ENSDARG000 | 10;10;10;10;1 | 1434956;1434 | 1435211;1435 | -;-;-;-;-;-;-; | 2233 | WDR70 | 312 | 178 | 252 | 133 | 248 | 230 |
| ENSDARG000 | 10;10;10 | 14542726;145 | 14543380;145 | -;-;- | 913 | ier3ip1 | 406 | 644 | 655 | 508 | 529 | 436 |
| ENSDARG000 | 10;10;10;10;1 | 4279521;4279 | 4279854;4279 | -;-;-;-;-;-;-; | 3273 | NA | 8 | 0 | 16 | 0 | 111 | 2 |
| ENSDARG000 | 10;10;10;10 | 31624143;316 | 31624759;316 | +;+;+;+ | 3145 | MSANTD2 | 122 | 74 | 128 | 52 | 157 | 98 |
| ENSDARG000 | 10 | 27063174 | 27063257 | + | 84 | dre-mir-194a | 0 | 0 | 0 | 0 | 0 | 0 |
| ENSDARG000 | 10;10;10;10 | 23520952;235 | 23521536;235 | +;+;+;+ | 1998 | NA | 0 | 0 | 2 | 0 | 0 | 8 |
| ENSDARG000 | 10;10;10;10;1 | 4046448;4046 | 4046467;4046 | +;+;+;+;+;+;+; | 867 | pitpnb | 254 | 211 | 300 | 90 | 329 | 212 |
| ENSDARG000 | 10;10;10;10 | 22097328;221 | 22098406;221 | -;-;-;- | 1503 | NA | 0 | 0 | 0 | 0 | 0 | 0 |
| ENSDARG000 | 10;10 | 22501763;225 | 22503956;225 | -;- | 2520 | zgc:172339 | 4 | 0 | 6 | 10 | 6 | 10 |
| ENSDARG000 | 10;10;10;10 | 3507861;3508 | 3507898;3508 | +;+;+;+ | 351 | NA | 0 | 2 | 6 | 0 | 2 | 25 |
| ENSDARG000 | 10 | 28923517 | 28923633 | - | 117 | 5S_rRNA | 0 | 0 | 0 | 0 | 0 | 0 |
| ENSDARG000 | 10;10 | 334627;34944 | 334645;34975 | +;+ | 330 | NA | 0 | 0 | 0 | 0 | 0 | 0 |
| ENSDARG000 | 10 | 9913063 | 9913177 | + | 115 | 5S_rRNA | 0 | 0 | 0 | 0 | 0 | 0 |
| ENSDARG000 | 10;10;10;10;1 | 12648430;126 | 12648599;126 | -;-;-;-;-;-;-; | 2189 | NA | 16 | 0 | 0 | 13 | 0 | 2 |
| ENSDARG000 | 10;10;10;10;1 | 42678520;426 | 42678563;426 | +;+;+;+;+;+;+; | 4730 | RHOBTB2 | 246 | 31 | 270 | 124 | 166 | 101 |
| ENSDARG000 | 10 | 6472706 | 6472832 | - | 127 | 5S_rRNA | 0 | 0 | 0 | 0 | 0 | 0 |
| ENSDARG000 | 10 | 12333253 | 12333369 | + | 117 | 5S_rRNA | 0 | 0 | 0 | 0 | 0 | 0 |
| ENSDARG000 | 10;10;10;10;1 | 19525839;195 | 19525907;195 | +;+;+;+;+;+;+; | 2879 | vsig8a | 6 | 4 | 31 | 0 | 10 | 0 |
| ENSDARG000 | 10 | 13607027 | 13607143 | - | 117 | 5S_rRNA | 0 | 0 | 0 | 0 | 0 | 0 |
| ENSDARG000 | 10 | 38548089 | 38548205 | + | 117 | 5S_rRNA | 26 | 22 | 58 | 9 | 28 | 46 |
| ENSDARG000 | 10 | 29473342 | 29473456 | + | 115 | 5S_rRNA | 0 | 0 | 0 | 0 | 0 | 0 |
| ENSDARG000 | 10;10;10;10;1 | 45345574;453 | 45345837;453 | +;+;+;+;+;+ | 858 | ppiab | 10312 | 13226 | 16477 | 12925 | 16203 | 11594 |
| ENSDARG000 | 10;10;10;10;1 | 36026576;360 | 36026635;360 | +;+;+;+;+;+;+; | 1938 | hmgb1a | 4649 | 2497 | 3736 | 2436 | 3982 | 3393 |
| ENSDARG000 | 10;10;10;10;1 | 8154874;8158 | 8154929;8158 | -;-;-;-;-;-;-; | 4140 | DHX29 | 404 | 342 | 518 | 220 | 584 | 238 |
| ENSDARG000 | 10;10;10 | 2669405;2670 | 2669460;2670 | +;+;+ | 352 | ccl27b | 49 | 4 | 7 | 0 | 0 | 3 |
| ENSDARG000 | 10;10;10;10;1 | 21861451;218 | 21861464;218 | -;-;-;-;-;-;-; | 519 | NA | 6 | 0 | 9 | 5 | 4 | 0 |
| ENSDARG000 | 10;10;10;10;1 | 39178450;391 | 39182914;391 | -;-;-;-;-;-;-;- | 5356 | NA | 86 | 6 | 115 | 84 | 63 | 42 |
| ENSDARG000 | 10;10;10;10;1 | 4367430;4372 | 4372452;4373 | -;-;-;-;-;-;- | 5800 | NA | 794 | 1699 | 454 | 1415 | 456 | 1330 |
| ENSDARG000 | 10 | 10329464 | 10329554 | - | 91 | dre-mir-219-1 | 4 | 0 | 0 | 0 | 0 | 0 |
| ENSDARG000 | 10 | 38376134 | 38376253 | - | 120 | 5S_rRNA | 0 | 0 | 0 | 0 | 0 | 0 |
| ENSDARG000 | 10 | 17695265 | 17695340 | + | 76 | dre-mir-218b | 0 | 0 | 0 | 0 | 0 | 0 |
| ENSDARG000 | 10 | 30736148 | 30736268 | + | 121 | 5S_rRNA | 0 | 0 | 0 | 0 | 0 | 0 |
| ENSDARG000 | 10 | 9725774 | 9725891 | + | 118 | 5S_rRNA | 0 | 0 | 0 | 0 | 2 | 0 |
| ENSDARG000 | 10;10;10;10;1 | 44296863;442 | 44297725;442 | -;-;-;-;-;-;-; | 2997 | ZSWIM6 | 292 | 144 | 158 | 155 | 181 | 158 |
| ENSDARG000 | 10;10;10 | 25355308;253 | 25355463;253 | +;+;+ | 1310 | NA | 0 | 4 | 0 | 2 | 3 | 0 |
| ENSDARG000 | 10;10;10;10;1 | 16857529;168 | 16857648;168 | -;-;-;-;-;-;-; | 1068 | stoml2 | 337 | 393 | 568 | 383 | 375 | 376 |
| ENSDARG000 | 10;10;10;10;1 | 29259882;292 | 29259917;292 | +;+;+;+;+;+;+; | 1094 | NA | 244 | 172 | 909 | 110 | 461 | 695 |
| ENSDARG000 | 10 | 38488708 | 38488829 | + | 122 | 5S_rRNA | 0 | 0 | 0 | 0 | 0 | 0 |
| ENSDARG000 | 10 | 45237956 | 45238042 | + | 87 | NA | 0 | 0 | 0 | 0 | 0 | 0 |
| ENSDARG000 | 10;10;10;10;1 | 44302220;443 | 44302287;443 | -;-;-;-;-;-;-; | 486 | CDK2AP1 | 56 | 151 | 206 | 78 | 107 | 112 |
| ENSDARG000 | 10 | 42911933 | 42912047 | - | 115 | 5S_rRNA | 0 | 0 | 0 | 0 | 0 | 0 |
| ENSDARG000 | 10;10;10;10;1 | 33382858;333 | 33383241;333 | +;+;+;+;+;+;+; | 4101 | mdh2 | 4959 | 5523 | 7024 | 6221 | 5296 | 5486 |
| ENSDARG000 | 10 | 28198894 | 28199041 | - | 148 | dre-mir-21-1 | 49 | 33 | 65 | 91 | 44 | 17 |
| ENSDARG000 | 10 | 35654245 | 35654354 | - | 110 | dre-mir-2184 | 0 | 0 | 0 | 0 | 0 | 0 |
| ENSDARG000 | 10;10;10;10;1 | 31646020;317 | 31646150;317 | +;+;+;+;+;+;+ | 1786 | esama | 94 | 3 | 17 | 20 | 12 | 44 |
| ENSDARG000 | 10 | 17123368 | 17123500 | + | 133 | U8 | 0 | 0 | 0 | 0 | 0 | 0 |
| ENSDARG000 | 10;10;10;10;1 | 40836378;408 | 40837014;408 | +;+;+;+;+;+ | 1651 | trim69 | 17 | 10 | 0 | 1 | 0 | 4 |
| ENSDARG000 | 10 | 22271789 | 22271903 | - | 115 | 5S_rRNA | 0 | 0 | 0 | 0 | 0 | 0 |
| ENSDARG000 | 10;10 | 16584382;165 | 16584911;165 | +;+ | 1249 | NA | 0 | 0 | 6 | 8 | 0 | 2 |
| ENSDARG000 | 10 | 17125340 | 17125466 | + | 127 | U8 | 0 | 0 | 0 | 0 | 0 | 0 |
| ENSDARG000 | 10 | 20483814 | 20483930 | - | 117 | 5S_rRNA | 1 | 0 | 0 | 0 | 0 | 0 |
| ENSDARG000 | 10 | 15954727 | 15954815 | + | 89 | dre-mir-27d | 0 | 0 | 0 | 0 | 0 | 0 |
| ENSDARG000 | 10 | 38074312 | 38074391 | - | 80 | dre-mir-99-2 | 0 | 0 | 0 | 0 | 0 | 0 |
| ENSDARG000 | 10 | 28252678 | 28252795 | + | 118 | 5S_rRNA | 0 | 0 | 0 | 0 | 0 | 0 |
| ENSDARG000 | 10;10 | 44513231;445 | 44513698;445 | -;- | 1804 | NA | 206 | 135 | 190 | 191 | 232 | 157 |
| ENSDARG000 | 10 | 9016721 | 9016837 | - | 117 | 5S_rRNA | 0 | 0 | 0 | 0 | 0 | 0 |
| ENSDARG000 | 10 | 44979533 | 44981295 | - | 1763 | purbb | 951 | 679 | 1046 | 426 | 1215 | 550 |
| ENSDARG000 | 10;10;10;10;1 | 36092564;360 | 36092815;360 | +;+;+;+;+;+;+; | 4032 | smyd4 | 100 | 135 | 177 | 72 | 98 | 131 |
| ENSDARG000 | 10;10;10;10;1 | 2573161;2582 | 2573166;2582 | -;-;-;-;-;-;- | 168 | NA | 2 | 0 | 0 | 2 | 0 | 0 |
| ENSDARG000 | 10;10 | 563252;56665 | 563522;56660 | +;+ | 1221 | BMP10 | 0 | 0 | 0 | 0 | 0 | 0 |
| ENSDARG000 | 10;10;10;10;1 | 31738501;317 | 31739394;317 | -;-;-;-;-;-;-; | 2143 | fez1 | 0 | 2 | 0 | 2 | 0 | 6 |
| ENSDARG000 | 10;10;10;10;1 | 44955106;449 | 44955405;449 | +;+;+;+;+;+;+; | 3351 | il1b | 7249 | 5814 | 5862 | 19560 | 3276 | 9499 |
| ENSDARG000 | 10;10;10;10;1 | 1052591;1056 | 1052771;1056 | +;+;+;+;+;+;+; | 2241 | UNC5C | 0 | 4 | 7 | 0 | 6 | 4 |
| ENSDARG000 | 10;10;10;10;1 | 3875716;3875 | 3875739;3875 | +;+;+;+;+;+;+; | 7238 | TTC28 | 897 | 477 | 614 | 645 | 922 | 472 |
| ENSDARG000 | 10;10;10;10;1 | 23537576;235 | 23537809;235 | +;+;+;+;+ | 1611 | NA | 4 | 0 | 8 | 0 | 4 | 0 |
| ENSDARG000 | 10 | 29852147 | 29852242 | + | 96 | SNORD14 | 7 | 0 | 5 | 5 | 28 | 0 |
| ENSDARG000 | 10;10;10;10 | 33999869;340 | 34000479;340 | -;-;-;- | 1406 | zar1l | 0 | 0 | 0 | 0 | 0 | 0 |
| ENSDARG000 | 10;10 | 26935155;269 | 26935236;269 | +;+ | 4257 | NA | 0 | 0 | 0 | 0 | 0 | 0 |
| ENSDARG000 | 10;10;10 | 44503819;445 | 44504021;445 | -;-;- | 615 | DUSP26 | 80 | 56 | 99 | 109 | 84 | 46 |
| ENSDARG000 | 10;10;10;10;1 | 22381802;224 | 22382512;224 | +;+;+;+;+;+;+; | 2807 | nlgn2b | 2 | 0 | 0 | 2 | 0 | 4 |
| ENSDARG000 | 10;10;10;10;1 | 1955475;1956 | 1956438;1956 | -;-;-;-;-;-;-; | 1790 | tdgf1 | 0 | 0 | 0 | 0 | 0 | 8 |
| ENSDARG000 | 10;10;10;10;1 | 42903091;429 | 42905837;429 | -;-;-;-;-;-;-; | 4319 | ACOT12 | 43 | 15 | 65 | 51 | 12 | 36 |
| ENSDARG000 | 10 | 35620333 | 35620449 | - | 117 | 5S_rRNA | 0 | 0 | 0 | 0 | 0 | 0 |
| ENSDARG000 | 10;10;10;10;1 | 5667401;5669 | 5668330;5670 | -;-;-;-;- | 2149 | st8sia4 | 335 | 157 | 434 | 264 | 168 | 454 |
| ENSDARG000 | 10;10;10;10 | 53587;55517; | 54003;55699; | -;-;-;- | 1642 | NA | 680 | 353 | 546 | 441 | 460 | 350 |
| ENSDARG000 | 10 | 11498693 | 11498809 | + | 117 | 5S_rRNA | 2 | 1 | 0 | 0 | 2 | 0 |
| ENSDARG000 | 10 | 27169956 | 27170072 | - | 117 | 5S_rRNA | 0 | 0 | 0 | 0 | 0 | 0 |
| ENSDARG000 | 10 | 28922135 | 28922251 | - | 117 | 5S_rRNA | 0 | 0 | 0 | 0 | 0 | 0 |
| ENSDARG000 | 10;10;10;10;1 | 14525184;145 | 14530043;145 | -;-;-;-;-;-;-; | 5593 | hdhd2 | 175 | 121 | 192 | 147 | 140 | 138 |
| ENSDARG000 | 10;10;10;10;1 | 20407888;204 | 20407955;204 | -;-;-;-;-;-;-; | 2214 | loxl2a | 52 | 124 | 127 | 80 | 162 | 8 |
| ENSDARG000 | 10;10 | 124995;13450 | 125302;13543 | -;- | 1233 | KCNJ6 | 0 | 2 | 0 | 0 | 0 | 0 |
| ENSDARG000 | 10 | 23417833 | 23417947 | + | 115 | 5S_rRNA | 0 | 0 | 0 | 0 | 0 | 0 |
| ENSDARG000 | 10 | 26306229 | 26306343 | + | 115 | 5S_rRNA | 0 | 0 | 0 | 0 | 0 | 0 |
| ENSDARG000 | 10 | 1664922 | 1665038 | + | 117 | 5S_rRNA | 0 | 0 | 0 | 0 | 0 | 0 |
| ENSDARG000 | 10 | 11115854 | 11115970 | + | 117 | 5S_rRNA | 0 | 0 | 1 | 1 | 0 | 0 |
| ENSDARG000 | 10 | 24383266 | 24383381 | - | 116 | 5S_rRNA | 0 | 0 | 0 | 1 | 0 | 0 |
| ENSDARG000 | 10;10;10;10;1 | 3111004;3111 | 3111655;3111 | +;+;+;+;+;+;+; | 2120 | ckmt2a | 0 | 0 | 0 | 0 | 0 | 0 |
| ENSDARG000 | 10;10 | 8656417;8656 | 8656517;8657 | +;+ | 1171 | pmaip1 | 234 | 50 | 89 | 80 | 154 | 244 |
| ENSDARG000 | 10;10;10;10;1 | 8146348;8148 | 8147438;8149 | -;-;-;-;- | 1743 | NA | 6 | 2 | 29 | 22 | 18 | 7 |
| ENSDARG000 | 10 | 3408783 | 3408900 | + | 118 | 5S_rRNA | 0 | 0 | 0 | 0 | 0 | 0 |
| ENSDARG000 | 10;10;10;10;1 | 2529037;2531 | 2529235;2531 | +;+;+;+;+;+;+; | 1923 | NA | 6 | 16 | 14 | 34 | 22 | 18 |
| ENSDARG000 | 10;10;10;10;1 | 323401;32473 | 324583;32479 | -;-;-;-;-;-;-; | 2384 | vaspa | 42 | 72 | 58 | 34 | 145 | 188 |
| ENSDARG000 | 10;10 | 21668606;216 | 21669422;216 | +;+ | 2520 | NA | 0 | 4 | 11 | 10 | 0 | 6 |
| ENSDARG000 | 10;10;10;10;1 | 29840725;298 | 29840789;298 | +;+;+;+;+;+;+; | 1697 | NA | 56 | 100 | 67 | 90 | 335 | 466 |
| ENSDARG000 | 10;10 | 17102902;171 | 17103281;171 | -;- | 708 | RNF208 | 159 | 90 | 268 | 123 | 182 | 84 |
| ENSDARG000 | 10;10 | 19595009;195 | 19596049;195 | +;+ | 1924 | NA | 0 | 0 | 0 | 0 | 0 | 0 |
| ENSDARG000 | 10;10;10;10;1 | 29269998;292 | 29270348;292 | -;-;-;-;-;-;-; | 915 | tmem126a | 239 | 321 | 512 | 277 | 392 | 264 |
| ENSDARG000 | 10;10;10;10;1 | 16911951;169 | 16911990;169 | +;+;+;+;+;+;+; | 4719 | UNC13B | 0 | 5 | 10 | 2 | 0 | 0 |
| ENSDARG000 | 10 | 30759929 | 30760044 | + | 116 | 5S_rRNA | 0 | 0 | 0 | 0 | 0 | 0 |
| ENSDARG000 | 10;10 | 19596214;195 | 19596635;195 | +;+ | 1430 | NA | 0 | 0 | 0 | 0 | 0 | 0 |
| ENSDARG000 | 10 | 43535960 | 43536095 | - | 136 | dre-mir-9-2 | 0 | 0 | 0 | 0 | 0 | 0 |
| ENSDARG000 | 10;10;10;10;1 | 25214270;252 | 25214401;252 | -;-;-;-;-;-;-; | 1738 | kpna7 | 30 | 4 | 0 | 0 | 4 | 28 |
| ENSDARG000 | 10 | 15960835 | 15960914 | + | 80 | dre-mir-24-1 | 0 | 0 | 0 | 0 | 0 | 0 |
| ENSDARG000 | 10;10;10;10 | 26987108;269 | 26987254;269 | -;-;-;- | 531 | MRPL49 | 162 | 271 | 257 | 144 | 186 | 172 |
| ENSDARG000 | 10 | 17890089 | 17890203 | + | 115 | 5S_rRNA | 0 | 0 | 0 | 0 | 0 | 0 |
| ENSDARG000 | 10;10;10;10 | 42169982;421 | 42170204;421 | +;+;+;+ | 1901 | NA | 4 | 0 | 0 | 0 | 0 | 0 |
| ENSDARG000 | 10;10;10;10;1 | 16473923;164 | 16478432;164 | -;-;-;-;-;-;-; | 6284 | NA | 130 | 84 | 141 | 216 | 195 | 295 |
| ENSDARG000 | 10 | 29850572 | 29850667 | + | 96 | SNORD14 | 1 | 3 | 4 | 3 | 3 | 1 |
| ENSDARG000 | 10 | 44940693 | 44942231 | + | 1539 | CNNM4 | 0 | 6 | 0 | 0 | 0 | 0 |
| ENSDARG000 | 10 | 22847272 | 22847404 | + | 133 | U8 | 0 | 0 | 0 | 0 | 0 | 0 |
| ENSDARG000 | 10 | 17454490 | 17454572 | + | 83 | dre-mir-301c | 0 | 0 | 0 | 0 | 0 | 0 |
| ENSDARG000 | 10;10 | 7671260;7672 | 7671536;7672 | +;+ | 418 | fam136a | 78 | 44 | 56 | 45 | 54 | 18 |
| ENSDARG000 | 10 | 24353750 | 24353866 | + | 117 | 5S_rRNA | 0 | 0 | 0 | 0 | 0 | 0 |
| ENSDARG000 | 10 | 33377130 | 33377214 | - | 85 | dre-mir-130a | 0 | 0 | 0 | 0 | 0 | 0 |
| ENSDARG000 | 10;10;10;10;1 | 42898103;429 | 42899024;429 | +;+;+;+;+;+ | 1701 | zcchc9 | 186 | 241 | 194 | 78 | 178 | 151 |
| ENSDARG000 | 10;10;10;10;1 | 161660;16376 | 162183;16381 | -;-;-;-;-;-;-; | 1281 | erg | 28 | 2 | 13 | 5 | 38 | 13 |
| ENSDARG000 | 10;10;10;10;1 | 573667;57585 | 573992;57616 | +;+;+;+;+;+;+; | 2788 | smad4a | 484 | 486 | 511 | 516 | 490 | 413 |
| ENSDARG000 | 10 | 2688565 | 2688681 | + | 117 | 5S_rRNA | 0 | 0 | 0 | 0 | 3 | 0 |
| ENSDARG000 | 10;10 | 17254502;172 | 17254846;172 | -;- | 1630 | GNAZ | 2085 | 1343 | 1610 | 1161 | 1797 | 1475 |
| ENSDARG000 | 10;10;10;10;1 | 2587234;2589 | 2587502;2590 | +;+;+;+;+;+ | 1115 | wu:fb59d01 | 10 | 0 | 29 | 79 | 15 | 31 |
| ENSDARG000 | 10;10;10;10;1 | 19573791;195 | 19573804;195 | +;+;+;+;+ | 1437 | NA | 0 | 0 | 0 | 0 | 0 | 0 |
| ENSDARG000 | 10;10;10 | 15187316;151 | 15187378;151 | +;+;+ | 438 | NA | 0 | 0 | 0 | 0 | 0 | 0 |
| ENSDARG000 | 10 | 7009374 | 7009490 | - | 117 | 5S_rRNA | 0 | 0 | 2 | 0 | 0 | 0 |
| ENSDARG000 | 10;10;10;10;1 | 3049636;3049 | 3049914;3049 | +;+;+;+;+;+;+; | 3864 | RASGRF2 | 0 | 1 | 2 | 0 | 8 | 0 |
| ENSDARG000 | 10 | 4493277 | 4493358 | + | 82 | dre-mir-430a- | 0 | 0 | 0 | 0 | 0 | 0 |
| ENSDARG000 | 10 | 38051652 | 38051766 | - | 115 | dre-mir-125b- | 0 | 0 | 0 | 0 | 0 | 0 |
| ENSDARG000 | 10 | 32395780 | 32395896 | + | 117 | 5S_rRNA | 0 | 0 | 1 | 0 | 0 | 0 |

| | | | | | | | | | | | | |
|---|---|---|---|---|---|---|---|---|---|---|---|---|
| ENSDARG000 | 10:10:10:10:1 | 20460551:204 | 20460619:204 | -:-:-:-:-:-:-:-: | 2274 | TACC1 | 3095 | 2661 | 2720 | 3064 | 3391 | 2873 |
| ENSDARG000 | 10:10:10:10:1 | 3187289:3189 | 3187340:3189 | -:-:-:-:-:-:-:-: | 6180 | pi4kaa | 1291 | 735 | 1225 | 580 | 1375 | 776 |
| ENSDARG000 | 10:10:10:10:1 | 41549819:415 | 41550525:415 | +:+:+:+:+:+:+:+: | 6452 | NA | 8 | 2 | 2 | 2 | 10 | 36 |
| ENSDARG000 | 10:10:10:10:1 | 3336772:3341 | 3337086:3341 | +:+:+:+:+:+:+:+: | 1984 | brap | 350 | 250 | 338 | 252 | 360 | 265 |
| ENSDARG000 | 10:10:10:10:1 | 44552548:445 | 44554719:445 | -:-:-:-:-:-:-:-: | 2767 | npm2b | 0 | 0 | 0 | 0 | 0 | 0 |
| ENSDARG000 | 10:10:10:10:1 | 2695464:2696 | 2695511:2697 | -:-:-:-:-:-:-:-: | 2321 | mier3a | 436 | 315 | 434 | 184 | 416 | 228 |
| ENSDARG000 | 10:10:10:10:1 | 44984946:449 | 44984976:449 | +:+:+:+:+:+:+:+: | 4282 | h2az2b | 5348 | 6418 | 7932 | 4973 | 6327 | 3495 |
| ENSDARG000 | 10 | 5950463 | 5950578 | - | 116 | 5S_rRNA | 0 | 0 | 0 | 0 | 0 | 0 |
| ENSDARG000 | 10 | 35644719 | 35644798 | + | 80 | NA | 0 | 0 | 0 | 0 | 0 | 0 |
| ENSDARG000 | 10 | 33376818 | 33376941 | - | 124 | dre-mir-454a | 0 | 4 | 8 | 0 | 0 | 0 |
| ENSDARG000 | 10 | 9141565 | 9141679 | - | 115 | 5S_rRNA | 0 | 0 | 0 | 0 | 0 | 0 |
| ENSDARG000 | 10:10:10:10:1 | 33393829:333 | 33394129:333 | +:+:+:+:+:+:+:+: | 4361 | zgc:153345 | 6 | 14 | 4 | 20 | 8 | 0 |
| ENSDARG000 | 10 | 25964284 | 25964399 | + | 116 | 5S_rRNA | 0 | 0 | 0 | 0 | 0 | 0 |
| ENSDARG000 | 10:10:10:10:1 | 33269371:332 | 33269385:332 | -:-:-:-:-:-:-:-: | 3078 | PRDM15 | 164 | 94 | 138 | 104 | 185 | 80 |
| ENSDARG000 | 10:10:10:10:1 | 3133098:3133 | 3133910:3133 | -;-;-;-;-;-;-;- | 1620 | ube2l3a | 1178 | 1244 | 1333 | 875 | 1288 | 1057 |
| ENSDARG000 | 10;10;10;10 | 8034688:8040 | 8037930:8040 | -;-;-;- | 3889 | NA | 151 | 110 | 190 | 80 | 146 | 204 |
| ENSDARG000 | 10 | 16756546 | 16757842 | + | 1297 | NA | 0 | 0 | 0 | 0 | 0 | 0 |
| ENSDARG000 | 10:10:10:10:1 | 9697097:9699 | 9697325:9699 | +:+:+:+:+:+:+:+: | 7089 | RABGAP1 | 2491 | 1793 | 2522 | 2780 | 2816 | 2376 |
| ENSDARG000 | 10 | 13931569 | 13931696 | - | 128 | 5S_rRNA | 0 | 0 | 0 | 0 | 0 | 0 |
| ENSDARG000 | 10:10:10:10:1 | 38643304:386 | 38643585:386 | +:+:+:+:+:+:+:+: | 2637 | mmp30 | 50 | 0 | 0 | 0 | 4 | 2 |
| ENSDARG000 | 10;10 | 14547941:145 | 14547981:145 | -;- | 768 | skor2 | 0 | 0 | 0 | 0 | 0 | 8 |
| ENSDARG000 | 10 | 33002319 | 33002433 | + | 115 | 5S_rRNA | 0 | 0 | 0 | 0 | 0 | 0 |
| ENSDARG000 | 10 | 17453363 | 17453503 | + | 141 | dre-mir-130c- | 0 | 0 | 2 | 0 | 0 | 0 |
| ENSDARG000 | 10:10:10:10:1 | 42762454:427 | 42762563:427 | -:-:-:-:-:-:-:-: | 1716 | wdr45 | 749 | 361 | 462 | 1243 | 443 | 622 |
| ENSDARG000 | 10 | 31692662 | 31692776 | + | 115 | 5S_rRNA | 0 | 0 | 0 | 0 | 0 | 0 |
| ENSDARG000 | 10:10:10:10:1 | 25204626:252 | 25205455:252 | +:+:+:+:+:+:+:+: | 1792 | chordc1b | 654 | 765 | 853 | 628 | 738 | 746 |
| ENSDARG000 | 10 | 16575078 | 16575175 | + | 98 | NA | 0 | 0 | 0 | 1 | 0 | 0 |
| ENSDARG000 | 10:10:10:10:1 | 42887204:428 | 42887528:428 | -;-;-;-;-;- | 828 | NA | 8 | 8 | 12 | 5 | 1 | 3 |
| ENSDARG000 | 10 | 21980547 | 21980665 | + | 119 | 5S_rRNA | 0 | 0 | 0 | 0 | 0 | 0 |
| ENSDARG000 | 10;10;10;10 | 16188761:161 | 16189028:161 | +;+;+;+ | 649 | CTXN3 | 4 | 30 | 14 | 42 | 11 | 19 |
| ENSDARG000 | 10 | 41839919 | 41840033 | + | 115 | 5S_rRNA | 0 | 0 | 0 | 0 | 0 | 0 |
| ENSDARG000 | 10:10:10:10:1 | 7255695:7255 | 7255882:7255 | +:+:+:+:+:+:+:+: | 1504 | fut10 | 111 | 58 | 121 | 59 | 104 | 86 |
| ENSDARG000 | 10:10:10:10:1 | 1082961:1084 | 1083086:1084 | -:-:-:-:-:-:-:-: | 2090 | bmpr1bb | 10 | 3 | 10 | 21 | 20 | 2 |
| ENSDARG000 | 10:10:10:10:1 | 31298972:312 | 31301384:313 | -:-:-:-:-:-:-:-: | 7082 | robo3 | 256 | 168 | 264 | 82 | 60 | 316 |
| ENSDARG000 | 10:10:10:10:1 | 8298984:8304 | 8299273:8304 | -;-;-;-;-;-;- | 1743 | CSGALNACT1 | 493 | 315 | 311 | 194 | 296 | 272 |
| ENSDARG000 | 10;10;10 | 7884128:7885 | 7884401:7885 | -;-;- | 414 | NA | 4 | 0 | 0 | 0 | 0 | 0 |
| ENSDARG000 | 10:10:10:10:1 | 33171501:331 | 33171919:331 | +;+;+;+;+;+;+ | 1107 | myl10 | 2 | 0 | 0 | 2 | 0 | 2 |
| ENSDARG000 | 10:10:10:10:1 | 29226180:292 | 29226453:292 | -:-:-:-:-:-:-:-: | 1489 | ccdc83 | 4 | 14 | 12 | 2 | 6 | 24 |
| ENSDARG000 | 10:10:10:10:1 | 44408675:444 | 44408885:444 | -;-;-;-;- | 885 | NA | 106 | 72 | 73 | 52 | 109 | 112 |
| ENSDARG000 | 10;10 | 34236168:342 | 34236276:342 | +;+ | 177 | NA | 0 | 0 | 0 | 0 | 0 | 0 |
| ENSDARG000 | 10 | 30531143 | 30531257 | - | 115 | 5S_rRNA | 0 | 0 | 0 | 0 | 0 | 0 |
| ENSDARG000 | 10 | 37147669 | 37147784 | + | 116 | 5S_rRNA | 0 | 0 | 0 | 0 | 0 | 0 |
| ENSDARG000 | 10:10:10:10:1 | 22960380:229 | 22960712:229 | -:-:-:-:-:-:-:-: | 1983 | NA | 64 | 80 | 108 | 60 | 130 | 72 |
| ENSDARG000 | 10:10:10:10:1 | 38708099:387 | 38708116:387 | +:+:+:+:+:+:+:+: | 1464 | tmprss2 | 78 | 23 | 22 | 21 | 70 | 79 |
| ENSDARG000 | 10 | 36960576 | 36960690 | + | 115 | 5S_rRNA | 0 | 0 | 0 | 0 | 0 | 0 |
| ENSDARG000 | 10 | 17119917 | 17120049 | + | 133 | U8 | 0 | 0 | 0 | 0 | 0 | 5 |
| ENSDARG000 | 10:10:10:10:1 | 39952995:400 | 39953695:400 | +;+;+;+;+;+;+ | 6337 | NA | 6 | 0 | 0 | 0 | 2 | 2 |
| ENSDARG000 | 10:10:10:10:1 | 36784756:367 | 36785066:367 | -;-;-;-;- | 1022 | dhrs13a.2 | 31 | 2 | 0 | 21 | 27 | 4 |
| ENSDARG000 | 10:10:10:10:1 | 3520256:3523 | 3520846:3523 | +;+;+;+;+;+;+ | 2315 | pla2g3 | 149 | 74 | 112 | 16 | 22 | 88 |
| ENSDARG000 | 10:10:10:10:1 | 15970:17325: | 16207:17457: | +:+:+:+:+:+:+:+: | 1833 | tiprl | 371 | 271 | 419 | 206 | 425 | 307 |
| ENSDARG000 | 10:10:10:10:1 | 311722:31191 | 311999:31199 | -:-:-:-:-:-:-:-: | 1819 | akt2l | 88 | 97 | 314 | 96 | 211 | 38 |
| ENSDARG000 | 10:10:10:10:1 | 7673446:7674 | 7674659:7674 | -:-:-:-:-:-:-:-: | 2780 | gmcl1 | 423 | 351 | 400 | 381 | 400 | 424 |
| ENSDARG000 | 10;10;10 | 4924388:4931 | 4925363:4931 | +;+;+ | 1221 | SLC46A2 | 12 | 20 | 4 | 4 | 32 | 14 |
| ENSDARG000 | 10;10 | 24684799:246 | 24684982:246 | +;+ | 1617 | NA | 0 | 0 | 0 | 0 | 0 | 0 |
| ENSDARG000 | 10:10:10:10:1 | 6907715:6911 | 6907786:6911 | +:+:+:+:+:+:+:+: | 1921 | slc38a9 | 54 | 51 | 47 | 37 | 129 | 26 |
| ENSDARG000 | 10 | 38434383 | 38434499 | - | 117 | 5S_rRNA | 0 | 0 | 0 | 0 | 0 | 0 |
| ENSDARG000 | 10 | 3768651 | 3768766 | - | 116 | 5S_rRNA | 0 | 0 | 0 | 0 | 0 | 0 |
| ENSDARG000 | 10 | 17342365 | 17342481 | - | 117 | 5S_rRNA | 0 | 0 | 0 | 0 | 0 | 0 |
| ENSDARG000 | 10 | 7512008 | 7512123 | - | 116 | 5S_rRNA | 0 | 0 | 0 | 0 | 0 | 0 |
| ENSDARG000 | 10 | 24455616 | 24455732 | - | 117 | 5S_rRNA | 0 | 0 | 0 | 0 | 0 | 0 |
| ENSDARG000 | 10;10;10 | 42733210:427 | 42733694:427 | +;+;+ | 3393 | NA | 371 | 106 | 360 | 398 | 95 | 275 |
| ENSDARG000 | 10:10:10:10:1 | 5644356:5644 | 5644517:5644 | +:+:+:+:+:+:+:+: | 825 | NA | 0 | 0 | 0 | 0 | 1 | 0 |
| ENSDARG000 | 10:10:10:10:1 | 44473851:444 | 44474397:444 | -:-:-:-:-:-:-:-: | 1523 | NA | 2 | 4 | 0 | 0 | 0 | 0 |
| ENSDARG000 | 10:10:10:10:1 | 39164638:391 | 39164870:391 | +;+;+;+;+;+ | 1434 | B3GAT1 | 0 | 0 | 5 | 4 | 0 | 0 |
| ENSDARG000 | 10:10:10:10:1 | 25369254:253 | 25369892:253 | +:+:+:+:+:+:+:+: | 6168 | bach1b | 462 | 301 | 575 | 453 | 574 | 364 |
| ENSDARG000 | 10:10:10:10:1 | 20033900:200 | 20034787:200 | -;-;-;-;- | 2198 | NA | 590 | 801 | 667 | 696 | 1638 | 611 |
| ENSDARG000 | 10:10:10:10:1 | 23088:23919: | 23239:24287: | +:+:+:+:+:+:+:+: | 2002 | riox2 | 212 | 197 | 409 | 138 | 268 | 101 |
| ENSDARG000 | 10:10:10:10:1 | 2600830:2601 | 2601036:2601 | +:+:+:+:+:+:+:+: | 3372 | NA | 495 | 259 | 286 | 125 | 177 | 106 |
| ENSDARG000 | 10;10 | 22494826:224 | 22495434:224 | -;- | 1011 | NA | 6 | 0 | 0 | 0 | 4 | 26 |
| ENSDARG000 | 10 | 13305065 | 13305183 | - | 119 | 5S_rRNA | 25 | 14 | 28 | 7 | 20 | 29 |
| ENSDARG000 | 10 | 211782 | 211947 | + | 166 | U1 | 0 | 0 | 0 | 0 | 0 | 0 |
| ENSDARG000 | 10;10;10;10 | 21511495:215 | 21514225:215 | +;+;+;+ | 3920 | NA | 0 | 2 | 0 | 0 | 0 | 0 |
| ENSDARG000 | 10 | 17627128 | 17627253 | - | 126 | 5S_rRNA | 0 | 0 | 0 | 0 | 0 | 0 |
| ENSDARG000 | 10;10;10 | 3145707:3153 | 3145806:3155 | +;+;+ | 2102 | hic2 | 156 | 102 | 260 | 119 | 202 | 121 |
| ENSDARG000 | 10 | 15954579 | 15954659 | + | 81 | dre-mir-23b | 0 | 0 | 0 | 0 | 0 | 0 |
| ENSDARG000 | 10 | 35645017 | 35645135 | + | 119 | dre-mir-132-2 | 0 | 0 | 0 | 0 | 0 | 0 |
| ENSDARG000 | 10 | 34327262 | 34327394 | - | 133 | 5S_rRNA | 0 | 0 | 4 | 5 | 5 | 0 |
| ENSDARG000 | 10 | 27063354 | 27063438 | + | 85 | dre-mir-192 | 0 | 0 | 0 | 0 | 0 | 0 |
| ENSDARG000 | 10:10:10:10:1 | 17026870:170 | 17027397:170 | +:+:+:+:+:+:+:+: | 4205 | NA | 129 | 2 | 14 | 4 | 14 | 17 |
| ENSDARG000 | 10;10;10;10 | 4806323:4807 | 4806334:4808 | +;+;+;+ | 3385 | NA | 4 | 2 | 2 | 0 | 6 | 14 |
| ENSDARG000 | 10:10:10:10:1 | 33305978:333 | 33306207:333 | -:-:-:-:-:-:-:-: | 2867 | NA | 2489 | 978 | 3907 | 1648 | 2810 | 3229 |
| ENSDARG000 | 10:10:10:10:1 | 44437080:444 | 44437179:444 | +:+:+:+:+:+:+:+: | 2262 | denr | 1220 | 1612 | 1686 | 1055 | 1536 | 1359 |
| ENSDARG000 | 10 | 39760754 | 39760879 | + | 126 | 5S_rRNA | 0 | 0 | 0 | 0 | 0 | 0 |
| ENSDARG000 | 10 | 13612133 | 13612248 | - | 116 | 5S_rRNA | 0 | 0 | 0 | 0 | 0 | 0 |
| ENSDARG000 | 10 | 36163179 | 36163294 | - | 116 | 5S_rRNA | 0 | 0 | 0 | 0 | 0 | 0 |
| ENSDARG000 | 10 | 40903121 | 40903237 | - | 117 | 5S_rRNA | 0 | 0 | 0 | 0 | 0 | 0 |
| ENSDARG000 | 10 | 17127316 | 17127448 | + | 133 | U8 | 0 | 0 | 0 | 0 | 0 | 0 |
| ENSDARG000 | 10 | 29851000 | 29851095 | + | 96 | SNORD14 | 3 | 1 | 7 | 10 | 11 | 0 |
| ENSDARG000 | 10 | 38683666 | 38683780 | + | 115 | 5S_rRNA | 0 | 0 | 0 | 0 | 0 | 0 |
| ENSDARG000 | 10;10 | 21522545:215 | 21525354:215 | +;+ | 2834 | NA | 0 | 2 | 4 | 0 | 0 | 10 |
| ENSDARG000 | 10:10:10:10:1 | 25325808:253 | 25329828:253 | -:-:-:-:-:-:-:-: | 6359 | cct8 | 6276 | 6074 | 9559 | 6798 | 6060 | 7831 |
| ENSDARG000 | 10;10;10 | 25272355:252 | 25272610:252 | +;+;+ | 690 | RAB38 | 33 | 4 | 36 | 14 | 3 | 22 |
| ENSDARG000 | 10 | 44239844 | 44241119 | - | 1276 | NA | 0 | 0 | 0 | 0 | 0 | 0 |
| ENSDARG000 | 10:10:10:10:1 | 38610741:386 | 38611117:386 | +:+:+:+:+:+:+:+: | 2556 | NA | 388 | 223 | 60 | 72 | 182 | 275 |
| ENSDARG000 | 10:10:10:10:1 | 23511816:235 | 23512121:235 | +;+;+;+;+;+ | 1420 | NA | 1 | 0 | 1 | 0 | 4 | 0 |
| ENSDARG000 | 10:10:10:10:1 | 4874542:4875 | 4875368:4875 | +:+:+:+:+:+:+:+: | 3296 | palm2akap2 | 805 | 347 | 791 | 498 | 1260 | 637 |
| ENSDARG000 | 10:10:10:10:1 | 1187653:1187 | 1187742:1187 | -:-:-:-:-:-:-:-: | 2373 | pdlim5b | 2372 | 533 | 514 | 614 | 714 | 312 |
| ENSDARG000 | 10 | 20684036 | 20684150 | + | 115 | 5S_rRNA | 0 | 0 | 0 | 0 | 0 | 0 |
| ENSDARG000 | 10 | 33377001 | 33377086 | - | 86 | dre-mir-301a | 0 | 0 | 0 | 0 | 0 | 0 |
| ENSDARG000 | 10 | 29708068 | 29708182 | - | 115 | 5S_rRNA | 83 | 2 | 0 | 0 | 2 | 16 |
| ENSDARG000 | 10:10:10:10:1 | 21359062:213 | 21359293:213 | -;-;-;-;-;-;-;- | 973 | avd | 0 | 0 | 0 | 0 | 0 | 0 |
| ENSDARG000 | 10 | 339912 | 340027 | - | 116 | 5S_rRNA | 0 | 0 | 0 | 0 | 0 | 0 |
| ENSDARG000 | 10 | 38072077 | 38072168 | - | 92 | dre-let-7c-2 | 0 | 0 | 0 | 0 | 0 | 0 |
| ENSDARG000 | 10;10;10;10 | 32901:38883: | 35256:39049: | -;-;-;- | 2965 | NA | 1417 | 756 | 1404 | 732 | 1633 | 1039 |
| ENSDARG000 | 10;10;10 | 1541263:1541 | 1541509:1542 | -;-;- | 1241 | NA | 64 | 61 | 43 | 18 | 58 | 38 |
| ENSDARG000 | 10:10:10:10:1 | 1544163:1545 | 1545638:1546 | -;-;-;-;- | 3087 | NA | 68 | 49 | 127 | 46 | 96 | 56 |
| ENSDARG000 | 10:10:10:10:1 | 1982550:1982 | 1983252:1982 | +;+;+;+;+ | 4181 | NA | 0 | 4 | 8 | 0 | 2 | 0 |
| ENSDARG000 | 10;10;10 | 10178101:101 | 10178267:101 | +;+;+ | 1392 | NA | 26 | 8 | 42 | 45 | 26 | 64 |
| ENSDARG000 | 10:10:10:10:1 | 15082410:150 | 15083370:150 | -;-;-;-;-;- | 1580 | NA | 140 | 45 | 12 | 18 | 56 | 10 |
| ENSDARG000 | 10:10:10:10:1 | 18798927:187 | 18799725:188 | +;+;+;+;+ | 3323 | NA | 0 | 0 | 0 | 0 | 0 | 0 |
| ENSDARG000 | 10:10:10:10:1 | 19583289:195 | 19583586:195 | +;+;+;+;+ | 5871 | NA | 0 | 0 | 0 | 0 | 0 | 0 |
| ENSDARG000 | 10:10:10:10:1 | 19602334:196 | 19602622:196 | +;+;+;+;+;+ | 663 | NA | 0 | 0 | 0 | 0 | 0 | 0 |
| ENSDARG000 | 10;10;10 | 22938231:229 | 22938236:229 | +;+;+ | 808 | NA | 6 | 12 | 7 | 5 | 6 | 10 |
| ENSDARG000 | 10;10;10 | 30978386:309 | 30978737:309 | +;+;+ | 1160 | NA | 0 | 0 | 0 | 0 | 0 | 0 |
| ENSDARG000 | 10;10;10;10 | 33361020:333 | 33361212:333 | +;+;+;+ | 1252 | NA | 6 | 4 | 0 | 15 | 2 | 10 |
| ENSDARG000 | 10:10:10:10:1 | 38040366:380 | 38040401:380 | -;-;-;-;-;- | 1938 | NA | 90 | 58 | 20 | 13 | 59 | 25 |
| ENSDARG000 | 10;10;10;10 | 38749266:387 | 38749625:387 | +;+;+;+ | 2765 | NA | 4 | 12 | 2 | 0 | 12 | 0 |
| ENSDARG000 | 10;10;10 | 42799115:428 | 42799380:428 | -;-;- | 675 | NA | 6 | 4 | 7 | 58 | 4 | 19 |
| ENSDARG000 | 10;10;10 | 44242236:442 | 44243282:442 | -;-;- | 1303 | NA | 0 | 6 | 5 | 0 | 4 | 0 |
| ENSDARG000 | 10:10:10:10:1 | 44268481:442 | 44268867:442 | -;-;-;-;- | 1933 | NA | 76 | 68 | 118 | 80 | 74 | 60 |
| ENSDARG000 | 10:10:10:10:1 | 44493919:444 | 44494844:444 | -:-:-:-:-:-:-:-: | 1955 | NA | 4 | 0 | 14 | 3 | 1 | 0 |
| ENSDARG000 | 24:24:24:24:2 | 17027310:170 | 17027743:170 | -;-;-;-;-;- | 1029 | ptgdsb.1 | 508 | 101 | 16 | 101 | 174 | 54 |
| ENSDARG000 | 24:24:24:24:2 | 7153628:7154 | 7153667:7154 | +;+;+;+;+ | 1088 | NA | 0 | 0 | 3 | 0 | 0 | 0 |
| ENSDARG000 | 24:24:24:24:2 | 29563767:295 | 29563811:295 | -;-;-;-;- | 505 | NA | 6 | 6 | 13 | 4 | 32 | 8 |
| ENSDARG000 | 24;24 | 38098896:380 | 38099515:380 | -;- | 708 | NA | 48 | 0 | 0 | 0 | 103 | 140 |
| ENSDARG000 | 24:24:24:24:2 | 38109799:381 | 38110510:381 | -;-;-;-;- | 845 | NA | 0 | 0 | 2 | 0 | 0 | 0 |
| ENSDARG000 | 24;24 | 38121403:381 | 38121770:381 | -;- | 420 | NA | 0 | 0 | 0 | 2 | 4 | 0 |
| ENSDARG000 | 24:24:24:24:2 | 25347503:253 | 25353565:253 | -;-;-;-;-;-;- | 8013 | PTCHD1 | 0 | 0 | 0 | 0 | 0 | 0 |
| ENSDARG000 | 24:24:24:24:2 | 25333832:253 | 25334110:253 | +;+;+;+;+;+;+ | 1025 | NA | 1 | 0 | 0 | 0 | 0 | 0 |
| ENSDARG000 | 24;24;24;24 | 33288455:332 | 33289141:332 | -;-;-;- | 1166 | si:ch1073-406 | 0 | 11 | 0 | 0 | 1 | 0 |
| ENSDARG000 | 24;24 | 38130604:381 | 38131229:381 | -;- | 678 | crp6 | 0 | 0 | 0 | 0 | 0 | 0 |
| ENSDARG000 | 24;24 | 33291541:332 | 33291786:332 | +;+ | 540 | NA | 0 | 0 | 0 | 0 | 0 | 0 |
| ENSDARG000 | 24:24:24:24:2 | 36303637:363 | 36305417:363 | -;-;-;-;-;-;- | 3659 | NAGLU | 464 | 222 | 332 | 155 | 309 | 353 |
| ENSDARG000 | 24:24:24:24:2 | 26997798:270 | 26998254:270 | +:+:+:+:+:+:+:+: | 2773 | larp4b | 417 | 366 | 781 | 482 | 567 | 433 |
| ENSDARG000 | 24;24 | 21368211:213 | 21368509:213 | -;- | 408 | NA | 0 | 0 | 0 | 0 | 0 | 0 |
| ENSDARG000 | 24:24:24:24:2 | 23840821:238 | 23841000:238 | +:+:+:+:+:+:+:+: | 5385 | pola1 | 165 | 910 | 989 | 362 | 910 | 490 |
| ENSDARG000 | 24:24:24:24:2 | 36317544:363 | 36317725:363 | +:+:+:+:+:+:+:+: | 2405 | pus3 | 90 | 130 | 112 | 54 | 76 | 140 |
| ENSDARG000 | 24:24:24:24:2 | 17031129:170 | 17031780:170 | -;-;-;-;-;-;- | 1364 | c8g | 0 | 0 | 16 | 0 | 0 | 4 |
| ENSDARG000 | 24:24:24:24:2 | 29622345:296 | 29622583:296 | -:-:-:-:-:-:-:-: | 1572 | NA | 5 | 8 | 19 | 20 | 57 | 10 |
| ENSDARG000 | 24:24:24:24:2 | 38155830:381 | 38155893:381 | +:+:+:+:+:+:+:+: | 6394 | NA | 3 | 0 | 4 | 0 | 4 | 0 |
| ENSDARG000 | 24:24:24:24:2 | 38137473:381 | 38138132:381 | -;-;-;-;-;- | 972 | crp7 | 0 | 4 | 0 | 0 | 2 | 0 |
| ENSDARG000 | 24:24:24:24:2 | 33301129:333 | 33301990:333 | -:-:-:-:-:-:-:-: | 4658 | slc4a2b | 216 | 314 | 285 | 129 | 273 | 646 |
| ENSDARG000 | 24:24:24:24:2 | 38194136:381 | 38194663:381 | -;-;-;-;- | 565 | NA | 0 | 0 | 2 | 5 | 8 | 4 |
| ENSDARG000 | 24:24:24:24:2 | 7294929:7295 | 7295023:7295 | -:-:-:-:-:-:-:-: | 1622 | actr3b | 0 | 4 | 0 | 0 | 0 | 45 |
| ENSDARG000 | 24;24 | 38197852:381 | 38197900:381 | +;+ | 405 | NA | 0 | 0 | 0 | 0 | 0 | 0 |
| ENSDARG000 | 24:24:24:24:2 | 17048078:170 | 17048420:170 | -:-:-:-:-:-:-:-: | 2757 | ARMC3 | 0 | 0 | 0 | 0 | 1 | 0 |
| ENSDARG000 | 24:24:24:24:2 | 17040211:170 | 17042912:170 | -:-:-:-:-:-:-:-: | 3997 | msrb2 | 0 | 0 | 2 | 0 | 0 | 0 |
| ENSDARG000 | 24:24:24:24:2 | 38199829:381 | 38199925:381 | +;+;+;+;+;+ | 585 | NA | 0 | 0 | 0 | 0 | 12 | 0 |
| ENSDARG000 | 24;24 | 38201089:382 | 38201137:382 | +;+ | 375 | NA | 0 | 0 | 0 | 0 | 0 | 0 |
| ENSDARG000 | 24:24:24:24:2 | 38202298:382 | 38202761:382 | -;-;-;-;- | 501 | NA | 0 | 0 | 0 | 4 | 5 | 0 |
| ENSDARG000 | 24:24:24:24:2 | 21371279:213 | 21372487:213 | -:-:-:-:-:-:-:-: | 7623 | ATP8A2 | 10 | 30 | 152 | 42 | 100 | 128 |
| ENSDARG000 | 24:24:24:24:2 | 25404358:254 | 25404857:254 | -:-:-:-:-:-:-:-: | 2999 | phex | 16 | 20 | 6 | 12 | 20 | 0 |
| ENSDARG000 | 24;24;24 | 36331032:363 | 36331388:363 | -;-;- | 1336 | si:ch211-40k2 | 70 | 90 | 66 | 58 | 53 | 97 |
| ENSDARG000 | 24;24 | 21501859:215 | 21502254:215 | +;+ | 419 | NA | 0 | 0 | 0 | 0 | 0 | 0 |
| ENSDARG000 | 24:24:24:24:2 | 21501842:215 | 21506055:215 | -;-;-;-;-;-;- | 4578 | rnf6 | 540 | 503 | 687 | 287 | 728 | 485 |
| ENSDARG000 | 24;24 | 38205798:382 | 38205846:382 | +;+ | 405 | NA | 0 | 0 | 0 | 0 | 0 | 0 |
| ENSDARG000 | 24:24:24:24:2 | 38211103:382 | 38211623:382 | -;-;-;-;- | 613 | NA | 0 | 0 | 2 | 7 | 11 | 0 |
| ENSDARG000 | 24:24:24:24:2 | 29783638:297 | 29786329:297 | -:-:-:-:-:-:-:-: | 6082 | AGL | 0 | 4 | 7 | 6 | 4 | 8 |
| ENSDARG000 | 24 | 29860769 | 29863372 | - | 2604 | NA | 0 | 0 | 0 | 0 | 0 | 0 |
| ENSDARG000 | 24:24:24:24:2 | 33392698:333 | 33392897:333 | +:+:+:+:+:+:+:+: | 4513 | NA | 649 | 484 | 711 | 481 | 613 | 576 |
| ENSDARG000 | 24:24:24:24:2 | 36339242:363 | 36339327:363 | +:+:+:+:+:+:+:+: | 3145 | grnb | 1802 | 854 | 1095 | 995 | 1265 | 1211 |
| ENSDARG000 | 24:24:24:24:2 | 27023274:270 | 27023694:270 | +:+:+:+:+:+:+:+: | 6066 | DIP2C | 1818 | 911 | 826 | 1216 | 1025 | 1253 |

| | | | | | | | | | | | | |
|---|---|---|---|---|---|---|---|---|---|---|---|---|
| ENSDARG000 | 24;24 | 10397865:104 | 10398285:104 | +;+ | 1746 | NA | 36 | 8 | 42 | 58 | 54 | 58 |
| ENSDARG000 | 24;24;24;24;2 | 10413484:104 | 10413711:104 | +;+;+;+;+ | 2100 | myca | 242 | 110 | 281 | 76 | 179 | 158 |
| ENSDARG000 | 24;24;24;24;2 | 7336411:7336 | 7337250:7337 | +;+;+;+;+;+;+; | 17730 | NA | 1834 | 706 | 1965 | 1150 | 1843 | 1417 |
| ENSDARG000 | 24;24 | 10430526:104 | 10430832:104 | +;+ | 825 | NA | 0 | 0 | 0 | 0 | 0 | 0 |
| ENSDARG000 | 24;24;24;24;2 | 10417365:104 | 10417450:104 | +;+;+;+;+ | 1093 | NA | 127 | 98 | 279 | 162 | 158 | 83 |
| ENSDARG000 | 24;24;24;24;2 | 3257376:3257 | 3257713:3257 | -;-;-;-;-;-;-;-;-; | 1574 | sugct | 16 | 8 | 43 | 48 | 24 | 14 |
| ENSDARG000 | 24;24;24;24;2 | 3328354:3330 | 3328577:3330 | +;+;+;+;+ | 1202 | bdh1 | 280 | 826 | 668 | 1003 | 747 | 685 |
| ENSDARG000 | 24;24;24;24;2 | 3307857:3309 | 3307906:3310 | +;+;+;+;+;+;+ | 1727 | gyg1b | 92 | 3306 | 406 | 3882 | 423 | 555 |
| ENSDARG000 | 24;24;24;24;2 | 10792773:107 | 10793677:107 | -;-;-;-;-;-;-;-;-; | 2220 | fam49bb | 245 | 267 | 414 | 191 | 400 | 305 |
| ENSDARG000 | 24;24;24;24;2 | 10898671:108 | 10898889:108 | +;+;+;+;+;+;+ | 1705 | eef1a1l3 | 0 | 0 | 0 | 0 | 0 | 0 |
| ENSDARG000 | 24;24;24;24;2 | 10897768:108 | 10902796:109 | -;-;-;-;-;-;-;-;-; | 9842 | ASAP1 | 184 | 329 | 441 | 244 | 342 | 529 |
| ENSDARG000 | 24;24;24;24;2 | 38301080:383 | 38301098:383 | +;+;+;+;+;+;+; | 3847 | mybpc2b | 176 | 699 | 355 | 353 | 332 | 2 |
| ENSDARG000 | 24;24;24;24;2 | 20188914:201 | 20189420:201 | -;-;-;-;- | 1485 | myd88 | 551 | 434 | 583 | 625 | 394 | 460 |
| ENSDARG000 | 24;24;24;24;2 | 14202327:142 | 14203108:142 | -;-;-;-;-;-;- | 1439 | xkr9 | 4 | 6 | 2 | 10 | 4 | 0 |
| ENSDARG000 | 24;24;24;24;2 | 29964976:299 | 29965545:299 | -;-;-;-;-;-;-;-;-; | 925 | NA | 0 | 0 | 0 | 8 | 12 | 0 |
| ENSDARG000 | 24;24;24;24;2 | 36373588:363 | 36374034:363 | +;+;+;+;+;+;+; | 5562 | NA | 36 | 22 | 61 | 38 | 52 | 29 |
| ENSDARG000 | 24;24;24;24;2 | 36392040:363 | 36392179:363 | +;+;+;+;+;+;+; | 1899 | arf2b | 2 | 4 | 1 | 0 | 2 | 8 |
| ENSDARG000 | 24;24;24;24;2 | 14214831:142 | 14215081:142 | +;+;+;+;+;+;+; | 2407 | tram1 | 1318 | 2140 | 2927 | 1740 | 1831 | 1414 |
| ENSDARG000 | 24;24;24;24;2 | 20194022:201 | 20194481:201 | -;-;-;-;-;-;-;-;-; | 2045 | cry-dash | 194 | 162 | 297 | 76 | 377 | 262 |
| ENSDARG000 | 24;24;24;24;2 | 17068724:170 | 17069754:170 | +;+;+;+;+;+;+; | 5883 | pip4k2aa | 514 | 705 | 791 | 705 | 721 | 582 |
| ENSDARG000 | 24;24;24;24;2 | 21514283:215 | 21514738:215 | +;+;+;+;+;+;+; | 2762 | cdk8 | 537 | 714 | 713 | 204 | 673 | 499 |
| ENSDARG000 | 24;24;24;24;2 | 30053651:300 | 30053847:300 | -;-;-;-;-;-;-;-;-; | 2426 | NA | 0 | 0 | 6 | 2 | 0 | 4 |
| ENSDARG000 | 24;24;24;24;2 | 25447911:254 | 25449553:254 | -;-;-;-;-;-;-;-;-; | 3026 | mbtps2 | 664 | 553 | 638 | 530 | 630 | 600 |
| ENSDARG000 | 24;24;24;24;2 | 30120815:301 | 30121181:301 | +;+;+;+;+;+ | 1309 | plppr5b | 18 | 121 | 169 | 64 | 441 | 135 |
| ENSDARG000 | 24;24;24;24;2 | 20214418:202 | 20214646:202 | -;-;-;-;-;-;-;-;-; | 2841 | map3k22 | 126 | 82 | 90 | 69 | 185 | 98 |
| ENSDARG000 | 24;24;24;24;2 | 30225967:302 | 30228837:302 | -;-;-;-;-;-;-;-;-; | 4734 | snx7 | 527 | 507 | 560 | 421 | 565 | 599 |
| ENSDARG000 | 24;24;24;24;2 | 41167269:411 | 41167460:411 | -;-;-;-;-;-;-;-;-; | 2072 | acvr2ba | 26 | 6 | 55 | 32 | 12 | 32 |
| ENSDARG000 | 24;24;24 | 30215475:302 | 30215588:302 | +;+;+ | 1665 | NA | 0 | 0 | 0 | 0 | 0 | 0 |
| ENSDARG000 | 24;24;24;24;2 | 23936829:239 | 23937136:239 | -;-;-;-;- | 1522 | arxa | 0 | 0 | 2 | 0 | 2 | 0 |
| ENSDARG000 | 24;24;24;24;2 | 33439243:334 | 33439306:334 | -;-;-;-;-;-;-;-;-; | 2346 | NA | 0 | 0 | 0 | 0 | 4 | 0 |
| ENSDARG000 | 24;24;24;24;2 | 20325312:203 | 20325604:203 | +;+;+;+;+;+;+; | 1781 | acaa1 | 222 | 173 | 305 | 161 | 331 | 291 |
| ENSDARG000 | 24;24;24;24;2 | 21540842:215 | 21540984:215 | +;+;+;+;+;+;+; | 2501 | wasf3b | 489 | 537 | 703 | 956 | 409 | 1083 |
| ENSDARG000 | 24;24;24;24;2 | 41273252:412 | 41273368:412 | +;+;+;+;+;+;+; | 2004 | NA | 10 | 6 | 4 | 8 | 10 | 8 |
| ENSDARG000 | 24;24;24;24;2 | 36414829:364 | 36414997:364 | +;+;+;+;+;+;+; | 1595 | GRB7 | 2 | 0 | 0 | 0 | 2 | 0 |
| ENSDARG000 | 24;24;24;24;2 | 36452924:364 | 36453115:364 | +;+;+;+;+;+;+; | 1750 | srcin1b | 0 | 0 | 0 | 0 | 0 | 0 |
| ENSDARG000 | 24 | 17136749 | 17139114 | - | 2366 | NA | 65 | 62 | 100 | 86 | 141 | 138 |
| ENSDARG000 | 24;24;24;24;2 | 14240196:142 | 14240285:142 | +;+;+;+;+;+;+; | 8148 | ncoa2 | 1164 | 1016 | 1500 | 1014 | 2089 | 922 |
| ENSDARG000 | 24;24 | 3413854:3419 | 3415367:3419 | -;- | 1798 | slc35g2b | 10 | 23 | 12 | 92 | 18 | 48 |
| ENSDARG000 | 24;24;24;24;2 | 25485264:255 | 25485305:255 | -;-;-;-;-;-;-;-;-; | 3458 | CNKSR2 | 0 | 0 | 0 | 0 | 0 | 0 |
| ENSDARG000 | 24;24;24;24;2 | 3347510:3348 | 3348640:3348 | -;-;-;-;-;-;-;-;-; | 3177 | NCK1 | 1649 | 3649 | 3601 | 2870 | 3903 | 3259 |
| ENSDARG000 | 24;24 | 36636208:366 | 36636652:366 | +;+ | 1151 | NA | 29 | 10 | 8 | 0 | 21 | 2 |
| ENSDARG000 | 24;24 | 36673686:366 | 36676129:366 | -;- | 2550 | ccr10 | 0 | 0 | 0 | 0 | 0 | 2 |
| ENSDARG000 | 24;24;24;24;2 | 33462800:334 | 33462990:334 | +;+;+;+;+;+;+; | 2416 | rmc1 | 888 | 397 | 492 | 393 | 522 | 600 |
| ENSDARG000 | 24;24;24;24;2 | 7554582:7558 | 7558781:7558 | -;-;-;-;-;-;-;-;-; | 6900 | galnt11 | 1122 | 1196 | 1135 | 809 | 1315 | 1038 |
| ENSDARG000 | 24;24 | 36705855:367 | 36705997:367 | +;+ | 395 | NA | 6 | 0 | 6 | 0 | 0 | 5 |
| ENSDARG000 | 24;24;24;24;2 | 25467465:254 | 25467748:254 | +;+;+;+;+;+;+; | 5502 | smpx | 0 | 0 | 2 | 0 | 0 | 0 |
| ENSDARG000 | 24;24;24 | 21584074:215 | 21587343:215 | -;-;- | 3518 | GPR12 | 0 | 0 | 0 | 0 | 0 | 0 |
| ENSDARG000 | 24;24;24 | 36721341:367 | 36723122:367 | -;-;- | 1817 | NA | 30 | 0 | 8 | 2 | 28 | 14 |
| ENSDARG000 | 24;24;24;24;2 | 3469373:3474 | 3471946:3474 | -;-;-;-;- | 3274 | idi1 | 774 | 971 | 1508 | 381 | 1166 | 1196 |
| ENSDARG000 | 24;24;24;24;2 | 21600320:216 | 21601099:216 | -;-;-;-;-;-;-;-;-; | 2050 | usp12a | 690 | 633 | 628 | 649 | 413 | 528 |
| ENSDARG000 | 24;24;24;24;2 | 21621654:216 | 21621739:216 | +;+;+;+;+;+;+; | 2385 | rpl21 | 10472 | 10137 | 15218 | 12188 | 9314 | 14507 |
| ENSDARG000 | 24;24;24;24;2 | 14451404:144 | 14451950:144 | +;+;+;+;+;+;+; | 1707 | prdm14 | 4 | 3 | 0 | 0 | 0 | 0 |
| ENSDARG000 | 24;24;24;24 | 14527935:145 | 14528142:145 | +;+;+;+ | 829 | NA | 0 | 0 | 0 | 0 | 0 | 0 |
| ENSDARG000 | 24;24;24;24;2 | 27223952:272 | 27226669:272 | -;-;-;-;-;-;-;-;-; | 7809 | zmynd11 | 2421 | 1933 | 2800 | 2467 | 2419 | 2634 |
| ENSDARG000 | 24;24;24;24;2 | 20332979:203 | 20333251:203 | -;-;-;-;-;-;-;-;-; | 5656 | NA | 80 | 122 | 105 | 188 | 203 | 97 |
| ENSDARG000 | 24;24;24;24;2 | 17140938:171 | 17142955:171 | +;+;+;+;+;+;+; | 11328 | NA | 1449 | 1083 | 1813 | 1611 | 1850 | 1502 |
| ENSDARG000 | 24;24 | 30349981:303 | 30350267:303 | +;+ | 2466 | NA | 14 | 4 | 8 | 2 | 4 | 24 |
| ENSDARG000 | 24;24;24;24 | 30329467:303 | 30329643:303 | -;-;-;- | 498 | NA | 0 | 0 | 0 | 0 | 0 | 0 |
| ENSDARG000 | 24;24 | 14533842:145 | 14533990:145 | -;- | 833 | NA | 0 | 0 | 0 | 0 | 0 | 0 |
| ENSDARG000 | 24;24;24 | 23973784:239 | 23973881:239 | +;+;+ | 303 | NA | 0 | 0 | 0 | 0 | 0 | 0 |
| ENSDARG000 | 24;24;24;24;2 | 23971056:239 | 23971222:239 | -;-;-;-;-;-;-;-;-; | 772 | ndufb4 | 0 | 0 | 0 | 0 | 0 | 0 |
| ENSDARG000 | 24;24;24;24;2 | 23959107:239 | 23959673:239 | +;+;+;+;+;+;+; | 6190 | hgd | 52 | 2 | 29 | 6 | 98 | 50 |
| ENSDARG000 | 24;24;24;24;2 | 30392834:304 | 30393236:304 | +;+;+;+;+;+;+; | 1142 | dpyda.1 | 203 | 103 | 240 | 109 | 183 | 217 |
| ENSDARG000 | 24;24;24;24 | 41305391:413 | 41305887:413 | -;-;-;- | 981 | crygn2 | 8 | 4 | 2 | 6 | 6 | 4 |
| ENSDARG000 | 24;24;24;24;2 | 38369351:383 | 38371288:383 | -;-;-;-;-;-;-;-;-; | 2753 | LRRC4B | 0 | 0 | 0 | 0 | 0 | 0 |
| ENSDARG000 | 24;24;24;24;2 | 21640635:216 | 21640880:216 | +;+;+;+;+;+;+; | 1215 | gtf3ab | 0 | 0 | 0 | 4 | 0 | 0 |
| ENSDARG000 | 24;24 | 3509377:3534 | 3509579:3534 | -;- | 328 | NA | 0 | 0 | 0 | 0 | 0 | 0 |
| ENSDARG000 | 24;24;24;24;2 | 3478871:3478 | 3479120:3479 | +;+;+;+;+;+;+; | 3153 | wdr37 | 296 | 529 | 640 | 357 | 558 | 445 |
| ENSDARG000 | 24;24;24;24;2 | 41690545:417 | 41690754:417 | +;+;+;+;+;+;+; | 9581 | lama1 | 3471 | 1304 | 2064 | 958 | 1267 | 916 |
| ENSDARG000 | 24;24;24;24;2 | 25683071:256 | 25683432:256 | -;-;-;-;-;-;-;-;-; | 2019 | chrnd | 0 | 0 | 0 | 0 | 0 | 0 |
| ENSDARG000 | 24;24 | 38396572:383 | 38397151:383 | -;- | 663 | NA | 0 | 0 | 0 | 6 | 0 | 0 |
| ENSDARG000 | 24;24;24;24;2 | 7596009:7602 | 7596396:7602 | -;-;-;-;-;-;-;-;-; | 3010 | atp6v0a1b | 4 | 0 | 38 | 12 | 17 | 10 |
| ENSDARG000 | 24;24;24;24;2 | 7631797:7634 | 7632180:7634 | +;+;+;+;+;+;+ | 2874 | cavin1b | 16853 | 11844 | 12469 | 11805 | 18232 | 9231 |
| ENSDARG000 | 24;24;24;24;2 | 21646317:216 | 21646865:216 | -;-;-;-;-;-;-;-;-; | 2870 | lnx2a | 2 | 8 | 0 | 20 | 11 | 14 |
| ENSDARG000 | 24;24 | 38446880:384 | 38447071:384 | -;- | 483 | NA | 0 | 0 | 0 | 0 | 0 | 0 |
| ENSDARG000 | 24;24;24 | 21676921:216 | 21677208:216 | +;+;+ | 1924 | si:ch211-140l | 209 | 194 | 198 | 198 | 173 | 233 |
| ENSDARG000 | 24;24;24;24;2 | 27268001:272 | 27268150:272 | +;+;+;+;+;+;+; | 1419 | vipr2 | 0 | 0 | 13 | 10 | 14 | 2 |
| ENSDARG000 | 24;24;24;24;2 | 30835929:308 | 30836293:308 | -;-;-;-;-;-;-;-;-; | 2063 | ptbp2a | 444 | 390 | 599 | 425 | 385 | 459 |
| ENSDARG000 | 24;24;24;24;2 | 38522254:385 | 38522415:385 | +;+;+;+;+;+;+; | 1432 | NA | 6 | 19 | 13 | 13 | 3 | 10 |
| ENSDARG000 | 24;24 | 21687276:216 | 21687805:216 | -;- | 759 | urad | 0 | 0 | 0 | 0 | 0 | 0 |
| ENSDARG000 | 24;24 | 21684189:216 | 21684666:216 | +;+ | 1117 | pdx1 | 0 | 6 | 6 | 4 | 30 | 28 |
| ENSDARG000 | 24;24 | 38523993:385 | 38524110:385 | -;- | 546 | NA | 0 | 0 | 0 | 0 | 0 | 0 |
| ENSDARG000 | 24;24;24;24;2 | 14541013:145 | 14541068:145 | +;+;+;+;+;+;+; | 3751 | ngdn | 330 | 365 | 700 | 356 | 456 | 500 |
| ENSDARG000 | 24;24;24;24 | 33690310:336 | 33690395:336 | -;-;-;- | 896 | NA | 0 | 0 | 0 | 0 | 0 | 0 |
| ENSDARG000 | 24;24;24;24;2 | 33589667:335 | 33589723:335 | +;+;+;+;+;+;+; | 13897 | NA | 0 | 0 | 10 | 16 | 18 | 24 |
| ENSDARG000 | 24;24;24;24;2 | 11070110:110 | 11070478:110 | -;-;-;-;- | 1212 | chmp4c | 4 | 0 | 2 | 0 | 6 | 0 |
| ENSDARG000 | 24;24;24;24 | 27381661:273 | 27381752:273 | +;+;+;+ | 577 | NA | 0 | 0 | 0 | 0 | 0 | 6 |
| ENSDARG000 | 24;24;24;24;2 | 20372497:203 | 20372675:203 | +;+;+;+;+;+;+; | 5461 | plcd1a | 674 | 1281 | 536 | 1086 | 1351 | 802 |
| ENSDARG000 | 24;24;24 | 38536928:385 | 38537025:385 | -;-;- | 630 | NA | 0 | 1 | 0 | 0 | 0 | 0 |
| ENSDARG000 | 24;24;24;24;2 | 38534550:385 | 38534918:385 | +;+;+;+;+;+;+ | 2081 | zgc:154125 | 281 | 410 | 782 | 605 | 528 | 191 |
| ENSDARG000 | 24;24;24;24;2 | 27396989:273 | 27397557:273 | -;-;-;-;-;-;-;-;-; | 1576 | ccl34b.1 | 0 | 0 | 10 | 4 | 4 | 6 |
| ENSDARG000 | 24;24;24;24;2 | 3537762:3539 | 3537925:3540 | -;-;-;-;-;-;-;-;-; | 2895 | ADARB2 | 0 | 14 | 2 | 10 | 4 | 4 |
| ENSDARG000 | 24;24;24;24;2 | 24014880:240 | 24015173:240 | +;+;+;+;+;+;+ | 1382 | chodl | 0 | 0 | 0 | 0 | 2 | 0 |
| ENSDARG000 | 24;24;24;24;2 | 38557989:385 | 38558414:385 | -;-;-;-;-;-;-;-;-; | 1903 | SLC17A7 | 0 | 6 | 3 | 4 | 0 | 0 |
| ENSDARG000 | 24;24;24;24;2 | 27407955:274 | 27408176:274 | -;-;-;-;- | 627 | NA | 0 | 0 | 0 | 0 | 12 | 2 |
| ENSDARG000 | 24;24;24;24;2 | 25760986:257 | 25765532:257 | -;-;-;-;-;-;-;-;-; | 6341 | KLHL24 | 1120 | 313 | 449 | 787 | 402 | 680 |
| ENSDARG000 | 24;24;24;24;2 | 21691939:216 | 21692998:216 | -;-;-;-;-;-;-;-;-; | 3849 | NA | 10 | 4 | 12 | 10 | 34 | 34 |
| ENSDARG000 | 24;24;24;24;2 | 37301852:373 | 37303395:373 | -;-;-;-;-;-;-;-;-; | 3213 | pdpk1b | 1325 | 1554 | 1848 | 1598 | 1529 | 1120 |
| ENSDARG000 | 24;24;24;24 | 27416592:274 | 27416887:274 | -;-;-;- | 704 | ccl34b.4 | 20 | 4 | 8 | 12 | 6 | 108 |
| ENSDARG000 | 24;24;24;24;2 | 14559328:145 | 14560043:145 | -;-;-;-;-;-;-;-;-; | 4419 | pabpn1 | 4977 | 4464 | 6044 | 4501 | 5299 | 4657 |
| ENSDARG000 | 24;24;24;24;2 | 3963684:3963 | 3964019:3964 | +;+;+;+;+;+;+; | 4477 | PFKP | 741 | 3350 | 8081 | 5144 | 3089 | 1374 |
| ENSDARG000 | 24;24;24;24;2 | 37338169:373 | 37338565:373 | +;+;+;+;+;+;+; | 5266 | gnptg | 499 | 263 | 421 | 486 | 399 | 428 |
| ENSDARG000 | 24;24;24;24;2 | 11083146:110 | 11083219:110 | +;+;+;+;+;+;+; | 1084 | zfand1 | 108 | 141 | 134 | 26 | 74 | 103 |
| ENSDARG000 | 24;24;24;24;2 | 37329119:373 | 37329667:373 | -;-;-;-;-;-;-;-;-; | 1832 | tsr3 | 330 | 274 | 378 | 200 | 191 | 228 |
| ENSDARG000 | 24;24;24;24;2 | 7686464:7686 | 7687374:7687 | -;-;-;-;-;-;-;-;-; | 2275 | syt5b | 0 | 6 | 27 | 4 | 2 | 3 |
| ENSDARG000 | 24;24;24;24;2 | 31081504:310 | 31083128:310 | -;-;-;-;-;-;-;-;-; | 2487 | hccsb | 578 | 854 | 856 | 482 | 706 | 502 |
| ENSDARG000 | 24;24;24;24;2 | 38618502:386 | 38619326:386 | -;-;-;-;-;-;-;-;-; | 2816 | slc6a16b | 144 | 306 | 500 | 134 | 332 | 310 |
| ENSDARG000 | 24;24;24;24 | 24035863:240 | 24036051:240 | -;-;-;- | 603 | lyz | 0 | 0 | 8 | 3 | 52 | 46 |
| ENSDARG000 | 24;24;24;24;2 | 38648885:386 | 38649677:386 | -;-;-;-;-;- | 1323 | NA | 10 | 0 | 4 | 6 | 0 | 0 |
| ENSDARG000 | 24;24;24;24;2 | 24036195:240 | 24040134:240 | -;-;-;-;-;-;-;-;-; | 6496 | NA | 0 | 0 | 4 | 0 | 29 | 24 |
| ENSDARG000 | 24;24;24;24;2 | 20450027:204 | 20450929:204 | -;-;-;-;-;-;-;-;-; | 2364 | ctdspla | 34 | 46 | 129 | 54 | 52 | 100 |
| ENSDARG000 | 24;24;24;24;2 | 25794318:258 | 25794624:258 | +;+;+;+;+;+;+; | 2652 | NA | 2674 | 2576 | 3997 | 3832 | 2885 | 2578 |
| ENSDARG000 | 24;24;24;24;2 | 27421076:274 | 27421767:274 | -;-;-;-;-;-;- | 1023 | NA | 2 | 2 | 4 | 22 | 14 | 0 |
| ENSDARG000 | 24;24;24;24;2 | 21750810:217 | 21751146:217 | -;-;-;-;-;-;-;-;-; | 5499 | flt1 | 42 | 32 | 50 | 68 | 38 | 38 |
| ENSDARG000 | 24;24;24;24;2 | 31098534:310 | 31098979:310 | -;-;-;-;-;-;-;-;-; | 1460 | TLCD4 | 70 | 36 | 77 | 58 | 61 | 67 |
| ENSDARG000 | 24;24;24;24;2 | 21720304:217 | 21720915:217 | +;+;+;+;+;+;+; | 5270 | pan3 | 1092 | 710 | 1051 | 641 | 1013 | 703 |
| ENSDARG000 | 24;24;24;24;2 | 31170362:311 | 31171551:311 | -;-;-;-;-;-;- | 2000 | cnn3a | 356 | 308 | 162 | 607 | 144 | 336 |
| ENSDARG000 | 24;24;24;24;2 | 17256793:172 | 17256970:172 | +;+;+;+;+;+;+; | 1174 | commd3 | 471 | 566 | 403 | 409 | 525 | 371 |
| ENSDARG000 | 24;24;24;24;2 | 7714014:7715 | 7714155:7715 | -;-;-;-;-;-;-;-;-; | 2338 | NA | 2 | 0 | 22 | 0 | 0 | 4 |
| ENSDARG000 | 24;24;24;24;2 | 27450085:274 | 27450273:274 | -;-;-;-;- | 533 | NA | 20 | 119 | 113 | 94 | 297 | 62 |
| ENSDARG000 | 24;24;24;24;2 | 31201195:312 | 31201462:312 | -;-;-;-;-;-;- | 965 | alg14 | 98 | 148 | 152 | 106 | 162 | 128 |
| ENSDARG000 | 24;24;24;24;2 | 24064562:240 | 24064688:240 | +;+;+;+;+;+;+ | 1237 | zgc:112408 | 0 | 0 | 0 | 0 | 2 | 0 |
| ENSDARG000 | 24;24;24;24;2 | 11105786:111 | 11105944:111 | +;+;+;+;+ | 551 | NA | 0 | 0 | 0 | 0 | 0 | 0 |
| ENSDARG000 | 24;24;24;24;2 | 37361189:373 | 37361291:373 | -;-;-;-;-;-;-;- | 1265 | NA | 0 | 0 | 0 | 0 | 0 | 0 |
| ENSDARG000 | 24;24;24;24;2 | 37361089:373 | 37361279:373 | +;+;+;+;+;+;+; | 1424 | SBK1 | 0 | 0 | 0 | 0 | 0 | 0 |
| ENSDARG000 | 24;24;24;24;2 | 20536056:205 | 20536352:205 | +;+;+;+;+;+;+; | 2622 | gars1 | 4484 | 4814 | 8376 | 4039 | 4233 | 4767 |
| ENSDARG000 | 24;24 | 38665082:386 | 38667295:386 | -;- | 2378 | NA | 12 | 0 | 0 | 2 | 8 | 0 |
| ENSDARG000 | 24;24;24;24;2 | 38658309:386 | 38658398:386 | +;+;+;+;+;+;+; | 2665 | NA | 30 | 0 | 0 | 13 | 6 | 2 |
| ENSDARG000 | 24;24;24;24;2 | 27459604:274 | 27459904:274 | -;-;-;-;- | 627 | ccl34b.9 | 0 | 22 | 0 | 14 | 22 | 20 |
| ENSDARG000 | 24;24;24;24;2 | 27470097:274 | 27470316:274 | -;-;-;-;- | 604 | cxl34b.11 | 16 | 0 | 0 | 0 | 0 | 0 |
| ENSDARG000 | 24;24;24;24;2 | 31277360:312 | 31277574:312 | +;+;+;+;+;+ | 1372 | f3a | 120 | 4 | 6 | 110 | 11 | 38 |
| ENSDARG000 | 24;24;24;24;2 | 17260001:172 | 17260085:172 | +;+;+;+;+;+;+; | 2655 | bmi1a | 290 | 401 | 388 | 326 | 357 | 387 |
| ENSDARG000 | 24;24;24;24;2 | 11112113:111 | 11116118:111 | -;-;-;-;-;-;-;-;-; | 7405 | myripb | 0 | 3 | 0 | 0 | 0 | 0 |
| ENSDARG000 | 24;24;24;24;2 | 24086249:240 | 24086269:240 | +;+;+;+;+;+;+; | 1762 | NA | 0 | 2 | 0 | 0 | 0 | 4 |
| ENSDARG000 | 24;24;24;24;2 | 25913162:259 | 25913566:259 | +;+;+;+;+;+;+; | 7593 | MAP3K15 | 90 | 56 | 56 | 10 | 78 | 100 |
| ENSDARG000 | 24;24;24;24;2 | 7741521:7742 | 7741653:7742 | +;+;+;+;+;+;+; | 1188 | zgc:56095 | 712 | 560 | 876 | 626 | 878 | 744 |
| ENSDARG000 | 24;24;24 | 27511734:275 | 27511858:275 | +;+;+ | 798 | cxcl32b.1 | 26 | 21 | 12 | 27 | 75 | 9 |
| ENSDARG000 | 24;24;24;24;2 | 11334733:113 | 11334919:113 | +;+;+;+;+;+;+; | 2009 | NA | 211 | 4 | 144 | 156 | 332 | 100 |
| ENSDARG000 | 24;24;24;24;2 | 37406535:374 | 37406621:374 | +;+;+;+;+;+;+; | 3046 | NA | 2266 | 620 | 1706 | 664 | 617 | 1005 |
| ENSDARG000 | 24;24;24;24;2 | 31294019:312 | 31294418:312 | -;-;-;-;-;- | 1274 | acp5b | 181 | 76 | 44 | 116 | 34 | 56 |
| ENSDARG000 | 24;24;24;24;2 | 42127983:421 | 42128200:421 | +;+;+;+;+;+;+; | 4213 | WWP1 | 2177 | 1268 | 1986 | 1626 | 1978 | 1459 |
| ENSDARG000 | 24;24;24;24;2 | 14581864:145 | 14581995:145 | +;+;+;+;+;+;+; | 3213 | thtpa | 142 | 133 | 160 | 81 | 184 | 121 |
| ENSDARG000 | 24;24;24;24;2 | 24101811:241 | 24102392:241 | -;-;-;-;-;-;-;-;-; | 3299 | zgc:112982 | 1966 | 1423 | 2258 | 2306 | 2365 | 2465 |
| ENSDARG000 | 24;24;24;24;2 | 20559009:205 | 20559153:205 | +;+;+;+;+;+;+; | 2949 | hhatlb | 2 | 0 | 6 | 0 | 8 | 0 |
| ENSDARG000 | 24;24;24;24;2 | 4141484:4141 | 4143819:4143 | -;-;-;-;-;-;-;- | 3369 | klf6a | 4724 | 8184 | 4952 | 7602 | 4302 | 4975 |
| ENSDARG000 | 24;24;24;24;2 | 24118193:241 | 24122043:241 | -;-;-;-;-;-;-;-;-; | 6261 | MAP7D2 | 1073 | 37 | 25 | 34 | 137 | 58 |
| ENSDARG000 | 24;24;24;24;2 | 7748751:7748 | 7749175:7749 | -;-;-;-;-;-;-;-;-; | 4881 | NA | 0 | 0 | 0 | 0 | 0 | 0 |
| ENSDARG000 | 24;24;24;24;2 | 27548474:275 | 27548698:275 | +;+;+;+;+;+;+; | 5007 | ek1 | 9 | 0 | 2 | 0 | 0 | 0 |
| ENSDARG000 | 24;24 | 4154959:4156 | 4155463:4156 | +;+ | 532 | NA | 0 | 0 | 0 | 0 | 0 | 0 |
| ENSDARG000 | 24;24;24;24;2 | 31310640:313 | 31311376:313 | -;-;-;-;-;-;-;-;-; | 2802 | abcd3a | 1442 | 1196 | 1306 | 842 | 1567 | 1039 |
| ENSDARG000 | 24;24;24;24;2 | 7782313:7794 | 7782621:7794 | +;+;+;+;+;+;+; | 3733 | NA | 0 | 0 | 6 | 0 | 2 | 0 |
| ENSDARG000 | 24;24;24;24;2 | 17269849:172 | 17270010:172 | +;+;+;+;+;+;+; | 4954 | spag6 | 0 | 4 | 2 | 0 | 0 | 0 |
| ENSDARG000 | 24;24;24;24;2 | 38726737:387 | 38728017:387 | -;-;-;-;-;-;-;-;-; | 3442 | ntn2 | 4 | 0 | 4 | 0 | 6 | 0 |
| ENSDARG000 | 24;24;24;24;2 | 37449893:374 | 37450846:374 | +;+;+;+;+;+;+; | 2352 | NLRC3 | 21 | 38 | 6 | 12 | 26 | 58 |
| ENSDARG000 | 24;24;24;24;2 | 4373033:4373 | 4373508:4373 | +;+;+;+;+;+;+; | 1982 | CCNY | 589 | 1024 | 1483 | 1086 | 965 | 901 |
| ENSDARG000 | 24;24;24;24;2 | 20575259:205 | 20576444:205 | +;+;+;+;+;+ | 1904 | klhl40b | 18 | 13 | 18 | 2 | 19 | 100 |
| ENSDARG000 | 24;24;24;24;2 | 14623831:146 | 14624170:146 | -;-;-;-;-;-;-;- | 3243 | jph1a | 14 | 0 | 0 | 0 | 16 | 0 |
| ENSDARG000 | 24;24;24 | 14623831:146 | 14624183:146 | +;+;+ | 704 | NA | 0 | 0 | 0 | 0 | 0 | 0 |
| ENSDARG000 | 24;24 | 33697130:337 | 33699959:337 | -;- | 3492 | cavin4b | 0 | 0 | 0 | 0 | 2 | 0 |
| ENSDARG000 | 24 | 38814734 | 38816725 | - | 1992 | nog2 | 10 | 0 | 4 | 0 | 2 | 0 |
| ENSDARG000 | 24;24 | 7820042:7825 | 7821071:7826 | -;- | 1881 | NA | 176 | 217 | 339 | 172 | 301 | 298 |
| ENSDARG000 | 24;24;24;24;2 | 17322644:173 | 17322672:173 | +;+;+;+;+;+;+; | 1409 | NA | 68 | 261 | 60 | 254 | 328 | 34 |

| | | | | | | | | | | | | |
|---|---|---|---|---|---|---|---|---|---|---|---|---|
| ENSDARG000 | 24;24;24;24;2 | 31334209;313 | 31334374;313 | +;+;+;+;+;+;+; | 2186 | fam168b | 947 | 981 | 1837 | 1218 | 1056 | 1034 |
| ENSDARG000 | 24;24 | 27783946;278 | 27784648;278 | -;- | 1224 | hs6st1b | 109 | 117 | 159 | 47 | 133 | 84 |
| ENSDARG000 | 24;24 | 28137210;282 | 28137729;282 | -;- | 1828 | bcl2a | 364 | 199 | 331 | 145 | 448 | 201 |
| ENSDARG000 | 24;24;24;24;2 | 39105051;391 | 39105461;391 | +;+;+;+;+;+;+; | 2246 | NA | 0 | 2 | 6 | 0 | 0 | 0 |
| ENSDARG000 | 24 | 4447635 | 4450238 | - | 2604 | fzd8a | 20 | 0 | 26 | 14 | 14 | 40 |
| ENSDARG000 | 24;24;24;24;2 | 33708804;337 | 33709627;337 | -;-;-;-;-;-;-;-; | 2138 | tmeff1b | 10 | 10 | 76 | 20 | 65 | 26 |
| ENSDARG000 | 24;24;24;24;2 | 37466770;374 | 37467284;374 | -;-;-;-;-;-;-;-; | 1986 | cluap1 | 138 | 94 | 96 | 141 | 116 | 174 |
| ENSDARG000 | 24;24;24;24;2 | 20589458;205 | 20589607;205 | -;-;-;-;-;-;-;-; | 2424 | ZBTB47 | 581 | 314 | 371 | 185 | 664 | 743 |
| ENSDARG000 | 24;24;24;24;2 | 14713776;147 | 14713964;147 | +;+;+;+;+;+ | 1493 | gdap1 | 32 | 6 | 0 | 0 | 12 | 10 |
| ENSDARG000 | 24;24;24;24;2 | 4755391;4756 | 4756473;4756 | -;-;-;-;-;-;-;-; | 2220 | cpb1 | 0 | 0 | 30 | 0 | 4 | 2 |
| ENSDARG000 | 24;24;24;24;2 | 7828097;7828 | 7828284;7828 | +;+;+;+;+;+;+; | 1287 | zgc:101569 | 87 | 138 | 248 | 234 | 160 | 90 |
| ENSDARG000 | 24 | 39117499 | 39118364 | + | 866 | NA | 0 | 6 | 4 | 0 | 0 | 0 |
| ENSDARG000 | 24;24;24;24;2 | 42141313;421 | 42141610;421 | -;-;-;-;-;-;-;-; | 1844 | rmdn1 | 385 | 302 | 407 | 334 | 327 | 371 |
| ENSDARG000 | 24;24;24;24;2 | 28231602;282 | 28231870;282 | -;-;-;-;-;-;-;- | 782 | cox4i1l | 293 | 1161 | 2327 | 1802 | 574 | 280 |
| ENSDARG000 | 24;24;24;24;2 | 7861373;7884 | 7862342;7885 | +;+;+;+;+;+;+; | 2491 | bmp6 | 2 | 2 | 0 | 0 | 12 | 12 |
| ENSDARG000 | 24;24;24;24;2 | 39129316;391 | 39129397;391 | +;+;+;+;+;+;+; | 6195 | TBC1D24 | 380 | 294 | 1033 | 264 | 536 | 684 |
| ENSDARG000 | 24;24;24;24;2 | 42148140;421 | 42148226;421 | +;+;+;+;+;+;+; | 1667 | serpinb1l3 | 744 | 952 | 708 | 628 | 933 | 423 |
| ENSDARG000 | 24;24;24;24;2 | 26006730;260 | 26006781;260 | +;+;+;+;+;+;+; | 979 | ccl20b | 6 | 0 | 0 | 0 | 0 | 8 |
| ENSDARG000 | 24;24;24;24;2 | 11350330;113 | 11350831;113 | -;-;-;-;-;-;-;-; | 3288 | acad11 | 101 | 154 | 189 | 140 | 123 | 102 |
| ENSDARG000 | 24;24;24;24;2 | 26017094;260 | 26017245;260 | +;+;+;+;+;+;+; | 3097 | tfr1b | 4961 | 10220 | 11817 | 8558 | 6062 | 19562 |
| ENSDARG000 | 24;24 | 11381400;113 | 11381453;113 | +;+ | 3687 | ackr4b | 30 | 36 | 98 | 315 | 186 | 39 |
| ENSDARG000 | 24;24;24;24;2 | 14801844;148 | 14801960;148 | +;+;+;+;+;+ | 1164 | pi15a | 6 | 4 | 0 | 0 | 4 | 2 |
| ENSDARG000 | 24;24;24 | 4767806;4767 | 4769067;4768 | -;-;- | 1393 | AGTR1 | 0 | 0 | 8 | 2 | 6 | 2 |
| ENSDARG000 | 24;24;24;24;2 | 28252520;282 | 28253872;282 | -;-;-;-;-;-;-;-; | 4468 | prkag2a | 700 | 587 | 656 | 610 | 450 | 366 |
| ENSDARG000 | 24;24;24 | 4970580;4971 | 4971444;4972 | -;-;- | 2262 | zic1 | 6 | 0 | 0 | 0 | 10 | 10 |
| ENSDARG000 | 24;24;24;24;2 | 20643658;206 | 20644234;206 | -;-;-;-;-;-;-;-; | 5053 | NKTR | 3778 | 1829 | 4739 | 6316 | 4682 | 8264 |
| ENSDARG000 | 24;24;24;24;2 | 7941373;7945 | 7943400;7945 | -;-;-;-;-;-;-;-; | 3224 | txndc5 | 4741 | 3487 | 5925 | 2674 | 5706 | 3738 |
| ENSDARG000 | 24;24;24;24 | 8244085;8248 | 8244651;8248 | -;-;-;- | 1015 | EEF1E1 | 477 | 451 | 724 | 470 | 909 | 683 |
| ENSDARG000 | 24;24;24;24;2 | 17334678;173 | 17334779;173 | +;+;+;+;+;+;+; | 2429 | pdia4 | 1714 | 4361 | 5490 | 1472 | 3589 | 1555 |
| ENSDARG000 | 24;24;24;24;2 | 33767771;337 | 33768738;337 | -;-;-;-;-;-;-;-; | 1899 | cdk5 | 90 | 104 | 70 | 47 | 114 | 119 |
| ENSDARG000 | 24;24;24;24;2 | 4977862;4978 | 4979207;4978 | +;+;+;+;+ | 3561 | zic4 | 0 | 0 | 0 | 0 | 4 | 0 |
| ENSDARG000 | 24;24;24;24;2 | 14859238;148 | 14859546;148 | +;+;+;+;+;+;+; | 2374 | CRISPLD1 | 8 | 4 | 53 | 9 | 353 | 4 |
| ENSDARG000 | 24;24;24;24 | 39178599;391 | 39178646;391 | +;+;+;+ | 542 | NA | 2 | 4 | 6 | 3 | 1 | 4 |
| ENSDARG000 | 24;24;24;24;2 | 39178348;391 | 39178655;391 | -;-;-;-;-;-;-;-; | 1057 | nubp2 | 12 | 12 | 7 | 4 | 10 | 7 |
| ENSDARG000 | 24;24;24 | 39158283;391 | 39158529;391 | +;+;+ | 1751 | atp6v0cb | 1670 | 1767 | 1799 | 1988 | 1774 | 1724 |
| ENSDARG000 | 24;24;24;24;2 | 8393290;8393 | 8393946;8393 | -;-;-;-;-;-;-;-; | 2387 | slc35b3 | 404 | 288 | 427 | 234 | 438 | 520 |
| ENSDARG000 | 24;24;24;24;2 | 24164227;241 | 24165122;241 | -;-;-;-;-;-;-;-; | 1994 | eif1axb | 2209 | 2054 | 2683 | 1446 | 2415 | 1924 |
| ENSDARG000 | 24;24;24;24;2 | 21863935;218 | 21865295;218 | -;-;-;-;-;-;-;-; | 5572 | NA | 2231 | 2160 | 5465 | 3576 | 4854 | 2648 |
| ENSDARG000 | 24;24;24;24;2 | 11451077;114 | 11453762;114 | -;-;-;-;-;-;-;-; | 3851 | PXDC1 | 369 | 252 | 170 | 328 | 148 | 288 |
| ENSDARG000 | 24;24;24;24;2 | 5208171;5208 | 5208237;5208 | +;+;+;+;+;+;+; | 1468 | NA | 23 | 62 | 24 | 12 | 102 | 54 |
| ENSDARG000 | 24;24;24;24;2 | 31348595;313 | 31348812;313 | +;+;+;+;+;+;+; | 1903 | cremb | 26 | 4 | 68 | 8 | 30 | 50 |
| ENSDARG000 | 24;24;24;24;2 | 37484661;374 | 37484890;374 | +;+;+;+;+;+;+; | 5827 | WDR90 | 39 | 145 | 183 | 87 | 184 | 65 |
| ENSDARG000 | 24;24;24;24;2 | 11469811;114 | 11474316;114 | -;-;-;-;-;-;-;-; | 7454 | prpf4bb | 797 | 839 | 987 | 464 | 902 | 768 |
| ENSDARG000 | 24;24;24;24;2 | 31402894;314 | 31403069;314 | -;-;-;-;-;-;-;-; | 2959 | NA | 0 | 0 | 1 | 0 | 0 | 0 |
| ENSDARG000 | 24;24;24;24;2 | 39186940;391 | 39187069;391 | +;+;+;+;+;+ | 884 | NA | 0 | 2 | 0 | 4 | 4 | 0 |
| ENSDARG000 | 24;24;24;24;2 | 31374324;313 | 31374621;313 | +;+;+;+;+;+;+; | 2608 | cpne3 | 712 | 740 | 735 | 588 | 720 | 680 |
| ENSDARG000 | 24;24;24;24;2 | 20658760;206 | 20659107;206 | +;+;+;+;+;+;+; | 1696 | SEC22C | 12 | 15 | 8 | 3 | 14 | 4 |
| ENSDARG000 | 24;24;24;24;2 | 17345521;173 | 17345893;173 | +;+;+;+;+;+;+; | 3762 | ezh2 | 236 | 693 | 648 | 164 | 859 | 345 |
| ENSDARG000 | 24;24;24;24;2 | 14937205;150 | 14937396;150 | +;+;+;+;+;+;+; | 1659 | dok6 | 53 | 67 | 48 | 28 | 74 | 28 |
| ENSDARG000 | 24;24;24;24;2 | 21917012;219 | 21918180;219 | -;-;-;-;-;-;-;-; | 7106 | tagln3b | 6808 | 6613 | 9570 | 8057 | 8239 | 6452 |
| ENSDARG000 | 24;24 | 24170914;241 | 24171375;241 | +;+ | 3501 | NA | 118 | 84 | 226 | 48 | 134 | 188 |
| ENSDARG000 | 24;24;24;24;2 | 26036395;260 | 26036547;260 | +;+;+;+;+;+;+; | 3660 | TNK2 | 10 | 4 | 21 | 9 | 22 | 0 |
| ENSDARG000 | 24;24;24;24;2 | 20708174;207 | 20708519;207 | -;-;-;-;-;-;-;-; | 1741 | vipr1b | 0 | 0 | 0 | 0 | 0 | 0 |
| ENSDARG000 | 24;24;24;24;2 | 21938741;219 | 21939090;219 | -;-;-;-;-;-;-;-; | 1873 | acot9.2 | 166 | 44 | 172 | 140 | 135 | 94 |
| ENSDARG000 | 24;24;24;24;2 | 33781485;337 | 33781916;337 | -;-;-;-;-;-;-;-; | 2628 | abcb8 | 382 | 319 | 352 | 237 | 306 | 243 |
| ENSDARG000 | 24 | 39353393 | 39354829 | - | 1437 | NA | 159 | 290 | 212 | 285 | 189 | 77 |
| ENSDARG000 | 24;24;24;24;2 | 5237709;5237 | 5238040;5238 | +;+;+;+;+;+;+; | 3966 | plod2 | 2022 | 1679 | 3390 | 3204 | 1384 | 39 |
| ENSDARG000 | 24;24;24;24;2 | 39227519;392 | 39227523;392 | +;+;+;+;+;+;+; | 5711 | NA | 672 | 372 | 527 | 641 | 544 | 739 |
| ENSDARG000 | 24;24;24;24;2 | 37533728;375 | 37533880;375 | +;+;+;+;+;+;+; | 2513 | rhot2 | 453 | 356 | 428 | 197 | 397 | 319 |
| ENSDARG000 | 24;24;24;24;2 | 24183752;241 | 24184727;241 | -;-;-;-;-;-;-;-; | 3190 | rps6ka3b | 241 | 158 | 298 | 262 | 202 | 349 |
| ENSDARG000 | 24;24;24;24;2 | 28383278;283 | 28383558;283 | +;+;+;+;+;+;+; | 2062 | sh3glb1a | 1025 | 852 | 1083 | 857 | 910 | 716 |
| ENSDARG000 | 24;24;24;24;2 | 854877;86500 | 854981;86530 | +;+;+;+;+;+;+; | 8595 | PIEZO2 | 397 | 173 | 330 | 128 | 606 | 141 |
| ENSDARG000 | 24;24;24;24;2 | 26134029;261 | 26134572;261 | +;+;+;+;+;+;+; | 3069 | tmtopsb | 317 | 260 | 97 | 161 | 110 | 0 |
| ENSDARG000 | 24;24;24;24;2 | 8827603;8827 | 8828757;8828 | -;-;-;-;-;-;-;-; | 2145 | gcm2 | 0 | 0 | 0 | 0 | 0 | 0 |
| ENSDARG000 | 24;24;24;24;2 | 20892782;208 | 20893462;208 | -;-;-;-;-;-;-;-; | 4567 | NA | 170 | 483 | 604 | 171 | 432 | 272 |
| ENSDARG000 | 24;24;24 | 26276805;262 | 26276868;262 | +;+;+ | 979 | adipoqa | 0 | 3 | 2 | 0 | 0 | 0 |
| ENSDARG000 | 24;24;24;24;2 | 15130979;151 | 15132315;151 | -;-;-;-;-;-;-;-; | 8188 | RTTN | 108 | 186 | 260 | 72 | 267 | 111 |
| ENSDARG000 | 24;24 | 11564896;115 | 11565136;115 | +;+ | 526 | NA | 0 | 0 | 0 | 0 | 0 | 0 |
| ENSDARG000 | 24 | 37567029 | 37568359 | - | 1331 | h1-0 | 5599 | 4758 | 6785 | 8402 | 3682 | 3788 |
| ENSDARG000 | 24;24;24;24;2 | 15114092;151 | 15114764;151 | -;-;-;-;-;-;-;-; | 2518 | cd226 | 8 | 13 | 2 | 2 | 20 | 21 |
| ENSDARG000 | 24;24;24;24;2 | 11622196;116 | 11622296;116 | +;+;+;+;+;+;+; | 1544 | NA | 0 | 0 | 0 | 0 | 0 | 0 |
| ENSDARG000 | 24;24;24;24;2 | 11528270;115 | 11532090;115 | -;-;-;-;-;-;-;-; | 6688 | gli3 | 60 | 100 | 180 | 38 | 90 | 20 |
| ENSDARG000 | 24;24;24;24;2 | 17364745;173 | 17365182;173 | -;-;-;-;-;-;-;-; | 3503 | cul1b | 727 | 733 | 832 | 920 | 892 | 628 |
| ENSDARG000 | 24;24;24;24;2 | 8861272;8865 | 8864251;8865 | -;-;-;-;-;-;-;- | 3840 | elovl2 | 30 | 169 | 199 | 43 | 58 | 26 |
| ENSDARG000 | 24;24;24 | 26284514;263 | 26284659;263 | -;-;- | 392 | cops9 | 220 | 186 | 138 | 68 | 177 | 183 |
| ENSDARG000 | 24;24;24;24;2 | 5653162;5653 | 5653707;5653 | -;-;-;-;-;-;- | 2086 | dipk2ab | 210 | 250 | 1648 | 487 | 983 | 345 |
| ENSDARG000 | 24 | 961874 | 962759 | - | 886 | NA | 0 | 8 | 0 | 6 | 4 | 2 |
| ENSDARG000 | 24;24;24;24;2 | 984822;98800 | 986061;98850 | -;-;-;-;-;-;-;-; | 5471 | NA | 624 | 1126 | 1677 | 684 | 1526 | 746 |
| ENSDARG000 | 24;24;24;24;2 | 33802528;338 | 33802628;338 | +;+;+;+;+;+;+; | 7882 | atg9b | 1310 | 485 | 723 | 849 | 775 | 1254 |
| ENSDARG000 | 24;24;24;24;2 | 947473;96888 | 949115;96909 | -;-;-;-;-;-;-;-; | 2526 | napga | 537 | 307 | 870 | 326 | 887 | 762 |
| ENSDARG000 | 24;24;24;24;2 | 26306362;263 | 26306942;263 | -;-;-;-;-;-;-;-; | 1059 | apodb | 6 | 4 | 6 | 8 | 8 | 24 |
| ENSDARG000 | 24;24;24;24;2 | 24273371;242 | 24274083;242 | -;-;-;-;-;-;-;-; | 2432 | pdha1b | 2 | 4 | 2 | 8 | 4 | 0 |
| ENSDARG000 | 24 | 5784264 | 5786759 | - | 2496 | CHST2 | 54 | 8 | 44 | 56 | 22 | 12 |
| ENSDARG000 | 24;24;24;24;2 | 31430890;314 | 31430922;314 | -;-;-;-;-;-;-;-; | 2361 | NA | 0 | 0 | 4 | 10 | 0 | 0 |
| ENSDARG000 | 24;24;24 | 28425900;284 | 28426836;284 | -;-;- | 1789 | fbxo45 | 310 | 350 | 304 | 178 | 263 | 293 |
| ENSDARG000 | 24;24;24 | 28410963;284 | 28413070;284 | -;-;- | 2351 | nrros | 122 | 110 | 149 | 132 | 555 | 623 |
| ENSDARG000 | 24;24;24;24;2 | 5789582;5789 | 5789637;5789 | +;+;+;+;+;+;+; | 1779 | NA | 537 | 201 | 1371 | 1943 | 2261 | 1386 |
| ENSDARG000 | 24;24;24;24;2 | 37640626;376 | 37640744;376 | +;+;+;+;+;+;+; | 2986 | wdr24 | 276 | 225 | 358 | 210 | 326 | 189 |
| ENSDARG000 | 24;24;24;24;2 | 37631632;376 | 37632039;376 | -;-;-;-;-;- | 1067 | zgc:112496 | 136 | 295 | 356 | 260 | 245 | 166 |
| ENSDARG000 | 24;24;24;24;2 | 12058002;120 | 12058031;120 | +;+;+;+;+;+ | 890 | NA | 0 | 0 | 0 | 0 | 0 | 0 |
| ENSDARG000 | 24;24;24;24;2 | 5799764;5800 | 5800661;5800 | +;+;+;+;+;+;+; | 2851 | NA | 77 | 32 | 20 | 28 | 223 | 94 |
| ENSDARG000 | 24;24;24;24;2 | 39509721;395 | 39511741;395 | -;-;-;-;-;-;-;-; | 2418 | lyrm1 | 49 | 61 | 67 | 41 | 45 | 23 |
| ENSDARG000 | 24;24;24;24;2 | 39443874;394 | 39444066;394 | +;+;+;+;+;+;+; | 12391 | DNAH3 | 3 | 36 | 6 | 32 | 10 | 2 |
| ENSDARG000 | 24;24;24;24;2 | 31688977;316 | 31689221;316 | +;+;+;+;+ | 1189 | NA | 0 | 0 | 0 | 0 | 0 | 6 |
| ENSDARG000 | 24;24;24;24;2 | 21959708;219 | 21961058;219 | -;-;-;-;-;-;-;-; | 2673 | acot9.1 | 833 | 588 | 427 | 613 | 598 | 483 |
| ENSDARG000 | 24;24;24;24;2 | 15642577;156 | 15646345;156 | -;-;-;-;-;-;- | 5116 | cbln2b | 0 | 0 | 4 | 0 | 0 | 0 |
| ENSDARG000 | 24;24;24;24;2 | 12074340;120 | 12074397;120 | +;+;+;+;+;+;+; | 3278 | CCR9 | 18 | 0 | 18 | 4 | 24 | 11 |
| ENSDARG000 | 24;24;24;24;2 | 20909922;209 | 20910482;209 | -;-;-;-;-;-;-;- | 1442 | NA | 15 | 40 | 35 | 58 | 12 | 37 |
| ENSDARG000 | 24;24;24;24;2 | 8904135;8904 | 8905073;8905 | +;+;+;+;+;+;+; | 1940 | gnal | 190 | 129 | 163 | 162 | 200 | 234 |
| ENSDARG000 | 24;24;24;24;2 | 8979452;8980 | 8980271;8980 | -;-;-;-;-;-;-;-; | 1971 | mppe1 | 116 | 149 | 152 | 122 | 121 | 194 |
| ENSDARG000 | 24;24;24;24;2 | 31832458;318 | 31834827;318 | -;-;-;-;-;-;-;-;- | 3752 | STEAP2 | 16 | 25 | 33 | 58 | 12 | 13 |
| ENSDARG000 | 24;24;24;24;2 | 31842621;318 | 31843115;318 | +;+;+;+;+;+;+; | 2687 | NA | 0 | 0 | 0 | 0 | 0 | 0 |
| ENSDARG000 | 24;24;24;24;2 | 5811350;5811 | 5811872;5811 | +;+;+;+;+;+;+; | 2612 | NA | 145 | 94 | 16 | 37 | 247 | 139 |
| ENSDARG000 | 24;24;24;24;2 | 33868109;338 | 33868321;338 | -;-;-;-;-;-;-;-; | 1603 | asic1c | 0 | 0 | 0 | 0 | 0 | 0 |
| ENSDARG000 | 24;24 | 28552567;285 | 28553333;285 | -;- | 901 | NA | 0 | 0 | 0 | 0 | 0 | 0 |
| ENSDARG000 | 24;24;24;24;2 | 28561038;285 | 28561131;285 | +;+;+;+;+;+;+; | 7818 | abca4a | 10 | 22 | 10 | 16 | 38 | 68 |
| ENSDARG000 | 24;24;24;24;2 | 21973929;219 | 21974156;219 | +;+;+;+;+;+ | 1340 | sat1b | 22 | 6 | 32 | 33 | 13 | 6 |
| ENSDARG000 | 24;24 | 17403470;174 | 17404605;174 | -;- | 1217 | NA | 8 | 28 | 37 | 46 | 26 | 32 |
| ENSDARG000 | 24;24;24;24;2 | 21978230;219 | 21979016;219 | -;-;-;-;-;-;-;-;- | 1464 | apoob | 165 | 142 | 177 | 205 | 174 | 147 |
| ENSDARG000 | 24;24;24;24;2 | 28644418;286 | 28645169;286 | -;-;-;-;-;-;-;-; | 1516 | NA | 0 | 0 | 5 | 0 | 0 | 0 |
| ENSDARG000 | 24;24;24;24;2 | 5826005;5829 | 5826242;5829 | +;+;+;+;+;+;+; | 1887 | NA | 9 | 0 | 10 | 1 | 22 | 12 |
| ENSDARG000 | 24;24;24;24;2 | 15655069;156 | 15655762;156 | +;+;+;+;+;+;+; | 2136 | NA | 7 | 16 | 36 | 28 | 14 | 18 |
| ENSDARG000 | 24;24;24;24;2 | 28448387;284 | 28448691;284 | +;+;+;+;+;+;+; | 5287 | NA | 52 | 24 | 54 | 6 | 104 | 84 |
| ENSDARG000 | 24;24 | 12128505;121 | 12128985;121 | -;- | 585 | NA | 0 | 0 | 2 | 0 | 0 | 0 |
| ENSDARG000 | 24;24 | 12129726;121 | 12129762;121 | +;+ | 312 | NA | 0 | 4 | 0 | 0 | 0 | 0 |
| ENSDARG000 | 24;24;24;24;2 | 5840258;5840 | 5841199;5841 | +;+;+;+;+;+;+; | 3396 | trpc1 | 0 | 3 | 0 | 0 | 14 | 0 |
| ENSDARG000 | 24;24;24;24;2 | 20927135;209 | 20927215;209 | +;+;+;+;+;+;+; | 870 | fam162a | 1515 | 1505 | 1124 | 1637 | 1291 | 995 |
| ENSDARG000 | 24;24;24;24;2 | 17409681;174 | 17410541;174 | -;-;-;-;- | 1129 | NA | 10 | 26 | 30 | 39 | 0 | 18 |
| ENSDARG000 | 24;24;24;24;2 | 26328787;263 | 26328875;263 | +;+;+;+;+;+;+; | 3044 | mynn | 286 | 492 | 586 | 162 | 468 | 352 |
| ENSDARG000 | 24;24 | 34069675;340 | 34069862;340 | +;+ | 587 | H2BK1 | 0 | 0 | 0 | 0 | 0 | 0 |
| ENSDARG000 | 24;24;24;24;2 | 37661140;376 | 37662074;376 | -;-;-;-;-;-;-;-; | 2891 | anks3 | 154 | 97 | 165 | 114 | 219 | 190 |
| ENSDARG000 | 24;24;24;24;2 | 34085940;340 | 34086351;340 | +;+;+;+;+;+;+; | 1937 | asb10 | 8 | 2 | 0 | 0 | 22 | 20 |
| ENSDARG000 | 24;24;24;24;2 | 15897717;158 | 15898651;159 | +;+;+;+;+;+;+; | 4029 | NETO1 | 0 | 20 | 28 | 12 | 34 | 0 |
| ENSDARG000 | 24;24 | 16074089;160 | 16074251;160 | -;- | 739 | NA | 0 | 0 | 0 | 0 | 0 | 0 |
| ENSDARG000 | 24;24;24;24;2 | 20941360;209 | 20942986;209 | -;-;-;-;-;-;-;-; | 3208 | KPNA1 | 1842 | 2376 | 1506 | 1646 | 2327 | 1288 |
| ENSDARG000 | 24;24;24 | 20920563;209 | 20920751;209 | +;+;+ | 721 | cst14a.2 | 2523 | 2518 | 1859 | 2094 | 1934 | 1687 |
| ENSDARG000 | 24;24;24;24;2 | 1011158;1015 | 1015519;1015 | -;-;-;-;-;-;-;-; | 5261 | ralaa | 263 | 751 | 464 | 288 | 483 | 389 |
| ENSDARG000 | 24;24;24;24;2 | 20922544;209 | 20922806;209 | -;-;-;-;- | 673 | mix23 | 90 | 152 | 147 | 88 | 73 | 98 |
| ENSDARG000 | 24;24;24;24;2 | 20916751;209 | 20916916;209 | +;+;+;+;+;+;+ | 1010 | cst14a.1 | 64 | 77 | 93 | 71 | 100 | 89 |
| ENSDARG000 | 24;24 | 34113424;341 | 34114005;341 | +;+ | 1824 | gbx1 | 0 | 0 | 0 | 0 | 2 | 6 |
| ENSDARG000 | 24;24 | 34112756;341 | 34113020;341 | -;- | 511 | NA | 0 | 0 | 0 | 0 | 0 | 0 |
| ENSDARG000 | 24;24;24;24 | 1042594;1054 | 1042735;1055 | +;+;+;+ | 1666 | NA | 0 | 0 | 8 | 0 | 2 | 0 |
| ENSDARG000 | 24;24;24;24;2 | 1033508;1104 | 1033564;1104 | -;-;-;-;-;-;-;- | 1053 | NA | 0 | 0 | 0 | 0 | 4 | 0 |
| ENSDARG000 | 24;24;24;24;2 | 1023839;1024 | 1024039;1024 | +;+;+;+;+;+;+; | 2076 | zgc:111976 | 2 | 8 | 1 | 0 | 4 | 8 |
| ENSDARG000 | 24;24;24;24;2 | 26345427;263 | 26345536;263 | +;+;+;+;+;+;+; | 3639 | NA | 10 | 6 | 16 | 0 | 10 | 4 |
| ENSDARG000 | 24;24;24;24;2 | 1106270;1106 | 1106453;1106 | +;+;+;+;+ | 1286 | NA | 0 | 0 | 0 | 0 | 0 | 0 |
| ENSDARG000 | 24;24;24;24;2 | 34488316;344 | 34489353;344 | -;-;-;-;-;-;-;-; | 3734 | ctnna1 | 4740 | 4081 | 4570 | 2836 | 4342 | 3004 |
| ENSDARG000 | 24;24;24;24;2 | 31896724;318 | 31897276;318 | -;-;-;-;-;-;-;-; | 1212 | dtx3la | 14 | 0 | 0 | 0 | 0 | 0 |
| ENSDARG000 | 24;24;24;24;2 | 12133814;121 | 12134075;121 | +;+;+;+;+;+;+; | 1908 | lztfl1 | 444 | 410 | 526 | 368 | 308 | 335 |
| ENSDARG000 | 24;24 | 34606966;346 | 34607604;346 | +;+ | 5226 | LRRTM2 | 2 | 0 | 16 | 0 | 4 | 22 |
| ENSDARG000 | 24;24 | 12155107;121 | 12155729;121 | -;- | 912 | NA | 0 | 0 | 0 | 0 | 0 | 0 |
| ENSDARG000 | 24;24;24;24;2 | 16140423;161 | 16140790;161 | +;+;+;+;+;+ | 1127 | timm21 | 148 | 130 | 257 | 110 | 134 | 206 |
| ENSDARG000 | 24;24;24;24;2 | 17417150;174 | 17421921;174 | -;-;-;-;-;-;-;-; | 8686 | cntnap2a | 214 | 336 | 256 | 133 | 893 | 40 |
| ENSDARG000 | 24;24;24;24;2 | 26354768;263 | 26355281;263 | -;-;-;-;-;-;-;-; | 1982 | NA | 8 | 4 | 20 | 2 | 35 | 58 |
| ENSDARG000 | 24;24;24;24;2 | 24285751;242 | 24286114;242 | +;+;+;+;+ | 3099 | ythdf3 | 534 | 417 | 629 | 378 | 782 | 414 |
| ENSDARG000 | 24 | 21990964 | 21991770 | - | 807 | si:rp71-1f1.4 | 11 | 26 | 10 | 15 | 14 | 18 |
| ENSDARG000 | 24 | 22002090 | 22003031 | - | 942 | NA | 2 | 31 | 10 | 12 | 0 | 1 |
| ENSDARG000 | 24;24;24;24;2 | 5889721;5910 | 5889878;5910 | -;-;-;-;-;-;-;-; | 1463 | NA | 8 | 0 | 0 | 0 | 0 | 3 |
| ENSDARG000 | 24;24;24;24;2 | 5871589;5872 | 5872182;5872 | -;-;-;-;-;-;-;-; | 2446 | pcolce2b | 366 | 597 | 249 | 285 | 621 | 161 |
| ENSDARG000 | 24;24;24;24;2 | 5893134;5894 | 5893572;5894 | +;+;+;+;+;+;+; | 3502 | mastl | 101 | 574 | 478 | 346 | 357 | 206 |
| ENSDARG000 | 24;24;24 | 9104613;9114 | 9104852;9114 | -;-;- | 679 | NA | 0 | 0 | 0 | 0 | 0 | 0 |
| ENSDARG000 | 24;24;24;24;2 | 37681082;376 | 37681609;376 | -;-;-;-;- | 1370 | zgc:112185 | 135 | 98 | 128 | 96 | 106 | 96 |
| ENSDARG000 | 24;24;24;24;2 | 9162512;9162 | 9163044;9163 | +;+;+;+;+;+;+; | 3105 | DLGAP1 | 21 | 25 | 61 | 29 | 15 | 5 |
| ENSDARG000 | 24;24;24 | 34806822;348 | 34807222;348 | +;+;+ | 1042 | NA | 1829 | 2229 | 2091 | 1943 | 2034 | 2319 |
| ENSDARG000 | 24 | 22014810 | 22015751 | - | 942 | NA | 5 | 26 | 2 | 5 | 6 | 0 |
| ENSDARG000 | 24;24;24;24;2 | 16149251;161 | 16150255;161 | +;+;+;+;+;+;+; | 4541 | NA | 1318 | 1310 | 1775 | 1341 | 1658 | 1905 |
| ENSDARG000 | 24;24;24;24;2 | 1110138;1112 | 1113351;1113 | -;-;-;-;-;-;-;-; | 6015 | itgb1a | 4199 | 3219 | 4813 | 3724 | 6193 | 1923 |
| ENSDARG000 | 24;24;24;24;2 | 22021675;220 | 22021830;220 | +;+;+;+;+;+;+; | 1192 | ropn1l | 10 | 20 | 12 | 29 | 34 | 17 |
| ENSDARG000 | 24;24;24;24;2 | 12629649;126 | 12630369;126 | -;-;-;-;-;- | 1278 | pdcd6 | 1358 | 1234 | 1102 | 1219 | 1084 | 1279 |
| ENSDARG000 | 24;24;24;24 | 37688729;376 | 37688817;376 | +;+;+;+ | 925 | h3f3d | 2781 | 1821 | 3858 | 2426 | 2362 | 1723 |
| ENSDARG000 | 24;24;24;24;2 | 24416875;244 | 24417498;244 | -;-;-;-;-;- | 1120 | NA | 0 | 0 | 0 | 0 | 0 | 0 |
| ENSDARG000 | 24;24 | 24461341;244 | 24463346;244 | +;+ | 2006 | bhlhe22 | 0 | 0 | 0 | 0 | 0 | 0 |

| | | | | | | | | | | | | |
|---|---|---|---|---|---|---|---|---|---|---|---|---|
| ENSDARG000 | 24;24;24;24;2 | 22039964:220 | 22040383:220 | +;+;+;+;+;+;+ | 1994 | ANKRD33B | 57 | 84 | 60 | 33 | 68 | 42 |
| ENSDARG000 | 24;24;24 | 35387517:353 | 35387727:353 | +;+;+ | 2992 | snai2 | 2609 | 1411 | 1159 | 2177 | 495 | 3063 |
| ENSDARG000 | 24;24;24;24;2 | 35679797:356 | 35679835:356 | -;-;-;-;-;-;-;-; | 1510 | mapre2 | 1434 | 1297 | 1182 | 1091 | 1221 | 1240 |
| ENSDARG000 | 24;24;24;24;2 | 26395205:263 | 26395486:263 | -;-;-;-;- | 1008 | zgc:194621 | 0 | 0 | 0 | 0 | 0 | 0 |
| ENSDARG000 | 24;24;24;24;2 | 26379441:264 | 26379538:264 | +;+;+;+;+;+;+; | 5378 | NA | 488 | 540 | 689 | 413 | 572 | 587 |
| ENSDARG000 | 24;24 | 9269949:928 | 9270102:928 | -;- | 386 | NA | 1 | 1 | 6 | 2 | 6 | 2 |
| ENSDARG000 | 24;24;24;24;2 | 12690163:126 | 12690374:126 | +;+;+;+;+;+;+; | 2314 | rec8b | 2 | 0 | 0 | 0 | 0 | 0 |
| ENSDARG000 | 24;24;24;24;2 | 18444189:184 | 18444350:184 | -;-;-;-;-;-;-;-; | 1170 | sec61g | 2 | 4 | 262 | 265 | 124 | 242 |
| ENSDARG000 | 24;24;24;24;2 | 26373341:263 | 26373614:263 | -;-;-;-;-;-;-;-; | 2193 | skilb | 393 | 307 | 508 | 356 | 447 | 319 |
| ENSDARG000 | 24;24;24;24;2 | 18286427:182 | 18286461:182 | +;+;+;+;+;+;+; | 2041 | tpk1 | 829 | 712 | 414 | 835 | 1037 | 812 |
| ENSDARG000 | 24;24 | 37695768:376 | 37696574:376 | -;- | 1194 | NA | 38 | 76 | 90 | 92 | 108 | 66 |
| ENSDARG000 | 24;24;24;24;2 | 24701780:247 | 24702710:247 | -;-;-;-;-;-;-;-; | 2082 | armc1 | 701 | 701 | 769 | 436 | 828 | 511 |
| ENSDARG000 | 24;24;24;24;2 | 22070290:220 | 22071797:220 | -;-;-;-;-;-;-;-; | 2290 | dap | 3239 | 3018 | 2615 | 3509 | 2772 | 3020 |
| ENSDARG000 | 24;24;24;24;2 | 16174199:161 | 16174327:161 | +;+;+;+;+;+;+; | 1252 | NA | 52 | 0 | 0 | 8 | 9 | 5 |
| ENSDARG000 | 24;24;24;24;2 | 37709191:377 | 37709314:377 | +;+;+;+;+;+;+; | 2307 | decr2 | 240 | 151 | 283 | 184 | 241 | 226 |
| ENSDARG000 | 24;24;24;24;2 | 9287409:9287 | 9289026:9289 | -;-;-;-;-;-;- | 3683 | tgif1 | 1159 | 773 | 1171 | 908 | 1141 | 1127 |
| ENSDARG000 | 24;24;24;24 | 39518774:395 | 39519147:395 | +;+;+;+ | 2904 | dcun1d3 | 311 | 237 | 278 | 305 | 332 | 304 |
| ENSDARG000 | 24;24;24;24;2 | 39537565:395 | 39537626:395 | +;+;+;+;+;+;+; | 2005 | NA | 108 | 43 | 66 | 99 | 88 | 76 |
| ENSDARG000 | 24;24;24;24;2 | 24726956:247 | 24727050:247 | +;+;+;+;+;+;+; | 2944 | mtfr1 | 185 | 187 | 247 | 169 | 200 | 204 |
| ENSDARG000 | 24;24;24;24;2 | 1162325:1162 | 1163353:1163 | -;-;-;-;-;-;-;-; | 4950 | nrp1a | 1108 | 675 | 2431 | 2329 | 866 | 4272 |
| ENSDARG000 | 24;24;24 | 39576071:395 | 39576564:395 | +;+;+ | 1517 | NA | 145 | 62 | 88 | 118 | 107 | 101 |
| ENSDARG000 | 24;24;24;24;2 | 5915895:5915 | 5917537:5917 | -;-;-;-;-;-;-;-; | 3480 | acbd5a | 122 | 144 | 258 | 232 | 190 | 227 |
| ENSDARG000 | 24;24 | 35746156:357 | 35746409:357 | +;+ | 354 | NA | 0 | 0 | 0 | 0 | 0 | 0 |
| ENSDARG000 | 24;24;24;24;2 | 35713659:357 | 35713908:357 | -;-;-;-;-;-;-;-; | 2415 | dtna | 0 | 0 | 3 | 0 | 0 | 0 |
| ENSDARG000 | 24;24;24;24;2 | 5912635:5912 | 5913214:5913 | +;+;+;+;+;+;+; | 1621 | NA | 0 | 0 | 2 | 0 | 2 | 2 |
| ENSDARG000 | 24;24;24;24;2 | 39586701:395 | 39586829:395 | +;+;+;+;+;+;+; | 1062 | NA | 95 | 64 | 99 | 109 | 87 | 139 |
| ENSDARG000 | 24;24;24;24 | 39606053:396 | 39606300:396 | -;-;-;- | 633 | cox6b1 | 2530 | 2983 | 3692 | 4297 | 1909 | 2652 |
| ENSDARG000 | 24;24;24;24;2 | 16207562:162 | 16207630:162 | +;+;+;+;+;+;+ | 394 | NA | 0 | 0 | 0 | 0 | 0 | 0 |
| ENSDARG000 | 24;24;24;24;2 | 18448108:184 | 18448456:184 | -;-;-;-;-;-;-;-; | 1958 | zgc:112332 | 44 | 26 | 50 | 237 | 10 | 74 |
| ENSDARG000 | 24;24;24;24;2 | 39620461:396 | 39620719:396 | -;-;-;-;-;-;-;-; | 1923 | NA | 302 | 53 | 177 | 117 | 286 | 284 |
| ENSDARG000 | 24;24;24;24;2 | 39625164:396 | 39625604:396 | +;+;+;+;+ | 1922 | NA | 0 | 0 | 0 | 0 | 0 | 1 |
| ENSDARG000 | 24 | 9604442 | 9605320 | + | 879 | NA | 24 | 13 | 28 | 4 | 7 | 9 |
| ENSDARG000 | 24;24;24;24;2 | 9372292:9372 | 9372589:9372 | +;+;+;+;+;+;+; | 8341 | NA | 451 | 87 | 586 | 514 | 600 | 500 |
| ENSDARG000 | 24;24;24;24;2 | 22485710:224 | 22485738:224 | +;+;+;+;+;+;+; | 1460 | NA | 113 | 315 | 71 | 304 | 134 | 25 |
| ENSDARG000 | 24;24;24;24;2 | 22118804:221 | 22124049:221 | -;-;-;-;-;-;-;-; | 9362 | NA | 216 | 100 | 241 | 352 | 220 | 66 |
| ENSDARG000 | 24;24;24 | 1294176:1295 | 1294307:1295 | +;+;+ | 337 | NA | 0 | 0 | 0 | 0 | 0 | 0 |
| ENSDARG000 | 24;24;24;24;2 | 9612218:9612 | 9612660:9612 | +;+;+;+;+;+;+; | 4790 | nphp3 | 357 | 225 | 279 | 256 | 372 | 267 |
| ENSDARG000 | 24;24;24 | 12747377:127 | 12748644:127 | -;-;- | 1568 | cox19 | 128 | 151 | 182 | 164 | 155 | 232 |
| ENSDARG000 | 24;24;24;24;2 | 24733307:247 | 24735011:247 | -;-;-;-;-;-;-;-; | 3958 | pde7a | 669 | 386 | 630 | 197 | 479 | 495 |
| ENSDARG000 | 24;24;24 | 12741936:127 | 12742448:127 | -;-;- | 2832 | NA | 123 | 36 | 87 | 117 | 144 | 95 |
| ENSDARG000 | 24;24;24 | 12740848:127 | 12741218:127 | +;+;+ | 714 | si:ch211-196f | 19 | 10 | 24 | 16 | 6 | 28 |
| ENSDARG000 | 24;24 | 22690983:227 | 22691394:227 | -;- | 547 | NA | 0 | 4 | 20 | 0 | 7 | 1 |
| ENSDARG000 | 24;24;24;24;2 | 9673227:9673 | 9673717:9673 | -;-;-;-;-;-;-;-; | 1835 | uba5 | 440 | 498 | 409 | 269 | 455 | 239 |
| ENSDARG000 | 24;24;24;24;2 | 9566799:9568 | 9566979:9568 | -;-;-;-;-;- | 609 | NA | 0 | 0 | 0 | 0 | 2 | 0 |
| ENSDARG000 | 24;24;24;24 | 9590188:9591 | 9590223:9591 | +;+;+;+ | 1949 | NA | 0 | 0 | 0 | 0 | 0 | 0 |
| ENSDARG000 | 24;24;24;24;2 | 37722048:377 | 37727432:377 | +;+;+;+;+;+;+; | 8313 | rab11fip3 | 121 | 50 | 82 | 44 | 56 | 44 |
| ENSDARG000 | 24;24;24;24 | 20960216:209 | 20960417:209 | +;+;+;+ | 1047 | NA | 0 | 2 | 4 | 2 | 14 | 0 |
| ENSDARG000 | 24;24;24;24;2 | 16393302:163 | 16393904:163 | +;+;+;+;+;+;+; | 5780 | sema5a | 6 | 6 | 8 | 20 | 6 | 0 |
| ENSDARG000 | 24;24;24;24;2 | 12754013:127 | 12754500:127 | -;-;-;-;-;-;-;-; | 4176 | NA | 1437 | 1523 | 2107 | 589 | 1837 | 892 |
| ENSDARG000 | 24;24;24;24;2 | 20968308:209 | 20968600:209 | +;+;+;+;+;+;+; | 2307 | drd3 | 2 | 0 | 2 | 0 | 0 | 4 |
| ENSDARG000 | 24;24;24;24;2 | 26432541:264 | 26432904:264 | +;+;+;+;+;+;+ | 2843 | cldn11b | 146 | 297 | 311 | 166 | 211 | 262 |
| ENSDARG000 | 24;24;24;24;2 | 5935377:5935 | 5935634:5935 | +;+;+;+;+;+;+; | 3788 | abi1a | 723 | 1376 | 1642 | 1699 | 1167 | 1113 |
| ENSDARG000 | 24;24;24;24;2 | 39638555:396 | 39638752:396 | +;+;+;+;+;+;+; | 2947 | luc7l | 1743 | 1421 | 2189 | 1499 | 2086 | 2018 |
| ENSDARG000 | 24;24;24;24;2 | 6015299:6016 | 6015784:6016 | -;-;-;-;-;-;-;-; | 1625 | pdss1 | 100 | 245 | 439 | 113 | 345 | 246 |
| ENSDARG000 | 24;24;24;24;2 | 26446321:264 | 26446631:264 | -;-;-;-;-;-;- | 2271 | NA | 0 | 0 | 0 | 0 | 4 | 1 |
| ENSDARG000 | 24;24;24;24;2 | 12920019:129 | 12921541:129 | -;-;-;-;-;-;-;-; | 3414 | pck2 | 508 | 798 | 1607 | 903 | 750 | 852 |
| ENSDARG000 | 24;24;24;24;2 | 37799818:378 | 37800460:378 | +;+;+;+;+;+;+; | 2981 | telo2 | 324 | 355 | 436 | 176 | 363 | 346 |
| ENSDARG000 | 24;24;24;24;2 | 12894282:128 | 12894426:128 | +;+;+;+;+;+;+; | 2271 | NA | 19 | 30 | 36 | 38 | 161 | 56 |
| ENSDARG000 | 24;24;24;24;2 | 24808955:248 | 24808986:248 | +;+;+;+;+;+;+; | 833 | DNAJC5B | 0 | 0 | 0 | 0 | 0 | 0 |
| ENSDARG000 | 24;24;24;24;2 | 21058779:210 | 21060548:210 | -;-;-;-;-;-;-;-; | 3577 | qtrt2 | 102 | 108 | 196 | 62 | 148 | 126 |
| ENSDARG000 | 24;24;24;24;2 | 26475182:264 | 26475440:264 | -;-;-;-;-;-;-;-; | 787 | rpl22l1 | 2087 | 2135 | 3457 | 2288 | 2676 | 4892 |
| ENSDARG000 | 24;24;24;24;2 | 35787629:357 | 35787768:357 | +;+;+;+;+;+;+; | 2069 | NA | 0 | 0 | 0 | 0 | 6 | 0 |
| ENSDARG000 | 24;24;24;24;2 | 16912248:169 | 16913181:169 | -;-;-;-;-;- | 1756 | cmbl | 117 | 127 | 132 | 241 | 129 | 413 |
| ENSDARG000 | 24;24;24;24;2 | 24831727:248 | 24832060:248 | +;+;+;+;+;+;+; | 1674 | trim55b | 0 | 0 | 0 | 0 | 0 | 0 |
| ENSDARG000 | 24;24;24;24;2 | 16905188:169 | 16905436:169 | +;+;+;+;+;+;+; | 1906 | cct5 | 5482 | 6422 | 8896 | 5847 | 6439 | 6344 |
| ENSDARG000 | 24;24;24;24;2 | 16851213:168 | 16856150:168 | -;-;-;-;-;-;-;-; | 8128 | MTRR | 325 | 312 | 443 | 241 | 437 | 286 |
| ENSDARG000 | 24;24;24 | 35827766:358 | 35827825:358 | +;+;+ | 807 | NA | 0 | 0 | 0 | 0 | 0 | 0 |
| ENSDARG000 | 24;24;24;24;2 | 18469766:184 | 18470907:184 | -;-;-;-;-;-;-;-; | 2057 | si:dkev-73n8.: | 20 | 22 | 1871 | 165 | 677 | 922 |
| ENSDARG000 | 24;24;24;24;2 | 21092156:210 | 21092363:210 | +;+;+;+;+;+;+; | 3418 | ccdc191 | 44 | 22 | 30 | 30 | 34 | 6 |
| ENSDARG000 | 24;24;24;24;2 | 35911020:359 | 35911331:359 | +;+;+;+;+;+;+; | 2637 | abcd4 | 352 | 230 | 420 | 249 | 340 | 461 |
| ENSDARG000 | 24;24;24;24;2 | 18485866:184 | 18487880:184 | -;-;-;-;-;-;-;-; | 3624 | NA | 158 | 89 | 215 | 120 | 306 | 145 |
| ENSDARG000 | 24;24;24;24 | 16726748:168 | 16726922:168 | +;+;+;+ | 671 | NA | 0 | 0 | 0 | 0 | 0 | 0 |
| ENSDARG000 | 24;24;24;24;2 | 24847683:248 | 24848633:248 | -;-;-;-;-;-;- | 1217 | crhb | 0 | 0 | 0 | 0 | 0 | 0 |
| ENSDARG000 | 24;24;24;24;2 | 35881124:358 | 35881304:358 | +;+;+;+;+;+;+; | 2028 | KLHL14 | 0 | 0 | 0 | 0 | 2 | 0 |
| ENSDARG000 | 24;24 | 35880330:359 | 35880723:359 | -;- | 431 | NA | 0 | 0 | 0 | 0 | 0 | 0 |
| ENSDARG000 | 24;24;24;24;2 | 21122836:211 | 21123222:211 | -;-;-;-;- | 1697 | zdhhc23b | 184 | 108 | 147 | 34 | 150 | 182 |
| ENSDARG000 | 24;24;24;24;2 | 24923166:249 | 24923307:249 | +;+;+;+;+;+;+; | 1262 | pcyt1ba | 14 | 0 | 41 | 10 | 26 | 2 |
| ENSDARG000 | 24;24;24;24;2 | 1370627:1371 | 1370698:1371 | +;+;+;+;+;+;+; | 1492 | NA | 59 | 4 | 9 | 76 | 33 | 30 |
| ENSDARG000 | 24;24;24;24;2 | 28651094:286 | 28651191:286 | +;+;+;+;+;+;+; | 1175 | NA | 2 | 0 | 1 | 0 | 0 | 0 |
| ENSDARG000 | 24;24;24;24;2 | 22731228:227 | 22731441:227 | +;+;+;+;+ | 1239 | NA | 73 | 151 | 401 | 95 | 392 | 393 |
| ENSDARG000 | 24;24;24;24;2 | 28764021:287 | 28765319:287 | -;-;-;-;-;-;-;-; | 6999 | col11a1a | 1403 | 35 | 121 | 20 | 134 | 40 |
| ENSDARG000 | 24;24;24;24;2 | 6025538:6026 | 6026218:6026 | -;-;-;-;- | 1733 | NA | 0 | 0 | 0 | 0 | 0 | 0 |
| ENSDARG000 | 24;24;24;24;2 | 26481300:264 | 26482775:264 | -;-;-;-;-;-;-;-; | 2301 | eif5a | 16584 | 17673 | 23807 | 18913 | 18956 | 25379 |
| ENSDARG000 | 24;24;24;24;2 | 24944352:249 | 24944633:249 | -;-;-;-;-;-;-;-; | 1607 | PDK3 | 260 | 160 | 514 | 322 | 262 | 178 |
| ENSDARG000 | 24;24;24;24;2 | 6040029:6044 | 6041867:6044 | -;-;-;-;-;-;-;-; | 3287 | apbb1ip | 68 | 44 | 96 | 38 | 218 | 186 |
| ENSDARG000 | 24;24;24;24;2 | 21131874:211 | 21134005:211 | -;-;-;-;-;-;-;-; | 4666 | GRAMD1C | 42 | 6 | 16 | 4 | 10 | 72 |
| ENSDARG000 | 24;24;24;24;2 | 6034990:6035 | 6035513:6035 | -;-;-;-;-;-;-;-; | 1692 | NA | 0 | 0 | 0 | 0 | 0 | 0 |
| ENSDARG000 | 24;24;24;24;2 | 22740543:227 | 22741567:227 | -;-;-;-;-;-;-;-; | 2009 | zc2hc1a | 709 | 696 | 916 | 783 | 614 | 994 |
| ENSDARG000 | 24;24;24;24;2 | 12945803:129 | 12946057:129 | +;+;+;+;+;+;+; | 1173 | psme1 | 2289 | 2208 | 2421 | 1109 | 1909 | 1766 |
| ENSDARG000 | 24;24;24;24;2 | 28686287:286 | 28688208:286 | -;-;-;-;-;- | 2910 | olfm3a | 0 | 0 | 0 | 0 | 0 | 0 |
| ENSDARG000 | 24;24;24;24;2 | 1431732:1431 | 1432211:1432 | +;+;+;+;+;+;+; | 2257 | NA | 0 | 2 | 2 | 6 | 10 | 0 |
| ENSDARG000 | 24;24;24;24;2 | 39656834:396 | 39657060:396 | +;+;+;+;+;+;+; | 1519 | stub1 | 494 | 426 | 480 | 296 | 492 | 489 |
| ENSDARG000 | 24;24;24;24;2 | 24989145:249 | 24989446:249 | -;-;-;-;-;-;-;-; | 1536 | zdhhc20b | 225 | 179 | 194 | 81 | 162 | 144 |
| ENSDARG000 | 24;24;24;24;2 | 6094164:6095 | 6094257:6095 | -;-;-;-;- | 917 | NA | 14 | 0 | 0 | 0 | 20 | 26 |
| ENSDARG000 | 24;24 | 24983135:249 | 24983244:249 | +;+ | 751 | NA | 0 | 0 | 1 | 0 | 3 | 0 |
| ENSDARG000 | 24;24;24;24;2 | 24961899:249 | 24962545:249 | -;-;-;-;-;-;-;-; | 1838 | slc51a | 28 | 8 | 0 | 2 | 4 | 0 |
| ENSDARG000 | 24;24;24;24;2 | 18520402:185 | 18521705:185 | -;-;-;-;-;-;-;-; | 5037 | relch | 939 | 839 | 896 | 970 | 1029 | 1138 |
| ENSDARG000 | 24;24;24;24;2 | 37825634:378 | 37825774:378 | +;+;+;+;+;+;+; | 4887 | IFT140 | 256 | 307 | 260 | 134 | 236 | 194 |
| ENSDARG000 | 24;24;24;24;2 | 37854002:378 | 37854616:378 | -;-;-;-;-;-;- | 2078 | tmem204 | 76 | 22 | 20 | 14 | 46 | 24 |
| ENSDARG000 | 24;24;24;24;2 | 6107901:6108 | 6108213:6108 | +;+;+;+;+ | 3525 | NA | 14 | 0 | 4 | 14 | 2 | 0 |
| ENSDARG000 | 24;24;24;24;2 | 32176155:321 | 32176798:321 | +;+;+;+;+;+;+; | 1861 | vim | 55 | 32 | 71 | 10 | 84 | 54 |
| ENSDARG000 | 24;24;24;24;2 | 35933869:359 | 35934084:359 | +;+;+;+;+;+;+; | 13260 | NA | 13 | 34 | 74 | 6 | 175 | 26 |
| ENSDARG000 | 24;24;24;24;2 | 32151629:321 | 32151831:321 | -;-;-;-;-;-;-;-; | 2223 | trdmt1 | 65 | 65 | 164 | 94 | 118 | 116 |
| ENSDARG000 | 24;24;24;24;2 | 21154120:211 | 21154760:211 | -;-;-;-;-;-;-;-; | 2555 | ATP6V1A | 5187 | 5650 | 5456 | 3835 | 5601 | 4694 |
| ENSDARG000 | 24;24;24;24;2 | 39740382:397 | 39740611:397 | -;-;-;-;-;- | 1657 | GFOD1 | 119 | 190 | 152 | 205 | 193 | 165 |
| ENSDARG000 | 24;24;24 | 1627457:1639 | 1628091:1640 | -;-;- | 916 | NA | 6 | 18 | 26 | 6 | 0 | 8 |
| ENSDARG000 | 24;24;24;24;2 | 36158745:361 | 36158890:361 | -;-;-;-;-;-;-;-; | 1326 | pak1ip1 | 409 | 682 | 1040 | 764 | 594 | 619 |
| ENSDARG000 | 24;24;24 | 22759451:227 | 22759775:227 | +;+;+ | 1479 | si:dkey-7n6.2 | 134 | 92 | 330 | 16 | 306 | 130 |
| ENSDARG000 | 24;24;24;24;2 | 6133147:6142 | 6137663:6142 | -;-;-;-;-;-;-;-; | 6525 | gad2 | 49 | 4 | 16 | 8 | 39 | 54 |
| ENSDARG000 | 24;24 | 23202545:232 | 23202707:232 | +;+ | 1297 | pex2 | 127 | 143 | 134 | 90 | 99 | 89 |
| ENSDARG000 | 24;24;24;24;2 | 32209778:322 | 32209862:322 | +;+;+;+;+;+ | 1046 | NA | 0 | 0 | 0 | 0 | 0 | 0 |
| ENSDARG000 | 24;24;24;24;2 | 37902687:379 | 37902924:379 | +;+;+;+;+;+;+; | 4541 | NA | 0 | 0 | 0 | 0 | 0 | 2 |
| ENSDARG000 | 24;24;24;24 | 32406894:324 | 32407026:324 | -;-;-;- | 353 | ppp1r17 | 0 | 0 | 0 | 0 | 0 | 0 |
| ENSDARG000 | 24;24;24;24;2 | 18689980:186 | 18690080:186 | +;+;+;+;+;+;+; | 4272 | CSPP1 | 48 | 72 | 83 | 63 | 68 | 39 |
| ENSDARG000 | 24;24;24;24;2 | 9693951:9696 | 9694215:9696 | +;+;+;+;+;+;+; | 7096 | topbp1 | 132 | 548 | 548 | 234 | 528 | 317 |
| ENSDARG000 | 24;24;24;24;2 | 39778648:397 | 39779656:397 | -;-;-;-;-;-;-;-; | 4818 | SLC12A7 | 1775 | 2337 | 2057 | 1021 | 2149 | 1476 |
| ENSDARG000 | 24;24;24;24;2 | 32414643:324 | 32414756:324 | -;-;-;-;-;-;-;-; | 2629 | NA | 77 | 6 | 4 | 0 | 19 | 10 |
| ENSDARG000 | 24;24;24;24;2 | 32411753:324 | 32411935:324 | +;+;+;+;+;+;+ | 3019 | neurod6a | 0 | 0 | 0 | 0 | 2 | 0 |
| ENSDARG000 | 24;24;24;24;2 | 23241444:232 | 23246667:232 | -;-;-;-;-;-;-;-; | 14894 | ZFHX4 | 1795 | 1686 | 7697 | 3941 | 9073 | 5306 |
| ENSDARG000 | 24;24;24;24;2 | 25008211:250 | 25008945:250 | -;-;-;-;-;-;-;-; | 3482 | nup58 | 476 | 467 | 960 | 428 | 617 | 500 |
| ENSDARG000 | 24;24;24;24;2 | 2318351:2318 | 2318449:2318 | +;+;+;+;+;+;+; | 4154 | NA | 207 | 361 | 652 | 320 | 577 | 473 |
| ENSDARG000 | 24;24;24;24;2 | 2283959:2284 | 2284679:2284 | -;-;-;-;-;-;-;-; | 3155 | CUL2 | 568 | 747 | 1120 | 548 | 993 | 706 |
| ENSDARG000 | 24;24;24;24;2 | 36177388:361 | 36178350:361 | -;-;-;-;-;-;-;-; | 3864 | NA | 360 | 400 | 612 | 264 | 483 | 390 |
| ENSDARG000 | 24;24;24;24;2 | 12989727:129 | 12989762:129 | +;+;+;+;+;+;+; | 3095 | ube2wb | 115 | 108 | 171 | 140 | 132 | 232 |
| ENSDARG000 | 24;24;24;24;2 | 12983434:129 | 12983512:129 | +;+;+;+;+;+;+; | 866 | eloca | 417 | 302 | 288 | 264 | 353 | 356 |
| ENSDARG000 | 24;24;24;24;2 | 6161037:6161 | 6163329:6163 | -;-;-;-;-;-;-;-; | 8031 | myo3a | 345 | 6 | 111 | 22 | 201 | 246 |
| ENSDARG000 | 24;24;24;24;2 | 12963119:129 | 12963430:129 | -;-;-;-;-;-;-;-; | 2117 | dcaf11 | 634 | 377 | 542 | 491 | 545 | 492 |
| ENSDARG000 | 24;24;24;24;2 | 9744012:9746 | 9744312:9746 | +;+;+;+;+;+;+; | 2378 | NA | 1375 | 1350 | 1132 | 399 | 597 | 351 |
| ENSDARG000 | 24;24 | 18876733:188 | 18877121:188 | +;+ | 1006 | NA | 1 | 0 | 0 | 0 | 0 | 1 |
| ENSDARG000 | 24;24;24;24;2 | 18736916:187 | 18738411:187 | -;-;-;-;-;-;-;-; | 7176 | ARFGEF1 | 1490 | 1581 | 1425 | 1581 | 2081 | 1345 |
| ENSDARG000 | 24;24;24;24;2 | 21174523:211 | 21174631:211 | +;+;+;+;+;+;+; | 2764 | naa50 | 966 | 2017 | 2657 | 1388 | 2207 | 2304 |
| ENSDARG000 | 24;24;24;24 | 37941618:379 | 37944026:379 | -;-;-;- | 3210 | metrn | 49 | 18 | 20 | 12 | 22 | 40 |
| ENSDARG000 | 24;24;24;24;2 | 32517915:325 | 32518303:325 | -;-;-;-;-;-;-;- | 896 | eif1b | 1810 | 1703 | 2066 | 1141 | 1570 | 1530 |
| ENSDARG000 | 24;24;24;24;2 | 9881219:9885 | 9881682:9885 | +;+;+;+;+ | 2171 | cdv3 | 2778 | 3432 | 3237 | 1138 | 2658 | 1819 |
| ENSDARG000 | 24;24;24;24;2 | 26491904:264 | 26494491:264 | -;-;-;-;-;-;-;-; | 6819 | NA | 1736 | 1673 | 2511 | 3265 | 2166 | 1351 |
| ENSDARG000 | 24;24;24;24;2 | 23646644:236 | 23648286:236 | -;-;-;-;-;-;-;-; | 3490 | hnf4g | 0 | 0 | 4 | 2 | 2 | 0 |
| ENSDARG000 | 24;24;24 | 37999090:380 | 37999803:380 | -;-;- | 2293 | NA | 0 | 0 | 0 | 0 | 0 | 0 |
| ENSDARG000 | 24;24 | 38073223:380 | 38073839:380 | -;- | 678 | NA | 38 | 0 | 6 | 0 | 4 | 2 |
| ENSDARG000 | 24;24;24;24 | 38082471:380 | 38083148:380 | -;-;-;- | 880 | crp2 | 0 | 0 | 2 | 0 | 0 | 0 |
| ENSDARG000 | 24;24 | 38085317:380 | 38085711:380 | -;- | 429 | NA | 0 | 0 | 0 | 0 | 0 | 0 |
| ENSDARG000 | 24;24;24;24;2 | 38092608:380 | 38093286:380 | -;-;-;-;-;-;- | 785 | crp3 | 0 | 0 | 24 | 0 | 2 | 0 |
| ENSDARG000 | 24;24;24 | 21187136:211 | 21187269:211 | +;+;+ | 748 | NA | 0 | 0 | 2 | 12 | 4 | 0 |
| ENSDARG000 | 24;24;24;24;2 | 36204028:362 | 36204462:362 | +;+;+;+;+;+;+; | 2532 | rbbp8 | 32 | 126 | 86 | 72 | 109 | 51 |
| ENSDARG000 | 24;24;24;24;2 | 21187027:211 | 21187269:211 | +;+;+;+;+;+;+; | 9400 | USF3 | 779 | 485 | 890 | 705 | 1027 | 845 |
| ENSDARG000 | 24;24;24;24;2 | 16962969:169 | 16967015:169 | -;-;-;-;-;-;-;-; | 5819 | klhl15 | 308 | 229 | 410 | 339 | 378 | 264 |
| ENSDARG000 | 24;24;24;24;2 | 6286599:6287 | 6286720:6288 | -;-;-;-;-;-;-;-; | 3441 | zgc:174877 | 65 | 119 | 136 | 105 | 152 | 89 |
| ENSDARG000 | 24;24;24;24;2 | 32525146:325 | 32525279:325 | +;+;+;+;+;+;+; | 3038 | YME1L1 | 767 | 1060 | 1826 | 1125 | 999 | 624 |
| ENSDARG000 | 24;24;24;24;2 | 23687410:236 | 23687967:236 | -;-;-;-;-;-;-;-; | 3477 | pign | 200 | 158 | 216 | 116 | 216 | 288 |
| ENSDARG000 | 24;24 | 39900674:399 | 39900730:399 | +;+ | 427 | NA | 0 | 0 | 0 | 0 | 0 | 0 |
| ENSDARG000 | 24;24;24;24;2 | 40001290:400 | 40001856:400 | -;-;-;-;- | 2639 | aoc1 | 0 | 0 | 6 | 4 | 8 | 0 |
| ENSDARG000 | 24;24;24;24;2 | 2346514:2346 | 2349693:2349 | -;-;-;-;-;-;-;-; | 8609 | RREB1 | 542 | 815 | 1390 | 1183 | 1498 | 772 |
| ENSDARG000 | 24;24;24;24;2 | 18880254:188 | 18884220:188 | -;-;-;-;-;-;-;-; | 5744 | CPA6 | 0 | 8 | 16 | 4 | 8 | 0 |
| ENSDARG000 | 24;24;24;24;2 | 6360423:6363 | 6363146:6363 | -;-;-;-;-;-;-;-; | 3639 | anxa13 | 1829 | 3422 | 5033 | 3302 | 3161 | 3341 |
| ENSDARG000 | 24;24;24;24;2 | 9895851:9896 | 9896309:9896 | -;-;-;-;-;-;-;-; | 3273 | vps41 | 1784 | 1309 | 2274 | 1503 | 3037 | 2106 |
| ENSDARG000 | 24;24;24;24;2 | 21206243:212 | 21207422:212 | -;-;-;-;-;-;-;-; | 4677 | boc | 315 | 182 | 337 | 392 | 258 | 176 |
| ENSDARG000 | 24;24;24;24;2 | 23716918:237 | 23717286:237 | +;+;+;+;+;+;+; | 1387 | cops5 | 896 | 746 | 736 | 442 | 804 | 560 |
| ENSDARG000 | 24;24;24;24;2 | 28934875:289 | 28935322:289 | +;+;+;+;+;+;+; | 6855 | rnpc3 | 296 | 271 | 524 | 426 | 396 | 380 |
| ENSDARG000 | 24;24;24;24;2 | 32567648:325 | 32567731:325 | +;+;+;+;+;+;+; | 1004 | NA | 11 | 0 | 0 | 0 | 0 | 0 |
| ENSDARG000 | 24;24;24;24;2 | 17005647:170 | 17005743:170 | +;+;+;+;+;+;+; | 5687 | NA | 1022 | 1089 | 1102 | 853 | 1156 | 910 |
| ENSDARG000 | 24;24;24;24;2 | 32592748:326 | 32592833:326 | +;+;+;+;+;+;+; | 1550 | NA | 431 | 210 | 316 | 243 | 285 | 340 |
| ENSDARG000 | 24;24;24;24;2 | 16983269:169 | 16983377:169 | +;+;+;+;+;+;+; | 2175 | eif2s3 | 5462 | 6395 | 10479 | 8238 | 5296 | 7395 |
| ENSDARG000 | 24;24;24;24;2 | 36227074:362 | 36227574:362 | -;-;-;-;-;-;-;-; | 1417 | TMEM241 | 69 | 42 | 74 | 37 | 62 | 28 |

| | | | | | | | | | | | | |
|---|---|---|---|---|---|---|---|---|---|---|---|---|
| ENSDARG000 | 24:24:24:24:2 | 6395385:6395 | 6397298:6397 | -;-;-;-;-;-;-;-;-; | 4095 | GPR158 | 0 | 17 | 0 | 0 | 6 | 2 |
| ENSDARG000 | 24:24:24:24:2 | 40016889:400 | 40016975:400 | +;+;+;+;+;+;+;+;+; | 14109 | NA | 17 | 0 | 0 | 0 | 21 | 18 |
| ENSDARG000 | 24:24:24:24:2 | 17021513:170 | 17022158:170 | -;-;-;-;-;- | 1237 | ptgdsb.2 | 0 | 0 | 0 | 0 | 0 | 0 |
| ENSDARG000 | 24:24:24:24:2 | 25032340:250 | 25032557:250 | +;+;+;+;+;+;+;+;+; | 2739 | mtmr6 | 902 | 641 | 820 | 750 | 882 | 1115 |
| ENSDARG000 | 24:24:24:24:2 | 2495197:2498 | 2495246:2498 | +;+;+;+;+;+;+;+;+; | 3541 | f13a1a.1 | 0 | 2 | 0 | 4 | 30 | 0 |
| ENSDARG000 | 24:24:24:24:2 | 9981933:9981 | 9982896:9984 | -;-;-;-;-;-;-;-;-; | 5933 | zgc:152652 | 0 | 0 | 0 | 0 | 0 | 0 |
| ENSDARG000 | 24:24:24:24:2 | 9976212:9976 | 9977854:9977 | -;-;-;-;-;-;-;-;-; | 3269 | zgc:171977 | 0 | 0 | 0 | 0 | 0 | 0 |
| ENSDARG000 | 24:24:24:24:2 | 23758510:237 | 23759819:237 | -;-;-;-;-;-;-;-;-; | 2936 | sgk3 | 1055 | 519 | 1046 | 653 | 787 | 817 |
| ENSDARG000 | 24:24:24:24:2 | 23730061:237 | 23730247:237 | +;+;+;+;+;+;+;+;+; | 1554 | ppp1r42 | 0 | 0 | 2 | 0 | 0 | 0 |
| ENSDARG000 | 24:24:24:24:2 | 6513843:6514 | 6515669:6515 | -;-;-;-;-;-;-;-;-; | 6892 | ARHGAP21 | 716 | 703 | 729 | 793 | 925 | 849 |
| ENSDARG000 | 24;24 | 32870701:330 | 32870963:330 | -;- | 574 | NA | 0 | 0 | 0 | 0 | 0 | 0 |
| ENSDARG000 | 24:24:24:24:2 | 33155509:332 | 33155742:332 | -;-;-;-;- | 1125 | NA | 0 | 0 | 0 | 0 | 0 | 0 |
| ENSDARG000 | 24:24:24:24:2 | 6673200:6673 | 6673509:6673 | -;-;-;-;- | 1286 | enkur | 9 | 13 | 33 | 22 | 8 | 38 |
| ENSDARG000 | 24:24:24:24:2 | 18948665:189 | 18948957:189 | +;+;+;+;+;+;+;+;+; | 6069 | PREX2 | 122 | 453 | 28 | 46 | 78 | 26 |
| ENSDARG000 | 24;24;24 | 2519761:2522 | 2520008:2522 | +;+;+ | 688 | nrn1a | 0 | 0 | 2 | 0 | 0 | 0 |
| ENSDARG000 | 24;24;24 | 9995077:9997 | 9997569:9997 | -;-;- | 2714 | NA | 0 | 0 | 0 | 0 | 0 | 0 |
| ENSDARG000 | 24:24:24:24:2 | 36239617:362 | 36239927:362 | +;+;+;+;+;+;+;+;+; | 1821 | riok3 | 1152 | 1196 | 2041 | 1817 | 861 | 1251 |
| ENSDARG000 | 24;24;24 | 10003426:100 | 10005895:100 | -;-;- | 2572 | zgc:171750 | 0 | 0 | 0 | 0 | 0 | 0 |
| ENSDARG000 | 24:24:24:24:2 | 10010630:100 | 10013003:100 | -;-;-;-;-;-;- | 3838 | NA | 0 | 0 | 0 | 0 | 0 | 0 |
| ENSDARG000 | 24:24:24:24:2 | 13017586:130 | 13017742:130 | +;+;+;+;+;+;+;+;+; | 3981 | stau2 | 5744 | 4881 | 6982 | 6032 | 7567 | 6102 |
| ENSDARG000 | 24;24;24 | 10018653:100 | 10021028:100 | -;-;- | 2537 | zgc:173856 | 0 | 0 | 0 | 0 | 0 | 0 |
| ENSDARG000 | 24 | 25069609 | 25073032 | + | 3424 | amer2 | 306 | 48 | 234 | 242 | 76 | 368 |
| ENSDARG000 | 24:24:24:24:2 | 33226331:332 | 33226485:332 | -;-;-;-;-;-;-;-;-; | 1665 | nrbp2b | 2 | 16 | 36 | 14 | 8 | 18 |
| ENSDARG000 | 24:24:24:24:2 | 2736815:2762 | 2736900:2762 | +;+;+;+;+;+;+ | 781 | LYRM4 | 22 | 10 | 60 | 10 | 34 | 12 |
| ENSDARG000 | 24:24:24:24:2 | 2557794:2637 | 2558378:2637 | -;-;-;-;-;-;- | 2075 | fars2 | 101 | 86 | 185 | 92 | 187 | 183 |
| ENSDARG000 | 24;24 | 26742226:267 | 26743015:267 | +;+ | 1098 | GHSR | 18 | 8 | 6 | 0 | 13 | 8 |
| ENSDARG000 | 24:24:24:24:2 | 10027902:100 | 10028081:100 | +;+;+;+;+;+;+ | 4743 | NA | 0 | 0 | 0 | 0 | 0 | 0 |
| ENSDARG000 | 24:24:24:24:2 | 23791758:237 | 23792286:237 | +;+;+;+;+;+;+;+;+; | 4604 | MYBL1 | 4 | 177 | 146 | 86 | 110 | 80 |
| ENSDARG000 | 24;24 | 28972084:289 | 28972564:289 | -;- | 539 | NA | 0 | 0 | 0 | 0 | 0 | 0 |
| ENSDARG000 | 24;24;24 | 28975027:289 | 28975193:289 | +;+;+ | 836 | NA | 0 | 0 | 0 | 0 | 0 | 0 |
| ENSDARG000 | 24;24 | 6771308:6772 | 6771673:6772 | -;- | 664 | NA | 0 | 0 | 0 | 0 | 0 | 0 |
| ENSDARG000 | 24:24:24:24:2 | 29027428:290 | 29027547:290 | -;-;-;-;-;-;-;-;-; | 1396 | NA | 2 | 0 | 0 | 0 | 4 | 0 |
| ENSDARG000 | 24:24:24:24:2 | 6687892:6694 | 6688041:6694 | -;-;-;-;-;-;-;-;-; | 3272 | DPP6 | 0 | 45 | 12 | 23 | 16 | 0 |
| ENSDARG000 | 24:24:24:24:2 | 13251480:132 | 13252890:132 | -;-;-;-;-;-;-;-;-; | 2399 | rdh10a | 128 | 101 | 113 | 10 | 280 | 609 |
| ENSDARG000 | 24:24:24:24:2 | 10039165:100 | 10039266:100 | +;+;+;+;+;+;+;+;+; | 4587 | POU6F2 | 0 | 0 | 0 | 0 | 8 | 0 |
| ENSDARG000 | 24:24:24:24:2 | 26753956:267 | 26754323:267 | -;-;-;-;-;-;-;-;-; | 4249 | FNDC3B | 726 | 709 | 936 | 741 | 1122 | 695 |
| ENSDARG000 | 24 | 23837519 | 23839647 | - | 2129 | rrs1 | 292 | 713 | 1081 | 304 | 628 | 432 |
| ENSDARG000 | 24:24:24:24:2 | 25144333:251 | 25144499:251 | +;+;+;+;+;+;+;+;+; | 2589 | NA | 0 | 0 | 0 | 0 | 0 | 0 |
| ENSDARG000 | 24:24:24:24:2 | 25173470:251 | 25173826:251 | -;-;-;-;-;- | 1702 | PLCXD2 | 0 | 0 | 0 | 0 | 0 | 0 |
| ENSDARG000 | 24:24:24:24:2 | 25092316:250 | 25094843:250 | -;-;-;-;-;-;-;-;-; | 7034 | PHLDB2 | 62 | 12 | 148 | 10 | 95 | 111 |
| ENSDARG000 | 24;24;24 | 36255949:362 | 36255979:362 | +;+;+ | 265 | NA | 1 | 0 | 2 | 3 | 0 | 9 |
| ENSDARG000 | 24:24:24:24:2 | 36253255:362 | 36253379:362 | -;-;-;-;-;-;-;-;-; | 1889 | si:ch211-40k2 | 76 | 92 | 132 | 91 | 86 | 117 |
| ENSDARG000 | 24:24:24:24:2 | 13277573:132 | 13277867:132 | +;+;+;+;+;+ | 768 | NA | 0 | 0 | 0 | 0 | 0 | 0 |
| ENSDARG000 | 24 | 2826210 | 2829049 | - | 2840 | NA | 0 | 2 | 6 | 0 | 0 | 0 |
| ENSDARG000 | 24:24:24:24:2 | 10302955:103 | 10303357:103 | +;+;+;+;+;+;+;+;+; | 3616 | NA | 179 | 196 | 250 | 98 | 282 | 184 |
| ENSDARG000 | 24;24;24 | 10303097:103 | 10303357:103 | +;+;+ | 931 | NA | 0 | 0 | 0 | 0 | 0 | 0 |
| ENSDARG000 | 24:24:24:24:2 | 10303071:103 | 10303357:103 | +;+;+;+;+ | 1268 | otulina | 147 | 87 | 155 | 69 | 131 | 102 |
| ENSDARG000 | 24:24:24:24:2 | 13336327:133 | 13337819:133 | -;-;-;-;-;-;-;-;-; | 2987 | NA | 44 | 235 | 208 | 98 | 110 | 162 |
| ENSDARG000 | 24:24:24:24:2 | 19210001:192 | 19210389:192 | +;+;+;+;+;+;+;+;+; | 5108 | C24H8orf34 | 0 | 0 | 0 | 0 | 8 | 0 |
| ENSDARG000 | 24:24:24:24:2 | 29381946:293 | 29381976:293 | +;+;+;+;+;+;+;+;+; | 2099 | NTNG1 | 2 | 0 | 2 | 2 | 0 | 0 |
| ENSDARG000 | 24:24:24:24:2 | 26895748:268 | 26895982:268 | +;+;+;+;+ | 1926 | nceh1b.1 | 34 | 1 | 33 | 20 | 18 | 4 |
| ENSDARG000 | 24:24:24:24:2 | 33279654:332 | 33280082:332 | -;-;-;-;-;-;-;-;-; | 1457 | zgc:195173 | 90 | 76 | 103 | 16 | 33 | 98 |
| ENSDARG000 | 24:24:24:24:2 | 2836307:2839 | 2837371:2839 | -;-;-;-;- | 1517 | cyb5a | 108 | 294 | 284 | 318 | 194 | 276 |
| ENSDARG000 | 24:24:24:24:2 | 2843268:2843 | 2843508:2844 | +;+;+;+;+;+;+;+;+; | 2177 | NA | 2 | 4 | 0 | 2 | 0 | 0 |
| ENSDARG000 | 24:24:24:24:2 | 26932472:269 | 26932657:269 | +;+;+;+;+;+;+;+;+; | 1879 | tnfsf10 | 689 | 76 | 171 | 71 | 162 | 49 |
| ENSDARG000 | 24:24:24:24:2 | 25191415:251 | 25191732:251 | +;+;+;+;+;+ | 1747 | si:ch73-264p1 | 12 | 30 | 34 | 18 | 10 | 20 |
| ENSDARG000 | 24:24:24:24:2 | 2889335:2892 | 2894249:2894 | -;-;-;-;-;-;- | 6251 | dipk1c | 26 | 4 | 8 | 0 | 18 | 6 |
| ENSDARG000 | 24:24:24:24:2 | 10349147:103 | 10350339:103 | -;-;-;-;-;-;-;-;-; | 3161 | ankha | 41 | 10 | 65 | 32 | 98 | 44 |
| ENSDARG000 | 24:24:24:24:2 | 19414561:194 | 19414727:194 | +;+;+;+;+;+;+;+;+; | 8402 | sulf1 | 144 | 662 | 442 | 52 | 145 | 63 |
| ENSDARG000 | 24:24:24:24:2 | 36275660:362 | 36276229:362 | -;-;-;-;-;-;-;-;-; | 2353 | coasy | 375 | 319 | 371 | 264 | 476 | 277 |
| ENSDARG000 | 24:24:24:24:2 | 25210015:252 | 25210429:252 | +;+;+;+;+;+;+;+;+; | 3142 | cip2a | 104 | 270 | 347 | 118 | 238 | 205 |
| ENSDARG000 | 24;24 | 26941197:269 | 26943623:269 | -;- | 2784 | msl2b | 357 | 366 | 479 | 305 | 389 | 434 |
| ENSDARG000 | 24:24:24:24:2 | 2933687:2935 | 2933785:2935 | +;+;+;+;+;+;+;+;+; | 2307 | eci2 | 364 | 946 | 1066 | 729 | 1070 | 826 |
| ENSDARG000 | 24;24;24;24 | 13361846:133 | 13363759:133 | -;-;-;- | 2536 | KCNB2 | 0 | 0 | 2 | 0 | 6 | 0 |
| ENSDARG000 | 24:24:24:24:2 | 2941000:2941 | 2942066:2942 | -;-;-;-;-;-;-;- | 1423 | tubb6 | 18 | 47 | 0 | 0 | 4 | 2 |
| ENSDARG000 | 24:24:24:24:2 | 2949937:2950 | 2950400:2950 | -;-;-;-;-;-;-;- | 1083 | NA | 0 | 0 | 0 | 0 | 0 | 0 |
| ENSDARG000 | 24:24:24:24:2 | 26948712:269 | 26949687:269 | -;-;-;-;-;-;-;-;-; | 4970 | STAG1 | 736 | 714 | 945 | 532 | 678 | 576 |
| ENSDARG000 | 24:24:24:24:2 | 13635108:136 | 13635442:136 | +;+;+;+;+;+;+;+;+; | 6032 | trpa1b | 0 | 10 | 26 | 12 | 32 | 4 |
| ENSDARG000 | 24:24:24:24:2 | 25236381:252 | 25236696:252 | -;-;-;-;-;-;-;- | 1478 | NA | 18 | 0 | 12 | 4 | 26 | 0 |
| ENSDARG000 | 24:24:24:24:2 | 25244946:252 | 25245496:252 | -;-;-;-;-;-;-;-;-; | 2158 | NA | 0 | 0 | 8 | 0 | 0 | 0 |
| ENSDARG000 | 24:24:24:24:2 | 25258904:252 | 25259293:252 | +;+;+;+;+;+;+;+;+; | 1672 | NA | 2 | 0 | 14 | 18 | 0 | 0 |
| ENSDARG000 | 24:24:24:24:2 | 13869092:139 | 13869183:139 | +;+;+;+;+;+;+;+;+; | 5078 | eya1 | 6 | 10 | 0 | 2 | 10 | 4 |
| ENSDARG000 | 24:24:24:24:2 | 19591893:195 | 19591964:195 | +;+;+;+;+;+;+;+;+; | 4699 | SLCO5A1 | 271 | 73 | 156 | 68 | 142 | 224 |
| ENSDARG000 | 24:24:24:24:2 | 19707609:197 | 19708617:197 | -;-;-;-;-;-;-;-;-; | 2040 | csrnp1b | 9173 | 3691 | 4433 | 4343 | 3301 | 4510 |
| ENSDARG000 | 24:24:24:24:2 | 20006711:200 | 20006968:200 | -;-;-;-;-;-;-;-;-; | 1940 | slc22a13b | 0 | 0 | 11 | 2 | 59 | 2 |
| ENSDARG000 | 24:24:24:24:2 | 20032331:200 | 20032596:200 | -;-;-;-;-;-;-;-;-; | 1667 | NA | 0 | 0 | 0 | 0 | 11 | 10 |
| ENSDARG000 | 24;24 | 20061677:200 | 20061708:200 | +;+ | 559 | NA | 0 | 0 | 0 | 0 | 2 | 1 |
| ENSDARG000 | 24:24:24:24:2 | 20064088:200 | 20066778:200 | -;-;-;-;-;-;-;-;-; | 4352 | oxsr1b | 780 | 1247 | 1859 | 1331 | 1837 | 1118 |
| ENSDARG000 | 24;24 | 20062044:200 | 20062156:200 | +;+ | 401 | NA | 0 | 0 | 4 | 0 | 0 | 16 |
| ENSDARG000 | 24;24;24;24 | 20060824:200 | 20061345:200 | -;-;-;- | 992 | NA | 44 | 14 | 18 | 0 | 10 | 6 |
| ENSDARG000 | 24;24 | 39695688:396 | 39695724:396 | +;+ | 363 | NA | 0 | 0 | 5 | 0 | 9 | 0 |
| ENSDARG000 | 24;24;24 | 39690607:396 | 39690642:396 | -;-;- | 327 | NA | 0 | 0 | 0 | 0 | 0 | 0 |
| ENSDARG000 | 24:24:24:24:2 | 42948:43089: | 42966:43149: | +;+;+;+;+ | 288 | tmx3b | 97 | 124 | 157 | 34 | 166 | 85 |
| ENSDARG000 | 24 | 35564668 | 35565927 | + | 1260 | cebpd | 813 | 387 | 344 | 521 | 290 | 221 |
| ENSDARG000 | 24:24:24:24:2 | 28410651:284 | 28410680:284 | +;+;+;+;+;+;+;+;+; | 282 | NA | 8 | 5 | 0 | 2 | 18 | 12 |
| ENSDARG000 | 24:24:24:24:2 | 40905100:409 | 40905414:409 | +;+;+;+;+;+;+;+;+; | 6471 | scn5lab | 0 | 2 | 0 | 0 | 0 | 2 |
| ENSDARG000 | 24;24;24;24 | 37080771:371 | 37081388:371 | +;+;+;+ | 1944 | kcnc3b | 2 | 2 | 0 | 0 | 2 | 0 |
| ENSDARG000 | 24:24:24:24:2 | 21813127:218 | 21813294:218 | -;-;-;-;- | 891 | NA | 0 | 0 | 0 | 0 | 0 | 0 |
| ENSDARG000 | 24;24 | 39683268:396 | 39683467:396 | -;- | 288 | NA | 0 | 0 | 0 | 0 | 0 | 0 |
| ENSDARG000 | 24;24;24 | 12200260:122 | 12200447:122 | +;+;+ | 642 | NA | 0 | 0 | 0 | 0 | 4 | 0 |
| ENSDARG000 | 24:24:24:24:2 | 195072:19659 | 195280:19672 | -;-;-;-;- | 1326 | NA | 0 | 0 | 6 | 0 | 8 | 0 |
| ENSDARG000 | 24:24:24:24:2 | 744713:76665 | 745131:76686 | +;+;+;+;+;+;+ | 3896 | stk17a | 2407 | 3645 | 921 | 2724 | 2178 | 1506 |
| ENSDARG000 | 24 | 21963052 | 21963168 | - | 117 | 5S_rRNA | 1 | 0 | 0 | 0 | 2 | 0 |
| ENSDARG000 | 24:24:24:24:2 | 36798558:368 | 36799100:368 | -;-;-;-;- | 1190 | dcakd | 97 | 87 | 143 | 52 | 74 | 112 |
| ENSDARG000 | 24;24 | 21346796:213 | 21347099:213 | +;+ | 885 | SHISA2 | 21 | 4 | 46 | 0 | 8 | 9 |
| ENSDARG000 | 24:24:24:24:2 | 8736497:8736 | 8736558:8736 | +;+;+;+;+ | 518 | tmem14ca | 262 | 260 | 291 | 192 | 263 | 343 |
| ENSDARG000 | 24:24:24:24:2 | 40763020:407 | 40763055:407 | -;-;-;-;-;-;-;-;-; | 5838 | NA | 2 | 0 | 0 | 0 | 0 | 0 |
| ENSDARG000 | 24:24:24:24:2 | 792429:79334 | 792488:79348 | +;+;+;+;+;+;+;+;+; | 828 | impa2 | 87 | 210 | 113 | 97 | 190 | 119 |
| ENSDARG000 | 24 | 36183099 | 36183216 | + | 118 | 5S_rRNA | 0 | 0 | 3 | 0 | 0 | 0 |
| ENSDARG000 | 24:24:24:24:2 | 34970680:349 | 34970750:349 | +;+;+;+;+;+;+;+;+; | 4869 | RB1CC1 | 1223 | 1066 | 1520 | 978 | 1223 | 1004 |
| ENSDARG000 | 24:24:24:24:2 | 13316737:133 | 13317358:133 | +;+;+;+;+ | 1182 | SBSPON | 0 | 0 | 0 | 0 | 0 | 0 |
| ENSDARG000 | 24 | 1802268 | 1802384 | + | 117 | 5S_rRNA | 0 | 0 | 0 | 0 | 0 | 0 |
| ENSDARG000 | 24 | 11048446 | 11048560 | - | 115 | 5S_rRNA | 0 | 0 | 0 | 0 | 0 | 0 |
| ENSDARG000 | 24 | 10948064 | 10948145 | - | 82 | dre-mir-7147 | 0 | 0 | 0 | 0 | 0 | 0 |
| ENSDARG000 | 24:24:24:24:2 | 6353394:6356 | 6354226:6356 | +;+;+;+;+;+ | 1880 | NA | 4 | 6 | 0 | 0 | 0 | 0 |
| ENSDARG000 | 24;24;24;24 | 21841238:218 | 21843384:218 | -;-;-;- | 3213 | NA | 0 | 18 | 2 | 0 | 4 | 4 |
| ENSDARG000 | 24:24:24:24:2 | 673161:67316 | 673657:67365 | +;+;+;+;+;+;+;+;+; | 6314 | HECW1 | 26 | 80 | 117 | 136 | 102 | 30 |
| ENSDARG000 | 24 | 41710688 | 41710804 | + | 117 | 5S_rRNA | 0 | 0 | 0 | 0 | 0 | 0 |
| ENSDARG000 | 24;24 | 23742690:237 | 23743118:237 | +;+ | 531 | TCF24 | 0 | 0 | 0 | 0 | 0 | 0 |
| ENSDARG000 | 24 | 28748101 | 28748217 | + | 117 | 5S_rRNA | 0 | 0 | 0 | 0 | 0 | 0 |
| ENSDARG000 | 24 | 10255914 | 10256030 | - | 117 | 5S_rRNA | 0 | 0 | 0 | 0 | 0 | 0 |
| ENSDARG000 | 24 | 20788168 | 20788284 | + | 117 | 5S_rRNA | 0 | 0 | 0 | 0 | 0 | 0 |
| ENSDARG000 | 24 | 41678257 | 41679356 | - | 1100 | lrrc30a | 0 | 0 | 0 | 0 | 0 | 0 |
| ENSDARG000 | 24:24:24:24:2 | 20916128:209 | 20916153:209 | -;-;-;-;-;-;- | 171 | NA | 12 | 6 | 12 | 8 | 13 | 7 |
| ENSDARG000 | 24:24:24:24:2 | 40712794:407 | 40712905:407 | -;-;-;-;-;-;-;-;-; | 5985 | smyhc3 | 0 | 0 | 0 | 0 | 0 | 0 |
| ENSDARG000 | 24:24:24:24:2 | 33552198:335 | 33552273:335 | -;-;-;-;-;-;-;-;-; | 1893 | NA | 0 | 0 | 0 | 0 | 0 | 0 |
| ENSDARG000 | 24 | 14635303 | 14635419 | + | 117 | 5S_rRNA | 0 | 0 | 0 | 0 | 0 | 0 |
| ENSDARG000 | 24:24:24:24:2 | 25285758:252 | 25285861:252 | -;-;-;-;-;-;-;-;-; | 3114 | NA | 14 | 4 | 0 | 16 | 2 | 0 |
| ENSDARG000 | 24 | 31640312 | 31640427 | - | 116 | 5S_rRNA | 0 | 0 | 0 | 0 | 0 | 0 |
| ENSDARG000 | 24:24:24:24:2 | 40656510:406 | 40656611:406 | -;-;-;-;-;-;-;-;-; | 6147 | smyhc1 | 2 | 0 | 0 | 0 | 0 | 0 |
| ENSDARG000 | 24 | 20187316 | 20187430 | - | 115 | 5S_rRNA | 0 | 0 | 0 | 0 | 0 | 0 |
| ENSDARG000 | 24;24;24 | 40820699:408 | 40821632:408 | -;-;- | 1186 | cebp1 | 6 | 6 | 4 | 0 | 15 | 10 |
| ENSDARG000 | 24:24:24:24:2 | 36018164:360 | 36018285:360 | +;+;+;+;+;+;+;+;+; | 1293 | NA | 720 | 437 | 499 | 457 | 671 | 613 |
| ENSDARG000 | 24:24:24:24:2 | 36812640:368 | 36814396:368 | -;-;-;-;-;-;- | 3169 | FAM171A2 | 2 | 0 | 0 | 0 | 18 | 6 |
| ENSDARG000 | 24;24;24;24 | 19986432:199 | 19986744:199 | +;+;+;+ | 5260 | LIMD2 | 8 | 3 | 23 | 8 | 31 | 0 |
| ENSDARG000 | 24;24 | 32560395:325 | 32562334:325 | -;- | 2004 | PTCHD3 | 16 | 8 | 22 | 14 | 20 | 0 |
| ENSDARG000 | 24 | 39120725 | 39121900 | - | 1176 | NA | 0 | 2 | 0 | 0 | 0 | 0 |
| ENSDARG000 | 24 | 6063671 | 6063785 | + | 115 | 5S_rRNA | 0 | 0 | 0 | 0 | 0 | 0 |
| ENSDARG000 | 24:24:24:24:2 | 11908480:119 | 11908584:119 | +;+;+;+;+;+;+;+;+; | 3320 | fen1 | 170 | 1207 | 1715 | 454 | 1231 | 703 |
| ENSDARG000 | 24 | 238247 | 238363 | + | 117 | 5S_rRNA | 0 | 0 | 0 | 0 | 0 | 0 |
| ENSDARG000 | 24:24:24:24:2 | 80653:80653: | 80689:80689: | +;+;+;+;+;+;+;+;+; | 4316 | RECK | 920 | 1316 | 2067 | 1098 | 1959 | 1289 |
| ENSDARG000 | 24:24:24:24:2 | 36903172:369 | 36903310:369 | -;-;-;-;-;-;-;-;-; | 1410 | NA | 0 | 0 | 0 | 0 | 0 | 0 |
| ENSDARG000 | 24:24:24:24:2 | 40860320:408 | 40860392:408 | +;+;+;+;+;+;+;+;+; | 1269 | gorasp1b | 6 | 0 | 0 | 0 | 0 | 18 |
| ENSDARG000 | 24:24:24:24:2 | 36995295:369 | 36995433:369 | -;-;-;-;-;-;-;-;-; | 1410 | dnaaf3 | 0 | 0 | 0 | 0 | 0 | 0 |
| ENSDARG000 | 24;24;24 | 39688072:396 | 39688108:396 | +;+;+ | 531 | NA | 2 | 0 | 3 | 0 | 8 | 2 |
| ENSDARG000 | 24:24:24:24:2 | 37257596:372 | 37258306:372 | -;-;-;-;-;-;- | 1252 | ube2ib | 1001 | 1259 | 1486 | 1124 | 1175 | 1164 |
| ENSDARG000 | 24 | 15670013 | 15671038 | + | 1026 | NA | 0 | 6 | 0 | 0 | 2 | 0 |
| ENSDARG000 | 24:24:24:24:2 | 38208823:382 | 38208828:382 | +;+;+;+;+;+;+;+;+; | 342 | NA | 0 | 0 | 0 | 0 | 0 | 0 |
| ENSDARG000 | 24;24 | 29961238:299 | 29962184:299 | -;- | 1083 | NA | 0 | 0 | 2 | 16 | 6 | 0 |
| ENSDARG000 | 24 | 17738519 | 17738635 | + | 117 | 5S_rRNA | 0 | 0 | 0 | 0 | 0 | 0 |
| ENSDARG000 | 24 | 26088143 | 26089609 | + | 1467 | NA | 0 | 0 | 0 | 0 | 0 | 0 |
| ENSDARG000 | 24 | 37552985 | 37553117 | + | 133 | 5S_rRNA | 0 | 0 | 0 | 0 | 0 | 0 |
| ENSDARG000 | 24 | 29473856 | 29473975 | - | 120 | 5S_rRNA | 0 | 0 | 0 | 0 | 0 | 0 |
| ENSDARG000 | 24 | 19288654 | 19288768 | - | 115 | 5S_rRNA | 0 | 0 | 0 | 0 | 0 | 0 |
| ENSDARG000 | 24 | 21272612 | 21272728 | + | 117 | 5S_rRNA | 0 | 0 | 0 | 0 | 0 | 0 |
| ENSDARG000 | 24:24:24:24:2 | 40680153:406 | 40680173:406 | -;-;-;-;-;-;-;-;-; | 5896 | smyhc2 | 8 | 2 | 0 | 0 | 0 | 0 |
| ENSDARG000 | 24 | 252805 | 252926 | + | 122 | 5S_rRNA | 0 | 0 | 0 | 0 | 0 | 0 |
| ENSDARG000 | 24:24:24:24:2 | 21603588:216 | 21603607:216 | -;-;-;-;-;-;- | 186 | NA | 0 | 0 | 0 | 0 | 0 | 0 |
| ENSDARG000 | 24:24:24:24:2 | 39027481:390 | 39027575:390 | +;+;+;+;+;+;+;+;+; | 3557 | CAPN15 | 1312 | 904 | 1110 | 746 | 1116 | 760 |
| ENSDARG000 | 24;24 | 25692802:256 | 25693189:256 | +;+ | 1716 | camk2n2 | 2 | 0 | 0 | 0 | 0 | 0 |
| ENSDARG000 | 24:24:24:24:2 | 6657312:6660 | 6657359:6661 | -;-;-;-;-;-;-;- | 1137 | NA | 171 | 401 | 541 | 268 | 155 | 875 |
| ENSDARG000 | 24 | 24455632 | 24455748 | + | 117 | dre-mir-124-3 | 0 | 0 | 0 | 0 | 0 | 0 |
| ENSDARG000 | 24:24:24:24:2 | 7988458:7991 | 7992588:7992 | -;-;-;-;-;-;-;-;-; | 4775 | bloc1s5 | 322 | 373 | 256 | 326 | 314 | 334 |
| ENSDARG000 | 24;24 | 26658132:266 | 26658384:266 | +;+ | 461 | NA | 0 | 0 | 0 | 0 | 3 | 0 |
| ENSDARG000 | 24 | 21629725 | 21629840 | + | 116 | 5S_rRNA | 0 | 0 | 0 | 0 | 0 | 0 |
| ENSDARG000 | 24:24:24:24:2 | 37700259:377 | 37700351:377 | +;+;+;+;+;+ | 777 | NA | 51 | 96 | 104 | 24 | 95 | 57 |
| ENSDARG000 | 24:24:24:24:2 | 2470061:2470 | 2470207:2470 | +;+;+;+;+;+;+;+;+; | 1688 | F13A1 | 0 | 0 | 0 | 0 | 0 | 0 |
| ENSDARG000 | 24:24:24:24:2 | 42094833:420 | 42094940:420 | +;+;+;+;+;+ | 1329 | ahrra | 45 | 4 | 2 | 6 | 13 | 18 |
| ENSDARG000 | 24:24:24:24:2 | 35536664:355 | 35537185:355 | -;-;-;-;-;-;-;-;-; | 3015 | mcm4 | 1139 | 3379 | 3843 | 1298 | 3295 | 2008 |
| ENSDARG000 | 24;24 | 39700552:397 | 39700877:397 | -;- | 375 | NA | 0 | 0 | 6 | 0 | 8 | 0 |
| ENSDARG000 | 24:24:24:24:2 | 7322116:7323 | 7322190:7324 | +;+;+;+;+;+ | 1639 | XRCC2 | 0 | 20 | 22 | 30 | 4 | 0 |
| ENSDARG000 | 24;24;24;24 | 41554613:415 | 41554705:415 | -;-;-;- | 718 | NA | 34 | 32 | 80 | 40 | 22 | 92 |

| | | | | | | | | | | | |
|---|---|---|---|---|---|---|---|---|---|---|---|
| ENSDARG000 | 24:24:24:24:2 | 29912509:29 | 29912698:29 | +:+:+:+:+:+:+: | 1626 | FRRS1 | 631 | 714 | 821 | 551 | 1012 | 931 |
| ENSDARG000 | 24 | 26185410 | 26185500 | + | 91 | dre-mir-727 | 0 | 0 | 0 | 0 | 0 | 0 |
| ENSDARG000 | 24;24;24 | 9298198:929 | 9298281:929 | +;+;+ | 1239 | NA | 254 | 150 | 241 | 180 | 178 | 170 |
| ENSDARG000 | 24:24:24:24:2 | 21905717:21 | 21909542:21 | -;-;-;-;-;-;-;- | 4946 | c1qtnf9 | 10 | 4 | 2 | 10 | 40 | 0 |
| ENSDARG000 | 24 | 260902 | 261906 | + | 1005 | NA | 6 | 58 | 33 | 14 | 84 | 63 |
| ENSDARG000 | 24:24:24:24:2 | 30695120:30 | 30695192:30 | +;+;+;+;+;+;+ | 804 | NA | 0 | 8 | 12 | 20 | 0 | 0 |
| ENSDARG000 | 24 | 22456844 | 22456961 | + | 118 | 5S_rRNA | 0 | 0 | 0 | 0 | 0 | 0 |
| ENSDARG000 | 24 | 33769287 | 33769403 | + | 117 | 5S_rRNA | 0 | 0 | 0 | 2 | 0 | 2 |
| ENSDARG000 | 24 | 29034624 | 29034740 | - | 117 | 5S_rRNA | 0 | 0 | 0 | 0 | 0 | 0 |
| ENSDARG000 | 24 | 11411822 | 11411941 | - | 120 | 5S_rRNA | 0 | 0 | 0 | 0 | 0 | 0 |
| ENSDARG000 | 24:24:24:24:2 | 35232716:35 | 35232898:35 | -:-:-:-:-:-:-:-: | 1601 | SNTG1 | 6 | 4 | 8 | 0 | 6 | 2 |
| ENSDARG000 | 24:24:24:24:2 | 21330753:21 | 21331069:21 | -:-:-:-:-:-:-:-: | 2498 | spice1 | 18 | 22 | 180 | 56 | 146 | 210 |
| ENSDARG000 | 24:24:24:24:2 | 119680:1388 | 119839:1390 | +:+:+:+:+:+:+: | 1918 | tgfbr1b | 115 | 106 | 282 | 125 | 285 | 191 |
| ENSDARG000 | 24:24:24:24:2 | 37195463:37 | 37195629:37 | -:-:-:-:-:-:-:-: | 2202 | cog7 | 409 | 390 | 684 | 256 | 614 | 460 |
| ENSDARG000 | 24:24:24:24:2 | 40529346:40 | 40529517:40 | +:+:+:+:+:+:+: | 3648 | NA | 251 | 82 | 109 | 122 | 281 | 373 |
| ENSDARG000 | 24:24:24:24:2 | 37274759:37 | 37277618:37 | -;-;-;-;- | 3686 | KCTD5 | 362 | 102 | 94 | 354 | 82 | 64 |
| ENSDARG000 | 24:24:24:24:2 | 31940504:31 | 31940683:31 | -;-;-;-;- | 804 | C1QL3 | 2 | 0 | 11 | 4 | 18 | 14 |
| ENSDARG000 | 24 | 18077543 | 18077659 | + | 117 | 5S_rRNA | 0 | 0 | 4 | 0 | 4 | 0 |
| ENSDARG000 | 24 | 39787473 | 39787564 | + | 92 | NA | 2 | 5 | 4 | 4 | 2 | 9 |
| ENSDARG000 | 24 | 31786463 | 31786577 | - | 115 | 5S_rRNA | 0 | 0 | 0 | 0 | 0 | 0 |
| ENSDARG000 | 24 | 264839 | 265858 | + | 1020 | NA | 14 | 76 | 38 | 12 | 81 | 130 |
| ENSDARG000 | 24;24 | 40855741:40 | 40858315:40 | -;- | 3606 | NA | 0 | 2 | 0 | 0 | 3 | 1 |
| ENSDARG000 | 24:24:24:24:2 | 32650227:32 | 32650346:32 | -;-;-;-;-;-;- | 1456 | ca2 | 117 | 12 | 15 | 11 | 73 | 60 |
| ENSDARG000 | 24;24 | 4434117:443 | 4434590:443 | +;+ | 1679 | NA | 0 | 2 | 8 | 2 | 0 | 2 |
| ENSDARG000 | 24;24;24;24 | 41316714:41 | 41317493:41 | -;-;-;- | 969 | NA | 84 | 100 | 60 | 86 | 47 | 27 |
| ENSDARG000 | 24;24;24;24 | 36129216:36 | 36129353:36 | -;-;-;- | 503 | tmem14cb | 136 | 109 | 131 | 144 | 99 | 123 |
| ENSDARG000 | 24 | 19863246 | 19868789 | + | 5544 | NA | 0 | 0 | 0 | 0 | 0 | 0 |
| ENSDARG000 | 24 | 18625252 | 18625367 | - | 116 | 5S_rRNA | 0 | 0 | 0 | 0 | 0 | 0 |
| ENSDARG000 | 24 | 15213871 | 15213985 | - | 115 | 5S_rRNA | 0 | 0 | 0 | 0 | 0 | 0 |
| ENSDARG000 | 24:24:24:24:2 | 41931585:41 | 41931652:41 | +:+:+:+:+:+:+: | 3418 | epb41l3a | 4 | 88 | 217 | 52 | 99 | 46 |
| ENSDARG000 | 24 | 30030534 | 30030650 | + | 117 | 5S_rRNA | 0 | 0 | 0 | 0 | 0 | 0 |
| ENSDARG000 | 24 | 37779010 | 37779127 | + | 118 | 5S_rRNA | 0 | 0 | 0 | 0 | 0 | 0 |
| ENSDARG000 | 24:24:24:24:2 | 23988970:23 | 23993247:23 | -;-;-;-;- | 5436 | NA | 66 | 39 | 51 | 32 | 54 | 153 |
| ENSDARG000 | 24:24:24:24:2 | 38269088:38 | 38269093:38 | -;-;-;-;-;-;- | 909 | zgc:91999 | 2 | 0 | 0 | 0 | 2 | 2 |
| ENSDARG000 | 24 | 6582052 | 6582168 | + | 117 | 5S_rRNA | 0 | 0 | 0 | 0 | 0 | 0 |
| ENSDARG000 | 24 | 15277216 | 15278775 | + | 1560 | SOCS6 | 136 | 104 | 163 | 92 | 119 | 85 |
| ENSDARG000 | 24 | 23273242 | 23273359 | - | 118 | 5S_rRNA | 0 | 0 | 0 | 0 | 0 | 0 |
| ENSDARG000 | 24 | 39990695 | 39992254 | + | 1560 | NA | 0 | 0 | 0 | 0 | 0 | 0 |
| ENSDARG000 | 24:24:24:24:2 | 40728978:40 | 40729010:40 | -;-;-;-;-;- | 762 | NA | 0 | 0 | 0 | 0 | 0 | 0 |
| ENSDARG000 | 24:24:24:24:2 | 8721676:872 | 8723079:872 | -:-:-:-:-:-:-:-: | 3092 | tfap2a | 9451 | 11677 | 11565 | 6772 | 15329 | 11274 |
| ENSDARG000 | 24:24:24:24:2 | 40876247:40 | 40877003:40 | -:-:-:-:-:-:-:-: | 2222 | wdr48a | 158 | 164 | 138 | 81 | 170 | 146 |
| ENSDARG000 | 24;24;24 | 41989108:41 | 41989511:41 | +;+;+ | 1946 | zbtb14 | 384 | 377 | 477 | 206 | 400 | 331 |
| ENSDARG000 | 24;24;24 | 39684134:39 | 39684177:39 | -;-;- | 315 | NA | 0 | 0 | 0 | 0 | 0 | 0 |
| ENSDARG000 | 24;24;24 | 31655939:31 | 31656314:31 | +;+;+ | 2509 | NA | 49 | 19 | 32 | 30 | 7 | 26 |
| ENSDARG000 | 24 | 41826706 | 41826822 | - | 117 | 5S_rRNA | 0 | 0 | 0 | 0 | 0 | 0 |
| ENSDARG000 | 24:24:24:24:2 | 42081728:42 | 42081836:42 | -;-;-;-;-;-;- | 885 | ssr1 | 1865 | 1930 | 4210 | 1696 | 2469 | 1490 |
| ENSDARG000 | 24 | 16807521 | 16807648 | + | 128 | 5S_rRNA | 0 | 0 | 0 | 0 | 0 | 0 |
| ENSDARG000 | 24;24;24;24 | 26302781:26 | 26303386:26 | -;-;-;- | 760 | otos | 6 | 2 | 0 | 0 | 6 | 0 |
| ENSDARG000 | 24;24;24;24 | 27428141:27 | 27428243:27 | -;-;-;- | 336 | NA | 9 | 18 | 2 | 8 | 4 | 6 |
| ENSDARG000 | 24;24;24;24 | 42102204:42 | 42102850:42 | -;-;-;- | 1619 | pigm | 172 | 173 | 218 | 100 | 237 | 102 |
| ENSDARG000 | 24:24:24:24:2 | 15602:17377 | 15808:17671 | +:+:+:+:+:+:+: | 4353 | NA | 2 | 0 | 6 | 0 | 10 | 2 |
| ENSDARG000 | 24:24:24:24:2 | 39211288:39 | 39211309:39 | +:+:+:+:+:+:+: | 1425 | NA | 155 | 104 | 84 | 91 | 102 | 40 |
| ENSDARG000 | 24:24:24:24:2 | 31493:34574 | 34171:34876 | -;-;-;-;-;-;-;- | 4771 | NA | 0 | 0 | 0 | 0 | 0 | 0 |
| ENSDARG000 | 24:24:24:24:2 | 34139146:34 | 34139364:34 | -:-:-:-:-:-:-:-: | 3405 | AGAP3 | 116 | 48 | 130 | 27 | 87 | 80 |
| ENSDARG000 | 24:24:24:24:2 | 42060967:42 | 42061522:42 | -:-:-:-:-:-:-:-: | 1536 | ppp4r1 | 607 | 555 | 704 | 574 | 713 | 476 |
| ENSDARG000 | 24 | 17546753 | 17546869 | - | 117 | 5S_rRNA | 0 | 0 | 0 | 0 | 0 | 0 |
| ENSDARG000 | 24 | 28337955 | 28338069 | + | 115 | 5S_rRNA | 0 | 0 | 0 | 0 | 0 | 0 |
| ENSDARG000 | 24 | 30576364 | 30576479 | + | 116 | 5S_rRNA | 0 | 0 | 0 | 0 | 0 | 0 |
| ENSDARG000 | 24:24:24:24:2 | 41233755:41 | 41233811:41 | -:-:-:-:-:-:-:-: | 1590 | xylb | 16 | 39 | 50 | 18 | 48 | 50 |
| ENSDARG000 | 24 | 41254874 | 41254989 | - | 116 | 5S_rRNA | 0 | 0 | 0 | 0 | 0 | 0 |
| ENSDARG000 | 24:24:24:24:2 | 166892:1697 | 167191:1700 | +:+:+:+:+:+:+: | 3271 | lancl2 | 186 | 514 | 437 | 338 | 489 | 397 |
| ENSDARG000 | 24 | 41915878 | 41917794 | + | 1917 | TMEM200C | 2 | 10 | 16 | 0 | 50 | 0 |
| ENSDARG000 | 24 | 9712495 | 9712610 | - | 116 | 5S_rRNA | 0 | 0 | 0 | 0 | 0 | 0 |
| ENSDARG000 | 24:24:24:24:2 | 23729908:23 | 23730171:23 | -:-:-:-:-:-:-:-: | 2608 | MCMDC2 | 35 | 33 | 178 | 95 | 81 | 121 |
| ENSDARG000 | 24 | 2689078 | 2689194 | - | 117 | 5S_rRNA | 0 | 0 | 0 | 0 | 0 | 0 |
| ENSDARG000 | 24 | 4059456 | 4059573 | + | 118 | 5S_rRNA | 0 | 0 | 0 | 0 | 0 | 0 |
| ENSDARG000 | 24 | 16554869 | 16554983 | - | 115 | 5S_rRNA | 0 | 0 | 0 | 0 | 0 | 0 |
| ENSDARG000 | 24:24:24:24:2 | 29895249:29 | 29896333:29 | -;-;-;-;- | 1784 | NA | 0 | 0 | 0 | 0 | 1 | 0 |
| ENSDARG000 | 24:24:24:24:2 | 8742279:874 | 8742439:874 | -:-:-:-:-:-:-:-: | 2503 | mak | 471 | 297 | 491 | 295 | 348 | 385 |
| ENSDARG000 | 24:24:24:24:2 | 11887885:11 | 11890464:11 | -:-:-:-:-:-:-:-: | 4168 | tm9sf1 | 609 | 363 | 615 | 371 | 539 | 362 |
| ENSDARG000 | 24:24:24:24:2 | 11058151:11 | 11062971:11 | -;-;-;-;- | 5773 | NA | 292 | 539 | 624 | 166 | 352 | 442 |
| ENSDARG000 | 24:24:24:24:2 | 8133726:814 | 8134225:814 | -;-;-;-;-;-;-;- | 1728 | NA | 0 | 0 | 0 | 0 | 0 | 0 |
| ENSDARG000 | 24 | 38177961 | 38178027 | - | 67 | dre-mir-735 | 0 | 0 | 0 | 0 | 0 | 0 |
| ENSDARG000 | 24 | 39614853 | 39616374 | + | 1522 | NA | 1461 | 257 | 327 | 395 | 1006 | 1848 |
| ENSDARG000 | 24:24:24:24:2 | 8849804:884 | 8849809:884 | +:+:+:+:+:+:+: | 343 | NA | 0 | 4 | 12 | 1 | 0 | 0 |
| ENSDARG000 | 24;24;24 | 38209946:38 | 38209994:38 | +;+;+ | 360 | NA | 0 | 0 | 0 | 0 | 0 | 0 |
| ENSDARG000 | 24:24:24:24:2 | 40734996:40 | 40735179:40 | -:-:-:-:-:-:-:-: | 4170 | NA | 0 | 0 | 0 | 0 | 0 | 0 |
| ENSDARG000 | 24 | 29352039 | 29353521 | + | 1483 | prmt6 | 130 | 217 | 233 | 92 | 225 | 114 |
| ENSDARG000 | 24 | 22308944 | 22309058 | - | 115 | 5S_rRNA | 0 | 0 | 0 | 0 | 0 | 0 |
| ENSDARG000 | 24 | 6405194 | 6405316 | + | 123 | 5S_rRNA | 0 | 0 | 0 | 0 | 0 | 0 |
| ENSDARG000 | 24 | 29052708 | 29052826 | - | 119 | 5S_rRNA | 0 | 0 | 0 | 0 | 0 | 0 |
| ENSDARG000 | 24;24 | 12954489:12 | 12955884:12 | -;- | 1754 | NA | 0 | 4 | 0 | 10 | 0 | 0 |
| ENSDARG000 | 24:24:24:24:2 | 36840652:36 | 36840792:36 | +:+:+:+:+:+:+: | 2370 | NA | 232 | 160 | 276 | 142 | 472 | 223 |
| ENSDARG000 | 24:24:24:24:2 | 35183595:35 | 35183670:35 | +:+:+:+:+:+:+: | 4122 | pcmtd1 | 618 | 344 | 368 | 370 | 355 | 552 |
| ENSDARG000 | 24 | 4318179 | 4318293 | - | 115 | 5S_rRNA | 0 | 0 | 0 | 0 | 0 | 0 |
| ENSDARG000 | 24;24 | 13735616:13 | 13736230:13 | +;+ | 3404 | MSC | 0 | 0 | 0 | 0 | 0 | 0 |
| ENSDARG000 | 24 | 2919405 | 2919521 | - | 117 | 5S_rRNA | 0 | 0 | 0 | 0 | 0 | 0 |
| ENSDARG000 | 24 | 39726846 | 39726978 | - | 133 | 5S_rRNA | 2 | 0 | 0 | 0 | 4 | 1 |
| ENSDARG000 | 24 | 22371986 | 22372100 | - | 115 | 5S_rRNA | 0 | 0 | 0 | 0 | 0 | 0 |
| ENSDARG000 | 24 | 11011436 | 11011551 | - | 116 | 5S_rRNA | 0 | 0 | 0 | 0 | 0 | 0 |
| ENSDARG000 | 24;24;24 | 39330654:39 | 39330658:39 | +;+;+ | 321 | NA | 0 | 0 | 0 | 0 | 0 | 0 |
| ENSDARG000 | 24:24:24:24:2 | 20418959:20 | 20419072:20 | -:-:-:-:-:-:-:-: | 2698 | VILL | 26 | 14 | 74 | 2 | 75 | 31 |
| ENSDARG000 | 24 | 41537062 | 41537191 | + | 130 | 5S_rRNA | 0 | 0 | 0 | 0 | 0 | 0 |
| ENSDARG000 | 24 | 41610068 | 41611201 | + | 1134 | NA | 0 | 0 | 0 | 0 | 0 | 0 |
| ENSDARG000 | 24 | 27412139 | 27412253 | + | 115 | 5S_rRNA | 0 | 0 | 0 | 0 | 0 | 0 |
| ENSDARG000 | 24 | 36905287 | 36905401 | + | 115 | 5S_rRNA | 0 | 0 | 0 | 0 | 0 | 0 |
| ENSDARG000 | 24 | 38177965 | 38178027 | - | 63 | dre-mir-735 | 0 | 0 | 0 | 0 | 0 | 0 |
| ENSDARG000 | 24 | 7967484 | 7967600 | - | 117 | 5S_rRNA | 0 | 0 | 0 | 0 | 0 | 0 |
| ENSDARG000 | 24:24:24:24:2 | 40473032:40 | 40473063:40 | +:+:+:+:+:+:+: | 2304 | KCNH2 | 0 | 0 | 0 | 0 | 0 | 0 |
| ENSDARG000 | 24:24:24:24:2 | 8900916:890 | 8900943:890 | -:-:-:-:-:-:-:-: | 468 | NA | 10 | 0 | 2 | 3 | 5 | 1 |
| ENSDARG000 | 24:24:24:24:2 | 402493:4039 | 402710:4040 | +;+;+;+;+ | 1141 | vstm2a | 0 | 0 | 0 | 0 | 0 | 0 |
| ENSDARG000 | 24:24:24:24:2 | 36581016:36 | 36581116:36 | -:-:-:-:-:-:-:-: | 1320 | NA | 9 | 0 | 0 | 0 | 0 | 0 |
| ENSDARG000 | 24 | 13609598 | 13609714 | - | 117 | 5S_rRNA | 0 | 0 | 0 | 0 | 0 | 0 |
| ENSDARG000 | 24 | 14350900 | 14351020 | + | 121 | 5S_rRNA | 0 | 0 | 0 | 0 | 0 | 0 |
| ENSDARG000 | 24:24:24:24:2 | 31459715:31 | 31459841:31 | +:+:+:+:+:+:+: | 3096 | NA | 0 | 0 | 0 | 11 | 4 | 0 |
| ENSDARG000 | 24 | 10905172 | 10905286 | - | 115 | 5S_rRNA | 0 | 0 | 0 | 0 | 0 | 1 |
| ENSDARG000 | 24 | 27484936 | 27485059 | + | 124 | U1 | 0 | 0 | 0 | 0 | 0 | 0 |
| ENSDARG000 | 24:24:24:24:2 | 41786274:41 | 41786615:41 | -;-;-;-;-;-;-;- | 1183 | NA | 87 | 48 | 27 | 27 | 39 | 23 |
| ENSDARG000 | 24:24:24:24:2 | 42117231:42 | 42117479:42 | +:+:+:+:+:+:+: | 885 | snx16 | 610 | 491 | 669 | 636 | 629 | 519 |
| ENSDARG000 | 24;24 | 3050020:305 | 3050434:306 | +;+ | 3863 | inhbaa | 158 | 60 | 66 | 50 | 26 | 72 |
| ENSDARG000 | 24:24:24:24:2 | 26659010:26 | 26664090:26 | -:-:-:-:-:-:-:-: | 8017 | NA | 458 | 370 | 198 | 465 | 154 | 204 |
| ENSDARG000 | 24:24:24:24:2 | 26887487:26 | 26887624:26 | +;+;+;+;+ | 1227 | NCEH1 | 49 | 22 | 27 | 64 | 8 | 16 |
| ENSDARG000 | 24;24 | 38216340:38 | 38216355:38 | +;+ | 324 | NA | 0 | 0 | 0 | 0 | 0 | 0 |
| ENSDARG000 | 24:24:24:24:2 | 32573480:32 | 32574322:32 | -:-:-:-:-:-:-:-: | 1885 | rdh12l | 68 | 37 | 20 | 24 | 26 | 42 |
| ENSDARG000 | 24:24:24:24:2 | 31952333:31 | 31952435:31 | -:-:-:-:-:-:-:-: | 1185 | RSU1 | 598 | 515 | 635 | 447 | 778 | 638 |
| ENSDARG000 | 24:24:24:24:2 | 2460178:246 | 2462241:246 | -:-:-:-:-:-:-:-: | 2669 | ly86 | 4 | 0 | 12 | 0 | 32 | 0 |
| ENSDARG000 | 24 | 26190552 | 26190634 | + | 83 | dre-mir-728 | 0 | 0 | 0 | 0 | 0 | 0 |
| ENSDARG000 | 24:24:24:24:2 | 41466841:41 | 41466920:41 | -;-;-;-;-;-;- | 915 | NA | 114 | 208 | 303 | 265 | 94 | 216 |
| ENSDARG000 | 24:24:24:24:2 | 42004640:42 | 42004761:42 | +:+:+:+:+:+:+: | 1650 | top1mt | 278 | 139 | 231 | 240 | 103 | 198 |
| ENSDARG000 | 24:24:24:24:2 | 211217:2112 | 211572:2115 | +:+:+:+:+:+:+: | 2336 | NA | 2 | 0 | 0 | 0 | 4 | 0 |
| ENSDARG000 | 24:24:24:24:2 | 29758272:29 | 29758796:29 | -:-:-:-:-:-:-:-: | 2359 | NA | 52 | 0 | 0 | 0 | 0 | 2 |
| ENSDARG000 | 24;24;24;24 | 27434876:27 | 27434992:27 | -;-;-;- | 342 | NA | 0 | 0 | 0 | 0 | 0 | 0 |
| ENSDARG000 | 24:24:24:24:2 | 26005629:26 | 26005654:26 | -;-;-;-;-;-;- | 147 | NA | 3 | 3 | 4 | 3 | 9 | 0 |
| ENSDARG000 | 24 | 994016 | 994130 | - | 115 | 5S_rRNA | 1 | 0 | 4 | 0 | 0 | 2 |
| ENSDARG000 | 24 | 10817960 | 10818085 | + | 126 | 5S_rRNA | 0 | 0 | 0 | 0 | 0 | 0 |
| ENSDARG000 | 24 | 15412989 | 15413105 | + | 117 | 5S_rRNA | 0 | 0 | 0 | 0 | 0 | 0 |
| ENSDARG000 | 24 | 18369702 | 18369817 | + | 116 | 5S_rRNA | 0 | 0 | 0 | 0 | 0 | 0 |
| ENSDARG000 | 24 | 527269 | 527383 | - | 115 | 5S_rRNA | 0 | 0 | 0 | 0 | 0 | 0 |
| ENSDARG000 | 24 | 30352608 | 30352735 | + | 128 | dre-mir-137-2 | 0 | 0 | 0 | 0 | 0 | 0 |
| ENSDARG000 | 24;24;24;24 | 36090794:36 | 36090934:36 | -;-;-;- | 408 | NA | 306 | 187 | 189 | 122 | 284 | 233 |
| ENSDARG000 | 24;24;24;24 | 35522301:35 | 35524089:35 | -;-;-;- | 2102 | ube2v2 | 3829 | 5267 | 5454 | 4035 | 4732 | 4538 |
| ENSDARG000 | 24:24:24:24:2 | 12970119:12 | 12970128:12 | -;-;-;-;-;- | 147 | NA | 0 | 1 | 7 | 7 | 4 | 7 |
| ENSDARG000 | 24;24;24;24 | 12835935:12 | 12836607:12 | +;+;+;+ | 2959 | nanog | 0 | 0 | 0 | 0 | 0 | 0 |
| ENSDARG000 | 24;24;24 | 38223618:38 | 38223666:38 | +;+;+ | 360 | NA | 0 | 0 | 0 | 0 | 4 | 0 |
| ENSDARG000 | 24:24:24:24:2 | 1845689:186 | 1845708:186 | -:-:-:-:-:-:-:-: | 1593 | NA | 22 | 2 | 26 | 10 | 8 | 4 |
| ENSDARG000 | 24 | 29103393 | 29103509 | + | 117 | 5S_rRNA | 0 | 0 | 0 | 0 | 0 | 0 |
| ENSDARG000 | 24:24:24:24:2 | 30521619:30 | 30521962:30 | +:+:+:+:+:+:+: | 2288 | DPYD | 490 | 371 | 625 | 468 | 512 | 620 |
| ENSDARG000 | 24:24:24:24:2 | 37598441:37 | 37598870:37 | +;+;+;+;+;+;+ | 1426 | rhbdl1 | 27 | 14 | 26 | 37 | 19 | 17 |
| ENSDARG000 | 24 | 40648256 | 40648372 | + | 117 | 5S_rRNA | 52 | 12 | 25 | 14 | 13 | 27 |
| ENSDARG000 | 24:24:24:24:2 | 32035759:32 | 32035773:32 | -:-:-:-:-:-:-:-: | 10878 | CUBN | 32 | 22 | 207 | 88 | 336 | 14 |
| ENSDARG000 | 24:24:24:24:2 | 26277390:26 | 26281476:26 | -;-;-;-;-;-;-;- | 4494 | and1 | 143 | 50 | 60 | 55 | 66 | 26 |
| ENSDARG000 | 24 | 11614257 | 11614373 | + | 117 | 5S_rRNA | 0 | 0 | 0 | 0 | 0 | 0 |
| ENSDARG000 | 24:24:24:24:2 | 9003998:900 | 9004133:900 | +:+:+:+:+:+:+: | 1840 | NA | 88 | 26 | 161 | 16 | 88 | 109 |
| ENSDARG000 | 24:24:24:24:2 | 7708652:770 | 7708711:770 | +;+;+;+;+ | 3539 | NA | 22 | 8 | 24 | 30 | 14 | 2 |
| ENSDARG000 | 24;24;24;24 | 42074143:42 | 42074200:42 | +;+;+;+ | 450 | NA | 681 | 559 | 921 | 577 | 709 | 453 |
| ENSDARG000 | 24 | 20470954 | 20471090 | - | 137 | dre-mir-26a-3 | 0 | 0 | 0 | 0 | 0 | 0 |
| ENSDARG000 | 24 | 23089389 | 23089505 | + | 117 | 5S_rRNA | 0 | 0 | 0 | 0 | 0 | 0 |
| ENSDARG000 | 24:24:24:24:2 | 1673250:170 | 1673780:170 | -:-:-:-:-:-:-:-: | 3724 | NA | 6 | 0 | 11 | 4 | 6 | 0 |
| ENSDARG000 | 24:24:24:24:2 | 3816698:381 | 3816947:381 | -;-;-;-;-;- | 808 | NA | 0 | 2 | 0 | 0 | 0 | 9 |
| ENSDARG000 | 24:24:24:24:2 | 3867036:387 | 3867690:387 | -;-;-;-;-;-;- | 1513 | NA | 0 | 0 | 0 | 0 | 0 | 1 |
| ENSDARG000 | 24;24;24 | 17318521:17 | 17318850:17 | +;+;+ | 1116 | NA | 0 | 0 | 0 | 0 | 0 | 0 |
| ENSDARG000 | 24;24;24 | 22969054:22 | 22969134:22 | -;-;- | 398 | NA | 0 | 4 | 0 | 0 | 0 | 0 |
| ENSDARG000 | 24;24;24;24 | 23024880:23 | 23026122:23 | -;-;- | 2099 | NA | 0 | 0 | 0 | 0 | 0 | 0 |
| ENSDARG000 | 24:24:24:24:2 | 23113614:23 | 23114036:23 | -;-;-;-;-;- | 1325 | NA | 0 | 0 | 0 | 0 | 0 | 0 |
| ENSDARG000 | 24:24:24:24:2 | 23589745:23 | 23589771:23 | -;-;-;-;-;-;- | 1381 | NA | 0 | 0 | 0 | 0 | 0 | 0 |
| ENSDARG000 | 24:24:24:24:2 | 27391512:27 | 27392732:27 | +;+;+;+;+;+;+ | 1670 | NA | 0 | 0 | 0 | 0 | 0 | 4 |
| ENSDARG000 | 24:24:24:24:2 | 29140066:29 | 29140347:29 | +;+;+;+;+ | 903 | NA | 0 | 0 | 0 | 0 | 0 | 0 |
| ENSDARG000 | 24;24;24 | 31055418:31 | 31055548:31 | -;-;- | 750 | NA | 0 | 0 | 0 | 0 | 0 | 0 |
| ENSDARG000 | 24:24:24:24:2 | 31846895:31 | 31846920:31 | +:+:+:+:+:+:+: | 2154 | NA | 0 | 0 | 0 | 0 | 0 | 0 |
| ENSDARG000 | 24;24;24 | 37012723:37 | 37013045:37 | +;+;+ | 927 | NA | 0 | 0 | 0 | 0 | 0 | 0 |
| ENSDARG000 | 22:22:22:22:2 | 1092479:110 | 1093223:110 | +;+;+;+;+;+;+ | 5202 | guca1e | 2 | 0 | 2 | 0 | 0 | 0 |
| ENSDARG000 | 22:22:22:22:2 | 5923190:592 | 5923965:592 | -;-;-;-;-;-;-;- | 2142 | NA | 748 | 1092 | 676 | 791 | 1124 | 912 |

| | | | | | | | | | | | | |
|---|---|---|---|---|---|---|---|---|---|---|---|---|
| ENSDARG000 | 22:22:22:22:2 | 12161738:121 | 12162664:121 | +:+:+:+:+:+:+: | 4340 | ccnt2b | 1190 | 1342 | 2394 | 1149 | 1667 | 1325 |
| ENSDARG000 | 22;22 | 1110855:112( | 1111083:112( | +;+ | 800 | NA | 0 | 0 | 0 | 0 | 0 | 0 |
| ENSDARG000 | 22:22:22:22:2 | 4524367:4526 | 4524741:4526 | -:-:-:-:-:-:-:-: | 8805 | fbn2b | 10848 | 2464 | 15221 | 1984 | 12857 | 1165 |
| ENSDARG000 | 22;22;22 | 14275823:142 | 14276351:142 | -;-;- | 686 | NA | 0 | 0 | 0 | 0 | 0 | 0 |
| ENSDARG000 | 22:22:22:22:2 | 1123110:1123 | 1123477:1123 | +;+;+;+;+ | 815 | NA | 4 | 0 | 0 | 0 | 0 | 1 |
| ENSDARG000 | 22:22:22:22:2 | 2364835:2367 | 2365234:237( | +:+:+:+:+:+:+: | 3255 | NA | 69 | 48 | 79 | 58 | 64 | 73 |
| ENSDARG000 | 22;22;22 | 23969080:239 | 23969140:239 | -;-;- | 234 | NA | 0 | 0 | 0 | 0 | 0 | 0 |
| ENSDARG000 | 22:22:22:22:2 | 2368904:2384 | 2368929:2384 | +:+:+:+:+:+:+: | 1356 | si:dkev-4c15.: | 100 | 82 | 57 | 38 | 62 | 78 |
| ENSDARG000 | 22;22 | 24157807:241 | 24158457:241 | +;+ | 2909 | b3galt2 | 234 | 354 | 246 | 252 | 346 | 292 |
| ENSDARG000 | 22;22;22 | 14304677:143 | 14305046:143 | -;-;- | 911 | NA | 0 | 0 | 0 | 0 | 0 | 0 |
| ENSDARG000 | 22;22;22;22 | 14338178:143 | 14338239:143 | -;-;-;- | 913 | NA | 0 | 0 | 0 | 0 | 0 | 12 |
| ENSDARG000 | 22;22 | 14350839:143 | 14351091:143 | -;- | 724 | NA | 0 | 0 | 0 | 0 | 0 | 0 |
| ENSDARG000 | 22:22:22:22:2 | 5940398:5942 | 5940441:5942 | -;-;-;-;-;- | 1237 | NA | 910 | 847 | 953 | 512 | 1258 | 867 |
| ENSDARG000 | 22:22:22:22:2 | 38001162:380( | 38001541:380( | -:-:-:-:-:-:-:-: | 3691 | NA | 107 | 136 | 163 | 107 | 185 | 210 |
| ENSDARG000 | 22:22:22:22:2 | 4707663:4711 | 4707947:4711 | +:+:+:+:+:+:+: | 1869 | cers4a | 24 | 18 | 27 | 16 | 30 | 2 |
| ENSDARG000 | 22;22;22;22 | 29911326:299 | 29912026:299 | -;-;-;- | 1522 | dusp5 | 1243 | 427 | 633 | 906 | 354 | 627 |
| ENSDARG000 | 22:22:22:22:2 | 1137997:1138 | 1138214:1138 | +:+:+:+:+:+:+: | 676 | GUCA1A | 0 | 2 | 0 | 0 | 0 | 4 |
| ENSDARG000 | 22:22:22:22:2 | 24214665:242 | 24214745:242 | +:+:+:+:+:+:+: | 1260 | glrx2 | 134 | 114 | 128 | 121 | 131 | 50 |
| ENSDARG000 | 22:22:22:22:2 | 158112:16054 | 158624:16066 | -:-:-:-:-:-:-:-: | 1428 | fblim1 | 40 | 24 | 8 | 48 | 26 | 2 |
| ENSDARG000 | 22:22:22:22:2 | 16293071:162 | 16293236:162 | +:+:+:+:+:+:+: | 2374 | dbt | 1080 | 831 | 1169 | 863 | 1309 | 1120 |
| ENSDARG000 | 22:22:22:22:2 | 35131890:351 | 35132029:351 | +:+:+:+:+:+:+: | 4818 | rnf13 | 6416 | 3123 | 1712 | 3529 | 2841 | 5825 |
| ENSDARG000 | 22:22:22:22:2 | 16314902:163 | 16314985:163 | -;-;-;-;- | 451 | NA | 0 | 0 | 0 | 0 | 0 | 0 |
| ENSDARG000 | 22:22:22:22:2 | 16311740:163 | 16312429:163 | -:-:-:-:-:-:-:-: | 2023 | trmt13 | 109 | 82 | 93 | 54 | 90 | 76 |
| ENSDARG000 | 22;22 | 10658018:106 | 10658295:106 | -;- | 499 | NA | 0 | 0 | 0 | 0 | 0 | 0 |
| ENSDARG000 | 22:22:22:22:2 | 14445713:144 | 14445870:144 | -:-:-:-:-:-:-:-: | 10386 | LRP1B | 8 | 2 | 2 | 4 | 58 | 10 |
| ENSDARG000 | 22:22:22:22:2 | 10645088:106 | 10645422:106 | +:+:+:+:+:+:+: | 2448 | rassf1 | 590 | 1041 | 1084 | 552 | 1450 | 807 |
| ENSDARG000 | 22:22:22:22:2 | 9472814:9472 | 9472894:9472 | +:+:+:+:+:+:+: | 3680 | CACNA2D2 | 4 | 26 | 24 | 28 | 0 | 38 |
| ENSDARG000 | 22:22:22:22:2 | 16308450:163 | 16308670:163 | +:+:+:+:+:+:+: | 1522 | lrrc39 | 11 | 65 | 12 | 16 | 114 | 42 |
| ENSDARG000 | 22:22:22:22:2 | 10660140:106 | 10660289:106 | +;+;+;+;+;+;+ | 1346 | tusc2b | 186 | 249 | 257 | 174 | 358 | 186 |
| ENSDARG000 | 22;22 | 6044172:6045 | 6044778:6046 | -;- | 1283 | NA | 12 | 0 | 0 | 3 | 0 | 1 |
| ENSDARG000 | 22;22;22;22 | 38930725:389 | 38930977:389 | -;-;-;- | 922 | ncbp2 | 337 | 413 | 410 | 224 | 306 | 202 |
| ENSDARG000 | 22:22:22:22:2 | 341990:34233 | 342743:34274 | -:-:-:-:-:-:-:-: | 1873 | necap2 | 638 | 676 | 904 | 475 | 824 | 466 |
| ENSDARG000 | 22:22:22:22:2 | 25066110:25( | 25067113:25( | -:-:-:-:-:-:-:-: | 3627 | kat14 | 530 | 471 | 638 | 375 | 646 | 444 |
| ENSDARG000 | 22;22 | 1158069:1159 | 1158351:1159 | -;- | 642 | NA | 0 | 0 | 0 | 0 | 0 | 2 |
| ENSDARG000 | 22:22:22:22:2 | 35179575:351 | 35181938:351 | -;-;-;-;-;- | 3249 | pfn2b | 19223 | 23090 | 17108 | 22197 | 17159 | 21442 |
| ENSDARG000 | 22:22:22:22:2 | 2390544:239( | 2390620:239( | +;+;+;+;+;+ | 612 | NA | 6 | 0 | 17 | 0 | 6 | 16 |
| ENSDARG000 | 22;22 | 6065987:6066 | 6066262:6066 | -;- | 389 | NA | 0 | 0 | 2 | 0 | 13 | 0 |
| ENSDARG000 | 22;22;22 | 349121:35136 | 349535:35157 | -;-;- | 1327 | TMEM240 | 0 | 0 | 0 | 0 | 0 | 0 |
| ENSDARG000 | 22:22:22:22:2 | 14489758:144 | 14490060:144 | +:+:+:+:+:+:+: | 2722 | NA | 4 | 8 | 4 | 0 | 4 | 2 |
| ENSDARG000 | 22:22:22:22:2 | 14363952:143 | 14364092:143 | -:-:-:-:-:-:-:-: | 4937 | NA | 22 | 14 | 13 | 19 | 17 | 17 |
| ENSDARG000 | 22:22:22:22:2 | 2403068:2403 | 2403109:2403 | +;+;+;+;+;+ | 733 | zgc:112977 | 25 | 32 | 33 | 27 | 2 | 48 |
| ENSDARG000 | 22:22:22:22:2 | 4745105:475( | 4747992:475( | -:-:-:-:-:-:-:-: | 4049 | rad23ab | 1075 | 862 | 917 | 671 | 1139 | 741 |
| ENSDARG000 | 22;22;22 | 1164656:1167 | 1165099:1167 | -;-;- | 538 | NA | 0 | 0 | 0 | 0 | 0 | 0 |
| ENSDARG000 | 22:22:22:22:2 | 19453822:194 | 19453902:194 | +;+;+;+;+ | 844 | NA | 9 | 8 | 0 | 22 | 24 | 2 |
| ENSDARG000 | 22:22:22:22:2 | 359273:36027 | 359705:36039 | -;-;-;-;- | 985 | ssu72 | 842 | 832 | 848 | 635 | 817 | 721 |
| ENSDARG000 | 22:22:22:22:2 | 29990448:299 | 29990524:299 | +:+:+:+:+:+:+: | 1080 | si:dkev-286i15 | 551 | 272 | 382 | 394 | 395 | 329 |
| ENSDARG000 | 22;22 | 370915:3725( | 372348:37272 | -;- | 1653 | FNDC10 | 4 | 6 | 16 | 2 | 10 | 12 |
| ENSDARG000 | 22;22;22;22 | 6096303:6098 | 6096378:6098 | +;+;+;+ | 1071 | zgc:171887 | 11 | 9 | 2 | 3 | 2 | 18 |
| ENSDARG000 | 22:22:22:22:2 | 6073628:6074 | 6074156:6075 | -:-:-:-:-:-:-:-: | 2130 | si:dkev-19a16 | 100 | 109 | 147 | 130 | 272 | 83 |
| ENSDARG000 | 22:22:22:22:2 | 26839241:268 | 26840121:268 | -;-;-;-;-;- | 2135 | gde1 | 58 | 108 | 119 | 111 | 165 | 125 |
| ENSDARG000 | 22:22:22:22:2 | 374478:37447 | 374565:37456 | +;+;+;+;+ | 1036 | ora4 | 0 | 6 | 0 | 0 | 0 | 0 |
| ENSDARG000 | 22;22 | 6084771:6085 | 6084819:6085 | +;+ | 399 | NA | 0 | 0 | 0 | 0 | 0 | 0 |
| ENSDARG000 | 22;22 | 6085307:6087 | 6085579:6087 | -;- | 475 | NA | 0 | 0 | 0 | 0 | 0 | 0 |
| ENSDARG000 | 22:22:22:22:2 | 26822332:268 | 26822710:268 | -;-;-;-;- | 1187 | mcrip2 | 18 | 85 | 144 | 64 | 47 | 112 |
| ENSDARG000 | 22:22:22:22:2 | 1170294:117( | 1170515:117( | +:+:+:+:+:+:+: | 2751 | irf6 | 70 | 2 | 4 | 22 | 24 | 4 |
| ENSDARG000 | 22;22 | 26813157:268 | 26813369:268 | +;+ | 406 | NA | 10 | 2 | 4 | 0 | 4 | 4 |
| ENSDARG000 | 22;22 | 26832849:268 | 26833307:268 | +;+ | 648 | NA | 0 | 0 | 0 | 0 | 0 | 0 |
| ENSDARG000 | 22:22:22:22:2 | 26798853:267 | 26799122:267 | +:+:+:+:+:+:+: | 5597 | NA | 166 | 262 | 343 | 87 | 304 | 150 |
| ENSDARG000 | 22;22;22 | 26804197:268 | 26804367:268 | +;+;+ | 505 | NA | 0 | 0 | 0 | 0 | 0 | 0 |
| ENSDARG000 | 22:22:22:22:2 | 35205017:352 | 35206891:352 | +:+:+:+:+:+:+: | 4683 | tsc22d2 | 698 | 1341 | 1284 | 1713 | 1021 | 1185 |
| ENSDARG000 | 22;22;22 | 465269:46545 | 465313:46565 | +;+;+ | 728 | NA | 32 | 21 | 68 | 70 | 69 | 31 |
| ENSDARG000 | 22:22:22:22:2 | 2409175:2411 | 2409232:2411 | +;+;+;+;+;+ | 1739 | zgc:113220 | 94 | 140 | 141 | 33 | 139 | 117 |
| ENSDARG000 | 22:22:22:22:2 | 413349:41373 | 413720:41685 | +:+:+:+:+:+:+: | 8877 | celsr2 | 419 | 334 | 854 | 573 | 766 | 415 |
| ENSDARG000 | 22:22:22:22:2 | 24220937:242 | 24221090:242 | +:+:+:+:+:+:+: | 1390 | uchl5 | 712 | 862 | 987 | 399 | 812 | 592 |
| ENSDARG000 | 22:22:22:22:2 | 38935060:389 | 38935164:389 | +:+:+:+:+:+:+: | 2694 | SIRT7 | 37 | 43 | 26 | 30 | 17 | 15 |
| ENSDARG000 | 22:22:22:22:2 | 19473804:194 | 19473890:194 | +:+:+:+:+:+:+: | 797 | NA | 6 | 2 | 4 | 0 | 6 | 8 |
| ENSDARG000 | 22:22:22:22:2 | 12217920:122 | 12219367:122 | -:-:-:-:-:-:-:-: | 5146 | ZRANB3 | 16 | 172 | 166 | 80 | 186 | 132 |
| ENSDARG000 | 22:22:22:22:2 | 38944615:389 | 38946943:389 | -:-:-:-:-:-:-:-: | 3423 | rbbp9 | 441 | 738 | 733 | 404 | 658 | 619 |
| ENSDARG000 | 22:22:22:22:2 | 9522971:9523 | 9523630:9523 | +:+:+:+:+:+:+: | 3320 | strip1 | 1116 | 966 | 1010 | 594 | 1196 | 866 |
| ENSDARG000 | 22;22;22 | 2574765:2576 | 2574949:2577 | +;+;+ | 857 | NA | 30 | 36 | 71 | 19 | 20 | 18 |
| ENSDARG000 | 22:22:22:22:2 | 30009926:30( | 30009992:30( | +:+:+:+:+:+:+: | 2630 | si:dkev-286i15 | 883 | 921 | 1402 | 719 | 1258 | 843 |
| ENSDARG000 | 22;22;22 | 2751887:2757 | 2751946:2757 | +;+;+ | 703 | NA | 15 | 6 | 8 | 14 | 9 | 14 |
| ENSDARG000 | 22:22:22:22:2 | 2844865:2845 | 2845082:2845 | +;+;+;+;+ | 1484 | NA | 12 | 8 | 40 | 18 | 30 | 17 |
| ENSDARG000 | 22;22 | 25136351:251 | 25136455:251 | +;+ | 141 | NA | 0 | 0 | 0 | 0 | 0 | 0 |
| ENSDARG000 | 22;22 | 2504524:2504 | 2504685:2505 | +;+ | 493 | NA | 28 | 22 | 19 | 4 | 35 | 36 |
| ENSDARG000 | 22:22:22:22:2 | 25086567:25( | 25086991:25( | +:+:+:+:+:+:+: | 5883 | rrbp1b | 273 | 385 | 2819 | 464 | 1430 | 293 |
| ENSDARG000 | 22:22:22:22:2 | 2685883:2686 | 2686063:2686 | +;+;+;+;+ | 1459 | NA | 36 | 16 | 14 | 22 | 13 | 42 |
| ENSDARG000 | 22;22;22 | 2769236:2769 | 2769285:277( | +;+;+ | 623 | NA | 21 | 11 | 36 | 40 | 33 | 13 |
| ENSDARG000 | 22;22;22 | 2819613:282( | 2819831:282( | +;+;+ | 915 | NA | 14 | 19 | 29 | 1 | 27 | 8 |
| ENSDARG000 | 22;22;22 | 2778600:2779 | 2778669:2779 | +;+;+ | 672 | NA | 9 | 16 | 16 | 8 | 10 | 26 |
| ENSDARG000 | 22:22:22:22:2 | 2660505:266( | 2660745:266( | +:+:+:+:+:+:+: | 1631 | znf1162 | 30 | 15 | 16 | 16 | 19 | 39 |
| ENSDARG000 | 22:22:22:22:2 | 38037530:38( | 38037699:38( | +;+;+;+;+ | 914 | commd2 | 575 | 543 | 686 | 677 | 608 | 619 |
| ENSDARG000 | 22:22:22:22:2 | 2860260:2861 | 2860275:2861 | +;+;+;+;+;+;+ | 1417 | NA | 67 | 80 | 96 | 88 | 68 | 78 |
| ENSDARG000 | 22:22:22:22:2 | 2510828:251( | 2510904:2511 | +:+:+:+:+:+:+: | 2095 | zgc:173726 | 56 | 50 | 140 | 48 | 124 | 97 |
| ENSDARG000 | 22:22:22:22:2 | 2549044:2549 | 2549098:2549 | +:+:+:+:+:+:+: | 1851 | zgc:175284 | 142 | 55 | 69 | 96 | 140 | 111 |
| ENSDARG000 | 22;22 | 2709467:2712 | 2709539:2712 | +;+ | 645 | NA | 53 | 29 | 74 | 49 | 34 | 61 |
| ENSDARG000 | 22;22;22;22 | 2801464:2804 | 2801532:2804 | +;+;+;+ | 899 | NA | 14 | 21 | 20 | 24 | 5 | 24 |
| ENSDARG000 | 22;22;22;22 | 2715815:2716 | 2715840:2716 | +;+;+;+ | 880 | NA | 28 | 15 | 29 | 22 | 22 | 4 |
| ENSDARG000 | 22;22;22;22 | 2533514:2535 | 2533557:2535 | +;+;+;+ | 761 | NA | 7 | 23 | 19 | 3 | 16 | 32 |
| ENSDARG000 | 22:22:22:22:2 | 2830703:283( | 2830886:283( | +:+:+:+:+:+:+: | 2280 | znf983 | 52 | 51 | 50 | 40 | 72 | 195 |
| ENSDARG000 | 22;22 | 2793382:2795 | 2793559:2795 | +;+ | 625 | NA | 14 | 31 | 38 | 6 | 31 | 2 |
| ENSDARG000 | 22;22;22;22 | 2524615:2525 | 2524681:2525 | +;+;+;+ | 1644 | znf1003 | 38 | 4 | 34 | 30 | 26 | 40 |
| ENSDARG000 | 22:22:22:22:2 | 10673151:106 | 10673243:106 | +:+:+:+:+:+:+: | 5569 | HYAL2 | 827 | 605 | 545 | 370 | 2192 | 885 |
| ENSDARG000 | 22;22;22 | 10673169:106 | 10673206:106 | -;-;- | 394 | NA | 2 | 1 | 4 | 0 | 1 | 2 |
| ENSDARG000 | 22:22:22:22:2 | 35275206:352 | 35275502:352 | +:+:+:+:+:+:+: | 3133 | NA | 288 | 347 | 235 | 230 | 272 | 301 |
| ENSDARG000 | 22:22:22:22:2 | 38049130:38( | 38049811:38( | +;+;+;+;+;+ | 1782 | wwtr1 | 2395 | 643 | 1498 | 949 | 1181 | 924 |
| ENSDARG000 | 22:22:22:22:2 | 21672656:216 | 21672763:216 | -:-:-:-:-:-:-:-: | 3522 | NA | 242 | 80 | 250 | 160 | 308 | 233 |
| ENSDARG000 | 22:22:22:22:2 | 12361317:123 | 12361357:123 | +:+:+:+:+:+:+: | 6340 | NA | 764 | 583 | 1291 | 834 | 922 | 662 |
| ENSDARG000 | 22:22:22:22:2 | 19478140:194 | 19478226:194 | +:+:+:+:+:+:+: | 745 | NA | 0 | 0 | 5 | 0 | 19 | 0 |
| ENSDARG000 | 22:22:22:22:2 | 9632831:9632 | 9635378:9635 | -:-:-:-:-:-:-:-: | 12665 | NA | 42 | 21 | 12 | 6 | 10 | 2130 |
| ENSDARG000 | 22:22:22:22:2 | 10698549:106 | 10698719:106 | +;+;+;+;+ | 1361 | abhd14a | 127 | 86 | 150 | 83 | 174 | 186 |
| ENSDARG000 | 22:22:22:22:2 | 6104892:6112 | 6107696:6112 | -;-;-;-;-;- | 3798 | si:dkev-19a16 | 36 | 10 | 32 | 66 | 6 | 56 |
| ENSDARG000 | 22:22:22:22:2 | 1186866:1189 | 1186999:1189 | +:+:+:+:+:+:+: | 2383 | NA | 60 | 62 | 95 | 73 | 81 | 54 |
| ENSDARG000 | 22:22:22:22:2 | 9553680:9554 | 9553853:9554 | -:-:-:-:-:-:-:-: | 5550 | NA | 10 | 8 | 9 | 0 | 3 | 156 |
| ENSDARG000 | 22:22:22:22:2 | 9607677:961( | 9607850:961( | -:-:-:-:-:-:-:-: | 3536 | NA | 17 | 6 | 6 | 6 | 18 | 6 |
| ENSDARG000 | 22:22:22:22:2 | 19486711:194 | 19486791:194 | +:+:+:+:+:+:+: | 773 | NA | 4 | 0 | 0 | 2 | 32 | 14 |
| ENSDARG000 | 22:22:22:22:2 | 472327:47326 | 473680:47368 | -:-:-:-:-:-:-:-: | 3842 | dstyk | 5749 | 3791 | 4331 | 6104 | 3196 | 3010 |
| ENSDARG000 | 22;22 | 35425946:354 | 35426471:354 | -;- | 700 | si:ch211-276k | 10 | 49 | 2 | 0 | 0 | 2 |
| ENSDARG000 | 22;22 | 35431169:354 | 35431559:354 | -;- | 539 | NA | 22 | 90 | 0 | 0 | 0 | 10 |
| ENSDARG000 | 22:22:22:22:2 | 6149121:6152 | 6149439:6152 | +;+;+;+;+ | 1188 | NA | 37 | 11 | 80 | 13 | 12 | 9 |
| ENSDARG000 | 22:22:22:22:2 | 24246213:242 | 24246827:242 | -;-;-;-;-;-;- | 1299 | rgs2 | 66 | 66 | 74 | 44 | 132 | 124 |
| ENSDARG000 | 22:22:22:22:2 | 19494996:194 | 19495076:194 | +:+:+:+:+:+:+: | 871 | NA | 0 | 0 | 0 | 0 | 0 | 0 |
| ENSDARG000 | 22;22 | 35454382:354 | 35454595:354 | +;+ | 427 | NA | 0 | 0 | 0 | 0 | 0 | 0 |
| ENSDARG000 | 22:22:22:22:2 | 38127467:381 | 38127963:381 | -;-;-;-;-;-;-;- | 2044 | tm4sf4 | 4 | 0 | 5 | 0 | 0 | 0 |
| ENSDARG000 | 22:22:22:22:2 | 19504185:195 | 19504355:195 | -;-;-;-;-;-;- | 784 | NA | 2 | 2 | 12 | 4 | 6 | 0 |
| ENSDARG000 | 22:22:22:22:2 | 12431608:124 | 12431940:124 | +;+;+;+;+ | 4501 | rnd3a | 268 | 138 | 419 | 369 | 291 | 382 |
| ENSDARG000 | 22:22:22:22:2 | 26853254:268 | 26858362:268 | -:-:-:-:-:-:-:-: | 6194 | hmox2a | 303 | 123 | 385 | 323 | 290 | 402 |
| ENSDARG000 | 22:22:22:22:2 | 16320076:163 | 16320360:163 | +:+:+:+:+:+:+: | 3299 | OSBPL1A | 1736 | 1707 | 1100 | 1639 | 1663 | 2132 |
| ENSDARG000 | 22;22;22;22 | 6170371:617( | 6170673:6171 | -;-;-;- | 926 | NA | 0 | 0 | 0 | 0 | 0 | 0 |
| ENSDARG000 | 22;22 | 24250187:242 | 24250350:242 | -;- | 297 | NA | 0 | 0 | 0 | 0 | 0 | 0 |
| ENSDARG000 | 22;22 | 12557663:125 | 12557736:125 | +;+ | 292 | NA | 0 | 0 | 0 | 0 | 0 | 0 |
| ENSDARG000 | 22:22:22:22:2 | 35472653:354 | 35472792:354 | +:+:+:+:+:+:+: | 993 | dynlt2b | 72 | 28 | 50 | 22 | 70 | 52 |
| ENSDARG000 | 22:22:22:22:2 | 24264633:242 | 24264902:242 | -;-;-;-;-;-;-;- | 889 | RGS13 | 36 | 12 | 9 | 4 | 10 | 16 |
| ENSDARG000 | 22:22:22:22:2 | 38159823:381 | 38160043:381 | +;+;+;+;+;+;+ | 1508 | tm4sf18 | 13438 | 5240 | 6689 | 3338 | 11445 | 13343 |
| ENSDARG000 | 22:22:22:22:2 | 26866198:268 | 26866371:268 | +;+;+;+;+ | 750 | NA | 0 | 0 | 0 | 0 | 0 | 0 |
| ENSDARG000 | 22:22:22:22:2 | 6182981:6182 | 6187611:6187 | -:-:-:-:-:-:-:-: | 5650 | NA | 6142 | 2607 | 3185 | 1760 | 0 | 6134 |
| ENSDARG000 | 22:22:22:22:2 | 19517610:195 | 19517754:195 | +;+;+;+;+;+ | 799 | si:dkev-78l4.1 | 0 | 0 | 0 | 0 | 0 | 0 |
| ENSDARG000 | 22:22:22:22:2 | 38968246:389 | 38968318:389 | +:+:+:+:+:+:+: | 1732 | arhgef3 | 367 | 220 | 236 | 147 | 297 | 171 |
| ENSDARG000 | 22:22:22:22:2 | 537717:53966 | 538066:53981 | -;-;-;-;-;-;-;- | 1257 | ccnd3 | 0 | 2 | 0 | 0 | 2 | 0 |
| ENSDARG000 | 22:22:22:22:2 | 508290:52049 | 508490:52061 | +;+;+;+;+;+ | 2611 | NUAK2 | 194 | 146 | 56 | 190 | 82 | 124 |
| ENSDARG000 | 22:22:22:22:2 | 4762478:4763 | 4762964:4763 | -;-;-;-;-;-;-;- | 1712 | calr3a | 3628 | 5725 | 8645 | 2215 | 5738 | 3348 |
| ENSDARG000 | 22:22:22:22:2 | 16350555:163 | 16352833:163 | -:-:-:-:-:-:-:-: | 4309 | ttc39c | 389 | 363 | 223 | 303 | 371 | 369 |
| ENSDARG000 | 22:22:22:22:2 | 9782369:9783 | 9783088:9783 | +:+:+:+:+:+:+: | 1725 | NA | 40 | 107 | 30 | 154 | 243 | 377 |
| ENSDARG000 | 22:22:22:22:2 | 24280772:242 | 24281672:242 | -:-:-:-:-:-:-:-: | 1842 | NA | 0 | 0 | 0 | 0 | 0 | 0 |
| ENSDARG000 | 22:22:22:22:2 | 12595144:125 | 12595224:125 | +:+:+:+:+:+:+: | 1574 | mmadhca | 238 | 232 | 350 | 166 | 264 | 173 |
| ENSDARG000 | 22;22;22 | 6198742:6199 | 6199059:6199 | +;+;+ | 734 | NA | 0 | 0 | 0 | 0 | 0 | 0 |
| ENSDARG000 | 22:22:22:22:2 | 12604302:126 | 12604469:126 | -:-:-:-:-:-:-:-: | 2647 | cep70 | 71 | 166 | 119 | 47 | 170 | 78 |
| ENSDARG000 | 22:22:22:22:2 | 10713713:107 | 10713977:107 | +:+:+:+:+:+:+: | 3426 | mfsd14a1 | 1442 | 1380 | 1432 | 1083 | 1591 | 1562 |
| ENSDARG000 | 22:22:22:22:2 | 19522982:195 | 19523500:195 | +:+:+:+:+:+:+: | 1995 | NA | 0 | 0 | 0 | 0 | 3 | 0 |
| ENSDARG000 | 22:22:22:22:2 | 39054012:39( | 39054708:39( | +:+:+:+:+:+:+: | 6625 | IP6K1 | 269 | 326 | 336 | 265 | 388 | 275 |
| ENSDARG000 | 22;22 | 35485100:354 | 35485410:354 | -;- | 1080 | NMUR1 | 0 | 0 | 0 | 0 | 0 | 0 |
| ENSDARG000 | 22:22:22:22:2 | 30046365:30( | 30047333:30( | +:+:+:+:+:+:+: | 6148 | add3a | 8702 | 5703 | 4120 | 4695 | 6251 | 6712 |
| ENSDARG000 | 22:22:22:22:2 | 6211114:6211 | 6216011:6212 | -:-:-:-:-:-:-:-: | 7248 | NA | 2 | 2 | 0 | 0 | 0 | 17 |
| ENSDARG000 | 22:22:22:22:2 | 16382502:163 | 16383363:163 | -:-:-:-:-:-:-:-: | 6018 | NA | 0 | 0 | 0 | 0 | 16 | 0 |
| ENSDARG000 | 22:22:22:22:2 | 36547670:365 | 36547930:365 | +:+:+:+:+:+:+: | 5448 | NA | 116 | 50 | 88 | 108 | 126 | 34 |
| ENSDARG000 | 22:22:22:22:2 | 19547476:195 | 19547919:195 | -;-;-;-;-;- | 1298 | micos13 | 736 | 709 | 827 | 564 | 975 | 722 |
| ENSDARG000 | 22:22:22:22:2 | 550828:55209 | 551283:5522( | -;-;-;-;-;-;- | 1504 | bysl | 1359 | 1279 | 1976 | 716 | 1416 | 924 |
| ENSDARG000 | 22:22:22:22:2 | 9806867:9806 | 9806994:9806 | +;+;+;+;+;+;+ | 966 | NA | 0 | 153 | 0 | 66 | 240 | 139 |
| ENSDARG000 | 22:22:22:22:2 | 17611836:176 | 17612294:176 | -:-:-:-:-:-:-:-: | 4387 | ARHGAP45 | 469 | 401 | 250 | 595 | 560 | 577 |
| ENSDARG000 | 22;22 | 9824521:9826 | 9825000:9826 | -;- | 574 | NA | 10 | 32 | 17 | 32 | 28 | 40 |
| ENSDARG000 | 22:22:22:22:2 | 6254194:6254 | 6254288:6254 | +;+;+;+;+ | 859 | rnasel4 | 0 | 0 | 0 | 0 | 0 | 0 |
| ENSDARG000 | 22:22:22:22:2 | 17632821:176 | 17633131:176 | -;-;-;-;-;- | 1483 | NA | 0 | 0 | 0 | 0 | 0 | 0 |
| ENSDARG000 | 22;22;22 | 6273581:6273 | 6273683:6274 | -;-;- | 664 | NA | 0 | 0 | 1 | 0 | 9 | 2 |
| ENSDARG000 | 22:22:22:22:2 | 1205787:1205 | 1205999:1205 | -:-:-:-:-:-:-:-: | 2546 | AMPD2 | 460 | 397 | 625 | 434 | 605 | 336 |
| ENSDARG000 | 22;22;22;22 | 25140904:251 | 25140974:251 | +;+;+;+ | 338 | NA | 0 | 0 | 0 | 0 | 0 | 0 |
| ENSDARG000 | 22;22;22;22 | 26929554:269 | 26932074:269 | -;-;-;- | 2844 | cxcl12b | 102 | 4 | 4 | 0 | 10 | 36 |
| ENSDARG000 | 22;22;22;22 | 36656680:366 | 36656868:366 | +;+;+;+ | 2276 | b3gnt7 | 117 | 42 | 125 | 76 | 28 | 71 |
| ENSDARG000 | 22:22:22:22:2 | 27065286:27( | 27065634:27( | -:-:-:-:-:-:-:-: | 1904 | NA | 4 | 4 | 4 | 2 | 9 | 6 |
| ENSDARG000 | 22:22:22:22:2 | 9827288:9827 | 9832177:9832 | -:-:-:-:-:-:-:-: | 7916 | NA | 228 | 194 | 414 | 235 | 178 | 493 |
| ENSDARG000 | 22:22:22:22:2 | 16418688:164 | 16418968:164 | +:+:+:+:+:+:+: | 1658 | ankrd29 | 6 | 25 | 16 | 49 | 12 | 26 |
| ENSDARG000 | 22:22:22:22:2 | 9862243:9862 | 9862616:9862 | +:+:+:+:+:+:+: | 1998 | si:dkev-253d2 | 176 | 77 | 232 | 103 | 0 | 3084 |
| ENSDARG000 | 22:22:22:22:2 | 16406920:164 | 16407295:164 | -:-:-:-:-:-:-:-: | 2471 | NA | 0 | 0 | 2 | 15 | 4 | 20 |

| | | | | | | | | | | | | |
|---|---|---|---|---|---|---|---|---|---|---|---|---|
| ENSDARG000 | 22;22;22 | 27090136:270 | 27090239:270 | +;+;+ | 539 | NA | 0 | 0 | 4 | 0 | 2 | 0 |
| ENSDARG000 | 22;22;22;22 | 4868448:4870 | 4868760:4870 | -;-;-;- | 1288 | NA | 0 | 0 | 0 | 0 | 0 | 0 |
| ENSDARG000 | 22;22;22;22 | 4908913:4909 | 4909236:4909 | -;-;-;- | 1174 | NA | 0 | 0 | 0 | 0 | 0 | 4 |
| ENSDARG000 | 22;22;22;22;2 | 4819410:4827 | 4819913:4828 | -;-;-;-;-;- | 1638 | NA | 0 | 0 | 0 | 0 | 0 | 0 |
| ENSDARG000 | 22;22;22;22;2 | 4819410:4827 | 4819913:4828 | -;-;-;-;-;-;-; | 3224 | NA | 2 | 4 | 0 | 0 | 3 | 17 |
| ENSDARG000 | 22;22;22;22 | 4926777:4929 | 4926928:4929 | -;-;-;- | 1020 | NA | 0 | 2 | 8 | 0 | 9 | 11 |
| ENSDARG000 | 22;22;22;22;2 | 4884980:4885 | 4886837:4885 | -;-;-;-;-;-;-;- | 2626 | NA | 0 | 0 | 0 | 0 | 1 | 12 |
| ENSDARG000 | 22;22;22;22;2 | 9871238:9871 | 9871630:9871 | +;+;+;+;+;+;+; | 4679 | NA | 534 | 251 | 960 | 589 | 0 | 6413 |
| ENSDARG000 | 22;22;22;22;2 | 39096911:390 | 39097217:390 | +;+;+;+;+;+;+; | 1453 | lmcd1 | 37 | 87 | 124 | 149 | 157 | 48 |
| ENSDARG000 | 22;22;22;22;2 | 19552987:195 | 19553096:195 | +;+;+;+;+;+;+; | 1611 | hsd11b1la | 614 | 144 | 678 | 200 | 3111 | 145 |
| ENSDARG000 | 22;22;22;22 | 6284038:6285 | 6284855:6285 | -;-;-;- | 1659 | NA | 0 | 17 | 16 | 8 | 16 | 0 |
| ENSDARG000 | 22;22;22;22;2 | 39084107:390 | 39085027:390 | +;+;+;+;+;+;+; | 5066 | gmppb | 1079 | 1753 | 2003 | 1105 | 2004 | 952 |
| ENSDARG000 | 22;22;22;22;2 | 9886903:9887 | 9887466:9888 | -;-;-;-;-;- | 2079 | NA | 198 | 183 | 114 | 209 | 144 | 201 |
| ENSDARG000 | 22;22 | 1250239:1251 | 1250351:1251 | +;+ | 410 | NA | 0 | 0 | 0 | 0 | 0 | 0 |
| ENSDARG000 | 22;22;22;22;2 | 1249788:1259 | 1250351:1262 | +;+;+;+;+ | 4043 | NA | 16 | 19 | 8 | 3 | 8 | 0 |
| ENSDARG000 | 22;22 | 6293563:6293 | 6293638:6294 | +;+ | 957 | rnasel2 | 0 | 6 | 24 | 8 | 14 | 6 |
| ENSDARG000 | 22;22;22;22;2 | 567733:56950 | 567816:57040 | +;+;+;+;+;+;+; | 5353 | usp49 | 205 | 172 | 247 | 110 | 188 | 127 |
| ENSDARG000 | 22;22;22;22;2 | 558287:56077 | 558300:56092 | +;+;+;+;+;+;+ | 4527 | med20 | 154 | 262 | 246 | 126 | 266 | 89 |
| ENSDARG000 | 22;22;22;22;2 | 24287774:242 | 24288272:242 | -;-;-;-;-;-;-;- | 994 | NA | 94 | 2 | 0 | 14 | 2 | 2 |
| ENSDARG000 | 22;22;22;22;2 | 36673312:366 | 36674340:366 | -;-;-;-;-;-;-; | 3574 | ncl | 12341 | 12597 | 20982 | 10316 | 14068 | 11478 |
| ENSDARG000 | 22;22;22 | 9893111:9894 | 9893781:9894 | -;-;- | 880 | znf990 | 26 | 11 | 9 | 2 | 8 | 40 |
| ENSDARG000 | 22;22;22;22;2 | 21794037:217 | 21795009:217 | -;-;-;-;-;-;- | 1692 | tle5 | 497 | 344 | 494 | 301 | 469 | 267 |
| ENSDARG000 | 22;22;22 | 1276680:1277 | 1276737:1277 | +;+;+ | 820 | si:ch73-138e1 | 44 | 42 | 51 | 16 | 46 | 36 |
| ENSDARG000 | 22;22;22;22;2 | 25184459:251 | 25184496:251 | +;+;+;+;+;+;+; | 637 | si:ch211-226l | 0 | 0 | 0 | 0 | 0 | 0 |
| ENSDARG000 | 22;22;22;22;2 | 10735909:107 | 10737154:107 | -;-;-;-;-;-;-; | 3346 | sass6 | 38 | 88 | 108 | 10 | 66 | 114 |
| ENSDARG000 | 22;22;22;22;2 | 38173960:381 | 38174156:381 | +;+;+;+;+;+;+; | 3590 | cp | 79 | 26 | 204 | 142 | 67 | 98 |
| ENSDARG000 | 22;22;22 | 1285182:1285 | 1285273:1285 | +;+;+ | 967 | NA | 18 | 18 | 14 | 24 | 27 | 21 |
| ENSDARG000 | 22;22;22;22;2 | 21852070:218 | 21852303:218 | -;-;-;-;-;-;-; | 1328 | gna11a | 820 | 825 | 915 | 395 | 628 | 773 |
| ENSDARG000 | 22;22;22;22;2 | 9898028:9899 | 9899880:9899 | -;-;-;-;- | 2049 | NA | 136 | 153 | 169 | 120 | 120 | 190 |
| ENSDARG000 | 22;22 | 24306145:243 | 24307393:243 | -;- | 1272 | NA | 0 | 0 | 0 | 0 | 0 | 0 |
| ENSDARG000 | 22;22;22;22;2 | 6300311:6301 | 6300575:6301 | -;-;-;-;-;-;-; | 2661 | NA | 397 | 151 | 209 | 77 | 339 | 292 |
| ENSDARG000 | 22;22;22;22 | 4939559:4940 | 4939710:4940 | -;-;-;- | 1038 | NA | 0 | 0 | 0 | 0 | 0 | 2 |
| ENSDARG000 | 22;22;22;22;2 | 4947923:4952 | 4948177:4952 | +;+;+;+;+ | 1251 | NA | 30 | 15 | 36 | 16 | 36 | 23 |
| ENSDARG000 | 22;22;22 | 1291651:1292 | 1291747:1292 | +;+;+ | 844 | NA | 20 | 18 | 30 | 48 | 29 | 12 |
| ENSDARG000 | 22;22;22;22;2 | 25219153:252 | 25219175:252 | +;+;+;+;+ | 400 | NA | 0 | 0 | 0 | 13 | 0 | 0 |
| ENSDARG000 | 22;22;22;22;2 | 24389135:244 | 24389874:244 | +;+;+;+;+;+;+; | 5927 | P3H2 | 517 | 820 | 653 | 700 | 1060 | 1022 |
| ENSDARG000 | 22;22 | 9904253:9906 | 9904908:9906 | -;- | 793 | NA | 4 | 0 | 2 | 2 | 0 | 8 |
| ENSDARG000 | 22;22;22;22;2 | 591809:59295 | 592494:59309 | -;-;-;-;- | 1970 | tmcc2 | 12 | 8 | 6 | 0 | 2 | 0 |
| ENSDARG000 | 22;22;22;22;2 | 6407445:6408 | 6407821:6408 | -;-;-;-;- | 1292 | zgc:171699 | 0 | 12 | 2 | 4 | 0 | 10 |
| ENSDARG000 | 22;22 | 1300587:1306 | 1300659:1307 | +;+ | 476 | NA | 4 | 13 | 0 | 1 | 1 | 4 |
| ENSDARG000 | 22;22 | 6320862:6322 | 6321182:6322 | +;+ | 612 | NA | 6362 | 1734 | 2739 | 2187 | 1 | 4216 |
| ENSDARG000 | 22;22;22;22;2 | 6393227:6393 | 6393892:6394 | -;-;-;-;-;-;-; | 1879 | si:rp71-1i20.2 | 45 | 0 | 0 | 0 | 10 | 4 |
| ENSDARG000 | 22;22;22;22;2 | 1313046:1317 | 1313105:1317 | +;+;+;+;+ | 6315 | NA | 259 | 210 | 336 | 151 | 271 | 191 |
| ENSDARG000 | 22;22 | 1323514:1326 | 1323690:1329 | +;+ | 2890 | NA | 618 | 356 | 618 | 331 | 721 | 588 |
| ENSDARG000 | 22;22;22 | 9932767:9933 | 9933179:9934 | -;-;- | 635 | NA | 5 | 86 | 26 | 282 | 733 | 18 |
| ENSDARG000 | 22;22;22;22;2 | 9926090:9926 | 9927321:9927 | -;-;-;-;- | 1505 | si:dkev-253d2 | 29 | 67 | 23 | 320 | 444 | 72 |
| ENSDARG000 | 22;22;22;22;2 | 6321038:6382 | 6321182:6382 | +;+;+;+;+;+;+; | 2177 | NA | 0 | 4 | 1 | 1 | 0 | 7 |
| ENSDARG000 | 22;22;22;22;2 | 30330574:303 | 30330853:303 | +;+;+;+;+;+;+; | 4028 | mxi1 | 1743 | 1190 | 1568 | 2370 | 1494 | 1240 |
| ENSDARG000 | 22;22;22;22;2 | 6342781:6342 | 6343097:6343 | -;-;-;-;-;-;-; | 1470 | zgc:113298 | 4160 | 626 | 1604 | 1164 | 200 | 2436 |
| ENSDARG000 | 22;22 | 30331414:303 | 30331657:303 | +;+ | 1438 | NA | 0 | 0 | 1 | 1 | 0 | 16 |
| ENSDARG000 | 22;22;22;22;2 | 19640309:196 | 19640483:196 | +;+;+;+;+;+;+ | 2052 | rgmd | 490 | 448 | 849 | 600 | 907 | 412 |
| ENSDARG000 | 22;22 | 21933125:219 | 21933437:219 | -;- | 774 | NA | 0 | 0 | 0 | 0 | 0 | 0 |
| ENSDARG000 | 22;22;22;22;2 | 603382:60371 | 603944:60394 | -;-;-;-;-;- | 1219 | cdkn1a | 337 | 723 | 1512 | 272 | 2016 | 531 |
| ENSDARG000 | 22;22;22;22;2 | 21950838:219 | 21951021:219 | +;+;+;+;+;+;+ | 1125 | gna15.4 | 0 | 2 | 4 | 0 | 0 | 0 |
| ENSDARG000 | 22;22;22;22 | 21965148:219 | 21965248:219 | +;+;+;+ | 501 | NA | 2 | 0 | 0 | 1 | 0 | 0 |
| ENSDARG000 | 22;22;22;22;2 | 25236657:252 | 25236741:252 | +;+;+;+;+;+;+; | 733 | zgc:172218 | 0 | 0 | 0 | 0 | 0 | 0 |
| ENSDARG000 | 22;22 | 1377331:1383 | 1377445:1384 | +;+ | 256 | NA | 0 | 0 | 0 | 0 | 0 | 0 |
| ENSDARG000 | 22;22;22;22;2 | 19665617:196 | 19665764:196 | -;-;-;-;-;-;-; | 1699 | celf5a | 0 | 0 | 8 | 0 | 8 | 26 |
| ENSDARG000 | 22;22;22;22;2 | 27092469:270 | 27092765:270 | -;-;-;-;-;-;-; | 2371 | xpnpep1 | 1317 | 1267 | 1382 | 1083 | 1157 | 1337 |
| ENSDARG000 | 22;22;22;22;2 | 9946793:9948 | 9947172:9948 | -;-;-;-;-;-;-; | 4600 | NA | 174 | 205 | 232 | 355 | 1175 | 90 |
| ENSDARG000 | 22;22;22;22;2 | 1462177:1462 | 1462203:1463 | +;+;+;+;+;+;+ | 957 | si:dkeyp-53d3 | 90 | 62 | 111 | 69 | 63 | 96 |
| ENSDARG000 | 22;22;22;22;2 | 16432404:164 | 16434511:164 | -;-;-;-;-;-;-; | 3065 | plekhb2 | 432 | 326 | 519 | 430 | 382 | 462 |
| ENSDARG000 | 22;22;22;22;2 | 30446987:304 | 30447204:304 | +;+;+;+;+;+;+; | 1509 | si:dkev-169i5. | 14 | 0 | 6 | 3 | 5 | 25 |
| ENSDARG000 | 22;22;22;22;2 | 21972824:219 | 21973004:219 | +;+;+;+;+;+;+; | 4148 | NA | 1 | 0 | 0 | 2 | 8 | 2 |
| ENSDARG000 | 22;22 | 1469685:1471 | 1469763:1472 | +;+ | 374 | NA | 18 | 6 | 9 | 6 | 12 | 2 |
| ENSDARG000 | 22;22;22;22;2 | 1501566:1502 | 1501688:1502 | +;+;+;+;+;+;+; | 1658 | zgc:173486 | 14 | 25 | 17 | 9 | 45 | 16 |
| ENSDARG000 | 22 | 20104706 | 20105969 | - | 1264 | rxfp3.2b | 0 | 0 | 0 | 0 | 0 | 0 |
| ENSDARG000 | 22;22;22;22;2 | 1480988:1481 | 1481066:1481 | +;+;+;+;+ | 1945 | NA | 39 | 53 | 45 | 36 | 52 | 41 |
| ENSDARG000 | 22;22;22 | 30463165:304 | 30463379:304 | +;+;+ | 709 | NA | 10 | 11 | 105 | 36 | 39 | 63 |
| ENSDARG000 | 22;22 | 1508545:1508 | 1508838:1509 | +;+ | 408 | NA | 0 | 0 | 0 | 0 | 0 | 0 |
| ENSDARG000 | 22;22 | 1491664:1492 | 1491762:1493 | +;+ | 493 | NA | 11 | 10 | 30 | 4 | 10 | 6 |
| ENSDARG000 | 22;22;22;22;2 | 4962370:4962 | 4962697:4962 | -;-;-;-;-;-;-; | 3336 | cherp | 834 | 970 | 1663 | 663 | 1247 | 756 |
| ENSDARG000 | 22;22;22;22;2 | 20112715:201 | 20113805:201 | -;-;-;-;-;-;-; | 2158 | creb3l3a | 0 | 0 | 0 | 0 | 0 | 0 |
| ENSDARG000 | 22;22;22;22;2 | 10044823:100 | 10045276:100 | -;-;-;-;-;-;- | 2027 | zgc:174564 | 53 | 14 | 50 | 20 | 28 | 52 |
| ENSDARG000 | 22;22;22;22;2 | 10752511:107 | 10752569:107 | +;+;+;+;+;+;+ | 909 | lsm7 | 462 | 926 | 686 | 540 | 1080 | 586 |
| ENSDARG000 | 22;22;22;22;2 | 613265:61326 | 613688:61368 | -;-;-;-;-;-;-; | 2636 | srsf3b | 2721 | 2885 | 4562 | 2647 | 4059 | 3493 |
| ENSDARG000 | 22;22;22 | 24491627:244 | 24491724:244 | +;+;+ | 533 | NA | 0 | 0 | 0 | 0 | 0 | 0 |
| ENSDARG000 | 22;22;22;22;2 | 25242322:252 | 25242406:252 | +;+;+;+;+;+;+; | 714 | NA | 0 | 0 | 0 | 0 | 0 | 0 |
| ENSDARG000 | 22;22;22;22;2 | 36692891:366 | 36693347:366 | +;+;+;+;+;+ | 1427 | trim35-34 | 0 | 6 | 0 | 24 | 0 | 6 |
| ENSDARG000 | 22;22;22;22;2 | 36692681:366 | 36693347:366 | +;+;+;+;+;+;+; | 5460 | NA | 5 | 8 | 6 | 0 | 12 | 0 |
| ENSDARG000 | 22;22;22;22;2 | 22004082:220 | 22004151:220 | +;+;+;+;+;+;+; | 1459 | NA | 2 | 0 | 0 | 0 | 0 | 0 |
| ENSDARG000 | 22;22;22;22;2 | 10757848:107 | 10757887:107 | +;+;+;+;+;+;+; | 4081 | NA | 0 | 0 | 2 | 0 | 7 | 0 |
| ENSDARG000 | 22;22 | 10054336:100 | 10055028:100 | -;- | 790 | NA | 17 | 10 | 19 | 9 | 23 | 34 |
| ENSDARG000 | 22;22;22;22;2 | 24673168:246 | 24673295:246 | +;+;+;+;+ | 554 | NA | 0 | 0 | 0 | 0 | 0 | 0 |
| ENSDARG000 | 22;22;22;22;2 | 30480418:304 | 30480721:304 | -;-;-;-;- | 954 | NA | 0 | 0 | 40 | 8 | 2 | 11 |
| ENSDARG000 | 22;22;22;22;2 | 17657612:176 | 17658060:176 | -;-;-;-;-;-;-; | 4575 | tjp3 | 143 | 10 | 2 | 30 | 34 | 20 |
| ENSDARG000 | 22;22;22;22;2 | 10066846:100 | 10067176:100 | +;+;+;+;+;+;+; | 1436 | si:ch211-222l | 232 | 142 | 153 | 141 | 119 | 610 |
| ENSDARG000 | 22;22;22;22;2 | 20135443:201 | 20135479:201 | +;+;+;+;+;+;+; | 3825 | eef2a.1 | 442 | 629 | 856 | 509 | 738 | 626 |
| ENSDARG000 | 22;22;22;22 | 10076578:100 | 10077789:100 | -;-;-;- | 1491 | NA | 28 | 4 | 22 | 10 | 4 | 24 |
| ENSDARG000 | 22;22;22;22;2 | 10090673:100 | 10090823:100 | +;+;+;+;+;+;+; | 5707 | si:dkey-102c8 | 50 | 0 | 0 | 0 | 14 | 18 |
| ENSDARG000 | 22;22 | 30497494:304 | 30497584:304 | +;+ | 354 | NA | 0 | 0 | 0 | 0 | 0 | 2 |
| ENSDARG000 | 22;22 | 10085572:100 | 10085865:100 | -;- | 641 | NA | 70 | 32 | 97 | 28 | 11 | 71 |
| ENSDARG000 | 22;22;22 | 10771115:107 | 10771972:107 | -;-;- | 1178 | timm13 | 430 | 495 | 606 | 209 | 450 | 344 |
| ENSDARG000 | 22;22;22;22;2 | 1517318:1519 | 1517384:1519 | +;+;+;+;+ | 548 | NA | 53 | 28 | 41 | 22 | 28 | 61 |
| ENSDARG000 | 22;22;22;22;2 | 22021936:220 | 22022321:220 | +;+;+;+;+;+;+; | 2872 | gna15.1 | 59 | 2 | 18 | 0 | 0 | 20 |
| ENSDARG000 | 22;22;22;22;2 | 16446052:164 | 16446209:164 | +;+;+;+;+;+;+; | 5608 | NA | 28 | 6 | 30 | 14 | 34 | 8 |
| ENSDARG000 | 22;22;22;22;2 | 24811396:248 | 24811634:248 | -;-;-;-;-;-;-; | 4408 | vtg6 | 0 | 0 | 0 | 0 | 0 | 0 |
| ENSDARG000 | 22;22;22;22 | 2874838:2877 | 2875060:2877 | -;-;-;- | 994 | aqp12 | 3 | 0 | 12 | 0 | 0 | 6 |
| ENSDARG000 | 22;22;22;22;2 | 20145036:201 | 20145094:201 | +;+;+;+;+;+;+; | 3143 | eef2a.2 | 253 | 26 | 299 | 46 | 157 | 176 |
| ENSDARG000 | 22;22;22;22;2 | 24784013:247 | 24784266:247 | -;-;-;-;-;-;-; | 4453 | vtg4 | 0 | 0 | 0 | 0 | 0 | 2 |
| ENSDARG000 | 22;22;22;22;2 | 30502555:305 | 30502710:305 | +;+;+;+;+;+;+; | 2429 | NA | 2895 | 2224 | 1903 | 2803 | 2303 | 4157 |
| ENSDARG000 | 22;22;22;22;2 | 30517227:305 | 30517384:305 | +;+;+;+;+ | 2846 | NA | 130 | 35 | 70 | 273 | 53 | 193 |
| ENSDARG000 | 22;22;22 | 30676491:306 | 30676848:306 | -;-;- | 519 | NA | 0 | 0 | 0 | 0 | 0 | 0 |
| ENSDARG000 | 22;22;22 | 1576863:1577 | 1577204:1577 | +;+;+ | 642 | NA | 40 | 33 | 51 | 49 | 42 | 63 |
| ENSDARG000 | 22;22;22 | 641342:64406 | 641928:64459 | -;-;- | 1389 | apobec2a | 4 | 2 | 2 | 0 | 0 | 0 |
| ENSDARG000 | 22;22 | 1568306:1568 | 1568357:1569 | +;+ | 147 | NA | 13 | 10 | 8 | 10 | 8 | 15 |
| ENSDARG000 | 22;22;22;22;2 | 1539373:1540 | 1539424:1540 | +;+;+;+;+;+;+; | 8074 | NA | 253 | 114 | 286 | 302 | 233 | 267 |
| ENSDARG000 | 22;22;22;22;2 | 6434672:6438 | 6434883:6438 | -;-;-;-;-;- | 1396 | NA | 20 | 2 | 11 | 28 | 2 | 14 |
| ENSDARG000 | 22;22;22;22;2 | 1526040:1526 | 1526093:1526 | +;+;+;+;+;+ | 971 | NA | 67 | 53 | 93 | 39 | 54 | 35 |
| ENSDARG000 | 22;22 | 1556948:1557 | 1556984:1557 | +;+ | 109 | NA | 0 | 0 | 2 | 5 | 3 | 0 |
| ENSDARG000 | 22;22;22;22;2 | 25248961:252 | 25249046:252 | +;+;+;+;+;+;+; | 731 | si:ch211-226l | 0 | 0 | 0 | 0 | 0 | 0 |
| ENSDARG000 | 22;22 | 1586060:1586 | 1586163:1586 | +;+ | 184 | NA | 4 | 0 | 5 | 4 | 0 | 1 |
| ENSDARG000 | 22;22;22;22 | 16535575:165 | 16535654:165 | +;+;+;+ | 2643 | tal1 | 28 | 4 | 10 | 2 | 28 | 6 |
| ENSDARG000 | 22;22;22;22;2 | 24715282:247 | 24715428:247 | +;+;+;+;+;+;+; | 1629 | SSX2IP | 4 | 0 | 2 | 0 | 7 | 0 |
| ENSDARG000 | 22;22;22;22;2 | 24751101:247 | 24751339:247 | -;-;-;-;-;-;-; | 4386 | vtg5 | 0 | 2 | 0 | 0 | 10 | 0 |
| ENSDARG000 | 22;22;22;22;2 | 6456003:6457 | 6456468:6457 | -;-;-;-;-;- | 1736 | zgc:172133 | 36 | 5 | 0 | 16 | 4 | 34 |
| ENSDARG000 | 22;22;22;22;2 | 2895511:2897 | 2898204:2898 | -;-;-;-;-;-;-; | 4527 | pak2b | 1845 | 2058 | 2062 | 1560 | 1834 | 1674 |
| ENSDARG000 | 22;22;22;22;2 | 12641580:126 | 12645297:126 | -;-;-;-;-;-;-; | 6006 | adarb1a | 39 | 185 | 40 | 83 | 2 | 103 |
| ENSDARG000 | 22;22;22;22;2 | 22141040:221 | 22142132:221 | -;-;-;-;-;-;-;- | 1838 | cdc34a | 118 | 67 | 90 | 24 | 160 | 72 |
| ENSDARG000 | 22;22;22;22;2 | 24798458:247 | 24798570:247 | -;-;-;-;-;-;-; | 3655 | NA | 0 | 0 | 0 | 0 | 0 | 0 |
| ENSDARG000 | 22;22 | 6471584:6475 | 6471806:6475 | +;+ | 521 | NA | 0 | 0 | 0 | 2 | 0 | 1 |
| ENSDARG000 | 22;22;22;22;2 | 38202614:382 | 38202911:382 | -;-;-;-;-;-;-; | 3340 | hps3 | 3842 | 3702 | 4041 | 3949 | 3872 | 4650 |
| ENSDARG000 | 22;22;22;22;2 | 661505:66171 | 661623:66185 | +;+;+;+;+;+;+; | 2386 | ELF3 | 1068 | 9 | 55 | 66 | 84 | 62 |
| ENSDARG000 | 22;22;22;22;2 | 24730965:247 | 24731403:247 | -;-;-;-;-;-;-; | 4595 | vtg1 | 0 | 0 | 2 | 0 | 8 | 0 |
| ENSDARG000 | 22;22;22 | 14764998:147 | 14766596:147 | -;-;- | 1773 | NA | 2 | 0 | 2 | 0 | 0 | 0 |
| ENSDARG000 | 22;22;22;22;2 | 10110358:101 | 10113537:101 | -;-;-;-;-;-;-; | 4102 | rdh5 | 4 | 4 | 34 | 0 | 12 | 42 |
| ENSDARG000 | 22;22;22;22;2 | 10098446:100 | 10098971:100 | -;-;-;-;-;-;-; | 2244 | gls2b | 2 | 0 | 24 | 0 | 0 | 4 |
| ENSDARG000 | 22;22;22;22;2 | 24770744:247 | 24770828:247 | +;+;+;+;+;+;+; | 713 | NA | 6 | 0 | 0 | 0 | 0 | 0 |
| ENSDARG000 | 22;22;22;22;2 | 24851227:248 | 24851459:248 | -;-;-;-;-;-;-; | 4203 | vtg7 | 2 | 0 | 0 | 0 | 0 | 0 |
| ENSDARG000 | 22;22;22;22 | 16555939:165 | 16556349:165 | +;+;+;+ | 4276 | pdzk1ip1 | 38 | 16 | 129 | 117 | 383 | 76 |
| ENSDARG000 | 22;22;22;22;2 | 30811775:308 | 30813071:308 | -;-;-;-;-;-;-; | 3682 | NA | 206 | 18 | 58 | 12 | 32 | 176 |
| ENSDARG000 | 22;22;22;22;2 | 20159067:201 | 20160093:201 | -;-;-;-;-;-;-; | 2709 | btbd2a | 577 | 411 | 454 | 390 | 626 | 607 |
| ENSDARG000 | 22;22;22 | 10781894:107 | 10782084:107 | +;+;+ | 3381 | enc3 | 127 | 209 | 420 | 306 | 348 | 232 |
| ENSDARG000 | 22;22;22;22;2 | 30839702:308 | 30839788:308 | +;+;+;+;+;+;+; | 1301 | NA | 16 | 0 | 0 | 0 | 0 | 0 |
| ENSDARG000 | 22;22;22;22;2 | 20169352:201 | 20169441:201 | +;+;+;+;+;+;+; | 4815 | hmg20b | 610 | 568 | 723 | 594 | 820 | 703 |
| ENSDARG000 | 22;22;22;22;2 | 22260210:222 | 22260479:222 | +;+;+;+;+;+;+; | 2038 | ubxn6 | 598 | 453 | 658 | 496 | 502 | 609 |
| ENSDARG000 | 22;22;22;22;2 | 36719821:367 | 36720032:367 | +;+;+;+;+;+;+ | 1600 | NA | 33 | 43 | 84 | 41 | 104 | 73 |
| ENSDARG000 | 22;22;22;22;2 | 6487048:6487 | 6491424:6489 | -;-;-;-;-;-;-; | 5131 | NA | 0 | 56 | 108 | 140 | 56 | 590 |
| ENSDARG000 | 22;22;22;22;2 | 38229321:382 | 38229589:382 | +;+;+;+;+;+;+; | 1444 | si:ch211-284e | 0 | 3 | 19 | 15 | 0 | 2 |
| ENSDARG000 | 22;22;22;22;2 | 20195280:201 | 20195349:201 | +;+;+;+;+;+;+; | 5563 | unc13a | 33 | 242 | 122 | 92 | 1350 | 126 |
| ENSDARG000 | 22;22;22;22 | 38239302:382 | 38239663:382 | -;-;-;- | 635 | NA | 0 | 0 | 1 | 0 | 0 | 0 |
| ENSDARG000 | 22;22 | 6515397:6519 | 6515472:6520 | -;- | 227 | NA | 0 | 0 | 0 | 0 | 0 | 0 |
| ENSDARG000 | 22;22;22;22;2 | 696931:69702 | 697512:69751 | +;+;+;+;+ | 1485 | GPR37L1 | 1 | 0 | 0 | 0 | 0 | 2 |
| ENSDARG000 | 22;22;22;22;2 | 25274712:252 | 25274797:252 | +;+;+;+;+;+ | 708 | NA | 0 | 0 | 0 | 0 | 0 | 0 |
| ENSDARG000 | 22;22;22;22;2 | 25289360:252 | 25289445:252 | +;+;+;+;+;+ | 586 | zgc:171717 | 0 | 0 | 0 | 6 | 0 | 0 |
| ENSDARG000 | 22;22;22;22;2 | 20246587:202 | 20246730:202 | -;-;-;-;-;-;-; | 9191 | mbd3b | 373 | 230 | 411 | 166 | 616 | 224 |
| ENSDARG000 | 22;22 | 6527414:6527 | 6527626:6529 | -;- | 1893 | NA | 2 | 114 | 161 | 72 | 218 | 296 |
| ENSDARG000 | 22;22;22;22;2 | 6546697:6547 | 6547079:6547 | -;-;-;-;- | 1429 | zgc:171490 | 2 | 10 | 9 | 18 | 11 | 34 |
| ENSDARG000 | 22;22;22;22;2 | 710759:71344 | 712885:71351 | -;-;-;-;-;-;- | 2734 | arl8a | 2469 | 3056 | 2950 | 2226 | 2500 | 2736 |
| ENSDARG000 | 22;22 | 12699121:127 | 12699510:127 | -;- | 523 | NA | 0 | 2 | 1 | 0 | 0 | 3 |
| ENSDARG000 | 22;22;22;22;2 | 12699121:127 | 12699510:127 | -;-;-;-;-;- | 1127 | slx9 | 215 | 188 | 247 | 160 | 144 | 149 |
| ENSDARG000 | 22;22;22;22;2 | 2931340:2931 | 2931590:2931 | -;-;-;-;-;-;-; | 902 | pigx | 183 | 174 | 218 | 200 | 275 | 153 |
| ENSDARG000 | 22;22;22;22;2 | 2937485:2937 | 2937746:2937 | +;+;+;+;+;+;+; | 2047 | cep19 | 70 | 47 | 47 | 70 | 77 | 107 |
| ENSDARG000 | 22;22;22;22;2 | 10814204:108 | 10814402:108 | -;-;-;-;-;-;-; | 3696 | ARHGEF18 | 96 | 125 | 142 | 85 | 212 | 84 |
| ENSDARG000 | 22;22;22 | 6571847:6574 | 6572159:6574 | +;+;+ | 738 | NA | 4 | 0 | 2 | 1 | 2 | 0 |
| ENSDARG000 | 22;22;22;22;2 | 38252686:382 | 38253502:382 | -;-;-;-;-;-;-; | 2539 | elavl2 | 0 | 0 | 2 | 0 | 0 | 0 |
| ENSDARG000 | 22;22 | 6587550:6601 | 6587867:6601 | -;- | 458 | NA | 0 | 0 | 0 | 0 | 0 | 0 |
| ENSDARG000 | 22;22;22;22;2 | 25331766:253 | 25331863:253 | +;+;+;+;+;+;+; | 1529 | NA | 0 | 6 | 0 | 0 | 0 | 0 |
| ENSDARG000 | 22;22;22;22;2 | 4997784:4999 | 5001288:5001 | -;-;-;-;-;-;-;- | 4765 | rx1 | 0 | 0 | 0 | 0 | 0 | 0 |
| ENSDARG000 | 22;22;22;22;2 | 12729329:127 | 12731129:127 | -;-;-;-;-;-;-; | 4009 | NA | 0 | 4 | 2 | 2 | 0 | 0 |
| ENSDARG000 | 22;22;22;22;2 | 724639:72743 | 725148:72760 | +;+;+;+;+;+;+ | 2415 | zgc:162255 | 802 | 500 | 841 | 687 | 745 | 619 |
| ENSDARG000 | 22;22;22;22;2 | 1610183:1610 | 1610232:1610 | +;+;+;+;+ | 897 | NA | 43 | 20 | 30 | 31 | 20 | 61 |

| | | | | | | | | | | | | |
|---|---|---|---|---|---|---|---|---|---|---|---|---|
| ENSDARG000 | 22;22;22 | 1615691:161( | 1615766:161( | +;+;+ | 231 | NA | 4 | 1 | 12 | 4 | 9 | 7 |
| ENSDARG000 | 22;22;22;22;2 | 1630686:163( | 1630753:163( | +;+;+;+;+;+ | 458 | NA | 13 | 8 | 11 | 9 | 23 | 1 |
| ENSDARG000 | 22;22;22;22;2 | 20388621:20: | 20390294:20: | -;-;-;-;-;-;-;-; | 3193 | map2k2a | 771 | 1334 | 865 | 503 | 1265 | 725 |
| ENSDARG000 | 22;22;22;22;2 | 6613020:667 | 6613353:667 | +;+;+;+;+;+;+; | 2023 | NA | 26 | 4 | 11 | 38 | 0 | 22 |
| ENSDARG000 | 22;22 | 6661712:666: | 6662070:666: | -;- | 646 | NA | 0 | 0 | 0 | 0 | 0 | 0 |
| ENSDARG000 | 22;22;22;22;2 | 6632702:663( | 6635157:663( | -;-;-;-;-;- | 3747 | NA | 301 | 144 | 247 | 128 | 0 | 555 |
| ENSDARG000 | 22;22;22;22;2 | 14836040:14 | 14836271:14 | +;+;+;+;+;+;+; | 5822 | gtpbp1l | 224 | 144 | 196 | 68 | 364 | 100 |
| ENSDARG000 | 22;22;22;22 | 10123729:10: | 10124024:10: | -;-;-;- | 563 | bloc1s1 | 305 | 272 | 246 | 342 | 240 | 278 |
| ENSDARG000 | 22;22;22;22;2 | 22276283:22: | 22277226:22: | -;-;-;-;-;-;-;-; | 3668 | chaf1a | 534 | 1698 | 2603 | 903 | 1448 | 973 |
| ENSDARG000 | 22;22;22;22;2 | 20372533:20: | 20373025:20: | -;-;-;-;-;-;-;-; | 1917 | NA | 240 | 245 | 228 | 247 | 412 | 228 |
| ENSDARG000 | 22;22;22;22;2 | 17691381:17( | 17695432:17( | -;-;-;-;-;-;-;-; | 7899 | SBNO2 | 3818 | 3550 | 3042 | 3924 | 2085 | 1275 |
| ENSDARG000 | 22;22;22;22;2 | 36755263:36: | 36756558:36: | -;-;-;-;-;-;-;-; | 2874 | acy1 | 870 | 776 | 1179 | 1067 | 1264 | 1098 |
| ENSDARG000 | 22;22;22;22;2 | 25335320:25: | 25335391:25: | +;+;+;+;+;+;+; | 1307 | si:ch211-226l | 0 | 0 | 0 | 0 | 0 | 0 |
| ENSDARG000 | 22;22;22;22;2 | 16759010:16: | 16759150:16: | +;+;+;+;+;+;+; | 1038 | tm2d1 | 229 | 157 | 238 | 189 | 195 | 231 |
| ENSDARG000 | 22;22;22 | 16657317:16( | 16657837:16( | -;-;- | 641 | NA | 0 | 0 | 0 | 0 | 0 | 0 |
| ENSDARG000 | 22;22;22;22;2 | 16569123:16! | 16570128:16! | -;-;-;-;-;-;-;-; | 7616 | PATJ | 1590 | 667 | 751 | 1110 | 1796 | 1634 |
| ENSDARG000 | 22;22;22;22;2 | 38276024:38: | 38276212:38: | +;+;+;+;+;+;+; | 1869 | rcor3 | 92 | 122 | 208 | 257 | 267 | 252 |
| ENSDARG000 | 22;22;22;22;2 | 2959879:295! | 2960538:296( | +;+;+;+;+;+;+; | 1569 | impact | 273 | 298 | 288 | 242 | 211 | 242 |
| ENSDARG000 | 22;22;22;22;2 | 5036179:503( | 5036355:503( | -;-;-;-;-;-;-;-; | 1347 | MATK | 17 | 45 | 83 | 81 | 103 | 54 |
| ENSDARG000 | 22;22;22;22;2 | 2949111:294! | 2949451:294! | -;-;-;-;-;-;-;-; | 1219 | ing5a | 315 | 239 | 200 | 251 | 149 | 444 |
| ENSDARG000 | 22;22;22;22;2 | 737211:7415: | 737240:7416( | +;+;+;+;+;+;+; | 1745 | znf76 | 169 | 133 | 201 | 238 | 191 | 279 |
| ENSDARG000 | 22;22;22 | 30945349:30! | 30945473:30! | -;-;- | 307 | NA | 0 | 0 | 0 | 0 | 0 | 0 |
| ENSDARG000 | 22;22 | 27117768:27: | 27118188:27: | -;- | 509 | NA | 4 | 2 | 25 | 8 | 7 | 7 |
| ENSDARG000 | 22;22;22;22;2 | 14995782:14! | 14995851:14! | +;+;+;+;+;+ | 286 | NA | 0 | 0 | 0 | 0 | 0 | 0 |
| ENSDARG000 | 22;22 | 27134809:27: | 27135307:27: | -;- | 662 | NA | 0 | 0 | 14 | 0 | 9 | 0 |
| ENSDARG000 | 22;22;22;22;2 | 10927370:10! | 10928225:10! | -;-;-;-;-;-;-;-; | 5644 | insrb | 690 | 537 | 1452 | 1058 | 952 | 821 |
| ENSDARG000 | 22 | 15009168 | 15010688 | - | 1521 | ELFN2 | 0 | 0 | 0 | 0 | 0 | 0 |
| ENSDARG000 | 22;22;22;22;2 | 30959733:30! | 30960043:30! | -;-;-;-;-;-;-;-; | 1339 | ssuh2.2 | 133 | 131 | 20 | 14 | 8 | 22 |
| ENSDARG000 | 22;22;22;22;2 | 15251527:15: | 15252138:15: | -;-;-;-;-;-;- | 1857 | mfng | 32 | 54 | 58 | 16 | 158 | 84 |
| ENSDARG000 | 22;22;22;22 | 25343200:25: | 25343392:25: | -;-;-;- | 421 | NA | 0 | 2 | 0 | 0 | 0 | 0 |
| ENSDARG000 | 22;22;22 | 1641547:164: | 1641747:164: | +;+;+ | 621 | si:dkev-1b17.: | 8 | 18 | 19 | 24 | 13 | 10 |
| ENSDARG000 | 22;22;22;22;2 | 12752286:12: | 12752832:12: | -;-;-;-;-;-;- | 1420 | cnot9 | 534 | 884 | 778 | 569 | 797 | 475 |
| ENSDARG000 | 22;22 | 27284462:27: | 27284668:27: | +;+ | 620 | NA | 2 | 8 | 1 | 4 | 0 | 6 |
| ENSDARG000 | 22;22;22;22 | 27295601:27: | 27296008:27: | -;-;-;- | 911 | NA | 0 | 0 | 0 | 0 | 0 | 0 |
| ENSDARG000 | 22;22;22;22;2 | 27189979:27: | 27190286:27: | -;-;-;-;-;-;-;-; | 1650 | NA | 167 | 335 | 209 | 205 | 256 | 366 |
| ENSDARG000 | 22;22;22;22 | 27238034:27: | 27238424:27: | -;-;-;- | 736 | NA | 26 | 80 | 180 | 49 | 30 | 34 |
| ENSDARG000 | 22;22;22;22;2 | 6788761:679: | 6788960:679: | -;-;-;-;- | 432 | NA | 1 | 6 | 4 | 4 | 0 | 74 |
| ENSDARG000 | 22;22;22 | 1650128:165: | 1650337:165: | +;+;+ | 651 | NA | 58 | 43 | 76 | 54 | 53 | 72 |
| ENSDARG000 | 22;22;22 | 6943403:694 | 6943683:694 | +;+;+ | 907 | NA | 3 | 0 | 1 | 4 | 0 | 11 |
| ENSDARG000 | 22;22;22;22;2 | 10129788:10: | 10130067:10: | +;+;+;+;+;+;+; | 1413 | tbc1d20 | 262 | 165 | 250 | 131 | 250 | 233 |
| ENSDARG000 | 22;22;22;22;2 | 6952107:695: | 6954499:695: | -;-;-;-;-;-;- | 3255 | NA | 0 | 18 | 39 | 62 | 64 | 49 |
| ENSDARG000 | 22;22;22;22;2 | 1660415:166( | 1660452:166: | +;+;+;+;+ | 1443 | NA | 48 | 26 | 70 | 69 | 42 | 39 |
| ENSDARG000 | 22;22;22;22;2 | 15308526:15: | 15309106:15: | +;+;+;+;+;+;+; | 3582 | NA | 0 | 0 | 0 | 0 | 0 | 0 |
| ENSDARG000 | 22;22;22 | 1668471:167( | 1668500:167( | +;+;+ | 592 | NA | 50 | 30 | 21 | 22 | 30 | 38 |
| ENSDARG000 | 22;22;22;22;2 | 30991365:30! | 30991464:30! | -;-;-;-;-;-;-;-; | 834 | ssuh2.3 | 28 | 23 | 12 | 29 | 29 | 28 |
| ENSDARG000 | 22;22;22;22;2 | 6962924:696: | 6963317:696: | -;-;-;-;- | 1168 | NA | 0 | 26 | 23 | 64 | 50 | 78 |
| ENSDARG000 | 22;22;22;22;2 | 5056288:505( | 5057799:505 | -;-;-;-;-;-;-;-; | 4856 | ZFR2 | 1730 | 1295 | 2325 | 1250 | 2156 | 1586 |
| ENSDARG000 | 22;22;22;22;2 | 3121086:312( | 3124347:312 | -;-;-;-;-;-;-;-; | 5330 | lmnb2 | 1294 | 2653 | 3182 | 1562 | 2142 | 1595 |
| ENSDARG000 | 22;22;22;22;2 | 6969477:697: | 6969983:697: | -;-;-;-;-;-;-;-; | 2284 | NA | 4 | 541 | 360 | 828 | 824 | 519 |
| ENSDARG000 | 22;22;22;22;2 | 1680628:168: | 1680663:168: | +;+;+;+;+;+ | 1788 | NA | 30 | 18 | 58 | 35 | 29 | 44 |
| ENSDARG000 | 22;22;22;22;2 | 22302614:22: | 22302637:22: | +;+;+;+;+;+;+; | 3957 | scamp4 | 5887 | 5694 | 7575 | 4520 | 5214 | 5002 |
| ENSDARG000 | 22;22;22;22 | 1691293:169: | 1691405:169: | +;+;+;+ | 980 | znf1160 | 77 | 41 | 32 | 30 | 59 | 58 |
| ENSDARG000 | 22;22;22;22;2 | 25415980:25 | 25416066:25 | -;-;-;-;-;- | 543 | si:ch73-189n: | 0 | 0 | 0 | 0 | 0 | 0 |
| ENSDARG000 | 22;22;22;22;2 | 38301365:38: | 38301453:38: | +;+;+;+;+;+;+; | 2040 | NA | 20 | 61 | 39 | 16 | 68 | 50 |
| ENSDARG000 | 22;22;22;22;2 | 10158502:10: | 10158553:10: | +;+;+;+;+;+;+; | 1169 | rpp14 | 178 | 243 | 312 | 104 | 154 | 242 |
| ENSDARG000 | 22;22;22;22;2 | 25432849:25 | 25433176:25 | -;-;-;-;-;-;-;-; | 1100 | NA | 0 | 0 | 0 | 0 | 0 | 0 |
| ENSDARG000 | 22;22;22;22;2 | 10145098:10: | 10146163:10: | -;-;-;-;-;-;-;-; | 3011 | rbck1 | 353 | 130 | 218 | 110 | 219 | 251 |
| ENSDARG000 | 22;22;22;22;2 | 25453334:25 | 25453406:25 | +;+;+;+;+;+;+; | 653 | zgc:152936 | 0 | 0 | 0 | 0 | 0 | 0 |
| ENSDARG000 | 22;22;22;22;2 | 770001:7700( | 770372:7703 | -;-;-;-;-;-;-;-; | 2069 | rbbp5 | 646 | 616 | 768 | 779 | 811 | 506 |
| ENSDARG000 | 22;22;22;22;2 | 11070842:11( | 11071964:11( | -;-;-;-;-;-;-;-; | 1836 | use1 | 188 | 224 | 246 | 222 | 348 | 220 |
| ENSDARG000 | 22;22;22 | 1700552:170: | 1700581:170: | +;+;+ | 1113 | NA | 22 | 25 | 26 | 30 | 33 | 18 |
| ENSDARG000 | 22;22;22;22;2 | 27529632:27! | 27529961:27! | -;-;-;-;-;-;-;-; | 1540 | NA | 2 | 0 | 8 | 14 | 1 | 6 |
| ENSDARG000 | 22;22 | 1708760:171: | 1708817:171: | +;+ | 168 | NA | 0 | 2 | 0 | 0 | 4 | 0 |
| ENSDARG000 | 22;22;22;22;2 | 17784414:17 | 17784877:17 | +;+;+;+;+;+;+; | 5595 | tm6sf2b | 1 | 8 | 2 | 0 | 0 | 0 |
| ENSDARG000 | 22;22;22;22;2 | 31004735:31( | 31004885:31( | -;-;-;-;-;-;-;-; | 1248 | ssuh2.4 | 172 | 93 | 51 | 165 | 83 | 204 |
| ENSDARG000 | 22;22 | 27594566:27! | 27594799:27! | -;- | 517 | NA | 0 | 0 | 0 | 0 | 0 | 4 |
| ENSDARG000 | 22;22;22;22;2 | 36794438:36 | 36796041:36 | -;-;-;-;-;-;-;-; | 5454 | MAP1S | 3688 | 2667 | 2601 | 3247 | 3056 | 2492 |
| ENSDARG000 | 22;22;22 | 1725568:172( | 1725752:172( | +;+;+ | 686 | NA | 20 | 5 | 17 | 9 | 21 | 9 |
| ENSDARG000 | 22;22;22;22 | 11082776:11( | 11082852:11( | +;+;+;+ | 1015 | NA | 25 | 109 | 86 | 71 | 43 | 106 |
| ENSDARG000 | 22;22;22;22;2 | 15315655:15: | 15315914:15: | +;+;+;+;+;+ | 1065 | sult3st3 | 0 | 0 | 0 | 0 | 0 | 0 |
| ENSDARG000 | 22;22;22;22;2 | 38394502:38: | 38394937:38: | -;-;-;-;-;-;-;-; | 4239 | ptk7a | 2138 | 1509 | 2981 | 1359 | 3668 | 1703 |
| ENSDARG000 | 22;22;22;22;2 | 7009062:700! | 7010222:701( | -;-;-;-;-;-;-;-; | 3049 | smarcd1 | 1059 | 1329 | 1859 | 774 | 1710 | 1017 |
| ENSDARG000 | 22;22;22;22;2 | 22316855:22: | 22317634:22: | -;-;-;-;-;-;-;-; | 2872 | si:ch211-129( | 61 | 2 | 9 | 11 | 13 | 10 |
| ENSDARG000 | 22;22;22;22 | 28207968:28: | 28208436:28: | -;-;-;- | 925 | si:dkey-222p3 | 0 | 26 | 9 | 14 | 7 | 8 |
| ENSDARG000 | 22;22;22;22;2 | 6996046:699( | 6996379:699( | -;-;-;-;-;-;-;-; | 1463 | gpd1b | 1852 | 2235 | 743 | 1309 | 3224 | 334 |
| ENSDARG000 | 22;22;22;22 | 25473300:25 | 25473401:25 | -;-;-;- | 463 | NA | 0 | 0 | 0 | 0 | 0 | 0 |
| ENSDARG000 | 22;22 | 1734981:173 | 1735090:173 | +;+ | 473 | NA | 7 | 9 | 17 | 27 | 29 | 9 |
| ENSDARG000 | 22;22;22 | 1751640:175: | 1751698:175: | +;+;+ | 915 | NA | 36 | 23 | 29 | 41 | 37 | 24 |
| ENSDARG000 | 22;22 | 1767959:177: | 1768022:177: | +;+ | 951 | NA | 42 | 30 | 26 | 21 | 11 | 31 |
| ENSDARG000 | 22;22;22 | 11090338:11( | 11090533:11( | -;-;- | 587 | NA | 6 | 431 | 44 | 124 | 973 | 370 |
| ENSDARG000 | 22;22;22;22;2 | 5103349:510( | 5103816:510( | +;+;+;+;+;+;+; | 2674 | ATCAY | 238 | 82 | 122 | 45 | 0 | 56 |
| ENSDARG000 | 22;22;22;22;2 | 36863599:36 | 36863667:36 | +;+;+;+;+;+ | 860 | NA | 6 | 13 | 3 | 1 | 16 | 58 |
| ENSDARG000 | 22;22;22;22;2 | 3153876:315( | 3153945:315( | +;+;+;+;+;+;+ | 1202 | rpl36 | 4641 | 4870 | 8246 | 7909 | 4855 | 7787 |
| ENSDARG000 | 22;22;22 | 36832648:36 | 36832894:36 | -;-;- | 472 | CARTPT | 0 | 0 | 0 | 0 | 0 | 0 |
| ENSDARG000 | 22;22;22;22;2 | 36875400:36 | 36875665:36 | +;+;+;+;+;+;+; | 2740 | mccc1 | 1189 | 792 | 983 | 696 | 1110 | 888 |
| ENSDARG000 | 22;22;22 | 1779401:178( | 1779453:178( | +;+;+ | 244 | NA | 17 | 9 | 9 | 18 | 6 | 9 |
| ENSDARG000 | 22;22;22 | 1796057:179( | 1796109:179( | +;+;+ | 1427 | NA | 35 | 29 | 32 | 13 | 33 | 38 |
| ENSDARG000 | 22;22;22;22;2 | 36839080:36 | 36839482:36 | -;-;-;-;-;-;-;-; | 1731 | kng1 | 0 | 14 | 24 | 6 | 10 | 6 |
| ENSDARG000 | 22;22;22 | 11097512:11( | 11097838:11( | +;+;+ | 641 | NA | 0 | 750 | 0 | 199 | 2055 | 690 |
| ENSDARG000 | 22;22;22;22;2 | 28236737:28: | 28236911:28: | +;+;+;+;+;+;+; | 2922 | senp7b | 374 | 285 | 509 | 256 | 554 | 292 |
| ENSDARG000 | 22;22;22;22;2 | 12770877:12: | 12770959:12: | +;+;+;+;+;+;+; | 1707 | ftcd | 0 | 0 | 4 | 26 | 44 | 0 |
| ENSDARG000 | 22;22;22;22;2 | 22345136:22: | 22345504:22: | -;-;-;-;-;-;-;-; | 4293 | camsap2a | 246 | 60 | 5 | 0 | 15 | 40 |
| ENSDARG000 | 22;22;22;22;2 | 36902352:36! | 36902898:36! | +;+;+;+;+;+ | 1656 | trim35-20 | 17 | 9 | 1 | 10 | 2 | 12 |
| ENSDARG000 | 22;22;22;22 | 25540945:25! | 25540998:25! | -;-;-;- | 462 | NA | 0 | 0 | 0 | 0 | 0 | 0 |
| ENSDARG000 | 22;22;22;22;2 | 16839467:16 | 16840669:16 | -;-;-;-;-;-;-;-; | 3522 | nfia | 57 | 69 | 48 | 2 | 53 | 35 |
| ENSDARG000 | 22;22;22;22;2 | 3160447:316: | 3161601:316: | -;-;-;-;-;-;-;-; | 4114 | LONP1 | 613 | 804 | 1144 | 409 | 1016 | 408 |
| ENSDARG000 | 22;22 | 1806235:180 | 1806344:180! | +;+ | 961 | NA | 81 | 18 | 80 | 18 | 42 | 52 |
| ENSDARG000 | 22;22 | 1815408:181 | 1815511:181 | +;+ | 1248 | NA | 208 | 61 | 97 | 155 | 150 | 231 |
| ENSDARG000 | 22;22;22;22 | 11102991:11: | 11103019:11: | +;+;+;+ | 988 | NA | 0 | 36 | 0 | 4 | 164 | 124 |
| ENSDARG000 | 22;22;22;22;2 | 11106391:11: | 11106527:11: | +;+;+;+;+ | 739 | NA | 0 | 0 | 0 | 0 | 2 | 0 |
| ENSDARG000 | 22;22;22;22 | 17828267:17 | 17828487:17 | +;+;+;+ | 1935 | hapln4 | 0 | 0 | 0 | 0 | 0 | 0 |
| ENSDARG000 | 22;22;22;22;2 | 15323930:15: | 15324197:15: | +;+;+;+;+;+ | 1318 | sult3st5 | 0 | 0 | 0 | 0 | 0 | 0 |
| ENSDARG000 | 22;22;22;22;2 | 20406410:20 | 20409253:20 | -;-;-;-;-;-;-;-; | 4101 | pias4a | 972 | 988 | 1202 | 576 | 1259 | 944 |
| ENSDARG000 | 22;22;22;22;2 | 38468036:38 | 38468686:38 | -;-;-;-;-;-;-;- | 1404 | soul4 | 287 | 300 | 362 | 306 | 238 | 381 |
| ENSDARG000 | 22;22;22 | 1821718:182: | 1821747:182: | +;+;+ | 792 | NA | 9 | 8 | 6 | 8 | 6 | 14 |
| ENSDARG000 | 22;22;22;22 | 31039091:31( | 31039134:31( | +;+;+;+ | 513 | rpl32 | 9849 | 10016 | 15716 | 14400 | 9592 | 15529 |
| ENSDARG000 | 22;22;22;22;2 | 786556:7897! | 786660:7899! | +;+;+;+;+;+;+; | 4275 | cry3b | 117 | 90 | 168 | 88 | 57 | 210 |
| ENSDARG000 | 22;22;22;22;2 | 25554232:25! | 25554466:25! | +;+;+;+;+;+;+; | 6245 | AARS2 | 206 | 138 | 302 | 124 | 232 | 182 |
| ENSDARG000 | 22;22;22;22;2 | 1836782:183 | 1836806:183 | +;+;+;+;+;+;+ | 841 | NA | 6 | 7 | 26 | 26 | 56 | 24 |
| ENSDARG000 | 22;22;22 | 1846147:184( | 1846173:184( | +;+;+ | 631 | NA | 31 | 15 | 27 | 3 | 17 | 30 |
| ENSDARG000 | 22;22;22 | 1853999:185 | 1854048:185 | +;+;+ | 1242 | znf1174 | 18 | 23 | 33 | 14 | 18 | 39 |
| ENSDARG000 | 22;22;22;22;2 | 5118361:511 | 5118513:511 | +;+;+;+;+;+;+; | 2287 | mibp | 92 | 767 | 1086 | 431 | 569 | 343 |
| ENSDARG000 | 22;22;22;22;2 | 1878147:187 | 1878201:187 | +;+;+;+;+;+;+; | 482 | NA | 0 | 0 | 0 | 0 | 0 | 0 |
| ENSDARG000 | 22;22;22;22;2 | 1868983:188: | 1869009:188: | +;+;+;+;+;+;+; | 4825 | NA | 0 | 1 | 0 | 0 | 0 | 0 |
| ENSDARG000 | 22;22;22;22;2 | 11123487:11: | 11124125:11: | -;-;-;-;-;-;-;-; | 2175 | atp6ap2 | 4242 | 5385 | 4364 | 4747 | 5903 | 4871 |
| ENSDARG000 | 22;22;22;22;2 | 22417751:22 | 22418497:22 | +;+;+;+;+;+;+; | 4696 | ddx59 | 383 | 153 | 314 | 196 | 223 | 250 |
| ENSDARG000 | 22;22;22;22;2 | 17118225:17: | 17118245:17: | +;+;+;+;+;+;+; | 6894 | frem1b | 108 | 6 | 4 | 46 | 10 | 11 |
| ENSDARG000 | 22;22;22;22;2 | 31045277:31( | 31045768:31( | -;-;-;-;-;-;-;-; | 4872 | cand2 | 1 | 0 | 0 | 0 | 0 | 6 |
| ENSDARG000 | 22;22;22 | 12790837:12 | 12791197:12 | +;+;+ | 1101 | mstna | 0 | 0 | 0 | 0 | 0 | 0 |
| ENSDARG000 | 22;22;22;22;2 | 28320794:28: | 28320902:28: | +;+;+;+;+;+;+; | 1561 | NA | 0 | 0 | 0 | 0 | 0 | 0 |
| ENSDARG000 | 22;22;22;22;2 | 28320792:28: | 28320902:28: | +;+;+;+;+;+;+; | 1814 | NA | 0 | 0 | 0 | 0 | 0 | 0 |
| ENSDARG000 | 22;22;22;22;2 | 31059919:31( | 31060081:31( | +;+;+;+;+;+;+; | 1473 | sec13 | 945 | 1252 | 1294 | 699 | 1417 | 733 |
| ENSDARG000 | 22;22 | 1899142:189! | 1899257:190: | +;+ | 1938 | NA | 62 | 44 | 79 | 48 | 65 | 111 |
| ENSDARG000 | 22;22;22;22;2 | 38484447:38 | 38484793:38 | -;-;-;-;-;-;-;-; | 2922 | klc4 | 363 | 444 | 406 | 264 | 305 | 185 |
| ENSDARG000 | 22;22;22;22;2 | 25611415:25( | 25611501:25( | -;-;-;-;-;- | 652 | si:ch211-12h: | 0 | 0 | 0 | 0 | 0 | 0 |
| ENSDARG000 | 22;22;22;22;2 | 15331214:15: | 15331476:15: | +;+;+;+;+;+ | 2161 | sult3st4 | 0 | 0 | 0 | 0 | 0 | 0 |
| ENSDARG000 | 22;22;22;22;2 | 10166407:10: | 10166520:10: | +;+;+;+;+;+;+; | 4698 | pxk | 4412 | 2112 | 2971 | 3430 | 2233 | 5826 |
| ENSDARG000 | 22;22;22;22;2 | 812848:8129: | 813448:8134 | -;-;-;-;-;- | 1063 | zgc:153675 | 295 | 258 | 335 | 364 | 263 | 274 |
| ENSDARG000 | 22;22;22 | 1904546:190! | 1904595:190! | +;+;+ | 220 | NA | 4 | 3 | 8 | 4 | 2 | 3 |
| ENSDARG000 | 22;22;22;22;2 | 1911269:191: | 1911319:191: | +;+;+;+;+;+ | 1269 | znf1156 | 22 | 17 | 41 | 24 | 10 | 14 |
| ENSDARG000 | 22;22;22;22;2 | 17864872:17 | 17865002:17 | -;-;-;-;-;-;-;-; | 4470 | NCAN | 2 | 4 | 15 | 0 | 4 | 4 |
| ENSDARG000 | 22;22 | 20427170:20 | 20427655:20 | +;+ | 957 | foxq2 | 0 | 0 | 0 | 6 | 0 | 0 |
| ENSDARG000 | 22;22;22 | 1922293:192: | 1922366:192: | +;+;+ | 1228 | NA | 30 | 2 | 44 | 14 | 8 | 40 |
| ENSDARG000 | 22;22 | 25672155:25( | 25672248:25( | +;+ | 941 | b3galt10.2 | 2 | 0 | 0 | 0 | 0 | 0 |
| ENSDARG000 | 22;22;22;22;2 | 3184500:318 | 3184726:318 | +;+;+;+;+;+;+; | 5256 | ftsj3 | 970 | 764 | 1298 | 361 | 1238 | 628 |
| ENSDARG000 | 22;22 | 28372015:28: | 28372414:28: | -;- | 578 | NA | 0 | 0 | 0 | 0 | 0 | 0 |
| ENSDARG000 | 22;22;22;22;2 | 1930589:193( | 1930655:193( | +;+;+;+;+;+ | 1508 | NA | 28 | 50 | 97 | 104 | 68 | 35 |
| ENSDARG000 | 22;22;22;22;2 | 22437561:22 | 22438965:22 | +;+;+;+;+;+;+; | 6428 | kif14 | 108 | 456 | 483 | 185 | 412 | 154 |
| ENSDARG000 | 22;22 | 20461488:20 | 20462546:20 | +;+ | 1339 | NA | 0 | 0 | 0 | 0 | 0 | 0 |
| ENSDARG000 | 22;22;22;22;2 | 11144153:11: | 11144779:11: | +;+;+;+;+;+;+; | 7187 | bcor | 2545 | 1255 | 2568 | 1814 | 1645 | 1910 |
| ENSDARG000 | 22;22;22;22;2 | 25681911:25( | 25681962:25( | +;+;+;+;+;+;+; | 717 | si:dkeyp-98a7 | 0 | 0 | 0 | 0 | 0 | 0 |
| ENSDARG000 | 22;22;22;22;2 | 17192767:17: | 17192979:17: | +;+;+;+;+;+;+; | 2142 | atpaf1 | 301 | 325 | 462 | 206 | 435 | 318 |
| ENSDARG000 | 22;22;22;22;2 | 20473229:20 | 20479633:20 | -;-;-;-;-;-;-;-; | 10062 | NA | 2 | 4 | 0 | 0 | 2 | 0 |
| ENSDARG000 | 22;22;22;22;2 | 25681912:25( | 25681962:25( | +;+;+;+;+;+ | 649 | NA | 0 | 0 | 0 | 0 | 0 | 0 |
| ENSDARG000 | 22;22;22;22;2 | 28446365:28 | 28446764:28 | +;+;+;+;+;+;+; | 4075 | abi3bpb | 527 | 22 | 358 | 33 | 1552 | 34 |
| ENSDARG000 | 22;22;22;22;2 | 25693295:25 | 25693346:25 | +;+;+;+;+;+;+; | 718 | si:dkeyp-98a7 | 0 | 0 | 0 | 0 | 0 | 0 |
| ENSDARG000 | 22;22;22;22;2 | 25704430:25 | 25704464:25 | +;+;+;+;+;+ | 640 | si:dkeyp-98a7 | 0 | 0 | 0 | 0 | 0 | 0 |
| ENSDARG000 | 22;22;22;22;2 | 25693297:25( | 25693346:25( | +;+;+;+;+;+ | 662 | NA | 0 | 0 | 0 | 0 | 0 | 0 |
| ENSDARG000 | 22;22;22;22;2 | 10201826:10: | 10201912:10: | +;+;+;+;+;+;+; | 1595 | pdhb | 2122 | 2075 | 2222 | 1739 | 1885 | 1833 |
| ENSDARG000 | 22;22;22 | 1940595:194: | 1940621:194: | +;+;+ | 693 | NA | 40 | 21 | 28 | 24 | 36 | 29 |
| ENSDARG000 | 22;22 | 5140530:514 | 5140991:514! | -;- | 508 | NA | 4 | 1 | 9 | 4 | 4 | 2 |
| ENSDARG000 | 22;22;22;22;2 | 5123479:512: | 5124322:512 | +;+;+;+;+;+;+; | 1650 | loxl5a | 0 | 0 | 0 | 4 | 0 | 0 |
| ENSDARG000 | 22;22;22;22;2 | 38520209:38! | 38526209:38! | -;-;-;-;-;-;-;-; | 8818 | NA | 624 | 670 | 1078 | 1000 | 1000 | 910 |
| ENSDARG000 | 22;22;22;22;2 | 5153461:515 | 5157125:515! | -;-;-;-;- | 4322 | tnfaip8l1 | 30 | 20 | 24 | 2 | 68 | 18 |
| ENSDARG000 | 22;22;22;22;2 | 5135193:513! | 5135421:513! | +;+;+;+;+;+;+; | 1516 | mydgf | 252 | 677 | 744 | 437 | 706 | 678 |
| ENSDARG000 | 22;22;22;22;2 | 12798569:12 | 12798703:12 | +;+;+;+;+;+;+; | 4430 | stat1a | 4950 | 4000 | 4487 | 3676 | 3543 | 4533 |
| ENSDARG000 | 22;22;22;22;2 | 15336752:15: | 15337022:15: | +;+;+;+;+;+ | 1278 | sult3st2 | 0 | 0 | 0 | 0 | 0 | 0 |
| ENSDARG000 | 22;22;22;22 | 11203429:11: | 11203628:11: | +;+;+;+ | 4708 | NA | 0 | 0 | 7 | 0 | 6 | 10 |
| ENSDARG000 | 22;22;22;22;2 | 25715925:25 | 25715976:25 | +;+;+;+;+;+ | 665 | NA | 0 | 0 | 0 | 0 | 0 | 0 |
| ENSDARG000 | 22;22;22;22;2 | 819656:8204 | 820167:8205! | -;-;-;-;-;-;-;-; | 2229 | cry4 | 360 | 764 | 581 | 891 | 355 | 482 |
| ENSDARG000 | 22;22 | 11436447:11 | 11438138:11 | -;- | 1851 | mid1ip1b | 765 | 1102 | 836 | 314 | 804 | 638 |
| ENSDARG000 | 22;22;22;22;2 | 1946676:194 | 1946775:194 | +;+;+;+;+;+;+ | 686 | NA | 29 | 17 | 73 | 54 | 45 | 23 |

| | | | | | | | | | | | | |
|---|---|---|---|---|---|---|---|---|---|---|---|---|
| ENSDARG000 | 22;22;22;22;2 | 7033381;703 | 7033446;703 | +;+;+;+;+ | 1020 | NA | 0 | 0 | 0 | 0 | 0 | 0 |
| ENSDARG000 | 22;22;22;22;2 | 7044625;704 | 7045610;704 | -;-;-;-;-;-;-;-; | 4044 | asic1b | 0 | 2 | 21 | 6 | 1 | 53 |
| ENSDARG000 | 22;22;22;22;2 | 10214994;10 | 10215144;10 | +;+;+;+;+;+ | 1304 | kctd6a | 115 | 88 | 216 | 149 | 114 | 257 |
| ENSDARG000 | 22;22;22;22;2 | 25720725;25 | 25720776;25 | +;+;+;+;+;+ | 667 | si:dkeyp-98a7 | 0 | 0 | 0 | 0 | 0 | 0 |
| ENSDARG000 | 22;22;22;22;2 | 18160330;18 | 18160431;18 | -;-;-;-;-;-;-;-; | 847 | rfxank | 105 | 81 | 78 | 37 | 82 | 72 |
| ENSDARG000 | 22;22;22;22 | 28588052;28 | 28588209;28 | -;-;-;- | 558 | NA | 0 | 0 | 8 | 0 | 0 | 0 |
| ENSDARG000 | 22;22;22;22;2 | 25733817;25 | 25734231;25 | +;+;+;+;+;+;+; | 2119 | si:dkeyp-98a7 | 0 | 0 | 0 | 0 | 0 | 0 |
| ENSDARG000 | 22;22 | 10232527;10 | 10232624;10 | +;+ | 2664 | ccdc71 | 586 | 536 | 620 | 471 | 652 | 475 |
| ENSDARG000 | 22;22;22;22;2 | 28610372;28 | 28610420;28 | -;-;-;-;-;-;-;-; | 1109 | NA | 1 | 0 | 0 | 0 | 0 | 0 |
| ENSDARG000 | 22;22;22;22;2 | 18156000;18 | 18156042;18 | +;+;+;+;+;+;+; | 721 | nr2c2ap | 63 | 78 | 64 | 54 | 50 | 30 |
| ENSDARG000 | 22;22;22;22;2 | 28639026;28 | 28643282;28 | -;-;-;-;- | 4430 | jagn1b | 736 | 791 | 950 | 483 | 842 | 716 |
| ENSDARG000 | 22;22;22;22;2 | 1953096;195 | 1953162;195 | +;+;+;+;+;+;+; | 3027 | NA | 20 | 31 | 38 | 32 | 34 | 16 |
| ENSDARG000 | 22;22;22 | 22533586;22 | 22533743;22 | -;-;- | 394 | NA | 0 | 0 | 0 | 0 | 0 | 0 |
| ENSDARG000 | 22;22;22;22;2 | 17203752;17 | 17203921;17 | +;+;+;+;+;+;+; | 5237 | NA | 2 | 2 | 0 | 2 | 4 | 4 |
| ENSDARG000 | 22;22;22;22;2 | 15343953;15 | 15344173;15 | +;+;+;+;+;+;+; | 1398 | rrp36 | 140 | 174 | 242 | 156 | 276 | 94 |
| ENSDARG000 | 22;22;22 | 22566174;22 | 22566214;22 | +;+;+ | 267 | NA | 0 | 0 | 0 | 0 | 0 | 0 |
| ENSDARG000 | 22;22 | 22721223;22 | 22721469;22 | +;+ | 1374 | NA | 4 | 0 | 0 | 0 | 0 | 6 |
| ENSDARG000 | 22 | 20546612 | 20547407 | + | 796 | NA | 0 | 2 | 0 | 0 | 12 | 0 |
| ENSDARG000 | 22;22 | 7359679;740 | 7359874;740 | -;- | 561 | NA | 0 | 0 | 17 | 0 | 0 | 21 |
| ENSDARG000 | 22;22;22;22;2 | 28647130;28 | 28649788;28 | -;-;-;-;-;-;-;- | 3998 | COL8A1 | 6802 | 302 | 272 | 3514 | 1872 | 494 |
| ENSDARG000 | 22 | 15384456 | 15385442 | - | 987 | tmem264 | 0 | 0 | 0 | 0 | 0 | 0 |
| ENSDARG000 | 22;22;22;22;2 | 15429099;15 | 15430015;15 | -;-;-;-;- | 1394 | rpp21 | 56 | 92 | 76 | 74 | 98 | 96 |
| ENSDARG000 | 22;22;22;22;2 | 7439255;743 | 7439589;743 | -;-;-;-;- | 662 | NA | 0 | 0 | 0 | 0 | 0 | 0 |
| ENSDARG000 | 22;22;22;22;2 | 7439186;743 | 7439570;743 | +;+;+;+;+;+;+; | 5031 | cela1.6 | 4 | 5 | 4 | 2 | 2 | 6 |
| ENSDARG000 | 22;22;22;22;2 | 22977268;22 | 22978160;22 | -;-;-;-;-;-;-;-; | 4286 | NA | 124 | 147 | 150 | 78 | 392 | 354 |
| ENSDARG000 | 22;22;22;22;2 | 25931782;25 | 25932034;25 | +;+;+;+;+;+;+; | 2136 | dnaja3b | 148 | 430 | 244 | 434 | 310 | 298 |
| ENSDARG000 | 22;22;22;22;2 | 11440705;11 | 11442773;11 | -;-;-;-;-;-;-;- | 2891 | tspan7b | 14 | 38 | 12 | 10 | 30 | 18 |
| ENSDARG000 | 22 | 20560041 | 20560787 | + | 747 | nfil3-4 | 0 | 1 | 4 | 0 | 12 | 10 |
| ENSDARG000 | 22;22;22;22;2 | 3223489;322 | 3223714;322 | +;+;+;+;+;+ | 865 | lim2.2 | 42 | 24 | 47 | 6 | 47 | 24 |
| ENSDARG000 | 22 | 20573862 | 20575679 | - | 1818 | lingo3a | 0 | 14 | 0 | 0 | 2 | 4 |
| ENSDARG000 | 22;22;22;22;2 | 28739668;28 | 28739973;28 | -;-;-;-;-;-;-;-; | 5575 | si:dkeyp-34c1 | 8 | 0 | 10 | 4 | 0 | 0 |
| ENSDARG000 | 22;22 | 3238474;323 | 3239055;323 | +;+ | 703 | NA | 0 | 0 | 0 | 0 | 0 | 0 |
| ENSDARG000 | 22;22;22;22;2 | 18166599;18 | 18166738;18 | +;+;+;+;+;+;+; | 922 | borcs8 | 115 | 148 | 152 | 144 | 146 | 138 |
| ENSDARG000 | 22;22;22;22;2 | 11497366;11 | 11500380;11 | -;-;-;-;-;-;-;-; | 5042 | slc26a11 | 3831 | 1142 | 945 | 1853 | 992 | 3192 |
| ENSDARG000 | 22;22;22 | 18174125;18 | 18174892;18 | -;-;- | 1361 | NA | 14 | 6 | 2 | 10 | 20 | 2 |
| ENSDARG000 | 22;22;22;22;2 | 20689958;20 | 20690159;20 | -;-;-;-;-;- | 893 | org | 0 | 0 | 0 | 0 | 0 | 0 |
| ENSDARG000 | 22;22;22;22;2 | 11520249;11 | 11520636;11 | +;+;+;+;+;+;+ | 1953 | sgsh | 791 | 347 | 489 | 591 | 580 | 849 |
| ENSDARG000 | 22;22;22;22;2 | 17214389;17 | 17214456;17 | +;+;+;+;+;+;+; | 4234 | nrd1b | 204 | 242 | 317 | 222 | 501 | 250 |
| ENSDARG000 | 22;22 | 2083693;208 | 2084962;208 | -;- | 1316 | NA | 31 | 23 | 55 | 26 | 56 | 26 |
| ENSDARG000 | 22;22;22 | 3252936;325 | 3255337;325 | -;-;- | 3130 | gpr35.1 | 21 | 19 | 89 | 63 | 121 | 25 |
| ENSDARG000 | 22;22;22;22;2 | 3262413;326 | 3263990;326 | -;-;-;-;-;-;-;-; | 3425 | NA | 7464 | 6071 | 5342 | 4780 | 6643 | 5908 |
| ENSDARG000 | 22;22;22 | 38614887;38 | 38615328;38 | -;-;- | 951 | nppc | 8 | 4 | 4 | 6 | 0 | 0 |
| ENSDARG000 | 22;22;22;22;2 | 15438513;15 | 15439034;15 | +;+;+;+;+;+;+; | 2201 | gpc5b | 0 | 0 | 6 | 0 | 0 | 0 |
| ENSDARG000 | 22;22;22;22;2 | 2111120;211 | 2111213;211 | +;+;+;+;+;+;+; | 1209 | NA | 26 | 23 | 19 | 18 | 19 | 18 |
| ENSDARG000 | 22;22;22;22;2 | 7462997;746 | 7463068;746 | +;+;+;+;+;+;+; | 1246 | cela1.1 | 0 | 0 | 0 | 0 | 0 | 0 |
| ENSDARG000 | 22;22;22;22;2 | 38778649;38 | 38778956;38 | +;+;+;+;+;+;+; | 4363 | alpi.2 | 4 | 0 | 2 | 0 | 0 | 0 |
| ENSDARG000 | 22;22;22;22;2 | 835728;8380 | 835858;8381 | +;+;+;+;+;+;+; | 1797 | dennd2db | 38 | 39 | 100 | 10 | 32 | 54 |
| ENSDARG000 | 22;22;22;22;2 | 856869;8594 | 857843;8595 | -;-;-;-;-;-;-;- | 1954 | dram2b | 1380 | 1259 | 1254 | 1085 | 1425 | 1459 |
| ENSDARG000 | 22;22;22;22;2 | 36914636;36 | 36915003;36 | +;+;+;+;+;+ | 1231 | NA | 2 | 0 | 0 | 0 | 0 | 0 |
| ENSDARG000 | 22;22;22;22 | 20710434;20 | 20710576;20 | +;+;+;+ | 850 | NA | 0 | 0 | 4 | 0 | 2 | 0 |
| ENSDARG000 | 22;22;22;22;2 | 38762693;38 | 38762958;38 | +;+;+;+;+;+;+; | 3509 | alpi.1 | 14 | 0 | 8 | 0 | 0 | 0 |
| ENSDARG000 | 22;22;22;22;2 | 25991130;25 | 25991609;25 | -;-;-;-;-;- | 1464 | pdgfaa | 0 | 2 | 0 | 0 | 0 | 0 |
| ENSDARG000 | 22;22;22;22;2 | 15552025;15 | 15552390;15 | -;-;-;-;-;-;-;-; | 2726 | NA | 0 | 0 | 1 | 0 | 6 | 2 |
| ENSDARG000 | 22;22;22;22;2 | 15507218;15 | 15507401;15 | +;+;+;+;+;+;+; | 1705 | gpc1a | 0 | 0 | 0 | 0 | 0 | 0 |
| ENSDARG000 | 22;22;22;22;2 | 38751797;38 | 38752096;38 | -;-;-;-;-;-;-;- | 1167 | NA | 0 | 0 | 0 | 0 | 0 | 0 |
| ENSDARG000 | 22;22;22;22;2 | 28803739;28 | 28804218;28 | +;+;+;+;+;+;+; | 5031 | tp53bp2b | 467 | 210 | 279 | 174 | 425 | 398 |
| ENSDARG000 | 22;22;22;22;2 | 23105059;23 | 23106646;23 | -;-;-;-;-;-;-;-; | 2640 | nek7 | 1258 | 1522 | 1760 | 1439 | 1436 | 1525 |
| ENSDARG000 | 22;22;22;22;2 | 5176255;517 | 5176938;517 | +;+;+;+;+;+;+; | 5942 | CERS1 | 61 | 60 | 91 | 74 | 293 | 108 |
| ENSDARG000 | 22;22;22;22;2 | 2120815;212 | 2120885;212 | +;+;+;+;+;+;+; | 2041 | NA | 43 | 39 | 45 | 86 | 65 | 40 |
| ENSDARG000 | 22;22;22;22;2 | 18187857;18 | 18188225;18 | +;+;+;+;+;+;+; | 4583 | mef2b | 18 | 22 | 8 | 18 | 20 | 80 |
| ENSDARG000 | 22;22;22;22;2 | 7472420;747 | 7472491;747 | +;+;+;+;+;+;+; | 924 | cela1.2 | 0 | 0 | 0 | 0 | 0 | 0 |
| ENSDARG000 | 22;22;22;22;2 | 36923240;36 | 36923868;36 | -;-;-;-;-;- | 1645 | si:dkev-37m8. | 23 | 6 | 15 | 4 | 4 | 10 |
| ENSDARG000 | 22;22 | 11535131;11 | 11535472;11 | +;+ | 1344 | npb | 18 | 6 | 6 | 12 | 6 | 23 |
| ENSDARG000 | 22;22;22;22;2 | 11538392;11 | 11539079;11 | -;-;-;-;-;-;-;- | 1587 | gnav1 | 13 | 12 | 18 | 21 | 14 | 2 |
| ENSDARG000 | 22;22;22;22;2 | 5233396;523 | 5234196;523 | -;-;-;-;-;-;-;-; | 2564 | ncln | 1678 | 2092 | 2588 | 1211 | 1887 | 1610 |
| ENSDARG000 | 22;22;22;22;2 | 20714624;20 | 20715246;20 | -;-;-;-;-;-;-;- | 1426 | oaz1a | 6283 | 6033 | 7447 | 8106 | 6859 | 6033 |
| ENSDARG000 | 22;22;22;22;2 | 3303671;330 | 3304117;330 | +;+;+;+;+;+;+ | 1754 | gipc3 | 38 | 72 | 42 | 92 | 34 | 44 |
| ENSDARG000 | 22;22;22;22;2 | 36931219;36 | 36931336;36 | -;-;-;-;- | 934 | NA | 0 | 0 | 0 | 0 | 0 | 0 |
| ENSDARG000 | 22;22;22;22;2 | 18230864;18 | 18233850;18 | -;-;-;-;-;-;-;-; | 4300 | tmem161a | 380 | 423 | 621 | 316 | 374 | 596 |
| ENSDARG000 | 22;22;22;22;2 | 31207226;31 | 31208194;31 | +;+;+;+;+;+;+; | 5001 | GRIP2 | 0 | 0 | 2 | 12 | 15 | 0 |
| ENSDARG000 | 22;22 | 3375262;337 | 3375350;337 | +;+ | 464 | NA | 0 | 0 | 0 | 0 | 0 | 0 |
| ENSDARG000 | 22;22;22;22;2 | 7480465;748 | 7480543;748 | +;+;+;+;+;+;+; | 924 | cela1.3 | 0 | 0 | 0 | 0 | 0 | 0 |
| ENSDARG000 | 22;22;22;22;2 | 15577827;15 | 15579438;15 | -;-;-;-;-;-;-;-; | 3460 | tpm4a | 249 | 151 | 765 | 113 | 326 | 246 |
| ENSDARG000 | 22;22;22;22;2 | 12831675;12 | 12831828;12 | -;-;-;-;-;-;-;-; | 1946 | glsa | 383 | 243 | 545 | 225 | 260 | 407 |
| ENSDARG000 | 22;22 | 3384732;338 | 3384964;338 | +;+ | 592 | NA | 2 | 0 | 2 | 0 | 15 | 0 |
| ENSDARG000 | 22;22;22;22;2 | 15570965;15 | 15572592;15 | -;-;-;-;-;-;-;-; | 2525 | hsh2d | 6 | 0 | 16 | 2 | 16 | 4 |
| ENSDARG000 | 22;22;22;22;2 | 20720808;20 | 20721000;20 | +;+;+;+;+ | 619 | NA | 0 | 0 | 0 | 0 | 0 | 0 |
| ENSDARG000 | 22;22;22;22;2 | 20729063;20 | 20729583;20 | -;-;-;-;- | 816 | NA | 2 | 0 | 2 | 0 | 0 | 0 |
| ENSDARG000 | 22;22 | 10300526;10 | 10300606;10 | +;+ | 204 | NA | 0 | 0 | 0 | 0 | 0 | 0 |
| ENSDARG000 | 22;22;22;22;2 | 10281420;10 | 10282199;10 | -;-;-;-;-;-;-;-; | 9122 | stab1 | 191 | 50 | 559 | 88 | 259 | 520 |
| ENSDARG000 | 22;22;22;22;2 | 23238790;23 | 23243661;23 | -;-;-;-;-;-;-;-; | 6522 | lhx9 | 14 | 13 | 0 | 0 | 4 | 14 |
| ENSDARG000 | 22;22;22;22;2 | 2129384;212 | 2129581;212 | +;+;+;+;+;+;+; | 590 | NA | 0 | 0 | 0 | 0 | 4 | 0 |
| ENSDARG000 | 22;22 | 5320415;532 | 5322129;532 | -;- | 1772 | s1pr4 | 25 | 4 | 4 | 26 | 18 | 70 |
| ENSDARG000 | 22;22;22;22;2 | 36944527;36 | 36948824;36 | -;-;-;-;-;-;-;-; | 7360 | ATP11B | 1429 | 1228 | 1350 | 725 | 1597 | 1159 |
| ENSDARG000 | 22;22;22;22;2 | 18247292;18 | 18247958;18 | +;+;+;+;+;+;+ | 1740 | NA | 0 | 0 | 0 | 0 | 2 | 0 |
| ENSDARG000 | 22;22;22;22;2 | 20747182;20 | 20749045;20 | -;-;-;-;-;-;- | 3243 | amh | 2 | 1 | 0 | 26 | 0 | 0 |
| ENSDARG000 | 22;22;22;22;2 | 28858225;28 | 28858499;28 | -;-;-;-;-;-;-;-; | 1904 | gtpbp2b | 149 | 110 | 260 | 148 | 244 | 198 |
| ENSDARG000 | 22;22;22;22;2 | 866681;8696 | 869998;8699 | -;-;-;-;-;-;-;-; | 4886 | cept1b | 578 | 694 | 831 | 517 | 1064 | 526 |
| ENSDARG000 | 22;22;22 | 23264622;23 | 23264833;23 | -;-;- | 715 | C22H1orf53 | 27 | 32 | 50 | 8 | 22 | 47 |
| ENSDARG000 | 22 | 18386253 | 18387059 | - | 807 | tssk6 | 0 | 0 | 0 | 0 | 0 | 0 |
| ENSDARG000 | 22;22;22;22;2 | 5331675;533 | 5332027;533 | +;+;+;+;+;+;+; | 6675 | MYO9B | 3316 | 1494 | 2372 | 1606 | 2055 | 2064 |
| ENSDARG000 | 22;22;22;22;2 | 18315490;18 | 18316545;18 | +;+;+;+;+;+;+; | 8585 | gatad2ab | 1414 | 1019 | 1575 | 1168 | 1630 | 1413 |
| ENSDARG000 | 22;22;22;22;2 | 7486867;748 | 7486934;748 | +;+;+;+;+;+;+; | 1373 | cela1.4 | 0 | 0 | 0 | 0 | 0 | 0 |
| ENSDARG000 | 22;22;22;22;2 | 26129250;26 | 26129960;26 | -;-;-;-;-;-;-;-; | 5732 | c3b.2 | 0 | 2 | 14 | 0 | 3 | 0 |
| ENSDARG000 | 22;22;22;22;2 | 31784419;31 | 31784476;31 | -;-;-;-;-;-;- | 1040 | NA | 1 | 0 | 0 | 0 | 0 | 0 |
| ENSDARG000 | 22;22;22;22;2 | 3389347;338 | 3389978;338 | -;-;-;-;-;-;-;-; | 6580 | PTPRS | 30 | 10 | 70 | 3 | 80 | 20 |
| ENSDARG000 | 22;22;22;22;2 | 11596432;11 | 11599250;11 | -;-;-;-;-;-;-;-; | 4295 | pcyt2 | 2522 | 3128 | 2993 | 2669 | 2085 | 2648 |
| ENSDARG000 | 22;22;22;22;2 | 7497319;749 | 7497386;749 | +;+;+;+;+;+;+; | 921 | cela1.5 | 0 | 0 | 0 | 0 | 0 | 0 |
| ENSDARG000 | 22;22 | 3595268;368 | 3595439;368 | -;- | 277 | NA | 0 | 0 | 0 | 0 | 0 | 0 |
| ENSDARG000 | 22;22;22;22;2 | 5667371;566 | 5667503;566 | -;-;-;-;-;-;-;-; | 990 | dnase1l4.1 | 0 | 0 | 4 | 0 | 30 | 0 |
| ENSDARG000 | 22;22;22 | 11614776;11 | 11615052;11 | +;+;+ | 743 | NA | 0 | 0 | 0 | 1 | 5 | 1 |
| ENSDARG000 | 22;22;22;22;2 | 20782915;20 | 20783375;20 | -;-;-;-;-;-;-;-; | 4908 | DOT1L | 362 | 526 | 936 | 626 | 596 | 229 |
| ENSDARG000 | 22;22;22;22;2 | 11617520;11 | 11617922;11 | -;-;-;-;-;-;-;-; | 2317 | gcga | 6 | 2 | 14 | 0 | 2 | 14 |
| ENSDARG000 | 22;22;22;22;2 | 23359369;23 | 23359505;23 | +;+;+;+;+;+;+; | 3070 | DENND1B | 76 | 25 | 40 | 34 | 68 | 43 |
| ENSDARG000 | 22;22;22;22 | 5663529;566 | 5663706;566 | +;+;+;+ | 1275 | tma7 | 812 | 1088 | 1131 | 405 | 774 | 829 |
| ENSDARG000 | 22;22;22;22;2 | 28888781;28 | 28888858;28 | +;+;+;+;+ | 756 | NA | 0 | 0 | 0 | 0 | 0 | 0 |
| ENSDARG000 | 22;22;22;22;2 | 12870275;12 | 12871033;12 | -;-;-;-;-;-;- | 2152 | nab1a | 24 | 24 | 36 | 22 | 49 | 54 |
| ENSDARG000 | 22;22;22;22;2 | 17237219;17 | 17237442;17 | +;+;+;+;+;+;+; | 3208 | axdnd1 | 18 | 18 | 68 | 70 | 63 | 120 |
| ENSDARG000 | 22;22;22;22;2 | 883678;8863 | 883887;8865 | +;+;+;+;+;+;+; | 1666 | STK38 | 344 | 347 | 416 | 319 | 504 | 475 |
| ENSDARG000 | 22;22;22;22;2 | 23493170;23 | 23493454;23 | -;-;-;-;-;-;-;-; | 4952 | crb1 | 2 | 2 | 6 | 0 | 6 | 0 |
| ENSDARG000 | 22;22;22;22 | 28969071;28 | 28969341;28 | +;+;+;+ | 679 | NA | 0 | 0 | 0 | 0 | 0 | 0 |
| ENSDARG000 | 22;22;22;22;2 | 2143107;214 | 2143175;214 | +;+;+;+;+;+;+; | 1618 | NA | 41 | 33 | 58 | 41 | 26 | 42 |
| ENSDARG000 | 22;22 | 23492326;23 | 23492625;23 | +;+ | 806 | NA | 2 | 0 | 2 | 0 | 0 | 0 |
| ENSDARG000 | 22;22;22;22;2 | 7614672;761 | 7615999;761 | -;-;-;-;-;- | 1596 | NA | 8 | 2 | 0 | 0 | 0 | 10 |
| ENSDARG000 | 22;22;22;22;2 | 12911236;12 | 12911585;12 | -;-;-;-;-;- | 2558 | mfsd6a | 329 | 94 | 114 | 115 | 204 | 142 |
| ENSDARG000 | 22;22;22;22;2 | 17251275;17 | 17251811;17 | -;-;-;-;-;-;-;-; | 1808 | nphs2 | 2 | 0 | 2 | 0 | 41 | 16 |
| ENSDARG000 | 22;22;22;22;2 | 15695746;15 | 15696208;15 | -;-;-;-;-;-;-;-; | 3216 | safb | 4378 | 3680 | 5323 | 2932 | 5049 | 3901 |
| ENSDARG000 | 22;22;22;22;2 | 12933485;12 | 12934553;12 | -;-;-;-;-;- | 1460 | NA | 10 | 5 | 5 | 0 | 0 | 0 |
| ENSDARG000 | 22;22;22;22;2 | 38796339;38 | 38796824;38 | -;-;-;-;-;-;-;-; | 1512 | si:ch211-262l | 234 | 415 | 445 | 302 | 490 | 351 |
| ENSDARG000 | 22;22;22;22;2 | 17261801;17 | 17262846;17 | +;+;+;+;+;+;+; | 4083 | tdrd5 | 0 | 2 | 4 | 10 | 4 | 2 |
| ENSDARG000 | 22;22;22;22;2 | 28995391;28 | 28995468;28 | +;+;+;+;+ | 756 | NA | 0 | 0 | 0 | 0 | 0 | 0 |
| ENSDARG000 | 22 | 37347896 | 37349967 | - | 2072 | sox2 | 760 | 1133 | 1253 | 845 | 1144 | 17 |
| ENSDARG000 | 22;22;22;22;2 | 7675388;768 | 7677770;768 | -;-;-;-;-;-;-;-; | 5141 | NA | 2 | 0 | 0 | 0 | 0 | 792 |
| ENSDARG000 | 22;22;22;22;2 | 37347440;37 | 37347542;37 | -;-;-;-;-;-;-;-; | 1113 | NA | 73 | 19 | 22 | 10 | 10 | 18 |
| ENSDARG000 | 22;22;22;22;2 | 15643174;15 | 15646723;15 | -;-;-;-;-;-;-;-; | 5905 | NA | 18 | 2 | 20 | 6 | 14 | 4 |
| ENSDARG000 | 22;22;22;22;2 | 31821815;31 | 31822262;31 | +;+;+;+;+;+;+; | 7518 | DOCK3 | 1351 | 907 | 3163 | 2331 | 3097 | 1651 |
| ENSDARG000 | 22;22;22;22;2 | 29009541;29 | 29009618;29 | +;+;+;+;+ | 756 | NA | 0 | 0 | 0 | 0 | 0 | 0 |
| ENSDARG000 | 22;22 | 7738966;773 | 7739117;773 | +;+ | 496 | NA | 0 | 0 | 0 | 0 | 0 | 2 |
| ENSDARG000 | 22;22;22;22;2 | 12979213;12 | 12980116;12 | -;-;-;-;-;-;-;-; | 4032 | ahr1b | 160 | 330 | 332 | 251 | 275 | 261 |
| ENSDARG000 | 22;22;22;22;2 | 18389271;18 | 18389431;18 | +;+;+;+;+;+ | 856 | yjefn3 | 3330 | 2749 | 2351 | 2598 | 2861 | 2435 |
| ENSDARG000 | 22;22;22;22;2 | 29027037;29 | 29027114;29 | +;+;+;+;+ | 756 | NA | 0 | 0 | 0 | 0 | 0 | 0 |
| ENSDARG000 | 22;22;22;22;2 | 20847580;20 | 20849758;20 | -;-;-;-;-;-;-;-; | 4151 | ell | 473 | 483 | 823 | 734 | 690 | 669 |
| ENSDARG000 | 22;22 | 2160039;216 | 2160093;216 | +;+ | 347 | NA | 13 | 17 | 16 | 6 | 5 | 19 |
| ENSDARG000 | 22;22;22;22;2 | 18469004;18 | 18469067;18 | +;+;+;+;+;+;+; | 3776 | CILP2 | 84 | 52 | 12 | 56 | 89 | 6 |
| ENSDARG000 | 22;22;22 | 17286841;17 | 17286976;17 | +;+;+ | 632 | fam163ab | 0 | 0 | 16 | 14 | 14 | 0 |
| ENSDARG000 | 22;22;22;22;2 | 7742211;774 | 7742273;774 | +;+;+;+;+;+;+; | 1648 | NA | 0 | 2 | 0 | 0 | 0 | 0 |
| ENSDARG000 | 22;22;22;22;2 | 17312111;17 | 17312248;17 | +;+;+;+;+;+;+; | 5077 | NA | 3145 | 2955 | 3314 | 3004 | 2956 | 3696 |
| ENSDARG000 | 22;22;22;22;2 | 37534094;37 | 37534228;37 | +;+;+;+;+;+ | 954 | dnajc19 | 101 | 95 | 90 | 115 | 73 | 110 |
| ENSDARG000 | 22 | 17359346 | 17360458 | + | 1113 | GPR52 | 2 | 0 | 0 | 2 | 6 | 0 |
| ENSDARG000 | 22;22;22 | 18488701;18 | 18489217;18 | -;-;- | 925 | si:ch211-212 | 187 | 245 | 263 | 233 | 286 | 232 |
| ENSDARG000 | 22;22 | 4311803;431 | 4311933;431 | -;- | 422 | NA | 0 | 0 | 0 | 0 | 0 | 0 |
| ENSDARG000 | 22;22;22;22;2 | 23546926;23 | 23547295;23 | +;+;+;+;+;+;+; | 10507 | aspm | 1000 | 1660 | 2347 | 954 | 2109 | 920 |
| ENSDARG000 | 22;22;22;22;2 | 11628438;11 | 11629574;11 | -;-;-;-;-;-;-;-; | 3362 | dpp4 | 10 | 18 | 26 | 20 | 43 | 49 |
| ENSDARG000 | 22;22;22;22;2 | 29067388;29 | 29067449;29 | +;+;+;+;+ | 750 | NA | 0 | 0 | 0 | 0 | 0 | 0 |
| ENSDARG000 | 22;22;22;22;2 | 13101736;13 | 13101909;13 | -;-;-;-;- | 623 | NA | 2 | 2 | 2 | 0 | 3 | 0 |
| ENSDARG000 | 22;22;22;22;2 | 13046270;13 | 13050126;13 | -;-;-;-;-;-;-;-; | 7511 | ahr2 | 5134 | 4261 | 6664 | 5129 | 6507 | 4146 |
| ENSDARG000 | 22;22;22;22;2 | 4313605;431 | 4317316;431 | -;-;-;-;-;-;-;-; | 6944 | kdm4b | 2380 | 1220 | 3212 | 2715 | 2204 | 1730 |
| ENSDARG000 | 22;22;22;22 | 38804330;38 | 38804643;38 | -;-;-;- | 671 | NA | 0 | 0 | 0 | 0 | 0 | 0 |
| ENSDARG000 | 22;22;22;22;2 | 2170887;217 | 2171109;217 | +;+;+;+;+;+;+; | 1181 | NA | 45 | 31 | 17 | 44 | 31 | 54 |
| ENSDARG000 | 22;22;22;22;2 | 29075042;29 | 29078817;29 | -;-;-;-;-;-;-;-; | 5708 | cbx6a | 764 | 616 | 826 | 327 | 739 | 649 |
| ENSDARG000 | 22;22;22 | 38807943;38 | 38808360;38 | +;+;+ | 1046 | NA | 4 | 2 | 1 | 16 | 0 | 0 |
| ENSDARG000 | 22;22;22;22 | 32228882;32 | 32229112;32 | +;+;+;+ | 1637 | manf | 685 | 1363 | 2295 | 365 | 1418 | 508 |
| ENSDARG000 | 22;22;22;22 | 7792791;779 | 7792906;779 | +;+;+;+ | 1470 | NA | 0 | 0 | 16 | 8 | 9 | 4 |
| ENSDARG000 | 22;22;22;22;2 | 7758462;775 | 7761900;776 | -;-;-;-;-;-;-;-; | 5936 | NA | 195 | 133 | 180 | 134 | 191 | 77 |
| ENSDARG000 | 22;22;22;22;2 | 7758462;780 | 7761900;780 | -;-;-;-;-;-;-;- | 4342 | NA | 132 | 81 | 186 | 76 | 152 | 42 |
| ENSDARG000 | 22;22;22;22;2 | 10361170;10 | 10362346;10 | -;-;-;-;-;-;-;-; | 5196 | NISCH | 2110 | 1829 | 2537 | 2668 | 3517 | 3130 |
| ENSDARG000 | 22;22;22 | 32235922;32 | 32241124;32 | +;+;+ | 5203 | rbm15b | 430 | 302 | 598 | 155 | 522 | 288 |
| ENSDARG000 | 22;22;22;22;2 | 20925849;20 | 20926768;20 | -;-;-;-;-;-;-;- | 2125 | fkbp8 | 633 | 792 | 639 | 660 | 652 | 605 |
| ENSDARG000 | 22;22;22;22;2 | 13276693;13 | 13282718;13 | -;-;-;-;-;-;-;-; | 6675 | NA | 2 | 0 | 7 | 0 | 110 | 4 |
| ENSDARG000 | 22;22;22;22;2 | 912450;9139 | 912975;9140 | -;-;-;-;-;-;-;-; | 5650 | cacna1sa | 6 | 0 | 0 | 0 | 0 | 0 |
| ENSDARG000 | 22;22;22;22;2 | 2175857;217 | 2176086;217 | +;+;+;+;+ | 549 | NA | 13 | 23 | 9 | 4 | 3 | 5 |
| ENSDARG000 | 22;22 | 32266906;32 | 32267058;32 | -;- | 564 | NA | 0 | 0 | 0 | 1 | 0 | 0 |

| | | | | | | | | | | | | |
|---|---|---|---|---|---|---|---|---|---|---|---|---|
| ENSDARG000 | 22:22:22:22:2 | 18530395:185 | 18530460:185 | +:+:+:+:+:+:+: | 3916 | NA | 30 | 42 | 14 | 30 | 60 | 72 |
| ENSDARG000 | 22;22;22;22 | 11657139:116 | 11657774:116 | -;-;-;- | 1058 | mettl21a | 110 | 124 | 126 | 54 | 102 | 92 |
| ENSDARG000 | 22:22:22:22:2 | 11661766:116 | 11661864:116 | +;+;+;+;+;+;+ | 1746 | NA | 0 | 0 | 3 | 0 | 4 | 0 |
| ENSDARG000 | 22;22 | 32278262:322 | 32278567:322 | -;- | 795 | NA | 32 | 0 | 2 | 6 | 0 | 2 |
| ENSDARG000 | 22:22:22:22:2 | 38812549:388 | 38813477:388 | -:-:-:-:-:-:-:-: | 2056 | si:ch211-262h | 0 | 0 | 6 | 20 | 8 | 0 |
| ENSDARG000 | 22:22:22:22:2 | 26180184:261 | 26181005:261 | -:-:-:-:-:-:-:-: | 6026 | c3b.1 | 6 | 0 | 31 | 0 | 11 | 0 |
| ENSDARG000 | 22:22:22:22:2 | 18542010:185 | 18544107:185 | -:-:-:-:-:-:-:-: | 3511 | cirbpb | 29409 | 31289 | 41487 | 37277 | 27631 | 35414 |
| ENSDARG000 | 22:22:22:22:2 | 2183110:2183 | 2183181:2183 | +:+:+:+:+:+:+: | 1089 | NA | 43 | 59 | 29 | 22 | 40 | 84 |
| ENSDARG000 | 22:22:22:22:2 | 29087308:290 | 29087500:290 | +:+:+:+:+:+:+: | 7687 | pla2g6 | 1073 | 1029 | 1226 | 845 | 1264 | 990 |
| ENSDARG000 | 22:22:22:22:2 | 37542741:375 | 37543670:375 | -:-:-:-:-:-:-:-: | 2939 | fxr1 | 1252 | 1281 | 1830 | 1501 | 1306 | 1035 |
| ENSDARG000 | 22:22:22:22:2 | 2189356:2189 | 2189584:2189 | +:+:+:+:+:+:+: | 928 | NA | 22 | 11 | 48 | 32 | 20 | 38 |
| ENSDARG000 | 22:22:22:22:2 | 5682635:5682 | 5682729:5682 | +:+:+:+:+:+:+: | 3425 | dnase1l4.2 | 76 | 56 | 66 | 38 | 225 | 108 |
| ENSDARG000 | 22:22:22:22:2 | 17455711:174 | 17456169:174 | -:-:-:-:-:-:-:-: | 1765 | ifi44f1 | 17 | 2 | 7 | 1 | 30 | 6 |
| ENSDARG000 | 22:22:22:22:2 | 32289567:322 | 32292450:322 | -:-:-:-:-:-:-:-: | 4219 | pcbp4 | 3771 | 2917 | 4986 | 4903 | 4280 | 3096 |
| ENSDARG000 | 22:22:22:22:2 | 8497039:8497 | 8497959:8497 | -;-;-;-;-;- | 2168 | NA | 132 | 76 | 150 | 135 | 61 | 86 |
| ENSDARG000 | 22:22:22:22:2 | 8506168:8507 | 8506524:8508 | -;-;-;-;-;- | 1439 | NA | 27 | 34 | 78 | 29 | 59 | 25 |
| ENSDARG000 | 22:22:22:22:2 | 4408643:4412 | 4411431:4412 | -:-:-:-:-:-:-:-: | 5079 | uhrf1 | 444 | 1344 | 1611 | 621 | 1582 | 1017 |
| ENSDARG000 | 22:22:22:22:2 | 13458735:134 | 13463231:134 | -:-:-:-:-:-:-:-: | 7892 | CNTNAP5 | 8 | 0 | 526 | 6 | 904 | 0 |
| ENSDARG000 | 22:22:22:22:2 | 10479965:104 | 10480838:104 | -:-:-:-:-:-:-:-: | 1973 | aspn | 247 | 12 | 8 | 2 | 22 | 10 |
| ENSDARG000 | 22:22:22:22:2 | 11710467:117 | 11710898:117 | -;-;-;-;-;-;- | 1085 | crygs1 | 0 | 0 | 0 | 0 | 0 | 0 |
| ENSDARG000 | 22;22;22;22 | 8596974:8598 | 8597072:8598 | +;+;+;+ | 815 | NA | 22 | 0 | 0 | 0 | 0 | 12 |
| ENSDARG000 | 22:22:22:22:2 | 8581752:8581 | 8586795:8586 | -:-:-:-:-:-:-:-: | 8460 | NA | 947 | 776 | 856 | 592 | 456 | 493 |
| ENSDARG000 | 22:22:22:22:2 | 10454603:104 | 10455317:104 | -;-;-;-;-;- | 1518 | ogna | 40 | 2 | 30 | 2 | 56 | 0 |
| ENSDARG000 | 22;22;22;22 | 13638586:136 | 13638680:136 | -;-;-;- | 537 | NA | 0 | 0 | 0 | 0 | 0 | 0 |
| ENSDARG000 | 22:22:22:22:2 | 8531200:8531 | 8531252:8532 | -;-;-;-;- | 716 | NA | 12 | 12 | 71 | 6 | 0 | 2 |
| ENSDARG000 | 22;22;22;22 | 8536838:8540 | 8536896:8540 | +;+;+;+ | 769 | si:ch73-27e22 | 4 | 31 | 149 | 43 | 0 | 4 |
| ENSDARG000 | 22:22:22:22:2 | 17460871:174 | 17461044:174 | -:-:-:-:-:-:-:-: | 1549 | NA | 24 | 4 | 12 | 7 | 11 | 10 |
| ENSDARG000 | 22:22:22:22:2 | 10440991:104 | 10441035:104 | +:+:+:+:+:+:+: | 1708 | cenpp | 92 | 240 | 265 | 104 | 165 | 148 |
| ENSDARG000 | 22:22:22:22:2 | 8589423:8713 | 8589830:8716 | -:-:-:-:-:-:-:-: | 5685 | NA | 3 | 220 | 2 | 107 | 347 | 457 |
| ENSDARG000 | 22:22:22:22:2 | 8695096:8695 | 8695146:8696 | +;+;+;+;+ | 1774 | NA | 0 | 0 | 0 | 0 | 1 | 13 |
| ENSDARG000 | 22;22 | 8517657:8546 | 8517799:8546 | -;- | 212 | NA | 0 | 2 | 13 | 0 | 0 | 2 |
| ENSDARG000 | 22;22;22 | 10463890:104 | 10464427:104 | -;-;- | 1861 | OMD | 8 | 0 | 2 | 0 | 0 | 0 |
| ENSDARG000 | 22:22:22:22:2 | 10424995:104 | 10425705:104 | -:-:-:-:-:-:-:-: | 4255 | nol8 | 283 | 349 | 589 | 250 | 362 | 298 |
| ENSDARG000 | 22:22:22:22:2 | 10490937:104 | 10491426:104 | -:-:-:-:-:-:-:-: | 2908 | NA | 33 | 4 | 28 | 14 | 25 | 11 |
| ENSDARG000 | 22:22:22:22:2 | 20951211:209 | 20955826:209 | -:-:-:-:-:-:-:-: | 5936 | ssbp4 | 590 | 472 | 783 | 434 | 918 | 722 |
| ENSDARG000 | 22:22:22:22:2 | 11717429:117 | 11720180:117 | -:-:-:-:-:-:-:-: | 3989 | KRT222 | 39 | 2 | 2 | 0 | 10 | 4 |
| ENSDARG000 | 22:22:22:22:2 | 13711434:137 | 13712880:137 | -:-:-:-:-:-:-:-: | 2866 | NA | 29 | 10 | 45 | 13 | 26 | 12 |
| ENSDARG000 | 22:22:22:22:2 | 2239254:2239 | 2239325:2239 | +:+:+:+:+:+:+: | 1268 | NA | 55 | 56 | 81 | 32 | 59 | 55 |
| ENSDARG000 | 22:22:22:22:2 | 2207502:2207 | 2207560:2207 | +:+:+:+:+:+:+: | 859 | NA | 38 | 13 | 30 | 12 | 26 | 16 |
| ENSDARG000 | 22:22:22:22:2 | 26239424:262 | 26240006:262 | -:-:-:-:-:-:-:-: | 1476 | yju2b | 178 | 150 | 251 | 207 | 164 | 116 |
| ENSDARG000 | 22;22;22 | 2254972:2255 | 2255030:2255 | +;+;+ | 396 | NA | 7 | 8 | 18 | 7 | 8 | 26 |
| ENSDARG000 | 22:22:22:22:2 | 2198246:2271 | 2198439:2271 | +;+;+;+;+;+;+ | 2308 | znf1157 | 19 | 6 | 29 | 33 | 8 | 30 |
| ENSDARG000 | 22:22:22:22:2 | 2198323:2198 | 2198467:2198 | +:+:+:+:+:+:+: | 684 | NA | 12 | 25 | 11 | 4 | 17 | 6 |
| ENSDARG000 | 22:22:22:22:2 | 24869690:248 | 24869885:248 | -:-:-:-:-:-:-:-: | 5002 | vtg2 | 0 | 4 | 6 | 0 | 0 | 4 |
| ENSDARG000 | 22;22;22 | 2260497:2261 | 2260556:2261 | +;+;+ | 557 | NA | 30 | 3 | 10 | 10 | 11 | 30 |
| ENSDARG000 | 22;22;22 | 2269395:2271 | 2269446:2271 | +;+;+ | 534 | NA | 4 | 8 | 9 | 10 | 14 | 5 |
| ENSDARG000 | 22;22 | 2195963:2206 | 2197169:2206 | +;+ | 1537 | NA | 42 | 48 | 26 | 18 | 22 | 53 |
| ENSDARG000 | 22:22:22:22:2 | 2217935:2217 | 2217980:2217 | +:+:+:+:+:+:+: | 986 | NA | 19 | 20 | 14 | 9 | 7 | 42 |
| ENSDARG000 | 22:22:22:22:2 | 2228919:2229 | 2228981:2230 | +;+;+;+;+ | 1545 | znf1161 | 84 | 37 | 80 | 38 | 93 | 55 |
| ENSDARG000 | 22:22:22:22:2 | 2247645:2247 | 2247716:2247 | +;+;+;+;+;+;+ | 646 | NA | 7 | 18 | 25 | 8 | 6 | 2 |
| ENSDARG000 | 22:22:22:22:2 | 26264543:262 | 26264819:262 | -;-;-;-;-;-;-;- | 1230 | wdr83 | 154 | 140 | 194 | 117 | 223 | 149 |
| ENSDARG000 | 22:22:22:22:2 | 11756040:117 | 11756509:117 | +:+:+:+:+:+:+: | 1503 | krt97 | 280 | 42 | 18 | 24 | 145 | 2 |
| ENSDARG000 | 22:22:22:22:2 | 38824012:388 | 38824223:388 | +:+:+:+:+:+:+: | 4367 | anos1b | 86 | 58 | 102 | 66 | 12 | 6 |
| ENSDARG000 | 22:22:22:22:2 | 13846130:138 | 13848159:138 | -;-;-;-;-;- | 2903 | s100b | 6 | 2 | 2 | 0 | 2 | 0 |
| ENSDARG000 | 22:22:22:22:2 | 38875921:388 | 38876797:388 | -;-;-;-;- | 1079 | NA | 0 | 0 | 0 | 0 | 0 | 2 |
| ENSDARG000 | 22:22:22:22:2 | 29165330:291 | 29169150:291 | -:-:-:-:-:-:-:-: | 5333 | pvalb7 | 12 | 4 | 6 | 0 | 0 | 8 |
| ENSDARG000 | 22:22:22:22:2 | 29204885:292 | 29204968:292 | +:+:+:+:+:+:+: | 2216 | ncf4 | 31 | 62 | 57 | 42 | 140 | 69 |
| ENSDARG000 | 22:22:22:22:2 | 17467591:174 | 17467785:174 | -:-:-:-:-:-:-:-: | 1606 | NA | 770 | 451 | 390 | 294 | 562 | 499 |
| ENSDARG000 | 22:22:22:22:2 | 18725101:187 | 18728162:187 | -:-:-:-:-:-:-:-: | 4686 | midn | 3947 | 2142 | 2451 | 2241 | 2593 | 3150 |
| ENSDARG000 | 22:22:22:22:2 | 13886821:138 | 13887005:138 | +:+:+:+:+:+:+: | 6497 | sh3bp4a | 294 | 358 | 498 | 318 | 226 | 274 |
| ENSDARG000 | 22;22;22 | 4442473:4443 | 4442688:4445 | +;+;+ | 2361 | ticam1 | 300 | 270 | 248 | 383 | 295 | 260 |
| ENSDARG000 | 22:22:22:22:2 | 29135846:291 | 29135899:291 | +:+:+:+:+:+:+: | 1551 | baiap2l2b | 0 | 2 | 0 | 0 | 0 | 0 |
| ENSDARG000 | 22 | 13977772 | 13979393 | + | 1622 | arl4ca | 30 | 2 | 22 | 10 | 4 | 10 |
| ENSDARG000 | 22:22:22:22:2 | 11778053:117 | 11781769:117 | -:-:-:-:-:-:-:-: | 6573 | ptpn4b | 30 | 36 | 97 | 38 | 148 | 78 |
| ENSDARG000 | 22:22:22:22:2 | 11775269:117 | 11775674:117 | +:+:+:+:+:+:+: | 1438 | krt96 | 0 | 6 | 9 | 0 | 7 | 0 |
| ENSDARG000 | 22:22:22:22:2 | 37588528:375 | 37591791:375 | -:-:-:-:-:-:-:-: | 5018 | ttc14 | 556 | 278 | 556 | 541 | 704 | 579 |
| ENSDARG000 | 22:22:22:22:2 | 17475034:174 | 17476621:174 | -:-:-:-:-:-:-:-: | 3403 | ifi44f4 | 625 | 395 | 235 | 112 | 115 | 271 |
| ENSDARG000 | 22:22:22:22:2 | 978867:98029 | 978971:98043 | +:+:+:+:+:+:+: | 4165 | DEF6 | 28 | 12 | 7 | 12 | 10 | 10 |
| ENSDARG000 | 22:22:22:22:2 | 15706062:157 | 15706697:157 | -:-:-:-:-:-:-:-: | 3148 | malt2 | 72 | 34 | 102 | 69 | 160 | 48 |
| ENSDARG000 | 22:22:22:22:2 | 18766061:187 | 18766191:187 | -;-;-;-;-;-;-;- | 1013 | atp5f1d | 1876 | 2891 | 2994 | 1994 | 2061 | 1481 |
| ENSDARG000 | 22:22:22:22:2 | 38914983:389 | 38915132:389 | +:+:+:+:+:+:+: | 2850 | senp5 | 145 | 140 | 193 | 76 | 207 | 86 |
| ENSDARG000 | 22:22:22:22:2 | 17484851:174 | 17485042:174 | -;-;-;-;-;-;- | 1353 | ifi44f5 | 180 | 129 | 16 | 13 | 9 | 48 |
| ENSDARG000 | 22;22 | 38914983:389 | 38915132:389 | +;+ | 801 | NA | 0 | 0 | 0 | 0 | 0 | 0 |
| ENSDARG000 | 22:22:22:22:2 | 21128113:211 | 21129747:211 | -;-;-;-;-;-;- | 2947 | tmem59l | 3023 | 3524 | 3529 | 5631 | 3655 | 3506 |
| ENSDARG000 | 22:22:22:22:2 | 23584797:235 | 23585186:235 | -:-:-:-:-:-:-:-: | 876 | NA | 0 | 0 | 0 | 0 | 0 | 0 |
| ENSDARG000 | 22:22:22:22:2 | 4447387:4447 | 4447790:4447 | +:+:+:+:+:+:+: | 5627 | timm44 | 508 | 481 | 657 | 273 | 566 | 398 |
| ENSDARG000 | 22;22 | 33132095:331 | 33132262:331 | -;- | 204 | NA | 0 | 0 | 0 | 0 | 0 | 0 |
| ENSDARG000 | 22:22:22:22:2 | 33131891:331 | 33132418:331 | +;+;+;+;+;+ | 5843 | dag1 | 1028 | 387 | 1148 | 552 | 922 | 746 |
| ENSDARG000 | 22:22:22:22:2 | 14051245:140 | 14051947:140 | +:+:+:+:+:+:+: | 4688 | NA | 59 | 32 | 228 | 38 | 349 | 109 |
| ENSDARG000 | 22:22:22:22:2 | 18786797:187 | 18787173:187 | +:+:+:+:+:+:+: | 3848 | CBARP | 282 | 172 | 434 | 70 | 228 | 66 |
| ENSDARG000 | 22:22:22:22:2 | 997838:10065 | 997897:10068 | +:+:+:+:+:+:+: | 1813 | PPARD | 56 | 73 | 61 | 54 | 70 | 49 |
| ENSDARG000 | 22:22:22:22:2 | 21152847:211 | 21152930:211 | +:+:+:+:+:+:+: | 1734 | crlf1b | 64 | 15 | 15 | 22 | 13 | 22 |
| ENSDARG000 | 22:22:22:22:2 | 8753092:8753 | 8753410:8753 | +:+:+:+:+:+:+: | 2336 | NA | 31 | 4 | 2 | 0 | 8 | 0 |
| ENSDARG000 | 22 | 1018601 | 1019152 | + | 552 | NA | 8 | 8 | 8 | 16 | 2 | 3 |
| ENSDARG000 | 22:22:22:22:2 | 17493170:174 | 17493655:174 | -;-;-;-;-;-;- | 1762 | NA | 0 | 4 | 0 | 0 | 0 | 0 |
| ENSDARG000 | 22:22:22:22:2 | 18884971:188 | 18885046:188 | +:+:+:+:+:+:+: | 5159 | fstl3 | 81 | 28 | 12 | 15 | 12 | 24 |
| ENSDARG000 | 22:22:22:22:2 | 17499905:174 | 17500025:175 | +:+:+:+:+:+:+: | 1930 | rab11ba | 1228 | 1195 | 1105 | 1156 | 1262 | 1117 |
| ENSDARG000 | 22:22:22:22:2 | 33362552:333 | 33362716:333 | +;+;+;+;+;+;+ | 1222 | nicn1 | 26 | 14 | 20 | 8 | 14 | 22 |
| ENSDARG000 | 22:22:22:22:2 | 23593192:235 | 23593298:235 | -;-;-;-;- | 921 | NA | 0 | 0 | 0 | 0 | 0 | 0 |
| ENSDARG000 | 22:22:22:22:2 | 8772363:8775 | 8772420:8775 | +;+;+;+;+;+ | 5311 | NA | 131 | 30 | 65 | 195 | 184 | 218 |
| ENSDARG000 | 22:22:22:22:2 | 24900284:249 | 24901578:249 | -:-:-:-:-:-:-:-: | 3349 | NA | 0 | 19 | 18 | 30 | 20 | 21 |
| ENSDARG000 | 22;22;22 | 8790643:8792 | 8791249:8793 | -;-;- | 1892 | NA | 0 | 0 | 8 | 0 | 0 | 18 |
| ENSDARG000 | 22;22;22 | 24936865:249 | 24937098:249 | -;-;- | 615 | NA | 4 | 0 | 2 | 0 | 6 | 0 |
| ENSDARG000 | 22:22:22:22:2 | 24888649:248 | 24888694:248 | -:-:-:-:-:-:-:-: | 4684 | NA | 0 | 0 | 9 | 4 | 14 | 0 |
| ENSDARG000 | 22:22:22:22:2 | 1028724:1032 | 1028845:1032 | +:+:+:+:+:+:+: | 1579 | NA | 273 | 186 | 200 | 138 | 254 | 201 |
| ENSDARG000 | 22;22 | 1039548:1039 | 1039597:1039 | +;+ | 126 | NA | 0 | 0 | 0 | 0 | 0 | 0 |
| ENSDARG000 | 22;22;22 | 15719347:157 | 15719703:157 | -;-;- | 699 | mrpl54 | 162 | 227 | 184 | 111 | 142 | 232 |
| ENSDARG000 | 22;22;22;22 | 15720381:157 | 15720692:157 | +;+;+;+ | 983 | fam32a | 702 | 648 | 783 | 858 | 875 | 992 |
| ENSDARG000 | 22:22:22:22:2 | 26277618:262 | 26278296:262 | -;-;-;-;-;-;- | 1925 | sde2 | 303 | 454 | 404 | 333 | 399 | 419 |
| ENSDARG000 | 22 | 33791073 | 33791336 | - | 264 | NA | 0 | 0 | 0 | 0 | 0 | 0 |
| ENSDARG000 | 22;22;22;22 | 4488454:4513 | 4488501:4514 | +;+;+;+ | 1142 | CTXN1 | 4 | 6 | 14 | 10 | 5 | 0 |
| ENSDARG000 | 22:22:22:22:2 | 8799855:8800 | 8800530:8800 | +:+:+:+:+:+:+: | 3022 | NA | 9 | 3 | 7 | 0 | 0 | 12 |
| ENSDARG000 | 22:22:22:22:2 | 23598093:236 | 23598371:236 | -;-;-;-;-;-;- | 1192 | cfhl5 | 0 | 14 | 2 | 0 | 8 | 2 |
| ENSDARG000 | 22:22:22:22:2 | 34350349:343 | 34350463:343 | -:-:-:-:-:-:-:-: | 3918 | RNF123 | 209 | 144 | 173 | 94 | 185 | 163 |
| ENSDARG000 | 22:22:22:22:2 | 23598036:236 | 23598371:236 | -:-:-:-:-:-:-:-: | 2438 | cfhl4 | 0 | 0 | 28 | 0 | 0 | 0 |
| ENSDARG000 | 22 | 34430310 | 34431770 | + | 1461 | AMIGO3 | 0 | 0 | 0 | 0 | 0 | 0 |
| ENSDARG000 | 22:22:22:22:2 | 5704559:5719 | 5705003:5720 | -;-;-;-;-;- | 2675 | NA | 18 | 28 | 112 | 16 | 134 | 37 |
| ENSDARG000 | 22;22 | 5710003:5713 | 5710503:5713 | -;- | 696 | NA | 4 | 10 | 50 | 6 | 2 | 4 |
| ENSDARG000 | 22:22:22:22:2 | 34598340:345 | 34599503:345 | -:-:-:-:-:-:-:-: | 2611 | terf2ip | 92 | 117 | 204 | 156 | 229 | 169 |
| ENSDARG000 | 22:22:22:22:2 | 18929412:189 | 18929642:189 | +:+:+:+:+:+:+: | 2304 | bsg | 9890 | 9087 | 17060 | 13518 | 11486 | 10280 |
| ENSDARG000 | 22;22 | 15732967:157 | 15733063:157 | +;+ | 540 | NA | 0 | 0 | 0 | 0 | 0 | 0 |
| ENSDARG000 | 22;22 | 15747337:157 | 15747449:157 | +;+ | 660 | NA | 0 | 0 | 0 | 0 | 3 | 0 |
| ENSDARG000 | 22:22:22:22:2 | 37631034:376 | 37631432:376 | +:+:+:+:+:+:+: | 3555 | psmd1 | 4973 | 5460 | 6798 | 2466 | 7161 | 3250 |
| ENSDARG000 | 22:22:22:22:2 | 34615591:346 | 34616624:346 | +:+:+:+:+:+:+: | 2715 | NA | 8 | 20 | 6 | 6 | 10 | 22 |
| ENSDARG000 | 22:22:22:22:2 | 21167096:211 | 21167326:211 | -:-:-:-:-:-:-:-: | 1314 | rex1bd | 306 | 246 | 362 | 237 | 336 | 285 |
| ENSDARG000 | 22;22;22 | 15898221:158 | 15898611:158 | +;+;+ | 2180 | klf2a | 467 | 70 | 34 | 96 | 282 | 92 |
| ENSDARG000 | 22 | 26290209 | 26290863 | + | 655 | mrps28 | 120 | 231 | 210 | 150 | 184 | 128 |
| ENSDARG000 | 22:22:22:22:2 | 2298419:2300 | 2298490:2300 | +;+;+;+;+;+;+ | 727 | NA | 29 | 35 | 29 | 27 | 23 | 49 |
| ENSDARG000 | 22:22:22:22:2 | 26299016:262 | 26299101:262 | -:-:-:-:-:-:-:-: | 2103 | capn1b | 411 | 301 | 299 | 609 | 320 | 516 |
| ENSDARG000 | 22:22:22:22:2 | 26328063:263 | 26328377:263 | -:-:-:-:-:-:-:-: | 2454 | capn2b | 4535 | 4194 | 1556 | 5801 | 3943 | 4321 |
| ENSDARG000 | 22:22:22:22:2 | 1049367:1055 | 1049487:1056 | +:+:+:+:+:+:+: | 2381 | camk1ga | 2 | 10 | 8 | 0 | 4 | 4 |
| ENSDARG000 | 22;22 | 34645215:346 | 34645284:346 | +;+ | 496 | NA | 4 | 0 | 0 | 0 | 0 | 0 |
| ENSDARG000 | 22:22:22:22:2 | 5727753:5731 | 5728117:5731 | +:+:+:+:+:+:+: | 1651 | NA | 4 | 0 | 0 | 0 | 4 | 4 |
| ENSDARG000 | 22:22:22:22:2 | 34661401:346 | 34661561:346 | +:+:+:+:+:+:+: | 4014 | NA | 0 | 0 | 0 | 0 | 0 | 0 |
| ENSDARG000 | 22:22:22:22:2 | 5742173:5742 | 5742786:5742 | -;-;-;-;- | 854 | BCDIN3D | 8 | 22 | 24 | 22 | 24 | 16 |
| ENSDARG000 | 22:22:22:22:2 | 5745303:5746 | 5745648:5747 | -:-:-:-:-:-:-:-: | 1570 | NA | 0 | 0 | 0 | 0 | 0 | 0 |
| ENSDARG000 | 22:22:22:22:2 | 18951096:189 | 18951103:189 | +:+:+:+:+:+:+: | 3334 | HCN2 | 0 | 3 | 0 | 8 | 16 | 6 |
| ENSDARG000 | 22:22:22:22:2 | 11857356:118 | 11857765:118 | +:+:+:+:+:+:+: | 4017 | MRAS | 264 | 164 | 288 | 189 | 376 | 359 |
| ENSDARG000 | 22:22:22:22:2 | 1073413:1073 | 1073499:1073 | -;-;-;-;-;- | 918 | NA | 95 | 24 | 0 | 2 | 0 | 29 |
| ENSDARG000 | 22:22:22:22:2 | 8979955:8982 | 8980098:8983 | +;+;+;+;+;+ | 2576 | NA | 684 | 1497 | 658 | 922 | 1190 | 4100 |
| ENSDARG000 | 22:22:22:22:2 | 10516332:105 | 10517387:105 | -:-:-:-:-:-:-:-: | 2412 | ippk | 279 | 298 | 357 | 249 | 396 | 254 |
| ENSDARG000 | 22:22:22:22:2 | 29312228:293 | 29312923:293 | -:-:-:-:-:-:-:-: | 2516 | NA | 62 | 23 | 71 | 36 | 170 | 62 |
| ENSDARG000 | 22:22:22:22:2 | 2306386:2306 | 2306458:2306 | +:+:+:+:+:+:+: | 835 | NA | 48 | 44 | 70 | 4 | 40 | 36 |
| ENSDARG000 | 22;22;22;22 | 2315996:2316 | 2316067:2316 | +;+;+;+ | 569 | NA | 63 | 26 | 69 | 59 | 67 | 37 |
| ENSDARG000 | 22:22:22:22:2 | 21252790:212 | 21252985:212 | +:+:+:+:+:+:+: | 7728 | CPAMD8 | 26 | 0 | 8 | 6 | 26 | 10 |
| ENSDARG000 | 22:22:22:22:2 | 11898661:118 | 11899230:118 | -:-:-:-:-:-:-:-: | 1997 | PTH2R | 0 | 0 | 0 | 0 | 16 | 0 |
| ENSDARG000 | 22:22:22:22:2 | 34710036:347 | 34710622:347 | +:+:+:+:+:+:+: | 1789 | hoga1 | 44 | 68 | 123 | 50 | 160 | 37 |
| ENSDARG000 | 22;22 | 29584800:295 | 29586078:295 | -;- | 1432 | adra2a | 6 | 0 | 0 | 0 | 26 | 0 |
| ENSDARG000 | 22:22:22:22:2 | 24954103:249 | 24954487:249 | -:-:-:-:-:-:-:-: | 1712 | NA | 0 | 0 | 0 | 0 | 0 | 0 |
| ENSDARG000 | 22:22:22:22:2 | 34691020:346 | 34691148:346 | +:+:+:+:+:+:+: | 1224 | HYKK | 4 | 6 | 0 | 0 | 4 | 0 |
| ENSDARG000 | 22;22;22;22 | 34701848:347 | 34702043:347 | +;+;+;+ | 901 | atpv0e2 | 214 | 290 | 262 | 318 | 220 | 374 |
| ENSDARG000 | 22:22:22:22:2 | 15912253:159 | 15912399:159 | -:-:-:-:-:-:-:-: | 5515 | EPS15L1 | 954 | 905 | 1510 | 1607 | 1062 | 879 |
| ENSDARG000 | 22:22:22:22:2 | 19018602:190 | 19018651:190 | -:-:-:-:-:-:-:-: | 3846 | POLRMT | 259 | 279 | 356 | 146 | 254 | 231 |
| ENSDARG000 | 22:22:22:22:2 | 2328299:2333 | 2328344:2333 | +;+;+;+;+;+;+ | 2336 | NA | 218 | 196 | 274 | 173 | 236 | 249 |
| ENSDARG000 | 22;22 | 5752257:5756 | 5752359:5756 | +;+ | 440 | cox14 | 110 | 136 | 144 | 72 | 82 | 78 |
| ENSDARG000 | 22;22;22;22 | 2345752:2347 | 2345806:2347 | +;+;+;+ | 564 | NA | 6 | 19 | 13 | 4 | 31 | 10 |
| ENSDARG000 | 22;22;22 | 19111444:191 | 19111768:191 | +;+;+ | 624 | fgf22 | 0 | 0 | 0 | 0 | 0 | 0 |
| ENSDARG000 | 22:22:22:22:2 | 17509422:175 | 17509574:175 | +;+;+;+;+ | 1701 | marchf2 | 277 | 220 | 263 | 241 | 283 | 209 |
| ENSDARG000 | 22:22:22:22:2 | 10540446:105 | 10541020:105 | -;-;-;-;- | 865 | si:dkev-42i9.4 | 344 | 412 | 1225 | 1481 | 329 | 490 |
| ENSDARG000 | 22:22:22:22:2 | 29596646:295 | 29596854:295 | -;-;-;-;-;-;-;- | 2133 | shoc2 | 280 | 482 | 430 | 327 | 560 | 310 |
| ENSDARG000 | 22;22;22 | 2357347:2358 | 2357418:2358 | +;+;+ | 455 | NA | 0 | 4 | 8 | 6 | 3 | 12 |
| ENSDARG000 | 22:22:22:22:2 | 24973865:249 | 24975836:249 | -;-;-;-;-;-;- | 2843 | dnal4b | 88 | 90 | 146 | 56 | 124 | 84 |
| ENSDARG000 | 22:22:22:22:2 | 9009496:9010 | 9009816:9010 | +:+:+:+:+:+:+: | 11125 | NA | 156 | 220 | 244 | 326 | 578 | 590 |
| ENSDARG000 | 22:22:22:22:2 | 19175651:191 | 19175723:191 | +;+;+;+;+ | 766 | NA | 0 | 10 | 0 | 1 | 6 | 6 |
| ENSDARG000 | 22;22;22 | 25007941:250 | 25008018:250 | -;-;- | 488 | NA | 0 | 0 | 0 | 0 | 0 | 0 |
| ENSDARG000 | 22:22:22:22:2 | 10543329:105 | 10543525:105 | +:+:+:+:+:+:+: | 2568 | atrip | 166 | 302 | 559 | 206 | 325 | 212 |
| ENSDARG000 | 22;22;22 | 5810877:5812 | 5811173:5812 | +;+;+ | 563 | NA | 0 | 0 | 0 | 0 | 0 | 0 |
| ENSDARG000 | 22:22:22:22:2 | 5758090:5761 | 5758746:5762 | -:-:-:-:-:-:-:-: | 2156 | cers5 | 840 | 891 | 817 | 730 | 597 | 892 |
| ENSDARG000 | 22;22;22 | 29648854:296 | 29648963:296 | +;+;+ | 696 | bbip1 | 196 | 76 | 236 | 190 | 140 | 158 |
| ENSDARG000 | 22:22:22:22:2 | 37733107:377 | 37733186:377 | -:-:-:-:-:-:-:-: | 3529 | acap2b | 975 | 523 | 601 | 636 | 745 | 582 |
| ENSDARG000 | 22:22:22:22:2 | 11972786:119 | 11972911:119 | +:+:+:+:+:+:+: | 5447 | mgat5 | 1599 | 1555 | 2655 | 1282 | 3215 | 1310 |
| ENSDARG000 | 22:22:22:22:2 | 21313332:213 | 21313817:213 | -;-;-;-;-;-;- | 792 | cks2 | 28 | 182 | 152 | 70 | 112 | 106 |

| | | | | | | | | | | | | |
|---|---|---|---|---|---|---|---|---|---|---|---|---|
| ENSDARG000 | 22:22:22:22:2 | 19277361:192 | 19278006:192 | +;+;+;+;+;+;+; | 1748 | NA | 16 | 4 | 2 | 10 | 72 | 49 |
| ENSDARG000 | 22:22:22:22:2 | 9060699:9061 | 9060906:9061 | +;+;+;+;+;+;+; | 5993 | NA | 2264 | 1633 | 3033 | 1548 | 2914 | 2715 |
| ENSDARG000 | 22:22:22:22:2 | 19262161:192 | 19262235:192 | +;+;+;+;+;+;+; | 927 | NA | 0 | 2 | 0 | 0 | 0 | 0 |
| ENSDARG000 | 22:22:22:22:2 | 25025136:250 | 25025369:250 | -;-;-;-;- | 1782 | nptxrb | 0 | 2 | 0 | 0 | 0 | 0 |
| ENSDARG000 | 22:22:22:22:2 | 19247255:192 | 19247323:192 | +;+;+;+;+ | 839 | NA | 98 | 2 | 18 | 30 | 206 | 114 |
| ENSDARG000 | 22:22:22:22:2 | 14102397:141 | 14102500:141 | +;+;+;+;+;+;+; | 1042 | he1.3 | 0 | 0 | 4 | 0 | 6 | 12 |
| ENSDARG000 | 22;22;22;22 | 19215022:192 | 19215425:192 | -;-;-;- | 652 | NA | 0 | 0 | 0 | 0 | 0 | 0 |
| ENSDARG000 | 22:22:22:22:2 | 19237031:192 | 19237111:192 | +;+;+;+;+ | 750 | NA | 0 | 0 | 0 | 0 | 0 | 0 |
| ENSDARG000 | 22:22:22:22:2 | 19283978:192 | 19284067:192 | +;+;+;+;+;+;+; | 861 | NA | 0 | 0 | 0 | 0 | 2 | 0 |
| ENSDARG000 | 22:22:22:22:2 | 14093306:140 | 14093428:140 | +;+;+;+;+;+;+; | 1060 | he1.1 | 0 | 2 | 12 | 0 | 5 | 8 |
| ENSDARG000 | 22:22:22:22:2 | 19188809:191 | 19188942:191 | +;+;+;+;+;+;+; | 1142 | NA | 36 | 6 | 0 | 4 | 56 | 40 |
| ENSDARG000 | 22:22:22:22:2 | 14083729:140 | 14084086:140 | -;-;-;-;-;-;-;-;-; | 4511 | aox5 | 20 | 18 | 1422 | 174 | 504 | 834 |
| ENSDARG000 | 22:22:22:22:2 | 19405517:194 | 19405606:194 | +;+;+;+;+;+;+; | 863 | NA | 12 | 70 | 0 | 4 | 80 | 78 |
| ENSDARG000 | 22:22:22:22:2 | 23641801:236 | 23642200:236 | -;-;-;-;-;-;-;-;-; | 3412 | cfh | 294 | 170 | 376 | 242 | 1639 | 500 |
| ENSDARG000 | 22:22:22:22:2 | 19264181:194 | 19264264:194 | +;+;+;+;+;+;+; | 1267 | NA | 22 | 0 | 0 | 0 | 12 | 0 |
| ENSDARG000 | 22:22:22:22:2 | 19365220:193 | 19365352:193 | +;+;+;+;+;+;+; | 1024 | NA | 20 | 0 | 18 | 30 | 96 | 96 |
| ENSDARG000 | 22:22:22:22:2 | 19218733:192 | 19219390:192 | +;+;+;+;+;+;+; | 1673 | NA | 6 | 4 | 0 | 0 | 0 | 12 |
| ENSDARG000 | 22;22 | 34802598:348 | 34802649:348 | +;+ | 255 | NA | 0 | 0 | 0 | 0 | 0 | 0 |
| ENSDARG000 | 22:22:22:22:2 | 19264188:192 | 19264264:192 | +;+;+;+;+;+;+; | 847 | NA | 0 | 2 | 0 | 0 | 0 | 0 |
| ENSDARG000 | 22:22:22:22:2 | 19311411:193 | 19311503:193 | +;+;+;+;+;+;+; | 880 | si:dkev-21e2. | 4 | 4 | 0 | 0 | 10 | 20 |
| ENSDARG000 | 22:22:22:22:2 | 14098028:140 | 14098150:140 | +;+;+;+;+ | 980 | he1.2 | 0 | 0 | 0 | 0 | 1 | 0 |
| ENSDARG000 | 22:22:22:22:2 | 19230434:192 | 19230520:192 | +;+;+;+;+ | 796 | NA | 0 | 0 | 0 | 0 | 0 | 0 |
| ENSDARG000 | 22;22;22;22 | 19206321:192 | 19206724:192 | -;-;-;- | 652 | NA | 0 | 0 | 0 | 0 | 0 | 0 |
| ENSDARG000 | 22:22:22:22:2 | 19397226:193 | 19397554:193 | -;-;-;-;-;-;-;-;-; | 883 | NA | 0 | 0 | 0 | 0 | 0 | 0 |
| ENSDARG000 | 22:22:22:22:2 | 19289970:192 | 19290056:192 | +;+;+;+;+;+;+; | 861 | NA | 4 | 0 | 0 | 0 | 4 | 4 |
| ENSDARG000 | 22:22:22:22:2 | 19415080:194 | 19415153:194 | +;+;+;+;+ | 811 | NA | 2 | 24 | 8 | 6 | 14 | 94 |
| ENSDARG000 | 22:22:22:22:2 | 29655981:296 | 29659667:296 | -;-;-;-;-;-;-;-;-; | 5157 | pdcd4b | 387 | 242 | 410 | 588 | 248 | 307 |
| ENSDARG000 | 22;22 | 5835693:5839 | 5835995:5839 | -;- | 636 | NA | 105 | 75 | 114 | 34 | 0 | 169 |
| ENSDARG000 | 22:22:22:22:2 | 9151581:9151 | 9152219:9152 | -;-;-;-;-;-;-;-;-; | 10105 | NA | 34 | 14 | 25 | 13 | 18 | 160 |
| ENSDARG000 | 22:22:22:22:2 | 21317597:213 | 21317749:213 | +;+;+;+;+;+;+; | 3929 | shc2 | 1079 | 447 | 1402 | 884 | 1323 | 1326 |
| ENSDARG000 | 22:22:22:22:2 | 9286489:9286 | 9286593:9286 | +;+;+;+;+;+;+; | 4715 | NA | 79 | 42 | 27 | 18 | 34 | 84 |
| ENSDARG000 | 22:22:22:22:2 | 9262800:9264 | 9262874:9264 | +;+;+;+;+;+;+; | 4375 | NA | 73 | 41 | 46 | 19 | 32 | 24 |
| ENSDARG000 | 22:22:22:22:2 | 25034740:250 | 25035068:250 | -;-;-;-;-;-;-;- | 1453 | polr3f | 162 | 137 | 426 | 148 | 231 | 164 |
| ENSDARG000 | 22:22:22:22:2 | 9239831:9240 | 9240026:9241 | +;+;+;+;+;+;+; | 3731 | NA | 82 | 48 | 58 | 12 | 50 | 14 |
| ENSDARG000 | 22:22:22:22:2 | 9389801:9390 | 9389905:9390 | +;+;+;+;+;+;+; | 17041 | NA | 57 | 41 | 33 | 12 | 34 | 566 |
| ENSDARG000 | 22:22:22:22:2 | 15973122:159 | 15973184:159 | +;+;+;+;+;+;+; | 3747 | rc3h1a | 403 | 320 | 443 | 175 | 479 | 348 |
| ENSDARG000 | 22:22:22:22:2 | 9110998:9113 | 9111111:9113 | +;+;+;+;+;+;+; | 5944 | NA | 19 | 25 | 14 | 11 | 17 | 6 |
| ENSDARG000 | 22:22:22:22:2 | 9359899:9360 | 9360094:9362 | +;+;+;+;+;+;+; | 6328 | NA | 10 | 8 | 1 | 0 | 4 | 11 |
| ENSDARG000 | 22:22:22:22:2 | 9126557:9206 | 9126807:9206 | +;+;+;+;+;+;+; | 4678 | NA | 50 | 36 | 34 | 7 | 29 | 21 |
| ENSDARG000 | 22:22:22:22:2 | 15959844:159 | 15960275:159 | +;+;+;+;+;+;+; | 5645 | stil | 42 | 103 | 174 | 38 | 128 | 72 |
| ENSDARG000 | 22:22:22:22:2 | 17543607:175 | 17543966:175 | -;-;-;-;-;-;-;-;-; | 3081 | NA | 229 | 82 | 137 | 88 | 190 | 109 |
| ENSDARG000 | 22:22:22:22:2 | 5871035:5871 | 5871048:5871 | -;-;-;-;-;-;-;-;-; | 614 | NA | 1 | 0 | 2 | 0 | 2 | 0 |
| ENSDARG000 | 22:22:22:22:2 | 5858467:5868 | 5858547:5868 | +;+;+;+;+;+;+; | 1266 | NA | 4 | 2 | 0 | 4 | 0 | 6 |
| ENSDARG000 | 22:22:22:22:2 | 17531020:175 | 17531251:175 | +;+;+;+;+;+;+; | 6051 | hnrnpm | 5920 | 5024 | 5720 | 3372 | 5448 | 4259 |
| ENSDARG000 | 22:22:22:22:2 | 37804781:378 | 37805503:378 | -;-;-;-;-;- | 1493 | ppp1r2 | 299 | 443 | 542 | 297 | 392 | 364 |
| ENSDARG000 | 22:22:22:22:2 | 10560686:105 | 10562047:105 | -;-;-;-;- | 2184 | shisa10.1 | 1349 | 1768 | 1633 | 909 | 1302 | 1185 |
| ENSDARG000 | 22:22:22:22:2 | 34843282:348 | 34843971:348 | -;-;-;-;-;-;-;-;-; | 5149 | slit1b | 2 | 14 | 39 | 6 | 10 | 32 |
| ENSDARG000 | 22:22:22:22:2 | 5908556:5910 | 5911888:5910 | -;-;-;-;-;-;-;- | 4814 | NA | 46 | 76 | 56 | 112 | 149 | 36 |
| ENSDARG000 | 22:22:22:22:2 | 21359164:213 | 21359602:213 | -;-;-;-;-;-;-;-;-; | 3637 | NA | 47 | 22 | 37 | 122 | 59 | 77 |
| ENSDARG000 | 22:22:22:22:2 | 10575060:105 | 10575487:105 | -;-;-;-;- | 963 | shisa10.2 | 51 | 172 | 19 | 0 | 10 | 0 |
| ENSDARG000 | 22;22;22;22 | 25045692:250 | 25046886:250 | -;-;-;- | 1905 | NA | 0 | 0 | 0 | 0 | 0 | 4 |
| ENSDARG000 | 22:22:22:22:2 | 10586296:105 | 10586677:105 | -;-;-;-;- | 992 | NA | 8 | 4 | 0 | 0 | 0 | 2 |
| ENSDARG000 | 22;22;22;22 | 10582222:105 | 10582404:105 | -;-;-;- | 840 | NA | 0 | 0 | 0 | 4 | 0 | 0 |
| ENSDARG000 | 22:22:22:22:2 | 37982164:379 | 37982257:379 | +;+;+;+;+;+;+; | 841 | si:ch73-197b | 18 | 5 | 12 | 6 | 19 | 20 |
| ENSDARG000 | 22:22:22:22:2 | 12094484:120 | 12094934:120 | -;-;-;-;-;-;-;-;-; | 2142 | TMEM163 | 2 | 4 | 0 | 16 | 0 | 0 |
| ENSDARG000 | 22:22:22:22:2 | 23693195:236 | 23693863:236 | -;-;-;-;-;-;-;-;-; | 2682 | cfhl1 | 0 | 2 | 0 | 0 | 0 | 0 |
| ENSDARG000 | 22:22:22:22:2 | 29692252:296 | 29692320:296 | -;-;-;-;-;-;-;-;-; | 3882 | NA | 4 | 2 | 0 | 0 | 2 | 6 |
| ENSDARG000 | 22:22:22:22:2 | 21398508:213 | 21398684:213 | +;+;+;+;+;+;+; | 2580 | shdb | 106 | 55 | 84 | 40 | 52 | 18 |
| ENSDARG000 | 22:22:22:22:2 | 25049563:250 | 25049676:250 | +;+;+;+;+;+;+; | 2586 | dzank1 | 0 | 0 | 0 | 0 | 0 | 0 |
| ENSDARG000 | 22:22:22:22:2 | 34960346:349 | 34960799:349 | -;-;-;-;-;-;-;-;-; | 1936 | ARHGAP19 | 24 | 173 | 137 | 100 | 118 | 66 |
| ENSDARG000 | 22:22:22:22:2 | 10591958:105 | 10592774:105 | -;-;-;-;-;-;-;-;-; | 3880 | bap1 | 452 | 372 | 487 | 356 | 529 | 336 |
| ENSDARG000 | 22:22:22:22:2 | 23734001:237 | 23734711:237 | -;-;-;-;-;-;-;-;-; | 2376 | cfhl2 | 0 | 0 | 0 | 0 | 4 | 0 |
| ENSDARG000 | 22;22 | 26665958:266 | 26666009:266 | -;- | 378 | NA | 0 | 0 | 0 | 0 | 0 | 0 |
| ENSDARG000 | 22;22 | 26649505:266 | 26649756:266 | -;- | 545 | NA | 0 | 0 | 0 | 0 | 0 | 4 |
| ENSDARG000 | 22;22 | 26644234:266 | 26644485:266 | -;- | 545 | NA | 1 | 0 | 0 | 0 | 0 | 2 |
| ENSDARG000 | 22 | 21475648 | 21476836 | - | 1189 | s1pr3a | 20 | 0 | 0 | 2 | 6 | 2 |
| ENSDARG000 | 22:22:22:22:2 | 26600834:266 | 26602598:266 | +;+;+;+;+;+;+; | 4527 | ADCY9 | 172 | 82 | 166 | 77 | 290 | 174 |
| ENSDARG000 | 22:22:22:22:2 | 16022211:160 | 16022314:160 | +;+;+;+;+;+;+; | 2424 | serpinc1 | 81 | 44 | 85 | 47 | 127 | 112 |
| ENSDARG000 | 22:22:22:22:2 | 34985533:349 | 34985580:349 | -;-;-;-;-;-;-;-;-; | 1263 | KCNIP2 | 0 | 2 | 0 | 0 | 0 | 0 |
| ENSDARG000 | 22:22:22:22:2 | 23752586:237 | 23752873:237 | -;-;-;-;- | 945 | NA | 0 | 0 | 0 | 0 | 0 | 0 |
| ENSDARG000 | 22;22;22 | 23757955:237 | 23758132:237 | -;-;- | 536 | NA | 0 | 0 | 0 | 0 | 0 | 0 |
| ENSDARG000 | 22:22:22:22:2 | 29858535:298 | 29859439:298 | -;-;-;-;-;-;-;-;-; | 4680 | smc3 | 2934 | 3019 | 4767 | 2410 | 4004 | 2825 |
| ENSDARG000 | 22:22:22:22:2 | 14117078:141 | 14117184:141 | +;+;+;+;+;+;+; | 2050 | bzw1a | 2562 | 2844 | 4775 | 1769 | 4775 | 2108 |
| ENSDARG000 | 22:22:22:22:2 | 10606573:106 | 10607163:106 | +;+;+;+;+;+;+; | 8876 | RAD54L2 | 859 | 703 | 883 | 608 | 981 | 693 |
| ENSDARG000 | 22;22;22 | 10639440:106 | 10640461:106 | -;-;- | 1300 | cyb561d2 | 15 | 6 | 10 | 5 | 19 | 4 |
| ENSDARG000 | 22:22:22:22:2 | 21516689:215 | 21516756:215 | +;+;+;+;+;+;+; | 5050 | mier2 | 639 | 393 | 572 | 491 | 635 | 544 |
| ENSDARG000 | 22:22:22:22:2 | 26703026:267 | 26703596:267 | +;+;+;+;+;+;+; | 9682 | CREBBP | 2419 | 1319 | 2525 | 1556 | 2445 | 1994 |
| ENSDARG000 | 22:22:22:22:2 | 26791306:267 | 26791351:267 | +;+;+;+;+;+;+; | 899 | NA | 1 | 2 | 0 | 0 | 8 | 0 |
| ENSDARG000 | 22:22:22:22:2 | 21549419:215 | 21549861:215 | +;+;+;+;+;+ | 1856 | plpp2b | 1006 | 475 | 1945 | 1714 | 1412 | 836 |
| ENSDARG000 | 22:22:22:22:2 | 35013443:350 | 35016282:350 | -;-;-;-;-;-;-;-;-; | 4895 | armh3 | 1010 | 724 | 983 | 613 | 1011 | 914 |
| ENSDARG000 | 22:22:22:22:2 | 23767557:237 | 23768463:237 | -;-;-;-;-;-;-;-;-; | 3399 | cfhl3 | 0 | 0 | 0 | 0 | 0 | 0 |
| ENSDARG000 | 22;22 | 14127472:141 | 14127845:141 | -;- | 1025 | NA | 0 | 0 | 0 | 0 | 0 | 0 |
| ENSDARG000 | 22:22:22:22:2 | 17606588:176 | 17606803:176 | +;+;+;+;+;+;+; | 1215 | polr2eb | 709 | 682 | 1051 | 570 | 732 | 522 |
| ENSDARG000 | 22:22:22:22:2 | 35068046:350 | 35068143:350 | +;+;+;+;+;+;+; | 2223 | NA | 78 | 6 | 10 | 12 | 20 | 12 |
| ENSDARG000 | 22;22;22 | 16032160:160 | 16032681:160 | -;-;- | 1566 | zbtb37 | 134 | 80 | 121 | 33 | 142 | 132 |
| ENSDARG000 | 22:22:22:22:2 | 17589608:175 | 17589903:175 | -;-;-;-;-;-;-;-;-; | 1491 | gpx4a | 804 | 443 | 674 | 535 | 703 | 719 |
| ENSDARG000 | 22:22:22:22:2 | 14145510:141 | 14145742:141 | -;-;-;-;-;-;-;- | 979 | NA | 2 | 0 | 0 | 0 | 2 | 0 |
| ENSDARG000 | 22;22 | 16038495:160 | 16040707:160 | -;- | 2352 | s1pr1 | 104 | 56 | 82 | 18 | 70 | 38 |
| ENSDARG000 | 22:22:22:22:2 | 21618121:216 | 21618744:216 | +;+;+;+;+;+;+; | 3848 | tle2a | 1376 | 398 | 1658 | 1511 | 1693 | 1867 |
| ENSDARG000 | 22:22:22:22:2 | 16097185:161 | 16097507:161 | +;+;+;+;+;+;+ | 1505 | dph5 | 282 | 500 | 548 | 179 | 523 | 313 |
| ENSDARG000 | 22:22:22:22:2 | 35089031:350 | 35089798:350 | +;+;+;+;+;+;+; | 3037 | srfa | 199 | 159 | 276 | 116 | 259 | 267 |
| ENSDARG000 | 22:22:22:22:2 | 16123090:161 | 16123768:161 | -;-;-;-;-;-;-;-;-; | 1942 | slc30a7 | 1946 | 2187 | 2615 | 1727 | 2633 | 2108 |
| ENSDARG000 | 22:22:22:22:2 | 16169447:161 | 16172129:161 | -;-;-;-;-;-;-;-;-; | 5771 | vcam1b | 132 | 82 | 202 | 218 | 262 | 264 |
| ENSDARG000 | 22:22:22:22:2 | 14178813:141 | 14179578:141 | -;-;-;-;-;-;-;-;-; | 5233 | NA | 2 | 0 | 2 | 0 | 14 | 2 |
| ENSDARG000 | 22:22:22:22:2 | 16190231:161 | 16193831:161 | -;-;-;-;-;-;-;-;-; | 5949 | cdc14ab | 1807 | 823 | 846 | 1351 | 1511 | 2108 |
| ENSDARG000 | 22:22:22:22:2 | 16282144:162 | 16282988:162 | -;-;-;-;-;-;-;-;-; | 2731 | rtca | 2003 | 2761 | 2317 | 2012 | 3046 | 1823 |
| ENSDARG000 | 22:22:22:22:2 | 37902976:379 | 37903048:379 | +;+;+;+;+;+;+; | 2871 | kcnh1b | 0 | 0 | 7 | 0 | 0 | 0 |
| ENSDARG000 | 22:22:22:22:2 | 6905210:6906 | 6905282:6907 | +;+;+;+;+;+ | 1766 | NA | 0 | 8 | 20 | 16 | 34 | 0 |
| ENSDARG000 | 22:22:22:22:2 | 6764980:6770 | 6769552:6771 | -;-;-;-;- | 5369 | NA | 568 | 264 | 87 | 90 | 64 | 395 |
| ENSDARG000 | 22:22:22:22:2 | 18497759:184 | 18497865:184 | +;+;+;+;+;+ | 774 | NA | 0 | 0 | 0 | 0 | 0 | 0 |
| ENSDARG000 | 22 | 3232925 | 3234494 | + | 1570 | NA | 3 | 0 | 2 | 1 | 9 | 24 |
| ENSDARG000 | 22;22;22;22 | 2459591:2623 | 2460607:2623 | +;+;+;+ | 4431 | NA | 50 | 31 | 60 | 49 | 76 | 40 |
| ENSDARG000 | 22;22 | 2495929:2497 | 2496112:2499 | +;+ | 1877 | NA | 50 | 35 | 85 | 38 | 36 | 25 |
| ENSDARG000 | 22;22;22;22 | 344763:34596 | 344788:34597 | +;+;+;+ | 111 | NA | 0 | 0 | 0 | 0 | 0 | 0 |
| ENSDARG000 | 22;22;22;22 | 30757562:307 | 30757623:307 | -;-;-;- | 714 | NA | 14 | 0 | 0 | 6 | 21 | 5 |
| ENSDARG000 | 22:22:22:22:2 | 225937:23331 | 226008:23334 | -;-;-;-;-;-;-;-;-; | 1698 | NA | 0 | 0 | 0 | 0 | 0 | 0 |
| ENSDARG000 | 22;22 | 44365;50487 | 48178;50878 | -;- | 4206 | NA | 2 | 8 | 0 | 0 | 0 | 0 |
| ENSDARG000 | 22:22:22:22:2 | 60264:66067: | 60281:66089: | +;+;+;+;+;+;+; | 207 | NA | 2 | 1 | 1 | 0 | 0 | 0 |
| ENSDARG000 | 22;22;22 | 8690263:8691 | 8690813:8691 | -;-;- | 1041 | NA | 0 | 40 | 0 | 4 | 77 | 0 |
| ENSDARG000 | 22 | 17425678 | 17425796 | - | 119 | 5S_rRNA | 0 | 0 | 0 | 0 | 0 | 0 |
| ENSDARG000 | 22 | 28193836 | 28193951 | + | 116 | 5S_rRNA | 0 | 0 | 0 | 0 | 0 | 0 |
| ENSDARG000 | 22;22 | 26853254:268 | 26854181:268 | +;+ | 1606 | tmem186 | 0 | 0 | 0 | 0 | 0 | 0 |
| ENSDARG000 | 22;22 | 1330477:1332 | 1330716:1333 | +;+ | 1524 | NA | 12 | 24 | 8 | 4 | 8 | 8 |
| ENSDARG000 | 22;22;22 | 38374543:383 | 38374680:383 | -;-;- | 369 | NA | 44 | 37 | 47 | 6 | 38 | 48 |
| ENSDARG000 | 22:22:22:22:2 | 36498980:364 | 36499060:364 | -;-;-;-;-;-;-;-;-; | 1545 | NA | 17 | 10 | 38 | 11 | 8 | 13 |
| ENSDARG000 | 22:22:22:22:2 | 19538626:195 | 19538683:195 | +;+;+;+;+ | 3033 | NA | 4 | 0 | 16 | 0 | 14 | 7 |
| ENSDARG000 | 22:22:22:22:2 | 35930526:359 | 35930651:359 | +;+;+;+;+;+;+; | 1506 | NA | 42 | 5 | 49 | 19 | 30 | 18 |
| ENSDARG000 | 22:22:22:22:2 | 26400519:264 | 26400812:264 | +;+;+;+;+;+;+; | 2341 | capn8 | 18 | 95 | 0 | 2 | 18 | 18 |
| ENSDARG000 | 22;22 | 635813:63727 | 635898:63755 | +;+ | 366 | pth4 | 0 | 0 | 0 | 0 | 0 | 0 |
| ENSDARG000 | 22 | 208644 | 209741 | - | 1098 | ora1 | 0 | 2 | 0 | 0 | 0 | 0 |
| ENSDARG000 | 22:22:22:22:2 | 5632029:5634 | 5634187:5634 | -;-;-;-;-;-;-;-;-; | 5678 | mcm2 | 977 | 3360 | 4176 | 1290 | 3573 | 1716 |
| ENSDARG000 | 22 | 34416730 | 34416844 | - | 115 | 5S_rRNA | 0 | 0 | 0 | 0 | 0 | 0 |
| ENSDARG000 | 22:22:22:22:2 | 5123914:5125 | 5123940:5125 | -;-;-;-;-;-;-;-;-; | 255 | NA | 0 | 0 | 1 | 0 | 0 | 0 |
| ENSDARG000 | 22 | 22825790 | 22825878 | - | 89 | dre-mir-181b- | 15 | 29 | 11 | 15 | 47 | 8 |
| ENSDARG000 | 22 | 37403125 | 37403314 | - | 190 | NA | 0 | 0 | 0 | 0 | 0 | 0 |
| ENSDARG000 | 22;22 | 9003090:9005 | 9003666:9009 | +;+ | 3963 | NA | 24 | 22 | 44 | 26 | 54 | 44 |
| ENSDARG000 | 22:22:22:22:2 | 25464880:254 | 25467000:254 | -;-;-;-;-;- | 3088 | NA | 0 | 0 | 0 | 0 | 0 | 0 |
| ENSDARG000 | 22;22;22 | 1396444:1398 | 1396466:1398 | +;+;+ | 1381 | zgc:113336 | 54 | 68 | 97 | 52 | 42 | 60 |
| ENSDARG000 | 22:22:22:22:2 | 36741346:367 | 36741512:367 | -;-;-;-;-;-;-;-;-; | 1286 | ACY1 | 232 | 224 | 244 | 137 | 338 | 216 |
| ENSDARG000 | 22 | 19771698 | 19771812 | + | 115 | 5S_rRNA | 0 | 0 | 0 | 0 | 0 | 0 |
| ENSDARG000 | 22 | 20151925 | 20151981 | + | 57 | NA | 0 | 0 | 0 | 0 | 0 | 0 |
| ENSDARG000 | 22:22:22:22:2 | 34674224:346 | 34674437:346 | +;+;+;+;+ | 3871 | NA | 0 | 0 | 0 | 0 | 0 | 0 |
| ENSDARG000 | 22;22 | 6881445:6884 | 6882637:6884 | -;- | 1512 | NA | 0 | 0 | 0 | 0 | 0 | 6 |
| ENSDARG000 | 22 | 21093851 | 21093965 | + | 115 | 5S_rRNA | 0 | 0 | 0 | 0 | 0 | 0 |
| ENSDARG000 | 22 | 28779715 | 28779812 | - | 98 | dre-mir-217 | 0 | 0 | 0 | 0 | 0 | 0 |
| ENSDARG000 | 22 | 28780257 | 28780333 | - | 77 | dre-mir-216b | 0 | 0 | 0 | 0 | 0 | 0 |
| ENSDARG000 | 22 | 32883761 | 32883877 | - | 117 | 5S_rRNA | 0 | 0 | 0 | 0 | 0 | 0 |
| ENSDARG000 | 22;22;22 | 25256801:252 | 25256831:252 | +;+;+ | 165 | NA | 0 | 0 | 0 | 0 | 0 | 0 |
| ENSDARG000 | 22 | 38581012 | 38582101 | + | 1090 | NA | 0 | 0 | 0 | 0 | 0 | 0 |
| ENSDARG000 | 22:22:22:22:2 | 6932299:6935 | 6933137:6935 | -;-;-;-;-;- | 1795 | zgc:171500 | 0 | 23 | 32 | 8 | 24 | 16 |
| ENSDARG000 | 22 | 8346075 | 8346200 | - | 126 | 5S_rRNA | 0 | 0 | 2 | 0 | 0 | 0 |
| ENSDARG000 | 22 | 14252414 | 14252530 | + | 117 | 5S_rRNA | 0 | 0 | 0 | 0 | 0 | 0 |
| ENSDARG000 | 22:22:22:22:2 | 81799:83026: | 81892:83133: | +;+;+;+;+;+;+; | 3634 | zgc:112334 | 4 | 22 | 2 | 22 | 10 | 9 |
| ENSDARG000 | 22;22;22;22 | 24378064:243 | 24378274:243 | +;+;+;+ | 618 | NA | 0 | 26 | 14 | 18 | 51 | 8 |
| ENSDARG000 | 22:22:22:22:2 | 6829466:6831 | 6829748:6831 | -;-;-;-;- | 780 | NA | 0 | 0 | 0 | 0 | 0 | 8 |
| ENSDARG000 | 22;22 | 31722434:317 | 31722583:317 | -;- | 564 | NA | 0 | 0 | 0 | 0 | 0 | 0 |
| ENSDARG000 | 22:22:22:22:2 | 15633013:156 | 15633111:156 | +;+;+;+;+;+;+; | 1964 | NA | 0 | 0 | 76 | 10 | 34 | 42 |
| ENSDARG000 | 22:22:22:22:2 | 85597:89989: | 90453:90284: | -;-;-;-;-;-;-;-;-; | 5523 | ndufaf3 | 583 | 530 | 715 | 294 | 633 | 318 |
| ENSDARG000 | 22:22:22:22:2 | 291091:29131 | 291270:29150 | -;-;-;-;-;-;-;-;-; | 1782 | NA | 0 | 4 | 2 | 8 | 8 | 2 |
| ENSDARG000 | 22:22:22:22:2 | 36780999:367 | 36781166:367 | -;-;-;-;-;-;-;- | 1408 | NA | 132 | 132 | 39 | 64 | 20 | 57 |
| ENSDARG000 | 22:22:22:22:2 | 8446780:8450 | 8448345:8450 | -;-;-;-;-;-;- | 3563 | NA | 2 | 8 | 4 | 4 | 0 | 2 |
| ENSDARG000 | 22;22 | 6711549:6711 | 6711834:6712 | +;+ | 561 | NA | 0 | 0 | 0 | 0 | 0 | 0 |
| ENSDARG000 | 22;22 | 2032023:2033 | 2032077:2034 | +;+ | 855 | NA | 81 | 72 | 83 | 62 | 65 | 43 |
| ENSDARG000 | 22:22:22:22:2 | 5549734:5550 | 5549853:5550 | -;-;-;-;-;-;- | 1486 | NA | 150 | 184 | 153 | 96 | 145 | 201 |
| ENSDARG000 | 22 | 5307485 | 5307574 | + | 90 | dre-mir-24-4 | 0 | 0 | 0 | 0 | 0 | 0 |
| ENSDARG000 | 22;22 | 191931:19236 | 191976:19323 | -;- | 918 | FBXO42 | 236 | 148 | 175 | 185 | 205 | 123 |
| ENSDARG000 | 22:22:22:22:2 | 1786230:1786 | 1786288:1786 | +;+;+;+;+;+ | 2894 | NA | 34 | 98 | 117 | 44 | 58 | 40 |
| ENSDARG000 | 22:22:22:22:2 | 7923713:7924 | 7923971:7924 | +;+;+;+;+;+ | 2931 | NA | 0 | 0 | 0 | 0 | 0 | 0 |
| ENSDARG000 | 22:22:22:22:2 | 6323600:6323 | 6323626:6323 | +;+;+;+;+;+;+; | 348 | NA | 957 | 176 | 460 | 292 | 108 | 669 |
| ENSDARG000 | 22:22:22:22:2 | 5655756:5657 | 5657913:5657 | -;-;-;-;-;- | 2841 | ccdc51 | 224 | 276 | 173 | 110 | 236 | 146 |
| ENSDARG000 | 22:22:22:22:2 | 1320991:1322 | 1321027:1322 | -;-;-;-;-;-;-;-;-; | 270 | NA | 52 | 60 | 63 | 31 | 47 | 54 |
| ENSDARG000 | 22:22:22:22:2 | 7421631:7424 | 7421642:7424 | -;-;-;-;-;-;-;-;-; | 294 | NA | 7 | 12 | 15 | 6 | 7 | 6 |

| | | | | | | | | | | | | |
|---|---|---|---|---|---|---|---|---|---|---|---|---|
| ENSDARG000 | 22 | 10763933 | 10764047 | - | 115 | 5S_rRNA | 0 | 0 | 0 | 0 | 0 | 0 |
| ENSDARG000 | 22:22:22:22:2 | 22172272:221 | 22172398:221 | -:-:-:-:-:-:-:-:-: | 3825 | ap3d1 | 4984 | 4391 | 5496 | 4538 | 4754 | 3719 |
| ENSDARG000 | 22:22:22:22:2 | 5959984:5959 | 5960213:5960 | +:+:+:+:+:+:+: | 2167 | NA | 183 | 39 | 71 | 44 | 8 | 73 |
| ENSDARG000 | 22 | 8265533 | 8265649 | + | 117 | 5S_rRNA | 0 | 0 | 0 | 0 | 0 | 0 |
| ENSDARG000 | 22:22:22:22:2 | 37874691:378 | 37874936:378 | +;+;+;+;+;+;+ | 1522 | ahsg1 | 0 | 0 | 12 | 0 | 8 | 0 |
| ENSDARG000 | 22:22:22:22:2 | 8522577:8637 | 8522947:8639 | -;-;-;-;-;-;-;- | 2887 | NA | 8 | 46 | 82 | 45 | 187 | 75 |
| ENSDARG000 | 22;22;22 | 28078927:280 | 28079081:280 | -;-;- | 522 | wu:fu71h07 | 867 | 401 | 1608 | 369 | 960 | 630 |
| ENSDARG000 | 22 | 36683988 | 36684056 | - | 69 | SNORD82 | 0 | 0 | 0 | 0 | 0 | 0 |
| ENSDARG000 | 22:22:22:22:2 | 7542472:7545 | 7542839:7545 | -;-;-;-;- | 1962 | NA | 0 | 0 | 0 | 0 | 0 | 226 |
| ENSDARG000 | 22 | 12643956 | 12644079 | + | 124 | 5S_rRNA | 0 | 0 | 0 | 0 | 0 | 0 |
| ENSDARG000 | 22;22 | 7460029:7461 | 7460418:7461 | -;- | 543 | NA | 91 | 80 | 32 | 38 | 53 | 67 |
| ENSDARG000 | 22:22:22:22:2 | 6922006:6922 | 6922145:6922 | -;-;-;-;-;- | 879 | NA | 64 | 14 | 9 | 9 | 2 | 18 |
| ENSDARG000 | 22 | 5293213 | 5293301 | + | 89 | dre-mir-23a-1 | 0 | 0 | 0 | 0 | 0 | 0 |
| ENSDARG000 | 22 | 33757766 | 33757880 | - | 115 | 5S_rRNA | 0 | 0 | 0 | 0 | 0 | 0 |
| ENSDARG000 | 22;22;22;22 | 8380715:8385 | 8382375:8385 | -;-;-;- | 2626 | NA | 10 | 0 | 0 | 0 | 2 | 8 |
| ENSDARG000 | 22:22:22:22:2 | 6384693:6386 | 6384723:6386 | +:+:+:+:+:+:+: | 351 | NA | 0 | 0 | 0 | 0 | 0 | 3 |
| ENSDARG000 | 22:22:22:22:2 | 2648931:2650 | 2648983:2650 | +;+;+;+;+;+;+ | 1439 | zgc:110821 | 103 | 74 | 98 | 83 | 70 | 93 |
| ENSDARG000 | 22;22;22 | 8328377:8331 | 8328854:8331 | +;+;+ | 956 | NA | 0 | 0 | 4 | 0 | 0 | 2 |
| ENSDARG000 | 22;22;22;22 | 381143:38143 | 381361:38168 | -;-;-;- | 996 | NA | 0 | 0 | 0 | 0 | 0 | 0 |
| ENSDARG000 | 22;22;22;22 | 4782287:4799 | 4783196:4800 | -;-;-;- | 2683 | NA | 108 | 37 | 17 | 78 | 0 | 67 |
| ENSDARG000 | 22;22;22;22 | 27599775:276 | 27599959:276 | -;-;-;- | 1423 | NA | 0 | 0 | 0 | 0 | 0 | 0 |
| ENSDARG000 | 22:22:22:22:2 | 3910610:3910 | 3910626:3910 | -:-:-:-:-:-:-:-:-: | 1443 | NA | 0 | 0 | 0 | 0 | 0 | 0 |
| ENSDARG000 | 22:22:22:22:2 | 26585096:265 | 26589866:265 | -;-;-;-;- | 5571 | SRL | 0 | 0 | 8 | 14 | 8 | 3 |
| ENSDARG000 | 22;22 | 9764731:9767 | 9765486:9768 | +;+ | 1548 | NA | 22 | 12 | 4 | 1 | 0 | 21 |
| ENSDARG000 | 22:22:22:22:2 | 8157374:8158 | 8157429:8158 | -;-;-;-;-;- | 906 | NA | 2496 | 2281 | 3164 | 1948 | 1593 | 3231 |
| ENSDARG000 | 22:22:22:22:2 | 8295097:8295 | 8295158:8295 | -;-;-;-;- | 873 | NA | 211 | 88 | 287 | 139 | 1 | 244 |
| ENSDARG000 | 22;22;22 | 6232552:6234 | 6232676:6234 | -;-;- | 753 | NA | 0 | 0 | 0 | 0 | 0 | 15 |
| ENSDARG000 | 22;22;22 | 4735436:4738 | 4735960:4739 | -;-;- | 2001 | NA | 0 | 0 | 2 | 0 | 0 | 0 |
| ENSDARG000 | 22 | 13945953 | 13946067 | - | 115 | 5S_rRNA | 0 | 0 | 0 | 0 | 0 | 0 |
| ENSDARG000 | 22 | 28779949 | 28780037 | - | 89 | dre-mir-216a | 0 | 0 | 0 | 0 | 0 | 0 |
| ENSDARG000 | 22 | 38692001 | 38692121 | + | 121 | 5S_rRNA | 0 | 0 | 0 | 0 | 0 | 0 |
| ENSDARG000 | 22 | 10667858 | 10667974 | - | 117 | 5S_rRNA | 0 | 0 | 0 | 0 | 0 | 0 |
| ENSDARG000 | 22:22:22:22:2 | 7896072:7897 | 7897205:7897 | -;-;-;-;-;-;-;- | 2814 | NA | 2050 | 1950 | 2024 | 1545 | 2107 | 1368 |
| ENSDARG000 | 22:22:22:22:2 | 25352530:253 | 25352921:253 | +:+:+:+:+:+:+: | 1156 | si:dkev-240e1 | 0 | 0 | 0 | 0 | 0 | 0 |
| ENSDARG000 | 22:22:22:22:2 | 3914318:3914 | 3914435:3914 | +:+:+:+:+:+:+: | 1389 | NA | 0 | 0 | 0 | 0 | 0 | 0 |
| ENSDARG000 | 22;22 | 3336723:3344 | 3337058:3344 | -;- | 609 | tbxa2r | 2 | 0 | 0 | 0 | 6 | 0 |
| ENSDARG000 | 22;22 | 2680895:2681 | 2681184:2682 | +;+ | 1456 | zgc:194221 | 0 | 0 | 0 | 0 | 0 | 0 |
| ENSDARG000 | 22 | 19429806 | 19429924 | - | 119 | 5S_rRNA | 0 | 0 | 0 | 0 | 0 | 0 |
| ENSDARG000 | 22;22 | 22888;23103 | 23012;24275 | +;+ | 1298 | NA | 105 | 46 | 87 | 81 | 77 | 32 |
| ENSDARG000 | 22;22;22 | 9981079:9982 | 9982188:9982 | +;+;+ | 1529 | NA | 10 | 0 | 4 | 8 | 30 | 1 |
| ENSDARG000 | 22:22:22:22:2 | 7522728:7523 | 7522849:7523 | -;-;-;-;-;-;-;- | 1641 | NA | 0 | 0 | 0 | 0 | 0 | 308 |
| ENSDARG000 | 22;22 | 2562101:2675 | 2562155:2676 | +;+ | 909 | NA | 20 | 22 | 17 | 10 | 26 | 10 |
| ENSDARG000 | 22 | 22129358 | 22130623 | - | 1266 | C2CD4C | 4 | 31 | 0 | 6 | 2 | 4 |
| ENSDARG000 | 22:22:22:22:2 | 8904046:8904 | 8905865:8904 | -:-:-:-:-:-:-:-:-: | 2669 | NA | 0 | 0 | 0 | 0 | 2 | 84 |
| ENSDARG000 | 22:22:22:22:2 | 5574952:5577 | 5575300:5577 | +:+:+:+:+:+:+: | 2320 | GGH | 88 | 79 | 16 | 16 | 37 | 2 |
| ENSDARG000 | 22:22:22:22:2 | 24318131:243 | 24318564:243 | +:+:+:+:+:+:+: | 6043 | NA | 30 | 83 | 46 | 30 | 76 | 50 |
| ENSDARG000 | 22:22:22:22:2 | 22234415:222 | 22234788:222 | -:-:-:-:-:-:-:-:-: | 2247 | hdgfl2 | 715 | 1261 | 1406 | 473 | 1309 | 846 |
| ENSDARG000 | 22:22:22:22:2 | 37888249:378 | 37888455:378 | +;+;+;+;+;+;+ | 3203 | NA | 0 | 0 | 281 | 18 | 2 | 0 |
| ENSDARG000 | 22:22:22:22:2 | 33535575:335 | 33535898:335 | -:-:-:-:-:-:-:-:-: | 2070 | GRM7 | 6 | 0 | 0 | 0 | 0 | 4 |
| ENSDARG000 | 22:22:22:22:2 | 26443235:264 | 26443486:264 | +:+:+:+:+:+:+: | 2597 | capn2l | 2 | 48 | 0 | 18 | 10 | 6 |
| ENSDARG000 | 22:22:22:22:2 | 24311193:243 | 24311421:243 | -;-;-;-;- | 604 | uts2d | 0 | 1 | 0 | 0 | 0 | 0 |
| ENSDARG000 | 22:22:22:22:2 | 2040454:2048 | 2040508:2048 | +:+:+:+:+:+:+: | 1566 | znf1153 | 23 | 22 | 41 | 76 | 51 | 44 |
| ENSDARG000 | 22;22 | 1589407:1593 | 1589689:1594 | +;+ | 1017 | NA | 102 | 58 | 67 | 48 | 81 | 63 |
| ENSDARG000 | 22;22 | 8827347:8828 | 8827852:8829 | -;- | 758 | NA | 26 | 18 | 42 | 0 | 0 | 18 |
| ENSDARG000 | 22 | 6476142 | 6476258 | - | 117 | 5S_rRNA | 0 | 0 | 0 | 0 | 0 | 0 |
| ENSDARG000 | 22 | 17838913 | 17844117 | - | 5205 | NA | 0 | 0 | 0 | 0 | 0 | 6 |
| ENSDARG000 | 22 | 36558553 | 36558667 | + | 115 | 5S_rRNA | 0 | 0 | 0 | 0 | 0 | 0 |
| ENSDARG000 | 22:22:22:22:2 | 24559947:245 | 24560355:245 | +:+:+:+:+:+:+: | 3749 | wdr47b | 0 | 7 | 6 | 2 | 14 | 16 |
| ENSDARG000 | 22:22:22:22:2 | 5483588:5486 | 5483905:5486 | -;-;-;-;-;-;-;- | 1844 | NA | 54 | 28 | 40 | 23 | 39 | 8 |
| ENSDARG000 | 22:22:22:22:2 | 7971101:7974 | 7973003:7974 | -;-;-;-;- | 3146 | NA | 2757 | 2522 | 2518 | 1813 | 2902 | 1372 |
| ENSDARG000 | 22;22;22;22 | 8272393:8272 | 8272543:8273 | +;+;+;+ | 1383 | NA | 4 | 8 | 12 | 0 | 18 | 0 |
| ENSDARG000 | 22 | 11894362 | 11894479 | + | 118 | 5S_rRNA | 0 | 0 | 0 | 0 | 0 | 0 |
| ENSDARG000 | 22:22:22:22:2 | 35505619:355 | 35505802:355 | +:+:+:+:+:+:+: | 6355 | hykk.2 | 70 | 93 | 138 | 92 | 116 | 56 |
| ENSDARG000 | 22:22:22:22:2 | 35912986:359 | 35913016:359 | -:-:-:-:-:-:-:-:-: | 1734 | NA | 35 | 3 | 8 | 17 | 9 | 16 |
| ENSDARG000 | 22:22:22:22:2 | 30520839:305 | 30520976:305 | -:-:-:-:-:-:-:-:-: | 2213 | NA | 4 | 6 | 2 | 26 | 18 | 18 |
| ENSDARG000 | 22 | 18254769 | 18254855 | + | 87 | dre-mir-9-4 | 0 | 0 | 0 | 0 | 0 | 0 |
| ENSDARG000 | 22;22 | 2632961:2637 | 2633390:2639 | +;+ | 1966 | NA | 45 | 8 | 26 | 19 | 14 | 22 |
| ENSDARG000 | 22:22:22:22:2 | 24726066:247 | 24726082:247 | +;+;+;+;+;+;+ | 147 | NA | 0 | 0 | 0 | 0 | 0 | 0 |
| ENSDARG000 | 22:22:22:22:2 | 35321110:353 | 35321336:353 | -;-;-;-;-;-;-;- | 1790 | NA | 0 | 0 | 0 | 0 | 4 | 0 |
| ENSDARG000 | 22:22:22:22:2 | 898131:90158 | 898156:90185 | -:-:-:-:-:-:-:-:-: | 1266 | KCTD20 | 2 | 10 | 2 | 0 | 0 | 0 |
| ENSDARG000 | 22;22;22;22 | 8012532:8014 | 8013553:8014 | +;+;+;+ | 2501 | NA | 42 | 24 | 70 | 58 | 65 | 26 |
| ENSDARG000 | 22 | 38185763 | 38185883 | + | 121 | 5S_rRNA | 1 | 0 | 0 | 0 | 1 | 4 |
| ENSDARG000 | 22 | 30832796 | 30832911 | - | 116 | 5S_rRNA | 0 | 0 | 0 | 0 | 0 | 0 |
| ENSDARG000 | 22;22 | 5478353:5478 | 5478386:5479 | +;+ | 894 | NA | 284 | 58 | 45 | 45 | 19 | 51 |
| ENSDARG000 | 22;22;22 | 2451732:2452 | 2452220:2452 | +;+;+ | 819 | NA | 0 | 0 | 18 | 8 | 30 | 2 |
| ENSDARG000 | 22:22:22:22:2 | 5532003:5534 | 5532022:5534 | +:+:+:+:+:+:+: | 681 | NA | 2 | 4 | 0 | 0 | 0 | 0 |
| ENSDARG000 | 22 | 7668434 | 7669624 | - | 1191 | NA | 0 | 0 | 0 | 0 | 0 | 0 |
| ENSDARG000 | 22;22 | 8489400:8490 | 8489690:8491 | +;+ | 586 | NA | 0 | 0 | 0 | 0 | 0 | 0 |
| ENSDARG000 | 22:22:22:22:2 | 34696880:346 | 34696913:346 | -:-:-:-:-:-:-:-:-: | 285 | NA | 0 | 0 | 0 | 0 | 0 | 0 |
| ENSDARG000 | 22 | 12584839 | 12584954 | - | 116 | 5S_rRNA | 0 | 0 | 0 | 0 | 0 | 0 |
| ENSDARG000 | 22;22;22 | 1352192:1353 | 1352378:1353 | +;+;+ | 2239 | znf45l | 63 | 24 | 48 | 14 | 61 | 21 |
| ENSDARG000 | 22;22;22 | 25814941:258 | 25815115:258 | +;+;+ | 4491 | vasna | 64 | 4 | 30 | 14 | 32 | 56 |
| ENSDARG000 | 22 | 33726550 | 33726664 | - | 115 | 5S_rRNA | 0 | 0 | 0 | 0 | 0 | 0 |
| ENSDARG000 | 22:22:22:22:2 | 5156:5415:55 | 5322:5484:56 | -;-;-;-;-;- | 663 | atad3 | 294 | 346 | 570 | 132 | 393 | 168 |
| ENSDARG000 | 22;22 | 6512469:6515 | 6512763:6515 | +;+ | 1097 | NA | 0 | 5 | 6 | 6 | 4 | 2 |
| ENSDARG000 | 22;22 | 26519299:265 | 26521777:265 | -;- | 2661 | b3galt10.1 | 0 | 2 | 0 | 0 | 0 | 2 |
| ENSDARG000 | 22;22;22;22 | 6843363:6849 | 6843695:6849 | +;+;+;+ | 1400 | NA | 0 | 0 | 0 | 0 | 4 | 74 |
| ENSDARG000 | 22;22;22;22 | 387073:38942 | 387330:38968 | -;-;-;- | 1035 | NA | 0 | 0 | 0 | 0 | 0 | 0 |
| ENSDARG000 | 22 | 24366193 | 24366272 | - | 80 | dre-mir-725 | 0 | 0 | 0 | 0 | 0 | 0 |
| ENSDARG000 | 22 | 12374786 | 12374883 | + | 98 | dre-mir-128-1 | 0 | 0 | 0 | 0 | 0 | 0 |
| ENSDARG000 | 22;22 | 6587280:6587 | 6587565:6587 | +;+ | 561 | NA | 0 | 2 | 0 | 0 | 0 | 0 |
| ENSDARG000 | 22;22 | 7655271:7661 | 7655537:7662 | +;+ | 1581 | NA | 0 | 0 | 0 | 0 | 0 | 121 |
| ENSDARG000 | 22;22 | 6757741:6758 | 6758206:6759 | +;+ | 1186 | NA | 0 | 0 | 6 | 0 | 0 | 11 |
| ENSDARG000 | 22 | 1440702 | 1441490 | + | 789 | si:dkeyp-53d3 | 12 | 40 | 28 | 10 | 18 | 6 |
| ENSDARG000 | 22:22:22:22:2 | 20312180:203 | 20316022:203 | -:-:-:-:-:-:-:-:-: | 6424 | tcf3b | 2758 | 2420 | 3654 | 2505 | 3922 | 2696 |
| ENSDARG000 | 22 | 22840558 | 22840675 | + | 118 | 5S_rRNA | 0 | 0 | 1 | 0 | 0 | 0 |
| ENSDARG000 | 22 | 56644 | 57177 | - | 534 | NA | 60 | 2 | 10 | 0 | 9 | 8 |
| ENSDARG000 | 22:22:22:22:2 | 8313513:8313 | 8313660:8313 | +;+;+;+;+;+;+ | 3164 | NA | 321 | 208 | 452 | 140 | 0 | 352 |
| ENSDARG000 | 22:22:22:22:2 | 9939901:9939 | 9940158:9940 | +:+:+:+:+:+:+: | 1546 | zgc:171686 | 828 | 106 | 970 | 542 | 121 | 1941 |
| ENSDARG000 | 22:22:22:22:2 | 21889822:218 | 21889835:218 | +:+:+:+:+:+:+: | 453 | NA | 8 | 2 | 0 | 0 | 0 | 2 |
| ENSDARG000 | 22:22:22:22:2 | 94692:96156: | 94862:96268: | +:+:+:+:+:+:+: | 2280 | NCKIPSD | 1579 | 872 | 1122 | 1071 | 1329 | 966 |
| ENSDARG000 | 22;22;22;22 | 8628064:8632 | 8628106:8632 | +;+;+;+ | 804 | NA | 30 | 0 | 7 | 0 | 4 | 6 |
| ENSDARG000 | 22;22;22;22 | 30742373:307 | 30742502:307 | +;+;+;+ | 1080 | NA | 0 | 2 | 0 | 0 | 16 | 6 |
| ENSDARG000 | 22;22 | 6697927:6698 | 6698212:6698 | +;+ | 561 | NA | 2 | 0 | 1 | 0 | 0 | 4 |
| ENSDARG000 | 22:22:22:22:2 | 22634007:226 | 22636534:226 | -;-;-;-;-;-;- | 4090 | nr5a2 | 26 | 0 | 0 | 4 | 12 | 30 |
| ENSDARG000 | 22;22 | 26263290:262 | 26263368:262 | +;+ | 480 | NA | 0 | 0 | 0 | 0 | 0 | 0 |
| ENSDARG000 | 22;22;22 | 30496360:304 | 30496423:304 | +;+;+ | 522 | NA | 0 | 0 | 7 | 4 | 0 | 2 |
| ENSDARG000 | 22 | 12117620 | 12117744 | + | 125 | 5S_rRNA | 0 | 0 | 0 | 0 | 0 | 0 |
| ENSDARG000 | 22;22 | 2431585:2443 | 2431639:2444 | +;+ | 876 | NA | 55 | 26 | 63 | 37 | 28 | 35 |
| ENSDARG000 | 22:22:22:22:2 | 254390:25956 | 254443:25960 | -:-:-:-:-:-:-:-:-: | 1686 | NA | 0 | 0 | 0 | 0 | 0 | 0 |
| ENSDARG000 | 22 | 17824459 | 17824573 | + | 115 | 5S_rRNA | 0 | 0 | 0 | 0 | 0 | 0 |
| ENSDARG000 | 22 | 39042630 | 39042744 | + | 115 | 5S_rRNA | 0 | 0 | 0 | 0 | 0 | 0 |
| ENSDARG000 | 22;22;22;22 | 5433093:5436 | 5433211:5436 | -;-;-;- | 1050 | zgc:194627 | 49 | 135 | 101 | 36 | 118 | 29 |
| ENSDARG000 | 22;22;22;22 | 8728969:8729 | 8729395:8729 | +;+;+;+ | 1499 | NA | 0 | 0 | 0 | 0 | 8 | 0 |
| ENSDARG000 | 22;22 | 6727266:6727 | 6727551:6727 | +;+ | 561 | NA | 548 | 203 | 286 | 197 | 4 | 429 |
| ENSDARG000 | 22:22:22:22:2 | 6026595:6026 | 6026641:6026 | -:-:-:-:-:-:-:-:-: | 1791 | NA | 345 | 260 | 583 | 240 | 608 | 471 |
| ENSDARG000 | 22 | 3760757 | 3760871 | + | 115 | 5S_rRNA | 0 | 0 | 0 | 0 | 0 | 0 |
| ENSDARG000 | 22 | 15781608 | 15781724 | - | 117 | 5S_rRNA | 0 | 0 | 0 | 0 | 0 | 0 |
| ENSDARG000 | 22 | 2631618 | 2631738 | + | 121 | 5S_rRNA | 0 | 0 | 0 | 0 | 0 | 0 |
| ENSDARG000 | 22;22 | 30656903:306 | 30662800:306 | -;- | 5940 | NA | 12 | 0 | 0 | 10 | 6 | 0 |
| ENSDARG000 | 22 | 26483105 | 26484268 | - | 1164 | NA | 4 | 0 | 0 | 0 | 0 | 0 |
| ENSDARG000 | 22:22:22:22:2 | 11395:12588: | 11457:12721: | +:+:+:+:+:+:+: | 3272 | sdhb | 1400 | 1305 | 1337 | 1103 | 1418 | 1083 |
| ENSDARG000 | 22 | 22944461 | 22944585 | - | 125 | 5S_rRNA | 0 | 0 | 0 | 0 | 0 | 0 |
| ENSDARG000 | 22;22;22 | 3045495:3055 | 3045816:3055 | +;+;+ | 1368 | NA | 0 | 2 | 0 | 0 | 0 | 0 |
| ENSDARG000 | 22;22;22;22 | 31660964:316 | 31662010:316 | -;-;-;- | 1650 | LSM3 | 122 | 185 | 321 | 130 | 213 | 152 |
| ENSDARG000 | 22:22:22:22:2 | 24645325:246 | 24645421:246 | +;+;+;+;+ | 2369 | lpar3 | 26 | 6 | 77 | 44 | 152 | 6 |
| ENSDARG000 | 22;22;22;22 | 5851122:5855 | 5851185:5855 | +;+;+;+ | 1895 | NA | 2 | 2 | 2 | 0 | 0 | 0 |
| ENSDARG000 | 22 | 16497670 | 16499153 | + | 1484 | ier5 | 3282 | 1835 | 2256 | 2406 | 1300 | 1926 |
| ENSDARG000 | 22:22:22:22:2 | 24623936:246 | 24624056:246 | +:+:+:+:+:+:+: | 1373 | mcoln2 | 0 | 4 | 8 | 4 | 40 | 2 |
| ENSDARG000 | 22;22 | 6858787:6859 | 6859069:6859 | +;+ | 564 | NA | 0 | 1 | 0 | 0 | 0 | 0 |
| ENSDARG000 | 22;22;22 | 4862637:4863 | 4863138:4863 | -;-;- | 1082 | NA | 0 | 0 | 0 | 0 | 0 | 0 |
| ENSDARG000 | 22 | 23095688 | 23095804 | + | 117 | 5S_rRNA | 0 | 0 | 0 | 0 | 0 | 0 |
| ENSDARG000 | 22:22:22:22:2 | 39007533:390 | 39007629:390 | +:+:+:+:+:+:+: | 6026 | NA | 657 | 604 | 898 | 463 | 995 | 646 |
| ENSDARG000 | 22 | 1229906 | 1230022 | - | 117 | 5S_rRNA | 0 | 0 | 1 | 0 | 0 | 0 |
| ENSDARG000 | 22 | 38901103 | 38901221 | + | 119 | 5S_rRNA | 0 | 0 | 0 | 0 | 0 | 0 |
| ENSDARG000 | 22:22:22:22:2 | 8878680:8879 | 8879236:8879 | -;-;-;-;- | 1379 | NA | 18 | 24 | 16 | 18 | 42 | 42 |
| ENSDARG000 | 22 | 35741654 | 35741770 | + | 117 | 5S_rRNA | 0 | 0 | 0 | 0 | 0 | 0 |
| ENSDARG000 | 22 | 23496815 | 23496929 | - | 115 | 5S_rRNA | 0 | 0 | 0 | 0 | 0 | 0 |
| ENSDARG000 | 22 | 16639310 | 16639425 | + | 116 | 5S_rRNA | 0 | 0 | 0 | 0 | 0 | 0 |
| ENSDARG000 | 22 | 21118902 | 21119016 | - | 115 | 5S_rRNA | 0 | 0 | 0 | 0 | 0 | 0 |
| ENSDARG000 | 22 | 5299951 | 5300056 | + | 106 | dre-mir-27e | 0 | 0 | 0 | 0 | 0 | 0 |
| ENSDARG000 | 22 | 15781894 | 15782009 | - | 116 | 5S_rRNA | 0 | 0 | 0 | 0 | 0 | 0 |
| ENSDARG000 | 22:22:22:22:2 | 110158:11570 | 110497:11573 | +:+:+:+:+:+:+: | 1979 | prkar2ab | 0 | 0 | 3 | 0 | 1 | 0 |
| ENSDARG000 | 22 | 8786167 | 8786293 | + | 127 | 5S_rRNA | 0 | 0 | 0 | 0 | 0 | 0 |
| ENSDARG000 | 22:22:22:22:2 | 9994833:9994 | 9995066:9995 | +:+:+:+:+:+:+: | 3890 | NA | 12 | 21 | 9 | 0 | 10 | 12 |
| ENSDARG000 | 22 | 36675801 | 36675931 | - | 131 | SNORA75 | 0 | 0 | 0 | 0 | 0 | 0 |
| ENSDARG000 | 22;22 | 9761930:9762 | 9762290:9763 | +;+ | 1937 | NA | 0 | 0 | 0 | 0 | 0 | 0 |
| ENSDARG000 | 22:22:22:22:2 | 25774750:257 | 25774807:257 | +;+;+;+;+;+ | 1704 | NA | 0 | 0 | 0 | 2 | 0 | 0 |
| ENSDARG000 | 22 | 34620589 | 34620751 | + | 163 | U1 | 0 | 0 | 0 | 0 | 0 | 0 |
| ENSDARG000 | 22;22;22;22 | 4797508:4797 | 4797769:4798 | -;-;-;- | 792 | zgc:195170 | 0 | 2 | 0 | 0 | 0 | 0 |
| ENSDARG000 | 22:22:22:22:2 | 396840:40360 | 396842:40369 | +:+:+:+:+:+:+: | 822 | capzb | 5222 | 5634 | 5610 | 5427 | 5648 | 3541 |
| ENSDARG000 | 22;22;22 | 8701238:8701 | 8701615:8701 | +;+;+ | 947 | NA | 0 | 56 | 0 | 37 | 50 | 2 |
| ENSDARG000 | 22;22;22 | 8612462:8744 | 8612795:8744 | +;+;+ | 1706 | NA | 0 | 5 | 4 | 11 | 0 | 0 |
| ENSDARG000 | 22:22:22:22:2 | 31493953:314 | 31494123:314 | -:-:-:-:-:-:-:-:-: | 1725 | slc6a6b | 91 | 107 | 120 | 62 | 530 | 43 |
| ENSDARG000 | 22:22:22:22:2 | 8337699:8345 | 8342590:8345 | -;-;-;-;-;-;- | 6477 | NA | 2410 | 1488 | 2494 | 1402 | 569 | 3133 |
| ENSDARG000 | 22:22:22:22:2 | 9917007:9918 | 9917037:9918 | +;+;+;+;+;+;+ | 1984 | blf | 1720 | 225 | 62 | 570 | 591 | 158 |
| ENSDARG000 | 22;22;22 | 10037276:100 | 10040967:100 | -;-;- | 4020 | NA | 10 | 30 | 42 | 12 | 48 | 94 |
| ENSDARG000 | 22:22:22:22:2 | 9751117:9752 | 9751245:9752 | +;+;+;+;+;+ | 1525 | NA | 0 | 0 | 0 | 0 | 0 | 0 |
| ENSDARG000 | 22:22:22:22:2 | 278790:27881 | 278805:27886 | -;-;-;-;- | 555 | NA | 0 | 0 | 0 | 0 | 0 | 0 |
| ENSDARG000 | 22;22;22 | 8365905:8366 | 8366097:8366 | +;+;+ | 840 | NA | 0 | 4 | 0 | 2 | 0 | 0 |
| ENSDARG000 | 22;22 | 1560991:1563 | 1561174:1567 | +;+ | 4474 | NA | 121 | 101 | 142 | 76 | 97 | 83 |
| ENSDARG000 | 22:22:22:22:2 | 30035719:300 | 30036205:300 | +;+;+;+;+ | 3153 | NA | 4 | 2 | 6 | 0 | 0 | 2 |

| | | | | | | | | | | | |
|---|---|---|---|---|---|---|---|---|---|---|---|
| ENSDARG000 | 22;22;22;22;2 | 268352:2685 | 268483:2686 | -;-;-;-;-;-;-;-; | 4893 | NA | 15 | 27 | 10 | 8 | 22 | 2 |
| ENSDARG000 | 22;22;22;22;2 | 12477996:124 | 12478062:124 | +;+;+;+;+;+;+ | 1065 | NA | 0 | 0 | 0 | 0 | 0 | 2 |
| ENSDARG000 | 22 | 25806441 | 25806557 | - | 117 | 5S_rRNA | 0 | 0 | 0 | 0 | 0 | 0 |
| ENSDARG000 | 22 | 36965621 | 36965735 | + | 115 | 5S_rRNA | 0 | 0 | 0 | 0 | 0 | 0 |
| ENSDARG000 | 22;22;22;22;2 | 25753972:257 | 25754077:257 | +;+;+;+;+ | 2168 | si:ch211-250 | 0 | 0 | 0 | 0 | 0 | 0 |
| ENSDARG000 | 22;22;22;22;2 | 105376:1059 | 105610:1060 | +;+;+;+;+;+;+; | 1394 | slc25a20 | 456 | 445 | 542 | 329 | 630 | 478 |
| ENSDARG000 | 22;22;22;22;2 | 3244950:3246 | 3245016:3246 | +;+;+;+;+;+;+; | 378 | NA | 39 | 36 | 22 | 11 | 19 | 12 |
| ENSDARG000 | 22;22 | 10015596:100 | 10017065:100 | -;- | 2336 | NA | 38 | 16 | 82 | 30 | 83 | 78 |
| ENSDARG000 | 22 | 14019604 | 14019722 | - | 119 | 5S_rRNA | 0 | 0 | 0 | 0 | 0 | 0 |
| ENSDARG000 | 22 | 35835035 | 35835169 | - | 135 | 5S_rRNA | 0 | 0 | 0 | 0 | 0 | 0 |
| ENSDARG000 | 22 | 31043509 | 31043647 | + | 139 | NA | 0 | 0 | 2 | 0 | 0 | 0 |
| ENSDARG000 | 22;22 | 8959490:8966 | 8960778:8966 | -;- | 1563 | NA | 0 | 0 | 0 | 0 | 0 | 5 |
| ENSDARG000 | 22;22;22;22 | 9772754:9774 | 9773897:9774 | +;+;+;+ | 2359 | NA | 0 | 0 | 0 | 0 | 0 | 0 |
| ENSDARG000 | 22;22 | 8929668:8930 | 8930577:8931 | -;- | 1008 | NA | 0 | 0 | 0 | 0 | 0 | 0 |
| ENSDARG000 | 22 | 36677247 | 36677370 | - | 124 | SNORA75 | 0 | 0 | 0 | 0 | 0 | 0 |
| ENSDARG000 | 22;22;22;22;2 | 24990337:249 | 24990893:249 | -;-;-;-;-;- | 966 | si:dkev-179i5. | 0 | 0 | 0 | 0 | 0 | 0 |
| ENSDARG000 | 22;22;22;22;2 | 130967:1327 | 131098:1329 | -;-;-;-;-;-;-;-; | 9420 | NA | 2083 | 1861 | 3049 | 1077 | 2898 | 1932 |
| ENSDARG000 | 22;22;22;22 | 9165:9730:98 | 9463:9807:10 | -;-;-;- | 650 | mrpl20 | 260 | 338 | 446 | 148 | 300 | 144 |
| ENSDARG000 | 22;22;22;22 | 30429681:304 | 30429787:304 | -;-;-;- | 429 | NA | 24 | 8 | 65 | 10 | 18 | 11 |
| ENSDARG000 | 22 | 26513454 | 26514386 | - | 933 | NA | 0 | 0 | 0 | 0 | 0 | 0 |
| ENSDARG000 | 22;22;22;22 | 8847831:8854 | 8848463:8854 | -;-;-;- | 1200 | NA | 0 | 1 | 0 | 5 | 11 | 22 |
| ENSDARG000 | 22;22;22;22;2 | 7445017:7446 | 7446571:7446 | -;-;-;-;-;-;- | 2982 | NA | 220 | 323 | 204 | 177 | 210 | 221 |
| ENSDARG000 | 22;22;22;22;2 | 31023205:310 | 31023454:310 | +;+;+;+;+ | 5571 | NA | 1088 | 1053 | 663 | 384 | 835 | 671 |
| ENSDARG000 | 22;22;22;22 | 21305682:213 | 21305797:213 | +;+;+;+ | 847 | odf3l2b | 0 | 2 | 0 | 0 | 0 | 0 |
| ENSDARG000 | 22 | 33283923 | 33284040 | - | 118 | 5S_rRNA | 0 | 0 | 0 | 0 | 0 | 0 |
| ENSDARG000 | 22 | 26781531 | 26781660 | + | 130 | NA | 0 | 0 | 5 | 2 | 4 | 0 |
| ENSDARG000 | 22;22;22;22 | 184670:18708 | 187115:18711 | -;-;-;- | 3016 | slc25a34 | 50 | 77 | 79 | 49 | 82 | 41 |
| ENSDARG000 | 22;22;22;22;2 | 38337270:383 | 38337612:383 | -;-;-;-;-;-;-;-; | 2031 | NA | 131 | 79 | 195 | 73 | 147 | 106 |
| ENSDARG000 | 22 | 19793119 | 19793233 | + | 115 | 5S_rRNA | 0 | 0 | 0 | 0 | 0 | 0 |
| ENSDARG000 | 22;22 | 2598998:2602 | 2599667:2604 | +;+ | 2709 | NA | 154 | 85 | 141 | 49 | 144 | 204 |
| ENSDARG000 | 22 | 30150790 | 30150914 | + | 125 | 5S_rRNA | 0 | 0 | 1 | 0 | 0 | 0 |
| ENSDARG000 | 22 | 20139525 | 20139592 | + | 68 | NA | 0 | 0 | 0 | 0 | 0 | 0 |
| ENSDARG000 | 22;22;22;22;2 | 26085343:260 | 26085816:260 | -;-;-;-;-;- | 1471 | PDGFA | 2 | 0 | 0 | 0 | 0 | 0 |
| ENSDARG000 | 22;22;22;22;2 | 36521139:365 | 36524511:365 | -;-;-;-;- | 4649 | polr2h | 206 | 404 | 315 | 153 | 278 | 294 |
| ENSDARG000 | 22;22;22 | 37671430:376 | 37672396:376 | -;-;- | 1469 | htr2b | 0 | 0 | 2 | 0 | 6 | 0 |
| ENSDARG000 | 22;22 | 8551813:8553 | 8552916:8553 | +;+ | 1449 | NA | 346 | 116 | 414 | 563 | 406 | 546 |
| ENSDARG000 | 22;22;22;22 | 5971404:5974 | 5971600:5974 | +;+;+;+ | 1087 | NA | 5 | 25 | 16 | 7 | 58 | 25 |
| ENSDARG000 | 22;22;22;22;2 | 24603930:246 | 24604496:246 | +;+;+;+;+;+;+; | 3716 | NA | 606 | 496 | 683 | 332 | 613 | 443 |
| ENSDARG000 | 22 | 30383999 | 30384127 | - | 129 | 5S_rRNA | 38 | 13 | 12 | 14 | 43 | 14 |
| ENSDARG000 | 22 | 1431743 | 1433028 | + | 1286 | NA | 34 | 84 | 120 | 88 | 114 | 28 |
| ENSDARG000 | 22 | 38627265 | 38627383 | + | 119 | 5S_rRNA | 0 | 0 | 0 | 0 | 0 | 2 |
| ENSDARG000 | 22 | 4677867 | 4677981 | + | 115 | 5S_rRNA | 0 | 0 | 0 | 0 | 1 | 0 |
| ENSDARG000 | 22;22 | 336256:33687 | 336397:33962 | +;+ | 2887 | btg2 | 18352 | 3212 | 13621 | 10624 | 4550 | 13248 |
| ENSDARG000 | 22;22;22;22;2 | 4772860:4776 | 4774301:4777 | -;-;-;-;-;-;-;-; | 3552 | NA | 0 | 1 | 4 | 0 | 0 | 4 |
| ENSDARG000 | 22;22;22;22;2 | 35377125:353 | 35377676:353 | -;-;-;-;- | 1317 | NA | 0 | 0 | 0 | 0 | 0 | 0 |
| ENSDARG000 | 22;22;22 | 34632156:346 | 34632204:346 | +;+;+ | 705 | NA | 10 | 0 | 0 | 0 | 0 | 16 |
| ENSDARG000 | 22 | 6290234 | 6290350 | + | 117 | 5S_rRNA | 0 | 0 | 0 | 0 | 0 | 0 |
| ENSDARG000 | 22;22;22;22 | 6892296:6892 | 6892559:6893 | +;+;+;+ | 1197 | NA | 0 | 0 | 0 | 0 | 0 | 2 |
| ENSDARG000 | 22;22;22;22;2 | 5518314:5522 | 5520989:5522 | -;-;-;-;-;- | 3826 | NA | 41 | 9 | 2 | 0 | 2 | 5 |
| ENSDARG000 | 22;22 | 1384104:1392 | 1384158:1397 | +;+ | 5270 | NA | 67 | 46 | 19 | 34 | 36 | 52 |
| ENSDARG000 | 22;22;22;22;2 | 34784075:347 | 34784151:347 | +;+;+;+;+;+ | 2045 | NA | 52 | 24 | 85 | 39 | 43 | 59 |
| ENSDARG000 | 22;22;22;22 | 2473222:2492 | 2473714:2492 | +;+;+;+ | 5954 | NA | 133 | 43 | 56 | 71 | 55 | 75 |
| ENSDARG000 | 22;22;22;22;2 | 76469:79525: | 79447:79574: | -;-;-;-;-;- | 3388 | NA | 260 | 273 | 470 | 274 | 554 | 282 |
| ENSDARG000 | 22;22;22;22 | 27703632:277 | 27705284:277 | -;-;-;- | 2898 | NA | 5 | 3 | 3 | 0 | 4 | 0 |
| ENSDARG000 | 22;22;22 | 9793923:9794 | 9793957:9794 | +;+;+ | 921 | NA | 4 | 1 | 9 | 0 | 0 | 9 |
| ENSDARG000 | 22;22;22;22;2 | 16482631:164 | 16486207:164 | -;-;-;-;-;-;-;- | 4271 | stx6 | 2223 | 1485 | 1685 | 1411 | 1928 | 1558 |
| ENSDARG000 | 22;22;22;22 | 8239451:8245 | 8243940:8246 | -;-;-;- | 5738 | NA | 343 | 283 | 449 | 163 | 210 | 288 |
| ENSDARG000 | 22 | 20490441 | 20490557 | - | 117 | 5S_rRNA | 0 | 0 | 0 | 0 | 0 | 0 |
| ENSDARG000 | 22;22;22 | 6740039:6740 | 6740738:6741 | +;+;+ | 1140 | NA | 5 | 3 | 2 | 1 | 2 | 5 |
| ENSDARG000 | 22 | 9714214 | 9714331 | - | 118 | 5S_rRNA | 0 | 0 | 0 | 0 | 0 | 0 |
| ENSDARG000 | 22 | 16277498 | 16277625 | + | 128 | U5 | 0 | 0 | 0 | 0 | 0 | 0 |
| ENSDARG000 | 22;22 | 10010292:100 | 10010595:100 | +;+ | 3707 | NA | 0 | 0 | 0 | 0 | 4 | 0 |
| ENSDARG000 | 22;22 | 1421212:1422 | 1421583:1424 | +;+ | 2088 | zgc:101130 | 238 | 147 | 219 | 161 | 279 | 210 |
| ENSDARG000 | 22;22 | 33915539:339 | 33916026:339 | -;- | 540 | NA | 8 | 0 | 0 | 0 | 0 | 0 |
| ENSDARG000 | 22;22 | 9854088:9854 | 9854505:9855 | +;+ | 935 | NA | 11 | 6 | 36 | 0 | 0 | 33 |
| ENSDARG000 | 22;22 | 8407707:8439 | 8408026:8440 | +;+ | 903 | NA | 0 | 0 | 2 | 0 | 2 | 12 |
| ENSDARG000 | 22 | 22955732 | 22955849 | + | 118 | 5S_rRNA | 0 | 0 | 0 | 0 | 0 | 0 |
| ENSDARG000 | 22 | 30631072 | 30635832 | + | 4761 | NA | 12 | 0 | 0 | 0 | 6 | 0 |
| ENSDARG000 | 22;22;22;22;2 | 7624599:7624 | 7629115:7629 | -;-;-;-;-;-;-;-;- | 5742 | NA | 0 | 0 | 0 | 0 | 0 | 188 |
| ENSDARG000 | 22;22;22;22;2 | 15624371:156 | 15624769:156 | +;+;+;+;+;+;+; | 3322 | lpl | 116 | 90 | 673 | 74 | 310 | 360 |
| ENSDARG000 | 22 | 14693112 | 14693228 | - | 117 | 5S_rRNA | 0 | 0 | 0 | 0 | 0 | 0 |
| ENSDARG000 | 22;22 | 25639255:256 | 25639318:256 | +;+ | 1248 | NA | 0 | 0 | 0 | 0 | 0 | 0 |
| ENSDARG000 | 22;22 | 4783644:4784 | 4784122:4784 | -;- | 809 | NA | 1 | 0 | 0 | 0 | 0 | 0 |
| ENSDARG000 | 22;22;22;22;2 | 26524760:265 | 26529514:265 | -;-;-;-;-;-;- | 5944 | NA | 226 | 641 | 637 | 1562 | 466 | 129 |
| ENSDARG000 | 22;22 | 8843870:8876 | 8844707:8876 | -;- | 966 | NA | 0 | 0 | 8 | 0 | 0 | 0 |
| ENSDARG000 | 22;22;22;22;2 | 6809366:6809 | 6810298:6809 | -;-;-;-;-;- | 1036 | NA | 0 | 0 | 0 | 0 | 0 | 0 |
| ENSDARG000 | 22;22;22 | 8939933:8950 | 8940995:8950 | -;-;- | 1968 | NA | 1 | 0 | 0 | 0 | 0 | 16 |
| ENSDARG000 | 22;22;22;22 | 9351372:9352 | 9351738:9353 | +;+;+;+ | 5735 | NA | 0 | 0 | 0 | 0 | 0 | 0 |
| ENSDARG000 | 22;22;22;22 | 10895963:108 | 10896719:108 | -;-;-;- | 1170 | NA | 1 | 0 | 0 | 6 | 1 | 0 |
| ENSDARG000 | 22;22;22 | 10903510:109 | 10904827:109 | -;-;- | 1963 | NA | 3 | 6 | 4 | 0 | 18 | 10 |
| ENSDARG000 | 22;22;22;22;2 | 13218865:132 | 13219146:132 | -;-;-;-;-;- | 1649 | NA | 0 | 0 | 0 | 0 | 0 | 0 |
| ENSDARG000 | 22;22;22;22;2 | 13422568:134 | 13423049:134 | -;-;-;-;-;-;- | 1243 | NA | 0 | 0 | 0 | 0 | 0 | 0 |
| ENSDARG000 | 22;22;22 | 15787981:158 | 15788840:158 | +;+;+ | 1188 | NA | 0 | 0 | 0 | 0 | 0 | 0 |
| ENSDARG000 | 22;22;22;22 | 18663099:187 | 18663680:187 | -;-;-;- | 1255 | NA | 0 | 0 | 0 | 0 | 2 | 0 |
| ENSDARG000 | 22;22;22;22;2 | 26562999:265 | 26563163:265 | +;+;+;+;+ | 3718 | NA | 4 | 2 | 0 | 0 | 0 | 0 |
| ENSDARG000 | 22;22;22;22;2 | 30723361:307 | 30723415:307 | +;+;+;+;+;+;+ | 1535 | NA | 0 | 0 | 0 | 0 | 0 | 0 |
| ENSDARG000 | 22;22;22;22;2 | 35543930:355 | 35544071:355 | +;+;+;+;+ | 798 | NA | 0 | 0 | 0 | 0 | 0 | 0 |
| ENSDARG000 | 22;22;22 | 36155966:361 | 36156193:361 | +;+;+ | 540 | NA | 0 | 0 | 0 | 0 | 0 | 0 |
| ENSDARG000 | 22;22;22 | 37275954:372 | 37278643:372 | -;-;- | 3027 | NA | 167 | 44 | 78 | 25 | 34 | 18 |
| ENSDARG000 | 22;22;22 | 37851344:378 | 37853189:378 | -;-;- | 2074 | NA | 0 | 0 | 0 | 0 | 0 | 0 |
| ENSDARG000 | 22;22;22 | 38245116:382 | 38245282:382 | +;+;+ | 1012 | NA | 0 | 0 | 0 | 0 | 0 | 0 |
| ENSDARG000 | 25;25;25;25;2 | 19356587:193 | 19361062:193 | -;-;-;-;-;-;-;-; | 16518 | NA | 242 | 864 | 637 | 376 | 1946 | 754 |
| ENSDARG000 | 25;25;25;25;2 | 21517551:215 | 21518848:215 | -;-;-;-;-;-;-;-; | 7653 | DOCK4 | 0 | 0 | 10 | 0 | 0 | 2 |
| ENSDARG000 | 25;25;25;25;2 | 15178225:151 | 15178812:151 | -;-;-;-;-;-;-;- | 1874 | wt1a | 0 | 2 | 2 | 0 | 0 | 0 |
| ENSDARG000 | 25;25 | 21517551:215 | 21518869:215 | +;+ | 5345 | NA | 2 | 4 | 16 | 2 | 0 | 2 |
| ENSDARG000 | 25;25;25;25;2 | 21716263:217 | 21716321:217 | +;+;+;+;+;+;+; | 1507 | znf277 | 558 | 235 | 678 | 456 | 608 | 514 |
| ENSDARG000 | 25;25;25;25;2 | 3224085:3225 | 3224591:3226 | -;-;-;-;-;-;-;-; | 1462 | cib1 | 211 | 167 | 175 | 208 | 182 | 166 |
| ENSDARG000 | 25;25;25;25;2 | 35913614:359 | 35913873:359 | +;+;+;+;+;+;+; | 2403 | gpia | 3774 | 8702 | 22213 | 15567 | 7287 | 5504 |
| ENSDARG000 | 25;25;25;25;2 | 3217419:3219 | 3217650:3219 | +;+;+;+;+;+;+; | 1452 | NA | 35 | 21 | 4 | 2 | 8 | 2 |
| ENSDARG000 | 25 | 8623709 | 8625601 | - | 1893 | gdpgp1 | 54 | 82 | 68 | 8 | 118 | 38 |
| ENSDARG000 | 25;25;25 | 19441920:194 | 19442271:194 | -;-;- | 684 | cart2 | 0 | 0 | 0 | 0 | 16 | 0 |
| ENSDARG000 | 25;25;25;25 | 6451038:6471 | 6453258:6471 | +;+;+;+ | 4452 | snx33 | 104 | 18 | 26 | 19 | 16 | 43 |
| ENSDARG000 | 25;25;25;25;2 | 6507141:6507 | 6508998:6508 | -;-;-;-;-;-;-;-; | 7091 | CSPG4 | 5225 | 2377 | 3058 | 4731 | 2117 | 2749 |
| ENSDARG000 | 25;25;25;25;2 | 8627748:8627 | 8627928:8627 | -;-;-;-;-;-;-;-; | 3337 | unc45a | 632 | 654 | 635 | 380 | 673 | 456 |
| ENSDARG000 | 25;25;25;25;2 | 15242410:152 | 15242786:152 | -;-;-;-;-;-;-;-; | 2391 | NA | 0 | 0 | 0 | 0 | 7 | 0 |
| ENSDARG000 | 25;25;25;25;2 | 37435335:374 | 37435482:374 | +;+;+;+;+;+;+; | 1436 | chmp1a | 774 | 658 | 550 | 483 | 590 | 523 |
| ENSDARG000 | 25;25 | 24250161:242 | 24250297:242 | -;- | 474 | NA | 28 | 11 | 17 | 22 | 24 | 22 |
| ENSDARG000 | 25;25;25 | 24250247:242 | 24250436:242 | +;+;+ | 1468 | tmem86a | 53 | 7 | 35 | 8 | 42 | 54 |
| ENSDARG000 | 25;25;25;25;2 | 31323970:313 | 31324122:313 | +;+;+;+;+;+;+; | 3959 | NA | 232 | 347 | 188 | 32 | 154 | 108 |
| ENSDARG000 | 25;25;25;25 | 31388899:313 | 31391198:313 | -;-;-;- | 3251 | NA | 16 | 36 | 17 | 2 | 4 | 26 |
| ENSDARG000 | 25;25;25;25;2 | 34811038:348 | 34811336:348 | +;+;+;+;+;+;+; | 1668 | adat1 | 60 | 43 | 79 | 36 | 83 | 115 |
| ENSDARG000 | 25;25;25;25;2 | 4635355:4637 | 4635416:4637 | +;+;+;+;+;+;+; | 2440 | pgghg | 140 | 120 | 368 | 325 | 467 | 260 |
| ENSDARG000 | 25;25 | 7229046:7229 | 7230905:7230 | +;+ | 1860 | LINGO1 | 0 | 2 | 0 | 0 | 2 | 0 |
| ENSDARG000 | 25;25;25;25;2 | 12924989:129 | 12925602:129 | -;-;-;-;- | 1092 | NA | 0 | 0 | 0 | 0 | 0 | 0 |
| ENSDARG000 | 25;25;25 | 12936778:129 | 12936896:129 | -;-;- | 325 | NA | 0 | 0 | 0 | 0 | 0 | 0 |
| ENSDARG000 | 25;25 | 37443194:374 | 37445449:374 | +;+ | 2256 | slc10a3 | 584 | 523 | 628 | 433 | 558 | 329 |
| ENSDARG000 | 25;25;25;25;2 | 15273370:152 | 15273509:152 | +;+;+;+;+;+;+; | 2655 | tcp11l1 | 275 | 252 | 203 | 214 | 208 | 279 |
| ENSDARG000 | 25;25;25;25;2 | 35942867:359 | 35943059:359 | +;+;+;+;+ | 1144 | hsd17b2 | 4 | 11 | 4 | 1 | 0 | 6 |
| ENSDARG000 | 25;25;25;25;2 | 24291156:243 | 24293001:243 | +;+;+;+;+;+;+; | 6590 | B4GALNT4 | 487 | 201 | 216 | 244 | 648 | 266 |
| ENSDARG000 | 25;25;25 | 24534449:245 | 24534843:245 | -;-;- | 865 | NA | 0 | 0 | 0 | 0 | 0 | 0 |
| ENSDARG000 | 25;25;25;25;2 | 3231812:3234 | 3232009:3235 | +;+;+;+;+;+;+; | 3314 | metap2b | 4124 | 4636 | 5664 | 3072 | 5108 | 4307 |
| ENSDARG000 | 25;25;25;25;2 | 31405266:314 | 31405334:314 | +;+;+;+;+;+;+; | 1387 | tnnt3a | 0 | 0 | 0 | 0 | 0 | 0 |
| ENSDARG000 | 25;25;25;25;2 | 35952036:359 | 35952888:359 | -;-;-;-;- | 1312 | mphosph6 | 273 | 267 | 346 | 213 | 322 | 260 |
| ENSDARG000 | 25;25;25;25;2 | 19485328:194 | 19485554:194 | -;-;-;-;-;-;-;- | 792 | zgc:193812 | 32 | 23 | 13 | 33 | 26 | 29 |
| ENSDARG000 | 25;25;25;25;2 | 17313568:173 | 17313675:173 | +;+;+;+;+;+;+; | 8596 | cnot1 | 4700 | 4300 | 6441 | 3565 | 6196 | 4241 |
| ENSDARG000 | 25;25;25;25 | 34835553:348 | 34836015:348 | -;-;-;- | 1079 | gabarapl2 | 751 | 758 | 713 | 784 | 803 | 761 |
| ENSDARG000 | 25;25;25;25;2 | 19485198:194 | 19485838:194 | +;+;+;+;+;+;+; | 3885 | NA | 11 | 1 | 11 | 16 | 12 | 0 |
| ENSDARG000 | 25;25;25;25;2 | 34845115:348 | 34845656:348 | +;+;+;+;+;+;+; | 1631 | tmem231 | 54 | 86 | 121 | 60 | 81 | 81 |
| ENSDARG000 | 25;25;25;25;2 | 12969653:129 | 12970109:129 | +;+;+;+;+;+;+ | 1340 | NA | 0 | 2 | 6 | 1 | 2 | 0 |
| ENSDARG000 | 25;25 | 12967795:129 | 12968305:129 | -;- | 732 | NA | 0 | 0 | 11 | 0 | 36 | 0 |
| ENSDARG000 | 25;25;25;25;2 | 8662469:8662 | 8662647:8662 | +;+;+;+;+;+;+; | 7815 | MAN2A2 | 1167 | 613 | 1053 | 632 | 1582 | 1098 |
| ENSDARG000 | 25;25;25;25;2 | 7241084:7241 | 7241468:7241 | +;+;+;+;+;+;+; | 2188 | hmg20a | 426 | 328 | 528 | 400 | 420 | 314 |
| ENSDARG000 | 25;25;25;25;2 | 15287036:152 | 15287400:152 | +;+;+;+;+;+;+; | 7933 | hipk3a | 1408 | 1225 | 2176 | 1729 | 2105 | 1410 |
| ENSDARG000 | 25;25;25 | 12981431:129 | 12981644:129 | -;-;- | 558 | NA | 0 | 4 | 14 | 12 | 112 | 30 |
| ENSDARG000 | 25;25;25 | 12995857:129 | 12995968:129 | -;-;- | 317 | NA | 14 | 0 | 4 | 6 | 10 | 8 |
| ENSDARG000 | 25;25;25;25;2 | 35957603:359 | 35958418:359 | -;-;-;-;-;-;-;-; | 1401 | SNX20 | 4 | 8 | 1 | 20 | 15 | 40 |
| ENSDARG000 | 25;25;25;25;2 | 235992:2359 | 236192:2361 | +;+;+;+;+;+;+; | 2380 | pus7 | 489 | 700 | 1261 | 555 | 1013 | 506 |
| ENSDARG000 | 25;25;25 | 31421253:314 | 31422222:314 | -;-;- | 1772 | myod1 | 22 | 16 | 14 | 6 | 6 | 17 |
| ENSDARG000 | 25;25;25 | 12997641:129 | 12997753:129 | -;-;- | 323 | ccl39.3 | 168 | 78 | 154 | 52 | 142 | 319 |
| ENSDARG000 | 25;25;25;25;2 | 28340171:283 | 28340933:283 | -;-;-;-;-;-;-;-; | 6135 | PTPRZ1 | 26 | 0 | 0 | 0 | 18 | 10 |
| ENSDARG000 | 25;25 | 3257932:3262 | 3258402:3262 | -;- | 578 | pthlhb | 0 | 0 | 0 | 0 | 0 | 0 |
| ENSDARG000 | 25;25;25 | 31424850:314 | 31427207:314 | -;-;- | 2859 | zgc:172145 | 126 | 179 | 197 | 88 | 138 | 132 |
| ENSDARG000 | 25;25;25;25;2 | 3294150:3298 | 3294768:3298 | +;+;+;+;+;+ | 2143 | tmpob | 390 | 723 | 811 | 287 | 687 | 406 |
| ENSDARG000 | 25;25;25;25;2 | 8755990:8764 | 8757057:8764 | -;-;-;-;-;-;-;-; | 3499 | furinb | 563 | 438 | 355 | 506 | 472 | 450 |
| ENSDARG000 | 25;25;25;25;2 | 24616717:246 | 24619335:246 | +;+;+;+;+;+;+; | 5504 | ABTB2 | 64 | 22 | 80 | 12 | 15 | 45 |
| ENSDARG000 | 25;25 | 31502082:315 | 31502346:315 | -;- | 461 | NA | 0 | 0 | 0 | 0 | 0 | 0 |
| ENSDARG000 | 25;25;25;25 | 21750386:217 | 21751447:217 | -;-;-;- | 3271 | TMEM168 | 164 | 74 | 217 | 34 | 205 | 134 |
| ENSDARG000 | 25;25;25;25;2 | 3306620:3306 | 3306816:3306 | +;+;+;+;+;+;+; | 2319 | slc25a3b | 10615 | 11592 | 13347 | 10216 | 11164 | 9060 |
| ENSDARG000 | 25;25;25;25;2 | 13017489:130 | 13017923:130 | -;-;-;-;-;-;-;-; | 1168 | NA | 8 | 20 | 80 | 64 | 314 | 262 |
| ENSDARG000 | 25;25;25;25;2 | 31546828:315 | 31546967:315 | -;-;-;-;-;-;-;-; | 5358 | lamb1a | 216 | 52 | 284 | 38 | 240 | 78 |
| ENSDARG000 | 25;25;25 | 13022907:130 | 13023037:130 | -;-;- | 419 | ccl39.1 | 0 | 20 | 55 | 22 | 235 | 54 |
| ENSDARG000 | 25;25;25;25;2 | 7346800:7398 | 7346971:7401 | +;+;+;+;+;+;+; | 5698 | PEAK1 | 976 | 552 | 1006 | 1030 | 1082 | 1146 |
| ENSDARG000 | 25;25;25;25;2 | 25085349:250 | 25085699:250 | +;+;+;+;+;+;+; | 1670 | NA | 0 | 0 | 0 | 0 | 0 | 94 |
| ENSDARG000 | 25;25 | 34862225:348 | 34862388:348 | +;+ | 1431 | chst6 | 28 | 10 | 18 | 15 | 14 | 10 |
| ENSDARG000 | 25;25;25;25;2 | 25092874:250 | 25092950:250 | +;+;+;+;+;+ | 1475 | NA | 0 | 0 | 0 | 0 | 0 | 6 |
| ENSDARG000 | 25;25;25 | 35986345:359 | 35987373:359 | -;-;- | 4574 | SALL1 | 14 | 12 | 4 | 14 | 2 | 8 |
| ENSDARG000 | 25;25;25;25;2 | 19522954:195 | 19523143:195 | +;+;+;+;+;+;+; | 3153 | sh3gl3b | 313 | 175 | 217 | 44 | 120 | 360 |
| ENSDARG000 | 25;25;25;25;2 | 34887999:348 | 34888164:348 | +;+;+;+;+;+;+; | 4034 | spire2 | 0 | 12 | 15 | 0 | 38 | 0 |
| ENSDARG000 | 25;25;25;25;2 | 8921425:8925 | 8922344:8926 | +;+;+;+;+;+;+; | 4361 | accs | 277 | 214 | 302 | 147 | 261 | 244 |
| ENSDARG000 | 25;25 | 21765404:217 | 21765648:217 | -;- | 439 | NA | 0 | 0 | 0 | 0 | 0 | 0 |
| ENSDARG000 | 25;25;25;25;2 | 21770962:217 | 21771768:217 | -;-;-;-;- | 1367 | bmt2 | 604 | 577 | 757 | 329 | 676 | 704 |
| ENSDARG000 | 25;25;25;25;2 | 28555584:285 | 28556131:285 | +;+;+;+;+;+;+; | 2382 | SLC15A5 | 0 | 0 | 0 | 0 | 0 | 0 |

| | | | | | | | | | | | |
|---|---|---|---|---|---|---|---|---|---|---|---|
| ENSDARG000 | 25:25:25:25:2 | 15354095:154 | 15354732:154 | +:+:+:+:+:+:+: | 6217 | NA | 2 | 0 | 0 | 0 | 0 | 0 |
| ENSDARG000 | 25:25:25:25:2 | 17349265:173 | 17349777:173 | -:-:-:-:-:-:-:-: | 1764 | setd6 | 170 | 191 | 323 | 142 | 184 | 166 |
| ENSDARG000 | 25;25;25;25 | 21787173:217 | 21791470:217 | -;-;-;- | 5785 | NA | 0 | 0 | 4 | 2 | 0 | 2 |
| ENSDARG000 | 25:25:25:25:2 | 31633322:316 | 31633672:316 | -:-:-:-:-:-:-:-: | 6090 | lamb4 | 2 | 0 | 2 | 10 | 6 | 5 |
| ENSDARG000 | 25:25:25:25:2 | 21800397:218 | 21800553:218 | -:-:-:-:-:-:-:-: | 1961 | ahcyl2a | 0 | 0 | 16 | 0 | 4 | 0 |
| ENSDARG000 | 25:25:25:25:2 | 4660639:4666 | 4661262:4666 | +:+:+:+:+:+:+: | 2795 | deaf1 | 162 | 112 | 246 | 94 | 270 | 167 |
| ENSDARG000 | 25:25:25:25:2 | 245018:24543 | 245151:24548 | +:+:+:+:+:+:+: | 3183 | vwa5a | 2373 | 1926 | 1849 | 1954 | 1678 | 2384 |
| ENSDARG000 | 25:25:25:25:2 | 25123385:251 | 25123484:251 | +:+:+:+:+:+:+: | 2182 | ldha | 3584 | 10360 | 24618 | 17548 | 7394 | 5526 |
| ENSDARG000 | 25:25:25:25:2 | 7423770:7427 | 7423809:7427 | +;+;+;+;+;+ | 1405 | NA | 42 | 40 | 327 | 212 | 45 | 49 |
| ENSDARG000 | 25:25:25:25:2 | 17358287:173 | 17363198:173 | -:-:-:-:-:-:-:-: | 6746 | cyp2x6 | 0 | 0 | 18 | 0 | 0 | 0 |
| ENSDARG000 | 25;25;25 | 13048201:130 | 13048744:130 | -;-;- | 803 | ccl35.1 | 17 | 46 | 91 | 106 | 133 | 83 |
| ENSDARG000 | 25;25;25;25 | 15487494:154 | 15487605:154 | +;+;+;+ | 471 | NA | 0 | 0 | 0 | 0 | 0 | 0 |
| ENSDARG000 | 25:25:25:25:2 | 36010815:360 | 36016106:360 | -:-:-:-:-:-:-:-: | 8770 | RBL2 | 865 | 474 | 730 | 495 | 1236 | 906 |
| ENSDARG000 | 25;25 | 7504314:7506 | 7504554:7506 | +;+ | 684 | ifitm5 | 0 | 4 | 0 | 0 | 10 | 0 |
| ENSDARG000 | 25:25:25:25:2 | 7492663:7494 | 7492998:7494 | +:+:+:+:+:+:+: | 4525 | cat | 611 | 544 | 832 | 774 | 909 | 783 |
| ENSDARG000 | 25;25 | 15490212:154 | 15490280:154 | +;+ | 175 | NA | 0 | 0 | 0 | 0 | 0 | 0 |
| ENSDARG000 | 25:25:25:25:2 | 314608:32062 | 317916:32145 | -;-;-;-;-;-;-;- | 5108 | prickle1a | 46 | 24 | 60 | 0 | 86 | 12 |
| ENSDARG000 | 25:25:25:25:2 | 3326885:3327 | 3327071:3327 | +:+:+:+:+:+:+: | 1781 | ldhbb | 8 | 0 | 27 | 0 | 10 | 10 |
| ENSDARG000 | 25;25;25;25 | 8955530:8955 | 8955697:8956 | +;+;+;+ | 1424 | NA | 2 | 10 | 2 | 2 | 30 | 27 |
| ENSDARG000 | 25:25:25:25:2 | 17372290:173 | 17372548:173 | -;-;-;-;-;-;-;-;- | 1728 | cyp2x7 | 30 | 66 | 18 | 16 | 70 | 38 |
| ENSDARG000 | 25:25:25:25:2 | 13068019:130 | 13068143:130 | -;-;-;-;-;-;- | 2078 | smpd3 | 2 | 4 | 2 | 4 | 14 | 20 |
| ENSDARG000 | 25;25;25 | 4696377:4699 | 4696594:4699 | +;+;+ | 440 | NA | 2 | 0 | 0 | 0 | 0 | 0 |
| ENSDARG000 | 25:25:25:25:2 | 25132703:251 | 25133215:251 | -:-:-:-:-:-:-:-: | 1898 | tsg101a | 1132 | 975 | 1249 | 828 | 1264 | 1125 |
| ENSDARG000 | 25:25:25:25:2 | 9003230:9003 | 9003708:9003 | +:+:+:+:+:+:+: | 5426 | rag1 | 0 | 0 | 24 | 0 | 10 | 28 |
| ENSDARG000 | 25:25:25:25:2 | 31714335:317 | 31715006:317 | -:-:-:-:-:-:-:-: | 1899 | acot19 | 2 | 0 | 0 | 12 | 4 | 0 |
| ENSDARG000 | 25:25:25:25:2 | 28596989:285 | 28597241:285 | -:-:-:-:-:-:-:-: | 3471 | agbl2 | 2 | 6 | 2 | 6 | 0 | 0 |
| ENSDARG000 | 25:25:25:25:2 | 19541019:195 | 19541203:195 | +:+:+:+:+:+:+: | 3427 | nedd1 | 348 | 667 | 524 | 389 | 738 | 552 |
| ENSDARG000 | 25:25:25:25:2 | 4707173:4707 | 4708746:4708 | -:-:-:-:-:-:-:-: | 2845 | drd4a | 0 | 0 | 0 | 0 | 0 | 0 |
| ENSDARG000 | 25:25:25:25:2 | 15494040:154 | 15494870:154 | -;-;-;-;-;- | 1657 | NA | 0 | 0 | 0 | 0 | 0 | 0 |
| ENSDARG000 | 25;25;25 | 4703200:4708 | 4703273:4708 | +;+;+ | 658 | NA | 0 | 0 | 0 | 0 | 0 | 0 |
| ENSDARG000 | 25:25:25:25:2 | 469855:47256 | 470001:47267 | +:+:+:+:+:+:+: | 1012 | rsl24d1 | 3267 | 2896 | 6462 | 6141 | 3190 | 6333 |
| ENSDARG000 | 25:25:25:25:2 | 36045072:360 | 36045139:360 | +:+:+:+:+:+:+: | 8684 | rpgrip1l | 176 | 101 | 157 | 76 | 166 | 108 |
| ENSDARG000 | 25:25:25:25:2 | 485353:48850 | 488923:48892 | -:-:-:-:-:-:-:-: | 10157 | myo9ab | 5091 | 4705 | 5492 | 5863 | 3387 | 3422 |
| ENSDARG000 | 25:25:25:25:2 | 34915576:349 | 34916302:349 | +:+:+:+:+:+:+: | 5990 | SNTB2 | 1191 | 1148 | 1688 | 1248 | 1293 | 1842 |
| ENSDARG000 | 25:25:25:25:2 | 28640459:286 | 28641430:286 | -:-:-:-:-:-:-:-: | 3518 | fnbp4 | 2121 | 1434 | 2302 | 1845 | 2009 | 2677 |
| ENSDARG000 | 25;25;25;25 | 15496732:154 | 15496891:154 | -;-;-;- | 846 | NA | 0 | 0 | 0 | 0 | 0 | 0 |
| ENSDARG000 | 25:25:25:25:2 | 15502659:155 | 15502944:155 | -;-;-;-;-;- | 1112 | NA | 0 | 0 | 0 | 0 | 0 | 0 |
| ENSDARG000 | 25;25;25;25 | 479997:48187 | 480505:48208 | -;-;-;- | 1331 | NA | 234 | 221 | 117 | 90 | 106 | 198 |
| ENSDARG000 | 25:25:25:25:2 | 418932:41986 | 419410:42015 | +:+:+:+:+:+:+: | 4251 | NA | 0 | 4 | 2 | 0 | 22 | 28 |
| ENSDARG000 | 25:25:25:25:2 | 17386763:173 | 17387036:173 | -;-;-;-;-;-;-;-;- | 1575 | cyp2x8 | 0 | 0 | 0 | 0 | 0 | 0 |
| ENSDARG000 | 25;25 | 9010425:9013 | 9012954:9013 | -;- | 2682 | rag2 | 0 | 6 | 14 | 0 | 14 | 16 |
| ENSDARG000 | 25:25:25:25:2 | 9013342:9013 | 9013422:9013 | +:+:+:+:+:+:+: | 2109 | NA | 74 | 70 | 87 | 52 | 115 | 84 |
| ENSDARG000 | 25:25:25:25:2 | 13157165:131 | 13157274:131 | +:+:+:+:+:+:+: | 1688 | NA | 7 | 0 | 0 | 0 | 0 | 0 |
| ENSDARG000 | 25:25:25:25:2 | 15504780:155 | 15504959:155 | -:-:-:-:-:-:-:-: | 1256 | NA | 0 | 0 | 0 | 0 | 0 | 0 |
| ENSDARG000 | 25:25:25:25:2 | 4750972:4751 | 4751399:4751 | +:+:+:+:+:+:+: | 1169 | NA | 8 | 0 | 0 | 0 | 0 | 0 |
| ENSDARG000 | 25:25:25:25:2 | 13180468:131 | 13184348:131 | -:-:-:-:-:-:-:-: | 4458 | NA | 535 | 253 | 450 | 228 | 506 | 303 |
| ENSDARG000 | 25;25 | 7519673:7520 | 7520186:7520 | -;- | 1160 | NA | 28 | 58 | 50 | 36 | 24 | 22 |
| ENSDARG000 | 25:25:25:25:2 | 4760489:4761 | 4760686:4761 | +:+:+:+:+:+:+: | 2281 | NA | 0 | 0 | 0 | 0 | 0 | 0 |
| ENSDARG000 | 25:25:25:25:2 | 7532627:7532 | 7533718:7533 | +:+:+:+:+:+:+: | 6952 | ptdss2 | 494 | 737 | 867 | 314 | 679 | 566 |
| ENSDARG000 | 25;25 | 25202213:252 | 25202329:252 | +;+ | 212 | NA | 0 | 0 | 0 | 0 | 0 | 0 |
| ENSDARG000 | 25:25:25:25:2 | 19560647:195 | 19560963:195 | -:-:-:-:-:-:-:-: | 1900 | si:ch211-59o9 | 68 | 120 | 108 | 62 | 65 | 70 |
| ENSDARG000 | 25:25:25:25:2 | 15510884:155 | 15511204:155 | -;-;-;-;-;- | 1147 | NA | 0 | 0 | 0 | 0 | 0 | 0 |
| ENSDARG000 | 25;25 | 25351512:253 | 25352089:253 | -;- | 834 | NA | 0 | 0 | 0 | 0 | 6 | 0 |
| ENSDARG000 | 25;25;25 | 19557628:195 | 19557709:195 | +;+;+ | 1323 | lyrm5b | 23 | 28 | 37 | 8 | 40 | 3 |
| ENSDARG000 | 25:25:25:25:2 | 3341774:3342 | 3342327:3342 | -;-;-;-;- | 1233 | golt1bb | 432 | 697 | 690 | 330 | 708 | 462 |
| ENSDARG000 | 25:25:25:25:2 | 34956368:349 | 34956898:349 | -:-:-:-:-:-:-:-: | 2108 | cdk10 | 160 | 139 | 267 | 126 | 235 | 173 |
| ENSDARG000 | 25:25:25:25:2 | 21829777:218 | 21830070:218 | +:+:+:+:+:+:+: | 2287 | ckmt1 | 4 | 4 | 24 | 0 | 2 | 0 |
| ENSDARG000 | 25:25:25:25:2 | 3347461:3347 | 3347709:3347 | +:+:+:+:+:+:+: | 2176 | slc35b4 | 499 | 505 | 389 | 260 | 809 | 370 |
| ENSDARG000 | 25:25:25:25:2 | 31743651:317 | 31744301:317 | -:-:-:-:-:-:-:-: | 1797 | ubl7a | 401 | 348 | 424 | 453 | 420 | 411 |
| ENSDARG000 | 25:25:25:25:2 | 7612606:7614 | 7612758:7614 | -:-:-:-:-:-:-:-: | 5554 | NA | 264 | 286 | 226 | 75 | 284 | 154 |
| ENSDARG000 | 25:25:25:25:2 | 25395590:254 | 25396304:254 | -;-;-;-;-;- | 1526 | hrasa | 332 | 688 | 626 | 704 | 398 | 494 |
| ENSDARG000 | 25;25;25;25 | 15513004:155 | 15513219:155 | -;-;-;- | 902 | NA | 0 | 0 | 0 | 0 | 0 | 0 |
| ENSDARG000 | 25;25 | 34976101:349 | 34976682:349 | -;- | 940 | NA | 0 | 2 | 12 | 6 | 0 | 0 |
| ENSDARG000 | 25:25:25:25:2 | 25381548:253 | 25382205:253 | -;-;-;-;-;- | 1200 | zgc:123278 | 2 | 0 | 0 | 0 | 0 | 0 |
| ENSDARG000 | 25:25:25:25:2 | 21860746:218 | 21861137:218 | -:-:-:-:-:-:-:-: | 2232 | FBXO31 | 122 | 128 | 100 | 107 | 110 | 84 |
| ENSDARG000 | 25:25:25:25:2 | 7653902:7654 | 7654694:7654 | -;-;-;-;-;- | 1222 | arpp19b | 992 | 1486 | 1489 | 877 | 1212 | 992 |
| ENSDARG000 | 25:25:25:25:2 | 4787431:4787 | 4787633:4787 | +:+:+:+:+:+:+: | 6123 | MYO5C | 16 | 20 | 2 | 46 | 16 | 0 |
| ENSDARG000 | 25:25:25:25:2 | 34984333:349 | 34984774:349 | +:+:+:+:+:+:+: | 3292 | NA | 0 | 6 | 0 | 0 | 0 | 0 |
| ENSDARG000 | 25:25:25:25:2 | 3358701:3359 | 3358892:3360 | +:+:+:+:+:+:+: | 2103 | chchd3b | 74 | 109 | 119 | 67 | 126 | 123 |
| ENSDARG000 | 25:25:25:25:2 | 7670683:7671 | 7670800:7672 | +:+:+:+:+:+:+: | 2055 | kcnj11l | 16 | 28 | 118 | 61 | 6 | 23 |
| ENSDARG000 | 25:25:25:25:2 | 15647750:156 | 15648289:156 | +:+:+:+:+:+:+: | 3819 | spon1b | 46 | 18 | 18 | 56 | 32 | 12 |
| ENSDARG000 | 25;25;25 | 36152215:361 | 36152559:361 | +;+;+ | 3129 | irx5b | 0 | 0 | 0 | 0 | 0 | 0 |
| ENSDARG000 | 25:25:25:25:2 | 7667634:7667 | 7668184:7668 | -:-:-:-:-:-:-:-: | 1668 | bet1l | 160 | 262 | 297 | 222 | 202 | 152 |
| ENSDARG000 | 25:25:25:25:2 | 7675629:7679 | 7679889:7679 | -:-:-:-:-:-:-:-: | 4597 | NA | 18 | 36 | 123 | 149 | 35 | 52 |
| ENSDARG000 | 25;25;25 | 3378135:3391 | 3383700:3392 | -;-;- | 6740 | GPR22 | 6 | 0 | 10 | 0 | 0 | 0 |
| ENSDARG000 | 25:25:25:25:2 | 31837228:318 | 31837885:318 | -;-;-;-;-;-;- | 1406 | scamp5a | 52 | 166 | 236 | 82 | 170 | 46 |
| ENSDARG000 | 25:25:25:25:2 | 19578048:195 | 19578312:195 | -:-:-:-:-:-:-:-: | 1281 | sycp3 | 0 | 0 | 2 | 4 | 0 | 2 |
| ENSDARG000 | 25;25;25;25 | 21895182:218 | 21895231:218 | +;+;+;+ | 452 | dynlrb2 | 0 | 0 | 6 | 0 | 0 | 2 |
| ENSDARG000 | 25:25:25:25:2 | 36197733:361 | 36200159:361 | -:-:-:-:-:-:-:-: | 4243 | amfra | 1829 | 1176 | 1559 | 1208 | 1652 | 1800 |
| ENSDARG000 | 25:25:25:25:2 | 19585318:195 | 19585812:195 | -:-:-:-:-:-:-:-: | 1781 | actr6 | 223 | 269 | 317 | 105 | 368 | 324 |
| ENSDARG000 | 25:25:25:25:2 | 3475384:3475 | 3475517:3475 | -:-:-:-:-:-:-:-: | 1595 | PRKAR2B | 0 | 4 | 8 | 0 | 23 | 0 |
| ENSDARG000 | 25:25:25:25:2 | 17495615:174 | 17496224:174 | -;-;-;-;-;-;-;- | 1544 | trim35-40 | 74 | 102 | 132 | 51 | 140 | 111 |
| ENSDARG000 | 25:25:25:25:2 | 17479264:174 | 17479787:174 | -;-;-;-;-;-;-;- | 2601 | NA | 524 | 297 | 655 | 378 | 460 | 388 |
| ENSDARG000 | 25:25:25:25:2 | 821725:82394 | 821803:82427 | -:-:-:-:-:-:-:-: | 5005 | znf609a | 422 | 334 | 815 | 440 | 824 | 438 |
| ENSDARG000 | 25:25:25:25:2 | 15841670:158 | 15842082:158 | +;+;+;+;+;+;+;+ | 2294 | syt9b | 2 | 0 | 4 | 6 | 0 | 8 |
| ENSDARG000 | 25:25:25:25:2 | 31868268:318 | 31868350:318 | +;+;+;+;+;+;+;+ | 1704 | parp16 | 770 | 652 | 831 | 852 | 656 | 772 |
| ENSDARG000 | 25:25:25:25:2 | 13191391:131 | 13191495:131 | +:+:+:+:+:+:+: | 1972 | NA | 626 | 477 | 570 | 210 | 688 | 370 |
| ENSDARG000 | 25:25:25:25:2 | 3507368:3510 | 3507516:3511 | +;+;+;+;+;+;+;+ | 1298 | ccdc34 | 9 | 60 | 9 | 16 | 3 | 10 |
| ENSDARG000 | 25;25;25 | 9853189:9931 | 9853529:9931 | -;-;- | 545 | NA | 0 | 0 | 0 | 0 | 0 | 0 |
| ENSDARG000 | 25:25:25:25:2 | 1035656:1035 | 1036031:1036 | +:+:+:+:+:+:+: | 1047 | NA | 0 | 0 | 0 | 0 | 0 | 0 |
| ENSDARG000 | 25:25:25:25:2 | 15933411:159 | 15933858:159 | +:+:+:+:+:+:+: | 11194 | ppfibp2b | 5043 | 2696 | 2930 | 4244 | 3602 | 4085 |
| ENSDARG000 | 25:25:25:25:2 | 36219436:362 | 36219908:362 | -:-:-:-:-:-:-:-: | 3663 | NA | 263 | 131 | 154 | 130 | 221 | 201 |
| ENSDARG000 | 25:25:25:25:2 | 22017182:220 | 22017336:220 | +:+:+:+:+:+:+: | 4072 | NA | 22 | 6 | 98 | 2 | 126 | 46 |
| ENSDARG000 | 25:25:25:25:2 | 4837915:4841 | 4838131:4841 | +:+:+:+:+:+:+: | 1443 | gnb5a | 2 | 20 | 0 | 2 | 0 | 8 |
| ENSDARG000 | 25:25:25:25:2 | 3549401:3549 | 3549545:3549 | +:+:+:+:+:+:+: | 2339 | ccdc77 | 72 | 122 | 132 | 42 | 116 | 48 |
| ENSDARG000 | 25:25:25:25:2 | 13198224:131 | 13199489:131 | -:-:-:-:-:-:-:-: | 1894 | NA | 58 | 81 | 16 | 3 | 10 | 10 |
| ENSDARG000 | 25:25:25:25:2 | 22107643:221 | 22107796:221 | +:+:+:+:+:+:+: | 3135 | sigirr | 8 | 0 | 20 | 0 | 20 | 0 |
| ENSDARG000 | 25:25:25:25:2 | 31890273:318 | 31890569:318 | -:-:-:-:-:-:-:-: | 1213 | EIF3J | 0 | 0 | 0 | 0 | 0 | 0 |
| ENSDARG000 | 25;25;25;25 | 31890256:318 | 31890569:318 | -;-;-;- | 1474 | NA | 2 | 0 | 12 | 22 | 6 | 4 |
| ENSDARG000 | 25:25:25:25:2 | 31900825:319 | 31901912:319 | -:-:-:-:-:-:-:-: | 2028 | eif3ja | 438 | 723 | 1120 | 502 | 718 | 470 |
| ENSDARG000 | 25:25:25:25:2 | 10410620:104 | 10410734:104 | +:+:+:+:+:+:+: | 1727 | ehf | 80 | 0 | 0 | 0 | 2 | 0 |
| ENSDARG000 | 25:25:25:25:2 | 13205878:132 | 13205944:132 | +:+:+:+:+:+:+: | 821 | NA | 6 | 0 | 0 | 0 | 0 | 12 |
| ENSDARG000 | 25:25:25:25:2 | 35019693:350 | 35020901:350 | +:+:+:+:+:+:+: | 9456 | FLNC | 0 | 2 | 2 | 6 | 4 | 0 |
| ENSDARG000 | 25:25:25:25:2 | 17508235:175 | 17508770:175 | -:-:-:-:-:-:-:-: | 4968 | NA | 12 | 0 | 6 | 6 | 16 | 28 |
| ENSDARG000 | 25:25:25:25:2 | 7692075:7692 | 7698053:7698 | -:-:-:-:-:-:-:-: | 7754 | PRDM11 | 618 | 450 | 622 | 410 | 662 | 572 |
| ENSDARG000 | 25;25;25 | 1131420:1131 | 1131452:1131 | +;+;+ | 1869 | ciao2a | 386 | 277 | 510 | 306 | 579 | 646 |
| ENSDARG000 | 25:25:25:25:2 | 25438165:254 | 25438469:254 | +:+:+:+:+:+:+: | 2783 | LRRC56 | 0 | 0 | 0 | 0 | 0 | 0 |
| ENSDARG000 | 25;25 | 28672539:286 | 28673418:286 | -;- | 973 | lrrc10 | 0 | 0 | 0 | 0 | 0 | 0 |
| ENSDARG000 | 25;25;25 | 13208362:132 | 13208540:132 | +;+;+ | 685 | NA | 0 | 0 | 0 | 0 | 0 | 0 |
| ENSDARG000 | 25:25:25:25:2 | 10458990:104 | 10459240:104 | +:+:+:+:+:+:+: | 2535 | ric8a | 502 | 255 | 360 | 659 | 236 | 268 |
| ENSDARG000 | 25:25:25:25:2 | 16044734:160 | 16047056:160 | -:-:-:-:-:-:-:-: | 3344 | cyb5r2 | 14 | 62 | 6 | 42 | 17 | 8 |
| ENSDARG000 | 25:25:25:25:2 | 28678851:286 | 28678930:286 | +:+:+:+:+:+:+: | 2505 | cnot2 | 655 | 745 | 1008 | 480 | 809 | 800 |
| ENSDARG000 | 25:25:25:25:2 | 4855549:4857 | 4855839:4858 | +:+:+:+:+:+:+: | 3989 | AP4E1 | 405 | 266 | 294 | 139 | 282 | 206 |
| ENSDARG000 | 25:25:25:25:2 | 7718967:7719 | 7720248:7720 | -:-:-:-:-:-:-:-: | 4071 | PHF21A | 381 | 195 | 479 | 333 | 559 | 327 |
| ENSDARG000 | 25:25:25:25:2 | 13313941:133 | 13314967:133 | -;-;-;-;-;-;-;-;- | 2173 | NA | 2 | 0 | 2 | 4 | 42 | 4 |
| ENSDARG000 | 25:25:25:25:2 | 22151271:221 | 22152461:221 | -:-:-:-:-:-:-:-: | 4452 | pkp3a | 490 | 32 | 6 | 49 | 126 | 64 |
| ENSDARG000 | 25:25:25:25:2 | 36249279:362 | 36250316:362 | -:-:-:-:-:-:-:-: | 2821 | DUS2 | 454 | 446 | 744 | 278 | 593 | 398 |
| ENSDARG000 | 25;25 | 1195282:1196 | 1195351:1196 | -;- | 157 | NA | 0 | 0 | 0 | 0 | 0 | 0 |
| ENSDARG000 | 25:25:25:25:2 | 17559399:175 | 17560315:175 | -:-:-:-:-:-:-:-: | 7468 | MMP15 | 878 | 394 | 844 | 472 | 679 | 925 |
| ENSDARG000 | 25:25:25:25:2 | 17589906:175 | 17590681:175 | +:+:+:+:+:+:+: | 5330 | vac14 | 1646 | 1358 | 1545 | 781 | 1193 | 1663 |
| ENSDARG000 | 25:25:25:25:2 | 4890767:4896 | 4893586:4896 | -:-:-:-:-:-:-:-: | 4167 | cyp19a1b | 19 | 10 | 7 | 21 | 4 | 2 |
| ENSDARG000 | 25 | 1304173 | 1305384 | + | 1212 | NA | 0 | 0 | 0 | 0 | 0 | 0 |
| ENSDARG000 | 25:25:25:25:2 | 1266945:1276 | 1267968:1276 | -:-:-:-:-:-:-:-: | 3991 | CALML4 | 0 | 0 | 0 | 0 | 0 | 0 |
| ENSDARG000 | 25:25:25:25:2 | 17581393:175 | 17581837:175 | -:-:-:-:-:-:-:-: | 1655 | zpd | 0 | 0 | 0 | 0 | 0 | 0 |
| ENSDARG000 | 25:25:25:25:2 | 13321307:133 | 13321427:133 | +;+;+;+;+ | 1433 | NA | 0 | 10 | 2 | 6 | 24 | 20 |
| ENSDARG000 | 25:25:25:25:2 | 19594160:195 | 19594655:195 | -:-:-:-:-:-:-:-: | 3354 | gtse1 | 100 | 352 | 418 | 236 | 236 | 134 |
| ENSDARG000 | 25:25:25:25:2 | 25453221:254 | 25453653:254 | +:+:+:+:+:+:+: | 1978 | RASSF7 | 0 | 6 | 8 | 0 | 30 | 2 |
| ENSDARG000 | 25:25:25:25:2 | 25508495:255 | 25508585:255 | +:+:+:+:+:+:+: | 6483 | NA | 1077 | 804 | 1192 | 606 | 1142 | 899 |
| ENSDARG000 | 25:25:25:25:2 | 31912153:319 | 31912476:319 | +:+:+:+:+:+:+: | 3302 | NA | 46 | 4 | 56 | 101 | 39 | 120 |
| ENSDARG000 | 25:25:25:25:2 | 16074350:160 | 16074493:160 | -;-;-;-;-;-;-;-;- | 1159 | OVCH2 | 8 | 8 | 38 | 2 | 9 | 30 |
| ENSDARG000 | 25:25:25:25:2 | 10485103:104 | 10485283:104 | +:+:+:+:+:+:+: | 1498 | psmd13 | 1964 | 2281 | 2564 | 1247 | 2165 | 1340 |
| ENSDARG000 | 25:25:25:25:2 | 13332280:133 | 13337390:133 | -:-:-:-:-:-:-:-: | 6419 | ndrg4 | 1454 | 417 | 3420 | 3603 | 2510 | 3195 |
| ENSDARG000 | 25:25:25:25:2 | 7784582:7814 | 7785239:7814 | +:+:+:+:+:+:+: | 6619 | DGKZ | 181 | 114 | 187 | 56 | 329 | 298 |
| ENSDARG000 | 25:25:25:25:2 | 16079913:160 | 16080310:160 | +:+:+:+:+:+:+: | 2787 | far1 | 1329 | 1252 | 1248 | 1432 | 1113 | 1261 |
| ENSDARG000 | 25:25:25:25:2 | 17689565:177 | 17690069:177 | +:+:+:+:+:+:+: | 2601 | galnt18a | 0 | 0 | 2 | 0 | 2 | 0 |
| ENSDARG000 | 25:25:25:25:2 | 28714527:287 | 28716366:287 | -:-:-:-:-:-:-:-: | 5366 | WASHC4 | 2610 | 2279 | 2540 | 1772 | 2885 | 2741 |
| ENSDARG000 | 25:25:25:25:2 | 22216723:222 | 22217124:222 | +:+:+:+:+:+:+: | 2090 | hexa | 620 | 814 | 547 | 559 | 878 | 665 |
| ENSDARG000 | 25:25:25:25:2 | 25546789:255 | 25547317:255 | -:-:-:-:-:-:-:-: | 1843 | irf7 | 422 | 830 | 656 | 934 | 119 | 176 |
| ENSDARG000 | 25:25:25:25:2 | 4954220:4954 | 4954464:4954 | +:+:+:+:+:+:+: | 1857 | NA | 2 | 22 | 4 | 9 | 1 | 6 |
| ENSDARG000 | 25;25;25 | 17847054:178 | 17847333:178 | +;+;+ | 810 | PYURF | 49 | 20 | 52 | 34 | 56 | 34 |
| ENSDARG000 | 25;25 | 10499642:105 | 10499863:105 | -;- | 600 | cox8b | 179 | 63 | 90 | 116 | 68 | 212 |
| ENSDARG000 | 25;25;25;25 | 17848923:178 | 17850583:178 | -;-;-;- | 1810 | NA | 135 | 130 | 283 | 273 | 148 | 48 |
| ENSDARG000 | 25:25:25:25:2 | 3573095:3573 | 3573870:3573 | -:-:-:-:-:-:-:-: | 6611 | NA | 200 | 209 | 345 | 185 | 301 | 178 |
| ENSDARG000 | 25:25:25:25:2 | 1325946:1326 | 1326454:1326 | -:-:-:-:-:-:-:-: | 1450 | CLN6 | 12 | 2 | 10 | 2 | 4 | 20 |
| ENSDARG000 | 25:25:25:25:2 | 10514209:105 | 10514404:105 | -:-:-:-:-:-:-:-: | 1499 | NA | 157 | 10 | 15 | 2 | 2 | 63 |
| ENSDARG000 | 25:25:25:25:2 | 25564558:255 | 25566372:255 | -;-;-;-;- | 2036 | hic1l | 123 | 78 | 147 | 104 | 79 | 214 |
| ENSDARG000 | 25;25 | 1335530:1338 | 1336466:1340 | +;+ | 3378 | NA | 345 | 283 | 318 | 261 | 335 | 377 |
| ENSDARG000 | 25:25:25:25:2 | 25592082:255 | 25592277:255 | -:-:-:-:-:-:-:-: | 2951 | TBC1D2B | 263 | 152 | 395 | 197 | 350 | 308 |
| ENSDARG000 | 25:25:25:25:2 | 1343161:1343 | 1343941:1343 | -:-:-:-:-:-:-:-: | 4484 | ITGA11 | 6 | 4 | 2 | 6 | 4 | 0 |
| ENSDARG000 | 25:25:25:25:2 | 10556274:105 | 10556540:105 | -:-:-:-:-:-:-:-: | 1602 | gal | 0 | 0 | 0 | 0 | 0 | 0 |
| ENSDARG000 | 25:25:25:25:2 | 25666392:256 | 25667171:256 | -;-;-;-;-;- | 1539 | cib2 | 10 | 21 | 44 | 6 | 36 | 10 |
| ENSDARG000 | 25:25:25:25:2 | 10547228:105 | 10547465:105 | +;+;+;+;+;+;+ | 1040 | zgc:110339 | 8697 | 1035 | 634 | 925 | 382 | 1265 |
| ENSDARG000 | 25:25:25:25:2 | 25737386:257 | 25737759:257 | +:+:+:+:+:+:+: | 1589 | lrrc61 | 87 | 54 | 112 | 34 | 69 | 74 |
| ENSDARG000 | 25:25:25:25:2 | 4960509:4967 | 4960542:4967 | +;+;+;+;+;+ | 1496 | NA | 84 | 149 | 36 | 69 | 75 | 39 |
| ENSDARG000 | 25;25 | 1464379:1470 | 1464532:1471 | -;- | 570 | NA | 1 | 0 | 6 | 0 | 2 | 0 |
| ENSDARG000 | 25:25:25:25:2 | 32031536:320 | 32031959:320 | +:+:+:+:+:+:+: | 8945 | NA | 1098 | 839 | 1338 | 944 | 1195 | 742 |
| ENSDARG000 | 25:25:25:25:2 | 16132580:161 | 16132846:161 | -;-;-;-;-;-;- | 1118 | dkk3b | 94 | 77 | 108 | 65 | 131 | 79 |
| ENSDARG000 | 25:25:25:25:2 | 17860798:178 | 17860841:178 | +;+;+;+;+;+ | 681 | pth1a | 0 | 0 | 0 | 8 | 4 | 0 |
| ENSDARG000 | 25:25:25:25:2 | 19609713:196 | 19610752:196 | -:-:-:-:-:-:-:-: | 5392 | atp2b1b | 2 | 22 | 10 | 20 | 40 | 32 |
| ENSDARG000 | 25:25:25:25:2 | 36292057:362 | 36292551:362 | +:+:+:+:+:+:+: | 3148 | bmb | 576 | 1046 | 712 | 756 | 1790 | 467 |
| ENSDARG000 | 25:25:25:25:2 | 25766033:257 | 25766159:257 | +:+:+:+:+:+:+: | 1438 | idh3a | 1294 | 1161 | 1440 | 1021 | 1583 | 1296 |
| ENSDARG000 | 25:25:25:25:2 | 28772632:287 | 28772894:287 | +:+:+:+:+:+:+: | 5666 | NA | 12 | 22 | 24 | 199 | 16 | 162 |

| | | | | | | | | | | | |
|---|---|---|---|---|---|---|---|---|---|---|---|
| ENSDARG000 | 25;25;25;25;2 | 16154688:161 | 16155963:161 | -;-;-;-;-;- | 2102 | NA | 0 | 0 | 0 | 0 | 0 | 0 |
| ENSDARG000 | 25;25 | 1549838:155 | 1551092:155 | +;+ | 1590 | avpr1aa | 12 | 0 | 0 | 2 | 14 | 2 |
| ENSDARG000 | 25;25;25;25;2 | 36266782:362 | 36267455:362 | -;-;-;-;-;-;-;-; | 2275 | SLC7A10 | 31 | 16 | 15 | 15 | 26 | 5 |
| ENSDARG000 | 25;25;25;25;2 | 4979446:497 | 4979698:497 | +;+;+;+;+;+;+; | 2165 | NA | 65 | 89 | 61 | 33 | 47 | 35 |
| ENSDARG000 | 25;25;25;25;2 | 36287745:362 | 36288132:362 | -;-;-;-;-;- | 1434 | faap24 | 6 | 73 | 51 | 7 | 26 | 20 |
| ENSDARG000 | 25;25 | 35044524:350 | 35045250:350 | -;- | 727 | NA | 80 | 47 | 134 | 51 | 60 | 12 |
| ENSDARG000 | 25 | 35045803 | 35046114 | - | 312 | NA | 16 | 17 | 26 | 10 | 20 | 12 |
| ENSDARG000 | 25 | 35047454 | 35047838 | - | 385 | NA | 0 | 0 | 12 | 0 | 0 | 0 |
| ENSDARG000 | 25;25;25;25;2 | 1591964:160 | 1592323:160 | +;+;+;+;+;+;+; | 2120 | ppm1h | 389 | 356 | 414 | 210 | 452 | 316 |
| ENSDARG000 | 25;25;25 | 10568514:105 | 10568895:105 | -;-;- | 628 | NA | 0 | 0 | 0 | 0 | 0 | 0 |
| ENSDARG000 | 25;25;25;25 | 7872135:787 | 7872277:787 | +;+;+;+ | 1100 | mdkb | 2049 | 2588 | 11108 | 2861 | 20822 | 8410 |
| ENSDARG000 | 25;25;25;25 | 16158490:161 | 16158718:161 | -;-;-;- | 791 | NA | 0 | 0 | 0 | 0 | 0 | 0 |
| ENSDARG000 | 25;25;25;25;2 | 10571090:105 | 10571352:105 | +;+;+;+;+;+;+; | 1199 | NA | 0 | 0 | 0 | 0 | 0 | 0 |
| ENSDARG000 | 25 | 35048945 | 35049318 | + | 374 | NA | 2 | 6 | 13 | 0 | 0 | 0 |
| ENSDARG000 | 25 | 35050385 | 35051005 | + | 621 | NA | 2 | 8 | 9 | 2 | 0 | 2 |
| ENSDARG000 | 25;25;25;25;2 | 17862338:178 | 17862498:178 | +;+;+;+;+;+;+; | 3579 | btbd10a | 1216 | 579 | 965 | 744 | 705 | 462 |
| ENSDARG000 | 25;25 | 25880883:258 | 25881003:258 | +;+ | 176 | NA | 0 | 0 | 0 | 0 | 0 | 0 |
| ENSDARG000 | 25 | 35051656 | 35052155 | + | 500 | hist2h3c | 6 | 39 | 56 | 21 | 20 | 0 |
| ENSDARG000 | 25 | 35054246 | 35054619 | + | 374 | NA | 0 | 0 | 0 | 0 | 0 | 0 |
| ENSDARG000 | 25 | 35056619 | 35056932 | + | 314 | NA | 16 | 18 | 10 | 6 | 22 | 0 |
| ENSDARG000 | 25;25 | 25896626:259 | 25896873:259 | -;- | 410 | NA | 0 | 0 | 0 | 0 | 0 | 0 |
| ENSDARG000 | 25;25;25;25;2 | 16172504:161 | 16172668:161 | -;-;-;-;-;- | 915 | NA | 0 | 6 | 2 | 6 | 2 | 6 |
| ENSDARG000 | 25;25 | 3677650:368 | 3677706:368 | +;+ | 3813 | prnprs3 | 547 | 617 | 200 | 136 | 173 | 348 |
| ENSDARG000 | 25;25 | 3704007:370 | 3704420:370 | +;+ | 616 | NA | 4 | 0 | 0 | 2 | 0 | 0 |
| ENSDARG000 | 25;25;25 | 5012791:501 | 5012937:501 | +;+;+ | 359 | NA | 0 | 1 | 0 | 1 | 0 | 0 |
| ENSDARG000 | 25;25;25;25;2 | 22274642:222 | 22275031:222 | +;+;+;+;+;+;+; | 2072 | nr2e3 | 0 | 2 | 0 | 12 | 0 | 0 |
| ENSDARG000 | 25 | 35058088 | 35058462 | + | 375 | zgc:112234 | 18 | 29 | 55 | 44 | 23 | 14 |
| ENSDARG000 | 25;25 | 25968971:259 | 25969254:259 | +;+ | 813 | NA | 0 | 1 | 0 | 0 | 0 | 0 |
| ENSDARG000 | 25;25;25;25 | 16186344:161 | 16186530:161 | +;+;+;+ | 1033 | NA | 0 | 0 | 0 | 0 | 0 | 0 |
| ENSDARG000 | 25;25 | 7882847:788 | 7884283:788 | -;- | 2058 | NA | 4 | 6 | 65 | 18 | 106 | 30 |
| ENSDARG000 | 25 | 35061044 | 35061658 | + | 615 | NA | 0 | 9 | 2 | 0 | 0 | 0 |
| ENSDARG000 | 25;25;25;25;2 | 1631828:163 | 1635916:163 | -;-;-;-;-;-;-;-; | 9135 | MON2 | 1760 | 1426 | 1852 | 1225 | 1910 | 1339 |
| ENSDARG000 | 25;25;25;25;2 | 28813157:288 | 28813584:288 | +;+;+;+;+;+;+; | 1461 | NA | 0 | 0 | 4 | 2 | 0 | 3 |
| ENSDARG000 | 25 | 35062353 | 35063658 | + | 1306 | zgc:113983 | 8 | 17 | 45 | 0 | 24 | 0 |
| ENSDARG000 | 25 | 35064642 | 35065028 | + | 387 | NA | 2 | 0 | 2 | 0 | 0 | 0 |
| ENSDARG000 | 25;25;25;25;2 | 22281441:222 | 22281771:222 | +;+;+;+;+;+;+ | 2347 | stoml1 | 198 | 209 | 256 | 132 | 253 | 296 |
| ENSDARG000 | 25;25;25;25;2 | 13406069:134 | 13406969:134 | +;+;+;+;+;+ | 2888 | znrf1 | 21 | 17 | 12 | 10 | 35 | 38 |
| ENSDARG000 | 25;25;25;25;2 | 13399967:134 | 13400371:134 | -;-;-;-;-;-;- | 1269 | gins3 | 42 | 152 | 156 | 69 | 98 | 105 |
| ENSDARG000 | 25 | 35065802 | 35066113 | + | 312 | zgc:165555 | 8 | 10 | 18 | 18 | 4 | 10 |
| ENSDARG000 | 25 | 35067318 | 35067692 | + | 375 | NA | 6 | 10 | 12 | 2 | 4 | 2 |
| ENSDARG000 | 25;25;25;25;2 | 7888019:788 | 7893377:789 | -;-;-;-;-;-;-;-; | 9320 | ambra1b | 763 | 410 | 692 | 420 | 936 | 617 |
| ENSDARG000 | 25;25;25;25;2 | 3708087:371 | 3708838:371 | -;-;-;-;-;-;-;-; | 4054 | morc2 | 1099 | 1137 | 1707 | 937 | 1538 | 1269 |
| ENSDARG000 | 25;25;25;25;2 | 25993240:259 | 25993316:259 | -;-;-;-;-;-;-;-; | 2250 | ACSBG1 | 0 | 0 | 0 | 0 | 0 | 0 |
| ENSDARG000 | 25;25 | 1973524:197 | 1973635:197 | -;- | 165 | NA | 0 | 0 | 0 | 0 | 0 | 0 |
| ENSDARG000 | 25;25;25;25;2 | 16189044:161 | 16189129:161 | +;+;+;+;+;+ | 2065 | NA | 0 | 0 | 2 | 0 | 0 | 0 |
| ENSDARG000 | 25;25;25;25;2 | 2002715:200 | 2004475:200 | -;-;-;-;-;-;-;- | 2955 | WNT7B | 0 | 4 | 0 | 0 | 2 | 0 |
| ENSDARG000 | 25;25 | 2012677:201 | 2012838:201 | -;- | 249 | NA | 0 | 0 | 0 | 0 | 0 | 0 |
| ENSDARG000 | 25;25 | 26044619:260 | 26044883:260 | +;+ | 528 | NA | 0 | 0 | 0 | 0 | 0 | 0 |
| ENSDARG000 | 25;25;25;25;2 | 13473938:134 | 13474441:134 | -;-;-;-;-;-;-;-; | 1900 | ldhd | 109 | 110 | 144 | 131 | 223 | 170 |
| ENSDARG000 | 25 | 26065997 | 26066384 | - | 388 | NA | 0 | 0 | 0 | 0 | 0 | 0 |
| ENSDARG000 | 25;25;25 | 26669680:266 | 26669756:266 | -;-;- | 943 | NA | 215 | 121 | 177 | 103 | 198 | 159 |
| ENSDARG000 | 25;25;25;25;2 | 22296666:222 | 22296946:222 | +;+;+;+;+;+;+; | 1794 | CYP11A1 | 0 | 0 | 0 | 0 | 2 | 0 |
| ENSDARG000 | 25;25;25;25;2 | 5015019:501 | 5015241:501 | +;+;+;+;+;+;+; | 2287 | hdac10 | 247 | 295 | 417 | 207 | 308 | 254 |
| ENSDARG000 | 25;25;25;25;2 | 13498188:135 | 13498452:135 | +;+;+;+;+;+ | 2817 | snrkb | 505 | 1599 | 716 | 850 | 2272 | 637 |
| ENSDARG000 | 25;25;25;25 | 19666621:196 | 19666742:196 | +;+;+;+ | 1304 | zgc:171459 | 44 | 54 | 62 | 55 | 70 | 76 |
| ENSDARG000 | 25;25;25;25;2 | 2126816:212 | 2127173:212 | -;-;-;-;-;- | 1102 | NA | 10 | 20 | 72 | 30 | 28 | 24 |
| ENSDARG000 | 25;25;25;25;2 | 17893784:178 | 17894248:178 | -;-;-;-;-;-;-;-; | 2413 | bmal1a | 328 | 464 | 339 | 304 | 328 | 306 |
| ENSDARG000 | 25;25;25;25;2 | 7918695:791 | 7920322:792 | -;-;-;-;-;-;- | 2660 | glcea | 398 | 329 | 352 | 328 | 502 | 317 |
| ENSDARG000 | 25;25;25;25;2 | 32294719:322 | 32296252:322 | -;-;-;-;-;-;-;-; | 3216 | slc27a2b | 50 | 40 | 66 | 26 | 132 | 52 |
| ENSDARG000 | 25;25 | 26727104:267 | 26727776:267 | -;- | 1208 | fbxl22 | 1 | 0 | 0 | 0 | 0 | 0 |
| ENSDARG000 | 25;25;25;25;2 | 19670273:196 | 19670437:196 | +;+;+;+;+;+ | 5057 | zgc:113426 | 88 | 60 | 65 | 108 | 46 | 100 |
| ENSDARG000 | 25;25;25;25;2 | 35068166:350 | 35068444:350 | -;-;-;-;-;-;-;-; | 4956 | NA | 133 | 730 | 891 | 267 | 684 | 357 |
| ENSDARG000 | 25;25;25;25;2 | 7927349:792 | 7927480:792 | -;-;-;-;-;-;-;-; | 2228 | NOX5 | 0 | 0 | 0 | 0 | 0 | 2 |
| ENSDARG000 | 25;25;25;25;2 | 16194450:161 | 16194494:161 | +;+;+;+;+;+;+; | 9748 | mical2b | 410 | 227 | 151 | 224 | 102 | 171 |
| ENSDARG000 | 25;25;25;25;2 | 2198781:222 | 2198952:222 | +;+;+;+;+;+;+; | 1590 | pparab | 81 | 34 | 89 | 38 | 114 | 154 |
| ENSDARG000 | 25 | 36310250 | 36310561 | - | 312 | NA | 0 | 0 | 0 | 0 | 0 | 0 |
| ENSDARG000 | 25 | 36311333 | 36311719 | + | 387 | NA | 0 | 0 | 1 | 0 | 0 | 0 |
| ENSDARG000 | 25 | 36312459 | 36312869 | + | 411 | NA | 0 | 3 | 3 | 0 | 0 | 2 |
| ENSDARG000 | 25 | 36315127 | 36315438 | + | 312 | NA | 0 | 0 | 0 | 0 | 0 | 0 |
| ENSDARG000 | 25;25;25 | 36316280:363 | 36316364:363 | +;+;+ | 747 | NA | 0 | 0 | 4 | 0 | 4 | 4 |
| ENSDARG000 | 25;25;25;25;2 | 3749902:375 | 3750708:375 | -;-;-;-;-;-;- | 1713 | zgc:158398 | 536 | 551 | 949 | 610 | 704 | 658 |
| ENSDARG000 | 25 | 36325793 | 36326104 | + | 312 | NA | 6 | 10 | 4 | 0 | 0 | 0 |
| ENSDARG000 | 25;25;25;25;2 | 13528257:135 | 13528875:135 | -;-;-;-;-;-;-;-; | 3107 | ano10b | 2680 | 1711 | 1137 | 2448 | 1305 | 1814 |
| ENSDARG000 | 25;25;25;25;2 | 10602193:106 | 10603334:106 | -;-;-;-;-;-;-;-; | 5219 | ppp6r3 | 2176 | 1925 | 2183 | 1053 | 1992 | 1442 |
| ENSDARG000 | 25;25;25;25;2 | 19680962:196 | 19681011:196 | +;+;+;+;+;+;+; | 1934 | NA | 0 | 6 | 12 | 0 | 6 | 0 |
| ENSDARG000 | 25;25;25;25;2 | 22319940:223 | 22319988:223 | +;+;+;+;+;+;+; | 2832 | cyp11a1.1 | 0 | 0 | 0 | 0 | 0 | 0 |
| ENSDARG000 | 25;25;25;25;2 | 26738450:267 | 26739316:267 | -;-;-;-;-;-;-;-; | 2825 | usp3 | 180 | 202 | 217 | 128 | 194 | 200 |
| ENSDARG000 | 25;25;25;25;2 | 5035343:503 | 5035646:503 | +;+;+;+;+;+;+; | 1836 | parvb | 56 | 28 | 86 | 62 | 150 | 94 |
| ENSDARG000 | 25;25;25;25;2 | 7954140:795 | 7954506:795 | -;-;-;-;-;-;-;-; | 2253 | hal | 0 | 0 | 14 | 2 | 6 | 0 |
| ENSDARG000 | 25 | 36327034 | 36327904 | + | 871 | zgc:110216 | 29 | 33 | 54 | 14 | 33 | 8 |
| ENSDARG000 | 25;25;25;25;2 | 17920361:179 | 17921191:179 | +;+;+;+;+;+;+; | 2112 | borcs5 | 154 | 101 | 196 | 88 | 132 | 92 |
| ENSDARG000 | 25 | 36328524 | 36329005 | + | 482 | hist1h2a4 | 1 | 0 | 0 | 0 | 0 | 0 |
| ENSDARG000 | 25 | 36329623 | 36330033 | + | 411 | zgc:173552 | 0 | 0 | 0 | 0 | 0 | 0 |
| ENSDARG000 | 25 | 36332782 | 36333093 | + | 312 | NA | 1 | 4 | 12 | 0 | 10 | 8 |
| ENSDARG000 | 25;25;25;25;2 | 32315319:323 | 32316296:323 | -;-;-;-;-;-;-;-; | 5302 | CEP152 | 128 | 282 | 297 | 251 | 164 | 153 |
| ENSDARG000 | 25 | 36333993 | 36334367 | + | 375 | NA | 10 | 2 | 6 | 0 | 0 | 6 |
| ENSDARG000 | 25 | 36336920 | 36337306 | + | 387 | NA | 2 | 3 | 3 | 0 | 2 | 0 |
| ENSDARG000 | 25 | 36338289 | 36338600 | + | 312 | NA | 2 | 6 | 0 | 0 | 0 | 0 |
| ENSDARG000 | 25 | 36339867 | 36340516 | + | 650 | NA | 5 | 0 | 1 | 0 | 2 | 0 |
| ENSDARG000 | 25;25;25;25;2 | 17934426:179 | 17934866:179 | -;-;-;-;-;-;-;-; | 7706 | cep290 | 425 | 415 | 488 | 339 | 456 | 443 |
| ENSDARG000 | 25 | 36341356 | 36341742 | + | 387 | NA | 0 | 0 | 0 | 0 | 0 | 2 |
| ENSDARG000 | 25;25;25;25;2 | 3759553:375 | 3759820:375 | +;+;+;+;+;+;+; | 4245 | thoc5 | 909 | 1059 | 1715 | 1005 | 993 | 964 |
| ENSDARG000 | 25;25;25;25;2 | 19695652:196 | 19695755:196 | +;+;+;+;+;+;+; | 2739 | mxg | 0 | 0 | 4 | 0 | 0 | 0 |
| ENSDARG000 | 25 | 36342875 | 36343285 | + | 411 | zgc:173552 | 0 | 2 | 0 | 0 | 0 | 0 |
| ENSDARG000 | 25;25;25;25;2 | 28823952:288 | 28824014:288 | +;+;+;+;+;+;+; | 1428 | nfybb | 603 | 801 | 481 | 973 | 745 | 1479 |
| ENSDARG000 | 25;25;25;25;2 | 18002368:180 | 18003274:180 | +;+;+;+;+;+;+; | 5589 | TMTC3 | 1749 | 1508 | 1846 | 1203 | 2057 | 1892 |
| ENSDARG000 | 25;25;25;25;2 | 2282265:228 | 2282915:228 | +;+;+;+;+;+ | 2075 | cdkn1bb | 918 | 716 | 1129 | 1530 | 699 | 855 |
| ENSDARG000 | 25;25;25;25;2 | 2263857:226 | 2264390:226 | +;+;+;+;+;+ | 1794 | yars2 | 95 | 94 | 165 | 43 | 124 | 105 |
| ENSDARG000 | 25;25;25;25;2 | 17925424:179 | 17925646:179 | +;+;+;+;+;+;+ | 1596 | C25H12orf29 | 140 | 162 | 247 | 124 | 183 | 132 |
| ENSDARG000 | 25;25;25;25 | 2276598:228 | 2276928:228 | -;-;-;- | 712 | NA | 9 | 0 | 1 | 0 | 8 | 0 |
| ENSDARG000 | 25;25;25;25;2 | 13558992:135 | 13562776:135 | -;-;-;-;-;-;-;-; | 5723 | fa2h | 106 | 50 | 53 | 68 | 98 | 160 |
| ENSDARG000 | 25;25;25;25;2 | 7982979:798 | 7983126:798 | +;+;+;+;+;+;+; | 1422 | ucmab | 0 | 0 | 0 | 0 | 0 | 0 |
| ENSDARG000 | 25;25;25;25;2 | 7987920:799 | 7990611:799 | -;-;-;-;-;-;-;-; | 4401 | camk1db | 12 | 8 | 0 | 8 | 26 | 22 |
| ENSDARG000 | 25 | 36345867 | 36346178 | + | 312 | NA | 0 | 0 | 0 | 0 | 0 | 2 |
| ENSDARG000 | 25 | 36347126 | 36347500 | + | 375 | NA | 6 | 1 | 6 | 0 | 4 | 8 |
| ENSDARG000 | 25 | 36348401 | 36349012 | + | 612 | hist1h2a3 | 0 | 5 | 3 | 0 | 6 | 0 |
| ENSDARG000 | 25;25;25;25;2 | 16312258:163 | 16312524:163 | +;+;+;+;+;+;+; | 2165 | parvaa | 673 | 888 | 697 | 756 | 740 | 691 |
| ENSDARG000 | 25;25 | 7984270:798 | 7984437:798 | +;+ | 459 | NA | 0 | 0 | 0 | 0 | 0 | 0 |
| ENSDARG000 | 25;25;25;25;2 | 13620555:136 | 13620669:136 | +;+;+;+;+ | 1281 | NA | 0 | 0 | 0 | 0 | 4 | 0 |
| ENSDARG000 | 25 | 36349574 | 36349984 | + | 411 | zgc:173552 | 5 | 12 | 7 | 9 | 11 | 10 |
| ENSDARG000 | 25;25;25;25;2 | 26754578:267 | 26754954:267 | -;-;-;-;-;- | 979 | si:dkeyp-73b1 | 0 | 2 | 0 | 0 | 4 | 0 |
| ENSDARG000 | 25 | 36361250 | 36361697 | - | 448 | NA | 0 | 0 | 0 | 6 | 2 | 2 |
| ENSDARG000 | 25;25;25;25;2 | 22335914:223 | 22336060:223 | -;-;-;-;-;-;-;-; | 2852 | NA | 0 | 0 | 0 | 0 | 0 | 0 |
| ENSDARG000 | 25;25;25;25;2 | 19718504:197 | 19718619:197 | +;+;+;+;+;+;+; | 2472 | NA | 6 | 1019 | 6 | 1048 | 1008 | 1756 |
| ENSDARG000 | 25 | 36362647 | 36362958 | - | 312 | NA | 0 | 0 | 0 | 0 | 0 | 0 |
| ENSDARG000 | 25 | 36365567 | 36365977 | - | 411 | zgc:173552 | 0 | 6 | 4 | 12 | 4 | 0 |
| ENSDARG000 | 25 | 36367137 | 36367523 | - | 387 | NA | 0 | 0 | 0 | 0 | 0 | 2 |
| ENSDARG000 | 25 | 36368458 | 36369057 | - | 600 | NA | 4 | 2 | 2 | 0 | 0 | 0 |
| ENSDARG000 | 25 | 36369921 | 36370292 | - | 372 | NA | 0 | 0 | 4 | 0 | 0 | 0 |
| ENSDARG000 | 25;25;25;25;2 | 13639232:136 | 13639760:136 | -;-;-;-;-;-;-;-; | 2547 | NA | 651 | 701 | 1146 | 557 | 975 | 745 |
| ENSDARG000 | 25;25;25;25 | 28862660:288 | 28862714:288 | +;+;+;+ | 5290 | NA | 3008 | 2954 | 2577 | 3033 | 5145 | 5425 |
| ENSDARG000 | 25;25;25;25;2 | 28893615:288 | 28893804:288 | +;+;+;+;+;+;+; | 1997 | amn1 | 63 | 97 | 109 | 120 | 935 | 790 |
| ENSDARG000 | 25;25;25;25;2 | 32366071:323 | 32366820:323 | -;-;-;-;-;-;-;-; | 2009 | cops2 | 1494 | 1743 | 2312 | 1419 | 1712 | 1623 |
| ENSDARG000 | 25;25;25;25;2 | 2546343:254 | 2546374:254 | -;-;-;-;-;-;-;-; | 3453 | NA | 4203 | 3288 | 3981 | 4107 | 4077 | 3761 |
| ENSDARG000 | 25;25;25;25;2 | 16356083:163 | 16356269:163 | +;+;+;+;+;+;+; | 2095 | tead1a | 123 | 86 | 100 | 138 | 117 | 143 |
| ENSDARG000 | 25;25;25;25;2 | 28907798:289 | 28908229:289 | -;-;-;-;-;- | 918 | etfbkmt | 64 | 43 | 97 | 51 | 77 | 117 |
| ENSDARG000 | 25;25;25;25;2 | 36405021:364 | 36405152:364 | +;+;+;+;+;+;+; | 2926 | ANKRD27 | 1520 | 937 | 877 | 832 | 917 | 1406 |
| ENSDARG000 | 25;25 | 22489689:224 | 22489728:224 | +;+ | 178 | NA | 0 | 0 | 0 | 0 | 0 | 0 |
| ENSDARG000 | 25;25;25;25;2 | 26781731:267 | 26782186:267 | +;+;+;+;+;+;+; | 2000 | NA | 0 | 0 | 0 | 0 | 0 | 0 |
| ENSDARG000 | 25;25 | 22507496:225 | 22507634:225 | -;- | 179 | NA | 0 | 0 | 0 | 0 | 0 | 0 |
| ENSDARG000 | 25;25;25;25;2 | 19733704:197 | 19733882:197 | +;+;+;+;+;+;+; | 8186 | zgc:101783 | 281 | 272 | 353 | 264 | 264 | 345 |
| ENSDARG000 | 25;25;25;25;2 | 5068442:507 | 5068580:507 | +;+;+;+;+;+;+; | 1492 | parvg | 80 | 44 | 72 | 16 | 140 | 184 |
| ENSDARG000 | 25;25 | 28989285:290 | 28989795:290 | +;+ | 645 | NA | 0 | 0 | 0 | 0 | 0 | 0 |
| ENSDARG000 | 25;25;25;25;2 | 28987957:289 | 28989940:289 | -;-;-;-;-;-;-;-; | 4075 | arid3b | 128 | 114 | 276 | 126 | 241 | 223 |
| ENSDARG000 | 25;25;25;25 | 35097162:350 | 35098847:350 | -;-;-;- | 2025 | zgc:162611 | 90 | 62 | 101 | 137 | 48 | 64 |
| ENSDARG000 | 25;25;25;25;2 | 32390794:323 | 32390963:323 | +;+;+;+;+;+;+; | 1973 | galk2 | 501 | 496 | 410 | 342 | 391 | 512 |
| ENSDARG000 | 25;25;25;25;2 | 3788074:378 | 3788183:378 | +;+;+;+;+;+;+; | 1639 | chid1 | 442 | 406 | 441 | 282 | 395 | 414 |
| ENSDARG000 | 25;25;25;25;2 | 2645488:264 | 2646184:264 | +;+;+;+;+;+;+; | 3084 | ADCK2 | 150 | 136 | 192 | 144 | 168 | 140 |
| ENSDARG000 | 25;25;25;25;2 | 29075145:290 | 29075302:290 | -;-;-;-;-;-;-;- | 690 | cox5aa | 1545 | 2312 | 2551 | 1530 | 1884 | 1647 |
| ENSDARG000 | 25;25 | 35102371:351 | 35102781:351 | -;- | 454 | NA | 2 | 9 | 35 | 4 | 6 | 6 |
| ENSDARG000 | 25;25;25;25;2 | 8031999:803 | 8032728:803 | -;-;-;-;-;-;-;-; | 3661 | dhtkd1 | 1135 | 707 | 1247 | 438 | 1378 | 1298 |
| ENSDARG000 | 25;25 | 35103413:351 | 35104129:351 | -;- | 717 | NA | 17 | 40 | 30 | 12 | 16 | 24 |
| ENSDARG000 | 25;25 | 35105627:351 | 35106751:351 | -;- | 1157 | zgc:194989 | 22 | 6 | 18 | 0 | 0 | 2 |
| ENSDARG000 | 25 | 35107480 | 35107791 | - | 312 | NA | 6 | 0 | 0 | 4 | 2 | 0 |
| ENSDARG000 | 25 | 35109126 | 35109536 | - | 411 | NA | 0 | 4 | 0 | 0 | 1 | 0 |
| ENSDARG000 | 25;25;25;25;2 | 22571475:225 | 22572424:225 | +;+;+;+;+;+;+; | 3848 | stra6 | 0 | 2 | 12 | 0 | 2 | 0 |
| ENSDARG000 | 25 | 35110309 | 35110695 | - | 387 | NA | 6 | 0 | 1 | 0 | 0 | 0 |
| ENSDARG000 | 25 | 35112092 | 35112466 | - | 375 | NA | 0 | 1 | 4 | 0 | 0 | 0 |
| ENSDARG000 | 25;25;25;25;2 | 26798673:268 | 26798942:268 | +;+;+;+;+;+;+; | 2072 | CA12 | 0 | 4 | 4 | 0 | 2 | 0 |
| ENSDARG000 | 25 | 35113580 | 35113891 | - | 312 | NA | 0 | 3 | 14 | 2 | 0 | 0 |
| ENSDARG000 | 25;25;25;25;2 | 32404938:324 | 32405605:324 | -;-;-;-;-;-;-;-; | 4080 | ATP8B4 | 16 | 20 | 101 | 171 | 26 | 13 |
| ENSDARG000 | 25;25;25;25;2 | 29099477:291 | 29104479:291 | -;-;-;-;-;-;-;-; | 7158 | PARP6 | 805 | 564 | 622 | 411 | 811 | 1051 |
| ENSDARG000 | 25;25;25;25;2 | 5098062:511 | 5098328:511 | -;-;-;-;-;- | 1884 | SHISAL1 | 0 | 0 | 0 | 0 | 0 | 0 |
| ENSDARG000 | 25 | 16516154 | 16516562 | + | 409 | NA | 0 | 0 | 0 | 0 | 0 | 0 |
| ENSDARG000 | 25;25;25;25;2 | 16467860:164 | 16468295:164 | -;-;-;-;-;-;-;-; | 2902 | NA | 20 | 82 | 172 | 42 | 64 | 47 |
| ENSDARG000 | 25;25;25;25;2 | 13662606:136 | 13662689:136 | +;+;+;+;+;+;+; | 1144 | ccdc113 | 4 | 9 | 56 | 2 | 8 | 46 |
| ENSDARG000 | 25 | 35120281 | 35120691 | - | 411 | NA | 0 | 0 | 6 | 0 | 2 | 0 |
| ENSDARG000 | 25;25;25;25;2 | 29160102:291 | 29160320:291 | +;+;+;+;+;+;+; | 2970 | pkmb | 0 | 0 | 2 | 4 | 8 | 2 |
| ENSDARG000 | 25 | 35121332 | 35121946 | - | 615 | NA | 0 | 4 | 2 | 0 | 0 | 0 |
| ENSDARG000 | 25 | 35124202 | 35124576 | - | 375 | NA | 1 | 6 | 2 | 0 | 0 | 0 |
| ENSDARG000 | 25 | 35125713 | 35126024 | - | 312 | NA | 0 | 0 | 0 | 0 | 0 | 0 |
| ENSDARG000 | 25;25;25 | 29167487:291 | 29167869:291 | -;-;- | 498 | NA | 34 | 0 | 14 | 10 | 4 | 24 |

| | | | | | | | | | | | | |
|---|---|---|---|---|---|---|---|---|---|---|---|---|
| ENSDARG000 | 25:25:25:25:2 | 18118906:181 | 18119413:181 | -:-:-:-:-:-:-:-: | 1613 | kitlga | 3812 | 3449 | 4801 | 2608 | 3416 | 4717 |
| ENSDARG000 | 25;25;25 | 2672491:2685 | 2672706:2685 | -;-;- | 501 | NA | 0 | 0 | 0 | 0 | 0 | 0 |
| ENSDARG000 | 25:25:25:25:2 | 2668881:2668 | 2668929:2668 | +:+:+:+:+:+:+: | 2545 | bbs4 | 60 | 104 | 56 | 46 | 90 | 93 |
| ENSDARG000 | 25:25:25:25:2 | 26811170:268 | 26812072:268 | +:+:+:+:+:+:+: | 4724 | MAN2C1 | 636 | 366 | 559 | 358 | 583 | 531 |
| ENSDARG000 | 25:25:25:25:2 | 5935938:5940 | 5940835:5941 | -;-;-;-;-;-;- | 6570 | NUAK1 | 488 | 343 | 592 | 310 | 635 | 529 |
| ENSDARG000 | 25 | 35127231 | 35127617 | - | 387 | NA | 0 | 4 | 0 | 14 | 0 | 2 |
| ENSDARG000 | 25 | 8061531 | 8061979 | + | 449 | NA | 0 | 0 | 0 | 0 | 0 | 0 |
| ENSDARG000 | 25;25 | 13671997:136 | 13674710:136 | -;- | 3233 | ZNF319 | 483 | 307 | 689 | 249 | 601 | 624 |
| ENSDARG000 | 25:25:25:25:2 | 5972690:5974 | 5972870:5974 | +:+:+:+:+:+:+: | 1826 | DEPDC7 | 53 | 29 | 9 | 105 | 18 | 19 |
| ENSDARG000 | 25:25:25:25:2 | 18159795:181 | 18159907:181 | -:-:-:-:-:-:-:-: | 2001 | NA | 566 | 257 | 775 | 353 | 982 | 574 |
| ENSDARG000 | 25:25:25:25:2 | 13677299:136 | 13677556:136 | +;+;+;+;+;+;+ | 1953 | usb1 | 188 | 164 | 257 | 145 | 196 | 245 |
| ENSDARG000 | 25:25:25:25:2 | 5983430:5984 | 5983742:5984 | +;+;+;+;+ | 1068 | ppib | 11404 | 12655 | 17131 | 11041 | 14946 | 10265 |
| ENSDARG000 | 25:25:25:25:2 | 3801227:3801 | 3801851:3801 | -:-:-:-:-:-:-:-: | 2895 | gatd1 | 585 | 357 | 483 | 452 | 444 | 424 |
| ENSDARG000 | 25;25;25;25 | 8063455:8089 | 8064021:8090 | +;+;+;+ | 1797 | kcnc1b | 2 | 2 | 0 | 2 | 5 | 0 |
| ENSDARG000 | 25:25:25:25:2 | 29211792:292 | 29213176:292 | -:-:-:-:-:-:-:-: | 3118 | HCN4 | 34 | 6 | 8 | 44 | 29 | 2 |
| ENSDARG000 | 25;25;25;25 | 16553209:165 | 16553349:165 | -;-;-;- | 569 | NA | 0 | 0 | 0 | 0 | 0 | 0 |
| ENSDARG000 | 25:25:25:25:2 | 32473277:324 | 32473488:324 | +:+:+:+:+:+:+: | 1998 | sqor | 896 | 622 | 1400 | 1944 | 727 | 721 |
| ENSDARG000 | 25:25:25:25:2 | 8086519:8105 | 8087126:8105 | -:-:-:-:-:-:-:-: | 2273 | SERGEF | 2 | 43 | 41 | 11 | 34 | 26 |
| ENSDARG000 | 25:25:25:25:2 | 18326298:183 | 18327943:183 | -;-;-;-;- | 2970 | dusp6 | 17001 | 15043 | 21625 | 22839 | 26254 | 29626 |
| ENSDARG000 | 25;25 | 22635166:226 | 22638334:226 | -;- | 3400 | islr2 | 2 | 2 | 0 | 0 | 2 | 0 |
| ENSDARG000 | 25 | 35132090 | 35132464 | + | 375 | NA | 2 | 1 | 6 | 0 | 6 | 0 |
| ENSDARG000 | 25 | 35133404 | 35133790 | + | 387 | NA | 2 | 6 | 0 | 0 | 2 | 0 |
| ENSDARG000 | 25 | 35134393 | 35134803 | + | 411 | NA | 0 | 0 | 4 | 0 | 0 | 0 |
| ENSDARG000 | 25;25 | 29393532:294 | 29394177:294 | +;+ | 1702 | NA | 0 | 0 | 0 | 0 | 0 | 0 |
| ENSDARG000 | 25 | 29393632 | 29395206 | - | 1575 | NA | 0 | 0 | 0 | 0 | 0 | 0 |
| ENSDARG000 | 25 | 29373572 | 29375149 | - | 1578 | NA | 4 | 0 | 0 | 0 | 0 | 0 |
| ENSDARG000 | 25;25 | 29407671:294 | 29409274:294 | -;- | 1960 | ugt5a1 | 0 | 0 | 0 | 0 | 0 | 0 |
| ENSDARG000 | 25;25 | 29401246:294 | 29403259:294 | -;- | 2357 | ugt5a2 | 0 | 0 | 0 | 0 | 0 | 0 |
| ENSDARG000 | 25 | 29381862 | 29383370 | - | 1509 | ugt5a4 | 0 | 0 | 0 | 0 | 0 | 0 |
| ENSDARG000 | 25;25 | 29373563:293 | 29373844:293 | +;+ | 856 | NA | 0 | 0 | 0 | 0 | 0 | 0 |
| ENSDARG000 | 25 | 35135676 | 35136267 | - | 592 | zgc:173585 | 0 | 2 | 4 | 0 | 2 | 0 |
| ENSDARG000 | 25 | 35137080 | 35137391 | - | 312 | NA | 0 | 0 | 0 | 0 | 0 | 0 |
| ENSDARG000 | 25 | 35139110 | 35139520 | - | 411 | NA | 0 | 0 | 0 | 0 | 0 | 0 |
| ENSDARG000 | 25:25:25:25:2 | 19753890:197 | 19757596:197 | +:+:+:+:+:+:+: | 14192 | NA | 108 | 4 | 69 | 30 | 48 | 30 |
| ENSDARG000 | 25 | 35140360 | 35140746 | - | 387 | NA | 0 | 0 | 3 | 0 | 0 | 0 |
| ENSDARG000 | 25:25:25:25:2 | 5992789:5992 | 5993292:5993 | -:-:-:-:-:-:-:-: | 1066 | snx22 | 881 | 579 | 573 | 938 | 415 | 692 |
| ENSDARG000 | 25:25:25:25:2 | 13689060:136 | 13690507:136 | -;-;-;-;-;- | 2345 | pla2g15 | 4496 | 2222 | 2155 | 2861 | 2553 | 4299 |
| ENSDARG000 | 25 | 35141528 | 35142127 | - | 600 | NA | 0 | 0 | 6 | 0 | 0 | 0 |
| ENSDARG000 | 25;25 | 36674715:366 | 36674738:366 | +;+ | 2217 | mafb | 12 | 2 | 4 | 0 | 0 | 0 |
| ENSDARG000 | 25 | 35143049 | 35143360 | - | 312 | NA | 0 | 0 | 0 | 0 | 0 | 0 |
| ENSDARG000 | 25 | 35144234 | 35144620 | - | 387 | NA | 2 | 1 | 0 | 0 | 0 | 2 |
| ENSDARG000 | 25:25:25:25:2 | 18366978:183 | 18367970:183 | -:-:-:-:-:-:-:-: | 2450 | poc1b | 108 | 170 | 131 | 130 | 102 | 76 |
| ENSDARG000 | 25:25:25:25:2 | 26827140:268 | 26827292:268 | -:-:-:-:-:-:-:-: | 1745 | neil1 | 106 | 197 | 275 | 134 | 142 | 190 |
| ENSDARG000 | 25:25:25:25:2 | 6016642:6017 | 6017528:6017 | -:-:-:-:-:-:-:-: | 2918 | snx1a | 11115 | 10158 | 13677 | 9805 | 11108 | 10900 |
| ENSDARG000 | 25 | 35145546 | 35145920 | - | 375 | NA | 0 | 0 | 0 | 4 | 0 | 0 |
| ENSDARG000 | 25:25:25:25:2 | 8139220:8142 | 8139847:8142 | -:-:-:-:-:-:-:-: | 1977 | tph1a | 2 | 0 | 0 | 0 | 4 | 0 |
| ENSDARG000 | 25:25:25:25:2 | 18448186:184 | 18448551:184 | -:-:-:-:-:-:-:-: | 1479 | cpa1 | 0 | 0 | 0 | 0 | 0 | 2 |
| ENSDARG000 | 25 | 35147387 | 35147698 | - | 312 | si:dkev-261m | 0 | 0 | 0 | 0 | 2 | 0 |
| ENSDARG000 | 25 | 35150523 | 35150933 | - | 411 | zgc:113984 | 6 | 20 | 4 | 6 | 10 | 6 |
| ENSDARG000 | 25:25:25:25:2 | 3809733:3810 | 3810333:3810 | -:-:-:-:-:-:-:-: | 1865 | cd151 | 522 | 388 | 364 | 1016 | 444 | 467 |
| ENSDARG000 | 25:25:25:25:2 | 18436301:184 | 18436372:184 | +:+:+:+:+:+:+: | 1229 | cep41 | 26 | 25 | 18 | 22 | 48 | 33 |
| ENSDARG000 | 25:25:25:25:2 | 10762016:107 | 10762220:107 | +:+:+:+:+:+:+: | 1741 | ap3s2 | 682 | 1012 | 782 | 713 | 811 | 1416 |
| ENSDARG000 | 25 | 35151841 | 35152227 | - | 387 | NA | 0 | 4 | 1 | 0 | 0 | 0 |
| ENSDARG000 | 25:25:25:25:2 | 29427852:294 | 29430341:294 | -:-:-:-:-:-:-:-: | 4088 | ube2q2 | 1229 | 1062 | 1415 | 1212 | 1358 | 1308 |
| ENSDARG000 | 25 | 35153283 | 35153985 | - | 703 | zgc:153405 | 0 | 0 | 4 | 0 | 0 | 0 |
| ENSDARG000 | 25;25 | 36734247:367 | 36734440:367 | -;- | 496 | NA | 0 | 0 | 0 | 0 | 0 | 2 |
| ENSDARG000 | 25:25:25:25:2 | 6049614:6049 | 6049759:6049 | +:+:+:+:+:+:+: | 1820 | NA | 768 | 584 | 628 | 731 | 774 | 719 |
| ENSDARG000 | 25:25:25:25:2 | 22643954:226 | 22644053:226 | +:+:+:+:+:+:+: | 2596 | ush1c | 0 | 0 | 22 | 2 | 0 | 10 |
| ENSDARG000 | 25:25:25:25:2 | 36960724:369 | 36960945:369 | -:-:-:-:-:-:-:-: | 1652 | wwox | 318 | 226 | 259 | 161 | 307 | 175 |
| ENSDARG000 | 25:25:25:25:2 | 19870603:198 | 19871124:198 | +:+:+:+:+:+:+: | 7697 | GRAMD4 | 970 | 1383 | 1275 | 1114 | 1279 | 1277 |
| ENSDARG000 | 25:25:25:25:2 | 3844582:3846 | 3845459:3846 | -:-:-:-:-:-:-:-: | 1991 | cracr2b | 0 | 0 | 0 | 2 | 6 | 0 |
| ENSDARG000 | 25:25:25:25:2 | 35155618:351 | 35157253:351 | -:-:-:-:-:-:-:-: | 4233 | NA | 12 | 24 | 64 | 2 | 76 | 4 |
| ENSDARG000 | 25:25:25:25:2 | 8180539:8180 | 8180825:8180 | -:-:-:-:-:-:-:-: | 1782 | saal1 | 200 | 239 | 236 | 118 | 207 | 230 |
| ENSDARG000 | 25;25;25;25 | 3831136:3833 | 3831523:3833 | -;-;-;- | 675 | NA | 0 | 0 | 0 | 0 | 0 | 0 |
| ENSDARG000 | 25:25:25:25:2 | 16554817:165 | 16555198:165 | -:-:-:-:-:-:-:-: | 4122 | NA | 0 | 0 | 0 | 0 | 0 | 0 |
| ENSDARG000 | 25 | 35155676 | 35156872 | + | 1197 | ano9a | 0 | 0 | 0 | 0 | 0 | 0 |
| ENSDARG000 | 25:25:25:25:2 | 26833879:268 | 26834147:268 | -:-:-:-:-:-:-:-: | 1186 | commd4 | 268 | 306 | 324 | 312 | 312 | 291 |
| ENSDARG000 | 25:25:25:25:2 | 2706722:2706 | 2708771:2707 | -:-:-:-:-:-:-:-: | 3429 | adpgk | 478 | 443 | 615 | 335 | 586 | 398 |
| ENSDARG000 | 25:25:25:25:2 | 32525131:325 | 32525331:325 | +:+:+:+:+:+:+: | 5116 | scaper | 589 | 587 | 907 | 493 | 834 | 777 |
| ENSDARG000 | 25:25:25:25:2 | 37073189:370 | 37073937:370 | -;-;-;-;-;-;-;-;- | 1554 | hprt1l | 238 | 249 | 476 | 197 | 173 | 372 |
| ENSDARG000 | 25:25:25:25:2 | 10811551:108 | 10812178:108 | +:+:+:+:+:+:+: | 3269 | anpepb.1 | 0 | 0 | 14 | 24 | 48 | 16 |
| ENSDARG000 | 25:25:25:25:2 | 32496723:324 | 32496901:324 | +:+:+:+:+:+:+: | 2561 | ctdspl2a | 423 | 586 | 670 | 445 | 666 | 615 |
| ENSDARG000 | 25:25:25:25:2 | 6065011:6067 | 6065147:6067 | -:-:-:-:-:-:-:-: | 1763 | cpeb1a | 151 | 373 | 472 | 209 | 388 | 190 |
| ENSDARG000 | 25:25:25:25:2 | 32518578:325 | 32518938:325 | -;-;-;-;- | 1262 | NA | 2 | 0 | 1 | 0 | 6 | 3 |
| ENSDARG000 | 25;25;25;25 | 2727223:2729 | 2727450:2729 | +;+;+;+ | 556 | NA | 0 | 0 | 0 | 0 | 0 | 0 |
| ENSDARG000 | 25:25:25:25:2 | 18459924:184 | 18460181:184 | -:-:-:-:-:-:-:-: | 1353 | cpa5 | 14 | 12 | 20 | 54 | 218 | 236 |
| ENSDARG000 | 25:25:25:25:2 | 10830269:108 | 10830820:108 | +:+:+:+:+:+:+: | 3730 | ANPEP | 2 | 0 | 18 | 0 | 0 | 0 |
| ENSDARG000 | 25:25:25:25:2 | 13716737:137 | 13717262:137 | -;-;-;-;-;- | 1275 | lcat | 14 | 8 | 6 | 12 | 0 | 26 |
| ENSDARG000 | 25:25:25:25:2 | 10843985:108 | 10844090:108 | +:+:+:+:+:+:+: | 3825 | NA | 0 | 0 | 0 | 0 | 0 | 0 |
| ENSDARG000 | 25;25;25 | 19947096:199 | 19948778:199 | +;+;+ | 2541 | kcna6a | 0 | 0 | 0 | 1 | 0 | 0 |
| ENSDARG000 | 25:25:25:25:2 | 2776511:2871 | 2777372:2871 | +:+:+:+:+:+:+: | 6769 | NEO1 | 508 | 196 | 385 | 260 | 448 | 304 |
| ENSDARG000 | 25:25:25:25:2 | 13731542:137 | 13731605:137 | +:+:+:+:+:+:+: | 3002 | DRC7 | 14 | 7 | 27 | 56 | 39 | 43 |
| ENSDARG000 | 25:25:25:25:2 | 10913725:109 | 10913841:109 | +;+;+;+;+;+;+ | 1932 | mhc1lja | 0 | 0 | 0 | 0 | 0 | 0 |
| ENSDARG000 | 25;25;25;25 | 10923100:109 | 10923212:109 | +;+;+;+ | 931 | NA | 0 | 0 | 0 | 0 | 0 | 0 |
| ENSDARG000 | 25:25:25:25:2 | 10947743:109 | 10947872:109 | +;+;+;+;+;+;+ | 2289 | mhc1lfa | 0 | 0 | 0 | 0 | 0 | 0 |
| ENSDARG000 | 25:25:25:25:2 | 37094476:370 | 37095642:370 | -:-:-:-:-:-:-:-: | 5382 | nfat5a | 26 | 32 | 48 | 48 | 4 | 38 |
| ENSDARG000 | 25;25 | 2776865:2871 | 2777372:2871 | +;+ | 828 | NA | 0 | 0 | 0 | 0 | 0 | 0 |
| ENSDARG000 | 25;25;25 | 19954576:199 | 19955062:199 | +;+;+ | 3380 | KCNA1 | 0 | 0 | 2 | 0 | 0 | 4 |
| ENSDARG000 | 25:25:25:25:2 | 20004835:200 | 20005154:200 | -:-:-:-:-:-:-:-: | 2619 | CNOT4 | 673 | 708 | 726 | 441 | 921 | 670 |
| ENSDARG000 | 25:25:25:25:2 | 20027175:200 | 20027605:200 | -:-:-:-:-:-:-:-: | 2724 | zgc:136858 | 0 | 2 | 4 | 0 | 6 | 0 |
| ENSDARG000 | 25:25:25:25:2 | 10953351:109 | 10955366:109 | +:+:+:+:+:+:+: | 3093 | mhc1lda | 0 | 0 | 0 | 0 | 0 | 0 |
| ENSDARG000 | 25:25:25:25:2 | 37126921:371 | 37127043:371 | +;+;+;+;+;+;+ | 3152 | NA | 0 | 6 | 8 | 0 | 6 | 2 |
| ENSDARG000 | 25:25:25:25:2 | 10972288:109 | 10972419:109 | +:+:+:+:+:+:+: | 1130 | mhc1lla | 0 | 0 | 0 | 0 | 0 | 0 |
| ENSDARG000 | 25:25:25:25:2 | 18475032:184 | 18475196:184 | +:+:+:+:+:+:+: | 2077 | tes | 486 | 574 | 224 | 310 | 330 | 362 |
| ENSDARG000 | 25:25:25:25:2 | 8316953:8317 | 8316984:8317 | +:+:+:+:+:+:+: | 11489 | NA | 0 | 0 | 0 | 0 | 0 | 0 |
| ENSDARG000 | 25:25:25:25:2 | 35189555:351 | 35189797:351 | +:+:+:+:+:+:+: | 2984 | ANO5 | 608 | 539 | 701 | 343 | 793 | 647 |
| ENSDARG000 | 25:25:25:25:2 | 10987817:109 | 10988268:109 | +;+;+;+;+;+ | 2984 | si:dkey-52p2.! | 0 | 0 | 0 | 0 | 0 | 0 |
| ENSDARG000 | 25:25:25:25:2 | 13745551:137 | 13745790:137 | -:-:-:-:-:-:-:-: | 6928 | ckap5 | 2362 | 3216 | 3928 | 1810 | 3509 | 1912 |
| ENSDARG000 | 25:25:25:25:2 | 6122823:6122 | 6123352:6123 | +:+:+:+:+:+:+: | 4834 | rbpms2a | 118 | 18 | 58 | 2 | 28 | 35 |
| ENSDARG000 | 25:25:25:25:2 | 11002640:110 | 11002751:110 | +:+:+:+:+:+:+: | 1657 | mhc1lga | 0 | 0 | 0 | 0 | 0 | 0 |
| ENSDARG000 | 25;25;25 | 18556588:185 | 18556820:185 | +;+;+ | 1705 | cav2 | 1670 | 911 | 743 | 2853 | 257 | 828 |
| ENSDARG000 | 25:25:25:25:2 | 35250976:352 | 35251365:352 | +:+:+:+:+:+:+: | 2608 | slc17a6a | 0 | 0 | 0 | 0 | 0 | 0 |
| ENSDARG000 | 25:25:25:25:2 | 37142582:371 | 37144323:371 | -:-:-:-:-:-:-:-: | 3979 | NA | 618 | 831 | 1222 | 186 | 922 | 511 |
| ENSDARG000 | 25:25:25:25:2 | 3868438:3870 | 3868624:3870 | +;+;+;+;+;+ | 4841 | tmem138 | 158 | 69 | 84 | 67 | 85 | 75 |
| ENSDARG000 | 25;25;25;25 | 3880906:3881 | 3881525:3882 | -;-;-;- | 1554 | NA | 0 | 0 | 0 | 0 | 0 | 0 |
| ENSDARG000 | 25:25:25:25:2 | 22719852:227 | 22720726:227 | +:+:+:+:+:+:+: | 5947 | abcc8 | 2 | 0 | 0 | 8 | 8 | 2 |
| ENSDARG000 | 25:25:25:25:2 | 11008419:110 | 11008537:110 | +;+;+;+;+ | 1078 | mhc1lia | 0 | 2 | 0 | 0 | 4 | 8 |
| ENSDARG000 | 25:25:25:25:2 | 18563476:185 | 18563719:185 | +;+;+;+;+ | 3907 | cav1 | 8939 | 4016 | 7070 | 13863 | 3496 | 6222 |
| ENSDARG000 | 25:25:25:25:2 | 26844028:268 | 26844309:268 | +:+:+:+:+:+:+: | 3232 | sema7a | 16 | 6 | 12 | 14 | 17 | 54 |
| ENSDARG000 | 25;25;25;25 | 3887270:3890 | 3887439:3890 | -;-;-;- | 518 | tmem258 | 342 | 428 | 625 | 339 | 529 | 464 |
| ENSDARG000 | 25:25:25:25:2 | 29472361:294 | 29472505:294 | +:+:+:+:+:+:+: | 1630 | NA | 0 | 0 | 2 | 0 | 4 | 0 |
| ENSDARG000 | 25:25:25:25:2 | 3897907:3901 | 3899267:3901 | -:-:-:-:-:-:-:-: | 5038 | MYRF | 0 | 0 | 0 | 8 | 12 | 0 |
| ENSDARG000 | 25 | 35294670 | 35296165 | - | 1496 | fancf | 12 | 16 | 6 | 8 | 8 | 18 |
| ENSDARG000 | 25:25:25:25:2 | 33033633:330 | 33033644:330 | +:+:+:+:+:+:+: | 6130 | TLN2 | 555 | 368 | 672 | 451 | 588 | 362 |
| ENSDARG000 | 25;25 | 3891909:3916 | 3892110:3916 | +;+ | 500 | NA | 0 | 3 | 0 | 0 | 0 | 0 |
| ENSDARG000 | 25;25 | 11015417:110 | 11015620:110 | -;- | 923 | mespab | 0 | 0 | 0 | 0 | 0 | 0 |
| ENSDARG000 | 25:25:25:25:2 | 18583877:185 | 18584071:185 | +:+:+:+:+:+:+: | 4382 | met | 36 | 26 | 26 | 42 | 7 | 26 |
| ENSDARG000 | 25:25:25:25:2 | 6186823:6186 | 6186970:6186 | +:+:+:+:+:+:+: | 4613 | oaz2a | 796 | 704 | 1364 | 970 | 953 | 1474 |
| ENSDARG000 | 25:25:25:25:2 | 20077225:200 | 20077285:200 | +:+:+:+:+:+:+: | 1164 | NA | 0 | 12 | 2 | 10 | 7 | 0 |
| ENSDARG000 | 25 | 29610121 | 29611476 | - | 1356 | NA | 0 | 0 | 0 | 0 | 0 | 0 |
| ENSDARG000 | 25;25 | 11025903:110 | 11025952:110 | -;- | 860 | mespbb | 0 | 0 | 0 | 0 | 0 | 0 |
| ENSDARG000 | 25:25:25:25:2 | 16582172:165 | 16582409:165 | -:-:-:-:-:-:-:-: | 1397 | cpa4 | 0 | 4 | 0 | 0 | 8 | 0 |
| ENSDARG000 | 25:25:25:25:2 | 35348607:353 | 35349616:353 | -;-;-;-;-;-;-;-;- | 1620 | svip | 240 | 135 | 299 | 210 | 175 | 252 |
| ENSDARG000 | 25:25:25:25:2 | 20089986:200 | 20090101:200 | +:+:+:+:+:+:+: | 4325 | tnni4b.2 | 2 | 0 | 0 | 0 | 0 | 0 |
| ENSDARG000 | 25;25 | 20094340:200 | 20094743:200 | -;- | 470 | NA | 1 | 0 | 0 | 0 | 0 | 0 |
| ENSDARG000 | 25:25:25:25:2 | 29662411:296 | 29662803:296 | +;+;+;+;+;+ | 2348 | pim3 | 300 | 338 | 230 | 138 | 345 | 114 |
| ENSDARG000 | 25:25:25:25:2 | 6211677:6213 | 6211950:6213 | -:-:-:-:-:-:-:-: | 1448 | psma4 | 3041 | 3242 | 3942 | 1854 | 3502 | 2314 |
| ENSDARG000 | 25:25:25:25:2 | 13791627:137 | 13791927:137 | +:+:+:+:+:+:+: | 1665 | cstpp1 | 204 | 229 | 270 | 189 | 283 | 270 |
| ENSDARG000 | 25:25:25:25:2 | 11034750:110 | 11038930:110 | -:-:-:-:-:-:-:-: | 6448 | SV2B | 6 | 0 | 0 | 0 | 0 | 0 |
| ENSDARG000 | 25:25:25:25:2 | 13801716:138 | 13804390:138 | -:-:-:-:-:-:-:-: | 4883 | mapk8ip1a | 680 | 1029 | 601 | 978 | 600 | 1191 |
| ENSDARG000 | 25:25:25:25:2 | 8356707:8357 | 8356932:8357 | +:+:+:+:+:+:+: | 16267 | NA | 6 | 8 | 0 | 44 | 8 | 8 |
| ENSDARG000 | 25:25:25:25:2 | 18711804:187 | 18712373:187 | +:+:+:+:+:+:+: | 2101 | NA | 144 | 137 | 208 | 90 | 107 | 180 |
| ENSDARG000 | 25:25:25:25:2 | 26877705:268 | 26877968:268 | -:-:-:-:-:-:-:-: | 1806 | dldh | 2631 | 2729 | 3167 | 2658 | 2259 | 2565 |
| ENSDARG000 | 25:25:25:25:2 | 18723618:187 | 18723845:187 | -;-;-;-;-;-;-;- | 974 | NA | 9 | 0 | 2 | 0 | 2 | 2 |
| ENSDARG000 | 25:25:25:25:2 | 3015676:3016 | 3016615:3016 | -:-:-:-:-:-:-:-: | 2206 | scamp2 | 2078 | 2171 | 2541 | 2201 | 2756 | 2334 |
| ENSDARG000 | 25 | 22853718 | 22854863 | + | 1146 | kcnj11 | 0 | 6 | 0 | 0 | 0 | 0 |
| ENSDARG000 | 25:25:25:25:2 | 6228496:6230 | 6228740:6230 | -:-:-:-:-:-:-:-: | 3091 | IREB2 | 1136 | 1157 | 1723 | 1003 | 1294 | 820 |
| ENSDARG000 | 25:25:25:25:2 | 18743954:187 | 18744303:187 | -:-:-:-:-:-:-:-: | 1384 | NA | 0 | 0 | 4 | 4 | 6 | 0 |
| ENSDARG000 | 25:25:25:25:2 | 33849647:339 | 33849815:339 | +:+:+:+:+:+:+: | 8351 | roraa | 160 | 205 | 355 | 304 | 338 | 178 |
| ENSDARG000 | 25:25:25:25:2 | 20116407:201 | 20116964:201 | +:+:+:+:+:+:+: | 2654 | bpgm | 166 | 149 | 270 | 230 | 140 | 146 |
| ENSDARG000 | 25:25:25:25:2 | 16590636:165 | 16590863:165 | -:-:-:-:-:-:-:-: | 1341 | cpa2 | 2 | 22 | 0 | 4 | 0 | 0 |
| ENSDARG000 | 25:25:25:25:2 | 37161532:371 | 37162209:371 | -:-:-:-:-:-:-:-: | 4995 | NA | 0 | 4 | 10 | 0 | 6 | 0 |
| ENSDARG000 | 25:25:25:25:2 | 29688671:296 | 29689043:296 | +:+:+:+:+:+:+: | 7467 | brd1b | 474 | 385 | 570 | 327 | 527 | 489 |
| ENSDARG000 | 25:25:25:25:2 | 20140926:201 | 20141027:201 | +:+:+:+:+:+:+: | 2349 | NA | 179 | 85 | 556 | 65 | 403 | 14 |
| ENSDARG000 | 25:25:25:25:2 | 13846883:138 | 13847600:138 | -:-:-:-:-:-:-:-: | 3101 | cry2 | 1180 | 875 | 1356 | 1342 | 1689 | 1169 |
| ENSDARG000 | 25:25:25:25:2 | 35375848:354 | 35376087:354 | +:+:+:+:+:+:+: | 5648 | ANO3 | 18 | 8 | 19 | 10 | 59 | 58 |
| ENSDARG000 | 25:25:25:25:2 | 18877675:188 | 18878582:188 | +:+:+:+:+:+:+: | 4783 | NA | 245 | 292 | 342 | 201 | 297 | 303 |
| ENSDARG000 | 25;25 | 18884678:188 | 18884764:188 | +;+ | 246 | NA | 2 | 1 | 2 | 7 | 8 | 0 |
| ENSDARG000 | 25:25:25:25:2 | 14087045:140 | 14087098:140 | +:+:+:+:+:+:+: | 1467 | acte1 | 12 | 0 | 0 | 0 | 2 | 0 |
| ENSDARG000 | 25:25:25:25:2 | 22863028:228 | 22863805:228 | -;-;-;-;- | 1339 | mob2a | 174 | 507 | 57 | 344 | 160 | 356 |
| ENSDARG000 | 25;25;25;25 | 14092871:140 | 14093178:140 | +;+;+;+ | 888 | guca1g | 0 | 0 | 0 | 0 | 0 | 9 |
| ENSDARG000 | 25:25:25:25:2 | 20215964:202 | 20216021:202 | +:+:+:+:+:+:+: | 968 | tnnt2d | 2 | 0 | 22 | 0 | 4 | 0 |
| ENSDARG000 | 25;25;25 | 14032845:140 | 14036581:140 | -;-;- | 3935 | zgc:101566 | 754 | 560 | 924 | 928 | 743 | 966 |
| ENSDARG000 | 25;25;25;25 | 14017609:140 | 14018077:140 | +;+;+;+ | 2769 | chst1 | 2 | 2 | 14 | 12 | 6 | 14 |
| ENSDARG000 | 25:25:25:25:2 | 22979743:229 | 22981922:229 | -;-;-;-;-;- | 3407 | dusp8a | 163 | 51 | 187 | 285 | 128 | 62 |
| ENSDARG000 | 25:25:25:25:2 | 3035384:3035 | 3035914:3035 | +:+:+:+:+:+:+: | 1855 | mpi | 191 | 138 | 223 | 84 | 216 | 195 |
| ENSDARG000 | 25:25:25:25:2 | 3047129:3053 | 3047568:3053 | -;-;-;-;-;- | 1362 | plex9.2 | 32 | 62 | 80 | 22 | 49 | 22 |
| ENSDARG000 | 25:25:25:25:2 | 26895962:268 | 26896126:268 | +:+:+:+:+:+:+: | 6146 | znf800a | 316 | 338 | 579 | 269 | 511 | 290 |
| ENSDARG000 | 25;25;25;25 | 26895675:268 | 26895791:268 | +;+;+;+ | 434 | NA | 0 | 0 | 0 | 0 | 0 | 0 |
| ENSDARG000 | 25;25;25 | 26895394:268 | 26895791:268 | +;+;+ | 1581 | NA | 0 | 0 | 0 | 2 | 0 | 4 |
| ENSDARG000 | 25:25:25:25:2 | 8407892:8408 | 8407992:8408 | +:+:+:+:+:+:+: | 18751 | NA | 2 | 2 | 8 | 0 | 10 | 0 |
| ENSDARG000 | 25:25:25:25:2 | 29911977:299 | 29914059:299 | -;-;-;-;-;- | 3364 | TAFA5 | 26 | 20 | 64 | 26 | 162 | 64 |

| | | | | | | | | | | | |
|---|---|---|---|---|---|---|---|---|---|---|---|
| ENSDARG000 | 25:25:25:25:2 | 33972019:339 | 33972483:339 | +:+:+:+:+:+:+: | 4463 | ICE2 | 242 | 183 | 329 | 152 | 255 | 193 |
| ENSDARG000 | 25:25:25:25:2 | 37193052:371 | 37193312:371 | -;-;-;-;-;- | 928 | NA | 6 | 11 | 12 | 8 | 9 | 7 |
| ENSDARG000 | 25:25:25:25:2 | 30074947:300 | 30075162:300 | +:+:+:+:+:+:+: | 1404 | NA | 6 | 4 | 0 | 0 | 8 | 14 |
| ENSDARG000 | 25:25:25:25:2 | 37209619:372 | 37209834:372 | +;+;+;+;+;+;+ | 1521 | si:dkev-234i1 | 14 | 2 | 6 | 0 | 2 | 1 |
| ENSDARG000 | 25:25:25:25:2 | 11110380:111 | 11110731:111 | +:+:+:+:+:+:+: | 12789 | NA | 2581 | 1691 | 2653 | 1742 | 3172 | 1831 |
| ENSDARG000 | 25:25:25:25:2 | 20229573:202 | 20229910:202 | -:-:-:-:-:-:-:-: | 2687 | dnm1l | 1302 | 1538 | 1458 | 739 | 1315 | 1058 |
| ENSDARG000 | 25:25:25:25:2 | 16601839:166 | 16602057:166 | +:+:+:+:+:+:+: | 1872 | atp6v1e1a | 1118 | 1246 | 1494 | 1392 | 1353 | 1116 |
| ENSDARG000 | 25:25:25:25:2 | 3058924:3061 | 3059323:3061 | +;+;+;+;+ | 1606 | fth1b | 502 | 438 | 995 | 832 | 740 | 937 |
| ENSDARG000 | 25:25:25:25:2 | 18909607:189 | 18909737:189 | -;-;-;-;-;-;-;- | 1166 | parietopsin | 10 | 0 | 0 | 2 | 0 | 0 |
| ENSDARG000 | 25:25:25:25:2 | 23280220:232 | 23280309:232 | +:+:+:+:+:+:+: | 1964 | NA | 0 | 0 | 8 | 8 | 9 | 14 |
| ENSDARG000 | 25:25:25:25:2 | 18889332:188 | 18889491:188 | +:+:+:+:+:+:+: | 2688 | NA | 204 | 193 | 162 | 137 | 201 | 360 |
| ENSDARG000 | 25:25:25:25:2 | 35474987:354 | 35475363:354 | -:-:-:-:-:-:-:-: | 2363 | slc5a12 | 8 | 11 | 82 | 68 | 185 | 2 |
| ENSDARG000 | 25:25:25:25:2 | 3069671:3070 | 3070589:3070 | -:-:-:-:-:-:-:-: | 2019 | BEST1 | 759 | 386 | 985 | 559 | 982 | 886 |
| ENSDARG000 | 25;25;25 | 20264117:202 | 20264556:202 | -;-;- | 693 | dnajb9a | 71 | 160 | 366 | 247 | 234 | 95 |
| ENSDARG000 | 25;25;25 | 35502552:355 | 35502602:355 | +;+;+ | 858 | FIBIN | 4 | 4 | 4 | 0 | 4 | 0 |
| ENSDARG000 | 25:25:25:25:2 | 16646113:166 | 16646449:166 | +:+:+:+:+:+:+: | 7530 | NA | 190 | 299 | 418 | 300 | 426 | 243 |
| ENSDARG000 | 25:25:25:25:2 | 16615178:166 | 16617687:166 | -:-:-:-:-:-:-:-: | 4403 | NA | 46 | 142 | 130 | 65 | 133 | 96 |
| ENSDARG000 | 25;25;25;25 | 6266009:6267 | 6266187:6268 | +;+;+;+ | 1324 | slc25a44a | 77 | 45 | 118 | 16 | 89 | 95 |
| ENSDARG000 | 25:25:25:25:2 | 26921480:269 | 26921891:269 | +:+:+:+:+:+:+: | 3939 | grm8b | 0 | 2 | 0 | 0 | 14 | 0 |
| ENSDARG000 | 25:25:25:25:2 | 37219187:372 | 37219848:372 | -:-:-:-:-:-:-:-: | 4552 | GLG1 | 8940 | 5919 | 7139 | 6943 | 8247 | 5998 |
| ENSDARG000 | 25:25:25:25:2 | 6280991:6281 | 6281080:6281 | -:-:-:-:-:-:-:-: | 1319 | skic8 | 148 | 229 | 251 | 148 | 178 | 92 |
| ENSDARG000 | 25:25:25:25:2 | 3099073:3104 | 3099288:3105 | +:+:+:+:+:+:+: | 2062 | RAB3IL1 | 3855 | 3601 | 4689 | 3569 | 5555 | 4100 |
| ENSDARG000 | 25:25:25:25:2 | 3994391:3994 | 3994530:3994 | +:+:+:+:+:+:+: | 3940 | EPS8L2 | 2 | 8 | 10 | 48 | 42 | 9 |
| ENSDARG000 | 25:25:25:25:2 | 20272145:202 | 20273384:202 | +:+:+:+:+:+:+: | 2946 | NA | 207 | 154 | 307 | 105 | 237 | 191 |
| ENSDARG000 | 25;25 | 18921825:189 | 18921949:189 | +;+ | 306 | NA | 13 | 19 | 6 | 9 | 4 | 39 |
| ENSDARG000 | 25:25:25:25:2 | 34013093:340 | 34013366:340 | +:+:+:+:+:+:+: | 1723 | anxa2a | 6785 | 18691 | 25405 | 71913 | 13771 | 6425 |
| ENSDARG000 | 25:25:25:25:2 | 23336310:233 | 23336392:233 | +:+:+:+:+:+:+: | 1042 | NA | 1 | 0 | 0 | 0 | 0 | 0 |
| ENSDARG000 | 25:25:25:25:2 | 30131055:301 | 30131365:301 | +:+:+:+:+:+:+: | 2649 | api5 | 1432 | 1702 | 2347 | 1088 | 1691 | 1201 |
| ENSDARG000 | 25:25:25:25:2 | 6306885:6306 | 6307083:6307 | +:+:+:+:+:+:+: | 872 | crabp1a | 4 | 0 | 4 | 0 | 22 | 0 |
| ENSDARG000 | 25:25:25:25:2 | 18928189:189 | 18928705:189 | -:-:-:-:-:-:-:-: | 2584 | NT5DC3 | 75 | 31 | 36 | 26 | 88 | 65 |
| ENSDARG000 | 25:25:25:25:2 | 14165447:141 | 14165631:141 | +:+:+:+:+:+:+: | 5411 | shank2b | 88 | 53 | 194 | 147 | 223 | 153 |
| ENSDARG000 | 25;25;25 | 18951554:189 | 18951879:189 | -;-;- | 591 | uqcc6 | 51 | 56 | 84 | 36 | 106 | 64 |
| ENSDARG000 | 25;25;25;25 | 11309594:113 | 11309738:113 | +;+;+;+ | 534 | NA | 0 | 0 | 0 | 0 | 0 | 0 |
| ENSDARG000 | 25:25:25:25:2 | 23358924:234 | 23359402:234 | +:+:+:+:+:+:+: | 5493 | OSBPL5 | 522 | 385 | 628 | 293 | 513 | 525 |
| ENSDARG000 | 25:25:25:25:2 | 11389220:114 | 11389486:114 | +:+:+:+:+:+:+: | 6721 | NA | 0 | 2 | 0 | 0 | 0 | 0 |
| ENSDARG000 | 25:25:25:25:2 | 3119810:3120 | 3120306:3120 | -:-:-:-:-:-:-:-: | 3261 | NA | 521 | 403 | 948 | 687 | 959 | 779 |
| ENSDARG000 | 25;25 | 3131809:3137 | 3132046:3137 | +;+ | 459 | NA | 0 | 0 | 0 | 0 | 0 | 0 |
| ENSDARG000 | 25;25;25;25 | 35533790:355 | 35534346:355 | -;-;-;- | 1370 | NA | 25 | 4 | 30 | 20 | 38 | 12 |
| ENSDARG000 | 25:25:25:25:2 | 18953575:189 | 18953861:189 | +:+:+:+:+:+:+: | 5660 | TDG | 150 | 236 | 176 | 130 | 108 | 161 |
| ENSDARG000 | 25:25:25:25:2 | 27405738:274 | 27405836:274 | +:+:+:+:+:+:+: | 3035 | pot1 | 49 | 135 | 142 | 99 | 104 | 102 |
| ENSDARG000 | 25:25:25:25:2 | 18964782:189 | 18964889:189 | +:+:+:+:+:+:+: | 2898 | tdg.1 | 582 | 549 | 666 | 330 | 832 | 541 |
| ENSDARG000 | 25:25:25:25:2 | 4133488:4133 | 4133777:4133 | -:-:-:-:-:-:-:-: | 1667 | fads2 | 1652 | 2067 | 557 | 550 | 354 | 137 |
| ENSDARG000 | 25:25:25:25:2 | 8477843:8482 | 8478983:8482 | -:-:-:-:-:-:-:-: | 4996 | POLG | 105 | 183 | 193 | 103 | 155 | 113 |
| ENSDARG000 | 25:25:25:25:2 | 6349892:6354 | 6351332:6354 | -:-:-:-:-:-:-:-: | 5230 | sin3aa | 668 | 485 | 904 | 474 | 836 | 636 |
| ENSDARG000 | 25:25:25:25:2 | 34035541:340 | 34035625:340 | +:+:+:+:+:+:+: | 1097 | NA | 38 | 10 | 0 | 0 | 0 | 26 |
| ENSDARG000 | 25:25:25:25:2 | 8447565:8447 | 8447674:8447 | +:+:+:+:+:+:+: | 4488 | fanci | 79 | 291 | 287 | 96 | 217 | 204 |
| ENSDARG000 | 25 | 34154171 | 34156152 | - | 1982 | foxb1a | 0 | 0 | 4 | 0 | 0 | 0 |
| ENSDARG000 | 25:25:25:25:2 | 30171955:301 | 30172002:301 | +:+:+:+:+:+:+: | 2617 | si:ch211-147l | 0 | 0 | 0 | 0 | 2 | 2 |
| ENSDARG000 | 25:25:25:25:2 | 11795683:118 | 11806434:118 | -:-:-:-:-:-:-:-: | 13341 | ntrk3a | 2 | 8 | 0 | 0 | 2 | 4 |
| ENSDARG000 | 25:25:25:25:2 | 14369493:143 | 14370039:143 | -:-:-:-:-:-:-:-: | 1472 | polr2c | 514 | 786 | 924 | 665 | 691 | 696 |
| ENSDARG000 | 25;25;25 | 23524796:235 | 23525133:235 | -;-;- | 1291 | phlda2 | 2033 | 1446 | 2109 | 1461 | 788 | 1336 |
| ENSDARG000 | 25:25:25:25:2 | 14381405:143 | 14381811:143 | -;-;-;-;-;-;-;- | 1387 | coq9 | 496 | 410 | 620 | 220 | 709 | 575 |
| ENSDARG000 | 25:25:25:25:2 | 35565590:355 | 35565603:355 | -:-:-:-:-:-:-:-: | 3272 | CLPX | 354 | 299 | 445 | 197 | 351 | 278 |
| ENSDARG000 | 25:25:25:25:2 | 20357731:203 | 20360116:203 | -:-:-:-:-:-:-:-: | 3946 | kctd15a | 3063 | 1317 | 4179 | 1841 | 5121 | 2482 |
| ENSDARG000 | 25:25:25:25:2 | 3148431:3152 | 3149089:3153 | -:-:-:-:-:-:-:-: | 4389 | hps5 | 2349 | 2192 | 3079 | 2915 | 4089 | 2960 |
| ENSDARG000 | 25;25 | 27493444:274 | 27494818:275 | +;+ | 2581 | GPR37 | 8 | 0 | 4 | 0 | 5 | 0 |
| ENSDARG000 | 25;25 | 35588165:355 | 35588334:355 | +;+ | 517 | NA | 0 | 0 | 0 | 0 | 0 | 0 |
| ENSDARG000 | 25:25:25:25:2 | 16689633:166 | 16689730:166 | +:+:+:+:+:+:+: | 2484 | ada2a | 1143 | 995 | 1170 | 1578 | 1411 | 1302 |
| ENSDARG000 | 25:25:25:25:2 | 35553542:355 | 35553733:355 | +;+;+;+;+ | 2287 | spi1a | 93 | 113 | 152 | 38 | 413 | 239 |
| ENSDARG000 | 25:25:25:25:2 | 27541796:275 | 27544534:275 | -;-;-;-;-;- | 4353 | HYAL4 | 4 | 0 | 0 | 0 | 20 | 20 |
| ENSDARG000 | 25:25:25:25:2 | 27531874:275 | 27536569:275 | -;-;-;-;-;- | 5755 | spam1 | 0 | 0 | 0 | 0 | 0 | 0 |
| ENSDARG000 | 25:25:25:25:2 | 14398650:144 | 14398751:144 | +:+:+:+:+:+:+: | 1289 | ciapin1 | 707 | 839 | 952 | 514 | 1037 | 549 |
| ENSDARG000 | 25:25:25:25:2 | 23553532:235 | 23556554:235 | -:-:-:-:-:-:-:-: | 4221 | nap1l4a | 2028 | 1677 | 2195 | 1290 | 1751 | 1672 |
| ENSDARG000 | 25:25:25:25:2 | 6403836:6404 | 6404736:6404 | -:-:-:-:-:-:-:-: | 3414 | PTPN9 | 385 | 404 | 490 | 281 | 485 | 325 |
| ENSDARG000 | 25:25:25:25:2 | 34246625:342 | 34247400:342 | +:+:+:+:+:+:+: | 2963 | bnip2 | 563 | 621 | 690 | 560 | 938 | 502 |
| ENSDARG000 | 25:25:25:25:2 | 12399046:123 | 12399433:123 | -:-:-:-:-:-:-:-: | 2057 | det1 | 136 | 124 | 184 | 68 | 154 | 116 |
| ENSDARG000 | 25:25:25:25:2 | 4180301:4180 | 4180365:4180 | -:-:-:-:-:-:-:-: | 1137 | NA | 0 | 8 | 0 | 0 | 0 | 0 |
| ENSDARG000 | 25;25 | 12446575:124 | 12446650:124 | +;+ | 134 | NA | 0 | 0 | 0 | 0 | 0 | 0 |
| ENSDARG000 | 25:25:25:25:2 | 16772392:167 | 16772801:167 | -:-:-:-:-:-:-:-: | 1802 | ndufa9a | 1274 | 1424 | 1844 | 882 | 1574 | 1114 |
| ENSDARG000 | 25:25:25:25:2 | 14408936:144 | 14409412:144 | -:-:-:-:-:-:-:-: | 2526 | prmt7 | 526 | 998 | 1288 | 461 | 924 | 534 |
| ENSDARG000 | 25;25;25;25 | 12494079:124 | 12494321:124 | +;+;+;+ | 1816 | map1lc3b | 2765 | 1536 | 1314 | 3397 | 1887 | 2737 |
| ENSDARG000 | 25:25:25:25:2 | 16708584:167 | 16709030:167 | -:-:-:-:-:-:-:-: | 5194 | GALNT8 | 0 | 0 | 0 | 0 | 0 | 0 |
| ENSDARG000 | 25:25:25:25:2 | 16747475:167 | 16747864:167 | -:-:-:-:-:-:-:-: | 1914 | ribc2 | 0 | 4 | 4 | 0 | 0 | 0 |
| ENSDARG000 | 25:25:25:25:2 | 6432578:6435 | 6436261:6436 | -:-:-:-:-:-:-:-: | 5312 | snupn | 184 | 260 | 290 | 176 | 222 | 268 |
| ENSDARG000 | 25:25:25:25:2 | 20567821:205 | 20568364:205 | -:-:-:-:-:-:-:-: | 1500 | adal | 44 | 97 | 66 | 24 | 36 | 80 |
| ENSDARG000 | 25:25:25:25:2 | 37249963:372 | 37250312:372 | -:-:-:-:-:-:-:-: | 2292 | NA | 129 | 191 | 222 | 68 | 187 | 75 |
| ENSDARG000 | 25:25:25:25:2 | 27596076:276 | 27596595:276 | -;-;-;-;-;-;-;- | 2092 | hyal6 | 0 | 0 | 0 | 4 | 0 | 0 |
| ENSDARG000 | 25:25:25:25:2 | 16706101:167 | 16706509:167 | -:-:-:-:-:-:-:-: | 2171 | GALNT8 | 4 | 0 | 0 | 0 | 2 | 1 |
| ENSDARG000 | 25;25;25 | 20577890:205 | 20578810:205 | -;-;- | 1215 | cplx3a | 0 | 0 | 0 | 0 | 0 | 4 |
| ENSDARG000 | 25:25:25:25:2 | 27629006:276 | 27629988:276 | -:-:-:-:-:-:-:-: | 4504 | NA | 796 | 651 | 645 | 500 | 657 | 768 |
| ENSDARG000 | 25:25:25:25:2 | 34274691:342 | 34274717:342 | -;-;-;-;-;-;- | 1635 | GCNT3 | 4 | 4 | 0 | 741 | 0 | 2 |
| ENSDARG000 | 25:25:25:25:2 | 18978648:189 | 18979002:189 | -:-:-:-:-:-:-:-: | 4693 | efcab6 | 2 | 6 | 6 | 2 | 14 | 10 |
| ENSDARG000 | 25;25;25 | 18990151:189 | 18990332:189 | -;-;- | 368 | NA | 0 | 0 | 0 | 0 | 0 | 0 |
| ENSDARG000 | 25:25:25:25:2 | 30196039:301 | 30196349:301 | +:+:+:+:+:+:+: | 2682 | hsd17b12a | 95 | 99 | 134 | 84 | 190 | 52 |
| ENSDARG000 | 25:25:25:25:2 | 20601653:206 | 20602853:206 | -:-:-:-:-:-:-:-: | 3125 | csk | 886 | 1133 | 1392 | 1173 | 1435 | 1038 |
| ENSDARG000 | 25:25:25:25:2 | 34407529:344 | 34407824:344 | +:+:+:+:+:+:+: | 3076 | OTUD7A | 4 | 8 | 4 | 0 | 2 | 0 |
| ENSDARG000 | 25:25:25:25:2 | 14425834:144 | 14426380:144 | -:-:-:-:-:-:-:-: | 2817 | exoc3l1 | 5515 | 7112 | 15663 | 10159 | 15451 | 5192 |
| ENSDARG000 | 25:25:25:25:2 | 4231238:4231 | 4231468:4231 | -:-:-:-:-:-:-:-: | 2126 | SYT7 | 30 | 147 | 102 | 121 | 39 | 94 |
| ENSDARG000 | 25;25 | 14431539:144 | 14431880:144 | +;+ | 1306 | NA | 8 | 30 | 71 | 46 | 86 | 41 |
| ENSDARG000 | 25:25:25:25:2 | 20668272:206 | 20672819:206 | -:-:-:-:-:-:-:-: | 6731 | edc3 | 957 | 540 | 1200 | 580 | 993 | 946 |
| ENSDARG000 | 25:25:25:25:2 | 3196855:3198 | 3197606:3198 | -:-:-:-:-:-:-:-: | 2461 | gtf2h1 | 246 | 236 | 296 | 145 | 325 | 232 |
| ENSDARG000 | 25:25:25:25:2 | 12510793:125 | 12510966:125 | -:-:-:-:-:-:-:-: | 1882 | NA | 0 | 0 | 0 | 0 | 0 | 0 |
| ENSDARG000 | 25;25;25;25 | 19041329:190 | 19041417:190 | +;+;+;+ | 1419 | NA | 0 | 0 | 0 | 0 | 0 | 0 |
| ENSDARG000 | 25:25:25:25:2 | 19008497:190 | 19008671:190 | +:+:+:+:+:+:+: | 1833 | samm50 | 635 | 827 | 879 | 342 | 809 | 436 |
| ENSDARG000 | 25:25:25:25:2 | 19023737:190 | 19023829:190 | -:-:-:-:-:-:-:-: | 6151 | NA | 0 | 2 | 4 | 0 | 4 | 0 |
| ENSDARG000 | 25:25:25:25:2 | 14510773:145 | 14512607:145 | -:-:-:-:-:-:-:-: | 8269 | NAV2 | 2148 | 899 | 1007 | 1231 | 1159 | 1374 |
| ENSDARG000 | 25;25;25;25 | 14507567:145 | 14508012:145 | +;+;+;+ | 1401 | dbx1b | 30 | 454 | 2346 | 568 | 4042 | 14 |
| ENSDARG000 | 25:25:25:25:2 | 4460039:4460 | 4460604:4460 | -:-:-:-:-:-:-:-: | 1471 | NA | 83 | 45 | 80 | 33 | 128 | 68 |
| ENSDARG000 | 25:25:25:25:2 | 23591990:235 | 23592155:235 | +:+:+:+:+:+:+: | 5280 | NA | 227 | 131 | 369 | 282 | 398 | 569 |
| ENSDARG000 | 25:25:25:25:2 | 35603107:356 | 35604580:356 | -:-:-:-:-:-:-:-: | 9170 | lrrk2 | 1013 | 655 | 1441 | 889 | 1428 | 1000 |
| ENSDARG000 | 25;25;25;25 | 20694177:206 | 20694517:206 | +;+;+;+ | 1119 | kxd1 | 359 | 262 | 401 | 345 | 247 | 428 |
| ENSDARG000 | 25:25:25:25:2 | 30233638:302 | 30233785:302 | +:+:+:+:+:+:+: | 2937 | alkbh3 | 156 | 102 | 186 | 134 | 171 | 188 |
| ENSDARG000 | 25:25:25:25:2 | 35663933:356 | 35664911:356 | +:+:+:+:+:+:+: | 5918 | slc2a13b | 697 | 493 | 464 | 509 | 556 | 611 |
| ENSDARG000 | 25:25:25:25:2 | 4507288:4507 | 4507540:4507 | -:-:-:-:-:-:-:-: | 3002 | NA | 154 | 158 | 211 | 194 | 242 | 222 |
| ENSDARG000 | 25:25:25:25:2 | 27714932:277 | 27715556:277 | -:-:-:-:-:-:-:-: | 2511 | zgc:153935 | 322 | 294 | 433 | 255 | 438 | 479 |
| ENSDARG000 | 25:25:25:25:2 | 20715950:207 | 20716736:207 | +:+:+:+:+:+:+: | 2243 | ergic2 | 1072 | 795 | 1294 | 745 | 1116 | 988 |
| ENSDARG000 | 25 | 34511205 | 34512477 | + | 1273 | NA | 0 | 6 | 4 | 30 | 17 | 10 |
| ENSDARG000 | 25;25 | 34487700:345 | 34489502:345 | -;- | 2550 | klf13 | 76 | 129 | 80 | 57 | 324 | 30 |
| ENSDARG000 | 25:25:25:25:2 | 37268900:372 | 37269507:372 | +;+;+;+;+ | 1534 | NA | 3 | 1 | 12 | 18 | 26 | 0 |
| ENSDARG000 | 25;25;25 | 37282088:372 | 37282540:372 | -;-;- | 863 | NA | 20 | 20 | 18 | 14 | 63 | 12 |
| ENSDARG000 | 25:25:25:25:2 | 12579655:125 | 12579974:125 | -:-:-:-:-:-:-:-: | 4239 | ZCCHC14 | 185 | 135 | 252 | 209 | 287 | 287 |
| ENSDARG000 | 25:25:25:25:2 | 12640211:126 | 12640592:126 | +;+;+;+;+ | 3839 | jph3b | 18 | 10 | 169 | 180 | 199 | 80 |
| ENSDARG000 | 25;25;25;25 | 37285737:372 | 37285982:372 | +;+;+;+ | 829 | NA | 0 | 0 | 0 | 0 | 0 | 0 |
| ENSDARG000 | 25:25:25:25:2 | 12707546:127 | 12707887:127 | -:-:-:-:-:-:-:-: | 2566 | klhdc4 | 888 | 556 | 1032 | 313 | 834 | 798 |
| ENSDARG000 | 25;25 | 37309093:373 | 37309895:373 | -;- | 1316 | NA | 0 | 0 | 4 | 0 | 0 | 0 |
| ENSDARG000 | 25;25;25 | 37294866:372 | 37295625:372 | -;-;- | 1096 | NA | 0 | 0 | 0 | 0 | 0 | 0 |
| ENSDARG000 | 25;25 | 37290206:372 | 37290326:372 | +;+ | 435 | NA | 0 | 0 | 0 | 0 | 0 | 0 |
| ENSDARG000 | 25:25:25:25:2 | 14694876:146 | 14695146:146 | +:+:+:+:+:+:+: | 2873 | mpped2 | 184 | 198 | 216 | 156 | 166 | 138 |
| ENSDARG000 | 25;25 | 37305086:373 | 37305418:373 | -;- | 565 | NA | 0 | 0 | 0 | 1 | 0 | 0 |
| ENSDARG000 | 25:25:25:25:2 | 37289088:372 | 37289133:372 | +:+:+:+:+:+:+: | 1270 | NA | 0 | 0 | 0 | 0 | 2 | 2 |
| ENSDARG000 | 25;25;25 | 20729668:207 | 20730380:207 | -;-;- | 950 | NA | 110 | 124 | 105 | 144 | 147 | 110 |
| ENSDARG000 | 25;25 | 37297659:372 | 37297926:372 | +;+ | 603 | NA | 0 | 0 | 0 | 0 | 0 | 0 |
| ENSDARG000 | 25;25 | 37299390:372 | 37299527:373 | +;+ | 633 | NA | 0 | 0 | 0 | 0 | 0 | 0 |
| ENSDARG000 | 25:25:25:25:2 | 16783584:167 | 16786567:167 | -:-:-:-:-:-:-:-: | 7285 | DYRK4 | 2 | 12 | 392 | 14 | 104 | 70 |
| ENSDARG000 | 25;25 | 37313713:373 | 37314411:373 | -;- | 893 | NA | 0 | 0 | 0 | 0 | 0 | 0 |
| ENSDARG000 | 25:25:25:25:2 | 8518356:8518 | 8518785:8518 | +:+:+:+:+:+:+: | 2169 | NA | 8 | 4 | 0 | 3 | 21 | 45 |
| ENSDARG000 | 25:25:25:25:2 | 20743118:207 | 20743174:207 | -:-:-:-:-:-:-:-: | 9326 | MAGI2 | 71 | 16 | 125 | 52 | 75 | 78 |
| ENSDARG000 | 25:25:25:25:2 | 34681018:346 | 34681130:346 | -:-:-:-:-:-:-:-: | 3577 | nup160 | 828 | 950 | 1581 | 674 | 1163 | 724 |
| ENSDARG000 | 25:25:25:25:2 | 14870043:148 | 14870125:148 | +:+:+:+:+:+:+: | 1879 | dnajc24 | 40 | 80 | 74 | 24 | 59 | 32 |
| ENSDARG000 | 25 | 30298377 | 30299662 | + | 1286 | C25H11orf96 | 165 | 10 | 42 | 50 | 23 | 38 |
| ENSDARG000 | 25:25:25:25:2 | 16853386:168 | 16853414:168 | +:+:+:+:+:+:+: | 2541 | C25H12orf4 | 232 | 140 | 156 | 92 | 202 | 188 |
| ENSDARG000 | 25:25:25:25:2 | 30316839:303 | 30317525:303 | +:+:+:+:+:+:+: | 5907 | ticrr | 112 | 216 | 332 | 122 | 282 | 164 |
| ENSDARG000 | 25:25:25:25:2 | 27744293:277 | 27744684:277 | +:+:+:+:+:+:+: | 2934 | wasla | 1295 | 1008 | 1747 | 1157 | 1386 | 1150 |
| ENSDARG000 | 25:25:25:25:2 | 12750340:127 | 12750395:127 | -:-:-:-:-:-:-:-: | 1708 | slc7a5 | 2129 | 1793 | 3986 | 1500 | 3119 | 2515 |
| ENSDARG000 | 25;25;25 | 8525440:8546 | 8525858:8547 | -;-;- | 645 | NA | 0 | 0 | 0 | 0 | 0 | 0 |
| ENSDARG000 | 25:25:25:25:2 | 20997671:210 | 21000017:210 | -;-;-;-;-;-;-;- | 3744 | gnaia | 753 | 742 | 836 | 538 | 768 | 755 |
| ENSDARG000 | 25;25 | 27816115:278 | 27816840:278 | -;- | 1058 | lmod2a | 0 | 2 | 0 | 0 | 0 | 0 |
| ENSDARG000 | 25:25:25:25:2 | 12798270:127 | 12798647:127 | -:-:-:-:-:-:-:-: | 2021 | ca5a | 610 | 540 | 480 | 266 | 663 | 503 |
| ENSDARG000 | 25:25:25:25:2 | 35683956:357 | 35684199:357 | +:+:+:+:+:+:+: | 8333 | KIF21A | 28 | 25 | 120 | 40 | 106 | 30 |
| ENSDARG000 | 25;25;25;25 | 19095231:190 | 19095350:190 | +;+;+;+ | 1514 | NA | 453 | 720 | 1040 | 817 | 735 | 448 |
| ENSDARG000 | 25:25:25:25:2 | 8581541:8583 | 8581759:8583 | -:-:-:-:-:-:-:-: | 1692 | rhcgb | 0 | 6 | 2 | 3 | 0 | 0 |
| ENSDARG000 | 25:25:25:25:2 | 23702410:237 | 23702712:237 | +:+:+:+:+:+:+: | 2731 | zgc:123060 | 2 | 0 | 0 | 0 | 0 | 0 |
| ENSDARG000 | 25:25:25:25:2 | 19105804:191 | 19105972:191 | +:+:+:+:+:+:+: | 2224 | rlbp1b | 7 | 8 | 10 | 65 | 2 | 18 |
| ENSDARG000 | 25:25:25:25:2 | 23656879:236 | 23658880:236 | -:-:-:-:-:-:-:-: | 4396 | NA | 6 | 6 | 6 | 0 | 6 | 44 |
| ENSDARG000 | 25:25:25:25:2 | 19095302:190 | 19095350:190 | +:+:+:+:+:+:+: | 794 | NA | 0 | 0 | 0 | 0 | 0 | 0 |
| ENSDARG000 | 25:25:25:25:2 | 16973945:169 | 16974305:169 | -:-:-:-:-:-:-:-: | 1551 | ccnd2a | 2312 | 2175 | 3818 | 1267 | 4967 | 5406 |
| ENSDARG000 | 25:25:25:25:2 | 27822188:278 | 27824591:278 | -:-:-:-:-:-:-:-: | 4906 | ASB15 | 4 | 2 | 3 | 8 | 0 | 0 |
| ENSDARG000 | 25:25:25:25:2 | 30343012:303 | 30343647:303 | -:-:-:-:-:-:-:-: | 1847 | pdia3 | 8955 | 12764 | 17447 | 9324 | 11625 | 8143 |
| ENSDARG000 | 25;25;25 | 16945348:169 | 16946095:169 | +;+;+ | 1611 | fgf6a | 0 | 0 | 0 | 0 | 0 | 0 |
| ENSDARG000 | 25:25:25:25:2 | 16880990:168 | 16881120:168 | +:+:+:+:+:+:+: | 3525 | zgc:77158 | 266 | 158 | 257 | 176 | 235 | 110 |
| ENSDARG000 | 25:25:25:25:2 | 16962972:169 | 16964550:169 | -;-;-;-;-;- | 2119 | tigara | 121 | 70 | 62 | 103 | 130 | 175 |
| ENSDARG000 | 25;25;25;25 | 16938706:169 | 16939070:169 | -;-;-;- | 1472 | NA | 0 | 2 | 0 | 0 | 0 | 0 |
| ENSDARG000 | 25:25:25:25:2 | 21046856:210 | 21047167:210 | +;+;+;+;+ | 6272 | CHRM2 | 4 | 0 | 0 | 0 | 0 | 0 |
| ENSDARG000 | 25:25:25:25:2 | 23777469:237 | 23777593:237 | +;+;+;+;+ | 1119 | mrpl23 | 848 | 801 | 1157 | 1045 | 843 | 922 |
| ENSDARG000 | 25:25:25:25:2 | 15029041:150 | 15030078:150 | -:-:-:-:-:-:-:-: | 3490 | pax6a | 0 | 0 | 4 | 0 | 4 | 0 |
| ENSDARG000 | 25;25;25;25 | 24039364:240 | 24042257:240 | -;-;-;- | 3385 | igf2b | 2422 | 1680 | 7733 | 5888 | 5466 | 2162 |
| ENSDARG000 | 25 | 21064019 | 21066136 | - | 2118 | fbxl14a | 196 | 218 | 252 | 200 | 314 | 184 |
| ENSDARG000 | 25:25:25:25:2 | 35774544:357 | 35774736:357 | +:+:+:+:+:+:+: | 3016 | NA | 0 | 0 | 0 | 0 | 0 | 0 |
| ENSDARG000 | 25:25:25:25:2 | 12811709:128 | 12812016:128 | -:-:-:-:-:-:-:-: | 2487 | uba2 | 644 | 848 | 1267 | 460 | 1074 | 805 |
| ENSDARG000 | 25;25;25 | 12841940:128 | 12842065:128 | +;+;+ | 490 | ccl33.3 | 10 | 17 | 14 | 3 | 24 | 26 |
| ENSDARG000 | 25:25:25:25:2 | 27873671:278 | 27874112:278 | +:+:+:+:+:+:+: | 2627 | IQUB | 0 | 6 | 8 | 0 | 8 | 10 |
| ENSDARG000 | 25:25:25:25:2 | 12853111:128 | 12854442:128 | -:-:-:-:-:-:-:-: | 3064 | septin15 | 762 | 570 | 611 | 400 | 612 | 725 |

| | | | | | | | | | | | |
|---|---|---|---|---|---|---|---|---|---|---|---|
| ENSDARG000 | 25;25 | 37318550:373 | 37318869:373 | -;- | 456 | NA | 0 | 0 | 0 | 0 | 0 | 0 |
| ENSDARG000 | 25:25:25:25:2 | 35889102:358 | 35889371:358 | +:+:+:+:+:+:+: | 3889 | lsm14aa | 823 | 562 | 920 | 396 | 847 | 735 |
| ENSDARG000 | 25:25:25:25:2 | 21075227:210 | 21075753:210 | -:-:-:-:-:-:-:-: | 2067 | PRR5 | 54 | 37 | 129 | 65 | 41 | 53 |
| ENSDARG000 | 25;25;25 | 12849609:128 | 12849759:128 | +;+;+ | 385 | ccl33.2 | 0 | 0 | 0 | 0 | 0 | 0 |
| ENSDARG000 | 25:25:25:25:2 | 24055341:240 | 24055500:240 | -:-:-:-:-:-:-:-: | 1470 | th | 0 | 0 | 0 | 0 | 0 | 0 |
| ENSDARG000 | 25;25;25 | 37320928:373 | 37321339:373 | -;-;- | 757 | NA | 0 | 0 | 0 | 0 | 0 | 0 |
| ENSDARG000 | 25;25;25 | 24051048:240 | 24051201:240 | +;+;+ | 372 | NA | 0 | 0 | 0 | 0 | 0 | 0 |
| ENSDARG000 | 25:25:25:25:2 | 17244532:172 | 17244679:172 | +:+:+:+:+:+:+: | 5063 | kdm7ab | 334 | 313 | 346 | 292 | 338 | 210 |
| ENSDARG000 | 25:25:25:25:2 | 19120529:191 | 19121358:191 | -:-:-:-:-:-:-:-: | 2174 | abhd2b | 0 | 0 | 10 | 8 | 6 | 0 |
| ENSDARG000 | 25:25:25:25:2 | 37325054:373 | 37325896:373 | -;-;-;-;-;- | 1860 | ldlrad3 | 597 | 334 | 510 | 436 | 390 | 533 |
| ENSDARG000 | 25;25 | 37331128:373 | 37331478:373 | +;+ | 3077 | NA | 44 | 32 | 30 | 80 | 65 | 38 |
| ENSDARG000 | 25:25:25:25:2 | 34727928:347 | 34729250:347 | -;-;-;-;-;-;- | 2364 | frs2b | 464 | 408 | 722 | 558 | 455 | 491 |
| ENSDARG000 | 25:25:25:25:2 | 37332636:373 | 37334511:373 | -;-;-;-;-;-;-;- | 3041 | trim44 | 270 | 211 | 221 | 138 | 157 | 383 |
| ENSDARG000 | 25:25:25:25:2 | 21098675:210 | 21099071:210 | +:+:+:+:+:+:+: | 2434 | rad52 | 19 | 66 | 96 | 39 | 42 | 25 |
| ENSDARG000 | 25;25;25 | 19180930:191 | 19181489:191 | -;-;- | 824 | NA | 6 | 2 | 4 | 7 | 9 | 16 |
| ENSDARG000 | 25:25:25:25:2 | 19149241:191 | 19149466:191 | +:+:+:+:+:+:+: | 5482 | mfge8b | 14920 | 3312 | 4255 | 8066 | 11754 | 6132 |
| ENSDARG000 | 25:25:25:25:2 | 30385691:303 | 30385722:303 | -:-:-:-:-:-:-:-: | 3286 | NA | 18 | 18 | 86 | 17 | 86 | 35 |
| ENSDARG000 | 25:25:25:25:2 | 21114671:211 | 21117009:211 | -:-:-:-:-:-:-:-: | 8187 | NA | 1317 | 1189 | 1869 | 989 | 1473 | 1317 |
| ENSDARG000 | 25:25:25:25:2 | 27923846:279 | 27924112:279 | +:+:+:+:+:+:+: | 2571 | slc13a1 | 0 | 0 | 8 | 8 | 40 | 0 |
| ENSDARG000 | 25:25:25:25:2 | 37340722:373 | 37340827:373 | +:+:+:+:+:+:+: | 4857 | PAMR1 | 141 | 50 | 73 | 108 | 47 | 46 |
| ENSDARG000 | 25:25:25:25:2 | 19201231:192 | 19201293:192 | +;+;+;+;+ | 1993 | hapln3 | 617 | 251 | 160 | 574 | 265 | 514 |
| ENSDARG000 | 25:25:25:25:2 | 37366698:373 | 37366955:373 | +:+:+:+:+:+:+: | 4291 | slc1a2b | 16 | 8 | 0 | 0 | 12 | 10 |
| ENSDARG000 | 25:25:25:25:2 | 24121611:241 | 24121721:241 | +:+:+:+:+:+:+: | 7042 | NA | 107 | 44 | 143 | 120 | 114 | 155 |
| ENSDARG000 | 25:25:25:25:2 | 19213872:192 | 19213957:192 | -:-:-:-:-:-:-:-: | 4262 | NA | 0 | 2 | 0 | 0 | 10 | 2 |
| ENSDARG000 | 25:25:25:25:2 | 27937585:279 | 27938191:279 | +:+:+:+:+:+:+: | 5070 | NA | 4 | 0 | 8 | 4 | 44 | 2 |
| ENSDARG000 | 25:25:25:25:2 | 37408792:374 | 37408994:374 | -:-:-:-:-:-:-:-: | 1334 | NA | 19 | 20 | 33 | 14 | 42 | 14 |
| ENSDARG000 | 25:25:25:25:2 | 24181067:241 | 24181767:241 | -:-:-:-:-:-:-:-: | 1991 | uevld | 178 | 180 | 192 | 126 | 206 | 208 |
| ENSDARG000 | 25:25:25:25:2 | 19238175:192 | 19238313:192 | +:+:+:+:+:+:+: | 4770 | NA | 344 | 233 | 322 | 242 | 331 | 266 |
| ENSDARG000 | 25:25:25:25:2 | 30442389:304 | 30442547:304 | +;+;+;+;+;+ | 1164 | pdcd2l | 172 | 189 | 337 | 109 | 242 | 204 |
| ENSDARG000 | 25:25:25:25:2 | 30460378:304 | 30460458:304 | +:+:+:+:+:+:+: | 2420 | banp | 174 | 204 | 222 | 176 | 194 | 114 |
| ENSDARG000 | 25:25:25:25:2 | 34748125:347 | 34748146:347 | +:+:+:+:+:+:+: | 7047 | wwp2 | 6689 | 3708 | 5182 | 3592 | 4626 | 3374 |
| ENSDARG000 | 25:25:25:25:2 | 24214393:242 | 24214596:242 | -;-;-;-;-;- | 1008 | NA | 0 | 0 | 0 | 0 | 0 | 0 |
| ENSDARG000 | 25;25;25;25 | 30915178:309 | 30915877:309 | +;+;+;+ | 13525 | NA | 98 | 98 | 264 | 74 | 402 | 136 |
| ENSDARG000 | 25:25:25:25:2 | 31222069:312 | 31222119:312 | +:+:+:+:+:+:+: | 831 | tnni2a.1 | 0 | 0 | 12 | 6 | 0 | 0 |
| ENSDARG000 | 25:25:25:25:2 | 31238606:312 | 31238701:312 | +;+;+;+;+;+;+ | 3676 | tnni2a.2 | 0 | 0 | 0 | 2 | 0 | 0 |
| ENSDARG000 | 25:25:25:25:2 | 31264155:312 | 31264914:312 | +:+:+:+:+:+:+: | 1676 | tnni2a.3 | 10 | 0 | 42 | 10 | 24 | 16 |
| ENSDARG000 | 25;25;25 | 24231968:242 | 24232082:242 | -;-;- | 493 | NA | 0 | 0 | 0 | 0 | 0 | 0 |
| ENSDARG000 | 25:25:25:25:2 | 24233010:242 | 24237924:242 | -:-:-:-:-:-:-:-: | 7486 | spty2d1 | 788 | 613 | 1078 | 713 | 822 | 774 |
| ENSDARG000 | 25;25;25;25 | 28272908:282 | 28273987:282 | +;+;+;+ | 2069 | fezf1 | 0 | 0 | 0 | 0 | 0 | 0 |
| ENSDARG000 | 25:25:25:25:2 | 37428187:374 | 37428764:374 | -:-:-:-:-:-:-:-: | 2026 | pdhx | 683 | 587 | 859 | 561 | 768 | 457 |
| ENSDARG000 | 25:25:25:25:2 | 21176937:211 | 21177496:211 | +:+:+:+:+:+:+: | 3874 | erc1a | 0 | 1 | 0 | 8 | 6 | 0 |
| ENSDARG000 | 25;25;25 | 21174391:211 | 21174584:211 | -;-;- | 766 | NA | 0 | 0 | 0 | 0 | 0 | 0 |
| ENSDARG000 | 25:25:25:25:2 | 31276842:312 | 31277520:312 | +:+:+:+:+:+:+: | 1433 | tnni2a.4 | 0 | 2 | 0 | 2 | 8 | 0 |
| ENSDARG000 | 25:25:25:25:2 | 28279577:282 | 28279806:282 | +:+:+:+:+:+:+: | 6520 | aass | 888 | 541 | 858 | 514 | 842 | 829 |
| ENSDARG000 | 25;25 | 21277478:212 | 21278511:212 | -;- | 1187 | tmem60 | 116 | 107 | 139 | 95 | 135 | 121 |
| ENSDARG000 | 25;25 | 21267579:212 | 21271282:212 | +;+ | 5280 | NA | 2 | 2 | 4 | 16 | 72 | 32 |
| ENSDARG000 | 25;25 | 21324588:213 | 21324856:213 | +;+ | 3264 | LRRN3 | 6 | 4 | 18 | 2 | 4 | 8 |
| ENSDARG000 | 25:25:25:25:2 | 21296588:212 | 21298548:212 | -:-:-:-:-:-:-:-: | 2619 | immp2l | 82 | 80 | 92 | 102 | 112 | 74 |
| ENSDARG000 | 25;25;25 | 186583:18710 | 186670:18728 | +;+;+ | 306 | NA | 98 | 304 | 331 | 82 | 173 | 173 |
| ENSDARG000 | 25:25:25:25:2 | 17396263:173 | 17397078:173 | -;-;-;-;-;-;-;- | 2146 | cyp2x9 | 0 | 0 | 0 | 0 | 6 | 0 |
| ENSDARG000 | 25;25;25;25 | 37187952:371 | 37188605:371 | -;-;-;- | 1360 | urahb | 0 | 2 | 9 | 0 | 0 | 0 |
| ENSDARG000 | 25:25:25:25:2 | 753364:75554 | 753388:75562 | +:+:+:+:+:+:+: | 1050 | twf1a | 750 | 870 | 843 | 737 | 820 | 723 |
| ENSDARG000 | 25:25:25:25:2 | 710139:71306 | 710163:71328 | -:-:-:-:-:-:-:-: | 1209 | TMEM117 | 4 | 2 | 0 | 1 | 15 | 10 |
| ENSDARG000 | 25:25:25:25:2 | 30380705:303 | 30380914:303 | -;-;-;-;- | 813 | NA | 12 | 6 | 34 | 6 | 53 | 9 |
| ENSDARG000 | 25:25:25:25:2 | 1088693:1088 | 1089066:1089 | -:-:-:-:-:-:-:-: | 7183 | spg11 | 2558 | 1467 | 2183 | 1976 | 2309 | 2160 |
| ENSDARG000 | 25:25:25:25:2 | 32755485:327 | 32755573:327 | +:+:+:+:+:+:+: | 1607 | etfa | 1583 | 1393 | 1730 | 1197 | 1599 | 1444 |
| ENSDARG000 | 25;25;25 | 3525494:3528 | 3525688:3528 | +;+;+ | 654 | RAB19 | 0 | 8 | 2 | 0 | 4 | 0 |
| ENSDARG000 | 25:25:25:25:2 | 265360:26662 | 265625:26675 | -;-;-;-;- | 972 | NA | 134 | 36 | 46 | 30 | 78 | 44 |
| ENSDARG000 | 25 | 3564220 | 3565263 | + | 1044 | zgc:194202 | 0 | 0 | 4 | 0 | 0 | 0 |
| ENSDARG000 | 25:25:25:25:2 | 2333258:2345 | 2333959:2345 | -;-;-;-;-;-;- | 1513 | mrps35 | 448 | 545 | 656 | 602 | 570 | 414 |
| ENSDARG000 | 25:25:25:25:2 | 5702094:5702 | 5702196:5702 | -:-:-:-:-:-:-:-: | 3030 | AP3B2 | 4 | 50 | 32 | 39 | 38 | 20 |
| ENSDARG000 | 25;25;25;25 | 28584162:285 | 28584295:285 | -;-;-;- | 777 | NA | 0 | 0 | 6 | 0 | 0 | 0 |
| ENSDARG000 | 25:25:25:25:2 | 34938317:349 | 34938504:349 | +:+:+:+:+:+:+: | 1508 | vps4a | 484 | 301 | 339 | 191 | 402 | 400 |
| ENSDARG000 | 25:25:25:25:2 | 7461624:7477 | 7462155:7477 | +:+:+:+:+:+:+: | 3461 | syt12 | 28 | 26 | 26 | 20 | 18 | 26 |
| ENSDARG000 | 25:25:25:25:2 | 37489929:374 | 37490125:374 | -;-;-;-;-;-;- | 951 | NA | 0 | 0 | 2 | 0 | 2 | 2 |
| ENSDARG000 | 25 | 15119597 | 15119730 | + | 134 | 5S_rRNA | 0 | 0 | 0 | 0 | 0 | 0 |
| ENSDARG000 | 25;25;25 | 12919595:129 | 12919819:129 | -;-;- | 431 | NA | 0 | 2 | 0 | 2 | 0 | 0 |
| ENSDARG000 | 25;25 | 10909850:109 | 10910418:109 | +;+ | 2545 | NA | 0 | 0 | 0 | 0 | 0 | 0 |
| ENSDARG000 | 25 | 32020881 | 32020995 | - | 115 | 5S_rRNA | 0 | 0 | 0 | 0 | 0 | 0 |
| ENSDARG000 | 25 | 12472467 | 12472545 | + | 79 | dre-mir-7a-1 | 0 | 0 | 0 | 0 | 0 | 0 |
| ENSDARG000 | 25 | 475542 | 475658 | + | 117 | 5S_rRNA | 0 | 0 | 0 | 0 | 0 | 0 |
| ENSDARG000 | 25 | 23489979 | 23495448 | + | 5470 | NA | 0 | 0 | 0 | 0 | 0 | 0 |
| ENSDARG000 | 25 | 26867723 | 26867839 | + | 117 | 5S_rRNA | 0 | 0 | 0 | 0 | 0 | 0 |
| ENSDARG000 | 25 | 23546579 | 23546696 | + | 118 | 5S_rRNA | 0 | 0 | 0 | 0 | 0 | 0 |
| ENSDARG000 | 25 | 8551240 | 8551340 | - | 101 | dre-mir-9-3 | 0 | 0 | 0 | 0 | 0 | 0 |
| ENSDARG000 | 25 | 6073932 | 6074050 | - | 119 | 5S_rRNA | 0 | 0 | 0 | 0 | 0 | 0 |
| ENSDARG000 | 25:25:25:25:2 | 16064495:160 | 16064692:160 | -;-;-;-;- | 936 | NA | 2 | 0 | 1 | 2 | 19 | 10 |
| ENSDARG000 | 25 | 7799946 | 7800062 | + | 117 | 5S_rRNA | 0 | 0 | 0 | 0 | 0 | 0 |
| ENSDARG000 | 25 | 31318012 | 31318128 | - | 117 | 5S_rRNA | 0 | 0 | 0 | 0 | 0 | 0 |
| ENSDARG000 | 25 | 27616938 | 27617054 | + | 117 | 5S_rRNA | 0 | 1 | 0 | 0 | 0 | 0 |
| ENSDARG000 | 25 | 195790 | 195914 | + | 125 | SNORA71 | 0 | 0 | 0 | 0 | 0 | 0 |
| ENSDARG000 | 25 | 36509674 | 36509789 | + | 116 | 5S_rRNA | 0 | 0 | 0 | 0 | 0 | 0 |
| ENSDARG000 | 25 | 29277005 | 29277126 | - | 122 | 5S_rRNA | 0 | 0 | 0 | 0 | 0 | 0 |
| ENSDARG000 | 25 | 23033488 | 23033619 | - | 132 | 5S_rRNA | 0 | 0 | 0 | 0 | 0 | 0 |
| ENSDARG000 | 25 | 35881415 | 35881531 | + | 117 | 5S_rRNA | 0 | 0 | 0 | 0 | 0 | 0 |
| ENSDARG000 | 25;25;25;25 | 371164:37317 | 372355:37327 | -;-;-;- | 1730 | szl | 8 | 2 | 2 | 0 | 0 | 0 |
| ENSDARG000 | 25:25:25:25:2 | 10640623:106 | 10641776:106 | -:-:-:-:-:-:-:-: | 10218 | lrp5 | 832 | 470 | 708 | 546 | 536 | 600 |
| ENSDARG000 | 25:25:25:25:2 | 17423515:174 | 17423849:174 | +:+:+:+:+:+:+: | 2582 | ELMO3 | 234 | 114 | 206 | 72 | 213 | 133 |
| ENSDARG000 | 25:25:25:25:2 | 25653066:256 | 25653087:256 | +:+:+:+:+:+:+: | 198 | NA | 0 | 0 | 0 | 0 | 0 | 0 |
| ENSDARG000 | 25:25:25:25:2 | 10882597:108 | 10884438:108 | -;-;-;-;-;-;- | 3003 | NA | 0 | 0 | 0 | 0 | 0 | 0 |
| ENSDARG000 | 25:25:25:25:2 | 28589669:285 | 28589783:285 | +;+;+;+;+;+ | 576 | tmem17 | 67 | 106 | 102 | 294 | 69 | 93 |
| ENSDARG000 | 25:25:25:25:2 | 35304903:353 | 35305051:353 | +:+:+:+:+:+:+: | 1090 | GAS2 | 99 | 47 | 85 | 48 | 203 | 784 |
| ENSDARG000 | 25:25:25:25:2 | 37446861:374 | 37446921:374 | +:+:+:+:+:+:+: | 3954 | NA | 43 | 208 | 346 | 164 | 247 | 183 |
| ENSDARG000 | 25:25:25:25:2 | 37483308:374 | 37485211:374 | -:-:-:-:-:-:-:-: | 2633 | psma1 | 2409 | 2472 | 2930 | 1622 | 2787 | 1792 |
| ENSDARG000 | 25:25:25:25:2 | 31165878:311 | 31166150:311 | -;-;-;-;-;- | 848 | NA | 204 | 143 | 208 | 190 | 183 | 226 |
| ENSDARG000 | 25;25;25;25 | 207422:20863 | 207619:20873 | -;-;-;- | 483 | NA | 30 | 7 | 6 | 30 | 17 | 13 |
| ENSDARG000 | 25:25:25:25:2 | 21259229:212 | 21259259:212 | +:+:+:+:+:+:+: | 267 | NA | 2 | 2 | 7 | 0 | 7 | 4 |
| ENSDARG000 | 25:25:25:25:2 | 5249513:5250 | 5249703:5250 | +:+:+:+:+:+:+: | 1272 | NA | 16 | 9 | 10 | 4 | 20 | 14 |
| ENSDARG000 | 25;25;25 | 30799236:307 | 30799742:307 | +;+;+ | 642 | NA | 0 | 0 | 0 | 0 | 0 | 0 |
| ENSDARG000 | 25;25;25;25 | 19999623:200 | 20000046:200 | +;+;+;+ | 1165 | tmem178ba | 6 | 86 | 0 | 2 | 89 | 133 |
| ENSDARG000 | 25:25:25:25:2 | 202323:20232 | 202507:20250 | -;-;-;-;-;-;- | 486 | NA | 119 | 23 | 105 | 117 | 83 | 70 |
| ENSDARG000 | 25 | 15217824 | 15218109 | + | 286 | NA | 0 | 0 | 0 | 0 | 0 | 0 |
| ENSDARG000 | 25 | 3209055 | 3209170 | + | 116 | 5S_rRNA | 0 | 0 | 0 | 0 | 0 | 10 |
| ENSDARG000 | 25;25;25;25 | 34641536:346 | 34641541:346 | +;+;+;+ | 788 | rps27l | 958 | 1750 | 2237 | 1839 | 1189 | 1059 |
| ENSDARG000 | 25:25:25:25:2 | 450343:45125 | 450370:45126 | -:-:-:-:-:-:-:-: | 321 | NA | 286 | 291 | 326 | 299 | 281 | 326 |
| ENSDARG000 | 25:25:25:25:2 | 26441054:264 | 26441130:264 | +:+:+:+:+:+:+: | 1107 | DAPK2 | 2 | 0 | 0 | 0 | 5 | 0 |
| ENSDARG000 | 25;25 | 32886143:328 | 32887674:328 | -;- | 1679 | c2cd4a | 10 | 0 | 252 | 82 | 50 | 28 |
| ENSDARG000 | 25 | 11553756 | 11553873 | + | 118 | 5S_rRNA | 0 | 0 | 0 | 0 | 0 | 0 |
| ENSDARG000 | 25 | 34791362 | 34791466 | + | 105 | mir140 | 6 | 0 | 7 | 0 | 2 | 0 |
| ENSDARG000 | 25 | 27961280 | 27961394 | + | 115 | 5S_rRNA | 0 | 0 | 0 | 0 | 0 | 0 |
| ENSDARG000 | 25:25:25:25:2 | 33002963:330 | 33003098:330 | +:+:+:+:+:+:+: | 936 | NA | 43 | 21 | 92 | 34 | 77 | 19 |
| ENSDARG000 | 25 | 33729284 | 33729398 | + | 115 | 5S_rRNA | 0 | 0 | 0 | 0 | 0 | 0 |
| ENSDARG000 | 25;25;25 | 9678421:9707 | 9680660:9707 | -;-;- | 2462 | lrrc4ca | 267 | 292 | 173 | 508 | 205 | 170 |
| ENSDARG000 | 25 | 24684964 | 24685078 | - | 115 | 5S_rRNA | 0 | 0 | 0 | 0 | 0 | 0 |
| ENSDARG000 | 25:25:25:25:2 | 192116:19256 | 192139:19271 | +;+;+;+;+ | 471 | RPS17 | 13592 | 13438 | 19167 | 17004 | 12608 | 15619 |
| ENSDARG000 | 25 | 10203220 | 10203338 | + | 119 | 5S_rRNA | 0 | 0 | 0 | 0 | 0 | 0 |
| ENSDARG000 | 25:25:25:25:2 | 135932:16690 | 136058:16712 | -;-;-;-;- | 981 | NA | 40 | 3 | 4 | 7 | 12 | 6 |
| ENSDARG000 | 25 | 25506483 | 25506555 | - | 73 | dre-mir-210 | 0 | 2 | 1 | 0 | 4 | 0 |
| ENSDARG000 | 25 | 25206625 | 25206753 | + | 129 | 5S_rRNA | 0 | 0 | 0 | 0 | 0 | 0 |
| ENSDARG000 | 25 | 68423 | 68538 | + | 116 | 5S_rRNA | 0 | 0 | 0 | 0 | 0 | 0 |
| ENSDARG000 | 25:25:25:25:2 | 1732838:1732 | 1732943:1732 | +:+:+:+:+:+:+: | 2094 | fbln1 | 623 | 83 | 162 | 146 | 21 | 296 |
| ENSDARG000 | 25 | 14940788 | 14940902 | + | 115 | 5S_rRNA | 0 | 0 | 0 | 0 | 0 | 0 |
| ENSDARG000 | 25 | 14684233 | 14684347 | + | 115 | 5S_rRNA | 0 | 0 | 0 | 0 | 0 | 0 |
| ENSDARG000 | 25 | 24367976 | 24368090 | + | 115 | 5S_rRNA | 0 | 0 | 0 | 0 | 0 | 0 |
| ENSDARG000 | 25 | 30176471 | 30176559 | + | 89 | dre-mir-129-4 | 0 | 0 | 0 | 0 | 0 | 0 |
| ENSDARG000 | 25:25:25:25:2 | 29315924:293 | 29319882:293 | -;-;-;-;-;-;-;- | 5541 | nptna | 22 | 12 | 4 | 0 | 2 | 2 |
| ENSDARG000 | 25 | 5340962 | 5341102 | - | 141 | U4 | 0 | 0 | 0 | 0 | 0 | 0 |
| ENSDARG000 | 25:25:25:25:2 | 1220279:1223 | 1220407:1223 | -:-:-:-:-:-:-:-: | 1482 | CORO2B | 41 | 0 | 0 | 2 | 5 | 4 |
| ENSDARG000 | 25:25:25:25:2 | 5304652:5309 | 5304930:5310 | +:+:+:+:+:+:+: | 2171 | NA | 6 | 8 | 0 | 0 | 4 | 4 |
| ENSDARG000 | 25 | 15628863 | 15628980 | - | 118 | 5S_rRNA | 0 | 0 | 0 | 0 | 0 | 0 |
| ENSDARG000 | 25:25:25:25:2 | 10776498:107 | 10779350:107 | -;-;-;-;- | 3524 | NA | 492 | 609 | 539 | 338 | 447 | 640 |
| ENSDARG000 | 25 | 14254501 | 14254615 | - | 115 | 5S_rRNA | 0 | 0 | 4 | 0 | 0 | 0 |
| ENSDARG000 | 25:25:25:25:2 | 5755314:5755 | 5755501:5755 | +:+:+:+:+:+:+: | 10087 | NA | 809 | 867 | 1042 | 503 | 1011 | 774 |
| ENSDARG000 | 25 | 19774559 | 19774675 | + | 117 | 5S_rRNA | 0 | 0 | 0 | 0 | 0 | 0 |
| ENSDARG000 | 25:25:25:25:2 | 3831090:3834 | 3831101:3834 | +:+:+:+:+:+:+: | 324 | NA | 0 | 0 | 0 | 0 | 0 | 0 |
| ENSDARG000 | 25 | 36456800 | 36456918 | + | 119 | 5S_rRNA | 0 | 0 | 0 | 0 | 6 | 4 |
| ENSDARG000 | 25:25:25:25:2 | 36279157:362 | 36279187:362 | +:+:+:+:+:+:+: | 165 | NA | 0 | 0 | 0 | 0 | 0 | 0 |
| ENSDARG000 | 25 | 25310063 | 25315481 | + | 5419 | NA | 0 | 12 | 2 | 0 | 0 | 0 |
| ENSDARG000 | 25:25:25:25:2 | 33192404:331 | 33193049:331 | +:+:+:+:+:+:+: | 9091 | tpm1 | 2779 | 2020 | 4255 | 4912 | 6647 | 3097 |
| ENSDARG000 | 25:25:25:25:2 | 7435291:7435 | 7435301:7435 | +:+:+:+:+:+:+: | 4624 | prc1a | 148 | 491 | 486 | 150 | 404 | 200 |
| ENSDARG000 | 25:25:25:25:2 | 37480285:374 | 37480393:374 | +;+;+;+;+;+ | 714 | NA | 8 | 13 | 6 | 28 | 0 | 15 |
| ENSDARG000 | 25 | 172331 | 173165 | - | 835 | NA | 8 | 4 | 6 | 0 | 10 | 6 |
| ENSDARG000 | 25:25:25:25:2 | 388258:39263 | 388358:39266 | +:+:+:+:+:+:+: | 9330 | RFX7 | 1315 | 617 | 1121 | 628 | 1469 | 730 |
| ENSDARG000 | 25:25:25:25:2 | 32787363:327 | 32788043:327 | -:-:-:-:-:-:-:-: | 2529 | tmem266 | 27 | 2 | 22 | 26 | 30 | 20 |
| ENSDARG000 | 25 | 22433726 | 22433844 | - | 119 | 5S_rRNA | 0 | 0 | 0 | 0 | 0 | 0 |
| ENSDARG000 | 25:25:25:25:2 | 25367853:253 | 25367870:253 | +:+:+:+:+:+:+: | 171 | NA | 0 | 0 | 0 | 1 | 0 | 3 |
| ENSDARG000 | 25:25:25:25:2 | 18766165:187 | 18766401:187 | -:-:-:-:-:-:-:-: | 2272 | NA | 0 | 1 | 4 | 2 | 0 | 6 |
| ENSDARG000 | 25 | 36058520 | 36058655 | - | 136 | 5S_rRNA | 0 | 0 | 0 | 0 | 0 | 0 |
| ENSDARG000 | 25 | 6276718 | 6276855 | - | 138 | 5S_rRNA | 13 | 13 | 13 | 14 | 16 | 3 |
| ENSDARG000 | 25:25:25:25:2 | 768014:76801 | 768111:76811 | -:-:-:-:-:-:-:-: | 1464 | irak4 | 292 | 175 | 257 | 207 | 198 | 178 |
| ENSDARG000 | 25 | 6842208 | 6842323 | - | 116 | 5S_rRNA | 0 | 0 | 0 | 0 | 0 | 0 |
| ENSDARG000 | 25 | 5928171 | 5928286 | - | 116 | 5S_rRNA | 0 | 0 | 0 | 0 | 0 | 0 |
| ENSDARG000 | 25 | 3316057 | 3316283 | + | 227 | NA | 0 | 4 | 0 | 0 | 0 | 0 |
| ENSDARG000 | 25:25:25:25:2 | 16757355:167 | 16757489:167 | -:-:-:-:-:-:-:-: | 1758 | GALNT8 | 0 | 0 | 0 | 0 | 0 | 0 |
| ENSDARG000 | 25;25;25;25 | 20385583:203 | 20390248:203 | -;-;-;- | 5201 | CHST8 | 2 | 0 | 2 | 0 | 0 | 0 |
| ENSDARG000 | 25 | 33780513 | 33780628 | + | 116 | 5S_rRNA | 0 | 0 | 0 | 0 | 0 | 0 |
| ENSDARG000 | 25:25:25:25:2 | 217686:23023 | 217715:23023 | -;-;-;-;-;-;- | 159 | NA | 31 | 7 | 17 | 19 | 27 | 9 |
| ENSDARG000 | 25:25:25:25:2 | 12012:135291 | 12057:136241 | +:+:+:+:+:+:+: | 3586 | NA | 78 | 86 | 85 | 91 | 170 | 56 |
| ENSDARG000 | 25:25:25:25:2 | 33256012:332 | 33256563:332 | +;+;+;+;+;+ | 2092 | lactb | 408 | 236 | 306 | 344 | 330 | 250 |
| ENSDARG000 | 25:25:25:25:2 | 5790536:5791 | 5790801:5791 | -:-:-:-:-:-:-:-: | 2157 | appl2 | 282 | 249 | 397 | 273 | 369 | 301 |
| ENSDARG000 | 25 | 31678079 | 31678193 | - | 115 | 5S_rRNA | 0 | 0 | 0 | 0 | 0 | 0 |
| ENSDARG000 | 25:25:25:25:2 | 8269947:8269 | 8270197:8270 | +:+:+:+:+:+:+: | 4530 | NA | 2 | 2 | 7 | 0 | 48 | 34 |

| | | | | | | | | | | | | |
|---|---|---|---|---|---|---|---|---|---|---|---|---|
| ENSDARG000 | 25;25 | 33266586:332 | 33271072:332 | -;- | 4889 | NA | 1 | 0 | 3 | 6 | 2 | 0 |
| ENSDARG000 | 25;25;25;25 | 19170500:191 | 19170753:191 | +;+;+;+ | 690 | NA | 0 | 0 | 0 | 0 | 0 | 0 |
| ENSDARG000 | 25:25:25:25:2 | 150570:15434 | 150648:15449 | +:+:+:+:+:+:+: | 1800 | NA | 769 | 355 | 525 | 607 | 671 | 573 |
| ENSDARG000 | 25 | 8324738 | 8324855 | - | 118 | 5S_rRNA | 0 | 0 | 0 | 0 | 0 | 0 |
| ENSDARG000 | 25 | 115173 | 115289 | + | 117 | 5S_rRNA | 0 | 0 | 0 | 0 | 0 | 0 |
| ENSDARG000 | 25:25:25:25:2 | 17405201:174 | 17405666:174 | +:+:+:+:+:+:+: | 3359 | e2f4 | 383 | 654 | 734 | 338 | 661 | 552 |
| ENSDARG000 | 25:25:25:25:2 | 3449959:3451 | 3451933:3451 | -:-:-:-:-:-:-:-: | 3455 | hbp1 | 1177 | 327 | 462 | 1038 | 518 | 973 |
| ENSDARG000 | 25 | 36844127 | 36844249 | + | 123 | 5S_rRNA | 0 | 0 | 0 | 0 | 0 | 0 |
| ENSDARG000 | 25 | 196605 | 196729 | + | 125 | SNORA71 | 0 | 0 | 1 | 3 | 0 | 0 |
| ENSDARG000 | 25 | 28503850 | 28503965 | + | 116 | 5S_rRNA | 0 | 0 | 0 | 0 | 0 | 0 |
| ENSDARG000 | 25 | 34255145 | 34255273 | - | 129 | 5S_rRNA | 0 | 0 | 0 | 0 | 0 | 0 |
| ENSDARG000 | 25 | 20592483 | 20592597 | + | 115 | 5S_rRNA | 0 | 0 | 0 | 0 | 0 | 0 |
| ENSDARG000 | 25 | 2192682 | 2192775 | + | 94 | NA | 0 | 12 | 19 | 13 | 35 | 30 |
| ENSDARG000 | 25 | 36826484 | 36827491 | - | 1008 | NA | 0 | 0 | 0 | 0 | 0 | 0 |
| ENSDARG000 | 25;25 | 29083447:290 | 29084220:290 | -;- | 917 | rpp25a | 0 | 0 | 0 | 0 | 4 | 0 |
| ENSDARG000 | 25:25:25:25:2 | 7571920:7573 | 7572302:7573 | +:+:+:+:+:+:+: | 3684 | NA | 422 | 280 | 371 | 203 | 372 | 261 |
| ENSDARG000 | 25:25:25:25:2 | 7591293:7596 | 7591930:7596 | +;+;+;+;+;+ | 2790 | NA | 0 | 4 | 0 | 4 | 0 | 0 |
| ENSDARG000 | 25:25:25:25:2 | 34623747:346 | 34623851:346 | -;-;-;-;-;- | 519 | NA | 100 | 129 | 176 | 62 | 160 | 49 |
| ENSDARG000 | 25 | 30981333 | 30981447 | + | 115 | 5S_rRNA | 0 | 0 | 0 | 0 | 0 | 0 |
| ENSDARG000 | 25 | 11973945 | 11974059 | + | 115 | 5S_rRNA | 0 | 0 | 0 | 0 | 0 | 0 |
| ENSDARG000 | 25:25:25:25:2 | 1695547:1698 | 1695729:1698 | -:-:-:-:-:-:-:-: | 1848 | SLC6A13 | 36 | 8 | 6 | 12 | 4 | 46 |
| ENSDARG000 | 25:25:25:25:2 | 23513713:235 | 23513734:235 | +:+:+:+:+:+:+: | 186 | NA | 2 | 0 | 1 | 0 | 2 | 0 |
| ENSDARG000 | 25;25;25;25 | 10929790:109 | 10929888:109 | +;+;+;+ | 1891 | NA | 6 | 0 | 0 | 0 | 0 | 0 |
| ENSDARG000 | 25 | 30839147 | 30839263 | - | 117 | 5S_rRNA | 0 | 0 | 0 | 0 | 0 | 0 |
| ENSDARG000 | 25;25;25 | 196313:20558 | 196335:20562 | -;-;- | 90 | NA | 39 | 15 | 30 | 43 | 32 | 25 |
| ENSDARG000 | 25:25:25:25:2 | 34662358:346 | 34662582:346 | -;-;-;-;-;-;-;- | 1134 | NA | 302 | 299 | 372 | 402 | 327 | 171 |
| ENSDARG000 | 25 | 19105149 | 19105267 | - | 119 | 5S_rRNA | 0 | 0 | 0 | 0 | 0 | 0 |
| ENSDARG000 | 25 | 16771791 | 16771908 | + | 118 | 5S_rRNA | 0 | 0 | 0 | 0 | 0 | 0 |
| ENSDARG000 | 25:25:25:25:2 | 34576067:345 | 34576166:345 | +:+:+:+:+:+:+: | 5653 | trpm1b | 13168 | 6353 | 9185 | 9084 | 9916 | 12303 |
| ENSDARG000 | 25 | 13271458 | 13280481 | + | 9024 | NA | 0 | 0 | 0 | 0 | 0 | 0 |
| ENSDARG000 | 25 | 1629655 | 1629749 | - | 95 | dre-let-7i | 0 | 7 | 4 | 2 | 3 | 2 |
| ENSDARG000 | 25:25:25:25:2 | 31122806:311 | 31123140:311 | +;+;+;+;+ | 1771 | RASSF8 | 454 | 399 | 505 | 306 | 635 | 406 |
| ENSDARG000 | 25 | 336503 | 337522 | + | 1020 | NA | 0 | 0 | 0 | 0 | 0 | 0 |
| ENSDARG000 | 25 | 13012917 | 13013052 | - | 136 | 5S_rRNA | 0 | 0 | 0 | 0 | 0 | 0 |
| ENSDARG000 | 25:25:25:25:2 | 3648497:3649 | 3648636:3649 | +:+:+:+:+:+:+: | 2448 | NA | 395 | 1234 | 1208 | 464 | 908 | 464 |
| ENSDARG000 | 25:25:25:25:2 | 3536238:3539 | 3536863:3539 | -:-:-:-:-:-:-:-: | 2537 | hdhd5 | 155 | 84 | 89 | 32 | 80 | 74 |
| ENSDARG000 | 25 | 28888529 | 28888645 | - | 117 | 5S_rRNA | 11 | 3 | 3 | 0 | 29 | 5 |
| ENSDARG000 | 25 | 13630820 | 13630938 | - | 119 | 5S_rRNA | 0 | 0 | 0 | 0 | 0 | 0 |
| ENSDARG000 | 25 | 3319164 | 3319390 | + | 227 | NA | 1 | 0 | 3 | 0 | 0 | 3 |
| ENSDARG000 | 25 | 20603896 | 20604012 | + | 117 | 5S_rRNA | 0 | 0 | 0 | 0 | 0 | 0 |
| ENSDARG000 | 25 | 4325616 | 4325731 | - | 116 | 5S_rRNA | 0 | 0 | 0 | 0 | 0 | 0 |
| ENSDARG000 | 25 | 31044110 | 31044207 | - | 98 | dre-mir-135a | 0 | 0 | 0 | 0 | 0 | 0 |
| ENSDARG000 | 25 | 22251689 | 22251804 | - | 116 | 5S_rRNA | 0 | 0 | 0 | 0 | 0 | 0 |
| ENSDARG000 | 25:25:25:25:2 | 31944165:319 | 31944663:319 | -;-;-;-;- | 2201 | DUOXA2 | 0 | 0 | 0 | 0 | 0 | 0 |
| ENSDARG000 | 25 | 20516089 | 20516212 | - | 124 | 5S_rRNA | 0 | 0 | 0 | 0 | 0 | 0 |
| ENSDARG000 | 25:25:25:25:2 | 1047627:1052 | 1050083:1053 | -:-:-:-:-:-:-:-: | 3540 | NA | 0 | 0 | 2 | 0 | 0 | 0 |
| ENSDARG000 | 25;25 | 18905472:189 | 18909737:189 | -;- | 6229 | NA | 215 | 112 | 154 | 163 | 75 | 249 |
| ENSDARG000 | 25 | 11376854 | 11378623 | - | 1770 | KLHL25 | 54 | 28 | 133 | 137 | 195 | 46 |
| ENSDARG000 | 25:25:25:25:2 | 2361721:2389 | 2362136:2389 | +:+:+:+:+:+:+: | 1623 | NA | 0 | 2 | 0 | 0 | 0 | 12 |
| ENSDARG000 | 25;25 | 16043246:160 | 16043267:160 | +;+ | 813 | sb:cb470 | 1 | 1 | 0 | 1 | 0 | 1 |
| ENSDARG000 | 25:25:25:25:2 | 4541211:4545 | 4541767:4546 | +:+:+:+:+:+:+: | 2935 | PNPLA2 | 119 | 38 | 98 | 38 | 95 | 114 |
| ENSDARG000 | 25:25:25:25:2 | 31958911:319 | 31958983:319 | +:+:+:+:+:+:+: | 4847 | NA | 14 | 0 | 6 | 0 | 6 | 0 |
| ENSDARG000 | 25;25 | 4581214:4596 | 4581734:4596 | +;+ | 656 | NA | 74 | 48 | 70 | 26 | 37 | 21 |
| ENSDARG000 | 25 | 14134590 | 14134704 | + | 115 | 5S_rRNA | 0 | 0 | 0 | 0 | 0 | 0 |
| ENSDARG000 | 25 | 23115845 | 23115960 | - | 116 | 5S_rRNA | 0 | 0 | 0 | 0 | 0 | 0 |
| ENSDARG000 | 25 | 33139075 | 33139200 | + | 126 | dre-mir-190a | 0 | 0 | 0 | 0 | 0 | 0 |
| ENSDARG000 | 25 | 254462 | 255971 | - | 1510 | NA | 2 | 0 | 0 | 0 | 2 | 0 |
| ENSDARG000 | 25 | 30971844 | 30971960 | + | 117 | 5S_rRNA | 0 | 0 | 0 | 0 | 0 | 0 |
| ENSDARG000 | 25 | 7133655 | 7133769 | + | 115 | 5S_rRNA | 0 | 0 | 0 | 0 | 0 | 0 |
| ENSDARG000 | 25:25:25:25:2 | 5604512:5612 | 5605621:5612 | +:+:+:+:+:+:+: | 5541 | plxnb2b | 47 | 63 | 324 | 188 | 375 | 198 |
| ENSDARG000 | 25:25:25:25:2 | 5288665:5292 | 5288796:5292 | +;+;+;+;+;+ | 1074 | nansb | 2 | 5 | 7 | 2 | 19 | 0 |
| ENSDARG000 | 25 | 18097747 | 18097861 | + | 115 | 5S_rRNA | 0 | 0 | 0 | 0 | 0 | 0 |
| ENSDARG000 | 25 | 27152633 | 27152747 | + | 115 | 5S_rRNA | 0 | 0 | 0 | 0 | 0 | 0 |
| ENSDARG000 | 25:25:25:25:2 | 581227:58236 | 581272:58249 | +:+:+:+:+:+:+: | 2442 | NELL2 | 8 | 21 | 70 | 29 | 36 | 35 |
| ENSDARG000 | 25:25:25:25:2 | 24721849:247 | 24723248:247 | -:-:-:-:-:-:-:-: | 8093 | BRSK2 | 410 | 121 | 433 | 162 | 146 | 224 |
| ENSDARG000 | 25:25:25:25:2 | 37460726:374 | 37461492:374 | -:-:-:-:-:-:-:-: | 2199 | znf276 | 616 | 417 | 645 | 395 | 538 | 657 |
| ENSDARG000 | 25;25 | 36488164:364 | 36489050:364 | -;- | 1216 | irx3b | 0 | 0 | 2 | 2 | 2 | 0 |
| ENSDARG000 | 25 | 30046223 | 30047477 | - | 1255 | NA | 0 | 2 | 0 | 0 | 0 | 0 |
| ENSDARG000 | 25:25:25:25:2 | 32746014:327 | 32746674:327 | -;-;-;-;-;- | 1686 | isl2a | 0 | 0 | 2 | 0 | 0 | 0 |
| ENSDARG000 | 25:25:25:25:2 | 31099908:311 | 31100024:311 | -:-:-:-:-:-:-:-: | 1107 | kras | 262 | 266 | 290 | 167 | 363 | 270 |
| ENSDARG000 | 25 | 37144968 | 37145083 | - | 116 | 5S_rRNA | 2 | 2 | 8 | 0 | 4 | 0 |
| ENSDARG000 | 25 | 15134602 | 15134718 | + | 117 | 5S_rRNA | 0 | 0 | 0 | 0 | 0 | 0 |
| ENSDARG000 | 25 | 27083330 | 27083444 | + | 115 | 5S_rRNA | 0 | 0 | 0 | 0 | 0 | 0 |
| ENSDARG000 | 25:25:25:25:2 | 16731802:167 | 16731811:167 | +:+:+:+:+:+:+: | 366 | NA | 0 | 0 | 0 | 0 | 1 | 4 |
| ENSDARG000 | 25 | 28301324 | 28301448 | + | 125 | 5S_rRNA | 0 | 0 | 2 | 0 | 0 | 0 |
| ENSDARG000 | 25 | 13281610 | 13281666 | + | 57 | NA | 0 | 0 | 0 | 0 | 0 | 0 |
| ENSDARG000 | 25 | 6546173 | 6546287 | + | 115 | 5S_rRNA | 0 | 0 | 0 | 0 | 0 | 0 |
| ENSDARG000 | 25;25;25 | 25172983:251 | 25173469:251 | +;+;+ | 1514 | ccl35.2 | 0 | 10 | 6 | 0 | 5 | 18 |
| ENSDARG000 | 25:25:25:25:2 | 222921:22413 | 223118:22424 | -;-;-;-;- | 483 | NA | 31 | 9 | 8 | 22 | 40 | 11 |
| ENSDARG000 | 25:25:25:25:2 | 5821910:5829 | 5822247:5829 | +;+;+;+;+ | 524 | NA | 447 | 438 | 272 | 337 | 413 | 474 |
| ENSDARG000 | 25 | 2189964 | 2190062 | + | 99 | NA | 0 | 0 | 4 | 0 | 0 | 1 |
| ENSDARG000 | 25 | 13196626 | 13196742 | + | 117 | 5S_rRNA | 4 | 0 | 0 | 0 | 0 | 2 |
| ENSDARG000 | 25 | 28429618 | 28429732 | - | 115 | 5S_rRNA | 0 | 0 | 0 | 0 | 0 | 0 |
| ENSDARG000 | 25:25:25:25:2 | 23202011:232 | 23202094:232 | +:+:+:+:+:+:+: | 1582 | NA | 0 | 6 | 0 | 0 | 0 | 0 |
| ENSDARG000 | 25:25:25:25:2 | 288661:28866 | 288815:28881 | -;-;-;-;- | 483 | NA | 0 | 2 | 1 | 0 | 4 | 10 |
| ENSDARG000 | 25;25;25 | 5883742:5902 | 5885274:5902 | -;-;- | 1969 | NA | 0 | 0 | 0 | 0 | 12 | 0 |
| ENSDARG000 | 25:25:25:25:2 | 6734961:6744 | 6735185:6744 | +;+;+;+;+ | 1328 | NA | 0 | 0 | 0 | 0 | 0 | 0 |
| ENSDARG000 | 25:25:25:25:2 | 9517201:9517 | 9517284:9517 | -;-;-;-;- | 374 | NA | 0 | 0 | 0 | 0 | 0 | 0 |
| ENSDARG000 | 25;25;25;25 | 9642095:9642 | 9642778:9642 | -;-;-;- | 1405 | NA | 0 | 0 | 0 | 0 | 0 | 0 |
| ENSDARG000 | 25:25:25:25:2 | 9968518:1001 | 9968674:1001 | +;+;+;+;+ | 2753 | NA | 0 | 0 | 0 | 0 | 0 | 0 |
| ENSDARG000 | 25;25;25;25 | 16174902:161 | 16175336:161 | +;+;+;+ | 3552 | NA | 2 | 0 | 0 | 0 | 0 | 0 |
| ENSDARG000 | 25:25:25:25:2 | 23064627:230 | 23064940:230 | +:+:+:+:+:+:+: | 1238 | NA | 0 | 0 | 0 | 0 | 0 | 2 |
| ENSDARG000 | 25;25;25;25 | 23128698:231 | 23128797:231 | +;+;+;+ | 498 | NA | 0 | 0 | 0 | 0 | 0 | 0 |
| ENSDARG000 | 25:25:25:25:2 | 23140524:231 | 23140792:231 | +;+;+;+;+;+;+ | 1282 | NA | 0 | 0 | 0 | 0 | 0 | 0 |
| ENSDARG000 | 25;25;25;25 | 23147602:231 | 23147783:231 | +;+;+;+ | 440 | NA | 0 | 0 | 0 | 0 | 0 | 0 |
| ENSDARG000 | 25;25;25;25 | 23157622:231 | 23157703:231 | +;+;+;+ | 834 | NA | 0 | 0 | 0 | 0 | 0 | 0 |
| ENSDARG000 | 25:25:25:25:2 | 23160076:231 | 23160167:231 | +;+;+;+;+;+ | 365 | NA | 0 | 0 | 0 | 0 | 0 | 0 |
| ENSDARG000 | 25;25;25;25 | 23163952:231 | 23164061:231 | +;+;+;+ | 431 | NA | 0 | 0 | 0 | 0 | 0 | 0 |
| ENSDARG000 | 25:25:25:25:2 | 23165191:231 | 23165292:231 | +;+;+;+;+;+;+ | 497 | NA | 0 | 0 | 0 | 0 | 0 | 0 |
| ENSDARG000 | 25:25:25:25:2 | 23170301:231 | 23170401:231 | +;+;+;+;+;+ | 449 | NA | 0 | 0 | 0 | 0 | 0 | 0 |
| ENSDARG000 | 25;25;25 | 23187376:231 | 23187796:231 | +;+;+ | 632 | NA | 0 | 0 | 0 | 0 | 0 | 0 |
| ENSDARG000 | 25;25;25 | 23200066:232 | 23200346:232 | +;+;+ | 585 | NA | 0 | 0 | 0 | 0 | 0 | 0 |
| ENSDARG000 | 25:25:25:25:2 | 23230158:232 | 23230265:232 | +:+:+:+:+:+:+: | 810 | NA | 0 | 0 | 0 | 0 | 0 | 0 |
| ENSDARG000 | 25:25:25:25:2 | 23252369:232 | 23253353:232 | +:+:+:+:+:+:+: | 1874 | NA | 0 | 0 | 2 | 0 | 2 | 0 |
| ENSDARG000 | 25;25;25 | 28941582:289 | 28942191:289 | +;+;+ | 865 | NA | 2 | 0 | 0 | 0 | 0 | 0 |
| ENSDARG000 | 25;25;25;25 | 28949389:289 | 28949769:289 | +;+;+;+ | 2220 | NA | 0 | 0 | 0 | 0 | 0 | 0 |
| ENSDARG000 | 25;25;25 | 36096364:360 | 36096495:360 | +;+;+ | 887 | NA | 0 | 0 | 0 | 0 | 0 | 0 |
| ENSDARG000 | 25:25:25:25:2 | 37466657:374 | 37467041:374 | +:+:+:+:+:+:+: | 3411 | NA | 96 | 128 | 223 | 52 | 290 | 197 |
| ENSDARG000 | MT | 951 | 1019 | + | 69 | NA | 0 | 0 | 0 | 0 | 0 | 6 |
| ENSDARG000 | MT | 1020 | 1971 | + | 952 | NA | 444 | 597 | 2229 | 1888 | 447 | 2324 |
| ENSDARG000 | MT | 1972 | 2042 | + | 71 | NA | 0 | 0 | 0 | 0 | 0 | 0 |
| ENSDARG000 | MT | 2043 | 3725 | + | 1683 | NA | 3498 | 2738 | 6725 | 2630 | 3283 | 6386 |
| ENSDARG000 | MT | 3726 | 3802 | + | 77 | NA | 0 | 0 | 0 | 0 | 0 | 0 |
| ENSDARG000 | MT | 3803 | 4777 | + | 975 | mt-nd1 | 20640 | 19285 | 20521 | 14024 | 22802 | 25345 |
| ENSDARG000 | MT | 4781 | 4854 | + | 74 | NA | 0 | 0 | 0 | 0 | 0 | 0 |
| ENSDARG000 | MT | 4852 | 4922 | - | 71 | NA | 0 | 0 | 0 | 0 | 0 | 0 |
| ENSDARG000 | MT | 4924 | 4992 | + | 69 | NA | 0 | 0 | 0 | 0 | 0 | 0 |
| ENSDARG000 | MT | 4993 | 6039 | + | 1047 | mt-nd2 | 18337 | 14450 | 16900 | 12195 | 21042 | 22767 |
| ENSDARG000 | MT | 6038 | 6107 | + | 70 | NA | 0 | 0 | 0 | 0 | 0 | 0 |
| ENSDARG000 | MT | 6109 | 6176 | - | 68 | NA | 0 | 0 | 0 | 0 | 0 | 0 |
| ENSDARG000 | MT | 6178 | 6250 | - | 73 | NA | 0 | 0 | 0 | 0 | 0 | 0 |
| ENSDARG000 | MT | 6283 | 6354 | - | 72 | NA | 0 | 0 | 0 | 0 | 0 | 0 |
| ENSDARG000 | MT | 6353 | 6423 | - | 71 | NA | 0 | 0 | 0 | 0 | 0 | 0 |
| ENSDARG000 | MT | 6425 | 7975 | + | 1551 | mt-co1 | 184334 | 166424 | 182238 | 186315 | 201028 | 207739 |
| ENSDARG000 | MT | 7976 | 8046 | - | 71 | NA | 0 | 0 | 0 | 0 | 0 | 0 |
| ENSDARG000 | MT | 8049 | 8117 | + | 69 | NA | 0 | 0 | 0 | 0 | 0 | 0 |
| ENSDARG000 | MT | 8130 | 8820 | + | 691 | mt-co2 | 46983 | 55277 | 58515 | 56805 | 60088 | 79058 |
| ENSDARG000 | MT | 8821 | 8893 | + | 73 | NA | 0 | 0 | 0 | 0 | 0 | 0 |
| ENSDARG000 | MT | 8894 | 9058 | + | 165 | mt-atp8 | 58 | 73 | 57 | 23 | 56 | 65 |
| ENSDARG000 | MT | 9052 | 9735 | + | 684 | mt-atp6 | 37027 | 44768 | 44797 | 27521 | 49708 | 52422 |
| ENSDARG000 | MT | 9735 | 10520 | + | 786 | mt-co3 | 48465 | 55032 | 53061 | 50389 | 63174 | 77930 |
| ENSDARG000 | MT | 10520 | 10589 | + | 70 | NA | 0 | 0 | 0 | 0 | 0 | 0 |
| ENSDARG000 | MT | 10590 | 10940 | + | 351 | mt-nd3 | 2310 | 2479 | 1684 | 1240 | 1912 | 2929 |
| ENSDARG000 | MT | 10939 | 11008 | + | 70 | NA | 0 | 0 | 0 | 0 | 0 | 0 |
| ENSDARG000 | MT | 11009 | 11305 | + | 297 | mt-nd4l | 2622 | 1593 | 2130 | 1541 | 2191 | 2446 |
| ENSDARG000 | MT | 11299 | 12679 | + | 1381 | mt-nd4 | 25731 | 21605 | 28051 | 24314 | 32139 | 35414 |
| ENSDARG000 | MT | 12681 | 12750 | + | 70 | NA | 0 | 0 | 0 | 0 | 0 | 0 |
| ENSDARG000 | MT | 12750 | 12819 | + | 70 | NA | 0 | 0 | 0 | 0 | 0 | 0 |
| ENSDARG000 | MT | 12823 | 12895 | + | 73 | NA | 0 | 0 | 0 | 0 | 0 | 0 |
| ENSDARG000 | MT | 12897 | 14717 | + | 1821 | mt-nd5 | 18204 | 15272 | 19276 | 12836 | 21942 | 21778 |
| ENSDARG000 | MT | 14714 | 15232 | - | 519 | mt-nd6 | 5528 | 3942 | 5688 | 4316 | 5892 | 5671 |
| ENSDARG000 | MT | 15233 | 15301 | - | 69 | NA | 0 | 0 | 0 | 0 | 0 | 0 |
| ENSDARG000 | MT | 15308 | 16448 | + | 1141 | mt-cyb | 33220 | 39868 | 48381 | 43561 | 43405 | 55988 |
| ENSDARG000 | MT | 16449 | 16520 | + | 72 | NA | 0 | 0 | 0 | 0 | 0 | 0 |
| ENSDARG000 | MT | 16527 | 16596 | - | 70 | NA | 0 | 0 | 0 | 0 | 0 | 0 |
| ENSDARG000 | KN149696.2:I | 82274:82860: | 82400:83383: | -:-:-:-:-:-:-:-: | 2988 | NA | 21 | 12 | 55 | 0 | 10 | 1 |
| ENSDARG000 | KN149696.2:I | 287891:29398 | 290252:29413 | -;- | 2517 | NA | 11 | 1 | 5 | 2 | 5 | 6 |
| ENSDARG000 | KN149696.2:I | 165289:16566 | 165578:16578 | -:-:-:-:-:-:-:-: | 1776 | zgc:112970 | 7 | 6 | 394 | 249 | 90 | 65 |
| ENSDARG000 | KN149696.2:I | 161124:16144 | 161195:16147 | +:+:+:+:+:+:+: | 1515 | zgc:174680 | 0 | 2 | 67 | 111 | 4 | 5 |
| ENSDARG000 | KN149696.2:I | 129959:12995 | 130160:13016 | -;-;-;-;-;-;-;- | 2438 | NA | 26 | 13 | 30 | 2 | 4 | 10 |
| ENSDARG000 | KN149696.2:I | 183761:18679 | 186323:18699 | -;-;- | 3171 | NA | 3 | 4 | 50 | 0 | 6 | 1 |
| ENSDARG000 | KN149696.2:I | 45311:45816: | 45344:47583: | +;+;+;+;+;+ | 2796 | NA | 3 | 6 | 8 | 0 | 6 | 4 |
| ENSDARG000 | KN147651.2 | 332150 | 332256 | + | 107 | U6 | 0 | 0 | 0 | 0 | 0 | 0 |
| ENSDARG000 | KN147651.2 | 332680 | 332753 | + | 74 | U6 | 0 | 0 | 0 | 0 | 0 | 0 |
| ENSDARG000 | KN147651.2:I | 277542:28282 | 277798:28291 | +;+;+;+;+;+;+ | 8657 | NA | 692 | 281 | 879 | 171 | 332 | 486 |
| ENSDARG000 | KN147651.2 | 337203 | 337309 | + | 107 | U6 | 0 | 0 | 0 | 0 | 0 | 0 |
| ENSDARG000 | KN147651.2 | 336149 | 336255 | + | 107 | U6 | 0 | 0 | 0 | 0 | 0 | 0 |
| ENSDARG000 | KN147651.2:I | 267828:27479 | 268822:27487 | -;- | 1080 | NA | 2 | 6 | 6 | 0 | 8 | 0 |
| ENSDARG000 | KN147651.2 | 336676 | 336782 | + | 107 | U6 | 0 | 0 | 0 | 0 | 0 | 0 |
| ENSDARG000 | KN147651.2:I | 199184:20126 | 201153:20151 | -;-;-;-;-;-;-;- | 3336 | NA | 34 | 58 | 44 | 20 | 37 | 2 |
| ENSDARG000 | KN147651.2 | 335622 | 335728 | + | 107 | U6 | 0 | 0 | 0 | 0 | 0 | 0 |
| ENSDARG000 | KN149690.1:I | 45579:45579: | 45761:45761: | -;-;-;-;-;-;- | 2707 | NA | 12 | 12 | 10 | 0 | 0 | 8 |

| | | | | | | | | | | | | |
|---|---|---|---|---|---|---|---|---|---|---|---|---|
| ENSDARG000 | KN149690.1:K | 294797:29732 | 295286:29751 | +;+;+;+;+ | 2858 | NA | 12 | 14 | 30 | 2 | 10 | 6 |
| ENSDARG000 | KN149690.1:K | 73094:85124: | 73228:85228: | -;-;- | 353 | NA | 0 | 0 | 0 | 0 | 0 | 0 |
| ENSDARG000 | KN149686.1:K | 931:1969:142 | 1084:2118:17 | +;+;+ | 3434 | NA | 17 | 34 | 20 | 0 | 14 | 6 |
| ENSDARG000 | KN147652.2 | 157259 | 157360 | + | 102 | U6 | 0 | 0 | 0 | 0 | 0 | 0 |
| ENSDARG000 | KN147652.2 | 157774 | 157880 | + | 107 | U6 | 0 | 0 | 0 | 0 | 0 | 0 |
| ENSDARG000 | KN147652.2 | 201314 | 201420 | + | 107 | U6 | 0 | 0 | 0 | 0 | 0 | 0 |
| ENSDARG000 | KN147652.2 | 202905 | 203011 | + | 107 | U6 | 0 | 0 | 0 | 0 | 0 | 0 |
| ENSDARG000 | KN147652.2 | 159880 | 159986 | + | 107 | U6 | 0 | 0 | 0 | 0 | 0 | 0 |
| ENSDARG000 | KN147652.2 | 205017 | 205123 | + | 107 | U6 | 0 | 0 | 0 | 0 | 0 | 0 |
| ENSDARG000 | KN147652.2 | 160407 | 160513 | + | 107 | U6 | 0 | 0 | 0 | 0 | 0 | 0 |
| ENSDARG000 | KN147652.2 | 153159 | 153307 | - | 149 | U1 | 0 | 1 | 1 | 0 | 0 | 0 |
| ENSDARG000 | KN147652.2 | 158830 | 158936 | + | 107 | U6 | 0 | 0 | 0 | 0 | 0 | 0 |
| ENSDARG000 | KN147652.2 | 190488 | 190644 | - | 157 | U1 | 2 | 0 | 0 | 0 | 0 | 0 |
| ENSDARG000 | KN147652.2 | 200784 | 200890 | + | 107 | U6 | 0 | 0 | 0 | 0 | 0 | 0 |
| ENSDARG000 | KN147652.2 | 203435 | 203541 | + | 107 | U6 | 0 | 0 | 0 | 0 | 0 | 0 |
| ENSDARG000 | KN147652.2 | 159360 | 159466 | + | 107 | U6 | 0 | 0 | 0 | 0 | 0 | 0 |
| ENSDARG000 | KN147652.2 | 201845 | 201951 | + | 107 | U6 | 0 | 0 | 0 | 0 | 0 | 0 |
| ENSDARG000 | KN147652.2 | 151853 | 152016 | - | 164 | U1 | 0 | 0 | 0 | 0 | 0 | 0 |
| ENSDARG000 | KN147652.2 | 192137 | 192300 | - | 164 | U1 | 0 | 0 | 0 | 0 | 0 | 0 |
| ENSDARG000 | KN147652.2 | 158303 | 158409 | + | 107 | U6 | 0 | 0 | 0 | 0 | 0 | 0 |
| ENSDARG000 | KN149688.2:K | 16364:16364: | 16510:16510: | +:+:+:+:+:+:+: | 9487 | NA | 33 | 4 | 45 | 12 | 16 | 11 |
| ENSDARG000 | KN149688.2 | 56952 | 57100 | + | 149 | U1 | 2 | 0 | 1 | 0 | 0 | 0 |
| ENSDARG000 | KN149688.2 | 98697 | 98848 | + | 152 | U1 | 0 | 0 | 1 | 0 | 0 | 0 |
| ENSDARG000 | KN149688.2 | 96087 | 96250 | + | 164 | U1 | 0 | 0 | 0 | 0 | 0 | 0 |
| ENSDARG000 | KN149688.2:K | 220107:23764 | 220278:23971 | +;+;+;+;+;+ | 5264 | NA | 26 | 6 | 32 | 2 | 0 | 10 |
| ENSDARG000 | KN149688.2 | 58242 | 58405 | + | 164 | U1 | 0 | 0 | 0 | 0 | 0 | 0 |
| ENSDARG000 | KN149688.2 | 109712 | 109853 | + | 142 | U1 | 0 | 0 | 0 | 0 | 0 | 0 |
| ENSDARG000 | KN149688.2 | 14841 | 14959 | + | 119 | 5S_rRNA | 0 | 0 | 0 | 0 | 0 | 0 |
| ENSDARG000 | KN149688.2 | 59547 | 59710 | + | 164 | U1 | 0 | 0 | 0 | 0 | 0 | 0 |
| ENSDARG000 | KN149688.2 | 97392 | 97555 | + | 164 | U1 | 0 | 0 | 0 | 0 | 0 | 0 |
| ENSDARG000 | KN147636.1 | 69767 | 69868 | + | 102 | U6 | 0 | 0 | 0 | 0 | 0 | 0 |
| ENSDARG000 | KN147636.1 | 55367 | 55530 | - | 164 | U1 | 0 | 0 | 0 | 0 | 0 | 0 |
| ENSDARG000 | KN147636.1 | 70811 | 70884 | + | 74 | U6 | 0 | 0 | 0 | 0 | 0 | 0 |
| ENSDARG000 | KN147636.1:K | 158783:16108 | 159800:16175 | +;+ | 1684 | NA | 0 | 0 | 0 | 0 | 0 | 0 |
| ENSDARG000 | KN147636.1:K | 139905:13990 | 142339:14233 | -:-:-:-:-:-:-:-: | 3312 | NA | 0 | 0 | 0 | 0 | 0 | 0 |
| ENSDARG000 | KN147636.1 | 70282 | 70388 | + | 107 | U6 | 0 | 0 | 0 | 0 | 0 | 0 |
| ENSDARG000 | KN147636.1 | 33723 | 33837 | - | 115 | 5S_rRNA | 0 | 0 | 0 | 0 | 0 | 0 |
| ENSDARG000 | KN147636.1 | 56677 | 56825 | - | 149 | U1 | 1 | 0 | 0 | 0 | 0 | 0 |
| ENSDARG000 | KN149689.2 | 114447 | 114561 | + | 115 | 5S_rRNA | 0 | 0 | 0 | 0 | 0 | 0 |
| ENSDARG000 | KN149689.2:K | 123981:12398 | 124282:12428 | +:+:+:+:+:+:+: | 8012 | NA | 36 | 11 | 0 | 0 | 17 | 10 |
| ENSDARG000 | KN149695.1:K | 52148:53398: | 52223:53430: | -;-;-;-;- | 822 | NA | 2 | 10 | 6 | 0 | 1 | 2 |
| ENSDARG000 | KN149695.1:K | 152954:15459 | 153023:15894 | +;+ | 4419 | NA | 2 | 0 | 0 | 0 | 0 | 6 |
| ENSDARG000 | KN150691.1 | 89732 | 89845 | - | 114 | 5S_rRNA | 0 | 0 | 0 | 0 | 0 | 0 |
| ENSDARG000 | KN150691.1 | 17078 | 17195 | + | 118 | 5S_rRNA | 0 | 0 | 0 | 0 | 0 | 0 |
| ENSDARG000 | KN150691.1:K | 84292:84292: | 84390:84390: | -;-;-;-;- | 490 | NA | 0 | 0 | 0 | 0 | 0 | 0 |
| ENSDARG000 | KN147632.2:K | 95132;99992 | 96135;10007 | -;- | 1089 | NA | 24 | 10 | 2 | 0 | 0 | 4 |
| ENSDARG000 | KN147632.2:K | 27234;27448 | 27411;28412 | -;- | 1143 | NA | 8 | 0 | 0 | 0 | 0 | 2 |
| ENSDARG000 | KN147637.2:K | 107459:11712 | 110935:11729 | -;-;- | 3817 | NA | 0 | 12 | 1 | 0 | 3 | 2 |
| ENSDARG000 | KN147637.2:K | 117568:12485 | 117780:12779 | -;-;-;- | 4085 | NA | 5 | 0 | 7 | 6 | 4 | 2 |
| ENSDARG000 | KN147637.2:K | 157645:15767 | 157674:15854 | -;-;- | 984 | NA | 14 | 5 | 11 | 0 | 2 | 5 |
| ENSDARG000 | KN147637.2:K | 37082:37801: | 37675:37980: | +:+:+:+:+:+:+: | 6926 | NA | 0 | 0 | 4 | 0 | 0 | 0 |
| ENSDARG000 | KN149685.1 | 52649 | 59470 | - | 6822 | NA | 16 | 14 | 30 | 26 | 20 | 16 |
| ENSDARG000 | KN150214.1:K | 2983:4249:48 | 3054:4290:48 | +:+:+:+:+:+:+: | 2369 | NA | 8 | 6 | 0 | 42 | 4 | 8 |
| ENSDARG000 | KN150214.1:K | 31556:31884: | 31749:31961: | -:-:-:-:-:-:-:-: | 4869 | NA | 18 | 0 | 15 | 0 | 32 | 0 |
| ENSDARG000 | KZ116066.1 | 46424 | 46539 | + | 116 | 5S_rRNA | 0 | 0 | 0 | 0 | 0 | 0 |
| ENSDARG000 | KN149684.1:K | 38615:46052: | 38735:46291: | +:+:+:+:+:+:+: | 2962 | NA | 18 | 2 | 4 | 0 | 10 | 8 |
| ENSDARG000 | KZ116064.1:K | 67457:73772: | 67719:74050: | +;+;+ | 2424 | NA | 0 | 0 | 0 | 0 | 0 | 0 |
| ENSDARG000 | KZ116064.1:K | 28518:31986: | 28964:32159: | +;+;+ | 796 | NA | 0 | 0 | 2 | 0 | 0 | 0 |
| ENSDARG000 | KN150230.1:K | 48970:63111: | 49258:63202: | +:+:+:+:+:+:+: | 4754 | FAM135B | 2 | 2 | 2 | 21 | 2 | 4 |
| ENSDARG000 | KN150608.1:K | 7296:9752:10 | 7533:9930:11 | +;+;+ | 1304 | NA | 2 | 1 | 0 | 0 | 10 | 0 |
| ENSDARG000 | KN150608.1:K | 59559:59559: | 59717:59717: | +:+:+:+:+:+:+: | 2805 | NA | 14 | 29 | 16 | 8 | 16 | 33 |
| ENSDARG000 | KN150608.1:K | 102275:12361 | 102510:12378 | +;+;+;+;+;+ | 2053 | NA | 61 | 32 | 8 | 4 | 13 | 28 |
| ENSDARG000 | KN150171.1:K | 22085:30898: | 26021:31623: | -:-:-:-:-:-:-:-: | 7455 | clcn2b | 243 | 86 | 442 | 282 | 444 | 345 |
| ENSDARG000 | KN149682.1:K | 61071;64549 | 61155;65540 | +;+ | 1077 | NA | 0 | 0 | 0 | 0 | 0 | 0 |
| ENSDARG000 | KN149847.1:K | 66755;74409 | 67932;74493 | -;- | 1263 | NA | 16 | 2 | 8 | 0 | 1 | 16 |
| ENSDARG000 | KN150487.1:K | 423:423:836: | 994:657:994: | +:+:+:+:+:+:+: | 4224 | KDM5A | 354 | 612 | 795 | 258 | 915 | 436 |
| ENSDARG000 | KN150487.1:K | 31702:31702: | 31778:31778: | +:+:+:+:+:+:+: | 3289 | NA | 31 | 30 | 98 | 31 | 21 | 44 |
| ENSDARG000 | KN150487.1:K | 23438:25097: | 23604:25203: | +:+:+:+:+:+:+: | 3838 | NA | 289 | 469 | 567 | 244 | 615 | 418 |
| ENSDARG000 | KN150487.1:K | 15455:15698: | 15543:15825: | +;+;+ | 432 | NA | 20 | 52 | 36 | 6 | 50 | 6 |
| ENSDARG000 | KN149948.1:K | 69090:73434: | 71301:73505: | -;-;-;-;-;-;- | 2975 | CCDC134 | 0 | 2 | 47 | 0 | 77 | 82 |
| ENSDARG000 | KN149948.1 | 55371 | 55485 | + | 115 | 5S_rRNA | 0 | 0 | 0 | 0 | 0 | 0 |
| ENSDARG000 | KN149948.1:K | 89256:92615: | 89340:92887: | +;+;+;+;+ | 1149 | mhc1zka | 3 | 8 | 13 | 0 | 38 | 52 |
| ENSDARG000 | KN149948.1:K | 3740:3740:82 | 4675:4675:83 | -:-:-:-:-:-:-:-: | 2581 | NA | 0 | 0 | 0 | 0 | 0 | 0 |
| ENSDARG000 | KN149790.1 | 1312 | 1418 | + | 107 | U6 | 0 | 0 | 0 | 0 | 0 | 0 |
| ENSDARG000 | KN149790.1:K | 22627:22627: | 25391:25391: | -:-:-:-:-:-:-:-: | 4796 | NA | 100 | 197 | 156 | 74 | 152 | 155 |
| ENSDARG000 | KN149790.1:K | 106986:10799 | 107541:10820 | -;-;-;- | 1166 | TMEM19 | 263 | 690 | 1947 | 421 | 1539 | 897 |
| ENSDARG000 | KN149790.1:K | 75795:76925: | 78809:78809: | -:-:-:-:-:-:-:-: | 4229 | NA | 188 | 225 | 278 | 126 | 210 | 234 |
| ENSDARG000 | KN149790.1:K | 39:39:1249:1 | 1041:1041:13 | -;-;-;-;-;-;-;- | 1646 | NA | 0 | 0 | 0 | 0 | 0 | 0 |
| ENSDARG000 | KN149778.1:K | 21;8025 | 758;8477 | -;- | 1191 | NA | 0 | 0 | 0 | 0 | 0 | 0 |
| ENSDARG000 | KN150425.1 | 87099 | 87218 | + | 120 | 5S_rRNA | 0 | 0 | 0 | 0 | 0 | 0 |
| ENSDARG000 | KN150425.1:K | 103570:10383 | 106640:10538 | +;+ | 3071 | cyp1c1 | 82 | 4 | 32 | 18 | 66 | 110 |
| ENSDARG000 | KN150425.1:K | 19107:20148: | 20075:20288: | +:+:+:+:+:+:+: | 3703 | NA | 10 | 0 | 0 | 0 | 0 | 0 |
| ENSDARG000 | KN150326.1:K | 26197:28430: | 26520:28564: | -:-:-:-:-:-:-:-: | 4065 | EIF2AK4 | 601 | 290 | 721 | 414 | 474 | 498 |
| ENSDARG000 | KN150326.1:K | 6045:15954:1 | 8164:16022:1 | -;-;-;-;- | 2480 | NA | 158 | 57 | 114 | 101 | 99 | 100 |
| ENSDARG000 | KN149698.1:K | 44067:44479: | 44111:44498: | +;+;+;+ | 807 | NA | 0 | 0 | 0 | 0 | 2 | 0 |
| ENSDARG000 | KN149698.1:K | 88501:91031: | 88994:91186: | +;+;+;+;+;+;+ | 2766 | NA | 270 | 290 | 514 | 308 | 465 | 378 |
| ENSDARG000 | KN149698.1 | 35592 | 36353 | + | 762 | NA | 0 | 0 | 0 | 0 | 0 | 0 |
| ENSDARG000 | KN149698.1:K | 55899:55899: | 56083:56083: | +:+:+:+:+:+:+: | 933 | NA | 0 | 0 | 0 | 0 | 0 | 0 |
| ENSDARG000 | KN150568.1 | 39414 | 39530 | - | 117 | 5S_rRNA | 0 | 0 | 0 | 0 | 0 | 0 |
| ENSDARG000 | KN150158.1:K | 121:4567:503 | 2290:4725:52 | -;-;-;-;-;-;- | 3147 | psmd2 | 1514 | 1500 | 1688 | 636 | 1517 | 852 |
| ENSDARG000 | KN150158.1 | 86505 | 86625 | - | 121 | 5S_rRNA | 0 | 0 | 0 | 0 | 0 | 0 |
| ENSDARG000 | KN150578.1:K | 14229:21963: | 14289:22124: | +;+;+;+;+;+;+ | 918 | DTX1 | 0 | 18 | 0 | 0 | 8 | 17 |
| ENSDARG000 | KN150578.1 | 87116 | 87234 | + | 119 | 5S_rRNA | 0 | 0 | 0 | 0 | 0 | 0 |
| ENSDARG000 | KN150605.1 | 59641 | 62078 | + | 2438 | NA | 41 | 24 | 68 | 44 | 31 | 0 |
| ENSDARG000 | KN150605.1:K | 65045:69904: | 65273:70182: | +:+:+:+:+:+:+: | 3258 | NA | 92 | 24 | 43 | 14 | 22 | 6 |
| ENSDARG000 | KN149797.1:K | 55555:60494: | 55985:60661: | -:-:-:-:-:-:-:-: | 2195 | NA | 42 | 28 | 4 | 8 | 10 | 22 |
| ENSDARG000 | KN149797.1:K | 26121:27109: | 26526:27174: | -:-:-:-:-:-:-:-: | 4136 | NA | 679 | 412 | 781 | 493 | 872 | 578 |
| ENSDARG000 | KN149702.1:K | 22079:26991: | 22287:27113: | -:-:-:-:-:-:-:-: | 1186 | KHDRBS3 | 0 | 0 | 9 | 0 | 0 | 0 |
| ENSDARG000 | KN150603.1 | 64973 | 65090 | + | 118 | 5S_rRNA | 0 | 0 | 0 | 0 | 0 | 0 |
| ENSDARG000 | KN149984.1:K | 32189:37715: | 33843:37795: | -:-:-:-:-:-:-:-: | 3429 | ARHGAP6 | 32 | 12 | 8 | 0 | 15 | 8 |
| ENSDARG000 | KN150258.1:K | 33006:35501: | 33023:35623: | -;-;- | 1008 | NA | 0 | 0 | 0 | 0 | 0 | 0 |
| ENSDARG000 | KN149895.1:K | 10606:11065: | 10978:11149: | +;+;+ | 935 | METTL23 | 0 | 0 | 0 | 0 | 0 | 0 |
| ENSDARG000 | KN149895.1:K | 56178:58914: | 56281:58997: | +;+;+;+ | 465 | NA | 16 | 0 | 0 | 0 | 13 | 0 |
| ENSDARG000 | KN150432.1:K | 6941:9581:13 | 7055:9700:13 | -:-:-:-:-:-:-:-: | 2433 | NCAM2 | 486 | 1623 | 412 | 731 | 552 | 707 |
| ENSDARG000 | KN149826.1 | 58199 | 58312 | - | 114 | 5S_rRNA | 0 | 0 | 0 | 0 | 0 | 0 |
| ENSDARG000 | KN149826.1 | 35565 | 35679 | + | 115 | 5S_rRNA | 0 | 5 | 0 | 0 | 0 | 0 |
| ENSDARG000 | KN150617.1:K | 66381;73787 | 66868;79211 | +;+ | 5913 | fzd4 | 590 | 300 | 638 | 434 | 444 | 958 |
| ENSDARG000 | KN149932.1:K | 72714:72808: | 72731:72909: | +:+:+:+:+:+:+: | 915 | NA | 3 | 0 | 2 | 0 | 0 | 6 |
| ENSDARG000 | KN149904.1 | 46459 | 46574 | + | 116 | 5S_rRNA | 0 | 0 | 0 | 0 | 0 | 0 |
| ENSDARG000 | KN150196.1:K | 54331;63004 | 54642;63490 | +;+ | 799 | KCTD15 | 34 | 10 | 18 | 10 | 31 | 14 |
| ENSDARG000 | KN150616.1:K | 78145;78839 | 78673;79086 | +;+ | 777 | NA | 12 | 2 | 53 | 26 | 34 | 20 |
| ENSDARG000 | KN150616.1 | 60073 | 61356 | + | 1284 | NA | 6 | 7 | 62 | 5 | 80 | 14 |
| ENSDARG000 | KN150663.1:K | 47682:47682: | 47840:47840: | -;-;-;-;-;-;-;- | 824 | SYNDIG1 | 0 | 0 | 0 | 0 | 12 | 0 |
| ENSDARG000 | KN150663.1:K | 58;368 | 287;936 | -;- | 799 | NA | 0 | 4 | 2 | 0 | 2 | 1 |
| ENSDARG000 | KN150630.1:K | 17799;21554 | 18704;24120 | +;+ | 3473 | NA | 36 | 2 | 8 | 12 | 60 | 0 |
| ENSDARG000 | KN150125.1:K | 14601:19333: | 15331:19487: | +:+:+:+:+:+:+: | 3209 | NA | 0 | 0 | 8 | 0 | 0 | 2 |
| ENSDARG000 | KN149968.1:K | 15950:18909: | 16370:18928: | +:+:+:+:+:+:+: | 1134 | NHERF1 | 56 | 28 | 61 | 20 | 97 | 38 |
| ENSDARG000 | KN150342.1:K | 49191:54256: | 54082:55964: | -;-;-;-;- | 7850 | BNC1 | 28 | 0 | 0 | 0 | 4 | 0 |
| ENSDARG000 | KN150342.1:K | 26584:28332: | 26951:28607: | -;-;-;- | 2031 | NA | 0 | 0 | 0 | 0 | 0 | 0 |
| ENSDARG000 | KN150529.1:K | 16574:32949: | 16714:33056: | +:+:+:+:+:+:+: | 3963 | RIC1 | 167 | 61 | 129 | 84 | 113 | 103 |
| ENSDARG000 | KN150646.1:K | 56022:62037: | 56228:62167: | +;+;+;+;+ | 822 | NA | 48 | 12 | 123 | 18 | 67 | 39 |
| ENSDARG000 | KN150670.1:K | 733:1234:293 | 1033:1348:30 | -:-:-:-:-:-:-:-: | 2886 | NA | 1191 | 1034 | 1181 | 556 | 1493 | 838 |
| ENSDARG000 | KN150670.1:K | 34341:40016: | 38044:40100: | -;-;-;-;-;-;- | 7177 | NA | 184 | 127 | 166 | 55 | 268 | 103 |
| ENSDARG000 | KN149912.1:K | 29879:37217: | 30155:37311: | +;+;+;+ | 1194 | SPHK1 | 6 | 2 | 6 | 0 | 0 | 0 |
| ENSDARG000 | KN149912.1 | 76624 | 77959 | + | 1336 | NA | 0 | 0 | 0 | 0 | 0 | 0 |
| ENSDARG000 | KN149912.1:K | 57102:57102: | 57367:60973: | -:-:-:-:-:-:-:-: | 6126 | NA | 141 | 4 | 4 | 18 | 18 | 2 |
| ENSDARG000 | KN150405.1:K | 1010:2326:84 | 5772:5772:85 | -:-:-:-:-:-:-:-: | 6266 | NA | 7485 | 4536 | 8004 | 7622 | 7920 | 7505 |
| ENSDARG000 | KN149998.1:K | 45287:49057: | 45450:49401: | -;-;-;- | 989 | NA | 0 | 0 | 0 | 0 | 0 | 0 |
| ENSDARG000 | KN149998.1:K | 683:4853:791 | 823:5197:825 | -;-;-;- | 1191 | NA | 8 | 4 | 0 | 0 | 2 | 0 |
| ENSDARG000 | KN150552.1:K | 10471:10537: | 10494:10638: | +:+:+:+:+:+:+: | 909 | ASS1 | 27 | 7 | 47 | 18 | 21 | 2 |
| ENSDARG000 | KN149784.1 | 55886 | 56002 | - | 117 | 5S_rRNA | 0 | 0 | 0 | 0 | 0 | 0 |
| ENSDARG000 | KN149784.1:K | 29639:29639: | 30256:30256: | +;+;+;+;+;+;+ | 1115 | NA | 0 | 0 | 0 | 0 | 0 | 0 |
| ENSDARG000 | KN150455.1 | 29252 | 30557 | - | 1306 | NA | 0 | 0 | 0 | 0 | 0 | 0 |
| ENSDARG000 | KN150700.2:K | 501:3255:403 | 1672:3428:42 | -:-:-:-:-:-:-:-: | 10509 | NA | 339 | 190 | 196 | 306 | 271 | 133 |
| ENSDARG000 | KN150131.1 | 36873 | 37008 | + | 136 | 5S_rRNA | 0 | 0 | 0 | 0 | 0 | 0 |
| ENSDARG000 | KN150131.1:K | 549:1209:179 | 666:1409:198 | +:+:+:+:+:+:+: | 1266 | acp7 | 396 | 158 | 152 | 180 | 374 | 194 |
| ENSDARG000 | KN150131.1:K | 60456:60659: | 60529:61089: | +;+;+;+;+;+ | 5473 | NA | 59 | 16 | 11 | 2 | 29 | 18 |
| ENSDARG000 | KN149725.1:K | 34665:34665: | 34811:34811: | +;+;+;+;+ | 1354 | NA | 0 | 0 | 0 | 0 | 0 | 0 |
| ENSDARG000 | KN150000.1 | 12517 | 12630 | + | 114 | 5S_rRNA | 0 | 0 | 0 | 0 | 0 | 0 |
| ENSDARG000 | KN150000.1:K | 7817:7859:78 | 8009:8009:80 | +:+:+:+:+:+:+: | 5292 | PTPRD | 38 | 28 | 38 | 53 | 66 | 18 |
| ENSDARG000 | KN150176.1 | 56608 | 56724 | + | 117 | 5S_rRNA | 0 | 0 | 0 | 0 | 0 | 0 |
| ENSDARG000 | KN150176.1 | 59159 | 59275 | - | 117 | 5S_rRNA | 0 | 0 | 0 | 0 | 0 | 0 |
| ENSDARG000 | KN150098.1:K | 40966:40966: | 41886:41886: | -:-:-:-:-:-:-:-: | 2586 | NA | 0 | 0 | 0 | 0 | 0 | 0 |
| ENSDARG000 | KN150677.1:K | 44475;46353 | 44798;48192 | -;- | 2164 | NA | 4 | 2 | 4 | 0 | 0 | 0 |
| ENSDARG000 | KN150041.2:K | 10631:14249: | 11207:14370: | +:+:+:+:+:+:+: | 8220 | DMXL1 | 1164 | 711 | 1357 | 887 | 1530 | 871 |
| ENSDARG000 | KN150709.1:K | 28312:30077: | 28527:30209: | -;-;- | 978 | NA | 2 | 0 | 0 | 0 | 0 | 0 |
| ENSDARG000 | KN149880.1:K | 51895:55523: | 52143:55701: | -;-;-;-;-;-;- | 1193 | sec14l1 | 194 | 88 | 198 | 225 | 199 | 169 |
| ENSDARG000 | KN149880.1:K | 4887:5109:12 | 4952:5348:12 | +:+:+:+:+:+:+: | 4545 | NA | 1433 | 57 | 1923 | 20 | 1763 | 1148 |
| ENSDARG000 | KN150449.1:K | 4696:5176:70 | 5085:5325:73 | -;-;-;-;-;-;- | 1460 | NA | 2 | 0 | 0 | 0 | 0 | 0 |
| ENSDARG000 | KN150449.1:K | 25237:26519: | 25265:26636: | +;+;+;+;+ | 501 | RAC1 | 190 | 165 | 166 | 89 | 122 | 172 |
| ENSDARG000 | KN150449.1:K | 38186:42447: | 42357:42594: | -;-;-;-;-;- | 4888 | STK19 | 388 | 324 | 365 | 477 | 323 | 403 |
| ENSDARG000 | KN150477.1:K | 42099:44997: | 42575:45170: | -;-;- | 2779 | NA | 10 | 8 | 25 | 6 | 10 | 21 |
| ENSDARG000 | KN150596.1:K | 17881:19940: | 18545:20053: | +;+;+;+;+ | 5883 | PPIC | 970 | 823 | 1063 | 1018 | 1927 | 1349 |
| ENSDARG000 | KN150241.1:K | 43860:48212: | 44068:48421: | +;+;+;+;+ | 1047 | MCHR2 | 0 | 0 | 0 | 0 | 0 | 0 |
| ENSDARG000 | KN150241.1:K | 21712:23544: | 21863:23641: | +:+:+:+:+:+:+: | 1408 | NA | 8 | 12 | 6 | 4 | 84 | 22 |
| ENSDARG000 | KN149959.1:K | 2273:3796:59 | 2975:3935:61 | +;+;+;+;+;+ | 3878 | NA | 0 | 0 | 0 | 0 | 0 | 0 |
| ENSDARG000 | KN149959.1:K | 21026:25293: | 21309:26282: | +;+;+ | 3230 | NA | 422 | 255 | 336 | 258 | 396 | 278 |
| ENSDARG000 | KN149959.1:K | 14158:16806: | 15151:16926: | -;-;-;- | 1342 | NA | 14 | 2 | 14 | 2 | 12 | 0 |
| ENSDARG000 | KN149813.1:K | 12050:14857: | 13007:14995: | -:-:-:-:-:-:-:-: | 2995 | NA | 2 | 0 | 5 | 0 | 6 | 10 |
| ENSDARG000 | KN149992.1:K | 51454;56413 | 52673;56497 | -;- | 1305 | NA | 0 | 0 | 2 | 0 | 0 | 0 |
| ENSDARG000 | KN150030.1:K | 3051:5042:66 | 3245:5091:66 | -;-;-;- | 507 | golim4a | 233 | 132 | 231 | 159 | 339 | 186 |
| ENSDARG000 | KN150030.1:K | 51413:51413: | 55230:55230: | -;-;-;-;- | 5408 | NA | 62 | 0 | 0 | 22 | 8 | 2 |

| | | | | | | | | | | | | |
|---|---|---|---|---|---|---|---|---|---|---|---|---|
| ENSDARG000 | KN150653.1:K | 48664:48838: | 48744:48920: | -;-;-;-;- | 795 | NA | 0 | 0 | 0 | 0 | 0 | 0 |
| ENSDARG000 | KN150653.1:K | 31269:32980: | 32856:33059: | -;-;-;-;-;- | 2619 | NA | 0 | 0 | 0 | 0 | 0 | 0 |
| ENSDARG000 | KN150653.1 | 20994 | 21110 | - | 117 | 5S_rRNA | 0 | 0 | 0 | 0 | 0 | 0 |
| ENSDARG000 | KN150170.1 | 35264 | 35370 | - | 107 | Y_RNA | 4 | 1 | 0 | 0 | 0 | 0 |
| ENSDARG000 | KN150170.1:K | 10890:11163: | 10941:11290: | +:+:+:+:+:+:+: | 855 | HUS1B | 54 | 165 | 234 | 156 | 234 | 85 |
| ENSDARG000 | KN150170.1:K | 36770:37563: | 36838:37657: | +;+;+ | 312 | NA | 0 | 0 | 0 | 0 | 0 | 0 |
| ENSDARG000 | KN150642.1 | 9441 | 10244 | + | 804 | atoh1b | 0 | 0 | 0 | 0 | 0 | 0 |
| ENSDARG000 | KN150642.1:K | 52178:52362: | 52187:52498: | -;-;-;-;- | 513 | NA | 185 | 106 | 156 | 109 | 129 | 141 |
| ENSDARG000 | KN150642.1:K | 16216:19033: | 17190:19202: | -;-;-;- | 1838 | NA | 8 | 5 | 4 | 0 | 6 | 8 |
| ENSDARG000 | KN150118.1 | 50723 | 50836 | - | 114 | 5S_rRNA | 0 | 0 | 0 | 0 | 0 | 0 |
| ENSDARG000 | KN150361.1:K | 21:1436:1588 | 3874:1458:17 | -:-:-:-:-:-:-:-:-: | 5775 | NA | 844 | 751 | 744 | 473 | 1165 | 858 |
| ENSDARG000 | KN150361.1:K | 20035:21416: | 20065:21426: | -:-:-:-:-:-:-:-:-: | 951 | CNNM4 | 0 | 5 | 0 | 0 | 13 | 8 |
| ENSDARG000 | KN149859.1:K | 8728:16744:2 | 8906:16908:2 | +:+:+:+:+:+:+: | 2638 | SLC44A2 | 1231 | 677 | 480 | 830 | 449 | 744 |
| ENSDARG000 | KN150104.1:K | 1741:1741:40 | 2504:2504:43 | +:+:+:+:+:+:+: | 5656 | THBS1 | 297 | 62 | 38 | 70 | 138 | 37 |
| ENSDARG000 | KN150104.1:K | 20111:20111: | 22461:22461: | +;+;+;+;+ | 4295 | NA | 11 | 7 | 4 | 11 | 0 | 0 |
| ENSDARG000 | KN149884.1:K | 9445:10343:1 | 10004:10369: | -:-:-:-:-:-:-:-:-: | 1467 | NA | 53 | 8 | 28 | 16 | 30 | 13 |
| ENSDARG000 | KN149884.1:K | 28781:35855: | 31924:36296: | -;-;-;- | 4540 | NA | 2 | 0 | 0 | 0 | 4 | 0 |
| ENSDARG000 | KZ116011.1:K | 20449;28609 | 20627;28768 | -;- | 339 | bmf1 | 24 | 12 | 16 | 0 | 2 | 2 |
| ENSDARG000 | KN149739.1:K | 39153:41108: | 40274:41251: | -;-;- | 1494 | SLC16A7 | 98 | 21 | 25 | 80 | 7 | 56 |
| ENSDARG000 | KN149739.1:K | 4228:11990:1 | 4407:12084:1 | +;+;+;+;+;+ | 4411 | NA | 0 | 4 | 6 | 2 | 0 | 0 |
| ENSDARG000 | KN150265.1:K | 23250:26707: | 23451:26941: | -;-;- | 1425 | TLX3 | 0 | 0 | 0 | 0 | 0 | 0 |
| ENSDARG000 | KN150265.1:K | 6124:8955:91 | 6232:9014:93 | +:+:+:+:+:+:+: | 826 | KCNIP1 | 0 | 0 | 5 | 0 | 0 | 0 |
| ENSDARG000 | KN149710.1:K | 3782:3782:14 | 4544:4544:15 | +:+:+:+:+:+:+: | 5433 | NA | 4502 | 2500 | 2826 | 3666 | 4680 | 3850 |
| ENSDARG000 | KN150173.1 | 14560 | 14650 | - | 91 | NA | 0 | 9 | 0 | 0 | 0 | 0 |
| ENSDARG000 | KN150173.1:K | 42072:43023: | 42914:43123: | -;-;-;-;- | 1538 | NA | 0 | 0 | 0 | 0 | 0 | 0 |
| ENSDARG000 | KN150173.1 | 15259 | 15346 | - | 88 | NA | 0 | 3 | 0 | 2 | 0 | 0 |
| ENSDARG000 | KN150173.1:K | 2840:3421:43 | 3106:3490:44 | -;-;-;-;-;-;- | 771 | MMD2 | 55 | 30 | 37 | 70 | 98 | 66 |
| ENSDARG000 | KN150173.1:K | 20647:21174: | 21102:21274: | -;-;-;-;-;-;- | 1471 | NA | 41 | 1140 | 31 | 31 | 21 | 105 |
| ENSDARG000 | KN150173.1 | 15936 | 16023 | - | 88 | NA | 0 | 1 | 0 | 0 | 0 | 0 |
| ENSDARG000 | KN150259.1:K | 16397:16397: | 16780:16780: | -:-:-:-:-:-:-:-:-: | 3906 | NA | 83 | 30 | 78 | 8 | 28 | 16 |
| ENSDARG000 | KN150259.1 | 45362 | 50078 | - | 4717 | NA | 0 | 0 | 0 | 0 | 0 | 0 |
| ENSDARG000 | KN150351.1 | 43446 | 43594 | + | 149 | U1 | 0 | 0 | 0 | 3 | 0 | 7 |
| ENSDARG000 | KN150351.1 | 46323 | 46486 | + | 164 | U1 | 0 | 0 | 0 | 0 | 0 | 0 |
| ENSDARG000 | KN150351.1 | 47650 | 47813 | + | 164 | U1 | 1 | 0 | 0 | 0 | 0 | 0 |
| ENSDARG000 | KN150351.1 | 44985 | 45148 | + | 164 | U1 | 0 | 0 | 0 | 0 | 0 | 0 |
| ENSDARG000 | KN150351.1 | 30189 | 30335 | - | 147 | U1 | 0 | 0 | 0 | 0 | 0 | 0 |
| ENSDARG000 | KN150336.1 | 21882 | 21999 | - | 118 | 5S_rRNA | 0 | 0 | 0 | 1 | 0 | 0 |
| ENSDARG000 | KN150479.1 | 10852 | 10965 | - | 114 | 5S_rRNA | 0 | 0 | 0 | 0 | 0 | 0 |
| ENSDARG000 | KN150703.1:K | 188:10007:14 | 5992:10148:1 | -;-;- | 6177 | NA | 14 | 30 | 34 | 28 | 42 | 38 |
| ENSDARG000 | KN150703.1:K | 43473:46582: | 44569:46711: | -;-;- | 1656 | NA | 22 | 18 | 30 | 20 | 46 | 39 |
| ENSDARG000 | KN150703.1:K | 36295;40060 | 38242;40144 | -;- | 2033 | NA | 24 | 42 | 40 | 69 | 11 | 42 |
| ENSDARG000 | KN150703.1:K | 23110:23110: | 25996:25996: | -;-;-;-;-;-;-;- | 3442 | NA | 66 | 74 | 150 | 62 | 100 | 131 |
| ENSDARG000 | KN150703.1:K | 15203:16491: | 15410:16614: | +;+;+ | 5099 | NA | 6 | 14 | 22 | 4 | 20 | 40 |
| ENSDARG000 | KN149717.1 | 39834 | 39948 | - | 115 | 5S_rRNA | 0 | 0 | 0 | 0 | 0 | 0 |
| ENSDARG000 | KN149840.1:K | 1246;2357 | 1383;2551 | +;+ | 333 | NA | 2 | 0 | 0 | 0 | 0 | 0 |
| ENSDARG000 | KN150335.1:K | 34020:37235: | 34212:37577: | -;-;- | 558 | NA | 0 | 0 | 0 | 0 | 0 | 58 |
| ENSDARG000 | KN150335.1:K | 11961:15526: | 12735:15644: | -;-;-;-;-;- | 2241 | NA | 0 | 63 | 135 | 34 | 242 | 275 |
| ENSDARG000 | KN150244.1:K | 18842:19219: | 19125:19343: | -;-;-;-;-;-;-;- | 1312 | NA | 0 | 0 | 0 | 0 | 0 | 0 |
| ENSDARG000 | KN150244.1:K | 12960:12960: | 14095:14095: | +;+;+;+;+;+;+ | 2506 | NA | 0 | 0 | 0 | 0 | 0 | 0 |
| ENSDARG000 | KN150708.1:K | 15753:20489: | 18987:21274: | -;-;-;-;-;- | 4921 | slc16a6a | 8 | 22 | 18 | 0 | 0 | 18 |
| ENSDARG000 | KN150708.1:K | 33580:33580: | 33874:33874: | +:+:+:+:+:+:+: | 2368 | NA | 14 | 52 | 8 | 4 | 16 | 58 |
| ENSDARG000 | KN150708.1:K | 36:4504:5174 | 664:4512:576 | -;-;- | 1226 | NA | 7 | 27 | 6 | 5 | 3 | 13 |
| ENSDARG000 | KN150191.1:K | 1722:11756:1 | 1799:11970:1 | +:+:+:+:+:+:+: | 3519 | PITPNM2 | 634 | 361 | 552 | 332 | 469 | 403 |
| ENSDARG000 | KN150570.1 | 13067 | 13540 | + | 474 | thrsp | 0 | 0 | 2 | 0 | 0 | 0 |
| ENSDARG000 | KN149800.1:K | 2019:16407:1 | 2981:16563:1 | -;-;- | 1227 | NA | 43 | 37 | 44 | 28 | 83 | 22 |
| ENSDARG000 | KN149697.1:K | 4287:7606:81 | 4316:8005:83 | +;+;+;+ | 777 | NA | 0 | 0 | 0 | 0 | 0 | 0 |
| ENSDARG000 | KN149697.1:K | 28917:30910: | 28962:31260: | +;+;+;+ | 768 | NA | 0 | 0 | 0 | 0 | 0 | 0 |
| ENSDARG000 | KN150293.1:K | 37108:37509: | 37277:37647: | -;-;-;- | 447 | NA | 0 | 0 | 0 | 0 | 0 | 0 |
| ENSDARG000 | KN150172.1:K | 19282:19282: | 19571:19571: | -:-:-:-:-:-:-:-:-: | 1280 | HAPLN1 | 0 | 0 | 0 | 0 | 0 | 0 |
| ENSDARG000 | KN150518.1:K | 17591:28933: | 18648:29058: | -:-:-:-:-:-:-:-:-: | 1815 | sema6a | 12 | 0 | 0 | 0 | 8 | 10 |
| ENSDARG000 | KN149749.1:K | 28504:29714: | 28575:29833: | -;-;-;- | 354 | NA | 39 | 45 | 35 | 47 | 48 | 67 |
| ENSDARG000 | KN149909.1:K | 4310:12590:1 | 4331:12610:1 | +;+;+;+;+;+;+ | 189 | NA | 0 | 0 | 0 | 0 | 0 | 0 |
| ENSDARG000 | KN149909.1:K | 22664:26266: | 22784:26334: | -:-:-:-:-:-:-:-:-: | 951 | NA | 232 | 237 | 341 | 210 | 189 | 224 |
| ENSDARG000 | KN150505.1:K | 18029:19019: | 18934:19083: | -;-;-;- | 1301 | NA | 255 | 276 | 586 | 359 | 639 | 498 |
| ENSDARG000 | KN150505.1:K | 5069:8633:94 | 5311:8791:96 | -;-;-;-;-;-;- | 1068 | NA | 535 | 507 | 981 | 872 | 1352 | 992 |
| ENSDARG000 | KN150189.1:K | 3:743:1020:1 | 608:927:1272 | +:+:+:+:+:+:+: | 2871 | NA | 0 | 0 | 9 | 0 | 78 | 34 |
| ENSDARG000 | KN149680.1:K | 2792:2792:31 | 3218:3032:32 | -;-;-;-;-;-;- | 2760 | NA | 16 | 2 | 4 | 0 | 0 | 0 |
| ENSDARG000 | KZ116065.1:K | 2333:2655:28 | 2562:2741:29 | +:+:+:+:+:+:+: | 1502 | KYAT3 | 2 | 0 | 0 | 0 | 0 | 0 |
| ENSDARG000 | KZ116044.1:K | 4289;12797 | 4464;14027 | +;+ | 1407 | NA | 0 | 0 | 4 | 0 | 0 | 2 |
| ENSDARG000 | KZ116044.1 | 30259 | 30377 | - | 119 | 5S_rRNA | 0 | 0 | 0 | 0 | 0 | 0 |
| ENSDARG000 | KN150137.1:K | 17751:18525: | 18132:18625: | +;+;+;+;+;+;+ | 2534 | BPHL | 276 | 184 | 222 | 168 | 169 | 188 |
| ENSDARG000 | KN150137.1 | 28074 | 28189 | - | 116 | 5S_rRNA | 5 | 0 | 1 | 5 | 0 | 2 |
| ENSDARG000 | KN150306.1:K | 15511:23675: | 18927:23885: | -;-;-;-;-;-;-;- | 4393 | NA | 3 | 0 | 0 | 0 | 4 | 0 |
| ENSDARG000 | KN150631.1:K | 15268:16119: | 15426:16386: | +;+;+;+;+;+ | 1059 | NA | 143 | 172 | 187 | 145 | 153 | 9 |
| ENSDARG000 | KN150631.1:K | 20777:22099: | 21749:22269: | +;+;+;+;+;+ | 3018 | ACADSB | 423 | 452 | 601 | 464 | 395 | 147 |
| ENSDARG000 | KN150631.1:K | 29810:31650: | 30519:31738: | +;+;+;+ | 1215 | NA | 6 | 1 | 1 | 0 | 0 | 0 |
| ENSDARG000 | KN149782.1 | 29662 | 32060 | - | 2399 | trim32 | 216 | 190 | 341 | 231 | 261 | 210 |
| ENSDARG000 | KN149782.1 | 27110 | 27223 | - | 114 | 5S_rRNA | 10 | 0 | 2 | 17 | 14 | 7 |
| ENSDARG000 | KN150062.1:K | 8580:8672:94 | 9594:8890:95 | +:+:+:+:+:+:+: | 1810 | NA | 319 | 138 | 215 | 182 | 353 | 115 |
| ENSDARG000 | KN150400.1 | 32941 | 33016 | + | 76 | U6 | 0 | 0 | 0 | 0 | 0 | 0 |
| ENSDARG000 | KN149795.1 | 17232 | 17300 | - | 69 | NA | 14 | 26 | 35 | 5 | 24 | 7 |
| ENSDARG000 | KN149795.1 | 21189 | 21265 | - | 77 | NA | 2 | 1 | 0 | 0 | 0 | 0 |
| ENSDARG000 | KN149795.1:K | 2783:2783:65 | 2976:2976:66 | +;+;+;+;+;+;+ | 569 | NA | 14 | 21 | 18 | 6 | 32 | 2 |
| ENSDARG000 | KZ116019.1:K | 24340:24736: | 24458:24855: | -;-;-;-;-;-;- | 1703 | NA | 84 | 58 | 57 | 89 | 53 | 41 |
| ENSDARG000 | KZ116019.1:K | 3917:5149:78 | 4957:5404:81 | -;-;-;-;- | 1945 | NA | 205 | 189 | 209 | 132 | 188 | 104 |
| ENSDARG000 | KN149989.1:K | 15290:19619: | 15577:21433: | +;+;+;+;+;+ | 2675 | NA | 5 | 2 | 1 | 0 | 0 | 0 |
| ENSDARG000 | KN149989.1:K | 142:142:1168 | 928:928:1281 | +;+;+;+;+;+;+ | 1297 | NA | 2 | 0 | 0 | 0 | 6 | 0 |
| ENSDARG000 | KN150102.1 | 11947 | 12065 | + | 119 | 5S_rRNA | 0 | 0 | 0 | 0 | 0 | 0 |
| ENSDARG000 | KN150102.1:K | 29:681:1635:1 | 595:722:1760 | +;+;+;+;+;+ | 2246 | NA | 0 | 0 | 2 | 4 | 0 | 0 |
| ENSDARG000 | KZ116034.1:K | 24596:25110: | 25024:25280: | -;-;-;- | 777 | NA | 263 | 200 | 209 | 148 | 128 | 94 |
| ENSDARG000 | KZ116034.1:K | 20255:20255: | 20446:20371: | +:+:+:+:+:+:+: | 1858 | NA | 131 | 135 | 153 | 170 | 194 | 142 |
| ENSDARG000 | KZ116034.1:K | 1250:2741:29 | 1288:2837:31 | +;+;+;+;+;+ | 3484 | dusp27 | 23 | 8 | 20 | 36 | 4 | 10 |
| ENSDARG000 | KZ116034.1:K | 26713;27283 | 26869;27407 | +;+ | 282 | NA | 1 | 2 | 2 | 12 | 3 | 4 |
| ENSDARG000 | KZ116034.1:K | 26611:30384: | 26673:30410: | -;-;-;-;-;-;- | 363 | cd247 | 6 | 0 | 0 | 0 | 2 | 0 |
| ENSDARG000 | KN149996.1:K | 28363;28946 | 28722;30899 | +;+ | 2314 | NA | 0 | 6 | 6 | 4 | 5 | 5 |
| ENSDARG000 | KN150015.1:K | 3538:4084:64 | 3658:4196:65 | +:+:+:+:+:+:+: | 1906 | PDCL3 | 301 | 148 | 211 | 343 | 812 | 281 |
| ENSDARG000 | KN150127.1 | 17150 | 17259 | + | 110 | dre-mir-2189 | 1 | 1 | 6 | 2 | 1 | 0 |
| ENSDARG000 | KN150127.1:K | 6611:12926:1 | 6633:13009:1 | -;-;- | 730 | NA | 18 | 13 | 26 | 28 | 9 | 5 |
| ENSDARG000 | KN150127.1:K | 13896:13896: | 16152:17026: | -;-;-;-;-;- | 4096 | NA | 73 | 17 | 103 | 82 | 28 | 35 |
| ENSDARG000 | KN150127.1:K | 27702:28886: | 28803:29544: | -;-;- | 1847 | NA | 290 | 36 | 397 | 263 | 240 | 128 |
| ENSDARG000 | KN149706.1 | 28026 | 28114 | - | 89 | dre-mir-723 | 0 | 0 | 0 | 0 | 0 | 0 |
| ENSDARG000 | KN149843.1:K | 2888:5512:56 | 3770:5558:57 | -;-;-;- | 1158 | si:dkey-71b5. | 1 | 0 | 0 | 0 | 0 | 0 |
| ENSDARG000 | KN150156.1:K | 8987;12844 | 10620;13887 | -;- | 2678 | NA | 12 | 6 | 59 | 3 | 6 | 4 |
| ENSDARG000 | KN149946.1:K | 20160:20160: | 25261:25261: | -;-;-;-;- | 6014 | NA | 118 | 78 | 122 | 158 | 52 | 122 |
| ENSDARG000 | KN149946.1:K | 21:11787:127 | 2458:12594:1 | -;-;- | 3298 | NA | 2597 | 2153 | 2608 | 1745 | 2463 | 3317 |
| ENSDARG000 | KN150165.1:K | 1367:3505:10 | 1883:3650:10 | +;+;+;+;+;+ | 1246 | ABCD2 | 2 | 2 | 8 | 0 | 16 | 0 |
| ENSDARG000 | KN150165.1 | 8239 | 8354 | + | 116 | 5S_rRNA | 0 | 0 | 0 | 0 | 0 | 0 |
| ENSDARG000 | KN150165.1 | 7444 | 7560 | - | 117 | 5S_rRNA | 0 | 0 | 0 | 0 | 0 | 0 |
| ENSDARG000 | KN150698.1:K | 2793;8019 | 3018;8412 | +;+ | 620 | NA | 4 | 0 | 0 | 0 | 0 | 0 |
| ENSDARG000 | KN150698.1:K | 28829:28829: | 29490:29250: | -;-;-;-;-;-;-;-;- | 2606 | NA | 14 | 2 | 20 | 6 | 6 | 0 |
| ENSDARG000 | KN150200.1:K | 8091:10607:1 | 8139:10930:1 | +;+;+;+ | 696 | NA | 0 | 0 | 8 | 0 | 2 | 0 |
| ENSDARG000 | KN150240.1:K | 21641:22383: | 21769:22547: | -;-;-;-;-;-;-;- | 1999 | NA | 39 | 79 | 92 | 54 | 108 | 38 |
| ENSDARG000 | KN150263.1:K | 232;1626 | 362;4079 | +;+ | 2585 | NA | 28 | 2 | 22 | 2 | 2 | 16 |
| ENSDARG000 | KN150042.1 | 7613 | 7719 | + | 107 | 5S_rRNA | 0 | 0 | 0 | 0 | 0 | 0 |
| ENSDARG000 | KN150620.1 | 23926 | 23991 | + | 66 | U6 | 0 | 0 | 0 | 0 | 0 | 0 |
| ENSDARG000 | KN149753.1 | 24186 | 24308 | + | 123 | 5S_rRNA | 0 | 0 | 0 | 0 | 0 | 19 |
| ENSDARG000 | KN149753.1:K | 11147:11147: | 16418:16644: | -;-;-;- | 5783 | NA | 1 | 2 | 4 | 0 | 0 | 10 |
| ENSDARG000 | KN150379.1:K | 9051:10351:1 | 9080:10379:1 | -;-;-;-;- | 738 | NA | 0 | 0 | 0 | 0 | 0 | 0 |
| ENSDARG000 | KN149943.1:K | 18756:22874: | 19012:23306: | -;-;-;-;- | 1292 | NA | 0 | 0 | 0 | 0 | 0 | 0 |
| ENSDARG000 | KN149943.1:K | 8050:10410:1 | 8370:10513:1 | -;-;-;- | 647 | fgf21 | 2 | 0 | 0 | 0 | 0 | 0 |
| ENSDARG000 | KN150032.1:K | 21:3785:6910 | 1329:4120:72 | -;-;-;- | 2027 | NA | 0 | 134 | 121 | 182 | 289 | 166 |
| ENSDARG000 | KN150032.1:K | 20270:20710: | 20318:21027: | +;+;+;+;+;+ | 2298 | NA | 4 | 2 | 2 | 0 | 0 | 78 |
| ENSDARG000 | KN150066.1 | 17631 | 17762 | - | 132 | 5S_rRNA | 0 | 0 | 0 | 0 | 0 | 0 |
| ENSDARG000 | KN149765.1 | 15024 | 15139 | + | 116 | 5S_rRNA | 0 | 0 | 0 | 0 | 0 | 0 |
| ENSDARG000 | KN150572.1:K | 9109:9109:91 | 9318:9318:93 | +:+:+:+:+:+:+: | 6898 | NA | 70 | 61 | 104 | 50 | 61 | 49 |
| ENSDARG000 | KN149846.1 | 4737 | 4833 | - | 97 | NA | 0 | 0 | 0 | 0 | 0 | 0 |
| ENSDARG000 | KN150120.1:K | 21111:25182: | 21146:25703: | +;+;+ | 1296 | NA | 14 | 9 | 33 | 17 | 17 | 19 |
| ENSDARG000 | KN150067.1:K | 3520:5663:88 | 5076:5691:89 | -:-:-:-:-:-:-:-:-: | 2665 | NA | 22 | 8 | 19 | 11 | 22 | 48 |
| ENSDARG000 | KN150343.1:K | 18176:18246: | 18226:18347: | +;+;+;+ | 378 | NA | 54 | 29 | 45 | 25 | 37 | 45 |
| ENSDARG000 | KN150489.1:K | 2126:6967:95 | 4190:7062:96 | -;-;-;-;- | 3272 | NA | 174 | 127 | 172 | 119 | 155 | 47 |
| ENSDARG000 | KN150699.1:K | 8780:8966:92 | 8795:9079:93 | +:+:+:+:+:+:+: | 1641 | NA | 221 | 168 | 294 | 158 | 219 | 197 |
| ENSDARG000 | KN150699.1:K | 14154;15053 | 14653;15108 | -;- | 556 | h2af1al | 0 | 0 | 0 | 0 | 6 | 4 |
| ENSDARG000 | KN150699.1:K | 15387:17489: | 16064:17608: | +;+;+;+ | 1494 | creg1 | 347 | 240 | 381 | 378 | 505 | 370 |
| ENSDARG000 | KN149818.1:K | 21:1124:3003 | 1047:1250:30 | -:-:-:-:-:-:-:-:-: | 2739 | PSMD2 | 2334 | 2149 | 2826 | 1082 | 2458 | 1357 |
| ENSDARG000 | KN150285.1 | 22571 | 22685 | - | 115 | 5S_rRNA | 0 | 0 | 0 | 0 | 0 | 0 |
| ENSDARG000 | KN150285.1:K | 12751;15973 | 13489;16472 | -;- | 1239 | NA | 0 | 0 | 0 | 0 | 0 | 0 |
| ENSDARG000 | KN150084.1:K | 231:231:2376 | 1266:1266:24 | +:+:+:+:+:+:+: | 4321 | NA | 4 | 2 | 14 | 2 | 0 | 6 |
| ENSDARG000 | KN150324.1:K | 8882:12432:1 | 9871:12605:1 | +;+;+;+ | 5903 | NA | 370 | 417 | 253 | 166 | 385 | 318 |
| ENSDARG000 | KN150297.1 | 87 | 5872 | - | 5786 | NA | 14 | 0 | 2 | 0 | 0 | 0 |
| ENSDARG000 | KN149855.1:K | 8111;9486 | 9143;10664 | +;+ | 2212 | NA | 0 | 0 | 0 | 0 | 4 | 0 |
| ENSDARG000 | KN149855.1:K | 17504:18797: | 17563:18899: | +;+;+;+;+ | 534 | NA | 544 | 721 | 72 | 68 | 132 | 86 |
| ENSDARG000 | KN150490.1:K | 19245:19442: | 19367:19602: | -;-;-;- | 627 | NA | 4 | 0 | 0 | 0 | 2 | 0 |
| ENSDARG000 | KN150490.1:K | 882:5170:531 | 907:5439:543 | +:+:+:+:+:+:+: | 1648 | UBLCP1 | 0 | 1 | 4 | 0 | 2 | 0 |
| ENSDARG000 | KN150562.1:K | 227:3951:472 | 1288:4135:48 | +;+;+;+;+;+ | 6103 | NA | 48 | 60 | 74 | 20 | 53 | 32 |
| ENSDARG000 | KN150332.1:K | 622:629:935: | 707:757:973: | +:+:+:+:+:+:+: | 2206 | NA | 0 | 6 | 1 | 0 | 2 | 2 |
| ENSDARG000 | KN150645.1:K | 9855;9974 | 9893;10129 | +;+ | 195 | NA | 4 | 2 | 27 | 23 | 6 | 2 |
| ENSDARG000 | KN149979.1 | 12828 | 13855 | + | 1028 | NA | 13 | 0 | 0 | 0 | 15 | 11 |
| ENSDARG000 | KN150334.1:K | 14450;16056 | 15742;16282 | -;- | 1520 | EMC4 | 38 | 101 | 132 | 100 | 130 | 114 |
| ENSDARG000 | KN150561.1:K | 10489:11547: | 10591:11623: | -;-;-;-;-;-;-;- | 588 | NA | 12 | 23 | 54 | 47 | 45 | 31 |
| ENSDARG000 | KN150561.1 | 5595 | 5709 | - | 115 | 5S_rRNA | 6 | 18 | 18 | 20 | 15 | 30 |
| ENSDARG000 | KZ115964.1 | 6769 | 6901 | - | 133 | 5S_rRNA | 0 | 0 | 0 | 0 | 0 | 0 |
| ENSDARG000 | KZ115982.1:K | 8065:8711:10 | 8198:8854:10 | +:+:+:+:+:+:+: | 2094 | lrig3 | 92 | 8 | 20 | 32 | 18 | 28 |
| ENSDARG000 | KN149779.1:K | 15622:17376: | 15711:17613: | +;+;+;+ | 591 | SYNGR1 | 2 | 0 | 1 | 0 | 0 | 0 |
| ENSDARG000 | KN149779.1:K | 7696:12650:1 | 12146:12759: | -;-;-;-;-;-;- | 4913 | ift27 | 246 | 205 | 220 | 272 | 226 | 293 |
| ENSDARG000 | KN149779.1:K | 312:8008:958 | 1340:8091:97 | +;+;+;+ | 2198 | cacng2b | 0 | 0 | 0 | 0 | 0 | 0 |
| ENSDARG000 | KN150532.1:K | 3963:4212:56 | 4110:4373:57 | +;+;+ | 393 | NA | 0 | 0 | 0 | 0 | 0 | 0 |
| ENSDARG000 | KN150521.1 | 9405 | 9520 | + | 116 | 5S_rRNA | 0 | 0 | 0 | 0 | 0 | 0 |
| ENSDARG000 | KN150411.1:K | 7221:8630:92 | 8627:9223:92 | -;-;-;-;-;-;- | 3564 | NA | 6 | 1 | 1 | 0 | 20 | 6 |
| ENSDARG000 | KN150527.1:K | 12064;16791 | 12148;17794 | +;+ | 1089 | NA | 0 | 1 | 2 | 0 | 0 | 4 |
| ENSDARG000 | KN149961.1:K | 18138;19360 | 18389;19567 | +;+ | 460 | NA | 0 | 0 | 0 | 0 | 0 | 0 |
| ENSDARG000 | KN150636.1:K | 18646;18850 | 18719;19078 | +;+ | 303 | thfp5 | 31 | 2 | 0 | 1 | 0 | 0 |
| ENSDARG000 | KN150636.1:K | 2551:6768:67 | 2599:7528:72 | +:+:+:+:+:+:+: | 3605 | NA | 0 | 0 | 0 | 0 | 1 | 6 |

| | | | | | | | | | | | |
|---|---|---|---|---|---|---|---|---|---|---|---|
| ENSDARG000 | KN150327.1: | 5738:8777:10 | 5947:8874:10 | +;+;+;+;+ | 641 | NA | 0 | 0 | 0 | 0 | 0 | 0 |
| ENSDARG000 | KN150019.2: | 4903;9532 | 5229;10929 | +;+ | 1725 | NA | 0 | 0 | 0 | 0 | 0 | 0 |
| ENSDARG000 | KN149764.1: | 4168:5209:54 | 4526:5323:55 | -;-;-;-;-;-;-;- | 1952 | NA | 585 | 284 | 494 | 253 | 593 | 466 |
| ENSDARG000 | KN149955.1: | 4425:7903:11 | 4528:8360:12 | +;+;+ | 1633 | NA | 190 | 449 | 559 | 729 | 177 | 528 |
| ENSDARG000 | KN150322.1: | 4302:7933:85 | 7625:8104:86 | -;-;-;-;-;-;- | 4522 | TULP1 | 8 | 10 | 13 | 26 | 44 | 0 |
| ENSDARG000 | KN150353.1: | 391:5979:767 | 4844:6131:79 | -;-;-;-;-;-;-;- | 6085 | POLR3D | 150 | 304 | 457 | 144 | 245 | 134 |
| ENSDARG000 | KN150633.1: | 21:2207:3675 | 583:2334:378 | +;+;+;+ | 988 | NA | 53 | 23 | 27 | 30 | 31 | 24 |
| ENSDARG000 | KZ116043.1:K | 21:1309:2646 | 952:1474:284 | +;+;+;+;+ | 5905 | FH | 654 | 524 | 650 | 322 | 584 | 587 |
| ENSDARG000 | KN149715.1 | 7841 | 8773 | + | 933 | NA | 0 | 0 | 0 | 0 | 0 | 0 |
| ENSDARG000 | KN150599.1 | 3839 | 4980 | + | 1142 | NA | 0 | 0 | 0 | 0 | 0 | 0 |
| ENSDARG000 | KN150589.1: | 3626:3912:42 | 3838:4048:43 | -;-;-;-;-;-;- | 849 | NA | 41 | 6 | 1 | 12 | 4 | 2 |
| ENSDARG000 | KN150212.1: | 7721:7819:81 | 7893:7893:81 | -;-;-;-;-;-;-;-; | 749 | NA | 2 | 0 | 0 | 0 | 0 | 0 |
| ENSDARG000 | KN150212.1 | 4956 | 5070 | - | 115 | 5S_rRNA | 0 | 0 | 0 | 0 | 0 | 0 |
| ENSDARG000 | KN150115.1: | 21;4529 | 172;5710 | -;- | 1334 | RAB19 | 2 | 6 | 2 | 0 | 0 | 2 |
| ENSDARG000 | KN150115.1: | 10553:14686: | 10787:14819: | -;-;-;-;-;- | 1008 | NA | 151 | 87 | 163 | 232 | 139 | 142 |
| ENSDARG000 | KN150574.1: | 6811:6811:97 | 8923:8923:99 | -;-;-;-;-;-;-;-; | 4867 | NA | 808 | 824 | 1334 | 1201 | 1452 | 979 |
| ENSDARG000 | KN150654.1: | 8142:11830:1 | 10004:11980: | -;-;-;-;-;- | 2723 | NA | 0 | 0 | 0 | 0 | 0 | 0 |
| ENSDARG000 | KN149914.1: | 21:212:388:3 | 292:292:565: | -;-;-;-;-;-;-;-; | 2135 | NA | 328 | 171 | 258 | 260 | 260 | 169 |
| ENSDARG000 | KZ115972.1:K | 6562:7493:84 | 6895:8405:85 | +;+;+ | 1308 | NA | 0 | 0 | 1 | 0 | 0 | 0 |
| ENSDARG000 | KZ115972.1:K | 17084:17084: | 17888:17278: | +;+;+;+;+ | 921 | NA | 0 | 0 | 0 | 0 | 0 | 0 |
| ENSDARG000 | KN150652.1: | 3488:4875:54 | 4627:4990:56 | +;+;+;+ | 4176 | NA | 538 | 351 | 505 | 491 | 654 | 557 |
| ENSDARG000 | KN150545.1: | 7895:8462:10 | 8110:8578:11 | -;-;-;-;-;-;-;-; | 1332 | CORO7 | 46 | 61 | 0 | 46 | 94 | 88 |
| ENSDARG000 | KN150328.1: | 3117:3218:52 | 3635:3635:53 | +;+;+;+;+;+;+; | 1334 | NA | 72 | 0 | 0 | 0 | 0 | 0 |
| ENSDARG000 | KZ116008.1:K | 16081;16737 | 16699;16987 | +;+ | 870 | NA | 17 | 4 | 6 | 3 | 3 | 4 |
| ENSDARG000 | KZ116008.1:K | 1272:2456:26 | 1296:2485:26 | -;-;-;-;-;-;-;-; | 1158 | NA | 0 | 1 | 0 | 0 | 0 | 0 |
| ENSDARG000 | KN150362.1: | 6063:7866:10 | 8315:8315:12 | -;-;-;-;- | 4833 | NA | 0 | 4 | 2 | 0 | 18 | 20 |
| ENSDARG000 | KN150141.1: | 16513;17058 | 16932;17105 | -;- | 468 | NA | 34 | 75 | 68 | 32 | 69 | 14 |
| ENSDARG000 | KN150141.1: | 426:1939:246 | 1863:2066:26 | +;+;+;+;+;+;+; | 3651 | NA | 1908 | 834 | 1561 | 804 | 1135 | 1168 |
| ENSDARG000 | KN150345.1 | 9178 | 9292 | + | 115 | 5S_rRNA | 0 | 0 | 0 | 0 | 0 | 0 |
| ENSDARG000 | KN150683.1: | 2789:3554:47 | 2885:3729:48 | +;+;+;+;+;+ | 1946 | NA | 99 | 0 | 2 | 0 | 227 | 0 |
| ENSDARG000 | KN149978.1: | 12292:14593: | 14305:14779: | -;-;- | 2326 | LEPROTL1 | 2404 | 2362 | 2881 | 2928 | 2623 | 2622 |
| ENSDARG000 | KN149978.1: | 3824:3824:75 | 7616:7287:76 | -;-;-;-;-;-;-;-; | 4590 | NA | 4 | 6 | 2 | 0 | 0 | 0 |
| ENSDARG000 | KZ116053.1:K | 7782;10763 | 8017;10784 | +;+ | 258 | LSM10 | 36 | 31 | 67 | 13 | 29 | 42 |
| ENSDARG000 | KN149936.1: | 9176:9381:97 | 9295:9498:10 | +;+;+ | 1172 | NA | 129 | 245 | 269 | 365 | 465 | 555 |
| ENSDARG000 | KN149936.1: | 10489:14068: | 10534:14182: | +;+;+;+ | 339 | NA | 1 | 6 | 0 | 6 | 10 | 4 |
| ENSDARG000 | KN149732.1: | 11675;12626 | 12598;12673 | -;- | 972 | NA | 2 | 0 | 0 | 0 | 0 | 2 |
| ENSDARG000 | KN149803.1: | 9945:9945:10 | 10104:10104: | +;+;+;+;+ | 449 | NA | 0 | 0 | 0 | 0 | 0 | 0 |
| ENSDARG000 | KN150311.1: | 155:807:1571 | 887:887:1714 | -;-;-;-;-;-;-;-; | 2849 | SNX1 | 122 | 120 | 123 | 86 | 93 | 119 |
| ENSDARG000 | KN150008.1: | 12653;13646 | 13519;14179 | -;- | 1401 | NA | 0 | 2 | 0 | 0 | 0 | 0 |
| ENSDARG000 | KN150387.1: | 3916:6027:97 | 4114:7027:10 | -;-;-;-;- | 1548 | NA | 1370 | 462 | 865 | 953 | 1301 | 1246 |
| ENSDARG000 | KN150071.1: | 41:725:11522 | 421:895:1490 | +;+;+ | 3936 | NA | 151 | 121 | 155 | 123 | 192 | 106 |
| ENSDARG000 | KN150647.1 | 11706 | 11883 | + | 178 | U1 | 1 | 0 | 0 | 0 | 0 | 0 |
| ENSDARG000 | KN150647.1 | 13027 | 13190 | + | 164 | U1 | 0 | 0 | 0 | 0 | 0 | 0 |
| ENSDARG000 | KZ116020.1:K | 6856:8686:10 | 6945:8796:10 | +;+;+ | 345 | NA | 0 | 0 | 0 | 0 | 0 | 0 |
| ENSDARG000 | KN150550.1: | 918:918:1110 | 978:978:1234 | +;+;+;+;+;+;+; | 4273 | fanca | 23 | 52 | 38 | 20 | 71 | 22 |
| ENSDARG000 | KN150317.1: | 4727;8769 | 5812;8822 | -;- | 1140 | NA | 81 | 61 | 23 | 6 | 40 | 15 |
| ENSDARG000 | KN149861.1 | 7181 | 7336 | - | 156 | 5_8S_rRNA | 0 | 0 | 0 | 0 | 0 | 0 |
| ENSDARG000 | KN150506.1: | 22:1079:2093 | 621:1238:221 | +;+;+;+;+;+;+ | 2405 | NA | 69 | 83 | 232 | 164 | 95 | 137 |
| ENSDARG000 | KZ116047.1:K | 3806:6743:91 | 4171:7005:92 | +;+;+;+;+;+ | 1263 | HPDL | 44 | 164 | 137 | 23 | 111 | 24 |
| ENSDARG000 | KN150679.1: | 255:1774:309 | 367:2007:318 | +;+;+;+;+;+;+; | 4347 | NA | 372 | 405 | 588 | 313 | 458 | 399 |
| ENSDARG000 | KN149897.1 | 6228 | 7239 | + | 1012 | NA | 0 | 0 | 0 | 0 | 0 | 0 |
| ENSDARG000 | KN149897.1 | 10052 | 11063 | + | 1012 | NA | 0 | 0 | 0 | 0 | 0 | 0 |
| ENSDARG000 | KN150680.1: | 8442:8643:88 | 8566:8836:89 | +;+;+;+;+ | 1134 | bglapl | 0 | 0 | 0 | 0 | 0 | 0 |
| ENSDARG000 | KN150072.1: | 1636:1636:19 | 1810:1810:22 | +;+;+;+;+;+;+; | 2482 | NA | 87 | 66 | 122 | 30 | 55 | 84 |
| ENSDARG000 | KN149793.1: | 8547:10095:1 | 8647:10238:1 | +;+;+ | 495 | NA | 332 | 366 | 600 | 399 | 323 | 420 |
| ENSDARG000 | KN150491.1: | 2817:4564:51 | 3024:4715:53 | -;-;-;- | 744 | NA | 154 | 93 | 184 | 89 | 123 | 113 |
| ENSDARG000 | KN150193.1 | 4511 | 5548 | - | 1038 | NA | 155 | 0 | 87 | 0 | 184 | 94 |
| ENSDARG000 | KN150314.1: | 1230;6916 | 6344;7448 | -;- | 5648 | FAM167A | 48 | 42 | 8 | 20 | 64 | 20 |
| ENSDARG000 | KN149883.1: | 2235:2235:24 | 2310:2310:24 | -;-;-;-;-;-;-;- | 263 | NA | 145 | 258 | 94 | 50 | 93 | 68 |
| ENSDARG000 | KN150541.1: | 9481:9481:10 | 9959:10151:1 | -;-;-;-;-;-;-;- | 1559 | NA | 0 | 0 | 0 | 0 | 0 | 0 |
| ENSDARG000 | KN150128.1: | 513:513:513: | 682:682:682: | +;+;+;+;+;+;+; | 4221 | WDR48 | 20 | 8 | 12 | 2 | 38 | 66 |
| ENSDARG000 | KN150380.1: | 4419:5556:64 | 4754:5775:65 | -;-;-;- | 875 | NA | 53 | 42 | 109 | 45 | 73 | 56 |
| ENSDARG000 | KN150090.1 | 9154 | 9268 | + | 115 | 5S_rRNA | 0 | 0 | 0 | 0 | 0 | 0 |
| ENSDARG000 | KN149995.1: | 4573:6918:72 | 4727:7005:74 | +;+;+ | 468 | CPLX3 | 0 | 0 | 0 | 0 | 0 | 0 |
| ENSDARG000 | KN150665.1 | 773 | 846 | + | 74 | U6 | 0 | 0 | 0 | 0 | 0 | 0 |
| ENSDARG000 | KN150665.1 | 6125 | 6198 | + | 74 | U6 | 0 | 0 | 0 | 0 | 0 | 0 |
| ENSDARG000 | KN150665.1 | 10591 | 10697 | + | 107 | U6 | 0 | 0 | 0 | 0 | 0 | 0 |
| ENSDARG000 | KN150665.1 | 9006 | 9112 | + | 107 | U6 | 0 | 0 | 0 | 0 | 0 | 0 |
| ENSDARG000 | KN150665.1 | 11106 | 11212 | + | 107 | U6 | 0 | 0 | 0 | 0 | 0 | 0 |
| ENSDARG000 | KN150665.1 | 11632 | 11705 | + | 74 | U6 | 0 | 0 | 0 | 0 | 0 | 0 |
| ENSDARG000 | KN150665.1 | 10061 | 10167 | + | 107 | U6 | 0 | 0 | 0 | 0 | 0 | 0 |
| ENSDARG000 | KN150665.1 | 243 | 349 | + | 107 | U6 | 0 | 0 | 0 | 0 | 0 | 0 |
| ENSDARG000 | KN150665.1 | 9536 | 9642 | + | 107 | U6 | 0 | 0 | 0 | 0 | 0 | 0 |
| ENSDARG000 | KN149976.1: | 21:2859:3732 | 1459:3046:37 | -;-;-;-;-;-;- | 2216 | NA | 22 | 6 | 0 | 12 | 2 | 20 |
| ENSDARG000 | KN150287.1: | 6535:6887:83 | 6790:6946:84 | -;-;-;- | 405 | NA | 2 | 0 | 0 | 0 | 0 | 0 |
| ENSDARG000 | KN149711.1: | 21:474:1479: | 139:508:1514 | -;-;-;-;-;-;-;- | 932 | NA | 21 | 23 | 42 | 57 | 37 | 38 |
| ENSDARG000 | KZ115980.1 | 81 | 192 | + | 112 | 5S_rRNA | 0 | 0 | 0 | 0 | 0 | 0 |
| ENSDARG000 | KZ115980.1:K | 559:3123:473 | 1393:3245:49 | +;+;+;+;+;+;+; | 5176 | NA | 0 | 0 | 0 | 2 | 2 | 0 |
| ENSDARG000 | KZ115989.1:K | 57:2952:3317 | 624:3054:416 | -;-;-;-;-;- | 1917 | NA | 105 | 45 | 102 | 26 | 23 | 42 |
| ENSDARG000 | KN150164.1: | 4431:4820:55 | 4669:4904:56 | -;-;-;-;-;-;-;-; | 1680 | NA | 719 | 508 | 671 | 333 | 737 | 437 |
| ENSDARG000 | KZ115969.1:K | 6244:6528:67 | 6444:6671:69 | +;+;+;+ | 555 | NA | 0 | 0 | 0 | 0 | 0 | 0 |
| ENSDARG000 | KN149900.1 | 5302 | 5420 | - | 119 | 5S_rRNA | 0 | 0 | 0 | 0 | 0 | 0 |
| ENSDARG000 | KZ116018.1:K | 8110:8523:87 | 8399:8624:89 | +;+;+;+;+;+;+ | 1493 | HYCC2 | 90 | 11 | 118 | 70 | 32 | 21 |
| ENSDARG000 | KN150162.1: | 4256;4694 | 4411;4885 | -;- | 348 | NA | 0 | 0 | 0 | 0 | 0 | 0 |
| ENSDARG000 | KN150365.1: | 4514;6237 | 4692;6428 | -;- | 371 | NA | 6 | 0 | 0 | 0 | 0 | 4 |
| ENSDARG000 | KN150365.1 | 900 | 1018 | - | 119 | 5S_rRNA | 8 | 0 | 0 | 0 | 0 | 0 |
| ENSDARG000 | KZ115966.1:K | 4009:5883:58 | 4022:5973:59 | +;+;+;+;+;+;+ | 332 | NA | 2 | 2 | 2 | 0 | 5 | 6 |
| ENSDARG000 | KN150155.1: | 1808:3620:47 | 2372:3720:85 | +;+;+ | 4508 | NA | 0 | 0 | 0 | 0 | 0 | 0 |
| ENSDARG000 | KN149857.1 | 2025 | 2880 | + | 856 | NA | 2 | 3 | 1 | 0 | 0 | 2 |
| ENSDARG000 | KN149816.1: | 2954;3124 | 3001;3294 | -;- | 219 | NA | 19 | 6 | 24 | 14 | 20 | 10 |
| ENSDARG000 | KN149816.1: | 8179:8933:95 | 8352:9364:97 | -;-;-;- | 879 | slc39a3 | 162 | 140 | 144 | 75 | 116 | 89 |
| ENSDARG000 | KN149713.1: | 686:888:1015 | 799:939:1098 | +;+;+;+;+;+;+; | 1796 | NA | 0 | 0 | 0 | 0 | 0 | 6 |
| ENSDARG000 | KN150412.1: | 21:2704:2894 | 2606:2807:29 | -;-;-;-;- | 3073 | NA | 42 | 18 | 52 | 20 | 54 | 50 |
| ENSDARG000 | KZ116002.1 | 3325 | 5436 | + | 2112 | NA | 0 | 0 | 0 | 0 | 0 | 0 |
| ENSDARG000 | KZ115962.1:K | 21:2488:3902 | 2700:2685:39 | -;-;-;-;-;-;-;- | 3292 | YBEY | 4 | 0 | 1 | 0 | 2 | 2 |
| ENSDARG000 | KN150350.1: | 486:1277:159 | 1049:1509:17 | +;+;+;+;+;+;+; | 2177 | USP30 | 30 | 6 | 10 | 13 | 25 | 0 |
| ENSDARG000 | KN150614.1: | 286:867:1818 | 442:1050:193 | +;+;+;+;+;+;+; | 1218 | GALNT10 | 563 | 357 | 363 | 300 | 529 | 374 |
| ENSDARG000 | KN150614.1: | 7174:8530:88 | 7359:8652:90 | +;+;+ | 438 | NA | 11 | 24 | 11 | 8 | 12 | 4 |
| ENSDARG000 | KN150185.1 | 5614 | 5949 | + | 336 | NA | 0 | 0 | 0 | 0 | 0 | 0 |
| ENSDARG000 | KN150185.1: | 182:1701:247 | 274:1786:268 | +;+;+;+ | 438 | NA | 0 | 0 | 0 | 0 | 0 | 0 |
| ENSDARG000 | KN150013.1: | 14:556:3758: | 35:752:3891: | +;+;+;+;+;+;+; | 1419 | wbp11 | 170 | 218 | 224 | 100 | 290 | 118 |
| ENSDARG000 | KZ116023.1 | 1152 | 2674 | - | 1523 | zgc:153137 | 0 | 0 | 0 | 0 | 0 | 0 |
| ENSDARG000 | KN149707.2: | 4029:4046:48 | 4221:4221:49 | +;+;+;+;+;+;+; | 1283 | cdo1 | 310 | 193 | 303 | 136 | 226 | 106 |
| ENSDARG000 | KZ116055.1:K | 291;751 | 635;1335 | -;- | 930 | NA | 0 | 0 | 0 | 0 | 0 | 0 |
| ENSDARG000 | KN150355.1: | 1608;2654 | 1851;2790 | -;- | 381 | NA | 0 | 0 | 0 | 0 | 0 | 0 |
| ENSDARG000 | KN150355.1: | 2275;4194 | 3761;4765 | -;- | 2059 | NA | 0 | 0 | 0 | 0 | 0 | 0 |
| ENSDARG000 | KN149997.1: | 4401:4623:48 | 4479:4732:49 | +;+;+;+;+ | 1343 | NA | 358 | 418 | 493 | 211 | 643 | 252 |
| ENSDARG000 | KN149950.1: | 74;2097 | 1199;5493 | +;+ | 4523 | NA | 0 | 16 | 0 | 0 | 0 | 1336 |
| ENSDARG000 | KZ116046.1:K | 21;4644 | 3100;5787 | -;- | 4224 | NA | 0 | 2 | 2 | 0 | 0 | 101 |
| ENSDARG000 | KN149735.1: | 127:944:1629 | 326:1121:175 | -;-;-;-;- | 681 | zgc:86738 | 0 | 0 | 0 | 0 | 0 | 0 |
| ENSDARG000 | KN149729.1: | 130;2697 | 381;2711 | -;- | 267 | RAMP3 | 47 | 49 | 77 | 30 | 56 | 37 |
| ENSDARG000 | KZ115963.1 | 1221 | 1388 | + | 168 | NA | 0 | 0 | 2 | 1 | 0 | 5 |
| ENSDARG000 | KN149874.1: | 11:436:630:1 | 101:521:674: | -;-;-;-;-;-;-;- | 804 | ttc8 | 91 | 80 | 136 | 39 | 157 | 24 |
| ENSDARG000 | KN149951.1: | 442:631:1067 | 538:675:1141 | +;+;+;+;+ | 345 | NA | 0 | 7 | 0 | 0 | 3 | 0 |
| ENSDARG000 | KN149916.1 | 3617 | 4292 | - | 676 | NA | 0 | 0 | 2 | 0 | 0 | 2 |
| ENSDARG000 | KZ115991.1:K | 1443:1539:18 | 1470:1747:19 | +;+;+;+;+ | 477 | PCNP | 0 | 125 | 4 | 1 | 0 | 0 |
| ENSDARG000 | KN150039.1: | 79;680 | 159;763 | -;- | 165 | NA | 14 | 34 | 31 | 18 | 15 | 14 |
| ENSDARG000 | KN150650.1: | 1563:2454:27 | 2176:2553:30 | +;+;+ | 965 | NA | 0 | 2 | 0 | 8 | 0 | 0 |
| ENSDARG000 | KZ115967.1 | 382 | 496 | - | 115 | 5S_rRNA | 0 | 0 | 0 | 0 | 0 | 0 |
| ENSDARG000 | KN150470.1: | 1:270:463:18 | 167:370:575: | +;+;+;+ | 510 | NA | 103 | 109 | 158 | 24 | 158 | 104 |
| ENSDARG000 | KN150051.1: | 86:86:478:47 | 273:273:487: | -;-;-;-;-;-;-;-; | 651 | LAMP5 | 0 | 0 | 0 | 0 | 0 | 0 |
| ENSDARG000 | KN150086.1: | 21;1305 | 302;1638 | -;- | 616 | RHO | 4 | 0 | 0 | 7 | 0 | 2 |
